## Supplementary material for "Lysine-specific demethylase 1a is obligatory for gene regulation during kidney development": Suppl Table-5

**Supplementary Table-5**

| Chr | start | end | KWT_1_IP | KWT_2_IP | KKO_1_IP | KKO_2_IP | logFC | logCPM | PValue |
|---|---|---|---|---|---|---|---|---|---|
| 2 | 37235000 | 37235399 | 9.354114 | 18.26523 | 1.591277 | 0.503359 | 4.149143 | -0.87402 | 0.002314 |
| Y | 89278400 | 89279799 | 23.04072 | 42.34629 | 6.449196 | 2.740593 | 3.112038 | 0.282769 | 0.00333 |
| 8 | 3080800 | 3083999 | 31.75927 | 55.14625 | 15.19385 | 6.668385 | 2.234585 | 0.826015 | 0.003552 |
| 13 | 45154800 | 45155999 | 2.716848 | 5.070275 | 42.99939 | 17.87302 | -2.59411 | 0.121146 | 0.004074 |
| 1 | 108872800 | 108876999 | 95.95085 | 151.9922 | 63.94609 | 26.59433 | 1.575745 | 2.3727 | 0.004333 |
| 4 | 122156200 | 122156599 | 11.12339 | 21.74562 | 4.605345 | 1.580892 | 2.889897 | -0.56032 | 0.004422 |
| 13 | 65555400 | 65555799 | 10.58269 | 20.68396 | 5.269298 | 1.819228 | 2.625467 | -0.54745 | 0.004543 |
| 14 | 113216600 | 113217599 | 12.54142 | 24.43438 | 6.460414 | 2.349956 | 2.514329 | -0.33954 | 0.004705 |
| 3 | 36562200 | 36562999 | 45.7132 | 76.0379 | 21.57133 | 9.362505 | 2.190341 | 1.27641 | 0.004815 |
| 12 | 22699800 | 22700799 | 10.47684 | 20.49544 | 2.495428 | 0.827864 | 3.669355 | -0.69151 | 0.004822 |
| 15 | 101270600 | 101271799 | 2.362194 | 4.534421 | 36.88069 | 14.45916 | -2.45696 | -0.10361 | 0.005132 |
| 4 | 131623400 | 131623999 | 103.4319 | 164.6496 | 34.587 | 14.45833 | 2.580865 | 2.234144 | 0.005358 |
| 1 | 145695200 | 145696799 | 22.3329 | 41.33653 | 8.260696 | 3.476795 | 2.737784 | 0.245141 | 0.005442 |
| 9 | 62029000 | 62030999 | 4.91568 | 8.741825 | 52.57093 | 23.18023 | -2.17316 | 0.450147 | 0.005626 |
| 15 | 80760000 | 80760999 | 2.676105 | 5.115466 | 38.45081 | 15.25763 | -2.36435 | -0.05696 | 0.005876 |
| X | 122685800 | 122686799 | 13.76981 | 26.82181 | 5.505155 | 2.008211 | 2.868891 | -0.31754 | 0.006443 |
| 1 | 145885600 | 145885999 | 9.349093 | 18.27626 | 3.277139 | 1.056924 | 3.181384 | -0.80094 | 0.006582 |
| 10 | 28228800 | 28229199 | 9.172852 | 17.9406 | 4.680391 | 1.535828 | 2.650602 | -0.74105 | 0.006821 |
| 17 | 39272000 | 39272599 | 11.96796 | 23.38061 | 4.171071 | 1.456032 | 3.103459 | -0.49303 | 0.007175 |
| 16 | 81578400 | 81578799 | 21.88823 | 39.83035 | 12.22517 | 5.262975 | 2.106375 | 0.342317 | 0.007255 |
| 7 | 21006000 | 21006399 | 9.985937 | 19.53523 | 4.359639 | 1.445521 | 2.859936 | -0.69437 | 0.007286 |
| 2 | 89217000 | 89218599 | 17.35331 | 32.77193 | 10.3181 | 4.201123 | 2.126849 | 0.087553 | 0.007345 |
| X | 32283800 | 32284399 | 13.96122 | 27.20466 | 5.492733 | 1.994045 | 2.894183 | -0.33 | 0.007492 |
| 18 | 37523200 | 37524399 | 14.45627 | 27.97456 | 8.567686 | 3.266468 | 2.24948 | -0.15269 | 0.007766 |
| 17 | 25452800 | 25455199 | 4.438165 | 8.028547 | 48.15128 | 20.88326 | -2.15185 | 0.330132 | 0.007866 |
| 12 | 50980600 | 50981399 | 10.40353 | 20.3399 | 4.926882 | 1.687919 | 2.700774 | -0.5767 | 0.008444 |
| 2 | 32613400 | 32616199 | 6.134708 | 10.81064 | 54.04737 | 23.92188 | -1.91676 | 0.548564 | 0.008894 |
| 6 | 144266800 | 144267599 | 1.717089 | 3.349379 | 28.42423 | 10.1794 | -2.38364 | -0.45427 | 0.008898 |
| 17 | 82408400 | 82409199 | 13.77766 | 26.79748 | 7.411458 | 2.740445 | 2.432238 | -0.26465 | 0.008973 |
| 12 | 3055000 | 3055599 | 37.31356 | 63.28878 | 24.03898 | 10.52981 | 1.771481 | 1.069326 | 0.008977 |
| 10 | 79905400 | 79907399 | 3.421965 | 6.430968 | 39.37902 | 16.17565 | -2.11918 | 0.072593 | 0.009229 |
| 11 | 79486200 | 79487199 | 1.716296 | 3.350465 | 28.12344 | 9.943846 | -2.35146 | -0.50212 | 0.009239 |
| 1 | 144729200 | 144730799 | 15.62402 | 30.01508 | 9.256562 | 3.620897 | 2.205281 | -0.05666 | 0.009317 |
| 15 | 92391000 | 92391399 | 7.940074 | 15.51813 | 4.7161 | 1.51488 | 2.459936 | -0.83547 | 0.009433 |
| 7 | 22198200 | 22199199 | 13.70376 | 26.49616 | 9.564116 | 3.660659 | 2.010439 | -0.14553 | 0.009544 |
| 13 | 66397400 | 66397799 | 8.88827 | 17.38383 | 5.364655 | 1.759607 | 2.418634 | -0.74427 | 0.009571 |
| 1 | 146161600 | 146161999 | 11.10723 | 21.72376 | 4.629809 | 1.566033 | 2.900433 | -0.60832 | 0.009809 |
| 7 | 7461600 | 7463599 | 30.83761 | 53.26146 | 21.90632 | 9.610328 | 1.660338 | 0.880658 | 0.009888 |
| 12 | 22464200 | 22466399 | 30.29138 | 52.61938 | 20.34676 | 8.932275 | 1.750128 | 0.822524 | 0.009941 |
| 12 | 24073200 | 24075199 | 31.25079 | 54.05002 | 21.78921 | 9.559358 | 1.689475 | 0.870898 | 0.010154 |
| 7 | 7721600 | 7721799 | 11.12849 | 21.73305 | 6.909401 | 2.426447 | 2.296609 | -0.47503 | 0.010297 |
| 7 | 67121000 | 67123799 | 5.188827 | 9.338867 | 48.0192 | 20.95885 | -1.93912 | 0.365441 | 0.010301 |
| 4 | 133887200 | 133888399 | 10.86104 | 21.22852 | 4.045568 | 1.395863 | 3.032911 | -0.55198 | 0.010411 |
| 6 | 60535800 | 60535999 | 7.505095 | 14.64884 | 4.039286 | 1.271897 | 2.613771 | -0.91446 | 0.010543 |
| 12 | 67831600 | 67832199 | 12.36409 | 24.05839 | 8.451701 | 3.114281 | 2.096857 | -0.29056 | 0.01066 |

| | | | | | | | | | |
|---|---|---|---|---|---|---|---|---|---|
| 15 | 23242600 | 23243599 | 11.13038 | 21.72847 | 7.155183 | 2.529289 | 2.240594 | -0.4525 | 0.010669 |
| 3 | 129623200 | 129625399 | 5.767232 | 10.32892 | 49.00149 | 21.47801 | -1.82879 | 0.403555 | 0.010728 |
| 5 | 41368000 | 41368399 | 8.494974 | 16.57166 | 3.313887 | 1.037311 | 3.064192 | -0.91593 | 0.010797 |
| 11 | 90139400 | 90141399 | 3.331654 | 6.218231 | 40.79362 | 16.95348 | -2.23138 | 0.120026 | 0.010806 |
| 4 | 133643600 | 133647199 | 86.83124 | 138.2728 | 50.473 | 21.1095 | 1.787295 | 2.209694 | 0.010846 |
| 16 | 79621400 | 79621999 | 9.73775 | 19.04794 | 4.979011 | 1.664216 | 2.62434 | -0.66849 | 0.010926 |
| 1 | 160896800 | 160898999 | 4.967683 | 9.045239 | 45.09673 | 19.3996 | -1.88038 | 0.289853 | 0.011005 |
| 12 | 22337200 | 22337799 | 10.1282 | 19.79113 | 6.208963 | 2.15632 | 2.329174 | -0.53155 | 0.011036 |
| 3 | 119277400 | 119278599 | 14.76707 | 28.4423 | 10.11899 | 3.928955 | 2.013244 | -0.08619 | 0.011037 |
| 17 | 24025400 | 24032399 | 88.76814 | 141.3842 | 55.81535 | 23.35002 | 1.67478 | 2.206392 | 0.011102 |
| 1 | 118055400 | 118056799 | 35.70731 | 60.66275 | 23.24801 | 10.18835 | 1.758667 | 1.048008 | 0.011131 |
| 3 | 117898400 | 117898999 | 1.394666 | 2.727266 | 25.60565 | 8.725985 | -2.45238 | -0.64638 | 0.011329 |
| 12 | 23978000 | 23978399 | 8.491584 | 16.58027 | 4.033676 | 1.274966 | 2.788492 | -0.87999 | 0.011502 |
| 1 | 100153800 | 100155599 | 32.02447 | 55.92536 | 15.42 | 6.791987 | 2.233569 | 0.752342 | 0.011797 |
| 10 | 31363400 | 31364399 | 27.91574 | 48.88925 | 18.47549 | 8.156364 | 1.782836 | 0.692117 | 0.011813 |
| 1 | 181440000 | 181440799 | 0.966477 | 1.889397 | 28.77441 | 9.879178 | -3.13414 | -0.61935 | 0.011921 |
| 14 | 113571400 | 113574399 | 87.17694 | 138.6251 | 66.24456 | 27.65186 | 1.397551 | 2.258484 | 0.01196 |
| 7 | 8654200 | 8654399 | 7.866176 | 15.35666 | 4.745968 | 1.497558 | 2.459571 | -0.89492 | 0.012023 |
| 1 | 117164200 | 117165799 | 27.63084 | 48.69622 | 14.99528 | 6.636917 | 2.075927 | 0.600297 | 0.012226 |
| 13 | 66274200 | 66275199 | 10.55807 | 20.64069 | 6.305058 | 2.163685 | 2.379951 | -0.56248 | 0.012459 |
| 10 | 37840400 | 37841799 | 20.39697 | 37.71612 | 11.54753 | 4.866685 | 2.129605 | 0.242728 | 0.012487 |
| 9 | 9866600 | 9867999 | 13.78564 | 26.77282 | 8.626083 | 3.228414 | 2.202093 | -0.22 | 0.012582 |
| X | 128050200 | 128050599 | 8.265436 | 16.16507 | 6.036569 | 1.976628 | 2.153601 | -0.75834 | 0.012589 |
| 1 | 108978800 | 108980199 | 19.35571 | 35.412 | 14.01355 | 5.992095 | 1.749876 | 0.296851 | 0.012626 |
| 11 | 96208400 | 96211799 | 8.200783 | 14.27518 | 61.09307 | 26.85494 | -1.6986 | 0.750426 | 0.012663 |
| 7 | 22208400 | 22209399 | 29.68186 | 51.60305 | 20.89858 | 9.179273 | 1.683956 | 0.809318 | 0.012678 |
| X | 33904800 | 33905999 | 11.9694 | 23.36269 | 6.88416 | 2.440149 | 2.393431 | -0.43182 | 0.012706 |
| X | 57383200 | 57384799 | 14.9291 | 29.00415 | 7.107961 | 2.652365 | 2.590766 | -0.2208 | 0.012794 |
| 1 | 110099000 | 110101399 | 41.23018 | 68.97595 | 25.75892 | 11.22104 | 1.795987 | 1.207335 | 0.01283 |
| 15 | 71669000 | 71670599 | 16.82053 | 31.62793 | 11.89534 | 4.881845 | 1.864726 | 0.113849 | 0.012832 |
| 3 | 15977200 | 15977999 | 13.29919 | 25.96392 | 4.863322 | 1.717368 | 3.030033 | -0.43281 | 0.012854 |
| X | 30764600 | 30765199 | 12.92916 | 25.13331 | 9.057041 | 3.36383 | 2.052055 | -0.25458 | 0.012874 |
| 7 | 21973200 | 21973599 | 8.317425 | 16.25712 | 5.057992 | 1.62725 | 2.431976 | -0.82074 | 0.013018 |
| 9 | 118449200 | 118455599 | 22.69072 | 36.5687 | 126.123 | 53.67216 | -1.42558 | 1.789486 | 0.013022 |
| 12 | 23634800 | 23635799 | 12.83254 | 24.96529 | 8.670719 | 3.199774 | 2.113185 | -0.27864 | 0.013133 |
| 9 | 66740200 | 66743199 | 9.769621 | 16.60054 | 76.09834 | 33.35032 | -1.805 | 1.012412 | 0.013161 |
| 18 | 69477200 | 69477599 | 0.653088 | 1.277382 | 22.06799 | 7.253566 | -3.21794 | -0.80094 | 0.013244 |
| 7 | 8891600 | 8892399 | 15.60988 | 30.14647 | 8.687243 | 3.3432 | 2.322723 | -0.1088 | 0.013349 |
| 4 | 80724200 | 80725599 | 14.00463 | 26.99225 | 10.54558 | 4.086553 | 1.881775 | -0.09947 | 0.013491 |
| 14 | 18409200 | 18410999 | 22.2772 | 40.13549 | 14.71274 | 6.413455 | 1.842223 | 0.405951 | 0.013518 |
| 18 | 89929800 | 89930599 | 27.94164 | 48.71105 | 21.20116 | 9.328246 | 1.58063 | 0.762739 | 0.013695 |
| 7 | 20188200 | 20189399 | 15.88144 | 30.24088 | 11.60708 | 4.644149 | 1.867014 | 0.022936 | 0.013784 |
| 7 | 17124200 | 17126799 | 34.78578 | 59.6041 | 23.14375 | 10.14965 | 1.741145 | 0.969631 | 0.014116 |
| 16 | 17551600 | 17552999 | 4.499952 | 8.304038 | 41.62356 | 17.5906 | -1.86883 | 0.200608 | 0.014159 |
| X | 32624400 | 32624799 | 11.13251 | 21.7233 | 8.156736 | 2.904747 | 2.045926 | -0.42335 | 0.014192 |
| 4 | 155162600 | 155163999 | 2.960118 | 5.717435 | 37.09715 | 14.26799 | -2.11449 | -0.16456 | 0.014306 |
| 7 | 22187000 | 22187399 | 9.607607 | 18.77989 | 6.295742 | 2.168455 | 2.241485 | -0.56661 | 0.014319 |

| | | | | | | | | | |
|---|---|---|---|---|---|---|---|---|---|
| 11 | 47306200 | 47306599 | 8.547743 | 16.72077 | 6.653455 | 2.21704 | 2.043895 | -0.69235 | 0.01432 |
| 5 | 6979200 | 6980799 | 20.42543 | 37.6676 | 12.50892 | 5.291131 | 2.011506 | 0.259877 | 0.014358 |
| 1 | 8103800 | 8105399 | 22.79082 | 41.30807 | 12.99115 | 5.62064 | 2.067301 | 0.373076 | 0.014481 |
| 7 | 85548000 | 85549599 | 21.38464 | 39.00816 | 13.048 | 5.600246 | 1.986131 | 0.327175 | 0.014549 |
| 7 | 22222800 | 22223199 | 10.09415 | 19.73454 | 6.975269 | 2.39118 | 2.176589 | -0.57048 | 0.014632 |
| 2 | 62665400 | 62666799 | 15.38964 | 29.83786 | 6.762629 | 2.555304 | 2.682404 | -0.17541 | 0.01473 |
| 10 | 38612800 | 38614799 | 26.00565 | 46.11869 | 14.42382 | 6.374876 | 2.056504 | 0.525268 | 0.014861 |
| 6 | 19601400 | 19602199 | 11.82523 | 22.9816 | 9.257102 | 3.445087 | 1.89096 | -0.26164 | 0.014957 |
| 12 | 22640000 | 22642599 | 65.11352 | 104.7866 | 50.34046 | 21.37789 | 1.408721 | 1.862606 | 0.014992 |
| 1 | 169290000 | 169290799 | 2.340233 | 4.560043 | 28.91452 | 10.50012 | -1.9972 | -0.39601 | 0.015006 |
| 5 | 94921000 | 94921799 | 26.35705 | 46.47854 | 16.74418 | 7.410116 | 1.852192 | 0.589763 | 0.015063 |
| X | 145401400 | 145401799 | 8.887483 | 17.3851 | 6.65064 | 2.218918 | 2.098719 | -0.68181 | 0.015299 |
| 7 | 22258600 | 22258999 | 12.53452 | 24.46385 | 5.897264 | 2.093081 | 2.670306 | -0.42271 | 0.015301 |
| 1 | 111922000 | 111926399 | 65.22656 | 104.9563 | 50.35912 | 21.38223 | 1.410298 | 1.865853 | 0.015318 |
| 12 | 23922600 | 23922999 | 8.773716 | 17.16267 | 5.003322 | 1.653378 | 2.482958 | -0.72365 | 0.015416 |
| 7 | 7951600 | 7953799 | 20.25983 | 37.74045 | 11.20607 | 4.672343 | 2.184962 | 0.200264 | 0.015422 |
| 6 | 98484400 | 98484799 | 2.116212 | 4.12544 | 28.98929 | 10.46356 | -2.13148 | -0.41854 | 0.015462 |
| 9 | 3103800 | 3106999 | 73.84202 | 118.1748 | 58.77931 | 24.75576 | 1.349533 | 2.046232 | 0.015587 |
| X | 33378600 | 33379999 | 45.93384 | 76.22875 | 29.80195 | 12.91421 | 1.730574 | 1.312341 | 0.015702 |
| 14 | 42518000 | 42519199 | 13.8033 | 26.68123 | 10.59731 | 4.061368 | 1.873365 | -0.13657 | 0.015731 |
| X | 26490800 | 26492399 | 18.6434 | 35.1342 | 10.90362 | 4.457878 | 2.143892 | 0.115128 | 0.015774 |
| 6 | 113789600 | 113799999 | 42.72013 | 67.06355 | 208.422 | 86.44313 | -1.31479 | 2.561045 | 0.015827 |
| 5 | 95162600 | 95163399 | 22.91047 | 41.07893 | 15.09328 | 6.609194 | 1.834707 | 0.43933 | 0.015965 |
| 8 | 123563000 | 123565199 | 5.290229 | 9.591221 | 44.43396 | 19.21752 | -1.78092 | 0.316448 | 0.015985 |
| 4 | 150190200 | 150194399 | 11.80186 | 19.92489 | 76.68625 | 33.54369 | -1.55508 | 1.07733 | 0.016058 |
| 7 | 9065000 | 9066399 | 19.97353 | 36.34171 | 14.70525 | 6.331623 | 1.712363 | 0.334132 | 0.016161 |
| X | 31361400 | 31361999 | 13.85219 | 27.00632 | 4.406761 | 1.588892 | 3.190769 | -0.35724 | 0.016168 |
| 18 | 32926800 | 32927399 | 8.888212 | 17.38393 | 5.727867 | 1.879753 | 2.324858 | -0.74018 | 0.016222 |
| 17 | 24804600 | 24806999 | 6.612869 | 11.7012 | 50.9167 | 22.51772 | -1.71615 | 0.498493 | 0.016232 |
| 12 | 48581200 | 48582399 | 46.70385 | 77.65065 | 25.52797 | 11.07634 | 1.979226 | 1.290141 | 0.016233 |
| 17 | 29533400 | 29536399 | 8.90005 | 15.28313 | 66.8361 | 29.31591 | -1.73401 | 0.902115 | 0.016247 |
| 14 | 100115600 | 100116399 | 14.75701 | 28.54486 | 9.571915 | 3.656308 | 2.119076 | -0.13949 | 0.016283 |
| 1 | 110258000 | 110259199 | 14.24656 | 27.25316 | 11.23611 | 4.449028 | 1.777409 | -0.02792 | 0.016292 |
| 7 | 85520200 | 85520799 | 10.20341 | 19.9288 | 8.205792 | 2.875853 | 1.934708 | -0.49042 | 0.016358 |
| 1 | 111885000 | 111885799 | 10.77223 | 21.0134 | 8.36016 | 2.992244 | 1.95821 | -0.41108 | 0.016361 |
| 4 | 132729600 | 132731199 | 2.343835 | 4.55584 | 29.57879 | 10.99346 | -2.06148 | -0.3088 | 0.016495 |
| 5 | 7468800 | 7471399 | 24.64276 | 43.45751 | 18.45947 | 8.169138 | 1.616507 | 0.581556 | 0.016495 |
| 11 | 15456000 | 15456399 | 8.716536 | 17.05122 | 6.398954 | 2.116781 | 2.13364 | -0.71398 | 0.016612 |
| 15 | 27925000 | 27926199 | 3.059512 | 5.919993 | 32.23413 | 12.29443 | -1.85157 | -0.20258 | 0.016624 |
| 12 | 23480600 | 23481399 | 12.54413 | 24.436 | 8.491472 | 3.094845 | 2.12711 | -0.32446 | 0.01664 |
| 15 | 90113200 | 90114399 | 2.028842 | 3.963467 | 27.95507 | 9.761802 | -2.09372 | -0.54529 | 0.01665 |
| 6 | 92755200 | 92756799 | 2.654892 | 5.139176 | 31.56657 | 12.01683 | -2.01839 | -0.21533 | 0.016714 |
| 4 | 137632200 | 137641599 | 38.22337 | 60.29664 | 192.9436 | 80.07406 | -1.35127 | 2.458372 | 0.016846 |
| 11 | 12191000 | 12191999 | 12.85889 | 24.89989 | 10.81212 | 4.109473 | 1.757838 | -0.17364 | 0.016896 |
| X | 32655200 | 32656199 | 15.47429 | 29.6175 | 10.83337 | 4.283022 | 1.951223 | -0.0206 | 0.017115 |
| Y | 90763200 | 90763999 | 38.33165 | 64.15494 | 30.34599 | 13.2229 | 1.45664 | 1.196633 | 0.017156 |
| 1 | 140856600 | 140857199 | 10.51031 | 20.52835 | 6.631946 | 2.324017 | 2.272559 | -0.49282 | 0.017189 |

| | | | | | | | | | |
|---|---|---|---|---|---|---|---|---|---|
| 12 | 24045600 | 24046599 | 10.58291 | 20.68357 | 5.973312 | 2.06443 | 2.446007 | -0.54001 | 0.01722 |
| 12 | 22420000 | 22420599 | 22.18462 | 40.3201 | 13.61348 | 5.869659 | 1.969298 | 0.354991 | 0.017227 |
| 4 | 147391200 | 147393799 | 65.04176 | 104.6486 | 51.39752 | 21.82001 | 1.376606 | 1.870334 | 0.017299 |
| 1 | 154712000 | 154713599 | 4.126688 | 7.823212 | 37.01336 | 14.94875 | -1.73219 | 0.012389 | 0.017326 |
| 12 | 20115600 | 20116799 | 37.17264 | 62.47953 | 30.3757 | 13.26887 | 1.41717 | 1.151381 | 0.017339 |
| 12 | 19442800 | 19444799 | 33.70158 | 57.47701 | 26.43981 | 11.59076 | 1.497261 | 1.009335 | 0.017347 |
| 4 | 126322400 | 126323599 | 2.652803 | 5.141454 | 31.08935 | 11.76733 | -1.98805 | -0.23674 | 0.017434 |
| 7 | 8561000 | 8561799 | 15.12206 | 29.17341 | 9.077132 | 3.508025 | 2.207267 | -0.09965 | 0.017435 |
| 1 | 144484400 | 144485599 | 2.255026 | 4.387684 | 34.48781 | 12.71296 | -2.32472 | -0.32979 | 0.017457 |
| 14 | 90074800 | 90075999 | 13.39936 | 26.04263 | 8.83101 | 3.284667 | 2.135641 | -0.24444 | 0.017524 |
| 1 | 104220200 | 104221399 | 21.34643 | 38.34729 | 16.15493 | 7.062772 | 1.641216 | 0.419384 | 0.017529 |
| 7 | 15188600 | 15190599 | 16.41268 | 31.00881 | 12.70764 | 5.169363 | 1.753775 | 0.082813 | 0.017555 |
| 15 | 34477000 | 34477199 | 12.17603 | 23.69127 | 9.711864 | 3.580097 | 1.879024 | -0.28682 | 0.017597 |
| 5 | 94432200 | 94433399 | 18.41072 | 34.36549 | 13.01582 | 5.408315 | 1.842194 | 0.176452 | 0.017598 |
| 1 | 116577400 | 116577999 | 12.18358 | 23.69512 | 7.902648 | 2.923941 | 2.16705 | -0.28192 | 0.017633 |
| 4 | 50378800 | 50379199 | 9.060972 | 17.72208 | 5.726346 | 1.880453 | 2.352036 | -0.73477 | 0.017714 |
| 11 | 114627800 | 114633199 | 14.54742 | 23.87105 | 95.82884 | 41.26588 | -1.62116 | 1.419067 | 0.017901 |
| 7 | 8068000 | 8068599 | 16.15958 | 30.40051 | 13.24925 | 5.432831 | 1.656437 | 0.108619 | 0.017939 |
| 7 | 22278400 | 22279399 | 26.15458 | 45.75016 | 21.17995 | 9.340854 | 1.492198 | 0.704843 | 0.018017 |
| 9 | 45895200 | 45896599 | 4.300847 | 7.843337 | 44.55756 | 19.13013 | -2.06432 | 0.273321 | 0.018065 |
| 16 | 17110400 | 17111999 | 3.365208 | 6.501817 | 33.38229 | 12.82118 | -1.7793 | -0.17318 | 0.018113 |
| 12 | 19678800 | 19679199 | 10.45316 | 20.4465 | 3.521232 | 1.160043 | 3.21301 | -0.70896 | 0.018115 |
| 17 | 8353800 | 8355799 | 3.06152 | 5.917338 | 32.09598 | 12.30509 | -1.85308 | -0.18467 | 0.018116 |
| 12 | 22430600 | 22431399 | 11.60104 | 22.62466 | 8.546052 | 3.068519 | 2.027034 | -0.3874 | 0.018143 |
| 9 | 124257400 | 124259799 | 11.31029 | 22.09591 | 6.644113 | 2.319093 | 2.380848 | -0.4931 | 0.018162 |
| 17 | 40141600 | 40142199 | 10.48908 | 20.47025 | 8.379057 | 2.977647 | 1.926835 | -0.43911 | 0.018182 |
| 4 | 130006000 | 130006999 | 2.029201 | 3.963016 | 25.74876 | 9.036632 | -1.98244 | -0.53291 | 0.018183 |
| 7 | 8148400 | 8148799 | 9.396052 | 18.36113 | 7.824895 | 2.715025 | 1.89815 | -0.5386 | 0.018216 |
| 13 | 106297200 | 106298399 | 10.79766 | 21.05242 | 8.533655 | 3.074492 | 1.921171 | -0.38791 | 0.01827 |
| 1 | 102959200 | 102967599 | 208.3928 | 311.9099 | 169.1436 | 69.75919 | 1.145807 | 3.513616 | 0.018305 |
| 9 | 45190600 | 45194399 | 12.88519 | 21.7035 | 76.14362 | 33.28293 | -1.42239 | 1.101525 | 0.018356 |
| 4 | 132734400 | 132735799 | 2.266331 | 4.377849 | 32.46725 | 12.48121 | -2.29761 | -0.18299 | 0.018462 |
| 3 | 32049000 | 32050599 | 16.93131 | 32.27182 | 10.82277 | 4.321314 | 2.061415 | 0.02493 | 0.018482 |
| 11 | 83319800 | 83321199 | 5.443363 | 9.782279 | 45.82393 | 20.02741 | -1.80677 | 0.374491 | 0.018488 |
| 5 | 143778200 | 143779199 | 23.38505 | 41.52827 | 16.25092 | 7.174089 | 1.737927 | 0.503357 | 0.018508 |
| 4 | 122497800 | 122499599 | 40.79411 | 67.75089 | 34.22706 | 14.83958 | 1.362433 | 1.290872 | 0.018537 |
| 5 | 129863800 | 129864999 | 2.982323 | 5.693594 | 34.81589 | 13.85405 | -2.0749 | -0.05282 | 0.018577 |
| 7 | 21097600 | 21099599 | 43.22274 | 71.35424 | 35.50173 | 15.3285 | 1.384014 | 1.367334 | 0.018579 |
| 12 | 111000200 | 111006999 | 21.39088 | 34.3217 | 141.6049 | 59.87089 | -1.69139 | 1.933826 | 0.018606 |
| 8 | 117704400 | 117707199 | 9.941088 | 17.17111 | 64.37971 | 28.24352 | -1.51183 | 0.84819 | 0.018612 |
| 12 | 19647800 | 19653999 | 175.5993 | 265.9909 | 147.3468 | 60.97007 | 1.130094 | 3.273428 | 0.018654 |
| 7 | 21937200 | 21937599 | 8.548169 | 16.72001 | 7.275498 | 2.444389 | 1.908337 | -0.66206 | 0.018735 |
| 4 | 128785200 | 128790799 | 15.68 | 25.65746 | 98.08873 | 42.18983 | -1.55297 | 1.459082 | 0.018735 |
| 1 | 6733600 | 6736199 | 106.2619 | 168.188 | 58.69185 | 24.39907 | 1.844018 | 2.400908 | 0.018753 |
| 11 | 112548000 | 112548799 | 1.948838 | 3.805652 | 29.12663 | 10.24134 | -2.21948 | -0.51971 | 0.018833 |
| 6 | 100647200 | 100648399 | 3.261226 | 6.290048 | 36.34398 | 14.05537 | -1.95808 | -0.14475 | 0.018908 |
| 12 | 22329800 | 22331599 | 36.98687 | 62.82229 | 25.78697 | 11.30026 | 1.660116 | 1.058337 | 0.018979 |

| | | | | | | | | | |
|---|---|---|---|---|---|---|---|---|---|
| 13 | 7573800 | 7574599 | 13.4088 | 26.01859 | 10.05735 | 3.78575 | 1.935647 | -0.2005 | 0.019024 |
| X | 28660600 | 28661199 | 11.59089 | 22.65103 | 5.243811 | 1.817578 | 2.751632 | -0.51723 | 0.019056 |
| 7 | 130867000 | 130867999 | 2.951833 | 5.725936 | 31.92468 | 12.03501 | -1.86938 | -0.24658 | 0.019151 |
| 7 | 137906600 | 137908399 | 4.249441 | 8.051971 | 37.58788 | 15.19655 | -1.71444 | 0.018917 | 0.019188 |
| 17 | 77884600 | 77885199 | 10.8203 | 21.16456 | 4.970815 | 1.667852 | 2.772875 | -0.63704 | 0.019231 |
| 4 | 59342000 | 59343399 | 4.110246 | 7.845674 | 36.83132 | 14.66098 | -1.70272 | -0.03744 | 0.019322 |
| 14 | 31382800 | 31392599 | 43.7069 | 68.56186 | 206.6492 | 85.7048 | -1.27149 | 2.57578 | 0.019336 |
| 6 | 132850000 | 132850799 | 13.36414 | 25.72948 | 10.89618 | 4.235767 | 1.763676 | -0.09588 | 0.019368 |
| 4 | 43695400 | 43698799 | 14.87379 | 24.71845 | 84.32116 | 36.5749 | -1.38286 | 1.256912 | 0.01954 |
| 15 | 45498800 | 45501599 | 52.40389 | 85.03238 | 45.23156 | 19.38295 | 1.274513 | 1.6295 | 0.019561 |
| 3 | 121056000 | 121059399 | 13.05527 | 21.98623 | 76.75967 | 33.55048 | -1.41549 | 1.103521 | 0.019605 |
| 12 | 23377200 | 23378599 | 18.19842 | 34.05465 | 12.87304 | 5.326281 | 1.84932 | 0.158857 | 0.01972 |
| 1 | 116539400 | 116540999 | 11.89778 | 23.16946 | 8.900108 | 3.257453 | 1.978178 | -0.31754 | 0.019746 |
| 12 | 24054800 | 24055799 | 11.89778 | 23.16946 | 8.900108 | 3.257453 | 1.978178 | -0.31754 | 0.019746 |
| 2 | 164698200 | 164700399 | 7.592801 | 13.28584 | 56.76764 | 25.0428 | -1.69516 | 0.662442 | 0.019806 |
| 10 | 81591200 | 81591599 | 10.48731 | 20.47381 | 8.605709 | 3.040188 | 1.895089 | -0.45775 | 0.019904 |
| X | 148895600 | 148896399 | 15.98626 | 30.42877 | 12.1986 | 4.8848 | 1.80452 | 0.028431 | 0.019974 |
| 2 | 22313600 | 22314399 | 13.98309 | 27.02838 | 9.767397 | 3.743548 | 2.005937 | -0.13706 | 0.019983 |
| 12 | 22560800 | 22562199 | 40.9243 | 67.98747 | 34.31185 | 14.87941 | 1.364015 | 1.289224 | 0.020037 |
| 7 | 9316200 | 9317399 | 22.23476 | 39.80452 | 16.97392 | 7.442602 | 1.621128 | 0.448616 | 0.020066 |
| 1 | 112552600 | 112554199 | 23.76236 | 42.01244 | 18.3274 | 8.106537 | 1.579409 | 0.548371 | 0.020082 |
| 13 | 52280800 | 52283999 | 14.75438 | 24.50176 | 87.24877 | 37.82534 | -1.44479 | 1.266368 | 0.020119 |
| 15 | 48530200 | 48532799 | 32.35164 | 55.99344 | 22.00345 | 9.653557 | 1.726254 | 0.870815 | 0.020142 |
| 12 | 23937400 | 23939999 | 29.06174 | 50.76799 | 19.7563 | 8.703762 | 1.741517 | 0.739977 | 0.02015 |
| 7 | 20593000 | 20594399 | 14.80819 | 28.34871 | 12.26641 | 4.846575 | 1.712304 | -0.02498 | 0.02019 |
| 4 | 38342200 | 38342599 | 8.892907 | 17.37446 | 3.571323 | 1.14058 | 3.00012 | -0.83958 | 0.020362 |
| 18 | 78183800 | 78186999 | 9.567537 | 16.319 | 69.44753 | 30.44489 | -1.69692 | 0.970475 | 0.020386 |
| 9 | 60004600 | 60005799 | 3.378672 | 6.484967 | 33.43928 | 13.11381 | -1.81361 | -0.10151 | 0.020389 |
| 14 | 104852400 | 104853199 | 11.42163 | 22.25534 | 8.925729 | 3.241686 | 1.929487 | -0.35014 | 0.020401 |
| 4 | 17370400 | 17373399 | 33.61832 | 57.91232 | 22.4564 | 9.850863 | 1.744129 | 0.917373 | 0.020411 |
| 13 | 66145200 | 66148799 | 140.3655 | 215.5903 | 122.2836 | 50.77254 | 1.111805 | 2.985408 | 0.020502 |
| 11 | 79727800 | 79734599 | 24.83639 | 39.87285 | 132.3881 | 56.02277 | -1.37794 | 1.917317 | 0.020508 |
| 7 | 40683200 | 40684599 | 13.89392 | 26.85482 | 10.40666 | 3.989403 | 1.90719 | -0.1353 | 0.020652 |
| 7 | 31317400 | 31321599 | 93.962 | 148.6759 | 79.26785 | 32.94785 | 1.233559 | 2.400731 | 0.02066 |
| 1 | 112948400 | 112950999 | 23.8631 | 41.98576 | 19.58362 | 8.670616 | 1.481273 | 0.602552 | 0.020686 |
| 15 | 76761600 | 76766199 | 14.27886 | 23.74478 | 86.68914 | 37.61848 | -1.4804 | 1.248329 | 0.020771 |
| 19 | 5322200 | 5323599 | 4.27545 | 8.00981 | 38.19534 | 15.77287 | -1.77045 | 0.098556 | 0.020828 |
| 10 | 74074200 | 74074799 | 13.04649 | 25.25725 | 11.24726 | 4.27953 | 1.720464 | -0.16636 | 0.020888 |
| 6 | 27044600 | 27046399 | 31.36739 | 54.01646 | 24.74421 | 10.85433 | 1.505098 | 0.912548 | 0.020963 |
| 11 | 78822200 | 78829799 | 31.34502 | 50.00383 | 150.5582 | 63.12396 | -1.24803 | 2.128758 | 0.020973 |
| 6 | 126916000 | 126917799 | 3.289015 | 6.261266 | 35.32752 | 14.14247 | -1.96963 | -0.02905 | 0.020994 |
| 12 | 22432600 | 22433399 | 13.71072 | 26.50936 | 9.959091 | 3.812013 | 1.955485 | -0.14393 | 0.021013 |
| 15 | 101915800 | 101917799 | 5.695313 | 10.43711 | 42.59962 | 18.17417 | -1.58699 | 0.25167 | 0.021034 |
| 14 | 96449600 | 96450399 | 11.89159 | 23.17987 | 7.616362 | 2.752397 | 2.213006 | -0.36526 | 0.021067 |
| 7 | 31900800 | 31902199 | 41.0536 | 68.05697 | 34.91804 | 15.12001 | 1.340151 | 1.313682 | 0.021085 |
| 12 | 23600600 | 23603999 | 55.32336 | 89.7191 | 44.15282 | 18.91283 | 1.385689 | 1.646834 | 0.021123 |
| 15 | 21902400 | 21903599 | 11.79958 | 22.98889 | 8.926229 | 3.247201 | 1.970775 | -0.33928 | 0.021136 |

| | | | | | | | | | |
|---|---|---|---|---|---|---|---|---|---|
| 3 | 61544600 | 61545799 | 19.59487 | 36.43364 | 12.8044 | 5.355678 | 1.941451 | 0.209961 | 0.021151 |
| 9 | 17279600 | 17280399 | 10.41654 | 20.31986 | 9.198983 | 3.291798 | 1.773902 | -0.41349 | 0.021216 |
| 10 | 81719600 | 81724799 | 90.59601 | 143.388 | 82.67033 | 34.36672 | 1.121203 | 2.39342 | 0.021257 |
| 1 | 189652600 | 189655199 | 11.63035 | 19.63268 | 73.77741 | 32.27 | -1.52052 | 1.07631 | 0.021305 |
| 17 | 78441800 | 78442799 | 15.15061 | 28.74611 | 11.99401 | 4.83546 | 1.737671 | 0.039758 | 0.021328 |
| 8 | 23669600 | 23672799 | 9.912857 | 17.127 | 62.69226 | 27.50346 | -1.47707 | 0.844851 | 0.021362 |
| 15 | 89367600 | 89369599 | 4.332956 | 8.135859 | 44.3085 | 18.22763 | -1.95807 | 0.093392 | 0.021435 |
| 16 | 61446200 | 61447599 | 18.73831 | 34.78945 | 13.97865 | 5.858961 | 1.748393 | 0.213539 | 0.021508 |
| 14 | 50086800 | 50087799 | 11.72029 | 22.80873 | 9.523408 | 3.505378 | 1.853138 | -0.29876 | 0.02152 |
| 17 | 17268000 | 17273599 | 214.4163 | 322.1376 | 144.0934 | 59.48008 | 1.428158 | 3.441823 | 0.021567 |
| 2 | 23909400 | 23909999 | 9.225541 | 18.028 | 8.236968 | 2.857712 | 1.79882 | -0.53984 | 0.021588 |
| 3 | 87061200 | 87062799 | 3.364793 | 6.502329 | 32.15446 | 12.33795 | -1.72409 | -0.17925 | 0.021659 |
| 1 | 176372200 | 176372999 | 14.62906 | 27.96315 | 12.40026 | 4.920945 | 1.671756 | -0.01284 | 0.021752 |
| 15 | 17156800 | 17158599 | 33.89912 | 58.60968 | 20.84686 | 9.145831 | 1.86878 | 0.885812 | 0.021779 |
| 7 | 20488800 | 20489199 | 9.738693 | 19.04592 | 6.726893 | 2.286392 | 2.185237 | -0.6054 | 0.021818 |
| 18 | 39076400 | 39078599 | 6.549906 | 11.64327 | 47.78516 | 21.07734 | -1.62831 | 0.455199 | 0.021864 |
| 10 | 6480800 | 6481599 | 9.740163 | 19.04302 | 7.388895 | 2.531132 | 2.044409 | -0.57679 | 0.021898 |
| 7 | 38578200 | 38580199 | 22.88166 | 40.5663 | 17.76003 | 7.851279 | 1.575407 | 0.512332 | 0.022041 |
| 1 | 102964800 | 102967599 | 104.2756 | 163.8153 | 88.82884 | 36.84531 | 1.201351 | 2.56288 | 0.022043 |
| 13 | 43059200 | 43060799 | 7.356254 | 13.03691 | 49.88763 | 22.05574 | -1.53173 | 0.483205 | 0.022112 |
| 6 | 64385000 | 64386199 | 13.97542 | 27.1563 | 8.8234 | 3.287656 | 2.194662 | -0.23134 | 0.022125 |
| 6 | 91953400 | 91954599 | 4.647032 | 8.679105 | 39.8641 | 16.5483 | -1.72499 | 0.122759 | 0.022148 |
| X | 4020800 | 4021399 | 12.43774 | 24.3249 | 3.472785 | 1.179295 | 3.44213 | -0.5778 | 0.022163 |
| 7 | 9330400 | 9330999 | 11.70923 | 22.82546 | 7.618692 | 2.751604 | 2.191216 | -0.36986 | 0.022164 |
| 3 | 5987800 | 5988399 | 10.84808 | 21.17662 | 8.601216 | 3.042297 | 1.942694 | -0.44818 | 0.022191 |
| 1 | 167770400 | 167771999 | 5.087142 | 9.451843 | 40.2458 | 16.85128 | -1.62721 | 0.161374 | 0.022314 |
| 4 | 90552000 | 90553199 | 19.29504 | 35.89263 | 12.63005 | 5.278646 | 1.939873 | 0.203918 | 0.022338 |
| 13 | 61035800 | 61038599 | 25.39123 | 44.89506 | 16.43244 | 7.268273 | 1.830217 | 0.554495 | 0.022387 |
| 13 | 65509600 | 65509999 | 7.779225 | 15.20368 | 5.064231 | 1.626549 | 2.32965 | -0.83681 | 0.022389 |
| 11 | 87999800 | 88002999 | 8.663839 | 15.13125 | 58.46328 | 25.75249 | -1.55177 | 0.699734 | 0.022436 |
| 3 | 122067600 | 122068999 | 2.960643 | 5.716859 | 31.97571 | 12.31422 | -1.90305 | -0.17112 | 0.022453 |
| 7 | 8778600 | 8783799 | 37.8953 | 63.58256 | 30.75008 | 13.42002 | 1.424654 | 1.170283 | 0.022521 |
| 18 | 80520000 | 80523199 | 10.706 | 18.4306 | 64.324 | 28.21621 | -1.41014 | 0.88313 | 0.022536 |
| 13 | 98554800 | 98555799 | 82.46309 | 131.7975 | 47.22718 | 19.85323 | 1.819377 | 2.093796 | 0.02255 |
| 12 | 23502400 | 23504399 | 30.31474 | 52.0966 | 26.22797 | 11.50449 | 1.369344 | 0.928496 | 0.022583 |
| 4 | 18226800 | 18227999 | 14.14228 | 27.09412 | 12.11837 | 4.777829 | 1.666865 | -0.03955 | 0.022655 |
| X | 122189400 | 122190799 | 23.48585 | 41.97902 | 16.39249 | 7.197074 | 1.745988 | 0.465787 | 0.022697 |
| X | 32956200 | 32958399 | 31.65974 | 55.31574 | 17.34445 | 7.642013 | 2.050625 | 0.748088 | 0.02275 |
| 17 | 74796400 | 74797199 | 4.108643 | 7.847875 | 35.045 | 13.93016 | -1.62912 | -0.04594 | 0.022788 |
| 18 | 40481200 | 40481799 | 9.738217 | 19.04692 | 6.044567 | 2.038126 | 2.34358 | -0.63463 | 0.02284 |
| 10 | 74780800 | 74781799 | 5.073758 | 9.467371 | 40.45047 | 16.8179 | -1.62411 | 0.13354 | 0.022852 |
| 7 | 140700800 | 140704199 | 9.313065 | 16.17898 | 63.39999 | 27.83401 | -1.57346 | 0.785572 | 0.022864 |
| 17 | 57585200 | 57587799 | 9.330321 | 15.95963 | 67.26598 | 29.49527 | -1.68196 | 0.942217 | 0.022904 |
| 15 | 3076800 | 3078399 | 62.58795 | 100.7974 | 52.45853 | 22.3015 | 1.294932 | 1.83779 | 0.022904 |
| 4 | 138531400 | 138537399 | 24.01838 | 38.64992 | 127.5894 | 54.17758 | -1.36541 | 1.83937 | 0.022914 |
| 12 | 22231800 | 22233999 | 38.02661 | 63.53493 | 33.66398 | 14.65275 | 1.293949 | 1.218109 | 0.022939 |
| 7 | 21173600 | 21176999 | 64.5068 | 103.9621 | 48.12945 | 20.48218 | 1.464459 | 1.816625 | 0.022945 |

| | | | | | | | | | |
|---|---|---|---|---|---|---|---|---|---|
| 12 | 22456800 | 22460999 | 61.35288 | 98.74414 | 55.16654 | 23.43065 | 1.191763 | 1.856205 | 0.022957 |
| 1 | 99334800 | 99335799 | 11.9992 | 23.31378 | 10.89245 | 4.062022 | 1.678759 | -0.24603 | 0.023004 |
| 7 | 135660200 | 135662399 | 6.10788 | 10.98728 | 46.42369 | 20.27272 | -1.65993 | 0.370765 | 0.023011 |
| 12 | 23602600 | 23603999 | 37.62527 | 63.28612 | 28.53596 | 12.46796 | 1.525459 | 1.146178 | 0.023052 |
| 15 | 55024000 | 55024599 | 17.3436 | 32.95557 | 11.40347 | 4.583058 | 2.009474 | 0.050868 | 0.023092 |
| 7 | 22910800 | 22911599 | 15.6597 | 30.00852 | 11.51113 | 4.535249 | 1.887332 | -0.02708 | 0.023097 |
| 6 | 120945000 | 120947799 | 8.993942 | 15.6519 | 60.62674 | 26.64573 | -1.5562 | 0.755757 | 0.023111 |
| 7 | 33343000 | 33344199 | 46.74258 | 76.40095 | 41.42614 | 17.81097 | 1.254638 | 1.502826 | 0.02322 |
| 15 | 71101400 | 71103599 | 25.7214 | 45.19153 | 20.02019 | 8.862376 | 1.554982 | 0.632825 | 0.023235 |
| 7 | 38624200 | 38626399 | 17.05461 | 32.34028 | 11.80936 | 4.766747 | 1.926972 | 0.062401 | 0.023238 |
| 2 | 131054200 | 131056799 | 9.442749 | 16.35669 | 61.0679 | 26.79294 | -1.5044 | 0.81724 | 0.023299 |
| 10 | 7492400 | 7493399 | 14.58733 | 27.69961 | 12.33633 | 4.965029 | 1.64755 | 0.029914 | 0.023318 |
| 7 | 126753000 | 126756599 | 13.58915 | 22.74588 | 79.99485 | 34.85871 | -1.42633 | 1.169198 | 0.023398 |
| 12 | 59113200 | 59114199 | 3.254609 | 6.296491 | 32.9262 | 12.56776 | -1.79692 | -0.19821 | 0.023431 |
| 12 | 52645000 | 52646399 | 24.13123 | 42.40453 | 20.70334 | 9.166043 | 1.415603 | 0.621488 | 0.023439 |
| 11 | 115290600 | 115292199 | 3.367013 | 6.49957 | 32.02323 | 12.34999 | -1.72574 | -0.16164 | 0.023531 |
| 14 | 58034400 | 58036599 | 8.142947 | 14.23714 | 55.68649 | 24.54932 | -1.56837 | 0.678875 | 0.023537 |
| 14 | 20402800 | 20403399 | 1.714532 | 3.352909 | 23.02499 | 7.837689 | -2.01442 | -0.65482 | 0.02355 |
| 17 | 18069800 | 18070999 | 6.168049 | 10.93349 | 48.79248 | 21.57064 | -1.75067 | 0.47243 | 0.023581 |
| 11 | 69300600 | 69303199 | 8.976941 | 15.58555 | 59.99056 | 26.32711 | -1.54725 | 0.79091 | 0.023597 |
| 3 | 88203600 | 88204999 | 5.675427 | 10.21392 | 46.96116 | 20.49888 | -1.78012 | 0.365822 | 0.023629 |
| 12 | 22353200 | 22354399 | 20.73969 | 37.34069 | 17.15148 | 7.482729 | 1.519536 | 0.40423 | 0.023631 |
| 5 | 114212000 | 114220599 | 37.37309 | 59.0925 | 183.2225 | 76.12889 | -1.30312 | 2.400163 | 0.023636 |
| 17 | 46958600 | 46966999 | 30.09729 | 47.94483 | 155.6735 | 65.13977 | -1.35888 | 2.175413 | 0.023695 |
| 17 | 12365000 | 12365999 | 2.947214 | 5.730645 | 31.15258 | 11.54601 | -1.81044 | -0.3103 | 0.023771 |
| 5 | 5868400 | 5868799 | 1.332533 | 2.602834 | 20.23163 | 6.436378 | -2.09263 | -0.91513 | 0.023803 |
| 10 | 81664400 | 81665799 | 19.5011 | 35.26915 | 16.05841 | 6.966505 | 1.536517 | 0.366358 | 0.023847 |
| 6 | 10236800 | 10237399 | 7.656378 | 14.97052 | 6.451276 | 2.091295 | 1.962392 | -0.80103 | 0.023921 |
| 18 | 4684200 | 4686799 | 7.475423 | 13.30781 | 50.00737 | 22.03052 | -1.50208 | 0.453247 | 0.02396 |
| 17 | 40081400 | 40082999 | 24.23419 | 42.87724 | 17.60804 | 7.787242 | 1.666039 | 0.542405 | 0.023969 |
| 13 | 120126600 | 120127199 | 13.97749 | 27.26068 | 5.866295 | 2.104915 | 2.819024 | -0.36661 | 0.024095 |
| 1 | 30004400 | 30005199 | 10.0305 | 19.57587 | 9.413312 | 3.343516 | 1.698771 | -0.4431 | 0.024121 |
| 1 | 110495800 | 110496599 | 10.85195 | 21.16682 | 9.383171 | 3.362755 | 1.803704 | -0.40097 | 0.024163 |
| 12 | 20230600 | 20232599 | 57.16217 | 92.20047 | 52.36917 | 22.30996 | 1.171616 | 1.78256 | 0.024209 |
| 7 | 126496000 | 126497799 | 4.627002 | 8.520759 | 40.44973 | 17.14295 | -1.7938 | 0.212375 | 0.024257 |
| 12 | 18247200 | 18251199 | 106.8048 | 167.496 | 93.49043 | 38.77492 | 1.157782 | 2.5981 | 0.024283 |
| 6 | 119025800 | 119027999 | 6.025154 | 10.96381 | 42.94541 | 18.48867 | -1.53736 | 0.296337 | 0.024341 |
| 9 | 99467400 | 99467999 | 1.791892 | 3.502739 | 26.88328 | 9.249517 | -2.1882 | -0.60221 | 0.024347 |
| 10 | 48098000 | 48098399 | 7.377798 | 14.41105 | 5.801747 | 1.845764 | 2.078875 | -0.87239 | 0.024349 |
| 7 | 72279600 | 72281599 | 63.69951 | 102.6518 | 49.77415 | 21.17949 | 1.397756 | 1.818848 | 0.024409 |
| 7 | 107464000 | 107464599 | 11.03078 | 21.53252 | 8.598968 | 3.043354 | 1.966196 | -0.4434 | 0.024438 |
| 8 | 89087600 | 89089199 | 4.804516 | 8.868758 | 39.99102 | 16.8937 | -1.71763 | 0.197585 | 0.024462 |
| 12 | 23591800 | 23596999 | 76.97397 | 122.5814 | 72.8656 | 30.47677 | 1.08555 | 2.204334 | 0.024531 |
| 10 | 11704000 | 11704599 | 9.677531 | 18.90313 | 8.640116 | 3.02406 | 1.787101 | -0.50138 | 0.024548 |
| 15 | 78284800 | 78291999 | 21.2401 | 34.08795 | 139.1923 | 58.8723 | -1.67604 | 1.925186 | 0.02456 |
| 3 | 150450200 | 150451399 | 13.3104 | 25.82946 | 9.861221 | 3.710718 | 1.952488 | -0.20345 | 0.024624 |
| 10 | 121837000 | 121839999 | 14.17917 | 23.78957 | 77.90691 | 33.99948 | -1.32394 | 1.143585 | 0.024646 |

| | | | | | | | | | |
|---|---|---|---|---|---|---|---|---|---|
| 12 | 19017000 | 19019399 | 76.90584 | 122.5711 | 70.65141 | 29.58345 | 1.131079 | 2.179184 | 0.024683 |
| 10 | 33194600 | 33196399 | 22.30303 | 39.76951 | 17.8069 | 7.83386 | 1.547951 | 0.474035 | 0.024693 |
| 11 | 70860000 | 70862199 | 6.489819 | 11.46516 | 50.35947 | 22.2784 | -1.73039 | 0.514744 | 0.024697 |
| 2 | 141450800 | 141451799 | 13.31201 | 25.80486 | 9.182218 | 3.472594 | 2.044391 | -0.18982 | 0.024733 |
| 8 | 76739000 | 76739599 | 9.45636 | 18.49402 | 7.416476 | 2.519713 | 2.008099 | -0.60842 | 0.024798 |
| 4 | 64627000 | 64628399 | 14.16209 | 27.46909 | 9.19583 | 3.468057 | 2.135947 | -0.18816 | 0.024882 |
| 15 | 25372000 | 25373399 | 4.737256 | 8.950597 | 38.12317 | 15.50672 | -1.5929 | 0.042815 | 0.024925 |
| 18 | 3058200 | 3058399 | 10.59316 | 20.6658 | 7.975162 | 2.851333 | 2.003003 | -0.41925 | 0.025022 |
| 7 | 9036800 | 9039399 | 32.55541 | 56.62265 | 19.08925 | 8.386778 | 1.946276 | 0.815426 | 0.025042 |
| 7 | 135754800 | 135755599 | 2.336962 | 4.563834 | 25.83625 | 9.075397 | -1.79156 | -0.52646 | 0.025054 |
| 7 | 32941200 | 32941999 | 26.93909 | 47.21472 | 20.35274 | 8.990886 | 1.59422 | 0.677974 | 0.025125 |
| 1 | 111890000 | 111891199 | 15.18709 | 29.37729 | 7.099948 | 2.711838 | 2.576187 | -0.14225 | 0.025171 |
| 19 | 23879200 | 23882599 | 13.17102 | 21.94023 | 82.47661 | 35.83127 | -1.52192 | 1.223228 | 0.025172 |
| 19 | 43346800 | 43347599 | 12.9776 | 25.03722 | 11.50302 | 4.445091 | 1.6559 | -0.11652 | 0.025177 |
| 15 | 80132000 | 80132799 | 1.795057 | 3.496652 | 27.96794 | 10.19796 | -2.3223 | -0.38612 | 0.025237 |
| 7 | 102668000 | 102669599 | 22.94764 | 41.27975 | 15.70286 | 6.857275 | 1.788138 | 0.422256 | 0.025244 |
| 1 | 9420000 | 9420399 | 8.26447 | 16.16662 | 7.713195 | 2.570442 | 1.787738 | -0.69151 | 0.025268 |
| 15 | 79430200 | 79433799 | 14.76648 | 24.49032 | 85.34873 | 36.96874 | -1.4139 | 1.280645 | 0.025276 |
| 7 | 138002400 | 138003799 | 3.984226 | 7.627065 | 34.59104 | 13.66886 | -1.64337 | -0.0689 | 0.025294 |
| 3 | 159498800 | 159500599 | 23.62953 | 42.47511 | 15.11088 | 6.602863 | 1.883235 | 0.430312 | 0.025302 |
| 12 | 19015400 | 19019399 | 99.20641 | 156.1461 | 91.99024 | 38.166 | 1.083967 | 2.522638 | 0.025342 |
| 5 | 69771400 | 69772599 | 12.53932 | 24.44988 | 7.631759 | 2.747057 | 2.291967 | -0.36923 | 0.025344 |
| 7 | 10869400 | 10870599 | 45.33874 | 74.55508 | 39.20254 | 16.8966 | 1.302726 | 1.423303 | 0.025394 |
| 12 | 69621600 | 69623199 | 4.215593 | 7.914911 | 39.48711 | 16.24621 | -1.83204 | 0.083125 | 0.025413 |
| 11 | 38374600 | 38375599 | 13.08575 | 25.19696 | 11.48156 | 4.461574 | 1.660117 | -0.0982 | 0.025452 |
| 1 | 131765000 | 131767999 | 7.408327 | 12.98413 | 53.52348 | 23.64408 | -1.64318 | 0.630909 | 0.025571 |
| X | 33584000 | 33585399 | 13.40945 | 26.06058 | 9.674377 | 3.600217 | 2.008745 | -0.2396 | 0.025572 |
| 16 | 81880800 | 81883399 | 30.20604 | 52.52556 | 21.30011 | 9.356887 | 1.681653 | 0.808708 | 0.025603 |
| 4 | 133126600 | 133127799 | 2.538047 | 4.906733 | 31.97066 | 12.24341 | -2.10761 | -0.19139 | 0.025632 |
| 14 | 27779600 | 27780799 | 3.374431 | 6.490339 | 32.37644 | 12.62125 | -1.758 | -0.12421 | 0.025633 |
| 14 | 19596000 | 19598999 | 33.17993 | 56.30334 | 29.44369 | 12.89902 | 1.312693 | 1.055666 | 0.02577 |
| 10 | 100228000 | 100229599 | 23.64034 | 42.50615 | 14.34299 | 6.265883 | 1.958352 | 0.426633 | 0.025788 |
| 12 | 22230800 | 22233999 | 46.89372 | 76.85198 | 40.78618 | 17.55405 | 1.287531 | 1.467705 | 0.025807 |
| 14 | 60928200 | 60930799 | 8.986431 | 15.57433 | 60.4939 | 26.54152 | -1.56077 | 0.80985 | 0.025827 |
| 4 | 46265800 | 46266799 | 1.951441 | 3.80329 | 26.93936 | 9.755086 | -2.14631 | -0.41679 | 0.025835 |
| X | 19204400 | 19204999 | 10.82313 | 21.15187 | 7.204992 | 2.493729 | 2.220004 | -0.52638 | 0.025848 |
| 17 | 30794200 | 30796999 | 8.66662 | 15.12914 | 57.35752 | 25.25811 | -1.52456 | 0.705213 | 0.02585 |
| 5 | 79768000 | 79768599 | 10.38595 | 20.26815 | 9.40925 | 3.346102 | 1.747691 | -0.43361 | 0.025853 |
| 7 | 21904000 | 21904399 | 12.34825 | 24.09075 | 6.966307 | 2.489284 | 2.406957 | -0.39867 | 0.025876 |
| X | 3356400 | 3357599 | 20.90798 | 38.60608 | 12.92295 | 5.456907 | 2.002132 | 0.258566 | 0.02588 |
| 6 | 69999200 | 69999999 | 12.57968 | 24.35869 | 11.86142 | 4.508744 | 1.59466 | -0.17298 | 0.025882 |
| 1 | 184885600 | 184886399 | 2.643691 | 5.151381 | 27.71176 | 10.06237 | -1.76534 | -0.39673 | 0.025922 |
| 11 | 101777000 | 101778199 | 3.3292 | 6.220671 | 38.00351 | 15.76375 | -2.12679 | 0.107058 | 0.025928 |
| 6 | 106873800 | 106874999 | 12.95309 | 25.07816 | 11.08043 | 4.214692 | 1.731138 | -0.16926 | 0.025928 |
| 4 | 147403400 | 147406599 | 57.64239 | 93.3304 | 45.25715 | 19.363 | 1.404841 | 1.683167 | 0.026002 |
| 9 | 19111200 | 19112799 | 33.82617 | 57.67936 | 26.82897 | 11.7612 | 1.481237 | 1.011477 | 0.026024 |
| 12 | 22355200 | 22356799 | 20.2617 | 36.81205 | 16.52991 | 7.130317 | 1.562014 | 0.349152 | 0.026114 |

| | | | | | | | | | |
|---|---|---|---|---|---|---|---|---|---|
| 10 | 47374000 | 47375199 | 15.903 | 30.19278 | 13.04764 | 5.253198 | 1.690297 | 0.04671 | 0.026114 |
| 7 | 134986000 | 134986599 | 1.641707 | 3.211354 | 23.77326 | 7.849018 | -2.08211 | -0.7658 | 0.026163 |
| 7 | 20751800 | 20753799 | 37.25089 | 62.33721 | 33.69684 | 14.68167 | 1.265165 | 1.199711 | 0.026172 |
| X | 148106200 | 148107199 | 14.19325 | 27.39091 | 11.62176 | 4.48704 | 1.770039 | -0.10494 | 0.026233 |
| 9 | 21673800 | 21674399 | 1.641729 | 3.211321 | 22.36566 | 7.376588 | -1.99331 | -0.77388 | 0.026237 |
| 1 | 114055800 | 114057199 | 13.14595 | 25.42209 | 10.41931 | 3.984722 | 1.829968 | -0.15191 | 0.026251 |
| 2 | 47592800 | 47593999 | 13.66167 | 26.22822 | 12.9013 | 5.056013 | 1.540158 | -0.0606 | 0.026277 |
| X | 3625000 | 3626199 | 26.41848 | 46.28481 | 20.84626 | 9.205963 | 1.531836 | 0.682659 | 0.026308 |
| 18 | 87381800 | 87382199 | 8.099106 | 15.84326 | 7.715071 | 2.569437 | 1.759246 | -0.69678 | 0.026321 |
| 13 | 96564200 | 96567199 | 7.295557 | 13.02493 | 47.35937 | 20.81355 | -1.45196 | 0.430285 | 0.026335 |
| 7 | 19805600 | 19806799 | 14.44156 | 27.60119 | 13.28197 | 5.272788 | 1.554338 | -0.01106 | 0.026387 |
| 3 | 108425000 | 108425599 | 12.18053 | 23.68059 | 10.75083 | 3.989421 | 1.72558 | -0.26082 | 0.02642 |
| 2 | 103857800 | 103858799 | 3.940677 | 7.338491 | 39.08096 | 16.30332 | -1.94021 | 0.135539 | 0.026436 |
| 7 | 21013000 | 21014799 | 45.81139 | 75.24181 | 40.25793 | 17.34266 | 1.277055 | 1.439347 | 0.026456 |
| 15 | 97750000 | 97751999 | 5.57217 | 10.01318 | 45.29043 | 19.79521 | -1.75661 | 0.375389 | 0.026485 |
| 4 | 53713600 | 53714799 | 5.686106 | 10.45121 | 41.02965 | 17.43384 | -1.5269 | 0.231385 | 0.026522 |
| 12 | 18205800 | 18207399 | 51.12689 | 83.01405 | 46.84023 | 20.0819 | 1.190569 | 1.615456 | 0.02659 |
| 5 | 71470600 | 71472599 | 24.0645 | 43.06695 | 15.50052 | 6.798309 | 1.863041 | 0.460174 | 0.02664 |
| 14 | 23219000 | 23219799 | 11.03452 | 21.52193 | 9.381159 | 3.364047 | 1.827112 | -0.3963 | 0.026664 |
| 7 | 22298800 | 22299199 | 11.60842 | 22.60575 | 10.3593 | 3.791993 | 1.730157 | -0.3152 | 0.026666 |
| 1 | 26881200 | 26882999 | 28.0237 | 48.54009 | 23.66717 | 10.38372 | 1.417138 | 0.844876 | 0.026667 |
| 13 | 110324800 | 110325599 | 14.01487 | 26.99738 | 12.20686 | 4.737681 | 1.672722 | -0.08897 | 0.026748 |
| 15 | 84075600 | 84077999 | 8.146248 | 14.3311 | 51.42033 | 22.76683 | -1.44417 | 0.573259 | 0.026788 |
| 6 | 115014400 | 115014999 | 2.64545 | 5.149463 | 28.05656 | 10.30798 | -1.79888 | -0.35239 | 0.026826 |
| 9 | 114581800 | 114582599 | 2.64545 | 5.149463 | 28.05656 | 10.30798 | -1.79888 | -0.35239 | 0.026826 |
| 2 | 119444200 | 119444399 | 7.420978 | 14.4597 | 4.399534 | 1.36024 | 2.498172 | -0.97326 | 0.026844 |
| 4 | 43668000 | 43670199 | 5.445493 | 9.944782 | 41.92882 | 17.96986 | -1.63786 | 0.268143 | 0.026849 |
| 12 | 23500400 | 23501199 | 13.03105 | 25.28951 | 10.50709 | 3.95219 | 1.833438 | -0.20592 | 0.026902 |
| 7 | 8494600 | 8496599 | 27.1402 | 47.49907 | 21.50583 | 9.488771 | 1.524166 | 0.703505 | 0.026975 |
| 2 | 165079800 | 165094999 | 83.839 | 126.0866 | 355.9808 | 146.9845 | -1.22649 | 3.409929 | 0.026987 |
| 4 | 132970000 | 132971199 | 4.434009 | 8.388181 | 35.77717 | 14.51383 | -1.59029 | 0.027084 | 0.027006 |
| 7 | 17817200 | 17818399 | 42.31203 | 69.91724 | 38.10036 | 16.4613 | 1.253375 | 1.356766 | 0.027075 |
| 4 | 140768000 | 140785599 | 69.74504 | 105.855 | 315.9812 | 130.8274 | -1.2953 | 3.220397 | 0.027112 |
| 12 | 23522400 | 23522999 | 11.50515 | 22.42774 | 8.941356 | 3.229097 | 1.945517 | -0.3678 | 0.027162 |
| 7 | 118454400 | 118455399 | 4.742056 | 8.944677 | 36.84249 | 15.03896 | -1.54922 | 0.052404 | 0.027185 |
| 10 | 81710800 | 81711199 | 13.05238 | 25.24501 | 10.54506 | 4.029931 | 1.806661 | -0.15465 | 0.027206 |
| 2 | 167694600 | 167695799 | 3.709662 | 7.026757 | 35.1544 | 14.22308 | -1.81263 | 0.010828 | 0.027238 |
| 5 | 59907400 | 59908599 | 21.64119 | 38.76357 | 17.68116 | 7.749058 | 1.525465 | 0.44294 | 0.027263 |
| 7 | 42409200 | 42411199 | 48.29366 | 78.8956 | 42.99369 | 18.48146 | 1.246952 | 1.515072 | 0.027312 |
| 13 | 12151400 | 12152999 | 21.21752 | 39.05681 | 13.3114 | 5.644608 | 1.972113 | 0.278457 | 0.027336 |
| 8 | 121476600 | 121484599 | 35.80017 | 56.81071 | 166.8298 | 69.49474 | -1.22014 | 2.301834 | 0.027371 |
| 4 | 123728200 | 123730199 | 5.028687 | 9.384688 | 45.17338 | 18.77672 | -1.79439 | 0.142685 | 0.027376 |
| 7 | 20921000 | 20925799 | 69.12999 | 111.1857 | 49.83012 | 21.14337 | 1.507717 | 1.887841 | 0.027432 |
| 4 | 138011000 | 138019799 | 48.81061 | 75.98797 | 223.5447 | 92.79229 | -1.24524 | 2.714933 | 0.027449 |
| 7 | 130360600 | 130362799 | 7.659265 | 13.49301 | 50.35183 | 22.28758 | -1.49907 | 0.554587 | 0.027461 |
| 16 | 76916000 | 76917199 | 13.03917 | 25.27252 | 11.10843 | 4.203742 | 1.745726 | -0.18443 | 0.02752 |
| 1 | 190334400 | 190335399 | 4.113505 | 7.841217 | 34.34038 | 13.70884 | -1.60657 | -0.03278 | 0.02757 |

| | | | | | | | | | |
|---|---|---|---|---|---|---|---|---|---|
| 17 | 58347800 | 58348599 | 11.07701 | 21.58501 | 8.936417 | 3.243252 | 1.881736 | -0.35752 | 0.027606 |
| 14 | 44581800 | 44582199 | 9.808434 | 19.18169 | 7.41223 | 2.521448 | 2.059645 | -0.5982 | 0.027606 |
| 7 | 9190000 | 9196799 | 39.44371 | 66.04713 | 29.78649 | 12.98295 | 1.525154 | 1.19597 | 0.027619 |
| X | 37268800 | 37269999 | 47.30658 | 78.47385 | 29.87256 | 12.94105 | 1.768994 | 1.322638 | 0.02764 |
| 7 | 60696400 | 60696999 | 12.07433 | 23.52169 | 9.569754 | 3.486128 | 1.904216 | -0.32828 | 0.027646 |
| 12 | 22218400 | 22219799 | 18.45397 | 34.85577 | 11.52335 | 4.690363 | 2.059736 | 0.100947 | 0.027647 |
| 7 | 31882200 | 31884199 | 34.92853 | 59.27066 | 29.71584 | 13.01826 | 1.373251 | 1.063202 | 0.027653 |
| 2 | 180363400 | 180365599 | 6.791452 | 12.30406 | 45.59996 | 19.73963 | -1.46435 | 0.331981 | 0.027654 |
| 15 | 70734600 | 70735599 | 11.63753 | 22.5978 | 11.47508 | 4.299635 | 1.553988 | -0.2326 | 0.02767 |
| 4 | 70058800 | 70059599 | 13.02131 | 25.31006 | 9.268016 | 3.444811 | 2.026823 | -0.24998 | 0.027763 |
| 13 | 118261200 | 118261599 | 8.888206 | 17.38314 | 8.30605 | 2.818102 | 1.764479 | -0.61832 | 0.02782 |
| 1 | 74963400 | 74969199 | 26.27638 | 42.19248 | 129.2923 | 54.72815 | -1.26224 | 1.911934 | 0.027841 |
| 2 | 57746800 | 57747199 | 1.332133 | 2.603284 | 19.38224 | 6.215966 | -2.0412 | -0.88885 | 0.027853 |
| 12 | 24039800 | 24041199 | 15.76159 | 30.14278 | 11.92177 | 4.72438 | 1.835837 | -0.00695 | 0.027868 |
| 13 | 79717800 | 79718399 | 48.0499 | 78.88691 | 32.97011 | 14.20075 | 1.631782 | 1.444288 | 0.027926 |
| 12 | 48554600 | 48555199 | 11.5269 | 22.43124 | 10.78791 | 3.972448 | 1.653184 | -0.29501 | 0.027998 |
| 15 | 70333800 | 70334599 | 11.5269 | 22.43124 | 10.78791 | 3.972448 | 1.653184 | -0.29501 | 0.027998 |
| 7 | 9438000 | 9439599 | 21.81137 | 40.03209 | 12.30164 | 5.237334 | 2.114489 | 0.29607 | 0.028173 |
| 11 | 112836600 | 112839799 | 13.34 | 22.54955 | 72.97795 | 31.93359 | -1.30632 | 1.067738 | 0.028213 |
| 7 | 79180800 | 79186799 | 33.00533 | 52.60757 | 155.365 | 65.06244 | -1.22164 | 2.157958 | 0.028215 |
| 7 | 60787800 | 60788999 | 20.52523 | 38.4833 | 10.4353 | 4.303513 | 2.325695 | 0.161978 | 0.028223 |
| 1 | 117627600 | 117631599 | 56.45194 | 91.19847 | 50.22685 | 21.44084 | 1.218474 | 1.734885 | 0.028325 |
| 12 | 23117000 | 23122199 | 122.2814 | 189.2397 | 115.4563 | 47.95763 | 1.014961 | 2.836661 | 0.028328 |
| 14 | 113261000 | 113261199 | 9.523834 | 18.62015 | 8.276562 | 2.834907 | 1.855503 | -0.57643 | 0.028331 |
| 12 | 22682800 | 22685799 | 33.88774 | 57.54805 | 30.26659 | 13.26258 | 1.304526 | 1.053148 | 0.02838 |
| 15 | 25527000 | 25528399 | 9.06281 | 15.90764 | 53.59538 | 23.70582 | -1.35498 | 0.614076 | 0.028398 |
| 12 | 22328400 | 22328999 | 12.25943 | 23.89289 | 8.94528 | 3.239741 | 2.028978 | -0.34762 | 0.028572 |
| 17 | 39883000 | 39883999 | 19.94879 | 36.3369 | 16.55492 | 7.118167 | 1.544183 | 0.33268 | 0.028608 |
| 6 | 149002000 | 149003399 | 7.762871 | 13.59061 | 56.79404 | 25.06563 | -1.66359 | 0.653621 | 0.028624 |
| 4 | 133645400 | 133645799 | 59.53429 | 96.27583 | 35.911 | 15.3498 | 1.78048 | 1.706209 | 0.02866 |
| 3 | 102745200 | 102745799 | 42.57244 | 71.65349 | 22.36387 | 9.77316 | 2.05259 | 1.150444 | 0.028679 |
| 5 | 113065400 | 113067799 | 5.915991 | 10.4911 | 53.10297 | 23.47056 | -1.93128 | 0.472611 | 0.028695 |
| 4 | 146306800 | 146309199 | 78.32865 | 124.7461 | 74.31997 | 31.08763 | 1.082398 | 2.204322 | 0.028708 |
| 15 | 38867600 | 38871399 | 12.11667 | 20.82709 | 66.67888 | 29.24782 | -1.28732 | 0.904241 | 0.028715 |
| 7 | 8129200 | 8129799 | 8.94651 | 17.48336 | 8.445194 | 2.927259 | 1.72226 | -0.54644 | 0.028717 |
| 16 | 61876600 | 61878399 | 24.1478 | 42.37541 | 21.82588 | 9.651819 | 1.339162 | 0.645155 | 0.028736 |
| 7 | 9412200 | 9412999 | 12.10696 | 23.47822 | 11.89651 | 4.490776 | 1.547185 | -0.20095 | 0.028759 |
| 1 | 78972800 | 78974199 | 13.97408 | 27.2781 | 4.449675 | 1.57469 | 3.215729 | -0.41685 | 0.028759 |
| 6 | 83606600 | 83614399 | 31.21997 | 49.80366 | 150.8089 | 63.22788 | -1.25625 | 2.128189 | 0.028771 |
| 12 | 23486000 | 23486399 | 12.188 | 23.66299 | 11.0276 | 4.1372 | 1.67528 | -0.22284 | 0.028795 |
| 18 | 3049400 | 3053799 | 106.9672 | 167.4363 | 100.8967 | 41.8547 | 1.045376 | 2.631228 | 0.028796 |
| X | 150361000 | 150362799 | 21.05478 | 38.57331 | 14.1145 | 6.024023 | 1.863731 | 0.304064 | 0.028808 |
| 5 | 137651600 | 137653999 | 6.479828 | 11.47286 | 48.78957 | 21.57435 | -1.68266 | 0.487071 | 0.028817 |
| 7 | 44886200 | 44890999 | 16.55675 | 27.24895 | 89.59145 | 38.62841 | -1.33265 | 1.3817 | 0.028819 |
| 17 | 33468400 | 33468799 | 7.533389 | 14.7301 | 7.129321 | 2.311574 | 1.800314 | -0.79882 | 0.028921 |
| 17 | 10691600 | 10694199 | 8.574812 | 15.08841 | 52.50387 | 23.24554 | -1.40024 | 0.575704 | 0.028961 |
| 1 | 146003400 | 146003799 | 19.77874 | 36.35735 | 12.28155 | 5.217847 | 1.982755 | 0.270757 | 0.029024 |

| | | | | | | | | | |
|---|---|---|---|---|---|---|---|---|---|
| 8 | 26365400 | 26365799 | 9.219912 | 18.03529 | 5.343267 | 1.783517 | 2.447729 | -0.6803 | 0.029073 |
| 18 | 26566800 | 26567799 | 9.746591 | 19.03041 | 9.009243 | 3.17506 | 1.7312 | -0.47533 | 0.029078 |
| 12 | 22690800 | 22691599 | 12.5657 | 24.39445 | 9.873571 | 3.706514 | 1.871823 | -0.22062 | 0.029113 |
| 7 | 13978800 | 13979799 | 15.0608 | 28.56643 | 13.0345 | 5.258553 | 1.609565 | 0.044143 | 0.029162 |
| 2 | 59487600 | 59488399 | 3.675671 | 7.067611 | 32.42268 | 12.65021 | -1.64096 | -0.11896 | 0.029206 |
| 14 | 41795200 | 41795799 | 10.19621 | 19.94436 | 4.579588 | 1.533163 | 2.798181 | -0.65618 | 0.029225 |
| 16 | 72398400 | 72399599 | 16.95033 | 31.75569 | 14.66548 | 6.056079 | 1.566124 | 0.145504 | 0.029237 |
| 1 | 59095200 | 59095799 | 16.69649 | 32.38266 | 6.7682 | 2.553335 | 2.801065 | -0.16512 | 0.029252 |
| 4 | 38406600 | 38408399 | 14.70589 | 28.20038 | 12.73899 | 5.008986 | 1.657344 | -0.03952 | 0.029295 |
| 5 | 17301800 | 17304199 | 21.84087 | 38.88604 | 18.94012 | 8.343128 | 1.426399 | 0.482532 | 0.029313 |
| 11 | 110368200 | 110370999 | 8.122171 | 14.3589 | 49.87612 | 22.06151 | -1.39484 | 0.515173 | 0.029319 |
| 12 | 18641200 | 18646999 | 104.5325 | 164.0188 | 97.4244 | 40.40433 | 1.068764 | 2.587332 | 0.029352 |
| 12 | 22714400 | 22716799 | 35.75844 | 60.12733 | 33.0603 | 14.44338 | 1.240513 | 1.14619 | 0.029428 |
| 7 | 20248200 | 20248599 | 8.331752 | 16.29715 | 4.718302 | 1.555069 | 2.502333 | -0.74432 | 0.029451 |
| 7 | 140750400 | 140752799 | 5.992929 | 11.01024 | 41.04139 | 17.44943 | -1.45386 | 0.235746 | 0.029486 |
| 10 | 51223200 | 51224399 | 11.1646 | 21.72939 | 10.79363 | 3.969848 | 1.608361 | -0.30393 | 0.029523 |
| 5 | 116150000 | 116154199 | 16.72746 | 27.52702 | 90.24893 | 38.91017 | -1.32869 | 1.383366 | 0.029551 |
| 12 | 19402800 | 19404399 | 36.94136 | 61.82381 | 34.23955 | 14.91883 | 1.230345 | 1.198608 | 0.029567 |
| 2 | 124029800 | 124030399 | 9.855469 | 19.23438 | 9.860569 | 3.502018 | 1.607412 | -0.44431 | 0.029621 |
| 17 | 90128800 | 90129399 | 9.345806 | 18.28298 | 6.093791 | 2.020328 | 2.295814 | -0.69613 | 0.029626 |
| 1 | 10599400 | 10600399 | 14.00381 | 27.06226 | 11.37205 | 4.363223 | 1.793145 | -0.12779 | 0.029661 |
| 6 | 117566200 | 117566399 | 7.091494 | 13.85817 | 6.348107 | 2.035387 | 1.893045 | -0.84978 | 0.029727 |
| 5 | 111370200 | 111381399 | 46.73499 | 72.74357 | 227.9202 | 94.61105 | -1.33622 | 2.717361 | 0.029756 |
| 3 | 95531000 | 95532799 | 5.379975 | 10.01573 | 39.80026 | 16.61205 | -1.52533 | 0.14954 | 0.029774 |
| 3 | 41474200 | 41475599 | 3.247163 | 6.303764 | 32.81004 | 12.31874 | -1.76772 | -0.25654 | 0.029826 |
| 1 | 171357800 | 171359399 | 6.773242 | 12.02398 | 55.95441 | 24.70886 | -1.81107 | 0.475094 | 0.029828 |
| 2 | 104323200 | 104325199 | 7.430187 | 12.96451 | 56.16854 | 24.72707 | -1.71487 | 0.706533 | 0.029836 |
| 12 | 22457600 | 22460799 | 53.30394 | 86.28008 | 50.1279 | 21.44219 | 1.144479 | 1.68297 | 0.029848 |
| 19 | 6677400 | 6686599 | 40.4984 | 63.90443 | 187.9701 | 78.02146 | -1.22965 | 2.451314 | 0.029877 |
| 19 | 18689400 | 18691999 | 8.450584 | 14.87673 | 51.44095 | 22.77283 | -1.39102 | 0.566686 | 0.0299 |
| 2 | 117328800 | 117329599 | 3.659628 | 7.086543 | 32.51922 | 12.35941 | -1.60579 | -0.20969 | 0.030001 |
| 8 | 20126200 | 20127599 | 31.38714 | 53.60113 | 28.66958 | 12.56961 | 1.281142 | 0.990657 | 0.030033 |
| 4 | 151595600 | 151596999 | 15.84087 | 29.94572 | 13.44415 | 5.462315 | 1.625595 | 0.075861 | 0.030036 |
| 18 | 54604000 | 54604999 | 16.53135 | 31.14445 | 13.84064 | 5.661589 | 1.632009 | 0.105587 | 0.030045 |
| 12 | 103217200 | 103219799 | 9.86483 | 17.19892 | 56.8549 | 25.01674 | -1.32851 | 0.729676 | 0.030062 |
| 4 | 121914800 | 121916399 | 16.22098 | 30.29055 | 14.90301 | 6.188178 | 1.46954 | 0.159322 | 0.030069 |
| 17 | 23464000 | 23464999 | 11.34548 | 19.49179 | 63.77626 | 27.9742 | -1.31832 | 0.903511 | 0.030116 |
| 8 | 95393200 | 95393999 | 2.643629 | 5.151447 | 27.0786 | 9.82781 | -1.73154 | -0.40011 | 0.030151 |
| 12 | 80178800 | 80180199 | 5.194202 | 9.721288 | 37.70573 | 15.59714 | -1.4794 | 0.109952 | 0.030192 |
| 13 | 41258600 | 41259399 | 2.642048 | 5.153165 | 26.73606 | 9.585834 | -1.69704 | -0.44609 | 0.030206 |
| 14 | 42603000 | 42605199 | 33.48251 | 56.98824 | 28.42169 | 12.45756 | 1.381255 | 1.03068 | 0.030217 |
| X | 4831000 | 4832599 | 19.32299 | 35.39213 | 16.17006 | 6.903573 | 1.54685 | 0.296743 | 0.03027 |
| 2 | 137735800 | 137736999 | 14.57831 | 28.0701 | 11.41352 | 4.435511 | 1.821354 | -0.08137 | 0.030309 |
| 9 | 64334800 | 64336799 | 6.062221 | 11.03059 | 43.50938 | 18.7329 | -1.54794 | 0.296328 | 0.030346 |
| 19 | 12853800 | 12854399 | 10.45882 | 20.42655 | 8.41344 | 2.951344 | 1.93509 | -0.48232 | 0.030349 |
| 17 | 31595400 | 31597599 | 5.847899 | 10.77409 | 40.57587 | 17.18551 | -1.46453 | 0.216749 | 0.030354 |
| 19 | 45488400 | 45489999 | 6.315639 | 11.5414 | 41.997 | 17.9844 | -1.42726 | 0.270653 | 0.030392 |

| | | | | | | | | | |
|---|---|---|---|---|---|---|---|---|---|
| 17 | 47388400 | 47390399 | 107.7376 | 170.8466 | 52.43954 | 21.82884 | 2.031004 | 2.356215 | 0.030405 |
| 7 | 20803200 | 20804399 | 36.89658 | 61.90644 | 33.52377 | 14.63046 | 1.262415 | 1.17088 | 0.030441 |
| 2 | 117418600 | 117419599 | 4.22058 | 7.910565 | 38.2435 | 15.78076 | -1.79022 | 0.092358 | 0.030503 |
| 11 | 72182200 | 72183199 | 3.054452 | 5.926696 | 29.44512 | 11.08539 | -1.70226 | -0.25489 | 0.030512 |
| 8 | 127507600 | 127509599 | 6.451461 | 11.79986 | 44.27283 | 18.93802 | -1.47036 | 0.270872 | 0.030518 |
| 7 | 122853400 | 122857999 | 15.62936 | 26.1071 | 81.98966 | 35.68165 | -1.26405 | 1.198624 | 0.030548 |
| X | 31585400 | 31586799 | 14.37371 | 27.87269 | 10.04325 | 3.792016 | 2.032331 | -0.17915 | 0.030581 |
| 1 | 113962400 | 113964399 | 33.11194 | 56.12318 | 28.75763 | 12.59406 | 1.341676 | 1.062368 | 0.030613 |
| 10 | 122790000 | 122791999 | 7.528555 | 13.33203 | 53.14916 | 23.5013 | -1.59143 | 0.49675 | 0.030661 |
| 1 | 108886200 | 108888999 | 77.67858 | 123.8645 | 69.03099 | 28.92493 | 1.180298 | 2.161949 | 0.030663 |
| 8 | 123506600 | 123519399 | 47.07332 | 73.04635 | 231.3163 | 96.0647 | -1.35497 | 2.771927 | 0.030677 |
| 6 | 126667600 | 126667799 | 7.131018 | 13.89414 | 5.147423 | 1.59077 | 2.228264 | -0.97685 | 0.030735 |
| 8 | 121881400 | 121887799 | 25.64223 | 41.18444 | 128.6506 | 54.47713 | -1.28942 | 1.902936 | 0.03083 |
| 4 | 148744800 | 148749199 | 18.39363 | 30.01833 | 97.0986 | 41.72372 | -1.31573 | 1.502711 | 0.03084 |
| 18 | 57218200 | 57220999 | 7.64704 | 13.59721 | 48.35877 | 21.32552 | -1.4243 | 0.458667 | 0.030857 |
| 9 | 66146400 | 66149199 | 12.98297 | 21.96878 | 71.76175 | 31.41127 | -1.31932 | 1.058188 | 0.030875 |
| 13 | 66288000 | 66288999 | 45.5132 | 74.55526 | 41.94729 | 18.05027 | 1.203389 | 1.471877 | 0.030881 |
| 15 | 36629000 | 36630599 | 6.989962 | 12.30075 | 54.62566 | 24.18452 | -1.7494 | 0.566778 | 0.030894 |
| 7 | 85780800 | 85782599 | 23.12459 | 41.25554 | 17.81069 | 7.832154 | 1.600847 | 0.477379 | 0.030913 |
| 6 | 38526800 | 38529399 | 9.239829 | 16.17883 | 54.14767 | 23.90077 | -1.3456 | 0.655636 | 0.03097 |
| 16 | 24049800 | 24051599 | 5.967047 | 10.93611 | 46.01411 | 19.62837 | -1.63087 | 0.260063 | 0.030983 |
| 3 | 75463600 | 75465399 | 10.21324 | 17.74396 | 59.58155 | 26.15872 | -1.35189 | 0.784503 | 0.030985 |
| 5 | 104023600 | 104025399 | 9.735176 | 16.9793 | 59.63437 | 26.24598 | -1.41571 | 0.725282 | 0.030999 |
| 13 | 77205000 | 77205799 | 12.19052 | 23.68388 | 10.55016 | 3.936539 | 1.744227 | -0.2433 | 0.031002 |
| 4 | 96096600 | 96098599 | 15.69663 | 30.20666 | 8.794502 | 3.423146 | 2.291042 | -0.07212 | 0.031045 |
| 13 | 18118000 | 18120399 | 11.78816 | 20.26978 | 65.49674 | 28.72961 | -1.30021 | 0.8997 | 0.031132 |
| 6 | 83142200 | 83143599 | 3.248809 | 6.30214 | 30.74244 | 11.58818 | -1.68001 | -0.24726 | 0.031198 |
| 7 | 19839800 | 19839999 | 7.231671 | 14.08886 | 3.626577 | 1.11959 | 2.721966 | -0.98746 | 0.031271 |
| 4 | 154248200 | 154258599 | 44.89098 | 70.2933 | 205.283 | 85.15386 | -1.22803 | 2.608236 | 0.031296 |
| 7 | 14313800 | 14318999 | 113.5354 | 176.7946 | 109.3626 | 45.39418 | 1.001891 | 2.724914 | 0.03136 |
| 1 | 80919600 | 80920599 | 10.78063 | 20.99717 | 10.98085 | 4.010918 | 1.544784 | -0.33135 | 0.031419 |
| 3 | 111626800 | 111627799 | 10.78063 | 20.99717 | 10.98085 | 4.010918 | 1.544784 | -0.33135 | 0.031419 |
| 17 | 28586800 | 28588399 | 4.991133 | 9.188882 | 40.4898 | 17.16612 | -1.68916 | 0.214321 | 0.031429 |
| 15 | 100365600 | 100367799 | 6.575063 | 11.73944 | 45.74469 | 20.10273 | -1.55052 | 0.421854 | 0.031441 |
| 4 | 62650200 | 62651599 | 3.576614 | 6.839762 | 33.84632 | 13.41395 | -1.77057 | -0.06451 | 0.031472 |
| 7 | 22902400 | 22902599 | 6.813625 | 13.28679 | 5.850079 | 1.823957 | 1.977343 | -0.95119 | 0.031473 |
| 7 | 33027800 | 33030799 | 65.48597 | 105.4265 | 51.49646 | 21.881 | 1.385489 | 1.852115 | 0.031554 |
| 6 | 46559200 | 46559599 | 8.773761 | 17.16277 | 6.077225 | 2.026352 | 2.201296 | -0.68869 | 0.03159 |
| 2 | 132785600 | 132788999 | 14.98964 | 25.06282 | 77.83435 | 33.89402 | -1.24773 | 1.183763 | 0.031615 |
| 8 | 123930800 | 123937599 | 20.99093 | 33.78808 | 123.4486 | 52.4415 | -1.51054 | 1.825259 | 0.031618 |
| 4 | 23480800 | 23481999 | 14.70445 | 27.83127 | 14.33689 | 5.810053 | 1.431663 | 0.058801 | 0.031636 |
| 12 | 59107400 | 59109399 | 7.519181 | 13.33993 | 50.46226 | 22.29522 | -1.51476 | 0.475251 | 0.031659 |
| 14 | 41858800 | 41863399 | 46.26785 | 75.69947 | 43.52669 | 18.72081 | 1.171229 | 1.489331 | 0.031686 |
| 7 | 9852000 | 9853199 | 38.51168 | 64.19449 | 35.75027 | 15.53781 | 1.222514 | 1.245639 | 0.03173 |
| 5 | 83178600 | 83180399 | 24.01039 | 42.2176 | 21.59043 | 9.560366 | 1.349297 | 0.614778 | 0.031752 |
| 3 | 101851000 | 101852599 | 4.281399 | 8.192159 | 35.23389 | 13.94138 | -1.5704 | -0.06092 | 0.031861 |
| 11 | 118558800 | 118561199 | 8.223711 | 14.35192 | 56.7795 | 24.99914 | -1.58544 | 0.706557 | 0.031869 |

| | | | | | | | | | |
|---|---|---|---|---|---|---|---|---|---|
| 15 | 95612000 | 95614399 | 10.79953 | 18.77885 | 62.97431 | 27.66336 | -1.35014 | 0.779293 | 0.031955 |
| 19 | 12573000 | 12574599 | 6.076915 | 10.68761 | 51.88459 | 22.9737 | -1.87594 | 0.566186 | 0.032083 |
| X | 111280400 | 111280999 | 207.3509 | 310.5978 | 159.4839 | 65.78546 | 1.225521 | 3.49823 | 0.032101 |
| 4 | 132556200 | 132557199 | 2.946762 | 5.731104 | 28.61098 | 10.58406 | -1.68568 | -0.32331 | 0.032128 |
| X | 32626000 | 32626599 | 18.92149 | 35.20716 | 13.728 | 5.73493 | 1.795478 | 0.202438 | 0.032178 |
| 7 | 59418400 | 59420199 | 49.1389 | 80.40968 | 37.66279 | 16.19969 | 1.466201 | 1.490964 | 0.032186 |
| 2 | 68178400 | 68181199 | 13.27247 | 22.27904 | 78.98916 | 34.48224 | -1.43797 | 1.135565 | 0.032251 |
| 15 | 7163600 | 7164799 | 3.966803 | 7.647243 | 35.20844 | 13.63503 | -1.63784 | -0.136 | 0.032275 |
| 6 | 51612600 | 51614399 | 4.94409 | 9.234346 | 38.7559 | 16.08698 | -1.59639 | 0.122197 | 0.032319 |
| 1 | 99299200 | 99300199 | 12.27578 | 23.86495 | 10.57808 | 3.926412 | 1.758418 | -0.25943 | 0.032383 |
| 4 | 149691400 | 149696199 | 14.06064 | 23.50919 | 77.68352 | 33.82803 | -1.33675 | 1.181178 | 0.032423 |
| 8 | 73245400 | 73246999 | 6.995549 | 12.5532 | 44.77751 | 19.5944 | -1.41997 | 0.390045 | 0.032436 |
| 18 | 18778200 | 18778799 | 13.30087 | 25.5011 | 13.02109 | 5.122781 | 1.481804 | -0.0521 | 0.032498 |
| 4 | 138965200 | 138966399 | 2.864885 | 5.524075 | 32.45827 | 12.56266 | -1.98033 | -0.15476 | 0.032555 |
| 5 | 145140800 | 145142399 | 4.716181 | 8.976925 | 36.17771 | 14.487 | -1.49371 | -0.01436 | 0.032567 |
| 10 | 33372800 | 33374399 | 20.30008 | 36.79727 | 16.80707 | 7.270704 | 1.534689 | 0.362167 | 0.032609 |
| 7 | 140934600 | 140942599 | 34.71675 | 55.07197 | 168.8953 | 70.33027 | -1.28312 | 2.311168 | 0.032678 |
| 1 | 14026200 | 14026799 | 9.461906 | 18.4826 | 9.030159 | 3.158663 | 1.695906 | -0.50462 | 0.032696 |
| 8 | 111397400 | 111400799 | 10.68176 | 18.20711 | 69.59565 | 30.50816 | -1.54359 | 0.979568 | 0.032727 |
| 11 | 115402800 | 115405399 | 7.321016 | 12.82733 | 56.77613 | 25.07471 | -1.74575 | 0.637817 | 0.032756 |
| 10 | 127994000 | 127994599 | 2.428514 | 4.744391 | 25.33913 | 8.843889 | -1.70006 | -0.54848 | 0.032781 |
| 7 | 144730200 | 144731399 | 3.878687 | 7.396031 | 34.93882 | 13.93807 | -1.71409 | -0.0377 | 0.032811 |
| 17 | 25487200 | 25490799 | 10.34687 | 18.01089 | 57.97262 | 25.48326 | -1.29082 | 0.75525 | 0.032813 |
| 13 | 65985600 | 65987199 | 23.06544 | 41.87604 | 12.60766 | 5.443283 | 2.13062 | 0.365136 | 0.032829 |
| X | 4077200 | 4077599 | 8.314122 | 16.26394 | 7.73565 | 2.558403 | 1.802427 | -0.71499 | 0.032836 |
| 2 | 146674000 | 146678599 | 26.27896 | 42.22104 | 127.9168 | 54.19271 | -1.24487 | 1.892669 | 0.032847 |
| 14 | 25555000 | 25557399 | 9.055646 | 15.91802 | 51.971 | 23.01437 | -1.30978 | 0.588147 | 0.032861 |
| 7 | 32847600 | 32850799 | 34.31083 | 58.28647 | 27.72303 | 12.14927 | 1.448798 | 1.048699 | 0.03288 |
| 12 | 114830200 | 114832399 | 61.52566 | 99.14141 | 53.62401 | 22.81524 | 1.24041 | 1.818932 | 0.032881 |
| 4 | 32358000 | 32364199 | 22.95508 | 37.14644 | 108.3813 | 46.35117 | -1.17806 | 1.666704 | 0.032935 |
| 4 | 60674000 | 60675999 | 24.77081 | 43.36948 | 23.30145 | 10.28447 | 1.2788 | 0.686756 | 0.032975 |
| 8 | 8357200 | 8358599 | 3.866219 | 7.407964 | 38.41248 | 15.13895 | -1.832 | -0.07196 | 0.032991 |
| 4 | 147448200 | 147448799 | 13.10962 | 25.5714 | 6.957323 | 2.492105 | 2.491302 | -0.3799 | 0.03302 |
| 7 | 14405800 | 14408799 | 91.65644 | 144.9792 | 86.81218 | 36.07755 | 1.066031 | 2.409334 | 0.033041 |
| 3 | 47540400 | 47543399 | 39.80958 | 66.33383 | 35.75403 | 15.53588 | 1.269575 | 1.254721 | 0.033066 |
| 5 | 4354400 | 4355599 | 18.03928 | 33.60406 | 13.69618 | 5.710566 | 1.732708 | 0.186794 | 0.033071 |
| 7 | 59579800 | 59580799 | 13.12807 | 25.4733 | 11.13707 | 4.192605 | 1.760659 | -0.19982 | 0.033084 |
| 3 | 51606400 | 51610799 | 13.51224 | 22.81881 | 73.55253 | 32.17566 | -1.30068 | 1.07683 | 0.033101 |
| 14 | 42466200 | 42467599 | 22.04987 | 39.55838 | 16.5461 | 7.241893 | 1.649785 | 0.43268 | 0.033104 |
| X | 3880600 | 3881799 | 11.69162 | 22.78849 | 9.787213 | 3.540153 | 1.837352 | -0.35636 | 0.033109 |
| 7 | 22132800 | 22133199 | 12.11488 | 23.46364 | 12.46976 | 4.735298 | 1.471497 | -0.17962 | 0.033145 |
| 16 | 3272200 | 3277399 | 116.6251 | 181.4334 | 109.8784 | 45.61477 | 1.031386 | 2.743244 | 0.033157 |
| 7 | 25087200 | 25088599 | 3.978754 | 7.633463 | 32.93875 | 12.93351 | -1.56332 | -0.09404 | 0.03316 |
| 17 | 18375400 | 18376199 | 10.23619 | 19.98462 | 8.389938 | 2.961249 | 1.895788 | -0.46576 | 0.033177 |
| 18 | 3051200 | 3053199 | 79.71883 | 126.7547 | 79.03004 | 32.98597 | 1.014515 | 2.258054 | 0.033202 |
| 7 | 11458600 | 11459399 | 29.86975 | 51.4109 | 28.10969 | 12.33045 | 1.251293 | 0.915684 | 0.033273 |
| 5 | 111204600 | 111207799 | 11.34952 | 19.53666 | 63.30761 | 27.77028 | -1.30394 | 0.883962 | 0.033284 |

| | | | | | | | | | |
|---|---|---|---|---|---|---|---|---|---|
| 7 | 14234200 | 14235399 | 45.56844 | 75.06154 | 35.09301 | 15.13635 | 1.47259 | 1.401202 | 0.033348 |
| 9 | 27124400 | 27124999 | 17.40337 | 32.63107 | 10.84842 | 4.473642 | 2.033204 | 0.133493 | 0.033364 |
| 12 | 111370000 | 111381599 | 49.97287 | 77.79963 | 223.8035 | 92.89929 | -1.21295 | 2.714583 | 0.033393 |
| 1 | 130466800 | 130467999 | 12.09569 | 23.49906 | 11.18494 | 4.174245 | 1.650384 | -0.24146 | 0.033439 |
| 5 | 12961000 | 12962199 | 14.81051 | 28.34351 | 13.16257 | 5.205741 | 1.609973 | -0.01925 | 0.033447 |
| 11 | 76346400 | 76348399 | 4.974556 | 9.204815 | 39.53451 | 16.64613 | -1.64525 | 0.182636 | 0.033456 |
| 10 | 128117200 | 128120999 | 10.61869 | 17.92249 | 75.37637 | 32.96801 | -1.6821 | 1.074382 | 0.033459 |
| 1 | 22224200 | 22225199 | 25.20883 | 44.64841 | 17.75677 | 7.851376 | 1.713052 | 0.539532 | 0.033494 |
| 8 | 70933200 | 70953999 | 115.2714 | 169.4139 | 474.555 | 195.4376 | -1.2459 | 3.826412 | 0.033548 |
| 10 | 90959000 | 90960199 | 4.876204 | 9.176569 | 36.16962 | 14.82274 | -1.4907 | 0.068259 | 0.033556 |
| 11 | 115205000 | 115231799 | 163.6233 | 234.6834 | 600.1217 | 247.8175 | -1.14894 | 4.200806 | 0.033588 |
| 9 | 66925000 | 66926599 | 4.715553 | 8.97772 | 35.58599 | 14.24337 | -1.46931 | -0.01716 | 0.033598 |
| 1 | 191448800 | 191451599 | 10.5226 | 18.27729 | 60.16121 | 26.40944 | -1.32349 | 0.789479 | 0.03363 |
| 14 | 26911400 | 26916399 | 19.17762 | 31.35858 | 97.16811 | 41.78312 | -1.2521 | 1.474773 | 0.033664 |
| X | 78400600 | 78401799 | 13.30526 | 25.87711 | 9.291502 | 3.437264 | 2.06139 | -0.26185 | 0.03367 |
| 4 | 156535200 | 156535599 | 8.054154 | 15.75417 | 6.892482 | 2.310126 | 1.90329 | -0.68162 | 0.033691 |
| 5 | 74217400 | 74217799 | 1.714685 | 3.35282 | 22.05759 | 7.159435 | -1.89126 | -0.82939 | 0.033698 |
| 7 | 42649800 | 42652199 | 25.17896 | 44.18405 | 22.05391 | 9.752495 | 1.384146 | 0.651986 | 0.033714 |
| 13 | 88014800 | 88015199 | 15.04708 | 28.59606 | 13.93775 | 5.599836 | 1.521999 | 0.034675 | 0.033753 |
| 14 | 10765800 | 10767799 | 7.189996 | 12.82248 | 46.80358 | 20.5882 | -1.45863 | 0.434061 | 0.033794 |
| 14 | 50870200 | 50872799 | 35.20722 | 60.38887 | 23.96417 | 10.51032 | 1.710155 | 0.963848 | 0.033798 |
| 15 | 64049600 | 64050599 | 2.739096 | 5.346271 | 27.73868 | 9.834769 | -1.68287 | -0.48189 | 0.033823 |
| 12 | 86528400 | 86540199 | 65.19252 | 100.3732 | 251.0385 | 104.2666 | -1.02391 | 2.924845 | 0.033841 |
| 13 | 23922600 | 23923999 | 4.724957 | 8.602842 | 41.81333 | 17.9883 | -1.84347 | 0.281563 | 0.033881 |
| 3 | 146204400 | 146206399 | 6.849218 | 12.31599 | 44.81821 | 19.57815 | -1.4469 | 0.375207 | 0.033904 |
| 12 | 22441600 | 22444999 | 47.73547 | 77.96996 | 44.06994 | 18.94289 | 1.19423 | 1.516196 | 0.033924 |
| 7 | 7467200 | 7470999 | 38.19321 | 64.02378 | 30.94714 | 13.49823 | 1.425612 | 1.182116 | 0.033936 |
| 6 | 115651400 | 115653999 | 11.0182 | 18.97324 | 62.15078 | 27.26313 | -1.31908 | 0.879374 | 0.033967 |
| 17 | 14004600 | 14006799 | 25.87205 | 45.45548 | 19.62971 | 8.689405 | 1.591086 | 0.632219 | 0.033968 |
| 19 | 39394000 | 39396199 | 41.68645 | 68.89232 | 39.18054 | 16.92949 | 1.192003 | 1.354762 | 0.033976 |
| 9 | 107335400 | 107343399 | 30.55699 | 48.78968 | 145.5013 | 61.07968 | -1.2328 | 2.098265 | 0.033998 |
| 10 | 81717000 | 81718599 | 28.5684 | 49.95 | 19.51894 | 8.605038 | 1.73466 | 0.722054 | 0.034009 |
| 15 | 77839600 | 77844999 | 22.22132 | 35.84719 | 115.9142 | 49.38993 | -1.33102 | 1.755769 | 0.034021 |
| 15 | 32242000 | 32246599 | 22.96694 | 37.13416 | 109.2723 | 46.69631 | -1.19155 | 1.687033 | 0.034044 |
| 1 | 113606800 | 113611799 | 68.7148 | 110.1797 | 58.08808 | 24.53496 | 1.268852 | 1.981502 | 0.034141 |
| 10 | 89725600 | 89726199 | 10.84431 | 21.1862 | 7.754952 | 2.705436 | 2.104581 | -0.49721 | 0.034175 |
| Y | 4900200 | 4902399 | 19.10404 | 34.70378 | 12.7847 | 5.515783 | 1.843678 | 0.330046 | 0.034186 |
| 12 | 63494600 | 63495399 | 12.64057 | 24.6082 | 9.330746 | 3.42487 | 1.99365 | -0.29626 | 0.034211 |
| 14 | 4340800 | 4341199 | 2.739574 | 5.345586 | 26.24009 | 9.352113 | -1.61014 | -0.46674 | 0.034223 |
| 4 | 107594200 | 107598399 | 12.30895 | 20.72134 | 74.36467 | 32.49984 | -1.45488 | 1.102365 | 0.034253 |
| 10 | 85036400 | 85037799 | 25.81415 | 45.14255 | 23.77024 | 10.48086 | 1.307876 | 0.712256 | 0.034255 |
| 4 | 122356200 | 122356599 | 10.5681 | 20.61378 | 9.996106 | 3.581009 | 1.67713 | -0.4037 | 0.034323 |
| 10 | 62099400 | 62102799 | 13.73699 | 22.9896 | 76.52979 | 33.34535 | -1.34726 | 1.169275 | 0.034347 |
| 12 | 22662400 | 22662999 | 11.6086 | 22.60527 | 10.81424 | 3.960431 | 1.668161 | -0.31187 | 0.034352 |
| 17 | 38009000 | 38010199 | 10.84821 | 21.17627 | 9.045725 | 3.20114 | 1.87025 | -0.44462 | 0.034362 |
| 7 | 6002600 | 6004599 | 17.32184 | 32.91528 | 11.01539 | 4.426787 | 2.055551 | 0.049301 | 0.03437 |
| 18 | 72403000 | 72404399 | 8.574912 | 15.15385 | 49.88179 | 22.06559 | -1.31817 | 0.521838 | 0.034383 |

| | | | | | | | | | |
|---|---|---|---|---|---|---|---|---|---|
| 14 | 42249600 | 42253199 | 29.1205 | 50.20562 | 27.55387 | 12.08725 | 1.246252 | 0.896304 | 0.034402 |
| 17 | 47884200 | 47886199 | 5.489826 | 10.2904 | 38.39823 | 15.84225 | -1.42172 | 0.102888 | 0.034403 |
| 2 | 165334800 | 165336399 | 5.084117 | 9.455356 | 38.01944 | 15.896 | -1.54331 | 0.15119 | 0.034429 |
| 2 | 165229800 | 165230399 | 1.641752 | 3.211363 | 21.37082 | 7.153771 | -1.94728 | -0.7235 | 0.034448 |
| 1 | 124689200 | 124689999 | 11.97609 | 23.34138 | 9.597651 | 3.474653 | 1.897195 | -0.35029 | 0.034453 |
| 7 | 51510800 | 51512599 | 11.37551 | 19.51011 | 64.30211 | 28.20173 | -1.3293 | 0.91925 | 0.034482 |
| 3 | 112603800 | 112604599 | 12.76651 | 24.71469 | 12.31952 | 4.687993 | 1.559878 | -0.16569 | 0.034514 |
| 7 | 60376200 | 60378199 | 29.11272 | 50.22967 | 27.70989 | 12.15598 | 1.239209 | 0.891208 | 0.034519 |
| 3 | 37826200 | 37829199 | 13.09986 | 22.36655 | 67.67818 | 29.67165 | -1.20811 | 0.970863 | 0.034543 |
| 13 | 62863000 | 62864999 | 59.95627 | 96.95517 | 45.82548 | 19.58684 | 1.440027 | 1.713135 | 0.034554 |
| 12 | 113784000 | 113784999 | 35.41361 | 59.50628 | 34.10829 | 14.8981 | 1.180738 | 1.151662 | 0.034603 |
| 8 | 46752200 | 46752799 | 2.338598 | 4.561939 | 25.39468 | 9.088276 | -1.7915 | -0.45923 | 0.034626 |
| 5 | 114347000 | 114359399 | 72.15765 | 110.4165 | 271.7834 | 112.7732 | -1.00779 | 3.05069 | 0.034635 |
| 10 | 126270400 | 126271599 | 15.17439 | 28.70037 | 14.45867 | 5.868201 | 1.46225 | 0.068959 | 0.034638 |
| 10 | 56031800 | 56032399 | 12.59517 | 24.32836 | 13.001 | 4.99944 | 1.447046 | -0.13145 | 0.034644 |
| 10 | 55290800 | 55291199 | 9.060353 | 17.72291 | 7.471715 | 2.49712 | 1.958412 | -0.66862 | 0.034694 |
| 1 | 36977800 | 36979199 | 6.322417 | 11.53213 | 41.39094 | 17.76651 | -1.40969 | 0.280919 | 0.034719 |
| 4 | 45548400 | 45551399 | 11.45462 | 19.83988 | 62.95139 | 27.61923 | -1.27192 | 0.826788 | 0.034731 |
| 15 | 102552000 | 102552799 | 10.8894 | 21.18117 | 11.08344 | 4.096082 | 1.529196 | -0.29006 | 0.034732 |
| 2 | 179691800 | 179697799 | 24.49459 | 39.41069 | 122.5534 | 52.02857 | -1.27962 | 1.843012 | 0.034749 |
| 7 | 22137200 | 22137599 | 15.22463 | 28.99112 | 13.82824 | 5.532497 | 1.557915 | 0.023004 | 0.034774 |
| 7 | 11275800 | 11276399 | 14.36302 | 27.39103 | 13.53249 | 5.397207 | 1.511711 | 0.00363 | 0.034794 |
| 7 | 10837800 | 10839799 | 43.18746 | 71.08568 | 39.79773 | 17.1601 | 1.213102 | 1.406758 | 0.034829 |
| 2 | 118218800 | 118219599 | 4.394892 | 8.439443 | 34.30936 | 13.43691 | -1.47585 | -0.09571 | 0.034881 |
| 18 | 65546000 | 65547199 | 21.67101 | 38.90027 | 16.55129 | 7.240778 | 1.62567 | 0.426425 | 0.034915 |
| 7 | 20121400 | 20121799 | 8.382891 | 16.39173 | 8.922434 | 3.04534 | 1.571969 | -0.60472 | 0.034931 |
| 19 | 20819400 | 20819999 | 8.382891 | 16.39173 | 8.922434 | 3.04534 | 1.571969 | -0.60472 | 0.034931 |
| 17 | 49754400 | 49754799 | 23.96819 | 42.29967 | 17.75703 | 7.857097 | 1.633862 | 0.568005 | 0.034935 |
| 13 | 60894400 | 60902199 | 42.11552 | 66.83066 | 166.8573 | 69.50435 | -0.98664 | 2.306243 | 0.034987 |
| 14 | 19183800 | 19185599 | 18.29849 | 33.80192 | 16.12304 | 6.804301 | 1.496734 | 0.242358 | 0.035041 |
| 12 | 23707800 | 23710399 | 60.81921 | 97.73417 | 58.85747 | 24.94239 | 1.081097 | 1.904148 | 0.035042 |
| 19 | 51151200 | 51152799 | 17.99081 | 33.33875 | 15.43726 | 6.486932 | 1.542936 | 0.22059 | 0.035057 |
| 11 | 106810400 | 106812199 | 6.425575 | 11.51812 | 45.28143 | 19.83274 | -1.55986 | 0.393629 | 0.035095 |
| 12 | 19644600 | 19646999 | 87.32087 | 138.1889 | 82.3137 | 34.21653 | 1.074116 | 2.393755 | 0.035108 |
| 4 | 122598600 | 122600599 | 22.00974 | 39.24479 | 18.82362 | 8.281458 | 1.449507 | 0.474509 | 0.035111 |
| 7 | 59838000 | 59839199 | 22.19871 | 39.29825 | 18.44395 | 8.155998 | 1.475739 | 0.52544 | 0.035119 |
| 1 | 134238600 | 134257999 | 75.3967 | 113.5118 | 358.2693 | 147.972 | -1.38575 | 3.387014 | 0.035121 |
| 15 | 81575600 | 81582399 | 30.04535 | 48.00084 | 141.7799 | 59.56687 | -1.21797 | 2.079134 | 0.035145 |
| X | 32723400 | 32724999 | 20.45338 | 38.24841 | 11.65215 | 4.826675 | 2.156051 | 0.180228 | 0.035183 |
| 13 | 58362800 | 58364199 | 5.333973 | 9.701177 | 42.36108 | 18.24865 | -1.69286 | 0.292357 | 0.035204 |
| 7 | 9408600 | 9411399 | 20.76225 | 38.04308 | 15.02448 | 6.41109 | 1.755819 | 0.303662 | 0.035269 |
| 7 | 23116800 | 23119399 | 76.19554 | 121.5806 | 69.65761 | 29.21449 | 1.141569 | 2.142491 | 0.035302 |
| 9 | 38305600 | 38307999 | 27.98124 | 48.71575 | 24.82531 | 10.91377 | 1.354715 | 0.787631 | 0.03532 |
| 1 | 135782400 | 135785599 | 10.91701 | 18.66278 | 65.23054 | 28.60141 | -1.41413 | 0.950188 | 0.035404 |
| 15 | 102418800 | 102420599 | 6.839961 | 12.32557 | 44.33434 | 19.3211 | -1.42795 | 0.361123 | 0.035448 |
| 4 | 121702400 | 121703599 | 11.53547 | 22.41412 | 11.96075 | 4.458248 | 1.490105 | -0.2497 | 0.035471 |
| 9 | 95407200 | 95408999 | 4.280514 | 8.193178 | 34.04084 | 13.45652 | -1.51947 | -0.06665 | 0.035472 |

| | | | | | | | | | |
|---|---|---|---|---|---|---|---|---|---|
| 7 | 11014400 | 11019199 | 106.4982 | 166.5528 | 105.2962 | 43.68378 | 0.975279 | 2.647097 | 0.03549 |
| 7 | 9557800 | 9558999 | 13.50071 | 26.23546 | 9.914427 | 3.692365 | 1.981158 | -0.23501 | 0.035534 |
| 4 | 155271800 | 155278199 | 19.85663 | 32.26949 | 100.1789 | 42.9779 | -1.26118 | 1.570937 | 0.035549 |
| 5 | 59052400 | 59053799 | 13.59774 | 26.00823 | 13.88822 | 5.501439 | 1.40858 | -0.02501 | 0.035551 |
| 11 | 89624200 | 89625199 | 4.734168 | 8.954475 | 35.23478 | 14.299 | -1.47619 | 0.029222 | 0.035572 |
| 7 | 115903400 | 115905599 | 6.483503 | 11.46994 | 48.25544 | 21.34166 | -1.66764 | 0.495713 | 0.035599 |
| 5 | 136158400 | 136160199 | 5.510381 | 10.13209 | 40.46434 | 17.18471 | -1.55024 | 0.227971 | 0.035632 |
| 8 | 29142600 | 29142999 | 8.948369 | 17.48081 | 9.469638 | 3.308013 | 1.549829 | -0.51561 | 0.035656 |
| 6 | 113437800 | 113439799 | 5.954401 | 11.06649 | 41.50623 | 17.37201 | -1.44649 | 0.168807 | 0.035713 |
| X | 153306800 | 153307199 | 11.02908 | 21.53737 | 8.405719 | 2.955552 | 2.005599 | -0.46582 | 0.035716 |
| 19 | 8824600 | 8825999 | 2.186241 | 4.216808 | 34.15338 | 13.20709 | -2.43185 | -0.16053 | 0.035734 |
| 17 | 47799400 | 47804599 | 18.76248 | 30.42572 | 104.0211 | 44.55542 | -1.40276 | 1.615799 | 0.035753 |
| 18 | 7531600 | 7532599 | 12.4703 | 24.20536 | 11.77239 | 4.424615 | 1.611245 | -0.21087 | 0.035768 |
| 12 | 97015600 | 97016999 | 24.40222 | 42.80821 | 22.83059 | 10.0934 | 1.289359 | 0.653909 | 0.035773 |
| 3 | 109235600 | 109238799 | 15.56804 | 26.12895 | 77.52247 | 33.83918 | -1.18221 | 1.14429 | 0.035775 |
| 7 | 115629600 | 115633999 | 16.16168 | 26.47702 | 93.64723 | 40.29883 | -1.44032 | 1.441311 | 0.035788 |
| 10 | 17197600 | 17197999 | 16.34111 | 30.7805 | 14.72777 | 6.026315 | 1.525901 | 0.10724 | 0.035793 |
| 14 | 42470000 | 42472399 | 16.34111 | 30.7805 | 14.72777 | 6.026315 | 1.525901 | 0.10724 | 0.035793 |
| X | 136865200 | 136866399 | 22.54921 | 41.75579 | 10.7204 | 4.509268 | 2.382434 | 0.251807 | 0.035839 |
| 11 | 50100800 | 50102199 | 5.05524 | 9.489003 | 36.67627 | 15.09614 | -1.46851 | 0.086637 | 0.03587 |
| 7 | 136024800 | 136026799 | 7.918377 | 14.04349 | 48.8314 | 21.58071 | -1.3939 | 0.480039 | 0.035899 |
| 8 | 13607400 | 13614599 | 33.87592 | 54.1934 | 138.8125 | 58.43126 | -1.01082 | 2.039987 | 0.035945 |
| 14 | 65580600 | 65582599 | 6.498539 | 11.82152 | 42.42306 | 18.27094 | -1.4131 | 0.299104 | 0.03595 |
| 10 | 44305400 | 44308399 | 11.09989 | 19.00277 | 66.45648 | 29.14233 | -1.41494 | 0.936793 | 0.03601 |
| 11 | 109541200 | 109542799 | 5.966595 | 10.76456 | 44.88117 | 19.53035 | -1.6374 | 0.348879 | 0.036022 |
| 12 | 74818600 | 74819399 | 10.8501 | 21.17148 | 9.66942 | 3.445516 | 1.767387 | -0.41909 | 0.036096 |
| 7 | 21977400 | 21977599 | 7.859249 | 15.37078 | 7.764318 | 2.543168 | 1.72911 | -0.75527 | 0.036137 |
| 7 | 103284600 | 103285199 | 10.31396 | 20.1187 | 10.28963 | 3.68465 | 1.6003 | -0.40806 | 0.036138 |
| 4 | 67678200 | 67678599 | 7.736566 | 15.12998 | 5.73848 | 1.874878 | 2.128819 | -0.77812 | 0.036143 |
| 2 | 130405200 | 130407199 | 5.869797 | 10.37889 | 48.24158 | 21.33773 | -1.80978 | 0.498914 | 0.036153 |
| 18 | 39048000 | 39054199 | 22.98102 | 37.15588 | 110.5761 | 47.25177 | -1.20786 | 1.687926 | 0.036198 |
| 11 | 104074600 | 104089399 | 73.59575 | 111.9914 | 296.4818 | 122.85 | -1.11919 | 3.16625 | 0.036221 |
| 14 | 42586000 | 42587399 | 21.96736 | 38.91247 | 20.2746 | 8.964423 | 1.326415 | 0.521612 | 0.036253 |
| 15 | 81400200 | 81400399 | 8.6568 | 16.93456 | 7.311281 | 2.419819 | 1.939805 | -0.70835 | 0.036255 |
| 17 | 27488200 | 27493999 | 124.1546 | 191.8184 | 123.3582 | 51.24033 | 0.937256 | 2.868173 | 0.036311 |
| X | 152899000 | 152899999 | 15.33006 | 29.60236 | 10.18198 | 3.920739 | 2.071017 | -0.10514 | 0.036311 |
| 7 | 132894600 | 132897399 | 11.64194 | 20.08571 | 63.97837 | 28.06646 | -1.27876 | 0.864222 | 0.036352 |
| 8 | 85762600 | 85765199 | 8.946327 | 15.62057 | 54.65206 | 24.06977 | -1.40918 | 0.701147 | 0.036358 |
| 3 | 87695200 | 87697399 | 8.652571 | 15.13835 | 54.12168 | 23.87304 | -1.44003 | 0.665815 | 0.036376 |
| 2 | 69036000 | 69036799 | 3.964657 | 7.649627 | 32.12846 | 12.41183 | -1.50265 | -0.15093 | 0.036377 |
| 19 | 44667200 | 44675999 | 40.86529 | 64.76158 | 174.2984 | 72.50849 | -1.09665 | 2.34511 | 0.036416 |
| 12 | 23507400 | 23508999 | 21.44065 | 38.55806 | 17.15687 | 7.494002 | 1.563367 | 0.415394 | 0.036445 |
| 5 | 121965800 | 121984599 | 82.69299 | 124.1606 | 370.0959 | 152.746 | -1.30683 | 3.441747 | 0.036449 |
| 19 | 34007800 | 34008799 | 14.42243 | 27.66058 | 13.34096 | 5.243628 | 1.564685 | -0.04179 | 0.036462 |
| 11 | 114657400 | 114661799 | 15.85301 | 26.27348 | 82.53415 | 35.7319 | -1.26482 | 1.290518 | 0.03652 |
| 12 | 23926400 | 23930999 | 81.9431 | 130.0876 | 83.18185 | 34.66453 | 0.976002 | 2.305145 | 0.036533 |
| 7 | 21988600 | 21988999 | 7.372717 | 14.41862 | 7.769989 | 2.540183 | 1.63888 | -0.77155 | 0.03656 |

| | | | | | | | | | |
|---|---|---|---|---|---|---|---|---|---|
| 12 | 20428600 | 20431199 | 76.19986 | 121.313 | 77.91086 | 32.57377 | 0.973747 | 2.214438 | 0.036585 |
| 5 | 122266400 | 122277799 | 47.20238 | 73.8875 | 207.9543 | 86.26497 | -1.17526 | 2.615113 | 0.036591 |
| 9 | 96057200 | 96057999 | 11.82674 | 22.93673 | 12.50274 | 4.717768 | 1.443912 | -0.20341 | 0.036629 |
| 12 | 56571800 | 56572599 | 11.82674 | 22.93673 | 12.50274 | 4.717768 | 1.443912 | -0.20341 | 0.036629 |
| 10 | 17334000 | 17334799 | 11.03091 | 21.53214 | 9.043058 | 3.202146 | 1.893798 | -0.43984 | 0.036651 |
| 2 | 141750200 | 141751199 | 3.777341 | 7.309808 | 35.70386 | 13.61039 | -1.69968 | -0.19009 | 0.036676 |
| 15 | 78660000 | 78661199 | 16.82294 | 27.6637 | 89.9827 | 38.78326 | -1.31762 | 1.39494 | 0.036749 |
| 1 | 116588600 | 116592399 | 64.86965 | 104.3488 | 49.8396 | 21.15188 | 1.416243 | 1.873187 | 0.036756 |
| 4 | 147377600 | 147378999 | 31.64645 | 55.08715 | 20.71017 | 9.103641 | 1.790447 | 0.802482 | 0.036757 |
| 11 | 88608400 | 88609999 | 4.935618 | 9.242627 | 37.72122 | 15.58841 | -1.55112 | 0.102721 | 0.036758 |
| 7 | 22985600 | 22991799 | 103.9725 | 162.9711 | 102.0566 | 42.32994 | 0.991484 | 2.606149 | 0.036811 |
| 2 | 142446400 | 142446999 | 2.944088 | 5.73387 | 27.63887 | 10.1015 | -1.61947 | -0.37008 | 0.036827 |
| 17 | 10029200 | 10029599 | 11.81171 | 22.96553 | 11.22034 | 4.160757 | 1.621494 | -0.26637 | 0.03683 |
| 17 | 17386400 | 17388799 | 61.34221 | 98.76452 | 52.40972 | 22.27203 | 1.266467 | 1.843376 | 0.036862 |
| 9 | 101402200 | 101402999 | 12.01043 | 23.28997 | 12.49959 | 4.719434 | 1.465386 | -0.19915 | 0.036884 |
| 16 | 3207600 | 3223199 | 81.61783 | 129.7231 | 70.64349 | 29.47395 | 1.208784 | 2.267378 | 0.036893 |
| 10 | 117545600 | 117548799 | 10.43515 | 17.97867 | 62.69801 | 27.50362 | -1.40881 | 0.8714 | 0.036946 |
| 1 | 51601800 | 51602999 | 3.246527 | 6.304339 | 29.64355 | 11.10408 | -1.61884 | -0.2724 | 0.03699 |
| 13 | 66003000 | 66006199 | 70.91392 | 113.3998 | 68.71659 | 28.90885 | 1.064177 | 2.069514 | 0.037015 |
| 15 | 17515000 | 17516399 | 18.33651 | 33.70588 | 16.7743 | 7.126708 | 1.429506 | 0.270965 | 0.037066 |
| 9 | 32420800 | 32421799 | 2.942401 | 5.735597 | 27.95013 | 10.09898 | -1.62029 | -0.41166 | 0.037066 |
| 5 | 132677800 | 132679399 | 4.589673 | 8.73853 | 36.1881 | 14.48273 | -1.5319 | -0.0182 | 0.037068 |
| 3 | 36916800 | 36918199 | 7.080726 | 12.55932 | 48.29345 | 21.34092 | -1.5379 | 0.470801 | 0.037072 |
| 11 | 88509600 | 88511399 | 4.323353 | 8.146153 | 35.59435 | 14.55669 | -1.63479 | 0.053424 | 0.037077 |
| X | 9304400 | 9305399 | 12.93419 | 25.13846 | 10.73947 | 3.994619 | 1.809737 | -0.24328 | 0.037081 |
| 13 | 48426400 | 48427999 | 4.863067 | 9.195796 | 35.22668 | 14.30389 | -1.43854 | 0.032971 | 0.037108 |
| 5 | 104690200 | 104691999 | 19.08594 | 34.79585 | 17.5833 | 7.552019 | 1.396876 | 0.323299 | 0.037111 |
| 5 | 124667000 | 124669799 | 8.231395 | 14.34761 | 56.76053 | 24.97141 | -1.58555 | 0.724808 | 0.037156 |
| X | 64752000 | 64754199 | 25.02968 | 43.78697 | 23.96163 | 10.56856 | 1.252711 | 0.702798 | 0.03717 |
| 11 | 15418000 | 15418799 | 9.847681 | 19.2473 | 7.796029 | 2.691877 | 1.972946 | -0.54712 | 0.037188 |
| 4 | 150456000 | 150456799 | 2.548968 | 4.969224 | 27.87543 | 10.05753 | -1.81697 | -0.42237 | 0.037199 |
| 14 | 42486400 | 42488399 | 19.17883 | 35.33982 | 13.97269 | 5.91686 | 1.762036 | 0.257862 | 0.037217 |
| 9 | 107033600 | 107035999 | 5.312202 | 9.993794 | 40.93817 | 16.78629 | -1.54791 | 0.083328 | 0.037218 |
| 1 | 191020400 | 191021999 | 6.986307 | 12.56444 | 43.74856 | 19.10839 | -1.38343 | 0.373846 | 0.037225 |
| 10 | 3202600 | 3202999 | 24.13918 | 42.49469 | 19.95871 | 8.835724 | 1.472194 | 0.600859 | 0.037251 |
| 5 | 95340800 | 95341399 | 9.462176 | 18.48203 | 9.917874 | 3.472708 | 1.560703 | -0.49736 | 0.037258 |
| 15 | 83671800 | 83676999 | 18.57693 | 30.28345 | 97.55375 | 41.90214 | -1.31099 | 1.519642 | 0.03726 |
| 11 | 116325000 | 116329199 | 13.71417 | 22.7115 | 85.5888 | 37.03497 | -1.52629 | 1.295628 | 0.037338 |
| 9 | 48771200 | 48772799 | 4.747168 | 8.938401 | 35.61281 | 14.58166 | -1.50501 | 0.061899 | 0.037363 |
| 13 | 76166200 | 76167599 | 6.024236 | 11.06701 | 41.61966 | 17.6969 | -1.46714 | 0.235755 | 0.03738 |
| 2 | 172917600 | 172935599 | 85.60388 | 129.2389 | 333.6347 | 137.9087 | -1.09255 | 3.332073 | 0.037423 |
| 7 | 13450200 | 13451399 | 34.08932 | 57.61495 | 32.98434 | 14.43259 | 1.182631 | 1.096941 | 0.037424 |
| 6 | 51406000 | 51407599 | 6.408431 | 11.53348 | 44.28783 | 19.33114 | -1.52269 | 0.365737 | 0.037424 |
| 4 | 138305400 | 138306799 | 5.179012 | 9.742369 | 36.19426 | 14.8433 | -1.40759 | 0.073018 | 0.037429 |
| 17 | 32414400 | 32418199 | 13.10972 | 21.99243 | 75.44771 | 32.92357 | -1.39044 | 1.139806 | 0.037443 |
| 16 | 18777000 | 18779199 | 113.6152 | 176.9911 | 109.7826 | 45.56574 | 0.998432 | 2.718982 | 0.037448 |
| 17 | 17236600 | 17245399 | 378.4944 | 541.9538 | 262.2569 | 108.3265 | 1.249287 | 4.23319 | 0.037476 |

| | | | | | | | | | |
|---|---|---|---|---|---|---|---|---|---|
| 14 | 98857600 | 98858999 | 5.805068 | 10.43886 | 43.72353 | 19.0993 | -1.64691 | 0.367189 | 0.03748 |
| 8 | 9169000 | 9169799 | 12.33274 | 23.79831 | 12.97849 | 5.01211 | 1.411964 | -0.12106 | 0.037488 |
| 12 | 22528600 | 22532999 | 66.8543 | 106.9695 | 67.24408 | 28.31246 | 1.012184 | 2.046191 | 0.037498 |
| 9 | 19642200 | 19642599 | 8.946979 | 17.49964 | 7.900949 | 2.660101 | 1.855703 | -0.64434 | 0.037502 |
| 19 | 44145400 | 44147199 | 7.185637 | 12.73215 | 46.69381 | 20.6445 | -1.47027 | 0.480482 | 0.037516 |
| 1 | 112416800 | 112417799 | 11.64438 | 22.58599 | 12.50588 | 4.716104 | 1.422256 | -0.20767 | 0.037535 |
| 1 | 74807800 | 74818599 | 52.60612 | 81.80421 | 223.1261 | 92.63418 | -1.1376 | 2.735812 | 0.03756 |
| 12 | 23593400 | 23596999 | 58.37468 | 93.96673 | 56.08868 | 23.82813 | 1.096844 | 1.846849 | 0.037647 |
| 3 | 142632800 | 142633999 | 6.433227 | 11.51161 | 45.25009 | 19.84836 | -1.56112 | 0.40515 | 0.0377 |
| 13 | 70774600 | 70774999 | 2.335101 | 4.565973 | 24.54648 | 8.379588 | -1.68038 | -0.63358 | 0.037704 |
| 2 | 28202600 | 28204399 | 5.265586 | 9.76156 | 39.62814 | 16.64317 | -1.5626 | 0.173546 | 0.037741 |
| 12 | 22409200 | 22410399 | 11.35288 | 22.06486 | 10.62824 | 3.953578 | 1.639035 | -0.26371 | 0.037779 |
| 12 | 20456600 | 20461599 | 117.771 | 182.8667 | 114.6041 | 47.59016 | 0.979931 | 2.775706 | 0.037784 |
| 12 | 19567200 | 19567599 | 11.90273 | 23.12438 | 11.95479 | 4.461237 | 1.534051 | -0.24099 | 0.037805 |
| 15 | 65601000 | 65602799 | 19.27621 | 35.13182 | 17.57989 | 7.553488 | 1.410606 | 0.3266 | 0.037823 |
| 16 | 90245800 | 90247599 | 5.302638 | 10.00951 | 38.13418 | 15.54175 | -1.43651 | 0.055101 | 0.037826 |
| 5 | 142312600 | 142313399 | 3.546501 | 6.869052 | 31.90128 | 12.11162 | -1.62136 | -0.217 | 0.037839 |
| 7 | 9074800 | 9075399 | 13.95889 | 26.7342 | 13.78204 | 5.438759 | 1.463786 | -0.0333 | 0.037868 |
| 1 | 101909800 | 101913199 | 65.83812 | 105.518 | 60.99653 | 25.74518 | 1.135587 | 1.993962 | 0.037878 |
| 7 | 5432200 | 5436599 | 112.8217 | 175.427 | 114.1414 | 47.38812 | 0.927471 | 2.750331 | 0.037964 |
| 1 | 88248800 | 88251199 | 7.148777 | 12.86198 | 47.0274 | 20.53332 | -1.45376 | 0.376533 | 0.037979 |
| 16 | 37920400 | 37921599 | 20.11075 | 36.07654 | 19.24333 | 8.423003 | 1.302499 | 0.426104 | 0.037983 |
| 4 | 124563200 | 124570199 | 36.86271 | 58.35708 | 174.2448 | 72.44362 | -1.24732 | 2.368832 | 0.037996 |
| 12 | 19425200 | 19426799 | 19.96813 | 35.7985 | 17.39751 | 7.619049 | 1.435144 | 0.424124 | 0.038002 |
| 10 | 78078600 | 78081999 | 9.914933 | 17.12393 | 60.47916 | 26.53225 | -1.42572 | 0.846258 | 0.038016 |
| 17 | 25584200 | 25597599 | 57.82513 | 89.06391 | 250.8929 | 104.2116 | -1.19485 | 2.914956 | 0.03808 |
| 19 | 16547400 | 16548399 | 12.10761 | 23.47701 | 12.36324 | 4.669268 | 1.491472 | -0.19782 | 0.038105 |
| 6 | 30873000 | 30873999 | 2.189152 | 4.214883 | 34.02156 | 13.2238 | -2.43381 | -0.14326 | 0.038108 |
| 16 | 22736600 | 22737999 | 6.431153 | 11.83288 | 41.12517 | 17.44979 | -1.35197 | 0.230438 | 0.038128 |
| 14 | 109434400 | 109435999 | 14.33712 | 27.44705 | 13.77408 | 5.442189 | 1.500782 | -0.02543 | 0.038144 |
| 12 | 23586200 | 23586399 | 9.74883 | 19.02604 | 10.484 | 3.724193 | 1.505229 | -0.44234 | 0.038164 |
| 2 | 88322200 | 88323199 | 14.70625 | 27.82848 | 14.79045 | 5.996682 | 1.386376 | 0.061526 | 0.038242 |
| 7 | 22989000 | 22991599 | 57.02044 | 92.00642 | 54.51704 | 23.23624 | 1.111427 | 1.774394 | 0.038308 |
| 3 | 95075800 | 95081799 | 31.89843 | 50.77754 | 154.4541 | 64.5653 | -1.26609 | 2.199805 | 0.038337 |
| 4 | 129494000 | 129494999 | 3.872003 | 7.402472 | 33.88657 | 13.44126 | -1.66165 | -0.05942 | 0.038387 |
| 1 | 104217000 | 104218199 | 11.96128 | 23.11909 | 11.9696 | 4.588765 | 1.495546 | -0.15276 | 0.038392 |
| 11 | 62385600 | 62387799 | 5.678621 | 10.56743 | 39.82068 | 16.63197 | -1.45062 | 0.154101 | 0.038395 |
| 7 | 31929400 | 31930799 | 18.44294 | 34.90408 | 11.78395 | 4.776359 | 2.034432 | 0.088781 | 0.03841 |
| 7 | 44461600 | 44466999 | 21.37179 | 34.56713 | 107.6927 | 46.0398 | -1.27295 | 1.674543 | 0.038426 |
| 7 | 141481800 | 141484199 | 5.837296 | 10.7873 | 38.99933 | 16.4496 | -1.40154 | 0.195988 | 0.038441 |
| 12 | 18849000 | 18855799 | 198.2399 | 297.1219 | 185.074 | 76.3497 | 0.947838 | 3.487408 | 0.038444 |
| 2 | 127886000 | 127886599 | 9.290932 | 18.14126 | 10.05551 | 3.542004 | 1.507626 | -0.47937 | 0.038462 |
| 12 | 111642000 | 111663399 | 73.39295 | 110.1933 | 372.9075 | 153.9051 | -1.48974 | 3.441181 | 0.038464 |
| X | 26866800 | 26867799 | 14.0896 | 27.32407 | 10.66984 | 4.026997 | 1.919031 | -0.1816 | 0.038488 |
| 7 | 31956400 | 31960599 | 84.68884 | 134.4181 | 82.82221 | 34.50734 | 1.029127 | 2.314269 | 0.038508 |
| 11 | 102203000 | 102205599 | 7.179845 | 12.7402 | 46.18351 | 20.40168 | -1.45253 | 0.467432 | 0.038513 |
| 16 | 94113600 | 94116199 | 8.433409 | 14.89563 | 48.80343 | 21.59093 | -1.31168 | 0.52529 | 0.038524 |

| | | | | | | | | | |
|---|---|---|---|---|---|---|---|---|---|
| 10 | 120470400 | 120473399 | 10.04489 | 17.51041 | 57.991 | 25.5144 | -1.33152 | 0.732098 | 0.038552 |
| 14 | 18554000 | 18554599 | 5.145604 | 9.385875 | 40.79621 | 17.51063 | -1.68205 | 0.271771 | 0.038604 |
| 6 | 19982800 | 19983599 | 12.12265 | 23.44934 | 13.03654 | 4.979557 | 1.3996 | -0.15862 | 0.038613 |
| 1 | 48900000 | 48901599 | 17.87124 | 33.20044 | 15.73147 | 6.588016 | 1.514173 | 0.207602 | 0.038619 |
| 6 | 83261400 | 83263399 | 6.856083 | 12.30887 | 44.25062 | 19.35597 | -1.43071 | 0.38466 | 0.038649 |
| 5 | 67865600 | 67865999 | 2.642753 | 5.152396 | 25.95888 | 9.358033 | -1.66196 | -0.42804 | 0.038656 |
| 8 | 39600000 | 39600999 | 2.642753 | 5.152396 | 25.95888 | 9.358033 | -1.66196 | -0.42804 | 0.038656 |
| 1 | 132952600 | 132954199 | 3.486097 | 6.619676 | 35.84168 | 14.42531 | -1.91916 | -0.00944 | 0.038688 |
| 19 | 15047800 | 15048599 | 16.52257 | 31.57724 | 11.09379 | 4.404494 | 2.001181 | 0.004247 | 0.038702 |
| 11 | 116055600 | 116057599 | 4.502346 | 8.474468 | 37.90206 | 15.52817 | -1.67146 | 0.065221 | 0.038703 |
| 5 | 60718000 | 60718799 | 11.43562 | 22.28098 | 9.379115 | 3.409588 | 1.856316 | -0.34501 | 0.038729 |
| 12 | 23615800 | 23619999 | 65.40195 | 104.7503 | 66.72076 | 28.13316 | 0.994848 | 2.014201 | 0.038731 |
| 9 | 118303600 | 118309799 | 42.62476 | 67.57264 | 167.9281 | 69.88194 | -0.98134 | 2.335702 | 0.038746 |
| 2 | 30811000 | 30820199 | 33.03676 | 52.58293 | 155.6188 | 65.04078 | -1.22686 | 2.204744 | 0.03877 |
| 3 | 91794200 | 91794599 | 9.675593 | 18.90596 | 8.453228 | 2.938755 | 1.82545 | -0.52464 | 0.038793 |
| 7 | 23083200 | 23086599 | 101.3892 | 158.7252 | 103.6306 | 42.98824 | 0.930014 | 2.625247 | 0.038844 |
| 2 | 62475800 | 62476799 | 4.270749 | 8.204131 | 33.71329 | 13.18924 | -1.48968 | -0.10263 | 0.038862 |
| 13 | 21815800 | 21816399 | 19.3149 | 35.0445 | 18.22438 | 7.874536 | 1.350629 | 0.353495 | 0.038896 |
| 6 | 83340200 | 83351399 | 35.56197 | 56.0127 | 193.4536 | 80.26977 | -1.46284 | 2.487521 | 0.038914 |
| 16 | 39069200 | 39070399 | 15.05803 | 29.03178 | 11.43633 | 4.427696 | 1.872027 | -0.08736 | 0.038962 |
| 7 | 9525200 | 9529199 | 36.11698 | 60.37492 | 34.74986 | 15.13024 | 1.174958 | 1.206694 | 0.038981 |
| 8 | 123101800 | 123103999 | 5.341414 | 9.694822 | 42.31116 | 18.27392 | -1.69448 | 0.30475 | 0.03904 |
| 7 | 20604800 | 20609199 | 37.13918 | 64.24923 | 17.89843 | 7.852449 | 2.218676 | 0.892976 | 0.039055 |
| 7 | 31743000 | 31745599 | 52.04154 | 84.75749 | 44.08458 | 18.92918 | 1.311007 | 1.567854 | 0.039079 |
| 3 | 152100000 | 152102599 | 10.03118 | 17.48039 | 55.77776 | 24.53524 | -1.27791 | 0.735695 | 0.039197 |
| X | 28854800 | 28855199 | 10.78083 | 20.99678 | 11.4468 | 4.183126 | 1.484792 | -0.32801 | 0.0392 |
| 14 | 57762200 | 57765399 | 13.03312 | 22.01136 | 70.62159 | 30.89422 | -1.29373 | 1.073125 | 0.0392 |
| 1 | 116860000 | 116860799 | 12.49181 | 24.15724 | 13.03022 | 4.983083 | 1.441363 | -0.15028 | 0.039236 |
| 14 | 114453200 | 114456599 | 40.97092 | 68.06125 | 37.37074 | 16.20531 | 1.242879 | 1.292643 | 0.039256 |
| 6 | 68905400 | 68905999 | 10.56276 | 20.62797 | 9.078336 | 3.188799 | 1.837318 | -0.47349 | 0.039272 |
| 16 | 8646600 | 8649799 | 12.6842 | 21.19923 | 78.00701 | 33.96023 | -1.49104 | 1.181971 | 0.039304 |
| 4 | 119959600 | 119962199 | 10.18133 | 17.77289 | 56.36501 | 24.82065 | -1.26907 | 0.710914 | 0.039324 |
| 7 | 14573400 | 14576999 | 100.5143 | 158.2158 | 88.64702 | 36.77926 | 1.156325 | 2.521519 | 0.03935 |
| 18 | 69566400 | 69567799 | 5.685768 | 10.5598 | 39.16384 | 16.4073 | -1.4311 | 0.165352 | 0.039355 |
| 16 | 79524600 | 79526799 | 13.19457 | 25.33594 | 13.95626 | 5.466685 | 1.379271 | -0.06397 | 0.039386 |
| 12 | 19534000 | 19537199 | 72.61426 | 116.1492 | 66.61259 | 28.03411 | 1.143836 | 2.063401 | 0.039392 |
| 3 | 100786800 | 100787999 | 14.38691 | 27.75376 | 12.07547 | 4.667463 | 1.732768 | -0.09786 | 0.039396 |
| 2 | 164808400 | 164818999 | 52.91113 | 82.00973 | 236.737 | 98.33296 | -1.22302 | 2.793893 | 0.039406 |
| 12 | 22209400 | 22214399 | 90.54053 | 143.0586 | 92.03305 | 38.22623 | 0.96149 | 2.434917 | 0.039411 |
| 13 | 16588000 | 16589199 | 14.52771 | 27.80605 | 13.7701 | 5.443905 | 1.519047 | -0.0215 | 0.039418 |
| 9 | 45049400 | 45050799 | 5.256447 | 9.77007 | 39.15342 | 16.38554 | -1.54036 | 0.157302 | 0.03944 |
| 9 | 12771200 | 12771599 | 11.52297 | 22.43905 | 10.65216 | 3.899879 | 1.67846 | -0.31493 | 0.039461 |
| X | 139907000 | 139909999 | 6.935137 | 12.44981 | 47.4726 | 20.76671 | -1.51486 | 0.394196 | 0.039474 |
| 13 | 34500000 | 34501999 | 7.625722 | 13.62 | 45.77136 | 20.10865 | -1.33912 | 0.425365 | 0.039507 |
| 13 | 97574800 | 97576399 | 5.563781 | 9.981995 | 46.34576 | 20.28402 | -1.79664 | 0.383864 | 0.039534 |
| 1 | 49042200 | 49042799 | 9.677694 | 18.90289 | 9.091808 | 3.183772 | 1.713852 | -0.49775 | 0.039539 |
| 5 | 102589600 | 102589999 | 8.080866 | 15.76431 | 5.118949 | 1.602835 | 2.399997 | -0.91267 | 0.039548 |

| | | | | | | | | | |
|---|---|---|---|---|---|---|---|---|---|
| 11 | 61834600 | 61836399 | 4.907885 | 9.26987 | 38.75567 | 15.77519 | -1.568 | 0.046626 | 0.039553 |
| 17 | 31052000 | 31053399 | 10.05774 | 17.5008 | 57.95747 | 25.47038 | -1.33172 | 0.758963 | 0.039571 |
| 8 | 121175000 | 121195599 | 103.5747 | 154.0941 | 401.9551 | 165.6121 | -1.12663 | 3.617071 | 0.039665 |
| 8 | 80315200 | 80316799 | 4.093617 | 7.868562 | 31.86884 | 12.44628 | -1.46529 | -0.11264 | 0.039677 |
| 12 | 19317600 | 19324599 | 194.9497 | 292.2306 | 186.0132 | 76.73919 | 0.916788 | 3.484185 | 0.0397 |
| 6 | 100886600 | 100887999 | 5.677807 | 10.5683 | 39.2561 | 16.38989 | -1.42955 | 0.151544 | 0.039709 |
| 6 | 27116400 | 27117199 | 12.36739 | 24.04068 | 11.21049 | 4.164458 | 1.686093 | -0.2532 | 0.039762 |
| 17 | 24642400 | 24646599 | 11.09847 | 18.93917 | 65.75367 | 28.82665 | -1.40503 | 0.967666 | 0.039766 |
| 14 | 3328200 | 3332199 | 64.75034 | 104.0703 | 59.16587 | 25.07968 | 1.164324 | 1.905316 | 0.039774 |
| 1 | 95388000 | 95389199 | 26.20265 | 46.05063 | 20.20726 | 8.947412 | 1.568442 | 0.631273 | 0.0398 |
| 2 | 172280800 | 172283999 | 9.27582 | 16.20201 | 55.77454 | 24.56998 | -1.38631 | 0.697029 | 0.0398 |
| 1 | 83137200 | 83138999 | 4.583054 | 8.745543 | 36.33313 | 14.47188 | -1.53051 | -0.03406 | 0.039819 |
| 6 | 90012600 | 90013399 | 19.99046 | 36.23223 | 18.09209 | 7.827626 | 1.407555 | 0.364124 | 0.039848 |
| 16 | 16688200 | 16689399 | 4.767441 | 8.913609 | 36.49414 | 15.12169 | -1.55782 | 0.110758 | 0.03988 |
| 9 | 9238200 | 9240599 | 10.30844 | 18.02694 | 55.27508 | 24.37213 | -1.22037 | 0.683918 | 0.039888 |
| 11 | 100232800 | 100233199 | 8.602185 | 16.82383 | 8.943302 | 3.033042 | 1.614555 | -0.62135 | 0.039917 |
| 3 | 102427000 | 102431399 | 21.46575 | 34.83474 | 101.4142 | 43.46211 | -1.17087 | 1.603627 | 0.039931 |
| 4 | 98418600 | 98420599 | 13.97824 | 23.44413 | 75.45246 | 32.92095 | -1.29866 | 1.147163 | 0.039962 |
| 1 | 112536800 | 112538599 | 25.0142 | 44.45762 | 16.29977 | 7.190709 | 1.831058 | 0.509381 | 0.039971 |
| 12 | 22546000 | 22555999 | 208.915 | 311.7694 | 198.049 | 81.63365 | 0.913901 | 3.569151 | 0.039985 |
| 5 | 93203600 | 93205199 | 21.97507 | 38.81004 | 21.15388 | 9.358785 | 1.260048 | 0.553967 | 0.039995 |
| 13 | 66825000 | 66830399 | 121.3991 | 188.4085 | 113.2189 | 47.01835 | 1.039947 | 2.786815 | 0.039997 |
| 12 | 20420200 | 20424599 | 139.054 | 212.938 | 137.4556 | 57.04519 | 0.922151 | 3.044326 | 0.040037 |
| 1 | 44971000 | 44972799 | 33.38763 | 56.75247 | 31.24529 | 13.69373 | 1.238939 | 1.043996 | 0.04006 |
| 7 | 10707400 | 10712999 | 131.6578 | 202.6667 | 129.8488 | 53.93012 | 0.93865 | 2.940053 | 0.040063 |
| 3 | 20718800 | 20720799 | 19.52624 | 35.41898 | 17.9773 | 7.770234 | 1.384739 | 0.35529 | 0.040084 |
| 16 | 94171000 | 94172799 | 6.622187 | 11.69192 | 48.24442 | 21.34531 | -1.64095 | 0.519895 | 0.040108 |
| 12 | 19528400 | 19532599 | 97.20385 | 152.8099 | 100.3725 | 41.6372 | 0.925925 | 2.545875 | 0.040125 |
| 5 | 30798000 | 30799999 | 6.389622 | 11.55056 | 43.32895 | 18.80836 | -1.48398 | 0.337273 | 0.040125 |
| 7 | 85554000 | 85554799 | 13.71388 | 26.50166 | 12.12103 | 4.650837 | 1.671176 | -0.12874 | 0.040126 |
| 16 | 13566800 | 13569399 | 15.59901 | 26.11148 | 75.67886 | 32.97912 | -1.1485 | 1.171614 | 0.04015 |
| 13 | 65703000 | 65705799 | 72.4853 | 115.6708 | 73.36209 | 30.77207 | 0.995357 | 2.136227 | 0.040155 |
| 10 | 79120800 | 79121599 | 16.76412 | 31.03903 | 16.84467 | 7.086579 | 1.315307 | 0.218654 | 0.040171 |
| 1 | 130643200 | 130645799 | 11.71717 | 19.71574 | 75.40573 | 32.95008 | -1.54625 | 1.105325 | 0.040173 |
| 10 | 75047200 | 75048599 | 5.938547 | 10.78651 | 43.4502 | 18.74874 | -1.58007 | 0.305453 | 0.040173 |
| 4 | 106893000 | 106893999 | 3.200539 | 6.053812 | 35.61029 | 14.45073 | -2.04632 | 0.016444 | 0.040176 |
| 9 | 46139400 | 46142399 | 10.83211 | 18.43791 | 67.86512 | 29.74625 | -1.48956 | 0.992069 | 0.040205 |
| 7 | 21823200 | 21823599 | 11.3354 | 22.09621 | 9.410409 | 3.399743 | 1.847811 | -0.36721 | 0.040216 |
| 5 | 140404200 | 140406799 | 7.760257 | 13.59112 | 54.08597 | 23.87799 | -1.59325 | 0.643954 | 0.040229 |
| 1 | 25068800 | 25069399 | 10.42816 | 20.30196 | 11.11014 | 4.07672 | 1.474516 | -0.3202 | 0.040263 |
| 8 | 85659600 | 85664999 | 33.7896 | 53.89847 | 147.9857 | 62.03726 | -1.11547 | 2.132211 | 0.040273 |
| 11 | 62977600 | 62978599 | 4.60835 | 8.718852 | 35.24287 | 14.29411 | -1.51395 | 0.025468 | 0.040295 |
| 4 | 3531000 | 3532599 | 18.77334 | 34.31542 | 17.6118 | 7.5396 | 1.377631 | 0.30655 | 0.040306 |
| 7 | 80884000 | 80885799 | 11.23517 | 19.25644 | 64.27436 | 28.18803 | -1.34761 | 0.924758 | 0.040308 |
| 6 | 29715200 | 29719399 | 16.21195 | 26.80437 | 84.78202 | 36.64472 | -1.27511 | 1.32046 | 0.040312 |
| 14 | 41923800 | 41930399 | 89.05115 | 141.5425 | 71.18282 | 29.69999 | 1.323451 | 2.274223 | 0.040321 |
| 12 | 22633600 | 22634199 | 19.54994 | 36.68185 | 12.11407 | 4.989572 | 2.047026 | 0.153231 | 0.040332 |

| | | | | | | | | | |
|---|---|---|---|---|---|---|---|---|---|
| 17 | 17224800 | 17226399 | 43.34329 | 71.11467 | 42.43779 | 18.27373 | 1.119508 | 1.445417 | 0.040335 |
| 15 | 60836800 | 60837799 | 10.67377 | 20.80819 | 10.88528 | 3.928378 | 1.559249 | -0.37407 | 0.040359 |
| 8 | 111654400 | 111658599 | 13.96129 | 23.28102 | 81.01104 | 35.21783 | -1.41114 | 1.211789 | 0.040373 |
| 15 | 100348800 | 100350599 | 7.694726 | 13.54764 | 49.29343 | 21.82162 | -1.4633 | 0.558666 | 0.040374 |
| 4 | 152352000 | 152373399 | 116.4242 | 170.8185 | 478.3885 | 197.0372 | -1.24793 | 3.853251 | 0.040375 |
| 4 | 118499000 | 118502199 | 11.28881 | 19.57134 | 59.05176 | 25.90906 | -1.19908 | 0.813858 | 0.040451 |
| 3 | 79707800 | 79708399 | 3.256114 | 6.294998 | 30.35345 | 11.62368 | -1.6849 | -0.19251 | 0.040451 |
| 2 | 164327600 | 164330399 | 9.453032 | 16.1725 | 67.91886 | 29.78199 | -1.67733 | 0.938591 | 0.040479 |
| 9 | 58133000 | 58135399 | 8.497568 | 14.83159 | 54.6264 | 24.05293 | -1.48266 | 0.704373 | 0.040489 |
| 7 | 30031400 | 30040199 | 38.96782 | 62.05295 | 157.6585 | 65.93715 | -1.00659 | 2.194007 | 0.040494 |
| 9 | 119007200 | 119010199 | 14.39911 | 24.01323 | 78.15052 | 33.97618 | -1.3149 | 1.210845 | 0.040496 |
| 6 | 113291800 | 113292399 | 2.24975 | 4.392173 | 25.03824 | 8.841654 | -1.8081 | -0.51198 | 0.040504 |
| 12 | 20062800 | 20073999 | 356.4808 | 508.3188 | 311.8682 | 128.9053 | 0.901077 | 4.292249 | 0.04051 |
| 9 | 83396000 | 83396599 | 13.11454 | 25.63109 | 4.503795 | 1.557114 | 3.139608 | -0.50481 | 0.040512 |
| 9 | 42098800 | 42100199 | 3.897004 | 7.542751 | 33.76461 | 12.86019 | -1.57351 | -0.19513 | 0.040525 |
| 5 | 135394000 | 135405399 | 44.22699 | 69.08271 | 220.6289 | 91.54279 | -1.35975 | 2.65302 | 0.04053 |
| 2 | 95306000 | 95306799 | 11.07334 | 21.52002 | 12.10223 | 4.50448 | 1.417517 | -0.2598 | 0.040548 |
| 13 | 11203800 | 11204799 | 25.16182 | 43.9585 | 24.98535 | 11.00797 | 1.198367 | 0.725831 | 0.040578 |
| 2 | 95483400 | 95484599 | 15.42867 | 29.29909 | 14.11564 | 5.68038 | 1.535841 | 0.043503 | 0.040592 |
| 13 | 65741600 | 65746399 | 119.9898 | 186.0861 | 116.914 | 48.55815 | 0.974986 | 2.797648 | 0.040601 |
| 11 | 107755800 | 107759399 | 11.56768 | 19.7664 | 65.87587 | 28.88335 | -1.34626 | 0.955833 | 0.040621 |
| 5 | 107008600 | 107009799 | 3.350529 | 6.520079 | 28.18732 | 10.37969 | -1.47586 | -0.33616 | 0.040714 |
| 3 | 21149800 | 21151399 | 12.96249 | 25.06244 | 12.13633 | 4.645589 | 1.592453 | -0.14533 | 0.040717 |
| 19 | 5795400 | 5801999 | 27.18145 | 43.45839 | 141.4837 | 59.50628 | -1.35676 | 2.052011 | 0.040744 |
| 11 | 85030200 | 85031399 | 4.107583 | 7.695694 | 39.95957 | 16.50006 | -1.89348 | 0.096612 | 0.040773 |
| 16 | 52647400 | 52648199 | 9.785595 | 19.10602 | 9.061706 | 3.19513 | 1.724817 | -0.47337 | 0.040815 |
| 10 | 100424800 | 100425199 | 1.272637 | 2.488931 | 22.94968 | 7.795388 | -2.42138 | -0.67181 | 0.040816 |
| 7 | 19976200 | 19976599 | 9.000358 | 17.60448 | 5.923916 | 1.990276 | 2.272368 | -0.66161 | 0.040818 |
| 2 | 135500600 | 135502199 | 5.5023 | 10.27391 | 37.07833 | 15.39834 | -1.38131 | 0.126212 | 0.040835 |
| 2 | 160063600 | 160065399 | 5.811712 | 10.81936 | 38.12583 | 15.91345 | -1.35362 | 0.149969 | 0.040852 |
| 17 | 94672200 | 94672799 | 9.91912 | 19.38073 | 9.117318 | 3.17419 | 1.753416 | -0.51278 | 0.040853 |
| 8 | 22139600 | 22139999 | 7.817825 | 15.2866 | 6.873051 | 2.230659 | 1.900841 | -0.79135 | 0.040871 |
| 2 | 167041400 | 167043999 | 8.303489 | 14.55905 | 51.42354 | 22.72438 | -1.42205 | 0.625845 | 0.040886 |
| 2 | 102903400 | 102904399 | 3.679239 | 7.06329 | 31.72664 | 12.43499 | -1.61677 | -0.10505 | 0.040893 |
| X | 90939600 | 90940399 | 12.1664 | 23.69962 | 10.20551 | 3.720682 | 1.823571 | -0.32183 | 0.040933 |
| 15 | 52561600 | 52562199 | 9.568825 | 18.69719 | 9.122765 | 3.172175 | 1.702658 | -0.52261 | 0.040994 |
| 4 | 27741200 | 27742399 | 15.55524 | 29.8067 | 11.784 | 4.643489 | 1.84323 | -0.02697 | 0.041006 |
| 7 | 115519600 | 115520399 | 2.335041 | 4.566039 | 23.19473 | 7.910015 | -1.59783 | -0.64119 | 0.041036 |
| 3 | 115236000 | 115237999 | 11.29956 | 19.58705 | 60.1761 | 26.40225 | -1.22521 | 0.81551 | 0.041043 |
| 7 | 85141600 | 85142799 | 15.75719 | 30.24058 | 11.98398 | 4.701295 | 1.846793 | -0.03773 | 0.041043 |
| 11 | 95962400 | 95969999 | 25.23032 | 40.74056 | 114.4128 | 48.81087 | -1.12673 | 1.730214 | 0.041051 |
| 11 | 120123400 | 120153599 | 113.4094 | 165.8228 | 508.7033 | 209.5684 | -1.38407 | 3.907704 | 0.041095 |
| 4 | 151782200 | 151806199 | 134.9903 | 196.633 | 510.2807 | 210.2942 | -1.14813 | 3.967394 | 0.041101 |
| 9 | 45340000 | 45348799 | 40.17463 | 63.43269 | 186.0442 | 77.24621 | -1.22477 | 2.435578 | 0.041124 |
| 7 | 141552000 | 141558799 | 24.77408 | 39.6957 | 138.7043 | 58.5226 | -1.45486 | 1.979774 | 0.041136 |
| 19 | 56049600 | 56051999 | 11.44934 | 19.84591 | 59.62513 | 26.16048 | -1.19315 | 0.816252 | 0.041209 |
| X | 122655200 | 122655799 | 12.73634 | 24.81951 | 9.147091 | 3.317811 | 2.051418 | -0.33533 | 0.041247 |

| | | | | | | | | | |
|---|---|---|---|---|---|---|---|---|---|
| 13 | 41243600 | 41245199 | 5.920356 | 10.80083 | 43.04147 | 18.47057 | -1.55949 | 0.278114 | 0.041262 |
| 7 | 5905200 | 5906599 | 41.73228 | 68.95171 | 40.2426 | 17.38555 | 1.154802 | 1.358759 | 0.041263 |
| 13 | 66064200 | 66066599 | 54.29548 | 87.96355 | 49.62671 | 21.24179 | 1.187883 | 1.666875 | 0.041287 |
| 19 | 4175400 | 4178399 | 11.51204 | 19.81526 | 63.33643 | 27.78288 | -1.28439 | 0.884563 | 0.041352 |
| 8 | 3064600 | 3068399 | 48.18106 | 79.07457 | 40.1877 | 17.30692 | 1.350586 | 1.454493 | 0.041354 |
| 17 | 19911000 | 19911999 | 36.14815 | 60.7656 | 33.40645 | 14.59336 | 1.240606 | 1.149151 | 0.041388 |
| 11 | 115077400 | 115080399 | 9.863553 | 17.24685 | 54.17735 | 23.88565 | -1.25497 | 0.681422 | 0.041426 |
| 3 | 85385800 | 85387199 | 3.463784 | 6.738004 | 31.40356 | 11.59829 | -1.58785 | -0.31533 | 0.041448 |
| 7 | 136102600 | 136107599 | 25.42293 | 40.98396 | 117.9575 | 50.21622 | -1.16486 | 1.780603 | 0.041503 |
| 9 | 38117800 | 38118799 | 21.36039 | 38.0392 | 19.95768 | 8.789682 | 1.320131 | 0.479964 | 0.041513 |
| 5 | 94336000 | 94337599 | 15.25388 | 29.403 | 11.43265 | 4.429023 | 1.889903 | -0.08329 | 0.041523 |
| 7 | 72810400 | 72812599 | 12.10092 | 20.87473 | 62.89466 | 27.59094 | -1.199 | 0.866285 | 0.041546 |
| 1 | 84126400 | 84127799 | 12.82102 | 24.64171 | 12.61484 | 4.927981 | 1.487532 | -0.08073 | 0.041596 |
| 17 | 45003400 | 45004799 | 3.77536 | 7.312731 | 30.03363 | 11.39955 | -1.44492 | -0.21786 | 0.041619 |
| 8 | 121604800 | 121644799 | 258.687 | 356.9343 | 891.7609 | 370.5446 | -1.17182 | 4.78259 | 0.041631 |
| 18 | 79274000 | 79274599 | 11.03486 | 21.52096 | 10.27834 | 3.689427 | 1.695458 | -0.38937 | 0.041707 |
| 12 | 19063000 | 19069799 | 162.8818 | 247.0826 | 160.5161 | 66.44886 | 0.902157 | 3.242516 | 0.041722 |
| 1 | 145190800 | 145191799 | 14.06559 | 26.8842 | 14.46451 | 5.741274 | 1.395748 | -0.01161 | 0.041756 |
| 1 | 180677800 | 180678999 | 2.949329 | 5.728466 | 28.35773 | 10.60376 | -1.68778 | -0.28446 | 0.04176 |
| 12 | 23992800 | 23993799 | 15.443 | 28.92054 | 15.94454 | 6.588597 | 1.312309 | 0.137538 | 0.041897 |
| 7 | 8137800 | 8139399 | 15.23672 | 29.45721 | 10.21343 | 3.911001 | 2.067247 | -0.12379 | 0.041937 |
| 4 | 45890200 | 45894999 | 14.64042 | 24.57471 | 73.40556 | 32.0443 | -1.19169 | 1.136647 | 0.04194 |
| 11 | 119008800 | 119044399 | 153.779 | 218.6307 | 675.9131 | 279.5161 | -1.43537 | 4.329868 | 0.04196 |
| 1 | 103035200 | 103036199 | 15.04426 | 28.27363 | 14.52668 | 5.966868 | 1.41975 | 0.107993 | 0.041973 |
| 3 | 122519400 | 122524399 | 27.35786 | 44.02756 | 119.9555 | 50.94242 | -1.08907 | 1.838335 | 0.041977 |
| 10 | 77344800 | 77348799 | 16.86883 | 27.78138 | 86.79607 | 37.43388 | -1.25885 | 1.372006 | 0.042001 |
| 12 | 22182200 | 22183799 | 41.96687 | 69.64232 | 37.35278 | 16.18574 | 1.276484 | 1.307285 | 0.042074 |
| 2 | 77355200 | 77356999 | 4.704608 | 8.991439 | 37.69825 | 14.96642 | -1.53967 | -0.04052 | 0.042075 |
| 5 | 118891000 | 118894999 | 10.43935 | 17.90543 | 63.09121 | 27.67077 | -1.42475 | 0.914989 | 0.042085 |
| 7 | 31202200 | 31208999 | 145.495 | 222.8133 | 137.6264 | 57.11677 | 0.985762 | 3.04523 | 0.042091 |
| 10 | 81694600 | 81697199 | 33.46076 | 57.60498 | 25.11719 | 11.01786 | 1.576053 | 0.926394 | 0.04211 |
| 6 | 115958600 | 115978199 | 83.96602 | 126.0458 | 362.9102 | 149.772 | -1.2571 | 3.445904 | 0.042153 |
| 13 | 66773600 | 66774999 | 46.49669 | 75.78148 | 45.72692 | 19.63954 | 1.09813 | 1.545923 | 0.042194 |
| 4 | 37981200 | 37981999 | 14.79885 | 27.99314 | 15.37138 | 6.237193 | 1.339111 | 0.066898 | 0.042207 |
| 9 | 36079400 | 36080199 | 12.37131 | 24.03043 | 11.80465 | 4.411577 | 1.604581 | -0.23094 | 0.04221 |
| 8 | 13484600 | 13487399 | 12.14034 | 20.83204 | 63.285 | 27.75674 | -1.21233 | 0.916305 | 0.042234 |
| 3 | 15703400 | 15704999 | 21.74059 | 38.84848 | 18.78255 | 8.248392 | 1.440406 | 0.46091 | 0.042251 |
| 14 | 46020600 | 46021199 | 9.745014 | 19.03348 | 9.745083 | 3.415327 | 1.625395 | -0.49088 | 0.042265 |
| 4 | 80822400 | 80824199 | 97.18081 | 153.3762 | 78.62824 | 32.63973 | 1.287209 | 2.464052 | 0.042271 |
| 18 | 41100400 | 41101399 | 10.49926 | 20.44976 | 11.58883 | 4.231369 | 1.43127 | -0.33397 | 0.042272 |
| 1 | 60527000 | 60529399 | 8.774198 | 15.40089 | 50.90497 | 22.51562 | -1.32731 | 0.608464 | 0.042291 |
| 13 | 54591600 | 54593599 | 8.774198 | 15.40089 | 50.90497 | 22.51562 | -1.32731 | 0.608464 | 0.042291 |
| 12 | 19808400 | 19813399 | 176.1689 | 266.495 | 163.0133 | 67.42406 | 0.985563 | 3.30174 | 0.042295 |
| 5 | 123054200 | 123057799 | 11.51872 | 19.80968 | 63.30897 | 27.7701 | -1.28442 | 0.892627 | 0.042305 |
| 12 | 23482200 | 23485199 | 48.47287 | 79.47296 | 40.73102 | 17.53339 | 1.337942 | 1.467281 | 0.042312 |
| 5 | 104780600 | 104781999 | 12.48517 | 24.17205 | 12.93233 | 4.919271 | 1.459844 | -0.16811 | 0.042318 |
| 11 | 115066600 | 115076399 | 37.16203 | 58.58214 | 190.6147 | 79.10043 | -1.37606 | 2.470689 | 0.042329 |

| | | | | | | | | | |
|---|---|---|---|---|---|---|---|---|---|
| 5 | 83160000 | 83160599 | 9.004791 | 17.59599 | 9.951463 | 3.45573 | 1.496396 | -0.53209 | 0.042392 |
| 10 | 8355600 | 8356199 | 10.67823 | 20.79818 | 11.12833 | 4.063649 | 1.513325 | -0.3328 | 0.042407 |
| 12 | 19586600 | 19589599 | 34.8936 | 59.12718 | 29.30941 | 12.83466 | 1.389506 | 1.074335 | 0.042412 |
| X | 34301200 | 34301599 | 8.947301 | 17.50005 | 6.833123 | 2.245581 | 2.086464 | -0.7285 | 0.042477 |
| 16 | 91132600 | 91135199 | 13.08331 | 22.38806 | 64.95136 | 28.48125 | -1.14706 | 0.947426 | 0.042481 |
| 3 | 122722000 | 122723399 | 14.91771 | 28.48493 | 13.59123 | 5.406129 | 1.563734 | 0.000712 | 0.042508 |
| 1 | 127416200 | 127419799 | 19.96658 | 32.64684 | 94.9405 | 40.8244 | -1.16093 | 1.475965 | 0.042515 |
| 12 | 22573400 | 22574999 | 20.63432 | 36.84297 | 20.2121 | 8.882014 | 1.258562 | 0.460253 | 0.042535 |
| 19 | 59454200 | 59461399 | 35.92626 | 57.20094 | 158.9527 | 66.46461 | -1.13579 | 2.196194 | 0.042539 |
| 8 | 71123000 | 71124799 | 5.114587 | 9.420251 | 38.85594 | 16.4637 | -1.59376 | 0.210348 | 0.042545 |
| 2 | 20335200 | 20336599 | 16.23149 | 30.64665 | 14.75839 | 6.011814 | 1.522149 | 0.091436 | 0.042585 |
| 19 | 41469800 | 41472399 | 11.15659 | 19.29031 | 59.53276 | 26.11797 | -1.23229 | 0.838454 | 0.042604 |
| 3 | 10903800 | 10907399 | 36.39347 | 61.30595 | 32.83925 | 14.35468 | 1.278025 | 1.132618 | 0.042604 |
| 8 | 5063400 | 5064799 | 13.04781 | 25.25449 | 12.16831 | 4.634527 | 1.606615 | -0.1602 | 0.042621 |
| 11 | 102353600 | 102359199 | 23.95863 | 38.66231 | 112.2965 | 47.86844 | -1.17615 | 1.747269 | 0.042638 |
| 7 | 9642400 | 9646599 | 62.45633 | 100.41 | 60.69055 | 25.73595 | 1.076654 | 1.894601 | 0.042673 |
| 18 | 67336000 | 67339599 | 14.7014 | 24.59441 | 80.21849 | 34.94435 | -1.31834 | 1.177274 | 0.042673 |
| 4 | 155493000 | 155494199 | 4.862472 | 9.196666 | 34.65852 | 14.06672 | -1.41447 | 0.030248 | 0.042688 |
| 1 | 132053000 | 132072199 | 117.8711 | 173.8302 | 447.1341 | 184.136 | -1.11831 | 3.770017 | 0.042693 |
| 17 | 87923800 | 87924599 | 2.856663 | 5.530558 | 30.42206 | 11.57326 | -1.86216 | -0.22019 | 0.042708 |
| 4 | 134136600 | 134137399 | 10.63623 | 20.69349 | 10.06887 | 3.714221 | 1.634488 | -0.30405 | 0.042711 |
| 4 | 126915200 | 126916799 | 6.165986 | 11.30716 | 39.89717 | 17.00586 | -1.37817 | 0.244821 | 0.042744 |
| 12 | 22443600 | 22444999 | 31.90419 | 54.15197 | 32.52502 | 14.25 | 1.113842 | 1.05226 | 0.042781 |
| 2 | 97767400 | 97767999 | 9.393276 | 18.36534 | 7.830686 | 2.680521 | 1.91083 | -0.58311 | 0.0428 |
| 12 | 42736000 | 42736599 | 9.879816 | 19.32066 | 8.082512 | 2.751776 | 1.948256 | -0.591 | 0.042802 |
| 9 | 72892400 | 72893399 | 5.369037 | 10.02794 | 37.08555 | 15.39391 | -1.41572 | 0.122628 | 0.042824 |
| 1 | 146705400 | 146706199 | 12.01777 | 23.27452 | 13.06647 | 4.962941 | 1.3934 | -0.17783 | 0.042826 |
| 7 | 38486600 | 38489199 | 38.1835 | 64.4021 | 30.65536 | 13.40516 | 1.447695 | 1.127368 | 0.042885 |
| 7 | 7498800 | 7500199 | 17.34644 | 32.94595 | 12.27503 | 4.9381 | 1.902726 | 0.056395 | 0.04289 |
| 11 | 56073800 | 56074599 | 6.329124 | 11.52299 | 40.80023 | 17.5536 | -1.39231 | 0.291095 | 0.042896 |
| 5 | 140329400 | 140334399 | 17.02044 | 27.69906 | 99.16429 | 42.56757 | -1.46436 | 1.543115 | 0.042905 |
| X | 31666400 | 31668799 | 61.05996 | 98.68213 | 43.14882 | 18.42885 | 1.551096 | 1.731538 | 0.042909 |
| 19 | 42076800 | 42079599 | 9.053398 | 15.92128 | 49.83587 | 22.06687 | -1.2489 | 0.580092 | 0.04292 |
| 17 | 92260200 | 92260799 | 9.921014 | 19.37654 | 9.742273 | 3.416432 | 1.650643 | -0.48602 | 0.042949 |
| 19 | 6116400 | 6118399 | 6.501024 | 11.4563 | 49.27924 | 21.81141 | -1.70168 | 0.541991 | 0.042955 |
| 2 | 28348000 | 28350799 | 6.940958 | 12.62015 | 41.87952 | 18.04789 | -1.30184 | 0.304234 | 0.042986 |
| 6 | 88901000 | 88905399 | 19.82925 | 32.38457 | 96.51531 | 41.48396 | -1.1974 | 1.493497 | 0.042988 |
| 4 | 91393800 | 91394199 | 8.432344 | 16.49322 | 8.788041 | 2.956308 | 1.620169 | -0.65211 | 0.042988 |
| 3 | 25689000 | 25690199 | 11.75865 | 22.75104 | 13.04287 | 4.976036 | 1.357132 | -0.16697 | 0.043017 |
| 10 | 80438200 | 80445999 | 28.38713 | 45.19016 | 156.2286 | 65.31128 | -1.45017 | 2.195761 | 0.043047 |
| 13 | 43876000 | 43877799 | 7.649628 | 13.35382 | 57.29588 | 25.23099 | -1.70154 | 0.699791 | 0.043055 |
| 7 | 21111800 | 21114399 | 61.56102 | 98.85628 | 63.27768 | 26.78754 | 0.992124 | 1.929263 | 0.043098 |
| 5 | 33612000 | 33623599 | 57.48736 | 88.56292 | 248.2324 | 103.1096 | -1.18736 | 2.910363 | 0.043105 |
| 18 | 75684800 | 75688399 | 16.50643 | 27.30539 | 83.09391 | 35.92792 | -1.21949 | 1.314411 | 0.04314 |
| 5 | 123677600 | 123678399 | 18.27081 | 33.49767 | 17.64727 | 7.524233 | 1.344252 | 0.286238 | 0.043166 |
| 2 | 82300000 | 82300799 | 11.63826 | 22.59655 | 12.40381 | 4.652036 | 1.441481 | -0.22625 | 0.043181 |
| 6 | 31089600 | 31091799 | 6.267978 | 11.3036 | 44.88319 | 19.54138 | -1.56837 | 0.352958 | 0.043183 |

| | | | | | | | | | |
|---|---|---|---|---|---|---|---|---|---|
| 1 | 12134600 | 12135799 | 5.477776 | 10.30639 | 40.3946 | 16.56041 | -1.485 | 0.081738 | 0.043188 |
| 5 | 57871800 | 57879599 | 50.46473 | 78.62302 | 226.0185 | 93.80844 | -1.21117 | 2.701814 | 0.043212 |
| 1 | 164282000 | 164289799 | 36.22881 | 57.71647 | 154.3851 | 64.60709 | -1.07995 | 2.176533 | 0.043216 |
| 12 | 23712000 | 23714999 | 58.53303 | 94.47927 | 53.04226 | 22.61759 | 1.189529 | 1.765054 | 0.043232 |
| 5 | 82143000 | 82143799 | 13.91219 | 26.84753 | 12.70364 | 4.905251 | 1.614754 | -0.10429 | 0.043234 |
| 7 | 33317800 | 33319799 | 54.14832 | 87.74697 | 49.6221 | 21.24371 | 1.184801 | 1.661885 | 0.043264 |
| 12 | 10962000 | 10962999 | 14.61081 | 27.6451 | 15.37547 | 6.235198 | 1.321498 | 0.063146 | 0.043283 |
| 6 | 17306000 | 17308399 | 13.19945 | 22.29107 | 70.6609 | 30.9109 | -1.27648 | 1.073648 | 0.043283 |
| 5 | 129213800 | 129249599 | 174.9668 | 248.7782 | 677.5065 | 280.1704 | -1.25238 | 4.329276 | 0.043317 |
| 5 | 61408200 | 61411199 | 64.64825 | 103.829 | 55.11118 | 23.33558 | 1.26184 | 1.925856 | 0.043339 |
| 14 | 70782400 | 70799599 | 83.93146 | 125.6422 | 385.669 | 159.0645 | -1.35306 | 3.500009 | 0.043375 |
| 8 | 122200800 | 122205199 | 17.69306 | 29.08342 | 89.163 | 38.42376 | -1.23297 | 1.398715 | 0.043388 |
| 3 | 145793800 | 145793999 | 7.858473 | 15.37224 | 8.389065 | 2.771579 | 1.609687 | -0.72351 | 0.043422 |
| 7 | 136534400 | 136540399 | 36.01144 | 57.49062 | 142.3745 | 59.75521 | -0.9656 | 2.107068 | 0.04345 |
| 2 | 24937000 | 24946399 | 40.12768 | 63.05013 | 195.804 | 81.21924 | -1.31262 | 2.539837 | 0.043455 |
| 1 | 193486200 | 193486999 | 12.66673 | 24.55442 | 11.58091 | 4.379403 | 1.645897 | -0.1898 | 0.043475 |
| 9 | 106562000 | 106563599 | 5.021204 | 9.16172 | 40.80066 | 17.50595 | -1.71644 | 0.26845 | 0.043496 |
| 5 | 106094600 | 106096799 | 13.1748 | 22.26985 | 69.53874 | 30.42848 | -1.25468 | 1.065203 | 0.043528 |
| 9 | 36946400 | 36946999 | 9.006303 | 17.59315 | 10.08202 | 3.524808 | 1.469802 | -0.50872 | 0.043532 |
| 7 | 80270400 | 80271999 | 6.923664 | 12.64161 | 42.01207 | 18.01028 | -1.29909 | 0.279189 | 0.043631 |
| 18 | 76573400 | 76575599 | 9.389868 | 16.47326 | 52.53868 | 23.22835 | -1.27655 | 0.621433 | 0.043656 |
| 7 | 103625600 | 103626799 | 15.25561 | 28.57798 | 15.94773 | 6.586642 | 1.295476 | 0.133921 | 0.04368 |
| 5 | 125465800 | 125467199 | 3.895697 | 7.379863 | 34.6254 | 14.00615 | -1.72175 | 0.009206 | 0.043684 |
| 11 | 102282200 | 102299599 | 74.82663 | 112.8348 | 345.3684 | 142.7057 | -1.33944 | 3.354642 | 0.04372 |
| 4 | 135964400 | 135978199 | 68.31176 | 104.1748 | 291.0126 | 120.6567 | -1.19408 | 3.120083 | 0.043722 |
| 14 | 86680200 | 86680999 | 11.15174 | 21.7501 | 8.770093 | 3.145784 | 1.93309 | -0.39641 | 0.04373 |
| 16 | 20366200 | 20368199 | 5.95538 | 10.77327 | 43.33359 | 18.79358 | -1.58267 | 0.329932 | 0.043738 |
| Y | 6955400 | 6956599 | 11.19283 | 21.80619 | 5.817418 | 2.11702 | 2.499718 | -0.36067 | 0.043754 |
| 7 | 14461000 | 14462999 | 60.63178 | 97.48235 | 59.93177 | 25.4163 | 1.052274 | 1.889978 | 0.043791 |
| 17 | 27273800 | 27286199 | 53.46126 | 83.00205 | 223.5632 | 92.83764 | -1.12126 | 2.764002 | 0.043797 |
| 1 | 44186600 | 44187599 | 6.13009 | 11.34949 | 39.08851 | 16.44552 | -1.32992 | 0.186979 | 0.043801 |
| X | 33680000 | 33686199 | 70.62529 | 113.3457 | 57.64336 | 24.38067 | 1.322235 | 1.95841 | 0.043822 |
| 15 | 59516800 | 59518399 | 5.663291 | 10.58386 | 40.04336 | 16.60485 | -1.4481 | 0.126067 | 0.043823 |
| 13 | 22129000 | 22129999 | 5.693344 | 10.55174 | 38.52849 | 16.18721 | -1.41163 | 0.176494 | 0.043857 |
| 14 | 124902800 | 124903599 | 11.2838 | 21.89225 | 12.51217 | 4.712781 | 1.3784 | -0.21621 | 0.043857 |
| 7 | 38695200 | 38697399 | 27.04908 | 46.92127 | 27.63223 | 12.1239 | 1.146753 | 0.833874 | 0.04386 |
| 11 | 119781600 | 119783399 | 4.240914 | 8.065732 | 33.65604 | 13.5004 | -1.54336 | -0.01595 | 0.043871 |
| 7 | 14177600 | 14179999 | 59.69279 | 96.15351 | 57.30155 | 24.36726 | 1.100382 | 1.830032 | 0.043886 |
| 7 | 33235200 | 33235999 | 46.37693 | 75.62793 | 46.79071 | 20.10047 | 1.062701 | 1.538991 | 0.043892 |
| 6 | 87907600 | 87908799 | 26.05635 | 45.61155 | 23.662 | 10.44192 | 1.329199 | 0.698227 | 0.0439 |
| 7 | 76630200 | 76630799 | 11.34987 | 22.07024 | 11.85453 | 4.391657 | 1.488111 | -0.27332 | 0.043903 |
| X | 24272000 | 24272999 | 14.91168 | 28.52896 | 13.76214 | 5.447342 | 1.55511 | -0.01365 | 0.043904 |
| 5 | 35642000 | 35643599 | 17.82827 | 32.63968 | 17.63037 | 7.531648 | 1.306281 | 0.289841 | 0.043945 |
| 13 | 65557800 | 65563599 | 84.59272 | 134.1675 | 85.86383 | 35.74846 | 0.973508 | 2.336363 | 0.043964 |
| 11 | 75037800 | 75039799 | 7.001202 | 12.54641 | 43.69018 | 19.14035 | -1.38651 | 0.397159 | 0.043967 |
| 5 | 73312800 | 73316399 | 13.5292 | 22.80239 | 71.75813 | 31.3719 | -1.26626 | 1.093301 | 0.04398 |
| 1 | 174836600 | 174837799 | 15.44052 | 29.81046 | 10.82573 | 4.172015 | 1.993688 | -0.09928 | 0.043986 |

| | | | | | | | | | |
|---|---|---|---|---|---|---|---|---|---|
| 7 | 22339400 | 22339799 | 9.447482 | 18.44029 | 9.235794 | 3.273549 | 1.641462 | -0.46054 | 0.044015 |
| 9 | 44404000 | 44413599 | 30.37078 | 48.24332 | 164.5334 | 68.60708 | -1.43398 | 2.26659 | 0.044015 |
| 11 | 92961600 | 92963999 | 32.53668 | 55.14895 | 32.97108 | 14.43935 | 1.120408 | 1.066471 | 0.044033 |
| 11 | 101623400 | 101624999 | 4.898298 | 9.279639 | 36.54528 | 14.78111 | -1.4741 | 0.020563 | 0.044083 |
| 1 | 112559400 | 112560399 | 17.83664 | 32.8804 | 17.25319 | 7.301854 | 1.357608 | 0.25262 | 0.044123 |
| 11 | 69118400 | 69123999 | 16.82899 | 27.49707 | 95.1555 | 40.90624 | -1.41141 | 1.481002 | 0.044124 |
| 10 | 108364400 | 108365399 | 5.617407 | 10.55713 | 36.77882 | 15.10703 | -1.31808 | 0.084096 | 0.044176 |
| 1 | 64729200 | 64731599 | 8.781481 | 15.39449 | 51.44439 | 22.73027 | -1.34304 | 0.630058 | 0.044211 |
| 12 | 20658600 | 20660799 | 32.42389 | 54.98296 | 31.8682 | 13.9583 | 1.16486 | 1.060069 | 0.044229 |
| 13 | 66329600 | 66332399 | 110.9737 | 172.4848 | 112.8506 | 46.85497 | 0.919036 | 2.755595 | 0.04424 |
| 14 | 113516400 | 113517199 | 12.71491 | 24.47172 | 13.99688 | 5.446124 | 1.334357 | -0.08993 | 0.044258 |
| 10 | 45453800 | 45454999 | 8.465355 | 14.86196 | 50.36379 | 22.28026 | -1.36282 | 0.602945 | 0.044271 |
| 7 | 130351200 | 130352799 | 4.165819 | 7.960079 | 34.51402 | 13.70818 | -1.58691 | -0.0515 | 0.044277 |
| 13 | 66823400 | 66824199 | 31.8693 | 54.61165 | 29.00425 | 12.71985 | 1.29202 | 0.963877 | 0.044293 |
| 16 | 80105800 | 80106199 | 8.335187 | 16.29413 | 8.900141 | 3.058559 | 1.557792 | -0.58341 | 0.044315 |
| 13 | 116445400 | 116447799 | 13.29819 | 22.59635 | 67.52058 | 29.5911 | -1.19117 | 1.016281 | 0.044353 |
| 12 | 22397200 | 22397799 | 11.36364 | 22.04563 | 12.63847 | 4.761881 | 1.374278 | -0.21327 | 0.044361 |
| 1 | 44516800 | 44517199 | 7.935948 | 15.52411 | 8.161631 | 2.717889 | 1.650078 | -0.69808 | 0.044365 |
| 5 | 139797200 | 139802599 | 21.60583 | 34.78956 | 118.8361 | 50.5067 | -1.41309 | 1.813908 | 0.044368 |
| 1 | 92249400 | 92255199 | 26.38358 | 42.50943 | 117.2852 | 49.89307 | -1.10501 | 1.798543 | 0.044375 |
| 6 | 38274800 | 38276599 | 6.174483 | 10.92807 | 47.20635 | 20.87585 | -1.70469 | 0.487732 | 0.044389 |
| 15 | 48381600 | 48383799 | 23.96546 | 42.30461 | 20.22925 | 8.950587 | 1.447249 | 0.570014 | 0.0444 |
| 15 | 83504000 | 83515999 | 74.66072 | 114.1869 | 269.6025 | 111.8547 | -0.94839 | 3.061765 | 0.044438 |
| 15 | 77723600 | 77726799 | 12.41955 | 20.63286 | 81.50353 | 35.34454 | -1.59307 | 1.252497 | 0.044522 |
| 11 | 73844800 | 73846399 | 20.54456 | 36.50332 | 20.29057 | 8.953425 | 1.235701 | 0.48902 | 0.044524 |
| 3 | 27654400 | 27655199 | 3.240548 | 6.310164 | 29.642 | 10.85806 | -1.58787 | -0.35414 | 0.044539 |
| 5 | 108590000 | 108593599 | 9.821253 | 16.97719 | 62.19715 | 27.28667 | -1.47841 | 0.836968 | 0.044544 |
| 15 | 84100600 | 84104199 | 12.70793 | 21.54482 | 69.0669 | 30.24935 | -1.29226 | 1.035611 | 0.044553 |
| 7 | 11450800 | 11452199 | 35.11509 | 59.34519 | 32.87788 | 14.38575 | 1.229779 | 1.101253 | 0.044568 |
| X | 26186200 | 26189799 | 31.06145 | 53.83182 | 24.55319 | 10.77264 | 1.513025 | 0.856909 | 0.044598 |
| 4 | 115278000 | 115278999 | 19.53601 | 35.22686 | 17.90699 | 7.797551 | 1.375554 | 0.38741 | 0.044602 |
| 5 | 94809000 | 94809399 | 8.38421 | 16.38977 | 9.078824 | 3.120923 | 1.539015 | -0.5801 | 0.044604 |
| 12 | 23669400 | 23669999 | 12.82939 | 25.05642 | 6.62868 | 2.325221 | 2.558213 | -0.45407 | 0.04463 |
| 17 | 23298800 | 23299599 | 10.31931 | 20.10911 | 11.48316 | 4.165994 | 1.427865 | -0.35901 | 0.044647 |
| 7 | 14563000 | 14566799 | 94.54987 | 148.9499 | 98.25575 | 40.77024 | 0.92207 | 2.50295 | 0.044682 |
| 3 | 102533000 | 102534599 | 15.27381 | 28.87205 | 15.24062 | 6.192983 | 1.393574 | 0.075485 | 0.04469 |
| 7 | 17858600 | 17859799 | 35.93099 | 60.10864 | 36.84164 | 16.04853 | 1.084677 | 1.20152 | 0.044703 |
| 17 | 37785000 | 37785799 | 12.53579 | 24.48433 | 6.320625 | 2.215086 | 2.589859 | -0.4622 | 0.044719 |
| 8 | 45408800 | 45414199 | 33.4335 | 53.41308 | 142.1886 | 59.73741 | -1.06847 | 2.081838 | 0.044724 |
| 9 | 108959200 | 108961799 | 23.28636 | 40.92525 | 22.60699 | 10.00996 | 1.238545 | 0.61746 | 0.044733 |
| 1 | 69739000 | 69739599 | 13.28494 | 25.5065 | 14.07498 | 5.515024 | 1.376813 | -0.06118 | 0.04477 |
| 14 | 82275800 | 82276399 | 10.94137 | 21.38898 | 5.64349 | 1.939095 | 2.576945 | -0.55405 | 0.044821 |
| 12 | 23291400 | 23297399 | 126.4811 | 194.839 | 126.8618 | 52.69476 | 0.916203 | 2.923632 | 0.044828 |
| 6 | 102807800 | 102808799 | 15.83046 | 29.97163 | 14.67005 | 5.94123 | 1.505721 | 0.069013 | 0.04483 |
| 17 | 92015800 | 92016999 | 16.26088 | 30.56531 | 14.42579 | 5.923712 | 1.54049 | 0.117648 | 0.044872 |
| X | 25778400 | 25778999 | 13.87585 | 26.94229 | 9.672741 | 3.626353 | 2.044263 | -0.20913 | 0.044888 |
| 12 | 24393200 | 24396399 | 81.85511 | 130.1831 | 79.08887 | 33.0195 | 1.052218 | 2.252623 | 0.044892 |

| | | | | | | | | | |
|---|---|---|---|---|---|---|---|---|---|
| 12 | 22363600 | 22364199 | 13.89026 | 26.50172 | 14.10921 | 5.621716 | 1.406262 | -0.00245 | 0.044946 |
| 8 | 24294600 | 24295599 | 5.147365 | 9.786713 | 37.40026 | 15.01199 | -1.42155 | 2.62E-05 | 0.044963 |
| 2 | 26813800 | 26817199 | 8.495907 | 14.90319 | 52.50867 | 23.2128 | -1.41902 | 0.618668 | 0.044964 |
| 9 | 19189600 | 19190599 | 16.34277 | 30.77624 | 15.1848 | 6.215845 | 1.481528 | 0.109909 | 0.044983 |
| 1 | 112862000 | 112863799 | 14.63587 | 27.96384 | 13.55149 | 5.383358 | 1.544511 | -0.00559 | 0.045009 |
| 12 | 23542600 | 23543599 | 29.62146 | 50.79272 | 30.3855 | 13.32629 | 1.121392 | 0.95097 | 0.045023 |
| 14 | 18547000 | 18547399 | 10.9645 | 21.33886 | 12.0269 | 4.425252 | 1.429082 | -0.30018 | 0.045042 |
| 3 | 44589800 | 44590799 | 13.2078 | 25.31216 | 14.51192 | 5.716114 | 1.314945 | -0.04463 | 0.045047 |
| 11 | 100985600 | 100986999 | 5.710962 | 10.53306 | 38.93712 | 16.46747 | -1.43617 | 0.205898 | 0.045075 |
| X | 33888000 | 33889399 | 21.3857 | 39.54067 | 12.99741 | 5.478436 | 2.029188 | 0.255739 | 0.045095 |
| 12 | 23654200 | 23656999 | 95.17499 | 150.1504 | 86.00856 | 35.69618 | 1.126943 | 2.47299 | 0.0451 |
| 7 | 98435600 | 98437399 | 6.468771 | 11.85421 | 40.44693 | 17.25642 | -1.33154 | 0.25157 | 0.045109 |
| 15 | 99487000 | 99502799 | 71.95015 | 108.6496 | 339.0649 | 140.1631 | -1.36561 | 3.325203 | 0.04511 |
| 14 | 99346000 | 99347199 | 5.148495 | 9.657457 | 37.19475 | 15.32725 | -1.46418 | 0.095503 | 0.045123 |
| 10 | 63173800 | 63175599 | 8.635714 | 15.14925 | 51.43968 | 22.74126 | -1.36595 | 0.617509 | 0.045124 |
| 17 | 8454800 | 8465199 | 53.08827 | 82.95192 | 205.5384 | 85.27853 | -0.99379 | 2.649798 | 0.045134 |
| 6 | 89768800 | 89771399 | 31.28504 | 53.54479 | 29.56649 | 12.96514 | 1.235849 | 0.972188 | 0.045135 |
| 8 | 48972000 | 48973799 | 7.803505 | 13.89468 | 48.94666 | 21.55954 | -1.40975 | 0.450554 | 0.045136 |
| 11 | 35931600 | 35942799 | 48.77049 | 76.49466 | 209.3196 | 86.81353 | -1.13253 | 2.579921 | 0.045138 |
| 4 | 140580600 | 140594799 | 90.17178 | 136.1746 | 324.1546 | 134.0023 | -0.9753 | 3.326738 | 0.045156 |
| 7 | 8787800 | 8788199 | 9.632909 | 18.83279 | 8.512211 | 2.91807 | 1.828691 | -0.57094 | 0.045165 |
| 9 | 56167600 | 56168999 | 78.44626 | 124.8961 | 68.52973 | 28.65376 | 1.200479 | 2.213648 | 0.045184 |
| 3 | 103183600 | 103187999 | 20.76062 | 33.933 | 95.69345 | 41.1428 | -1.11716 | 1.483048 | 0.045202 |
| 9 | 45848600 | 45855599 | 23.03317 | 36.88259 | 139.2544 | 58.69749 | -1.5673 | 1.998986 | 0.04521 |
| 5 | 103571600 | 103573799 | 9.600999 | 16.72318 | 56.85051 | 25 | -1.36891 | 0.740546 | 0.045219 |
| 17 | 67557200 | 67559199 | 5.610521 | 10.56789 | 36.88235 | 15.09206 | -1.31629 | 0.069393 | 0.04523 |
| 12 | 22223400 | 22224799 | 26.92061 | 47.07657 | 23.67006 | 10.43651 | 1.374194 | 0.71906 | 0.045247 |
| 19 | 5599800 | 5607199 | 31.6908 | 50.57331 | 144.1564 | 60.46987 | -1.1686 | 2.115478 | 0.045254 |
| 7 | 15282400 | 15282799 | 15.40699 | 28.98354 | 15.76099 | 6.463195 | 1.340276 | 0.108755 | 0.045275 |
| 3 | 50516800 | 50517599 | 13.99012 | 26.67673 | 14.90162 | 5.944749 | 1.335572 | 0.004333 | 0.045324 |
| 18 | 83247400 | 83249599 | 5.900552 | 10.81646 | 42.09707 | 17.95247 | -1.51936 | 0.247779 | 0.045333 |
| 4 | 152404400 | 152493399 | 540.2318 | 696.3192 | 1887.831 | 791.5065 | -1.38844 | 5.873601 | 0.045352 |
| 17 | 24954200 | 24956199 | 6.992889 | 12.55641 | 43.19053 | 18.88981 | -1.36719 | 0.383307 | 0.045407 |
| 17 | 55174200 | 55174799 | 11.24247 | 21.89346 | 11.29355 | 4.133116 | 1.561259 | -0.31755 | 0.045473 |
| 4 | 122186200 | 122186599 | 7.816131 | 15.28931 | 8.191203 | 2.705386 | 1.634128 | -0.72699 | 0.045541 |
| 4 | 151846600 | 151849599 | 10.37793 | 17.97577 | 58.41651 | 25.62963 | -1.30558 | 0.817842 | 0.045549 |
| 5 | 116141200 | 116146399 | 27.98659 | 44.75881 | 140.9089 | 59.28919 | -1.30804 | 2.043059 | 0.045551 |
| 2 | 31229200 | 31230799 | 7.734141 | 13.86208 | 44.76516 | 19.60864 | -1.27876 | 0.403423 | 0.045555 |
| 9 | 103604000 | 103605399 | 4.480384 | 8.493553 | 36.52117 | 14.75064 | -1.59731 | 0.011885 | 0.045559 |
| 7 | 8499800 | 8502999 | 51.58648 | 84.01739 | 45.17857 | 19.39901 | 1.263188 | 1.567736 | 0.04556 |
| 1 | 132020800 | 132021999 | 3.658455 | 7.087907 | 29.43387 | 11.16 | -1.45914 | -0.22519 | 0.045575 |
| 14 | 18563600 | 18564399 | 9.404408 | 18.34846 | 10.6135 | 3.789361 | 1.429964 | -0.42989 | 0.045578 |
| 8 | 35660800 | 35663799 | 13.29103 | 22.60439 | 66.99641 | 29.36349 | -1.17925 | 1.007341 | 0.045578 |
| 19 | 53332200 | 53333799 | 5.289214 | 10.03155 | 37.2383 | 15.02544 | -1.38657 | 0.019317 | 0.045585 |
| 11 | 107561000 | 107562199 | 3.866842 | 7.407544 | 32.83098 | 12.94729 | -1.60742 | -0.08146 | 0.045606 |
| 17 | 22770000 | 22770799 | 36.80852 | 61.31348 | 36.41436 | 15.81959 | 1.130102 | 1.247575 | 0.045636 |
| 7 | 101898800 | 101900599 | 7.085259 | 12.55584 | 47.22119 | 20.87716 | -1.50664 | 0.477436 | 0.045646 |

| | | | | | | | | | |
|---|---|---|---|---|---|---|---|---|---|
| 16 | 36694200 | 36696199 | 7.49289 | 13.36323 | 45.74215 | 20.12056 | -1.36669 | 0.433585 | 0.045648 |
| 17 | 27365200 | 27380199 | 66.56195 | 101.7156 | 280.5053 | 116.3557 | -1.17313 | 3.076983 | 0.045651 |
| 8 | 52663400 | 52664199 | 11.88052 | 23.17901 | 8.098668 | 2.892458 | 2.141202 | -0.40262 | 0.045699 |
| 19 | 49563400 | 49564399 | 12.2647 | 23.87702 | 10.63928 | 3.904451 | 1.766216 | -0.29697 | 0.045741 |
| 14 | 108313000 | 108313599 | 11.10422 | 21.55116 | 11.60667 | 4.364772 | 1.465411 | -0.22679 | 0.045742 |
| 5 | 35855800 | 35864999 | 34.67894 | 54.83292 | 182.0794 | 75.65412 | -1.40177 | 2.397702 | 0.045755 |
| 4 | 121534200 | 121535999 | 43.71182 | 71.96528 | 42.38611 | 18.27793 | 1.14032 | 1.407197 | 0.045797 |
| 5 | 94522800 | 94524399 | 58.56713 | 94.21373 | 58.78398 | 24.94906 | 1.031933 | 1.869021 | 0.045847 |
| 15 | 99775200 | 99778599 | 12.40364 | 21.39794 | 61.23322 | 26.86213 | -1.12492 | 0.866026 | 0.045894 |
| 5 | 62618400 | 62620399 | 26.26065 | 45.87697 | 24.8949 | 10.96794 | 1.264941 | 0.732597 | 0.045896 |
| 6 | 83430000 | 83437999 | 29.15451 | 46.54576 | 148.9515 | 62.51926 | -1.33447 | 2.100183 | 0.045897 |
| 7 | 51071800 | 51072399 | 9.119927 | 17.80746 | 10.51974 | 3.705151 | 1.416679 | -0.4806 | 0.0459 |
| 18 | 50047400 | 50048599 | 14.7642 | 28.05803 | 14.72944 | 5.918106 | 1.415237 | 0.033713 | 0.045957 |
| X | 33287000 | 33288999 | 47.66504 | 78.08397 | 41.92169 | 18.03963 | 1.270855 | 1.478003 | 0.045959 |
| 16 | 4747200 | 4750799 | 14.39514 | 24.06649 | 77.78582 | 33.87082 | -1.30511 | 1.182143 | 0.045969 |
| 4 | 94937400 | 94942199 | 23.31861 | 37.65916 | 110.7061 | 47.23846 | -1.19209 | 1.722049 | 0.04597 |
| 7 | 9858000 | 9858599 | 26.6652 | 46.35755 | 27.14425 | 11.9224 | 1.155367 | 0.804419 | 0.046013 |
| 1 | 100966600 | 100968599 | 23.61074 | 41.46709 | 22.0352 | 9.751167 | 1.294553 | 0.630358 | 0.046018 |
| 7 | 8581000 | 8584399 | 48.49801 | 79.26157 | 41.39813 | 17.79822 | 1.308397 | 1.50939 | 0.04602 |
| 2 | 92874800 | 92877399 | 11.22027 | 19.27006 | 62.67763 | 27.49202 | -1.30994 | 0.903975 | 0.04603 |
| 17 | 68148400 | 68148999 | 2.250318 | 4.391696 | 24.90264 | 8.848696 | -1.8085 | -0.48948 | 0.046054 |
| 11 | 101332600 | 101334799 | 6.836924 | 12.32868 | 42.73973 | 18.60996 | -1.37412 | 0.354275 | 0.046065 |
| 15 | 72929800 | 72930399 | 2.739485 | 5.345715 | 24.96225 | 8.888032 | -1.53723 | -0.47376 | 0.046085 |
| 5 | 93965200 | 93966599 | 14.78531 | 28.40081 | 12.46512 | 4.87749 | 1.703913 | -0.05477 | 0.046094 |
| 3 | 108460600 | 108462199 | 4.516265 | 8.690459 | 32.62746 | 12.7026 | -1.35377 | -0.11735 | 0.046128 |
| 5 | 36013000 | 36021399 | 29.77555 | 47.55409 | 145.6009 | 61.14354 | -1.2703 | 2.089018 | 0.046131 |
| 8 | 44041400 | 44043399 | 22.32825 | 40.02265 | 17.57163 | 7.696443 | 1.580052 | 0.44252 | 0.046135 |
| 7 | 62195000 | 62196199 | 40.4555 | 67.1402 | 39.00577 | 16.90305 | 1.161679 | 1.302994 | 0.04615 |
| 12 | 60806200 | 60806799 | 9.857849 | 19.23044 | 10.92798 | 3.909367 | 1.452038 | -0.41548 | 0.046154 |
| 2 | 126774400 | 126775399 | 3.241792 | 6.308931 | 28.19431 | 10.37626 | -1.5225 | -0.34061 | 0.046193 |
| 17 | 48012200 | 48024399 | 51.04422 | 78.9579 | 242.6077 | 100.7733 | -1.31516 | 2.830975 | 0.046195 |
| 9 | 59443600 | 59448199 | 17.05703 | 28.08616 | 88.68593 | 38.24597 | -1.27437 | 1.374796 | 0.0462 |
| 14 | 81940800 | 81942599 | 28.72074 | 50.19866 | 21.15217 | 9.322283 | 1.627047 | 0.731682 | 0.046205 |
| 10 | 19416000 | 19417799 | 23.00464 | 40.38339 | 23.37821 | 10.34133 | 1.171589 | 0.635688 | 0.046206 |
| 15 | 96931400 | 96931799 | 1.715644 | 3.35158 | 21.04625 | 6.709712 | -1.80151 | -0.89753 | 0.046213 |
| 4 | 150991600 | 150995399 | 16.33462 | 26.81627 | 94.71944 | 40.79624 | -1.43732 | 1.414337 | 0.046223 |
| 10 | 22105400 | 22105999 | 9.232048 | 18.01936 | 10.49155 | 3.720158 | 1.428546 | -0.4566 | 0.04626 |
| X | 33906800 | 33911199 | 53.26496 | 87.24447 | 36.30075 | 15.61984 | 1.637 | 1.488887 | 0.046262 |
| 10 | 66782000 | 66784199 | 18.32058 | 33.74593 | 17.12504 | 7.255385 | 1.404266 | 0.260427 | 0.046263 |
| 15 | 76184000 | 76194399 | 43.35 | 67.7433 | 209.1532 | 86.77732 | -1.31042 | 2.643967 | 0.046265 |
| 7 | 23087400 | 23089199 | 50.53934 | 82.28011 | 46.81828 | 20.10024 | 1.181475 | 1.571256 | 0.04628 |
| 1 | 150230000 | 150231599 | 18.35068 | 33.72501 | 16.97363 | 7.213402 | 1.412603 | 0.271916 | 0.046286 |
| 7 | 42028000 | 42030199 | 59.29911 | 95.74467 | 50.86117 | 21.696 | 1.269519 | 1.75626 | 0.046314 |
| 15 | 72922400 | 72928999 | 22.16608 | 35.64388 | 127.1088 | 53.91789 | -1.47764 | 1.859355 | 0.046318 |
| 13 | 99363600 | 99366199 | 15.01974 | 25.07354 | 77.19628 | 33.5827 | -1.23528 | 1.200218 | 0.046328 |
| 19 | 5686800 | 5692799 | 23.15805 | 37.3827 | 110.9719 | 47.32388 | -1.20688 | 1.735907 | 0.046386 |
| 3 | 83986000 | 83987399 | 22.17397 | 39.18753 | 21.37566 | 9.455639 | 1.259673 | 0.549948 | 0.046404 |

| | | | | | | | | | |
|---|---|---|---|---|---|---|---|---|---|
| 7 | 21139800 | 21142999 | 21.18313 | 37.47744 | 20.38895 | 9.017192 | 1.263744 | 0.528911 | 0.046409 |
| 12 | 69735600 | 69736799 | 12.74671 | 24.7576 | 11.00684 | 4.116125 | 1.744536 | -0.22708 | 0.046431 |
| 14 | 49408600 | 49410999 | 11.11036 | 18.99416 | 65.2917 | 28.62833 | -1.39043 | 0.949014 | 0.046451 |
| 4 | 45228800 | 45231399 | 9.829769 | 17.25271 | 51.46415 | 22.76187 | -1.18053 | 0.616006 | 0.046462 |
| 16 | 68555600 | 68556199 | 14.28891 | 27.60228 | 12.10992 | 4.654923 | 1.728471 | -0.11635 | 0.046465 |
| 11 | 96908200 | 96913799 | 23.90375 | 38.21535 | 150.6334 | 63.3485 | -1.63205 | 2.052518 | 0.046486 |
| 16 | 88085400 | 88088199 | 13.58454 | 23.15348 | 67.12391 | 29.42449 | -1.14713 | 0.987792 | 0.046523 |
| 2 | 172172800 | 172174199 | 18.50451 | 33.54311 | 18.66297 | 8.073378 | 1.252515 | 0.352613 | 0.046542 |
| 1 | 8411400 | 8411999 | 16.68564 | 31.56681 | 13.29494 | 5.392961 | 1.717402 | 0.078136 | 0.046569 |
| 9 | 12897000 | 12898199 | 16.68564 | 31.56681 | 13.29494 | 5.392961 | 1.717402 | 0.078136 | 0.046569 |
| 1 | 174203800 | 174206799 | 23.12095 | 40.58599 | 23.27932 | 10.29725 | 1.184759 | 0.636417 | 0.046573 |
| 10 | 22921400 | 22921999 | 9.790407 | 19.10004 | 10.29808 | 3.68108 | 1.52688 | -0.42211 | 0.046579 |
| 13 | 35999200 | 36001599 | 9.393152 | 16.47064 | 52.00151 | 22.98097 | -1.26198 | 0.629252 | 0.046582 |
| 8 | 95065200 | 95068599 | 15.04154 | 25.00194 | 79.14304 | 34.33333 | -1.27526 | 1.251379 | 0.046601 |
| 14 | 14466000 | 14466999 | 4.402589 | 8.429295 | 32.27191 | 12.74657 | -1.40107 | -0.07133 | 0.046695 |
| 7 | 17443400 | 17446199 | 77.03698 | 122.6875 | 75.43339 | 31.55284 | 1.037044 | 2.205381 | 0.046751 |
| X | 29860800 | 29861199 | 7.863546 | 15.36209 | 6.208361 | 1.979996 | 2.071224 | -0.85109 | 0.046777 |
| 13 | 108116600 | 108118599 | 7.614827 | 13.36085 | 54.08997 | 23.91671 | -1.6179 | 0.61199 | 0.04678 |
| 5 | 110097000 | 110098999 | 54.79789 | 89.03302 | 44.65345 | 19.15362 | 1.360902 | 1.610765 | 0.046784 |
| 8 | 106937000 | 106937999 | 3.998546 | 7.610607 | 33.10107 | 13.25627 | -1.60046 | -0.02642 | 0.046797 |
| 7 | 20144600 | 20145599 | 12.15582 | 23.73327 | 7.957625 | 2.806907 | 2.218767 | -0.44047 | 0.046824 |
| 2 | 157731600 | 157743799 | 48.35812 | 75.26661 | 219.7774 | 91.23153 | -1.23473 | 2.717983 | 0.046844 |
| 9 | 43446200 | 43451199 | 22.1114 | 35.90829 | 100.0875 | 42.91624 | -1.10735 | 1.588499 | 0.046894 |
| 7 | 8497400 | 8503199 | 75.65702 | 120.6767 | 74.30746 | 31.1486 | 1.03723 | 2.154218 | 0.046932 |
| 14 | 65966800 | 65970199 | 16.89256 | 28.18168 | 78.03761 | 33.93481 | -1.08315 | 1.213153 | 0.046941 |
| 13 | 48519800 | 48522199 | 8.60135 | 15.17565 | 49.33202 | 21.82956 | -1.30127 | 0.538618 | 0.046952 |
| 18 | 82595000 | 82604999 | 62.01774 | 95.72413 | 240.7666 | 100.0097 | -1.02924 | 2.875261 | 0.046954 |
| 3 | 8980800 | 8981799 | 3.844841 | 7.428233 | 32.2778 | 12.39907 | -1.54358 | -0.17256 | 0.046955 |
| 17 | 4970400 | 4973399 | 9.346048 | 16.48628 | 51.4998 | 22.78971 | -1.24462 | 0.548843 | 0.046966 |
| 8 | 118040800 | 118053199 | 47.00625 | 73.29668 | 215.7298 | 89.52686 | -1.24407 | 2.685045 | 0.046988 |
| 11 | 117930400 | 117943799 | 69.29941 | 105.1842 | 308.2175 | 127.615 | -1.26853 | 3.218633 | 0.046992 |
| 6 | 101114600 | 101116999 | 21.79219 | 39.08373 | 18.24794 | 7.988886 | 1.493367 | 0.43829 | 0.047027 |
| X | 139901800 | 139905999 | 13.10145 | 22.36443 | 64.86664 | 28.43851 | -1.14722 | 0.970433 | 0.047039 |
| 19 | 8690600 | 8691599 | 5.032085 | 9.516335 | 34.69152 | 14.08366 | -1.36796 | 0.031357 | 0.047058 |
| 13 | 40098600 | 40100399 | 20.20039 | 36.60347 | 17.83742 | 7.719885 | 1.441775 | 0.365908 | 0.047079 |
| 5 | 71533000 | 71534999 | 21.78973 | 38.71768 | 19.81529 | 8.743326 | 1.35378 | 0.495602 | 0.047093 |
| 17 | 39802800 | 39804399 | 32.07818 | 55.11086 | 27.87529 | 12.22695 | 1.362578 | 0.941276 | 0.047097 |
| 7 | 9307400 | 9310399 | 32.92684 | 56.592 | 26.73931 | 11.72882 | 1.460421 | 0.940135 | 0.047124 |
| 16 | 33004200 | 33005999 | 6.129145 | 11.35063 | 38.53459 | 16.20686 | -1.30891 | 0.184468 | 0.047129 |
| 5 | 133123800 | 133127399 | 19.39123 | 31.95225 | 88.70044 | 38.26367 | -1.089 | 1.371127 | 0.047138 |
| 9 | 65291200 | 65292399 | 4.28419 | 8.189102 | 32.7536 | 12.99818 | -1.47006 | -0.05613 | 0.047139 |
| 5 | 90371400 | 90372399 | 6.350831 | 11.58601 | 41.40944 | 17.77051 | -1.40384 | 0.278532 | 0.047148 |
| 12 | 114659600 | 114660799 | 33.42079 | 56.50183 | 34.03123 | 14.892 | 1.109729 | 1.092947 | 0.047182 |
| 2 | 30528000 | 30535399 | 29.5052 | 47.01305 | 152.719 | 63.92498 | -1.35915 | 2.16538 | 0.047193 |
| 8 | 127314600 | 127323999 | 33.13184 | 52.59819 | 168.7418 | 70.32039 | -1.34689 | 2.286364 | 0.047203 |
| 16 | 80087600 | 80089199 | 22.6114 | 40.34705 | 19.08386 | 8.390781 | 1.470666 | 0.476205 | 0.047223 |
| 11 | 77353000 | 77365599 | 56.90496 | 87.5947 | 253.1216 | 105.1287 | -1.23221 | 2.926327 | 0.047228 |

| | | | | | | | | | |
|---|---|---|---|---|---|---|---|---|---|
| 11 | 12515800 | 12517399 | 17.09942 | 31.48098 | 17.81023 | 7.550773 | 1.24799 | 0.254477 | 0.047238 |
| 10 | 127932600 | 127933799 | 4.58708 | 8.741343 | 33.83854 | 13.51704 | -1.43266 | -0.02942 | 0.047266 |
| 5 | 112699000 | 112701599 | 9.683527 | 16.66694 | 62.04378 | 27.2158 | -1.50243 | 0.877511 | 0.047293 |
| 11 | 113506400 | 113513199 | 31.39028 | 50.06571 | 146.7596 | 61.51379 | -1.20978 | 2.133641 | 0.047354 |
| 13 | 55475000 | 55480399 | 15.7351 | 25.68991 | 95.89071 | 41.2107 | -1.52083 | 1.491054 | 0.04737 |
| 17 | 24903400 | 24906999 | 14.9215 | 25.14374 | 71.36088 | 31.19625 | -1.11806 | 1.099241 | 0.047393 |
| 4 | 150078000 | 150090999 | 57.61883 | 89.29242 | 228.6365 | 94.96914 | -1.05046 | 2.797918 | 0.04741 |
| X | 5024600 | 5025999 | 21.9565 | 40.05694 | 15.26123 | 6.549015 | 1.801732 | 0.337104 | 0.04741 |
| 4 | 61996400 | 61996999 | 8.666789 | 16.93632 | 9.956463 | 3.453221 | 1.442475 | -0.54196 | 0.047432 |
| 9 | 108773200 | 108800599 | 134.0818 | 194.4596 | 543.1926 | 223.9596 | -1.2604 | 4.03387 | 0.047443 |
| 17 | 49454200 | 49455999 | 6.449763 | 11.87526 | 40.04752 | 16.97671 | -1.3081 | 0.223075 | 0.047446 |
| 12 | 23443800 | 23446799 | 27.10416 | 47.64334 | 19.88838 | 8.807206 | 1.640334 | 0.634793 | 0.047452 |
| 11 | 101823000 | 101824999 | 4.890676 | 9.287397 | 36.08976 | 14.52319 | -1.4485 | 0.002398 | 0.047474 |
| 6 | 85833200 | 85834399 | 7.664482 | 13.57853 | 46.17116 | 20.41405 | -1.36252 | 0.48295 | 0.047475 |
| 3 | 16637600 | 16639399 | 14.44484 | 27.59485 | 14.20817 | 5.645956 | 1.456444 | -0.00539 | 0.047504 |
| 17 | 17229200 | 17231399 | 111.3448 | 173.9333 | 99.36547 | 41.22818 | 1.120011 | 2.670373 | 0.047515 |
| 4 | 151172600 | 151175599 | 11.77591 | 20.01412 | 68.49212 | 30.01742 | -1.38568 | 1.009462 | 0.047543 |
| 7 | 98998000 | 99003199 | 18.69047 | 30.25108 | 107.1476 | 45.82996 | -1.45632 | 1.655736 | 0.047551 |
| 7 | 32199800 | 32201399 | 22.14989 | 39.94241 | 17.18891 | 7.484333 | 1.614423 | 0.406141 | 0.047554 |
| 1 | 148462400 | 148463799 | 17.99916 | 34.3337 | 10.41892 | 4.153273 | 2.204809 | 0.02975 | 0.047558 |
| 5 | 65005600 | 65006599 | 11.35428 | 22.06234 | 12.44049 | 4.636577 | 1.41137 | -0.25085 | 0.047572 |
| 17 | 46747600 | 46759999 | 64.85506 | 99.26454 | 274.0555 | 113.7219 | -1.17293 | 3.04372 | 0.047586 |
| 7 | 28661200 | 28661999 | 3.597485 | 6.81907 | 34.03433 | 13.74943 | -1.80711 | 0.001557 | 0.047634 |
| 4 | 3145000 | 3145999 | 15.93527 | 30.10811 | 13.89122 | 5.650324 | 1.583768 | 0.080822 | 0.047651 |
| 4 | 52228200 | 52229799 | 14.1296 | 27.11306 | 12.44419 | 4.883859 | 1.63556 | -0.05302 | 0.047668 |
| 18 | 79945000 | 79945799 | 4.713041 | 8.980902 | 33.25211 | 13.28428 | -1.36901 | -0.02837 | 0.047683 |
| 8 | 84721200 | 84732399 | 46.24198 | 72.34398 | 204.6186 | 84.88609 | -1.18297 | 2.624376 | 0.047693 |
| 4 | 91684200 | 91684599 | 7.695519 | 15.04752 | 7.570726 | 2.4576 | 1.743893 | -0.78914 | 0.0477 |
| 7 | 9078000 | 9079999 | 16.91448 | 31.45672 | 16.65692 | 6.964495 | 1.356563 | 0.195081 | 0.047711 |
| 1 | 143585800 | 143587999 | 19.23913 | 34.61092 | 19.70882 | 8.606161 | 1.211002 | 0.403342 | 0.047744 |
| 9 | 106047400 | 106049199 | 7.249029 | 12.76884 | 48.76456 | 21.5855 | -1.5321 | 0.55012 | 0.04778 |
| 7 | 110908600 | 110909599 | 4.073804 | 7.895924 | 32.11467 | 12.15255 | -1.42845 | -0.22133 | 0.047786 |
| 13 | 54921200 | 54922799 | 23.87432 | 41.81067 | 24.50228 | 10.81684 | 1.154363 | 0.680323 | 0.04779 |
| 7 | 20760000 | 20762199 | 19.96846 | 36.7015 | 15.79219 | 6.710489 | 1.636965 | 0.28223 | 0.047791 |
| 8 | 122327800 | 122340399 | 51.39063 | 79.7116 | 227.5138 | 94.4916 | -1.20583 | 2.780004 | 0.047809 |
| 7 | 9009800 | 9010999 | 27.53068 | 48.09166 | 21.4232 | 9.437912 | 1.547859 | 0.734808 | 0.047838 |
| 19 | 7460000 | 7460799 | 3.243267 | 6.307479 | 28.05428 | 10.382 | -1.52299 | -0.32075 | 0.04784 |
| 17 | 38806600 | 38807799 | 26.68547 | 46.69954 | 23.30513 | 10.28196 | 1.385017 | 0.706443 | 0.047841 |
| 1 | 179908600 | 179909599 | 14.39813 | 27.31557 | 15.06737 | 6.075825 | 1.340801 | 0.043017 | 0.047848 |
| 18 | 79897400 | 79898799 | 18.14947 | 33.35574 | 17.67609 | 7.511772 | 1.339255 | 0.272447 | 0.047884 |
| 3 | 15814800 | 15816599 | 19.81207 | 37.17702 | 10.63519 | 4.379786 | 2.249672 | 0.151121 | 0.047917 |
| 5 | 25291200 | 25293599 | 9.219391 | 16.20026 | 49.84023 | 22.06533 | -1.22455 | 0.590915 | 0.047918 |
| 5 | 65190000 | 65191199 | 3.15338 | 6.149082 | 28.11038 | 10.10882 | -1.52294 | -0.42371 | 0.047928 |
| 11 | 75544000 | 75553599 | 30.56115 | 48.53764 | 167.2537 | 69.72975 | -1.44911 | 2.271804 | 0.047939 |
| 6 | 108804400 | 108806999 | 12.29065 | 20.79414 | 73.49096 | 32.1667 | -1.43266 | 1.055659 | 0.047957 |
| 12 | 22480200 | 22483999 | 104.5836 | 163.3461 | 108.7252 | 45.11237 | 0.899664 | 2.669375 | 0.047967 |
| 5 | 36497000 | 36498599 | 7.398634 | 13.41175 | 42.3523 | 18.32067 | -1.23462 | 0.32632 | 0.04797 |

| | | | | | | | | | |
|---|---|---|---|---|---|---|---|---|---|
| 3 | 37231000 | 37231599 | 5.018112 | 9.533085 | 34.3296 | 13.80356 | -1.33828 | -0.00211 | 0.047977 |
| 14 | 30718400 | 30719799 | 4.081315 | 7.885532 | 30.43952 | 11.68993 | -1.37361 | -0.17339 | 0.047989 |
| 4 | 137265400 | 137281399 | 74.68745 | 113.0228 | 318.924 | 131.9255 | -1.21784 | 3.28182 | 0.047991 |
| 3 | 18457800 | 18459999 | 7.873187 | 14.18779 | 44.90841 | 19.57067 | -1.24538 | 0.369212 | 0.048005 |
| 4 | 145606000 | 145606999 | 32.75527 | 55.98062 | 29.77409 | 13.05466 | 1.289701 | 0.993585 | 0.048005 |
| 2 | 179321600 | 179330599 | 41.77905 | 66.33467 | 160.0775 | 66.71841 | -0.93679 | 2.287667 | 0.048013 |
| 11 | 13311200 | 13313399 | 27.70639 | 48.07911 | 27.47889 | 12.05675 | 1.189787 | 0.833528 | 0.048018 |
| 3 | 31308600 | 31309999 | 6.727915 | 12.40147 | 41.2271 | 17.44975 | -1.28612 | 0.221661 | 0.048018 |
| 18 | 34756600 | 34758399 | 5.478995 | 10.30478 | 35.65176 | 14.62538 | -1.30652 | 0.075107 | 0.048026 |
| 12 | 23927800 | 23930799 | 68.07824 | 108.7967 | 72.82562 | 30.61023 | 0.919938 | 2.086131 | 0.048043 |
| 11 | 83855000 | 83855599 | 1.947258 | 3.807011 | 22.32556 | 7.651818 | -1.80446 | -0.63181 | 0.048057 |
| 6 | 51105800 | 51106799 | 3.89369 | 7.548833 | 30.78183 | 11.63372 | -1.42901 | -0.2289 | 0.048087 |
| 19 | 46290200 | 46324999 | 171.7287 | 243.1308 | 694.7692 | 287.5147 | -1.32799 | 4.392124 | 0.048089 |
| 15 | 100566600 | 100567999 | 10.39628 | 17.95468 | 59.4415 | 26.077 | -1.33314 | 0.846497 | 0.048096 |
| 9 | 88278200 | 88279599 | 15.00372 | 28.70232 | 13.62471 | 5.39441 | 1.577279 | -0.01248 | 0.0481 |
| 1 | 136693600 | 136696399 | 17.79631 | 29.34333 | 84.77684 | 36.5812 | -1.14576 | 1.357988 | 0.048111 |
| 4 | 8711600 | 8720999 | 70.84491 | 108.9305 | 253.1527 | 105.1255 | -0.9196 | 2.952767 | 0.048144 |
| 14 | 28239800 | 28240799 | 4.178412 | 7.948491 | 34.28243 | 13.7457 | -1.59125 | -0.01968 | 0.048152 |
| 18 | 19469400 | 19470399 | 16.36028 | 30.73119 | 15.74166 | 6.471528 | 1.423257 | 0.127074 | 0.048169 |
| 13 | 25197000 | 25198799 | 19.80373 | 36.27214 | 14.7508 | 6.29776 | 1.713336 | 0.297167 | 0.048179 |
| 3 | 44955200 | 44956999 | 19.46897 | 35.46341 | 18.02718 | 7.751062 | 1.387495 | 0.332259 | 0.048189 |
| 1 | 187383200 | 187385599 | 6.794164 | 12.00728 | 47.72934 | 21.1133 | -1.58721 | 0.508022 | 0.048193 |
| 12 | 23563200 | 23569599 | 90.57223 | 142.922 | 96.7961 | 40.17803 | 0.885957 | 2.46632 | 0.048196 |
| 5 | 95798600 | 95798999 | 11.11154 | 21.70645 | 8.432416 | 2.946037 | 2.020753 | -0.4854 | 0.048206 |
| 1 | 108912800 | 108917999 | 103.6976 | 162.4858 | 94.01185 | 38.99456 | 1.105048 | 2.608491 | 0.048219 |
| 2 | 39602600 | 39606599 | 57.16215 | 92.24036 | 56.25222 | 23.97759 | 1.070083 | 1.77338 | 0.048245 |
| 2 | 91020200 | 91021199 | 3.551349 | 6.864529 | 30.40389 | 11.65285 | -1.5662 | -0.18708 | 0.048257 |
| 4 | 155789200 | 155790399 | 4.419778 | 8.406741 | 33.12413 | 13.28426 | -1.46198 | -0.0176 | 0.04827 |
| 4 | 140471400 | 140489199 | 79.11326 | 119.8638 | 311.1495 | 128.7582 | -1.09611 | 3.25778 | 0.048274 |
| 15 | 80262200 | 80293599 | 193.9155 | 273.9366 | 700.6492 | 290.0538 | -1.17135 | 4.425561 | 0.048288 |
| 15 | 97990600 | 97991599 | 3.154819 | 6.092069 | 31.43721 | 12.10437 | -1.78955 | -0.17286 | 0.048299 |
| 17 | 31547000 | 31560199 | 66.83975 | 102.5729 | 259.4463 | 107.7104 | -1.04377 | 2.990392 | 0.048306 |
| 6 | 116169400 | 116171599 | 7.819274 | 13.80868 | 47.20955 | 20.88647 | -1.37216 | 0.522168 | 0.048327 |
| 5 | 101224400 | 101225399 | 12.58098 | 24.35611 | 12.79904 | 4.869927 | 1.484517 | -0.16681 | 0.048337 |
| 2 | 105240800 | 105241999 | 4.511788 | 8.697491 | 32.77045 | 12.69247 | -1.35193 | -0.13446 | 0.048341 |
| 10 | 128132400 | 128134999 | 6.854226 | 12.31075 | 43.19402 | 18.88705 | -1.39539 | 0.38016 | 0.048357 |
| 11 | 96807000 | 96807599 | 2.642691 | 5.152462 | 25.3379 | 9.129813 | -1.62658 | -0.43147 | 0.048426 |
| 2 | 126547400 | 126549599 | 8.363529 | 14.94302 | 46.32766 | 20.34701 | -1.22358 | 0.429409 | 0.048427 |
| 2 | 11617000 | 11618199 | 4.073312 | 7.896606 | 30.85278 | 11.66375 | -1.36947 | -0.22757 | 0.048489 |
| 10 | 100470600 | 100471799 | 4.073312 | 7.896606 | 30.85278 | 11.66375 | -1.36947 | -0.22757 | 0.048489 |
| X | 29999600 | 30000999 | 18.23647 | 33.51793 | 17.34005 | 7.368279 | 1.374044 | 0.272449 | 0.048495 |
| 2 | 163665800 | 163671199 | 23.16278 | 37.37843 | 111.4111 | 47.49163 | -1.21326 | 1.745613 | 0.048501 |
| 12 | 15624800 | 15625599 | 2.547951 | 4.970037 | 26.10186 | 9.348505 | -1.71291 | -0.45422 | 0.048537 |
| 14 | 42612000 | 42614399 | 25.92853 | 45.62389 | 19.49768 | 8.632412 | 1.60651 | 0.614373 | 0.048538 |
| 8 | 3101600 | 3103399 | 22.93098 | 40.33677 | 23.26965 | 10.30306 | 1.176947 | 0.607233 | 0.048553 |
| 17 | 38011400 | 38017599 | 114.3982 | 178.238 | 111.0709 | 46.09941 | 0.991907 | 2.716563 | 0.048565 |
| 18 | 81484000 | 81485199 | 14.97187 | 28.37486 | 15.2825 | 6.175992 | 1.371687 | 0.055563 | 0.048567 |

| | | | | | | | | | |
|---|---|---|---|---|---|---|---|---|---|
| 14 | 50694800 | 50695599 | 11.06548 | 21.53463 | 11.89193 | 4.376823 | 1.457138 | -0.29876 | 0.048578 |
| 18 | 58633600 | 58635399 | 8.919261 | 15.69949 | 50.92063 | 22.54315 | -1.29967 | 0.569031 | 0.048581 |
| 4 | 154222000 | 154228199 | 30.05731 | 48.0665 | 139.1003 | 58.5213 | -1.18701 | 2.047209 | 0.048607 |
| 4 | 117468800 | 117470999 | 5.501434 | 10.02288 | 42.97198 | 18.47225 | -1.66532 | 0.283145 | 0.04861 |
| 12 | 73115800 | 73117799 | 7.15616 | 12.77338 | 44.16351 | 19.41245 | -1.3796 | 0.425336 | 0.04861 |
| 17 | 29927200 | 29936399 | 42.85354 | 67.76768 | 172.7087 | 71.76565 | -1.02061 | 2.398269 | 0.048615 |
| 10 | 105608400 | 105609199 | 2.117616 | 4.123074 | 26.69699 | 9.788139 | -2.03426 | -0.36636 | 0.048634 |
| 4 | 112021200 | 112022199 | 27.0375 | 46.97391 | 26.29408 | 11.54422 | 1.220019 | 0.81458 | 0.048668 |
| 3 | 87078800 | 87082799 | 17.01501 | 28.11863 | 86.15069 | 37.22396 | -1.22956 | 1.328097 | 0.048678 |
| 10 | 6473600 | 6474599 | 13.59119 | 26.481 | 8.017451 | 2.914366 | 2.323714 | -0.32061 | 0.048688 |
| 10 | 114635800 | 114639199 | 25.43449 | 40.9757 | 116.9021 | 49.72214 | -1.15333 | 1.800653 | 0.048698 |
| 12 | 92175400 | 92177599 | 24.41537 | 43.07654 | 20.80224 | 9.205062 | 1.433111 | 0.58 | 0.048724 |
| 2 | 12654600 | 12656199 | 25.83638 | 45.38096 | 21.24419 | 9.402004 | 1.475822 | 0.639517 | 0.048725 |
| 13 | 66183400 | 66183799 | 8.265279 | 16.16506 | 9.410533 | 3.189186 | 1.485608 | -0.62775 | 0.048725 |
| 5 | 23504800 | 23506199 | 15.61348 | 29.70799 | 14.27776 | 5.72194 | 1.545148 | 0.033472 | 0.048745 |
| 3 | 127509200 | 127510199 | 3.952437 | 7.663246 | 30.86168 | 11.6602 | -1.41197 | -0.23181 | 0.048792 |
| 11 | 84409200 | 84421799 | 73.61695 | 112.1521 | 290.8564 | 120.5604 | -1.08816 | 3.141967 | 0.048799 |
| 3 | 152044600 | 152049999 | 19.27151 | 31.37284 | 103.7086 | 44.52364 | -1.34981 | 1.542651 | 0.048821 |
| 6 | 72858400 | 72860199 | 5.926932 | 11.10706 | 37.27147 | 15.38364 | -1.27059 | 0.105948 | 0.048826 |
| 3 | 69281800 | 69282599 | 2.942188 | 5.735807 | 26.02712 | 9.390679 | -1.51612 | -0.42192 | 0.048843 |
| 13 | 92744000 | 92748799 | 27.47203 | 44.3227 | 115.2787 | 49.12644 | -1.01782 | 1.758978 | 0.048857 |
| 5 | 105231400 | 105232599 | 10.17646 | 19.83312 | 10.23589 | 3.707633 | 1.572083 | -0.37199 | 0.048876 |
| 4 | 137568400 | 137591199 | 117.9224 | 174.4286 | 424.9933 | 175.0237 | -1.03606 | 3.719748 | 0.048888 |
| 19 | 6291200 | 6318599 | 122.6158 | 178.8831 | 503.3801 | 207.4171 | -1.26273 | 3.942933 | 0.0489 |
| 7 | 3088400 | 3092599 | 71.75526 | 114.8613 | 65.93631 | 27.77897 | 1.14365 | 2.04089 | 0.048937 |
| 6 | 71228000 | 71228799 | 3.044344 | 5.940136 | 25.67137 | 9.152962 | -1.43103 | -0.46399 | 0.048945 |
| 1 | 113410200 | 113412799 | 34.27907 | 58.34486 | 29.42659 | 12.89803 | 1.364872 | 1.033 | 0.048963 |
| 4 | 146640400 | 146641799 | 10.59421 | 20.66396 | 10.11459 | 3.62519 | 1.660737 | -0.40186 | 0.048967 |
| 16 | 34992400 | 34994999 | 5.442156 | 9.94821 | 39.7817 | 17.02924 | -1.5607 | 0.25855 | 0.048994 |
| 16 | 83318400 | 83319399 | 11.90304 | 23.12361 | 12.4302 | 4.640871 | 1.477671 | -0.23779 | 0.049009 |
| 19 | 45718400 | 45737999 | 105.6123 | 156.9919 | 418.9287 | 172.5905 | -1.16056 | 3.632341 | 0.04901 |
| 17 | 46907400 | 46909799 | 11.38547 | 19.4995 | 63.15066 | 27.69356 | -1.30438 | 0.931641 | 0.049017 |
| 6 | 100131600 | 100135199 | 14.15065 | 23.85818 | 71.87709 | 31.42726 | -1.20374 | 1.091662 | 0.049042 |
| 12 | 18798400 | 18805199 | 296.5001 | 429.8022 | 268.7674 | 110.8207 | 0.897127 | 4.048515 | 0.04906 |
| 10 | 117848400 | 117853999 | 28.65389 | 46.07917 | 121.8673 | 51.69883 | -1.04768 | 1.863884 | 0.04906 |
| 6 | 42068400 | 42070399 | 23.84505 | 41.95629 | 22.713 | 10.05604 | 1.268345 | 0.609374 | 0.049066 |
| 1 | 155585200 | 155592599 | 38.11345 | 60.43878 | 166.2632 | 69.20961 | -1.12717 | 2.323208 | 0.049074 |
| 11 | 114787000 | 114789199 | 4.522123 | 8.457833 | 37.06049 | 15.34748 | -1.65438 | 0.106162 | 0.049118 |
| 1 | 180489200 | 180489599 | 41.60441 | 70.23148 | 25.42107 | 11.11914 | 1.840431 | 1.126771 | 0.049129 |
| 19 | 3678000 | 3683199 | 17.65127 | 28.68203 | 99.54376 | 42.70161 | -1.42143 | 1.569336 | 0.04914 |
| 6 | 71215400 | 71218199 | 14.29297 | 24.11872 | 70.78561 | 30.95805 | -1.16607 | 1.083839 | 0.049175 |
| 7 | 22152200 | 22152599 | 14.00154 | 26.65575 | 13.64909 | 5.463564 | 1.456141 | 0.011608 | 0.04919 |
| 15 | 83088200 | 83092599 | 19.88411 | 32.68454 | 87.81365 | 37.84204 | -1.04334 | 1.403424 | 0.049214 |
| 13 | 64838800 | 64839399 | 10.43296 | 20.29458 | 12.13657 | 4.483094 | 1.339521 | -0.29379 | 0.049219 |
| 9 | 3150200 | 3164599 | 307.217 | 443.8312 | 282.4571 | 116.5176 | 0.867007 | 4.099047 | 0.049236 |
| 13 | 66599000 | 66600399 | 38.79219 | 64.46638 | 39.5599 | 17.1591 | 1.083104 | 1.283411 | 0.049255 |
| 7 | 31674800 | 31678199 | 57.08011 | 92.17808 | 54.11268 | 23.08825 | 1.126092 | 1.749798 | 0.049259 |

| | | | | | | | | | |
|---|---|---|---|---|---|---|---|---|---|
| 8 | 25338400 | 25338999 | 10.59168 | 20.66834 | 9.484786 | 3.376643 | 1.760032 | -0.42699 | 0.04928 |
| 2 | 88009200 | 88010799 | 19.79517 | 35.45999 | 19.55966 | 8.571732 | 1.252973 | 0.432125 | 0.049295 |
| 13 | 54267000 | 54268799 | 6.340875 | 11.59516 | 40.92912 | 17.51266 | -1.38321 | 0.263449 | 0.049343 |
| 13 | 51642800 | 51644999 | 10.14269 | 17.78765 | 52.01784 | 22.99116 | -1.15224 | 0.631299 | 0.049416 |
| 10 | 80498200 | 80507999 | 29.68683 | 47.2746 | 155.4449 | 65.01296 | -1.37758 | 2.185345 | 0.049417 |
| 8 | 91571000 | 91572599 | 5.787049 | 10.85055 | 36.71243 | 15.13959 | -1.28131 | 0.099688 | 0.049424 |
| 1 | 3515800 | 3516799 | 11.18288 | 21.70004 | 12.18679 | 4.586666 | 1.405142 | -0.22069 | 0.04943 |
| 19 | 25346000 | 25350399 | 15.75427 | 26.3934 | 74.05288 | 32.28755 | -1.10187 | 1.161223 | 0.049435 |
| 13 | 100969200 | 100970199 | 5.835537 | 10.7895 | 37.90532 | 15.97717 | -1.35965 | 0.190994 | 0.04944 |
| X | 125690200 | 125690799 | 11.76548 | 23.00396 | 6.313057 | 2.159589 | 2.537988 | -0.5532 | 0.04949 |
| 8 | 45785600 | 45787399 | 5.430715 | 9.77042 | 49.14152 | 21.45722 | -1.90925 | 0.368554 | 0.049495 |
| 10 | 80988200 | 81001599 | 67.67052 | 103.1438 | 290.2101 | 120.3097 | -1.20501 | 3.130472 | 0.049509 |
| 4 | 94927400 | 94927999 | 1.643291 | 3.20998 | 23.147 | 8.095509 | -2.11907 | -0.5587 | 0.049529 |
| 9 | 3179000 | 3182599 | 70.7677 | 113.073 | 73.15067 | 30.73848 | 0.968747 | 2.095427 | 0.04955 |
| 11 | 107950800 | 107954399 | 17.3775 | 28.94663 | 78.59488 | 34.14435 | -1.05491 | 1.231659 | 0.049591 |
| 16 | 15412000 | 15413799 | 8.013116 | 14.14813 | 52.57014 | 23.25905 | -1.49262 | 0.531065 | 0.049591 |
| 7 | 23191800 | 23193799 | 57.94372 | 93.60557 | 52.46709 | 22.39626 | 1.193199 | 1.740972 | 0.0496 |
| 7 | 62116000 | 62118199 | 26.91659 | 46.97927 | 24.62111 | 10.84099 | 1.314534 | 0.747087 | 0.049615 |
| 18 | 3150200 | 3152799 | 70.47034 | 113.1595 | 56.56381 | 23.94417 | 1.347979 | 1.941197 | 0.049631 |
| 9 | 82856200 | 82858199 | 9.339542 | 16.15921 | 59.41168 | 26.06537 | -1.48275 | 0.824552 | 0.049632 |
| 4 | 38912000 | 38912999 | 10.99084 | 21.37519 | 11.86128 | 4.388996 | 1.44292 | -0.28214 | 0.049649 |
| 9 | 63604400 | 63606799 | 10.25121 | 17.70702 | 59.44083 | 26.07684 | -1.35301 | 0.844041 | 0.049649 |
| 7 | 45838200 | 45840199 | 5.391833 | 10.00258 | 37.41757 | 15.69985 | -1.44421 | 0.166944 | 0.049702 |
| 15 | 11063200 | 11065999 | 9.5859 | 16.73199 | 55.24395 | 24.32562 | -1.32663 | 0.70773 | 0.049718 |
| 1 | 162946600 | 162947199 | 2.334628 | 4.566484 | 22.86906 | 7.682266 | -1.55779 | -0.69643 | 0.049726 |
| 14 | 46433400 | 46434199 | 3.140847 | 6.102504 | 31.70966 | 11.82496 | -1.75614 | -0.28627 | 0.049736 |
| 8 | 52210600 | 52211799 | 12.30756 | 23.83398 | 12.18377 | 4.629274 | 1.524954 | -0.17701 | 0.049766 |
| 17 | 31239200 | 31240999 | 6.328261 | 11.52416 | 40.26731 | 17.31916 | -1.37299 | 0.288727 | 0.049771 |
| 8 | 88670000 | 88671199 | 4.450606 | 8.366763 | 34.95783 | 14.35187 | -1.57511 | 0.069218 | 0.049777 |
| 4 | 45818200 | 45827399 | 45.76713 | 71.60557 | 205.9364 | 85.43225 | -1.20693 | 2.622994 | 0.049795 |
| 19 | 5752000 | 5756999 | 14.11088 | 23.28901 | 85.15389 | 36.76022 | -1.48312 | 1.339494 | 0.049807 |
| 15 | 98334800 | 98336599 | 8.635208 | 15.14951 | 50.90672 | 22.50709 | -1.35095 | 0.615538 | 0.049819 |
| 11 | 53915600 | 53920599 | 18.64555 | 30.3347 | 101.5113 | 43.57029 | -1.36792 | 1.55026 | 0.049832 |
| 12 | 22636400 | 22642799 | 83.7449 | 133.3927 | 70.94751 | 29.6833 | 1.245995 | 2.204775 | 0.049836 |
| 8 | 89014000 | 89023399 | 43.28736 | 68.18947 | 184.7953 | 76.67843 | -1.11357 | 2.487879 | 0.049841 |
| 18 | 29375000 | 29375999 | 12.38224 | 24.0022 | 12.96636 | 4.904025 | 1.453914 | -0.18752 | 0.04988 |
| 8 | 59919400 | 59920599 | 14.26775 | 27.78198 | 7.980967 | 2.924287 | 2.388661 | -0.28544 | 0.049883 |
| 4 | 112178200 | 112181399 | 73.89569 | 117.8714 | 76.04015 | 31.87647 | 0.970212 | 2.15079 | 0.049886 |
| 14 | 31288200 | 31295199 | 25.86659 | 41.75723 | 110.6738 | 47.20008 | -1.04384 | 1.737667 | 0.049891 |
| 13 | 13131200 | 13131599 | 11.43997 | 22.21306 | 12.66224 | 4.74638 | 1.389503 | -0.22895 | 0.049897 |
| 12 | 70294600 | 70295399 | 3.241062 | 6.309671 | 32.30081 | 11.85472 | -1.71378 | -0.34088 | 0.049922 |
| 5 | 99793200 | 99793999 | 11.91285 | 23.09971 | 13.09706 | 4.946069 | 1.387207 | -0.19734 | 0.049958 |
| 3 | 41812600 | 41813399 | 9.172937 | 17.9397 | 8.549808 | 2.905068 | 1.76452 | -0.6076 | 0.050007 |
| 19 | 41808400 | 41810199 | 5.659183 | 10.5883 | 37.19199 | 15.39259 | -1.33913 | 0.113048 | 0.050009 |
| 7 | 138655800 | 138660399 | 20.78054 | 33.88234 | 94.53235 | 40.60726 | -1.10418 | 1.519429 | 0.05001 |
| 6 | 52766800 | 52768799 | 9.697041 | 16.97092 | 52.01048 | 22.94772 | -1.21927 | 0.663624 | 0.050018 |
| 7 | 8474400 | 8477199 | 60.2416 | 97.01397 | 53.44776 | 22.72045 | 1.212924 | 1.835826 | 0.050021 |

| | | | | | | | | | |
|---|---|---|---|---|---|---|---|---|---|
| X | 31624600 | 31626199 | 17.24461 | 32.83176 | 12.31015 | 4.926986 | 1.900149 | 0.039878 | 0.050039 |
| 16 | 30426000 | 30428199 | 8.051456 | 14.39995 | 44.73037 | 19.6288 | -1.22512 | 0.418854 | 0.050058 |
| 15 | 25932600 | 25933799 | 4.155271 | 7.970051 | 33.585 | 13.1971 | -1.53155 | -0.08993 | 0.050066 |
| X | 25245400 | 25247199 | 20.82808 | 38.33518 | 14.60737 | 6.195202 | 1.812355 | 0.278984 | 0.050076 |
| 5 | 95348800 | 95349199 | 7.858347 | 15.37232 | 8.552347 | 2.847949 | 1.573493 | -0.69648 | 0.050089 |
| 2 | 169580000 | 169584399 | 27.80538 | 44.81334 | 114.0685 | 48.54479 | -0.98852 | 1.790661 | 0.050098 |
| 19 | 5198600 | 5200199 | 4.861877 | 9.197537 | 34.09364 | 13.83111 | -1.39015 | 0.027522 | 0.050104 |
| 7 | 49243600 | 49259199 | 62.21644 | 95.32659 | 273.1771 | 113.3851 | -1.22542 | 3.021241 | 0.050107 |
| 10 | 81215800 | 81218399 | 8.429812 | 14.83403 | 48.77149 | 21.59284 | -1.31851 | 0.568841 | 0.050125 |
| 8 | 106247200 | 106250199 | 9.702677 | 16.65336 | 63.06087 | 27.659 | -1.52778 | 0.904933 | 0.050126 |
| 7 | 24795600 | 24797599 | 9.570257 | 16.62754 | 56.77302 | 24.92628 | -1.3752 | 0.779765 | 0.050127 |
| 19 | 3078400 | 3079599 | 20.69051 | 37.81618 | 15.57279 | 6.667492 | 1.691849 | 0.317179 | 0.05014 |
| 10 | 75411400 | 75416599 | 27.65129 | 44.49695 | 117.8162 | 50.02931 | -1.04797 | 1.841496 | 0.050144 |
| 13 | 16219800 | 16220799 | 15.70648 | 29.45931 | 16.42006 | 6.767417 | 1.299821 | 0.132088 | 0.050146 |
| 17 | 24126800 | 24127799 | 4.842794 | 9.225613 | 33.24444 | 13.28895 | -1.33095 | -0.02452 | 0.050183 |
| 3 | 27659600 | 27661799 | 6.718514 | 12.0677 | 43.7251 | 19.11833 | -1.44148 | 0.379263 | 0.050184 |
| 8 | 111696000 | 111697999 | 9.2025 | 16.10358 | 53.58219 | 23.6389 | -1.33683 | 0.667268 | 0.050215 |
| 8 | 57675600 | 57676799 | 14.50337 | 27.86306 | 13.09394 | 5.121524 | 1.607377 | -0.05706 | 0.050225 |
| 15 | 78056000 | 78062999 | 27.23003 | 43.75172 | 122.5513 | 51.92414 | -1.13181 | 1.890384 | 0.050271 |
| 4 | 51071600 | 51073799 | 21.66246 | 39.0828 | 17.78104 | 7.73736 | 1.53548 | 0.401178 | 0.050271 |
| 7 | 63016000 | 63018399 | 36.36516 | 61.36277 | 32.21558 | 14.08934 | 1.306897 | 1.117375 | 0.050276 |
| 19 | 43587000 | 43595799 | 58.37446 | 91.0055 | 213.5575 | 88.62791 | -0.91805 | 2.692965 | 0.050329 |
| 17 | 29141200 | 29151599 | 46.98354 | 73.7064 | 200.825 | 83.28876 | -1.12606 | 2.574672 | 0.050329 |
| 8 | 126721800 | 126724999 | 12.69163 | 21.55989 | 67.45429 | 29.56084 | -1.25715 | 1.016376 | 0.050368 |
| 5 | 94798200 | 94801599 | 34.6092 | 58.81396 | 28.52234 | 12.5001 | 1.420903 | 1.047295 | 0.050376 |
| 7 | 13308800 | 13310199 | 39.48864 | 65.89816 | 33.86366 | 14.72845 | 1.338224 | 1.238261 | 0.050385 |
| 9 | 19928000 | 19930399 | 60.07606 | 96.51461 | 63.78782 | 27.02147 | 0.946814 | 1.913717 | 0.050393 |
| 3 | 89130200 | 89136199 | 22.45937 | 36.07788 | 125.6202 | 53.20603 | -1.4451 | 1.893735 | 0.050413 |
| 5 | 4488800 | 4490599 | 25.02226 | 43.90288 | 23.28295 | 10.29476 | 1.297552 | 0.656521 | 0.050422 |
| 6 | 59635400 | 59635999 | 9.74897 | 19.02576 | 10.95999 | 3.895223 | 1.441194 | -0.43881 | 0.050426 |
| 6 | 87851000 | 87851599 | 9.74897 | 19.02576 | 10.95999 | 3.895223 | 1.441194 | -0.43881 | 0.050426 |
| 4 | 140438800 | 140470199 | 139.4781 | 201.2057 | 570.1053 | 235.2223 | -1.28858 | 4.114121 | 0.050464 |
| 11 | 5104800 | 5129199 | 125.3967 | 183.7768 | 478.114 | 196.938 | -1.14325 | 3.871505 | 0.050475 |
| 10 | 57252400 | 57253399 | 12.81177 | 24.65575 | 13.87409 | 5.399849 | 1.356838 | -0.08871 | 0.050484 |
| 8 | 88191400 | 88201199 | 65.26887 | 100.0722 | 265.5538 | 110.2323 | -1.11387 | 3.008017 | 0.050497 |
| 9 | 62150000 | 62152799 | 10.87983 | 18.71031 | 61.01088 | 26.76203 | -1.31285 | 0.889757 | 0.050504 |
| 9 | 100039800 | 100041199 | 3.155417 | 6.145564 | 33.77729 | 12.34203 | -1.80793 | -0.35142 | 0.050518 |
| 12 | 23526000 | 23526399 | 8.661324 | 16.92297 | 5.459092 | 1.745155 | 2.383315 | -0.83076 | 0.05055 |
| 13 | 38817400 | 38819199 | 7.699918 | 13.6217 | 48.8097 | 21.58686 | -1.43893 | 0.502123 | 0.050552 |
| 7 | 142461600 | 142464199 | 8.137018 | 14.40504 | 48.2894 | 21.35376 | -1.34321 | 0.496012 | 0.050558 |
| 7 | 31991000 | 31993599 | 61.16972 | 98.28296 | 64.05115 | 27.13755 | 0.967203 | 1.911059 | 0.050561 |
| 4 | 117585600 | 117587599 | 8.870173 | 15.71115 | 47.23228 | 20.88412 | -1.18685 | 0.493701 | 0.050567 |
| 19 | 41818200 | 41824599 | 29.86214 | 47.80209 | 133.3186 | 56.16824 | -1.13219 | 2.014832 | 0.0506 |
| 9 | 87164200 | 87164599 | 6.632728 | 12.96021 | 6.351377 | 2.032586 | 1.798721 | -0.86657 | 0.050603 |
| 1 | 178007000 | 178008199 | 10.54034 | 20.48861 | 12.11012 | 4.49954 | 1.348405 | -0.27292 | 0.050638 |
| 14 | 29850800 | 29861599 | 48.94594 | 76.71814 | 200.1578 | 83.01947 | -1.06457 | 2.591231 | 0.050639 |
| 13 | 19240000 | 19241599 | 17.60182 | 32.29838 | 18.23136 | 7.764358 | 1.246639 | 0.277601 | 0.050671 |

| | | | | | | | | | |
|---|---|---|---|---|---|---|---|---|---|
| 14 | 42343800 | 42346199 | 31.90825 | 54.14549 | 33.51417 | 14.68227 | 1.070624 | 1.055346 | 0.05069 |
| 13 | 66666200 | 66672799 | 145.0707 | 222.5537 | 125.9495 | 52.28618 | 1.113772 | 3.010278 | 0.050697 |
| 15 | 78933800 | 78940999 | 25.91119 | 41.46724 | 139.3655 | 58.69512 | -1.40096 | 2.01976 | 0.050777 |
| X | 122907800 | 122908199 | 10.77842 | 21.00142 | 11.3341 | 4.117933 | 1.506006 | -0.34841 | 0.050795 |
| 19 | 41853400 | 41855199 | 4.514664 | 8.464102 | 36.58256 | 15.08832 | -1.63001 | 0.089158 | 0.050803 |
| 7 | 134305600 | 134315999 | 55.02111 | 86.00249 | 201.3062 | 83.51956 | -0.91139 | 2.644113 | 0.050811 |
| 1 | 49270800 | 49272999 | 31.46294 | 53.62807 | 32.40728 | 14.20614 | 1.105764 | 1.013924 | 0.050867 |
| 19 | 7634200 | 7635999 | 16.93597 | 31.41386 | 17.20785 | 7.221611 | 1.304343 | 0.211197 | 0.05087 |
| 2 | 27116400 | 27120999 | 13.36456 | 22.13197 | 82.53022 | 35.71087 | -1.51088 | 1.294149 | 0.050905 |
| 12 | 20469800 | 20474199 | 129.7176 | 199.4549 | 136.7316 | 56.77997 | 0.839969 | 2.961381 | 0.05091 |
| 12 | 19087600 | 19092799 | 115.7476 | 180.2715 | 111.1765 | 46.14589 | 1.00646 | 2.723807 | 0.050915 |
| 7 | 33053800 | 33054199 | 8.615594 | 16.83191 | 9.473769 | 3.305428 | 1.496515 | -0.52537 | 0.050933 |
| 7 | 22036200 | 22036599 | 7.531522 | 14.73311 | 9.005688 | 2.996715 | 1.43987 | -0.70306 | 0.050937 |
| 18 | 25747800 | 25751199 | 13.63326 | 23.13042 | 70.36414 | 30.8243 | -1.21724 | 1.032449 | 0.050954 |
| 7 | 28762600 | 28779999 | 108.6266 | 161.6208 | 394.9257 | 162.717 | -1.03237 | 3.616195 | 0.050979 |
| 6 | 29396000 | 29399599 | 12.83967 | 21.81051 | 66.90144 | 29.31821 | -1.22871 | 1.01705 | 0.050985 |
| 10 | 3584000 | 3585199 | 5.471029 | 10.31539 | 35.17552 | 14.36944 | -1.28089 | 0.057642 | 0.051 |
| 14 | 30608400 | 30624399 | 93.70665 | 141.0945 | 337.4611 | 139.3821 | -0.98586 | 3.386964 | 0.051003 |
| 8 | 11361600 | 11369799 | 38.42695 | 60.929 | 164.0588 | 68.28425 | -1.09646 | 2.326852 | 0.051035 |
| 15 | 78550800 | 78586199 | 169.7934 | 239.0069 | 741.2434 | 307.0396 | -1.45443 | 4.477465 | 0.051035 |
| 14 | 14463400 | 14463799 | 7.373499 | 14.41744 | 7.575145 | 2.455887 | 1.683408 | -0.80014 | 0.05105 |
| 17 | 22754200 | 22754999 | 24.54851 | 42.94549 | 25.43812 | 11.21984 | 1.139105 | 0.702422 | 0.05109 |
| 5 | 59353200 | 59358599 | 34.78207 | 58.88605 | 29.92973 | 13.10278 | 1.353613 | 1.082444 | 0.051118 |
| 10 | 78671600 | 78672399 | 10.58944 | 20.67223 | 8.840264 | 3.125562 | 1.867772 | -0.45261 | 0.051128 |
| 13 | 120644600 | 120645199 | 10.58944 | 20.67223 | 8.840264 | 3.125562 | 1.867772 | -0.45261 | 0.051128 |
| 14 | 118068000 | 118070199 | 12.02388 | 20.54568 | 64.25334 | 28.17192 | -1.25495 | 0.956172 | 0.051146 |
| 4 | 129179600 | 129188399 | 43.72842 | 68.96339 | 179.2524 | 74.38957 | -1.05214 | 2.464508 | 0.051178 |
| 1 | 27680000 | 27680599 | 8.218325 | 16.07 | 9.383027 | 3.202222 | 1.471892 | -0.60598 | 0.051179 |
| 4 | 121963200 | 121963799 | 11.2922 | 21.88056 | 13.07924 | 4.955883 | 1.30662 | -0.19472 | 0.051185 |
| 16 | 90204400 | 90205999 | 5.9328 | 11.00948 | 39.11653 | 16.40595 | -1.37036 | 0.174876 | 0.051192 |
| 18 | 3054600 | 3056399 | 17.73603 | 33.5923 | 12.05418 | 4.877875 | 1.948397 | 0.077567 | 0.051234 |
| 4 | 135152200 | 135153999 | 4.444099 | 8.375126 | 34.47339 | 14.09409 | -1.54883 | 0.051755 | 0.051293 |
| 4 | 112178400 | 112181399 | 73.39786 | 117.0969 | 76.03115 | 31.88005 | 0.96114 | 2.144989 | 0.051333 |
| 10 | 117618200 | 117618999 | 3.661017 | 7.084892 | 29.3178 | 11.17461 | -1.46125 | -0.20697 | 0.051334 |
| 7 | 60292600 | 60294199 | 31.15537 | 53.11517 | 32.46213 | 14.23043 | 1.089599 | 1.011103 | 0.051359 |
| 4 | 28321200 | 28321599 | 10.78853 | 20.98194 | 12.60543 | 4.664005 | 1.330958 | -0.28168 | 0.051361 |
| 18 | 75405800 | 75412599 | 28.5807 | 45.78699 | 132.9668 | 56.08672 | -1.189 | 1.987932 | 0.051366 |
| 14 | 121595600 | 121600199 | 19.45737 | 31.91176 | 91.10727 | 39.22431 | -1.13206 | 1.431584 | 0.051379 |
| 17 | 71521400 | 71522199 | 3.648903 | 7.099126 | 29.53549 | 10.89578 | -1.42518 | -0.32374 | 0.051403 |
| 8 | 95868600 | 95871399 | 10.18911 | 17.72147 | 55.75506 | 24.49562 | -1.25793 | 0.765041 | 0.051411 |
| 12 | 57927000 | 57929199 | 28.91134 | 50.2073 | 26.2659 | 11.52865 | 1.316802 | 0.828655 | 0.051434 |
| 14 | 13704000 | 13705599 | 15.97737 | 30.39895 | 12.7611 | 5.114631 | 1.736371 | 0.032437 | 0.051436 |
| 16 | 3287600 | 3293399 | 136.7772 | 209.8878 | 139.8286 | 58.05004 | 0.878987 | 3.003033 | 0.051443 |
| 12 | 21873200 | 21875399 | 75.66521 | 120.4536 | 79.2756 | 33.14123 | 0.938423 | 2.217491 | 0.051453 |
| X | 97952200 | 97953999 | 18.41203 | 33.908 | 17.24258 | 7.307054 | 1.401344 | 0.262881 | 0.051461 |
| 14 | 60016600 | 60017999 | 15.39531 | 28.70331 | 16.46418 | 6.854747 | 1.247061 | 0.165704 | 0.051484 |
| 9 | 58489600 | 58490399 | 3.780146 | 7.305704 | 29.79642 | 11.42788 | -1.44913 | -0.18184 | 0.051503 |

| | | | | | | | | | |
|---|---|---|---|---|---|---|---|---|---|
| 6 | 123687400 | 123688799 | 12.39459 | 23.97065 | 13.15614 | 5.029853 | 1.417421 | -0.15156 | 0.051513 |
| 8 | 112074600 | 112086199 | 72.19396 | 111.223 | 243.9388 | 101.3266 | -0.83398 | 2.91434 | 0.051518 |
| 6 | 79766600 | 79768399 | 14.51457 | 27.46025 | 15.95634 | 6.472135 | 1.258874 | 0.064766 | 0.051519 |
| 1 | 102775400 | 102776199 | 11.61274 | 22.59469 | 11.88168 | 4.380837 | 1.524986 | -0.28541 | 0.051522 |
| 4 | 129822200 | 129823999 | 6.074771 | 11.01851 | 41.25006 | 17.83056 | -1.47651 | 0.311673 | 0.051524 |
| 16 | 52947000 | 52947599 | 9.574904 | 18.68671 | 10.96294 | 3.893928 | 1.415792 | -0.44356 | 0.051525 |
| 4 | 129839800 | 129843599 | 13.54791 | 22.72963 | 72.63795 | 31.69932 | -1.28845 | 1.138808 | 0.051529 |
| 11 | 29642000 | 29644599 | 15.45405 | 25.87895 | 74.58256 | 32.5094 | -1.1404 | 1.165147 | 0.051559 |
| 5 | 33977000 | 33978999 | 9.604669 | 16.72081 | 56.2976 | 24.74871 | -1.35512 | 0.747665 | 0.051563 |
| 12 | 22062800 | 22067999 | 155.5741 | 236.5688 | 158.9095 | 65.83147 | 0.856809 | 3.191817 | 0.051566 |
| 5 | 137100800 | 137107199 | 27.3905 | 44.04167 | 123.942 | 52.56896 | -1.13729 | 1.866856 | 0.051582 |
| 14 | 8768000 | 8770199 | 19.479 | 34.97036 | 19.57805 | 8.563913 | 1.233486 | 0.41667 | 0.051611 |
| 6 | 114663800 | 114666199 | 5.926169 | 11.10819 | 36.70426 | 15.14361 | -1.24796 | 0.103321 | 0.051622 |
| 3 | 42191800 | 42193199 | 16.60806 | 30.96646 | 16.69412 | 6.94969 | 1.335872 | 0.176761 | 0.051634 |
| 2 | 153434200 | 153435999 | 7.457337 | 13.33184 | 43.65973 | 19.16568 | -1.3008 | 0.415702 | 0.051648 |
| 5 | 110917600 | 110918999 | 7.457337 | 13.33184 | 43.65973 | 19.16568 | -1.3008 | 0.415702 | 0.051648 |
| 7 | 8221000 | 8221999 | 13.11981 | 25.14643 | 14.3967 | 5.66885 | 1.316912 | -0.0474 | 0.05165 |
| 2 | 90677400 | 90678199 | 4.695566 | 9.002969 | 32.42208 | 12.75395 | -1.30892 | -0.08263 | 0.051675 |
| 5 | 28612600 | 28613999 | 4.695566 | 9.002969 | 32.42208 | 12.75395 | -1.30892 | -0.08263 | 0.051675 |
| 5 | 14838800 | 14839799 | 11.83497 | 22.92104 | 13.54662 | 5.144746 | 1.319994 | -0.17895 | 0.0517 |
| 8 | 127551200 | 127553999 | 10.47454 | 18.01066 | 64.32576 | 28.21595 | -1.44384 | 0.890855 | 0.051725 |
| 19 | 31695000 | 31697199 | 10.29207 | 18.05569 | 53.10164 | 23.47685 | -1.16046 | 0.628228 | 0.05174 |
| 5 | 140421400 | 140439199 | 87.82172 | 132.404 | 331.6038 | 137.0101 | -1.05059 | 3.360508 | 0.051745 |
| 5 | 76965400 | 76965799 | 1.015621 | 1.98195 | 16.97868 | 5.335012 | -2.19659 | -0.97617 | 0.051762 |
| 4 | 116483800 | 116485199 | 6.476618 | 11.84552 | 39.84088 | 17.0429 | -1.31363 | 0.261985 | 0.051767 |
| 5 | 142755600 | 142781799 | 159.5022 | 229.8353 | 557.1315 | 229.9033 | -1.06517 | 4.13187 | 0.051769 |
| 4 | 133540200 | 133542199 | 7.076616 | 12.5623 | 46.19011 | 20.39738 | -1.47284 | 0.46222 | 0.051772 |
| 1 | 144619600 | 144621199 | 26.6047 | 47.02663 | 17.29846 | 7.651781 | 1.823322 | 0.567922 | 0.051773 |
| 11 | 101977800 | 101980199 | 6.082902 | 10.68511 | 50.82797 | 22.49599 | -1.84698 | 0.582356 | 0.051778 |
| 8 | 121257200 | 121314799 | 250.3921 | 341.3212 | 1042.103 | 433.8434 | -1.47874 | 4.97157 | 0.051783 |
| 18 | 12861800 | 12866999 | 24.16809 | 38.93192 | 117.0323 | 49.77088 | -1.22865 | 1.80201 | 0.051807 |
| 9 | 14736200 | 14736799 | 1.641772 | 3.211256 | 19.70525 | 6.485622 | -1.80915 | -0.79007 | 0.051823 |
| 12 | 20440400 | 20446399 | 123.4435 | 190.5104 | 127.213 | 52.84185 | 0.881863 | 2.889113 | 0.051835 |
| 5 | 143267600 | 143289599 | 137.301 | 200.1982 | 509.3601 | 209.893 | -1.11823 | 3.952157 | 0.051842 |
| 8 | 88993400 | 88995399 | 7.740054 | 13.85114 | 44.20391 | 19.38817 | -1.26304 | 0.412691 | 0.051847 |
| 14 | 22028000 | 22033799 | 25.29058 | 40.74566 | 120.7849 | 51.37689 | -1.20844 | 1.799748 | 0.051904 |
| Y | 4035800 | 4037399 | 44.4548 | 73.19625 | 35.84973 | 15.45998 | 1.405192 | 1.399392 | 0.051922 |
| 14 | 103336000 | 103338999 | 11.2926 | 19.26681 | 66.92123 | 29.33806 | -1.40599 | 0.969709 | 0.051939 |
| 7 | 41924000 | 41924799 | 25.96549 | 45.27387 | 26.40771 | 11.62033 | 1.161042 | 0.75886 | 0.051939 |
| 18 | 67581400 | 67582599 | 4.963642 | 9.215367 | 37.41009 | 15.68017 | -1.55953 | 0.15892 | 0.051948 |
| 6 | 29692400 | 29696399 | 14.84379 | 24.84789 | 75.63074 | 32.95864 | -1.21884 | 1.167945 | 0.051957 |
| 1 | 12605200 | 12606199 | 14.1484 | 26.79511 | 14.80247 | 5.991028 | 1.333066 | 0.050243 | 0.051969 |
| 8 | 71041400 | 71065199 | 121.7434 | 178.1734 | 480.8284 | 198.0734 | -1.19797 | 3.893491 | 0.051983 |
| 5 | 18209600 | 18210799 | 4.82153 | 9.257107 | 33.13797 | 12.98459 | -1.29518 | -0.09235 | 0.051984 |
| 7 | 5992200 | 5994199 | 60.21554 | 96.85168 | 62.39707 | 26.47708 | 0.985585 | 1.876958 | 0.051999 |
| 4 | 37953000 | 37956599 | 50.13543 | 81.3403 | 51.75649 | 22.17681 | 1.016694 | 1.631317 | 0.052032 |
| 17 | 29557200 | 29563399 | 22.3275 | 36.09119 | 106.4329 | 45.46815 | -1.19497 | 1.692782 | 0.052034 |

| | | | | | | | | | |
|---|---|---|---|---|---|---|---|---|---|
| 15 | 89186200 | 89187999 | 6.360336 | 11.48074 | 41.68725 | 18.12899 | -1.43844 | 0.344727 | 0.052038 |
| 9 | 44416000 | 44423599 | 29.48929 | 47.02198 | 148.9636 | 62.41598 | -1.32187 | 2.140558 | 0.052052 |
| 10 | 55722800 | 55723999 | 18.04776 | 33.59499 | 15.35479 | 6.410137 | 1.568366 | 0.195968 | 0.052061 |
| 12 | 19004400 | 19008599 | 90.11264 | 142.8901 | 82.57325 | 34.37072 | 1.120194 | 2.347053 | 0.052072 |
| 4 | 130661000 | 130665799 | 22.96764 | 37.16721 | 105.3637 | 45.06107 | -1.13658 | 1.664869 | 0.052077 |
| 6 | 91655800 | 91656999 | 3.244713 | 6.306055 | 27.92644 | 10.3909 | -1.52393 | -0.3012 | 0.052115 |
| 9 | 59601200 | 59604199 | 10.47872 | 17.94294 | 63.61331 | 27.896 | -1.43409 | 0.930561 | 0.052135 |
| 10 | 85176800 | 85177999 | 12.94773 | 25.09127 | 12.48384 | 4.727784 | 1.569999 | -0.17791 | 0.052135 |
| 11 | 75836800 | 75839199 | 7.500342 | 13.35645 | 45.18601 | 19.90111 | -1.35107 | 0.442615 | 0.052136 |
| 4 | 130669800 | 130689999 | 113.5569 | 167.5232 | 443.0077 | 182.4373 | -1.15776 | 3.763947 | 0.052137 |
| 12 | 40845400 | 40847399 | 5.207073 | 9.816551 | 36.90866 | 15.08045 | -1.42116 | 0.058295 | 0.052157 |
| 8 | 38248200 | 38248999 | 13.83435 | 26.27382 | 14.62908 | 5.885879 | 1.330037 | 0.028857 | 0.052162 |
| 12 | 112106000 | 112168999 | 331.1164 | 443.9112 | 1217.397 | 508.1061 | -1.34818 | 5.236426 | 0.052169 |
| 14 | 41194400 | 41199399 | 52.94525 | 85.96179 | 48.48277 | 20.78527 | 1.191182 | 1.622661 | 0.052176 |
| 7 | 22959400 | 22959999 | 13.94988 | 26.43145 | 14.60443 | 5.905046 | 1.33474 | 0.045284 | 0.052178 |
| 17 | 47153600 | 47159199 | 23.59871 | 38.20864 | 105.7774 | 45.25561 | -1.10181 | 1.654762 | 0.052203 |
| 12 | 76662200 | 76671199 | 49.76636 | 77.91587 | 197.4952 | 81.92466 | -1.02421 | 2.612222 | 0.052205 |
| 7 | 5900400 | 5906599 | 155.6028 | 237.0639 | 152.4225 | 63.18011 | 0.922124 | 3.152959 | 0.052239 |
| 10 | 126728800 | 126732999 | 22.42055 | 36.4381 | 98.5914 | 42.2985 | -1.0636 | 1.57248 | 0.052248 |
| 1 | 189788200 | 189789399 | 4.274112 | 8.200405 | 31.80417 | 12.48679 | -1.41141 | -0.09467 | 0.052278 |
| 7 | 22928400 | 22929999 | 11.32111 | 22.06666 | 10.71865 | 3.876261 | 1.661786 | -0.35875 | 0.052296 |
| 8 | 35389800 | 35390199 | 1.946409 | 3.807607 | 20.93937 | 6.95914 | -1.6724 | -0.74727 | 0.052316 |
| 6 | 55464200 | 55464999 | 2.844146 | 5.541895 | 26.44211 | 9.603279 | -1.59401 | -0.39438 | 0.052348 |
| 12 | 24060400 | 24062599 | 23.75419 | 41.88245 | 21.9438 | 9.711485 | 1.316015 | 0.58494 | 0.05235 |
| 11 | 95895200 | 95899399 | 14.36247 | 23.74502 | 84.98494 | 36.73084 | -1.45248 | 1.316797 | 0.05236 |
| 12 | 20637800 | 20650599 | 171.7425 | 259.7561 | 170.6885 | 70.5949 | 0.882125 | 3.30411 | 0.05237 |
| 4 | 33940600 | 33945199 | 23.41691 | 37.89958 | 103.5575 | 44.2932 | -1.08341 | 1.662895 | 0.052384 |
| 7 | 14533800 | 14537799 | 88.23429 | 139.8966 | 85.32565 | 35.51259 | 1.042331 | 2.348731 | 0.05239 |
| 3 | 15806600 | 15808399 | 23.02503 | 40.84207 | 19.67172 | 8.692626 | 1.439409 | 0.514433 | 0.052401 |
| 18 | 3156600 | 3158599 | 57.55101 | 92.96674 | 53.51553 | 22.8424 | 1.154848 | 1.743174 | 0.052404 |
| 16 | 85591400 | 85593799 | 7.866659 | 14.19767 | 46.06856 | 20.03798 | -1.27928 | 0.36189 | 0.052434 |
| 4 | 141065200 | 141081599 | 74.75534 | 113.3476 | 307.0686 | 127.0998 | -1.15668 | 3.24124 | 0.052439 |
| 5 | 119804000 | 119810999 | 37.60756 | 59.69518 | 162.615 | 67.75761 | -1.11164 | 2.292832 | 0.052458 |
| 19 | 39895800 | 39896599 | 19.16898 | 34.62159 | 19.75237 | 8.583551 | 1.212846 | 0.381223 | 0.052464 |
| 5 | 95440200 | 95441999 | 23.03911 | 40.8131 | 20.24256 | 8.950164 | 1.396694 | 0.527298 | 0.052469 |
| 12 | 107749800 | 107750799 | 2.949216 | 5.728575 | 27.75621 | 10.37403 | -1.65636 | -0.28765 | 0.052478 |
| 15 | 74300000 | 74302399 | 9.911836 | 17.2516 | 56.86281 | 24.99358 | -1.32483 | 0.75274 | 0.052492 |
| 11 | 68274200 | 68280599 | 29.4044 | 47.06853 | 137.198 | 57.80121 | -1.19581 | 2.015202 | 0.052493 |
| 4 | 61542000 | 61543199 | 19.09005 | 34.48521 | 19.64485 | 8.534914 | 1.215029 | 0.378901 | 0.052531 |
| 15 | 36282600 | 36283599 | 4.536074 | 8.446158 | 37.46574 | 15.63297 | -1.68068 | 0.136993 | 0.052549 |
| 2 | 173355800 | 173368599 | 77.09682 | 117.0379 | 304.6682 | 126.1514 | -1.09779 | 3.216513 | 0.052565 |
| 11 | 115877200 | 115877999 | 2.631316 | 5.116983 | 29.70061 | 10.99784 | -1.89691 | -0.31521 | 0.052574 |
| 4 | 61426200 | 61427399 | 17.16466 | 32.05373 | 16.1286 | 6.696726 | 1.437532 | 0.17098 | 0.052585 |
| 14 | 31347400 | 31348999 | 4.787556 | 9.029921 | 37.46599 | 15.29555 | -1.56066 | 0.052478 | 0.052585 |
| 2 | 118501400 | 118508799 | 37.04476 | 58.97511 | 150.4775 | 62.91093 | -1.01303 | 2.199602 | 0.052589 |
| 8 | 44249800 | 44251599 | 14.08626 | 26.85668 | 14.78079 | 5.898117 | 1.356123 | 0.005482 | 0.052607 |
| 2 | 35144800 | 35150199 | 19.42715 | 31.61155 | 100.5571 | 43.16338 | -1.29493 | 1.54866 | 0.052638 |

| | | | | | | | | | |
|---|---|---|---|---|---|---|---|---|---|
| 4 | 135300400 | 135304199 | 12.70795 | 21.25308 | 78.09539 | 34.013 | -1.48913 | 1.173901 | 0.052639 |
| 16 | 78948600 | 78948999 | 1.643839 | 3.208944 | 19.11004 | 6.025603 | -1.71126 | -0.94737 | 0.052647 |
| 7 | 11298200 | 11301799 | 113.4101 | 176.069 | 120.7144 | 50.12861 | 0.85035 | 2.777879 | 0.052674 |
| 8 | 127514200 | 127514999 | 2.941243 | 5.736771 | 25.5329 | 9.153555 | -1.48011 | -0.44683 | 0.052691 |
| 5 | 98196000 | 98197399 | 15.92418 | 30.14537 | 14.53573 | 5.88836 | 1.526424 | 0.070116 | 0.052704 |
| 8 | 121954200 | 121959999 | 21.75009 | 35.12909 | 111.5464 | 47.60236 | -1.3028 | 1.717181 | 0.052718 |
| 12 | 84230800 | 84243599 | 58.14609 | 89.68667 | 244.6015 | 101.6031 | -1.14662 | 2.886965 | 0.052719 |
| 2 | 119121200 | 119125399 | 15.48814 | 25.52495 | 87.23517 | 37.63414 | -1.38737 | 1.359654 | 0.052747 |
| 5 | 33681600 | 33687599 | 30.04819 | 48.07231 | 136.8974 | 57.62916 | -1.16317 | 2.033359 | 0.052777 |
| 14 | 73055400 | 73056399 | 14.71805 | 27.7773 | 16.03076 | 6.535996 | 1.262437 | 0.083436 | 0.052804 |
| 5 | 33688800 | 33695399 | 39.31079 | 62.538 | 151.0541 | 63.08974 | -0.93521 | 2.225457 | 0.052837 |
| 8 | 96071400 | 96081999 | 43.81146 | 68.32526 | 223.4115 | 92.71266 | -1.39547 | 2.681452 | 0.052843 |
| 19 | 56409400 | 56414599 | 17.00566 | 27.77526 | 95.16085 | 40.90291 | -1.39744 | 1.486616 | 0.052862 |
| 2 | 36958200 | 36959199 | 14.70809 | 28.19492 | 13.66758 | 5.379403 | 1.555227 | -0.03375 | 0.052878 |
| 12 | 61787200 | 61788199 | 14.70809 | 28.19492 | 13.66758 | 5.379403 | 1.555227 | -0.03375 | 0.052878 |
| 13 | 17703200 | 17704199 | 14.70809 | 28.19492 | 13.66758 | 5.379403 | 1.555227 | -0.03375 | 0.052878 |
| 7 | 34815400 | 34824399 | 40.65911 | 64.18899 | 181.3221 | 75.28041 | -1.17081 | 2.43849 | 0.052897 |
| 6 | 149089800 | 149091399 | 9.835002 | 17.27323 | 51.46741 | 22.77594 | -1.17907 | 0.60403 | 0.052906 |
| 11 | 69791200 | 69802399 | 38.64862 | 60.87171 | 191.9342 | 79.63885 | -1.33181 | 2.489383 | 0.052907 |
| 1 | 152081000 | 152082599 | 15.91115 | 29.50234 | 16.97344 | 7.126725 | 1.233731 | 0.203725 | 0.052933 |
| 2 | 31522400 | 31535399 | 88.2479 | 132.6261 | 350.9492 | 144.8803 | -1.13398 | 3.423349 | 0.052942 |
| 9 | 97913600 | 97915799 | 14.0962 | 23.53156 | 77.0457 | 33.51697 | -1.32353 | 1.197647 | 0.052947 |
| 14 | 117162600 | 117164599 | 9.882295 | 16.85273 | 65.59973 | 28.75764 | -1.56889 | 0.969697 | 0.052952 |
| 12 | 23374600 | 23376199 | 54.58253 | 88.00966 | 56.03096 | 23.85946 | 1.006918 | 1.790233 | 0.05302 |
| 1 | 190383200 | 190402599 | 110.8403 | 164.426 | 416.3786 | 171.5042 | -1.08779 | 3.669186 | 0.053091 |
| 1 | 49373600 | 49374599 | 10.48761 | 20.47319 | 9.518844 | 3.366174 | 1.750151 | -0.45063 | 0.053092 |
| 12 | 23382000 | 23383399 | 19.41462 | 36.07132 | 14.87084 | 6.226741 | 1.713009 | 0.218089 | 0.053108 |
| 15 | 35937200 | 35939599 | 5.518064 | 10.01187 | 42.85481 | 18.517 | -1.66773 | 0.308001 | 0.053108 |
| 9 | 61195800 | 61198199 | 8.600005 | 15.17678 | 48.26865 | 21.35803 | -1.2697 | 0.534485 | 0.053154 |
| 4 | 128984800 | 128992599 | 36.03632 | 57.46796 | 143.2842 | 60.04565 | -0.97705 | 2.141661 | 0.053165 |
| 11 | 57819600 | 57821399 | 7.094786 | 12.54906 | 47.21219 | 20.88104 | -1.50817 | 0.499006 | 0.053168 |
| 8 | 122984200 | 123030599 | 210.9745 | 292.1083 | 866.6829 | 359.9261 | -1.41466 | 4.728293 | 0.053194 |
| 17 | 6875800 | 6877799 | 4.1455 | 7.979305 | 33.8719 | 13.1762 | -1.52852 | -0.12356 | 0.053279 |
| X | 148591000 | 148591399 | 8.774625 | 17.15668 | 9.094529 | 3.108911 | 1.609679 | -0.59124 | 0.053288 |
| 11 | 72094400 | 72098799 | 19.63803 | 32.13633 | 91.37622 | 39.30366 | -1.12761 | 1.461442 | 0.05329 |
| 2 | 168527800 | 168534199 | 31.39236 | 50.08725 | 145.9521 | 61.20618 | -1.20065 | 2.121262 | 0.053301 |
| 3 | 143546200 | 143547799 | 15.75027 | 29.75258 | 15.10438 | 6.146406 | 1.447391 | 0.084046 | 0.053328 |
| 4 | 154341600 | 154387599 | 256.3976 | 352.255 | 923.5616 | 384.0208 | -1.24767 | 4.849076 | 0.053365 |
| 2 | 128657400 | 128661799 | 12.47897 | 21.36483 | 65.50564 | 28.72549 | -1.22635 | 0.939733 | 0.053371 |
| 17 | 18789000 | 18791799 | 73.74026 | 117.8554 | 69.5666 | 29.24397 | 1.101133 | 2.089342 | 0.053372 |
| 15 | 89845000 | 89845799 | 12.85114 | 24.59605 | 14.60432 | 5.774406 | 1.259866 | -0.03627 | 0.0534 |
| 6 | 34077000 | 34081399 | 27.37393 | 43.98892 | 120.6941 | 51.14738 | -1.10178 | 1.886113 | 0.053402 |
| 4 | 138808000 | 138816999 | 37.48734 | 59.41805 | 168.066 | 69.92725 | -1.16789 | 2.337013 | 0.053406 |
| 5 | 73547400 | 73552199 | 24.34604 | 39.5032 | 101.0411 | 43.2976 | -0.98503 | 1.610513 | 0.053408 |
| 7 | 143152600 | 143154199 | 6.82628 | 12.33973 | 41.72794 | 18.11372 | -1.33555 | 0.337685 | 0.053417 |
| 5 | 35500400 | 35509199 | 37.62662 | 59.69611 | 166.1256 | 69.18635 | -1.14309 | 2.307893 | 0.053417 |
| X | 51283800 | 51284399 | 24.22371 | 44.06982 | 13.43876 | 5.78714 | 2.116378 | 0.365663 | 0.053421 |

| | | | | | | | | | |
|---|---|---|---|---|---|---|---|---|---|
| 5 | 64125000 | 64128599 | 13.97018 | 23.32749 | 84.09111 | 36.58815 | -1.4621 | 1.198013 | 0.053466 |
| 19 | 7271800 | 7273399 | 5.463627 | 10.04793 | 41.62329 | 17.67257 | -1.60361 | 0.224589 | 0.053466 |
| 10 | 80143400 | 80174599 | 162.3604 | 231.2162 | 648.3272 | 268.0331 | -1.29215 | 4.305517 | 0.053474 |
| 2 | 72033400 | 72035399 | 5.615981 | 10.26221 | 40.32645 | 17.27032 | -1.53672 | 0.261892 | 0.053479 |
| 10 | 104974600 | 104975999 | 5.031419 | 9.517141 | 34.12457 | 13.84713 | -1.34359 | 0.028633 | 0.05349 |
| 1 | 108616400 | 108618799 | 29.81442 | 51.68633 | 24.45599 | 10.73009 | 1.46016 | 0.846652 | 0.053512 |
| 13 | 63306000 | 63306799 | 23.314 | 40.93882 | 23.92717 | 10.58686 | 1.158058 | 0.634756 | 0.053531 |
| 7 | 15003000 | 15003599 | 9.52235 | 18.6245 | 7.887488 | 2.662053 | 1.939561 | -0.62651 | 0.053542 |
| 12 | 20027400 | 20030199 | 69.62628 | 111.7978 | 58.23104 | 24.64782 | 1.288669 | 1.942187 | 0.053545 |
| 12 | 23696400 | 23705599 | 192.7353 | 288.6523 | 192.0922 | 79.23327 | 0.851503 | 3.500535 | 0.053546 |
| 12 | 114008800 | 114010399 | 27.63778 | 48.01956 | 27.47099 | 12.06137 | 1.188641 | 0.817658 | 0.053551 |
| 15 | 13079400 | 13081599 | 9.832185 | 17.27598 | 52.54763 | 23.26273 | -1.20873 | 0.598044 | 0.05356 |
| 11 | 100093800 | 100104799 | 69.12035 | 106.4986 | 244.3297 | 101.4903 | -0.89871 | 2.910779 | 0.053564 |
| 18 | 71218600 | 71219199 | 9.172559 | 17.90408 | 9.870259 | 3.497043 | 1.506282 | -0.46333 | 0.053573 |
| 10 | 80180600 | 80233599 | 301.7462 | 408.3802 | 1094.428 | 456.1321 | -1.30118 | 5.08578 | 0.053586 |
| 5 | 106604800 | 106608999 | 33.26734 | 53.1548 | 138.2614 | 58.09859 | -1.03487 | 2.076752 | 0.053586 |
| 7 | 11771800 | 11772999 | 21.33071 | 38.67064 | 17.25727 | 7.464208 | 1.568443 | 0.370962 | 0.053599 |
| 10 | 5754600 | 5756399 | 10.07381 | 17.48871 | 57.36555 | 25.17403 | -1.31825 | 0.792163 | 0.053608 |
| 14 | 32967400 | 32970999 | 13.7621 | 23.38162 | 66.44486 | 29.11936 | -1.11909 | 1.018174 | 0.053619 |
| X | 81007600 | 81008799 | 20.69341 | 37.73496 | 16.26753 | 6.984925 | 1.625106 | 0.332597 | 0.05363 |
| 13 | 49331200 | 49361599 | 154.8618 | 222.5567 | 585.6893 | 241.7827 | -1.18747 | 4.170877 | 0.05364 |
| 12 | 23341000 | 23342599 | 43.75043 | 71.87834 | 44.57514 | 19.20489 | 1.065068 | 1.434304 | 0.053652 |
| 10 | 61561400 | 61562399 | 3.794661 | 7.284587 | 31.14318 | 12.20768 | -1.54618 | -0.10398 | 0.053701 |
| 13 | 103422400 | 103424399 | 22.74172 | 40.17884 | 20.80974 | 9.205853 | 1.333198 | 0.553832 | 0.053701 |
| 10 | 99963600 | 99965199 | 8.708982 | 15.4302 | 46.1719 | 20.41406 | -1.17972 | 0.488652 | 0.053724 |
| 7 | 17231400 | 17233399 | 20.23705 | 36.41694 | 19.71103 | 8.604938 | 1.284243 | 0.409772 | 0.053727 |
| 8 | 71192800 | 71204199 | 52.95221 | 82.32118 | 219.6614 | 91.19925 | -1.10626 | 2.739932 | 0.053736 |
| 7 | 85009000 | 85010199 | 22.0816 | 39.736 | 17.60981 | 7.687702 | 1.569906 | 0.417785 | 0.053782 |
| 4 | 125796600 | 125797399 | 3.650186 | 7.097604 | 28.11047 | 10.41758 | -1.36057 | -0.31054 | 0.053784 |
| 5 | 148299000 | 148302599 | 17.03021 | 28.06494 | 85.17298 | 36.74372 | -1.21676 | 1.362951 | 0.053792 |
| 4 | 78958000 | 78959999 | 17.22598 | 31.62903 | 18.23915 | 7.760895 | 1.216833 | 0.270842 | 0.053803 |
| 11 | 115936200 | 115975799 | 182.2921 | 257.0102 | 710.6099 | 294.273 | -1.28653 | 4.454225 | 0.053803 |
| 12 | 110338200 | 110348599 | 58.79288 | 91.03474 | 228.8384 | 95.05341 | -1.02484 | 2.814102 | 0.053812 |
| 12 | 23580600 | 23582199 | 24.63497 | 43.49817 | 20.59211 | 9.110649 | 1.462414 | 0.574019 | 0.053822 |
| 14 | 50268600 | 50270199 | 25.51647 | 44.61787 | 25.26594 | 11.13948 | 1.203629 | 0.714564 | 0.053878 |
| X | 26233600 | 26235799 | 23.45406 | 41.53005 | 20.59865 | 9.108401 | 1.396967 | 0.536795 | 0.053881 |
| 6 | 98147400 | 98148199 | 11.12893 | 21.73197 | 8.169942 | 2.873521 | 2.056356 | -0.46422 | 0.05389 |
| 1 | 113215400 | 113216799 | 13.08745 | 25.17366 | 14.46626 | 5.63844 | 1.325493 | -0.07892 | 0.053909 |
| 4 | 99082000 | 99082799 | 2.24759 | 4.393938 | 23.0433 | 7.909022 | -1.65163 | -0.62131 | 0.053934 |
| 17 | 27203400 | 27209599 | 39.92683 | 63.608 | 147.698 | 61.8089 | -0.87622 | 2.179086 | 0.053951 |
| 7 | 7988000 | 7992999 | 95.48361 | 150.021 | 102.905 | 42.6846 | 0.862886 | 2.556976 | 0.053963 |
| 9 | 86349000 | 86349799 | 3.553954 | 6.862109 | 30.29219 | 11.66909 | -1.56835 | -0.1694 | 0.053998 |
| 7 | 28321000 | 28322999 | 8.241958 | 14.69118 | 48.99295 | 21.56036 | -1.33053 | 0.44632 | 0.053998 |
| 4 | 126184800 | 126198399 | 62.24023 | 95.6277 | 258.2838 | 107.2452 | -1.137 | 2.965274 | 0.054004 |
| 1 | 155944400 | 155955999 | 63.69512 | 97.88096 | 255.901 | 106.2584 | -1.08989 | 2.961952 | 0.054009 |
| 18 | 61309400 | 61312599 | 11.60562 | 20.1207 | 58.54966 | 25.68877 | -1.14726 | 0.813362 | 0.05402 |
| 12 | 22948600 | 22953199 | 136.9093 | 209.6619 | 142.3518 | 59.07765 | 0.849412 | 3.043488 | 0.054024 |

| | | | | | | | | | |
|---|---|---|---|---|---|---|---|---|---|
| 11 | 50512200 | 50523999 | 68.54687 | 104.7572 | 277.7868 | 115.2301 | -1.11655 | 3.075556 | 0.054033 |
| 16 | 64814400 | 64815599 | 9.753307 | 19.01734 | 11.65151 | 4.194585 | 1.338025 | -0.39214 | 0.054039 |
| 1 | 7546600 | 7547799 | 13.82075 | 26.66902 | 13.31723 | 5.143309 | 1.538112 | -0.10255 | 0.054046 |
| 11 | 101854000 | 101861199 | 31.13636 | 49.8544 | 131.4719 | 55.41083 | -1.0512 | 2.007655 | 0.054049 |
| 7 | 42493800 | 42494199 | 19.58495 | 35.53436 | 17.47343 | 7.550091 | 1.431078 | 0.354444 | 0.054061 |
| 1 | 108883000 | 108892799 | 190.7435 | 286.7314 | 183.3355 | 75.69004 | 0.913889 | 3.428796 | 0.054074 |
| 13 | 65700400 | 65705999 | 109.9122 | 171.8964 | 105.2967 | 43.68386 | 1.020813 | 2.649333 | 0.054084 |
| 11 | 63361200 | 63361799 | 13.91012 | 26.4611 | 15.56405 | 6.240371 | 1.256011 | 0.021625 | 0.054086 |
| 12 | 22593000 | 22595799 | 24.2226 | 42.97247 | 19.22174 | 8.492763 | 1.545813 | 0.52167 | 0.054089 |
| 15 | 83946600 | 83966599 | 128.3446 | 189.5158 | 426.3466 | 175.5778 | -0.92341 | 3.749702 | 0.054121 |
| 8 | 121231000 | 121248999 | 73.74628 | 111.4325 | 331.119 | 136.9058 | -1.2942 | 3.312163 | 0.054125 |
| 19 | 6054000 | 6056199 | 7.630924 | 13.61429 | 44.15898 | 19.41831 | -1.28907 | 0.430083 | 0.054127 |
| 5 | 13448800 | 13449599 | 2.943042 | 5.734933 | 25.8974 | 9.3982 | -1.51666 | -0.40084 | 0.054154 |
| 7 | 130452600 | 130453399 | 2.641144 | 5.154137 | 24.34103 | 8.65755 | -1.55179 | -0.48199 | 0.054163 |
| X | 39570600 | 39571399 | 12.65577 | 24.56459 | 11.60835 | 4.363619 | 1.653768 | -0.20812 | 0.054188 |
| 2 | 176988400 | 176989399 | 13.42424 | 26.00637 | 12.33857 | 4.67976 | 1.635437 | -0.16814 | 0.054189 |
| 11 | 29122000 | 29123599 | 6.35851 | 11.24134 | 47.72662 | 21.11059 | -1.68099 | 0.501346 | 0.054194 |
| 4 | 106672600 | 106676399 | 19.30103 | 31.63392 | 89.72789 | 38.61938 | -1.12301 | 1.439613 | 0.054199 |
| 11 | 4407600 | 4408399 | 3.354641 | 6.514949 | 27.80499 | 10.40679 | -1.47888 | -0.27764 | 0.054204 |
| 9 | 121603800 | 121634399 | 165.5065 | 235.6139 | 649.2664 | 268.4372 | -1.26759 | 4.310793 | 0.05421 |
| 17 | 27482400 | 27484799 | 55.4128 | 89.3617 | 58.9592 | 25.1112 | 0.955793 | 1.788422 | 0.05424 |
| 17 | 38348000 | 38349799 | 27.34266 | 47.77481 | 24.04413 | 10.59417 | 1.372773 | 0.734223 | 0.054272 |
| 12 | 88123600 | 88124999 | 21.32293 | 37.64673 | 20.93527 | 9.262639 | 1.23126 | 0.552807 | 0.054287 |
| 5 | 75968800 | 75970999 | 6.60378 | 11.81553 | 45.2543 | 19.85201 | -1.52438 | 0.405999 | 0.054317 |
| 13 | 53842600 | 53844399 | 6.358651 | 11.57879 | 40.81441 | 17.55582 | -1.38608 | 0.28872 | 0.054335 |
| 1 | 85294400 | 85294799 | 14.76611 | 27.73643 | 16.43796 | 6.757891 | 1.214562 | 0.113862 | 0.054345 |
| 8 | 102656200 | 102657399 | 13.15 | 25.41618 | 13.3653 | 5.124297 | 1.47395 | -0.1335 | 0.05437 |
| 11 | 78049800 | 78058999 | 33.93206 | 53.83992 | 165.6492 | 68.99482 | -1.28739 | 2.302568 | 0.054413 |
| 18 | 3090400 | 3093199 | 76.03829 | 121.2583 | 75.38372 | 31.59018 | 1.023128 | 2.16219 | 0.054443 |
| 18 | 38985600 | 38987399 | 6.000647 | 11.09048 | 38.46867 | 16.22317 | -1.34246 | 0.194481 | 0.05445 |
| 1 | 100561600 | 100564799 | 44.04909 | 72.0322 | 45.70123 | 19.65238 | 1.029271 | 1.493804 | 0.05445 |
| 1 | 144995200 | 144996399 | 23.62712 | 41.42973 | 24.49049 | 10.8231 | 1.141903 | 0.659731 | 0.054466 |
| 14 | 25841000 | 25843799 | 9.183074 | 16.24272 | 48.29401 | 21.35952 | -1.17205 | 0.512406 | 0.054474 |
| 10 | 9954600 | 9956199 | 13.99548 | 27.0408 | 11.95749 | 4.592096 | 1.717525 | -0.12334 | 0.054476 |
| 15 | 7119400 | 7120399 | 3.951915 | 7.663819 | 29.61853 | 11.17971 | -1.35153 | -0.23807 | 0.054488 |
| 11 | 23255600 | 23257199 | 7.278024 | 12.8413 | 49.83957 | 22.054 | -1.55472 | 0.531359 | 0.05453 |
| 10 | 53266200 | 53267199 | 12.32518 | 23.80906 | 13.34844 | 5.130836 | 1.378383 | -0.13541 | 0.054537 |
| 4 | 154743400 | 154750199 | 38.68644 | 61.49567 | 153.9323 | 64.23169 | -0.98777 | 2.249998 | 0.054574 |
| 14 | 50172400 | 50173199 | 12.59646 | 24.32583 | 13.94844 | 5.368945 | 1.345034 | -0.12542 | 0.054586 |
| 13 | 106878800 | 106879199 | 9.121956 | 17.80427 | 11.10782 | 3.939483 | 1.330608 | -0.45462 | 0.054612 |
| 19 | 41079600 | 41080199 | 2.334982 | 4.566104 | 21.8863 | 7.456162 | -1.51324 | -0.64881 | 0.054616 |
| 2 | 92841600 | 92849599 | 34.96164 | 55.6652 | 151.7511 | 63.45351 | -1.10812 | 2.194177 | 0.054625 |
| 1 | 116582600 | 116583599 | 14.19548 | 27.38559 | 12.55096 | 4.850685 | 1.658634 | -0.09896 | 0.054638 |
| 2 | 96860200 | 96861799 | 14.19548 | 27.38559 | 12.55096 | 4.850685 | 1.658634 | -0.09896 | 0.054638 |
| 13 | 66142600 | 66148799 | 160.2934 | 243.9861 | 151.9265 | 62.95724 | 0.967236 | 3.173122 | 0.054658 |
| 7 | 20765200 | 20767199 | 24.61809 | 44.07954 | 16.52687 | 7.245419 | 1.805483 | 0.464005 | 0.054659 |
| 6 | 27122000 | 27123399 | 12.78212 | 24.68124 | 13.94449 | 5.37061 | 1.365454 | -0.1213 | 0.05468 |

| | | | | | | | | | |
|---|---|---|---|---|---|---|---|---|---|
| 7 | 11153000 | 11155599 | 56.11963 | 90.66537 | 54.63977 | 23.32587 | 1.089031 | 1.736649 | 0.054681 |
| 4 | 147240800 | 147242599 | 13.27484 | 25.52867 | 14.46211 | 5.640271 | 1.345189 | -0.0749 | 0.054685 |
| X | 25663200 | 25664399 | 29.24359 | 50.42543 | 29.2863 | 12.84729 | 1.165329 | 0.899952 | 0.05469 |
| 13 | 97841000 | 97841599 | 10.97203 | 21.3226 | 12.69408 | 4.725729 | 1.336438 | -0.25796 | 0.054751 |
| 12 | 22540000 | 22543999 | 65.85118 | 105.3987 | 71.69225 | 30.1992 | 0.899211 | 2.037857 | 0.054765 |
| 7 | 125798600 | 125800599 | 5.543231 | 10.3254 | 38.14195 | 15.9049 | -1.42003 | 0.142875 | 0.054768 |
| 6 | 113830800 | 113831999 | 2.947829 | 5.729992 | 27.26362 | 10.13105 | -1.62265 | -0.31019 | 0.054786 |
| 17 | 48586400 | 48588399 | 8.708148 | 15.4691 | 47.28233 | 20.86772 | -1.20849 | 0.46891 | 0.054789 |
| 9 | 7441400 | 7442599 | 14.44804 | 27.59381 | 14.92492 | 5.934029 | 1.386205 | -0.00099 | 0.054812 |
| 7 | 27051000 | 27052799 | 6.136823 | 11.34142 | 37.91492 | 15.99106 | -1.28967 | 0.195447 | 0.054827 |
| 7 | 19142800 | 19173199 | 190.0975 | 270.1977 | 645.1072 | 266.7879 | -1.06312 | 4.334815 | 0.05488 |
| 16 | 5348200 | 5365399 | 97.15376 | 146.3396 | 333.4288 | 137.7305 | -0.91545 | 3.38005 | 0.054886 |
| 7 | 8946400 | 8947199 | 14.72033 | 28.164 | 14.23803 | 5.635057 | 1.488191 | -0.01472 | 0.054917 |
| 5 | 122509600 | 122511199 | 6.236112 | 11.32755 | 42.37609 | 18.28414 | -1.47435 | 0.304944 | 0.054929 |
| 7 | 61627600 | 61628999 | 30.40562 | 52.14402 | 30.68234 | 13.45662 | 1.145087 | 0.953016 | 0.05493 |
| 18 | 60539400 | 60540799 | 17.53505 | 32.80651 | 15.56252 | 6.441319 | 1.525349 | 0.161548 | 0.054952 |
| 6 | 72331800 | 72336599 | 17.54313 | 28.79015 | 88.26008 | 38.00851 | -1.23376 | 1.419178 | 0.054953 |
| 17 | 39096800 | 39099399 | 61.59659 | 99.10493 | 60.8104 | 25.81586 | 1.056328 | 1.869048 | 0.054988 |
| 16 | 22295800 | 22299599 | 12.50735 | 21.34138 | 65.94324 | 28.90954 | -1.23823 | 0.972062 | 0.054995 |
| 17 | 65010200 | 65011599 | 5.913009 | 11.12777 | 38.06862 | 15.60091 | -1.29009 | 0.079508 | 0.055027 |
| 5 | 138267600 | 138269799 | 9.511489 | 16.75435 | 49.33007 | 21.83575 | -1.16052 | 0.562055 | 0.05505 |
| 15 | 61461200 | 61462199 | 14.09828 | 26.81152 | 15.56025 | 6.24255 | 1.274507 | 0.025455 | 0.055085 |
| 17 | 33771800 | 33773199 | 5.286081 | 10.03666 | 34.3123 | 13.81259 | -1.26536 | 0.005512 | 0.055091 |
| 12 | 21896800 | 21898999 | 47.52188 | 77.19174 | 51.12527 | 21.92554 | 0.960605 | 1.602655 | 0.055098 |
| 1 | 142961800 | 142962599 | 11.00552 | 21.35415 | 13.11319 | 4.93722 | 1.276786 | -0.21868 | 0.055101 |
| 2 | 98861600 | 98863199 | 21.19222 | 38.26467 | 17.65811 | 7.676264 | 1.515679 | 0.39248 | 0.055108 |
| 7 | 33246800 | 33248199 | 41.84506 | 69.59078 | 36.73534 | 15.94202 | 1.29955 | 1.282647 | 0.05511 |
| 2 | 92867800 | 92872199 | 21.24554 | 34.34843 | 107.5373 | 45.95061 | -1.28069 | 1.685488 | 0.055137 |
| 2 | 160487200 | 160488999 | 7.907976 | 14.06604 | 45.69552 | 20.14519 | -1.29373 | 0.456134 | 0.055145 |
| 1 | 51142200 | 51142599 | 7.65442 | 14.97347 | 8.827956 | 2.93853 | 1.48922 | -0.70121 | 0.055157 |
| 12 | 22694200 | 22697399 | 28.64087 | 50.07778 | 22.42573 | 9.886697 | 1.540207 | 0.728822 | 0.05516 |
| 15 | 79706600 | 79719199 | 50.67322 | 78.47954 | 235.075 | 97.64383 | -1.27709 | 2.807449 | 0.055164 |
| 2 | 169928800 | 169950399 | 122.6293 | 181.2568 | 430.3059 | 177.2102 | -0.99962 | 3.732455 | 0.055171 |
| 4 | 140334200 | 140346199 | 41.13536 | 64.26655 | 212.7042 | 88.25289 | -1.41082 | 2.647441 | 0.055194 |
| 12 | 71061800 | 71064199 | 12.99637 | 22.09279 | 66.96456 | 29.34842 | -1.21154 | 1.010238 | 0.055221 |
| 11 | 92134200 | 92136599 | 25.8854 | 45.25404 | 25.26746 | 11.13829 | 1.223913 | 0.719923 | 0.055226 |
| 14 | 56609800 | 56610999 | 4.253031 | 8.223377 | 32.48185 | 12.43129 | -1.40313 | -0.18073 | 0.055235 |
| 6 | 71416600 | 71419799 | 12.78991 | 21.86229 | 63.78924 | 27.96922 | -1.15543 | 0.954954 | 0.055235 |
| 9 | 119651600 | 119659599 | 37.5922 | 59.70647 | 160.0149 | 66.71409 | -1.0873 | 2.274515 | 0.055261 |
| 2 | 158526200 | 158532199 | 29.78326 | 47.86902 | 122.5319 | 51.93936 | -1.00169 | 1.882954 | 0.055261 |
| 12 | 112863800 | 112930799 | 327.1841 | 437.6433 | 1272.103 | 531.1162 | -1.43539 | 5.272583 | 0.055279 |
| 16 | 18965200 | 18966199 | 12.65577 | 24.26949 | 14.52313 | 5.71019 | 1.255868 | -0.05651 | 0.055279 |
| 16 | 91063000 | 91068599 | 34.48665 | 54.89959 | 152.6627 | 63.81967 | -1.13695 | 2.199534 | 0.055287 |
| 3 | 113505600 | 113507799 | 26.17794 | 45.93508 | 22.51031 | 9.953958 | 1.410706 | 0.660154 | 0.0553 |
| 2 | 180752400 | 180756599 | 15.36552 | 25.56945 | 78.19285 | 33.9477 | -1.22561 | 1.237137 | 0.055305 |
| 15 | 86280200 | 86281799 | 4.26896 | 8.206205 | 31.32864 | 12.23271 | -1.38138 | -0.11435 | 0.05533 |
| 1 | 174870000 | 174870799 | 11.50582 | 22.42567 | 10.71541 | 3.877396 | 1.684608 | -0.35415 | 0.055397 |

| | | | | | | | | | |
|---|---|---|---|---|---|---|---|---|---|
| 2 | 33018800 | 33024199 | 21.44474 | 34.64694 | 112.9247 | 48.21037 | -1.33984 | 1.707367 | 0.055402 |
| 13 | 98490400 | 98492599 | 12.53287 | 21.21813 | 67.82971 | 29.69481 | -1.28809 | 1.04808 | 0.055411 |
| 11 | 120880400 | 120884999 | 15.72667 | 26.09723 | 82.74101 | 35.8529 | -1.27805 | 1.273547 | 0.05543 |
| 11 | 87482000 | 87483799 | 4.82505 | 8.999498 | 37.55697 | 15.62604 | -1.59081 | 0.127565 | 0.05544 |
| 4 | 155200600 | 155231199 | 148.8157 | 214.2754 | 577.9423 | 238.5147 | -1.2202 | 4.142423 | 0.055444 |
| 6 | 116605600 | 116607599 | 6.302431 | 11.55948 | 38.81234 | 16.53935 | -1.30669 | 0.243339 | 0.055448 |
| 2 | 3820000 | 3821799 | 7.224673 | 13.1887 | 41.47091 | 17.78253 | -1.22028 | 0.281059 | 0.055488 |
| 1 | 102960400 | 102962599 | 58.29886 | 93.81166 | 53.86737 | 22.87277 | 1.151985 | 1.856068 | 0.055517 |
| X | 61800400 | 61801399 | 13.9817 | 27.23939 | 8.011941 | 2.915851 | 2.36369 | -0.31149 | 0.05552 |
| 13 | 6959000 | 6960799 | 20.7871 | 37.97631 | 16.1682 | 6.926159 | 1.644096 | 0.321965 | 0.055522 |
| 6 | 92299400 | 92299799 | 11.99162 | 23.32994 | 11.08974 | 4.091582 | 1.66664 | -0.28143 | 0.055528 |
| 9 | 95067000 | 95067999 | 11.99162 | 23.32994 | 11.08974 | 4.091582 | 1.66664 | -0.28143 | 0.055528 |
| 13 | 66285800 | 66289199 | 62.43127 | 100.3501 | 63.00513 | 26.70985 | 1.021583 | 1.901129 | 0.055536 |
| 7 | 9671200 | 9672999 | 15.36361 | 29.60581 | 12.05643 | 4.67462 | 1.823625 | -0.07749 | 0.05554 |
| 9 | 44317400 | 44320999 | 12.7657 | 21.54682 | 71.74451 | 31.37976 | -1.34718 | 1.077212 | 0.055544 |
| 13 | 52961600 | 52964799 | 13.90903 | 23.66689 | 66.53805 | 29.16374 | -1.10343 | 1.00396 | 0.055591 |
| 19 | 5034400 | 5041999 | 39.20393 | 62.23526 | 158.6262 | 66.10209 | -1.01572 | 2.290745 | 0.055592 |
| 5 | 134516600 | 134547599 | 177.3123 | 253.1692 | 608.8209 | 251.5899 | -1.06691 | 4.267798 | 0.055622 |
| 1 | 34989400 | 34989999 | 11.95591 | 23.07258 | 14.15624 | 5.465965 | 1.244561 | -0.12167 | 0.055633 |
| 15 | 85927000 | 85927999 | 4.255489 | 8.220756 | 31.10031 | 11.95253 | -1.34692 | -0.16908 | 0.05564 |
| 7 | 10753400 | 10754599 | 28.93566 | 49.90709 | 29.63465 | 13.00021 | 1.133436 | 0.89559 | 0.055654 |
| 6 | 56904800 | 56906399 | 16.92132 | 32.43935 | 11.15155 | 4.38845 | 2.044488 | -0.0184 | 0.055668 |
| X | 32461800 | 32463999 | 34.83641 | 59.50806 | 28.36464 | 12.43619 | 1.446787 | 1.003977 | 0.055691 |
| 4 | 59581200 | 59582199 | 5.343971 | 10.05608 | 35.09546 | 14.38445 | -1.31788 | 0.068762 | 0.055706 |
| 9 | 54528200 | 54529999 | 4.428171 | 8.221862 | 39.60487 | 16.59835 | -1.80386 | 0.157517 | 0.055709 |
| 16 | 18775400 | 18779399 | 127.4052 | 196.633 | 128.8787 | 53.53374 | 0.908754 | 2.889936 | 0.055717 |
| 9 | 43549600 | 43553399 | 24.51422 | 39.72338 | 101.7911 | 43.57636 | -0.98945 | 1.639157 | 0.055734 |
| 4 | 141271600 | 141275199 | 11.70217 | 20.0705 | 63.80905 | 27.9855 | -1.27777 | 0.918767 | 0.055736 |
| 9 | 77891600 | 77894999 | 11.59322 | 20.13253 | 58.03503 | 25.47262 | -1.13332 | 0.794413 | 0.055759 |
| 13 | 66222200 | 66226599 | 68.25462 | 109.6653 | 56.66154 | 24.00632 | 1.301273 | 1.918504 | 0.055796 |
| 11 | 6831200 | 6831999 | 14.8853 | 27.8941 | 16.40921 | 6.773204 | 1.220526 | 0.129295 | 0.055808 |
| 12 | 7367800 | 7368799 | 11.92513 | 23.1237 | 12.8487 | 4.852077 | 1.414856 | -0.19895 | 0.055838 |
| 16 | 55894400 | 55896999 | 11.27061 | 19.61503 | 56.39681 | 24.78754 | -1.12911 | 0.759519 | 0.05585 |
| 8 | 86308600 | 86310999 | 26.01639 | 45.94573 | 19.06614 | 8.435202 | 1.650752 | 0.57715 | 0.055869 |
| 5 | 33403000 | 33403599 | 3.960636 | 7.654091 | 29.84875 | 11.451 | -1.38663 | -0.18055 | 0.055873 |
| 13 | 41211200 | 41212199 | 3.960636 | 7.654091 | 29.84875 | 11.451 | -1.38663 | -0.18055 | 0.055873 |
| 8 | 121470200 | 121472999 | 6.147954 | 10.95089 | 44.659 | 19.66485 | -1.61711 | 0.43305 | 0.055896 |
| 10 | 15672800 | 15673199 | 2.737162 | 5.349064 | 23.6108 | 8.180938 | -1.42128 | -0.57665 | 0.055897 |
| 5 | 137015600 | 137032599 | 94.31672 | 140.9982 | 380.5474 | 156.911 | -1.16938 | 3.526544 | 0.055905 |
| 19 | 45647800 | 45650399 | 8.042781 | 14.13159 | 50.88444 | 22.5261 | -1.44988 | 0.586515 | 0.05591 |
| 6 | 67858000 | 67858799 | 16.11214 | 30.12917 | 16.73622 | 6.93303 | 1.298601 | 0.154606 | 0.055912 |
| 1 | 138304600 | 138305599 | 3.65171 | 7.09581 | 27.9783 | 10.42487 | -1.36133 | -0.2911 | 0.055915 |
| 5 | 31069600 | 31079599 | 45.30191 | 71.20745 | 191.814 | 79.56819 | -1.10737 | 2.533683 | 0.05593 |
| 13 | 29045600 | 29046399 | 3.043081 | 5.94182 | 26.6769 | 9.397634 | -1.47013 | -0.50522 | 0.055953 |
| 12 | 69355400 | 69358199 | 29.56643 | 50.69118 | 31.29921 | 13.72687 | 1.075882 | 0.952487 | 0.055973 |
| 15 | 99432200 | 99434799 | 8.146946 | 14.39729 | 47.74378 | 21.1199 | -1.3286 | 0.515257 | 0.055984 |
| 19 | 42108200 | 42110399 | 7.985737 | 13.8762 | 58.86431 | 25.84638 | -1.68624 | 0.77144 | 0.05599 |

| | | | | | | | | | |
|---|---|---|---|---|---|---|---|---|---|
| 11 | 104332200 | 104333999 | 7.481137 | 13.37373 | 44.1884 | 19.39857 | -1.31393 | 0.415662 | 0.056011 |
| 19 | 6915000 | 6923799 | 34.90463 | 55.37081 | 166.58 | 69.3671 | -1.25544 | 2.309879 | 0.056016 |
| 17 | 10838600 | 10840799 | 8.037757 | 14.46102 | 43.7793 | 19.11555 | -1.18347 | 0.377351 | 0.056039 |
| 7 | 38677200 | 38679999 | 18.74872 | 34.77038 | 16.03669 | 6.731659 | 1.550341 | 0.225123 | 0.056132 |
| 14 | 111555400 | 111555999 | 9.971124 | 19.42511 | 10.69466 | 3.894303 | 1.475792 | -0.35427 | 0.056138 |
| X | 31835400 | 31837599 | 21.54968 | 39.42472 | 15.44042 | 6.602491 | 1.765853 | 0.319452 | 0.05614 |
| 12 | 23524400 | 23525199 | 11.58706 | 22.39432 | 13.60708 | 5.218131 | 1.267276 | -0.15035 | 0.056146 |
| 2 | 153643200 | 153647999 | 22.96313 | 37.17111 | 104.2372 | 44.58921 | -1.12055 | 1.658157 | 0.056158 |
| 5 | 123160200 | 123165799 | 26.73633 | 43.02194 | 119.7638 | 50.85179 | -1.12023 | 1.840959 | 0.056161 |
| 9 | 111926400 | 111928599 | 10.92739 | 19.10853 | 53.67599 | 23.66619 | -1.09479 | 0.68499 | 0.056162 |
| 8 | 121815200 | 121838199 | 110.9578 | 162.9325 | 476.9523 | 196.4356 | -1.31059 | 3.839474 | 0.056188 |
| 1 | 87140600 | 87142199 | 5.551108 | 10.3183 | 38.05743 | 15.92571 | -1.42187 | 0.15675 | 0.056193 |
| 14 | 60977000 | 60980399 | 12.01315 | 20.59449 | 64.94664 | 28.48339 | -1.26638 | 0.92501 | 0.056195 |
| 2 | 168115200 | 168124599 | 48.2854 | 75.35199 | 211.3096 | 87.68444 | -1.1735 | 2.670596 | 0.056196 |
| 12 | 109911600 | 109920799 | 53.09576 | 82.91254 | 202.5086 | 84.02677 | -0.97375 | 2.660823 | 0.056212 |
| X | 67078200 | 67079599 | 23.58388 | 41.66679 | 21.60443 | 9.55741 | 1.332224 | 0.562815 | 0.056217 |
| 17 | 54091600 | 54092999 | 14.21293 | 26.95965 | 15.99552 | 6.452413 | 1.235954 | 0.04473 | 0.056226 |
| 1 | 190188000 | 190188599 | 8.833512 | 17.26635 | 9.822067 | 3.385148 | 1.496637 | -0.56091 | 0.056236 |
| 7 | 51456600 | 51457599 | 7.980239 | 15.60911 | 9.443794 | 3.173566 | 1.441739 | -0.66039 | 0.056262 |
| 11 | 75851600 | 75852599 | 13.11138 | 25.12548 | 15.09572 | 5.947353 | 1.248013 | -0.04294 | 0.056272 |
| 9 | 43557000 | 43557999 | 3.891085 | 7.553618 | 29.685 | 11.14781 | -1.36734 | -0.25383 | 0.056273 |
| 17 | 39704200 | 39705199 | 30.89403 | 52.95337 | 30.80862 | 13.51142 | 1.16135 | 0.961233 | 0.056289 |
| 3 | 34906600 | 34908199 | 13.59045 | 25.99811 | 13.69558 | 5.422837 | 1.429505 | -0.02629 | 0.056302 |
| X | 30078600 | 30079199 | 9.813902 | 19.16711 | 10.38761 | 3.643492 | 1.54467 | -0.48402 | 0.05631 |
| 11 | 35183200 | 35186599 | 13.35995 | 22.96373 | 62.89673 | 27.58882 | -1.06339 | 0.904447 | 0.056322 |
| 1 | 3305000 | 3306799 | 31.85678 | 55.32115 | 23.3968 | 10.2692 | 1.621788 | 0.839897 | 0.056336 |
| 10 | 111501800 | 111503399 | 7.527529 | 13.70103 | 41.93155 | 18.0466 | -1.18498 | 0.30259 | 0.056356 |
| 18 | 81843600 | 81849399 | 29.75128 | 47.89051 | 119.0893 | 50.59167 | -0.95726 | 1.833405 | 0.056369 |
| 4 | 117350400 | 117354399 | 20.716 | 33.35584 | 121.1799 | 51.50081 | -1.5018 | 1.813489 | 0.056388 |
| 6 | 108654000 | 108656399 | 8.937072 | 15.68738 | 50.37996 | 22.28412 | -1.28615 | 0.607217 | 0.056389 |
| 13 | 42718800 | 42721199 | 8.713387 | 15.42334 | 46.16895 | 20.41663 | -1.18079 | 0.499436 | 0.056393 |
| 1 | 180553200 | 180554399 | 6.348791 | 11.58783 | 40.33056 | 17.29717 | -1.36512 | 0.27376 | 0.056416 |
| 1 | 115381000 | 115381799 | 9.523984 | 18.61969 | 9.184633 | 3.14924 | 1.705796 | -0.56887 | 0.056434 |
| 18 | 12261400 | 12262399 | 3.247908 | 6.302996 | 28.30056 | 10.64715 | -1.55838 | -0.25984 | 0.056484 |
| 6 | 121038400 | 121046199 | 32.66584 | 52.12572 | 144.014 | 60.40429 | -1.1238 | 2.117973 | 0.056514 |
| 12 | 107146200 | 107152799 | 37.38541 | 59.47127 | 152.3412 | 63.62177 | -1.01991 | 2.225657 | 0.056576 |
| 1 | 99177000 | 99177799 | 10.85248 | 21.16545 | 10.75977 | 3.861838 | 1.606426 | -0.39058 | 0.05662 |
| X | 32539800 | 32542399 | 77.16675 | 123.7948 | 51.68861 | 21.84887 | 1.605312 | 1.9775 | 0.056642 |
| 7 | 100159800 | 100161999 | 7.702946 | 13.95748 | 42.35212 | 18.33029 | -1.17821 | 0.330467 | 0.056661 |
| 1 | 138487200 | 138489799 | 10.62283 | 18.58093 | 53.12921 | 23.43042 | -1.12024 | 0.67769 | 0.056669 |
| 5 | 125252200 | 125257999 | 20.34767 | 32.85904 | 113.5163 | 48.43361 | -1.42426 | 1.720235 | 0.056698 |
| 1 | 16203200 | 16205399 | 6.428512 | 11.8996 | 38.54668 | 16.222 | -1.24275 | 0.188944 | 0.0567 |
| 11 | 121872200 | 121873399 | 32.39293 | 55.21543 | 27.0008 | 11.83618 | 1.409296 | 1.008254 | 0.056713 |
| 13 | 30143600 | 30154599 | 80.04453 | 122.1416 | 274.2175 | 113.7093 | -0.87843 | 3.109844 | 0.056732 |
| 17 | 13718400 | 13720599 | 11.41331 | 19.88712 | 55.86757 | 24.57336 | -1.09562 | 0.742419 | 0.056752 |
| 4 | 125798200 | 125813399 | 75.43505 | 114.9222 | 287.8121 | 119.2986 | -1.03783 | 3.142107 | 0.056772 |
| 7 | 128456000 | 128457199 | 5.777823 | 10.86228 | 35.67305 | 14.64444 | -1.23328 | 0.079854 | 0.056774 |

| | | | | | | | | | |
|---|---|---|---|---|---|---|---|---|---|
| 11 | 102918800 | 102919599 | 3.464421 | 6.737013 | 27.42513 | 10.15604 | -1.3972 | -0.3152 | 0.056791 |
| 5 | 125564200 | 125618999 | 351.7665 | 472.3758 | 1173.233 | 489.5535 | -1.20285 | 5.210312 | 0.056804 |
| 14 | 25586000 | 25588799 | 10.08626 | 17.47857 | 57.87292 | 25.39149 | -1.33224 | 0.811088 | 0.056813 |
| 14 | 54923400 | 54924599 | 4.170927 | 7.955414 | 33.23468 | 13.25074 | -1.53832 | -0.04113 | 0.056818 |
| 15 | 82238000 | 82240799 | 13.42008 | 22.88784 | 64.31723 | 28.19561 | -1.10202 | 0.979262 | 0.056821 |
| 19 | 6530400 | 6573399 | 210.368 | 292.0498 | 848.1922 | 352.0987 | -1.37994 | 4.687238 | 0.056827 |
| 1 | 133641800 | 133648999 | 31.17504 | 49.83877 | 136.6077 | 57.44786 | -1.10934 | 2.057427 | 0.056878 |
| 2 | 90066000 | 90066599 | 8.663955 | 16.94036 | 9.198569 | 3.144176 | 1.572135 | -0.59413 | 0.056884 |
| 3 | 108497800 | 108499599 | 7.28308 | 12.83857 | 49.83014 | 22.05383 | -1.55525 | 0.541768 | 0.056895 |
| 10 | 8741800 | 8743599 | 7.852246 | 13.85387 | 47.73158 | 21.12139 | -1.38406 | 0.532591 | 0.056897 |
| 17 | 38002400 | 38003399 | 15.8551 | 29.53855 | 16.57043 | 6.907684 | 1.277883 | 0.175346 | 0.056899 |
| 9 | 102699400 | 102701399 | 6.398719 | 11.81602 | 42.96428 | 18.14034 | -1.41138 | 0.213869 | 0.056899 |
| 12 | 80689400 | 80691599 | 7.104731 | 12.54202 | 47.72767 | 21.1173 | -1.52567 | 0.522298 | 0.05691 |
| 7 | 74989600 | 74991399 | 6.75025 | 12.41981 | 40.61054 | 17.2318 | -1.26527 | 0.229917 | 0.056921 |
| 9 | 42762800 | 42764399 | 6.093734 | 11.39339 | 37.20263 | 15.41519 | -1.23642 | 0.121299 | 0.056928 |
| 12 | 108170000 | 108196599 | 118.0788 | 172.1363 | 513.0562 | 211.4204 | -1.34664 | 3.953841 | 0.056942 |
| 17 | 22804000 | 22804999 | 35.85949 | 60.2956 | 35.61863 | 15.56093 | 1.137512 | 1.14993 | 0.056954 |
| 5 | 139587200 | 139618399 | 234.4669 | 329.3013 | 706.5065 | 292.7544 | -0.92636 | 4.513157 | 0.05696 |
| 4 | 155289800 | 155296199 | 27.71495 | 44.51137 | 122.3762 | 51.81651 | -1.10576 | 1.904996 | 0.056987 |
| 12 | 112387800 | 112406399 | 105.1717 | 156.1244 | 407.4033 | 167.8191 | -1.13012 | 3.656296 | 0.05699 |
| 7 | 25702800 | 25704199 | 32.47085 | 54.83604 | 35.21876 | 15.40657 | 1.01747 | 1.100291 | 0.057008 |
| 8 | 115291000 | 115297199 | 34.37167 | 54.96117 | 134.8325 | 56.71895 | -0.94939 | 2.053335 | 0.057009 |
| 7 | 14309000 | 14310799 | 43.2737 | 71.13393 | 37.29417 | 16.07165 | 1.306744 | 1.420458 | 0.05701 |
| 1 | 114229000 | 114230799 | 16.91976 | 31.47512 | 15.94529 | 6.663873 | 1.419786 | 0.191027 | 0.057018 |
| 3 | 47449400 | 47450799 | 14.04288 | 26.93481 | 14.29084 | 5.615673 | 1.428511 | -0.04394 | 0.057027 |
| 12 | 45156000 | 45157999 | 8.031809 | 14.46851 | 45.45668 | 19.81838 | -1.23539 | 0.372364 | 0.057049 |
| 8 | 20183200 | 20183599 | 1.946726 | 3.80718 | 20.5822 | 6.730467 | -1.62693 | -0.80693 | 0.057086 |
| 12 | 23822000 | 23826399 | 118.8583 | 183.9584 | 125.61 | 52.17519 | 0.85283 | 2.835207 | 0.057088 |
| 7 | 14061200 | 14064399 | 82.57616 | 130.7653 | 89.95241 | 37.40415 | 0.867503 | 2.375278 | 0.057091 |
| 1 | 85721000 | 85725399 | 19.39951 | 31.96334 | 89.95394 | 38.8031 | -1.10878 | 1.372246 | 0.057095 |
| 14 | 41967200 | 41970199 | 24.15331 | 42.62466 | 20.68381 | 9.152209 | 1.426108 | 0.574974 | 0.057113 |
| 10 | 3450600 | 3451999 | 25.08374 | 44.0652 | 22.71469 | 10.054 | 1.338198 | 0.637022 | 0.057115 |
| 8 | 104902400 | 104904199 | 6.743722 | 12.04681 | 44.17262 | 19.40341 | -1.46313 | 0.41609 | 0.057116 |
| 9 | 27090000 | 27092999 | 13.02442 | 22.46464 | 64.09956 | 28.11939 | -1.12112 | 0.872457 | 0.057164 |
| 19 | 8865800 | 8866599 | 3.767249 | 7.324705 | 28.10356 | 10.42101 | -1.31596 | -0.30615 | 0.057176 |
| 13 | 66248800 | 66250799 | 26.002 | 45.569 | 23.65209 | 10.44774 | 1.328469 | 0.680565 | 0.05719 |
| 5 | 25971600 | 25972799 | 14.49874 | 27.49073 | 15.88168 | 6.41194 | 1.272784 | 0.049633 | 0.057225 |
| 8 | 122020200 | 122027999 | 30.96697 | 49.267 | 154.7461 | 64.63691 | -1.31314 | 2.217424 | 0.057262 |
| X | 106068000 | 106069399 | 15.28535 | 28.8305 | 15.56623 | 6.354727 | 1.356378 | 0.091886 | 0.057264 |
| 7 | 139379600 | 139382599 | 17.03368 | 28.4709 | 77.04904 | 33.54493 | -1.05021 | 1.191539 | 0.057281 |
| 14 | 63481600 | 63495399 | 97.25241 | 146.1371 | 351.8367 | 145.2409 | -0.998 | 3.427557 | 0.057412 |
| 7 | 101144200 | 101154599 | 49.3031 | 76.79519 | 213.5892 | 88.65283 | -1.16376 | 2.704674 | 0.057427 |
| 19 | 25165000 | 25166399 | 11.603 | 20.07534 | 58.46405 | 25.64962 | -1.14879 | 0.834016 | 0.057429 |
| 2 | 151526800 | 151528399 | 6.109506 | 11.37428 | 37.02784 | 15.45393 | -1.24038 | 0.149415 | 0.057434 |
| 5 | 135101200 | 135103399 | 7.873576 | 13.83802 | 49.28955 | 21.82502 | -1.43395 | 0.579748 | 0.057461 |
| 7 | 9356800 | 9359799 | 22.17252 | 39.49677 | 20.41323 | 8.98814 | 1.342128 | 0.485475 | 0.057477 |
| 5 | 144482000 | 144485199 | 22.15578 | 35.86326 | 103.2999 | 44.18735 | -1.15902 | 1.657638 | 0.057484 |

| | | | | | | | | | |
|---|---|---|---|---|---|---|---|---|---|
| 17 | 46164400 | 46165599 | 4.252467 | 8.223963 | 31.23989 | 11.94459 | -1.34576 | -0.18684 | 0.057486 |
| 7 | 112026800 | 112027599 | 3.566735 | 6.849856 | 31.15479 | 12.19782 | -1.63308 | -0.11196 | 0.057497 |
| 11 | 44556000 | 44557399 | 3.774921 | 7.31338 | 28.82653 | 10.9309 | -1.3846 | -0.22407 | 0.057519 |
| 2 | 157208600 | 157210999 | 8.375849 | 14.92273 | 44.6675 | 19.66475 | -1.1754 | 0.445352 | 0.057522 |
| 6 | 92003200 | 92008999 | 31.14452 | 49.82893 | 132.619 | 55.83233 | -1.06568 | 2.032543 | 0.05755 |
| 7 | 38895400 | 38896399 | 10.13796 | 19.77554 | 10.77287 | 3.8573 | 1.510312 | -0.40926 | 0.057567 |
| 18 | 37726800 | 37728799 | 42.22334 | 69.32479 | 45.66409 | 19.6687 | 0.977979 | 1.438608 | 0.057587 |
| 10 | 26249400 | 26253399 | 19.54324 | 31.87804 | 96.22778 | 41.34174 | -1.21701 | 1.510803 | 0.057588 |
| 2 | 22473800 | 22475199 | 12.85347 | 24.90768 | 12.82866 | 4.859201 | 1.519651 | -0.17772 | 0.057596 |
| 18 | 18076800 | 18077599 | 10.31687 | 20.11349 | 11.37568 | 4.102499 | 1.448557 | -0.37988 | 0.057627 |
| 14 | 41430400 | 41433399 | 40.62408 | 68.12338 | 30.42611 | 13.274 | 1.538918 | 1.185366 | 0.057629 |
| 8 | 71207200 | 71213999 | 25.35149 | 40.64341 | 128.9188 | 54.43973 | -1.31453 | 1.96095 | 0.057638 |
| 5 | 113661800 | 113666399 | 23.34271 | 37.67484 | 111.6791 | 47.61578 | -1.20513 | 1.740448 | 0.057638 |
| 14 | 61283000 | 61284399 | 5.262027 | 9.76489 | 37.42331 | 15.69544 | -1.47841 | 0.163439 | 0.057646 |
| 1 | 42890000 | 42890999 | 4.592977 | 8.735118 | 33.15953 | 13.30187 | -1.40988 | -0.01645 | 0.057649 |
| 13 | 99982000 | 99983199 | 3.841127 | 7.431503 | 30.58065 | 11.67019 | -1.45643 | -0.19953 | 0.05765 |
| 11 | 99024600 | 99026399 | 4.90659 | 9.132441 | 36.3591 | 15.18138 | -1.52661 | 0.142504 | 0.057653 |
| 7 | 10707600 | 10713399 | 130.696 | 201.1044 | 136.1821 | 56.55736 | 0.858434 | 2.948243 | 0.057661 |
| 9 | 106166200 | 106169199 | 7.726312 | 13.60447 | 49.29843 | 21.82344 | -1.45765 | 0.556559 | 0.057687 |
| 8 | 122265800 | 122302799 | 201.2166 | 282.2218 | 737.2492 | 305.5572 | -1.21222 | 4.532229 | 0.057727 |
| 3 | 129372200 | 129375799 | 16.59826 | 27.67956 | 82.72844 | 35.96549 | -1.19306 | 1.218729 | 0.057743 |
| 5 | 76287200 | 76288799 | 6.768277 | 12.40063 | 39.90838 | 17.03119 | -1.24835 | 0.253465 | 0.057745 |
| 7 | 89056200 | 89057799 | 6.659786 | 12.11909 | 41.35407 | 17.80273 | -1.34075 | 0.295302 | 0.057746 |
| 3 | 94097400 | 94099399 | 16.80234 | 31.32813 | 15.98037 | 6.652762 | 1.414418 | 0.176234 | 0.057753 |
| 14 | 95346000 | 95347599 | 18.66579 | 34.20165 | 17.39614 | 7.423117 | 1.393346 | 0.291483 | 0.057764 |
| 1 | 67621000 | 67621999 | 11.84856 | 22.89525 | 13.71772 | 5.265356 | 1.286723 | -0.14334 | 0.057813 |
| 2 | 91287200 | 91290799 | 14.82918 | 24.86008 | 73.99439 | 32.27556 | -1.18669 | 1.150404 | 0.057816 |
| 1 | 91627800 | 91632799 | 21.38745 | 34.97926 | 94.79373 | 40.76519 | -1.05931 | 1.474356 | 0.057848 |
| 15 | 88772400 | 88773799 | 5.205474 | 9.818082 | 35.76029 | 14.59946 | -1.37459 | 0.052915 | 0.057868 |
| 14 | 112566200 | 112566799 | 11.0292 | 21.53698 | 8.864726 | 3.118537 | 1.929311 | -0.46222 | 0.057875 |
| 3 | 100394400 | 100397999 | 15.8736 | 26.34461 | 81.00666 | 35.10466 | -1.23398 | 1.271465 | 0.057876 |
| 7 | 45511400 | 45520999 | 44.52487 | 69.91613 | 194.2964 | 80.58743 | -1.15336 | 2.553929 | 0.05788 |
| 7 | 76865200 | 76866799 | 21.23445 | 38.96905 | 14.40662 | 6.134411 | 1.85116 | 0.297121 | 0.057886 |
| X | 31782200 | 31782999 | 16.27991 | 30.5078 | 16.40933 | 6.773147 | 1.349272 | 0.142975 | 0.057894 |
| 8 | 23620400 | 23636999 | 77.81715 | 117.905 | 309.118 | 127.9192 | -1.11035 | 3.256481 | 0.057895 |
| 2 | 93026200 | 93045199 | 90.42453 | 135.5882 | 362.5019 | 149.5584 | -1.15176 | 3.468807 | 0.0579 |
| 8 | 36931800 | 36933799 | 36.24574 | 60.63756 | 37.92822 | 16.52226 | 1.055591 | 1.204538 | 0.057916 |
| 14 | 118248600 | 118250199 | 9.966551 | 17.54214 | 50.93587 | 22.55009 | -1.14112 | 0.576754 | 0.057928 |
| 15 | 97676800 | 97678999 | 6.249997 | 11.63254 | 37.02027 | 15.45819 | -1.20849 | 0.152957 | 0.057932 |
| 8 | 108273200 | 108285399 | 59.39668 | 91.403 | 252.2349 | 104.7562 | -1.16618 | 2.932692 | 0.057936 |
| 6 | 52185400 | 52197399 | 43.2557 | 67.92376 | 200.7306 | 83.25621 | -1.24195 | 2.553539 | 0.057969 |
| 4 | 148127600 | 148131399 | 14.15556 | 23.85367 | 70.71992 | 30.91628 | -1.18068 | 1.095625 | 0.05797 |
| 7 | 31454000 | 31457199 | 48.79412 | 79.67348 | 45.70153 | 19.6419 | 1.172751 | 1.524626 | 0.057988 |
| 13 | 65214200 | 65214799 | 9.291214 | 18.14075 | 10.99885 | 3.878198 | 1.378282 | -0.47221 | 0.057989 |
| 15 | 79759400 | 79764999 | 26.29218 | 42.57328 | 107.7738 | 46.11269 | -0.97311 | 1.658497 | 0.057995 |
| 7 | 5588800 | 5590799 | 56.99854 | 91.81851 | 60.14485 | 25.57834 | 0.964335 | 1.822976 | 0.058009 |
| 2 | 71307200 | 71308399 | 17.77114 | 32.37409 | 19.28521 | 8.290905 | 1.160564 | 0.321178 | 0.058036 |

| | | | | | | | | | |
|---|---|---|---|---|---|---|---|---|---|
| 9 | 120512000 | 120517399 | 21.69056 | 35.48007 | 91.15272 | 39.20145 | -0.98229 | 1.472014 | 0.058036 |
| 10 | 92604400 | 92607999 | 41.82024 | 69.11745 | 41.77113 | 18.04958 | 1.104634 | 1.355809 | 0.058041 |
| 19 | 3056200 | 3058199 | 60.39084 | 97.13938 | 60.48016 | 25.66587 | 1.034912 | 1.876572 | 0.058043 |
| 9 | 70069600 | 70073599 | 20.64988 | 34.01916 | 88.25027 | 38.06614 | -0.9917 | 1.377361 | 0.058077 |
| 4 | 82777400 | 82778399 | 3.442908 | 6.6544 | 31.61773 | 12.12249 | -1.66737 | -0.18514 | 0.058124 |
| X | 51017800 | 51018199 | 8.65669 | 16.93475 | 8.179763 | 2.710153 | 1.778545 | -0.70029 | 0.058127 |
| 1 | 113115400 | 113118199 | 33.11876 | 56.61284 | 28.11347 | 12.32674 | 1.388107 | 0.99045 | 0.058133 |
| 8 | 126001200 | 126002599 | 24.66961 | 43.20553 | 24.96245 | 11.02043 | 1.174886 | 0.685669 | 0.058136 |
| 10 | 116199600 | 116200799 | 5.449079 | 10.34489 | 34.35019 | 13.83139 | -1.22456 | 0.006644 | 0.058143 |
| 12 | 110575400 | 110576599 | 5.198218 | 9.82508 | 35.87195 | 14.57991 | -1.37243 | 0.037903 | 0.058167 |
| 5 | 115066200 | 115077199 | 62.58262 | 97.20357 | 218.9748 | 90.92664 | -0.86342 | 2.759712 | 0.058175 |
| 4 | 45515600 | 45518199 | 6.988926 | 12.30053 | 49.80775 | 22.0509 | -1.61654 | 0.558719 | 0.058181 |
| 12 | 20171800 | 20174599 | 47.05613 | 77.03347 | 36.23373 | 15.58746 | 1.460489 | 1.478675 | 0.058183 |
| 12 | 116520400 | 116522199 | 14.5903 | 27.66003 | 15.98713 | 6.456636 | 1.272055 | 0.052302 | 0.058186 |
| 5 | 131660000 | 131660599 | 2.737605 | 5.348423 | 23.46506 | 8.184874 | -1.42125 | -0.55291 | 0.058192 |
| 9 | 99750200 | 99755999 | 47.55349 | 74.73913 | 187.1484 | 77.63179 | -1.00229 | 2.536703 | 0.058198 |
| 3 | 125898000 | 125899199 | 15.95907 | 30.73867 | 11.59133 | 4.500418 | 1.931881 | -0.06828 | 0.058213 |
| 2 | 173389200 | 173414199 | 139.4871 | 202.5973 | 529.5051 | 218.2824 | -1.16243 | 4.011541 | 0.058215 |
| 5 | 101720600 | 101721799 | 8.210337 | 14.6887 | 43.67381 | 19.1601 | -1.16227 | 0.415272 | 0.058228 |
| 5 | 93131400 | 93139199 | 39.45535 | 62.81806 | 153.1599 | 64.03986 | -0.94751 | 2.19891 | 0.058236 |
| 18 | 61826600 | 61833599 | 35.48015 | 56.38154 | 156.1085 | 65.11959 | -1.13339 | 2.25712 | 0.058247 |
| 1 | 48543600 | 48544399 | 9.796362 | 19.09261 | 11.49201 | 4.161854 | 1.354612 | -0.37275 | 0.058256 |
| 1 | 86595200 | 86603999 | 47.59415 | 74.72637 | 192.3363 | 79.77316 | -1.04309 | 2.557825 | 0.058262 |
| 4 | 107697400 | 107714599 | 68.28814 | 103.7822 | 299.9289 | 124.2308 | -1.24705 | 3.191386 | 0.058298 |
| 9 | 72162600 | 72163399 | 18.24614 | 33.94502 | 16.07489 | 6.717203 | 1.51728 | 0.203561 | 0.058345 |
| 11 | 6362800 | 6366999 | 16.56364 | 27.34173 | 85.33832 | 36.84412 | -1.25626 | 1.340156 | 0.058353 |
| 12 | 102612000 | 102625599 | 62.24457 | 95.34705 | 272.8593 | 113.247 | -1.2237 | 3.025979 | 0.058362 |
| 7 | 80261000 | 80263199 | 6.556502 | 12.12732 | 39.06542 | 16.45729 | -1.23551 | 0.197403 | 0.058362 |
| 13 | 13101800 | 13103199 | 22.87134 | 40.28132 | 22.02769 | 9.750764 | 1.254455 | 0.592418 | 0.058368 |
| 15 | 95965000 | 95968399 | 14.55153 | 24.68511 | 67.04582 | 29.36947 | -1.05434 | 1.034785 | 0.058372 |
| 7 | 44649200 | 44663599 | 55.75275 | 85.79405 | 250.3803 | 103.9857 | -1.24677 | 2.932173 | 0.058386 |
| 8 | 95127200 | 95129599 | 9.013528 | 15.9442 | 47.75344 | 21.12007 | -1.18206 | 0.511678 | 0.058404 |
| 9 | 69196600 | 69198799 | 9.131784 | 15.95621 | 54.14796 | 23.85887 | -1.36561 | 0.688697 | 0.058412 |
| 5 | 83733800 | 83735599 | 16.06054 | 29.90005 | 17.25532 | 7.201852 | 1.235449 | 0.18249 | 0.058416 |
| 11 | 5204200 | 5208799 | 16.77654 | 27.91008 | 79.03883 | 34.3087 | -1.11545 | 1.243933 | 0.058423 |
| 4 | 12158800 | 12159599 | 13.71451 | 26.50012 | 12.59778 | 4.836119 | 1.615379 | -0.12572 | 0.058439 |
| 11 | 48575600 | 48577799 | 9.242895 | 16.22073 | 50.9234 | 22.52044 | -1.2538 | 0.612725 | 0.058461 |
| 2 | 26083400 | 26099599 | 107.7158 | 160.3032 | 394.5995 | 162.5864 | -1.04266 | 3.611769 | 0.058462 |
| 15 | 83039600 | 83048399 | 41.1211 | 64.92946 | 176.0985 | 73.11805 | -1.11196 | 2.434311 | 0.058463 |
| 7 | 121766600 | 121774599 | 37.74196 | 59.95343 | 156.7858 | 65.37773 | -1.05176 | 2.269858 | 0.05849 |
| 4 | 24256600 | 24257199 | 12.22888 | 23.62196 | 13.95653 | 5.365696 | 1.303667 | -0.13365 | 0.058495 |
| 3 | 13122800 | 13126399 | 94.39111 | 148.7277 | 100.2359 | 41.59288 | 0.891388 | 2.499515 | 0.058509 |
| 7 | 59083200 | 59085999 | 25.7572 | 45.08715 | 24.69322 | 10.89685 | 1.251487 | 0.698646 | 0.058514 |
| 7 | 141026000 | 141028799 | 10.4508 | 18.32689 | 51.49028 | 22.75682 | -1.09487 | 0.632832 | 0.058549 |
| 12 | 20792400 | 20795599 | 78.28517 | 124.4973 | 84.24937 | 35.17081 | 0.8968 | 2.253127 | 0.058574 |
| 2 | 168579600 | 168587599 | 55.91147 | 87.22215 | 206.6962 | 85.77318 | -0.93141 | 2.679875 | 0.058591 |
| 5 | 105646600 | 105652599 | 38.6294 | 61.55668 | 144.8684 | 60.64613 | -0.89514 | 2.168882 | 0.058593 |

| | | | | | | | | | |
|---|---|---|---|---|---|---|---|---|---|
| 7 | 8230000 | 8231999 | 29.87728 | 51.89798 | 25.02999 | 10.98795 | 1.433364 | 0.82791 | 0.058594 |
| 4 | 146347200 | 146350599 | 21.44438 | 38.2762 | 18.38967 | 8.083506 | 1.447827 | 0.463147 | 0.058605 |
| 7 | 5900400 | 5904199 | 97.73972 | 153.711 | 102.563 | 42.548 | 0.903722 | 2.539377 | 0.058623 |
| 7 | 8014600 | 8015599 | 16.32807 | 31.20706 | 11.55004 | 4.5851 | 1.926962 | 0.003184 | 0.058625 |
| 1 | 155034200 | 155034999 | 3.537058 | 6.87772 | 28.74504 | 10.65165 | -1.43747 | -0.31169 | 0.058633 |
| 2 | 31358600 | 31393999 | 181.6608 | 257.9428 | 647.8531 | 267.9694 | -1.13765 | 4.349444 | 0.058648 |
| 13 | 20670000 | 20671799 | 17.37849 | 32.8375 | 13.89268 | 5.649983 | 1.708574 | 0.094716 | 0.058716 |
| 6 | 57773400 | 57773799 | 2.24712 | 4.394306 | 22.54034 | 7.678219 | -1.61008 | -0.65007 | 0.058723 |
| 18 | 36824400 | 36825599 | 3.648501 | 7.099597 | 27.6243 | 10.17621 | -1.32721 | -0.33358 | 0.05874 |
| 17 | 6704000 | 6705799 | 3.960989 | 7.43224 | 35.39675 | 14.57873 | -1.76384 | 0.080691 | 0.05874 |
| 7 | 38910600 | 38911799 | 12.36226 | 23.75605 | 14.56027 | 5.690683 | 1.229654 | -0.07825 | 0.058749 |
| 5 | 137717400 | 137720399 | 9.424195 | 16.49143 | 54.71705 | 24.13945 | -1.3335 | 0.665784 | 0.058796 |
| 14 | 85261800 | 85262199 | 2.938973 | 5.739071 | 25.34626 | 8.914716 | -1.44431 | -0.5172 | 0.0588 |
| 13 | 46084800 | 46086399 | 18.97838 | 34.15736 | 19.59985 | 8.554707 | 1.200243 | 0.397883 | 0.058801 |
| 7 | 5427400 | 5429799 | 55.97485 | 90.22262 | 60.58207 | 25.78563 | 0.929723 | 1.805147 | 0.058804 |
| 4 | 49596800 | 49598999 | 9.754112 | 17.01005 | 56.88966 | 25.03588 | -1.34548 | 0.723146 | 0.058805 |
| 9 | 19827200 | 19827799 | 10.42557 | 20.30595 | 11.93667 | 4.359147 | 1.37785 | -0.3337 | 0.05881 |
| 12 | 95542800 | 95544999 | 10.46749 | 18.27487 | 53.63846 | 23.62199 | -1.15764 | 0.708825 | 0.058816 |
| 7 | 9611800 | 9614199 | 20.44214 | 36.22785 | 19.84037 | 8.771604 | 1.254895 | 0.504223 | 0.058823 |
| 11 | 118602400 | 118632399 | 163.7558 | 234.1791 | 609.9942 | 252.0112 | -1.17984 | 4.244081 | 0.058838 |
| 1 | 189071600 | 189078799 | 36.58682 | 58.22196 | 158.746 | 66.32976 | -1.10927 | 2.213617 | 0.058841 |
| 6 | 121186200 | 121189799 | 20.67373 | 33.62279 | 97.07403 | 41.66053 | -1.15604 | 1.55743 | 0.058848 |
| 14 | 18376800 | 18377999 | 10.3658 | 20.15079 | 12.11274 | 4.497897 | 1.325002 | -0.2773 | 0.058853 |
| 17 | 81999400 | 82000399 | 10.3658 | 20.15079 | 12.11274 | 4.497897 | 1.325002 | -0.2773 | 0.058853 |
| 5 | 136269000 | 136291999 | 98.23384 | 146.1901 | 398.8846 | 164.3518 | -1.19109 | 3.611 | 0.05887 |
| 7 | 68648600 | 68649199 | 9.639031 | 18.81853 | 10.99287 | 3.880797 | 1.430104 | -0.4626 | 0.058875 |
| 7 | 35961200 | 35962799 | 5.54239 | 10.32616 | 37.58406 | 15.66627 | -1.39831 | 0.140308 | 0.058876 |
| 17 | 34781600 | 34797199 | 77.19057 | 116.3587 | 340.4646 | 140.6656 | -1.27495 | 3.361722 | 0.058878 |
| 4 | 128600000 | 128607399 | 31.22672 | 49.79813 | 144.5016 | 60.55603 | -1.19534 | 2.13764 | 0.058881 |
| 6 | 97124800 | 97126199 | 5.587666 | 10.28563 | 38.88028 | 16.48701 | -1.47084 | 0.215715 | 0.058887 |
| 19 | 9392800 | 9394199 | 16.85727 | 31.57173 | 15.44758 | 6.382339 | 1.482522 | 0.148157 | 0.058903 |
| 13 | 51994400 | 52007999 | 54.9381 | 84.9804 | 234.3695 | 97.35142 | -1.1595 | 2.831842 | 0.058915 |
| 16 | 3259200 | 3261799 | 27.87182 | 48.84632 | 21.13515 | 9.33601 | 1.58892 | 0.685994 | 0.05892 |
| 5 | 123241600 | 123245399 | 14.79532 | 24.85918 | 71.22668 | 31.11031 | -1.13185 | 1.124552 | 0.058928 |
| 3 | 13054400 | 13055999 | 17.83172 | 33.84418 | 12.08723 | 4.86968 | 1.960842 | 0.065113 | 0.058941 |
| 12 | 19525800 | 19527599 | 34.70629 | 58.36014 | 32.83107 | 14.34338 | 1.207781 | 1.143187 | 0.058959 |
| 9 | 17413200 | 17413999 | 3.241537 | 6.309176 | 26.94619 | 9.907506 | -1.45621 | -0.3472 | 0.058982 |
| 9 | 65797400 | 65798199 | 3.241537 | 6.309176 | 26.94619 | 9.907506 | -1.45621 | -0.3472 | 0.058982 |
| 16 | 37908400 | 37910599 | 11.32611 | 19.24387 | 67.87038 | 29.74421 | -1.42887 | 1.010239 | 0.058992 |
| 11 | 61820600 | 61822199 | 6.740779 | 12.38158 | 38.86913 | 16.5306 | -1.20873 | 0.238081 | 0.058999 |
| 12 | 62283600 | 62284799 | 17.18134 | 31.70107 | 17.60995 | 7.443679 | 1.276841 | 0.242332 | 0.059028 |
| X | 49287000 | 49288199 | 13.89983 | 27.02576 | 10.1268 | 3.754924 | 2.000679 | -0.24374 | 0.059046 |
| 14 | 3392400 | 3392799 | 9.060687 | 17.72177 | 8.582899 | 2.8937 | 1.751924 | -0.63475 | 0.059056 |
| 1 | 77279000 | 77280199 | 4.075035 | 7.894221 | 35.34553 | 13.40755 | -1.56967 | -0.20578 | 0.059058 |
| 12 | 23452600 | 23453799 | 21.72201 | 39.8427 | 13.77988 | 5.871769 | 1.94536 | 0.302383 | 0.059059 |
| 13 | 65971200 | 65973399 | 24.04539 | 42.23173 | 23.82992 | 10.54564 | 1.208532 | 0.636104 | 0.059081 |
| 1 | 101836800 | 101837199 | 9.00833 | 17.58935 | 11.13703 | 3.923368 | 1.31838 | -0.47861 | 0.059089 |

| | | | | | | | | | |
|---|---|---|---|---|---|---|---|---|---|
| 15 | 101615000 | 101623599 | 41.03045 | 65.01801 | 166.8601 | 69.40846 | -1.02822 | 2.346347 | 0.059091 |
| 9 | 96599200 | 96602399 | 13.92239 | 23.6527 | 66.47728 | 29.13345 | -1.10331 | 1.018719 | 0.059094 |
| 1 | 145996400 | 145997199 | 19.94761 | 35.82668 | 17.27196 | 7.552501 | 1.447965 | 0.412598 | 0.059094 |
| 17 | 14008000 | 14008599 | 11.08277 | 21.50255 | 13.22693 | 4.981646 | 1.274526 | -0.21574 | 0.059111 |
| 13 | 10206800 | 10208199 | 4.646918 | 8.940001 | 32.61849 | 12.70672 | -1.31371 | -0.11332 | 0.059165 |
| 14 | 11295600 | 11296799 | 13.86646 | 26.55066 | 14.85927 | 5.867996 | 1.345834 | -0.02874 | 0.05917 |
| 1 | 131661600 | 131668599 | 35.88646 | 57.21224 | 144.3353 | 60.46166 | -0.99446 | 2.150959 | 0.059182 |
| 1 | 153082600 | 153083799 | 4.681542 | 9.020783 | 34.06767 | 13.21149 | -1.35797 | -0.12726 | 0.059195 |
| 11 | 49324600 | 49325199 | 12.37162 | 24.0296 | 12.28717 | 4.594105 | 1.546722 | -0.22775 | 0.059232 |
| 7 | 14807000 | 14809999 | 80.12209 | 127.5223 | 78.73534 | 32.90597 | 1.030051 | 2.226601 | 0.059292 |
| 17 | 27495200 | 27499399 | 149.5388 | 227.6835 | 159.3955 | 66.05819 | 0.798727 | 3.163869 | 0.059294 |
| 15 | 61443400 | 61444799 | 7.41689 | 13.43452 | 44.53917 | 19.28518 | -1.30572 | 0.333275 | 0.059298 |
| 12 | 83466600 | 83469199 | 11.72723 | 20.41349 | 57.52976 | 25.28844 | -1.10053 | 0.76011 | 0.059301 |
| 19 | 3755800 | 3758599 | 9.22913 | 16.22902 | 50.37998 | 22.30816 | -1.23745 | 0.580767 | 0.059308 |
| 7 | 31378000 | 31380199 | 32.1996 | 55.34939 | 28.44009 | 12.47489 | 1.340246 | 0.938615 | 0.059348 |
| 4 | 139685600 | 139698599 | 99.30424 | 149.174 | 343.1178 | 141.63 | -0.93256 | 3.433962 | 0.05935 |
| 7 | 8663600 | 8665399 | 23.2319 | 40.9514 | 21.04777 | 9.315398 | 1.343821 | 0.58389 | 0.05935 |
| 2 | 172745000 | 172754599 | 43.56291 | 68.32743 | 194.8653 | 80.81865 | -1.19193 | 2.576331 | 0.059377 |
| 13 | 33636800 | 33637999 | 16.64757 | 30.88531 | 17.78917 | 7.46347 | 1.232939 | 0.209224 | 0.059383 |
| 17 | 87675000 | 87692399 | 73.42532 | 110.7554 | 344.1723 | 142.2251 | -1.36086 | 3.347875 | 0.059408 |
| 8 | 124783600 | 124784399 | 2.339209 | 4.561227 | 24.66626 | 8.875962 | -1.757 | -0.44104 | 0.059427 |
| 9 | 30936400 | 30937199 | 11.19201 | 21.68544 | 13.67117 | 5.181264 | 1.230751 | -0.19293 | 0.059449 |
| 3 | 148758000 | 148758599 | 46.39788 | 76.67075 | 34.45052 | 14.88465 | 1.530299 | 1.36279 | 0.059495 |
| 6 | 17710200 | 17712599 | 16.21225 | 27.20685 | 73.67355 | 32.15662 | -1.051 | 1.142843 | 0.05951 |
| 2 | 79670000 | 79671199 | 4.818574 | 9.261512 | 35.74792 | 13.96352 | -1.39914 | -0.0974 | 0.059511 |
| 11 | 95466000 | 95473199 | 28.25982 | 45.04738 | 155.49 | 65.12227 | -1.44588 | 2.150638 | 0.059519 |
| 16 | 46594600 | 46595199 | 8.219802 | 16.06812 | 9.991023 | 3.435926 | 1.37339 | -0.5776 | 0.059524 |
| 12 | 23694400 | 23718399 | 450.9043 | 629.4759 | 399.6955 | 165.7768 | 0.820856 | 4.607684 | 0.059529 |
| 9 | 111251200 | 111252399 | 5.22939 | 9.795464 | 36.05862 | 14.89436 | -1.4033 | 0.099533 | 0.059542 |
| 9 | 8887000 | 8888999 | 23.86921 | 41.81979 | 25.33702 | 11.18943 | 1.106531 | 0.674652 | 0.059551 |
| 14 | 3141400 | 3144799 | 82.03224 | 130.0992 | 90.43914 | 37.65126 | 0.854308 | 2.336067 | 0.059557 |
| X | 19517200 | 19518399 | 15.76395 | 30.13702 | 12.84014 | 5.093287 | 1.728358 | -0.00126 | 0.059574 |
| 18 | 70554600 | 70555999 | 5.016765 | 9.534724 | 33.17969 | 13.32878 | -1.28792 | -0.00766 | 0.059574 |
| 14 | 112108600 | 112109799 | 15.832 | 29.96782 | 15.13998 | 6.134454 | 1.459867 | 0.071739 | 0.059616 |
| 16 | 90186200 | 90190399 | 28.69109 | 46.04801 | 124.1752 | 52.52432 | -1.07914 | 1.927815 | 0.059618 |
| 12 | 81826400 | 81828999 | 15.45971 | 25.90126 | 75.80204 | 33.05127 | -1.16257 | 1.159831 | 0.059628 |
| 17 | 34884800 | 34909399 | 182.7281 | 260.2582 | 623.264 | 257.6653 | -1.06449 | 4.304246 | 0.059636 |
| 17 | 91615000 | 91616599 | 18.85985 | 34.22949 | 19.01962 | 8.214959 | 1.255791 | 0.349293 | 0.059637 |
| 5 | 87083600 | 87085999 | 25.82747 | 44.86879 | 27.62516 | 12.12813 | 1.083072 | 0.809559 | 0.059638 |
| 3 | 150046000 | 150046799 | 16.45831 | 30.54424 | 17.79308 | 7.461772 | 1.21713 | 0.205725 | 0.059641 |
| 5 | 93404400 | 93405399 | 17.10082 | 32.18314 | 13.67667 | 5.604577 | 1.69225 | 0.116224 | 0.059646 |
| 7 | 15441600 | 15445199 | 104.274 | 162.6697 | 114.9856 | 47.71576 | 0.811712 | 2.690424 | 0.059674 |
| 7 | 46699600 | 46707199 | 31.37919 | 50.11446 | 144.4269 | 60.64923 | -1.18323 | 2.090337 | 0.059694 |
| 7 | 31953800 | 31955399 | 38.57177 | 63.93247 | 40.2623 | 17.43223 | 1.045718 | 1.31408 | 0.059701 |
| 5 | 93849800 | 93852599 | 75.11648 | 119.891 | 74.00299 | 31.04876 | 1.034704 | 2.133541 | 0.059722 |
| 7 | 125150400 | 125153799 | 15.08026 | 25.41587 | 70.27577 | 30.72357 | -1.08059 | 1.096778 | 0.059723 |
| 19 | 45050600 | 45057199 | 36.63723 | 58.54236 | 137.0212 | 57.58182 | -0.88287 | 2.078165 | 0.059739 |

| | | | | | | | | | |
|---|---|---|---|---|---|---|---|---|---|
| 12 | 20398400 | 20401599 | 106.1264 | 165.3695 | 110.8786 | 46.01878 | 0.886553 | 2.710471 | 0.059773 |
| 12 | 21738000 | 21741799 | 71.54327 | 114.247 | 74.97863 | 31.48112 | 0.947195 | 2.113511 | 0.059795 |
| 12 | 100072000 | 100073199 | 4.382492 | 8.455785 | 30.36783 | 11.73181 | -1.27947 | -0.15049 | 0.059841 |
| 5 | 135277400 | 135300999 | 161.5275 | 234.802 | 515.0999 | 212.3259 | -0.90882 | 4.000615 | 0.059857 |
| 18 | 74672800 | 74678799 | 38.45559 | 61.31547 | 142.7597 | 59.81157 | -0.87876 | 2.149405 | 0.059857 |
| 5 | 37118400 | 37147399 | 136.3843 | 197.4627 | 552.973 | 228.0434 | -1.26646 | 4.059259 | 0.059895 |
| 9 | 118335200 | 118336799 | 6.796787 | 12.37056 | 40.27538 | 17.33457 | -1.27314 | 0.293868 | 0.059932 |
| 7 | 18947600 | 18949799 | 7.128991 | 12.88155 | 41.72506 | 18.12194 | -1.27468 | 0.341793 | 0.05994 |
| 18 | 9117600 | 9118399 | 5.27286 | 10.05845 | 33.96455 | 13.53813 | -1.23517 | -0.02869 | 0.059941 |
| 19 | 41582800 | 41609399 | 157.9883 | 228.7901 | 526.3473 | 217.057 | -0.98269 | 4.057389 | 0.059965 |
| 17 | 39933200 | 39934799 | 29.43144 | 51.44476 | 21.73385 | 9.579263 | 1.623614 | 0.736322 | 0.059967 |
| 10 | 82365600 | 82367199 | 16.72116 | 31.94021 | 11.54141 | 4.587239 | 1.959865 | 0.01097 | 0.059981 |
| 6 | 92457800 | 92458599 | 3.04505 | 5.939196 | 24.89884 | 8.92714 | -1.39463 | -0.44564 | 0.059988 |
| 12 | 23114800 | 23115599 | 22.14794 | 38.98137 | 23.46554 | 10.38862 | 1.116297 | 0.597201 | 0.059992 |
| 17 | 13580400 | 13630999 | 1771.611 | 2111.665 | 3954.532 | 1673.496 | -0.96916 | 7.127192 | 0.059997 |
| 1 | 132998800 | 133008799 | 48.72116 | 76.14466 | 205.7198 | 85.35192 | -1.11813 | 2.643622 | 0.06001 |
| 17 | 19064400 | 19065799 | 7.19448 | 13.19501 | 40.50323 | 17.26098 | -1.17923 | 0.253111 | 0.06004 |
| 5 | 85457200 | 85458199 | 13.8555 | 26.23833 | 16.0772 | 6.508035 | 1.185559 | 0.052383 | 0.060091 |
| 2 | 25175000 | 25196799 | 122.0169 | 179.0782 | 469.2502 | 193.2697 | -1.15164 | 3.848548 | 0.0601 |
| 7 | 85734800 | 85736799 | 27.22867 | 47.57352 | 24.61397 | 10.84494 | 1.333206 | 0.735416 | 0.06012 |
| 11 | 58794600 | 58799399 | 22.48335 | 36.4415 | 104.6198 | 44.79546 | -1.15261 | 1.630175 | 0.060148 |
| 18 | 84275400 | 84277999 | 13.64457 | 23.12066 | 68.62261 | 30.04998 | -1.18183 | 1.042297 | 0.060171 |
| 19 | 4032400 | 4043199 | 49.66389 | 77.27509 | 215.968 | 89.65436 | -1.17195 | 2.72354 | 0.060178 |
| X | 19702800 | 19703999 | 15.73902 | 30.4534 | 10.2357 | 3.904459 | 2.117258 | -0.13017 | 0.060186 |
| 11 | 74787200 | 74789799 | 9.958321 | 16.89216 | 68.74021 | 30.11112 | -1.63388 | 1.022007 | 0.060207 |
| 11 | 68743000 | 68744199 | 4.111383 | 7.844126 | 32.05011 | 12.77058 | -1.50453 | -0.04406 | 0.06021 |
| 15 | 23160200 | 23161599 | 15.07296 | 28.23089 | 16.86891 | 6.969176 | 1.19719 | 0.135533 | 0.060211 |
| 13 | 87768600 | 87769599 | 16.4746 | 31.38615 | 13.5569 | 5.418529 | 1.700617 | 0.031654 | 0.060292 |
| 6 | 69738200 | 69738799 | 12.30785 | 23.79958 | 13.99014 | 5.352192 | 1.317833 | -0.14834 | 0.060311 |
| 1 | 133852000 | 133855199 | 17.50631 | 29.23325 | 76.39852 | 33.24189 | -0.99997 | 1.204192 | 0.060319 |
| 12 | 10792800 | 10794599 | 21.19965 | 37.52009 | 22.37438 | 9.894553 | 1.132614 | 0.530235 | 0.060327 |
| 7 | 3057600 | 3059999 | 48.25519 | 78.72608 | 47.88976 | 20.57595 | 1.087513 | 1.537965 | 0.060329 |
| 8 | 121350600 | 121364999 | 72.99796 | 111.0066 | 291.9815 | 120.9607 | -1.11066 | 3.179702 | 0.060331 |
| 13 | 96026400 | 96031199 | 21.3207 | 34.68296 | 100.4475 | 43.11189 | -1.16044 | 1.555829 | 0.060331 |
| 8 | 96574600 | 96576199 | 6.135424 | 11.25465 | 38.80198 | 16.53198 | -1.34396 | 0.242384 | 0.060336 |
| 13 | 65876000 | 65876599 | 9.952314 | 19.46695 | 7.460868 | 2.501463 | 2.090914 | -0.64239 | 0.060339 |
| 15 | 75998400 | 76007399 | 35.13594 | 55.62558 | 175.7734 | 73.08059 | -1.32887 | 2.366342 | 0.060353 |
| 9 | 106768600 | 106770799 | 6.253992 | 11.31401 | 42.27044 | 18.33138 | -1.47717 | 0.32943 | 0.060355 |
| 7 | 132319000 | 132320599 | 6.825263 | 12.34077 | 41.2001 | 17.87937 | -1.31686 | 0.335378 | 0.060359 |
| 14 | 36465600 | 36467199 | 26.67549 | 46.33579 | 28.65892 | 12.58092 | 1.07669 | 0.818419 | 0.06038 |
| 2 | 149490000 | 149491199 | 13.89777 | 26.88415 | 11.9965 | 4.581386 | 1.712268 | -0.14218 | 0.060381 |
| 2 | 180552600 | 180553799 | 3.740734 | 7.203893 | 33.82787 | 13.13975 | -1.66987 | -0.13288 | 0.060383 |
| 18 | 39261200 | 39262199 | 2.545749 | 4.971774 | 24.61889 | 8.641767 | -1.60252 | -0.53206 | 0.060396 |
| 7 | 32944200 | 32946799 | 67.35067 | 107.911 | 68.72839 | 28.99589 | 0.995613 | 2.007073 | 0.0604 |
| 8 | 118245800 | 118256999 | 48.62723 | 76.21875 | 198.6896 | 82.4105 | -1.06336 | 2.590744 | 0.060424 |
| 4 | 129320400 | 129322599 | 8.604535 | 15.17268 | 47.73452 | 21.12505 | -1.25452 | 0.542839 | 0.060425 |
| 15 | 78481400 | 78492999 | 66.36187 | 101.8473 | 256.6174 | 106.537 | -1.03811 | 2.988432 | 0.060454 |

| | | | | | | | | | |
|---|---|---|---|---|---|---|---|---|---|
| 4 | 59189000 | 59192999 | 13.19228 | 22.71569 | 63.50997 | 27.85931 | -1.09243 | 0.887646 | 0.060458 |
| 1 | 33507000 | 33510399 | 18.70011 | 30.97249 | 81.34663 | 35.20245 | -1.00786 | 1.303147 | 0.060473 |
| 4 | 135581400 | 135584599 | 16.37273 | 27.45857 | 73.07644 | 31.88312 | -1.02603 | 1.15017 | 0.060511 |
| 13 | 47426200 | 47427399 | 4.252186 | 8.224256 | 30.62586 | 11.70425 | -1.31656 | -0.18989 | 0.060534 |
| 1 | 144607800 | 144610199 | 27.25325 | 47.42377 | 26.81882 | 11.78633 | 1.205237 | 0.793376 | 0.060537 |
| 2 | 172891600 | 172900999 | 41.82437 | 65.54549 | 199.2843 | 82.65854 | -1.28504 | 2.590891 | 0.060541 |
| 5 | 145979400 | 145982799 | 79.09797 | 125.533 | 88.12528 | 36.71355 | 0.841139 | 2.313455 | 0.060544 |
| 7 | 100978000 | 100980599 | 7.521291 | 13.24755 | 47.72121 | 21.12394 | -1.44829 | 0.551787 | 0.060544 |
| 5 | 134212000 | 134214799 | 10.82936 | 18.77458 | 56.83202 | 24.93514 | -1.20348 | 0.809689 | 0.06055 |
| 11 | 117027800 | 117037799 | 40.31941 | 63.2972 | 194.9291 | 80.84733 | -1.30148 | 2.557158 | 0.060555 |
| 4 | 154715600 | 154728599 | 74.95905 | 113.8898 | 302.6116 | 125.3319 | -1.12631 | 3.198143 | 0.060581 |
| 12 | 22653400 | 22655199 | 46.23381 | 75.29021 | 50.5756 | 21.71542 | 0.94302 | 1.560415 | 0.060583 |
| 3 | 90349800 | 90356399 | 28.76212 | 46.3711 | 113.3923 | 48.27826 | -0.93026 | 1.781238 | 0.060585 |
| 11 | 57597400 | 57607599 | 61.9443 | 95.81611 | 231.4445 | 96.13646 | -0.96856 | 2.834339 | 0.0606 |
| 19 | 3168800 | 3170199 | 17.8797 | 33.6087 | 13.65942 | 5.608871 | 1.75401 | 0.1309 | 0.060614 |
| 3 | 74467600 | 74468999 | 15.43614 | 29.263 | 12.70257 | 5.130334 | 1.678439 | 0.050157 | 0.060614 |
| 18 | 65550200 | 65551799 | 8.396829 | 14.93702 | 44.64374 | 19.6799 | -1.17475 | 0.454269 | 0.060625 |
| 7 | 20979200 | 20982799 | 92.08592 | 145.334 | 96.61472 | 40.10601 | 0.912936 | 2.462459 | 0.060644 |
| 2 | 31687200 | 31737199 | 224.8323 | 309.6086 | 916.7159 | 381.034 | -1.41964 | 4.812202 | 0.060682 |
| 12 | 58020200 | 58021399 | 12.14374 | 23.46154 | 13.35241 | 5.129323 | 1.357654 | -0.13954 | 0.060695 |
| 2 | 170190000 | 170191799 | 6.427506 | 11.90077 | 37.99403 | 15.98389 | -1.2215 | 0.186435 | 0.060702 |
| 3 | 155471800 | 155473199 | 19.96143 | 37.24715 | 12.92756 | 5.374792 | 1.965418 | 0.192165 | 0.060741 |
| 18 | 77325000 | 77326999 | 9.033491 | 15.98429 | 47.22777 | 20.88633 | -1.16273 | 0.505244 | 0.060756 |
| 7 | 14848200 | 14849799 | 44.1222 | 73.14007 | 36.71052 | 15.8956 | 1.37171 | 1.324894 | 0.060791 |
| 7 | 3141200 | 3143599 | 40.69124 | 67.31685 | 42.3174 | 18.29789 | 1.04811 | 1.342507 | 0.06081 |
| 4 | 91778200 | 91778799 | 10.38059 | 20.28084 | 8.908334 | 3.106217 | 1.847742 | -0.50133 | 0.060822 |
| 1 | 92623000 | 92628199 | 23.49029 | 37.955 | 110.5256 | 47.19128 | -1.17771 | 1.706889 | 0.060829 |
| 11 | 120055000 | 120057399 | 9.084308 | 15.9361 | 49.85634 | 22.04115 | -1.24855 | 0.61789 | 0.060832 |
| 17 | 17367600 | 17367999 | 11.02733 | 21.54233 | 8.202724 | 2.864895 | 2.047317 | -0.48867 | 0.060836 |
| 17 | 72237800 | 72238199 | 10.13624 | 19.82712 | 6.108825 | 2.014863 | 2.415573 | -0.69834 | 0.060839 |
| 10 | 81549000 | 81549599 | 16.26973 | 30.20418 | 17.79699 | 7.460075 | 1.20119 | 0.202222 | 0.060843 |
| 12 | 44836400 | 44837799 | 17.56802 | 32.70858 | 16.67305 | 6.958094 | 1.41348 | 0.194501 | 0.06085 |
| 9 | 19737600 | 19738799 | 16.62623 | 31.94134 | 11.19128 | 4.377336 | 2.025443 | -0.03992 | 0.060864 |
| 18 | 25507200 | 25508599 | 8.713806 | 15.46273 | 45.66054 | 20.17 | -1.15978 | 0.473439 | 0.060874 |
| 17 | 60463600 | 60465199 | 20.50996 | 36.67449 | 21.12928 | 9.273624 | 1.189607 | 0.452567 | 0.060883 |
| 1 | 132794200 | 132795799 | 6.100828 | 11.38479 | 36.55324 | 15.1954 | -1.21594 | 0.132853 | 0.060892 |
| 11 | 97405400 | 97419999 | 60.2326 | 92.32905 | 267.8473 | 111.1809 | -1.24253 | 3.011261 | 0.060893 |
| 2 | 11607400 | 11608199 | 5.578208 | 10.29405 | 38.39903 | 16.22737 | -1.44823 | 0.199976 | 0.060902 |
| 14 | 20751800 | 20753799 | 5.525611 | 10.34135 | 37.20013 | 15.38849 | -1.37267 | 0.109439 | 0.060916 |
| 1 | 75118000 | 75121399 | 12.17958 | 20.82996 | 63.75913 | 27.9572 | -1.22401 | 0.947428 | 0.06093 |
| 1 | 52085000 | 52086799 | 9.03212 | 15.95245 | 46.68849 | 20.65544 | -1.14991 | 0.526529 | 0.060964 |
| 19 | 5429800 | 5436199 | 18.88216 | 30.45614 | 113.3864 | 48.30398 | -1.53359 | 1.755142 | 0.060968 |
| 17 | 41030200 | 41030799 | 13.29487 | 25.48474 | 13.72078 | 5.401008 | 1.406235 | -0.04763 | 0.060973 |
| 9 | 74540600 | 74541199 | 10.50642 | 20.43545 | 12.72664 | 4.704779 | 1.28122 | -0.2875 | 0.060987 |
| 11 | 27441600 | 27441999 | 10.50642 | 20.43545 | 12.72664 | 4.704779 | 1.28122 | -0.2875 | 0.060987 |
| 14 | 38065400 | 38066799 | 19.85581 | 35.4746 | 21.33081 | 9.368792 | 1.127724 | 0.453749 | 0.061009 |
| 11 | 13772200 | 13774599 | 29.05661 | 50.11267 | 29.56649 | 12.97028 | 1.142578 | 0.896194 | 0.061012 |

| | | | | | | | | | |
|---|---|---|---|---|---|---|---|---|---|
| 3 | 68622000 | 68629399 | 40.49264 | 64.3003 | 167.4829 | 69.8137 | -1.04664 | 2.283322 | 0.061017 |
| 12 | 95124200 | 95125599 | 8.064747 | 14.3768 | 44.14839 | 19.42671 | -1.2113 | 0.439283 | 0.06102 |
| 7 | 8990800 | 8991799 | 16.26532 | 30.54948 | 16.32534 | 6.712722 | 1.362749 | 0.128657 | 0.061024 |
| 3 | 38489200 | 38489799 | 12.86027 | 24.88925 | 13.40687 | 5.108423 | 1.448027 | -0.15664 | 0.061052 |
| 8 | 115022600 | 115023599 | 4.144059 | 7.98058 | 32.05173 | 12.45 | -1.44702 | -0.13244 | 0.061063 |
| 4 | 105011400 | 105016999 | 25.69833 | 41.53276 | 110.7217 | 47.28975 | -1.05035 | 1.70305 | 0.061065 |
| 8 | 121029000 | 121032199 | 11.32855 | 19.51538 | 59.4494 | 26.07846 | -1.21469 | 0.873914 | 0.061089 |
| 11 | 59888400 | 59890999 | 6.956738 | 12.31827 | 47.21658 | 20.87844 | -1.53431 | 0.485291 | 0.061096 |
| 7 | 136407800 | 136414399 | 37.65302 | 60.02967 | 143.7974 | 60.2378 | -0.91987 | 2.151788 | 0.061096 |
| 14 | 32944200 | 32944799 | 2.027274 | 3.965424 | 19.76936 | 6.516824 | -1.52148 | -0.77749 | 0.061111 |
| 6 | 10051200 | 10052799 | 34.76484 | 58.22363 | 37.88482 | 16.51473 | 0.998701 | 1.18667 | 0.061147 |
| 17 | 21132400 | 21133599 | 14.73387 | 27.74449 | 16.11498 | 6.597381 | 1.248588 | 0.098178 | 0.061149 |
| 7 | 11177600 | 11185399 | 202.2319 | 302.0647 | 204.8622 | 84.45661 | 0.820713 | 3.55228 | 0.061174 |
| 8 | 112129200 | 112130799 | 9.337853 | 16.46667 | 48.26611 | 21.36007 | -1.15307 | 0.548998 | 0.061175 |
| 13 | 60346000 | 60347399 | 22.16806 | 39.2005 | 22.62574 | 10.00743 | 1.178882 | 0.545692 | 0.061179 |
| 5 | 100484400 | 100485999 | 4.528915 | 8.670822 | 31.68656 | 12.51679 | -1.33453 | -0.07027 | 0.061179 |
| 10 | 3050200 | 3053999 | 95.28923 | 150.975 | 80.70516 | 33.56768 | 1.23132 | 2.373819 | 0.06118 |
| 4 | 155874800 | 155878399 | 14.4856 | 24.32703 | 71.15257 | 31.07061 | -1.16138 | 1.12873 | 0.061185 |
| 9 | 11553400 | 11557799 | 71.37376 | 113.8831 | 77.62115 | 32.55291 | 0.891469 | 2.138682 | 0.061199 |
| 13 | 55606400 | 55615199 | 27.9355 | 44.76073 | 130.2365 | 54.94885 | -1.19143 | 1.981466 | 0.06121 |
| 12 | 85747400 | 85759799 | 65.41511 | 101.0035 | 231.8462 | 96.3042 | -0.89709 | 2.869362 | 0.061219 |
| 4 | 119887400 | 119897199 | 47.0319 | 73.6696 | 200.4911 | 83.16337 | -1.12609 | 2.602043 | 0.061222 |
| 7 | 15371200 | 15372999 | 50.83673 | 82.35 | 54.50059 | 23.32586 | 0.958107 | 1.665148 | 0.061235 |
| 2 | 167585600 | 167627999 | 190.5945 | 266.9337 | 764.5617 | 316.949 | -1.34711 | 4.553735 | 0.061246 |
| 12 | 63782400 | 63783599 | 17.33217 | 32.99429 | 12.13433 | 4.857922 | 1.926738 | 0.041011 | 0.061264 |
| 15 | 60677400 | 60678599 | 15.09787 | 28.47194 | 16.48607 | 6.732446 | 1.255813 | 0.093549 | 0.061266 |
| 9 | 118673000 | 118674999 | 6.084851 | 11.4042 | 36.73272 | 15.15856 | -1.2121 | 0.104378 | 0.061267 |
| 10 | 79292200 | 79293999 | 22.96144 | 40.54131 | 22.51815 | 9.962889 | 1.233092 | 0.566469 | 0.061271 |
| 13 | 92735600 | 92736799 | 3.764421 | 7.32888 | 29.05167 | 10.65829 | -1.34865 | -0.34257 | 0.061301 |
| 4 | 6010200 | 6011799 | 15.28917 | 29.29019 | 13.65449 | 5.384082 | 1.608878 | -0.0219 | 0.061317 |
| 2 | 80056000 | 80057399 | 16.95681 | 31.07068 | 18.8902 | 8.060494 | 1.138469 | 0.281771 | 0.061339 |
| 1 | 91839600 | 91861799 | 104.2998 | 155.1452 | 399.0666 | 164.419 | -1.10667 | 3.619867 | 0.061354 |
| 19 | 13602800 | 13605999 | 39.9474 | 66.93519 | 33.28182 | 14.51255 | 1.385074 | 1.195075 | 0.061359 |
| 4 | 146188000 | 146188399 | 6.632468 | 12.96052 | 7.162907 | 2.294752 | 1.626069 | -0.85804 | 0.061364 |
| 14 | 15441600 | 15443399 | 8.179572 | 14.37383 | 49.29638 | 21.8255 | -1.37991 | 0.583327 | 0.061383 |
| 7 | 71167600 | 71168399 | 10.56289 | 20.62761 | 9.550711 | 3.356435 | 1.764433 | -0.46988 | 0.061396 |
| 9 | 66954400 | 66958399 | 17.59406 | 28.84271 | 91.20375 | 39.25832 | -1.27907 | 1.433448 | 0.061406 |
| 9 | 109144000 | 109144599 | 2.428909 | 4.743779 | 23.26936 | 8.162429 | -1.58451 | -0.53607 | 0.061409 |
| 18 | 72482000 | 72485199 | 16.48099 | 27.40108 | 81.30579 | 35.27733 | -1.18268 | 1.251665 | 0.061415 |
| 9 | 107393600 | 107409199 | 73.81199 | 112.1308 | 302.9577 | 125.4698 | -1.15055 | 3.200931 | 0.061428 |
| 1 | 154331600 | 154333799 | 24.58284 | 43.13656 | 24.10855 | 10.66079 | 1.222145 | 0.652844 | 0.061431 |
| 13 | 102667800 | 102668399 | 10.67639 | 20.8023 | 11.96497 | 4.348023 | 1.415671 | -0.3465 | 0.061432 |
| 17 | 21349600 | 21350399 | 10.90359 | 21.15763 | 13.22978 | 4.979713 | 1.251832 | -0.22002 | 0.061435 |
| 16 | 9742800 | 9744199 | 16.62398 | 31.30105 | 14.30523 | 5.857304 | 1.589587 | 0.110456 | 0.061466 |
| 7 | 31518800 | 31519599 | 22.33042 | 40.01783 | 18.02072 | 7.894595 | 1.543588 | 0.444739 | 0.061481 |
| 12 | 19072800 | 19075199 | 48.9373 | 79.38828 | 52.74738 | 22.59967 | 0.954385 | 1.632988 | 0.061494 |
| 11 | 98206000 | 98231599 | 179.3437 | 256.4918 | 600.1508 | 247.9413 | -1.02504 | 4.243541 | 0.061497 |

| | | | | | | | | | |
|---|---|---|---|---|---|---|---|---|---|
| 2 | 65849200 | 65850199 | 4.835441 | 9.236482 | 32.19516 | 12.79631 | -1.27602 | -0.04611 | 0.061507 |
| 7 | 20654600 | 20656599 | 25.26875 | 44.59967 | 20.10093 | 8.894034 | 1.532139 | 0.57999 | 0.061515 |
| 8 | 70779200 | 70791999 | 64.43036 | 98.75748 | 260.7809 | 108.2467 | -1.10709 | 3.013519 | 0.061518 |
| 8 | 70860400 | 70864399 | 14.63008 | 24.29682 | 80.82107 | 35.04017 | -1.34672 | 1.260434 | 0.061519 |
| 14 | 124576400 | 124577599 | 9.455145 | 18.49681 | 5.758729 | 1.909874 | 2.386694 | -0.69427 | 0.061519 |
| 6 | 10196400 | 10199199 | 49.39633 | 80.18478 | 51.22035 | 21.95588 | 1.011766 | 1.617916 | 0.061534 |
| 14 | 41711000 | 41713799 | 27.02092 | 46.96637 | 27.39091 | 12.0292 | 1.160903 | 0.808143 | 0.061542 |
| 8 | 125474400 | 125479399 | 21.44025 | 34.95627 | 97.8926 | 42.04993 | -1.10976 | 1.521571 | 0.061558 |
| 1 | 117397400 | 117402399 | 85.31573 | 134.9804 | 94.91963 | 39.45141 | 0.834764 | 2.400209 | 0.06157 |
| 2 | 128809000 | 128810199 | 4.68971 | 9.010428 | 31.35997 | 12.26283 | -1.25188 | -0.10511 | 0.061571 |
| 2 | 157236600 | 157240799 | 26.49842 | 42.78407 | 107.809 | 45.98781 | -0.97086 | 1.732986 | 0.061576 |
| 2 | 169636200 | 169639799 | 16.61121 | 27.64296 | 77.26041 | 33.54338 | -1.09644 | 1.239381 | 0.06159 |
| 9 | 108717000 | 108726799 | 45.9018 | 71.53292 | 215.1717 | 89.303 | -1.27606 | 2.694559 | 0.061596 |
| 7 | 72891000 | 72892599 | 8.232977 | 14.69973 | 44.70537 | 19.6458 | -1.19663 | 0.428806 | 0.061597 |
| 17 | 29047000 | 29087599 | 318.4153 | 434.3587 | 957.4838 | 398.5708 | -1.00792 | 4.961306 | 0.061615 |
| 7 | 80741800 | 80745999 | 19.94088 | 32.2356 | 106.3316 | 45.42899 | -1.35579 | 1.687321 | 0.061622 |
| 6 | 77630000 | 77630999 | 4.249393 | 8.227229 | 31.39163 | 11.93995 | -1.34499 | -0.20485 | 0.061629 |
| 2 | 173466600 | 173468799 | 7.413439 | 13.43778 | 42.90786 | 18.56238 | -1.25091 | 0.326302 | 0.061637 |
| 2 | 35096800 | 35099199 | 5.424829 | 10.07649 | 40.30382 | 16.88188 | -1.53923 | 0.163397 | 0.061647 |
| 14 | 42192000 | 42193999 | 14.24541 | 27.26476 | 14.85022 | 5.871515 | 1.383195 | -0.02088 | 0.061656 |
| 15 | 6928600 | 6929999 | 21.0862 | 37.25065 | 22.8235 | 10.09688 | 1.092683 | 0.549103 | 0.061664 |
| 10 | 80313600 | 80342199 | 186.0972 | 264.7234 | 639.7836 | 264.5507 | -1.07952 | 4.322819 | 0.061677 |
| 1 | 118059200 | 118060599 | 36.06747 | 60.34124 | 38.44089 | 16.74592 | 1.029243 | 1.203918 | 0.061692 |
| 4 | 5781200 | 5781999 | 3.460001 | 6.744002 | 26.71947 | 9.664555 | -1.32757 | -0.40321 | 0.061699 |
| 1 | 52158000 | 52159999 | 7.76609 | 13.92677 | 44.24891 | 19.3738 | -1.25547 | 0.396648 | 0.061704 |
| 18 | 23494200 | 23494799 | 8.722484 | 17.03913 | 10.69197 | 3.736542 | 1.341669 | -0.51158 | 0.061708 |
| 8 | 44994600 | 44995199 | 2.941171 | 5.736842 | 24.90414 | 8.923846 | -1.4437 | -0.4503 | 0.061713 |
| 15 | 76334200 | 76359799 | 86.18122 | 127.886 | 434.7197 | 179.0645 | -1.51175 | 3.661125 | 0.061772 |
| 1 | 184736400 | 184738599 | 7.000759 | 12.62259 | 42.74301 | 18.61419 | -1.34093 | 0.355157 | 0.061774 |
| 2 | 27682600 | 27685399 | 15.05647 | 25.44115 | 69.2444 | 30.2963 | -1.05775 | 1.072786 | 0.0618 |
| 1 | 88632800 | 88634599 | 7.576791 | 13.62251 | 42.15512 | 18.41684 | -1.21479 | 0.377416 | 0.061827 |
| 6 | 83039800 | 83044599 | 24.84405 | 40.26885 | 101.5132 | 43.46618 | -0.96557 | 1.632865 | 0.061835 |
| 15 | 70209800 | 70210599 | 4.811853 | 9.27151 | 33.43702 | 12.96839 | -1.29207 | -0.12617 | 0.061842 |
| 14 | 42121600 | 42123199 | 20.53158 | 37.33609 | 17.88288 | 7.705221 | 1.470802 | 0.349584 | 0.061844 |
| 15 | 96524600 | 96528399 | 26.00512 | 42.01219 | 106.0484 | 45.27098 | -0.97234 | 1.714588 | 0.061849 |
| 18 | 32085600 | 32110399 | 147.6814 | 214.4313 | 511.3523 | 210.806 | -1.03072 | 4.016871 | 0.061855 |
| 5 | 106221200 | 106221999 | 3.764548 | 7.328691 | 29.70472 | 10.90311 | -1.38121 | -0.33926 | 0.061879 |
| 8 | 13957200 | 13958199 | 4.207892 | 8.119683 | 30.31803 | 11.70934 | -1.33376 | -0.15175 | 0.061899 |
| 10 | 79950800 | 79993599 | 178.0208 | 250.2988 | 746.8805 | 309.4313 | -1.40048 | 4.494849 | 0.061908 |
| 10 | 81560600 | 81562799 | 36.83349 | 62.69459 | 27.05234 | 11.8574 | 1.588746 | 1.040596 | 0.061939 |
| 12 | 22448600 | 22451799 | 52.49652 | 85.0905 | 52.36519 | 22.42199 | 1.063501 | 1.65533 | 0.061949 |
| 4 | 152165800 | 152179799 | 66.788 | 102.2317 | 265.3422 | 110.1084 | -1.08382 | 3.042104 | 0.061976 |
| 7 | 9101600 | 9105199 | 43.05678 | 72.12798 | 29.93061 | 13.04929 | 1.644759 | 1.205613 | 0.061985 |
| 5 | 141023800 | 141030599 | 39.1339 | 62.30338 | 147.3943 | 61.62516 | -0.90412 | 2.19964 | 0.061988 |
| 3 | 93350000 | 93355399 | 44.21007 | 69.75479 | 176.8324 | 73.39504 | -1.01562 | 2.45409 | 0.061997 |
| 3 | 100876200 | 100877599 | 5.777024 | 10.8633 | 35.10839 | 14.40696 | -1.20978 | 0.077193 | 0.062013 |
| 7 | 8668400 | 8670599 | 20.67078 | 37.85352 | 16.19938 | 6.917891 | 1.63981 | 0.308461 | 0.062015 |

| | | | | | | | | | |
|---|---|---|---|---|---|---|---|---|---|
| 9 | 119251000 | 119260599 | 41.21428 | 64.8919 | 185.9724 | 77.16193 | -1.19462 | 2.496708 | 0.062021 |
| 12 | 22587800 | 22591999 | 49.71751 | 80.78198 | 46.83129 | 20.08839 | 1.152506 | 1.597846 | 0.06206 |
| 14 | 121531400 | 121533399 | 5.918639 | 11.03934 | 37.61368 | 15.64935 | -1.3012 | 0.139521 | 0.06212 |
| 7 | 138641200 | 138646199 | 27.79771 | 44.83358 | 111.4356 | 47.4688 | -0.95304 | 1.767688 | 0.062128 |
| 1 | 109061200 | 109063199 | 43.493 | 71.12724 | 47.85018 | 20.57694 | 0.945029 | 1.492067 | 0.062138 |
| 18 | 50981200 | 50981999 | 10.57291 | 20.60111 | 12.0923 | 4.393047 | 1.387783 | -0.34797 | 0.062156 |
| 9 | 43192000 | 43192599 | 1.871792 | 3.657276 | 25.26923 | 8.79578 | -2.05945 | -0.56966 | 0.06217 |
| 7 | 9070800 | 9073399 | 24.21655 | 43.46485 | 16.39358 | 7.172446 | 1.798602 | 0.446423 | 0.062184 |
| 5 | 115747200 | 115760199 | 55.67576 | 85.27773 | 272.6694 | 113.166 | -1.38355 | 3.026149 | 0.062185 |
| 17 | 20753400 | 20754199 | 22.34112 | 39.23816 | 22.57815 | 9.991626 | 1.180268 | 0.622599 | 0.062186 |
| 1 | 177274800 | 177276199 | 6.135883 | 11.34255 | 37.37053 | 15.75604 | -1.26839 | 0.192951 | 0.06219 |
| 4 | 57372800 | 57374999 | 10.91741 | 19.12372 | 53.67802 | 23.70128 | -1.09374 | 0.656523 | 0.062194 |
| 9 | 120912800 | 120932199 | 139.7972 | 204.8369 | 460.7017 | 189.7684 | -0.93364 | 3.875989 | 0.062199 |
| 8 | 86243400 | 86244799 | 3.557555 | 6.858732 | 30.18966 | 11.68926 | -1.5711 | -0.15197 | 0.062203 |
| 7 | 144621000 | 144622999 | 7.926675 | 14.16407 | 43.65465 | 19.17124 | -1.21477 | 0.42048 | 0.062204 |
| 13 | 61550400 | 61553799 | 67.68988 | 108.3775 | 70.90845 | 29.88335 | 0.955689 | 2.029953 | 0.062233 |
| 10 | 22586200 | 22586999 | 2.93963 | 5.738403 | 24.54045 | 8.682017 | -1.40572 | -0.49805 | 0.062249 |
| 17 | 19655400 | 19656999 | 35.55 | 59.78078 | 36.16834 | 15.80153 | 1.103208 | 1.148791 | 0.06226 |
| 17 | 5141600 | 5143399 | 62.90569 | 101.146 | 47.00137 | 19.93488 | 1.455125 | 1.884405 | 0.062267 |
| 15 | 76380200 | 76384999 | 17.53819 | 28.74144 | 88.58917 | 38.12746 | -1.24232 | 1.438085 | 0.062267 |
| 8 | 112435200 | 112448999 | 60.05136 | 92.34913 | 250.633 | 104.0814 | -1.14289 | 2.94612 | 0.062272 |
| 12 | 23984400 | 23986999 | 56.63921 | 91.19105 | 62.72067 | 26.65318 | 0.893227 | 1.841333 | 0.062277 |
| 1 | 137652200 | 137653799 | 20.60069 | 37.12382 | 19.61476 | 8.547771 | 1.320397 | 0.404415 | 0.06228 |
| 3 | 20892600 | 20893999 | 17.10711 | 31.79781 | 17.11909 | 7.163923 | 1.331928 | 0.201141 | 0.062284 |
| 13 | 66235000 | 66236399 | 18.79066 | 35.18415 | 14.37331 | 5.940945 | 1.740405 | 0.164917 | 0.062291 |
| 9 | 67471200 | 67474199 | 11.89706 | 20.66599 | 59.17853 | 25.97912 | -1.12391 | 0.792503 | 0.062304 |
| 12 | 112064400 | 112104799 | 212.518 | 296.646 | 777.9656 | 322.6731 | -1.22484 | 4.604491 | 0.062312 |
| 4 | 40970000 | 40972199 | 7.507725 | 13.34975 | 44.63988 | 19.68523 | -1.33554 | 0.451576 | 0.062372 |
| 4 | 118036600 | 118037599 | 4.159142 | 7.966445 | 32.28638 | 12.73732 | -1.48092 | -0.0792 | 0.062393 |
| X | 53910200 | 53911199 | 12.29086 | 23.7987 | 13.18268 | 5.011705 | 1.41202 | -0.17071 | 0.062415 |
| 10 | 85221400 | 85227199 | 30.58892 | 49.14741 | 121.9395 | 51.66311 | -0.9574 | 1.894463 | 0.062439 |
| 13 | 65630400 | 65635399 | 76.45128 | 122.0041 | 72.61754 | 30.46134 | 1.086879 | 2.140368 | 0.062449 |
| 10 | 55625200 | 55628199 | 34.23426 | 57.98453 | 33.31774 | 14.58798 | 1.177389 | 1.079448 | 0.062498 |
| 19 | 38384000 | 38385999 | 7.019747 | 12.60666 | 43.19594 | 18.88963 | -1.36183 | 0.381025 | 0.062501 |
| 2 | 173127600 | 173138799 | 59.74473 | 92.58812 | 224.4086 | 93.21301 | -0.97134 | 2.797953 | 0.062517 |
| 4 | 30640200 | 30641399 | 18.37579 | 33.67705 | 17.98657 | 7.672958 | 1.323888 | 0.289594 | 0.062521 |
| 7 | 56282600 | 56283599 | 11.08337 | 21.50144 | 13.70263 | 5.163358 | 1.223242 | -0.2126 | 0.062536 |
| 6 | 83045400 | 83052599 | 40.08316 | 63.14041 | 199.0352 | 82.58684 | -1.33134 | 2.48819 | 0.06255 |
| 7 | 98952200 | 98954799 | 9.304977 | 16.18336 | 55.719 | 24.47941 | -1.38709 | 0.760361 | 0.062552 |
| 14 | 42149600 | 42150199 | 9.235398 | 18.06335 | 8.580189 | 2.894616 | 1.778926 | -0.62957 | 0.062563 |
| 4 | 149958000 | 149986199 | 138.1294 | 200.6736 | 514.6371 | 212.1479 | -1.13477 | 4.007747 | 0.062581 |
| 7 | 127156400 | 127160199 | 13.15862 | 22.33202 | 66.35684 | 29.07154 | -1.18313 | 1.025579 | 0.0626 |
| 6 | 5823600 | 5825599 | 9.330881 | 16.50748 | 46.69566 | 20.65177 | -1.10035 | 0.509056 | 0.062603 |
| 18 | 77983600 | 77983999 | 1.714381 | 3.353139 | 20.61228 | 6.951792 | -1.84359 | -0.6959 | 0.062611 |
| 2 | 167509200 | 167510799 | 5.172664 | 9.356272 | 41.17754 | 17.86105 | -1.70967 | 0.323665 | 0.062614 |
| 12 | 19228800 | 19233999 | 159.4272 | 241.7792 | 169.3343 | 70.09389 | 0.793493 | 3.247461 | 0.062651 |
| 3 | 97231400 | 97233799 | 10.61801 | 18.5858 | 52.04734 | 22.9672 | -1.09029 | 0.664386 | 0.062652 |

| | | | | | | | | | |
|---|---|---|---|---|---|---|---|---|---|
| 5 | 106245200 | 106246599 | 5.665283 | 10.58175 | 35.99104 | 14.9384 | -1.29616 | 0.122067 | 0.062654 |
| 10 | 60694400 | 60701999 | 30.76885 | 49.07502 | 146.6357 | 61.46263 | -1.23731 | 2.132355 | 0.062656 |
| 12 | 27897400 | 27900599 | 11.72221 | 20.41892 | 56.98762 | 25.06105 | -1.08643 | 0.749364 | 0.062694 |
| 2 | 25855600 | 25857199 | 4.954902 | 9.223847 | 36.38265 | 15.18708 | -1.51368 | 0.139984 | 0.0627 |
| 4 | 128126600 | 128128199 | 9.702099 | 17.00278 | 50.94356 | 22.50271 | -1.18694 | 0.638542 | 0.062715 |
| 7 | 116934800 | 116936399 | 20.90001 | 36.94044 | 21.48291 | 9.502858 | 1.167754 | 0.539935 | 0.062732 |
| 7 | 13817400 | 13818599 | 32.67876 | 55.0845 | 35.80988 | 15.65631 | 1.000024 | 1.117799 | 0.062746 |
| 9 | 3136400 | 3138999 | 73.27486 | 116.6539 | 82.65515 | 34.55634 | 0.832209 | 2.213752 | 0.062749 |
| 6 | 76054600 | 76056199 | 21.69779 | 38.75546 | 19.74609 | 8.674657 | 1.364406 | 0.463828 | 0.062775 |
| 7 | 135009400 | 135016799 | 35.76628 | 56.9609 | 155.3189 | 64.96833 | -1.10812 | 2.186557 | 0.062795 |
| 13 | 112600200 | 112601199 | 3.54526 | 6.87017 | 28.83851 | 10.92261 | -1.47296 | -0.23252 | 0.062802 |
| 6 | 23890800 | 23891799 | 11.95703 | 23.07073 | 14.633 | 5.652871 | 1.196391 | -0.11868 | 0.062809 |
| 12 | 119680600 | 119680999 | 7.737366 | 15.12402 | 7.598834 | 2.44654 | 1.756899 | -0.8147 | 0.062812 |
| 11 | 94022000 | 94030399 | 27.18931 | 43.48816 | 137.4106 | 57.82558 | -1.31307 | 2.0383 | 0.062827 |
| 3 | 106726600 | 106727399 | 11.88858 | 23.15916 | 10.50691 | 3.828988 | 1.748345 | -0.3261 | 0.062832 |
| 12 | 112554000 | 112584799 | 152.5288 | 217.0778 | 659.7125 | 272.769 | -1.40915 | 4.314346 | 0.062834 |
| 8 | 21538600 | 21539199 | 7.775875 | 15.2085 | 8.096466 | 2.734435 | 1.617257 | -0.65479 | 0.062847 |
| 2 | 24839200 | 24841399 | 8.276037 | 14.65754 | 45.64512 | 20.18269 | -1.23707 | 0.488397 | 0.062877 |
| 7 | 126987800 | 126993399 | 28.21905 | 45.29912 | 124.8849 | 52.83992 | -1.11061 | 1.920975 | 0.062881 |
| 6 | 136626800 | 136633799 | 40.92637 | 65.09995 | 156.0943 | 65.18344 | -0.92493 | 2.23072 | 0.062885 |
| 9 | 118691200 | 118698599 | 36.69917 | 58.48371 | 144.8834 | 60.65653 | -0.96892 | 2.165282 | 0.062911 |
| 7 | 10740800 | 10749399 | 119.6581 | 186.528 | 105.2709 | 43.68883 | 1.135247 | 2.711173 | 0.062958 |
| 7 | 59410400 | 59414799 | 47.94392 | 77.94161 | 47.3349 | 20.31131 | 1.086398 | 1.587255 | 0.062973 |
| 11 | 76886200 | 76889999 | 16.96214 | 27.80401 | 90.40585 | 38.91323 | -1.31911 | 1.433356 | 0.062976 |
| X | 26640200 | 26641199 | 13.51455 | 26.34433 | 8.524014 | 3.079064 | 2.241427 | -0.34102 | 0.062981 |
| 10 | 40558400 | 40561199 | 11.11005 | 19.3629 | 55.86398 | 24.57514 | -1.13383 | 0.737196 | 0.062991 |
| 15 | 79166800 | 79179399 | 48.39385 | 75.32876 | 220.2354 | 91.42047 | -1.23645 | 2.716087 | 0.06302 |
| 5 | 91549400 | 91550999 | 7.451831 | 13.40077 | 42.69045 | 18.64726 | -1.25653 | 0.374224 | 0.063028 |
| X | 34459200 | 34460799 | 18.72375 | 34.89946 | 15.19002 | 6.327159 | 1.641864 | 0.194353 | 0.063039 |
| 7 | 75879200 | 75879799 | 4.373303 | 8.467803 | 30.14256 | 11.4606 | -1.24497 | -0.20682 | 0.063059 |
| 8 | 120716600 | 120731799 | 63.70435 | 97.62255 | 269.6221 | 111.9131 | -1.17206 | 3.017898 | 0.063062 |
| 7 | 5381000 | 5381999 | 29.78788 | 51.38448 | 29.21703 | 12.81706 | 1.195906 | 0.899101 | 0.063084 |
| 15 | 95735600 | 95737199 | 4.81093 | 9.272897 | 32.21033 | 12.48039 | -1.23688 | -0.13212 | 0.063085 |
| 12 | 22598800 | 22600199 | 39.63567 | 65.67486 | 41.28471 | 17.87095 | 1.048326 | 1.322134 | 0.063107 |
| 14 | 5727400 | 5729599 | 7.422104 | 13.42937 | 42.30724 | 18.34366 | -1.23362 | 0.33614 | 0.063115 |
| 1 | 134259000 | 134268199 | 39.61762 | 62.72175 | 172.014 | 71.51811 | -1.12489 | 2.368173 | 0.063126 |
| 11 | 102815400 | 102816399 | 2.770551 | 5.341487 | 33.66769 | 13.03562 | -2.08142 | -0.15286 | 0.063162 |
| 8 | 56264200 | 56265399 | 16.79626 | 31.69582 | 13.71806 | 5.594439 | 1.671748 | 0.0968 | 0.063166 |
| 5 | 142884200 | 142885399 | 4.083654 | 7.882303 | 29.73571 | 11.47219 | -1.34688 | -0.15884 | 0.063184 |
| 10 | 110200000 | 110200799 | 5.009288 | 9.543733 | 32.70656 | 13.0712 | -1.25945 | -0.02617 | 0.063193 |
| 8 | 85676400 | 85677999 | 7.174888 | 12.83644 | 43.14336 | 18.92713 | -1.33734 | 0.40729 | 0.063254 |
| 10 | 39381000 | 39388599 | 41.92558 | 66.6044 | 160.4554 | 66.9132 | -0.93358 | 2.270662 | 0.063263 |
| 8 | 22116600 | 22118599 | 24.3632 | 42.69997 | 24.95556 | 11.02417 | 1.158356 | 0.672684 | 0.063266 |
| 12 | 22669200 | 22670199 | 12.51975 | 24.09662 | 14.69502 | 5.717423 | 1.243542 | -0.08943 | 0.063272 |
| 7 | 105896600 | 105898799 | 22.26405 | 39.73414 | 19.73686 | 8.676625 | 1.400255 | 0.473032 | 0.06328 |
| 5 | 146900200 | 146901199 | 2.943878 | 5.734077 | 25.77996 | 9.408838 | -1.51768 | -0.38011 | 0.063319 |
| 2 | 168228400 | 168239199 | 74.91705 | 114.4334 | 269.7011 | 111.8627 | -0.94726 | 3.087521 | 0.063323 |

| | | | | | | | | | |
|---|---|---|---|---|---|---|---|---|---|
| 5 | 35339600 | 35341599 | 9.657969 | 16.99502 | 48.78568 | 21.59918 | -1.12452 | 0.575207 | 0.063331 |
| 13 | 105853200 | 105854399 | 14.55771 | 28.18672 | 10.71895 | 4.075915 | 1.94478 | -0.15226 | 0.063334 |
| 8 | 121767000 | 121794599 | 190.7909 | 271.9899 | 620.6301 | 256.5371 | -0.99348 | 4.291157 | 0.06335 |
| 13 | 65113400 | 65115599 | 30.92315 | 52.90867 | 31.73653 | 13.91636 | 1.117069 | 0.976966 | 0.063352 |
| 13 | 102910000 | 102911799 | 4.994334 | 9.561771 | 36.69591 | 14.49608 | -1.40695 | -0.05731 | 0.063367 |
| 12 | 18386200 | 18391199 | 155.2014 | 236.1154 | 160.6153 | 66.54768 | 0.839221 | 3.182333 | 0.063395 |
| 10 | 62013400 | 62033199 | 104.4704 | 156.1306 | 375.8614 | 154.9698 | -1.00497 | 3.533342 | 0.063406 |
| 7 | 103750000 | 103751999 | 20.77521 | 37.60128 | 18.27986 | 7.922889 | 1.444026 | 0.377903 | 0.063411 |
| 4 | 130501200 | 130504199 | 10.79723 | 18.83691 | 54.21676 | 23.8658 | -1.13006 | 0.719261 | 0.063413 |
| 14 | 121507600 | 121518799 | 50.98014 | 79.27628 | 215.9626 | 89.66003 | -1.13575 | 2.734387 | 0.06342 |
| 1 | 190020800 | 190022199 | 7.439655 | 13.35561 | 42.135 | 18.43167 | -1.24371 | 0.385962 | 0.063422 |
| 8 | 127268000 | 127270799 | 10.78356 | 18.848 | 54.22918 | 23.90045 | -1.12934 | 0.691567 | 0.063426 |
| 15 | 3145000 | 3146799 | 24.58977 | 43.11942 | 24.67847 | 10.90654 | 1.188159 | 0.664494 | 0.063451 |
| 14 | 108402200 | 108403999 | 18.52075 | 34.52458 | 15.37637 | 6.403685 | 1.608408 | 0.191839 | 0.063458 |
| X | 26347400 | 26348199 | 11.96744 | 23.34634 | 9.652905 | 3.45214 | 1.905447 | -0.39083 | 0.063496 |
| 14 | 105043200 | 105043599 | 1.643942 | 3.208827 | 17.74607 | 5.589632 | -1.60395 | -0.95611 | 0.063516 |
| 4 | 129891800 | 129902199 | 59.85333 | 92.06803 | 247.974 | 102.9808 | -1.13162 | 2.940997 | 0.063519 |
| 3 | 95724400 | 95727599 | 12.21839 | 21.17657 | 57.477 | 25.2179 | -1.04741 | 0.817984 | 0.06355 |
| 12 | 19613800 | 19615999 | 54.85214 | 88.50879 | 59.47277 | 25.34887 | 0.93048 | 1.772206 | 0.063554 |
| 12 | 20152200 | 20152799 | 12.44684 | 23.91172 | 15.14126 | 5.92263 | 1.182372 | -0.07254 | 0.063559 |
| 5 | 125142600 | 125238199 | 549.1548 | 708.2432 | 1820.169 | 763.0941 | -1.31044 | 5.863828 | 0.063578 |
| 6 | 41891800 | 41892399 | 8.497843 | 16.61597 | 9.201352 | 3.143166 | 1.544782 | -0.59919 | 0.063587 |
| 5 | 35755200 | 35756599 | 14.42507 | 24.36871 | 67.98686 | 29.74404 | -1.09308 | 1.073084 | 0.063605 |
| 15 | 75752800 | 75766599 | 71.09284 | 108.4043 | 296.7143 | 123.0176 | -1.16484 | 3.123334 | 0.063618 |
| 6 | 90710400 | 90714799 | 20.61589 | 33.67347 | 92.51174 | 39.76495 | -1.08019 | 1.491171 | 0.063624 |
| 14 | 28645200 | 28645799 | 2.737132 | 5.349109 | 22.96272 | 7.952377 | -1.38071 | -0.58035 | 0.063624 |
| 6 | 48371800 | 48374599 | 11.63245 | 20.09446 | 58.46706 | 25.6497 | -1.14815 | 0.847032 | 0.063673 |
| 9 | 43170200 | 43175799 | 26.15599 | 42.29094 | 104.711 | 44.74788 | -0.94319 | 1.689422 | 0.063673 |
| 17 | 22837200 | 22843799 | 234.1526 | 345.5391 | 237.6391 | 97.86372 | 0.784554 | 3.765987 | 0.063705 |
| 12 | 18424600 | 18429999 | 183.469 | 276.8243 | 173.6969 | 71.78456 | 0.945744 | 3.355603 | 0.063723 |
| 1 | 133189000 | 133204199 | 81.68388 | 124.3202 | 283.6205 | 117.5267 | -0.90457 | 3.163723 | 0.063736 |
| 4 | 135898400 | 135905599 | 23.75318 | 38.17845 | 123.5477 | 52.37245 | -1.33879 | 1.874997 | 0.063742 |
| 15 | 66756600 | 66760799 | 18.0705 | 29.94298 | 80.69808 | 34.93251 | -1.04485 | 1.295682 | 0.063745 |
| 7 | 45954200 | 45961599 | 39.46671 | 62.80343 | 148.2102 | 61.92562 | -0.9013 | 2.215809 | 0.063747 |
| 8 | 99859200 | 99860399 | 14.0825 | 27.27246 | 11.38494 | 4.32483 | 1.81393 | -0.15889 | 0.063755 |
| 2 | 86156400 | 86157799 | 18.09747 | 33.19226 | 17.40981 | 7.41922 | 1.350517 | 0.28139 | 0.063764 |
| 1 | 75212800 | 75218199 | 23.79043 | 38.45495 | 107.0484 | 45.72958 | -1.11219 | 1.694764 | 0.063766 |
| 5 | 70038200 | 70039799 | 17.37947 | 32.76835 | 13.70365 | 5.598001 | 1.718999 | 0.107922 | 0.063784 |
| 5 | 102992000 | 102994199 | 6.214664 | 11.34374 | 40.87525 | 17.52938 | -1.41482 | 0.272823 | 0.063806 |
| 5 | 114127200 | 114129399 | 6.214664 | 11.34374 | 40.87525 | 17.52938 | -1.41482 | 0.272823 | 0.063806 |
| 12 | 20102000 | 20106399 | 103.6384 | 161.9882 | 107.2651 | 44.50323 | 0.907931 | 2.656587 | 0.063806 |
| 2 | 152764000 | 152776399 | 47.36427 | 73.83877 | 215.2192 | 89.31786 | -1.23029 | 2.688622 | 0.063807 |
| 15 | 82061200 | 82073999 | 93.76087 | 141.4846 | 322.8396 | 133.4259 | -0.91527 | 3.342894 | 0.063816 |
| 14 | 42068200 | 42069799 | 26.59459 | 46.89499 | 19.64065 | 8.692032 | 1.636972 | 0.598162 | 0.063818 |
| 3 | 147539000 | 147539599 | 2.841003 | 5.547822 | 24.601 | 8.650917 | -1.44727 | -0.5176 | 0.063836 |
| 1 | 86375200 | 86376399 | 3.585363 | 6.954516 | 28.8843 | 10.89129 | -1.45024 | -0.24388 | 0.063838 |
| 5 | 38540800 | 38542199 | 5.909151 | 11.13353 | 35.19041 | 14.39284 | -1.17414 | 0.066134 | 0.063845 |

| | | | | | | | | | |
|---|---|---|---|---|---|---|---|---|---|
| 13 | 58991200 | 58992999 | 4.916866 | 9.261053 | 34.5214 | 14.12669 | -1.40962 | 0.057628 | 0.063846 |
| 3 | 12206000 | 12208799 | 37.37884 | 62.13578 | 40.10192 | 17.39772 | 1.010588 | 1.275362 | 0.063862 |
| 2 | 23606800 | 23607799 | 16.8314 | 30.91491 | 18.91697 | 8.045633 | 1.132638 | 0.267986 | 0.063864 |
| 4 | 124426800 | 124428999 | 8.209374 | 14.69007 | 43.14718 | 18.92571 | -1.14452 | 0.413056 | 0.063865 |
| 7 | 20576200 | 20578599 | 36.59664 | 62.57314 | 25.39941 | 11.13694 | 1.677205 | 0.999145 | 0.063876 |
| 1 | 156869400 | 156870199 | 3.846892 | 7.42639 | 30.33757 | 11.69639 | -1.45992 | -0.16403 | 0.06388 |
| 2 | 172482600 | 172491599 | 53.18554 | 83.70533 | 187.7054 | 77.87668 | -0.8419 | 2.511933 | 0.063896 |
| 13 | 65969800 | 65970199 | 8.502488 | 16.61026 | 10.55642 | 3.685791 | 1.323404 | -0.51971 | 0.063904 |
| 14 | 4359600 | 4360399 | 21.77354 | 38.36455 | 23.82904 | 10.54723 | 1.071835 | 0.583348 | 0.063919 |
| 8 | 115438400 | 115439199 | 6.409439 | 11.86815 | 37.42112 | 15.74137 | -1.20312 | 0.18641 | 0.063923 |
| 7 | 29135400 | 29142799 | 42.72135 | 67.48918 | 172.7907 | 71.76236 | -1.02841 | 2.42294 | 0.063925 |
| 11 | 79494400 | 79496999 | 9.712264 | 16.8852 | 55.7082 | 24.46873 | -1.32582 | 0.76997 | 0.063936 |
| 7 | 41512600 | 41522599 | 33.81909 | 57.28734 | 31.32043 | 13.71391 | 1.248747 | 1.074435 | 0.063941 |
| 1 | 185405800 | 185409599 | 20.98721 | 34.51395 | 86.90057 | 37.45599 | -0.94977 | 1.399592 | 0.063948 |
| 10 | 39335200 | 39339599 | 16.41567 | 27.40884 | 74.54771 | 32.43421 | -1.05722 | 1.200301 | 0.063952 |
| Y | 4049400 | 4051199 | 11.21293 | 21.83663 | 5.211387 | 1.906407 | 2.647893 | -0.34521 | 0.063953 |
| 15 | 86049000 | 86058199 | 39.93066 | 63.21167 | 168.4927 | 70.05087 | -1.08397 | 2.370373 | 0.063957 |
| 8 | 34793200 | 34795999 | 12.54934 | 21.30683 | 66.87251 | 29.30115 | -1.26166 | 1.019916 | 0.063969 |
| 14 | 61135800 | 61137199 | 5.3108 | 10.09398 | 38.05663 | 15.28658 | -1.40381 | 0.003944 | 0.063978 |
| 17 | 13649800 | 13655199 | 104.9004 | 157.5416 | 340.1324 | 140.3884 | -0.84162 | 3.439725 | 0.064005 |
| 1 | 135726800 | 135728999 | 10.05895 | 17.49958 | 55.22966 | 24.2687 | -1.26248 | 0.758739 | 0.064013 |
| 4 | 130956400 | 130958199 | 11.24943 | 19.63539 | 57.54352 | 25.33798 | -1.15619 | 0.718116 | 0.064024 |
| 13 | 66125400 | 66131999 | 108.3812 | 169.1307 | 103.7267 | 43.04225 | 1.016637 | 2.685616 | 0.064029 |
| 6 | 87857800 | 87860199 | 6.707254 | 12.07743 | 42.17608 | 18.39766 | -1.3862 | 0.360699 | 0.064036 |
| 6 | 130990200 | 130993799 | 93.05019 | 147.2367 | 88.48348 | 36.7788 | 1.061242 | 2.402794 | 0.064037 |
| 12 | 55482000 | 55487199 | 19.27002 | 31.6645 | 86.51269 | 37.27611 | -1.06762 | 1.404909 | 0.064039 |
| 14 | 42184200 | 42185199 | 9.689961 | 18.88495 | 12.1075 | 4.385641 | 1.265 | -0.37083 | 0.06408 |
| 7 | 70458800 | 70461399 | 11.24751 | 19.63702 | 55.88731 | 24.61263 | -1.11403 | 0.712504 | 0.064095 |
| 2 | 21952600 | 21955399 | 16.15654 | 27.26574 | 71.02219 | 31.06232 | -0.99489 | 1.08914 | 0.064124 |
| X | 169326200 | 169326599 | 14.87707 | 28.21315 | 13.95365 | 5.631387 | 1.4939 | 0.045854 | 0.064126 |
| 14 | 35139400 | 35139799 | 9.993496 | 19.51147 | 9.18663 | 3.240536 | 1.739361 | -0.46731 | 0.064132 |
| 4 | 73581200 | 73581799 | 10.42922 | 20.30034 | 12.52347 | 4.60203 | 1.3014 | -0.31028 | 0.064132 |
| 4 | 140358200 | 140381199 | 105.5322 | 156.5689 | 413.0567 | 170.1378 | -1.14667 | 3.666508 | 0.064144 |
| 12 | 23616600 | 23619799 | 57.06882 | 91.80669 | 63.81846 | 27.08717 | 0.876476 | 1.867281 | 0.064151 |
| 4 | 44755800 | 44757199 | 5.541548 | 10.32693 | 37.02875 | 15.4289 | -1.37638 | 0.137738 | 0.064165 |
| 17 | 46328800 | 46344999 | 71.15278 | 108.1311 | 300.5915 | 124.5036 | -1.19114 | 3.19265 | 0.064176 |
| 11 | 117306800 | 117317999 | 53.18602 | 83.2611 | 194.5157 | 80.68956 | -0.90687 | 2.61519 | 0.064179 |
| 12 | 20075800 | 20082199 | 178.3392 | 269.0971 | 178.2818 | 73.68051 | 0.867435 | 3.353713 | 0.064183 |
| 11 | 120994800 | 120996199 | 4.020317 | 7.783645 | 31.24534 | 11.88482 | -1.41547 | -0.20326 | 0.064198 |
| 15 | 52575600 | 52576599 | 6.249193 | 11.63368 | 36.46773 | 15.22164 | -1.1863 | 0.150399 | 0.064206 |
| 19 | 55196000 | 55196999 | 3.350168 | 6.520528 | 26.32724 | 9.680923 | -1.37591 | -0.34605 | 0.064222 |
| 17 | 19213400 | 19214199 | 15.76885 | 29.70674 | 16.36546 | 6.694834 | 1.325005 | 0.105724 | 0.064232 |
| 12 | 19618000 | 19627799 | 255.4422 | 374.3947 | 254.2796 | 104.7388 | 0.793337 | 3.877336 | 0.064248 |
| 17 | 46232400 | 46246999 | 61.92676 | 94.83266 | 271.3896 | 112.6294 | -1.22404 | 3.031764 | 0.064283 |
| 19 | 46429400 | 46432399 | 10.14098 | 17.79009 | 49.87261 | 22.04868 | -1.09148 | 0.623437 | 0.064286 |
| 19 | 27575600 | 27576399 | 4.195651 | 8.140505 | 32.28526 | 12.1582 | -1.38584 | -0.23558 | 0.064287 |
| 7 | 99051400 | 99055799 | 18.23299 | 29.83265 | 91.01936 | 39.14805 | -1.22897 | 1.460553 | 0.064295 |

| | | | | | | | | | |
|---|---|---|---|---|---|---|---|---|---|
| 15 | 13354400 | 13355199 | 3.764167 | 7.329257 | 27.76407 | 10.17614 | -1.28227 | -0.3492 | 0.064326 |
| 12 | 22475800 | 22476799 | 11.11805 | 21.68188 | 10.90752 | 3.918425 | 1.621448 | -0.38277 | 0.064342 |
| 12 | 24042600 | 24043199 | 11.25865 | 21.85998 | 13.62414 | 5.108831 | 1.261342 | -0.22692 | 0.064347 |
| 7 | 112475400 | 112476399 | 3.981789 | 7.629967 | 31.09284 | 12.25157 | -1.48586 | -0.08617 | 0.064347 |
| 2 | 174512200 | 174516399 | 20.88192 | 34.23713 | 92.66132 | 39.88794 | -1.05557 | 1.438956 | 0.064353 |
| 6 | 140229600 | 140244399 | 95.79257 | 144.8816 | 322.6639 | 133.4496 | -0.87747 | 3.296643 | 0.064356 |
| 4 | 140067600 | 140130599 | 465.6355 | 614.8779 | 1370.291 | 573.0707 | -1.07077 | 5.480002 | 0.064395 |
| 1 | 70260200 | 70261399 | 14.28953 | 27.6006 | 12.58547 | 4.840078 | 1.672755 | -0.11333 | 0.064432 |
| 7 | 11676200 | 11679799 | 97.75439 | 153.6835 | 104.2512 | 43.24651 | 0.879588 | 2.546127 | 0.06444 |
| 15 | 72652600 | 72653799 | 4.599432 | 8.728302 | 33.06855 | 13.32682 | -1.41287 | -0.00086 | 0.06445 |
| 8 | 25980200 | 25981999 | 10.61303 | 18.57736 | 51.50129 | 22.72652 | -1.07565 | 0.664476 | 0.064462 |
| 11 | 113179800 | 113196799 | 97.71994 | 146.4741 | 357.0093 | 147.2815 | -1.01892 | 3.476571 | 0.064488 |
| 9 | 110209400 | 110219199 | 39.55485 | 62.37018 | 181.1706 | 75.18389 | -1.21235 | 2.46568 | 0.064495 |
| 7 | 12382400 | 12385999 | 74.30632 | 118.59 | 75.61337 | 31.72138 | 0.987899 | 2.135267 | 0.064504 |
| 7 | 80724200 | 80731399 | 41.56403 | 65.73962 | 170.333 | 70.78453 | -1.04419 | 2.392819 | 0.064506 |
| 4 | 43044000 | 43046399 | 10.5267 | 18.27334 | 56.30555 | 24.7075 | -1.22858 | 0.794302 | 0.064508 |
| 7 | 59250200 | 59251399 | 25.09669 | 43.64391 | 27.29484 | 11.99083 | 1.060823 | 0.792055 | 0.064524 |
| 10 | 41352000 | 41367399 | 68.47974 | 103.903 | 313.2863 | 129.6998 | -1.31013 | 3.225587 | 0.064542 |
| 15 | 23985800 | 23986999 | 13.1067 | 25.16581 | 14.31316 | 5.607378 | 1.332805 | -0.06384 | 0.064552 |
| 19 | 42297200 | 42298999 | 7.062819 | 12.9073 | 39.29397 | 16.82611 | -1.17212 | 0.270551 | 0.064555 |
| X | 31860400 | 31861999 | 19.68134 | 35.74528 | 19.02847 | 8.211642 | 1.318251 | 0.352503 | 0.064558 |
| 8 | 40964000 | 40965399 | 8.489232 | 14.90593 | 49.83849 | 22.05043 | -1.34371 | 0.598889 | 0.064565 |
| 7 | 4876600 | 4881599 | 26.45842 | 42.82929 | 105.2677 | 45.03019 | -0.93102 | 1.663867 | 0.064568 |
| 5 | 144095400 | 144097199 | 7.35025 | 13.47893 | 40.51791 | 17.27033 | -1.14973 | 0.254063 | 0.064592 |
| 7 | 8040200 | 8041799 | 16.16363 | 30.94149 | 13.02143 | 5.14588 | 1.751912 | -0.00784 | 0.064593 |
| 17 | 38987000 | 38989999 | 44.06114 | 73.1516 | 36.16343 | 15.67539 | 1.393685 | 1.308067 | 0.064595 |
| 1 | 91727200 | 91730799 | 19.66553 | 31.84879 | 105.937 | 45.33267 | -1.36525 | 1.643083 | 0.06467 |
| 9 | 73020200 | 73022999 | 11.65396 | 20.07362 | 59.49489 | 26.09832 | -1.17555 | 0.875124 | 0.064683 |
| 1 | 19513800 | 19515599 | 17.73871 | 32.6875 | 18.2824 | 7.74142 | 1.265882 | 0.256578 | 0.064688 |
| 5 | 109138800 | 109139199 | 30.21629 | 51.60819 | 31.11807 | 13.64323 | 1.109418 | 0.990283 | 0.064694 |
| 2 | 130121000 | 130123399 | 7.72877 | 13.92357 | 41.66322 | 18.16294 | -1.16483 | 0.364221 | 0.064698 |
| 8 | 87471600 | 87472999 | 7.879189 | 14.21436 | 42.77469 | 18.61499 | -1.17167 | 0.357772 | 0.064702 |
| 19 | 40819200 | 40820799 | 5.551258 | 9.989997 | 43.20469 | 18.86025 | -1.69204 | 0.358586 | 0.064705 |
| 5 | 134237200 | 134248399 | 75.90764 | 115.9585 | 269.1691 | 111.6448 | -0.92522 | 3.085551 | 0.06472 |
| 4 | 5644000 | 5645199 | 6.092044 | 11.39545 | 36.07829 | 14.93771 | -1.19118 | 0.116089 | 0.064739 |
| 6 | 94200600 | 94202399 | 6.092044 | 11.39545 | 36.07829 | 14.93771 | -1.19118 | 0.116089 | 0.064739 |
| 7 | 125675200 | 125685399 | 52.87807 | 82.00585 | 224.1602 | 93.10095 | -1.14374 | 2.782797 | 0.06476 |
| 2 | 26907600 | 26918399 | 42.50915 | 66.29422 | 217.7155 | 90.34892 | -1.40168 | 2.680469 | 0.064772 |
| 5 | 100276800 | 100278599 | 9.493638 | 16.77477 | 47.75417 | 21.1251 | -1.11038 | 0.524646 | 0.064785 |
| 15 | 21909400 | 21910399 | 15.68767 | 29.15854 | 17.55668 | 7.347345 | 1.172203 | 0.190659 | 0.06479 |
| 7 | 81141600 | 81143799 | 10.34695 | 18.0448 | 54.19313 | 23.84996 | -1.19101 | 0.722433 | 0.064791 |
| 17 | 29564400 | 29567199 | 13.24519 | 22.2991 | 71.12818 | 31.08609 | -1.28567 | 1.101388 | 0.064797 |
| 10 | 81667200 | 81667799 | 15.10792 | 28.17915 | 16.58197 | 6.898951 | 1.211458 | 0.161118 | 0.064808 |
| 19 | 8902800 | 8908999 | 24.17601 | 38.89134 | 117.8726 | 50.02875 | -1.24288 | 1.84692 | 0.064818 |
| 9 | 46668800 | 46669599 | 3.135733 | 6.10633 | 29.65065 | 10.85503 | -1.6345 | -0.35863 | 0.064844 |
| 1 | 153267400 | 153271799 | 41.08832 | 67.80647 | 43.99409 | 18.99055 | 1.002492 | 1.37326 | 0.064869 |
| 7 | 21866400 | 21866799 | 9.170277 | 17.90631 | 9.724797 | 3.423412 | 1.534804 | -0.48581 | 0.064896 |

| | | | | | | | | | |
|---|---|---|---|---|---|---|---|---|---|
| 6 | 125071800 | 125079999 | 31.08309 | 49.47944 | 149.9991 | 62.70266 | -1.26117 | 2.198733 | 0.064899 |
| 14 | 50692600 | 50693799 | 21.19131 | 37.53719 | 22.73056 | 10.05036 | 1.110994 | 0.521674 | 0.0649 |
| 7 | 10747400 | 10748799 | 38.33805 | 63.93788 | 37.9929 | 16.51785 | 1.129524 | 1.244742 | 0.064909 |
| 5 | 114979400 | 114993399 | 73.27472 | 111.5203 | 286.0608 | 118.5374 | -1.07349 | 3.162326 | 0.064937 |
| 17 | 20124400 | 20126799 | 59.87279 | 96.7141 | 51.941 | 22.16939 | 1.254598 | 1.747371 | 0.06494 |
| 5 | 120518200 | 120525399 | 53.00073 | 83.49574 | 180.7339 | 74.99381 | -0.78988 | 2.491949 | 0.064944 |
| 11 | 78162200 | 78164799 | 7.719938 | 13.60854 | 48.25204 | 21.35543 | -1.42575 | 0.542157 | 0.064956 |
| 3 | 149505400 | 149505999 | 11.07815 | 21.51108 | 13.6277 | 5.107203 | 1.238643 | -0.23123 | 0.064967 |
| 14 | 46758800 | 46759999 | 15.6491 | 30.14644 | 11.88707 | 4.613138 | 1.867833 | -0.0722 | 0.064972 |
| 12 | 23982800 | 23986999 | 66.06355 | 105.7921 | 71.84631 | 30.28661 | 0.902308 | 2.023193 | 0.064998 |
| 17 | 29401000 | 29411599 | 45.84197 | 71.26028 | 223.9267 | 92.97152 | -1.34187 | 2.739134 | 0.065021 |
| 14 | 78961200 | 78964799 | 22.32608 | 36.52485 | 91.36729 | 39.29578 | -0.94384 | 1.470938 | 0.06504 |
| 11 | 99662800 | 99663799 | 5.023472 | 9.526675 | 33.08581 | 13.35251 | -1.29087 | 0.007851 | 0.065066 |
| 14 | 41756600 | 41758999 | 20.99571 | 37.51832 | 21.01582 | 9.228959 | 1.229336 | 0.459567 | 0.065087 |
| 7 | 62324400 | 62326199 | 32.23042 | 54.98554 | 32.35066 | 14.18234 | 1.144415 | 1.009787 | 0.065089 |
| 15 | 89252400 | 89263999 | 53.56149 | 82.67176 | 241.5586 | 100.3387 | -1.24512 | 2.872429 | 0.065095 |
| 4 | 117746600 | 117750399 | 12.94601 | 21.76442 | 71.57715 | 31.26882 | -1.32958 | 1.113961 | 0.065096 |
| 7 | 125554400 | 125561399 | 39.47708 | 62.79381 | 149.7849 | 62.54755 | -0.91739 | 2.22952 | 0.065153 |
| 13 | 38528200 | 38532199 | 22.75101 | 36.91876 | 98.16282 | 42.06726 | -1.04052 | 1.604843 | 0.065162 |
| 5 | 121735200 | 121738599 | 8.695326 | 15.10833 | 56.19602 | 24.67389 | -1.49771 | 0.773309 | 0.065164 |
| 11 | 114160600 | 114170199 | 44.77452 | 70.49708 | 188.1481 | 78.06458 | -1.09213 | 2.498549 | 0.065171 |
| 12 | 22338600 | 22341399 | 32.53585 | 54.88984 | 35.8058 | 15.65853 | 0.995144 | 1.110374 | 0.06519 |
| 1 | 44293200 | 44294999 | 19.94389 | 36.39956 | 16.61359 | 7.12583 | 1.543088 | 0.321163 | 0.065199 |
| 4 | 43794400 | 43796199 | 23.18571 | 40.76557 | 24.28473 | 10.75352 | 1.130757 | 0.616174 | 0.065206 |
| 8 | 107937200 | 107941399 | 16.27995 | 27.16213 | 76.83902 | 33.41551 | -1.11382 | 1.208751 | 0.065235 |
| 12 | 108987200 | 109028799 | 302.4729 | 410.6924 | 1031.948 | 429.8777 | -1.20354 | 5.034422 | 0.065236 |
| 2 | 73285000 | 73285999 | 4.830005 | 9.244528 | 31.71924 | 12.54038 | -1.24634 | -0.06515 | 0.065239 |
| 6 | 117860000 | 117861599 | 6.856776 | 12.69683 | 40.85119 | 17.18358 | -1.23282 | 0.195912 | 0.06524 |
| 7 | 104087600 | 104088199 | 8.33687 | 16.29238 | 9.961458 | 3.450718 | 1.387758 | -0.55184 | 0.065253 |
| 7 | 8176600 | 8177399 | 12.7058 | 24.4495 | 14.69155 | 5.719568 | 1.26393 | -0.0854 | 0.065263 |
| 4 | 148254000 | 148275999 | 136.2337 | 198.8446 | 488.6486 | 201.337 | -1.06672 | 3.936321 | 0.065268 |
| 16 | 62186000 | 62187599 | 20.49167 | 36.4394 | 22.20217 | 9.790124 | 1.104956 | 0.488065 | 0.0653 |
| 2 | 4656400 | 4657199 | 4.068327 | 7.90352 | 28.64899 | 10.69485 | -1.2449 | -0.27782 | 0.065305 |
| 7 | 125346200 | 125347599 | 4.068327 | 7.90352 | 28.64899 | 10.69485 | -1.2449 | -0.27782 | 0.065305 |
| 12 | 114773000 | 114773799 | 23.87579 | 42.21393 | 21.38446 | 9.458689 | 1.365818 | 0.556061 | 0.065312 |
| 11 | 59461400 | 59463799 | 10.51823 | 18.31272 | 56.91704 | 25.02229 | -1.24076 | 0.750333 | 0.065316 |
| 17 | 28128000 | 28131399 | 11.55608 | 19.77672 | 62.60657 | 27.45329 | -1.27192 | 0.938365 | 0.065322 |
| 2 | 154915800 | 154916399 | 3.648367 | 7.099754 | 27.00017 | 9.941565 | -1.29376 | -0.33687 | 0.065336 |
| 14 | 70420000 | 70423399 | 12.68873 | 21.5625 | 65.79696 | 28.83591 | -1.22111 | 1.011689 | 0.065358 |
| 7 | 45438800 | 45469999 | 148.8703 | 215.0285 | 543.5232 | 224.2168 | -1.12212 | 4.095419 | 0.065397 |
| 7 | 20682800 | 20687799 | 55.41551 | 89.3662 | 59.35664 | 25.28053 | 0.946253 | 1.790434 | 0.065427 |
| 6 | 71201800 | 71202599 | 3.046736 | 5.93695 | 25.27777 | 9.174253 | -1.43258 | -0.39966 | 0.065434 |
| 16 | 4273200 | 4276399 | 13.01028 | 22.48285 | 59.67612 | 26.18042 | -1.01651 | 0.850465 | 0.065437 |
| 12 | 22257400 | 22258199 | 20.76487 | 37.15398 | 19.0322 | 8.348734 | 1.3588 | 0.44689 | 0.065452 |
| 8 | 121531000 | 121563999 | 196.0045 | 277.1306 | 679.5358 | 281.2726 | -1.10923 | 4.41249 | 0.065491 |
| 7 | 9089400 | 9090799 | 18.37914 | 34.00757 | 17.06084 | 7.183753 | 1.426937 | 0.236548 | 0.065498 |
| 11 | 6036400 | 6038799 | 11.42055 | 19.87877 | 54.764 | 24.072 | -1.06766 | 0.75664 | 0.065507 |

| | | | | | | | | | |
|---|---|---|---|---|---|---|---|---|---|
| 2 | 28757000 | 28758799 | 4.456129 | 8.5152 | 33.37793 | 13.25566 | -1.44315 | -0.05211 | 0.065511 |
| 11 | 116700400 | 116706199 | 21.33427 | 34.64112 | 98.65957 | 42.29935 | -1.1385 | 1.589257 | 0.065527 |
| 8 | 71256200 | 71281399 | 159.9964 | 230.7247 | 547.3295 | 225.8355 | -1.03291 | 4.120223 | 0.065528 |
| 4 | 132457000 | 132461999 | 29.12508 | 46.85391 | 117.2059 | 49.74752 | -0.96686 | 1.851444 | 0.065553 |
| 2 | 105495800 | 105514399 | 145.7141 | 212.4234 | 497.0583 | 204.828 | -0.9978 | 3.955569 | 0.065565 |
| 12 | 104409600 | 104415799 | 36.21734 | 57.72517 | 145.6301 | 60.98271 | -0.99488 | 2.159448 | 0.065567 |
| 10 | 61804800 | 61823999 | 116.3451 | 172.8462 | 397.9739 | 163.9479 | -0.94865 | 3.643395 | 0.065568 |
| 2 | 30409800 | 30417799 | 35.38313 | 56.10677 | 164.1283 | 68.31802 | -1.21564 | 2.322414 | 0.065573 |
| 14 | 110429400 | 110430999 | 18.28047 | 33.82972 | 17.18038 | 7.23271 | 1.409865 | 0.235566 | 0.065579 |
| 16 | 90497400 | 90504999 | 46.38103 | 73.41377 | 167.4931 | 69.63094 | -0.86014 | 2.37732 | 0.065587 |
| 2 | 23640000 | 23641199 | 14.91637 | 28.13858 | 16.2746 | 6.641897 | 1.257528 | 0.088298 | 0.065587 |
| 1 | 113875800 | 113877999 | 28.87405 | 50.08919 | 27.96176 | 12.2685 | 1.223638 | 0.84371 | 0.065608 |
| 7 | 39000400 | 39001999 | 21.30584 | 37.73774 | 22.6306 | 10.00629 | 1.124822 | 0.522468 | 0.065621 |
| 18 | 38329000 | 38339599 | 55.64557 | 86.32362 | 223.5034 | 92.82403 | -1.06528 | 2.778359 | 0.065621 |
| 5 | 120249400 | 120277599 | 160.1226 | 229.9477 | 573.8843 | 236.9368 | -1.11206 | 4.181901 | 0.06566 |
| 15 | 68797000 | 68802399 | 23.95095 | 38.72472 | 107.9024 | 46.11022 | -1.11316 | 1.687099 | 0.065662 |
| 7 | 38452800 | 38453799 | 19.33087 | 35.37056 | 17.96361 | 7.679927 | 1.393978 | 0.306353 | 0.065682 |
| 17 | 17499000 | 17500199 | 16.11922 | 30.07507 | 17.0372 | 7.084501 | 1.266896 | 0.170132 | 0.065706 |
| 4 | 139413000 | 139440599 | 167.5985 | 240.6589 | 570.8089 | 235.6708 | -1.03882 | 4.183778 | 0.065736 |
| 3 | 112686200 | 112687999 | 21.39324 | 38.96397 | 15.98102 | 6.872288 | 1.693774 | 0.342823 | 0.065797 |
| 12 | 22068800 | 22077599 | 257.6307 | 377.1371 | 258.52 | 106.4936 | 0.778324 | 3.896629 | 0.065801 |
| 8 | 41315000 | 41315999 | 6.756626 | 12.41295 | 38.88794 | 16.53449 | -1.20588 | 0.235647 | 0.06583 |
| 2 | 107681600 | 107682599 | 11.92571 | 23.06871 | 13.74154 | 5.248505 | 1.301953 | -0.15813 | 0.065847 |
| 1 | 75459000 | 75470599 | 61.27622 | 94.39309 | 241.4822 | 100.3047 | -1.05587 | 2.912221 | 0.065873 |
| X | 33285800 | 33288999 | 55.19099 | 89.45661 | 49.59715 | 21.23654 | 1.213658 | 1.66167 | 0.065881 |
| 6 | 105157400 | 105158599 | 15.26124 | 29.29965 | 12.29324 | 4.817477 | 1.765754 | -0.04558 | 0.065885 |
| 3 | 103328400 | 103334999 | 38.33429 | 61.02327 | 149.0713 | 62.31631 | -0.95049 | 2.202666 | 0.065888 |
| 6 | 30633600 | 30646199 | 66.32812 | 101.9384 | 253.6432 | 105.3225 | -1.01848 | 2.960171 | 0.065934 |
| 1 | 7689400 | 7691799 | 32.0082 | 54.14971 | 35.71145 | 15.63042 | 0.979489 | 1.084547 | 0.065935 |
| 18 | 56914400 | 56918599 | 17.75163 | 29.12281 | 91.99543 | 39.61164 | -1.27724 | 1.424174 | 0.065951 |
| 6 | 67012800 | 67014799 | 9.884923 | 17.26654 | 53.64336 | 23.63167 | -1.23926 | 0.696206 | 0.065962 |
| 13 | 84491400 | 84497799 | 41.22914 | 65.63197 | 158.4832 | 66.25139 | -0.93391 | 2.206444 | 0.06598 |
| 8 | 115500200 | 115502799 | 8.927456 | 15.69318 | 48.77623 | 21.5995 | -1.23909 | 0.581236 | 0.065996 |
| 17 | 56555200 | 56567199 | 52.92928 | 82.71417 | 200.9213 | 83.36178 | -0.96503 | 2.646704 | 0.066001 |
| 5 | 10962400 | 10965199 | 31.79035 | 54.92378 | 26.82864 | 11.76987 | 1.414223 | 0.891511 | 0.066045 |
| 7 | 127047600 | 127049199 | 5.76848 | 10.8742 | 34.63088 | 14.15139 | -1.18383 | 0.059749 | 0.066054 |
| 10 | 130377400 | 130378399 | 5.76848 | 10.8742 | 34.63088 | 14.15139 | -1.18383 | 0.059749 | 0.066054 |
| 5 | 115677800 | 115694399 | 63.9431 | 97.41882 | 290.6057 | 120.4581 | -1.28976 | 3.138771 | 0.066058 |
| 1 | 89843200 | 89865999 | 168.4231 | 243.1866 | 548.0843 | 226.1087 | -0.95724 | 4.101693 | 0.066073 |
| 4 | 128614000 | 128619799 | 19.21322 | 31.32703 | 93.56861 | 40.19336 | -1.20207 | 1.515688 | 0.0661 |
| 8 | 92560400 | 92565599 | 30.49298 | 48.87159 | 131.6555 | 55.57007 | -1.08023 | 1.975322 | 0.066101 |
| 1 | 160465800 | 160467399 | 17.14865 | 32.095 | 16.03873 | 6.634434 | 1.451432 | 0.156944 | 0.066111 |
| 11 | 74719400 | 74745999 | 125.9856 | 184.0297 | 498.239 | 205.272 | -1.20528 | 3.923723 | 0.066126 |
| X | 142740400 | 142741599 | 11.39754 | 22.26395 | 8.196873 | 2.866402 | 2.094014 | -0.47889 | 0.066138 |
| 15 | 45500800 | 45501999 | 29.3576 | 50.19977 | 32.56668 | 14.27969 | 1.0045 | 0.977735 | 0.066143 |
| 10 | 48805400 | 48807199 | 17.03182 | 32.51158 | 12.17742 | 4.846967 | 1.907862 | 0.020407 | 0.066145 |
| 8 | 123537000 | 123549399 | 50.5001 | 78.21455 | 230.7987 | 95.86756 | -1.25545 | 2.80634 | 0.066149 |

| | | | | | | | | | |
|---|---|---|---|---|---|---|---|---|---|
| 18 | 3624800 | 3625999 | 12.79863 | 24.67577 | 13.78709 | 5.33776 | 1.373658 | -0.10568 | 0.066152 |
| 9 | 83429800 | 83432199 | 15.48097 | 25.85204 | 80.78553 | 35.15145 | -1.25684 | 1.200248 | 0.066168 |
| 17 | 17215400 | 17218999 | 126.0912 | 194.8084 | 126.8378 | 52.68567 | 0.91947 | 2.870986 | 0.066169 |
| 12 | 60790800 | 60791399 | 9.745157 | 19.03319 | 10.23069 | 3.587342 | 1.555439 | -0.48726 | 0.066185 |
| 2 | 167739000 | 167740999 | 6.925261 | 12.24139 | 46.16868 | 20.42212 | -1.51162 | 0.504685 | 0.066185 |
| 9 | 107975400 | 107977399 | 7.201075 | 12.81183 | 44.12621 | 19.4493 | -1.37708 | 0.44555 | 0.066203 |
| 5 | 104993000 | 104996799 | 97.44954 | 153.4358 | 100.4435 | 41.67511 | 0.93252 | 2.516219 | 0.066211 |
| 10 | 75720200 | 75727599 | 37.15547 | 59.27583 | 141.0455 | 59.13941 | -0.90919 | 2.129004 | 0.06622 |
| 9 | 59894000 | 59896799 | 12.16978 | 20.79911 | 62.61048 | 27.45204 | -1.20003 | 0.953629 | 0.066232 |
| 4 | 9706200 | 9707999 | 8.008391 | 14.49795 | 43.43352 | 18.81939 | -1.16128 | 0.339243 | 0.066239 |
| 10 | 115180800 | 115183799 | 10.4606 | 18.3137 | 50.96361 | 22.49253 | -1.08107 | 0.659827 | 0.066241 |
| 5 | 99590600 | 99591799 | 4.639057 | 8.954766 | 33.53574 | 12.93543 | -1.33782 | -0.14469 | 0.066249 |
| 1 | 178806200 | 178809999 | 19.46746 | 32.29708 | 83.93076 | 36.36245 | -0.99252 | 1.287899 | 0.06625 |
| 5 | 104037800 | 104038599 | 2.845478 | 5.539286 | 26.89078 | 9.873018 | -1.63538 | -0.35756 | 0.066252 |
| 15 | 62565400 | 62565799 | 2.845478 | 5.539286 | 26.89078 | 9.873018 | -1.63538 | -0.35756 | 0.066252 |
| 10 | 63604000 | 63605199 | 4.987537 | 9.570107 | 35.02052 | 13.74582 | -1.32995 | -0.08241 | 0.066269 |
| 8 | 6209400 | 6210399 | 11.37246 | 22.02994 | 14.1522 | 5.369416 | 1.202437 | -0.1856 | 0.066273 |
| 15 | 99117400 | 99150199 | 177.467 | 252.1455 | 649.4517 | 268.6024 | -1.17308 | 4.339988 | 0.066291 |
| 14 | 26691800 | 26692799 | 4.124009 | 7.684143 | 37.45406 | 15.61002 | -1.81327 | 0.128863 | 0.066315 |
| 2 | 179724200 | 179732999 | 41.46689 | 65.47158 | 179.2748 | 74.43464 | -1.12584 | 2.435746 | 0.066331 |
| 5 | 142557200 | 142576399 | 115.5908 | 172.5254 | 374.3476 | 154.3066 | -0.85693 | 3.559367 | 0.066334 |
| 4 | 45943000 | 45957999 | 83.03782 | 125.6794 | 311.6969 | 128.9425 | -1.03165 | 3.279399 | 0.066345 |
| 5 | 114554800 | 114563999 | 33.5144 | 53.04738 | 172.4423 | 71.68804 | -1.3698 | 2.370319 | 0.066346 |
| 4 | 136311400 | 136312199 | 2.940328 | 5.737698 | 24.40553 | 8.688072 | -1.40601 | -0.47571 | 0.066346 |
| 3 | 52569400 | 52573399 | 19.16822 | 31.75829 | 80.87506 | 35.00637 | -0.96347 | 1.30009 | 0.06635 |
| 2 | 85846400 | 85848199 | 19.40033 | 36.12559 | 14.75439 | 6.155776 | 1.730114 | 0.204475 | 0.066351 |
| 10 | 79889200 | 79894599 | 22.89917 | 36.89759 | 114.8959 | 48.88105 | -1.27878 | 1.791155 | 0.066355 |
| 11 | 51222800 | 51224199 | 13.55748 | 26.06361 | 14.56771 | 5.686835 | 1.363734 | -0.06807 | 0.066372 |
| 17 | 83193000 | 83193599 | 11.04006 | 21.58209 | 4.869123 | 1.672673 | 2.79104 | -0.55612 | 0.066408 |
| 7 | 139695600 | 139696599 | 5.335009 | 10.0662 | 34.05533 | 13.89229 | -1.26768 | 0.048543 | 0.066426 |
| 10 | 95540400 | 95541199 | 3.542261 | 6.872954 | 28.35248 | 10.67592 | -1.44 | -0.25422 | 0.066428 |
| 15 | 74710000 | 74710999 | 3.542261 | 6.872954 | 28.35248 | 10.67592 | -1.44 | -0.25422 | 0.066428 |
| 7 | 21144000 | 21147399 | 14.12474 | 27.15608 | 12.47635 | 4.869426 | 1.643595 | -0.06934 | 0.066432 |
| 1 | 95741600 | 95742199 | 9.581883 | 18.67472 | 12.22706 | 4.427694 | 1.236042 | -0.37232 | 0.066438 |
| 2 | 180184200 | 180188799 | 23.64254 | 38.17325 | 108.2099 | 46.15923 | -1.14014 | 1.728612 | 0.066443 |
| 7 | 30227400 | 30235399 | 34.32492 | 54.66785 | 151.9791 | 63.57525 | -1.13585 | 2.183431 | 0.066446 |
| 19 | 13764400 | 13766599 | 31.05903 | 52.90621 | 33.40428 | 14.64244 | 1.042597 | 1.018285 | 0.066451 |
| 7 | 15450800 | 15452199 | 42.94053 | 70.78838 | 42.87829 | 18.50045 | 1.100668 | 1.388532 | 0.066456 |
| 5 | 139255800 | 139285399 | 160.9656 | 231.112 | 585.6022 | 241.7823 | -1.13423 | 4.18484 | 0.066462 |
| 11 | 84303600 | 84310399 | 37.50458 | 59.76264 | 144.5001 | 60.48962 | -0.93409 | 2.168351 | 0.06647 |
| 5 | 70208400 | 70211199 | 31.59055 | 53.77818 | 33.55647 | 14.70841 | 1.059595 | 1.027907 | 0.066473 |
| 2 | 25312000 | 25339399 | 113.3408 | 165.3413 | 507.4025 | 209.0759 | -1.38777 | 3.943056 | 0.066492 |
| 12 | 82110600 | 82114399 | 16.55828 | 27.69701 | 74.1306 | 32.28956 | -1.0342 | 1.179931 | 0.066501 |
| 4 | 83443200 | 83444199 | 6.153799 | 11.32122 | 37.79738 | 16.04039 | -1.29415 | 0.222011 | 0.066501 |
| 6 | 147389200 | 147390599 | 5.119498 | 9.82623 | 33.18192 | 13.01307 | -1.21357 | -0.08715 | 0.066515 |
| 7 | 45742200 | 45763199 | 106.6796 | 157.4925 | 443.0284 | 182.4472 | -1.24584 | 3.750907 | 0.066515 |
| 13 | 101185400 | 101186599 | 8.652738 | 15.51843 | 44.80097 | 19.61375 | -1.1183 | 0.403818 | 0.06654 |

| | | | | | | | | | |
|---|---|---|---|---|---|---|---|---|---|
| 4 | 120894200 | 120903599 | 57.2545 | 88.86166 | 221.574 | 92.0162 | -1.01051 | 2.770509 | 0.066541 |
| 9 | 108446800 | 108461599 | 58.4437 | 89.25468 | 284.336 | 117.9278 | -1.38165 | 3.087695 | 0.066544 |
| 5 | 67803400 | 67805199 | 8.306124 | 14.68457 | 49.89137 | 22.06811 | -1.36353 | 0.513804 | 0.066547 |
| 5 | 95453400 | 95455799 | 25.66562 | 44.81069 | 26.56086 | 11.69752 | 1.138325 | 0.740695 | 0.066548 |
| 14 | 110421200 | 110421799 | 9.858051 | 19.23011 | 11.41833 | 4.086803 | 1.388755 | -0.412 | 0.066551 |
| 8 | 15167400 | 15167999 | 11.84979 | 22.89291 | 14.6673 | 5.635267 | 1.189504 | -0.13729 | 0.066564 |
| X | 31701200 | 31703399 | 37.82825 | 63.18087 | 37.9465 | 16.51305 | 1.114079 | 1.227593 | 0.066568 |
| 5 | 142827600 | 142832999 | 30.93671 | 49.63821 | 124.2119 | 52.51744 | -0.97206 | 1.938966 | 0.066581 |
| 11 | 117842000 | 117843399 | 4.133822 | 7.813521 | 33.41828 | 13.58174 | -1.59507 | 0.026509 | 0.066583 |
| 4 | 147140800 | 147141199 | 8.432365 | 16.49315 | 9.267305 | 3.119185 | 1.543854 | -0.6482 | 0.066599 |
| 8 | 100537200 | 100537599 | 8.432365 | 16.49315 | 9.267305 | 3.119185 | 1.543854 | -0.6482 | 0.066599 |
| 11 | 106162200 | 106164599 | 11.54896 | 19.78313 | 62.08518 | 27.22672 | -1.25922 | 0.928938 | 0.066605 |
| 4 | 152336600 | 152339399 | 10.03865 | 17.24043 | 63.68952 | 27.93585 | -1.49249 | 0.899823 | 0.066639 |
| 5 | 33581400 | 33595999 | 97.50958 | 145.8615 | 365.9202 | 150.8979 | -1.06317 | 3.515053 | 0.066645 |
| 5 | 113761800 | 113774399 | 62.53809 | 96.15906 | 251.3885 | 104.3931 | -1.08916 | 2.949621 | 0.066645 |
| 4 | 130067200 | 130071799 | 17.26877 | 28.66211 | 78.93018 | 34.20646 | -1.07554 | 1.272944 | 0.066656 |
| 9 | 72714000 | 72715399 | 7.41476 | 13.07475 | 47.72599 | 21.12126 | -1.4669 | 0.536436 | 0.066693 |
| 2 | 40618600 | 40619999 | 18.68607 | 33.63753 | 20.16816 | 8.800704 | 1.137628 | 0.396113 | 0.066694 |
| 4 | 124274600 | 124276199 | 8.097324 | 14.38822 | 44.10707 | 19.46157 | -1.21172 | 0.459325 | 0.066701 |
| 15 | 79631800 | 79636999 | 23.04642 | 37.44408 | 96.1418 | 41.23847 | -0.9886 | 1.579402 | 0.066737 |
| 12 | 21853800 | 21856399 | 43.08491 | 70.53117 | 45.69374 | 19.65694 | 1.000261 | 1.476937 | 0.066744 |
| 3 | 16108000 | 16109399 | 38.11649 | 63.25113 | 41.74221 | 18.08745 | 0.978435 | 1.299281 | 0.066763 |
| 6 | 75208600 | 75209599 | 13.35322 | 25.41735 | 16.04808 | 6.426073 | 1.156025 | 0.012906 | 0.066792 |
| 7 | 45015000 | 45020199 | 22.12786 | 35.65642 | 117.2317 | 49.87809 | -1.35689 | 1.789064 | 0.066796 |
| 5 | 36380200 | 36386599 | 30.57 | 49.16411 | 120.1 | 50.95448 | -0.9333 | 1.863137 | 0.066798 |
| X | 25509000 | 25509599 | 11.7027 | 22.75156 | 12.58996 | 4.672009 | 1.445024 | -0.25966 | 0.066798 |
| 2 | 181653200 | 181654999 | 67.72225 | 109.0978 | 43.14227 | 18.34861 | 1.690197 | 1.830357 | 0.066805 |
| 13 | 21624800 | 21625599 | 4.370326 | 8.471705 | 30.28813 | 11.45468 | -1.24394 | -0.22512 | 0.066809 |
| 4 | 18378200 | 18379599 | 15.69207 | 29.49208 | 16.80968 | 6.904226 | 1.27158 | 0.120331 | 0.066813 |
| 10 | 127343000 | 127400799 | 345.5496 | 463.9992 | 1173.045 | 489.4791 | -1.22851 | 5.211141 | 0.066819 |
| 1 | 25176800 | 25177999 | 19.92512 | 35.935 | 20.19093 | 8.790099 | 1.233216 | 0.396278 | 0.066825 |
| 4 | 52236400 | 52237999 | 19.92512 | 35.935 | 20.19093 | 8.790099 | 1.233216 | 0.396278 | 0.066825 |
| 6 | 52249800 | 52251799 | 13.13612 | 22.35469 | 64.78477 | 28.39558 | -1.14676 | 0.998793 | 0.066858 |
| 1 | 131524600 | 131528599 | 20.3356 | 33.51016 | 84.71676 | 36.54602 | -0.95441 | 1.370436 | 0.06687 |
| 18 | 63439600 | 63440199 | 11.22121 | 21.87161 | 11.35895 | 4.108657 | 1.567061 | -0.35683 | 0.066883 |
| 7 | 59175200 | 59177599 | 20.27009 | 36.79142 | 18.31952 | 7.911923 | 1.412941 | 0.358484 | 0.066885 |
| 2 | 36797000 | 36797799 | 18.2436 | 33.13799 | 19.46612 | 8.39926 | 1.177646 | 0.342426 | 0.066886 |
| 6 | 60921000 | 60922199 | 5.897112 | 11.15154 | 37.16321 | 15.089 | -1.24113 | 0.04432 | 0.066894 |
| 8 | 31334400 | 31335199 | 10.82324 | 21.01964 | 13.02915 | 4.876245 | 1.270852 | -0.24169 | 0.066935 |
| 16 | 93442000 | 93444999 | 14.24941 | 24.14177 | 66.40348 | 29.07684 | -1.07246 | 1.044956 | 0.066939 |
| 12 | 111229400 | 111247399 | 123.9732 | 182.5863 | 437.09 | 179.9971 | -1.0165 | 3.792466 | 0.066941 |
| 2 | 167877000 | 167893599 | 80.24744 | 121.5146 | 313.3688 | 129.6539 | -1.08733 | 3.268998 | 0.066954 |
| 13 | 21363400 | 21366599 | 14.34653 | 24.06493 | 71.64956 | 31.26425 | -1.18693 | 1.141564 | 0.066964 |
| 10 | 115286800 | 115290999 | 27.92012 | 45.1142 | 107.8704 | 46.05965 | -0.89413 | 1.713184 | 0.06697 |
| 4 | 44425000 | 44425999 | 2.969644 | 5.70751 | 30.50871 | 11.9187 | -1.85719 | -0.12534 | 0.066988 |
| 1 | 153629800 | 153631399 | 6.269796 | 11.66338 | 38.7274 | 16.18514 | -1.27087 | 0.158102 | 0.066994 |
| 12 | 23943000 | 23944599 | 38.70421 | 64.02492 | 43.43552 | 18.7804 | 0.938531 | 1.338676 | 0.066997 |

| | | | | | | | | | |
|---|---|---|---|---|---|---|---|---|---|
| 6 | 79129600 | 79131199 | 24.3053 | 42.87474 | 21.83489 | 9.662358 | 1.356978 | 0.583338 | 0.067011 |
| 1 | 63673600 | 63674199 | 3.645856 | 7.102718 | 27.28557 | 9.93395 | -1.29351 | -0.37749 | 0.067029 |
| 1 | 128731000 | 128731999 | 6.927405 | 12.6369 | 39.25054 | 16.84566 | -1.20295 | 0.279885 | 0.067036 |
| 17 | 39502400 | 39502999 | 21.72981 | 38.22125 | 23.91898 | 10.5907 | 1.06015 | 0.601515 | 0.06704 |
| 2 | 129533600 | 129543599 | 50.47286 | 79.33617 | 189.189 | 78.47939 | -0.93185 | 2.535682 | 0.067045 |
| 5 | 144298400 | 144300999 | 12.08579 | 20.89073 | 57.96425 | 25.42961 | -1.07984 | 0.842516 | 0.067052 |
| X | 150426400 | 150428599 | 8.187747 | 14.68667 | 50.29251 | 22.01516 | -1.36296 | 0.409779 | 0.067061 |
| 6 | 30031400 | 30031999 | 2.027509 | 3.965127 | 19.2513 | 6.29255 | -1.47261 | -0.80973 | 0.067067 |
| 19 | 16516800 | 16517199 | 2.027509 | 3.965127 | 19.2513 | 6.29255 | -1.47261 | -0.80973 | 0.067067 |
| 8 | 78166800 | 78187199 | 250.4724 | 351.4012 | 700.6753 | 290.3869 | -0.8223 | 4.52982 | 0.067071 |
| X | 147073600 | 147073999 | 7.939129 | 15.52002 | 10.1438 | 3.485231 | 1.304472 | -0.58431 | 0.067076 |
| 5 | 129167400 | 129176999 | 52.43018 | 82.36978 | 187.366 | 77.71786 | -0.86443 | 2.547309 | 0.067101 |
| 12 | 23389600 | 23392599 | 40.51104 | 67.3645 | 38.9263 | 16.89149 | 1.169584 | 1.281602 | 0.067104 |
| 7 | 31778200 | 31782199 | 119.2554 | 184.2701 | 130.7976 | 54.33048 | 0.795156 | 2.865659 | 0.067108 |
| 7 | 9742400 | 9745199 | 83.11104 | 132.015 | 85.24451 | 35.54385 | 0.962701 | 2.29192 | 0.067114 |
| 9 | 21001400 | 21005999 | 17.09868 | 28.40008 | 77.76838 | 33.71959 | -1.06728 | 1.263688 | 0.067139 |
| 5 | 95409000 | 95409399 | 9.467493 | 18.47121 | 10.75557 | 3.841255 | 1.421126 | -0.42673 | 0.067142 |
| 6 | 108910800 | 108911799 | 2.952756 | 5.745302 | 38.25926 | 14.1028 | -2.09219 | -0.30961 | 0.067143 |
| 10 | 81808400 | 81810999 | 23.43928 | 41.12268 | 25.32435 | 11.19675 | 1.083085 | 0.651183 | 0.067162 |
| 6 | 29836800 | 29837799 | 17.26266 | 32.2238 | 16.71071 | 6.943146 | 1.393921 | 0.176108 | 0.067164 |
| 1 | 64769400 | 64778399 | 40.53356 | 63.93924 | 183.1072 | 75.99157 | -1.19147 | 2.457757 | 0.067164 |
| 16 | 40543600 | 40544799 | 14.81002 | 28.38215 | 14.27297 | 5.622326 | 1.502195 | -0.0281 | 0.067218 |
| 12 | 22507400 | 22517999 | 45.25422 | 74.5488 | 38.64624 | 16.66939 | 1.324147 | 1.402106 | 0.067234 |
| 4 | 56801800 | 56802999 | 7.9008 | 14.14621 | 42.62807 | 18.68875 | -1.18047 | 0.406879 | 0.067239 |
| 9 | 36301800 | 36304199 | 37.01611 | 62.32065 | 33.24181 | 14.52823 | 1.284041 | 1.139335 | 0.067241 |
| 5 | 100968200 | 100968999 | 5.001706 | 9.552893 | 32.23293 | 12.81468 | -1.23054 | -0.04494 | 0.067244 |
| 12 | 18779800 | 18784399 | 125.1855 | 193.4828 | 128.195 | 53.24932 | 0.894705 | 2.863018 | 0.067248 |
| 12 | 23896000 | 23896999 | 27.15269 | 47.14839 | 26.04831 | 11.43241 | 1.238501 | 0.820674 | 0.067262 |
| 7 | 9354600 | 9355799 | 11.69635 | 22.77276 | 11.91297 | 4.368452 | 1.539865 | -0.30217 | 0.067263 |
| 18 | 78085800 | 78101799 | 110.7294 | 165.2249 | 380.9037 | 157.0008 | -0.94464 | 3.563789 | 0.067266 |
| 2 | 71757200 | 71758599 | 13.07575 | 25.19732 | 14.39059 | 5.579495 | 1.341049 | -0.09581 | 0.067271 |
| 2 | 31578000 | 31579399 | 3.086539 | 5.87773 | 35.9229 | 14.37091 | -2.08313 | -0.03394 | 0.067275 |
| 17 | 70377800 | 70378399 | 10.15285 | 19.75181 | 12.26412 | 4.526295 | 1.287526 | -0.29963 | 0.067296 |
| 15 | 78516000 | 78525799 | 39.75772 | 62.63167 | 185.1701 | 76.83434 | -1.23884 | 2.484967 | 0.067301 |
| 2 | 166022200 | 166044999 | 114.3451 | 169.3042 | 417.771 | 172.0512 | -1.05299 | 3.702573 | 0.067306 |
| 17 | 21926200 | 21927599 | 6.60802 | 12.16502 | 40.06558 | 16.98744 | -1.27476 | 0.224045 | 0.067329 |
| 12 | 117807000 | 117808999 | 9.349906 | 16.50796 | 47.74807 | 21.12568 | -1.13362 | 0.532241 | 0.067339 |
| 8 | 38679800 | 38680999 | 14.12885 | 27.14521 | 14.32668 | 5.602713 | 1.442674 | -0.05763 | 0.067347 |
| 4 | 149796200 | 149807599 | 57.77351 | 89.49195 | 222.2752 | 92.32707 | -1.00706 | 2.806092 | 0.067369 |
| 14 | 6235600 | 6236199 | 11.8356 | 22.91985 | 14.03694 | 5.333571 | 1.268434 | -0.17586 | 0.067391 |
| 6 | 29433200 | 29437199 | 14.37381 | 23.50676 | 95.94267 | 41.25821 | -1.64741 | 1.462594 | 0.067435 |
| 14 | 67507000 | 67509799 | 9.674165 | 17.02432 | 53.66226 | 23.75799 | -1.25934 | 0.579071 | 0.067453 |
| 18 | 56684200 | 56685999 | 5.44065 | 10.3563 | 33.30413 | 13.33523 | -1.1715 | -0.01452 | 0.06746 |
| Y | 89966800 | 89967599 | 8.615628 | 16.85116 | 4.067041 | 1.372014 | 2.723564 | -0.66611 | 0.067475 |
| 3 | 156063800 | 156064599 | 5.044733 | 9.501347 | 33.95982 | 13.89824 | -1.34951 | 0.058643 | 0.067499 |
| 15 | 82828600 | 82843999 | 72.71704 | 110.5495 | 292.835 | 121.3044 | -1.12114 | 3.185121 | 0.067539 |
| 17 | 86548600 | 86551799 | 15.68655 | 26.04386 | 78.02672 | 33.82197 | -1.19638 | 1.265795 | 0.067564 |

| | | | | | | | | | |
|---|---|---|---|---|---|---|---|---|---|
| 17 | 17773800 | 17775599 | 11.38956 | 19.49513 | 62.03555 | 27.20327 | -1.27918 | 0.936155 | 0.067571 |
| 7 | 131755200 | 131759799 | 20.66682 | 33.66273 | 96.25006 | 41.32982 | -1.14054 | 1.533864 | 0.067593 |
| 12 | 19070800 | 19071799 | 23.21767 | 40.65067 | 24.68486 | 10.89517 | 1.103416 | 0.680946 | 0.067598 |
| 3 | 10867200 | 10867999 | 12.21683 | 23.59601 | 14.65983 | 5.639077 | 1.232038 | -0.12904 | 0.067599 |
| 19 | 39730000 | 39730799 | 12.21683 | 23.59601 | 14.65983 | 5.639077 | 1.232038 | -0.12904 | 0.067599 |
| 18 | 3878600 | 3880599 | 42.83482 | 70.14769 | 47.84349 | 20.58496 | 0.926456 | 1.471986 | 0.067606 |
| 14 | 56353000 | 56356999 | 28.45585 | 45.84875 | 112.9603 | 48.054 | -0.94212 | 1.799451 | 0.06761 |
| 9 | 58100400 | 58112599 | 72.6283 | 110.6693 | 283.6334 | 117.5739 | -1.07083 | 3.136945 | 0.067617 |
| 2 | 24984000 | 24990799 | 30.40803 | 48.87639 | 118.615 | 50.28386 | -0.92477 | 1.882035 | 0.067624 |
| 12 | 20099400 | 20106399 | 136.7609 | 209.4062 | 143.6465 | 59.61308 | 0.834456 | 3.046713 | 0.067629 |
| 12 | 81055600 | 81058399 | 10.12531 | 17.81275 | 48.79109 | 21.6027 | -1.05771 | 0.579566 | 0.067651 |
| 12 | 110675600 | 110676599 | 5.833778 | 10.7917 | 36.82151 | 15.50967 | -1.31696 | 0.185991 | 0.067661 |
| 4 | 132737000 | 132741599 | 19.18955 | 31.3857 | 91.82874 | 39.49063 | -1.16924 | 1.468081 | 0.067669 |
| 7 | 139598400 | 139599799 | 18.51522 | 33.41521 | 19.51555 | 8.488766 | 1.178572 | 0.380338 | 0.067682 |
| 4 | 141014200 | 141031799 | 74.30008 | 112.6788 | 303.1875 | 125.5005 | -1.14664 | 3.236853 | 0.067684 |
| 15 | 12822600 | 12826999 | 16.78939 | 27.92426 | 81.40978 | 35.33157 | -1.15734 | 1.247875 | 0.067687 |
| 7 | 74644000 | 74644799 | 3.897831 | 7.541295 | 29.30586 | 11.1831 | -1.37305 | -0.19861 | 0.067691 |
| 11 | 3125400 | 3131599 | 371.2435 | 526.6457 | 346.5028 | 143.3452 | 0.792822 | 4.368601 | 0.067711 |
| 13 | 35149600 | 35150199 | 2.526021 | 4.931169 | 24.38992 | 8.62375 | -1.60699 | -0.50123 | 0.06773 |
| 8 | 27599200 | 27613199 | 63.19165 | 96.73654 | 266.2421 | 110.4844 | -1.16818 | 3.03909 | 0.067731 |
| 17 | 6156600 | 6157999 | 5.998697 | 11.09248 | 37.37659 | 15.75178 | -1.3001 | 0.189486 | 0.067732 |
| 12 | 20592600 | 20595599 | 40.04804 | 66.35677 | 38.55716 | 16.69004 | 1.161268 | 1.31892 | 0.067742 |
| 19 | 58284400 | 58285999 | 6.278094 | 11.65516 | 38.07872 | 15.96151 | -1.25082 | 0.169325 | 0.067758 |
| 7 | 5239000 | 5240599 | 32.14926 | 54.51956 | 35.06827 | 15.36012 | 1.01543 | 1.063841 | 0.067771 |
| 3 | 96127200 | 96128999 | 6.6053 | 12.11261 | 38.27961 | 16.31669 | -1.22055 | 0.245241 | 0.067778 |
| 16 | 3327000 | 3329199 | 37.30142 | 63.04463 | 31.58382 | 13.81983 | 1.374259 | 1.107392 | 0.067783 |
| 15 | 76177600 | 76183199 | 25.13055 | 40.19494 | 138.4781 | 58.27421 | -1.43746 | 2.036828 | 0.067798 |
| 5 | 104116600 | 104117599 | 3.53405 | 6.880551 | 30.38359 | 11.14464 | -1.50293 | -0.3449 | 0.067803 |
| 2 | 113157600 | 113158399 | 4.373034 | 8.468153 | 29.5303 | 11.22243 | -1.2148 | -0.20991 | 0.067814 |
| 7 | 101646600 | 101651599 | 17.82399 | 29.08289 | 95.84097 | 41.18031 | -1.34269 | 1.502684 | 0.067833 |
| 7 | 45621400 | 45637799 | 93.94491 | 141.4175 | 334.3948 | 138.1063 | -0.96973 | 3.391661 | 0.067839 |
| 5 | 27991200 | 27992199 | 13.61073 | 25.95535 | 16.07603 | 6.412193 | 1.188619 | 0.00379 | 0.06784 |
| X | 148319800 | 148320599 | 10.74823 | 20.97185 | 10.79535 | 3.849439 | 1.597141 | -0.41362 | 0.067859 |
| 1 | 131065400 | 131066399 | 3.55086 | 6.864969 | 29.21802 | 11.18779 | -1.5077 | -0.19317 | 0.067864 |
| 5 | 145138000 | 145138799 | 3.55086 | 6.864969 | 29.21802 | 11.18779 | -1.5077 | -0.19317 | 0.067864 |
| X | 4752000 | 4752599 | 11.78137 | 22.96116 | 10.86791 | 3.936139 | 1.698192 | -0.34644 | 0.067877 |
| 2 | 120306200 | 120308399 | 9.560677 | 16.74568 | 51.47947 | 22.72882 | -1.22384 | 0.647388 | 0.067877 |
| 5 | 107276800 | 107277399 | 2.247086 | 4.394327 | 21.89118 | 7.453269 | -1.56751 | -0.65388 | 0.0679 |
| 4 | 140824600 | 140848399 | 146.55 | 212.3066 | 522.1334 | 215.3084 | -1.07836 | 4.050981 | 0.067937 |
| 10 | 51237800 | 51238599 | 14.1627 | 27.05377 | 15.51677 | 6.166969 | 1.303058 | -0.00321 | 0.06795 |
| X | 117569800 | 117571599 | 17.58523 | 33.82784 | 9.217755 | 3.591928 | 2.385498 | -0.04679 | 0.06796 |
| 11 | 69284600 | 69292999 | 30.65209 | 48.83702 | 152.7202 | 63.91921 | -1.30454 | 2.168025 | 0.067971 |
| 18 | 35997400 | 35999999 | 10.64291 | 18.56002 | 53.11913 | 23.37657 | -1.12182 | 0.723981 | 0.067973 |
| 12 | 19096200 | 19101999 | 188.2807 | 282.4755 | 190.3718 | 78.55381 | 0.835555 | 3.467386 | 0.06798 |
| 3 | 38582000 | 38583799 | 17.10385 | 31.47303 | 18.73355 | 7.947754 | 1.174527 | 0.25938 | 0.068006 |
| 11 | 68296200 | 68298999 | 10.97416 | 19.07077 | 55.26397 | 24.26699 | -1.14039 | 0.77909 | 0.068009 |
| 7 | 95705200 | 95706799 | 6.542088 | 12.19699 | 37.118 | 15.45424 | -1.14147 | 0.143598 | 0.068026 |

| | | | | | | | | | |
|---|---|---|---|---|---|---|---|---|---|
| 5 | 98178000 | 98180199 | 11.8671 | 20.30467 | 62.63398 | 27.46484 | -1.23487 | 0.941271 | 0.068054 |
| 12 | 18889800 | 18896799 | 161.4668 | 244.7801 | 171.5114 | 70.98703 | 0.792419 | 3.255338 | 0.068084 |
| 6 | 134704200 | 134706199 | 8.935995 | 15.68788 | 49.31899 | 21.81765 | -1.2555 | 0.603248 | 0.068107 |
| 6 | 125295600 | 125300799 | 23.99353 | 38.69818 | 112.1406 | 47.7694 | -1.17372 | 1.76225 | 0.068109 |
| 7 | 113112000 | 113113799 | 7.543836 | 13.71115 | 40.80439 | 17.59284 | -1.14667 | 0.307917 | 0.068113 |
| 1 | 155611200 | 155619799 | 57.17168 | 88.55419 | 227.2067 | 94.37552 | -1.05395 | 2.806874 | 0.068125 |
| 3 | 17158600 | 17160999 | 26.41489 | 45.73823 | 29.76865 | 13.06055 | 1.002747 | 0.849652 | 0.068136 |
| 15 | 86209800 | 86211999 | 9.108043 | 15.96578 | 51.4633 | 22.73719 | -1.29166 | 0.631502 | 0.068137 |
| 12 | 83809000 | 83831599 | 105.0639 | 155.2394 | 436.1803 | 179.6287 | -1.24299 | 3.736473 | 0.068156 |
| 16 | 48186600 | 48187599 | 13.80716 | 26.70766 | 12.63358 | 4.825709 | 1.629452 | -0.14041 | 0.068158 |
| 4 | 143838000 | 143838399 | 11.46474 | 22.23572 | 13.47398 | 5.083133 | 1.292765 | -0.20567 | 0.06816 |
| 11 | 18094000 | 18094799 | 11.46474 | 22.23572 | 13.47398 | 5.083133 | 1.292765 | -0.20567 | 0.06816 |
| 8 | 120942200 | 120997399 | 320.4276 | 433.9008 | 1062.541 | 442.8049 | -1.17031 | 5.077261 | 0.068186 |
| 1 | 66446600 | 66447199 | 2.93886 | 5.739179 | 24.04196 | 8.447601 | -1.36725 | -0.5244 | 0.068195 |
| 13 | 21677600 | 21678199 | 2.93886 | 5.739179 | 24.04196 | 8.447601 | -1.36725 | -0.5244 | 0.068195 |
| 5 | 124894200 | 124903199 | 67.12845 | 103.326 | 239.4207 | 99.43758 | -0.91412 | 2.930313 | 0.06821 |
| 5 | 136594000 | 136597999 | 19.16268 | 31.33269 | 89.85733 | 38.63877 | -1.14065 | 1.47247 | 0.068234 |
| 12 | 32583400 | 32586599 | 21.63863 | 35.17036 | 97.24008 | 41.72142 | -1.09451 | 1.568343 | 0.068235 |
| 7 | 45431400 | 45437999 | 28.92083 | 46.26276 | 133.5992 | 56.23046 | -1.1834 | 2.035952 | 0.068255 |
| 5 | 25046600 | 25048799 | 9.994986 | 17.54444 | 50.9414 | 22.53256 | -1.14207 | 0.612697 | 0.068255 |
| 10 | 5368000 | 5368599 | 2.524101 | 4.935303 | 24.53178 | 8.384848 | -1.57017 | -0.62348 | 0.068305 |
| 15 | 89019000 | 89034599 | 69.37571 | 105.2516 | 320.3932 | 132.6379 | -1.32402 | 3.228014 | 0.068307 |
| 5 | 123279200 | 123283799 | 14.63715 | 24.04365 | 92.57497 | 39.8825 | -1.56063 | 1.402094 | 0.068323 |
| X | 54357200 | 54358399 | 15.15103 | 29.28713 | 11.32566 | 4.339963 | 1.911748 | -0.11899 | 0.068325 |
| 12 | 20234400 | 20238999 | 123.8366 | 190.8491 | 136.589 | 56.73687 | 0.780412 | 2.915552 | 0.068342 |
| 5 | 97960800 | 97962199 | 5.119005 | 9.826935 | 32.58038 | 12.77099 | -1.18655 | -0.09006 | 0.068368 |
| 8 | 70754600 | 70756599 | 5.805861 | 10.55614 | 41.84156 | 18.03186 | -1.55559 | 0.295122 | 0.068369 |
| 5 | 124247600 | 124269399 | 93.96261 | 139.3412 | 417.1144 | 171.8013 | -1.32937 | 3.673472 | 0.068372 |
| 11 | 73134800 | 73135599 | 3.136974 | 6.105378 | 28.20129 | 10.37284 | -1.56898 | -0.34506 | 0.068373 |
| 2 | 158242400 | 158244999 | 10.16831 | 17.78397 | 51.49956 | 22.70965 | -1.1382 | 0.675289 | 0.068378 |
| 15 | 76692000 | 76694399 | 10.16831 | 17.78397 | 51.49956 | 22.70965 | -1.1382 | 0.675289 | 0.068378 |
| 10 | 130336000 | 130336599 | 9.747157 | 19.02929 | 10.84874 | 3.831059 | 1.46343 | -0.46102 | 0.068386 |
| 6 | 148732200 | 148733199 | 4.872313 | 9.30626 | 32.82073 | 13.04689 | -1.29379 | -0.04599 | 0.068395 |
| 16 | 20354000 | 20357999 | 15.1015 | 25.3934 | 70.18031 | 30.66052 | -1.08001 | 1.117421 | 0.0684 |
| 13 | 34827600 | 34833599 | 28.51464 | 45.8125 | 120.5017 | 51.05048 | -1.04146 | 1.893213 | 0.068417 |
| 8 | 97582600 | 97583599 | 18.27512 | 33.48854 | 18.56853 | 7.922511 | 1.27029 | 0.291066 | 0.06842 |
| 18 | 65780000 | 65781999 | 8.35481 | 14.66047 | 50.90256 | 22.50899 | -1.39789 | 0.609964 | 0.068426 |
| X | 94842600 | 94843199 | 10.21249 | 19.90969 | 12.0078 | 4.331345 | 1.357417 | -0.37795 | 0.068427 |
| 18 | 37723400 | 37728999 | 69.56979 | 111.3984 | 69.35379 | 29.23252 | 1.027355 | 2.029291 | 0.068434 |
| 2 | 152777200 | 152779399 | 8.702558 | 15.47515 | 44.63709 | 19.68311 | -1.1243 | 0.458125 | 0.068438 |
| 8 | 122462400 | 122474199 | 58.86537 | 90.65326 | 245.388 | 101.9231 | -1.13755 | 2.917247 | 0.068444 |
| 10 | 3083600 | 3085799 | 53.26719 | 86.62325 | 46.83285 | 20.09822 | 1.253918 | 1.593472 | 0.068455 |
| 18 | 65342600 | 65344599 | 6.766138 | 12.40285 | 38.82572 | 16.55799 | -1.20787 | 0.248621 | 0.068477 |
| 19 | 41045600 | 41047399 | 6.766138 | 12.40285 | 38.82572 | 16.55799 | -1.20787 | 0.248621 | 0.068477 |
| 12 | 69514000 | 69515199 | 5.016092 | 9.535544 | 32.61017 | 13.09389 | -1.26235 | -0.01044 | 0.068488 |
| 12 | 23450400 | 23450799 | 8.077944 | 15.77342 | 6.232387 | 1.971652 | 2.114395 | -0.8724 | 0.068546 |
| 4 | 139854600 | 139879799 | 170.845 | 246.1533 | 542.7753 | 223.9823 | -0.92878 | 4.133924 | 0.068563 |

| | | | | | | | | | |
|---|---|---|---|---|---|---|---|---|---|
| 4 | 136943800 | 136946199 | 11.08925 | 19.37136 | 52.60397 | 23.17466 | -1.0463 | 0.700548 | 0.068568 |
| X | 148738600 | 148739999 | 13.01145 | 24.98104 | 14.91792 | 5.845571 | 1.263237 | -0.06213 | 0.068571 |
| 1 | 43865600 | 43867799 | 13.26822 | 22.6528 | 64.3732 | 28.22255 | -1.11794 | 0.967659 | 0.068581 |
| 7 | 14822000 | 14825599 | 89.99569 | 142.166 | 97.06219 | 40.31054 | 0.875603 | 2.44144 | 0.068586 |
| 6 | 142646400 | 142649599 | 10.1236 | 17.50688 | 59.48429 | 26.09682 | -1.37017 | 0.831379 | 0.068598 |
| 17 | 8219400 | 8222599 | 11.67602 | 20.08889 | 61.13464 | 26.8167 | -1.21401 | 0.886493 | 0.068605 |
| 7 | 23028600 | 23036599 | 214.9712 | 320.1495 | 212.7772 | 87.67673 | 0.846047 | 3.607909 | 0.068635 |
| 2 | 166221200 | 166255799 | 188.2127 | 266.9836 | 652.7981 | 270.0594 | -1.10037 | 4.364011 | 0.068645 |
| 12 | 18897600 | 18900199 | 71.6792 | 114.2716 | 81.15559 | 33.99441 | 0.830969 | 2.168415 | 0.068663 |
| X | 31333400 | 31341999 | 41.78699 | 69.60677 | 33.19562 | 14.42132 | 1.445554 | 1.263322 | 0.068676 |
| 2 | 148896200 | 148898199 | 19.69769 | 36.10952 | 17.40229 | 7.420957 | 1.471571 | 0.29798 | 0.068694 |
| 17 | 73922000 | 73923599 | 7.851395 | 14.18759 | 42.80827 | 18.59122 | -1.17311 | 0.350439 | 0.0687 |
| 6 | 117652200 | 117653999 | 6.958884 | 12.65798 | 41.35621 | 17.81313 | -1.27931 | 0.29951 | 0.068703 |
| 19 | 45061800 | 45075999 | 70.10579 | 107.0027 | 279.1941 | 115.7797 | -1.09474 | 3.102057 | 0.068732 |
| X | 70878200 | 70880199 | 23.4248 | 41.99038 | 17.98393 | 7.876685 | 1.616593 | 0.452657 | 0.068736 |
| 16 | 35304800 | 35306799 | 9.889619 | 17.26388 | 53.63959 | 23.62067 | -1.23946 | 0.705448 | 0.06875 |
| 8 | 85775600 | 85777199 | 5.447679 | 10.34679 | 33.19787 | 13.35495 | -1.17412 | 0.001106 | 0.068752 |
| 7 | 106019200 | 106020199 | 14.53337 | 27.42419 | 16.97118 | 6.921852 | 1.162011 | 0.085105 | 0.068767 |
| 17 | 49279600 | 49280799 | 5.901255 | 11.14534 | 34.72118 | 14.13509 | -1.14766 | 0.048524 | 0.068796 |
| 7 | 5602400 | 5602999 | 23.51141 | 41.22201 | 25.5029 | 11.26941 | 1.076646 | 0.662914 | 0.068827 |
| 19 | 3050600 | 3060399 | 215.9799 | 321.7334 | 193.6599 | 79.80135 | 0.989207 | 3.602919 | 0.068834 |
| 6 | 132535000 | 132535799 | 13.62547 | 25.92424 | 16.14185 | 6.469585 | 1.175213 | 0.019442 | 0.068852 |
| 6 | 125289000 | 125292599 | 12.67262 | 21.27308 | 73.74759 | 32.19496 | -1.4054 | 1.129192 | 0.068854 |
| 15 | 10950400 | 10952999 | 13.8836 | 23.69387 | 63.8508 | 27.99261 | -1.04183 | 0.973503 | 0.068854 |
| 5 | 57155000 | 57156599 | 24.41506 | 43.29297 | 19.66703 | 8.693051 | 1.523301 | 0.526786 | 0.06886 |
| 1 | 98695600 | 98698999 | 45.29067 | 73.79691 | 50.04571 | 21.49243 | 0.929872 | 1.548497 | 0.068873 |
| 2 | 116949600 | 116952599 | 13.26552 | 22.63307 | 63.21652 | 27.71395 | -1.09317 | 0.973775 | 0.068874 |
| 10 | 80070800 | 80079599 | 31.4293 | 50.06457 | 148.6661 | 62.20392 | -1.23037 | 2.175452 | 0.068876 |
| 5 | 77403600 | 77414799 | 51.56878 | 80.78732 | 212.5789 | 88.17608 | -1.07752 | 2.602522 | 0.068889 |
| 8 | 36491000 | 36495799 | 25.51514 | 41.28657 | 104.9362 | 44.87574 | -0.97975 | 1.669687 | 0.068901 |
| X | 73729400 | 73733199 | 36.99216 | 62.8013 | 30.06478 | 13.17296 | 1.439711 | 1.069108 | 0.068903 |
| 11 | 114431200 | 114437199 | 46.35229 | 73.47015 | 161.2309 | 67.08425 | -0.80244 | 2.342215 | 0.068908 |
| 8 | 120736200 | 120752999 | 94.31962 | 141.3405 | 361.5374 | 149.142 | -1.08885 | 3.480974 | 0.068932 |
| 14 | 77732800 | 77733799 | 13.23781 | 25.25845 | 16.08457 | 6.407944 | 1.150278 | -0.00395 | 0.068934 |
| 1 | 65823600 | 65825599 | 18.52221 | 34.47121 | 13.97253 | 5.835082 | 1.7385 | 0.197433 | 0.068936 |
| 2 | 18051600 | 18052999 | 5.126756 | 9.538333 | 37.49721 | 15.66742 | -1.51081 | 0.146122 | 0.068936 |
| 14 | 14388600 | 14389599 | 3.153895 | 6.148213 | 25.39083 | 9.167322 | -1.38233 | -0.41613 | 0.068938 |
| 14 | 57982800 | 57983999 | 6.464748 | 11.85858 | 38.2854 | 16.31221 | -1.25069 | 0.241864 | 0.068939 |
| 14 | 41963800 | 41966199 | 38.77816 | 64.48838 | 40.64067 | 17.63644 | 1.044999 | 1.277847 | 0.068987 |
| 2 | 163042200 | 163058999 | 117.8638 | 175.8888 | 370.7704 | 152.8272 | -0.81547 | 3.562786 | 0.069008 |
| 4 | 138689000 | 138689999 | 4.654897 | 8.925471 | 31.78238 | 12.50039 | -1.29166 | -0.08276 | 0.069027 |
| 4 | 50123800 | 50124199 | 3.763264 | 7.330594 | 28.57438 | 10.41905 | -1.31655 | -0.36624 | 0.06904 |
| 13 | 63367600 | 63367799 | 15.3893 | 28.71198 | 15.15631 | 6.302826 | 1.36739 | 0.157985 | 0.069043 |
| 6 | 17693800 | 17694999 | 4.769966 | 9.044905 | 34.81305 | 14.0529 | -1.43854 | 0.008539 | 0.069045 |
| 5 | 76137600 | 76143599 | 36.65237 | 58.52688 | 138.5099 | 58.15158 | -0.8999 | 2.09912 | 0.069073 |
| 4 | 124184200 | 124187399 | 11.14835 | 19.35672 | 61.35718 | 26.92792 | -1.26985 | 0.797508 | 0.069091 |
| 13 | 66165400 | 66171999 | 112.7403 | 175.7176 | 108.7388 | 45.12909 | 1.002351 | 2.709557 | 0.069096 |

| | | | | | | | | | |
|---|---|---|---|---|---|---|---|---|---|
| 10 | 86893600 | 86895199 | 9.537161 | 16.72786 | 49.33667 | 21.80849 | -1.1641 | 0.622207 | 0.069096 |
| 8 | 84454400 | 84458799 | 16.56736 | 27.339 | 84.12071 | 36.31046 | -1.23575 | 1.343392 | 0.06911 |
| 1 | 111147400 | 111147999 | 8.504478 | 16.60778 | 10.67104 | 3.750505 | 1.300073 | -0.49666 | 0.069113 |
| X | 32668200 | 32669199 | 12.7366 | 24.81815 | 10.04664 | 3.647688 | 1.916389 | -0.32857 | 0.069136 |
| 9 | 96077800 | 96079199 | 6.170878 | 11.05038 | 44.72963 | 19.60777 | -1.60245 | 0.396707 | 0.069161 |
| 15 | 100604000 | 100614799 | 51.40346 | 80.08815 | 208.8678 | 86.68967 | -1.07078 | 2.700407 | 0.069179 |
| 17 | 17649600 | 17651999 | 23.27705 | 40.76649 | 26.06295 | 11.50624 | 1.029559 | 0.679238 | 0.069184 |
| 7 | 11977000 | 11981599 | 85.22521 | 135.0255 | 92.96353 | 38.66568 | 0.86683 | 2.368019 | 0.06919 |
| 12 | 67876400 | 67876999 | 16.40319 | 31.03101 | 14.38631 | 5.836237 | 1.58053 | 0.078724 | 0.069207 |
| 10 | 17822800 | 17824999 | 9.016518 | 15.97548 | 45.1181 | 19.94804 | -1.09691 | 0.488189 | 0.069223 |
| 18 | 80087200 | 80088799 | 4.148592 | 7.976474 | 31.33593 | 12.22828 | -1.42154 | -0.11836 | 0.069248 |
| 2 | 167411400 | 167427199 | 67.03843 | 101.9013 | 300.5965 | 124.514 | -1.27639 | 3.187145 | 0.069264 |
| 3 | 29125600 | 29126399 | 4.24655 | 8.230259 | 32.1929 | 12.18551 | -1.37401 | -0.22 | 0.069272 |
| 5 | 77606200 | 77613599 | 47.51539 | 75.22499 | 164.5043 | 68.39567 | -0.79885 | 2.373331 | 0.069286 |
| 11 | 16209200 | 16211599 | 13.84766 | 23.73232 | 63.42924 | 27.81723 | -1.02907 | 0.933364 | 0.069292 |
| 10 | 75789000 | 75790799 | 7.067655 | 12.9434 | 39.91242 | 17.0432 | -1.18806 | 0.257776 | 0.069292 |
| 16 | 92214600 | 92225399 | 54.06417 | 84.51603 | 203.5878 | 84.46521 | -0.95263 | 2.641128 | 0.069319 |
| 12 | 5200800 | 5202999 | 10.63819 | 18.58483 | 54.22402 | 23.88951 | -1.14941 | 0.698196 | 0.069321 |
| 2 | 5610800 | 5614599 | 16.57727 | 27.67754 | 74.59572 | 32.45426 | -1.04417 | 1.200778 | 0.06935 |
| 2 | 131417000 | 131418799 | 15.03665 | 28.62169 | 15.23781 | 6.101615 | 1.399653 | 0.027536 | 0.069352 |
| 7 | 61209400 | 61210399 | 15.17297 | 29.12797 | 12.9316 | 5.068748 | 1.685495 | -0.04387 | 0.069358 |
| 4 | 55281600 | 55285999 | 20.51475 | 33.73946 | 85.07468 | 36.67014 | -0.95178 | 1.396065 | 0.069392 |
| 1 | 108876200 | 108876799 | 20.28301 | 35.93396 | 21.48493 | 9.499304 | 1.129243 | 0.511174 | 0.069415 |
| 6 | 71807800 | 71808999 | 4.824892 | 9.252107 | 31.24271 | 12.28553 | -1.21617 | -0.08445 | 0.069421 |
| 17 | 30396400 | 30397799 | 6.861254 | 12.7205 | 38.6334 | 16.22397 | -1.14857 | 0.183363 | 0.069425 |
| 8 | 72081600 | 72121599 | 273.4918 | 375.5119 | 909.4826 | 378.203 | -1.1342 | 4.858753 | 0.069428 |
| 14 | 50279200 | 50280999 | 26.79629 | 46.91189 | 24.02826 | 10.60452 | 1.347521 | 0.701565 | 0.069434 |
| 12 | 19635000 | 19637999 | 70.51394 | 112.4449 | 80.46364 | 33.7174 | 0.820508 | 2.158285 | 0.069443 |
| 12 | 26996400 | 26996999 | 11.72306 | 22.75765 | 13.50509 | 5.071448 | 1.32917 | -0.21696 | 0.069445 |
| 14 | 122177600 | 122178599 | 11.72306 | 22.75765 | 13.50509 | 5.071448 | 1.32917 | -0.21696 | 0.069445 |
| 7 | 9082800 | 9084599 | 10.8894 | 21.18117 | 11.08344 | 4.096082 | 1.529196 | -0.29006 | 0.069464 |
| 5 | 113332600 | 113348199 | 63.2042 | 96.50447 | 281.2761 | 116.653 | -1.2538 | 3.092963 | 0.069469 |
| 2 | 165957400 | 165959199 | 9.527088 | 16.69821 | 49.86794 | 22.02997 | -1.18187 | 0.635958 | 0.069471 |
| 5 | 135744200 | 135768199 | 125.3007 | 183.5627 | 466.173 | 192.0243 | -1.109 | 3.877663 | 0.069479 |
| 3 | 144437000 | 144437799 | 12.8524 | 24.90802 | 12.61619 | 4.776557 | 1.54542 | -0.17922 | 0.069492 |
| 10 | 124248200 | 124250599 | 44.9638 | 73.36961 | 48.37905 | 20.78755 | 0.971742 | 1.528791 | 0.069539 |
| 10 | 127453400 | 127458199 | 18.48567 | 30.32985 | 86.65156 | 37.31212 | -1.13259 | 1.423393 | 0.069547 |
| 15 | 89846800 | 89848799 | 24.39717 | 42.94271 | 23.06967 | 10.21324 | 1.279523 | 0.609862 | 0.069557 |
| 3 | 133625000 | 133626599 | 6.219249 | 11.62283 | 37.15944 | 15.40903 | -1.20666 | 0.127448 | 0.069562 |
| 19 | 39196800 | 39199799 | 58.15457 | 93.57647 | 64.47252 | 27.37458 | 0.88974 | 1.862464 | 0.069582 |
| X | 54365200 | 54366999 | 26.79822 | 47.14763 | 20.7864 | 9.203238 | 1.562521 | 0.62527 | 0.069584 |
| 15 | 102263600 | 102264999 | 5.148667 | 9.784891 | 32.60309 | 13.09862 | -1.22583 | -0.00662 | 0.069585 |
| 2 | 152274800 | 152276199 | 3.372878 | 6.429075 | 38.47681 | 15.36189 | -2.05275 | -0.03356 | 0.069587 |
| 7 | 31610600 | 31622599 | 60.9677 | 97.95974 | 58.86248 | 24.9397 | 1.084056 | 1.908328 | 0.069611 |
| 17 | 76230200 | 76231799 | 14.93639 | 28.1052 | 16.827 | 6.897451 | 1.203224 | 0.105655 | 0.069643 |
| 18 | 14762400 | 14763199 | 2.740063 | 5.344887 | 24.22068 | 8.67231 | -1.50139 | -0.45531 | 0.069644 |
| 13 | 51635800 | 51637999 | 8.859857 | 15.72422 | 44.60835 | 19.70275 | -1.10209 | 0.47216 | 0.069646 |

| | | | | | | | | | |
|---|---|---|---|---|---|---|---|---|---|
| 7 | 13542000 | 13543199 | 20.65348 | 37.46413 | 18.31077 | 7.914255 | 1.438959 | 0.364982 | 0.069648 |
| 5 | 106022000 | 106025199 | 15.49678 | 25.8717 | 75.58833 | 32.88464 | -1.16006 | 1.198989 | 0.06965 |
| 5 | 5553000 | 5553599 | 2.736256 | 5.350381 | 22.60739 | 7.717591 | -1.33934 | -0.63314 | 0.069655 |
| 1 | 175663200 | 175663999 | 13.9282 | 26.42426 | 16.57751 | 6.684665 | 1.156786 | 0.042573 | 0.06966 |
| 17 | 19125000 | 19129799 | 93.08021 | 146.7395 | 101.389 | 42.07425 | 0.856103 | 2.488664 | 0.069686 |
| 18 | 24197600 | 24198599 | 11.86641 | 22.9276 | 13.96463 | 5.362457 | 1.261618 | -0.14191 | 0.069701 |
| 4 | 133637400 | 133638399 | 3.247683 | 6.30321 | 27.70478 | 10.41797 | -1.52712 | -0.263 | 0.06972 |
| 13 | 63905000 | 63907399 | 32.38335 | 54.71602 | 36.26075 | 15.86483 | 0.972495 | 1.097422 | 0.069752 |
| 15 | 84270400 | 84343599 | 477.5757 | 627.1618 | 1459.678 | 610.8375 | -1.14136 | 5.569498 | 0.069787 |
| 1 | 58462000 | 58465999 | 19.99507 | 33.02509 | 86.7552 | 37.47079 | -1.00871 | 1.341793 | 0.069787 |
| 8 | 127777400 | 127778999 | 5.220181 | 9.804161 | 35.02218 | 14.40148 | -1.35496 | 0.079764 | 0.069789 |
| 10 | 126913000 | 126938199 | 154.8711 | 222.1343 | 594.483 | 245.4883 | -1.21453 | 4.199591 | 0.069815 |
| 4 | 45527600 | 45531399 | 15.66672 | 26.1049 | 76.04555 | 33.04323 | -1.1558 | 1.221511 | 0.069822 |
| 4 | 67593600 | 67594799 | 19.00783 | 34.55005 | 19.61497 | 8.456163 | 1.227014 | 0.344143 | 0.069823 |
| 17 | 84920800 | 84922599 | 13.15755 | 22.29195 | 66.27365 | 29.02 | -1.18392 | 1.041751 | 0.069846 |
| 15 | 79574200 | 79575799 | 5.413763 | 10.08482 | 38.71084 | 16.14056 | -1.47495 | 0.14189 | 0.069847 |
| 7 | 9635600 | 9637799 | 33.00787 | 55.56409 | 37.32903 | 16.31381 | 0.952721 | 1.131722 | 0.069864 |
| 7 | 60390400 | 60390999 | 11.17455 | 21.71339 | 12.93646 | 4.820919 | 1.332803 | -0.25199 | 0.069871 |
| 12 | 23358600 | 23361199 | 35.96793 | 61.28231 | 28.78099 | 12.6161 | 1.467527 | 1.031254 | 0.069874 |
| 1 | 154568600 | 154569199 | 10.39087 | 20.25653 | 12.00435 | 4.332675 | 1.381838 | -0.37333 | 0.069876 |
| 18 | 85024600 | 85026599 | 19.54176 | 35.65774 | 18.51025 | 7.941536 | 1.359531 | 0.324603 | 0.069905 |
| 12 | 23623400 | 23625399 | 28.41937 | 49.16292 | 29.70273 | 13.03125 | 1.109494 | 0.868504 | 0.069906 |
| 6 | 87872400 | 87873199 | 2.844316 | 5.540192 | 26.39748 | 9.63325 | -1.60067 | -0.38127 | 0.069908 |
| 8 | 21804600 | 21806999 | 55.41109 | 89.49978 | 57.32194 | 24.46353 | 1.001133 | 1.744482 | 0.069915 |
| 17 | 30286600 | 30287599 | 3.764424 | 7.184896 | 33.20768 | 13.22117 | -1.68002 | -0.05004 | 0.069915 |
| 19 | 5774000 | 5775799 | 6.27151 | 11.30079 | 42.18539 | 18.38548 | -1.48055 | 0.353457 | 0.069922 |
| 1 | 109061600 | 109065799 | 85.57552 | 135.5779 | 86.51972 | 35.98521 | 0.976125 | 2.36648 | 0.069931 |
| 13 | 65862600 | 65864999 | 54.42631 | 88.18829 | 51.82766 | 22.18624 | 1.129378 | 1.666632 | 0.069959 |
| 7 | 22284800 | 22285199 | 8.603034 | 16.82306 | 6.582267 | 2.142498 | 2.091641 | -0.76745 | 0.069969 |
| 2 | 156265400 | 156266399 | 13.25344 | 25.23056 | 16.1496 | 6.465022 | 1.137071 | 0.011766 | 0.069972 |
| 2 | 168588800 | 168604599 | 132.4825 | 194.8564 | 437.1952 | 180.0431 | -0.92491 | 3.815377 | 0.069977 |
| 1 | 75448800 | 75456999 | 40.96409 | 64.7112 | 175.484 | 72.8798 | -1.11125 | 2.422561 | 0.069986 |
| 7 | 17545400 | 17551599 | 151.7196 | 232.0576 | 141.6696 | 58.77786 | 1.001239 | 3.073132 | 0.069998 |
| 5 | 97259000 | 97259999 | 5.32729 | 10.07506 | 33.58219 | 13.63201 | -1.24012 | 0.030769 | 0.070018 |
| 9 | 107499200 | 107512799 | 47.57494 | 73.739 | 231.6587 | 96.22357 | -1.34484 | 2.791342 | 0.070028 |
| 15 | 75611200 | 75634399 | 126.3193 | 184.936 | 471.8253 | 194.3603 | -1.11652 | 3.887869 | 0.070029 |
| 4 | 125175400 | 125189399 | 87.98521 | 133.6825 | 286.3729 | 118.6067 | -0.81587 | 3.200591 | 0.070041 |
| 4 | 129134200 | 129137199 | 11.35844 | 19.56851 | 60.03863 | 26.337 | -1.22517 | 0.871983 | 0.070071 |
| 7 | 9230200 | 9230999 | 14.67192 | 27.87353 | 15.80566 | 6.354914 | 1.304447 | 0.038126 | 0.070072 |
| 6 | 8258800 | 8259999 | 7.270241 | 12.84524 | 47.73454 | 21.11671 | -1.49146 | 0.512532 | 0.070082 |
| 13 | 66492800 | 66498999 | 114.0054 | 177.4133 | 107.6638 | 44.69333 | 1.028781 | 2.736861 | 0.070083 |
| 17 | 27051800 | 27064799 | 85.6315 | 129.7844 | 306.7567 | 126.9543 | -0.9606 | 3.251368 | 0.070098 |
| 13 | 97381000 | 97387799 | 27.0633 | 43.53183 | 118.361 | 50.22898 | -1.08698 | 1.852862 | 0.070105 |
| 11 | 73948400 | 73949199 | 13.37848 | 25.67432 | 15.12198 | 5.93308 | 1.282226 | -0.05258 | 0.070109 |
| X | 160965000 | 160966999 | 25.02567 | 44.08111 | 21.70615 | 9.60819 | 1.404926 | 0.604361 | 0.070111 |
| 1 | 99106200 | 99108599 | 45.72212 | 75.42378 | 40.10553 | 17.30895 | 1.288346 | 1.387297 | 0.070117 |
| 11 | 118332000 | 118335799 | 13.60677 | 22.78684 | 73.74479 | 32.14539 | -1.3067 | 1.158524 | 0.070138 |

| | | | | | | | | | |
|---|---|---|---|---|---|---|---|---|---|
| 15 | 101030800 | 101036199 | 20.56329 | 33.3441 | 100.4336 | 43.01699 | -1.2208 | 1.616974 | 0.070149 |
| 18 | 83376600 | 83377999 | 6.990229 | 12.55962 | 41.62108 | 18.19365 | -1.31309 | 0.376551 | 0.070159 |
| 10 | 75756000 | 75757799 | 4.424939 | 8.400012 | 32.47779 | 13.07937 | -1.44006 | -0.00476 | 0.070167 |
| 18 | 50125000 | 50127199 | 8.880268 | 15.73232 | 46.70266 | 20.64908 | -1.16877 | 0.489397 | 0.07018 |
| 11 | 78830800 | 78832999 | 11.87189 | 20.34109 | 64.36391 | 28.22656 | -1.27131 | 0.928852 | 0.070187 |
| 2 | 94124600 | 94128199 | 18.20541 | 30.21914 | 78.51808 | 34.02976 | -0.99204 | 1.274127 | 0.070202 |
| 2 | 105147400 | 105149199 | 6.735139 | 12.43619 | 43.06501 | 18.185 | -1.3427 | 0.213362 | 0.07023 |
| 3 | 6118000 | 6121799 | 67.34327 | 107.5507 | 77.01381 | 32.34019 | 0.821835 | 2.107365 | 0.070232 |
| 11 | 69701200 | 69710399 | 77.69077 | 118.7684 | 265.3861 | 110.0941 | -0.86949 | 3.070934 | 0.070293 |
| 1 | 183078000 | 183079999 | 10.89569 | 18.69014 | 60.94596 | 26.73013 | -1.31352 | 0.913676 | 0.070293 |
| 9 | 118962200 | 118988799 | 151.6614 | 219.1646 | 539.3222 | 222.4698 | -1.08278 | 4.088375 | 0.07032 |
| 14 | 56056800 | 56058399 | 5.312277 | 10.09238 | 33.20595 | 13.35047 | -1.20937 | -0.00271 | 0.070332 |
| 3 | 83748200 | 83749199 | 6.10337 | 11.30944 | 36.8668 | 15.49321 | -1.24929 | 0.179411 | 0.070343 |
| 11 | 101066800 | 101072199 | 20.76773 | 33.89705 | 91.12585 | 39.15896 | -1.04964 | 1.502221 | 0.07036 |
| 4 | 146461200 | 146466799 | 12.41071 | 23.97511 | 13.00413 | 4.997682 | 1.426511 | -0.13557 | 0.070386 |
| 5 | 114494600 | 114514399 | 78.06605 | 117.6808 | 338.4438 | 139.8314 | -1.25006 | 3.361247 | 0.070391 |
| 8 | 36684200 | 36685999 | 7.183738 | 12.73476 | 44.61561 | 19.72382 | -1.40425 | 0.47195 | 0.070404 |
| 10 | 61153000 | 61155199 | 9.688413 | 17.01255 | 49.32755 | 21.82549 | -1.13981 | 0.603204 | 0.070452 |
| 7 | 21163000 | 21170599 | 92.33353 | 146.102 | 89.73165 | 37.29752 | 1.029914 | 2.402186 | 0.070469 |
| 17 | 44732400 | 44740399 | 38.02593 | 60.51731 | 157.5857 | 65.85286 | -1.04292 | 2.211523 | 0.070483 |
| 7 | 9209600 | 9212399 | 45.40216 | 74.56579 | 41.36052 | 17.81725 | 1.225266 | 1.440578 | 0.070501 |
| 16 | 75754200 | 75755399 | 13.52218 | 25.79053 | 15.97848 | 6.371174 | 1.188252 | 0.001074 | 0.070509 |
| 16 | 17611800 | 17623199 | 64.92526 | 99.95209 | 243.5584 | 101.1585 | -0.98646 | 2.925553 | 0.070512 |
| 14 | 42114200 | 42115799 | 17.8347 | 32.54588 | 19.76935 | 8.480754 | 1.134683 | 0.313481 | 0.070519 |
| 17 | 78611200 | 78612399 | 4.440014 | 8.529921 | 33.1663 | 12.97249 | -1.41106 | -0.10434 | 0.070524 |
| 8 | 75759800 | 75761399 | 9.221456 | 16.19793 | 48.26066 | 21.35938 | -1.17843 | 0.594926 | 0.070529 |
| 3 | 146154000 | 146154999 | 3.851168 | 7.422548 | 30.23046 | 11.71525 | -1.46253 | -0.14665 | 0.07054 |
| 15 | 73436000 | 73448999 | 76.38114 | 116.0222 | 294.0312 | 121.7693 | -1.05839 | 3.203334 | 0.070547 |
| 2 | 87517600 | 87518599 | 12.38852 | 24.01458 | 12.70554 | 4.805179 | 1.482408 | -0.18913 | 0.070549 |
| 8 | 52959400 | 52960399 | 12.38852 | 24.01458 | 12.70554 | 4.805179 | 1.482408 | -0.18913 | 0.070549 |
| 15 | 79192800 | 79194199 | 31.4311 | 53.42327 | 34.15051 | 14.96691 | 1.024587 | 1.041907 | 0.070571 |
| 18 | 78044000 | 78045799 | 6.879053 | 12.69759 | 37.92414 | 16.0216 | -1.13071 | 0.207778 | 0.070585 |
| 12 | 22484800 | 22489399 | 135.9387 | 208.0803 | 137.2527 | 56.95544 | 0.890544 | 3.051391 | 0.070594 |
| 14 | 5884600 | 5886799 | 23.89914 | 41.76616 | 27.10397 | 11.94413 | 1.008257 | 0.717972 | 0.070606 |
| 10 | 128793400 | 128797799 | 24.72428 | 39.99845 | 103.9966 | 44.467 | -1.01264 | 1.672563 | 0.070623 |
| 12 | 22110000 | 22114999 | 105.0977 | 163.8101 | 119.4169 | 49.56056 | 0.766313 | 2.706076 | 0.070633 |
| 2 | 172638400 | 172651399 | 106.1717 | 158.8774 | 363.6381 | 149.9675 | -0.93048 | 3.509151 | 0.070645 |
| 7 | 66151800 | 66157999 | 34.96195 | 56.05156 | 127.5317 | 53.855 | -0.8366 | 1.970171 | 0.070649 |
| 17 | 35814400 | 35824399 | 43.25398 | 68.24725 | 178.6477 | 74.14903 | -1.06179 | 2.453003 | 0.070649 |
| X | 32448200 | 32450399 | 22.41864 | 41.06195 | 14.38042 | 6.139551 | 1.926233 | 0.317284 | 0.070697 |
| 7 | 38257400 | 38258399 | 30.33713 | 51.63661 | 32.19675 | 14.11225 | 1.060575 | 1.022212 | 0.070699 |
| 13 | 51935000 | 51952399 | 96.33459 | 144.9149 | 338.0022 | 139.5705 | -0.95087 | 3.405633 | 0.070701 |
| 14 | 61374800 | 61377399 | 8.807208 | 15.43278 | 50.92187 | 22.49102 | -1.32497 | 0.635776 | 0.070712 |
| 15 | 79620000 | 79624599 | 17.30935 | 28.58712 | 81.33873 | 35.12959 | -1.12308 | 1.334628 | 0.070716 |
| 4 | 57307800 | 57310799 | 12.41769 | 21.39745 | 59.01696 | 25.88898 | -1.07233 | 0.87375 | 0.070722 |
| 15 | 79128400 | 79153799 | 148.5634 | 215.5017 | 514.1052 | 211.9624 | -1.03264 | 4.031707 | 0.07073 |
| 5 | 80803600 | 80805399 | 14.31322 | 27.49887 | 13.58056 | 5.311243 | 1.536545 | -0.05814 | 0.07074 |

| | | | | | | | | | |
|---|---|---|---|---|---|---|---|---|---|
| 3 | 90128000 | 90128399 | 1.946971 | 3.806868 | 19.39334 | 6.283373 | -1.52989 | -0.84407 | 0.070752 |
| 5 | 78963600 | 78964199 | 9.706457 | 18.95093 | 10.20379 | 3.599705 | 1.548517 | -0.46654 | 0.070758 |
| 7 | 11736400 | 11737399 | 13.61159 | 26.39713 | 12.22659 | 4.614502 | 1.675987 | -0.18206 | 0.070795 |
| 17 | 7094800 | 7095799 | 4.077966 | 7.890159 | 28.7641 | 10.9741 | -1.28251 | -0.20038 | 0.070807 |
| 3 | 42756600 | 42757599 | 15.5545 | 29.03331 | 17.9725 | 7.468241 | 1.140509 | 0.164576 | 0.070817 |
| 3 | 96062000 | 96063599 | 4.082152 | 7.714103 | 35.13328 | 14.28562 | -1.68694 | 0.028288 | 0.070819 |
| 3 | 120964400 | 120965599 | 8.097631 | 14.43702 | 44.67919 | 19.65844 | -1.22285 | 0.437006 | 0.070824 |
| 7 | 38484800 | 38485599 | 11.23466 | 21.90966 | 9.920748 | 3.563143 | 1.768238 | -0.38635 | 0.070833 |
| 5 | 134754400 | 134758799 | 20.31623 | 33.10118 | 93.06493 | 39.96631 | -1.1159 | 1.528703 | 0.070851 |
| 5 | 119125200 | 119134599 | 51.76604 | 80.97475 | 198.7395 | 82.44835 | -0.97882 | 2.628469 | 0.070901 |
| 5 | 113928600 | 113938999 | 44.93805 | 70.40449 | 201.0645 | 83.39943 | -1.19525 | 2.597221 | 0.070902 |
| 16 | 89630800 | 89634599 | 18.2077 | 30.23078 | 80.9342 | 35.08318 | -1.03513 | 1.271897 | 0.070927 |
| 1 | 147585800 | 147586999 | 14.7532 | 27.70456 | 17.10956 | 7.037267 | 1.155599 | 0.118037 | 0.070954 |
| 3 | 87885400 | 87917599 | 187.7965 | 265.7864 | 679.6322 | 281.2668 | -1.16827 | 4.397627 | 0.070966 |
| 9 | 118628200 | 118633199 | 30.57147 | 49.15759 | 117.3412 | 49.77099 | -0.90033 | 1.868476 | 0.07097 |
| 3 | 116051000 | 116053399 | 8.823743 | 15.7709 | 44.19403 | 19.40409 | -1.07859 | 0.425362 | 0.070982 |
| 11 | 21320600 | 21321999 | 5.924148 | 10.68235 | 41.14286 | 17.91541 | -1.52348 | 0.347125 | 0.070994 |
| 14 | 42436600 | 42440999 | 87.04859 | 137.3318 | 100.2687 | 41.61483 | 0.777454 | 2.469065 | 0.070998 |
| 4 | 143735800 | 143736999 | 24.30232 | 42.665 | 22.28988 | 9.86061 | 1.318307 | 0.638703 | 0.071006 |
| 12 | 22844800 | 22853399 | 217.4322 | 323.1608 | 221.1806 | 91.1207 | 0.801027 | 3.643486 | 0.071008 |
| 11 | 6353000 | 6357799 | 19.38167 | 31.60091 | 93.64309 | 40.22499 | -1.19076 | 1.516065 | 0.071028 |
| 4 | 139469000 | 139476799 | 44.73884 | 70.88441 | 163.0386 | 67.813 | -0.87063 | 2.351903 | 0.071043 |
| 12 | 15673800 | 15674399 | 17.43364 | 33.78918 | 7.833752 | 2.9648 | 2.652926 | -0.14318 | 0.071068 |
| 11 | 43895800 | 43897999 | 21.90904 | 38.88209 | 22.08024 | 9.75277 | 1.204356 | 0.50996 | 0.07107 |
| 8 | 35457400 | 35459599 | 9.026005 | 15.99186 | 45.64267 | 20.18007 | -1.11225 | 0.488132 | 0.071076 |
| 5 | 137345400 | 137354999 | 34.39842 | 54.66804 | 164.2246 | 68.51205 | -1.25074 | 2.253634 | 0.071082 |
| 2 | 26276800 | 26279999 | 10.37551 | 18.0248 | 55.77257 | 24.48549 | -1.23444 | 0.781314 | 0.071091 |
| X | 69172400 | 69173199 | 11.99978 | 23.26666 | 13.21265 | 4.991336 | 1.385112 | -0.19441 | 0.071124 |
| 17 | 29958600 | 29969999 | 59.55459 | 92.36795 | 220.5465 | 91.5989 | -0.94884 | 2.783323 | 0.071125 |
| 8 | 22136200 | 22137999 | 13.79121 | 26.83464 | 9.770682 | 3.601051 | 2.047056 | -0.26617 | 0.071126 |
| 11 | 69124800 | 69128399 | 18.22086 | 29.84195 | 89.38908 | 38.46178 | -1.20174 | 1.446271 | 0.071133 |
| 14 | 60614800 | 60617799 | 20.19569 | 32.82102 | 96.11001 | 41.23279 | -1.17713 | 1.568795 | 0.071144 |
| 16 | 15703600 | 15706599 | 15.48971 | 25.87729 | 75.05603 | 32.66691 | -1.14959 | 1.191138 | 0.071147 |
| 13 | 93439800 | 93442399 | 13.39593 | 22.58625 | 71.23467 | 31.1466 | -1.26944 | 1.088042 | 0.071165 |
| 15 | 58744400 | 58745999 | 5.102093 | 9.557567 | 36.62491 | 15.1268 | -1.46077 | 0.098533 | 0.071192 |
| 16 | 4944600 | 4967599 | 123.3695 | 182.1865 | 433.8454 | 178.6661 | -1.00528 | 3.74737 | 0.071194 |
| 2 | 30428600 | 30434199 | 20.79417 | 33.90074 | 92.76226 | 39.84527 | -1.07632 | 1.519797 | 0.071199 |
| 8 | 27465800 | 27472599 | 30.63457 | 48.81856 | 148.8852 | 62.33081 | -1.26813 | 2.161126 | 0.071199 |
| 11 | 13845600 | 13846599 | 13.99951 | 27.07624 | 12.62921 | 4.826952 | 1.648804 | -0.13624 | 0.071211 |
| 11 | 115737600 | 115759399 | 117.5558 | 172.6297 | 460.2797 | 189.5683 | -1.17588 | 3.838988 | 0.07124 |
| 1 | 114198200 | 114200199 | 22.9549 | 41.25639 | 17.45537 | 7.628254 | 1.635238 | 0.431243 | 0.071247 |
| 5 | 25964400 | 25965599 | 6.011668 | 10.98317 | 38.71657 | 16.58481 | -1.38118 | 0.264708 | 0.071282 |
| X | 106645000 | 106646199 | 13.87428 | 26.51903 | 15.15768 | 6.01609 | 1.309994 | -0.0111 | 0.071289 |
| 1 | 8454400 | 8455999 | 25.46941 | 44.52773 | 26.38733 | 11.63215 | 1.138595 | 0.719648 | 0.071295 |
| 17 | 31508400 | 31510399 | 6.492717 | 11.82779 | 39.21251 | 16.85809 | -1.29757 | 0.28491 | 0.071314 |
| 10 | 81219200 | 81272999 | 253.7088 | 347.8123 | 945.0247 | 393.0786 | -1.30225 | 4.882435 | 0.071332 |
| 17 | 56444200 | 56457199 | 71.152 | 108.3981 | 283.6661 | 117.5804 | -1.10111 | 3.140761 | 0.071347 |

| | | | | | | | | | |
|---|---|---|---|---|---|---|---|---|---|
| 5 | 115739400 | 115744999 | 22.37092 | 36.13417 | 107.4089 | 45.84056 | -1.20769 | 1.714844 | 0.071347 |
| 14 | 92336400 | 92337599 | 12.47379 | 24.21234 | 11.45883 | 4.306595 | 1.648491 | -0.21248 | 0.07135 |
| 7 | 32216400 | 32218199 | 12.36347 | 23.75431 | 15.04021 | 5.881151 | 1.182503 | -0.07533 | 0.071355 |
| 14 | 106732600 | 106733999 | 12.36347 | 23.75431 | 15.04021 | 5.881151 | 1.182503 | -0.07533 | 0.071355 |
| 10 | 59057000 | 59059199 | 14.21532 | 23.84311 | 73.37609 | 32.01615 | -1.23457 | 1.142243 | 0.071388 |
| 14 | 104343800 | 104345399 | 14.83957 | 27.69759 | 16.02818 | 6.660815 | 1.236394 | 0.15243 | 0.071411 |
| 13 | 94837200 | 94841799 | 25.3642 | 41.01215 | 103.9338 | 44.41772 | -0.97656 | 1.68655 | 0.071446 |
| 15 | 100624600 | 100629199 | 26.06358 | 41.99264 | 112.1101 | 47.68961 | -1.05763 | 1.797475 | 0.071451 |
| 1 | 128396200 | 128398199 | 8.529016 | 15.2427 | 45.80571 | 20.11332 | -1.179 | 0.425799 | 0.071452 |
| 7 | 8468200 | 8470399 | 33.23025 | 57.903 | 19.98046 | 8.788505 | 1.913414 | 0.796833 | 0.071459 |
| 14 | 101948800 | 101949999 | 3.73158 | 7.210307 | 32.28654 | 12.39524 | -1.58578 | -0.17669 | 0.071473 |
| 5 | 112484800 | 112486399 | 4.924066 | 9.253976 | 34.44959 | 14.15059 | -1.41206 | 0.072377 | 0.071516 |
| 4 | 147224600 | 147227599 | 55.52215 | 89.46834 | 61.20356 | 26.04045 | 0.902457 | 1.811417 | 0.071517 |
| 1 | 42332000 | 42334199 | 17.29391 | 31.52052 | 19.85534 | 8.530657 | 1.081179 | 0.316026 | 0.071547 |
| 9 | 71333000 | 71334399 | 13.25035 | 25.23606 | 15.22423 | 6.08876 | 1.223072 | 0.006173 | 0.071548 |
| 12 | 92360600 | 92361399 | 12.8652 | 24.6235 | 14.36901 | 5.68102 | 1.283911 | -0.03784 | 0.07155 |
| 5 | 37102000 | 37112999 | 50.0119 | 78.16942 | 201.7076 | 83.68644 | -1.05184 | 2.642052 | 0.071566 |
| 2 | 68605800 | 68606799 | 15.69522 | 29.50615 | 16.10776 | 6.61283 | 1.333198 | 0.116145 | 0.071579 |
| 2 | 157261200 | 157285599 | 128.2427 | 188.7508 | 443.0998 | 182.4707 | -0.98921 | 3.803658 | 0.071582 |
| 19 | 45138200 | 45159999 | 144.4308 | 210.571 | 488.8095 | 201.427 | -0.98617 | 3.95406 | 0.071601 |
| 5 | 18235000 | 18235799 | 18.33617 | 33.0785 | 20.85861 | 9.077623 | 1.068129 | 0.383994 | 0.071605 |
| 3 | 4752600 | 4753399 | 11.6016 | 22.62319 | 9.913702 | 3.564886 | 1.813699 | -0.37704 | 0.071614 |
| 9 | 43861400 | 43862799 | 6.134944 | 11.34368 | 36.82875 | 15.52228 | -1.24691 | 0.190453 | 0.071616 |
| 4 | 44554000 | 44565199 | 64.42773 | 98.9049 | 257.629 | 106.9609 | -1.08573 | 2.982444 | 0.07163 |
| 3 | 95255800 | 95256999 | 12.96236 | 25.05412 | 13.85985 | 5.31332 | 1.401601 | -0.13461 | 0.071655 |
| 17 | 56165800 | 56166799 | 26.92381 | 46.75497 | 29.21302 | 12.82207 | 1.062186 | 0.824611 | 0.071701 |
| 5 | 14075200 | 14076399 | 9.519996 | 16.74509 | 47.72825 | 21.13282 | -1.11463 | 0.576366 | 0.071705 |
| 11 | 60667600 | 60671199 | 17.20125 | 28.37542 | 84.8335 | 36.61289 | -1.1947 | 1.349215 | 0.071711 |
| 6 | 54139400 | 54140399 | 4.678624 | 9.024395 | 30.40998 | 11.75842 | -1.19038 | -0.14515 | 0.071711 |
| 13 | 86024600 | 86025399 | 12.1819 | 23.64486 | 13.12182 | 4.932494 | 1.42445 | -0.20858 | 0.071713 |
| 3 | 98289200 | 98290999 | 11.88232 | 20.67725 | 54.78394 | 24.0769 | -1.01178 | 0.762465 | 0.071751 |
| 11 | 121107400 | 121111999 | 15.73171 | 26.4391 | 70.82446 | 30.93697 | -1.03531 | 1.124113 | 0.071783 |
| 12 | 18792400 | 18796999 | 129.1339 | 198.3739 | 136.8072 | 56.80416 | 0.830334 | 2.979735 | 0.071797 |
| 9 | 110685000 | 110688799 | 16.29852 | 27.14651 | 76.72297 | 33.32894 | -1.11242 | 1.227788 | 0.071801 |
| 6 | 82238800 | 82240199 | 6.454771 | 11.86962 | 37.8019 | 16.05206 | -1.22761 | 0.226388 | 0.071807 |
| 1 | 130782000 | 130783199 | 4.815597 | 9.00715 | 35.97218 | 14.89559 | -1.52216 | 0.105597 | 0.071818 |
| 1 | 160477200 | 160478199 | 4.302214 | 8.169256 | 32.48343 | 13.07413 | -1.47942 | -0.00857 | 0.071834 |
| 1 | 147683400 | 147684399 | 16.66844 | 31.21131 | 16.64306 | 6.87879 | 1.359668 | 0.151294 | 0.071838 |
| 6 | 143560400 | 143562199 | 10.65273 | 18.5753 | 54.21018 | 23.85523 | -1.15002 | 0.725737 | 0.071852 |
| 3 | 40348400 | 40350399 | 42.80306 | 70.20502 | 47.28316 | 20.35729 | 0.945497 | 1.454311 | 0.071856 |
| 11 | 87651200 | 87657799 | 35.20976 | 55.8474 | 161.6727 | 67.31466 | -1.20019 | 2.313679 | 0.07186 |
| 12 | 22209400 | 22217599 | 151.2573 | 230.5895 | 157.4827 | 65.2889 | 0.835874 | 3.13939 | 0.071884 |
| 4 | 155169000 | 155172199 | 12.20471 | 20.84823 | 65.97209 | 28.92478 | -1.2722 | 0.959847 | 0.071908 |
| 12 | 19619000 | 19627199 | 234.2394 | 345.4618 | 240.5557 | 99.06421 | 0.765843 | 3.776069 | 0.071918 |
| 1 | 187076600 | 187081399 | 32.42023 | 51.8924 | 129.0003 | 54.3421 | -0.96677 | 2.019389 | 0.071923 |
| 3 | 6308400 | 6309199 | 10.15352 | 19.75075 | 13.21517 | 4.881998 | 1.179376 | -0.29308 | 0.071923 |
| 5 | 70373800 | 70374799 | 10.15352 | 19.75075 | 13.21517 | 4.881998 | 1.179376 | -0.29308 | 0.071923 |

| | | | | | | | | | |
|---|---|---|---|---|---|---|---|---|---|
| 8 | 106362600 | 106364399 | 6.499954 | 11.5683 | 46.22494 | 20.36859 | -1.58958 | 0.441235 | 0.071924 |
| 6 | 114373400 | 114379399 | 39.60236 | 63.09063 | 143.7396 | 60.1518 | -0.84885 | 2.177818 | 0.071928 |
| 7 | 112805200 | 112807999 | 10.81799 | 18.78904 | 55.23866 | 24.24749 | -1.16105 | 0.787149 | 0.071937 |
| 4 | 136512600 | 136516999 | 17.31091 | 28.62203 | 80.91691 | 34.9747 | -1.11375 | 1.320179 | 0.071941 |
| 7 | 7696200 | 7697199 | 16.55123 | 31.73179 | 12.42749 | 4.889714 | 1.860111 | -0.01896 | 0.071947 |
| 17 | 90786400 | 90786999 | 9.520871 | 18.54864 | 11.7326 | 4.270185 | 1.278589 | -0.35809 | 0.071948 |
| 7 | 11423200 | 11429399 | 195.1958 | 291.9537 | 206.6873 | 85.23163 | 0.760622 | 3.525733 | 0.071955 |
| 8 | 88499400 | 88501799 | 9.652856 | 17.00517 | 47.73363 | 21.12864 | -1.09169 | 0.56092 | 0.071966 |
| Y | 8956400 | 8957999 | 12.13525 | 23.53082 | 6.939982 | 2.620932 | 2.312893 | -0.21863 | 0.071976 |
| 3 | 36260200 | 36260999 | 2.464673 | 4.790897 | 29.6618 | 11.021 | -1.99669 | -0.31178 | 0.071992 |
| 10 | 79725800 | 79759399 | 161.4288 | 229.9238 | 632.645 | 261.5432 | -1.2647 | 4.303126 | 0.071993 |
| 5 | 71014000 | 71015599 | 17.30182 | 32.15911 | 16.4039 | 6.864811 | 1.408607 | 0.200612 | 0.071997 |
| 15 | 79381600 | 79396599 | 78.8556 | 119.441 | 307.1025 | 127.0726 | -1.08265 | 3.262838 | 0.072001 |
| 7 | 19035600 | 19046399 | 57.94245 | 89.82766 | 221.817 | 92.13267 | -0.99779 | 2.790596 | 0.072003 |
| 18 | 24993400 | 24996199 | 12.31857 | 21.12116 | 63.31506 | 27.76816 | -1.19356 | 0.922999 | 0.072011 |
| 4 | 152196400 | 152221999 | 142.7645 | 206.4012 | 548.2062 | 226.1209 | -1.19221 | 4.081087 | 0.07202 |
| 8 | 123177800 | 123197799 | 91.32427 | 136.4226 | 385.543 | 158.9477 | -1.23671 | 3.540141 | 0.072027 |
| 12 | 111234600 | 111242599 | 58.94749 | 90.9561 | 237.2629 | 98.55453 | -1.08206 | 2.879782 | 0.07204 |
| 7 | 7989000 | 7992599 | 85.63365 | 135.2912 | 97.32983 | 40.42411 | 0.800362 | 2.435165 | 0.072055 |
| 6 | 88170200 | 88184399 | 82.99681 | 125.2526 | 333.9535 | 138.023 | -1.13957 | 3.339258 | 0.072065 |
| 5 | 35122200 | 35124399 | 8.525185 | 14.80463 | 56.16675 | 24.65258 | -1.52595 | 0.781373 | 0.072075 |
| 18 | 38286400 | 38288399 | 7.689563 | 13.97512 | 40.26142 | 17.36012 | -1.10016 | 0.308844 | 0.072083 |
| 19 | 56743400 | 56745599 | 10.52611 | 18.2738 | 55.76542 | 24.47172 | -1.21466 | 0.792523 | 0.072113 |
| 10 | 79540000 | 79549999 | 44.07434 | 69.11089 | 194.3559 | 80.60963 | -1.17205 | 2.581064 | 0.072116 |
| 19 | 25104800 | 25106199 | 4.699476 | 8.998029 | 31.1499 | 12.3022 | -1.25756 | -0.07195 | 0.07213 |
| 18 | 32452800 | 32455399 | 31.08021 | 53.26532 | 31.11602 | 13.64609 | 1.155356 | 0.962258 | 0.072136 |
| 4 | 140223200 | 140227199 | 32.78099 | 52.39371 | 134.4605 | 56.52589 | -1.01497 | 2.065821 | 0.072158 |
| 2 | 7131600 | 7133599 | 7.199241 | 13.21918 | 43.40746 | 18.46928 | -1.27478 | 0.249904 | 0.07216 |
| 12 | 20447600 | 20455599 | 193.3991 | 289.9678 | 198.676 | 81.9666 | 0.810927 | 3.480642 | 0.072164 |
| 8 | 110673600 | 110675999 | 13.33391 | 22.99599 | 59.65416 | 26.16918 | -0.98423 | 0.87003 | 0.072177 |
| 2 | 25238000 | 25240799 | 7.010004 | 12.29423 | 50.86502 | 22.4824 | -1.64814 | 0.612807 | 0.072204 |
| 7 | 5375800 | 5378399 | 54.66222 | 88.62328 | 50.20382 | 21.50054 | 1.182975 | 1.653585 | 0.072205 |
| 11 | 28184200 | 28185799 | 21.63451 | 38.77996 | 19.78393 | 8.665286 | 1.365567 | 0.442642 | 0.072215 |
| 9 | 85670600 | 85673999 | 20.17796 | 33.24143 | 83.42566 | 35.98495 | -0.944 | 1.373183 | 0.072215 |
| 8 | 121850800 | 121857999 | 34.60615 | 55.16444 | 142.976 | 59.88225 | -1.03346 | 2.153569 | 0.072228 |
| 5 | 147255200 | 147259399 | 30.05789 | 48.41743 | 113.2666 | 48.16617 | -0.86807 | 1.810202 | 0.072243 |
| 12 | 113090400 | 113132599 | 215.7115 | 299.7897 | 817.8997 | 339.4671 | -1.28821 | 4.670929 | 0.072254 |
| 12 | 31573600 | 31574999 | 19.42736 | 35.2185 | 19.46925 | 8.421406 | 1.26193 | 0.361361 | 0.072258 |
| 9 | 69005800 | 69008599 | 11.12786 | 19.3486 | 55.28909 | 24.28648 | -1.12032 | 0.770982 | 0.072264 |
| 15 | 42514000 | 42515599 | 12.67833 | 21.52795 | 65.19161 | 28.5655 | -1.21011 | 1.01916 | 0.072288 |
| 2 | 75265000 | 75265599 | 5.008611 | 9.54456 | 32.13402 | 12.83636 | -1.23337 | -0.02898 | 0.072327 |
| 9 | 56766400 | 56769199 | 10.35504 | 18.00158 | 54.68807 | 24.01973 | -1.20782 | 0.77088 | 0.072332 |
| 10 | 24993200 | 24995999 | 10.82355 | 18.83995 | 57.48881 | 25.27 | -1.21455 | 0.755409 | 0.072342 |
| 4 | 128745400 | 128753399 | 47.81675 | 75.75958 | 163.6159 | 68.05241 | -0.77998 | 2.354934 | 0.072351 |
| 6 | 68170200 | 68171799 | 38.19984 | 63.7055 | 39.46495 | 17.15752 | 1.069564 | 1.244317 | 0.072357 |
| 1 | 153109200 | 153110399 | 5.42669 | 10.37525 | 33.5468 | 13.3045 | -1.16716 | -0.04638 | 0.072365 |
| 9 | 57270400 | 57271799 | 5.42669 | 10.37525 | 33.5468 | 13.3045 | -1.16716 | -0.04638 | 0.072365 |

| | | | | | | | | | |
|---|---|---|---|---|---|---|---|---|---|
| 8 | 71214800 | 71231199 | 70.57568 | 107.0032 | 307.5722 | 127.3057 | -1.24211 | 3.241441 | 0.07239 |
| 5 | 147309200 | 147309799 | 3.344434 | 6.527684 | 25.9106 | 9.189295 | -1.30458 | -0.47986 | 0.072403 |
| 6 | 53853800 | 53854799 | 3.344434 | 6.527684 | 25.9106 | 9.189295 | -1.30458 | -0.47986 | 0.072403 |
| 16 | 32427400 | 32428399 | 4.834831 | 9.237382 | 31.62213 | 12.56262 | -1.24949 | -0.04895 | 0.072429 |
| 2 | 143829400 | 143831999 | 6.978345 | 12.64146 | 41.24822 | 17.85794 | -1.28217 | 0.324134 | 0.07243 |
| 3 | 71436800 | 71437999 | 26.35141 | 46.14515 | 23.92733 | 10.56232 | 1.329739 | 0.694121 | 0.072436 |
| 1 | 35757200 | 35758399 | 11.97001 | 23.04928 | 15.18383 | 5.899709 | 1.134721 | -0.09861 | 0.072442 |
| 7 | 33541600 | 33544599 | 77.94422 | 123.9391 | 86.62178 | 36.15643 | 0.850159 | 2.256927 | 0.072485 |
| 6 | 38410200 | 38411799 | 8.345234 | 14.66326 | 49.8305 | 22.05983 | -1.36703 | 0.586078 | 0.07249 |
| 9 | 48564000 | 48574999 | 66.57051 | 102.8299 | 231.3979 | 96.11779 | -0.86803 | 2.86163 | 0.072516 |
| 15 | 31659800 | 31660799 | 4.469753 | 8.50309 | 33.16666 | 13.29704 | -1.44799 | -0.02028 | 0.07252 |
| 1 | 46463800 | 46465999 | 10.19034 | 17.7644 | 53.09924 | 23.36254 | -1.18407 | 0.727135 | 0.072522 |
| 7 | 32289400 | 32291999 | 54.94591 | 88.64008 | 59.34425 | 25.28672 | 0.93539 | 1.779868 | 0.072525 |
| 4 | 141340400 | 141370199 | 137.2057 | 198.1246 | 551.3002 | 227.4335 | -1.26104 | 4.098905 | 0.072545 |
| 2 | 69552400 | 69553999 | 5.024873 | 9.294926 | 40.67298 | 17.133 | -1.67321 | 0.185166 | 0.072553 |
| 13 | 12948400 | 12949399 | 13.78719 | 26.76787 | 10.80917 | 4.055185 | 1.877176 | -0.20413 | 0.072554 |
| 2 | 155462800 | 155478799 | 92.06511 | 137.5215 | 379.0125 | 156.2536 | -1.20058 | 3.541265 | 0.072562 |
| 16 | 33348200 | 33356999 | 62.12562 | 96.83362 | 206.9562 | 85.89095 | -0.7836 | 2.697555 | 0.072585 |
| 17 | 8056800 | 8057799 | 11.26457 | 21.84782 | 14.188 | 5.352127 | 1.194904 | -0.20519 | 0.072612 |
| 2 | 23850400 | 23851399 | 15.87935 | 29.4897 | 18.47471 | 7.7405 | 1.114219 | 0.199242 | 0.072619 |
| 15 | 98788800 | 98807599 | 97.33432 | 146.1532 | 340.6662 | 140.6022 | -0.95225 | 3.441858 | 0.072631 |
| 16 | 51002400 | 51002999 | 9.521082 | 18.54837 | 12.20174 | 4.443067 | 1.221895 | -0.35473 | 0.072642 |
| 15 | 97567600 | 97570799 | 13.22795 | 22.31207 | 69.51904 | 30.40019 | -1.2518 | 1.083036 | 0.072656 |
| 12 | 13318200 | 13320199 | 9.52189 | 16.76842 | 50.94782 | 22.55298 | -1.20601 | 0.566059 | 0.072661 |
| 3 | 35142400 | 35145199 | 13.55571 | 23.18499 | 62.22017 | 27.28254 | -1.03525 | 0.950485 | 0.072662 |
| 7 | 8568800 | 8570199 | 18.4377 | 34.29283 | 15.81045 | 6.609497 | 1.554522 | 0.20557 | 0.072705 |
| 19 | 25213000 | 25215799 | 17.66812 | 29.14097 | 84.31809 | 36.38801 | -1.14778 | 1.352051 | 0.07271 |
| 7 | 136741400 | 136742399 | 5.74566 | 10.90363 | 34.37839 | 13.85431 | -1.152 | 0.011574 | 0.072737 |
| 7 | 14698800 | 14709399 | 86.01511 | 136.1965 | 87.85352 | 36.52939 | 0.960016 | 2.381194 | 0.072741 |
| 19 | 31033200 | 31036999 | 24.47442 | 39.80623 | 96.89413 | 41.58414 | -0.91071 | 1.563059 | 0.072745 |
| 16 | 52732400 | 52735599 | 22.38364 | 39.61984 | 21.85116 | 9.662968 | 1.24429 | 0.534591 | 0.072771 |
| X | 30959400 | 30960799 | 24.76387 | 43.62035 | 22.50658 | 9.96248 | 1.33778 | 0.603447 | 0.072784 |
| 17 | 91521200 | 91523199 | 21.96378 | 38.62834 | 23.72899 | 10.50682 | 1.086552 | 0.603008 | 0.072794 |
| 5 | 140488200 | 140494999 | 34.36024 | 54.64908 | 154.911 | 64.67885 | -1.166 | 2.229223 | 0.072796 |
| 15 | 102934600 | 102936799 | 10.86535 | 18.81661 | 58.48356 | 25.65891 | -1.24203 | 0.819731 | 0.072805 |
| 16 | 38533000 | 38533999 | 4.418628 | 8.408254 | 31.99638 | 12.82004 | -1.41079 | -0.02318 | 0.072805 |
| 11 | 53520800 | 53523399 | 9.391051 | 16.20284 | 60.45627 | 26.52139 | -1.50494 | 0.851188 | 0.072824 |
| 12 | 23425000 | 23425599 | 17.88005 | 32.77559 | 19.25658 | 8.212322 | 1.188136 | 0.288236 | 0.072844 |
| 11 | 112671800 | 112676399 | 22.15672 | 35.64676 | 120.5201 | 51.16035 | -1.39994 | 1.841072 | 0.072848 |
| 14 | 52597200 | 52599199 | 24.34137 | 42.94488 | 21.1434 | 9.356074 | 1.405244 | 0.579923 | 0.072865 |
| 4 | 129635200 | 129637799 | 13.97864 | 23.62535 | 68.57401 | 30.00508 | -1.14999 | 1.066446 | 0.072866 |
| 5 | 43977600 | 43982399 | 24.95715 | 40.56812 | 96.82575 | 41.54613 | -0.883 | 1.572871 | 0.072883 |
| 3 | 67062000 | 67062399 | 9.065858 | 17.70813 | 11.16437 | 3.908386 | 1.332899 | -0.4982 | 0.072884 |
| 10 | 68163200 | 68164599 | 5.728879 | 10.86773 | 34.91202 | 14.08047 | -1.17935 | 0.017002 | 0.072888 |
| 1 | 92665200 | 92670999 | 24.66754 | 40.02077 | 98.04996 | 42.0127 | -0.92318 | 1.611801 | 0.072888 |
| 15 | 78662000 | 78666799 | 14.09955 | 23.18773 | 87.17971 | 37.57679 | -1.52539 | 1.384138 | 0.072903 |
| 2 | 172691400 | 172726999 | 209.1753 | 293.2469 | 731.6495 | 303.2585 | -1.14664 | 4.53933 | 0.072919 |

| | | | | | | | | | |
|---|---|---|---|---|---|---|---|---|---|
| 15 | 80343000 | 80343999 | 3.236188 | 6.314356 | 25.75266 | 9.184581 | -1.35055 | -0.46243 | 0.072929 |
| 2 | 180953400 | 180956399 | 15.11197 | 25.40772 | 71.34206 | 31.16684 | -1.1029 | 1.118755 | 0.07294 |
| 17 | 30410200 | 30411999 | 6.591237 | 11.82375 | 43.17516 | 18.89852 | -1.45361 | 0.385537 | 0.072963 |
| 17 | 13596000 | 13630999 | 1225.131 | 1507.072 | 2792.063 | 1176.489 | -0.90728 | 6.628183 | 0.072991 |
| 13 | 105358400 | 105359599 | 5.277942 | 10.05004 | 32.69131 | 13.08032 | -1.18629 | -0.01847 | 0.073003 |
| 6 | 20898400 | 20900599 | 28.94547 | 49.88687 | 29.4971 | 12.93957 | 1.139177 | 0.900668 | 0.073005 |
| 6 | 72735200 | 72739799 | 27.62673 | 44.59126 | 108.7077 | 46.35165 | -0.92389 | 1.744026 | 0.073061 |
| 11 | 104071400 | 104071999 | 2.840963 | 5.547903 | 23.96028 | 8.421435 | -1.40876 | -0.5212 | 0.073068 |
| 3 | 52174200 | 52177799 | 18.11288 | 29.56854 | 92.96389 | 39.95114 | -1.2745 | 1.495074 | 0.073069 |
| 18 | 73705000 | 73707599 | 14.48935 | 24.05343 | 80.77292 | 35.00959 | -1.36036 | 1.26463 | 0.073082 |
| X | 28806600 | 28807199 | 14.68688 | 28.31301 | 13.16458 | 5.098463 | 1.636176 | -0.08498 | 0.073094 |
| 15 | 76087200 | 76097199 | 40.21358 | 63.40634 | 180.9473 | 75.09085 | -1.1869 | 2.466542 | 0.073095 |
| 2 | 93461600 | 93483999 | 135.881 | 198.6934 | 490.5562 | 202.0916 | -1.0709 | 3.907815 | 0.073097 |
| 2 | 180562000 | 180562999 | 3.967487 | 7.646436 | 29.64311 | 11.48893 | -1.392 | -0.14556 | 0.073099 |
| 7 | 20036800 | 20036999 | 6.829237 | 13.34599 | 7.594747 | 2.436087 | 1.588638 | -0.8453 | 0.073108 |
| 11 | 74543400 | 74544799 | 4.968635 | 9.488278 | 32.18798 | 12.80095 | -1.23797 | -0.04222 | 0.073121 |
| 2 | 152787200 | 152794599 | 30.74627 | 49.03787 | 142.4772 | 59.71768 | -1.19696 | 2.132788 | 0.073131 |
| 1 | 87047600 | 87049399 | 7.523712 | 13.7072 | 39.77303 | 17.09713 | -1.10726 | 0.293123 | 0.073165 |
| 7 | 23265000 | 23267599 | 75.1691 | 119.6761 | 84.184 | 35.1982 | 0.842749 | 2.214771 | 0.073166 |
| 19 | 5581800 | 5586599 | 14.56295 | 24.3407 | 75.5178 | 32.87784 | -1.24628 | 1.183483 | 0.07318 |
| 13 | 26921200 | 26921599 | 8.602222 | 16.8237 | 9.8918 | 3.35824 | 1.469466 | -0.61365 | 0.073182 |
| 12 | 62040800 | 62042999 | 40.09557 | 66.22924 | 44.5215 | 19.23046 | 0.951598 | 1.360547 | 0.073221 |
| 10 | 44432400 | 44433399 | 18.32925 | 33.6649 | 18.35173 | 7.805598 | 1.298348 | 0.279756 | 0.073228 |
| 7 | 98502200 | 98520399 | 81.29994 | 122.1772 | 347.5518 | 143.4757 | -1.23827 | 3.423261 | 0.073234 |
| 7 | 15240600 | 15245399 | 139.3445 | 212.9901 | 145.2185 | 60.24116 | 0.841258 | 3.082354 | 0.073243 |
| 1 | 100247400 | 100249799 | 26.64731 | 46.16472 | 30.25614 | 13.27464 | 0.992945 | 0.847342 | 0.073259 |
| 12 | 22622600 | 22624399 | 39.8538 | 66.23587 | 34.97323 | 15.17047 | 1.299196 | 1.282951 | 0.073281 |
| 1 | 8282800 | 8283799 | 15.3878 | 28.99928 | 16.8496 | 6.888705 | 1.249361 | 0.10101 | 0.073291 |
| 4 | 137198000 | 137204999 | 27.05456 | 43.53727 | 116.786 | 49.59281 | -1.0667 | 1.837933 | 0.073298 |
| 11 | 96159000 | 96163799 | 19.2203 | 31.36386 | 93.19038 | 40.04266 | -1.19376 | 1.502961 | 0.073311 |
| 8 | 122440200 | 122446999 | 26.49346 | 42.48847 | 127.285 | 53.77723 | -1.23178 | 1.950189 | 0.073317 |
| 12 | 57584800 | 57590799 | 44.35402 | 70.45797 | 153.7395 | 64.10858 | -0.79106 | 2.273923 | 0.073337 |
| 2 | 99093200 | 99094399 | 14.35361 | 27.41149 | 15.29537 | 6.082249 | 1.341036 | -0.00089 | 0.07334 |
| 10 | 48468400 | 48469399 | 16.11859 | 30.49805 | 15.00599 | 6.085337 | 1.496301 | 0.076594 | 0.073341 |
| 18 | 83420400 | 83422199 | 6.533108 | 12.20916 | 36.64707 | 15.19703 | -1.11707 | 0.126932 | 0.073353 |
| X | 24523400 | 24524799 | 29.85232 | 51.49013 | 29.28617 | 12.84735 | 1.195498 | 0.900922 | 0.073356 |
| X | 148433800 | 148435799 | 43.97185 | 71.92543 | 48.9212 | 21.03911 | 0.929519 | 1.49263 | 0.073387 |
| 8 | 120380400 | 120398199 | 90.61953 | 135.9698 | 356.742 | 147.2079 | -1.12377 | 3.45554 | 0.07339 |
| 8 | 122004000 | 122019199 | 74.70311 | 113.7228 | 290.3844 | 120.3381 | -1.06665 | 3.157096 | 0.073391 |
| 1 | 170395200 | 170395399 | 6.076892 | 11.85847 | 7.218267 | 2.26736 | 1.514055 | -0.93546 | 0.073391 |
| 9 | 49186000 | 49186599 | 10.27998 | 20.06739 | 10.83867 | 3.834489 | 1.538596 | -0.44666 | 0.073403 |
| 17 | 29073400 | 29078199 | 71.02404 | 108.9644 | 249.4225 | 103.5451 | -0.90017 | 2.997052 | 0.073405 |
| 6 | 4600200 | 4601799 | 8.225256 | 14.66747 | 43.10226 | 18.96143 | -1.1481 | 0.435743 | 0.073405 |
| 11 | 117087600 | 117097599 | 46.46726 | 72.6116 | 205.0309 | 85.06734 | -1.18187 | 2.64508 | 0.073418 |
| 8 | 56013800 | 56015199 | 17.55835 | 31.98683 | 19.94936 | 8.576297 | 1.09512 | 0.321708 | 0.073437 |
| 3 | 94839800 | 94848199 | 30.38935 | 48.57271 | 136.1593 | 57.24264 | -1.14195 | 2.062782 | 0.073448 |
| 13 | 70886400 | 70887199 | 2.739397 | 5.345844 | 23.71921 | 8.437226 | -1.46266 | -0.48079 | 0.07345 |

| | | | | | | | | | |
|---|---|---|---|---|---|---|---|---|---|
| 17 | 87940400 | 87948799 | 44.07922 | 69.8738 | 164.2001 | 68.32612 | -0.90088 | 2.337233 | 0.073457 |
| 17 | 70018600 | 70018999 | 9.355503 | 18.25862 | 11.71201 | 4.159615 | 1.290192 | -0.44315 | 0.07346 |
| 19 | 48720000 | 48720599 | 3.041705 | 5.943656 | 25.70975 | 8.929781 | -1.39841 | -0.55899 | 0.073478 |
| 9 | 71651000 | 71655599 | 25.81699 | 41.80677 | 102.1472 | 43.70749 | -0.92192 | 1.653644 | 0.073492 |
| 11 | 53838400 | 53839399 | 4.367295 | 8.475683 | 30.44653 | 11.45218 | -1.24333 | -0.24368 | 0.073501 |
| 9 | 3151600 | 3160199 | 219.0721 | 324.7432 | 226.3016 | 93.20548 | 0.771586 | 3.68969 | 0.073501 |
| 4 | 153476200 | 153533799 | 453.3232 | 598.633 | 1372.456 | 573.9746 | -1.11163 | 5.47948 | 0.073532 |
| 2 | 179392400 | 179393799 | 5.373903 | 10.02252 | 35.36251 | 14.71469 | -1.35095 | 0.128982 | 0.073572 |
| 4 | 148134400 | 148157599 | 105.2868 | 155.1182 | 457.4597 | 188.3862 | -1.31673 | 3.785066 | 0.073573 |
| 7 | 38317200 | 38319799 | 29.08277 | 49.82506 | 32.077 | 14.06714 | 1.015916 | 0.959585 | 0.073587 |
| 6 | 84153400 | 84153999 | 11.09064 | 21.48806 | 14.25845 | 5.404976 | 1.157693 | -0.19113 | 0.073602 |
| 5 | 114008400 | 114024399 | 88.01941 | 132.4528 | 336.8425 | 139.1046 | -1.07506 | 3.397446 | 0.073603 |
| 1 | 156802000 | 156803399 | 8.365648 | 14.93961 | 43.11957 | 18.94572 | -1.12112 | 0.427533 | 0.073616 |
| 1 | 166444600 | 166446199 | 19.74462 | 35.27228 | 21.88394 | 9.612577 | 1.082842 | 0.45511 | 0.073618 |
| 3 | 97973600 | 97976999 | 12.20701 | 21.19227 | 58.08418 | 25.48954 | -1.06111 | 0.800756 | 0.073627 |
| 11 | 119087400 | 119109799 | 108.6843 | 160.7037 | 427.6617 | 176.1219 | -1.16372 | 3.724738 | 0.073636 |
| 13 | 68540200 | 68545399 | 134.9764 | 207.0453 | 146.1915 | 60.68844 | 0.794788 | 3.010104 | 0.073639 |
| 1 | 120541200 | 120546999 | 42.90666 | 68.12814 | 155.4343 | 64.78615 | -0.8559 | 2.286001 | 0.073644 |
| 12 | 80155800 | 80164199 | 31.87276 | 50.88392 | 144.3382 | 60.58027 | -1.16101 | 2.102134 | 0.073646 |
| 2 | 25787600 | 25808199 | 102.7672 | 152.8467 | 393.117 | 161.9657 | -1.10669 | 3.621742 | 0.073646 |
| 12 | 71713200 | 71715399 | 24.52596 | 42.75976 | 28.14885 | 12.38574 | 0.987495 | 0.755685 | 0.073661 |
| 2 | 81958600 | 81959999 | 15.54258 | 29.43281 | 15.02179 | 6.080965 | 1.445949 | 0.06532 | 0.073671 |
| 7 | 20253800 | 20254799 | 12.64366 | 24.60963 | 11.11218 | 4.084898 | 1.745381 | -0.28471 | 0.073672 |
| 17 | 31998600 | 32016199 | 85.75312 | 129.1338 | 340.4172 | 140.6075 | -1.12595 | 3.383045 | 0.073682 |
| 10 | 79365200 | 79399599 | 244.7764 | 341.9245 | 742.603 | 307.9709 | -0.95184 | 4.594235 | 0.073685 |
| 5 | 122107800 | 122158199 | 375.7877 | 500.932 | 1247.862 | 521.2402 | -1.21779 | 5.327932 | 0.073691 |
| 10 | 126817800 | 126829799 | 48.36458 | 75.6605 | 199.7401 | 82.86271 | -1.08371 | 2.625483 | 0.073697 |
| 3 | 41481400 | 41482999 | 10.31694 | 18.04353 | 50.96778 | 22.47473 | -1.10248 | 0.67608 | 0.073697 |
| 2 | 34493000 | 34494599 | 6.483255 | 11.83817 | 38.72356 | 16.59932 | -1.27558 | 0.269915 | 0.07374 |
| 3 | 24299800 | 24300399 | 9.467993 | 18.47019 | 12.17303 | 4.353941 | 1.242209 | -0.41626 | 0.073771 |
| 3 | 72558000 | 72558599 | 9.467993 | 18.47019 | 12.17303 | 4.353941 | 1.242209 | -0.41626 | 0.073771 |
| 17 | 7990000 | 7993399 | 19.01678 | 31.48197 | 79.5656 | 34.42039 | -0.95259 | 1.308964 | 0.073795 |
| 8 | 41960400 | 41961599 | 11.09553 | 21.56195 | 12.31385 | 4.585285 | 1.393483 | -0.25819 | 0.073798 |
| X | 29634200 | 29635399 | 19.67636 | 35.65919 | 19.33694 | 8.367318 | 1.289766 | 0.366081 | 0.073799 |
| 17 | 35282000 | 35288799 | 28.27347 | 45.6085 | 109.171 | 46.51414 | -0.89854 | 1.762652 | 0.073804 |
| 6 | 144766200 | 144772999 | 42.09553 | 66.85479 | 153.0953 | 63.82513 | -0.8609 | 2.279014 | 0.073817 |
| 4 | 144547800 | 144550799 | 41.76401 | 68.59689 | 47.27347 | 20.3652 | 0.913214 | 1.434224 | 0.073817 |
| 4 | 140535600 | 140567199 | 278.2722 | 380.4777 | 938.1272 | 390.3727 | -1.1661 | 4.924235 | 0.073822 |
| 12 | 111382600 | 111399399 | 87.88659 | 132.0822 | 341.8915 | 141.1408 | -1.1022 | 3.42315 | 0.073825 |
| 14 | 61681200 | 61683599 | 17.37182 | 31.65679 | 19.95256 | 8.574363 | 1.080343 | 0.318409 | 0.073827 |
| 5 | 104011800 | 104015799 | 15.40782 | 25.92579 | 69.69251 | 30.45357 | -1.04018 | 1.111727 | 0.073855 |
| 5 | 66774400 | 66776799 | 9.354194 | 16.21965 | 58.39026 | 25.61894 | -1.45198 | 0.80219 | 0.073862 |
| 16 | 40440800 | 40441999 | 12.78276 | 24.67993 | 14.43684 | 5.56292 | 1.315133 | -0.1183 | 0.073865 |
| 13 | 66987600 | 66990799 | 59.33414 | 95.75207 | 55.84459 | 23.80602 | 1.134428 | 1.77489 | 0.07387 |
| 6 | 38157600 | 38162199 | 24.68555 | 39.99917 | 99.47982 | 42.58548 | -0.94651 | 1.638715 | 0.073873 |
| 12 | 23080000 | 23083599 | 90.32268 | 142.4978 | 101.3981 | 42.08367 | 0.814539 | 2.472459 | 0.073881 |
| 6 | 132929400 | 132929999 | 4.069747 | 7.90155 | 27.90411 | 10.46722 | -1.21385 | -0.26198 | 0.073882 |

| | | | | | | | | | |
|---|---|---|---|---|---|---|---|---|---|
| 6 | 52219400 | 52224199 | 22.78321 | 36.94375 | 99.89181 | 42.78494 | -1.06572 | 1.619971 | 0.073902 |
| 4 | 152229400 | 152251999 | 129.9943 | 190.4058 | 471.9968 | 194.4253 | -1.07438 | 3.880616 | 0.073906 |
| X | 33032000 | 33032999 | 12.64223 | 24.63482 | 10.22608 | 3.711772 | 1.882098 | -0.33003 | 0.073916 |
| 5 | 139453800 | 139466399 | 56.10362 | 87.46796 | 206.2396 | 85.59056 | -0.92486 | 2.690524 | 0.073942 |
| 2 | 67693600 | 67695399 | 23.29245 | 40.98812 | 24.10007 | 10.66994 | 1.149822 | 0.606888 | 0.073949 |
| 3 | 101918600 | 101920799 | 15.83549 | 26.33747 | 76.49695 | 33.19919 | -1.15162 | 1.243677 | 0.073967 |
| 14 | 48739800 | 48741799 | 7.934297 | 14.15615 | 43.10883 | 18.95582 | -1.19874 | 0.429606 | 0.074015 |
| 14 | 102131400 | 102131999 | 3.532599 | 6.881879 | 28.59166 | 10.41335 | -1.40623 | -0.3753 | 0.074022 |
| 5 | 124230000 | 124234399 | 17.12876 | 27.69811 | 106.9035 | 45.6913 | -1.58088 | 1.673814 | 0.074038 |
| 10 | 47918400 | 47919399 | 14.54286 | 27.4491 | 16.28451 | 6.638209 | 1.222487 | 0.080869 | 0.074047 |
| 7 | 80771600 | 80790199 | 109.6121 | 163.3731 | 379.6949 | 156.4695 | -0.95778 | 3.584753 | 0.074049 |
| 9 | 63043600 | 63076999 | 248.1807 | 344.1386 | 815.7759 | 338.7041 | -1.08879 | 4.705885 | 0.07408 |
| 15 | 85448200 | 85467999 | 116.6282 | 171.0001 | 472.0371 | 194.4296 | -1.22807 | 3.863678 | 0.074096 |
| 7 | 127534600 | 127538399 | 17.1809 | 28.7296 | 73.00381 | 31.79257 | -0.95954 | 1.185244 | 0.074108 |
| 5 | 118465000 | 118472799 | 43.656 | 69.06545 | 170.5503 | 70.8837 | -0.97471 | 2.387757 | 0.074114 |
| 7 | 84666800 | 84668199 | 22.74355 | 40.11412 | 23.73744 | 10.50457 | 1.141982 | 0.579378 | 0.074117 |
| 1 | 133361800 | 133367999 | 28.65337 | 46.09263 | 117.7254 | 49.96423 | -0.9969 | 1.852309 | 0.074118 |
| 2 | 30802200 | 30805199 | 12.07319 | 20.89079 | 56.8761 | 24.95301 | -1.05218 | 0.832516 | 0.074122 |
| 5 | 30971800 | 30974199 | 9.054011 | 15.96328 | 46.68354 | 20.66045 | -1.14991 | 0.545285 | 0.074131 |
| 11 | 50114800 | 50131999 | 79.46611 | 119.9413 | 328.8031 | 135.9004 | -1.17943 | 3.335724 | 0.074148 |
| 4 | 26237800 | 26238599 | 9.898421 | 19.30061 | 10.74455 | 3.867223 | 1.472269 | -0.3957 | 0.07416 |
| 14 | 12289400 | 12289799 | 9.898421 | 19.30061 | 10.74455 | 3.867223 | 1.472269 | -0.3957 | 0.07416 |
| 8 | 58104600 | 58106799 | 10.47482 | 18.31761 | 51.51137 | 22.7123 | -1.09623 | 0.678686 | 0.074171 |
| 7 | 23050200 | 23054999 | 116.9676 | 181.3439 | 126.5595 | 52.56661 | 0.823243 | 2.803587 | 0.074188 |
| 3 | 44462800 | 44464399 | 16.10126 | 30.53987 | 14.43517 | 5.823645 | 1.559818 | 0.058763 | 0.074191 |
| 5 | 3065200 | 3069599 | 65.24227 | 104.4185 | 74.46691 | 31.36557 | 0.831017 | 2.040022 | 0.074195 |
| 19 | 27835400 | 27835999 | 4.916118 | 9.261795 | 33.96538 | 13.8937 | -1.38572 | 0.054947 | 0.074196 |
| 8 | 84934400 | 84940799 | 22.57752 | 36.3826 | 113.5813 | 48.32807 | -1.28224 | 1.787497 | 0.074217 |
| 9 | 54922600 | 54924199 | 10.9794 | 19.09093 | 55.83044 | 24.52466 | -1.15358 | 0.770224 | 0.074218 |
| 16 | 69073000 | 69073999 | 10.91112 | 21.14517 | 13.78325 | 5.219155 | 1.184588 | -0.19847 | 0.074219 |
| 14 | 100695800 | 100697599 | 4.667106 | 8.799177 | 39.22486 | 16.02501 | -1.66447 | 0.056842 | 0.074221 |
| 4 | 8785200 | 8786199 | 15.2899 | 28.81854 | 16.9539 | 6.929811 | 1.23209 | 0.099933 | 0.074244 |
| 11 | 94145800 | 94152999 | 42.45519 | 67.32392 | 158.0516 | 65.79158 | -0.89887 | 2.325213 | 0.074251 |
| 1 | 75739600 | 75740599 | 4.499165 | 8.71764 | 32.17146 | 12.19184 | -1.29238 | -0.21154 | 0.074254 |
| 6 | 118847000 | 118848999 | 7.886633 | 14.20625 | 41.66243 | 18.16533 | -1.13637 | 0.365097 | 0.074258 |
| 18 | 75019600 | 75020799 | 5.74774 | 10.83333 | 34.59599 | 14.14107 | -1.1876 | 0.062364 | 0.07427 |
| 19 | 42184000 | 42191399 | 46.00574 | 72.56722 | 181.2527 | 75.22137 | -0.99463 | 2.461548 | 0.074277 |
| 16 | 59770600 | 59771599 | 24.84605 | 43.47002 | 23.9607 | 10.56903 | 1.242316 | 0.700101 | 0.074283 |
| 7 | 11082600 | 11085199 | 27.61125 | 48.2495 | 22.6766 | 9.992392 | 1.470487 | 0.730537 | 0.074289 |
| 4 | 112831800 | 112832399 | 21.48446 | 37.82841 | 24.01291 | 10.63012 | 1.040361 | 0.589908 | 0.074308 |
| 3 | 139637200 | 139637599 | 10.32553 | 20.098 | 13.13794 | 4.828821 | 1.219028 | -0.3084 | 0.074309 |
| 4 | 33246000 | 33248999 | 13.32954 | 22.6306 | 68.67587 | 30.09097 | -1.21363 | 1.023352 | 0.07434 |
| 15 | 102856000 | 102860999 | 23.61381 | 38.5824 | 95.71365 | 41.146 | -0.9333 | 1.494154 | 0.074341 |
| 10 | 5633600 | 5635799 | 16.06251 | 29.89682 | 17.72687 | 7.401554 | 1.196254 | 0.185042 | 0.074347 |
| 5 | 113004600 | 113020399 | 82.84821 | 125.5602 | 307.0933 | 127.0917 | -1.00992 | 3.251711 | 0.074366 |
| 2 | 16595200 | 16597799 | 40.31351 | 66.28624 | 45.65104 | 19.67528 | 0.914693 | 1.41493 | 0.07437 |
| 2 | 40258200 | 40259599 | 20.98352 | 38.31355 | 16.61887 | 7.124238 | 1.616842 | 0.327665 | 0.074382 |

| | | | | | | | | | |
|---|---|---|---|---|---|---|---|---|---|
| 7 | 20691600 | 20693999 | 31.20403 | 53.45977 | 31.73532 | 13.91732 | 1.13234 | 0.967411 | 0.074395 |
| 7 | 117029200 | 117029999 | 3.532688 | 6.881802 | 29.25072 | 10.65844 | -1.43955 | -0.37194 | 0.07441 |
| 14 | 60317000 | 60317999 | 3.532688 | 6.881802 | 29.25072 | 10.65844 | -1.43955 | -0.37194 | 0.07441 |
| 3 | 79454200 | 79457599 | 20.95131 | 34.57333 | 85.60365 | 36.95059 | -0.92373 | 1.354709 | 0.074415 |
| 18 | 76163000 | 76171199 | 33.85339 | 53.90447 | 151.4906 | 63.35075 | -1.1518 | 2.190489 | 0.074433 |
| 7 | 59134400 | 59141799 | 96.62037 | 151.6354 | 110.4623 | 45.81261 | 0.774972 | 2.580534 | 0.074439 |
| 3 | 5770000 | 5772599 | 29.40487 | 50.53381 | 31.84935 | 13.97022 | 1.047048 | 0.93298 | 0.07444 |
| 11 | 68681600 | 68682999 | 5.655899 | 10.59185 | 34.95828 | 14.4457 | -1.24787 | 0.102586 | 0.074441 |
| 14 | 89496400 | 89496999 | 10.45927 | 20.42502 | 10.21756 | 3.591532 | 1.655348 | -0.46788 | 0.074451 |
| 11 | 79771200 | 79774999 | 19.92409 | 32.68622 | 87.05595 | 37.48446 | -1.03195 | 1.427 | 0.074464 |
| 9 | 118910000 | 118927399 | 75.24462 | 113.9379 | 324.1875 | 134.1296 | -1.22905 | 3.268423 | 0.074486 |
| 14 | 42520400 | 42523199 | 37.02531 | 61.66988 | 41.14524 | 17.87398 | 0.963011 | 1.254924 | 0.074487 |
| 13 | 73656800 | 73658399 | 5.884172 | 10.82929 | 38.34721 | 16.26533 | -1.37799 | 0.217643 | 0.0745 |
| 3 | 17445800 | 17450599 | 55.42741 | 89.43315 | 59.61315 | 25.40728 | 0.941942 | 1.773464 | 0.0745 |
| 2 | 169041200 | 169044599 | 14.6419 | 25.0348 | 63.40654 | 27.80203 | -0.95238 | 0.95877 | 0.074504 |
| 11 | 112666000 | 112670999 | 20.66118 | 33.69577 | 98.40729 | 42.27546 | -1.16981 | 1.515667 | 0.074504 |
| 12 | 19376400 | 19379399 | 62.24066 | 100.2801 | 52.58123 | 22.36721 | 1.284814 | 1.822685 | 0.074551 |
| 5 | 136896200 | 136916799 | 118.4514 | 176.1716 | 390.27 | 160.7925 | -0.89151 | 3.623709 | 0.074553 |
| 2 | 6167400 | 6169999 | 12.03117 | 20.95317 | 60.90035 | 26.77857 | -1.14492 | 0.758441 | 0.074558 |
| 17 | 34944600 | 34956799 | 63.69857 | 97.84365 | 250.1408 | 103.8604 | -1.05809 | 2.970291 | 0.074567 |
| 14 | 58308000 | 58316199 | 41.64918 | 65.70538 | 178.9628 | 74.27421 | -1.11914 | 2.455767 | 0.074588 |
| 6 | 64310800 | 64311999 | 10.77596 | 21.00616 | 11.2218 | 4.052976 | 1.527395 | -0.36916 | 0.074594 |
| 2 | 89527800 | 89529599 | 37.5618 | 62.54042 | 41.14991 | 17.87185 | 0.983036 | 1.260826 | 0.074595 |
| 9 | 3134200 | 3135599 | 34.64611 | 58.38327 | 36.23555 | 15.84098 | 1.066686 | 1.128932 | 0.074596 |
| 13 | 84246600 | 84247799 | 14.90196 | 28.55561 | 14.14517 | 5.573514 | 1.522948 | -0.02692 | 0.074601 |
| 11 | 88060400 | 88060999 | 2.546365 | 4.971282 | 23.84545 | 8.419607 | -1.56446 | -0.51275 | 0.074607 |
| 18 | 64649200 | 64649999 | 2.546365 | 4.971282 | 23.84545 | 8.419607 | -1.56446 | -0.51275 | 0.074607 |
| 16 | 38573000 | 38576199 | 12.05068 | 20.92847 | 55.86812 | 24.52266 | -1.02299 | 0.79299 | 0.074617 |
| 5 | 143512600 | 143514399 | 7.234464 | 12.86533 | 45.69192 | 20.1472 | -1.42191 | 0.449835 | 0.07462 |
| 8 | 106474200 | 106478399 | 16.38916 | 27.45528 | 73.05596 | 31.8516 | -1.02582 | 1.161888 | 0.074626 |
| 5 | 120591200 | 120605599 | 84.56524 | 128.194 | 298.9758 | 123.7419 | -0.94107 | 3.246998 | 0.07463 |
| 8 | 23457600 | 23466399 | 47.48107 | 74.46906 | 195.4014 | 81.04141 | -1.07204 | 2.578427 | 0.074674 |
| 3 | 72671200 | 72671799 | 10.95363 | 21.36223 | 10.59701 | 3.80453 | 1.639894 | -0.38906 | 0.074689 |
| 16 | 50661200 | 50662399 | 5.180486 | 9.842527 | 33.78951 | 13.58356 | -1.27014 | -0.00375 | 0.074702 |
| 16 | 91720000 | 91721599 | 5.112231 | 9.422959 | 37.25063 | 15.76746 | -1.53165 | 0.202936 | 0.074721 |
| X | 34664600 | 34665399 | 11.13802 | 21.70996 | 11.21436 | 4.055109 | 1.574098 | -0.35994 | 0.074726 |
| 11 | 85201400 | 85203199 | 6.892888 | 12.3576 | 43.69883 | 19.13722 | -1.40849 | 0.391765 | 0.074758 |
| 6 | 51148800 | 51151999 | 10.17538 | 17.80345 | 53.12304 | 23.43345 | -1.18142 | 0.669574 | 0.074762 |
| 1 | 189988400 | 189993999 | 32.35979 | 51.96142 | 120.0277 | 50.80761 | -0.85556 | 1.915844 | 0.074779 |
| 8 | 121756400 | 121766199 | 61.61623 | 95.67521 | 213.9339 | 88.8373 | -0.853 | 2.763796 | 0.074799 |
| 10 | 41979200 | 41988199 | 59.93523 | 93.20759 | 211.5531 | 87.82857 | -0.87279 | 2.736182 | 0.07481 |
| 5 | 114780200 | 114781799 | 5.420974 | 10.07941 | 38.06475 | 15.92143 | -1.45518 | 0.153224 | 0.07485 |
| 7 | 67652400 | 67654399 | 18.32733 | 30.04328 | 87.04461 | 37.46724 | -1.15323 | 1.435096 | 0.074877 |
| 3 | 133005200 | 133006199 | 16.20954 | 29.93817 | 18.9791 | 8.011421 | 1.089704 | 0.233062 | 0.074882 |
| 10 | 107753800 | 107754399 | 10.99015 | 21.49833 | 6.07663 | 2.026367 | 2.524625 | -0.64836 | 0.074889 |
| 11 | 94861400 | 94864399 | 11.84916 | 20.27674 | 61.51894 | 26.97619 | -1.21087 | 0.939763 | 0.074918 |
| 1 | 186508600 | 186509799 | 12.07897 | 23.22671 | 13.75978 | 5.367809 | 1.281181 | -0.08935 | 0.074935 |

| | | | | | | | | | |
|---|---|---|---|---|---|---|---|---|---|
| 3 | 48393800 | 48395199 | 21.98003 | 39.54027 | 18.65189 | 8.144896 | 1.480475 | 0.421431 | 0.074939 |
| 1 | 191068400 | 191082199 | 75.23586 | 114.349 | 292.6619 | 121.2233 | -1.07191 | 3.19085 | 0.07497 |
| 9 | 30997000 | 31011799 | 102.5199 | 153.8019 | 342.4359 | 141.2987 | -0.88734 | 3.460335 | 0.074973 |
| 4 | 61131200 | 61132399 | 13.409 | 25.63473 | 16.01892 | 6.354343 | 1.182608 | -0.01596 | 0.074974 |
| 10 | 81286200 | 81332399 | 344.5058 | 463.161 | 1147.475 | 478.7231 | -1.19757 | 5.191709 | 0.074995 |
| 16 | 11071200 | 11072999 | 6.552459 | 11.85177 | 41.73947 | 18.10529 | -1.39306 | 0.33124 | 0.074997 |
| 1 | 114819800 | 114820599 | 12.28557 | 23.84482 | 12.74793 | 4.793127 | 1.475563 | -0.20885 | 0.075002 |
| 2 | 167526800 | 167538799 | 60.90084 | 93.68444 | 249.8945 | 103.7791 | -1.11762 | 2.939846 | 0.075003 |
| 2 | 38388000 | 38392799 | 22.13308 | 36.30918 | 89.96706 | 38.73749 | -0.9283 | 1.433369 | 0.075007 |
| 19 | 41833200 | 41837999 | 19.95186 | 32.62897 | 88.89512 | 38.22753 | -1.06674 | 1.470746 | 0.075011 |
| 4 | 148710800 | 148711799 | 5.426167 | 10.37597 | 32.95464 | 13.06344 | -1.14085 | -0.04923 | 0.075018 |
| 12 | 114170600 | 114171399 | 21.86651 | 38.40183 | 24.47555 | 10.83064 | 1.033621 | 0.624201 | 0.075019 |
| 4 | 152152000 | 152164599 | 58.8627 | 90.5827 | 244.654 | 101.608 | -1.13515 | 2.931973 | 0.075048 |
| 2 | 180417000 | 180422199 | 31.30127 | 50.12915 | 129.2892 | 54.50863 | -1.01883 | 1.999948 | 0.075072 |
| 12 | 18647800 | 18651599 | 64.563 | 103.6761 | 63.8583 | 27.033 | 1.047449 | 1.932342 | 0.075083 |
| 7 | 31943800 | 31945199 | 14.87769 | 28.62206 | 12.97877 | 5.0562 | 1.663439 | -0.06567 | 0.075091 |
| 7 | 44484600 | 44519599 | 164.5178 | 234.9358 | 613.5814 | 253.5508 | -1.18569 | 4.264817 | 0.075101 |
| 13 | 66063000 | 66066599 | 65.38367 | 105.0985 | 61.0492 | 25.88171 | 1.133546 | 1.903459 | 0.075111 |
| 11 | 69894600 | 69897399 | 9.56892 | 16.74112 | 51.48367 | 22.70889 | -1.22432 | 0.666475 | 0.075112 |
| 8 | 107408200 | 107410399 | 12.50437 | 21.74646 | 56.44668 | 24.79815 | -0.98259 | 0.776272 | 0.075121 |
| 11 | 102642600 | 102646599 | 12.06868 | 20.21549 | 74.07684 | 32.29459 | -1.48481 | 1.152261 | 0.075127 |
| 4 | 149012600 | 149038599 | 126.1162 | 183.9487 | 500.2923 | 206.1495 | -1.21391 | 3.946193 | 0.075152 |
| 5 | 37958800 | 37996999 | 293.8321 | 405.3835 | 841.7506 | 349.7704 | -0.90518 | 4.784772 | 0.075153 |
| 11 | 73986800 | 73987399 | 15.70857 | 29.11968 | 18.09119 | 7.598539 | 1.123747 | 0.206767 | 0.075163 |
| X | 25784800 | 25785399 | 11.78314 | 22.95337 | 11.45951 | 4.177092 | 1.614529 | -0.32257 | 0.075177 |
| 4 | 152374200 | 152403599 | 152.2321 | 219.1615 | 568.3653 | 234.5669 | -1.16381 | 4.144627 | 0.07519 |
| 8 | 85510800 | 85528599 | 102.9048 | 153.1601 | 389.4355 | 160.4605 | -1.08922 | 3.609076 | 0.075193 |
| X | 169203400 | 169203799 | 13.7736 | 26.7448 | 9.454012 | 3.543246 | 2.065141 | -0.21209 | 0.075197 |
| 10 | 113518800 | 113520599 | 19.26183 | 35.16436 | 18.40081 | 7.889691 | 1.348352 | 0.318896 | 0.075217 |
| 7 | 9971400 | 9975199 | 62.60002 | 101.0538 | 50.35835 | 21.47589 | 1.361164 | 1.768775 | 0.075228 |
| 9 | 45770200 | 45771799 | 5.468885 | 10.31825 | 33.48954 | 13.66302 | -1.20836 | 0.049562 | 0.075232 |
| 9 | 64247400 | 64248799 | 4.640887 | 8.685386 | 34.85605 | 14.41931 | -1.52705 | 0.099322 | 0.075248 |
| 14 | 30850400 | 30865799 | 77.59055 | 117.6308 | 300.1638 | 124.2368 | -1.0706 | 3.244132 | 0.075288 |
| 12 | 30961000 | 30962399 | 8.783861 | 15.44301 | 48.77635 | 21.59892 | -1.26206 | 0.578398 | 0.075298 |
| 2 | 149316600 | 149318999 | 23.60886 | 41.63392 | 21.72269 | 9.613273 | 1.321806 | 0.580695 | 0.075303 |
| 6 | 58571000 | 58572199 | 28.85665 | 49.85252 | 30.12669 | 13.21672 | 1.108726 | 0.88217 | 0.075319 |
| 13 | 28412400 | 28414799 | 35.52939 | 59.91767 | 34.35841 | 15.02385 | 1.180175 | 1.121177 | 0.07532 |
| 5 | 73332600 | 73333399 | 9.692021 | 18.88193 | 11.2781 | 4.105187 | 1.36051 | -0.35691 | 0.075321 |
| 7 | 32919800 | 32922999 | 75.5497 | 120.3148 | 82.89669 | 34.67302 | 0.873 | 2.205203 | 0.075325 |
| 4 | 3186600 | 3187999 | 12.72931 | 24.8173 | 8.509276 | 3.06781 | 2.156783 | -0.35773 | 0.075331 |
| 19 | 45023800 | 45030599 | 28.13869 | 45.00533 | 134.5157 | 56.60501 | -1.23311 | 2.039533 | 0.075345 |
| 9 | 118889200 | 118909199 | 101.6625 | 150.9367 | 406.6205 | 167.499 | -1.17587 | 3.653116 | 0.075353 |
| 5 | 42251200 | 42253799 | 34.18588 | 58.03748 | 33.30557 | 14.59295 | 1.17937 | 1.06076 | 0.075369 |
| 3 | 38602600 | 38604999 | 27.90245 | 48.15595 | 30.68384 | 13.46073 | 1.032342 | 0.886247 | 0.075375 |
| 16 | 70185800 | 70187999 | 25.27285 | 44.243 | 25.72987 | 11.35511 | 1.165405 | 0.696353 | 0.075383 |
| 1 | 156605600 | 156609999 | 22.31826 | 36.53442 | 89.06211 | 38.31302 | -0.90607 | 1.460862 | 0.07539 |
| 3 | 3112400 | 3113199 | 19.27962 | 36.07354 | 14.2128 | 5.881818 | 1.790483 | 0.173161 | 0.075412 |

| | | | | | | | | | |
|---|---|---|---|---|---|---|---|---|---|
| 4 | 156311400 | 156322399 | 46.33833 | 71.74806 | 237.2327 | 98.54022 | -1.41967 | 2.810537 | 0.075442 |
| 7 | 32863600 | 32864799 | 29.98871 | 51.1959 | 34.00241 | 14.90727 | 0.970401 | 0.995872 | 0.075455 |
| 4 | 119670000 | 119672999 | 13.0304 | 22.06209 | 66.82864 | 29.25721 | -1.21091 | 1.046614 | 0.075491 |
| 8 | 21944200 | 21946399 | 18.20711 | 32.91674 | 20.88341 | 9.063879 | 1.061923 | 0.371218 | 0.075496 |
| 17 | 45393200 | 45394399 | 4.98449 | 9.573911 | 31.3977 | 12.28821 | -1.16869 | -0.09989 | 0.075501 |
| 1 | 112491200 | 112493599 | 38.29931 | 63.4059 | 43.40565 | 18.77787 | 0.925578 | 1.327582 | 0.075504 |
| 13 | 3251800 | 3256599 | 83.80119 | 133.0178 | 88.01907 | 36.67496 | 0.926513 | 2.313446 | 0.075512 |
| 9 | 99276600 | 99281199 | 16.73493 | 27.61258 | 84.766 | 36.58735 | -1.23255 | 1.345098 | 0.075567 |
| 10 | 42021000 | 42029199 | 45.48398 | 71.82158 | 173.9292 | 72.21899 | -0.94882 | 2.434937 | 0.07557 |
| 8 | 107060200 | 107064999 | 27.47383 | 44.33192 | 109.1519 | 46.52404 | -0.93862 | 1.752243 | 0.07558 |
| 7 | 13461600 | 13463799 | 29.683 | 50.67981 | 31.17783 | 13.66905 | 1.080506 | 0.990971 | 0.075594 |
| 14 | 25606600 | 25609199 | 11.1887 | 19.33259 | 59.55742 | 26.12824 | -1.23015 | 0.84311 | 0.075604 |
| 3 | 121977400 | 121979199 | 9.546255 | 16.75338 | 49.86457 | 22.05248 | -1.17739 | 0.612293 | 0.07562 |
| 15 | 97827200 | 97838199 | 62.46698 | 96.53882 | 229.0406 | 95.13838 | -0.94367 | 2.850896 | 0.075658 |
| 9 | 77952400 | 77954999 | 14.39597 | 24.10719 | 75.62305 | 32.963 | -1.26218 | 1.161788 | 0.075666 |
| 6 | 90523400 | 90531999 | 50.71357 | 79.52669 | 192.7801 | 79.95551 | -0.95812 | 2.582548 | 0.075675 |
| 4 | 81893600 | 81894599 | 11.97414 | 23.3475 | 9.906653 | 3.566632 | 1.858393 | -0.36776 | 0.075675 |
| 2 | 28908600 | 28913999 | 27.04024 | 43.54621 | 114.7719 | 48.78946 | -1.04004 | 1.813652 | 0.075685 |
| 18 | 47499400 | 47504399 | 26.49084 | 42.51744 | 126.7388 | 53.61009 | -1.22329 | 1.924731 | 0.075686 |
| 12 | 19555400 | 19564599 | 285.2142 | 413.5768 | 277.7603 | 114.5238 | 0.79464 | 4.042817 | 0.07569 |
| 2 | 158217400 | 158220399 | 14.73852 | 24.92105 | 66.97358 | 29.30887 | -1.03934 | 1.064029 | 0.075709 |
| 1 | 177364200 | 177367199 | 23.67927 | 41.91353 | 21.27514 | 9.408223 | 1.363392 | 0.542575 | 0.07571 |
| 7 | 69980000 | 69985199 | 21.7604 | 35.47063 | 95.50907 | 41.02288 | -1.05349 | 1.524121 | 0.075714 |
| 8 | 85180000 | 85194799 | 98.76438 | 149.1468 | 311.649 | 128.8315 | -0.78746 | 3.328314 | 0.075716 |
| 11 | 4161800 | 4173799 | 57.37199 | 88.4652 | 242.4935 | 100.7276 | -1.15424 | 2.892125 | 0.075721 |
| 11 | 98354000 | 98363999 | 46.62027 | 73.58803 | 171.7079 | 71.28302 | -0.89573 | 2.444919 | 0.075755 |
| 16 | 67924400 | 67927199 | 30.65176 | 52.36983 | 32.2375 | 14.13442 | 1.079523 | 0.988037 | 0.075771 |
| 1 | 11724800 | 11725599 | 13.24881 | 25.28595 | 15.39559 | 6.130118 | 1.214431 | -0.00834 | 0.075774 |
| 5 | 140548200 | 140554999 | 31.54225 | 50.33965 | 140.3601 | 58.88306 | -1.13671 | 2.112015 | 0.075785 |
| 5 | 121798200 | 121799399 | 3.734276 | 7.20843 | 31.5401 | 12.16439 | -1.55884 | -0.16212 | 0.075799 |
| 7 | 8403400 | 8404999 | 45.80266 | 74.61071 | 51.64289 | 22.17613 | 0.900266 | 1.556563 | 0.075801 |
| 11 | 58667800 | 58669999 | 12.94197 | 22.08605 | 62.61485 | 27.45068 | -1.11434 | 0.971046 | 0.07581 |
| 18 | 44422600 | 44423599 | 5.166044 | 9.856734 | 34.6213 | 13.79386 | -1.29171 | -0.03245 | 0.075853 |
| 12 | 85654600 | 85660399 | 17.12677 | 28.12883 | 92.63893 | 39.90742 | -1.33652 | 1.40869 | 0.075899 |
| 11 | 53870000 | 53871599 | 10.19462 | 17.79364 | 53.65557 | 23.62548 | -1.19668 | 0.708783 | 0.075916 |
| 18 | 75548600 | 75549199 | 2.64418 | 5.150842 | 25.12209 | 9.156721 | -1.62958 | -0.38944 | 0.075923 |
| 15 | 34659200 | 34660199 | 13.90107 | 26.50505 | 15.26723 | 6.091522 | 1.29105 | 0.004429 | 0.075926 |
| X | 79693200 | 79694399 | 18.96251 | 35.05236 | 17.0479 | 7.187959 | 1.470123 | 0.246927 | 0.075944 |
| 2 | 61275000 | 61275399 | 2.737101 | 5.349155 | 22.32513 | 7.727678 | -1.33967 | -0.58405 | 0.075956 |
| 5 | 61205400 | 61207799 | 28.4976 | 48.79809 | 33.05542 | 14.49564 | 0.942668 | 0.961829 | 0.075963 |
| 8 | 84135200 | 84135799 | 3.763954 | 7.329573 | 26.50298 | 9.704676 | -1.21426 | -0.35585 | 0.075977 |
| 11 | 3122000 | 3124199 | 326.7408 | 472.335 | 266.1259 | 109.7715 | 1.043539 | 4.092087 | 0.075999 |
| 17 | 28542800 | 28550199 | 37.53884 | 59.74547 | 149.2216 | 62.36244 | -0.9827 | 2.208189 | 0.076009 |
| 10 | 76958600 | 76964199 | 24.55566 | 39.75748 | 103.0204 | 44.0756 | -1.00668 | 1.655118 | 0.076033 |
| 15 | 86321000 | 86325399 | 19.05288 | 31.47139 | 82.39711 | 35.59042 | -1.00371 | 1.33614 | 0.076035 |
| 1 | 101905800 | 101908599 | 30.65571 | 52.1309 | 35.10196 | 15.38451 | 0.949931 | 1.034402 | 0.076038 |
| 7 | 10578600 | 10581599 | 58.45423 | 93.88008 | 64.85863 | 27.46249 | 0.882451 | 1.919871 | 0.076039 |

| | | | | | | | | | |
|---|---|---|---|---|---|---|---|---|---|
| 10 | 60719600 | 60726799 | 42.44131 | 67.34139 | 156.2306 | 65.07177 | -0.88098 | 2.307175 | 0.076048 |
| 11 | 59850400 | 59859999 | 38.65343 | 61.22028 | 172.1375 | 71.58427 | -1.16033 | 2.357432 | 0.076051 |
| 2 | 35608400 | 35613999 | 27.07146 | 43.50423 | 118.8498 | 50.36806 | -1.09549 | 1.883585 | 0.076055 |
| 3 | 104499600 | 104499999 | 22.03591 | 39.04484 | 22.53008 | 9.961128 | 1.180265 | 0.523657 | 0.076062 |
| 17 | 8614600 | 8618599 | 19.05749 | 31.44863 | 81.65751 | 35.24746 | -0.99184 | 1.347895 | 0.076074 |
| 8 | 102731200 | 102732199 | 10.39319 | 20.25109 | 12.5916 | 4.573969 | 1.305544 | -0.34915 | 0.076077 |
| 5 | 137110400 | 137110999 | 10.22018 | 19.89366 | 13.24419 | 4.8665 | 1.193866 | -0.30983 | 0.076089 |
| 7 | 15109000 | 15109599 | 10.22018 | 19.89366 | 13.24419 | 4.8665 | 1.193866 | -0.30983 | 0.076089 |
| 3 | 111829200 | 111830799 | 18.70973 | 34.36791 | 17.01197 | 7.234551 | 1.436821 | 0.279233 | 0.076089 |
| 4 | 112578000 | 112580199 | 25.60283 | 44.63849 | 24.6312 | 10.83813 | 1.240703 | 0.755248 | 0.076097 |
| 4 | 59039000 | 59039599 | 3.458141 | 6.746964 | 25.73866 | 9.190388 | -1.2566 | -0.45304 | 0.076107 |
| 9 | 61866200 | 61869199 | 12.44627 | 21.36787 | 60.02433 | 26.32769 | -1.09972 | 0.909298 | 0.076111 |
| 2 | 163215400 | 163237999 | 163.6423 | 237.9178 | 497.1164 | 204.9093 | -0.83832 | 3.997826 | 0.076155 |
| 16 | 54322200 | 54323399 | 13.73905 | 26.45537 | 14.48746 | 5.629337 | 1.39895 | -0.08082 | 0.076163 |
| 17 | 46080000 | 46085399 | 24.63311 | 39.72708 | 111.7237 | 47.58772 | -1.13075 | 1.764769 | 0.076165 |
| 15 | 57997400 | 57998799 | 5.403233 | 9.784321 | 43.34241 | 18.77366 | -1.71919 | 0.319334 | 0.076191 |
| 16 | 71719600 | 71722599 | 15.78746 | 26.81473 | 68.35878 | 29.9558 | -0.96347 | 1.028768 | 0.076195 |
| 4 | 138257400 | 138268799 | 68.98101 | 106.6125 | 226.297 | 93.9987 | -0.78323 | 2.852037 | 0.07621 |
| 16 | 5088600 | 5090599 | 11.10623 | 19.35109 | 53.13267 | 23.3697 | -1.06252 | 0.738939 | 0.076211 |
| 9 | 101970800 | 101974999 | 19.97614 | 33.04077 | 90.46668 | 39.11221 | -1.06817 | 1.327568 | 0.076214 |
| 17 | 50274200 | 50275799 | 8.200567 | 14.73199 | 42.67967 | 18.65882 | -1.12174 | 0.385426 | 0.076217 |
| 11 | 77872600 | 77876199 | 13.9509 | 23.03285 | 86.55873 | 37.37552 | -1.52279 | 1.332652 | 0.076222 |
| 5 | 149441400 | 149445399 | 21.16443 | 34.766 | 85.67935 | 36.91194 | -0.91962 | 1.412808 | 0.076228 |
| 5 | 94358200 | 94359799 | 18.9028 | 34.44731 | 18.9617 | 8.148375 | 1.273861 | 0.327096 | 0.076254 |
| 4 | 44973000 | 44974599 | 7.572195 | 13.679 | 40.65864 | 17.66424 | -1.15195 | 0.344506 | 0.076255 |
| 2 | 181233600 | 181234799 | 4.694162 | 9.004779 | 30.6704 | 12.04758 | -1.22694 | -0.09132 | 0.076255 |
| 16 | 8555200 | 8561399 | 26.67615 | 43.06185 | 110.4362 | 47.09554 | -0.99663 | 1.739408 | 0.076261 |
| 8 | 72572000 | 72574399 | 48.63286 | 78.9656 | 53.87525 | 23.09827 | 0.917457 | 1.615346 | 0.076304 |
| 6 | 134684200 | 134690599 | 34.42093 | 54.92257 | 138.687 | 58.16424 | -0.99438 | 2.120867 | 0.076309 |
| 3 | 142888600 | 142889599 | 3.459863 | 6.744225 | 25.45311 | 9.197701 | -1.25663 | -0.41003 | 0.076337 |
| 1 | 142252200 | 142252999 | 8.783551 | 17.15136 | 11.00781 | 3.874311 | 1.29902 | -0.48664 | 0.076359 |
| 4 | 74332800 | 74333799 | 10.8908 | 21.17884 | 12.97916 | 4.805943 | 1.300942 | -0.27703 | 0.076384 |
| 8 | 123133600 | 123135599 | 4.088294 | 7.7095 | 35.04613 | 14.31281 | -1.68973 | 0.043328 | 0.076392 |
| 7 | 47080200 | 47081599 | 15.59983 | 29.32364 | 16.70391 | 6.85883 | 1.272414 | 0.117732 | 0.076394 |
| 16 | 17223200 | 17224399 | 5.187928 | 9.835224 | 33.68493 | 13.60403 | -1.27256 | 0.011729 | 0.076426 |
| 9 | 48604600 | 48606599 | 8.535135 | 15.23671 | 44.17147 | 19.41389 | -1.12826 | 0.430525 | 0.076429 |
| 16 | 60668200 | 60669399 | 13.28815 | 25.49957 | 15.50795 | 6.085482 | 1.235761 | -0.05246 | 0.076437 |
| 5 | 73366000 | 73368399 | 10.91595 | 19.12813 | 51.5145 | 22.75299 | -1.03436 | 0.646731 | 0.076442 |
| 2 | 137242200 | 137242399 | 6.522319 | 12.7306 | 7.675634 | 2.416887 | 1.525267 | -0.91378 | 0.076451 |
| 17 | 40011400 | 40012999 | 19.75486 | 35.16712 | 22.21233 | 9.785689 | 1.054101 | 0.475939 | 0.076451 |
| 10 | 79700800 | 79703599 | 11.52928 | 19.75213 | 60.97379 | 26.73963 | -1.23535 | 0.927417 | 0.07646 |
| 8 | 70990800 | 70993599 | 16.47471 | 27.37603 | 77.13519 | 33.45546 | -1.1082 | 1.256054 | 0.076467 |
| 8 | 125540200 | 125562599 | 143.353 | 209.1731 | 475.1705 | 195.7898 | -0.9538 | 3.941392 | 0.076467 |
| 3 | 71709200 | 71710199 | 13.22314 | 25.28469 | 16.02359 | 6.352383 | 1.163255 | -0.01987 | 0.076498 |
| 13 | 66046200 | 66048399 | 68.44924 | 109.9017 | 60.63861 | 25.66469 | 1.205618 | 1.941451 | 0.0765 |
| 11 | 101315000 | 101316399 | 5.31063 | 9.996346 | 33.94868 | 13.90733 | -1.2775 | 0.066022 | 0.076501 |
| 17 | 56857800 | 56864399 | 38.86365 | 61.74801 | 152.9316 | 63.78376 | -0.97315 | 2.263734 | 0.076528 |

| | | | | | | | | | |
|---|---|---|---|---|---|---|---|---|---|
| 18 | 34137400 | 34140599 | 15.13314 | 25.39023 | 71.81195 | 31.33964 | -1.11335 | 1.14051 | 0.076536 |
| 2 | 158113400 | 158119999 | 24.12052 | 38.96318 | 106.8204 | 45.59391 | -1.09132 | 1.714459 | 0.076538 |
| 1 | 59801800 | 59804199 | 9.941869 | 17.28013 | 58.50066 | 25.6913 | -1.36363 | 0.773911 | 0.076538 |
| 13 | 101828800 | 101832999 | 12.5066 | 21.3742 | 66.5921 | 29.19775 | -1.24986 | 0.95668 | 0.076546 |
| 3 | 62246000 | 62247999 | 9.210709 | 15.98408 | 59.49974 | 26.10639 | -1.49981 | 0.79463 | 0.076571 |
| 15 | 99408200 | 99427199 | 92.82717 | 139.3768 | 346.7352 | 143.1046 | -1.04621 | 3.442303 | 0.076573 |
| 4 | 8760600 | 8762999 | 12.50906 | 21.73014 | 56.44669 | 24.78062 | -0.9839 | 0.791828 | 0.076591 |
| 5 | 137634200 | 137645799 | 54.36779 | 83.96824 | 237.5025 | 98.65356 | -1.19761 | 2.860645 | 0.076597 |
| 11 | 85711200 | 85735999 | 107.8653 | 158.4967 | 481.1729 | 198.1656 | -1.36217 | 3.828539 | 0.076598 |
| 10 | 80343000 | 80375799 | 199.0403 | 280.8757 | 687.492 | 284.6573 | -1.10948 | 4.441246 | 0.076601 |
| 16 | 3294600 | 3297999 | 73.27829 | 117.1099 | 72.44603 | 30.45175 | 1.033529 | 2.091692 | 0.076611 |
| 4 | 154401400 | 154445399 | 293.9987 | 401.7862 | 948.5359 | 394.7341 | -1.10413 | 4.931949 | 0.076628 |
| 8 | 106876000 | 106877999 | 8.514945 | 15.23433 | 42.61604 | 18.69536 | -1.07492 | 0.41479 | 0.076632 |
| 15 | 79999600 | 80003599 | 19.17122 | 31.05708 | 102.7744 | 43.98854 | -1.35735 | 1.635529 | 0.076636 |
| 7 | 10373600 | 10375799 | 60.30525 | 96.78345 | 69.26997 | 29.29836 | 0.830429 | 1.947943 | 0.076663 |
| 1 | 137158400 | 137159799 | 17.77561 | 32.58852 | 19.35929 | 8.254615 | 1.172668 | 0.287293 | 0.07667 |
| 6 | 20539400 | 20539999 | 10.91174 | 21.14415 | 14.26179 | 5.403001 | 1.135021 | -0.19536 | 0.076701 |
| 13 | 56910200 | 56925599 | 92.17617 | 139.359 | 310.7452 | 128.5048 | -0.87968 | 3.303171 | 0.076706 |
| 1 | 72624800 | 72625399 | 1.870938 | 3.657744 | 23.62053 | 8.098957 | -1.94295 | -0.62928 | 0.076716 |
| 3 | 27331200 | 27334999 | 21.97471 | 36.04535 | 87.46552 | 37.65861 | -0.89822 | 1.433554 | 0.076731 |
| 12 | 19747600 | 19748199 | 21.08645 | 37.31308 | 23.37922 | 10.33929 | 1.061666 | 0.533648 | 0.076756 |
| X | 32937400 | 32940199 | 36.33885 | 61.01456 | 36.76496 | 16.05227 | 1.109129 | 1.166912 | 0.076767 |
| 10 | 60050400 | 60057599 | 37.68437 | 60.00421 | 145.7301 | 60.94138 | -0.94166 | 2.193132 | 0.076772 |
| 1 | 98671400 | 98672399 | 18.18886 | 32.95164 | 20.81078 | 9.009085 | 1.070715 | 0.359204 | 0.076793 |
| 2 | 91066800 | 91067999 | 5.952451 | 11.14008 | 35.5275 | 14.69683 | -1.20053 | 0.109871 | 0.076814 |
| 5 | 138689400 | 138692799 | 16.16114 | 27.26151 | 69.8983 | 30.56669 | -0.97222 | 1.091493 | 0.076837 |
| 14 | 3341800 | 3342799 | 22.92074 | 40.12707 | 26.14794 | 11.54003 | 1.002221 | 0.683287 | 0.076859 |
| 12 | 76234800 | 76238199 | 13.20352 | 22.33095 | 67.9418 | 29.73477 | -1.21742 | 1.057346 | 0.076893 |
| 2 | 156453400 | 156473999 | 116.9092 | 173.153 | 407.7126 | 167.9094 | -0.98503 | 3.697693 | 0.076899 |
| X | 18769800 | 18770999 | 14.85501 | 28.79053 | 10.74934 | 4.068817 | 1.977521 | -0.1631 | 0.076905 |
| X | 32426800 | 32430199 | 40.82606 | 68.4398 | 33.32778 | 14.53693 | 1.415143 | 1.19486 | 0.076929 |
| 6 | 130851200 | 130852199 | 42.35112 | 69.77074 | 44.55549 | 19.21874 | 1.024455 | 1.398666 | 0.076947 |
| 7 | 85435800 | 85437599 | 30.29421 | 51.88482 | 32.457 | 14.23344 | 1.057139 | 0.968051 | 0.076948 |
| 9 | 64408400 | 64409799 | 4.989791 | 9.567426 | 31.28372 | 12.30494 | -1.17116 | -0.08325 | 0.076965 |
| 12 | 52022400 | 52023799 | 4.989791 | 9.567426 | 31.28372 | 12.30494 | -1.17116 | -0.08325 | 0.076965 |
| 16 | 41795200 | 41796199 | 6.878029 | 12.69891 | 37.37783 | 15.78538 | -1.10935 | 0.205294 | 0.076966 |
| 3 | 68710200 | 68711199 | 4.366766 | 8.476374 | 29.19005 | 10.9689 | -1.1814 | -0.25 | 0.076969 |
| 8 | 126159200 | 126164599 | 25.92557 | 41.7741 | 115.4407 | 49.1129 | -1.10718 | 1.794497 | 0.076971 |
| 5 | 117456600 | 117469399 | 66.73985 | 102.6962 | 242.392 | 100.6674 | -0.94105 | 2.935587 | 0.076971 |
| 3 | 77375800 | 77377199 | 21.2936 | 37.52802 | 24.38271 | 10.79177 | 1.007421 | 0.578879 | 0.076977 |
| 3 | 88135400 | 88156399 | 96.13726 | 143.4218 | 394.185 | 162.4686 | -1.19831 | 3.565858 | 0.076981 |
| 4 | 43024600 | 43026999 | 8.492614 | 14.90421 | 49.3166 | 21.81007 | -1.32872 | 0.606822 | 0.077008 |
| 5 | 18563800 | 18565999 | 15.03079 | 25.44797 | 66.44211 | 29.08764 | -0.9977 | 1.054272 | 0.077016 |
| 3 | 9472400 | 9474599 | 16.00065 | 26.99903 | 68.13931 | 29.80024 | -0.94933 | 1.088114 | 0.077021 |
| 15 | 39036600 | 39038199 | 5.118018 | 9.828344 | 31.38943 | 12.29229 | -1.13158 | -0.0959 | 0.077025 |
| 15 | 16461000 | 16464999 | 111.3308 | 174.7505 | 95.21566 | 39.49014 | 1.193522 | 2.585072 | 0.07703 |
| 4 | 132972200 | 132986799 | 75.59783 | 114.7848 | 295.5666 | 122.39 | -1.08184 | 3.211713 | 0.077035 |

| | | | | | | | | | |
|---|---|---|---|---|---|---|---|---|---|
| 15 | 98618400 | 98627799 | 38.59105 | 61.20425 | 159.932 | 66.583 | -1.05292 | 2.31856 | 0.077042 |
| 4 | 127872800 | 127889999 | 119.4506 | 176.8538 | 417.144 | 171.7929 | -0.98801 | 3.703895 | 0.077046 |
| 16 | 91518000 | 91519599 | 6.296187 | 11.63732 | 37.38447 | 15.76481 | -1.23278 | 0.19394 | 0.07705 |
| 18 | 66216600 | 66219999 | 21.81598 | 35.78845 | 87.46383 | 37.65942 | -0.90843 | 1.431794 | 0.077063 |
| 1 | 187106600 | 187107199 | 3.945319 | 7.671119 | 27.54806 | 10.21298 | -1.2222 | -0.30799 | 0.077069 |
| 9 | 71395400 | 71396399 | 3.945319 | 7.671119 | 27.54806 | 10.21298 | -1.2222 | -0.30799 | 0.077069 |
| 4 | 147130600 | 147132399 | 24.55243 | 43.38755 | 21.26813 | 9.408218 | 1.412645 | 0.56562 | 0.077073 |
| 5 | 87866400 | 87867399 | 15.21561 | 28.61049 | 17.40216 | 7.141697 | 1.179636 | 0.11459 | 0.077087 |
| 5 | 54155000 | 54156999 | 11.08432 | 19.37715 | 51.52722 | 22.71304 | -1.01609 | 0.687455 | 0.077091 |
| 2 | 129138600 | 129139999 | 6.132733 | 11.07323 | 42.7592 | 18.58819 | -1.52676 | 0.34059 | 0.077104 |
| 2 | 18771600 | 18772399 | 3.532422 | 6.882033 | 27.29301 | 9.930944 | -1.33828 | -0.38203 | 0.077107 |
| 18 | 76463200 | 76473199 | 59.9144 | 92.54708 | 236.9352 | 98.41786 | -1.05391 | 2.859737 | 0.077126 |
| 19 | 49061200 | 49062999 | 22.75209 | 40.57838 | 19.60546 | 8.62365 | 1.439332 | 0.479974 | 0.077132 |
| 13 | 27351400 | 27352999 | 18.70984 | 33.72423 | 21.30892 | 9.283354 | 1.064325 | 0.392641 | 0.077166 |
| 4 | 118116200 | 118123399 | 27.64636 | 44.28296 | 131.9078 | 55.62631 | -1.22581 | 1.991488 | 0.077178 |
| 17 | 88840200 | 88845799 | 23.22438 | 37.75646 | 100.2622 | 43.02269 | -1.03639 | 1.567923 | 0.077185 |
| 2 | 130748000 | 130749199 | 6.035784 | 11.05571 | 38.23966 | 16.3248 | -1.351 | 0.247089 | 0.077191 |
| 7 | 21089800 | 21092799 | 30.86909 | 53.3077 | 28.36783 | 12.44495 | 1.29113 | 0.892633 | 0.077204 |
| 11 | 5711800 | 5712599 | 4.245121 | 8.231747 | 29.05828 | 10.97245 | -1.22342 | -0.23561 | 0.07722 |
| 7 | 28836000 | 28841599 | 26.63862 | 43.03159 | 104.8474 | 44.75286 | -0.9216 | 1.718485 | 0.077226 |
| 12 | 42585200 | 42586399 | 18.90944 | 34.80917 | 16.13365 | 6.841086 | 1.532157 | 0.265229 | 0.077253 |
| 12 | 47986000 | 47988199 | 28.00853 | 48.64501 | 28.44577 | 12.48634 | 1.157169 | 0.826273 | 0.077256 |
| 18 | 80262800 | 80268999 | 34.65649 | 55.51355 | 126.7361 | 53.44832 | -0.84295 | 1.996999 | 0.077293 |
| 11 | 99211200 | 99214399 | 12.35086 | 21.47244 | 55.34219 | 24.30792 | -0.97241 | 0.778874 | 0.077295 |
| 11 | 79530000 | 79533399 | 9.940739 | 17.23362 | 56.7919 | 24.91752 | -1.32523 | 0.805058 | 0.077295 |
| 7 | 10166600 | 10168999 | 42.38153 | 70.49055 | 37.22327 | 16.15497 | 1.299075 | 1.283689 | 0.077331 |
| 4 | 53825000 | 53826799 | 6.250579 | 11.68252 | 37.79728 | 15.67094 | -1.2244 | 0.124942 | 0.077337 |
| 5 | 79380600 | 79381199 | 10.37095 | 20.14482 | 13.15294 | 4.915548 | 1.198837 | -0.25171 | 0.077349 |
| 4 | 100512800 | 100514199 | 12.14797 | 20.78157 | 61.52001 | 26.97598 | -1.17566 | 0.944419 | 0.077353 |
| 1 | 112076600 | 112077799 | 15.608 | 29.00074 | 18.38301 | 7.696666 | 1.097829 | 0.193219 | 0.077365 |
| 8 | 108496000 | 108500199 | 46.2812 | 75.20777 | 52.15262 | 22.373 | 0.895105 | 1.593399 | 0.077378 |
| 19 | 57012200 | 57021999 | 55.62182 | 87.08969 | 192.0406 | 79.66141 | -0.82347 | 2.612674 | 0.077395 |
| 4 | 61694600 | 61696599 | 23.80265 | 41.87052 | 23.8247 | 10.5488 | 1.196938 | 0.614748 | 0.077427 |
| 4 | 141316400 | 141328999 | 60.04326 | 92.45838 | 246.1716 | 102.2472 | -1.11385 | 2.919306 | 0.077427 |
| 14 | 41828000 | 41830199 | 33.75273 | 57.8537 | 28.20691 | 12.37043 | 1.414826 | 0.968934 | 0.077436 |
| X | 31820200 | 31823599 | 35.88036 | 60.94965 | 31.24564 | 13.69279 | 1.341516 | 1.061738 | 0.077442 |
| 6 | 56210600 | 56210799 | 17.95841 | 33.70997 | 11.07364 | 4.557757 | 2.052043 | 0.132708 | 0.077444 |
| 15 | 38395200 | 38396999 | 7.000717 | 12.99893 | 39.84692 | 16.69878 | -1.15984 | 0.178286 | 0.077447 |
| 6 | 35234600 | 35235799 | 4.194441 | 8.142609 | 28.50821 | 10.70486 | -1.20299 | -0.25451 | 0.07745 |
| 9 | 119340000 | 119349199 | 67.56867 | 103.266 | 270.9497 | 112.3949 | -1.10124 | 3.074119 | 0.077477 |
| 19 | 4311800 | 4313999 | 6.648851 | 12.05602 | 39.66073 | 17.15026 | -1.29192 | 0.315342 | 0.077486 |
| 6 | 99497400 | 99500599 | 16.64584 | 27.6406 | 78.29235 | 33.94056 | -1.11593 | 1.265352 | 0.077504 |
| 5 | 31684000 | 31685399 | 4.35241 | 8.274836 | 35.48674 | 14.24548 | -1.58567 | -0.01296 | 0.077506 |
| 5 | 83715200 | 83715799 | 7.937812 | 15.52133 | 6.591701 | 2.140064 | 1.977454 | -0.7894 | 0.077515 |
| 3 | 13373200 | 13374199 | 12.8471 | 24.92492 | 12.73453 | 4.796863 | 1.53817 | -0.19601 | 0.077518 |
| 6 | 29916200 | 29919399 | 12.3612 | 21.46002 | 56.39983 | 24.75044 | -1.00058 | 0.801722 | 0.077526 |
| 4 | 62558600 | 62562799 | 17.8433 | 29.28545 | 85.93404 | 37.00826 | -1.17073 | 1.41783 | 0.077528 |

| | | | | | | | | | |
|---|---|---|---|---|---|---|---|---|---|
| 5 | 111695800 | 111706599 | 56.32243 | 88.06678 | 198.0287 | 82.15686 | -0.85319 | 2.637587 | 0.07753 |
| 17 | 20667400 | 20668999 | 36.26773 | 60.42296 | 38.65237 | 16.79366 | 1.023542 | 1.249004 | 0.077537 |
| 9 | 65283600 | 65286199 | 12.90135 | 22.17806 | 60.66215 | 26.60874 | -1.06129 | 0.898584 | 0.077538 |
| 10 | 5177400 | 5179599 | 10.96356 | 19.10111 | 54.76478 | 24.08645 | -1.12483 | 0.739909 | 0.077542 |
| 4 | 152180600 | 152183199 | 8.9689 | 15.66763 | 51.99282 | 22.90562 | -1.33321 | 0.68976 | 0.07755 |
| 5 | 95624200 | 95626799 | 26.63589 | 46.36798 | 26.73487 | 11.75238 | 1.177317 | 0.783736 | 0.077561 |
| 6 | 86381000 | 86381799 | 2.165734 | 4.229616 | 26.51369 | 9.306896 | -1.93639 | -0.53251 | 0.077574 |
| 7 | 12275600 | 12276599 | 10.60001 | 20.65395 | 11.83487 | 4.29943 | 1.420535 | -0.34959 | 0.077582 |
| 4 | 142571200 | 142571999 | 6.686595 | 12.46569 | 36.55097 | 15.21965 | -1.08826 | 0.144586 | 0.077594 |
| 16 | 28736600 | 28740599 | 12.52141 | 21.36502 | 66.53068 | 29.16703 | -1.24949 | 0.971937 | 0.077599 |
| 4 | 131600200 | 131601999 | 8.132914 | 14.40791 | 45.12229 | 19.95044 | -1.24499 | 0.483291 | 0.077611 |
| 7 | 12655400 | 12656599 | 6.355607 | 11.58151 | 39.21727 | 16.85379 | -1.32754 | 0.281604 | 0.077635 |
| 6 | 26035800 | 26036399 | 9.926986 | 19.3634 | 12.04825 | 4.315718 | 1.321918 | -0.40537 | 0.077672 |
| 13 | 31399000 | 31402199 | 15.68451 | 26.08926 | 76.49471 | 33.20077 | -1.16517 | 1.241716 | 0.077673 |
| 8 | 14960600 | 14980399 | 93.98029 | 140.3518 | 381.4727 | 157.2613 | -1.18084 | 3.545528 | 0.077685 |
| 15 | 76110200 | 76117999 | 43.21638 | 68.26344 | 172.424 | 71.60592 | -1.00906 | 2.425552 | 0.07769 |
| 16 | 37359600 | 37360599 | 4.550936 | 8.779946 | 30.41778 | 11.75434 | -1.22923 | -0.14923 | 0.077693 |
| 16 | 55849600 | 55850599 | 6.11568 | 11.36684 | 35.85885 | 15.00727 | -1.19836 | 0.158158 | 0.077694 |
| 11 | 112595000 | 112598199 | 15.82921 | 26.7587 | 68.2293 | 29.85618 | -0.96403 | 1.070249 | 0.0777 |
| 2 | 172992600 | 173005599 | 62.11441 | 95.32511 | 254.7457 | 105.7594 | -1.12293 | 2.987804 | 0.077728 |
| 19 | 38348600 | 38349399 | 11.33657 | 22.02522 | 13.65548 | 5.094456 | 1.275783 | -0.24282 | 0.077769 |
| 4 | 99993800 | 99995399 | 6.214739 | 11.68249 | 39.8586 | 16.36551 | -1.28907 | 0.092241 | 0.077771 |
| 8 | 119574000 | 119583999 | 62.86248 | 97.78071 | 215.6667 | 89.53351 | -0.83132 | 2.73389 | 0.077775 |
| 3 | 52817200 | 52817799 | 24.46086 | 42.67569 | 27.73972 | 12.21102 | 1.005864 | 0.746093 | 0.077776 |
| 5 | 123251800 | 123261999 | 49.40386 | 77.13739 | 204.3077 | 84.77443 | -1.09063 | 2.66098 | 0.077799 |
| 9 | 86346600 | 86347199 | 3.461597 | 6.741468 | 25.82696 | 9.444713 | -1.29354 | -0.36513 | 0.077802 |
| 14 | 58127000 | 58128399 | 7.572418 | 13.37637 | 47.73684 | 21.1186 | -1.43383 | 0.516249 | 0.077812 |
| 3 | 87949400 | 87950799 | 4.386771 | 8.450187 | 29.67383 | 11.51775 | -1.25354 | -0.1362 | 0.077825 |
| 13 | 54928600 | 54935599 | 40.01125 | 63.52583 | 156.0576 | 65.04089 | -0.96247 | 2.28617 | 0.077833 |
| 18 | 88911800 | 88912799 | 6.057852 | 11.43725 | 35.32604 | 14.39154 | -1.13679 | 0.052327 | 0.077835 |
| 2 | 164914200 | 164924399 | 45.9459 | 72.19924 | 185.4817 | 76.93859 | -1.03938 | 2.539255 | 0.077838 |
| 12 | 23677000 | 23679199 | 51.82245 | 83.92305 | 51.74314 | 22.14108 | 1.059652 | 1.671329 | 0.077848 |
| 7 | 44837400 | 44843799 | 29.38776 | 47.00847 | 131.7308 | 55.44207 | -1.14014 | 2.037114 | 0.077854 |
| 12 | 23301200 | 23302599 | 39.8689 | 65.76516 | 45.61752 | 19.68745 | 0.906181 | 1.375666 | 0.077861 |
| 7 | 31681000 | 31681599 | 9.73709 | 18.96079 | 11.57521 | 4.239428 | 1.320294 | -0.33482 | 0.077887 |
| 10 | 82404200 | 82411199 | 164.1981 | 248.6122 | 177.1447 | 73.28993 | 0.766716 | 3.281328 | 0.07789 |
| 19 | 56995200 | 56998599 | 12.98211 | 21.73701 | 73.05492 | 31.84122 | -1.36076 | 1.158612 | 0.077912 |
| 14 | 70310600 | 70318999 | 45.81813 | 71.56549 | 203.9852 | 84.63736 | -1.19588 | 2.652891 | 0.077913 |
| 11 | 73700000 | 73701399 | 20.37806 | 36.88527 | 19.44718 | 8.428031 | 1.328248 | 0.377518 | 0.077914 |
| 10 | 79797400 | 79853999 | 348.1835 | 467.2858 | 1144.893 | 477.7699 | -1.18407 | 5.219687 | 0.077927 |
| 7 | 20900600 | 20903599 | 31.03323 | 54.09078 | 22.09519 | 9.720486 | 1.671646 | 0.783803 | 0.077927 |
| 6 | 92760000 | 92761999 | 6.71462 | 12.45903 | 37.52723 | 15.74157 | -1.13541 | 0.174814 | 0.077934 |
| 16 | 95381200 | 95382999 | 7.875727 | 14.21806 | 41.17983 | 17.90525 | -1.11587 | 0.350899 | 0.077934 |
| 15 | 76103400 | 76105199 | 6.181641 | 11.36884 | 38.8958 | 16.51365 | -1.33017 | 0.224479 | 0.077941 |
| 17 | 25759600 | 25764199 | 24.66292 | 39.68035 | 116.33 | 49.38439 | -1.19479 | 1.842034 | 0.077942 |
| 4 | 101565200 | 101567799 | 17.06919 | 31.58251 | 16.94026 | 7.133349 | 1.330393 | 0.22411 | 0.07797 |
| 2 | 90780800 | 90781399 | 2.639754 | 5.15563 | 22.71635 | 7.966846 | -1.43415 | -0.53806 | 0.077979 |

| | | | | | | | | | |
|---|---|---|---|---|---|---|---|---|---|
| 18 | 27269400 | 27270599 | 13.72178 | 25.88687 | 15.56885 | 6.35295 | 1.20193 | 0.074377 | 0.078002 |
| 10 | 117574200 | 117578399 | 16.63268 | 27.62124 | 77.0957 | 33.42388 | -1.09471 | 1.264149 | 0.078009 |
| 17 | 64887600 | 64891199 | 12.36093 | 20.81065 | 75.0163 | 32.7855 | -1.46142 | 1.100616 | 0.078011 |
| 10 | 118787400 | 118788999 | 7.458258 | 13.05403 | 51.41002 | 22.68461 | -1.57777 | 0.650023 | 0.078014 |
| 2 | 34764400 | 34765399 | 3.236125 | 6.314412 | 25.10704 | 8.949967 | -1.3135 | -0.46594 | 0.078031 |
| 8 | 4224400 | 4224999 | 3.236125 | 6.314412 | 25.10704 | 8.949967 | -1.3135 | -0.46594 | 0.078031 |
| 7 | 24695200 | 24719999 | 151.1722 | 218.8854 | 525.0803 | 216.539 | -1.04324 | 4.059069 | 0.078038 |
| 7 | 24940200 | 24950999 | 39.34597 | 61.82891 | 190.9289 | 79.19822 | -1.30431 | 2.536169 | 0.078046 |
| 3 | 21265800 | 21266599 | 22.66155 | 39.75565 | 24.66529 | 10.90477 | 1.072556 | 0.647225 | 0.078051 |
| 14 | 41982200 | 41983799 | 12.30097 | 23.81351 | 13.91565 | 5.295322 | 1.333185 | -0.16615 | 0.078067 |
| 6 | 120642200 | 120644399 | 8.824158 | 15.42534 | 51.99119 | 22.9167 | -1.35549 | 0.677741 | 0.078076 |
| 7 | 98610200 | 98613199 | 14.89883 | 25.18773 | 67.5647 | 29.566 | -1.03676 | 1.066073 | 0.078077 |
| 4 | 43035600 | 43037999 | 8.74706 | 15.42703 | 46.16237 | 20.42849 | -1.18241 | 0.539561 | 0.078089 |
| 7 | 14829200 | 14832399 | 73.41378 | 117.2355 | 75.51148 | 31.70545 | 0.974241 | 2.11714 | 0.078093 |
| 15 | 61347200 | 61347799 | 11.11628 | 21.68846 | 10.78871 | 3.851727 | 1.644659 | -0.40419 | 0.078096 |
| 1 | 88627800 | 88630399 | 12.86439 | 21.7848 | 66.2448 | 29.00315 | -1.21636 | 1.044549 | 0.078103 |
| 7 | 78976800 | 78978799 | 6.368698 | 11.9348 | 35.5876 | 14.68888 | -1.10171 | 0.106382 | 0.078109 |
| 2 | 148283400 | 148286399 | 14.36777 | 24.45038 | 63.77271 | 27.95196 | -0.99554 | 1.002845 | 0.078115 |
| 17 | 58625600 | 58626999 | 18.00269 | 32.62433 | 20.8146 | 9.0073 | 1.056459 | 0.355967 | 0.078118 |
| 7 | 98894200 | 98895999 | 14.82404 | 27.64834 | 17.59528 | 7.321136 | 1.099335 | 0.161676 | 0.078123 |
| 7 | 13746000 | 13763799 | 35.29598 | 59.64849 | 32.37895 | 14.16727 | 1.259075 | 1.101907 | 0.078124 |
| 12 | 36091000 | 36092799 | 7.243501 | 12.85984 | 45.66727 | 20.1648 | -1.42313 | 0.460886 | 0.078125 |
| 13 | 108283600 | 108284799 | 7.488074 | 13.76498 | 39.48829 | 16.77607 | -1.07924 | 0.239602 | 0.078136 |
| 6 | 72781000 | 72783999 | 15.68161 | 26.48935 | 68.76071 | 30.08188 | -0.98977 | 1.076653 | 0.078141 |
| X | 130894200 | 130894599 | 8.840101 | 17.25554 | 10.78193 | 3.818574 | 1.331049 | -0.4645 | 0.078145 |
| 14 | 9214000 | 9214999 | 18.88386 | 34.82188 | 17.48487 | 7.396721 | 1.420957 | 0.260088 | 0.078154 |
| 9 | 11825800 | 11828199 | 26.92881 | 47.04876 | 25.65867 | 11.30511 | 1.257724 | 0.735826 | 0.07817 |
| 8 | 83954200 | 83958199 | 16.23346 | 27.18436 | 71.84784 | 31.31755 | -1.01601 | 1.165145 | 0.078171 |
| 14 | 25137600 | 25139999 | 10.00226 | 17.53925 | 49.88057 | 22.04427 | -1.1122 | 0.628396 | 0.078173 |
| 11 | 65501000 | 65502799 | 16.36358 | 30.72275 | 16.68523 | 6.864967 | 1.338663 | 0.132352 | 0.078184 |
| 12 | 103173800 | 103183799 | 54.54563 | 85.22575 | 196.4634 | 81.5136 | -0.88979 | 2.650395 | 0.078187 |
| 10 | 21868800 | 21870599 | 7.952474 | 14.13746 | 43.59191 | 19.22666 | -1.21964 | 0.454138 | 0.078216 |
| 16 | 62207000 | 62207999 | 12.88382 | 24.82799 | 15.10013 | 5.854234 | 1.252024 | -0.09531 | 0.078222 |
| 11 | 87876400 | 87879399 | 16.29039 | 27.15329 | 75.61958 | 32.86515 | -1.09123 | 1.218711 | 0.078222 |
| 2 | 171941200 | 171946399 | 34.96117 | 56.04745 | 126.284 | 53.324 | -0.82264 | 1.971144 | 0.078222 |
| 3 | 42960800 | 42962199 | 14.1966 | 27.38292 | 13.03642 | 5.040849 | 1.603674 | -0.09597 | 0.078224 |
| 5 | 19535600 | 19537199 | 15.41078 | 28.97688 | 16.70858 | 6.857298 | 1.255502 | 0.114067 | 0.078239 |
| 3 | 14377200 | 14379799 | 46.53212 | 75.62784 | 53.32078 | 22.87688 | 0.871428 | 1.590196 | 0.078241 |
| 17 | 26027200 | 26040399 | 55.58287 | 85.46044 | 259.3789 | 107.7105 | -1.30423 | 2.948794 | 0.078247 |
| 12 | 23339600 | 23348399 | 191.5601 | 286.9627 | 199.9737 | 82.48814 | 0.785398 | 3.496722 | 0.078266 |
| 11 | 76281000 | 76287399 | 40.73087 | 64.88764 | 143.5967 | 60.09095 | -0.80701 | 2.179021 | 0.078268 |
| X | 81160600 | 81161799 | 19.49322 | 36.294 | 14.60212 | 6.093548 | 1.750968 | 0.205483 | 0.078275 |
| 1 | 133253000 | 133255199 | 7.045388 | 12.58624 | 43.12666 | 18.94326 | -1.3662 | 0.41561 | 0.078279 |
| 7 | 9340600 | 9345599 | 22.28818 | 39.38701 | 21.26623 | 9.407346 | 1.274169 | 0.550726 | 0.078304 |
| 7 | 31623600 | 31626599 | 60.36632 | 97.06662 | 62.57875 | 26.54325 | 0.984048 | 1.884156 | 0.078357 |
| 6 | 10221800 | 10222599 | 10.51059 | 20.42707 | 13.77175 | 5.124436 | 1.159583 | -0.26163 | 0.078367 |
| 15 | 100913400 | 100917999 | 19.32379 | 31.57113 | 89.72729 | 38.57172 | -1.12847 | 1.484687 | 0.078377 |

| | | | | | | | | | |
|---|---|---|---|---|---|---|---|---|---|
| 11 | 69376800 | 69392199 | 79.71739 | 120.9858 | 294.2237 | 121.8176 | -0.99999 | 3.221795 | 0.078381 |
| 7 | 44524200 | 44539199 | 68.77759 | 104.6246 | 291.1513 | 120.6291 | -1.19146 | 3.171362 | 0.078383 |
| 12 | 22533800 | 22562199 | 569.979 | 778.4079 | 510.4046 | 212.4282 | 0.742538 | 4.946545 | 0.078409 |
| 5 | 35525400 | 35557599 | 257.197 | 357.5648 | 780.4758 | 323.9176 | -0.96603 | 4.666844 | 0.07841 |
| 5 | 130056000 | 130058399 | 11.59783 | 20.14188 | 58.05591 | 25.48286 | -1.13326 | 0.792522 | 0.078434 |
| 17 | 79244200 | 79245199 | 4.711159 | 8.983289 | 31.54047 | 12.58276 | -1.29095 | -0.03682 | 0.078453 |
| 7 | 32137800 | 32142999 | 108.1077 | 168.4416 | 120.7454 | 50.11457 | 0.790157 | 2.713988 | 0.07846 |
| 1 | 171329400 | 171331799 | 11.08394 | 19.38272 | 53.70217 | 23.6782 | -1.07526 | 0.683691 | 0.078478 |
| 6 | 127332400 | 127336999 | 19.83254 | 32.76526 | 81.35148 | 35.14324 | -0.92741 | 1.335606 | 0.078479 |
| 4 | 152529000 | 152563999 | 238.3653 | 332.3795 | 757.9683 | 314.4298 | -1.02479 | 4.620861 | 0.078496 |
| 17 | 25827400 | 25833399 | 25.63668 | 41.24615 | 118.0465 | 50.1115 | -1.16031 | 1.843785 | 0.078497 |
| 18 | 47241000 | 47241999 | 16.54265 | 30.46984 | 18.74653 | 7.94279 | 1.128346 | 0.249143 | 0.078497 |
| 18 | 19054800 | 19055799 | 16.41758 | 30.31408 | 18.77928 | 7.930038 | 1.121995 | 0.235015 | 0.078498 |
| 10 | 40025000 | 40026199 | 5.311555 | 10.09323 | 32.6346 | 13.11465 | -1.18374 | -0.00548 | 0.0785 |
| 4 | 137097000 | 137103399 | 29.97333 | 48.11999 | 123.1353 | 52.10755 | -1.00324 | 1.918985 | 0.078507 |
| 9 | 109704200 | 109706399 | 10.18903 | 17.79655 | 53.1195 | 23.39992 | -1.1819 | 0.697677 | 0.078507 |
| 6 | 93511000 | 93513199 | 9.292374 | 16.56209 | 45.21243 | 19.89812 | -1.04364 | 0.448002 | 0.078516 |
| 5 | 111327800 | 111333199 | 19.76451 | 32.41389 | 86.34257 | 37.17219 | -1.03229 | 1.430833 | 0.078518 |
| 11 | 115126400 | 115154199 | 178.0896 | 252.8998 | 648.7961 | 268.3545 | -1.16806 | 4.347478 | 0.07852 |
| 16 | 17966000 | 17973199 | 30.97174 | 49.32249 | 150.2156 | 62.82896 | -1.26722 | 2.183926 | 0.078533 |
| Y | 9869400 | 9870599 | 10.94671 | 21.3188 | 5.810926 | 2.124924 | 2.46245 | -0.34782 | 0.078535 |
| 5 | 18998800 | 19000599 | 22.46163 | 39.4745 | 25.30886 | 11.20598 | 1.025144 | 0.618852 | 0.078547 |
| 6 | 83459600 | 83460399 | 10.15738 | 19.74463 | 13.29225 | 4.937892 | 1.163685 | -0.27361 | 0.078549 |
| 4 | 35112400 | 35114399 | 9.665674 | 17.03082 | 47.7374 | 21.12955 | -1.08984 | 0.556691 | 0.078555 |
| 3 | 108052800 | 108054799 | 4.761376 | 9.052296 | 33.76912 | 13.55619 | -1.38666 | -0.01252 | 0.078559 |
| 11 | 96885600 | 96895999 | 54.09965 | 84.47391 | 201.4254 | 83.57796 | -0.93925 | 2.661754 | 0.078586 |
| 14 | 18624000 | 18625199 | 13.71637 | 26.47442 | 12.40908 | 4.788367 | 1.627644 | -0.10998 | 0.078594 |
| 8 | 71962000 | 71966599 | 22.79686 | 36.9016 | 101.0781 | 43.23752 | -1.08667 | 1.657173 | 0.078602 |
| 19 | 47719200 | 47720399 | 8.087183 | 14.40045 | 43.08755 | 18.97805 | -1.17514 | 0.443891 | 0.078607 |
| 12 | 105460400 | 105469199 | 41.6705 | 66.04293 | 162.7193 | 67.69699 | -0.96926 | 2.341173 | 0.078611 |
| 10 | 69066200 | 69066799 | 11.10926 | 21.71533 | 7.550467 | 2.600052 | 2.192406 | -0.53613 | 0.078616 |
| 2 | 59955000 | 59956199 | 5.112827 | 9.835752 | 31.5135 | 12.27838 | -1.12933 | -0.11273 | 0.078619 |
| 12 | 66462000 | 66462999 | 13.62536 | 26.29756 | 13.05055 | 5.036955 | 1.546539 | -0.10824 | 0.07862 |
| 7 | 85031600 | 85032399 | 17.18004 | 31.99614 | 17.36931 | 7.24486 | 1.324411 | 0.191491 | 0.078632 |
| 10 | 21608800 | 21609799 | 11.38659 | 22.00489 | 14.77455 | 5.668211 | 1.124729 | -0.14683 | 0.078639 |
| 15 | 35786600 | 35789199 | 25.25586 | 44.00907 | 28.63381 | 12.59495 | 1.004329 | 0.767954 | 0.078657 |
| 6 | 21851000 | 21853799 | 13.3465 | 22.98027 | 59.0624 | 25.90831 | -0.97134 | 0.884624 | 0.078664 |
| 10 | 41685400 | 41687999 | 13.74403 | 23.42186 | 63.79727 | 27.96589 | -1.05752 | 0.986317 | 0.078674 |
| 8 | 122544400 | 122548799 | 18.21546 | 30.16762 | 78.81257 | 34.10417 | -1.00005 | 1.302539 | 0.078675 |
| 5 | 137514800 | 137531999 | 66.84927 | 101.688 | 295.0221 | 122.2317 | -1.25156 | 3.171663 | 0.078685 |
| 10 | 77511600 | 77514599 | 14.64672 | 24.56695 | 70.5355 | 30.77692 | -1.13471 | 1.142829 | 0.078697 |
| 12 | 35584200 | 35586199 | 10.91945 | 19.12467 | 50.97887 | 22.50688 | -1.01957 | 0.654411 | 0.078725 |
| 15 | 100571400 | 100574399 | 8.896413 | 15.71835 | 46.16316 | 20.42158 | -1.15482 | 0.519369 | 0.078732 |
| 13 | 53729200 | 53731199 | 6.736048 | 12.13277 | 46.51101 | 20.2813 | -1.52031 | 0.364745 | 0.078737 |
| X | 25417800 | 25419799 | 28.7886 | 50.35519 | 22.99491 | 10.14088 | 1.512352 | 0.725431 | 0.078743 |
| 5 | 53840400 | 53841199 | 4.67145 | 9.033305 | 30.05785 | 11.49823 | -1.15762 | -0.18333 | 0.078743 |
| 2 | 157728600 | 157730799 | 7.007909 | 12.61651 | 41.64709 | 18.17105 | -1.30602 | 0.362478 | 0.078753 |

| | | | | | | | | | |
|---|---|---|---|---|---|---|---|---|---|
| 7 | 8671400 | 8672799 | 18.33334 | 34.68643 | 12.496 | 5.068133 | 1.940288 | 0.091559 | 0.078755 |
| 1 | 124621200 | 124621999 | 10.74873 | 20.87696 | 12.42076 | 4.644926 | 1.329732 | -0.24785 | 0.078758 |
| 11 | 76404800 | 76406999 | 8.172733 | 14.37763 | 47.72313 | 21.12988 | -1.33242 | 0.567162 | 0.078767 |
| 11 | 15800400 | 15801599 | 15.25907 | 28.90108 | 15.63575 | 6.327122 | 1.363373 | 0.063179 | 0.078807 |
| 12 | 29298000 | 29298999 | 12.7688 | 24.38038 | 16.19076 | 6.440845 | 1.092226 | -0.01287 | 0.078822 |
| 2 | 10116600 | 10117399 | 12.21454 | 23.60134 | 12.81429 | 4.919671 | 1.427528 | -0.14113 | 0.078825 |
| 2 | 165991000 | 166006599 | 77.14786 | 117.4482 | 285.488 | 118.3101 | -0.99565 | 3.156815 | 0.078843 |
| 9 | 44515200 | 44516799 | 7.771958 | 13.92148 | 42.63312 | 18.68421 | -1.20347 | 0.401502 | 0.078849 |
| 17 | 38015400 | 38017599 | 50.62635 | 82.07514 | 55.02226 | 23.56309 | 0.940689 | 1.650485 | 0.078862 |
| 12 | 20784400 | 20791199 | 188.5746 | 283.3524 | 193.4011 | 79.83669 | 0.819225 | 3.43721 | 0.078864 |
| 16 | 13129400 | 13131599 | 17.79884 | 29.3761 | 81.41238 | 35.14536 | -1.08539 | 1.343155 | 0.078868 |
| 7 | 34704400 | 34709199 | 28.11383 | 45.35381 | 108.5586 | 46.25692 | -0.89839 | 1.760221 | 0.078869 |
| 4 | 154084000 | 154085199 | 6.83661 | 12.73303 | 38.2248 | 15.94273 | -1.12409 | 0.155845 | 0.078902 |
| 14 | 118015400 | 118016999 | 7.632206 | 13.66907 | 43.69011 | 19.1499 | -1.26504 | 0.402838 | 0.078903 |
| 4 | 106297200 | 106308799 | 55.11198 | 85.24221 | 230.2462 | 95.63873 | -1.12958 | 2.833043 | 0.078916 |
| 4 | 38157000 | 38157999 | 15.91716 | 29.75817 | 17.89765 | 7.416863 | 1.184875 | 0.157728 | 0.078917 |
| 1 | 128412000 | 128412999 | 12.85403 | 24.90613 | 13.32244 | 5.048729 | 1.464966 | -0.17462 | 0.078918 |
| 1 | 117024200 | 117025399 | 12.09183 | 23.50623 | 11.54263 | 4.286055 | 1.611815 | -0.25735 | 0.078946 |
| 12 | 23334200 | 23336999 | 37.79268 | 64.54062 | 25.98106 | 11.39088 | 1.689136 | 1.016228 | 0.078948 |
| 4 | 134138600 | 134142799 | 16.85992 | 28.22177 | 71.89849 | 31.33146 | -0.96324 | 1.171786 | 0.078998 |
| 12 | 23368200 | 23376399 | 147.3707 | 224.4385 | 164.2071 | 68.05732 | 0.735467 | 3.158548 | 0.079006 |
| 9 | 73489800 | 73491399 | 19.5613 | 34.94687 | 22.34162 | 9.813124 | 1.039887 | 0.454226 | 0.079008 |
| 18 | 9307400 | 9308599 | 19.91954 | 35.51254 | 22.1266 | 9.735829 | 1.074817 | 0.467899 | 0.079049 |
| 9 | 96512600 | 96514799 | 11.01737 | 19.06433 | 57.37732 | 25.17294 | -1.19602 | 0.827167 | 0.079069 |
| 9 | 19017000 | 19017599 | 12.34803 | 23.7201 | 15.72194 | 6.151101 | 1.11706 | -0.07083 | 0.079095 |
| 3 | 3986800 | 3987799 | 20.85629 | 36.84017 | 23.92642 | 10.58505 | 1.009214 | 0.551701 | 0.079096 |
| 2 | 171603600 | 171606199 | 11.22066 | 19.66813 | 52.05726 | 22.99871 | -1.00949 | 0.644518 | 0.079108 |
| 2 | 93073800 | 93077199 | 11.71309 | 19.95528 | 63.54431 | 27.85438 | -1.28128 | 0.983794 | 0.079124 |
| 14 | 65562800 | 65564799 | 6.784677 | 12.38325 | 38.72361 | 16.60825 | -1.21195 | 0.274177 | 0.079175 |
| 7 | 22260400 | 22260799 | 10.73796 | 21.00346 | 4.937278 | 1.656973 | 2.763182 | -0.63654 | 0.079183 |
| 18 | 75578400 | 75583599 | 25.39115 | 41.00791 | 106.493 | 45.44308 | -1.01369 | 1.721815 | 0.079195 |
| 6 | 117226200 | 117228999 | 11.88293 | 20.67799 | 55.34402 | 24.32282 | -1.02646 | 0.762361 | 0.079212 |
| 10 | 93370200 | 93370599 | 2.335776 | 4.565195 | 21.64438 | 7.477005 | -1.51561 | -0.59934 | 0.079221 |
| 4 | 139548200 | 139577199 | 205.0518 | 290.1438 | 658.2482 | 272.4269 | -0.99604 | 4.401447 | 0.079221 |
| 13 | 68179000 | 68181599 | 70.39274 | 112.1681 | 82.15826 | 34.39209 | 0.785797 | 2.181035 | 0.079232 |
| 11 | 58962600 | 58973799 | 63.98435 | 97.73645 | 274.7263 | 113.9477 | -1.20108 | 3.084294 | 0.079249 |
| 4 | 115683000 | 115691399 | 38.50572 | 61.277 | 150.2537 | 62.78342 | -0.95639 | 2.21288 | 0.079249 |
| 7 | 24671200 | 24675399 | 16.60182 | 27.65246 | 75.01682 | 32.58837 | -1.05351 | 1.227625 | 0.079269 |
| 2 | 156153800 | 156154999 | 6.463742 | 11.85967 | 37.75072 | 16.07898 | -1.22999 | 0.239431 | 0.07927 |
| 6 | 93810600 | 93812599 | 9.524772 | 16.7661 | 48.79349 | 21.60123 | -1.14432 | 0.568145 | 0.079272 |
| 7 | 9662800 | 9664599 | 23.15334 | 40.57205 | 26.52652 | 11.71632 | 0.997392 | 0.670915 | 0.079283 |
| 1 | 180395800 | 180403599 | 55.20456 | 86.24951 | 196.9053 | 81.69752 | -0.87591 | 2.65171 | 0.079289 |
| 11 | 14864000 | 14864999 | 18.73127 | 33.58052 | 21.25624 | 9.309072 | 1.056598 | 0.423792 | 0.079302 |
| 4 | 138004800 | 138009799 | 22.00368 | 35.61527 | 101.4278 | 43.38493 | -1.14272 | 1.657566 | 0.079305 |
| 10 | 40224600 | 40226999 | 12.55154 | 21.68688 | 58.5539 | 25.68797 | -1.04099 | 0.847946 | 0.07931 |
| 13 | 91492600 | 91494599 | 26.86058 | 46.64448 | 29.63475 | 13.00705 | 1.038141 | 0.824498 | 0.079328 |
| 11 | 119646800 | 119649199 | 11.23782 | 19.64359 | 51.52923 | 22.71207 | -0.99655 | 0.69014 | 0.079329 |

| | | | | | | | | | |
|---|---|---|---|---|---|---|---|---|---|
| 9 | 107165200 | 107170399 | 22.04342 | 35.6595 | 104.4305 | 44.65079 | -1.18376 | 1.669627 | 0.079335 |
| 12 | 27391200 | 27394999 | 20.83278 | 33.89984 | 94.8513 | 40.71466 | -1.11036 | 1.548318 | 0.07935 |
| 12 | 18051600 | 18056599 | 142.5874 | 217.607 | 159.3475 | 66.07847 | 0.736595 | 3.115386 | 0.079357 |
| 13 | 48810600 | 48837199 | 175.985 | 252.6554 | 584.0703 | 241.1529 | -1.00207 | 4.186756 | 0.079364 |
| 15 | 76681600 | 76683999 | 10.42404 | 18.36331 | 49.88394 | 22.08055 | -1.04527 | 0.566855 | 0.079384 |
| 2 | 112095800 | 112097599 | 7.591602 | 13.65715 | 41.10912 | 17.95096 | -1.1748 | 0.370597 | 0.079385 |
| X | 53907000 | 53908599 | 14.52479 | 27.83932 | 14.88139 | 5.859701 | 1.415767 | -0.03036 | 0.079393 |
| 1 | 75397600 | 75409199 | 60.01784 | 92.44447 | 243.0057 | 100.936 | -1.09511 | 2.913755 | 0.079401 |
| 7 | 31524400 | 31525399 | 16.18937 | 30.32889 | 17.24305 | 7.120949 | 1.269019 | 0.145632 | 0.079432 |
| 14 | 58266800 | 58268199 | 7.718609 | 13.60931 | 47.2053 | 20.8912 | -1.394 | 0.538046 | 0.079456 |
| 10 | 69937800 | 69953999 | 108.6636 | 162.2965 | 369.7365 | 152.4261 | -0.92626 | 3.545348 | 0.079466 |
| 9 | 20488600 | 20489199 | 2.427223 | 4.746466 | 21.24513 | 7.238445 | -1.41547 | -0.64756 | 0.079468 |
| 12 | 5299200 | 5301999 | 10.65214 | 18.22828 | 62.52654 | 27.41802 | -1.38691 | 0.93565 | 0.079481 |
| 10 | 45406600 | 45408399 | 24.17165 | 42.12492 | 28.22307 | 12.41523 | 0.962197 | 0.759437 | 0.079482 |
| 9 | 94418800 | 94421199 | 13.79024 | 23.37781 | 65.84644 | 28.83816 | -1.10649 | 1.036804 | 0.079496 |
| 11 | 32404600 | 32406399 | 7.891389 | 14.16036 | 41.60415 | 18.20617 | -1.14227 | 0.390859 | 0.079496 |
| 17 | 83812000 | 83815399 | 14.36116 | 24.47417 | 65.54857 | 28.73364 | -1.03348 | 0.989242 | 0.079496 |
| 8 | 92809200 | 92810199 | 12.83879 | 24.56361 | 15.74545 | 6.232007 | 1.149657 | -0.02898 | 0.079506 |
| 11 | 70209600 | 70215999 | 38.4927 | 61.2796 | 145.8707 | 60.98408 | -0.91307 | 2.200038 | 0.079509 |
| 6 | 125515000 | 125518599 | 19.29245 | 32.06228 | 79.28365 | 34.39026 | -0.92078 | 1.264517 | 0.079548 |
| Y | 1786400 | 1787399 | 10.75383 | 20.94253 | 6.988993 | 2.556736 | 2.178335 | -0.34406 | 0.079569 |
| 15 | 57507800 | 57508599 | 3.135399 | 6.106555 | 27.08434 | 9.896887 | -1.50212 | -0.37196 | 0.079585 |
| 4 | 92457000 | 92457799 | 12.53249 | 24.06935 | 15.718 | 6.153295 | 1.137586 | -0.06683 | 0.079592 |
| 2 | 19460800 | 19462999 | 9.65284 | 17.04315 | 50.9914 | 22.56058 | -1.1825 | 0.537899 | 0.079595 |
| 11 | 4669000 | 4709399 | 250.6319 | 347.7016 | 807.6349 | 335.2995 | -1.05878 | 4.698748 | 0.079602 |
| 8 | 91634400 | 91636399 | 10.42923 | 18.03362 | 58.96365 | 25.86833 | -1.31513 | 0.832737 | 0.079622 |
| 4 | 146121400 | 146121999 | 15.64009 | 29.195 | 17.60484 | 7.314641 | 1.17846 | 0.164542 | 0.079626 |
| 7 | 17063200 | 17064599 | 17.51428 | 32.91529 | 15.62596 | 6.418529 | 1.533152 | 0.134601 | 0.079645 |
| 18 | 60051800 | 60052999 | 15.1642 | 28.71752 | 16.44885 | 6.658043 | 1.282369 | 0.06636 | 0.079645 |
| 11 | 115892600 | 115894799 | 15.08372 | 25.38904 | 68.50074 | 29.93588 | -1.04529 | 1.107755 | 0.079649 |
| 7 | 10005800 | 10006599 | 13.34682 | 25.75915 | 14.42367 | 5.567753 | 1.375455 | -0.10599 | 0.079659 |
| 17 | 31295400 | 31299599 | 14.06008 | 23.55977 | 72.14182 | 31.45826 | -1.22723 | 1.152208 | 0.079667 |
| 1 | 134487800 | 134489599 | 7.227422 | 12.7877 | 45.1329 | 19.96015 | -1.41623 | 0.49354 | 0.079676 |
| 3 | 23131400 | 23132999 | 13.8676 | 26.54833 | 15.34902 | 6.064362 | 1.298697 | -0.02588 | 0.079683 |
| 1 | 117015800 | 117020399 | 72.96063 | 116.0784 | 80.72779 | 33.72031 | 0.858055 | 2.232346 | 0.079727 |
| 7 | 11269000 | 11271999 | 57.99996 | 93.36489 | 64.54413 | 27.42082 | 0.885583 | 1.85004 | 0.079742 |
| 6 | 140184400 | 140185599 | 5.42235 | 10.38116 | 34.89393 | 13.78445 | -1.21758 | -0.05687 | 0.079749 |
| 10 | 4640800 | 4641999 | 17.04956 | 31.5185 | 18.79496 | 7.923806 | 1.178511 | 0.234585 | 0.079796 |
| 2 | 42991600 | 42992999 | 20.99768 | 37.51377 | 21.47757 | 9.433444 | 1.197841 | 0.461761 | 0.07981 |
| 5 | 120701000 | 120705999 | 14.34054 | 23.7592 | 80.52771 | 34.85551 | -1.37382 | 1.288454 | 0.079825 |
| X | 32143000 | 32145599 | 53.17918 | 86.22615 | 52.32398 | 22.40993 | 1.084218 | 1.651989 | 0.079843 |
| 5 | 122068000 | 122093399 | 123.3071 | 181.3841 | 440.4566 | 181.3847 | -1.03874 | 3.812403 | 0.079874 |
| 17 | 32292400 | 32303199 | 63.46602 | 97.66521 | 242.666 | 100.7821 | -1.01497 | 2.933287 | 0.079892 |
| 10 | 48001600 | 48003399 | 17.41262 | 32.25376 | 18.24288 | 7.670266 | 1.256962 | 0.225689 | 0.079899 |
| 6 | 58844600 | 58845599 | 4.068052 | 7.903901 | 27.41446 | 10.22441 | -1.18039 | -0.28423 | 0.0799 |
| 18 | 62311600 | 62315399 | 19.08318 | 31.44357 | 83.29694 | 35.93218 | -1.02158 | 1.36751 | 0.079911 |
| 9 | 31524600 | 31526199 | 8.545264 | 15.14641 | 44.60244 | 19.71852 | -1.15618 | 0.481459 | 0.079928 |

| | | | | | | | | | |
|---|---|---|---|---|---|---|---|---|---|
| 15 | 82013400 | 82016199 | 9.530948 | 16.76205 | 48.78666 | 21.60205 | -1.14495 | 0.57831 | 0.079937 |
| 6 | 23735400 | 23737999 | 13.53228 | 23.2108 | 60.64256 | 26.59687 | -0.99609 | 0.922184 | 0.07994 |
| 11 | 76216200 | 76217799 | 7.731448 | 13.95847 | 41.71645 | 18.13714 | -1.16082 | 0.349989 | 0.07996 |
| 3 | 130152000 | 130152999 | 3.889372 | 7.556747 | 35.27515 | 13.15601 | -1.60522 | -0.2663 | 0.079981 |
| 16 | 95092000 | 95098599 | 41.03209 | 65.45805 | 142.3839 | 59.69764 | -0.78005 | 2.134597 | 0.079995 |
| 3 | 158871400 | 158872599 | 11.00058 | 21.36125 | 13.52089 | 5.065616 | 1.239675 | -0.23419 | 0.079997 |
| 10 | 27950400 | 27952399 | 15.64535 | 26.12368 | 73.88789 | 32.14793 | -1.11338 | 1.196805 | 0.08 |
| 12 | 33092200 | 33095999 | 20.75102 | 33.96418 | 89.96207 | 38.6981 | -1.02552 | 1.459293 | 0.080017 |
| 14 | 39269800 | 39270799 | 10.59185 | 20.66806 | 9.969545 | 3.550997 | 1.688449 | -0.42348 | 0.080049 |
| 4 | 138152400 | 138155799 | 11.56753 | 19.81457 | 62.65765 | 27.47773 | -1.27025 | 0.928817 | 0.080057 |
| 4 | 132927200 | 132931999 | 19.96336 | 32.64961 | 88.96388 | 38.25789 | -1.06692 | 1.469419 | 0.080062 |
| 8 | 124059000 | 124061399 | 10.54854 | 18.29573 | 56.85003 | 24.94337 | -1.24101 | 0.802793 | 0.08007 |
| 16 | 94582600 | 94591199 | 58.91652 | 91.66571 | 206.1898 | 85.59603 | -0.85927 | 2.727884 | 0.080075 |
| 12 | 22530600 | 22537999 | 168.3447 | 253.9456 | 183.9498 | 76.01628 | 0.738432 | 3.356911 | 0.080082 |
| 7 | 109292800 | 109295599 | 11.72295 | 20.41788 | 53.70271 | 23.61478 | -1.00111 | 0.747319 | 0.080084 |
| 1 | 192084000 | 192086799 | 15.32229 | 26.01503 | 72.33036 | 31.69265 | -1.08827 | 1.032969 | 0.080094 |
| 10 | 43653600 | 43654599 | 7.346088 | 13.48474 | 38.33483 | 16.31766 | -1.06811 | 0.244339 | 0.080094 |
| 5 | 21397000 | 21398999 | 40.94036 | 67.77469 | 42.84094 | 18.5329 | 1.040278 | 1.336808 | 0.080107 |
| 9 | 106241600 | 106266999 | 130.8776 | 191.6498 | 464.8312 | 191.4767 | -1.04329 | 3.88475 | 0.08011 |
| 9 | 8014400 | 8016599 | 9.26146 | 16.2481 | 53.63191 | 23.71183 | -1.32607 | 0.620539 | 0.080154 |
| 12 | 22434200 | 22437999 | 52.45461 | 84.66427 | 54.32122 | 23.16164 | 0.997387 | 1.757072 | 0.080155 |
| 12 | 22434400 | 22437999 | 52.45461 | 84.66427 | 54.32122 | 23.16164 | 0.997387 | 1.757072 | 0.080155 |
| 7 | 135139200 | 135142599 | 19.66832 | 32.14362 | 90.59179 | 38.94759 | -1.11622 | 1.480076 | 0.080157 |
| 18 | 7380600 | 7381999 | 20.43752 | 36.83423 | 19.85065 | 8.649262 | 1.291809 | 0.402069 | 0.080161 |
| 18 | 61144400 | 61153599 | 44.9762 | 70.73082 | 188.3083 | 78.119 | -1.08991 | 2.522607 | 0.080163 |
| 12 | 23547400 | 23550799 | 49.56957 | 80.30753 | 56.64574 | 24.24624 | 0.866615 | 1.66185 | 0.080168 |
| 7 | 96956000 | 96984199 | 174.2696 | 250.1878 | 564.8898 | 233.2343 | -0.96809 | 4.186916 | 0.08017 |
| 4 | 38638400 | 38639199 | 11.18547 | 21.6959 | 14.08899 | 5.313062 | 1.194815 | -0.20813 | 0.080174 |
| 12 | 22115800 | 22116599 | 21.34299 | 37.6686 | 22.12904 | 9.791516 | 1.152736 | 0.56044 | 0.080202 |
| 1 | 97319800 | 97321599 | 23.7074 | 41.82788 | 22.97886 | 10.16821 | 1.248586 | 0.580131 | 0.080206 |
| 4 | 154139800 | 154147399 | 30.12766 | 48.0498 | 143.6197 | 60.19406 | -1.23781 | 2.133057 | 0.080224 |
| 5 | 119823200 | 119827399 | 21.43316 | 34.93721 | 92.32193 | 39.65394 | -1.02636 | 1.522732 | 0.080234 |
| 9 | 15520800 | 15521599 | 17.37389 | 31.65284 | 20.41244 | 8.774665 | 1.047222 | 0.320775 | 0.080237 |
| 4 | 106235600 | 106236599 | 4.262752 | 8.213002 | 29.67992 | 11.51322 | -1.29376 | -0.14025 | 0.080238 |
| 1 | 95914400 | 95915599 | 18.00471 | 32.91735 | 19.69234 | 8.428071 | 1.1586 | 0.304256 | 0.080238 |
| 7 | 113725600 | 113726599 | 4.865446 | 9.313413 | 31.76632 | 12.55492 | -1.23812 | -0.06789 | 0.080265 |
| 15 | 14172600 | 14173999 | 14.19894 | 26.98738 | 16.41073 | 6.59178 | 1.205879 | 0.032139 | 0.080269 |
| 15 | 78526600 | 78529599 | 12.41998 | 20.92721 | 68.75286 | 30.05556 | -1.3276 | 1.091262 | 0.080283 |
| 3 | 148254000 | 148254799 | 7.365379 | 13.42043 | 39.76721 | 17.09194 | -1.13692 | 0.292198 | 0.080285 |
| 4 | 104768800 | 104769599 | 2.937709 | 5.740352 | 27.11133 | 9.420404 | -1.52475 | -0.55654 | 0.080291 |
| 19 | 21813400 | 21814999 | 16.42974 | 30.55427 | 18.39466 | 7.691475 | 1.173375 | 0.19616 | 0.080312 |
| 2 | 23005200 | 23006399 | 13.47915 | 25.86557 | 14.79444 | 5.805827 | 1.322642 | -0.05297 | 0.080323 |
| 9 | 102591600 | 102593199 | 5.326576 | 10.0759 | 33.02046 | 13.39783 | -1.21519 | 0.028044 | 0.080335 |
| 1 | 155911000 | 155912399 | 6.531311 | 11.86715 | 40.77144 | 17.58549 | -1.35223 | 0.302085 | 0.080337 |
| 7 | 17308800 | 17310999 | 40.05676 | 66.47335 | 37.06162 | 16.05971 | 1.220893 | 1.302379 | 0.080337 |
| 2 | 179796000 | 179798199 | 5.37946 | 9.884684 | 37.7313 | 16.03875 | -1.48602 | 0.225417 | 0.080359 |
| 7 | 139477800 | 139483399 | 28.15972 | 45.3244 | 114.102 | 48.46649 | -0.97501 | 1.832564 | 0.080398 |

| | | | | | | | | | |
|---|---|---|---|---|---|---|---|---|---|
| 4 | 141185200 | 141208799 | 207.7855 | 291.8243 | 699.3458 | 289.7878 | -1.08591 | 4.512443 | 0.080405 |
| 5 | 95871800 | 95871999 | 6.932224 | 13.55915 | 8.152054 | 2.682521 | 1.47795 | -0.75625 | 0.080411 |
| 5 | 93911200 | 93912399 | 17.90031 | 33.55467 | 14.71067 | 6.068847 | 1.640879 | 0.150485 | 0.080415 |
| 17 | 6025600 | 6030199 | 18.90886 | 30.88223 | 93.32549 | 40.11331 | -1.21723 | 1.48921 | 0.080426 |
| 12 | 24032600 | 24035199 | 23.83678 | 41.68303 | 26.70495 | 11.77446 | 1.026764 | 0.7081 | 0.080443 |
| 1 | 39026200 | 39030999 | 23.54557 | 38.25299 | 97.04923 | 41.62604 | -0.97153 | 1.580339 | 0.080446 |
| 1 | 191205200 | 191210799 | 37.7848 | 60.31609 | 136.6379 | 57.34 | -0.83745 | 2.109788 | 0.080452 |
| 12 | 114029000 | 114029799 | 24.41392 | 42.53951 | 28.29481 | 12.44538 | 0.972685 | 0.76398 | 0.080465 |
| 2 | 170572600 | 170575199 | 15.59824 | 26.16536 | 71.29569 | 31.11044 | -1.0598 | 1.14372 | 0.080472 |
| X | 4254800 | 4256199 | 16.92507 | 32.63145 | 10.60322 | 4.105138 | 2.145924 | -0.06942 | 0.080472 |
| 4 | 16333800 | 16334999 | 16.5563 | 31.07535 | 16.68056 | 6.866503 | 1.354879 | 0.135997 | 0.080477 |
| 6 | 74682800 | 74684199 | 12.03326 | 23.24201 | 15.15597 | 5.829279 | 1.162884 | -0.13015 | 0.080485 |
| 17 | 57037000 | 57039399 | 7.601324 | 13.36269 | 49.28691 | 21.82542 | -1.48391 | 0.574076 | 0.080491 |
| 4 | 150773400 | 150775999 | 12.44847 | 21.38937 | 60.61454 | 26.58747 | -1.11221 | 0.901215 | 0.080503 |
| 11 | 116101800 | 116126799 | 147.0276 | 212.3041 | 541.4351 | 223.3642 | -1.13537 | 4.099582 | 0.080504 |
| 7 | 9926600 | 9927599 | 34.16511 | 57.54206 | 36.8192 | 16.09353 | 1.022871 | 1.132506 | 0.080505 |
| 5 | 18239400 | 18240199 | 11.3011 | 22.05707 | 10.1682 | 3.607525 | 1.760336 | -0.42478 | 0.080541 |
| 8 | 84792600 | 84794799 | 7.999084 | 14.15625 | 46.16832 | 20.41714 | -1.30377 | 0.49534 | 0.08055 |
| 3 | 90131600 | 90136599 | 22.30045 | 36.56173 | 91.1672 | 39.24424 | -0.93778 | 1.441623 | 0.080554 |
| 14 | 8846000 | 8846799 | 11.48085 | 22.21077 | 14.60989 | 5.577141 | 1.160158 | -0.16325 | 0.080555 |
| 8 | 123585400 | 123600199 | 137.2515 | 199.6055 | 496.2253 | 204.5359 | -1.08847 | 3.991752 | 0.080606 |
| 7 | 30670200 | 30677799 | 32.50775 | 51.89252 | 141.179 | 59.24599 | -1.10103 | 2.10527 | 0.080609 |
| 9 | 118954400 | 118959999 | 17.68468 | 28.78566 | 95.85624 | 41.15047 | -1.3601 | 1.539082 | 0.080611 |
| 15 | 59697600 | 59700999 | 16.26239 | 27.17678 | 74.05712 | 32.23862 | -1.06 | 1.18888 | 0.080612 |
| 8 | 124439800 | 124447599 | 30.49723 | 48.85444 | 127.9118 | 53.94965 | -1.03999 | 1.990344 | 0.080616 |
| 15 | 33738600 | 33740399 | 18.90739 | 34.78569 | 17.33075 | 7.354357 | 1.428621 | 0.271614 | 0.080621 |
| 1 | 189229400 | 189232399 | 13.02944 | 22.46127 | 59.08268 | 25.91815 | -1.00429 | 0.871648 | 0.080626 |
| 7 | 10986000 | 10988999 | 62.69872 | 100.7458 | 65.40111 | 27.71198 | 0.972928 | 1.912089 | 0.080633 |
| 11 | 50021600 | 50024799 | 10.68696 | 18.55232 | 55.78393 | 24.47925 | -1.19356 | 0.793164 | 0.080643 |
| 2 | 180594800 | 180596799 | 5.095 | 9.255259 | 41.78961 | 18.02993 | -1.74211 | 0.295866 | 0.080651 |
| 17 | 31409800 | 31419399 | 44.26726 | 69.75602 | 182.6715 | 75.80043 | -1.06385 | 2.480763 | 0.080657 |
| 8 | 48295200 | 48296199 | 5.296117 | 10.11115 | 32.26255 | 12.83733 | -1.15225 | -0.03988 | 0.080666 |
| 16 | 93341800 | 93343199 | 5.296117 | 10.11115 | 32.26255 | 12.83733 | -1.15225 | -0.03988 | 0.080666 |
| X | 29552800 | 29554799 | 21.88068 | 39.20175 | 20.12617 | 8.818873 | 1.356638 | 0.449443 | 0.080681 |
| 13 | 54616000 | 54617999 | 8.26787 | 14.6652 | 44.08439 | 19.4807 | -1.18534 | 0.47113 | 0.080683 |
| 9 | 107978200 | 107999799 | 144.1477 | 208.0212 | 539.9696 | 222.7772 | -1.16169 | 4.108554 | 0.080684 |
| 13 | 51866800 | 51893799 | 143.0295 | 209.6237 | 456.275 | 187.9421 | -0.88609 | 3.872151 | 0.080684 |
| 12 | 22480200 | 22489599 | 247.1224 | 362.4649 | 247.0491 | 101.7559 | 0.789233 | 3.864677 | 0.080687 |
| 8 | 125457600 | 125468999 | 54.89088 | 85.72606 | 197.6271 | 82.00028 | -0.89039 | 2.658657 | 0.08069 |
| 17 | 76826000 | 76827399 | 14.18736 | 27.40516 | 12.43743 | 4.781596 | 1.67916 | -0.1162 | 0.080697 |
| X | 33975600 | 33977599 | 24.29296 | 44.39595 | 12.47147 | 5.340332 | 2.237249 | 0.339573 | 0.080706 |
| 7 | 20871000 | 20872199 | 24.14555 | 42.18616 | 27.65516 | 12.18479 | 0.993811 | 0.724501 | 0.08072 |
| 1 | 128528400 | 128531999 | 12.85818 | 22.21265 | 57.44184 | 25.19996 | -0.97899 | 0.851248 | 0.080723 |
| 11 | 106651000 | 106656199 | 20.43955 | 33.46488 | 92.39504 | 39.74973 | -1.08503 | 1.455266 | 0.080725 |
| 1 | 102595400 | 102596999 | 19.29908 | 34.98001 | 20.25256 | 8.762089 | 1.195981 | 0.364239 | 0.08073 |
| 7 | 107647800 | 107649199 | 14.42561 | 27.65244 | 14.76977 | 5.813802 | 1.416874 | -0.03313 | 0.080742 |
| 4 | 73977400 | 73977799 | 7.159363 | 13.95121 | 6.298421 | 1.949205 | 1.952447 | -0.96404 | 0.080748 |

| | | | | | | | | | |
|---|---|---|---|---|---|---|---|---|---|
| 5 | 33838200 | 33847999 | 45.74168 | 72.01378 | 183.0476 | 75.94748 | -1.02196 | 2.500176 | 0.080761 |
| 4 | 149723800 | 149748599 | 171.4169 | 245.6678 | 571.4339 | 236.0001 | -1.01348 | 4.212291 | 0.080774 |
| 1 | 23622800 | 23623799 | 7.650615 | 14.02697 | 39.9734 | 17.04282 | -1.07372 | 0.255961 | 0.080786 |
| 3 | 11835400 | 11837399 | 23.11424 | 40.81484 | 21.73108 | 9.614482 | 1.293417 | 0.563806 | 0.080793 |
| 7 | 9137400 | 9139199 | 20.18131 | 38.21078 | 9.410489 | 3.810792 | 2.481072 | 0.096348 | 0.08081 |
| 2 | 33512000 | 33538199 | 171.3732 | 245.6131 | 582.8353 | 240.7077 | -1.04224 | 4.211699 | 0.080832 |
| 18 | 81685600 | 81688599 | 11.39622 | 19.90626 | 52.07815 | 22.94204 | -0.99287 | 0.70202 | 0.08085 |
| 9 | 108933200 | 108936199 | 12.32779 | 20.70567 | 70.86972 | 30.94941 | -1.38676 | 1.120894 | 0.080856 |
| 7 | 14109200 | 14113599 | 104.2434 | 163.0958 | 113.6216 | 47.13564 | 0.835915 | 2.640083 | 0.08086 |
| 1 | 72878400 | 72880399 | 7.965776 | 14.17854 | 45.18153 | 19.90712 | -1.26619 | 0.447315 | 0.080881 |
| 1 | 175422400 | 175423999 | 16.68818 | 31.18016 | 16.49055 | 6.840987 | 1.366789 | 0.163851 | 0.080897 |
| 13 | 54936400 | 54948799 | 67.13751 | 102.5598 | 275.6615 | 114.3373 | -1.13651 | 3.083523 | 0.080897 |
| 2 | 156558000 | 156561199 | 10.98884 | 19.08477 | 55.81323 | 24.49888 | -1.15376 | 0.787687 | 0.080899 |
| 2 | 172872200 | 172890399 | 91.97895 | 137.3785 | 380.3854 | 156.8165 | -1.20743 | 3.543112 | 0.080905 |
| 12 | 74435000 | 74436999 | 21.06822 | 37.35452 | 23.19458 | 10.25315 | 1.075569 | 0.512249 | 0.080906 |
| 7 | 21061800 | 21062999 | 19.23486 | 36.26597 | 12.43665 | 5.080582 | 2.002006 | 0.122301 | 0.080919 |
| 18 | 69575800 | 69583199 | 61.03098 | 95.44794 | 199.0901 | 82.59551 | -0.74474 | 2.636806 | 0.08092 |
| 8 | 92366600 | 92371399 | 18.94689 | 31.57749 | 77.16933 | 33.53619 | -0.90367 | 1.228681 | 0.080928 |
| 5 | 105462800 | 105464399 | 21.43538 | 38.84599 | 17.57529 | 7.605286 | 1.54815 | 0.374168 | 0.08093 |
| 4 | 83959400 | 83960799 | 17.1092 | 31.79332 | 17.59332 | 7.365018 | 1.292257 | 0.203675 | 0.080933 |
| 14 | 94742800 | 94744399 | 17.1092 | 31.79332 | 17.59332 | 7.365018 | 1.292257 | 0.203675 | 0.080933 |
| 5 | 113021200 | 113027999 | 36.74046 | 58.45687 | 148.3614 | 61.97668 | -1.00622 | 2.217775 | 0.080935 |
| 10 | 98749400 | 98752199 | 18.58419 | 30.27661 | 93.21026 | 40.02749 | -1.24634 | 1.526887 | 0.080963 |
| 18 | 75648000 | 75655599 | 36.19901 | 57.42852 | 164.1573 | 68.3635 | -1.18172 | 2.307628 | 0.080972 |
| 1 | 26166600 | 26167799 | 15.9939 | 30.40864 | 14.47963 | 5.812205 | 1.555653 | 0.042338 | 0.080977 |
| 3 | 55046000 | 55050599 | 30.21014 | 48.68492 | 111.0662 | 47.26174 | -0.83107 | 1.795233 | 0.080979 |
| 11 | 117691800 | 117707199 | 55.7611 | 85.45711 | 276.4393 | 114.744 | -1.39969 | 3.014311 | 0.080996 |
| 16 | 45282000 | 45283199 | 14.97595 | 28.03961 | 17.91955 | 7.407519 | 1.100528 | 0.139714 | 0.081008 |
| 6 | 90190800 | 90192999 | 29.97558 | 51.68636 | 30.18975 | 13.24361 | 1.157192 | 0.904918 | 0.081009 |
| 2 | 67116200 | 67117199 | 18.19376 | 33.25259 | 19.68813 | 8.429561 | 1.173108 | 0.307589 | 0.081031 |
| 2 | 34042600 | 34046999 | 16.417 | 27.09587 | 83.5377 | 36.06324 | -1.23855 | 1.340261 | 0.081048 |
| 6 | 98239200 | 98240999 | 28.82545 | 50.42086 | 22.77748 | 10.0452 | 1.527609 | 0.724126 | 0.081063 |
| 14 | 41935200 | 41938599 | 29.11577 | 50.00587 | 32.28997 | 14.16323 | 1.011961 | 0.936599 | 0.081079 |
| 16 | 31093400 | 31094199 | 4.131015 | 7.991709 | 30.02433 | 11.46221 | -1.32768 | -0.19712 | 0.081095 |
| X | 20937400 | 20937799 | 10.96512 | 21.33749 | 13.01493 | 4.793462 | 1.314957 | -0.2936 | 0.081105 |
| 15 | 12561000 | 12563999 | 14.15656 | 24.2594 | 62.90218 | 27.58588 | -0.9856 | 0.934648 | 0.081128 |
| 1 | 79583000 | 79583799 | 11.50821 | 22.41852 | 11.81594 | 4.305243 | 1.536651 | -0.32684 | 0.08113 |
| 11 | 107021800 | 107023599 | 7.885484 | 14.20748 | 41.13624 | 17.9307 | -1.11771 | 0.36282 | 0.081143 |
| 17 | 37251800 | 37255399 | 25.6384 | 41.57216 | 100.7101 | 43.13434 | -0.90785 | 1.625435 | 0.081151 |
| 8 | 23422200 | 23422999 | 4.215983 | 8.106173 | 30.1161 | 11.75174 | -1.34038 | -0.11744 | 0.081176 |
| 7 | 74203400 | 74205999 | 12.11481 | 20.88235 | 59.01612 | 25.88886 | -1.1072 | 0.8689 | 0.081178 |
| 17 | 88936800 | 88937799 | 3.834061 | 7.437746 | 28.40372 | 10.70449 | -1.33212 | -0.24861 | 0.081185 |
| X | 4343400 | 4351399 | 119.9082 | 185.6147 | 126.8853 | 52.70488 | 0.851385 | 2.833485 | 0.081196 |
| 9 | 25251000 | 25252199 | 8.055172 | 14.39355 | 42.60476 | 18.71031 | -1.15651 | 0.421405 | 0.08127 |
| 9 | 46420600 | 46425199 | 26.62033 | 43.08955 | 102.2459 | 43.7362 | -0.88042 | 1.66356 | 0.081284 |
| 5 | 108220400 | 108222399 | 19.19216 | 34.30611 | 22.34629 | 9.810659 | 1.013438 | 0.448063 | 0.081288 |
| 7 | 29315200 | 29315399 | 2.637348 | 5.158162 | 21.44904 | 7.257773 | -1.30471 | -0.67127 | 0.081292 |

| | | | | | | | | | |
|---|---|---|---|---|---|---|---|---|---|
| 7 | 17368000 | 17368999 | 30.93267 | 52.85354 | 33.45197 | 14.66698 | 1.039863 | 0.99234 | 0.081305 |
| 7 | 11937000 | 11939199 | 50.62738 | 82.20396 | 52.85539 | 22.66098 | 1.00277 | 1.618904 | 0.081315 |
| 4 | 114417200 | 114418799 | 6.614657 | 12.15899 | 38.35215 | 16.29418 | -1.21477 | 0.229803 | 0.081318 |
| X | 153144800 | 153146399 | 35.53498 | 59.94252 | 35.13821 | 15.36603 | 1.148798 | 1.122856 | 0.081322 |
| 1 | 190235000 | 190241999 | 47.41816 | 74.87912 | 171.5075 | 71.21504 | -0.86853 | 2.434755 | 0.081331 |
| 7 | 59347600 | 59349399 | 16.45937 | 30.47955 | 17.62954 | 7.415195 | 1.224889 | 0.218194 | 0.081342 |
| 17 | 37320400 | 37321399 | 3.828127 | 7.442917 | 28.81139 | 10.68528 | -1.33022 | -0.3059 | 0.081364 |
| 11 | 52369600 | 52372199 | 10.51175 | 17.98872 | 63.0699 | 27.65636 | -1.41834 | 0.934951 | 0.081372 |
| 8 | 41457200 | 41459199 | 8.170551 | 14.7633 | 42.83166 | 18.60297 | -1.11766 | 0.349891 | 0.08138 |
| 4 | 143654800 | 143658799 | 92.36135 | 146.2318 | 88.7386 | 36.89571 | 1.047887 | 2.388659 | 0.081384 |
| 7 | 15315600 | 15318799 | 83.38327 | 132.7508 | 76.24464 | 31.87764 | 1.134642 | 2.220968 | 0.081388 |
| 7 | 96154600 | 96155599 | 5.259992 | 10.07967 | 31.85346 | 12.54989 | -1.12468 | -0.07245 | 0.081395 |
| 4 | 152613400 | 152662999 | 361.6197 | 484.3327 | 1177.36 | 491.4668 | -1.17567 | 5.252132 | 0.081399 |
| 1 | 151630000 | 151631799 | 9.843818 | 17.2647 | 48.81451 | 21.57659 | -1.10363 | 0.623744 | 0.081401 |
| 6 | 73248600 | 73250599 | 12.71127 | 21.94248 | 57.98745 | 25.43875 | -1.01045 | 0.856962 | 0.081404 |
| 5 | 121581800 | 121583199 | 4.995013 | 9.561052 | 31.17993 | 12.32456 | -1.17396 | -0.06683 | 0.081411 |
| 4 | 97760000 | 97760599 | 2.735962 | 5.350804 | 25.56182 | 8.681526 | -1.50856 | -0.64306 | 0.081429 |
| 2 | 176504600 | 176505599 | 13.69382 | 26.60558 | 10.84799 | 4.046355 | 1.871068 | -0.22424 | 0.081441 |
| 7 | 9748000 | 9754599 | 160.5004 | 243.6097 | 170.2105 | 70.47415 | 0.797907 | 3.231216 | 0.081449 |
| 17 | 13650800 | 13655199 | 101.2699 | 152.4431 | 323.4109 | 133.5697 | -0.81338 | 3.392928 | 0.081461 |
| 12 | 21864600 | 21867399 | 43.79223 | 71.78429 | 48.38486 | 20.82633 | 0.94403 | 1.464497 | 0.081464 |
| 10 | 127011800 | 127017399 | 22.82326 | 36.87669 | 104.5692 | 44.64717 | -1.13931 | 1.705312 | 0.081477 |
| 1 | 188746800 | 188748799 | 6.867632 | 12.71228 | 36.89699 | 15.52802 | -1.08564 | 0.18937 | 0.08148 |
| 14 | 83103200 | 83104599 | 15.78243 | 30.09105 | 13.90582 | 5.548786 | 1.605365 | 0.020172 | 0.081482 |
| 2 | 127318400 | 127327999 | 50.10824 | 78.13645 | 208.8173 | 86.65773 | -1.10502 | 2.684606 | 0.081485 |
| X | 4245200 | 4247399 | 29.7956 | 52.17032 | 20.42079 | 9.013524 | 1.732593 | 0.709999 | 0.08152 |
| 3 | 47582400 | 47583399 | 12.49361 | 24.15318 | 14.48322 | 5.546834 | 1.288029 | -0.14114 | 0.081523 |
| 7 | 20998000 | 21002999 | 49.71554 | 81.99266 | 36.21431 | 15.62805 | 1.554923 | 1.399201 | 0.081526 |
| 1 | 158957800 | 158958399 | 10.57815 | 20.58743 | 13.23779 | 4.869909 | 1.242175 | -0.30089 | 0.081527 |
| 18 | 33861600 | 33862799 | 12.2174 | 23.59467 | 15.15176 | 5.831115 | 1.18409 | -0.12603 | 0.081532 |
| 11 | 84566800 | 84568199 | 6.383623 | 11.55589 | 40.15913 | 17.40189 | -1.37248 | 0.323395 | 0.081547 |
| 8 | 121018400 | 121024199 | 22.96208 | 36.90906 | 121.3955 | 51.46445 | -1.36196 | 1.871592 | 0.081575 |
| 7 | 127522800 | 127530799 | 29.49321 | 47.01965 | 144.8093 | 60.66003 | -1.28146 | 2.145716 | 0.081581 |
| 19 | 57672200 | 57672799 | 2.627608 | 5.127025 | 26.14869 | 9.322338 | -1.66206 | -0.46359 | 0.081582 |
| 12 | 35600800 | 35602199 | 22.53958 | 40.60078 | 18.07917 | 7.882722 | 1.564412 | 0.417022 | 0.081585 |
| 10 | 79612400 | 79621199 | 39.77942 | 62.95545 | 166.7947 | 69.33556 | -1.07553 | 2.376381 | 0.081597 |
| 5 | 73069800 | 73072999 | 15.39093 | 25.95674 | 70.38605 | 30.778 | -1.05266 | 1.090858 | 0.081661 |
| 6 | 52166200 | 52169399 | 9.450011 | 16.48025 | 53.61366 | 23.59465 | -1.3054 | 0.717806 | 0.081674 |
| 7 | 112194400 | 112196599 | 10.50481 | 18.31936 | 54.20838 | 23.85606 | -1.16987 | 0.723109 | 0.08168 |
| 15 | 5127800 | 5129999 | 16.1258 | 26.88487 | 74.43336 | 32.35411 | -1.0827 | 1.215475 | 0.081689 |
| 13 | 54456600 | 54458599 | 7.726688 | 13.92627 | 40.61502 | 17.69571 | -1.12738 | 0.359662 | 0.081696 |
| 7 | 12239200 | 12243399 | 55.08498 | 89.14609 | 54.06621 | 23.12358 | 1.082491 | 1.692975 | 0.081708 |
| 14 | 51472000 | 51475399 | 37.56801 | 62.23769 | 43.95305 | 19.02365 | 0.880935 | 1.319313 | 0.08172 |
| 1 | 7724200 | 7725199 | 13.18762 | 25.04713 | 16.59458 | 6.6758 | 1.081416 | 0.027371 | 0.08176 |
| 7 | 42653200 | 42657199 | 42.59019 | 69.63587 | 48.90092 | 21.02714 | 0.883139 | 1.494703 | 0.08177 |
| 4 | 18522400 | 18524399 | 22.28668 | 40.42653 | 17.12746 | 7.402838 | 1.643522 | 0.375112 | 0.081778 |
| 7 | 79101200 | 79102199 | 4.567105 | 8.76285 | 30.67709 | 12.04309 | -1.26543 | -0.09529 | 0.081781 |

| | | | | | | | | | |
|---|---|---|---|---|---|---|---|---|---|
| 5 | 112921200 | 112925999 | 28.74763 | 46.39098 | 109.6052 | 46.72202 | -0.87917 | 1.751797 | 0.081785 |
| 7 | 10673200 | 10674799 | 50.65745 | 82.00204 | 56.58168 | 24.20403 | 0.897234 | 1.681487 | 0.081789 |
| 4 | 47767200 | 47768399 | 13.6122 | 25.95225 | 16.56127 | 6.608918 | 1.145286 | 0.006601 | 0.081813 |
| 6 | 121052200 | 121059799 | 49.98879 | 78.5389 | 182.3488 | 75.63735 | -0.89382 | 2.544824 | 0.081833 |
| 7 | 33221800 | 33225799 | 105.6513 | 165.9785 | 100.0092 | 41.48287 | 1.049524 | 2.565304 | 0.081848 |
| 2 | 27771200 | 27782199 | 49.88321 | 77.9353 | 204.8485 | 84.99325 | -1.07891 | 2.649554 | 0.08185 |
| 1 | 60081600 | 60083599 | 10.83707 | 18.83274 | 56.36402 | 24.7502 | -1.18688 | 0.77823 | 0.081851 |
| 4 | 63231200 | 63237599 | 26.76585 | 43.02638 | 119.7454 | 50.77033 | -1.12166 | 1.872877 | 0.081861 |
| 12 | 112279600 | 112290799 | 73.2714 | 112.3618 | 250.8499 | 104.1312 | -0.86467 | 3.005834 | 0.081861 |
| 12 | 20135600 | 20149799 | 351.6298 | 500.3145 | 343.4901 | 142.0275 | 0.735777 | 4.324071 | 0.081872 |
| 15 | 97571600 | 97572799 | 6.215235 | 11.68179 | 34.55684 | 14.19203 | -1.08439 | 0.082947 | 0.081881 |
| 6 | 114845600 | 114850199 | 17.70633 | 29.02226 | 86.91164 | 37.40835 | -1.20075 | 1.43486 | 0.081889 |
| 4 | 130882200 | 130888599 | 32.37484 | 51.63154 | 144.7194 | 60.65133 | -1.14545 | 2.136156 | 0.081897 |
| 3 | 87709000 | 87711799 | 12.52844 | 21.27941 | 64.64301 | 28.32753 | -1.2146 | 1.015371 | 0.081904 |
| 1 | 35668400 | 35669999 | 17.18798 | 31.32382 | 20.41598 | 8.7728 | 1.032315 | 0.317475 | 0.081906 |
| 15 | 20185000 | 20186199 | 16.59297 | 31.01472 | 15.89208 | 6.588236 | 1.412449 | 0.158707 | 0.081915 |
| 15 | 25704600 | 25707799 | 25.17152 | 40.79224 | 100.4007 | 42.98405 | -0.9314 | 1.636123 | 0.081924 |
| 13 | 65642400 | 65647599 | 122.3176 | 190.0656 | 114.7897 | 47.66202 | 1.034066 | 2.766471 | 0.081925 |
| X | 3724800 | 3726399 | 18.47135 | 34.93292 | 13.44532 | 5.457988 | 1.846588 | 0.098486 | 0.081926 |
| 17 | 48773200 | 48776999 | 25.44069 | 41.34011 | 96.23892 | 41.28455 | -0.84758 | 1.580265 | 0.081932 |
| 18 | 3160400 | 3162799 | 41.51064 | 68.4544 | 45.06857 | 19.44827 | 0.981058 | 1.382774 | 0.081932 |
| 1 | 134483000 | 134484999 | 8.095295 | 14.43885 | 43.62807 | 19.18923 | -1.18809 | 0.432632 | 0.081937 |
| 18 | 80538200 | 80541199 | 11.51092 | 19.81571 | 58.9316 | 25.85077 | -1.18059 | 0.879137 | 0.081938 |
| 18 | 83723400 | 83724399 | 10.79687 | 20.96586 | 14.23129 | 5.331398 | 1.140887 | -0.23373 | 0.081946 |
| 11 | 119381800 | 119382799 | 2.463194 | 4.791802 | 28.55515 | 10.5424 | -1.93343 | -0.33797 | 0.081948 |
| X | 148130600 | 148131799 | 13.42527 | 25.60308 | 16.56598 | 6.606854 | 1.126178 | 0.002736 | 0.081963 |
| 7 | 4472200 | 4482599 | 67.83777 | 104.8698 | 225.4801 | 93.65927 | -0.80151 | 2.847063 | 0.082 |
| 17 | 26076000 | 26080999 | 19.05521 | 31.08813 | 91.872 | 39.47324 | -1.18611 | 1.505421 | 0.082 |
| 17 | 28487800 | 28515399 | 147.6279 | 213.6828 | 522.4916 | 215.4808 | -1.07127 | 4.063823 | 0.082022 |
| 15 | 98698600 | 98709799 | 61.3805 | 94.49978 | 248.9538 | 103.4001 | -1.09881 | 2.923442 | 0.082024 |
| 17 | 18213400 | 18219199 | 90.64015 | 143.1818 | 98.60885 | 40.95256 | 0.86307 | 2.44234 | 0.082042 |
| 12 | 102279000 | 102280199 | 5.568826 | 10.63339 | 36.54563 | 14.53695 | -1.25877 | -0.02828 | 0.082048 |
| 13 | 87160800 | 87162199 | 16.01674 | 29.94055 | 17.79627 | 7.376434 | 1.201291 | 0.158762 | 0.082063 |
| 6 | 127267200 | 127269599 | 12.02211 | 20.95809 | 54.81778 | 24.11897 | -0.99316 | 0.736284 | 0.082073 |
| 12 | 57313600 | 57314199 | 3.343625 | 6.52869 | 25.42094 | 8.954285 | -1.26816 | -0.50598 | 0.082082 |
| 12 | 22300000 | 22301399 | 20.43899 | 37.62412 | 15.52128 | 6.581591 | 1.698921 | 0.277065 | 0.082094 |
| 4 | 129853000 | 129856199 | 12.51679 | 21.33347 | 63.6832 | 27.9162 | -1.18881 | 0.980783 | 0.082096 |
| 11 | 59617200 | 59618799 | 2.460012 | 4.793719 | 28.36597 | 10.29298 | -1.90075 | -0.40246 | 0.082101 |
| 10 | 90749200 | 90751999 | 10.30303 | 18.07339 | 50.41849 | 22.28901 | -1.08473 | 0.624082 | 0.082105 |
| 11 | 120488200 | 120489599 | 7.075318 | 12.93555 | 38.77035 | 16.5932 | -1.14952 | 0.26575 | 0.08211 |
| 13 | 37918000 | 37922199 | 27.44117 | 44.34806 | 103.6183 | 44.25404 | -0.86056 | 1.704859 | 0.082118 |
| 10 | 80003200 | 80004599 | 4.70425 | 8.991985 | 31.05947 | 12.32605 | -1.26086 | -0.05569 | 0.082118 |
| 7 | 11282800 | 11287399 | 71.64304 | 114.3412 | 73.29199 | 30.74815 | 0.980277 | 2.13138 | 0.082122 |
| 18 | 84105400 | 84107799 | 8.54425 | 14.9439 | 53.59133 | 23.63251 | -1.44459 | 0.668654 | 0.082139 |
| 11 | 114761200 | 114768799 | 31.98956 | 51.12288 | 134.7507 | 56.63864 | -1.05359 | 2.06851 | 0.082152 |
| 1 | 37198800 | 37200399 | 39.50777 | 65.79931 | 39.43238 | 17.12901 | 1.117275 | 1.261854 | 0.082177 |
| 5 | 121358000 | 121359799 | 6.102561 | 11.31082 | 36.32515 | 15.2603 | -1.22748 | 0.176892 | 0.082188 |

| | | | | | | | | | |
|---|---|---|---|---|---|---|---|---|---|
| 15 | 82086000 | 82112599 | 135.4313 | 196.7154 | 521.1179 | 214.8238 | -1.18181 | 4.010323 | 0.082195 |
| 10 | 76544400 | 76556199 | 75.54593 | 114.9157 | 288.4112 | 119.4923 | -1.04271 | 3.173449 | 0.082207 |
| 6 | 69463600 | 69466399 | 34.14378 | 57.51419 | 37.18569 | 16.25442 | 1.007812 | 1.129651 | 0.082208 |
| 2 | 123921600 | 123922599 | 5.741917 | 10.90845 | 37.48051 | 15.06955 | -1.27253 | 0.009935 | 0.082256 |
| 5 | 117824600 | 117834799 | 51.73709 | 81.00027 | 194.2218 | 80.56691 | -0.94424 | 2.612284 | 0.082266 |
| 4 | 48539400 | 48543199 | 12.99949 | 22.12265 | 65.9186 | 28.89259 | -1.18678 | 0.997986 | 0.082266 |
| 3 | 120890600 | 120891599 | 7.826717 | 14.24036 | 41.37424 | 17.81492 | -1.11086 | 0.30681 | 0.082281 |
| 8 | 110505000 | 110506199 | 5.349861 | 10.04946 | 33.9093 | 13.93977 | -1.27286 | 0.07812 | 0.082286 |
| 5 | 113124200 | 113132599 | 57.24592 | 89.63631 | 190.8001 | 79.1465 | -0.77257 | 2.612691 | 0.082296 |
| 4 | 117168000 | 117170599 | 16.09007 | 26.89511 | 72.27028 | 31.46542 | -1.03972 | 1.185719 | 0.082311 |
| 14 | 42100000 | 42100599 | 11.14668 | 21.6892 | 13.01093 | 4.794832 | 1.338074 | -0.28915 | 0.08233 |
| 7 | 23166400 | 23170599 | 97.17938 | 152.5355 | 111.0954 | 46.07609 | 0.775437 | 2.578563 | 0.082331 |
| 15 | 102994200 | 102995599 | 7.428402 | 13.42322 | 40.66437 | 17.66018 | -1.17886 | 0.341293 | 0.082332 |
| 6 | 31997800 | 31999399 | 10.28554 | 18.06747 | 48.266 | 21.36217 | -1.02223 | 0.600505 | 0.08235 |
| 1 | 179786600 | 179787999 | 6.453761 | 11.87072 | 37.26524 | 15.81881 | -1.20658 | 0.223934 | 0.082369 |
| 12 | 19550800 | 19564599 | 364.3828 | 517.4314 | 348.8937 | 144.3099 | 0.758904 | 4.353909 | 0.082378 |
| 9 | 63013200 | 63016999 | 20.62814 | 34.06087 | 82.65284 | 35.69182 | -0.89462 | 1.344613 | 0.082388 |
| 7 | 17277000 | 17279199 | 22.10982 | 39.05457 | 23.5529 | 10.4198 | 1.115311 | 0.55818 | 0.082393 |
| 7 | 15287200 | 15290599 | 75.10078 | 120.0565 | 70.15165 | 29.49917 | 1.116098 | 2.086009 | 0.082397 |
| 19 | 41337000 | 41337799 | 2.844241 | 5.540245 | 25.78469 | 9.40521 | -1.56635 | -0.38462 | 0.082415 |
| 8 | 118149800 | 118158199 | 35.79813 | 56.89862 | 151.8781 | 63.36806 | -1.08038 | 2.250343 | 0.08242 |
| 9 | 119438200 | 119439999 | 9.568341 | 16.74135 | 50.95098 | 22.47534 | -1.20934 | 0.664559 | 0.082448 |
| 19 | 4007600 | 4025399 | 86.09761 | 130.3702 | 300.873 | 124.4829 | -0.92732 | 3.270327 | 0.082465 |
| 10 | 60591400 | 60605399 | 73.95927 | 113.3488 | 253.0499 | 105.0358 | -0.86531 | 3.017981 | 0.08248 |
| 16 | 65169000 | 65170399 | 18.81267 | 34.12364 | 20.3621 | 8.801319 | 1.153645 | 0.356858 | 0.082486 |
| 5 | 35285800 | 35291999 | 40.42647 | 64.35121 | 145.228 | 60.70637 | -0.83648 | 2.205095 | 0.08249 |
| 6 | 23245400 | 23250799 | 45.6393 | 72.07906 | 171.4554 | 71.19792 | -0.92282 | 2.430659 | 0.082494 |
| 2 | 165580200 | 165582599 | 11.07209 | 19.39158 | 50.4438 | 22.27023 | -0.98451 | 0.655212 | 0.082499 |
| 1 | 93677400 | 93678599 | 5.496667 | 10.36777 | 34.64593 | 14.14318 | -1.25155 | 0.052332 | 0.082515 |
| 9 | 61935600 | 61942399 | 36.35505 | 58.001 | 140.9216 | 59.09044 | -0.93916 | 2.126565 | 0.082516 |
| 10 | 84316600 | 84319199 | 12.66698 | 21.98707 | 57.55803 | 25.25595 | -0.99526 | 0.804667 | 0.08252 |
| 1 | 191566000 | 191575799 | 44.73061 | 70.16942 | 194.0759 | 80.48999 | -1.14763 | 2.573389 | 0.082533 |
| X | 26219200 | 26221199 | 25.94287 | 45.57846 | 22.96247 | 10.16443 | 1.370995 | 0.641098 | 0.082542 |
| 12 | 23350800 | 23351399 | 10.80281 | 21.12329 | 3.855791 | 1.251369 | 3.151556 | -0.74818 | 0.082551 |
| 19 | 58439800 | 58445199 | 26.50142 | 42.51316 | 126.9677 | 53.6661 | -1.22689 | 1.940786 | 0.082559 |
| 17 | 56402000 | 56411999 | 37.04491 | 58.60866 | 168.2543 | 69.93077 | -1.19143 | 2.383094 | 0.082563 |
| 14 | 105663800 | 105665199 | 8.074991 | 14.4558 | 43.15392 | 18.92182 | -1.16765 | 0.40769 | 0.082569 |
| 17 | 20670400 | 20673399 | 55.31703 | 89.59574 | 49.08616 | 21.00647 | 1.229759 | 1.673852 | 0.082574 |
| 1 | 171345600 | 171346599 | 7.818587 | 14.27112 | 44.307 | 19.00158 | -1.20274 | 0.291281 | 0.082609 |
| 1 | 120737200 | 120738799 | 18.38332 | 33.58856 | 19.68392 | 8.431052 | 1.187501 | 0.310918 | 0.082613 |
| 2 | 44761600 | 44762199 | 9.291354 | 18.14049 | 11.50023 | 4.057029 | 1.314018 | -0.46863 | 0.082621 |
| 18 | 42108000 | 42108599 | 9.291354 | 18.14049 | 11.50023 | 4.057029 | 1.314018 | -0.46863 | 0.082621 |
| 4 | 138060400 | 138076799 | 69.69381 | 106.2849 | 282.4345 | 117.1008 | -1.12203 | 3.11909 | 0.082635 |
| 11 | 65410400 | 65411199 | 13.5467 | 26.0911 | 14.26276 | 5.538307 | 1.401325 | -0.08646 | 0.082645 |
| 9 | 61775800 | 61778399 | 51.9252 | 83.94136 | 58.86838 | 25.15133 | 0.871947 | 1.714479 | 0.082651 |
| 7 | 10543000 | 10554799 | 304.4281 | 438.6681 | 307.6988 | 126.9735 | 0.723049 | 4.136834 | 0.082658 |
| 5 | 9384800 | 9386999 | 20.83559 | 37.44812 | 20.3928 | 8.91181 | 1.274323 | 0.422301 | 0.082664 |

| | | | | | | | | | |
|---|---|---|---|---|---|---|---|---|---|
| 6 | 85826600 | 85831799 | 28.10345 | 45.37026 | 109.6685 | 46.774 | -0.91132 | 1.738406 | 0.082664 |
| 4 | 100683400 | 100684799 | 15.40129 | 28.6247 | 18.57875 | 7.775649 | 1.064741 | 0.191209 | 0.082693 |
| 7 | 53173400 | 53177399 | 38.77924 | 64.68037 | 39.46297 | 17.1581 | 1.091503 | 1.24549 | 0.082703 |
| 3 | 158378200 | 158378999 | 10.36657 | 20.14991 | 13.07042 | 4.858155 | 1.214935 | -0.27084 | 0.082709 |
| 2 | 30435000 | 30436199 | 5.834986 | 10.87182 | 36.48912 | 15.20866 | -1.28248 | 0.139752 | 0.082711 |
| 1 | 19269200 | 19270799 | 9.316543 | 16.54103 | 46.74097 | 20.63806 | -1.09664 | 0.474421 | 0.082718 |
| 7 | 13765400 | 13769999 | 106.1043 | 165.5431 | 119.5954 | 49.62814 | 0.780435 | 2.690119 | 0.082722 |
| 7 | 100571800 | 100582199 | 35.38117 | 55.88497 | 181.4053 | 75.33427 | -1.37029 | 2.422785 | 0.082745 |
| 8 | 30045800 | 30047399 | 15.89694 | 29.52379 | 18.2773 | 7.657289 | 1.130882 | 0.197735 | 0.082751 |
| 3 | 6118800 | 6121799 | 62.42143 | 99.97249 | 73.06257 | 30.80533 | 0.796891 | 2.015134 | 0.082751 |
| 2 | 152739800 | 152744799 | 22.99895 | 37.50906 | 90.90898 | 39.05447 | -0.90152 | 1.518064 | 0.082756 |
| 12 | 22365600 | 22367799 | 25.40584 | 44.15232 | 29.82081 | 13.09506 | 0.950454 | 0.807385 | 0.082787 |
| X | 32912400 | 32916399 | 51.81103 | 84.01225 | 53.96292 | 23.11278 | 1.002485 | 1.648644 | 0.082811 |
| 14 | 25401200 | 25404199 | 10.99096 | 18.79487 | 65.32169 | 28.64208 | -1.40629 | 0.944947 | 0.082814 |
| Y | 89575400 | 89576799 | 11.31306 | 22.01678 | 5.204111 | 1.915134 | 2.653738 | -0.32368 | 0.082823 |
| 14 | 118943000 | 118945599 | 13.84939 | 23.72739 | 60.6217 | 26.58526 | -0.96423 | 0.934729 | 0.082826 |
| 6 | 129944600 | 129945599 | 19.34327 | 34.98078 | 20.58713 | 8.932101 | 1.169717 | 0.377104 | 0.082845 |
| 5 | 134228000 | 134236399 | 47.95719 | 75.60422 | 181.0334 | 75.12502 | -0.93446 | 2.473537 | 0.08285 |
| 11 | 35500600 | 35503399 | 15.51812 | 26.23801 | 68.78797 | 30.10245 | -1.00399 | 1.067418 | 0.082855 |
| 4 | 102616400 | 102617999 | 9.609017 | 16.76304 | 55.25273 | 24.321 | -1.32442 | 0.714936 | 0.082877 |
| 14 | 39033600 | 39034999 | 17.67248 | 32.47955 | 19.19106 | 8.155438 | 1.18349 | 0.272021 | 0.082895 |
| 14 | 44047200 | 44050799 | 60.15991 | 96.54887 | 69.79864 | 29.52135 | 0.815972 | 1.94855 | 0.082897 |
| 17 | 34319200 | 34320999 | 6.469037 | 11.91293 | 40.07401 | 16.9837 | -1.30462 | 0.220626 | 0.082909 |
| 4 | 129470800 | 129473199 | 8.90193 | 15.7136 | 46.16159 | 20.42465 | -1.15571 | 0.529894 | 0.082942 |
| 4 | 21727400 | 21728599 | 7.355326 | 13.4718 | 38.27577 | 16.34172 | -1.07039 | 0.25725 | 0.082946 |
| 10 | 88214400 | 88216199 | 6.976307 | 12.30733 | 47.20366 | 20.88816 | -1.53721 | 0.527797 | 0.082951 |
| 15 | 57897200 | 57897999 | 4.668268 | 9.037263 | 30.19589 | 11.49077 | -1.15641 | -0.20131 | 0.082953 |
| 8 | 80681400 | 80681999 | 5.823925 | 10.80403 | 35.79622 | 15.01981 | -1.27077 | 0.167371 | 0.082964 |
| 1 | 112128800 | 112130799 | 26.90047 | 46.89912 | 27.04457 | 11.89977 | 1.177741 | 0.76802 | 0.082965 |
| 9 | 102557800 | 102559199 | 3.531364 | 6.883005 | 27.45595 | 9.932718 | -1.33914 | -0.4029 | 0.082968 |
| 12 | 48752400 | 48753199 | 10.26634 | 19.94346 | 12.79734 | 4.779089 | 1.225222 | -0.25627 | 0.082974 |
| 14 | 22034800 | 22041799 | 36.12891 | 57.43527 | 153.15 | 63.91085 | -1.07863 | 2.244595 | 0.082979 |
| 8 | 85318600 | 85328599 | 49.27058 | 77.16572 | 191.4469 | 79.41296 | -0.99293 | 2.605223 | 0.082985 |
| 18 | 35783800 | 35790999 | 46.86709 | 73.79503 | 182.456 | 75.70331 | -0.98198 | 2.498051 | 0.082992 |
| 7 | 23044400 | 23045399 | 36.78334 | 60.96783 | 41.313 | 17.88708 | 0.940339 | 1.308198 | 0.083046 |
| 4 | 106863400 | 106864999 | 4.372723 | 8.26001 | 35.15934 | 14.30789 | -1.59211 | 0.033128 | 0.083049 |
| 3 | 137680600 | 137681999 | 6.150252 | 11.3947 | 38.00855 | 15.97609 | -1.2834 | 0.179466 | 0.083057 |
| 7 | 108328000 | 108330799 | 29.08425 | 50.33095 | 29.5692 | 12.97281 | 1.14974 | 0.86821 | 0.08307 |
| 12 | 19829400 | 19833199 | 136.2694 | 208.6498 | 150.6118 | 62.50342 | 0.760984 | 3.047595 | 0.083101 |
| 7 | 15927800 | 15942799 | 70.37462 | 107.1116 | 285.4116 | 118.2701 | -1.12828 | 3.160378 | 0.083103 |
| 3 | 114463200 | 114463599 | 2.525988 | 4.931245 | 23.76708 | 8.399405 | -1.56922 | -0.50479 | 0.083106 |
| 7 | 43120200 | 43128799 | 47.07096 | 74.00025 | 184.0165 | 76.33524 | -0.99198 | 2.530332 | 0.083106 |
| 14 | 41949600 | 41949999 | 10.04726 | 19.54472 | 13.32017 | 4.919419 | 1.153992 | -0.29449 | 0.083108 |
| 1 | 105439800 | 105441199 | 21.10112 | 37.2289 | 24.29599 | 10.75108 | 1.001543 | 0.565703 | 0.08312 |
| 16 | 64472800 | 64474199 | 4.365363 | 8.478212 | 30.61475 | 11.45437 | -1.24371 | -0.26254 | 0.083123 |
| 15 | 61993600 | 61995799 | 9.151942 | 16.26471 | 44.10096 | 19.45632 | -1.03606 | 0.465054 | 0.083162 |
| 7 | 4494000 | 4496599 | 9.166242 | 16.24022 | 44.59414 | 19.71651 | -1.05657 | 0.489049 | 0.083172 |

| | | | | | | | | | |
|---|---|---|---|---|---|---|---|---|---|
| 11 | 77576600 | 77578799 | 13.06914 | 22.40649 | 60.02808 | 26.32643 | -1.03203 | 0.924873 | 0.083173 |
| 11 | 96766400 | 96769599 | 8.752216 | 15.46409 | 46.68977 | 20.65441 | -1.1945 | 0.518522 | 0.083181 |
| 5 | 122278800 | 122285399 | 36.3051 | 58.03793 | 132.9095 | 55.88977 | -0.85073 | 2.062266 | 0.083207 |
| X | 90322600 | 90325199 | 31.91961 | 55.87906 | 18.16255 | 8.015514 | 1.998589 | 0.721441 | 0.083209 |
| 7 | 20910800 | 20919999 | 147.7999 | 225.8477 | 152.2425 | 63.15065 | 0.857345 | 3.091529 | 0.083238 |
| 2 | 168410800 | 168427399 | 116.3272 | 172.2956 | 408.4015 | 168.1931 | -0.99459 | 3.697074 | 0.083242 |
| 11 | 73117800 | 73132999 | 84.92957 | 128.8033 | 298.4491 | 123.5404 | -0.93117 | 3.237664 | 0.083264 |
| 12 | 21984200 | 21985799 | 57.33841 | 92.14279 | 62.50675 | 26.48975 | 0.909764 | 1.899517 | 0.083265 |
| 4 | 150209400 | 150215199 | 27.86492 | 44.807 | 117.156 | 49.69474 | -1.03129 | 1.863672 | 0.083274 |
| X | 27685600 | 27686799 | 9.175629 | 17.93385 | 10.3137 | 3.560844 | 1.479173 | -0.5472 | 0.083278 |
| 6 | 45648000 | 45648599 | 12.65534 | 24.21361 | 16.22478 | 6.420987 | 1.086273 | -0.03004 | 0.083279 |
| 16 | 57426800 | 57427999 | 5.751582 | 10.89598 | 33.12242 | 13.39716 | -1.10418 | 0.02144 | 0.083283 |
| 7 | 74628400 | 74629999 | 10.62826 | 18.57516 | 50.98117 | 22.46752 | -1.0613 | 0.688844 | 0.083283 |
| 13 | 29592200 | 29592799 | 3.942018 | 7.674832 | 28.4857 | 10.44855 | -1.25542 | -0.34445 | 0.083289 |
| 4 | 149183600 | 149184399 | 3.439105 | 6.657101 | 29.32973 | 11.16613 | -1.5493 | -0.21535 | 0.083342 |
| 4 | 111497000 | 111498199 | 5.107551 | 9.843289 | 31.6488 | 12.26758 | -1.12743 | -0.12979 | 0.083359 |
| 10 | 18324400 | 18325999 | 6.243604 | 11.68948 | 39.05916 | 16.14723 | -1.26738 | 0.115915 | 0.083381 |
| 16 | 3337400 | 3340999 | 70.80476 | 112.8389 | 83.96731 | 35.15551 | 0.763225 | 2.178599 | 0.083418 |
| 5 | 26649400 | 26650199 | 10.4948 | 20.45869 | 11.87476 | 4.287132 | 1.410338 | -0.37184 | 0.083421 |
| 12 | 112627000 | 112655999 | 138.6043 | 199.9884 | 556.7567 | 229.7081 | -1.26285 | 4.110483 | 0.083423 |
| 4 | 107871400 | 107873999 | 8.553785 | 14.94235 | 54.12245 | 23.84371 | -1.45895 | 0.689295 | 0.083464 |
| 2 | 25059200 | 25072399 | 63.65559 | 97.88137 | 244.2363 | 101.4236 | -1.02193 | 2.948807 | 0.083467 |
| 11 | 58884200 | 58894999 | 58.83424 | 90.99627 | 225.1712 | 93.53071 | -1.0034 | 2.834809 | 0.083473 |
| 8 | 79786800 | 79787799 | 6.134004 | 11.34481 | 36.28967 | 15.28982 | -1.22522 | 0.187952 | 0.083476 |
| 15 | 99224600 | 99228399 | 16.33577 | 27.50558 | 71.00611 | 31.03388 | -0.98218 | 1.108835 | 0.083484 |
| 5 | 105087600 | 105088599 | 11.37308 | 22.02883 | 14.64937 | 5.560768 | 1.15233 | -0.1825 | 0.083495 |
| 6 | 149393800 | 149394799 | 12.18999 | 23.65823 | 13.37935 | 5.03161 | 1.39566 | -0.20704 | 0.083503 |
| 17 | 8647400 | 8658199 | 74.25149 | 113.865 | 249.9913 | 103.7748 | -0.84057 | 3.006047 | 0.083509 |
| 9 | 64343200 | 64345599 | 7.933237 | 14.1572 | 42.58895 | 18.72394 | -1.18101 | 0.427413 | 0.083509 |
| 3 | 119822200 | 119822799 | 11.15752 | 21.66309 | 14.22402 | 5.334858 | 1.187031 | -0.2251 | 0.083523 |
| 14 | 24845400 | 24847399 | 7.054183 | 12.95706 | 38.34894 | 16.3097 | -1.12495 | 0.23756 | 0.08354 |
| 15 | 27508200 | 27509399 | 5.26482 | 10.0717 | 31.74331 | 12.56799 | -1.12745 | -0.05612 | 0.083549 |
| 18 | 17591800 | 17593599 | 12.80604 | 24.6315 | 15.55949 | 6.065164 | 1.19035 | -0.07829 | 0.083563 |
| 18 | 81580200 | 81584999 | 19.26534 | 31.69352 | 84.94016 | 36.6165 | -1.03925 | 1.388651 | 0.083568 |
| X | 25988400 | 25989599 | 17.34679 | 32.44725 | 16.74045 | 6.931413 | 1.405353 | 0.164676 | 0.083574 |
| 9 | 120591000 | 120597199 | 38.43472 | 61.33205 | 139.3241 | 58.43822 | -0.84203 | 2.121501 | 0.083587 |
| 6 | 54524400 | 54525799 | 6.515687 | 12.23287 | 36.27417 | 14.92542 | -1.0906 | 0.095625 | 0.083595 |
| 11 | 50442000 | 50443999 | 27.37086 | 47.1941 | 29.08458 | 12.75878 | 1.080173 | 0.891494 | 0.083596 |
| 7 | 116755200 | 116756599 | 5.917922 | 11.17609 | 35.3361 | 14.3874 | -1.16956 | 0.0486 | 0.083633 |
| 3 | 94861200 | 94867799 | 26.51027 | 42.78765 | 106.441 | 45.38299 | -0.95296 | 1.742942 | 0.083646 |
| 7 | 9679600 | 9682799 | 41.90018 | 70.80165 | 26.3965 | 11.5493 | 1.798145 | 1.121305 | 0.083657 |
| 14 | 67999400 | 68001199 | 9.087632 | 15.97385 | 48.77893 | 21.60093 | -1.21395 | 0.581985 | 0.083665 |
| 7 | 141270400 | 141303999 | 255.9018 | 354.9614 | 793.9281 | 329.6163 | -1.0045 | 4.701066 | 0.083672 |
| 7 | 41432400 | 41433999 | 43.81804 | 72.20784 | 43.9867 | 18.97567 | 1.092429 | 1.398268 | 0.083672 |
| 3 | 25614800 | 25615599 | 2.840865 | 5.548033 | 31.73794 | 11.14397 | -1.81056 | -0.50449 | 0.083694 |
| 8 | 84698200 | 84701199 | 9.110491 | 15.91458 | 51.46531 | 22.67287 | -1.29612 | 0.690544 | 0.083709 |
| 12 | 54529000 | 54529999 | 13.23937 | 25.25567 | 16.5707 | 6.604792 | 1.106904 | -0.00113 | 0.083727 |

| | | | | | | | | | |
|---|---|---|---|---|---|---|---|---|---|
| 14 | 59316600 | 59317199 | 3.643663 | 7.105293 | 26.3098 | 9.458488 | -1.22441 | -0.42638 | 0.083728 |
| 14 | 42541600 | 42544199 | 34.62146 | 58.79173 | 31.50037 | 13.80304 | 1.277729 | 1.056593 | 0.083736 |
| 17 | 23803800 | 23805599 | 7.27756 | 13.17795 | 41.24573 | 17.86625 | -1.2233 | 0.328274 | 0.08374 |
| 10 | 84355600 | 84359999 | 21.31874 | 35.03619 | 85.21716 | 36.72022 | -0.90045 | 1.40645 | 0.083747 |
| 12 | 19009800 | 19013599 | 90.64299 | 142.5456 | 106.6504 | 44.24343 | 0.738745 | 2.536235 | 0.083764 |
| 13 | 62584200 | 62585399 | 38.41727 | 63.46928 | 42.96436 | 18.56034 | 0.9416 | 1.352602 | 0.083769 |
| X | 101080800 | 101081599 | 3.702937 | 7.193739 | 29.13359 | 10.875 | -1.40081 | -0.27737 | 0.083772 |
| 7 | 34896600 | 34908799 | 80.98755 | 122.979 | 288.9786 | 119.6651 | -0.94988 | 3.210893 | 0.083777 |
| 17 | 7385200 | 7390399 | 20.27377 | 33.58887 | 80.46943 | 34.83 | -0.87573 | 1.303178 | 0.083778 |
| 18 | 37948000 | 37951199 | 26.66148 | 43.01426 | 106.7104 | 45.47409 | -0.94965 | 1.756319 | 0.083793 |
| 12 | 21848000 | 21852999 | 98.49588 | 154.1944 | 115.2706 | 47.81684 | 0.734934 | 2.625857 | 0.083799 |
| 8 | 55387200 | 55389199 | 25.83409 | 45.18232 | 26.30074 | 11.59767 | 1.164231 | 0.715724 | 0.083808 |
| 11 | 79633800 | 79635199 | 25.42403 | 44.42528 | 26.16809 | 11.53047 | 1.147489 | 0.728298 | 0.08381 |
| 19 | 45801800 | 45809599 | 35.06936 | 55.95427 | 139.0483 | 58.31143 | -0.97145 | 2.122744 | 0.083811 |
| 5 | 129671200 | 129673999 | 16.7019 | 27.55209 | 81.694 | 35.25814 | -1.18259 | 1.347328 | 0.083818 |
| 12 | 23867800 | 23871799 | 84.6485 | 133.8798 | 99.03685 | 41.15565 | 0.761447 | 2.409514 | 0.08382 |
| 12 | 22375600 | 22380599 | 50.68558 | 82.18735 | 54.43686 | 23.31578 | 0.958323 | 1.647143 | 0.083827 |
| 11 | 117821600 | 117826199 | 16.36595 | 27.09094 | 79.6868 | 34.47052 | -1.1705 | 1.303232 | 0.08383 |
| 2 | 58246200 | 58248999 | 13.86569 | 23.71293 | 61.69723 | 27.05303 | -0.99086 | 0.951753 | 0.083843 |
| 3 | 25197600 | 25200799 | 15.79101 | 26.80161 | 68.36641 | 29.95265 | -0.96442 | 1.034375 | 0.083858 |
| 11 | 67746000 | 67747399 | 7.374263 | 13.44543 | 38.71173 | 16.62465 | -1.09511 | 0.285064 | 0.083866 |
| 10 | 42064000 | 42075799 | 89.23625 | 134.2693 | 336.1845 | 138.8288 | -1.05275 | 3.399604 | 0.083867 |
| 9 | 51913200 | 51915199 | 11.55511 | 20.16684 | 53.16946 | 23.41042 | -1.00425 | 0.715667 | 0.083876 |
| 12 | 20651400 | 20653999 | 40.61186 | 67.04353 | 41.317 | 17.83924 | 1.076439 | 1.364706 | 0.083896 |
| 12 | 18979000 | 18993599 | 367.3527 | 520.8209 | 360.0673 | 148.9713 | 0.720557 | 4.37684 | 0.083899 |
| 13 | 92491000 | 92492199 | 7.571713 | 13.37667 | 47.21147 | 20.8857 | -1.41782 | 0.514164 | 0.083899 |
| 12 | 23621000 | 23621799 | 12.09585 | 23.20201 | 15.69499 | 6.166168 | 1.082097 | -0.06105 | 0.083902 |
| 18 | 44960400 | 44962199 | 6.371617 | 11.96586 | 36.28326 | 14.92183 | -1.12201 | 0.091972 | 0.083937 |
| 9 | 44951600 | 44955999 | 21.94742 | 35.63603 | 95.69736 | 41.02876 | -1.0539 | 1.590155 | 0.083943 |
| 18 | 40389000 | 40390399 | 8.228486 | 14.70401 | 42.60071 | 18.70771 | -1.12619 | 0.42 | 0.083946 |
| 10 | 18097400 | 18098999 | 7.010012 | 12.9594 | 38.51601 | 16.23735 | -1.12123 | 0.202826 | 0.083972 |
| 9 | 5930200 | 5931399 | 19.27421 | 34.33373 | 22.31679 | 9.82634 | 1.013558 | 0.469034 | 0.083988 |
| 8 | 38890800 | 38891399 | 16.87976 | 31.13335 | 19.33924 | 8.178669 | 1.116898 | 0.246832 | 0.083997 |
| 12 | 71319600 | 71322799 | 19.80077 | 32.80211 | 79.83407 | 34.55273 | -0.8984 | 1.302144 | 0.084001 |
| 8 | 112426800 | 112434399 | 33.83744 | 53.87892 | 146.6548 | 61.32828 | -1.10572 | 2.190148 | 0.084004 |
| 17 | 8529600 | 8530799 | 4.893464 | 9.284675 | 32.5295 | 13.11439 | -1.30203 | 0.001301 | 0.08401 |
| 2 | 27364400 | 27366999 | 14.43186 | 24.41024 | 66.97081 | 29.31047 | -1.06901 | 1.059672 | 0.084011 |
| 8 | 121742000 | 121755399 | 83.62267 | 127.0961 | 286.5467 | 118.6904 | -0.88917 | 3.192322 | 0.084013 |
| 8 | 129713800 | 129714999 | 20.40218 | 36.47011 | 22.15317 | 9.725629 | 1.113617 | 0.456131 | 0.084014 |
| 6 | 42748200 | 42750599 | 22.69797 | 40.24115 | 21.85678 | 9.66233 | 1.267234 | 0.520542 | 0.084016 |
| 12 | 111812200 | 111816999 | 26.17218 | 42.27236 | 104.1657 | 44.45411 | -0.93805 | 1.720731 | 0.084018 |
| 6 | 147947600 | 147949399 | 7.720014 | 13.96909 | 41.23544 | 17.87785 | -1.14046 | 0.335637 | 0.08402 |
| 12 | 52225400 | 52227199 | 19.69629 | 35.37699 | 21.63593 | 9.460351 | 1.107618 | 0.420925 | 0.084021 |
| 2 | 58372800 | 58376999 | 18.56601 | 30.71244 | 80.68072 | 34.88454 | -1.00831 | 1.316333 | 0.084032 |
| 4 | 128993800 | 129008199 | 70.99441 | 108.7729 | 253.342 | 105.1516 | -0.92667 | 3.023312 | 0.084032 |
| 14 | 45651800 | 45653799 | 8.001686 | 14.50635 | 40.20738 | 17.39068 | -1.04787 | 0.325308 | 0.084039 |
| 4 | 130309200 | 130311999 | 10.60771 | 18.59617 | 49.35936 | 21.8092 | -1.01326 | 0.635457 | 0.084048 |

| | | | | | | | | | |
|---|---|---|---|---|---|---|---|---|---|
| 12 | 75611200 | 75612799 | 5.878435 | 11.17961 | 33.90591 | 13.60192 | -1.09061 | -0.00292 | 0.084054 |
| 11 | 118747400 | 118756599 | 53.31744 | 83.18119 | 202.714 | 84.12009 | -0.97161 | 2.676943 | 0.084072 |
| 6 | 10233800 | 10235199 | 17.68898 | 32.13682 | 20.84611 | 8.992513 | 1.035615 | 0.339647 | 0.084074 |
| 12 | 85559000 | 85561799 | 10.50981 | 18.31679 | 54.2034 | 23.84464 | -1.17002 | 0.732174 | 0.08408 |
| 12 | 50684200 | 50685399 | 16.8955 | 31.17199 | 18.66633 | 7.890782 | 1.169416 | 0.242869 | 0.084087 |
| 19 | 5501600 | 5507799 | 23.48387 | 37.95853 | 105.7563 | 45.17624 | -1.11342 | 1.694207 | 0.084091 |
| 7 | 13495800 | 13497999 | 65.26526 | 104.3739 | 76.48727 | 32.181 | 0.790658 | 2.064383 | 0.084097 |
| 13 | 3071600 | 3073599 | 31.04848 | 53.11359 | 32.17957 | 14.11043 | 1.102638 | 0.979153 | 0.084103 |
| 3 | 102568800 | 102571999 | 13.42626 | 22.92138 | 64.4029 | 28.23544 | -1.10176 | 0.968224 | 0.084111 |
| 6 | 117511200 | 117512399 | 10.13266 | 17.80207 | 47.73583 | 21.13068 | -1.02757 | 0.595689 | 0.08413 |
| 5 | 134313200 | 134333399 | 133.9341 | 197.1494 | 435.766 | 179.4508 | -0.90224 | 3.802156 | 0.08413 |
| 14 | 47020800 | 47022999 | 6.990232 | 13.01459 | 38.81683 | 16.19925 | -1.11585 | 0.159426 | 0.084139 |
| 10 | 80508800 | 80516399 | 31.68986 | 50.63043 | 138.4275 | 58.16212 | -1.10674 | 2.076918 | 0.084146 |
| 7 | 43898600 | 43904199 | 20.19963 | 32.76704 | 97.89525 | 41.94173 | -1.20844 | 1.608399 | 0.084147 |
| 10 | 76539600 | 76540799 | 17.68545 | 32.83152 | 17.57965 | 7.369244 | 1.33815 | 0.214201 | 0.084164 |
| 5 | 148573800 | 148590399 | 102.7631 | 154.623 | 331.2845 | 136.8055 | -0.82816 | 3.401653 | 0.084165 |
| 9 | 6750800 | 6751399 | 9.456004 | 18.4948 | 7.642547 | 2.578584 | 1.971891 | -0.62991 | 0.084184 |
| 7 | 80957600 | 80962999 | 29.10141 | 46.87488 | 111.7593 | 47.5225 | -0.89542 | 1.809281 | 0.084209 |
| 10 | 81068600 | 81105399 | 239.1956 | 332.0364 | 797.1459 | 330.9138 | -1.10553 | 4.689302 | 0.084214 |
| 7 | 89338000 | 89339199 | 16.39907 | 30.27057 | 19.44355 | 8.213983 | 1.070111 | 0.238987 | 0.084232 |
| 8 | 115569000 | 115572999 | 16.56993 | 27.70242 | 74.14111 | 32.28379 | -1.03415 | 1.184896 | 0.084237 |
| 15 | 93391200 | 93396999 | 26.97423 | 43.58154 | 105.9665 | 45.241 | -0.91839 | 1.71224 | 0.084238 |
| 7 | 9704600 | 9707199 | 53.32618 | 86.23887 | 53.41562 | 22.83282 | 1.051234 | 1.702849 | 0.084246 |
| 18 | 82897800 | 82905199 | 44.03954 | 69.91062 | 156.4926 | 65.21202 | -0.82883 | 2.293621 | 0.084257 |
| X | 45179800 | 45181999 | 23.24015 | 42.30271 | 15.52125 | 6.679525 | 1.853056 | 0.36013 | 0.084262 |
| 9 | 51534000 | 51537199 | 18.0493 | 29.97853 | 79.18876 | 34.33506 | -1.01574 | 1.265934 | 0.084277 |
| 13 | 54442600 | 54444599 | 8.661044 | 15.52138 | 42.64017 | 18.67987 | -1.04779 | 0.40416 | 0.084312 |
| 8 | 120595000 | 120614399 | 137.8954 | 202.0599 | 453.1688 | 186.6649 | -0.92948 | 3.87481 | 0.084336 |
| 7 | 135197600 | 135200599 | 11.12317 | 19.33082 | 54.18826 | 23.79548 | -1.09267 | 0.778048 | 0.084342 |
| X | 54345600 | 54346599 | 33.64719 | 56.59493 | 37.90787 | 16.56275 | 0.957091 | 1.142727 | 0.084354 |
| 2 | 166930400 | 166952399 | 150.3801 | 219.1521 | 476.2737 | 196.2699 | -0.89171 | 3.960917 | 0.084359 |
| 13 | 115140400 | 115142599 | 7.478087 | 13.78132 | 39.01572 | 16.51856 | -1.05669 | 0.22402 | 0.084391 |
| 5 | 30276600 | 30278799 | 9.54076 | 16.75623 | 48.79459 | 21.59289 | -1.14596 | 0.598403 | 0.084393 |
| 11 | 15958800 | 15959599 | 14.85765 | 27.88396 | 17.95499 | 7.392509 | 1.094378 | 0.124371 | 0.084397 |
| 13 | 65751600 | 65761599 | 283.6221 | 411.2358 | 289.6432 | 119.4245 | 0.725941 | 4.044008 | 0.084398 |
| 13 | 60606200 | 60607599 | 7.032908 | 12.95118 | 37.31025 | 15.81263 | -1.08223 | 0.221993 | 0.084399 |
| 17 | 45743800 | 45745199 | 6.980731 | 12.57119 | 40.60426 | 17.70969 | -1.27408 | 0.360231 | 0.084403 |
| 17 | 78968800 | 78970399 | 6.383375 | 11.91068 | 35.43171 | 14.73205 | -1.10674 | 0.134833 | 0.084408 |
| 11 | 78079000 | 78080599 | 7.589076 | 13.36782 | 48.24738 | 21.35657 | -1.4516 | 0.549605 | 0.084414 |
| 9 | 114445600 | 114446599 | 2.9429 | 5.735072 | 24.67573 | 8.946381 | -1.44606 | -0.40761 | 0.08442 |
| 19 | 57093800 | 57095599 | 10.10266 | 17.84598 | 46.68952 | 20.65526 | -0.98947 | 0.530052 | 0.084425 |
| X | 4743400 | 4747199 | 35.36179 | 60.63904 | 27.22329 | 11.93948 | 1.533541 | 0.972409 | 0.084426 |
| 5 | 45677400 | 45679199 | 4.490163 | 8.485059 | 33.47155 | 13.61018 | -1.48196 | 0.028724 | 0.08443 |
| 13 | 7486600 | 7489799 | 31.21484 | 53.24974 | 33.29986 | 14.59839 | 1.056652 | 1.005637 | 0.084433 |
| 14 | 99654400 | 99655599 | 6.032568 | 11.43175 | 36.70407 | 14.83634 | -1.18212 | 0.03016 | 0.084438 |
| 9 | 56979200 | 56984799 | 27.10554 | 43.50545 | 122.4135 | 51.78308 | -1.14015 | 1.923717 | 0.084441 |
| 14 | 25471000 | 25472799 | 5.007934 | 9.545387 | 31.56546 | 12.60332 | -1.20702 | -0.03179 | 0.084443 |

| | | | | | | | | | |
|---|---|---|---|---|---|---|---|---|---|
| 9 | 107424000 | 107429999 | 26.26881 | 42.16528 | 121.278 | 51.30794 | -1.17156 | 1.921432 | 0.084453 |
| 5 | 135117200 | 135123199 | 31.1281 | 49.88746 | 126.6631 | 53.45952 | -0.995 | 1.976173 | 0.08446 |
| 3 | 148692000 | 148697199 | 21.9518 | 35.19768 | 133.718 | 56.4777 | -1.57381 | 1.953194 | 0.084462 |
| 1 | 110254600 | 110257199 | 22.01613 | 38.80936 | 24.76044 | 10.95851 | 1.033684 | 0.582194 | 0.084467 |
| 12 | 77279400 | 77279999 | 10.66734 | 20.8227 | 10.00527 | 3.542133 | 1.702098 | -0.44229 | 0.084471 |
| 15 | 77626600 | 77627199 | 10.66734 | 20.8227 | 10.00527 | 3.542133 | 1.702098 | -0.44229 | 0.084471 |
| 12 | 45814600 | 45815799 | 13.35821 | 25.72862 | 14.98766 | 5.81943 | 1.311414 | -0.08596 | 0.084476 |
| 10 | 76995600 | 76997999 | 7.609348 | 13.63743 | 41.59182 | 18.21971 | -1.19678 | 0.396188 | 0.084497 |
| 5 | 24060000 | 24061799 | 7.631035 | 13.66985 | 43.16247 | 18.91527 | -1.24731 | 0.400612 | 0.084513 |
| 5 | 129081000 | 129086399 | 22.42306 | 36.1417 | 112.4523 | 47.86197 | -1.27702 | 1.779919 | 0.084516 |
| 17 | 52855000 | 52856199 | 16.75061 | 31.064 | 18.15472 | 7.620999 | 1.21131 | 0.212737 | 0.084523 |
| 13 | 66380800 | 66381199 | 8.027932 | 15.69288 | 7.357749 | 2.371221 | 1.851196 | -0.80714 | 0.084526 |
| 2 | 87011800 | 87012799 | 14.71839 | 28.2044 | 14.87682 | 5.861472 | 1.434097 | -0.0264 | 0.084547 |
| 4 | 154267400 | 154271999 | 22.79363 | 36.93536 | 99.66903 | 42.66797 | -1.06375 | 1.634144 | 0.084563 |
| 16 | 88942000 | 88942599 | 10.26705 | 19.9425 | 13.74668 | 5.138481 | 1.121538 | -0.24989 | 0.084573 |
| 7 | 65071200 | 65075599 | 24.94674 | 40.58776 | 95.25016 | 40.8835 | -0.85778 | 1.558353 | 0.084575 |
| 4 | 118901000 | 118903999 | 13.64949 | 23.08253 | 66.28765 | 29.0098 | -1.13441 | 1.059323 | 0.084583 |
| 5 | 94904800 | 94905399 | 9.349496 | 18.27323 | 10.31034 | 3.561896 | 1.505716 | -0.54222 | 0.084596 |
| 13 | 66021800 | 66023399 | 23.0533 | 40.78378 | 21.73876 | 9.614266 | 1.292896 | 0.544231 | 0.08461 |
| 7 | 109066600 | 109069199 | 11.07832 | 19.38675 | 52.61332 | 23.21212 | -1.04552 | 0.670434 | 0.084612 |
| 10 | 46918200 | 46919799 | 9.45335 | 16.81918 | 44.64449 | 19.67755 | -1.00485 | 0.46001 | 0.084612 |
| 1 | 112370200 | 112373599 | 52.5294 | 84.72631 | 61.4698 | 26.18629 | 0.819431 | 1.780771 | 0.084613 |
| 2 | 30654600 | 30673399 | 88.72956 | 133.8446 | 322.6211 | 133.3206 | -0.99473 | 3.349042 | 0.084629 |
| 4 | 146239400 | 146242599 | 30.20147 | 52.47125 | 26.08705 | 11.45338 | 1.389983 | 0.829196 | 0.084642 |
| 11 | 114730400 | 114738599 | 55.9877 | 88.01156 | 182.0102 | 75.50222 | -0.72655 | 2.536927 | 0.084642 |
| 12 | 114810200 | 114811999 | 39.93624 | 66.23853 | 41.68648 | 18.05722 | 1.04665 | 1.3099 | 0.084657 |
| 3 | 18361000 | 18361799 | 9.753671 | 19.01662 | 12.63919 | 4.554616 | 1.220481 | -0.38527 | 0.084663 |
| 17 | 66029600 | 66033999 | 25.48002 | 41.31762 | 100.7081 | 43.13525 | -0.91658 | 1.623867 | 0.084668 |
| 14 | 15824000 | 15825199 | 8.495762 | 15.26052 | 41.61418 | 18.1948 | -1.03497 | 0.387221 | 0.084681 |
| 12 | 102048800 | 102052599 | 12.84574 | 21.87647 | 67.05914 | 29.39417 | -1.22739 | 0.991428 | 0.084681 |
| 11 | 77824600 | 77831399 | 35.14222 | 56.27638 | 131.4853 | 55.42841 | -0.8768 | 2.006367 | 0.084691 |
| 12 | 28812800 | 28814799 | 10.27712 | 18.07348 | 49.31839 | 21.8377 | -1.05238 | 0.586525 | 0.084692 |
| 1 | 178354800 | 178358799 | 21.58285 | 34.93141 | 106.3895 | 45.50466 | -1.23955 | 1.658998 | 0.084713 |
| 12 | 86463600 | 86474799 | 64.21357 | 99.03469 | 237.0908 | 98.48329 | -0.95896 | 2.89004 | 0.084714 |
| 10 | 81722800 | 81724799 | 50.47214 | 81.55131 | 54.86264 | 23.42467 | 0.930927 | 1.723126 | 0.084728 |
| 6 | 137635600 | 137641199 | 34.21043 | 54.71326 | 131.1614 | 55.18878 | -0.91585 | 2.046246 | 0.084729 |
| 4 | 141655200 | 141664599 | 47.0225 | 74.0332 | 178.7581 | 74.16813 | -0.94791 | 2.499878 | 0.08473 |
| 2 | 180990800 | 180994799 | 11.66589 | 20.06199 | 58.91333 | 25.84195 | -1.16271 | 0.889618 | 0.084735 |
| 5 | 92284000 | 92286399 | 10.84198 | 18.83018 | 56.35446 | 24.73684 | -1.18691 | 0.786935 | 0.08474 |
| 14 | 76761400 | 76766599 | 24.62033 | 39.73468 | 109.0187 | 46.47811 | -1.09399 | 1.742561 | 0.084742 |
| 15 | 100317400 | 100324799 | 31.08271 | 49.90399 | 119.6275 | 50.62773 | -0.90912 | 1.917919 | 0.084747 |
| 4 | 104118400 | 104120399 | 10.76791 | 18.85201 | 49.9059 | 22.02601 | -1.00946 | 0.659406 | 0.084749 |
| 2 | 86983400 | 86984399 | 10.53198 | 20.49966 | 12.39822 | 4.557288 | 1.328645 | -0.30885 | 0.084781 |
| 5 | 139915400 | 139921399 | 29.36438 | 47.07254 | 125.4837 | 52.98177 | -1.06463 | 1.966366 | 0.084805 |
| 3 | 134257200 | 134259199 | 21.40433 | 38.22956 | 20.7898 | 9.133517 | 1.271119 | 0.464305 | 0.084822 |
| 1 | 75445800 | 75447199 | 4.096588 | 7.864467 | 30.03466 | 11.77199 | -1.38572 | -0.10461 | 0.084829 |
| 14 | 108385800 | 108386999 | 17.10281 | 32.17918 | 14.14495 | 5.798827 | 1.643554 | 0.118883 | 0.084831 |

| | | | | | | | | | |
|---|---|---|---|---|---|---|---|---|---|
| 12 | 84695000 | 84697999 | 21.39397 | 34.92831 | 89.50406 | 38.4657 | -0.98045 | 1.498999 | 0.084835 |
| 8 | 79764600 | 79765599 | 3.23802 | 6.312582 | 24.84231 | 8.962319 | -1.31418 | -0.42265 | 0.084849 |
| 3 | 8802600 | 8806999 | 17.67672 | 29.16601 | 85.60666 | 36.94992 | -1.16821 | 1.347417 | 0.084871 |
| 5 | 33849600 | 33860599 | 51.63333 | 80.34607 | 211.9053 | 87.96677 | -1.08836 | 2.722155 | 0.084875 |
| 1 | 128798000 | 128801599 | 16.03659 | 26.58043 | 78.57435 | 34.01995 | -1.17741 | 1.286261 | 0.084878 |
| 9 | 87965000 | 87966799 | 9.464113 | 16.809 | 45.14419 | 19.92705 | -1.02332 | 0.470992 | 0.084879 |
| 13 | 37866600 | 37866999 | 1.574313 | 3.076808 | 20.85299 | 6.696868 | -1.91968 | -0.87835 | 0.084905 |
| 9 | 119310200 | 119339199 | 196.2281 | 279.0898 | 606.7887 | 250.9141 | -0.92718 | 4.325569 | 0.084905 |
| 4 | 58400400 | 58401799 | 6.049261 | 11.4479 | 34.86598 | 14.13369 | -1.11035 | 0.034528 | 0.084907 |
| 7 | 10633200 | 10634799 | 29.16557 | 49.73764 | 34.64235 | 15.18654 | 0.901909 | 1.002819 | 0.084917 |
| 14 | 52545600 | 52555199 | 75.12136 | 115.4161 | 241.473 | 100.2646 | -0.76903 | 2.968794 | 0.084923 |
| 15 | 87427200 | 87443599 | 108.6857 | 162.8967 | 349.049 | 143.9951 | -0.83335 | 3.479471 | 0.084928 |
| 10 | 84877800 | 84880999 | 14.7865 | 24.89613 | 69.09086 | 30.19648 | -1.08584 | 1.103371 | 0.084929 |
| 10 | 59961800 | 59964199 | 10.15385 | 17.77161 | 49.89452 | 22.0154 | -1.09354 | 0.662078 | 0.084949 |
| 5 | 136259200 | 136268199 | 46.26334 | 72.7562 | 186.2665 | 77.27214 | -1.03352 | 2.522703 | 0.084987 |
| 19 | 8942400 | 8945799 | 16.31491 | 27.52633 | 68.21206 | 29.83108 | -0.92316 | 1.08915 | 0.084994 |
| 4 | 109951600 | 109954199 | 20.16815 | 33.26486 | 81.84779 | 35.32214 | -0.91484 | 1.356546 | 0.085013 |
| 4 | 155032600 | 155049799 | 95.01822 | 141.7831 | 381.2366 | 157.1384 | -1.16642 | 3.561416 | 0.085022 |
| 15 | 25621600 | 25623999 | 12.90971 | 22.13441 | 60.00528 | 26.31655 | -1.04894 | 0.92429 | 0.085046 |
| 15 | 90887800 | 90890399 | 10.42149 | 18.37176 | 50.42898 | 22.31866 | -1.05973 | 0.560735 | 0.085049 |
| 1 | 97211600 | 97213399 | 28.48026 | 49.52004 | 27.22844 | 11.96044 | 1.245537 | 0.811175 | 0.08505 |
| 2 | 72900400 | 72905199 | 19.11126 | 31.13622 | 96.53169 | 41.45427 | -1.25661 | 1.52736 | 0.085055 |
| 13 | 57022200 | 57023199 | 2.171819 | 4.225906 | 27.26506 | 10.04816 | -2.04073 | -0.35038 | 0.085056 |
| 5 | 94352400 | 94356599 | 45.84544 | 75.03236 | 47.87949 | 20.59534 | 1.021797 | 1.487079 | 0.085086 |
| 4 | 123941800 | 123947799 | 38.39816 | 61.37251 | 137.5374 | 57.82057 | -0.81994 | 2.071053 | 0.085104 |
| 8 | 80663600 | 80663999 | 2.544919 | 4.972416 | 22.84006 | 7.953365 | -1.48444 | -0.56634 | 0.085111 |
| 16 | 75733800 | 75734799 | 17.16918 | 31.35695 | 20.34708 | 8.719696 | 1.041047 | 0.304977 | 0.085114 |
| 13 | 110567000 | 110567599 | 3.531275 | 6.883083 | 26.8058 | 9.692878 | -1.30413 | -0.4063 | 0.085117 |
| 6 | 50845000 | 50846199 | 4.365515 | 8.478016 | 31.26071 | 11.70153 | -1.27433 | -0.25935 | 0.085121 |
| 11 | 72686200 | 72687799 | 5.100451 | 9.558868 | 35.5089 | 14.6545 | -1.41523 | 0.093281 | 0.085126 |
| 3 | 94849600 | 94852599 | 13.33099 | 22.99956 | 58.00903 | 25.44778 | -0.94366 | 0.864901 | 0.085142 |
| 9 | 28104400 | 28105399 | 4.240617 | 8.236521 | 30.6245 | 11.45146 | -1.28479 | -0.26686 | 0.085174 |
| 8 | 55547600 | 55548799 | 16.39973 | 31.08532 | 15.03982 | 6.076114 | 1.525297 | 0.067959 | 0.085188 |
| 16 | 60849800 | 60851599 | 21.95092 | 39.25212 | 20.22125 | 8.874901 | 1.349631 | 0.460119 | 0.085196 |
| 8 | 126485600 | 126495999 | 73.70697 | 112.4881 | 271.9543 | 112.7748 | -0.98494 | 3.104576 | 0.08522 |
| 1 | 37418600 | 37420799 | 9.8141 | 17.29375 | 46.68077 | 20.66153 | -1.03563 | 0.55548 | 0.085222 |
| 11 | 103024600 | 103027799 | 9.8141 | 17.29375 | 46.68077 | 20.66153 | -1.03563 | 0.55548 | 0.085222 |
| 13 | 20036200 | 20037999 | 8.7893 | 15.81608 | 44.88541 | 19.59546 | -1.09101 | 0.383659 | 0.085244 |
| 8 | 72159200 | 72183399 | 145.9251 | 212.6223 | 482.6362 | 198.8955 | -0.9547 | 3.96319 | 0.085257 |
| 11 | 117044000 | 117062399 | 101.2974 | 151.2816 | 367.083 | 151.3296 | -1.01729 | 3.546033 | 0.085263 |
| 2 | 172772000 | 172789999 | 101.9837 | 152.7037 | 355.1633 | 146.4898 | -0.95279 | 3.496809 | 0.085263 |
| 14 | 79951000 | 79952599 | 11.71394 | 20.42937 | 52.61779 | 23.15707 | -0.97067 | 0.727532 | 0.085267 |
| 7 | 38940200 | 38942199 | 16.17209 | 30.01142 | 18.84601 | 7.90392 | 1.109674 | 0.206272 | 0.085275 |
| 13 | 43697400 | 43699799 | 9.398086 | 16.46646 | 49.34725 | 21.79363 | -1.18697 | 0.638905 | 0.085277 |
| 1 | 135188800 | 135191599 | 14.47016 | 24.40287 | 70.2697 | 30.72674 | -1.13886 | 1.088212 | 0.085282 |
| 3 | 128532000 | 128533399 | 18.73454 | 34.81981 | 16.40314 | 6.865114 | 1.522867 | 0.214142 | 0.085308 |
| 8 | 36560800 | 36567199 | 27.0007 | 43.57207 | 109.7554 | 46.78624 | -0.9713 | 1.748883 | 0.085311 |

| | | | | | | | | | |
|---|---|---|---|---|---|---|---|---|---|
| 14 | 27149400 | 27152599 | 27.07848 | 43.87845 | 102.4592 | 43.86229 | -0.85584 | 1.643903 | 0.085317 |
| 6 | 130421000 | 130422399 | 22.86726 | 40.4609 | 21.73955 | 9.614322 | 1.281498 | 0.541283 | 0.085318 |
| 4 | 6286200 | 6291599 | 42.36294 | 69.9677 | 40.73497 | 17.59374 | 1.158291 | 1.364598 | 0.08532 |
| 2 | 49180800 | 49181599 | 11.69421 | 22.77982 | 11.81216 | 4.306409 | 1.559288 | -0.3223 | 0.085328 |
| 4 | 17690000 | 17691199 | 15.13632 | 28.80033 | 15.83338 | 6.345261 | 1.353122 | 0.033024 | 0.085352 |
| 16 | 92257800 | 92266799 | 42.61537 | 67.58812 | 158.7077 | 66.07467 | -0.89901 | 2.32012 | 0.085357 |
| 19 | 36903000 | 36904199 | 6.570997 | 11.83828 | 41.65806 | 18.15979 | -1.39648 | 0.355241 | 0.085378 |
| 7 | 23255800 | 23256599 | 16.49874 | 30.18704 | 19.08091 | 8.158136 | 1.080889 | 0.286856 | 0.085381 |
| 2 | 90712200 | 90720999 | 53.25723 | 83.62633 | 184.4519 | 76.50403 | -0.8206 | 2.557709 | 0.085385 |
| 7 | 127236200 | 127237199 | 3.53486 | 6.879766 | 26.38877 | 9.706221 | -1.30451 | -0.34461 | 0.085387 |
| 16 | 23042600 | 23045599 | 12.80439 | 21.87668 | 62.69168 | 27.48693 | -1.12969 | 0.957648 | 0.085399 |
| 17 | 54931400 | 54932199 | 15.51539 | 29.51254 | 15.10633 | 6.057823 | 1.453824 | 0.036337 | 0.0854 |
| 18 | 90116600 | 90118999 | 23.95083 | 41.89856 | 27.49423 | 12.12639 | 0.992234 | 0.703254 | 0.085406 |
| 15 | 92995800 | 92996399 | 10.64854 | 20.75742 | 12.12139 | 4.378862 | 1.402671 | -0.3656 | 0.085408 |
| 12 | 18709400 | 18716599 | 293.5452 | 424.5648 | 294.8562 | 121.614 | 0.742685 | 4.081657 | 0.085415 |
| 7 | 20649200 | 20650999 | 13.90857 | 26.89913 | 13.25791 | 5.068325 | 1.570396 | -0.13446 | 0.085418 |
| 12 | 15916800 | 15917199 | 8.314456 | 16.26249 | 9.931066 | 3.34329 | 1.426487 | -0.64608 | 0.085426 |
| 7 | 38883400 | 38885199 | 21.74239 | 38.96895 | 19.66273 | 8.613016 | 1.381088 | 0.443486 | 0.085426 |
| 4 | 31987000 | 31987999 | 7.84351 | 14.25313 | 40.26577 | 17.3647 | -1.07247 | 0.309757 | 0.085449 |
| 5 | 41753800 | 41754999 | 5.163787 | 9.859033 | 32.85284 | 13.07073 | -1.2143 | -0.04093 | 0.085451 |
| 19 | 17359600 | 17361199 | 5.163787 | 9.859033 | 32.85284 | 13.07073 | -1.2143 | -0.04093 | 0.085451 |
| 5 | 116354600 | 116367999 | 81.44579 | 124.5429 | 259.5968 | 107.6992 | -0.76897 | 3.066059 | 0.085466 |
| 1 | 75227200 | 75232599 | 21.37318 | 34.60999 | 100.8208 | 43.13931 | -1.17464 | 1.64692 | 0.085476 |
| 17 | 17912200 | 17914199 | 22.76856 | 39.98065 | 25.78857 | 11.41037 | 1.016524 | 0.634286 | 0.085507 |
| 1 | 115814400 | 115814999 | 11.27787 | 21.82065 | 14.80595 | 5.648873 | 1.117331 | -0.16583 | 0.085529 |
| 4 | 16882200 | 16882599 | 7.371944 | 14.41985 | 8.714412 | 2.918923 | 1.448771 | -0.68569 | 0.08554 |
| 13 | 35426600 | 35427999 | 19.77288 | 35.50869 | 21.73255 | 9.503921 | 1.106574 | 0.423208 | 0.085546 |
| 7 | 132225400 | 132228799 | 14.33056 | 24.11604 | 69.57684 | 30.40413 | -1.14163 | 1.106976 | 0.085552 |
| 5 | 130681400 | 130683199 | 6.6701 | 12.49158 | 36.16721 | 14.94441 | -1.06126 | 0.113681 | 0.085555 |
| 19 | 45350800 | 45352999 | 10.62191 | 18.56296 | 50.9745 | 22.46345 | -1.06194 | 0.69083 | 0.085557 |
| 17 | 35116800 | 35118799 | 5.858692 | 10.52264 | 42.65949 | 18.65339 | -1.60151 | 0.372135 | 0.085593 |
| 7 | 6389800 | 6390999 | 18.78553 | 33.8546 | 21.40327 | 9.326544 | 1.063471 | 0.394949 | 0.085608 |
| 5 | 137575200 | 137578999 | 18.90959 | 31.20769 | 82.29596 | 35.5255 | -1.01413 | 1.345904 | 0.085618 |
| X | 4786600 | 4789599 | 38.08193 | 63.77314 | 36.57433 | 15.94253 | 1.180088 | 1.197302 | 0.085622 |
| 8 | 36978800 | 36979799 | 5.655078 | 10.59274 | 34.40735 | 14.21253 | -1.22448 | 0.099965 | 0.085637 |
| 10 | 59732200 | 59739799 | 30.89107 | 49.3187 | 137.8171 | 57.84664 | -1.13925 | 2.099327 | 0.08564 |
| 11 | 98543800 | 98544199 | 17.58796 | 32.24666 | 19.83017 | 8.454627 | 1.123178 | 0.286344 | 0.085653 |
| 16 | 78145600 | 78147199 | 18.40391 | 33.96547 | 17.37821 | 7.343172 | 1.39485 | 0.250759 | 0.085657 |
| 7 | 19570600 | 19571599 | 11.09719 | 21.47603 | 14.32918 | 5.461064 | 1.142803 | -0.17308 | 0.085663 |
| 14 | 3226800 | 3228599 | 35.30431 | 58.93947 | 39.64773 | 17.24858 | 0.95118 | 1.225435 | 0.085666 |
| 16 | 12137200 | 12138399 | 20.68299 | 37.03227 | 21.50133 | 9.426524 | 1.179519 | 0.446552 | 0.085699 |
| 7 | 57726200 | 57728599 | 23.78879 | 41.92334 | 23.63824 | 10.46238 | 1.21087 | 0.594538 | 0.08571 |
| 10 | 7117800 | 7122799 | 32.67062 | 52.49008 | 118.122 | 50.04294 | -0.81698 | 1.897306 | 0.085723 |
| 8 | 121668800 | 121681999 | 63.37891 | 97.73993 | 234.1723 | 97.27104 | -0.96015 | 2.891214 | 0.085736 |
| 11 | 5063400 | 5066799 | 13.01976 | 22.47044 | 57.4435 | 25.19996 | -0.96281 | 0.860135 | 0.085737 |
| 18 | 84788200 | 84788799 | 2.428341 | 4.744675 | 22.1478 | 7.710779 | -1.50368 | -0.56677 | 0.085745 |
| 17 | 85748000 | 85753399 | 31.56907 | 50.67391 | 120.9834 | 51.18474 | -0.90371 | 1.925399 | 0.085755 |

| | | | | | | | | | |
|---|---|---|---|---|---|---|---|---|---|
| 8 | 119163600 | 119165599 | 7.825411 | 14.27303 | 41.47861 | 17.80503 | -1.10883 | 0.289559 | 0.085758 |
| 13 | 56891800 | 56893799 | 6.295151 | 11.63834 | 36.8411 | 15.53034 | -1.21126 | 0.191443 | 0.085767 |
| 16 | 22926000 | 22927399 | 6.295151 | 11.63834 | 36.8411 | 15.53034 | -1.21126 | 0.191443 | 0.085767 |
| 11 | 69557800 | 69560999 | 15.50896 | 26.24648 | 68.25412 | 29.87711 | -0.99227 | 1.058785 | 0.085825 |
| 6 | 23245600 | 23250999 | 45.63861 | 72.07962 | 170.8035 | 70.92802 | -0.91729 | 2.430017 | 0.085827 |
| 9 | 21221800 | 21227199 | 25.53609 | 41.22834 | 104.9544 | 44.76739 | -0.98553 | 1.732365 | 0.085829 |
| 3 | 8309600 | 8311799 | 76.08989 | 121.0156 | 88.28481 | 36.85946 | 0.788686 | 2.249048 | 0.085835 |
| 13 | 30167600 | 30170999 | 20.96357 | 34.16632 | 92.17771 | 39.58971 | -1.05633 | 1.524121 | 0.085855 |
| 3 | 90519000 | 90526399 | 36.94506 | 59.06209 | 137.9169 | 57.99696 | -0.87882 | 2.062998 | 0.085866 |
| 7 | 125836200 | 125837999 | 9.957668 | 17.5585 | 46.68317 | 20.65982 | -1.01341 | 0.550158 | 0.085877 |
| 4 | 155467000 | 155473999 | 38.8449 | 61.7648 | 149.1757 | 62.2643 | -0.93587 | 2.240539 | 0.085879 |
| 13 | 111834800 | 111836399 | 22.53964 | 39.55781 | 26.042 | 11.51737 | 0.98745 | 0.642641 | 0.085883 |
| 4 | 58482600 | 58483799 | 23.0872 | 40.71299 | 23.44662 | 10.37624 | 1.180545 | 0.581665 | 0.085909 |
| 5 | 30519800 | 30544199 | 144.22 | 209.6191 | 498.5949 | 205.525 | -1.02558 | 4.000803 | 0.085946 |
| 11 | 94813600 | 94815599 | 10.18026 | 17.77333 | 51.50223 | 22.68115 | -1.13923 | 0.70326 | 0.085954 |
| 5 | 119386200 | 119394199 | 46.70355 | 73.91799 | 167.1207 | 69.47314 | -0.84718 | 2.379044 | 0.085977 |
| 8 | 78167000 | 78181999 | 217.3079 | 309.0121 | 600.9341 | 248.5011 | -0.76662 | 4.329329 | 0.085983 |
| 4 | 140697200 | 140715199 | 99.52746 | 149.3389 | 339.4796 | 140.0855 | -0.91706 | 3.456299 | 0.085997 |
| 4 | 63318800 | 63324999 | 23.63995 | 37.96184 | 126.0626 | 53.36692 | -1.37762 | 1.904522 | 0.086 |
| 17 | 91918600 | 91922999 | 55.0628 | 89.06611 | 55.18733 | 23.59446 | 1.05102 | 1.702951 | 0.086005 |
| 16 | 17742000 | 17750199 | 35.92468 | 57.19036 | 146.0227 | 61.04618 | -1.01399 | 2.198603 | 0.086016 |
| 4 | 123576200 | 123578599 | 9.14606 | 16.28787 | 43.6011 | 19.20232 | -1.01609 | 0.449584 | 0.086019 |
| 7 | 10568600 | 10572199 | 70.53115 | 112.4135 | 83.80332 | 35.09146 | 0.760754 | 2.17581 | 0.086035 |
| 13 | 55641800 | 55643399 | 6.325339 | 11.60936 | 37.7561 | 16.07443 | -1.26031 | 0.236048 | 0.086036 |
| 2 | 118990400 | 118992599 | 8.10472 | 14.43098 | 43.60586 | 19.20712 | -1.18956 | 0.443843 | 0.08605 |
| 9 | 88320400 | 88321599 | 8.10472 | 14.43098 | 43.60586 | 19.20712 | -1.18956 | 0.443843 | 0.08605 |
| 7 | 33203200 | 33204999 | 53.86224 | 87.10033 | 56.1224 | 23.98792 | 0.994388 | 1.709043 | 0.086064 |
| 12 | 105650400 | 105652399 | 10.73013 | 18.51961 | 57.84342 | 25.37543 | -1.2499 | 0.850849 | 0.086077 |
| 3 | 157272400 | 157276399 | 21.42127 | 34.97039 | 92.07512 | 39.56962 | -1.01964 | 1.499733 | 0.086098 |
| 11 | 117903200 | 117911199 | 35.02986 | 55.63709 | 156.1261 | 65.09313 | -1.15342 | 2.271817 | 0.086116 |
| 1 | 88687800 | 88688999 | 6.247787 | 11.68533 | 36.10151 | 14.95046 | -1.15677 | 0.117135 | 0.086135 |
| 10 | 56598400 | 56600199 | 16.37378 | 30.02195 | 19.55633 | 8.337327 | 1.040765 | 0.27557 | 0.086146 |
| 14 | 70819200 | 70826599 | 33.9437 | 54.17851 | 139.0184 | 58.3298 | -1.01696 | 2.109814 | 0.086158 |
| 9 | 53639000 | 53639599 | 12.50076 | 24.13744 | 15.04876 | 5.796488 | 1.224878 | -0.12065 | 0.086162 |
| 19 | 23928600 | 23928999 | 4.06961 | 7.901739 | 27.29942 | 10.23564 | -1.18176 | -0.26515 | 0.086169 |
| X | 32969800 | 32971999 | 21.11334 | 38.3488 | 18.32891 | 7.909046 | 1.472653 | 0.361719 | 0.086169 |
| 17 | 12336800 | 12338799 | 10.62165 | 18.59469 | 51.51649 | 22.73554 | -1.07498 | 0.660447 | 0.086171 |
| 18 | 37020600 | 37026799 | 194.6172 | 290.8428 | 212.8583 | 87.762 | 0.711614 | 3.541486 | 0.086172 |
| 2 | 157239200 | 157240799 | 15.96305 | 27.05557 | 66.04 | 28.92108 | -0.90106 | 1.046565 | 0.086181 |
| 7 | 14390400 | 14393399 | 74.43387 | 118.7593 | 78.05303 | 32.73152 | 0.943786 | 2.144325 | 0.086187 |
| 7 | 6916600 | 6917599 | 4.514154 | 8.693752 | 29.66774 | 11.52228 | -1.21342 | -0.13215 | 0.086207 |
| 6 | 89327000 | 89343199 | 80.56704 | 121.9337 | 307.1256 | 127.0495 | -1.054 | 3.279727 | 0.086209 |
| 3 | 35061600 | 35064399 | 9.943344 | 17.58568 | 46.16371 | 20.41774 | -0.9936 | 0.514443 | 0.086223 |
| 6 | 108357600 | 108358999 | 9.281277 | 16.57796 | 44.1838 | 19.41016 | -1.0074 | 0.432374 | 0.086234 |
| 5 | 4916200 | 4917799 | 4.424204 | 8.5436 | 33.01223 | 12.7087 | -1.38063 | -0.15875 | 0.086255 |
| 11 | 35683200 | 35684599 | 5.873101 | 10.83881 | 37.32444 | 15.77212 | -1.334 | 0.199437 | 0.086259 |
| 6 | 103114800 | 103115599 | 9.799466 | 19.08874 | 12.56939 | 4.583097 | 1.218198 | -0.34538 | 0.086263 |

| | | | | | | | | | |
|---|---|---|---|---|---|---|---|---|---|
| 5 | 5351600 | 5352199 | 2.543414 | 4.973539 | 22.61941 | 7.712249 | -1.44281 | -0.64326 | 0.086263 |
| 14 | 74269000 | 74269599 | 9.297559 | 18.12939 | 12.75292 | 4.588838 | 1.141787 | -0.39604 | 0.086276 |
| 17 | 23182800 | 23184399 | 29.12218 | 49.96008 | 33.00179 | 14.47428 | 0.979249 | 0.949071 | 0.086292 |
| 14 | 33035600 | 33038999 | 17.81355 | 29.78045 | 72.53172 | 31.58225 | -0.89865 | 1.188858 | 0.086293 |
| 2 | 94900000 | 94900799 | 11.38536 | 22.00706 | 13.83542 | 5.302806 | 1.220185 | -0.1529 | 0.086293 |
| 5 | 94423600 | 94424599 | 16.32255 | 30.38678 | 16.67547 | 6.960252 | 1.308314 | 0.184964 | 0.086295 |
| 1 | 139909800 | 139911799 | 20.95193 | 37.07751 | 23.65439 | 10.46084 | 1.035597 | 0.531414 | 0.086299 |
| 4 | 154161000 | 154178399 | 93.89876 | 141.1426 | 336.5661 | 138.949 | -0.98373 | 3.420249 | 0.086311 |
| 1 | 36101400 | 36139399 | 215.1046 | 302.6499 | 700.1472 | 290.0383 | -1.03242 | 4.485353 | 0.086315 |
| 4 | 140864000 | 140868599 | 15.09187 | 25.42444 | 69.1606 | 30.23207 | -1.05715 | 1.099107 | 0.086322 |
| 5 | 137498400 | 137501599 | 15.09187 | 25.42444 | 69.1606 | 30.23207 | -1.05715 | 1.099107 | 0.086322 |
| 14 | 82207000 | 82207999 | 13.51782 | 26.15316 | 12.49486 | 4.768302 | 1.615623 | -0.14747 | 0.086323 |
| 1 | 165651200 | 165652799 | 7.833332 | 14.26429 | 40.32317 | 17.34426 | -1.07099 | 0.297337 | 0.086326 |
| 1 | 39095600 | 39096799 | 6.903062 | 12.70565 | 39.52189 | 16.76145 | -1.19342 | 0.225971 | 0.086326 |
| 5 | 104770600 | 104770999 | 10.49734 | 20.45359 | 12.47371 | 4.532431 | 1.332123 | -0.34768 | 0.08633 |
| 2 | 115060600 | 115062399 | 17.92627 | 32.49168 | 20.71934 | 8.964066 | 1.057227 | 0.353622 | 0.086353 |
| 5 | 86827600 | 86828799 | 13.30701 | 25.49121 | 15.36398 | 6.059288 | 1.241386 | -0.03769 | 0.086356 |
| 8 | 15004600 | 15021799 | 73.21664 | 111.7037 | 272.3258 | 112.9205 | -0.99736 | 3.111159 | 0.086356 |
| 6 | 72727200 | 72733199 | 28.80649 | 46.34869 | 114.7392 | 48.71167 | -0.95154 | 1.844971 | 0.08637 |
| 6 | 144247400 | 144248999 | 14.95894 | 28.39997 | 16.39053 | 6.599326 | 1.277926 | 0.047379 | 0.086379 |
| 9 | 30557800 | 30558199 | 11.63642 | 22.53446 | 14.73784 | 5.599514 | 1.175503 | -0.17538 | 0.086381 |
| X | 88625000 | 88625399 | 11.63642 | 22.53446 | 14.73784 | 5.599514 | 1.175503 | -0.17538 | 0.086381 |
| 7 | 97826000 | 97827999 | 6.665383 | 12.03475 | 40.12082 | 17.44166 | -1.31595 | 0.341932 | 0.086387 |
| 14 | 107134000 | 107134199 | 19.39542 | 36.50131 | 10.43894 | 4.277823 | 2.254466 | 0.129437 | 0.086395 |
| 6 | 84539400 | 84571999 | 195.7541 | 276.0125 | 692.0319 | 286.5758 | -1.14535 | 4.453149 | 0.086403 |
| 9 | 42836600 | 42844199 | 40.25395 | 63.75673 | 168.9272 | 70.2501 | -1.07474 | 2.35736 | 0.086404 |
| 7 | 15780000 | 15783199 | 14.99062 | 24.78829 | 81.64984 | 35.29411 | -1.33315 | 1.313625 | 0.086409 |
| 19 | 55575400 | 55578799 | 16.36997 | 27.44791 | 70.18334 | 30.61661 | -0.96839 | 1.15121 | 0.086427 |
| 5 | 93638000 | 93641799 | 34.3338 | 58.4356 | 32.12349 | 14.08007 | 1.241406 | 1.038032 | 0.08643 |
| 6 | 3305000 | 3308399 | 27.34145 | 47.08395 | 31.29802 | 13.72925 | 0.970924 | 0.903215 | 0.086435 |
| 7 | 22143200 | 22144199 | 24.66815 | 43.10245 | 22.62621 | 9.969518 | 1.312281 | 0.716366 | 0.086452 |
| 3 | 46049400 | 46051199 | 21.88121 | 38.65081 | 24.20902 | 10.71006 | 1.061181 | 0.558704 | 0.086463 |
| 4 | 131162600 | 131167399 | 32.18129 | 51.72794 | 116.2359 | 49.27709 | -0.81399 | 1.881944 | 0.086464 |
| 16 | 97908200 | 97908999 | 195.869 | 295.1186 | 166.839 | 68.92218 | 1.094297 | 3.383962 | 0.086471 |
| 10 | 85449800 | 85455399 | 24.01196 | 39.04282 | 103.0018 | 44.20074 | -1.02691 | 1.568105 | 0.086475 |
| 14 | 23572600 | 23573399 | 8.955011 | 17.47166 | 12.18234 | 4.349482 | 1.163708 | -0.43035 | 0.086504 |
| 17 | 39467600 | 39468399 | 17.74673 | 33.67174 | 13.18559 | 5.315901 | 1.829222 | 0.0695 | 0.086504 |
| 17 | 38029600 | 38030999 | 14.59323 | 27.65319 | 16.94756 | 6.851017 | 1.18702 | 0.057784 | 0.086513 |
| 5 | 47642400 | 47644199 | 6.831584 | 12.7421 | 40.04681 | 16.66176 | -1.18715 | 0.14961 | 0.086517 |
| 10 | 60540000 | 60566599 | 226.3284 | 318.5966 | 670.6083 | 277.7797 | -0.89549 | 4.478651 | 0.086528 |
| 1 | 24501600 | 24503599 | 46.75776 | 76.13318 | 52.80133 | 22.66996 | 0.896961 | 1.564852 | 0.086533 |
| 6 | 133646400 | 133647599 | 27.11759 | 47.04137 | 30.25608 | 13.27515 | 1.020424 | 0.83918 | 0.086534 |
| 10 | 65424600 | 65426999 | 32.71115 | 55.19395 | 37.79366 | 16.52862 | 0.925437 | 1.111549 | 0.086542 |
| 5 | 120278400 | 120341199 | 336.9908 | 451.985 | 1168.677 | 487.74 | -1.26264 | 5.229323 | 0.086544 |
| 4 | 129965000 | 129986199 | 82.3658 | 123.8892 | 343.8871 | 141.9966 | -1.2018 | 3.405436 | 0.086547 |
| 10 | 81584000 | 81585399 | 48.95059 | 79.99976 | 43.50349 | 18.70325 | 1.250355 | 1.511327 | 0.086557 |
| 2 | 77645400 | 77648599 | 16.56235 | 27.70864 | 73.60527 | 32.06426 | -1.02341 | 1.176958 | 0.086559 |

| | | | | | | | | | |
|---|---|---|---|---|---|---|---|---|---|
| 1 | 92603000 | 92605799 | 13.28115 | 22.58338 | 63.63392 | 27.88935 | -1.10644 | 1.007014 | 0.086574 |
| 3 | 87698400 | 87708199 | 54.16657 | 84.46249 | 208.4326 | 86.49841 | -0.99028 | 2.686559 | 0.086574 |
| 15 | 84601800 | 84608799 | 29.33602 | 47.09135 | 121.4903 | 51.40235 | -1.01513 | 1.921689 | 0.086577 |
| 7 | 26852000 | 26868799 | 92.2911 | 138.4964 | 351.8333 | 145.1875 | -1.07709 | 3.452911 | 0.086579 |
| 17 | 37704600 | 37705999 | 16.31596 | 30.12281 | 19.35245 | 8.173413 | 1.069877 | 0.236518 | 0.086585 |
| 7 | 38508000 | 38510599 | 54.01497 | 87.26383 | 59.02338 | 25.19676 | 0.922969 | 1.735724 | 0.086588 |
| 1 | 86509800 | 86510799 | 5.473159 | 10.3899 | 34.9816 | 14.08765 | -1.24522 | 0.006737 | 0.086599 |
| 7 | 20742200 | 20747199 | 43.58807 | 72.27877 | 38.91438 | 16.85375 | 1.271037 | 1.322427 | 0.086616 |
| 6 | 6478400 | 6479599 | 14.11817 | 27.17423 | 14.248 | 5.542823 | 1.458729 | -0.07433 | 0.086621 |
| 7 | 21878600 | 21879599 | 13.70681 | 25.83987 | 17.15339 | 7.008986 | 1.059507 | 0.083976 | 0.086622 |
| 6 | 95425600 | 95426799 | 4.54064 | 8.790394 | 30.82551 | 11.72909 | -1.22552 | -0.20242 | 0.086626 |
| 10 | 22520800 | 22526999 | 44.97087 | 71.50733 | 155.3679 | 64.85069 | -0.7836 | 2.245496 | 0.086633 |
| 14 | 91172800 | 91173599 | 13.07856 | 25.20759 | 13.65305 | 5.290606 | 1.416472 | -0.10037 | 0.086633 |
| 1 | 67631000 | 67631999 | 19.17196 | 35.87818 | 14.05967 | 5.816777 | 1.798018 | 0.170593 | 0.086634 |
| 8 | 71469600 | 71470799 | 19.17196 | 35.87818 | 14.05967 | 5.816777 | 1.798018 | 0.170593 | 0.086634 |
| 9 | 31191000 | 31193199 | 10.13248 | 17.81517 | 47.72867 | 21.13472 | -1.02629 | 0.583554 | 0.086636 |
| 3 | 20723000 | 20723999 | 15.46742 | 29.20941 | 16.88455 | 6.875211 | 1.261777 | 0.088879 | 0.086647 |
| 18 | 3200600 | 3203599 | 56.43145 | 90.99066 | 61.71323 | 26.27904 | 0.915907 | 1.797539 | 0.086655 |
| 4 | 133933000 | 133939599 | 30.44601 | 48.87976 | 120.0421 | 50.80007 | -0.94401 | 1.918444 | 0.086673 |
| 1 | 107517200 | 107518199 | 4.037539 | 7.74841 | 32.40631 | 12.71425 | -1.51853 | -0.09972 | 0.086683 |
| 5 | 66519800 | 66522599 | 8.922256 | 15.73457 | 50.95663 | 22.55053 | -1.29661 | 0.546343 | 0.086692 |
| 15 | 85808200 | 85816799 | 32.58298 | 51.87463 | 150.7406 | 63.02541 | -1.20008 | 2.193753 | 0.0867 |
| 5 | 123405800 | 123407799 | 16.32237 | 27.51753 | 68.18339 | 29.81152 | -0.92306 | 1.096113 | 0.086701 |
| 17 | 14413200 | 14426999 | 77.06374 | 117.513 | 276.8111 | 114.7667 | -0.94844 | 3.122517 | 0.086708 |
| 11 | 5914400 | 5955199 | 223.5885 | 311.1565 | 783.1685 | 324.9838 | -1.16999 | 4.650535 | 0.086709 |
| 2 | 27404200 | 27406999 | 11.46179 | 19.87676 | 58.57206 | 25.69881 | -1.16534 | 0.80898 | 0.086716 |
| 6 | 136637000 | 136644399 | 46.43107 | 73.36729 | 171.2495 | 71.13096 | -0.895 | 2.418467 | 0.086717 |
| 4 | 153837600 | 153846999 | 62.55604 | 96.9053 | 225.1234 | 93.51009 | -0.91073 | 2.806196 | 0.08675 |
| 19 | 42647800 | 42653399 | 31.77607 | 50.90796 | 127.9382 | 53.9687 | -0.9809 | 1.988269 | 0.086751 |
| 3 | 147741600 | 147742799 | 4.940125 | 9.538047 | 31.12453 | 11.99286 | -1.13981 | -0.15135 | 0.086768 |
| 14 | 58148000 | 58149199 | 4.066326 | 7.906299 | 26.92376 | 9.982558 | -1.14626 | -0.30685 | 0.086776 |
| 8 | 5185800 | 5187999 | 29.1141 | 50.56688 | 27.88088 | 12.23864 | 1.24165 | 0.831078 | 0.0868 |
| 1 | 112478400 | 112480199 | 16.82735 | 32.31509 | 11.65027 | 4.561978 | 1.983383 | -0.03301 | 0.086801 |
| 2 | 16360400 | 16362599 | 11.99732 | 20.31639 | 70.67478 | 30.94263 | -1.40974 | 1.043635 | 0.086803 |
| 11 | 98727000 | 98730799 | 15.35009 | 25.53911 | 75.25918 | 32.66992 | -1.17223 | 1.238066 | 0.086808 |
| 13 | 12319000 | 12320999 | 8.011261 | 14.49426 | 40.16124 | 17.41488 | -1.04977 | 0.337468 | 0.086836 |
| 7 | 27325600 | 27328999 | 17.56096 | 29.16542 | 76.0599 | 32.97687 | -0.99709 | 1.265471 | 0.086838 |
| 2 | 104956000 | 104965799 | 53.98579 | 84.22865 | 208.3793 | 86.47051 | -0.99326 | 2.676281 | 0.086852 |
| 2 | 167101400 | 167123799 | 130.3023 | 190.7924 | 471.4298 | 194.1958 | -1.07019 | 3.885846 | 0.086853 |
| 5 | 143109800 | 143125999 | 129.6186 | 191.7352 | 411.177 | 169.3338 | -0.85199 | 3.719871 | 0.086854 |
| 2 | 22929200 | 22930599 | 6.400228 | 11.62422 | 42.38618 | 18.29132 | -1.43817 | 0.305856 | 0.086857 |
| 4 | 30517200 | 30518199 | 11.03117 | 21.53138 | 9.998041 | 3.543889 | 1.749576 | -0.43275 | 0.086857 |
| 4 | 137943400 | 137945199 | 7.476907 | 13.37736 | 42.097 | 18.46741 | -1.24313 | 0.406808 | 0.086887 |
| 17 | 57475000 | 57476399 | 3.730735 | 7.210858 | 30.46056 | 11.67768 | -1.50015 | -0.18579 | 0.086892 |
| 6 | 53771200 | 53772999 | 5.052396 | 9.597565 | 34.96045 | 14.05924 | -1.3556 | -0.00201 | 0.086913 |
| 13 | 66701800 | 66704999 | 84.43962 | 133.9441 | 90.97259 | 37.88095 | 0.888079 | 2.333041 | 0.086924 |
| 6 | 47571000 | 47572799 | 9.352239 | 16.22038 | 57.30841 | 25.14447 | -1.42493 | 0.798654 | 0.086935 |

| | | | | | | | | | |
|---|---|---|---|---|---|---|---|---|---|
| 7 | 35149000 | 35150599 | 8.151525 | 14.77357 | 40.20087 | 17.39438 | -1.02183 | 0.328555 | 0.086936 |
| 12 | 110444000 | 110447399 | 13.9252 | 23.64964 | 64.24285 | 28.1514 | -1.05431 | 1.019814 | 0.086949 |
| 7 | 10029200 | 10033599 | 113.4895 | 176.069 | 128.293 | 53.28133 | 0.761845 | 2.791123 | 0.086951 |
| 7 | 7543400 | 7548999 | 83.07136 | 131.7278 | 86.67086 | 36.07715 | 0.933322 | 2.340884 | 0.086971 |
| 1 | 116702200 | 116703399 | 15.61216 | 29.26541 | 17.93731 | 7.400011 | 1.162948 | 0.138912 | 0.086971 |
| 19 | 45207800 | 45227799 | 127.6742 | 187.1375 | 476.0902 | 196.1029 | -1.11084 | 3.86934 | 0.086985 |
| 12 | 20605600 | 20605799 | 7.495967 | 14.66111 | 8.696708 | 2.935934 | 1.464856 | -0.65784 | 0.08699 |
| 12 | 23631000 | 23632799 | 32.09882 | 54.42849 | 32.58363 | 14.2714 | 1.118573 | 1.061675 | 0.087003 |
| X | 109372400 | 109373999 | 18.16199 | 34.52544 | 12.90942 | 5.183601 | 1.90112 | 0.060674 | 0.087005 |
| 4 | 118228800 | 118233999 | 22.61976 | 36.40329 | 117.1385 | 49.74776 | -1.32818 | 1.830358 | 0.087013 |
| 1 | 16771800 | 16773999 | 23.31089 | 40.70287 | 27.18692 | 11.97386 | 0.96672 | 0.726425 | 0.087027 |
| 6 | 65648000 | 65650199 | 13.78614 | 23.35251 | 65.24027 | 28.56587 | -1.0947 | 1.044149 | 0.087047 |
| 9 | 124241200 | 124241999 | 13.18051 | 25.34096 | 15.55008 | 6.068917 | 1.230341 | -0.07024 | 0.087058 |
| 13 | 117830400 | 117831199 | 12.54284 | 24.453 | 9.896075 | 3.569257 | 1.923986 | -0.35386 | 0.087059 |
| 4 | 33176200 | 33177599 | 6.19206 | 11.71495 | 34.85589 | 14.13774 | -1.07764 | 0.038285 | 0.08708 |
| 2 | 87134600 | 87135599 | 13.45691 | 25.54179 | 17.16042 | 6.913044 | 1.060043 | 0.036617 | 0.087089 |
| 4 | 122363200 | 122364399 | 9.357213 | 18.25448 | 12.28516 | 4.39277 | 1.213423 | -0.41779 | 0.087092 |
| 2 | 32340000 | 32343199 | 23.72833 | 38.48159 | 97.74432 | 41.86937 | -0.97561 | 1.617883 | 0.087099 |
| 9 | 102320000 | 102322399 | 26.31159 | 45.5027 | 31.35689 | 13.75684 | 0.920423 | 0.862586 | 0.0871 |
| 17 | 45461000 | 45466999 | 39.74369 | 63.35791 | 143.5116 | 60.11166 | -0.83946 | 2.155442 | 0.087116 |
| 5 | 45307400 | 45308999 | 23.96403 | 41.8701 | 28.13184 | 12.39502 | 0.958467 | 0.723717 | 0.087121 |
| 3 | 133249000 | 133251199 | 10.65614 | 18.57289 | 53.12729 | 23.3721 | -1.12124 | 0.7311 | 0.087148 |
| 1 | 45467400 | 45469199 | 19.55563 | 35.36447 | 19.90332 | 8.635525 | 1.233435 | 0.376573 | 0.087156 |
| 2 | 101664800 | 101667399 | 11.0659 | 19.40351 | 49.89241 | 22.04867 | -0.96774 | 0.63598 | 0.08716 |
| X | 81704400 | 81705199 | 14.81221 | 27.93185 | 17.53891 | 7.162998 | 1.139862 | 0.09413 | 0.08716 |
| 2 | 136774200 | 136774999 | 10.10672 | 19.70317 | 12.63212 | 4.557471 | 1.270625 | -0.37602 | 0.087166 |
| 1 | 155751800 | 155752799 | 16.29072 | 30.09023 | 18.61438 | 7.85671 | 1.12509 | 0.233052 | 0.087177 |
| 19 | 10458000 | 10460199 | 8.188602 | 14.42763 | 49.31563 | 21.82829 | -1.37403 | 0.550675 | 0.087191 |
| 11 | 120192200 | 120202799 | 45.71726 | 72.03221 | 179.8318 | 74.6209 | -0.99517 | 2.483721 | 0.087196 |
| 9 | 106305000 | 106317999 | 56.20363 | 87.37758 | 210.4636 | 87.38026 | -0.95872 | 2.739966 | 0.087196 |
| 11 | 98253800 | 98255399 | 4.915371 | 9.262536 | 33.41283 | 13.6623 | -1.36159 | 0.052264 | 0.087227 |
| 12 | 74167600 | 74168399 | 4.127496 | 7.994691 | 29.54552 | 11.21502 | -1.29625 | -0.21833 | 0.087266 |
| 12 | 23274800 | 23276599 | 25.22561 | 44.07637 | 27.35344 | 12.0524 | 1.07233 | 0.727996 | 0.087286 |
| 1 | 74928000 | 74933399 | 34.49949 | 55.25227 | 126.6418 | 53.39415 | -0.84898 | 2.002021 | 0.087306 |
| 7 | 38385200 | 38388399 | 60.4871 | 97.14184 | 69.50721 | 29.42935 | 0.831978 | 1.927403 | 0.087324 |
| X | 32782200 | 32784199 | 22.57695 | 40.93283 | 17.2577 | 7.463743 | 1.650257 | 0.380701 | 0.087336 |
| 12 | 72121200 | 72122199 | 13.05453 | 24.9101 | 16.57542 | 6.602731 | 1.087464 | -0.00501 | 0.087369 |
| 7 | 17571800 | 17572999 | 44.23164 | 72.43393 | 46.75524 | 20.11659 | 1.00578 | 1.477742 | 0.087371 |
| 7 | 10812000 | 10823399 | 318.4314 | 457.7585 | 317.5461 | 131.0827 | 0.735662 | 4.17299 | 0.087388 |
| 16 | 58687400 | 58688999 | 13.89791 | 26.48615 | 16.45937 | 6.573931 | 1.181947 | 0.011608 | 0.087406 |
| 2 | 167830800 | 167855799 | 137.8876 | 200.1991 | 507.3311 | 209.1495 | -1.11842 | 4.016915 | 0.087418 |
| 17 | 46005800 | 46006999 | 4.137295 | 7.986376 | 29.78936 | 11.4893 | -1.33132 | -0.16167 | 0.087418 |
| 14 | 78228200 | 78229799 | 24.46267 | 42.67156 | 28.69426 | 12.62984 | 0.957134 | 0.749799 | 0.087434 |
| 6 | 50542400 | 50544199 | 7.545185 | 13.39053 | 45.66238 | 20.16886 | -1.36558 | 0.464714 | 0.087442 |
| 7 | 102135600 | 102138199 | 13.15248 | 22.75921 | 57.5199 | 25.2355 | -0.9457 | 0.835789 | 0.087444 |
| 8 | 48356400 | 48357399 | 5.110053 | 9.839708 | 34.73627 | 13.49738 | -1.26488 | -0.11495 | 0.087468 |
| X | 26544800 | 26545799 | 26.63928 | 46.18686 | 30.37894 | 13.32883 | 0.988178 | 0.842016 | 0.087493 |

| | | | | | | | | | |
|---|---|---|---|---|---|---|---|---|---|
| 5 | 55038800 | 55039599 | 11.62523 | 22.61908 | 11.59239 | 4.274048 | 1.559979 | -0.28703 | 0.08751 |
| 16 | 29507000 | 29508599 | 8.198338 | 14.73427 | 41.62207 | 18.18992 | -1.0851 | 0.380918 | 0.087519 |
| 2 | 90972600 | 90974199 | 4.483012 | 8.491373 | 32.99252 | 13.34923 | -1.45396 | 0.010764 | 0.087525 |
| 11 | 76775600 | 76776399 | 4.378574 | 8.4609 | 28.703 | 11.01677 | -1.18879 | -0.17707 | 0.087526 |
| 2 | 87145200 | 87145999 | 25.51794 | 44.37488 | 25.9917 | 11.41809 | 1.155023 | 0.793678 | 0.087533 |
| 1 | 179437400 | 179438199 | 3.846358 | 7.426847 | 29.16259 | 11.23286 | -1.40184 | -0.17005 | 0.087533 |
| 17 | 18738800 | 18740399 | 43.28163 | 71.12111 | 46.19448 | 19.90411 | 0.998558 | 1.430321 | 0.087547 |
| 8 | 60296800 | 60298199 | 17.53234 | 32.38533 | 18.68308 | 7.885136 | 1.22472 | 0.242467 | 0.087548 |
| 11 | 99383400 | 99385399 | 8.96348 | 16.0547 | 42.63145 | 18.68455 | -0.99951 | 0.410396 | 0.087555 |
| 4 | 130550000 | 130557799 | 44.07697 | 69.47818 | 175.7162 | 72.9173 | -1.01328 | 2.473981 | 0.087563 |
| 11 | 69898200 | 69905999 | 27.22119 | 43.47218 | 139.1299 | 58.4216 | -1.33394 | 2.087277 | 0.087564 |
| 18 | 23857200 | 23858599 | 17.98316 | 32.87624 | 19.88977 | 8.513119 | 1.142809 | 0.305729 | 0.087565 |
| 2 | 145465400 | 145467399 | 20.23782 | 35.83558 | 23.47343 | 10.37956 | 0.99829 | 0.521015 | 0.087568 |
| 12 | 86873000 | 86874199 | 3.462393 | 6.639373 | 31.07021 | 12.2179 | -1.67966 | -0.09924 | 0.087583 |
| 1 | 91656000 | 91663399 | 45.28446 | 71.60073 | 171.5039 | 71.2594 | -0.9315 | 2.402454 | 0.087583 |
| 6 | 113419400 | 113420399 | 4.272779 | 8.201947 | 30.07123 | 11.78948 | -1.32875 | -0.1034 | 0.087588 |
| 10 | 128657200 | 128658199 | 4.272779 | 8.201947 | 30.07123 | 11.78948 | -1.32875 | -0.1034 | 0.087588 |
| 6 | 112837400 | 112840599 | 19.65444 | 32.53059 | 79.7423 | 34.49171 | -0.90879 | 1.312172 | 0.087601 |
| 13 | 66754800 | 66760399 | 135.8482 | 208.9798 | 137.1575 | 56.9605 | 0.903242 | 2.95522 | 0.087614 |
| 7 | 10077600 | 10078399 | 26.96577 | 46.64926 | 30.92568 | 13.5678 | 0.976275 | 0.863632 | 0.087634 |
| 7 | 122798000 | 122800199 | 10.61158 | 18.59218 | 49.36611 | 21.80151 | -1.01375 | 0.645138 | 0.087659 |
| 18 | 11790000 | 11791399 | 6.530197 | 11.86793 | 40.23949 | 17.35092 | -1.33298 | 0.299734 | 0.087671 |
| 11 | 61374000 | 61382999 | 39.65292 | 62.42953 | 190.2635 | 78.94169 | -1.28328 | 2.497007 | 0.087702 |
| 4 | 14851200 | 14851999 | 11.15046 | 21.68007 | 13.59127 | 5.03955 | 1.267517 | -0.26637 | 0.087712 |
| 13 | 98537800 | 98541399 | 24.44488 | 39.83997 | 93.65946 | 40.22604 | -0.85851 | 1.531192 | 0.087714 |
| 2 | 65701200 | 65703399 | 11.1475 | 19.35643 | 56.92157 | 24.98223 | -1.16189 | 0.791899 | 0.087784 |
| 12 | 22048400 | 22052599 | 106.0889 | 165.9738 | 112.7939 | 46.79254 | 0.871591 | 2.642169 | 0.087798 |
| 13 | 66583000 | 66589599 | 169.082 | 256.6752 | 161.7098 | 66.95765 | 0.947246 | 3.230051 | 0.087812 |
| 7 | 26248400 | 26248799 | 13.32106 | 25.19225 | 16.16424 | 6.561966 | 1.116032 | 0.056213 | 0.087823 |
| 14 | 52995200 | 52997799 | 29.41926 | 50.1647 | 34.55194 | 15.14676 | 0.917858 | 1.003931 | 0.087832 |
| 17 | 84997600 | 85008399 | 45.0409 | 70.68735 | 192.0393 | 79.64556 | -1.12134 | 2.564844 | 0.087837 |
| 1 | 98336400 | 98337399 | 14.08644 | 26.83787 | 16.45428 | 6.575824 | 1.200583 | 0.015463 | 0.087841 |
| X | 26579200 | 26580799 | 16.61637 | 31.9813 | 11.03568 | 4.293686 | 2.053415 | -0.05652 | 0.087842 |
| 4 | 87949400 | 87952599 | 14.22029 | 23.56844 | 82.93025 | 35.90434 | -1.42775 | 1.284349 | 0.087843 |
| 12 | 71877400 | 71878999 | 5.949497 | 10.77771 | 40.16622 | 17.38941 | -1.47133 | 0.316015 | 0.087847 |
| 4 | 135318200 | 135327199 | 58.18357 | 90.75752 | 198.5718 | 82.40261 | -0.81648 | 2.68216 | 0.087848 |
| 10 | 120832800 | 120833199 | 10.89949 | 21.16443 | 14.13279 | 5.295939 | 1.163452 | -0.23237 | 0.087854 |
| 4 | 96099600 | 96101399 | 18.80121 | 33.95363 | 21.13894 | 9.187355 | 1.087501 | 0.381688 | 0.087861 |
| 12 | 4905600 | 4922599 | 91.53576 | 138.6699 | 296.9657 | 122.8845 | -0.81895 | 3.26156 | 0.087875 |
| X | 24128400 | 24131599 | 40.15852 | 67.28837 | 34.96803 | 15.24774 | 1.32181 | 1.19905 | 0.087887 |
| 8 | 9428800 | 9429399 | 11.00827 | 21.51776 | 8.268254 | 2.847825 | 2.052864 | -0.53406 | 0.087892 |
| 1 | 92770600 | 92790199 | 89.88603 | 134.8875 | 347.664 | 143.4671 | -1.09795 | 3.452568 | 0.087898 |
| 11 | 78944000 | 78946599 | 10.30829 | 18.05286 | 49.36549 | 21.79315 | -1.05583 | 0.651303 | 0.087919 |
| 2 | 105293800 | 105294399 | 3.645607 | 7.103004 | 25.39247 | 9.231576 | -1.18846 | -0.3876 | 0.087924 |
| 9 | 108066400 | 108067999 | 7.36441 | 13.45913 | 38.22417 | 16.36698 | -1.07271 | 0.270031 | 0.087929 |
| 4 | 61923800 | 61924799 | 12.46206 | 23.94143 | 14.93161 | 5.84029 | 1.203333 | -0.07412 | 0.087985 |
| 11 | 19457400 | 19458999 | 15.45441 | 28.8434 | 18.54098 | 7.705864 | 1.0865 | 0.166128 | 0.088012 |

| | | | | | | | | | |
|---|---|---|---|---|---|---|---|---|---|
| 3 | 86391200 | 86392999 | 16.42181 | 30.61387 | 17.64709 | 7.34857 | 1.239645 | 0.178336 | 0.088014 |
| 7 | 5472800 | 5476999 | 31.72307 | 53.85337 | 32.52428 | 14.25038 | 1.105924 | 1.050045 | 0.088016 |
| 11 | 69921600 | 69922799 | 3.188182 | 6.064282 | 32.41557 | 12.99952 | -1.89392 | -0.03109 | 0.088024 |
| 6 | 119201200 | 119204999 | 14.65592 | 24.28017 | 80.01524 | 34.63218 | -1.33353 | 1.289484 | 0.088038 |
| 7 | 5941600 | 5944399 | 71.53813 | 114.0248 | 84.03826 | 35.19255 | 0.777306 | 2.175223 | 0.088043 |
| 18 | 61239000 | 61254599 | 90.85537 | 137.3698 | 307.1801 | 127.0327 | -0.88369 | 3.30162 | 0.088049 |
| 15 | 8473200 | 8475599 | 14.24603 | 24.16672 | 64.80136 | 28.38624 | -1.03582 | 1.031427 | 0.088069 |
| 9 | 27036000 | 27039999 | 25.26131 | 41.10934 | 94.72568 | 40.6619 | -0.83122 | 1.555516 | 0.088079 |
| 14 | 51326200 | 51328999 | 12.27414 | 21.1505 | 60.13375 | 26.37889 | -1.11605 | 0.872911 | 0.088082 |
| Y | 41230200 | 41231199 | 8.456972 | 16.54251 | 3.437362 | 1.149575 | 2.937697 | -0.70138 | 0.088094 |
| 8 | 109029200 | 109030999 | 8.99554 | 16.02408 | 43.60537 | 19.19989 | -1.03943 | 0.446534 | 0.088097 |
| 17 | 40055800 | 40059199 | 83.59487 | 132.0887 | 100.2319 | 41.63236 | 0.723684 | 2.4304 | 0.088108 |
| 1 | 111405600 | 111407399 | 22.65877 | 39.93611 | 23.44457 | 10.37644 | 1.152646 | 0.584027 | 0.08811 |
| 8 | 121099800 | 121121999 | 98.36743 | 146.5798 | 391.7036 | 161.4144 | -1.15932 | 3.587393 | 0.088126 |
| X | 154227200 | 154228999 | 19.6153 | 36.41266 | 15.78277 | 6.616366 | 1.640258 | 0.226641 | 0.088135 |
| 4 | 56090600 | 56092799 | 8.02572 | 14.49913 | 43.9213 | 19.08032 | -1.17985 | 0.351279 | 0.088143 |
| 4 | 17295400 | 17296599 | 12.3925 | 23.77234 | 14.41299 | 5.661602 | 1.237836 | -0.06325 | 0.088149 |
| 8 | 23897400 | 23902599 | 30.40991 | 48.90388 | 114.873 | 48.73286 | -0.87692 | 1.863425 | 0.088152 |
| 1 | 36640600 | 36644599 | 15.23543 | 25.68807 | 68.0745 | 29.76496 | -1.0195 | 1.091327 | 0.088153 |
| 4 | 156112400 | 156120999 | 43.8447 | 68.95836 | 186.4164 | 77.3352 | -1.11178 | 2.519779 | 0.088154 |
| 19 | 7317800 | 7319199 | 6.925425 | 12.63934 | 38.18874 | 16.38015 | -1.16271 | 0.275114 | 0.088165 |
| 19 | 6574600 | 6588999 | 66.04154 | 100.6156 | 295.4443 | 122.4636 | -1.26707 | 3.13901 | 0.088183 |
| 17 | 37683000 | 37684399 | 17.62241 | 32.54556 | 18.78175 | 7.928814 | 1.224216 | 0.24494 | 0.088189 |
| 1 | 48739400 | 48740999 | 25.65002 | 45.00796 | 24.20369 | 10.70261 | 1.277701 | 0.661102 | 0.088193 |
| 7 | 11294000 | 11295399 | 51.47572 | 83.17082 | 58.1359 | 24.82146 | 0.875855 | 1.729124 | 0.088197 |
| 7 | 43543000 | 43544599 | 5.431397 | 10.36886 | 31.67726 | 12.60587 | -1.09008 | -0.03883 | 0.088223 |
| 12 | 23559800 | 23561199 | 14.71404 | 28.15257 | 13.97665 | 5.531265 | 1.513169 | -0.01629 | 0.088234 |
| 10 | 8472200 | 8474999 | 36.60814 | 61.9245 | 33.27054 | 14.56156 | 1.273949 | 1.100665 | 0.088242 |
| 6 | 34287800 | 34291399 | 16.6194 | 27.63458 | 75.46838 | 32.74734 | -1.06317 | 1.247755 | 0.088269 |
| 11 | 43394800 | 43396199 | 6.169807 | 11.37841 | 37.87231 | 16.0189 | -1.28676 | 0.20634 | 0.08829 |
| 12 | 23087600 | 23089399 | 47.10645 | 76.47716 | 55.48334 | 23.78542 | 0.828887 | 1.615176 | 0.088296 |
| 8 | 8429800 | 8431399 | 8.634395 | 15.5519 | 43.83012 | 19.11247 | -1.08006 | 0.373649 | 0.088297 |
| 6 | 134399800 | 134401399 | 8.871408 | 15.70949 | 44.07607 | 19.48953 | -1.08769 | 0.491771 | 0.088308 |
| 13 | 68145800 | 68149199 | 59.76883 | 96.08132 | 67.4291 | 28.58996 | 0.861536 | 1.895175 | 0.088317 |
| 10 | 121851400 | 121862199 | 82.77683 | 126.0722 | 272.8219 | 113.0758 | -0.82761 | 3.149055 | 0.088323 |
| 1 | 181655400 | 181658799 | 15.04258 | 25.45598 | 65.92492 | 28.85711 | -0.98613 | 1.056576 | 0.088325 |
| 17 | 11129800 | 11131799 | 8.156215 | 14.76676 | 47.35856 | 20.51799 | -1.25922 | 0.346577 | 0.088327 |
| 3 | 100069400 | 100070199 | 4.605164 | 8.72228 | 32.43129 | 13.12362 | -1.39106 | 0.011793 | 0.088336 |
| 2 | 128540000 | 128541599 | 5.996747 | 11.09448 | 36.29496 | 15.28543 | -1.25694 | 0.184481 | 0.088352 |
| 8 | 121042000 | 121076999 | 180.6155 | 256.5416 | 635.7804 | 262.9615 | -1.11791 | 4.344367 | 0.088365 |
| 8 | 121841600 | 121849999 | 41.74263 | 66.00972 | 170.4785 | 70.83825 | -1.03975 | 2.396938 | 0.088368 |
| 13 | 30920400 | 30920799 | 1.573476 | 3.077851 | 20.73479 | 6.933016 | -1.96308 | -0.73265 | 0.088375 |
| 6 | 38637200 | 38641399 | 21.98509 | 35.69553 | 98.30466 | 42.14514 | -1.09034 | 1.590827 | 0.088398 |
| 19 | 44733800 | 44756199 | 109.8487 | 163.3173 | 398.5621 | 164.2028 | -1.03153 | 3.629051 | 0.088398 |
| 10 | 80267400 | 80298799 | 171.5173 | 246.1964 | 564.6919 | 233.1586 | -0.99101 | 4.189103 | 0.088402 |
| 9 | 20940600 | 20953799 | 82.41692 | 125.3838 | 281.4521 | 116.6136 | -0.88172 | 3.172027 | 0.088405 |
| 17 | 89415600 | 89416199 | 8.948836 | 17.48017 | 10.90016 | 3.813544 | 1.347145 | -0.50469 | 0.088426 |

| | | | | | | | | | |
|---|---|---|---|---|---|---|---|---|---|
| 4 | 125594200 | 125608199 | 83.73088 | 126.9062 | 295.7828 | 122.4134 | -0.94036 | 3.250433 | 0.088436 |
| 10 | 59121600 | 59132599 | 63.59041 | 97.66495 | 251.6296 | 104.4767 | -1.06942 | 2.972544 | 0.088442 |
| 1 | 117328400 | 117330399 | 26.46569 | 46.07618 | 26.50074 | 11.65017 | 1.18111 | 0.782533 | 0.088453 |
| 1 | 74640000 | 74640999 | 4.443778 | 8.526558 | 31.26436 | 12.27225 | -1.33163 | -0.09637 | 0.088454 |
| 15 | 91556800 | 91557999 | 4.443778 | 8.526558 | 31.26436 | 12.27225 | -1.33163 | -0.09637 | 0.088454 |
| 3 | 101511000 | 101512599 | 7.636077 | 13.60864 | 42.58171 | 18.74194 | -1.23827 | 0.434773 | 0.088478 |
| 10 | 127154800 | 127179399 | 128.5396 | 188.63 | 457.7798 | 188.5468 | -1.04119 | 3.850491 | 0.088482 |
| 3 | 12317400 | 12318999 | 19.4164 | 34.81661 | 22.27987 | 9.755514 | 1.04149 | 0.430682 | 0.088496 |
| 7 | 63802800 | 63812199 | 51.42881 | 80.87631 | 184.136 | 76.38788 | -0.86463 | 2.526025 | 0.088509 |
| 14 | 117269400 | 117269999 | 3.343507 | 6.528834 | 24.10646 | 8.482852 | -1.19073 | -0.51316 | 0.088528 |
| 4 | 106462000 | 106468999 | 53.6127 | 84.09597 | 185.2664 | 76.83704 | -0.82005 | 2.578835 | 0.08854 |
| 5 | 120695200 | 120699399 | 14.48664 | 24.05484 | 79.05413 | 34.27041 | -1.32928 | 1.260602 | 0.088541 |
| 2 | 82164600 | 82165799 | 15.04211 | 29.0786 | 11.1461 | 4.269765 | 1.923737 | -0.12187 | 0.088564 |
| 9 | 109912800 | 109917599 | 18.06058 | 29.56028 | 88.0247 | 37.86456 | -1.1936 | 1.454199 | 0.088565 |
| 14 | 42875000 | 42876599 | 23.54609 | 41.12542 | 28.1347 | 12.39354 | 0.932444 | 0.72562 | 0.088567 |
| 7 | 29747800 | 29752199 | 17.20577 | 28.72278 | 73.55774 | 31.99978 | -0.9708 | 1.202958 | 0.08857 |
| 7 | 52489400 | 52490799 | 15.30364 | 29.31226 | 14.3838 | 5.674798 | 1.535812 | -0.01744 | 0.088577 |
| 8 | 101999000 | 101999599 | 15.34084 | 28.4418 | 19.02198 | 7.988032 | 1.018212 | 0.204816 | 0.088583 |
| 7 | 11793800 | 11795799 | 44.88819 | 73.39522 | 50.05929 | 21.52545 | 0.925109 | 1.499943 | 0.088587 |
| 14 | 72946200 | 72948199 | 10.22431 | 17.77739 | 54.70079 | 24.02793 | -1.22624 | 0.766409 | 0.088606 |
| 1 | 112363600 | 112373799 | 118.4442 | 183.4492 | 130.0079 | 54.00177 | 0.800034 | 2.82172 | 0.088609 |
| 2 | 4510800 | 4511999 | 6.072441 | 11.41937 | 34.00417 | 13.9527 | -1.09275 | 0.076618 | 0.08861 |
| 7 | 5432200 | 5444999 | 274.309 | 399.5384 | 281.4172 | 115.9772 | 0.732201 | 3.97535 | 0.08862 |
| 2 | 28340200 | 28343999 | 18.21822 | 29.84392 | 87.63546 | 37.71061 | -1.173 | 1.442666 | 0.088626 |
| 2 | 30603400 | 30610199 | 27.03346 | 43.51398 | 112.3996 | 47.7471 | -1.01186 | 1.829421 | 0.088627 |
| 7 | 9528000 | 9529199 | 18.32001 | 32.86661 | 21.01172 | 9.196463 | 1.043174 | 0.416734 | 0.088661 |
| X | 147358000 | 147358599 | 9.640928 | 18.81414 | 12.08981 | 4.299795 | 1.285174 | -0.43332 | 0.088668 |
| 1 | 102075200 | 102075799 | 10.84354 | 20.99362 | 13.29675 | 5.058046 | 1.219286 | -0.18633 | 0.088698 |
| 5 | 120857000 | 120862999 | 54.1741 | 85.2305 | 177.9192 | 73.81305 | -0.73912 | 2.518873 | 0.088709 |
| 8 | 73072400 | 73077399 | 24.87774 | 40.25244 | 100.9902 | 43.18605 | -0.96109 | 1.668859 | 0.08874 |
| 19 | 6939400 | 6954599 | 58.5718 | 89.92301 | 258.6033 | 107.3666 | -1.22822 | 2.979014 | 0.088743 |
| 12 | 23607400 | 23610799 | 42.6454 | 69.76036 | 49.98017 | 21.49505 | 0.854818 | 1.489203 | 0.088759 |
| 5 | 146519800 | 146523799 | 25.36533 | 41.01104 | 102.0503 | 43.60877 | -0.95037 | 1.687975 | 0.088766 |
| 4 | 139724200 | 139734599 | 72.86971 | 111.2854 | 270.0723 | 112.0118 | -0.98965 | 3.090358 | 0.088768 |
| 4 | 56738800 | 56741999 | 27.01707 | 43.54608 | 109.608 | 46.65101 | -0.97221 | 1.785198 | 0.088777 |
| 13 | 107769400 | 107770199 | 14.78315 | 28.00526 | 16.94243 | 6.852986 | 1.204881 | 0.061556 | 0.088805 |
| 16 | 61530200 | 61532399 | 16.5656 | 31.54827 | 13.88537 | 5.554004 | 1.672375 | 0.035586 | 0.088824 |
| 12 | 19500600 | 19508399 | 136.6756 | 209.5415 | 137.6903 | 57.15523 | 0.897824 | 3.020743 | 0.088828 |
| 10 | 79235800 | 79240999 | 72.51634 | 116.0428 | 69.67399 | 29.34049 | 1.078521 | 2.051657 | 0.088833 |
| 5 | 31960800 | 31961999 | 16.76226 | 31.38787 | 16.51196 | 6.824274 | 1.380007 | 0.152787 | 0.088835 |
| 6 | 124113200 | 124116399 | 27.96986 | 48.74462 | 26.06952 | 11.46795 | 1.285409 | 0.775376 | 0.088848 |
| 8 | 73677600 | 73678799 | 17.76638 | 32.27161 | 20.93762 | 9.034023 | 1.035298 | 0.342008 | 0.088867 |
| 13 | 62040800 | 62042999 | 47.42285 | 76.89208 | 53.21258 | 22.79038 | 0.895109 | 1.640104 | 0.088875 |
| X | 3290000 | 3290799 | 12.74835 | 24.75628 | 12.77655 | 4.784979 | 1.531581 | -0.21592 | 0.088884 |
| 1 | 97107600 | 97109399 | 28.27167 | 48.75699 | 31.8493 | 13.97169 | 0.996608 | 0.897552 | 0.088907 |
| 6 | 114453600 | 114459199 | 22.35034 | 36.14842 | 102.558 | 43.84336 | -1.13839 | 1.674109 | 0.088947 |
| 15 | 95740400 | 95741799 | 5.122639 | 9.821762 | 30.6947 | 12.07462 | -1.10648 | -0.08218 | 0.08895 |

| | | | | | | | | | |
|---|---|---|---|---|---|---|---|---|---|
| 1 | 190634800 | 190635599 | 10.7109 | 18.48679 | 57.28646 | 25.13112 | -1.2384 | 0.850963 | 0.088972 |
| 11 | 120154400 | 120191399 | 176.824 | 250.0766 | 671.1764 | 277.8005 | -1.2391 | 4.407794 | 0.088972 |
| 16 | 35874400 | 35877199 | 14.80094 | 25.29069 | 62.30159 | 27.31615 | -0.91266 | 0.96377 | 0.088995 |
| 11 | 100630400 | 100632799 | 6.808376 | 12.41681 | 40.8833 | 17.55164 | -1.2873 | 0.281328 | 0.089008 |
| 9 | 107466800 | 107471799 | 19.07362 | 31.02282 | 94.18127 | 40.42019 | -1.22796 | 1.5529 | 0.089009 |
| 14 | 41417200 | 41417999 | 15.96695 | 30.92639 | 10.45808 | 3.970053 | 2.116549 | -0.14213 | 0.089019 |
| 11 | 59916000 | 59919599 | 14.83098 | 24.77547 | 72.57645 | 31.58664 | -1.16352 | 1.19007 | 0.089046 |
| 4 | 128838800 | 128844599 | 26.05556 | 41.97626 | 110.1087 | 46.83267 | -1.03238 | 1.799864 | 0.089052 |
| 5 | 31651800 | 31652399 | 2.638596 | 5.156865 | 21.70272 | 7.503932 | -1.34995 | -0.59271 | 0.089066 |
| 15 | 102573400 | 102574399 | 2.638596 | 5.156865 | 21.70272 | 7.503932 | -1.34995 | -0.59271 | 0.089066 |
| 12 | 70147800 | 70151599 | 14.01497 | 23.9837 | 62.2979 | 27.31784 | -0.98842 | 0.94623 | 0.089067 |
| 14 | 61090600 | 61092999 | 12.99792 | 22.49883 | 56.408 | 24.74792 | -0.93398 | 0.831677 | 0.089092 |
| 7 | 62073000 | 62074399 | 14.41205 | 27.73067 | 14.35872 | 5.59123 | 1.475981 | -0.06748 | 0.089108 |
| 10 | 88489400 | 88489999 | 13.19808 | 25.32966 | 15.40876 | 6.044547 | 1.235384 | -0.05524 | 0.089123 |
| 9 | 4387000 | 4388199 | 26.6898 | 47.15554 | 18.78952 | 8.312095 | 1.708931 | 0.576287 | 0.089125 |
| 12 | 16594400 | 16596399 | 8.485976 | 15.27404 | 41.11488 | 17.94394 | -1.01459 | 0.373228 | 0.089162 |
| X | 127440800 | 127442199 | 18.25999 | 34.62042 | 13.33107 | 5.382812 | 1.852114 | 0.079744 | 0.089178 |
| 9 | 4952800 | 4954199 | 15.29143 | 28.81452 | 17.43387 | 7.12889 | 1.191464 | 0.102609 | 0.089189 |
| 10 | 80011800 | 80033199 | 88.65625 | 132.4387 | 378.5765 | 156.076 | -1.25311 | 3.53936 | 0.089193 |
| 7 | 38493400 | 38499199 | 115.1043 | 179.1926 | 117.8678 | 48.92638 | 0.913137 | 2.731472 | 0.089195 |
| 19 | 58782800 | 58783999 | 6.086784 | 11.33783 | 35.35591 | 14.74325 | -1.17715 | 0.14425 | 0.089209 |
| 3 | 65947600 | 65949599 | 8.299385 | 15.05268 | 42.96254 | 18.57212 | -1.08985 | 0.328893 | 0.089224 |
| 13 | 54483000 | 54487599 | 28.91041 | 46.63171 | 107.2627 | 45.6957 | -0.84135 | 1.76658 | 0.08923 |
| 18 | 65756000 | 65758599 | 18.2466 | 30.03251 | 84.48322 | 36.42717 | -1.10833 | 1.383318 | 0.089256 |
| 3 | 3102000 | 3103399 | 32.59102 | 55.63718 | 32.12261 | 14.08311 | 1.171482 | 1.003953 | 0.089261 |
| 12 | 21857600 | 21863599 | 112.204 | 173.7719 | 129.8 | 53.91545 | 0.724163 | 2.820754 | 0.089263 |
| 18 | 11046200 | 11056399 | 69.67579 | 107.2341 | 238.1244 | 98.89755 | -0.85295 | 2.932439 | 0.089287 |
| 17 | 30082400 | 30115599 | 279.3602 | 385.7544 | 826.2615 | 343.2864 | -0.94869 | 4.770911 | 0.089293 |
| 2 | 28094200 | 28102999 | 40.80182 | 64.4643 | 172.2279 | 71.53284 | -1.08959 | 2.41838 | 0.089295 |
| 8 | 122875600 | 122884999 | 45.28822 | 71.21834 | 183.4379 | 76.09811 | -1.04229 | 2.523376 | 0.0893 |
| 2 | 27589200 | 27594799 | 22.50863 | 36.39042 | 103.0037 | 44.01325 | -1.13566 | 1.685748 | 0.089302 |
| 16 | 18422600 | 18455799 | 227.8665 | 318.4969 | 743.6749 | 308.3863 | -1.05539 | 4.584507 | 0.089334 |
| 17 | 26042600 | 26065999 | 90.95983 | 135.4803 | 390.6494 | 160.9726 | -1.26952 | 3.594655 | 0.089345 |
| 14 | 72294200 | 72295399 | 7.953569 | 14.18689 | 44.15677 | 19.42083 | -1.2306 | 0.431753 | 0.089346 |
| 6 | 48148400 | 48149999 | 6.694191 | 12.48189 | 36.5798 | 15.22745 | -1.08756 | 0.142053 | 0.089376 |
| 6 | 146370000 | 146371399 | 6.694191 | 12.48189 | 36.5798 | 15.22745 | -1.08756 | 0.142053 | 0.089376 |
| 6 | 55435400 | 55443799 | 44.97554 | 71.06839 | 167.536 | 69.58898 | -0.90921 | 2.417224 | 0.089398 |
| 3 | 127346600 | 127347599 | 5.178998 | 9.844033 | 32.64215 | 13.1101 | -1.21913 | -0.0093 | 0.089403 |
| 7 | 10843200 | 10847599 | 95.2724 | 149.5838 | 111.7284 | 46.34028 | 0.739396 | 2.571319 | 0.089407 |
| 6 | 145241200 | 145243999 | 10.34822 | 18.06928 | 54.76014 | 24.11818 | -1.204 | 0.704004 | 0.08943 |
| 7 | 46061400 | 46093399 | 216.3464 | 303.5182 | 711.5119 | 294.8788 | -1.05571 | 4.528676 | 0.089436 |
| 5 | 35418400 | 35419599 | 5.646401 | 10.60211 | 33.92364 | 13.95543 | -1.19816 | 0.082846 | 0.089447 |
| 5 | 42763000 | 42764199 | 16.7073 | 31.51776 | 14.81424 | 6.045614 | 1.553499 | 0.101092 | 0.089454 |
| 5 | 134245000 | 134246599 | 18.30879 | 30.10753 | 83.40204 | 35.94719 | -1.08685 | 1.392112 | 0.089456 |
| 15 | 95126000 | 95127199 | 6.544735 | 12.14945 | 36.38615 | 15.25327 | -1.12572 | 0.171248 | 0.08947 |
| 9 | 116886000 | 116886399 | 2.027376 | 3.965302 | 20.05613 | 6.766984 | -1.57229 | -0.69284 | 0.08947 |
| X | 33969000 | 33973399 | 73.81152 | 118.0508 | 64.658 | 27.2071 | 1.209948 | 2.069435 | 0.089493 |

| | | | | | | | | | |
|---|---|---|---|---|---|---|---|---|---|
| 7 | 9534400 | 9534799 | 10.17253 | 19.86553 | 10.87639 | 3.821562 | 1.52821 | -0.47071 | 0.089516 |
| 12 | 90818000 | 90818999 | 14.11464 | 26.52704 | 17.49284 | 7.184988 | 1.06281 | 0.107117 | 0.089516 |
| 9 | 63635000 | 63636999 | 7.913631 | 14.17742 | 41.58483 | 18.22244 | -1.14144 | 0.400164 | 0.089536 |
| 8 | 115543000 | 115544799 | 6.28459 | 11.64876 | 36.35732 | 15.27282 | -1.18737 | 0.175401 | 0.089547 |
| 1 | 192475200 | 192485399 | 59.76201 | 93.37625 | 200.535 | 83.20294 | -0.78782 | 2.652402 | 0.089563 |
| 6 | 144342200 | 144344199 | 8.8477 | 15.76164 | 44.13574 | 19.43247 | -1.08044 | 0.445658 | 0.089563 |
| 11 | 99276800 | 99277599 | 3.459794 | 6.744337 | 24.83182 | 8.968902 | -1.22053 | -0.41345 | 0.089565 |
| 16 | 90625200 | 90629399 | 20.3102 | 33.17382 | 90.67128 | 38.97371 | -1.07276 | 1.489482 | 0.089579 |
| 3 | 89388400 | 89390799 | 12.86685 | 22.20695 | 58.55528 | 25.68771 | -1.0074 | 0.859292 | 0.08958 |
| X | 55008000 | 55008999 | 21.39172 | 38.34298 | 20.69337 | 9.06397 | 1.285488 | 0.444598 | 0.089584 |
| 12 | 22969400 | 22977399 | 197.4516 | 295.0543 | 212.5504 | 87.63365 | 0.734319 | 3.542863 | 0.089589 |
| 7 | 130247400 | 130248999 | 5.840531 | 10.53406 | 41.67084 | 18.13995 | -1.56203 | 0.343907 | 0.08959 |
| 3 | 52800000 | 52800599 | 2.334922 | 4.566168 | 20.6247 | 7.019173 | -1.42677 | -0.65645 | 0.089592 |
| 13 | 53812600 | 53814199 | 6.23183 | 11.7013 | 37.44257 | 15.40315 | -1.19955 | 0.09361 | 0.089606 |
| 1 | 32042600 | 32043799 | 11.44879 | 22.26048 | 12.20698 | 4.52567 | 1.456819 | -0.26868 | 0.089607 |
| 3 | 148914200 | 148917799 | 14.82422 | 25.26384 | 62.75242 | 27.50798 | -0.92507 | 0.98983 | 0.08961 |
| 7 | 17491600 | 17494799 | 74.26198 | 118.6938 | 72.37706 | 30.42703 | 1.054423 | 2.091101 | 0.089616 |
| 3 | 97202400 | 97205799 | 16.08764 | 26.94073 | 73.01892 | 31.82842 | -1.05226 | 1.164382 | 0.089621 |
| 5 | 25257000 | 25267999 | 51.52716 | 80.13748 | 219.157 | 90.98444 | -1.14123 | 2.731774 | 0.089628 |
| 2 | 98238000 | 98238799 | 12.04208 | 23.22405 | 15.70879 | 6.076555 | 1.103045 | -0.10986 | 0.089638 |
| 2 | 162992800 | 162997399 | 27.30843 | 44.08421 | 106.8759 | 45.57955 | -0.91561 | 1.738103 | 0.089659 |
| 11 | 116010800 | 116012999 | 7.311308 | 13.5339 | 40.3266 | 16.97075 | -1.12388 | 0.198869 | 0.089683 |
| 4 | 146377200 | 146379199 | 26.67253 | 46.75241 | 23.91864 | 10.56707 | 1.349083 | 0.681877 | 0.089686 |
| 17 | 86254200 | 86255199 | 5.539026 | 10.32922 | 35.37933 | 14.72472 | -1.30926 | 0.130015 | 0.089699 |
| 13 | 108024800 | 108028799 | 17.63805 | 28.6214 | 105.4481 | 45.18884 | -1.50944 | 1.598094 | 0.089707 |
| 8 | 126732800 | 126735999 | 14.09644 | 23.92712 | 67.07378 | 29.38694 | -1.09977 | 1.026448 | 0.089708 |
| 7 | 8075400 | 8076199 | 13.21442 | 25.75913 | 9.20045 | 3.32368 | 2.098681 | -0.34193 | 0.08974 |
| 10 | 42866800 | 42877399 | 62.5209 | 96.86616 | 217.2331 | 90.23262 | -0.8597 | 2.80307 | 0.089744 |
| 2 | 177533800 | 177534999 | 7.718808 | 13.97019 | 40.70346 | 17.64203 | -1.12141 | 0.333323 | 0.089748 |
| 2 | 130125200 | 130126399 | 4.254371 | 8.221918 | 28.70887 | 11.01263 | -1.22927 | -0.18121 | 0.089749 |
| 18 | 32538800 | 32550599 | 63.94763 | 98.82971 | 227.3118 | 94.42024 | -0.89893 | 2.85057 | 0.089775 |
| 11 | 78905000 | 78925799 | 89.09444 | 133.5237 | 360.7314 | 148.8124 | -1.1675 | 3.478161 | 0.089776 |
| 19 | 56698000 | 56699399 | 6.232771 | 11.70035 | 38.02155 | 15.64748 | -1.22217 | 0.096254 | 0.089785 |
| 15 | 96326600 | 96327199 | 8.454809 | 16.51108 | 10.52991 | 3.699772 | 1.309995 | -0.49988 | 0.089789 |
| 1 | 117633400 | 117636999 | 82.4519 | 130.696 | 82.87871 | 34.48555 | 0.985885 | 2.349695 | 0.089791 |
| 12 | 55608200 | 55613399 | 42.20987 | 67.19879 | 144.4381 | 60.38705 | -0.76607 | 2.203128 | 0.089791 |
| 4 | 118331000 | 118334599 | 17.74454 | 29.45122 | 77.89175 | 33.75599 | -1.01752 | 1.273101 | 0.089793 |
| 17 | 30962600 | 30968799 | 44.29236 | 70.07751 | 163.1307 | 67.80257 | -0.88956 | 2.385946 | 0.089794 |
| 2 | 103799600 | 103801799 | 9.198037 | 15.98675 | 57.34643 | 25.1839 | -1.4463 | 0.770104 | 0.089795 |
| 17 | 81087000 | 81090799 | 11.43901 | 19.88851 | 59.18603 | 25.99736 | -1.17902 | 0.776136 | 0.089802 |
| 16 | 94000800 | 94002999 | 9.836968 | 17.24162 | 48.82024 | 21.56743 | -1.10556 | 0.635531 | 0.089805 |
| 2 | 10084600 | 10085999 | 6.034801 | 11.05664 | 37.70986 | 16.09324 | -1.33048 | 0.244668 | 0.08981 |
| 7 | 7404400 | 7406399 | 30.00621 | 51.58933 | 31.67558 | 13.89418 | 1.084631 | 0.930989 | 0.089812 |
| 4 | 101272800 | 101277799 | 24.48948 | 39.80006 | 95.56018 | 40.99766 | -0.89181 | 1.574378 | 0.089839 |
| 13 | 4448800 | 4449199 | 18.32709 | 33.26752 | 20.46841 | 8.839075 | 1.11068 | 0.349455 | 0.089854 |
| 17 | 5141400 | 5148799 | 120.0447 | 185.3758 | 115.5486 | 47.99649 | 0.981704 | 2.874861 | 0.089855 |
| 2 | 62241400 | 62245199 | 19.31663 | 32.03789 | 81.07909 | 35.1202 | -0.95444 | 1.287028 | 0.089865 |

| | | | | | | | | | |
|---|---|---|---|---|---|---|---|---|---|
| 11 | 3444400 | 3445999 | 9.354241 | 16.47478 | 46.15982 | 20.43309 | -1.08905 | 0.559318 | 0.089883 |
| 6 | 83590000 | 83592599 | 12.8714 | 22.19843 | 57.42528 | 25.19142 | -0.98005 | 0.865921 | 0.089895 |
| 7 | 132212200 | 132213599 | 4.864932 | 9.313969 | 31.18707 | 12.32013 | -1.21097 | -0.07076 | 0.089898 |
| 14 | 52341000 | 52342199 | 4.864932 | 9.313969 | 31.18707 | 12.32013 | -1.21097 | -0.07076 | 0.089898 |
| 1 | 151985800 | 151989399 | 14.61215 | 25.09495 | 61.84162 | 27.12531 | -0.91195 | 0.916575 | 0.08991 |
| 7 | 20323800 | 20324599 | 13.281 | 25.48968 | 15.22545 | 5.972035 | 1.262861 | -0.05378 | 0.089913 |
| 12 | 24397600 | 24399399 | 48.20181 | 78.14148 | 56.63287 | 24.25214 | 0.82842 | 1.645981 | 0.089914 |
| 11 | 120236400 | 120261599 | 117.6132 | 172.5382 | 461.9828 | 190.2818 | -1.18337 | 3.85494 | 0.089917 |
| 12 | 114554000 | 114554999 | 27.18646 | 47.11254 | 30.68696 | 13.46378 | 1.001926 | 0.848141 | 0.089921 |
| 5 | 84116600 | 84117599 | 14.80382 | 27.98387 | 16.80165 | 6.827179 | 1.209583 | 0.075042 | 0.089923 |
| 14 | 76060000 | 76062599 | 16.14084 | 27.28663 | 66.55422 | 29.12355 | -0.90018 | 1.070048 | 0.089925 |
| 15 | 82957600 | 82959399 | 7.735353 | 13.91495 | 40.58448 | 17.72274 | -1.12959 | 0.371498 | 0.089929 |
| 8 | 95735800 | 95742199 | 26.80458 | 43.00674 | 123.9725 | 52.41353 | -1.1756 | 1.935184 | 0.089932 |
| 5 | 120657600 | 120686799 | 142.7953 | 206.5496 | 528.8294 | 218.1139 | -1.13855 | 4.073449 | 0.08997 |
| 3 | 131479600 | 131481399 | 8.30057 | 15.05125 | 43.50916 | 18.81401 | -1.10843 | 0.331223 | 0.089973 |
| 1 | 99939000 | 99940199 | 13.06472 | 25.21964 | 14.3181 | 5.521943 | 1.356314 | -0.11294 | 0.089977 |
| 10 | 105650000 | 105651999 | 20.88846 | 37.31474 | 22.04683 | 9.684327 | 1.152753 | 0.463121 | 0.089985 |
| 12 | 87940000 | 87941599 | 16.41473 | 30.59228 | 18.33106 | 7.636928 | 1.184159 | 0.182418 | 0.089987 |
| 9 | 124100800 | 124101999 | 10.44679 | 20.27342 | 14.30073 | 5.38012 | 1.080437 | -0.22362 | 0.089992 |
| 12 | 110362400 | 110362999 | 10.44679 | 20.27342 | 14.30073 | 5.38012 | 1.080437 | -0.22362 | 0.089992 |
| 12 | 110949200 | 110953399 | 17.70162 | 29.11628 | 84.55554 | 36.44817 | -1.15461 | 1.387401 | 0.090001 |
| 8 | 80021000 | 80022199 | 8.206485 | 14.69418 | 41.57868 | 18.22801 | -1.09036 | 0.406395 | 0.09002 |
| 11 | 116533800 | 116556399 | 88.63709 | 132.5819 | 366.5237 | 151.1451 | -1.20319 | 3.514763 | 0.090034 |
| 11 | 115666600 | 115671799 | 17.25245 | 28.34075 | 86.05322 | 37.07345 | -1.21939 | 1.40384 | 0.09005 |
| 16 | 30783400 | 30792599 | 53.93961 | 84.21022 | 199.0054 | 82.57523 | -0.92651 | 2.664461 | 0.090059 |
| 2 | 166070400 | 166094399 | 150.1923 | 218.6859 | 482.6827 | 198.9293 | -0.91528 | 3.974186 | 0.090074 |
| 7 | 41926000 | 41929399 | 51.89197 | 84.92977 | 41.21942 | 17.73063 | 1.415292 | 1.49769 | 0.090082 |
| 12 | 19982800 | 19985999 | 71.85204 | 114.3764 | 83.91675 | 35.07853 | 0.781844 | 2.216718 | 0.090083 |
| 1 | 28577200 | 28578399 | 13.32081 | 25.46907 | 15.92224 | 6.3139 | 1.182042 | -0.0187 | 0.09009 |
| 6 | 138247600 | 138248799 | 11.82268 | 22.73355 | 14.57157 | 5.684755 | 1.167816 | -0.09029 | 0.090096 |
| 2 | 155667800 | 155669799 | 8.763653 | 15.45591 | 46.68418 | 20.65951 | -1.19609 | 0.539493 | 0.090103 |
| 9 | 90225000 | 90227199 | 32.48445 | 56.27239 | 24.83227 | 10.89493 | 1.560309 | 0.868278 | 0.090106 |
| 4 | 92885200 | 92885999 | 11.80893 | 22.92202 | 14.1126 | 5.303753 | 1.276148 | -0.21085 | 0.090106 |
| X | 168662000 | 168665199 | 9.632758 | 17.04257 | 53.76633 | 23.77858 | -1.25733 | 0.520976 | 0.090111 |
| 4 | 154630400 | 154642799 | 59.17658 | 91.29605 | 241.8651 | 100.4663 | -1.10455 | 2.88244 | 0.090119 |
| 4 | 124978400 | 124979799 | 6.114754 | 11.36796 | 35.3162 | 14.77493 | -1.17593 | 0.155612 | 0.090131 |
| 11 | 40623800 | 40624999 | 6.114754 | 11.36796 | 35.3162 | 14.77493 | -1.17593 | 0.155612 | 0.090131 |
| 15 | 77815000 | 77816599 | 4.962807 | 9.498335 | 31.13656 | 12.3114 | -1.18113 | -0.06408 | 0.090135 |
| 2 | 169662600 | 169665399 | 19.68576 | 32.49711 | 80.67136 | 34.83045 | -0.92722 | 1.3441 | 0.090138 |
| 16 | 94582600 | 94584799 | 22.70502 | 36.95101 | 92.19873 | 39.57785 | -0.94549 | 1.549803 | 0.090152 |
| X | 54996000 | 54997399 | 21.66489 | 38.96952 | 19.93586 | 8.706302 | 1.364721 | 0.423356 | 0.090169 |
| 6 | 126825400 | 126826599 | 4.147677 | 7.977393 | 30.16897 | 11.76156 | -1.36558 | -0.12425 | 0.09018 |
| 5 | 146594000 | 146602399 | 50.92714 | 80.12801 | 183.1055 | 75.96541 | -0.86937 | 2.515338 | 0.090184 |
| 14 | 111219400 | 111220399 | 10.93751 | 21.30883 | 12.58109 | 4.57825 | 1.377492 | -0.33548 | 0.090203 |
| 6 | 120768800 | 120771199 | 8.331427 | 14.66899 | 47.72686 | 21.12802 | -1.30352 | 0.557707 | 0.090204 |
| 14 | 73910600 | 73912199 | 5.871102 | 11.19068 | 34.03234 | 13.58818 | -1.08857 | -0.01864 | 0.090206 |
| 8 | 38221800 | 38223599 | 17.52146 | 32.84713 | 15.94133 | 6.575666 | 1.496376 | 0.150007 | 0.090209 |

| | | | | | | | | | |
|---|---|---|---|---|---|---|---|---|---|
| 1 | 190978200 | 190979599 | 21.81379 | 38.28555 | 24.94225 | 11.03138 | 1.00215 | 0.633964 | 0.090216 |
| 11 | 53352400 | 53358799 | 31.59067 | 50.30773 | 153.656 | 64.26718 | -1.2714 | 2.185774 | 0.090243 |
| 14 | 15995200 | 15998599 | 27.90205 | 47.80625 | 33.00333 | 14.47348 | 0.915465 | 0.95307 | 0.090253 |
| X | 32042000 | 32042999 | 14.5974 | 28.50102 | 5.560752 | 1.960565 | 2.976319 | -0.41577 | 0.090259 |
| 9 | 116019600 | 116023399 | 15.33154 | 26.00505 | 65.47749 | 28.68137 | -0.94561 | 1.03619 | 0.090264 |
| 9 | 110656000 | 110658599 | 9.52636 | 16.69946 | 48.81441 | 21.56731 | -1.15106 | 0.632051 | 0.090274 |
| 6 | 30047600 | 30049199 | 7.879479 | 13.90317 | 47.73393 | 21.12205 | -1.37922 | 0.530458 | 0.090286 |
| 18 | 25366800 | 25368199 | 13.31205 | 25.44746 | 16.60713 | 6.589093 | 1.120532 | -0.01429 | 0.09029 |
| 19 | 6821000 | 6834999 | 71.70354 | 109.1368 | 283.553 | 117.5008 | -1.09184 | 3.160016 | 0.090304 |
| 8 | 126604000 | 126607999 | 17.6769 | 29.50087 | 73.60973 | 32.01664 | -0.93343 | 1.207449 | 0.090317 |
| X | 61147800 | 61151999 | 60.04891 | 96.96034 | 54.09364 | 23.07643 | 1.199276 | 1.76147 | 0.090318 |
| 2 | 89526400 | 89529999 | 60.12456 | 96.84262 | 61.45586 | 26.13136 | 1.00981 | 1.833486 | 0.090322 |
| 4 | 67236000 | 67238199 | 21.29613 | 38.03233 | 21.36695 | 9.387902 | 1.224518 | 0.465665 | 0.090329 |
| 13 | 7263600 | 7265399 | 19.39022 | 34.99575 | 20.45307 | 8.896123 | 1.177181 | 0.387551 | 0.090344 |
| 7 | 20672800 | 20678799 | 86.92105 | 137.5551 | 97.07708 | 40.35328 | 0.829913 | 2.396818 | 0.090361 |
| 11 | 98685400 | 98687799 | 7.453839 | 13.05561 | 50.35844 | 22.234 | -1.54789 | 0.636556 | 0.09039 |
| 13 | 99062000 | 99063399 | 6.220685 | 11.33906 | 39.22204 | 16.84949 | -1.35767 | 0.278295 | 0.090392 |
| 5 | 147245400 | 147256199 | 68.4004 | 105.2689 | 243.2887 | 101.0421 | -0.91055 | 2.932512 | 0.090399 |
| 11 | 115155000 | 115164999 | 54.48181 | 84.20732 | 230.1877 | 95.61478 | -1.14759 | 2.846245 | 0.090421 |
| 1 | 90555200 | 90558599 | 15.22666 | 25.69622 | 67.54559 | 29.5417 | -1.00778 | 1.08285 | 0.090433 |
| 16 | 4866800 | 4868999 | 10.98277 | 19.08858 | 54.73904 | 24.03873 | -1.12541 | 0.77539 | 0.090453 |
| 19 | 5088600 | 5090199 | 6.78364 | 12.38432 | 38.19351 | 16.37599 | -1.19173 | 0.271788 | 0.090459 |
| 7 | 38512200 | 38515199 | 55.1057 | 89.03287 | 57.45926 | 24.5315 | 0.990595 | 1.734001 | 0.090494 |
| 3 | 7065200 | 7065799 | 10.91913 | 21.13195 | 14.8127 | 5.64472 | 1.072277 | -0.1742 | 0.090505 |
| 15 | 79003600 | 79012999 | 44.80157 | 70.1213 | 198.4109 | 82.30712 | -1.183 | 2.614726 | 0.090509 |
| 10 | 77843600 | 77846999 | 15.88626 | 26.37405 | 79.33263 | 34.38679 | -1.20232 | 1.265967 | 0.090526 |
| 13 | 30463200 | 30464199 | 19.63102 | 35.40282 | 21.21759 | 9.237631 | 1.140787 | 0.396773 | 0.090527 |
| 2 | 69691000 | 69693199 | 9.96974 | 17.56298 | 47.73814 | 21.13074 | -1.04597 | 0.560331 | 0.090528 |
| 8 | 71667000 | 71668199 | 5.194535 | 9.828778 | 33.02744 | 13.39301 | -1.25032 | 0.024289 | 0.090541 |
| X | 32838000 | 32839399 | 24.80476 | 43.91298 | 20.69669 | 9.152118 | 1.469869 | 0.547617 | 0.090551 |
| 12 | 109449200 | 109450999 | 7.97927 | 14.52653 | 39.76543 | 17.10951 | -1.02491 | 0.300644 | 0.090551 |
| X | 26657400 | 26661199 | 38.83902 | 65.54314 | 32.15566 | 14.06352 | 1.404422 | 1.127689 | 0.090556 |
| 16 | 35936000 | 35937399 | 20.62942 | 36.3845 | 23.56059 | 10.42635 | 1.012645 | 0.565368 | 0.090558 |
| 3 | 39762600 | 39763999 | 9.992515 | 17.5251 | 48.28935 | 21.34406 | -1.0666 | 0.624554 | 0.090566 |
| 11 | 59730000 | 59731199 | 3.72433 | 7.21528 | 30.11144 | 11.41878 | -1.46785 | -0.22498 | 0.090569 |
| 13 | 66337800 | 66339599 | 25.12108 | 44.15212 | 23.3246 | 10.32817 | 1.302712 | 0.629762 | 0.090575 |
| 11 | 100533400 | 100539599 | 28.19341 | 45.3155 | 117.6 | 49.8516 | -1.02129 | 1.878132 | 0.090582 |
| 4 | 24897800 | 24898599 | 4.418053 | 8.40901 | 31.43857 | 12.59068 | -1.38482 | -0.02598 | 0.090583 |
| 11 | 59382200 | 59384799 | 10.18763 | 17.7971 | 52.04493 | 22.92898 | -1.15244 | 0.693909 | 0.090598 |
| 12 | 20211600 | 20217799 | 164.7202 | 248.8023 | 185.0184 | 76.48934 | 0.702183 | 3.330283 | 0.090603 |
| 14 | 62997000 | 62999599 | 12.07298 | 20.54971 | 64.1479 | 28.1169 | -1.25322 | 0.992263 | 0.090617 |
| 12 | 87600200 | 87601599 | 29.40729 | 50.02863 | 34.64547 | 15.18494 | 0.90962 | 1.024501 | 0.090621 |
| 8 | 88445600 | 88446399 | 3.544764 | 6.870616 | 27.65221 | 10.46327 | -1.41125 | -0.23875 | 0.090621 |
| 7 | 122003800 | 122006999 | 12.76436 | 21.90883 | 60.61685 | 26.58708 | -1.07817 | 0.912138 | 0.090626 |
| 17 | 19415600 | 19417399 | 39.61644 | 65.40604 | 46.14664 | 19.92573 | 0.882025 | 1.36603 | 0.090641 |
| 9 | 61548000 | 61555399 | 48.55474 | 76.65714 | 175.4896 | 72.86049 | -0.86808 | 2.440798 | 0.090641 |
| 15 | 12203600 | 12205799 | 11.14167 | 19.35925 | 56.38545 | 24.75734 | -1.14798 | 0.781427 | 0.090645 |

| | | | | | | | | | |
|---|---|---|---|---|---|---|---|---|---|
| 4 | 155362800 | 155375599 | 55.38517 | 86.05157 | 210.2766 | 87.31073 | -0.98016 | 2.750909 | 0.090656 |
| 2 | 29600000 | 29600599 | 2.247949 | 4.393642 | 21.64817 | 7.473818 | -1.56976 | -0.6043 | 0.090664 |
| 2 | 18527000 | 18527599 | 7.937917 | 15.52163 | 10.51833 | 3.590058 | 1.260546 | -0.60517 | 0.090667 |
| 11 | 69886200 | 69891799 | 23.1635 | 37.44451 | 104.4624 | 44.62938 | -1.11511 | 1.690314 | 0.090685 |
| 11 | 76288400 | 76290199 | 7.832155 | 14.26556 | 39.78396 | 17.10721 | -1.05124 | 0.29497 | 0.090691 |
| 7 | 16646200 | 16660799 | 74.93122 | 114.2269 | 276.1993 | 114.5051 | -0.98646 | 3.128196 | 0.090713 |
| 10 | 19181000 | 19181399 | 2.334803 | 4.566189 | 19.31836 | 6.320442 | -1.2819 | -0.8024 | 0.090722 |
| 14 | 28272200 | 28272599 | 2.334803 | 4.566189 | 19.31836 | 6.320442 | -1.2819 | -0.8024 | 0.090722 |
| 19 | 43654000 | 43656399 | 7.207715 | 13.20885 | 37.74611 | 16.1033 | -1.07782 | 0.251465 | 0.09073 |
| 2 | 97633800 | 97636599 | 26.06621 | 45.93185 | 20.42251 | 9.039282 | 1.550662 | 0.602434 | 0.090748 |
| 5 | 54152600 | 54153599 | 3.733717 | 7.208795 | 30.344 | 11.69213 | -1.50198 | -0.16813 | 0.090767 |
| 1 | 190494400 | 190505999 | 70.63302 | 108.0453 | 261.8088 | 108.6254 | -0.98557 | 3.056187 | 0.09077 |
| 5 | 115990400 | 115993599 | 15.14445 | 25.38093 | 71.1883 | 31.04566 | -1.10129 | 1.152427 | 0.090773 |
| 13 | 47346400 | 47349999 | 15.36412 | 25.9709 | 66.48857 | 29.09392 | -0.96972 | 1.067913 | 0.090777 |
| 13 | 73889800 | 73893799 | 20.39157 | 33.48136 | 86.13969 | 37.10318 | -0.98173 | 1.416175 | 0.090778 |
| 8 | 125793400 | 125796199 | 12.70207 | 21.95163 | 56.9167 | 24.96988 | -0.98271 | 0.84522 | 0.0908 |
| 7 | 22968400 | 22969399 | 10.20796 | 19.91919 | 11.30999 | 4.027665 | 1.45858 | -0.42427 | 0.090802 |
| 8 | 61802400 | 61802999 | 7.325213 | 13.51412 | 37.37593 | 15.80114 | -1.02153 | 0.213165 | 0.090843 |
| X | 32140800 | 32145799 | 74.13433 | 118.55 | 70.67312 | 29.73209 | 1.087761 | 2.075587 | 0.090844 |
| 4 | 9269200 | 9270799 | 7.324029 | 13.50263 | 38.45009 | 16.2705 | -1.06415 | 0.220575 | 0.090863 |
| 6 | 51505400 | 51507999 | 10.94465 | 19.10002 | 51.52732 | 22.68635 | -1.03692 | 0.712579 | 0.090866 |
| 11 | 15163600 | 15164799 | 13.5238 | 25.78773 | 16.46956 | 6.570148 | 1.144188 | 0.003886 | 0.090868 |
| 11 | 61407200 | 61410599 | 10.84127 | 18.83047 | 55.81143 | 24.49972 | -1.17295 | 0.785148 | 0.090888 |
| 17 | 38751200 | 38755599 | 67.52572 | 108.5588 | 55.45183 | 23.51409 | 1.318787 | 1.89966 | 0.09089 |
| 3 | 97588000 | 97589999 | 7.074214 | 12.93665 | 38.23522 | 16.35924 | -1.12915 | 0.263346 | 0.090894 |
| 6 | 11194800 | 11196599 | 20.11997 | 35.90783 | 22.79438 | 10.02072 | 1.049258 | 0.465632 | 0.090895 |
| 14 | 47178400 | 47179799 | 3.946852 | 7.669406 | 26.81251 | 9.989537 | -1.19022 | -0.29179 | 0.090908 |
| 11 | 94942000 | 94951799 | 37.79102 | 59.90994 | 159.5333 | 66.38873 | -1.08067 | 2.330952 | 0.09091 |
| 1 | 100786600 | 100788399 | 44.23793 | 72.29062 | 51.13458 | 21.98324 | 0.872165 | 1.507229 | 0.090916 |
| 4 | 135681800 | 135686399 | 16.04193 | 26.57594 | 78.52895 | 33.98852 | -1.17688 | 1.292279 | 0.090927 |
| 1 | 113111600 | 113113399 | 16.81696 | 32.36035 | 11.03134 | 4.294677 | 2.070066 | -0.05249 | 0.090933 |
| 17 | 16049800 | 16052599 | 19.93824 | 32.69764 | 87.06159 | 37.48117 | -1.03177 | 1.431092 | 0.090941 |
| 1 | 171332800 | 171339599 | 66.3075 | 103.0683 | 211.6152 | 87.86059 | -0.72888 | 2.747096 | 0.090969 |
| 11 | 5084000 | 5097999 | 76.05141 | 115.7673 | 277.48 | 114.988 | -0.97551 | 3.158782 | 0.090981 |
| 8 | 121383000 | 121387799 | 27.50859 | 44.3024 | 111.1094 | 47.23555 | -0.96847 | 1.812109 | 0.090986 |
| 10 | 22219200 | 22221799 | 13.72921 | 23.09312 | 70.61788 | 30.85462 | -1.22523 | 1.109925 | 0.091 |
| 9 | 64363200 | 64365799 | 10.9565 | 19.10532 | 53.14536 | 23.38691 | -1.08122 | 0.725332 | 0.091002 |
| 4 | 39954600 | 39954999 | 7.938938 | 15.52026 | 9.251156 | 3.175256 | 1.437454 | -0.59184 | 0.091004 |
| 6 | 38510400 | 38512599 | 9.586548 | 16.73109 | 52.01723 | 22.90304 | -1.24014 | 0.7057 | 0.091012 |
| 3 | 88125800 | 88134599 | 58.87428 | 92.18706 | 189.4113 | 78.57028 | -0.72161 | 2.613383 | 0.091018 |
| X | 157353200 | 157353799 | 2.523805 | 4.936046 | 22.01879 | 7.456641 | -1.40302 | -0.66406 | 0.091027 |
| 13 | 28071200 | 28073399 | 21.61809 | 38.82171 | 19.68518 | 8.608705 | 1.375649 | 0.431187 | 0.091028 |
| 5 | 112390200 | 112397999 | 46.67228 | 73.96372 | 158.7853 | 66.05454 | -0.77104 | 2.346822 | 0.091056 |
| 4 | 125371200 | 125387199 | 102.8459 | 153.9774 | 354.8386 | 146.3527 | -0.93965 | 3.498961 | 0.091057 |
| 16 | 43709800 | 43710599 | 3.427421 | 6.665397 | 28.12429 | 10.41074 | -1.44961 | -0.31934 | 0.091091 |
| 7 | 80247000 | 80248799 | 8.845914 | 15.67086 | 45.64625 | 20.18215 | -1.14095 | 0.491695 | 0.091097 |
| 4 | 44701400 | 44715599 | 101.691 | 152.9441 | 334.8513 | 138.2625 | -0.85989 | 3.40988 | 0.091109 |

| | | | | | | | | | |
|---|---|---|---|---|---|---|---|---|---|
| 9 | 106156600 | 106160999 | 13.44847 | 22.90445 | 64.32077 | 28.19401 | -1.10149 | 0.99086 | 0.091112 |
| 9 | 92494200 | 92495199 | 4.572123 | 8.757428 | 30.58396 | 12.06586 | -1.26858 | -0.07876 | 0.091121 |
| 12 | 20220200 | 20223399 | 84.42122 | 133.3547 | 102.2113 | 42.44813 | 0.708918 | 2.438236 | 0.091125 |
| 10 | 44841400 | 44843799 | 25.64393 | 44.37887 | 30.9253 | 13.56778 | 0.904606 | 0.852828 | 0.091128 |
| 10 | 61162800 | 61197199 | 208.4722 | 293.4467 | 688.0708 | 285.0153 | -1.05122 | 4.478633 | 0.091138 |
| 10 | 80085000 | 80087399 | 7.171817 | 12.90422 | 42.69837 | 18.64219 | -1.31046 | 0.367893 | 0.091144 |
| 17 | 54317000 | 54318599 | 23.57215 | 41.29547 | 26.84654 | 11.85513 | 1.005446 | 0.677008 | 0.091146 |
| 18 | 3059400 | 3070599 | 250.1299 | 368.8958 | 219.1 | 90.22844 | 0.995161 | 3.778036 | 0.09115 |
| 19 | 53587800 | 53589799 | 10.93964 | 19.08999 | 51.51934 | 22.68187 | -1.03735 | 0.714543 | 0.091152 |
| 15 | 89218800 | 89222999 | 25.97252 | 42.05361 | 98.98631 | 42.35129 | -0.86832 | 1.655737 | 0.091157 |
| 7 | 98803000 | 98809199 | 23.44141 | 37.961 | 100.2839 | 42.91119 | -1.03409 | 1.648346 | 0.091162 |
| 19 | 32525200 | 32528199 | 10.11594 | 17.83179 | 52.6189 | 23.28821 | -1.16382 | 0.561037 | 0.091185 |
| 14 | 122004000 | 122008599 | 20.94125 | 34.2312 | 95.9214 | 41.24211 | -1.10799 | 1.480794 | 0.09119 |
| 4 | 128508200 | 128512799 | 16.85459 | 27.59354 | 91.45377 | 39.34368 | -1.34737 | 1.448847 | 0.091191 |
| 15 | 76213400 | 76220399 | 26.44089 | 42.49104 | 118.0362 | 50.02352 | -1.11947 | 1.882307 | 0.091208 |
| 10 | 78946000 | 78946999 | 5.277847 | 10.13262 | 32.0308 | 12.55289 | -1.1186 | -0.09177 | 0.091211 |
| 11 | 71357200 | 71359599 | 14.66629 | 25.00107 | 62.77862 | 27.52003 | -0.94069 | 0.985837 | 0.091231 |
| 8 | 102922400 | 102923999 | 13.3083 | 23.02793 | 56.97267 | 24.99538 | -0.9151 | 0.836526 | 0.09125 |
| 11 | 94412000 | 94413199 | 4.074806 | 7.894533 | 27.68482 | 10.49693 | -1.21834 | -0.22474 | 0.091254 |
| 17 | 16631000 | 16631999 | 9.406013 | 16.46038 | 49.8847 | 22.00844 | -1.20305 | 0.660015 | 0.091255 |
| 16 | 38341800 | 38343399 | 7.207167 | 13.17343 | 38.21976 | 16.3625 | -1.10284 | 0.269092 | 0.091269 |
| 17 | 8840800 | 8842399 | 5.133516 | 9.806308 | 31.07925 | 12.35157 | -1.14028 | -0.04672 | 0.091271 |
| 5 | 78565400 | 78565799 | 7.499039 | 14.65664 | 6.628471 | 2.130336 | 1.900784 | -0.83167 | 0.091276 |
| 11 | 4124600 | 4125599 | 3.827818 | 7.443185 | 27.55492 | 10.20966 | -1.26494 | -0.31239 | 0.091278 |
| 7 | 28798000 | 28804799 | 43.9744 | 69.5695 | 164.4275 | 68.33912 | -0.91153 | 2.38729 | 0.091283 |
| 8 | 63489200 | 63490599 | 12.56049 | 24.07679 | 16.07895 | 6.329442 | 1.097642 | -0.04903 | 0.091297 |
| 16 | 11083800 | 11084599 | 12.56049 | 24.07679 | 16.07895 | 6.329442 | 1.097642 | -0.04903 | 0.091297 |
| 12 | 22254600 | 22256599 | 40.53191 | 66.99229 | 39.24711 | 16.95965 | 1.149387 | 1.35042 | 0.091298 |
| 7 | 31725600 | 31726999 | 18.11438 | 33.438 | 17.97743 | 7.59425 | 1.324397 | 0.248823 | 0.091313 |
| 3 | 87679400 | 87691599 | 60.93765 | 93.8246 | 237.4441 | 98.62062 | -1.0408 | 2.921983 | 0.091346 |
| 11 | 7047000 | 7052999 | 27.14551 | 43.824 | 105.5726 | 45.0274 | -0.90634 | 1.735776 | 0.091353 |
| 10 | 125223800 | 125225999 | 11.386 | 19.54892 | 59.95512 | 26.29748 | -1.22559 | 0.904192 | 0.091358 |
| 19 | 32733000 | 32736199 | 19.23065 | 31.39532 | 92.74185 | 39.85667 | -1.18493 | 1.49532 | 0.091366 |
| 5 | 144664800 | 144681799 | 126.3712 | 188.1145 | 387.222 | 159.5508 | -0.7845 | 3.60924 | 0.091368 |
| 7 | 57738800 | 57740599 | 22.03701 | 39.04206 | 22.99632 | 10.16751 | 1.150734 | 0.525773 | 0.091371 |
| 18 | 35835400 | 35836599 | 5.022134 | 9.528301 | 31.9639 | 12.88781 | -1.23991 | 0.002336 | 0.091392 |
| 17 | 64638200 | 64639799 | 11.9111 | 20.64031 | 55.27073 | 24.24977 | -1.02788 | 0.81632 | 0.091407 |
| 2 | 104233800 | 104234999 | 6.041364 | 11.45774 | 34.98999 | 14.11914 | -1.10844 | 0.019251 | 0.091479 |
| 4 | 79308000 | 79309199 | 13.94414 | 26.24007 | 15.42435 | 6.321708 | 1.229208 | 0.091284 | 0.091481 |
| 15 | 99349200 | 99357199 | 29.57471 | 47.34901 | 133.0198 | 56.05437 | -1.14245 | 2.008604 | 0.091487 |
| 8 | 115406000 | 115411999 | 34.31976 | 55.0167 | 122.7405 | 51.82418 | -0.80841 | 1.971143 | 0.091492 |
| 7 | 13494200 | 13497999 | 76.99894 | 122.3809 | 90.34158 | 37.69209 | 0.770768 | 2.270203 | 0.091503 |
| 19 | 7239400 | 7249399 | 57.71538 | 89.26101 | 226.0338 | 93.88907 | -1.0367 | 2.835272 | 0.091506 |
| 9 | 78335000 | 78336999 | 9.812568 | 16.98133 | 57.29567 | 25.13719 | -1.35961 | 0.821305 | 0.091525 |
| 1 | 144474800 | 144475999 | 11.76072 | 22.74795 | 14.5016 | 5.540632 | 1.203421 | -0.15779 | 0.091534 |
| 13 | 27486400 | 27487799 | 14.48385 | 27.96943 | 13.07046 | 5.032059 | 1.636391 | -0.1062 | 0.091544 |
| 16 | 55658000 | 55659599 | 8.020739 | 14.48231 | 40.12027 | 17.44084 | -1.05181 | 0.349514 | 0.091563 |

| | | | | | | | | | |
|---|---|---|---|---|---|---|---|---|---|
| 12 | 76579600 | 76585599 | 28.86092 | 46.30768 | 121.5168 | 51.37806 | -1.04035 | 1.935945 | 0.091567 |
| 17 | 47200200 | 47203599 | 8.811661 | 15.12813 | 61.3565 | 26.91203 | -1.62575 | 0.897213 | 0.091591 |
| 2 | 27918200 | 27924799 | 31.46297 | 50.37839 | 127.2152 | 53.61849 | -0.98871 | 2.005873 | 0.091615 |
| 18 | 85775200 | 85776199 | 14.50483 | 27.50317 | 15.6572 | 6.320873 | 1.293163 | 0.048079 | 0.091623 |
| 1 | 172308000 | 172340799 | 156.5004 | 225.0171 | 577.8408 | 238.5252 | -1.15149 | 4.163673 | 0.091637 |
| X | 4570800 | 4571399 | 12.93499 | 25.13549 | 12.17291 | 4.53438 | 1.628945 | -0.23363 | 0.091644 |
| 18 | 42644200 | 42645399 | 6.62563 | 11.80101 | 43.61989 | 19.2083 | -1.477 | 0.433406 | 0.091653 |
| X | 120414800 | 120415599 | 15.86883 | 30.43545 | 13.54245 | 5.322405 | 1.679695 | -0.02626 | 0.091658 |
| 17 | 50276800 | 50277999 | 16.71307 | 31.5882 | 15.71396 | 6.387241 | 1.478688 | 0.092093 | 0.091667 |
| 13 | 41216800 | 41217199 | 7.654891 | 14.97287 | 9.942924 | 3.338863 | 1.309626 | -0.66688 | 0.091671 |
| 6 | 83629200 | 83633999 | 32.66945 | 52.48939 | 118.2215 | 50.08673 | -0.81818 | 1.896263 | 0.091673 |
| 3 | 17508400 | 17509399 | 12.08172 | 23.50148 | 12.23236 | 4.518968 | 1.536577 | -0.27186 | 0.091683 |
| 17 | 35857200 | 35872199 | 95.6506 | 144.2418 | 317.5919 | 131.23 | -0.86462 | 3.356205 | 0.091691 |
| 18 | 44593800 | 44595399 | 17.93684 | 32.64024 | 20.86645 | 8.982968 | 1.058483 | 0.333006 | 0.091692 |
| 8 | 5976600 | 5977399 | 10.51438 | 20.41947 | 14.32921 | 5.363516 | 1.094825 | -0.2394 | 0.091698 |
| 12 | 80012200 | 80014599 | 6.08001 | 11.01345 | 39.64187 | 17.16375 | -1.42153 | 0.316932 | 0.091702 |
| 17 | 63824200 | 63825799 | 17.08806 | 31.42626 | 19.87157 | 8.437029 | 1.087516 | 0.265759 | 0.091705 |
| 6 | 145808000 | 145815999 | 46.46051 | 73.34833 | 174.3599 | 72.3902 | -0.92234 | 2.440244 | 0.09171 |
| 13 | 91246800 | 91247399 | 8.221342 | 16.06616 | 11.08282 | 3.841922 | 1.215794 | -0.54611 | 0.091722 |
| 14 | 70337000 | 70338399 | 4.26762 | 8.207761 | 29.58701 | 11.53597 | -1.29703 | -0.12317 | 0.091725 |
| 2 | 101062200 | 101063199 | 18.92053 | 34.63833 | 18.42603 | 7.871868 | 1.329124 | 0.302618 | 0.091737 |
| 6 | 16989200 | 16989799 | 3.265352 | 6.368081 | 25.51532 | 9.163252 | -1.3323 | -0.43287 | 0.091742 |
| 11 | 17415400 | 17417999 | 13.97782 | 24.03337 | 59.06468 | 25.90819 | -0.90752 | 0.900482 | 0.091757 |
| 4 | 138684600 | 138688199 | 19.06694 | 31.43808 | 80.9707 | 34.9387 | -0.98063 | 1.358138 | 0.091762 |
| 13 | 71734400 | 71737199 | 25.13801 | 43.77872 | 29.05609 | 12.77611 | 0.975605 | 0.774811 | 0.091776 |
| 15 | 85715600 | 85721599 | 33.52229 | 53.72417 | 123.5785 | 52.15715 | -0.85288 | 1.978606 | 0.091784 |
| 9 | 14291200 | 14293799 | 12.2321 | 21.16938 | 56.3712 | 24.73163 | -1.02046 | 0.829541 | 0.091793 |
| 11 | 106846800 | 106851599 | 19.952 | 32.71074 | 90.07501 | 38.77374 | -1.08039 | 1.435796 | 0.091796 |
| 1 | 173003600 | 173004999 | 13.75969 | 26.39018 | 15.59581 | 6.129455 | 1.275269 | -0.04173 | 0.091813 |
| 5 | 136161000 | 136184799 | 133.5772 | 195.4722 | 465.7031 | 191.8439 | -1.01844 | 3.895415 | 0.091815 |
| 9 | 35574200 | 35575799 | 38.90818 | 64.26476 | 46.13993 | 19.92883 | 0.856938 | 1.358637 | 0.091817 |
| 16 | 70206000 | 70206999 | 6.055471 | 11.4402 | 33.61294 | 13.67479 | -1.06328 | 0.044206 | 0.09183 |
| 8 | 93332400 | 93338599 | 34.61751 | 55.55245 | 123.8163 | 52.36216 | -0.80526 | 1.937988 | 0.091833 |
| 4 | 16744600 | 16745399 | 14.69978 | 28.17958 | 13.38853 | 5.269256 | 1.582858 | -0.03533 | 0.091838 |
| 4 | 135807000 | 135824399 | 90.99263 | 137.5322 | 305.4943 | 126.3225 | -0.87446 | 3.308422 | 0.091875 |
| 8 | 100785200 | 100787599 | 16.09133 | 27.35322 | 67.29871 | 29.4948 | -0.91232 | 1.021096 | 0.091879 |
| 18 | 57057800 | 57059599 | 10.36427 | 18.03254 | 53.63566 | 23.56974 | -1.17742 | 0.756525 | 0.091897 |
| 15 | 103050600 | 103052799 | 9.699908 | 17.00408 | 48.82001 | 21.57021 | -1.1256 | 0.630733 | 0.091904 |
| 17 | 36477800 | 36481199 | 24.7092 | 39.99241 | 99.7853 | 42.68078 | -0.95269 | 1.661825 | 0.091905 |
| 11 | 108778000 | 108782999 | 29.23479 | 47.1551 | 108.2332 | 46.10938 | -0.8383 | 1.765944 | 0.091915 |
| 2 | 179852400 | 179871199 | 109.3317 | 162.6022 | 386.1353 | 159.0884 | -0.99174 | 3.623076 | 0.091934 |
| 17 | 69997600 | 70000799 | 39.97341 | 67.09844 | 33.15398 | 14.4731 | 1.393991 | 1.176855 | 0.091935 |
| 1 | 36483800 | 36484599 | 1.871295 | 3.657538 | 22.83419 | 7.879351 | -1.90273 | -0.60846 | 0.091936 |
| 12 | 23541400 | 23552199 | 144.7048 | 220.4053 | 165.3002 | 68.5129 | 0.699909 | 3.155919 | 0.091936 |
| 7 | 31402800 | 31405999 | 72.495 | 115.858 | 74.66352 | 31.3838 | 0.974658 | 2.0918 | 0.091937 |
| 10 | 62876800 | 62878799 | 6.129043 | 11.41251 | 37.05623 | 15.46154 | -1.23677 | 0.146892 | 0.091939 |
| 13 | 61932200 | 61932999 | 12.78777 | 24.6924 | 13.70195 | 5.276942 | 1.390145 | -0.1229 | 0.091948 |

| | | | | | | | | | |
|---|---|---|---|---|---|---|---|---|---|
| 7 | 59895800 | 59899799 | 45.53262 | 74.80545 | 45.09424 | 19.42845 | 1.10578 | 1.438587 | 0.09195 |
| 7 | 8210800 | 8211999 | 13.61644 | 26.33937 | 13.13338 | 5.01606 | 1.554313 | -0.14152 | 0.091958 |
| 4 | 108332800 | 108334399 | 15.19936 | 28.64585 | 17.33912 | 7.088126 | 1.190884 | 0.09999 | 0.091959 |
| 11 | 59020400 | 59033199 | 75.75999 | 114.9422 | 303.8458 | 125.7892 | -1.12061 | 3.22765 | 0.091967 |
| 15 | 19684400 | 19684999 | 14.46634 | 27.54609 | 16.44409 | 6.579613 | 1.237371 | 0.02316 | 0.091978 |
| 10 | 80666400 | 80689599 | 148.1856 | 214.1021 | 527.549 | 217.6186 | -1.0849 | 4.090862 | 0.091981 |
| 4 | 138978400 | 138979799 | 5.671306 | 10.57529 | 34.82996 | 14.49748 | -1.25318 | 0.131002 | 0.091982 |
| 10 | 100133400 | 100134599 | 7.988088 | 14.49256 | 42.38451 | 18.31517 | -1.1237 | 0.327104 | 0.091992 |
| 7 | 113492000 | 113492599 | 11.79011 | 22.70376 | 15.67775 | 6.090924 | 1.067392 | -0.0997 | 0.091997 |
| X | 120345400 | 120346999 | 17.11485 | 33.08909 | 9.321673 | 3.569077 | 2.361799 | -0.10566 | 0.092019 |
| 15 | 88941600 | 88958599 | 99.36147 | 148.1692 | 378.7288 | 156.0853 | -1.09424 | 3.573943 | 0.09202 |
| 9 | 65143200 | 65146399 | 10.94572 | 19.11197 | 52.61079 | 23.17106 | -1.066 | 0.705156 | 0.092024 |
| 19 | 4446400 | 4448599 | 9.65794 | 16.74678 | 57.32992 | 25.15373 | -1.37986 | 0.801804 | 0.092036 |
| 1 | 152302000 | 152303399 | 15.5928 | 28.76523 | 18.96589 | 8.018665 | 1.03157 | 0.233496 | 0.092058 |
| 3 | 153837600 | 153841199 | 16.26823 | 27.58203 | 67.81496 | 29.70141 | -0.91162 | 1.04504 | 0.09206 |
| 13 | 89133800 | 89135799 | 19.93891 | 36.36159 | 18.26587 | 7.842301 | 1.405556 | 0.329749 | 0.092062 |
| 7 | 11277200 | 11279599 | 27.40501 | 47.60308 | 29.56645 | 12.97933 | 1.070709 | 0.823499 | 0.092066 |
| 6 | 87681800 | 87684399 | 12.96827 | 22.53737 | 55.92922 | 24.55997 | -0.9183 | 0.787077 | 0.092068 |
| 12 | 101924000 | 101924799 | 4.677652 | 9.025598 | 29.22693 | 11.28991 | -1.13191 | -0.15113 | 0.092075 |
| 6 | 3111000 | 3111599 | 9.702716 | 18.96267 | 10.30362 | 3.564003 | 1.558141 | -0.53227 | 0.092088 |
| 5 | 23992200 | 23994199 | 9.49837 | 16.78661 | 47.2239 | 20.88966 | -1.09335 | 0.520408 | 0.092101 |
| X | 24289800 | 24292399 | 33.93164 | 57.90985 | 31.61804 | 13.86159 | 1.251783 | 1.012545 | 0.092132 |
| 8 | 39438400 | 39439999 | 21.34056 | 37.67297 | 24.76603 | 10.95781 | 0.99145 | 0.562439 | 0.09216 |
| 5 | 120746200 | 120774399 | 170.7045 | 243.8011 | 600.9479 | 248.325 | -1.1021 | 4.263098 | 0.092163 |
| 14 | 41467600 | 41469999 | 20.16321 | 35.73721 | 23.75188 | 10.5007 | 0.977755 | 0.510218 | 0.092167 |
| 5 | 137124000 | 137124599 | 3.344993 | 6.526984 | 23.83634 | 8.493714 | -1.19119 | -0.46824 | 0.092168 |
| 16 | 70110200 | 70111799 | 7.963353 | 14.18009 | 44.13167 | 19.43777 | -1.23189 | 0.442969 | 0.092175 |
| 7 | 55103800 | 55105999 | 13.43229 | 25.95965 | 12.74658 | 4.888968 | 1.570887 | -0.13112 | 0.092176 |
| 2 | 30572200 | 30591199 | 81.69391 | 123.76 | 301.7013 | 124.8441 | -1.00569 | 3.259683 | 0.092193 |
| 3 | 100832200 | 100834399 | 9.516412 | 16.77222 | 47.20893 | 20.89454 | -1.09563 | 0.551761 | 0.0922 |
| 4 | 126007200 | 126009999 | 9.981998 | 17.55316 | 47.72906 | 21.13427 | -1.04748 | 0.580714 | 0.092227 |
| 11 | 114556400 | 114569399 | 58.03653 | 89.84856 | 228.0223 | 94.71455 | -1.03877 | 2.816245 | 0.092236 |
| 12 | 22272800 | 22275999 | 53.73152 | 86.58668 | 63.65873 | 27.08677 | 0.798863 | 1.810362 | 0.092247 |
| 10 | 13467200 | 13468799 | 22.22759 | 39.06318 | 25.94692 | 11.48849 | 0.974536 | 0.619371 | 0.092254 |
| 10 | 60346800 | 60365799 | 101.5287 | 152.06 | 347.7914 | 143.4561 | -0.92831 | 3.492097 | 0.092261 |
| 19 | 37607000 | 37607999 | 11.72323 | 22.54062 | 15.15668 | 5.9143 | 1.099395 | -0.08858 | 0.092264 |
| 19 | 4179600 | 4182199 | 6.341471 | 11.5062 | 39.14258 | 16.91232 | -1.33871 | 0.308841 | 0.092264 |
| 7 | 5922600 | 5929199 | 162.2188 | 245.9374 | 174.7516 | 72.32966 | 0.772331 | 3.255289 | 0.092279 |
| 4 | 148532200 | 148533599 | 5.283576 | 10.04075 | 32.03133 | 12.87037 | -1.16363 | -0.00558 | 0.092285 |
| 2 | 179494000 | 179502999 | 36.40321 | 57.69197 | 169.7448 | 70.61629 | -1.22485 | 2.339753 | 0.092288 |
| 7 | 121332200 | 121334399 | 10.06465 | 17.53472 | 54.17932 | 23.82872 | -1.23192 | 0.735299 | 0.092294 |
| 4 | 140231400 | 140253599 | 150.486 | 219.7939 | 461.1846 | 190.0079 | -0.83796 | 3.927013 | 0.092303 |
| X | 37267200 | 37270999 | 66.52047 | 107.1216 | 54.80825 | 23.29753 | 1.319379 | 1.849678 | 0.092311 |
| 10 | 108740600 | 108741999 | 7.498592 | 13.76286 | 38.34386 | 16.32443 | -1.03965 | 0.245295 | 0.092312 |
| 7 | 12136600 | 12137799 | 27.31978 | 46.90392 | 33.05209 | 14.49724 | 0.886436 | 0.933334 | 0.092313 |
| 1 | 182168400 | 182169399 | 3.432881 | 6.661569 | 28.35849 | 10.67188 | -1.48418 | -0.2585 | 0.092324 |
| 6 | 132078000 | 132078399 | 24.23066 | 42.22755 | 24.90231 | 10.9544 | 1.145299 | 0.754262 | 0.092326 |

| | | | | | | | | | |
|---|---|---|---|---|---|---|---|---|---|
| 10 | 20501200 | 20505199 | 32.7188 | 52.43492 | 123.1546 | 51.9759 | -0.88291 | 1.978955 | 0.092341 |
| 11 | 49788200 | 49794599 | 39.92308 | 63.60551 | 140.6744 | 58.8741 | -0.806 | 2.175377 | 0.092343 |
| 19 | 46560800 | 46562799 | 10.43839 | 18.34744 | 48.25862 | 21.37019 | -1.00022 | 0.589149 | 0.09237 |
| 1 | 27406000 | 27406599 | 9.17859 | 17.92748 | 11.5394 | 4.042956 | 1.301354 | -0.49295 | 0.092377 |
| 2 | 99113200 | 99114199 | 16.05234 | 29.86284 | 18.8809 | 7.890126 | 1.103808 | 0.191649 | 0.09238 |
| 4 | 139373400 | 139399999 | 207.8986 | 294.781 | 629.5776 | 260.4721 | -0.90595 | 4.370964 | 0.09238 |
| 2 | 76096600 | 76097599 | 11.52398 | 22.37138 | 14.21674 | 5.338323 | 1.232395 | -0.21649 | 0.092388 |
| 14 | 122284600 | 122286599 | 9.50912 | 16.72825 | 46.6817 | 20.66887 | -1.08396 | 0.574419 | 0.09239 |
| 4 | 41493000 | 41508599 | 99.35381 | 149.3511 | 332.3995 | 137.231 | -0.88422 | 3.41979 | 0.092427 |
| 8 | 53477000 | 53480399 | 31.71852 | 53.60208 | 37.81821 | 16.54713 | 0.882506 | 1.094939 | 0.092439 |
| 3 | 89054200 | 89058999 | 23.77589 | 38.43875 | 102.1473 | 43.64661 | -1.04496 | 1.687687 | 0.092445 |
| 4 | 125388000 | 125399399 | 58.3534 | 90.6663 | 212.5272 | 88.24478 | -0.92024 | 2.750992 | 0.092446 |
| 10 | 66559400 | 66560399 | 17.85933 | 32.59252 | 19.56467 | 8.392401 | 1.151342 | 0.312013 | 0.092471 |
| 9 | 19001400 | 19002799 | 27.00656 | 47.14244 | 26.79611 | 11.79963 | 1.198396 | 0.750657 | 0.092497 |
| 4 | 128322600 | 128326199 | 22.85906 | 37.2304 | 91.22881 | 39.1725 | -0.91867 | 1.537867 | 0.092509 |
| 11 | 62918800 | 62920799 | 8.883066 | 15.72972 | 44.59233 | 19.71791 | -1.10278 | 0.491713 | 0.092511 |
| 4 | 71091400 | 71092599 | 4.363319 | 8.480881 | 29.49987 | 10.96767 | -1.18165 | -0.28813 | 0.092527 |
| 9 | 78105000 | 78105799 | 4.363319 | 8.480881 | 29.49987 | 10.96767 | -1.18165 | -0.28813 | 0.092527 |
| 8 | 85707800 | 85710199 | 9.328693 | 16.52856 | 44.58649 | 19.713 | -1.0314 | 0.487668 | 0.092535 |
| 19 | 4524600 | 4528999 | 14.78329 | 24.53352 | 77.874 | 33.74563 | -1.27938 | 1.26791 | 0.092536 |
| 5 | 36359800 | 36374999 | 116.4583 | 173.5249 | 367.6351 | 151.4951 | -0.82474 | 3.59085 | 0.092539 |
| 1 | 31865000 | 31866599 | 20.1507 | 36.64827 | 18.81738 | 8.106542 | 1.371319 | 0.347725 | 0.09254 |
| 11 | 98672600 | 98674999 | 7.457433 | 13.39535 | 41.09271 | 17.96688 | -1.20323 | 0.379181 | 0.092574 |
| 4 | 136916600 | 136942399 | 149.931 | 217.8599 | 497.6378 | 205.1363 | -0.9676 | 4.005287 | 0.092577 |
| 10 | 25172400 | 25173199 | 3.833786 | 7.437981 | 27.80125 | 10.47237 | -1.30064 | -0.25175 | 0.092583 |
| 9 | 77965200 | 77967199 | 6.571161 | 12.19988 | 36.98087 | 15.5002 | -1.14364 | 0.168776 | 0.092585 |
| 3 | 35052600 | 35054599 | 8.611219 | 15.58438 | 41.79346 | 18.11588 | -1.00295 | 0.342888 | 0.092591 |
| 2 | 172727800 | 172735599 | 45.14537 | 71.29418 | 167.8882 | 69.71423 | -0.90834 | 2.432446 | 0.092591 |
| 11 | 50752200 | 50756999 | 37.46604 | 59.79913 | 135.7816 | 56.96969 | -0.841 | 2.113367 | 0.092605 |
| 7 | 8268600 | 8269199 | 12.25289 | 23.91265 | 7.862692 | 2.793778 | 2.23213 | -0.41489 | 0.092609 |
| 7 | 14194000 | 14196199 | 58.22973 | 93.83555 | 61.43372 | 26.14009 | 0.965744 | 1.815051 | 0.092628 |
| 7 | 142726200 | 142730399 | 19.98422 | 33.03622 | 80.29403 | 34.7012 | -0.89669 | 1.327599 | 0.092629 |
| 2 | 161640600 | 161642199 | 7.493423 | 13.73806 | 39.93978 | 17.02967 | -1.10187 | 0.257433 | 0.092634 |
| 7 | 54639000 | 54640399 | 20.39979 | 36.1948 | 23.76038 | 10.49704 | 0.996358 | 0.504593 | 0.092636 |
| 17 | 18293600 | 18294399 | 11.34609 | 22.07701 | 12.2496 | 4.514606 | 1.447954 | -0.28961 | 0.092647 |
| 11 | 117018000 | 117024399 | 25.61842 | 41.2765 | 115.3487 | 49.06913 | -1.12338 | 1.795863 | 0.092674 |
| 12 | 48916200 | 48917399 | 12.07784 | 23.51211 | 11.62217 | 4.266803 | 1.617679 | -0.29484 | 0.09268 |
| 15 | 69610800 | 69611199 | 4.133888 | 7.989258 | 29.30535 | 11.24023 | -1.29972 | -0.18231 | 0.092684 |
| 18 | 26976800 | 26977799 | 4.667682 | 9.037985 | 28.97066 | 11.01394 | -1.09554 | -0.20748 | 0.092717 |
| 1 | 17967800 | 17968999 | 16.84151 | 31.21851 | 18.73378 | 7.868901 | 1.173089 | 0.217748 | 0.09272 |
| 12 | 20099400 | 20101199 | 28.89947 | 49.32958 | 34.68578 | 15.20671 | 0.888507 | 0.994185 | 0.092721 |
| 2 | 26043400 | 26048999 | 23.75893 | 38.45385 | 100.1365 | 42.83057 | -1.01416 | 1.660684 | 0.092763 |
| 15 | 102862000 | 102866999 | 20.12201 | 32.6103 | 100.7986 | 43.15555 | -1.25877 | 1.627557 | 0.092764 |
| 3 | 129853600 | 129855199 | 8.661217 | 15.18951 | 49.85558 | 22.03544 | -1.31734 | 0.619363 | 0.092769 |
| 19 | 10665600 | 10670399 | 19.28264 | 31.70329 | 86.70355 | 37.36735 | -1.06876 | 1.396588 | 0.092779 |
| 3 | 46694800 | 46695199 | 8.661199 | 16.92327 | 5.87904 | 1.880432 | 2.277872 | -0.82647 | 0.092779 |
| 6 | 62821600 | 62822199 | 11.01596 | 21.48245 | 11.53146 | 4.143417 | 1.529476 | -0.38083 | 0.092821 |

| | | | | | | | | | |
|---|---|---|---|---|---|---|---|---|---|
| 9 | 11418400 | 11419799 | 21.00593 | 36.99048 | 25.01796 | 11.07476 | 0.949488 | 0.587491 | 0.092836 |
| 2 | 65231000 | 65231799 | 4.983476 | 9.575179 | 30.22309 | 11.81709 | -1.11246 | -0.10574 | 0.092845 |
| 17 | 25403000 | 25403999 | 4.983476 | 9.575179 | 30.22309 | 11.81709 | -1.11246 | -0.10574 | 0.092845 |
| 5 | 102995400 | 102996599 | 6.040565 | 11.45875 | 34.40553 | 13.87683 | -1.08354 | 0.016496 | 0.092857 |
| 19 | 3650600 | 3652599 | 8.688956 | 15.49015 | 42.5687 | 18.73099 | -1.05259 | 0.438286 | 0.092862 |
| 18 | 80379800 | 80383799 | 22.34071 | 36.53106 | 90.19011 | 38.78264 | -0.92545 | 1.476487 | 0.09288 |
| 7 | 31952000 | 31960999 | 144.9677 | 221.5921 | 154.8344 | 64.23059 | 0.805965 | 3.084374 | 0.092899 |
| 4 | 81623800 | 81624999 | 4.968103 | 9.593681 | 30.58044 | 11.77394 | -1.10578 | -0.15721 | 0.092964 |
| 12 | 85543000 | 85545199 | 8.396127 | 14.88949 | 43.56885 | 19.25639 | -1.14663 | 0.474188 | 0.092974 |
| 6 | 57226200 | 57227399 | 13.21129 | 25.30593 | 15.96691 | 6.298434 | 1.175974 | -0.036 | 0.092997 |
| X | 26451600 | 26452999 | 16.40505 | 31.64645 | 10.40696 | 4.02272 | 2.129494 | -0.08041 | 0.093 |
| 9 | 29555400 | 29570799 | 86.49583 | 131.0433 | 301.5291 | 124.7754 | -0.92238 | 3.259135 | 0.093001 |
| 13 | 88815600 | 88815999 | 9.879945 | 19.32012 | 9.021464 | 3.07471 | 1.790522 | -0.58335 | 0.093038 |
| 11 | 12890400 | 12891799 | 13.088 | 24.85543 | 17.16911 | 6.908401 | 1.021686 | 0.029011 | 0.093057 |
| 14 | 21548600 | 21553799 | 24.6503 | 39.7434 | 112.2547 | 47.79624 | -1.13748 | 1.773162 | 0.093068 |
| 8 | 122448000 | 122451599 | 19.45344 | 31.84962 | 86.5483 | 37.234 | -1.06187 | 1.452834 | 0.093073 |
| 4 | 135010000 | 135011999 | 8.447493 | 15.32761 | 41.32473 | 17.85493 | -1.00756 | 0.325149 | 0.093073 |
| 17 | 78671000 | 78677399 | 46.70139 | 73.91977 | 165.1635 | 68.66198 | -0.83009 | 2.37705 | 0.093075 |
| 17 | 27681000 | 27751399 | 433.2284 | 572.9423 | 1336.769 | 558.9481 | -1.13477 | 5.455501 | 0.093086 |
| 7 | 59252200 | 59253599 | 15.93385 | 30.14237 | 16.21698 | 6.579251 | 1.368708 | 0.079834 | 0.09309 |
| 5 | 68573000 | 68573399 | 9.000093 | 17.60499 | 7.684789 | 2.568912 | 1.905682 | -0.6681 | 0.093095 |
| 10 | 25253200 | 25256599 | 19.64052 | 32.5455 | 78.67006 | 34.05534 | -0.88851 | 1.297605 | 0.093103 |
| 5 | 95835600 | 95836199 | 12.71914 | 24.41846 | 16.20267 | 6.350009 | 1.113317 | -0.05992 | 0.093133 |
| 3 | 99915200 | 99915999 | 3.530374 | 6.88392 | 28.30212 | 10.18354 | -1.37547 | -0.42068 | 0.093154 |
| 7 | 33923600 | 33927199 | 12.68666 | 21.60227 | 64.23374 | 28.15536 | -1.18351 | 0.99043 | 0.093168 |
| 1 | 184529000 | 184529999 | 16.09279 | 30.14469 | 17.82951 | 7.364495 | 1.212455 | 0.147202 | 0.093171 |
| 3 | 73206400 | 73207199 | 13.71497 | 26.08458 | 16.7085 | 6.70211 | 1.13396 | 0.024922 | 0.093189 |
| 2 | 166380000 | 166422599 | 258.6566 | 357.5966 | 832.1499 | 345.6665 | -1.06648 | 4.751857 | 0.093202 |
| 12 | 82265200 | 82266399 | 4.733448 | 9.076627 | 32.4779 | 12.77176 | -1.30003 | -0.08539 | 0.093216 |
| 2 | 67352400 | 67353999 | 19.62502 | 34.78222 | 22.02997 | 9.739506 | 1.046823 | 0.503548 | 0.093218 |
| 9 | 54440400 | 54441799 | 5.750838 | 10.89695 | 32.55652 | 13.16218 | -1.07872 | 0.018696 | 0.093223 |
| 11 | 13043800 | 13044399 | 8.387483 | 16.38491 | 11.67326 | 4.078644 | 1.160384 | -0.51405 | 0.093233 |
| 3 | 72069000 | 72069599 | 9.823044 | 19.14343 | 12.33645 | 4.471295 | 1.258445 | -0.36463 | 0.093233 |
| 2 | 113538200 | 113539599 | 12.87079 | 24.56642 | 16.58013 | 6.600672 | 1.067857 | -0.00888 | 0.093233 |
| 4 | 130087200 | 130099199 | 85.77018 | 131.077 | 266.9182 | 110.7213 | -0.73607 | 3.079632 | 0.093237 |
| 7 | 79032400 | 79034199 | 6.60248 | 12.17008 | 37.32926 | 15.80027 | -1.17074 | 0.211713 | 0.093241 |
| 2 | 167434800 | 167449799 | 69.6594 | 106.5469 | 262.5039 | 108.9117 | -1.00961 | 3.05769 | 0.093246 |
| 18 | 88743800 | 88744399 | 6.029084 | 11.43774 | 33.78062 | 13.62285 | -1.05923 | 0.016407 | 0.093254 |
| 1 | 189517400 | 189521799 | 16.66065 | 27.59319 | 78.01956 | 33.76151 | -1.1136 | 1.297111 | 0.093302 |
| 8 | 81930000 | 81933799 | 77.37819 | 123.0346 | 88.5581 | 36.96331 | 0.807713 | 2.257242 | 0.09331 |
| 13 | 53622000 | 53631399 | 56.98046 | 89.08741 | 199.9442 | 82.95233 | -0.85057 | 2.63937 | 0.093315 |
| 5 | 114286600 | 114295599 | 47.34479 | 74.56011 | 179.4685 | 74.46541 | -0.94314 | 2.494435 | 0.093343 |
| 8 | 4935800 | 4938399 | 33.8111 | 57.04517 | 34.63534 | 15.14698 | 1.098189 | 1.112706 | 0.093373 |
| 16 | 95960200 | 95965599 | 20.6253 | 33.43016 | 105.3963 | 45.12817 | -1.28722 | 1.626667 | 0.093381 |
| 10 | 123805200 | 123806799 | 23.2629 | 41.81409 | 16.09002 | 7.03099 | 1.770247 | 0.429322 | 0.093406 |
| 1 | 91597800 | 91599999 | 14.48739 | 24.76607 | 61.17479 | 26.82307 | -0.91635 | 0.957654 | 0.093418 |
| 1 | 67018800 | 67019599 | 13.55064 | 25.78669 | 15.83537 | 6.344243 | 1.194467 | 0.015185 | 0.093432 |

| | | | | | | | | | |
|---|---|---|---|---|---|---|---|---|---|
| 3 | 34692600 | 34698399 | 37.24165 | 59.62533 | 127.9565 | 53.92285 | -0.75477 | 2.015445 | 0.093447 |
| X | 26198200 | 26200199 | 21.68616 | 39.4584 | 17.03283 | 7.333673 | 1.619921 | 0.353183 | 0.093456 |
| 6 | 113158000 | 113163199 | 23.4395 | 38.34439 | 92.32819 | 39.7083 | -0.88912 | 1.473549 | 0.09346 |
| 11 | 80753200 | 80754999 | 8.363839 | 14.94263 | 42.07788 | 18.4815 | -1.08533 | 0.42313 | 0.093489 |
| 10 | 61887400 | 61899599 | 60.95667 | 94.38126 | 226.3966 | 94.0392 | -0.95742 | 2.814789 | 0.093499 |
| 15 | 98855600 | 98857799 | 7.629631 | 14.05516 | 39.58618 | 16.7689 | -1.05016 | 0.227458 | 0.093502 |
| 7 | 79142000 | 79162799 | 150.7409 | 219.9204 | 472.622 | 194.7427 | -0.87402 | 3.943922 | 0.093504 |
| 7 | 33371600 | 33375199 | 90.00161 | 141.9975 | 103.2731 | 42.86283 | 0.783114 | 2.470204 | 0.093506 |
| 13 | 13371800 | 13373599 | 28.53745 | 48.73292 | 34.19055 | 14.99012 | 0.891752 | 0.988004 | 0.09352 |
| 3 | 4990600 | 4991599 | 16.30577 | 30.46803 | 17.68715 | 7.336417 | 1.234111 | 0.163382 | 0.09352 |
| 1 | 129264600 | 129265199 | 5.641994 | 10.51894 | 34.81029 | 14.49326 | -1.25982 | 0.133535 | 0.093533 |
| 5 | 117306200 | 117310399 | 25.76493 | 41.85728 | 95.72685 | 41.05786 | -0.82236 | 1.584768 | 0.093545 |
| 7 | 120450000 | 120451999 | 8.392587 | 14.94182 | 42.56126 | 18.74882 | -1.10492 | 0.445592 | 0.093557 |
| 13 | 110038600 | 110040599 | 9.563069 | 16.74409 | 49.88637 | 22.01902 | -1.17875 | 0.651179 | 0.093567 |
| 14 | 30942600 | 30956999 | 57.26076 | 88.19327 | 242.7739 | 100.839 | -1.1616 | 2.91382 | 0.093579 |
| 5 | 134461200 | 134482399 | 157.1386 | 228.5464 | 481.3377 | 198.3978 | -0.84922 | 3.991653 | 0.093586 |
| 5 | 74390600 | 74391199 | 2.637336 | 5.158169 | 20.80189 | 7.035177 | -1.26013 | -0.67513 | 0.093597 |
| 12 | 107182600 | 107184199 | 7.873419 | 14.22052 | 40.12598 | 17.43685 | -1.07782 | 0.346306 | 0.093602 |
| 14 | 108352800 | 108354999 | 19.00703 | 34.63278 | 18.84613 | 8.100018 | 1.289978 | 0.328012 | 0.093606 |
| 7 | 102673800 | 102674399 | 20.2939 | 35.97081 | 23.02192 | 10.17787 | 1.032145 | 0.510343 | 0.093611 |
| 7 | 24232800 | 24239799 | 42.84738 | 67.79576 | 166.23 | 69.09304 | -0.96428 | 2.383218 | 0.093631 |
| 4 | 140793000 | 140809199 | 77.59114 | 117.7306 | 293.0672 | 121.3301 | -1.03389 | 3.226243 | 0.093633 |
| 5 | 136227400 | 136228199 | 3.760973 | 7.333997 | 25.65671 | 9.22463 | -1.14321 | -0.42519 | 0.093669 |
| X | 15945800 | 15948399 | 19.77242 | 36.32111 | 17.67011 | 7.51411 | 1.461518 | 0.289118 | 0.093673 |
| 9 | 20299400 | 20301399 | 36.36 | 60.43655 | 43.34715 | 18.80441 | 0.858934 | 1.277286 | 0.093674 |
| 6 | 85487400 | 85490999 | 27.21395 | 43.78536 | 114.1578 | 48.46325 | -1.02608 | 1.84366 | 0.09368 |
| 17 | 21771600 | 21771999 | 2.027765 | 3.964802 | 18.08841 | 5.858027 | -1.37154 | -0.84693 | 0.093682 |
| 8 | 49099200 | 49099999 | 5.325861 | 10.07673 | 32.46232 | 13.16533 | -1.19001 | 0.025316 | 0.093686 |
| 18 | 81028600 | 81033399 | 36.242 | 57.72482 | 146.3423 | 61.22274 | -1.00299 | 2.181636 | 0.09369 |
| 18 | 26954000 | 26955799 | 20.88485 | 37.03158 | 23.66147 | 10.45901 | 1.034557 | 0.511385 | 0.093696 |
| 10 | 105831400 | 105831999 | 9.122248 | 17.80381 | 12.10039 | 4.295788 | 1.20709 | -0.44755 | 0.093698 |
| X | 19846600 | 19847399 | 12.3461 | 24.12259 | 7.500738 | 2.611822 | 2.337996 | -0.4811 | 0.093713 |
| 8 | 31604400 | 31605199 | 11.6542 | 22.5691 | 14.54489 | 5.525963 | 1.195651 | -0.17698 | 0.093716 |
| 12 | 23662200 | 23664199 | 29.07427 | 50.36062 | 29.5026 | 12.94396 | 1.154047 | 0.859868 | 0.093723 |
| 17 | 54949800 | 54950999 | 5.908218 | 11.18648 | 34.29784 | 13.88895 | -1.11853 | 0.028034 | 0.093734 |
| 4 | 59233200 | 59237399 | 30.52705 | 49.20213 | 115.2216 | 49.03851 | -0.86858 | 1.792149 | 0.093743 |
| 5 | 115775200 | 115794999 | 84.21819 | 126.5734 | 343.7121 | 141.8937 | -1.17121 | 3.421633 | 0.093769 |
| 10 | 81189400 | 81211599 | 101.1722 | 150.205 | 400.389 | 164.9317 | -1.16063 | 3.653293 | 0.093779 |
| 1 | 184601800 | 184608799 | 32.4024 | 51.94088 | 126.7817 | 53.52704 | -0.93796 | 1.970377 | 0.093783 |
| 14 | 51494600 | 51495599 | 13.43116 | 25.98134 | 13.8883 | 5.304043 | 1.456147 | -0.14103 | 0.093813 |
| 11 | 101536800 | 101538199 | 5.282987 | 10.12657 | 31.31659 | 12.32845 | -1.09315 | -0.07809 | 0.093819 |
| 3 | 106839800 | 106840199 | 9.931647 | 19.35322 | 13.21503 | 4.795751 | 1.173422 | -0.35638 | 0.093825 |
| X | 124900800 | 124902599 | 20.5115 | 36.87009 | 21.17763 | 9.25384 | 1.198409 | 0.421975 | 0.093831 |
| 3 | 108940200 | 108941999 | 23.34316 | 40.87035 | 27.55985 | 12.16409 | 0.953156 | 0.687028 | 0.093846 |
| 4 | 139477800 | 139501399 | 150.3391 | 218.9829 | 483.6884 | 199.3359 | -0.9158 | 3.968324 | 0.093849 |
| 13 | 66592400 | 66596799 | 99.82492 | 157.138 | 101.4861 | 42.1064 | 0.951737 | 2.522971 | 0.093854 |
| 6 | 95644800 | 95646599 | 10.08839 | 17.86709 | 47.77904 | 21.1263 | -1.01952 | 0.502329 | 0.093862 |

| | | | | | | | | | |
|---|---|---|---|---|---|---|---|---|---|
| 16 | 4372600 | 4379199 | 38.28983 | 61.06361 | 137.0337 | 57.42877 | -0.82538 | 2.14131 | 0.093872 |
| 10 | 37604400 | 37607799 | 24.39053 | 42.96566 | 23.43891 | 10.37503 | 1.257881 | 0.60178 | 0.093874 |
| 5 | 34668600 | 34671799 | 13.69493 | 23.46886 | 60.64474 | 26.59592 | -0.98052 | 0.930618 | 0.093903 |
| 13 | 65948800 | 65949599 | 10.63613 | 20.80213 | 6.881181 | 2.330058 | 2.280964 | -0.60104 | 0.093927 |
| 5 | 117913400 | 117931799 | 113.9731 | 169.3851 | 392.0937 | 161.5316 | -0.95585 | 3.636146 | 0.09393 |
| 13 | 66771000 | 66776399 | 80.28808 | 127.6648 | 86.51511 | 36.11155 | 0.894581 | 2.259483 | 0.093938 |
| 18 | 75739800 | 75741999 | 15.36556 | 25.94758 | 66.39767 | 29.0454 | -0.96899 | 1.078256 | 0.093946 |
| 5 | 6879200 | 6879999 | 13.80772 | 26.70594 | 13.12849 | 5.017218 | 1.573868 | -0.13735 | 0.093971 |
| 12 | 55071200 | 55072399 | 13.80772 | 26.70594 | 13.12849 | 5.017218 | 1.573868 | -0.13735 | 0.093971 |
| 15 | 10705400 | 10710999 | 29.36323 | 47.45926 | 107.2581 | 45.81472 | -0.8126 | 1.706506 | 0.093978 |
| 17 | 20504000 | 20504799 | 17.39579 | 32.30053 | 18.17211 | 7.615435 | 1.267823 | 0.212224 | 0.094 |
| 16 | 45543000 | 45545599 | 14.12439 | 24.30495 | 59.10693 | 25.92736 | -0.89224 | 0.893043 | 0.094008 |
| 7 | 59132600 | 59133599 | 12.69729 | 24.46953 | 15.60261 | 6.048339 | 1.184487 | -0.09641 | 0.094033 |
| 17 | 81546200 | 81547799 | 14.27755 | 27.20132 | 15.74871 | 6.294164 | 1.281806 | 0.014949 | 0.094043 |
| 15 | 80595000 | 80596399 | 4.41165 | 8.417393 | 30.95424 | 12.33273 | -1.35458 | -0.04467 | 0.094056 |
| 14 | 36387400 | 36388199 | 3.643494 | 7.105487 | 25.0254 | 8.988063 | -1.15134 | -0.43328 | 0.094069 |
| 16 | 29824000 | 29825399 | 5.589609 | 10.66481 | 32.87478 | 13.09822 | -1.10512 | -0.03201 | 0.094072 |
| 1 | 100456400 | 100459599 | 61.6424 | 98.79839 | 73.73619 | 31.12459 | 0.767928 | 1.991878 | 0.094094 |
| 8 | 114708800 | 114710399 | 10.18445 | 17.76955 | 51.50208 | 22.67218 | -1.13961 | 0.712451 | 0.094095 |
| 15 | 75164400 | 75166199 | 12.6541 | 22.01322 | 54.27728 | 23.84944 | -0.90875 | 0.76969 | 0.094095 |
| 17 | 20473000 | 20473599 | 17.8193 | 32.29407 | 21.28453 | 9.209913 | 1.009913 | 0.355047 | 0.094107 |
| 8 | 23903400 | 23903999 | 2.941028 | 5.736983 | 23.67259 | 8.474383 | -1.36966 | -0.45724 | 0.094139 |
| 2 | 153996000 | 153996799 | 12.66765 | 24.19028 | 16.2855 | 6.476805 | 1.073297 | -0.01403 | 0.09414 |
| X | 33474800 | 33476199 | 44.51859 | 73.6302 | 40.02594 | 17.30409 | 1.256949 | 1.354156 | 0.094145 |
| 5 | 145478800 | 145480599 | 21.06959 | 37.35137 | 23.65976 | 10.45954 | 1.046873 | 0.514375 | 0.094163 |
| 7 | 9178000 | 9182199 | 47.65443 | 78.55374 | 38.92295 | 16.79336 | 1.389341 | 1.399633 | 0.094165 |
| 10 | 115757000 | 115757999 | 10.55711 | 20.4665 | 13.32271 | 5.036262 | 1.188702 | -0.21007 | 0.094179 |
| 8 | 99864800 | 99865199 | 9.352354 | 18.26622 | 11.53589 | 4.044201 | 1.327843 | -0.4881 | 0.094182 |
| X | 26603800 | 26608399 | 65.78896 | 106.4933 | 40.76299 | 17.43689 | 1.74461 | 1.703893 | 0.094184 |
| 1 | 85308800 | 85313399 | 180.8105 | 272.2198 | 192.4612 | 79.49044 | 0.77096 | 3.397487 | 0.094191 |
| 9 | 110840400 | 110841799 | 6.900677 | 12.70764 | 38.4229 | 16.28424 | -1.15197 | 0.221049 | 0.0942 |
| 6 | 121047000 | 121051399 | 16.53934 | 27.72754 | 71.99605 | 31.40427 | -0.99066 | 1.152871 | 0.094217 |
| 1 | 90943200 | 90945599 | 11.44303 | 19.87181 | 54.75218 | 24.0318 | -1.06825 | 0.789832 | 0.094226 |
| 6 | 90535600 | 90537199 | 6.639079 | 12.06863 | 38.6474 | 16.65955 | -1.2503 | 0.298316 | 0.094236 |
| 15 | 7010000 | 7011799 | 9.676665 | 16.99611 | 47.21696 | 20.89599 | -1.07816 | 0.597306 | 0.094248 |
| 11 | 72391800 | 72400799 | 37.94836 | 60.20685 | 158.2459 | 65.90503 | -1.06079 | 2.306603 | 0.094257 |
| X | 34600400 | 34601799 | 17.27265 | 32.17822 | 16.56998 | 6.907957 | 1.400874 | 0.18891 | 0.094261 |
| 1 | 103838200 | 103839999 | 25.50016 | 44.77796 | 23.90558 | 10.57736 | 1.287798 | 0.647227 | 0.094262 |
| 17 | 27329000 | 27351799 | 115.6252 | 169.5651 | 463.1877 | 190.7821 | -1.21264 | 3.860074 | 0.094277 |
| 3 | 19855200 | 19857799 | 34.92285 | 59.24213 | 33.81659 | 14.81333 | 1.187034 | 1.071126 | 0.09428 |
| 13 | 89887000 | 89891199 | 27.64571 | 44.59463 | 110.3917 | 47.03258 | -0.94696 | 1.762027 | 0.094293 |
| 15 | 73504600 | 73508999 | 16.43706 | 27.43312 | 74.60311 | 32.44977 | -1.05709 | 1.203653 | 0.094295 |
| 6 | 31059200 | 31060399 | 5.564027 | 10.64093 | 31.77361 | 12.59071 | -1.05187 | -0.05102 | 0.094299 |
| 19 | 5910400 | 5924199 | 75.38108 | 115.1349 | 261.8673 | 108.6119 | -0.89603 | 3.088658 | 0.094308 |
| 14 | 113070400 | 113070799 | 10.02695 | 19.58252 | 10.05667 | 3.529553 | 1.618111 | -0.4808 | 0.094309 |
| X | 147512200 | 147514399 | 25.11771 | 43.91778 | 26.62808 | 11.73939 | 1.105959 | 0.717581 | 0.094322 |
| 6 | 91591200 | 91600599 | 50.57938 | 79.63885 | 178.1377 | 73.91142 | -0.83773 | 2.49999 | 0.094323 |

| | | | | | | | | | |
|---|---|---|---|---|---|---|---|---|---|
| 17 | 27301800 | 27312599 | 47.15413 | 73.65195 | 205.8356 | 85.40514 | -1.16754 | 2.653137 | 0.094342 |
| 9 | 118315600 | 118318999 | 20.14587 | 32.93337 | 90.21383 | 38.78909 | -1.07507 | 1.476125 | 0.094363 |
| 4 | 59832800 | 59836799 | 28.10689 | 45.36724 | 107.0633 | 45.65173 | -0.8771 | 1.743125 | 0.094379 |
| 17 | 40371600 | 40372199 | 12.08926 | 23.48088 | 13.42192 | 5.018878 | 1.388042 | -0.22707 | 0.09441 |
| 4 | 16110400 | 16111599 | 14.03008 | 26.31229 | 17.65804 | 7.278929 | 1.033983 | 0.1205 | 0.09444 |
| 7 | 130285800 | 130286399 | 4.238452 | 8.238768 | 29.50906 | 10.96477 | -1.22277 | -0.2925 | 0.094442 |
| 4 | 138205600 | 138207599 | 7.303104 | 13.09628 | 41.08075 | 17.98692 | -1.23627 | 0.389969 | 0.094464 |
| 15 | 8871400 | 8872199 | 4.3252 | 8.385399 | 28.98369 | 10.95422 | -1.19356 | -0.22845 | 0.094467 |
| 8 | 85935200 | 85937199 | 21.23317 | 38.47058 | 18.75973 | 8.120189 | 1.441046 | 0.377014 | 0.094469 |
| 3 | 116823400 | 116823999 | 4.470703 | 8.60472 | 31.37217 | 12.206 | -1.30983 | -0.12046 | 0.0945 |
| 10 | 77520200 | 77527999 | 43.73824 | 69.01729 | 175.0143 | 72.64413 | -1.01592 | 2.450521 | 0.094505 |
| 17 | 13980200 | 13982199 | 11.70631 | 20.02146 | 60.95448 | 26.72751 | -1.21591 | 0.943549 | 0.094509 |
| 5 | 139097600 | 139100199 | 12.53847 | 21.69516 | 55.83056 | 24.49429 | -0.97143 | 0.832759 | 0.094512 |
| 7 | 32807800 | 32819999 | 175.8075 | 265.9628 | 173.5876 | 71.79893 | 0.892165 | 3.30139 | 0.094516 |
| 11 | 101013800 | 101022999 | 53.40015 | 83.90128 | 181.0876 | 75.11455 | -0.78867 | 2.545234 | 0.094527 |
| 8 | 110003000 | 110004799 | 5.884217 | 11.1709 | 32.64194 | 13.14377 | -1.04187 | 0.007084 | 0.09453 |
| 11 | 106186200 | 106194399 | 80.07853 | 122.564 | 253.3665 | 105.1367 | -0.75599 | 3.046789 | 0.094534 |
| 2 | 89094200 | 89096199 | 51.30988 | 82.80711 | 60.95686 | 25.98712 | 0.799431 | 1.761111 | 0.094535 |
| 4 | 128162800 | 128171599 | 50.41188 | 78.99846 | 190.4355 | 78.98877 | -0.95086 | 2.594662 | 0.094536 |
| 8 | 71823800 | 71843999 | 115.1879 | 170.806 | 396.9594 | 163.4988 | -0.96458 | 3.676337 | 0.094546 |
| 5 | 7014600 | 7016399 | 19.09699 | 34.57067 | 19.33549 | 8.378351 | 1.243252 | 0.365496 | 0.094555 |
| 4 | 150597000 | 150598599 | 5.932981 | 11.16035 | 34.01097 | 13.94864 | -1.12533 | 0.072943 | 0.094572 |
| 7 | 28600200 | 28607199 | 25.3586 | 40.71191 | 121.9818 | 51.62044 | -1.23006 | 1.913452 | 0.094575 |
| 17 | 83376800 | 83381399 | 21.48092 | 35.30149 | 89.55091 | 38.58707 | -0.9611 | 1.408573 | 0.094578 |
| 14 | 66074600 | 66075399 | 4.24455 | 8.232342 | 27.84195 | 10.503 | -1.16063 | -0.24188 | 0.094579 |
| 15 | 100331200 | 100336999 | 32.3772 | 51.94656 | 120.1946 | 50.81451 | -0.85939 | 1.941374 | 0.094583 |
| 3 | 31238800 | 31240199 | 15.35186 | 28.71985 | 18.39272 | 7.613692 | 1.09605 | 0.149535 | 0.094584 |
| 13 | 22080600 | 22082599 | 9.629771 | 17.06465 | 47.24437 | 20.88749 | -1.06893 | 0.491261 | 0.094586 |
| 14 | 64822600 | 64833399 | 63.97698 | 98.84759 | 230.6266 | 95.79729 | -0.91985 | 2.855794 | 0.094588 |
| 2 | 128200000 | 128204399 | 22.72141 | 36.99416 | 93.07756 | 39.96158 | -0.95708 | 1.542102 | 0.09459 |
| 2 | 129516800 | 129518199 | 6.702084 | 12.47304 | 35.94337 | 15.01041 | -1.06704 | 0.153423 | 0.094596 |
| 7 | 92307000 | 92307999 | 5.3028 | 10.1034 | 31.58945 | 12.62352 | -1.12847 | -0.02678 | 0.094598 |
| 12 | 100978200 | 100997199 | 102.6077 | 153.573 | 359.965 | 148.4581 | -0.96453 | 3.504788 | 0.094617 |
| 7 | 24786200 | 24793199 | 28.60342 | 45.81985 | 129.8531 | 54.7669 | -1.15398 | 1.988986 | 0.094631 |
| 11 | 52228400 | 52234599 | 29.64997 | 47.59556 | 119.6412 | 50.62037 | -0.9777 | 1.921626 | 0.094634 |
| 2 | 143239200 | 143240199 | 11.72338 | 22.75693 | 14.01244 | 5.264468 | 1.275821 | -0.21381 | 0.094664 |
| 12 | 3050000 | 3055999 | 182.8232 | 276.2333 | 160.9336 | 66.53897 | 1.054362 | 3.327509 | 0.094665 |
| 11 | 3989200 | 3990399 | 5.264349 | 10.07247 | 31.16564 | 12.33341 | -1.10033 | -0.05897 | 0.094692 |
| 18 | 22476200 | 22478399 | 12.89809 | 22.18062 | 59.02089 | 25.88912 | -1.02155 | 0.89355 | 0.094697 |
| 7 | 70410600 | 70410999 | 7.818957 | 15.28467 | 11.24319 | 3.865756 | 1.135824 | -0.5785 | 0.094722 |
| 11 | 58870400 | 58873999 | 11.26344 | 19.28653 | 61.51679 | 26.97661 | -1.28256 | 0.930422 | 0.094765 |
| 11 | 103655400 | 103679199 | 135.4075 | 197.552 | 486.6308 | 200.5148 | -1.07077 | 3.942632 | 0.094767 |
| 10 | 48278000 | 48278799 | 16.68778 | 31.16353 | 17.67701 | 7.339509 | 1.266216 | 0.170544 | 0.09477 |
| 5 | 136847600 | 136879999 | 248.4223 | 348.2137 | 697.4811 | 289.1032 | -0.83015 | 4.542779 | 0.094779 |
| 8 | 95563800 | 95565599 | 8.628097 | 15.20185 | 47.73198 | 21.12736 | -1.25232 | 0.551064 | 0.09478 |
| 11 | 57557400 | 57572799 | 93.11476 | 140.8452 | 301.5888 | 124.7371 | -0.82068 | 3.293321 | 0.094797 |
| 11 | 75051600 | 75064799 | 72.29051 | 110.4663 | 262.7101 | 108.9733 | -0.95977 | 3.078017 | 0.094798 |

| | | | | | | | | | |
|---|---|---|---|---|---|---|---|---|---|
| 15 | 71282200 | 71283799 | 14.75297 | 28.55803 | 11.19276 | 4.260786 | 1.900502 | -0.14498 | 0.094798 |
| 2 | 160442600 | 160447199 | 23.62446 | 38.56832 | 89.4816 | 38.45549 | -0.83768 | 1.503271 | 0.0948 |
| 5 | 142709400 | 142731599 | 146.5833 | 213.3944 | 488.9164 | 201.502 | -0.96934 | 3.976462 | 0.094801 |
| 6 | 124262200 | 124263399 | 20.47732 | 36.29032 | 23.0218 | 10.17813 | 1.044781 | 0.513329 | 0.09482 |
| 2 | 48093800 | 48095199 | 5.824757 | 10.88076 | 35.45858 | 14.71478 | -1.23511 | 0.120513 | 0.094827 |
| 6 | 52616800 | 52618399 | 5.824757 | 10.88076 | 35.45858 | 14.71478 | -1.23511 | 0.120513 | 0.094827 |
| X | 4347000 | 4350599 | 72.6145 | 115.6793 | 78.80351 | 32.9763 | 0.889876 | 2.192711 | 0.094834 |
| 17 | 34828600 | 34838399 | 40.39745 | 63.65661 | 179.7323 | 74.58058 | -1.17221 | 2.479026 | 0.094875 |
| 17 | 46513200 | 46514599 | 5.752507 | 10.5903 | 38.35322 | 16.26092 | -1.40974 | 0.214225 | 0.094883 |
| 11 | 9402800 | 9403399 | 13.57473 | 25.69973 | 17.60632 | 7.131071 | 1.025275 | 0.055705 | 0.094888 |
| 11 | 98093600 | 98095199 | 5.210057 | 9.813759 | 33.42958 | 13.67879 | -1.28101 | 0.057049 | 0.094896 |
| 4 | 138545200 | 138546599 | 6.334598 | 11.60079 | 37.7105 | 16.10317 | -1.26273 | 0.248983 | 0.094901 |
| 5 | 134933600 | 134936599 | 12.09066 | 20.86507 | 56.82647 | 24.92921 | -1.05347 | 0.857498 | 0.094916 |
| 5 | 37050600 | 37060999 | 62.5138 | 97.27885 | 204.6235 | 84.94367 | -0.76249 | 2.725766 | 0.094916 |
| 4 | 127273400 | 127280599 | 44.64212 | 70.58461 | 171.2249 | 71.14321 | -0.94974 | 2.402081 | 0.09493 |
| 17 | 27313400 | 27315399 | 8.912809 | 15.70424 | 46.16039 | 20.43154 | -1.15752 | 0.550688 | 0.094971 |
| 5 | 56446800 | 56448199 | 16.40484 | 31.02717 | 14.86674 | 6.034 | 1.532844 | 0.08144 | 0.094975 |
| 11 | 101150200 | 101156399 | 30.70946 | 49.10518 | 133.5701 | 56.18784 | -1.09795 | 2.048775 | 0.09498 |
| 9 | 60417400 | 60418599 | 3.887305 | 7.387916 | 31.90962 | 12.81926 | -1.5942 | -0.02011 | 0.094982 |
| 5 | 122375600 | 122379999 | 19.02817 | 31.10972 | 88.69023 | 38.13555 | -1.13222 | 1.472345 | 0.094983 |
| 2 | 38330400 | 38332799 | 10.60678 | 18.6037 | 50.42762 | 22.29047 | -1.04361 | 0.627585 | 0.094984 |
| 4 | 139886600 | 139896599 | 69.78845 | 107.4557 | 232.9656 | 96.76098 | -0.8179 | 2.923582 | 0.095 |
| 2 | 167744200 | 167748399 | 22.82558 | 36.94122 | 101.7629 | 43.52445 | -1.09514 | 1.660454 | 0.095006 |
| 13 | 30731800 | 30735399 | 15.65245 | 26.51903 | 66.04344 | 28.91925 | -0.92998 | 1.047694 | 0.095016 |
| 12 | 86728400 | 86750599 | 130.5013 | 191.7833 | 443.4278 | 182.6192 | -0.96939 | 3.827897 | 0.095049 |
| 12 | 82179600 | 82182799 | 38.76133 | 64.05639 | 46.13161 | 19.93259 | 0.852584 | 1.352408 | 0.095066 |
| 19 | 27191000 | 27197399 | 33.96275 | 54.56022 | 118.3334 | 50.12472 | -0.76408 | 1.902977 | 0.095073 |
| 7 | 28587000 | 28598599 | 56.68388 | 87.80264 | 221.4637 | 91.98998 | -1.02926 | 2.805317 | 0.095091 |
| 7 | 22585600 | 22592799 | 248.0507 | 365.3457 | 247.5975 | 101.9626 | 0.803149 | 3.799787 | 0.095135 |
| 3 | 87447600 | 87460799 | 66.92355 | 102.5682 | 253.9075 | 105.3921 | -1.01415 | 3.015601 | 0.095157 |
| 14 | 14750000 | 14750599 | 3.541761 | 6.873404 | 27.16049 | 10.21722 | -1.37694 | -0.26052 | 0.095159 |
| 1 | 162484400 | 162485599 | 5.914263 | 11.1259 | 33.42756 | 13.70778 | -1.10432 | 0.072877 | 0.095169 |
| 19 | 46488200 | 46497999 | 41.3454 | 65.18124 | 177.3276 | 73.58727 | -1.1182 | 2.469643 | 0.095171 |
| 5 | 107868000 | 107876799 | 65.18524 | 100.8704 | 219.8765 | 91.33121 | -0.82014 | 2.826969 | 0.095171 |
| 6 | 66799200 | 66799399 | 9.958848 | 19.44247 | 10.01972 | 3.538631 | 1.60477 | -0.46142 | 0.095177 |
| 4 | 18026800 | 18027999 | 13.33828 | 25.44116 | 16.47465 | 6.568259 | 1.125065 | 1.91E-05 | 0.095193 |
| 8 | 72037000 | 72041399 | 18.81772 | 30.90796 | 85.2212 | 36.71295 | -1.08096 | 1.40817 | 0.095195 |
| 8 | 110898000 | 110904599 | 44.19269 | 70.18254 | 151.5647 | 63.18443 | -0.77655 | 2.280985 | 0.095214 |
| 7 | 20509800 | 20509999 | 6.517902 | 12.73782 | 8.967931 | 2.877548 | 1.284365 | -0.84372 | 0.095216 |
| 15 | 85086400 | 85115999 | 238.7984 | 335.176 | 687.2208 | 284.7905 | -0.86185 | 4.522296 | 0.095218 |
| 17 | 81512600 | 81515599 | 33.18555 | 56.44783 | 31.85832 | 13.96318 | 1.20355 | 1.038851 | 0.095219 |
| 2 | 15460600 | 15461799 | 17.58635 | 32.32731 | 19.09165 | 8.111125 | 1.184236 | 0.269581 | 0.095221 |
| 11 | 61468600 | 61471799 | 17.9279 | 30.09686 | 72.7897 | 31.77809 | -0.88871 | 1.143091 | 0.095239 |
| 10 | 35139800 | 35141799 | 16.21163 | 30.67031 | 14.87241 | 6.032736 | 1.516422 | 0.077692 | 0.095241 |
| 11 | 26325400 | 26325999 | 10.87005 | 21.12136 | 14.26341 | 5.316121 | 1.155206 | -0.24969 | 0.095249 |
| 8 | 111154200 | 111157399 | 23.76977 | 38.84599 | 89.08746 | 38.30064 | -0.81996 | 1.488184 | 0.09525 |
| 4 | 150038600 | 150040199 | 7.062174 | 12.57318 | 43.09634 | 18.98457 | -1.3695 | 0.438161 | 0.095262 |

| | | | | | | | | | |
|---|---|---|---|---|---|---|---|---|---|
| 15 | 14891400 | 14892799 | 17.32798 | 32.11303 | 17.44188 | 7.328501 | 1.314903 | 0.219184 | 0.095264 |
| 4 | 125741800 | 125747199 | 33.39162 | 53.44683 | 127.6823 | 53.78633 | -0.90954 | 2.019407 | 0.095269 |
| 8 | 12257200 | 12267799 | 59.30955 | 92.19385 | 210.1349 | 87.24553 | -0.87934 | 2.743262 | 0.09527 |
| 5 | 48793200 | 48796599 | 34.78987 | 58.1853 | 41.13155 | 17.91446 | 0.879751 | 1.205858 | 0.095284 |
| 6 | 125469800 | 125475999 | 25.43638 | 41.01939 | 111.4 | 47.44515 | -1.08069 | 1.767598 | 0.095298 |
| 5 | 132980400 | 132981199 | 3.234417 | 6.316034 | 24.1129 | 8.480065 | -1.2377 | -0.51797 | 0.095301 |
| 9 | 3277600 | 3284399 | 149.302 | 226.9996 | 171.4728 | 71.03306 | 0.687459 | 3.193726 | 0.095306 |
| 5 | 137291400 | 137293599 | 9.892287 | 17.26211 | 52.03004 | 22.90653 | -1.19583 | 0.70903 | 0.095329 |
| 7 | 135467400 | 135472399 | 23.97278 | 39.0665 | 93.58613 | 40.19303 | -0.88569 | 1.531287 | 0.095347 |
| 11 | 33556000 | 33559399 | 16.30178 | 27.1127 | 74.84873 | 32.48363 | -1.07863 | 1.243978 | 0.095355 |
| 14 | 57621800 | 57622799 | 3.575332 | 6.974578 | 27.71033 | 9.914714 | -1.31772 | -0.43789 | 0.095361 |
| 18 | 77577600 | 77584799 | 31.79263 | 50.89743 | 128.5839 | 54.18115 | -0.98969 | 2.011679 | 0.095363 |
| 17 | 20048200 | 20049199 | 21.25422 | 37.67283 | 23.6592 | 10.45962 | 1.059143 | 0.517361 | 0.095371 |
| 4 | 118382200 | 118406199 | 256.0611 | 356.2381 | 755.6745 | 313.5898 | -0.92378 | 4.656001 | 0.095385 |
| 7 | 7996600 | 8003999 | 88.23427 | 140.7293 | 62.84893 | 26.34309 | 1.499504 | 2.167131 | 0.095386 |
| 17 | 87979400 | 87980799 | 6.654402 | 12.1237 | 38.68792 | 16.63021 | -1.24304 | 0.283463 | 0.095398 |
| 9 | 119460400 | 119468999 | 46.1151 | 72.49886 | 185.8352 | 77.08987 | -1.03566 | 2.529282 | 0.095409 |
| 11 | 4056000 | 4077799 | 141.1508 | 207.8299 | 421.0116 | 173.3754 | -0.77613 | 3.79823 | 0.09541 |
| 7 | 22481000 | 22484399 | 109.7258 | 170.9287 | 121.673 | 50.50104 | 0.800044 | 2.719441 | 0.095411 |
| 3 | 151870000 | 151874999 | 27.44239 | 44.35359 | 102.4934 | 43.77812 | -0.84454 | 1.701642 | 0.095412 |
| 5 | 28360200 | 28369199 | 54.87037 | 85.80358 | 201.183 | 83.46474 | -0.91337 | 2.635216 | 0.095419 |
| 7 | 32166400 | 32178199 | 196.7851 | 294.3441 | 208.2256 | 85.86675 | 0.761751 | 3.526034 | 0.095428 |
| 9 | 65536400 | 65539799 | 18.11333 | 29.9269 | 80.47428 | 34.76585 | -1.04196 | 1.329798 | 0.095433 |
| 8 | 10671000 | 10672399 | 7.156307 | 13.27162 | 36.97492 | 15.52096 | -1.02463 | 0.180294 | 0.095447 |
| 12 | 116663400 | 116664799 | 14.65765 | 27.90248 | 16.439 | 6.581509 | 1.255523 | 0.027003 | 0.09545 |
| 14 | 51744200 | 51745199 | 9.046765 | 15.97058 | 45.11783 | 19.9625 | -1.09951 | 0.528664 | 0.095456 |
| 17 | 17906800 | 17907399 | 3.765282 | 7.327611 | 25.76769 | 9.484227 | -1.18108 | -0.339 | 0.095463 |
| 8 | 127921400 | 127922799 | 4.85902 | 9.320157 | 30.70903 | 12.06597 | -1.18058 | -0.09011 | 0.095466 |
| 15 | 83984200 | 83996399 | 67.724 | 104.2462 | 238.8285 | 99.19211 | -0.89775 | 2.928655 | 0.095477 |
| 1 | 82150400 | 82152199 | 13.30652 | 23.04631 | 56.42583 | 24.75617 | -0.89974 | 0.828214 | 0.095477 |
| 4 | 129392400 | 129394999 | 10.0233 | 17.52484 | 50.43528 | 22.23443 | -1.12921 | 0.678192 | 0.095482 |
| 11 | 65724200 | 65725399 | 5.789028 | 10.91218 | 35.23636 | 14.40051 | -1.20434 | 0.05982 | 0.095488 |
| 6 | 126955000 | 126957799 | 16.77841 | 28.32336 | 67.71842 | 29.61996 | -0.87171 | 1.08548 | 0.095492 |
| 1 | 194482000 | 194483599 | 4.727814 | 9.081619 | 32.60854 | 12.75921 | -1.29828 | -0.10204 | 0.095495 |
| 1 | 152149000 | 152151599 | 11.21647 | 19.6873 | 50.41893 | 22.30027 | -0.96203 | 0.621142 | 0.0955 |
| 16 | 53817800 | 53818399 | 5.288586 | 10.11998 | 31.209 | 12.34656 | -1.09575 | -0.06171 | 0.095506 |
| 13 | 66705800 | 66709399 | 88.52544 | 140.6769 | 79.65788 | 33.22899 | 1.152439 | 2.283718 | 0.095515 |
| 9 | 66702200 | 66703399 | 7.25684 | 13.15047 | 39.12842 | 16.93292 | -1.14937 | 0.32132 | 0.095515 |
| 10 | 110290600 | 110292199 | 8.15468 | 14.39111 | 45.64524 | 20.19716 | -1.26533 | 0.528004 | 0.095522 |
| 14 | 90903000 | 90905999 | 33.60303 | 56.33666 | 39.44971 | 17.20823 | 0.893124 | 1.17312 | 0.095531 |
| 17 | 56314200 | 56344199 | 147.3506 | 213.0833 | 532.538 | 219.6511 | -1.10411 | 4.077489 | 0.095532 |
| 8 | 89755000 | 89759999 | 30.14831 | 48.34627 | 122.1726 | 51.61141 | -0.98709 | 1.955359 | 0.095546 |
| 4 | 154116800 | 154122799 | 25.14448 | 40.43792 | 115.5512 | 49.0235 | -1.15868 | 1.856201 | 0.095546 |
| 8 | 120144600 | 120200799 | 321.6405 | 434.7346 | 1067.017 | 444.7971 | -1.17631 | 5.106588 | 0.095562 |
| 5 | 24620400 | 24625999 | 26.35492 | 42.53336 | 105.7532 | 45.08487 | -0.95234 | 1.744834 | 0.095571 |
| 1 | 151514200 | 151516999 | 28.68365 | 49.09087 | 34.54787 | 15.14947 | 0.887728 | 0.969035 | 0.095584 |
| X | 114598000 | 114598799 | 12.30399 | 23.76028 | 15.24304 | 5.868127 | 1.185517 | -0.12317 | 0.095624 |

| | | | | | | | | | |
|---|---|---|---|---|---|---|---|---|---|
| 14 | 7506800 | 7507799 | 12.07488 | 23.52014 | 10.99908 | 4.012677 | 1.703979 | -0.31822 | 0.095628 |
| 11 | 69562000 | 69563199 | 5.397136 | 9.99672 | 35.74723 | 15.03105 | -1.38163 | 0.173016 | 0.095629 |
| 4 | 151451200 | 151472999 | 126.2668 | 186.8096 | 406.3884 | 167.3621 | -0.87238 | 3.716517 | 0.095635 |
| 7 | 67397200 | 67400999 | 19.13764 | 31.79368 | 75.79007 | 32.86679 | -0.86806 | 1.265303 | 0.095636 |
| X | 17371400 | 17372799 | 20.95095 | 38.42283 | 16.94588 | 7.222267 | 1.599477 | 0.305454 | 0.095636 |
| 4 | 60203600 | 60205999 | 18.39739 | 33.23477 | 21.80484 | 9.473058 | 1.012944 | 0.379017 | 0.095663 |
| 12 | 23941000 | 23944599 | 71.10945 | 113.4533 | 76.14643 | 31.93124 | 0.913544 | 2.140847 | 0.095664 |
| 13 | 120670200 | 120672999 | 23.60864 | 41.22824 | 28.68518 | 12.63479 | 0.908269 | 0.729465 | 0.095668 |
| 4 | 6231400 | 6234399 | 28.84801 | 49.24089 | 35.14083 | 15.40619 | 0.867147 | 0.994057 | 0.095674 |
| 6 | 84243400 | 84248999 | 34.98969 | 56.01434 | 126.4569 | 53.2831 | -0.82786 | 2.015561 | 0.095675 |
| 2 | 132657400 | 132658399 | 5.83202 | 10.7939 | 35.74867 | 15.04743 | -1.27346 | 0.180979 | 0.095677 |
| 6 | 93486800 | 93488199 | 7.041711 | 12.96976 | 37.32636 | 15.81552 | -1.08084 | 0.219543 | 0.09569 |
| 4 | 138027000 | 138048199 | 120.7084 | 178.9285 | 396.0882 | 163.134 | -0.89492 | 3.68321 | 0.095693 |
| 16 | 77366400 | 77367799 | 16.81639 | 30.94241 | 19.78548 | 8.394648 | 1.071974 | 0.259911 | 0.095708 |
| 7 | 83734200 | 83736199 | 12.8489 | 21.84106 | 64.18069 | 28.12823 | -1.1669 | 1.007539 | 0.095732 |
| 4 | 16347400 | 16348199 | 14.69066 | 27.83439 | 16.84772 | 6.812432 | 1.204224 | 0.058893 | 0.095739 |
| 14 | 96148000 | 96149799 | 22.22342 | 39.71329 | 20.79503 | 9.131785 | 1.325968 | 0.467644 | 0.095742 |
| 13 | 53615600 | 53617599 | 12.38431 | 21.43195 | 55.83009 | 24.4942 | -0.98889 | 0.830279 | 0.095747 |
| 17 | 57538600 | 57545199 | 33.99829 | 54.15983 | 145.4605 | 60.86737 | -1.08576 | 2.174536 | 0.095749 |
| 1 | 26132400 | 26133599 | 13.00314 | 25.01921 | 13.61385 | 5.301507 | 1.403075 | -0.08626 | 0.095754 |
| 2 | 154905200 | 154910999 | 27.90046 | 44.76029 | 121.6221 | 51.41155 | -1.09071 | 1.939685 | 0.095755 |
| 6 | 60313400 | 60314199 | 14.51502 | 27.87041 | 14.80934 | 5.801453 | 1.430875 | -0.04673 | 0.095765 |
| 8 | 118739400 | 118742999 | 20.80329 | 34.30335 | 81.97454 | 35.37288 | -0.87307 | 1.361069 | 0.095774 |
| 18 | 68243600 | 68247999 | 20.80329 | 34.30335 | 81.97454 | 35.37288 | -0.87307 | 1.361069 | 0.095774 |
| 17 | 20648000 | 20651199 | 58.05786 | 93.58367 | 56.68385 | 24.12761 | 1.078158 | 1.806958 | 0.09578 |
| 5 | 113260200 | 113268199 | 42.54843 | 67.23792 | 166.943 | 69.34517 | -0.98378 | 2.414239 | 0.095789 |
| 2 | 120049200 | 120051199 | 35.88065 | 59.89992 | 41.64222 | 18.11595 | 0.903798 | 1.22804 | 0.095794 |
| 5 | 138255400 | 138262799 | 30.75232 | 49.12425 | 141.2415 | 59.33029 | -1.17946 | 2.081615 | 0.095797 |
| 11 | 115772000 | 115797399 | 118.4347 | 173.3316 | 476.0577 | 196.1081 | -1.2233 | 3.892597 | 0.095804 |
| 11 | 75789000 | 75792599 | 15.02815 | 25.1056 | 73.27011 | 31.88914 | -1.15837 | 1.188869 | 0.09586 |
| 11 | 94767200 | 94770999 | 13.76662 | 23.42031 | 64.8755 | 28.43462 | -1.08216 | 1.002701 | 0.095868 |
| 15 | 86082000 | 86089799 | 35.77351 | 56.95676 | 147.6612 | 61.74174 | -1.03579 | 2.194109 | 0.095868 |
| 1 | 108883200 | 108885199 | 43.69669 | 71.18355 | 51.06751 | 21.92943 | 0.848733 | 1.549453 | 0.095877 |
| 2 | 71503400 | 71504799 | 7.12468 | 12.88578 | 39.62259 | 17.18871 | -1.19881 | 0.332566 | 0.095878 |
| 3 | 43768400 | 43769199 | 13.34162 | 25.38301 | 17.19741 | 6.893332 | 1.054407 | 0.020044 | 0.095884 |
| 9 | 90192000 | 90192799 | 4.536934 | 8.79415 | 30.35574 | 11.48342 | -1.19492 | -0.22379 | 0.095891 |
| 3 | 9136600 | 9140199 | 18.90759 | 31.20932 | 81.13287 | 35.02724 | -0.99355 | 1.343344 | 0.09591 |
| 3 | 90374800 | 90376399 | 9.827181 | 17.29687 | 48.26399 | 21.3672 | -1.08411 | 0.569733 | 0.095912 |
| 11 | 72307600 | 72310599 | 10.23776 | 17.80677 | 56.89365 | 24.99645 | -1.28051 | 0.762919 | 0.095914 |
| 15 | 35292400 | 35294399 | 14.96045 | 25.12866 | 69.00018 | 30.13386 | -1.0707 | 1.125991 | 0.095916 |
| 4 | 129228400 | 129230599 | 8.537933 | 14.94457 | 51.98609 | 22.93983 | -1.4008 | 0.653404 | 0.095951 |
| 17 | 20098400 | 20101599 | 71.46674 | 114.3944 | 70.62148 | 29.75097 | 1.038922 | 2.043492 | 0.095954 |
| 7 | 33518800 | 33521199 | 74.82778 | 118.8718 | 89.48162 | 37.31498 | 0.741958 | 2.282122 | 0.095974 |
| 7 | 32205800 | 32210799 | 60.2641 | 97.08169 | 60.9452 | 25.91977 | 1.025615 | 1.828712 | 0.096012 |
| 1 | 167246000 | 167247399 | 7.459729 | 13.81126 | 39.18851 | 16.48754 | -1.05341 | 0.197333 | 0.096028 |
| 7 | 31713400 | 31714399 | 12.10671 | 23.18621 | 15.29269 | 6.035404 | 1.112455 | -0.04756 | 0.096031 |
| 11 | 40529800 | 40530399 | 10.8289 | 21.12827 | 11.43474 | 4.080741 | 1.525731 | -0.40694 | 0.096039 |

| | | | | | | | | | |
|---|---|---|---|---|---|---|---|---|---|
| 5 | 135081000 | 135085999 | 19.54142 | 31.87909 | 91.43133 | 39.28288 | -1.1432 | 1.506702 | 0.096047 |
| 8 | 120202200 | 120209799 | 40.28192 | 63.70072 | 166.7826 | 69.30303 | -1.05939 | 2.395684 | 0.096053 |
| 5 | 38553000 | 38558399 | 24.32613 | 39.55065 | 95.03184 | 40.7795 | -0.89241 | 1.566716 | 0.096054 |
| 3 | 3523800 | 3525599 | 18.36605 | 34.56797 | 14.15066 | 5.79776 | 1.746613 | 0.129034 | 0.096088 |
| 7 | 42699200 | 42701599 | 56.06868 | 90.44109 | 60.81363 | 25.90903 | 0.928897 | 1.783211 | 0.096094 |
| 10 | 56261200 | 56264999 | 33.20969 | 53.20562 | 124.526 | 52.53065 | -0.8784 | 1.988798 | 0.096104 |
| 3 | 105034000 | 105035999 | 9.125168 | 16.3138 | 42.60122 | 18.70073 | -0.97725 | 0.422675 | 0.096123 |
| 3 | 149222400 | 149225799 | 13.64064 | 23.51925 | 60.25386 | 26.43206 | -0.96652 | 0.872952 | 0.096128 |
| 7 | 14437000 | 14439199 | 47.59019 | 77.63884 | 50.64539 | 21.75968 | 0.987018 | 1.538328 | 0.096134 |
| 3 | 14053400 | 14054399 | 16.03533 | 29.57972 | 17.70636 | 7.486787 | 1.170285 | 0.235413 | 0.096134 |
| 1 | 7984800 | 7986199 | 20.83071 | 36.9723 | 21.59672 | 9.538349 | 1.164262 | 0.502114 | 0.096147 |
| 4 | 141091600 | 141115599 | 119.3063 | 176.1094 | 427.6861 | 176.1288 | -1.03415 | 3.753942 | 0.096166 |
| 3 | 7256200 | 7259799 | 28.27149 | 49.17552 | 25.99411 | 11.42095 | 1.301833 | 0.801861 | 0.096175 |
| 11 | 86469800 | 86471199 | 5.605586 | 10.57561 | 32.9282 | 13.43552 | -1.14896 | 0.050581 | 0.096176 |
| 11 | 98274000 | 98282199 | 33.4133 | 53.08112 | 154.1862 | 64.29834 | -1.20276 | 2.264059 | 0.096179 |
| 4 | 130695600 | 130709399 | 95.24372 | 143.7066 | 311.9809 | 128.9338 | -0.84361 | 3.344927 | 0.096207 |
| 6 | 88191600 | 88219999 | 180.6598 | 258.6521 | 576.7591 | 238.2373 | -0.95406 | 4.227436 | 0.096213 |
| 5 | 19430200 | 19433399 | 15.3836 | 25.59654 | 75.92191 | 32.95911 | -1.18172 | 1.236423 | 0.096232 |
| 12 | 107929400 | 107936599 | 42.78457 | 67.4469 | 175.1523 | 72.68413 | -1.05127 | 2.471394 | 0.096232 |
| 4 | 134619000 | 134621199 | 10.82995 | 18.83617 | 54.74402 | 24.05151 | -1.14455 | 0.764041 | 0.096243 |
| 4 | 112097200 | 112099599 | 45.09166 | 73.31153 | 54.34614 | 23.3206 | 0.799661 | 1.582903 | 0.096249 |
| 4 | 156026000 | 156028999 | 15.93293 | 27.10117 | 64.47134 | 28.2571 | -0.86367 | 1.012716 | 0.096258 |
| X | 122857600 | 122859399 | 20.54458 | 37.78973 | 16.8568 | 7.155303 | 1.587085 | 0.285893 | 0.096263 |
| 5 | 125270000 | 125284199 | 89.24455 | 134.7353 | 310.9182 | 128.5188 | -0.93085 | 3.331707 | 0.096263 |
| 7 | 136798000 | 136804199 | 38.21937 | 60.74109 | 151.1225 | 63.04664 | -0.97929 | 2.254343 | 0.096284 |
| 11 | 113347000 | 113364999 | 136.3821 | 201.592 | 405.2459 | 166.8899 | -0.75976 | 3.732769 | 0.096305 |
| 4 | 151291200 | 151330799 | 242.2677 | 337.8246 | 747.7888 | 310.2065 | -0.98184 | 4.620685 | 0.096306 |
| 12 | 106727400 | 106729999 | 22.74271 | 39.81023 | 27.56233 | 12.16291 | 0.915214 | 0.686328 | 0.096322 |
| 7 | 136914000 | 136925399 | 85.15811 | 129.0267 | 299.3319 | 123.8693 | -0.93409 | 3.257244 | 0.096341 |
| 7 | 61867600 | 61870199 | 33.15447 | 56.81663 | 30.55301 | 13.39911 | 1.274408 | 0.972597 | 0.09636 |
| 17 | 87627200 | 87640599 | 91.71313 | 138.3113 | 318.9149 | 131.7789 | -0.93106 | 3.354408 | 0.096365 |
| 1 | 10427000 | 10431799 | 49.64969 | 80.54278 | 55.10292 | 23.60865 | 0.91224 | 1.634302 | 0.096379 |
| 1 | 8084200 | 8085199 | 12.90346 | 24.81647 | 15.45956 | 6.027884 | 1.209529 | -0.07707 | 0.096383 |
| 7 | 14643000 | 14645199 | 56.25612 | 90.98904 | 54.6678 | 23.37096 | 1.095264 | 1.706777 | 0.096404 |
| 14 | 95289200 | 95290399 | 11.1657 | 21.72761 | 12.2539 | 4.513518 | 1.425335 | -0.29406 | 0.096412 |
| 7 | 33214200 | 33220999 | 145.3346 | 222.0175 | 157.5769 | 65.3591 | 0.782736 | 3.096775 | 0.096431 |
| 11 | 63809200 | 63818199 | 67.6101 | 103.9368 | 242.1318 | 100.5463 | -0.92325 | 2.954482 | 0.096445 |
| X | 91105600 | 91105999 | 7.247322 | 14.17096 | 8.748568 | 2.838442 | 1.455499 | -0.79376 | 0.096454 |
| 9 | 19068200 | 19069399 | 30.25697 | 51.33516 | 35.81856 | 15.69104 | 0.898648 | 1.05447 | 0.096476 |
| 7 | 31800200 | 31802199 | 48.94731 | 79.48271 | 55.03552 | 23.59715 | 0.896294 | 1.614662 | 0.096488 |
| 16 | 4824000 | 4833399 | 39.62144 | 62.71924 | 165.7672 | 68.91611 | -1.07179 | 2.370512 | 0.096521 |
| 3 | 90712600 | 90714399 | 16.87979 | 31.51284 | 17.67195 | 7.341056 | 1.28207 | 0.174119 | 0.096533 |
| 2 | 151731800 | 151741799 | 60.80632 | 94.43363 | 211.1436 | 87.67639 | -0.8527 | 2.760009 | 0.096544 |
| 15 | 88701800 | 88710599 | 33.41736 | 53.1468 | 154.5268 | 64.51395 | -1.20261 | 2.229761 | 0.096552 |
| 8 | 122452400 | 122461399 | 76.01107 | 116.2918 | 256.0086 | 106.2231 | -0.8471 | 3.053921 | 0.096566 |
| 2 | 144300200 | 144302599 | 13.66288 | 23.49902 | 59.63395 | 26.15814 | -0.95359 | 0.895576 | 0.096567 |
| 11 | 98265200 | 98273199 | 26.70858 | 42.73437 | 132.395 | 55.74285 | -1.28408 | 2.025441 | 0.096588 |

| | | | | | | | | | |
|---|---|---|---|---|---|---|---|---|---|
| 15 | 75767400 | 75770599 | 11.30723 | 19.60443 | 55.80779 | 24.48581 | -1.11555 | 0.808064 | 0.096592 |
| X | 25557400 | 25559199 | 28.64267 | 50.43372 | 18.76256 | 8.305822 | 1.805753 | 0.624157 | 0.096593 |
| 18 | 61775800 | 61794399 | 87.75243 | 132.5047 | 320.5802 | 132.5195 | -0.99884 | 3.327903 | 0.096597 |
| X | 118800200 | 118801599 | 17.31869 | 33.4788 | 9.318165 | 3.569915 | 2.378324 | -0.10151 | 0.096598 |
| 8 | 122085600 | 122105799 | 116.8752 | 173.2072 | 404.2177 | 166.4784 | -0.97136 | 3.686798 | 0.096612 |
| 8 | 92591000 | 92594199 | 15.62113 | 26.16951 | 72.4031 | 31.5676 | -1.08177 | 1.158346 | 0.096623 |
| 2 | 118531200 | 118551599 | 117.7852 | 173.3334 | 456.8559 | 188.1352 | -1.15634 | 3.805058 | 0.096623 |
| 7 | 73207400 | 73208799 | 10.17968 | 17.7738 | 50.97141 | 22.44856 | -1.12427 | 0.701383 | 0.096639 |
| 11 | 63690200 | 63691999 | 15.16444 | 28.37753 | 18.39751 | 7.611891 | 1.079023 | 0.145931 | 0.096643 |
| 5 | 130415200 | 130416999 | 11.24706 | 19.6205 | 52.05667 | 22.91408 | -1.01299 | 0.721688 | 0.09665 |
| 11 | 52411000 | 52412199 | 3.840044 | 7.432431 | 28.18541 | 10.73568 | -1.33656 | -0.21182 | 0.096655 |
| 2 | 20524200 | 20527599 | 14.18826 | 24.23068 | 63.93558 | 28.03135 | -1.01154 | 0.967026 | 0.096674 |
| 9 | 60325600 | 60333199 | 57.92495 | 90.24357 | 205.2107 | 85.17256 | -0.87345 | 2.703227 | 0.096689 |
| 10 | 110148200 | 110149999 | 18.54615 | 33.35771 | 22.30854 | 9.742984 | 0.980677 | 0.405716 | 0.096691 |
| 9 | 118829000 | 118831999 | 27.47648 | 44.28932 | 107.6966 | 45.84114 | -0.92239 | 1.784544 | 0.096697 |
| 8 | 73447800 | 73450999 | 21.05136 | 34.47769 | 86.49734 | 37.21752 | -0.94698 | 1.450232 | 0.096703 |
| 5 | 83409800 | 83410399 | 9.582261 | 18.67407 | 13.22221 | 4.79271 | 1.122922 | -0.36553 | 0.096726 |
| 7 | 17196400 | 17198399 | 20.60564 | 36.89984 | 21.3992 | 9.380447 | 1.181184 | 0.44429 | 0.09673 |
| 1 | 162269200 | 162270399 | 17.68573 | 33.38844 | 14.96385 | 6.097032 | 1.624454 | 0.106475 | 0.096736 |
| X | 109128800 | 109131599 | 32.06721 | 55.55157 | 25.99757 | 11.40621 | 1.476202 | 0.868039 | 0.096741 |
| 9 | 3089200 | 3091799 | 107.7874 | 167.6122 | 117.584 | 48.81682 | 0.81909 | 2.74798 | 0.096744 |
| 12 | 84280000 | 84283399 | 19.64484 | 32.54447 | 81.05317 | 35.08086 | -0.93158 | 1.301418 | 0.096766 |
| 19 | 39831800 | 39832199 | 7.779687 | 15.20428 | 10.72931 | 3.711784 | 1.186772 | -0.56138 | 0.096768 |
| 17 | 10465400 | 10466399 | 4.322081 | 8.391887 | 33.12185 | 12.41942 | -1.37326 | -0.24686 | 0.096777 |
| 13 | 12932000 | 12933199 | 11.9839 | 23.02641 | 16.2202 | 6.34146 | 1.030769 | -0.07591 | 0.096795 |
| 10 | 54323800 | 54323999 | 10.5034 | 20.44149 | 13.6441 | 5.02027 | 1.187917 | -0.30064 | 0.096854 |
| 3 | 97535200 | 97536999 | 6.377565 | 11.95889 | 34.49849 | 14.22429 | -1.05334 | 0.098549 | 0.096857 |
| 10 | 58090600 | 58091799 | 6.377565 | 11.95889 | 34.49849 | 14.22429 | -1.05334 | 0.098549 | 0.096857 |
| 5 | 109595600 | 109596799 | 19.3154 | 35.87477 | 15.65265 | 6.557134 | 1.631093 | 0.22063 | 0.096859 |
| 1 | 133982800 | 133984599 | 6.168742 | 11.3793 | 37.3305 | 15.78434 | -1.26559 | 0.203862 | 0.096863 |
| 11 | 120468400 | 120473399 | 20.84525 | 33.95242 | 96.99979 | 41.65133 | -1.13947 | 1.534094 | 0.096871 |
| 15 | 27466200 | 27467999 | 8.42799 | 14.94766 | 44.07859 | 19.47985 | -1.15809 | 0.47194 | 0.096873 |
| 12 | 94959000 | 94960999 | 26.58793 | 46.32468 | 28.42228 | 12.50096 | 1.088532 | 0.775961 | 0.09688 |
| 11 | 60073000 | 60087999 | 90.95343 | 136.9026 | 326.3451 | 134.7787 | -0.98146 | 3.392593 | 0.09689 |
| 16 | 18398400 | 18416399 | 123.9454 | 182.2738 | 443.0838 | 182.4697 | -1.04064 | 3.816479 | 0.096906 |
| 3 | 14531400 | 14531999 | 10.82058 | 21.1627 | 6.878455 | 2.330615 | 2.305353 | -0.59589 | 0.096919 |
| 16 | 30053000 | 30066399 | 86.45319 | 131.1654 | 302.486 | 125.2261 | -0.92389 | 3.230576 | 0.096926 |
| 4 | 99536800 | 99546599 | 83.88852 | 128.0196 | 267.4211 | 110.8935 | -0.77441 | 3.107946 | 0.096926 |
| 8 | 71360600 | 71372799 | 68.23752 | 104.6341 | 252.9813 | 105.0148 | -0.97961 | 3.005822 | 0.096929 |
| 5 | 114325400 | 114345999 | 127.1131 | 187.5961 | 416.5326 | 171.5352 | -0.90523 | 3.75799 | 0.096955 |
| 16 | 68947600 | 68949199 | 16.13483 | 30.08443 | 18.24074 | 7.593407 | 1.167838 | 0.176326 | 0.096962 |
| 15 | 28833800 | 28834399 | 9.642849 | 18.80973 | 12.67727 | 4.539248 | 1.208961 | -0.4081 | 0.096966 |
| 8 | 22042200 | 22044799 | 25.83875 | 45.37434 | 22.64381 | 10.01942 | 1.384201 | 0.645458 | 0.096969 |
| 5 | 139314200 | 139320399 | 23.23716 | 37.43926 | 113.1744 | 48.14319 | -1.23625 | 1.793062 | 0.096988 |
| 8 | 126361200 | 126361799 | 10.68253 | 20.78858 | 13.64003 | 5.021745 | 1.211721 | -0.29618 | 0.096995 |
| 7 | 58878000 | 58879599 | 11.14404 | 19.30549 | 55.22151 | 24.22796 | -1.12244 | 0.816059 | 0.096996 |
| 12 | 22686800 | 22689599 | 38.679 | 64.20767 | 40.1626 | 17.40731 | 1.055538 | 1.296473 | 0.097009 |

| | | | | | | | | | |
|---|---|---|---|---|---|---|---|---|---|
| 12 | 21879400 | 21883199 | 115.0976 | 177.9216 | 136.3428 | 56.63368 | 0.685266 | 2.856544 | 0.097017 |
| 15 | 83162800 | 83166599 | 22.2789 | 36.58184 | 84.64604 | 36.4605 | -0.82921 | 1.42005 | 0.09703 |
| 7 | 45052600 | 45082399 | 144.3977 | 208.6751 | 537.742 | 221.8169 | -1.14921 | 4.087186 | 0.097038 |
| 1 | 72587000 | 72587799 | 5.634737 | 10.53017 | 34.32425 | 14.23403 | -1.2335 | 0.116824 | 0.097046 |
| 12 | 24459000 | 24460399 | 25.38459 | 43.98174 | 30.49951 | 13.38146 | 0.912001 | 0.841444 | 0.097047 |
| 14 | 112063000 | 112063999 | 15.28345 | 28.46424 | 19.07832 | 7.957549 | 1.022602 | 0.179014 | 0.097048 |
| 14 | 41810600 | 41812799 | 29.86797 | 51.70918 | 28.86612 | 12.66452 | 1.223092 | 0.866799 | 0.097053 |
| 5 | 120687800 | 120694399 | 30.40559 | 48.58467 | 135.3217 | 56.86726 | -1.13317 | 2.071371 | 0.097061 |
| 1 | 154904400 | 154909399 | 26.66856 | 43.07857 | 109.6286 | 46.79175 | -0.9844 | 1.718658 | 0.097061 |
| 8 | 89105400 | 89117999 | 89.91831 | 136.803 | 274.984 | 113.9355 | -0.72253 | 3.172706 | 0.097065 |
| 17 | 15365400 | 15367199 | 8.809343 | 15.78957 | 41.58284 | 18.21431 | -0.98701 | 0.402821 | 0.097072 |
| 8 | 59363400 | 59366599 | 19.12512 | 31.81105 | 75.34218 | 32.69966 | -0.85865 | 1.250099 | 0.097073 |
| 14 | 61921400 | 61925399 | 16.88554 | 27.89352 | 83.1872 | 35.93358 | -1.19079 | 1.330048 | 0.097074 |
| 10 | 3199800 | 3202999 | 92.01003 | 146.6704 | 60.39704 | 25.29603 | 1.615516 | 2.189501 | 0.097077 |
| 9 | 96001200 | 96002799 | 7.5394 | 13.71612 | 38.6713 | 16.65333 | -1.06788 | 0.298496 | 0.097081 |
| 4 | 129194800 | 129196999 | 22.24993 | 38.99748 | 24.76193 | 10.93895 | 1.039322 | 0.659027 | 0.097101 |
| 13 | 61639200 | 61640599 | 23.10822 | 41.17099 | 20.75982 | 9.139363 | 1.377725 | 0.491819 | 0.097106 |
| 2 | 173087000 | 173098599 | 48.11636 | 75.15958 | 203.6526 | 84.49884 | -1.12292 | 2.652257 | 0.097115 |
| 7 | 20750600 | 20757799 | 104.9069 | 163.2128 | 120.621 | 50.07397 | 0.744561 | 2.737993 | 0.09714 |
| 10 | 81752200 | 81755999 | 73.86599 | 118.0144 | 70.423 | 29.58906 | 1.084939 | 2.101503 | 0.097149 |
| 12 | 75864800 | 75866599 | 9.249683 | 16.16813 | 50.41152 | 22.21917 | -1.24366 | 0.680105 | 0.097167 |
| 7 | 17759000 | 17761999 | 40.85616 | 67.65711 | 39.56267 | 17.11853 | 1.152094 | 1.328092 | 0.097172 |
| 7 | 26951000 | 26974799 | 141.4419 | 206.8391 | 461.5906 | 190.158 | -0.92511 | 3.90685 | 0.097178 |
| 5 | 134145600 | 134156199 | 71.08043 | 109.1812 | 243.4327 | 101.075 | -0.86096 | 2.972033 | 0.097188 |
| 6 | 143226400 | 143228199 | 15.98654 | 29.38985 | 19.95899 | 8.478319 | 0.984585 | 0.258337 | 0.09722 |
| 10 | 80429000 | 80431199 | 6.555805 | 12.17848 | 35.31727 | 14.79116 | -1.07885 | 0.16367 | 0.097232 |
| 5 | 125620000 | 125627799 | 57.26475 | 89.62629 | 195.7072 | 81.18571 | -0.80983 | 2.620317 | 0.097234 |
| 12 | 33102800 | 33104199 | 7.294297 | 13.10653 | 40.57839 | 17.7372 | -1.21581 | 0.37606 | 0.097236 |
| 17 | 32120600 | 32141599 | 115.9342 | 171.4067 | 411.6963 | 169.5468 | -1.01605 | 3.726832 | 0.09726 |
| 2 | 172610200 | 172620399 | 66.03583 | 101.7733 | 234.9452 | 97.5924 | -0.90734 | 2.903671 | 0.097268 |
| 7 | 100718000 | 100721199 | 14.46485 | 24.79477 | 60.14302 | 26.37644 | -0.88964 | 0.93112 | 0.097268 |
| 16 | 22388800 | 22394799 | 31.82315 | 50.89221 | 132.8785 | 55.9066 | -1.03886 | 2.04573 | 0.097271 |
| 17 | 29773600 | 29776199 | 11.48668 | 19.83574 | 56.82379 | 24.92853 | -1.12583 | 0.847705 | 0.097275 |
| 11 | 86248000 | 86249399 | 6.413176 | 11.91735 | 35.32295 | 14.78707 | -1.10967 | 0.16015 | 0.097278 |
| 1 | 135976600 | 135982799 | 22.03401 | 35.48033 | 123.8263 | 52.63044 | -1.44414 | 1.812693 | 0.097282 |
| 13 | 92844600 | 92847399 | 10.13976 | 17.82056 | 50.39893 | 22.3159 | -1.10443 | 0.591518 | 0.097283 |
| 13 | 66987400 | 66990599 | 60.33858 | 97.28597 | 58.69422 | 24.99195 | 1.084258 | 1.803151 | 0.097305 |
| 7 | 126086200 | 126089799 | 23.80689 | 38.44329 | 105.2836 | 44.91799 | -1.09033 | 1.723893 | 0.097306 |
| 10 | 45598200 | 45599399 | 9.947388 | 17.58047 | 45.1068 | 19.95308 | -0.96111 | 0.520866 | 0.097307 |
| 9 | 30566400 | 30566999 | 3.575204 | 6.974848 | 26.38359 | 9.430809 | -1.24605 | -0.44486 | 0.097312 |
| 12 | 22100800 | 22105599 | 108.9751 | 169.9201 | 120.5789 | 50.03986 | 0.805578 | 2.702385 | 0.097317 |
| 8 | 46501800 | 46503199 | 5.352911 | 10.13127 | 37.80256 | 15.32036 | -1.4007 | 0.032032 | 0.097351 |
| X | 25693800 | 25695799 | 20.21448 | 37.39841 | 15.59472 | 6.56765 | 1.69066 | 0.248931 | 0.097361 |
| 1 | 118591200 | 118592799 | 6.102893 | 11.43461 | 37.9152 | 15.65443 | -1.2548 | 0.107057 | 0.097375 |
| 14 | 96716600 | 96717599 | 10.74197 | 20.79172 | 14.81608 | 5.642646 | 1.049469 | -0.17839 | 0.097379 |
| 15 | 37788400 | 37794399 | 29.14592 | 46.86025 | 116.9501 | 49.59726 | -0.96458 | 1.869317 | 0.097387 |
| 14 | 118872600 | 118881799 | 58.26331 | 90.70344 | 203.5004 | 84.47204 | -0.85489 | 2.716489 | 0.0974 |

| | | | | | | | | | |
|---|---|---|---|---|---|---|---|---|---|
| 12 | 100258800 | 100268599 | 66.4065 | 102.5659 | 224.8169 | 93.38423 | -0.83025 | 2.863038 | 0.097405 |
| 1 | 85438400 | 85444199 | 121.3133 | 187.1162 | 141.2218 | 58.66098 | 0.704792 | 2.899706 | 0.097429 |
| 18 | 5766200 | 5768199 | 14.47892 | 24.77143 | 61.77288 | 27.08706 | -0.92993 | 0.949828 | 0.097435 |
| 14 | 41454000 | 41458199 | 33.59898 | 57.21431 | 32.63081 | 14.30303 | 1.18862 | 1.030613 | 0.097436 |
| 7 | 24620600 | 24648599 | 184.926 | 263.2746 | 612.8593 | 253.3823 | -1.02422 | 4.310343 | 0.097466 |
| 4 | 115747400 | 115751799 | 20.04552 | 32.97257 | 82.67931 | 35.63986 | -0.94363 | 1.391292 | 0.097479 |
| 14 | 46624400 | 46629399 | 28.81959 | 46.35118 | 115.7973 | 49.13231 | -0.96541 | 1.857842 | 0.097488 |
| 7 | 104055400 | 104056399 | 11.99538 | 23.00757 | 16.2714 | 6.394005 | 1.018454 | -0.05935 | 0.0975 |
| 14 | 66544800 | 66547599 | 9.96988 | 17.55188 | 46.15489 | 20.43257 | -0.9986 | 0.566574 | 0.097514 |
| 1 | 109293200 | 109293999 | 16.98871 | 32.10455 | 14.84974 | 6.037798 | 1.581291 | 0.092658 | 0.097527 |
| 12 | 114840600 | 114841799 | 23.46866 | 41.0616 | 28.04622 | 12.37151 | 0.934756 | 0.699068 | 0.09754 |
| 19 | 47416200 | 47421199 | 22.90349 | 37.22301 | 93.91605 | 40.29267 | -0.96301 | 1.570842 | 0.097558 |
| 11 | 19811000 | 19811999 | 5.767173 | 10.79786 | 34.37762 | 14.21106 | -1.19616 | 0.106151 | 0.097558 |
| 7 | 140679400 | 140685599 | 40.0859 | 63.85817 | 140.683 | 58.86915 | -0.80054 | 2.178803 | 0.097588 |
| 18 | 35014600 | 35022199 | 32.05368 | 51.44306 | 121.1009 | 51.22122 | -0.88373 | 1.931069 | 0.09759 |
| 2 | 27232800 | 27246999 | 88.83706 | 134.4868 | 301.8955 | 124.8967 | -0.8877 | 3.275323 | 0.097591 |
| 15 | 64736200 | 64737199 | 12.66815 | 24.55197 | 12.55919 | 4.754048 | 1.528777 | -0.18357 | 0.097595 |
| 11 | 5003800 | 5022399 | 82.59798 | 124.7407 | 318.835 | 131.8045 | -1.07779 | 3.323881 | 0.097599 |
| 3 | 84078800 | 84080599 | 6.591426 | 12.18057 | 36.84685 | 15.54233 | -1.14725 | 0.195894 | 0.097611 |
| 7 | 26108400 | 26114599 | 138.1521 | 211.2841 | 158.4337 | 65.73165 | 0.704769 | 3.071713 | 0.097627 |
| Y | 1722200 | 1725399 | 19.05187 | 34.42678 | 11.83641 | 5.140483 | 1.935763 | 0.355664 | 0.097627 |
| 12 | 110955800 | 110980399 | 175.4247 | 253.8394 | 504.0753 | 207.8942 | -0.77166 | 4.070018 | 0.097644 |
| 14 | 42437400 | 42440999 | 79.72277 | 126.226 | 96.23639 | 40.01383 | 0.71918 | 2.378953 | 0.097649 |
| 1 | 89954800 | 89959199 | 22.14907 | 35.93609 | 97.49418 | 41.77567 | -1.06971 | 1.605788 | 0.09765 |
| 2 | 30224800 | 30227799 | 9.748488 | 17.0116 | 52.56165 | 23.14094 | -1.23143 | 0.708252 | 0.097651 |
| X | 31573200 | 31574799 | 12.59896 | 24.28241 | 15.69565 | 6.082696 | 1.165732 | -0.09764 | 0.097663 |
| 8 | 104276200 | 104279799 | 15.92877 | 27.12792 | 65.61274 | 28.76025 | -0.8872 | 1.00273 | 0.09768 |
| 10 | 18720200 | 18721399 | 6.76293 | 12.40619 | 37.21877 | 15.85664 | -1.14569 | 0.241336 | 0.097688 |
| 3 | 75925600 | 75926799 | 15.57569 | 29.82478 | 13.99682 | 5.526674 | 1.597132 | -0.01391 | 0.097704 |
| 2 | 153266200 | 153268199 | 6.559579 | 11.84652 | 40.64833 | 17.66575 | -1.35739 | 0.338702 | 0.097705 |
| 16 | 91006000 | 91036199 | 244.3554 | 342.5682 | 713.2243 | 295.6213 | -0.88568 | 4.540183 | 0.097719 |
| 1 | 90530800 | 90531599 | 4.549917 | 8.780975 | 29.23357 | 11.28568 | -1.17074 | -0.15522 | 0.097738 |
| 2 | 29623600 | 29631999 | 30.88453 | 49.32303 | 135.5223 | 56.90759 | -1.11445 | 2.089377 | 0.097743 |
| 8 | 26466200 | 26467399 | 6.676319 | 12.4818 | 34.9633 | 14.48897 | -1.01709 | 0.122727 | 0.097754 |
| 17 | 18830600 | 18834399 | 79.29474 | 126.323 | 79.97723 | 33.46805 | 0.995345 | 2.197983 | 0.097755 |
| 11 | 5210200 | 5213599 | 12.80974 | 21.89801 | 64.39946 | 28.23697 | -1.16692 | 0.953086 | 0.097762 |
| 8 | 44695600 | 44696799 | 17.13665 | 31.31129 | 19.61982 | 8.403411 | 1.091971 | 0.301672 | 0.09777 |
| 7 | 118615000 | 118628599 | 85.22737 | 129.6755 | 280.063 | 116.0424 | -0.82594 | 3.169912 | 0.097781 |
| 6 | 78122600 | 78123199 | 9.295295 | 18.13343 | 12.68442 | 4.536388 | 1.157175 | -0.41747 | 0.097797 |
| 5 | 140652800 | 140680799 | 169.4252 | 243.5716 | 547.8096 | 226.1329 | -0.96045 | 4.166082 | 0.097801 |
| 7 | 73524600 | 73526399 | 13.4583 | 23.30776 | 56.44446 | 24.7643 | -0.88411 | 0.828842 | 0.097814 |
| 15 | 101758600 | 101760599 | 8.75136 | 15.42242 | 45.64374 | 20.20217 | -1.16702 | 0.547874 | 0.097828 |
| 14 | 3628000 | 3628999 | 13.38043 | 25.66925 | 16.09739 | 6.322005 | 1.191189 | -0.04678 | 0.097846 |
| 2 | 153277000 | 153279799 | 11.81597 | 20.04321 | 66.78096 | 29.25286 | -1.34744 | 1.025079 | 0.097852 |
| 15 | 18664600 | 18665799 | 12.96297 | 25.05268 | 14.36322 | 5.508973 | 1.349903 | -0.13158 | 0.097863 |
| 13 | 66592200 | 66602599 | 180.9909 | 273.5437 | 173.2459 | 71.63604 | 0.934359 | 3.321807 | 0.097885 |
| 3 | 79793400 | 79795999 | 13.54966 | 23.20157 | 61.1705 | 26.82479 | -1.00956 | 0.93772 | 0.097895 |

| | | | | | | | | | |
|---|---|---|---|---|---|---|---|---|---|
| 1 | 38768400 | 38770799 | 8.520708 | 15.25029 | 42.09196 | 18.45968 | -1.05552 | 0.410316 | 0.097896 |
| 15 | 101009000 | 101027199 | 116.9256 | 174.074 | 378.2725 | 155.8651 | -0.86244 | 3.605676 | 0.097899 |
| 7 | 132030800 | 132033799 | 22.58588 | 37.12447 | 85.41156 | 36.80642 | -0.82041 | 1.408271 | 0.097904 |
| 13 | 95471000 | 95473799 | 22.09617 | 36.35427 | 83.7003 | 36.08358 | -0.82088 | 1.393591 | 0.09791 |
| 4 | 156272800 | 156282799 | 48.85071 | 76.45128 | 196.5402 | 81.53193 | -1.04502 | 2.617936 | 0.097924 |
| 9 | 114231800 | 114250599 | 210.5708 | 298.6465 | 618.8278 | 256.0136 | -0.86195 | 4.36733 | 0.09793 |
| 5 | 113467400 | 113498799 | 291.2419 | 402.1562 | 805.1563 | 334.5185 | -0.85134 | 4.77141 | 0.097935 |
| 2 | 163047000 | 163049599 | 18.63896 | 31.04442 | 75.98388 | 33.00769 | -0.90587 | 1.233058 | 0.097942 |
| 16 | 4559400 | 4561399 | 26.42645 | 46.15393 | 26.05748 | 11.47815 | 1.207825 | 0.737601 | 0.097943 |
| 1 | 8115800 | 8118199 | 79.35932 | 126.061 | 91.07429 | 37.97625 | 0.801031 | 2.289089 | 0.09795 |
| 8 | 49439800 | 49440199 | 9.012135 | 17.58223 | 12.79026 | 4.570183 | 1.102928 | -0.42369 | 0.097968 |
| 5 | 122660800 | 122666199 | 28.63303 | 46.11567 | 114.2158 | 48.55978 | -0.95044 | 1.811963 | 0.097973 |
| X | 148814000 | 148815399 | 31.09558 | 53.21235 | 33.3488 | 14.62353 | 1.054291 | 0.97941 | 0.097975 |
| 12 | 108267400 | 108280399 | 81.4328 | 123.771 | 281.3657 | 116.5452 | -0.90103 | 3.191279 | 0.097981 |
| 17 | 66681400 | 66689199 | 46.76267 | 73.50298 | 184.528 | 76.54573 | -1.00588 | 2.533406 | 0.097986 |
| 15 | 38454200 | 38460999 | 32.43415 | 51.93 | 131.0076 | 55.21132 | -0.98753 | 2.009043 | 0.097993 |
| 14 | 51433200 | 51435399 | 11.6859 | 20.47415 | 53.71168 | 23.72033 | -0.99694 | 0.656568 | 0.098 |
| 4 | 148540200 | 148541999 | 8.541196 | 15.23078 | 42.57223 | 18.72834 | -1.07671 | 0.435222 | 0.098007 |
| 13 | 72268800 | 72273399 | 23.54298 | 38.25509 | 94.66494 | 40.60832 | -0.9354 | 1.575907 | 0.098011 |
| 6 | 104332000 | 104334399 | 19.09697 | 34.473 | 21.03281 | 9.14612 | 1.115886 | 0.385764 | 0.098016 |
| 17 | 17302400 | 17304399 | 36.1408 | 61.3604 | 30.29008 | 13.27197 | 1.395345 | 1.062846 | 0.098022 |
| 19 | 3901000 | 3904999 | 16.79374 | 27.9209 | 76.70971 | 33.28304 | -1.07194 | 1.249431 | 0.098022 |
| 9 | 56500600 | 56502799 | 10.65 | 18.57674 | 52.05746 | 22.91273 | -1.09158 | 0.71831 | 0.098044 |
| 10 | 128360200 | 128362199 | 8.800865 | 15.43524 | 48.80412 | 21.57169 | -1.26366 | 0.618207 | 0.098049 |
| 5 | 134583000 | 134596599 | 90.14444 | 136.5688 | 302.5441 | 125.1948 | -0.86773 | 3.260141 | 0.098057 |
| 4 | 155844800 | 155846599 | 5.593329 | 10.28072 | 37.22317 | 15.81635 | -1.41101 | 0.221498 | 0.098058 |
| 1 | 132864600 | 132870999 | 47.19334 | 74.69775 | 160.212 | 66.60321 | -0.77114 | 2.377685 | 0.098071 |
| 2 | 104033600 | 104035599 | 10.60784 | 18.5859 | 48.83386 | 21.56673 | -0.99852 | 0.645225 | 0.098075 |
| 11 | 81039000 | 81045799 | 36.33584 | 58.01207 | 135.0097 | 56.66912 | -0.876 | 2.102918 | 0.098075 |
| 9 | 54466000 | 54468199 | 8.148811 | 14.39561 | 45.12568 | 19.96314 | -1.24785 | 0.515403 | 0.098076 |
| 19 | 13203400 | 13207999 | 27.85071 | 48.2674 | 28.18442 | 12.36583 | 1.15885 | 0.84749 | 0.0981 |
| 12 | 40033200 | 40036599 | 16.36511 | 27.48649 | 72.65007 | 31.72674 | -1.01627 | 1.132106 | 0.0981 |
| 1 | 191820400 | 191823199 | 11.21683 | 19.3157 | 58.9303 | 25.85045 | -1.21715 | 0.874311 | 0.098103 |
| 15 | 25270600 | 25270999 | 2.246839 | 4.394148 | 19.98852 | 6.539077 | -1.38545 | -0.8037 | 0.098106 |
| X | 3296200 | 3298199 | 35.17286 | 58.71155 | 40.69636 | 17.70318 | 0.908107 | 1.227743 | 0.098125 |
| 7 | 100355200 | 100357199 | 10.43051 | 18.35429 | 46.67671 | 20.6655 | -0.95102 | 0.575078 | 0.098139 |
| 11 | 33139600 | 33162599 | 195.2785 | 277.6295 | 620.1168 | 256.4424 | -0.96666 | 4.331045 | 0.098147 |
| 4 | 151208800 | 151230999 | 140.4719 | 204.5951 | 483.4686 | 199.2468 | -1.01322 | 3.968796 | 0.098156 |
| 19 | 39850000 | 39851799 | 22.62636 | 39.82697 | 25.13857 | 11.1291 | 1.048243 | 0.601633 | 0.098174 |
| 19 | 58500200 | 58500999 | 6.264269 | 11.6689 | 35.39242 | 14.75746 | -1.13817 | 0.142748 | 0.098191 |
| X | 147536000 | 147538599 | 52.23303 | 84.71212 | 54.04304 | 23.15024 | 1.012498 | 1.644973 | 0.098219 |
| 8 | 85721000 | 85725799 | 21.5957 | 35.20543 | 91.21522 | 39.17992 | -0.99792 | 1.520814 | 0.098222 |
| 11 | 82453000 | 82460399 | 40.8225 | 64.792 | 153.247 | 63.84794 | -0.90833 | 2.296543 | 0.098232 |
| 7 | 38128800 | 38129199 | 11.36674 | 22.10313 | 11.55923 | 4.282126 | 1.524514 | -0.27494 | 0.098232 |
| 17 | 25154200 | 25156199 | 8.762916 | 15.45638 | 46.16092 | 20.42748 | -1.17977 | 0.537433 | 0.098241 |
| 3 | 11861200 | 11862399 | 12.58229 | 24.35353 | 13.79295 | 5.253248 | 1.376263 | -0.16066 | 0.09826 |
| 4 | 146711800 | 146712599 | 12.58229 | 24.35353 | 13.79295 | 5.253248 | 1.376263 | -0.16066 | 0.09826 |

| | | | | | | | | | |
|---|---|---|---|---|---|---|---|---|---|
| 13 | 52039400 | 52044199 | 17.13336 | 28.44528 | 82.56232 | 35.78742 | -1.15128 | 1.26998 | 0.09827 |
| 7 | 14744600 | 14746199 | 49.47109 | 80.77862 | 47.90268 | 20.5879 | 1.125 | 1.529222 | 0.098282 |
| 11 | 98871200 | 98879999 | 48.73166 | 76.5133 | 185.5584 | 76.96209 | -0.95734 | 2.557091 | 0.098284 |
| 15 | 97863800 | 97866599 | 10.01871 | 17.52792 | 49.90155 | 22.01092 | -1.11367 | 0.66685 | 0.098289 |
| 6 | 146826200 | 146827399 | 4.808625 | 9.276365 | 29.2203 | 11.29415 | -1.09319 | -0.14705 | 0.098296 |
| 9 | 57967800 | 57975199 | 34.73603 | 55.44202 | 135.2396 | 56.74262 | -0.9441 | 2.11088 | 0.09832 |
| 15 | 74957200 | 74958799 | 1505.722 | 1897.089 | 1182.701 | 496.9907 | 0.698841 | 6.24562 | 0.098325 |
| 11 | 105857600 | 105861399 | 18.58202 | 30.37148 | 90.37665 | 38.85658 | -1.19417 | 1.476018 | 0.098333 |
| 6 | 47854200 | 47855799 | 8.123306 | 14.4155 | 43.57071 | 19.24614 | -1.19272 | 0.465978 | 0.098339 |
| 6 | 140549200 | 140550799 | 5.156066 | 9.866788 | 31.79808 | 12.57819 | -1.15872 | -0.06277 | 0.098343 |
| 8 | 117343600 | 117359199 | 104.1913 | 156.6643 | 333.6907 | 137.7738 | -0.82057 | 3.415276 | 0.098371 |
| 10 | 66239000 | 66239799 | 14.0151 | 27.02869 | 14.2936 | 5.529409 | 1.454101 | -0.09249 | 0.09838 |
| 10 | 85810400 | 85810799 | 14.0151 | 27.02869 | 14.2936 | 5.529409 | 1.454101 | -0.09249 | 0.09838 |
| 1 | 128733000 | 128738799 | 27.93084 | 45.1262 | 106.9295 | 45.65095 | -0.88138 | 1.714247 | 0.098389 |
| 14 | 60933600 | 60935599 | 5.697658 | 10.31798 | 42.28087 | 18.31274 | -1.60793 | 0.318829 | 0.098392 |
| 11 | 49058800 | 49061199 | 15.77928 | 26.82604 | 63.88927 | 28.00083 | -0.86539 | 1.016281 | 0.098405 |
| 7 | 20318600 | 20319799 | 10.72368 | 20.93168 | 10.97491 | 3.888627 | 1.580362 | -0.43389 | 0.098411 |
| 11 | 19669600 | 19671599 | 17.78669 | 29.81773 | 70.38744 | 30.702 | -0.85372 | 1.156874 | 0.09842 |
| 2 | 180869800 | 180878599 | 40.78409 | 64.12255 | 185.56 | 76.97644 | -1.21018 | 2.525405 | 0.098432 |
| 1 | 101524000 | 101527599 | 49.29381 | 80.20492 | 48.46166 | 20.8013 | 1.094504 | 1.57682 | 0.098438 |
| 7 | 101465800 | 101475999 | 56.10957 | 86.7545 | 227.6052 | 94.5419 | -1.08801 | 2.839695 | 0.098444 |
| 17 | 24573200 | 24576399 | 11.86198 | 20.70311 | 51.55388 | 22.69957 | -0.92218 | 0.715356 | 0.098446 |
| 16 | 94013600 | 94016199 | 29.72883 | 50.7876 | 35.05326 | 15.36885 | 0.915359 | 0.990393 | 0.098455 |
| 3 | 43449600 | 43452199 | 28.77418 | 50.05859 | 26.56888 | 11.67476 | 1.296202 | 0.803878 | 0.098456 |
| 7 | 11146000 | 11155599 | 166.5136 | 252.4966 | 171.7532 | 71.09266 | 0.835443 | 3.252249 | 0.098457 |
| X | 33715200 | 33718399 | 66.56893 | 106.6427 | 71.76538 | 30.27035 | 0.916144 | 2.013531 | 0.098458 |
| X | 31528800 | 31530999 | 23.37233 | 41.41478 | 21.74494 | 9.613907 | 1.315499 | 0.530281 | 0.098459 |
| 4 | 140540800 | 140555999 | 178.6059 | 254.4925 | 598.6034 | 247.4531 | -1.03796 | 4.297815 | 0.09846 |
| 4 | 136037800 | 136038999 | 5.388404 | 10.00639 | 35.25732 | 14.77299 | -1.35678 | 0.15681 | 0.098468 |
| 4 | 11297000 | 11298599 | 7.146153 | 13.28425 | 37.05326 | 15.50374 | -1.02298 | 0.166603 | 0.098472 |
| 12 | 75642000 | 75643799 | 12.87448 | 21.85908 | 66.38539 | 29.0878 | -1.21455 | 1.019446 | 0.098475 |
| 7 | 15124000 | 15124599 | 19.99474 | 35.62566 | 23.15198 | 10.19205 | 1.014086 | 0.474473 | 0.098478 |
| 7 | 119346400 | 119348199 | 25.01982 | 43.571 | 29.61244 | 13.02039 | 0.941598 | 0.775975 | 0.098478 |
| 13 | 60908200 | 60911599 | 23.32793 | 38.01753 | 90.25874 | 38.7647 | -0.87258 | 1.529152 | 0.098515 |
| 1 | 108489800 | 108490999 | 16.62926 | 30.60797 | 19.79025 | 8.392983 | 1.056467 | 0.256503 | 0.098516 |
| 15 | 31872400 | 31874399 | 16.62926 | 30.60797 | 19.79025 | 8.392983 | 1.056467 | 0.256503 | 0.098516 |
| 7 | 46228800 | 46229799 | 4.841653 | 9.337958 | 30.45954 | 11.78082 | -1.14494 | -0.14389 | 0.098531 |
| 19 | 10071000 | 10072199 | 4.841653 | 9.337958 | 30.45954 | 11.78082 | -1.14494 | -0.14389 | 0.098531 |
| 1 | 35008400 | 35009799 | 6.828879 | 12.76251 | 35.53882 | 14.73627 | -1.00972 | 0.126369 | 0.098535 |
| 9 | 55844600 | 55845599 | 4.130424 | 7.992192 | 28.82057 | 10.9922 | -1.26756 | -0.20326 | 0.098538 |
| 17 | 86680400 | 86689799 | 49.11056 | 77.35929 | 175.7456 | 72.92295 | -0.85959 | 2.490311 | 0.098556 |
| 13 | 95659000 | 95663199 | 21.79041 | 35.82127 | 85.40948 | 36.8074 | -0.87172 | 1.403504 | 0.098582 |
| 3 | 111175400 | 111176199 | 8.588014 | 15.06068 | 48.79681 | 21.56694 | -1.29808 | 0.621575 | 0.098584 |
| 2 | 44655200 | 44656799 | 18.72945 | 34.3619 | 19.06112 | 8.119541 | 1.271521 | 0.289879 | 0.098586 |
| 7 | 90091800 | 90093399 | 9.414209 | 16.49447 | 49.87934 | 22.02835 | -1.20011 | 0.63882 | 0.09859 |
| 14 | 52999800 | 53003599 | 85.76038 | 135.8082 | 83.8662 | 34.87343 | 1.022932 | 2.378359 | 0.098594 |
| 6 | 24954600 | 24956599 | 15.65621 | 26.11405 | 73.25106 | 31.84949 | -1.10142 | 1.208235 | 0.098613 |

| | | | | | | | | | |
|---|---|---|---|---|---|---|---|---|---|
| 7 | 59700000 | 59700599 | 17.43725 | 31.82306 | 20.90276 | 8.966096 | 1.022948 | 0.313074 | 0.098622 |
| 2 | 79283400 | 79290799 | 50.69005 | 79.94144 | 173.792 | 72.12341 | -0.79489 | 2.467069 | 0.098637 |
| 8 | 106001400 | 106012399 | 61.35114 | 94.81836 | 228.0717 | 94.73587 | -0.96341 | 2.849309 | 0.098642 |
| 7 | 47099800 | 47101599 | 29.18537 | 50.10926 | 30.25544 | 13.27062 | 1.10823 | 0.936653 | 0.098652 |
| 4 | 139685600 | 139690599 | 47.13508 | 74.75894 | 156.0421 | 64.96714 | -0.72937 | 2.322089 | 0.098658 |
| 5 | 77000000 | 77007199 | 51.16825 | 79.90934 | 197.8266 | 82.08337 | -0.99308 | 2.65741 | 0.098668 |
| 2 | 8942800 | 8943999 | 12.50932 | 24.15671 | 14.33233 | 5.517602 | 1.295494 | -0.12526 | 0.098681 |
| 9 | 109148400 | 109149999 | 12.50932 | 24.15671 | 14.33233 | 5.517602 | 1.295494 | -0.12526 | 0.098681 |
| X | 31273000 | 31278399 | 75.50049 | 120.4369 | 78.23817 | 32.80001 | 0.960306 | 2.153859 | 0.098682 |
| 4 | 144276600 | 144278799 | 26.41774 | 46.17145 | 25.47823 | 11.22848 | 1.2406 | 0.72654 | 0.098686 |
| 1 | 175228400 | 175230999 | 19.84498 | 35.49566 | 22.62894 | 9.929407 | 1.044728 | 0.449071 | 0.098689 |
| 6 | 74242200 | 74244799 | 35.13862 | 59.29415 | 35.98948 | 15.73993 | 1.098623 | 1.117195 | 0.09869 |
| 19 | 56415800 | 56419199 | 24.03932 | 39.00589 | 96.18075 | 41.21516 | -0.93225 | 1.607005 | 0.098697 |
| 5 | 123037600 | 123039199 | 6.684101 | 12.09759 | 39.63307 | 17.1779 | -1.28847 | 0.32521 | 0.098707 |
| 17 | 79955200 | 79956999 | 6.354038 | 11.98657 | 36.50769 | 14.89887 | -1.11945 | 0.062535 | 0.098716 |
| 12 | 16797800 | 16797999 | 49.39802 | 81.39049 | 34.34717 | 14.81625 | 1.619477 | 1.404432 | 0.098716 |
| 5 | 138114000 | 138115599 | 4.319847 | 8.299391 | 33.78131 | 13.21155 | -1.47651 | -0.10541 | 0.098739 |
| 10 | 80802000 | 80808999 | 38.54124 | 61.24417 | 150.3629 | 62.72106 | -0.96035 | 2.258757 | 0.098761 |
| 7 | 38635200 | 38636399 | 14.16555 | 27.05381 | 15.79581 | 6.28052 | 1.2763 | -0.00195 | 0.098762 |
| 12 | 100029600 | 100031999 | 12.35983 | 21.46288 | 54.76973 | 24.03877 | -0.95816 | 0.796382 | 0.098776 |
| 7 | 59108400 | 59111199 | 41.41142 | 68.63723 | 41.10937 | 17.79864 | 1.117748 | 1.32133 | 0.098794 |
| 12 | 102092800 | 102099999 | 42.1105 | 66.84153 | 159.0709 | 66.27963 | -0.91717 | 2.295169 | 0.0988 |
| 9 | 18966600 | 18968799 | 48.36352 | 78.53553 | 55.05857 | 23.60721 | 0.878424 | 1.612232 | 0.09882 |
| 15 | 89006400 | 89008599 | 4.652648 | 8.356034 | 42.13404 | 18.42449 | -1.91128 | 0.365537 | 0.098836 |
| 14 | 26827000 | 26828599 | 6.93406 | 12.62861 | 38.1526 | 16.40758 | -1.16498 | 0.287698 | 0.09886 |
| 17 | 24518800 | 24519399 | 3.532157 | 6.882264 | 25.3982 | 9.228356 | -1.23318 | -0.39214 | 0.098867 |
| 1 | 144554000 | 144556799 | 20.42946 | 36.09626 | 24.30005 | 10.74975 | 0.957707 | 0.545785 | 0.098872 |
| 4 | 141133800 | 141154199 | 119.6276 | 176.7236 | 417.4412 | 171.9112 | -0.9931 | 3.740703 | 0.098872 |
| 9 | 5533800 | 5534999 | 16.48555 | 30.20758 | 20.39037 | 8.700458 | 0.988695 | 0.281348 | 0.098876 |
| 2 | 90947800 | 90951799 | 19.07707 | 31.11475 | 91.8758 | 39.47069 | -1.18527 | 1.509265 | 0.098884 |
| 5 | 115982200 | 115989399 | 41.58016 | 66.11856 | 146.4781 | 61.14262 | -0.81143 | 2.240128 | 0.098885 |
| 7 | 79973800 | 79987199 | 60.79535 | 93.40771 | 249.1939 | 103.4708 | -1.11918 | 2.964041 | 0.098888 |
| 17 | 69882400 | 69883199 | 3.132123 | 6.108942 | 25.60817 | 9.183138 | -1.39684 | -0.4453 | 0.098898 |
| 19 | 39258400 | 39260599 | 26.25717 | 45.6026 | 30.74247 | 13.49436 | 0.953096 | 0.821726 | 0.0989 |
| 6 | 97558600 | 97561199 | 7.497437 | 13.76417 | 37.80214 | 16.08835 | -1.01872 | 0.24286 | 0.098913 |
| 10 | 130246800 | 130249399 | 26.30441 | 42.55783 | 99.44516 | 42.52317 | -0.85889 | 1.672478 | 0.09892 |
| 11 | 31029600 | 31030399 | 13.40649 | 25.33312 | 17.66281 | 7.182704 | 0.99518 | 0.066996 | 0.098928 |
| 8 | 93088200 | 93095599 | 53.19606 | 83.31151 | 197.1678 | 81.78622 | -0.92507 | 2.606905 | 0.098931 |
| 2 | 11781800 | 11783799 | 13.56116 | 22.81958 | 68.9098 | 30.11197 | -1.20702 | 1.104989 | 0.098944 |
| 2 | 131027600 | 131030199 | 10.62486 | 18.59253 | 50.44682 | 22.25561 | -1.04496 | 0.666116 | 0.098946 |
| 3 | 58179800 | 58180199 | 8.497951 | 16.61584 | 9.694729 | 3.313433 | 1.469753 | -0.59539 | 0.098947 |
| 11 | 76736800 | 76737999 | 6.442757 | 11.88322 | 36.24426 | 15.32777 | -1.16127 | 0.205785 | 0.098952 |
| 2 | 125427000 | 125430599 | 13.29039 | 22.65664 | 64.34759 | 28.20791 | -1.11751 | 0.981079 | 0.098961 |
| 10 | 67868000 | 67872799 | 34.77913 | 55.8046 | 123.8285 | 52.357 | -0.79911 | 1.94205 | 0.098987 |
| 7 | 37821000 | 37831599 | 57.54796 | 89.75997 | 199.2291 | 82.67599 | -0.83725 | 2.682647 | 0.098992 |
| 2 | 27157000 | 27162599 | 30.96457 | 49.58267 | 125.5359 | 52.91446 | -0.99237 | 2.004372 | 0.09901 |
| 17 | 20044800 | 20046399 | 36.10572 | 60.87069 | 36.03461 | 15.75532 | 1.134587 | 1.129485 | 0.099017 |

| | | | | | | | | | |
|---|---|---|---|---|---|---|---|---|---|
| 8 | 13208200 | 13209999 | 7.246841 | 13.16228 | 38.63863 | 16.67342 | -1.12735 | 0.306698 | 0.099026 |
| 11 | 113407000 | 113430799 | 180.7736 | 259.9481 | 539.6312 | 222.7486 | -0.84459 | 4.163201 | 0.099033 |
| 13 | 52778800 | 52783399 | 20.7277 | 33.98282 | 86.61542 | 37.28469 | -0.96915 | 1.435948 | 0.099035 |
| 13 | 48931400 | 48937999 | 38.44926 | 61.31748 | 139.4128 | 58.42532 | -0.84424 | 2.141168 | 0.099036 |
| 4 | 105128200 | 105128999 | 4.06184 | 7.912545 | 26.70577 | 9.732132 | -1.11078 | -0.37032 | 0.099043 |
| 14 | 108916400 | 108917199 | 10.32585 | 20.09742 | 13.64817 | 5.018793 | 1.163916 | -0.3051 | 0.099065 |
| 4 | 119279000 | 119280399 | 6.099619 | 11.09342 | 40.24472 | 17.3375 | -1.42864 | 0.292252 | 0.09907 |
| 12 | 22709000 | 22709999 | 12.83015 | 24.58151 | 16.65864 | 6.56697 | 1.075187 | -0.03944 | 0.099076 |
| 7 | 138516000 | 138517399 | 8.013404 | 14.46653 | 40.11741 | 17.44342 | -1.05324 | 0.351833 | 0.09908 |
| 7 | 59416800 | 59417399 | 15.26192 | 28.89254 | 16.83111 | 6.817776 | 1.256997 | 0.070192 | 0.099081 |
| X | 145063400 | 145064999 | 23.53446 | 42.15447 | 19.19144 | 8.411013 | 1.528315 | 0.46014 | 0.099108 |
| 11 | 62429200 | 62430399 | 5.115924 | 9.546785 | 35.37342 | 14.70512 | -1.41991 | 0.12184 | 0.099118 |
| 5 | 139949600 | 139968399 | 107.5802 | 159.9898 | 387.9734 | 159.845 | -1.02199 | 3.623621 | 0.09912 |
| 17 | 44918200 | 44921199 | 12.05066 | 20.93154 | 55.33964 | 24.29296 | -1.00918 | 0.789311 | 0.099135 |
| 9 | 118828800 | 118836399 | 47.64344 | 74.68288 | 189.126 | 78.44315 | -1.02146 | 2.587975 | 0.099146 |
| 4 | 109917400 | 109919799 | 38.05678 | 63.33541 | 42.19919 | 18.32211 | 0.964999 | 1.267927 | 0.09915 |
| 19 | 39687400 | 39688199 | 10.86319 | 21.13861 | 13.63595 | 5.023217 | 1.235326 | -0.29173 | 0.09916 |
| 7 | 17755600 | 17757599 | 33.85279 | 57.87616 | 31.11635 | 13.64356 | 1.274301 | 0.997039 | 0.099163 |
| 3 | 75703800 | 75704799 | 11.60946 | 22.60307 | 12.28455 | 4.505519 | 1.484266 | -0.30192 | 0.099175 |
| 5 | 139523400 | 139524599 | 6.245926 | 11.6872 | 34.98478 | 14.47676 | -1.11051 | 0.111918 | 0.099181 |
| 5 | 120614000 | 120635599 | 127.0764 | 187.0075 | 432.26 | 178.0065 | -0.96706 | 3.805667 | 0.099185 |
| 1 | 9368400 | 9370999 | 20.12161 | 33.31745 | 78.68892 | 34.04569 | -0.85526 | 1.307826 | 0.099203 |
| 14 | 41864400 | 41867199 | 27.45809 | 47.75419 | 28.99675 | 12.73912 | 1.103328 | 0.806323 | 0.099204 |
| 6 | 136560000 | 136561999 | 9.553557 | 16.74916 | 48.81799 | 21.5711 | -1.14721 | 0.627969 | 0.099208 |
| 6 | 91351600 | 91354799 | 10.92182 | 19.12219 | 49.37961 | 21.79443 | -0.97391 | 0.658187 | 0.099213 |
| 5 | 64315200 | 64319399 | 19.88289 | 32.4157 | 93.86733 | 40.32022 | -1.15776 | 1.517195 | 0.099216 |
| 6 | 128781600 | 128782599 | 10.26741 | 19.94202 | 14.24219 | 5.326222 | 1.07024 | -0.2467 | 0.099221 |
| 3 | 55363600 | 55364999 | 5.55612 | 10.65346 | 35.04557 | 13.78401 | -1.18085 | -0.0693 | 0.099224 |
| 5 | 112430000 | 112441399 | 46.55947 | 72.92549 | 194.7911 | 80.79947 | -1.0992 | 2.601989 | 0.099241 |
| 1 | 130824200 | 130826799 | 11.25588 | 19.63662 | 53.16324 | 23.40171 | -1.04237 | 0.719532 | 0.099248 |
| 19 | 5114600 | 5123399 | 42.6991 | 67.54322 | 168.6987 | 70.10974 | -0.99121 | 2.389607 | 0.099279 |
| 17 | 48613200 | 48615599 | 11.42711 | 19.87118 | 53.13669 | 23.34267 | -1.02493 | 0.768885 | 0.099293 |
| 2 | 111034800 | 111035199 | 8.557671 | 16.70366 | 12.32648 | 4.368221 | 1.093149 | -0.46019 | 0.099293 |
| 11 | 77743800 | 77759599 | 62.53035 | 95.58648 | 271.3533 | 112.568 | -1.21446 | 3.068235 | 0.099299 |
| 17 | 86965600 | 86966999 | 8.983507 | 16.0369 | 42.58067 | 18.71494 | -1.00233 | 0.430928 | 0.099303 |
| 17 | 66786000 | 66787799 | 10.63428 | 18.5868 | 50.98792 | 22.47163 | -1.06071 | 0.686856 | 0.099311 |
| 16 | 93877600 | 93896399 | 126.4796 | 187.2868 | 407.9558 | 168.0092 | -0.87308 | 3.702037 | 0.099313 |
| 1 | 187336200 | 187344799 | 44.58667 | 70.26713 | 178.8911 | 74.23279 | -1.02308 | 2.478141 | 0.099317 |
| 19 | 10221800 | 10251999 | 143.3334 | 206.8804 | 544.4402 | 224.6167 | -1.18128 | 4.105647 | 0.099322 |
| 2 | 22018800 | 22019399 | 11.32736 | 21.83159 | 14.81847 | 5.74278 | 1.095579 | -0.11498 | 0.099326 |
| 2 | 76366400 | 76368999 | 11.06675 | 19.39783 | 48.8356 | 21.57489 | -0.93747 | 0.639743 | 0.099335 |
| 7 | 55901800 | 55902599 | 3.716617 | 7.220531 | 30.57251 | 11.40938 | -1.46702 | -0.28124 | 0.099337 |
| 16 | 69858200 | 69859199 | 11.94131 | 23.03197 | 15.79764 | 6.109208 | 1.083814 | -0.11111 | 0.099359 |
| 4 | 123200200 | 123221599 | 126.0044 | 185.7037 | 426.628 | 175.69 | -0.95624 | 3.780881 | 0.099362 |
| 3 | 91202200 | 91203399 | 18.65352 | 33.92879 | 19.40783 | 8.359843 | 1.21654 | 0.336134 | 0.099364 |
| 9 | 65040200 | 65044199 | 15.53593 | 25.60715 | 87.36588 | 37.69267 | -1.38492 | 1.356837 | 0.099368 |
| 12 | 80965400 | 80967599 | 13.44469 | 23.32709 | 56.48079 | 24.78163 | -0.88331 | 0.811835 | 0.09937 |

| | | | | | | | | | |
|---|---|---|---|---|---|---|---|---|---|
| 2 | 67973800 | 67975399 | 15.59713 | 29.30349 | 17.87777 | 7.347229 | 1.173802 | 0.124563 | 0.099374 |
| 3 | 107992200 | 107994199 | 8.850332 | 15.46974 | 53.06851 | 23.38982 | -1.38105 | 0.679548 | 0.099384 |
| X | 117170400 | 117172199 | 24.90618 | 44.54621 | 17.99744 | 7.897462 | 1.698002 | 0.475954 | 0.099407 |
| 1 | 74133600 | 74139599 | 24.45588 | 39.49229 | 106.8379 | 45.58264 | -1.07272 | 1.72395 | 0.099415 |
| 7 | 125688600 | 125702399 | 83.25914 | 126.5384 | 281.0138 | 116.3972 | -0.86744 | 3.192969 | 0.099431 |
| 2 | 104301400 | 104302799 | 5.588828 | 10.6657 | 32.29118 | 12.85961 | -1.07869 | -0.03483 | 0.09945 |
| 15 | 72834800 | 72835999 | 5.588828 | 10.6657 | 32.29118 | 12.85961 | -1.07869 | -0.03483 | 0.09945 |
| 9 | 95117400 | 95119399 | 24.29179 | 42.59288 | 26.19474 | 11.57573 | 1.085055 | 0.666869 | 0.099469 |
| 11 | 102947000 | 102948999 | 7.962142 | 14.18086 | 43.60937 | 19.20438 | -1.21452 | 0.440793 | 0.099476 |
| 15 | 5863800 | 5865199 | 19.85736 | 36.1218 | 19.40863 | 8.359221 | 1.306656 | 0.345889 | 0.099482 |
| 4 | 60779000 | 60781399 | 23.39762 | 41.2086 | 23.91268 | 10.58515 | 1.169015 | 0.597533 | 0.099487 |
| 1 | 4473200 | 4475199 | 17.81828 | 32.57864 | 20.17053 | 8.632023 | 1.109446 | 0.303313 | 0.099494 |
| 7 | 141977000 | 141977999 | 15.08086 | 28.15609 | 19.02734 | 7.903309 | 1.01545 | 0.16156 | 0.09951 |
| 2 | 168767400 | 168791799 | 175.61 | 252.5739 | 544.4864 | 224.7456 | -0.89871 | 4.159886 | 0.09951 |
| 12 | 73784800 | 73787199 | 10.59739 | 18.60304 | 47.75133 | 21.12265 | -0.96501 | 0.611995 | 0.099512 |
| 4 | 57859200 | 57860599 | 6.352344 | 11.98858 | 35.34807 | 14.41238 | -1.07169 | 0.057144 | 0.099513 |
| 4 | 69124400 | 69124999 | 11.58079 | 22.40279 | 14.5062 | 5.539085 | 1.181832 | -0.16196 | 0.099526 |
| 15 | 78669800 | 78683599 | 68.06176 | 104.2691 | 252.9331 | 104.9795 | -0.98538 | 3.024447 | 0.099556 |
| 11 | 100769800 | 100772999 | 19.38832 | 31.59562 | 91.74237 | 39.4012 | -1.16198 | 1.522822 | 0.099558 |
| 17 | 19337800 | 19340999 | 68.23063 | 109.2767 | 72.26291 | 30.46833 | 0.940867 | 2.021981 | 0.09957 |
| 12 | 112302400 | 112317599 | 94.93455 | 143.0479 | 322.684 | 133.3048 | -0.90055 | 3.37185 | 0.099578 |
| 7 | 10716200 | 10730999 | 109.3143 | 170.0416 | 116.6358 | 48.4208 | 0.851839 | 2.741888 | 0.099579 |
| 7 | 31167800 | 31168599 | 28.49169 | 48.6374 | 32.72714 | 14.34811 | 0.951689 | 0.989032 | 0.099583 |
| 8 | 22062400 | 22062799 | 8.22486 | 16.06164 | 11.75845 | 4.133118 | 1.114322 | -0.49589 | 0.099592 |
| 14 | 34303000 | 34325599 | 123.8156 | 182.2803 | 435.1522 | 179.1983 | -1.01306 | 3.798788 | 0.099595 |
| 3 | 99327600 | 99328199 | 20.61436 | 36.44744 | 24.47723 | 10.82668 | 0.961803 | 0.541731 | 0.099652 |
| 11 | 93516600 | 93517799 | 14.73717 | 28.1215 | 15.77951 | 6.285376 | 1.331098 | 0.009691 | 0.099669 |
| 15 | 72953600 | 72955399 | 12.72176 | 24.17876 | 16.69881 | 6.708083 | 1.0239 | 0.018615 | 0.099681 |
| 15 | 11384000 | 11384999 | 4.977575 | 9.582429 | 29.74354 | 11.56494 | -1.08092 | -0.12562 | 0.099693 |
| X | 3217400 | 3218799 | 42.48425 | 70.18832 | 43.94825 | 18.98537 | 1.053753 | 1.364991 | 0.099702 |
| 2 | 15606000 | 15607999 | 18.6705 | 34.19289 | 18.3321 | 7.827914 | 1.31728 | 0.2963 | 0.099704 |
| 13 | 53141000 | 53144599 | 14.53327 | 24.71665 | 63.25269 | 27.72264 | -0.96844 | 1.007533 | 0.099706 |
| 7 | 33206400 | 33210999 | 85.09698 | 134.7494 | 97.59133 | 40.57938 | 0.793301 | 2.381542 | 0.099723 |
| 10 | 80827600 | 80838999 | 47.57296 | 74.07731 | 211.2173 | 87.67237 | -1.1999 | 2.708659 | 0.099724 |
| 5 | 4012400 | 4013799 | 4.703787 | 8.992578 | 30.49454 | 12.09615 | -1.23377 | -0.05854 | 0.099726 |
| 5 | 37730800 | 37731599 | 4.703787 | 8.992578 | 30.49454 | 12.09615 | -1.23377 | -0.05854 | 0.099726 |
| 2 | 166647200 | 166648199 | 4.751958 | 9.060457 | 32.14823 | 12.8271 | -1.30705 | -0.0367 | 0.099735 |
| 7 | 96474400 | 96475799 | 3.054505 | 5.899085 | 31.44371 | 12.09991 | -1.83509 | -0.17697 | 0.099749 |
| 9 | 34848800 | 34851399 | 10.57804 | 18.6274 | 47.20621 | 20.8969 | -0.94578 | 0.567668 | 0.099751 |
| 6 | 86824800 | 86827399 | 12.80231 | 21.87829 | 61.59781 | 27.00784 | -1.1042 | 0.954414 | 0.099752 |
| 17 | 38624400 | 38625799 | 28.62039 | 48.95129 | 33.69697 | 14.77559 | 0.919438 | 0.974643 | 0.099758 |
| 10 | 77854600 | 77857399 | 14.50852 | 24.73927 | 61.6543 | 27.02779 | -0.92968 | 0.98209 | 0.099761 |
| 7 | 28450600 | 28461799 | 52.63752 | 81.52495 | 224.6109 | 93.29732 | -1.15658 | 2.806319 | 0.099765 |
| 7 | 22377000 | 22377999 | 10.74315 | 20.98705 | 9.391033 | 3.280271 | 1.819608 | -0.48789 | 0.099774 |
| 12 | 23988600 | 23990599 | 38.74612 | 64.38788 | 39.11376 | 16.96524 | 1.097733 | 1.284586 | 0.099779 |
| 9 | 17032000 | 17032599 | 11.78534 | 22.94373 | 12.53109 | 4.598817 | 1.478077 | -0.2958 | 0.099784 |
| 17 | 24420800 | 24432599 | 45.15876 | 70.68074 | 203.0946 | 84.24999 | -1.2052 | 2.614787 | 0.099787 |

| | | | | | | | | | |
|---|---|---|---|---|---|---|---|---|---|
| 15 | 75652800 | 75657399 | 15.40072 | 25.53801 | 77.32129 | 33.48786 | -1.21161 | 1.278882 | 0.099794 |
| 4 | 125167800 | 125174599 | 37.09869 | 59.34797 | 127.7639 | 53.77908 | -0.7606 | 2.040897 | 0.099794 |
| X | 90812400 | 90813799 | 19.17508 | 35.33145 | 18.0622 | 7.649031 | 1.394495 | 0.26843 | 0.099801 |
| 2 | 76736000 | 76736999 | 4.964723 | 9.597682 | 30.71986 | 11.76712 | -1.1047 | -0.17485 | 0.099821 |
| 8 | 89285600 | 89287999 | 10.21838 | 17.78059 | 53.63387 | 23.57053 | -1.19755 | 0.753946 | 0.099825 |
| 17 | 48228000 | 48230799 | 10.21838 | 17.78059 | 53.63387 | 23.57053 | -1.19755 | 0.753946 | 0.099825 |
| 1 | 178792600 | 178793599 | 5.8644 | 11.20082 | 34.76989 | 13.82265 | -1.11259 | -0.03173 | 0.09983 |
| 13 | 55967600 | 55976999 | 60.7424 | 94.13692 | 219.0633 | 90.9927 | -0.91266 | 2.796959 | 0.099835 |
| 5 | 52784800 | 52787599 | 10.88481 | 18.8044 | 57.34161 | 25.15602 | -1.2153 | 0.841444 | 0.099841 |
| 3 | 100314800 | 100316199 | 10.05655 | 17.50112 | 53.08644 | 23.33174 | -1.20537 | 0.751465 | 0.099851 |
| 13 | 23889800 | 23894799 | 21.64493 | 35.55198 | 85.87955 | 36.99672 | -0.89095 | 1.41403 | 0.099852 |
| 14 | 84033000 | 84034199 | 14.12411 | 27.13665 | 14.08048 | 5.506116 | 1.466059 | -0.05922 | 0.099876 |
| 13 | 33546400 | 33547999 | 18.30795 | 32.81408 | 19.46957 | 8.528256 | 1.149046 | 0.417856 | 0.099887 |
| 4 | 80107800 | 80109799 | 23.14122 | 40.7717 | 23.34182 | 10.33171 | 1.188409 | 0.590528 | 0.099892 |
| 1 | 135705400 | 135707199 | 4.618335 | 8.708444 | 32.84794 | 13.41659 | -1.42336 | 0.044779 | 0.099895 |
| 2 | 179735000 | 179748399 | 75.10597 | 114.4136 | 275.4194 | 114.1596 | -0.98084 | 3.142648 | 0.099898 |
| 9 | 103927600 | 103928599 | 3.135231 | 6.106667 | 25.84264 | 9.434291 | -1.43354 | -0.37864 | 0.099906 |
| 18 | 36858000 | 36859599 | 7.759438 | 13.93253 | 41.08579 | 17.9696 | -1.1472 | 0.383193 | 0.099927 |
| 17 | 24673800 | 24674999 | 4.988783 | 9.568683 | 30.12454 | 11.83766 | -1.11547 | -0.08904 | 0.099938 |
| 4 | 154125000 | 154127599 | 16.72151 | 27.92541 | 71.7246 | 31.2101 | -0.97482 | 1.197332 | 0.09994 |
| 18 | 61731400 | 61732399 | 21.57912 | 37.87071 | 26.0422 | 11.5171 | 0.924741 | 0.636452 | 0.099955 |
| 5 | 120784000 | 120824999 | 254.0618 | 352.6534 | 786.5464 | 326.5164 | -0.99943 | 4.690375 | 0.099956 |
| 11 | 117472400 | 117473999 | 7.645795 | 14.02605 | 38.32073 | 16.32564 | -1.01209 | 0.251104 | 0.099961 |
| 17 | 31531600 | 31534999 | 12.15159 | 20.85359 | 59.46046 | 26.07953 | -1.1212 | 0.910825 | 0.099975 |
| 12 | 78338600 | 78340799 | 29.01971 | 50.21816 | 29.99787 | 13.16087 | 1.125785 | 0.868093 | 0.099976 |
| 19 | 11923200 | 11924799 | 16.87767 | 30.83053 | 20.91452 | 8.960693 | 0.977689 | 0.303109 | 0.099981 |
| 4 | 114674400 | 114681199 | 35.33499 | 56.50873 | 133.7363 | 56.26794 | -0.8975 | 2.048906 | 0.099983 |
| 10 | 31396600 | 31399399 | 15.41707 | 26.36957 | 62.93405 | 27.59565 | -0.86703 | 0.957552 | 0.099983 |
| 17 | 69146800 | 69150199 | 22.03428 | 35.97032 | 88.47364 | 38.02156 | -0.92164 | 1.500812 | 0.099986 |
| 15 | 48123800 | 48125199 | 15.78308 | 30.15485 | 14.57059 | 5.788562 | 1.547821 | 0.008818 | 0.099997 |
| X | 32861200 | 32863999 | 39.81773 | 67.70146 | 27.138 | 11.89392 | 1.694681 | 1.064577 | 0.100007 |
| 5 | 66442200 | 66444599 | 30.43761 | 51.89329 | 34.9671 | 15.32857 | 0.949348 | 1.009954 | 0.100016 |
| 16 | 32248800 | 32251199 | 10.28273 | 18.08081 | 46.67638 | 20.66842 | -0.97296 | 0.582365 | 0.100019 |
| 13 | 44455400 | 44455799 | 2.16404 | 4.230535 | 24.37381 | 8.369102 | -1.78678 | -0.6188 | 0.100025 |
| 17 | 24469000 | 24472199 | 11.86499 | 20.69336 | 52.6401 | 23.16729 | -0.95304 | 0.726253 | 0.10003 |
| 6 | 126138400 | 126144399 | 26.35255 | 42.55051 | 105.4753 | 44.99503 | -0.94716 | 1.729587 | 0.100053 |
| 19 | 59544600 | 59546199 | 15.87938 | 29.19677 | 20.05093 | 8.515809 | 0.968898 | 0.257379 | 0.100059 |
| 2 | 84867400 | 84868599 | 7.022046 | 12.96723 | 36.28688 | 15.32075 | -1.03658 | 0.203771 | 0.100072 |
| 4 | 140274200 | 140297799 | 163.0366 | 234.8692 | 536.4145 | 221.3616 | -0.97964 | 4.135342 | 0.100087 |
| 9 | 123787600 | 123788399 | 12.97025 | 25.03615 | 14.93654 | 5.761666 | 1.285618 | -0.11119 | 0.100091 |
| 1 | 174545000 | 174545599 | 10.16116 | 19.73864 | 13.37251 | 4.99489 | 1.147822 | -0.25444 | 0.100095 |
| 10 | 66836600 | 66837399 | 13.8451 | 26.20027 | 17.6859 | 7.169848 | 1.045633 | 0.062118 | 0.100097 |
| 11 | 97610200 | 97620599 | 55.78351 | 86.51236 | 214.5288 | 89.10061 | -1.00337 | 2.783144 | 0.100105 |
| 14 | 123651000 | 123652999 | 12.52284 | 24.09008 | 16.15946 | 6.296556 | 1.105235 | -0.08059 | 0.100116 |
| 2 | 163153800 | 163156599 | 11.43228 | 19.87988 | 53.6857 | 23.58364 | -1.03919 | 0.76878 | 0.100129 |
| 14 | 24396800 | 24400799 | 24.90882 | 40.63323 | 93.34735 | 40.10536 | -0.82444 | 1.518455 | 0.100139 |
| 2 | 162840800 | 162842399 | 5.178254 | 9.844786 | 32.07404 | 12.87593 | -1.19322 | -0.01208 | 0.100145 |

| | | | | | | | | | |
|---|---|---|---|---|---|---|---|---|---|
| 14 | 52481600 | 52482599 | 5.178254 | 9.844786 | 32.07404 | 12.87593 | -1.19322 | -0.01208 | 0.100145 |
| 2 | 96032800 | 96033599 | 16.52386 | 31.16539 | 16.04282 | 6.549795 | 1.423741 | 0.104301 | 0.100153 |
| 8 | 81082400 | 81083199 | 15.16374 | 28.31269 | 18.63676 | 7.740202 | 1.053475 | 0.161526 | 0.100165 |
| 3 | 44768200 | 44769599 | 15.59225 | 28.95494 | 19.51811 | 8.178266 | 1.009355 | 0.19979 | 0.100178 |
| 12 | 112291600 | 112301599 | 49.73439 | 77.66241 | 201.3196 | 83.53355 | -1.0595 | 2.658547 | 0.100183 |
| 2 | 102004600 | 102005799 | 5.923508 | 11.17025 | 33.52968 | 13.69351 | -1.09875 | 0.055461 | 0.100188 |
| 8 | 112026000 | 112027199 | 5.923508 | 11.17025 | 33.52968 | 13.69351 | -1.09875 | 0.055461 | 0.100188 |
| 12 | 54359000 | 54360399 | 5.923508 | 11.17025 | 33.52968 | 13.69351 | -1.09875 | 0.055461 | 0.100188 |
| 9 | 39412000 | 39413999 | 16.14222 | 30.45467 | 16.76327 | 6.839618 | 1.329222 | 0.101117 | 0.100193 |
| 8 | 121921400 | 121950799 | 164.5544 | 236.6199 | 560.0268 | 231.1683 | -1.0337 | 4.162563 | 0.100198 |
| 9 | 22203400 | 22203999 | 9.921286 | 19.37593 | 10.73414 | 3.7681 | 1.511189 | -0.47879 | 0.100245 |
| 5 | 139367400 | 139372199 | 18.67165 | 30.65562 | 85.74325 | 36.93151 | -1.10177 | 1.413137 | 0.100256 |
| 17 | 7244200 | 7246599 | 6.72765 | 12.13732 | 42.75589 | 18.60607 | -1.39686 | 0.348768 | 0.100262 |
| 10 | 72227200 | 72228199 | 10.93162 | 21.3283 | 11.29473 | 4.031967 | 1.555443 | -0.4054 | 0.100269 |
| 5 | 76917800 | 76920199 | 9.759712 | 17.36139 | 44.10915 | 19.44428 | -0.94229 | 0.461703 | 0.100276 |
| 14 | 67104400 | 67107599 | 9.633781 | 16.75526 | 55.21648 | 24.25868 | -1.32454 | 0.759861 | 0.100282 |
| 7 | 44372000 | 44374399 | 9.614894 | 17.07662 | 44.08784 | 19.46094 | -0.96648 | 0.471913 | 0.100286 |
| 4 | 63326000 | 63329199 | 13.68514 | 23.1212 | 67.98562 | 29.74446 | -1.16859 | 1.067667 | 0.100293 |
| 10 | 50150400 | 50153399 | 44.78907 | 73.58016 | 45.64492 | 19.66528 | 1.064516 | 1.436835 | 0.100298 |
| 12 | 22903000 | 22908399 | 243.9208 | 357.9069 | 252.3042 | 103.9179 | 0.741169 | 3.858846 | 0.100302 |
| 13 | 30382000 | 30383399 | 5.46272 | 10.3999 | 33.35073 | 13.34957 | -1.16762 | -0.01722 | 0.100317 |
| X | 54796400 | 54797399 | 14.01616 | 27.07172 | 13.95364 | 5.367046 | 1.499161 | -0.11108 | 0.10032 |
| 19 | 33616200 | 33616999 | 11.27214 | 21.83233 | 15.2482 | 5.789303 | 1.082354 | -0.18071 | 0.10033 |
| 8 | 61058200 | 61059999 | 17.14671 | 32.71052 | 13.28287 | 5.293936 | 1.791697 | 0.028762 | 0.10033 |
| 11 | 23121200 | 23122799 | 9.502883 | 16.78291 | 46.15968 | 20.42174 | -1.06127 | 0.526791 | 0.100342 |
| 12 | 23200600 | 23210599 | 268.1246 | 393.6591 | 245.8249 | 101.2454 | 0.916802 | 3.852201 | 0.100348 |
| 11 | 107533800 | 107535599 | 6.848742 | 12.38828 | 40.68055 | 17.64489 | -1.29296 | 0.330737 | 0.10035 |
| 12 | 108997400 | 109001199 | 48.06522 | 75.4733 | 182.8769 | 75.85074 | -0.95596 | 2.555315 | 0.100357 |
| 3 | 91175600 | 91176399 | 10.91975 | 21.13094 | 15.3054 | 5.835305 | 1.024726 | -0.17112 | 0.100366 |
| 18 | 42748400 | 42750199 | 8.934698 | 16.09826 | 41.67397 | 18.16455 | -0.95765 | 0.37091 | 0.100375 |
| 7 | 32853200 | 32855199 | 29.05804 | 50.62082 | 24.82096 | 10.91597 | 1.409786 | 0.784082 | 0.100385 |
| 13 | 118713000 | 118713599 | 9.819297 | 19.15304 | 12.67369 | 4.540681 | 1.234535 | -0.40342 | 0.100395 |
| 11 | 77106400 | 77107799 | 3.980978 | 7.630935 | 29.96142 | 11.79455 | -1.43116 | -0.09194 | 0.100405 |
| 13 | 117431200 | 117432799 | 24.75686 | 43.18078 | 25.17428 | 11.07988 | 1.161788 | 0.745944 | 0.100408 |
| 13 | 64517600 | 64520199 | 13.04992 | 22.12447 | 69.17584 | 30.297 | -1.25663 | 1.035013 | 0.100415 |
| 12 | 20129000 | 20131799 | 74.96844 | 120.1127 | 63.0244 | 26.58818 | 1.274486 | 2.01698 | 0.100429 |
| 7 | 31133400 | 31137999 | 137.264 | 210.9175 | 141.0953 | 58.58669 | 0.874519 | 2.978694 | 0.100436 |
| 18 | 56337200 | 56338399 | 23.20112 | 40.91146 | 23.34666 | 10.33212 | 1.193518 | 0.582426 | 0.10044 |
| 7 | 25340800 | 25342799 | 6.105582 | 10.98923 | 40.60217 | 17.72079 | -1.4665 | 0.357624 | 0.100452 |
| 8 | 45187600 | 45188199 | 3.3442 | 6.52797 | 23.33363 | 8.259115 | -1.15176 | -0.49405 | 0.100461 |
| 8 | 122859200 | 122870199 | 57.20776 | 88.49812 | 223.2433 | 92.72984 | -1.03088 | 2.83028 | 0.100469 |
| 4 | 144473600 | 144474599 | 12.31907 | 23.55589 | 14.25368 | 5.650777 | 1.229341 | -0.03475 | 0.100472 |
| 9 | 112978200 | 112979999 | 9.098846 | 16.34678 | 43.76898 | 19.13401 | -1.00917 | 0.392581 | 0.100478 |
| 9 | 114228600 | 114250399 | 230.1969 | 323.7204 | 673.7761 | 279.136 | -0.88073 | 4.493422 | 0.10048 |
| 12 | 23719200 | 23726599 | 127.3542 | 196.0599 | 132.2373 | 54.92283 | 0.864719 | 2.936887 | 0.100482 |
| 10 | 127069400 | 127080399 | 62.22562 | 95.7353 | 245.5188 | 101.9638 | -1.06082 | 2.937026 | 0.100487 |
| 16 | 30899000 | 30900999 | 6.283549 | 11.64979 | 35.81473 | 15.03969 | -1.16528 | 0.17288 | 0.100496 |

| | | | | | | | | | |
|---|---|---|---|---|---|---|---|---|---|
| 10 | 16298200 | 16304199 | 29.86422 | 48.19532 | 115.0806 | 49.05927 | -0.89444 | 1.753528 | 0.1005 |
| 11 | 35725000 | 35726399 | 5.294657 | 10.11287 | 31.11129 | 12.36754 | -1.09864 | -0.04555 | 0.100512 |
| 13 | 46225000 | 46226999 | 18.18938 | 33.64992 | 18.01081 | 7.585757 | 1.333939 | 0.238069 | 0.100512 |
| 3 | 108258200 | 108260199 | 9.483479 | 16.78629 | 44.06443 | 19.48602 | -0.99281 | 0.499311 | 0.100518 |
| 2 | 22373400 | 22374399 | 14.71237 | 27.74721 | 17.63253 | 7.199638 | 1.123227 | 0.093048 | 0.100537 |
| 6 | 127126000 | 127132999 | 35.93935 | 57.17888 | 146.5895 | 61.23148 | -1.0208 | 2.218794 | 0.100549 |
| 6 | 91855000 | 91855799 | 3.360864 | 6.5072 | 27.51777 | 10.46466 | -1.4872 | -0.22172 | 0.100551 |
| 2 | 5899800 | 5900799 | 5.162284 | 9.860567 | 31.69248 | 12.59716 | -1.16125 | -0.0466 | 0.100565 |
| 6 | 5600200 | 5601199 | 4.936821 | 9.544071 | 30.04163 | 11.50548 | -1.07976 | -0.17502 | 0.100565 |
| 10 | 60657800 | 60673399 | 76.52621 | 116.2631 | 297.5496 | 123.2329 | -1.07235 | 3.199229 | 0.100582 |
| 7 | 5344000 | 5345799 | 47.62852 | 77.21102 | 57.6457 | 24.68561 | 0.785691 | 1.646739 | 0.100584 |
| 17 | 86175200 | 86181799 | 40.58696 | 64.60132 | 146.1249 | 61.07427 | -0.83991 | 2.207459 | 0.100585 |
| 12 | 20614800 | 20619999 | 110.0003 | 170.5426 | 132.0798 | 54.85943 | 0.673228 | 2.800654 | 0.100622 |
| 7 | 99808200 | 99809599 | 5.901371 | 10.81554 | 37.71442 | 16.08838 | -1.36179 | 0.2413 | 0.100639 |
| 14 | 124635200 | 124636599 | 16.00034 | 29.97655 | 17.73302 | 7.322563 | 1.212255 | 0.144628 | 0.100669 |
| 12 | 18770400 | 18778999 | 190.06 | 284.9397 | 207.0197 | 85.40763 | 0.726308 | 3.481918 | 0.100671 |
| 7 | 34838000 | 34854199 | 87.7872 | 132.8793 | 298.3541 | 123.4263 | -0.88818 | 3.278001 | 0.100675 |
| 7 | 46351400 | 46353399 | 6.238437 | 11.64886 | 34.34301 | 14.26149 | -1.09228 | 0.126082 | 0.100693 |
| X | 151443000 | 151444599 | 6.529642 | 12.21388 | 34.41745 | 14.25031 | -1.02456 | 0.116521 | 0.100709 |
| 6 | 79234800 | 79236199 | 14.44763 | 27.26061 | 17.35945 | 7.081203 | 1.120584 | 0.085167 | 0.100709 |
| 16 | 10728200 | 10733799 | 27.66536 | 44.55552 | 109.894 | 46.71983 | -0.9444 | 1.811595 | 0.100711 |
| 8 | 83403000 | 83404599 | 15.74637 | 29.36921 | 18.92334 | 7.873433 | 1.081732 | 0.173293 | 0.100713 |
| 17 | 77530400 | 77532199 | 15.74637 | 29.36921 | 18.92334 | 7.873433 | 1.081732 | 0.173293 | 0.100713 |
| 7 | 140549200 | 140553799 | 19.17427 | 31.72621 | 78.31468 | 33.8671 | -0.91869 | 1.315622 | 0.100715 |
| 6 | 135703000 | 135712399 | 90.36522 | 137.2904 | 281.1307 | 116.4334 | -0.75093 | 3.201686 | 0.100719 |
| 9 | 120378600 | 120380199 | 8.20653 | 14.72575 | 41.07477 | 17.97462 | -1.0682 | 0.390337 | 0.100732 |
| 5 | 113222600 | 113236599 | 86.02271 | 130.4886 | 289.3361 | 119.7757 | -0.86748 | 3.233301 | 0.100738 |
| 2 | 66086600 | 66087999 | 6.157944 | 11.3883 | 36.84713 | 15.52616 | -1.24216 | 0.187976 | 0.100739 |
| 2 | 124900400 | 124905199 | 32.20127 | 51.37353 | 139.7549 | 58.60043 | -1.10179 | 2.123087 | 0.10074 |
| 11 | 26309200 | 26310799 | 22.05923 | 39.12764 | 22.73693 | 10.04737 | 1.171187 | 0.516481 | 0.100743 |
| 6 | 31632000 | 31635599 | 15.99853 | 27.00741 | 67.09004 | 29.3453 | -0.92661 | 1.081844 | 0.100743 |
| 7 | 45887000 | 45888399 | 7.705688 | 13.95382 | 39.11676 | 16.9431 | -1.06508 | 0.328705 | 0.100755 |
| 11 | 62084000 | 62085399 | 14.33907 | 27.13059 | 16.22049 | 6.57806 | 1.217604 | 0.062187 | 0.100759 |
| 4 | 130293400 | 130295199 | 7.559544 | 13.69326 | 39.1218 | 16.93896 | -1.09193 | 0.325465 | 0.100765 |
| 7 | 44578600 | 44579599 | 4.738472 | 9.072216 | 31.77399 | 12.55058 | -1.27524 | -0.07182 | 0.100773 |
| 1 | 86312200 | 86313199 | 14.35433 | 27.06515 | 14.80437 | 6.047429 | 1.337029 | 0.082752 | 0.100774 |
| 6 | 124966200 | 124984199 | 131.0775 | 193.5053 | 412.0858 | 169.7045 | -0.84463 | 3.753303 | 0.100785 |
| 9 | 119956000 | 119960599 | 18.52732 | 30.4042 | 86.84907 | 37.4003 | -1.13234 | 1.419306 | 0.100791 |
| 4 | 139037200 | 139039199 | 9.567182 | 16.74181 | 49.89108 | 22.01057 | -1.17907 | 0.660721 | 0.1008 |
| 15 | 3076600 | 3087999 | 246.3918 | 362.5017 | 255.9743 | 105.4118 | 0.742417 | 3.817778 | 0.100804 |
| 6 | 26861000 | 26863599 | 20.03088 | 36.51361 | 18.85189 | 8.098256 | 1.36608 | 0.334501 | 0.100807 |
| 11 | 33817600 | 33823799 | 37.23967 | 59.61796 | 127.3353 | 53.65537 | -0.74806 | 2.017217 | 0.100808 |
| 11 | 46198400 | 46204599 | 32.06723 | 51.44321 | 127.3633 | 53.83561 | -0.9571 | 1.945431 | 0.100811 |
| 14 | 21731800 | 21737199 | 20.70791 | 33.99886 | 85.62141 | 36.87906 | -0.95104 | 1.41713 | 0.100825 |
| 17 | 18686000 | 18689399 | 58.46212 | 94.6052 | 51.36762 | 21.96835 | 1.241571 | 1.69973 | 0.100848 |
| 14 | 79520800 | 79521399 | 9.923195 | 19.37172 | 11.35596 | 4.014638 | 1.422458 | -0.45266 | 0.100852 |
| 11 | 74326200 | 74333799 | 45.23563 | 71.64079 | 158.6298 | 65.96489 | -0.81633 | 2.362084 | 0.100861 |

| | | | | | | | | | |
|---|---|---|---|---|---|---|---|---|---|
| 8 | 107329600 | 107333799 | 21.64684 | 35.19284 | 94.94031 | 40.73951 | -1.05886 | 1.562622 | 0.100874 |
| 8 | 80832000 | 80832999 | 11.45385 | 22.18144 | 15.2439 | 5.791146 | 1.104705 | -0.1765 | 0.100876 |
| 6 | 71470400 | 71471999 | 7.512965 | 13.34489 | 43.07494 | 19.01213 | -1.28562 | 0.456163 | 0.100876 |
| 11 | 103126200 | 103130999 | 25.10485 | 40.49086 | 108.1767 | 46.08132 | -1.05677 | 1.76205 | 0.100879 |
| 12 | 19551600 | 19553599 | 51.83056 | 84.2656 | 50.68749 | 21.75042 | 1.100264 | 1.596625 | 0.100893 |
| 5 | 132827600 | 132828399 | 13.40098 | 25.61451 | 16.69294 | 6.625333 | 1.122423 | -0.01155 | 0.100897 |
| X | 9239200 | 9243599 | 45.49324 | 75.93203 | 31.49937 | 13.70167 | 1.645522 | 1.252649 | 0.100898 |
| 1 | 33758800 | 33761999 | 14.37158 | 24.46461 | 63.27782 | 27.73575 | -0.98354 | 0.997856 | 0.100928 |
| 8 | 59823600 | 59827399 | 60.76515 | 97.36164 | 74.15508 | 31.28639 | 0.738141 | 2.00008 | 0.10093 |
| 2 | 173595000 | 173596999 | 5.687835 | 10.55763 | 35.26413 | 14.785 | -1.28147 | 0.161333 | 0.100933 |
| 10 | 63592400 | 63593799 | 8.323718 | 15.02309 | 40.11142 | 17.44611 | -0.99964 | 0.353603 | 0.10094 |
| 1 | 138533000 | 138533599 | 3.760895 | 7.334115 | 25.01907 | 8.991033 | -1.10647 | -0.42864 | 0.100946 |
| 18 | 68737200 | 68737999 | 11.40872 | 22.22436 | 12.45427 | 4.538747 | 1.449672 | -0.32493 | 0.100953 |
| 15 | 82844800 | 82866599 | 116.8236 | 173.3145 | 399.3793 | 164.5035 | -0.9517 | 3.667865 | 0.100961 |
| 10 | 117417000 | 117418599 | 6.878366 | 12.72802 | 38.02112 | 16.00817 | -1.12776 | 0.1919 | 0.100972 |
| 19 | 24653800 | 24655999 | 6.878366 | 12.72802 | 38.02112 | 16.00817 | -1.12776 | 0.1919 | 0.100972 |
| 10 | 11659000 | 11659799 | 12.59778 | 24.32335 | 14.94625 | 5.75857 | 1.244841 | -0.1194 | 0.100977 |
| 7 | 28175600 | 28187999 | 76.44999 | 117.4123 | 239.4827 | 99.433 | -0.73281 | 2.976634 | 0.100987 |
| 8 | 70818200 | 70819799 | 5.19545 | 9.487489 | 38.6948 | 16.58525 | -1.58994 | 0.259822 | 0.100987 |
| 9 | 120830800 | 120834399 | 28.45659 | 45.83745 | 108.9489 | 46.32919 | -0.89086 | 1.807285 | 0.100996 |
| 5 | 74670600 | 74671799 | 5.865828 | 11.15144 | 34.44104 | 13.82757 | -1.11724 | 0.002553 | 0.101009 |
| 6 | 83084400 | 83086799 | 10.00052 | 17.5402 | 48.28992 | 21.3454 | -1.06552 | 0.622505 | 0.10101 |
| 2 | 150548000 | 150548799 | 8.557379 | 15.17697 | 43.04739 | 19.0243 | -1.10203 | 0.472858 | 0.101012 |
| 3 | 59563200 | 59564799 | 19.70434 | 35.68591 | 20.50609 | 8.880677 | 1.205622 | 0.371569 | 0.10103 |
| 7 | 42454000 | 42457799 | 38.0839 | 62.99146 | 46.11818 | 19.93871 | 0.828966 | 1.340542 | 0.101033 |
| 17 | 53676800 | 53677799 | 17.77792 | 32.36277 | 20.68281 | 8.899883 | 1.05898 | 0.328258 | 0.10104 |
| 2 | 125760000 | 125760399 | 4.189601 | 8.150971 | 27.0458 | 9.978412 | -1.10266 | -0.32213 | 0.101053 |
| 8 | 27649800 | 27651999 | 8.835271 | 15.47172 | 51.99804 | 22.9535 | -1.3516 | 0.647271 | 0.101073 |
| 16 | 93282800 | 93287199 | 39.44147 | 62.8286 | 143.2332 | 59.93237 | -0.84991 | 2.179683 | 0.101075 |
| X | 33467400 | 33467999 | 18.93472 | 34.57888 | 18.9607 | 8.126319 | 1.282686 | 0.317597 | 0.101085 |
| 2 | 165730200 | 165732599 | 5.569164 | 10.18711 | 37.66845 | 16.11081 | -1.44744 | 0.252272 | 0.101096 |
| 17 | 34809000 | 34814999 | 24.1482 | 38.9723 | 107.6303 | 45.88557 | -1.10339 | 1.74168 | 0.101101 |
| 3 | 122121800 | 122122999 | 7.363698 | 13.42379 | 38.70307 | 16.62462 | -1.09702 | 0.287448 | 0.101103 |
| 13 | 66174200 | 66182399 | 133.7049 | 205.4896 | 136.5659 | 56.70749 | 0.884165 | 2.971749 | 0.101105 |
| 7 | 94377800 | 94378399 | 9.682511 | 18.89585 | 11.32142 | 4.024451 | 1.383806 | -0.43847 | 0.101115 |
| 5 | 139057800 | 139067399 | 53.4032 | 83.89796 | 180.2901 | 74.78279 | -0.78248 | 2.547097 | 0.101119 |
| 12 | 48920800 | 48922599 | 21.96474 | 38.91766 | 22.89692 | 10.12338 | 1.152287 | 0.523592 | 0.101126 |
| 7 | 26413800 | 26415999 | 47.97979 | 77.86651 | 56.61055 | 24.2621 | 0.825389 | 1.625267 | 0.101136 |
| 5 | 135988600 | 135997199 | 42.24715 | 67.11945 | 149.0968 | 62.18033 | -0.81658 | 2.267454 | 0.101137 |
| 2 | 179846200 | 179850599 | 19.57276 | 31.85739 | 93.37036 | 40.08259 | -1.17671 | 1.541758 | 0.101149 |
| 15 | 84051800 | 84054399 | 12.91586 | 21.83414 | 67.91049 | 29.71682 | -1.24909 | 1.0601 | 0.10117 |
| 10 | 86230400 | 86231199 | 11.43348 | 22.22797 | 13.47865 | 5.002128 | 1.313455 | -0.26058 | 0.101184 |
| 16 | 27832200 | 27834599 | 18.50432 | 33.81535 | 18.89059 | 8.089543 | 1.255686 | 0.307986 | 0.101209 |
| 10 | 127107600 | 127110799 | 9.758872 | 17.00733 | 53.09032 | 23.35437 | -1.24636 | 0.728331 | 0.10122 |
| X | 97148800 | 97149799 | 12.58366 | 24.32306 | 15.0868 | 5.782674 | 1.239095 | -0.13523 | 0.101232 |
| X | 110717600 | 110718799 | 15.08654 | 28.49961 | 17.38314 | 7.07326 | 1.185686 | 0.084125 | 0.101239 |
| 1 | 28927200 | 28930599 | 50.42208 | 81.43491 | 59.17182 | 25.25035 | 0.819397 | 1.738775 | 0.10124 |

| | | | | | | | | | |
|---|---|---|---|---|---|---|---|---|---|
| 3 | 17985200 | 17987199 | 22.52852 | 39.58332 | 26.34016 | 11.66044 | 0.971894 | 0.624921 | 0.101241 |
| 10 | 81391000 | 81437799 | 296.0659 | 404.878 | 919.6992 | 382.6938 | -1.04758 | 4.921586 | 0.101241 |
| 17 | 13394800 | 13461399 | 2967.907 | 3399.029 | 6012.407 | 2560.372 | -0.94567 | 7.755539 | 0.101293 |
| 8 | 125622200 | 125626599 | 22.13195 | 35.91964 | 95.70916 | 41.02033 | -1.04331 | 1.598943 | 0.101306 |
| 6 | 43765200 | 43769199 | 37.43412 | 61.88221 | 45.59644 | 19.70556 | 0.819738 | 1.344913 | 0.101308 |
| 2 | 92261400 | 92264999 | 15.25102 | 25.69118 | 69.19852 | 30.24815 | -1.043 | 1.09962 | 0.101313 |
| 4 | 138604200 | 138611799 | 33.47826 | 53.42094 | 139.9459 | 58.69612 | -1.04724 | 2.117923 | 0.101319 |
| 4 | 15147600 | 15149999 | 14.63772 | 25.02678 | 64.57836 | 28.31576 | -0.97931 | 0.95852 | 0.101329 |
| 18 | 17597800 | 17599999 | 46.55451 | 75.87714 | 52.20402 | 22.42158 | 0.909513 | 1.551769 | 0.101334 |
| X | 63660000 | 63660999 | 13.81145 | 26.74385 | 13.29986 | 5.055498 | 1.565454 | -0.15352 | 0.101338 |
| 2 | 169671000 | 169675399 | 31.22306 | 50.18481 | 117.4726 | 49.79725 | -0.87294 | 1.881517 | 0.101344 |
| 13 | 82697800 | 82698999 | 11.25125 | 21.87528 | 13.48311 | 5.000837 | 1.290806 | -0.26498 | 0.10135 |
| 11 | 13902200 | 13902599 | 8.551276 | 16.71461 | 10.983 | 3.785659 | 1.292024 | -0.56011 | 0.101361 |
| 12 | 34015800 | 34018999 | 22.10817 | 35.9645 | 93.97494 | 40.33411 | -1.01236 | 1.554078 | 0.10137 |
| 6 | 115999000 | 115999599 | 92.24987 | 146.0093 | 71.63415 | 29.77927 | 1.353674 | 2.393418 | 0.101372 |
| 14 | 114752200 | 114756999 | 40.38169 | 64.40638 | 136.8295 | 57.35148 | -0.74637 | 2.139338 | 0.101377 |
| 14 | 115277400 | 115279999 | 14.25907 | 23.85699 | 74.41497 | 32.41846 | -1.25396 | 1.17005 | 0.101384 |
| 7 | 8094800 | 8100399 | 58.90256 | 94.6755 | 68.82746 | 29.17695 | 0.810556 | 1.900115 | 0.10139 |
| 5 | 116056400 | 116058399 | 8.75648 | 15.46093 | 45.63909 | 20.1924 | -1.16244 | 0.524901 | 0.101409 |
| 5 | 135215200 | 135216999 | 8.75648 | 15.46093 | 45.63909 | 20.1924 | -1.16244 | 0.524901 | 0.101409 |
| 7 | 32170800 | 32174599 | 89.21795 | 140.4264 | 106.4336 | 44.15849 | 0.721056 | 2.518902 | 0.10142 |
| 14 | 42655400 | 42657999 | 51.61714 | 83.93009 | 50.69892 | 21.75641 | 1.094317 | 1.593759 | 0.10142 |
| 7 | 79191800 | 79197399 | 35.20681 | 56.21861 | 133.2748 | 55.9537 | -0.90229 | 2.096839 | 0.101434 |
| 11 | 97305200 | 97310399 | 15.59387 | 25.52477 | 89.86393 | 38.65673 | -1.43406 | 1.448649 | 0.101442 |
| X | 3180000 | 3184599 | 124.9444 | 194.1875 | 112.9814 | 46.90973 | 1.088101 | 2.76534 | 0.101447 |
| 7 | 18052000 | 18062599 | 275.8432 | 401.0827 | 291.7698 | 120.2652 | 0.683273 | 4.001269 | 0.10145 |
| 3 | 51322800 | 51324799 | 10.97666 | 19.09241 | 53.66781 | 23.57953 | -1.09659 | 0.762989 | 0.101475 |
| 14 | 46742800 | 46742999 | 7.407073 | 14.48644 | 9.390068 | 3.075709 | 1.375617 | -0.75572 | 0.101476 |
| 8 | 85007400 | 85011599 | 16.25021 | 27.16666 | 71.18913 | 30.99749 | -1.00364 | 1.183379 | 0.101488 |
| 8 | 96686600 | 96687799 | 7.706039 | 13.98205 | 39.68862 | 17.14837 | -1.08101 | 0.316466 | 0.101489 |
| 15 | 78703800 | 78705399 | 7.706039 | 13.98205 | 39.68862 | 17.14837 | -1.08101 | 0.316466 | 0.101489 |
| 14 | 11509200 | 11510199 | 5.899219 | 11.19615 | 33.83384 | 13.632 | -1.09136 | 0.009941 | 0.101489 |
| 4 | 151835400 | 151839399 | 18.70815 | 30.6652 | 89.13892 | 38.36747 | -1.15822 | 1.435467 | 0.10149 |
| 7 | 81622800 | 81624999 | 14.44553 | 24.39788 | 66.363 | 29.03166 | -1.05669 | 1.072322 | 0.101493 |
| 2 | 4886000 | 4887599 | 8.1129 | 14.42411 | 43.06915 | 18.99448 | -1.17366 | 0.452797 | 0.101497 |
| 3 | 88549400 | 88551799 | 8.1129 | 14.42411 | 43.06915 | 18.99448 | -1.17366 | 0.452797 | 0.101497 |
| 7 | 14076200 | 14079999 | 76.99053 | 122.8068 | 77.6751 | 32.56139 | 0.998653 | 2.155588 | 0.101498 |
| 1 | 92827000 | 92828399 | 5.035954 | 9.511757 | 32.37457 | 13.17784 | -1.27269 | 0.035608 | 0.101517 |
| 13 | 17992800 | 17996399 | 38.1256 | 60.81069 | 135.7597 | 56.90649 | -0.81767 | 2.136238 | 0.101525 |
| 18 | 77880400 | 77887799 | 38.03282 | 60.51312 | 149.248 | 62.34704 | -0.96507 | 2.218196 | 0.101525 |
| X | 90869000 | 90869799 | 13.31262 | 25.90501 | 10.93275 | 4.027326 | 1.838415 | -0.26992 | 0.101527 |
| 1 | 143093400 | 143094599 | 13.15826 | 25.39584 | 14.93169 | 5.763216 | 1.305752 | -0.10709 | 0.101527 |
| 13 | 3469000 | 3471999 | 109.3151 | 169.4942 | 131.4409 | 54.59341 | 0.671417 | 2.798606 | 0.101532 |
| 4 | 150123200 | 150128399 | 24.39608 | 39.50906 | 100.0125 | 42.79667 | -0.97268 | 1.648288 | 0.101545 |
| 8 | 85752600 | 85754199 | 5.380217 | 10.01549 | 34.76889 | 14.51371 | -1.33129 | 0.140412 | 0.101548 |
| 9 | 102931000 | 102935599 | 24.64014 | 40.07048 | 95.18114 | 40.84702 | -0.87567 | 1.563758 | 0.10155 |
| 7 | 14021600 | 14022599 | 32.54711 | 54.86935 | 38.85939 | 16.99021 | 0.876967 | 1.1193 | 0.101558 |

| | | | | | | | | | |
|---|---|---|---|---|---|---|---|---|---|
| 15 | 96182200 | 96186799 | 23.95239 | 38.72364 | 102.9148 | 43.97387 | -1.04517 | 1.687878 | 0.101563 |
| 18 | 50261200 | 50261999 | 2.553697 | 4.965315 | 26.03145 | 9.650577 | -1.75523 | -0.32683 | 0.101567 |
| 18 | 52771200 | 52772599 | 9.551141 | 16.71399 | 49.36076 | 21.77687 | -1.16593 | 0.66081 | 0.101568 |
| Y | 20806600 | 20807199 | 8.128355 | 15.89037 | 8.711626 | 2.991012 | 1.551735 | -0.59173 | 0.101587 |
| 19 | 53242800 | 53256199 | 84.07977 | 127.7529 | 280.8565 | 116.3231 | -0.85317 | 3.198494 | 0.10162 |
| 11 | 60029800 | 60032399 | 6.774182 | 12.02209 | 43.59597 | 19.25632 | -1.45261 | 0.458646 | 0.101631 |
| 3 | 4355200 | 4356599 | 20.25243 | 36.21429 | 21.95779 | 9.637189 | 1.1162 | 0.451631 | 0.10164 |
| 15 | 81741800 | 81744199 | 8.026871 | 14.13885 | 46.68341 | 20.66432 | -1.32406 | 0.549993 | 0.101648 |
| 7 | 7427400 | 7429999 | 28.52951 | 49.86963 | 24.41491 | 10.7612 | 1.412504 | 0.735675 | 0.10165 |
| 16 | 46746400 | 46747199 | 3.23794 | 6.312656 | 24.2283 | 8.736631 | -1.27764 | -0.42608 | 0.101652 |
| 12 | 16903600 | 16904599 | 4.536312 | 8.794766 | 29.11082 | 11.00183 | -1.13341 | -0.23003 | 0.101654 |
| 13 | 92620400 | 92621199 | 4.536312 | 8.794766 | 29.11082 | 11.00183 | -1.13341 | -0.23003 | 0.101654 |
| 7 | 61534600 | 61536599 | 26.81328 | 46.29788 | 32.34679 | 14.19046 | 0.900244 | 0.879803 | 0.10166 |
| 12 | 80078800 | 80079599 | 4.853019 | 9.326454 | 30.23028 | 11.81287 | -1.14967 | -0.10975 | 0.101681 |
| 11 | 113063200 | 113083999 | 114.3887 | 168.9143 | 423.7129 | 174.4932 | -1.08031 | 3.74712 | 0.101684 |
| 5 | 116009200 | 116010799 | 6.673948 | 12.10649 | 39.14279 | 16.91761 | -1.26692 | 0.310658 | 0.101702 |
| 7 | 62119200 | 62120999 | 24.14496 | 43.13569 | 19.03432 | 8.360459 | 1.570808 | 0.479213 | 0.101704 |
| 7 | 23224200 | 23227199 | 86.22196 | 136.3525 | 99.76348 | 41.45516 | 0.777157 | 2.414189 | 0.101706 |
| 5 | 138977600 | 138994199 | 134.119 | 198.0605 | 412.464 | 169.8608 | -0.81197 | 3.74807 | 0.101733 |
| 12 | 39459800 | 39460599 | 10.97675 | 21.3124 | 14.73468 | 5.525804 | 1.113319 | -0.22624 | 0.101734 |
| 5 | 114946600 | 114962599 | 131.9874 | 196.0184 | 378.5718 | 155.9497 | -0.69567 | 3.651295 | 0.101742 |
| 9 | 33271000 | 33271599 | 4.667389 | 9.038346 | 28.36615 | 10.77895 | -1.06456 | -0.21057 | 0.10176 |
| 16 | 31844400 | 31845599 | 4.667389 | 9.038346 | 28.36615 | 10.77895 | -1.06456 | -0.21057 | 0.10176 |
| 12 | 103275400 | 103314599 | 247.8854 | 344.6016 | 777 | 322.4776 | -1.01292 | 4.667224 | 0.10176 |
| 3 | 154076400 | 154077199 | 4.664143 | 9.042388 | 28.48888 | 10.76794 | -1.06283 | -0.22891 | 0.101777 |
| 10 | 84190200 | 84191199 | 4.664143 | 9.042388 | 28.48888 | 10.76794 | -1.06283 | -0.22891 | 0.101777 |
| 3 | 124439200 | 124441199 | 15.39244 | 29.00912 | 16.63692 | 6.80132 | 1.267486 | 0.099467 | 0.101781 |
| 14 | 104910800 | 104911399 | 10.10081 | 19.71776 | 11.3521 | 4.015715 | 1.447556 | -0.44788 | 0.101784 |
| 15 | 62019200 | 62022799 | 20.46866 | 33.4456 | 90.22048 | 38.78542 | -1.05348 | 1.48347 | 0.101802 |
| 17 | 19431200 | 19432599 | 42.25212 | 69.39796 | 48.92048 | 21.07464 | 0.880647 | 1.436358 | 0.101802 |
| 11 | 68803800 | 68804799 | 12.30192 | 23.50513 | 16.29263 | 6.471923 | 1.032919 | -0.02181 | 0.10182 |
| 15 | 88727600 | 88758599 | 165.0126 | 237.5873 | 555.6044 | 229.2971 | -1.01451 | 4.143408 | 0.101822 |
| 2 | 130544000 | 130545199 | 5.868904 | 11.19399 | 32.27392 | 12.86782 | -1.01018 | -0.02709 | 0.101835 |
| 4 | 154919200 | 154924799 | 27.61724 | 44.2952 | 124.7197 | 52.70157 | -1.14241 | 1.947006 | 0.101847 |
| 2 | 59209400 | 59209999 | 4.123615 | 7.997972 | 28.45426 | 10.73283 | -1.23303 | -0.243 | 0.101847 |
| 2 | 166550800 | 166562399 | 57.59023 | 89.75438 | 204.8541 | 85.01998 | -0.87837 | 2.696688 | 0.10186 |
| 9 | 115698000 | 115706799 | 61.72952 | 96.40748 | 195.2441 | 81.01142 | -0.70375 | 2.661055 | 0.101867 |
| 1 | 118752000 | 118757599 | 34.08245 | 54.42387 | 133.4396 | 56.02387 | -0.95073 | 2.095592 | 0.101872 |
| 5 | 112401600 | 112408799 | 28.40689 | 45.56952 | 123.895 | 52.36668 | -1.09177 | 1.942136 | 0.101876 |
| 7 | 23202000 | 23204999 | 68.17036 | 109.0578 | 75.81183 | 31.91257 | 0.867194 | 2.057862 | 0.101891 |
| 7 | 30143400 | 30163399 | 145.3928 | 212.6351 | 456.4178 | 188.0259 | -0.86889 | 3.905771 | 0.101893 |
| 1 | 191173200 | 191174999 | 11.05851 | 19.40761 | 48.28635 | 21.35476 | -0.92051 | 0.618234 | 0.101898 |
| 18 | 19196200 | 19197599 | 15.45397 | 29.24795 | 16.82557 | 6.81956 | 1.274285 | 0.07395 | 0.101973 |
| 3 | 52036200 | 52037599 | 7.978853 | 14.5353 | 38.70563 | 16.63987 | -0.98469 | 0.293532 | 0.101978 |
| X | 28087200 | 28087599 | 9.187759 | 17.96392 | 6.604957 | 2.136344 | 2.189333 | -0.77604 | 0.101985 |
| 1 | 80966400 | 80967199 | 11.51788 | 22.44921 | 10.37369 | 3.753667 | 1.729895 | -0.35579 | 0.101994 |
| 2 | 72115800 | 72116799 | 4.482357 | 8.491973 | 32.43687 | 13.11839 | -1.42888 | 0.00802 | 0.10201 |

| | | | | | | | | | |
|---|---|---|---|---|---|---|---|---|---|
| 4 | 133479000 | 133482199 | 14.2375 | 23.86788 | 73.38581 | 32.00988 | -1.23331 | 1.147347 | 0.102019 |
| 13 | 44650800 | 44651799 | 5.77975 | 10.92047 | 34.77092 | 14.14269 | -1.17824 | 0.042158 | 0.102023 |
| 17 | 43251000 | 43253799 | 12.4994 | 21.73128 | 55.3757 | 24.32306 | -0.95624 | 0.777618 | 0.102025 |
| 12 | 111909400 | 111915999 | 27.2645 | 43.75069 | 120.992 | 51.16416 | -1.11562 | 1.930858 | 0.102026 |
| 10 | 10563800 | 10565799 | 11.70184 | 20.44047 | 51.01464 | 22.47379 | -0.92528 | 0.701608 | 0.102026 |
| 19 | 13451400 | 13452399 | 14.35703 | 27.83463 | 10.56696 | 3.99358 | 1.954356 | -0.17452 | 0.102032 |
| 11 | 63846400 | 63848999 | 13.38321 | 22.9451 | 59.52883 | 26.10749 | -0.98605 | 0.92439 | 0.10204 |
| 9 | 63864600 | 63868199 | 12.79419 | 21.88479 | 61.07511 | 26.78067 | -1.09131 | 0.945087 | 0.102051 |
| 17 | 66383200 | 66384799 | 7.983665 | 14.10631 | 44.61369 | 19.73596 | -1.26022 | 0.512532 | 0.102053 |
| 13 | 66827000 | 66829199 | 70.94494 | 113.0919 | 84.39979 | 35.34938 | 0.759472 | 2.171832 | 0.102056 |
| 11 | 117544400 | 117546999 | 13.7094 | 23.45492 | 60.60142 | 26.57339 | -0.9807 | 0.946177 | 0.102091 |
| 17 | 66627600 | 66629599 | 9.034783 | 15.98276 | 44.07392 | 19.49248 | -1.06362 | 0.503306 | 0.102103 |
| 1 | 171330800 | 171336199 | 42.36705 | 67.40111 | 150.405 | 62.81991 | -0.82121 | 2.225747 | 0.102118 |
| 17 | 87060000 | 87065399 | 24.05515 | 39.0103 | 97.9042 | 41.93682 | -0.95842 | 1.618582 | 0.102121 |
| 7 | 14597000 | 14602599 | 117.5348 | 183.0103 | 116.1735 | 48.22171 | 0.964592 | 2.730402 | 0.102136 |
| 7 | 141641200 | 141652799 | 50.92019 | 79.41723 | 205.4675 | 85.26536 | -1.05809 | 2.681209 | 0.102136 |
| 5 | 27108800 | 27111399 | 19.8127 | 32.39819 | 87.72824 | 37.72439 | -1.05805 | 1.470622 | 0.102162 |
| 3 | 37718400 | 37721199 | 16.07649 | 27.39664 | 66.23508 | 29.03451 | -0.88653 | 0.9974 | 0.102171 |
| 9 | 56498200 | 56499199 | 5.418667 | 10.38624 | 30.72476 | 12.09683 | -1.02966 | -0.07702 | 0.102177 |
| 4 | 153903000 | 153931799 | 220.2672 | 310.6862 | 664.532 | 275.1717 | -0.91571 | 4.448657 | 0.102187 |
| 6 | 48039600 | 48041999 | 12.84032 | 21.80971 | 63.58189 | 27.86275 | -1.15552 | 1.015028 | 0.102191 |
| 5 | 95668800 | 95669399 | 11.56553 | 22.35479 | 13.38227 | 5.129689 | 1.290215 | -0.15179 | 0.102195 |
| 17 | 31187200 | 31191199 | 21.69486 | 35.50623 | 87.16175 | 37.49245 | -0.91626 | 1.461032 | 0.1022 |
| 4 | 113422200 | 113423199 | 17.1725 | 31.57564 | 19.96109 | 8.477196 | 1.087805 | 0.268203 | 0.102212 |
| 17 | 5239800 | 5240799 | 4.653998 | 8.808459 | 34.71568 | 14.06932 | -1.47745 | 0.020093 | 0.102218 |
| 8 | 23879400 | 23881199 | 8.770063 | 15.84161 | 41.20229 | 17.90229 | -0.96141 | 0.353514 | 0.102222 |
| 4 | 124550000 | 124562399 | 80.50106 | 122.4762 | 283.6447 | 117.5241 | -0.92665 | 3.170202 | 0.102248 |
| 13 | 56060000 | 56062799 | 12.66274 | 21.58646 | 62.04718 | 27.19947 | -1.13439 | 0.97828 | 0.102252 |
| 13 | 24130200 | 24133999 | 12.81959 | 22.2721 | 54.26815 | 23.82591 | -0.8919 | 0.789755 | 0.102268 |
| 6 | 71277400 | 71279799 | 13.32119 | 23.01173 | 56.39306 | 24.73983 | -0.90189 | 0.851487 | 0.102274 |
| 3 | 79192000 | 79195199 | 13.85943 | 23.38514 | 70.22598 | 30.71249 | -1.19908 | 1.081248 | 0.102275 |
| 11 | 51746400 | 51752999 | 31.61117 | 50.64973 | 124.7091 | 52.61597 | -0.95118 | 1.983725 | 0.102281 |
| 7 | 31835000 | 31836399 | 37.51155 | 62.13712 | 43.9781 | 19.033 | 0.877842 | 1.321781 | 0.102281 |
| 9 | 91279000 | 91279999 | 12.15453 | 23.39152 | 16.16907 | 6.292599 | 1.063783 | -0.08866 | 0.102314 |
| X | 81694200 | 81695599 | 21.0415 | 37.57579 | 22.24124 | 9.77243 | 1.150175 | 0.467607 | 0.102315 |
| 2 | 115890200 | 115895399 | 28.82783 | 46.34605 | 116.142 | 49.25025 | -0.97056 | 1.870728 | 0.102318 |
| 1 | 9696800 | 9702199 | 22.57019 | 36.75681 | 95.60061 | 41.04843 | -1.00464 | 1.538682 | 0.102321 |
| 10 | 20187600 | 20189399 | 8.936688 | 15.72649 | 48.25599 | 21.36448 | -1.2202 | 0.566981 | 0.102327 |
| 10 | 118702400 | 118706599 | 18.98271 | 30.88565 | 96.83499 | 41.56434 | -1.2741 | 1.546977 | 0.102331 |
| 4 | 45745200 | 45746999 | 6.580248 | 12.19122 | 36.36402 | 15.28511 | -1.12343 | 0.179892 | 0.102339 |
| 7 | 60772800 | 60774199 | 19.53406 | 35.30654 | 21.05217 | 9.140102 | 1.150408 | 0.382612 | 0.102343 |
| 7 | 79674000 | 79675399 | 4.175351 | 7.95141 | 31.44892 | 12.58028 | -1.46387 | -0.03367 | 0.102352 |
| 4 | 133356600 | 133361999 | 19.31111 | 31.6239 | 86.0308 | 37.01499 | -1.06335 | 1.448559 | 0.102364 |
| 1 | 76121400 | 76123599 | 12.98248 | 22.55718 | 56.44322 | 24.7823 | -0.93008 | 0.792669 | 0.102366 |
| 5 | 123493400 | 123494999 | 6.633258 | 12.14238 | 37.71009 | 16.11266 | -1.19833 | 0.25329 | 0.102367 |
| 5 | 73495400 | 73497399 | 8.993177 | 16.02653 | 42.55905 | 18.73262 | -1.00395 | 0.442176 | 0.102369 |
| 14 | 19818200 | 19820199 | 12.01793 | 20.97299 | 51.55561 | 22.69876 | -0.90366 | 0.718002 | 0.102374 |

| | | | | | | | | | |
|---|---|---|---|---|---|---|---|---|---|
| 12 | 111424200 | 111430999 | 27.49987 | 44.32291 | 109.1872 | 46.47009 | -0.9413 | 1.786264 | 0.102376 |
| 9 | 45316800 | 45322199 | 26.88421 | 43.27716 | 111.8818 | 47.53254 | -1.01293 | 1.826008 | 0.102392 |
| 9 | 122304000 | 122306199 | 9.494915 | 16.77315 | 44.59068 | 19.72657 | -1.01209 | 0.522694 | 0.102395 |
| 11 | 3106800 | 3109999 | 100.6342 | 157.3149 | 120.1904 | 49.86505 | 0.701994 | 2.653783 | 0.102412 |
| 2 | 168594000 | 168604199 | 105.5582 | 158.4492 | 330.385 | 136.3467 | -0.79206 | 3.449501 | 0.10243 |
| 1 | 152124200 | 152124999 | 11.52093 | 22.38047 | 14.15608 | 5.286835 | 1.245883 | -0.23527 | 0.10243 |
| 4 | 134458800 | 134467599 | 38.67665 | 61.52286 | 147.0977 | 61.42852 | -0.92069 | 2.226499 | 0.102436 |
| 10 | 75515200 | 75516999 | 7.121083 | 12.82313 | 40.08394 | 17.48413 | -1.22718 | 0.361135 | 0.102444 |
| 5 | 65030600 | 65032599 | 24.87703 | 43.6697 | 24.38582 | 10.78693 | 1.223223 | 0.648561 | 0.102457 |
| 6 | 133163400 | 133170799 | 148.2788 | 225.5735 | 171.6481 | 71.11793 | 0.677574 | 3.181539 | 0.10246 |
| 11 | 100858400 | 100864199 | 26.0973 | 42.01429 | 112.9966 | 48.01257 | -1.06965 | 1.822061 | 0.102481 |
| 2 | 130198800 | 130205999 | 34.4481 | 54.94802 | 141.9951 | 59.52464 | -1.02822 | 2.131064 | 0.102494 |
| 1 | 13872800 | 13873999 | 4.795866 | 9.295617 | 29.09443 | 11.00866 | -1.05497 | -0.22155 | 0.102498 |
| 2 | 61637400 | 61637999 | 4.795866 | 9.295617 | 29.09443 | 11.00866 | -1.05497 | -0.22155 | 0.102498 |
| 8 | 3189200 | 3193799 | 60.53686 | 97.13656 | 72.09163 | 30.48293 | 0.777889 | 1.955667 | 0.102499 |
| 8 | 26062000 | 26064999 | 14.23064 | 23.83256 | 71.59207 | 31.20779 | -1.19973 | 1.157895 | 0.10251 |
| 18 | 73116400 | 73117799 | 16.77777 | 30.72914 | 20.29185 | 8.663748 | 1.019424 | 0.285648 | 0.102519 |
| 8 | 70801000 | 70806999 | 21.98099 | 35.45219 | 114.1003 | 48.6041 | -1.32463 | 1.759432 | 0.102526 |
| 15 | 88784600 | 88788999 | 22.36964 | 36.4715 | 89.52293 | 38.45471 | -0.92002 | 1.520602 | 0.102537 |
| 2 | 93157400 | 93161999 | 21.39782 | 34.96947 | 88.01128 | 37.83791 | -0.95357 | 1.482437 | 0.102537 |
| X | 25634400 | 25635399 | 23.50779 | 41.23067 | 26.83489 | 11.86164 | 1.003827 | 0.659107 | 0.102542 |
| 2 | 174411400 | 174415399 | 21.67156 | 35.53686 | 89.31191 | 38.44844 | -0.94896 | 1.436402 | 0.102546 |
| 5 | 105350400 | 105352199 | 31.99114 | 54.82194 | 31.55632 | 13.83909 | 1.176578 | 0.967829 | 0.102562 |
| 11 | 105311400 | 105316199 | 25.78235 | 41.83626 | 95.37794 | 40.87269 | -0.81939 | 1.60896 | 0.102593 |
| 3 | 64105000 | 64105599 | 10.79718 | 20.96525 | 14.73879 | 5.524143 | 1.090119 | -0.23055 | 0.102605 |
| 14 | 9540200 | 9540999 | 10.79718 | 20.96525 | 14.73879 | 5.524143 | 1.090119 | -0.23055 | 0.102605 |
| 8 | 60925800 | 60928199 | 18.0968 | 30.37439 | 72.14025 | 31.49061 | -0.86255 | 1.146901 | 0.102613 |
| 7 | 71981800 | 71983199 | 6.22559 | 11.70756 | 39.32323 | 16.13479 | -1.26619 | 0.086986 | 0.102617 |
| 9 | 111633800 | 111634599 | 5.111338 | 9.837885 | 29.73633 | 11.56907 | -1.04372 | -0.12158 | 0.102625 |
| 9 | 28038400 | 28040199 | 7.130021 | 13.30547 | 37.77612 | 15.70356 | -1.04065 | 0.143894 | 0.10263 |
| 11 | 85586600 | 85587599 | 4.93424 | 9.548768 | 30.79744 | 11.73913 | -1.10823 | -0.18985 | 0.102632 |
| 7 | 26047000 | 26048799 | 50.61669 | 81.86723 | 59.95332 | 25.63044 | 0.81046 | 1.699034 | 0.102633 |
| 7 | 65518000 | 65523799 | 30.57031 | 49.16365 | 113.742 | 48.25563 | -0.85499 | 1.861865 | 0.102633 |
| 4 | 147443600 | 147443999 | 9.062885 | 17.71801 | 5.052605 | 1.63148 | 2.537434 | -0.79161 | 0.102664 |
| 13 | 62394000 | 62396199 | 46.13356 | 75.0934 | 54.41964 | 23.36212 | 0.833453 | 1.569204 | 0.102707 |
| 15 | 74396800 | 74421199 | 140.5641 | 204.3862 | 502.5656 | 207.1532 | -1.07291 | 3.994327 | 0.102711 |
| 8 | 95497600 | 95509399 | 58.84829 | 91.39855 | 210.0947 | 87.23999 | -0.8925 | 2.758105 | 0.102717 |
| 5 | 145066000 | 145071199 | 21.60063 | 35.20136 | 91.14784 | 39.14607 | -0.99737 | 1.525904 | 0.102723 |
| 4 | 21573200 | 21575799 | 32.08778 | 54.57851 | 35.52158 | 15.5687 | 0.99888 | 1.039204 | 0.102723 |
| 2 | 32344600 | 32348399 | 17.87651 | 29.72524 | 75.14933 | 32.60915 | -0.95241 | 1.249854 | 0.102724 |
| 17 | 45565600 | 45572599 | 48.12512 | 76.24716 | 162.8596 | 67.7374 | -0.76409 | 2.354429 | 0.102732 |
| X | 29445600 | 29449399 | 40.80955 | 68.21101 | 36.59783 | 15.93364 | 1.276066 | 1.227715 | 0.102734 |
| 2 | 127964200 | 127970999 | 31.24525 | 49.6346 | 164.1803 | 68.46286 | -1.38996 | 2.264579 | 0.102749 |
| 8 | 12576600 | 12578399 | 10.25358 | 18.11572 | 45.62956 | 20.18564 | -0.93496 | 0.526669 | 0.102761 |
| 6 | 133492200 | 133498799 | 108.9079 | 170.4534 | 110.0755 | 45.6629 | 0.945563 | 2.636423 | 0.10277 |
| 17 | 29544800 | 29556399 | 46.63972 | 72.58867 | 215.064 | 89.27558 | -1.2557 | 2.717002 | 0.102775 |
| 12 | 73449000 | 73451599 | 13.3815 | 22.9554 | 61.21554 | 26.84847 | -1.02555 | 0.919702 | 0.102778 |

| | | | | | | | | | |
|---|---|---|---|---|---|---|---|---|---|
| 3 | 108351600 | 108352799 | 6.721257 | 12.45165 | 35.8141 | 15.0553 | -1.07135 | 0.180864 | 0.102781 |
| 9 | 113108400 | 113109599 | 6.721257 | 12.45165 | 35.8141 | 15.0553 | -1.07135 | 0.180864 | 0.102781 |
| 7 | 22456200 | 22457199 | 15.10811 | 28.8839 | 15.20459 | 6.030774 | 1.427654 | -0.0012 | 0.102793 |
| 19 | 57160800 | 57161999 | 5.604549 | 10.64813 | 32.08852 | 12.89996 | -1.08388 | -0.00331 | 0.102804 |
| 4 | 3322600 | 3323999 | 12.46175 | 23.88572 | 16.66823 | 6.562828 | 1.034761 | -0.04734 | 0.102812 |
| 11 | 93396600 | 93397599 | 12.46175 | 23.88572 | 16.66823 | 6.562828 | 1.034761 | -0.04734 | 0.102812 |
| 8 | 117590000 | 117591399 | 9.124029 | 16.31519 | 42.07564 | 18.46674 | -0.95911 | 0.420466 | 0.102825 |
| X | 25671600 | 25673399 | 29.21688 | 50.49756 | 30.55889 | 13.40651 | 1.106863 | 0.880554 | 0.102838 |
| 3 | 153322600 | 153328799 | 57.08917 | 89.38124 | 190.2876 | 78.93476 | -0.77292 | 2.613939 | 0.102855 |
| X | 102873400 | 102873999 | 16.59729 | 31.52084 | 15.19834 | 6.115038 | 1.535924 | 0.058112 | 0.102856 |
| 12 | 18966400 | 18977799 | 308.6763 | 444.7956 | 316.9501 | 130.7909 | 0.70036 | 4.137506 | 0.102859 |
| 15 | 27743800 | 27746599 | 9.905899 | 17.2062 | 54.12087 | 23.75796 | -1.25763 | 0.780809 | 0.102863 |
| X | 33043800 | 33046999 | 36.39713 | 62.54694 | 25.3996 | 11.14112 | 1.677784 | 0.956425 | 0.102864 |
| 7 | 27290200 | 27298599 | 38.83954 | 61.77454 | 146.9812 | 61.36967 | -0.91386 | 2.230652 | 0.102864 |
| 11 | 71920000 | 71923599 | 17.82473 | 29.76938 | 71.94187 | 31.30568 | -0.88743 | 1.198846 | 0.102866 |
| 1 | 96065600 | 96068799 | 33.86199 | 57.12908 | 37.69401 | 16.48445 | 0.978731 | 1.116349 | 0.102867 |
| 12 | 34316200 | 34316799 | 10.75974 | 20.93803 | 13.73518 | 5.058347 | 1.212252 | -0.29315 | 0.102868 |
| 11 | 70024600 | 70030599 | 26.26491 | 42.29704 | 115.9002 | 49.26814 | -1.09603 | 1.812718 | 0.102889 |
| 19 | 32689000 | 32690399 | 6.816262 | 12.41114 | 39.75327 | 17.10099 | -1.24954 | 0.289129 | 0.102892 |
| 5 | 66526400 | 66529399 | 11.36956 | 19.93598 | 49.36776 | 21.81682 | -0.9139 | 0.635467 | 0.102901 |
| 7 | 108771400 | 108778999 | 40.59298 | 64.59493 | 148.5843 | 62.08095 | -0.86449 | 2.215787 | 0.102908 |
| 5 | 119793600 | 119799399 | 31.3695 | 50.44946 | 114.0336 | 48.38011 | -0.82152 | 1.86252 | 0.102915 |
| 4 | 139441600 | 139456999 | 97.90666 | 147.5259 | 321.0169 | 132.616 | -0.84866 | 3.37234 | 0.102948 |
| 9 | 7370800 | 7373999 | 29.09042 | 49.52915 | 32.25044 | 14.13612 | 0.998316 | 1.013844 | 0.102954 |
| 6 | 3402400 | 3405999 | 37.23994 | 61.89165 | 43.87137 | 19.03023 | 0.875872 | 1.282204 | 0.102957 |
| 4 | 121793200 | 121793599 | 7.371639 | 14.42028 | 9.98364 | 3.323579 | 1.261573 | -0.70036 | 0.10296 |
| 14 | 55869800 | 55874199 | 15.214 | 25.36429 | 74.90292 | 32.55785 | -1.17537 | 1.212776 | 0.102967 |
| 2 | 126405400 | 126407799 | 10.09156 | 17.85732 | 47.77375 | 21.1276 | -1.02075 | 0.510877 | 0.102986 |
| 9 | 20118200 | 20118799 | 10.03848 | 19.56099 | 13.11226 | 4.759807 | 1.198841 | -0.3549 | 0.102987 |
| 7 | 15253400 | 15258399 | 27.8395 | 48.29331 | 28.11117 | 12.33404 | 1.163575 | 0.83904 | 0.102989 |
| 17 | 69957200 | 69959399 | 9.482188 | 16.7997 | 48.86393 | 21.6044 | -1.13996 | 0.494652 | 0.102992 |
| 7 | 133238400 | 133240199 | 9.737284 | 17.01608 | 51.49659 | 22.69252 | -1.20142 | 0.685979 | 0.103004 |
| 2 | 74591200 | 74594799 | 20.90708 | 34.21483 | 86.95858 | 37.40421 | -0.96619 | 1.460391 | 0.103005 |
| 11 | 99606200 | 99607399 | 58.7213 | 94.94776 | 46.36364 | 19.81408 | 1.393119 | 1.716858 | 0.103015 |
| 12 | 23834400 | 23841799 | 182.2851 | 274.3355 | 192.351 | 79.43686 | 0.782468 | 3.406454 | 0.103016 |
| 9 | 109760200 | 109762999 | 13.57284 | 23.18167 | 61.64983 | 27.02939 | -1.02257 | 0.962475 | 0.10303 |
| 7 | 38859600 | 38862599 | 56.25293 | 90.49386 | 66.94647 | 28.41469 | 0.786909 | 1.869224 | 0.10305 |
| 7 | 17996000 | 17999399 | 94.54945 | 148.2313 | 101.8919 | 42.26074 | 0.857437 | 2.595698 | 0.103051 |
| 5 | 30738000 | 30752399 | 72.95369 | 111.872 | 247.2054 | 102.618 | -0.84989 | 3.005884 | 0.103062 |
| 2 | 165233400 | 165251399 | 120.8214 | 178.7093 | 407.8514 | 167.9641 | -0.9418 | 3.719997 | 0.103077 |
| 15 | 87542200 | 87574599 | 239.9188 | 337.8203 | 668.4287 | 276.8643 | -0.80591 | 4.474906 | 0.103097 |
| 14 | 75723400 | 75724799 | 14.16335 | 26.66755 | 18.17957 | 7.443852 | 1.019412 | 0.099334 | 0.103098 |
| 6 | 84123400 | 84130399 | 38.81683 | 61.7923 | 144.0877 | 60.23421 | -0.88345 | 2.201396 | 0.103099 |
| 13 | 44234000 | 44236999 | 14.78094 | 25.30326 | 65.75927 | 28.83646 | -0.98934 | 0.948668 | 0.103141 |
| 10 | 22666600 | 22668999 | 20.6106 | 34.08175 | 79.24382 | 34.2543 | -0.83294 | 1.325181 | 0.103149 |
| 2 | 125514600 | 125515399 | 4.069473 | 7.901928 | 26.7012 | 10.0067 | -1.14931 | -0.26833 | 0.103149 |
| 2 | 92851000 | 92866999 | 99.32127 | 148.9331 | 348.8776 | 143.9386 | -0.96119 | 3.468733 | 0.10318 |

| | | | | | | | | | |
|---|---|---|---|---|---|---|---|---|---|
| 4 | 106946600 | 106956599 | 37.53964 | 59.4022 | 166.215 | 69.08956 | -1.1543 | 2.378286 | 0.103181 |
| 3 | 74410800 | 74412799 | 24.7463 | 43.003 | 28.99401 | 12.73142 | 0.952987 | 0.8042 | 0.103183 |
| 4 | 137754600 | 137756199 | 5.38278 | 10.10828 | 35.04682 | 14.41467 | -1.31304 | 0.080834 | 0.103191 |
| 11 | 118369800 | 118412799 | 216.6273 | 300.5027 | 856.5311 | 355.6038 | -1.35406 | 4.699398 | 0.103196 |
| 12 | 23587600 | 23588599 | 16.1283 | 29.53539 | 18.64125 | 7.960567 | 1.08419 | 0.277746 | 0.103201 |
| 18 | 13172400 | 13175399 | 107.1871 | 169.9578 | 64.1018 | 26.68097 | 1.734466 | 2.361287 | 0.103211 |
| 10 | 16702000 | 16703799 | 23.00823 | 40.52383 | 24.8452 | 10.99788 | 1.090572 | 0.598658 | 0.103234 |
| 1 | 163458600 | 163458799 | 2.030741 | 3.961051 | 17.9634 | 5.616283 | -1.3181 | -0.97466 | 0.103239 |
| 1 | 30911800 | 30913399 | 8.071136 | 14.18228 | 50.89048 | 22.52993 | -1.44514 | 0.584435 | 0.10325 |
| 18 | 3044400 | 3048199 | 138.8713 | 212.8393 | 150.0988 | 62.30323 | 0.795642 | 3.029489 | 0.103254 |
| 3 | 138967200 | 138967999 | 21.64092 | 37.97386 | 26.591 | 11.75842 | 0.898809 | 0.640477 | 0.103265 |
| 13 | 3297000 | 3298599 | 29.50999 | 50.51074 | 34.09431 | 14.95072 | 0.947818 | 0.972512 | 0.103269 |
| 1 | 62250600 | 62250999 | 2.427367 | 4.746524 | 20.83889 | 6.773876 | -1.32674 | -0.81737 | 0.103272 |
| 11 | 116805600 | 116811999 | 31.63716 | 50.64425 | 128.4795 | 54.13067 | -0.99583 | 2.014031 | 0.103273 |
| 13 | 85761200 | 85761999 | 15.35205 | 28.65594 | 18.63302 | 7.742472 | 1.070502 | 0.165101 | 0.103287 |
| 4 | 45268200 | 45271399 | 17.57637 | 29.63954 | 69.45202 | 30.36866 | -0.84291 | 1.09908 | 0.103311 |
| 3 | 138564000 | 138565799 | 7.589544 | 13.65943 | 40.06905 | 17.48927 | -1.13732 | 0.366061 | 0.103347 |
| 1 | 90036800 | 90041199 | 28.57199 | 46.1476 | 103.7409 | 44.27177 | -0.80609 | 1.723956 | 0.103368 |
| 16 | 89936800 | 89942599 | 29.64073 | 47.62563 | 119.2579 | 50.52958 | -0.97056 | 1.890667 | 0.10339 |
| 4 | 141303600 | 141310599 | 40.3953 | 64.37798 | 140.1058 | 58.66154 | -0.78231 | 2.164927 | 0.103429 |
| 18 | 80352600 | 80362199 | 70.15371 | 108.403 | 224.2553 | 93.15073 | -0.7464 | 2.855621 | 0.103431 |
| 1 | 8511800 | 8513399 | 17.44292 | 31.91257 | 20.18041 | 8.628962 | 1.079908 | 0.296634 | 0.103442 |
| 12 | 23384200 | 23385799 | 15.53578 | 28.60659 | 18.48386 | 7.834739 | 1.05783 | 0.241726 | 0.103466 |
| 12 | 104080600 | 104082799 | 9.921507 | 17.61384 | 44.60308 | 19.6953 | -0.9394 | 0.475687 | 0.103469 |
| 4 | 150972400 | 150978399 | 27.86804 | 44.765 | 116.841 | 49.48577 | -1.0306 | 1.898195 | 0.10348 |
| X | 58894800 | 58895399 | 10.11375 | 19.68599 | 13.84013 | 5.086514 | 1.116206 | -0.30795 | 0.103495 |
| 13 | 68460200 | 68468199 | 169.3471 | 255.9268 | 185.5356 | 76.71598 | 0.73951 | 3.320278 | 0.103495 |
| 11 | 116569200 | 116577999 | 32.23708 | 51.35431 | 144.5972 | 60.50632 | -1.1537 | 2.173077 | 0.103498 |
| 10 | 127443400 | 127452399 | 44.40938 | 70.37858 | 157.8469 | 65.6651 | -0.83401 | 2.344484 | 0.1035 |
| 8 | 112400800 | 112405999 | 33.32119 | 53.52738 | 117.3142 | 49.69004 | -0.77917 | 1.903784 | 0.1035 |
| 17 | 49491400 | 49494199 | 10.79611 | 18.8528 | 52.60904 | 23.17188 | -1.08554 | 0.702497 | 0.103505 |
| 17 | 57016000 | 57019999 | 14.68089 | 24.0265 | 91.39868 | 39.31464 | -1.54543 | 1.449866 | 0.103512 |
| 11 | 75321000 | 75336599 | 89.75136 | 135.6604 | 306.676 | 126.8124 | -0.89974 | 3.307295 | 0.103517 |
| 7 | 66537400 | 66542799 | 28.2844 | 45.59691 | 106.9332 | 45.52305 | -0.87013 | 1.781029 | 0.103519 |
| 1 | 160619400 | 160620799 | 6.757336 | 12.12039 | 43.17622 | 18.90072 | -1.41857 | 0.386397 | 0.10352 |
| 6 | 53173000 | 53174999 | 11.95249 | 20.61399 | 56.82775 | 24.92931 | -1.07114 | 0.861463 | 0.103525 |
| 15 | 73516600 | 73518599 | 11.95249 | 20.61399 | 56.82775 | 24.92931 | -1.07114 | 0.861463 | 0.103525 |
| 15 | 37420400 | 37425999 | 39.61132 | 63.07667 | 144.9312 | 60.61293 | -0.86178 | 2.191691 | 0.103527 |
| 12 | 56746400 | 56747399 | 6.512184 | 12.23766 | 34.01165 | 13.97248 | -0.99572 | 0.085028 | 0.103537 |
| X | 33055000 | 33055399 | 11.11333 | 21.69958 | 10.03471 | 3.53484 | 1.763753 | -0.4518 | 0.103544 |
| 17 | 18834800 | 18839599 | 122.7795 | 189.2943 | 141.3347 | 58.70801 | 0.719831 | 2.907566 | 0.103574 |
| 5 | 139665800 | 139681799 | 100.5215 | 150.8094 | 335.3967 | 138.3955 | -0.88567 | 3.459061 | 0.10358 |
| 17 | 48399400 | 48416799 | 96.87724 | 146.1265 | 319.4806 | 132.0204 | -0.85417 | 3.351333 | 0.10359 |
| 2 | 93470200 | 93476799 | 48.16688 | 75.81823 | 178.6769 | 74.13234 | -0.9132 | 2.504524 | 0.103602 |
| 7 | 20510800 | 20518999 | 218.0723 | 322.8942 | 242.2205 | 99.75401 | 0.66383 | 3.709048 | 0.103635 |
| 3 | 31187800 | 31192799 | 29.87234 | 48.18849 | 106.5826 | 45.41229 | -0.78481 | 1.763331 | 0.103642 |
| 4 | 89131600 | 89136799 | 38.55048 | 61.67738 | 129.0519 | 54.32988 | -0.71946 | 2.039216 | 0.103645 |

| | | | | | | | | | |
|---|---|---|---|---|---|---|---|---|---|
| 19 | 42589600 | 42597199 | 32.23479 | 51.15241 | 168.7354 | 70.28792 | -1.38755 | 2.297659 | 0.103654 |
| 1 | 93731200 | 93734599 | 13.92366 | 23.67085 | 63.74684 | 27.93836 | -1.04177 | 1.009206 | 0.103659 |
| 12 | 31284600 | 31285399 | 4.062775 | 7.911242 | 25.93882 | 9.50165 | -1.07591 | -0.35324 | 0.103686 |
| 7 | 5622400 | 5626599 | 91.149 | 144.4386 | 89.25247 | 37.12652 | 1.022814 | 2.368209 | 0.103692 |
| 11 | 101442800 | 101443999 | 4.036611 | 7.749209 | 31.23544 | 12.24315 | -1.46424 | -0.10554 | 0.103697 |
| 4 | 40265000 | 40269999 | 20.98825 | 34.2043 | 94.09781 | 40.41341 | -1.08454 | 1.524979 | 0.103717 |
| 4 | 134449200 | 134453799 | 22.16723 | 36.25741 | 86.44935 | 37.17782 | -0.87492 | 1.471653 | 0.103728 |
| 7 | 48096800 | 48097799 | 17.86437 | 32.81321 | 19.66277 | 8.362482 | 1.162592 | 0.277844 | 0.103733 |
| 14 | 78313400 | 78314199 | 14.38912 | 27.335 | 16.90015 | 6.795645 | 1.180877 | 0.038724 | 0.103735 |
| 8 | 13618000 | 13635999 | 116.0376 | 172.4573 | 383.014 | 157.7914 | -0.89612 | 3.635874 | 0.103773 |
| 12 | 23037600 | 23039599 | 57.72466 | 93.85785 | 43.4969 | 18.66558 | 1.475379 | 1.604175 | 0.103773 |
| 11 | 78066400 | 78070399 | 16.02233 | 26.59235 | 76.3823 | 33.10145 | -1.13589 | 1.268795 | 0.103778 |
| 10 | 40386400 | 40394199 | 52.90061 | 83.64128 | 171.9703 | 71.45249 | -0.71161 | 2.408798 | 0.10378 |
| X | 28654600 | 28654999 | 14.28291 | 26.82169 | 18.6272 | 7.661353 | 0.987621 | 0.117598 | 0.103783 |
| 17 | 27852400 | 27860399 | 37.31601 | 59.20782 | 159.5279 | 66.44367 | -1.09641 | 2.30389 | 0.103789 |
| 10 | 116488600 | 116490599 | 13.62672 | 23.587 | 57.51966 | 25.23563 | -0.89424 | 0.836682 | 0.103804 |
| 7 | 33256000 | 33257599 | 75.82821 | 120.4063 | 86.90096 | 36.22234 | 0.801953 | 2.295738 | 0.10381 |
| 14 | 122715800 | 122716599 | 7.040728 | 12.93966 | 36.72169 | 15.60562 | -1.06347 | 0.232662 | 0.103831 |
| 19 | 58175000 | 58176399 | 7.040728 | 12.93966 | 36.72169 | 15.60562 | -1.06347 | 0.232662 | 0.103831 |
| 5 | 130118200 | 130119799 | 6.611864 | 12.10401 | 37.17308 | 15.8818 | -1.18181 | 0.253287 | 0.103862 |
| 3 | 62034000 | 62036199 | 38.48743 | 64.46376 | 37.09547 | 16.17152 | 1.175274 | 1.197837 | 0.103865 |
| 6 | 41428200 | 41429199 | 13.78921 | 26.35004 | 15.80667 | 6.277287 | 1.238973 | -0.00974 | 0.103866 |
| X | 25462400 | 25464199 | 40.24382 | 66.52093 | 44.52225 | 19.24013 | 0.958029 | 1.354655 | 0.103884 |
| 7 | 78543800 | 78546199 | 10.62153 | 18.2494 | 59.36794 | 26.04064 | -1.30974 | 0.894195 | 0.103893 |
| 10 | 127000600 | 127004199 | 18.1054 | 29.93323 | 79.36698 | 34.30286 | -1.02164 | 1.321341 | 0.103898 |
| 10 | 77000600 | 77004399 | 16.56154 | 27.3821 | 81.43414 | 35.19745 | -1.18658 | 1.317992 | 0.103915 |
| 7 | 15307800 | 15311799 | 73.78634 | 117.8635 | 71.68616 | 30.11122 | 1.057181 | 2.107091 | 0.103928 |
| 8 | 85097600 | 85102599 | 25.54709 | 41.27717 | 103.7177 | 44.28289 | -0.96555 | 1.708185 | 0.10393 |
| 6 | 108189400 | 108191599 | 8.720703 | 15.48696 | 44.08953 | 19.4633 | -1.10673 | 0.464797 | 0.103947 |
| 8 | 70808000 | 70816999 | 39.3267 | 62.53192 | 146.748 | 61.252 | -0.89444 | 2.239995 | 0.10395 |
| 8 | 70954800 | 70959799 | 18.79309 | 30.37746 | 108.6418 | 46.41071 | -1.47228 | 1.687275 | 0.10396 |
| 9 | 98445200 | 98446399 | 9.052002 | 15.96523 | 45.11996 | 19.96709 | -1.10056 | 0.539112 | 0.103966 |
| 17 | 43262200 | 43264999 | 11.53009 | 20.18871 | 50.99976 | 22.51097 | -0.94265 | 0.661296 | 0.103968 |
| 15 | 15032200 | 15033799 | 23.44888 | 41.377 | 23.35024 | 10.33228 | 1.209974 | 0.57716 | 0.103982 |
| 1 | 180158000 | 180161799 | 16.25252 | 26.86183 | 81.32276 | 35.14087 | -1.21217 | 1.321569 | 0.103993 |
| 17 | 47828200 | 47844999 | 80.56227 | 122.7198 | 274.6653 | 113.8454 | -0.87596 | 3.144651 | 0.104 |
| 4 | 113068400 | 113069599 | 29.64042 | 50.97089 | 32.17809 | 14.11469 | 1.044738 | 0.927894 | 0.104007 |
| 11 | 118896000 | 118901399 | 38.10885 | 60.82962 | 133.3696 | 55.96357 | -0.79046 | 2.111508 | 0.104019 |
| 11 | 100740800 | 100743199 | 8.69341 | 15.18017 | 51.98774 | 22.8975 | -1.37846 | 0.693707 | 0.104027 |
| 15 | 100537600 | 100544399 | 29.82058 | 47.83711 | 120.4042 | 50.8909 | -0.98073 | 1.943674 | 0.104053 |
| 13 | 59732800 | 59734199 | 8.384989 | 14.98207 | 42.59686 | 18.70771 | -1.0995 | 0.420842 | 0.104054 |
| 11 | 3111000 | 3113399 | 378.0823 | 539.972 | 303.1628 | 125.2739 | 1.031308 | 4.272021 | 0.104055 |
| 6 | 94120200 | 94120999 | 10.14993 | 19.75647 | 13.65223 | 5.017318 | 1.13972 | -0.30957 | 0.104079 |
| 15 | 102839000 | 102851999 | 79.85876 | 121.5595 | 288.1515 | 119.4093 | -0.95962 | 3.159821 | 0.104079 |
| 16 | 4235800 | 4238799 | 18.80944 | 30.88344 | 83.93722 | 36.15441 | -1.06041 | 1.412603 | 0.10409 |
| 9 | 88209000 | 88210599 | 10.58499 | 18.62337 | 46.67639 | 20.66597 | -0.93022 | 0.577931 | 0.104103 |
| 5 | 134915600 | 134931799 | 87.4454 | 132.0287 | 320.2393 | 132.3747 | -1.0026 | 3.329677 | 0.104108 |

| | | | | | | | | | |
|---|---|---|---|---|---|---|---|---|---|
| 2 | 71355000 | 71363599 | 56.45379 | 87.63219 | 215.9859 | 89.69352 | -0.99359 | 2.766783 | 0.10411 |
| X | 4534200 | 4535799 | 41.24302 | 67.90226 | 46.17565 | 19.91226 | 0.933742 | 1.405173 | 0.104111 |
| 1 | 53138200 | 53140599 | 11.72244 | 20.42007 | 53.17391 | 23.38447 | -0.9868 | 0.743548 | 0.104113 |
| 6 | 137932200 | 137933199 | 4.025866 | 7.757204 | 30.86675 | 11.97091 | -1.43145 | -0.14243 | 0.104114 |
| 1 | 73594000 | 73594599 | 5.563521 | 10.64172 | 31.19376 | 12.35507 | -1.02467 | -0.05387 | 0.104116 |
| 7 | 80072800 | 80081199 | 49.77109 | 78.36041 | 174.674 | 72.4736 | -0.83281 | 2.500973 | 0.104156 |
| 5 | 35979000 | 35983199 | 15.77788 | 26.41797 | 71.2462 | 31.05271 | -1.04499 | 1.164151 | 0.104159 |
| 1 | 62804600 | 62811999 | 36.69232 | 58.50998 | 144.0244 | 60.34996 | -0.95875 | 2.143182 | 0.104172 |
| 4 | 141083400 | 141090599 | 37.37225 | 59.48289 | 143.7451 | 60.0773 | -0.93508 | 2.206016 | 0.104182 |
| 2 | 57320000 | 57321199 | 18.78192 | 33.98857 | 21.07137 | 9.134478 | 1.095831 | 0.369752 | 0.104189 |
| 1 | 104588200 | 104588999 | 11.98155 | 23.32407 | 12.27586 | 4.507724 | 1.528763 | -0.29295 | 0.104193 |
| 18 | 37541800 | 37543799 | 36.14426 | 61.33359 | 32.01159 | 14.024 | 1.315519 | 1.070281 | 0.104206 |
| 1 | 110696600 | 110697999 | 15.82902 | 29.59138 | 18.28648 | 7.578864 | 1.145662 | 0.157764 | 0.104214 |
| 5 | 54935400 | 54936999 | 15.82902 | 29.59138 | 18.28648 | 7.578864 | 1.145662 | 0.157764 | 0.104214 |
| 6 | 24893400 | 24894999 | 8.234145 | 14.41689 | 50.91264 | 22.47151 | -1.42215 | 0.646108 | 0.104223 |
| 17 | 15467000 | 15470599 | 12.85585 | 22.21607 | 56.93579 | 24.97821 | -0.96609 | 0.845839 | 0.104229 |
| 9 | 60816600 | 60819199 | 26.13619 | 45.70092 | 26.70183 | 11.77242 | 1.158857 | 0.72099 | 0.104248 |
| 5 | 74311400 | 74313799 | 13.37942 | 22.93903 | 58.96468 | 25.86011 | -0.97265 | 0.924511 | 0.104259 |
| 2 | 153413800 | 153417999 | 21.11019 | 34.42247 | 89.89462 | 38.61616 | -1.00896 | 1.516095 | 0.104262 |
| 9 | 96418800 | 96419999 | 3.956837 | 7.658308 | 27.5959 | 10.51227 | -1.26354 | -0.21071 | 0.104265 |
| 10 | 84285400 | 84286799 | 3.956837 | 7.658308 | 27.5959 | 10.51227 | -1.26354 | -0.21071 | 0.104265 |
| 1 | 55818800 | 55820399 | 21.28435 | 37.54242 | 21.2623 | 9.409143 | 1.204877 | 0.564545 | 0.104276 |
| 2 | 103577200 | 103579399 | 9.160385 | 16.28489 | 44.10047 | 19.45089 | -1.03434 | 0.460635 | 0.104278 |
| 3 | 96586600 | 96592999 | 34.72158 | 55.45467 | 132.633 | 55.69976 | -0.91471 | 2.09006 | 0.104279 |
| 7 | 30107200 | 30122999 | 110.9063 | 165.8124 | 349.2542 | 144.0102 | -0.81188 | 3.526556 | 0.1043 |
| 2 | 127727600 | 127730399 | 11.38133 | 19.91733 | 51.54187 | 22.73972 | -0.97727 | 0.670015 | 0.1043 |
| 2 | 31971200 | 31981199 | 58.46236 | 90.97149 | 207.5977 | 86.17873 | -0.87989 | 2.72445 | 0.104316 |
| 7 | 7577800 | 7581999 | 73.57989 | 117.4499 | 78.16914 | 32.80122 | 0.926376 | 2.129559 | 0.104328 |
| 5 | 122367800 | 122374799 | 31.64612 | 50.639 | 129.4218 | 54.49779 | -1.00712 | 2.026402 | 0.104328 |
| 8 | 11954800 | 11956199 | 6.099997 | 11.43709 | 36.20149 | 14.92937 | -1.18669 | 0.099175 | 0.104364 |
| 7 | 37085800 | 37090199 | 21.85104 | 35.76106 | 86.57603 | 37.2402 | -0.89629 | 1.461568 | 0.104364 |
| 10 | 77634200 | 77638199 | 22.52273 | 36.76273 | 89.27228 | 38.36288 | -0.90344 | 1.503118 | 0.104374 |
| 2 | 29855200 | 29856199 | 4.546344 | 8.784587 | 28.74789 | 11.03694 | -1.13855 | -0.17579 | 0.104386 |
| 17 | 48314800 | 48321799 | 26.43993 | 42.53753 | 116.5077 | 49.45666 | -1.09714 | 1.844602 | 0.1044 |
| 12 | 80738400 | 80742599 | 24.9934 | 40.54524 | 95.74841 | 41.02339 | -0.87041 | 1.612945 | 0.104404 |
| 3 | 122301600 | 122305199 | 17.21731 | 28.72513 | 74.137 | 32.2398 | -0.98199 | 1.209224 | 0.104435 |
| 15 | 79784200 | 79785399 | 4.698546 | 8.999225 | 30.01027 | 11.84087 | -1.20257 | -0.0777 | 0.104465 |
| 7 | 110223000 | 110225999 | 10.8684 | 18.81438 | 56.29635 | 24.699 | -1.18747 | 0.821188 | 0.104469 |
| 6 | 88877200 | 88890599 | 59.92213 | 92.25628 | 244.6895 | 101.6292 | -1.10849 | 2.922228 | 0.10448 |
| 7 | 62159400 | 62161999 | 70.15101 | 112.0153 | 80.60372 | 33.83935 | 0.814537 | 2.116477 | 0.104482 |
| 6 | 27559600 | 27562999 | 25.69427 | 44.6976 | 30.11333 | 13.23171 | 0.95408 | 0.794974 | 0.104489 |
| 11 | 12828400 | 12829199 | 6.848736 | 12.71121 | 35.84881 | 15.03216 | -1.04026 | 0.173231 | 0.104514 |
| 7 | 14505200 | 14506799 | 40.8274 | 68.14603 | 37.73444 | 16.41408 | 1.230849 | 1.243231 | 0.104514 |
| 6 | 24716400 | 24718399 | 17.1929 | 28.41334 | 82.7205 | 35.74248 | -1.15614 | 1.325199 | 0.104527 |
| 16 | 30036200 | 30037399 | 4.513736 | 8.694418 | 29.09013 | 11.29246 | -1.18442 | -0.13511 | 0.104527 |
| 2 | 22316000 | 22320599 | 46.37992 | 76.10659 | 46.20657 | 19.89763 | 1.094978 | 1.457595 | 0.104532 |
| 17 | 13580800 | 13595199 | 489.1574 | 651.0738 | 1167.512 | 487.778 | -0.74585 | 5.35315 | 0.104538 |

| | | | | | | | | | |
|---|---|---|---|---|---|---|---|---|---|
| 7 | 34954400 | 34969799 | 94.62951 | 143.4855 | 290.322 | 120.169 | -0.73603 | 3.243517 | 0.104539 |
| 7 | 70017400 | 70020599 | 12.62121 | 21.64841 | 60.5947 | 26.57591 | -1.09496 | 0.917679 | 0.10454 |
| 7 | 10563400 | 10572199 | 134.9124 | 206.657 | 155.9479 | 64.72406 | 0.697436 | 3.038728 | 0.104541 |
| 13 | 54827800 | 54836399 | 36.52063 | 57.93789 | 160.2905 | 66.75216 | -1.13468 | 2.307601 | 0.104546 |
| 19 | 5344600 | 5345799 | 3.043456 | 5.905282 | 30.75801 | 11.58036 | -1.77213 | -0.25579 | 0.104551 |
| 3 | 137770800 | 137777999 | 64.80375 | 99.37836 | 253.2353 | 105.1217 | -1.0552 | 3.002957 | 0.104561 |
| 4 | 117291000 | 117292599 | 8.018113 | 14.5058 | 40.69788 | 17.64889 | -1.068 | 0.337447 | 0.104572 |
| 7 | 35281000 | 35287399 | 39.12168 | 62.31643 | 142.3539 | 59.56077 | -0.85289 | 2.180769 | 0.104578 |
| 12 | 47769800 | 47770999 | 16.36279 | 30.3474 | 19.32312 | 8.110528 | 1.08904 | 0.212288 | 0.104586 |
| 11 | 100829600 | 100831399 | 8.363079 | 14.9761 | 41.05378 | 17.99634 | -1.04499 | 0.405022 | 0.104587 |
| 8 | 86294600 | 86296599 | 16.03581 | 29.42256 | 19.36934 | 8.249867 | 1.02615 | 0.267346 | 0.104588 |
| 10 | 80967400 | 80987399 | 101.711 | 151.0909 | 391.1821 | 161.1484 | -1.11785 | 3.643155 | 0.104595 |
| 13 | 53370800 | 53412199 | 363.2547 | 493.9998 | 942.7348 | 392.6185 | -0.80445 | 5.010382 | 0.104615 |
| 6 | 143112000 | 143113799 | 8.050948 | 14.47638 | 41.1 | 17.95443 | -1.09187 | 0.375502 | 0.104615 |
| 7 | 31845800 | 31848399 | 54.24565 | 87.53518 | 62.83236 | 26.78279 | 0.835591 | 1.771454 | 0.104624 |
| 11 | 82338800 | 82343199 | 17.0344 | 28.48125 | 71.92032 | 31.31837 | -0.95057 | 1.183793 | 0.104631 |
| 7 | 53986400 | 53987999 | 15.65987 | 29.56566 | 16.67174 | 6.792048 | 1.296082 | 0.091053 | 0.104639 |
| 6 | 5612000 | 5612399 | 9.57316 | 18.68971 | 11.36368 | 4.012488 | 1.371643 | -0.46224 | 0.104655 |
| 2 | 117609800 | 117611799 | 9.848198 | 17.28488 | 48.27907 | 21.35297 | -1.08625 | 0.609882 | 0.104658 |
| 11 | 117177600 | 117183599 | 33.30945 | 53.11731 | 138.1701 | 57.89941 | -1.03801 | 2.139425 | 0.104661 |
| 2 | 150909000 | 150910999 | 23.06608 | 40.75747 | 22.78269 | 10.07838 | 1.224205 | 0.5589 | 0.104673 |
| X | 30888400 | 30889999 | 31.64041 | 54.46908 | 27.63243 | 12.12117 | 1.359105 | 0.923168 | 0.10468 |
| 2 | 172946000 | 172974799 | 192.7532 | 274.3809 | 620.1133 | 256.3873 | -0.98181 | 4.312602 | 0.104687 |
| 12 | 99930600 | 99940799 | 73.56845 | 113.0028 | 246.6494 | 102.4106 | -0.83032 | 2.972653 | 0.104697 |
| 6 | 116437400 | 116439999 | 14.13725 | 24.28822 | 58.5202 | 25.66874 | -0.87934 | 0.907393 | 0.104706 |
| Y | 9712200 | 9715199 | 14.21779 | 27.2554 | 5.12146 | 2.015674 | 2.892225 | -0.06863 | 0.104712 |
| 8 | 36125200 | 36125999 | 10.6973 | 20.75547 | 14.38871 | 5.418231 | 1.104611 | -0.2164 | 0.104734 |
| 1 | 126525000 | 126527199 | 10.4133 | 18.3713 | 47.75079 | 21.13014 | -0.98112 | 0.546206 | 0.104746 |
| 15 | 82337800 | 82343799 | 40.7146 | 64.90644 | 137.7799 | 57.71181 | -0.74595 | 2.15531 | 0.104747 |
| 1 | 75437200 | 75442799 | 25.53476 | 41.25114 | 102.2692 | 43.65609 | -0.94641 | 1.712917 | 0.10475 |
| 4 | 79538400 | 79539399 | 10.05141 | 19.537 | 14.3679 | 5.341113 | 1.036857 | -0.26854 | 0.104756 |
| 8 | 89767600 | 89774999 | 39.04901 | 62.00477 | 153.8614 | 64.13249 | -0.97678 | 2.281591 | 0.10477 |
| 10 | 98942600 | 98948199 | 53.8828 | 84.24139 | 201.8188 | 83.73035 | -0.94461 | 2.639537 | 0.104775 |
| 17 | 83548600 | 83551199 | 14.26232 | 24.58068 | 59.72431 | 26.19942 | -0.8906 | 0.879133 | 0.10478 |
| 2 | 132846600 | 132847399 | 10.88669 | 21.18569 | 12.9018 | 4.75032 | 1.316881 | -0.29664 | 0.104789 |
| 12 | 23331800 | 23332999 | 13.30126 | 25.47099 | 16.55951 | 6.537557 | 1.132439 | -0.03037 | 0.104817 |
| X | 32883000 | 32883999 | 21.27247 | 38.80041 | 17.92147 | 7.692731 | 1.526582 | 0.339585 | 0.104834 |
| 9 | 83155000 | 83156599 | 9.49602 | 16.78833 | 45.63438 | 20.18512 | -1.04378 | 0.514135 | 0.104864 |
| 2 | 128785400 | 128786199 | 3.657517 | 7.088996 | 27.06389 | 10.24176 | -1.33579 | -0.23763 | 0.104868 |
| 16 | 36648000 | 36649199 | 5.823857 | 10.88155 | 34.90803 | 14.48073 | -1.21208 | 0.117916 | 0.104877 |
| 2 | 11710000 | 11713599 | 24.79443 | 40.30742 | 92.50936 | 39.69541 | -0.82648 | 1.569637 | 0.104886 |
| 12 | 23270800 | 23273999 | 73.41177 | 116.7568 | 90.22181 | 37.66843 | 0.705833 | 2.246524 | 0.104898 |
| 5 | 36545800 | 36558799 | 56.06308 | 86.50055 | 236.5326 | 98.2512 | -1.15002 | 2.879474 | 0.104899 |
| X | 33222200 | 33228799 | 76.076 | 121.9801 | 59.05925 | 24.94428 | 1.391467 | 1.994971 | 0.104928 |
| 7 | 38476600 | 38478999 | 19.16898 | 34.62159 | 19.75237 | 8.583551 | 1.212846 | 0.381223 | 0.104929 |
| 12 | 48584000 | 48586399 | 20.07772 | 36.375 | 18.65678 | 8.076215 | 1.3689 | 0.368784 | 0.104929 |
| 10 | 83446400 | 83458799 | 81.88749 | 125.0268 | 262.9014 | 109.0316 | -0.78336 | 3.09727 | 0.104943 |

| | | | | | | | | | |
|---|---|---|---|---|---|---|---|---|---|
| 10 | 62977600 | 62982199 | 22.63536 | 37.06174 | 86.83992 | 37.36057 | -0.84899 | 1.456871 | 0.104944 |
| 4 | 92362600 | 92364199 | 14.72437 | 27.76421 | 17.93661 | 7.326309 | 1.098718 | 0.094189 | 0.104965 |
| 18 | 90434800 | 90436599 | 23.08841 | 40.71033 | 23.91987 | 10.58593 | 1.151739 | 0.58371 | 0.104965 |
| 6 | 10475000 | 10475799 | 11.43737 | 22.21899 | 14.06026 | 5.249423 | 1.245056 | -0.23825 | 0.104966 |
| 1 | 64483200 | 64486799 | 14.15229 | 24.26019 | 65.75634 | 28.83835 | -1.0494 | 0.933282 | 0.104995 |
| 14 | 104759400 | 104760399 | 5.356912 | 10.12822 | 34.75722 | 14.11742 | -1.28329 | 0.033573 | 0.104997 |
| 1 | 23960600 | 23961599 | 7.695557 | 13.96717 | 38.62775 | 16.68381 | -1.043 | 0.314142 | 0.104999 |
| 15 | 76800200 | 76802399 | 10.46399 | 18.32624 | 48.84109 | 21.56234 | -1.01903 | 0.65008 | 0.105003 |
| 19 | 13576800 | 13578599 | 61.74348 | 99.02295 | 73.54225 | 31.07257 | 0.776035 | 1.971829 | 0.105017 |
| 4 | 154966000 | 154986599 | 113.1544 | 167.7304 | 399.947 | 164.7233 | -1.00207 | 3.68239 | 0.105031 |
| 12 | 23671000 | 23676199 | 88.40652 | 139.4863 | 104.4368 | 43.34665 | 0.741332 | 2.470268 | 0.10504 |
| 4 | 121627200 | 121627599 | 7.339757 | 14.3538 | 9.714202 | 3.304935 | 1.267423 | -0.62979 | 0.10504 |
| 2 | 17909200 | 17911399 | 27.38932 | 47.19134 | 32.29126 | 14.16517 | 0.929508 | 0.900334 | 0.105046 |
| 6 | 25336400 | 25338599 | 23.46242 | 41.07446 | 27.96824 | 12.34315 | 0.939138 | 0.68965 | 0.105047 |
| 4 | 100573800 | 100576199 | 13.81561 | 23.78228 | 57.97253 | 25.43003 | -0.89535 | 0.888471 | 0.105048 |
| 4 | 125869800 | 125871999 | 10.22821 | 17.7751 | 54.15783 | 23.7818 | -1.21214 | 0.773381 | 0.105088 |
| 2 | 152748200 | 152763199 | 78.7806 | 119.5958 | 290.3458 | 120.2216 | -0.9972 | 3.215645 | 0.10509 |
| 1 | 36544600 | 36547199 | 10.1662 | 17.78557 | 49.37529 | 21.77839 | -1.07749 | 0.667643 | 0.105091 |
| 8 | 73612800 | 73614199 | 11.57274 | 20.13297 | 53.13088 | 23.34635 | -1.00583 | 0.764551 | 0.1051 |
| 7 | 82428800 | 82431199 | 9.369253 | 16.51529 | 47.20945 | 20.89416 | -1.11771 | 0.548875 | 0.105101 |
| 2 | 169884400 | 169891799 | 57.97115 | 91.0137 | 183.999 | 76.3146 | -0.69532 | 2.563264 | 0.105112 |
| 13 | 58117000 | 58121199 | 18.55788 | 30.73433 | 80.24822 | 34.72511 | -0.99946 | 1.301684 | 0.10512 |
| 10 | 59838400 | 59839399 | 6.236118 | 11.69703 | 34.50568 | 14.22031 | -1.08479 | 0.094911 | 0.105128 |
| 10 | 129290400 | 129290999 | 6.236118 | 11.69703 | 34.50568 | 14.22031 | -1.08479 | 0.094911 | 0.105128 |
| 10 | 65419200 | 65420199 | 11.48152 | 22.20974 | 15.1149 | 5.772726 | 1.110849 | -0.16019 | 0.105149 |
| 13 | 66850000 | 66852999 | 45.67134 | 75.57147 | 36.20942 | 15.65909 | 1.438342 | 1.349469 | 0.105164 |
| 16 | 46316400 | 46318599 | 15.9169 | 26.35617 | 79.68008 | 34.47534 | -1.20986 | 1.297546 | 0.105174 |
| 4 | 140032400 | 140065399 | 339.6385 | 463.8937 | 889.5703 | 370.2283 | -0.80509 | 4.941751 | 0.105183 |
| 4 | 132776000 | 132783799 | 31.62818 | 50.63786 | 126.6009 | 53.35145 | -0.97452 | 2.008891 | 0.1052 |
| 17 | 40890600 | 40891999 | 19.65121 | 35.025 | 23.42839 | 10.31089 | 0.973237 | 0.472007 | 0.105219 |
| 14 | 35014000 | 35014599 | 4.779724 | 9.198253 | 30.82324 | 11.99706 | -1.19056 | -0.11814 | 0.105244 |
| 2 | 99868600 | 99870399 | 18.35757 | 33.71178 | 18.37705 | 7.817988 | 1.297291 | 0.279216 | 0.105249 |
| X | 61151000 | 61151599 | 20.85695 | 37.23816 | 19.68062 | 8.648917 | 1.312519 | 0.462598 | 0.105256 |
| 3 | 7718600 | 7719999 | 20.08506 | 36.35309 | 20.49648 | 8.88334 | 1.232209 | 0.378019 | 0.10526 |
| 14 | 15837200 | 15839799 | 26.99101 | 46.42457 | 33.49656 | 14.69312 | 0.852973 | 0.916171 | 0.105261 |
| 8 | 22524200 | 22531199 | 43.81629 | 69.72847 | 145.8665 | 60.95068 | -0.72769 | 2.216371 | 0.105265 |
| 5 | 95567800 | 95569799 | 21.493 | 38.46084 | 20.11249 | 8.821834 | 1.328495 | 0.451686 | 0.10527 |
| 14 | 120928600 | 120930599 | 14.62778 | 25.06751 | 60.12075 | 26.36612 | -0.87339 | 0.935235 | 0.105282 |
| 5 | 112424200 | 112428999 | 17.53245 | 28.8452 | 82.756 | 35.67623 | -1.1369 | 1.383247 | 0.105291 |
| 9 | 102686600 | 102691399 | 38.71201 | 61.91419 | 129.599 | 54.53326 | -0.7206 | 2.049883 | 0.105297 |
| 7 | 7926800 | 7929199 | 25.77908 | 45.0243 | 27.84215 | 12.26446 | 1.077471 | 0.737842 | 0.105312 |
| 4 | 139769600 | 139773199 | 19.67706 | 32.50634 | 78.98048 | 34.11749 | -0.89624 | 1.334426 | 0.105314 |
| 2 | 147022000 | 147038399 | 119.959 | 177.3546 | 424.406 | 174.7809 | -1.01075 | 3.727376 | 0.105328 |
| 2 | 128123600 | 128130999 | 29.58467 | 47.64688 | 110.642 | 47.03823 | -0.8577 | 1.813517 | 0.105353 |
| 11 | 60066000 | 60069799 | 17.31903 | 28.61425 | 79.09498 | 34.16972 | -1.08143 | 1.32804 | 0.105363 |
| 17 | 33109000 | 33111199 | 9.946709 | 17.58132 | 44.58283 | 19.72084 | -0.94417 | 0.518776 | 0.105365 |
| 6 | 42590400 | 42591599 | 5.932097 | 11.16127 | 33.45389 | 13.71477 | -1.10104 | 0.070276 | 0.105372 |

| | | | | | | | | | |
|---|---|---|---|---|---|---|---|---|---|
| 16 | 93934600 | 93936199 | 5.932097 | 11.16127 | 33.45389 | 13.71477 | -1.10104 | 0.070276 | 0.105372 |
| 3 | 31112600 | 31118599 | 31.68204 | 50.97042 | 113.7823 | 48.29867 | -0.80293 | 1.851203 | 0.105374 |
| 12 | 20868800 | 20880799 | 300.6317 | 435.3826 | 297.0346 | 122.4913 | 0.77021 | 4.062978 | 0.105392 |
| 7 | 126938200 | 126946199 | 36.71058 | 58.4731 | 141.0131 | 58.99596 | -0.93093 | 2.180691 | 0.105392 |
| 15 | 77260800 | 77273999 | 95.00061 | 144.1728 | 283.3577 | 117.3168 | -0.69312 | 3.225857 | 0.105395 |
| 6 | 48409600 | 48426799 | 92.04231 | 138.614 | 327.5759 | 135.3059 | -0.96832 | 3.382163 | 0.105401 |
| 3 | 99153400 | 99165999 | 90.03649 | 136.653 | 295.1834 | 122.2166 | -0.82916 | 3.222529 | 0.105407 |
| 1 | 182865800 | 182871999 | 69.20534 | 105.8851 | 255.7081 | 106.0997 | -0.98041 | 3.051537 | 0.105408 |
| 7 | 9872000 | 9874199 | 58.51694 | 93.99887 | 71.03318 | 30.08563 | 0.75375 | 1.918454 | 0.105411 |
| 5 | 117479800 | 117488999 | 58.67283 | 91.12904 | 208.48 | 86.56907 | -0.88559 | 2.757534 | 0.105423 |
| 5 | 73556800 | 73560999 | 20.7525 | 34.35822 | 81.18833 | 35.12437 | -0.85612 | 1.311781 | 0.105436 |
| 13 | 97649800 | 97651599 | 7.635186 | 14.04764 | 37.31475 | 15.8339 | -0.9678 | 0.230763 | 0.105438 |
| 8 | 109133400 | 109135599 | 8.198739 | 14.42304 | 48.24786 | 21.36231 | -1.34385 | 0.567089 | 0.105451 |
| 17 | 15127200 | 15129999 | 12.8968 | 22.19162 | 60.70974 | 26.6304 | -1.06133 | 0.888746 | 0.105453 |
| 8 | 95074800 | 95085599 | 66.28076 | 102.3276 | 227.1588 | 94.35723 | -0.84902 | 2.870932 | 0.105467 |
| 11 | 20257600 | 20259199 | 8.453873 | 15.31867 | 39.6636 | 17.16593 | -0.95098 | 0.330376 | 0.10548 |
| 11 | 74513800 | 74515799 | 8.477941 | 15.29245 | 41.16512 | 17.91757 | -1.01232 | 0.359064 | 0.105492 |
| 19 | 40413800 | 40415999 | 8.477941 | 15.29245 | 41.16512 | 17.91757 | -1.01232 | 0.359064 | 0.105492 |
| 18 | 14203400 | 14204799 | 7.179652 | 13.22054 | 36.7655 | 15.58314 | -1.0317 | 0.222951 | 0.105499 |
| 14 | 56905200 | 56906399 | 5.028782 | 9.292508 | 37.85853 | 15.96997 | -1.57198 | 0.186142 | 0.105502 |
| 12 | 92911000 | 92911599 | 3.641722 | 7.107568 | 25.99118 | 9.226017 | -1.19003 | -0.47355 | 0.105505 |
| 2 | 44208600 | 44211199 | 11.05199 | 19.42892 | 53.12743 | 23.52606 | -1.0562 | 0.598455 | 0.105515 |
| 3 | 30720600 | 30721799 | 17.2815 | 32.16608 | 18.21022 | 7.603212 | 1.262951 | 0.197603 | 0.105523 |
| 7 | 130175600 | 130176999 | 6.113923 | 11.36896 | 34.77683 | 14.54375 | -1.15325 | 0.153064 | 0.105529 |
| 19 | 10611400 | 10615599 | 14.25041 | 23.81917 | 72.6112 | 31.61311 | -1.22083 | 1.180419 | 0.105537 |
| 2 | 181404800 | 181406799 | 10.93402 | 19.1188 | 50.99584 | 22.48319 | -1.02056 | 0.680872 | 0.105552 |
| 1 | 89029000 | 89036999 | 41.19316 | 65.28303 | 160.5135 | 66.77585 | -0.96633 | 2.3418 | 0.105557 |
| 17 | 86694000 | 86695799 | 16.31812 | 30.11902 | 19.83102 | 8.378473 | 1.034346 | 0.238995 | 0.105566 |
| 5 | 101171600 | 101173199 | 5.149738 | 9.87319 | 31.32478 | 12.32438 | -1.129 | -0.08205 | 0.105566 |
| 2 | 156482200 | 156491599 | 58.70812 | 91.13278 | 211.4658 | 87.81623 | -0.90666 | 2.767316 | 0.105573 |
| 1 | 135651600 | 135652799 | 4.133595 | 7.989498 | 28.71474 | 11.0085 | -1.2698 | -0.18535 | 0.105576 |
| 12 | 23954800 | 23957999 | 49.94029 | 80.72399 | 55.8774 | 23.87016 | 0.890571 | 1.712817 | 0.105577 |
| 8 | 21784400 | 21787799 | 34.85416 | 59.45281 | 30.92635 | 13.55777 | 1.321107 | 1.019444 | 0.105578 |
| 13 | 66729800 | 66730799 | 29.75243 | 51.203 | 31.4155 | 13.78051 | 1.08616 | 0.923176 | 0.105583 |
| 17 | 56739000 | 56827799 | 866.1938 | 1094.628 | 2083.31 | 875.1694 | -0.90674 | 6.195453 | 0.105585 |
| 5 | 24791400 | 24811599 | 100.3414 | 149.5037 | 382.4577 | 157.6024 | -1.09657 | 3.589481 | 0.105586 |
| 2 | 102934600 | 102935399 | 3.241154 | 6.309543 | 25.12751 | 9.225672 | -1.35397 | -0.35711 | 0.105591 |
| 5 | 33174000 | 33177199 | 11.4204 | 19.5703 | 61.56488 | 26.99937 | -1.26266 | 0.923181 | 0.105602 |
| 4 | 152396800 | 152400599 | 27.21869 | 43.80054 | 113.7298 | 48.29363 | -1.01987 | 1.837593 | 0.105607 |
| 8 | 76071000 | 76072599 | 16.15751 | 29.97506 | 19.50591 | 8.184129 | 1.058608 | 0.210299 | 0.105611 |
| 11 | 31314600 | 31316199 | 7.798249 | 14.30289 | 38.8897 | 16.57251 | -1.00603 | 0.252061 | 0.105611 |
| X | 31555400 | 31557199 | 20.16897 | 37.02932 | 17.17115 | 7.307362 | 1.528499 | 0.292902 | 0.105614 |
| 2 | 181334600 | 181340399 | 33.68231 | 53.98253 | 122.4403 | 51.67951 | -0.83259 | 1.977153 | 0.105616 |
| 2 | 168154600 | 168167799 | 85.65415 | 129.4503 | 309.5374 | 127.9903 | -0.98081 | 3.309295 | 0.105622 |
| 12 | 56082200 | 56083999 | 17.37803 | 31.64495 | 21.34938 | 9.183024 | 0.98198 | 0.325498 | 0.105628 |
| 5 | 8848000 | 8848999 | 7.008597 | 13.0052 | 36.43027 | 15.28032 | -1.03156 | 0.174266 | 0.105635 |
| 9 | 120100800 | 120105999 | 32.01961 | 51.4692 | 114.4883 | 48.53776 | -0.79935 | 1.879696 | 0.105642 |

| | | | | | | | | | |
|---|---|---|---|---|---|---|---|---|---|
| 12 | 31161000 | 31162999 | 10.14254 | 17.78782 | 47.76611 | 21.11239 | -1.02958 | 0.625363 | 0.105643 |
| 2 | 163757400 | 163767199 | 54.88492 | 85.46281 | 212.9491 | 88.39123 | -1.0058 | 2.711334 | 0.105646 |
| X | 31740200 | 31742199 | 17.45962 | 32.56661 | 17.65675 | 7.345701 | 1.328844 | 0.18482 | 0.105654 |
| 1 | 72300800 | 72302199 | 6.229988 | 11.66082 | 33.85865 | 14.00376 | -1.06577 | 0.109253 | 0.105657 |
| 17 | 18743600 | 18745799 | 49.21806 | 79.85897 | 56.18109 | 24.07424 | 0.872388 | 1.630665 | 0.105659 |
| 18 | 36006400 | 36016799 | 65.71045 | 102.0227 | 213.7032 | 88.74274 | -0.75902 | 2.766562 | 0.105663 |
| X | 96883200 | 96883999 | 12.93391 | 25.17019 | 11.75033 | 4.325566 | 1.697176 | -0.27391 | 0.105664 |
| 7 | 31785800 | 31788399 | 40.72939 | 67.24034 | 46.14518 | 19.92445 | 0.921815 | 1.368927 | 0.105667 |
| 11 | 103680400 | 103708399 | 229.3269 | 321.9025 | 686.0244 | 284.2919 | -0.91751 | 4.520966 | 0.10568 |
| 18 | 53666800 | 53669199 | 12.97084 | 22.53399 | 54.2837 | 23.83215 | -0.8756 | 0.790401 | 0.105691 |
| 10 | 3196800 | 3198399 | 41.37905 | 69.3267 | 34.36078 | 14.98203 | 1.389862 | 1.203853 | 0.105694 |
| 7 | 121439400 | 121441999 | 14.0276 | 23.97181 | 61.15102 | 26.81191 | -0.96269 | 0.95839 | 0.105695 |
| 16 | 3219600 | 3222599 | 70.61387 | 112.5117 | 84.63076 | 35.42326 | 0.747419 | 2.186384 | 0.105707 |
| 4 | 60323400 | 60324799 | 13.71 | 26.54927 | 13.17436 | 5.006164 | 1.568321 | -0.15643 | 0.105717 |
| 12 | 22390600 | 22393399 | 27.58306 | 47.93621 | 29.49586 | 12.9524 | 1.084192 | 0.817424 | 0.105732 |
| 9 | 69360800 | 69361799 | 4.833697 | 9.239057 | 30.48889 | 12.10098 | -1.19561 | -0.05463 | 0.105738 |
| 1 | 167033400 | 167036799 | 21.14261 | 34.80351 | 84.27734 | 36.34035 | -0.89316 | 1.384003 | 0.105747 |
| 7 | 15412400 | 15414599 | 64.30958 | 102.8391 | 78.00014 | 32.81461 | 0.741036 | 2.06595 | 0.105751 |
| 1 | 92328000 | 92339799 | 75.79264 | 116.106 | 249.8846 | 103.7128 | -0.81304 | 3.027336 | 0.105763 |
| 17 | 3131000 | 3136399 | 791.21 | 1049.598 | 662.5173 | 276.9156 | 0.755405 | 5.40853 | 0.105786 |
| 13 | 112937000 | 112938599 | 9.028705 | 15.98895 | 43.55136 | 19.25729 | -1.04533 | 0.490452 | 0.105788 |
| 3 | 49081400 | 49082399 | 16.64015 | 30.84582 | 19.4068 | 8.151391 | 1.105729 | 0.218277 | 0.105794 |
| 4 | 135990000 | 135993199 | 12.34072 | 21.47622 | 53.18905 | 23.37685 | -0.91502 | 0.760907 | 0.105805 |
| 11 | 116360600 | 116361999 | 4.47511 | 8.498395 | 31.95487 | 12.85872 | -1.39997 | -0.0102 | 0.105806 |
| 2 | 172936400 | 172940599 | 22.9656 | 37.19807 | 99.00528 | 42.37043 | -1.04452 | 1.642791 | 0.105813 |
| 4 | 100965800 | 100968199 | 9.626691 | 17.06775 | 49.4206 | 21.84702 | -1.13314 | 0.488935 | 0.105815 |
| 2 | 130263800 | 130266799 | 15.06253 | 25.44912 | 66.45934 | 29.07698 | -0.99827 | 1.070645 | 0.105827 |
| 7 | 42372400 | 42373999 | 36.18447 | 60.14652 | 43.87862 | 19.03535 | 0.834493 | 1.27669 | 0.105836 |
| 19 | 44719800 | 44731399 | 67.52851 | 104.7732 | 213.6928 | 88.74758 | -0.72139 | 2.779385 | 0.105838 |
| 8 | 18895800 | 18898799 | 13.01034 | 22.48262 | 55.83346 | 24.4946 | -0.92076 | 0.846685 | 0.105845 |
| 17 | 50920600 | 50922799 | 23.75553 | 41.60938 | 27.33031 | 12.06696 | 0.990716 | 0.681669 | 0.105861 |
| 13 | 11224800 | 11225799 | 12.47214 | 24.20119 | 13.25137 | 4.987947 | 1.440094 | -0.20142 | 0.105861 |
| 13 | 100627800 | 100630199 | 9.6925 | 17.03226 | 48.78969 | 21.60094 | -1.12254 | 0.589141 | 0.105867 |
| 17 | 83521000 | 83535599 | 67.03499 | 102.4037 | 275.9867 | 114.4724 | -1.1404 | 3.083081 | 0.105868 |
| 15 | 95748600 | 95755599 | 52.91206 | 83.58226 | 170.5121 | 70.81354 | -0.70142 | 2.430091 | 0.10587 |
| 2 | 121668000 | 121669799 | 30.36389 | 51.98745 | 33.85398 | 14.84568 | 0.999299 | 0.971752 | 0.105911 |
| 11 | 50831000 | 50835799 | 24.49671 | 39.80386 | 95.56662 | 40.99439 | -0.89205 | 1.578033 | 0.10592 |
| 16 | 78284000 | 78286399 | 10.91129 | 19.13225 | 51.51845 | 22.76709 | -1.03429 | 0.635002 | 0.105925 |
| 10 | 8286200 | 8287399 | 16.45572 | 30.15137 | 20.56855 | 8.777108 | 0.97381 | 0.282824 | 0.105987 |
| 12 | 115224800 | 115226799 | 32.63697 | 54.85194 | 39.42256 | 17.22038 | 0.855673 | 1.145346 | 0.105992 |
| 6 | 99702800 | 99704599 | 8.573164 | 15.15604 | 44.08972 | 19.50207 | -1.14027 | 0.509436 | 0.106 |
| 4 | 100479600 | 100481999 | 8.912109 | 15.7048 | 45.64059 | 20.20101 | -1.14112 | 0.54864 | 0.106005 |
| 6 | 21183800 | 21184199 | 10.54321 | 20.55331 | 11.76472 | 4.246705 | 1.432969 | -0.37051 | 0.106036 |
| 4 | 135851800 | 135855399 | 17.61991 | 29.56533 | 69.28457 | 30.2385 | -0.84323 | 1.145236 | 0.106056 |
| 17 | 5847800 | 5852599 | 22.07134 | 35.94965 | 90.49609 | 38.85875 | -0.95731 | 1.534798 | 0.106073 |
| 4 | 44691400 | 44691999 | 3.533196 | 6.88131 | 25.28109 | 9.238 | -1.23407 | -0.37151 | 0.106079 |
| 1 | 10798800 | 10800199 | 17.30715 | 31.26571 | 21.43113 | 9.310527 | 0.950069 | 0.369411 | 0.106081 |

| | | | | | | | | | |
|---|---|---|---|---|---|---|---|---|---|
| 13 | 58919400 | 58921199 | 6.741688 | 12.42898 | 36.24628 | 15.338 | -1.09801 | 0.2102 | 0.106085 |
| 12 | 98544000 | 98545399 | 5.898395 | 11.19706 | 33.25535 | 13.39268 | -1.06589 | 0.007177 | 0.106098 |
| 17 | 84123000 | 84133399 | 54.96031 | 85.7506 | 204.8659 | 85.01238 | -0.94294 | 2.675916 | 0.106109 |
| 12 | 72874800 | 72879799 | 27.12974 | 43.83825 | 102.3362 | 43.6981 | -0.85952 | 1.707605 | 0.106111 |
| 17 | 18485600 | 18488399 | 41.75109 | 68.6213 | 48.91366 | 21.07772 | 0.865031 | 1.426909 | 0.106111 |
| 17 | 12368200 | 12369199 | 5.898395 | 11.14961 | 32.46517 | 13.1924 | -1.0483 | 0.037673 | 0.106117 |
| 6 | 29417200 | 29419199 | 8.490385 | 15.25165 | 41.08904 | 17.96441 | -1.0173 | 0.387255 | 0.10614 |
| 7 | 101266600 | 101267799 | 4.841103 | 9.338489 | 29.86231 | 11.54418 | -1.11577 | -0.14688 | 0.106145 |
| X | 33635600 | 33637199 | 25.22553 | 44.75839 | 19.4792 | 8.605438 | 1.585805 | 0.529454 | 0.106173 |
| 11 | 107605200 | 107610599 | 36.73238 | 58.44169 | 144.8085 | 60.48914 | -0.97171 | 2.21891 | 0.106178 |
| 11 | 74067800 | 74068399 | 25.48141 | 44.2226 | 30.49771 | 13.38134 | 0.920455 | 0.828233 | 0.106222 |
| 8 | 123848600 | 123858599 | 53.93047 | 83.87026 | 209.06 | 86.79339 | -1.00773 | 2.733186 | 0.106223 |
| X | 17858000 | 17859799 | 24.86179 | 44.30708 | 18.27214 | 8.042366 | 1.665348 | 0.4967 | 0.106225 |
| 18 | 25955800 | 25957399 | 9.090665 | 16.3706 | 42.19813 | 18.40521 | -0.95218 | 0.376362 | 0.106225 |
| 11 | 120373600 | 120376799 | 12.00203 | 20.52325 | 60.41582 | 26.4909 | -1.16764 | 0.946568 | 0.106228 |
| 6 | 51183800 | 51185399 | 6.819217 | 12.77514 | 35.06345 | 14.48008 | -0.98427 | 0.109493 | 0.106246 |
| 8 | 92337600 | 92339999 | 8.778957 | 15.82973 | 40.62206 | 17.68916 | -0.94414 | 0.363164 | 0.106261 |
| 17 | 38581600 | 38584399 | 26.41631 | 46.29047 | 22.98292 | 10.1513 | 1.391997 | 0.682787 | 0.106294 |
| 17 | 27841200 | 27842999 | 7.66859 | 13.64277 | 43.08089 | 18.98955 | -1.25313 | 0.445902 | 0.106297 |
| 5 | 80840000 | 80841199 | 21.63162 | 39.65163 | 15.30339 | 6.526612 | 1.788249 | 0.308681 | 0.106302 |
| 5 | 151658000 | 151658199 | 10.09578 | 19.73032 | 9.443077 | 3.268712 | 1.735128 | -0.52769 | 0.106307 |
| 18 | 56650800 | 56653599 | 15.93294 | 27.07283 | 66.73816 | 29.24816 | -0.91522 | 1.022949 | 0.10631 |
| X | 90884200 | 90885199 | 12.66623 | 24.52693 | 11.85395 | 4.505347 | 1.607683 | -0.17155 | 0.106346 |
| 19 | 3109800 | 3110399 | 10.45938 | 20.42464 | 10.72278 | 3.771043 | 1.585938 | -0.46428 | 0.106353 |
| 7 | 31239000 | 31252999 | 255.897 | 377.7903 | 225.2193 | 92.74947 | 0.991192 | 3.761342 | 0.106362 |
| 11 | 64682600 | 64684199 | 13.10517 | 24.82753 | 17.70339 | 7.16015 | 0.969889 | 0.047061 | 0.106411 |
| 2 | 87538200 | 87539599 | 13.86875 | 26.54601 | 15.84929 | 6.265077 | 1.252071 | -0.02301 | 0.106422 |
| X | 26443800 | 26445599 | 15.0036 | 28.27752 | 18.01879 | 7.365701 | 1.117832 | 0.100512 | 0.106459 |
| 5 | 45295600 | 45296999 | 6.839939 | 12.74809 | 42.41978 | 17.67156 | -1.27098 | 0.157362 | 0.106469 |
| 1 | 79899800 | 79901599 | 8.232084 | 14.73233 | 43.68223 | 19.15862 | -1.15845 | 0.410759 | 0.106479 |
| 19 | 10790000 | 10791799 | 7.694356 | 13.99296 | 39.20349 | 16.89017 | -1.05955 | 0.301772 | 0.106479 |
| 19 | 53932200 | 53934799 | 14.08558 | 23.62597 | 74.54904 | 32.52835 | -1.2706 | 1.141475 | 0.106485 |
| 7 | 20892000 | 20893199 | 11.62085 | 22.57379 | 14.05576 | 5.250814 | 1.267476 | -0.23391 | 0.106495 |
| 18 | 12885200 | 12885799 | 3.043437 | 5.941345 | 22.65971 | 8.009579 | -1.24057 | -0.50396 | 0.1065 |
| 10 | 127126200 | 127127799 | 6.353638 | 11.65187 | 39.93076 | 17.019 | -1.33707 | 0.243353 | 0.106512 |
| 11 | 77803000 | 77807199 | 23.05425 | 37.51498 | 94.94167 | 40.75356 | -0.96614 | 1.551238 | 0.106527 |
| 9 | 70118400 | 70119599 | 7.068309 | 12.89929 | 37.65699 | 16.15335 | -1.11347 | 0.27605 | 0.106529 |
| 5 | 144114200 | 144122999 | 48.84206 | 76.79883 | 178.9954 | 74.25415 | -0.89845 | 2.526894 | 0.106558 |
| 17 | 25551200 | 25553399 | 10.44151 | 18.3447 | 47.20932 | 20.89762 | -0.96881 | 0.595174 | 0.106559 |
| 10 | 59778400 | 59791599 | 65.89507 | 101.8521 | 222.2714 | 92.32651 | -0.82307 | 2.848578 | 0.106561 |
| 11 | 75170000 | 75175199 | 23.52091 | 38.27414 | 92.56854 | 39.73482 | -0.90087 | 1.551642 | 0.106608 |
| 5 | 41371400 | 41373399 | 24.24431 | 42.20814 | 30.2916 | 13.31656 | 0.86346 | 0.777579 | 0.106612 |
| 15 | 96130000 | 96131799 | 6.180468 | 11.70107 | 38.47797 | 15.58419 | -1.21909 | 0.042145 | 0.106636 |
| 11 | 110675000 | 110676399 | 14.42667 | 27.64973 | 15.26871 | 6.013163 | 1.368603 | -0.03025 | 0.106648 |
| 10 | 95717400 | 95718799 | 5.57616 | 10.67999 | 33.13261 | 13.07334 | -1.10152 | -0.0643 | 0.106649 |
| 8 | 21549200 | 21553199 | 27.81534 | 48.60055 | 25.74515 | 11.34364 | 1.299692 | 0.741244 | 0.106657 |
| 17 | 58545000 | 58546999 | 20.16449 | 35.73518 | 24.21961 | 10.70774 | 0.949639 | 0.512341 | 0.10667 |

| | | | | | | | | | |
|---|---|---|---|---|---|---|---|---|---|
| 14 | 104302400 | 104303999 | 20.44195 | 36.08016 | 24.85207 | 10.99628 | 0.924334 | 0.558289 | 0.106673 |
| 2 | 41771000 | 41772399 | 16.04404 | 30.27415 | 16.65995 | 6.795219 | 1.329634 | 0.098487 | 0.106679 |
| 6 | 29735200 | 29738799 | 16.86 | 28.23207 | 70.24712 | 30.61879 | -0.92931 | 1.165945 | 0.106717 |
| 2 | 122125000 | 122139799 | 96.67159 | 145.6177 | 326.9639 | 135.0611 | -0.89431 | 3.378521 | 0.10674 |
| 14 | 86026200 | 86026999 | 10.78 | 20.84626 | 13.73475 | 5.253223 | 1.155204 | -0.16823 | 0.106742 |
| 11 | 31949600 | 31953999 | 19.00728 | 31.52085 | 79.86331 | 34.59057 | -0.95609 | 1.286587 | 0.106743 |
| 16 | 47828400 | 47829199 | 11.97224 | 23.04561 | 16.17386 | 6.290622 | 1.042792 | -0.0927 | 0.106745 |
| 18 | 39621800 | 39624999 | 21.83259 | 35.41841 | 96.74808 | 41.45234 | -1.07965 | 1.607559 | 0.106747 |
| 1 | 155416600 | 155421999 | 62.63559 | 100.2599 | 76.96772 | 32.42506 | 0.725158 | 2.033951 | 0.106755 |
| 5 | 136303400 | 136304399 | 8.38029 | 14.85664 | 44.08814 | 19.49045 | -1.16687 | 0.478518 | 0.106755 |
| 19 | 16845400 | 16856199 | 83.51119 | 127.2649 | 273.9093 | 113.5463 | -0.81909 | 3.136622 | 0.106757 |
| 1 | 100156800 | 100158199 | 12.65373 | 24.57191 | 13.4087 | 5.022758 | 1.452316 | -0.21409 | 0.106759 |
| 14 | 31820800 | 31822599 | 18.00183 | 32.82283 | 20.89101 | 8.971507 | 1.067127 | 0.323005 | 0.106768 |
| X | 100427000 | 100430199 | 13.98704 | 24.02613 | 59.02365 | 25.88943 | -0.90717 | 0.910258 | 0.106772 |
| 7 | 47130200 | 47133399 | 38.87082 | 65.48641 | 33.12979 | 14.48217 | 1.360182 | 1.141643 | 0.106773 |
| 8 | 71125600 | 71133599 | 34.14856 | 54.41363 | 142.5173 | 59.65768 | -1.04916 | 2.165448 | 0.106774 |
| 17 | 37182200 | 37183799 | 6.968263 | 13.04758 | 35.61791 | 14.722 | -0.97759 | 0.115732 | 0.106778 |
| 4 | 57141000 | 57143799 | 15.11744 | 25.42538 | 70.25638 | 30.69565 | -1.07986 | 1.114209 | 0.106782 |
| 12 | 69491600 | 69493599 | 7.019005 | 12.99402 | 36.36299 | 15.30082 | -1.03343 | 0.187856 | 0.106801 |
| 6 | 135802600 | 135804399 | 8.768878 | 15.84318 | 40.66979 | 17.66573 | -0.94229 | 0.351216 | 0.106803 |
| 4 | 52459400 | 52459999 | 12.60488 | 24.00821 | 17.21495 | 6.884022 | 0.975979 | 0.004678 | 0.106807 |
| 3 | 63683400 | 63685599 | 18.57339 | 33.23266 | 22.25985 | 9.764624 | 0.973982 | 0.436536 | 0.10681 |
| 2 | 28167200 | 28170799 | 12.76412 | 21.54769 | 67.266 | 29.42241 | -1.25432 | 1.072199 | 0.106815 |
| 5 | 52940200 | 52940999 | 5.469979 | 10.39302 | 32.66602 | 13.13064 | -1.14408 | -0.00434 | 0.106818 |
| 1 | 88544600 | 88546199 | 18.59344 | 33.8717 | 20.69297 | 8.89624 | 1.124646 | 0.331438 | 0.106819 |
| 5 | 143228400 | 143251799 | 163.7826 | 235.1707 | 554.7561 | 229.0445 | -1.03099 | 4.18386 | 0.106822 |
| 5 | 113306000 | 113331799 | 132.0387 | 192.4744 | 485.0007 | 199.8607 | -1.10465 | 3.95524 | 0.106831 |
| 13 | 66044600 | 66048599 | 108.8747 | 169.9563 | 111.9191 | 46.44021 | 0.91451 | 2.681162 | 0.106831 |
| 18 | 79151200 | 79153799 | 12.26171 | 21.14259 | 56.84681 | 24.9375 | -1.03543 | 0.864506 | 0.106839 |
| 16 | 12587400 | 12588799 | 24.50578 | 42.79358 | 26.09046 | 11.49168 | 1.097725 | 0.730109 | 0.106862 |
| 14 | 13021000 | 13021999 | 17.26929 | 31.45097 | 21.4389 | 9.220054 | 0.967445 | 0.324583 | 0.106873 |
| 6 | 90382200 | 90384199 | 6.527969 | 11.86949 | 39.18199 | 16.88495 | -1.29397 | 0.295025 | 0.10689 |
| 12 | 107937400 | 107940799 | 16.85109 | 28.24809 | 71.47488 | 31.17549 | -0.95351 | 1.153994 | 0.106918 |
| 1 | 171264800 | 171265799 | 4.368974 | 8.473468 | 27.23684 | 10.27584 | -1.08794 | -0.24076 | 0.106923 |
| 7 | 48791800 | 48796599 | 21.13327 | 34.36642 | 92.2837 | 39.60312 | -1.05181 | 1.558314 | 0.106925 |
| 18 | 41621800 | 41623599 | 17.50605 | 32.0943 | 20.21283 | 8.618567 | 1.08872 | 0.286338 | 0.106934 |
| 7 | 142606200 | 142613199 | 26.77191 | 43.0232 | 117.6241 | 49.84921 | -1.09655 | 1.881969 | 0.106937 |
| 17 | 46829800 | 46837799 | 46.79806 | 73.50468 | 188.6854 | 78.26282 | -1.03878 | 2.548491 | 0.106937 |
| 7 | 49263400 | 49267199 | 15.45477 | 25.92766 | 71.89727 | 31.3752 | -1.08506 | 1.141505 | 0.106947 |
| 16 | 16043600 | 16044199 | 4.272335 | 8.202461 | 29.50308 | 11.5612 | -1.30063 | -0.10631 | 0.106954 |
| 10 | 35728000 | 35731399 | 35.81505 | 60.19854 | 37.64733 | 16.44537 | 1.055562 | 1.154167 | 0.106957 |
| 7 | 43999400 | 44016599 | 107.4744 | 160.3415 | 368.2219 | 151.7688 | -0.93927 | 3.565928 | 0.106968 |
| X | 126295000 | 126295999 | 16.09942 | 30.66678 | 15.25078 | 6.097493 | 1.499567 | 0.033916 | 0.106986 |
| 11 | 49915600 | 49916799 | 7.273993 | 13.18077 | 39.65721 | 17.16303 | -1.16572 | 0.321304 | 0.106988 |
| 12 | 8127400 | 8127999 | 3.531008 | 6.883315 | 24.90048 | 8.991015 | -1.19648 | -0.41654 | 0.10699 |
| 17 | 34254600 | 34267799 | 99.00088 | 148.6941 | 330.31 | 136.3373 | -0.88256 | 3.436573 | 0.106996 |
| 9 | 53778000 | 53779199 | 14.6053 | 27.28752 | 19.12305 | 7.933543 | 0.964847 | 0.153368 | 0.106996 |

| | | | | | | | | | |
|---|---|---|---|---|---|---|---|---|---|
| 16 | 10613800 | 10617599 | 19.17654 | 31.75792 | 77.36781 | 33.47797 | -0.89973 | 1.302702 | 0.107011 |
| 11 | 116016200 | 116026399 | 45.21773 | 70.94128 | 190.4381 | 78.98041 | -1.10448 | 2.571095 | 0.107013 |
| 11 | 81854800 | 81861599 | 36.54396 | 58.24592 | 142.6938 | 59.75338 | -0.95261 | 2.158431 | 0.107013 |
| 13 | 51798400 | 51800199 | 10.16185 | 17.80944 | 49.36016 | 21.80429 | -1.0753 | 0.636886 | 0.10702 |
| 19 | 32463000 | 32464999 | 10.16185 | 17.80944 | 49.36016 | 21.80429 | -1.0753 | 0.636886 | 0.10702 |
| 10 | 79405800 | 79424599 | 137.0345 | 201.2339 | 444.6484 | 183.1305 | -0.90504 | 3.840166 | 0.107038 |
| 10 | 79563400 | 79582399 | 340.3972 | 459.5409 | 1032.193 | 430.3585 | -1.05014 | 5.125774 | 0.107039 |
| 11 | 114692400 | 114702599 | 62.08137 | 96.10376 | 218.8156 | 90.89032 | -0.88248 | 2.818546 | 0.107045 |
| 15 | 101057800 | 101064199 | 21.41718 | 34.99243 | 91.68971 | 39.41578 | -1.01189 | 1.486847 | 0.107047 |
| 7 | 32856000 | 32865599 | 188.2171 | 281.4859 | 210.1263 | 86.648 | 0.684053 | 3.526795 | 0.107049 |
| 4 | 152009400 | 152010199 | 4.836955 | 9.342531 | 29.98303 | 11.53178 | -1.11399 | -0.16434 | 0.107088 |
| 1 | 189939200 | 189940399 | 4.721026 | 9.087717 | 31.54008 | 12.26666 | -1.24134 | -0.12483 | 0.107093 |
| 19 | 35252800 | 35254199 | 16.00779 | 30.42019 | 15.70019 | 6.308654 | 1.440088 | 0.049447 | 0.107094 |
| 2 | 89092800 | 89096399 | 67.5167 | 107.9089 | 79.86821 | 33.57463 | 0.775394 | 2.085482 | 0.10711 |
| 14 | 23022400 | 23024599 | 19.74172 | 35.58955 | 21.58515 | 9.40192 | 1.123173 | 0.399917 | 0.107113 |
| 1 | 140818600 | 140819199 | 9.863563 | 19.221 | 13.11625 | 4.75845 | 1.174012 | -0.35947 | 0.107118 |
| 11 | 55254400 | 55255799 | 5.778912 | 10.92126 | 34.19876 | 13.90359 | -1.15372 | 0.039445 | 0.107118 |
| 16 | 20805000 | 20808199 | 46.44204 | 75.31722 | 57.05045 | 24.43779 | 0.7654 | 1.635952 | 0.107118 |
| 5 | 102232200 | 102233999 | 7.447318 | 13.83157 | 37.59558 | 15.75004 | -0.98728 | 0.176181 | 0.107119 |
| 7 | 8351000 | 8353199 | 56.0115 | 90.39905 | 60.60564 | 25.83893 | 0.934136 | 1.770175 | 0.107129 |
| 7 | 15482200 | 15486999 | 102.5294 | 160.5664 | 115.2447 | 47.80432 | 0.794012 | 2.622551 | 0.107131 |
| 14 | 72824000 | 72824799 | 5.548191 | 10.666 | 32.14774 | 12.55005 | -1.04518 | -0.10052 | 0.107159 |
| 14 | 3806400 | 3808799 | 60.76935 | 97.62355 | 64.98676 | 27.52737 | 0.936377 | 1.916929 | 0.107173 |
| 8 | 12180400 | 12186399 | 62.55142 | 96.11447 | 243.8528 | 101.2543 | -1.04669 | 2.962625 | 0.107173 |
| 1 | 12921200 | 12922599 | 7.653559 | 14.03144 | 40.5466 | 17.28876 | -1.09406 | 0.255964 | 0.107187 |
| 4 | 138529400 | 138530599 | 6.394591 | 11.93898 | 34.35714 | 14.27033 | -1.05828 | 0.127119 | 0.107195 |
| 12 | 18641000 | 18657199 | 246.1802 | 362.4333 | 254.081 | 104.6321 | 0.753754 | 3.806525 | 0.107201 |
| 2 | 173099400 | 173126799 | 137.6281 | 200.7559 | 488.8254 | 201.4229 | -1.05432 | 3.945418 | 0.107211 |
| 13 | 118251600 | 118252399 | 10.89116 | 21.17826 | 13.49202 | 4.998258 | 1.244941 | -0.27378 | 0.107217 |
| 19 | 5540200 | 5541799 | 6.298856 | 11.56438 | 36.68432 | 15.61145 | -1.22364 | 0.2336 | 0.10722 |
| 19 | 55594800 | 55595999 | 4.781905 | 9.03482 | 32.93673 | 13.39755 | -1.37019 | 0.030852 | 0.107227 |
| 17 | 22167000 | 22169399 | 56.75804 | 91.61269 | 60.20864 | 25.67284 | 0.962917 | 1.767584 | 0.107232 |
| 6 | 29522400 | 29531599 | 44.55886 | 70.2953 | 177.4048 | 73.62741 | -1.00934 | 2.456729 | 0.107262 |
| 5 | 61394400 | 61396199 | 18.24566 | 33.49191 | 19.65252 | 8.365473 | 1.191852 | 0.284608 | 0.107281 |
| 1 | 82460000 | 82461399 | 5.079452 | 9.797502 | 34.83869 | 13.47096 | -1.268 | -0.12927 | 0.107284 |
| 9 | 80617800 | 80618799 | 12.01347 | 20.96665 | 51.56369 | 22.7033 | -0.9044 | 0.716037 | 0.107305 |
| 2 | 126516400 | 126518799 | 11.15861 | 19.34998 | 55.80724 | 24.48569 | -1.13424 | 0.805554 | 0.107329 |
| 12 | 70157200 | 70160799 | 18.74677 | 30.91881 | 79.75643 | 34.41882 | -0.98263 | 1.353334 | 0.107348 |
| 11 | 116561800 | 116563399 | 6.332292 | 11.66479 | 40.08241 | 16.97996 | -1.33463 | 0.217203 | 0.107352 |
| 17 | 6879200 | 6885999 | 27.11991 | 43.49643 | 122.4514 | 51.74106 | -1.14215 | 1.94731 | 0.107354 |
| 11 | 117352800 | 117360999 | 39.04172 | 62.00181 | 151.9308 | 63.3366 | -0.95845 | 2.277331 | 0.107355 |
| 2 | 167228400 | 167230199 | 8.855614 | 15.72961 | 42.53388 | 18.77348 | -1.03245 | 0.463565 | 0.107359 |
| 17 | 28330200 | 28335599 | 30.05088 | 48.0902 | 129.4831 | 54.5298 | -1.08196 | 2.022408 | 0.10736 |
| 19 | 5941800 | 5946999 | 21.9621 | 35.73333 | 97.09504 | 41.676 | -1.06851 | 1.551816 | 0.107367 |
| 5 | 24950000 | 24965599 | 92.50081 | 140.0026 | 301.7368 | 124.8225 | -0.82928 | 3.280355 | 0.107368 |
| 7 | 73421000 | 73426999 | 34.63827 | 55.52511 | 125.0412 | 52.79242 | -0.82202 | 1.972281 | 0.107378 |
| 9 | 55307200 | 55312199 | 19.74652 | 32.12676 | 93.96342 | 40.33103 | -1.17417 | 1.548259 | 0.107382 |

| | | | | | | | | | |
|---|---|---|---|---|---|---|---|---|---|
| 2 | 145725400 | 145727599 | 8.916609 | 15.7373 | 47.21414 | 20.89095 | -1.18622 | 0.531899 | 0.107383 |
| 6 | 59848600 | 59850199 | 24.2102 | 42.43742 | 24.97358 | 11.03336 | 1.147976 | 0.667204 | 0.107398 |
| 7 | 38649800 | 38650999 | 18.9812 | 33.81028 | 23.33316 | 10.27425 | 0.92769 | 0.469507 | 0.107403 |
| 6 | 117985200 | 117997799 | 71.01522 | 109.2866 | 238.3944 | 99.00861 | -0.82736 | 2.934151 | 0.107413 |
| 10 | 43644000 | 43646799 | 29.15701 | 49.59075 | 36.23169 | 15.87986 | 0.832677 | 1.028107 | 0.107414 |
| 17 | 36131400 | 36136599 | 19.58345 | 31.927 | 95.6425 | 41.08249 | -1.2066 | 1.517305 | 0.10743 |
| X | 64556600 | 64557199 | 10.28178 | 20.06231 | 11.95781 | 4.261814 | 1.389374 | -0.41759 | 0.107448 |
| 7 | 33422600 | 33424599 | 48.19031 | 78.73298 | 48.46037 | 20.832 | 1.071987 | 1.517287 | 0.10745 |
| 8 | 115284400 | 115289599 | 38.23888 | 61.11113 | 131.5027 | 55.27489 | -0.76161 | 2.072896 | 0.107452 |
| 1 | 114910600 | 114911599 | 13.60743 | 25.64366 | 18.19092 | 7.437283 | 0.964026 | 0.088257 | 0.107456 |
| 8 | 3395600 | 3396799 | 13.60743 | 25.64366 | 18.19092 | 7.437283 | 0.964026 | 0.088257 | 0.107456 |
| 15 | 60964400 | 60964999 | 9.30212 | 18.12125 | 13.38617 | 4.876319 | 1.056848 | -0.35093 | 0.10746 |
| 15 | 3084000 | 3084799 | 24.28242 | 42.3546 | 29.66358 | 13.05601 | 0.898708 | 0.750874 | 0.107487 |
| 14 | 120832800 | 120834199 | 6.350475 | 11.96492 | 33.5092 | 13.70545 | -1.00117 | 0.06657 | 0.107489 |
| 8 | 88481400 | 88486399 | 25.33076 | 41.0562 | 98.08135 | 41.99554 | -0.88828 | 1.632188 | 0.107498 |
| 10 | 8829000 | 8831799 | 9.297885 | 16.55994 | 43.07182 | 18.96759 | -0.97486 | 0.448236 | 0.107519 |
| 16 | 40283400 | 40284399 | 4.971498 | 9.589645 | 29.26284 | 11.31371 | -1.04883 | -0.14579 | 0.10754 |
| 4 | 143375600 | 143377599 | 20.12317 | 35.6262 | 24.47967 | 10.82489 | 0.929654 | 0.524441 | 0.107543 |
| 12 | 107963000 | 107973199 | 56.10306 | 87.1218 | 212.6947 | 88.3216 | -0.97943 | 2.759692 | 0.10756 |
| 14 | 66534000 | 66536599 | 11.45488 | 19.86218 | 54.73117 | 24.0135 | -1.06874 | 0.807043 | 0.107561 |
| 7 | 79457000 | 79458599 | 9.920321 | 17.61537 | 44.07699 | 19.45961 | -0.92205 | 0.47354 | 0.107569 |
| 13 | 97199400 | 97204599 | 33.6696 | 53.99383 | 124.3814 | 52.54568 | -0.85396 | 1.958353 | 0.107571 |
| 18 | 65020200 | 65022799 | 13.66063 | 23.50094 | 58.5385 | 25.67782 | -0.92673 | 0.892212 | 0.107576 |
| 7 | 139615000 | 139617599 | 11.89585 | 20.32529 | 62.06625 | 27.21281 | -1.2207 | 0.953321 | 0.10758 |
| 7 | 121820800 | 121822199 | 6.213578 | 11.71969 | 37.69061 | 15.38694 | -1.19795 | 0.064208 | 0.107594 |
| 5 | 143353200 | 143368599 | 107.371 | 160.472 | 353.6053 | 145.7946 | -0.87737 | 3.532421 | 0.10761 |
| 5 | 74204800 | 74206799 | 7.839998 | 14.25692 | 38.66629 | 16.66005 | -1.01301 | 0.302699 | 0.107617 |
| 7 | 8630600 | 8630999 | 7.779787 | 15.20417 | 11.21401 | 3.881447 | 1.122982 | -0.55767 | 0.107622 |
| 5 | 135325800 | 135327599 | 6.602429 | 12.11635 | 36.68421 | 15.62084 | -1.15788 | 0.237946 | 0.107635 |
| 1 | 170985800 | 170988599 | 12.24474 | 21.16907 | 57.46006 | 25.20854 | -1.04839 | 0.839557 | 0.107647 |
| 9 | 59166800 | 59168599 | 6.466465 | 11.58506 | 42.63462 | 18.68211 | -1.46578 | 0.391766 | 0.107651 |
| 3 | 51593000 | 51602399 | 60.00831 | 93.53778 | 198.3501 | 82.31871 | -0.77221 | 2.69484 | 0.107679 |
| 1 | 112495400 | 112497999 | 30.52375 | 52.1221 | 31.05842 | 13.61684 | 1.126335 | 0.993137 | 0.107688 |
| 12 | 16805800 | 16806799 | 12.41866 | 23.67618 | 15.7979 | 6.304591 | 1.081544 | -0.0076 | 0.107694 |
| 10 | 78275200 | 78278399 | 15.79381 | 26.37797 | 71.63651 | 31.17314 | -1.05492 | 1.193217 | 0.107698 |
| 9 | 64286600 | 64297399 | 86.45311 | 131.3708 | 282.3284 | 116.9359 | -0.82036 | 3.19686 | 0.107701 |
| 9 | 11134400 | 11135799 | 20.14514 | 36.30545 | 20.90284 | 9.108289 | 1.197159 | 0.402563 | 0.107701 |
| 14 | 6974200 | 6975799 | 20.14514 | 36.30545 | 20.90284 | 9.108289 | 1.197159 | 0.402563 | 0.107701 |
| 7 | 126345000 | 126346599 | 6.046083 | 11.4519 | 32.57498 | 13.18057 | -1.00991 | 0.023589 | 0.107717 |
| 9 | 119985200 | 119986199 | 5.266219 | 10.14637 | 32.31607 | 12.53582 | -1.11561 | -0.12557 | 0.107729 |
| 8 | 122476200 | 122506799 | 216.8445 | 305.7296 | 663.8162 | 274.8944 | -0.93796 | 4.454734 | 0.107738 |
| 1 | 157370600 | 157371599 | 5.58098 | 10.67456 | 31.82023 | 12.60509 | -1.04954 | -0.05375 | 0.107742 |
| 6 | 45036200 | 45037599 | 7.645092 | 14.03433 | 37.25616 | 15.85739 | -0.97005 | 0.24381 | 0.107743 |
| 4 | 152052200 | 152060199 | 49.01759 | 77.02813 | 179.5927 | 74.49607 | -0.89962 | 2.539604 | 0.107744 |
| Y | 1957000 | 1959799 | 13.79527 | 26.39506 | 7.438495 | 2.943669 | 2.316817 | -0.04833 | 0.107744 |
| 3 | 157107200 | 157110199 | 35.22106 | 59.31159 | 37.67039 | 16.46483 | 1.03346 | 1.136999 | 0.107744 |
| 7 | 16149600 | 16151599 | 7.383056 | 13.46707 | 38.18874 | 16.39151 | -1.07305 | 0.280298 | 0.107771 |

| | | | | | | | | | |
|---|---|---|---|---|---|---|---|---|---|
| 7 | 18803600 | 18823799 | 100.4054 | 150.2782 | 353.2148 | 145.6744 | -0.96848 | 3.504152 | 0.107776 |
| 7 | 31975800 | 31978599 | 56.25171 | 90.52483 | 67.3255 | 28.59063 | 0.77987 | 1.85613 | 0.107783 |
| X | 61616200 | 61617199 | 17.34734 | 32.46231 | 16.05346 | 6.6421 | 1.46742 | 0.161007 | 0.107786 |
| 1 | 131639000 | 131640399 | 5.391184 | 10.1019 | 34.97095 | 14.43692 | -1.3151 | 0.09534 | 0.107797 |
| 9 | 38223000 | 38229799 | 121.2543 | 187.2185 | 127.7307 | 53.05676 | 0.851397 | 2.879559 | 0.107803 |
| 1 | 119377000 | 119379799 | 11.55244 | 20.16898 | 51.01419 | 22.46622 | -0.94453 | 0.708183 | 0.107803 |
| 17 | 54323200 | 54324599 | 23.66472 | 41.621 | 25.04406 | 11.08906 | 1.116467 | 0.61659 | 0.107827 |
| 3 | 56505400 | 56506399 | 17.12573 | 31.33688 | 20.93652 | 8.950476 | 1.001755 | 0.29623 | 0.107831 |
| 2 | 124505800 | 124507399 | 10.63366 | 18.58709 | 50.45458 | 22.23796 | -1.04557 | 0.68496 | 0.107843 |
| 13 | 3352600 | 3355799 | 40.46064 | 66.94164 | 45.0191 | 19.46735 | 0.951157 | 1.343694 | 0.107844 |
| 1 | 103947000 | 103948199 | 16.71362 | 30.74961 | 20.35783 | 8.638901 | 1.022154 | 0.2614 | 0.107857 |
| 11 | 117953800 | 117988799 | 229.4927 | 319.4696 | 764.7215 | 317.3142 | -1.09715 | 4.645985 | 0.107859 |
| 3 | 20951800 | 20953199 | 15.37164 | 29.0256 | 17.46908 | 7.114743 | 1.204015 | 0.090489 | 0.107869 |
| 7 | 16753200 | 16763199 | 47.40091 | 74.51604 | 181.5465 | 75.31023 | -0.96255 | 2.530769 | 0.10787 |
| 18 | 66638600 | 66638999 | 2.246853 | 4.394123 | 19.32312 | 6.318071 | -1.33629 | -0.8078 | 0.107872 |
| 11 | 93658000 | 93659599 | 17.90209 | 33.55002 | 15.18758 | 6.26812 | 1.594656 | 0.153102 | 0.10788 |
| 10 | 62888000 | 62890399 | 13.2301 | 22.68621 | 59.52838 | 26.10762 | -1.0023 | 0.922034 | 0.107884 |
| 7 | 33309000 | 33316599 | 180.5204 | 272.8052 | 175.9091 | 72.73497 | 0.90835 | 3.323629 | 0.107917 |
| 9 | 108910800 | 108912399 | 9.776193 | 17.33574 | 43.53837 | 19.24396 | -0.92831 | 0.481521 | 0.107921 |
| 16 | 94890800 | 94893599 | 14.10021 | 24.33641 | 57.52219 | 25.23452 | -0.85041 | 0.863503 | 0.107922 |
| 4 | 12195000 | 12198199 | 35.44061 | 59.45993 | 38.77926 | 16.92341 | 0.995275 | 1.171376 | 0.107925 |
| 3 | 38891400 | 38893199 | 6.99003 | 13.02523 | 37.15353 | 15.48293 | -1.05042 | 0.149209 | 0.107932 |
| 14 | 55747600 | 55755399 | 37.42584 | 59.15267 | 170.9648 | 71.02172 | -1.20232 | 2.407063 | 0.107932 |
| 13 | 97594200 | 97596199 | 10.40773 | 18.39867 | 46.16163 | 20.41866 | -0.92896 | 0.52328 | 0.107934 |
| 3 | 94015800 | 94018599 | 55.9326 | 90.10413 | 65.18104 | 27.72135 | 0.821563 | 1.823135 | 0.107937 |
| 1 | 152935200 | 152936399 | 4.677166 | 9.0262 | 28.64309 | 11.05897 | -1.10219 | -0.15413 | 0.107945 |
| 5 | 60912000 | 60916399 | 36.25788 | 60.1593 | 45.00014 | 19.49745 | 0.798388 | 1.299106 | 0.107956 |
| 1 | 184455000 | 184460399 | 39.5915 | 63.08832 | 146.5044 | 61.329 | -0.87601 | 2.169269 | 0.107967 |
| 5 | 3090000 | 3093799 | 49.07488 | 79.52495 | 58.33622 | 24.97419 | 0.810525 | 1.656944 | 0.107992 |
| 11 | 60523600 | 60524399 | 2.335747 | 4.565227 | 21.03513 | 7.26289 | -1.47399 | -0.60306 | 0.107992 |
| 19 | 42074200 | 42074999 | 3.132037 | 6.109 | 24.97384 | 8.951352 | -1.36023 | -0.44876 | 0.107998 |
| 1 | 36386800 | 36388199 | 4.064427 | 7.908941 | 25.81824 | 9.510946 | -1.07708 | -0.33313 | 0.108 |
| 4 | 107374200 | 107380199 | 32.07339 | 51.43442 | 123.1197 | 52.01317 | -0.90913 | 1.95619 | 0.10801 |
| 16 | 87520400 | 87523799 | 14.86413 | 24.86829 | 72.21081 | 31.4574 | -1.15109 | 1.170983 | 0.108014 |
| 15 | 49056000 | 49057999 | 19.29593 | 34.35773 | 23.0523 | 10.15385 | 0.968061 | 0.474827 | 0.108024 |
| 15 | 46280000 | 46281799 | 21.20546 | 37.50793 | 24.68525 | 10.91779 | 0.990856 | 0.54072 | 0.108026 |
| 1 | 108956800 | 108957599 | 12.65408 | 24.57415 | 12.6465 | 4.734798 | 1.535263 | -0.21889 | 0.108032 |
| 14 | 55149400 | 55151199 | 8.570629 | 15.20287 | 43.04199 | 19.01939 | -1.09947 | 0.470663 | 0.108047 |
| 14 | 41902200 | 41906199 | 50.50237 | 82.09669 | 49.02097 | 21.03441 | 1.110655 | 1.592932 | 0.108059 |
| 15 | 11902400 | 11907399 | 24.10013 | 38.74936 | 121.5786 | 51.56331 | -1.29373 | 1.862905 | 0.10807 |
| 6 | 10002200 | 10003199 | 12.46705 | 24.21269 | 12.65122 | 4.733794 | 1.51423 | -0.22321 | 0.108091 |
| 15 | 78475200 | 78480199 | 35.71369 | 56.97696 | 136.6491 | 57.29778 | -0.92045 | 2.125709 | 0.108136 |
| 6 | 4734600 | 4738199 | 12.18106 | 21.21354 | 53.19062 | 23.38805 | -0.93264 | 0.749414 | 0.108163 |
| 17 | 87753400 | 87754999 | 6.942132 | 12.67207 | 38.19672 | 16.37995 | -1.15941 | 0.272722 | 0.108186 |
| 3 | 144824000 | 144824999 | 4.81351 | 9.269031 | 29.12235 | 11.31482 | -1.09647 | -0.12982 | 0.108194 |
| 2 | 24423200 | 24424999 | 15.67567 | 28.98178 | 18.79677 | 7.923415 | 1.057905 | 0.221204 | 0.108223 |
| 2 | 156562200 | 156565199 | 17.23847 | 28.70857 | 74.59776 | 32.40325 | -0.9917 | 1.229629 | 0.108248 |

| | | | | | | | | | |
|---|---|---|---|---|---|---|---|---|---|
| 6 | 31912200 | 31916199 | 16.7563 | 27.63926 | 84.18983 | 36.33388 | -1.22155 | 1.348178 | 0.108254 |
| 5 | 115591200 | 115593199 | 7.446446 | 13.40582 | 40.07368 | 17.48488 | -1.164 | 0.362887 | 0.108264 |
| 10 | 126790800 | 126803599 | 68.06122 | 104.8165 | 240.889 | 100.0546 | -0.90174 | 2.91887 | 0.108273 |
| 2 | 152184800 | 152189399 | 16.86665 | 27.93362 | 87.06372 | 37.67451 | -1.25407 | 1.300951 | 0.108275 |
| 17 | 29331600 | 29338599 | 33.70714 | 53.95949 | 125.2901 | 52.78922 | -0.86832 | 2.014937 | 0.108281 |
| 8 | 122170000 | 122182999 | 61.9173 | 95.48347 | 231.2636 | 96.06284 | -0.9758 | 2.89093 | 0.10829 |
| 5 | 31301800 | 31303399 | 5.949825 | 11.14282 | 33.87039 | 13.99503 | -1.13021 | 0.102019 | 0.108291 |
| 14 | 29390800 | 29392399 | 7.615087 | 14.07492 | 37.45151 | 15.79328 | -0.96383 | 0.204259 | 0.108295 |
| 10 | 80816600 | 80826599 | 38.69066 | 61.2183 | 170.4138 | 70.83178 | -1.14702 | 2.381015 | 0.108296 |
| 7 | 6328800 | 6331799 | 9.938535 | 17.59127 | 47.23738 | 20.88926 | -1.0259 | 0.505808 | 0.108302 |
| 8 | 88601400 | 88606399 | 19.14561 | 31.38671 | 84.98729 | 36.58271 | -1.05599 | 1.433609 | 0.108317 |
| 2 | 27485200 | 27494599 | 59.24675 | 91.82115 | 214.4086 | 89.05925 | -0.91755 | 2.796022 | 0.108329 |
| 17 | 43872400 | 43874599 | 36.18998 | 60.76251 | 38.21368 | 16.68444 | 1.047543 | 1.166496 | 0.108335 |
| 12 | 23577000 | 23578199 | 13.0388 | 24.66275 | 16.72943 | 6.788897 | 1.037069 | 0.054081 | 0.108337 |
| 8 | 4249400 | 4261399 | 73.4652 | 112.6871 | 248.3046 | 103.0781 | -0.84553 | 3.000399 | 0.108342 |
| 11 | 66967400 | 66978399 | 55.94856 | 87.61223 | 191.4592 | 79.41918 | -0.81035 | 2.609843 | 0.108345 |
| 14 | 98588600 | 98590399 | 16.85902 | 30.86443 | 20.85595 | 8.909907 | 0.986002 | 0.29047 | 0.108348 |
| 5 | 120737400 | 120741599 | 17.52496 | 28.55556 | 93.06055 | 39.96554 | -1.32795 | 1.521634 | 0.108353 |
| 4 | 123213800 | 123218799 | 42.0881 | 66.49323 | 168.7553 | 70.08876 | -1.01568 | 2.419858 | 0.108359 |
| 3 | 10125000 | 10126199 | 6.673311 | 12.5054 | 35.07152 | 14.47634 | -1.01461 | 0.105867 | 0.108374 |
| 8 | 58678400 | 58679799 | 15.51371 | 29.07771 | 18.51316 | 7.638974 | 1.108486 | 0.140485 | 0.108376 |
| 4 | 141493400 | 141495999 | 10.49124 | 18.30363 | 50.98281 | 22.44266 | -1.08268 | 0.713908 | 0.108387 |
| 13 | 39904400 | 39904999 | 4.364451 | 8.479391 | 26.85538 | 10.01963 | -1.0518 | -0.28171 | 0.108404 |
| 7 | 8393800 | 8397599 | 87.40056 | 138.1246 | 102.1715 | 42.44153 | 0.760635 | 2.429654 | 0.108426 |
| 11 | 113932600 | 113937999 | 33.29138 | 53.55152 | 114.0866 | 48.41732 | -0.73596 | 1.858675 | 0.108448 |
| 5 | 137285000 | 137290599 | 31.91368 | 51.16782 | 121.4749 | 51.30234 | -0.89756 | 1.962634 | 0.108454 |
| 5 | 125434000 | 125441999 | 47.48461 | 75.21579 | 161.6728 | 67.23631 | -0.77341 | 2.359002 | 0.108465 |
| 2 | 49606400 | 49608799 | 11.39879 | 19.90322 | 50.47218 | 22.22928 | -0.94808 | 0.705623 | 0.108478 |
| 17 | 41148000 | 41149999 | 17.04614 | 31.28498 | 19.65051 | 8.366466 | 1.093763 | 0.274747 | 0.1085 |
| 3 | 107422800 | 107425999 | 16.25385 | 27.20175 | 73.03175 | 31.82123 | -1.03869 | 1.171451 | 0.108519 |
| 3 | 9506200 | 9515999 | 51.05038 | 80.41493 | 176.2264 | 73.12247 | -0.80774 | 2.491166 | 0.108523 |
| 7 | 44984200 | 44993399 | 48.65259 | 76.19868 | 190.9384 | 79.20192 | -1.00725 | 2.603936 | 0.108533 |
| 7 | 37205600 | 37207799 | 9.70777 | 16.99834 | 48.83413 | 21.55554 | -1.12647 | 0.650046 | 0.108537 |
| 5 | 95581600 | 95584999 | 37.3066 | 63.03024 | 33.1159 | 14.4886 | 1.30591 | 1.111893 | 0.108547 |
| 16 | 35019200 | 35023199 | 17.25019 | 28.67972 | 74.43674 | 32.29678 | -0.9901 | 1.249871 | 0.108548 |
| 3 | 135431800 | 135436799 | 26.98023 | 43.56541 | 111.8226 | 47.70471 | -0.99738 | 1.732362 | 0.10855 |
| 11 | 76836200 | 76836999 | 3.23688 | 6.313678 | 23.72614 | 8.500899 | -1.2391 | -0.45104 | 0.108561 |
| 2 | 170097800 | 170101799 | 22.90061 | 37.24513 | 93.53668 | 40.14269 | -0.95557 | 1.558581 | 0.108567 |
| 2 | 86174000 | 86174999 | 13.42057 | 25.31103 | 17.25263 | 7.044452 | 1.022646 | 0.079186 | 0.10859 |
| 7 | 78420200 | 78423399 | 18.52116 | 30.75115 | 76.32377 | 33.08459 | -0.92618 | 1.269674 | 0.108592 |
| 6 | 125023000 | 125028599 | 22.83633 | 36.93414 | 101.5323 | 43.40464 | -1.09304 | 1.674214 | 0.108593 |
| 3 | 68398000 | 68403199 | 23.51962 | 37.96793 | 107.6162 | 45.89529 | -1.14031 | 1.732616 | 0.108596 |
| 3 | 140742200 | 140743599 | 7.619153 | 14.07626 | 37.98103 | 16.02615 | -0.98423 | 0.209224 | 0.1086 |
| X | 3271600 | 3274199 | 33.46224 | 57.37908 | 30.12368 | 13.21148 | 1.309097 | 0.969088 | 0.108603 |
| 5 | 24522200 | 24545599 | 114.0089 | 167.9694 | 446.6713 | 183.9434 | -1.16758 | 3.785308 | 0.108623 |
| 12 | 116465600 | 116468999 | 20.74309 | 34.39846 | 78.81765 | 34.137 | -0.81153 | 1.292916 | 0.108652 |
| 11 | 89234400 | 89236199 | 7.772406 | 14.34285 | 40.22839 | 17.00107 | -1.04182 | 0.220141 | 0.108657 |

| | | | | | | | | | |
|---|---|---|---|---|---|---|---|---|---|
| 18 | 12776200 | 12779599 | 17.79676 | 29.8012 | 70.36887 | 30.6722 | -0.85415 | 1.168543 | 0.10866 |
| 13 | 13564600 | 13565999 | 5.713879 | 10.94516 | 31.93001 | 12.59422 | -1.01283 | -0.06611 | 0.108676 |
| 3 | 92645000 | 92645799 | 4.846899 | 9.332806 | 29.75068 | 11.56079 | -1.11824 | -0.12966 | 0.108699 |
| 1 | 143879400 | 143880799 | 5.683463 | 10.64033 | 36.10123 | 14.92546 | -1.28861 | 0.105259 | 0.108708 |
| 12 | 22727200 | 22729399 | 37.63021 | 62.93706 | 34.38042 | 14.97427 | 1.250271 | 1.209354 | 0.108762 |
| 1 | 46933600 | 46936199 | 12.00952 | 21.00721 | 51.54212 | 22.73119 | -0.90071 | 0.682848 | 0.108768 |
| 1 | 126810800 | 126811999 | 12.86139 | 24.88619 | 14.41011 | 5.496084 | 1.34344 | -0.1505 | 0.108778 |
| 9 | 65659800 | 65660799 | 12.86139 | 24.88619 | 14.41011 | 5.496084 | 1.34344 | -0.1505 | 0.108778 |
| 8 | 41869600 | 41872399 | 14.27523 | 24.56611 | 58.57991 | 25.69623 | -0.86405 | 0.891957 | 0.108804 |
| 14 | 28785600 | 28788399 | 12.16279 | 21.22897 | 54.84476 | 24.14917 | -0.97536 | 0.718759 | 0.108816 |
| 7 | 44105200 | 44125199 | 128.8315 | 189.0803 | 447.2296 | 184.2001 | -1.00397 | 3.848486 | 0.108825 |
| 12 | 112685800 | 112712799 | 133.9479 | 194.3587 | 512.7309 | 211.3892 | -1.17724 | 4.025907 | 0.108827 |
| 16 | 94522600 | 94525999 | 13.44339 | 22.55557 | 70.45036 | 30.74577 | -1.25561 | 1.134473 | 0.108839 |
| 1 | 30153800 | 30154799 | 12.99376 | 24.98291 | 16.05904 | 6.26683 | 1.164011 | -0.07133 | 0.108845 |
| 14 | 87183000 | 87184199 | 13.17252 | 25.06909 | 17.02694 | 6.820506 | 1.05103 | 0.014649 | 0.108848 |
| 2 | 35090200 | 35091399 | 6.067524 | 11.11313 | 38.78595 | 16.55949 | -1.36448 | 0.247095 | 0.108848 |
| 5 | 65835600 | 65836599 | 13.94748 | 26.38493 | 18.08638 | 7.337542 | 1.022479 | 0.065943 | 0.108867 |
| 15 | 78715200 | 78722199 | 34.79197 | 55.43505 | 142.7665 | 59.75633 | -1.02503 | 2.168155 | 0.108877 |
| 2 | 158134600 | 158142399 | 49.45964 | 78.27283 | 160.651 | 66.78051 | -0.70795 | 2.382518 | 0.108879 |
| 8 | 26007600 | 26011799 | 20.20645 | 33.22187 | 82.04461 | 35.35932 | -0.92197 | 1.397436 | 0.108885 |
| 2 | 172605200 | 172609199 | 19.30735 | 32.04378 | 75.81233 | 32.85391 | -0.85738 | 1.27651 | 0.108889 |
| 12 | 9397800 | 9398999 | 13.35702 | 25.34963 | 17.7315 | 7.144624 | 1.002327 | 0.038254 | 0.108896 |
| 13 | 73621400 | 73627999 | 31.58716 | 50.6665 | 121.1377 | 51.19401 | -0.90698 | 1.947992 | 0.108918 |
| 19 | 24976600 | 24979799 | 12.62863 | 21.64314 | 60.57313 | 26.56467 | -1.09497 | 0.925543 | 0.108962 |
| 9 | 96749200 | 96758799 | 66.91519 | 103.7264 | 214.3067 | 89.01475 | -0.74102 | 2.795084 | 0.108968 |
| 4 | 140254400 | 140271799 | 85.78491 | 129.3511 | 319.7027 | 132.098 | -1.03133 | 3.356546 | 0.108979 |
| 6 | 125579200 | 125582199 | 12.52219 | 21.36429 | 63.18869 | 27.70169 | -1.17539 | 0.969905 | 0.108993 |
| 7 | 130582800 | 130587399 | 16.23171 | 26.8736 | 79.73974 | 34.49774 | -1.18306 | 1.299885 | 0.109004 |
| 14 | 86441800 | 86443999 | 12.18291 | 21.23041 | 52.09351 | 22.91491 | -0.90126 | 0.740645 | 0.109021 |
| 4 | 133908000 | 133909399 | 5.497904 | 10.27972 | 33.78884 | 13.99985 | -1.24436 | 0.110625 | 0.109028 |
| 17 | 35452200 | 35456999 | 19.5823 | 32.18975 | 82.50151 | 35.55346 | -0.97542 | 1.398458 | 0.109055 |
| 15 | 60664000 | 60667999 | 54.87663 | 88.46003 | 65.05989 | 27.69459 | 0.798857 | 1.803223 | 0.109061 |
| 6 | 147907000 | 147907999 | 4.492878 | 8.727779 | 27.46004 | 10.25588 | -1.04383 | -0.27417 | 0.109066 |
| 16 | 31245800 | 31249199 | 21.51845 | 35.26941 | 86.16828 | 37.08783 | -0.90846 | 1.440792 | 0.109066 |
| 10 | 105852000 | 105852799 | 10.96364 | 21.17086 | 14.75374 | 5.681082 | 1.066119 | -0.14043 | 0.109069 |
| 5 | 57516200 | 57517199 | 12.10154 | 23.4882 | 13.26138 | 4.985628 | 1.397748 | -0.20999 | 0.10907 |
| 4 | 155440600 | 155456799 | 87.25344 | 131.8402 | 306.1643 | 126.5874 | -0.9389 | 3.313846 | 0.109074 |
| 15 | 35296400 | 35301999 | 37.87583 | 60.25922 | 149.1268 | 62.29052 | -0.97005 | 2.220207 | 0.109088 |
| 6 | 84491600 | 84492999 | 5.302074 | 10.10426 | 31.02297 | 12.39133 | -1.10177 | -0.02959 | 0.109094 |
| 17 | 26751000 | 26752599 | 5.920043 | 10.80079 | 38.16898 | 16.37862 | -1.38674 | 0.269283 | 0.109095 |
| 2 | 147294200 | 147295399 | 5.411861 | 10.39551 | 35.85954 | 14.0316 | -1.24194 | -0.08696 | 0.109097 |
| 4 | 148277000 | 148297199 | 135.9034 | 200.2356 | 422.526 | 174.0003 | -0.83408 | 3.786451 | 0.109103 |
| 4 | 130620600 | 130621599 | 7.372256 | 13.44821 | 37.65335 | 16.16045 | -1.05442 | 0.280302 | 0.109108 |
| 1 | 87050200 | 87057199 | 47.35565 | 74.54339 | 176.8034 | 73.3553 | -0.92239 | 2.503475 | 0.109125 |
| 4 | 131336400 | 131337399 | 3.711314 | 7.172919 | 29.15352 | 11.17953 | -1.4424 | -0.18202 | 0.109139 |
| 11 | 94544400 | 94544999 | 2.427404 | 4.746482 | 19.45848 | 6.318426 | -1.22724 | -0.82568 | 0.109143 |
| 4 | 4654800 | 4655399 | 12.49421 | 24.15183 | 14.99271 | 5.744756 | 1.237884 | -0.1381 | 0.109156 |

| | | | | | | | | | |
|---|---|---|---|---|---|---|---|---|---|
| 15 | 70766200 | 70767399 | 12.32183 | 21.51362 | 51.56674 | 22.71049 | -0.86746 | 0.712153 | 0.109198 |
| 5 | 140863400 | 140869799 | 48.28746 | 76.10009 | 173.1323 | 71.84319 | -0.86108 | 2.479585 | 0.109198 |
| 16 | 38065800 | 38067199 | 5.022875 | 9.62232 | 34.92873 | 13.77368 | -1.32529 | -0.06869 | 0.109219 |
| 9 | 50339600 | 50343599 | 23.92032 | 38.74497 | 99.32851 | 42.50917 | -0.99065 | 1.643794 | 0.109243 |
| 7 | 44880600 | 44885399 | 20.75515 | 33.64327 | 99.25581 | 42.50143 | -1.19149 | 1.623433 | 0.109244 |
| 18 | 14915000 | 14917999 | 13.34263 | 22.98499 | 56.88615 | 24.95405 | -0.91688 | 0.877846 | 0.109245 |
| 18 | 35160000 | 35161199 | 6.078629 | 11.35185 | 34.32867 | 14.25258 | -1.12799 | 0.125043 | 0.109247 |
| 4 | 129082400 | 129084199 | 9.57126 | 16.73956 | 49.89551 | 22.00209 | -1.17937 | 0.670194 | 0.109253 |
| 9 | 22299800 | 22300999 | 7.538291 | 13.71737 | 38.1435 | 16.42116 | -1.04772 | 0.296135 | 0.109264 |
| 11 | 115426200 | 115428599 | 15.20286 | 25.70889 | 64.79612 | 28.35688 | -0.9471 | 1.062835 | 0.109282 |
| 15 | 59545800 | 59547199 | 5.612248 | 10.63956 | 32.00126 | 12.92428 | -1.08693 | 0.012168 | 0.109284 |
| 4 | 91754200 | 91755199 | 11.00095 | 21.36072 | 14.03022 | 5.25893 | 1.186197 | -0.23102 | 0.109296 |
| 19 | 39371400 | 39371799 | 10.75468 | 20.9528 | 13.09632 | 4.765249 | 1.296144 | -0.33666 | 0.109307 |
| 10 | 71073200 | 71074599 | 9.813033 | 17.27871 | 45.63645 | 20.20237 | -1.00461 | 0.563781 | 0.109375 |
| 6 | 145195000 | 145196799 | 11.83313 | 20.33355 | 58.37644 | 25.60583 | -1.13056 | 0.89656 | 0.109376 |
| 13 | 93908000 | 93909399 | 6.991072 | 12.99844 | 36.45586 | 15.24909 | -1.02982 | 0.165558 | 0.10938 |
| X | 31767000 | 31769599 | 39.19133 | 65.33388 | 40.54604 | 17.62325 | 1.067107 | 1.253723 | 0.109382 |
| 1 | 135440600 | 135445599 | 15.58605 | 26.155 | 68.46055 | 29.88057 | -1.00192 | 1.138061 | 0.109399 |
| X | 46499600 | 46500199 | 11.78163 | 22.95994 | 11.84794 | 4.295312 | 1.573685 | -0.33965 | 0.109409 |
| X | 49477800 | 49482399 | 52.4879 | 85.68074 | 44.01614 | 18.9156 | 1.331016 | 1.536179 | 0.109438 |
| 11 | 20581400 | 20583599 | 11.4316 | 19.88032 | 53.1482 | 23.34872 | -1.02467 | 0.766973 | 0.109439 |
| 1 | 132109800 | 132120799 | 54.09141 | 84.88307 | 188.037 | 77.98465 | -0.82757 | 2.570533 | 0.109452 |
| 10 | 62721200 | 62723199 | 9.132429 | 15.95533 | 50.94454 | 22.44559 | -1.27797 | 0.686675 | 0.10947 |
| 11 | 67345400 | 67347799 | 9.132429 | 15.95533 | 50.94454 | 22.44559 | -1.27797 | 0.686675 | 0.10947 |
| 1 | 133492000 | 133494199 | 12.66288 | 21.99066 | 54.27451 | 23.82246 | -0.91045 | 0.793993 | 0.10948 |
| 4 | 22486200 | 22492999 | 51.09651 | 79.95979 | 200.0599 | 82.99264 | -1.00607 | 2.621291 | 0.109505 |
| 16 | 58588200 | 58590199 | 28.55636 | 49.73671 | 27.29536 | 12.00264 | 1.248647 | 0.790014 | 0.10952 |
| 2 | 93432400 | 93435599 | 14.64874 | 24.63342 | 68.47182 | 29.91461 | -1.08819 | 1.113506 | 0.109532 |
| 3 | 104934200 | 104939599 | 79.82767 | 121.6233 | 269.1564 | 111.5683 | -0.85946 | 3.140738 | 0.109543 |
| 17 | 6052400 | 6054199 | 9.363588 | 16.46194 | 46.16418 | 20.44228 | -1.09128 | 0.579684 | 0.109546 |
| 9 | 56165000 | 56168799 | 91.38615 | 143.693 | 101.2651 | 42.00858 | 0.824782 | 2.539596 | 0.109561 |
| 3 | 56064600 | 56066199 | 8.948357 | 15.72229 | 48.25703 | 21.36246 | -1.22124 | 0.587207 | 0.109575 |
| 7 | 22897600 | 22901199 | 26.29364 | 45.78245 | 29.54755 | 12.99063 | 1.015898 | 0.785221 | 0.109581 |
| 17 | 31426800 | 31431599 | 35.21444 | 56.21865 | 133.8029 | 56.15825 | -0.90833 | 2.103451 | 0.109607 |
| 7 | 20914400 | 20916999 | 61.98176 | 99.26928 | 76.47768 | 32.24571 | 0.720983 | 2.015813 | 0.109638 |
| 5 | 75329600 | 75331799 | 10.75094 | 18.87332 | 47.2248 | 20.89042 | -0.92849 | 0.610746 | 0.109645 |
| 8 | 45387800 | 45391999 | 27.5211 | 44.27319 | 112.1493 | 47.60067 | -0.98481 | 1.8487 | 0.109645 |
| 3 | 70928800 | 70931199 | 30.8023 | 52.9255 | 31.55628 | 13.84159 | 1.126558 | 0.939288 | 0.109656 |
| 7 | 17186600 | 17189399 | 33.60472 | 57.4603 | 31.55653 | 13.83673 | 1.243734 | 0.994608 | 0.109658 |
| 14 | 3147200 | 3154999 | 229.8579 | 339.7686 | 246.9335 | 101.6931 | 0.707213 | 3.738767 | 0.109674 |
| 11 | 71960800 | 71965999 | 27.24279 | 43.74085 | 118.2003 | 50.02712 | -1.08133 | 1.912137 | 0.109676 |
| 9 | 63651200 | 63652599 | 5.595967 | 10.65773 | 31.61244 | 12.64333 | -1.05465 | -0.02177 | 0.109679 |
| 2 | 152498800 | 152503999 | 30.56398 | 49.16081 | 111.8016 | 47.44268 | -0.83 | 1.857206 | 0.109687 |
| 18 | 6008400 | 6009599 | 12.11748 | 23.45883 | 14.43045 | 5.490609 | 1.259913 | -0.16727 | 0.10969 |
| 10 | 34709800 | 34711999 | 19.57198 | 35.21498 | 22.12572 | 9.660321 | 1.070411 | 0.410714 | 0.109691 |
| 1 | 179224800 | 179226799 | 20.94179 | 36.91018 | 25.10349 | 11.11066 | 0.942071 | 0.578641 | 0.109704 |
| 5 | 116011600 | 116014199 | 9.022567 | 15.99521 | 43.02839 | 19.02197 | -1.02682 | 0.477477 | 0.109705 |

| | | | | | | | | | |
|---|---|---|---|---|---|---|---|---|---|
| 8 | 87790200 | 87792799 | 15.80601 | 26.78461 | 64.37841 | 28.19188 | -0.8789 | 1.045287 | 0.10971 |
| 3 | 92684400 | 92685199 | 4.120572 | 8.000528 | 27.9689 | 10.48836 | -1.2 | -0.26492 | 0.109713 |
| 17 | 64579200 | 64583199 | 21.65038 | 35.55304 | 88.3103 | 38.04031 | -0.93125 | 1.417568 | 0.109727 |
| 18 | 36867600 | 36868999 | 5.116539 | 9.830458 | 29.63613 | 11.5889 | -1.0468 | -0.10466 | 0.109733 |
| 19 | 10632400 | 10634199 | 7.818316 | 14.28071 | 38.22602 | 16.37921 | -0.98903 | 0.27524 | 0.109743 |
| 17 | 22836600 | 22850799 | 344.4467 | 492.6721 | 335.912 | 138.7734 | 0.753345 | 4.247709 | 0.109749 |
| 7 | 126219400 | 126221399 | 8.280785 | 15.07527 | 39.23671 | 16.88228 | -0.95298 | 0.30012 | 0.109761 |
| 19 | 6590000 | 6600999 | 60.90299 | 94.01217 | 228.7258 | 95.00818 | -0.98117 | 2.871872 | 0.109762 |
| 7 | 76869400 | 76869599 | 6.802049 | 13.30507 | 10.16528 | 3.351008 | 1.135193 | -0.7419 | 0.109773 |
| 18 | 83721000 | 83721399 | 10.94256 | 21.29224 | 13.73151 | 5.059992 | 1.235919 | -0.28869 | 0.109789 |
| 8 | 125881000 | 125888599 | 33.57608 | 53.68823 | 129.4741 | 54.46377 | -0.92471 | 2.051216 | 0.109793 |
| 2 | 163260800 | 163263199 | 9.055533 | 15.75007 | 57.35589 | 25.19911 | -1.46785 | 0.758739 | 0.109798 |
| 11 | 53893000 | 53896399 | 12.93752 | 22.16512 | 61.14443 | 26.8145 | -1.0744 | 0.930117 | 0.109798 |
| 17 | 68372800 | 68376599 | 20.68234 | 33.99655 | 82.10973 | 35.38398 | -0.89012 | 1.401431 | 0.109864 |
| 10 | 17767600 | 17768799 | 21.9472 | 40.09812 | 16.48576 | 7.061665 | 1.694257 | 0.331792 | 0.109865 |
| X | 33371200 | 33371799 | 21.9472 | 40.09812 | 16.48576 | 7.061665 | 1.694257 | 0.331792 | 0.109865 |
| 14 | 42174600 | 42175599 | 15.31621 | 28.78908 | 17.79006 | 7.305429 | 1.156046 | 0.118306 | 0.109873 |
| 1 | 180307200 | 180311999 | 27.62858 | 44.58939 | 104.2957 | 44.46419 | -0.86445 | 1.745733 | 0.109876 |
| 13 | 99890200 | 99892599 | 9.420312 | 16.86114 | 43.15292 | 18.92426 | -0.94762 | 0.417424 | 0.109902 |
| 15 | 51903200 | 51907199 | 20.70946 | 34.01103 | 87.51255 | 37.69813 | -0.98185 | 1.413932 | 0.109909 |
| 15 | 99315600 | 99326199 | 42.76778 | 67.15105 | 187.8906 | 77.92965 | -1.16324 | 2.55387 | 0.10991 |
| 2 | 168340400 | 168354399 | 94.71572 | 143.0937 | 305.0119 | 126.1048 | -0.81561 | 3.318198 | 0.109911 |
| 2 | 165439200 | 165440799 | 5.796073 | 10.90597 | 34.01777 | 13.94458 | -1.15808 | 0.069264 | 0.109911 |
| 4 | 134476200 | 134478199 | 6.847577 | 12.38906 | 40.1552 | 17.41194 | -1.27391 | 0.328424 | 0.109915 |
| 7 | 127207600 | 127214999 | 54.81343 | 85.02651 | 213.7236 | 88.76328 | -1.02266 | 2.778704 | 0.109919 |
| 18 | 77031600 | 77032799 | 4.792736 | 9.300356 | 29.23818 | 11.00313 | -1.05389 | -0.24008 | 0.109929 |
| 3 | 138907800 | 138911999 | 31.49108 | 50.76551 | 122.433 | 52.1204 | -0.9107 | 1.791391 | 0.109935 |
| 17 | 36452000 | 36458599 | 37.35248 | 59.52329 | 151.0046 | 63.21696 | -1.00327 | 2.166435 | 0.109959 |
| 9 | 37321000 | 37323599 | 10.28201 | 18.08159 | 46.15312 | 20.43626 | -0.95662 | 0.580349 | 0.109968 |
| 2 | 80410600 | 80412599 | 9.656251 | 17.03949 | 45.10927 | 19.96036 | -1.00687 | 0.536043 | 0.109969 |
| 10 | 81140200 | 81177999 | 194.5743 | 273.1442 | 711.5592 | 294.8697 | -1.20694 | 4.518541 | 0.109974 |
| 17 | 30173400 | 30175599 | 5.976048 | 11.11571 | 34.78222 | 14.53906 | -1.18519 | 0.149532 | 0.109977 |
| 5 | 90291400 | 90294399 | 15.74872 | 26.86862 | 62.33173 | 27.32596 | -0.82684 | 0.981691 | 0.109977 |
| 18 | 49969600 | 49971399 | 28.67234 | 49.28138 | 33.49657 | 14.69281 | 0.938698 | 0.931458 | 0.109984 |
| 6 | 117403400 | 117406999 | 17.20157 | 28.73744 | 73.06719 | 31.80151 | -0.96049 | 1.193577 | 0.109984 |
| 7 | 110366600 | 110368999 | 10.18111 | 17.80012 | 50.4429 | 22.23711 | -1.1072 | 0.678889 | 0.110029 |
| 18 | 84684200 | 84690999 | 31.84293 | 51.23002 | 111.8444 | 47.47707 | -0.77083 | 1.850838 | 0.110031 |
| 16 | 21800000 | 21806199 | 44.00471 | 70.02098 | 144.6078 | 60.41597 | -0.70932 | 2.221027 | 0.110054 |
| 4 | 37623400 | 37624799 | 36.16159 | 60.3047 | 42.75965 | 18.58958 | 0.875524 | 1.242189 | 0.110055 |
| 11 | 113318200 | 113342399 | 218.1061 | 310.116 | 597.7178 | 247.1752 | -0.75388 | 4.330524 | 0.110056 |
| 6 | 134663000 | 134663799 | 3.343388 | 6.528979 | 22.82915 | 8.025421 | -1.11136 | -0.52035 | 0.110063 |
| 17 | 56716400 | 56722199 | 33.23649 | 53.18189 | 126.6416 | 53.32432 | -0.90538 | 2.028986 | 0.110107 |
| 15 | 73819600 | 73820799 | 3.737472 | 7.206135 | 29.65427 | 11.47953 | -1.47577 | -0.1537 | 0.110117 |
| 6 | 46655400 | 46658599 | 37.92028 | 62.56704 | 47.2283 | 20.38486 | 0.784749 | 1.368823 | 0.110129 |
| 9 | 96415600 | 96417799 | 9.80023 | 17.30891 | 44.58529 | 19.725 | -0.96708 | 0.526403 | 0.110132 |
| 11 | 107079000 | 107080399 | 5.408556 | 10.08878 | 35.38519 | 14.7201 | -1.34268 | 0.126447 | 0.110136 |
| 4 | 151499400 | 151500999 | 9.1591 | 16.28597 | 43.57617 | 19.2163 | -1.01688 | 0.458475 | 0.110139 |

| | | | | | | | | | |
|---|---|---|---|---|---|---|---|---|---|
| 7 | 89280200 | 89282399 | 9.1591 | 16.28597 | 43.57617 | 19.2163 | -1.01688 | 0.458475 | 0.110139 |
| 12 | 84720200 | 84725999 | 31.56006 | 50.67134 | 116.5487 | 49.32618 | -0.84956 | 1.917378 | 0.110145 |
| 9 | 72196400 | 72197399 | 5.868172 | 11.1951 | 31.69526 | 12.63113 | -0.98346 | -0.02991 | 0.110157 |
| 15 | 59044000 | 59046399 | 14.01035 | 23.9894 | 64.01679 | 28.07286 | -1.02718 | 0.941616 | 0.110187 |
| 5 | 142284600 | 142285799 | 5.890097 | 11.20599 | 33.36937 | 13.37606 | -1.06378 | -0.00839 | 0.110208 |
| 9 | 47619200 | 47620599 | 5.890097 | 11.20599 | 33.36937 | 13.37606 | -1.06378 | -0.00839 | 0.110208 |
| 4 | 154581400 | 154591799 | 49.34122 | 77.21865 | 195.5219 | 81.10952 | -1.02316 | 2.617978 | 0.110215 |
| X | 4772400 | 4779399 | 64.35413 | 103.7027 | 56.47148 | 24.02794 | 1.230861 | 1.826455 | 0.110219 |
| 18 | 11297600 | 11298999 | 22.46808 | 40.28754 | 20.25228 | 8.867935 | 1.387201 | 0.44819 | 0.11022 |
| 2 | 168258200 | 168309999 | 431.3225 | 575.4756 | 1155.193 | 482.4727 | -0.90511 | 5.314093 | 0.110221 |
| 5 | 124848600 | 124866999 | 154.5181 | 225.6681 | 456.8857 | 188.238 | -0.78652 | 3.928113 | 0.110231 |
| 2 | 25872200 | 25874599 | 8.34707 | 14.66224 | 47.73401 | 21.12655 | -1.3053 | 0.588099 | 0.110236 |
| 8 | 76984200 | 76984999 | 10.04096 | 19.55639 | 13.6936 | 5.002264 | 1.12887 | -0.33109 | 0.110249 |
| 5 | 37445200 | 37450199 | 25.63008 | 41.57217 | 95.23576 | 40.79946 | -0.8269 | 1.616941 | 0.11027 |
| 3 | 9015400 | 9023199 | 53.33289 | 83.96973 | 175.348 | 72.75343 | -0.73878 | 2.502312 | 0.110277 |
| 13 | 9045600 | 9046599 | 5.477928 | 10.38547 | 32.57147 | 13.15309 | -1.14677 | 0.011135 | 0.110286 |
| 1 | 109831400 | 109832399 | 13.30357 | 25.14169 | 17.75157 | 7.216955 | 0.977691 | 0.0659 | 0.110288 |
| 2 | 25497800 | 25500799 | 11.91853 | 20.31059 | 63.08222 | 27.6524 | -1.24567 | 0.979329 | 0.110299 |
| 17 | 8582800 | 8586199 | 11.09924 | 19.37064 | 51.00303 | 22.45613 | -1.00212 | 0.711393 | 0.110311 |
| 7 | 44226000 | 44266599 | 309.2751 | 424.033 | 866.6056 | 360.4546 | -0.89139 | 4.881703 | 0.110313 |
| 17 | 59889200 | 59890999 | 12.65749 | 24.20956 | 17.20571 | 6.815787 | 1.0007 | -0.02434 | 0.110314 |
| 14 | 34431400 | 34432399 | 3.529361 | 6.884856 | 29.19434 | 10.44657 | -1.41261 | -0.43875 | 0.110358 |
| 4 | 61498600 | 61500399 | 33.18042 | 56.04309 | 35.17523 | 15.38825 | 1.050563 | 1.101922 | 0.110364 |
| 11 | 53036200 | 53042799 | 37.08161 | 59.36247 | 125.2404 | 52.76954 | -0.7304 | 2.018017 | 0.110386 |
| 15 | 59829000 | 59830199 | 12.74484 | 24.42088 | 16.57505 | 6.531854 | 1.073081 | -0.0422 | 0.110401 |
| 17 | 17292400 | 17296999 | 74.69273 | 119.5399 | 63.9855 | 26.94958 | 1.244029 | 2.04926 | 0.110404 |
| 12 | 52549800 | 52552199 | 12.78181 | 21.91474 | 62.26669 | 27.3083 | -1.11673 | 0.923221 | 0.110411 |
| 17 | 92874800 | 92875799 | 13.76141 | 26.04014 | 18.09161 | 7.335368 | 1.003913 | 0.06219 | 0.110433 |
| 7 | 40449200 | 40451599 | 27.19339 | 47.11119 | 31.35652 | 13.75742 | 0.970978 | 0.852989 | 0.110438 |
| 14 | 83681600 | 83682799 | 15.04545 | 28.16391 | 18.19698 | 7.524829 | 1.085187 | 0.143809 | 0.110443 |
| 7 | 103870200 | 103872399 | 7.704827 | 13.98316 | 39.15894 | 16.91451 | -1.06131 | 0.314128 | 0.110453 |
| 5 | 24928600 | 24938399 | 46.37981 | 73.03798 | 177.5932 | 73.687 | -0.95773 | 2.494291 | 0.110457 |
| X | 29508200 | 29509199 | 14.73876 | 27.42745 | 18.22493 | 7.612099 | 1.033438 | 0.177074 | 0.110471 |
| 2 | 172562400 | 172583199 | 136.2841 | 200.519 | 442.1006 | 182.0591 | -0.89923 | 3.809563 | 0.110474 |
| 6 | 46295000 | 46296199 | 20.80695 | 36.74314 | 25.02244 | 11.07053 | 0.94121 | 0.557213 | 0.110476 |
| 4 | 36621400 | 36622599 | 12.04281 | 23.02299 | 15.73266 | 6.24072 | 1.054738 | -0.03129 | 0.110491 |
| 9 | 43007000 | 43007399 | 2.735764 | 5.351128 | 20.28285 | 6.809909 | -1.16202 | -0.69991 | 0.110495 |
| 9 | 78863800 | 78865599 | 20.71369 | 37.2961 | 20.88692 | 9.111724 | 1.235897 | 0.412074 | 0.110498 |
| 15 | 89973200 | 89973999 | 4.141623 | 7.982637 | 29.10676 | 11.27865 | -1.305 | -0.14729 | 0.1105 |
| 19 | 7198000 | 7207599 | 52.6186 | 82.59948 | 181.7676 | 75.38783 | -0.81759 | 2.562876 | 0.110503 |
| 5 | 37397600 | 37407399 | 73.07742 | 112.2291 | 243.8669 | 101.2529 | -0.82406 | 2.975829 | 0.110528 |
| 11 | 7239200 | 7242599 | 18.56778 | 31.14324 | 75.72194 | 33.03959 | -0.89645 | 1.157062 | 0.110528 |
| 6 | 28369200 | 28369999 | 5.412645 | 10.39449 | 30.24739 | 11.84388 | -0.99868 | -0.09653 | 0.110533 |
| 3 | 30470200 | 30471399 | 5.401925 | 10.4092 | 31.08579 | 12.05404 | -1.02278 | -0.12745 | 0.110537 |
| 9 | 55135000 | 55136799 | 6.961556 | 13.0577 | 36.85773 | 15.18968 | -1.02223 | 0.106615 | 0.110546 |
| 4 | 135513000 | 135517399 | 21.20563 | 34.71508 | 85.20432 | 36.65436 | -0.91565 | 1.45627 | 0.110558 |
| 7 | 23038400 | 23043199 | 106.9355 | 166.7995 | 122.3911 | 50.78955 | 0.757769 | 2.695025 | 0.110563 |

| | | | | | | | | | |
|---|---|---|---|---|---|---|---|---|---|
| 3 | 10547000 | 10547999 | 25.91187 | 44.7316 | 32.55692 | 14.28254 | 0.841555 | 0.87886 | 0.110576 |
| 5 | 151419000 | 151420599 | 23.32674 | 41.20296 | 23.35647 | 10.33296 | 1.204108 | 0.566037 | 0.110579 |
| 3 | 151493800 | 151494599 | 12.04311 | 23.22199 | 16.21312 | 6.274778 | 1.05704 | -0.10688 | 0.110584 |
| 8 | 21857000 | 21859999 | 29.32416 | 50.73322 | 29.93109 | 13.13148 | 1.143537 | 0.871175 | 0.110587 |
| 9 | 120409800 | 120415599 | 17.75655 | 29.15778 | 88.59307 | 38.16154 | -1.22075 | 1.409983 | 0.110593 |
| 15 | 76008200 | 76009199 | 5.520515 | 10.34602 | 33.86512 | 13.97635 | -1.2344 | 0.093713 | 0.110606 |
| 6 | 54747600 | 54749999 | 13.91837 | 24.12381 | 59.26444 | 26.00216 | -0.90466 | 0.826296 | 0.110628 |
| 15 | 52152200 | 52153599 | 12.98743 | 24.66194 | 17.74033 | 7.139766 | 0.963608 | 0.03065 | 0.110629 |
| 16 | 19472800 | 19473599 | 12.98743 | 24.66194 | 17.74033 | 7.139766 | 0.963608 | 0.03065 | 0.110629 |
| 17 | 35789800 | 35790999 | 4.431574 | 8.53746 | 29.7224 | 11.53284 | -1.24163 | -0.13901 | 0.110636 |
| 18 | 68585200 | 68587799 | 25.14045 | 44.09126 | 25.5383 | 11.28788 | 1.171023 | 0.667023 | 0.110639 |
| 14 | 19634600 | 19635999 | 7.595806 | 13.69681 | 40.62334 | 17.68895 | -1.15119 | 0.354163 | 0.11064 |
| 10 | 69204600 | 69213799 | 65.88941 | 102.227 | 209.0319 | 86.81229 | -0.72508 | 2.779157 | 0.110643 |
| 7 | 14769400 | 14771399 | 53.69894 | 86.75039 | 60.71247 | 25.91671 | 0.873828 | 1.73696 | 0.110654 |
| 4 | 155232000 | 155238399 | 25.75197 | 41.49283 | 108.9835 | 46.36325 | -1.03401 | 1.794014 | 0.110656 |
| 1 | 185082000 | 185083399 | 3.438587 | 6.657453 | 28.14799 | 10.70606 | -1.48888 | -0.22152 | 0.110666 |
| 4 | 42862200 | 42863999 | 8.341698 | 15.01792 | 41.65294 | 18.17336 | -1.05745 | 0.370025 | 0.11067 |
| 3 | 21513200 | 21515799 | 42.93272 | 70.20937 | 52.20636 | 22.45003 | 0.801044 | 1.494692 | 0.110675 |
| 5 | 34963600 | 34964199 | 3.344935 | 6.527055 | 23.21777 | 8.26927 | -1.15283 | -0.47175 | 0.110683 |
| 1 | 194621000 | 194621999 | 4.01052 | 7.7675 | 30.78634 | 11.69036 | -1.39706 | -0.21635 | 0.110688 |
| 6 | 6858800 | 6861799 | 19.19486 | 31.75125 | 79.06447 | 34.18463 | -0.93142 | 1.317062 | 0.110689 |
| 15 | 19937600 | 19938799 | 13.69441 | 26.16622 | 16.41262 | 6.519336 | 1.17523 | -0.00806 | 0.110689 |
| 7 | 90442400 | 90442799 | 2.247107 | 4.393811 | 19.46923 | 6.313988 | -1.33661 | -0.83662 | 0.110689 |
| 7 | 10606600 | 10608399 | 44.85554 | 73.06944 | 54.34915 | 23.33841 | 0.796795 | 1.55618 | 0.110692 |
| 14 | 57521400 | 57526799 | 28.24625 | 45.64293 | 105.8797 | 45.21191 | -0.85054 | 1.709567 | 0.110703 |
| 1 | 151375800 | 151376999 | 7.240619 | 13.2081 | 38.72671 | 16.6211 | -1.12091 | 0.279363 | 0.110704 |
| 7 | 9108600 | 9113799 | 82.70999 | 131.1057 | 96.63922 | 40.21146 | 0.769211 | 2.354251 | 0.110708 |
| 15 | 85876600 | 85879199 | 9.381707 | 16.5076 | 47.20315 | 20.89886 | -1.11917 | 0.569395 | 0.110708 |
| 13 | 76738800 | 76739199 | 6.730999 | 12.44083 | 35.76008 | 15.08007 | -1.07368 | 0.194365 | 0.110714 |
| 17 | 19241800 | 19245199 | 98.78059 | 156.3919 | 83.07259 | 34.53954 | 1.239633 | 2.393802 | 0.110717 |
| 17 | 34337200 | 34341999 | 21.58558 | 34.89343 | 103.3522 | 44.15453 | -1.20114 | 1.687963 | 0.110723 |
| 12 | 19076000 | 19086399 | 248.8562 | 364.3584 | 272.7929 | 112.3719 | 0.651383 | 3.893372 | 0.110736 |
| 16 | 4463800 | 4474999 | 45.60334 | 71.41036 | 194.7108 | 80.76823 | -1.12911 | 2.605898 | 0.110753 |
| 8 | 3076400 | 3078399 | 17.64155 | 32.48776 | 19.79736 | 8.390175 | 1.142364 | 0.262986 | 0.110761 |
| 16 | 88950600 | 88953199 | 31.5123 | 53.53204 | 36.58725 | 16.03387 | 0.928315 | 1.04853 | 0.110765 |
| 7 | 103109600 | 103110399 | 13.54336 | 25.69613 | 17.72708 | 7.147056 | 1.021435 | 0.042049 | 0.110778 |
| 12 | 22175200 | 22178599 | 43.17912 | 71.94 | 36.60264 | 15.90344 | 1.352539 | 1.268424 | 0.110801 |
| 17 | 27849200 | 27851399 | 6.395606 | 11.62681 | 40.24494 | 17.34677 | -1.36224 | 0.296454 | 0.110809 |
| 12 | 57258200 | 57259599 | 5.765392 | 10.87816 | 32.39644 | 13.21581 | -1.08545 | 0.048979 | 0.110812 |
| 11 | 118793600 | 118803799 | 67.26685 | 103.1331 | 251.5272 | 104.4095 | -0.99225 | 3.007841 | 0.110845 |
| 1 | 134567200 | 134580199 | 63.50036 | 98.40961 | 214.1773 | 88.96318 | -0.81622 | 2.798028 | 0.110847 |
| 12 | 40271600 | 40273599 | 10.51263 | 18.31506 | 52.58671 | 23.12816 | -1.12664 | 0.735666 | 0.110858 |
| 1 | 75334000 | 75340399 | 31.60531 | 50.67093 | 126.0365 | 53.22485 | -0.96482 | 1.964162 | 0.11086 |
| 1 | 191604400 | 191615199 | 65.08735 | 100.3221 | 238.4194 | 99.03547 | -0.94908 | 2.901266 | 0.110866 |
| 17 | 37964600 | 37966999 | 53.81848 | 86.94761 | 60.60174 | 25.87102 | 0.879835 | 1.737163 | 0.110908 |
| 1 | 119177200 | 119184999 | 61.57746 | 96.1446 | 198.6922 | 82.44436 | -0.73322 | 2.665557 | 0.110926 |
| 13 | 51406000 | 51416999 | 77.13976 | 118.8592 | 231.4109 | 96.12443 | -0.66215 | 2.911696 | 0.110926 |

| | | | | | | | | | |
|---|---|---|---|---|---|---|---|---|---|
| 8 | 96050400 | 96056799 | 39.68361 | 63.03064 | 152.5689 | 63.61217 | -0.9406 | 2.273074 | 0.110928 |
| 3 | 130706600 | 130707599 | 4.960967 | 9.602128 | 30.24843 | 11.52068 | -1.07404 | -0.19581 | 0.110943 |
| 18 | 4564800 | 4565999 | 4.960967 | 9.602128 | 30.24843 | 11.52068 | -1.07404 | -0.19581 | 0.110943 |
| 2 | 137523600 | 137526999 | 55.70783 | 89.65103 | 66.74139 | 28.34326 | 0.778466 | 1.854052 | 0.110945 |
| 18 | 73573800 | 73574999 | 5.105546 | 9.846146 | 29.25567 | 11.31777 | -1.01151 | -0.14171 | 0.110958 |
| 2 | 117940400 | 117943999 | 13.0555 | 22.12151 | 68.03091 | 29.79048 | -1.23282 | 1.039159 | 0.11096 |
| 11 | 98764200 | 98765399 | 6.023621 | 11.47997 | 34.07551 | 13.60897 | -1.05462 | -0.01749 | 0.110963 |
| 3 | 137849000 | 137849999 | 4.663848 | 9.042752 | 27.88086 | 10.53303 | -1.03115 | -0.23203 | 0.11097 |
| 3 | 20469600 | 20470599 | 14.41295 | 27.72756 | 14.85479 | 5.787296 | 1.426615 | -0.06453 | 0.110971 |
| 9 | 120727800 | 120734999 | 41.85051 | 66.6666 | 144.3468 | 60.39894 | -0.77571 | 2.181523 | 0.110978 |
| 2 | 87714200 | 87716999 | 32.85592 | 55.75994 | 35.99896 | 15.77138 | 1.01016 | 1.06135 | 0.110984 |
| 15 | 81952400 | 81958199 | 46.83316 | 74.22381 | 155.5469 | 64.71183 | -0.73619 | 2.344429 | 0.110985 |
| 12 | 3079400 | 3082599 | 58.59537 | 94.16764 | 70.17017 | 29.73969 | 0.774707 | 1.903196 | 0.110987 |
| 3 | 105365000 | 105367799 | 20.0635 | 32.9731 | 83.20965 | 35.8551 | -0.95334 | 1.402306 | 0.111022 |
| 19 | 60849600 | 60851199 | 5.098244 | 9.856577 | 29.48844 | 11.29128 | -1.00738 | -0.17679 | 0.111033 |
| 5 | 91663600 | 91667199 | 19.01981 | 31.50179 | 77.36535 | 33.47942 | -0.91132 | 1.300804 | 0.111061 |
| 11 | 100648200 | 100650999 | 16.71969 | 27.94608 | 70.67469 | 30.76971 | -0.95259 | 1.186509 | 0.111075 |
| 9 | 4024000 | 4029199 | 75.20513 | 120.3148 | 69.08825 | 29.08332 | 1.142366 | 2.063905 | 0.111081 |
| 2 | 90288400 | 90289399 | 6.485046 | 11.90092 | 37.76359 | 16.08049 | -1.22562 | 0.237006 | 0.111088 |
| 12 | 22809400 | 22819999 | 268.5278 | 392.0023 | 279.2617 | 115.0618 | 0.718978 | 3.940389 | 0.111102 |
| 18 | 80089800 | 80093399 | 21.2692 | 34.69494 | 89.44508 | 38.42838 | -0.99005 | 1.510194 | 0.111105 |
| 12 | 29403200 | 29408599 | 24.43442 | 39.52336 | 105.8987 | 45.27512 | -1.05618 | 1.674854 | 0.111106 |
| 17 | 19986400 | 19987399 | 24.07564 | 42.13284 | 28.04358 | 12.37264 | 0.971946 | 0.699788 | 0.111106 |
| 7 | 13371600 | 13377199 | 128.36 | 196.9119 | 151.8922 | 63.05566 | 0.667417 | 3.006399 | 0.11111 |
| 12 | 108250200 | 108266399 | 91.50255 | 138.1656 | 311.2758 | 128.6728 | -0.89609 | 3.328565 | 0.111114 |
| 2 | 96125400 | 96125999 | 3.644436 | 7.104377 | 24.27629 | 8.766515 | -1.11499 | -0.41536 | 0.11112 |
| 7 | 116974600 | 116975599 | 3.644436 | 7.104377 | 24.27629 | 8.766515 | -1.11499 | -0.41536 | 0.11112 |
| 10 | 114215000 | 114215399 | 3.644436 | 7.104377 | 24.27629 | 8.766515 | -1.11499 | -0.41536 | 0.11112 |
| 2 | 177136000 | 177136599 | 12.22748 | 23.57245 | 16.20827 | 6.276764 | 1.078142 | -0.10281 | 0.111133 |
| 8 | 53059000 | 53060799 | 19.10545 | 34.21045 | 22.64135 | 9.925663 | 0.991835 | 0.43668 | 0.111134 |
| 2 | 156852600 | 156854399 | 8.801911 | 15.80359 | 42.13861 | 18.43082 | -1.00357 | 0.391195 | 0.111138 |
| 6 | 76456200 | 76457799 | 21.29819 | 37.75198 | 23.47036 | 10.37606 | 1.073446 | 0.515972 | 0.111143 |
| 4 | 127324200 | 127329199 | 27.9726 | 45.10271 | 109.0156 | 46.42078 | -0.91331 | 1.774629 | 0.111161 |
| 12 | 106274800 | 106284799 | 73.60109 | 113.2973 | 233.8055 | 97.10642 | -0.7471 | 2.929409 | 0.111166 |
| 14 | 44294400 | 44295599 | 12.72013 | 24.41618 | 16.70149 | 6.548726 | 1.06913 | -0.05702 | 0.111171 |
| 2 | 20933600 | 20935599 | 17.16478 | 31.22241 | 20.19763 | 8.699534 | 1.040491 | 0.326023 | 0.111178 |
| 18 | 56844600 | 56847999 | 14.48419 | 24.76605 | 60.6503 | 26.59358 | -0.90397 | 0.954288 | 0.111184 |
| 11 | 80944400 | 80947799 | 12.67216 | 22.01174 | 55.32262 | 24.28595 | -0.93633 | 0.794583 | 0.111192 |
| 14 | 41451400 | 41452599 | 14.78949 | 28.44521 | 14.10753 | 5.499901 | 1.535146 | -0.06101 | 0.111193 |
| 13 | 23076200 | 23078799 | 26.3917 | 46.62767 | 19.84617 | 8.779579 | 1.614608 | 0.576739 | 0.1112 |
| 5 | 116396000 | 116429399 | 234.7284 | 329.9645 | 683.5139 | 283.1874 | -0.87444 | 4.499271 | 0.111265 |
| 2 | 49341000 | 49342799 | 11.4561 | 19.87958 | 55.84188 | 24.50141 | -1.09627 | 0.800132 | 0.111265 |
| 17 | 27770000 | 27781599 | 41.48438 | 65.46063 | 173.101 | 71.84989 | -1.07624 | 2.449144 | 0.11127 |
| 8 | 23900000 | 23902599 | 19.00915 | 31.50278 | 76.20784 | 32.992 | -0.88948 | 1.29363 | 0.111277 |
| 7 | 25057200 | 25076799 | 91.57617 | 137.2407 | 352.2418 | 145.3097 | -1.09361 | 3.478527 | 0.111281 |
| 1 | 35926200 | 35930999 | 38.89687 | 62.12796 | 131.8363 | 55.37125 | -0.74238 | 2.091174 | 0.111285 |
| 5 | 87492400 | 87494799 | 16.72515 | 28.41599 | 67.33368 | 29.5089 | -0.85845 | 1.028344 | 0.111286 |

| | | | | | | | | | |
|---|---|---|---|---|---|---|---|---|---|
| 6 | 31527800 | 31530599 | 10.9242 | 18.52503 | 67.78213 | 29.68875 | -1.48176 | 1.025504 | 0.111305 |
| 2 | 131012800 | 131015199 | 10.32313 | 18.0367 | 49.91419 | 21.99247 | -1.07297 | 0.691012 | 0.111308 |
| 2 | 173293200 | 173302999 | 86.47117 | 131.9586 | 260.6943 | 108.1037 | -0.69402 | 3.108919 | 0.111319 |
| 6 | 8549400 | 8549799 | 8.340946 | 16.28805 | 11.6412 | 4.093126 | 1.147396 | -0.49429 | 0.111335 |
| 10 | 8586200 | 8586599 | 2.935937 | 5.742051 | 22.18756 | 7.518479 | -1.2044 | -0.66052 | 0.111339 |
| 13 | 66202800 | 66206999 | 83.54527 | 132.4452 | 95.67464 | 39.81473 | 0.798463 | 2.351439 | 0.111341 |
| 3 | 105594200 | 105595599 | 6.793198 | 12.42742 | 38.78295 | 16.5851 | -1.20661 | 0.259057 | 0.111362 |
| 2 | 132002600 | 132004399 | 9.368449 | 16.51577 | 46.68215 | 20.66031 | -1.10146 | 0.546823 | 0.111363 |
| 12 | 85582600 | 85586799 | 24.61154 | 40.09247 | 91.21624 | 39.16988 | -0.81185 | 1.537127 | 0.111378 |
| 12 | 3940800 | 3948199 | 28.48055 | 45.56139 | 132.4599 | 55.756 | -1.19296 | 2.032256 | 0.111382 |
| 12 | 42872600 | 42876399 | 41.33264 | 68.6655 | 41.10776 | 17.82642 | 1.118726 | 1.297686 | 0.111391 |
| 9 | 62682800 | 62686799 | 13.7257 | 23.09523 | 68.9472 | 30.12783 | -1.19062 | 1.105498 | 0.111395 |
| X | 117671200 | 117672599 | 25.77806 | 45.4103 | 22.50579 | 9.961978 | 1.395732 | 0.609735 | 0.111403 |
| 18 | 82981800 | 82991399 | 43.84774 | 69.28742 | 175.2067 | 72.77576 | -1.01026 | 2.41713 | 0.111407 |
| 3 | 117813800 | 117815599 | 20.88614 | 37.68085 | 20.33782 | 8.850325 | 1.290962 | 0.401259 | 0.111407 |
| 9 | 42285400 | 42286999 | 8.647039 | 15.53716 | 40.56048 | 17.72937 | -0.97228 | 0.383581 | 0.111417 |
| 15 | 82793400 | 82816199 | 102.4564 | 152.8751 | 367.2152 | 151.3577 | -1.00388 | 3.562854 | 0.11142 |
| 6 | 34252800 | 34254399 | 6.855778 | 12.74889 | 37.63048 | 15.73625 | -1.10348 | 0.162096 | 0.11143 |
| 16 | 92579000 | 92583599 | 23.58779 | 38.23967 | 97.67569 | 41.82869 | -0.98418 | 1.624585 | 0.111448 |
| X | 3744400 | 3745599 | 15.55646 | 30.20529 | 10.13911 | 3.801384 | 2.141501 | -0.18707 | 0.111488 |
| 12 | 23639200 | 23650399 | 297.3917 | 428.7568 | 305.265 | 125.9603 | 0.702298 | 4.128836 | 0.11149 |
| 12 | 85240600 | 85250199 | 47.67131 | 75.05959 | 175.9224 | 72.99219 | -0.90497 | 2.498268 | 0.111501 |
| 7 | 13472400 | 13475599 | 64.2624 | 103.7042 | 52.06431 | 22.19427 | 1.350043 | 1.782725 | 0.111508 |
| 8 | 115662600 | 115665399 | 12.4763 | 21.7929 | 52.1089 | 22.95559 | -0.86393 | 0.707467 | 0.111512 |
| 7 | 17788600 | 17789199 | 18.49138 | 33.32285 | 22.42172 | 9.770942 | 0.974141 | 0.395443 | 0.111528 |
| 18 | 80689600 | 80701199 | 55.08862 | 86.00032 | 197.772 | 82.06381 | -0.8873 | 2.665254 | 0.111532 |
| 4 | 44569400 | 44572199 | 13.62732 | 23.13516 | 63.6694 | 27.89696 | -1.07301 | 1.021119 | 0.111545 |
| 12 | 22139200 | 22143599 | 106.0994 | 164.7276 | 127.6259 | 52.99833 | 0.674231 | 2.773811 | 0.111545 |
| 5 | 136624600 | 136628199 | 17.93856 | 30.09246 | 69.28947 | 30.23585 | -0.81793 | 1.149395 | 0.111579 |
| 1 | 192910000 | 192910399 | 13.6515 | 26.24739 | 13.4603 | 5.252793 | 1.484674 | -0.07297 | 0.111584 |
| 3 | 155870400 | 155871599 | 18.15799 | 32.74363 | 22.25229 | 9.689117 | 0.960316 | 0.387643 | 0.111591 |
| 14 | 32671600 | 32679999 | 36.09693 | 57.48429 | 150.9568 | 63.13885 | -1.054 | 2.187215 | 0.111594 |
| 6 | 140211600 | 140215999 | 27.77278 | 44.86507 | 103.0397 | 43.98302 | -0.83655 | 1.7167 | 0.111597 |
| 2 | 26334600 | 26338199 | 11.64933 | 19.81535 | 65.7626 | 28.82309 | -1.3414 | 0.997668 | 0.11161 |
| 2 | 74478000 | 74479399 | 9.691827 | 17.03239 | 48.25845 | 21.36713 | -1.10678 | 0.587135 | 0.111611 |
| 12 | 112831800 | 112846199 | 86.1663 | 130.6183 | 297.115 | 122.9709 | -0.9051 | 3.247137 | 0.111619 |
| 1 | 72972000 | 72975199 | 14.1466 | 24.26493 | 66.91683 | 29.34871 | -1.07416 | 0.928735 | 0.111619 |
| 7 | 19174000 | 19182999 | 41.46701 | 65.81284 | 151.456 | 63.09977 | -0.86893 | 2.297752 | 0.111647 |
| 13 | 40881600 | 40899599 | 152.1405 | 222.21 | 465.0019 | 191.5807 | -0.83409 | 3.926856 | 0.111652 |
| 7 | 22986000 | 22987599 | 30.18658 | 51.28707 | 37.21017 | 16.30731 | 0.842522 | 1.042934 | 0.111654 |
| 7 | 86950400 | 86952599 | 14.90104 | 28.16422 | 16.69531 | 6.785717 | 1.227255 | 0.076137 | 0.111663 |
| 10 | 32042800 | 32043799 | 18.67736 | 34.01839 | 20.78479 | 8.937749 | 1.124462 | 0.333817 | 0.111669 |
| 9 | 115967600 | 115970599 | 18.3264 | 30.52236 | 73.60393 | 31.97286 | -0.8844 | 1.231413 | 0.111669 |
| 1 | 91269200 | 91276799 | 40.45359 | 63.99673 | 168.3783 | 69.97968 | -1.06601 | 2.386 | 0.111681 |
| 13 | 6597000 | 6598799 | 11.77131 | 20.39798 | 56.93133 | 24.97835 | -1.08761 | 0.815358 | 0.111693 |
| 5 | 95471000 | 95474799 | 54.29823 | 87.63165 | 58.36306 | 24.88198 | 0.94354 | 1.764756 | 0.111709 |
| 15 | 82152400 | 82157799 | 19.48396 | 31.6246 | 98.11588 | 42.05437 | -1.26197 | 1.5932 | 0.111717 |

| | | | | | | | | | |
|---|---|---|---|---|---|---|---|---|---|
| 5 | 138750200 | 138755799 | 26.8717 | 43.3024 | 109.5286 | 46.60358 | -0.97962 | 1.791092 | 0.11174 |
| 4 | 155733800 | 155749599 | 73.8258 | 112.3907 | 276.8236 | 114.7194 | -1.01463 | 3.15577 | 0.111741 |
| 1 | 29894400 | 29895599 | 12.86807 | 24.86815 | 14.98248 | 5.747702 | 1.279188 | -0.12982 | 0.111748 |
| 5 | 35709000 | 35715999 | 29.06965 | 46.5632 | 126.0244 | 53.14608 | -1.08783 | 1.991264 | 0.111766 |
| 1 | 107967800 | 107968799 | 17.51726 | 31.96328 | 20.98987 | 9.005503 | 1.023251 | 0.315466 | 0.111766 |
| 11 | 94985200 | 94994599 | 51.31497 | 80.59337 | 186.9319 | 77.53478 | -0.89283 | 2.552006 | 0.11177 |
| 8 | 60958800 | 60959599 | 4.267172 | 8.208277 | 29.01637 | 11.30803 | -1.26833 | -0.12611 | 0.111786 |
| 5 | 95158400 | 95159799 | 23.87295 | 42.48677 | 20.74655 | 9.141981 | 1.42297 | 0.503957 | 0.111788 |
| 5 | 146955400 | 146961599 | 32.32872 | 51.62345 | 134.1151 | 56.30744 | -1.03402 | 2.093836 | 0.111803 |
| 15 | 24895600 | 24897199 | 13.73033 | 26.48319 | 14.91648 | 5.768058 | 1.364905 | -0.09483 | 0.111814 |
| 8 | 21873200 | 21874199 | 16.69405 | 31.77953 | 14.75679 | 5.90781 | 1.596168 | 0.042657 | 0.111814 |
| 4 | 117991600 | 117992599 | 4.542752 | 8.788221 | 28.2613 | 10.78926 | -1.10577 | -0.19668 | 0.111841 |
| 6 | 32238200 | 32239199 | 4.542752 | 8.788221 | 28.2613 | 10.78926 | -1.10577 | -0.19668 | 0.111841 |
| 2 | 143386400 | 143391199 | 28.94005 | 46.62206 | 108.7028 | 46.23298 | -0.86299 | 1.803077 | 0.111843 |
| 17 | 18691600 | 18695199 | 50.13491 | 82.85315 | 34.35667 | 14.84498 | 1.645843 | 1.37521 | 0.111856 |
| 4 | 102976400 | 102977199 | 5.161532 | 9.861334 | 31.11837 | 12.36312 | -1.13429 | -0.04944 | 0.111902 |
| 5 | 65649000 | 65650999 | 10.73954 | 18.55647 | 57.34093 | 25.15585 | -1.23428 | 0.838983 | 0.111903 |
| 9 | 63822200 | 63824599 | 7.843717 | 13.92094 | 44.61326 | 19.70475 | -1.27668 | 0.464247 | 0.111904 |
| X | 3590000 | 3591199 | 14.36951 | 27.88916 | 11.28549 | 4.243505 | 1.871702 | -0.18808 | 0.111909 |
| 14 | 64300400 | 64302799 | 11.13354 | 19.36294 | 54.2214 | 23.82094 | -1.09128 | 0.765455 | 0.11192 |
| 8 | 71995000 | 72003799 | 43.46738 | 68.82521 | 159.7685 | 66.43103 | -0.88463 | 2.365964 | 0.111924 |
| 1 | 74935000 | 74945399 | 64.81446 | 99.75426 | 239.9297 | 99.64763 | -0.96798 | 2.930272 | 0.111928 |
| 9 | 56098600 | 56103199 | 17.59332 | 28.88442 | 86.70854 | 37.34687 | -1.20339 | 1.411409 | 0.111936 |
| 7 | 14212400 | 14213999 | 27.03116 | 46.74644 | 32.29397 | 14.16795 | 0.916892 | 0.867591 | 0.111964 |
| X | 35067000 | 35068999 | 19.63803 | 36.34527 | 17.06333 | 7.182559 | 1.522639 | 0.246537 | 0.111978 |
| 10 | 65779400 | 65781399 | 17.00705 | 30.98853 | 21.35786 | 9.179581 | 0.952003 | 0.318907 | 0.111988 |
| X | 117149800 | 117150599 | 11.50838 | 22.41801 | 12.32843 | 4.494161 | 1.47545 | -0.32348 | 0.111994 |
| 4 | 129514400 | 129518599 | 8.387808 | 14.64996 | 51.44745 | 22.66286 | -1.41419 | 0.68847 | 0.112006 |
| 11 | 113737800 | 113751799 | 95.39143 | 144.1418 | 305.3494 | 126.2518 | -0.80646 | 3.314394 | 0.112013 |
| 13 | 35829200 | 35830999 | 20.91638 | 36.96476 | 25.32612 | 11.20316 | 0.932812 | 0.549612 | 0.112039 |
| 13 | 46580400 | 46583799 | 23.27172 | 38.09427 | 86.32397 | 37.13472 | -0.80104 | 1.461266 | 0.112056 |
| X | 33865200 | 33869399 | 41.58551 | 69.51562 | 36.07417 | 15.70674 | 1.324047 | 1.229504 | 0.112057 |
| 2 | 59474800 | 59479599 | 20.8614 | 34.25492 | 83.86776 | 36.12553 | -0.91038 | 1.415228 | 0.112081 |
| 1 | 4640800 | 4642599 | 7.506505 | 13.75371 | 37.21001 | 15.87882 | -0.99989 | 0.253361 | 0.112082 |
| 7 | 10346600 | 10348799 | 33.44973 | 57.76437 | 26.47829 | 11.616 | 1.505202 | 0.902225 | 0.112085 |
| 12 | 3337200 | 3338799 | 22.88637 | 40.1828 | 26.18373 | 11.58406 | 1.001878 | 0.636995 | 0.112089 |
| 3 | 97134000 | 97136599 | 14.25063 | 24.17998 | 64.2792 | 28.15943 | -1.02343 | 1.0282 | 0.112092 |
| 8 | 110666000 | 110667799 | 7.687784 | 13.62906 | 43.57108 | 19.26184 | -1.27373 | 0.470103 | 0.112101 |
| 2 | 166126600 | 166130999 | 19.21033 | 31.40661 | 89.4141 | 38.44639 | -1.13036 | 1.472591 | 0.112111 |
| 12 | 20479600 | 20480999 | 39.10504 | 65.15657 | 41.06375 | 17.84249 | 1.044941 | 1.258329 | 0.112115 |
| 3 | 11446800 | 11448199 | 16.61357 | 30.95512 | 18.12748 | 7.555318 | 1.216123 | 0.18445 | 0.11212 |
| 4 | 100724400 | 100727199 | 21.05294 | 37.12958 | 25.87453 | 11.45047 | 0.907379 | 0.573226 | 0.112131 |
| 11 | 102246200 | 102257799 | 54.9936 | 85.33053 | 214.4759 | 89.07192 | -1.02227 | 2.773463 | 0.112133 |
| 1 | 10643600 | 10644199 | 4.660587 | 9.046819 | 28.00604 | 10.52297 | -1.02952 | -0.25068 | 0.112135 |
| 13 | 16537000 | 16537999 | 7.998335 | 14.51056 | 38.62062 | 16.68956 | -0.98877 | 0.31831 | 0.112141 |
| 9 | 53881200 | 53883799 | 15.43862 | 25.93763 | 70.82763 | 30.93513 | -1.06296 | 1.125102 | 0.112189 |
| 2 | 87282000 | 87283799 | 26.51112 | 45.9988 | 31.35659 | 13.75809 | 0.937054 | 0.83643 | 0.112196 |

| | | | | | | | | | |
|---|---|---|---|---|---|---|---|---|---|
| 16 | 60315200 | 60315799 | 9.634371 | 18.82933 | 10.14521 | 3.508119 | 1.569081 | -0.53558 | 0.112196 |
| 19 | 40022800 | 40024399 | 21.65297 | 37.95901 | 25.58798 | 11.30583 | 0.953317 | 0.652626 | 0.112217 |
| 1 | 168467400 | 168471399 | 38.73399 | 65.31148 | 33.12532 | 14.48396 | 1.356575 | 1.134294 | 0.112218 |
| 12 | 20051400 | 20061999 | 263.2294 | 383.3509 | 275.6085 | 113.5848 | 0.702449 | 3.975748 | 0.11222 |
| 3 | 152364200 | 152365399 | 15.7132 | 29.4441 | 18.32786 | 7.565793 | 1.139909 | 0.142578 | 0.112221 |
| 6 | 47615000 | 47616799 | 19.80865 | 35.56704 | 22.00621 | 9.624938 | 1.090591 | 0.424019 | 0.112249 |
| 1 | 18557600 | 18558799 | 12.16695 | 23.36918 | 16.71603 | 6.542493 | 1.007416 | -0.06899 | 0.112261 |
| 17 | 78924000 | 78935199 | 70.5275 | 108.8972 | 227.7784 | 94.61454 | -0.76317 | 2.870016 | 0.112265 |
| 11 | 115171000 | 115180999 | 46.27315 | 72.73598 | 181.4841 | 75.28337 | -0.99697 | 2.53224 | 0.112281 |
| 6 | 117382800 | 117386599 | 16.89694 | 28.22511 | 75.34569 | 32.7907 | -1.03055 | 1.195151 | 0.112333 |
| 10 | 71062200 | 71063399 | 12.24083 | 23.60273 | 15.5217 | 6.007561 | 1.141836 | -0.10738 | 0.112338 |
| 6 | 87725400 | 87728799 | 17.73336 | 29.44257 | 74.98074 | 32.50199 | -0.96303 | 1.267884 | 0.112363 |
| 8 | 114702600 | 114707399 | 28.86845 | 46.69535 | 100.7137 | 43.05824 | -0.74404 | 1.681593 | 0.112364 |
| 10 | 47200600 | 47201999 | 21.82429 | 38.50837 | 25.15132 | 11.12935 | 1.000485 | 0.571072 | 0.112369 |
| 2 | 36664400 | 36667999 | 94.5672 | 148.6608 | 111.7192 | 46.34395 | 0.731971 | 2.547751 | 0.112384 |
| 11 | 50103000 | 50104599 | 4.957465 | 9.507539 | 30.08312 | 11.82522 | -1.12241 | -0.08628 | 0.112392 |
| 1 | 36744000 | 36749399 | 23.76116 | 38.49796 | 99.40108 | 42.54902 | -1.00052 | 1.637565 | 0.112398 |
| 4 | 106710600 | 106714799 | 23.90223 | 38.7379 | 97.01094 | 41.5349 | -0.95612 | 1.631357 | 0.112417 |
| X | 32443000 | 32443799 | 16.37508 | 31.25896 | 14.55475 | 5.792818 | 1.598739 | 0.020475 | 0.112423 |
| 17 | 69845600 | 69846999 | 56.31298 | 91.91951 | 40.68958 | 17.48567 | 1.545087 | 1.546553 | 0.112424 |
| 5 | 85087400 | 85088199 | 21.52066 | 37.76724 | 26.20259 | 11.58781 | 0.912163 | 0.637782 | 0.112426 |
| 7 | 17040000 | 17044199 | 112.9247 | 174.5593 | 136.4896 | 56.69466 | 0.656196 | 2.855552 | 0.112426 |
| 19 | 3362200 | 3365399 | 14.81134 | 25.2731 | 61.72831 | 27.06174 | -0.90071 | 0.975654 | 0.11243 |
| 7 | 85506800 | 85507399 | 9.812383 | 19.17106 | 10.7774 | 3.756788 | 1.499515 | -0.50342 | 0.112436 |
| 2 | 31280800 | 31283199 | 12.41082 | 21.41726 | 56.88171 | 24.95343 | -1.01763 | 0.856893 | 0.112446 |
| 11 | 118876400 | 118892799 | 84.83134 | 128.1032 | 310.3851 | 128.3081 | -1.00082 | 3.325824 | 0.112467 |
| 4 | 121469600 | 121470999 | 20.90393 | 37.62721 | 20.88161 | 9.112871 | 1.248615 | 0.415237 | 0.112468 |
| 7 | 24615600 | 24619799 | 17.67727 | 29.09763 | 81.66113 | 35.21149 | -1.10492 | 1.376919 | 0.112472 |
| 7 | 19024200 | 19025999 | 7.184994 | 13.23612 | 36.2414 | 15.35193 | -1.00907 | 0.218035 | 0.11248 |
| 16 | 78091800 | 78093599 | 7.184994 | 13.23612 | 36.2414 | 15.35193 | -1.00907 | 0.218035 | 0.11248 |
| 4 | 140716000 | 140725399 | 46.76717 | 73.48668 | 181.9962 | 75.49237 | -0.98664 | 2.539733 | 0.112482 |
| 1 | 124893600 | 124894399 | 9.975668 | 19.4187 | 13.65629 | 5.015844 | 1.115331 | -0.31404 | 0.112484 |
| 1 | 100908200 | 100911399 | 33.8477 | 58.57375 | 24.1347 | 10.58855 | 1.658326 | 0.882599 | 0.112498 |
| 12 | 70952600 | 70953799 | 18.46537 | 33.09519 | 22.55215 | 9.878779 | 0.951128 | 0.427966 | 0.112508 |
| 5 | 129942800 | 129948999 | 34.63484 | 55.53882 | 119.9293 | 50.66082 | -0.76092 | 1.960922 | 0.112518 |
| 18 | 86275200 | 86276599 | 18.86305 | 34.12106 | 21.63879 | 9.385389 | 1.063014 | 0.374389 | 0.112522 |
| X | 62089400 | 62090799 | 19.23092 | 35.70224 | 16.98606 | 7.119962 | 1.507604 | 0.226195 | 0.112531 |
| 3 | 3212000 | 3214799 | 59.039 | 94.9298 | 69.0211 | 29.27468 | 0.811013 | 1.8886 | 0.112535 |
| 11 | 59340800 | 59341799 | 5.890636 | 11.16125 | 31.98733 | 12.93353 | -1.01922 | 0.019728 | 0.112547 |
| 4 | 146215800 | 146217999 | 26.14125 | 45.5549 | 25.76521 | 11.33363 | 1.20502 | 0.76537 | 0.112548 |
| 14 | 42829800 | 42832799 | 36.40778 | 60.93331 | 39.33982 | 17.14156 | 1.009929 | 1.200139 | 0.112556 |
| 10 | 79303200 | 79303799 | 2.54269 | 4.973896 | 22.46994 | 7.489728 | -1.4037 | -0.7251 | 0.112571 |
| 13 | 62859400 | 62865199 | 89.24147 | 141.5229 | 83.80069 | 34.88514 | 1.085232 | 2.343376 | 0.112592 |
| 18 | 67263800 | 67267399 | 23.19611 | 37.77724 | 92.70377 | 39.80502 | -0.92093 | 1.538324 | 0.11261 |
| 10 | 69289000 | 69289999 | 3.599691 | 6.926545 | 29.47778 | 11.47726 | -1.52875 | -0.1309 | 0.112648 |
| 6 | 5672000 | 5674599 | 35.11063 | 58.39014 | 43.90856 | 19.05478 | 0.790994 | 1.26903 | 0.11265 |
| 2 | 73103200 | 73105399 | 18.43108 | 30.42062 | 79.22948 | 34.20381 | -0.9961 | 1.342591 | 0.112657 |

| | | | | | | | | | |
|---|---|---|---|---|---|---|---|---|---|
| 8 | 70316000 | 70352799 | 270.3533 | 373.0316 | 823.643 | 342.2402 | -0.99359 | 4.782372 | 0.112661 |
| 12 | 22874000 | 22874799 | 36.46332 | 60.52918 | 40.83282 | 17.69771 | 0.947011 | 1.291856 | 0.112679 |
| 10 | 60082200 | 60083999 | 6.984062 | 13.02383 | 34.89972 | 14.52531 | -0.95891 | 0.141454 | 0.11268 |
| 15 | 41651400 | 41653399 | 7.761721 | 13.65745 | 48.76999 | 21.59247 | -1.43719 | 0.562651 | 0.112683 |
| 18 | 73307400 | 73308999 | 18.02276 | 33.27516 | 17.8669 | 7.54562 | 1.326226 | 0.246354 | 0.112721 |
| 11 | 115399800 | 115401599 | 4.869738 | 9.308991 | 30.52484 | 12.11134 | -1.1868 | -0.05736 | 0.112738 |
| X | 9497400 | 9499599 | 28.06018 | 49.21367 | 22.86677 | 10.10738 | 1.487268 | 0.680096 | 0.112755 |
| 7 | 28328600 | 28333999 | 28.16861 | 45.33139 | 112.1858 | 47.64137 | -0.95069 | 1.836803 | 0.112756 |
| 13 | 69524400 | 69527199 | 12.48134 | 21.76213 | 52.11637 | 22.93269 | -0.86648 | 0.732954 | 0.11276 |
| 2 | 119151000 | 119163999 | 65.71716 | 100.6371 | 259.1468 | 107.5485 | -1.07191 | 3.03178 | 0.112765 |
| 11 | 98039800 | 98041399 | 5.342683 | 9.94402 | 34.75229 | 14.51074 | -1.34057 | 0.142932 | 0.112777 |
| 12 | 23109600 | 23113999 | 90.73096 | 142.7598 | 111.1607 | 46.11772 | 0.681831 | 2.527289 | 0.112797 |
| 19 | 6962000 | 6965599 | 12.66822 | 21.61516 | 62.09604 | 27.22285 | -1.1335 | 0.96899 | 0.112801 |
| 14 | 84680400 | 84686199 | 40.69597 | 64.21958 | 176.3472 | 73.19906 | -1.13053 | 2.44763 | 0.112806 |
| 4 | 90736200 | 90736999 | 11.52108 | 22.1486 | 16.31493 | 6.369014 | 0.968991 | -0.08515 | 0.112811 |
| 7 | 31056600 | 31064799 | 183.995 | 276.2196 | 206.9793 | 85.42102 | 0.6835 | 3.45469 | 0.112823 |
| 7 | 110755200 | 110756199 | 3.038191 | 5.908111 | 30.60375 | 11.33383 | -1.74233 | -0.31654 | 0.112834 |
| 16 | 64590800 | 64592399 | 7.882038 | 14.21115 | 39.56939 | 17.23274 | -1.06071 | 0.355977 | 0.112844 |
| 11 | 109045200 | 109055599 | 64.44408 | 99.96627 | 212.5414 | 88.27222 | -0.7815 | 2.78165 | 0.112853 |
| 17 | 92548000 | 92549199 | 13.54783 | 26.08873 | 14.76822 | 5.737451 | 1.350789 | -0.08351 | 0.112862 |
| 4 | 121493000 | 121496799 | 83.05104 | 131.3232 | 102.6976 | 42.67195 | 0.681115 | 2.413306 | 0.112865 |
| 15 | 99772400 | 99773599 | 4.312791 | 8.304734 | 32.10682 | 12.4748 | -1.39379 | -0.13119 | 0.112879 |
| 7 | 9643600 | 9645599 | 34.60021 | 57.62531 | 43.35332 | 18.83211 | 0.790379 | 1.25087 | 0.112883 |
| 6 | 38852000 | 38853799 | 9.071498 | 15.98293 | 46.15834 | 20.43087 | -1.13243 | 0.551453 | 0.112883 |
| 19 | 12998200 | 13001199 | 36.1581 | 61.32323 | 31.38086 | 13.74538 | 1.343698 | 1.07311 | 0.112887 |
| 9 | 115337800 | 115339999 | 15.42177 | 26.34345 | 61.80223 | 27.09654 | -0.84266 | 0.967902 | 0.112926 |
| 7 | 131659400 | 131659999 | 3.423781 | 6.668022 | 26.52179 | 9.694944 | -1.34832 | -0.36934 | 0.112938 |
| 14 | 20529600 | 20530399 | 3.423781 | 6.668022 | 26.52179 | 9.694944 | -1.34832 | -0.36934 | 0.112938 |
| 7 | 32213200 | 32215599 | 40.89142 | 67.7395 | 40.08681 | 17.34957 | 1.134976 | 1.324921 | 0.112953 |
| 11 | 22461600 | 22462999 | 6.886666 | 12.7204 | 36.85165 | 15.55393 | -1.08626 | 0.200338 | 0.112969 |
| 12 | 76572800 | 76574599 | 6.886666 | 12.7204 | 36.85165 | 15.55393 | -1.08626 | 0.200338 | 0.112969 |
| 13 | 99025600 | 99027599 | 6.323236 | 11.61125 | 36.69335 | 15.61144 | -1.21835 | 0.23117 | 0.112979 |
| 4 | 144976000 | 144977999 | 7.073167 | 12.97165 | 42.22583 | 17.99674 | -1.26423 | 0.25217 | 0.112987 |
| 19 | 36092400 | 36094399 | 6.818291 | 12.77635 | 34.50342 | 14.24324 | -0.96055 | 0.106869 | 0.113021 |
| 16 | 30989000 | 30990199 | 5.875523 | 11.18399 | 31.59738 | 12.65204 | -0.98642 | -0.01408 | 0.113023 |
| 11 | 88590400 | 88592799 | 7.124182 | 12.93648 | 40.23639 | 17.36796 | -1.21094 | 0.308107 | 0.113025 |
| 1 | 71810800 | 71812599 | 30.85802 | 52.46834 | 35.47982 | 15.54996 | 0.943688 | 1.035358 | 0.11303 |
| 6 | 6870200 | 6876399 | 40.69419 | 64.91042 | 137.9344 | 57.82152 | -0.74661 | 2.136404 | 0.113034 |
| 2 | 95249800 | 95251999 | 17.34197 | 31.44458 | 21.85558 | 9.451152 | 0.934374 | 0.349987 | 0.113058 |
| 3 | 22055400 | 22057999 | 22.82269 | 40.54679 | 20.48974 | 9.042114 | 1.371792 | 0.504864 | 0.113068 |
| 13 | 66607800 | 66609999 | 31.52932 | 54.19324 | 30.99001 | 13.59335 | 1.186671 | 0.93922 | 0.11307 |
| 15 | 74613200 | 74626599 | 78.51242 | 119.4421 | 279.9844 | 116.0051 | -0.94415 | 3.171154 | 0.11307 |
| 2 | 151966400 | 151967399 | 4.720469 | 9.088184 | 30.9414 | 12.02798 | -1.2131 | -0.12779 | 0.113076 |
| 9 | 98830800 | 98848199 | 128.5092 | 189.8606 | 421.7127 | 173.6705 | -0.9043 | 3.740021 | 0.113125 |
| 17 | 39332400 | 39332999 | 14.77794 | 28.4807 | 13.51952 | 5.240061 | 1.605039 | -0.08078 | 0.113125 |
| 17 | 44499000 | 44503399 | 21.68057 | 35.5192 | 85.52338 | 36.80328 | -0.88775 | 1.446728 | 0.113132 |
| 18 | 36661200 | 36668999 | 38.30828 | 61.04081 | 138.0384 | 57.7814 | -0.83777 | 2.169551 | 0.113134 |

| | | | | | | | | | |
|---|---|---|---|---|---|---|---|---|---|
| X | 89231600 | 89232799 | 18.28357 | 34.48811 | 14.94806 | 6.101431 | 1.670286 | 0.11761 | 0.113138 |
| 8 | 61084200 | 61087999 | 22.72838 | 37.02182 | 96.94656 | 41.62937 | -1.01426 | 1.536284 | 0.11314 |
| 9 | 83637400 | 83638199 | 4.007423 | 7.769578 | 31.57635 | 11.93222 | -1.4264 | -0.2316 | 0.113143 |
| 13 | 69203200 | 69204599 | 12.01406 | 23.32158 | 12.62003 | 4.7407 | 1.458592 | -0.2161 | 0.113147 |
| 1 | 15812800 | 15814599 | 5.858283 | 11.21009 | 34.92558 | 13.81701 | -1.11137 | -0.04783 | 0.113159 |
| 16 | 35501800 | 35503599 | 7.828626 | 14.26937 | 38.17908 | 16.40237 | -0.99085 | 0.287853 | 0.113164 |
| 5 | 113872000 | 113887799 | 89.42316 | 134.4229 | 331.5535 | 136.8817 | -1.03231 | 3.418227 | 0.113166 |
| 8 | 87523800 | 87526999 | 16.90759 | 28.61106 | 67.27283 | 29.44682 | -0.84761 | 1.063302 | 0.113174 |
| 17 | 38431600 | 38437799 | 107.4701 | 167.0469 | 129.6307 | 53.82163 | 0.673207 | 2.756342 | 0.11318 |
| 12 | 20107200 | 20112199 | 33.30891 | 56.44749 | 33.97097 | 14.87641 | 1.110994 | 1.072823 | 0.113181 |
| 9 | 87833400 | 87835599 | 29.5533 | 47.67161 | 105.8884 | 45.11382 | -0.79099 | 1.763753 | 0.113192 |
| 12 | 112767000 | 112779199 | 60.50482 | 92.91164 | 249.5842 | 103.6253 | -1.1297 | 2.974525 | 0.113201 |
| 16 | 11335200 | 11341199 | 45.5634 | 72.14192 | 160.1999 | 66.60943 | -0.82079 | 2.367841 | 0.113208 |
| 4 | 49385600 | 49386399 | 17.36277 | 32.39139 | 17.55282 | 7.30039 | 1.329606 | 0.182277 | 0.113213 |
| 5 | 37062000 | 37076799 | 80.99779 | 123.1396 | 280.6751 | 116.2677 | -0.90457 | 3.185798 | 0.113232 |
| 11 | 57717800 | 57729599 | 72.24587 | 110.4192 | 270.3341 | 112.1408 | -1.00142 | 3.073757 | 0.113234 |
| 3 | 132664800 | 132666599 | 8.130197 | 14.80561 | 39.24365 | 16.87887 | -0.97876 | 0.296823 | 0.113265 |
| 5 | 130697200 | 130698199 | 12.00778 | 22.98727 | 16.80894 | 6.641655 | 0.963467 | -0.04014 | 0.113281 |
| 11 | 21942600 | 21955599 | 106.9169 | 159.8294 | 363.6041 | 149.9238 | -0.92307 | 3.52797 | 0.113282 |
| 17 | 56637000 | 56644399 | 28.71519 | 46.02777 | 122.4891 | 51.71041 | -1.06225 | 1.968231 | 0.113292 |
| 11 | 51863400 | 51865199 | 7.815336 | 14.27616 | 37.68388 | 16.14564 | -0.96865 | 0.275243 | 0.1133 |
| 9 | 70114400 | 70117599 | 12.78497 | 21.89215 | 60.0084 | 26.31518 | -1.06524 | 0.934061 | 0.113303 |
| 1 | 83071000 | 83072799 | 13.70898 | 25.90106 | 16.89321 | 6.871792 | 1.089575 | 0.067576 | 0.113304 |
| 13 | 78885200 | 78885999 | 13.70898 | 25.90106 | 16.89321 | 6.871792 | 1.089575 | 0.067576 | 0.113304 |
| 5 | 146908800 | 146930799 | 139.3518 | 205.5092 | 419.0528 | 172.5717 | -0.78344 | 3.771112 | 0.113314 |
| 8 | 109473600 | 109476199 | 10.41702 | 18.3674 | 46.15173 | 20.42487 | -0.93276 | 0.550408 | 0.113315 |
| 9 | 119402200 | 119409999 | 49.31913 | 77.23314 | 193.4885 | 80.26074 | -1.00711 | 2.60633 | 0.113325 |
| 16 | 47060400 | 47061599 | 14.97673 | 28.365 | 16.7389 | 6.774187 | 1.239233 | 0.06377 | 0.113326 |
| 6 | 68622000 | 68624799 | 38.72217 | 65.35189 | 32.55007 | 14.23642 | 1.382621 | 1.125926 | 0.113326 |
| 17 | 31114600 | 31134399 | 106.5626 | 159.4546 | 350.9559 | 144.7359 | -0.87412 | 3.509542 | 0.113331 |
| 15 | 95726800 | 95729599 | 10.34437 | 18.07018 | 52.05591 | 22.93322 | -1.13102 | 0.694598 | 0.113341 |
| 14 | 54167000 | 54168599 | 12.59859 | 24.01791 | 15.34059 | 6.122792 | 1.144052 | -0.00654 | 0.113341 |
| 5 | 122894400 | 122895599 | 8.343024 | 14.99982 | 40.04781 | 17.4965 | -1.00377 | 0.377301 | 0.113351 |
| 8 | 118058400 | 118062399 | 22.92981 | 37.19998 | 95.20588 | 40.79286 | -0.98591 | 1.608118 | 0.113352 |
| 6 | 112459400 | 112468199 | 48.15048 | 75.45529 | 189.2699 | 78.50525 | -1.00808 | 2.593246 | 0.113353 |
| 17 | 17502800 | 17507399 | 40.79703 | 67.19661 | 44.0178 | 18.98516 | 0.987631 | 1.394236 | 0.113356 |
| 15 | 34034400 | 34038199 | 23.56414 | 38.6416 | 88.47522 | 38.08879 | -0.81498 | 1.440119 | 0.11336 |
| 9 | 74763200 | 74776199 | 100.5306 | 151.6052 | 318.4971 | 131.6052 | -0.79694 | 3.356193 | 0.113366 |
| 1 | 90372200 | 90373799 | 6.367073 | 11.97123 | 33.46059 | 13.73394 | -1.00277 | 0.078703 | 0.113375 |
| 2 | 89031000 | 89032799 | 15.86229 | 29.22503 | 20.46499 | 8.667527 | 0.943748 | 0.246785 | 0.113394 |
| 7 | 38315000 | 38315799 | 16.61138 | 31.39387 | 16.08532 | 6.53912 | 1.435907 | 0.092153 | 0.113396 |
| 2 | 93784200 | 93789799 | 35.84433 | 57.2585 | 135.5483 | 56.93907 | -0.8997 | 2.084925 | 0.113396 |
| 4 | 89307600 | 89309399 | 16.0077 | 29.62958 | 19.87249 | 8.363812 | 1.011834 | 0.221268 | 0.113421 |
| 12 | 84363400 | 84364599 | 5.79075 | 10.56557 | 38.69796 | 16.6043 | -1.43799 | 0.268348 | 0.113425 |
| 1 | 80112600 | 80113599 | 12.778 | 24.66537 | 15.64158 | 6.033258 | 1.199205 | -0.1107 | 0.113435 |
| 7 | 32095200 | 32097399 | 20.1761 | 35.71678 | 24.77477 | 10.95557 | 0.915869 | 0.525166 | 0.113465 |
| 1 | 36346800 | 36367199 | 95.84271 | 143.7989 | 346.784 | 143.0966 | -1.00234 | 3.456822 | 0.113469 |

| | | | | | | | | | |
|---|---|---|---|---|---|---|---|---|---|
| 4 | 125625200 | 125626599 | 7.786148 | 14.31629 | 38.41522 | 16.31495 | -0.98355 | 0.236632 | 0.113475 |
| 13 | 74361800 | 74363599 | 7.786148 | 14.31629 | 38.41522 | 16.31495 | -0.98355 | 0.236632 | 0.113475 |
| 9 | 36165000 | 36167199 | 33.4442 | 56.84779 | 34.27069 | 15.01964 | 1.108651 | 1.047727 | 0.113478 |
| 2 | 136076000 | 136076999 | 7.604884 | 14.0889 | 37.5298 | 15.77612 | -0.96213 | 0.190796 | 0.113484 |
| 12 | 10498800 | 10499999 | 7.604884 | 14.0889 | 37.5298 | 15.77612 | -0.96213 | 0.190796 | 0.113484 |
| 17 | 49282200 | 49285799 | 14.70592 | 24.96637 | 63.7827 | 27.94663 | -0.96633 | 1.024342 | 0.113487 |
| 7 | 69989400 | 69990399 | 5.628099 | 10.62197 | 32.40282 | 13.2109 | -1.11913 | 0.04526 | 0.113497 |
| 16 | 90731000 | 90732199 | 5.628099 | 10.62197 | 32.40282 | 13.2109 | -1.11913 | 0.04526 | 0.113497 |
| 17 | 23876200 | 23903799 | 165.4041 | 239.0428 | 518.2316 | 213.7677 | -0.89997 | 4.087609 | 0.113506 |
| 12 | 40237600 | 40239599 | 18.28057 | 33.17478 | 21.11539 | 9.121318 | 1.061314 | 0.350306 | 0.113508 |
| 8 | 82184200 | 82185599 | 15.33504 | 28.75245 | 18.33823 | 7.562507 | 1.106101 | 0.135326 | 0.113509 |
| 7 | 79840200 | 79842599 | 8.48193 | 14.90884 | 46.68487 | 20.67359 | -1.24944 | 0.576859 | 0.113509 |
| 4 | 139815000 | 139824399 | 53.81881 | 83.92504 | 198.9826 | 82.57601 | -0.93252 | 2.684925 | 0.113518 |
| 4 | 124873000 | 124885799 | 68.99046 | 105.6763 | 258.2004 | 107.162 | -0.99596 | 3.027341 | 0.113531 |
| 11 | 88949400 | 88955199 | 29.53897 | 47.68589 | 105.0704 | 44.81062 | -0.77809 | 1.740125 | 0.113536 |
| 17 | 29948400 | 29950799 | 9.921248 | 17.24555 | 53.60754 | 23.5435 | -1.24068 | 0.768374 | 0.113545 |
| 17 | 13210200 | 13211999 | 10.60383 | 18.59247 | 47.77144 | 21.11056 | -0.9664 | 0.631628 | 0.113554 |
| 1 | 6159800 | 6162399 | 38.07088 | 63.07048 | 40.22864 | 17.4116 | 1.027257 | 1.316389 | 0.113556 |
| 4 | 63460600 | 63463599 | 11.39951 | 19.90417 | 51.01245 | 22.467 | -0.96343 | 0.705527 | 0.113556 |
| 7 | 23405600 | 23406199 | 15.62593 | 30.10238 | 11.53592 | 4.476616 | 1.906836 | -0.0738 | 0.113591 |
| 12 | 37709200 | 37710999 | 32.83613 | 55.85283 | 33.7074 | 14.7736 | 1.107139 | 1.03778 | 0.113594 |
| 14 | 75991400 | 75992199 | 6.528776 | 12.21505 | 33.86758 | 14.0172 | -1.00084 | 0.113912 | 0.113594 |
| 18 | 15049000 | 15052399 | 16.09544 | 26.91368 | 70.62614 | 30.75651 | -1.00566 | 1.179933 | 0.113596 |
| 1 | 182879200 | 182882199 | 13.978 | 24.01804 | 59.08693 | 25.91743 | -0.9093 | 0.904891 | 0.113601 |
| 6 | 139639000 | 139641599 | 15.59994 | 26.57281 | 63.4156 | 27.79759 | -0.86794 | 0.995911 | 0.113606 |
| 16 | 17050800 | 17052999 | 10.19114 | 17.79503 | 50.97831 | 22.45256 | -1.12288 | 0.69941 | 0.113625 |
| 14 | 88400600 | 88402799 | 43.65124 | 71.88567 | 46.17191 | 19.91282 | 1.01593 | 1.407085 | 0.113629 |
| 2 | 88969000 | 88969999 | 13.96027 | 26.41588 | 17.41654 | 7.061975 | 1.078317 | 0.061673 | 0.113653 |
| 5 | 63649800 | 63651199 | 13.96027 | 26.41588 | 17.41654 | 7.061975 | 1.078317 | 0.061673 | 0.113653 |
| 13 | 41492200 | 41493999 | 8.625792 | 15.56179 | 40.10159 | 17.45095 | -0.94947 | 0.357685 | 0.113661 |
| 2 | 128753800 | 128754399 | 2.523992 | 4.935582 | 21.24082 | 7.241402 | -1.36026 | -0.64249 | 0.113661 |
| 7 | 113853200 | 113853999 | 4.977054 | 9.583046 | 29.16016 | 11.3326 | -1.05173 | -0.12858 | 0.113665 |
| 11 | 51496400 | 51498199 | 8.516652 | 14.9473 | 49.31949 | 21.81246 | -1.32486 | 0.604762 | 0.113667 |
| 10 | 62858600 | 62860599 | 11.10943 | 19.36281 | 51.53686 | 22.67232 | -1.01785 | 0.731467 | 0.113671 |
| 15 | 85893600 | 85919399 | 190.9241 | 273.1448 | 561.3792 | 231.9114 | -0.83753 | 4.238849 | 0.113678 |
| 7 | 131152800 | 131159399 | 49.49335 | 78.21536 | 166.1144 | 69.00212 | -0.75884 | 2.416759 | 0.113684 |
| 2 | 114327800 | 114328599 | 4.322497 | 8.391095 | 27.89161 | 10.47441 | -1.12878 | -0.25338 | 0.113687 |
| 1 | 160498200 | 160498799 | 9.014462 | 17.57789 | 13.36025 | 4.8055 | 1.03218 | -0.39875 | 0.1137 |
| 2 | 163691200 | 163697799 | 48.01729 | 75.9323 | 167.6689 | 69.67088 | -0.81418 | 2.400095 | 0.113701 |
| 2 | 32601200 | 32605399 | 15.08505 | 25.06447 | 75.74611 | 32.85083 | -1.20866 | 1.252645 | 0.113705 |
| 7 | 13373200 | 13376799 | 103.6282 | 161.2466 | 123.8522 | 51.41426 | 0.689153 | 2.733923 | 0.113707 |
| 4 | 61288200 | 61294999 | 75.91621 | 120.6662 | 92.28403 | 38.50448 | 0.719866 | 2.267341 | 0.113725 |
| 2 | 100982000 | 100982799 | 10.91416 | 21.2256 | 12.21975 | 4.522422 | 1.389317 | -0.2819 | 0.113749 |
| 15 | 45641800 | 45643199 | 16.44711 | 30.93781 | 17.2063 | 7.056807 | 1.308429 | 0.123166 | 0.113752 |
| 17 | 39325400 | 39330599 | 139.0646 | 212.393 | 164.01 | 68.02393 | 0.660939 | 3.099634 | 0.113758 |
| 3 | 151450200 | 151451799 | 14.84915 | 27.61047 | 19.53464 | 8.17034 | 0.941756 | 0.185723 | 0.113765 |
| 15 | 86374800 | 86383999 | 53.65961 | 84.05484 | 186.0649 | 77.17905 | -0.82849 | 2.603003 | 0.113771 |

| | | | | | | | | | |
|---|---|---|---|---|---|---|---|---|---|
| 8 | 81740400 | 81742199 | 7.581022 | 13.37174 | 46.16608 | 20.42976 | -1.38702 | 0.531026 | 0.113778 |
| 19 | 42471200 | 42474599 | 13.33365 | 22.99202 | 56.92066 | 24.96993 | -0.9171 | 0.867864 | 0.113786 |
| 8 | 117929200 | 117942199 | 76.7157 | 117.7386 | 246.3103 | 102.2581 | -0.77011 | 2.990117 | 0.113806 |
| 1 | 57445200 | 57446399 | 7.484786 | 13.77874 | 36.78028 | 15.59506 | -0.97397 | 0.224892 | 0.113814 |
| 15 | 30991400 | 30993799 | 9.33023 | 16.27479 | 57.43215 | 25.27847 | -1.42251 | 0.717106 | 0.113838 |
| 1 | 116693800 | 116699799 | 74.57432 | 119.3312 | 67.72617 | 28.51859 | 1.159474 | 2.053913 | 0.113862 |
| 8 | 126832400 | 126833399 | 13.50745 | 25.81617 | 16.41815 | 6.517614 | 1.156167 | -0.01196 | 0.113863 |
| 5 | 127791600 | 127795799 | 27.21107 | 44.17527 | 97.7855 | 41.91887 | -0.77646 | 1.602344 | 0.113873 |
| 7 | 125607400 | 125610399 | 11.71163 | 19.71887 | 68.68475 | 30.01896 | -1.41159 | 1.094449 | 0.113875 |
| 4 | 27171400 | 27172399 | 16.79688 | 30.97832 | 19.72702 | 8.343461 | 1.081001 | 0.246838 | 0.113881 |
| 9 | 62670000 | 62681599 | 66.42064 | 101.6772 | 265.0195 | 109.9765 | -1.08981 | 3.04064 | 0.113882 |
| 15 | 36440600 | 36462399 | 148.0439 | 216.8609 | 454.5484 | 187.237 | -0.83239 | 3.880395 | 0.113884 |
| 10 | 35407600 | 35410199 | 28.5071 | 49.7683 | 24.80876 | 10.9254 | 1.38639 | 0.752563 | 0.113893 |
| 5 | 3153000 | 3157199 | 74.42651 | 118.3222 | 90.84911 | 37.91342 | 0.71448 | 2.261419 | 0.113913 |
| 15 | 75862800 | 75890799 | 167.8634 | 240.15 | 576.9731 | 238.3539 | -1.06269 | 4.237905 | 0.113917 |
| 11 | 71450600 | 71452999 | 24.78975 | 43.39757 | 27.25749 | 12.02934 | 1.054998 | 0.695975 | 0.113932 |
| 13 | 66227800 | 66228999 | 11.56203 | 22.36163 | 15.71327 | 6.006103 | 1.064453 | -0.15425 | 0.113936 |
| 9 | 38724200 | 38724799 | 10.60292 | 20.64893 | 12.95073 | 4.736801 | 1.283538 | -0.32251 | 0.113944 |
| 2 | 157531800 | 157540399 | 40.84002 | 64.7932 | 155.7102 | 64.84756 | -0.93188 | 2.308554 | 0.113948 |
| 3 | 10979200 | 10979999 | 11.79125 | 22.70204 | 16.17866 | 6.288647 | 1.021624 | -0.09674 | 0.113954 |
| 12 | 19417000 | 19423399 | 78.15215 | 124.4764 | 81.83614 | 34.23591 | 0.940713 | 2.202747 | 0.113954 |
| 18 | 20878200 | 20880599 | 13.26081 | 22.63832 | 60.49009 | 26.51994 | -1.02926 | 0.965749 | 0.113959 |
| 10 | 85020200 | 85021999 | 9.876189 | 17.27057 | 49.90403 | 22.00339 | -1.13495 | 0.673593 | 0.113966 |
| 15 | 85493600 | 85501999 | 43.45565 | 68.49352 | 176.3673 | 73.18677 | -1.03926 | 2.474613 | 0.113995 |
| 19 | 24870000 | 24879199 | 52.69425 | 82.93214 | 183.8922 | 76.29465 | -0.8257 | 2.510254 | 0.114001 |
| 4 | 115297200 | 115297999 | 12.11969 | 23.16733 | 15.82788 | 6.280552 | 1.054979 | -0.02855 | 0.114019 |
| 1 | 41221600 | 41223399 | 11.99167 | 20.98888 | 52.09835 | 22.98843 | -0.9176 | 0.671297 | 0.114028 |
| 17 | 84853800 | 84854199 | 3.042717 | 5.942304 | 22.15302 | 7.776961 | -1.19907 | -0.53039 | 0.114046 |
| 2 | 27015200 | 27020399 | 26.62465 | 43.0763 | 98.33166 | 42.04729 | -0.82525 | 1.672247 | 0.114055 |
| 3 | 37946800 | 37947199 | 9.12229 | 17.84367 | 9.110231 | 3.050581 | 1.686172 | -0.65333 | 0.114057 |
| 4 | 91426400 | 91428199 | 20.2012 | 36.17798 | 22.54807 | 9.883507 | 1.07817 | 0.44393 | 0.114062 |
| 2 | 150205800 | 150206599 | 12.94174 | 24.74575 | 17.11681 | 6.786458 | 1.03809 | -0.01926 | 0.114064 |
| 19 | 9969200 | 9970999 | 6.729733 | 12.05785 | 41.08038 | 17.99963 | -1.355 | 0.391315 | 0.114067 |
| X | 4363000 | 4364199 | 17.49113 | 32.46284 | 18.75134 | 7.863023 | 1.229698 | 0.217253 | 0.114075 |
| 6 | 115566000 | 115570399 | 22.77031 | 37.34483 | 87.15116 | 37.5209 | -0.84227 | 1.435703 | 0.114087 |
| 7 | 27031800 | 27040799 | 50.96248 | 80.09844 | 180.5102 | 74.87833 | -0.8505 | 2.536561 | 0.114103 |
| 12 | 23606000 | 23606599 | 7.658676 | 14.96766 | 11.33236 | 3.921212 | 1.086922 | -0.55934 | 0.114105 |
| 11 | 97720200 | 97725599 | 20.52848 | 33.43834 | 93.88049 | 40.31274 | -1.11425 | 1.53143 | 0.114107 |
| 7 | 52928200 | 52929799 | 18.59985 | 33.25859 | 23.45619 | 10.295 | 0.899866 | 0.444618 | 0.114109 |
| 11 | 118585400 | 118591199 | 27.40313 | 44.05326 | 116.263 | 49.29781 | -1.04513 | 1.870631 | 0.114118 |
| 5 | 134334600 | 134344599 | 45.34681 | 71.56495 | 172.7367 | 71.70351 | -0.94471 | 2.447727 | 0.114131 |
| 9 | 69674400 | 69678999 | 24.51543 | 39.80104 | 97.82719 | 41.93826 | -0.92696 | 1.595803 | 0.114153 |
| 4 | 64265800 | 64266399 | 2.335325 | 4.565709 | 20.52473 | 7.034573 | -1.42906 | -0.63137 | 0.114167 |
| 19 | 36760000 | 36762199 | 10.42301 | 18.36131 | 46.15009 | 20.42785 | -0.93368 | 0.56072 | 0.114168 |
| 1 | 111945000 | 111947599 | 46.72532 | 75.78868 | 57.69199 | 24.7155 | 0.75848 | 1.63299 | 0.114169 |
| X | 32778600 | 32781399 | 36.57948 | 63.2259 | 22.68794 | 9.953456 | 1.856575 | 0.905381 | 0.11417 |
| 17 | 85347600 | 85351399 | 15.87535 | 26.72862 | 69.84033 | 30.52376 | -0.99953 | 1.106384 | 0.114172 |

| | | | | | | | | | |
|---|---|---|---|---|---|---|---|---|---|
| 4 | 6041200 | 6041799 | 4.626508 | 8.979261 | 28.55475 | 10.7377 | -1.06836 | -0.24458 | 0.114173 |
| 19 | 47736400 | 47738399 | 10.59021 | 18.61356 | 48.26081 | 21.37046 | -0.97974 | 0.589896 | 0.114176 |
| 12 | 22404200 | 22408199 | 62.25699 | 99.60461 | 77.38254 | 32.57697 | 0.707199 | 2.048096 | 0.114177 |
| 8 | 93236800 | 93239199 | 38.12571 | 62.91985 | 47.76511 | 20.61891 | 0.776686 | 1.367608 | 0.114181 |
| 12 | 113847800 | 113849399 | 31.62901 | 53.30747 | 39.48671 | 17.26311 | 0.81249 | 1.119749 | 0.114182 |
| 3 | 139155200 | 139157399 | 20.36684 | 36.26127 | 23.61611 | 10.40299 | 1.010426 | 0.482725 | 0.114189 |
| 1 | 57018600 | 57021999 | 16.18319 | 27.24934 | 72.1798 | 31.54708 | -1.01927 | 1.108056 | 0.114196 |
| 13 | 3123800 | 3124599 | 20.87087 | 36.81404 | 25.87513 | 11.45039 | 0.895158 | 0.570335 | 0.114224 |
| 9 | 11411200 | 11411999 | 10.09449 | 19.7336 | 8.772682 | 3.013665 | 1.848055 | -0.5554 | 0.114224 |
| 1 | 117331200 | 117334999 | 44.61459 | 74.09065 | 37.80332 | 16.3893 | 1.348452 | 1.309709 | 0.114225 |
| 4 | 129355000 | 129356399 | 5.860903 | 10.84934 | 35.76321 | 15.04981 | -1.26692 | 0.178497 | 0.114228 |
| 1 | 142308200 | 142309999 | 18.45478 | 33.77934 | 20.18723 | 8.626482 | 1.161814 | 0.303117 | 0.114229 |
| X | 148900200 | 148902599 | 63.95445 | 102.761 | 66.38007 | 28.12576 | 0.980069 | 1.917471 | 0.114242 |
| 11 | 98989800 | 98995799 | 33.97705 | 54.51675 | 121.0217 | 51.16898 | -0.7997 | 1.940639 | 0.114253 |
| 10 | 69158800 | 69164399 | 26.58396 | 42.77074 | 120.9058 | 51.32621 | -1.14258 | 1.84551 | 0.114254 |
| 15 | 79878200 | 79892999 | 76.63208 | 116.6016 | 276.6578 | 114.6327 | -0.96139 | 3.167312 | 0.114259 |
| 11 | 97420800 | 97429399 | 51.49876 | 80.46374 | 197.5488 | 81.9641 | -0.98057 | 2.648484 | 0.114327 |
| 2 | 93017600 | 93025399 | 32.59084 | 51.84863 | 148.3867 | 61.97749 | -1.17926 | 2.218108 | 0.114327 |
| 5 | 88822600 | 88824399 | 11.71924 | 20.09715 | 61.06807 | 26.78307 | -1.21279 | 0.916607 | 0.114328 |
| X | 3199000 | 3200599 | 34.22989 | 58.45471 | 31.49568 | 13.80863 | 1.270864 | 1.007877 | 0.114329 |
| 11 | 119855600 | 119858999 | 12.84687 | 21.84263 | 63.089 | 27.65029 | -1.14205 | 1.004402 | 0.114329 |
| 7 | 8450200 | 8454399 | 91.70944 | 143.9695 | 114.6045 | 47.53232 | 0.647511 | 2.572937 | 0.114337 |
| 16 | 39024600 | 39026799 | 21.77879 | 38.35589 | 25.70468 | 11.37849 | 0.962573 | 0.591481 | 0.114343 |
| 15 | 98262600 | 98264599 | 6.654576 | 12.50126 | 39.2342 | 16.12669 | -1.17066 | 0.100576 | 0.114347 |
| 2 | 115890800 | 115894999 | 26.68935 | 43.05226 | 106.2939 | 45.28712 | -0.94306 | 1.759422 | 0.114348 |
| 4 | 141252400 | 141259999 | 49.44761 | 78.28663 | 159.0936 | 66.14719 | -0.69318 | 2.372208 | 0.114349 |
| 10 | 85511400 | 85511999 | 3.457245 | 6.748407 | 23.32835 | 8.262133 | -1.10475 | -0.48929 | 0.114351 |
| 8 | 48441200 | 48445599 | 21.01555 | 34.18878 | 94.3395 | 40.49132 | -1.09066 | 1.551281 | 0.114359 |
| X | 128480400 | 128482599 | 25.18625 | 44.56901 | 21.15759 | 9.356741 | 1.459408 | 0.555616 | 0.114372 |
| 8 | 117363200 | 117369999 | 36.41287 | 58.33912 | 126.5591 | 53.39057 | -0.76916 | 1.991234 | 0.114383 |
| 17 | 38353200 | 38354399 | 17.02224 | 31.53334 | 19.39785 | 8.154928 | 1.137137 | 0.225245 | 0.114384 |
| 12 | 18916200 | 18917599 | 40.33941 | 66.24477 | 49.95372 | 21.51816 | 0.783641 | 1.435343 | 0.114402 |
| 5 | 30981800 | 30984199 | 9.067386 | 15.90101 | 47.763 | 21.10963 | -1.18983 | 0.621873 | 0.114421 |
| 17 | 31453800 | 31458599 | 39.56129 | 63.13812 | 133.2168 | 55.88701 | -0.73545 | 2.117488 | 0.114423 |
| 5 | 8436800 | 8438399 | 19.41505 | 34.95304 | 21.46453 | 9.365308 | 1.103505 | 0.403859 | 0.114434 |
| 5 | 18759200 | 18759999 | 11.92791 | 23.06336 | 15.70374 | 6.009631 | 1.10804 | -0.14592 | 0.114441 |
| 1 | 47991000 | 47992799 | 19.06086 | 34.78074 | 20.04115 | 8.598191 | 1.210171 | 0.324181 | 0.114444 |
| 5 | 129300200 | 129317399 | 115.3466 | 171.6625 | 377.1509 | 155.3974 | -0.87874 | 3.611603 | 0.114467 |
| 11 | 36144400 | 36145199 | 15.07009 | 29.26449 | 10.81974 | 4.052594 | 2.006081 | -0.19388 | 0.114473 |
| 4 | 139258800 | 139266999 | 42.09278 | 66.85585 | 148.7295 | 62.01009 | -0.81908 | 2.275361 | 0.114479 |
| 17 | 29352600 | 29389599 | 262.5622 | 363.178 | 796.2515 | 330.7283 | -0.97981 | 4.744238 | 0.114481 |
| 2 | 35856400 | 35858999 | 9.034102 | 15.98343 | 43.55621 | 19.26307 | -1.0465 | 0.501201 | 0.114486 |
| 3 | 69412000 | 69415999 | 21.05459 | 34.90364 | 81.32679 | 35.21581 | -0.83579 | 1.297638 | 0.114487 |
| 18 | 82182400 | 82183999 | 16.53774 | 30.66028 | 19.49777 | 8.188042 | 1.090752 | 0.217286 | 0.114544 |
| 13 | 41022200 | 41024599 | 9.603321 | 17.09046 | 44.11498 | 19.43988 | -0.96483 | 0.456449 | 0.114548 |
| 17 | 19701000 | 19702799 | 48.78699 | 78.87128 | 60.295 | 25.76215 | 0.747896 | 1.708405 | 0.114552 |
| 4 | 85579000 | 85580199 | 4.667096 | 9.038707 | 27.76736 | 10.54635 | -1.03323 | -0.21367 | 0.114575 |

| | | | | | | | | | |
|---|---|---|---|---|---|---|---|---|---|
| 12 | 76884200 | 76884999 | 4.667096 | 9.038707 | 27.76736 | 10.54635 | -1.03323 | -0.21367 | 0.114575 |
| 14 | 41850600 | 41853599 | 31.20864 | 53.44795 | 33.24485 | 14.57887 | 1.065219 | 0.9723 | 0.114577 |
| 8 | 66559200 | 66559599 | 17.83811 | 32.62473 | 19.49245 | 8.338575 | 1.160457 | 0.299445 | 0.114577 |
| 14 | 25558800 | 25566799 | 39.44874 | 62.82415 | 140.0982 | 58.5999 | -0.81851 | 2.187957 | 0.114584 |
| 15 | 75579800 | 75601199 | 117.9918 | 173.397 | 440.0159 | 181.2142 | -1.10355 | 3.827042 | 0.114588 |
| 6 | 133727000 | 133727599 | 14.10087 | 27.15617 | 12.19632 | 4.737032 | 1.679206 | -0.08637 | 0.114592 |
| 13 | 38514400 | 38515799 | 5.281906 | 10.12788 | 30.15268 | 11.85889 | -1.03729 | -0.08387 | 0.114599 |
| 17 | 30223000 | 30224999 | 12.65232 | 22.01558 | 52.65264 | 23.13916 | -0.86485 | 0.764249 | 0.114607 |
| 16 | 18315600 | 18321199 | 21.84438 | 35.46896 | 98.97853 | 42.43572 | -1.10929 | 1.589087 | 0.114612 |
| 2 | 150046600 | 150046999 | 9.126824 | 17.79661 | 13.26755 | 4.773527 | 1.05882 | -0.39724 | 0.114612 |
| 8 | 75780000 | 75785999 | 26.15217 | 42.30768 | 98.85036 | 42.266 | -0.85887 | 1.673302 | 0.11462 |
| 5 | 31854600 | 31855999 | 6.096627 | 11.3899 | 33.80543 | 14.02607 | -1.10081 | 0.119902 | 0.11462 |
| 16 | 5071000 | 5074399 | 18.23103 | 30.20985 | 76.67548 | 33.19187 | -0.9585 | 1.292891 | 0.114631 |
| 8 | 123791000 | 123792799 | 5.750447 | 10.59179 | 37.27745 | 15.7941 | -1.36795 | 0.209295 | 0.114633 |
| 15 | 101208800 | 101215999 | 43.70664 | 69.4206 | 148.0873 | 61.74363 | -0.75862 | 2.275963 | 0.114647 |
| 14 | 42263600 | 42265599 | 28.00211 | 48.60533 | 28.8011 | 12.63765 | 1.137949 | 0.833453 | 0.114648 |
| 13 | 37620200 | 37622599 | 10.24985 | 18.12869 | 44.5804 | 19.71726 | -0.89977 | 0.514048 | 0.114652 |
| 2 | 58164200 | 58165799 | 8.602734 | 15.58882 | 44.05835 | 19.06729 | -1.07711 | 0.337695 | 0.114701 |
| 10 | 47675600 | 47676599 | 7.496097 | 14.66101 | 10.56126 | 3.57285 | 1.184852 | -0.64234 | 0.114708 |
| 6 | 40374000 | 40376399 | 33.59221 | 56.36011 | 41.05092 | 17.91503 | 0.836596 | 1.168402 | 0.114711 |
| 17 | 63718600 | 63720199 | 30.03792 | 51.29836 | 35.51827 | 15.57231 | 0.910715 | 0.994852 | 0.114712 |
| 7 | 47317400 | 47323999 | 99.98363 | 156.9005 | 112.6251 | 46.70911 | 0.796131 | 2.58489 | 0.114718 |
| 10 | 4335800 | 4338999 | 16.38954 | 27.45478 | 69.64074 | 30.36181 | -0.95692 | 1.159912 | 0.114728 |
| 1 | 7741600 | 7742999 | 20.23181 | 35.84683 | 24.77738 | 10.9548 | 0.921454 | 0.516645 | 0.114729 |
| 2 | 162663000 | 162667599 | 26.4175 | 42.5464 | 112.8387 | 47.97269 | -1.04886 | 1.809336 | 0.11473 |
| 1 | 146071400 | 146080199 | 212.0265 | 315.5966 | 185.0631 | 76.25279 | 1.025848 | 3.614598 | 0.114731 |
| X | 16735000 | 16735599 | 10.65055 | 20.75037 | 13.20067 | 4.801853 | 1.271919 | -0.33813 | 0.114744 |
| 3 | 92924600 | 92926399 | 11.69637 | 20.44623 | 49.94113 | 22.0131 | -0.89425 | 0.688523 | 0.114749 |
| 1 | 152698400 | 152700599 | 14.75911 | 25.34241 | 59.10918 | 25.92661 | -0.833 | 0.914993 | 0.114763 |
| 14 | 41961200 | 41966199 | 67.90351 | 108.6734 | 71.28553 | 30.02388 | 0.95132 | 2.043636 | 0.114778 |
| 19 | 53011200 | 53020799 | 59.46712 | 92.82794 | 193.4749 | 80.28145 | -0.74569 | 2.668717 | 0.114802 |
| 11 | 92713200 | 92716799 | 43.02424 | 70.52541 | 50.55279 | 21.75947 | 0.855567 | 1.464631 | 0.114808 |
| 8 | 36924400 | 36925599 | 7.873598 | 14.24525 | 41.23788 | 17.88164 | -1.11286 | 0.336532 | 0.114809 |
| 7 | 41196200 | 41198599 | 64.13053 | 103.2207 | 60.94626 | 25.88767 | 1.112387 | 1.863212 | 0.114827 |
| 7 | 27880800 | 27887199 | 28.24327 | 45.30509 | 121.8375 | 51.49366 | -1.07608 | 1.942992 | 0.114838 |
| 1 | 118960000 | 118965999 | 30.7567 | 49.40854 | 116.9105 | 49.52016 | -0.88942 | 1.898153 | 0.114842 |
| 7 | 7163200 | 7163999 | 16.92104 | 31.04216 | 20.88652 | 8.897865 | 0.995116 | 0.28013 | 0.114844 |
| 7 | 29114800 | 29115399 | 2.638562 | 5.156899 | 21.0868 | 7.287305 | -1.308 | -0.59642 | 0.114855 |
| 4 | 115253800 | 115255399 | 14.61051 | 27.61289 | 17.98254 | 7.31014 | 1.093253 | 0.078271 | 0.114859 |
| 17 | 40063000 | 40065799 | 73.67212 | 117.2735 | 89.37359 | 37.35951 | 0.727143 | 2.219588 | 0.11486 |
| 1 | 184461600 | 184469399 | 45.82066 | 72.70457 | 156.2768 | 65.09207 | -0.77108 | 2.308122 | 0.114861 |
| 10 | 50776400 | 50781599 | 29.50878 | 47.37398 | 123.1355 | 52.10729 | -1.02574 | 1.917867 | 0.114862 |
| 15 | 38518400 | 38520199 | 10.54448 | 18.298 | 54.69636 | 23.99881 | -1.18512 | 0.795705 | 0.114864 |
| 4 | 128359800 | 128360599 | 2.737478 | 5.348609 | 21.59264 | 7.520483 | -1.3 | -0.56388 | 0.114881 |
| 9 | 43022600 | 43023599 | 3.944878 | 7.671606 | 25.71473 | 9.519832 | -1.12142 | -0.31774 | 0.114893 |
| 15 | 18756400 | 18757399 | 12.18591 | 23.66781 | 13.30304 | 4.975793 | 1.411118 | -0.22571 | 0.114904 |
| 19 | 23461800 | 23465199 | 18.29237 | 30.55436 | 77.89592 | 33.89937 | -0.96459 | 1.201401 | 0.114909 |

| | | | | | | | | | |
|---|---|---|---|---|---|---|---|---|---|
| 1 | 192610000 | 192611399 | 14.23444 | 26.91705 | 17.99384 | 7.30616 | 1.057176 | 0.07079 | 0.114925 |
| 17 | 40128000 | 40128999 | 12.63671 | 24.63545 | 9.696568 | 3.498319 | 1.961737 | -0.35237 | 0.114925 |
| 1 | 42732400 | 42738999 | 42.83977 | 67.80104 | 162.4728 | 67.54012 | -0.93092 | 2.376477 | 0.114931 |
| 15 | 35298800 | 35301599 | 24.41579 | 39.47267 | 100.5247 | 42.95243 | -0.98377 | 1.688263 | 0.114935 |
| 19 | 45012000 | 45017599 | 20.93602 | 33.90886 | 100.897 | 43.17829 | -1.20491 | 1.641083 | 0.114959 |
| 18 | 11021200 | 11023799 | 18.04106 | 33.00378 | 18.31451 | 7.832126 | 1.266188 | 0.296517 | 0.114961 |
| 7 | 122074200 | 122075199 | 23.07294 | 40.55244 | 26.2555 | 11.626 | 1.011228 | 0.623438 | 0.114971 |
| 1 | 86052200 | 86052799 | 8.385971 | 16.38716 | 11.59387 | 4.023609 | 1.178389 | -0.53748 | 0.114976 |
| 2 | 136800400 | 136801399 | 5.711989 | 10.89849 | 31.66205 | 12.61459 | -1.01957 | -0.03107 | 0.114987 |
| 14 | 62498200 | 62499599 | 11.62091 | 20.04242 | 56.77346 | 24.9055 | -1.10984 | 0.862068 | 0.115005 |
| 10 | 79558400 | 79577999 | 353.6376 | 476.3906 | 1048.019 | 437.1011 | -1.02327 | 5.159988 | 0.115007 |
| 4 | 154179400 | 154192599 | 71.53162 | 108.9537 | 276.6477 | 114.6641 | -1.05787 | 3.144564 | 0.115013 |
| 17 | 66608600 | 66610199 | 24.40817 | 42.90281 | 25.03955 | 11.08837 | 1.160019 | 0.6279 | 0.115027 |
| 6 | 95458400 | 95461799 | 18.91015 | 31.61545 | 72.75308 | 31.67973 | -0.81722 | 1.190376 | 0.115042 |
| 10 | 62173200 | 62179799 | 31.25092 | 50.16485 | 118.408 | 50.09764 | -0.88718 | 1.923051 | 0.115048 |
| 7 | 97437200 | 97439599 | 7.917548 | 14.21186 | 41.60517 | 18.20169 | -1.1373 | 0.386303 | 0.115049 |
| 12 | 22357600 | 22359999 | 44.61631 | 72.83362 | 47.86468 | 20.56973 | 0.976903 | 1.519467 | 0.115058 |
| 14 | 54827200 | 54837599 | 64.42159 | 99.99717 | 212.793 | 88.36801 | -0.78203 | 2.770083 | 0.115062 |
| 18 | 76365400 | 76367399 | 27.96661 | 48.00447 | 34.58948 | 15.17131 | 0.854509 | 0.941077 | 0.115075 |
| 4 | 155685600 | 155686799 | 4.673713 | 9.030484 | 28.15455 | 10.81031 | -1.06925 | -0.17469 | 0.115085 |
| 7 | 102090800 | 102091599 | 4.673713 | 9.030484 | 28.15455 | 10.81031 | -1.06925 | -0.17469 | 0.115085 |
| X | 91151000 | 91151799 | 26.55489 | 46.57237 | 24.3902 | 10.78051 | 1.31555 | 0.673509 | 0.115098 |
| 9 | 14371600 | 14373399 | 7.123717 | 13.31636 | 39.03718 | 16.18108 | -1.08332 | 0.134982 | 0.115101 |
| 4 | 63142200 | 63145799 | 15.44038 | 25.60338 | 78.56556 | 34.02639 | -1.23102 | 1.278601 | 0.115108 |
| 13 | 42251400 | 42252799 | 17.81019 | 32.0712 | 22.3226 | 9.736885 | 0.924431 | 0.393061 | 0.115111 |
| 4 | 15930800 | 15931999 | 16.09437 | 30.14053 | 18.31749 | 7.569084 | 1.173151 | 0.149814 | 0.11512 |
| 7 | 30404800 | 30408399 | 16.79853 | 28.29805 | 66.51875 | 29.07677 | -0.84737 | 1.10336 | 0.115137 |
| 11 | 101164800 | 101169999 | 22.53087 | 36.44258 | 102.1268 | 43.66099 | -1.12063 | 1.672219 | 0.115137 |
| 7 | 10349800 | 10350199 | 14.2362 | 26.61343 | 19.1313 | 7.929137 | 0.929437 | 0.146188 | 0.115152 |
| X | 4102600 | 4105999 | 28.93032 | 50.75596 | 21.70774 | 9.597662 | 1.605962 | 0.678321 | 0.115155 |
| 12 | 44256000 | 44257399 | 8.047398 | 14.12941 | 47.73983 | 21.12101 | -1.35833 | 0.59453 | 0.115171 |
| 12 | 20402400 | 20405799 | 72.04767 | 114.6076 | 86.47862 | 36.11447 | 0.740386 | 2.240032 | 0.115177 |
| 2 | 170892400 | 170894199 | 17.38337 | 31.94651 | 20.25094 | 8.606469 | 1.082835 | 0.27255 | 0.11521 |
| 12 | 56578000 | 56585599 | 68.23273 | 105.3468 | 224.1041 | 93.08832 | -0.78755 | 2.870413 | 0.115211 |
| 11 | 101795400 | 101803799 | 39.43702 | 62.83368 | 139.401 | 58.34451 | -0.81041 | 2.172827 | 0.115216 |
| 4 | 147728200 | 147728999 | 9.990878 | 19.5191 | 10.77356 | 3.757772 | 1.525031 | -0.49852 | 0.115218 |
| 11 | 97593800 | 97595599 | 5.000563 | 9.310347 | 36.40681 | 15.19302 | -1.50162 | 0.137463 | 0.115224 |
| 19 | 29155600 | 29157199 | 24.10894 | 42.02654 | 30.21998 | 13.29583 | 0.860808 | 0.759103 | 0.115231 |
| 12 | 113312800 | 113314799 | 21.32422 | 38.39223 | 19.00365 | 8.291023 | 1.41202 | 0.411914 | 0.115238 |
| 4 | 136497200 | 136502999 | 32.36259 | 51.96268 | 117.0479 | 49.54155 | -0.81944 | 1.915968 | 0.115241 |
| 7 | 90220200 | 90221799 | 4.320836 | 8.39455 | 34.00472 | 12.68829 | -1.40406 | -0.26264 | 0.11526 |
| 12 | 23397800 | 23399399 | 28.25895 | 48.58725 | 30.86839 | 13.54013 | 1.035576 | 0.921516 | 0.115268 |
| 13 | 3534800 | 3535999 | 46.67372 | 75.6828 | 56.99865 | 24.41319 | 0.773524 | 1.639951 | 0.115272 |
| 13 | 12744200 | 12746599 | 27.14647 | 46.7667 | 33.39783 | 14.65053 | 0.868176 | 0.902515 | 0.115274 |
| 13 | 44882800 | 44885799 | 12.42704 | 21.40509 | 57.38924 | 25.17464 | -1.03177 | 0.874984 | 0.115278 |
| 13 | 45888200 | 45889999 | 8.446296 | 15.32451 | 41.88109 | 18.09668 | -1.02734 | 0.325132 | 0.11528 |
| 10 | 122324400 | 122325199 | 5.238834 | 10.11498 | 29.95896 | 11.54262 | -1.00153 | -0.152 | 0.115293 |

| | | | | | | | | | |
|---|---|---|---|---|---|---|---|---|---|
| 19 | 8309200 | 8311599 | 40.09502 | 66.38528 | 45.00225 | 19.4699 | 0.939742 | 1.334053 | 0.1153 |
| 14 | 15266800 | 15268199 | 4.808164 | 9.277058 | 28.63701 | 11.06329 | -1.06347 | -0.15004 | 0.115303 |
| 7 | 143626600 | 143629999 | 17.91324 | 30.11514 | 68.84454 | 30.07505 | -0.80761 | 1.126114 | 0.115311 |
| 17 | 46739600 | 46742599 | 12.40401 | 21.0591 | 64.10061 | 28.08621 | -1.21752 | 1.017226 | 0.115325 |
| 2 | 98802600 | 98802999 | 9.815494 | 19.06878 | 13.85292 | 5.170804 | 1.048674 | -0.25993 | 0.115327 |
| 4 | 111989000 | 111989999 | 20.13204 | 35.60769 | 24.56371 | 10.86413 | 0.923532 | 0.535064 | 0.115328 |
| 14 | 14853600 | 14854599 | 13.00553 | 24.95633 | 16.61106 | 6.519019 | 1.10679 | -0.05183 | 0.115333 |
| 5 | 115080000 | 115092399 | 57.87188 | 89.26729 | 235.1666 | 97.68393 | -1.09662 | 2.884921 | 0.115347 |
| 1 | 84815600 | 84818599 | 17.00855 | 28.12331 | 78.56519 | 33.95922 | -1.09654 | 1.317125 | 0.11535 |
| 10 | 52160400 | 52161999 | 11.22611 | 21.63109 | 15.86523 | 6.152819 | 0.984054 | -0.11028 | 0.115356 |
| 8 | 66371600 | 66373399 | 20.35349 | 36.43871 | 22.73181 | 9.966813 | 1.076836 | 0.448453 | 0.115365 |
| 6 | 121072600 | 121077599 | 25.527 | 41.29121 | 100.566 | 42.99564 | -0.9188 | 1.674922 | 0.115369 |
| 8 | 21924200 | 21925199 | 13.51259 | 25.83527 | 15.20558 | 6.030034 | 1.267514 | -0.01921 | 0.11538 |
| 4 | 139980200 | 139984199 | 23.21667 | 37.74497 | 92.28664 | 39.60078 | -0.9174 | 1.564549 | 0.115392 |
| 14 | 91217200 | 91218399 | 16.44306 | 30.57327 | 18.68117 | 7.813773 | 1.151326 | 0.197188 | 0.1154 |
| 15 | 70359400 | 70360999 | 16.44306 | 30.57327 | 18.68117 | 7.813773 | 1.151326 | 0.197188 | 0.1154 |
| 18 | 7954000 | 7955199 | 22.01841 | 38.80427 | 25.70755 | 11.37821 | 0.979524 | 0.586274 | 0.115405 |
| 10 | 80644200 | 80646599 | 11.25391 | 19.29299 | 59.91464 | 26.27653 | -1.24385 | 0.917651 | 0.115412 |
| 2 | 27564200 | 27579599 | 62.65448 | 95.84312 | 264.3078 | 109.6629 | -1.17188 | 3.053551 | 0.115413 |
| 4 | 61302600 | 61303799 | 12.37223 | 24.02794 | 13.29821 | 4.976901 | 1.43254 | -0.22138 | 0.115428 |
| 6 | 3369400 | 3370599 | 26.56934 | 45.68369 | 32.66457 | 14.32802 | 0.865987 | 0.917229 | 0.11543 |
| 7 | 139341400 | 139344399 | 11.83891 | 20.36047 | 58.95566 | 25.86055 | -1.14272 | 0.888416 | 0.115431 |
| 1 | 187326800 | 187327799 | 5.236636 | 10.11874 | 31.94118 | 12.26096 | -1.0879 | -0.16049 | 0.115434 |
| 16 | 84232200 | 84233599 | 22.00104 | 39.47869 | 20.14575 | 8.815897 | 1.366024 | 0.439678 | 0.115444 |
| 10 | 77508800 | 77510599 | 8.891285 | 15.68448 | 44.60804 | 19.73987 | -1.10932 | 0.536277 | 0.115444 |
| 2 | 178068800 | 178071599 | 12.02695 | 20.61796 | 61.63987 | 27.03313 | -1.18963 | 0.9212 | 0.115446 |
| 4 | 113346800 | 113348799 | 32.57348 | 54.79957 | 38.84244 | 16.97208 | 0.875599 | 1.135507 | 0.115452 |
| 3 | 17072800 | 17074399 | 15.01359 | 28.6512 | 14.41831 | 5.742228 | 1.485688 | 0.00751 | 0.115462 |
| 10 | 46687800 | 46688799 | 8.151511 | 14.78313 | 39.14636 | 16.92437 | -0.98216 | 0.32155 | 0.115479 |
| 6 | 146303200 | 146306599 | 21.07939 | 34.39839 | 88.13259 | 37.86974 | -0.98065 | 1.504952 | 0.115505 |
| 7 | 10694400 | 10697799 | 74.08204 | 118.0348 | 85.32019 | 35.71064 | 0.804693 | 2.189094 | 0.115509 |
| 14 | 95332400 | 95335199 | 71.91915 | 114.5856 | 79.23422 | 33.16215 | 0.868499 | 2.186398 | 0.115514 |
| 10 | 16359800 | 16360999 | 15.56539 | 29.38431 | 17.27215 | 7.038158 | 1.236677 | 0.092662 | 0.115518 |
| X | 32766200 | 32769599 | 56.73457 | 92.52791 | 41.22487 | 17.71014 | 1.534906 | 1.558956 | 0.115519 |
| 3 | 85600200 | 85603599 | 9.530743 | 16.7836 | 51.49844 | 22.79683 | -1.22033 | 0.565986 | 0.115527 |
| 5 | 8417000 | 8419399 | 17.82992 | 29.76345 | 71.34363 | 31.03575 | -0.87568 | 1.203878 | 0.115555 |
| 15 | 101554600 | 101555799 | 3.135148 | 6.106723 | 25.23316 | 9.207468 | -1.39867 | -0.38199 | 0.115557 |
| 18 | 32043200 | 32044399 | 7.693138 | 13.99407 | 38.67209 | 16.65627 | -1.03955 | 0.299414 | 0.115587 |
| 3 | 17708000 | 17710999 | 35.98034 | 60.45012 | 36.49499 | 15.9399 | 1.106085 | 1.155794 | 0.115631 |
| 9 | 65965600 | 65967999 | 8.473945 | 14.96133 | 47.21414 | 20.88947 | -1.25837 | 0.525289 | 0.115632 |
| 10 | 129830800 | 129832599 | 24.69238 | 42.89446 | 31.3492 | 13.7626 | 0.837092 | 0.81123 | 0.115632 |
| 16 | 50249400 | 50250799 | 5.682531 | 10.64102 | 35.54189 | 14.68852 | -1.26564 | 0.102641 | 0.115655 |
| 12 | 42182400 | 42183199 | 17.8479 | 32.02008 | 22.46889 | 9.834125 | 0.909659 | 0.416326 | 0.115661 |
| 16 | 94367000 | 94371999 | 21.39765 | 34.70211 | 100.9181 | 43.22919 | -1.17025 | 1.613964 | 0.115667 |
| 19 | 23993200 | 23995399 | 9.811381 | 17.30824 | 46.1554 | 20.42444 | -1.01768 | 0.540945 | 0.115671 |
| 1 | 117015400 | 117019599 | 69.1596 | 110.1694 | 80.06906 | 33.50335 | 0.796518 | 2.188337 | 0.115688 |
| 8 | 45927200 | 45927999 | 11.3954 | 21.98934 | 16.3093 | 6.298594 | 0.973513 | -0.12038 | 0.115692 |

| | | | | | | | | | |
|---|---|---|---|---|---|---|---|---|---|
| 5 | 108650200 | 108651399 | 6.037369 | 11.46278 | 32.101 | 12.92324 | -0.98112 | 0.005449 | 0.115695 |
| 15 | 56278200 | 56278999 | 6.037369 | 11.46278 | 32.101 | 12.92324 | -0.98112 | 0.005449 | 0.115695 |
| 3 | 37776200 | 37787399 | 50.82337 | 79.84871 | 194.9191 | 80.85127 | -0.96613 | 2.545392 | 0.115706 |
| 15 | 97514000 | 97518599 | 26.9077 | 43.63861 | 95.47251 | 40.89683 | -0.7609 | 1.62232 | 0.115709 |
| 8 | 92511800 | 92520799 | 51.72219 | 81.85744 | 171.4548 | 71.27328 | -0.73722 | 2.384263 | 0.115724 |
| 4 | 155571200 | 155576999 | 22.38287 | 36.19586 | 103.2455 | 44.13033 | -1.14639 | 1.677344 | 0.11575 |
| 14 | 84020600 | 84022199 | 17.11601 | 32.17962 | 15.86617 | 6.512809 | 1.478945 | 0.128384 | 0.115782 |
| 16 | 92491600 | 92496999 | 32.23444 | 51.68121 | 120.9628 | 51.0847 | -0.87715 | 1.96331 | 0.115827 |
| 17 | 15883800 | 15884599 | 17.52307 | 32.84233 | 16.42554 | 6.778131 | 1.452935 | 0.152625 | 0.11583 |
| 2 | 157938200 | 157944999 | 29.70554 | 47.59106 | 125.2042 | 52.81585 | -1.04669 | 1.9853 | 0.115838 |
| 1 | 53303800 | 53305199 | 23.41793 | 40.94836 | 28.26389 | 12.46123 | 0.91989 | 0.709718 | 0.115842 |
| 17 | 35301000 | 35304599 | 13.60546 | 23.17326 | 63.79857 | 27.96525 | -1.07305 | 0.98983 | 0.115843 |
| 1 | 98389000 | 98390399 | 16.17523 | 29.92842 | 20.03928 | 8.438325 | 1.01412 | 0.226238 | 0.115849 |
| 12 | 87915600 | 87916799 | 14.13091 | 27.13977 | 15.32321 | 5.99838 | 1.344971 | -0.0518 | 0.115853 |
| 11 | 95689200 | 95691199 | 12.36275 | 21.45711 | 54.22365 | 23.79154 | -0.9442 | 0.803223 | 0.115858 |
| 5 | 120350600 | 120364399 | 96.00285 | 144.7797 | 312.334 | 129.0593 | -0.83513 | 3.355012 | 0.11588 |
| 8 | 104365000 | 104366999 | 15.76466 | 26.83115 | 62.83664 | 27.54197 | -0.84102 | 1.004043 | 0.115908 |
| 3 | 27737000 | 27745199 | 46.66623 | 73.96276 | 155.6764 | 64.76878 | -0.7424 | 2.342628 | 0.115909 |
| 17 | 29606000 | 29609799 | 18.31046 | 30.53652 | 71.92512 | 31.26991 | -0.85052 | 1.216139 | 0.115915 |
| 12 | 113057400 | 113086999 | 155.4594 | 224.6241 | 522.1386 | 215.3852 | -1.00079 | 4.090116 | 0.115917 |
| 15 | 79398200 | 79401399 | 21.64455 | 35.54639 | 82.82837 | 35.68017 | -0.83915 | 1.414777 | 0.115928 |
| X | 25280200 | 25280799 | 16.34844 | 30.78458 | 15.66538 | 6.413395 | 1.438518 | 0.112996 | 0.11593 |
| 6 | 105208800 | 105209799 | 5.786321 | 10.9146 | 33.53706 | 13.68922 | -1.13158 | 0.05175 | 0.115937 |
| 12 | 32924400 | 32931999 | 45.96658 | 72.98611 | 152.3585 | 63.50356 | -0.72795 | 2.287482 | 0.115939 |
| 5 | 143309800 | 143312399 | 12.23726 | 21.16457 | 55.27544 | 24.25041 | -0.99273 | 0.834461 | 0.115955 |
| 12 | 111503800 | 111506599 | 11.23677 | 18.97421 | 67.64027 | 29.58843 | -1.44456 | 1.065691 | 0.11598 |
| 5 | 92433000 | 92433999 | 5.403536 | 10.33666 | 31.1585 | 12.33784 | -1.06375 | -0.05505 | 0.115986 |
| 11 | 108819200 | 108831799 | 71.63803 | 109.5883 | 257.7796 | 106.955 | -0.94272 | 3.054969 | 0.115988 |
| 13 | 53618600 | 53620599 | 13.72165 | 23.45352 | 61.69854 | 27.05239 | -1.00687 | 0.955384 | 0.116004 |
| 1 | 143088000 | 143088999 | 21.20327 | 37.51285 | 20.30368 | 8.979489 | 1.27154 | 0.530355 | 0.116008 |
| 7 | 83308000 | 83310999 | 19.12372 | 31.81431 | 73.5798 | 31.93862 | -0.82438 | 1.24752 | 0.116033 |
| 5 | 119437600 | 119439999 | 8.707921 | 15.49675 | 43.06767 | 18.97859 | -1.07038 | 0.449414 | 0.116043 |
| 6 | 85639000 | 85640199 | 13.95173 | 26.74879 | 15.89278 | 6.252682 | 1.265526 | -0.03646 | 0.116043 |
| 1 | 59709200 | 59710599 | 9.511341 | 16.75433 | 45.11799 | 19.97123 | -1.03216 | 0.555999 | 0.116056 |
| 18 | 19479400 | 19480999 | 15.14713 | 28.40852 | 18.34341 | 7.560865 | 1.088985 | 0.131694 | 0.116074 |
| 3 | 85907400 | 85910199 | 18.8223 | 31.28141 | 75.8834 | 32.91571 | -0.8932 | 1.258436 | 0.116074 |
| 7 | 22430000 | 22430599 | 16.11046 | 29.73623 | 20.4898 | 8.656604 | 0.969536 | 0.239524 | 0.116074 |
| 14 | 73562200 | 73563999 | 11.92742 | 20.65153 | 55.82866 | 24.49395 | -1.04197 | 0.822823 | 0.116076 |
| 15 | 37344400 | 37348999 | 28.76336 | 46.38628 | 108.7625 | 46.32402 | -0.8693 | 1.770047 | 0.116079 |
| 8 | 48482200 | 48483799 | 21.1259 | 37.4113 | 24.03559 | 10.62795 | 1.02586 | 0.525832 | 0.116092 |
| 14 | 59696600 | 59697399 | 16.79904 | 31.28297 | 18.80473 | 7.845102 | 1.178121 | 0.192068 | 0.116097 |
| 18 | 27061000 | 27062599 | 16.79904 | 31.28297 | 18.80473 | 7.845102 | 1.178121 | 0.192068 | 0.116097 |
| 11 | 24930200 | 24930799 | 3.641525 | 7.107791 | 24.02854 | 8.516772 | -1.07551 | -0.48416 | 0.116105 |
| 17 | 84458800 | 84476799 | 98.86812 | 148.7101 | 325.939 | 134.5854 | -0.86135 | 3.407545 | 0.116107 |
| 16 | 71853000 | 71854999 | 21.22044 | 38.20098 | 19.59731 | 8.551258 | 1.360878 | 0.413309 | 0.116115 |
| 17 | 48232400 | 48237199 | 21.72728 | 35.14193 | 101.1992 | 43.26361 | -1.15972 | 1.671701 | 0.116121 |
| 13 | 100658800 | 100660399 | 8.520337 | 15.22696 | 41.03652 | 18.01939 | -1.02223 | 0.419562 | 0.116133 |

| | | | | | | | | | |
|---|---|---|---|---|---|---|---|---|---|
| 7 | 107604400 | 107607799 | 32.15681 | 54.50549 | 37.11521 | 16.25427 | 0.933537 | 1.069974 | 0.116139 |
| 7 | 129288800 | 129295799 | 32.9133 | 52.69326 | 127.8854 | 53.88944 | -0.93191 | 2.011158 | 0.116139 |
| 7 | 57483600 | 57484799 | 12.63413 | 24.25393 | 16.62153 | 6.515202 | 1.066534 | -0.0598 | 0.116146 |
| 9 | 29849600 | 29855399 | 30.34612 | 48.9623 | 112.015 | 47.73892 | -0.8332 | 1.759285 | 0.116167 |
| 18 | 23944000 | 23947999 | 13.93893 | 24.05245 | 59.74775 | 26.21067 | -0.92202 | 0.866087 | 0.116187 |
| 1 | 102527200 | 102528599 | 12.87487 | 24.85091 | 15.54867 | 5.999053 | 1.217705 | -0.10944 | 0.116191 |
| 10 | 84585800 | 84599599 | 100.3662 | 151.0126 | 326.4819 | 134.8213 | -0.84119 | 3.401305 | 0.116196 |
| 7 | 9486600 | 9487199 | 11.67821 | 22.52803 | 16.22281 | 6.27081 | 1.014286 | -0.11504 | 0.116214 |
| 8 | 3726200 | 3727799 | 8.729568 | 15.48003 | 43.55035 | 19.24902 | -1.09089 | 0.473614 | 0.116218 |
| 7 | 37474000 | 37476999 | 13.61218 | 23.54535 | 62.03243 | 27.21494 | -1.00585 | 0.845042 | 0.116223 |
| 9 | 120000000 | 120005799 | 47.17686 | 74.72582 | 156.1987 | 64.95865 | -0.7332 | 2.359853 | 0.116291 |
| 5 | 134960000 | 134983599 | 133.634 | 196.5227 | 431.59 | 177.7363 | -0.89423 | 3.817049 | 0.116292 |
| 17 | 57185600 | 57223199 | 304.2978 | 417.7782 | 852.4846 | 354.5048 | -0.88714 | 4.860327 | 0.116295 |
| 2 | 115375200 | 115375599 | 2.524981 | 4.933384 | 22.144 | 7.714069 | -1.44861 | -0.56187 | 0.116329 |
| 17 | 18473800 | 18475599 | 24.06677 | 42.5297 | 22.78178 | 10.07778 | 1.285548 | 0.565224 | 0.116329 |
| 5 | 15108600 | 15112599 | 118.3935 | 183.0166 | 139.1276 | 57.79042 | 0.696822 | 2.858009 | 0.116333 |
| 7 | 108104000 | 108105799 | 15.92017 | 30.18342 | 16.15233 | 6.522332 | 1.382019 | 0.064773 | 0.11635 |
| 6 | 124278200 | 124281399 | 27.18815 | 43.8174 | 109.0992 | 46.42837 | -0.95676 | 1.787641 | 0.116356 |
| 8 | 60947400 | 60948799 | 10.58373 | 18.61815 | 48.26542 | 21.3702 | -0.97909 | 0.579802 | 0.116366 |
| 12 | 51904000 | 51905799 | 8.070271 | 14.45952 | 41.05992 | 17.99089 | -1.09508 | 0.398789 | 0.116377 |
| 11 | 59472000 | 59472799 | 11.3415 | 21.81169 | 15.83288 | 6.174055 | 0.991403 | -0.0921 | 0.116379 |
| 9 | 101980600 | 101983199 | 12.95471 | 22.53251 | 54.85535 | 24.10098 | -0.89069 | 0.772809 | 0.116448 |
| 17 | 68112600 | 68113199 | 4.060625 | 7.914238 | 25.57688 | 9.25832 | -1.03983 | -0.39786 | 0.11645 |
| 18 | 82569800 | 82574399 | 24.7768 | 39.9267 | 108.0085 | 45.95702 | -1.07613 | 1.78899 | 0.116457 |
| 4 | 148332400 | 148364199 | 265.3138 | 369.8575 | 730.3768 | 302.9966 | -0.81762 | 4.625475 | 0.116457 |
| X | 114781600 | 114782999 | 20.73119 | 37.56872 | 20.04175 | 8.672906 | 1.313084 | 0.375739 | 0.116471 |
| 7 | 41441200 | 41442599 | 21.16828 | 37.58427 | 23.48946 | 10.3713 | 1.067228 | 0.504358 | 0.116474 |
| 18 | 19441000 | 19442199 | 16.33111 | 29.99487 | 20.59616 | 8.760262 | 0.967836 | 0.269043 | 0.116477 |
| 11 | 103731000 | 103733799 | 12.83079 | 22.24483 | 54.2556 | 23.80474 | -0.89385 | 0.813691 | 0.11648 |
| 15 | 97872600 | 97906399 | 249.0868 | 346.8972 | 746.4172 | 309.698 | -0.9428 | 4.639825 | 0.116483 |
| 12 | 23757600 | 23757999 | 8.145968 | 15.93448 | 9.817669 | 3.277701 | 1.423256 | -0.67745 | 0.116486 |
| X | 152791400 | 152791799 | 8.145968 | 15.93448 | 9.817669 | 3.277701 | 1.423256 | -0.67745 | 0.116486 |
| 4 | 154259400 | 154266399 | 39.99739 | 63.53687 | 149.3601 | 62.28196 | -0.89827 | 2.269463 | 0.116494 |
| 7 | 112967800 | 112969399 | 9.280704 | 16.24243 | 50.41601 | 22.24438 | -1.23757 | 0.657209 | 0.1165 |
| 7 | 21086600 | 21088199 | 18.16088 | 34.34533 | 14.23777 | 5.779833 | 1.73992 | 0.097512 | 0.116511 |
| 6 | 88978800 | 88979599 | 2.845117 | 5.53956 | 25.08316 | 9.19625 | -1.53356 | -0.3675 | 0.116512 |
| 10 | 38463600 | 38464999 | 12.19776 | 23.64015 | 14.47354 | 5.47887 | 1.273827 | -0.18242 | 0.116514 |
| 4 | 121315400 | 121316199 | 22.06536 | 38.70246 | 27.61421 | 12.20657 | 0.872031 | 0.652002 | 0.116515 |
| 7 | 14945800 | 14946199 | 8.14661 | 15.93352 | 8.500088 | 2.789866 | 1.646785 | -0.73976 | 0.116516 |
| 18 | 65713400 | 65715399 | 6.54307 | 12.15258 | 35.30016 | 14.78771 | -1.08109 | 0.166176 | 0.116527 |
| 6 | 116525200 | 116529599 | 57.09386 | 89.36679 | 186.1291 | 77.21169 | -0.74157 | 2.61847 | 0.116528 |
| 6 | 95591400 | 95593999 | 11.52054 | 20.21369 | 48.2925 | 21.35322 | -0.86239 | 0.624538 | 0.11653 |
| 4 | 86363200 | 86365399 | 12.2037 | 21.19482 | 55.90386 | 24.53878 | -1.00593 | 0.791741 | 0.116532 |
| 15 | 43340200 | 43340999 | 6.751252 | 12.41855 | 36.19985 | 15.36548 | -1.10055 | 0.223402 | 0.116536 |
| 16 | 19403800 | 19406599 | 46.5062 | 75.87611 | 51.76346 | 22.24067 | 0.922613 | 1.53442 | 0.11655 |
| 7 | 29682600 | 29683799 | 7.284195 | 13.11831 | 39.56391 | 17.2549 | -1.17598 | 0.359736 | 0.116556 |
| 4 | 110276000 | 110276799 | 18.40873 | 32.95402 | 21.64247 | 9.489916 | 1.003011 | 0.432685 | 0.116564 |

| | | | | | | | | | |
|---|---|---|---|---|---|---|---|---|---|
| 8 | 122534600 | 122536799 | 7.949326 | 14.14058 | 42.04805 | 18.53608 | -1.16697 | 0.447646 | 0.11657 |
| 13 | 60359400 | 60360799 | 25.74243 | 44.60497 | 32.40063 | 14.21559 | 0.845384 | 0.844995 | 0.116573 |
| 16 | 91386200 | 91390199 | 19.26303 | 31.67131 | 81.85951 | 35.2798 | -0.98737 | 1.395014 | 0.116591 |
| 2 | 128720600 | 128723999 | 18.59949 | 31.08256 | 73.28772 | 31.90378 | -0.85218 | 1.194213 | 0.116602 |
| 15 | 45391000 | 45392599 | 26.61111 | 45.68896 | 32.12827 | 14.09207 | 0.889442 | 0.928442 | 0.11661 |
| 5 | 129654800 | 129663999 | 49.70696 | 77.71618 | 198.8665 | 82.50516 | -1.03944 | 2.635606 | 0.116613 |
| 1 | 194326200 | 194327199 | 4.241456 | 8.235604 | 26.75088 | 10.02798 | -1.09419 | -0.26696 | 0.116615 |
| 8 | 80406800 | 80407599 | 4.241456 | 8.235604 | 26.75088 | 10.02798 | -1.09419 | -0.26696 | 0.116615 |
| 6 | 140779200 | 140779599 | 3.040109 | 5.945782 | 21.51465 | 7.291175 | -1.11059 | -0.65926 | 0.116619 |
| 15 | 86503800 | 86506799 | 19.26881 | 32.09242 | 73.1294 | 31.76704 | -0.80302 | 1.234105 | 0.116623 |
| 3 | 113468200 | 113469799 | 17.31073 | 32.53596 | 15.86084 | 6.51392 | 1.494668 | 0.132046 | 0.116631 |
| 3 | 61579800 | 61582999 | 34.36167 | 58.36062 | 34.27205 | 15.01834 | 1.14627 | 1.058837 | 0.116633 |
| 9 | 106698400 | 106700799 | 10.93793 | 19.11641 | 50.46424 | 22.23974 | -1.00565 | 0.688336 | 0.116638 |
| 5 | 119272600 | 119280999 | 43.48555 | 68.82377 | 164.3109 | 68.30455 | -0.92558 | 2.37662 | 0.116647 |
| 17 | 84338200 | 84363199 | 194.9819 | 280.6042 | 536.1793 | 221.2959 | -0.72393 | 4.152124 | 0.116648 |
| 4 | 19189800 | 19190399 | 4.840553 | 9.339021 | 29.27 | 11.30967 | -1.08627 | -0.14987 | 0.116648 |
| 2 | 174932000 | 174933799 | 19.75222 | 35.25569 | 23.73421 | 10.43283 | 0.965074 | 0.46387 | 0.116683 |
| 8 | 120211000 | 120216599 | 27.8992 | 44.76126 | 119.7768 | 50.63668 | -1.06853 | 1.937041 | 0.11669 |
| 1 | 35324000 | 35325399 | 14.79987 | 27.963 | 17.9769 | 7.312132 | 1.11106 | 0.082006 | 0.116691 |
| 12 | 23505400 | 23506599 | 16.54714 | 30.37573 | 20.89611 | 8.894178 | 0.964178 | 0.273376 | 0.116724 |
| X | 13290200 | 13290799 | 11.18072 | 21.86868 | 7.188994 | 2.420396 | 2.301995 | -0.60927 | 0.116751 |
| 15 | 78592800 | 78605199 | 77.7699 | 118.1324 | 286.124 | 118.4948 | -0.99311 | 3.202767 | 0.116764 |
| 11 | 61384000 | 61385599 | 6.71389 | 12.14512 | 41.21368 | 17.87472 | -1.34001 | 0.329823 | 0.116768 |
| 8 | 121249800 | 121256199 | 26.61353 | 42.75345 | 117.2466 | 49.66359 | -1.10158 | 1.893432 | 0.116787 |
| 3 | 68580000 | 68581399 | 18.3285 | 33.72572 | 19.00374 | 8.062645 | 1.252968 | 0.27023 | 0.116794 |
| 4 | 54215600 | 54216399 | 12.18183 | 23.67745 | 12.70234 | 4.722716 | 1.484947 | -0.2479 | 0.116794 |
| 3 | 30568200 | 30571199 | 15.03929 | 25.09744 | 72.63653 | 31.59213 | -1.1463 | 1.200356 | 0.1168 |
| 18 | 84702000 | 84704199 | 7.852136 | 13.91632 | 44.60151 | 19.71854 | -1.27784 | 0.475182 | 0.116803 |
| 17 | 87250800 | 87252799 | 10.13137 | 17.80393 | 46.16294 | 20.43843 | -0.97925 | 0.58968 | 0.116817 |
| 12 | 18836600 | 18848199 | 314.8335 | 452.3521 | 331.8456 | 136.9966 | 0.654268 | 4.180227 | 0.116824 |
| 6 | 25606200 | 25606799 | 8.062542 | 15.74521 | 11.76117 | 4.131576 | 1.086237 | -0.50072 | 0.116829 |
| 10 | 79854800 | 79869999 | 71.93385 | 109.4525 | 281.8924 | 116.8015 | -1.07959 | 3.166332 | 0.116833 |
| 7 | 18954000 | 18957399 | 15.2726 | 25.65348 | 67.92799 | 29.66599 | -1.01852 | 1.12567 | 0.116835 |
| 19 | 12942800 | 12944799 | 53.59441 | 86.54682 | 62.30599 | 26.5832 | 0.832532 | 1.749583 | 0.116839 |
| 2 | 58990600 | 58992599 | 8.101233 | 14.43373 | 42.0509 | 18.51251 | -1.13656 | 0.437311 | 0.11686 |
| 16 | 12416600 | 12419599 | 21.24447 | 37.39749 | 25.78618 | 11.41563 | 0.921455 | 0.593051 | 0.116882 |
| 3 | 12994200 | 12996399 | 23.72141 | 41.44658 | 29.03087 | 12.79121 | 0.898624 | 0.722833 | 0.116895 |
| 8 | 117454200 | 117464199 | 81.78151 | 125.5112 | 248.329 | 103.0959 | -0.68977 | 2.992251 | 0.116896 |
| 12 | 20574800 | 20579199 | 107.8238 | 167.8263 | 112.533 | 46.71323 | 0.885221 | 2.730513 | 0.116903 |
| 6 | 30302800 | 30304799 | 8.225118 | 14.70722 | 41.04213 | 18.01373 | -1.07177 | 0.413374 | 0.116903 |
| 8 | 122032000 | 122075799 | 255.2181 | 353.9281 | 775.8882 | 322.1387 | -0.9759 | 4.704551 | 0.116911 |
| 6 | 52690600 | 52691599 | 6.312847 | 11.62097 | 36.20444 | 15.35152 | -1.19439 | 0.215604 | 0.116928 |
| 1 | 92104400 | 92108799 | 20.07169 | 32.98399 | 84.43854 | 36.38344 | -0.97405 | 1.403398 | 0.116962 |
| 3 | 87003000 | 87007599 | 24.37285 | 39.52585 | 96.79149 | 41.46046 | -0.92307 | 1.618344 | 0.116982 |
| 8 | 105105600 | 105108399 | 10.3386 | 18.07213 | 51.51903 | 22.70797 | -1.11592 | 0.683385 | 0.116992 |
| 6 | 90730200 | 90734799 | 18.92386 | 31.19563 | 80.39686 | 34.68802 | -0.98167 | 1.359356 | 0.116994 |
| 7 | 87198200 | 87199999 | 15.5418 | 29.05983 | 18.87683 | 7.821076 | 1.074734 | 0.155731 | 0.117 |

| | | | | | | | | | |
|---|---|---|---|---|---|---|---|---|---|
| X | 77151200 | 77152999 | 19.50802 | 34.89705 | 23.373 | 10.25488 | 0.974348 | 0.448762 | 0.117001 |
| 11 | 72188800 | 72194399 | 31.23078 | 50.17491 | 115.1609 | 48.78583 | -0.84537 | 1.895518 | 0.117006 |
| 10 | 82306800 | 82308399 | 33.16662 | 55.58937 | 41.14567 | 17.94484 | 0.813581 | 1.178602 | 0.117021 |
| 15 | 19452400 | 19453399 | 19.99735 | 35.44001 | 24.48161 | 10.82351 | 0.922605 | 0.51293 | 0.117021 |
| 14 | 7824600 | 7825999 | 5.236073 | 10.11971 | 30.70136 | 11.77381 | -1.02964 | -0.16655 | 0.11707 |
| 15 | 74530800 | 74539999 | 53.73526 | 84.39953 | 178.8408 | 74.17936 | -0.76252 | 2.551868 | 0.117076 |
| 15 | 10823800 | 10824999 | 13.37077 | 25.31978 | 17.78059 | 7.197938 | 0.990961 | 0.053489 | 0.117079 |
| 11 | 61065000 | 61066399 | 9.247615 | 16.21743 | 47.76397 | 21.11047 | -1.16194 | 0.620553 | 0.117086 |
| 7 | 125494200 | 125501399 | 45.98269 | 72.96252 | 154.9816 | 64.55335 | -0.75397 | 2.307757 | 0.117102 |
| 8 | 85439200 | 85441599 | 9.871322 | 17.27302 | 49.37124 | 21.78015 | -1.11933 | 0.662222 | 0.117117 |
| 11 | 75082400 | 75087399 | 25.05621 | 40.50413 | 100.6953 | 43.01995 | -0.9493 | 1.691935 | 0.117123 |
| 12 | 19509600 | 19515399 | 164.126 | 247.5632 | 183.7978 | 75.95168 | 0.702834 | 3.357038 | 0.117159 |
| 4 | 73114800 | 73116799 | 15.20105 | 28.64217 | 17.83252 | 7.292783 | 1.150087 | 0.102665 | 0.117168 |
| 14 | 48888800 | 48890199 | 12.0956 | 20.85783 | 56.27409 | 24.68631 | -1.04013 | 0.864013 | 0.117168 |
| 10 | 80380800 | 80398399 | 68.62268 | 104.3873 | 283.2864 | 117.37 | -1.15527 | 3.171194 | 0.117194 |
| 4 | 43010000 | 43011799 | 4.629681 | 8.825827 | 33.30741 | 13.29435 | -1.39594 | -0.03494 | 0.117194 |
| 7 | 140921600 | 140923999 | 9.398263 | 16.50033 | 47.74633 | 21.12291 | -1.1368 | 0.601565 | 0.117201 |
| 3 | 153367200 | 153370399 | 11.10471 | 19.37663 | 52.62105 | 23.16585 | -1.04673 | 0.715029 | 0.117202 |
| 1 | 110128600 | 110129199 | 10.37172 | 20.14395 | 14.14489 | 5.291272 | 1.093599 | -0.24533 | 0.117204 |
| 2 | 93805000 | 93809799 | 18.16469 | 30.25725 | 72.37666 | 31.44252 | -0.87267 | 1.229712 | 0.117208 |
| 12 | 37427600 | 37428799 | 18.93346 | 33.7163 | 23.97151 | 10.55769 | 0.884979 | 0.473077 | 0.117246 |
| 8 | 124217800 | 124219599 | 5.822958 | 10.88235 | 34.36055 | 14.24814 | -1.18882 | 0.115318 | 0.117249 |
| 9 | 121995200 | 121997799 | 12.6835 | 21.6044 | 62.59308 | 27.43699 | -1.14605 | 0.985673 | 0.11726 |
| 11 | 101933600 | 101945599 | 68.00722 | 104.3861 | 243.9455 | 101.278 | -0.92949 | 2.985291 | 0.117267 |
| 1 | 184376800 | 184379199 | 8.905429 | 16.14294 | 41.85472 | 18.10575 | -0.95288 | 0.334886 | 0.117276 |
| 16 | 17644400 | 17660599 | 86.68324 | 131.2134 | 297.3414 | 123.0089 | -0.90141 | 3.276745 | 0.117299 |
| 5 | 119384600 | 119385399 | 4.425665 | 8.542347 | 29.23997 | 11.2816 | -1.2097 | -0.15932 | 0.117305 |
| 4 | 92646400 | 92647999 | 20.79478 | 36.72189 | 25.24039 | 11.16694 | 0.927658 | 0.555773 | 0.117315 |
| 1 | 156500200 | 156502799 | 10.58001 | 18.31659 | 57.3756 | 25.17256 | -1.25326 | 0.819711 | 0.117321 |
| 3 | 69137200 | 69138399 | 11.26572 | 21.84544 | 15.20421 | 5.741143 | 1.094501 | -0.19893 | 0.117325 |
| 17 | 76331400 | 76332599 | 9.247998 | 16.6451 | 42.72325 | 18.6466 | -0.94673 | 0.381803 | 0.117332 |
| 18 | 12987800 | 12989399 | 9.011806 | 16.02499 | 44.10496 | 19.44853 | -1.05734 | 0.457604 | 0.117336 |
| 17 | 94129000 | 94130799 | 17.73465 | 33.14664 | 16.98393 | 7.041607 | 1.413237 | 0.172923 | 0.117341 |
| 13 | 65638000 | 65639799 | 31.72138 | 53.77571 | 37.7008 | 16.51149 | 0.891724 | 1.068248 | 0.117341 |
| 16 | 94627200 | 94629599 | 15.18541 | 25.7401 | 66.03901 | 28.92151 | -0.97269 | 1.041086 | 0.117342 |
| 19 | 38854400 | 38855599 | 5.587266 | 10.66749 | 31.13572 | 12.38775 | -1.02493 | -0.04049 | 0.117349 |
| 18 | 9957400 | 9958799 | 10.91208 | 19.13151 | 47.77412 | 21.1104 | -0.92571 | 0.633074 | 0.117378 |
| 17 | 45044600 | 45050799 | 34.50187 | 54.93639 | 147.2791 | 61.58858 | -1.08391 | 2.190192 | 0.117379 |
| 14 | 51662200 | 51664399 | 16.64126 | 31.29365 | 17.20116 | 7.058278 | 1.324703 | 0.126835 | 0.117389 |
| X | 3652600 | 3653799 | 16.64126 | 31.29365 | 17.20116 | 7.058278 | 1.324703 | 0.126835 | 0.117389 |
| 9 | 91232200 | 91238199 | 44.79646 | 71.23496 | 148.8996 | 62.15512 | -0.72773 | 2.241566 | 0.117397 |
| 11 | 103022800 | 103023799 | 5.347228 | 10.13601 | 33.71742 | 13.62061 | -1.23169 | 0.012853 | 0.117437 |
| 7 | 11186400 | 11204599 | 492.0417 | 679.9283 | 476.7368 | 198.0467 | 0.663009 | 4.763198 | 0.117443 |
| 3 | 79425200 | 79426799 | 6.429888 | 11.94449 | 36.43631 | 15.25996 | -1.15183 | 0.162725 | 0.117478 |
| 11 | 23458600 | 23465799 | 47.13976 | 74.71741 | 161.0147 | 66.9935 | -0.7763 | 2.341528 | 0.11748 |
| 12 | 4023000 | 4032599 | 45.47046 | 71.82772 | 165.4652 | 68.71785 | -0.87635 | 2.42468 | 0.117494 |
| X | 32806000 | 32807599 | 23.44105 | 41.89722 | 20.33111 | 8.926617 | 1.435445 | 0.47496 | 0.117495 |

| | | | | | | | | | |
|---|---|---|---|---|---|---|---|---|---|
| 17 | 25645800 | 25657799 | 68.65772 | 105.3769 | 243.4394 | 101.0669 | -0.91296 | 2.98696 | 0.117501 |
| 14 | 55328400 | 55330199 | 10.90329 | 19.14003 | 47.22474 | 20.88972 | -0.90847 | 0.611481 | 0.11751 |
| 7 | 31129800 | 31132599 | 71.80717 | 114.4714 | 85.16528 | 35.67217 | 0.764003 | 2.171771 | 0.117531 |
| 9 | 60634600 | 60635799 | 30.96318 | 52.64425 | 31.66213 | 13.87669 | 1.112147 | 1.03358 | 0.117535 |
| 14 | 42355400 | 42356999 | 14.30017 | 27.56976 | 14.16735 | 5.48554 | 1.493501 | -0.08718 | 0.117535 |
| 11 | 38194200 | 38194999 | 11.03069 | 21.3181 | 15.76972 | 6.048672 | 0.986248 | -0.14904 | 0.117551 |
| 15 | 8581800 | 8583599 | 33.54598 | 56.01444 | 42.24747 | 18.38176 | 0.786627 | 1.218731 | 0.117553 |
| 10 | 99097400 | 99100799 | 17.49674 | 28.90665 | 79.78333 | 34.4661 | -1.07937 | 1.328331 | 0.117578 |
| 17 | 33576800 | 33578199 | 11.06956 | 19.39271 | 48.84528 | 21.56973 | -0.93822 | 0.647372 | 0.117588 |
| 6 | 126079000 | 126083799 | 26.92262 | 43.62093 | 96.46024 | 41.29072 | -0.7776 | 1.642118 | 0.117601 |
| 19 | 53742800 | 53747199 | 28.23027 | 45.655 | 99.95235 | 42.72657 | -0.76578 | 1.683596 | 0.117607 |
| X | 16210200 | 16212599 | 30.72561 | 53.65083 | 23.36611 | 10.29122 | 1.580401 | 0.761798 | 0.117617 |
| 14 | 88964800 | 88966399 | 8.79037 | 15.81474 | 41.63837 | 18.18108 | -0.98373 | 0.377237 | 0.117628 |
| 4 | 140995200 | 141006799 | 64.37254 | 99.30376 | 229.9527 | 95.51815 | -0.91072 | 2.884955 | 0.117635 |
| 7 | 47073000 | 47077599 | 54.30863 | 87.6144 | 63.50486 | 27.06052 | 0.821128 | 1.777947 | 0.117639 |
| 7 | 51042200 | 51043199 | 26.68772 | 45.90744 | 34.04308 | 14.9329 | 0.813684 | 0.914863 | 0.117641 |
| 4 | 9387000 | 9388599 | 11.06573 | 19.39898 | 47.77606 | 21.10951 | -0.90584 | 0.635837 | 0.117657 |
| X | 32984800 | 32987399 | 22.85232 | 40.95363 | 20.24546 | 8.869125 | 1.4108 | 0.454413 | 0.117676 |
| 11 | 49468000 | 49469999 | 10.315 | 18.04549 | 48.85095 | 21.54671 | -1.04132 | 0.668437 | 0.11768 |
| 7 | 42549400 | 42556799 | 138.0765 | 211.4102 | 156.897 | 65.1113 | 0.720968 | 3.048492 | 0.117698 |
| 9 | 22040000 | 22047199 | 31.02865 | 49.58068 | 131.573 | 55.29259 | -1.06347 | 2.071271 | 0.117698 |
| 7 | 97093600 | 97094999 | 8.464507 | 15.30597 | 40.15281 | 17.42305 | -0.9724 | 0.342479 | 0.117701 |
| 1 | 178985400 | 178986799 | 12.99976 | 22.48971 | 55.32384 | 24.27186 | -0.90687 | 0.834743 | 0.117711 |
| 5 | 36347200 | 36355599 | 40.12781 | 63.81746 | 146.2167 | 61.04244 | -0.85967 | 2.234384 | 0.117722 |
| 13 | 29764600 | 29767399 | 15.37467 | 25.97867 | 69.35628 | 30.3452 | -1.03018 | 1.072189 | 0.117729 |
| 9 | 21202000 | 21203999 | 8.962714 | 15.71851 | 48.80801 | 21.57272 | -1.23787 | 0.618934 | 0.117739 |
| 12 | 44035600 | 44036199 | 10.33398 | 20.0829 | 14.78752 | 5.504454 | 1.032954 | -0.25956 | 0.117742 |
| 9 | 20936000 | 20939399 | 13.97623 | 23.65363 | 66.41206 | 29.07117 | -1.10216 | 1.051593 | 0.117743 |
| 5 | 140589400 | 140604999 | 83.1399 | 125.9643 | 293.4418 | 121.4305 | -0.94009 | 3.25746 | 0.117753 |
| 3 | 94626200 | 94627199 | 4.658724 | 9.049122 | 28.14014 | 10.51763 | -1.02902 | -0.2696 | 0.11777 |
| 1 | 85583600 | 85586799 | 41.75252 | 68.61858 | 49.46528 | 21.31475 | 0.848805 | 1.428137 | 0.117777 |
| X | 157936200 | 157937599 | 16.84782 | 31.13801 | 19.93857 | 8.408716 | 1.076507 | 0.237602 | 0.117777 |
| 6 | 39045200 | 39049799 | 24.19024 | 39.26412 | 93.8768 | 40.23796 | -0.88739 | 1.59942 | 0.117821 |
| 14 | 32172800 | 32179199 | 36.14599 | 57.77984 | 128.9193 | 54.20687 | -0.81332 | 2.063106 | 0.117833 |
| 1 | 54809600 | 54811199 | 26.2856 | 45.89216 | 27.99241 | 12.32781 | 1.097305 | 0.747074 | 0.117848 |
| 9 | 18875000 | 18875999 | 14.3404 | 26.80486 | 19.04518 | 7.895125 | 0.945734 | 0.147225 | 0.117849 |
| 5 | 142120000 | 142155599 | 304.6024 | 421.7878 | 787.8953 | 327.2007 | -0.74725 | 4.728495 | 0.117867 |
| 7 | 101484400 | 101488399 | 16.04846 | 26.64673 | 78.23971 | 33.91605 | -1.16759 | 1.263736 | 0.117882 |
| 10 | 87469600 | 87470599 | 13.15137 | 25.41062 | 15.07286 | 5.787708 | 1.300759 | -0.12281 | 0.117901 |
| X | 33173000 | 33173999 | 13.9004 | 27.02195 | 11.54198 | 4.285953 | 1.812089 | -0.23403 | 0.117903 |
| 5 | 117965600 | 117966399 | 11.47254 | 22.13947 | 15.89092 | 6.135999 | 1.02107 | -0.12053 | 0.11792 |
| 18 | 4054200 | 4057199 | 31.31283 | 53.18699 | 37.21107 | 16.30668 | 0.89482 | 1.050685 | 0.117924 |
| 7 | 31139400 | 31146399 | 173.0157 | 260.7708 | 197.4923 | 81.5924 | 0.673216 | 3.375437 | 0.117936 |
| 11 | 101782200 | 101794599 | 64.01072 | 99.19119 | 217.4241 | 90.31185 | -0.82662 | 2.799885 | 0.11796 |
| 7 | 20813000 | 20815799 | 75.85993 | 120.6865 | 85.15366 | 35.56714 | 0.837265 | 2.240894 | 0.11796 |
| 3 | 3260800 | 3268199 | 123.2045 | 191.1457 | 122.4481 | 50.85369 | 0.947455 | 2.793954 | 0.117972 |
| 9 | 73098000 | 73100999 | 17.59398 | 29.58759 | 68.84147 | 30.07654 | -0.83293 | 1.121898 | 0.117972 |

| | | | | | | | | | |
|---|---|---|---|---|---|---|---|---|---|
| 13 | 59800400 | 59814799 | 87.58982 | 132.847 | 290.5088 | 120.2561 | -0.84767 | 3.236235 | 0.117973 |
| X | 31604000 | 31606199 | 35.35597 | 60.57095 | 28.04521 | 12.29901 | 1.488892 | 0.980441 | 0.117979 |
| 9 | 23898200 | 23899399 | 15.08257 | 28.15271 | 19.51261 | 8.108068 | 0.978752 | 0.164141 | 0.117984 |
| 3 | 159644800 | 159645399 | 135.488 | 211.6895 | 64.67527 | 26.83417 | 2.021313 | 2.672738 | 0.117986 |
| 8 | 95160200 | 95162399 | 10.47658 | 18.31589 | 49.38811 | 21.77131 | -1.03581 | 0.680467 | 0.117994 |
| 19 | 29892400 | 29893799 | 13.15902 | 25.40019 | 14.19921 | 5.477581 | 1.377571 | -0.11169 | 0.118003 |
| X | 11600000 | 11602999 | 9.603239 | 17.10427 | 45.19709 | 19.90303 | -0.99739 | 0.456335 | 0.118014 |
| 12 | 110198600 | 110206799 | 41.39083 | 65.52718 | 164.2668 | 68.2961 | -0.99552 | 2.368413 | 0.118019 |
| 8 | 69832200 | 69832599 | 15.92041 | 29.75028 | 18.86643 | 7.82454 | 1.108112 | 0.162912 | 0.11802 |
| 2 | 144582800 | 144584599 | 6.567837 | 12.20308 | 35.3349 | 14.79484 | -1.07677 | 0.161163 | 0.118031 |
| 2 | 121125200 | 121138799 | 63.90627 | 97.69742 | 273.4755 | 113.4504 | -1.19415 | 3.066733 | 0.118038 |
| 8 | 123586600 | 123594199 | 95.39217 | 142.7159 | 349.9423 | 144.3132 | -1.03001 | 3.511584 | 0.118051 |
| 14 | 54985600 | 54987199 | 11.37478 | 19.92172 | 49.92085 | 22.03937 | -0.93102 | 0.654701 | 0.118055 |
| 1 | 179989000 | 179991199 | 9.022479 | 16.01699 | 44.08015 | 19.4652 | -1.05864 | 0.468627 | 0.118068 |
| 10 | 62413600 | 62415199 | 7.142391 | 13.28418 | 35.37212 | 14.78749 | -0.95499 | 0.161477 | 0.118077 |
| 19 | 27933400 | 27934799 | 9.306675 | 16.55084 | 42.52919 | 18.75224 | -0.95865 | 0.457184 | 0.118077 |
| 7 | 30146200 | 30147799 | 25.36757 | 43.82506 | 31.58221 | 13.85526 | 0.855982 | 0.868093 | 0.118087 |
| 1 | 185498000 | 185500999 | 25.4426 | 41.34684 | 92.74034 | 39.7863 | -0.79393 | 1.576007 | 0.118091 |
| 19 | 52998200 | 53008799 | 56.1961 | 87.38435 | 205.2051 | 85.19434 | -0.9219 | 2.735772 | 0.118097 |
| 11 | 55019400 | 55022399 | 13.81425 | 23.76722 | 57.44916 | 25.20004 | -0.88349 | 0.891245 | 0.118098 |
| X | 4012000 | 4013599 | 22.4004 | 39.92783 | 22.02214 | 9.691416 | 1.249854 | 0.48761 | 0.118099 |
| 6 | 83677800 | 83692999 | 111.3858 | 166.1435 | 369.1987 | 152.1654 | -0.89212 | 3.570099 | 0.118108 |
| 9 | 26012000 | 26012999 | 19.61506 | 35.00531 | 20.8738 | 9.176711 | 1.138675 | 0.457467 | 0.118111 |
| 7 | 22992800 | 23001799 | 217.5377 | 322.3999 | 243.874 | 100.4407 | 0.653046 | 3.693515 | 0.118114 |
| 5 | 116268200 | 116277199 | 47.97016 | 75.59415 | 174.4365 | 72.38367 | -0.88159 | 2.480831 | 0.118123 |
| 2 | 156681400 | 156687199 | 26.45334 | 42.55276 | 117.7435 | 49.97044 | -1.1121 | 1.849432 | 0.118134 |
| 15 | 79295000 | 79298799 | 19.5116 | 31.89969 | 87.66223 | 37.69335 | -1.07945 | 1.472473 | 0.118145 |
| 9 | 19412400 | 19413599 | 13.77414 | 26.36292 | 16.45595 | 6.506293 | 1.188503 | -0.02133 | 0.118145 |
| 10 | 81649400 | 81650999 | 13.77414 | 26.36292 | 16.45595 | 6.506293 | 1.188503 | -0.02133 | 0.118145 |
| 10 | 95506400 | 95506999 | 3.534577 | 6.880023 | 25.1752 | 9.251058 | -1.23565 | -0.35121 | 0.118164 |
| 12 | 50454000 | 50455999 | 23.74219 | 42.62572 | 17.14792 | 7.500658 | 1.706214 | 0.442351 | 0.118168 |
| 10 | 11523800 | 11526199 | 16.72671 | 28.38897 | 65.07018 | 28.50827 | -0.81071 | 1.035034 | 0.118182 |
| 15 | 100930600 | 100965599 | 225.9225 | 318.0611 | 660.4274 | 273.5575 | -0.87569 | 4.476722 | 0.118195 |
| 10 | 129763200 | 129763999 | 14.60024 | 27.59869 | 17.66842 | 7.179574 | 1.118765 | 0.077111 | 0.118202 |
| 19 | 46120200 | 46126399 | 33.06509 | 52.9381 | 124.5516 | 52.48739 | -0.88711 | 2.010131 | 0.118211 |
| 15 | 79823400 | 79858999 | 246.5513 | 344.7542 | 706.1161 | 292.7928 | -0.86581 | 4.578834 | 0.118223 |
| 10 | 74756600 | 74778399 | 161.9881 | 234.2988 | 510.2619 | 210.4573 | -0.90535 | 4.075125 | 0.118231 |
| 17 | 21398000 | 21400999 | 15.73637 | 26.91364 | 61.82504 | 27.1092 | -0.81212 | 0.958944 | 0.118237 |
| 19 | 49171800 | 49173199 | 15.45762 | 28.83596 | 19.50288 | 8.112032 | 1.012775 | 0.171279 | 0.11826 |
| 4 | 36949600 | 36953999 | 34.30357 | 55.02406 | 121.5642 | 51.37482 | -0.79334 | 1.950614 | 0.118268 |
| 10 | 77728600 | 77739599 | 60.13055 | 93.06681 | 218.1151 | 90.59947 | -0.92432 | 2.82125 | 0.118276 |
| 12 | 77625800 | 77626999 | 11.51007 | 22.24245 | 14.46838 | 5.551998 | 1.168375 | -0.14715 | 0.11829 |
| 8 | 81919000 | 81921599 | 24.24648 | 42.88719 | 22.20696 | 9.821677 | 1.334614 | 0.555604 | 0.118301 |
| 16 | 49978800 | 49981599 | 10.88707 | 19.17032 | 46.14864 | 20.42901 | -0.87201 | 0.569332 | 0.118321 |
| 3 | 94744600 | 94746199 | 4.926517 | 9.358944 | 35.55672 | 14.29741 | -1.4158 | -0.00305 | 0.118326 |
| 11 | 71566800 | 71571199 | 20.58881 | 34.10202 | 78.84911 | 34.12258 | -0.82474 | 1.304638 | 0.118333 |
| 14 | 63504600 | 63509799 | 37.24572 | 59.61094 | 125.1501 | 52.71364 | -0.72373 | 2.025415 | 0.118333 |

| | | | | | | | | | |
|---|---|---|---|---|---|---|---|---|---|
| 12 | 21786400 | 21787199 | 15.73728 | 29.06631 | 20.49875 | 8.652738 | 0.937086 | 0.232634 | 0.118336 |
| 11 | 76619000 | 76621599 | 11.06429 | 19.40662 | 47.76668 | 21.1147 | -0.9049 | 0.628134 | 0.118341 |
| 7 | 31548000 | 31550199 | 20.32584 | 37.0349 | 18.72874 | 8.049807 | 1.395117 | 0.338711 | 0.118347 |
| 7 | 28866800 | 28878599 | 58.97221 | 91.26182 | 217.7184 | 90.43473 | -0.95005 | 2.82359 | 0.11835 |
| 7 | 14471800 | 14473399 | 35.10597 | 59.06063 | 38.78265 | 16.94601 | 0.985557 | 1.146231 | 0.118353 |
| 16 | 70734200 | 70737799 | 43.7228 | 71.94034 | 46.73143 | 20.14665 | 0.99918 | 1.417133 | 0.118363 |
| 10 | 103707600 | 103708599 | 11.78492 | 22.99048 | 9.75405 | 3.487896 | 1.865771 | -0.39369 | 0.11837 |
| 4 | 46687800 | 46698999 | 70.68503 | 108.7351 | 234.5025 | 97.38693 | -0.81135 | 2.941912 | 0.118375 |
| 4 | 132662000 | 132671599 | 46.1402 | 72.82306 | 167.3978 | 69.49023 | -0.87426 | 2.446201 | 0.118377 |
| 17 | 20618600 | 20621199 | 82.18938 | 129.9698 | 103.2307 | 42.89498 | 0.658828 | 2.412258 | 0.118383 |
| 11 | 114790600 | 114794799 | 26.41948 | 42.87556 | 95.68408 | 41.00772 | -0.78858 | 1.606803 | 0.118385 |
| 5 | 144758600 | 144762199 | 15.13145 | 25.79368 | 61.15749 | 26.80935 | -0.85826 | 0.986173 | 0.118402 |
| X | 80688600 | 80692199 | 39.10666 | 65.19003 | 40.58866 | 17.64132 | 1.062479 | 1.254773 | 0.118425 |
| 12 | 20240200 | 20242799 | 61.45293 | 98.58756 | 72.04405 | 30.45393 | 0.799815 | 1.960174 | 0.118452 |
| 4 | 135672400 | 135674599 | 8.960291 | 16.06177 | 41.05366 | 17.98096 | -0.94419 | 0.401443 | 0.118471 |
| 7 | 130120800 | 130124399 | 22.67336 | 37.04444 | 90.79508 | 39.03105 | -0.91588 | 1.488934 | 0.118471 |
| 5 | 21310800 | 21313999 | 41.56183 | 68.12702 | 51.10523 | 21.99709 | 0.790097 | 1.459947 | 0.118474 |
| 10 | 20462000 | 20463799 | 17.62046 | 31.86057 | 22.46465 | 9.748478 | 0.910854 | 0.369918 | 0.118506 |
| 4 | 83112200 | 83114799 | 13.32773 | 22.993 | 59.14917 | 25.94792 | -0.97223 | 0.86467 | 0.118526 |
| X | 33127000 | 33130199 | 46.3453 | 76.4637 | 41.24531 | 17.80203 | 1.267996 | 1.39055 | 0.118539 |
| 1 | 110981400 | 110982399 | 14.32297 | 27.50277 | 15.31773 | 5.999913 | 1.363738 | -0.04781 | 0.118548 |
| 2 | 177559200 | 177560199 | 14.32297 | 27.50277 | 15.31773 | 5.999913 | 1.363738 | -0.04781 | 0.118548 |
| 6 | 29624000 | 29624799 | 3.714115 | 7.222181 | 28.18745 | 10.44295 | -1.3405 | -0.31354 | 0.11855 |
| 14 | 52226000 | 52228399 | 11.53057 | 19.83695 | 58.39275 | 25.6139 | -1.16612 | 0.883733 | 0.118562 |
| 5 | 116218400 | 116221599 | 19.8225 | 32.78294 | 77.9731 | 33.70812 | -0.86548 | 1.320534 | 0.118568 |
| 12 | 22169400 | 22172199 | 40.40023 | 66.68874 | 46.68865 | 20.1574 | 0.893204 | 1.371762 | 0.118573 |
| 9 | 80190800 | 80192199 | 14.59912 | 27.66526 | 16.75081 | 6.771012 | 1.203822 | 0.056229 | 0.118585 |
| 6 | 100409200 | 100415599 | 37.65383 | 60.03305 | 137.4483 | 57.58091 | -0.85467 | 2.148801 | 0.118586 |
| X | 122901400 | 122906799 | 72.24516 | 115.3646 | 77.96618 | 32.7342 | 0.90498 | 2.11915 | 0.118589 |
| 7 | 108623600 | 108627799 | 25.2222 | 40.78329 | 102.2138 | 43.68369 | -0.96057 | 1.684114 | 0.118592 |
| 7 | 5358000 | 5358999 | 22.69242 | 39.69235 | 28.73652 | 12.67302 | 0.851273 | 0.700861 | 0.118599 |
| 17 | 3484400 | 3486599 | 8.254579 | 15.12556 | 38.8648 | 16.58319 | -0.92628 | 0.262174 | 0.118606 |
| 18 | 58746800 | 58748799 | 9.881027 | 17.29565 | 51.50116 | 22.72603 | -1.17835 | 0.658383 | 0.118608 |
| 4 | 150365600 | 150366999 | 5.416657 | 10.08269 | 35.32532 | 14.74757 | -1.3452 | 0.140453 | 0.11862 |
| 2 | 118997600 | 119006799 | 45.41884 | 71.46403 | 176.4119 | 73.1888 | -0.98041 | 2.511525 | 0.118632 |
| 7 | 17847000 | 17848399 | 31.23298 | 53.38657 | 34.36342 | 15.06706 | 1.015698 | 0.990831 | 0.118649 |
| 16 | 89403800 | 89407599 | 29.20349 | 47.18683 | 102.9379 | 43.95222 | -0.76219 | 1.712234 | 0.118666 |
| 7 | 19781800 | 19788399 | 78.34542 | 124.6575 | 87.62503 | 36.60621 | 0.842876 | 2.237905 | 0.118693 |
| 17 | 20050000 | 20050999 | 21.62706 | 38.01604 | 26.57368 | 11.76765 | 0.901398 | 0.614244 | 0.118699 |
| 12 | 87490000 | 87490799 | 4.126602 | 7.99542 | 27.73829 | 10.5139 | -1.20356 | -0.22764 | 0.118708 |
| 1 | 81839600 | 81841399 | 18.43223 | 32.87291 | 22.4228 | 9.863133 | 0.94537 | 0.455994 | 0.118731 |
| 18 | 18573000 | 18576199 | 53.2028 | 85.68739 | 66.99152 | 28.4821 | 0.709411 | 1.827398 | 0.118749 |
| X | 31283200 | 31283999 | 9.291495 | 18.14024 | 12.02099 | 4.24288 | 1.250187 | -0.46505 | 0.118753 |
| 11 | 64391000 | 64392399 | 7.758323 | 14.36486 | 38.07864 | 16.01943 | -0.956 | 0.19679 | 0.118753 |
| 15 | 82029600 | 82032399 | 9.059802 | 15.74944 | 56.80242 | 24.94793 | -1.45402 | 0.765743 | 0.118756 |
| 2 | 118644400 | 118646199 | 4.997701 | 9.438082 | 32.34868 | 13.1715 | -1.28238 | 0.038237 | 0.118766 |
| 1 | 175701600 | 175703599 | 9.57242 | 17.1564 | 43.17808 | 18.91388 | -0.9223 | 0.411328 | 0.118769 |

| | | | | | | | | | |
|---|---|---|---|---|---|---|---|---|---|
| 5 | 112398800 | 112400599 | 8.038245 | 14.48769 | 40.08514 | 17.46504 | -1.05234 | 0.35914 | 0.118771 |
| 14 | 47805200 | 47806999 | 8.038245 | 14.48769 | 40.08514 | 17.46504 | -1.05234 | 0.35914 | 0.118771 |
| 18 | 11205000 | 11206599 | 6.521189 | 12.24806 | 35.74012 | 14.69581 | -1.06711 | 0.090391 | 0.118772 |
| 5 | 12793000 | 12794599 | 16.8213 | 31.70391 | 16.63811 | 6.801116 | 1.395184 | 0.113305 | 0.118774 |
| X | 32915200 | 32916199 | 23.80723 | 41.50691 | 30.27925 | 13.32318 | 0.840181 | 0.755823 | 0.118783 |
| 12 | 23640000 | 23650199 | 289.0842 | 417.6753 | 302.2566 | 124.6836 | 0.681863 | 4.096695 | 0.118806 |
| 15 | 96986800 | 96987999 | 5.154965 | 9.867945 | 30.63946 | 12.10837 | -1.10398 | -0.06851 | 0.118808 |
| 1 | 165591200 | 165593799 | 12.2939 | 21.13752 | 58.43589 | 25.63236 | -1.07636 | 0.892124 | 0.118808 |
| 15 | 100713400 | 100716999 | 22.88736 | 37.25514 | 91.89913 | 39.45209 | -0.92889 | 1.545227 | 0.118815 |
| 9 | 101610800 | 101612799 | 8.921635 | 16.11815 | 40.13239 | 17.43368 | -0.89858 | 0.352128 | 0.118817 |
| 18 | 36506600 | 36511199 | 22.24794 | 36.62733 | 81.41738 | 35.10888 | -0.76994 | 1.384782 | 0.118821 |
| 17 | 27660000 | 27679999 | 110.7806 | 163.8424 | 412.6073 | 169.9226 | -1.08384 | 3.72046 | 0.118841 |
| 2 | 145083600 | 145084599 | 9.073558 | 16.37909 | 41.72964 | 18.14627 | -0.93278 | 0.359898 | 0.118841 |
| 1 | 136067800 | 136077999 | 42.51565 | 67.29246 | 161.6894 | 67.21667 | -0.93472 | 2.37463 | 0.118845 |
| 8 | 122821800 | 122837599 | 89.58517 | 135.2864 | 305.5062 | 126.2926 | -0.89932 | 3.325895 | 0.118847 |
| 8 | 14529000 | 14531599 | 14.20794 | 24.21454 | 62.20307 | 27.26726 | -0.9734 | 0.984935 | 0.118847 |
| 12 | 91708200 | 91710999 | 14.431 | 24.89029 | 58.59908 | 25.70641 | -0.84492 | 0.873628 | 0.118857 |
| 5 | 113939800 | 113958399 | 105.0858 | 157.185 | 351.0588 | 144.7676 | -0.89571 | 3.516496 | 0.118864 |
| 16 | 20030000 | 20031599 | 7.315837 | 13.53392 | 36.91619 | 15.5486 | -0.99807 | 0.19483 | 0.118867 |
| 2 | 166997200 | 167011799 | 95.66042 | 144.2817 | 310.9736 | 128.5024 | -0.83365 | 3.352307 | 0.118881 |
| 11 | 115062000 | 115065399 | 11.56248 | 19.81775 | 59.93778 | 26.28623 | -1.20603 | 0.92059 | 0.118884 |
| 14 | 32180400 | 32192999 | 94.24968 | 142.9899 | 283.3781 | 117.3146 | -0.70545 | 3.231615 | 0.118885 |
| 3 | 101357400 | 101359399 | 10.12647 | 17.55193 | 57.91493 | 25.41009 | -1.32747 | 0.807347 | 0.118893 |
| 12 | 25288400 | 25292799 | 21.89711 | 36.1445 | 80.51004 | 34.75875 | -0.77152 | 1.350031 | 0.118896 |
| 19 | 14844000 | 14844399 | 8.779741 | 17.14261 | 12.35202 | 4.353173 | 1.134791 | -0.47462 | 0.118903 |
| 8 | 74970400 | 74971599 | 15.16935 | 29.22238 | 13.50905 | 5.242527 | 1.641396 | -0.07266 | 0.118905 |
| 4 | 7786800 | 7787399 | 12.51199 | 24.1134 | 16.11706 | 6.24704 | 1.117207 | -0.0975 | 0.118912 |
| 5 | 95160600 | 95161599 | 12.17938 | 23.65361 | 14.04227 | 5.254997 | 1.333587 | -0.22091 | 0.118918 |
| 7 | 92247200 | 92249199 | 20.38539 | 36.22374 | 24.15793 | 10.65914 | 0.974881 | 0.495854 | 0.118922 |
| 11 | 117612000 | 117614399 | 11.00537 | 19.07284 | 54.7009 | 23.99984 | -1.1262 | 0.809932 | 0.118923 |
| 11 | 84671000 | 84672399 | 7.643988 | 14.0358 | 36.71963 | 15.62372 | -0.94871 | 0.241373 | 0.118934 |
| 4 | 155564200 | 155565999 | 7.205693 | 13.2369 | 37.81621 | 16.08056 | -1.07435 | 0.236079 | 0.118945 |
| 14 | 37977800 | 37979799 | 22.41047 | 40.43445 | 19.28244 | 8.390146 | 1.468618 | 0.410428 | 0.118946 |
| 6 | 135143400 | 135146599 | 11.12161 | 19.36869 | 53.15208 | 23.37126 | -1.06202 | 0.744043 | 0.118975 |
| 6 | 105243600 | 105244999 | 17.59042 | 32.6425 | 18.65089 | 7.822406 | 1.244901 | 0.218248 | 0.118979 |
| 12 | 20115600 | 20126799 | 289.9636 | 418.8426 | 308.5507 | 127.2841 | 0.65582 | 4.100482 | 0.118981 |
| 11 | 102346400 | 102352599 | 29.54282 | 47.3071 | 125.21 | 52.77884 | -1.05618 | 2.0014 | 0.119004 |
| 7 | 3050000 | 3056599 | 104.3159 | 163.8833 | 98.97209 | 41.05278 | 1.046253 | 2.563985 | 0.119019 |
| 19 | 6897400 | 6900799 | 15.50968 | 26.24566 | 64.88225 | 28.40046 | -0.9194 | 1.056771 | 0.119021 |
| 9 | 44790400 | 44794399 | 17.06935 | 28.11846 | 82.2871 | 35.49288 | -1.16468 | 1.365232 | 0.119029 |
| 15 | 72843800 | 72851199 | 44.66384 | 70.95772 | 148.3264 | 61.85798 | -0.72909 | 2.269263 | 0.11904 |
| 1 | 46014200 | 46015599 | 17.02542 | 32.41727 | 12.43355 | 4.975461 | 1.866524 | 0.037253 | 0.119049 |
| 10 | 7326800 | 7327999 | 15.49973 | 29.11733 | 18.46092 | 7.588283 | 1.118734 | 0.126147 | 0.11905 |
| 8 | 22922800 | 22926399 | 17.66176 | 29.52049 | 72.01392 | 31.35264 | -0.901 | 1.188816 | 0.119064 |
| 10 | 18466400 | 18467999 | 12.84583 | 22.22756 | 54.76819 | 24.02822 | -0.90905 | 0.832377 | 0.119082 |
| 2 | 74655200 | 74656999 | 8.462434 | 15.30672 | 39.09421 | 16.95748 | -0.93339 | 0.340192 | 0.119122 |
| 19 | 44057400 | 44058999 | 5.958091 | 11.13425 | 33.81085 | 14.02156 | -1.13299 | 0.116311 | 0.119132 |

| | | | | | | | | | |
|---|---|---|---|---|---|---|---|---|---|
| 11 | 12697400 | 12698399 | 16.11472 | 30.54367 | 16.1464 | 6.523822 | 1.398838 | 0.068552 | 0.119199 |
| 17 | 42453600 | 42454799 | 13.09474 | 25.12461 | 16.6918 | 6.552889 | 1.109386 | -0.04905 | 0.119199 |
| 12 | 20007400 | 20022799 | 501.7306 | 692.1575 | 482.4681 | 200.4811 | 0.669048 | 4.789501 | 0.119201 |
| 1 | 49578800 | 49580399 | 20.58062 | 36.27605 | 26.03201 | 11.52146 | 0.865072 | 0.576968 | 0.119218 |
| 12 | 99528400 | 99530199 | 7.817133 | 14.28199 | 37.6915 | 16.14531 | -0.96839 | 0.272845 | 0.119218 |
| 10 | 3869800 | 3872999 | 28.07366 | 48.0561 | 35.64018 | 15.6287 | 0.81248 | 0.966498 | 0.119228 |
| 4 | 151363600 | 151375799 | 74.87652 | 114.8887 | 240.789 | 99.9626 | -0.77298 | 2.994397 | 0.119253 |
| 7 | 21130000 | 21132799 | 34.37094 | 58.91425 | 29.2045 | 12.80795 | 1.39112 | 0.973236 | 0.119255 |
| 4 | 149668400 | 149672399 | 18.12912 | 29.60549 | 90.10487 | 38.72746 | -1.22739 | 1.487933 | 0.119289 |
| 5 | 84906600 | 84908799 | 24.32294 | 42.29087 | 30.79225 | 13.52577 | 0.84253 | 0.795683 | 0.119292 |
| 12 | 57592000 | 57598999 | 50.06306 | 78.90337 | 172.1594 | 71.44007 | -0.8008 | 2.479014 | 0.119293 |
| 11 | 8634000 | 8641199 | 41.23296 | 65.62376 | 142.9389 | 59.73537 | -0.78567 | 2.210375 | 0.119301 |
| 18 | 84026600 | 84034599 | 66.61374 | 103.1515 | 212.3491 | 88.2042 | -0.73695 | 2.814414 | 0.119319 |
| 12 | 22601800 | 22602799 | 15.5273 | 28.84868 | 18.76626 | 7.857909 | 1.060405 | 0.193242 | 0.119322 |
| 11 | 61652600 | 61653599 | 6.978566 | 13.03766 | 36.12073 | 14.98651 | -1.00349 | 0.130005 | 0.119326 |
| 5 | 150587200 | 150590199 | 14.08972 | 23.91406 | 63.15292 | 27.66852 | -1.0138 | 1.024557 | 0.119331 |
| 8 | 43509400 | 43510999 | 15.86256 | 29.31024 | 19.26945 | 8.129101 | 1.037716 | 0.22715 | 0.119333 |
| 12 | 20580000 | 20586999 | 103.3978 | 161.7496 | 99.71845 | 41.36857 | 1.011839 | 2.640282 | 0.119339 |
| 13 | 36227200 | 36230399 | 37.47847 | 62.06139 | 46.63945 | 20.18009 | 0.791545 | 1.326872 | 0.119365 |
| 7 | 61610800 | 61612599 | 46.69782 | 75.66211 | 57.62748 | 24.66816 | 0.756249 | 1.654072 | 0.119387 |
| 2 | 118692200 | 118692999 | 3.76178 | 7.332801 | 24.2711 | 8.769712 | -1.0702 | -0.41076 | 0.119388 |
| 3 | 154981600 | 154982399 | 13.66724 | 25.64082 | 18.1337 | 7.470556 | 0.959461 | 0.115856 | 0.11939 |
| 17 | 30029600 | 30033799 | 25.39944 | 41.0014 | 103.7612 | 44.24965 | -0.97728 | 1.735357 | 0.119402 |
| 1 | 49924200 | 49925399 | 21.48407 | 37.875 | 25.9532 | 11.48619 | 0.931473 | 0.581119 | 0.119414 |
| 14 | 41093000 | 41094399 | 19.72823 | 36.06739 | 18.18368 | 7.782654 | 1.403027 | 0.313944 | 0.119419 |
| 7 | 25380000 | 25392799 | 75.09761 | 114.7628 | 257.6418 | 106.8725 | -0.87664 | 3.077029 | 0.119428 |
| 5 | 34708400 | 34710599 | 7.455353 | 13.39729 | 40.05682 | 17.5077 | -1.16597 | 0.374671 | 0.119439 |
| 10 | 61046600 | 61049599 | 14.63129 | 25.04011 | 59.57733 | 26.125 | -0.86229 | 0.945638 | 0.119475 |
| 10 | 65474000 | 65475799 | 30.81707 | 52.1851 | 38.38117 | 16.80385 | 0.822905 | 1.075888 | 0.119475 |
| 3 | 89549000 | 89551399 | 9.854425 | 17.25475 | 48.31193 | 21.32464 | -1.08953 | 0.650845 | 0.11948 |
| 1 | 136454600 | 136455599 | 4.560561 | 8.769997 | 29.04542 | 11.32914 | -1.1771 | -0.12086 | 0.119481 |
| 6 | 87729800 | 87735599 | 30.7303 | 49.42278 | 112.612 | 47.77878 | -0.83298 | 1.860303 | 0.119488 |
| 18 | 59270600 | 59272599 | 16.6902 | 30.38163 | 20.3061 | 8.73816 | 0.994614 | 0.31854 | 0.119502 |
| X | 118735800 | 118736799 | 14.29971 | 27.75216 | 12.24877 | 4.606894 | 1.750107 | -0.18464 | 0.119502 |
| 4 | 58942800 | 58944199 | 7.805135 | 14.29193 | 37.1977 | 15.88709 | -0.94529 | 0.260086 | 0.119505 |
| X | 3421600 | 3424599 | 39.79799 | 65.68269 | 47.78059 | 20.62625 | 0.838032 | 1.371694 | 0.119513 |
| 15 | 103010400 | 103012399 | 8.603469 | 15.58782 | 39.66287 | 17.16826 | -0.92632 | 0.331276 | 0.119525 |
| 1 | 114138600 | 114140999 | 49.32341 | 79.77933 | 59.98662 | 25.64558 | 0.772501 | 1.693988 | 0.119543 |
| 15 | 101930800 | 101941399 | 53.45195 | 83.49352 | 196.971 | 81.7266 | -0.92341 | 2.653952 | 0.119547 |
| 14 | 42096200 | 42099199 | 27.81438 | 47.89049 | 29.96607 | 13.1449 | 1.057879 | 0.906579 | 0.119548 |
| 19 | 54945200 | 54946199 | 12.4502 | 23.90587 | 16.62676 | 6.513296 | 1.046148 | -0.06379 | 0.119548 |
| 10 | 69234000 | 69238999 | 24.14715 | 38.99763 | 106.2294 | 45.32883 | -1.08246 | 1.719989 | 0.119551 |
| 9 | 21152800 | 21163599 | 64.18961 | 99.35858 | 214.8026 | 89.22347 | -0.80792 | 2.820858 | 0.119564 |
| 2 | 90898400 | 90901799 | 14.87583 | 25.21001 | 63.73982 | 27.9128 | -0.95149 | 1.041076 | 0.119566 |
| 7 | 10713800 | 10714999 | 16.84705 | 31.60006 | 17.0492 | 7.026156 | 1.346088 | 0.143315 | 0.119569 |
| 4 | 134446000 | 134448199 | 9.077334 | 15.97891 | 46.15841 | 20.43456 | -1.13328 | 0.56172 | 0.119585 |
| 7 | 112383200 | 112383799 | 10.69835 | 20.75312 | 15.37925 | 5.796949 | 1.007832 | -0.21011 | 0.119597 |

| | | | | | | | | | |
|---|---|---|---|---|---|---|---|---|---|
| 5 | 147651800 | 147664599 | 80.38616 | 122.592 | 273.0536 | 113.2116 | -0.86765 | 3.120682 | 0.119615 |
| 4 | 14192800 | 14194999 | 23.73008 | 42.26689 | 19.33163 | 8.512652 | 1.516635 | 0.493797 | 0.11962 |
| 12 | 100489200 | 100490199 | 14.31411 | 26.76086 | 18.73996 | 7.766065 | 0.967331 | 0.146149 | 0.119635 |
| 11 | 54677800 | 54679799 | 9.488391 | 16.76205 | 44.07742 | 19.49941 | -0.99622 | 0.522758 | 0.11964 |
| 4 | 29734400 | 29735199 | 12.32688 | 23.76175 | 16.12229 | 6.245224 | 1.09648 | -0.10157 | 0.119648 |
| 6 | 111073400 | 111074799 | 13.43165 | 25.97955 | 14.39486 | 5.500203 | 1.404214 | -0.13797 | 0.119654 |
| 3 | 26505600 | 26507999 | 15.55589 | 26.64322 | 64.13365 | 28.12473 | -0.87913 | 0.945919 | 0.119668 |
| X | 30896800 | 30897599 | 18.38655 | 34.58848 | 15.38676 | 6.309789 | 1.627699 | 0.135893 | 0.119675 |
| 2 | 6083200 | 6096399 | 95.85242 | 145.1771 | 290.7158 | 120.2899 | -0.72246 | 3.266555 | 0.119694 |
| 15 | 53052000 | 53053199 | 15.33207 | 29.59696 | 11.58586 | 4.467859 | 1.884887 | -0.09614 | 0.119706 |
| 9 | 119833000 | 119835399 | 10.09945 | 17.85071 | 44.06049 | 19.48992 | -0.90544 | 0.519626 | 0.119714 |
| 9 | 82562800 | 82563599 | 7.736796 | 13.9534 | 39.57437 | 17.22855 | -1.08679 | 0.352784 | 0.119725 |
| 7 | 18098000 | 18105399 | 118.6492 | 185.1105 | 108.376 | 44.97259 | 1.083297 | 2.695447 | 0.119726 |
| 3 | 143245600 | 143246399 | 17.33119 | 31.62582 | 21.46376 | 9.20807 | 0.976071 | 0.314528 | 0.119732 |
| 16 | 61340600 | 61341599 | 12.20481 | 23.62378 | 15.04576 | 5.729384 | 1.209708 | -0.16129 | 0.119734 |
| 12 | 59258400 | 59259799 | 43.3002 | 70.63317 | 54.35735 | 23.35368 | 0.749139 | 1.532435 | 0.119755 |
| 4 | 148827000 | 148828799 | 5.45829 | 9.756415 | 43.13342 | 18.93731 | -1.72915 | 0.401998 | 0.11976 |
| 8 | 105187800 | 105191199 | 28.08704 | 48.72319 | 30.56489 | 13.41064 | 1.056223 | 0.843735 | 0.119763 |
| 7 | 75632800 | 75633599 | 5.26989 | 10.14209 | 29.77689 | 11.58801 | -1.00307 | -0.12034 | 0.119765 |
| X | 147578600 | 147579399 | 14.96617 | 28.99771 | 11.89236 | 4.507461 | 1.843059 | -0.15396 | 0.119765 |
| 12 | 72276600 | 72277999 | 5.259996 | 10.15345 | 31.85581 | 12.28699 | -1.08632 | -0.14582 | 0.119777 |
| 3 | 85892000 | 85893399 | 5.169203 | 9.85375 | 31.02952 | 12.38674 | -1.13729 | -0.03345 | 0.119787 |
| 16 | 93177000 | 93178599 | 5.169203 | 9.85375 | 31.02952 | 12.38674 | -1.13729 | -0.03345 | 0.119787 |
| 4 | 11395600 | 11400999 | 33.27587 | 53.57136 | 112.7536 | 47.90965 | -0.71701 | 1.831544 | 0.119795 |
| 18 | 45670600 | 45672399 | 14.84822 | 28.25815 | 16.23219 | 6.502167 | 1.290474 | 0.029274 | 0.119797 |
| 14 | 119090000 | 119092999 | 15.90083 | 26.71076 | 69.74639 | 30.46161 | -0.99866 | 1.126881 | 0.119809 |
| 14 | 3888000 | 3888599 | 12.6983 | 24.46721 | 16.11182 | 6.248858 | 1.137756 | -0.09344 | 0.11983 |
| 7 | 29312400 | 29313199 | 25.99464 | 44.98781 | 32.50383 | 14.26035 | 0.852846 | 0.857719 | 0.11983 |
| 6 | 145292400 | 145294999 | 13.01783 | 22.47181 | 57.48234 | 25.21716 | -0.96378 | 0.856471 | 0.119831 |
| 1 | 114039200 | 114044999 | 52.56532 | 84.93931 | 49.53231 | 21.15068 | 1.136641 | 1.722014 | 0.119834 |
| 8 | 126471200 | 126484799 | 74.66403 | 114.1346 | 262.7609 | 109.0036 | -0.91257 | 3.070578 | 0.119836 |
| 7 | 108121600 | 108123199 | 17.38306 | 32.75929 | 14.65568 | 5.991811 | 1.621797 | 0.113273 | 0.119836 |
| 10 | 12911800 | 12914399 | 34.58084 | 57.79683 | 42.71578 | 18.59656 | 0.815793 | 1.212958 | 0.119854 |
| 5 | 139114600 | 139120599 | 31.73924 | 50.92555 | 118.8531 | 50.25121 | -0.87171 | 1.938224 | 0.119861 |
| 17 | 41291200 | 41291999 | 11.71144 | 22.55489 | 15.58772 | 6.053935 | 1.066231 | -0.10252 | 0.119866 |
| 2 | 172492800 | 172519799 | 203.2296 | 289.0822 | 597.2332 | 246.9579 | -0.85339 | 4.323744 | 0.119877 |
| 7 | 10690600 | 10692999 | 25.2169 | 44.2681 | 23.12389 | 10.22906 | 1.318624 | 0.646797 | 0.119905 |
| 9 | 108024800 | 108036399 | 68.91979 | 106.3698 | 227.723 | 94.59174 | -0.79712 | 2.877267 | 0.119907 |
| X | 4309000 | 4312999 | 25.38317 | 44.86142 | 21.73143 | 9.612917 | 1.429907 | 0.571109 | 0.119908 |
| 14 | 32618400 | 32623599 | 28.59533 | 46.12848 | 105.013 | 44.74354 | -0.82628 | 1.760663 | 0.119919 |
| 3 | 56452200 | 56453599 | 12.1662 | 21.2598 | 50.48181 | 22.24459 | -0.85372 | 0.698455 | 0.119952 |
| 7 | 10582600 | 10590199 | 165.9828 | 250.6696 | 190.685 | 78.82806 | 0.669287 | 3.334599 | 0.119953 |
| 19 | 56977000 | 56980399 | 15.28608 | 26.0528 | 61.73123 | 27.06048 | -0.85742 | 0.988322 | 0.119954 |
| 2 | 173012000 | 173014999 | 15.23269 | 25.3525 | 74.78678 | 32.47255 | -1.17376 | 1.23172 | 0.11996 |
| 16 | 97652000 | 97656999 | 34.9979 | 56.00818 | 128.36 | 54.0586 | -0.8501 | 2.025873 | 0.119974 |
| 5 | 113814200 | 113843399 | 209.5582 | 298.259 | 586.6194 | 242.5451 | -0.78162 | 4.315492 | 0.119991 |
| 6 | 94737200 | 94738199 | 5.40653 | 10.40289 | 29.76919 | 11.59196 | -0.96716 | -0.11631 | 0.120012 |

| | | | | | | | | | |
|---|---|---|---|---|---|---|---|---|---|
| 3 | 3181400 | 3182799 | 23.03756 | 40.65262 | 24.49514 | 10.83881 | 1.116188 | 0.576843 | 0.120032 |
| Y | 89142400 | 89143399 | 9.5042 | 18.51798 | 10.38829 | 3.776054 | 1.448935 | -0.36952 | 0.120032 |
| 13 | 49497800 | 49500399 | 12.81849 | 22.27382 | 53.18701 | 23.35365 | -0.86281 | 0.786172 | 0.120034 |
| 8 | 120560600 | 120564799 | 17.26449 | 28.68838 | 75.01374 | 32.5353 | -1.00087 | 1.255916 | 0.12004 |
| 13 | 38558600 | 38559599 | 13.72918 | 26.10063 | 17.51106 | 7.030278 | 1.065984 | 0.028919 | 0.120043 |
| 13 | 61119400 | 61120599 | 13.72918 | 26.10063 | 17.51106 | 7.030278 | 1.065984 | 0.028919 | 0.120043 |
| 9 | 98013000 | 98013999 | 6.320441 | 12.01479 | 33.34048 | 13.38787 | -0.96495 | 0.00309 | 0.120057 |
| 18 | 38316400 | 38326999 | 55.6149 | 86.42896 | 214.4158 | 89.02624 | -1.00171 | 2.746057 | 0.120064 |
| 7 | 125184400 | 125194599 | 70.41129 | 108.98 | 218.4303 | 90.73039 | -0.699 | 2.824643 | 0.120065 |
| 6 | 143381400 | 143383599 | 10.44963 | 18.34489 | 48.81128 | 21.59275 | -1.01688 | 0.609034 | 0.120091 |
| 12 | 112200800 | 112224599 | 144.7548 | 211.0911 | 467.6871 | 192.7184 | -0.91859 | 3.95025 | 0.120093 |
| 2 | 31656200 | 31658199 | 7.039752 | 12.94109 | 36.18989 | 15.3744 | -1.04199 | 0.230214 | 0.120108 |
| 5 | 105631600 | 105634199 | 10.58283 | 18.61869 | 47.73317 | 21.13406 | -0.96304 | 0.577781 | 0.120119 |
| 7 | 31541400 | 31543999 | 26.68752 | 46.33295 | 31.29754 | 13.73482 | 0.950266 | 0.830427 | 0.120123 |
| 5 | 139526000 | 139535799 | 56.65158 | 87.8726 | 215.8833 | 89.66104 | -0.98981 | 2.77999 | 0.120124 |
| 13 | 62735600 | 62737599 | 49.29274 | 79.82401 | 59.96021 | 25.65664 | 0.775543 | 1.672138 | 0.120143 |
| 2 | 59309000 | 59309599 | 4.247408 | 8.229295 | 27.14233 | 10.28912 | -1.13128 | -0.22652 | 0.120164 |
| 10 | 34032400 | 34033799 | 4.247408 | 8.229295 | 27.14233 | 10.28912 | -1.13128 | -0.22652 | 0.120164 |
| 8 | 36054000 | 36060199 | 32.17369 | 51.37185 | 134.0048 | 56.25488 | -1.03999 | 2.096044 | 0.120167 |
| 5 | 73901200 | 73902799 | 6.340669 | 12.00255 | 33.16181 | 13.43536 | -0.97032 | 0.031207 | 0.120171 |
| 5 | 112381000 | 112388999 | 47.43756 | 74.87045 | 173.3459 | 71.95928 | -0.88468 | 2.446898 | 0.12019 |
| 18 | 58118600 | 58120599 | 10.50942 | 18.3385 | 54.77098 | 24.11283 | -1.18327 | 0.713882 | 0.120198 |
| 7 | 17247200 | 17248799 | 20.44666 | 37.60698 | 17.19999 | 7.301676 | 1.550925 | 0.285886 | 0.120202 |
| 6 | 143218000 | 143218799 | 2.458819 | 4.794389 | 25.34749 | 9.122737 | -1.72854 | -0.44071 | 0.120214 |
| 1 | 152885000 | 152885599 | 4.234036 | 8.24347 | 27.26607 | 10.0002 | -1.09131 | -0.34516 | 0.120223 |
| 15 | 20797200 | 20798399 | 4.234036 | 8.24347 | 27.26607 | 10.0002 | -1.09131 | -0.34516 | 0.120223 |
| 9 | 62072200 | 62075399 | 10.53326 | 18.30406 | 53.63976 | 23.55336 | -1.15636 | 0.774785 | 0.120228 |
| 7 | 10599400 | 10600399 | 11.88172 | 23.14318 | 12.31984 | 4.496351 | 1.5206 | -0.31442 | 0.120229 |
| 9 | 21278600 | 21280999 | 11.57014 | 20.14695 | 51.5416 | 22.66109 | -0.96113 | 0.748274 | 0.120248 |
| 14 | 42458800 | 42463399 | 51.46277 | 83.1084 | 62.2651 | 26.56734 | 0.775162 | 1.742821 | 0.120253 |
| 4 | 20198400 | 20199799 | 15.02301 | 28.65514 | 15.66266 | 6.244266 | 1.367245 | 0.014712 | 0.120261 |
| 8 | 127392200 | 127402199 | 49.19278 | 77.68943 | 165.0377 | 68.53215 | -0.75994 | 2.432443 | 0.120274 |
| 12 | 116540200 | 116542799 | 8.304435 | 14.68497 | 44.60136 | 19.72731 | -1.20219 | 0.503479 | 0.120291 |
| 5 | 115617600 | 115621199 | 14.67021 | 24.99665 | 61.10995 | 26.78758 | -0.90227 | 0.988577 | 0.120305 |
| 12 | 76591800 | 76597599 | 26.14004 | 42.03999 | 117.4596 | 49.83488 | -1.12643 | 1.85581 | 0.120307 |
| 19 | 6797800 | 6813199 | 101.6921 | 153.1702 | 315.0986 | 130.1575 | -0.76821 | 3.380519 | 0.120324 |
| 11 | 90595400 | 90595799 | 10.67676 | 20.80146 | 12.9914 | 4.725622 | 1.296914 | -0.33975 | 0.12033 |
| 11 | 50733800 | 50751399 | 133.845 | 198.2278 | 393.3768 | 162.0053 | -0.73852 | 3.699735 | 0.120334 |
| 4 | 154218800 | 154219999 | 6.690395 | 12.48616 | 34.36441 | 14.2832 | -0.99554 | 0.131733 | 0.120357 |
| 3 | 11492200 | 11493399 | 12.38319 | 23.65134 | 16.85299 | 6.699941 | 0.992684 | -0.01625 | 0.120374 |
| 6 | 74619200 | 74620799 | 23.76982 | 41.36734 | 30.41414 | 13.36744 | 0.828775 | 0.780594 | 0.120376 |
| 7 | 7663000 | 7666999 | 53.7224 | 87.4095 | 48.49338 | 20.81483 | 1.217378 | 1.589079 | 0.120384 |
| 7 | 116362000 | 116364799 | 18.26551 | 30.61181 | 69.83044 | 30.45363 | -0.80451 | 1.161635 | 0.120386 |
| 17 | 56969400 | 56972399 | 14.47342 | 24.79227 | 59.00912 | 25.87771 | -0.86248 | 0.937414 | 0.120414 |
| 8 | 71282200 | 71286399 | 16.98185 | 28.11164 | 76.89116 | 33.26216 | -1.06597 | 1.302795 | 0.120415 |
| 16 | 39260400 | 39262199 | 21.5488 | 38.52726 | 21.85299 | 9.592271 | 1.21185 | 0.461962 | 0.120426 |
| 15 | 98728600 | 98729599 | 4.120402 | 8.000657 | 27.35952 | 10.25509 | -1.16776 | -0.2681 | 0.120437 |

| | | | | | | | | | |
|---|---|---|---|---|---|---|---|---|---|
| 9 | 91243000 | 91248199 | 42.16729 | 66.8064 | 160.0169 | 66.55432 | -0.92905 | 2.350396 | 0.120438 |
| 10 | 98319800 | 98320799 | 15.6463 | 29.17943 | 19.49802 | 8.114016 | 1.029573 | 0.174842 | 0.120451 |
| 1 | 154420000 | 154421199 | 19.69424 | 35.22049 | 23.37039 | 10.25612 | 0.987542 | 0.451853 | 0.120473 |
| 16 | 17387800 | 17396599 | 49.19558 | 77.29048 | 180.3897 | 74.82448 | -0.90136 | 2.543894 | 0.120487 |
| 17 | 10695000 | 10698399 | 18.57661 | 31.13057 | 69.91171 | 30.48745 | -0.78209 | 1.162617 | 0.120489 |
| 6 | 53044400 | 53045999 | 9.287973 | 16.19373 | 51.47398 | 22.64956 | -1.27184 | 0.718812 | 0.12049 |
| 4 | 135112600 | 135117999 | 30.70938 | 49.43881 | 109.4498 | 46.50485 | -0.78969 | 1.827408 | 0.120491 |
| 13 | 65515600 | 65517199 | 29.47755 | 50.62323 | 31.29775 | 13.72798 | 1.074411 | 0.938017 | 0.120493 |
| 3 | 33135600 | 33137399 | 7.454234 | 13.82757 | 39.8174 | 16.70849 | -1.07163 | 0.188802 | 0.120504 |
| X | 22250000 | 22258399 | 93.08687 | 146.8809 | 103.2621 | 42.86057 | 0.832114 | 2.470673 | 0.120509 |
| 10 | 95158000 | 95159799 | 6.573589 | 12.1546 | 35.21765 | 14.8435 | -1.08416 | 0.190878 | 0.120512 |
| 17 | 8384000 | 8385599 | 4.418547 | 8.548137 | 28.8644 | 11.01832 | -1.17557 | -0.19778 | 0.120529 |
| 7 | 64266800 | 64267999 | 10.40408 | 18.38055 | 45.62689 | 20.1847 | -0.91422 | 0.527451 | 0.12053 |
| 12 | 19778200 | 19785599 | 207.8792 | 311.108 | 202.346 | 83.451 | 0.883615 | 3.517456 | 0.120537 |
| 13 | 103910400 | 103912399 | 10.55264 | 18.67574 | 45.10243 | 19.94568 | -0.87366 | 0.509229 | 0.120537 |
| 15 | 36164200 | 36165799 | 5.330763 | 10.14922 | 33.94464 | 13.58576 | -1.22777 | -0.01826 | 0.120541 |
| 4 | 134679400 | 134686199 | 30.98451 | 49.62671 | 124.8192 | 52.6319 | -0.98246 | 1.995139 | 0.120553 |
| 2 | 29635000 | 29639599 | 21.54503 | 35.21155 | 86.19114 | 37.05597 | -0.91349 | 1.481934 | 0.120559 |
| 13 | 66948400 | 66952399 | 85.80434 | 135.7173 | 102.0466 | 42.40789 | 0.738041 | 2.409194 | 0.120567 |
| 13 | 74610400 | 74611599 | 8.575396 | 15.23292 | 44.0935 | 19.46074 | -1.13036 | 0.461777 | 0.120583 |
| 7 | 38260600 | 38264799 | 74.60251 | 118.9474 | 82.68347 | 34.64062 | 0.862055 | 2.167062 | 0.120589 |
| 15 | 87180800 | 87183999 | 51.49767 | 83.32486 | 59.03479 | 25.24507 | 0.858554 | 1.691058 | 0.120613 |
| 5 | 45023800 | 45026599 | 29.2473 | 50.40745 | 31.55914 | 13.84414 | 1.057197 | 0.905395 | 0.120617 |
| 7 | 7675200 | 7678999 | 27.55306 | 47.73669 | 26.92628 | 11.81371 | 1.20854 | 0.847547 | 0.120634 |
| 11 | 68025400 | 68026799 | 5.79516 | 10.90679 | 33.46024 | 13.71065 | -1.13378 | 0.066595 | 0.120635 |
| 4 | 61335800 | 61338199 | 32.8461 | 55.57992 | 35.12298 | 15.37392 | 1.040825 | 1.085184 | 0.12064 |
| X | 122903800 | 122906599 | 43.95619 | 71.9614 | 49.44013 | 21.26909 | 0.915915 | 1.482907 | 0.12072 |
| 8 | 90209000 | 90210199 | 7.473237 | 13.79232 | 36.2976 | 15.33752 | -0.95002 | 0.209184 | 0.120727 |
| 15 | 98537400 | 98550799 | 57.8085 | 89.18606 | 229.4252 | 95.29896 | -1.06209 | 2.881455 | 0.120748 |
| X | 4236400 | 4239399 | 25.21822 | 45.09897 | 17.09863 | 7.503805 | 1.788188 | 0.47625 | 0.120752 |
| 3 | 128508800 | 128509599 | 12.54605 | 24.04044 | 17.24845 | 6.796846 | 0.994277 | -0.04169 | 0.120754 |
| 5 | 139890400 | 139908999 | 121.0705 | 178.8828 | 423.4802 | 174.3985 | -0.9961 | 3.738079 | 0.120758 |
| 12 | 26795800 | 26797799 | 17.96678 | 32.6885 | 21.15521 | 9.109447 | 1.040382 | 0.333896 | 0.120762 |
| 14 | 18216200 | 18217999 | 20.34003 | 36.14115 | 24.33168 | 10.7363 | 0.961485 | 0.49725 | 0.120764 |
| 16 | 77611400 | 77612399 | 8.283317 | 14.69792 | 43.5641 | 19.24491 | -1.16498 | 0.466791 | 0.12077 |
| 10 | 46456400 | 46457599 | 10.23125 | 19.87055 | 14.91707 | 5.581452 | 0.999184 | -0.2419 | 0.120776 |
| 5 | 102083000 | 102086599 | 14.22654 | 24.18674 | 62.10628 | 27.21944 | -0.97327 | 1.00899 | 0.120776 |
| 15 | 3213200 | 3214999 | 28.75121 | 49.36386 | 34.45339 | 15.11193 | 0.900288 | 0.94147 | 0.120781 |
| 11 | 27657600 | 27660599 | 60.94462 | 97.89749 | 66.14892 | 28.01648 | 0.914765 | 1.920047 | 0.120781 |
| 12 | 105925400 | 105925999 | 6.357363 | 11.98267 | 32.97796 | 13.47907 | -0.9757 | 0.061273 | 0.120803 |
| 17 | 25369200 | 25370599 | 6.357363 | 11.98267 | 32.97796 | 13.47907 | -0.9757 | 0.061273 | 0.120803 |
| 6 | 115979000 | 115985599 | 27.55766 | 44.28397 | 115.2831 | 48.85399 | -1.02612 | 1.88395 | 0.120806 |
| 3 | 27207400 | 27208399 | 3.825559 | 7.445194 | 25.82978 | 9.504201 | -1.16261 | -0.34202 | 0.120813 |
| 12 | 51313800 | 51315599 | 23.59556 | 41.25215 | 29.02446 | 12.79478 | 0.892201 | 0.712894 | 0.120814 |
| 7 | 60535000 | 60535799 | 12.36434 | 24.04861 | 12.08596 | 4.468797 | 1.584781 | -0.26604 | 0.120822 |
| 4 | 52887200 | 52889399 | 15.77347 | 29.69408 | 17.81688 | 7.29747 | 1.201406 | 0.113729 | 0.120827 |
| 17 | 63988600 | 63990799 | 10.76363 | 18.86708 | 50.44017 | 22.28432 | -1.02389 | 0.638046 | 0.120829 |

| | | | | | | | | | |
|---|---|---|---|---|---|---|---|---|---|
| 13 | 37623400 | 37626799 | 12.04847 | 20.93285 | 53.71321 | 23.5826 | -0.96614 | 0.783942 | 0.120829 |
| 4 | 125075200 | 125080999 | 33.29751 | 53.54367 | 112.59 | 47.75926 | -0.71774 | 1.868814 | 0.120838 |
| 4 | 68055400 | 68056199 | 12.68655 | 24.15457 | 17.78244 | 7.116689 | 0.937529 | 0.010179 | 0.12084 |
| 5 | 147054000 | 147056799 | 16.0482 | 27.41162 | 63.5221 | 27.85044 | -0.82511 | 0.972583 | 0.120853 |
| 13 | 49183200 | 49185399 | 27.08542 | 46.45015 | 34.6792 | 15.20956 | 0.803298 | 0.944969 | 0.120859 |
| 9 | 19845000 | 19847799 | 37.42617 | 63.20334 | 34.253 | 14.98401 | 1.261452 | 1.118153 | 0.120864 |
| 7 | 79943400 | 79944799 | 6.244066 | 11.68907 | 33.88013 | 14.00879 | -1.0633 | 0.10669 | 0.12087 |
| X | 62516800 | 62517799 | 12.54299 | 24.45235 | 10.39177 | 3.749907 | 1.853795 | -0.35043 | 0.120871 |
| 1 | 60594400 | 60596399 | 18.54933 | 33.63796 | 21.67819 | 9.372984 | 1.042859 | 0.358233 | 0.120876 |
| 4 | 107528200 | 107531599 | 15.72365 | 26.447 | 66.91547 | 29.23667 | -0.95312 | 1.113831 | 0.120883 |
| 7 | 26050400 | 26062399 | 354.4763 | 503.6195 | 369.1777 | 152.6869 | 0.639128 | 4.346072 | 0.120885 |
| 7 | 100623200 | 100624999 | 5.64492 | 10.34433 | 39.32373 | 16.77755 | -1.48582 | 0.241833 | 0.120907 |
| 3 | 42369800 | 42372199 | 42.19603 | 69.12741 | 51.62098 | 22.21376 | 0.796368 | 1.47039 | 0.120925 |
| 3 | 9941200 | 9942399 | 6.818274 | 12.75151 | 38.39726 | 15.90289 | -1.12007 | 0.130299 | 0.120941 |
| 1 | 90213200 | 90214599 | 9.30356 | 16.54609 | 42.01637 | 18.52538 | -0.94142 | 0.457234 | 0.120951 |
| 11 | 14618600 | 14619999 | 24.4832 | 42.89202 | 27.25057 | 12.03366 | 1.038512 | 0.683071 | 0.120966 |
| 1 | 91765200 | 91773799 | 54.3744 | 85.02739 | 190.9326 | 79.21222 | -0.85108 | 2.634447 | 0.120966 |
| 12 | 51498600 | 51500799 | 26.82929 | 46.27203 | 33.39858 | 14.65136 | 0.853185 | 0.891391 | 0.121014 |
| 3 | 68175800 | 68176999 | 18.84952 | 33.63096 | 23.81732 | 10.47291 | 0.891988 | 0.459723 | 0.121019 |
| 3 | 104934400 | 104939799 | 81.14553 | 123.622 | 266.9695 | 110.6593 | -0.8243 | 3.142467 | 0.121049 |
| 14 | 48902200 | 48909199 | 37.87183 | 60.27242 | 153.0721 | 63.96723 | -1.00688 | 2.209753 | 0.121073 |
| 3 | 100587400 | 100587999 | 2.454566 | 4.796767 | 26.32542 | 9.123397 | -1.73325 | -0.57666 | 0.121078 |
| 19 | 35496000 | 35497799 | 18.88745 | 34.81041 | 18.43612 | 7.804653 | 1.343954 | 0.26501 | 0.12109 |
| 4 | 120103400 | 120148999 | 291.9455 | 400.5311 | 893.0011 | 371.3958 | -1.01596 | 4.871184 | 0.121098 |
| 6 | 82915600 | 82917999 | 13.19553 | 22.71508 | 57.42894 | 25.19159 | -0.94779 | 0.88343 | 0.121113 |
| 3 | 10974600 | 10975199 | 9.016568 | 17.57398 | 13.41857 | 4.855417 | 1.018316 | -0.37766 | 0.121115 |
| 13 | 34987400 | 34987999 | 11.03763 | 21.51314 | 12.38199 | 4.48044 | 1.419872 | -0.35462 | 0.121116 |
| 6 | 107527000 | 107527999 | 13.30905 | 25.20763 | 17.52584 | 7.091998 | 1.00494 | 0.049282 | 0.121124 |
| 5 | 17310800 | 17312599 | 22.00614 | 38.58431 | 26.91736 | 11.89481 | 0.904067 | 0.654325 | 0.121127 |
| 11 | 30444000 | 30444999 | 22.00614 | 38.58431 | 26.91736 | 11.89481 | 0.904067 | 0.654325 | 0.121127 |
| 7 | 16012000 | 16017199 | 21.92474 | 35.68946 | 90.36111 | 38.79231 | -0.96622 | 1.543217 | 0.12114 |
| 1 | 160021400 | 160023199 | 7.716961 | 13.67399 | 46.17174 | 20.41357 | -1.35289 | 0.478583 | 0.121152 |
| 2 | 39115600 | 39116599 | 7.681327 | 14.00512 | 38.18491 | 16.39866 | -1.01751 | 0.284542 | 0.121153 |
| 4 | 119616800 | 119618399 | 7.681327 | 14.00512 | 38.18491 | 16.39866 | -1.01751 | 0.284542 | 0.121153 |
| 9 | 77323200 | 77324199 | 5.400842 | 10.41066 | 29.8839 | 11.57664 | -0.96466 | -0.13341 | 0.121155 |
| 9 | 117354200 | 117355999 | 9.237635 | 16.64101 | 41.13384 | 17.93326 | -0.89174 | 0.37269 | 0.121162 |
| 10 | 86507200 | 86507799 | 3.422599 | 6.668845 | 26.02859 | 9.456457 | -1.31327 | -0.39332 | 0.12117 |
| 19 | 37236600 | 37239999 | 41.58991 | 66.10822 | 145.3586 | 60.65156 | -0.80114 | 2.251565 | 0.12118 |
| 11 | 22133200 | 22134599 | 15.05605 | 28.57432 | 16.78335 | 6.762496 | 1.251594 | 0.051255 | 0.121188 |
| 13 | 55312400 | 55318599 | 37.63908 | 60.04446 | 135.4696 | 56.79769 | -0.83261 | 2.129304 | 0.121196 |
| 5 | 73591800 | 73592399 | 3.342559 | 6.530011 | 22.32274 | 7.792616 | -1.06993 | -0.54717 | 0.121216 |
| 2 | 103166800 | 103167599 | 6.485793 | 12.27391 | 33.15285 | 13.43947 | -0.93864 | 0.034967 | 0.121219 |
| 16 | 90181600 | 90185199 | 18.72828 | 30.95936 | 77.70534 | 33.58195 | -0.94277 | 1.323505 | 0.121225 |
| 16 | 92768600 | 92771199 | 15.79603 | 26.7958 | 63.29546 | 27.72647 | -0.85384 | 1.034973 | 0.121228 |
| 3 | 6220800 | 6221199 | 8.060571 | 15.7474 | 11.67949 | 4.075853 | 1.104282 | -0.52381 | 0.121229 |
| 14 | 76168600 | 76168999 | 3.24421 | 6.306537 | 25.53626 | 9.483831 | -1.39292 | -0.31412 | 0.121235 |
| 12 | 109870200 | 109871199 | 12.75198 | 24.34199 | 16.84572 | 6.705043 | 1.033122 | -0.00848 | 0.121235 |

| | | | | | | | | | |
|---|---|---|---|---|---|---|---|---|---|
| 7 | 72512400 | 72512799 | 7.987726 | 15.59605 | 5.118412 | 1.617376 | 2.364613 | -0.88407 | 0.121243 |
| 15 | 93503600 | 93511999 | 53.68347 | 84.44574 | 178.8624 | 74.20289 | -0.76024 | 2.520724 | 0.121277 |
| 5 | 112662000 | 112687999 | 201.9503 | 288.7301 | 566.7005 | 234.1337 | -0.77207 | 4.248913 | 0.121277 |
| 14 | 104931600 | 104932199 | 3.938084 | 7.679201 | 25.73162 | 9.254186 | -1.08321 | -0.42367 | 0.121281 |
| 12 | 19853800 | 19870399 | 634.5351 | 856.9382 | 595.3262 | 248.1994 | 0.642951 | 5.123253 | 0.121291 |
| 4 | 61639200 | 61640399 | 19.33478 | 34.25465 | 24.39473 | 10.7858 | 0.87849 | 0.510378 | 0.121301 |
| 6 | 145245600 | 145254399 | 62.94661 | 98.10535 | 197.8506 | 82.11294 | -0.70004 | 2.69807 | 0.121309 |
| 4 | 105310400 | 105312599 | 9.327847 | 16.5428 | 45.64844 | 20.17446 | -1.06355 | 0.478874 | 0.121326 |
| 17 | 62703600 | 62705599 | 13.00008 | 24.99987 | 15.35607 | 5.989194 | 1.228908 | -0.07585 | 0.121329 |
| 9 | 118133200 | 118135199 | 7.360095 | 13.29219 | 44.93184 | 19.52745 | -1.3359 | 0.356973 | 0.121336 |
| 4 | 117848800 | 117858799 | 43.76694 | 69.02197 | 175.9076 | 73.0012 | -1.02383 | 2.463369 | 0.121358 |
| 2 | 71368800 | 71377599 | 62.37698 | 96.06606 | 243.3967 | 101.0967 | -1.04222 | 2.91639 | 0.121361 |
| 7 | 99462400 | 99463599 | 6.470615 | 11.85203 | 36.65226 | 15.64742 | -1.19087 | 0.247509 | 0.121371 |
| 10 | 14975200 | 14976799 | 21.72156 | 38.90414 | 21.2941 | 9.333174 | 1.264294 | 0.451557 | 0.121373 |
| 6 | 120456200 | 120460599 | 24.5621 | 39.75224 | 98.59457 | 42.16985 | -0.94415 | 1.66161 | 0.121374 |
| 4 | 148887000 | 148891799 | 25.94916 | 41.80312 | 115.4564 | 49.10428 | -1.10678 | 1.800653 | 0.12138 |
| 19 | 45438000 | 45439599 | 5.309388 | 10.09578 | 30.94373 | 12.41779 | -1.10521 | -0.01383 | 0.121394 |
| 14 | 14534800 | 14536599 | 6.769171 | 12.44447 | 37.82107 | 16.06598 | -1.16164 | 0.228319 | 0.121399 |
| 14 | 16181600 | 16182999 | 12.42725 | 21.38735 | 56.81635 | 24.92269 | -1.01868 | 0.883214 | 0.1214 |
| 4 | 126625200 | 126630799 | 37.61294 | 59.73956 | 156.4146 | 65.21321 | -1.05372 | 2.272883 | 0.121406 |
| 5 | 129734200 | 129735399 | 5.644752 | 10.6039 | 32.8275 | 13.49396 | -1.14984 | 0.077546 | 0.121413 |
| 2 | 128734200 | 128737999 | 28.75735 | 46.38147 | 104.9346 | 44.69974 | -0.81764 | 1.766392 | 0.121417 |
| 4 | 60593400 | 60594799 | 35.99306 | 59.69338 | 45.02192 | 19.5009 | 0.786509 | 1.304241 | 0.121424 |
| 13 | 36151600 | 36154799 | 21.28453 | 34.68204 | 89.83331 | 38.57981 | -0.99792 | 1.526662 | 0.121425 |
| 5 | 36253400 | 36255799 | 12.23484 | 20.82892 | 63.08509 | 27.65145 | -1.20985 | 0.989654 | 0.121442 |
| 12 | 115393400 | 115397799 | 152.837 | 231.6589 | 175.0662 | 72.45454 | 0.683227 | 3.257456 | 0.121446 |
| 16 | 32484600 | 32487399 | 8.426783 | 14.70495 | 53.57588 | 23.58725 | -1.46746 | 0.703249 | 0.121451 |
| 2 | 176435800 | 176437599 | 17.68924 | 32.84016 | 17.35267 | 7.273615 | 1.35666 | 0.21269 | 0.121451 |
| 7 | 130148200 | 130152599 | 29.03742 | 46.93707 | 101.1861 | 43.22632 | -0.74444 | 1.699885 | 0.121457 |
| 13 | 34834400 | 34837999 | 18.08394 | 29.66618 | 88.53974 | 38.1227 | -1.19562 | 1.423139 | 0.12146 |
| 5 | 72804600 | 72806399 | 9.119696 | 15.9989 | 50.39715 | 22.28109 | -1.25876 | 0.615755 | 0.121495 |
| 4 | 139459000 | 139463199 | 24.42229 | 39.86623 | 87.85567 | 37.75514 | -0.76382 | 1.505606 | 0.121496 |
| 7 | 33047600 | 33048999 | 18.15176 | 32.60481 | 22.56758 | 9.867206 | 0.930551 | 0.412466 | 0.121502 |
| 17 | 37632600 | 37638599 | 70.04663 | 112.0158 | 75.36527 | 31.70711 | 0.913584 | 2.069029 | 0.121517 |
| 12 | 77185600 | 77187799 | 10.92281 | 19.12509 | 49.38274 | 21.79692 | -0.97389 | 0.656168 | 0.121519 |
| 12 | 23408800 | 23410799 | 40.33502 | 66.13628 | 48.86047 | 21.03361 | 0.812268 | 1.453294 | 0.121526 |
| 17 | 17724000 | 17729399 | 68.10666 | 105.3497 | 214.7453 | 89.20001 | -0.72393 | 2.835622 | 0.121527 |
| 2 | 48703600 | 48705199 | 8.941383 | 15.72439 | 47.20231 | 20.90073 | -1.18901 | 0.573099 | 0.121533 |
| 9 | 120956200 | 120965199 | 46.85557 | 74.18736 | 158.0163 | 65.70129 | -0.76076 | 2.368705 | 0.121536 |
| 17 | 26134800 | 26139999 | 19.48128 | 31.89052 | 85.45581 | 36.76173 | -1.04187 | 1.453116 | 0.121566 |
| 2 | 168826400 | 168832199 | 47.87841 | 75.66688 | 163.2028 | 67.79433 | -0.781 | 2.414441 | 0.121567 |
| 1 | 127700800 | 127702599 | 12.54511 | 21.69213 | 55.29223 | 24.25761 | -0.95796 | 0.837555 | 0.12157 |
| 1 | 186539000 | 186541999 | 16.25116 | 27.60241 | 63.36834 | 27.76894 | -0.81287 | 1.025385 | 0.121585 |
| 1 | 16573200 | 16574399 | 6.225248 | 11.70796 | 33.46735 | 13.72997 | -1.03435 | 0.075031 | 0.121609 |
| 8 | 105493800 | 105494799 | 10.99518 | 21.27324 | 15.88541 | 6.061324 | 0.98044 | -0.16517 | 0.121613 |
| 13 | 21412400 | 21414199 | 21.09081 | 37.30294 | 25.24547 | 11.1657 | 0.950856 | 0.542028 | 0.121615 |
| 4 | 155480000 | 155483199 | 19.80585 | 32.80151 | 79.31864 | 34.32313 | -0.88918 | 1.303894 | 0.12163 |

| | | | | | | | | | |
|---|---|---|---|---|---|---|---|---|---|
| 17 | 64395200 | 64395999 | 44.59536 | 75.05539 | 24.25314 | 10.59866 | 2.00315 | 1.161845 | 0.121631 |
| 1 | 182691600 | 182703999 | 145.1485 | 213.4726 | 428.1541 | 176.3205 | -0.76332 | 3.816832 | 0.121633 |
| 16 | 90712200 | 90725799 | 88.969 | 134.6592 | 294.814 | 121.9593 | -0.85188 | 3.279327 | 0.121636 |
| 7 | 44277800 | 44293999 | 123.6906 | 183.2263 | 389.3588 | 160.3515 | -0.83688 | 3.695294 | 0.121688 |
| 18 | 82708800 | 82711799 | 17.81559 | 29.77972 | 70.27913 | 30.59874 | -0.8533 | 1.188163 | 0.121698 |
| 11 | 117400400 | 117421199 | 146.56 | 214.8458 | 444.5516 | 183.1106 | -0.81277 | 3.868521 | 0.121704 |
| 17 | 19844000 | 19850599 | 164.7438 | 248.7905 | 193.7934 | 80.11223 | 0.635089 | 3.334456 | 0.121705 |
| 7 | 32030200 | 32036399 | 170.6982 | 257.5273 | 193.8253 | 80.10034 | 0.683398 | 3.356251 | 0.12172 |
| 2 | 168366400 | 168372599 | 32.58118 | 52.17396 | 123.9165 | 52.2359 | -0.90031 | 2.002944 | 0.121738 |
| 1 | 131024400 | 131025799 | 5.36942 | 9.801618 | 39.22762 | 16.82126 | -1.56395 | 0.263153 | 0.121741 |
| X | 32187400 | 32189799 | 29.33483 | 50.84429 | 29.21691 | 12.81887 | 1.182153 | 0.857185 | 0.121746 |
| X | 18688800 | 18692199 | 49.80794 | 81.40872 | 47.35192 | 20.3585 | 1.153771 | 1.515949 | 0.121763 |
| 4 | 151435600 | 151450199 | 75.05525 | 114.8015 | 253.7263 | 105.2702 | -0.85305 | 3.058359 | 0.121767 |
| 8 | 36356200 | 36359199 | 15.26983 | 26.06996 | 61.22714 | 26.84324 | -0.84431 | 0.97161 | 0.12177 |
| 2 | 144120800 | 144122999 | 11.21277 | 19.6794 | 47.22857 | 20.88792 | -0.8687 | 0.617071 | 0.12178 |
| 1 | 85551000 | 85553599 | 127.068 | 195.3281 | 148.6383 | 61.72213 | 0.689214 | 2.966005 | 0.121781 |
| 11 | 49531000 | 49538999 | 52.01946 | 81.95072 | 171.14 | 71.01299 | -0.73816 | 2.489149 | 0.121785 |
| 14 | 26627200 | 26628199 | 18.48305 | 33.00204 | 23.35569 | 10.26338 | 0.893547 | 0.451398 | 0.121801 |
| 7 | 32917400 | 32918799 | 37.18478 | 62.21006 | 40.49797 | 17.64196 | 0.998219 | 1.210329 | 0.121808 |
| 2 | 119496200 | 119497399 | 2.640998 | 5.154288 | 22.51688 | 7.997079 | -1.4381 | -0.49256 | 0.12182 |
| 19 | 13185800 | 13188199 | 39.13888 | 64.58151 | 48.31967 | 20.85645 | 0.797527 | 1.372761 | 0.121824 |
| 15 | 63629200 | 63630799 | 9.091979 | 16.35545 | 40.57065 | 17.71662 | -0.89828 | 0.37904 | 0.12183 |
| 12 | 102906800 | 102907799 | 14.48445 | 27.96767 | 13.57308 | 5.228115 | 1.581809 | -0.1032 | 0.121832 |
| Y | 10548200 | 10551799 | 19.81357 | 35.8148 | 8.95614 | 3.888021 | 2.389001 | 0.351839 | 0.121837 |
| 11 | 34952800 | 34953399 | 2.427943 | 4.745314 | 21.03185 | 7.266129 | -1.41935 | -0.59811 | 0.121838 |
| 16 | 21331400 | 21333399 | 13.30679 | 23.0964 | 58.08779 | 25.48755 | -0.93763 | 0.811705 | 0.12184 |
| 2 | 70195000 | 70195599 | 9.591224 | 18.65873 | 14.40747 | 5.318387 | 0.976397 | -0.29822 | 0.12185 |
| 3 | 47535600 | 47539199 | 39.69536 | 65.74313 | 44.99247 | 19.46966 | 0.92607 | 1.328833 | 0.121873 |
| 12 | 62279600 | 62280799 | 19.56954 | 34.69159 | 24.86363 | 10.99178 | 0.869765 | 0.506894 | 0.121879 |
| 13 | 110206600 | 110207999 | 9.124982 | 16.31608 | 42.60191 | 18.69851 | -0.97711 | 0.420424 | 0.121894 |
| 1 | 55769200 | 55771199 | 22.88727 | 39.98294 | 29.16498 | 12.84904 | 0.840392 | 0.720712 | 0.121905 |
| 8 | 3097800 | 3100599 | 24.7226 | 43.05209 | 30.34944 | 13.34419 | 0.88931 | 0.777751 | 0.121911 |
| 1 | 98990000 | 98992199 | 21.40871 | 39.43715 | 15.38023 | 6.515494 | 1.780293 | 0.281153 | 0.121912 |
| 2 | 173950200 | 173976199 | 210.4631 | 298.5024 | 622.2281 | 257.4191 | -0.87051 | 4.36665 | 0.121914 |
| 12 | 70080600 | 70082199 | 5.712755 | 10.94666 | 30.75388 | 12.1187 | -0.95747 | -0.07187 | 0.121928 |
| 3 | 113463200 | 113463999 | 7.453639 | 13.82121 | 35.86018 | 15.05572 | -0.9227 | 0.182243 | 0.121944 |
| 4 | 52189000 | 52191399 | 45.05277 | 73.30012 | 55.4043 | 23.7808 | 0.772447 | 1.575016 | 0.121969 |
| 7 | 98304800 | 98306399 | 8.972284 | 15.716 | 49.34695 | 21.78834 | -1.25386 | 0.640321 | 0.12197 |
| 7 | 45347600 | 45358199 | 51.66818 | 81.05407 | 182.5823 | 75.72304 | -0.85196 | 2.575316 | 0.121974 |
| 13 | 111545400 | 111547199 | 17.27124 | 31.44685 | 21.91253 | 9.426619 | 0.935651 | 0.326943 | 0.12199 |
| 9 | 37319200 | 37319799 | 2.937901 | 5.740137 | 21.6488 | 7.542288 | -1.20568 | -0.56223 | 0.122 |
| 3 | 91359600 | 91360999 | 22.7446 | 39.80746 | 28.52634 | 12.58669 | 0.865776 | 0.69016 | 0.122004 |
| 6 | 73191800 | 73192199 | 5.259321 | 10.1542 | 30.62571 | 11.80137 | -1.0284 | -0.15183 | 0.12201 |
| 15 | 54862800 | 54864399 | 10.01757 | 17.52855 | 48.8469 | 21.54844 | -1.0829 | 0.663016 | 0.122017 |
| 6 | 138187800 | 138189399 | 29.75144 | 50.5552 | 37.78936 | 16.56134 | 0.799746 | 1.04118 | 0.122018 |
| 4 | 62433400 | 62438599 | 23.14175 | 37.81181 | 87.84166 | 37.76219 | -0.83871 | 1.488175 | 0.122031 |
| 9 | 57444200 | 57445399 | 5.444882 | 10.35059 | 30.9377 | 12.42255 | -1.06998 | -0.01 | 0.122052 |

| | | | | | | | | | |
|---|---|---|---|---|---|---|---|---|---|
| 2 | 85704800 | 85706199 | 17.78367 | 32.9902 | 18.64568 | 7.823743 | 1.260061 | 0.221744 | 0.122072 |
| 4 | 103462800 | 103466999 | 32.87388 | 55.35175 | 39.90575 | 17.44116 | 0.851311 | 1.131751 | 0.122074 |
| 2 | 172212000 | 172213799 | 7.56274 | 13.38096 | 44.61007 | 19.72893 | -1.33503 | 0.493023 | 0.12208 |
| X | 91405600 | 91406999 | 16.38005 | 30.66133 | 18.40238 | 7.610127 | 1.190475 | 0.155996 | 0.122088 |
| 7 | 7631200 | 7636799 | 82.61722 | 131.3995 | 84.90133 | 35.44782 | 0.963588 | 2.255496 | 0.122094 |
| 1 | 133303400 | 133308399 | 22.18556 | 35.94302 | 98.91799 | 42.34037 | -1.09228 | 1.636248 | 0.122108 |
| 9 | 85564000 | 85564799 | 11.83685 | 22.91746 | 15.05611 | 5.726435 | 1.166793 | -0.16969 | 0.122116 |
| 7 | 88281800 | 88283199 | 19.88717 | 35.69435 | 22.57419 | 9.876545 | 1.059024 | 0.428535 | 0.122119 |
| 7 | 7421000 | 7423399 | 45.33029 | 73.93525 | 49.49942 | 21.26556 | 0.949361 | 1.534734 | 0.122141 |
| 6 | 105398200 | 105399799 | 23.57974 | 41.67943 | 23.35969 | 10.33292 | 1.220858 | 0.560676 | 0.122142 |
| 3 | 105231800 | 105233399 | 10.28981 | 18.08137 | 46.68065 | 20.66669 | -0.97336 | 0.590348 | 0.122155 |
| 13 | 39630600 | 39631799 | 5.384159 | 10.37378 | 31.02571 | 12.0347 | -1.02475 | -0.12464 | 0.122156 |
| 2 | 86412600 | 86416599 | 77.57227 | 122.923 | 98.07381 | 40.79551 | 0.654694 | 2.358044 | 0.122175 |
| 2 | 71109200 | 71111599 | 11.21822 | 19.65781 | 50.99161 | 22.52205 | -0.98069 | 0.646219 | 0.12218 |
| 11 | 81884800 | 81886399 | 14.02057 | 26.57748 | 18.12514 | 7.321544 | 1.03539 | 0.053488 | 0.122204 |
| 2 | 127764000 | 127764799 | 3.134076 | 6.107504 | 24.73374 | 8.96989 | -1.36186 | -0.40612 | 0.122206 |
| 12 | 77066400 | 77067199 | 3.134076 | 6.107504 | 24.73374 | 8.96989 | -1.36186 | -0.40612 | 0.122206 |
| 11 | 77717800 | 77722199 | 20.86216 | 34.20536 | 83.506 | 35.94713 | -0.90701 | 1.43571 | 0.122209 |
| 11 | 69324400 | 69330399 | 34.67819 | 55.49475 | 126.1562 | 53.12806 | -0.83844 | 2.026406 | 0.122213 |
| 10 | 124211800 | 124213799 | 27.76174 | 48.27095 | 29.42453 | 12.92515 | 1.097698 | 0.811315 | 0.12222 |
| 19 | 55758800 | 55759399 | 3.04485 | 5.939462 | 23.06672 | 8.258318 | -1.28304 | -0.45605 | 0.122226 |
| 18 | 66856800 | 66859799 | 23.26274 | 40.85348 | 26.82216 | 11.86893 | 0.991307 | 0.638133 | 0.122231 |
| 13 | 52983200 | 52988999 | 34.19676 | 54.72066 | 125.1015 | 52.67887 | -0.84665 | 2.028347 | 0.122233 |
| 9 | 67495200 | 67497999 | 19.39275 | 32.42681 | 72.74992 | 31.68113 | -0.78071 | 1.191521 | 0.122265 |
| 4 | 47545200 | 47546999 | 19.07549 | 34.81785 | 19.3531 | 8.299891 | 1.26169 | 0.320335 | 0.122273 |
| 5 | 27918400 | 27921599 | 17.80965 | 29.78703 | 70.91421 | 30.88652 | -0.86594 | 1.181518 | 0.122279 |
| 7 | 105369000 | 105371199 | 11.08342 | 19.38248 | 49.39704 | 21.78063 | -0.95509 | 0.677853 | 0.122282 |
| 8 | 22154200 | 22156599 | 18.82365 | 35.04833 | 16.9145 | 7.058178 | 1.491795 | 0.20558 | 0.12229 |
| 6 | 128177000 | 128180199 | 19.63563 | 32.55065 | 75.78657 | 32.81633 | -0.83448 | 1.290996 | 0.122295 |
| 15 | 27473600 | 27478199 | 35.26707 | 56.57882 | 118.6553 | 50.15796 | -0.71805 | 1.946424 | 0.12233 |
| 17 | 56889800 | 56892799 | 11.12536 | 19.03975 | 60.95224 | 26.72787 | -1.28789 | 0.934235 | 0.122333 |
| 5 | 43512000 | 43513799 | 8.030067 | 14.19414 | 47.74737 | 21.12048 | -1.34917 | 0.510164 | 0.122338 |
| 7 | 25002400 | 25003599 | 6.365568 | 11.93995 | 33.36791 | 13.75119 | -1.00678 | 0.095858 | 0.122366 |
| 4 | 83415000 | 83415999 | 4.829484 | 9.349776 | 29.63891 | 11.27648 | -1.08149 | -0.20309 | 0.122372 |
| X | 24103800 | 24104799 | 27.09357 | 47.18459 | 29.0545 | 12.77544 | 1.083491 | 0.78683 | 0.122374 |
| 7 | 143959000 | 143972599 | 74.13362 | 112.9574 | 271.9062 | 112.7109 | -0.98053 | 3.137537 | 0.122387 |
| 5 | 128613600 | 128615199 | 6.2781 | 11.38256 | 42.33497 | 18.30908 | -1.46856 | 0.314899 | 0.122425 |
| 19 | 16416000 | 16417799 | 7.510503 | 13.72856 | 37.15778 | 15.90685 | -1.00345 | 0.268571 | 0.122465 |
| 10 | 75465200 | 75489199 | 156.5285 | 226.6368 | 506.2625 | 208.7792 | -0.94046 | 4.059302 | 0.12247 |
| 5 | 52748000 | 52754199 | 44.83038 | 71.22167 | 146.9161 | 61.26927 | -0.70998 | 2.269722 | 0.122485 |
| 2 | 37591800 | 37593999 | 11.47683 | 19.8438 | 55.22749 | 24.22917 | -1.08388 | 0.834179 | 0.122494 |
| 11 | 18224400 | 18225999 | 16.63458 | 30.50018 | 18.75356 | 7.995079 | 1.123747 | 0.276522 | 0.122495 |
| 19 | 23221200 | 23222399 | 5.850447 | 11.22198 | 32.65079 | 12.82843 | -1.00387 | -0.07572 | 0.122495 |
| 7 | 11291600 | 11291999 | 17.53356 | 31.52563 | 22.48608 | 9.823352 | 0.888061 | 0.400733 | 0.122512 |
| 7 | 64214600 | 64215399 | 4.411507 | 8.553882 | 29.11902 | 10.99841 | -1.17275 | -0.23428 | 0.122518 |
| 8 | 18096200 | 18097599 | 18.36665 | 33.53975 | 20.82008 | 8.923794 | 1.104724 | 0.317159 | 0.122524 |
| 9 | 62366200 | 62369799 | 11.39881 | 19.90461 | 50.47801 | 22.2328 | -0.94824 | 0.70365 | 0.122526 |

| | | | | | | | | | |
|---|---|---|---|---|---|---|---|---|---|
| 7 | 61674600 | 61677799 | 52.93227 | 85.70951 | 57.39107 | 24.55557 | 0.940795 | 1.679608 | 0.122535 |
| 12 | 22994200 | 22999999 | 140.3985 | 213.8916 | 163.7834 | 67.88852 | 0.6702 | 3.150239 | 0.122537 |
| 9 | 52079200 | 52080599 | 8.382524 | 14.98393 | 41.55348 | 18.24304 | -1.06332 | 0.416429 | 0.122557 |
| X | 24624600 | 24625199 | 9.708371 | 18.94486 | 12.71278 | 4.525014 | 1.223136 | -0.42662 | 0.122558 |
| 3 | 106540400 | 106541799 | 25.632 | 44.61173 | 30.66491 | 13.47834 | 0.925401 | 0.789275 | 0.122559 |
| 4 | 139626400 | 139638199 | 92.87899 | 140.5128 | 301.0764 | 124.5319 | -0.82143 | 3.289321 | 0.122575 |
| 16 | 16713800 | 16716599 | 12.14578 | 20.88365 | 58.97777 | 25.87008 | -1.10698 | 0.891409 | 0.12258 |
| 2 | 74526200 | 74531399 | 20.85236 | 33.97548 | 97.74681 | 41.97727 | -1.14923 | 1.528891 | 0.122594 |
| 2 | 63071000 | 63071799 | 3.93912 | 7.678044 | 24.95211 | 9.022616 | -1.04632 | -0.40585 | 0.122594 |
| 12 | 11505600 | 11506599 | 3.344289 | 6.448074 | 31.37639 | 12.14603 | -1.71448 | -0.15414 | 0.122595 |
| 2 | 63611200 | 63612399 | 8.970166 | 16.05112 | 41.03387 | 17.99905 | -0.9459 | 0.412939 | 0.122615 |
| 18 | 58629200 | 58631199 | 25.17057 | 43.99787 | 28.39746 | 12.51649 | 1.01611 | 0.723835 | 0.122644 |
| 17 | 21949000 | 21950799 | 14.28053 | 24.59762 | 57.46656 | 25.20856 | -0.83421 | 0.884104 | 0.122652 |
| 11 | 119472200 | 119476999 | 24.74785 | 40.3768 | 88.4523 | 38.00438 | -0.7558 | 1.513977 | 0.122673 |
| 3 | 42251000 | 42253999 | 26.66723 | 46.62316 | 25.9616 | 11.44475 | 1.227668 | 0.723843 | 0.122686 |
| 10 | 74960800 | 74973999 | 80.42833 | 122.8301 | 261.3955 | 108.4134 | -0.80034 | 3.09139 | 0.122694 |
| 8 | 4143200 | 4145199 | 46.97505 | 76.57446 | 53.3862 | 22.93062 | 0.890584 | 1.550115 | 0.122704 |
| 1 | 36637200 | 36639799 | 11.74229 | 20.40553 | 53.15931 | 23.34274 | -0.98774 | 0.778953 | 0.122721 |
| 9 | 30862400 | 30864999 | 11.74229 | 20.40553 | 53.15931 | 23.34274 | -0.98774 | 0.778953 | 0.122721 |
| X | 32816600 | 32819999 | 66.27647 | 106.5195 | 63.81223 | 27.04788 | 1.089004 | 1.911787 | 0.122733 |
| 4 | 52008800 | 52009999 | 11.64699 | 22.58149 | 14.48898 | 5.474759 | 1.209014 | -0.19515 | 0.122738 |
| 11 | 32534200 | 32547399 | 76.87484 | 117.2245 | 266.6232 | 110.5427 | -0.89792 | 3.122111 | 0.122748 |
| 1 | 27104000 | 27106199 | 23.01238 | 40.42046 | 26.74206 | 11.83513 | 0.980189 | 0.633262 | 0.12275 |
| 8 | 48141800 | 48145199 | 30.05096 | 48.42649 | 106.7116 | 45.40618 | -0.78119 | 1.794332 | 0.122787 |
| 5 | 135177600 | 135179999 | 11.11555 | 19.37157 | 52.6169 | 23.14602 | -1.04716 | 0.733212 | 0.122798 |
| X | 27804400 | 27805399 | 14.11691 | 27.21705 | 14.90631 | 5.771298 | 1.403455 | -0.08667 | 0.12281 |
| 6 | 102750000 | 102751799 | 11.09938 | 19.35913 | 50.46682 | 22.21157 | -0.98774 | 0.720636 | 0.122815 |
| 15 | 84166600 | 84170399 | 19.82599 | 32.77276 | 77.29889 | 33.40544 | -0.85345 | 1.326561 | 0.122821 |
| 13 | 99557400 | 99558399 | 4.798387 | 9.291807 | 27.76093 | 10.55012 | -0.99419 | -0.20946 | 0.122846 |
| 8 | 34467800 | 34469199 | 5.323184 | 10.1553 | 34.66887 | 13.81643 | -1.25184 | -0.03129 | 0.122857 |
| 2 | 19105000 | 19107999 | 19.00157 | 31.51129 | 75.10516 | 32.52965 | -0.86803 | 1.284948 | 0.122869 |
| 16 | 10806200 | 10827199 | 137.2542 | 202.8908 | 403.7749 | 166.2842 | -0.74519 | 3.731746 | 0.12288 |
| 12 | 22182200 | 22189599 | 156.8277 | 237.5968 | 184.9354 | 76.52782 | 0.640093 | 3.268408 | 0.122883 |
| 6 | 35060800 | 35062399 | 7.62597 | 14.06358 | 36.81782 | 15.57332 | -0.94336 | 0.217566 | 0.122885 |
| 11 | 27176800 | 27177399 | 12.01813 | 23.21015 | 15.8297 | 6.09164 | 1.098768 | -0.12553 | 0.12289 |
| 17 | 85241400 | 85243399 | 11.67773 | 20.48745 | 48.29441 | 21.35227 | -0.84314 | 0.62732 | 0.122893 |
| 7 | 126527600 | 126528799 | 4.013255 | 7.765653 | 29.42677 | 11.2222 | -1.33844 | -0.20437 | 0.122897 |
| 8 | 71159000 | 71160199 | 14.29725 | 26.61942 | 18.60058 | 7.762284 | 0.962021 | 0.170026 | 0.122899 |
| 17 | 36010600 | 36017199 | 30.02577 | 48.10327 | 124.6938 | 52.59918 | -1.02543 | 1.985966 | 0.122901 |
| 8 | 84816000 | 84826599 | 42.26852 | 66.754 | 169.9252 | 70.56168 | -1.02043 | 2.428607 | 0.122902 |
| 6 | 124721800 | 124724999 | 16.14664 | 26.89599 | 73.27331 | 31.83223 | -1.05952 | 1.223861 | 0.122903 |
| 6 | 144609000 | 144615799 | 47.69197 | 75.43305 | 161.3974 | 67.07176 | -0.7685 | 2.394566 | 0.122904 |
| 5 | 75355200 | 75356199 | 7.006329 | 13.00765 | 35.33381 | 14.8101 | -0.98666 | 0.169203 | 0.122916 |
| 8 | 93142600 | 93146599 | 19.38977 | 31.98729 | 80.62449 | 34.7971 | -0.94935 | 1.35055 | 0.122929 |
| 13 | 10248000 | 10248599 | 10.82244 | 21.1546 | 8.758634 | 3.016415 | 1.946817 | -0.53533 | 0.122932 |
| 11 | 3238600 | 3244999 | 34.41446 | 54.92833 | 133.1258 | 55.8551 | -0.9348 | 2.110452 | 0.122932 |
| 5 | 119650000 | 119650999 | 5.275672 | 10.13525 | 29.67221 | 11.60614 | -1.00582 | -0.10345 | 0.122938 |

| | | | | | | | | | |
|---|---|---|---|---|---|---|---|---|---|
| 9 | 37200400 | 37201599 | 13.24185 | 25.19064 | 17.76906 | 7.123997 | 0.996666 | 0.021693 | 0.12294 |
| 12 | 76374600 | 76377199 | 12.38438 | 21.43705 | 55.30609 | 24.26448 | -0.975 | 0.826676 | 0.122957 |
| 15 | 95027600 | 95029999 | 12.94817 | 22.56341 | 58.191 | 25.59667 | -0.97332 | 0.750125 | 0.12296 |
| 13 | 56397400 | 56411599 | 100.8088 | 151.695 | 321.0357 | 132.5761 | -0.81029 | 3.39918 | 0.122976 |
| 12 | 4867000 | 4867999 | 4.983128 | 9.321478 | 35.44353 | 14.67409 | -1.45225 | 0.104037 | 0.122986 |
| 17 | 5521000 | 5522999 | 11.85223 | 20.70405 | 50.48676 | 22.24184 | -0.89201 | 0.700413 | 0.122999 |
| 10 | 79762000 | 79778199 | 61.2179 | 93.64166 | 266.3303 | 110.501 | -1.21639 | 3.054008 | 0.123008 |
| 5 | 44993400 | 44993999 | 10.48997 | 20.46842 | 11.15917 | 3.97753 | 1.514486 | -0.41801 | 0.12302 |
| 11 | 29772600 | 29776999 | 31.4777 | 50.38116 | 126.2948 | 53.1985 | -0.97885 | 2.018406 | 0.123023 |
| 5 | 66498000 | 66501399 | 12.20833 | 21.19216 | 55.34966 | 24.2878 | -0.99184 | 0.798598 | 0.123025 |
| X | 30904400 | 30909799 | 124.032 | 192.2564 | 114.9686 | 47.7538 | 1.045623 | 2.810201 | 0.12303 |
| 11 | 82815800 | 82825799 | 64.36805 | 99.6588 | 217.0263 | 90.14703 | -0.81816 | 2.816165 | 0.123034 |
| X | 125081400 | 125084999 | 34.4961 | 58.32183 | 36.49775 | 15.97152 | 1.054794 | 1.099424 | 0.123036 |
| 7 | 141351600 | 141367599 | 109.9538 | 164.7601 | 335.1867 | 138.2703 | -0.75882 | 3.483581 | 0.123045 |
| 10 | 79876600 | 79878999 | 11.73908 | 20.40025 | 52.61289 | 23.10301 | -0.97315 | 0.779059 | 0.123054 |
| 13 | 31994600 | 31996199 | 11.73908 | 20.40025 | 52.61289 | 23.10301 | -0.97315 | 0.779059 | 0.123054 |
| 15 | 45821400 | 45821999 | 9.693041 | 18.88045 | 13.70166 | 4.999361 | 1.079074 | -0.34013 | 0.123056 |
| 4 | 123231200 | 123241799 | 61.84649 | 96.28516 | 199.3573 | 82.75181 | -0.73918 | 2.716834 | 0.123061 |
| 3 | 86973600 | 86975999 | 11.98503 | 20.61348 | 57.87465 | 25.38644 | -1.09833 | 0.887449 | 0.123084 |
| 15 | 35635000 | 35635799 | 5.71854 | 10.93904 | 30.65064 | 12.13738 | -0.96024 | -0.05555 | 0.123087 |
| 7 | 64923000 | 64931599 | 55.41806 | 86.88224 | 182.4212 | 75.6611 | -0.75139 | 2.588445 | 0.123122 |
| 8 | 103510200 | 103513999 | 24.25546 | 39.2484 | 99.71052 | 42.6411 | -0.97896 | 1.665253 | 0.123131 |
| 2 | 128959800 | 128961799 | 13.43293 | 23.31682 | 54.85648 | 24.07068 | -0.84192 | 0.804638 | 0.123134 |
| 1 | 20298600 | 20300199 | 14.48592 | 27.51546 | 16.80123 | 6.757644 | 1.198078 | 0.039848 | 0.123149 |
| 15 | 31316000 | 31322399 | 36.59758 | 58.57147 | 127.0303 | 53.50281 | -0.7706 | 2.02561 | 0.123156 |
| 3 | 96415000 | 96416199 | 11.20162 | 21.67009 | 15.7228 | 6.002581 | 1.020125 | -0.16259 | 0.123158 |
| 7 | 106147800 | 106148599 | 11.20162 | 21.67009 | 15.7228 | 6.002581 | 1.020125 | -0.16259 | 0.123158 |
| 10 | 82733000 | 82743399 | 63.7097 | 98.61265 | 215.3112 | 89.43475 | -0.82222 | 2.821165 | 0.12316 |
| 7 | 110832000 | 110833399 | 5.020041 | 9.62491 | 31.92253 | 12.55814 | -1.19246 | -0.08312 | 0.123169 |
| 7 | 32880600 | 32884399 | 43.23069 | 71.14198 | 47.27947 | 20.38429 | 0.966458 | 1.414372 | 0.123179 |
| 7 | 125074200 | 125078199 | 20.32332 | 33.53217 | 79.00709 | 34.10111 | -0.85231 | 1.350601 | 0.123193 |
| 8 | 88684400 | 88690199 | 31.83061 | 50.90173 | 131.6298 | 55.3775 | -1.02506 | 2.046631 | 0.1232 |
| 5 | 119742200 | 119744999 | 16.77044 | 28.32291 | 66.07259 | 28.90411 | -0.83634 | 1.079329 | 0.1232 |
| 4 | 119965000 | 119991599 | 188.3363 | 269.7668 | 554.8714 | 229.1797 | -0.83693 | 4.220902 | 0.12324 |
| 10 | 15002400 | 15003599 | 16.23151 | 30.62727 | 15.93319 | 6.49907 | 1.409321 | 0.097963 | 0.123249 |
| 18 | 85285800 | 85286199 | 2.735621 | 5.351353 | 19.76596 | 6.582817 | -1.11445 | -0.73048 | 0.123258 |
| 8 | 110215000 | 110216799 | 8.797013 | 15.80571 | 40.03462 | 17.50037 | -0.929 | 0.384479 | 0.123264 |
| 13 | 49915400 | 49916399 | 11.52736 | 20.22441 | 48.29213 | 21.35195 | -0.86152 | 0.626593 | 0.123266 |
| 2 | 146413000 | 146414599 | 9.078181 | 16.39223 | 40.65544 | 17.67268 | -0.89364 | 0.357632 | 0.123268 |
| 7 | 37225400 | 37230399 | 23.96112 | 39.07717 | 89.49707 | 38.44751 | -0.82019 | 1.518167 | 0.123273 |
| 5 | 140533800 | 140537999 | 20.38798 | 33.49944 | 84.50522 | 36.41018 | -0.95298 | 1.405593 | 0.12328 |
| 17 | 56499800 | 56506599 | 43.19484 | 68.22916 | 165.1187 | 68.57197 | -0.94737 | 2.425579 | 0.123287 |
| 5 | 94391400 | 94391599 | 10.18251 | 19.83263 | 12.8889 | 4.70664 | 1.237286 | -0.33163 | 0.123291 |
| 3 | 52547200 | 52552999 | 26.01268 | 42.0634 | 101.8633 | 43.53644 | -0.91111 | 1.683738 | 0.123299 |
| 8 | 84868600 | 84875399 | 31.97844 | 51.16769 | 131.638 | 55.42627 | -1.01675 | 2.028453 | 0.123327 |
| 5 | 131498800 | 131499799 | 7.891704 | 14.20069 | 39.5417 | 17.26278 | -1.06302 | 0.367837 | 0.123343 |
| 6 | 49194400 | 49195599 | 6.229176 | 11.66197 | 33.31359 | 13.77299 | -1.04186 | 0.106637 | 0.12335 |

| | | | | | | | | | |
|---|---|---|---|---|---|---|---|---|---|
| 10 | 11115200 | 11116599 | 6.977557 | 13.03877 | 35.55859 | 14.74756 | -0.98039 | 0.12741 | 0.123374 |
| 7 | 18769600 | 18771999 | 8.781625 | 15.39398 | 47.76773 | 21.10489 | -1.23642 | 0.626099 | 0.123376 |
| X | 156127200 | 156129599 | 24.40455 | 43.33085 | 20.47978 | 9.04234 | 1.467512 | 0.520266 | 0.123381 |
| 17 | 18973800 | 18974399 | 12.93526 | 25.1345 | 12.6832 | 4.72675 | 1.569706 | -0.23042 | 0.123393 |
| 7 | 21076200 | 21079799 | 35.74883 | 61.52548 | 25.20351 | 11.05562 | 1.665451 | 0.939614 | 0.123399 |
| 5 | 29259000 | 29262599 | 39.80756 | 65.66258 | 48.3286 | 20.8573 | 0.820952 | 1.378687 | 0.123424 |
| 9 | 17699800 | 17700199 | 7.658766 | 14.96754 | 11.8271 | 4.094495 | 1.025179 | -0.55563 | 0.123433 |
| 15 | 21472400 | 21473799 | 21.37415 | 38.97107 | 18.26202 | 7.842633 | 1.505291 | 0.342865 | 0.123436 |
| 3 | 12873200 | 12874399 | 15.095 | 28.44499 | 18.10412 | 7.402694 | 1.119465 | 0.103132 | 0.123456 |
| 9 | 8241400 | 8241999 | 9.357525 | 18.25371 | 13.2994 | 4.760136 | 1.098727 | -0.41083 | 0.123463 |
| 6 | 56683800 | 56685399 | 17.11563 | 31.1824 | 21.26723 | 9.142094 | 0.966757 | 0.319825 | 0.12347 |
| 11 | 114309200 | 114320599 | 85.13255 | 130.0935 | 257.3901 | 106.7672 | -0.69459 | 3.080139 | 0.123479 |
| 15 | 59017600 | 59021199 | 19.26529 | 32.10519 | 72.55025 | 31.527 | -0.79099 | 1.227944 | 0.123486 |
| 2 | 127236000 | 127239599 | 18.91467 | 31.61021 | 71.58108 | 31.16143 | -0.79407 | 1.194043 | 0.123494 |
| 4 | 70296000 | 70297599 | 268.6662 | 398.9338 | 170.3787 | 70.18494 | 1.479664 | 3.667313 | 0.123498 |
| 10 | 77686400 | 77702599 | 113.4054 | 169.8921 | 341.0799 | 140.6949 | -0.74005 | 3.488404 | 0.123509 |
| 5 | 125522000 | 125525199 | 10.08196 | 17.56171 | 55.8193 | 24.54722 | -1.27277 | 0.738875 | 0.123515 |
| 10 | 13198800 | 13200399 | 5.645893 | 10.66812 | 35.35619 | 14.37912 | -1.23556 | 0.041135 | 0.123527 |
| 18 | 65437200 | 65442399 | 25.69342 | 41.5503 | 101.1747 | 43.24637 | -0.91886 | 1.680855 | 0.123533 |
| 15 | 98747000 | 98755999 | 37.94224 | 60.18298 | 154.3963 | 64.28663 | -1.02619 | 2.313464 | 0.123542 |
| 11 | 98317600 | 98319799 | 9.238354 | 16.22268 | 46.70079 | 20.6637 | -1.12914 | 0.596847 | 0.123551 |
| 1 | 72466400 | 72470399 | 19.3347 | 32.02129 | 76.84723 | 33.25375 | -0.87816 | 1.30183 | 0.123553 |
| 18 | 88586800 | 88588799 | 25.64047 | 45.2741 | 20.90564 | 9.249346 | 1.497603 | 0.579585 | 0.123558 |
| 2 | 32183400 | 32186799 | 10.65885 | 18.57087 | 51.52293 | 22.66112 | -1.07733 | 0.734594 | 0.123576 |
| 11 | 4665600 | 4666999 | 5.70531 | 10.9107 | 31.18614 | 12.35932 | -0.9894 | -0.04996 | 0.123577 |
| 12 | 91745200 | 91746399 | 6.419145 | 11.95325 | 35.95839 | 15.00227 | -1.12752 | 0.146337 | 0.123578 |
| 2 | 92317600 | 92319199 | 8.831461 | 15.77629 | 41.54 | 18.24699 | -0.98968 | 0.423449 | 0.123594 |
| 1 | 176363000 | 176363599 | 9.13157 | 17.78914 | 13.88986 | 5.059284 | 0.977621 | -0.35211 | 0.123601 |
| 17 | 31069800 | 31072999 | 21.74149 | 35.87693 | 79.87631 | 34.47826 | -0.77087 | 1.352684 | 0.123613 |
| 5 | 121082600 | 121095199 | 76.50935 | 116.8157 | 263.5071 | 109.2829 | -0.88456 | 3.095652 | 0.123635 |
| 9 | 26179000 | 26180799 | 20.67414 | 37.81891 | 18.31215 | 7.831612 | 1.462144 | 0.319531 | 0.123638 |
| 1 | 109393600 | 109393799 | 8.391361 | 16.37914 | 12.83509 | 4.547996 | 1.00736 | -0.46141 | 0.12364 |
| 1 | 46457200 | 46458999 | 8.07946 | 14.87987 | 38.47477 | 16.304 | -0.92813 | 0.230346 | 0.123642 |
| 8 | 102098600 | 102100399 | 19.25244 | 35.23551 | 17.59752 | 7.520957 | 1.417548 | 0.302465 | 0.123647 |
| 18 | 37044400 | 37048999 | 102.2535 | 161.2615 | 94.50109 | 39.21784 | 1.093797 | 2.486902 | 0.123651 |
| 11 | 103051600 | 103053199 | 13.14461 | 22.75664 | 55.89937 | 24.52489 | -0.90468 | 0.828698 | 0.123652 |
| 2 | 153281800 | 153287599 | 22.44064 | 36.46975 | 93.25136 | 40.00666 | -0.98222 | 1.569799 | 0.123655 |
| 13 | 53052200 | 53055599 | 12.1951 | 20.85397 | 61.02713 | 26.75901 | -1.15944 | 0.945269 | 0.123657 |
| 10 | 129793400 | 129796199 | 27.9254 | 48.34756 | 31.68276 | 13.90028 | 0.993033 | 0.861556 | 0.123669 |
| 6 | 86094600 | 86097399 | 13.2363 | 22.68069 | 58.96598 | 25.85949 | -0.98914 | 0.928234 | 0.12369 |
| 7 | 20703200 | 20706999 | 83.72543 | 132.2698 | 97.69323 | 40.57401 | 0.762409 | 2.43569 | 0.123697 |
| X | 149042000 | 149043399 | 32.02945 | 54.42035 | 36.66265 | 16.06742 | 0.949151 | 1.05076 | 0.123703 |
| 18 | 3086000 | 3094599 | 157.1545 | 239.8211 | 153.3543 | 63.59216 | 0.931884 | 3.121406 | 0.123705 |
| 1 | 134075600 | 134084399 | 45.48162 | 72.20063 | 156.1237 | 65.05899 | -0.77902 | 2.293661 | 0.123705 |
| 19 | 44654800 | 44666199 | 54.8541 | 85.48122 | 206.9547 | 85.89312 | -0.96345 | 2.697191 | 0.123712 |
| 12 | 22563000 | 22568199 | 48.86787 | 79.2044 | 53.84769 | 23.05581 | 0.920142 | 1.650342 | 0.123733 |
| 17 | 20022600 | 20026199 | 45.96973 | 75.70893 | 43.95741 | 18.95831 | 1.161125 | 1.410596 | 0.123767 |

| | | | | | | | | | |
|---|---|---|---|---|---|---|---|---|---|
| 15 | 80185400 | 80192199 | 33.98116 | 54.13871 | 140.1485 | 58.65383 | -1.03252 | 2.170237 | 0.123771 |
| 4 | 140347200 | 140357399 | 38.50843 | 60.99432 | 169.6728 | 70.56147 | -1.14481 | 2.354974 | 0.123772 |
| 4 | 117341200 | 117347599 | 35.88986 | 57.23225 | 138.3387 | 57.97012 | -0.93243 | 2.140543 | 0.123781 |
| 4 | 147387800 | 147393799 | 117.4256 | 182.5793 | 122.8246 | 50.99337 | 0.879399 | 2.755283 | 0.123798 |
| 5 | 58763800 | 58765799 | 11.20089 | 19.67989 | 52.59958 | 23.28471 | -1.0237 | 0.605122 | 0.123803 |
| 19 | 25428400 | 25430999 | 12.64619 | 22.00639 | 52.66314 | 23.14476 | -0.86581 | 0.762328 | 0.123803 |
| 7 | 27740400 | 27752599 | 64.02009 | 98.85858 | 229.3481 | 95.26651 | -0.9123 | 2.865955 | 0.123806 |
| 7 | 105385400 | 105386399 | 5.99249 | 11.16392 | 36.50866 | 15.21993 | -1.24594 | 0.140778 | 0.123818 |
| 9 | 115138200 | 115140199 | 13.12137 | 22.41577 | 62.74587 | 27.51045 | -1.09614 | 0.958785 | 0.123822 |
| 9 | 19970000 | 19974399 | 64.71433 | 103.4396 | 81.76247 | 34.37545 | 0.680875 | 2.08007 | 0.123848 |
| 2 | 173246000 | 173263599 | 108.9863 | 163.1742 | 343.8581 | 141.8244 | -0.81068 | 3.499021 | 0.12385 |
| 4 | 4133800 | 4134399 | 4.502973 | 8.711567 | 27.62309 | 10.54391 | -1.08445 | -0.19687 | 0.12386 |
| 17 | 20546600 | 20548399 | 21.91264 | 39.02589 | 21.69748 | 9.555249 | 1.237162 | 0.486707 | 0.123872 |
| 18 | 12658000 | 12659199 | 3.645441 | 7.103195 | 24.16937 | 8.778587 | -1.11637 | -0.39435 | 0.123886 |
| 5 | 68650200 | 68654999 | 55.46093 | 89.12998 | 69.6784 | 29.5311 | 0.705673 | 1.898769 | 0.123891 |
| 14 | 108905600 | 108907199 | 23.14953 | 40.57995 | 27.88221 | 12.31885 | 0.92608 | 0.667141 | 0.123908 |
| 17 | 56170200 | 56171399 | 40.71702 | 67.03152 | 48.87477 | 21.07557 | 0.833424 | 1.405992 | 0.123918 |
| 8 | 120910600 | 120920399 | 53.49382 | 83.46321 | 199.0674 | 82.60617 | -0.94048 | 2.67477 | 0.123925 |
| 3 | 144649600 | 144650999 | 21.58242 | 38.43396 | 22.95485 | 10.10982 | 1.134777 | 0.488583 | 0.123931 |
| 14 | 30306800 | 30310799 | 15.05944 | 25.46353 | 67.07521 | 29.3533 | -1.01074 | 1.063432 | 0.12395 |
| 9 | 12802400 | 12804199 | 21.22476 | 37.74359 | 23.50894 | 10.36622 | 1.073446 | 0.49566 | 0.123978 |
| 9 | 58762400 | 58772399 | 52.34316 | 81.63228 | 197.446 | 81.93708 | -0.96111 | 2.682587 | 0.123993 |
| 2 | 82089000 | 82089799 | 7.125772 | 13.31279 | 34.96623 | 14.50731 | -0.92659 | 0.130954 | 0.123998 |
| 8 | 61745200 | 61746199 | 7.125772 | 13.31279 | 34.96623 | 14.50731 | -0.92659 | 0.130954 | 0.123998 |
| 14 | 16588600 | 16590599 | 11.11707 | 19.33802 | 52.05635 | 22.87254 | -1.03427 | 0.762041 | 0.124007 |
| 3 | 50988600 | 50995199 | 44.34478 | 70.42931 | 150.6611 | 62.81241 | -0.76279 | 2.278124 | 0.124013 |
| 1 | 91586600 | 91589199 | 14.59533 | 24.67473 | 65.31739 | 28.58262 | -1.01763 | 1.062748 | 0.124017 |
| 18 | 60800600 | 60809199 | 48.96916 | 77.0928 | 175.1853 | 72.68512 | -0.86059 | 2.501402 | 0.124019 |
| 3 | 80017200 | 80018999 | 17.61452 | 32.61008 | 18.51106 | 7.788759 | 1.25065 | 0.230161 | 0.124063 |
| 6 | 92098600 | 92099599 | 8.267976 | 15.09093 | 38.21612 | 16.39005 | -0.91063 | 0.282828 | 0.124067 |
| 18 | 25681000 | 25695399 | 80.78077 | 123.0863 | 277.3081 | 114.9503 | -0.88515 | 3.138796 | 0.124095 |
| 15 | 76596800 | 76598599 | 7.111433 | 12.83695 | 39.07752 | 16.99335 | -1.18627 | 0.344639 | 0.124099 |
| 5 | 60664000 | 60666999 | 41.18174 | 67.83805 | 47.81368 | 20.62335 | 0.882436 | 1.39675 | 0.124114 |
| 8 | 87959800 | 87965399 | 26.16579 | 42.31859 | 100.7363 | 43.06405 | -0.8861 | 1.678627 | 0.124116 |
| 19 | 47288800 | 47292599 | 17.08843 | 28.4335 | 73.33926 | 31.83817 | -0.98102 | 1.238967 | 0.124119 |
| 16 | 21151200 | 21152399 | 4.663554 | 9.043116 | 27.27881 | 10.3006 | -0.99911 | -0.23516 | 0.124131 |
| X | 147402000 | 147404199 | 22.27409 | 40.48011 | 17.48288 | 7.538618 | 1.618015 | 0.365333 | 0.124141 |
| 3 | 89679200 | 89682999 | 16.36556 | 27.48594 | 69.22851 | 30.23161 | -0.94686 | 1.130117 | 0.124144 |
| 1 | 149052400 | 149056599 | 69.30437 | 110.6112 | 72.91061 | 30.58913 | 0.940065 | 2.12792 | 0.124145 |
| 3 | 107490000 | 107491399 | 11.40951 | 19.89677 | 51.01449 | 22.45028 | -0.96419 | 0.723846 | 0.124173 |
| 11 | 96190000 | 96196199 | 22.65701 | 36.70368 | 98.40683 | 42.11892 | -1.05483 | 1.63863 | 0.124186 |
| 14 | 95387400 | 95390799 | 66.8694 | 106.9997 | 77.80802 | 32.7629 | 0.802654 | 2.04898 | 0.124196 |
| 10 | 47975800 | 47977799 | 27.18812 | 47.48066 | 27.19821 | 11.98007 | 1.18731 | 0.746064 | 0.124205 |
| 8 | 57764400 | 57767799 | 22.55925 | 37.14017 | 85.18789 | 36.73494 | -0.81518 | 1.386091 | 0.12421 |
| 11 | 67565000 | 67566799 | 6.155806 | 11.39009 | 35.76773 | 15.06098 | -1.1985 | 0.182969 | 0.12421 |
| 2 | 158450400 | 158461999 | 77.14141 | 117.528 | 273.8617 | 113.5179 | -0.93383 | 3.140243 | 0.124213 |
| 9 | 63516600 | 63520999 | 17.53034 | 29.23407 | 71.83716 | 31.2307 | -0.91138 | 1.214046 | 0.124218 |

| | | | | | | | | | |
|---|---|---|---|---|---|---|---|---|---|
| 7 | 47260000 | 47265999 | 82.15011 | 130.8462 | 77.66312 | 32.49087 | 1.087963 | 2.205368 | 0.124223 |
| 18 | 89786600 | 89788199 | 11.66033 | 20.09907 | 56.8107 | 24.92141 | -1.10719 | 0.864801 | 0.124247 |
| 3 | 37696600 | 37697999 | 7.725447 | 13.96392 | 39.0853 | 16.96842 | -1.06526 | 0.338488 | 0.124251 |
| 2 | 24908400 | 24909799 | 3.889108 | 7.557355 | 26.15588 | 9.743277 | -1.17426 | -0.29203 | 0.124259 |
| 18 | 53482200 | 53483999 | 24.63552 | 42.77244 | 31.03406 | 13.6196 | 0.847581 | 0.8193 | 0.124264 |
| 7 | 22032800 | 22033399 | 15.81024 | 29.31169 | 17.53359 | 7.363128 | 1.177983 | 0.205246 | 0.124286 |
| 11 | 100034600 | 100038799 | 16.3176 | 27.15951 | 74.41495 | 32.31222 | -1.06779 | 1.233361 | 0.124288 |
| 2 | 164787600 | 164805799 | 126.6295 | 187.0628 | 413.2793 | 170.1972 | -0.89672 | 3.741515 | 0.12431 |
| 17 | 72207600 | 72215599 | 50.73397 | 79.90022 | 177.3959 | 73.60577 | -0.82675 | 2.494951 | 0.124313 |
| 7 | 73199800 | 73200199 | 3.149328 | 6.156169 | 21.87689 | 7.52847 | -1.10478 | -0.60041 | 0.124333 |
| 4 | 83362800 | 83363599 | 9.12037 | 17.80677 | 12.02496 | 4.241707 | 1.223886 | -0.46985 | 0.124335 |
| 12 | 23489400 | 23493799 | 61.15971 | 98.3033 | 64.52266 | 27.35201 | 0.957631 | 1.901299 | 0.124365 |
| 7 | 14866600 | 14876599 | 109.8521 | 172.2972 | 104.0435 | 43.15119 | 1.044607 | 2.599393 | 0.124369 |
| 12 | 3960400 | 3962599 | 7.735641 | 13.66705 | 46.68822 | 20.65398 | -1.37122 | 0.512863 | 0.124375 |
| 1 | 62699800 | 62701799 | 13.67466 | 23.48812 | 57.39693 | 25.17586 | -0.89956 | 0.906669 | 0.124387 |
| 5 | 95475600 | 95478599 | 44.54584 | 73.56567 | 41.11778 | 17.75786 | 1.216816 | 1.370256 | 0.124398 |
| 13 | 31130800 | 31131999 | 6.83921 | 12.72832 | 34.82127 | 14.54165 | -0.9921 | 0.15437 | 0.124398 |
| 4 | 3199800 | 3201399 | 30.48617 | 52.97465 | 27.31033 | 11.9917 | 1.338263 | 0.831742 | 0.124422 |
| 11 | 119243000 | 119250399 | 32.98399 | 52.65457 | 134.8775 | 56.60431 | -1.0142 | 2.103278 | 0.124423 |
| 10 | 76365800 | 76368599 | 8.77342 | 15.44882 | 45.64246 | 20.20486 | -1.16516 | 0.556058 | 0.124423 |
| 18 | 77575200 | 77576799 | 7.894408 | 14.22934 | 40.61465 | 17.69435 | -1.09692 | 0.358234 | 0.124432 |
| 9 | 105007000 | 105009199 | 10.4541 | 18.33424 | 47.24319 | 20.88127 | -0.9706 | 0.624851 | 0.124438 |
| 12 | 79075000 | 79076599 | 7.067367 | 12.90066 | 37.13249 | 15.92359 | -1.09288 | 0.273661 | 0.124438 |
| 16 | 10723200 | 10727199 | 27.9045 | 45.14442 | 98.0594 | 41.92911 | -0.75391 | 1.675244 | 0.124447 |
| 8 | 121009200 | 121013399 | 16.62461 | 27.68169 | 74.50415 | 32.35921 | -1.04219 | 1.229521 | 0.124456 |
| 1 | 3196200 | 3197599 | 15.16048 | 28.10199 | 19.97859 | 8.391952 | 0.930378 | 0.206356 | 0.124457 |
| 1 | 4927600 | 4928999 | 8.774322 | 15.48669 | 47.74428 | 21.12538 | -1.22528 | 0.531056 | 0.124458 |
| 15 | 76130600 | 76134999 | 15.52182 | 25.56117 | 83.07045 | 35.826 | -1.31536 | 1.366515 | 0.124476 |
| 7 | 44744800 | 44749799 | 21.77954 | 35.83022 | 81.18441 | 34.99769 | -0.79782 | 1.393388 | 0.12451 |
| 1 | 134914600 | 134916399 | 10.82205 | 18.83992 | 52.59788 | 23.12211 | -1.08661 | 0.747894 | 0.124522 |
| 17 | 71972200 | 71990399 | 126.2338 | 187.0573 | 393.6996 | 162.1448 | -0.82259 | 3.689315 | 0.124532 |
| 7 | 10840800 | 10842399 | 34.66602 | 57.78533 | 40.77028 | 17.71982 | 0.882701 | 1.240764 | 0.124551 |
| 10 | 62983200 | 62984199 | 3.337371 | 6.451871 | 31.01866 | 11.88031 | -1.68269 | -0.19233 | 0.124552 |
| 9 | 15619600 | 15623799 | 35.58593 | 57.09491 | 118.8549 | 50.24845 | -0.70728 | 1.944218 | 0.124553 |
| 2 | 34561000 | 34562799 | 7.746521 | 13.59501 | 47.74513 | 21.1153 | -1.41351 | 0.600923 | 0.124576 |
| 4 | 124597200 | 124604399 | 28.4137 | 45.5663 | 122.3667 | 51.69634 | -1.07451 | 1.951649 | 0.124582 |
| 5 | 140720600 | 140727799 | 49.18939 | 77.30368 | 180.163 | 74.73331 | -0.89898 | 2.537662 | 0.124583 |
| 3 | 12032000 | 12033199 | 11.7269 | 22.79878 | 12.05826 | 4.474411 | 1.506604 | -0.2629 | 0.124595 |
| 14 | 21881000 | 21882599 | 9.513329 | 16.77447 | 45.11126 | 19.96429 | -1.02983 | 0.543551 | 0.1246 |
| 9 | 103528800 | 103529999 | 8.130109 | 14.80374 | 38.69163 | 16.64385 | -0.95828 | 0.299209 | 0.124608 |
| 8 | 4840400 | 4841799 | 14.57729 | 28.37621 | 9.581637 | 3.519292 | 2.158191 | -0.26795 | 0.124613 |
| 8 | 13438600 | 13440999 | 17.98652 | 30.02394 | 70.78182 | 30.79075 | -0.85178 | 1.204492 | 0.124619 |
| 16 | 32302200 | 32305199 | 11.88365 | 20.677 | 52.09494 | 22.89334 | -0.93951 | 0.760316 | 0.124619 |
| 19 | 15038400 | 15038999 | 10.72236 | 20.82075 | 14.65239 | 5.49007 | 1.088501 | -0.23351 | 0.124627 |
| 10 | 93475600 | 93479399 | 20.1244 | 33.31372 | 76.28471 | 32.99977 | -0.81075 | 1.309989 | 0.12463 |
| 8 | 18511600 | 18512399 | 3.262371 | 6.374801 | 24.33839 | 8.455163 | -1.22012 | -0.55652 | 0.124636 |
| 12 | 105556000 | 105573199 | 102.3967 | 152.4434 | 379.5046 | 156.3728 | -1.05839 | 3.603824 | 0.124641 |

| | | | | | | | | | |
|---|---|---|---|---|---|---|---|---|---|
| 8 | 115637200 | 115642199 | 29.06231 | 46.9066 | 102.7454 | 43.80725 | -0.76987 | 1.746018 | 0.124647 |
| 3 | 17963800 | 17965399 | 13.67553 | 25.81902 | 18.70628 | 7.621064 | 0.938179 | 0.078769 | 0.124649 |
| 17 | 82697600 | 82699399 | 19.1592 | 34.64183 | 21.53721 | 9.346362 | 1.090884 | 0.378487 | 0.124663 |
| 5 | 28689200 | 28690999 | 31.54086 | 50.69083 | 112.787 | 47.80445 | -0.8 | 1.884883 | 0.124667 |
| 19 | 47917000 | 47919799 | 10.16056 | 17.80987 | 48.30118 | 21.33922 | -1.04408 | 0.632983 | 0.124675 |
| 19 | 55137800 | 55138799 | 17.67583 | 32.38522 | 20.86735 | 8.905253 | 1.055486 | 0.293593 | 0.124684 |
| 1 | 18576000 | 18577199 | 17.19122 | 31.52183 | 20.96483 | 8.9374 | 1.011231 | 0.285926 | 0.124693 |
| 9 | 28216200 | 28219199 | 19.74191 | 32.87081 | 73.79954 | 32.05226 | -0.78175 | 1.238381 | 0.124715 |
| 16 | 59413800 | 59414999 | 14.43403 | 27.61586 | 15.10909 | 5.980031 | 1.374638 | -0.01568 | 0.124724 |
| 17 | 59042600 | 59043999 | 14.43403 | 27.61586 | 15.10909 | 5.980031 | 1.374638 | -0.01568 | 0.124724 |
| 1 | 99444600 | 99445199 | 13.81521 | 26.68451 | 14.22696 | 5.471295 | 1.449957 | -0.11379 | 0.124738 |
| 13 | 45211000 | 45213199 | 8.949147 | 16.07386 | 40.5512 | 17.73102 | -0.92376 | 0.387595 | 0.124741 |
| 1 | 31199400 | 31200999 | 16.69526 | 30.78718 | 20.30536 | 8.589479 | 1.030741 | 0.248336 | 0.124742 |
| 4 | 116842800 | 116845199 | 10.28897 | 18.04039 | 48.29288 | 21.34098 | -1.025 | 0.63214 | 0.124758 |
| 1 | 101289800 | 101294399 | 47.15757 | 76.98467 | 51.74225 | 22.23637 | 0.944719 | 1.529281 | 0.124761 |
| 3 | 89404400 | 89405999 | 6.657747 | 12.17743 | 39.86783 | 17.05324 | -1.27564 | 0.260523 | 0.124772 |
| X | 30910800 | 30912199 | 13.49581 | 26.33303 | 7.098046 | 2.530839 | 2.507715 | -0.38845 | 0.124779 |
| 6 | 72697400 | 72699599 | 11.9893 | 20.99105 | 49.93284 | 22.03851 | -0.8564 | 0.663601 | 0.124812 |
| 12 | 22151200 | 22157599 | 186.8433 | 279.3306 | 209.0355 | 86.19215 | 0.680001 | 3.532962 | 0.12482 |
| 5 | 3225600 | 3227199 | 18.09567 | 33.97052 | 15.83948 | 6.518369 | 1.556261 | 0.14665 | 0.124825 |
| 6 | 45832600 | 45834599 | 19.9969 | 36.10583 | 21.60968 | 9.394266 | 1.144089 | 0.393627 | 0.124851 |
| 6 | 119824600 | 119825799 | 4.579285 | 8.593203 | 33.26553 | 13.69874 | -1.46912 | 0.079548 | 0.124857 |
| 8 | 65185800 | 65186799 | 6.983181 | 13.02515 | 34.35111 | 14.29147 | -0.93556 | 0.138877 | 0.124862 |
| 1 | 90346600 | 90359599 | 95.7359 | 145.0157 | 293.411 | 121.4091 | -0.73725 | 3.264295 | 0.124863 |
| 3 | 35707000 | 35708399 | 13.46648 | 25.87827 | 15.95396 | 6.235182 | 1.221558 | -0.06209 | 0.124869 |
| 12 | 33821200 | 33822599 | 13.46648 | 25.87827 | 15.95396 | 6.235182 | 1.221558 | -0.06209 | 0.124869 |
| 10 | 21034000 | 21037799 | 19.53778 | 31.91594 | 89.35506 | 38.4096 | -1.10713 | 1.485183 | 0.12487 |
| 1 | 89144200 | 89145199 | 3.441367 | 6.65545 | 28.0585 | 10.72747 | -1.4918 | -0.20343 | 0.124896 |
| X | 30011600 | 30011999 | 11.83957 | 22.83796 | 15.88359 | 6.140719 | 1.064657 | -0.11236 | 0.124897 |
| 13 | 39067800 | 39069799 | 18.71823 | 34.10482 | 19.91503 | 8.559976 | 1.189064 | 0.328525 | 0.124904 |
| 7 | 55053200 | 55055199 | 19.04588 | 35.27689 | 17.31443 | 7.280514 | 1.459429 | 0.237114 | 0.124912 |
| 7 | 7535800 | 7542399 | 81.57456 | 129.7912 | 84.21842 | 35.18147 | 0.958038 | 2.240503 | 0.12493 |
| 1 | 152498600 | 152501199 | 9.811289 | 17.29673 | 44.59128 | 19.73488 | -0.96928 | 0.547281 | 0.12494 |
| 6 | 147311600 | 147312999 | 16.3443 | 30.38638 | 19.26792 | 8.060655 | 1.098257 | 0.198721 | 0.124952 |
| 7 | 38955200 | 38955599 | 10.44802 | 20.27153 | 15.30676 | 5.764288 | 0.981716 | -0.21733 | 0.124956 |
| 17 | 20232200 | 20237199 | 127.8004 | 196.3581 | 152.6671 | 63.39084 | 0.657727 | 2.977154 | 0.124957 |
| 15 | 98829600 | 98833599 | 11.89906 | 20.6636 | 53.1569 | 23.33149 | -0.96972 | 0.790192 | 0.124965 |
| 6 | 67170800 | 67172199 | 6.852686 | 12.75178 | 35.95913 | 15.02008 | -1.03656 | 0.154442 | 0.124969 |
| 3 | 107975800 | 107985199 | 40.65322 | 64.5456 | 149.6441 | 62.36838 | -0.87905 | 2.285405 | 0.124975 |
| 4 | 45671800 | 45679799 | 37.71673 | 60.01919 | 154.2408 | 64.4461 | -1.02407 | 2.213257 | 0.124978 |
| 12 | 22576000 | 22581999 | 67.90847 | 108.2904 | 86.5648 | 36.2742 | 0.661056 | 2.157259 | 0.124989 |
| 9 | 46125200 | 46126399 | 5.184969 | 9.83821 | 31.43213 | 12.67077 | -1.17037 | 0.000705 | 0.124994 |
| 7 | 11809000 | 11810399 | 50.07949 | 81.07728 | 58.73621 | 25.12815 | 0.827343 | 1.679742 | 0.124996 |
| 11 | 58868000 | 58869599 | 8.341391 | 14.6639 | 46.68424 | 20.67284 | -1.2731 | 0.574021 | 0.125003 |
| 15 | 83901200 | 83901999 | 11.29201 | 21.79197 | 16.38829 | 6.327321 | 0.954396 | -0.12163 | 0.125014 |
| 14 | 57928600 | 57930399 | 7.537452 | 13.68514 | 38.10696 | 16.45146 | -1.05204 | 0.310891 | 0.125016 |
| 18 | 76269600 | 76275399 | 34.47232 | 54.93623 | 141.608 | 59.28706 | -1.02601 | 2.161171 | 0.125021 |

| | | | | | | | | | |
|---|---|---|---|---|---|---|---|---|---|
| 17 | 10607600 | 10617399 | 46.83845 | 73.83538 | 179.2925 | 74.40194 | -0.95471 | 2.473588 | 0.125021 |
| 2 | 29506200 | 29510599 | 14.69584 | 24.97523 | 62.70594 | 27.48142 | -0.94121 | 1.013892 | 0.125023 |
| 8 | 115836800 | 115839799 | 14.69584 | 24.97523 | 62.70594 | 27.48142 | -0.94121 | 1.013892 | 0.125023 |
| 15 | 77780200 | 77780399 | 12.74118 | 24.79544 | 12.25841 | 4.51212 | 1.615427 | -0.27508 | 0.125036 |
| 17 | 9641800 | 9642599 | 14.34503 | 27.43824 | 16.43921 | 6.511481 | 1.244928 | -0.00957 | 0.125036 |
| 8 | 25134000 | 25135199 | 15.26335 | 28.88876 | 17.32818 | 7.022456 | 1.214581 | 0.072917 | 0.125049 |
| 13 | 46581600 | 46583799 | 19.43159 | 32.34388 | 73.14168 | 31.76019 | -0.79205 | 1.24084 | 0.125065 |
| 6 | 127905800 | 127917199 | 69.18447 | 106.5738 | 234.1953 | 97.27776 | -0.83687 | 2.913696 | 0.125071 |
| 6 | 143346200 | 143347599 | 6.634139 | 12.54848 | 33.71569 | 13.68183 | -0.93244 | 0.041449 | 0.125074 |
| 7 | 132614600 | 132617199 | 9.716008 | 17.02351 | 48.82915 | 21.56484 | -1.12436 | 0.638395 | 0.125078 |
| 19 | 37683400 | 37692199 | 63.4074 | 98.515 | 209.2887 | 86.90063 | -0.77856 | 2.753063 | 0.12509 |
| 11 | 21945600 | 21954999 | 75.51174 | 115.3987 | 264.2659 | 109.6209 | -0.90525 | 3.076415 | 0.125145 |
| 2 | 180343200 | 180345199 | 7.37054 | 13.47923 | 37.16907 | 15.90197 | -1.02984 | 0.262824 | 0.12515 |
| 6 | 9874600 | 9875799 | 13.23774 | 25.61075 | 13.65341 | 5.210957 | 1.459613 | -0.14681 | 0.125166 |
| 4 | 130511400 | 130520999 | 58.98862 | 92.06847 | 193.9843 | 80.49397 | -0.76147 | 2.670676 | 0.125178 |
| 15 | 99474000 | 99479999 | 27.64035 | 44.56622 | 104.9345 | 44.68198 | -0.87554 | 1.774621 | 0.12519 |
| 11 | 64399000 | 64400799 | 8.305293 | 15.0495 | 40.74565 | 17.63414 | -1.01528 | 0.329438 | 0.12519 |
| 7 | 19816400 | 19820199 | 79.5948 | 126.5667 | 88.99303 | 37.14709 | 0.841518 | 2.258114 | 0.125195 |
| 5 | 141832800 | 141834599 | 6.968356 | 13.04562 | 35.58705 | 14.71332 | -0.97656 | 0.118299 | 0.125198 |
| 10 | 52512400 | 52514199 | 45.46992 | 74.13222 | 54.44694 | 23.38753 | 0.815797 | 1.54511 | 0.1252 |
| 11 | 54703200 | 54705199 | 8.463224 | 15.30725 | 39.62345 | 17.18831 | -0.95294 | 0.340171 | 0.125211 |
| 17 | 7222600 | 7224399 | 8.463224 | 15.30725 | 39.62345 | 17.18831 | -0.95294 | 0.340171 | 0.125211 |
| 11 | 17206800 | 17208199 | 5.904945 | 11.19006 | 32.01837 | 12.94191 | -1.01697 | 0.017071 | 0.125226 |
| 10 | 61913800 | 61914599 | 19.08638 | 33.86139 | 24.48807 | 10.81935 | 0.857497 | 0.497969 | 0.125226 |
| 11 | 62622200 | 62622999 | 4.431047 | 8.537887 | 29.14051 | 11.30157 | -1.21252 | -0.14198 | 0.125232 |
| 16 | 37318600 | 37320199 | 9.071064 | 16.38227 | 40.6612 | 17.67129 | -0.89468 | 0.355311 | 0.125235 |
| X | 33266200 | 33271999 | 96.25442 | 151.6894 | 104.0215 | 43.16468 | 0.866604 | 2.49957 | 0.125239 |
| 9 | 102598800 | 102602599 | 15.71168 | 26.46992 | 67.00479 | 29.29015 | -0.95376 | 1.098454 | 0.125253 |
| 1 | 49224600 | 49225599 | 16.80156 | 30.57174 | 20.67951 | 8.903437 | 0.977628 | 0.324745 | 0.125262 |
| 14 | 87680200 | 87682399 | 9.919259 | 17.60901 | 45.13268 | 19.92995 | -0.95694 | 0.475633 | 0.12527 |
| 1 | 163978400 | 163979799 | 5.92183 | 11.17202 | 32.41536 | 13.22682 | -1.0489 | 0.050074 | 0.125313 |
| 4 | 35285400 | 35287799 | 27.81883 | 48.34204 | 29.36154 | 12.89275 | 1.102732 | 0.818706 | 0.125343 |
| 11 | 72025400 | 72035799 | 46.47335 | 73.32997 | 168.0547 | 69.75383 | -0.87021 | 2.452475 | 0.125344 |
| 2 | 119097000 | 119097799 | 3.235091 | 6.315383 | 22.7172 | 8.032233 | -1.15928 | -0.50237 | 0.125361 |
| 4 | 18848600 | 18848999 | 3.235091 | 6.315383 | 22.7172 | 8.032233 | -1.15928 | -0.50237 | 0.125361 |
| 7 | 59433600 | 59434399 | 15.50245 | 28.55105 | 18.24085 | 7.729588 | 1.074713 | 0.240752 | 0.125363 |
| 9 | 5256400 | 5257199 | 11.71564 | 22.77406 | 13.36025 | 4.962489 | 1.359091 | -0.25481 | 0.125368 |
| 17 | 38432000 | 38435199 | 72.24348 | 114.916 | 85.79471 | 35.82751 | 0.755654 | 2.24123 | 0.125369 |
| 7 | 106276800 | 106277799 | 12.52574 | 23.86119 | 17.13213 | 6.848653 | 0.974176 | 0.001973 | 0.12537 |
| X | 148548200 | 148550999 | 46.77017 | 76.8474 | 42.36781 | 18.25467 | 1.234255 | 1.439396 | 0.125383 |
| 2 | 59699800 | 59702799 | 10.7679 | 18.86507 | 49.9088 | 22.0402 | -1.00879 | 0.645767 | 0.125384 |
| 7 | 3099600 | 3110199 | 318.9848 | 456.883 | 339.427 | 140.1953 | 0.631558 | 4.226237 | 0.125391 |
| 8 | 87611200 | 87612599 | 7.52605 | 13.75193 | 38.77607 | 16.60725 | -1.06322 | 0.270967 | 0.125397 |
| 2 | 179402000 | 179406799 | 23.64564 | 38.20072 | 102.4409 | 43.7315 | -1.05923 | 1.708485 | 0.125403 |
| 7 | 59781800 | 59783399 | 38.42405 | 63.33989 | 48.86197 | 21.08178 | 0.75299 | 1.382414 | 0.125405 |
| 11 | 18957200 | 18959199 | 10.80515 | 18.84818 | 51.53351 | 22.68273 | -1.05631 | 0.717128 | 0.125408 |
| 2 | 124450000 | 124453199 | 16.25058 | 27.20381 | 71.33389 | 31.08688 | -1.00472 | 1.167179 | 0.12541 |

| | | | | | | | | | |
|---|---|---|---|---|---|---|---|---|---|
| 7 | 24587000 | 24588599 | 5.61148 | 10.64043 | 31.44238 | 12.6927 | -1.06093 | 0.009416 | 0.125418 |
| 6 | 112703800 | 112707399 | 16.50732 | 27.74934 | 66.98931 | 29.26388 | -0.88576 | 1.121176 | 0.125433 |
| 8 | 17911800 | 17913999 | 19.72516 | 35.63249 | 21.52174 | 9.350353 | 1.131382 | 0.388107 | 0.125434 |
| 6 | 135202000 | 135202799 | 4.239482 | 8.237657 | 26.25627 | 9.785931 | -1.05943 | -0.28927 | 0.12544 |
| 18 | 84124600 | 84125399 | 4.239482 | 8.237657 | 26.25627 | 9.785931 | -1.05943 | -0.28927 | 0.12544 |
| 2 | 116797200 | 116799999 | 11.08214 | 19.38653 | 51.54058 | 22.73201 | -1.016 | 0.674061 | 0.125441 |
| 9 | 26423600 | 26424199 | 6.970007 | 13.04696 | 36.78757 | 15.21312 | -1.02495 | 0.118397 | 0.125447 |
| 12 | 18994400 | 19003599 | 212.1199 | 315.2329 | 235.4929 | 97.01598 | 0.674685 | 3.645402 | 0.125486 |
| 17 | 15349600 | 15352399 | 12.21583 | 21.18742 | 55.33587 | 24.27902 | -0.99202 | 0.807184 | 0.125502 |
| 3 | 149550200 | 149551399 | 4.836298 | 9.343151 | 28.78606 | 11.06096 | -1.05413 | -0.17039 | 0.125507 |
| 14 | 3093600 | 3095199 | 29.21412 | 49.96112 | 35.18102 | 15.42618 | 0.886884 | 0.980528 | 0.125508 |
| 18 | 37705200 | 37706599 | 31.28507 | 53.0442 | 38.28078 | 16.76628 | 0.850151 | 1.06652 | 0.125525 |
| 1 | 92594200 | 92595599 | 6.528708 | 12.24078 | 34.51864 | 14.23561 | -1.02141 | 0.099609 | 0.125528 |
| 7 | 10367600 | 10369999 | 46.40203 | 75.53319 | 54.93402 | 23.58326 | 0.828396 | 1.570793 | 0.125551 |
| 7 | 17860800 | 17865199 | 89.37042 | 141.3876 | 98.43944 | 40.91319 | 0.849091 | 2.407041 | 0.125551 |
| 2 | 172257800 | 172258999 | 5.363674 | 9.910095 | 35.69539 | 15.06007 | -1.39489 | 0.189012 | 0.125593 |
| 7 | 13385000 | 13391399 | 125.0575 | 191.8943 | 152.2558 | 63.20878 | 0.627048 | 3.000943 | 0.125597 |
| 16 | 89899400 | 89922999 | 166.6189 | 241.9524 | 487.0787 | 200.7971 | -0.78635 | 4.015987 | 0.125599 |
| 1 | 184811800 | 184813599 | 18.08567 | 32.47736 | 23.40936 | 10.23758 | 0.872055 | 0.414668 | 0.125603 |
| 1 | 148336200 | 148336599 | 9.003743 | 17.59796 | 10.83747 | 3.741296 | 1.381655 | -0.54845 | 0.125611 |
| 11 | 35997200 | 35997999 | 7.314646 | 13.53516 | 36.36744 | 15.31218 | -0.97606 | 0.192327 | 0.125613 |
| 14 | 71066800 | 71067999 | 7.314646 | 13.53516 | 36.36744 | 15.31218 | -0.97606 | 0.192327 | 0.125613 |
| 10 | 16409000 | 16409799 | 15.29758 | 28.75869 | 17.68555 | 7.260535 | 1.164291 | 0.11873 | 0.125614 |
| 3 | 45890400 | 45890799 | 8.504937 | 16.60722 | 12.15035 | 4.276879 | 1.112697 | -0.48587 | 0.125624 |
| 1 | 147292600 | 147294799 | 38.58019 | 63.88674 | 36.07805 | 15.61052 | 1.202141 | 1.318106 | 0.125637 |
| 4 | 49520600 | 49521799 | 7.324597 | 13.13969 | 40.56226 | 17.75436 | -1.21324 | 0.385455 | 0.125647 |
| X | 33641000 | 33643399 | 34.28771 | 58.0053 | 34.07593 | 14.91451 | 1.145734 | 1.092478 | 0.125659 |
| 5 | 135036400 | 135042999 | 31.17319 | 49.88785 | 128.7902 | 54.24049 | -1.0214 | 2.021894 | 0.125669 |
| 12 | 22778600 | 22783399 | 105.0194 | 164.7573 | 109.2565 | 45.31214 | 0.909977 | 2.592947 | 0.12567 |
| 5 | 11904800 | 11905999 | 8.204312 | 14.72801 | 40.03515 | 17.51342 | -1.03078 | 0.385848 | 0.125672 |
| 3 | 132834800 | 132836199 | 15.27661 | 28.17901 | 20.173 | 8.530222 | 0.913983 | 0.235522 | 0.125673 |
| 4 | 144548600 | 144554399 | 69.92352 | 111.5777 | 83.46388 | 35.00707 | 0.757649 | 2.136486 | 0.125688 |
| 1 | 49358800 | 49359399 | 19.68644 | 34.89668 | 24.77896 | 10.95449 | 0.882998 | 0.507694 | 0.125691 |
| 19 | 24860600 | 24861399 | 2.166755 | 4.228997 | 23.67182 | 8.398271 | -1.78768 | -0.50127 | 0.125692 |
| 5 | 27045000 | 27046799 | 19.07691 | 34.03648 | 23.16726 | 10.18711 | 0.948662 | 0.459196 | 0.125714 |
| 1 | 44392200 | 44393799 | 20.85711 | 37.39343 | 21.888 | 9.585352 | 1.168919 | 0.440495 | 0.125723 |
| 3 | 16470400 | 16470999 | 9.407818 | 18.34329 | 13.16647 | 4.741324 | 1.111308 | -0.3911 | 0.125764 |
| 16 | 92921400 | 92923999 | 14.95381 | 25.56308 | 59.55833 | 26.11428 | -0.83238 | 0.957994 | 0.125767 |
| 4 | 131039200 | 131040399 | 13.49057 | 25.47683 | 18.71114 | 7.618607 | 0.919382 | 0.075044 | 0.125773 |
| 7 | 3057400 | 3072999 | 252.5334 | 372.2472 | 242.9506 | 100.0499 | 0.858537 | 3.787624 | 0.125774 |
| 19 | 42365000 | 42374599 | 50.48033 | 79.7401 | 163.9042 | 68.06892 | -0.71224 | 2.427836 | 0.125777 |
| 10 | 22606200 | 22607999 | 12.72497 | 21.61391 | 65.28017 | 28.60702 | -1.20656 | 1.014099 | 0.125793 |
| 7 | 7920400 | 7925599 | 75.32458 | 119.5021 | 94.93386 | 39.53604 | 0.66255 | 2.322034 | 0.125795 |
| 15 | 86943000 | 86946799 | 28.92369 | 46.64009 | 106.8531 | 45.50481 | -0.8361 | 1.772801 | 0.125807 |
| 5 | 37344200 | 37357199 | 81.80876 | 124.4384 | 276.7701 | 114.6682 | -0.86863 | 3.17475 | 0.125817 |
| 7 | 10694000 | 10697799 | 77.23471 | 122.877 | 88.87174 | 37.12151 | 0.801633 | 2.239558 | 0.125825 |
| 10 | 106747600 | 106750199 | 18.94122 | 34.64279 | 20.07722 | 8.588543 | 1.204647 | 0.310733 | 0.125833 |

| | | | | | | | | | |
|---|---|---|---|---|---|---|---|---|---|
| 4 | 134281000 | 134312799 | 192.168 | 273.6776 | 593.5643 | 245.391 | -0.92171 | 4.305512 | 0.125834 |
| 4 | 150828600 | 150837999 | 37.04657 | 58.91626 | 145.3585 | 60.68282 | -0.96618 | 2.234339 | 0.125838 |
| X | 34759600 | 34762799 | 56.29201 | 91.18448 | 52.93827 | 22.65675 | 1.146633 | 1.678479 | 0.12584 |
| 2 | 181303200 | 181305799 | 10.40941 | 18.37507 | 45.09867 | 19.95425 | -0.89834 | 0.535874 | 0.125842 |
| 7 | 20782000 | 20783199 | 26.21315 | 45.79273 | 25.07525 | 11.0472 | 1.252118 | 0.733892 | 0.125861 |
| 15 | 79570600 | 79572599 | 8.190563 | 14.42636 | 46.6826 | 20.6642 | -1.29542 | 0.550757 | 0.125868 |
| 5 | 9315800 | 9316799 | 6.125242 | 11.41574 | 34.85239 | 14.51976 | -1.14651 | 0.136625 | 0.125881 |
| 16 | 28734200 | 28735599 | 6.125242 | 11.41574 | 34.85239 | 14.51976 | -1.14651 | 0.136625 | 0.125881 |
| 4 | 8016800 | 8019399 | 35.57297 | 59.23493 | 44.38579 | 19.27844 | 0.79614 | 1.256159 | 0.125891 |
| 2 | 22428600 | 22431599 | 30.92462 | 52.27249 | 39.41692 | 17.24784 | 0.786932 | 1.091932 | 0.125894 |
| X | 29510200 | 29511199 | 19.56642 | 35.06649 | 22.92313 | 10.04179 | 1.010751 | 0.437446 | 0.125903 |
| 17 | 88089000 | 88107999 | 159.6875 | 233.4064 | 454.0719 | 187.0663 | -0.72788 | 3.915821 | 0.125911 |
| 18 | 38502400 | 38503199 | 12.07243 | 23.23622 | 16.08497 | 6.257787 | 1.06175 | -0.09147 | 0.125918 |
| 1 | 57008400 | 57013199 | 32.32683 | 51.65404 | 132.9962 | 55.88916 | -1.02009 | 2.072522 | 0.125942 |
| 11 | 104138200 | 104142799 | 22.49906 | 36.77417 | 85.87477 | 36.92121 | -0.84572 | 1.481789 | 0.125943 |
| 4 | 154738600 | 154742599 | 30.37156 | 48.94104 | 106.8052 | 45.44319 | -0.76733 | 1.796053 | 0.125947 |
| 15 | 17014200 | 17015199 | 16.94852 | 32.33423 | 13.77496 | 5.489416 | 1.723333 | 0.027704 | 0.125967 |
| 16 | 29881200 | 29886399 | 21.20201 | 34.44612 | 95.91108 | 41.13836 | -1.1052 | 1.574765 | 0.12597 |
| 3 | 17242200 | 17245599 | 84.81827 | 134.0054 | 106.0795 | 44.05851 | 0.662628 | 2.43559 | 0.125982 |
| 11 | 85707800 | 85709999 | 9.816793 | 17.30402 | 45.62922 | 20.19481 | -1.00198 | 0.549264 | 0.125984 |
| 6 | 6109000 | 6109999 | 5.438967 | 10.42285 | 31.94849 | 12.58659 | -1.08209 | -0.07401 | 0.125993 |
| 9 | 31201400 | 31202799 | 5.438967 | 10.42285 | 31.94849 | 12.58659 | -1.08209 | -0.07401 | 0.125993 |
| 11 | 103438200 | 103439399 | 5.438967 | 10.42285 | 31.94849 | 12.58659 | -1.08209 | -0.07401 | 0.125993 |
| 4 | 96052400 | 96053599 | 19.51397 | 36.22187 | 16.12356 | 6.759913 | 1.601745 | 0.226653 | 0.125996 |
| 10 | 20827800 | 20828999 | 6.106277 | 11.43179 | 34.43759 | 14.2379 | -1.11851 | 0.105692 | 0.126006 |
| 11 | 98996600 | 99002399 | 30.87513 | 49.69292 | 113.9446 | 48.39672 | -0.84077 | 1.836241 | 0.126007 |
| 3 | 18877800 | 18878599 | 12.08582 | 23.49024 | 13.35047 | 4.964709 | 1.40302 | -0.24605 | 0.126009 |
| 10 | 61681400 | 61706199 | 194.8727 | 277.7749 | 588.8148 | 243.3913 | -0.88742 | 4.292632 | 0.12601 |
| 9 | 62421400 | 62425399 | 15.07047 | 25.11597 | 74.29699 | 32.28716 | -1.1778 | 1.215752 | 0.126012 |
| 5 | 30018800 | 30019799 | 18.67967 | 33.64401 | 22.88432 | 9.979264 | 0.958019 | 0.400885 | 0.126014 |
| 19 | 47316200 | 47318399 | 8.684569 | 15.51478 | 42.06519 | 18.47679 | -1.03167 | 0.422566 | 0.126027 |
| 16 | 18893000 | 18895199 | 22.65105 | 40.20765 | 22.91398 | 10.11919 | 1.199514 | 0.515404 | 0.126028 |
| 17 | 87094400 | 87097399 | 12.25039 | 21.16525 | 56.35629 | 24.72382 | -1.02093 | 0.844435 | 0.126029 |
| 16 | 70276800 | 70277599 | 5.920831 | 11.11612 | 32.81738 | 13.50297 | -1.08298 | 0.084859 | 0.126029 |
| 11 | 61745000 | 61746999 | 10.46412 | 18.33811 | 48.83557 | 21.57147 | -1.01808 | 0.63841 | 0.126044 |
| 13 | 60760800 | 60767999 | 43.66379 | 69.45851 | 143.3542 | 59.86835 | -0.70885 | 2.228548 | 0.126045 |
| 7 | 29012400 | 29026799 | 79.296 | 120.645 | 274.7836 | 113.8533 | -0.90257 | 3.169358 | 0.126045 |
| 17 | 78289200 | 78289999 | 4.956826 | 9.607032 | 29.15387 | 11.03419 | -1.01182 | -0.22022 | 0.126054 |
| 1 | 109037600 | 109041399 | 38.79648 | 65.19767 | 34.75161 | 15.17959 | 1.285099 | 1.163125 | 0.126055 |
| 11 | 6228400 | 6232199 | 43.37654 | 70.76815 | 53.86068 | 23.14154 | 0.765202 | 1.529074 | 0.126059 |
| 13 | 89887400 | 89891199 | 26.97119 | 43.59257 | 101.8905 | 43.52133 | -0.86093 | 1.699445 | 0.126063 |
| 12 | 3115200 | 3119999 | 103.1138 | 160.9144 | 120.9295 | 50.18056 | 0.723858 | 2.686267 | 0.126065 |
| 1 | 92545400 | 92549599 | 21.35363 | 35.01027 | 83.76208 | 36.05697 | -0.87808 | 1.435557 | 0.126087 |
| 5 | 21504200 | 21504799 | 4.114622 | 8.0055 | 27.75873 | 10.23728 | -1.16621 | -0.3263 | 0.126094 |
| 16 | 49127800 | 49128399 | 4.114622 | 8.0055 | 27.75873 | 10.23728 | -1.16621 | -0.3263 | 0.126094 |
| 5 | 104292400 | 104294799 | 9.246259 | 16.62845 | 40.56647 | 17.71886 | -0.87462 | 0.382203 | 0.126109 |
| 12 | 84819400 | 84821399 | 10.44036 | 18.34551 | 46.15783 | 20.43484 | -0.93636 | 0.59117 | 0.126115 |

| | | | | | | | | | |
|---|---|---|---|---|---|---|---|---|---|
| 17 | 81034800 | 81036799 | 10.44036 | 18.34551 | 46.15783 | 20.43484 | -0.93636 | 0.59117 | 0.126115 |
| 6 | 18108600 | 18109399 | 7.997218 | 14.51196 | 38.09609 | 16.45803 | -0.9687 | 0.315973 | 0.126117 |
| 17 | 46760800 | 46762199 | 5.921364 | 11.03354 | 34.80022 | 14.50298 | -1.19217 | 0.140655 | 0.126129 |
| 4 | 151660600 | 151661599 | 4.311695 | 8.305433 | 30.9087 | 11.99762 | -1.33776 | -0.13712 | 0.12613 |
| X | 100773200 | 100777799 | 17.40048 | 29.44719 | 71.22956 | 31.17943 | -0.88839 | 1.069584 | 0.126144 |
| 12 | 5361200 | 5364199 | 19.89453 | 33.15909 | 73.25378 | 31.83593 | -0.75851 | 1.22791 | 0.126147 |
| 15 | 92941000 | 92941799 | 12.52387 | 24.0879 | 16.66897 | 6.49831 | 1.060032 | -0.07765 | 0.126151 |
| 2 | 156610800 | 156613199 | 12.94813 | 22.53842 | 53.76705 | 23.63427 | -0.86139 | 0.760338 | 0.126154 |
| 19 | 43188600 | 43190399 | 17.46623 | 32.17699 | 19.61376 | 8.308117 | 1.142043 | 0.258078 | 0.126165 |
| 1 | 183052000 | 183052799 | 10.55974 | 20.46304 | 15.26541 | 5.781948 | 0.991054 | -0.19761 | 0.126179 |
| 6 | 6216400 | 6217599 | 7.653791 | 14.02266 | 36.67181 | 15.65054 | -0.9513 | 0.254306 | 0.126181 |
| X | 146699000 | 146700799 | 30.18579 | 52.18517 | 29.86134 | 13.10059 | 1.18721 | 0.881623 | 0.126206 |
| X | 3635400 | 3637399 | 22.86961 | 42.27063 | 13.06215 | 5.50815 | 2.116566 | 0.270178 | 0.126207 |
| 13 | 75329000 | 75330399 | 18.21021 | 33.5911 | 19.04498 | 8.052974 | 1.247641 | 0.25623 | 0.126208 |
| 15 | 37168000 | 37169799 | 8.761527 | 15.86663 | 39.65063 | 17.17318 | -0.90083 | 0.336857 | 0.126208 |
| 11 | 104020800 | 104027799 | 46.48977 | 73.71072 | 154.889 | 64.46539 | -0.73948 | 2.330901 | 0.126216 |
| 4 | 94444400 | 94445199 | 4.963462 | 9.599156 | 28.2938 | 10.81721 | -0.9838 | -0.18705 | 0.126224 |
| 7 | 108216000 | 108217799 | 18.03748 | 33.19699 | 19.59708 | 8.312069 | 1.186739 | 0.268314 | 0.126268 |
| 7 | 133341800 | 133342999 | 5.105046 | 9.84686 | 28.66963 | 11.08562 | -0.9817 | -0.1447 | 0.126282 |
| 9 | 120164400 | 120175999 | 65.14812 | 100.2012 | 252.1061 | 104.6952 | -1.03364 | 2.944642 | 0.126295 |
| 5 | 118065600 | 118067599 | 12.00541 | 21.01671 | 49.92444 | 22.03255 | -0.85422 | 0.667633 | 0.1263 |
| 2 | 23443200 | 23444599 | 8.09712 | 14.84073 | 39.43852 | 16.8223 | -0.97446 | 0.258808 | 0.126314 |
| 4 | 138117200 | 138124999 | 68.75786 | 106.4702 | 214.6642 | 89.16582 | -0.70698 | 2.815121 | 0.126328 |
| 11 | 61857800 | 61860999 | 12.05352 | 20.92931 | 53.70816 | 23.57167 | -0.96632 | 0.792618 | 0.12634 |
| 5 | 125518200 | 125519399 | 6.729938 | 12.44201 | 35.22408 | 14.84894 | -1.05149 | 0.191868 | 0.126362 |
| 13 | 92991800 | 92993199 | 12.80112 | 24.67196 | 14.78988 | 5.731722 | 1.271824 | -0.09973 | 0.126365 |
| 1 | 140743600 | 140745199 | 13.54697 | 25.46968 | 18.64376 | 7.652868 | 0.914361 | 0.10296 | 0.126369 |
| 7 | 27967000 | 27967999 | 7.794723 | 14.30807 | 36.71344 | 15.62778 | -0.92138 | 0.244761 | 0.126375 |
| 11 | 113700200 | 113704199 | 19.26216 | 31.67217 | 81.2825 | 35.03219 | -0.9771 | 1.393774 | 0.126381 |
| 8 | 3849800 | 3850599 | 29.16057 | 49.65789 | 36.80819 | 16.13413 | 0.812301 | 1.019473 | 0.126386 |
| 7 | 89158800 | 89160199 | 7.122524 | 12.8879 | 38.58407 | 16.72842 | -1.15985 | 0.327939 | 0.126395 |
| 2 | 122002600 | 122004399 | 7.145871 | 13.26933 | 43.94613 | 18.41499 | -1.27066 | 0.188371 | 0.126405 |
| 1 | 144735800 | 144737799 | 27.8971 | 48.16458 | 26.02026 | 11.41499 | 1.269456 | 0.877337 | 0.126409 |
| 6 | 44676800 | 44677599 | 8.457326 | 16.50886 | 11.60699 | 4.108691 | 1.161936 | -0.47012 | 0.126428 |
| 13 | 101191200 | 101192999 | 15.09953 | 25.83188 | 59.60025 | 26.13427 | -0.81826 | 0.950866 | 0.126449 |
| 6 | 34833800 | 34834799 | 16.23551 | 29.61771 | 21.54471 | 9.246207 | 0.876173 | 0.307147 | 0.126452 |
| 16 | 94775400 | 94785999 | 67.24421 | 104.2426 | 212.6244 | 88.3152 | -0.72245 | 2.793863 | 0.126453 |
| 14 | 100051200 | 100052999 | 15.3655 | 29.43423 | 13.22147 | 5.214453 | 1.659909 | -0.0223 | 0.126477 |
| 16 | 51333400 | 51334599 | 15.18147 | 28.15688 | 18.83839 | 7.906905 | 1.017742 | 0.199821 | 0.12648 |
| 7 | 31381600 | 31382199 | 11.89608 | 23.14068 | 12.75476 | 4.711456 | 1.454894 | -0.27302 | 0.126501 |
| 17 | 66239800 | 66242199 | 11.87414 | 20.68756 | 51.02051 | 22.43966 | -0.90853 | 0.740743 | 0.126504 |
| 6 | 3169200 | 3170599 | 14.89814 | 28.14526 | 17.88588 | 7.276984 | 1.127068 | 0.083097 | 0.126515 |
| 8 | 117971200 | 117972799 | 7.299804 | 13.55053 | 39.9002 | 16.72739 | -1.10317 | 0.180296 | 0.126529 |
| 19 | 22964000 | 22965999 | 11.86784 | 20.6907 | 51.02676 | 22.45204 | -0.90849 | 0.729772 | 0.12654 |
| 5 | 107864800 | 107866199 | 8.172388 | 14.76123 | 39.07005 | 16.97696 | -0.98616 | 0.345806 | 0.126569 |
| 4 | 3937400 | 3939599 | 9.930235 | 17.2829 | 53.63028 | 23.57212 | -1.23816 | 0.748778 | 0.126574 |
| 12 | 39004600 | 39006199 | 12.79216 | 22.31497 | 51.58028 | 22.70377 | -0.81553 | 0.727275 | 0.126591 |

| | | | | | | | | | |
|---|---|---|---|---|---|---|---|---|---|
| 8 | 91925800 | 91928199 | 13.42295 | 23.32703 | 54.86939 | 24.09423 | -0.84148 | 0.787332 | 0.126592 |
| 4 | 46833200 | 46847199 | 100.1815 | 151.0681 | 312.2003 | 129.0005 | -0.77334 | 3.357382 | 0.126598 |
| 12 | 17385800 | 17386999 | 22.42935 | 39.32995 | 28.09383 | 12.41566 | 0.870448 | 0.659484 | 0.1266 |
| 15 | 85463800 | 85468199 | 29.76407 | 47.60305 | 132.1917 | 55.62385 | -1.12739 | 2.040833 | 0.126606 |
| 1 | 119881000 | 119882199 | 6.95476 | 13.06797 | 38.13928 | 15.67096 | -1.06676 | 0.09741 | 0.126607 |
| 14 | 59869400 | 59872399 | 11.83782 | 20.71604 | 49.93678 | 22.03145 | -0.87543 | 0.670391 | 0.126627 |
| 15 | 83497800 | 83503199 | 23.34635 | 37.63683 | 107.2078 | 45.64073 | -1.14963 | 1.774337 | 0.126631 |
| 19 | 33675000 | 33676199 | 8.910723 | 16.13483 | 39.64918 | 17.17521 | -0.87707 | 0.337754 | 0.126641 |
| 9 | 38350000 | 38351999 | 20.68682 | 37.0231 | 22.44358 | 9.84321 | 1.117346 | 0.450971 | 0.126655 |
| 7 | 29065400 | 29069399 | 18.98646 | 31.19974 | 85.4102 | 36.80167 | -1.07044 | 1.401664 | 0.126677 |
| 12 | 19341600 | 19348399 | 197.5245 | 294.8712 | 223.5434 | 92.14825 | 0.659441 | 3.562742 | 0.126679 |
| 13 | 66049400 | 66052399 | 24.07933 | 42.08329 | 26.16567 | 11.53116 | 1.069703 | 0.716873 | 0.126679 |
| 3 | 21566000 | 21567999 | 34.77031 | 57.92438 | 44.39638 | 19.28851 | 0.763627 | 1.249323 | 0.126682 |
| 7 | 26683600 | 26684799 | 27.55782 | 47.11709 | 35.72256 | 15.66333 | 0.78057 | 0.97489 | 0.126684 |
| 6 | 138955400 | 138958199 | 23.83226 | 41.99144 | 24.39201 | 10.79644 | 1.167878 | 0.597248 | 0.126689 |
| 2 | 152157000 | 152161399 | 18.47749 | 30.43495 | 82.01848 | 35.37546 | -1.04638 | 1.369077 | 0.126729 |
| 8 | 8488600 | 8491399 | 13.32383 | 23.00111 | 55.85222 | 24.50207 | -0.88894 | 0.85621 | 0.126752 |
| 10 | 122173200 | 122174399 | 15.6138 | 28.91046 | 20.53306 | 8.63778 | 0.930539 | 0.218322 | 0.126761 |
| 13 | 56926800 | 56939999 | 80.85974 | 122.7563 | 297.0676 | 123.0061 | -0.99257 | 3.215983 | 0.126762 |
| 7 | 122866400 | 122869399 | 11.07843 | 19.38611 | 48.86017 | 21.55551 | -0.93912 | 0.666491 | 0.126766 |
| 1 | 117878000 | 117879599 | 34.81194 | 57.98898 | 44.40582 | 19.29161 | 0.76495 | 1.250833 | 0.126769 |
| 6 | 130979600 | 130981399 | 20.0869 | 36.34761 | 20.97055 | 9.091556 | 1.198961 | 0.380322 | 0.126773 |
| 10 | 25867400 | 25870399 | 24.26889 | 39.60248 | 89.70147 | 38.54363 | -0.80368 | 1.509955 | 0.126782 |
| X | 104696000 | 104697799 | 35.53071 | 59.1243 | 44.42792 | 19.28817 | 0.792152 | 1.26492 | 0.126795 |
| 15 | 76399200 | 76411399 | 81.24904 | 123.3852 | 284.2629 | 117.7149 | -0.92132 | 3.208706 | 0.126795 |
| 12 | 20261000 | 20266599 | 201.1321 | 299.7133 | 230.378 | 94.9349 | 0.637136 | 3.596388 | 0.126829 |
| 7 | 33142000 | 33149599 | 183.715 | 277.1575 | 183.5382 | 75.84844 | 0.867874 | 3.358253 | 0.126841 |
| 2 | 179088400 | 179155999 | 911.7066 | 1158.564 | 1888.202 | 792.76 | -0.67492 | 6.105738 | 0.126855 |
| 12 | 23066400 | 23071799 | 114.0867 | 176.2741 | 140.7178 | 58.4511 | 0.625812 | 2.86456 | 0.126857 |
| 9 | 4794200 | 4796599 | 13.5843 | 23.63167 | 54.86818 | 24.10923 | -0.82261 | 0.774281 | 0.126861 |
| 9 | 58374400 | 58376799 | 8.1864 | 14.72299 | 39.53607 | 17.26822 | -1.0115 | 0.374171 | 0.126879 |
| 7 | 13771200 | 13772599 | 28.15324 | 48.53714 | 33.39748 | 14.65071 | 0.921896 | 0.902041 | 0.126907 |
| 15 | 25825200 | 25828999 | 22.55409 | 37.14555 | 85.83751 | 37.02092 | -0.8257 | 1.381697 | 0.12691 |
| 2 | 59629200 | 59631599 | 10.65342 | 18.57434 | 50.99229 | 22.43754 | -1.06208 | 0.723702 | 0.12691 |
| 16 | 33533800 | 33537799 | 18.36489 | 30.48465 | 74.5172 | 32.29635 | -0.9042 | 1.271314 | 0.126923 |
| 9 | 21705200 | 21710999 | 29.06278 | 46.56694 | 123.7459 | 52.21031 | -1.06089 | 1.980542 | 0.126929 |
| 5 | 102126000 | 102127199 | 11.21736 | 19.66406 | 47.77739 | 21.10943 | -0.88651 | 0.636558 | 0.126937 |
| 3 | 35278000 | 35279999 | 12.66151 | 21.99185 | 53.19307 | 23.35018 | -0.8814 | 0.790423 | 0.126957 |
| 11 | 117912200 | 117929399 | 120.4156 | 177.832 | 408.0734 | 168.0528 | -0.95178 | 3.745845 | 0.126961 |
| 4 | 88668200 | 88668599 | 3.341409 | 6.531434 | 22.60109 | 7.784026 | -1.06986 | -0.59497 | 0.126962 |
| 11 | 40582200 | 40583799 | 6.75708 | 12.45339 | 37.33983 | 15.80761 | -1.13869 | 0.212689 | 0.126964 |
| 5 | 111746000 | 111751199 | 18.89142 | 30.92572 | 89.01992 | 38.29536 | -1.14489 | 1.452019 | 0.126966 |
| 5 | 144779200 | 144799599 | 165.2763 | 239.2256 | 496.6868 | 204.8389 | -0.83544 | 4.064738 | 0.126966 |
| X | 89341400 | 89343999 | 53.63317 | 86.95191 | 53.9659 | 23.11112 | 1.051717 | 1.657251 | 0.126973 |
| 4 | 98283800 | 98284799 | 4.368704 | 8.473821 | 26.64452 | 10.04752 | -1.05567 | -0.24389 | 0.126977 |
| 12 | 22512600 | 22513599 | 24.18407 | 42.09854 | 30.89731 | 13.58188 | 0.831393 | 0.780664 | 0.126979 |
| 2 | 33731000 | 33734399 | 15.32577 | 26.01101 | 62.71478 | 27.47626 | -0.88321 | 1.028448 | 0.126991 |

| | | | | | | | | | |
|---|---|---|---|---|---|---|---|---|---|
| 11 | 119118800 | 119140399 | 154.3006 | 224.5132 | 474.9345 | 195.75 | -0.85499 | 3.984663 | 0.126999 |
| 7 | 134859000 | 134860999 | 12.78794 | 22.32161 | 51.57987 | 22.71254 | -0.81501 | 0.718138 | 0.127005 |
| 15 | 19786000 | 19789199 | 47.53264 | 77.34276 | 55.00503 | 23.61031 | 0.860191 | 1.578408 | 0.127017 |
| 6 | 145736800 | 145738799 | 7.848379 | 14.2645 | 40.27716 | 17.36577 | -1.07176 | 0.307394 | 0.127027 |
| 17 | 43933400 | 43942799 | 116.5628 | 174.6375 | 333.9357 | 137.7503 | -0.66969 | 3.487377 | 0.127033 |
| 9 | 119368600 | 119385399 | 112.691 | 167.1732 | 387.7965 | 159.7316 | -0.96134 | 3.668377 | 0.127037 |
| 10 | 101275000 | 101275399 | 9.536434 | 18.58542 | 12.9213 | 4.681463 | 1.150937 | -0.3673 | 0.127049 |
| 2 | 134445600 | 134447199 | 24.85797 | 43.51096 | 27.82192 | 12.277 | 1.029422 | 0.6999 | 0.127058 |
| 16 | 9808600 | 9813799 | 45.50169 | 71.78471 | 167.7414 | 69.61788 | -0.89837 | 2.456467 | 0.127078 |
| 1 | 116291000 | 116292999 | 24.25995 | 42.44791 | 28.52823 | 12.58459 | 0.958072 | 0.704398 | 0.127093 |
| 17 | 49308800 | 49313199 | 29.05488 | 46.91775 | 104.5337 | 44.59809 | -0.79361 | 1.73034 | 0.127099 |
| 6 | 10284200 | 10285799 | 22.26478 | 38.97686 | 28.5925 | 12.6184 | 0.832192 | 0.684032 | 0.127106 |
| 9 | 116855600 | 116856599 | 6.519197 | 12.25001 | 34.60382 | 14.21739 | -1.01956 | 0.085089 | 0.127108 |
| 19 | 33012200 | 33013399 | 6.519197 | 12.25001 | 34.60382 | 14.21739 | -1.01956 | 0.085089 | 0.127108 |
| 6 | 143861400 | 143863399 | 10.21525 | 18.18742 | 44.65651 | 19.67126 | -0.89278 | 0.459681 | 0.127113 |
| 8 | 71913600 | 71918199 | 25.28035 | 40.75024 | 107.9351 | 45.94394 | -1.04534 | 1.779399 | 0.127116 |
| 18 | 74194400 | 74198999 | 31.85742 | 51.21362 | 113.3514 | 48.06276 | -0.79197 | 1.875463 | 0.127123 |
| 8 | 73213800 | 73218999 | 20.25325 | 32.86312 | 95.76543 | 41.0376 | -1.17225 | 1.60036 | 0.127129 |
| X | 3708800 | 3710599 | 32.40331 | 56.45497 | 23.26856 | 10.23398 | 1.659266 | 0.799961 | 0.127132 |
| 5 | 129970200 | 129972399 | 12.26771 | 21.13711 | 56.29349 | 24.6942 | -1.02194 | 0.870982 | 0.127135 |
| 12 | 23909000 | 23909599 | 11.58807 | 22.31132 | 16.84679 | 6.54795 | 0.939584 | -0.09625 | 0.127178 |
| 17 | 18174000 | 18175799 | 42.74188 | 69.95441 | 52.23134 | 22.46746 | 0.795923 | 1.482444 | 0.127183 |
| 4 | 121024000 | 121026999 | 44.5786 | 72.99964 | 50.62406 | 21.78085 | 0.902537 | 1.479268 | 0.127193 |
| 8 | 120138800 | 120143399 | 18.25165 | 30.2111 | 77.78808 | 33.64819 | -0.97933 | 1.306028 | 0.127195 |
| 7 | 137055000 | 137056199 | 3.715787 | 7.221009 | 27.43326 | 10.21393 | -1.30836 | -0.29729 | 0.127203 |
| 17 | 66885200 | 66889999 | 37.06294 | 59.37432 | 122.7071 | 51.75722 | -0.69949 | 1.994765 | 0.127211 |
| 11 | 74299600 | 74301799 | 8.611484 | 15.16626 | 45.13058 | 19.97758 | -1.17513 | 0.553256 | 0.12722 |
| 16 | 17728600 | 17733599 | 26.55157 | 42.81975 | 110.645 | 47.12638 | -1.00907 | 1.766918 | 0.127223 |
| 10 | 122794400 | 122799799 | 49.81497 | 78.74698 | 160.9051 | 66.84828 | -0.70282 | 2.409609 | 0.127229 |
| 7 | 60468600 | 60472399 | 41.67594 | 69.90605 | 33.67946 | 14.69583 | 1.430455 | 1.191848 | 0.12723 |
| 1 | 191553000 | 191554199 | 3.845824 | 7.427303 | 28.0092 | 10.77851 | -1.34253 | -0.17608 | 0.127242 |
| 2 | 38897800 | 38898999 | 9.011076 | 16.00736 | 42.01015 | 18.54074 | -0.98963 | 0.462228 | 0.127281 |
| 7 | 41725400 | 41728199 | 56.08372 | 90.31406 | 67.82626 | 28.83085 | 0.767045 | 1.837468 | 0.127286 |
| 7 | 78237200 | 78237599 | 2.935908 | 5.742064 | 20.86565 | 7.063262 | -1.11504 | -0.66823 | 0.127296 |
| 11 | 43657800 | 43658199 | 2.935908 | 5.742064 | 20.86565 | 7.063262 | -1.11504 | -0.66823 | 0.127296 |
| 11 | 68702400 | 68706799 | 22.9945 | 37.51516 | 87.38109 | 37.54376 | -0.84392 | 1.511143 | 0.1273 |
| 12 | 38508400 | 38510199 | 16.13886 | 30.44938 | 16.54783 | 6.751221 | 1.346937 | 0.099569 | 0.12733 |
| 1 | 148502400 | 148502999 | 8.343117 | 16.28573 | 12.22597 | 4.328681 | 1.068856 | -0.46815 | 0.127334 |
| 2 | 16942000 | 16942999 | 11.38842 | 22.00163 | 16.27142 | 6.25151 | 0.984469 | -0.13774 | 0.12735 |
| 15 | 79398000 | 79404399 | 34.17695 | 54.73989 | 122.0647 | 51.47037 | -0.80921 | 1.998195 | 0.127356 |
| 14 | 70410400 | 70418999 | 34.02883 | 53.94771 | 171.902 | 71.53849 | -1.33914 | 2.327728 | 0.127364 |
| 16 | 4410800 | 4414599 | 17.97007 | 29.6692 | 79.22288 | 34.20851 | -1.0319 | 1.33703 | 0.12737 |
| 7 | 6047600 | 6054799 | 142.9649 | 217.5363 | 172.5244 | 71.48559 | 0.618189 | 3.176133 | 0.127377 |
| 8 | 89593000 | 89598199 | 32.78912 | 52.78223 | 122.2721 | 51.94623 | -0.85531 | 1.836408 | 0.127384 |
| 7 | 7982600 | 7984599 | 19.02781 | 35.33868 | 16.74846 | 7.016712 | 1.513004 | 0.221177 | 0.127411 |
| X | 5181800 | 5182599 | 13.81188 | 26.74186 | 13.80481 | 5.250041 | 1.511479 | -0.15044 | 0.127415 |
| 7 | 126037600 | 126039999 | 8.812933 | 15.42983 | 48.82431 | 21.54926 | -1.26471 | 0.647283 | 0.127417 |

| | | | | | | | | | |
|---|---|---|---|---|---|---|---|---|---|
| 8 | 25591000 | 25593999 | 15.27164 | 25.65432 | 67.37381 | 29.42566 | -1.00668 | 1.124208 | 0.127418 |
| 17 | 54803200 | 54804599 | 11.6256 | 22.61844 | 12.10489 | 4.465179 | 1.49767 | -0.28376 | 0.127423 |
| 2 | 36743000 | 36743399 | 13.60074 | 25.65509 | 16.342 | 6.670318 | 1.120169 | 0.077518 | 0.127444 |
| 5 | 137835400 | 137836799 | 11.41999 | 19.88898 | 51.54802 | 22.66641 | -0.97988 | 0.743743 | 0.127451 |
| 6 | 118568000 | 118573399 | 28.87642 | 46.67724 | 102.4785 | 43.77892 | -0.7707 | 1.699273 | 0.127455 |
| 14 | 70763800 | 70766799 | 10.17029 | 17.80517 | 48.84421 | 21.55687 | -1.0605 | 0.654197 | 0.127458 |
| 12 | 102693400 | 102694999 | 9.897302 | 17.6455 | 42.55227 | 18.7227 | -0.86565 | 0.44487 | 0.127461 |
| X | 24542400 | 24546799 | 67.31352 | 108.5564 | 50.99071 | 21.71812 | 1.444746 | 1.806302 | 0.127467 |
| 1 | 119146800 | 119152799 | 38.39411 | 61.40649 | 126.1253 | 53.07248 | -0.69328 | 2.048759 | 0.12747 |
| 17 | 56296800 | 56308799 | 84.12437 | 128.188 | 265.2151 | 109.9393 | -0.76227 | 3.1384 | 0.127509 |
| 4 | 125707800 | 125708599 | 3.712085 | 7.223596 | 27.70266 | 10.20198 | -1.3074 | -0.3366 | 0.127527 |
| 7 | 38517800 | 38518199 | 17.06805 | 30.88982 | 22.47627 | 9.742423 | 0.866701 | 0.360258 | 0.127528 |
| 7 | 134746000 | 134750199 | 23.90405 | 39.13888 | 85.27905 | 36.689 | -0.74448 | 1.459125 | 0.127533 |
| 8 | 70824200 | 70832999 | 64.77746 | 99.75382 | 233.762 | 97.09361 | -0.92982 | 2.918675 | 0.127536 |
| 9 | 3139800 | 3146399 | 164.9917 | 248.8661 | 196.0663 | 81.02113 | 0.617249 | 3.358876 | 0.127555 |
| 6 | 99732800 | 99733599 | 4.829151 | 9.350091 | 29.02526 | 11.03771 | -1.05076 | -0.20617 | 0.127562 |
| 17 | 13543600 | 13579599 | 1357.686 | 1655.313 | 3109.338 | 1311.672 | -0.94028 | 6.769849 | 0.127567 |
| 1 | 177623400 | 177624599 | 8.685963 | 15.23037 | 51.45712 | 22.7401 | -1.35914 | 0.623207 | 0.127568 |
| 3 | 80707400 | 80710399 | 20.31953 | 33.54158 | 79.69181 | 34.40628 | -0.86423 | 1.344675 | 0.127568 |
| 5 | 37077800 | 37097799 | 127.0855 | 188.2697 | 387.7888 | 159.7062 | -0.79181 | 3.694167 | 0.127577 |
| 3 | 145924800 | 145927599 | 13.99221 | 23.64338 | 66.35908 | 29.03409 | -1.10171 | 1.065821 | 0.127585 |
| 16 | 36748000 | 36749999 | 6.510433 | 12.24006 | 32.89937 | 13.50491 | -0.94678 | 0.079714 | 0.12759 |
| 11 | 111521000 | 111522599 | 20.31217 | 36.37266 | 22.45086 | 9.841822 | 1.091856 | 0.444772 | 0.127591 |
| 2 | 150746000 | 150746999 | 4.698081 | 8.999823 | 29.44751 | 11.61331 | -1.17465 | -0.08058 | 0.127598 |
| 9 | 34581600 | 34583599 | 8.097353 | 14.85324 | 37.24758 | 15.87017 | -0.88976 | 0.25153 | 0.127604 |
| 5 | 117227600 | 117244599 | 81.11043 | 123.0468 | 287.3442 | 118.9539 | -0.94204 | 3.23043 | 0.12761 |
| 7 | 11406000 | 11406599 | 24.82305 | 43.09044 | 31.85029 | 13.97626 | 0.820763 | 0.822231 | 0.127616 |
| 12 | 73439400 | 73443199 | 19.76676 | 32.83675 | 75.42248 | 32.70594 | -0.81478 | 1.265775 | 0.127621 |
| 3 | 75188200 | 75191199 | 29.58287 | 50.6193 | 35.5165 | 15.57396 | 0.89205 | 0.976199 | 0.127623 |
| 10 | 29167800 | 29168999 | 12.17343 | 23.69736 | 11.46658 | 4.210884 | 1.646786 | -0.29341 | 0.127634 |
| 11 | 57882800 | 57885199 | 12.6305 | 22.02131 | 52.12735 | 22.93694 | -0.84988 | 0.733627 | 0.127638 |
| 9 | 120791000 | 120798999 | 41.86317 | 66.28542 | 160.5207 | 66.74393 | -0.94553 | 2.364575 | 0.127666 |
| 17 | 17425200 | 17429999 | 179.8657 | 273.7009 | 139.1194 | 57.64521 | 1.25948 | 3.184272 | 0.127671 |
| 10 | 93723400 | 93726199 | 12.57501 | 21.67757 | 57.42579 | 25.1913 | -1.01436 | 0.86743 | 0.127671 |
| 3 | 29773400 | 29774199 | 7.487081 | 13.42406 | 41.59405 | 18.21411 | -1.21936 | 0.390792 | 0.127688 |
| 5 | 56587800 | 56589199 | 15.77411 | 29.70353 | 17.12121 | 7.009236 | 1.258815 | 0.109521 | 0.127714 |
| 2 | 15323200 | 15323999 | 18.34942 | 33.20658 | 22.39059 | 9.706111 | 0.97594 | 0.370814 | 0.127715 |
| 13 | 96396400 | 96396999 | 3.455357 | 6.751426 | 26.49367 | 9.220657 | -1.2639 | -0.54384 | 0.127716 |
| 6 | 42002600 | 42004799 | 22.12064 | 39.13427 | 24.59755 | 10.87858 | 1.057024 | 0.544886 | 0.127729 |
| 2 | 169892600 | 169903599 | 103.8489 | 156.6557 | 304.7426 | 125.9335 | -0.68568 | 3.350679 | 0.127749 |
| 19 | 47842400 | 47843999 | 9.440698 | 16.83438 | 44.15215 | 19.42378 | -0.98606 | 0.442236 | 0.127756 |
| X | 3307400 | 3309799 | 23.50993 | 41.30942 | 26.33898 | 11.66033 | 1.033359 | 0.631151 | 0.127758 |
| X | 16532400 | 16533999 | 20.60121 | 37.50675 | 19.5296 | 8.40239 | 1.353047 | 0.348203 | 0.127765 |
| 13 | 65651400 | 65652799 | 34.37415 | 57.17007 | 43.38196 | 18.82732 | 0.777947 | 1.264552 | 0.127771 |
| 2 | 66313400 | 66315199 | 14.56845 | 25.13795 | 56.99436 | 25.00277 | -0.79074 | 0.867282 | 0.127772 |
| 10 | 94259600 | 94260799 | 16.22414 | 29.74358 | 20.30601 | 8.658517 | 0.972927 | 0.275563 | 0.127797 |
| 19 | 58894200 | 58896999 | 13.11839 | 22.82073 | 52.66825 | 23.14231 | -0.81382 | 0.770054 | 0.127814 |

| | | | | | | | | | |
|---|---|---|---|---|---|---|---|---|---|
| 2 | 165028600 | 165044399 | 114.4716 | 170.7452 | 361.5436 | 149.0102 | -0.82253 | 3.57052 | 0.127816 |
| 1 | 189594400 | 189596199 | 7.88089 | 14.21238 | 39.05125 | 17.00215 | -1.04137 | 0.353691 | 0.127824 |
| 4 | 63244000 | 63249799 | 37.47732 | 59.78922 | 134.0879 | 56.22204 | -0.82389 | 2.127502 | 0.127828 |
| 8 | 71314200 | 71331199 | 78.41112 | 118.3883 | 320.1571 | 132.3407 | -1.15937 | 3.32801 | 0.12783 |
| 14 | 84582200 | 84583399 | 13.4844 | 25.31112 | 17.66808 | 7.272243 | 0.979163 | 0.109576 | 0.127832 |
| 4 | 111295400 | 111296999 | 8.767093 | 15.45329 | 45.12284 | 19.9708 | -1.14772 | 0.543704 | 0.127841 |
| 18 | 37504200 | 37505799 | 43.04999 | 70.46245 | 52.18232 | 22.44681 | 0.807796 | 1.484196 | 0.127849 |
| 13 | 43633400 | 43635199 | 9.525969 | 16.76469 | 45.63534 | 20.20394 | -1.04832 | 0.566169 | 0.127856 |
| 12 | 54537000 | 54541599 | 23.43501 | 38.34888 | 85.74147 | 36.87965 | -0.78206 | 1.465792 | 0.12787 |
| 14 | 120775200 | 120777199 | 10.92472 | 19.11927 | 48.32841 | 21.32293 | -0.94316 | 0.663866 | 0.127875 |
| 12 | 40183200 | 40185999 | 16.43144 | 27.81863 | 67.26802 | 29.44899 | -0.88784 | 1.056742 | 0.127895 |
| 1 | 46451400 | 46452799 | 12.9046 | 24.81437 | 15.9707 | 6.230275 | 1.162247 | -0.07413 | 0.127897 |
| 16 | 26038200 | 26039599 | 12.9046 | 24.81437 | 15.9707 | 6.230275 | 1.162247 | -0.07413 | 0.127897 |
| 4 | 131775400 | 131776599 | 5.975186 | 11.1166 | 34.2455 | 14.30927 | -1.16229 | 0.146978 | 0.127914 |
| 6 | 129176600 | 129178999 | 40.63973 | 67.00127 | 47.25817 | 20.39096 | 0.88183 | 1.38438 | 0.127918 |
| 4 | 23137600 | 23142199 | 59.26112 | 95.61227 | 59.79034 | 25.48051 | 1.03376 | 1.783735 | 0.12792 |
| 5 | 150418000 | 150420199 | 14.6145 | 25.05893 | 58.52428 | 25.66716 | -0.83515 | 0.926758 | 0.127922 |
| 9 | 62354600 | 62356799 | 11.5817 | 20.1425 | 52.08022 | 22.88036 | -0.97666 | 0.765922 | 0.127936 |
| 17 | 67276200 | 67276999 | 4.656825 | 9.051471 | 28.28679 | 10.51564 | -1.02897 | -0.28882 | 0.127941 |
| 1 | 183371200 | 183372999 | 9.450299 | 16.8261 | 43.59221 | 19.20184 | -0.96967 | 0.451235 | 0.127942 |
| 4 | 140322600 | 140333399 | 93.22998 | 141.3722 | 281.91 | 116.6893 | -0.71495 | 3.24192 | 0.127963 |
| 7 | 13353200 | 13355399 | 57.35325 | 92.16455 | 72.18946 | 30.59218 | 0.702799 | 1.905315 | 0.127971 |
| 2 | 180809400 | 180811799 | 26.88876 | 46.3638 | 33.94955 | 14.89295 | 0.832515 | 0.89491 | 0.12798 |
| 7 | 25245000 | 25246599 | 16.15959 | 29.62965 | 20.20057 | 8.611814 | 0.975771 | 0.276098 | 0.127991 |
| 12 | 18253200 | 18266999 | 507.8799 | 698.4911 | 503.1567 | 209.18 | 0.617154 | 4.836417 | 0.127995 |
| 7 | 35744600 | 35750399 | 30.89669 | 49.67638 | 113.1466 | 47.98601 | -0.83295 | 1.869495 | 0.12801 |
| 7 | 12023000 | 12034199 | 31.55822 | 53.63642 | 34.10312 | 14.94608 | 1.032258 | 1.042449 | 0.128011 |
| 13 | 64753400 | 64755399 | 26.65134 | 45.91893 | 34.08875 | 14.95367 | 0.812663 | 0.901807 | 0.128015 |
| 4 | 4849400 | 4850999 | 27.39844 | 47.35951 | 32.79566 | 14.38785 | 0.913365 | 0.874232 | 0.128016 |
| 7 | 4510400 | 4514999 | 25.31142 | 41.06317 | 94.043 | 40.29417 | -0.82621 | 1.61162 | 0.128016 |
| 4 | 79741800 | 79743399 | 16.00208 | 29.97273 | 18.22602 | 7.529149 | 1.172378 | 0.147242 | 0.128018 |
| 6 | 30158000 | 30187599 | 204.0088 | 288.9125 | 641.3531 | 265.3975 | -0.96344 | 4.388552 | 0.128032 |
| 2 | 35504200 | 35510799 | 31.42084 | 50.41782 | 118.8485 | 50.25401 | -0.88597 | 1.935636 | 0.128049 |
| 1 | 117993000 | 117994999 | 38.98883 | 64.12596 | 49.93861 | 21.52573 | 0.738164 | 1.412817 | 0.12805 |
| 2 | 127517400 | 127518199 | 10.35437 | 20.21932 | 11.38761 | 4.005686 | 1.48662 | -0.46228 | 0.128051 |
| 17 | 15327200 | 15332399 | 28.09253 | 45.37511 | 101.157 | 43.1714 | -0.79396 | 1.721045 | 0.128054 |
| 13 | 22537000 | 22537999 | 9.561723 | 17.15095 | 41.58723 | 18.20473 | -0.86828 | 0.402358 | 0.128054 |
| 3 | 96079600 | 96082799 | 13.92874 | 23.66657 | 62.6267 | 27.44623 | -1.01663 | 1.013444 | 0.128062 |
| 19 | 42598000 | 42609799 | 87.39665 | 133.475 | 254.7576 | 105.6609 | -0.64346 | 3.093109 | 0.12807 |
| 2 | 179982800 | 179986999 | 15.99597 | 26.67354 | 75.09209 | 32.64439 | -1.10682 | 1.211817 | 0.128088 |
| 8 | 35842400 | 35843999 | 8.334757 | 14.6668 | 46.16419 | 20.43849 | -1.25616 | 0.561838 | 0.128095 |
| 7 | 115733600 | 115737199 | 16.60969 | 27.64435 | 72.67887 | 31.55734 | -1.00834 | 1.234706 | 0.1281 |
| 15 | 8891800 | 8893799 | 25.0606 | 43.72148 | 29.467 | 12.97126 | 0.953794 | 0.751058 | 0.128106 |
| 5 | 70450400 | 70452599 | 29.12805 | 49.94551 | 35.00592 | 15.35266 | 0.89404 | 0.955646 | 0.128111 |
| 7 | 42183600 | 42183999 | 17.60164 | 31.64286 | 22.57322 | 9.862786 | 0.887838 | 0.403024 | 0.128128 |
| 7 | 44878000 | 44879399 | 4.773507 | 9.042012 | 31.90303 | 12.90718 | -1.31644 | 0.010171 | 0.128128 |
| 15 | 28377000 | 28377999 | 12.61057 | 24.25184 | 16.74524 | 6.530223 | 1.063128 | -0.07485 | 0.128139 |

| | | | | | | | | | |
|---|---|---|---|---|---|---|---|---|---|
| 7 | 96582600 | 96584599 | 21.11885 | 37.23803 | 26.42704 | 11.69546 | 0.881228 | 0.577398 | 0.128158 |
| 11 | 60426000 | 60430199 | 22.49725 | 36.78642 | 86.00297 | 36.9819 | -0.84699 | 1.475205 | 0.128164 |
| 18 | 50706000 | 50707399 | 6.222261 | 11.61677 | 34.32323 | 14.2572 | -1.0953 | 0.128626 | 0.128165 |
| 17 | 27864200 | 27887399 | 152.6807 | 222.3272 | 474.3721 | 195.5025 | -0.86632 | 3.972758 | 0.128165 |
| 6 | 146052200 | 146053399 | 4.542446 | 8.788523 | 27.67086 | 10.55883 | -1.07476 | -0.19974 | 0.128167 |
| 2 | 155557400 | 155560199 | 9.190432 | 16.25905 | 43.54919 | 19.26019 | -1.02195 | 0.501994 | 0.128178 |
| 4 | 146638200 | 146638799 | 14.15568 | 26.5455 | 17.36201 | 7.157963 | 1.069913 | 0.120028 | 0.128194 |
| 19 | 7172600 | 7177399 | 19.79661 | 32.14113 | 96.49351 | 41.3676 | -1.21431 | 1.58702 | 0.128222 |
| 14 | 29292400 | 29293999 | 15.42745 | 28.57474 | 20.53751 | 8.635811 | 0.913908 | 0.214844 | 0.128227 |
| 17 | 5044200 | 5047199 | 16.1286 | 26.60674 | 81.69328 | 35.25806 | -1.23278 | 1.344305 | 0.128228 |
| 14 | 67757400 | 67758999 | 6.986015 | 13.02959 | 34.9216 | 14.53041 | -0.95917 | 0.138918 | 0.12823 |
| 13 | 39144000 | 39144799 | 4.494344 | 8.725409 | 26.73956 | 10.03511 | -1.01245 | -0.25836 | 0.128254 |
| 2 | 32382000 | 32384199 | 8.574075 | 15.2337 | 43.57215 | 19.22731 | -1.11302 | 0.459623 | 0.128259 |
| 5 | 94345600 | 94346599 | 11.40275 | 22.24526 | 11.13808 | 3.982037 | 1.632722 | -0.39447 | 0.128269 |
| 9 | 54606000 | 54608199 | 8.873616 | 15.73778 | 42.51579 | 18.78943 | -1.03252 | 0.472338 | 0.128272 |
| 12 | 77189000 | 77190599 | 8.634952 | 15.55113 | 39.54071 | 17.2445 | -0.93223 | 0.367287 | 0.128305 |
| 2 | 12579800 | 12581799 | 21.06297 | 37.67549 | 22.43631 | 9.844602 | 1.142466 | 0.457156 | 0.12831 |
| 1 | 156189600 | 156190999 | 12.03298 | 20.95047 | 58.65213 | 25.78547 | -1.09103 | 0.760042 | 0.128317 |
| 16 | 42389200 | 42391399 | 7.914557 | 14.61805 | 38.0108 | 16.04462 | -0.93211 | 0.21119 | 0.128329 |
| 14 | 116779000 | 116781599 | 14.68717 | 24.98284 | 62.1822 | 27.25385 | -0.92851 | 1.004939 | 0.12834 |
| 7 | 31471000 | 31472799 | 19.74362 | 35.58448 | 22.05881 | 9.611104 | 1.091588 | 0.402188 | 0.128343 |
| 2 | 65265600 | 65267799 | 7.503124 | 13.77026 | 38.90632 | 16.5654 | -1.06016 | 0.245302 | 0.128357 |
| 8 | 118425600 | 118437799 | 73.79341 | 113.5737 | 234.6275 | 97.44553 | -0.74884 | 2.932645 | 0.128362 |
| 14 | 115614400 | 115614999 | 3.128493 | 6.11153 | 24.94511 | 8.71502 | -1.32487 | -0.54234 | 0.128363 |
| 13 | 74510600 | 74512199 | 7.301229 | 13.09844 | 40.05279 | 17.53061 | -1.19925 | 0.385488 | 0.128364 |
| 11 | 49666800 | 49670799 | 27.41641 | 44.38345 | 98.28115 | 42.04395 | -0.78077 | 1.660277 | 0.128365 |
| 17 | 45983400 | 45984999 | 4.72985 | 8.959907 | 31.32806 | 12.67279 | -1.30245 | 0.01008 | 0.128365 |
| 6 | 122949600 | 122949999 | 8.102486 | 15.83803 | 11.63912 | 4.007678 | 1.134492 | -0.56805 | 0.128378 |
| X | 24003800 | 24004599 | 26.11788 | 45.44554 | 30.60679 | 13.45059 | 0.954668 | 0.795067 | 0.128381 |
| 13 | 68150000 | 68155599 | 145.0941 | 221.8719 | 155.9205 | 64.68698 | 0.798145 | 3.075649 | 0.12839 |
| 7 | 144786800 | 144789399 | 10.81207 | 18.82437 | 51.5239 | 22.65137 | -1.05799 | 0.746168 | 0.128414 |
| 12 | 24023800 | 24024999 | 13.34834 | 25.46692 | 15.7756 | 6.286125 | 1.188488 | -0.00436 | 0.128429 |
| 5 | 37983000 | 37990399 | 64.17854 | 99.79427 | 202.7843 | 84.18962 | -0.71355 | 2.739014 | 0.128441 |
| 10 | 127573200 | 127575999 | 17.8011 | 29.79621 | 69.21359 | 30.16082 | -0.83055 | 1.17227 | 0.128442 |
| 12 | 84636600 | 84641599 | 25.33791 | 40.79687 | 118.0721 | 50.17834 | -1.17504 | 1.817517 | 0.128445 |
| 7 | 111733000 | 111736399 | 15.94096 | 26.60534 | 71.00342 | 30.88418 | -1.02974 | 1.201805 | 0.128451 |
| 10 | 42445800 | 42465199 | 166.9275 | 242.8225 | 479.0272 | 197.4422 | -0.7547 | 3.989501 | 0.128452 |
| 5 | 47623800 | 47625799 | 29.09936 | 49.59283 | 37.2418 | 16.32524 | 0.793695 | 1.011913 | 0.128456 |
| 7 | 86160600 | 86161599 | 13.41632 | 25.57358 | 17.72408 | 7.075996 | 1.027104 | 0.009922 | 0.128458 |
| 5 | 32767000 | 32769199 | 10.60819 | 18.60236 | 47.77023 | 21.11219 | -0.96584 | 0.627539 | 0.128489 |
| 7 | 17777400 | 17779599 | 48.87412 | 79.64039 | 52.3029 | 22.46226 | 0.976205 | 1.559692 | 0.128492 |
| X | 113566000 | 113566599 | 12.38238 | 23.96418 | 15.22237 | 5.800387 | 1.213568 | -0.15549 | 0.128493 |
| 4 | 94969200 | 94970999 | 6.873705 | 12.73234 | 35.82459 | 15.06249 | -1.04034 | 0.181863 | 0.1285 |
| 5 | 123507400 | 123511799 | 22.42411 | 36.43438 | 91.86001 | 39.40296 | -0.96224 | 1.574997 | 0.128503 |
| 8 | 21710000 | 21711999 | 32.12951 | 54.55629 | 36.51237 | 15.9989 | 0.958438 | 1.054044 | 0.128513 |
| 13 | 96644600 | 96646799 | 8.105865 | 14.83136 | 38.27851 | 16.36555 | -0.93487 | 0.266796 | 0.128513 |
| 3 | 22777400 | 22778399 | 14.5859 | 27.13947 | 18.9753 | 7.927243 | 0.959777 | 0.177131 | 0.128521 |

| | | | | | | | | | |
|---|---|---|---|---|---|---|---|---|---|
| X | 31287600 | 31289799 | 70.14863 | 112.4901 | 67.43755 | 28.48831 | 1.084184 | 1.985335 | 0.12854 |
| 10 | 97716800 | 97717999 | 14.5322 | 27.79572 | 15.72582 | 6.229245 | 1.326127 | -0.01008 | 0.12854 |
| 11 | 87295000 | 87297199 | 13.00846 | 22.48503 | 54.75571 | 24.02196 | -0.89249 | 0.843224 | 0.128563 |
| 4 | 133743400 | 133747599 | 21.98047 | 35.69822 | 94.14869 | 40.3719 | -1.02763 | 1.583124 | 0.128563 |
| 5 | 120733800 | 120735799 | 9.661253 | 17.03474 | 44.59149 | 19.73439 | -0.9911 | 0.544389 | 0.128565 |
| 9 | 7613600 | 7614199 | 9.70171 | 18.96591 | 10.18582 | 3.498388 | 1.582819 | -0.55629 | 0.128571 |
| 7 | 102859200 | 102860399 | 16.83255 | 31.18405 | 19.88525 | 8.359069 | 1.085684 | 0.22427 | 0.128583 |
| 1 | 133310400 | 133317199 | 27.30895 | 43.77776 | 123.5027 | 52.14567 | -1.14607 | 1.963244 | 0.128588 |
| 1 | 157575800 | 157577199 | 8.766505 | 15.83797 | 41.21033 | 17.90184 | -0.96199 | 0.351191 | 0.128606 |
| 18 | 83004600 | 83015999 | 65.92479 | 101.463 | 233.3065 | 96.89686 | -0.90316 | 2.930461 | 0.128619 |
| 10 | 80175400 | 80179799 | 23.63963 | 38.20453 | 101.4001 | 43.30695 | -1.04379 | 1.6975 | 0.128621 |
| 13 | 107080800 | 107083999 | 15.22138 | 26.14853 | 59.15167 | 25.94565 | -0.78897 | 0.912393 | 0.128623 |
| 5 | 130352400 | 130354799 | 49.0404 | 79.22829 | 62.01606 | 26.4748 | 0.712786 | 1.727621 | 0.128624 |
| 8 | 104972600 | 104975399 | 13.4185 | 22.92705 | 60.55762 | 26.55137 | -1.01263 | 0.956939 | 0.128626 |
| 3 | 6790200 | 6791199 | 13.80302 | 26.29664 | 17.55392 | 7.015983 | 1.078961 | 0.016068 | 0.128629 |
| 12 | 100165200 | 100167399 | 9.027253 | 15.69945 | 53.5718 | 23.53523 | -1.3741 | 0.756808 | 0.128634 |
| 17 | 9017400 | 9022799 | 42.53535 | 67.65922 | 143.499 | 59.92369 | -0.74818 | 2.229596 | 0.128639 |
| 3 | 153499000 | 153502399 | 18.12018 | 29.94516 | 79.37583 | 34.29679 | -1.0213 | 1.32578 | 0.128642 |
| 14 | 24381000 | 24381799 | 10.24183 | 19.97694 | 11.12238 | 3.985574 | 1.477243 | -0.40427 | 0.128643 |
| 4 | 136505800 | 136509999 | 20.32903 | 33.52637 | 78.95568 | 34.07238 | -0.85188 | 1.356359 | 0.128646 |
| 5 | 99767800 | 99770199 | 10.23933 | 18.13315 | 43.52494 | 19.24554 | -0.86441 | 0.496951 | 0.128654 |
| 1 | 162667000 | 162668799 | 23.24252 | 40.61639 | 29.29182 | 12.90766 | 0.856954 | 0.720092 | 0.128686 |
| 13 | 9233600 | 9240199 | 51.31877 | 80.99398 | 165.1992 | 68.57651 | -0.70211 | 2.449634 | 0.128693 |
| 6 | 10696800 | 10697999 | 13.47832 | 25.85014 | 16.51143 | 6.489586 | 1.16357 | -0.04269 | 0.128694 |
| 8 | 105061800 | 105063799 | 13.13057 | 22.78325 | 53.74367 | 23.58072 | -0.84571 | 0.806521 | 0.128716 |
| 15 | 89380000 | 89392599 | 82.3235 | 124.885 | 289.8536 | 119.9921 | -0.93323 | 3.23088 | 0.128716 |
| 1 | 88663600 | 88666599 | 13.29055 | 23.03231 | 54.82765 | 24.05543 | -0.85925 | 0.819219 | 0.128729 |
| 7 | 19542400 | 19561399 | 120.2622 | 179.1273 | 361.7182 | 149.0513 | -0.75602 | 3.597361 | 0.128732 |
| X | 3684000 | 3688599 | 69.54408 | 111.3452 | 66.68605 | 28.10346 | 1.08298 | 2.033129 | 0.128749 |
| 3 | 116301200 | 116301999 | 3.537961 | 6.876855 | 25.57908 | 9.507696 | -1.27424 | -0.30835 | 0.128752 |
| 19 | 18770600 | 18771399 | 3.537961 | 6.876855 | 25.57908 | 9.507696 | -1.27424 | -0.30835 | 0.128752 |
| 12 | 38564800 | 38565799 | 6.961102 | 13.06077 | 35.67827 | 14.69613 | -0.97428 | 0.103925 | 0.128754 |
| 2 | 33524400 | 33530799 | 46.82893 | 74.20598 | 156.336 | 65.03035 | -0.74405 | 2.348382 | 0.128757 |
| 15 | 98431000 | 98432399 | 5.208453 | 9.815294 | 32.33192 | 13.21925 | -1.23192 | 0.051683 | 0.128771 |
| 6 | 82068600 | 82069399 | 5.04002 | 9.60801 | 31.60358 | 12.61448 | -1.19963 | -0.0345 | 0.128773 |
| 12 | 69727200 | 69729199 | 9.725772 | 17.01998 | 49.36922 | 21.78103 | -1.14045 | 0.659507 | 0.128779 |
| 15 | 90762200 | 90763199 | 13.89157 | 26.90048 | 12.37866 | 4.70412 | 1.674478 | -0.15692 | 0.12879 |
| 15 | 100765800 | 100770399 | 26.69516 | 43.06272 | 106.3818 | 45.32605 | -0.94387 | 1.758294 | 0.128793 |
| 12 | 22923400 | 22937399 | 412.035 | 577.8502 | 420.0831 | 174.1087 | 0.632325 | 4.538722 | 0.128798 |
| 16 | 90493200 | 90496599 | 22.40247 | 36.88305 | 81.53933 | 35.162 | -0.76211 | 1.383804 | 0.128811 |
| 4 | 28054600 | 28055399 | 9.129316 | 17.79269 | 13.83984 | 5.011702 | 0.990236 | -0.37278 | 0.128812 |
| 10 | 80743200 | 80747399 | 13.36306 | 22.27832 | 74.00888 | 32.16912 | -1.34415 | 1.207798 | 0.128815 |
| 8 | 67580200 | 67583399 | 45.02896 | 73.13498 | 56.43959 | 24.20283 | 0.740582 | 1.604091 | 0.128837 |
| 6 | 103746200 | 103749199 | 41.80116 | 68.65332 | 50.55638 | 21.77885 | 0.817811 | 1.437445 | 0.128842 |
| 8 | 56910200 | 56910799 | 14.36788 | 26.97039 | 19.1897 | 7.898097 | 0.952377 | 0.122828 | 0.128876 |
| 8 | 84625400 | 84632999 | 32.48975 | 51.90147 | 132.7847 | 55.78065 | -1.01131 | 2.080555 | 0.128878 |
| 5 | 66919400 | 66920799 | 9.044462 | 16.00134 | 44.06843 | 19.4897 | -1.06189 | 0.501141 | 0.128891 |

| | | | | | | | | | |
|---|---|---|---|---|---|---|---|---|---|
| 10 | 100081400 | 100082999 | 17.49237 | 31.69826 | 22.49259 | 9.733916 | 0.904311 | 0.357001 | 0.128911 |
| 5 | 60018600 | 60019799 | 11.74746 | 22.51135 | 14.8418 | 5.847627 | 1.115165 | -0.05833 | 0.128924 |
| 6 | 38058600 | 38060799 | 7.40885 | 13.47545 | 40.28654 | 17.35403 | -1.15259 | 0.299916 | 0.128953 |
| 1 | 85979800 | 85981999 | 11.0571 | 19.41998 | 46.69029 | 20.66226 | -0.87117 | 0.604443 | 0.128954 |
| 14 | 110322200 | 110324999 | 32.27749 | 54.45072 | 39.91995 | 17.45741 | 0.827322 | 1.114344 | 0.128977 |
| 3 | 83033600 | 83037999 | 36.20995 | 57.75363 | 142.8527 | 59.87739 | -0.96537 | 2.134931 | 0.128981 |
| 15 | 66510400 | 66511199 | 13.84644 | 26.59708 | 15.94278 | 6.238458 | 1.260252 | -0.05409 | 0.128984 |
| 2 | 34923600 | 34925199 | 6.967978 | 13.04918 | 35.64948 | 14.73096 | -0.97867 | 0.113166 | 0.128993 |
| 8 | 3978800 | 3980399 | 19.1332 | 34.98281 | 20.07202 | 8.589871 | 1.218651 | 0.314068 | 0.12902 |
| 9 | 31822400 | 31826599 | 24.93733 | 40.27062 | 105.2046 | 44.88702 | -1.02244 | 1.72304 | 0.129033 |
| X | 118567400 | 118571199 | 43.81419 | 72.92452 | 37.11007 | 16.11363 | 1.352317 | 1.280818 | 0.129041 |
| 12 | 110209000 | 110214599 | 36.28091 | 58.06945 | 123.4651 | 52.0073 | -0.74183 | 2.022846 | 0.129045 |
| 8 | 127034000 | 127039599 | 41.52115 | 66.19097 | 135.4844 | 56.7488 | -0.69356 | 2.156099 | 0.129048 |
| 17 | 25752600 | 25758799 | 30.2335 | 48.66382 | 109.1818 | 46.37872 | -0.80921 | 1.832523 | 0.129051 |
| 10 | 111892400 | 111894799 | 21.55585 | 37.89906 | 26.25757 | 11.62717 | 0.913922 | 0.608728 | 0.129063 |
| 2 | 7024000 | 7025599 | 15.81215 | 29.62571 | 18.23157 | 7.527673 | 1.155788 | 0.143624 | 0.129086 |
| 11 | 114754400 | 114760199 | 27.99 | 45.0807 | 108.2577 | 46.02762 | -0.90586 | 1.809331 | 0.12909 |
| 2 | 28162000 | 28165999 | 17.62637 | 29.5613 | 68.1 | 29.71206 | -0.81856 | 1.150617 | 0.129103 |
| 18 | 47821400 | 47822999 | 13.79492 | 26.04792 | 17.96403 | 7.316592 | 1.008438 | 0.075618 | 0.129117 |
| 7 | 3068800 | 3069799 | 29.96145 | 51.43915 | 33.85182 | 14.84801 | 0.984754 | 0.946714 | 0.129119 |
| 1 | 105591000 | 105592599 | 12.59221 | 21.66691 | 57.93284 | 25.41229 | -1.02826 | 0.885384 | 0.129124 |
| 2 | 4923600 | 4925999 | 10.39345 | 18.05096 | 54.16694 | 23.77644 | -1.19049 | 0.782747 | 0.129135 |
| 7 | 14839000 | 14844399 | 149.6408 | 227.5369 | 176.417 | 73.08337 | 0.650008 | 3.191751 | 0.129147 |
| 7 | 58979000 | 58979799 | 9.476907 | 18.4522 | 13.45097 | 4.930942 | 1.068103 | -0.32613 | 0.12915 |
| 8 | 3094400 | 3096799 | 23.13383 | 40.69103 | 26.25771 | 11.62623 | 1.016414 | 0.615542 | 0.129159 |
| 19 | 26581600 | 26585399 | 23.33902 | 37.70714 | 103.0358 | 43.98811 | -1.0862 | 1.706531 | 0.129161 |
| 17 | 36470200 | 36473599 | 22.72404 | 37.40505 | 81.54256 | 35.16044 | -0.74206 | 1.387424 | 0.129161 |
| 2 | 12305200 | 12306999 | 11.18232 | 19.30222 | 55.71821 | 24.44321 | -1.13671 | 0.849482 | 0.129202 |
| 4 | 131635000 | 131636399 | 21.70435 | 38.07394 | 28.08966 | 12.41803 | 0.823982 | 0.648423 | 0.129217 |
| 4 | 140596200 | 140602799 | 75.43501 | 115.5359 | 245.46 | 101.8719 | -0.7946 | 3.031204 | 0.129229 |
| 8 | 84320600 | 84321799 | 6.104491 | 11.38037 | 33.75313 | 14.05489 | -1.10392 | 0.134008 | 0.129233 |
| 19 | 42507000 | 42507799 | 5.080449 | 9.574622 | 32.88402 | 13.44367 | -1.29168 | 0.050668 | 0.129235 |
| 18 | 41073000 | 41073599 | 10.68282 | 20.7879 | 14.16406 | 5.217215 | 1.157186 | -0.2929 | 0.129241 |
| 2 | 157391200 | 157406799 | 201.6092 | 287.2394 | 576.41 | 238.2829 | -0.80909 | 4.300162 | 0.129244 |
| 4 | 63794200 | 63795199 | 4.958426 | 9.605158 | 32.31494 | 12.26024 | -1.16313 | -0.20468 | 0.129269 |
| 5 | 148329600 | 148332999 | 17.37477 | 28.97815 | 71.83494 | 31.23212 | -0.92397 | 1.212049 | 0.129272 |
| 4 | 45766800 | 45769599 | 11.10335 | 19.36737 | 50.47262 | 22.21507 | -0.9874 | 0.718679 | 0.129277 |
| 7 | 62248800 | 62253399 | 93.20419 | 146.9524 | 105.6062 | 43.82499 | 0.799617 | 2.486698 | 0.129282 |
| 10 | 75871800 | 75880599 | 87.92816 | 134.1065 | 259.2348 | 107.4844 | -0.6633 | 3.121063 | 0.129292 |
| 19 | 40363600 | 40368199 | 18.33805 | 30.51085 | 72.96311 | 31.67352 | -0.87238 | 1.242574 | 0.129301 |
| 7 | 9737400 | 9738799 | 33.22107 | 55.82007 | 37.04259 | 16.17958 | 0.970497 | 1.150222 | 0.129302 |
| 3 | 141170400 | 141172999 | 15.18439 | 25.41233 | 73.46259 | 32.00925 | -1.14485 | 1.168236 | 0.129304 |
| 17 | 19383400 | 19385199 | 38.01716 | 62.76419 | 48.29521 | 20.85313 | 0.757312 | 1.364122 | 0.129336 |
| 3 | 4225000 | 4228999 | 68.35657 | 108.9893 | 87.34771 | 36.59498 | 0.657129 | 2.161985 | 0.129346 |
| 2 | 163915800 | 163917399 | 6.220618 | 11.67413 | 32.82668 | 13.51626 | -1.01451 | 0.089575 | 0.129359 |
| 5 | 149334600 | 149338999 | 24.70306 | 40.046 | 98.67346 | 42.25466 | -0.93255 | 1.627014 | 0.129382 |
| Y | 42467800 | 42468799 | 19.92006 | 35.80021 | 19.70629 | 8.612289 | 1.259577 | 0.416563 | 0.129385 |

| | | | | | | | | | |
|---|---|---|---|---|---|---|---|---|---|
| 13 | 98339800 | 98343399 | 15.46467 | 26.28742 | 62.78579 | 27.51689 | -0.86953 | 1.01434 | 0.129406 |
| 2 | 105515400 | 105521799 | 42.36396 | 67.39742 | 147.1885 | 61.478 | -0.79011 | 2.224305 | 0.129408 |
| 7 | 24809000 | 24823999 | 91.1548 | 137.6715 | 305.8524 | 126.4399 | -0.87563 | 3.323683 | 0.129408 |
| 1 | 3191000 | 3192399 | 14.19447 | 26.59927 | 18.76733 | 7.74669 | 0.961172 | 0.130907 | 0.129425 |
| 10 | 51105000 | 51105599 | 3.039352 | 5.907429 | 28.55348 | 10.61708 | -1.64828 | -0.30671 | 0.129427 |
| 17 | 35223200 | 35224999 | 5.424245 | 9.773234 | 41.13175 | 17.92821 | -1.65136 | 0.346301 | 0.12944 |
| 4 | 112124000 | 112124599 | 16.16332 | 29.62153 | 21.12603 | 9.012545 | 0.910441 | 0.280934 | 0.129464 |
| 11 | 10457200 | 10459799 | 36.32398 | 61.39436 | 34.76746 | 15.213 | 1.198286 | 1.106743 | 0.129488 |
| 13 | 53422800 | 53424199 | 60.79486 | 97.86588 | 62.77732 | 26.67097 | 0.99328 | 1.851056 | 0.129506 |
| 1 | 165125800 | 165128999 | 12.14642 | 21.2421 | 53.75371 | 23.70223 | -0.94544 | 0.687171 | 0.129509 |
| 17 | 32671200 | 32674599 | 17.68145 | 29.51056 | 74.27075 | 32.30561 | -0.9459 | 1.205778 | 0.129531 |
| 11 | 114641000 | 114644399 | 15.69755 | 26.4731 | 65.33708 | 28.56996 | -0.91722 | 1.08866 | 0.129553 |
| 16 | 84043000 | 84047799 | 44.61188 | 71.02325 | 142.7309 | 59.67379 | -0.66927 | 2.203159 | 0.129565 |
| 7 | 7665000 | 7665599 | 18.86659 | 33.9704 | 22.8801 | 9.981191 | 0.971809 | 0.404055 | 0.129576 |
| 9 | 98483200 | 98483999 | 3.132982 | 6.108301 | 24.23301 | 8.733285 | -1.32423 | -0.43069 | 0.129585 |
| 10 | 76073000 | 76075199 | 10.16947 | 17.78265 | 48.85539 | 21.54023 | -1.06248 | 0.675173 | 0.129589 |
| X | 5411200 | 5413199 | 31.80824 | 54.96839 | 28.93322 | 12.69325 | 1.307252 | 0.893229 | 0.1296 |
| 7 | 23334200 | 23335599 | 34.31433 | 57.35085 | 43.27184 | 18.83853 | 0.786099 | 1.213407 | 0.129604 |
| 16 | 20547200 | 20560999 | 85.02507 | 128.9351 | 294.5709 | 121.9313 | -0.91096 | 3.239173 | 0.12962 |
| 5 | 142021600 | 142029599 | 60.70577 | 94.51725 | 198.9154 | 82.56734 | -0.76258 | 2.714752 | 0.129632 |
| 1 | 193087000 | 193092199 | 27.64361 | 44.6047 | 107.5112 | 45.82317 | -0.90806 | 1.751637 | 0.129649 |
| 6 | 50499600 | 50506799 | 37.11455 | 59.3047 | 136.4588 | 57.34497 | -0.85837 | 2.07698 | 0.129649 |
| 14 | 76724000 | 76727399 | 14.57719 | 24.68903 | 64.26421 | 28.1373 | -0.99331 | 1.045412 | 0.129652 |
| 10 | 60834400 | 60843999 | 69.81346 | 107.8614 | 217.2343 | 90.23443 | -0.70792 | 2.858053 | 0.12966 |
| 15 | 81588200 | 81591799 | 14.23343 | 24.17962 | 62.08308 | 27.20719 | -0.97331 | 1.016353 | 0.129668 |
| 10 | 9153000 | 9153599 | 3.577459 | 6.97032 | 24.71492 | 8.98367 | -1.17492 | -0.3879 | 0.129675 |
| 10 | 81637800 | 81638199 | 10.28006 | 19.9248 | 15.35356 | 5.810232 | 0.946207 | -0.20317 | 0.129681 |
| 7 | 38592000 | 38592799 | 17.7991 | 32.5258 | 21.31129 | 9.126449 | 1.027995 | 0.309568 | 0.129688 |
| 8 | 58883000 | 58884599 | 17.7991 | 32.5258 | 21.31129 | 9.126449 | 1.027995 | 0.309568 | 0.129688 |
| 5 | 110772600 | 110773999 | 6.599203 | 12.1732 | 35.71725 | 15.10248 | -1.10588 | 0.204286 | 0.12971 |
| 7 | 86359600 | 86361799 | 25.26409 | 44.9005 | 19.30103 | 8.51493 | 1.604015 | 0.517988 | 0.129715 |
| 4 | 62585400 | 62588399 | 16.80595 | 28.28474 | 68.23561 | 29.819 | -0.88485 | 1.111458 | 0.12973 |
| 15 | 3079200 | 3083199 | 105.5846 | 164.3367 | 130.2504 | 54.0681 | 0.644257 | 2.732502 | 0.129739 |
| 11 | 82841200 | 82843999 | 12.20754 | 21.19251 | 54.80569 | 24.05032 | -0.9776 | 0.79682 | 0.129752 |
| 1 | 182454800 | 182458399 | 23.3154 | 38.06725 | 92.06649 | 39.56769 | -0.89747 | 1.500629 | 0.129756 |
| 10 | 61057400 | 61063199 | 39.29182 | 62.56376 | 139.1131 | 58.17435 | -0.81457 | 2.193299 | 0.12977 |
| 4 | 102427200 | 102428199 | 4.731333 | 9.078621 | 30.13783 | 11.82889 | -1.18977 | -0.09699 | 0.1298 |
| 7 | 59072600 | 59075199 | 56.68096 | 91.31299 | 66.34692 | 28.21867 | 0.815316 | 1.824555 | 0.129824 |
| 10 | 59974400 | 59979199 | 35.63514 | 57.0338 | 126.0704 | 53.102 | -0.79789 | 2.02291 | 0.129853 |
| 5 | 139640200 | 139662599 | 133.1969 | 195.9209 | 429.2023 | 176.7507 | -0.89036 | 3.813568 | 0.129854 |
| 1 | 162303800 | 162305799 | 16.00152 | 27.00138 | 64.81312 | 28.34584 | -0.87721 | 1.084476 | 0.129863 |
| 4 | 122150000 | 122150399 | 8.727991 | 17.02834 | 13.39861 | 4.785672 | 0.991824 | -0.42644 | 0.129866 |
| 17 | 71489800 | 71491799 | 9.387928 | 16.474 | 46.71449 | 20.65775 | -1.10749 | 0.609563 | 0.129868 |
| 12 | 9042200 | 9043599 | 5.126214 | 9.896436 | 31.86617 | 12.28384 | -1.1227 | -0.14992 | 0.129876 |
| X | 120133600 | 120134799 | 18.07311 | 33.50966 | 18.73683 | 7.867572 | 1.275014 | 0.227735 | 0.129885 |
| 4 | 32578200 | 32580599 | 10.21532 | 18.16319 | 44.09529 | 19.446 | -0.87775 | 0.466374 | 0.129905 |
| 2 | 86815200 | 86816199 | 12.15561 | 23.38957 | 16.67954 | 6.494481 | 1.018535 | -0.08571 | 0.129919 |

| | | | | | | | | | |
|---|---|---|---|---|---|---|---|---|---|
| 4 | 49383000 | 49383799 | 12.15561 | 23.38957 | 16.67954 | 6.494481 | 1.018535 | -0.08571 | 0.129919 |
| 12 | 5254800 | 5262199 | 46.18129 | 73.19448 | 154.1209 | 64.12194 | -0.74295 | 2.341894 | 0.129942 |
| 17 | 57020800 | 57036199 | 83.92297 | 127.0639 | 297.8414 | 123.2245 | -0.94986 | 3.271415 | 0.129951 |
| 7 | 32914600 | 32918399 | 62.21481 | 100.3782 | 56.4526 | 24.05823 | 1.186445 | 1.788328 | 0.129956 |
| X | 60979600 | 60980799 | 15.66229 | 29.13942 | 20.03102 | 8.369671 | 0.984434 | 0.191192 | 0.129973 |
| 3 | 83033800 | 83037999 | 35.40064 | 56.48802 | 141.6623 | 59.41535 | -0.9845 | 2.119525 | 0.129974 |
| 19 | 39129800 | 39131399 | 20.2095 | 36.58455 | 19.69627 | 8.534703 | 1.298029 | 0.37515 | 0.129978 |
| 2 | 24646800 | 24654799 | 52.82475 | 82.83741 | 187.378 | 77.71497 | -0.85842 | 2.582419 | 0.129986 |
| 14 | 29688200 | 29690799 | 11.9887 | 20.99159 | 49.39583 | 21.80288 | -0.84081 | 0.661673 | 0.129987 |
| 2 | 168792800 | 168794799 | 10.57818 | 18.6345 | 45.10482 | 19.9652 | -0.87943 | 0.561626 | 0.129987 |
| 12 | 20641400 | 20644799 | 63.92097 | 102.69 | 67.82317 | 28.73025 | 0.947785 | 1.921073 | 0.129993 |
| 7 | 33432200 | 33447199 | 406.3598 | 570.837 | 413.3991 | 171.2943 | 0.640248 | 4.513916 | 0.130022 |
| 6 | 59571600 | 59576399 | 78.90684 | 125.0138 | 98.92747 | 41.14601 | 0.6663 | 2.365488 | 0.130026 |
| 17 | 36295800 | 36301799 | 29.4177 | 47.39754 | 107.7553 | 45.83837 | -0.82643 | 1.798996 | 0.130027 |
| 11 | 84020000 | 84020999 | 4.362678 | 8.267467 | 32.44246 | 13.11316 | -1.46676 | 0.004244 | 0.130033 |
| 5 | 6565800 | 6567799 | 15.01407 | 28.04368 | 18.27128 | 7.583631 | 1.06793 | 0.154908 | 0.130052 |
| 11 | 22361800 | 22362999 | 12.77976 | 24.70458 | 13.6196 | 5.217854 | 1.40612 | -0.14036 | 0.130057 |
| 1 | 101861800 | 101864399 | 39.33766 | 64.91305 | 41.91453 | 18.09218 | 1.009355 | 1.366997 | 0.130072 |
| 7 | 134015200 | 134016399 | 6.012457 | 11.49406 | 31.84041 | 12.63128 | -0.94693 | -0.04476 | 0.130076 |
| 13 | 113990000 | 113991399 | 6.012457 | 11.49406 | 31.84041 | 12.63128 | -0.94693 | -0.04476 | 0.130076 |
| 12 | 104383800 | 104385999 | 9.545798 | 16.77743 | 48.25649 | 21.36807 | -1.12837 | 0.584304 | 0.130077 |
| 2 | 50046000 | 50046599 | 6.164225 | 11.75519 | 31.71984 | 12.65164 | -0.91619 | -0.02488 | 0.130097 |
| 14 | 120713800 | 120716799 | 20.09674 | 33.35807 | 74.71784 | 32.3824 | -0.77865 | 1.277384 | 0.130097 |
| 6 | 53314000 | 53314799 | 3.82378 | 7.446777 | 25.33333 | 9.2649 | -1.12647 | -0.36555 | 0.130097 |
| 13 | 37834800 | 37835599 | 3.82378 | 7.446777 | 25.33333 | 9.2649 | -1.12647 | -0.36555 | 0.130097 |
| 1 | 10952600 | 10954199 | 18.04855 | 32.82332 | 21.72313 | 9.358965 | 1.008 | 0.338615 | 0.1301 |
| 7 | 26322600 | 26325999 | 48.17319 | 78.10468 | 59.36569 | 25.42473 | 0.760142 | 1.646437 | 0.130116 |
| 2 | 174185800 | 174191199 | 39.08808 | 62.35931 | 132.5489 | 55.57719 | -0.74666 | 2.129127 | 0.130121 |
| 9 | 30852200 | 30856999 | 26.63062 | 43.10146 | 102.4185 | 43.80648 | -0.88264 | 1.664591 | 0.130134 |
| 3 | 5394400 | 5395799 | 5.263386 | 10.14957 | 29.29777 | 11.33718 | -0.97112 | -0.14046 | 0.130136 |
| 1 | 172341600 | 172343399 | 7.570947 | 13.71632 | 39.12498 | 16.93663 | -1.08969 | 0.323124 | 0.13014 |
| 7 | 129921600 | 129923399 | 7.570947 | 13.71632 | 39.12498 | 16.93663 | -1.08969 | 0.323124 | 0.13014 |
| 9 | 99021200 | 99022799 | 9.31227 | 16.5337 | 42.00414 | 18.54631 | -0.94347 | 0.468265 | 0.13015 |
| 18 | 82535200 | 82556599 | 117.4978 | 174.4583 | 389.3562 | 160.3884 | -0.90447 | 3.652996 | 0.130153 |
| 8 | 127223800 | 127228999 | 19.6332 | 32.21949 | 87.1857 | 37.5455 | -1.05464 | 1.420747 | 0.130157 |
| 17 | 40341600 | 40342799 | 19.12472 | 35.06624 | 18.83498 | 8.0297 | 1.316516 | 0.295131 | 0.130179 |
| X | 3691600 | 3692799 | 16.23454 | 30.70365 | 16.05939 | 6.5187 | 1.409484 | 0.085074 | 0.130184 |
| 10 | 59755200 | 59761999 | 33.70284 | 53.96344 | 122.9005 | 51.79784 | -0.84014 | 2.007979 | 0.130207 |
| 7 | 4590800 | 4595399 | 72.9414 | 116.5322 | 77.17958 | 32.42596 | 0.934773 | 2.102566 | 0.130242 |
| 7 | 28140400 | 28141999 | 6.408689 | 11.9618 | 35.48147 | 14.74462 | -1.10274 | 0.129752 | 0.130268 |
| 8 | 126449600 | 126451399 | 9.34422 | 16.51323 | 43.55426 | 19.2666 | -1.00046 | 0.515588 | 0.130277 |
| 12 | 19814400 | 19828599 | 511.9942 | 703.7655 | 504.7979 | 209.8804 | 0.622513 | 4.845122 | 0.130287 |
| 4 | 124585400 | 124587599 | 8.551446 | 15.24832 | 43.09336 | 18.95837 | -1.09278 | 0.435177 | 0.130308 |
| 8 | 78528600 | 78530999 | 13.4179 | 23.33221 | 54.8755 | 24.10597 | -0.84124 | 0.778591 | 0.130333 |
| 5 | 139780400 | 139796399 | 77.65356 | 117.8195 | 288.3661 | 119.3821 | -1.00954 | 3.226782 | 0.130338 |
| 13 | 51805400 | 51807599 | 6.176243 | 11.37308 | 36.20929 | 15.34701 | -1.22502 | 0.212184 | 0.130343 |
| 12 | 105738600 | 105740399 | 29.8344 | 50.53159 | 38.34931 | 16.79096 | 0.77773 | 1.06876 | 0.130345 |

| | | | | | | | | | |
|---|---|---|---|---|---|---|---|---|---|
| 5 | 134430600 | 134434199 | 20.17452 | 33.25762 | 78.84601 | 34.01902 | -0.86166 | 1.36167 | 0.130346 |
| 10 | 24745600 | 24746399 | 3.42512 | 6.667051 | 25.7835 | 9.473621 | -1.31474 | -0.35237 | 0.130347 |
| 1 | 170206000 | 170207399 | 5.281366 | 10.12853 | 29.57761 | 11.62715 | -1.00891 | -0.08677 | 0.130369 |
| 17 | 32551000 | 32556399 | 23.24481 | 37.66936 | 95.43991 | 40.85982 | -0.97287 | 1.63254 | 0.130371 |
| 12 | 23947800 | 23951399 | 73.5689 | 117.2248 | 78.63199 | 32.91427 | 0.912443 | 2.186438 | 0.130372 |
| 12 | 105527000 | 105534999 | 68.31074 | 105.6438 | 215.1969 | 89.38766 | -0.72318 | 2.838836 | 0.130377 |
| 7 | 25094600 | 25097799 | 16.13545 | 27.2938 | 63.78347 | 27.91615 | -0.83857 | 1.062457 | 0.130408 |
| 12 | 23145800 | 23152399 | 191.5553 | 286.0869 | 220.7749 | 91.01489 | 0.63438 | 3.553389 | 0.130412 |
| 6 | 138428600 | 138430399 | 9.060372 | 16.39604 | 40.71604 | 17.65049 | -0.89307 | 0.343279 | 0.13042 |
| 7 | 23547800 | 23549799 | 32.42 | 54.8271 | 38.78234 | 16.9727 | 0.878843 | 1.092561 | 0.13046 |
| 2 | 70768400 | 70770199 | 10.06908 | 17.88132 | 43.54578 | 19.22344 | -0.88317 | 0.472205 | 0.130468 |
| 12 | 88108400 | 88109799 | 30.00944 | 51.15873 | 36.58388 | 16.03721 | 0.863963 | 1.011334 | 0.130471 |
| 4 | 128106600 | 128119399 | 103.2412 | 156.1206 | 299.0683 | 123.6841 | -0.66051 | 3.300408 | 0.130481 |
| 2 | 30566000 | 30568199 | 8.605693 | 15.1714 | 44.61242 | 19.74422 | -1.15748 | 0.540881 | 0.130486 |
| 6 | 43703200 | 43704399 | 14.60145 | 27.59483 | 18.1516 | 7.37939 | 1.079332 | 0.079821 | 0.130493 |
| 16 | 57703000 | 57705599 | 16.13145 | 27.30591 | 63.77807 | 27.91964 | -0.83773 | 1.056983 | 0.130498 |
| 13 | 12679800 | 12683399 | 16.84679 | 28.25144 | 69.21842 | 30.1984 | -0.90717 | 1.14823 | 0.130501 |
| 12 | 22405000 | 22408199 | 57.81606 | 92.74397 | 72.86268 | 30.79541 | 0.695448 | 1.960392 | 0.130503 |
| 12 | 5269800 | 5272799 | 17.24193 | 29.10843 | 66.10893 | 28.91676 | -0.79787 | 1.084167 | 0.130516 |
| 7 | 9184400 | 9185199 | 13.10514 | 25.09591 | 17.23379 | 6.803384 | 1.054741 | -0.02986 | 0.130516 |
| 11 | 116628400 | 116635599 | 43.53632 | 69.17158 | 145.7632 | 60.79353 | -0.74056 | 2.266089 | 0.130521 |
| 17 | 18430600 | 18432399 | 35.39474 | 59.08942 | 43.24939 | 18.81527 | 0.829781 | 1.227733 | 0.13054 |
| 4 | 138919800 | 138922799 | 11.67581 | 20.0885 | 57.3166 | 25.14191 | -1.12123 | 0.882764 | 0.130546 |
| 18 | 64742000 | 64746999 | 42.91539 | 68.12289 | 156.2171 | 65.09415 | -0.86364 | 2.294949 | 0.13055 |
| 16 | 10070400 | 10072799 | 12.91765 | 22.1787 | 59.00279 | 25.87978 | -1.02176 | 0.907739 | 0.130557 |
| 2 | 69413800 | 69419399 | 26.56781 | 42.79771 | 109.8792 | 46.72905 | -1.00171 | 1.801787 | 0.130562 |
| 7 | 14811000 | 14819199 | 172.0284 | 260.7269 | 172.4531 | 71.37123 | 0.875214 | 3.263548 | 0.130584 |
| 4 | 45039000 | 45042599 | 16.82115 | 28.27105 | 67.02578 | 29.27797 | -0.85982 | 1.123786 | 0.130589 |
| 15 | 11963000 | 11965999 | 13.25852 | 23.08971 | 53.22874 | 23.40465 | -0.81215 | 0.75479 | 0.130595 |
| 15 | 72788400 | 72790999 | 11.25554 | 19.64304 | 55.36337 | 24.37564 | -1.10029 | 0.71588 | 0.130603 |
| 2 | 74533000 | 74536399 | 13.1944 | 22.71593 | 56.88716 | 24.95403 | -0.93406 | 0.881739 | 0.13061 |
| 14 | 45456400 | 45458599 | 11.66947 | 20.05837 | 57.28339 | 25.12661 | -1.12274 | 0.892614 | 0.130618 |
| 14 | 18527000 | 18528599 | 14.7054 | 27.22234 | 19.65512 | 8.27098 | 0.906538 | 0.209355 | 0.130626 |
| 11 | 95007400 | 95009599 | 8.408651 | 14.9232 | 42.02674 | 18.56274 | -1.09115 | 0.465563 | 0.130643 |
| 6 | 72364800 | 72370199 | 23.33154 | 38.04736 | 89.35644 | 38.38598 | -0.8564 | 1.517894 | 0.130653 |
| X | 33803400 | 33804999 | 19.19221 | 35.88895 | 15.77009 | 6.532763 | 1.633849 | 0.179912 | 0.130671 |
| 6 | 28974200 | 28988799 | 94.67035 | 143.5226 | 286.6182 | 118.6298 | -0.71735 | 3.246748 | 0.130681 |
| 6 | 117883000 | 117885199 | 7.636571 | 13.42289 | 50.34635 | 22.29494 | -1.50837 | 0.576015 | 0.130694 |
| 16 | 27535600 | 27539399 | 39.12729 | 64.91241 | 44.96865 | 19.48218 | 0.908744 | 1.309163 | 0.1307 |
| 15 | 85553200 | 85556399 | 13.85323 | 23.38843 | 67.43737 | 29.49819 | -1.14052 | 1.073731 | 0.13075 |
| 1 | 133333800 | 133341199 | 39.12472 | 62.31386 | 138.8239 | 58.07426 | -0.81694 | 2.184237 | 0.130753 |
| 2 | 31100200 | 31118199 | 115.4749 | 172.7008 | 343.5835 | 141.6812 | -0.72916 | 3.520447 | 0.130756 |
| 9 | 116715800 | 116716999 | 5.533647 | 10.63322 | 33.29808 | 13.01607 | -1.10127 | -0.09186 | 0.130773 |
| 13 | 65995200 | 65999799 | 61.58658 | 99.01253 | 67.73152 | 28.72201 | 0.898515 | 1.894946 | 0.130798 |
| X | 5313000 | 5314799 | 38.07348 | 63.74628 | 39.3865 | 17.16618 | 1.073264 | 1.204693 | 0.130802 |
| 6 | 126509000 | 126512799 | 22.05727 | 36.00127 | 88.61518 | 38.07985 | -0.92295 | 1.502071 | 0.130806 |
| 19 | 30000200 | 30001199 | 20.82429 | 36.70093 | 25.6462 | 11.35099 | 0.90326 | 0.579922 | 0.130809 |

| | | | | | | | | | |
|---|---|---|---|---|---|---|---|---|---|
| 15 | 99013200 | 99013599 | 9.887324 | 19.2934 | 12.70916 | 4.526455 | 1.248906 | -0.42189 | 0.130816 |
| 5 | 70026400 | 70027199 | 4.762528 | 9.238211 | 31.05206 | 11.70659 | -1.15152 | -0.22771 | 0.130816 |
| 10 | 6540200 | 6540799 | 11.0011 | 21.26072 | 15.92993 | 6.110409 | 0.968633 | -0.14745 | 0.130825 |
| 5 | 140926000 | 140934399 | 46.03749 | 72.90943 | 160.2185 | 66.62493 | -0.80546 | 2.362191 | 0.130834 |
| 3 | 16719200 | 16719999 | 9.700606 | 18.9695 | 9.533414 | 3.248861 | 1.685961 | -0.58462 | 0.13084 |
| 5 | 92110200 | 92111199 | 7.098969 | 12.95362 | 38.73255 | 16.61732 | -1.14866 | 0.276043 | 0.130842 |
| 4 | 147412200 | 147414399 | 44.05088 | 72.75916 | 40.66238 | 17.563 | 1.217095 | 1.368698 | 0.130845 |
| 9 | 23132600 | 23135399 | 15.55136 | 26.20736 | 65.86211 | 28.79315 | -0.94327 | 1.094965 | 0.130848 |
| 9 | 120684600 | 120689399 | 31.04713 | 49.93885 | 111.5723 | 47.34662 | -0.80445 | 1.85609 | 0.130852 |
| 14 | 67824800 | 67825199 | 5.097647 | 9.85743 | 28.28688 | 10.82088 | -0.94625 | -0.18288 | 0.130858 |
| 13 | 65334400 | 65335399 | 18.40994 | 33.8587 | 19.77815 | 8.396888 | 1.201319 | 0.27661 | 0.130859 |
| 11 | 41653000 | 41653799 | 5.400301 | 10.4114 | 29.29007 | 11.34104 | -0.93509 | -0.13639 | 0.130862 |
| 10 | 91154800 | 91156199 | 8.10007 | 14.43464 | 41.53649 | 18.28286 | -1.11861 | 0.435129 | 0.130865 |
| 15 | 10469800 | 10470999 | 8.10007 | 14.43464 | 41.53649 | 18.28286 | -1.11861 | 0.435129 | 0.130865 |
| 17 | 11117400 | 11118199 | 4.235614 | 8.241772 | 25.87457 | 9.534319 | -1.02264 | -0.33171 | 0.130882 |
| 5 | 146055000 | 146057399 | 50.64167 | 81.75419 | 63.26759 | 26.98308 | 0.727971 | 1.751747 | 0.130883 |
| 7 | 34160600 | 34163599 | 24.53237 | 40.2117 | 87.29809 | 37.57506 | -0.73866 | 1.448205 | 0.130883 |
| 19 | 37672600 | 37687999 | 113.8418 | 169.6047 | 372.0791 | 153.3244 | -0.87518 | 3.592201 | 0.130907 |
| 17 | 38452000 | 38455399 | 48.1714 | 78.63539 | 50.1391 | 21.54698 | 1.020386 | 1.52976 | 0.130912 |
| 11 | 33002000 | 33003399 | 7.358822 | 13.49068 | 36.6833 | 15.64212 | -1.00629 | 0.247568 | 0.130914 |
| 6 | 10568000 | 10569599 | 16.08904 | 29.48342 | 21.51975 | 9.181346 | 0.877007 | 0.280929 | 0.130917 |
| 2 | 150954800 | 150960799 | 100.5583 | 157.105 | 123.6208 | 51.29135 | 0.65887 | 2.663339 | 0.130917 |
| 11 | 100660800 | 100662399 | 6.950913 | 12.66455 | 37.63285 | 16.17568 | -1.14104 | 0.282948 | 0.130923 |
| 9 | 25979600 | 25981399 | 20.41565 | 36.16261 | 25.25076 | 11.16471 | 0.906767 | 0.521736 | 0.130931 |
| 13 | 66627800 | 66632999 | 119.5637 | 185.3855 | 129.4119 | 53.75065 | 0.822992 | 2.802937 | 0.130932 |
| 3 | 43084200 | 43085999 | 27.31699 | 47.54475 | 28.78088 | 12.65029 | 1.107644 | 0.793754 | 0.130935 |
| 6 | 135532200 | 135532999 | 14.26867 | 26.84961 | 19.07001 | 7.816461 | 0.959286 | 0.105737 | 0.130969 |
| 7 | 20672600 | 20674799 | 43.87264 | 71.48661 | 55.41348 | 23.79768 | 0.737644 | 1.551116 | 0.130987 |
| 18 | 79955200 | 79956999 | 25.65534 | 44.97601 | 26.19446 | 11.57393 | 1.163397 | 0.678557 | 0.130997 |
| 15 | 78931000 | 78932599 | 5.708657 | 10.62195 | 35.33361 | 14.76107 | -1.2722 | 0.145015 | 0.131019 |
| 6 | 77956000 | 77957999 | 9.562108 | 17.17604 | 42.14652 | 18.42747 | -0.88406 | 0.395274 | 0.13103 |
| 6 | 98905400 | 98908399 | 20.4138 | 33.48178 | 84.85603 | 36.53361 | -0.96072 | 1.428798 | 0.131034 |
| 14 | 57122800 | 57125599 | 15.82938 | 26.75977 | 64.89066 | 28.39548 | -0.8918 | 1.06659 | 0.131035 |
| 18 | 81575000 | 81579199 | 18.39063 | 30.88176 | 70.02723 | 30.5771 | -0.79609 | 1.13891 | 0.131048 |
| 17 | 57040200 | 57050599 | 66.48862 | 102.5967 | 227.5147 | 94.50531 | -0.84805 | 2.880139 | 0.131051 |
| 15 | 69267800 | 69270399 | 19.87933 | 33.20511 | 73.9163 | 32.16748 | -0.76951 | 1.20532 | 0.131054 |
| X | 147424000 | 147428799 | 51.8695 | 84.30988 | 52.35881 | 22.46559 | 1.054145 | 1.601196 | 0.13107 |
| 6 | 10652800 | 10655799 | 29.20402 | 50.25885 | 33.39688 | 14.64965 | 0.971541 | 0.922705 | 0.131093 |
| 5 | 39009400 | 39011199 | 21.60995 | 38.55085 | 22.69544 | 9.980738 | 1.157161 | 0.478983 | 0.131109 |
| 8 | 4770400 | 4772199 | 21.35377 | 37.6478 | 26.27041 | 11.62642 | 0.905257 | 0.578922 | 0.131114 |
| 7 | 68009600 | 68014399 | 30.18995 | 48.70364 | 104.9359 | 44.71398 | -0.747 | 1.760996 | 0.13115 |
| 8 | 50059400 | 50059999 | 8.835011 | 15.75617 | 41.01347 | 18.04121 | -0.97438 | 0.434791 | 0.13115 |
| 3 | 15333800 | 15334799 | 22.22973 | 39.05769 | 27.37552 | 12.11861 | 0.897432 | 0.625356 | 0.131167 |
| 15 | 30103000 | 30104799 | 22.22973 | 39.05769 | 27.37552 | 12.11861 | 0.897432 | 0.625356 | 0.131167 |
| 6 | 113021800 | 113024599 | 8.716888 | 15.48973 | 42.53168 | 18.76584 | -1.05425 | 0.458335 | 0.131176 |
| 6 | 97182000 | 97182999 | 5.860037 | 11.20742 | 30.64307 | 12.14153 | -0.92597 | -0.05164 | 0.131177 |
| 5 | 21772400 | 21774999 | 11.19944 | 19.69866 | 46.14485 | 20.43183 | -0.8336 | 0.583111 | 0.131188 |

| | | | | | | | | | |
|---|---|---|---|---|---|---|---|---|---|
| X | 24396400 | 24400199 | 61.68559 | 100.0083 | 44.64386 | 19.11914 | 1.525571 | 1.666603 | 0.131189 |
| 19 | 43597800 | 43608799 | 86.00968 | 130.0766 | 304.6254 | 125.9864 | -0.94996 | 3.29494 | 0.131196 |
| 7 | 47401200 | 47403399 | 62.6276 | 100.6968 | 66.6433 | 28.26487 | 0.946303 | 1.893279 | 0.131201 |
| 7 | 113204400 | 113205799 | 23.95322 | 41.89316 | 28.95269 | 12.76719 | 0.917875 | 0.708955 | 0.131212 |
| 5 | 91690400 | 91691999 | 14.9041 | 28.5499 | 15.1433 | 5.972683 | 1.423965 | -0.02117 | 0.131216 |
| 12 | 111258000 | 111266799 | 36.99686 | 58.39967 | 174.0038 | 72.23726 | -1.24764 | 2.438153 | 0.131219 |
| 8 | 120796800 | 120804399 | 33.98208 | 54.15395 | 139.2329 | 58.29117 | -1.02229 | 2.161402 | 0.131223 |
| 5 | 123681400 | 123683799 | 87.77106 | 140.4099 | 53.61298 | 22.56702 | 1.729468 | 2.077517 | 0.131228 |
| 15 | 75989400 | 75997399 | 48.15207 | 75.45406 | 187.165 | 77.63254 | -0.99204 | 2.59389 | 0.131237 |
| 2 | 18838000 | 18840599 | 15.89415 | 27.11158 | 63.49691 | 27.83634 | -0.84069 | 0.983724 | 0.13124 |
| 11 | 15631200 | 15632399 | 17.03535 | 31.09966 | 21.55331 | 9.241405 | 0.946652 | 0.310286 | 0.131267 |
| 5 | 110796800 | 110799999 | 11.35966 | 19.47193 | 58.79941 | 25.78718 | -1.20337 | 0.920247 | 0.131269 |
| 7 | 129862200 | 129864399 | 8.377888 | 14.91946 | 41.53789 | 18.29408 | -1.07165 | 0.443464 | 0.131296 |
| 4 | 92440200 | 92441399 | 11.66257 | 22.55474 | 15.62662 | 5.974628 | 1.083906 | -0.15296 | 0.131312 |
| 5 | 145321600 | 145322799 | 26.00542 | 45.22008 | 31.2271 | 13.71751 | 0.918718 | 0.80495 | 0.131314 |
| 7 | 29181600 | 29184399 | 11.86221 | 20.69547 | 50.49121 | 22.22623 | -0.8929 | 0.718816 | 0.131319 |
| 9 | 3120800 | 3131399 | 263.5309 | 384.7391 | 287.5988 | 118.4958 | 0.649709 | 3.939509 | 0.131323 |
| 7 | 22904400 | 22906999 | 28.04703 | 48.30858 | 30.92839 | 13.56718 | 1.02511 | 0.906114 | 0.131323 |
| 1 | 154973800 | 154975599 | 8.631149 | 15.15161 | 46.71012 | 20.66229 | -1.22702 | 0.599705 | 0.131354 |
| 9 | 12361000 | 12362399 | 16.93437 | 31.3683 | 19.79557 | 8.322955 | 1.100236 | 0.225258 | 0.131358 |
| 7 | 20363800 | 20363999 | 6.932509 | 13.55903 | 10.81922 | 3.653249 | 1.043326 | -0.65619 | 0.131372 |
| 7 | 33249000 | 33254799 | 169.879 | 256.8667 | 186.9973 | 77.33329 | 0.734136 | 3.310693 | 0.131379 |
| 1 | 109333000 | 109334799 | 21.59892 | 38.75973 | 21.31835 | 9.328505 | 1.258953 | 0.439315 | 0.131388 |
| 7 | 79308400 | 79311199 | 11.99241 | 20.60857 | 57.8578 | 25.37839 | -1.0985 | 0.89548 | 0.131389 |
| 9 | 19026200 | 19026799 | 12.45776 | 24.22792 | 12.74022 | 4.714489 | 1.520068 | -0.25975 | 0.131399 |
| 15 | 87106800 | 87109199 | 28.84067 | 49.0906 | 37.24342 | 16.32431 | 0.778694 | 1.021081 | 0.131406 |
| 5 | 7303000 | 7304799 | 15.47016 | 29.20159 | 17.86769 | 7.28239 | 1.179228 | 0.09427 | 0.131424 |
| 3 | 114287400 | 114287999 | 3.642278 | 7.106909 | 23.27052 | 8.295305 | -1.0371 | -0.46549 | 0.131425 |
| 5 | 137828200 | 137829999 | 11.53642 | 20.1829 | 48.86808 | 21.5548 | -0.88164 | 0.672615 | 0.131433 |
| 12 | 39593400 | 39594999 | 17.36113 | 32.39724 | 16.83745 | 6.999595 | 1.389331 | 0.178183 | 0.131439 |
| 17 | 54002600 | 54005399 | 38.46294 | 63.65354 | 47.19188 | 20.4107 | 0.810978 | 1.336578 | 0.131441 |
| X | 33182800 | 33183599 | 14.68544 | 28.4197 | 13.14964 | 5.011992 | 1.664606 | -0.13543 | 0.131445 |
| 19 | 52754400 | 52755199 | 16.40115 | 30.62678 | 18.76265 | 7.790908 | 1.156109 | 0.171103 | 0.131465 |
| 16 | 21937000 | 21938599 | 6.195743 | 11.70967 | 31.92295 | 12.97679 | -0.95484 | 0.039803 | 0.131467 |
| 12 | 99063000 | 99065199 | 11.11873 | 19.35543 | 51.53558 | 22.65441 | -1.01852 | 0.749422 | 0.131479 |
| 9 | 20881400 | 20885199 | 14.51291 | 24.37348 | 68.41089 | 29.87384 | -1.10219 | 1.125014 | 0.131489 |
| 7 | 44700600 | 44704399 | 16.25549 | 26.86002 | 79.54967 | 34.36876 | -1.18055 | 1.323557 | 0.1315 |
| 13 | 74739200 | 74741399 | 21.52999 | 35.70988 | 79.03571 | 34.23574 | -0.76176 | 1.292048 | 0.131527 |
| 10 | 65158600 | 65159799 | 13.15176 | 25.41164 | 14.87352 | 5.710891 | 1.318994 | -0.12441 | 0.131527 |
| 11 | 80793600 | 80796599 | 12.05752 | 20.93089 | 55.8935 | 24.52682 | -1.02371 | 0.79785 | 0.131531 |
| 2 | 130911600 | 130913599 | 7.009576 | 12.6747 | 41.21012 | 17.88252 | -1.27951 | 0.333943 | 0.131531 |
| 11 | 57705800 | 57708199 | 7.009576 | 12.6747 | 41.21012 | 17.88252 | -1.27951 | 0.333943 | 0.131531 |
| 18 | 19585200 | 19590399 | 54.82917 | 88.66821 | 57.50948 | 24.5823 | 0.984945 | 1.707742 | 0.13154 |
| 18 | 76023000 | 76039399 | 95.67533 | 143.9486 | 320.7979 | 132.4726 | -0.88496 | 3.401327 | 0.13154 |
| 7 | 114172200 | 114175399 | 48.67848 | 78.75953 | 61.47435 | 26.28737 | 0.719183 | 1.689889 | 0.131543 |
| 3 | 96742000 | 96748199 | 35.95931 | 57.54254 | 126.7094 | 53.3579 | -0.79267 | 2.028784 | 0.131556 |
| 7 | 98217400 | 98225999 | 54.58194 | 86.10889 | 166.8332 | 69.24049 | -0.62857 | 2.46186 | 0.131562 |

| | | | | | | | | | |
|---|---|---|---|---|---|---|---|---|---|
| 7 | 5511000 | 5513399 | 55.73293 | 89.87872 | 64.63959 | 27.53083 | 0.831788 | 1.792487 | 0.131581 |
| 18 | 35413600 | 35419799 | 38.04213 | 60.89566 | 125.0577 | 52.68706 | -0.69172 | 2.020683 | 0.131594 |
| 5 | 119024000 | 119027199 | 22.74726 | 37.37464 | 81.90748 | 35.29038 | -0.7507 | 1.410988 | 0.131596 |
| 12 | 10544400 | 10545799 | 15.82923 | 29.95687 | 15.37558 | 6.232432 | 1.435983 | 0.072908 | 0.131619 |
| 7 | 131765400 | 131766399 | 5.025418 | 9.620335 | 31.22425 | 12.33808 | -1.16736 | -0.06957 | 0.131628 |
| 3 | 18613200 | 18614599 | 9.415498 | 16.8668 | 41.0396 | 17.98465 | -0.87426 | 0.408537 | 0.131628 |
| 6 | 124985400 | 124987199 | 7.989195 | 14.16209 | 43.04994 | 19.03181 | -1.20172 | 0.471775 | 0.131642 |
| 14 | 39631200 | 39633199 | 24.5997 | 43.09447 | 27.17297 | 11.99901 | 1.049298 | 0.683778 | 0.131676 |
| 3 | 105444200 | 105448599 | 22.94015 | 37.59288 | 86.50778 | 37.23113 | -0.82256 | 1.445721 | 0.131694 |
| 7 | 89166200 | 89167599 | 7.786635 | 13.95294 | 42.11014 | 18.44925 | -1.18242 | 0.397007 | 0.131694 |
| 7 | 73061200 | 73071599 | 54.19709 | 83.8563 | 228.7236 | 95.00611 | -1.14325 | 2.825402 | 0.1317 |
| 1 | 5581400 | 5582599 | 12.95632 | 25.06838 | 14.29887 | 5.455175 | 1.364074 | -0.14922 | 0.1317 |
| 7 | 8811400 | 8812399 | 12.95632 | 25.06838 | 14.29887 | 5.455175 | 1.364074 | -0.14922 | 0.1317 |
| 2 | 158453000 | 158460599 | 56.59072 | 87.93135 | 211.2668 | 87.72089 | -0.95574 | 2.74898 | 0.131713 |
| 4 | 133566600 | 133568799 | 9.242561 | 16.61736 | 41.62696 | 18.18616 | -0.91305 | 0.384429 | 0.131715 |
| 9 | 106520400 | 106526399 | 37.99652 | 60.5248 | 138.4525 | 57.92926 | -0.85481 | 2.179102 | 0.131756 |
| 14 | 55153000 | 55166199 | 62.77219 | 97.12771 | 218.8158 | 90.89064 | -0.86774 | 2.827205 | 0.131756 |
| 13 | 77282800 | 77283999 | 6.643697 | 12.53901 | 34.21777 | 13.9502 | -0.96095 | 0.056561 | 0.131758 |
| 4 | 104770600 | 104772399 | 9.144822 | 16.29843 | 42.0312 | 18.4987 | -0.96196 | 0.440824 | 0.131765 |
| X | 77993800 | 77995399 | 23.97059 | 42.26679 | 24.48911 | 10.8378 | 1.172117 | 0.591286 | 0.131766 |
| 1 | 89849000 | 89860999 | 89.26262 | 135.0056 | 300.2367 | 124.1741 | -0.87535 | 3.294065 | 0.131768 |
| 17 | 37659600 | 37661999 | 29.98073 | 51.66885 | 32.17849 | 14.11581 | 1.064964 | 0.911688 | 0.1318 |
| 12 | 72541400 | 72543599 | 10.58772 | 18.61498 | 46.67219 | 20.66703 | -0.93136 | 0.583864 | 0.131806 |
| 3 | 43939000 | 43939999 | 14.3019 | 26.78463 | 19.64595 | 8.117546 | 0.904214 | 0.137189 | 0.131812 |
| 16 | 17761200 | 17761999 | 3.529744 | 6.884467 | 23.77943 | 8.527364 | -1.12128 | -0.44485 | 0.131816 |
| 14 | 52542000 | 52542799 | 3.717618 | 7.219814 | 27.31652 | 10.22466 | -1.3095 | -0.27808 | 0.131826 |
| 8 | 124558400 | 124563199 | 24.59143 | 40.11324 | 87.79691 | 37.71992 | -0.75472 | 1.516279 | 0.131831 |
| 7 | 117941400 | 117942199 | 4.801411 | 9.287237 | 27.65916 | 10.56669 | -0.99678 | -0.1914 | 0.131846 |
| 6 | 32614000 | 32616599 | 10.72473 | 18.92432 | 45.10137 | 19.957 | -0.85627 | 0.543831 | 0.131857 |
| 2 | 168750400 | 168756399 | 40.68806 | 64.54136 | 156.232 | 65.05402 | -0.94254 | 2.313213 | 0.131867 |
| 14 | 30972200 | 30973799 | 7.386098 | 13.49056 | 40.40959 | 17.31591 | -1.15051 | 0.274818 | 0.131873 |
| 7 | 31520400 | 31521599 | 19.97177 | 35.49848 | 24.80734 | 10.94314 | 0.908197 | 0.493364 | 0.131874 |
| 17 | 52907800 | 52911199 | 27.72004 | 44.91234 | 98.65785 | 42.23101 | -0.76813 | 1.64356 | 0.131876 |
| 1 | 49340800 | 49341399 | 13.5292 | 25.49441 | 18.59477 | 7.603304 | 0.923814 | 0.088318 | 0.131882 |
| 15 | 85618200 | 85621399 | 19.07201 | 31.47192 | 79.40726 | 34.27583 | -0.95061 | 1.345724 | 0.131907 |
| 18 | 53594000 | 53598599 | 35.53394 | 56.7375 | 132.2543 | 55.52046 | -0.8781 | 2.098189 | 0.131912 |
| 9 | 108902400 | 108904399 | 6.338164 | 11.66098 | 37.82406 | 16.05063 | -1.25353 | 0.220537 | 0.131941 |
| 14 | 42135400 | 42139199 | 72.52416 | 115.2654 | 93.56844 | 39.03728 | 0.633903 | 2.267592 | 0.131949 |
| 3 | 139874800 | 139876799 | 11.41319 | 19.89956 | 52.63295 | 23.15988 | -1.00903 | 0.727454 | 0.131952 |
| 12 | 79927600 | 79929599 | 11.38791 | 19.91234 | 49.40428 | 21.78138 | -0.91669 | 0.681246 | 0.131956 |
| 9 | 73317000 | 73317799 | 12.28091 | 23.82793 | 14.51766 | 5.466931 | 1.288098 | -0.1978 | 0.131958 |
| 17 | 34218000 | 34221999 | 28.93866 | 46.62315 | 106.8791 | 45.46202 | -0.83845 | 1.80019 | 0.131962 |
| 2 | 128020800 | 128024999 | 26.875 | 43.67845 | 93.11138 | 39.94805 | -0.72069 | 1.576959 | 0.131964 |
| 11 | 27349200 | 27349799 | 9.532891 | 18.53316 | 14.37773 | 5.33544 | 0.962505 | -0.28169 | 0.13197 |
| 14 | 22622400 | 22626799 | 24.7844 | 40.33434 | 89.83314 | 38.56212 | -0.78219 | 1.551574 | 0.131974 |
| 5 | 42418000 | 42418799 | 3.696362 | 7.210354 | 27.70148 | 9.917387 | -1.27056 | -0.43321 | 0.13199 |
| 7 | 32247400 | 32251599 | 53.42933 | 86.95445 | 48.51182 | 20.82464 | 1.209517 | 1.583671 | 0.132017 |

| | | | | | | | | | |
|---|---|---|---|---|---|---|---|---|---|
| 7 | 34580600 | 34589999 | 60.30337 | 94.1025 | 200.4383 | 83.17368 | -0.77739 | 2.674753 | 0.132031 |
| 12 | 80285000 | 80287399 | 10.4789 | 18.3313 | 49.38582 | 21.78168 | -1.03468 | 0.669049 | 0.132033 |
| 1 | 42542800 | 42543999 | 4.660055 | 9.047507 | 33.30815 | 12.49599 | -1.27616 | -0.24415 | 0.132037 |
| 4 | 32069600 | 32071599 | 33.35686 | 56.46523 | 37.70871 | 16.5076 | 0.961586 | 1.087931 | 0.132043 |
| 6 | 87954600 | 87959999 | 27.81308 | 44.8506 | 107.1866 | 45.64863 | -0.89681 | 1.770257 | 0.132048 |
| 11 | 111546600 | 111547799 | 14.55531 | 27.31414 | 19.1855 | 7.900317 | 0.970304 | 0.126473 | 0.13205 |
| 7 | 141034800 | 141050799 | 65.85361 | 100.3529 | 280.1891 | 116.1484 | -1.19412 | 3.133286 | 0.132066 |
| 16 | 4292000 | 4294799 | 10.38781 | 18.05322 | 53.63798 | 23.55414 | -1.17611 | 0.772233 | 0.132067 |
| 4 | 41696400 | 41700399 | 12.98706 | 21.8429 | 70.53963 | 30.81962 | -1.30352 | 1.107806 | 0.132079 |
| 17 | 93456600 | 93459599 | 60.4457 | 97.02653 | 67.6235 | 28.60989 | 0.868968 | 1.941777 | 0.132107 |
| 10 | 121435200 | 121447799 | 76.55091 | 117.0893 | 252.6808 | 104.8365 | -0.81866 | 3.058511 | 0.132117 |
| 3 | 131173600 | 131174599 | 4.976531 | 9.583662 | 28.58187 | 11.10246 | -1.02223 | -0.13154 | 0.132124 |
| 13 | 120660400 | 120662599 | 16.25192 | 29.59171 | 21.14342 | 9.095448 | 0.899784 | 0.317236 | 0.132129 |
| 2 | 121119600 | 121120999 | 5.510665 | 10.355 | 32.83256 | 13.48946 | -1.18351 | 0.073884 | 0.132134 |
| 11 | 94934200 | 94937999 | 14.0272 | 23.58564 | 67.26159 | 29.38489 | -1.12484 | 1.112162 | 0.13214 |
| 4 | 151331800 | 151339399 | 42.48027 | 67.31355 | 156.1456 | 64.94977 | -0.88266 | 2.346302 | 0.132148 |
| 7 | 113714000 | 113717199 | 14.72932 | 24.94572 | 63.71606 | 27.89697 | -0.96613 | 1.046076 | 0.132151 |
| 8 | 9034400 | 9038999 | 35.43098 | 56.82773 | 118.5721 | 50.10397 | -0.71115 | 1.954209 | 0.132152 |
| 14 | 37598800 | 37602199 | 74.32227 | 117.9716 | 95.71579 | 39.88184 | 0.632866 | 2.306576 | 0.132167 |
| 15 | 41042800 | 41044199 | 17.45529 | 31.49965 | 22.99439 | 10.00303 | 0.858856 | 0.380918 | 0.132173 |
| 4 | 150806200 | 150813999 | 34.765 | 55.46832 | 139.9204 | 58.67666 | -0.99304 | 2.121807 | 0.132187 |
| 7 | 24731200 | 24733199 | 8.138444 | 14.79685 | 38.13054 | 16.43308 | -0.94012 | 0.304543 | 0.132187 |
| 15 | 67951400 | 67954799 | 14.92407 | 25.56835 | 61.87293 | 27.13396 | -0.88659 | 0.937791 | 0.132201 |
| 11 | 107899200 | 107900799 | 5.403082 | 10.3375 | 30.58568 | 12.10526 | -1.03635 | -0.05791 | 0.132207 |
| 17 | 46644400 | 46645199 | 14.99586 | 27.92729 | 20.12626 | 8.395327 | 0.918732 | 0.179568 | 0.132225 |
| 19 | 5608000 | 5623599 | 78.47174 | 119.1999 | 285.4428 | 118.2131 | -0.9767 | 3.202437 | 0.132228 |
| 2 | 106122600 | 106124199 | 19.3905 | 36.10034 | 14.92292 | 6.228118 | 1.71171 | 0.205502 | 0.132235 |
| 7 | 80524600 | 80527599 | 19.28859 | 31.69916 | 83.08356 | 35.80057 | -1.00779 | 1.400299 | 0.13226 |
| 5 | 95465600 | 95467999 | 55.54252 | 89.44315 | 57.62635 | 24.49542 | 0.987603 | 1.830511 | 0.132262 |
| 3 | 52054000 | 52055199 | 7.771474 | 14.33262 | 36.84337 | 15.58308 | -0.91763 | 0.216082 | 0.132263 |
| 17 | 72044600 | 72063199 | 124.8825 | 185.9298 | 380.1409 | 156.6358 | -0.7745 | 3.605506 | 0.132268 |
| 10 | 130298000 | 130302399 | 32.56953 | 54.6763 | 41.58715 | 18.15575 | 0.7743 | 1.158971 | 0.132272 |
| 7 | 18319000 | 18325199 | 105.6108 | 165.2954 | 116.109 | 48.16574 | 0.824429 | 2.63523 | 0.132274 |
| 3 | 12300200 | 12301999 | 12.97088 | 25.03478 | 15.4524 | 5.963531 | 1.236371 | -0.1082 | 0.132274 |
| 11 | 35999000 | 36000799 | 7.180976 | 13.25802 | 36.8555 | 15.56514 | -1.02802 | 0.204774 | 0.132291 |
| 1 | 79194800 | 79195599 | 5.91303 | 11.1814 | 31.93675 | 12.96756 | -1.02013 | 0.032315 | 0.132304 |
| 12 | 44166000 | 44167599 | 15.11899 | 25.83571 | 60.08227 | 26.34321 | -0.82977 | 0.963665 | 0.132304 |
| 10 | 117245400 | 117249599 | 17.43256 | 29.32751 | 67.71553 | 29.58811 | -0.82186 | 1.116844 | 0.132314 |
| 14 | 15900400 | 15901999 | 12.40401 | 23.94709 | 16.16515 | 6.230816 | 1.110662 | -0.1159 | 0.132316 |
| 14 | 102926800 | 102927999 | 12.40401 | 23.94709 | 16.16515 | 6.230816 | 1.110662 | -0.1159 | 0.132316 |
| 11 | 69751400 | 69752799 | 5.603005 | 10.64989 | 30.96128 | 12.43521 | -1.03112 | -0.00886 | 0.132331 |
| 5 | 142941600 | 142968199 | 152.5359 | 220.3583 | 543.2823 | 224.1127 | -1.08595 | 4.093201 | 0.132335 |
| 17 | 40168800 | 40169799 | 12.37209 | 24.00318 | 14.61073 | 5.503768 | 1.289549 | -0.19485 | 0.132348 |
| 2 | 169359600 | 169360599 | 5.258646 | 10.15495 | 29.41421 | 11.32382 | -0.96907 | -0.15786 | 0.132355 |
| 13 | 65392800 | 65395399 | 60.10801 | 96.65242 | 68.74874 | 29.1608 | 0.842641 | 1.887863 | 0.132356 |
| 7 | 25824400 | 25826199 | 50.46884 | 81.44428 | 62.95504 | 26.83665 | 0.729063 | 1.764316 | 0.132359 |
| 6 | 56184000 | 56184799 | 18.4709 | 33.08373 | 23.93986 | 10.49237 | 0.864446 | 0.43463 | 0.132371 |

| | | | | | | | | | |
|---|---|---|---|---|---|---|---|---|---|
| 17 | 56425600 | 56443199 | 147.8481 | 216.4406 | 438.8745 | 180.7876 | -0.78543 | 3.890138 | 0.132372 |
| 4 | 114913400 | 114918199 | 30.9385 | 49.65446 | 118.2171 | 50.00329 | -0.89988 | 1.928079 | 0.13239 |
| 5 | 37628600 | 37629799 | 14.19354 | 26.42351 | 19.15982 | 7.997097 | 0.908971 | 0.171565 | 0.1324 |
| 8 | 84362800 | 84363399 | 14.13594 | 26.7281 | 18.57823 | 7.545053 | 1.001293 | 0.072407 | 0.132402 |
| 12 | 22583800 | 22586599 | 74.28939 | 118.4256 | 79.94607 | 33.48497 | 0.903914 | 2.172329 | 0.132402 |
| 19 | 47846200 | 47847599 | 6.661233 | 12.51908 | 33.47265 | 13.74858 | -0.94036 | 0.083444 | 0.132406 |
| 7 | 103066000 | 103066599 | 8.793196 | 17.13977 | 13.33247 | 4.819968 | 0.992139 | -0.38536 | 0.132437 |
| 13 | 95739200 | 95756799 | 171.3527 | 248.9112 | 476.2423 | 196.3229 | -0.71254 | 4.011157 | 0.132443 |
| 3 | 52252400 | 52253599 | 7.244758 | 13.16472 | 37.5951 | 16.21353 | -1.08718 | 0.302 | 0.132451 |
| 9 | 67665200 | 67667999 | 13.73595 | 23.44284 | 61.10546 | 26.78912 | -0.99394 | 0.96905 | 0.132455 |
| 16 | 30046200 | 30048999 | 18.96185 | 31.55619 | 72.48516 | 31.47117 | -0.8145 | 1.241326 | 0.132461 |
| 10 | 11024800 | 11026399 | 11.06686 | 19.39301 | 51.5316 | 22.76098 | -1.01531 | 0.645416 | 0.132467 |
| 14 | 38556600 | 38557799 | 16.65944 | 30.79192 | 20.426 | 8.613494 | 1.025941 | 0.236633 | 0.132478 |
| 3 | 83708800 | 83709999 | 17.40915 | 31.77012 | 21.07912 | 9.042336 | 1.008759 | 0.314543 | 0.132487 |
| 1 | 71236800 | 71237799 | 12.55814 | 24.38566 | 14.03327 | 5.257792 | 1.376698 | -0.21226 | 0.132487 |
| 4 | 8745800 | 8755799 | 49.73122 | 77.94218 | 191.6723 | 79.50069 | -0.97942 | 2.593008 | 0.132504 |
| 2 | 170334800 | 170340199 | 40.05868 | 63.88043 | 134.7071 | 56.44873 | -0.73591 | 2.14366 | 0.13252 |
| 7 | 125217000 | 125232999 | 102.5713 | 154.4179 | 315.2763 | 130.2137 | -0.75796 | 3.390329 | 0.132525 |
| 7 | 8855000 | 8855399 | 8.150808 | 15.92359 | 11.41528 | 3.984072 | 1.154681 | -0.52096 | 0.13254 |
| 12 | 114842600 | 114843999 | 22.11154 | 38.85155 | 27.44948 | 12.15175 | 0.886028 | 0.62462 | 0.132541 |
| 13 | 120597200 | 120600399 | 35.38829 | 58.71397 | 45.53817 | 19.72998 | 0.746301 | 1.29689 | 0.132547 |
| X | 8302000 | 8304199 | 47.93542 | 78.4439 | 48.42615 | 20.82962 | 1.069419 | 1.496634 | 0.132554 |
| 1 | 155651600 | 155655799 | 21.76178 | 35.46934 | 89.45626 | 38.42169 | -0.95935 | 1.522845 | 0.13256 |
| 12 | 22144600 | 22149999 | 134.7441 | 205.8729 | 155.1049 | 64.33617 | 0.696885 | 3.090178 | 0.132566 |
| 3 | 83011600 | 83014999 | 42.13229 | 69.42252 | 47.81032 | 20.62424 | 0.915944 | 1.397335 | 0.132573 |
| 4 | 135209400 | 135214399 | 19.63911 | 32.19281 | 85.18467 | 36.66626 | -1.02301 | 1.43481 | 0.132574 |
| 2 | 178217400 | 178220199 | 24.14885 | 39.29491 | 89.62126 | 38.46934 | -0.81644 | 1.552557 | 0.132579 |
| 17 | 18308400 | 18309399 | 12.46459 | 24.20494 | 13.93143 | 5.218012 | 1.37627 | -0.21523 | 0.132592 |
| 19 | 25266000 | 25267799 | 7.847053 | 14.26548 | 39.74023 | 17.12917 | -1.0521 | 0.305042 | 0.132598 |
| 4 | 152261600 | 152265999 | 19.27346 | 31.66093 | 81.7481 | 35.21883 | -0.98637 | 1.406131 | 0.132599 |
| X | 102235800 | 102237799 | 56.92677 | 91.81108 | 61.62635 | 26.25007 | 0.931283 | 1.793276 | 0.132606 |
| 5 | 71232000 | 71233199 | 11.4742 | 22.22105 | 15.06647 | 5.723492 | 1.123152 | -0.1781 | 0.132617 |
| 2 | 170509600 | 170512199 | 12.58305 | 21.67254 | 57.40841 | 25.183 | -1.01451 | 0.875589 | 0.132629 |
| 2 | 59229000 | 59238799 | 63.35946 | 98.56095 | 208.8243 | 86.69194 | -0.77332 | 2.731618 | 0.132638 |
| 16 | 92603200 | 92609399 | 28.79106 | 46.37636 | 110.6303 | 47.04444 | -0.89615 | 1.80652 | 0.132647 |
| 4 | 34391600 | 34392599 | 17.98831 | 32.86128 | 21.30665 | 9.128029 | 1.042692 | 0.312895 | 0.132649 |
| 4 | 131183600 | 131191599 | 43.18047 | 68.67185 | 145.686 | 60.82034 | -0.7489 | 2.236599 | 0.132653 |
| 1 | 97176400 | 97178399 | 25.40305 | 44.72361 | 22.53941 | 9.978019 | 1.370845 | 0.611326 | 0.132684 |
| 11 | 114675400 | 114683599 | 46.89097 | 73.36567 | 190.7123 | 79.11634 | -1.06122 | 2.621489 | 0.132694 |
| 3 | 142266000 | 142267599 | 5.743223 | 10.9541 | 33.50385 | 13.35866 | -1.09541 | -0.02803 | 0.132706 |
| 2 | 72809000 | 72811399 | 11.32211 | 19.30417 | 63.05326 | 27.64086 | -1.31777 | 0.971928 | 0.132723 |
| 15 | 85912200 | 85918399 | 68.45523 | 104.9728 | 243.2512 | 100.9765 | -0.91835 | 3.005107 | 0.132727 |
| 3 | 105354600 | 105362799 | 45.09592 | 71.36744 | 172.8668 | 71.8587 | -0.94658 | 2.379967 | 0.132742 |
| 13 | 64299800 | 64302199 | 16.49129 | 27.76743 | 67.07894 | 29.31746 | -0.88671 | 1.105867 | 0.132743 |
| 9 | 49400000 | 49401799 | 6.851655 | 12.75274 | 35.40731 | 14.78393 | -1.0138 | 0.151886 | 0.132756 |
| 6 | 34836200 | 34837199 | 3.941311 | 7.675607 | 24.71803 | 9.040783 | -1.04803 | -0.36439 | 0.132757 |
| 12 | 111330400 | 111335799 | 24.42653 | 39.86124 | 88.51898 | 38.03605 | -0.77517 | 1.50932 | 0.132763 |

| | | | | | | | | | |
|---|---|---|---|---|---|---|---|---|---|
| 1 | 18812400 | 18813799 | 17.97157 | 32.27608 | 23.49392 | 10.27359 | 0.858158 | 0.413812 | 0.132789 |
| 10 | 23224000 | 23226799 | 17.58339 | 29.18423 | 74.37064 | 32.23041 | -0.96397 | 1.270725 | 0.132791 |
| 7 | 126391000 | 126391799 | 4.418227 | 8.548387 | 28.2677 | 10.78541 | -1.1449 | -0.20085 | 0.132792 |
| 7 | 28550800 | 28552599 | 20.46176 | 36.06864 | 26.11027 | 11.55596 | 0.852639 | 0.576207 | 0.132798 |
| 7 | 14477400 | 14485399 | 228.2037 | 337.2528 | 253.2396 | 104.2899 | 0.659858 | 3.742198 | 0.132811 |
| 10 | 35648800 | 35651599 | 38.41176 | 63.61122 | 46.63762 | 20.18026 | 0.827046 | 1.327924 | 0.132816 |
| 9 | 4090000 | 4093199 | 41.92224 | 68.78918 | 51.12062 | 22.01338 | 0.804165 | 1.448598 | 0.132832 |
| 5 | 142169200 | 142176399 | 54.68833 | 85.54569 | 188.7503 | 78.3043 | -0.8254 | 2.628572 | 0.132832 |
| 19 | 4146600 | 4158799 | 59.9242 | 92.51355 | 225.6285 | 93.72156 | -0.98452 | 2.868771 | 0.132833 |
| 5 | 123195200 | 123196399 | 6.18934 | 11.74472 | 33.27847 | 13.41319 | -0.99996 | 0.012106 | 0.132833 |
| 12 | 73056600 | 73058199 | 10.0563 | 17.94106 | 44.16168 | 19.41979 | -0.8945 | 0.445508 | 0.132836 |
| 1 | 153377400 | 153378599 | 7.642884 | 14.03727 | 36.18561 | 15.39129 | -0.92716 | 0.238934 | 0.132852 |
| 15 | 83624800 | 83632599 | 42.3407 | 67.43918 | 139.0048 | 58.15378 | -0.70497 | 2.18518 | 0.13286 |
| 4 | 89257000 | 89258399 | 6.323748 | 12.02292 | 33.38597 | 13.40179 | -0.96591 | 0.000405 | 0.13287 |
| 15 | 10806600 | 10807799 | 15.77989 | 29.36259 | 19.2845 | 8.05572 | 1.04948 | 0.188143 | 0.132911 |
| 16 | 95452800 | 95455199 | 13.77419 | 23.85599 | 54.81312 | 24.04861 | -0.80852 | 0.82857 | 0.132913 |
| 14 | 21707000 | 21709599 | 10.1399 | 17.81973 | 46.67873 | 20.66758 | -0.99422 | 0.58752 | 0.132916 |
| 10 | 80696800 | 80710599 | 63.1545 | 96.8819 | 247.9559 | 102.9349 | -1.0611 | 2.995429 | 0.132923 |
| 17 | 80272200 | 80280599 | 45.48592 | 72.19743 | 153.9288 | 64.13543 | -0.75888 | 2.297429 | 0.132935 |
| 4 | 55049600 | 55051599 | 8.331277 | 15.01394 | 39.03439 | 17.00991 | -0.96315 | 0.360951 | 0.132938 |
| 16 | 30865000 | 30867599 | 10.0216 | 17.52577 | 48.85336 | 21.54108 | -1.0833 | 0.672476 | 0.132943 |
| 7 | 61802400 | 61803599 | 20.31557 | 36.10553 | 24.07111 | 10.61938 | 0.975371 | 0.493647 | 0.132945 |
| 15 | 80509200 | 80513999 | 33.15352 | 53.25876 | 113.8204 | 48.21116 | -0.74286 | 1.900652 | 0.132945 |
| 2 | 30023600 | 30025599 | 5.899318 | 10.8171 | 36.66159 | 15.62878 | -1.3202 | 0.236445 | 0.132951 |
| 7 | 3132200 | 3132999 | 16.30684 | 29.74972 | 21.16233 | 9.081384 | 0.908463 | 0.307159 | 0.132955 |
| 1 | 157177200 | 157178999 | 19.18087 | 34.4499 | 23.32592 | 10.19963 | 0.96198 | 0.421943 | 0.132968 |
| 19 | 6617200 | 6632199 | 78.51196 | 119.518 | 275.9128 | 114.34 | -0.92136 | 3.157517 | 0.132969 |
| 8 | 40393200 | 40394199 | 4.790151 | 9.304288 | 27.51129 | 10.28366 | -0.95694 | -0.26846 | 0.132979 |
| 12 | 118810400 | 118814199 | 27.96293 | 45.10065 | 104.757 | 44.62467 | -0.85546 | 1.764608 | 0.132979 |
| 3 | 154032800 | 154036399 | 32.0067 | 51.48242 | 110.5411 | 46.90919 | -0.74727 | 1.857313 | 0.132981 |
| 15 | 89436000 | 89445799 | 60.52766 | 93.56595 | 221.6699 | 92.07662 | -0.94128 | 2.844447 | 0.132981 |
| 13 | 66350000 | 66352399 | 39.87574 | 65.88356 | 43.97666 | 18.99859 | 0.961773 | 1.355985 | 0.132983 |
| 2 | 152217200 | 152218999 | 8.041515 | 14.18989 | 47.20991 | 20.88994 | -1.33427 | 0.529166 | 0.132986 |
| 10 | 81593800 | 81596999 | 57.91491 | 93.34935 | 64.16725 | 27.31148 | 0.896044 | 1.810843 | 0.132994 |
| 10 | 118714400 | 118717399 | 15.55389 | 26.62039 | 64.13534 | 28.12383 | -0.8806 | 0.951913 | 0.133004 |
| 2 | 46723000 | 46726999 | 52.03335 | 84.13667 | 60.70976 | 25.9462 | 0.831347 | 1.707745 | 0.133009 |
| 13 | 17869200 | 17870599 | 10.14802 | 17.80121 | 46.71857 | 20.65309 | -0.99675 | 0.61938 | 0.133011 |
| 17 | 44818800 | 44825199 | 53.08426 | 83.83131 | 164.0871 | 68.13583 | -0.64209 | 2.435925 | 0.133012 |
| 7 | 135997800 | 136001599 | 16.12407 | 27.2954 | 64.97126 | 28.4426 | -0.86509 | 1.055062 | 0.133015 |
| 17 | 66467200 | 66470999 | 90.73411 | 143.0467 | 108.5486 | 45.04559 | 0.721151 | 2.488451 | 0.133021 |
| 7 | 70153800 | 70157399 | 15.46773 | 26.28006 | 65.58737 | 28.74362 | -0.93291 | 1.020462 | 0.133028 |
| 1 | 66901000 | 66902799 | 7.776652 | 13.91717 | 40.53969 | 17.78044 | -1.13228 | 0.404106 | 0.133033 |
| 18 | 13975600 | 13980399 | 30.2826 | 48.61966 | 115.5368 | 48.89694 | -0.89648 | 1.915745 | 0.133038 |
| 4 | 153456000 | 153460999 | 26.20043 | 42.30977 | 104.3498 | 44.5215 | -0.9397 | 1.72495 | 0.133041 |
| 4 | 29414600 | 29415399 | 5.712193 | 10.9474 | 30.17229 | 11.88383 | -0.92932 | -0.07476 | 0.133042 |
| X | 32559400 | 32561399 | 52.07277 | 84.89499 | 48.47667 | 20.82282 | 1.177868 | 1.557864 | 0.133045 |
| 19 | 45003200 | 45010199 | 26.13244 | 42.04218 | 115.1881 | 48.89541 | -1.09761 | 1.844035 | 0.133051 |

| | | | | | | | | | |
|---|---|---|---|---|---|---|---|---|---|
| 1 | 22379800 | 22380799 | 5.130036 | 9.892815 | 29.90049 | 11.5691 | -1.03688 | -0.14155 | 0.133052 |
| 13 | 57394200 | 57395199 | 5.130036 | 9.892815 | 29.90049 | 11.5691 | -1.03688 | -0.14155 | 0.133052 |
| 1 | 38803200 | 38806599 | 16.04805 | 27.44413 | 61.29751 | 26.87768 | -0.77181 | 0.96021 | 0.133063 |
| 4 | 83869000 | 83870599 | 7.858415 | 14.25677 | 39.68958 | 17.15143 | -1.05353 | 0.317373 | 0.133071 |
| 16 | 36919800 | 36920999 | 13.31602 | 25.89009 | 12.06229 | 4.473341 | 1.689564 | -0.24398 | 0.133071 |
| 17 | 73414200 | 73416599 | 8.669404 | 15.23051 | 50.37576 | 22.29832 | -1.32863 | 0.589569 | 0.133074 |
| 12 | 23269000 | 23269599 | 20.54089 | 36.41808 | 24.86399 | 10.99148 | 0.939669 | 0.513239 | 0.133088 |
| 15 | 59585600 | 59589199 | 16.53542 | 28.15826 | 65.17775 | 28.56935 | -0.82426 | 1.00336 | 0.133091 |
| 17 | 35608800 | 35615199 | 47.6198 | 75.51871 | 152.1737 | 63.34899 | -0.6788 | 2.325147 | 0.133099 |
| 1 | 125838200 | 125842599 | 28.48695 | 45.85666 | 111.3286 | 47.29677 | -0.92258 | 1.827355 | 0.133101 |
| 6 | 121222600 | 121222999 | 3.232197 | 6.318106 | 22.60758 | 7.781523 | -1.11694 | -0.59995 | 0.133103 |
| 17 | 78055400 | 78055999 | 3.232197 | 6.318106 | 22.60758 | 7.781523 | -1.11694 | -0.59995 | 0.133103 |
| 10 | 39371000 | 39372999 | 11.87272 | 20.68648 | 51.02754 | 22.44371 | -0.90889 | 0.738803 | 0.133131 |
| 1 | 183401000 | 183402999 | 11.70333 | 20.43329 | 50.48735 | 22.23481 | -0.91099 | 0.706993 | 0.133133 |
| 6 | 72744800 | 72747399 | 16.11594 | 27.31051 | 63.30393 | 27.72157 | -0.82679 | 1.04495 | 0.133152 |
| 1 | 96601600 | 96603399 | 24.68557 | 43.15696 | 28.95613 | 12.76444 | 0.96046 | 0.719686 | 0.133158 |
| 4 | 146645000 | 146645399 | 11.54818 | 22.3889 | 15.11053 | 5.71083 | 1.136929 | -0.19335 | 0.133173 |
| 1 | 154025800 | 154028599 | 10.5678 | 18.63005 | 46.67978 | 20.65928 | -0.92896 | 0.553228 | 0.133177 |
| 9 | 57397600 | 57398799 | 23.72185 | 41.46473 | 29.64519 | 13.06622 | 0.869354 | 0.718759 | 0.133181 |
| 3 | 94456000 | 94457399 | 13.63572 | 23.53475 | 55.31264 | 24.26521 | -0.8421 | 0.859384 | 0.133181 |
| 7 | 23486400 | 23488799 | 64.16454 | 102.6958 | 79.57057 | 33.51627 | 0.711671 | 2.040235 | 0.133188 |
| 6 | 79605200 | 79606799 | 38.5086 | 64.50411 | 35.69599 | 15.56217 | 1.23156 | 1.197774 | 0.133195 |
| 11 | 22191000 | 22191599 | 4.658391 | 9.049526 | 26.90335 | 10.04596 | -0.96323 | -0.27599 | 0.133195 |
| 11 | 117284800 | 117305999 | 127.0387 | 187.8938 | 398.2596 | 164.0139 | -0.83531 | 3.721366 | 0.133239 |
| 8 | 126213000 | 126215999 | 22.26747 | 36.59729 | 81.25975 | 35.01574 | -0.76931 | 1.405776 | 0.133244 |
| 7 | 27351200 | 27362999 | 63.8768 | 97.9582 | 253.077 | 105.0562 | -1.07502 | 3.002061 | 0.133245 |
| 12 | 19390600 | 19390999 | 12.51696 | 23.79006 | 17.35194 | 6.970113 | 0.946322 | 0.019125 | 0.133249 |
| 7 | 59455400 | 59456799 | 10.9428 | 21.29141 | 14.24617 | 5.252205 | 1.182633 | -0.28541 | 0.133255 |
| 14 | 98827400 | 98828399 | 4.79503 | 9.296885 | 27.27236 | 10.30427 | -0.95998 | -0.23091 | 0.133264 |
| 17 | 83306800 | 83307999 | 4.79503 | 9.296885 | 27.27236 | 10.30427 | -0.95998 | -0.23091 | 0.133264 |
| 16 | 58962600 | 58965199 | 31.22238 | 53.41321 | 34.31547 | 15.04705 | 1.018568 | 0.983214 | 0.133269 |
| 2 | 60213000 | 60214199 | 5.020796 | 9.529928 | 30.85783 | 12.43035 | -1.18792 | -0.00319 | 0.133271 |
| 14 | 10912800 | 10918199 | 35.54305 | 56.73137 | 133.1902 | 55.88447 | -0.88899 | 2.109856 | 0.133285 |
| 11 | 50527600 | 50532599 | 32.96531 | 53.02073 | 113.4801 | 48.15162 | -0.74284 | 1.860912 | 0.133291 |
| 17 | 37123200 | 37125799 | 15.54089 | 26.66244 | 60.25699 | 26.42828 | -0.78801 | 0.926583 | 0.133294 |
| 19 | 20858600 | 20858999 | 7.088171 | 13.8633 | 11.318 | 3.826048 | 1.009259 | -0.64716 | 0.133294 |
| 7 | 122568400 | 122569999 | 9.486726 | 16.79553 | 43.53742 | 19.25216 | -0.97476 | 0.495025 | 0.133301 |
| 14 | 14176600 | 14179999 | 43.84554 | 71.79639 | 47.85446 | 20.58889 | 0.959497 | 1.4757 | 0.13331 |
| 6 | 7978200 | 7978799 | 10.74847 | 20.9711 | 11.82105 | 4.219412 | 1.466456 | -0.40662 | 0.133315 |
| 2 | 31543800 | 31550999 | 61.69384 | 96.45456 | 187.281 | 77.69882 | -0.64183 | 2.641451 | 0.133325 |
| 2 | 109938400 | 109940999 | 24.57956 | 43.28187 | 24.38651 | 10.79532 | 1.211397 | 0.60869 | 0.133327 |
| 10 | 128244000 | 128245999 | 8.226926 | 14.7355 | 41.57588 | 18.22186 | -1.0864 | 0.40187 | 0.133334 |
| 15 | 14706600 | 14707199 | 13.58139 | 25.99451 | 17.14193 | 6.777145 | 1.110602 | -0.02082 | 0.133336 |
| X | 117435800 | 117437399 | 20.57866 | 37.65134 | 18.19619 | 7.780026 | 1.464887 | 0.317122 | 0.133339 |
| 16 | 96965800 | 96967399 | 7.621854 | 14.06568 | 35.75028 | 15.1076 | -0.90016 | 0.210147 | 0.133367 |
| 2 | 98841000 | 98842199 | 14.50679 | 27.85517 | 14.57603 | 5.709725 | 1.451914 | -0.04832 | 0.13337 |
| 2 | 49706400 | 49707999 | 9.459809 | 16.81792 | 43.04267 | 18.98365 | -0.95336 | 0.460165 | 0.133385 |

| | | | | | | | | | |
|---|---|---|---|---|---|---|---|---|---|
| 2 | 153984800 | 153986999 | 9.459809 | 16.81792 | 43.04267 | 18.98365 | -0.95336 | 0.460165 | 0.133385 |
| 7 | 31426000 | 31430599 | 45.06538 | 73.67247 | 52.22502 | 22.45533 | 0.869518 | 1.505999 | 0.13339 |
| 3 | 81216000 | 81217999 | 21.93618 | 39.28826 | 20.59679 | 9.027595 | 1.325787 | 0.451025 | 0.133399 |
| 1 | 74984600 | 74993399 | 58.12776 | 90.13866 | 214.3561 | 89.03025 | -0.9432 | 2.784914 | 0.133402 |
| 5 | 101781800 | 101784199 | 9.383567 | 16.47731 | 46.18377 | 20.43455 | -1.09079 | 0.597665 | 0.13342 |
| 7 | 13561800 | 13566399 | 94.21511 | 149.2895 | 87.10172 | 36.2302 | 1.10544 | 2.370996 | 0.133421 |
| 8 | 107589200 | 107592399 | 15.51641 | 26.23939 | 64.30614 | 28.14253 | -0.90693 | 1.06237 | 0.133423 |
| 4 | 117691000 | 117693999 | 12.46572 | 21.37539 | 58.38328 | 25.6054 | -1.05966 | 0.918361 | 0.133431 |
| 5 | 73647200 | 73649999 | 12.46572 | 21.37539 | 58.38328 | 25.6054 | -1.05966 | 0.918361 | 0.133431 |
| 13 | 57868600 | 57873999 | 34.15633 | 54.75724 | 121.0849 | 51.12875 | -0.79559 | 1.967984 | 0.133441 |
| 11 | 94392600 | 94404599 | 75.6965 | 116.6069 | 228.4945 | 94.90999 | -0.6717 | 2.915423 | 0.133446 |
| X | 25576400 | 25576999 | 13.71376 | 26.71512 | 10.50807 | 3.828547 | 1.953654 | -0.30241 | 0.133447 |
| 6 | 36060600 | 36060999 | 7.775983 | 15.20791 | 7.778981 | 2.547788 | 1.70566 | -0.75818 | 0.133453 |
| 9 | 8811000 | 8811599 | 2.112536 | 4.13216 | 18.50468 | 6.097339 | -1.36746 | -0.78024 | 0.133456 |
| 7 | 138220600 | 138222999 | 11.0364 | 19.43436 | 45.61937 | 20.19448 | -0.83588 | 0.568056 | 0.133475 |
| 14 | 8600400 | 8600799 | 8.951148 | 17.47699 | 12.02893 | 4.240535 | 1.197384 | -0.47465 | 0.133483 |
| 15 | 73699800 | 73718999 | 135.2007 | 199.4382 | 406.2519 | 167.3006 | -0.78158 | 3.766302 | 0.133484 |
| 3 | 96869200 | 96869999 | 10.93415 | 21.10723 | 16.39617 | 6.322873 | 0.909502 | -0.12983 | 0.133516 |
| 11 | 114525800 | 114550399 | 189.1968 | 272.1426 | 523.1395 | 215.9309 | -0.73329 | 4.159031 | 0.133535 |
| 2 | 143756600 | 143757999 | 10.4446 | 18.34198 | 46.17148 | 20.43055 | -0.93704 | 0.601163 | 0.133557 |
| 6 | 55330400 | 55334999 | 19.02929 | 31.50404 | 77.37476 | 33.47362 | -0.91146 | 1.30532 | 0.13359 |
| 11 | 5420600 | 5425399 | 30.98937 | 49.98486 | 110.1943 | 46.94443 | -0.78037 | 1.768425 | 0.13359 |
| Y | 2316400 | 2318199 | 11.06192 | 21.49703 | 6.96175 | 2.591961 | 2.197839 | -0.28055 | 0.133613 |
| 15 | 14942400 | 14944199 | 25.22399 | 44.58847 | 21.5058 | 9.512752 | 1.435908 | 0.566808 | 0.133614 |
| 5 | 44383200 | 44386799 | 13.29585 | 23.09214 | 54.27337 | 23.82342 | -0.84016 | 0.797372 | 0.133634 |
| 2 | 177857800 | 177858999 | 16.69918 | 30.68241 | 18.7686 | 7.978858 | 1.134135 | 0.266079 | 0.133655 |
| 2 | 120098400 | 120102399 | 18.76188 | 30.95021 | 79.29548 | 34.22293 | -0.97277 | 1.349227 | 0.133656 |
| 6 | 51446000 | 51448399 | 9.270946 | 15.96252 | 58.77816 | 25.7836 | -1.48647 | 0.868234 | 0.13367 |
| 13 | 65787200 | 65793799 | 164.3176 | 249.3449 | 174.9546 | 72.4333 | 0.791556 | 3.237276 | 0.133701 |
| 7 | 60307800 | 60310999 | 62.0679 | 99.35611 | 79.49066 | 33.49024 | 0.665741 | 2.03168 | 0.13371 |
| 16 | 50957800 | 50960399 | 18.67364 | 31.01107 | 74.03904 | 32.09906 | -0.87031 | 1.266241 | 0.133711 |
| 8 | 45610600 | 45611799 | 26.93571 | 46.48158 | 33.94983 | 14.89344 | 0.836399 | 0.888589 | 0.133713 |
| 1 | 37730400 | 37733999 | 17.52932 | 29.23502 | 71.27301 | 30.98721 | -0.89998 | 1.212659 | 0.133715 |
| 9 | 37436600 | 37440399 | 21.92391 | 36.10327 | 82.11953 | 35.41556 | -0.80292 | 1.379616 | 0.13372 |
| 12 | 77008200 | 77009599 | 7.151539 | 12.85894 | 39.5517 | 17.27831 | -1.20572 | 0.368384 | 0.133733 |
| 11 | 118764600 | 118778399 | 95.14636 | 143.5591 | 306.562 | 126.6941 | -0.81983 | 3.344534 | 0.133737 |
| 13 | 56251600 | 56253599 | 6.52574 | 11.87105 | 38.13355 | 16.42343 | -1.25425 | 0.290307 | 0.133737 |
| 8 | 11749600 | 11758199 | 49.53888 | 78.17748 | 165.6755 | 68.77095 | -0.75734 | 2.450727 | 0.133739 |
| 4 | 105061600 | 105064799 | 16.25404 | 27.58429 | 63.92929 | 28.00768 | -0.82658 | 1.032551 | 0.133745 |
| 17 | 70968800 | 70970199 | 12.66164 | 22.04033 | 53.17288 | 23.37349 | -0.87717 | 0.761048 | 0.133748 |
| 12 | 101048400 | 101050399 | 10.91286 | 18.78707 | 57.27645 | 25.12476 | -1.21599 | 0.874336 | 0.133749 |
| 11 | 120462400 | 120464399 | 9.019821 | 16.01885 | 43.03808 | 18.99845 | -1.02374 | 0.464329 | 0.133752 |
| 18 | 46873800 | 46875599 | 9.673402 | 17.0236 | 45.1168 | 19.9745 | -1.00992 | 0.566998 | 0.133763 |
| 7 | 100256400 | 100257999 | 10.8264 | 18.81253 | 52.57814 | 23.08648 | -1.08829 | 0.776294 | 0.133766 |
| 5 | 24678000 | 24688599 | 53.1869 | 82.90756 | 198.2437 | 82.27415 | -0.94515 | 2.691238 | 0.133767 |
| 16 | 85441000 | 85442999 | 13.11352 | 22.82796 | 52.13035 | 22.91587 | -0.79851 | 0.759383 | 0.133772 |
| 10 | 33500000 | 33500799 | 6.708349 | 12.43177 | 34.72359 | 14.59042 | -1.0284 | 0.178319 | 0.133772 |

| | | | | | | | | | |
|---|---|---|---|---|---|---|---|---|---|
| 3 | 18490400 | 18491799 | 14.53824 | 27.41488 | 18.43929 | 7.530875 | 1.041089 | 0.093167 | 0.133781 |
| 8 | 34131600 | 34133399 | 4.916177 | 9.366191 | 33.9222 | 13.55748 | -1.33944 | -0.02711 | 0.133795 |
| 2 | 102058000 | 102060999 | 14.59577 | 24.69163 | 66.48156 | 29.09783 | -1.0421 | 1.057024 | 0.1338 |
| 19 | 6079400 | 6081999 | 10.41133 | 18.38106 | 44.0573 | 19.49293 | -0.86409 | 0.533869 | 0.133804 |
| 14 | 120543600 | 120545999 | 9.727305 | 17.41227 | 41.55637 | 18.22265 | -0.84722 | 0.416983 | 0.133816 |
| 19 | 3112200 | 3116999 | 42.98117 | 71.37109 | 39.35232 | 17.05971 | 1.236831 | 1.304525 | 0.133843 |
| 1 | 100958200 | 100958999 | 10.03124 | 19.57448 | 11.8384 | 4.215408 | 1.36861 | -0.42549 | 0.133853 |
| 5 | 106072600 | 106081199 | 77.75593 | 119.9125 | 225.3271 | 93.59689 | -0.61007 | 2.895379 | 0.133854 |
| 16 | 3763800 | 3766399 | 10.90687 | 19.13648 | 46.71023 | 20.65316 | -0.89297 | 0.619364 | 0.133858 |
| 5 | 140883600 | 140903799 | 183.0511 | 265.0402 | 489.706 | 201.9512 | -0.66681 | 4.060988 | 0.133874 |
| 7 | 27186200 | 27187999 | 9.395381 | 16.89113 | 41.09081 | 17.95281 | -0.87103 | 0.385329 | 0.133878 |
| 10 | 47756000 | 47757199 | 12.45424 | 24.23978 | 12.12876 | 4.460323 | 1.598422 | -0.28259 | 0.133884 |
| 19 | 59007800 | 59008799 | 19.90012 | 35.16015 | 25.95916 | 11.48375 | 0.825453 | 0.546761 | 0.13389 |
| 7 | 136910600 | 136913199 | 16.56276 | 27.69224 | 69.55289 | 30.28716 | -0.94268 | 1.181656 | 0.133911 |
| 10 | 42070800 | 42075199 | 57.65425 | 89.62217 | 202.5791 | 84.10826 | -0.86728 | 2.742191 | 0.133914 |
| 3 | 32771000 | 32771799 | 2.84215 | 5.541836 | 23.57827 | 8.485999 | -1.42029 | -0.44026 | 0.133941 |
| 12 | 93206000 | 93206599 | 5.698561 | 10.86147 | 33.93672 | 13.55086 | -1.12634 | -0.01709 | 0.133963 |
| 4 | 38060600 | 38061599 | 12.0847 | 23.45011 | 14.70949 | 5.536006 | 1.248171 | -0.20046 | 0.133976 |
| 12 | 81497000 | 81498599 | 49.32261 | 80.11168 | 56.18389 | 24.09386 | 0.87826 | 1.61163 | 0.13398 |
| 7 | 38504400 | 38506999 | 49.442 | 79.86046 | 63.65374 | 27.16662 | 0.686395 | 1.735916 | 0.133984 |
| 2 | 29779000 | 29784399 | 23.73223 | 38.47795 | 94.63192 | 40.52844 | -0.92948 | 1.621989 | 0.133985 |
| 11 | 119190200 | 119191599 | 6.315906 | 11.6759 | 37.40502 | 15.76955 | -1.22895 | 0.191467 | 0.133986 |
| 7 | 94566800 | 94570199 | 71.98048 | 114.5671 | 91.49477 | 38.23991 | 0.659548 | 2.221848 | 0.133988 |
| 8 | 106067000 | 106068399 | 4.957027 | 9.508283 | 29.514 | 11.59595 | -1.09423 | -0.08918 | 0.133992 |
| 15 | 31926400 | 31926799 | 7.816868 | 15.28807 | 11.07796 | 3.754292 | 1.174495 | -0.6282 | 0.134007 |
| 18 | 8016800 | 8020999 | 51.69751 | 83.55964 | 57.74357 | 24.66556 | 0.892863 | 1.717547 | 0.13401 |
| 14 | 41979600 | 41980999 | 22.08513 | 38.66619 | 28.5915 | 12.61901 | 0.820732 | 0.681315 | 0.134021 |
| 4 | 76945200 | 76945799 | 9.758618 | 19.00704 | 14.31342 | 5.226681 | 1.025839 | -0.33371 | 0.134042 |
| 13 | 29494800 | 29495599 | 9.758618 | 19.00704 | 14.31342 | 5.226681 | 1.025839 | -0.33371 | 0.134042 |
| 17 | 84224400 | 84226199 | 7.316737 | 12.90348 | 48.25272 | 21.35385 | -1.5018 | 0.533472 | 0.134042 |
| 9 | 72569000 | 72572199 | 10.60297 | 18.30729 | 57.32321 | 25.14752 | -1.25345 | 0.844844 | 0.134043 |
| 14 | 120728800 | 120729799 | 6.35645 | 11.98375 | 32.42073 | 13.24607 | -0.95064 | 0.058586 | 0.134055 |
| 12 | 87344000 | 87349199 | 23.21107 | 37.50594 | 108.7932 | 46.4549 | -1.17201 | 1.702983 | 0.134074 |
| 12 | 84312800 | 84315399 | 11.03503 | 19.05107 | 55.71732 | 24.44294 | -1.15544 | 0.847032 | 0.134077 |
| 5 | 33814600 | 33816399 | 11.83665 | 20.36156 | 57.87397 | 25.38626 | -1.11592 | 0.885049 | 0.134081 |
| 12 | 105972000 | 105977199 | 32.51291 | 52.2275 | 114.953 | 48.68796 | -0.78534 | 1.901053 | 0.134084 |
| 7 | 31597200 | 31602599 | 60.99188 | 98.26 | 62.14911 | 26.43408 | 1.014952 | 1.829041 | 0.134088 |
| 2 | 67244400 | 67244999 | 4.232591 | 8.244999 | 26.77885 | 9.761459 | -1.0572 | -0.36885 | 0.134102 |
| 12 | 79473400 | 79474599 | 13.05534 | 24.64108 | 16.80054 | 6.848785 | 1.024203 | 0.069065 | 0.134134 |
| 7 | 60049200 | 60049999 | 12.94894 | 25.08587 | 13.71002 | 5.199647 | 1.432702 | -0.17021 | 0.134135 |
| 17 | 54025200 | 54027199 | 14.62448 | 27.25143 | 20.13532 | 8.390871 | 0.88403 | 0.172472 | 0.13414 |
| 17 | 26203600 | 26205599 | 8.53767 | 15.234 | 41.01905 | 18.03558 | -1.02246 | 0.428652 | 0.134148 |
| 15 | 46413400 | 46417399 | 65.85945 | 105.5777 | 72.92873 | 30.79279 | 0.879639 | 1.990122 | 0.134149 |
| X | 5688400 | 5689999 | 18.91532 | 35.38484 | 16.15371 | 6.687304 | 1.580891 | 0.176822 | 0.13415 |
| 10 | 48386800 | 48388399 | 17.47039 | 32.09219 | 20.8204 | 8.853824 | 1.049196 | 0.277422 | 0.134164 |
| X | 20939400 | 20940399 | 21.46559 | 38.04141 | 23.47504 | 10.37853 | 1.084583 | 0.521798 | 0.134165 |
| 12 | 115751000 | 115753199 | 24.98806 | 44.75732 | 18.4694 | 8.094692 | 1.668979 | 0.469167 | 0.134171 |

| | | | | | | | | | |
|---|---|---|---|---|---|---|---|---|---|
| 17 | 18595400 | 18596199 | 18.73953 | 33.82235 | 22.44141 | 9.760611 | 0.996097 | 0.389143 | 0.134178 |
| 12 | 35903000 | 35905999 | 23.39194 | 38.40172 | 83.15123 | 35.81358 | -0.73393 | 1.423316 | 0.134191 |
| 15 | 23296400 | 23297999 | 15.75382 | 26.84335 | 61.2074 | 26.83032 | -0.80231 | 0.991835 | 0.134206 |
| 4 | 99536600 | 99543199 | 58.79048 | 92.26697 | 182.173 | 75.55375 | -0.66164 | 2.57034 | 0.13424 |
| 11 | 6230600 | 6231799 | 27.21525 | 46.65869 | 35.63841 | 15.62987 | 0.770437 | 0.948836 | 0.13425 |
| 6 | 93832800 | 93833999 | 6.224309 | 11.70892 | 32.91201 | 13.49683 | -1.00975 | 0.072366 | 0.134256 |
| 5 | 26340000 | 26341599 | 22.66149 | 40.6141 | 20.71727 | 9.075352 | 1.36589 | 0.453521 | 0.134259 |
| 8 | 70559800 | 70575999 | 98.51162 | 148.4462 | 310.4879 | 128.2684 | -0.79154 | 3.371027 | 0.134268 |
| 2 | 165448000 | 165469999 | 127.3499 | 188.6252 | 389.1662 | 160.2715 | -0.79446 | 3.697407 | 0.134285 |
| 5 | 128847400 | 128868799 | 196.6031 | 282.3626 | 531.9601 | 219.6245 | -0.70646 | 4.181962 | 0.134285 |
| 1 | 164617200 | 164619399 | 14.38891 | 24.92947 | 59.28772 | 26.01206 | -0.85822 | 0.831917 | 0.134304 |
| 7 | 31923400 | 31925399 | 29.26645 | 49.98958 | 31.11766 | 13.64313 | 1.063586 | 0.984652 | 0.134315 |
| 15 | 81196800 | 81198199 | 6.689447 | 12.48723 | 33.818 | 14.05068 | -0.97195 | 0.129147 | 0.134324 |
| 7 | 128443800 | 128445799 | 9.100438 | 16.34464 | 40.02405 | 17.50138 | -0.88104 | 0.388492 | 0.13433 |
| 2 | 38589000 | 38605399 | 113.7071 | 169.39 | 364.6456 | 150.26 | -0.84801 | 3.593686 | 0.134331 |
| 18 | 4430600 | 4433399 | 53.63938 | 86.68053 | 56.06716 | 23.94313 | 0.987731 | 1.727565 | 0.13436 |
| 7 | 11303200 | 11311199 | 202.4652 | 301.9861 | 227.3954 | 93.7204 | 0.668061 | 3.579445 | 0.134363 |
| 5 | 121986200 | 121996399 | 58.8114 | 90.93388 | 218.1862 | 90.62946 | -0.95928 | 2.840145 | 0.134364 |
| 10 | 22727800 | 22730599 | 13.58026 | 23.57398 | 58.19686 | 25.53738 | -0.91127 | 0.807331 | 0.134367 |
| X | 29560400 | 29560999 | 14.38208 | 26.93662 | 19.23816 | 7.948134 | 0.942734 | 0.137146 | 0.134374 |
| 6 | 136436800 | 136448599 | 87.31926 | 132.0669 | 302.7678 | 125.2211 | -0.91916 | 3.293603 | 0.134388 |
| 2 | 73111000 | 73111999 | 7.702404 | 13.98537 | 38.10618 | 16.45005 | -1.02137 | 0.309445 | 0.134391 |
| 16 | 17348000 | 17349399 | 7.702404 | 13.98537 | 38.10618 | 16.45005 | -1.02137 | 0.309445 | 0.134391 |
| 7 | 32168400 | 32169999 | 32.06561 | 54.08974 | 36.98928 | 16.17547 | 0.927448 | 1.112927 | 0.134394 |
| 1 | 93405600 | 93407599 | 10.61017 | 18.58212 | 47.79037 | 21.09834 | -0.96774 | 0.650968 | 0.134403 |
| 7 | 93449200 | 93450999 | 19.77027 | 35.643 | 21.38866 | 9.315684 | 1.138205 | 0.398478 | 0.13443 |
| 1 | 60448200 | 60449399 | 6.207223 | 11.34916 | 37.15103 | 15.8989 | -1.27489 | 0.256045 | 0.134433 |
| 17 | 71923000 | 71923999 | 6.207223 | 11.34916 | 37.15103 | 15.8989 | -1.27489 | 0.256045 | 0.134433 |
| 8 | 121499600 | 121512399 | 70.5766 | 108.2139 | 249.7458 | 103.6708 | -0.91266 | 3.007156 | 0.134437 |
| 6 | 135000000 | 135001799 | 7.124923 | 13.31425 | 34.41265 | 14.2721 | -0.90306 | 0.128358 | 0.134442 |
| 3 | 112868800 | 112870799 | 20.18251 | 36.06213 | 23.74568 | 10.4285 | 0.997807 | 0.461052 | 0.13445 |
| 4 | 126643600 | 126648199 | 18.72067 | 30.58144 | 87.2644 | 37.51105 | -1.13431 | 1.483653 | 0.134451 |
| 13 | 111686000 | 111692199 | 44.82323 | 71.19328 | 151.0457 | 62.97639 | -0.75085 | 2.276627 | 0.134452 |
| 15 | 10466000 | 10466399 | 2.027254 | 3.965452 | 18.93562 | 6.335276 | -1.47887 | -0.727 | 0.134458 |
| 14 | 21434800 | 21436799 | 24.35219 | 42.43181 | 28.9849 | 12.74889 | 0.934175 | 0.762496 | 0.134484 |
| 3 | 112585800 | 112586999 | 12.39329 | 23.7119 | 17.13247 | 6.780468 | 0.977922 | -0.031 | 0.134493 |
| 12 | 40933200 | 40936799 | 23.73827 | 38.89053 | 84.09722 | 36.18964 | -0.73319 | 1.450973 | 0.134497 |
| 6 | 38329400 | 38332999 | 23.06449 | 37.88801 | 83.20456 | 35.84609 | -0.75386 | 1.414264 | 0.134522 |
| 16 | 89690200 | 89695399 | 49.08169 | 77.38112 | 167.4201 | 69.47631 | -0.78826 | 2.472309 | 0.134532 |
| 11 | 107821600 | 107828399 | 50.26432 | 79.54642 | 158.4605 | 65.87001 | -0.66493 | 2.382423 | 0.13454 |
| 12 | 20140600 | 20149799 | 268.6733 | 390.1961 | 290.9891 | 119.9581 | 0.645804 | 4.018199 | 0.134544 |
| 7 | 30427200 | 30433199 | 29.46969 | 47.35985 | 113.5246 | 48.11367 | -0.90727 | 1.883227 | 0.134545 |
| 18 | 51617200 | 51617999 | 9.294167 | 16.56364 | 41.5081 | 18.26936 | -0.92086 | 0.441701 | 0.134549 |
| 5 | 122900000 | 122920999 | 108.7898 | 162.0584 | 384.0711 | 158.264 | -0.98668 | 3.594536 | 0.13456 |
| 10 | 80006000 | 80009599 | 17.08895 | 28.38354 | 73.69691 | 31.95395 | -0.99066 | 1.261899 | 0.134568 |
| 16 | 85349800 | 85351199 | 8.360928 | 14.97855 | 40.02003 | 17.53696 | -1.00774 | 0.400569 | 0.134601 |
| 2 | 9863000 | 9864199 | 5.293197 | 10.1146 | 29.97708 | 11.90541 | -1.04388 | -0.05123 | 0.134609 |

| | | | | | | | | | |
|---|---|---|---|---|---|---|---|---|---|
| 10 | 117938800 | 117939599 | 5.293197 | 10.1146 | 29.97708 | 11.90541 | -1.04388 | -0.05123 | 0.134609 |
| 7 | 105400800 | 105408399 | 32.69038 | 52.10881 | 138.2294 | 57.89153 | -1.06685 | 2.151956 | 0.134613 |
| 1 | 88557600 | 88560199 | 16.08217 | 27.34755 | 62.28716 | 27.2992 | -0.80122 | 1.012675 | 0.134618 |
| 14 | 3181600 | 3187799 | 111.4382 | 172.3378 | 140.0892 | 58.18971 | 0.600714 | 2.847506 | 0.134693 |
| 18 | 80171000 | 80174199 | 15.81998 | 26.76825 | 64.36115 | 28.17155 | -0.87951 | 1.057961 | 0.134703 |
| 2 | 18703400 | 18704799 | 10.58084 | 18.62411 | 45.10296 | 19.96761 | -0.88073 | 0.569751 | 0.134716 |
| 2 | 158360400 | 158367999 | 42.16513 | 66.80775 | 158.0796 | 65.75093 | -0.91138 | 2.348367 | 0.134718 |
| 3 | 45330000 | 45337199 | 50.87878 | 79.81662 | 190.3748 | 78.95477 | -0.93436 | 2.574651 | 0.134719 |
| X | 31602600 | 31602999 | 10.13655 | 19.82078 | 9.701376 | 3.311329 | 1.723698 | -0.57108 | 0.134745 |
| 5 | 149032600 | 149033799 | 4.596871 | 8.731069 | 30.8461 | 12.40821 | -1.31015 | -0.01194 | 0.134747 |
| 5 | 122166400 | 122184799 | 140.8794 | 207.2212 | 420.1682 | 173.0303 | -0.77884 | 3.814266 | 0.134749 |
| 2 | 173398800 | 173400599 | 21.02581 | 34.96099 | 77.96637 | 33.83091 | -0.77234 | 1.260605 | 0.134767 |
| 7 | 83677600 | 83679399 | 7.101761 | 12.90874 | 37.59911 | 16.20918 | -1.11512 | 0.298722 | 0.134775 |
| 17 | 47445200 | 47453399 | 47.06969 | 74.3908 | 163.7135 | 68.00741 | -0.80996 | 2.412995 | 0.134776 |
| 10 | 77803800 | 77811599 | 54.23567 | 84.76784 | 191.8027 | 79.57733 | -0.86261 | 2.643273 | 0.134784 |
| 11 | 58844600 | 58846199 | 7.515472 | 13.74339 | 36.6353 | 15.67497 | -0.98133 | 0.263766 | 0.13481 |
| 4 | 106478600 | 106489399 | 49.54326 | 77.47166 | 196.2409 | 81.41459 | -1.02465 | 2.63253 | 0.134819 |
| 6 | 129510000 | 129510799 | 5.355675 | 10.04293 | 32.76594 | 13.50939 | -1.22788 | 0.087389 | 0.134828 |
| 8 | 105868400 | 105870199 | 9.337902 | 16.51939 | 43.03218 | 19.03166 | -0.98202 | 0.502848 | 0.134829 |
| 10 | 81714800 | 81715999 | 15.58328 | 30.34734 | 8.430178 | 3.084359 | 2.439769 | -0.2719 | 0.134839 |
| 12 | 73445000 | 73448199 | 18.19612 | 30.70288 | 68.42846 | 29.92649 | -0.77092 | 1.094453 | 0.134863 |
| 1 | 34552400 | 34556799 | 31.15318 | 50.24791 | 105.734 | 45.04292 | -0.71333 | 1.768374 | 0.134865 |
| 13 | 3533400 | 3536599 | 69.82386 | 111.3561 | 86.19599 | 36.12471 | 0.707577 | 2.15684 | 0.134882 |
| 1 | 112517400 | 112529399 | 379.958 | 535.4313 | 373.8652 | 154.8352 | 0.69742 | 4.465815 | 0.134896 |
| 13 | 29162800 | 29163799 | 15.66145 | 29.56206 | 17.16934 | 6.998065 | 1.253263 | 0.093747 | 0.134901 |
| 14 | 84683400 | 84685999 | 25.80661 | 41.51026 | 113.5939 | 48.20571 | -1.09615 | 1.849642 | 0.13491 |
| 11 | 94249200 | 94252799 | 17.11352 | 28.41165 | 74.31659 | 32.21268 | -1.00142 | 1.266408 | 0.134932 |
| 11 | 118931200 | 118950399 | 118.6344 | 175.4642 | 412.9954 | 170.0825 | -0.98637 | 3.720455 | 0.134933 |
| 8 | 108394200 | 108395199 | 4.660257 | 9.047219 | 26.78858 | 10.05618 | -0.96446 | -0.257 | 0.134938 |
| 13 | 14127800 | 14129999 | 13.92056 | 23.68915 | 63.25089 | 27.72322 | -1.02927 | 0.998525 | 0.134949 |
| 5 | 46687200 | 46687799 | 16.7704 | 30.3714 | 22.10036 | 9.571493 | 0.867473 | 0.353825 | 0.134963 |
| 18 | 36119000 | 36123999 | 21.19821 | 34.41714 | 94.44342 | 40.48906 | -1.08506 | 1.58843 | 0.134975 |
| 13 | 52933400 | 52935399 | 11.35864 | 19.94806 | 46.68743 | 20.65965 | -0.83279 | 0.60592 | 0.134981 |
| 12 | 67837400 | 67838799 | 14.91872 | 27.60963 | 19.47581 | 8.198596 | 0.939189 | 0.211332 | 0.134986 |
| 3 | 69970600 | 69973199 | 27.34937 | 47.46929 | 31.06828 | 13.63192 | 0.995419 | 0.835473 | 0.134991 |
| 2 | 84493200 | 84493799 | 8.273746 | 16.14984 | 12.92752 | 4.579398 | 0.978015 | -0.46298 | 0.135 |
| 14 | 119954200 | 119958999 | 27.56273 | 44.65341 | 96.70083 | 41.39052 | -0.74771 | 1.6447 | 0.135002 |
| 3 | 37727000 | 37729999 | 14.74549 | 24.94691 | 65.37417 | 28.61384 | -1.00314 | 1.056161 | 0.135012 |
| 3 | 118227000 | 118228999 | 14.91644 | 25.59258 | 58.56347 | 25.68563 | -0.80576 | 0.92183 | 0.135013 |
| 8 | 5296800 | 5298199 | 12.65077 | 24.5829 | 13.3358 | 4.968047 | 1.467509 | -0.23294 | 0.13502 |
| 3 | 15983000 | 15984799 | 41.72784 | 68.28198 | 52.66992 | 22.65457 | 0.748961 | 1.484858 | 0.13504 |
| 8 | 43681200 | 43682199 | 12.87197 | 24.4952 | 18.27258 | 7.320681 | 0.917346 | 0.016811 | 0.135044 |
| 1 | 135690800 | 135696999 | 24.74258 | 40.02279 | 100.4001 | 42.92507 | -0.96128 | 1.673078 | 0.135079 |
| 13 | 18923800 | 18925399 | 25.71066 | 44.46103 | 33.4486 | 14.67449 | 0.794454 | 0.863343 | 0.135096 |
| 8 | 80838400 | 80839599 | 7.113875 | 13.32471 | 34.51452 | 14.2578 | -0.90192 | 0.111547 | 0.135102 |
| 19 | 22010800 | 22011999 | 17.01711 | 30.74024 | 22.45249 | 9.754849 | 0.859038 | 0.36991 | 0.135103 |
| 5 | 36905200 | 36907399 | 10.39987 | 18.39723 | 43.52822 | 19.25122 | -0.84437 | 0.510637 | 0.135121 |

| | | | | | | | | | |
|---|---|---|---|---|---|---|---|---|---|
| 6 | 19671000 | 19671399 | 9.703009 | 18.86588 | 12.9357 | 4.796237 | 1.140403 | -0.28702 | 0.135141 |
| 19 | 57498600 | 57501999 | 16.66428 | 28.01173 | 67.0244 | 29.27874 | -0.87303 | 1.12168 | 0.135191 |
| 9 | 42830400 | 42835799 | 23.25342 | 37.73552 | 94.246 | 40.38989 | -0.95042 | 1.601848 | 0.135198 |
| 1 | 89918800 | 89920999 | 17.51012 | 29.23732 | 70.1326 | 30.51401 | -0.87661 | 1.200131 | 0.135198 |
| 19 | 7227800 | 7232999 | 31.13991 | 50.26189 | 106.7453 | 45.51508 | -0.72548 | 1.747804 | 0.135201 |
| 17 | 26806600 | 26808199 | 7.005195 | 13.00887 | 34.78985 | 14.57702 | -0.96388 | 0.166668 | 0.135207 |
| X | 144064200 | 144065399 | 12.76138 | 24.72572 | 13.71552 | 5.198579 | 1.412195 | -0.17444 | 0.135211 |
| 4 | 39129800 | 39133199 | 36.04128 | 60.0727 | 43.8078 | 19.03899 | 0.835121 | 1.246684 | 0.135251 |
| 1 | 22674800 | 22677999 | 19.99762 | 35.88856 | 22.47881 | 9.835748 | 1.07267 | 0.429386 | 0.135263 |
| 11 | 116169400 | 116182399 | 73.08691 | 111.3115 | 282.4985 | 117.0856 | -1.05731 | 3.147764 | 0.135272 |
| 12 | 20199800 | 20203599 | 95.48738 | 150.0002 | 113.708 | 47.16456 | 0.718697 | 2.563296 | 0.135291 |
| 12 | 20200000 | 20203599 | 95.48738 | 150.0002 | 113.708 | 47.16456 | 0.718697 | 2.563296 | 0.135291 |
| 7 | 101216800 | 101220199 | 32.31179 | 52.04563 | 108.1428 | 45.98425 | -0.6975 | 1.813745 | 0.135298 |
| 15 | 98039600 | 98043799 | 11.89039 | 20.67436 | 53.17665 | 23.35849 | -0.96943 | 0.770845 | 0.135316 |
| 7 | 137012200 | 137013799 | 9.68967 | 17.03289 | 46.677 | 20.66993 | -1.05879 | 0.581107 | 0.135319 |
| 15 | 93124800 | 93127399 | 15.08623 | 25.82995 | 60.19186 | 26.3954 | -0.83246 | 0.942999 | 0.135323 |
| 16 | 92226400 | 92230799 | 37.41539 | 59.8614 | 124.2256 | 52.29816 | -0.70756 | 2.036441 | 0.135324 |
| 16 | 25462400 | 25464199 | 25.70781 | 45.23271 | 23.90257 | 10.58276 | 1.303102 | 0.623749 | 0.135334 |
| 3 | 84165800 | 84167799 | 7.109775 | 12.94622 | 38.68765 | 16.64155 | -1.15023 | 0.288615 | 0.135353 |
| 17 | 18335200 | 18343799 | 175.8188 | 265.3611 | 191.283 | 79.05995 | 0.746132 | 3.348846 | 0.135368 |
| 7 | 60914600 | 60917199 | 69.97442 | 111.4296 | 90.14399 | 37.70132 | 0.641731 | 2.203574 | 0.135401 |
| 7 | 8266800 | 8267599 | 15.67917 | 29.52153 | 17.72435 | 7.257499 | 1.200545 | 0.111117 | 0.135402 |
| 9 | 9870400 | 9871599 | 27.08279 | 46.86784 | 32.79697 | 14.38893 | 0.898644 | 0.862859 | 0.135405 |
| 16 | 8696800 | 8707999 | 83.97752 | 128.3327 | 253.9138 | 105.326 | -0.69459 | 3.078988 | 0.135411 |
| X | 111657600 | 111660199 | 32.0873 | 54.80027 | 34.45747 | 15.10727 | 1.049287 | 1.003477 | 0.135414 |
| X | 8844200 | 8845799 | 21.28604 | 38.1329 | 22.17467 | 9.717231 | 1.178182 | 0.450407 | 0.135416 |
| 8 | 114107600 | 114109599 | 12.32518 | 21.47859 | 52.66416 | 23.1661 | -0.9005 | 0.73927 | 0.135424 |
| 10 | 78055800 | 78057399 | 6.670435 | 12.50867 | 33.39941 | 13.77 | -0.94272 | 0.09798 | 0.135431 |
| 2 | 33935600 | 33951199 | 127.8204 | 190.2763 | 359.6242 | 148.1798 | -0.66137 | 3.608565 | 0.135441 |
| 8 | 48428000 | 48436199 | 50.28486 | 79.50592 | 162.3628 | 67.46081 | -0.70178 | 2.404657 | 0.135446 |
| 16 | 3263000 | 3268199 | 44.11378 | 72.98169 | 42.22292 | 18.25752 | 1.167515 | 1.352189 | 0.135452 |
| 15 | 37573000 | 37573799 | 6.489956 | 12.25376 | 33.02468 | 13.45237 | -0.94115 | 0.05269 | 0.135472 |
| 14 | 104702400 | 104713799 | 85.07767 | 129.8642 | 264.8467 | 109.8318 | -0.73959 | 3.102994 | 0.135486 |
| 7 | 8953600 | 8954399 | 11.58813 | 22.61543 | 10.49595 | 3.729086 | 1.747927 | -0.41493 | 0.135513 |
| 16 | 30662600 | 30664199 | 8.683278 | 15.51572 | 41.54355 | 18.24444 | -1.01347 | 0.420362 | 0.135518 |
| 13 | 79168600 | 79169599 | 17.2717 | 31.17776 | 22.99826 | 10.00094 | 0.844204 | 0.377722 | 0.13552 |
| X | 127557800 | 127558999 | 14.32098 | 27.54851 | 15.46518 | 6.026426 | 1.35989 | -0.06281 | 0.135526 |
| 15 | 78887600 | 78890999 | 13.04882 | 22.08855 | 64.63689 | 28.29585 | -1.16128 | 1.046004 | 0.135533 |
| 12 | 104533000 | 104538999 | 34.92175 | 56.08225 | 116.1895 | 49.19222 | -0.69875 | 1.912691 | 0.135537 |
| 5 | 139373200 | 139379199 | 24.10249 | 38.95034 | 100.2125 | 42.79658 | -0.9991 | 1.699935 | 0.13554 |
| 9 | 25377400 | 25379399 | 23.472 | 41.08457 | 28.66158 | 12.64741 | 0.904558 | 0.694888 | 0.135546 |
| 11 | 24907600 | 24909799 | 20.95462 | 36.88175 | 27.04765 | 11.97444 | 0.833211 | 0.596944 | 0.135546 |
| X | 3716400 | 3717999 | 19.79276 | 35.67314 | 21.86664 | 9.527337 | 1.108112 | 0.403665 | 0.135561 |
| 14 | 122718600 | 122722999 | 32.71572 | 52.45264 | 119.9358 | 50.65072 | -0.84356 | 1.963389 | 0.135564 |
| 7 | 83280600 | 83281399 | 3.713791 | 7.222395 | 26.94265 | 9.972372 | -1.2744 | -0.32004 | 0.135569 |
| 5 | 94894600 | 94896399 | 31.28569 | 52.96809 | 35.2065 | 15.413 | 0.968236 | 1.074314 | 0.135583 |
| 10 | 110526000 | 110527599 | 20.01821 | 35.36669 | 25.8803 | 11.449 | 0.838116 | 0.547538 | 0.135585 |

| | | | | | | | | | |
|---|---|---|---|---|---|---|---|---|---|
| 14 | 61410000 | 61410999 | 8.182669 | 14.75048 | 39.03936 | 17.0058 | -0.98837 | 0.357764 | 0.135601 |
| 9 | 107070000 | 107072799 | 16.44183 | 27.89878 | 68.86046 | 30.16666 | -0.91702 | 1.042566 | 0.135602 |
| 6 | 100518800 | 100520599 | 6.085787 | 11.10179 | 38.17188 | 16.38243 | -1.34805 | 0.270216 | 0.135614 |
| 15 | 99757800 | 99764599 | 38.73032 | 61.89071 | 127.5142 | 53.59125 | -0.69912 | 2.075983 | 0.135629 |
| 2 | 70993000 | 70994199 | 9.815988 | 17.30459 | 45.10654 | 19.96302 | -0.9853 | 0.547212 | 0.135631 |
| 3 | 152723600 | 152726999 | 18.06143 | 29.99571 | 81.04188 | 35.13615 | -1.0483 | 1.266997 | 0.135637 |
| 12 | 18289600 | 18291999 | 69.64135 | 111.0073 | 87.72359 | 36.73853 | 0.676969 | 2.173314 | 0.135638 |
| 18 | 38383000 | 38384199 | 5.573273 | 10.68337 | 30.17988 | 11.87978 | -0.96384 | -0.07871 | 0.135642 |
| 12 | 112619600 | 112626199 | 27.26418 | 43.79714 | 116.9371 | 49.52997 | -1.06318 | 1.894708 | 0.135644 |
| 19 | 46593000 | 46597199 | 21.32428 | 35.04482 | 82.94373 | 35.7404 | -0.86121 | 1.404308 | 0.135646 |
| 17 | 14249200 | 14250599 | 8.308778 | 15.03416 | 38.57581 | 16.72514 | -0.93934 | 0.336986 | 0.135646 |
| 1 | 156137000 | 156138999 | 11.40106 | 19.90487 | 52.09594 | 22.94275 | -0.99373 | 0.707306 | 0.135646 |
| 3 | 135309400 | 135310999 | 6.832218 | 12.77084 | 35.56705 | 14.7439 | -1.00992 | 0.123817 | 0.135653 |
| 9 | 122779600 | 122780599 | 5.031349 | 9.615272 | 31.12546 | 12.3587 | -1.17006 | -0.05334 | 0.13566 |
| 11 | 120355800 | 120358799 | 11.57127 | 20.15195 | 51.0175 | 22.43278 | -0.94615 | 0.74451 | 0.135665 |
| 4 | 125917000 | 125918799 | 9.655283 | 17.05304 | 45.63058 | 20.18742 | -1.0219 | 0.525492 | 0.135674 |
| 11 | 68582000 | 68584199 | 12.5202 | 21.71685 | 53.16702 | 23.32747 | -0.89896 | 0.807016 | 0.135674 |
| 5 | 122229000 | 122234999 | 43.37175 | 68.90519 | 147.1739 | 61.37753 | -0.7601 | 2.267788 | 0.135678 |
| 2 | 48577600 | 48579599 | 15.64443 | 29.60113 | 16.61328 | 6.737019 | 1.308399 | 0.07615 | 0.135708 |
| 18 | 32509800 | 32513799 | 21.31167 | 34.68876 | 90.87594 | 39.0122 | -1.01537 | 1.542791 | 0.13572 |
| 6 | 90438000 | 90456399 | 89.58613 | 133.9866 | 368.6424 | 152.0155 | -1.19647 | 3.5165 | 0.135727 |
| 4 | 153783600 | 153827199 | 350.8103 | 478.4989 | 895.3946 | 372.7311 | -0.77178 | 4.963205 | 0.135727 |
| 17 | 83831200 | 83838399 | 31.44136 | 50.42114 | 123.0133 | 51.96863 | -0.93651 | 1.954811 | 0.135736 |
| 12 | 55489600 | 55494799 | 30.89211 | 49.68254 | 113.4187 | 48.12039 | -0.83576 | 1.860459 | 0.135751 |
| 11 | 69706600 | 69709799 | 44.55889 | 70.65154 | 151.6428 | 63.11634 | -0.76993 | 2.328868 | 0.135755 |
| 6 | 32749600 | 32751399 | 7.291574 | 13.55914 | 35.95801 | 15.03713 | -0.95017 | 0.162525 | 0.135759 |
| 17 | 22178800 | 22181999 | 15.06159 | 25.94343 | 58.62121 | 25.71507 | -0.78644 | 0.88783 | 0.13576 |
| 13 | 52113800 | 52134999 | 123.3545 | 182.7505 | 396.5077 | 163.2977 | -0.86671 | 3.692821 | 0.135763 |
| 3 | 6575400 | 6576999 | 20.89807 | 37.00344 | 25.16577 | 11.12795 | 0.944349 | 0.528466 | 0.135804 |
| 8 | 15707400 | 15710999 | 26.05356 | 45.33676 | 30.54574 | 13.42434 | 0.954037 | 0.793151 | 0.135806 |
| 18 | 33516200 | 33519199 | 20.13603 | 33.29803 | 83.84275 | 36.24303 | -0.94755 | 1.326934 | 0.135808 |
| 5 | 120825800 | 120852799 | 142.797 | 207.8361 | 478.2468 | 197.1091 | -0.97588 | 3.97932 | 0.135819 |
| 7 | 46444800 | 46445799 | 4.366013 | 8.477347 | 26.14995 | 9.803724 | -1.02034 | -0.26583 | 0.135822 |
| 5 | 114951000 | 114961199 | 101.8042 | 154.2463 | 285.6108 | 118.1908 | -0.60915 | 3.261028 | 0.135823 |
| 3 | 31209800 | 31213399 | 16.73737 | 27.96127 | 71.28639 | 31.02598 | -0.96428 | 1.191307 | 0.135827 |
| 7 | 14451400 | 14457799 | 110.0102 | 171.4211 | 124.9541 | 51.86068 | 0.766167 | 2.714693 | 0.13585 |
| 16 | 20925600 | 20926799 | 22.27286 | 38.96558 | 29.14991 | 12.8573 | 0.804121 | 0.695372 | 0.135857 |
| 3 | 85528000 | 85532599 | 44.74638 | 72.84446 | 56.07809 | 24.07517 | 0.746616 | 1.564475 | 0.135861 |
| 7 | 7936800 | 7941599 | 46.60903 | 77.16731 | 38.26559 | 16.55486 | 1.389108 | 1.347874 | 0.135867 |
| 1 | 55170200 | 55170999 | 6.906512 | 12.70247 | 36.20441 | 15.37018 | -1.06885 | 0.224369 | 0.135869 |
| 7 | 40810400 | 40814999 | 82.08743 | 130.6397 | 83.20614 | 34.77055 | 0.985213 | 2.233031 | 0.135876 |
| 2 | 163130600 | 163136799 | 40.36401 | 64.43872 | 130.5033 | 54.77211 | -0.67597 | 2.108394 | 0.135877 |
| 13 | 72775000 | 72781599 | 51.99969 | 81.56377 | 182.072 | 75.51239 | -0.83905 | 2.577483 | 0.135902 |
| X | 153226600 | 153228199 | 38.34413 | 64.14862 | 39.89904 | 17.38088 | 1.063727 | 1.213549 | 0.13592 |
| 16 | 51465200 | 51465999 | 13.43737 | 25.28476 | 17.32391 | 7.101039 | 1.010967 | 0.09391 | 0.135955 |
| 5 | 84965800 | 84968199 | 27.99295 | 48.17066 | 34.49876 | 15.13295 | 0.863913 | 0.919098 | 0.135965 |
| 4 | 129589200 | 129590999 | 7.412949 | 13.43803 | 38.08359 | 16.47361 | -1.07935 | 0.317566 | 0.135978 |

| | | | | | | | | | |
|---|---|---|---|---|---|---|---|---|---|
| 6 | 72704800 | 72707799 | 15.24473 | 25.69561 | 66.41268 | 29.03556 | -0.98357 | 1.092178 | 0.135983 |
| 7 | 18716800 | 18722799 | 34.8112 | 55.77429 | 120.71 | 50.92488 | -0.76563 | 1.98775 | 0.135987 |
| 17 | 25862200 | 25901799 | 259.5496 | 359.1928 | 787.8615 | 327.2171 | -0.97971 | 4.736245 | 0.135988 |
| 3 | 126955600 | 126956999 | 7.535865 | 13.74407 | 38.19074 | 16.39495 | -1.04435 | 0.281228 | 0.136024 |
| 5 | 112956600 | 112964199 | 54.05982 | 84.90949 | 176.0867 | 73.03703 | -0.73147 | 2.551953 | 0.136031 |
| 12 | 24880600 | 24882399 | 9.715353 | 17.02364 | 48.30276 | 21.33372 | -1.10876 | 0.636448 | 0.136035 |
| 12 | 110039800 | 110044999 | 45.94879 | 73.01523 | 146.509 | 61.11354 | -0.66986 | 2.263329 | 0.136045 |
| 19 | 39496600 | 39497399 | 19.15001 | 33.96881 | 24.57109 | 10.8575 | 0.857161 | 0.500164 | 0.13605 |
| 15 | 68050200 | 68053999 | 23.222 | 37.75787 | 91.76886 | 39.3802 | -0.90871 | 1.562171 | 0.136061 |
| 4 | 141198200 | 141208199 | 123.2673 | 181.8264 | 407.0144 | 167.6145 | -0.91762 | 3.766604 | 0.136061 |
| 13 | 19121000 | 19124399 | 58.80979 | 95.40329 | 47.44852 | 20.32928 | 1.370971 | 1.650323 | 0.13608 |
| 9 | 14523600 | 14529199 | 27.25873 | 44.12251 | 96.9285 | 41.46137 | -0.76945 | 1.662449 | 0.136091 |
| 11 | 114807800 | 114810799 | 35.98846 | 59.70175 | 46.61206 | 20.1934 | 0.736815 | 1.302194 | 0.136096 |
| 14 | 82733000 | 82735599 | 30.76295 | 51.80175 | 40.05482 | 17.50676 | 0.750718 | 1.12434 | 0.136102 |
| 17 | 24963600 | 24967199 | 13.63152 | 23.52703 | 55.86942 | 24.50945 | -0.85705 | 0.859268 | 0.136106 |
| 3 | 33480600 | 33483199 | 57.47652 | 92.26611 | 74.30408 | 31.43923 | 0.661022 | 1.937939 | 0.136106 |
| 7 | 33571200 | 33573599 | 56.5187 | 91.0119 | 67.97959 | 28.89481 | 0.774809 | 1.838368 | 0.136121 |
| X | 8852600 | 8854599 | 19.84807 | 35.93994 | 20.44278 | 8.855196 | 1.220678 | 0.375391 | 0.136124 |
| 15 | 29934200 | 29935799 | 22.53704 | 39.35462 | 29.23357 | 12.87587 | 0.814229 | 0.724591 | 0.136153 |
| 15 | 12565400 | 12568199 | 12.18397 | 21.2054 | 53.74159 | 23.62107 | -0.94818 | 0.758221 | 0.136171 |
| 17 | 32304000 | 32315799 | 94.67392 | 143.7746 | 275.7351 | 114.1841 | -0.65699 | 3.211195 | 0.136184 |
| 7 | 60385800 | 60387399 | 20.36885 | 36.25715 | 24.09077 | 10.61398 | 0.981557 | 0.484887 | 0.136187 |
| 8 | 74999600 | 75001999 | 17.8853 | 32.18553 | 23.34639 | 10.19173 | 0.864624 | 0.399887 | 0.136191 |
| 12 | 23536000 | 23539399 | 36.62648 | 60.582 | 47.7303 | 20.63596 | 0.723485 | 1.337761 | 0.136207 |
| 1 | 182868200 | 182871999 | 58.32075 | 90.18198 | 218.0357 | 90.56692 | -0.97016 | 2.838545 | 0.136216 |
| 15 | 29159200 | 29159999 | 4.185711 | 8.157672 | 25.57004 | 9.261216 | -0.99688 | -0.39328 | 0.136257 |
| 10 | 123494600 | 123496599 | 21.44683 | 37.69663 | 27.51951 | 12.18631 | 0.839049 | 0.615422 | 0.136257 |
| 17 | 38745000 | 38747799 | 64.00816 | 102.2921 | 80.58215 | 33.87041 | 0.685484 | 2.085814 | 0.136262 |
| 18 | 34992800 | 34999599 | 40.16102 | 63.79246 | 148.0866 | 61.74671 | -0.88025 | 2.271241 | 0.136274 |
| 6 | 100089800 | 100105399 | 100.7606 | 151.3137 | 347.4292 | 143.3974 | -0.93045 | 3.440287 | 0.136274 |
| 6 | 68346400 | 68347999 | 12.15696 | 21.23315 | 50.49602 | 22.2438 | -0.85616 | 0.703766 | 0.136296 |
| 7 | 134853000 | 134856199 | 21.22178 | 35.14621 | 77.1715 | 33.39381 | -0.75045 | 1.306828 | 0.13631 |
| 1 | 104652200 | 104653599 | 17.8706 | 31.98953 | 23.46556 | 10.28964 | 0.84428 | 0.432233 | 0.136317 |
| 7 | 19183800 | 19192999 | 59.78337 | 92.93512 | 201.9407 | 83.84276 | -0.81041 | 2.742076 | 0.136318 |
| 17 | 35559200 | 35564199 | 29.39369 | 47.41155 | 103.2083 | 43.96881 | -0.762 | 1.764908 | 0.13632 |
| 11 | 66839600 | 66841599 | 18.07588 | 33.51644 | 18.53662 | 7.782983 | 1.29031 | 0.226226 | 0.136329 |
| 11 | 45742600 | 45743399 | 6.58233 | 12.14291 | 35.17663 | 14.8725 | -1.08708 | 0.204267 | 0.13634 |
| 13 | 13190200 | 13191799 | 17.74832 | 32.25587 | 19.98005 | 8.615154 | 1.101453 | 0.334393 | 0.136344 |
| 8 | 9435000 | 9437199 | 9.802668 | 17.31478 | 44.58052 | 19.72254 | -0.96653 | 0.524257 | 0.13635 |
| 11 | 112733400 | 112735799 | 9.802668 | 17.31478 | 44.58052 | 19.72254 | -0.96653 | 0.524257 | 0.13635 |
| 11 | 73071200 | 73072199 | 8.277215 | 15.07959 | 37.63661 | 16.17928 | -0.89191 | 0.293017 | 0.136352 |
| 1 | 18844000 | 18844999 | 11.66099 | 20.48374 | 48.27456 | 21.36611 | -0.84294 | 0.605496 | 0.136357 |
| 14 | 52150400 | 52152399 | 9.982624 | 17.26347 | 56.22618 | 24.66745 | -1.3092 | 0.826864 | 0.136369 |
| 7 | 99102000 | 99113399 | 59.27157 | 91.86319 | 212.735 | 88.36402 | -0.90555 | 2.794761 | 0.136375 |
| 19 | 16757400 | 16758799 | 9.400266 | 16.46457 | 47.78391 | 21.09702 | -1.14082 | 0.642724 | 0.1364 |
| 4 | 11485600 | 11486599 | 4.565665 | 8.764466 | 28.96538 | 11.3548 | -1.18079 | -0.10402 | 0.136406 |
| 6 | 3210800 | 3214799 | 80.08627 | 126.8013 | 101.3278 | 42.13075 | 0.651492 | 2.381879 | 0.136409 |

| | | | | | | | | | |
|---|---|---|---|---|---|---|---|---|---|
| 10 | 52387600 | 52388399 | 5.39332 | 10.4209 | 32.00626 | 12.28925 | -1.04966 | -0.15921 | 0.136431 |
| 18 | 36451200 | 36462199 | 51.93027 | 81.29034 | 195.4681 | 81.08515 | -0.94849 | 2.614332 | 0.136431 |
| 15 | 21424200 | 21425599 | 16.06245 | 30.2251 | 17.7132 | 7.260285 | 1.234117 | 0.118476 | 0.136469 |
| 18 | 77650600 | 77653999 | 16.41869 | 27.46276 | 72.47701 | 31.56875 | -1.01401 | 1.177066 | 0.136473 |
| 15 | 82263000 | 82264399 | 4.250497 | 8.226012 | 27.05252 | 10.30917 | -1.13427 | -0.2083 | 0.136492 |
| 7 | 32252800 | 32253999 | 17.39297 | 32.2767 | 19.33148 | 8.107643 | 1.177844 | 0.218779 | 0.136496 |
| 12 | 22938400 | 22947599 | 260.8608 | 379.984 | 293.7973 | 121.0781 | 0.597884 | 3.972773 | 0.136524 |
| 14 | 60871600 | 60874199 | 12.85967 | 22.21167 | 54.74253 | 24.01565 | -0.90992 | 0.849085 | 0.136562 |
| 13 | 116588400 | 116588999 | 3.639831 | 7.109772 | 24.34062 | 8.517444 | -1.07703 | -0.52976 | 0.136583 |
| 15 | 102098400 | 102100799 | 8.314006 | 15.04184 | 39.62989 | 17.18469 | -0.97791 | 0.336945 | 0.136594 |
| 6 | 76749400 | 76751599 | 10.20515 | 18.20466 | 44.15972 | 19.42069 | -0.87374 | 0.446339 | 0.136594 |
| 8 | 27930600 | 27932799 | 21.3108 | 37.72837 | 24.51216 | 10.83928 | 1.009694 | 0.530904 | 0.136612 |
| 3 | 96233200 | 96236599 | 20.96603 | 34.56245 | 80.87558 | 34.89471 | -0.84302 | 1.363272 | 0.136612 |
| 6 | 135349600 | 135351799 | 12.63242 | 22.04244 | 50.50343 | 22.23366 | -0.80279 | 0.720916 | 0.13662 |
| 4 | 116130200 | 116133399 | 41.89735 | 68.86343 | 49.98584 | 21.53997 | 0.83899 | 1.429347 | 0.136628 |
| 6 | 109334200 | 109335799 | 17.36555 | 31.98766 | 20.19786 | 8.556938 | 1.091565 | 0.259572 | 0.136628 |
| 18 | 16415400 | 16417399 | 19.5824 | 35.31501 | 21.39438 | 9.314639 | 1.124884 | 0.395296 | 0.136632 |
| 8 | 26743400 | 26745999 | 33.78949 | 56.98328 | 39.31995 | 17.19336 | 0.914418 | 1.121923 | 0.136634 |
| 14 | 88050600 | 88052799 | 22.05925 | 38.98714 | 25.07259 | 11.09086 | 1.023609 | 0.555329 | 0.136643 |
| 4 | 11620400 | 11621399 | 11.3543 | 21.79373 | 16.36959 | 6.419233 | 0.935171 | -0.07248 | 0.136645 |
| 8 | 122133600 | 122135399 | 14.58092 | 27.05287 | 20.15479 | 8.45525 | 0.864821 | 0.197374 | 0.136655 |
| 1 | 140982200 | 140983999 | 17.36564 | 31.90411 | 20.9079 | 8.889391 | 1.035095 | 0.276471 | 0.136656 |
| 13 | 62126800 | 62128399 | 43.74927 | 71.20276 | 56.00631 | 24.04208 | 0.715399 | 1.56589 | 0.136662 |
| 6 | 4972800 | 4974799 | 6.944245 | 13.08392 | 37.09641 | 15.1721 | -1.01985 | 0.077422 | 0.13668 |
| 5 | 22430200 | 22432599 | 19.8107 | 35.56297 | 22.48249 | 9.835055 | 1.059564 | 0.426256 | 0.136683 |
| 5 | 139237000 | 139241799 | 19.4079 | 31.97402 | 80.46475 | 34.70834 | -0.94786 | 1.367738 | 0.136685 |
| 2 | 153673200 | 153676599 | 17.98099 | 30.03431 | 71.46156 | 31.09926 | -0.86513 | 1.196382 | 0.136703 |
| 8 | 46616200 | 46616999 | 6.011862 | 11.49482 | 31.25566 | 12.39341 | -0.91959 | -0.04761 | 0.136721 |
| 2 | 49523000 | 49526399 | 14.08213 | 24.43615 | 57.04983 | 25.0314 | -0.831 | 0.815108 | 0.136721 |
| 7 | 138958800 | 138960399 | 5.866708 | 11.19731 | 30.55013 | 12.16326 | -0.92915 | -0.03556 | 0.13673 |
| 5 | 147524800 | 147528399 | 26.31817 | 42.58484 | 100.3696 | 42.92199 | -0.87121 | 1.668774 | 0.136732 |
| 18 | 49063600 | 49064999 | 32.43389 | 54.89416 | 37.84276 | 16.56549 | 0.915991 | 1.086901 | 0.136741 |
| 14 | 120197400 | 120199799 | 7.817603 | 15.28687 | 10.1704 | 3.468403 | 1.289062 | -0.61071 | 0.136754 |
| 13 | 65632400 | 65635199 | 47.81397 | 77.93559 | 52.21923 | 22.42871 | 0.947709 | 1.554541 | 0.136755 |
| 9 | 69109200 | 69109799 | 9.819822 | 19.06343 | 14.4034 | 5.407685 | 0.985141 | -0.23784 | 0.136761 |
| 8 | 101247200 | 101248399 | 15.69557 | 29.05639 | 20.60973 | 8.672251 | 0.93234 | 0.22081 | 0.136767 |
| 5 | 4484600 | 4485999 | 15.37044 | 28.6083 | 20.1172 | 8.399789 | 0.952853 | 0.186649 | 0.136776 |
| 2 | 89739200 | 89740399 | 19.12023 | 34.02534 | 24.90215 | 10.97397 | 0.842884 | 0.480444 | 0.136778 |
| 6 | 52340200 | 52341199 | 11.46101 | 22.16524 | 16.31123 | 6.235978 | 0.998482 | -0.15247 | 0.136782 |
| 18 | 80754600 | 80760199 | 27.04353 | 43.54391 | 109.0238 | 46.33421 | -0.96641 | 1.817454 | 0.136782 |
| 13 | 34527800 | 34529999 | 14.42507 | 24.41621 | 63.66313 | 27.86888 | -0.99572 | 1.050522 | 0.136788 |
| 10 | 8638800 | 8641799 | 15.20662 | 25.33026 | 72.52909 | 31.50898 | -1.13086 | 1.221448 | 0.136798 |
| 5 | 138276200 | 138280399 | 44.67161 | 70.95852 | 146.031 | 60.89079 | -0.70682 | 2.273932 | 0.13681 |
| 2 | 41963400 | 41963799 | 2.83741 | 5.545363 | 21.76587 | 7.528034 | -1.25329 | -0.59102 | 0.136817 |
| 4 | 48378800 | 48380199 | 17.93705 | 33.50573 | 17.6819 | 7.338107 | 1.370538 | 0.18062 | 0.136831 |
| 2 | 75807400 | 75809199 | 15.11908 | 25.82057 | 59.53905 | 26.10379 | -0.81772 | 0.967953 | 0.136836 |
| 4 | 43521200 | 43524799 | 18.36393 | 30.48556 | 73.94853 | 32.0517 | -0.89312 | 1.269975 | 0.136841 |

| | | | | | | | | | |
|---|---|---|---|---|---|---|---|---|---|
| 1 | 149375400 | 149376199 | 12.68799 | 24.15186 | 18.27749 | 7.318309 | 0.897442 | 0.012972 | 0.136851 |
| 17 | 19770800 | 19779199 | 154.0733 | 235.5041 | 150.0549 | 62.24752 | 0.938889 | 3.087283 | 0.136864 |
| 13 | 95009000 | 95010599 | 9.751249 | 17.01013 | 50.96726 | 22.43378 | -1.18728 | 0.71182 | 0.136876 |
| 9 | 19184200 | 19186199 | 38.7682 | 63.91871 | 49.41458 | 21.32183 | 0.750004 | 1.38164 | 0.136885 |
| 2 | 173427200 | 173438399 | 78.61446 | 120.4114 | 246.6459 | 102.3652 | -0.74194 | 3.030581 | 0.136885 |
| 13 | 89607200 | 89608399 | 15.05701 | 27.90683 | 20.54639 | 8.631874 | 0.880229 | 0.207876 | 0.136894 |
| 14 | 75183000 | 75185599 | 14.17905 | 24.23994 | 63.43517 | 27.81427 | -0.99952 | 0.956017 | 0.136903 |
| 1 | 116593400 | 116597399 | 28.67326 | 49.32888 | 30.68194 | 13.45859 | 1.06609 | 0.917115 | 0.136905 |
| 2 | 163416000 | 163419599 | 16.8751 | 28.1862 | 70.04496 | 30.48215 | -0.92735 | 1.195145 | 0.136922 |
| 8 | 117739800 | 117746399 | 45.99406 | 72.56306 | 167.6452 | 69.57868 | -0.882 | 2.455659 | 0.13693 |
| 19 | 29790000 | 29790599 | 3.455139 | 6.751794 | 22.4484 | 7.789128 | -1.0224 | -0.56591 | 0.136933 |
| 2 | 80456400 | 80457599 | 11.62502 | 22.34246 | 16.73056 | 6.536273 | 0.944118 | -0.081 | 0.136935 |
| 2 | 40317600 | 40319199 | 22.10787 | 39.6659 | 20.02656 | 8.764577 | 1.381055 | 0.440526 | 0.13694 |
| 6 | 127936000 | 127945399 | 47.45146 | 74.4903 | 182.7356 | 75.78958 | -0.97402 | 2.559466 | 0.136953 |
| 14 | 86130800 | 86133799 | 32.38168 | 54.9103 | 37.63071 | 16.48218 | 0.924401 | 1.068209 | 0.136956 |
| 9 | 120835200 | 120837199 | 7.823967 | 14.28325 | 39.85824 | 17.09023 | -1.04973 | 0.280018 | 0.136961 |
| 15 | 19148400 | 19150399 | 16.88454 | 31.97507 | 15.39043 | 6.229781 | 1.530095 | 0.079337 | 0.136966 |
| 17 | 80596400 | 80597799 | 5.459527 | 10.40304 | 31.0509 | 12.4057 | -1.06222 | -0.02843 | 0.13697 |
| 2 | 179680400 | 179686399 | 27.05379 | 43.5706 | 111.2022 | 47.27598 | -0.99363 | 1.80987 | 0.136982 |
| 7 | 47324800 | 47328199 | 58.77543 | 94.51315 | 70.08391 | 29.72872 | 0.782832 | 1.886961 | 0.136984 |
| 2 | 35991800 | 35999999 | 56.28058 | 88.11693 | 188.8261 | 78.32837 | -0.78231 | 2.612593 | 0.136999 |
| 18 | 36078400 | 36082199 | 18.30139 | 30.54704 | 70.86845 | 30.82685 | -0.82878 | 1.205441 | 0.137003 |
| 11 | 62480200 | 62484799 | 25.62234 | 41.58731 | 93.07099 | 39.89165 | -0.79239 | 1.601781 | 0.137008 |
| X | 40490400 | 40491199 | 11.90193 | 23.06855 | 15.27195 | 5.779074 | 1.163854 | -0.18325 | 0.137014 |
| 6 | 82717800 | 82729199 | 57.16164 | 89.33422 | 194.2179 | 80.58 | -0.80516 | 2.645879 | 0.137015 |
| 8 | 84762600 | 84775999 | 69.87272 | 107.5183 | 232.665 | 96.62787 | -0.81589 | 2.935258 | 0.13702 |
| 4 | 139275200 | 139283999 | 64.0191 | 99.50697 | 200.4272 | 83.21595 | -0.70128 | 2.746099 | 0.13704 |
| 2 | 28863000 | 28865599 | 14.04008 | 23.9601 | 60.01761 | 26.31204 | -0.93676 | 0.970428 | 0.137043 |
| 5 | 113981000 | 113990199 | 47.21303 | 74.27713 | 175.4511 | 72.78891 | -0.9171 | 2.514566 | 0.137052 |
| 14 | 43047400 | 43048399 | 11.73798 | 22.72501 | 15.67029 | 5.961294 | 1.097709 | -0.1679 | 0.137056 |
| 14 | 31406000 | 31408999 | 13.74722 | 23.83618 | 54.85888 | 24.06951 | -0.81071 | 0.816348 | 0.137063 |
| 10 | 92912400 | 92915599 | 16.52717 | 28.19979 | 62.96707 | 27.60473 | -0.77202 | 0.983716 | 0.137069 |
| 6 | 18513600 | 18515399 | 8.910849 | 15.66051 | 45.65777 | 20.2163 | -1.14691 | 0.581341 | 0.13707 |
| 7 | 115906400 | 115909799 | 16.60775 | 27.69252 | 72.87619 | 31.68024 | -1.00986 | 1.212732 | 0.137071 |
| 17 | 53323600 | 53325199 | 21.1126 | 37.69872 | 22.31482 | 9.805645 | 1.149535 | 0.46697 | 0.137075 |
| 8 | 49200800 | 49202199 | 8.733203 | 15.89108 | 38.68871 | 16.65674 | -0.85798 | 0.305261 | 0.137084 |
| 9 | 106800400 | 106801799 | 12.9303 | 24.76684 | 17.07998 | 6.738183 | 1.048889 | -0.03538 | 0.137092 |
| 13 | 44061600 | 44063799 | 10.75402 | 18.87015 | 46.7083 | 20.65403 | -0.91302 | 0.616578 | 0.137102 |
| 9 | 15826200 | 15827599 | 7.967526 | 14.5514 | 38.22932 | 16.38322 | -0.96261 | 0.276175 | 0.137104 |
| 5 | 102190000 | 102191799 | 19.40986 | 34.34318 | 25.49214 | 11.27392 | 0.818524 | 0.527458 | 0.137111 |
| 8 | 13336800 | 13338399 | 10.29427 | 18.06796 | 46.18719 | 20.42793 | -0.95898 | 0.610345 | 0.137113 |
| 7 | 98987400 | 98991799 | 21.55289 | 35.64798 | 78.30668 | 33.85448 | -0.75125 | 1.323628 | 0.137139 |
| 19 | 40312800 | 40315999 | 21.47762 | 35.29868 | 81.13761 | 34.96291 | -0.81908 | 1.403768 | 0.137152 |
| 15 | 42466800 | 42468199 | 15.69714 | 29.48057 | 18.27664 | 7.515825 | 1.149915 | 0.128264 | 0.13716 |
| 5 | 147227200 | 147231399 | 32.35978 | 51.96686 | 117.3379 | 49.67695 | -0.82262 | 1.91005 | 0.137175 |
| 7 | 134517200 | 134520799 | 18.72039 | 30.94901 | 76.50768 | 33.06634 | -0.92084 | 1.32235 | 0.137202 |
| 18 | 74854600 | 74863799 | 61.44072 | 95.52249 | 212.0952 | 88.05723 | -0.84146 | 2.740708 | 0.137212 |

| | | | | | | | | | |
|---|---|---|---|---|---|---|---|---|---|
| 5 | 66607800 | 66609799 | 12.9415 | 22.54438 | 52.68143 | 23.16837 | -0.83156 | 0.74776 | 0.13723 |
| 5 | 19706800 | 19711399 | 32.69939 | 52.46217 | 117.2151 | 49.55299 | -0.80907 | 1.941298 | 0.137246 |
| X | 147330600 | 147331799 | 13.14622 | 25.42959 | 15.01987 | 5.736779 | 1.313797 | -0.14037 | 0.137249 |
| 18 | 37598400 | 37598799 | 15.89531 | 29.36234 | 19.72063 | 8.322684 | 1.007662 | 0.232613 | 0.137251 |
| 11 | 97077800 | 97080999 | 12.75007 | 21.93187 | 59.04181 | 25.8985 | -1.03825 | 0.889353 | 0.137271 |
| 6 | 127337800 | 127340999 | 17.74039 | 29.8659 | 66.62609 | 29.11815 | -0.77234 | 1.111273 | 0.137287 |
| 12 | 32585200 | 32586599 | 14.8521 | 24.84063 | 69.90235 | 30.44594 | -1.10585 | 1.173411 | 0.137298 |
| 15 | 4986000 | 4987599 | 19.60419 | 35.27477 | 21.93468 | 9.576491 | 1.085045 | 0.409281 | 0.137304 |
| 12 | 69243400 | 69245199 | 8.90458 | 15.71115 | 44.08758 | 19.50875 | -1.09007 | 0.53208 | 0.137331 |
| 18 | 81998200 | 81999999 | 10.37082 | 18.4386 | 43.56238 | 19.21193 | -0.83873 | 0.467245 | 0.137357 |
| 14 | 120375200 | 120378599 | 17.05322 | 28.4493 | 70.62186 | 30.70977 | -0.92585 | 1.208325 | 0.137361 |
| 12 | 82239000 | 82240799 | 6.076456 | 11.45721 | 33.5532 | 13.70608 | -1.0641 | 0.056551 | 0.137365 |
| 4 | 43414400 | 43419399 | 29.01768 | 46.55845 | 117.7323 | 49.77672 | -0.98702 | 1.935729 | 0.137382 |
| X | 5534800 | 5537599 | 53.24517 | 86.53785 | 51.26494 | 21.99472 | 1.121926 | 1.605722 | 0.137384 |
| X | 52949000 | 52950199 | 28.60023 | 49.53251 | 31.24069 | 13.70647 | 1.048209 | 0.862606 | 0.137387 |
| 8 | 64915600 | 64919199 | 28.06147 | 48.73023 | 29.29822 | 12.8594 | 1.117163 | 0.828569 | 0.137405 |
| 5 | 81571600 | 81572599 | 14.14739 | 26.7566 | 17.91014 | 7.269791 | 1.055408 | 0.068143 | 0.137414 |
| 2 | 149382200 | 149383799 | 14.01446 | 26.63218 | 17.41554 | 6.99792 | 1.101393 | 0.033867 | 0.137414 |
| 6 | 18328600 | 18329999 | 14.01446 | 26.63218 | 17.41554 | 6.99792 | 1.101393 | 0.033867 | 0.137414 |
| 7 | 20479400 | 20480399 | 14.01799 | 26.33013 | 19.03982 | 7.828072 | 0.929887 | 0.113996 | 0.137415 |
| 16 | 87467600 | 87468999 | 10.21199 | 18.19302 | 43.0689 | 18.96189 | -0.83978 | 0.453163 | 0.137427 |
| 1 | 173675000 | 173676399 | 6.958278 | 13.05976 | 35.74842 | 14.71683 | -0.97719 | 0.098759 | 0.137429 |
| 1 | 108220400 | 108221599 | 17.47259 | 31.46741 | 23.05248 | 10.05361 | 0.851511 | 0.392721 | 0.137434 |
| 1 | 101182600 | 101183199 | 4.357586 | 8.488419 | 26.13132 | 9.526388 | -0.98061 | -0.36768 | 0.137438 |
| 16 | 14337400 | 14340799 | 25.8739 | 45.01649 | 28.64607 | 12.58812 | 1.03603 | 0.792123 | 0.137446 |
| 7 | 105650400 | 105650999 | 6.006702 | 11.44164 | 34.55924 | 13.81768 | -1.08018 | -0.00927 | 0.137451 |
| 2 | 70491000 | 70492799 | 8.291118 | 15.06181 | 39.72862 | 17.13975 | -0.97485 | 0.312507 | 0.137452 |
| 4 | 45580200 | 45582199 | 10.18974 | 17.79558 | 49.91996 | 21.98875 | -1.09263 | 0.69565 | 0.137486 |
| 5 | 147513800 | 147518199 | 28.9053 | 46.65163 | 102.9199 | 43.87976 | -0.78025 | 1.745494 | 0.137488 |
| 13 | 52630400 | 52634599 | 27.95421 | 45.09976 | 111.0841 | 47.33774 | -0.93947 | 1.758431 | 0.137525 |
| 4 | 41457800 | 41458799 | 12.45437 | 23.82552 | 17.36579 | 6.875502 | 0.965803 | -0.02668 | 0.137525 |
| 1 | 50573200 | 50574799 | 17.00861 | 30.85176 | 22.11749 | 9.559899 | 0.890797 | 0.347293 | 0.137532 |
| X | 33950800 | 33955399 | 92.06771 | 145.4242 | 101.8612 | 42.30141 | 0.838629 | 2.446647 | 0.137538 |
| 15 | 64544400 | 64545599 | 18.28159 | 33.47486 | 19.99869 | 8.541566 | 1.162407 | 0.298291 | 0.137542 |
| 13 | 53824200 | 53826599 | 13.05828 | 22.4516 | 60.16038 | 26.38878 | -1.03183 | 0.895707 | 0.137548 |
| 4 | 82521800 | 82524199 | 12.31003 | 21.48871 | 53.76212 | 23.67436 | -0.92929 | 0.715702 | 0.137549 |
| 18 | 39926400 | 39927999 | 22.84984 | 40.05706 | 28.43934 | 12.56565 | 0.879109 | 0.665621 | 0.137558 |
| 14 | 67147200 | 67149799 | 21.31543 | 34.65613 | 91.18795 | 39.1302 | -1.02277 | 1.560135 | 0.137562 |
| 19 | 39218800 | 39221399 | 39.03544 | 65.3442 | 38.15935 | 16.6293 | 1.154201 | 1.206789 | 0.137564 |
| 7 | 94566200 | 94570199 | 79.11444 | 125.5041 | 98.57298 | 41.04812 | 0.678842 | 2.328134 | 0.137567 |
| 7 | 126242400 | 126242599 | 6.3906 | 12.451 | 7.524874 | 2.324522 | 1.543799 | -0.98085 | 0.137569 |
| 4 | 127251000 | 127259999 | 51.0484 | 80.04503 | 185.5608 | 76.96197 | -0.89385 | 2.582996 | 0.137599 |
| 6 | 47021400 | 47023399 | 12.48928 | 21.75282 | 51.04142 | 22.44467 | -0.83726 | 0.747284 | 0.137614 |
| 9 | 54940800 | 54944999 | 24.0945 | 39.35853 | 86.1517 | 37.03227 | -0.75335 | 1.493541 | 0.137623 |
| 6 | 6873400 | 6876799 | 29.55608 | 47.67038 | 106.593 | 45.40697 | -0.8008 | 1.766774 | 0.137638 |
| 3 | 30905600 | 30906999 | 6.65707 | 12.51212 | 32.89305 | 13.50895 | -0.91559 | 0.083382 | 0.137649 |
| 12 | 23966800 | 23968799 | 40.53526 | 66.72434 | 42.47705 | 18.31585 | 1.028435 | 1.4009 | 0.137662 |

| | | | | | | | | | |
|---|---|---|---|---|---|---|---|---|---|
| 8 | 121569200 | 121592399 | 158.7827 | 231.0845 | 468.4422 | 193.0698 | -0.79325 | 3.981641 | 0.137684 |
| X | 31323200 | 31323999 | 15.0539 | 28.58778 | 15.75402 | 6.339513 | 1.34575 | 0.046101 | 0.137694 |
| 14 | 121147800 | 121149199 | 7.818695 | 13.93482 | 42.57334 | 18.73742 | -1.20448 | 0.433377 | 0.137706 |
| 4 | 154666800 | 154674199 | 38.82772 | 61.79316 | 145.3843 | 60.75381 | -0.89676 | 2.209662 | 0.137716 |
| 13 | 38820200 | 38823799 | 12.13971 | 21.2472 | 52.10916 | 22.99248 | -0.90042 | 0.672001 | 0.137718 |
| 7 | 3119400 | 3121999 | 58.36332 | 93.5976 | 73.33908 | 30.98455 | 0.698833 | 1.969835 | 0.137719 |
| 4 | 155425800 | 155431199 | 25.03086 | 40.23249 | 115.8972 | 49.13015 | -1.17135 | 1.873551 | 0.13772 |
| 12 | 20253800 | 20259999 | 183.4426 | 276.1174 | 197.5727 | 81.59647 | 0.753376 | 3.404048 | 0.137731 |
| 15 | 43145600 | 43147199 | 10.093 | 17.86024 | 43.01656 | 19.02368 | -0.86956 | 0.504781 | 0.137732 |
| 11 | 113084800 | 113092799 | 41.85457 | 66.28413 | 157.9387 | 65.67653 | -0.92197 | 2.359884 | 0.137738 |
| 1 | 21083400 | 21085399 | 9.587545 | 17.10596 | 42.03646 | 18.48416 | -0.89204 | 0.436552 | 0.137765 |
| 1 | 101038200 | 101039599 | 23.11143 | 41.15902 | 21.69541 | 9.554707 | 1.31381 | 0.496145 | 0.137774 |
| 5 | 37301600 | 37309399 | 44.79653 | 71.27427 | 141.279 | 59.01756 | -0.65038 | 2.223065 | 0.13778 |
| 14 | 45225800 | 45227399 | 13.15952 | 22.7443 | 55.32553 | 24.27189 | -0.8911 | 0.84373 | 0.137783 |
| 2 | 173483200 | 173484999 | 9.177057 | 16.27003 | 42.50272 | 18.78937 | -0.98485 | 0.476149 | 0.137783 |
| 14 | 14790000 | 14792399 | 24.81455 | 43.65031 | 24.28347 | 10.75193 | 1.228962 | 0.620233 | 0.137785 |
| 4 | 136195800 | 136198199 | 13.04389 | 22.43918 | 56.29904 | 24.69545 | -0.93679 | 0.889338 | 0.137789 |
| 8 | 75080200 | 75081599 | 19.36872 | 34.4599 | 24.51267 | 10.80435 | 0.883657 | 0.483533 | 0.137791 |
| 4 | 101593600 | 101594799 | 14.45425 | 27.57948 | 17.07496 | 6.802507 | 1.191103 | 0.011872 | 0.13782 |
| 18 | 53001600 | 53004599 | 12.1411 | 21.2702 | 48.85441 | 21.57399 | -0.80605 | 0.652892 | 0.13783 |
| 13 | 31999600 | 32000599 | 6.708188 | 12.46623 | 34.24681 | 14.33553 | -1.001 | 0.159599 | 0.137841 |
| 7 | 143148600 | 143150599 | 7.55581 | 13.38439 | 43.57626 | 19.26721 | -1.3003 | 0.477997 | 0.137842 |
| 8 | 33236400 | 33236999 | 5.125155 | 9.897393 | 30.02609 | 11.55813 | -1.03526 | -0.15895 | 0.137848 |
| 18 | 83568000 | 83568999 | 6.067606 | 11.46487 | 33.64824 | 13.68476 | -1.06172 | 0.041572 | 0.13785 |
| 8 | 127574200 | 127576999 | 13.83628 | 23.7443 | 57.39899 | 25.17533 | -0.88431 | 0.915195 | 0.137861 |
| 3 | 82555200 | 82556199 | 13.76526 | 25.75127 | 19.22297 | 7.957588 | 0.876881 | 0.137952 | 0.137861 |
| 4 | 145524200 | 145524999 | 17.00436 | 31.18128 | 21.44802 | 9.142556 | 0.963062 | 0.284976 | 0.137865 |
| 13 | 66251600 | 66252399 | 22.72453 | 39.8372 | 25.5041 | 11.26869 | 1.027416 | 0.659501 | 0.137873 |
| 17 | 38883600 | 38888399 | 108.8912 | 169.5285 | 119.063 | 49.42213 | 0.818773 | 2.728505 | 0.137906 |
| 5 | 36226800 | 36230399 | 15.3235 | 25.63889 | 70.01746 | 30.50345 | -1.06289 | 1.169872 | 0.137912 |
| 5 | 116499000 | 116499799 | 15.12095 | 28.4159 | 18.96673 | 7.791492 | 1.045704 | 0.121478 | 0.137922 |
| 15 | 57683000 | 57691999 | 54.58833 | 86.06468 | 168.5215 | 69.93357 | -0.64453 | 2.474518 | 0.137937 |
| X | 29338200 | 29341599 | 46.61467 | 77.62343 | 33.71998 | 14.64614 | 1.579606 | 1.28245 | 0.137942 |
| 2 | 83157400 | 83158999 | 15.50718 | 29.13246 | 18.28224 | 7.514344 | 1.132992 | 0.124611 | 0.137958 |
| 5 | 98711200 | 98712999 | 7.804152 | 14.29627 | 36.67319 | 15.65392 | -0.92409 | 0.255255 | 0.13796 |
| 14 | 78104000 | 78105399 | 6.799088 | 12.80104 | 33.51695 | 13.72515 | -0.90653 | 0.075039 | 0.137965 |
| 16 | 53960600 | 53961999 | 30.92644 | 52.26923 | 39.93464 | 17.47374 | 0.768082 | 1.093439 | 0.137966 |
| 12 | 117075200 | 117076799 | 7.904986 | 14.63067 | 38.65742 | 16.27161 | -0.95219 | 0.200288 | 0.137968 |
| 4 | 114726600 | 114728399 | 9.761445 | 17.35217 | 41.99388 | 18.52289 | -0.87314 | 0.461905 | 0.137974 |
| 1 | 172415200 | 172416599 | 8.348455 | 15.012 | 40.04506 | 17.49292 | -1.00258 | 0.37501 | 0.137988 |
| 9 | 47077600 | 47079399 | 8.348455 | 15.012 | 40.04506 | 17.49292 | -1.00258 | 0.37501 | 0.137988 |
| 17 | 34109000 | 34109999 | 4.07223 | 7.72173 | 31.87083 | 12.86562 | -1.5364 | -0.00342 | 0.13799 |
| 7 | 11819800 | 11820399 | 16.90404 | 30.53974 | 22.53905 | 9.790947 | 0.844391 | 0.369027 | 0.137996 |
| 17 | 8524400 | 8525799 | 8.786477 | 15.81845 | 40.05618 | 17.477 | -0.927 | 0.370434 | 0.138011 |
| 3 | 83667000 | 83674599 | 42.41188 | 67.36448 | 149.2403 | 62.22475 | -0.81308 | 2.274129 | 0.138018 |
| 3 | 108315800 | 108324599 | 98.11318 | 146.9904 | 337.3147 | 139.1432 | -0.93267 | 3.485557 | 0.138022 |
| 10 | 108849600 | 108849999 | 10.54509 | 20.54535 | 13.30041 | 4.83676 | 1.247924 | -0.33959 | 0.138036 |

| | | | | | | | | | |
|---|---|---|---|---|---|---|---|---|---|
| 11 | 77821400 | 77823799 | 16.70142 | 27.97778 | 68.56391 | 29.89271 | -0.90741 | 1.159597 | 0.13804 |
| 1 | 145961200 | 145962599 | 16.48803 | 30.40554 | 20.96377 | 8.867796 | 0.967471 | 0.248866 | 0.138058 |
| 18 | 32248400 | 32250399 | 15.78372 | 29.28554 | 20.48277 | 8.592422 | 0.955357 | 0.208343 | 0.138062 |
| 7 | 21111200 | 21121199 | 136.5061 | 209.99 | 141.5672 | 58.79174 | 0.864584 | 2.955881 | 0.138065 |
| 13 | 55885200 | 55895199 | 67.26804 | 104.2318 | 212.8311 | 88.4041 | -0.7245 | 2.801995 | 0.138072 |
| 6 | 137952000 | 137952999 | 11.67015 | 22.54174 | 16.18565 | 6.223934 | 1.025276 | -0.13235 | 0.138083 |
| 17 | 35878200 | 35883399 | 33.77941 | 53.94072 | 133.6691 | 56.12131 | -0.96604 | 2.093852 | 0.138089 |
| 3 | 144252600 | 144256199 | 16.05737 | 27.39912 | 60.70289 | 26.61219 | -0.76066 | 0.97974 | 0.138105 |
| 13 | 111822200 | 111824399 | 13.66497 | 23.49683 | 56.89009 | 24.95437 | -0.88602 | 0.895208 | 0.138106 |
| 6 | 34981400 | 34981999 | 8.223109 | 16.0639 | 12.19447 | 4.260324 | 1.069789 | -0.51529 | 0.138125 |
| 3 | 94488800 | 94490599 | 11.56821 | 20.15862 | 51.55847 | 22.67913 | -0.96087 | 0.735423 | 0.138135 |
| 15 | 92293200 | 92296799 | 26.44835 | 42.85769 | 98.41049 | 42.1293 | -0.83169 | 1.638242 | 0.138144 |
| X | 5330200 | 5333799 | 59.42481 | 95.77559 | 63.13218 | 26.86821 | 0.956256 | 1.815219 | 0.138157 |
| 1 | 87073200 | 87087199 | 101.8434 | 152.403 | 350.5148 | 144.5559 | -0.93742 | 3.50742 | 0.138159 |
| X | 126864200 | 126865199 | 35.1531 | 58.75599 | 43.24512 | 18.82756 | 0.821787 | 1.215521 | 0.138167 |
| 16 | 11726800 | 11728799 | 8.914755 | 16.11234 | 39.60705 | 17.19664 | -0.87986 | 0.347502 | 0.13817 |
| X | 54402000 | 54404399 | 43.43014 | 72.23572 | 38.84067 | 16.85709 | 1.273229 | 1.289653 | 0.138176 |
| X | 96532400 | 96532999 | 10.76323 | 20.92764 | 14.81638 | 5.492825 | 1.09483 | -0.26706 | 0.138177 |
| 5 | 27540400 | 27541599 | 6.939771 | 13.09071 | 34.19784 | 13.95805 | -0.89987 | 0.063993 | 0.13818 |
| 9 | 83325200 | 83326599 | 5.758717 | 10.58595 | 36.70105 | 15.59202 | -1.34914 | 0.220014 | 0.138203 |
| 7 | 60128400 | 60131799 | 41.11652 | 68.07104 | 44.99344 | 19.465 | 0.976082 | 1.338001 | 0.138207 |
| 6 | 125351400 | 125353399 | 6.710765 | 12.07424 | 39.56246 | 17.27355 | -1.29547 | 0.361178 | 0.138232 |
| 12 | 104518800 | 104529799 | 69.0775 | 106.6016 | 222.6289 | 92.47577 | -0.76148 | 2.87909 | 0.138232 |
| 4 | 133623800 | 133625399 | 14.88187 | 27.5614 | 18.28116 | 7.688109 | 1.028826 | 0.205315 | 0.138236 |
| 19 | 3461200 | 3470399 | 54.7177 | 85.52943 | 190.7844 | 79.15406 | -0.84194 | 2.641682 | 0.138238 |
| 11 | 51988400 | 51990399 | 9.416726 | 16.49315 | 48.30582 | 21.32718 | -1.15411 | 0.642638 | 0.138244 |
| 13 | 84070200 | 84071199 | 11.48218 | 22.20872 | 15.63183 | 5.973055 | 1.062051 | -0.15713 | 0.138267 |
| 14 | 22848800 | 22849599 | 4.763629 | 9.235516 | 34.29079 | 12.95922 | -1.29766 | -0.21206 | 0.138271 |
| 17 | 39059800 | 39062799 | 51.12929 | 83.46105 | 47.92512 | 20.59529 | 1.171005 | 1.536964 | 0.138285 |
| 11 | 68041800 | 68043199 | 6.181417 | 11.75294 | 33.9834 | 13.64006 | -1.02377 | -0.00064 | 0.138287 |
| 5 | 105888400 | 105896399 | 48.25069 | 76.55429 | 151.7675 | 63.20885 | -0.6547 | 2.311774 | 0.138296 |
| 7 | 33133000 | 33139999 | 158.3957 | 241.3725 | 160.5829 | 66.5671 | 0.873191 | 3.150588 | 0.138303 |
| 18 | 20893600 | 20894199 | 4.232491 | 8.245102 | 26.13866 | 9.523574 | -1.02187 | -0.37221 | 0.138304 |
| 19 | 17394600 | 17395199 | 4.232491 | 8.245102 | 26.13866 | 9.523574 | -1.02187 | -0.37221 | 0.138304 |
| 8 | 21771200 | 21772199 | 12.53573 | 24.11534 | 15.98187 | 6.227008 | 1.121862 | -0.08218 | 0.138304 |
| 2 | 43128200 | 43129799 | 15.33248 | 29.15202 | 16.82251 | 6.752202 | 1.281941 | 0.04241 | 0.138306 |
| 6 | 26014000 | 26015999 | 21.64827 | 35.54225 | 81.60913 | 35.15067 | -0.81812 | 1.41783 | 0.138314 |
| 2 | 159767800 | 159769399 | 10.85782 | 19.18868 | 46.69229 | 20.65461 | -0.88559 | 0.52772 | 0.138317 |
| 8 | 14405800 | 14406799 | 8.399038 | 15.39591 | 37.78514 | 16.11446 | -0.8601 | 0.258281 | 0.138331 |
| 4 | 63913400 | 63914799 | 6.330569 | 12.01472 | 32.11993 | 12.94167 | -0.91608 | 0.010381 | 0.138336 |
| 15 | 95717200 | 95723999 | 47.92717 | 76.04516 | 150.9876 | 62.88749 | -0.6568 | 2.309844 | 0.138345 |
| 6 | 86033000 | 86034599 | 8.578654 | 15.66382 | 40.4142 | 17.32468 | -0.93695 | 0.288803 | 0.138352 |
| 9 | 78752400 | 78753799 | 10.03563 | 17.94599 | 43.13499 | 18.93106 | -0.85849 | 0.427569 | 0.138352 |
| 16 | 21754800 | 21757799 | 14.88224 | 25.20372 | 63.16928 | 27.65725 | -0.93894 | 1.04674 | 0.138354 |
| X | 32899000 | 32902199 | 42.99705 | 71.43992 | 40.01936 | 17.356 | 1.214316 | 1.301701 | 0.138355 |
| 3 | 126066400 | 126068199 | 12.6465 | 24.28231 | 15.93148 | 6.241289 | 1.12885 | -0.06429 | 0.138359 |
| 6 | 17197600 | 17199199 | 5.750202 | 10.94771 | 32.21753 | 12.89472 | -1.04483 | -0.01782 | 0.138365 |

| | | | | | | | | | |
|---|---|---|---|---|---|---|---|---|---|
| 7 | 144303000 | 144314999 | 63.46621 | 98.39654 | 207.4461 | 86.16294 | -0.76992 | 2.79066 | 0.13837 |
| 5 | 116059200 | 116072999 | 90.27245 | 136.7922 | 288.6899 | 119.47 | -0.79752 | 3.255209 | 0.138376 |
| 19 | 3669000 | 3675799 | 40.86588 | 65.16911 | 133.677 | 56.01878 | -0.69607 | 2.143633 | 0.138377 |
| 12 | 3562000 | 3564799 | 38.39838 | 63.26832 | 49.41266 | 21.31521 | 0.734976 | 1.387941 | 0.138384 |
| 16 | 94508600 | 94511399 | 14.4479 | 24.79155 | 60.21003 | 26.40805 | -0.89132 | 0.919708 | 0.138384 |
| 1 | 160610000 | 160611599 | 9.542091 | 17.17829 | 41.6548 | 18.17516 | -0.86518 | 0.379011 | 0.138387 |
| X | 29383600 | 29385799 | 25.81556 | 44.67391 | 33.49606 | 14.69563 | 0.799557 | 0.858719 | 0.138395 |
| 6 | 147164200 | 147166599 | 14.71141 | 24.96147 | 62.66322 | 27.45138 | -0.94115 | 1.02853 | 0.138398 |
| 5 | 37873600 | 37876199 | 15.72488 | 26.42947 | 66.30206 | 28.96214 | -0.94081 | 1.120791 | 0.138405 |
| X | 31226400 | 31228799 | 30.34202 | 53.02295 | 23.7322 | 10.4578 | 1.541181 | 0.749264 | 0.138435 |
| X | 66258400 | 66259399 | 15.1428 | 28.7986 | 17.01442 | 6.825603 | 1.249477 | 0.040153 | 0.138449 |
| 19 | 58847800 | 58856599 | 61.71328 | 96.42346 | 195.6226 | 81.1649 | -0.70584 | 2.652667 | 0.138452 |
| 1 | 72909400 | 72914399 | 25.10343 | 40.51356 | 104.8867 | 44.71539 | -1.01047 | 1.742016 | 0.138458 |
| 5 | 122560000 | 122562399 | 8.060255 | 14.18308 | 47.72396 | 21.12925 | -1.35207 | 0.562209 | 0.13846 |
| 3 | 72895600 | 72897399 | 21.04394 | 38.12094 | 19.70744 | 8.532306 | 1.357176 | 0.378405 | 0.138461 |
| 19 | 25771200 | 25774599 | 21.55629 | 35.68425 | 77.65085 | 33.59087 | -0.73752 | 1.314829 | 0.138478 |
| 8 | 42510600 | 42511399 | 14.88802 | 27.7427 | 19.25653 | 8.024564 | 0.973854 | 0.173431 | 0.138481 |
| 5 | 112196000 | 112205799 | 63.90975 | 99.6437 | 194.6391 | 80.77559 | -0.65362 | 2.691712 | 0.138487 |
| 13 | 111454400 | 111455199 | 6.006303 | 11.50186 | 31.96273 | 12.61796 | -0.94488 | -0.06098 | 0.138518 |
| 19 | 7468600 | 7469799 | 6.006303 | 11.50186 | 31.96273 | 12.61796 | -0.94488 | -0.06098 | 0.138518 |
| 6 | 114875400 | 114877399 | 10.73243 | 18.88771 | 45.62106 | 20.19767 | -0.8772 | 0.572536 | 0.138523 |
| 7 | 11788800 | 11792799 | 74.43044 | 118.8539 | 78.75323 | 33.06456 | 0.933403 | 2.120461 | 0.138525 |
| 5 | 106136200 | 106141799 | 37.58345 | 60.1135 | 124.6517 | 52.4568 | -0.70689 | 2.045522 | 0.138533 |
| 14 | 19978400 | 19981199 | 19.36824 | 32.46973 | 71.72601 | 31.28628 | -0.75847 | 1.162409 | 0.138534 |
| 1 | 14823600 | 14828399 | 34.82945 | 55.758 | 126.0511 | 53.11238 | -0.82984 | 2.014155 | 0.138537 |
| 4 | 27065400 | 27066999 | 19.8544 | 35.6259 | 23.2291 | 10.1654 | 1.015292 | 0.432194 | 0.13854 |
| 12 | 109418000 | 109431999 | 78.66268 | 119.9841 | 261.8961 | 108.5898 | -0.83829 | 3.116603 | 0.138543 |
| 11 | 99993000 | 99996599 | 18.73316 | 30.95472 | 77.08697 | 33.30526 | -0.93152 | 1.328079 | 0.13857 |
| 5 | 38290400 | 38326999 | 279.1861 | 387.6438 | 746.7327 | 309.9743 | -0.78764 | 4.686335 | 0.138583 |
| 7 | 10756800 | 10758399 | 23.71968 | 41.68158 | 25.43468 | 11.26082 | 1.095926 | 0.627211 | 0.138597 |
| 9 | 123896400 | 123896999 | 9.816203 | 19.06792 | 14.84479 | 5.546281 | 0.948429 | -0.25353 | 0.138602 |
| 5 | 63244600 | 63245199 | 5.717885 | 10.88768 | 31.00332 | 12.40513 | -0.99619 | -0.01803 | 0.13863 |
| 11 | 59312400 | 59314599 | 8.7332 | 15.44164 | 43.05607 | 19.0442 | -1.0795 | 0.506032 | 0.138646 |
| 1 | 69495600 | 69496199 | 46.00072 | 76.5013 | 30.17942 | 13.09688 | 1.717353 | 1.28718 | 0.138659 |
| 14 | 42426200 | 42426599 | 8.19661 | 16.02372 | 7.848344 | 2.532309 | 1.788086 | -0.79735 | 0.138663 |
| 17 | 23750600 | 23750999 | 11.17975 | 21.70506 | 14.54826 | 5.45886 | 1.155976 | -0.22354 | 0.138667 |
| 7 | 41089400 | 41093999 | 69.35074 | 111.111 | 71.464 | 30.14813 | 0.981436 | 2.011614 | 0.138672 |
| 13 | 47082200 | 47084799 | 12.17732 | 20.86515 | 59.44279 | 26.0685 | -1.12039 | 0.924809 | 0.138681 |
| 18 | 67602600 | 67606999 | 21.88035 | 35.73246 | 86.20095 | 37.04995 | -0.89334 | 1.493153 | 0.138688 |
| 17 | 75312200 | 75317199 | 22.70409 | 37.00858 | 88.49392 | 38.00963 | -0.88263 | 1.522867 | 0.138696 |
| 14 | 75157200 | 75160599 | 14.62727 | 25.03578 | 59.59903 | 26.13473 | -0.8631 | 0.943883 | 0.138715 |
| 18 | 77269800 | 77277199 | 45.1404 | 71.34009 | 166.2721 | 69.06949 | -0.89264 | 2.412422 | 0.138751 |
| 12 | 87434000 | 87435599 | 8.727037 | 15.48187 | 42.51878 | 18.7866 | -1.05592 | 0.469332 | 0.138762 |
| 18 | 67907600 | 67909399 | 8.727037 | 15.48187 | 42.51878 | 18.7866 | -1.05592 | 0.469332 | 0.138762 |
| 11 | 95474000 | 95480399 | 32.21309 | 51.70425 | 118.8242 | 50.26621 | -0.84894 | 1.926774 | 0.138763 |
| 3 | 150716600 | 150717399 | 11.55547 | 22.37454 | 15.67556 | 5.959713 | 1.07574 | -0.17211 | 0.13878 |
| 10 | 43060400 | 43060999 | 11.55547 | 22.37454 | 15.67556 | 5.959713 | 1.07574 | -0.17211 | 0.13878 |

| | | | | | | | | | |
|---|---|---|---|---|---|---|---|---|---|
| X | 31481400 | 31482799 | 24.05479 | 43.56165 | 14.31836 | 6.201503 | 2.002925 | 0.390588 | 0.138784 |
| 7 | 7383400 | 7386399 | 31.59887 | 54.26085 | 31.43402 | 13.78771 | 1.167702 | 0.947554 | 0.138789 |
| 13 | 9854800 | 9855399 | 3.821378 | 7.448873 | 24.95837 | 9.019654 | -1.08928 | -0.41044 | 0.138791 |
| 17 | 38620800 | 38621999 | 29.80577 | 51.09374 | 34.31483 | 15.04924 | 0.955155 | 0.958657 | 0.138793 |
| 2 | 92390400 | 92391599 | 7.66816 | 14.01745 | 37.16486 | 15.90887 | -0.97434 | 0.267109 | 0.138794 |
| 18 | 9230600 | 9231799 | 7.66816 | 14.01745 | 37.16486 | 15.90887 | -0.97434 | 0.267109 | 0.138794 |
| 12 | 4596400 | 4597799 | 3.04739 | 5.936078 | 23.98762 | 8.747357 | -1.36372 | -0.38564 | 0.138795 |
| 5 | 138738000 | 138748799 | 54.82847 | 85.06558 | 210.7424 | 87.52272 | -1.00154 | 2.774933 | 0.138796 |
| 5 | 148724200 | 148729799 | 28.42981 | 45.87345 | 102.5869 | 43.72456 | -0.80015 | 1.752074 | 0.138804 |
| 6 | 102396400 | 102398399 | 20.52729 | 36.45634 | 24.79823 | 10.9466 | 0.946327 | 0.502406 | 0.138805 |
| 15 | 36126200 | 36127599 | 9.492885 | 16.79062 | 43.54179 | 19.25772 | -0.97584 | 0.505743 | 0.138852 |
| 15 | 8608200 | 8609399 | 11.59628 | 20.14206 | 53.15341 | 23.33306 | -1.00629 | 0.785117 | 0.138858 |
| 12 | 99105800 | 99108199 | 14.12542 | 24.31924 | 56.36498 | 24.72499 | -0.82284 | 0.886411 | 0.138867 |
| 4 | 23043600 | 23045999 | 27.8799 | 48.1923 | 31.56619 | 13.84847 | 0.993098 | 0.873241 | 0.138867 |
| 11 | 121592600 | 121595199 | 14.71205 | 24.93723 | 62.59076 | 27.4112 | -0.94082 | 1.039131 | 0.138872 |
| 10 | 3586600 | 3587999 | 11.43327 | 22.02217 | 16.18826 | 6.2847 | 0.978766 | -0.10484 | 0.138875 |
| 4 | 153563600 | 153569399 | 46.16292 | 72.82744 | 170.0077 | 70.55823 | -0.89698 | 2.456254 | 0.138891 |
| 7 | 133466600 | 133467999 | 11.08131 | 19.38052 | 48.33831 | 21.31541 | -0.92396 | 0.676038 | 0.138907 |
| 4 | 66396800 | 66398199 | 14.22415 | 27.3162 | 15.21354 | 5.957102 | 1.363735 | -0.0506 | 0.138908 |
| 5 | 82769200 | 82770399 | 14.22415 | 27.3162 | 15.21354 | 5.957102 | 1.363735 | -0.0506 | 0.138908 |
| 17 | 36877000 | 36878399 | 14.22415 | 27.3162 | 15.21354 | 5.957102 | 1.363735 | -0.0506 | 0.138908 |
| 13 | 14019400 | 14020399 | 4.953901 | 9.610517 | 30.57243 | 11.51969 | -1.07339 | -0.23245 | 0.138911 |
| 11 | 89225000 | 89227199 | 23.3261 | 40.90975 | 27.80363 | 12.28928 | 0.941707 | 0.660253 | 0.138918 |
| 1 | 24044200 | 24044999 | 3.880498 | 7.573283 | 25.31332 | 8.991639 | -1.06234 | -0.4676 | 0.138923 |
| 9 | 39076400 | 39076799 | 8.026583 | 15.694 | 11.78485 | 4.029029 | 1.114251 | -0.59089 | 0.138928 |
| 7 | 104992800 | 104994199 | 7.906167 | 14.6291 | 39.2208 | 16.5145 | -0.97349 | 0.202791 | 0.138938 |
| 2 | 34062800 | 34063999 | 4.00303 | 7.772406 | 28.59174 | 10.72083 | -1.27288 | -0.26605 | 0.138987 |
| 4 | 124981200 | 124989999 | 46.10428 | 72.86427 | 165.3859 | 68.70179 | -0.85448 | 2.412651 | 0.138991 |
| 17 | 13269800 | 13271199 | 7.193689 | 13.22562 | 35.663 | 15.14815 | -0.9899 | 0.228729 | 0.139012 |
| 4 | 31898400 | 31898999 | 10.21903 | 19.78914 | 14.34382 | 5.450408 | 1.027994 | -0.19407 | 0.139022 |
| 3 | 21989800 | 21991399 | 7.724247 | 13.96501 | 38.56534 | 16.73779 | -1.04563 | 0.336181 | 0.13904 |
| 2 | 147885200 | 147892199 | 57.84881 | 89.89099 | 203.8871 | 84.65606 | -0.87267 | 2.748545 | 0.139049 |
| Y | 2040600 | 2045199 | 39.32687 | 64.91266 | 35.0008 | 15.11059 | 1.268151 | 1.353378 | 0.139049 |
| X | 31251600 | 31252599 | 19.48015 | 35.20522 | 21.43578 | 9.307936 | 1.12085 | 0.385504 | 0.139059 |
| 9 | 90184200 | 90184999 | 9.631124 | 18.83706 | 8.12826 | 2.745658 | 1.909395 | -0.62086 | 0.139076 |
| 8 | 125505400 | 125513999 | 51.09248 | 80.38016 | 175.6161 | 72.85733 | -0.80476 | 2.516371 | 0.139079 |
| 5 | 149601400 | 149602999 | 11.22583 | 19.65625 | 47.79688 | 21.09749 | -0.88763 | 0.655835 | 0.139107 |
| 5 | 93992600 | 93993199 | 9.986648 | 19.53213 | 8.121421 | 2.746827 | 1.960912 | -0.61053 | 0.13911 |
| 5 | 27929800 | 27932999 | 20.88256 | 34.66883 | 74.91456 | 32.47179 | -0.72708 | 1.276499 | 0.13912 |
| 16 | 55494400 | 55495599 | 16.76621 | 30.37833 | 21.18747 | 9.170642 | 0.928854 | 0.34919 | 0.139139 |
| 8 | 120779200 | 120795599 | 144.3472 | 212.6783 | 409.8923 | 168.7976 | -0.70338 | 3.786914 | 0.139154 |
| 18 | 7112600 | 7114999 | 15.89818 | 27.15416 | 60.15169 | 26.37263 | -0.76018 | 0.969976 | 0.139169 |
| 1 | 151754000 | 151759799 | 30.1922 | 48.70147 | 104.2484 | 44.41424 | -0.73779 | 1.764303 | 0.139175 |
| 10 | 11285800 | 11286999 | 5.275129 | 10.13591 | 29.09422 | 11.37458 | -0.97684 | -0.10637 | 0.139186 |
| 11 | 103844000 | 103845199 | 5.275129 | 10.13591 | 29.09422 | 11.37458 | -0.97684 | -0.10637 | 0.139186 |
| 8 | 109260400 | 109265399 | 35.1253 | 56.28962 | 121.1148 | 51.11235 | -0.75687 | 1.980959 | 0.139189 |
| 19 | 38030800 | 38032199 | 6.01161 | 11.46813 | 31.11304 | 12.40051 | -0.92249 | -0.02885 | 0.139191 |

| | | | | | | | | | |
|---|---|---|---|---|---|---|---|---|---|
| 1 | 19751000 | 19751799 | 12.04413 | 23.21993 | 16.7281 | 6.477323 | 1.011511 | -0.1039 | 0.139197 |
| 11 | 54536800 | 54537399 | 12.10698 | 23.43336 | 15.65976 | 5.964459 | 1.141085 | -0.15951 | 0.139209 |
| 7 | 12051200 | 12054199 | 42.96723 | 70.66531 | 48.92919 | 21.09012 | 0.907165 | 1.422818 | 0.139214 |
| 17 | 86785800 | 86792199 | 28.66488 | 46.10914 | 113.4359 | 48.14064 | -0.94301 | 1.851694 | 0.139218 |
| 6 | 49171000 | 49177199 | 47.25875 | 75.14117 | 145.9459 | 60.91463 | -0.62215 | 2.247881 | 0.139228 |
| 14 | 60987800 | 60992399 | 23.83389 | 38.756 | 89.93282 | 38.59348 | -0.84196 | 1.56289 | 0.139251 |
| 13 | 87893000 | 87894199 | 24.05831 | 42.38971 | 23.71757 | 10.49792 | 1.221431 | 0.596724 | 0.139254 |
| 4 | 112511200 | 112512399 | 27.64036 | 47.53178 | 35.04784 | 15.37349 | 0.821853 | 0.923969 | 0.139265 |
| 18 | 14102600 | 14106199 | 16.61175 | 27.66796 | 72.1968 | 31.36256 | -0.99758 | 1.22554 | 0.139279 |
| 8 | 21804000 | 21807399 | 63.38174 | 101.7879 | 70.43108 | 29.81931 | 0.880202 | 1.933161 | 0.139283 |
| 3 | 68951600 | 68953399 | 10.60755 | 18.60266 | 47.24337 | 20.88068 | -0.94987 | 0.625577 | 0.139291 |
| 11 | 100186400 | 100191799 | 20.79994 | 33.97522 | 89.70837 | 38.56043 | -1.02308 | 1.488295 | 0.139294 |
| 15 | 85220000 | 85222999 | 12.13226 | 20.86847 | 56.7999 | 24.91507 | -1.05368 | 0.886482 | 0.139301 |
| 14 | 118891200 | 118892599 | 7.461902 | 13.80771 | 35.25817 | 14.84504 | -0.90271 | 0.193273 | 0.139302 |
| 2 | 153954800 | 153955599 | 4.114282 | 8.005781 | 26.49989 | 9.763731 | -1.09833 | -0.33286 | 0.139365 |
| 7 | 9161400 | 9162199 | 24.9662 | 43.45252 | 26.35993 | 11.58607 | 1.104663 | 0.779007 | 0.139369 |
| 5 | 104787400 | 104788399 | 23.6036 | 43.23581 | 14.66416 | 6.260004 | 1.972244 | 0.327174 | 0.13937 |
| 7 | 85003200 | 85005199 | 44.50166 | 72.41217 | 57.10326 | 24.51111 | 0.711552 | 1.572227 | 0.139381 |
| 15 | 79980200 | 79988799 | 45.52765 | 71.81077 | 171.5429 | 71.19144 | -0.9304 | 2.460386 | 0.139394 |
| 4 | 107389400 | 107396599 | 48.07611 | 76.29151 | 154.0885 | 64.18746 | -0.68126 | 2.3062 | 0.139405 |
| 13 | 31984600 | 31992999 | 52.29024 | 81.74073 | 192.3839 | 79.81709 | -0.91915 | 2.639402 | 0.139406 |
| 9 | 14175600 | 14176399 | 12.16803 | 23.36725 | 17.22617 | 6.745475 | 0.96362 | -0.06607 | 0.13941 |
| 19 | 38919000 | 38920199 | 4.453072 | 8.518127 | 29.94449 | 11.85852 | -1.28298 | -0.06921 | 0.139424 |
| 3 | 67541400 | 67542399 | 13.54827 | 25.54344 | 17.44173 | 7.125647 | 1.0187 | 0.081419 | 0.139433 |
| 16 | 28796600 | 28797999 | 26.54023 | 45.7314 | 34.4994 | 15.13384 | 0.78951 | 0.899874 | 0.139446 |
| 1 | 179521600 | 179525199 | 25.48908 | 41.30938 | 95.02793 | 40.68567 | -0.83399 | 1.633102 | 0.139455 |
| 11 | 29501600 | 29502999 | 16.28248 | 29.90506 | 20.81818 | 8.8549 | 0.947638 | 0.26756 | 0.13946 |
| 13 | 101124000 | 101126199 | 11.35387 | 19.93629 | 48.28111 | 21.36179 | -0.88201 | 0.609817 | 0.139474 |
| 10 | 110382400 | 110383399 | 12.85718 | 24.58689 | 17.03576 | 6.752091 | 1.03576 | -0.022 | 0.139484 |
| 6 | 10657000 | 10659199 | 43.13026 | 70.6915 | 51.68915 | 22.24757 | 0.826891 | 1.466082 | 0.139486 |
| 4 | 123981200 | 123982599 | 5.996148 | 11.49519 | 33.85336 | 13.32172 | -1.02346 | -0.06596 | 0.139486 |
| 16 | 39462800 | 39463399 | 5.996148 | 11.49519 | 33.85336 | 13.32172 | -1.02346 | -0.06596 | 0.139486 |
| 10 | 82013800 | 82014999 | 9.78909 | 17.32517 | 44.05292 | 19.48139 | -0.94747 | 0.500903 | 0.139494 |
| Y | 87093600 | 87094199 | 7.65922 | 14.98337 | 4.712745 | 1.561141 | 2.376518 | -0.74165 | 0.139498 |
| 4 | 20832800 | 20834599 | 15.24691 | 28.592 | 18.2482 | 7.523251 | 1.105184 | 0.132746 | 0.139505 |
| 7 | 17801800 | 17806399 | 87.73454 | 139.5921 | 78.2743 | 32.69831 | 1.16828 | 2.246332 | 0.139516 |
| 19 | 13572000 | 13579599 | 128.3003 | 197.8647 | 140.7851 | 58.47963 | 0.789602 | 2.905523 | 0.139519 |
| 5 | 100821600 | 100822399 | 5.561028 | 10.69733 | 30.40285 | 11.84769 | -0.95899 | -0.11213 | 0.139528 |
| 4 | 125542200 | 125547799 | 26.34642 | 42.55486 | 101.3343 | 43.24716 | -0.88877 | 1.718104 | 0.139538 |
| 15 | 71654600 | 71655199 | 3.527403 | 6.886621 | 28.26284 | 9.97804 | -1.34827 | -0.49008 | 0.139551 |
| 4 | 153182000 | 153183999 | 22.92158 | 37.6195 | 82.31934 | 35.45128 | -0.74914 | 1.426361 | 0.139553 |
| 10 | 47141000 | 47141599 | 11.00164 | 21.25957 | 16.42844 | 6.304666 | 0.923781 | -0.14442 | 0.139555 |
| 5 | 89963200 | 89965599 | 11.02696 | 19.4259 | 48.28115 | 21.37099 | -0.91792 | 0.565892 | 0.139565 |
| 5 | 11393000 | 11394999 | 14.13911 | 26.49477 | 19.00045 | 7.843301 | 0.937112 | 0.129426 | 0.139565 |
| 4 | 153534800 | 153604399 | 714.1356 | 917.6554 | 1719.41 | 721.1061 | -0.85994 | 5.923996 | 0.139585 |
| 4 | 119590600 | 119591599 | 6.301472 | 11.63214 | 35.18561 | 14.86351 | -1.14811 | 0.197376 | 0.139659 |
| 13 | 47077400 | 47081199 | 19.91928 | 33.11214 | 73.12321 | 31.72347 | -0.7581 | 1.257653 | 0.139665 |

| | | | | | | | | | |
|---|---|---|---|---|---|---|---|---|---|
| 10 | 79550800 | 79562599 | 75.3221 | 114.9809 | 259.2378 | 107.5075 | -0.88404 | 3.099331 | 0.139665 |
| 14 | 123209000 | 123209799 | 5.232128 | 10.1265 | 29.00094 | 11.04795 | -0.93781 | -0.19359 | 0.139676 |
| X | 46878000 | 46879399 | 13.80645 | 26.2426 | 17.75568 | 7.131321 | 1.05424 | 0.033171 | 0.139677 |
| 4 | 89558800 | 89560999 | 18.79767 | 33.70856 | 20.46051 | 8.958567 | 1.117136 | 0.421607 | 0.139682 |
| 5 | 129517200 | 129518199 | 11.39644 | 21.98753 | 16.81956 | 6.498902 | 0.928638 | -0.11739 | 0.139685 |
| X | 45398600 | 45401199 | 30.86694 | 53.93706 | 22.44074 | 9.88821 | 1.645585 | 0.750213 | 0.139695 |
| 15 | 36979000 | 36980399 | 7.263961 | 13.60191 | 34.50649 | 14.26145 | -0.87265 | 0.115166 | 0.139751 |
| 14 | 9346600 | 9348199 | 18.11133 | 33.40465 | 19.63311 | 8.30303 | 1.196138 | 0.257729 | 0.139779 |
| 18 | 55029400 | 55033399 | 18.72079 | 30.99504 | 78.52323 | 33.97179 | -0.95609 | 1.304155 | 0.139781 |
| 5 | 147849800 | 147852599 | 12.80175 | 21.90253 | 60.55394 | 26.55262 | -1.07784 | 0.941707 | 0.139782 |
| 2 | 49350600 | 49353199 | 45.03025 | 73.95171 | 48.39675 | 20.84792 | 0.987514 | 1.445661 | 0.139789 |
| 10 | 29237400 | 29238399 | 7.16856 | 13.26908 | 36.37461 | 15.30853 | -1.00435 | 0.188851 | 0.139795 |
| 4 | 37746000 | 37747399 | 30.51471 | 52.63166 | 30.93043 | 13.56865 | 1.148347 | 0.904435 | 0.139814 |
| 2 | 27338000 | 27340799 | 8.559835 | 14.94049 | 51.45688 | 22.65755 | -1.38658 | 0.698414 | 0.139815 |
| 8 | 128267600 | 128268999 | 5.437784 | 10.42406 | 30.77014 | 12.11076 | -1.02668 | -0.07978 | 0.139821 |
| 4 | 148750000 | 148754399 | 24.3124 | 39.5446 | 89.55607 | 38.43553 | -0.80671 | 1.559168 | 0.139846 |
| 10 | 8519000 | 8520599 | 25.35766 | 44.02686 | 32.29453 | 14.17289 | 0.831788 | 0.823003 | 0.139847 |
| 10 | 92931800 | 92934599 | 13.27418 | 23.04744 | 53.76557 | 23.59597 | -0.82968 | 0.798558 | 0.13986 |
| 8 | 120686000 | 120702399 | 121.9794 | 181.2278 | 376.3073 | 155.0232 | -0.79959 | 3.642094 | 0.13987 |
| 11 | 7362800 | 7364999 | 27.73264 | 47.60382 | 35.55532 | 15.59496 | 0.80285 | 0.940245 | 0.139874 |
| 1 | 75219000 | 75225999 | 27.6888 | 44.31948 | 130.1571 | 54.82993 | -1.20658 | 2.013098 | 0.139883 |
| 12 | 106271600 | 106273999 | 12.16231 | 21.24465 | 49.42247 | 21.77227 | -0.82437 | 0.70198 | 0.139902 |
| 4 | 137308800 | 137312399 | 19.55322 | 32.69714 | 71.6115 | 31.18658 | -0.74607 | 1.190729 | 0.139905 |
| 9 | 106097000 | 106105999 | 47.53169 | 74.81466 | 177.882 | 73.80204 | -0.92603 | 2.504632 | 0.139908 |
| 9 | 110946800 | 110947799 | 4.83597 | 9.343461 | 28.19555 | 10.82894 | -1.02369 | -0.17341 | 0.139928 |
| 10 | 34157800 | 34159399 | 6.78053 | 12.38754 | 36.61793 | 15.68634 | -1.12998 | 0.264606 | 0.139932 |
| 10 | 38128600 | 38130199 | 19.07213 | 34.40159 | 22.64582 | 9.857047 | 1.006413 | 0.394047 | 0.139933 |
| 15 | 50476200 | 50477999 | 19.07213 | 34.40159 | 22.64582 | 9.857047 | 1.006413 | 0.394047 | 0.139933 |
| 4 | 132648800 | 132651199 | 8.429642 | 15.35263 | 37.6308 | 16.18291 | -0.86631 | 0.296318 | 0.139949 |
| 18 | 46487200 | 46490399 | 31.88265 | 51.19301 | 115.0023 | 48.6771 | -0.81548 | 1.914224 | 0.139961 |
| 4 | 132536400 | 132540999 | 65.07873 | 104.1248 | 80.40882 | 33.85362 | 0.715959 | 2.050625 | 0.139962 |
| 3 | 91067600 | 91069399 | 55.30056 | 88.95116 | 69.63786 | 29.5519 | 0.705219 | 1.87291 | 0.139978 |
| 7 | 16961200 | 16962199 | 30.00176 | 51.37145 | 34.87422 | 15.29317 | 0.939522 | 0.970355 | 0.139986 |
| 12 | 24376200 | 24378999 | 69.90609 | 111.3677 | 90.11777 | 37.71233 | 0.642006 | 2.191036 | 0.139987 |
| 2 | 165834000 | 165835999 | 10.76328 | 18.86296 | 47.25611 | 20.87434 | -0.9303 | 0.63807 | 0.139991 |
| 8 | 107235800 | 107239599 | 16.4621 | 27.38743 | 72.63976 | 31.53138 | -1.02102 | 1.238992 | 0.139995 |
| 3 | 129007200 | 129012799 | 43.69381 | 68.7551 | 183.8634 | 76.28098 | -1.09561 | 2.508676 | 0.139996 |
| 7 | 20008400 | 20009199 | 12.1594 | 23.38268 | 13.44083 | 5.242544 | 1.325062 | -0.08949 | 0.14001 |
| 14 | 45730200 | 45736999 | 73.55853 | 117.0573 | 92.12422 | 38.49322 | 0.680377 | 2.230122 | 0.140013 |
| 5 | 115650200 | 115659999 | 56.43196 | 87.91764 | 196.2088 | 81.4363 | -0.84638 | 2.702499 | 0.140018 |
| 11 | 106580600 | 106581999 | 10.40529 | 18.38957 | 43.53269 | 19.2568 | -0.84563 | 0.52126 | 0.140022 |
| 16 | 9295000 | 9296599 | 12.47554 | 21.83136 | 51.01152 | 22.50501 | -0.83069 | 0.677611 | 0.140047 |
| 15 | 73857000 | 73862999 | 32.89556 | 52.69196 | 121.7403 | 51.3401 | -0.8601 | 1.993813 | 0.14005 |
| 2 | 172384800 | 172398199 | 92.49639 | 139.5632 | 317.2358 | 131.106 | -0.90984 | 3.343626 | 0.14005 |
| 12 | 108517200 | 108521999 | 25.44988 | 41.34863 | 94.01172 | 40.32451 | -0.81383 | 1.580622 | 0.14005 |
| 11 | 20198400 | 20199999 | 7.690703 | 13.99629 | 37.61611 | 16.19182 | -0.99898 | 0.294692 | 0.140061 |
| 8 | 116083200 | 116085599 | 10.11888 | 17.83285 | 45.62543 | 20.19396 | -0.95892 | 0.552916 | 0.140077 |

| | | | | | | | | | |
|---|---|---|---|---|---|---|---|---|---|
| 7 | 137449600 | 137450399 | 4.407634 | 8.557006 | 28.02059 | 10.5165 | -1.10843 | -0.25926 | 0.140089 |
| 17 | 7832000 | 7835199 | 26.45981 | 45.44163 | 34.22784 | 15.0132 | 0.790913 | 0.929573 | 0.140093 |
| 17 | 18339000 | 18345999 | 147.2887 | 224.5187 | 170.1341 | 70.53239 | 0.685946 | 3.141284 | 0.140109 |
| 11 | 121176200 | 121177999 | 7.770266 | 14.33402 | 36.2985 | 15.34735 | -0.89573 | 0.213606 | 0.140112 |
| 2 | 119448600 | 119449599 | 4.421659 | 8.545589 | 28.1662 | 10.80225 | -1.14724 | -0.18296 | 0.14012 |
| 3 | 142663600 | 142664399 | 4.421659 | 8.545589 | 28.1662 | 10.80225 | -1.14724 | -0.18296 | 0.14012 |
| X | 115304200 | 115304999 | 12.56211 | 24.36864 | 14.60597 | 5.50534 | 1.310868 | -0.19057 | 0.140133 |
| 12 | 114285400 | 114288999 | 43.40599 | 71.59833 | 45.60136 | 19.68032 | 1.028898 | 1.385659 | 0.140139 |
| 4 | 121355000 | 121358799 | 77.66829 | 123.1101 | 101.1726 | 42.09645 | 0.612482 | 2.35128 | 0.140145 |
| 3 | 108061400 | 108063399 | 7.631858 | 14.05209 | 35.69903 | 15.13278 | -0.90268 | 0.223388 | 0.140146 |
| 15 | 84174200 | 84175999 | 7.915057 | 13.89026 | 47.74676 | 21.11535 | -1.383 | 0.601658 | 0.14015 |
| 6 | 121060600 | 121068999 | 51.45858 | 80.85821 | 180.0016 | 74.66497 | -0.83311 | 2.541434 | 0.140157 |
| 1 | 136012000 | 136017599 | 21.89369 | 35.44732 | 99.88116 | 42.73236 | -1.12707 | 1.649276 | 0.140162 |
| 6 | 139951000 | 139952399 | 18.88492 | 34.07426 | 22.65118 | 9.855639 | 0.992683 | 0.390859 | 0.140164 |
| 2 | 94980400 | 94982599 | 22.02984 | 38.7796 | 26.74379 | 11.83959 | 0.921347 | 0.600494 | 0.140166 |
| 7 | 9594800 | 9596599 | 16.63372 | 31.35105 | 17.3402 | 7.087845 | 1.320607 | 0.113832 | 0.140179 |
| 4 | 115752600 | 115756999 | 25.51321 | 41.32406 | 102.3911 | 43.81909 | -0.94181 | 1.646954 | 0.14018 |
| 14 | 102123600 | 102125399 | 21.09577 | 34.86135 | 77.41588 | 33.44912 | -0.76686 | 1.334212 | 0.14019 |
| 16 | 20807000 | 20808199 | 27.14965 | 46.51609 | 34.76787 | 15.24728 | 0.801538 | 0.954522 | 0.140205 |
| 5 | 136635600 | 136637199 | 7.621994 | 14.06142 | 36.85088 | 15.57951 | -0.94486 | 0.212644 | 0.140208 |
| 10 | 88002000 | 88002599 | 9.47951 | 18.53722 | 10.38455 | 3.538527 | 1.534485 | -0.58427 | 0.140226 |
| X | 33349800 | 33351999 | 22.5789 | 39.84807 | 25.32099 | 11.20397 | 1.040338 | 0.575836 | 0.140227 |
| 1 | 131452400 | 131456199 | 19.58166 | 31.89179 | 91.1651 | 39.14482 | -1.14013 | 1.530908 | 0.140235 |
| 3 | 31132200 | 31133999 | 7.154326 | 12.91589 | 39.59339 | 17.21517 | -1.1968 | 0.342299 | 0.140235 |
| 11 | 119402800 | 119415799 | 90.02375 | 136.4273 | 284.6543 | 117.8031 | -0.78096 | 3.253547 | 0.14024 |
| X | 118081200 | 118083399 | 46.42524 | 75.94647 | 50.60858 | 21.76563 | 0.959145 | 1.499104 | 0.14024 |
| 12 | 69175000 | 69178399 | 19.55156 | 32.64176 | 72.21481 | 31.41738 | -0.76063 | 1.206876 | 0.14025 |
| 14 | 123219200 | 123221599 | 10.77716 | 18.85222 | 48.33435 | 21.31713 | -0.96351 | 0.670634 | 0.14027 |
| 9 | 39130000 | 39130999 | 11.09955 | 21.47168 | 16.32092 | 6.232241 | 0.953637 | -0.16081 | 0.140272 |
| 1 | 182831400 | 182836999 | 43.46976 | 69.24337 | 138.2098 | 57.82798 | -0.65856 | 2.183326 | 0.140303 |
| 6 | 132530200 | 132530999 | 10.71853 | 20.95763 | 8.800577 | 3.008007 | 1.936667 | -0.56129 | 0.140306 |
| X | 3720000 | 3723599 | 46.16095 | 76.19259 | 42.30019 | 18.26183 | 1.226703 | 1.385546 | 0.14031 |
| 7 | 65024600 | 65025399 | 4.532087 | 8.799015 | 26.79474 | 10.05279 | -1.00384 | -0.2613 | 0.140312 |
| 12 | 108758200 | 108760799 | 12.72837 | 21.92513 | 55.77522 | 24.46666 | -0.95624 | 0.874729 | 0.140317 |
| X | 147670200 | 147671399 | 25.27192 | 43.99044 | 30.84018 | 13.55642 | 0.897234 | 0.787314 | 0.140319 |
| 18 | 34043200 | 34045399 | 13.43078 | 23.34672 | 52.6797 | 23.13648 | -0.78158 | 0.782049 | 0.140335 |
| 10 | 117790600 | 117794799 | 28.10908 | 45.3767 | 105.4129 | 44.95559 | -0.85422 | 1.73784 | 0.140342 |
| X | 31949200 | 31952599 | 27.70291 | 48.08514 | 29.87321 | 13.10793 | 1.070088 | 0.833839 | 0.14035 |
| 9 | 7539800 | 7540399 | 5.071456 | 9.81509 | 29.55619 | 11.26181 | -1.00934 | -0.19181 | 0.140357 |
| 1 | 93022000 | 93031599 | 63.1349 | 97.64675 | 222.1189 | 92.26289 | -0.88214 | 2.835392 | 0.140379 |
| 9 | 115990400 | 115995599 | 46.10929 | 73.36563 | 142.9271 | 59.70906 | -0.62539 | 2.223925 | 0.140392 |
| 17 | 55641400 | 55642199 | 10.98302 | 21.29903 | 15.81379 | 5.968979 | 1.002922 | -0.20139 | 0.140399 |
| 5 | 150129800 | 150132199 | 12.81251 | 22.24211 | 57.04703 | 25.03345 | -0.96601 | 0.801646 | 0.140402 |
| 9 | 120368000 | 120369399 | 8.495211 | 15.275 | 39.52118 | 17.26493 | -0.95879 | 0.37591 | 0.14041 |
| 9 | 119618200 | 119621999 | 28.6148 | 46.1116 | 105.6565 | 44.95642 | -0.83735 | 1.792354 | 0.140421 |
| 7 | 13212600 | 13217599 | 77.0787 | 122.5732 | 91.6544 | 38.26032 | 0.752983 | 2.254347 | 0.140422 |
| 15 | 22012400 | 22014199 | 25.29405 | 43.71339 | 33.59053 | 14.73648 | 0.763922 | 0.867941 | 0.140457 |

| | | | | | | | | | |
|---|---|---|---|---|---|---|---|---|---|
| 4 | 46745600 | 46748399 | 12.93286 | 22.14983 | 58.38668 | 25.60442 | -1.00897 | 0.931483 | 0.140472 |
| 3 | 133653400 | 133654599 | 13.57045 | 23.61308 | 53.78663 | 23.63746 | -0.79514 | 0.768738 | 0.140473 |
| 15 | 63792800 | 63796999 | 16.36594 | 27.16048 | 77.10853 | 33.41432 | -1.11928 | 1.269558 | 0.140497 |
| 2 | 37009000 | 37010599 | 38.13028 | 62.79747 | 49.93554 | 21.53654 | 0.708865 | 1.394007 | 0.140508 |
| 12 | 22503000 | 22505999 | 50.0483 | 80.9254 | 62.25165 | 26.60454 | 0.739221 | 1.705327 | 0.14051 |
| 3 | 118223200 | 118225399 | 9.418478 | 16.85344 | 44.22682 | 19.39949 | -0.98398 | 0.419599 | 0.140516 |
| 14 | 31691000 | 31693199 | 11.92546 | 20.66384 | 56.95119 | 24.98702 | -1.06962 | 0.815989 | 0.14052 |
| 9 | 48391400 | 48392199 | 3.822543 | 7.447854 | 24.83462 | 9.026981 | -1.08981 | -0.38949 | 0.140549 |
| 16 | 64929600 | 64930599 | 11.4748 | 22.27958 | 12.63404 | 4.737713 | 1.393681 | -0.229 | 0.140556 |
| 12 | 112975200 | 112978999 | 19.90517 | 32.72319 | 81.46773 | 35.10918 | -0.93364 | 1.396804 | 0.140566 |
| 11 | 94429400 | 94430799 | 5.411587 | 10.39595 | 29.08761 | 11.37876 | -0.94105 | -0.10237 | 0.140581 |
| 14 | 70252400 | 70253799 | 5.784631 | 10.91611 | 32.42219 | 13.22209 | -1.08167 | 0.046357 | 0.140591 |
| 14 | 51513200 | 51514999 | 16.40495 | 30.25801 | 20.88572 | 8.832522 | 0.965924 | 0.246404 | 0.140592 |
| 7 | 9815800 | 9817999 | 30.44634 | 52.02417 | 31.59732 | 13.85385 | 1.099122 | 0.988338 | 0.140611 |
| 7 | 132414000 | 132415999 | 9.082613 | 15.97584 | 45.64211 | 20.20911 | -1.11781 | 0.569885 | 0.140618 |
| 6 | 122461800 | 122463999 | 10.25578 | 18.10867 | 44.57435 | 19.72121 | -0.90234 | 0.530884 | 0.140641 |
| 18 | 54599000 | 54599999 | 13.50218 | 25.79364 | 17.60908 | 6.997612 | 1.054082 | -0.00472 | 0.140649 |
| 5 | 26709400 | 26710999 | 24.31447 | 42.28967 | 28.58393 | 12.55827 | 0.949585 | 0.78985 | 0.140651 |
| X | 107159400 | 107161599 | 23.36876 | 41.21822 | 25.14739 | 11.12844 | 1.098315 | 0.586043 | 0.14067 |
| 4 | 46597400 | 46621199 | 119.5015 | 175.9097 | 430.8031 | 177.4071 | -1.05 | 3.799917 | 0.140679 |
| 7 | 13666400 | 13668399 | 51.99035 | 83.8063 | 65.14071 | 27.72853 | 0.719287 | 1.800171 | 0.140686 |
| 3 | 142235800 | 142236999 | 7.281731 | 13.56942 | 36.04157 | 15.01834 | -0.94831 | 0.148653 | 0.140696 |
| 13 | 62618200 | 62621799 | 50.5037 | 82.07045 | 54.56682 | 23.40889 | 0.95568 | 1.603749 | 0.140712 |
| 13 | 55977800 | 55997399 | 159.1718 | 231.8246 | 471.8251 | 194.4489 | -0.79797 | 3.970133 | 0.140713 |
| 4 | 156153200 | 156156799 | 18.29366 | 30.55469 | 70.33513 | 30.60785 | -0.81756 | 1.197614 | 0.140728 |
| 3 | 159108000 | 159109199 | 12.7753 | 22.34159 | 51.03254 | 22.50209 | -0.79827 | 0.68843 | 0.140741 |
| 15 | 46600200 | 46601799 | 21.24242 | 38.15165 | 20.63446 | 9.022928 | 1.283289 | 0.429392 | 0.140753 |
| 13 | 68654200 | 68656399 | 29.30699 | 51.31114 | 22.4307 | 9.900083 | 1.574408 | 0.711533 | 0.140784 |
| 2 | 134267000 | 134267399 | 4.78836 | 9.307015 | 27.64731 | 10.27897 | -0.95647 | -0.28767 | 0.140798 |
| 11 | 86739400 | 86740999 | 28.01265 | 47.90357 | 36.26266 | 15.90038 | 0.782917 | 0.97697 | 0.140841 |
| 18 | 58011000 | 58013999 | 14.23815 | 24.2029 | 63.80379 | 27.96308 | -1.01127 | 1.010301 | 0.140863 |
| X | 24560200 | 24560999 | 23.77482 | 41.60572 | 29.08013 | 12.82979 | 0.901812 | 0.700206 | 0.140877 |
| 16 | 94999400 | 95003199 | 21.02203 | 34.94483 | 76.81545 | 33.32501 | -0.75164 | 1.262583 | 0.14089 |
| 1 | 5756000 | 5756799 | 10.03867 | 19.56063 | 13.64199 | 4.954509 | 1.14165 | -0.35152 | 0.140898 |
| 3 | 54761800 | 54764199 | 18.47379 | 33.72268 | 21.20687 | 9.093869 | 1.085575 | 0.320462 | 0.140909 |
| 4 | 135446800 | 135458599 | 100.2156 | 151.5961 | 291.2707 | 120.4721 | -0.66439 | 3.292816 | 0.140931 |
| 14 | 67454200 | 67477999 | 196.3416 | 281.4272 | 544.9379 | 225.0526 | -0.74887 | 4.211054 | 0.140939 |
| 6 | 130838600 | 130843799 | 45.7411 | 74.50254 | 47.31948 | 20.31881 | 1.02382 | 1.554983 | 0.14096 |
| 9 | 106021200 | 106035599 | 90.07603 | 136.4624 | 287.0228 | 118.7712 | -0.79294 | 3.260091 | 0.140992 |
| 8 | 4827800 | 4829799 | 11.36871 | 19.95264 | 46.70355 | 20.65532 | -0.83284 | 0.617885 | 0.141003 |
| 2 | 78783400 | 78784799 | 5.735284 | 10.96142 | 34.24008 | 13.59308 | -1.12019 | -0.04119 | 0.141005 |
| 9 | 78346800 | 78351599 | 27.3695 | 44.43217 | 94.18995 | 40.37732 | -0.71418 | 1.60096 | 0.14101 |
| 15 | 48163200 | 48164199 | 13.25964 | 25.21956 | 18.11238 | 7.26463 | 0.969858 | 0.022916 | 0.141031 |
| 18 | 29149000 | 29150399 | 13.25964 | 25.21956 | 18.11238 | 7.26463 | 0.969858 | 0.022916 | 0.141031 |
| 12 | 20952600 | 20956999 | 148.2831 | 227.49 | 140.1511 | 58.18205 | 0.991495 | 3.013048 | 0.141035 |
| 8 | 121795600 | 121814399 | 152.1218 | 222.0302 | 455.2701 | 187.5845 | -0.80571 | 3.937136 | 0.141048 |
| 11 | 34660000 | 34662599 | 16.72984 | 28.38494 | 62.82545 | 27.52207 | -0.76041 | 1.036107 | 0.141065 |

| | | | | | | | | | |
|---|---|---|---|---|---|---|---|---|---|
| 12 | 82215800 | 82217599 | 10.35349 | 18.06648 | 50.98793 | 22.44741 | -1.10163 | 0.709303 | 0.141091 |
| 9 | 59623800 | 59627799 | 22.82807 | 37.28151 | 85.81886 | 36.8862 | -0.82592 | 1.494326 | 0.141113 |
| 1 | 43484600 | 43486399 | 9.505805 | 16.78037 | 44.06946 | 19.49878 | -0.99511 | 0.52899 | 0.141129 |
| 7 | 10041000 | 10043599 | 46.18784 | 76.33508 | 41.11981 | 17.76801 | 1.270086 | 1.369084 | 0.141137 |
| 2 | 23263600 | 23264999 | 21.13202 | 37.64834 | 23.3346 | 10.27328 | 1.082385 | 0.485312 | 0.141138 |
| 18 | 78797400 | 78800199 | 20.10297 | 32.91376 | 88.57111 | 38.10459 | -1.04796 | 1.453989 | 0.141154 |
| 13 | 66711600 | 66715799 | 130.7734 | 201.1357 | 147.462 | 61.23831 | 0.743516 | 2.958686 | 0.141156 |
| 15 | 33198800 | 33200399 | 18.22044 | 34.03011 | 16.85441 | 6.996426 | 1.460287 | 0.181113 | 0.141159 |
| 1 | 117591000 | 117593999 | 54.51037 | 87.84396 | 67.35547 | 28.66077 | 0.738399 | 1.81153 | 0.141164 |
| 10 | 121539000 | 121543999 | 28.99608 | 47.00591 | 97.75734 | 41.86269 | -0.68863 | 1.635083 | 0.141166 |
| 1 | 3095800 | 3097399 | 19.27227 | 34.16786 | 25.41675 | 11.2361 | 0.816482 | 0.505229 | 0.141168 |
| 1 | 76502200 | 76503199 | 14.69373 | 27.50342 | 19.47935 | 8.056161 | 0.952967 | 0.142888 | 0.14117 |
| 11 | 45916400 | 45918599 | 11.38921 | 19.90989 | 51.55485 | 22.72497 | -0.97824 | 0.686854 | 0.141172 |
| 17 | 8544000 | 8549999 | 33.21309 | 53.19735 | 121.0212 | 51.03217 | -0.83792 | 1.99644 | 0.141184 |
| 1 | 91135600 | 91140199 | 31.2582 | 50.16669 | 117.7048 | 49.78524 | -0.87889 | 1.92877 | 0.141186 |
| 11 | 110705600 | 110706599 | 10.65484 | 20.6725 | 14.61267 | 5.503292 | 1.075104 | -0.2178 | 0.141205 |
| 8 | 96578000 | 96580199 | 11.0483 | 19.43614 | 45.62444 | 20.20071 | -0.83634 | 0.58034 | 0.141213 |
| 10 | 64044600 | 64046799 | 31.74917 | 54.70928 | 30.35737 | 13.31688 | 1.230566 | 0.917824 | 0.141217 |
| 5 | 93361800 | 93363999 | 10.53815 | 18.67404 | 44.0416 | 19.47156 | -0.83899 | 0.492056 | 0.141225 |
| 7 | 14434600 | 14439999 | 101.7643 | 160.0928 | 104.5608 | 43.37862 | 0.934909 | 2.536139 | 0.141226 |
| 1 | 121515800 | 121516799 | 5.741514 | 10.95573 | 32.32792 | 12.87761 | -1.04271 | -0.03367 | 0.141261 |
| 16 | 18947600 | 18948399 | 14.45368 | 27.57925 | 16.89854 | 6.731791 | 1.205407 | 0.010299 | 0.14127 |
| 5 | 28370000 | 28376399 | 47.56057 | 74.79651 | 179.6463 | 74.52582 | -0.94154 | 2.522137 | 0.141274 |
| 10 | 94128200 | 94128799 | 3.936754 | 7.680674 | 24.59134 | 8.782365 | -1.00889 | -0.45224 | 0.141296 |
| 7 | 25014000 | 25017399 | 22.39385 | 36.46513 | 88.74379 | 38.10258 | -0.90897 | 1.538453 | 0.141298 |
| 18 | 72709000 | 72710999 | 25.91012 | 42.13324 | 90.57878 | 38.87107 | -0.73217 | 1.565293 | 0.141303 |
| 12 | 22119600 | 22124599 | 125.9439 | 193.6701 | 152.7899 | 63.44923 | 0.637632 | 2.960201 | 0.141315 |
| 7 | 22982400 | 22983599 | 34.46405 | 57.27587 | 43.94428 | 19.06132 | 0.762071 | 1.275043 | 0.141316 |
| 7 | 26198400 | 26207599 | 187.6447 | 281.0368 | 218.8213 | 90.25891 | 0.625192 | 3.500561 | 0.14133 |
| 4 | 135079800 | 135081199 | 5.689491 | 10.63594 | 34.3664 | 14.24369 | -1.22132 | 0.111723 | 0.14133 |
| 4 | 134770800 | 134799799 | 184.2141 | 264.1398 | 547.3545 | 226.0378 | -0.84614 | 4.205077 | 0.141346 |
| 17 | 39792400 | 39801799 | 51.97776 | 84.23732 | 53.47334 | 22.89401 | 1.018681 | 1.657479 | 0.141371 |
| 8 | 88385200 | 88395599 | 48.28478 | 75.45984 | 199.7985 | 82.89551 | -1.08909 | 2.642641 | 0.141377 |
| 1 | 115946600 | 115949399 | 33.90477 | 57.97654 | 30.79595 | 13.50329 | 1.291415 | 0.99295 | 0.141379 |
| 12 | 101741200 | 101744599 | 15.50131 | 26.25516 | 64.93579 | 28.43166 | -0.92007 | 1.047951 | 0.141382 |
| 11 | 11332400 | 11334199 | 22.11268 | 39.99975 | 19.08556 | 8.277846 | 1.470384 | 0.393089 | 0.141389 |
| 11 | 113197600 | 113247399 | 475.2591 | 634.9513 | 1103.356 | 460.7376 | -0.69508 | 5.292992 | 0.141391 |
| 14 | 34114600 | 34121999 | 59.02541 | 92.44725 | 183.4232 | 76.08434 | -0.67096 | 2.607486 | 0.141392 |
| 4 | 136979800 | 136983199 | 37.74616 | 60.34356 | 127.8166 | 53.75056 | -0.73832 | 2.060979 | 0.141394 |
| 11 | 49519800 | 49529799 | 56.497 | 87.96229 | 201.7416 | 83.74074 | -0.88647 | 2.713277 | 0.141406 |
| 4 | 139985200 | 140031599 | 484.5951 | 645.5424 | 1133.134 | 473.3607 | -0.71384 | 5.339575 | 0.141413 |
| 15 | 93094400 | 93095599 | 13.70777 | 23.43658 | 58.91003 | 25.83034 | -0.94109 | 0.952514 | 0.141428 |
| 7 | 11175200 | 11176399 | 26.69553 | 46.07973 | 33.90345 | 14.87321 | 0.825904 | 0.884328 | 0.141437 |
| 7 | 115020000 | 115022999 | 15.05775 | 25.45311 | 64.24974 | 28.11429 | -0.94935 | 1.064589 | 0.141442 |
| 3 | 120769000 | 120771199 | 9.089655 | 16.3584 | 39.52139 | 17.25139 | -0.85996 | 0.374509 | 0.141457 |
| 19 | 22859000 | 22863599 | 15.47127 | 25.93938 | 72.47334 | 31.61432 | -1.09593 | 1.148058 | 0.141473 |
| 5 | 32870600 | 32871799 | 4.963147 | 9.599524 | 27.70075 | 10.58543 | -0.95269 | -0.1901 | 0.141478 |

| | | | | | | | | | |
|---|---|---|---|---|---|---|---|---|---|
| 13 | 52009000 | 52011399 | 9.458467 | 16.81901 | 42.5211 | 18.75034 | -0.93557 | 0.458004 | 0.141519 |
| 7 | 30607600 | 30611399 | 17.65197 | 29.12074 | 78.43597 | 33.8527 | -1.04449 | 1.346484 | 0.141525 |
| 6 | 130482600 | 130485399 | 53.02681 | 86.35364 | 47.95544 | 20.59053 | 1.216719 | 1.573059 | 0.141531 |
| 2 | 160772800 | 160781399 | 45.37064 | 71.91119 | 154.6319 | 64.3359 | -0.77317 | 2.340395 | 0.141546 |
| 11 | 11556400 | 11559999 | 87.93087 | 138.3865 | 112.9137 | 46.84641 | 0.614722 | 2.521267 | 0.141548 |
| 2 | 34394200 | 34396399 | 10.56198 | 18.64593 | 44.05058 | 19.49019 | -0.84349 | 0.534646 | 0.141558 |
| 12 | 13161400 | 13162999 | 26.54759 | 45.91667 | 33.3517 | 14.63206 | 0.844937 | 0.865508 | 0.141561 |
| 8 | 108018800 | 108021399 | 9.958909 | 17.31373 | 57.42576 | 25.22288 | -1.33427 | 0.768401 | 0.141566 |
| 17 | 29777400 | 29780199 | 19.5035 | 32.25806 | 81.38251 | 35.18406 | -0.95024 | 1.321565 | 0.141581 |
| X | 65802000 | 65804999 | 28.82226 | 49.42597 | 35.55603 | 15.59405 | 0.856658 | 0.954364 | 0.141582 |
| 8 | 62328600 | 62331799 | 18.88992 | 31.65625 | 68.82467 | 30.01379 | -0.7355 | 1.162302 | 0.141599 |
| 13 | 23134800 | 23137199 | 36.73139 | 62.90248 | 28.04397 | 12.29807 | 1.543247 | 0.99078 | 0.141601 |
| 11 | 6338600 | 6342799 | 21.18397 | 34.75541 | 82.01214 | 35.3137 | -0.85778 | 1.42607 | 0.141601 |
| 7 | 115716800 | 115721199 | 32.04159 | 51.4466 | 114.2928 | 48.37449 | -0.79949 | 1.915671 | 0.141611 |
| 8 | 89751400 | 89752999 | 7.433869 | 13.14116 | 45.12905 | 19.96203 | -1.37808 | 0.502862 | 0.141616 |
| 8 | 30118400 | 30120599 | 25.0299 | 43.78587 | 27.67773 | 12.20724 | 1.045779 | 0.708665 | 0.141628 |
| 17 | 20770600 | 20772399 | 20.98471 | 37.26055 | 23.96825 | 10.58218 | 1.025914 | 0.503491 | 0.141632 |
| 2 | 168705600 | 168709799 | 26.92267 | 43.6261 | 96.01725 | 41.10462 | -0.77067 | 1.638635 | 0.141649 |
| 5 | 95433200 | 95433599 | 9.861136 | 19.22501 | 13.0554 | 4.708257 | 1.188122 | -0.38034 | 0.141655 |
| 7 | 89036200 | 89037399 | 4.79811 | 9.292225 | 27.16865 | 10.32007 | -0.96252 | -0.21255 | 0.141661 |
| 11 | 98664600 | 98667199 | 10.3706 | 18.06008 | 52.04948 | 22.88588 | -1.13205 | 0.740209 | 0.141667 |
| 3 | 58483000 | 58484799 | 9.322302 | 16.54655 | 43.54733 | 19.23254 | -0.9948 | 0.470307 | 0.141668 |
| 18 | 77374800 | 77378399 | 13.69231 | 23.47522 | 59.0276 | 25.88768 | -0.94121 | 0.923907 | 0.141671 |
| 6 | 59572200 | 59579399 | 94.84432 | 148.6959 | 118.5879 | 49.1855 | 0.643387 | 2.600402 | 0.141674 |
| 17 | 38875600 | 38878199 | 48.86865 | 79.6523 | 51.81765 | 22.25598 | 0.990018 | 1.553807 | 0.141685 |
| 1 | 182389400 | 182392999 | 15.55017 | 26.20841 | 65.31017 | 28.55287 | -0.93109 | 1.093475 | 0.141691 |
| 7 | 99219600 | 99223399 | 15.55017 | 26.20841 | 65.31017 | 28.55287 | -0.93109 | 1.093475 | 0.141691 |
| 1 | 110431800 | 110432599 | 11.86106 | 22.87188 | 16.73344 | 6.475402 | 0.990203 | -0.10798 | 0.141704 |
| 7 | 10602000 | 10608399 | 88.08153 | 139.7079 | 91.43404 | 38.0695 | 0.941368 | 2.339989 | 0.141715 |
| 4 | 126495400 | 126497199 | 8.449649 | 15.32096 | 38.60954 | 16.69619 | -0.91151 | 0.323375 | 0.141717 |
| 11 | 5462400 | 5464399 | 8.449649 | 15.32096 | 38.60954 | 16.69619 | -0.91151 | 0.323375 | 0.141717 |
| 8 | 55999000 | 56000199 | 17.53283 | 31.65807 | 21.70571 | 9.435527 | 0.948895 | 0.373466 | 0.141722 |
| 5 | 32152200 | 32153799 | 14.28402 | 26.8859 | 18.40661 | 7.541023 | 1.011979 | 0.101537 | 0.141724 |
| 3 | 4458800 | 4462999 | 62.12872 | 99.59505 | 76.95781 | 32.49299 | 0.718213 | 1.987951 | 0.14174 |
| 3 | 41140600 | 41142199 | 16.09595 | 30.13637 | 18.81289 | 7.7769 | 1.1343 | 0.152424 | 0.141765 |
| 6 | 137987000 | 137987199 | 1.575778 | 3.075444 | 17.0949 | 5.378379 | -1.60955 | -0.9662 | 0.141782 |
| 7 | 25818600 | 25820799 | 58.02565 | 93.12096 | 75.3765 | 31.8791 | 0.6532 | 1.948411 | 0.141784 |
| 1 | 47592600 | 47593599 | 16.73999 | 30.33062 | 21.37415 | 9.251518 | 0.913567 | 0.347719 | 0.141792 |
| 7 | 45785200 | 45786599 | 6.121591 | 11.35973 | 34.19827 | 14.34333 | -1.13335 | 0.164292 | 0.141793 |
| 7 | 79835800 | 79837599 | 7.603736 | 13.69051 | 39.55073 | 17.25431 | -1.11513 | 0.36149 | 0.141808 |
| 9 | 118989800 | 118992399 | 18.41964 | 30.89077 | 68.73426 | 29.98822 | -0.76871 | 1.154177 | 0.141811 |
| 13 | 52998800 | 53009999 | 62.51935 | 96.86768 | 209.6873 | 87.09829 | -0.80867 | 2.801641 | 0.141815 |
| 1 | 116813400 | 116813999 | 12.63975 | 24.17509 | 17.3617 | 6.878005 | 0.986242 | -0.02277 | 0.141818 |
| 9 | 117218600 | 117219999 | 14.2407 | 27.28916 | 16.48889 | 6.496533 | 1.240035 | -0.02686 | 0.141823 |
| 4 | 74289400 | 74290799 | 17.42014 | 31.56546 | 22.88777 | 9.90569 | 0.873105 | 0.356975 | 0.141835 |
| 10 | 60925000 | 60925999 | 4.414412 | 8.551487 | 27.7802 | 10.53884 | -1.11159 | -0.2221 | 0.141836 |
| 12 | 110586600 | 110587799 | 4.414412 | 8.551487 | 27.7802 | 10.53884 | -1.11159 | -0.2221 | 0.141836 |

| | | | | | | | | | |
|---|---|---|---|---|---|---|---|---|---|
| 8 | 93162800 | 93165199 | 11.85083 | 20.70512 | 49.41905 | 21.77378 | -0.86117 | 0.696643 | 0.141838 |
| 16 | 21710600 | 21713199 | 11.40024 | 19.90512 | 51.55685 | 22.70654 | -0.97873 | 0.705431 | 0.141845 |
| 1 | 189656200 | 189656999 | 5.527714 | 10.33949 | 33.26825 | 13.77507 | -1.21353 | 0.105463 | 0.141849 |
| 4 | 57383400 | 57387199 | 18.05505 | 30.38572 | 67.22284 | 29.37604 | -0.76042 | 1.115379 | 0.141875 |
| 6 | 95694000 | 95695999 | 9.96415 | 17.57469 | 46.68272 | 20.6585 | -1.01233 | 0.543778 | 0.141876 |
| 13 | 69926200 | 69927599 | 8.19379 | 14.67528 | 40.54784 | 17.77218 | -1.05568 | 0.404209 | 0.141877 |
| 1 | 44897600 | 44901199 | 61.76321 | 98.90434 | 79.53998 | 33.52919 | 0.658889 | 2.020436 | 0.141884 |
| 7 | 22315600 | 22315999 | 8.720092 | 17.04384 | 11.51001 | 3.971828 | 1.251724 | -0.55139 | 0.141898 |
| 10 | 118636000 | 118641999 | 43.70793 | 69.39511 | 151.4517 | 63.12134 | -0.79209 | 2.287216 | 0.141898 |
| 9 | 35797600 | 35799199 | 20.67519 | 36.57351 | 25.8838 | 11.44767 | 0.886912 | 0.539608 | 0.1419 |
| 14 | 50700200 | 50701399 | 14.96019 | 27.56484 | 19.14514 | 8.107784 | 0.955487 | 0.235411 | 0.141904 |
| 7 | 45103800 | 45111799 | 35.67156 | 56.67729 | 149.7286 | 62.44901 | -1.06596 | 2.259687 | 0.141932 |
| 4 | 127217600 | 127220599 | 16.03064 | 26.65557 | 75.468 | 32.74715 | -1.11504 | 1.244641 | 0.141933 |
| 3 | 152464200 | 152465599 | 4.024346 | 7.758209 | 29.1123 | 11.27406 | -1.34528 | -0.15136 | 0.141937 |
| 13 | 47201400 | 47205999 | 25.6261 | 41.58371 | 93.00264 | 39.8555 | -0.79176 | 1.606589 | 0.141944 |
| 12 | 105893200 | 105919799 | 211.4962 | 301.8706 | 566.3207 | 234.041 | -0.70936 | 4.273931 | 0.141946 |
| 2 | 105496800 | 105500999 | 36.54587 | 58.23842 | 137.7912 | 57.68597 | -0.90269 | 2.163651 | 0.141956 |
| 4 | 22763200 | 22766799 | 51.77932 | 83.71342 | 57.14327 | 24.41724 | 0.91099 | 1.711071 | 0.141965 |
| 2 | 178259800 | 178265999 | 42.65793 | 67.966 | 136.7978 | 57.25559 | -0.67021 | 2.174817 | 0.141967 |
| 11 | 96082400 | 96091599 | 56.5449 | 88.31075 | 198.0853 | 82.19028 | -0.85094 | 2.657871 | 0.14197 |
| 15 | 81862800 | 81866799 | 26.78521 | 43.35788 | 99.40231 | 42.52087 | -0.83086 | 1.660866 | 0.141983 |
| 5 | 112490400 | 112491599 | 5.853916 | 11.2167 | 30.16469 | 11.88787 | -0.89496 | -0.07081 | 0.141984 |
| 3 | 52948400 | 52949799 | 8.785365 | 15.48158 | 46.68187 | 20.66185 | -1.19427 | 0.547736 | 0.141988 |
| 1 | 6379800 | 6380199 | 8.228809 | 16.05655 | 12.8991 | 4.596759 | 0.964832 | -0.44406 | 0.141993 |
| 15 | 78836400 | 78838599 | 9.608838 | 17.08623 | 41.48746 | 18.2947 | -0.87744 | 0.458926 | 0.141994 |
| 9 | 121760800 | 121761599 | 10.29851 | 20.01627 | 13.96362 | 5.192971 | 1.112439 | -0.26439 | 0.141998 |
| 13 | 8228600 | 8229999 | 15.3466 | 29.10641 | 17.37312 | 7.00997 | 1.227238 | 0.060472 | 0.142011 |
| 7 | 19525800 | 19532399 | 41.39197 | 65.89389 | 138.4682 | 57.88774 | -0.73356 | 2.201746 | 0.142019 |
| 2 | 156566800 | 156568999 | 8.633729 | 15.55253 | 39.02119 | 17.01299 | -0.91282 | 0.365012 | 0.14202 |
| 12 | 20031600 | 20040799 | 298.4819 | 429.4568 | 324.7302 | 134.0298 | 0.612628 | 4.158727 | 0.142021 |
| 10 | 108059000 | 108060599 | 10.85824 | 19.2148 | 44.56888 | 19.7101 | -0.81589 | 0.510876 | 0.142024 |
| 10 | 75008600 | 75021799 | 80.98391 | 123.556 | 261.9776 | 108.63 | -0.79618 | 3.11174 | 0.142032 |
| 5 | 139520800 | 139522599 | 10.63181 | 18.58797 | 48.86575 | 21.54172 | -0.99948 | 0.679263 | 0.142038 |
| 6 | 120977800 | 120978799 | 4.823075 | 9.254794 | 28.95501 | 11.36416 | -1.10398 | -0.09606 | 0.142048 |
| 15 | 25762400 | 25766399 | 16.00873 | 26.66683 | 74.44857 | 32.34323 | -1.09477 | 1.223116 | 0.142053 |
| X | 6084000 | 6085199 | 14.18543 | 26.95999 | 16.7897 | 6.744907 | 1.172795 | 0.035262 | 0.142056 |
| 2 | 174935200 | 174936599 | 11.84004 | 22.71483 | 16.09778 | 6.3216 | 1.015597 | -0.0649 | 0.142058 |
| 1 | 136116400 | 136120199 | 14.60494 | 25.06964 | 57.45343 | 25.19971 | -0.80773 | 0.915561 | 0.142077 |
| 18 | 5132200 | 5134999 | 15.23442 | 26.10748 | 59.11365 | 25.92449 | -0.7909 | 0.934269 | 0.142083 |
| 9 | 21634400 | 21653399 | 108.574 | 162.2487 | 346.9545 | 143.0488 | -0.83429 | 3.534745 | 0.142087 |
| 4 | 151598400 | 151607999 | 70.2346 | 108.3593 | 222.9334 | 92.60239 | -0.74018 | 2.88428 | 0.14211 |
| 4 | 152061000 | 152069199 | 35.15191 | 55.96336 | 147.541 | 61.68579 | -1.06005 | 2.195133 | 0.142113 |
| 6 | 92679200 | 92683399 | 18.98175 | 31.53699 | 73.55906 | 31.90091 | -0.83675 | 1.261147 | 0.142114 |
| 15 | 20657400 | 20658999 | 16.86235 | 30.94974 | 20.13767 | 8.573103 | 1.043307 | 0.273796 | 0.142122 |
| 13 | 12114400 | 12117999 | 17.43361 | 29.31862 | 70.00716 | 30.58606 | -0.87026 | 1.121122 | 0.142122 |
| 8 | 118991600 | 118994399 | 15.58368 | 26.19545 | 67.47407 | 29.47523 | -0.9789 | 1.118415 | 0.142124 |
| 5 | 121997200 | 122008599 | 84.19882 | 128.036 | 270.5841 | 112.0953 | -0.79531 | 3.181551 | 0.142126 |

| | | | | | | | | | |
|---|---|---|---|---|---|---|---|---|---|
| 11 | 116723800 | 116731799 | 37.28101 | 59.56709 | 127.5898 | 53.61332 | -0.75526 | 2.078272 | 0.142139 |
| 18 | 74573200 | 74590599 | 141.7393 | 209.814 | 393.0921 | 161.8872 | -0.65626 | 3.708889 | 0.142147 |
| 17 | 6921000 | 6923999 | 10.58664 | 18.61775 | 45.1068 | 19.97286 | -0.88183 | 0.579909 | 0.14215 |
| 2 | 138071000 | 138071599 | 3.262785 | 6.37387 | 22.88433 | 7.995071 | -1.13934 | -0.5403 | 0.142151 |
| 13 | 55325400 | 55331399 | 22.94439 | 37.58924 | 85.26938 | 36.69378 | -0.80209 | 1.448701 | 0.142155 |
| 11 | 54413200 | 54414599 | 5.544171 | 10.40285 | 36.76715 | 15.14317 | -1.34284 | 0.089832 | 0.142155 |
| 14 | 27149200 | 27150799 | 21.04099 | 34.95867 | 75.52051 | 32.75179 | -0.7266 | 1.269019 | 0.142155 |
| 11 | 73631000 | 73631999 | 13.53655 | 25.712 | 15.41224 | 6.199066 | 1.225763 | 0.028243 | 0.142156 |
| 5 | 144491600 | 144493599 | 11.09415 | 19.36531 | 49.40815 | 21.75643 | -0.95669 | 0.707748 | 0.142172 |
| 12 | 52832000 | 52832399 | 2.543802 | 4.973235 | 20.57909 | 7.055356 | -1.31452 | -0.62964 | 0.142174 |
| 16 | 35758600 | 35759799 | 5.870653 | 11.22713 | 31.26425 | 12.38952 | -0.95294 | -0.05152 | 0.142177 |
| 5 | 38825400 | 38851999 | 321.4597 | 443.6399 | 775.7806 | 322.3418 | -0.65694 | 4.780427 | 0.142179 |
| 5 | 65853800 | 65856599 | 12.93304 | 22.16806 | 58.96501 | 25.85972 | -1.02188 | 0.923534 | 0.142184 |
| 14 | 11272200 | 11272799 | 10.17746 | 19.84909 | 13.18946 | 4.733475 | 1.226595 | -0.39061 | 0.142199 |
| 2 | 126380000 | 126381399 | 3.624656 | 6.997688 | 31.54764 | 12.1603 | -1.60086 | -0.16622 | 0.142239 |
| 11 | 73618200 | 73621199 | 26.52577 | 46.70645 | 21.94751 | 9.715735 | 1.471322 | 0.616336 | 0.142241 |
| 4 | 125519800 | 125521999 | 12.81201 | 22.25239 | 53.20656 | 23.35552 | -0.86494 | 0.791067 | 0.142251 |
| 14 | 23542800 | 23543599 | 4.358659 | 8.487006 | 25.37274 | 9.297832 | -0.9453 | -0.35063 | 0.142259 |
| 9 | 65452200 | 65455599 | 15.22581 | 25.70956 | 65.3585 | 28.59014 | -0.95968 | 1.075198 | 0.142264 |
| 6 | 81961200 | 81966399 | 31.66635 | 50.98073 | 109.9168 | 46.705 | -0.75161 | 1.82728 | 0.142291 |
| 6 | 34090000 | 34094399 | 21.02212 | 34.96655 | 75.64385 | 32.83072 | -0.72853 | 1.255184 | 0.142292 |
| 5 | 130066600 | 130067599 | 5.286975 | 10.12192 | 29.49262 | 11.65093 | -1.01233 | -0.07031 | 0.142297 |
| 8 | 61143600 | 61143799 | 8.36729 | 16.35519 | 11.34841 | 3.910387 | 1.217738 | -0.56018 | 0.142307 |
| 4 | 135305000 | 135307999 | 21.76511 | 35.85125 | 79.03105 | 34.08742 | -0.75758 | 1.375714 | 0.142317 |
| 3 | 98000200 | 98002199 | 6.341653 | 11.59422 | 36.62542 | 15.6752 | -1.22388 | 0.256985 | 0.142335 |
| 12 | 104978400 | 104981199 | 11.42966 | 19.89184 | 52.62578 | 23.1301 | -1.00963 | 0.754392 | 0.142338 |
| 11 | 82180800 | 82182199 | 4.528119 | 8.802962 | 27.03731 | 10.03588 | -1.00199 | -0.29966 | 0.142339 |
| 3 | 3250000 | 3254399 | 56.41402 | 92.03703 | 42.32522 | 18.18515 | 1.489748 | 1.554153 | 0.142352 |
| 4 | 134336000 | 134356199 | 137.1761 | 201.9592 | 422.4766 | 173.9785 | -0.82257 | 3.798783 | 0.142355 |
| 2 | 5849400 | 5850399 | 7.523497 | 13.75388 | 37.70322 | 16.13804 | -1.02214 | 0.266169 | 0.142361 |
| 10 | 83642800 | 83645799 | 17.45321 | 29.31865 | 66.47536 | 29.03309 | -0.79569 | 1.130728 | 0.142387 |
| 14 | 6320600 | 6321399 | 4.927095 | 9.561856 | 27.99086 | 10.52974 | -0.95115 | -0.2421 | 0.142388 |
| 4 | 139897400 | 139901999 | 21.36042 | 34.97252 | 83.46179 | 35.90648 | -0.87521 | 1.454416 | 0.142393 |
| 13 | 12784200 | 12784999 | 14.42353 | 27.261 | 18.48948 | 7.515347 | 1.03519 | 0.077243 | 0.142397 |
| 16 | 82339400 | 82340199 | 14.42353 | 27.261 | 18.48948 | 7.515347 | 1.03519 | 0.077243 | 0.142397 |
| 13 | 66445400 | 66454199 | 198.974 | 298.6123 | 199.1599 | 82.187 | 0.851 | 3.463789 | 0.142416 |
| 16 | 21649600 | 21651599 | 7.305963 | 13.53285 | 38.0248 | 15.98784 | -1.03805 | 0.193807 | 0.142431 |
| 2 | 68140200 | 68141399 | 21.21928 | 38.51738 | 19.14605 | 8.268047 | 1.415617 | 0.367275 | 0.142432 |
| 16 | 61057000 | 61061399 | 38.11442 | 63.56041 | 39.51504 | 17.17885 | 1.064517 | 1.24537 | 0.142441 |
| 6 | 121167000 | 121172799 | 30.6081 | 49.14847 | 115.9269 | 49.07142 | -0.88558 | 1.911428 | 0.142453 |
| 1 | 26869200 | 26870999 | 24.43636 | 43.21112 | 22.68045 | 10.03165 | 1.315056 | 0.560626 | 0.142454 |
| 15 | 78175800 | 78178599 | 13.08314 | 22.42448 | 58.42438 | 25.62255 | -0.99212 | 0.924238 | 0.142461 |
| 2 | 169487600 | 169489199 | 10.4207 | 18.36355 | 44.57905 | 19.73109 | -0.88352 | 0.554588 | 0.142462 |
| 4 | 148307000 | 148309199 | 11.20115 | 19.66705 | 49.35307 | 21.83507 | -0.9331 | 0.610987 | 0.142492 |
| 7 | 129954800 | 129956199 | 9.159854 | 16.27071 | 41.4943 | 18.31396 | -0.94852 | 0.463096 | 0.142501 |
| 11 | 45893600 | 45894999 | 16.06718 | 29.2714 | 20.68993 | 8.894256 | 0.915873 | 0.311576 | 0.142503 |
| X | 32719000 | 32720199 | 8.196363 | 16.02115 | 11.84883 | 4.082406 | 1.126631 | -0.56143 | 0.142508 |

| | | | | | | | | | |
|---|---|---|---|---|---|---|---|---|---|
| 6 | 131711000 | 131712999 | 41.94584 | 68.53453 | 54.36333 | 23.37001 | 0.707185 | 1.506149 | 0.14251 |
| 5 | 120120400 | 120125199 | 28.28065 | 45.62832 | 107.6981 | 45.8973 | -0.87809 | 1.755168 | 0.142517 |
| 19 | 10737400 | 10741599 | 17.03756 | 28.4975 | 74.31654 | 32.36969 | -0.99702 | 1.179828 | 0.14253 |
| 4 | 133958800 | 133966199 | 30.17206 | 48.35925 | 121.2767 | 51.19255 | -0.977 | 1.971057 | 0.142539 |
| 4 | 111095800 | 111096399 | 10.22822 | 19.87687 | 15.38906 | 5.729534 | 0.961358 | -0.25772 | 0.142539 |
| 10 | 83764800 | 83765999 | 5.393968 | 10.42002 | 33.26948 | 12.78647 | -1.10664 | -0.15316 | 0.142543 |
| X | 134183600 | 134184799 | 16.56089 | 31.06055 | 18.34383 | 7.560824 | 1.217261 | 0.14542 | 0.142547 |
| 18 | 17366200 | 17370199 | 28.22005 | 48.42191 | 35.51427 | 15.57647 | 0.828902 | 0.945543 | 0.142561 |
| 7 | 3619800 | 3621799 | 6.666031 | 12.17253 | 38.73919 | 16.60497 | -1.23689 | 0.268458 | 0.142565 |
| 2 | 32045800 | 32055799 | 56.22948 | 87.72752 | 192.4453 | 79.85856 | -0.81999 | 2.67675 | 0.142571 |
| 3 | 112533800 | 112535399 | 19.65652 | 35.01215 | 24.83074 | 10.93395 | 0.888799 | 0.478519 | 0.142573 |
| 14 | 42169000 | 42170199 | 13.7074 | 26.62705 | 12.66406 | 4.730802 | 1.651545 | -0.21302 | 0.142588 |
| 4 | 74857800 | 74858799 | 11.22358 | 19.66387 | 47.26197 | 20.87176 | -0.8708 | 0.646319 | 0.142599 |
| 4 | 119057600 | 119060599 | 11.90718 | 20.66471 | 53.70059 | 23.56197 | -0.98447 | 0.798717 | 0.1426 |
| 6 | 140610000 | 140612399 | 14.63517 | 25.02876 | 59.57903 | 26.12424 | -0.86319 | 0.951613 | 0.142602 |
| 13 | 21564400 | 21565399 | 7.29052 | 13.56024 | 35.40531 | 14.80033 | -0.92736 | 0.159976 | 0.142605 |
| 11 | 48503800 | 48504799 | 11.72499 | 22.44892 | 17.38216 | 6.865513 | 0.882248 | -0.04239 | 0.142612 |
| 16 | 3430800 | 3433599 | 57.53947 | 92.32704 | 74.60623 | 31.54622 | 0.655414 | 1.952219 | 0.142621 |
| 9 | 114913400 | 114930599 | 160.8127 | 234.9307 | 451.933 | 186.1918 | -0.71239 | 3.923808 | 0.142639 |
| 14 | 30798800 | 30801599 | 51.02907 | 82.25306 | 62.91411 | 26.77918 | 0.742307 | 1.797384 | 0.14264 |
| 6 | 132187400 | 132187999 | 12.48024 | 24.18302 | 14.36222 | 5.44156 | 1.315705 | -0.17687 | 0.142643 |
| 19 | 9426400 | 9428199 | 18.30454 | 33.13314 | 22.13217 | 9.591275 | 0.989262 | 0.36703 | 0.142658 |
| 7 | 38177600 | 38178599 | 26.82956 | 46.47855 | 32.84868 | 14.41208 | 0.884754 | 0.853256 | 0.142659 |
| 11 | 70559200 | 70562799 | 14.40777 | 24.4318 | 62.61413 | 27.42496 | -0.97081 | 1.03304 | 0.142666 |
| 5 | 34148800 | 34150799 | 28.92098 | 49.29073 | 37.75249 | 16.54915 | 0.765341 | 1.01123 | 0.142674 |
| 9 | 73569600 | 73570599 | 11.83806 | 22.63197 | 16.39036 | 6.501268 | 0.97224 | -0.03078 | 0.142683 |
| 16 | 91834800 | 91839399 | 36.7259 | 58.87877 | 119.3319 | 50.39132 | -0.67008 | 1.969723 | 0.142695 |
| 7 | 33240200 | 33244599 | 142.581 | 217.0439 | 173.0282 | 71.70346 | 0.611219 | 3.168058 | 0.142699 |
| 11 | 73667600 | 73669199 | 13.69091 | 26.14674 | 17.60335 | 6.999547 | 1.073313 | -0.00083 | 0.142707 |
| 14 | 42347000 | 42349999 | 26.82472 | 46.48336 | 32.18895 | 14.12275 | 0.913964 | 0.848408 | 0.142709 |
| 9 | 83844400 | 83846199 | 9.374244 | 16.94986 | 41.2365 | 17.8948 | -0.86466 | 0.352035 | 0.14273 |
| 15 | 100998600 | 100999999 | 9.374244 | 16.94986 | 41.2365 | 17.8948 | -0.86466 | 0.352035 | 0.14273 |
| 11 | 74432000 | 74432599 | 4.6487 | 8.936626 | 29.00066 | 11.32209 | -1.14838 | -0.11405 | 0.142734 |
| 16 | 4279400 | 4280199 | 4.6487 | 8.936626 | 29.00066 | 11.32209 | -1.14838 | -0.11405 | 0.142734 |
| X | 114468400 | 114469199 | 18.90687 | 33.85503 | 23.09203 | 10.12319 | 0.949169 | 0.438673 | 0.14274 |
| 12 | 99381600 | 99382799 | 6.005704 | 11.50263 | 31.37032 | 12.37816 | -0.91728 | -0.06385 | 0.142748 |
| 7 | 24305600 | 24312399 | 35.54938 | 56.74407 | 137.4354 | 57.66947 | -0.93382 | 2.108495 | 0.142751 |
| 7 | 35757000 | 35758799 | 11.38728 | 19.9127 | 48.87323 | 21.5486 | -0.90112 | 0.679342 | 0.142764 |
| 9 | 37524400 | 37525599 | 3.357629 | 6.439794 | 31.07551 | 12.21282 | -1.72278 | -0.10321 | 0.142767 |
| 6 | 118601000 | 118602399 | 6.994369 | 13.02056 | 34.30566 | 14.31986 | -0.93828 | 0.150311 | 0.14277 |
| 6 | 11035800 | 11038599 | 25.43088 | 41.36135 | 91.16774 | 39.12637 | -0.76792 | 1.561532 | 0.142771 |
| 19 | 27100800 | 27101999 | 5.696003 | 10.96876 | 32.35183 | 12.56832 | -1.00814 | -0.11622 | 0.142773 |
| 1 | 19079000 | 19080399 | 29.2934 | 49.69134 | 36.43081 | 15.95837 | 0.827549 | 1.053321 | 0.142774 |
| 7 | 96231000 | 96232199 | 5.619881 | 10.6883 | 35.75451 | 14.34234 | -1.23157 | -0.00491 | 0.142783 |
| 1 | 63484200 | 63485999 | 11.16903 | 19.36763 | 56.3668 | 24.73118 | -1.14739 | 0.805446 | 0.142792 |
| 8 | 72993000 | 72994999 | 11.35802 | 19.93325 | 47.75783 | 21.12103 | -0.86654 | 0.617721 | 0.142796 |
| 6 | 71934000 | 71934399 | 11.78117 | 22.63346 | 17.37992 | 6.797811 | 0.907287 | -0.07249 | 0.142812 |

| | | | | | | | | | |
|---|---|---|---|---|---|---|---|---|---|
| 9 | 97565800 | 97569199 | 12.29313 | 21.5546 | 50.45868 | 22.29822 | -0.83347 | 0.642162 | 0.142832 |
| 8 | 75956200 | 75956799 | 3.761703 | 7.332917 | 23.66178 | 8.545463 | -1.03309 | -0.41418 | 0.142845 |
| 10 | 7095600 | 7100199 | 20.80478 | 34.30533 | 79.0739 | 34.12102 | -0.82115 | 1.359044 | 0.142851 |
| 2 | 93436600 | 93438799 | 11.52645 | 20.19148 | 47.79063 | 21.10255 | -0.849 | 0.64969 | 0.142857 |
| 1 | 153992800 | 153994399 | 8.500267 | 15.25429 | 39.51306 | 17.28981 | -0.96185 | 0.389883 | 0.142862 |
| 13 | 108106600 | 108108799 | 8.500267 | 15.25429 | 39.51306 | 17.28981 | -0.96185 | 0.389883 | 0.142862 |
| 5 | 134779400 | 134781399 | 10.25635 | 18.11224 | 43.5381 | 19.2622 | -0.86803 | 0.528874 | 0.142868 |
| 14 | 68214000 | 68215199 | 17.92565 | 32.76258 | 20.55398 | 8.800275 | 1.089958 | 0.306635 | 0.142875 |
| 7 | 7369000 | 7375399 | 78.77479 | 125.3195 | 89.23242 | 37.26897 | 0.824081 | 2.244817 | 0.142878 |
| 1 | 86604800 | 86606799 | 10.4764 | 18.29251 | 49.39387 | 21.75425 | -1.03777 | 0.701068 | 0.142885 |
| 14 | 60610200 | 60613599 | 19.07513 | 31.8613 | 71.02558 | 30.90916 | -0.7715 | 1.199576 | 0.142889 |
| 3 | 95858000 | 95859399 | 6.417134 | 11.9549 | 34.85938 | 14.53263 | -1.08185 | 0.141207 | 0.142895 |
| 6 | 39110800 | 39113199 | 24.9679 | 43.86961 | 25.52148 | 11.29807 | 1.164739 | 0.638338 | 0.142899 |
| 18 | 87092600 | 87093599 | 10.99198 | 21.37358 | 13.3798 | 4.958063 | 1.269015 | -0.27239 | 0.142914 |
| 6 | 138662000 | 138665799 | 51.8896 | 83.80528 | 57.06313 | 24.3506 | 0.912905 | 1.740143 | 0.142934 |
| 16 | 4917800 | 4928599 | 58.27458 | 90.06694 | 223.7514 | 92.94131 | -1.00986 | 2.847014 | 0.142945 |
| 7 | 7410800 | 7415799 | 71.04899 | 113.2344 | 88.85277 | 37.20385 | 0.686923 | 2.17884 | 0.142945 |
| 11 | 87602600 | 87610399 | 48.87083 | 77.17075 | 165.692 | 68.79925 | -0.77545 | 2.435003 | 0.142955 |
| 15 | 76420000 | 76427199 | 26.67373 | 42.71881 | 124.8509 | 52.63869 | -1.19852 | 1.993958 | 0.142957 |
| 4 | 136970200 | 136995999 | 178.672 | 257.0033 | 532.9067 | 219.9625 | -0.8425 | 4.158273 | 0.142985 |
| 2 | 122120000 | 122124199 | 17.29358 | 28.68998 | 76.08135 | 32.96129 | -1.02134 | 1.275495 | 0.142999 |
| 15 | 101832800 | 101833599 | 4.560079 | 8.770541 | 28.47737 | 11.10222 | -1.148 | -0.1238 | 0.143015 |
| 12 | 70926400 | 70931599 | 29.2894 | 47.09013 | 111.0655 | 47.10046 | -0.88316 | 1.869518 | 0.143017 |
| 3 | 110589800 | 110590999 | 7.996102 | 14.51337 | 37.57388 | 16.22763 | -0.94845 | 0.313634 | 0.143022 |
| 1 | 147429200 | 147430799 | 17.49423 | 31.69403 | 22.96553 | 9.941577 | 0.874003 | 0.359318 | 0.143029 |
| 4 | 89220400 | 89222199 | 10.08677 | 17.85679 | 46.16704 | 20.41522 | -0.97149 | 0.502381 | 0.143032 |
| 6 | 26505600 | 26506399 | 14.59659 | 28.07535 | 14.62308 | 5.700225 | 1.465281 | -0.0621 | 0.143032 |
| 10 | 20585800 | 20587399 | 17.17109 | 32.02687 | 18.74041 | 7.797178 | 1.219738 | 0.185377 | 0.143036 |
| 1 | 103340600 | 103342999 | 30.39125 | 52.16239 | 33.75342 | 14.80452 | 1.008875 | 0.950236 | 0.14305 |
| 8 | 74508400 | 74509599 | 13.60425 | 25.99804 | 16.31906 | 6.480018 | 1.174208 | -0.0108 | 0.143057 |
| 6 | 83808200 | 83809399 | 5.262819 | 10.15021 | 28.70838 | 11.10366 | -0.9412 | -0.14345 | 0.143057 |
| 7 | 59688400 | 59690799 | 24.27932 | 42.75716 | 24.9604 | 11.04916 | 1.160657 | 0.607108 | 0.143063 |
| 3 | 13677800 | 13679399 | 20.06175 | 35.50801 | 25.80663 | 11.41227 | 0.848905 | 0.528585 | 0.143067 |
| 2 | 165278400 | 165281999 | 14.92378 | 25.19855 | 67.01952 | 29.31637 | -1.02465 | 1.075397 | 0.143067 |
| 2 | 164302800 | 164303799 | 3.336835 | 6.452136 | 29.8202 | 11.41053 | -1.62476 | -0.19842 | 0.143079 |
| 14 | 79927800 | 79928999 | 12.66074 | 24.26254 | 16.48939 | 6.495943 | 1.071249 | -0.04496 | 0.143095 |
| 1 | 115541000 | 115542599 | 16.97064 | 31.2709 | 20.86993 | 8.837559 | 1.012946 | 0.256684 | 0.143096 |
| 10 | 62159800 | 62171399 | 60.48881 | 93.94694 | 205.7253 | 85.42559 | -0.82259 | 2.757947 | 0.143105 |
| 9 | 123044200 | 123045799 | 11.5605 | 20.15988 | 49.94858 | 21.98107 | -0.91486 | 0.724718 | 0.143113 |
| 13 | 11957600 | 11958599 | 5.694856 | 10.9703 | 31.12715 | 12.0807 | -0.95123 | -0.12215 | 0.143117 |
| 10 | 62096800 | 62098399 | 5.788678 | 10.56681 | 37.64987 | 16.14498 | -1.3978 | 0.26357 | 0.143121 |
| 12 | 61767200 | 61768999 | 14.39195 | 27.6979 | 13.34384 | 5.192859 | 1.577545 | -0.07361 | 0.143139 |
| 5 | 30934600 | 30935599 | 3.4441 | 6.653481 | 27.97882 | 10.75161 | -1.49508 | -0.1856 | 0.143148 |
| X | 54213200 | 54216799 | 32.76727 | 56.82279 | 26.48942 | 11.62237 | 1.482208 | 0.864214 | 0.14315 |
| 12 | 103645600 | 103646999 | 7.112895 | 13.32595 | 33.956 | 14.0216 | -0.87791 | 0.108925 | 0.143153 |
| 14 | 64401600 | 64402399 | 3.643241 | 7.105779 | 23.16196 | 8.306771 | -1.03843 | -0.44366 | 0.143165 |
| 10 | 67120200 | 67121199 | 4.780502 | 9.03606 | 31.83834 | 12.93896 | -1.32012 | 0.025419 | 0.143179 |

| | | | | | | | | | |
|---|---|---|---|---|---|---|---|---|---|
| 9 | 61064000 | 61067599 | 16.17227 | 27.2565 | 67.66287 | 29.58131 | -0.92582 | 1.096177 | 0.143207 |
| 7 | 19306800 | 19308199 | 6.641902 | 12.54107 | 33.07888 | 13.4735 | -0.91095 | 0.051158 | 0.143212 |
| 16 | 43581800 | 43582999 | 6.641902 | 12.54107 | 33.07888 | 13.4735 | -0.91095 | 0.051158 | 0.143212 |
| 1 | 61417400 | 61419799 | 16.04889 | 27.38418 | 61.2779 | 26.86424 | -0.77506 | 0.979614 | 0.143221 |
| 5 | 148916800 | 148932199 | 167.2569 | 244.7742 | 438.8424 | 180.7781 | -0.60841 | 3.896958 | 0.143222 |
| 11 | 113595400 | 113602999 | 31.77363 | 50.90929 | 121.6146 | 51.30923 | -0.90765 | 1.98337 | 0.143228 |
| 2 | 178499000 | 178503399 | 21.4062 | 35.36545 | 80.60523 | 34.82111 | -0.80449 | 1.337228 | 0.14324 |
| 11 | 3578800 | 3580399 | 12.09021 | 20.91326 | 55.83063 | 24.49409 | -1.0243 | 0.831874 | 0.143251 |
| 12 | 17600400 | 17609999 | 59.91597 | 94.06035 | 183.844 | 76.24961 | -0.64676 | 2.565241 | 0.143259 |
| 7 | 99327400 | 99330599 | 13.86157 | 23.72241 | 58.99018 | 25.8675 | -0.92563 | 0.941861 | 0.143259 |
| X | 3958000 | 3960799 | 24.69859 | 43.11925 | 30.08384 | 13.24846 | 0.904345 | 0.741815 | 0.143274 |
| 5 | 67252400 | 67267199 | 112.8883 | 169.8865 | 322.7238 | 133.2763 | -0.6545 | 3.399538 | 0.143286 |
| 16 | 17423400 | 17425199 | 11.84643 | 20.72278 | 48.34181 | 21.32053 | -0.82817 | 0.676113 | 0.143289 |
| 2 | 107721000 | 107722799 | 21.51692 | 38.71829 | 19.46826 | 8.499332 | 1.389005 | 0.417331 | 0.143295 |
| 11 | 113057400 | 113062399 | 26.51522 | 42.79836 | 102.2262 | 43.58951 | -0.89431 | 1.739159 | 0.143304 |
| 12 | 72714600 | 72717999 | 40.14535 | 65.77464 | 52.67445 | 22.66808 | 0.695817 | 1.463995 | 0.143307 |
| X | 125721600 | 125722799 | 17.51619 | 32.97811 | 16.72909 | 6.850765 | 1.442205 | 0.12699 | 0.143314 |
| 6 | 79250200 | 79251399 | 17.90142 | 32.41368 | 22.79447 | 9.874396 | 0.916086 | 0.364334 | 0.143328 |
| 2 | 179159400 | 179171999 | 80.24253 | 122.2747 | 267.6106 | 110.9328 | -0.84328 | 3.136944 | 0.143351 |
| 3 | 152840400 | 152842999 | 13.49295 | 23.24786 | 56.37997 | 24.73213 | -0.88778 | 0.874938 | 0.143357 |
| 19 | 24104400 | 24110399 | 73.9684 | 114.5676 | 214.071 | 88.91929 | -0.5971 | 2.812674 | 0.143367 |
| 2 | 91196600 | 91197399 | 14.64543 | 27.93716 | 16.89251 | 6.733441 | 1.223678 | 0.014171 | 0.143376 |
| 13 | 97064800 | 97069399 | 31.19481 | 50.20484 | 110.175 | 46.79212 | -0.77765 | 1.837415 | 0.143383 |
| 5 | 56732400 | 56733999 | 18.61397 | 33.67709 | 22.00326 | 9.54113 | 1.021027 | 0.374066 | 0.143415 |
| 17 | 19943800 | 19944999 | 29.41744 | 50.01647 | 37.17391 | 16.29237 | 0.807942 | 1.032323 | 0.143416 |
| 14 | 56075800 | 56078599 | 11.69933 | 20.443 | 48.88391 | 21.53971 | -0.86369 | 0.694064 | 0.143419 |
| 12 | 26518200 | 26520199 | 7.957294 | 14.18396 | 41.53903 | 18.2799 | -1.14359 | 0.432069 | 0.143421 |
| 8 | 17104000 | 17105999 | 29.925 | 50.54535 | 39.50123 | 17.28085 | 0.735544 | 1.094637 | 0.143426 |
| 7 | 142926200 | 142932999 | 35.58865 | 56.69996 | 138.3495 | 57.90124 | -0.94739 | 2.170747 | 0.143462 |
| 13 | 38562600 | 38564199 | 20.05919 | 36.05945 | 22.49976 | 9.831012 | 1.079583 | 0.420132 | 0.143483 |
| 6 | 48143200 | 48144999 | 17.03574 | 31.55165 | 19.70379 | 8.285944 | 1.114874 | 0.226244 | 0.143485 |
| 19 | 15591000 | 15592799 | 8.431609 | 14.98144 | 44.09747 | 19.45818 | -1.15415 | 0.458754 | 0.143491 |
| 5 | 29773800 | 29776599 | 13.60524 | 23.55083 | 54.82913 | 24.05524 | -0.82771 | 0.8308 | 0.143496 |
| 7 | 80039000 | 80043799 | 15.44153 | 25.55642 | 77.19273 | 33.38777 | -1.20843 | 1.301456 | 0.143502 |
| 2 | 58946600 | 58949999 | 23.06316 | 37.44808 | 91.8033 | 39.35957 | -0.92278 | 1.589927 | 0.143502 |
| 14 | 79482000 | 79489599 | 54.92769 | 86.54669 | 170.2083 | 70.62792 | -0.65152 | 2.487056 | 0.143502 |
| 18 | 69796800 | 69798599 | 6.966831 | 13.04878 | 34.46974 | 14.24024 | -0.92949 | 0.113063 | 0.143521 |
| 12 | 35491200 | 35491999 | 8.578178 | 15.61746 | 38.16123 | 16.41851 | -0.86253 | 0.299606 | 0.143526 |
| 5 | 146546000 | 146548399 | 10.40786 | 18.37659 | 44.05101 | 19.4898 | -0.8643 | 0.531729 | 0.143527 |
| 14 | 41751600 | 41752399 | 10.76238 | 21.03255 | 7.240654 | 2.507898 | 2.194828 | -0.5239 | 0.143528 |
| X | 148173600 | 148174799 | 21.21959 | 37.55994 | 25.33069 | 11.20165 | 0.956441 | 0.535771 | 0.143531 |
| 17 | 57929400 | 57930999 | 20.34663 | 36.97498 | 19.76293 | 8.522421 | 1.312981 | 0.355677 | 0.143538 |
| 16 | 44491200 | 44493199 | 10.22171 | 18.15507 | 42.49582 | 18.75758 | -0.82616 | 0.470923 | 0.143539 |
| 12 | 22215400 | 22217599 | 48.49098 | 78.73252 | 57.81394 | 24.7863 | 0.811652 | 1.617825 | 0.143541 |
| 16 | 75101400 | 75104799 | 13.08483 | 22.36018 | 59.38648 | 26.03819 | -1.02016 | 0.956049 | 0.143542 |
| 12 | 84662800 | 84665199 | 10.12529 | 17.82809 | 45.6258 | 20.19776 | -0.9598 | 0.563193 | 0.143547 |
| 6 | 73243800 | 73247399 | 13.88799 | 24.12909 | 54.8896 | 24.09927 | -0.79356 | 0.793068 | 0.143547 |

| | | | | | | | | | |
|---|---|---|---|---|---|---|---|---|---|
| 3 | 55337200 | 55338199 | 17.64141 | 31.81836 | 23.4609 | 10.21286 | 0.843894 | 0.38639 | 0.143551 |
| 2 | 76005200 | 76008399 | 34.88553 | 58.47956 | 41.52886 | 18.11375 | 0.873036 | 1.181634 | 0.143555 |
| X | 31275400 | 31278199 | 51.28221 | 82.95141 | 59.84994 | 25.58801 | 0.832053 | 1.700397 | 0.143595 |
| 7 | 31287400 | 31290999 | 82.90331 | 131.2894 | 101.6913 | 42.28641 | 0.696669 | 2.377752 | 0.143597 |
| 15 | 88447800 | 88461399 | 109.5409 | 164.6954 | 325.7498 | 134.4924 | -0.71388 | 3.417165 | 0.143602 |
| 7 | 107226600 | 107227599 | 12.19459 | 23.6013 | 15.73887 | 5.996528 | 1.144103 | -0.15661 | 0.143614 |
| 12 | 102709400 | 102710399 | 6.035773 | 11.4648 | 30.97084 | 12.45665 | -0.92823 | -9.15E-05 | 0.143621 |
| X | 21730200 | 21731599 | 25.31967 | 44.16112 | 29.72723 | 13.08351 | 0.95578 | 0.758873 | 0.143622 |
| 17 | 31981200 | 31984999 | 23.1731 | 37.79428 | 88.7658 | 38.1342 | -0.85631 | 1.515071 | 0.143625 |
| 6 | 149173600 | 149173999 | 8.393234 | 16.37636 | 13.40516 | 4.782304 | 0.936686 | -0.43584 | 0.143626 |
| 11 | 69065200 | 69067399 | 8.301285 | 15.0529 | 39.14505 | 16.92642 | -0.95654 | 0.322454 | 0.143645 |
| 8 | 105896800 | 105898399 | 10.16448 | 17.80783 | 47.78394 | 21.10178 | -1.02874 | 0.640718 | 0.143645 |
| 1 | 109366600 | 109369799 | 32.75312 | 55.99125 | 32.51692 | 14.25757 | 1.163313 | 0.992735 | 0.143662 |
| 1 | 167167000 | 167169399 | 9.12812 | 15.99753 | 49.34625 | 21.79789 | -1.22855 | 0.631367 | 0.14367 |
| 12 | 52500000 | 52502799 | 47.26826 | 76.77791 | 57.80542 | 24.78994 | 0.776194 | 1.606679 | 0.143674 |
| 9 | 57237400 | 57241999 | 43.45086 | 69.28048 | 135.904 | 56.93962 | -0.63205 | 2.151435 | 0.143676 |
| 16 | 13636400 | 13638999 | 16.22421 | 27.63479 | 61.2328 | 26.84067 | -0.76128 | 0.996927 | 0.143677 |
| 8 | 11858400 | 11861599 | 17.39822 | 29.39818 | 64.43318 | 28.1913 | -0.74661 | 1.076416 | 0.14368 |
| 9 | 117683800 | 117693999 | 73.28343 | 113.1998 | 220.8329 | 91.72933 | -0.66223 | 2.862668 | 0.143681 |
| 5 | 100717200 | 100718999 | 8.537956 | 15.25694 | 42.06899 | 18.4743 | -1.05561 | 0.419479 | 0.143683 |
| 5 | 103981400 | 103984799 | 15.03834 | 25.4355 | 63.64027 | 27.85265 | -0.93644 | 1.06092 | 0.143683 |
| 1 | 182542600 | 182544599 | 10.85341 | 19.22229 | 45.10002 | 19.94166 | -0.83191 | 0.502265 | 0.143692 |
| 4 | 56648000 | 56648599 | 3.458724 | 6.746044 | 23.11031 | 8.2859 | -1.10761 | -0.44519 | 0.143693 |
| 19 | 23734200 | 23737799 | 14.64057 | 24.38358 | 77.08909 | 33.4867 | -1.27342 | 1.223014 | 0.143709 |
| 1 | 117092200 | 117096199 | 40.00206 | 65.6558 | 52.13753 | 22.4538 | 0.708947 | 1.442144 | 0.143714 |
| 7 | 13959800 | 13962799 | 48.76747 | 79.10006 | 58.9468 | 25.25283 | 0.789062 | 1.638534 | 0.143716 |
| 18 | 73753000 | 73757799 | 25.32763 | 41.06869 | 95.83997 | 41.0488 | -0.85399 | 1.621945 | 0.143719 |
| 19 | 16109600 | 16111199 | 9.698336 | 16.78605 | 57.85421 | 25.38235 | -1.39027 | 0.818709 | 0.143723 |
| 3 | 31217000 | 31217599 | 5.308666 | 10.09663 | 30.38837 | 12.18925 | -1.07849 | -0.01661 | 0.143739 |
| 4 | 154884800 | 154889799 | 20.21737 | 32.95193 | 92.76555 | 39.84295 | -1.11749 | 1.520752 | 0.143749 |
| 13 | 54899200 | 54919999 | 152.8885 | 223.7577 | 442.4054 | 182.2452 | -0.74943 | 3.894677 | 0.143782 |
| 17 | 46763000 | 46775199 | 73.03751 | 112.2154 | 234.5623 | 97.39528 | -0.76768 | 2.966977 | 0.143785 |
| 5 | 137371800 | 137378199 | 32.05441 | 51.44613 | 117.3915 | 49.65513 | -0.83877 | 1.928531 | 0.143798 |
| 19 | 57311400 | 57313799 | 15.51521 | 26.24048 | 63.75554 | 27.90258 | -0.89449 | 1.06085 | 0.143803 |
| 4 | 139954000 | 139977999 | 159.3026 | 231.206 | 482.0177 | 198.7216 | -0.83752 | 4.023477 | 0.143808 |
| 6 | 6721600 | 6722399 | 6.999953 | 13.00007 | 34.23627 | 14.34367 | -0.9414 | 0.166636 | 0.143828 |
| 5 | 93909000 | 93910199 | 41.41302 | 67.76618 | 50.46302 | 21.70526 | 0.799631 | 1.483296 | 0.143831 |
| 14 | 65904200 | 65907799 | 16.1492 | 27.27319 | 64.89901 | 28.39055 | -0.86478 | 1.076344 | 0.143833 |
| 1 | 140698200 | 140700799 | 23.36063 | 41.03054 | 26.01201 | 11.51166 | 1.04118 | 0.633476 | 0.14384 |
| 12 | 23368200 | 23373599 | 81.2668 | 128.6585 | 103.5012 | 43.0324 | 0.641749 | 2.384031 | 0.14384 |
| 1 | 146277800 | 146279999 | 18.32333 | 32.66153 | 23.89446 | 10.51513 | 0.844607 | 0.461706 | 0.143841 |
| 14 | 92056400 | 92060599 | 99.52554 | 155.3242 | 127.7095 | 52.99363 | 0.594368 | 2.682932 | 0.143843 |
| 13 | 51461400 | 51471599 | 61.5769 | 96.14512 | 191.8008 | 79.58475 | -0.68232 | 2.66457 | 0.143868 |
| 10 | 118652600 | 118657199 | 24.11276 | 39.34606 | 91.55441 | 39.33921 | -0.84253 | 1.510446 | 0.143868 |
| 9 | 35955400 | 35956399 | 11.79239 | 22.70032 | 16.6901 | 6.490658 | 0.97633 | -0.09379 | 0.143883 |
| 12 | 23073400 | 23086599 | 288.6479 | 416.6632 | 321.2929 | 132.5523 | 0.588996 | 4.110279 | 0.143885 |
| 1 | 75238600 | 75241399 | 18.28702 | 30.55835 | 71.53971 | 31.14154 | -0.84188 | 1.192419 | 0.143888 |

| | | | | | | | | | |
|---|---|---|---|---|---|---|---|---|---|
| 16 | 90130600 | 90137399 | 47.27746 | 74.9947 | 153.9736 | 64.1144 | -0.70543 | 2.317106 | 0.143891 |
| X | 34748400 | 34749999 | 36.39699 | 60.63255 | 44.38805 | 19.28454 | 0.829663 | 1.256077 | 0.143899 |
| 8 | 126737400 | 126740799 | 14.97928 | 25.5205 | 61.15875 | 26.80874 | -0.87375 | 0.98969 | 0.1439 |
| 13 | 111729800 | 111730199 | 3.641394 | 7.10794 | 22.76506 | 8.061013 | -0.99676 | -0.49126 | 0.143904 |
| 2 | 163895600 | 163897199 | 7.290358 | 13.56387 | 34.29247 | 14.32811 | -0.88026 | 0.15739 | 0.143911 |
| 4 | 42771200 | 42773999 | 10.43362 | 18.35132 | 45.10679 | 19.97227 | -0.90243 | 0.577064 | 0.143925 |
| 2 | 31202200 | 31209599 | 44.3653 | 70.41276 | 149.8018 | 62.41057 | -0.75586 | 2.298773 | 0.143932 |
| 7 | 8183200 | 8184599 | 16.28755 | 30.4861 | 18.80728 | 7.778472 | 1.150727 | 0.156032 | 0.143941 |
| 18 | 20000600 | 20002599 | 16.28755 | 30.4861 | 18.80728 | 7.778472 | 1.150727 | 0.156032 | 0.143941 |
| Y | 44605000 | 44605399 | 7.274516 | 14.23073 | 5.942758 | 1.968173 | 1.981666 | -0.74266 | 0.143941 |
| 7 | 7558200 | 7559799 | 18.05083 | 32.53894 | 23.28956 | 10.14523 | 0.885711 | 0.391321 | 0.143947 |
| 2 | 23045400 | 23047799 | 12.03014 | 20.94542 | 51.57138 | 22.67247 | -0.90646 | 0.750264 | 0.143954 |
| 7 | 9711400 | 9713999 | 33.87453 | 57.46967 | 36.55536 | 16.01354 | 1.031625 | 1.07007 | 0.143955 |
| 4 | 14867000 | 14869599 | 35.1721 | 58.7299 | 43.23376 | 18.81093 | 0.821486 | 1.224199 | 0.143972 |
| 16 | 94038400 | 94042399 | 29.10243 | 46.88652 | 107.668 | 45.79691 | -0.84098 | 1.800357 | 0.143978 |
| 5 | 68689600 | 68690799 | 12.94286 | 24.74371 | 17.62622 | 6.991816 | 0.995358 | -0.01641 | 0.143988 |
| X | 3242600 | 3244599 | 51.30852 | 83.37647 | 53.39879 | 22.90744 | 1.00954 | 1.606837 | 0.143991 |
| 9 | 60991600 | 60995799 | 17.97987 | 30.0352 | 70.8954 | 30.85465 | -0.85364 | 1.194979 | 0.143996 |
| 8 | 86549800 | 86551199 | 24.44376 | 42.48453 | 32.38826 | 14.22338 | 0.776595 | 0.804871 | 0.14401 |
| 12 | 77284600 | 77286599 | 12.89265 | 22.18488 | 56.31456 | 24.70246 | -0.95355 | 0.885132 | 0.14401 |
| 12 | 84701400 | 84707799 | 38.40673 | 61.35701 | 128.6762 | 54.0589 | -0.7251 | 2.083624 | 0.144018 |
| 11 | 79595000 | 79597199 | 11.67276 | 20.47107 | 47.2419 | 20.88033 | -0.81269 | 0.631119 | 0.144031 |
| 5 | 110416600 | 110419399 | 11.57347 | 20.15547 | 51.55797 | 22.67021 | -0.96116 | 0.744409 | 0.144045 |
| 5 | 26169000 | 26169999 | 12.85787 | 24.52428 | 18.24225 | 7.27333 | 0.927345 | 0.001138 | 0.144045 |
| 4 | 49731400 | 49732799 | 10.37073 | 17.9838 | 53.59215 | 23.51381 | -1.18058 | 0.801991 | 0.144046 |
| 11 | 85779000 | 85784199 | 41.42502 | 65.84825 | 149.1899 | 62.25219 | -0.84428 | 2.249891 | 0.144052 |
| 7 | 11086000 | 11089799 | 68.8776 | 109.9702 | 84.49189 | 35.46631 | 0.720008 | 2.119738 | 0.144083 |
| 7 | 136855000 | 136858399 | 18.73317 | 30.97218 | 77.18506 | 33.36061 | -0.93252 | 1.320769 | 0.144096 |
| 5 | 26807200 | 26810199 | 21.8943 | 38.61922 | 26.19266 | 11.59091 | 0.94634 | 0.577307 | 0.1441 |
| 6 | 120793800 | 120796399 | 20.0538 | 33.41012 | 72.12451 | 31.3366 | -0.72532 | 1.231934 | 0.144102 |
| 9 | 40483600 | 40484999 | 6.467809 | 12.29888 | 32.21385 | 12.92668 | -0.88164 | -0.00134 | 0.144104 |
| 15 | 32395600 | 32398799 | 14.85329 | 25.23224 | 67.72604 | 29.68002 | -1.03742 | 1.022835 | 0.14411 |
| 9 | 7524200 | 7524999 | 13.04438 | 25.26632 | 14.34542 | 5.445093 | 1.377805 | -0.1642 | 0.144114 |
| 7 | 44301200 | 44318999 | 126.6311 | 187.6516 | 384.2067 | 158.2357 | -0.78278 | 3.688728 | 0.144144 |
| 6 | 125459600 | 125461599 | 8.103309 | 14.83403 | 37.20233 | 15.89472 | -0.89294 | 0.261977 | 0.144147 |
| 2 | 171836400 | 171847199 | 75.82865 | 116.5781 | 239.147 | 99.30761 | -0.73994 | 2.955029 | 0.144151 |
| 13 | 89501600 | 89502599 | 11.11293 | 21.44722 | 16.89716 | 6.523037 | 0.887891 | -0.12273 | 0.144153 |
| 14 | 56497000 | 56499999 | 18.80367 | 31.29437 | 71.82856 | 31.18707 | -0.81331 | 1.241004 | 0.144163 |
| 13 | 20902600 | 20904599 | 10.21157 | 18.16784 | 42.51682 | 18.74005 | -0.82446 | 0.459889 | 0.144163 |
| 4 | 100471000 | 100471799 | 11.30327 | 21.76972 | 16.92561 | 6.572743 | 0.899159 | -0.10153 | 0.144172 |
| 11 | 64759800 | 64760399 | 11.30327 | 21.76972 | 16.92561 | 6.572743 | 0.899159 | -0.10153 | 0.144172 |
| 14 | 42871200 | 42873199 | 27.04764 | 46.94866 | 31.62985 | 13.87889 | 0.953902 | 0.834094 | 0.144184 |
| 2 | 143861000 | 143861999 | 6.66027 | 12.52018 | 32.91605 | 13.51464 | -0.9157 | 0.080787 | 0.14419 |
| 7 | 10575600 | 10577199 | 42.905 | 69.98569 | 54.8263 | 23.55481 | 0.723532 | 1.53375 | 0.144198 |
| 1 | 120045000 | 120049599 | 19.10745 | 31.46665 | 81.56522 | 35.17717 | -0.99064 | 1.372297 | 0.144199 |
| 5 | 122009600 | 122031199 | 166.6776 | 240.1055 | 528.8645 | 218.2471 | -0.92746 | 4.135698 | 0.144199 |
| 10 | 126535000 | 126537599 | 20.59189 | 34.10406 | 75.28694 | 32.57901 | -0.75808 | 1.304181 | 0.144201 |

| | | | | | | | | | |
|---|---|---|---|---|---|---|---|---|---|
| 13 | 45425800 | 45428199 | 9.872257 | 17.71117 | 44.28721 | 19.38368 | -0.91245 | 0.408242 | 0.144208 |
| 17 | 67800400 | 67804799 | 24.80728 | 40.31728 | 90.81722 | 38.96333 | -0.79991 | 1.572378 | 0.144213 |
| X | 98108600 | 98111999 | 14.89978 | 25.71165 | 58.64173 | 25.72551 | -0.79925 | 0.870907 | 0.144214 |
| 12 | 109708200 | 109717799 | 64.12109 | 99.10585 | 219.3797 | 91.12538 | -0.84362 | 2.848251 | 0.144222 |
| 9 | 73300000 | 73301199 | 13.71008 | 26.6126 | 13.2644 | 4.984684 | 1.577409 | -0.19121 | 0.144223 |
| 12 | 60823400 | 60825999 | 33.83061 | 56.69687 | 42.67868 | 18.61231 | 0.789349 | 1.182959 | 0.144238 |
| 7 | 40974800 | 40978199 | 53.29646 | 85.8829 | 68.17487 | 29.00707 | 0.688424 | 1.812979 | 0.144238 |
| 14 | 21710400 | 21711199 | 4.118156 | 8.002475 | 26.26194 | 9.782466 | -1.10043 | -0.29362 | 0.144239 |
| 12 | 20585400 | 20586999 | 49.58144 | 80.10306 | 55.81486 | 23.82775 | 0.880238 | 1.726482 | 0.144243 |
| 12 | 22627200 | 22632599 | 47.62201 | 77.32668 | 51.07714 | 21.89912 | 0.96405 | 1.611206 | 0.144245 |
| 7 | 28751200 | 28756799 | 37.18495 | 59.25271 | 136.5355 | 57.15476 | -0.86475 | 2.166613 | 0.144252 |
| 14 | 42418400 | 42424599 | 63.26068 | 101.3966 | 76.56594 | 32.32122 | 0.751155 | 1.992937 | 0.144254 |
| 11 | 106866200 | 106869999 | 19.63432 | 32.55354 | 75.26289 | 32.59307 | -0.82438 | 1.288205 | 0.144262 |
| 7 | 128341000 | 128343399 | 11.98751 | 20.99268 | 48.32729 | 21.33396 | -0.80929 | 0.657812 | 0.144266 |
| 3 | 24714000 | 24715599 | 21.37917 | 38.14975 | 22.8848 | 10.06165 | 1.130082 | 0.474458 | 0.144272 |
| 8 | 108287200 | 108288999 | 7.263643 | 13.14242 | 38.06461 | 16.50451 | -1.11242 | 0.328854 | 0.144273 |
| 11 | 58990200 | 58992199 | 10.59234 | 18.61185 | 46.15526 | 20.43303 | -0.91571 | 0.591914 | 0.144281 |
| 17 | 65862200 | 65862599 | 2.836244 | 5.546178 | 21.38328 | 7.290022 | -1.20899 | -0.64407 | 0.144284 |
| 11 | 115037400 | 115038999 | 8.091983 | 14.8461 | 37.25936 | 15.87124 | -0.89086 | 0.249105 | 0.144299 |
| 1 | 81477800 | 81478199 | 9.552621 | 18.65811 | 5.100176 | 1.621061 | 2.619165 | -0.83257 | 0.144308 |
| X | 32209200 | 32212199 | 41.60681 | 69.77363 | 34.82068 | 15.19153 | 1.37997 | 1.195791 | 0.144309 |
| 8 | 123263600 | 123266799 | 16.247 | 27.1895 | 69.02094 | 30.07311 | -0.95798 | 1.169618 | 0.14432 |
| 7 | 59205200 | 59205599 | 9.713261 | 18.92989 | 12.44613 | 4.509768 | 1.230756 | -0.36611 | 0.14433 |
| 10 | 4203000 | 4204999 | 7.080564 | 12.93013 | 36.61283 | 15.69257 | -1.06918 | 0.268877 | 0.144337 |
| 4 | 35155400 | 35158199 | 16.04109 | 26.61624 | 75.25596 | 32.6072 | -1.11328 | 1.270744 | 0.144346 |
| 6 | 53263200 | 53265999 | 17.67721 | 29.50042 | 70.1829 | 30.52551 | -0.86482 | 1.205492 | 0.144373 |
| 5 | 35375000 | 35376199 | 21.31603 | 37.81836 | 24.61511 | 10.87516 | 1.008639 | 0.513022 | 0.144378 |
| 7 | 49556800 | 49571199 | 121.9362 | 182.3204 | 341.3123 | 140.7381 | -0.64178 | 3.525853 | 0.144383 |
| 15 | 55251200 | 55253199 | 23.35431 | 40.65248 | 30.84039 | 13.55651 | 0.783685 | 0.775332 | 0.144396 |
| 12 | 39091200 | 39092199 | 14.94909 | 28.03103 | 19.51132 | 8.045202 | 0.981437 | 0.134845 | 0.14441 |
| 11 | 78935600 | 78942999 | 60.35012 | 94.46716 | 183.1307 | 75.96756 | -0.63818 | 2.618928 | 0.144426 |
| 17 | 20002000 | 20004799 | 46.62344 | 77.71559 | 31.91362 | 13.87029 | 1.660126 | 1.269608 | 0.14443 |
| 15 | 36627200 | 36627799 | 9.134094 | 17.78518 | 14.448 | 5.295335 | 0.913256 | -0.32846 | 0.144441 |
| 4 | 152942600 | 152978199 | 391.3162 | 531.179 | 939.7148 | 391.4726 | -0.69786 | 5.039769 | 0.144442 |
| 7 | 61625600 | 61630799 | 64.28629 | 103.2068 | 70.62114 | 29.88492 | 0.895689 | 1.943494 | 0.144455 |
| 17 | 20120600 | 20127799 | 129.3881 | 200.9422 | 113.6699 | 47.20089 | 1.12778 | 2.779621 | 0.144465 |
| 17 | 20594200 | 20595799 | 21.53355 | 38.56972 | 21.77317 | 9.542667 | 1.220063 | 0.450667 | 0.144496 |
| 1 | 93463600 | 93468599 | 59.75692 | 95.86899 | 75.39388 | 31.8711 | 0.694225 | 1.961291 | 0.144523 |
| 10 | 126770400 | 126775399 | 51.41502 | 80.88322 | 171.797 | 71.27241 | -0.76413 | 2.517999 | 0.144532 |
| 7 | 41001400 | 41004999 | 49.95263 | 81.14588 | 55.68763 | 23.88341 | 0.909643 | 1.609603 | 0.144532 |
| 15 | 97314400 | 97316599 | 17.44121 | 29.31257 | 69.4106 | 30.31951 | -0.85827 | 1.126463 | 0.144533 |
| 10 | 85857400 | 85858399 | 7.31042 | 13.53517 | 34.72583 | 14.60948 | -0.9086 | 0.187288 | 0.144541 |
| 12 | 104558600 | 104574999 | 111.4093 | 167.2779 | 326.0839 | 134.5815 | -0.69466 | 3.444207 | 0.144545 |
| 19 | 35979000 | 35981599 | 18.12077 | 32.37273 | 23.36083 | 10.26095 | 0.866007 | 0.445244 | 0.144562 |
| 6 | 98491200 | 98492399 | 5.124802 | 9.897712 | 29.42218 | 11.32031 | -1.0054 | -0.16197 | 0.144565 |
| 2 | 172175000 | 172201599 | 219.8159 | 312.1398 | 594.0818 | 245.7255 | -0.73759 | 4.349487 | 0.144594 |
| 8 | 22270000 | 22271999 | 19.81468 | 35.29943 | 24.10145 | 10.6114 | 0.943111 | 0.475767 | 0.144596 |

| | | | | | | | | | |
|---|---|---|---|---|---|---|---|---|---|
| 5 | 109314800 | 109318399 | 95.85232 | 149.8785 | 123.1325 | 51.08431 | 0.597663 | 2.647204 | 0.144596 |
| 10 | 41809200 | 41811199 | 10.45745 | 18.31782 | 47.26594 | 20.86985 | -0.97242 | 0.646291 | 0.144598 |
| 11 | 51279600 | 51280999 | 5.160114 | 9.862789 | 29.98315 | 11.90086 | -1.07951 | -0.05513 | 0.144601 |
| 18 | 44586600 | 44588199 | 8.633368 | 15.54916 | 39.02446 | 17.01814 | -0.91326 | 0.367316 | 0.144604 |
| 13 | 63115200 | 63116799 | 10.97936 | 19.09042 | 52.06173 | 22.86883 | -1.05307 | 0.766377 | 0.144625 |
| 4 | 104709000 | 104710999 | 15.1684 | 28.71355 | 17.23583 | 6.982722 | 1.213608 | 0.070261 | 0.144628 |
| 17 | 5336200 | 5337799 | 5.731353 | 10.86313 | 31.40808 | 12.69022 | -1.03033 | 0.015861 | 0.144631 |
| 8 | 80124400 | 80126399 | 9.835704 | 17.29098 | 45.63483 | 20.2081 | -1.0047 | 0.579919 | 0.144642 |
| 16 | 92402400 | 92404199 | 9.835704 | 17.29098 | 45.63483 | 20.2081 | -1.0047 | 0.579919 | 0.144642 |
| 18 | 83987600 | 83991199 | 17.88638 | 30.12969 | 67.24887 | 29.39566 | -0.7731 | 1.106378 | 0.144732 |
| 6 | 21068800 | 21070599 | 20.81633 | 37.17601 | 22.89639 | 10.05976 | 1.092864 | 0.465259 | 0.144737 |
| X | 126532800 | 126533799 | 20.77801 | 37.25777 | 21.7889 | 9.540626 | 1.170192 | 0.438224 | 0.144741 |
| 2 | 59671200 | 59674599 | 21.91534 | 35.72097 | 88.28586 | 37.91952 | -0.93018 | 1.522217 | 0.144748 |
| 14 | 69482000 | 69482999 | 4.122715 | 7.998707 | 26.65527 | 10.0397 | -1.13719 | -0.25243 | 0.144758 |
| 12 | 18690400 | 18695799 | 167.8377 | 254.4259 | 177.5843 | 73.49943 | 0.797931 | 3.259461 | 0.14478 |
| 7 | 59947400 | 59948999 | 20.539 | 37.31296 | 19.7574 | 8.52335 | 1.326102 | 0.358939 | 0.144787 |
| 16 | 8485200 | 8492599 | 37.60877 | 60.06503 | 131.499 | 55.22244 | -0.78745 | 2.091948 | 0.144796 |
| 4 | 152714200 | 152715599 | 7.928157 | 14.61794 | 36.2774 | 15.34891 | -0.86725 | 0.219503 | 0.144806 |
| 5 | 68223200 | 68225399 | 19.22809 | 34.67045 | 22.81424 | 9.934819 | 1.006817 | 0.398671 | 0.144809 |
| 4 | 49631800 | 49632599 | 6.509558 | 12.24126 | 32.34843 | 13.27353 | -0.92193 | 0.077053 | 0.144814 |
| 16 | 90883600 | 90884999 | 6.912623 | 12.65524 | 36.12148 | 15.43144 | -1.07705 | 0.252763 | 0.144819 |
| 12 | 36190000 | 36191399 | 7.477018 | 13.79138 | 37.95719 | 16.05132 | -1.01542 | 0.214159 | 0.144827 |
| 14 | 42072000 | 42073199 | 16.8085 | 31.75039 | 16.57502 | 6.74539 | 1.407923 | 0.098585 | 0.144844 |
| 11 | 50454800 | 50458799 | 21.95993 | 36.06749 | 81.14721 | 34.95726 | -0.78866 | 1.413278 | 0.14485 |
| 5 | 50442000 | 50444799 | 40.4571 | 66.51446 | 51.08654 | 22.01684 | 0.757845 | 1.421698 | 0.144857 |
| 12 | 9668000 | 9669199 | 13.83412 | 26.22548 | 18.6321 | 7.525457 | 0.976897 | 0.052451 | 0.144857 |
| 10 | 42637000 | 42638799 | 9.674638 | 17.04058 | 45.63013 | 20.19826 | -1.02443 | 0.556691 | 0.14487 |
| 7 | 34185800 | 34187199 | 12.45924 | 21.82161 | 49.3944 | 21.80743 | -0.78511 | 0.660305 | 0.144873 |
| 12 | 48677000 | 48678799 | 23.55211 | 41.76839 | 22.11438 | 9.773696 | 1.304027 | 0.524658 | 0.144889 |
| 5 | 60690200 | 60691599 | 19.66362 | 35.1658 | 23.58117 | 10.3486 | 0.972092 | 0.450387 | 0.144893 |
| 8 | 22029800 | 22035799 | 49.45338 | 80.8328 | 48.46902 | 20.83912 | 1.110067 | 1.514895 | 0.144896 |
| 7 | 9283600 | 9288399 | 91.02453 | 143.3961 | 99.25976 | 41.18684 | 0.852695 | 2.501743 | 0.144904 |
| 11 | 81463400 | 81464599 | 5.83051 | 10.87575 | 33.7631 | 14.04557 | -1.1682 | 0.126877 | 0.144904 |
| 16 | 89712400 | 89731999 | 158.9859 | 231.9101 | 457.7413 | 188.6145 | -0.75159 | 3.946704 | 0.144922 |
| 18 | 63824800 | 63825799 | 6.872909 | 12.7055 | 34.68617 | 14.62348 | -0.99945 | 0.192835 | 0.144935 |
| 5 | 130462400 | 130463399 | 15.37571 | 29.02783 | 17.77929 | 7.243971 | 1.177898 | 0.091637 | 0.144942 |
| 16 | 42422800 | 42423999 | 22.76429 | 41.3273 | 18.06975 | 7.801982 | 1.599594 | 0.376535 | 0.144954 |
| X | 148459600 | 148467799 | 108.8824 | 170.5985 | 103.6176 | 42.97918 | 1.035082 | 2.616839 | 0.144957 |
| 4 | 28636000 | 28637399 | 12.68075 | 24.47656 | 16.23833 | 6.264757 | 1.134715 | -0.10895 | 0.144958 |
| 5 | 149221200 | 149224799 | 18.50251 | 30.76725 | 72.97433 | 31.66615 | -0.86066 | 1.249256 | 0.144961 |
| 3 | 21049200 | 21050399 | 14.76151 | 27.68747 | 19.51663 | 8.043372 | 0.963909 | 0.131212 | 0.144977 |
| 6 | 104216600 | 104217799 | 14.76151 | 27.68747 | 19.51663 | 8.043372 | 0.963909 | 0.131212 | 0.144977 |
| 14 | 55193000 | 55196199 | 13.56011 | 23.19245 | 58.95005 | 25.84864 | -0.95717 | 0.946697 | 0.144993 |
| 5 | 119145800 | 119154999 | 59.55803 | 93.16265 | 187.2848 | 77.69634 | -0.69132 | 2.630638 | 0.145016 |
| 8 | 47188600 | 47195199 | 47.75591 | 75.7935 | 149.9396 | 62.46753 | -0.65117 | 2.30159 | 0.145025 |
| 7 | 96537800 | 96540399 | 15.72139 | 26.90688 | 59.09548 | 25.9138 | -0.74741 | 0.94906 | 0.145036 |
| 10 | 30764000 | 30765399 | 16.65585 | 30.86837 | 19.71516 | 8.283306 | 1.083513 | 0.219283 | 0.145042 |

| | | | | | | | | | |
|---|---|---|---|---|---|---|---|---|---|
| 8 | 80365200 | 80366999 | 9.418281 | 16.25955 | 58.31652 | 25.58318 | -1.44805 | 0.842478 | 0.145055 |
| 4 | 149658600 | 149663599 | 25.68081 | 41.54591 | 97.5769 | 41.72207 | -0.8663 | 1.671225 | 0.145061 |
| 9 | 3317000 | 3318599 | 27.61978 | 47.36645 | 36.14399 | 15.85198 | 0.771937 | 0.949142 | 0.145066 |
| 18 | 75959200 | 75962799 | 14.61733 | 24.63272 | 65.18469 | 28.49679 | -1.01725 | 1.092458 | 0.145081 |
| 7 | 10143600 | 10146599 | 58.54831 | 94.52134 | 59.28802 | 25.28684 | 1.030381 | 1.764056 | 0.145124 |
| 11 | 117361800 | 117384799 | 154.453 | 224.9056 | 470.6556 | 193.971 | -0.83836 | 3.972909 | 0.145131 |
| 13 | 112730200 | 112732599 | 8.995662 | 16.03685 | 42.03498 | 18.49622 | -0.98508 | 0.437764 | 0.145132 |
| 11 | 19873800 | 19877199 | 21.91863 | 35.7385 | 89.00579 | 38.23348 | -0.94085 | 1.516972 | 0.145158 |
| 12 | 22501800 | 22505999 | 61.89927 | 99.3298 | 73.81769 | 31.21618 | 0.776069 | 1.954396 | 0.145161 |
| 13 | 75671200 | 75672999 | 9.786092 | 17.32439 | 42.49976 | 18.79356 | -0.89564 | 0.496738 | 0.145161 |
| X | 25840000 | 25841799 | 28.25999 | 49.18323 | 25.65168 | 11.27449 | 1.321775 | 0.797129 | 0.145166 |
| 9 | 75276600 | 75283999 | 73.67187 | 113.986 | 212.6213 | 88.31761 | -0.59582 | 2.832866 | 0.145167 |
| 9 | 20030200 | 20035999 | 80.57436 | 127.8647 | 96.87275 | 40.35418 | 0.731241 | 2.32012 | 0.145187 |
| 3 | 38110800 | 38111599 | 13.77947 | 26.00637 | 19.12262 | 7.796308 | 0.915894 | 0.082554 | 0.145187 |
| 5 | 115660800 | 115674199 | 53.19094 | 82.32312 | 222.8392 | 92.56173 | -1.13193 | 2.819389 | 0.145199 |
| 8 | 13001000 | 13008599 | 39.96157 | 63.56661 | 143.1383 | 59.77489 | -0.83434 | 2.226581 | 0.14521 |
| 7 | 14210400 | 14213199 | 44.13094 | 72.72514 | 46.16391 | 19.91511 | 1.033235 | 1.399874 | 0.145226 |
| X | 30118400 | 30119599 | 15.17912 | 29.54106 | 9.569845 | 3.521373 | 2.215306 | -0.25454 | 0.14523 |
| 7 | 92119600 | 92119999 | 3.040087 | 5.94581 | 20.21855 | 6.844886 | -1.02021 | -0.66697 | 0.145239 |
| 11 | 68687200 | 68689999 | 12.3556 | 21.46569 | 52.09622 | 22.87216 | -0.886 | 0.785555 | 0.145247 |
| 2 | 120439400 | 120440199 | 5.450542 | 10.41166 | 30.57123 | 12.1503 | -1.03209 | -0.04722 | 0.145251 |
| 15 | 82920400 | 82924599 | 19.5788 | 32.61197 | 72.08339 | 31.31463 | -0.75925 | 1.232291 | 0.145255 |
| 15 | 33584400 | 33585799 | 21.64483 | 38.26063 | 25.72105 | 11.37736 | 0.960213 | 0.552925 | 0.145258 |
| 3 | 136321200 | 136322399 | 17.75219 | 33.09064 | 18.03279 | 7.510465 | 1.320108 | 0.191958 | 0.145258 |
| 3 | 115626800 | 115628799 | 23.15575 | 41.16788 | 21.56252 | 9.509806 | 1.321346 | 0.505688 | 0.145264 |
| 6 | 115677400 | 115683599 | 43.14019 | 68.70349 | 142.0585 | 59.42046 | -0.70976 | 2.190218 | 0.145265 |
| 10 | 81750400 | 81750799 | 10.34299 | 20.06684 | 15.90178 | 5.989628 | 0.912753 | -0.21552 | 0.14527 |
| 7 | 81412200 | 81413599 | 8.046761 | 14.18789 | 46.68387 | 20.66044 | -1.31879 | 0.53753 | 0.14528 |
| 1 | 34406400 | 34408399 | 12.54297 | 21.69388 | 54.2174 | 23.78625 | -0.92953 | 0.83408 | 0.145294 |
| 15 | 72702200 | 72705999 | 19.84334 | 32.77943 | 80.31222 | 34.6916 | -0.90837 | 1.334574 | 0.1453 |
| 8 | 72122400 | 72124999 | 13.40004 | 22.94105 | 58.96824 | 25.85865 | -0.97306 | 0.936617 | 0.145309 |
| 11 | 16186400 | 16187999 | 9.373751 | 16.91784 | 41.18576 | 17.91392 | -0.86789 | 0.361691 | 0.145328 |
| 4 | 43017400 | 43023599 | 24.02252 | 38.76517 | 108.9345 | 46.43492 | -1.12863 | 1.744289 | 0.145331 |
| 12 | 110412600 | 110414199 | 5.86909 | 10.84229 | 35.18898 | 14.84945 | -1.24747 | 0.189484 | 0.145332 |
| 16 | 95070800 | 95071799 | 6.482366 | 12.26824 | 32.55114 | 13.1937 | -0.91244 | 0.034886 | 0.145335 |
| 14 | 113825600 | 113826199 | 10.05202 | 19.53587 | 15.39329 | 5.727769 | 0.936908 | -0.26209 | 0.145335 |
| 5 | 28385200 | 28397599 | 68.60436 | 105.8886 | 225.7342 | 93.76558 | -0.79094 | 2.875783 | 0.145344 |
| 7 | 46844800 | 46847599 | 48.08509 | 77.70467 | 59.64582 | 25.47129 | 0.74135 | 1.718124 | 0.145347 |
| 7 | 23028600 | 23046999 | 427.6921 | 599.1098 | 423.6417 | 175.6155 | 0.67058 | 4.556773 | 0.145349 |
| 1 | 174329200 | 174332399 | 16.35091 | 27.16781 | 76.05032 | 32.98281 | -1.09887 | 1.254597 | 0.145368 |
| 5 | 31743400 | 31747999 | 26.58424 | 43.12746 | 93.16079 | 39.91574 | -0.74214 | 1.613618 | 0.14537 |
| 7 | 24118800 | 24138799 | 154.9297 | 225.8493 | 456.6518 | 188.178 | -0.78722 | 3.956241 | 0.145372 |
| 5 | 116078400 | 116102599 | 207.6722 | 297.0897 | 543.805 | 224.6532 | -0.67057 | 4.240427 | 0.145372 |
| 12 | 22473200 | 22473599 | 11.41522 | 21.95471 | 15.44827 | 6.023107 | 1.036591 | -0.09224 | 0.145385 |
| 8 | 123249000 | 123262399 | 71.81095 | 109.6092 | 265.9571 | 110.2913 | -0.98999 | 3.100563 | 0.14539 |
| 6 | 83101200 | 83102199 | 4.447483 | 8.523253 | 29.45786 | 11.60376 | -1.25142 | -0.08848 | 0.14539 |
| 12 | 20047000 | 20082199 | 972.2409 | 1266.089 | 876.3048 | 367.0366 | 0.595889 | 5.708029 | 0.145395 |

| | | | | | | | | | |
|---|---|---|---|---|---|---|---|---|---|
| 12 | 19104000 | 19113799 | 314.7782 | 451.0757 | 345.7454 | 142.7942 | 0.587209 | 4.219016 | 0.145401 |
| 17 | 29724400 | 29730599 | 28.26952 | 45.62973 | 104.1221 | 44.39761 | -0.82869 | 1.741661 | 0.14541 |
| 12 | 100864000 | 100870999 | 37.45799 | 59.80521 | 130.3233 | 54.70406 | -0.78124 | 2.101969 | 0.145413 |
| 2 | 68581600 | 68583799 | 25.97779 | 45.94255 | 20.91622 | 9.251091 | 1.518875 | 0.565995 | 0.145415 |
| 13 | 9228800 | 9231599 | 13.50179 | 23.24334 | 55.80937 | 24.48119 | -0.8736 | 0.883177 | 0.145415 |
| 11 | 61845800 | 61849999 | 20.12176 | 33.31718 | 75.18871 | 32.53093 | -0.78977 | 1.305916 | 0.14542 |
| 12 | 115836200 | 115839599 | 40.80335 | 67.55695 | 45.54265 | 19.70347 | 0.947755 | 1.336985 | 0.145424 |
| 7 | 43509200 | 43514599 | 36.78619 | 58.80128 | 129.3719 | 54.39465 | -0.79322 | 2.064114 | 0.145424 |
| 13 | 25219600 | 25220599 | 11.29042 | 19.61522 | 52.60439 | 23.09533 | -1.02928 | 0.780138 | 0.145439 |
| 7 | 88536400 | 88537799 | 15.0974 | 25.83441 | 58.50908 | 25.6563 | -0.79148 | 0.947609 | 0.145443 |
| 8 | 70735000 | 70744399 | 46.17888 | 72.80178 | 179.2321 | 74.37768 | -0.97441 | 2.472352 | 0.145459 |
| 12 | 19036400 | 19042199 | 94.66689 | 148.6769 | 101.0553 | 41.91513 | 0.8756 | 2.56432 | 0.145475 |
| 10 | 129228400 | 129228599 | 10.2432 | 20.03673 | 6.210633 | 2.050235 | 2.400998 | -0.69241 | 0.145485 |
| 14 | 26396600 | 26398199 | 16.12691 | 29.4162 | 22.09878 | 9.485366 | 0.829969 | 0.308599 | 0.145488 |
| 2 | 85968400 | 85970799 | 22.30528 | 39.10038 | 27.32395 | 12.07229 | 0.901588 | 0.659746 | 0.145493 |
| 2 | 171347200 | 171349399 | 13.76274 | 23.84475 | 53.74633 | 23.58082 | -0.78088 | 0.8232 | 0.145506 |
| 7 | 106346400 | 106347799 | 15.18536 | 28.67642 | 17.78578 | 7.242366 | 1.160619 | 0.087914 | 0.145517 |
| 10 | 120734200 | 120735399 | 5.184229 | 9.838957 | 30.87872 | 12.44193 | -1.14417 | -0.00206 | 0.145523 |
| 3 | 138229800 | 138233999 | 23.33543 | 38.05358 | 90.01732 | 38.66983 | -0.8668 | 1.517751 | 0.145555 |
| 5 | 100393000 | 100394999 | 9.575548 | 17.11788 | 41.53493 | 18.23365 | -0.87218 | 0.423037 | 0.14556 |
| 19 | 59436600 | 59452799 | 83.73965 | 127.6407 | 276.5565 | 114.6503 | -0.82846 | 3.131874 | 0.145569 |
| 15 | 92465200 | 92466999 | 7.773834 | 13.96062 | 41.08519 | 17.96691 | -1.1444 | 0.380911 | 0.145569 |
| 6 | 52286600 | 52289599 | 18.87642 | 31.67505 | 68.31227 | 29.81495 | -0.72392 | 1.147534 | 0.145573 |
| 5 | 117663000 | 117694999 | 266.2528 | 371.3536 | 713.113 | 295.8115 | -0.77656 | 4.617723 | 0.145573 |
| 7 | 20698000 | 20708199 | 146.5555 | 223.1538 | 161.2909 | 66.8448 | 0.75279 | 3.162222 | 0.145578 |
| 11 | 67951600 | 67953599 | 10.40955 | 18.36846 | 47.21821 | 20.89361 | -0.96522 | 0.542011 | 0.145589 |
| 6 | 142383400 | 142385399 | 24.52923 | 43.08038 | 26.01003 | 11.51018 | 1.111426 | 0.642499 | 0.145597 |
| 2 | 157578400 | 157580999 | 9.400941 | 16.90564 | 40.03667 | 17.48832 | -0.83206 | 0.383104 | 0.145598 |
| 7 | 66338600 | 66339999 | 7.856457 | 13.91405 | 43.57 | 19.26525 | -1.24482 | 0.481783 | 0.1456 |
| 17 | 19616200 | 19622999 | 17.03314 | 31.22052 | 18.0995 | 7.71974 | 1.207309 | 0.27936 | 0.145601 |
| 6 | 129349200 | 129350199 | 10.69028 | 20.77111 | 15.30451 | 5.706283 | 1.030001 | -0.24763 | 0.145607 |
| 1 | 135094200 | 135095599 | 6.710273 | 12.14708 | 39.63662 | 17.17569 | -1.28288 | 0.322875 | 0.145611 |
| 9 | 18897400 | 18899999 | 27.39324 | 47.58329 | 29.86753 | 13.11162 | 1.055383 | 0.822133 | 0.145632 |
| 6 | 6004800 | 6006799 | 7.792152 | 14.30953 | 36.18802 | 15.39508 | -0.90015 | 0.239893 | 0.145632 |
| X | 3386400 | 3388199 | 19.07675 | 35.84965 | 15.23608 | 6.258992 | 1.690819 | 0.147937 | 0.145646 |
| 5 | 148318600 | 148326999 | 55.02395 | 86.06701 | 191.3006 | 79.36263 | -0.83605 | 2.62926 | 0.145653 |
| 9 | 44837600 | 44839999 | 22.8077 | 39.90937 | 29.57025 | 13.04494 | 0.817937 | 0.69599 | 0.145657 |
| 9 | 70903000 | 70904199 | 17.57023 | 31.69563 | 23.38291 | 10.17663 | 0.843202 | 0.384077 | 0.145669 |
| 17 | 89540000 | 89541199 | 14.99143 | 28.31079 | 18.47105 | 7.521076 | 1.08867 | 0.088436 | 0.145679 |
| 11 | 46516000 | 46518199 | 10.54943 | 18.66069 | 43.52078 | 19.24816 | -0.82388 | 0.511428 | 0.145684 |
| 14 | 49941600 | 49942799 | 13.07802 | 24.61147 | 18.25858 | 7.482671 | 0.897138 | 0.091881 | 0.145689 |
| 17 | 54048400 | 54051799 | 53.03765 | 85.56094 | 65.38026 | 27.8584 | 0.745107 | 1.78019 | 0.14569 |
| 11 | 13268200 | 13270799 | 42.66631 | 70.41575 | 45.60953 | 19.68887 | 1.005056 | 1.380244 | 0.145691 |
| 7 | 3061800 | 3064799 | 46.03586 | 74.88414 | 57.16039 | 24.53271 | 0.758102 | 1.580655 | 0.145699 |
| 12 | 31409800 | 31410599 | 12.40874 | 23.69008 | 17.67463 | 7.032908 | 0.925147 | -0.01216 | 0.145699 |
| 3 | 115254200 | 115254999 | 4.23239 | 8.245204 | 25.50582 | 9.288592 | -0.98609 | -0.37557 | 0.145699 |
| 10 | 43878600 | 43880799 | 14.14509 | 24.26562 | 58.52304 | 25.6676 | -0.88123 | 0.919683 | 0.145699 |

| | | | | | | | | | |
|---|---|---|---|---|---|---|---|---|---|
| 3 | 101607000 | 101609399 | 12.68867 | 21.96482 | 53.69387 | 23.55718 | -0.8975 | 0.826404 | 0.145708 |
| 2 | 179703600 | 179709599 | 40.47153 | 64.30663 | 145.1358 | 60.53609 | -0.83923 | 2.261629 | 0.145725 |
| 6 | 82749200 | 82752799 | 20.51175 | 33.77598 | 80.65149 | 34.78165 | -0.87272 | 1.376277 | 0.145726 |
| 9 | 27210200 | 27223799 | 84.7326 | 129.0008 | 269.952 | 111.8733 | -0.77973 | 3.156573 | 0.145727 |
| 3 | 66830600 | 66833599 | 46.81835 | 75.91515 | 60.4327 | 25.88349 | 0.693703 | 1.64202 | 0.145745 |
| 10 | 71156000 | 71164399 | 45.65387 | 72.43636 | 157.0614 | 65.41562 | -0.78369 | 2.309071 | 0.145757 |
| 3 | 4285000 | 4285599 | 9.074443 | 17.67826 | 13.80536 | 5.026982 | 0.97716 | -0.35519 | 0.145771 |
| 1 | 111446400 | 111447199 | 12.54707 | 24.0383 | 17.75719 | 7.000733 | 0.951887 | -0.03882 | 0.145773 |
| 13 | 66889800 | 66894199 | 50.33581 | 81.62506 | 58.41159 | 25.01929 | 0.847124 | 1.646816 | 0.145775 |
| 5 | 61217600 | 61220599 | 28.37517 | 49.00873 | 32.68913 | 14.34076 | 0.966955 | 0.885246 | 0.145776 |
| 4 | 123145200 | 123148799 | 13.04891 | 22.44969 | 56.87069 | 24.94627 | -0.95078 | 0.887418 | 0.145803 |
| 16 | 34635000 | 34635999 | 4.23932 | 8.237819 | 25.65787 | 9.558434 | -1.0257 | -0.29248 | 0.145803 |
| 2 | 84500400 | 84501799 | 6.292018 | 11.69214 | 36.44317 | 15.25615 | -1.18225 | 0.159206 | 0.145814 |
| 19 | 16932000 | 16935599 | 18.96272 | 31.55445 | 74.28768 | 32.25153 | -0.85001 | 1.242411 | 0.145821 |
| 2 | 37052400 | 37054999 | 55.3535 | 89.17577 | 67.95643 | 28.90384 | 0.74675 | 1.822566 | 0.145826 |
| 7 | 108280400 | 108282799 | 25.57937 | 44.13374 | 32.94903 | 14.45432 | 0.804729 | 0.880842 | 0.145828 |
| 14 | 103968800 | 103971999 | 27.90643 | 47.85798 | 34.72314 | 15.22883 | 0.844491 | 0.950096 | 0.145835 |
| 8 | 36775200 | 36775999 | 14.31901 | 26.75135 | 20.17588 | 8.371013 | 0.859662 | 0.153929 | 0.145863 |
| 4 | 143932000 | 143933999 | 50.41808 | 81.5327 | 61.48667 | 26.28151 | 0.768025 | 1.704998 | 0.145864 |
| X | 25669200 | 25670799 | 36.23284 | 60.71688 | 41.02437 | 17.88976 | 0.944687 | 1.191146 | 0.145878 |
| 4 | 32243200 | 32246399 | 15.37722 | 26.39548 | 58.60333 | 25.70408 | -0.76232 | 0.919249 | 0.145902 |
| 7 | 135782800 | 135784399 | 7.900265 | 14.63691 | 36.42676 | 15.31134 | -0.86475 | 0.190251 | 0.145904 |
| 12 | 75886600 | 75891999 | 26.88421 | 43.66019 | 94.85203 | 40.67204 | -0.749 | 1.592429 | 0.145929 |
| 11 | 31541600 | 31542599 | 6.052312 | 11.44417 | 31.37617 | 12.74137 | -0.96157 | 0.03334 | 0.145931 |
| 6 | 149310200 | 149311199 | 5.055503 | 9.595161 | 31.43814 | 12.66591 | -1.20574 | -0.00309 | 0.145939 |
| 16 | 83135000 | 83137599 | 57.66913 | 92.59089 | 74.94844 | 31.71998 | 0.653939 | 1.933529 | 0.14594 |
| 4 | 134271400 | 134279599 | 56.91832 | 88.73295 | 194.9121 | 80.89028 | -0.82281 | 2.690706 | 0.145949 |
| 10 | 41440600 | 41442999 | 14.12093 | 24.29549 | 56.38293 | 24.73236 | -0.82502 | 0.890876 | 0.145955 |
| 7 | 144852600 | 144858599 | 25.65593 | 41.28572 | 113.9531 | 48.38841 | -1.10775 | 1.835067 | 0.145961 |
| 7 | 11855800 | 11857999 | 49.05858 | 80.04556 | 50.70617 | 21.7869 | 1.029332 | 1.539272 | 0.145975 |
| 2 | 17536000 | 17538199 | 9.752737 | 17.35453 | 44.11109 | 19.4399 | -0.94294 | 0.457271 | 0.145982 |
| 2 | 97595000 | 97596799 | 18.50288 | 34.11509 | 18.93379 | 8.011028 | 1.2774 | 0.260624 | 0.146006 |
| 7 | 30598800 | 30602199 | 17.82601 | 29.75752 | 70.13674 | 30.51111 | -0.85134 | 1.204155 | 0.146009 |
| 12 | 46486200 | 46487199 | 13.88063 | 26.50152 | 17.59763 | 7.001484 | 1.092373 | 0.003056 | 0.146021 |
| 5 | 147233400 | 147244599 | 89.90049 | 136.4315 | 285.7406 | 118.303 | -0.78475 | 3.22419 | 0.146025 |
| 1 | 88415800 | 88418399 | 16.56058 | 28.12868 | 62.29131 | 27.29719 | -0.7611 | 1.025043 | 0.146026 |
| 5 | 115594800 | 115598799 | 16.81391 | 27.93222 | 75.5365 | 32.75601 | -1.04929 | 1.257594 | 0.146032 |
| 14 | 118820600 | 118822599 | 13.09928 | 22.84919 | 51.05214 | 22.47036 | -0.76685 | 0.728727 | 0.14604 |
| 7 | 23187800 | 23197399 | 183.5032 | 277.4557 | 172.8034 | 71.46246 | 0.95902 | 3.313383 | 0.146041 |
| 17 | 20760800 | 20761799 | 14.0977 | 27.29004 | 13.67703 | 5.206079 | 1.552087 | -0.14494 | 0.146063 |
| 11 | 66378600 | 66381199 | 12.92863 | 22.55637 | 54.88268 | 24.15961 | -0.88948 | 0.728039 | 0.146068 |
| 5 | 110240800 | 110256799 | 189.3186 | 271.209 | 550.6588 | 227.4352 | -0.81806 | 4.218854 | 0.146071 |
| 12 | 22854200 | 22866599 | 301.8875 | 434.15 | 335.1553 | 138.343 | 0.582051 | 4.166124 | 0.146082 |
| 6 | 112364400 | 112366599 | 11.86637 | 20.68782 | 51.57032 | 22.69099 | -0.92403 | 0.729673 | 0.146083 |
| 13 | 52386800 | 52388999 | 12.46194 | 21.7607 | 51.04411 | 22.4819 | -0.83657 | 0.70899 | 0.146084 |
| 12 | 103045000 | 103046399 | 4.360472 | 8.484615 | 25.25768 | 9.307961 | -0.94662 | -0.33055 | 0.146086 |
| 15 | 84935000 | 84935599 | 4.360472 | 8.484615 | 25.25768 | 9.307961 | -0.94662 | -0.33055 | 0.146086 |

| | | | | | | | | | |
|---|---|---|---|---|---|---|---|---|---|
| 16 | 95722800 | 95731599 | 49.84712 | 78.68712 | 169.3764 | 70.31764 | -0.7795 | 2.443496 | 0.146088 |
| 17 | 41533600 | 41534199 | 9.85177 | 19.24051 | 10.5797 | 3.712737 | 1.520866 | -0.48203 | 0.146106 |
| 7 | 59595200 | 59596199 | 20.43625 | 36.71676 | 22.49235 | 9.832403 | 1.105561 | 0.42642 | 0.146123 |
| 5 | 108816800 | 108820599 | 25.19117 | 40.78926 | 96.52217 | 41.29758 | -0.87591 | 1.651893 | 0.146133 |
| 3 | 83675600 | 83680999 | 25.65334 | 41.58542 | 99.59827 | 42.64986 | -0.8918 | 1.629095 | 0.14614 |
| 12 | 112725800 | 112737199 | 79.55295 | 121.5915 | 253.1738 | 105.0231 | -0.76795 | 3.074271 | 0.146141 |
| 6 | 14974600 | 14974999 | 8.668915 | 16.93329 | 12.07786 | 4.226011 | 1.156198 | -0.50391 | 0.146142 |
| 4 | 46518200 | 46524399 | 49.48821 | 78.24787 | 158.174 | 65.72105 | -0.68704 | 2.403894 | 0.146147 |
| 10 | 122101800 | 122102599 | 12.08819 | 23.13077 | 17.86908 | 7.069713 | 0.883366 | -0.03167 | 0.146204 |
| 7 | 27756400 | 27756599 | 2.337248 | 4.563179 | 17.88062 | 5.646197 | -1.12661 | -0.93478 | 0.146209 |
| X | 80015400 | 80016799 | 17.80755 | 33.51213 | 16.61256 | 6.808043 | 1.474138 | 0.131733 | 0.146211 |
| 15 | 82128200 | 82129599 | 6.973034 | 12.69501 | 39.75837 | 17.10586 | -1.21776 | 0.290052 | 0.146216 |
| 1 | 61983200 | 61984399 | 18.98818 | 33.85296 | 24.92113 | 10.96521 | 0.836057 | 0.46855 | 0.146229 |
| 5 | 36133600 | 36135599 | 7.357625 | 13.49183 | 36.15295 | 15.41078 | -0.98489 | 0.245141 | 0.146237 |
| 5 | 30668000 | 30668799 | 2.748599 | 5.354179 | 25.78943 | 9.401586 | -1.61471 | -0.38914 | 0.146251 |
| 9 | 106079800 | 106085399 | 25.87001 | 41.75367 | 101.8887 | 43.44125 | -0.92519 | 1.741098 | 0.146262 |
| 10 | 80128200 | 80142599 | 79.87248 | 121.1726 | 288.5209 | 119.4421 | -0.97002 | 3.229254 | 0.146266 |
| 2 | 17229600 | 17231399 | 17.79225 | 32.09585 | 22.73534 | 9.894915 | 0.901238 | 0.383548 | 0.146268 |
| 8 | 16200600 | 16202199 | 17.79225 | 32.09585 | 22.73534 | 9.894915 | 0.901238 | 0.383548 | 0.146268 |
| 7 | 10535200 | 10558799 | 522.2908 | 716.8418 | 524.7783 | 218.2393 | 0.590894 | 4.868534 | 0.146275 |
| 2 | 178879200 | 178882199 | 20.44551 | 33.82874 | 75.77572 | 32.76853 | -0.77916 | 1.315494 | 0.146285 |
| 10 | 95133000 | 95136199 | 16.33146 | 27.50941 | 65.3787 | 28.57785 | -0.86311 | 1.100907 | 0.146288 |
| 1 | 131898400 | 131903199 | 20.49966 | 33.45312 | 89.43924 | 38.43151 | -1.04196 | 1.501795 | 0.14629 |
| 4 | 152252800 | 152258399 | 44.31594 | 70.47236 | 142.6183 | 59.53442 | -0.68108 | 2.239616 | 0.146294 |
| 6 | 96567400 | 96568999 | 8.941597 | 16.07814 | 42.70648 | 18.65594 | -0.99706 | 0.382629 | 0.146318 |
| 17 | 28322400 | 28329399 | 40.12524 | 63.8193 | 143.0233 | 59.71691 | -0.82766 | 2.230734 | 0.146322 |
| 4 | 119831200 | 119833999 | 8.971011 | 15.71612 | 48.29883 | 21.32825 | -1.22297 | 0.636436 | 0.146324 |
| 8 | 14744200 | 14746799 | 12.64126 | 22.03048 | 50.50768 | 22.21817 | -0.80385 | 0.739069 | 0.146332 |
| 2 | 26241400 | 26243199 | 6.640687 | 12.13573 | 36.62162 | 15.68198 | -1.1592 | 0.261268 | 0.146335 |
| 10 | 128744000 | 128750399 | 36.81639 | 58.78283 | 133.5794 | 56.07351 | -0.84158 | 2.100347 | 0.146338 |
| 11 | 100137200 | 100141799 | 25.03033 | 40.54013 | 96.59304 | 41.3364 | -0.88541 | 1.645698 | 0.146352 |
| 5 | 36687400 | 36689999 | 11.71578 | 20.4241 | 52.64686 | 23.16382 | -0.97218 | 0.73075 | 0.146366 |
| 19 | 13631400 | 13632799 | 20.68046 | 36.51355 | 26.27744 | 11.62616 | 0.861713 | 0.559136 | 0.14637 |
| X | 32475200 | 32478599 | 31.84989 | 54.11096 | 37.68817 | 16.51674 | 0.901292 | 1.05164 | 0.146384 |
| 18 | 37499200 | 37500399 | 29.8455 | 50.54952 | 37.46719 | 16.40465 | 0.811818 | 1.070157 | 0.146407 |
| 13 | 56534200 | 56546799 | 88.13378 | 133.9527 | 277.0803 | 114.7694 | -0.76501 | 3.192385 | 0.146414 |
| 14 | 16131600 | 16132799 | 17.67082 | 31.69744 | 23.39906 | 10.24259 | 0.836801 | 0.417665 | 0.146425 |
| X | 29709000 | 29710799 | 14.67353 | 28.28281 | 13.38454 | 5.18599 | 1.609175 | -0.08361 | 0.146429 |
| 7 | 32862000 | 32862599 | 29.04389 | 49.35194 | 33.89787 | 14.85585 | 0.921301 | 1.034417 | 0.146433 |
| 17 | 87641600 | 87652599 | 66.19137 | 102.0892 | 227.4111 | 94.4625 | -0.85507 | 2.88887 | 0.146439 |
| 3 | 43193000 | 43194199 | 12.43846 | 23.9264 | 16.59081 | 6.46568 | 1.057193 | -0.08046 | 0.14647 |
| 17 | 47214800 | 47224399 | 61.41734 | 95.88728 | 190.85 | 79.191 | -0.67912 | 2.666117 | 0.146479 |
| X | 26632200 | 26636199 | 57.37302 | 92.30235 | 68.09901 | 28.90687 | 0.790978 | 1.870424 | 0.146492 |
| 8 | 71957200 | 71961199 | 21.26584 | 34.69767 | 87.10645 | 37.42686 | -0.95154 | 1.505569 | 0.146501 |
| 10 | 106366200 | 106366799 | 10.56128 | 20.63188 | 9.907909 | 3.460457 | 1.719154 | -0.48913 | 0.146509 |
| 19 | 3087200 | 3090599 | 50.36709 | 81.31523 | 62.3751 | 26.60426 | 0.740824 | 1.752281 | 0.14652 |
| 16 | 45245600 | 45249799 | 34.34524 | 54.99017 | 121.8774 | 51.36402 | -0.80104 | 2.009292 | 0.14655 |

| | | | | | | | | | |
|---|---|---|---|---|---|---|---|---|---|
| 3 | 84687400 | 84689199 | 17.05529 | 31.50455 | 20.24567 | 8.544469 | 1.07031 | 0.242038 | 0.146551 |
| 7 | 22979600 | 22983599 | 57.34987 | 92.08209 | 74.88352 | 31.69449 | 0.647324 | 1.931945 | 0.146556 |
| 17 | 39797800 | 39801799 | 98.34528 | 153.5583 | 127.0723 | 52.72656 | 0.585648 | 2.673509 | 0.146572 |
| 12 | 84302800 | 84306199 | 24.60107 | 40.10335 | 87.73675 | 37.68531 | -0.75475 | 1.525204 | 0.146576 |
| 5 | 91079000 | 91079799 | 5.104545 | 9.847574 | 28.08889 | 10.85575 | -0.95156 | -0.14769 | 0.14658 |
| 12 | 54304800 | 54305799 | 13.84587 | 26.25448 | 17.96736 | 7.253029 | 1.030644 | 0.048151 | 0.146582 |
| 5 | 142865600 | 142874799 | 43.06539 | 67.67675 | 180.9322 | 75.052 | -1.09666 | 2.535513 | 0.146586 |
| 7 | 17694600 | 17696999 | 49.87536 | 80.94542 | 56.82372 | 24.35403 | 0.87579 | 1.627299 | 0.146611 |
| 10 | 95734400 | 95735799 | 11.23009 | 19.65078 | 48.8642 | 21.55662 | -0.91988 | 0.667196 | 0.146615 |
| 7 | 10647400 | 10656799 | 164.3885 | 248.3977 | 195.1329 | 80.6807 | 0.62359 | 3.322602 | 0.146628 |
| 3 | 40379400 | 40381599 | 8.742425 | 15.87863 | 38.11031 | 16.4455 | -0.83959 | 0.315282 | 0.146629 |
| 9 | 37295200 | 37297799 | 16.30649 | 27.53082 | 66.04253 | 28.88824 | -0.87646 | 1.079877 | 0.146629 |
| 10 | 55358800 | 55360199 | 15.75879 | 29.73454 | 17.76631 | 7.247185 | 1.212017 | 0.099069 | 0.146648 |
| 2 | 51442800 | 51445999 | 49.69839 | 78.44541 | 162.9393 | 67.64225 | -0.72816 | 2.446104 | 0.146648 |
| 14 | 28393800 | 28394799 | 12.11333 | 23.41646 | 16.21839 | 6.213311 | 1.082258 | -0.13872 | 0.146652 |
| 7 | 107634000 | 107635199 | 21.11976 | 37.02079 | 27.69257 | 12.23476 | 0.804126 | 0.655515 | 0.146669 |
| 11 | 96970600 | 96981999 | 94.35448 | 143.2906 | 274.2249 | 113.559 | -0.65392 | 3.210825 | 0.14668 |
| 10 | 85487200 | 85488599 | 6.565621 | 12.20521 | 34.25208 | 14.33147 | -1.03109 | 0.156075 | 0.146686 |
| 7 | 11108000 | 11123199 | 305.5395 | 438.9726 | 338.7262 | 139.8384 | 0.581292 | 4.180381 | 0.146696 |
| 7 | 11108200 | 11123199 | 305.5395 | 438.9726 | 338.7262 | 139.8384 | 0.581292 | 4.180381 | 0.146696 |
| 4 | 143115800 | 143116399 | 4.235452 | 8.241942 | 25.26339 | 9.304685 | -0.98769 | -0.33499 | 0.146697 |
| 7 | 57138200 | 57138999 | 4.235452 | 8.241942 | 25.26339 | 9.304685 | -0.98769 | -0.33499 | 0.146697 |
| 18 | 6515800 | 6516599 | 5.874068 | 11.18619 | 30.46651 | 12.18778 | -0.93262 | -0.01968 | 0.146709 |
| 8 | 12018200 | 12026999 | 42.50292 | 67.30692 | 158.3408 | 65.84175 | -0.90366 | 2.361482 | 0.146721 |
| 1 | 13059800 | 13065199 | 35.4524 | 56.44617 | 141.5747 | 59.19774 | -0.98807 | 2.192349 | 0.146725 |
| 5 | 106713800 | 106715199 | 8.192766 | 14.73995 | 39.01475 | 17.03515 | -0.99068 | 0.369611 | 0.146741 |
| 1 | 112521600 | 112529399 | 317.5581 | 455.0373 | 298.0003 | 123.0762 | 0.814036 | 4.218482 | 0.146744 |
| 2 | 28866600 | 28868999 | 11.90096 | 20.66827 | 53.1686 | 23.3376 | -0.96979 | 0.7883 | 0.14675 |
| 5 | 145907600 | 145908999 | 35.81962 | 59.28088 | 46.64747 | 20.17567 | 0.725331 | 1.328662 | 0.146753 |
| 9 | 21898200 | 21909799 | 75.38065 | 115.3443 | 249.1863 | 103.3959 | -0.81978 | 3.050055 | 0.146785 |
| X | 44224600 | 44226199 | 22.32985 | 40.10736 | 20.96087 | 9.165145 | 1.333235 | 0.439561 | 0.146788 |
| 3 | 30361000 | 30362799 | 7.467867 | 13.79925 | 39.15471 | 16.51732 | -1.0567 | 0.205804 | 0.146809 |
| 15 | 102141800 | 102144799 | 11.13645 | 19.36122 | 52.60588 | 23.10611 | -1.04794 | 0.768835 | 0.146813 |
| 15 | 75958600 | 75966399 | 50.86904 | 79.39631 | 194.3275 | 80.63629 | -0.97727 | 2.66714 | 0.146815 |
| 7 | 92285200 | 92286799 | 9.555988 | 16.74754 | 47.25546 | 20.87447 | -1.10055 | 0.63183 | 0.146842 |
| 9 | 107411400 | 107412599 | 4.473793 | 8.499607 | 30.85075 | 12.40296 | -1.34809 | -0.01575 | 0.146861 |
| 17 | 26286800 | 26288799 | 12.91679 | 22.1644 | 57.34448 | 25.15124 | -0.98171 | 0.912465 | 0.146861 |
| 3 | 89321200 | 89323999 | 12.63181 | 22.04324 | 49.96853 | 21.99934 | -0.78741 | 0.719052 | 0.146873 |
| 13 | 86453000 | 86454199 | 22.49691 | 39.4226 | 28.24016 | 12.47343 | 0.866507 | 0.671143 | 0.146889 |
| 8 | 3058600 | 3063199 | 47.46335 | 77.1922 | 55.4779 | 23.80903 | 0.84462 | 1.586613 | 0.146891 |
| 10 | 39862200 | 39863799 | 11.68592 | 20.45712 | 47.80226 | 21.09573 | -0.83057 | 0.661997 | 0.146892 |
| 12 | 84730200 | 84737199 | 34.74026 | 55.46402 | 132.5443 | 55.63341 | -0.9141 | 2.100805 | 0.146894 |
| 11 | 114098800 | 114127399 | 213.1986 | 304.2395 | 568.4517 | 234.9296 | -0.7038 | 4.276911 | 0.1469 |
| 2 | 25509800 | 25512199 | 9.055859 | 15.96123 | 44.09335 | 19.51542 | -1.06819 | 0.545377 | 0.1469 |
| 9 | 35197800 | 35202599 | 23.08149 | 37.86952 | 82.44986 | 35.50255 | -0.74214 | 1.429506 | 0.146927 |
| 2 | 131864400 | 131866199 | 6.325336 | 11.66953 | 36.80122 | 15.55806 | -1.2092 | 0.202363 | 0.146935 |
| 18 | 3458200 | 3461199 | 29.5334 | 49.98901 | 36.29016 | 15.88622 | 0.841413 | 1.06975 | 0.146945 |

| | | | | | | | | | |
|---|---|---|---|---|---|---|---|---|---|
| 6 | 117783400 | 117789799 | 34.43492 | 55.32306 | 113.0753 | 47.90285 | -0.6785 | 1.898536 | 0.146961 |
| X | 122295200 | 122295999 | 15.36049 | 29.06155 | 17.91835 | 7.267479 | 1.174497 | 0.078292 | 0.146967 |
| 2 | 90335000 | 90335599 | 3.130894 | 6.109809 | 23.22721 | 8.262936 | -1.24633 | -0.48118 | 0.146967 |
| 16 | 91974600 | 91981199 | 46.67369 | 73.94886 | 154.0077 | 64.05622 | -0.7276 | 2.351795 | 0.146973 |
| 12 | 23303600 | 23306999 | 72.58814 | 115.3789 | 94.70252 | 39.51445 | 0.618118 | 2.265527 | 0.146988 |
| 6 | 84179200 | 84188999 | 72.68704 | 112.0781 | 220.9502 | 91.7789 | -0.67931 | 2.896152 | 0.146989 |
| 1 | 77865600 | 77867399 | 17.78292 | 32.56877 | 21.26096 | 9.078028 | 1.036128 | 0.29694 | 0.146992 |
| 7 | 14192800 | 14199799 | 122.2702 | 189.5705 | 128.601 | 53.41433 | 0.864158 | 2.805685 | 0.147004 |
| 6 | 124726000 | 124726999 | 4.658225 | 9.049727 | 26.29467 | 9.814077 | -0.92974 | -0.27919 | 0.14701 |
| 8 | 58315200 | 58317199 | 18.73314 | 34.35107 | 20.02419 | 8.53573 | 1.199784 | 0.294715 | 0.147015 |
| 5 | 149436800 | 149438199 | 6.813349 | 12.78849 | 36.33183 | 14.95553 | -1.03041 | 0.097753 | 0.147031 |
| 12 | 58632600 | 58633599 | 6.377583 | 11.98738 | 34.61193 | 14.21374 | -1.05039 | 0.081425 | 0.147035 |
| 11 | 52259600 | 52262599 | 15.15255 | 25.76476 | 62.78496 | 27.5172 | -0.8983 | 1.009855 | 0.147038 |
| 7 | 11261600 | 11263399 | 19.58207 | 35.01623 | 23.92422 | 10.50009 | 0.945615 | 0.453219 | 0.147053 |
| 5 | 93704400 | 93707599 | 70.82457 | 112.8861 | 82.14408 | 34.39878 | 0.795654 | 2.173991 | 0.147058 |
| 13 | 30850400 | 30851199 | 3.380644 | 6.598204 | 25.15852 | 8.926159 | -1.24577 | -0.47548 | 0.14707 |
| 3 | 47881800 | 47883599 | 28.74323 | 49.6277 | 32.68811 | 14.34017 | 0.984959 | 0.890125 | 0.147072 |
| 10 | 86613600 | 86616199 | 15.00172 | 25.15393 | 69.54598 | 30.33872 | -1.08066 | 1.144459 | 0.147076 |
| 9 | 61538800 | 61540599 | 7.943007 | 14.57374 | 37.80869 | 16.10381 | -0.93835 | 0.248156 | 0.147083 |
| 4 | 134998800 | 135000999 | 11.11934 | 19.36954 | 51.54455 | 22.66795 | -1.01774 | 0.738538 | 0.147084 |
| 7 | 14581000 | 14588199 | 170.3978 | 258.2021 | 177.059 | 73.27267 | 0.822941 | 3.267373 | 0.14709 |
| 19 | 18608000 | 18609599 | 7.534596 | 13.74504 | 37.65955 | 16.16208 | -1.02384 | 0.278846 | 0.147111 |
| 9 | 12296000 | 12296799 | 10.9442 | 21.09076 | 16.43855 | 6.372661 | 0.897805 | -0.11261 | 0.147131 |
| 7 | 73722200 | 73726199 | 32.44257 | 52.2933 | 110.2442 | 46.92457 | -0.7172 | 1.790671 | 0.147131 |
| 14 | 121042800 | 121052999 | 78.63275 | 120.4212 | 246.4918 | 102.2977 | -0.74109 | 3.033551 | 0.147133 |
| 2 | 144052200 | 144055599 | 22.52992 | 37.18441 | 79.46692 | 34.30544 | -0.71201 | 1.352025 | 0.147155 |
| 5 | 88545400 | 88546999 | 9.564751 | 17.12867 | 41.55973 | 18.2175 | -0.87068 | 0.411612 | 0.147156 |
| 13 | 49304200 | 49305599 | 6.93329 | 13.10058 | 35.47335 | 14.42706 | -0.94687 | 0.054447 | 0.147159 |
| 9 | 108224600 | 108225599 | 4.058312 | 7.732334 | 30.89911 | 12.34747 | -1.4772 | -0.04032 | 0.147166 |
| 11 | 109012600 | 109013799 | 14.66036 | 27.89493 | 17.44328 | 6.990319 | 1.169104 | 0.032577 | 0.147171 |
| 7 | 85571400 | 85572999 | 46.94613 | 75.99831 | 61.4626 | 26.29297 | 0.668795 | 1.674659 | 0.147176 |
| 5 | 68825000 | 68826799 | 22.70115 | 40.2321 | 23.27354 | 10.28939 | 1.176709 | 0.526904 | 0.1472 |
| 7 | 24155400 | 24172599 | 113.7307 | 170.1484 | 343.0684 | 141.4779 | -0.74803 | 3.513768 | 0.1472 |
| 9 | 36797600 | 36798199 | 11.43285 | 22.14055 | 15.41008 | 5.854532 | 1.08733 | -0.17489 | 0.147215 |
| 12 | 60043600 | 60044399 | 9.93393 | 19.2629 | 14.37488 | 5.428054 | 0.994944 | -0.21791 | 0.147218 |
| 2 | 56998000 | 57002999 | 38.45305 | 61.31348 | 135.3884 | 56.72591 | -0.80239 | 2.145436 | 0.147222 |
| 12 | 113987400 | 113988199 | 23.86937 | 41.60529 | 31.31915 | 13.77868 | 0.795126 | 0.761449 | 0.147231 |
| 4 | 122516600 | 122518999 | 38.85528 | 65.27408 | 35.84715 | 15.6565 | 1.242239 | 1.167617 | 0.147253 |
| 2 | 77794400 | 77795599 | 17.24549 | 31.84545 | 20.24001 | 8.54588 | 1.085716 | 0.245483 | 0.147274 |
| 12 | 20211600 | 20227599 | 338.9778 | 482.5866 | 369.1687 | 152.6306 | 0.580607 | 4.315523 | 0.147295 |
| 1 | 115229800 | 115231799 | 25.24942 | 44.3074 | 25.35408 | 11.21181 | 1.18826 | 0.658056 | 0.147296 |
| X | 122604800 | 122608599 | 41.75576 | 69.99225 | 34.16055 | 14.89935 | 1.411834 | 1.198765 | 0.147296 |
| 7 | 5298400 | 5299799 | 32.78933 | 55.62344 | 37.6281 | 16.483 | 0.943101 | 1.067105 | 0.147297 |
| 6 | 134627200 | 134628399 | 12.33409 | 23.45428 | 16.88542 | 6.775518 | 0.966298 | 0.012523 | 0.147316 |
| 13 | 91274400 | 91274999 | 3.23334 | 6.317044 | 21.70112 | 7.567296 | -1.07542 | -0.5557 | 0.147316 |
| 12 | 78795000 | 78796399 | 6.284605 | 11.37934 | 40.69442 | 17.6286 | -1.41367 | 0.320142 | 0.14732 |
| 1 | 167340600 | 167342199 | 21.49897 | 38.6669 | 20.6527 | 9.020179 | 1.302506 | 0.42327 | 0.147323 |

| | | | | | | | | | |
|---|---|---|---|---|---|---|---|---|---|
| X | 115631600 | 115634599 | 49.87838 | 81.01683 | 56.18298 | 24.094 | 0.894508 | 1.612825 | 0.147329 |
| 11 | 115494200 | 115497999 | 20.21495 | 33.24509 | 81.00654 | 34.91605 | -0.90252 | 1.392187 | 0.147335 |
| 11 | 72431000 | 72433199 | 12.61808 | 22.03278 | 50.50691 | 22.24494 | -0.80351 | 0.709772 | 0.147351 |
| 15 | 98505600 | 98508399 | 14.66001 | 25.00797 | 59.49854 | 26.08363 | -0.86292 | 0.97626 | 0.147355 |
| 16 | 95765800 | 95766999 | 4.906496 | 9.372983 | 32.87019 | 13.06233 | -1.28598 | -0.04871 | 0.147359 |
| 13 | 56066400 | 56071599 | 26.61091 | 43.09314 | 94.97337 | 40.63843 | -0.77343 | 1.652098 | 0.147375 |
| 9 | 22351800 | 22357199 | 34.78698 | 55.87294 | 118.1469 | 50.03064 | -0.72793 | 1.911658 | 0.147378 |
| 4 | 144061400 | 144065599 | 46.34247 | 75.57471 | 53.44004 | 22.95723 | 0.870589 | 1.541282 | 0.147381 |
| 15 | 36608000 | 36611199 | 18.0788 | 30.35894 | 67.14159 | 29.3176 | -0.76003 | 1.135784 | 0.147388 |
| 5 | 3058600 | 3059799 | 43.2801 | 71.34914 | 46.71237 | 20.15484 | 0.988978 | 1.393441 | 0.147411 |
| 5 | 111023200 | 111025199 | 7.150828 | 13.27845 | 34.24327 | 14.34928 | -0.91183 | 0.167646 | 0.147412 |
| 17 | 26597400 | 26598999 | 7.150828 | 13.27845 | 34.24327 | 14.34928 | -0.91183 | 0.167646 | 0.147412 |
| 1 | 78389000 | 78390399 | 7.293346 | 13.20885 | 41.25598 | 17.86677 | -1.22036 | 0.325935 | 0.14742 |
| X | 24652600 | 24656199 | 53.35194 | 87.12586 | 45.12999 | 19.39709 | 1.319622 | 1.535023 | 0.147426 |
| 7 | 117842200 | 117844999 | 20.22304 | 33.25339 | 82.80928 | 35.6908 | -0.93397 | 1.394528 | 0.147427 |
| 11 | 32943200 | 32944999 | 8.179459 | 14.76969 | 40.12082 | 17.4399 | -1.02403 | 0.348081 | 0.14745 |
| 10 | 9276200 | 9277199 | 3.717453 | 7.219908 | 26.7167 | 9.995517 | -1.277 | -0.28127 | 0.147455 |
| 3 | 81787000 | 81787399 | 10.04115 | 19.55604 | 14.22437 | 5.198655 | 1.073908 | -0.32775 | 0.147456 |
| 8 | 71885000 | 71885999 | 34.19997 | 58.72996 | 26.19511 | 11.48981 | 1.542522 | 0.95074 | 0.147456 |
| 15 | 83569600 | 83588199 | 106.112 | 158.0557 | 390.5425 | 160.9293 | -1.04694 | 3.595685 | 0.147456 |
| 18 | 25856800 | 25859399 | 12.6645 | 21.99914 | 52.1091 | 22.87572 | -0.85121 | 0.788743 | 0.147461 |
| 10 | 62322000 | 62323199 | 5.552384 | 10.39791 | 36.10887 | 14.92138 | -1.32138 | 0.101642 | 0.147475 |
| 10 | 80638000 | 80643399 | 20.51143 | 33.34548 | 90.90143 | 39.00578 | -1.07371 | 1.560742 | 0.147499 |
| 7 | 112552200 | 112554999 | 20.05271 | 32.96507 | 80.25536 | 34.58641 | -0.90144 | 1.397501 | 0.147507 |
| 10 | 127577200 | 127583399 | 43.09285 | 68.35851 | 150.4634 | 62.65435 | -0.80554 | 2.313383 | 0.147522 |
| 14 | 42686600 | 42688399 | 27.03789 | 46.73329 | 30.86684 | 13.54162 | 0.981275 | 0.870695 | 0.147538 |
| 7 | 11700200 | 11706399 | 117.3394 | 181.3243 | 141.2361 | 58.66637 | 0.661386 | 2.865382 | 0.147546 |
| 15 | 77070200 | 77070799 | 11.12039 | 21.67313 | 13.02426 | 4.716617 | 1.358119 | -0.3481 | 0.147548 |
| 4 | 151473800 | 151476599 | 12.49073 | 21.76903 | 50.50013 | 22.21542 | -0.82069 | 0.738399 | 0.147554 |
| 15 | 6901200 | 6904599 | 13.83805 | 23.08079 | 73.58202 | 31.99284 | -1.28513 | 1.202972 | 0.147556 |
| X | 24667000 | 24667799 | 12.89957 | 24.43853 | 18.34958 | 7.421162 | 0.896429 | 0.047575 | 0.147576 |
| 14 | 43887600 | 43888599 | 6.018774 | 11.48613 | 30.57792 | 12.17751 | -0.89478 | -0.03439 | 0.147581 |
| 9 | 69665800 | 69672799 | 52.80877 | 82.84874 | 183.7352 | 76.20069 | -0.82944 | 2.574123 | 0.147601 |
| 16 | 91854600 | 91856399 | 8.012683 | 14.14748 | 44.09983 | 19.51142 | -1.24001 | 0.518864 | 0.147616 |
| 5 | 89462200 | 89463999 | 19.51551 | 35.0391 | 23.05938 | 10.08595 | 1.002192 | 0.424506 | 0.147633 |
| 9 | 64491800 | 64496999 | 29.50768 | 47.7408 | 97.86286 | 41.84683 | -0.67052 | 1.676709 | 0.147638 |
| 2 | 179396400 | 179400599 | 20.98391 | 34.52833 | 79.32976 | 34.20111 | -0.81762 | 1.387634 | 0.14764 |
| 18 | 43437600 | 43438399 | 13.6185 | 25.88677 | 18.25293 | 7.330186 | 0.99523 | 0.032125 | 0.147649 |
| 14 | 15909600 | 15910999 | 15.91172 | 27.13347 | 60.11014 | 26.35099 | -0.76063 | 0.985133 | 0.147651 |
| 2 | 111584600 | 111585399 | 10.84166 | 21.19304 | 7.274158 | 2.502543 | 2.20816 | -0.54447 | 0.147663 |
| 14 | 52243800 | 52249199 | 23.55413 | 38.26283 | 92.2324 | 39.55692 | -0.898 | 1.580018 | 0.147685 |
| 14 | 7449400 | 7450599 | 32.54557 | 54.55659 | 42.21673 | 18.41369 | 0.749697 | 1.176368 | 0.147699 |
| 1 | 89340400 | 89343599 | 16.12151 | 27.29761 | 63.86213 | 27.95909 | -0.84018 | 1.052004 | 0.147701 |
| 7 | 49280000 | 49292399 | 104.1674 | 157.312 | 299.4593 | 123.7939 | -0.65307 | 3.327536 | 0.147736 |
| 14 | 43006800 | 43007799 | 15.98054 | 29.16683 | 21.46633 | 9.20694 | 0.85977 | 0.30144 | 0.147743 |
| 7 | 15712200 | 15716599 | 17.55771 | 28.82419 | 82.47247 | 35.5214 | -1.13423 | 1.411073 | 0.147744 |
| 3 | 98277400 | 98279999 | 27.74268 | 47.81481 | 33.24715 | 14.58463 | 0.906556 | 0.900459 | 0.147744 |

| | | | | | | | | | |
|---|---|---|---|---|---|---|---|---|---|
| 13 | 12723200 | 12725799 | 15.03477 | 25.9091 | 57.56235 | 25.25094 | -0.76209 | 0.88081 | 0.14775 |
| 14 | 82784400 | 82786399 | 22.68621 | 40.27228 | 22.71288 | 10.02991 | 1.214259 | 0.514008 | 0.14776 |
| 3 | 9197000 | 9199599 | 13.09901 | 22.41206 | 58.9282 | 25.83983 | -1.00564 | 0.941472 | 0.147762 |
| 8 | 21515600 | 21517599 | 23.54033 | 41.14838 | 30.13486 | 13.28231 | 0.834819 | 0.717972 | 0.147778 |
| 2 | 47018600 | 47018999 | 8.965025 | 17.45785 | 13.94762 | 5.106499 | 0.93843 | -0.33628 | 0.147782 |
| 7 | 41988200 | 41997999 | 171.8021 | 261.1767 | 157.7189 | 65.33308 | 1.009995 | 3.201848 | 0.147801 |
| 19 | 24104600 | 24110399 | 73.96193 | 114.5758 | 212.8953 | 88.43088 | -0.58889 | 2.809633 | 0.147814 |
| 3 | 94881200 | 94882799 | 7.382135 | 13.49307 | 38.78269 | 16.60376 | -1.09035 | 0.267629 | 0.147819 |
| 12 | 16943200 | 16945799 | 11.31302 | 19.62154 | 54.74591 | 24.02017 | -1.08665 | 0.802655 | 0.147832 |
| 7 | 86107800 | 86108599 | 13.75084 | 26.41652 | 15.84995 | 6.200053 | 1.258878 | -0.05688 | 0.147834 |
| 4 | 78051600 | 78054199 | 41.50726 | 68.60542 | 45.56781 | 19.69119 | 0.968895 | 1.35671 | 0.147837 |
| 6 | 34123400 | 34124999 | 26.31444 | 45.26205 | 35.17462 | 15.42921 | 0.746451 | 0.916434 | 0.147845 |
| 17 | 71155600 | 71161199 | 28.15523 | 45.09761 | 129.1274 | 54.45301 | -1.16894 | 1.99068 | 0.147867 |
| 7 | 10247800 | 10248599 | 16.51201 | 30.1665 | 21.87816 | 9.372036 | 0.881994 | 0.301229 | 0.147868 |
| 6 | 123124800 | 123125399 | 9.010575 | 17.58514 | 13.2586 | 4.710283 | 1.058774 | -0.44205 | 0.147871 |
| 9 | 107775800 | 107780399 | 15.98029 | 26.36666 | 81.11714 | 35.01197 | -1.23552 | 1.341181 | 0.147874 |
| 6 | 135012000 | 135013399 | 8.753187 | 15.90655 | 40.81807 | 17.60238 | -0.93473 | 0.319409 | 0.147874 |
| 2 | 121185200 | 121186199 | 3.823684 | 7.446856 | 24.72316 | 9.037503 | -1.09087 | -0.36889 | 0.147878 |
| 2 | 174261200 | 174262599 | 7.131303 | 13.30283 | 33.81614 | 14.0664 | -0.88289 | 0.137329 | 0.147889 |
| 6 | 118914600 | 118915799 | 7.131303 | 13.30283 | 33.81614 | 14.0664 | -0.88289 | 0.137329 | 0.147889 |
| 4 | 152682000 | 152694799 | 101.8727 | 153.9049 | 300.6293 | 124.2925 | -0.68978 | 3.319269 | 0.147894 |
| 2 | 128823200 | 128824799 | 18.32655 | 32.9486 | 23.89546 | 10.43982 | 0.864295 | 0.410751 | 0.147904 |
| 1 | 13659200 | 13660599 | 10.47548 | 18.31667 | 48.33699 | 21.31061 | -1.00481 | 0.676662 | 0.147914 |
| 19 | 3059000 | 3059799 | 23.76365 | 41.37741 | 27.57879 | 12.1251 | 0.969728 | 0.769686 | 0.147918 |
| 11 | 103303000 | 103303999 | 5.388557 | 10.10392 | 33.3254 | 13.74161 | -1.24423 | 0.087449 | 0.14794 |
| X | 34761000 | 34761799 | 22.72187 | 39.937 | 26.96881 | 11.94226 | 0.951018 | 0.62322 | 0.147959 |
| 10 | 117964600 | 117965599 | 7.620747 | 14.06719 | 35.21244 | 14.8752 | -0.87787 | 0.207665 | 0.147969 |
| 11 | 101324200 | 101326199 | 7.956206 | 14.5676 | 36.66032 | 15.65897 | -0.8971 | 0.26103 | 0.147976 |
| 15 | 77388600 | 77389399 | 7.956206 | 14.5676 | 36.66032 | 15.65897 | -0.8971 | 0.26103 | 0.147976 |
| 8 | 71287400 | 71298199 | 45.34245 | 71.25712 | 184.6066 | 76.57645 | -1.0514 | 2.535925 | 0.147987 |
| 13 | 81804600 | 81806399 | 8.999229 | 16.01994 | 40.99346 | 18.06001 | -0.95155 | 0.446818 | 0.147991 |
| 15 | 49813000 | 49814399 | 17.67523 | 33.43474 | 15.36545 | 6.234514 | 1.593551 | 0.094354 | 0.148 |
| 5 | 135095000 | 135097799 | 19.15867 | 31.77588 | 74.01569 | 32.06059 | -0.83499 | 1.282738 | 0.148016 |
| 5 | 79170200 | 79172599 | 51.28815 | 82.6862 | 65.40598 | 27.84658 | 0.694243 | 1.79235 | 0.148031 |
| 6 | 114867800 | 114873999 | 37.39972 | 59.87037 | 122.2312 | 51.49996 | -0.68314 | 2.01792 | 0.148032 |
| 2 | 44454800 | 44455799 | 5.711631 | 10.94815 | 29.59526 | 11.65098 | -0.90085 | -0.07765 | 0.148041 |
| 7 | 30265600 | 30267399 | 4.913137 | 9.264776 | 31.77733 | 12.97824 | -1.28779 | 0.044197 | 0.148046 |
| 16 | 34995800 | 35000999 | 23.28416 | 37.73738 | 96.33821 | 41.24783 | -0.98369 | 1.628761 | 0.148049 |
| 13 | 94623000 | 94624399 | 7.589052 | 14.11069 | 34.84051 | 14.57265 | -0.8479 | 0.164487 | 0.148065 |
| 4 | 26284400 | 26286799 | 15.59862 | 26.68934 | 66.82083 | 29.30076 | -0.93573 | 0.961093 | 0.148094 |
| 2 | 51849600 | 51851199 | 19.10158 | 34.46486 | 21.98456 | 9.565699 | 1.05159 | 0.390328 | 0.148103 |
| 14 | 76414200 | 76419599 | 34.32211 | 55.01088 | 118.9313 | 50.20255 | -0.76345 | 1.975048 | 0.148129 |
| 3 | 5628600 | 5629399 | 11.77099 | 22.65478 | 17.35152 | 6.751594 | 0.917768 | -0.08925 | 0.148155 |
| 1 | 25401800 | 25404399 | 22.83697 | 40.08569 | 27.79554 | 12.29446 | 0.912865 | 0.644252 | 0.148166 |
| 6 | 125703000 | 125708399 | 35.64949 | 57.02304 | 126.925 | 53.41723 | -0.80883 | 2.041306 | 0.14819 |
| 8 | 51455400 | 51456999 | 25.8194 | 45.23243 | 26.5192 | 11.71059 | 1.153731 | 0.689425 | 0.148193 |
| 7 | 48085200 | 48086999 | 28.5616 | 48.69357 | 37.75217 | 16.54943 | 0.747882 | 1.006693 | 0.148202 |

| | | | | | | | | | |
|---|---|---|---|---|---|---|---|---|---|
| 10 | 60868800 | 60902399 | 225.045 | 318.1485 | 625.4983 | 258.9111 | -0.79087 | 4.415017 | 0.148209 |
| 3 | 107502000 | 107509999 | 50.40376 | 78.73561 | 200.5368 | 83.2057 | -1.03373 | 2.651682 | 0.148227 |
| 13 | 61781600 | 61784799 | 95.33269 | 149.4843 | 119.5726 | 49.59313 | 0.639263 | 2.598107 | 0.148247 |
| Y | 41243000 | 41243799 | 8.353676 | 16.32768 | 6.487118 | 2.238481 | 2.002056 | -0.58272 | 0.148251 |
| 2 | 28623200 | 28629799 | 29.24186 | 46.84272 | 124.3789 | 52.45837 | -1.06016 | 1.987987 | 0.148255 |
| 12 | 103024600 | 103027799 | 18.19152 | 30.70972 | 66.17556 | 28.94538 | -0.7224 | 1.088457 | 0.14826 |
| 17 | 17724400 | 17729999 | 71.59867 | 110.5987 | 215.4055 | 89.47488 | -0.65983 | 2.862542 | 0.148261 |
| 13 | 54651200 | 54652399 | 5.751586 | 10.82627 | 32.3132 | 13.23755 | -1.09304 | 0.066457 | 0.148263 |
| 5 | 110100200 | 110103399 | 41.17165 | 67.62722 | 51.63652 | 22.24524 | 0.765822 | 1.43554 | 0.148264 |
| 5 | 122203600 | 122222599 | 108.9698 | 162.5953 | 356.6133 | 146.9881 | -0.87277 | 3.563092 | 0.148272 |
| X | 166962800 | 166964399 | 24.14204 | 42.23996 | 29.08211 | 12.8285 | 0.923477 | 0.705608 | 0.148281 |
| 12 | 65074800 | 65076399 | 20.1175 | 35.63789 | 25.80949 | 11.41154 | 0.854506 | 0.520083 | 0.148281 |
| 4 | 71595600 | 71597999 | 26.10857 | 42.35136 | 93.01305 | 39.84938 | -0.76611 | 1.614888 | 0.148284 |
| 14 | 23459600 | 23460599 | 8.805288 | 15.47484 | 47.2049 | 20.90116 | -1.2123 | 0.580367 | 0.148293 |
| 15 | 82926800 | 82936599 | 75.57018 | 116.324 | 227.9392 | 94.66951 | -0.67252 | 2.930269 | 0.148296 |
| 11 | 46854600 | 46855199 | 10.15452 | 19.74916 | 14.75165 | 5.457501 | 1.020155 | -0.2833 | 0.148311 |
| 17 | 37959800 | 37960999 | 32.5536 | 54.71123 | 41.61468 | 18.17787 | 0.774347 | 1.149813 | 0.148327 |
| 7 | 15267200 | 15267999 | 25.19978 | 43.88698 | 28.63832 | 12.59254 | 1.000015 | 0.774917 | 0.148331 |
| 3 | 98574400 | 98575599 | 19.65382 | 34.76217 | 26.03919 | 11.5167 | 0.805378 | 0.534668 | 0.148332 |
| 13 | 3541000 | 3546399 | 48.04716 | 77.83696 | 60.55673 | 25.91926 | 0.725501 | 1.661225 | 0.148339 |
| 19 | 41065000 | 41068399 | 14.90651 | 25.60342 | 57.48926 | 25.21634 | -0.77827 | 0.910588 | 0.148344 |
| 15 | 54451000 | 54452199 | 4.486285 | 8.73842 | 26.76299 | 9.766933 | -0.97471 | -0.35981 | 0.148355 |
| 10 | 124417200 | 124418199 | 12.63523 | 24.25187 | 17.13657 | 6.72039 | 1.022102 | -0.05689 | 0.148357 |
| 19 | 8222200 | 8223799 | 37.00296 | 61.37484 | 47.15337 | 20.42563 | 0.759949 | 1.307505 | 0.148362 |
| 6 | 142510800 | 142513999 | 22.26573 | 36.59667 | 81.96385 | 35.32006 | -0.78176 | 1.404256 | 0.148363 |
| 7 | 81644400 | 81646599 | 12.63345 | 22.07246 | 49.95693 | 22.01077 | -0.78494 | 0.702601 | 0.148373 |
| 9 | 118777600 | 118788399 | 72.70965 | 112.5146 | 216.0248 | 89.73128 | -0.63727 | 2.828917 | 0.148406 |
| 3 | 36867200 | 36868599 | 7.094727 | 13.36667 | 36.50033 | 14.92978 | -0.96608 | 0.078439 | 0.148414 |
| 5 | 41911200 | 41922199 | 90.50585 | 137.4502 | 281.9837 | 116.773 | -0.75409 | 3.209171 | 0.148425 |
| 2 | 31462000 | 31463199 | 6.491004 | 12.27759 | 34.92414 | 14.1726 | -1.01465 | 0.0405 | 0.148427 |
| 12 | 22610400 | 22613799 | 59.73718 | 95.81199 | 76.52956 | 32.33819 | 0.671395 | 1.969502 | 0.148427 |
| X | 52912800 | 52915399 | 56.16717 | 90.59195 | 64.58861 | 27.51419 | 0.844559 | 1.790214 | 0.148429 |
| X | 31713600 | 31714199 | 11.29985 | 22.06233 | 10.54613 | 3.719182 | 1.715396 | -0.4434 | 0.14843 |
| X | 53813000 | 53813599 | 11.29985 | 22.06233 | 10.54613 | 3.719182 | 1.715396 | -0.4434 | 0.14843 |
| 5 | 143313200 | 143316199 | 12.75724 | 21.9281 | 58.47761 | 25.65044 | -1.02489 | 0.895707 | 0.148447 |
| 7 | 7813800 | 7815199 | 33.53796 | 56.30076 | 38.43179 | 16.77797 | 0.929791 | 1.159601 | 0.148454 |
| 11 | 7288000 | 7288999 | 33.0809 | 55.62062 | 41.07215 | 17.94316 | 0.816939 | 1.147894 | 0.14847 |
| 14 | 31832000 | 31836199 | 43.95562 | 70.01437 | 137.6979 | 57.61046 | -0.63734 | 2.185023 | 0.148481 |
| 7 | 41432400 | 41440399 | 166.7836 | 253.1666 | 172.022 | 71.22588 | 0.838216 | 3.231244 | 0.148485 |
| 6 | 29069400 | 29075199 | 32.17045 | 51.73234 | 111.1702 | 47.15836 | -0.74895 | 1.865469 | 0.148494 |
| 8 | 85580400 | 85581199 | 5.585705 | 10.66928 | 29.99705 | 11.92344 | -0.97001 | -0.04616 | 0.148515 |
| 18 | 74529800 | 74531199 | 5.585705 | 10.66928 | 29.99705 | 11.92344 | -0.97001 | -0.04616 | 0.148515 |
| 5 | 104834600 | 104835399 | 12.45532 | 23.82331 | 17.86001 | 7.074598 | 0.924827 | -0.02383 | 0.148518 |
| 18 | 61098800 | 61114799 | 88.1365 | 133.4614 | 289.0727 | 119.6018 | -0.83582 | 3.26932 | 0.148519 |
| 9 | 114328800 | 114347599 | 280.702 | 390.927 | 709.7188 | 294.4713 | -0.69776 | 4.640701 | 0.14852 |
| 7 | 10828200 | 10834399 | 111.0296 | 172.4042 | 133.1382 | 55.28631 | 0.679088 | 2.774255 | 0.148525 |
| 9 | 46030400 | 46031599 | 14.03744 | 26.30144 | 19.57607 | 8.082381 | 0.883859 | 0.130997 | 0.148545 |

| | | | | | | | | | |
|---|---|---|---|---|---|---|---|---|---|
| 5 | 113349000 | 113369399 | 115.4012 | 171.3427 | 375.9668 | 154.8726 | -0.88 | 3.653055 | 0.148571 |
| 15 | 18874400 | 18876199 | 21.77229 | 38.75323 | 23.42676 | 10.3219 | 1.117103 | 0.493778 | 0.148576 |
| 7 | 52497600 | 52500999 | 58.0118 | 93.33974 | 69.73649 | 29.60641 | 0.773004 | 1.866538 | 0.148587 |
| 11 | 64324600 | 64327599 | 22.12584 | 36.32054 | 81.73504 | 35.20253 | -0.78933 | 1.420413 | 0.148608 |
| 10 | 121349600 | 121350999 | 11.06033 | 19.4053 | 46.18826 | 20.42198 | -0.85679 | 0.620176 | 0.148633 |
| 1 | 111622800 | 111624399 | 15.35373 | 28.17899 | 21.06112 | 8.967009 | 0.845023 | 0.265022 | 0.148655 |
| 11 | 62678800 | 62680399 | 8.961266 | 16.0627 | 41.57815 | 18.21146 | -0.96256 | 0.401406 | 0.148658 |
| 5 | 97673400 | 97674799 | 9.647004 | 17.05823 | 44.57935 | 19.71777 | -0.98732 | 0.510732 | 0.148662 |
| 8 | 56746400 | 56748599 | 9.647004 | 17.05823 | 44.57935 | 19.71777 | -0.98732 | 0.510732 | 0.148662 |
| 6 | 125487400 | 125488799 | 8.055468 | 14.91583 | 36.97239 | 15.55291 | -0.85987 | 0.196246 | 0.148667 |
| 5 | 53581200 | 53583399 | 10.40867 | 18.36904 | 46.68471 | 20.65707 | -0.94878 | 0.539945 | 0.148675 |
| 9 | 72801000 | 72802999 | 10.54478 | 18.68852 | 42.99692 | 19.00913 | -0.80316 | 0.489986 | 0.148679 |
| 9 | 117311000 | 117314799 | 26.01149 | 42.4904 | 88.92538 | 38.22468 | -0.68888 | 1.499009 | 0.14868 |
| 5 | 5429600 | 5433799 | 18.76535 | 31.33041 | 71.55755 | 31.13222 | -0.80638 | 1.203381 | 0.148684 |
| 2 | 169139800 | 169145199 | 53.76764 | 84.77143 | 170.4823 | 70.74733 | -0.68301 | 2.473625 | 0.148691 |
| 5 | 148193600 | 148205799 | 92.01764 | 140.2997 | 265.8297 | 110.2109 | -0.63481 | 3.127843 | 0.148692 |
| 4 | 27859800 | 27864199 | 45.60426 | 75.15124 | 44.49573 | 19.19525 | 1.133301 | 1.40261 | 0.148702 |
| 3 | 108315400 | 108324599 | 101.5484 | 151.9101 | 341.5512 | 140.8499 | -0.90517 | 3.51414 | 0.148703 |
| 10 | 108229200 | 108233199 | 22.17629 | 36.71186 | 80.25753 | 34.71932 | -0.74432 | 1.315879 | 0.148706 |
| 15 | 54975600 | 54979999 | 21.5863 | 35.60246 | 82.27943 | 35.5078 | -0.82493 | 1.358863 | 0.148732 |
| 15 | 84502800 | 84507199 | 40.21208 | 64.15555 | 131.2363 | 55.03049 | -0.69138 | 2.127886 | 0.148738 |
| 7 | 86146000 | 86147799 | 27.54051 | 47.96789 | 27.55286 | 12.11595 | 1.182836 | 0.782615 | 0.148743 |
| 3 | 146856200 | 146858399 | 13.42211 | 23.40127 | 52.67947 | 23.17975 | -0.77776 | 0.739537 | 0.148747 |
| 7 | 3147200 | 3149199 | 47.9803 | 77.57209 | 58.51867 | 25.00523 | 0.76714 | 1.704655 | 0.148748 |
| 4 | 135523600 | 135525999 | 9.299275 | 16.23733 | 50.42713 | 22.20946 | -1.23839 | 0.694765 | 0.148756 |
| 6 | 142056200 | 142057999 | 12.7839 | 22.2741 | 52.1375 | 22.93192 | -0.83393 | 0.743318 | 0.148765 |
| 12 | 82387800 | 82389399 | 14.34823 | 24.47344 | 59.97465 | 26.29212 | -0.90527 | 0.978462 | 0.148782 |
| 6 | 19962400 | 19963599 | 14.30128 | 26.61472 | 19.54795 | 8.16397 | 0.889699 | 0.175133 | 0.148784 |
| 3 | 97674200 | 97675799 | 10.6335 | 18.59969 | 49.92287 | 22.01649 | -1.02949 | 0.671663 | 0.148787 |
| 9 | 60742800 | 60745999 | 13.57881 | 23.60312 | 52.69202 | 23.14132 | -0.7663 | 0.782699 | 0.148809 |
| 4 | 80917600 | 80920399 | 54.67999 | 87.95229 | 71.5531 | 30.36432 | 0.649861 | 1.87136 | 0.148824 |
| 15 | 8888600 | 8889599 | 14.97735 | 28.94266 | 13.60585 | 5.221095 | 1.632804 | -0.1094 | 0.148831 |
| 5 | 122139000 | 122148999 | 121.8708 | 180.2793 | 390.4327 | 160.7908 | -0.86611 | 3.718383 | 0.148852 |
| 4 | 97953400 | 97954799 | 5.234947 | 10.12166 | 28.27997 | 10.82457 | -0.90885 | -0.17873 | 0.148866 |
| 3 | 69060200 | 69061799 | 8.759474 | 15.87041 | 38.59525 | 16.70621 | -0.86117 | 0.332214 | 0.148869 |
| 1 | 156208600 | 156209399 | 13.30813 | 25.13366 | 19.20981 | 7.820753 | 0.862378 | 0.074019 | 0.148895 |
| 11 | 98306400 | 98314799 | 35.58737 | 56.67804 | 138.4022 | 57.89518 | -0.94903 | 2.182833 | 0.148896 |
| 16 | 17737600 | 17740399 | 13.39128 | 22.94772 | 58.44498 | 25.63114 | -0.95961 | 0.927171 | 0.148901 |
| 12 | 56000400 | 56001399 | 13.51749 | 25.75732 | 18.1502 | 7.251536 | 1.002162 | 0.013872 | 0.148902 |
| X | 4765200 | 4771399 | 70.24193 | 112.6406 | 68.52279 | 28.94708 | 1.063162 | 1.986366 | 0.148904 |
| 10 | 6125200 | 6126399 | 11.78926 | 22.61651 | 16.44338 | 6.45807 | 0.980191 | -0.06175 | 0.148914 |
| 18 | 66277600 | 66278999 | 14.64176 | 27.21903 | 20.66264 | 8.645222 | 0.84077 | 0.188789 | 0.148935 |
| 4 | 142888200 | 142889399 | 6.731688 | 12.47326 | 35.83098 | 15.05863 | -1.06961 | 0.178375 | 0.148944 |
| 10 | 72393000 | 72393799 | 12.35636 | 24.06497 | 12.17717 | 4.450812 | 1.590519 | -0.30393 | 0.148947 |
| 3 | 107603000 | 107604599 | 9.693169 | 17.00861 | 46.19636 | 20.42759 | -1.04579 | 0.611109 | 0.14896 |
| 19 | 43951000 | 43952999 | 9.864982 | 17.68817 | 41.57813 | 18.2066 | -0.82419 | 0.406342 | 0.148963 |
| 16 | 17986400 | 17989799 | 14.21654 | 24.19701 | 60.49846 | 26.51718 | -0.93467 | 0.996855 | 0.148976 |

| | | | | | | | | | |
|---|---|---|---|---|---|---|---|---|---|
| 8 | 93399600 | 93408999 | 74.65272 | 114.4706 | 244.4725 | 101.4827 | -0.80085 | 3.00682 | 0.148988 |
| 14 | 16481400 | 16482999 | 8.83394 | 15.75754 | 40.50031 | 17.81235 | -0.95598 | 0.432602 | 0.149001 |
| 10 | 32912200 | 32913599 | 7.571571 | 14.13486 | 35.55154 | 14.76924 | -0.86663 | 0.1392 | 0.149006 |
| 5 | 119553600 | 119555599 | 8.556147 | 15.64347 | 38.26738 | 16.37718 | -0.85924 | 0.274428 | 0.149007 |
| 3 | 136630800 | 136632399 | 15.49566 | 28.56065 | 20.9563 | 8.870911 | 0.877264 | 0.2424 | 0.149011 |
| 5 | 61202400 | 61203199 | 10.28765 | 19.91447 | 15.90107 | 6.052864 | 0.887565 | -0.18192 | 0.149026 |
| 6 | 122787600 | 122788599 | 4.66613 | 8.800001 | 32.8734 | 13.42329 | -1.40996 | 0.042164 | 0.149028 |
| 4 | 137187600 | 137191799 | 26.66045 | 43.06123 | 100.1393 | 42.73606 | -0.8547 | 1.719034 | 0.149029 |
| 2 | 162814400 | 162818599 | 19.01207 | 31.49943 | 74.45407 | 32.22696 | -0.85614 | 1.295676 | 0.149061 |
| 5 | 112472600 | 112483799 | 49.6133 | 77.72132 | 184.8435 | 76.67194 | -0.93174 | 2.59958 | 0.149061 |
| 4 | 109521400 | 109524599 | 17.73523 | 29.85678 | 67.21984 | 29.37751 | -0.78559 | 1.111138 | 0.149072 |
| 14 | 52734200 | 52737399 | 41.04039 | 68.27127 | 41.01025 | 17.80056 | 1.113896 | 1.280543 | 0.149078 |
| 5 | 53687600 | 53689599 | 9.382228 | 16.90733 | 40.07189 | 17.46273 | -0.83114 | 0.369014 | 0.149086 |
| 3 | 108817600 | 108819199 | 8.057142 | 14.47079 | 39.52145 | 17.28168 | -1.03708 | 0.380453 | 0.149086 |
| X | 33483000 | 33486999 | 64.79315 | 104.3333 | 60.53277 | 25.72849 | 1.138376 | 1.850469 | 0.149088 |
| 16 | 94043400 | 94050199 | 31.46337 | 50.40145 | 122.6534 | 51.73077 | -0.9347 | 1.989876 | 0.149095 |
| 9 | 48493400 | 48494199 | 11.07915 | 21.50922 | 15.17098 | 5.693674 | 1.083336 | -0.22172 | 0.149103 |
| 10 | 82352600 | 82356199 | 68.47749 | 109.3449 | 84.0309 | 35.27899 | 0.719857 | 2.114779 | 0.149115 |
| 1 | 142080200 | 142080999 | 4.704903 | 9.10046 | 28.90742 | 11.04416 | -1.09002 | -0.19232 | 0.149131 |
| 6 | 68836200 | 68837999 | 17.11774 | 32.17515 | 16.36029 | 6.718361 | 1.434467 | 0.131032 | 0.149139 |
| 5 | 108279800 | 108280999 | 7.817435 | 13.93546 | 42.05645 | 18.5067 | -1.18668 | 0.431192 | 0.14914 |
| 8 | 125589400 | 125590999 | 6.984061 | 12.68884 | 39.71056 | 17.12951 | -1.21912 | 0.302489 | 0.149144 |
| 9 | 121533400 | 121535399 | 6.984061 | 12.68884 | 39.71056 | 17.12951 | -1.21912 | 0.302489 | 0.149144 |
| 12 | 113219200 | 113242999 | 204.9626 | 291.9021 | 580.7304 | 240.0848 | -0.79722 | 4.305795 | 0.149147 |
| 3 | 153480200 | 153483199 | 19.52965 | 32.66578 | 71.73574 | 31.24555 | -0.75004 | 1.186075 | 0.149151 |
| 4 | 124170600 | 124174199 | 20.07957 | 33.35698 | 74.32677 | 32.2312 | -0.77113 | 1.265478 | 0.149159 |
| 6 | 72415800 | 72417999 | 8.41223 | 14.96001 | 41.51558 | 18.30548 | -1.06854 | 0.450163 | 0.149178 |
| 8 | 26359800 | 26361599 | 18.28922 | 32.76025 | 24.41452 | 10.69987 | 0.822213 | 0.433739 | 0.14918 |
| 5 | 89331000 | 89331599 | 6.32972 | 12.01576 | 31.55093 | 12.70649 | -0.8897 | 0.007617 | 0.149181 |
| 8 | 98854600 | 98855399 | 12.75889 | 24.39711 | 17.63193 | 6.989889 | 0.975446 | -0.02032 | 0.14919 |
| 14 | 70710600 | 70712999 | 10.09668 | 17.85159 | 43.0143 | 19.02591 | -0.87077 | 0.513284 | 0.149239 |
| 15 | 97125600 | 97141999 | 145.1432 | 215.2105 | 383.0185 | 157.7557 | -0.57999 | 3.680887 | 0.149241 |
| 14 | 54701600 | 54703999 | 9.403051 | 16.88171 | 40.01285 | 17.50211 | -0.83475 | 0.392499 | 0.14925 |
| 10 | 52293400 | 52294799 | 10.14748 | 17.80162 | 46.19816 | 20.42434 | -0.98061 | 0.61741 | 0.149253 |
| 10 | 23751600 | 23752199 | 14.49468 | 27.10947 | 17.36737 | 7.192292 | 1.095735 | 0.141948 | 0.14926 |
| 19 | 21151200 | 21153199 | 10.04173 | 17.9368 | 41.52196 | 18.24034 | -0.80537 | 0.432299 | 0.149261 |
| 12 | 106275400 | 106278399 | 23.33506 | 38.04445 | 88.11383 | 37.84904 | -0.83657 | 1.520704 | 0.149294 |
| 10 | 77251000 | 77255199 | 24.06598 | 39.39207 | 84.11149 | 36.18213 | -0.71556 | 1.462469 | 0.149319 |
| 2 | 169384200 | 169388799 | 20.12691 | 32.98936 | 88.96979 | 38.29399 | -1.0506 | 1.437844 | 0.149322 |
| 9 | 100955200 | 100957799 | 15.29881 | 26.03932 | 60.59137 | 26.55793 | -0.83163 | 1.000102 | 0.149326 |
| 13 | 53438400 | 53442599 | 35.932 | 57.59752 | 118.1018 | 49.86071 | -0.68705 | 1.973909 | 0.14936 |
| 17 | 54681000 | 54682799 | 23.32827 | 41.54499 | 20.99502 | 9.246292 | 1.373931 | 0.495614 | 0.149378 |
| 4 | 9829600 | 9831999 | 12.77368 | 22.28202 | 52.68324 | 23.18879 | -0.84825 | 0.727077 | 0.149385 |
| 11 | 68949200 | 68952399 | 13.4819 | 22.87892 | 62.55551 | 27.40551 | -1.06362 | 1.023091 | 0.149388 |
| 2 | 165854600 | 165857599 | 10.69377 | 18.94918 | 43.51376 | 19.23832 | -0.80069 | 0.492917 | 0.149432 |
| 4 | 67981400 | 67982799 | 15.7596 | 29.02491 | 21.5134 | 9.116223 | 0.861919 | 0.250738 | 0.149433 |
| 14 | 121701600 | 121703199 | 7.836486 | 14.2607 | 37.08697 | 15.96532 | -0.95187 | 0.295622 | 0.149439 |

| | | | | | | | | | |
|---|---|---|---|---|---|---|---|---|---|
| 11 | 114171000 | 114176599 | 26.70735 | 43.06996 | 106.9125 | 45.53521 | -0.95129 | 1.766728 | 0.149448 |
| 17 | 45995800 | 45998799 | 13.26602 | 22.67274 | 60.01343 | 26.31335 | -1.0156 | 0.95301 | 0.149473 |
| 11 | 48581800 | 48582999 | 11.99545 | 21.00689 | 47.79637 | 21.09985 | -0.79234 | 0.657901 | 0.149474 |
| 1 | 73090000 | 73091999 | 13.06587 | 22.43777 | 57.37849 | 25.16762 | -0.96486 | 0.905117 | 0.149484 |
| 1 | 101274600 | 101275399 | 12.67477 | 24.24313 | 17.04137 | 6.75027 | 1.015876 | -0.02591 | 0.149497 |
| 1 | 65746000 | 65750399 | 84.89304 | 134.005 | 109.6653 | 45.52746 | 0.613731 | 2.457457 | 0.149504 |
| 7 | 82642600 | 82646199 | 24.38544 | 39.90818 | 84.76565 | 36.46127 | -0.70818 | 1.46578 | 0.149504 |
| 7 | 140133400 | 140134199 | 7.758023 | 14.34811 | 35.81636 | 15.09085 | -0.87151 | 0.197757 | 0.149514 |
| X | 39953600 | 39956399 | 41.88792 | 69.761 | 39.9149 | 17.33823 | 1.183937 | 1.27161 | 0.149535 |
| 15 | 20472800 | 20473999 | 16.77092 | 30.50092 | 21.81126 | 9.396504 | 0.896885 | 0.327974 | 0.149543 |
| 8 | 57797600 | 57804199 | 48.40853 | 76.76676 | 160.3741 | 66.76033 | -0.7309 | 2.327199 | 0.149553 |
| 6 | 52290400 | 52297599 | 63.00521 | 98.05756 | 198.4807 | 82.39191 | -0.70691 | 2.722476 | 0.149556 |
| 3 | 18848400 | 18850399 | 16.54986 | 30.45971 | 20.1842 | 8.560865 | 1.020978 | 0.256506 | 0.149558 |
| 9 | 74977800 | 74979399 | 10.14399 | 17.81761 | 46.1676 | 20.43226 | -0.97854 | 0.595538 | 0.14956 |
| 18 | 80278400 | 80283799 | 30.18852 | 48.71234 | 101.8127 | 43.39582 | -0.70283 | 1.753992 | 0.149567 |
| 14 | 112319200 | 112321199 | 31.71427 | 53.79017 | 35.65746 | 15.61892 | 0.972195 | 1.062109 | 0.149573 |
| 14 | 51634800 | 51636999 | 22.17385 | 39.18732 | 24.98558 | 11.05251 | 1.035828 | 0.556106 | 0.149578 |
| 16 | 37233800 | 37235999 | 20.66812 | 36.77587 | 23.99668 | 10.57591 | 1.007166 | 0.488735 | 0.149588 |
| 2 | 104518600 | 104520399 | 5.903201 | 10.89362 | 37.88511 | 16.01054 | -1.34874 | 0.199458 | 0.149588 |
| 16 | 23805600 | 23807799 | 8.610915 | 15.21144 | 44.08261 | 19.50183 | -1.1356 | 0.515733 | 0.149594 |
| 6 | 143257200 | 143258399 | 6.787324 | 12.81705 | 37.84422 | 15.41093 | -1.07274 | 0.061542 | 0.149604 |
| 19 | 18463000 | 18463799 | 16.40586 | 29.72924 | 22.10683 | 9.567031 | 0.837049 | 0.347351 | 0.14965 |
| 11 | 69999800 | 70001399 | 16.09109 | 29.47974 | 22.00002 | 9.389481 | 0.844833 | 0.283343 | 0.149655 |
| 6 | 93219600 | 93220199 | 9.531273 | 18.59939 | 13.24621 | 4.714398 | 1.138189 | -0.42783 | 0.149666 |
| 8 | 105406000 | 105407399 | 12.44283 | 23.64225 | 18.24563 | 7.333645 | 0.864385 | 0.021987 | 0.149669 |
| 6 | 3171800 | 3175399 | 36.74416 | 61.10077 | 45.47659 | 19.73391 | 0.805762 | 1.275767 | 0.149676 |
| 5 | 121849000 | 121851599 | 12.265 | 21.15573 | 55.78698 | 24.47283 | -1.00744 | 0.859262 | 0.14968 |
| 4 | 149065400 | 149070799 | 21.21415 | 34.72428 | 83.46498 | 35.9041 | -0.88569 | 1.456692 | 0.149683 |
| 1 | 119764400 | 119766199 | 7.069702 | 12.94113 | 36.12067 | 15.43382 | -1.04551 | 0.25371 | 0.149689 |
| 7 | 113105600 | 113106199 | 2.246631 | 4.394553 | 18.98483 | 6.353739 | -1.34094 | -0.72517 | 0.149692 |
| 9 | 124144600 | 124147599 | 33.66882 | 56.97382 | 38.19597 | 16.71917 | 0.955994 | 1.092146 | 0.149695 |
| 4 | 46545400 | 46552599 | 44.18912 | 70.16718 | 148.3608 | 61.84025 | -0.74628 | 2.283854 | 0.149695 |
| 1 | 92400000 | 92402199 | 10.94158 | 19.12252 | 51.00558 | 22.47804 | -1.02065 | 0.688243 | 0.149703 |
| 19 | 3634200 | 3638999 | 23.63265 | 38.57046 | 86.54678 | 37.19027 | -0.78991 | 1.505194 | 0.149707 |
| 10 | 60778000 | 60792399 | 103.382 | 156.0657 | 300.9953 | 124.4138 | -0.67238 | 3.33522 | 0.149707 |
| 18 | 64187800 | 64189399 | 16.49991 | 30.8073 | 18.66688 | 7.752803 | 1.171399 | 0.172129 | 0.149721 |
| 6 | 64425000 | 64425999 | 5.532335 | 10.69143 | 30.13433 | 11.56073 | -0.92533 | -0.16417 | 0.149745 |
| 18 | 76841800 | 76850599 | 49.49765 | 78.21702 | 160.5114 | 66.67249 | -0.70943 | 2.417725 | 0.149766 |
| 6 | 136089200 | 136090599 | 8.378978 | 15.4245 | 38.47832 | 16.31554 | -0.87759 | 0.234732 | 0.149792 |
| 3 | 18027600 | 18029999 | 24.2224 | 42.24563 | 30.58869 | 13.46232 | 0.850842 | 0.75262 | 0.149822 |
| 18 | 90104000 | 90106199 | 27.88736 | 48.78933 | 24.69724 | 10.8936 | 1.364311 | 0.718272 | 0.149849 |
| 10 | 40758800 | 40760799 | 10.69107 | 18.57908 | 53.11959 | 23.32557 | -1.12105 | 0.773637 | 0.149849 |
| 13 | 77739600 | 77742399 | 20.4089 | 36.33812 | 23.43246 | 10.32162 | 1.024476 | 0.481323 | 0.149854 |
| 17 | 35826600 | 35829599 | 12.61576 | 21.65209 | 57.88148 | 25.38716 | -1.02866 | 0.909388 | 0.149861 |
| 18 | 36692400 | 36694399 | 13.75173 | 23.41434 | 60.47428 | 26.50736 | -0.98118 | 0.991779 | 0.149862 |
| 7 | 89756600 | 89759199 | 40.30929 | 66.30319 | 51.0813 | 22.01904 | 0.753689 | 1.415746 | 0.149868 |
| 17 | 17236600 | 17289399 | 180.7106 | 275.8846 | 111.3841 | 46.19269 | 1.595007 | 3.117955 | 0.149871 |

| | | | | | | | | | |
|---|---|---|---|---|---|---|---|---|---|
| 9 | 106760000 | 106765399 | 20.16376 | 32.67566 | 99.69463 | 42.68067 | -1.24017 | 1.627702 | 0.149875 |
| 10 | 43268800 | 43269399 | 4.231284 | 8.246378 | 26.2899 | 9.523642 | -1.02246 | -0.39297 | 0.149901 |
| 10 | 54725800 | 54726399 | 4.231284 | 8.246378 | 26.2899 | 9.523642 | -1.02246 | -0.39297 | 0.149901 |
| 10 | 40185200 | 40186999 | 11.35736 | 19.93363 | 47.2265 | 20.88741 | -0.85043 | 0.615743 | 0.149913 |
| 7 | 119575200 | 119576399 | 8.504989 | 15.26521 | 39.50708 | 17.28485 | -0.96068 | 0.387605 | 0.149921 |
| 15 | 31175400 | 31175999 | 6.321187 | 12.02605 | 31.64683 | 12.68579 | -0.88695 | -0.00795 | 0.14993 |
| 12 | 10424400 | 10425799 | 12.46242 | 21.81629 | 48.86634 | 21.5652 | -0.76999 | 0.66792 | 0.149939 |
| 2 | 170702600 | 170710599 | 49.94141 | 78.99708 | 162.8924 | 67.69567 | -0.71523 | 2.393765 | 0.149954 |
| 17 | 19009800 | 19014599 | 135.1916 | 207.3377 | 156.3505 | 64.90412 | 0.699814 | 3.015659 | 0.149954 |
| 5 | 112721800 | 112730199 | 60.28637 | 93.74325 | 203.7853 | 84.60506 | -0.81074 | 2.736906 | 0.149956 |
| 1 | 172344400 | 172346199 | 21.76755 | 38.61913 | 23.28632 | 10.28816 | 1.117875 | 0.511967 | 0.149957 |
| 7 | 64160400 | 64161399 | 5.572696 | 10.68404 | 29.60233 | 11.64686 | -0.93536 | -0.0816 | 0.149976 |
| 18 | 3085600 | 3088599 | 47.68925 | 77.82116 | 51.80708 | 22.26096 | 0.958011 | 1.534609 | 0.149978 |
| 7 | 41052000 | 41053999 | 31.78294 | 54.11668 | 35.90928 | 15.73995 | 0.971176 | 1.027776 | 0.149978 |
| 4 | 61560000 | 61560599 | 11.97375 | 22.9578 | 17.40994 | 6.848485 | 0.917747 | -0.05199 | 0.149988 |
| 9 | 70865600 | 70867399 | 12.46871 | 21.77655 | 49.42936 | 21.77324 | -0.78918 | 0.705331 | 0.150024 |
| 1 | 177269000 | 177271399 | 12.20839 | 20.84549 | 60.44271 | 26.49956 | -1.14649 | 0.959008 | 0.150041 |
| 14 | 13054000 | 13055799 | 27.71635 | 48.30331 | 26.96918 | 11.86378 | 1.223586 | 0.7745 | 0.150041 |
| 11 | 51370600 | 51372199 | 12.62836 | 22.02312 | 50.51153 | 22.22946 | -0.80443 | 0.728071 | 0.150056 |
| 8 | 129083600 | 129086599 | 15.77283 | 26.42173 | 68.45036 | 29.84285 | -0.98717 | 1.156998 | 0.150061 |
| 7 | 108547800 | 108550799 | 16.27777 | 27.56112 | 63.31098 | 27.71746 | -0.81391 | 1.052666 | 0.150083 |
| 6 | 99967600 | 99969599 | 8.753515 | 15.46283 | 43.56782 | 19.27422 | -1.09522 | 0.516579 | 0.150091 |
| 7 | 60943400 | 60944599 | 15.30842 | 29.18752 | 15.54115 | 6.201225 | 1.40332 | 0.019723 | 0.150099 |
| 13 | 67007200 | 67010599 | 72.38807 | 115.0075 | 95.05943 | 39.64497 | 0.607409 | 2.278502 | 0.150119 |
| 1 | 5017200 | 5017999 | 9.268097 | 16.6054 | 40.52778 | 17.75796 | -0.87816 | 0.407724 | 0.150121 |
| 8 | 12038400 | 12041999 | 18.66614 | 30.99428 | 73.33086 | 31.78895 | -0.85717 | 1.269347 | 0.150144 |
| 9 | 108539000 | 108540199 | 4.955866 | 9.608156 | 27.32 | 10.32511 | -0.91645 | -0.22958 | 0.150154 |
| 8 | 65301200 | 65303599 | 11.21619 | 19.665 | 46.72321 | 20.64631 | -0.85436 | 0.632646 | 0.15016 |
| 19 | 57908200 | 57909999 | 27.0983 | 47.09412 | 31.05726 | 13.63811 | 0.984718 | 0.817062 | 0.150168 |
| 15 | 98633800 | 98642999 | 47.38914 | 74.53511 | 176.1035 | 73.05714 | -0.91775 | 2.519648 | 0.150194 |
| 7 | 28979600 | 28985199 | 30.04326 | 48.43384 | 104.533 | 44.503 | -0.75059 | 1.780902 | 0.150196 |
| 9 | 44238000 | 44239399 | 5.757926 | 10.94062 | 31.54887 | 12.68036 | -1.02094 | -0.00494 | 0.150223 |
| 13 | 93328400 | 93329199 | 5.757926 | 10.94062 | 31.54887 | 12.68036 | -1.02094 | -0.00494 | 0.150223 |
| 18 | 75240000 | 75243799 | 27.37019 | 44.43702 | 92.32521 | 39.5804 | -0.6851 | 1.598913 | 0.150253 |
| 12 | 23198400 | 23208999 | 275.3164 | 403.363 | 259.6477 | 106.9526 | 0.87014 | 3.886341 | 0.15031 |
| 19 | 3310000 | 3326999 | 95.73079 | 144.5974 | 298.086 | 123.2333 | -0.76768 | 3.322232 | 0.150311 |
| 7 | 45522200 | 45523799 | 6.885463 | 12.36251 | 40.05924 | 17.52152 | -1.28167 | 0.376065 | 0.150316 |
| X | 33952400 | 33954799 | 61.43483 | 98.68127 | 71.91212 | 30.45575 | 0.805971 | 1.925262 | 0.150317 |
| X | 33081200 | 33083599 | 29.11743 | 51.14214 | 21.45997 | 9.497202 | 1.633045 | 0.66168 | 0.150318 |
| 2 | 178090000 | 178093399 | 22.5213 | 36.74959 | 91.56383 | 39.34341 | -0.94073 | 1.509058 | 0.150329 |
| 13 | 69844400 | 69847799 | 21.18774 | 35.18527 | 76.21217 | 33.04059 | -0.73056 | 1.273974 | 0.150337 |
| 3 | 92775600 | 92776799 | 7.201939 | 13.2399 | 36.20293 | 15.37889 | -1.01033 | 0.228741 | 0.150357 |
| 7 | 11964000 | 11964599 | 22.03795 | 38.53547 | 29.34806 | 12.93927 | 0.77862 | 0.70334 | 0.150357 |
| 16 | 14500800 | 14503399 | 43.52572 | 71.62721 | 48.37547 | 20.85679 | 0.943334 | 1.417232 | 0.150369 |
| X | 120854200 | 120856399 | 33.35664 | 58.16798 | 22.08806 | 9.72024 | 1.776785 | 0.791567 | 0.150375 |
| 8 | 24086400 | 24089199 | 13.89934 | 24.15311 | 53.23767 | 23.3766 | -0.74808 | 0.789594 | 0.150381 |
| 9 | 39743200 | 39747999 | 48.24293 | 78.48872 | 55.12042 | 23.65876 | 0.87817 | 1.580226 | 0.150389 |

| | | | | | | | | | |
|---|---|---|---|---|---|---|---|---|---|
| 19 | 7122800 | 7135799 | 71.09602 | 108.4437 | 263.5578 | 109.279 | -0.9931 | 3.114786 | 0.150395 |
| 13 | 34009000 | 34010399 | 22.23044 | 39.05586 | 27.85978 | 12.33216 | 0.872205 | 0.627348 | 0.150395 |
| 3 | 12051400 | 12051999 | 9.573308 | 18.68945 | 11.89509 | 4.202248 | 1.305859 | -0.45867 | 0.150402 |
| 11 | 21719600 | 21721799 | 15.84501 | 29.25371 | 20.90152 | 8.827497 | 0.917713 | 0.23609 | 0.150406 |
| 7 | 5419200 | 5420599 | 20.50557 | 37.45269 | 17.43167 | 7.469726 | 1.515999 | 0.323388 | 0.150409 |
| 7 | 38064800 | 38070399 | 108.2308 | 167.6763 | 139.1803 | 57.81166 | 0.572537 | 2.813675 | 0.150444 |
| 1 | 112943600 | 112945199 | 26.46253 | 45.50393 | 29.75367 | 13.05122 | 0.994362 | 0.91158 | 0.150476 |
| 2 | 129296200 | 129297399 | 12.66014 | 21.99305 | 52.11859 | 22.88085 | -0.85195 | 0.786847 | 0.150479 |
| 7 | 4504000 | 4505999 | 9.227344 | 16.25844 | 45.63412 | 20.1986 | -1.09162 | 0.550172 | 0.150501 |
| 11 | 29827600 | 29829599 | 18.60065 | 33.20133 | 23.73513 | 10.43295 | 0.87871 | 0.454419 | 0.150514 |
| 7 | 24240600 | 24242599 | 9.175296 | 16.25183 | 42.00629 | 18.57335 | -0.9699 | 0.487067 | 0.150517 |
| 7 | 36577000 | 36578599 | 7.599296 | 14.09659 | 34.77966 | 14.59462 | -0.85022 | 0.178197 | 0.150522 |
| 8 | 46783400 | 46785999 | 17.85276 | 30.22341 | 65.09455 | 28.49703 | -0.72146 | 1.06153 | 0.150522 |
| 9 | 32387400 | 32388799 | 5.226855 | 9.887138 | 34.19458 | 13.87459 | -1.29306 | 0.027101 | 0.150529 |
| 14 | 18449800 | 18451599 | 12.71457 | 21.94475 | 55.28065 | 24.25117 | -0.94165 | 0.8548 | 0.150539 |
| 7 | 89629200 | 89630599 | 9.984014 | 17.55114 | 45.11687 | 19.97887 | -0.96688 | 0.580744 | 0.150543 |
| 4 | 154058600 | 154060599 | 8.536494 | 15.23507 | 40.50541 | 17.80661 | -1.00407 | 0.426458 | 0.150551 |
| 19 | 57413400 | 57414599 | 8.536494 | 15.23507 | 40.50541 | 17.80661 | -1.00407 | 0.426458 | 0.150551 |
| 6 | 97155600 | 97156799 | 6.218546 | 11.41052 | 39.39411 | 16.77748 | -1.34683 | 0.237542 | 0.150551 |
| 12 | 21781200 | 21785399 | 90.02511 | 141.7274 | 114.1416 | 47.35782 | 0.633855 | 2.516078 | 0.150554 |
| 8 | 23744400 | 23747399 | 15.58528 | 26.58777 | 60.1317 | 26.36132 | -0.79029 | 0.978841 | 0.150555 |
| 13 | 40633400 | 40634799 | 6.783291 | 12.82992 | 33.13495 | 13.4477 | -0.87586 | 0.042477 | 0.150576 |
| 17 | 82714800 | 82719599 | 28.82177 | 46.74884 | 101.6925 | 43.56536 | -0.75264 | 1.624553 | 0.150581 |
| 1 | 25248600 | 25249799 | 14.47366 | 26.8576 | 20.23582 | 8.487636 | 0.849044 | 0.196375 | 0.150593 |
| 7 | 50885000 | 50885599 | 14.47366 | 26.8576 | 20.23582 | 8.487636 | 0.849044 | 0.196375 | 0.150593 |
| 4 | 28006600 | 28007999 | 21.13457 | 38.26883 | 20.29223 | 8.790169 | 1.320568 | 0.383052 | 0.150604 |
| 10 | 83864200 | 83865799 | 21.13457 | 38.26883 | 20.29223 | 8.790169 | 1.320568 | 0.383052 | 0.150604 |
| 2 | 55093000 | 55094799 | 39.84215 | 65.24454 | 50.99605 | 21.94105 | 0.730492 | 1.468965 | 0.150604 |
| 11 | 12631800 | 12632999 | 13.41903 | 25.65014 | 16.32498 | 6.47848 | 1.155129 | -0.01469 | 0.15061 |
| 15 | 72957400 | 72960199 | 23.99828 | 41.63707 | 30.07829 | 13.1972 | 0.853628 | 0.821551 | 0.150627 |
| 17 | 58817200 | 58818599 | 15.32354 | 28.37884 | 21.1025 | 8.874847 | 0.865137 | 0.21636 | 0.150635 |
| 14 | 70385000 | 70386599 | 12.926 | 24.48397 | 18.69049 | 7.562072 | 0.871729 | 0.048763 | 0.150659 |
| 15 | 10095600 | 10096799 | 15.71289 | 29.01232 | 21.13764 | 8.926156 | 0.890333 | 0.236609 | 0.150675 |
| 17 | 89668200 | 89669399 | 16.77459 | 30.36447 | 23.03387 | 9.981756 | 0.807259 | 0.35845 | 0.15068 |
| 4 | 126891800 | 126896599 | 26.95533 | 43.60971 | 100.5401 | 42.99048 | -0.83965 | 1.672495 | 0.150688 |
| 4 | 119537000 | 119549799 | 94.6094 | 143.2978 | 298.6617 | 123.5786 | -0.78009 | 3.266624 | 0.150701 |
| 2 | 174218200 | 174218999 | 11.89697 | 23.08638 | 15.23108 | 5.731672 | 1.176077 | -0.2016 | 0.150704 |
| 12 | 78907600 | 78909199 | 9.337187 | 16.52006 | 42.51755 | 18.80362 | -0.9646 | 0.500742 | 0.150708 |
| 2 | 97539400 | 97541399 | 21.79347 | 38.56862 | 24.42014 | 10.79938 | 1.046451 | 0.537607 | 0.150721 |
| 5 | 103832400 | 103833799 | 14.70975 | 27.21635 | 20.60276 | 8.675836 | 0.838011 | 0.214357 | 0.150722 |
| 3 | 3256800 | 3257799 | 15.41008 | 28.9469 | 18.87539 | 7.759569 | 1.078151 | 0.126194 | 0.150731 |
| 19 | 6731400 | 6733399 | 8.468167 | 14.91773 | 44.61727 | 19.74933 | -1.18222 | 0.548337 | 0.150741 |
| 5 | 34192400 | 34194799 | 32.51159 | 54.93377 | 39.2809 | 17.18631 | 0.863201 | 1.099962 | 0.150752 |
| 14 | 9060600 | 9062199 | 14.99745 | 27.92379 | 20.61402 | 8.602152 | 0.88379 | 0.182124 | 0.150753 |
| X | 122185000 | 122185799 | 14.99745 | 27.92379 | 20.61402 | 8.602152 | 0.88379 | 0.182124 | 0.150753 |
| 5 | 16709200 | 16709799 | 4.114112 | 8.005922 | 25.88084 | 9.531116 | -1.06377 | -0.33615 | 0.150771 |
| 12 | 104611000 | 104611599 | 4.114112 | 8.005922 | 25.88084 | 9.531116 | -1.06377 | -0.33615 | 0.150771 |

| | | | | | | | | | |
|---|---|---|---|---|---|---|---|---|---|
| 7 | 131391400 | 131393999 | 11.07927 | 19.3873 | 49.39428 | 21.79088 | -0.95477 | 0.666401 | 0.150772 |
| 5 | 47099600 | 47100199 | 30.17762 | 51.30198 | 34.69144 | 15.2042 | 0.94377 | 1.035151 | 0.150779 |
| 6 | 44708000 | 44709799 | 12.45281 | 21.83242 | 50.4606 | 22.2972 | -0.81514 | 0.644924 | 0.150792 |
| 11 | 51698600 | 51700599 | 56.43562 | 90.64998 | 72.91598 | 30.88029 | 0.663771 | 1.918861 | 0.150794 |
| 17 | 30116800 | 30123799 | 41.22734 | 65.61984 | 141.9049 | 59.30954 | -0.77518 | 2.207161 | 0.150797 |
| 11 | 27551000 | 27552599 | 20.6923 | 36.65817 | 23.94515 | 10.58711 | 1.002137 | 0.517711 | 0.150809 |
| 14 | 3131200 | 3131799 | 23.38875 | 40.76652 | 29.50446 | 12.98011 | 0.851647 | 0.757176 | 0.150824 |
| 14 | 101845000 | 101846799 | 10.62173 | 18.594 | 47.79867 | 21.09399 | -0.9673 | 0.656501 | 0.150839 |
| 2 | 177756800 | 177768799 | 76.5819 | 117.1241 | 252.017 | 104.5585 | -0.81457 | 3.06049 | 0.150846 |
| 7 | 117261400 | 117261799 | 7.411263 | 14.47853 | 7.902978 | 2.520487 | 1.648259 | -0.85153 | 0.150847 |
| 8 | 25583400 | 25584599 | 4.966539 | 9.595508 | 27.60486 | 10.60323 | -0.95534 | -0.17231 | 0.150851 |
| X | 120392600 | 120395199 | 30.67704 | 53.10356 | 29.86305 | 13.10186 | 1.212636 | 0.873681 | 0.150861 |
| 12 | 45117000 | 45121999 | 26.74964 | 43.38142 | 95.0319 | 40.70749 | -0.7628 | 1.620756 | 0.150866 |
| 3 | 101088600 | 101090599 | 20.03799 | 35.54962 | 25.17961 | 11.1249 | 0.887223 | 0.504996 | 0.15088 |
| 3 | 18819600 | 18820399 | 6.355538 | 11.98484 | 31.86715 | 13.01474 | -0.92532 | 0.055897 | 0.150882 |
| 14 | 20302800 | 20309399 | 48.90863 | 77.13931 | 165.1016 | 68.51777 | -0.77222 | 2.463685 | 0.150884 |
| 7 | 144485400 | 144488999 | 12.80519 | 21.87584 | 59.40812 | 26.04706 | -1.05234 | 0.955615 | 0.150888 |
| 6 | 144617200 | 144619199 | 11.33551 | 19.97503 | 45.61518 | 20.18981 | -0.79609 | 0.561434 | 0.150919 |
| 14 | 32125200 | 32126399 | 10.93476 | 21.10624 | 16.90154 | 6.520875 | 0.865325 | -0.12682 | 0.150948 |
| 17 | 89005200 | 89007199 | 22.31268 | 39.10098 | 28.99971 | 12.80925 | 0.81642 | 0.668929 | 0.150954 |
| 5 | 42659400 | 42660199 | 9.472971 | 18.46011 | 14.36057 | 5.20892 | 0.988124 | -0.36022 | 0.150956 |
| 14 | 101245000 | 101245799 | 5.405466 | 10.40436 | 28.60332 | 11.12715 | -0.9083 | -0.12222 | 0.150971 |
| 6 | 99972800 | 99979199 | 38.20908 | 61.15095 | 122.9808 | 51.79776 | -0.66188 | 2.02653 | 0.150981 |
| 10 | 19227800 | 19238999 | 90.25185 | 137.5911 | 270.5645 | 112.1708 | -0.6885 | 3.129824 | 0.150993 |
| 11 | 115190200 | 115195399 | 25.18209 | 41.20442 | 87.32949 | 37.56158 | -0.70533 | 1.471179 | 0.151001 |
| 2 | 157810600 | 157813199 | 12.96895 | 22.50609 | 54.29881 | 23.82441 | -0.8781 | 0.803893 | 0.151005 |
| 9 | 19114800 | 19118599 | 73.15013 | 116.5461 | 88.68631 | 37.11784 | 0.730652 | 2.190477 | 0.151005 |
| 7 | 24977800 | 24980199 | 10.22022 | 17.81094 | 52.04933 | 22.89611 | -1.15189 | 0.728599 | 0.151026 |
| 15 | 77863400 | 77864399 | 4.129537 | 7.992915 | 27.05734 | 10.305 | -1.17486 | -0.21248 | 0.151034 |
| 11 | 59154400 | 59160799 | 40.55406 | 64.65334 | 133.1128 | 55.76027 | -0.70184 | 2.152939 | 0.151042 |
| 18 | 76826800 | 76829599 | 13.65571 | 23.5007 | 57.46746 | 25.20828 | -0.90014 | 0.887038 | 0.151049 |
| 5 | 70257600 | 70258799 | 26.98262 | 46.27212 | 35.5975 | 15.61226 | 0.760133 | 0.94472 | 0.15107 |
| 7 | 142848400 | 142852599 | 54.38128 | 85.42513 | 174.2719 | 72.28544 | -0.70768 | 2.54948 | 0.151079 |
| 14 | 5073400 | 5074599 | 16.57166 | 30.55403 | 21.04083 | 8.902667 | 0.96911 | 0.251328 | 0.151081 |
| 14 | 49815000 | 49818599 | 43.46241 | 70.93366 | 55.51511 | 23.85549 | 0.725429 | 1.526185 | 0.151086 |
| 5 | 147547400 | 147548599 | 247.6775 | 368.5728 | 180.2612 | 74.27238 | 1.287147 | 3.623854 | 0.151108 |
| 8 | 109294000 | 109295799 | 20.78334 | 36.79813 | 25.7276 | 11.37652 | 0.904743 | 0.529783 | 0.151113 |
| 1 | 71691800 | 71693399 | 8.463695 | 14.96571 | 44.59689 | 19.72556 | -1.17512 | 0.50427 | 0.151134 |
| 1 | 90185200 | 90186799 | 8.463695 | 14.96571 | 44.59689 | 19.72556 | -1.17512 | 0.50427 | 0.151134 |
| 2 | 154579600 | 154580799 | 6.984008 | 13.03178 | 33.82275 | 14.06236 | -0.91211 | 0.133758 | 0.151153 |
| 5 | 31046000 | 31048399 | 9.779641 | 17.33149 | 41.9758 | 18.5579 | -0.87664 | 0.483809 | 0.151161 |
| 16 | 64553400 | 64554799 | 8.945734 | 15.72252 | 46.16472 | 20.44154 | -1.15748 | 0.579181 | 0.151164 |
| 9 | 124077400 | 124078399 | 12.1574 | 23.7239 | 10.48351 | 3.731509 | 1.815906 | -0.4007 | 0.151171 |
| 7 | 3192800 | 3194599 | 32.66449 | 54.57637 | 42.2399 | 18.38565 | 0.749434 | 1.208672 | 0.151189 |
| 5 | 64494200 | 64496399 | 10.87382 | 19.19073 | 44.04971 | 19.49097 | -0.80231 | 0.540472 | 0.15123 |
| 1 | 182808600 | 182825599 | 110.8173 | 166.0433 | 337.281 | 139.1325 | -0.7567 | 3.48448 | 0.15123 |
| 15 | 66986800 | 66989599 | 12.46722 | 21.39798 | 58.42059 | 25.6237 | -1.05895 | 0.90868 | 0.151236 |

| | | | | | | | | | |
|---|---|---|---|---|---|---|---|---|---|
| 7 | 44971200 | 44977599 | 58.27299 | 91.54929 | 175.3706 | 72.7423 | -0.61689 | 2.548703 | 0.151242 |
| 7 | 138680800 | 138686999 | 30.68463 | 49.10138 | 125.1448 | 52.6982 | -1.00302 | 2.024057 | 0.151251 |
| 12 | 23999000 | 24000999 | 37.37492 | 61.63765 | 48.28576 | 20.83706 | 0.731171 | 1.375392 | 0.151253 |
| 11 | 99083200 | 99084199 | 16.1415 | 27.2857 | 63.2114 | 27.66013 | -0.82606 | 1.068037 | 0.151257 |
| 9 | 30713600 | 30714399 | 3.8272 | 7.44372 | 25.12266 | 9.290928 | -1.12975 | -0.32541 | 0.151273 |
| 17 | 48112400 | 48127399 | 85.98424 | 130.7499 | 276.2277 | 114.4331 | -0.79487 | 3.18164 | 0.151283 |
| 2 | 117752400 | 117753399 | 7.747964 | 14.41041 | 38.23344 | 15.98383 | -0.95046 | 0.172035 | 0.151291 |
| 4 | 51498800 | 51500599 | 18.26964 | 33.87507 | 17.82525 | 7.484486 | 1.36107 | 0.225694 | 0.1513 |
| X | 4037000 | 4037599 | 21.00731 | 37.50464 | 22.89023 | 10.05987 | 1.106226 | 0.471207 | 0.151331 |
| 5 | 72654400 | 72656999 | 14.59157 | 25.06749 | 58.03455 | 25.45568 | -0.82218 | 0.907498 | 0.151342 |
| 8 | 45351800 | 45354199 | 10.84922 | 19.19834 | 45.09548 | 19.94309 | -0.83414 | 0.510814 | 0.151361 |
| 2 | 9341000 | 9341399 | 2.835706 | 5.546468 | 23.77287 | 8.000021 | -1.34374 | -0.68374 | 0.151362 |
| 13 | 58549000 | 58551999 | 20.62098 | 33.69559 | 92.65231 | 39.83133 | -1.08104 | 1.484632 | 0.151367 |
| 14 | 34395200 | 34396799 | 11.11322 | 19.37234 | 51.01171 | 22.44342 | -1.0025 | 0.727673 | 0.151368 |
| 5 | 145501000 | 145503599 | 29.58021 | 51.4942 | 26.97263 | 11.857 | 1.315289 | 0.800366 | 0.151371 |
| 6 | 13212200 | 13214399 | 26.26928 | 45.57403 | 32.79997 | 14.39135 | 0.859 | 0.837332 | 0.151372 |
| 3 | 125758800 | 125759799 | 6.194979 | 11.71077 | 31.36976 | 12.74606 | -0.92902 | 0.037086 | 0.151381 |
| 2 | 173843200 | 173862199 | 159.9989 | 234.7485 | 428.1463 | 176.3438 | -0.62963 | 3.856012 | 0.151387 |
| 3 | 111313400 | 111314799 | 13.96449 | 26.71708 | 16.96388 | 6.714455 | 1.16273 | -0.01455 | 0.15139 |
| 16 | 3247400 | 3250999 | 70.53992 | 112.1715 | 91.52808 | 38.20637 | 0.627203 | 2.250002 | 0.15139 |
| 3 | 130800800 | 130802999 | 30.21077 | 51.09254 | 39.93104 | 17.47558 | 0.735478 | 1.084774 | 0.151393 |
| 1 | 134093800 | 134095199 | 16.24215 | 29.51852 | 20.94371 | 9.030683 | 0.907305 | 0.325764 | 0.151396 |
| 7 | 10647200 | 10660399 | 194.4326 | 290.7424 | 223.0013 | 91.95475 | 0.644796 | 3.530482 | 0.151399 |
| 12 | 105063600 | 105068799 | 28.87465 | 46.35019 | 118.3108 | 50.05617 | -0.99979 | 1.919944 | 0.151419 |
| 4 | 79623800 | 79625399 | 16.6925 | 30.36981 | 21.22349 | 9.13896 | 0.930886 | 0.326187 | 0.15144 |
| 5 | 92363600 | 92369399 | 30.01233 | 48.45692 | 104.8067 | 44.70636 | -0.75119 | 1.735711 | 0.151447 |
| 5 | 139107400 | 139113599 | 34.76696 | 55.80819 | 116.5423 | 49.30733 | -0.71095 | 1.926602 | 0.151454 |
| 4 | 33606600 | 33607199 | 9.575345 | 18.68595 | 12.50686 | 4.448977 | 1.225887 | -0.43298 | 0.151476 |
| 7 | 26197800 | 26204999 | 158.3428 | 239.3666 | 194.9141 | 80.60133 | 0.572351 | 3.313033 | 0.151493 |
| 3 | 92882200 | 92884999 | 20.65226 | 37.10457 | 21.815 | 9.536362 | 1.164197 | 0.425888 | 0.151494 |
| 10 | 70278600 | 70283199 | 22.0122 | 36.45014 | 79.0823 | 34.21735 | -0.73334 | 1.31137 | 0.151516 |
| 7 | 42065200 | 42069199 | 73.76577 | 117.5454 | 87.97265 | 36.82714 | 0.754838 | 2.187028 | 0.151523 |
| 7 | 9198000 | 9199999 | 38.51862 | 64.46908 | 36.76463 | 16.01994 | 1.188447 | 1.20712 | 0.151524 |
| 12 | 15241000 | 15242599 | 11.50721 | 20.20821 | 46.68599 | 20.65834 | -0.81425 | 0.606658 | 0.151527 |
| 15 | 84496400 | 84508199 | 72.24249 | 110.9851 | 235.5792 | 97.81646 | -0.78989 | 2.968177 | 0.151533 |
| 5 | 142898600 | 142918799 | 119.5133 | 177.4253 | 376.6286 | 155.1432 | -0.83242 | 3.655662 | 0.151533 |
| 7 | 109117800 | 109120999 | 13.84818 | 23.73542 | 60.14313 | 26.37629 | -0.95244 | 0.927786 | 0.151534 |
| 17 | 53531800 | 53532399 | 11.54583 | 22.11051 | 17.38625 | 6.863021 | 0.860912 | -0.04633 | 0.151544 |
| 7 | 43574200 | 43575999 | 11.23235 | 19.65266 | 48.33159 | 21.31975 | -0.90366 | 0.671301 | 0.151545 |
| 9 | 35017200 | 35017999 | 3.533021 | 6.881462 | 24.07812 | 8.79016 | -1.16287 | -0.37819 | 0.151551 |
| 1 | 65650200 | 65651399 | 12.69857 | 24.41822 | 16.81972 | 6.561461 | 1.06647 | -0.07204 | 0.151563 |
| 10 | 40176800 | 40178799 | 11.8178 | 20.37184 | 56.29136 | 24.69363 | -1.07469 | 0.863704 | 0.151563 |
| 1 | 74544600 | 74545999 | 9.175798 | 16.27108 | 41.98909 | 18.55918 | -0.96711 | 0.474012 | 0.151575 |
| 18 | 32372400 | 32374799 | 9.175798 | 16.27108 | 41.98909 | 18.55918 | -0.96711 | 0.474012 | 0.151575 |
| 9 | 38228400 | 38228999 | 16.68905 | 30.26562 | 19.76089 | 8.54359 | 1.024365 | 0.339873 | 0.151602 |
| X | 31795800 | 31796199 | 11.09417 | 21.68452 | 9.46371 | 3.264037 | 1.872421 | -0.52326 | 0.151636 |
| 7 | 107193000 | 107194999 | 14.33276 | 24.50015 | 58.43184 | 25.62025 | -0.86586 | 0.954922 | 0.151646 |

| | | | | | | | | | |
|---|---|---|---|---|---|---|---|---|---|
| 7 | 127602800 | 127605799 | 14.33276 | 24.50015 | 58.43184 | 25.62025 | -0.86586 | 0.954922 | 0.151646 |
| 2 | 131870600 | 131874199 | 18.56622 | 30.74247 | 78.52066 | 33.97322 | -0.96779 | 1.30226 | 0.151684 |
| 11 | 47351800 | 47354599 | 22.91637 | 40.4747 | 24.98135 | 11.05199 | 1.082213 | 0.567785 | 0.151687 |
| 7 | 3968600 | 3970999 | 26.94605 | 46.99366 | 28.7656 | 12.65959 | 1.091953 | 0.765135 | 0.151687 |
| 7 | 8091400 | 8093999 | 26.58114 | 46.53668 | 24.69516 | 10.89914 | 1.296414 | 0.69936 | 0.151698 |
| 7 | 31199000 | 31200599 | 34.34938 | 57.62085 | 42.1043 | 18.3728 | 0.832034 | 1.174496 | 0.151715 |
| 17 | 42083400 | 42085399 | 16.3844 | 30.00351 | 21.91461 | 9.358784 | 0.874928 | 0.28764 | 0.151715 |
| 6 | 52447200 | 52448599 | 7.178532 | 13.26021 | 35.7693 | 15.09602 | -0.98409 | 0.199798 | 0.151718 |
| 5 | 135438200 | 135446599 | 53.41149 | 83.53279 | 190.9333 | 79.21145 | -0.87649 | 2.631094 | 0.151761 |
| 17 | 31540400 | 31546199 | 25.15575 | 40.81611 | 93.52237 | 40.07507 | -0.82673 | 1.607725 | 0.151767 |
| 15 | 88283000 | 88286599 | 23.43889 | 38.34589 | 85.15843 | 36.62562 | -0.77261 | 1.468581 | 0.151773 |
| 17 | 94605600 | 94607199 | 27.99997 | 48.9562 | 21.21336 | 9.352044 | 1.587927 | 0.725096 | 0.151778 |
| 2 | 71617000 | 71618599 | 10.18904 | 17.79586 | 49.39362 | 21.75806 | -1.07735 | 0.693768 | 0.151788 |
| 14 | 76082200 | 76083599 | 7.342595 | 13.12596 | 40.5441 | 17.79885 | -1.21698 | 0.408506 | 0.151808 |
| 16 | 17269400 | 17270599 | 6.536131 | 12.23361 | 33.3398 | 13.78961 | -0.9757 | 0.108739 | 0.151811 |
| 2 | 164947200 | 164960399 | 86.53349 | 131.4523 | 279.318 | 115.6783 | -0.80438 | 3.202755 | 0.151814 |
| 1 | 179148800 | 179149799 | 4.290711 | 8.318991 | 30.37457 | 11.47695 | -1.2737 | -0.23228 | 0.151819 |
| 3 | 152854400 | 152854999 | 4.112162 | 8.007594 | 26.00761 | 9.524006 | -1.06305 | -0.35628 | 0.151826 |
| X | 32965000 | 32968999 | 40.23382 | 67.28114 | 38.7642 | 16.88195 | 1.173753 | 1.223455 | 0.151837 |
| 4 | 93739000 | 93739999 | 13.58695 | 26.00571 | 16.97604 | 6.71115 | 1.124579 | -0.0224 | 0.151837 |
| 6 | 61553400 | 61554599 | 13.58695 | 26.00571 | 16.97604 | 6.71115 | 1.124579 | -0.0224 | 0.151837 |
| 6 | 134025400 | 134031199 | 40.3587 | 64.41219 | 131.1556 | 55.02407 | -0.6842 | 2.116466 | 0.151839 |
| 17 | 49109400 | 49110199 | 4.527945 | 8.803136 | 26.41805 | 9.801394 | -0.9681 | -0.3029 | 0.151841 |
| 7 | 28585200 | 28586199 | 4.441813 | 8.528469 | 28.97057 | 11.35013 | -1.21931 | -0.10801 | 0.151859 |
| X | 34574200 | 34575799 | 17.87442 | 33.80199 | 15.35921 | 6.235699 | 1.609077 | 0.098098 | 0.151888 |
| 16 | 31421800 | 31422399 | 3.82009 | 7.449996 | 24.45955 | 8.783085 | -1.05185 | -0.43521 | 0.151888 |
| 1 | 132466800 | 132467599 | 4.40539 | 8.558754 | 27.53321 | 10.27422 | -1.07534 | -0.28147 | 0.151891 |
| 15 | 25887600 | 25888999 | 16.02433 | 29.49743 | 21.58217 | 9.151708 | 0.879983 | 0.256603 | 0.151894 |
| 11 | 54264800 | 54265799 | 4.410531 | 8.554644 | 27.29169 | 10.29329 | -1.07764 | -0.24368 | 0.1519 |
| 14 | 64677600 | 64678399 | 4.410531 | 8.554644 | 27.29169 | 10.29329 | -1.07764 | -0.24368 | 0.1519 |
| 11 | 106238600 | 106250399 | 72.83908 | 112.3543 | 219.5743 | 91.20718 | -0.66635 | 2.889096 | 0.151904 |
| 7 | 29309600 | 29314399 | 62.58742 | 100.4482 | 73.33373 | 31.01814 | 0.801898 | 1.951134 | 0.151914 |
| 16 | 56941600 | 56942799 | 5.641594 | 10.34587 | 37.72356 | 16.07867 | -1.42486 | 0.234552 | 0.151925 |
| 4 | 76335400 | 76336199 | 10.28665 | 20.02438 | 12.27527 | 4.508034 | 1.309575 | -0.31636 | 0.151944 |
| 5 | 145076200 | 145077799 | 5.570407 | 10.38708 | 35.95137 | 14.96443 | -1.32494 | 0.130059 | 0.151947 |
| 10 | 77283400 | 77286799 | 13.6296 | 22.86678 | 71.0876 | 31.02283 | -1.24902 | 1.134836 | 0.15195 |
| 14 | 67051600 | 67054999 | 17.96301 | 30.05091 | 68.66369 | 29.91095 | -0.80685 | 1.177889 | 0.151993 |
| 5 | 95479400 | 95480799 | 25.58679 | 44.66513 | 28.19082 | 12.41397 | 1.04786 | 0.742927 | 0.152005 |
| 10 | 11156200 | 11157799 | 10.32643 | 18.0578 | 48.33497 | 21.31146 | -1.02518 | 0.673967 | 0.152024 |
| 10 | 66401000 | 66401399 | 3.127628 | 6.112105 | 23.11793 | 8.010958 | -1.20509 | -0.577 | 0.152025 |
| 2 | 162895400 | 162899199 | 16.87354 | 27.93024 | 79.91538 | 34.56862 | -1.13094 | 1.303102 | 0.152027 |
| 4 | 127179800 | 127186599 | 36.98132 | 59.03414 | 131.3618 | 55.12675 | -0.81163 | 2.106779 | 0.15205 |
| 5 | 147660600 | 147663399 | 22.99413 | 37.53855 | 85.82384 | 36.88328 | -0.81652 | 1.499899 | 0.15205 |
| 5 | 116713800 | 116732599 | 152.7993 | 224.6368 | 418.1713 | 172.2137 | -0.65635 | 3.823551 | 0.152057 |
| 1 | 89759400 | 89763599 | 23.30184 | 38.06593 | 85.43153 | 36.72348 | -0.78922 | 1.490411 | 0.152069 |
| X | 108406400 | 108408399 | 23.62929 | 42.3882 | 19.42437 | 8.502283 | 1.519896 | 0.451843 | 0.152073 |
| 4 | 32237400 | 32240799 | 19.06254 | 31.47896 | 77.72646 | 33.5668 | -0.9194 | 1.336073 | 0.152101 |

| | | | | | | | | | |
|---|---|---|---|---|---|---|---|---|---|
| 7 | 23138600 | 23144599 | 163.6295 | 246.9231 | 200.4995 | 82.86493 | 0.574273 | 3.349706 | 0.152104 |
| 4 | 153651000 | 153657999 | 46.10173 | 73.28609 | 143.6563 | 59.94637 | -0.63566 | 2.25174 | 0.152108 |
| 12 | 20637800 | 20640399 | 33.12694 | 55.85445 | 35.84853 | 15.67444 | 1.01851 | 1.118409 | 0.152108 |
| X | 8508400 | 8510199 | 36.60461 | 61.54509 | 38.78549 | 16.94417 | 1.044742 | 1.15787 | 0.152119 |
| 9 | 7386200 | 7387599 | 18.0308 | 32.86123 | 21.67028 | 9.311263 | 1.015518 | 0.326273 | 0.152122 |
| 15 | 52914800 | 52916599 | 23.95402 | 41.89136 | 29.44645 | 12.98408 | 0.893546 | 0.710853 | 0.152124 |
| 15 | 18960800 | 18962599 | 19.50691 | 34.57963 | 25.25379 | 11.16433 | 0.84267 | 0.506832 | 0.152135 |
| 10 | 60064200 | 60070999 | 40.26967 | 64.08965 | 139.0268 | 58.09923 | -0.77975 | 2.209819 | 0.152141 |
| 1 | 104385800 | 104386999 | 13.36063 | 25.72929 | 15.2912 | 5.939938 | 1.281331 | -0.0846 | 0.152152 |
| 17 | 70115600 | 70116399 | 12.13869 | 23.35252 | 17.34156 | 6.755974 | 0.960485 | -0.08118 | 0.152179 |
| 9 | 110601400 | 110605399 | 20.62389 | 34.06682 | 76.85064 | 33.19498 | -0.78955 | 1.336151 | 0.152188 |
| 2 | 111738400 | 111739199 | 11.87321 | 22.84931 | 17.27987 | 6.725585 | 0.935267 | -0.08802 | 0.152189 |
| 17 | 20811400 | 20813399 | 28.15578 | 48.82429 | 31.62788 | 13.87688 | 1.010078 | 0.849117 | 0.152195 |
| 2 | 3729600 | 3733599 | 20.61073 | 34.08152 | 75.73575 | 32.73759 | -0.76776 | 1.323279 | 0.152199 |
| 4 | 128548200 | 128549199 | 4.839162 | 9.230968 | 29.86206 | 11.90335 | -1.17246 | -0.04141 | 0.152215 |
| 3 | 84995000 | 84996199 | 10.69007 | 18.98822 | 44.60829 | 19.68891 | -0.83149 | 0.477598 | 0.152219 |
| 5 | 117631000 | 117636599 | 32.64854 | 52.50509 | 110.7444 | 46.98283 | -0.72196 | 1.863552 | 0.15222 |
| 1 | 166876600 | 166878199 | 7.248326 | 13.2334 | 42.06337 | 18.03399 | -1.23623 | 0.278662 | 0.152222 |
| 9 | 83905800 | 83908199 | 15.1 | 25.80451 | 61.82696 | 27.10828 | -0.8729 | 0.96155 | 0.152223 |
| 17 | 39227400 | 39230199 | 74.58332 | 118.6305 | 93.83163 | 39.18029 | 0.672419 | 2.24715 | 0.152234 |
| 5 | 25063600 | 25064599 | 11.0082 | 21.35032 | 15.12624 | 5.706407 | 1.06974 | -0.20612 | 0.152241 |
| 4 | 20007000 | 20009199 | 10.05464 | 17.89712 | 41.99851 | 18.50421 | -0.82798 | 0.454681 | 0.152264 |
| 2 | 58405800 | 58406399 | 3.419008 | 6.671337 | 24.54049 | 8.746259 | -1.20349 | -0.46789 | 0.152268 |
| 1 | 155418600 | 155421199 | 42.85981 | 69.85815 | 56.50057 | 24.2673 | 0.67684 | 1.544755 | 0.152276 |
| 5 | 32307600 | 32313199 | 26.37583 | 42.53347 | 104.1497 | 44.35602 | -0.93164 | 1.767565 | 0.152285 |
| 10 | 95250000 | 95254799 | 25.46508 | 41.00663 | 109.2519 | 46.43811 | -1.05557 | 1.811876 | 0.152292 |
| 4 | 37194400 | 37196199 | 21.5975 | 38.26959 | 23.85146 | 10.54526 | 1.06964 | 0.521871 | 0.152295 |
| 7 | 12768200 | 12769999 | 11.42888 | 19.89211 | 52.08976 | 22.89569 | -0.99487 | 0.75257 | 0.152305 |
| 4 | 45408000 | 45410399 | 9.344788 | 16.53115 | 43.54208 | 19.25727 | -0.9981 | 0.502787 | 0.152309 |
| 1 | 140968200 | 140971799 | 33.6411 | 56.56577 | 40.39005 | 17.64548 | 0.865066 | 1.145742 | 0.152346 |
| 9 | 120188600 | 120191999 | 15.79257 | 26.79965 | 61.64755 | 27.00759 | -0.81565 | 1.030329 | 0.152353 |
| 3 | 131131000 | 131132599 | 14.59724 | 25.1219 | 56.38286 | 24.73254 | -0.77684 | 0.891825 | 0.152359 |
| 4 | 18430000 | 18430599 | 9.423364 | 18.31977 | 14.96329 | 5.553861 | 0.889014 | -0.27984 | 0.152362 |
| 14 | 56030600 | 56032999 | 22.94735 | 40.089 | 29.03263 | 12.79095 | 0.8506 | 0.719419 | 0.152366 |
| 8 | 61940600 | 61941199 | 3.82128 | 7.448954 | 24.33436 | 8.789943 | -1.05233 | -0.41386 | 0.152374 |
| 16 | 21487000 | 21487799 | 12.85458 | 24.89904 | 15.07386 | 5.7212 | 1.287138 | -0.16371 | 0.152391 |
| 18 | 12220400 | 12224799 | 46.44594 | 73.81836 | 143.2927 | 59.78255 | -0.62186 | 2.258572 | 0.152395 |
| 6 | 26098200 | 26099199 | 10.78162 | 20.99524 | 13.47881 | 4.935215 | 1.249015 | -0.31468 | 0.152403 |
| 5 | 8131200 | 8133799 | 10.52455 | 18.69053 | 46.21581 | 20.40182 | -0.90533 | 0.478823 | 0.152403 |
| 7 | 38239800 | 38242599 | 59.93951 | 96.09316 | 77.88133 | 32.88682 | 0.649623 | 1.982599 | 0.152415 |
| 18 | 75828200 | 75832199 | 18.44911 | 30.11969 | 88.36268 | 37.97482 | -1.17483 | 1.491711 | 0.152451 |
| 17 | 94836200 | 94839199 | 34.57572 | 57.96456 | 42.07782 | 18.35406 | 0.841455 | 1.180518 | 0.152464 |
| 2 | 59181800 | 59184399 | 13.4048 | 23.34544 | 52.697 | 23.17147 | -0.78195 | 0.753623 | 0.152478 |
| 8 | 11053400 | 11060199 | 37.00543 | 58.70799 | 155.2872 | 64.66964 | -1.06995 | 2.306396 | 0.152479 |
| 7 | 46230600 | 46237799 | 34.11142 | 54.78962 | 116.1763 | 49.19852 | -0.73188 | 1.906102 | 0.152489 |
| 4 | 147244600 | 147246199 | 19.84098 | 35.66745 | 23.16258 | 10.12056 | 1.022516 | 0.420664 | 0.152489 |
| 2 | 58869000 | 58870399 | 10.87906 | 19.16452 | 44.57219 | 19.72944 | -0.82235 | 0.561109 | 0.1525 |

| | | | | | | | | | |
|---|---|---|---|---|---|---|---|---|---|
| 5 | 6406600 | 6412599 | 92.5588 | 145.6156 | 113.1327 | 46.93498 | 0.684871 | 2.530055 | 0.152506 |
| 10 | 60982000 | 60983999 | 11.27814 | 19.60735 | 51.53518 | 22.63567 | -1.00012 | 0.769696 | 0.152511 |
| 5 | 135872000 | 135876999 | 26.92039 | 43.62597 | 94.15124 | 40.30822 | -0.74228 | 1.63664 | 0.152518 |
| 9 | 118604000 | 118608199 | 19.12912 | 31.80555 | 76.02585 | 32.98768 | -0.87203 | 1.253168 | 0.152518 |
| 8 | 106244000 | 106245799 | 7.072908 | 12.97143 | 37.22849 | 15.86643 | -1.08321 | 0.243246 | 0.152521 |
| 9 | 73875800 | 73876399 | 3.04567 | 5.938368 | 22.98013 | 8.276323 | -1.28556 | -0.43445 | 0.152521 |
| 5 | 110039200 | 110042599 | 148.1557 | 224.75 | 183.4384 | 75.93863 | 0.57323 | 3.239794 | 0.152528 |
| 3 | 127252400 | 127253799 | 5.302415 | 10.17239 | 35.14433 | 13.80185 | -1.24953 | -0.08 | 0.152546 |
| 16 | 30099000 | 30105399 | 32.72552 | 52.45492 | 121.8112 | 51.42268 | -0.8663 | 1.971397 | 0.15255 |
| 2 | 33871000 | 33873799 | 15.10017 | 25.43639 | 66.38421 | 29.01773 | -0.99755 | 1.096976 | 0.152552 |
| 4 | 148833200 | 148845599 | 76.54074 | 117.8433 | 226.7141 | 94.16346 | -0.64582 | 2.926271 | 0.152565 |
| 15 | 58397800 | 58404799 | 34.92632 | 56.06095 | 117.8865 | 49.87191 | -0.72106 | 1.928659 | 0.152594 |
| 10 | 80839800 | 80841199 | 6.17964 | 11.70246 | 31.9734 | 12.94279 | -0.95227 | 0.027274 | 0.152596 |
| 9 | 74718000 | 74720599 | 13.09103 | 22.79824 | 54.33371 | 23.89066 | -0.86002 | 0.75392 | 0.152606 |
| 14 | 85022800 | 85024599 | 18.66618 | 33.6799 | 23.30438 | 10.13973 | 0.935461 | 0.39139 | 0.152608 |
| 10 | 127214200 | 127217199 | 14.11783 | 24.2867 | 57.48709 | 25.21659 | -0.85355 | 0.892441 | 0.152616 |
| 2 | 69652800 | 69653799 | 6.103656 | 11.38138 | 33.21822 | 13.82686 | -1.0804 | 0.131432 | 0.152628 |
| 11 | 49840800 | 49846999 | 39.8587 | 63.65684 | 129.6697 | 54.45231 | -0.68371 | 2.094705 | 0.152631 |
| 13 | 114530200 | 114531599 | 9.567517 | 17.16589 | 40.53565 | 17.74045 | -0.82985 | 0.400158 | 0.152638 |
| 12 | 22053600 | 22061799 | 230.0809 | 338.8554 | 267.3965 | 110.116 | 0.583607 | 3.799446 | 0.152642 |
| 10 | 121340200 | 121347199 | 29.51067 | 47.37304 | 118.6131 | 50.18714 | -0.972 | 1.919286 | 0.152649 |
| 6 | 8529200 | 8530799 | 21.68501 | 38.14005 | 27.52102 | 12.18575 | 0.856151 | 0.610336 | 0.152654 |
| 7 | 47326200 | 47328199 | 48.51245 | 78.46819 | 63.15182 | 26.99892 | 0.674722 | 1.696034 | 0.152659 |
| 13 | 20274200 | 20279399 | 28.15561 | 45.34 | 115.7776 | 49.21317 | -0.99466 | 1.816961 | 0.152676 |
| 1 | 81775000 | 81775799 | 7.416164 | 13.8829 | 34.49846 | 14.26511 | -0.84358 | 0.118781 | 0.152676 |
| 15 | 97006200 | 97008999 | 12.93395 | 22.62284 | 53.21413 | 23.46736 | -0.84052 | 0.689356 | 0.15268 |
| 6 | 145903400 | 145905799 | 9.30142 | 16.54913 | 40.9879 | 18.06562 | -0.90519 | 0.452904 | 0.152682 |
| 7 | 10575600 | 10592399 | 328.5926 | 469.4657 | 360.0152 | 148.7573 | 0.582211 | 4.26356 | 0.152687 |
| 11 | 60306800 | 60309199 | 9.378276 | 16.50905 | 45.11657 | 19.97323 | -1.05378 | 0.561276 | 0.152688 |
| 8 | 21748000 | 21749999 | 18.10618 | 32.44128 | 24.41738 | 10.69865 | 0.80821 | 0.430629 | 0.152695 |
| 3 | 94570200 | 94572399 | 12.13999 | 21.27148 | 47.78761 | 21.1056 | -0.77422 | 0.649008 | 0.152699 |
| 10 | 81633800 | 81634799 | 14.20989 | 26.40225 | 18.76067 | 7.855079 | 0.934604 | 0.182767 | 0.152704 |
| 12 | 103762200 | 103762799 | 17.49075 | 31.43342 | 23.5814 | 10.30608 | 0.815474 | 0.406664 | 0.152711 |
| 13 | 61267600 | 61275199 | 26.05623 | 45.30542 | 28.51044 | 12.52361 | 1.051809 | 0.800246 | 0.152712 |
| 17 | 72260000 | 72263199 | 26.53072 | 42.81592 | 104.1227 | 44.38821 | -0.92051 | 1.74423 | 0.152717 |
| 12 | 23971400 | 23976799 | 71.62431 | 114.198 | 81.9832 | 34.34688 | 0.815608 | 2.164161 | 0.152722 |
| 18 | 72734200 | 72736599 | 26.11196 | 45.00063 | 35.09035 | 15.39315 | 0.742072 | 0.897952 | 0.152722 |
| 9 | 3082600 | 3108599 | 486.8827 | 677.2756 | 428.3198 | 177.7477 | 0.821472 | 4.663484 | 0.152743 |
| 5 | 60614600 | 60616199 | 21.8665 | 38.4243 | 27.85807 | 12.33352 | 0.848846 | 0.62172 | 0.152744 |
| 15 | 85066200 | 85069199 | 10.06057 | 17.90852 | 41.48389 | 18.27672 | -0.80917 | 0.454731 | 0.152744 |
| 1 | 58397800 | 58398999 | 7.802938 | 14.29759 | 36.14216 | 15.42205 | -0.90265 | 0.252835 | 0.152746 |
| 19 | 6240000 | 6242199 | 9.558262 | 17.1557 | 40.00898 | 17.50442 | -0.81175 | 0.395639 | 0.152759 |
| 2 | 105489800 | 105494799 | 38.01445 | 60.53125 | 144.4375 | 60.40289 | -0.91622 | 2.191005 | 0.15276 |
| 6 | 124553800 | 124555199 | 9.538334 | 17.18354 | 40.06479 | 17.46611 | -0.80799 | 0.372196 | 0.152761 |
| 14 | 60370000 | 60371999 | 11.3353 | 20.00953 | 45.62031 | 20.18504 | -0.79272 | 0.542834 | 0.152793 |
| 7 | 127895800 | 127897399 | 7.66693 | 14.01858 | 36.63483 | 15.67726 | -0.9533 | 0.264707 | 0.152795 |
| 4 | 152716400 | 152729199 | 87.64487 | 133.0574 | 276.6756 | 114.5625 | -0.77392 | 3.216146 | 0.152796 |

| | | | | | | | | | |
|---|---|---|---|---|---|---|---|---|---|
| 2 | 8601600 | 8602799 | 17.10687 | 31.07611 | 22.92539 | 9.890714 | 0.851265 | 0.340664 | 0.152799 |
| 6 | 132439200 | 132440799 | 27.91045 | 48.9161 | 24.19572 | 10.68833 | 1.397463 | 0.691417 | 0.152816 |
| 15 | 97340200 | 97344999 | 36.20255 | 58.14846 | 117.4019 | 49.71763 | -0.66137 | 1.909429 | 0.152821 |
| 5 | 96099600 | 96100399 | 4.95229 | 9.612407 | 27.44043 | 10.31437 | -0.91471 | -0.2482 | 0.152839 |
| 6 | 142920400 | 142921199 | 4.95229 | 9.612407 | 27.44043 | 10.31437 | -0.91471 | -0.2482 | 0.152839 |
| 6 | 22703200 | 22704599 | 12.09896 | 20.89416 | 54.71275 | 24.00246 | -0.9969 | 0.84695 | 0.152841 |
| X | 126553600 | 126554799 | 19.63086 | 35.87623 | 19.26424 | 8.248915 | 1.312599 | 0.317792 | 0.152848 |
| 1 | 38604400 | 38606399 | 8.565963 | 15.63185 | 37.67522 | 16.16192 | -0.84008 | 0.284696 | 0.152852 |
| 3 | 87729200 | 87731199 | 11.51558 | 20.24878 | 46.14394 | 20.43262 | -0.79421 | 0.588796 | 0.15286 |
| 4 | 148001600 | 148027799 | 169.9236 | 245.2791 | 516.4406 | 213.0628 | -0.85955 | 4.105526 | 0.152864 |
| 7 | 31486200 | 31486799 | 12.45131 | 23.90393 | 17.1423 | 6.718554 | 1.001693 | -0.06088 | 0.152868 |
| 7 | 10628000 | 10636799 | 102.8737 | 161.4432 | 112.6589 | 46.72339 | 0.836882 | 2.586284 | 0.152876 |
| 11 | 76436200 | 76437599 | 6.587037 | 12.18477 | 34.69588 | 14.61492 | -1.05891 | 0.185899 | 0.152885 |
| 10 | 81359200 | 81361599 | 13.13183 | 22.76686 | 53.74488 | 23.58067 | -0.84708 | 0.813196 | 0.152903 |
| 8 | 72127400 | 72148799 | 193.1449 | 275.0175 | 591.1688 | 244.4083 | -0.90911 | 4.308097 | 0.152906 |
| 2 | 76998200 | 76998799 | 3.526834 | 6.887095 | 22.89335 | 8.050357 | -1.04099 | -0.51874 | 0.152914 |
| 6 | 115780600 | 115781199 | 3.526834 | 6.887095 | 22.89335 | 8.050357 | -1.04099 | -0.51874 | 0.152914 |
| 2 | 147436800 | 147437599 | 14.15477 | 27.07541 | 16.9578 | 6.716108 | 1.181562 | -0.01063 | 0.152918 |
| 9 | 44242800 | 44247999 | 20.74192 | 33.98923 | 84.22227 | 36.2469 | -0.92884 | 1.440591 | 0.153052 |
| X | 6879800 | 6881799 | 50.20924 | 82.24751 | 45.66668 | 19.65072 | 1.222502 | 1.486999 | 0.153052 |
| 9 | 119478600 | 119480999 | 15.20217 | 26.17365 | 56.99617 | 25.00188 | -0.73354 | 0.889642 | 0.153074 |
| 12 | 23123000 | 23144999 | 646.0511 | 869.9044 | 631.3118 | 263.3214 | 0.575654 | 5.170095 | 0.153089 |
| 7 | 11731200 | 11734799 | 53.68604 | 87.15023 | 53.53636 | 22.94904 | 1.068136 | 1.630441 | 0.153097 |
| 14 | 27057200 | 27059399 | 10.03795 | 17.54435 | 49.38734 | 21.76605 | -1.09754 | 0.681776 | 0.153098 |
| 10 | 46880800 | 46882399 | 25.98645 | 45.27509 | 30.53529 | 13.43002 | 0.952769 | 0.776761 | 0.153108 |
| 2 | 93180200 | 93184999 | 22.64171 | 36.71268 | 95.06631 | 40.71678 | -1.0035 | 1.618051 | 0.153119 |
| 17 | 85772200 | 85783199 | 97.5163 | 147.8195 | 286.2673 | 118.4802 | -0.67322 | 3.249547 | 0.153126 |
| 10 | 71690800 | 71691799 | 8.546032 | 15.65549 | 38.88108 | 16.59608 | -0.8783 | 0.264067 | 0.153182 |
| 18 | 10269200 | 10271599 | 35.5174 | 58.86995 | 47.1599 | 20.41868 | 0.699769 | 1.311265 | 0.153189 |
| 6 | 37274200 | 37278799 | 27.96548 | 45.10709 | 104.2392 | 44.40703 | -0.84804 | 1.762497 | 0.153193 |
| 19 | 43600600 | 43607199 | 55.80697 | 87.00151 | 199.6521 | 82.85727 | -0.88582 | 2.689809 | 0.1532 |
| 8 | 68728400 | 68730399 | 13.33836 | 22.98525 | 56.37955 | 24.73207 | -0.90404 | 0.872516 | 0.1532 |
| 17 | 81660400 | 81664199 | 79.84911 | 126.5054 | 103.2759 | 42.95442 | 0.621408 | 2.366714 | 0.153203 |
| 9 | 40597000 | 40598999 | 12.97199 | 22.51647 | 52.66593 | 23.10933 | -0.83338 | 0.800453 | 0.153203 |
| 5 | 24997000 | 24998399 | 8.090699 | 14.84745 | 36.72024 | 15.63629 | -0.86944 | 0.246673 | 0.153225 |
| 11 | 44994000 | 44995199 | 8.090699 | 14.84745 | 36.72024 | 15.63629 | -0.86944 | 0.246673 | 0.153225 |
| 13 | 38503200 | 38504999 | 8.090699 | 14.84745 | 36.72024 | 15.63629 | -0.86944 | 0.246673 | 0.153225 |
| 17 | 86658000 | 86659999 | 8.090699 | 14.84745 | 36.72024 | 15.63629 | -0.86944 | 0.246673 | 0.153225 |
| 7 | 132722400 | 132726999 | 23.47478 | 38.01084 | 100.4512 | 42.97924 | -1.03481 | 1.649395 | 0.153249 |
| 7 | 115842600 | 115849199 | 37.11438 | 59.31823 | 125.8074 | 52.88627 | -0.74055 | 2.06872 | 0.153255 |
| 17 | 66641000 | 66654199 | 107.3433 | 160.8756 | 330.5453 | 136.3602 | -0.77288 | 3.479713 | 0.153262 |
| 17 | 17430800 | 17435999 | 173.1411 | 262.3791 | 174.2504 | 72.1121 | 0.869221 | 3.267432 | 0.153262 |
| 8 | 12377000 | 12383199 | 29.62232 | 47.62667 | 111.4449 | 47.26321 | -0.87176 | 1.867983 | 0.153263 |
| 7 | 45944200 | 45950999 | 36.78206 | 58.81281 | 124.8602 | 52.52061 | -0.74129 | 2.054294 | 0.153296 |
| 5 | 137111800 | 137118199 | 51.49857 | 81.22105 | 165.3172 | 68.60687 | -0.69991 | 2.465412 | 0.153303 |
| 4 | 112263200 | 112263799 | 16.9611 | 30.95451 | 22.42985 | 9.620964 | 0.882284 | 0.312652 | 0.153315 |
| 9 | 22010200 | 22025199 | 83.10729 | 126.517 | 270.3177 | 112.0223 | -0.80979 | 3.157675 | 0.153319 |

| | | | | | | | | | |
|---|---|---|---|---|---|---|---|---|---|
| 2 | 177388800 | 177390999 | 25.72141 | 45.40186 | 21.96901 | 9.720463 | 1.43057 | 0.585837 | 0.153319 |
| 19 | 3624800 | 3626199 | 6.641598 | 12.18662 | 37.77082 | 16.08643 | -1.19243 | 0.237963 | 0.153322 |
| 5 | 8516400 | 8521599 | 89.30399 | 140.854 | 110.1438 | 45.71018 | 0.678354 | 2.478269 | 0.153327 |
| 2 | 174875800 | 174879799 | 37.76866 | 62.79425 | 43.81727 | 19.01176 | 0.898422 | 1.277886 | 0.153356 |
| 7 | 42698800 | 42708599 | 221.774 | 327.4734 | 261.2688 | 107.5955 | 0.571282 | 3.755337 | 0.153357 |
| 12 | 23188400 | 23192599 | 158.3387 | 239.3991 | 195.3607 | 80.7901 | 0.569446 | 3.309727 | 0.153358 |
| 18 | 37914200 | 37918399 | 54.25361 | 87.41178 | 68.91857 | 29.31694 | 0.697927 | 1.819067 | 0.153376 |
| 17 | 73172600 | 73174199 | 6.79672 | 12.80468 | 32.4253 | 13.2639 | -0.85767 | 0.067076 | 0.153376 |
| 9 | 70410400 | 70411199 | 5.458729 | 10.40383 | 30.48603 | 12.17431 | -1.03516 | -0.03124 | 0.153379 |
| 7 | 18090200 | 18096199 | 123.4698 | 190.6838 | 141.3234 | 58.7028 | 0.732376 | 2.878459 | 0.153387 |
| 14 | 34090600 | 34101199 | 51.35802 | 79.85152 | 206.1089 | 85.5711 | -1.0582 | 2.735684 | 0.153407 |
| 1 | 90938600 | 90941999 | 15.68065 | 26.49481 | 64.31282 | 28.13827 | -0.89325 | 1.069981 | 0.153413 |
| 2 | 129641800 | 129644599 | 14.1879 | 23.8932 | 67.38845 | 29.45428 | -1.10899 | 1.097311 | 0.153439 |
| X | 25675800 | 25678799 | 53.73052 | 87.16993 | 54.62396 | 23.40486 | 1.038762 | 1.643334 | 0.153449 |
| 13 | 65664000 | 65669799 | 170.1315 | 256.0474 | 207.2482 | 85.58931 | 0.575488 | 3.403927 | 0.153463 |
| 2 | 115411800 | 115412999 | 6.500206 | 12.25407 | 31.86126 | 13.01876 | -0.8938 | 0.059604 | 0.153467 |
| 15 | 84730600 | 84733199 | 10.30754 | 18.06983 | 46.72927 | 20.64701 | -0.9757 | 0.629879 | 0.153477 |
| 17 | 86870800 | 86873199 | 10.56124 | 18.64678 | 43.52983 | 19.25934 | -0.82627 | 0.532573 | 0.15348 |
| 2 | 39418400 | 39419599 | 19.79198 | 34.9342 | 26.5812 | 11.76128 | 0.781798 | 0.558491 | 0.153487 |
| 5 | 105918600 | 105921399 | 17.91503 | 30.11282 | 66.01604 | 28.83786 | -0.74734 | 1.125586 | 0.153491 |
| 12 | 102207000 | 102211599 | 17.30292 | 28.62943 | 75.76618 | 32.76468 | -1.01861 | 1.308456 | 0.153494 |
| 17 | 86711200 | 86713799 | 11.40242 | 19.90177 | 49.41877 | 21.75998 | -0.918 | 0.709114 | 0.153496 |
| 4 | 8744800 | 8749399 | 44.48019 | 70.75451 | 140.1673 | 58.53435 | -0.64989 | 2.23028 | 0.153499 |
| 18 | 11992400 | 11997199 | 20.9382 | 34.25092 | 89.4516 | 38.47048 | -1.00589 | 1.466466 | 0.15351 |
| 5 | 113172200 | 113179999 | 46.91837 | 74.55844 | 144.9616 | 60.47009 | -0.6244 | 2.262834 | 0.153514 |
| 16 | 33145400 | 33158599 | 85.19107 | 129.9744 | 273.1285 | 113.2536 | -0.7833 | 3.113961 | 0.153516 |
| 6 | 6578000 | 6578599 | 3.529654 | 6.884545 | 23.16663 | 8.30361 | -1.0832 | -0.44831 | 0.15352 |
| 13 | 44936400 | 44937399 | 5.250963 | 10.16364 | 29.06283 | 11.06803 | -0.9358 | -0.19647 | 0.153521 |
| 3 | 4826400 | 4827199 | 13.66067 | 25.91054 | 17.97344 | 7.251209 | 1.011963 | 0.044365 | 0.153529 |
| 2 | 4496400 | 4498999 | 13.8961 | 24.08975 | 55.42154 | 24.31624 | -0.81031 | 0.818742 | 0.153529 |
| 7 | 6064400 | 6070799 | 121.35 | 187.0926 | 148.6132 | 61.73139 | 0.630626 | 2.908408 | 0.153548 |
| 2 | 154278000 | 154278999 | 7.203376 | 13.21398 | 35.6258 | 15.17825 | -0.99279 | 0.241757 | 0.153551 |
| 1 | 154467600 | 154468799 | 6.162662 | 11.75746 | 30.57041 | 12.18166 | -0.86173 | -0.03052 | 0.153558 |
| 4 | 122095400 | 122096599 | 12.83226 | 24.5772 | 17.67681 | 6.975147 | 0.988736 | -0.03369 | 0.153564 |
| 11 | 67718000 | 67718999 | 9.86633 | 19.21643 | 14.2289 | 5.197273 | 1.049075 | -0.33227 | 0.153565 |
| 2 | 181111800 | 181114599 | 12.15362 | 21.22159 | 52.66766 | 23.19649 | -0.91763 | 0.709304 | 0.153577 |
| 7 | 141561000 | 141564999 | 17.2473 | 28.70155 | 72.25664 | 31.37098 | -0.94615 | 1.236699 | 0.153581 |
| 7 | 21750200 | 21750799 | 19.05778 | 34.14532 | 21.45245 | 9.399763 | 1.067182 | 0.431444 | 0.153581 |
| 6 | 59736000 | 59736399 | 9.536442 | 18.52861 | 13.95141 | 5.203143 | 0.998963 | -0.26565 | 0.153591 |
| 8 | 120284000 | 120286199 | 8.14783 | 14.78694 | 37.56884 | 16.22796 | -0.92186 | 0.314542 | 0.153593 |
| 7 | 11160800 | 11163599 | 26.1366 | 45.86766 | 25.44136 | 11.25111 | 1.233223 | 0.663546 | 0.153601 |
| 18 | 84264000 | 84264799 | 4.821073 | 9.357933 | 28.06377 | 10.54752 | -0.98519 | -0.24933 | 0.153607 |
| 4 | 12277800 | 12278999 | 6.314995 | 12.01061 | 37.59493 | 15.07984 | -1.13613 | 0.00968 | 0.153609 |
| 7 | 31778000 | 31794199 | 306.4896 | 443.1183 | 311.0115 | 128.2837 | 0.726914 | 4.088763 | 0.153614 |
| 12 | 107513200 | 107518199 | 34.28634 | 55.0566 | 113.2356 | 47.93469 | -0.68836 | 1.914653 | 0.153615 |
| 12 | 14438600 | 14439799 | 17.2688 | 32.20504 | 18.64318 | 7.758381 | 1.234681 | 0.186395 | 0.153616 |
| 4 | 57514600 | 57515199 | 3.457875 | 6.7474 | 22.60451 | 8.051661 | -1.06713 | -0.47055 | 0.153621 |

| | | | | | | | | | |
|---|---|---|---|---|---|---|---|---|---|
| 5 | 145642200 | 145644799 | 31.11906 | 53.74597 | 30.36177 | 13.31973 | 1.205483 | 0.895564 | 0.153628 |
| X | 32746200 | 32749199 | 32.64296 | 56.70121 | 25.90442 | 11.3717 | 1.511444 | 0.844907 | 0.15363 |
| 7 | 30409200 | 30420199 | 75.8037 | 116.1522 | 242.8769 | 100.8102 | -0.77118 | 3.021525 | 0.15363 |
| 13 | 41026400 | 41027799 | 17.02659 | 30.81312 | 22.6411 | 9.814532 | 0.852848 | 0.361704 | 0.15364 |
| 6 | 89161000 | 89163399 | 11.39221 | 19.91051 | 48.3509 | 21.30852 | -0.88571 | 0.688803 | 0.15364 |
| 13 | 99758600 | 99759599 | 12.88675 | 24.82068 | 16.62592 | 6.455454 | 1.111936 | -0.08642 | 0.153648 |
| 7 | 111364200 | 111365999 | 8.795774 | 15.8094 | 39.50995 | 17.26557 | -0.90958 | 0.379937 | 0.153653 |
| 4 | 132672800 | 132674599 | 5.898291 | 10.81788 | 36.13913 | 15.4009 | -1.29912 | 0.234015 | 0.153654 |
| 13 | 47258400 | 47259999 | 7.522221 | 13.75486 | 37.17034 | 15.90516 | -1.0013 | 0.263766 | 0.153659 |
| 9 | 44364200 | 44367799 | 15.75961 | 26.44951 | 68.56979 | 29.93057 | -0.98827 | 1.135385 | 0.15367 |
| 19 | 9498200 | 9499399 | 16.72649 | 31.06847 | 19.75202 | 8.273663 | 1.093307 | 0.208284 | 0.153682 |
| 4 | 33922600 | 33925399 | 18.65121 | 31.03243 | 71.87854 | 31.20309 | -0.82648 | 1.242274 | 0.153684 |
| 7 | 23670200 | 23674999 | 66.86771 | 106.6662 | 87.92964 | 36.86304 | 0.617261 | 2.145443 | 0.153685 |
| 9 | 43775000 | 43776199 | 4.619694 | 8.833148 | 31.10569 | 12.33322 | -1.28805 | -0.06232 | 0.153688 |
| 6 | 132120200 | 132124799 | 68.06783 | 109.3565 | 63.60272 | 26.94405 | 1.130852 | 1.924124 | 0.15369 |
| 16 | 10148800 | 10158199 | 68.23572 | 105.7636 | 211.0897 | 87.67973 | -0.69131 | 2.7965 | 0.153697 |
| 9 | 68809000 | 68810599 | 7.88937 | 14.65134 | 36.50442 | 15.29468 | -0.86308 | 0.176639 | 0.1537 |
| 2 | 83976800 | 83980999 | 43.9543 | 71.7228 | 55.5406 | 23.86478 | 0.740495 | 1.52991 | 0.153721 |
| 10 | 98097000 | 98097399 | 10.4647 | 20.40791 | 13.08507 | 4.700192 | 1.275895 | -0.38444 | 0.153733 |
| 11 | 6400600 | 6401799 | 22.8499 | 39.83374 | 30.45772 | 13.40084 | 0.772713 | 0.753022 | 0.153741 |
| 6 | 89655600 | 89658599 | 14.75428 | 24.95545 | 67.07231 | 29.35484 | -1.03963 | 1.059075 | 0.153747 |
| 6 | 26392400 | 26395199 | 39.09758 | 64.52753 | 44.02548 | 19.00538 | 0.930211 | 1.36691 | 0.153788 |
| 8 | 3112200 | 3120599 | 160.4846 | 242.2228 | 197.2829 | 81.54035 | 0.570109 | 3.345038 | 0.153803 |
| 7 | 41106200 | 41107599 | 29.60472 | 50.37377 | 32.41279 | 14.20658 | 1.015441 | 1.023454 | 0.153804 |
| 7 | 41843600 | 41851799 | 137.1644 | 212.0398 | 121.7779 | 50.58377 | 1.10091 | 2.864338 | 0.153816 |
| 9 | 8771600 | 8774199 | 32.17502 | 54.47164 | 38.71603 | 16.94923 | 0.87194 | 1.081672 | 0.153832 |
| 5 | 122958400 | 122960199 | 7.30009 | 13.55028 | 34.79245 | 14.58721 | -0.90649 | 0.171188 | 0.153833 |
| 1 | 119963200 | 119965399 | 6.009624 | 11.15189 | 35.28557 | 14.79948 | -1.20521 | 0.16335 | 0.153838 |
| 12 | 13482200 | 13487799 | 74.49577 | 115.25 | 212.2607 | 88.1679 | -0.57759 | 2.834857 | 0.153866 |
| 13 | 100244200 | 100244999 | 3.552741 | 6.863201 | 27.40985 | 10.53409 | -1.42135 | -0.1845 | 0.153881 |
| 1 | 174966400 | 174968399 | 20.66136 | 36.34125 | 23.83739 | 10.55462 | 0.992897 | 0.595966 | 0.15389 |
| 2 | 28664200 | 28666199 | 8.79485 | 15.80857 | 38.997 | 17.04067 | -0.89065 | 0.379968 | 0.15389 |
| 15 | 29994800 | 29998599 | 64.26834 | 103.2519 | 68.46885 | 29.00533 | 0.942072 | 1.920746 | 0.153892 |
| 12 | 22788000 | 22793399 | 115.0032 | 178.0792 | 139.6344 | 58.00042 | 0.65404 | 2.825492 | 0.153893 |
| 3 | 69479600 | 69480599 | 7.459304 | 13.80895 | 34.73148 | 14.61456 | -0.88061 | 0.188283 | 0.1539 |
| 7 | 10727600 | 10730999 | 91.80338 | 143.9571 | 105.3004 | 43.67346 | 0.768128 | 2.59135 | 0.153905 |
| 5 | 110006000 | 110008599 | 24.74483 | 43.23288 | 29.38976 | 12.94899 | 0.941561 | 0.729501 | 0.153908 |
| 4 | 141786600 | 141795399 | 49.85984 | 78.29495 | 178.1539 | 73.89227 | -0.86535 | 2.554025 | 0.153914 |
| 13 | 25949200 | 25950399 | 7.953416 | 14.56419 | 37.20928 | 15.89093 | -0.91909 | 0.258619 | 0.153933 |
| 6 | 146519000 | 146522199 | 21.7755 | 35.83581 | 80.84152 | 34.85491 | -0.79139 | 1.386584 | 0.153945 |
| 7 | 12762800 | 12767199 | 19.16529 | 31.78116 | 76.40433 | 33.09117 | -0.88056 | 1.285114 | 0.153945 |
| X | 32651800 | 32654199 | 25.72134 | 44.64496 | 32.80041 | 14.39221 | 0.829483 | 0.829806 | 0.153947 |
| 10 | 7465400 | 7468199 | 16.3129 | 27.52709 | 64.32342 | 28.13176 | -0.83868 | 1.084014 | 0.153961 |
| 14 | 21570000 | 21572999 | 7.797311 | 14.30379 | 38.90794 | 16.57616 | -1.00659 | 0.249637 | 0.153962 |
| 15 | 39806200 | 39807999 | 10.00676 | 17.55488 | 47.23729 | 20.88386 | -1.03276 | 0.616517 | 0.153974 |
| 17 | 21670400 | 21671599 | 7.570576 | 14.13624 | 34.99172 | 14.53103 | -0.84325 | 0.136613 | 0.153994 |
| 7 | 9368400 | 9374999 | 93.3462 | 146.698 | 101.6706 | 42.1739 | 0.848265 | 2.550909 | 0.154023 |

| | | | | | | | | | |
|---|---|---|---|---|---|---|---|---|---|
| 8 | 124062400 | 124069999 | 40.90155 | 65.10977 | 142.655 | 59.63377 | -0.79382 | 2.202585 | 0.154028 |
| 3 | 135013000 | 135014399 | 9.574238 | 17.11914 | 41.01107 | 18.00048 | -0.85366 | 0.420828 | 0.154039 |
| 6 | 134083600 | 134090599 | 48.53814 | 77.0828 | 157.9535 | 65.84717 | -0.70105 | 2.284843 | 0.154046 |
| 9 | 62579600 | 62586599 | 38.15313 | 60.78322 | 136.0383 | 56.93076 | -0.8231 | 2.173997 | 0.15406 |
| 17 | 45866400 | 45871599 | 25.07725 | 40.4875 | 101.2773 | 43.19891 | -0.96026 | 1.729553 | 0.154061 |
| 19 | 39453200 | 39454999 | 22.91734 | 40.36308 | 25.44764 | 11.2646 | 1.05011 | 0.597143 | 0.154067 |
| 2 | 127939200 | 127941199 | 11.02925 | 19.42352 | 45.61603 | 20.19264 | -0.83663 | 0.565947 | 0.154073 |
| 4 | 137682400 | 137692999 | 70.49541 | 108.2415 | 240.1562 | 99.70937 | -0.8543 | 2.978664 | 0.154073 |
| 12 | 112754400 | 112766199 | 61.75131 | 94.97972 | 237.146 | 98.48296 | -1.0225 | 2.941643 | 0.154076 |
| 1 | 188079800 | 188080599 | 8.840795 | 17.2544 | 13.26273 | 4.708915 | 1.031877 | -0.44679 | 0.154088 |
| 13 | 65829000 | 65830599 | 43.54542 | 70.79011 | 56.39843 | 24.20033 | 0.695907 | 1.583627 | 0.154088 |
| 12 | 16850200 | 16851799 | 9.272602 | 16.59037 | 39.99004 | 17.5429 | -0.86157 | 0.414713 | 0.15409 |
| 8 | 94536400 | 94540399 | 30.05737 | 48.42006 | 105.2524 | 44.7628 | -0.76216 | 1.804944 | 0.154098 |
| 9 | 88134000 | 88135999 | 11.1683 | 19.71259 | 47.22137 | 20.89389 | -0.86393 | 0.54399 | 0.154114 |
| 9 | 75008600 | 75010799 | 8.079241 | 14.8597 | 36.77909 | 15.61365 | -0.86741 | 0.23365 | 0.154119 |
| 12 | 81700000 | 81700799 | 2.745597 | 5.355845 | 24.91474 | 8.91727 | -1.54104 | -0.45963 | 0.154126 |
| 10 | 53578800 | 53579799 | 13.20513 | 25.2807 | 17.66527 | 6.979022 | 1.028564 | -0.02582 | 0.154138 |
| 1 | 8442000 | 8443599 | 17.8485 | 32.85139 | 20.09196 | 8.521195 | 1.13609 | 0.267356 | 0.154139 |
| 12 | 112264600 | 112271799 | 41.65757 | 66.05689 | 152.7786 | 63.59451 | -0.87739 | 2.323715 | 0.154141 |
| 1 | 5586400 | 5587199 | 12.20484 | 23.30136 | 18.32982 | 7.293234 | 0.849907 | -0.01165 | 0.154146 |
| 14 | 41373400 | 41373999 | 12.20484 | 23.30136 | 18.32982 | 7.293234 | 0.849907 | -0.01165 | 0.154146 |
| 6 | 111647800 | 111650799 | 23.71115 | 41.49367 | 29.50429 | 13.01676 | 0.877083 | 0.700109 | 0.154147 |
| 8 | 105168800 | 105169799 | 4.316318 | 8.302358 | 29.63107 | 11.5494 | -1.28324 | -0.12596 | 0.15415 |
| 15 | 20853000 | 20854399 | 14.12166 | 26.75981 | 18.54729 | 7.497575 | 1.011072 | 0.05734 | 0.15415 |
| 5 | 119931200 | 119950399 | 140.5195 | 207.6263 | 409.9502 | 168.8266 | -0.73442 | 3.739218 | 0.154154 |
| 11 | 31318600 | 31324399 | 43.66621 | 69.44664 | 141.8473 | 59.22051 | -0.69422 | 2.235447 | 0.154155 |
| 16 | 97461200 | 97461999 | 5.124449 | 9.898031 | 28.82336 | 11.08468 | -0.9752 | -0.16499 | 0.154158 |
| 10 | 15790400 | 15793599 | 52.69613 | 85.24783 | 60.65896 | 25.93599 | 0.852149 | 1.700079 | 0.154168 |
| 9 | 118177000 | 118186799 | 78.65894 | 121.1197 | 226.6125 | 94.12288 | -0.60552 | 2.924562 | 0.154172 |
| 15 | 81613000 | 81614399 | 6.242207 | 11.69095 | 32.78853 | 13.54692 | -1.01513 | 0.101452 | 0.154174 |
| 7 | 3902200 | 3903799 | 15.24388 | 28.23661 | 21.03034 | 8.842237 | 0.862924 | 0.21387 | 0.154191 |
| 1 | 118567600 | 118572199 | 23.33417 | 38.49496 | 83.11216 | 35.87101 | -0.72697 | 1.361712 | 0.154219 |
| 7 | 21039800 | 21041199 | 56.73022 | 91.29107 | 63.98423 | 27.17008 | 0.865279 | 1.860589 | 0.154219 |
| 15 | 84590000 | 84599799 | 56.17038 | 88.23603 | 173.8146 | 72.09581 | -0.65726 | 2.55025 | 0.154226 |
| 14 | 23278400 | 23279599 | 19.04762 | 35.27082 | 17.80126 | 7.487906 | 1.419164 | 0.239613 | 0.154228 |
| 13 | 66007600 | 66008599 | 20.34028 | 35.85271 | 26.90371 | 11.90725 | 0.800851 | 0.576056 | 0.154241 |
| 18 | 46417200 | 46417999 | 4.059297 | 7.916088 | 23.82104 | 8.559653 | -0.92801 | -0.42937 | 0.154244 |
| 2 | 40596600 | 40597599 | 11.88484 | 22.66723 | 16.74801 | 6.678277 | 0.936407 | -0.01357 | 0.154247 |
| 16 | 64218000 | 64218799 | 11.69712 | 22.7701 | 13.454 | 4.940839 | 1.364179 | -0.29224 | 0.154259 |
| 2 | 139710600 | 139711399 | 7.10253 | 13.33903 | 33.4761 | 13.76646 | -0.85131 | 0.091834 | 0.154283 |
| 12 | 85615400 | 85619799 | 22.00057 | 36.03392 | 83.1959 | 35.79769 | -0.8275 | 1.449438 | 0.154297 |
| 9 | 101738600 | 101740999 | 11.25633 | 19.64203 | 52.09336 | 22.93416 | -1.0128 | 0.713844 | 0.15432 |
| 8 | 26481800 | 26484599 | 11.70185 | 20.43416 | 49.42168 | 21.76779 | -0.88023 | 0.70324 | 0.154339 |
| 17 | 82200400 | 82201199 | 6.6474 | 12.51704 | 33.49554 | 13.71342 | -0.93713 | 0.073958 | 0.154341 |
| 5 | 72658800 | 72662199 | 17.94322 | 29.7094 | 77.20938 | 33.40472 | -0.99262 | 1.30093 | 0.154361 |
| 18 | 73622600 | 73623399 | 5.228688 | 10.13242 | 28.52058 | 10.80208 | -0.90507 | -0.21486 | 0.154362 |
| 2 | 162772800 | 162774399 | 7.988602 | 14.53225 | 37.60847 | 16.19716 | -0.94555 | 0.298902 | 0.154368 |

| | | | | | | | | | |
|---|---|---|---|---|---|---|---|---|---|
| 17 | 7525600 | 7529999 | 23.43754 | 38.34618 | 83.92241 | 36.09488 | -0.75146 | 1.467537 | 0.154378 |
| 19 | 43428600 | 43430399 | 11.78276 | 20.39117 | 54.73625 | 24.01431 | -1.03189 | 0.825283 | 0.154385 |
| 5 | 108836800 | 108854799 | 99.5663 | 149.7473 | 319.1459 | 131.7773 | -0.82107 | 3.408992 | 0.154405 |
| 15 | 99023800 | 99025199 | 7.025925 | 12.98662 | 34.69028 | 14.62789 | -0.96877 | 0.193824 | 0.154446 |
| 7 | 80738200 | 80740399 | 13.57282 | 23.65039 | 52.14671 | 22.93702 | -0.74808 | 0.747372 | 0.154451 |
| 16 | 47866200 | 47869599 | 49.61669 | 80.13313 | 65.37894 | 27.89865 | 0.652621 | 1.740413 | 0.154451 |
| 8 | 118120800 | 118127999 | 47.64393 | 75.08782 | 169.6016 | 70.37836 | -0.85064 | 2.477951 | 0.154453 |
| 9 | 25163000 | 25164799 | 13.21742 | 22.69714 | 56.84062 | 24.93166 | -0.93481 | 0.905828 | 0.154466 |
| 2 | 37664800 | 37668199 | 18.83831 | 30.95034 | 84.1841 | 36.27041 | -1.06127 | 1.40296 | 0.15447 |
| 5 | 10471800 | 10475399 | 34.79417 | 57.76145 | 46.60228 | 20.19904 | 0.689652 | 1.290231 | 0.154481 |
| 17 | 87332000 | 87333799 | 8.398016 | 14.93545 | 41.01406 | 18.08362 | -1.05318 | 0.450212 | 0.15449 |
| 4 | 105420800 | 105423399 | 17.83649 | 30.21239 | 66.83622 | 29.26436 | -0.76 | 1.057304 | 0.154503 |
| 11 | 115337400 | 115350999 | 86.64653 | 131.967 | 263.869 | 109.3653 | -0.71364 | 3.148745 | 0.154503 |
| 12 | 37614200 | 37614999 | 5.86368 | 11.23481 | 30.79065 | 12.13643 | -0.9228 | -0.07068 | 0.154511 |
| 8 | 125528400 | 125530199 | 8.879758 | 16.15186 | 39.2267 | 16.89607 | -0.85509 | 0.308535 | 0.154522 |
| 7 | 9536200 | 9538599 | 22.47107 | 40.27526 | 21.19486 | 9.284093 | 1.321255 | 0.452629 | 0.154525 |
| 2 | 64624000 | 64625799 | 16.21144 | 29.83223 | 21.57767 | 9.15382 | 0.896042 | 0.260013 | 0.154544 |
| 2 | 120047000 | 120051199 | 55.03038 | 88.58055 | 70.61979 | 29.99999 | 0.680296 | 1.849138 | 0.154551 |
| 7 | 81124600 | 81127799 | 15.56071 | 26.18345 | 65.22763 | 28.50268 | -0.93067 | 1.1089 | 0.154562 |
| 15 | 15603600 | 15604799 | 15.04539 | 28.61133 | 17.43114 | 6.993819 | 1.205012 | 0.040235 | 0.154563 |
| 15 | 64253800 | 64254999 | 6.765475 | 12.44698 | 36.20979 | 15.36588 | -1.09774 | 0.22096 | 0.154573 |
| 7 | 44633000 | 44637199 | 15.93106 | 26.38792 | 78.01677 | 33.763 | -1.17777 | 1.29214 | 0.154578 |
| 10 | 89823000 | 89831199 | 84.48315 | 129.8216 | 236.7867 | 98.32108 | -0.57111 | 2.967749 | 0.154584 |
| 8 | 75749200 | 75750999 | 19.7382 | 35.28582 | 24.14532 | 10.59956 | 0.942895 | 0.454839 | 0.154593 |
| 1 | 152564600 | 152565999 | 6.358922 | 12.00282 | 35.39111 | 14.42529 | -1.0717 | 0.054518 | 0.154595 |
| 3 | 75168200 | 75169399 | 12.6314 | 24.19707 | 17.82463 | 7.029656 | 0.955753 | -0.03605 | 0.154597 |
| 12 | 54119200 | 54122799 | 17.83879 | 30.24492 | 65.70209 | 28.77604 | -0.73374 | 1.048714 | 0.154605 |
| 15 | 63752000 | 63753799 | 6.687165 | 12.16032 | 38.65749 | 16.65658 | -1.24012 | 0.293586 | 0.15463 |
| 8 | 74303000 | 74304399 | 29.51109 | 50.77575 | 29.96697 | 13.14501 | 1.141816 | 0.921941 | 0.154633 |
| 13 | 7642600 | 7643999 | 16.34625 | 30.38228 | 19.76293 | 8.27078 | 1.061357 | 0.201257 | 0.154643 |
| 12 | 33977400 | 33980599 | 24.50567 | 40.21935 | 85.17931 | 36.68294 | -0.70231 | 1.429786 | 0.154653 |
| 18 | 61743000 | 61763999 | 147.7345 | 217.9703 | 412.5828 | 169.9081 | -0.67552 | 3.764192 | 0.154656 |
| 12 | 19114600 | 19121999 | 180.5783 | 271.4336 | 207.969 | 85.85856 | 0.653038 | 3.43039 | 0.154696 |
| X | 4800000 | 4801599 | 17.44248 | 33.397 | 13.21032 | 5.217177 | 1.840813 | 0.003738 | 0.154698 |
| 6 | 115706200 | 115713999 | 59.32343 | 92.21087 | 203.4008 | 84.45023 | -0.83218 | 2.743203 | 0.154714 |
| 11 | 82276000 | 82284999 | 62.75019 | 97.10691 | 213.2443 | 88.57633 | -0.83071 | 2.824428 | 0.154715 |
| 2 | 51442400 | 51445199 | 48.55755 | 76.65995 | 162.014 | 67.26487 | -0.75288 | 2.44038 | 0.154738 |
| 9 | 15832800 | 15834999 | 9.088419 | 16.35997 | 39.00135 | 17.01946 | -0.84052 | 0.372241 | 0.154743 |
| 1 | 88763800 | 88768399 | 30.54243 | 49.18894 | 108.5766 | 46.15956 | -0.78477 | 1.8137 | 0.154752 |
| 7 | 3074200 | 3095199 | 49.17438 | 79.76543 | 52.16724 | 22.34957 | 0.977023 | 1.635922 | 0.154755 |
| 3 | 17212400 | 17213999 | 20.50826 | 37.91402 | 15.91733 | 6.710232 | 1.679718 | 0.255849 | 0.154759 |
| 17 | 20714600 | 20717599 | 48.9786 | 80.16148 | 47.90343 | 20.60576 | 1.116248 | 1.495557 | 0.154763 |
| 11 | 110300000 | 110305199 | 39.19005 | 62.6474 | 128.3833 | 53.98346 | -0.69088 | 2.064512 | 0.154775 |
| 2 | 179213800 | 179220799 | 47.78172 | 75.77933 | 149.5945 | 62.2784 | -0.64913 | 2.323409 | 0.15479 |
| 2 | 27471400 | 27475799 | 29.218 | 47.17171 | 101.3514 | 43.22922 | -0.74164 | 1.73555 | 0.154795 |
| 3 | 83682400 | 83684599 | 9.931411 | 17.59733 | 42.48323 | 18.7829 | -0.87232 | 0.486742 | 0.154813 |
| 2 | 31420400 | 31421599 | 6.312533 | 12.03652 | 31.75245 | 12.66797 | -0.88452 | -0.02372 | 0.154818 |

| | | | | | | | | | |
|---|---|---|---|---|---|---|---|---|---|
| 6 | 126587600 | 126588999 | 6.312533 | 12.03652 | 31.75245 | 12.66797 | -0.88452 | -0.02372 | 0.154818 |
| X | 93769000 | 93769199 | 8.772831 | 17.15929 | 8.871599 | 2.919181 | 1.693195 | -0.71654 | 0.154844 |
| 9 | 100353000 | 100353999 | 5.848573 | 10.86003 | 34.20661 | 14.33436 | -1.19723 | 0.157272 | 0.15487 |
| 19 | 40356400 | 40357799 | 5.848573 | 10.86003 | 34.20661 | 14.33436 | -1.19723 | 0.157272 | 0.15487 |
| 4 | 149952600 | 149955999 | 17.11459 | 28.81204 | 65.44784 | 28.6031 | -0.79825 | 1.108311 | 0.15488 |
| X | 98948600 | 98953599 | 44.28483 | 72.43636 | 52.74208 | 22.68221 | 0.831572 | 1.497379 | 0.154891 |
| 3 | 135506000 | 135507199 | 13.60624 | 25.92237 | 18.22084 | 7.282171 | 1.00562 | 0.016582 | 0.154915 |
| 6 | 57511200 | 57513199 | 12.17594 | 21.20957 | 52.12021 | 22.92075 | -0.90372 | 0.743813 | 0.154916 |
| 4 | 154148200 | 154157399 | 49.76278 | 78.00509 | 183.7526 | 76.21444 | -0.91725 | 2.587847 | 0.154988 |
| X | 66055400 | 66057999 | 24.75781 | 43.33689 | 27.63581 | 12.19518 | 1.03358 | 0.700609 | 0.155001 |
| X | 88626400 | 88628799 | 33.15816 | 56.84201 | 32.17694 | 14.11167 | 1.200893 | 0.972718 | 0.155009 |
| 9 | 36435600 | 36436999 | 23.48216 | 41.04883 | 29.76988 | 13.12201 | 0.848954 | 0.717383 | 0.155013 |
| 1 | 130199400 | 130200999 | 8.554878 | 15.64496 | 37.72705 | 16.14095 | -0.83838 | 0.272027 | 0.155041 |
| X | 96528600 | 96531399 | 30.97507 | 53.28593 | 32.62852 | 14.31243 | 1.088779 | 0.932151 | 0.155047 |
| 18 | 11177400 | 11179199 | 21.001 | 37.33794 | 24.64178 | 10.86764 | 0.990479 | 0.498339 | 0.155051 |
| 4 | 60582600 | 60585399 | 27.4248 | 47.30086 | 34.4095 | 15.09485 | 0.842022 | 0.895753 | 0.155054 |
| 1 | 15584800 | 15586599 | 16.18284 | 27.72198 | 64.75574 | 28.39795 | -0.836 | 0.947375 | 0.155058 |
| 18 | 17611600 | 17613799 | 23.24339 | 41.09891 | 23.16143 | 10.24452 | 1.212858 | 0.553437 | 0.155063 |
| 5 | 64839600 | 64840599 | 5.965391 | 11.12671 | 33.22279 | 13.82213 | -1.11246 | 0.127865 | 0.155074 |
| 5 | 131936800 | 131937799 | 10.90018 | 21.16328 | 15.17603 | 5.692268 | 1.060357 | -0.22603 | 0.155081 |
| 1 | 120013200 | 120015199 | 12.3166 | 21.49968 | 49.43209 | 21.76731 | -0.8077 | 0.71189 | 0.155102 |
| 2 | 33915600 | 33925799 | 61.05499 | 94.98191 | 197.2221 | 81.87465 | -0.74412 | 2.729443 | 0.155103 |
| 5 | 54251400 | 54252999 | 7.808062 | 14.29541 | 38.29412 | 16.35829 | -0.98734 | 0.260089 | 0.155109 |
| 11 | 30727200 | 30727799 | 2.936224 | 5.741777 | 20.11129 | 6.853539 | -1.07101 | -0.64662 | 0.155123 |
| 6 | 84969800 | 84971399 | 25.20031 | 44.48631 | 23.24819 | 10.28625 | 1.320617 | 0.584757 | 0.155131 |
| 7 | 7736800 | 7740599 | 39.25919 | 65.33574 | 43.28191 | 18.79205 | 0.973386 | 1.274499 | 0.155132 |
| X | 32869600 | 32872799 | 28.15822 | 49.52335 | 21.33993 | 9.448961 | 1.594985 | 0.637775 | 0.155145 |
| 17 | 46854200 | 46860999 | 28.12898 | 45.34852 | 103.9137 | 44.24035 | -0.83666 | 1.777846 | 0.15515 |
| 7 | 42710000 | 42711599 | 40.53896 | 66.36437 | 53.74994 | 23.12278 | 0.679144 | 1.476513 | 0.15515 |
| 17 | 39715200 | 39718399 | 31.86049 | 55.215 | 28.29615 | 12.41484 | 1.346368 | 0.868504 | 0.155154 |
| 5 | 138932800 | 138943799 | 53.32 | 83.5729 | 181.2721 | 75.18636 | -0.79848 | 2.591276 | 0.155175 |
| 3 | 24549600 | 24550999 | 14.68973 | 27.24389 | 20.55528 | 8.627942 | 0.845989 | 0.200893 | 0.155192 |
| 13 | 59168000 | 59176399 | 72.62547 | 112.586 | 209.0736 | 86.84017 | -0.58722 | 2.795681 | 0.155192 |
| 11 | 59195800 | 59205799 | 55.18873 | 85.98199 | 202.044 | 83.85804 | -0.92069 | 2.700993 | 0.155195 |
| X | 90595400 | 90596599 | 21.71068 | 38.7871 | 22.99492 | 10.10001 | 1.148151 | 0.470844 | 0.155198 |
| 11 | 71946800 | 71959199 | 86.45161 | 131.2945 | 274.489 | 113.6697 | -0.78126 | 3.208242 | 0.155198 |
| 1 | 128518600 | 128521599 | 16.40657 | 27.45541 | 69.06057 | 30.08958 | -0.94486 | 1.170104 | 0.155209 |
| 5 | 137820400 | 137822599 | 11.99168 | 21.04816 | 47.7553 | 21.12232 | -0.78826 | 0.621093 | 0.15521 |
| 7 | 59053000 | 59060799 | 152.6089 | 232.0612 | 177.1918 | 73.40537 | 0.672131 | 3.191643 | 0.155227 |
| 9 | 123177000 | 123181799 | 70.86242 | 112.9019 | 90.54805 | 37.89735 | 0.654992 | 2.188701 | 0.155249 |
| 8 | 36613400 | 36615599 | 21.56259 | 35.19064 | 85.94165 | 36.92974 | -0.91159 | 1.502757 | 0.155252 |
| 15 | 86103200 | 86111399 | 59.93554 | 93.59854 | 187.5767 | 77.83067 | -0.68868 | 2.661392 | 0.155252 |
| 11 | 82574400 | 82576599 | 8.224347 | 14.73708 | 40.53448 | 17.75913 | -1.04943 | 0.397413 | 0.155271 |
| 7 | 16041400 | 16048599 | 37.82833 | 60.28209 | 137.1264 | 57.40761 | -0.84609 | 2.164331 | 0.155274 |
| 7 | 25510800 | 25516399 | 31.70514 | 50.95273 | 113.1785 | 47.96747 | -0.79769 | 1.884905 | 0.155282 |
| 3 | 97795200 | 97796399 | 9.206363 | 16.26973 | 44.58669 | 19.72327 | -1.05561 | 0.514676 | 0.155293 |
| X | 120458800 | 120461999 | 47.64544 | 77.45109 | 56.01751 | 24.0364 | 0.835038 | 1.594243 | 0.155305 |

| | | | | | | | | | |
|---|---|---|---|---|---|---|---|---|---|
| 2 | 32251600 | 32254999 | 13.95136 | 23.64771 | 62.56371 | 27.39931 | -1.01646 | 1.035325 | 0.155308 |
| 9 | 21929600 | 21950599 | 151.9447 | 223.1452 | 421.9187 | 173.7684 | -0.68026 | 3.840037 | 0.155318 |
| 9 | 4174000 | 4176199 | 28.35568 | 48.54998 | 36.69142 | 16.08998 | 0.785527 | 0.96788 | 0.155322 |
| 13 | 10826600 | 10829599 | 72.11639 | 114.6365 | 93.90044 | 39.18233 | 0.621163 | 2.263312 | 0.155322 |
| 2 | 164889200 | 164898799 | 67.71041 | 105.0213 | 206.6255 | 85.81666 | -0.66987 | 2.784235 | 0.155327 |
| 12 | 22723200 | 22724599 | 29.73426 | 50.47665 | 37.92455 | 16.61644 | 0.792418 | 1.051272 | 0.155336 |
| 13 | 54737600 | 54738799 | 5.757666 | 10.58664 | 36.17399 | 15.36293 | -1.32791 | 0.217563 | 0.155339 |
| 3 | 91261200 | 91263399 | 46.69901 | 75.73965 | 59.42336 | 25.45565 | 0.714882 | 1.635002 | 0.15535 |
| 5 | 28500800 | 28501799 | 12.00601 | 23.29923 | 14.42711 | 5.427752 | 1.265567 | -0.20503 | 0.155351 |
| 17 | 53316000 | 53318999 | 42.87347 | 70.48721 | 44.5871 | 19.21578 | 1.037028 | 1.424989 | 0.155355 |
| 4 | 139748200 | 139768799 | 172.4928 | 250.0826 | 479.6349 | 197.7635 | -0.71872 | 4.040451 | 0.155362 |
| 2 | 34054200 | 34054799 | 3.639006 | 7.110732 | 23.84379 | 8.284157 | -1.03801 | -0.55688 | 0.155394 |
| 5 | 7558600 | 7559599 | 13.66294 | 25.84548 | 19.17049 | 7.778118 | 0.909584 | 0.066531 | 0.155396 |
| 3 | 143893400 | 143893999 | 3.527723 | 6.886293 | 22.77028 | 8.058095 | -1.04163 | -0.49602 | 0.155409 |
| 17 | 84612000 | 84612599 | 3.232108 | 6.31818 | 21.30871 | 7.326994 | -1.0308 | -0.60745 | 0.155439 |
| X | 116815200 | 116818399 | 36.42663 | 62.55607 | 27.62885 | 12.1187 | 1.5574 | 0.966008 | 0.155441 |
| 8 | 67430800 | 67431399 | 13.21675 | 24.91533 | 18.83752 | 7.697497 | 0.873696 | 0.085074 | 0.155444 |
| 16 | 60733600 | 60734599 | 14.34601 | 27.43544 | 16.95171 | 6.717762 | 1.20023 | -0.00672 | 0.155447 |
| 5 | 28623000 | 28623999 | 3.845557 | 7.427532 | 27.44111 | 10.55496 | -1.31242 | -0.1791 | 0.155453 |
| 6 | 146960600 | 146961999 | 6.945379 | 13.0739 | 34.13493 | 13.98287 | -0.90359 | 0.076184 | 0.155463 |
| 14 | 42680800 | 42684999 | 43.8204 | 72.09817 | 47.29082 | 20.38748 | 0.985169 | 1.417399 | 0.155479 |
| 18 | 53593400 | 53598399 | 37.64568 | 60.04034 | 133.9921 | 56.15927 | -0.81728 | 2.13672 | 0.155489 |
| 5 | 118876200 | 118888199 | 65.65976 | 101.2826 | 222.028 | 92.22639 | -0.83181 | 2.886267 | 0.155494 |
| 7 | 132532400 | 132533999 | 9.081776 | 15.97617 | 45.12583 | 19.98006 | -1.10135 | 0.567862 | 0.155526 |
| 1 | 8674800 | 8676199 | 15.42722 | 28.90679 | 19.41817 | 8.016542 | 1.030867 | 0.143145 | 0.155538 |
| 7 | 53510400 | 53512599 | 25.67165 | 44.74163 | 29.90762 | 13.15674 | 0.965451 | 0.768455 | 0.155567 |
| 13 | 27556000 | 27557399 | 13.20042 | 25.32547 | 16.43137 | 6.452043 | 1.141983 | -0.04944 | 0.155569 |
| 2 | 38240400 | 38253399 | 97.26874 | 146.6776 | 309.9761 | 128.082 | -0.80553 | 3.356322 | 0.155571 |
| 3 | 51482400 | 51483599 | 13.84035 | 25.89348 | 18.83234 | 7.794996 | 0.914558 | 0.13791 | 0.155571 |
| 11 | 39194400 | 39197599 | 27.45097 | 47.48274 | 33.30188 | 14.61024 | 0.895322 | 0.869523 | 0.155571 |
| 15 | 102316600 | 102318199 | 5.2436 | 9.87626 | 34.01403 | 13.91956 | -1.29753 | 0.057121 | 0.155574 |
| 1 | 89739600 | 89741399 | 7.140339 | 13.29153 | 33.75692 | 14.09218 | -0.88571 | 0.151295 | 0.15558 |
| 11 | 3449600 | 3455399 | 29.59817 | 47.64241 | 108.4794 | 46.08205 | -0.83039 | 1.830138 | 0.155594 |
| 4 | 87513800 | 87514599 | 6.165833 | 11.73013 | 31.58232 | 12.66075 | -0.91896 | -0.00641 | 0.155602 |
| 17 | 27141800 | 27145799 | 18.75291 | 30.957 | 77.62165 | 33.51359 | -0.9415 | 1.339603 | 0.155607 |
| 18 | 35976800 | 35979999 | 11.14157 | 19.35867 | 52.60263 | 23.0961 | -1.04811 | 0.777591 | 0.155609 |
| 4 | 137541000 | 137543199 | 10.61663 | 18.59706 | 47.26465 | 20.86969 | -0.95092 | 0.644979 | 0.155624 |
| 7 | 14374400 | 14377599 | 66.08761 | 105.8441 | 77.35438 | 32.61486 | 0.796893 | 2.020978 | 0.155624 |
| 1 | 123746600 | 123748599 | 13.46074 | 23.28693 | 53.72216 | 23.56915 | -0.81465 | 0.835098 | 0.155624 |
| 11 | 95528800 | 95536999 | 51.66729 | 81.48164 | 164.394 | 68.22316 | -0.68729 | 2.467191 | 0.155628 |
| 4 | 99328600 | 99330599 | 12.78014 | 22.30116 | 49.97491 | 22.00158 | -0.77098 | 0.719731 | 0.155633 |
| 5 | 88985400 | 88987399 | 11.53522 | 20.18381 | 47.80974 | 21.09065 | -0.85009 | 0.668785 | 0.155635 |
| 4 | 132788000 | 132793799 | 27.55872 | 44.30258 | 112.5098 | 47.70513 | -0.98976 | 1.870424 | 0.155654 |
| 18 | 74864600 | 74872799 | 48.03043 | 75.92564 | 160.3635 | 66.62313 | -0.75051 | 2.407918 | 0.155658 |
| 5 | 22909000 | 22911799 | 26.37993 | 45.80259 | 32.18815 | 14.12637 | 0.89332 | 0.827682 | 0.155666 |
| 11 | 5048200 | 5049999 | 10.5917 | 18.61215 | 45.63128 | 20.20238 | -0.89927 | 0.589911 | 0.155676 |
| 3 | 30975000 | 30975999 | 16.43898 | 30.98586 | 17.83894 | 7.290966 | 1.26332 | 0.11281 | 0.155702 |

| | | | | | | | | | |
|---|---|---|---|---|---|---|---|---|---|
| 7 | 18888600 | 18891799 | 14.05455 | 23.94661 | 59.97715 | 26.29093 | -0.93693 | 0.985528 | 0.155718 |
| 8 | 88522400 | 88524399 | 8.730799 | 15.89431 | 37.6229 | 16.18819 | -0.817 | 0.300537 | 0.155721 |
| 14 | 63286800 | 63288399 | 10.27451 | 18.0923 | 44.58288 | 19.73575 | -0.9054 | 0.561989 | 0.155755 |
| 7 | 139485200 | 139488399 | 13.61832 | 23.55565 | 53.72265 | 23.56925 | -0.79824 | 0.837573 | 0.155758 |
| 3 | 95188600 | 95190999 | 8.709521 | 15.22955 | 50.93666 | 22.45762 | -1.34453 | 0.669248 | 0.155762 |
| 6 | 40518400 | 40519399 | 7.726008 | 14.41572 | 36.10477 | 15.01218 | -0.86163 | 0.145357 | 0.155763 |
| 10 | 82204400 | 82206999 | 55.35207 | 89.1616 | 68.24342 | 29.02031 | 0.740261 | 1.828691 | 0.155769 |
| 2 | 4570600 | 4572399 | 8.772528 | 15.83198 | 39.04662 | 16.98452 | -0.88625 | 0.353989 | 0.155769 |
| 14 | 75820200 | 75821799 | 7.402447 | 13.1541 | 43.07317 | 19.00463 | -1.30612 | 0.450932 | 0.155772 |
| 9 | 90425600 | 90426399 | 5.388169 | 10.4279 | 29.05418 | 11.07125 | -0.89943 | -0.19228 | 0.155775 |
| 12 | 112460000 | 112480199 | 118.6673 | 175.0857 | 430.4765 | 177.277 | -1.05255 | 3.761779 | 0.155793 |
| 12 | 26830200 | 26832799 | 26.90411 | 45.99143 | 35.30687 | 15.48084 | 0.762629 | 0.973422 | 0.155793 |
| 19 | 21449600 | 21450199 | 4.801137 | 9.287653 | 27.07574 | 10.33889 | -0.96546 | -0.19446 | 0.155801 |
| X | 32849800 | 32854799 | 76.02392 | 121.9891 | 53.83694 | 22.76481 | 1.526037 | 1.971346 | 0.155801 |
| 7 | 79643000 | 79645199 | 11.56392 | 20.15039 | 50.47403 | 22.2016 | -0.93058 | 0.737566 | 0.155804 |
| 5 | 145880200 | 145882199 | 59.65437 | 95.61078 | 78.8187 | 33.26928 | 0.624715 | 1.991136 | 0.155818 |
| 15 | 102663000 | 102664799 | 7.662388 | 14.01116 | 36.10496 | 15.44929 | -0.9328 | 0.264707 | 0.155828 |
| 2 | 164991000 | 164995999 | 25.39772 | 41.05408 | 102.8324 | 43.93009 | -0.96029 | 1.693345 | 0.155852 |
| Y | 6989400 | 6992399 | 18.28423 | 33.31074 | 10.07106 | 4.329329 | 2.127792 | 0.298957 | 0.155853 |
| 11 | 60466000 | 60466999 | 5.405955 | 10.09076 | 33.76158 | 14.02858 | -1.27359 | 0.118693 | 0.155861 |
| 1 | 58818600 | 58819199 | 2.835653 | 5.546464 | 21.68659 | 7.286578 | -1.21015 | -0.69546 | 0.15588 |
| 19 | 13387600 | 13388999 | 18.40463 | 33.92981 | 19.52833 | 8.26395 | 1.225365 | 0.262121 | 0.155884 |
| 7 | 97751400 | 97756799 | 22.39665 | 36.50076 | 88.42661 | 37.97818 | -0.90159 | 1.524629 | 0.155889 |
| X | 24577000 | 24579199 | 44.90666 | 74.11819 | 44.48903 | 19.20607 | 1.114453 | 1.383488 | 0.155897 |
| 8 | 120790000 | 120794999 | 39.68245 | 63.44707 | 125.2293 | 52.67199 | -0.63646 | 2.058982 | 0.155897 |
| 4 | 141583600 | 141603799 | 121.1132 | 179.6057 | 386.9041 | 159.3584 | -0.85507 | 3.674992 | 0.155897 |
| 6 | 134036200 | 134037199 | 4.015941 | 7.76384 | 28.13501 | 10.77492 | -1.28009 | -0.19254 | 0.155908 |
| 3 | 38939000 | 38947399 | 71.29501 | 109.4847 | 237.9682 | 98.80285 | -0.8245 | 2.976043 | 0.155913 |
| 2 | 167810400 | 167810999 | 2.426889 | 4.747081 | 18.97845 | 6.359426 | -1.23193 | -0.71476 | 0.15593 |
| 2 | 179313000 | 179315599 | 14.44294 | 24.43986 | 65.92515 | 28.85673 | -1.04471 | 1.053315 | 0.155931 |
| 15 | 102639000 | 102646199 | 38.66202 | 61.5368 | 140.6761 | 58.79606 | -0.85512 | 2.205109 | 0.155935 |
| 11 | 75854400 | 75858199 | 18.40204 | 30.46653 | 75.50096 | 32.66369 | -0.92424 | 1.302677 | 0.155941 |
| 8 | 53246000 | 53247199 | 12.36838 | 24.01749 | 14.56156 | 5.455142 | 1.30223 | -0.21341 | 0.155947 |
| 10 | 78248000 | 78249199 | 7.689485 | 13.99741 | 37.09168 | 15.96133 | -0.97842 | 0.292328 | 0.155951 |
| 10 | 122726800 | 122728999 | 26.16734 | 45.0913 | 35.1321 | 15.41142 | 0.743271 | 0.899765 | 0.155976 |
| 13 | 29051600 | 29060599 | 56.20558 | 88.59804 | 172.1251 | 71.42727 | -0.63346 | 2.479234 | 0.155987 |
| 18 | 74938800 | 74940599 | 8.544734 | 15.65702 | 38.33461 | 16.35728 | -0.85747 | 0.26165 | 0.15599 |
| 4 | 39943200 | 39945399 | 15.93035 | 29.77006 | 18.68467 | 7.748626 | 1.122546 | 0.161388 | 0.155992 |
| 2 | 84668000 | 84671599 | 19.95603 | 33.0654 | 75.27719 | 32.58462 | -0.80232 | 1.296578 | 0.155993 |
| 4 | 136775600 | 136782799 | 63.9869 | 99.99837 | 187.638 | 77.85025 | -0.59307 | 2.649467 | 0.156 |
| 4 | 45857200 | 45860999 | 27.37539 | 44.442 | 92.17874 | 39.51533 | -0.68274 | 1.601388 | 0.156006 |
| 6 | 118969600 | 118971999 | 16.01113 | 27.46218 | 59.72771 | 26.1956 | -0.73296 | 0.930232 | 0.15601 |
| 7 | 124133000 | 124135399 | 33.29786 | 56.68653 | 34.73861 | 15.22661 | 1.085396 | 1.034098 | 0.156011 |
| 14 | 33734800 | 33742199 | 38.87783 | 62.10731 | 137.7071 | 57.84988 | -0.80542 | 2.086131 | 0.156017 |
| 12 | 76856000 | 76860799 | 25.43168 | 41.36252 | 91.83391 | 39.41223 | -0.77838 | 1.561389 | 0.156021 |
| 13 | 5555400 | 5555999 | 9.828258 | 19.13005 | 14.94999 | 5.494956 | 0.965193 | -0.30724 | 0.156059 |
| 7 | 23526400 | 23527999 | 43.37345 | 71.44943 | 47.81684 | 20.62483 | 0.957028 | 1.404388 | 0.156063 |

| | | | | | | | | | |
|---|---|---|---|---|---|---|---|---|---|
| 11 | 7094200 | 7095599 | 5.766384 | 10.93285 | 31.46451 | 12.70486 | -1.02392 | 0.010541 | 0.156085 |
| 18 | 66079400 | 66079999 | 3.708725 | 7.225885 | 26.09111 | 9.487239 | -1.20438 | -0.38729 | 0.156086 |
| 10 | 84937600 | 84939799 | 12.84142 | 22.2279 | 53.70728 | 23.56302 | -0.88086 | 0.827031 | 0.156094 |
| 3 | 54961600 | 54965199 | 14.94948 | 25.18352 | 66.93672 | 29.25958 | -1.02384 | 1.096329 | 0.1561 |
| 14 | 84670400 | 84671999 | 21.17947 | 38.29299 | 18.95549 | 8.226443 | 1.416675 | 0.387482 | 0.156104 |
| 8 | 60119600 | 60121399 | 18.63654 | 35.3074 | 13.47928 | 5.451449 | 1.861437 | 0.088457 | 0.156105 |
| 5 | 27702000 | 27704199 | 16.32805 | 29.83968 | 21.8828 | 9.370386 | 0.866397 | 0.297898 | 0.156126 |
| 5 | 142793400 | 142820999 | 192.5582 | 276.0434 | 541.4881 | 223.6229 | -0.76735 | 4.208663 | 0.156137 |
| 7 | 48814800 | 48816399 | 8.25979 | 15.11674 | 36.65478 | 15.66606 | -0.8447 | 0.265331 | 0.156142 |
| 4 | 141395400 | 141401799 | 31.20275 | 50.20014 | 113.8784 | 48.34056 | -0.82605 | 1.848995 | 0.156156 |
| 16 | 32778600 | 32780399 | 6.617984 | 12.20089 | 37.34657 | 15.80367 | -1.16786 | 0.209258 | 0.156162 |
| 16 | 12413800 | 12414599 | 3.769597 | 7.321236 | 25.48807 | 9.535267 | -1.18814 | -0.28037 | 0.156165 |
| 13 | 40883000 | 40888599 | 42.76261 | 67.85435 | 149.4241 | 62.23968 | -0.80581 | 2.304133 | 0.156182 |
| 12 | 28743600 | 28747199 | 14.47014 | 24.77371 | 59.05226 | 25.89554 | -0.86489 | 0.939916 | 0.156184 |
| 11 | 27366000 | 27368399 | 44.55783 | 72.51611 | 57.63582 | 24.74126 | 0.700452 | 1.570919 | 0.156185 |
| 16 | 90304000 | 90306999 | 22.33246 | 36.98498 | 80.30393 | 34.74885 | -0.73445 | 1.311824 | 0.1562 |
| X | 14854200 | 14855799 | 17.65359 | 33.72442 | 13.80172 | 5.484548 | 1.784668 | 0.026368 | 0.156212 |
| 2 | 105305400 | 105307799 | 10.88963 | 19.14407 | 46.14291 | 20.43303 | -0.8749 | 0.585438 | 0.156216 |
| 7 | 101858800 | 101859999 | 7.772601 | 13.96136 | 40.56361 | 17.73565 | -1.12578 | 0.378659 | 0.15624 |
| 11 | 57384200 | 57385999 | 8.254507 | 15.09391 | 39.37119 | 16.84688 | -0.95123 | 0.272546 | 0.156255 |
| 17 | 20485200 | 20490999 | 93.50523 | 147.8042 | 99.1614 | 41.19566 | 0.901545 | 2.42962 | 0.156283 |
| 12 | 41262200 | 41263599 | 15.61688 | 29.25352 | 19.41249 | 8.018207 | 1.047838 | 0.146765 | 0.156308 |
| 17 | 53428800 | 53431399 | 27.3239 | 47.60765 | 29.3408 | 12.90507 | 1.082141 | 0.781111 | 0.15631 |
| 14 | 13336600 | 13340199 | 72.18281 | 114.6521 | 92.43587 | 38.54094 | 0.642902 | 2.284853 | 0.156312 |
| 6 | 89709800 | 89712199 | 27.11294 | 47.05312 | 32.19046 | 14.12406 | 0.931888 | 0.837736 | 0.156314 |
| 7 | 34884200 | 34895599 | 70.90529 | 108.9237 | 233.6812 | 97.02753 | -0.80531 | 2.969252 | 0.156316 |
| 6 | 88846200 | 88854799 | 51.61723 | 81.49836 | 165.6357 | 68.77196 | -0.69651 | 2.439205 | 0.156321 |
| 10 | 40076000 | 40077199 | 5.865976 | 11.19842 | 29.984 | 11.93222 | -0.90155 | -0.03839 | 0.156329 |
| X | 31770600 | 31772599 | 21.36094 | 38.03162 | 24.0928 | 10.61287 | 1.050344 | 0.491274 | 0.156338 |
| 4 | 136947600 | 136963199 | 106.0576 | 160.0162 | 302.0368 | 124.8225 | -0.64201 | 3.347024 | 0.15634 |
| 7 | 102835400 | 102836999 | 20.04134 | 35.33316 | 26.97863 | 11.9399 | 0.776072 | 0.572412 | 0.156354 |
| 1 | 119343800 | 119354799 | 84.14333 | 128.0018 | 273.0385 | 113.1254 | -0.80826 | 3.172863 | 0.156364 |
| 9 | 117567600 | 117569799 | 10.86621 | 19.17903 | 44.04328 | 19.48777 | -0.80304 | 0.538336 | 0.156365 |
| 13 | 112355200 | 112364999 | 90.71247 | 138.6183 | 254.9151 | 105.7402 | -0.58921 | 3.081818 | 0.156376 |
| 12 | 22566600 | 22567399 | 29.82047 | 50.43088 | 37.50742 | 16.41474 | 0.806831 | 1.082696 | 0.156379 |
| 17 | 69377600 | 69379599 | 10.40095 | 18.37453 | 46.15637 | 20.41904 | -0.93141 | 0.527386 | 0.156392 |
| 6 | 72832400 | 72833999 | 17.10929 | 31.75844 | 19.74112 | 8.276548 | 1.124728 | 0.215294 | 0.156392 |
| 17 | 17360000 | 17361599 | 35.96311 | 60.98859 | 34.74854 | 15.22019 | 1.189724 | 1.078558 | 0.156399 |
| 7 | 10889400 | 10907399 | 313.523 | 453.0865 | 307.448 | 126.8213 | 0.774985 | 4.095736 | 0.156402 |
| 3 | 100919200 | 100920599 | 7.723047 | 13.9661 | 38.04764 | 16.50829 | -1.02582 | 0.333872 | 0.156411 |
| 4 | 128926000 | 128927999 | 7.723047 | 13.9661 | 38.04764 | 16.50829 | -1.02582 | 0.333872 | 0.156411 |
| 7 | 128410600 | 128411999 | 8.593207 | 15.61327 | 37.5868 | 16.21225 | -0.84326 | 0.312024 | 0.156423 |
| 8 | 69246200 | 69247399 | 8.793879 | 15.43787 | 46.19242 | 20.43395 | -1.18434 | 0.598456 | 0.156429 |
| 6 | 121284800 | 121286599 | 8.386097 | 15.41432 | 36.76417 | 15.62076 | -0.81518 | 0.240457 | 0.156431 |
| 13 | 99332800 | 99333399 | 4.237322 | 8.23994 | 25.1595 | 9.317769 | -0.98942 | -0.31519 | 0.156436 |
| 14 | 20807200 | 20807999 | 4.237322 | 8.23994 | 25.1595 | 9.317769 | -0.98942 | -0.31519 | 0.156436 |
| 5 | 34493000 | 34494799 | 11.40624 | 19.8947 | 49.41924 | 21.75075 | -0.91851 | 0.72023 | 0.156451 |

| | | | | | | | | | |
|---|---|---|---|---|---|---|---|---|---|
| 10 | 60621800 | 60638199 | 76.61779 | 116.5008 | 278.0283 | 115.1771 | -0.97057 | 3.180838 | 0.156458 |
| 4 | 71016400 | 71018599 | 28.65442 | 49.58359 | 31.50861 | 13.82366 | 1.03695 | 0.868354 | 0.156462 |
| 2 | 164284400 | 164285799 | 6.618987 | 12.15496 | 35.63854 | 15.1616 | -1.11117 | 0.230641 | 0.15647 |
| 3 | 122777400 | 122778799 | 8.417912 | 14.98857 | 43.07517 | 18.97344 | -1.11801 | 0.443323 | 0.156477 |
| 5 | 29911000 | 29912399 | 6.929931 | 13.1057 | 33.16098 | 13.46167 | -0.84718 | 0.043583 | 0.156499 |
| 1 | 78171600 | 78186399 | 115.5272 | 173.3002 | 334.0026 | 137.8156 | -0.67946 | 3.463085 | 0.156563 |
| 17 | 27935400 | 27963999 | 153.2099 | 220.5409 | 560.8648 | 231.4758 | -1.13578 | 4.146577 | 0.156567 |
| 7 | 5577000 | 5578799 | 36.5286 | 61.28387 | 40.41295 | 17.6361 | 0.97955 | 1.179076 | 0.156572 |
| 4 | 14239800 | 14241199 | 16.67009 | 30.08481 | 21.60509 | 9.397829 | 0.88223 | 0.370841 | 0.156576 |
| X | 32775200 | 32776599 | 19.20679 | 35.81923 | 16.82564 | 7.00202 | 1.533481 | 0.198943 | 0.156597 |
| 16 | 61452800 | 61454599 | 21.61606 | 38.70267 | 22.35026 | 9.798758 | 1.187354 | 0.455139 | 0.156605 |
| 9 | 21111800 | 21117599 | 32.84998 | 52.72704 | 115.1275 | 48.70314 | -0.77519 | 1.92665 | 0.156613 |
| 4 | 152979000 | 152981399 | 17.31245 | 29.03201 | 67.58088 | 29.48377 | -0.83358 | 1.148543 | 0.156631 |
| 8 | 58408000 | 58409599 | 19.36029 | 34.22197 | 25.96036 | 11.48324 | 0.786757 | 0.53797 | 0.156632 |
| 12 | 35096800 | 35097999 | 7.644193 | 13.66001 | 40.53361 | 17.79984 | -1.16002 | 0.412444 | 0.15664 |
| 12 | 104795800 | 104797799 | 7.644193 | 13.66001 | 40.53361 | 17.79984 | -1.16002 | 0.412444 | 0.15664 |
| 17 | 68963400 | 68965399 | 20.97046 | 38.35523 | 17.45508 | 7.466581 | 1.550003 | 0.319923 | 0.156647 |
| 17 | 35268600 | 35272199 | 15.72019 | 26.45039 | 65.25975 | 28.51626 | -0.91686 | 1.109408 | 0.156652 |
| 6 | 137773200 | 137781799 | 84.79641 | 129.507 | 250.8109 | 104.0246 | -0.66441 | 3.091423 | 0.156657 |
| 13 | 85094000 | 85096399 | 23.98181 | 42.42449 | 22.57743 | 9.985273 | 1.295254 | 0.552591 | 0.156667 |
| 15 | 25469200 | 25469999 | 4.36225 | 8.482273 | 25.15433 | 9.321205 | -0.94842 | -0.31079 | 0.15669 |
| 3 | 129973800 | 129975599 | 10.52861 | 18.333 | 52.59436 | 23.12368 | -1.12564 | 0.742706 | 0.156691 |
| 4 | 133244400 | 133248399 | 19.92808 | 33.0949 | 73.09616 | 31.69066 | -0.75845 | 1.268471 | 0.156692 |
| 13 | 90785200 | 90786199 | 6.169847 | 11.76509 | 32.34539 | 12.90306 | -0.94362 | -0.02477 | 0.156694 |
| 14 | 81330400 | 81333799 | 31.99535 | 53.71596 | 42.68678 | 18.6366 | 0.711359 | 1.158298 | 0.156695 |
| 8 | 61275400 | 61275999 | 17.1795 | 31.20617 | 18.9108 | 8.159312 | 1.132616 | 0.331285 | 0.156696 |
| 13 | 47580400 | 47581599 | 9.188932 | 16.26019 | 42.51763 | 18.80309 | -0.98726 | 0.49778 | 0.156697 |
| 10 | 82230400 | 82232399 | 51.6276 | 83.38802 | 65.01925 | 27.74825 | 0.718017 | 1.741585 | 0.156698 |
| 1 | 25156400 | 25157199 | 9.649542 | 18.7342 | 14.40624 | 5.405649 | 0.960618 | -0.24214 | 0.156701 |
| 7 | 13312000 | 13313999 | 46.30661 | 75.37711 | 56.69476 | 24.33905 | 0.779985 | 1.570361 | 0.156703 |
| 2 | 122441200 | 122442599 | 9.567881 | 17.14241 | 39.99113 | 17.52291 | -0.81369 | 0.407201 | 0.15671 |
| 8 | 26475800 | 26477799 | 7.545663 | 13.70894 | 37.06819 | 15.99003 | -1.00934 | 0.30384 | 0.156715 |
| 9 | 85235400 | 85236199 | 11.49218 | 22.40687 | 12.415 | 4.471989 | 1.480534 | -0.36314 | 0.156717 |
| 7 | 118592800 | 118597599 | 30.0663 | 48.4223 | 111.0194 | 47.1983 | -0.83942 | 1.813332 | 0.156733 |
| 2 | 45211400 | 45211799 | 8.264964 | 16.16565 | 10.30588 | 3.470239 | 1.363403 | -0.64558 | 0.156746 |
| 4 | 61261600 | 61261999 | 8.264964 | 16.16565 | 10.30588 | 3.470239 | 1.363403 | -0.64558 | 0.156746 |
| 14 | 112556600 | 112556999 | 8.264964 | 16.16565 | 10.30588 | 3.470239 | 1.363403 | -0.64558 | 0.156746 |
| 8 | 15075000 | 15079599 | 25.88442 | 41.77906 | 102.0565 | 43.51565 | -0.92663 | 1.738819 | 0.15675 |
| 3 | 112673600 | 112675799 | 21.03524 | 37.16375 | 26.20141 | 11.59104 | 0.891538 | 0.554543 | 0.156756 |
| 14 | 122699200 | 122703399 | 36.28218 | 58.05535 | 123.485 | 51.99522 | -0.74286 | 2.030612 | 0.156758 |
| 8 | 11284200 | 11286399 | 6.936358 | 12.71553 | 38.85134 | 16.56778 | -1.1737 | 0.247153 | 0.156788 |
| 13 | 78311200 | 78312799 | 6.225906 | 11.12436 | 46.29304 | 20.33088 | -1.64451 | 0.412558 | 0.15679 |
| 6 | 7566400 | 7568599 | 16.14844 | 26.92128 | 72.22632 | 31.39378 | -1.03751 | 1.213247 | 0.156822 |
| 17 | 24291800 | 24292599 | 4.357396 | 8.488666 | 24.87448 | 9.059589 | -0.90864 | -0.3744 | 0.156824 |
| 9 | 91243200 | 91253999 | 87.01189 | 132.5071 | 266.2063 | 110.33 | -0.72062 | 3.151295 | 0.156831 |
| 2 | 70391000 | 70404999 | 82.5141 | 125.0893 | 302.8573 | 125.3499 | -0.99496 | 3.244949 | 0.156841 |
| 12 | 18368600 | 18385399 | 583.7619 | 793.3698 | 581.984 | 242.4013 | 0.573653 | 5.02304 | 0.156851 |

| | | | | | | | | | |
|---|---|---|---|---|---|---|---|---|---|
| 12 | 63460400 | 63463399 | 30.80501 | 52.65027 | 35.98841 | 15.77944 | 0.929412 | 0.992229 | 0.15687 |
| 17 | 31823400 | 31826599 | 15.62142 | 26.55944 | 60.57106 | 26.54669 | -0.80297 | 1.012012 | 0.156873 |
| 2 | 74408600 | 74409799 | 6.40117 | 11.93132 | 33.22107 | 13.83753 | -1.01414 | 0.136025 | 0.156874 |
| 9 | 75139200 | 75139999 | 6.40117 | 11.93132 | 33.22107 | 13.83753 | -1.01414 | 0.136025 | 0.156874 |
| 8 | 19322000 | 19325999 | 51.27962 | 82.73857 | 67.10698 | 28.60025 | 0.659548 | 1.771907 | 0.156874 |
| 13 | 65842400 | 65844599 | 40.03403 | 66.77526 | 41.00802 | 17.83021 | 1.082161 | 1.250922 | 0.156889 |
| 13 | 66634400 | 66640399 | 61.38676 | 98.56547 | 67.08146 | 28.39299 | 0.903713 | 1.935231 | 0.156905 |
| 13 | 66372600 | 66374199 | 36.22751 | 61.30347 | 35.93145 | 15.72793 | 1.149055 | 1.099623 | 0.156921 |
| 2 | 172317000 | 172319799 | 9.704081 | 17.0275 | 46.7126 | 20.6556 | -1.06033 | 0.611028 | 0.156923 |
| 9 | 30287000 | 30287999 | 5.699209 | 10.96457 | 29.213 | 11.37977 | -0.86585 | -0.114 | 0.156923 |
| 7 | 17808200 | 17812799 | 128.4708 | 197.4271 | 154.7528 | 64.25861 | 0.646217 | 2.97423 | 0.156925 |
| 1 | 7661600 | 7662199 | 10.46283 | 20.2489 | 16.40762 | 6.252436 | 0.86511 | -0.17465 | 0.156929 |
| 1 | 68388400 | 68389399 | 10.46283 | 20.2489 | 16.40762 | 6.252436 | 0.86511 | -0.17465 | 0.156929 |
| 9 | 105414000 | 105414999 | 5.39138 | 10.42352 | 28.3244 | 10.8445 | -0.87001 | -0.17744 | 0.156947 |
| 13 | 111399600 | 111400799 | 5.39138 | 10.42352 | 28.3244 | 10.8445 | -0.87001 | -0.17744 | 0.156947 |
| 6 | 99609200 | 99611399 | 9.943996 | 17.58473 | 42.50734 | 18.80101 | -0.8751 | 0.510399 | 0.156953 |
| 10 | 76489000 | 76491199 | 16.10191 | 27.31493 | 62.79966 | 27.50937 | -0.81504 | 1.034504 | 0.156955 |
| 1 | 142152400 | 142153599 | 12.01452 | 23.32092 | 13.13863 | 4.937877 | 1.400512 | -0.21294 | 0.156957 |
| 13 | 117714600 | 117716199 | 18.40968 | 32.73963 | 24.97642 | 11.01272 | 0.782773 | 0.479332 | 0.156964 |
| 10 | 41627400 | 41629599 | 14.55726 | 25.1537 | 54.83165 | 24.05503 | -0.73379 | 0.852142 | 0.156966 |
| 2 | 72221000 | 72227599 | 41.81724 | 66.69407 | 135.1256 | 56.63501 | -0.67813 | 2.140187 | 0.156981 |
| 13 | 54526000 | 54530199 | 22.45545 | 36.8229 | 81.62657 | 35.14018 | -0.76825 | 1.434859 | 0.156982 |
| 7 | 59281000 | 59281399 | 11.24044 | 21.60833 | 17.39356 | 6.790777 | 0.842083 | -0.08452 | 0.156991 |
| 9 | 38002400 | 38003199 | 11.24044 | 21.60833 | 17.39356 | 6.790777 | 0.842083 | -0.08452 | 0.156991 |
| 4 | 41649600 | 41651599 | 10.67707 | 18.9721 | 44.0794 | 19.44945 | -0.81537 | 0.473242 | 0.157013 |
| 2 | 173616600 | 173620799 | 24.25229 | 39.61749 | 86.28633 | 37.09055 | -0.74617 | 1.492612 | 0.157016 |
| 2 | 23393000 | 23394599 | 15.05124 | 28.14835 | 20.11871 | 8.332725 | 0.938728 | 0.154223 | 0.157019 |
| 4 | 46338000 | 46345799 | 60.54487 | 94.28344 | 197.2405 | 81.86995 | -0.75376 | 2.711094 | 0.157045 |
| 16 | 90309200 | 90310799 | 14.11876 | 26.44322 | 20.14354 | 8.322317 | 0.85007 | 0.136186 | 0.157061 |
| 5 | 124169000 | 124171599 | 8.259837 | 14.46988 | 51.44788 | 22.71837 | -1.43215 | 0.636398 | 0.15707 |
| 10 | 124947800 | 124949199 | 16.07718 | 30.168 | 18.94024 | 7.800224 | 1.130754 | 0.139725 | 0.15708 |
| 5 | 21283600 | 21285399 | 18.86542 | 34.52122 | 19.28788 | 8.245335 | 1.257133 | 0.304452 | 0.157089 |
| 2 | 74967200 | 74969399 | 8.311347 | 15.04409 | 38.57665 | 16.71816 | -0.93844 | 0.332316 | 0.157094 |
| 13 | 65781200 | 65783199 | 43.2436 | 70.84413 | 52.26928 | 22.49281 | 0.81366 | 1.471682 | 0.157096 |
| 1 | 97244400 | 97246399 | 22.41915 | 39.35627 | 28.4249 | 12.5739 | 0.854473 | 0.641983 | 0.157101 |
| 1 | 81218600 | 81219999 | 11.89241 | 20.66925 | 51.55873 | 22.64187 | -0.9253 | 0.776113 | 0.157111 |
| 7 | 132723400 | 132726599 | 19.9993 | 32.70814 | 87.34734 | 37.56261 | -1.03802 | 1.467175 | 0.157115 |
| 17 | 34238200 | 34253599 | 119.4046 | 177.4304 | 376.085 | 154.9341 | -0.82905 | 3.638836 | 0.157136 |
| 13 | 68538000 | 68539399 | 33.80474 | 56.43737 | 44.89863 | 19.53327 | 0.709989 | 1.224375 | 0.157146 |
| 9 | 50514800 | 50515599 | 4.327535 | 8.293706 | 30.02726 | 11.82672 | -1.31785 | -0.08954 | 0.157149 |
| 2 | 156746600 | 156748399 | 6.963922 | 13.05363 | 33.40776 | 13.78309 | -0.8831 | 0.102672 | 0.157152 |
| 3 | 143863600 | 143864199 | 3.639631 | 7.11 | 22.38098 | 7.819964 | -0.95474 | -0.54066 | 0.157153 |
| 10 | 127477800 | 127481399 | 18.12758 | 30.30462 | 68.63721 | 29.88596 | -0.79419 | 1.186443 | 0.157166 |
| 1 | 178879600 | 178881399 | 8.587773 | 15.60615 | 37.5883 | 16.20931 | -0.84396 | 0.309662 | 0.157176 |
| 18 | 20273600 | 20274199 | 3.757321 | 7.33944 | 23.0179 | 8.051366 | -0.95098 | -0.53216 | 0.157177 |
| 11 | 55363000 | 55364999 | 12.60964 | 22.0732 | 48.87012 | 21.56619 | -0.75337 | 0.668626 | 0.157177 |
| 14 | 63110400 | 63110999 | 4.232289 | 8.245307 | 24.88067 | 9.056628 | -0.94987 | -0.37893 | 0.157187 |

| | | | | | | | | | |
|---|---|---|---|---|---|---|---|---|---|
| 15 | 37313200 | 37313999 | 6.348102 | 11.96885 | 31.8457 | 13.0096 | -0.92619 | 0.058514 | 0.157191 |
| 4 | 149568600 | 149570599 | 9.16429 | 16.28117 | 41.4859 | 18.30781 | -0.94734 | 0.46089 | 0.157215 |
| 7 | 59659200 | 59663599 | 55.49727 | 90.00777 | 51.95327 | 22.25536 | 1.156453 | 1.649279 | 0.157236 |
| 1 | 56841200 | 56842599 | 6.79558 | 12.80938 | 35.85263 | 14.65262 | -0.99974 | 0.073591 | 0.157236 |
| 7 | 43847200 | 43847799 | 3.818915 | 7.451022 | 24.59822 | 8.779658 | -1.05196 | -0.45693 | 0.157249 |
| 5 | 137551600 | 137554599 | 23.41865 | 38.36602 | 82.92923 | 35.68668 | -0.73261 | 1.449119 | 0.157266 |
| 3 | 126232000 | 126233199 | 8.413305 | 15.35794 | 37.6818 | 16.15859 | -0.86533 | 0.281373 | 0.15728 |
| 6 | 85357200 | 85358999 | 8.413305 | 15.35794 | 37.6818 | 16.15859 | -0.86533 | 0.281373 | 0.15728 |
| X | 57789600 | 57791799 | 18.42602 | 33.79297 | 20.79438 | 8.86223 | 1.122919 | 0.294307 | 0.157297 |
| 7 | 17849200 | 17849799 | 17.55801 | 31.4918 | 23.9529 | 10.48594 | 0.793925 | 0.419045 | 0.157312 |
| X | 61698400 | 61700599 | 37.6527 | 63.59465 | 35.42346 | 15.49668 | 1.222221 | 1.120728 | 0.157317 |
| 16 | 93381200 | 93388799 | 39.31898 | 62.24053 | 162.5392 | 67.57417 | -1.05448 | 2.3692 | 0.157348 |
| 17 | 85811800 | 85814599 | 13.95266 | 23.66855 | 63.15675 | 27.66563 | -1.0288 | 1.027942 | 0.157354 |
| 14 | 9135600 | 9136999 | 8.407139 | 15.38441 | 36.65467 | 15.6688 | -0.81994 | 0.266274 | 0.157366 |
| 9 | 108700400 | 108707199 | 60.02294 | 92.80321 | 217.4185 | 90.3106 | -0.92495 | 2.8405 | 0.157375 |
| 4 | 152695600 | 152713399 | 143.782 | 211.8469 | 403.4544 | 166.1464 | -0.68619 | 3.786671 | 0.157378 |
| 7 | 110445600 | 110446199 | 13.76243 | 26.38184 | 17.10219 | 6.731433 | 1.140931 | -0.03307 | 0.157381 |
| 17 | 69538400 | 69539799 | 7.923868 | 14.207 | 40.02775 | 17.52843 | -1.08317 | 0.391198 | 0.157385 |
| 9 | 114633600 | 114645599 | 95.63976 | 145.4267 | 272.4117 | 112.8509 | -0.62154 | 3.184706 | 0.15739 |
| 3 | 20809200 | 20810799 | 16.65941 | 30.16695 | 22.64761 | 9.809905 | 0.822686 | 0.355246 | 0.157394 |
| 18 | 11955400 | 11956399 | 16.74577 | 31.02102 | 20.29433 | 8.531872 | 1.048704 | 0.224283 | 0.157396 |
| 7 | 11593600 | 11595599 | 45.17919 | 73.56691 | 57.17719 | 24.5492 | 0.73317 | 1.563842 | 0.15741 |
| 16 | 50485200 | 50487399 | 16.36684 | 27.89843 | 61.28026 | 26.86308 | -0.74864 | 0.989993 | 0.157415 |
| 7 | 5871000 | 5875399 | 105.0858 | 163.4954 | 133.8625 | 55.57079 | 0.597003 | 2.739162 | 0.157423 |
| 7 | 87737200 | 87738399 | 15.28262 | 28.16845 | 21.6002 | 9.143273 | 0.814376 | 0.242928 | 0.157438 |
| 18 | 38904400 | 38905599 | 6.9542 | 13.06424 | 33.48371 | 13.76282 | -0.88089 | 0.088176 | 0.157457 |
| 3 | 158582400 | 158584399 | 21.15048 | 37.15843 | 27.93264 | 12.36439 | 0.796897 | 0.620394 | 0.157473 |
| 15 | 44562600 | 44563599 | 6.048048 | 11.48228 | 32.71063 | 13.17055 | -1.00631 | 0.005542 | 0.157485 |
| 7 | 11421600 | 11437399 | 312.0617 | 449.9263 | 323.1932 | 133.3572 | 0.68957 | 4.130003 | 0.157489 |
| 3 | 12395400 | 12396799 | 16.71766 | 30.7413 | 21.33038 | 9.057915 | 0.954191 | 0.266289 | 0.15749 |
| 13 | 66241800 | 66242799 | 16.71766 | 30.7413 | 21.33038 | 9.057915 | 0.954191 | 0.266289 | 0.15749 |
| 15 | 49826000 | 49826799 | 9.748863 | 17.36651 | 40.97338 | 18.04056 | -0.83484 | 0.446461 | 0.15749 |
| 9 | 65099000 | 65101599 | 10.91042 | 19.13295 | 46.19952 | 20.41855 | -0.87741 | 0.627195 | 0.157491 |
| 12 | 20154000 | 20159199 | 149.0049 | 225.9363 | 184.1837 | 76.23689 | 0.574445 | 3.249365 | 0.157494 |
| 1 | 128010400 | 128011399 | 5.683394 | 10.95071 | 29.86732 | 11.58419 | -0.8931 | -0.12528 | 0.157494 |
| 5 | 28662800 | 28676399 | 100.5387 | 151.1316 | 327.6684 | 135.2776 | -0.84647 | 3.419328 | 0.157503 |
| 12 | 102626400 | 102630799 | 27.59122 | 44.61704 | 96.56226 | 41.27209 | -0.7493 | 1.68498 | 0.157507 |
| 8 | 123582200 | 123584399 | 9.430307 | 16.48788 | 48.84913 | 21.53498 | -1.17061 | 0.673106 | 0.157529 |
| 16 | 42473600 | 42475799 | 38.35154 | 63.58044 | 43.37527 | 18.78269 | 0.930686 | 1.31264 | 0.157547 |
| 10 | 81708000 | 81708799 | 20.24596 | 35.98765 | 23.98337 | 10.57922 | 0.97593 | 0.491439 | 0.157566 |
| X | 122198600 | 122199999 | 19.63562 | 35.69941 | 21.23623 | 9.152459 | 1.160909 | 0.352575 | 0.157568 |
| 1 | 31166400 | 31167399 | 15.44528 | 28.27828 | 21.19766 | 9.055005 | 0.837657 | 0.280261 | 0.157573 |
| 3 | 82904200 | 82905599 | 7.836509 | 14.21959 | 38.09824 | 16.46004 | -0.99752 | 0.317418 | 0.157577 |
| 1 | 182149600 | 182150599 | 10.76731 | 20.75429 | 16.44225 | 6.37021 | 0.875221 | -0.11668 | 0.15758 |
| 17 | 66522200 | 66523399 | 18.98281 | 33.62915 | 25.57389 | 11.30728 | 0.784533 | 0.512223 | 0.157582 |
| 1 | 96481600 | 96482599 | 13.69307 | 25.78233 | 19.72964 | 8.078418 | 0.85396 | 0.098894 | 0.157596 |
| 4 | 141762600 | 141785599 | 129.7184 | 191.684 | 398.5826 | 164.1453 | -0.80892 | 3.736502 | 0.157597 |

| | | | | | | | | | |
|---|---|---|---|---|---|---|---|---|---|
| 7 | 73292800 | 73293999 | 6.396077 | 11.97216 | 33.89554 | 14.0181 | -1.03029 | 0.107741 | 0.157604 |
| 4 | 133633200 | 133633999 | 5.535022 | 10.68708 | 28.80115 | 11.09546 | -0.86667 | -0.15265 | 0.157619 |
| 10 | 72932400 | 72936199 | 32.18268 | 51.72401 | 110.6328 | 46.8936 | -0.74315 | 1.883428 | 0.15762 |
| 8 | 19643000 | 19645999 | 59.14694 | 94.8536 | 77.79404 | 32.86633 | 0.6331 | 1.971846 | 0.157621 |
| 3 | 23492000 | 23492799 | 12.69152 | 24.22007 | 17.5879 | 7.004436 | 0.962577 | -0.00712 | 0.157628 |
| 6 | 49862200 | 49864999 | 23.68712 | 41.5595 | 27.72489 | 12.2586 | 0.968375 | 0.656093 | 0.157629 |
| 8 | 120827400 | 120828799 | 7.630749 | 14.05358 | 35.16573 | 14.90163 | -0.88055 | 0.220924 | 0.15764 |
| 1 | 102242000 | 102243199 | 13.87858 | 26.12387 | 19.725 | 8.080578 | 0.872616 | 0.102578 | 0.157646 |
| 13 | 107507400 | 107508799 | 7.437223 | 13.83544 | 34.30136 | 14.33477 | -0.85274 | 0.158407 | 0.157648 |
| 8 | 7017200 | 7017799 | 10.54521 | 20.54482 | 13.81044 | 5.024684 | 1.193467 | -0.33622 | 0.157649 |
| 10 | 84721800 | 84724399 | 12.93931 | 22.5462 | 51.05987 | 22.45876 | -0.78644 | 0.742248 | 0.15767 |
| 5 | 135052200 | 135075399 | 139.9892 | 206.0197 | 416.4516 | 171.4967 | -0.77369 | 3.805512 | 0.157671 |
| 7 | 14486400 | 14488199 | 43.999 | 72.69663 | 44.46451 | 19.20945 | 1.087456 | 1.368526 | 0.157684 |
| 7 | 26408000 | 26411799 | 50.1178 | 81.3211 | 57.91592 | 24.81814 | 0.854801 | 1.633492 | 0.15769 |
| 4 | 138799000 | 138805599 | 37.57051 | 59.71166 | 144.7556 | 60.39223 | -0.94181 | 2.252189 | 0.157696 |
| 15 | 63928200 | 63933399 | 23.91888 | 39.12178 | 83.86186 | 36.06483 | -0.72191 | 1.474024 | 0.157699 |
| 17 | 15813000 | 15814399 | 16.45748 | 30.00763 | 21.84905 | 9.38269 | 0.873983 | 0.311367 | 0.157706 |
| 9 | 120922600 | 120926399 | 41.50003 | 66.23009 | 130.0152 | 54.54136 | -0.63163 | 2.11966 | 0.157714 |
| 4 | 79381800 | 79383399 | 20.26467 | 35.78491 | 26.91031 | 11.90581 | 0.798842 | 0.556747 | 0.157725 |
| 18 | 58143200 | 58147999 | 22.0547 | 36.40012 | 77.61213 | 33.50476 | -0.70864 | 1.349848 | 0.157726 |
| 12 | 73090000 | 73095199 | 34.56315 | 55.23221 | 134.2554 | 56.42533 | -0.93718 | 2.071346 | 0.157741 |
| X | 5479800 | 5484599 | 155.8326 | 236.4479 | 181.9488 | 75.32695 | 0.658391 | 3.23825 | 0.157754 |
| 14 | 112248800 | 112249999 | 7.888185 | 14.65291 | 35.94799 | 15.05626 | -0.8405 | 0.174103 | 0.157771 |
| 8 | 105551600 | 105552399 | 6.707116 | 12.46743 | 33.71046 | 14.1061 | -0.97783 | 0.157055 | 0.157774 |
| 11 | 109316200 | 109317199 | 6.707116 | 12.46743 | 33.71046 | 14.1061 | -0.97783 | 0.157055 | 0.157774 |
| 11 | 106939200 | 106943199 | 13.4546 | 22.87476 | 60.94811 | 26.71188 | -1.02614 | 1.005411 | 0.157775 |
| 9 | 26008800 | 26009599 | 11.20226 | 21.66908 | 16.24484 | 6.204897 | 0.972688 | -0.15953 | 0.157777 |
| 17 | 53716600 | 53717199 | 4.788047 | 9.307495 | 26.40453 | 9.807652 | -0.88917 | -0.29413 | 0.157779 |
| 3 | 40636600 | 40638399 | 11.33474 | 19.97597 | 45.08778 | 19.95592 | -0.77924 | 0.559389 | 0.15778 |
| 4 | 65042600 | 65045999 | 18.92243 | 31.5856 | 72.74345 | 31.6437 | -0.8184 | 1.206648 | 0.157787 |
| 6 | 58421600 | 58422799 | 24.24551 | 42.1955 | 31.93314 | 14.03512 | 0.787368 | 0.786214 | 0.157788 |
| 4 | 45307400 | 45308999 | 6.517921 | 12.22984 | 32.28972 | 13.29928 | -0.92497 | 0.091645 | 0.157796 |
| 4 | 147276400 | 147278999 | 57.67496 | 92.8688 | 68.13544 | 28.95879 | 0.800493 | 1.841799 | 0.157801 |
| 4 | 153624600 | 153635999 | 81.92727 | 125.4368 | 245.9174 | 102.0537 | -0.6792 | 3.039202 | 0.157819 |
| 8 | 30147000 | 30149199 | 20.44083 | 36.91758 | 19.5391 | 8.491197 | 1.319617 | 0.388833 | 0.157819 |
| 13 | 19539600 | 19542799 | 13.55902 | 23.19334 | 58.40994 | 25.61205 | -0.94384 | 0.94507 | 0.157841 |
| 12 | 103348600 | 103351399 | 15.57623 | 26.64119 | 58.51036 | 25.65588 | -0.74743 | 0.954581 | 0.157859 |
| 13 | 67212400 | 67214399 | 43.38409 | 70.93181 | 54.39882 | 23.39095 | 0.756419 | 1.501527 | 0.157878 |
| X | 29481800 | 29484199 | 23.35671 | 40.86066 | 28.8017 | 12.70343 | 0.889823 | 0.703585 | 0.157889 |
| 17 | 83425800 | 83429399 | 18.20757 | 30.64076 | 67.25843 | 29.39112 | -0.74921 | 1.115887 | 0.157895 |
| 7 | 10535600 | 10540999 | 124.2939 | 191.1524 | 154.7307 | 64.25608 | 0.600693 | 2.957979 | 0.157896 |
| 4 | 135296200 | 135298999 | 11.73298 | 20.40676 | 51.01952 | 22.41516 | -0.92834 | 0.764871 | 0.157904 |
| 13 | 118787800 | 118789199 | 16.32959 | 29.70618 | 22.6205 | 9.741895 | 0.807587 | 0.32668 | 0.157907 |
| X | 25737000 | 25737799 | 22.67044 | 39.52523 | 30.45665 | 13.4013 | 0.761596 | 0.750406 | 0.157908 |
| 2 | 164041400 | 164043599 | 17.67391 | 30.01615 | 65.71628 | 28.79704 | -0.74484 | 1.033673 | 0.157917 |
| 7 | 62060800 | 62063399 | 26.74354 | 46.89107 | 25.35757 | 11.20564 | 1.269602 | 0.680156 | 0.15792 |
| 10 | 78187600 | 78189799 | 9.03596 | 16.00701 | 42.51001 | 18.79561 | -1.00891 | 0.484027 | 0.157921 |

| | | | | | | | | | |
|---|---|---|---|---|---|---|---|---|---|
| 17 | 38952000 | 38953999 | 35.96466 | 61.01607 | 33.55803 | 14.70055 | 1.240314 | 1.072687 | 0.157925 |
| 7 | 78307600 | 78308799 | 5.024292 | 9.621363 | 30.07333 | 11.87202 | -1.112 | -0.07531 | 0.157928 |
| 17 | 31089600 | 31094599 | 37.08125 | 59.3955 | 120.3216 | 50.74042 | -0.6709 | 1.999778 | 0.157943 |
| 8 | 25757600 | 25760599 | 18.78828 | 31.31504 | 70.24666 | 30.52891 | -0.78036 | 1.222791 | 0.157948 |
| 17 | 24479400 | 24485199 | 40.17218 | 64.29312 | 125.9201 | 53.03613 | -0.62376 | 2.030347 | 0.157956 |
| 8 | 74281600 | 74281999 | 8.265353 | 16.16491 | 10.94314 | 3.714416 | 1.268608 | -0.6162 | 0.157958 |
| 5 | 6403200 | 6404199 | 14.77572 | 27.72174 | 18.85219 | 7.765919 | 1.015363 | 0.127051 | 0.157963 |
| 11 | 72998400 | 73002999 | 22.90036 | 37.24517 | 89.92967 | 38.59485 | -0.89892 | 1.556819 | 0.157966 |
| 7 | 38897200 | 38899399 | 21.17725 | 37.36801 | 26.90714 | 11.90621 | 0.860829 | 0.571277 | 0.157969 |
| 9 | 113867600 | 113883199 | 132.6943 | 196.8375 | 375.2547 | 154.5648 | -0.67853 | 3.67116 | 0.15797 |
| X | 79580800 | 79586199 | 32.15924 | 54.62629 | 34.15193 | 14.96623 | 1.056465 | 1.050808 | 0.157972 |
| 7 | 140145600 | 140146399 | 21.27865 | 37.35334 | 28.63008 | 12.66444 | 0.769234 | 0.636387 | 0.157986 |
| 13 | 67550600 | 67552999 | 29.6639 | 50.53258 | 37.6521 | 16.5045 | 0.804762 | 1.017411 | 0.158021 |
| 8 | 14377400 | 14378799 | 7.6195 | 14.06392 | 35.76296 | 15.10918 | -0.90084 | 0.207683 | 0.158023 |
| 1 | 59461200 | 59464399 | 13.03663 | 22.45393 | 55.25593 | 24.23879 | -0.90866 | 0.876054 | 0.158034 |
| 4 | 38125000 | 38127799 | 35.59503 | 59.67529 | 41.50732 | 18.10558 | 0.902895 | 1.183274 | 0.158035 |
| 2 | 12207600 | 12209199 | 5.85337 | 11.21753 | 29.5882 | 11.65509 | -0.86651 | -0.07369 | 0.15804 |
| 5 | 41725000 | 41726599 | 14.85616 | 28.68133 | 12.76909 | 4.921689 | 1.702812 | -0.09959 | 0.158045 |
| 11 | 7555200 | 7558199 | 16.50831 | 28.22698 | 60.79373 | 26.65646 | -0.71965 | 0.962046 | 0.158075 |
| 9 | 106213000 | 106217399 | 22.83026 | 37.2894 | 85.30075 | 36.66506 | -0.8168 | 1.491846 | 0.158097 |
| 10 | 4464600 | 4466599 | 20.48408 | 36.57902 | 24.13087 | 10.60302 | 0.994587 | 0.467148 | 0.158109 |
| 1 | 37141800 | 37146799 | 28.74928 | 46.3892 | 101.4708 | 43.24916 | -0.76831 | 1.751854 | 0.158115 |
| 6 | 149258600 | 149267799 | 140.7447 | 207.7433 | 394.817 | 162.5926 | -0.68076 | 3.75655 | 0.158116 |
| 9 | 101188800 | 101189399 | 9.820525 | 19.06258 | 15.39887 | 5.786859 | 0.888201 | -0.23152 | 0.158142 |
| 14 | 51347000 | 51348799 | 22.5726 | 39.75359 | 26.66738 | 11.80466 | 0.96132 | 0.598941 | 0.158159 |
| 5 | 123329200 | 123331799 | 14.10806 | 24.29392 | 56.95949 | 24.98591 | -0.83957 | 0.882674 | 0.158168 |
| 11 | 89035200 | 89036799 | 7.418047 | 13.85858 | 34.44271 | 14.29055 | -0.84796 | 0.130437 | 0.158176 |
| 3 | 128068200 | 128068799 | 4.78984 | 9.304764 | 26.28849 | 9.817378 | -0.8904 | -0.27485 | 0.158188 |
| 14 | 103946600 | 103947399 | 4.78984 | 9.304764 | 26.28849 | 9.817378 | -0.8904 | -0.27485 | 0.158188 |
| 2 | 163444000 | 163444999 | 4.73637 | 9.074191 | 29.48178 | 11.62307 | -1.16485 | -0.08333 | 0.158201 |
| 5 | 39451600 | 39453199 | 15.56672 | 29.37273 | 18.45261 | 7.526815 | 1.140769 | 0.099593 | 0.158203 |
| 16 | 94054600 | 94058199 | 15.8349 | 26.43051 | 73.42206 | 31.93787 | -1.08762 | 1.199243 | 0.158216 |
| X | 78185800 | 78189999 | 75.84284 | 120.6227 | 92.50835 | 38.62284 | 0.716735 | 2.251841 | 0.158226 |
| 8 | 18697200 | 18699199 | 8.92792 | 16.10858 | 38.52141 | 16.76107 | -0.84205 | 0.357214 | 0.158235 |
| 1 | 193183200 | 193185999 | 10.06986 | 17.89464 | 41.46957 | 18.2966 | -0.81119 | 0.465801 | 0.158239 |
| 6 | 24603600 | 24605599 | 14.7387 | 25.3441 | 56.9614 | 24.98568 | -0.77929 | 0.898545 | 0.158262 |
| 3 | 36213200 | 36217199 | 61.34603 | 98.24369 | 80.52171 | 33.94692 | 0.631716 | 2.018719 | 0.158268 |
| 7 | 84712200 | 84714399 | 27.65827 | 48.01847 | 31.62713 | 13.87949 | 0.986483 | 0.834996 | 0.158277 |
| 5 | 23570200 | 23573599 | 65.92361 | 105.1913 | 87.745 | 36.80086 | 0.6007 | 2.135608 | 0.15828 |
| 7 | 17350600 | 17357199 | 135.4637 | 208.0732 | 150.6066 | 62.53282 | 0.760534 | 2.98612 | 0.158281 |
| 17 | 46018000 | 46019799 | 9.640865 | 17.05378 | 42.50766 | 18.80003 | -0.91892 | 0.504493 | 0.158282 |
| 15 | 38142200 | 38143999 | 28.62053 | 48.95079 | 37.16744 | 16.29731 | 0.778556 | 0.978316 | 0.158283 |
| 10 | 38481000 | 38483799 | 44.23683 | 72.54334 | 50.63265 | 21.79769 | 0.894057 | 1.461093 | 0.158288 |
| 14 | 42124800 | 42126799 | 24.08966 | 42.68954 | 22.79417 | 10.07762 | 1.291772 | 0.537114 | 0.158291 |
| 15 | 21148000 | 21148799 | 21.63897 | 38.27772 | 22.9237 | 10.13853 | 1.126399 | 0.540401 | 0.158292 |
| 5 | 42195800 | 42196999 | 14.83292 | 27.87988 | 19.55459 | 8.030416 | 0.975236 | 0.119403 | 0.158297 |
| 12 | 59789800 | 59791199 | 6.610427 | 12.58628 | 34.08469 | 13.64019 | -0.92618 | -0.00478 | 0.158312 |

| | | | | | | | | | |
|---|---|---|---|---|---|---|---|---|---|
| 18 | 68432600 | 68434199 | 11.77355 | 20.39624 | 54.21242 | 23.78524 | -1.0174 | 0.81507 | 0.158317 |
| 18 | 18607200 | 18611599 | 54.07616 | 86.96217 | 71.86507 | 30.48712 | 0.626932 | 1.878023 | 0.15832 |
| 2 | 53398800 | 53399599 | 4.697331 | 9.106402 | 29.17137 | 11.02775 | -1.08768 | -0.2287 | 0.158321 |
| 6 | 115880800 | 115884399 | 13.77735 | 23.80492 | 54.26237 | 23.80525 | -0.79788 | 0.848274 | 0.158333 |
| 16 | 11195200 | 11197199 | 11.69875 | 20.44356 | 48.35582 | 21.30831 | -0.84803 | 0.692174 | 0.158336 |
| 1 | 99083600 | 99084399 | 4.758969 | 9.246832 | 30.7646 | 11.46851 | -1.12179 | -0.2686 | 0.158364 |
| 2 | 50681200 | 50682399 | 23.9752 | 41.66318 | 31.84095 | 13.98155 | 0.772895 | 0.802865 | 0.158377 |
| 17 | 34415600 | 34456799 | 465.5651 | 622.1773 | 1086.448 | 453.6598 | -0.70171 | 5.288289 | 0.158377 |
| 17 | 8105800 | 8110799 | 24.20438 | 39.6615 | 86.18929 | 37.09283 | -0.7403 | 1.449044 | 0.158387 |
| 13 | 34800200 | 34801599 | 14.89966 | 28.14186 | 18.391 | 7.486017 | 1.086455 | 0.085799 | 0.158387 |
| 19 | 18221000 | 18222999 | 14.89966 | 28.14186 | 18.391 | 7.486017 | 1.086455 | 0.085799 | 0.158387 |
| X | 113483200 | 113484399 | 31.96174 | 54.01488 | 38.96089 | 17.04727 | 0.850905 | 1.099142 | 0.158391 |
| 10 | 63615600 | 63617999 | 21.86706 | 36.16824 | 77.20398 | 33.37815 | -0.71002 | 1.323342 | 0.158396 |
| 9 | 53447800 | 53449199 | 9.066861 | 15.95031 | 44.61823 | 19.75522 | -1.08722 | 0.567935 | 0.158397 |
| 6 | 63053800 | 63057399 | 38.53322 | 63.64821 | 49.38878 | 21.33272 | 0.745278 | 1.360677 | 0.158397 |
| 7 | 11086800 | 11089599 | 63.90824 | 102.3053 | 80.45021 | 33.89604 | 0.69064 | 2.033398 | 0.158414 |
| 2 | 162319400 | 162321599 | 19.31105 | 34.60159 | 24.25671 | 10.62786 | 0.910193 | 0.438728 | 0.158419 |
| 3 | 49935600 | 49937799 | 20.31775 | 36.10179 | 24.54875 | 10.83202 | 0.946877 | 0.495794 | 0.158423 |
| 13 | 97274200 | 97274799 | 2.843055 | 5.541139 | 23.49093 | 8.503893 | -1.42264 | -0.41898 | 0.158426 |
| 10 | 125735600 | 125736799 | 19.0275 | 33.74829 | 25.34077 | 11.19851 | 0.803272 | 0.500043 | 0.158436 |
| 1 | 117361000 | 117363599 | 45.85263 | 74.74936 | 49.99865 | 21.47607 | 0.950264 | 1.545641 | 0.158441 |
| 17 | 23643600 | 23647999 | 45.54748 | 74.62195 | 50.10516 | 21.56196 | 0.949222 | 1.475133 | 0.158446 |
| X | 28477200 | 28477799 | 11.50405 | 22.4309 | 11.61982 | 4.183404 | 1.575197 | -0.36822 | 0.158451 |
| 5 | 32372200 | 32376199 | 21.28484 | 35.07343 | 77.8168 | 33.56906 | -0.76659 | 1.369129 | 0.158454 |
| 10 | 14419200 | 14423599 | 24.71634 | 40.39869 | 90.17616 | 38.77071 | -0.78103 | 1.486438 | 0.158461 |
| 6 | 64166400 | 64167399 | 12.5584 | 24.01417 | 18.29437 | 7.252117 | 0.900586 | -0.01981 | 0.158463 |
| 18 | 88366200 | 88367199 | 11.82216 | 22.94544 | 14.43278 | 5.426575 | 1.243873 | -0.20931 | 0.158467 |
| 2 | 18068800 | 18070799 | 13.2729 | 22.64841 | 59.41154 | 26.04611 | -1.0029 | 0.968411 | 0.158475 |
| 12 | 34016800 | 34018999 | 19.88343 | 32.74014 | 79.32009 | 34.20694 | -0.89351 | 1.374959 | 0.158476 |
| 7 | 17470600 | 17472599 | 54.32955 | 87.33543 | 66.63569 | 28.25262 | 0.741151 | 1.888004 | 0.158478 |
| 9 | 37400000 | 37401599 | 10.28796 | 18.0823 | 45.11249 | 19.97634 | -0.92416 | 0.584327 | 0.158478 |
| 8 | 104939200 | 104955999 | 125.045 | 186.841 | 344.7833 | 142.1504 | -0.62196 | 3.539014 | 0.158507 |
| 13 | 61359600 | 61359999 | 10.23183 | 19.86933 | 15.94361 | 5.971209 | 0.90262 | -0.23554 | 0.158515 |
| 16 | 20927600 | 20928599 | 13.52379 | 25.41314 | 19.7706 | 8.122428 | 0.826537 | 0.109796 | 0.158531 |
| 12 | 49887400 | 49888599 | 7.212986 | 13.23074 | 36.15828 | 15.40649 | -1.01267 | 0.241764 | 0.15855 |
| 10 | 81593000 | 81596999 | 64.69407 | 103.8605 | 71.16087 | 30.11288 | 0.893819 | 1.945223 | 0.158551 |
| 8 | 92409200 | 92419399 | 76.42605 | 117.1977 | 239.7805 | 99.53594 | -0.73891 | 3.005891 | 0.158565 |
| 4 | 146237200 | 146238599 | 27.50685 | 47.84841 | 27.87101 | 12.24607 | 1.162876 | 0.801463 | 0.158565 |
| 7 | 85246600 | 85249599 | 59.46243 | 95.54883 | 72.60361 | 30.76532 | 0.74638 | 1.909565 | 0.158579 |
| 17 | 49846200 | 49850999 | 28.76203 | 46.39135 | 105.8039 | 45.07294 | -0.82916 | 1.764004 | 0.15858 |
| 16 | 53625200 | 53625799 | 4.057433 | 7.918688 | 24.73631 | 8.784906 | -0.96644 | -0.46956 | 0.158603 |
| 17 | 80804000 | 80805199 | 6.324178 | 11.67032 | 36.26308 | 15.32532 | -1.18759 | 0.199882 | 0.158604 |
| 17 | 93667400 | 93668199 | 10.40949 | 20.21289 | 15.93935 | 5.973073 | 0.926832 | -0.23122 | 0.158611 |
| 7 | 129979200 | 129979799 | 3.572972 | 6.979457 | 22.94792 | 8.024496 | -1.01664 | -0.53379 | 0.158625 |
| 4 | 103410200 | 103414599 | 26.18966 | 42.68486 | 91.79396 | 39.42524 | -0.73038 | 1.532087 | 0.158639 |
| 6 | 91746000 | 91755999 | 70.34162 | 109.003 | 205.5111 | 85.36352 | -0.60947 | 2.801944 | 0.158646 |
| 6 | 43766600 | 43769199 | 29.06374 | 49.18196 | 37.97181 | 16.62113 | 0.753057 | 1.07386 | 0.158649 |

| | | | | | | | | | |
|---|---|---|---|---|---|---|---|---|---|
| 7 | 30795600 | 30800599 | 22.77689 | 37.33955 | 84.15165 | 36.22294 | -0.7923 | 1.439147 | 0.15865 |
| 4 | 138149600 | 138150999 | 19.70032 | 34.90851 | 25.96654 | 11.48034 | 0.816409 | 0.515513 | 0.15865 |
| 14 | 10636600 | 10640599 | 18.6925 | 31.41413 | 68.39667 | 29.87621 | -0.73763 | 1.130367 | 0.158651 |
| 2 | 24272400 | 24278599 | 37.28611 | 59.5557 | 129.4923 | 54.38749 | -0.77739 | 2.088123 | 0.158685 |
| 17 | 39264200 | 39266599 | 18.85812 | 33.85087 | 21.82564 | 9.54821 | 1.031452 | 0.419069 | 0.158706 |
| 13 | 36300000 | 36302199 | 13.80833 | 23.76535 | 56.38211 | 24.73225 | -0.85659 | 0.886068 | 0.15871 |
| 6 | 127238400 | 127250999 | 80.69925 | 123.4379 | 252.8732 | 104.917 | -0.74365 | 3.058623 | 0.158714 |
| 12 | 78016600 | 78017199 | 10.1587 | 19.74253 | 15.31851 | 5.701511 | 0.958134 | -0.26071 | 0.158726 |
| 6 | 53709200 | 53710399 | 17.02844 | 30.80911 | 23.10705 | 10.01951 | 0.823165 | 0.364011 | 0.158744 |
| 5 | 104358600 | 104361399 | 17.13263 | 28.79152 | 67.09632 | 29.30785 | -0.8352 | 1.124794 | 0.158745 |
| 11 | 68961000 | 68962399 | 5.903309 | 11.19185 | 30.90075 | 12.47863 | -0.96456 | 0.011574 | 0.158756 |
| 1 | 104291400 | 104296799 | 57.8276 | 92.89059 | 74.5767 | 31.58599 | 0.66657 | 1.91858 | 0.158759 |
| 6 | 44852800 | 44854999 | 24.1944 | 42.53877 | 24.76548 | 10.96642 | 1.163401 | 0.621646 | 0.158761 |
| 18 | 76098800 | 76099599 | 13.1569 | 24.74412 | 19.29217 | 7.914801 | 0.824939 | 0.099809 | 0.158771 |
| 16 | 28697800 | 28721199 | 172.8339 | 252.1835 | 450.9841 | 185.8245 | -0.6089 | 3.943437 | 0.158784 |
| 7 | 14250200 | 14253799 | 72.21593 | 115.3631 | 80.4878 | 33.81165 | 0.859708 | 2.10613 | 0.158786 |
| 2 | 68945200 | 68949199 | 23.50431 | 38.70996 | 86.0367 | 37.1066 | -0.76975 | 1.384139 | 0.158794 |
| 9 | 51531200 | 51532199 | 9.291545 | 16.56322 | 40.48393 | 17.81473 | -0.8846 | 0.439565 | 0.158811 |
| 8 | 5065800 | 5066399 | 9.63575 | 18.82607 | 11.31577 | 3.94405 | 1.404215 | -0.50467 | 0.158816 |
| 4 | 18449200 | 18451799 | 16.36853 | 30.71011 | 18.16287 | 7.481968 | 1.21531 | 0.140249 | 0.158834 |
| 3 | 104771400 | 104772799 | 8.642781 | 15.1941 | 45.64969 | 20.21462 | -1.19011 | 0.573582 | 0.158834 |
| 12 | 8934600 | 8935999 | 5.063091 | 9.588894 | 31.36846 | 12.69552 | -1.20922 | 0.012321 | 0.158835 |
| 12 | 12791200 | 12792799 | 8.466945 | 15.30589 | 41.22616 | 17.89537 | -1.01086 | 0.344763 | 0.158843 |
| 16 | 96970000 | 96973999 | 26.88204 | 43.66206 | 93.05707 | 39.906 | -0.72126 | 1.589126 | 0.158855 |
| 7 | 60417200 | 60420199 | 73.27846 | 116.3817 | 94.18239 | 39.26554 | 0.637395 | 2.289795 | 0.158881 |
| 10 | 50989800 | 50991399 | 31.74523 | 54.41829 | 33.80307 | 14.8248 | 1.067469 | 0.968429 | 0.158886 |
| 16 | 78436200 | 78437399 | 5.549093 | 10.71063 | 31.28557 | 12.07157 | -0.98495 | -0.14347 | 0.158889 |
| 8 | 104217000 | 104218599 | 13.93633 | 25.96235 | 18.60793 | 7.757796 | 0.926673 | 0.162935 | 0.158892 |
| 12 | 80150200 | 80154399 | 37.21052 | 59.68239 | 117.1247 | 49.49292 | -0.6229 | 1.956178 | 0.158902 |
| 7 | 98492800 | 98493799 | 4.835642 | 9.343771 | 27.61065 | 10.5993 | -0.9929 | -0.17644 | 0.158904 |
| 10 | 80238200 | 80244999 | 30.61612 | 49.14348 | 115.6518 | 48.9279 | -0.88293 | 1.922942 | 0.158912 |
| 2 | 168071600 | 168082399 | 96.48091 | 146.6318 | 272.447 | 112.8485 | -0.61042 | 3.19533 | 0.158914 |
| X | 25707000 | 25709799 | 102.4018 | 161.4321 | 97.75573 | 40.56645 | 1.046156 | 2.497295 | 0.158915 |
| 6 | 146121000 | 146121799 | 4.487278 | 8.736818 | 25.36645 | 9.300947 | -0.90422 | -0.34615 | 0.158935 |
| 6 | 89152000 | 89155199 | 21.55988 | 35.65071 | 75.89545 | 32.80643 | -0.70611 | 1.325763 | 0.158947 |
| 11 | 75182400 | 75183999 | 7.578881 | 14.12474 | 34.36092 | 14.31531 | -0.82201 | 0.148059 | 0.15895 |
| 19 | 21132400 | 21133799 | 12.42671 | 23.66481 | 18.21231 | 7.285494 | 0.874201 | 0.006427 | 0.158953 |
| 3 | 41653400 | 41654399 | 5.226029 | 9.887721 | 33.62621 | 13.63771 | -1.26832 | 0.024374 | 0.158954 |
| 15 | 81637200 | 81641799 | 32.34286 | 51.98097 | 110.7289 | 46.93372 | -0.73729 | 1.883716 | 0.158962 |
| 2 | 180474600 | 180475399 | 6.305483 | 12.04504 | 33.05353 | 13.13623 | -0.93629 | -0.03401 | 0.158962 |
| 3 | 100153200 | 100154799 | 9.074337 | 16.36219 | 40.0861 | 17.45597 | -0.87797 | 0.362641 | 0.158972 |
| 19 | 25582000 | 25584799 | 26.27505 | 42.60906 | 94.28423 | 40.38535 | -0.77734 | 1.620938 | 0.158975 |
| 3 | 96308800 | 96319199 | 100.5947 | 152.5716 | 285.0288 | 117.9876 | -0.62069 | 3.238674 | 0.159006 |
| 15 | 82255400 | 82258999 | 16.91031 | 28.20039 | 70.62757 | 30.70505 | -0.93863 | 1.211178 | 0.15901 |
| 5 | 102171400 | 102175599 | 33.25671 | 53.58118 | 107.6402 | 45.78729 | -0.64854 | 1.805337 | 0.159033 |
| 13 | 86194000 | 86194999 | 8.390067 | 14.97772 | 40.50796 | 17.80374 | -1.02836 | 0.423381 | 0.159037 |
| 8 | 37953200 | 37955799 | 34.9643 | 59.1151 | 37.57788 | 16.44433 | 1.032391 | 1.103034 | 0.159037 |

| | | | | | | | | | |
|---|---|---|---|---|---|---|---|---|---|
| 3 | 6526800 | 6527199 | 14.77949 | 27.24062 | 19.14805 | 8.105481 | 0.938721 | 0.231985 | 0.159055 |
| 8 | 21506600 | 21508799 | 39.52318 | 66.02752 | 40.4277 | 17.59544 | 1.086374 | 1.232404 | 0.159065 |
| 4 | 145618600 | 145619199 | 13.44568 | 25.93321 | 16.05175 | 6.207699 | 1.230231 | -0.09419 | 0.159069 |
| 11 | 21521000 | 21522599 | 10.75369 | 18.86872 | 46.19312 | 20.42544 | -0.89694 | 0.618717 | 0.15907 |
| X | 154287400 | 154288999 | 17.8202 | 32.79594 | 20.44954 | 8.674059 | 1.108954 | 0.270338 | 0.159072 |
| 4 | 134430400 | 134436199 | 23.16961 | 37.49429 | 100.1405 | 42.82744 | -1.05081 | 1.661396 | 0.159098 |
| 10 | 3549600 | 3550999 | 18.61513 | 34.21789 | 20.06802 | 8.526116 | 1.194538 | 0.280954 | 0.159103 |
| 11 | 95294200 | 95296599 | 10.11962 | 17.82101 | 44.07919 | 19.51246 | -0.91043 | 0.561301 | 0.159106 |
| 7 | 4868000 | 4869599 | 10.92434 | 19.12748 | 49.92077 | 22.03413 | -0.9894 | 0.65608 | 0.159109 |
| 15 | 54953000 | 54954799 | 8.524332 | 15.26565 | 41.04458 | 17.99152 | -1.01756 | 0.403611 | 0.159112 |
| 9 | 77660000 | 77662599 | 12.47319 | 21.7982 | 48.88892 | 21.54432 | -0.7718 | 0.696152 | 0.159116 |
| 1 | 89723400 | 89726799 | 19.63213 | 32.55306 | 73.45357 | 31.81166 | -0.78931 | 1.28716 | 0.159122 |
| 16 | 31195000 | 31199199 | 50.26556 | 81.1324 | 66.51223 | 28.3603 | 0.644742 | 1.759052 | 0.159129 |
| 7 | 29215000 | 29219599 | 21.4233 | 35.35054 | 78.71648 | 33.97709 | -0.77111 | 1.350793 | 0.159137 |
| 17 | 18526000 | 18527399 | 19.24751 | 35.61798 | 18.50281 | 7.789837 | 1.377526 | 0.247084 | 0.159153 |
| 7 | 21007200 | 21015199 | 123.8649 | 192.4531 | 120.2906 | 49.9467 | 0.984529 | 2.769684 | 0.159155 |
| 9 | 105635400 | 105636199 | 4.824476 | 9.35461 | 27.32697 | 10.32164 | -0.95429 | -0.23383 | 0.159157 |
| 15 | 61065800 | 61068399 | 21.62194 | 38.02819 | 27.92949 | 12.36667 | 0.830695 | 0.610254 | 0.159163 |
| 7 | 16970600 | 16973199 | 62.44368 | 99.85555 | 83.24424 | 35.02094 | 0.604944 | 2.064519 | 0.159166 |
| 6 | 52298800 | 52302599 | 26.71838 | 43.40721 | 93.72059 | 40.19803 | -0.73958 | 1.58375 | 0.159171 |
| 7 | 7739200 | 7739799 | 17.12468 | 30.7992 | 23.12256 | 10.09899 | 0.815006 | 0.398104 | 0.159178 |
| 2 | 121696600 | 121698399 | 9.548702 | 16.77635 | 46.68661 | 20.66648 | -1.08089 | 0.588336 | 0.159201 |
| 7 | 9145000 | 9158199 | 204.6108 | 304.7586 | 236.2468 | 97.34885 | 0.624337 | 3.604677 | 0.159204 |
| 9 | 39202200 | 39204199 | 24.97361 | 43.44415 | 32.27688 | 14.18241 | 0.813721 | 0.795113 | 0.159215 |
| 17 | 18935800 | 18937599 | 42.35328 | 69.76423 | 48.91224 | 21.09672 | 0.890085 | 1.403047 | 0.159221 |
| 7 | 109558600 | 109559999 | 7.141519 | 12.92453 | 38.58381 | 16.72465 | -1.15586 | 0.325605 | 0.159223 |
| 12 | 107459000 | 107469799 | 66.95674 | 103.7455 | 214.3044 | 89.01609 | -0.74123 | 2.802359 | 0.159226 |
| 7 | 41594800 | 41597799 | 55.87914 | 89.99728 | 69.09428 | 29.37583 | 0.735564 | 1.832786 | 0.159227 |
| 19 | 39195200 | 39201199 | 78.9062 | 125.4845 | 91.65157 | 38.26078 | 0.786916 | 2.256251 | 0.159239 |
| 7 | 11549600 | 11558199 | 238.1754 | 349.9216 | 277.0903 | 114.1176 | 0.575272 | 3.839125 | 0.159241 |
| 9 | 57188200 | 57190799 | 9.8695 | 17.27369 | 47.79805 | 21.09011 | -1.0727 | 0.656457 | 0.159251 |
| 12 | 100958400 | 100966999 | 49.02595 | 77.4255 | 166.1007 | 68.97338 | -0.77411 | 2.431779 | 0.159252 |
| 11 | 94535400 | 94539199 | 18.12857 | 30.30008 | 70.96966 | 30.89758 | -0.84257 | 1.188987 | 0.159254 |
| 13 | 46642400 | 46643199 | 11.45496 | 22.17891 | 16.28399 | 6.192344 | 1.008819 | -0.17032 | 0.159287 |
| 7 | 18014800 | 18018799 | 114.623 | 177.0389 | 146.2785 | 60.76101 | 0.575809 | 2.872208 | 0.159302 |
| X | 131330600 | 131332999 | 34.78463 | 59.02159 | 35.99917 | 15.77053 | 1.091979 | 1.072372 | 0.159315 |
| 4 | 55349400 | 55350799 | 11.75378 | 20.38404 | 52.5983 | 23.07782 | -0.97428 | 0.804999 | 0.159349 |
| 13 | 68142800 | 68143399 | 21.2359 | 37.21979 | 27.83554 | 12.29665 | 0.804593 | 0.659586 | 0.159378 |
| 5 | 112911800 | 112913999 | 12.4932 | 21.74821 | 50.50928 | 22.20333 | -0.82255 | 0.754384 | 0.159384 |
| 17 | 29603200 | 29604999 | 6.98142 | 13.0278 | 33.26323 | 13.82821 | -0.88815 | 0.133714 | 0.159397 |
| 12 | 18907000 | 18914399 | 208.0343 | 308.5968 | 245.8035 | 101.2448 | 0.579803 | 3.676958 | 0.159399 |
| 1 | 76106800 | 76107399 | 4.358518 | 8.487191 | 24.75912 | 9.068639 | -0.90952 | -0.35395 | 0.159401 |
| 14 | 57661200 | 57661799 | 4.358518 | 8.487191 | 24.75912 | 9.068639 | -0.90952 | -0.35395 | 0.159401 |
| 16 | 89880800 | 89896599 | 104.6577 | 157.2129 | 324.637 | 134.0009 | -0.77711 | 3.433841 | 0.15942 |
| 7 | 59106200 | 59106999 | 23.42958 | 40.72431 | 31.99671 | 14.05197 | 0.733297 | 0.798386 | 0.159431 |
| 1 | 14312000 | 14313799 | 13.16018 | 25.02787 | 18.69839 | 7.501276 | 0.913315 | 0.024563 | 0.159435 |
| 14 | 124103400 | 124104199 | 13.16018 | 25.02787 | 18.69839 | 7.501276 | 0.913315 | 0.024563 | 0.159435 |

| | | | | | | | | | |
|---|---|---|---|---|---|---|---|---|---|
| 7 | 133833600 | 133835399 | 9.924 | 17.61047 | 41.46152 | 18.31492 | -0.83504 | 0.473769 | 0.159443 |
| 7 | 69657600 | 69658999 | 15.23915 | 28.97159 | 17.42506 | 6.995571 | 1.222727 | 0.044057 | 0.159447 |
| 5 | 107304800 | 107306599 | 8.741229 | 15.88024 | 37.58253 | 16.21293 | -0.81913 | 0.312936 | 0.159456 |
| 13 | 54367600 | 54370399 | 16.58448 | 28.10073 | 62.23277 | 27.25139 | -0.76129 | 1.046814 | 0.159459 |
| 3 | 14927800 | 14929799 | 9.423035 | 16.8577 | 39.97985 | 17.54003 | -0.83854 | 0.415559 | 0.159469 |
| 15 | 84806800 | 84808399 | 8.149553 | 14.44549 | 43.04248 | 19.02874 | -1.17331 | 0.472583 | 0.159489 |
| 6 | 121086200 | 121087599 | 5.369847 | 10.11816 | 32.34934 | 13.22921 | -1.19 | 0.052772 | 0.159501 |
| 12 | 112062600 | 112063599 | 5.369847 | 10.11816 | 32.34934 | 13.22921 | -1.19 | 0.052772 | 0.159501 |
| 5 | 141413600 | 141414999 | 7.166105 | 13.27129 | 35.28433 | 14.83941 | -0.95968 | 0.183831 | 0.159541 |
| 4 | 151415200 | 151417799 | 17.43993 | 28.96368 | 75.64396 | 32.79596 | -0.99932 | 1.262387 | 0.159542 |
| 11 | 4008200 | 4009199 | 5.093765 | 9.862986 | 27.20487 | 10.34273 | -0.88102 | -0.20703 | 0.159557 |
| 18 | 88858600 | 88859399 | 5.093765 | 9.862986 | 27.20487 | 10.34273 | -0.88102 | -0.20703 | 0.159557 |
| 5 | 33535000 | 33536399 | 5.535664 | 10.33229 | 33.22274 | 13.80602 | -1.21671 | 0.119682 | 0.159558 |
| 4 | 123004200 | 123017199 | 83.81397 | 126.9694 | 302.8834 | 125.3325 | -0.97446 | 3.260623 | 0.159562 |
| 17 | 39878400 | 39881399 | 47.94523 | 78.06221 | 55.1129 | 23.66193 | 0.871276 | 1.56958 | 0.15957 |
| 10 | 77115600 | 77116999 | 5.998672 | 11.15959 | 34.79892 | 14.54208 | -1.18035 | 0.147017 | 0.159572 |
| 1 | 185749000 | 185750399 | 6.09421 | 11.44206 | 32.85256 | 13.51658 | -1.04394 | 0.083342 | 0.159574 |
| 12 | 75781600 | 75782799 | 6.09421 | 11.44206 | 32.85256 | 13.51658 | -1.04394 | 0.083342 | 0.159574 |
| 2 | 165351400 | 165353999 | 12.79743 | 22.24951 | 54.8625 | 24.09769 | -0.90916 | 0.777117 | 0.159575 |
| 16 | 38218800 | 38222199 | 16.60717 | 28.06495 | 63.84941 | 27.93692 | -0.8002 | 1.071125 | 0.159577 |
| 7 | 31505800 | 31508599 | 23.69318 | 41.87237 | 23.74039 | 10.50169 | 1.20412 | 0.563528 | 0.159586 |
| 12 | 22526600 | 22527799 | 24.05548 | 42.19723 | 27.72632 | 12.25714 | 0.990207 | 0.661632 | 0.159602 |
| 7 | 103628400 | 103629399 | 14.21355 | 26.55766 | 20.25093 | 8.400366 | 0.844339 | 0.152896 | 0.159623 |
| 11 | 44120800 | 44125399 | 31.28525 | 50.54037 | 103.4697 | 44.16783 | -0.67119 | 1.721296 | 0.159637 |
| 1 | 147208000 | 147210999 | 48.10417 | 78.13529 | 57.86315 | 24.81461 | 0.799971 | 1.608584 | 0.159641 |
| 5 | 110987200 | 110989599 | 13.90307 | 24.11163 | 53.2316 | 23.35569 | -0.75083 | 0.813564 | 0.159647 |
| 3 | 45685400 | 45686399 | 19.55135 | 34.51786 | 26.50701 | 11.72788 | 0.768623 | 0.553445 | 0.15965 |
| 19 | 17084000 | 17084999 | 5.748504 | 10.94933 | 31.07126 | 12.42428 | -0.99141 | -0.02342 | 0.159665 |
| 19 | 44391400 | 44396999 | 24.64244 | 40.06823 | 89.70281 | 38.49389 | -0.79052 | 1.563593 | 0.15967 |
| 2 | 82712400 | 82715599 | 36.02502 | 60.43226 | 40.36763 | 17.61508 | 0.960957 | 1.176839 | 0.159682 |
| 7 | 4795800 | 4797799 | 20.42473 | 33.37053 | 85.21692 | 36.63309 | -0.97454 | 1.484582 | 0.15969 |
| X | 126856600 | 126859599 | 87.38075 | 138.011 | 103.0557 | 42.79592 | 0.746387 | 2.446253 | 0.159711 |
| 19 | 3329600 | 3333399 | 18.24241 | 30.23314 | 76.77845 | 33.23982 | -0.95938 | 1.289966 | 0.159733 |
| 12 | 20225200 | 20227399 | 44.24173 | 71.89397 | 58.07154 | 24.91226 | 0.675643 | 1.5934 | 0.159745 |
| 6 | 85454400 | 85462599 | 51.57406 | 80.77364 | 185.1977 | 76.82072 | -0.8793 | 2.604441 | 0.159746 |
| 6 | 54986800 | 54988999 | 7.279244 | 13.21642 | 40.23993 | 17.37217 | -1.1808 | 0.309018 | 0.159747 |
| 6 | 10798200 | 10800199 | 33.53855 | 56.15471 | 43.20748 | 18.83178 | 0.7578 | 1.190764 | 0.159757 |
| 1 | 59520600 | 59525199 | 27.3665 | 44.07086 | 108.0815 | 45.94417 | -0.93634 | 1.812024 | 0.159768 |
| 19 | 56941400 | 56944799 | 16.09435 | 26.95308 | 69.64509 | 30.35859 | -0.98355 | 1.160842 | 0.159799 |
| X | 115901600 | 115902199 | 18.77992 | 33.69237 | 23.10561 | 10.11275 | 0.943029 | 0.426388 | 0.159807 |
| X | 20872400 | 20873599 | 15.28668 | 29.53451 | 13.72454 | 5.271552 | 1.648781 | -0.10239 | 0.159817 |
| 2 | 9598200 | 9601399 | 16.69547 | 28.40093 | 62.36502 | 27.33414 | -0.74871 | 1.010876 | 0.15982 |
| 7 | 27178800 | 27181999 | 12.93164 | 22.62731 | 50.4886 | 22.27249 | -0.76463 | 0.679809 | 0.159832 |
| 2 | 177743400 | 177745399 | 10.18059 | 18.20708 | 42.59955 | 18.69444 | -0.81987 | 0.426202 | 0.159834 |
| 13 | 51553400 | 51556799 | 16.84804 | 28.25587 | 69.25546 | 30.21614 | -0.90776 | 1.14665 | 0.159841 |
| 11 | 19417000 | 19418199 | 20.92518 | 36.90671 | 26.19376 | 11.59158 | 0.880992 | 0.571025 | 0.159841 |
| 16 | 85597000 | 85601599 | 27.0439 | 43.55984 | 107.4658 | 45.6954 | -0.94453 | 1.805219 | 0.159862 |

| | | | | | | | | | |
|---|---|---|---|---|---|---|---|---|---|
| 7 | 45677200 | 45679999 | 15.03833 | 25.46971 | 62.64996 | 27.43041 | -0.91203 | 1.045739 | 0.159876 |
| 14 | 20295400 | 20298599 | 16.17838 | 27.25222 | 66.51887 | 29.07645 | -0.90153 | 1.100126 | 0.159879 |
| 6 | 56774000 | 56775399 | 9.814379 | 17.30572 | 44.06715 | 19.50212 | -0.95156 | 0.543102 | 0.159879 |
| 7 | 126733800 | 126737799 | 18.25472 | 30.13793 | 76.82986 | 33.17426 | -0.96514 | 1.338457 | 0.159891 |
| 13 | 113130000 | 113137399 | 50.1422 | 79.23099 | 160.9201 | 66.84029 | -0.69462 | 2.419264 | 0.1599 |
| 3 | 103479000 | 103486999 | 61.0547 | 95.41669 | 186.1868 | 77.2481 | -0.64943 | 2.647725 | 0.159912 |
| 15 | 9026800 | 9027799 | 10.33136 | 19.98491 | 13.8503 | 5.28973 | 1.085134 | -0.17785 | 0.159913 |
| 3 | 90895200 | 90896199 | 26.07566 | 45.30393 | 32.33958 | 14.19866 | 0.871095 | 0.81908 | 0.159926 |
| 2 | 84695200 | 84703799 | 56.77367 | 88.87914 | 192.0446 | 79.66436 | -0.7944 | 2.614701 | 0.159952 |
| 10 | 70275200 | 70277399 | 10.64658 | 18.57855 | 49.40659 | 21.75168 | -1.0162 | 0.708991 | 0.159964 |
| 5 | 92701800 | 92704599 | 16.67318 | 28.45746 | 61.32683 | 26.8863 | -0.72095 | 0.979475 | 0.159967 |
| 5 | 107112000 | 107112999 | 7.192599 | 13.22691 | 35.13507 | 14.91884 | -0.96798 | 0.226278 | 0.159967 |
| X | 139920000 | 139928199 | 37.38176 | 59.97129 | 118.4477 | 50.07007 | -0.63172 | 1.949977 | 0.15997 |
| X | 64321800 | 64324399 | 23.12337 | 42.26028 | 16.34488 | 6.998577 | 1.782124 | 0.339262 | 0.159973 |
| 9 | 19224600 | 19226799 | 32.5204 | 54.61054 | 42.66761 | 18.63105 | 0.735753 | 1.15785 | 0.159977 |
| 17 | 86106400 | 86108999 | 16.77199 | 27.93623 | 71.6797 | 31.13891 | -0.97341 | 1.224632 | 0.159978 |
| 12 | 114707400 | 114708999 | 36.2909 | 60.53661 | 44.33301 | 19.27687 | 0.829059 | 1.240481 | 0.159985 |
| 3 | 129581800 | 129583399 | 7.107285 | 12.94784 | 37.63222 | 16.17793 | -1.10975 | 0.283874 | 0.160009 |
| 4 | 124147800 | 124149999 | 10.12743 | 17.83319 | 49.88765 | 22.08407 | -1.08789 | 0.567133 | 0.16001 |
| X | 136726600 | 136729399 | 63.40139 | 101.4884 | 80.54902 | 33.93503 | 0.677268 | 2.036097 | 0.160012 |
| 17 | 24703800 | 24710599 | 32.52668 | 52.21772 | 115.2141 | 48.75342 | -0.78993 | 1.920216 | 0.160029 |
| 12 | 23028400 | 23031399 | 95.14024 | 148.9765 | 124.0874 | 51.47299 | 0.579415 | 2.622032 | 0.160046 |
| 3 | 104865800 | 104875599 | 57.99106 | 90.56949 | 188.7412 | 78.31318 | -0.7449 | 2.658163 | 0.160052 |
| 7 | 24269200 | 24278999 | 62.64113 | 97.56962 | 193.3809 | 80.26525 | -0.67567 | 2.708267 | 0.16006 |
| X | 24642600 | 24644799 | 28.13411 | 49.39685 | 23.22172 | 10.27368 | 1.470431 | 0.664751 | 0.160062 |
| 6 | 54195200 | 54198399 | 17.42017 | 29.36633 | 64.36221 | 28.14024 | -0.74669 | 1.097404 | 0.160064 |
| 5 | 9749800 | 9750599 | 16.11424 | 29.72789 | 21.4631 | 9.074176 | 0.90191 | 0.24447 | 0.160093 |
| 4 | 48584400 | 48587999 | 19.7479 | 32.84753 | 74.395 | 32.28898 | -0.79445 | 1.249221 | 0.160095 |
| 15 | 22677000 | 22678399 | 15.87024 | 29.88066 | 17.01245 | 6.966251 | 1.275912 | 0.110592 | 0.160107 |
| 2 | 60547800 | 60549799 | 10.01395 | 17.97878 | 42.13768 | 18.43128 | -0.81895 | 0.400139 | 0.160108 |
| 13 | 118523000 | 118525199 | 12.15782 | 21.21915 | 51.58177 | 22.71165 | -0.88788 | 0.714753 | 0.160109 |
| 9 | 50627400 | 50632799 | 23.83135 | 38.77373 | 88.31035 | 37.90502 | -0.81453 | 1.55329 | 0.160133 |
| 16 | 31265000 | 31272199 | 33.11858 | 52.93899 | 129.8977 | 54.61606 | -0.95019 | 2.060442 | 0.160134 |
| 4 | 109837000 | 109846599 | 81.4144 | 124.4345 | 257.6019 | 106.8596 | -0.75966 | 3.0749 | 0.160137 |
| 19 | 57481000 | 57486599 | 37.52555 | 60.22731 | 117.6834 | 49.7788 | -0.61555 | 1.936131 | 0.160137 |
| 7 | 8571400 | 8572999 | 16.3017 | 30.06395 | 21.4578 | 9.075945 | 0.917957 | 0.247903 | 0.160139 |
| 1 | 111502200 | 111504399 | 25.41446 | 44.91278 | 21.97907 | 9.722629 | 1.415005 | 0.571807 | 0.160145 |
| 15 | 60110400 | 60110799 | 2.935697 | 5.742211 | 19.70531 | 6.614655 | -1.0221 | -0.70201 | 0.16015 |
| 10 | 81757000 | 81758399 | 17.82794 | 32.13746 | 23.93132 | 10.42467 | 0.829139 | 0.391851 | 0.160155 |
| X | 42819800 | 42823999 | 79.86081 | 126.9493 | 91.22011 | 38.06458 | 0.809862 | 2.270144 | 0.160159 |
| 4 | 153442600 | 153449399 | 40.02354 | 63.89478 | 132.1103 | 55.4461 | -0.70584 | 2.107324 | 0.160169 |
| X | 15599200 | 15600799 | 16.94573 | 32.52152 | 13.27145 | 5.206391 | 1.805124 | -0.02152 | 0.160194 |
| 5 | 146846600 | 146851599 | 19.29855 | 31.73596 | 86.24704 | 37.17384 | -1.05962 | 1.392581 | 0.160198 |
| 7 | 124091200 | 124094399 | 17.56857 | 29.61448 | 64.97784 | 28.40881 | -0.74838 | 1.097785 | 0.160198 |
| 12 | 76688600 | 76703799 | 132.4583 | 196.6813 | 371.4111 | 152.997 | -0.66352 | 3.653627 | 0.160214 |
| 13 | 31796200 | 31805799 | 77.604 | 119.0078 | 243.6928 | 101.1604 | -0.74012 | 3.005509 | 0.160215 |
| 18 | 69571600 | 69572999 | 7.586161 | 14.09737 | 36.52132 | 15.28829 | -0.91808 | 0.169606 | 0.160227 |

| | | | | | | | | | |
|---|---|---|---|---|---|---|---|---|---|
| 12 | 17815200 | 17818599 | 20.23583 | 33.61256 | 74.95237 | 32.49981 | -0.77226 | 1.267278 | 0.160237 |
| 14 | 14537400 | 14538999 | 7.503058 | 13.75762 | 35.6203 | 15.18507 | -0.9357 | 0.246089 | 0.160252 |
| 13 | 76444800 | 76445999 | 16.00382 | 29.96891 | 18.72644 | 7.738965 | 1.132982 | 0.149853 | 0.160255 |
| 7 | 85020800 | 85025399 | 65.88046 | 105.2757 | 83.88973 | 35.2566 | 0.668844 | 2.09168 | 0.160266 |
| 10 | 81703400 | 81704799 | 13.82477 | 26.65801 | 15.32856 | 5.932081 | 1.333926 | -0.09061 | 0.160277 |
| 9 | 85521000 | 85522199 | 15.05893 | 28.56594 | 17.97619 | 7.250555 | 1.152256 | 0.058319 | 0.160286 |
| 1 | 109805000 | 109808799 | 43.92992 | 72.93185 | 39.89603 | 17.29301 | 1.248251 | 1.310886 | 0.160288 |
| 9 | 77437200 | 77442799 | 26.27268 | 42.60701 | 93.02016 | 39.84514 | -0.7579 | 1.620001 | 0.160293 |
| 5 | 116545400 | 116553999 | 40.09128 | 63.83829 | 143.3918 | 59.96563 | -0.82919 | 2.192414 | 0.160302 |
| 3 | 89986600 | 89988799 | 10.78597 | 18.85681 | 48.86995 | 21.54291 | -0.97911 | 0.67996 | 0.160306 |
| 9 | 121141600 | 121144799 | 10.92964 | 19.12589 | 49.9262 | 22.02595 | -0.98967 | 0.665607 | 0.160307 |
| 4 | 155854000 | 155857799 | 14.57769 | 24.37088 | 71.01976 | 30.92398 | -1.15615 | 1.177147 | 0.160325 |
| 8 | 6207400 | 6208399 | 12.74535 | 24.76582 | 13.21426 | 4.924139 | 1.490168 | -0.23157 | 0.160327 |
| 12 | 84686400 | 84691999 | 36.91955 | 59.0761 | 125.167 | 52.70457 | -0.73726 | 2.030728 | 0.160351 |
| 19 | 41790000 | 41790599 | 3.455668 | 6.750942 | 21.69168 | 7.573076 | -0.98141 | -0.54595 | 0.160354 |
| 8 | 51841400 | 51842399 | 12.14292 | 23.13374 | 18.29223 | 7.311203 | 0.83669 | 0.001432 | 0.160355 |
| 11 | 5491400 | 5492399 | 5.262253 | 10.15085 | 28.12423 | 10.8724 | -0.91094 | -0.14644 | 0.160366 |
| 3 | 36864600 | 36866799 | 22.52795 | 39.35004 | 30.81002 | 13.57266 | 0.738836 | 0.724292 | 0.160388 |
| 19 | 5637400 | 5639599 | 15.04435 | 25.93261 | 55.90469 | 24.52398 | -0.71862 | 0.877527 | 0.160422 |
| 7 | 24244200 | 24247399 | 15.97327 | 27.02936 | 63.83837 | 27.94328 | -0.85387 | 1.056965 | 0.160429 |
| 8 | 120024600 | 120042199 | 165.378 | 241.6797 | 439.3322 | 180.996 | -0.63032 | 3.919039 | 0.160433 |
| 13 | 20289600 | 20293399 | 16.33321 | 27.97154 | 61.95174 | 27.16766 | -0.75953 | 0.947538 | 0.160436 |
| 6 | 145561200 | 145562999 | 18.54582 | 30.69722 | 78.88075 | 34.12022 | -0.97641 | 1.309381 | 0.160461 |
| 14 | 104080400 | 104082999 | 14.25249 | 24.59135 | 54.80412 | 24.04189 | -0.76596 | 0.863776 | 0.160463 |
| 7 | 45085400 | 45088999 | 14.52154 | 24.73292 | 60.56612 | 26.54829 | -0.90478 | 0.990498 | 0.160471 |
| 8 | 35851000 | 35852799 | 13.09821 | 22.39285 | 58.35229 | 25.58565 | -0.99283 | 0.94932 | 0.160474 |
| 13 | 95011400 | 95014399 | 22.39868 | 36.21978 | 101.4565 | 43.36312 | -1.12036 | 1.677598 | 0.160505 |
| 10 | 7440400 | 7444199 | 21.53783 | 35.26638 | 84.29866 | 36.26799 | -0.87758 | 1.451988 | 0.16051 |
| 5 | 96521200 | 96526999 | 33.49073 | 53.74672 | 123.1879 | 52.10005 | -0.84543 | 1.93358 | 0.160511 |
| 14 | 20184400 | 20198399 | 91.61297 | 138.8441 | 285.81 | 118.2837 | -0.76143 | 3.252189 | 0.160537 |
| 15 | 13115600 | 13117599 | 27.02503 | 47.00273 | 30.42564 | 13.36718 | 1.011416 | 0.804698 | 0.160546 |
| 1 | 15916800 | 15917599 | 11.5364 | 22.33506 | 16.35195 | 6.220196 | 1.012877 | -0.16742 | 0.160584 |
| 2 | 52327800 | 52330599 | 13.4015 | 23.38837 | 54.89255 | 24.1685 | -0.83769 | 0.726834 | 0.160594 |
| 7 | 24986400 | 24992999 | 33.31035 | 53.13347 | 133.3742 | 55.91113 | -0.98625 | 2.129018 | 0.1606 |
| 19 | 5093800 | 5111399 | 79.83748 | 120.6711 | 307.3185 | 127.0767 | -1.07153 | 3.305769 | 0.160603 |
| 8 | 82938000 | 82939199 | 12.89787 | 24.79238 | 17.17699 | 6.707888 | 1.056162 | -0.06668 | 0.160606 |
| 18 | 66069400 | 66076799 | 43.94553 | 69.98206 | 141.0311 | 58.98609 | -0.67284 | 2.192491 | 0.160612 |
| 14 | 70767800 | 70772399 | 24.07958 | 39.01159 | 95.83455 | 41.02058 | -0.92892 | 1.637752 | 0.160617 |
| 15 | 41310000 | 41313199 | 22.30833 | 36.56406 | 84.02501 | 36.16534 | -0.82041 | 1.440658 | 0.16062 |
| 14 | 89865800 | 89866399 | 4.652591 | 9.056758 | 27.32137 | 10.03498 | -0.96236 | -0.33477 | 0.160635 |
| 2 | 42867200 | 42868199 | 11.3374 | 22.02292 | 15.21009 | 5.682596 | 1.119664 | -0.23322 | 0.160645 |
| 7 | 20550800 | 20552999 | 24.37826 | 43.17162 | 20.80458 | 9.199243 | 1.437647 | 0.540243 | 0.160656 |
| 6 | 112511800 | 112512599 | 9.937653 | 17.59165 | 42.49005 | 18.78949 | -0.87353 | 0.497535 | 0.160664 |
| 18 | 35904000 | 35905799 | 9.937653 | 17.59165 | 42.49005 | 18.78949 | -0.87353 | 0.497535 | 0.160664 |
| 5 | 140495800 | 140516999 | 204.2225 | 291.9039 | 548.4475 | 226.6015 | -0.70946 | 4.252666 | 0.160664 |
| 3 | 89958800 | 89960599 | 8.302199 | 14.68613 | 43.05588 | 19.04241 | -1.15117 | 0.497172 | 0.160669 |
| 13 | 9325800 | 9327799 | 12.05423 | 20.93195 | 53.72101 | 23.5783 | -0.96656 | 0.790728 | 0.16067 |

| | | | | | | | | | |
|---|---|---|---|---|---|---|---|---|---|
| 7 | 49673600 | 49674999 | 18.54517 | 33.92117 | 21.2387 | 9.084395 | 1.094288 | 0.310323 | 0.16068 |
| 14 | 83761200 | 83765199 | 50.21821 | 81.3246 | 61.10122 | 26.14532 | 0.775281 | 1.675078 | 0.160683 |
| 5 | 40936200 | 40936799 | 8.554358 | 16.70932 | 12.72334 | 4.452782 | 1.063465 | -0.5019 | 0.160702 |
| 14 | 102891600 | 102892199 | 8.554358 | 16.70932 | 12.72334 | 4.452782 | 1.063465 | -0.5019 | 0.160702 |
| 7 | 60417000 | 60420199 | 73.7808 | 117.1435 | 95.32867 | 39.73122 | 0.628944 | 2.299395 | 0.160714 |
| 9 | 37371000 | 37372199 | 7.462426 | 13.44002 | 39.55534 | 17.24999 | -1.14143 | 0.358311 | 0.160754 |
| 18 | 28666400 | 28668399 | 19.29846 | 34.11984 | 25.40963 | 11.23916 | 0.813248 | 0.533728 | 0.160773 |
| 4 | 100632600 | 100635599 | 14.5531 | 24.41417 | 72.37619 | 31.58354 | -1.18098 | 1.138759 | 0.160787 |
| 11 | 113294800 | 113303799 | 61.90344 | 96.65336 | 192.899 | 80.0406 | -0.68297 | 2.664792 | 0.160788 |
| 4 | 33030200 | 33031999 | 9.517714 | 16.77088 | 44.0795 | 19.51031 | -0.99724 | 0.549795 | 0.160788 |
| 14 | 32817000 | 32818999 | 11.33476 | 19.95061 | 47.20326 | 20.89755 | -0.84779 | 0.575665 | 0.160792 |
| 18 | 57016200 | 57017599 | 8.925144 | 16.12922 | 39.07703 | 16.969 | -0.85877 | 0.349851 | 0.160795 |
| 4 | 10096400 | 10098199 | 20.73437 | 37.23776 | 22.39211 | 9.791825 | 1.131761 | 0.430411 | 0.160797 |
| 7 | 38324200 | 38327599 | 46.50691 | 75.57157 | 53.32463 | 22.87512 | 0.869977 | 1.59383 | 0.160804 |
| 13 | 68729800 | 68735199 | 25.35953 | 41.0595 | 97.87354 | 41.87399 | -0.88655 | 1.653149 | 0.160809 |
| 5 | 42193400 | 42194799 | 13.61227 | 25.47939 | 18.58166 | 7.684777 | 0.910904 | 0.130218 | 0.160811 |
| 8 | 69805600 | 69807799 | 37.96442 | 62.73391 | 49.3898 | 21.33909 | 0.724443 | 1.353687 | 0.160818 |
| 16 | 13546400 | 13548599 | 14.88833 | 25.62253 | 56.43364 | 24.7548 | -0.74996 | 0.891218 | 0.160828 |
| 2 | 16085200 | 16087599 | 24.15081 | 42.17349 | 30.57791 | 13.46816 | 0.849096 | 0.735743 | 0.160836 |
| 7 | 115714000 | 115715999 | 8.635192 | 15.55428 | 39.53939 | 17.24016 | -0.93188 | 0.364984 | 0.160837 |
| 8 | 17846400 | 17846999 | 9.580131 | 18.67772 | 13.69853 | 4.938268 | 1.079278 | -0.38304 | 0.160845 |
| 2 | 76455200 | 76457999 | 72.95666 | 116.0666 | 94.00281 | 39.26276 | 0.638616 | 2.237982 | 0.160847 |
| X | 147441400 | 147448599 | 69.24979 | 110.8932 | 75.08259 | 31.65032 | 0.906702 | 2.030062 | 0.160864 |
| 6 | 66473600 | 66474199 | 9.940713 | 19.33351 | 15.43561 | 5.710113 | 0.925942 | -0.28284 | 0.160878 |
| 18 | 18960800 | 18965399 | 72.87686 | 115.7226 | 97.36033 | 40.58299 | 0.581152 | 2.294671 | 0.16089 |
| 17 | 26643000 | 26655399 | 67.20686 | 103.518 | 226.3961 | 94.03703 | -0.83006 | 2.914551 | 0.160894 |
| 11 | 47777400 | 47779399 | 8.212619 | 14.74529 | 40.03113 | 17.50929 | -1.02904 | 0.383567 | 0.160913 |
| 10 | 51307400 | 51308399 | 15.44738 | 28.27503 | 21.66488 | 9.257748 | 0.805876 | 0.282668 | 0.160915 |
| 17 | 8760600 | 8768999 | 51.14515 | 80.71782 | 167.7063 | 69.61632 | -0.72878 | 2.449494 | 0.160925 |
| 10 | 20247600 | 20248199 | 12.46397 | 23.88186 | 17.68834 | 6.971276 | 0.948222 | -0.04158 | 0.160964 |
| 8 | 77947400 | 77948999 | 10.73073 | 18.88902 | 44.57635 | 19.73425 | -0.84367 | 0.568478 | 0.160968 |
| 16 | 20359200 | 20362999 | 25.00595 | 40.54566 | 93.2392 | 39.93383 | -0.83282 | 1.62168 | 0.160976 |
| 12 | 81800000 | 81802199 | 12.13104 | 21.28228 | 47.23384 | 20.88298 | -0.75676 | 0.627618 | 0.160989 |
| 9 | 51944600 | 51945599 | 11.83747 | 22.91626 | 15.58413 | 5.930169 | 1.116804 | -0.16661 | 0.16099 |
| 11 | 83642800 | 83645199 | 12.60199 | 22.08359 | 49.39169 | 21.81622 | -0.768 | 0.651411 | 0.160998 |
| X | 24278200 | 24280599 | 28.63265 | 50.16021 | 24.39567 | 10.77274 | 1.422012 | 0.703661 | 0.161007 |
| 2 | 26800000 | 26803599 | 31.75406 | 50.91425 | 117.6712 | 49.69941 | -0.8588 | 1.959866 | 0.161022 |
| 18 | 31580000 | 31580599 | 2.836199 | 5.5462 | 20.74319 | 7.068176 | -1.16476 | -0.64788 | 0.161033 |
| 2 | 72348200 | 72349999 | 8.23056 | 15.13702 | 36.23074 | 15.38223 | -0.81941 | 0.234605 | 0.161037 |
| 13 | 105037000 | 105037999 | 8.23056 | 15.13702 | 36.23074 | 15.38223 | -0.81941 | 0.234605 | 0.161037 |
| 18 | 21028800 | 21029999 | 8.108272 | 14.42774 | 41.01816 | 18.0774 | -1.10247 | 0.444133 | 0.161045 |
| 1 | 180263600 | 180265999 | 9.61772 | 17.07595 | 41.46801 | 18.30966 | -0.87904 | 0.467745 | 0.161046 |
| 4 | 43566000 | 43571399 | 24.75409 | 40.03412 | 100.4056 | 42.92161 | -0.96121 | 1.676459 | 0.16106 |
| 4 | 62431000 | 62432599 | 5.956351 | 11.13607 | 32.73317 | 13.56419 | -1.08533 | 0.111108 | 0.16107 |
| 7 | 30238000 | 30241599 | 10.46175 | 18.32783 | 46.7427 | 20.64118 | -0.95576 | 0.642322 | 0.161075 |
| 6 | 3057200 | 3060399 | 84.44698 | 133.3488 | 108.9164 | 45.22476 | 0.616946 | 2.449089 | 0.161081 |
| 8 | 71845000 | 71851399 | 28.81469 | 46.354 | 109.6828 | 46.55356 | -0.88676 | 1.848533 | 0.16109 |

| | | | | | | | | | |
|---|---|---|---|---|---|---|---|---|---|
| 6 | 60469000 | 60471799 | 36.92589 | 61.53939 | 44.3381 | 19.26793 | 0.852537 | 1.251671 | 0.161104 |
| 17 | 66236000 | 66237399 | 6.06003 | 11.4346 | 31.31012 | 12.77145 | -0.96537 | 0.048396 | 0.16111 |
| 7 | 13830200 | 13841799 | 23.45515 | 41.0901 | 23.78223 | 10.50188 | 1.172349 | 0.672316 | 0.161119 |
| 7 | 18034000 | 18041199 | 171.8018 | 258.5744 | 209.1598 | 86.37995 | 0.576476 | 3.403222 | 0.161137 |
| 6 | 42325600 | 42329399 | 22.80054 | 37.00367 | 96.09642 | 41.18543 | -1.0065 | 1.599126 | 0.161145 |
| 2 | 32772400 | 32777199 | 19.54154 | 32.21291 | 77.53042 | 33.45085 | -0.88328 | 1.36035 | 0.161151 |
| 7 | 131162000 | 131162999 | 4.819111 | 9.359806 | 28.8352 | 10.78536 | -1.01742 | -0.26504 | 0.161155 |
| 7 | 7972600 | 7979199 | 84.6134 | 133.9967 | 101.3853 | 42.15902 | 0.730428 | 2.380993 | 0.161159 |
| 7 | 140617200 | 140622999 | 29.63744 | 48.03871 | 97.9234 | 41.92935 | -0.66001 | 1.638894 | 0.161162 |
| 6 | 118395400 | 118397199 | 5.396873 | 10.09763 | 33.27286 | 13.77034 | -1.24707 | 0.101856 | 0.16118 |
| 4 | 43757000 | 43758399 | 7.569581 | 14.13762 | 34.43475 | 14.29419 | -0.81962 | 0.134023 | 0.161187 |
| 9 | 48181000 | 48184599 | 18.47575 | 30.79331 | 77.27183 | 33.58144 | -0.94182 | 1.224488 | 0.161207 |
| 4 | 43837600 | 43838599 | 5.566037 | 10.69165 | 29.1201 | 11.39455 | -0.9034 | -0.10115 | 0.161211 |
| 19 | 6386600 | 6399999 | 62.34579 | 96.28367 | 219.0912 | 91.00593 | -0.88423 | 2.863015 | 0.161232 |
| 5 | 31101800 | 31102999 | 5.147523 | 9.875526 | 29.01346 | 11.39318 | -1.01604 | -0.09364 | 0.161236 |
| 14 | 20595800 | 20596799 | 5.147523 | 9.875526 | 29.01346 | 11.39318 | -1.01604 | -0.09364 | 0.161236 |
| 17 | 6272600 | 6273199 | 5.147523 | 9.875526 | 29.01346 | 11.39318 | -1.01604 | -0.09364 | 0.161236 |
| X | 159116600 | 159118999 | 24.46456 | 44.11406 | 17.82029 | 7.759762 | 1.705179 | 0.425393 | 0.161252 |
| 18 | 74979800 | 74982399 | 13.87304 | 23.71941 | 60.08046 | 26.34357 | -0.95243 | 0.951092 | 0.161258 |
| 7 | 41428000 | 41431599 | 69.75742 | 111.3126 | 86.73545 | 36.37922 | 0.698882 | 2.138631 | 0.161274 |
| X | 14319400 | 14321399 | 27.62812 | 48.69816 | 21.47852 | 9.506142 | 1.562938 | 0.61262 | 0.161305 |
| 4 | 37910800 | 37913799 | 34.88706 | 58.47654 | 42.06252 | 18.34541 | 0.854613 | 1.183064 | 0.161327 |
| 16 | 4752600 | 4755999 | 15.93364 | 27.08469 | 60.59484 | 26.55667 | -0.77549 | 1.0148 | 0.161341 |
| 5 | 92424600 | 92427399 | 17.58893 | 29.61986 | 64.89226 | 28.36299 | -0.74619 | 1.107937 | 0.161348 |
| 13 | 111649000 | 111651799 | 28.30852 | 48.06405 | 36.85075 | 16.14675 | 0.763138 | 1.040441 | 0.161361 |
| 7 | 62242800 | 62244599 | 43.601 | 70.98983 | 56.43458 | 24.22946 | 0.7006 | 1.562666 | 0.161366 |
| 5 | 138809600 | 138811999 | 11.4043 | 19.90308 | 50.48601 | 22.22852 | -0.94875 | 0.710887 | 0.161386 |
| 5 | 147839400 | 147840599 | 6.480914 | 12.28774 | 34.46771 | 13.91709 | -0.98822 | 0.022531 | 0.161394 |
| 15 | 71108200 | 71110999 | 30.46262 | 51.98289 | 37.0979 | 16.26384 | 0.867152 | 1.005943 | 0.161404 |
| 11 | 3114200 | 3124599 | 771.6195 | 1031.554 | 628.4065 | 262.386 | 0.817774 | 5.286059 | 0.161409 |
| 2 | 133732800 | 133735399 | 33.39127 | 56.47247 | 38.69127 | 16.93309 | 0.924693 | 1.095237 | 0.161412 |
| 12 | 23859200 | 23865799 | 145.8672 | 221.6645 | 183.3053 | 75.92316 | 0.556397 | 3.203885 | 0.161415 |
| X | 32431800 | 32434599 | 28.74011 | 50.07419 | 28.19926 | 12.40283 | 1.211625 | 0.787401 | 0.161418 |
| 8 | 22965600 | 22967599 | 9.005127 | 16.02967 | 41.50051 | 18.28414 | -0.96856 | 0.44677 | 0.161434 |
| 14 | 41941400 | 41942599 | 13.4348 | 25.95367 | 14.75995 | 5.672236 | 1.358563 | -0.11902 | 0.161435 |
| 2 | 88475000 | 88476199 | 15.60443 | 29.80045 | 16.30582 | 6.48394 | 1.369722 | 0.01499 | 0.161442 |
| 7 | 30261400 | 30264799 | 13.19315 | 22.40304 | 65.34364 | 28.63247 | -1.15661 | 1.017419 | 0.161443 |
| 14 | 29440800 | 29446799 | 32.88957 | 52.71013 | 122.4427 | 51.67775 | -0.86702 | 1.976215 | 0.161444 |
| 3 | 151821800 | 151825599 | 19.71846 | 32.88391 | 71.62538 | 31.13915 | -0.7382 | 1.21873 | 0.161475 |
| 5 | 94856000 | 94858599 | 36.4728 | 61.60981 | 34.10431 | 14.92017 | 1.230814 | 1.109981 | 0.161478 |
| 5 | 65690200 | 65691599 | 6.770883 | 12.83896 | 32.70289 | 13.20135 | -0.84958 | 0.021849 | 0.16148 |
| 7 | 65182600 | 65184799 | 36.41482 | 60.26258 | 47.17967 | 20.40507 | 0.732679 | 1.332595 | 0.161496 |
| 1 | 10584600 | 10585199 | 5.857221 | 11.24192 | 30.90868 | 12.12242 | -0.92068 | -0.0872 | 0.161501 |
| 9 | 71675400 | 71676999 | 6.23238 | 11.70081 | 32.30057 | 13.29077 | -0.98767 | 0.084348 | 0.16152 |
| 11 | 59838800 | 59840399 | 6.23238 | 11.70081 | 32.30057 | 13.29077 | -0.98767 | 0.084348 | 0.16152 |
| 11 | 113637000 | 113644999 | 33.8121 | 53.93704 | 135.9498 | 56.99558 | -0.99215 | 2.126964 | 0.161522 |
| 8 | 48028400 | 48032599 | 46.09591 | 74.7634 | 60.46126 | 25.90074 | 0.671349 | 1.634544 | 0.161545 |

| | | | | | | | | | |
|---|---|---|---|---|---|---|---|---|---|
| 11 | 63875000 | 63877599 | 9.226512 | 16.25883 | 45.11395 | 19.96791 | -1.07504 | 0.548124 | 0.161557 |
| 3 | 4082000 | 4084399 | 26.00019 | 45.23498 | 31.66585 | 13.91489 | 0.899193 | 0.798028 | 0.161574 |
| 1 | 191529200 | 191548799 | 137.4397 | 202.8529 | 395.3717 | 162.8209 | -0.71721 | 3.757174 | 0.161579 |
| 1 | 85151200 | 85155399 | 92.90428 | 145.7755 | 121.0888 | 50.21856 | 0.585656 | 2.584157 | 0.161588 |
| 10 | 84883400 | 84884399 | 15.24019 | 28.49336 | 20.11374 | 8.33481 | 0.956007 | 0.157818 | 0.161589 |
| 15 | 89234400 | 89240999 | 28.13437 | 45.08139 | 124.253 | 52.42786 | -1.11336 | 1.977519 | 0.161594 |
| 7 | 26988200 | 26991399 | 29.93653 | 50.84302 | 39.33433 | 17.23649 | 0.750274 | 1.048061 | 0.161597 |
| 11 | 79716000 | 79723799 | 49.05829 | 77.8053 | 153.2202 | 63.789 | -0.64564 | 2.32369 | 0.161616 |
| X | 120979400 | 120981199 | 25.12596 | 44.71699 | 20.5095 | 9.037488 | 1.512717 | 0.511995 | 0.161621 |
| 17 | 28589200 | 28595599 | 32.10231 | 51.41623 | 121.4297 | 51.2014 | -0.89186 | 1.995821 | 0.161622 |
| 2 | 32765400 | 32767999 | 11.0065 | 19.09911 | 53.65826 | 23.5433 | -1.09628 | 0.795956 | 0.161623 |
| 3 | 85914200 | 85916399 | 28.04783 | 48.03337 | 36.72772 | 16.10618 | 0.768799 | 0.96498 | 0.161639 |
| 9 | 68575000 | 68575799 | 4.820848 | 9.358139 | 27.44779 | 10.31117 | -0.95269 | -0.25249 | 0.161643 |
| 4 | 128101400 | 128105199 | 19.1598 | 31.7801 | 74.06356 | 32.08296 | -0.83575 | 1.281267 | 0.161652 |
| 11 | 120393800 | 120397199 | 12.46584 | 21.75747 | 49.97158 | 22.00314 | -0.80629 | 0.714442 | 0.161657 |
| 5 | 36953200 | 36954599 | 7.278555 | 13.57276 | 34.37586 | 14.30771 | -0.87865 | 0.140937 | 0.161706 |
| 8 | 34613400 | 34614399 | 7.278555 | 13.57276 | 34.37586 | 14.30771 | -0.87865 | 0.140937 | 0.161706 |
| 11 | 104862000 | 104863599 | 7.278555 | 13.57276 | 34.37586 | 14.30771 | -0.87865 | 0.140937 | 0.161706 |
| 10 | 83999800 | 84001199 | 6.300441 | 11.63316 | 34.65807 | 14.63566 | -1.12592 | 0.194889 | 0.16171 |
| 8 | 19670400 | 19671399 | 14.1374 | 26.7249 | 19.08302 | 7.753707 | 0.962149 | 0.075119 | 0.161737 |
| 7 | 13333800 | 13335399 | 43.37014 | 70.96537 | 53.87263 | 23.17141 | 0.771868 | 1.489855 | 0.16174 |
| 8 | 125194200 | 125199599 | 39.07644 | 62.35624 | 130.4771 | 54.72764 | -0.72365 | 2.119948 | 0.161741 |
| 7 | 120516800 | 120517599 | 4.310052 | 8.306479 | 29.14656 | 11.29716 | -1.25134 | -0.14604 | 0.161744 |
| 1 | 151714800 | 151715999 | 11.72689 | 22.53609 | 16.64768 | 6.505682 | 0.962894 | -0.07979 | 0.161755 |
| 18 | 73948000 | 73948999 | 6.235675 | 11.32758 | 37.59595 | 16.22016 | -1.30221 | 0.296184 | 0.161756 |
| 7 | 24339400 | 24340399 | 7.047349 | 12.96388 | 35.13963 | 14.9144 | -0.9965 | 0.222869 | 0.161792 |
| X | 57308000 | 57309599 | 15.44248 | 28.79983 | 20.14846 | 8.384438 | 0.964173 | 0.175369 | 0.16182 |
| X | 54463800 | 54467999 | 79.90096 | 127.0363 | 91.75446 | 38.29456 | 0.802675 | 2.26398 | 0.161844 |
| 1 | 20472600 | 20474199 | 7.116345 | 13.32157 | 41.57487 | 17.19137 | -1.17074 | 0.128645 | 0.161847 |
| 19 | 15239600 | 15240399 | 10.20907 | 18.17095 | 41.47355 | 18.27381 | -0.78816 | 0.455555 | 0.16185 |
| 5 | 99870200 | 99871199 | 4.962832 | 9.599892 | 27.11354 | 10.35611 | -0.92124 | -0.19316 | 0.161859 |
| 6 | 32723600 | 32726799 | 16.35724 | 27.97694 | 59.66704 | 26.16306 | -0.70541 | 0.958232 | 0.161871 |
| 19 | 13933800 | 13935799 | 24.82758 | 43.33837 | 30.20794 | 13.30194 | 0.905828 | 0.745746 | 0.161898 |
| 11 | 27773400 | 27773799 | 10.05578 | 19.52887 | 15.94788 | 5.969348 | 0.878207 | -0.23986 | 0.16191 |
| 4 | 147128600 | 147129599 | 26.88059 | 46.37914 | 31.47181 | 13.80629 | 0.94202 | 0.8864 | 0.16191 |
| 18 | 82589200 | 82591799 | 16.39067 | 27.87284 | 61.21229 | 26.82816 | -0.74887 | 1.012331 | 0.161917 |
| 7 | 15920000 | 15922199 | 10.41315 | 18.37116 | 43.53561 | 19.26486 | -0.84824 | 0.540114 | 0.161922 |
| 7 | 32183000 | 32184799 | 32.04822 | 54.25338 | 39.26942 | 17.19115 | 0.845799 | 1.082666 | 0.161929 |
| 13 | 33440800 | 33442999 | 23.87021 | 41.60323 | 31.81412 | 13.99569 | 0.772527 | 0.763286 | 0.161934 |
| 2 | 170029200 | 170030999 | 6.859431 | 12.74554 | 34.25453 | 14.34144 | -0.9701 | 0.160612 | 0.161953 |
| 19 | 39149800 | 39151599 | 23.44807 | 41.16827 | 27.79208 | 12.29543 | 0.951382 | 0.644885 | 0.161965 |
| 5 | 100125600 | 100128799 | 17.22398 | 28.73245 | 71.87538 | 31.25369 | -0.93706 | 1.208521 | 0.162016 |
| 3 | 64939800 | 64940999 | 16.02193 | 29.9292 | 19.27325 | 7.998147 | 1.085267 | 0.166528 | 0.162017 |
| 2 | 168125600 | 168140199 | 109.9122 | 165.7033 | 305.4692 | 126.2109 | -0.60893 | 3.363522 | 0.162029 |
| 8 | 46452400 | 46453599 | 6.607979 | 12.1649 | 35.14726 | 14.90219 | -1.08657 | 0.21507 | 0.162039 |
| 17 | 84156600 | 84158999 | 9.05638 | 16.01827 | 45.63604 | 20.18404 | -1.11094 | 0.505288 | 0.162048 |
| 7 | 112278000 | 112279399 | 6.260587 | 11.67259 | 33.22602 | 13.83296 | -1.04521 | 0.132463 | 0.162048 |

| | | | | | | | | | |
|---|---|---|---|---|---|---|---|---|---|
| 11 | 94080400 | 94084199 | 18.76403 | 31.36616 | 69.19805 | 30.12869 | -0.75638 | 1.191024 | 0.162056 |
| 2 | 113045400 | 113051199 | 40.62682 | 64.98933 | 128.8161 | 54.21882 | -0.64181 | 2.044729 | 0.162075 |
| 12 | 22657600 | 22660999 | 46.13907 | 75.3388 | 52.35121 | 22.49991 | 0.896949 | 1.521974 | 0.162079 |
| 5 | 74226600 | 74227799 | 4.589253 | 8.854877 | 30.47622 | 11.77308 | -1.22016 | -0.15206 | 0.162082 |
| 11 | 49540600 | 49542799 | 9.988543 | 17.54794 | 45.1312 | 19.97683 | -0.96764 | 0.590799 | 0.162082 |
| 2 | 148901600 | 148902799 | 13.81582 | 26.68291 | 14.74861 | 5.674672 | 1.397814 | -0.11078 | 0.162086 |
| 4 | 115792000 | 115803799 | 94.56822 | 143.3072 | 284.4512 | 117.7173 | -0.70909 | 3.255014 | 0.16211 |
| X | 24216200 | 24219199 | 64.87447 | 103.8653 | 79.76567 | 33.6139 | 0.724964 | 2.031888 | 0.162112 |
| 12 | 109793400 | 109806999 | 92.65203 | 140.6216 | 278.1423 | 115.1607 | -0.70217 | 3.222476 | 0.16212 |
| 1 | 47557000 | 47559399 | 27.76334 | 48.26561 | 30.42806 | 13.36455 | 1.049387 | 0.814898 | 0.162122 |
| 13 | 119564400 | 119566199 | 9.63506 | 16.75435 | 52.543 | 23.08141 | -1.25324 | 0.759637 | 0.162156 |
| 8 | 79329400 | 79331399 | 12.01356 | 20.99852 | 48.87896 | 21.54301 | -0.82498 | 0.692088 | 0.162165 |
| 8 | 107157600 | 107159999 | 12.3404 | 21.46211 | 54.29619 | 23.85398 | -0.94585 | 0.769534 | 0.162172 |
| X | 3963400 | 3963999 | 10.61368 | 20.75783 | 8.843842 | 2.999409 | 1.926291 | -0.58782 | 0.162181 |
| 10 | 60524600 | 60529399 | 30.06935 | 48.41814 | 108.3114 | 46.03454 | -0.8043 | 1.818867 | 0.162191 |
| 13 | 49120200 | 49122399 | 10.15862 | 17.81051 | 46.7273 | 20.64784 | -0.99639 | 0.627114 | 0.162196 |
| 6 | 107035000 | 107035799 | 16.11126 | 29.44389 | 22.52647 | 9.64585 | 0.805961 | 0.298427 | 0.162198 |
| 12 | 69418400 | 69420199 | 7.816175 | 13.9361 | 41.54156 | 18.27694 | -1.16874 | 0.429005 | 0.162213 |
| 6 | 118855000 | 118857399 | 12.29799 | 21.59071 | 49.35792 | 21.83536 | -0.7993 | 0.620696 | 0.162219 |
| 9 | 62248000 | 62253399 | 23.37825 | 37.74604 | 105.2898 | 44.91286 | -1.11699 | 1.725981 | 0.16222 |
| 5 | 95531200 | 95531999 | 12.37253 | 24.00147 | 15.13444 | 5.703897 | 1.238447 | -0.19171 | 0.162234 |
| 8 | 122554400 | 122557199 | 13.89055 | 23.7082 | 60.58388 | 26.56059 | -0.96549 | 0.968009 | 0.162262 |
| 4 | 34997000 | 34997999 | 5.231561 | 10.12748 | 27.79355 | 10.57783 | -0.87533 | -0.19975 | 0.162271 |
| 15 | 98990800 | 99009799 | 111.4841 | 166.8654 | 344.7782 | 142.1959 | -0.78267 | 3.504475 | 0.162272 |
| X | 75624600 | 75625799 | 16.73545 | 31.53743 | 17.43481 | 7.128567 | 1.321271 | 0.116454 | 0.162273 |
| 1 | 111995000 | 111996599 | 13.61253 | 25.63497 | 19.66143 | 8.04821 | 0.850802 | 0.096282 | 0.162276 |
| 6 | 21081000 | 21081999 | 13.61253 | 25.63497 | 19.66143 | 8.04821 | 0.850802 | 0.096282 | 0.162276 |
| X | 31749600 | 31751599 | 28.13614 | 49.36606 | 23.80674 | 10.52631 | 1.433982 | 0.67641 | 0.162276 |
| 10 | 4610600 | 4612199 | 8.539831 | 15.69973 | 40.69645 | 17.2873 | -0.93463 | 0.247945 | 0.162277 |
| 7 | 21046200 | 21047999 | 25.89599 | 45.33002 | 27.66929 | 12.21013 | 1.096094 | 0.704733 | 0.162291 |
| 6 | 125137200 | 125147199 | 178.206 | 255.1359 | 551.229 | 227.6922 | -0.90851 | 4.227179 | 0.162295 |
| 19 | 11888800 | 11890599 | 11.67896 | 20.11579 | 56.81277 | 24.92174 | -1.10636 | 0.871151 | 0.162297 |
| 1 | 29318000 | 29319199 | 14.29933 | 27.16253 | 17.3165 | 6.964087 | 1.136491 | 0.038805 | 0.1623 |
| 11 | 102457800 | 102459399 | 9.950737 | 17.57982 | 43.0211 | 19.03189 | -0.89343 | 0.520938 | 0.162317 |
| 19 | 16119200 | 16122599 | 21.87107 | 36.17804 | 75.994 | 32.85702 | -0.68685 | 1.322183 | 0.162318 |
| 17 | 20258600 | 20259799 | 19.70509 | 36.08611 | 19.30207 | 8.242566 | 1.320951 | 0.307611 | 0.162324 |
| 11 | 100002000 | 100005999 | 21.20647 | 34.74785 | 82.48338 | 35.49739 | -0.8671 | 1.442214 | 0.162326 |
| 2 | 30765000 | 30784199 | 126.4102 | 187.405 | 392.8206 | 161.7904 | -0.81607 | 3.681876 | 0.162335 |
| 13 | 22152800 | 22154799 | 19.65829 | 35.00787 | 25.30677 | 11.14555 | 0.86121 | 0.480685 | 0.162336 |
| 8 | 129960200 | 129961199 | 7.979541 | 14.2141 | 43.60932 | 19.20179 | -1.21125 | 0.438569 | 0.162336 |
| 7 | 31605600 | 31609599 | 56.84771 | 91.84749 | 59.28706 | 25.31189 | 0.990358 | 1.738576 | 0.162337 |
| 1 | 145720000 | 145721199 | 12.03837 | 22.93741 | 18.3648 | 7.338273 | 0.81938 | 0.000285 | 0.162342 |
| 1 | 141732600 | 141734399 | 23.39796 | 41.84078 | 19.25406 | 8.450225 | 1.510947 | 0.467681 | 0.162355 |
| X | 33607000 | 33609999 | 51.35139 | 84.22903 | 43.42922 | 18.69832 | 1.329713 | 1.47396 | 0.162374 |
| 14 | 41907000 | 41908799 | 22.96955 | 40.67401 | 21.99525 | 9.725868 | 1.272487 | 0.53356 | 0.162384 |
| 18 | 48891400 | 48891999 | 4.718195 | 9.089954 | 28.59389 | 11.09385 | -1.09694 | -0.13965 | 0.162389 |
| 5 | 119293200 | 119295599 | 25.17187 | 43.99346 | 29.38579 | 12.95044 | 0.966874 | 0.727602 | 0.162398 |

| | | | | | | | | | |
|---|---|---|---|---|---|---|---|---|---|
| 7 | 33566600 | 33569399 | 78.15376 | 123.7644 | 104.5509 | 43.47492 | 0.571663 | 2.375591 | 0.162413 |
| 7 | 11099400 | 11105199 | 124.0996 | 191.3199 | 137.8322 | 57.25315 | 0.771276 | 2.90865 | 0.162417 |
| 6 | 17670600 | 17671999 | 6.168965 | 11.76603 | 31.76152 | 12.66416 | -0.91675 | -0.02759 | 0.162431 |
| 8 | 115731000 | 115734799 | 18.44851 | 30.81488 | 69.81311 | 30.38165 | -0.79468 | 1.19669 | 0.162436 |
| 9 | 123507400 | 123508599 | 7.677656 | 14.00849 | 36.59972 | 15.70379 | -0.9554 | 0.277397 | 0.162458 |
| 6 | 112076800 | 112077399 | 5.618317 | 10.63285 | 30.82218 | 12.49323 | -1.03847 | 0.021995 | 0.162469 |
| 2 | 170897000 | 170903199 | 45.47138 | 72.27462 | 158.3822 | 66.08272 | -0.79648 | 2.258366 | 0.162474 |
| 3 | 91830200 | 91832599 | 24.46151 | 43.57775 | 20.40305 | 8.983051 | 1.483713 | 0.500683 | 0.1625 |
| 6 | 125261600 | 125263599 | 19.13027 | 31.80461 | 71.89388 | 31.19288 | -0.79157 | 1.252867 | 0.162504 |
| 2 | 130020400 | 130022399 | 8.741692 | 15.50081 | 44.06639 | 19.47787 | -1.10613 | 0.47356 | 0.162505 |
| 3 | 33898000 | 33898599 | 3.234161 | 6.316261 | 21.59631 | 7.580002 | -1.07705 | -0.53238 | 0.162509 |
| 7 | 89052400 | 89054599 | 14.7417 | 24.94942 | 63.7212 | 27.89337 | -0.96612 | 1.051584 | 0.162512 |
| 11 | 40567000 | 40568999 | 26.41362 | 45.34463 | 36.18131 | 15.86967 | 0.707954 | 0.934424 | 0.162514 |
| X | 31938600 | 31940599 | 18.70499 | 34.98656 | 16.88328 | 6.990959 | 1.500592 | 0.176881 | 0.162524 |
| 10 | 116850000 | 116855999 | 48.32641 | 76.46198 | 156.7487 | 65.15085 | -0.70649 | 2.386149 | 0.162534 |
| 12 | 40501600 | 40502999 | 6.038975 | 11.49038 | 32.8218 | 13.1538 | -1.00429 | -0.01004 | 0.162543 |
| 16 | 19922200 | 19924599 | 61.00027 | 98.31369 | 62.63117 | 26.65461 | 1.005368 | 1.818589 | 0.162546 |
| 3 | 38874400 | 38877199 | 10.14581 | 17.82708 | 48.2651 | 21.36689 | -1.04181 | 0.591452 | 0.162551 |
| 14 | 15579200 | 15580599 | 8.923175 | 16.10284 | 38.5186 | 16.75617 | -0.84243 | 0.354898 | 0.162554 |
| 1 | 117407000 | 117408599 | 14.52259 | 27.44487 | 18.40415 | 7.482591 | 1.050888 | 0.078337 | 0.162559 |
| 6 | 40383400 | 40385999 | 10.45663 | 18.34117 | 46.71654 | 20.65088 | -0.95407 | 0.620837 | 0.162561 |
| 11 | 109470000 | 109476999 | 34.83251 | 55.75493 | 121.8964 | 51.35136 | -0.78184 | 2.018218 | 0.162566 |
| 5 | 127094400 | 127098199 | 39.34071 | 64.95608 | 49.41192 | 21.33684 | 0.77381 | 1.367075 | 0.162572 |
| 17 | 71389200 | 71389999 | 5.423978 | 10.43765 | 29.81754 | 11.60699 | -0.96482 | -0.11911 | 0.162595 |
| 6 | 25071200 | 25071999 | 10.85771 | 21.15228 | 13.52427 | 4.924756 | 1.262178 | -0.33184 | 0.162596 |
| 5 | 113081800 | 113094799 | 81.67895 | 124.8577 | 249.7746 | 103.6163 | -0.71011 | 3.071856 | 0.162624 |
| 7 | 43138000 | 43141399 | 22.26362 | 36.5985 | 80.79777 | 34.81971 | -0.76097 | 1.401783 | 0.162627 |
| 15 | 94415400 | 94416999 | 12.64153 | 22.09635 | 50.48346 | 22.25036 | -0.79838 | 0.697179 | 0.162629 |
| 8 | 55912200 | 55918199 | 99.22651 | 154.6198 | 130.4496 | 54.14139 | 0.555513 | 2.709254 | 0.162633 |
| 2 | 32848400 | 32853799 | 27.32477 | 44.05767 | 103.0448 | 43.8777 | -0.86587 | 1.773077 | 0.162644 |
| 12 | 22468000 | 22468399 | 10.22726 | 19.78062 | 14.88861 | 5.690199 | 0.96643 | -0.17303 | 0.162652 |
| 8 | 21622400 | 21628399 | 132.946 | 203.6189 | 163.881 | 68.01459 | 0.604436 | 3.041716 | 0.162678 |
| 9 | 27891000 | 27891999 | 42.89923 | 70.52086 | 46.73458 | 20.14022 | 0.96993 | 1.426194 | 0.162681 |
| 1 | 107090800 | 107091599 | 10.82935 | 20.90398 | 16.97143 | 6.546078 | 0.846309 | -0.12808 | 0.162695 |
| 14 | 57593200 | 57595599 | 10.56043 | 18.66315 | 43.01339 | 19.02722 | -0.80721 | 0.522108 | 0.162702 |
| 1 | 78169000 | 78170799 | 10.82475 | 18.85732 | 52.61099 | 23.13767 | -1.08567 | 0.737002 | 0.162712 |
| 7 | 21065800 | 21066599 | 18.9689 | 35.02749 | 18.96044 | 8.005743 | 1.315498 | 0.256844 | 0.162712 |
| 3 | 133093000 | 133094399 | 14.03103 | 26.07164 | 19.22417 | 8.048419 | 0.881614 | 0.181786 | 0.162719 |
| 12 | 82524600 | 82525799 | 7.818609 | 14.28722 | 37.69959 | 16.14519 | -0.9682 | 0.270447 | 0.162724 |
| 5 | 115478600 | 115479799 | 4.84526 | 9.334387 | 28.01434 | 10.87022 | -1.0297 | -0.13856 | 0.162726 |
| 11 | 97689000 | 97693799 | 17.96404 | 29.6974 | 77.64881 | 33.5586 | -1.00154 | 1.320021 | 0.162733 |
| 17 | 20246200 | 20248999 | 32.66453 | 56.01047 | 32.73678 | 14.3575 | 1.155022 | 0.969104 | 0.16274 |
| 17 | 73311800 | 73316799 | 27.20248 | 44.19188 | 90.60164 | 38.85909 | -0.66509 | 1.585542 | 0.162762 |
| 18 | 38993000 | 38994399 | 12.84402 | 24.55292 | 18.21782 | 7.228042 | 0.936999 | -0.01474 | 0.162783 |
| 10 | 112105800 | 112108399 | 14.32565 | 24.14833 | 65.76209 | 28.7518 | -1.05852 | 1.088046 | 0.162789 |
| 6 | 132297600 | 132299199 | 19.75735 | 37.01546 | 14.4009 | 5.946566 | 1.810431 | 0.169942 | 0.162793 |
| 18 | 11379000 | 11382399 | 34.37066 | 58.99224 | 30.28537 | 13.28339 | 1.341246 | 0.963954 | 0.162801 |

| | | | | | | | | | |
|---|---|---|---|---|---|---|---|---|---|
| X | 25525200 | 25527799 | 23.29833 | 41.30744 | 23.47004 | 10.37516 | 1.203076 | 0.528703 | 0.162802 |
| 7 | 23748000 | 23751199 | 37.12777 | 62.7153 | 34.67612 | 15.17027 | 1.232593 | 1.114236 | 0.162808 |
| 6 | 49560800 | 49561799 | 5.548369 | 10.7114 | 30.05916 | 11.5874 | -0.92617 | -0.14947 | 0.162808 |
| 5 | 100297200 | 100298799 | 8.381595 | 15.46186 | 38.53933 | 16.29464 | -0.87323 | 0.224007 | 0.16281 |
| 14 | 3224800 | 3229999 | 72.32403 | 115.5515 | 80.32975 | 33.75177 | 0.865116 | 2.103069 | 0.162812 |
| 8 | 111391200 | 111393599 | 11.50025 | 20.27899 | 45.61823 | 20.18844 | -0.77405 | 0.556128 | 0.162823 |
| 11 | 74132200 | 74132999 | 11.97173 | 21.04299 | 46.66455 | 20.66465 | -0.75552 | 0.595136 | 0.162826 |
| 4 | 46629800 | 46631599 | 9.291687 | 16.56611 | 40.47544 | 17.80861 | -0.88408 | 0.437334 | 0.162842 |
| 4 | 41601400 | 41605199 | 15.9103 | 26.68818 | 66.84389 | 29.16741 | -0.9386 | 1.141421 | 0.162845 |
| 18 | 36611000 | 36612999 | 25.72205 | 44.26952 | 31.75437 | 13.92918 | 0.861681 | 0.903709 | 0.162849 |
| 5 | 132700400 | 132705799 | 28.85463 | 46.69226 | 100.5119 | 42.98503 | -0.74074 | 1.671237 | 0.162851 |
| 3 | 119853200 | 119853999 | 5.085051 | 9.875494 | 28.18658 | 10.54988 | -0.90891 | -0.25958 | 0.162854 |
| 8 | 34947200 | 34947999 | 5.085051 | 9.875494 | 28.18658 | 10.54988 | -0.90891 | -0.25958 | 0.162854 |
| 6 | 26859600 | 26860199 | 11.49324 | 22.09464 | 16.47566 | 6.43212 | 0.952059 | -0.08357 | 0.162862 |
| 8 | 59641400 | 59643199 | 9.882625 | 17.66478 | 40.48157 | 17.77297 | -0.79024 | 0.424806 | 0.162869 |
| 13 | 50996000 | 50998399 | 22.79467 | 40.30076 | 24.9868 | 11.05215 | 1.076292 | 0.556573 | 0.162885 |
| 4 | 58088200 | 58089599 | 5.651797 | 10.5963 | 32.2372 | 13.29547 | -1.12862 | 0.089453 | 0.162906 |
| 5 | 10125800 | 10126599 | 11.82239 | 22.88445 | 16.27329 | 6.195723 | 1.053061 | -0.16192 | 0.162908 |
| 18 | 54813800 | 54814999 | 7.94034 | 14.57611 | 36.72724 | 15.63258 | -0.89572 | 0.243289 | 0.162921 |
| 17 | 31003000 | 31005199 | 10.26744 | 18.09867 | 43.55007 | 19.27455 | -0.8705 | 0.549723 | 0.162924 |
| 7 | 31755600 | 31756399 | 36.66988 | 60.51322 | 46.68327 | 20.15098 | 0.753564 | 1.364237 | 0.162931 |
| 13 | 95690400 | 95694599 | 31.64645 | 50.99582 | 106.7419 | 45.41678 | -0.70732 | 1.796667 | 0.162931 |
| 2 | 150479200 | 150480599 | 8.219406 | 15.15087 | 36.29093 | 15.36025 | -0.81729 | 0.221453 | 0.162931 |
| 6 | 131145800 | 131147999 | 19.66912 | 36.84611 | 15.03863 | 6.211064 | 1.742239 | 0.171512 | 0.162933 |
| 12 | 19325600 | 19335799 | 242.1379 | 356.7602 | 247.4506 | 101.9025 | 0.770181 | 3.793046 | 0.162936 |
| 4 | 90195600 | 90196399 | 11.07189 | 21.41796 | 16.46142 | 6.286177 | 0.938463 | -0.15922 | 0.162945 |
| 18 | 19454000 | 19456999 | 29.25935 | 51.04962 | 25.79196 | 11.35392 | 1.367164 | 0.766794 | 0.162946 |
| 10 | 75542400 | 75546999 | 17.63333 | 29.21252 | 78.39797 | 33.93199 | -1.0388 | 1.293776 | 0.162975 |
| 16 | 94924400 | 94925599 | 8.730284 | 15.92402 | 37.66227 | 16.16613 | -0.8134 | 0.290404 | 0.162976 |
| 15 | 23810600 | 23812399 | 16.72146 | 31.51345 | 17.03149 | 6.962765 | 1.352676 | 0.113357 | 0.162979 |
| X | 61896000 | 61898799 | 46.59322 | 76.72524 | 45.06313 | 19.43406 | 1.144567 | 1.415682 | 0.162992 |
| 5 | 7225000 | 7226399 | 13.43161 | 25.53291 | 18.75937 | 7.532692 | 0.936423 | 0.03108 | 0.163002 |
| 14 | 33454200 | 33456999 | 13.27814 | 23.04378 | 52.67938 | 23.11389 | -0.80062 | 0.803614 | 0.163003 |
| 7 | 17149200 | 17158799 | 210.8907 | 312.3344 | 252.7774 | 104.102 | 0.554701 | 3.704901 | 0.163016 |
| 5 | 136385400 | 136386999 | 5.802118 | 10.90067 | 32.29821 | 13.27402 | -1.08724 | 0.075965 | 0.163026 |
| 6 | 70246600 | 70248599 | 21.4901 | 38.34349 | 22.78596 | 10.01913 | 1.143351 | 0.475283 | 0.163029 |
| 12 | 82835800 | 82836799 | 11.37166 | 19.94758 | 46.19212 | 20.42028 | -0.81735 | 0.625738 | 0.163031 |
| 11 | 44509000 | 44513399 | 28.15573 | 45.35039 | 107.2265 | 45.59325 | -0.8834 | 1.806623 | 0.163038 |
| 11 | 29607800 | 29608599 | 12.97556 | 24.68367 | 18.70417 | 7.499146 | 0.893735 | 0.020734 | 0.163059 |
| 17 | 74422000 | 74423599 | 12.97556 | 24.68367 | 18.70417 | 7.499146 | 0.893735 | 0.020734 | 0.163059 |
| 9 | 57061600 | 57066399 | 15.99037 | 26.67625 | 72.25931 | 31.42186 | -1.05133 | 1.204871 | 0.163068 |
| X | 122642200 | 122643199 | 16.94823 | 31.76936 | 18.14497 | 7.485901 | 1.263703 | 0.151141 | 0.163085 |
| 2 | 9875800 | 9884999 | 61.12548 | 95.36414 | 197.2826 | 81.86607 | -0.73559 | 2.678607 | 0.163095 |
| 7 | 44180600 | 44203399 | 134.4665 | 196.9374 | 441.3436 | 181.7997 | -0.92905 | 3.881762 | 0.163096 |
| 15 | 41326600 | 41328599 | 18.1549 | 32.35623 | 24.60818 | 10.83065 | 0.789077 | 0.464461 | 0.163113 |
| 15 | 37911800 | 37913199 | 7.288414 | 13.56245 | 34.30835 | 14.33081 | -0.88104 | 0.154872 | 0.163114 |
| 12 | 87312000 | 87314199 | 9.295742 | 16.56564 | 42.0235 | 18.49697 | -0.93862 | 0.441654 | 0.163115 |

| | | | | | | | | | |
|---|---|---|---|---|---|---|---|---|---|
| 7 | 111835800 | 111838399 | 14.39576 | 24.88189 | 54.28996 | 23.8176 | -0.73502 | 0.847953 | 0.163123 |
| 2 | 60523800 | 60528799 | 25.11695 | 40.84587 | 90.50544 | 38.84044 | -0.77558 | 1.562115 | 0.163136 |
| 15 | 36307800 | 36311599 | 24.52061 | 40.20069 | 83.23731 | 35.83032 | -0.67038 | 1.442461 | 0.163141 |
| 17 | 19827400 | 19830599 | 41.77182 | 69.47414 | 41.60718 | 18.05718 | 1.118341 | 1.287554 | 0.163141 |
| 18 | 68070800 | 68074399 | 18.23084 | 30.60394 | 69.46874 | 30.3273 | -0.79758 | 1.140511 | 0.163144 |
| 1 | 55066000 | 55066999 | 8.307871 | 15.04222 | 37.53498 | 16.26074 | -0.89872 | 0.330017 | 0.163153 |
| 17 | 18416000 | 18421399 | 85.65172 | 136.2172 | 84.04076 | 35.08546 | 1.030007 | 2.259057 | 0.163161 |
| 11 | 32106600 | 32108199 | 8.07795 | 14.86106 | 36.23778 | 15.37859 | -0.84562 | 0.231197 | 0.16317 |
| 14 | 14518000 | 14519999 | 8.07795 | 14.86106 | 36.23778 | 15.37859 | -0.84562 | 0.231197 | 0.16317 |
| 17 | 18880000 | 18881999 | 38.42542 | 63.67714 | 47.16789 | 20.41929 | 0.812425 | 1.32273 | 0.163172 |
| 7 | 137958600 | 137960399 | 7.734344 | 14.3755 | 35.40185 | 14.81579 | -0.84508 | 0.168044 | 0.16319 |
| 13 | 12056800 | 12058399 | 8.438417 | 15.34029 | 37.06782 | 15.97762 | -0.84794 | 0.306397 | 0.163192 |
| 2 | 117309600 | 117310999 | 7.036073 | 12.9979 | 36.31782 | 15.32615 | -1.0345 | 0.198833 | 0.163196 |
| 18 | 5333800 | 5335199 | 7.036073 | 12.9979 | 36.31782 | 15.32615 | -1.0345 | 0.198833 | 0.163196 |
| 10 | 82017000 | 82025999 | 172.6138 | 261.4205 | 178.3307 | 73.7865 | 0.82983 | 3.278276 | 0.163225 |
| 19 | 6926200 | 6930599 | 15.18579 | 25.38179 | 69.9522 | 30.45482 | -1.076 | 1.1809 | 0.163234 |
| 8 | 22183800 | 22185399 | 26.43619 | 45.92478 | 32.18312 | 14.1289 | 0.897465 | 0.820997 | 0.163237 |
| 10 | 5612400 | 5613599 | 22.1189 | 38.80934 | 25.57969 | 11.31042 | 0.985668 | 0.642827 | 0.163244 |
| 10 | 111000200 | 111002799 | 13.32482 | 22.99038 | 59.1718 | 25.95793 | -0.97298 | 0.862842 | 0.163254 |
| 1 | 119958000 | 119959399 | 6.491352 | 12.26624 | 31.37979 | 12.76064 | -0.86455 | 0.041931 | 0.163256 |
| 10 | 81333400 | 81339999 | 38.34472 | 61.02802 | 141.192 | 59.00665 | -0.87244 | 2.20679 | 0.163269 |
| 17 | 83330400 | 83333199 | 12.53507 | 21.70205 | 54.24338 | 23.79838 | -0.92947 | 0.823806 | 0.163277 |
| 7 | 47769600 | 47770999 | 15.17222 | 27.85474 | 21.06516 | 8.96484 | 0.828568 | 0.26164 | 0.163291 |
| 7 | 3765800 | 3768799 | 40.17942 | 66.05311 | 44.07006 | 18.99235 | 0.96023 | 1.416361 | 0.163297 |
| 4 | 61332000 | 61334799 | 25.42065 | 44.55361 | 25.27327 | 11.16458 | 1.200699 | 0.676739 | 0.163301 |
| 6 | 124731400 | 124733999 | 12.03458 | 20.94222 | 51.03719 | 22.43026 | -0.89177 | 0.757339 | 0.163307 |
| 6 | 86464200 | 86466199 | 9.577409 | 17.12927 | 39.97653 | 17.54255 | -0.81572 | 0.418659 | 0.163314 |
| 4 | 146189600 | 146190199 | 9.756442 | 19.01125 | 14.27617 | 5.182824 | 1.037029 | -0.35418 | 0.163333 |
| 10 | 14464800 | 14465399 | 4.190735 | 8.149041 | 25.10857 | 9.30172 | -1.00177 | -0.31219 | 0.163345 |
| 3 | 77682200 | 77682999 | 11.06663 | 21.43235 | 16.43165 | 6.241262 | 0.94912 | -0.17714 | 0.163347 |
| 5 | 75915200 | 75918599 | 17.01607 | 28.94077 | 61.81249 | 27.09155 | -0.70889 | 1.0138 | 0.163359 |
| 3 | 99145200 | 99152399 | 52.38326 | 82.8062 | 165.4255 | 68.72597 | -0.67039 | 2.411052 | 0.163362 |
| 2 | 138155000 | 138156999 | 8.718446 | 15.88513 | 40.93315 | 17.59404 | -0.93821 | 0.299886 | 0.163366 |
| 9 | 42002800 | 42005599 | 13.08153 | 22.87577 | 50.50107 | 22.26663 | -0.74936 | 0.689916 | 0.163368 |
| 12 | 113052000 | 113056399 | 29.69139 | 47.94873 | 100.3865 | 42.83237 | -0.70392 | 1.728017 | 0.16339 |
| 1 | 149381400 | 149381999 | 12.74478 | 24.14739 | 18.69579 | 7.559791 | 0.852183 | 0.044992 | 0.163405 |
| 9 | 63108000 | 63114399 | 46.41204 | 73.8267 | 141.157 | 58.94675 | -0.59885 | 2.231936 | 0.163408 |
| 4 | 136012000 | 136013799 | 7.854457 | 14.25969 | 38.10319 | 16.4513 | -0.99379 | 0.310356 | 0.16342 |
| 11 | 53781800 | 53782999 | 7.854457 | 14.25969 | 38.10319 | 16.4513 | -0.99379 | 0.310356 | 0.16342 |
| 3 | 112465600 | 112466999 | 14.63539 | 27.93895 | 14.93349 | 5.943137 | 1.400323 | 0.002575 | 0.163422 |
| 8 | 3243800 | 3244799 | 22.49134 | 39.2173 | 30.45558 | 13.40175 | 0.750414 | 0.747788 | 0.163422 |
| 18 | 75920000 | 75922799 | 16.64293 | 28.03148 | 64.85924 | 28.35 | -0.82466 | 1.102065 | 0.163445 |
| 7 | 10879600 | 10881999 | 58.98576 | 94.73401 | 74.71177 | 31.63464 | 0.691954 | 1.925718 | 0.163468 |
| 14 | 42325200 | 42328199 | 22.36472 | 39.63357 | 24.51539 | 10.83855 | 1.080937 | 0.528755 | 0.163472 |
| 4 | 127330600 | 127333799 | 15.62817 | 26.53279 | 65.03667 | 28.49629 | -0.90712 | 1.026766 | 0.163483 |
| 5 | 124485800 | 124493799 | 32.15385 | 51.4222 | 129.3505 | 54.42392 | -0.98516 | 2.04385 | 0.163485 |
| 18 | 81052000 | 81054999 | 17.15428 | 28.77099 | 66.39827 | 28.97051 | -0.82107 | 1.146741 | 0.163497 |

| | | | | | | | | | |
|---|---|---|---|---|---|---|---|---|---|
| 2 | 169724200 | 169728599 | 24.22239 | 39.25289 | 94.05507 | 40.26642 | -0.8927 | 1.632618 | 0.163504 |
| 17 | 95012400 | 95013799 | 8.165075 | 14.78027 | 39.10829 | 16.94721 | -0.9834 | 0.3314 | 0.163505 |
| 5 | 30756000 | 30769599 | 77.8736 | 119.0529 | 248.8884 | 103.2511 | -0.77347 | 3.068941 | 0.163513 |
| 7 | 9667000 | 9667799 | 10.17274 | 19.86477 | 11.9267 | 4.194882 | 1.395483 | -0.46351 | 0.163522 |
| 10 | 60002000 | 60012199 | 96.43761 | 146.6454 | 269.2092 | 111.5255 | -0.59241 | 3.184096 | 0.163527 |
| 4 | 115823600 | 115830199 | 40.06193 | 63.85275 | 141.8596 | 59.40364 | -0.8119 | 2.160646 | 0.163534 |
| 10 | 121311000 | 121312999 | 15.46895 | 26.28336 | 61.1173 | 26.78472 | -0.83109 | 1.017005 | 0.163552 |
| 6 | 84201000 | 84208399 | 35.59474 | 56.70633 | 137.7235 | 57.63453 | -0.94079 | 2.172305 | 0.163572 |
| 12 | 105822400 | 105852999 | 328.7282 | 450.3016 | 842.5917 | 350.5158 | -0.76547 | 4.895821 | 0.163582 |
| 12 | 53292600 | 53295999 | 23.71445 | 38.49404 | 91.37056 | 39.16537 | -0.87681 | 1.596334 | 0.163589 |
| 18 | 79997800 | 79999199 | 11.1968 | 19.68279 | 45.0949 | 19.96921 | -0.80208 | 0.587087 | 0.16359 |
| 19 | 47567000 | 47569199 | 11.4673 | 19.87271 | 53.66267 | 23.5443 | -1.03988 | 0.810179 | 0.163614 |
| 1 | 117958600 | 117961399 | 42.22908 | 69.44251 | 50.04297 | 21.56903 | 0.849589 | 1.422297 | 0.163618 |
| 4 | 153461800 | 153474799 | 118.5939 | 177.113 | 340.4528 | 140.3513 | -0.68146 | 3.547759 | 0.163647 |
| 11 | 56105200 | 56107199 | 23.67468 | 41.32293 | 31.19531 | 13.7361 | 0.791131 | 0.740764 | 0.163649 |
| 5 | 65225800 | 65227599 | 8.598508 | 15.59356 | 37.55274 | 16.23568 | -0.84614 | 0.321964 | 0.163656 |
| 9 | 48127200 | 48128599 | 8.598508 | 15.59356 | 37.55274 | 16.23568 | -0.84614 | 0.321964 | 0.163656 |
| 10 | 49302400 | 49303999 | 23.09783 | 41.20791 | 21.61116 | 9.503367 | 1.322444 | 0.485104 | 0.163662 |
| 18 | 54003800 | 54006199 | 22.07001 | 38.95983 | 26.11681 | 11.55552 | 0.963498 | 0.569882 | 0.163665 |
| 2 | 29462800 | 29464599 | 9.342798 | 16.51455 | 42.52602 | 18.81088 | -0.96587 | 0.511405 | 0.16367 |
| 4 | 66883000 | 66884599 | 9.342798 | 16.51455 | 42.52602 | 18.81088 | -0.96587 | 0.511405 | 0.16367 |
| 3 | 19951400 | 19955399 | 53.20616 | 85.76803 | 68.41163 | 29.12169 | 0.68208 | 1.801294 | 0.163675 |
| X | 26457400 | 26459599 | 55.90753 | 90.35371 | 60.19028 | 25.70677 | 0.945522 | 1.731802 | 0.163682 |
| 3 | 109371000 | 109371999 | 9.516866 | 17.21375 | 40.16419 | 17.42186 | -0.80439 | 0.348313 | 0.163684 |
| 2 | 164504000 | 164511399 | 59.35055 | 93.43492 | 173.3114 | 71.9075 | -0.56822 | 2.507662 | 0.163699 |
| 3 | 102059400 | 102062399 | 15.68932 | 26.95207 | 57.52634 | 25.23272 | -0.70545 | 0.912738 | 0.163704 |
| 5 | 115612400 | 115614799 | 13.25893 | 23.1271 | 50.51596 | 22.234 | -0.73431 | 0.729502 | 0.163709 |
| 13 | 60776200 | 60777599 | 7.432941 | 13.8755 | 35.5067 | 14.75161 | -0.89089 | 0.14071 | 0.16371 |
| 12 | 14901000 | 14902799 | 19.43095 | 35.4145 | 21.18201 | 9.10099 | 1.155797 | 0.336997 | 0.163712 |
| 4 | 147249000 | 147250999 | 25.13044 | 44.01818 | 26.43895 | 11.67353 | 1.118835 | 0.686697 | 0.16373 |
| 13 | 31488200 | 31489399 | 6.303755 | 12.04717 | 31.86823 | 12.6531 | -0.88241 | -0.03968 | 0.163745 |
| 13 | 107806200 | 107813599 | 46.48603 | 73.70633 | 151.663 | 63.12405 | -0.70921 | 2.329533 | 0.163746 |
| 3 | 68432200 | 68434999 | 20.03837 | 32.99442 | 78.74604 | 33.95902 | -0.87194 | 1.37552 | 0.163755 |
| 18 | 59756800 | 59757599 | 11.61093 | 22.26771 | 17.91823 | 7.043369 | 0.833693 | -0.05703 | 0.163764 |
| 10 | 82197600 | 82203399 | 163.496 | 247.6832 | 186.2672 | 77.07733 | 0.689603 | 3.270729 | 0.163768 |
| 7 | 13943400 | 13944399 | 23.848 | 41.68014 | 29.77214 | 13.12082 | 0.870799 | 0.722729 | 0.163796 |
| 13 | 77768400 | 77769599 | 19.84319 | 35.32755 | 25.30346 | 11.14665 | 0.87424 | 0.483724 | 0.163798 |
| 1 | 98197600 | 98198999 | 6.143176 | 11.78586 | 30.89025 | 12.12965 | -0.85376 | -0.07924 | 0.163798 |
| 2 | 136732600 | 136734399 | 10.41281 | 18.36588 | 45.09544 | 19.95612 | -0.89953 | 0.544168 | 0.163803 |
| 7 | 133377800 | 133379799 | 10.41281 | 18.36588 | 45.09544 | 19.95612 | -0.89953 | 0.544168 | 0.163803 |
| X | 25268000 | 25270199 | 20.77548 | 37.98948 | 18.67084 | 7.989188 | 1.440111 | 0.322849 | 0.163803 |
| 14 | 30601200 | 30605799 | 16.59743 | 27.15843 | 91.38912 | 39.30807 | -1.36969 | 1.454739 | 0.163818 |
| 1 | 152262000 | 152262599 | 4.012329 | 7.766231 | 27.64448 | 10.52748 | -1.2467 | -0.21359 | 0.163818 |
| 12 | 109550200 | 109562999 | 106.7068 | 159.7537 | 337.3147 | 139.1243 | -0.81357 | 3.499277 | 0.163818 |
| 11 | 106377600 | 106379799 | 10.19393 | 17.79325 | 49.39748 | 21.7512 | -1.07775 | 0.703011 | 0.16382 |
| 11 | 69308200 | 69312799 | 20.45457 | 33.47646 | 85.78984 | 36.90236 | -0.97803 | 1.456817 | 0.163842 |
| 15 | 65609400 | 65610399 | 10.13366 | 17.81362 | 44.60108 | 19.7506 | -0.92907 | 0.581568 | 0.163847 |

| | | | | | | | | | |
|---|---|---|---|---|---|---|---|---|---|
| 17 | 10554000 | 10555799 | 5.890626 | 10.90085 | 36.8592 | 15.51782 | -1.30441 | 0.181032 | 0.163855 |
| 10 | 80576000 | 80578599 | 10.78549 | 18.84569 | 48.34927 | 21.30377 | -0.96443 | 0.689401 | 0.163857 |
| 8 | 124425400 | 124429799 | 20.10682 | 33.33412 | 73.555 | 31.8516 | -0.75728 | 1.289967 | 0.163863 |
| 17 | 47627800 | 47628999 | 5.678373 | 10.96028 | 29.99185 | 11.57183 | -0.89078 | -0.14253 | 0.163865 |
| 15 | 75402600 | 75406999 | 22.58135 | 37.11332 | 81.44384 | 35.09513 | -0.75249 | 1.405218 | 0.163893 |
| 15 | 63222800 | 63224399 | 9.734335 | 17.42574 | 41.03443 | 17.99301 | -0.82776 | 0.417024 | 0.163913 |
| X | 61012600 | 61014599 | 34.73019 | 57.80162 | 46.02893 | 19.98549 | 0.708741 | 1.262201 | 0.163915 |
| 3 | 122306000 | 122307999 | 9.955364 | 17.5702 | 43.03732 | 19.04292 | -0.89517 | 0.53359 | 0.163917 |
| 17 | 36580800 | 36584599 | 20.96182 | 34.56574 | 78.55359 | 33.89711 | -0.80077 | 1.358198 | 0.163948 |
| 6 | 133728400 | 133729999 | 22.35051 | 39.05498 | 29.64964 | 13.06462 | 0.782994 | 0.714598 | 0.16398 |
| 7 | 19653800 | 19655599 | 14.92508 | 25.56727 | 58.54675 | 25.67506 | -0.80716 | 0.935748 | 0.163982 |
| 1 | 85028400 | 85030199 | 21.84945 | 39.04125 | 22.22206 | 9.759232 | 1.206279 | 0.468068 | 0.164019 |
| 4 | 101121600 | 101122799 | 8.035416 | 14.51051 | 40.65593 | 17.67111 | -1.06839 | 0.34717 | 0.164022 |
| 7 | 35501800 | 35504799 | 14.79704 | 25.28702 | 58.47225 | 25.63742 | -0.82161 | 0.958374 | 0.164037 |
| 13 | 55062800 | 55076199 | 105.2425 | 158.8966 | 293.8806 | 121.4783 | -0.61181 | 3.332868 | 0.164037 |
| 9 | 17721000 | 17721999 | 14.79607 | 27.61661 | 20.58439 | 8.554123 | 0.87464 | 0.164699 | 0.164041 |
| 12 | 108602800 | 108604199 | 4.00523 | 7.770946 | 27.25461 | 10.26434 | -1.2103 | -0.25355 | 0.164064 |
| 11 | 88217400 | 88221999 | 28.89241 | 46.66103 | 99.28261 | 42.36438 | -0.72702 | 1.726542 | 0.16407 |
| 10 | 63648200 | 63649799 | 21.21203 | 37.26131 | 28.48773 | 12.6086 | 0.772709 | 0.624459 | 0.164088 |
| 4 | 121366400 | 121368199 | 52.80411 | 85.00221 | 69.54903 | 29.54775 | 0.643058 | 1.844607 | 0.164102 |
| 5 | 93859200 | 93862199 | 65.25405 | 104.4917 | 78.97228 | 33.28857 | 0.748293 | 2.025476 | 0.164104 |
| 4 | 110208000 | 110209399 | 8.535273 | 15.25853 | 41.0272 | 18.01045 | -1.01907 | 0.415063 | 0.16411 |
| 1 | 117900000 | 117901199 | 22.81519 | 39.89616 | 29.8277 | 13.15119 | 0.805264 | 0.708651 | 0.164113 |
| 15 | 79590600 | 79593399 | 15.61534 | 26.54826 | 61.71209 | 27.04671 | -0.83053 | 1.011755 | 0.164118 |
| 7 | 37567200 | 37572599 | 28.70964 | 46.42535 | 98.32085 | 42.02343 | -0.71821 | 1.686025 | 0.164122 |
| 17 | 88955800 | 88956999 | 14.16583 | 27.03408 | 17.66766 | 7.035924 | 1.114364 | 0.009663 | 0.164133 |
| 2 | 17059200 | 17060199 | 11.56243 | 22.18523 | 16.69221 | 6.552501 | 0.93072 | -0.06717 | 0.164147 |
| 3 | 50788600 | 50789599 | 11.56243 | 22.18523 | 16.69221 | 6.552501 | 0.93072 | -0.06717 | 0.164147 |
| 2 | 119990000 | 119992599 | 12.63676 | 22.06611 | 49.42758 | 21.77001 | -0.77011 | 0.709965 | 0.164158 |
| 6 | 149262800 | 149267799 | 90.13473 | 136.0258 | 301.4662 | 124.5963 | -0.87308 | 3.339322 | 0.16416 |
| 8 | 81369400 | 81371799 | 33.82822 | 56.88725 | 41.5118 | 18.1362 | 0.833952 | 1.148297 | 0.164166 |
| 8 | 110749200 | 110753199 | 20.58015 | 34.10405 | 76.00402 | 32.90225 | -0.77172 | 1.296592 | 0.164167 |
| 11 | 70894000 | 70896199 | 6.993698 | 12.6834 | 39.14195 | 16.92244 | -1.20109 | 0.312473 | 0.164171 |
| 2 | 76586000 | 76587599 | 38.60777 | 63.50215 | 51.59372 | 22.23959 | 0.677204 | 1.411565 | 0.164185 |
| 12 | 8448200 | 8449399 | 6.505562 | 12.26331 | 32.43881 | 13.25632 | -0.91903 | 0.059675 | 0.164208 |
| 10 | 56980000 | 56981799 | 17.02702 | 31.31344 | 20.07759 | 8.524706 | 1.067 | 0.264273 | 0.164213 |
| 10 | 81045200 | 81050799 | 53.81245 | 84.3283 | 180.6188 | 74.91692 | -0.78054 | 2.594955 | 0.16422 |
| 17 | 27888400 | 27914199 | 178.131 | 256.903 | 505.0988 | 208.4076 | -0.762 | 4.118999 | 0.164256 |
| 5 | 141976800 | 141989199 | 89.82185 | 137.0946 | 257.3951 | 106.7396 | -0.62045 | 3.105581 | 0.164257 |
| 8 | 60553400 | 60556199 | 33.88518 | 56.6128 | 44.32885 | 19.29423 | 0.732739 | 1.215522 | 0.164258 |
| 4 | 133627200 | 133631999 | 22.76722 | 37.348 | 81.16487 | 34.94691 | -0.73953 | 1.430363 | 0.164266 |
| 14 | 61918200 | 61920599 | 10.40927 | 18.07817 | 54.71778 | 24.01802 | -1.20301 | 0.782641 | 0.16429 |
| X | 28805200 | 28805399 | 10.42753 | 20.39584 | 8.153619 | 2.741081 | 2.025491 | -0.62229 | 0.164294 |
| 5 | 110500600 | 110502999 | 10.49165 | 18.34172 | 50.45827 | 22.24252 | -1.06487 | 0.680283 | 0.164315 |
| 11 | 83433400 | 83434599 | 14.32351 | 26.58892 | 20.08017 | 8.418061 | 0.845822 | 0.191368 | 0.164326 |
| 17 | 57781400 | 57782199 | 11.47278 | 21.97314 | 17.31369 | 6.832128 | 0.858092 | -0.04908 | 0.164346 |
| 8 | 5756600 | 5758799 | 21.47158 | 38.38724 | 22.23038 | 9.758344 | 1.181909 | 0.46191 | 0.164361 |

| | | | | | | | | | |
|---|---|---|---|---|---|---|---|---|---|
| 4 | 91395400 | 91396999 | 15.78559 | 29.28136 | 20.97656 | 8.8027 | 0.920658 | 0.210863 | 0.164368 |
| 10 | 7435600 | 7436599 | 17.34753 | 32.02945 | 20.14889 | 8.509011 | 1.100077 | 0.246456 | 0.164379 |
| 10 | 30718200 | 30720799 | 17.38321 | 28.69872 | 83.32523 | 35.97856 | -1.15244 | 1.337171 | 0.164387 |
| 5 | 124076400 | 124079799 | 20.73832 | 34.36519 | 75.4301 | 32.65317 | -0.74984 | 1.297177 | 0.164388 |
| 1 | 76099800 | 76100799 | 6.623166 | 12.56592 | 31.53508 | 12.71482 | -0.82631 | 0.015215 | 0.164397 |
| 11 | 21926600 | 21927399 | 6.778619 | 12.82865 | 32.03257 | 12.98408 | -0.82618 | 0.034377 | 0.164437 |
| 7 | 31747600 | 31751799 | 52.12689 | 84.81866 | 50.21218 | 21.55307 | 1.123969 | 1.582999 | 0.164445 |
| 5 | 95429000 | 95429599 | 14.99558 | 28.72241 | 15.02451 | 5.927457 | 1.443048 | -0.01999 | 0.164449 |
| 9 | 46245600 | 46250999 | 29.538 | 47.69661 | 99.9721 | 42.65046 | -0.70566 | 1.732289 | 0.16445 |
| 5 | 103150000 | 103151799 | 8.886263 | 16.17242 | 37.61673 | 16.19167 | -0.79224 | 0.30383 | 0.164455 |
| X | 77977600 | 77978599 | 13.99656 | 27.16344 | 13.13083 | 4.93896 | 1.619358 | -0.18559 | 0.164457 |
| 17 | 36115600 | 36116799 | 6.631322 | 12.55331 | 32.04094 | 12.98 | -0.85692 | 0.030616 | 0.164476 |
| 10 | 43290600 | 43291799 | 12.03373 | 23.03357 | 17.14291 | 6.776481 | 0.936986 | -0.03885 | 0.164481 |
| 6 | 149125000 | 149126399 | 14.80513 | 27.31669 | 21.20471 | 8.963203 | 0.798379 | 0.232622 | 0.164486 |
| 2 | 177618800 | 177621799 | 24.62758 | 43.02612 | 29.95275 | 13.19667 | 0.907493 | 0.730694 | 0.164486 |
| 11 | 116619000 | 116623399 | 35.38274 | 56.91211 | 111.5943 | 47.35535 | -0.61668 | 1.862363 | 0.164488 |
| 11 | 94362800 | 94367399 | 23.79315 | 38.50103 | 98.43783 | 42.0974 | -0.98809 | 1.663172 | 0.164488 |
| 3 | 97808400 | 97808999 | 3.93661 | 7.680827 | 23.32303 | 8.321295 | -0.93168 | -0.45923 | 0.164491 |
| 11 | 73847800 | 73848999 | 19.74221 | 34.81308 | 27.05229 | 11.972 | 0.751038 | 0.568762 | 0.164492 |
| 8 | 91875200 | 91897399 | 203.9277 | 293.9046 | 506.1948 | 208.8789 | -0.57209 | 4.131062 | 0.164505 |
| 13 | 114730200 | 114731599 | 28.01288 | 47.6703 | 37.82052 | 16.57694 | 0.714362 | 1.020494 | 0.164509 |
| 7 | 55610800 | 55612599 | 8.792135 | 15.8137 | 42.71184 | 18.65399 | -1.02081 | 0.379469 | 0.164515 |
| 1 | 167076800 | 167080199 | 14.51252 | 24.74004 | 60.04286 | 26.32086 | -0.89169 | 0.981401 | 0.164523 |
| 5 | 91449200 | 91451399 | 11.21339 | 19.65979 | 47.24754 | 20.87773 | -0.87089 | 0.632562 | 0.164557 |
| 10 | 94235400 | 94239999 | 33.57747 | 54.09436 | 108.3501 | 46.08477 | -0.64442 | 1.808009 | 0.164559 |
| 14 | 105622600 | 105623999 | 5.72561 | 10.9707 | 31.38868 | 12.37074 | -0.98434 | -0.07174 | 0.164572 |
| 11 | 78764600 | 78765399 | 4.952044 | 9.612691 | 26.83128 | 10.08061 | -0.88182 | -0.25136 | 0.164573 |
| 2 | 168889200 | 168901599 | 113.2667 | 170.0162 | 323.7171 | 133.5941 | -0.66113 | 3.4502 | 0.164578 |
| X | 25851200 | 25856599 | 94.4049 | 149.1396 | 98.80536 | 41.03566 | 0.919042 | 2.44548 | 0.164582 |
| 13 | 68498800 | 68503399 | 96.42065 | 150.9802 | 124.1613 | 51.50368 | 0.597797 | 2.622493 | 0.164587 |
| 7 | 25973000 | 25973999 | 25.27614 | 43.96385 | 32.22585 | 14.15865 | 0.833025 | 0.798323 | 0.164592 |
| 4 | 145225200 | 145228999 | 57.12402 | 92.09265 | 64.8789 | 27.62137 | 0.86095 | 1.802829 | 0.164599 |
| 5 | 93618600 | 93620799 | 33.03382 | 55.68242 | 37.79913 | 16.52597 | 0.937851 | 1.121104 | 0.164621 |
| 4 | 11469200 | 11469799 | 2.940106 | 5.737911 | 21.97099 | 7.806214 | -1.25269 | -0.48979 | 0.164639 |
| 12 | 69462000 | 69466799 | 26.39585 | 42.90403 | 89.33363 | 38.3301 | -0.68661 | 1.572863 | 0.164672 |
| 6 | 70538800 | 70539599 | 11.94419 | 23.02549 | 17.32385 | 6.70947 | 0.949423 | -0.10217 | 0.164676 |
| 12 | 44846600 | 44847199 | 11.94419 | 23.02549 | 17.32385 | 6.70947 | 0.949423 | -0.10217 | 0.164676 |
| 7 | 10318800 | 10322199 | 51.1446 | 83.07653 | 51.7641 | 22.19956 | 1.048492 | 1.611857 | 0.164678 |
| 8 | 71719400 | 71722199 | 16.74653 | 28.34763 | 62.79028 | 27.48738 | -0.76164 | 1.05605 | 0.164686 |
| 3 | 18842400 | 18844799 | 22.19404 | 38.89154 | 29.55446 | 13.05386 | 0.781567 | 0.670156 | 0.164692 |
| 17 | 3164600 | 3166799 | 13.90744 | 24.14062 | 52.68989 | 23.12008 | -0.73409 | 0.805107 | 0.164705 |
| 1 | 166338200 | 166339199 | 10.83112 | 21.00958 | 15.13148 | 5.704936 | 1.046977 | -0.21039 | 0.164706 |
| 19 | 44271000 | 44273599 | 11.21561 | 19.66547 | 46.19912 | 20.41603 | -0.8381 | 0.630688 | 0.164722 |
| 16 | 58343000 | 58344199 | 9.758967 | 17.35494 | 40.96125 | 18.06112 | -0.83677 | 0.457584 | 0.164727 |
| 7 | 62493200 | 62496199 | 38.06988 | 63.01557 | 44.49645 | 19.24755 | 0.881054 | 1.330763 | 0.164739 |
| 5 | 106941200 | 106942599 | 6.882029 | 12.72469 | 34.69549 | 14.62376 | -0.99773 | 0.19036 | 0.164747 |
| 4 | 133983200 | 133996599 | 79.98234 | 122.7416 | 235.2657 | 97.67166 | -0.64409 | 2.992143 | 0.164765 |

| | | | | | | | | | |
|---|---|---|---|---|---|---|---|---|---|
| 12 | 9720400 | 9721799 | 15.98227 | 30.51251 | 15.57099 | 6.19583 | 1.467939 | 0.018355 | 0.164793 |
| 7 | 48028600 | 48031999 | 48.12596 | 78.62522 | 51.24544 | 22.02897 | 0.989667 | 1.520168 | 0.164817 |
| 12 | 22317600 | 22317999 | 9.018462 | 17.57046 | 13.003 | 4.730375 | 1.056504 | -0.3603 | 0.164817 |
| 7 | 94275600 | 94278199 | 32.40047 | 55.3465 | 33.6028 | 14.7328 | 1.099363 | 0.998925 | 0.16483 |
| 7 | 60883600 | 60888799 | 94.01881 | 147.0581 | 123.4173 | 51.20083 | 0.567311 | 2.640627 | 0.164836 |
| 15 | 31408800 | 31419599 | 77.23255 | 118.8727 | 230.5 | 95.73196 | -0.6575 | 2.932316 | 0.164845 |
| 17 | 41293600 | 41294599 | 11.965 | 22.97783 | 17.88392 | 7.002632 | 0.886553 | -0.06557 | 0.164846 |
| 1 | 184936000 | 184936799 | 4.673136 | 9.031193 | 26.99785 | 10.3564 | -1.00763 | -0.18076 | 0.164857 |
| 8 | 72620400 | 72621799 | 14.50579 | 27.08609 | 20.66306 | 8.580758 | 0.842311 | 0.160096 | 0.164863 |
| 14 | 53835600 | 53836999 | 8.066365 | 14.8735 | 36.29843 | 15.35678 | -0.84363 | 0.21802 | 0.164868 |
| 16 | 13353400 | 13357799 | 23.29117 | 38.08 | 85.03475 | 36.56395 | -0.78116 | 1.477434 | 0.164871 |
| 5 | 66018000 | 66022399 | 20.20788 | 33.63657 | 75.09917 | 32.61053 | -0.77384 | 1.242506 | 0.164872 |
| 11 | 18758000 | 18758799 | 29.31754 | 49.64913 | 39.99888 | 17.51246 | 0.691969 | 1.071222 | 0.164873 |
| 7 | 104304800 | 104307999 | 51.94464 | 84.2112 | 57.84711 | 24.77258 | 0.90566 | 1.656281 | 0.164885 |
| 12 | 22913600 | 22914599 | 23.58262 | 41.40257 | 27.4571 | 12.14675 | 0.977095 | 0.647036 | 0.164893 |
| 14 | 49839200 | 49839999 | 5.225476 | 10.13795 | 28.65901 | 10.7958 | -0.90383 | -0.23333 | 0.164896 |
| 7 | 14672200 | 14674199 | 45.89768 | 74.52398 | 59.83605 | 25.65375 | 0.683192 | 1.613801 | 0.164898 |
| 5 | 129828400 | 129830399 | 7.936272 | 14.60334 | 35.68743 | 15.1409 | -0.84797 | 0.23022 | 0.164904 |
| 18 | 34195200 | 34197599 | 12.28295 | 21.53204 | 48.31305 | 21.34609 | -0.77256 | 0.642008 | 0.164904 |
| 1 | 87717000 | 87719999 | 13.40114 | 22.92727 | 57.85245 | 25.36796 | -0.9465 | 0.942867 | 0.164905 |
| 3 | 16682400 | 16683999 | 25.01887 | 43.74847 | 27.03181 | 11.91819 | 1.078253 | 0.712789 | 0.164915 |
| 12 | 109347200 | 109347999 | 24.02538 | 41.76928 | 32.33478 | 14.20244 | 0.754533 | 0.797873 | 0.164915 |
| 7 | 36134200 | 36137599 | 15.99821 | 27.47993 | 60.88188 | 26.70448 | -0.75923 | 0.917757 | 0.164923 |
| 6 | 68829200 | 68830999 | 17.67197 | 32.8674 | 18.48772 | 7.729209 | 1.270326 | 0.205687 | 0.164923 |
| 13 | 99480800 | 99481599 | 4.566385 | 8.613016 | 31.74559 | 12.95939 | -1.3886 | 0.042008 | 0.16497 |
| 12 | 88644800 | 88646199 | 14.27612 | 27.17214 | 18.12718 | 7.259681 | 1.07775 | 0.029258 | 0.164973 |
| 2 | 96347200 | 96347999 | 11.88377 | 23.17096 | 10.22185 | 3.680797 | 1.801315 | -0.36746 | 0.164978 |
| 7 | 11147800 | 11151399 | 66.20198 | 106.0132 | 78.29465 | 33.00464 | 0.781593 | 2.026559 | 0.164987 |
| 19 | 55918400 | 55920399 | 19.80298 | 32.79939 | 73.99312 | 32.02043 | -0.78915 | 1.30095 | 0.164995 |
| 6 | 72163200 | 72166799 | 12.83486 | 22.24003 | 52.63618 | 23.09385 | -0.85071 | 0.816923 | 0.165017 |
| 4 | 153677600 | 153709399 | 262.6849 | 368.8514 | 651.0038 | 269.765 | -0.64511 | 4.516785 | 0.165038 |
| 16 | 20780000 | 20780399 | 2.735746 | 5.351158 | 19.02405 | 6.380646 | -1.06884 | -0.70775 | 0.165039 |
| 13 | 95148200 | 95149799 | 6.697365 | 12.47835 | 33.22289 | 13.84779 | -0.95115 | 0.14061 | 0.165041 |
| 16 | 17539400 | 17540799 | 6.697365 | 12.47835 | 33.22289 | 13.84779 | -0.95115 | 0.14061 | 0.165041 |
| 6 | 29167200 | 29171799 | 23.42449 | 38.01396 | 94.39736 | 40.4554 | -0.94217 | 1.600969 | 0.165056 |
| 12 | 12989000 | 12993199 | 23.42449 | 38.01396 | 94.39736 | 40.4554 | -0.94217 | 1.600969 | 0.165056 |
| 14 | 82584600 | 82585799 | 16.03981 | 29.89013 | 19.81564 | 8.257019 | 1.039093 | 0.183 | 0.16506 |
| 17 | 57005400 | 57012399 | 30.61878 | 49.14183 | 115.5597 | 48.88001 | -0.88203 | 1.926759 | 0.16508 |
| 12 | 56288200 | 56289399 | 13.18017 | 24.98797 | 15.94241 | 6.437906 | 1.131489 | 0.038753 | 0.165081 |
| 1 | 35848200 | 35851399 | 15.37274 | 25.97966 | 64.86231 | 28.38038 | -0.9337 | 1.067155 | 0.165098 |
| 18 | 3196400 | 3196999 | 20.2862 | 35.74416 | 24.8153 | 10.98375 | 0.912488 | 0.577449 | 0.165117 |
| 5 | 21906200 | 21907399 | 5.466804 | 10.39613 | 30.40983 | 12.20099 | -1.03853 | -0.01547 | 0.165122 |
| 13 | 38124000 | 38127799 | 12.87729 | 22.20723 | 59.12694 | 25.93786 | -1.02174 | 0.865574 | 0.165132 |
| 16 | 28895600 | 28896199 | 9.459466 | 18.48757 | 10.68122 | 3.693086 | 1.469732 | -0.53683 | 0.16514 |
| 2 | 85849200 | 85853199 | 23.69743 | 38.92331 | 82.16206 | 35.39527 | -0.69703 | 1.416179 | 0.16515 |
| 18 | 12953600 | 12954599 | 12.1567 | 23.38763 | 17.20013 | 6.700505 | 0.973788 | -0.08277 | 0.165153 |
| 10 | 115422000 | 115425199 | 15.83375 | 26.43098 | 72.85627 | 31.69357 | -1.07646 | 1.197846 | 0.165172 |

| | | | | | | | | | |
|---|---|---|---|---|---|---|---|---|---|
| X | 91137400 | 91139399 | 18.93573 | 35.154 | 17.84661 | 7.480735 | 1.414416 | 0.225227 | 0.165177 |
| 15 | 96919400 | 96920399 | 10.58237 | 20.57624 | 15.35084 | 5.690365 | 1.020017 | -0.26822 | 0.165183 |
| 19 | 29505000 | 29507399 | 26.12307 | 45.47446 | 31.21724 | 13.72263 | 0.927422 | 0.791282 | 0.165194 |
| 2 | 93089800 | 93092799 | 13.94275 | 24.06532 | 54.25087 | 23.79944 | -0.78225 | 0.859008 | 0.165205 |
| 15 | 83357000 | 83361999 | 24.17591 | 39.30128 | 90.81136 | 38.96658 | -0.83623 | 1.5659 | 0.165209 |
| 2 | 129427600 | 129428399 | 3.321664 | 6.460186 | 28.13119 | 10.40733 | -1.49394 | -0.32375 | 0.165215 |
| 9 | 62664000 | 62669199 | 28.42619 | 45.88581 | 101.7063 | 43.37138 | -0.78671 | 1.739118 | 0.165222 |
| 18 | 62041200 | 62045199 | 17.96949 | 30.04371 | 69.23406 | 30.14798 | -0.81912 | 1.184274 | 0.165231 |
| 9 | 26296400 | 26297599 | 12.26311 | 23.30724 | 18.25049 | 7.331286 | 0.844252 | 0.018171 | 0.165246 |
| 1 | 184609600 | 184611799 | 12.44145 | 21.80774 | 48.31491 | 21.34513 | -0.75435 | 0.644765 | 0.165246 |
| 7 | 51876000 | 51876599 | 14.82868 | 27.55024 | 21.14653 | 8.854789 | 0.824138 | 0.194844 | 0.165253 |
| 8 | 113722000 | 113723799 | 15.59804 | 28.94297 | 20.98175 | 8.800938 | 0.904067 | 0.207365 | 0.165256 |
| X | 78406800 | 78409199 | 23.06013 | 41.21239 | 21.74184 | 9.546796 | 1.315574 | 0.475382 | 0.165261 |
| 2 | 32177400 | 32178799 | 5.77473 | 10.9252 | 31.38897 | 12.73201 | -1.02718 | 0.025838 | 0.165268 |
| 9 | 121661000 | 121668599 | 71.76043 | 110.9095 | 211.8407 | 87.99386 | -0.63113 | 2.85163 | 0.165296 |
| 2 | 27556800 | 27563199 | 30.73787 | 49.421 | 110.6077 | 46.90924 | -0.80768 | 1.868433 | 0.165315 |
| X | 55372800 | 55374799 | 35.17369 | 59.19214 | 39.41441 | 17.2236 | 0.965683 | 1.146825 | 0.165346 |
| 18 | 80330600 | 80339799 | 62.76131 | 97.49548 | 202.4594 | 84.067 | -0.7459 | 2.755032 | 0.165365 |
| 6 | 23271600 | 23273399 | 8.556834 | 15.67198 | 43.9665 | 18.77214 | -1.05353 | 0.275412 | 0.165383 |
| 13 | 75089400 | 75091799 | 34.32007 | 58.22569 | 35.83393 | 15.69752 | 1.078878 | 1.068993 | 0.165384 |
| 7 | 9149200 | 9156799 | 143.874 | 218.7334 | 181.6035 | 75.22834 | 0.551176 | 3.193778 | 0.165391 |
| 2 | 84877000 | 84882799 | 23.86395 | 38.49699 | 106.1802 | 45.24223 | -1.10223 | 1.752972 | 0.165395 |
| 2 | 44499600 | 44502399 | 25.52785 | 41.74105 | 87.06545 | 37.43849 | -0.68301 | 1.483873 | 0.165406 |
| 12 | 109407200 | 109410799 | 17.19398 | 28.74318 | 69.11715 | 30.09448 | -0.88022 | 1.183722 | 0.165411 |
| 6 | 86269200 | 86271799 | 16.3495 | 27.91707 | 60.77283 | 26.64435 | -0.73535 | 0.973281 | 0.165428 |
| 2 | 153187200 | 153188599 | 7.200688 | 13.2409 | 35.67028 | 15.14748 | -0.98857 | 0.22629 | 0.165431 |
| 9 | 63608600 | 63610799 | 11.71812 | 20.42273 | 49.96258 | 21.97909 | -0.89694 | 0.732505 | 0.165433 |
| 8 | 3461200 | 3466799 | 28.23997 | 45.6476 | 98.6159 | 42.1278 | -0.74748 | 1.697267 | 0.165433 |
| 12 | 60323800 | 60325599 | 17.12758 | 32.17696 | 17.58221 | 7.226487 | 1.331197 | 0.13786 | 0.165435 |
| 8 | 49295600 | 49298399 | 19.94196 | 32.69454 | 82.28758 | 35.42168 | -0.95085 | 1.432265 | 0.165442 |
| 6 | 38098000 | 38103999 | 29.85377 | 48.19812 | 103.5085 | 44.14927 | -0.74103 | 1.736614 | 0.165454 |
| 13 | 117561600 | 117563199 | 15.65172 | 26.51842 | 62.72831 | 27.46781 | -0.85594 | 1.044011 | 0.165467 |
| 8 | 107858000 | 107860799 | 10.41953 | 18.36104 | 45.09648 | 19.9602 | -0.90044 | 0.554517 | 0.165473 |
| 12 | 21888600 | 21892399 | 101.8374 | 158.4497 | 129.4704 | 53.74706 | 0.599962 | 2.736939 | 0.165476 |
| 12 | 30633200 | 30635599 | 11.86577 | 20.69227 | 49.43165 | 21.75361 | -0.86267 | 0.724214 | 0.165478 |
| 10 | 76923000 | 76934999 | 80.2102 | 122.8142 | 240.7052 | 99.89186 | -0.67869 | 3.038066 | 0.165484 |
| 15 | 22268200 | 22269999 | 25.28339 | 44.55172 | 23.02687 | 10.19204 | 1.335155 | 0.602487 | 0.165497 |
| 19 | 44806400 | 44808799 | 9.807337 | 17.31108 | 43.54445 | 19.26684 | -0.9334 | 0.53061 | 0.165508 |
| 14 | 124877800 | 124878999 | 15.2521 | 28.92423 | 17.97 | 7.252372 | 1.169901 | 0.062105 | 0.165513 |
| 2 | 61916400 | 61917599 | 8.618585 | 16.82885 | 12.03686 | 4.238196 | 1.143798 | -0.48427 | 0.165528 |
| 9 | 96679400 | 96680399 | 4.52777 | 8.803311 | 25.80567 | 9.569677 | -0.9338 | -0.30614 | 0.165532 |
| 13 | 59916200 | 59919399 | 14.46894 | 24.77462 | 58.50737 | 25.65685 | -0.85147 | 0.93828 | 0.165538 |
| 2 | 151599000 | 151601799 | 15.90305 | 27.10249 | 60.13417 | 26.36025 | -0.76306 | 0.989214 | 0.165541 |
| 3 | 121072600 | 121074199 | 8.711547 | 15.95871 | 37.76945 | 16.12158 | -0.80893 | 0.265013 | 0.165545 |
| 19 | 6789800 | 6796799 | 151.5761 | 221.7667 | 431.4163 | 177.7219 | -0.72648 | 3.899921 | 0.165558 |
| 13 | 66876600 | 66878799 | 40.27005 | 66.02897 | 53.27327 | 22.93313 | 0.685511 | 1.454846 | 0.165575 |
| 4 | 114434600 | 114442199 | 38.37794 | 61.36526 | 127.7326 | 53.72976 | -0.71292 | 2.05517 | 0.165593 |

| | | | | | | | | | |
|---|---|---|---|---|---|---|---|---|---|
| 5 | 138984600 | 138993999 | 91.08087 | 139.1077 | 254.9597 | 105.7458 | -0.58499 | 3.092547 | 0.1656 |
| 6 | 123834600 | 123835199 | 10.49519 | 20.45791 | 12.9326 | 4.673631 | 1.287355 | -0.36501 | 0.165614 |
| 15 | 4731000 | 4732599 | 21.37593 | 37.63857 | 27.52447 | 12.18455 | 0.837584 | 0.596658 | 0.165626 |
| 10 | 47250400 | 47254599 | 52.59259 | 85.00638 | 62.96435 | 26.89565 | 0.793035 | 1.721981 | 0.165631 |
| 4 | 156286600 | 156291199 | 27.2944 | 44.10969 | 100.7789 | 43.0399 | -0.82839 | 1.702536 | 0.165633 |
| 12 | 19431400 | 19433599 | 32.54336 | 54.87618 | 37.8299 | 16.5413 | 0.915742 | 1.116329 | 0.165643 |
| 15 | 82867600 | 82869399 | 6.564633 | 12.20616 | 33.71565 | 14.10172 | -1.00787 | 0.153528 | 0.165646 |
| 17 | 92561000 | 92562999 | 24.85343 | 43.52516 | 28.23433 | 12.46434 | 1.008715 | 0.692377 | 0.165658 |
| 17 | 20204200 | 20205399 | 17.92087 | 33.0556 | 20.13074 | 8.5127 | 1.145304 | 0.256765 | 0.165684 |
| 7 | 31908600 | 31911199 | 52.04811 | 84.10983 | 63.51323 | 27.12325 | 0.764952 | 1.726274 | 0.165685 |
| 11 | 114271200 | 114276199 | 20.09893 | 33.33473 | 75.40959 | 32.66396 | -0.79311 | 1.284969 | 0.165699 |
| 7 | 65098800 | 65099399 | 14.35099 | 26.68948 | 20.74241 | 8.672577 | 0.808004 | 0.184257 | 0.1657 |
| X | 33855600 | 33857999 | 26.09075 | 45.8463 | 24.94782 | 11.04489 | 1.260652 | 0.643331 | 0.165712 |
| 6 | 3267200 | 3270599 | 50.50689 | 81.46443 | 66.83426 | 28.4717 | 0.642538 | 1.78092 | 0.165712 |
| X | 43649600 | 43651399 | 19.60473 | 36.53969 | 16.81414 | 7.00426 | 1.561924 | 0.206047 | 0.165715 |
| 6 | 38603000 | 38605399 | 12.93335 | 22.58214 | 49.43787 | 21.77341 | -0.73739 | 0.711334 | 0.165735 |
| 2 | 150382000 | 150382599 | 9.291636 | 18.13998 | 12.56067 | 4.4356 | 1.186903 | -0.46148 | 0.165737 |
| 15 | 7152000 | 7152599 | 9.291636 | 18.13998 | 12.56067 | 4.4356 | 1.186903 | -0.46148 | 0.165737 |
| 8 | 109440400 | 109442999 | 8.097744 | 14.43647 | 40.5139 | 17.82654 | -1.08224 | 0.43076 | 0.165741 |
| 10 | 40017200 | 40018599 | 8.097744 | 14.43647 | 40.5139 | 17.82654 | -1.08224 | 0.43076 | 0.165741 |
| 7 | 138446800 | 138449199 | 11.79005 | 20.36777 | 54.16468 | 23.76227 | -1.01885 | 0.84215 | 0.165753 |
| 17 | 19574000 | 19577999 | 115.9026 | 178.7197 | 149.6424 | 62.15883 | 0.554851 | 2.903981 | 0.165765 |
| 8 | 105863600 | 105864999 | 8.042695 | 14.93509 | 35.93996 | 15.05954 | -0.81329 | 0.177617 | 0.165768 |
| 12 | 23396200 | 23396799 | 9.475213 | 18.45559 | 14.92735 | 5.448774 | 0.924623 | -0.33634 | 0.165769 |
| 15 | 100697000 | 100703199 | 35.32377 | 56.51552 | 124.4121 | 52.37737 | -0.7925 | 2.032995 | 0.165778 |
| 4 | 135232600 | 135239599 | 29.1233 | 46.8815 | 109.0818 | 46.34291 | -0.86141 | 1.826936 | 0.165786 |
| 2 | 167521200 | 167525799 | 21.27048 | 35.08876 | 77.4517 | 33.42729 | -0.75861 | 1.353583 | 0.165788 |
| 2 | 119167200 | 119168999 | 7.095216 | 12.95607 | 37.14109 | 15.92023 | -1.08726 | 0.268876 | 0.16579 |
| 7 | 48002400 | 48003799 | 36.20513 | 59.91191 | 48.79407 | 21.10241 | 0.675865 | 1.333174 | 0.165806 |
| 7 | 92705400 | 92706199 | 4.969921 | 9.591507 | 27.51917 | 10.62395 | -0.95838 | -0.15478 | 0.165809 |
| 6 | 73033600 | 73035199 | 43.59823 | 71.07361 | 56.62803 | 24.32353 | 0.69855 | 1.542844 | 0.165811 |
| 8 | 106019000 | 106020599 | 7.23743 | 13.12224 | 37.57285 | 16.24898 | -1.09323 | 0.316676 | 0.165819 |
| X | 24241400 | 24243599 | 39.65982 | 66.02446 | 42.74156 | 18.5615 | 1.006556 | 1.272369 | 0.16582 |
| 11 | 73137800 | 73139199 | 16.18538 | 29.57399 | 22.59896 | 9.67908 | 0.807617 | 0.300828 | 0.165848 |
| 3 | 54124000 | 54124999 | 14.25826 | 26.57827 | 20.63375 | 8.593511 | 0.813745 | 0.167929 | 0.165871 |
| 18 | 20836800 | 20838199 | 6.620715 | 12.57585 | 33.90393 | 13.63643 | -0.92573 | 0.013499 | 0.165888 |
| 5 | 13714000 | 13715399 | 17.68769 | 32.79948 | 19.7242 | 8.280643 | 1.170905 | 0.225781 | 0.165909 |
| 5 | 123633400 | 123639799 | 34.90228 | 55.68891 | 131.0413 | 54.9553 | -0.89278 | 2.120643 | 0.165909 |
| 7 | 22105800 | 22106799 | 13.97691 | 27.15104 | 11.16192 | 4.169097 | 1.856754 | -0.21533 | 0.165918 |
| 9 | 41067600 | 41072999 | 31.49159 | 50.7334 | 107.1869 | 45.58935 | -0.72119 | 1.80458 | 0.165919 |
| 9 | 54857800 | 54858799 | 4.730277 | 9.079619 | 28.99552 | 11.36965 | -1.13284 | -0.10281 | 0.165921 |
| 9 | 21980800 | 21982599 | 9.491317 | 16.79176 | 42.50783 | 18.79954 | -0.94106 | 0.501536 | 0.165927 |
| 4 | 40826200 | 40830399 | 23.64595 | 38.56513 | 87.07406 | 37.4073 | -0.79958 | 1.515337 | 0.165944 |
| 9 | 21665400 | 21669399 | 21.03003 | 34.51094 | 81.49228 | 35.09333 | -0.85867 | 1.421724 | 0.165947 |
| 8 | 92608200 | 92610199 | 7.389658 | 13.46056 | 36.57854 | 15.72745 | -1.01342 | 0.285737 | 0.165949 |
| 7 | 23154800 | 23156799 | 43.07368 | 70.54577 | 53.30381 | 22.93525 | 0.779304 | 1.477297 | 0.165956 |
| 6 | 34153200 | 34154199 | 6.385358 | 11.98099 | 33.41282 | 13.76208 | -1.00392 | 0.090699 | 0.165962 |

| | | | | | | | | | |
|---|---|---|---|---|---|---|---|---|---|
| 7 | 31356000 | 31357999 | 30.80939 | 53.3712 | 29.93092 | 13.13192 | 1.216896 | 0.869062 | 0.165988 |
| 13 | 79779400 | 79780199 | 14.20109 | 27.41995 | 14.73727 | 5.67711 | 1.436402 | -0.10255 | 0.165993 |
| 11 | 52273200 | 52276399 | 12.60702 | 21.67338 | 58.47715 | 25.65036 | -1.04162 | 0.893316 | 0.165998 |
| 4 | 17972200 | 17972799 | 9.816557 | 19.06749 | 15.36036 | 5.741626 | 0.898941 | -0.25034 | 0.165998 |
| 8 | 93686400 | 93689399 | 20.9797 | 34.5464 | 78.34364 | 33.78631 | -0.7985 | 1.376703 | 0.166006 |
| 12 | 102020800 | 102023999 | 12.78778 | 22.27093 | 51.05905 | 22.45128 | -0.80413 | 0.748613 | 0.16601 |
| 2 | 52747600 | 52749799 | 11.29767 | 19.62754 | 53.14991 | 23.33463 | -1.04332 | 0.780032 | 0.166018 |
| 17 | 57765200 | 57766999 | 26.14627 | 45.13591 | 35.0487 | 15.37567 | 0.748576 | 0.883486 | 0.166018 |
| 12 | 81036200 | 81040199 | 24.16219 | 39.27934 | 88.8361 | 38.11913 | -0.8053 | 1.566328 | 0.166023 |
| 5 | 28328200 | 28340599 | 72.78923 | 112.1577 | 234.3209 | 97.33321 | -0.76377 | 2.909017 | 0.166052 |
| 12 | 9748200 | 9749599 | 12.9434 | 24.45826 | 19.22047 | 7.815995 | 0.823935 | 0.066556 | 0.166053 |
| 1 | 36178400 | 36183999 | 24.66057 | 40.05254 | 90.61777 | 38.85895 | -0.8071 | 1.585451 | 0.166062 |
| 9 | 70374600 | 70376399 | 22.65852 | 40.02917 | 24.8954 | 11.01343 | 1.071275 | 0.562725 | 0.166072 |
| 19 | 42685600 | 42689599 | 17.67768 | 29.19249 | 79.85052 | 34.48445 | -1.06656 | 1.333173 | 0.16609 |
| 15 | 75650800 | 75651399 | 4.250216 | 8.226304 | 26.4775 | 10.08525 | -1.10273 | -0.21137 | 0.16609 |
| 2 | 166340000 | 166356999 | 145.6778 | 214.8047 | 401.617 | 165.3911 | -0.65858 | 3.773958 | 0.166094 |
| 18 | 3398000 | 3403399 | 69.28121 | 110.8062 | 79.72741 | 33.54852 | 0.817186 | 2.067424 | 0.166104 |
| 17 | 18620400 | 18620999 | 16.37047 | 29.90289 | 22.59495 | 9.681124 | 0.823401 | 0.304157 | 0.166108 |
| X | 116117600 | 116120799 | 30.21366 | 51.84065 | 32.29024 | 14.16238 | 1.063575 | 0.950758 | 0.166109 |
| 8 | 122114200 | 122116399 | 8.04113 | 14.1311 | 45.66443 | 20.21815 | -1.29425 | 0.576511 | 0.166126 |
| 6 | 40693600 | 40695199 | 19.07409 | 34.39693 | 23.12722 | 10.06959 | 0.975794 | 0.396323 | 0.166147 |
| 18 | 5986600 | 5990199 | 25.29346 | 41.09211 | 92.15649 | 39.52706 | -0.79416 | 1.580584 | 0.166148 |
| 8 | 31579600 | 31584399 | 27.0247 | 43.58199 | 105.7931 | 45.05878 | -0.91913 | 1.766511 | 0.166171 |
| 3 | 8249000 | 8251199 | 26.27533 | 45.14884 | 36.18077 | 15.87006 | 0.702016 | 0.925937 | 0.166171 |
| 18 | 64788800 | 64795399 | 38.34076 | 61.44702 | 119.6741 | 50.50995 | -0.61333 | 1.982081 | 0.166173 |
| 9 | 35121200 | 35122999 | 6.946671 | 12.70966 | 38.25291 | 16.35585 | -1.15479 | 0.257593 | 0.166213 |
| 15 | 38562600 | 38564999 | 18.10884 | 32.22349 | 24.59573 | 10.83951 | 0.782557 | 0.473324 | 0.166227 |
| 3 | 48144800 | 48146399 | 16.95365 | 31.42352 | 18.42676 | 7.740898 | 1.205333 | 0.217234 | 0.166243 |
| 15 | 97142800 | 97145799 | 22.04494 | 35.72134 | 97.80622 | 41.8698 | -1.08469 | 1.631348 | 0.166247 |
| 14 | 63307400 | 63309799 | 10.55042 | 18.64449 | 44.04137 | 19.48215 | -0.84287 | 0.52198 | 0.166249 |
| 14 | 32577000 | 32583799 | 37.17234 | 59.27011 | 134.1712 | 56.21422 | -0.83819 | 2.144799 | 0.166262 |
| X | 93349000 | 93350599 | 6.457797 | 12.31284 | 31.1645 | 12.43481 | -0.82532 | -0.02267 | 0.166271 |
| 7 | 37452000 | 37452599 | 4.658059 | 9.049929 | 25.69282 | 9.584961 | -0.89587 | -0.2824 | 0.166284 |
| 12 | 19160200 | 19182799 | 775.2079 | 1028.743 | 741.283 | 309.8239 | 0.564948 | 5.402849 | 0.166286 |
| 4 | 8891400 | 8893799 | 16.32987 | 27.97595 | 60.28339 | 26.43681 | -0.71994 | 0.942627 | 0.166312 |
| 7 | 23496200 | 23500999 | 110.1852 | 170.7471 | 141.086 | 58.60317 | 0.57937 | 2.809902 | 0.166321 |
| 11 | 49426200 | 49427799 | 6.340174 | 11.982 | 31.36336 | 12.75075 | -0.89667 | 0.040828 | 0.166324 |
| 2 | 30358600 | 30365599 | 29.26427 | 46.87125 | 126.975 | 53.54052 | -1.08934 | 1.993065 | 0.166327 |
| 16 | 48706200 | 48707399 | 14.37088 | 26.96327 | 20.17375 | 8.309753 | 0.879434 | 0.128098 | 0.166334 |
| 17 | 69687400 | 69691799 | 31.30979 | 50.48999 | 108.3716 | 46.15414 | -0.74228 | 1.774643 | 0.166336 |
| 1 | 38981400 | 38985599 | 18.52685 | 31.17565 | 66.13956 | 28.90151 | -0.70019 | 1.117173 | 0.166344 |
| 11 | 93607200 | 93608199 | 14.16265 | 26.36492 | 19.32879 | 8.068187 | 0.893523 | 0.173074 | 0.166346 |
| 15 | 48854200 | 48855599 | 19.5847 | 35.31073 | 21.8749 | 9.52669 | 1.092618 | 0.397571 | 0.166355 |
| 13 | 73401800 | 73417799 | 131.9074 | 195.1471 | 397.3361 | 163.634 | -0.777 | 3.717095 | 0.166356 |
| 5 | 105570000 | 105570799 | 12.4046 | 23.94575 | 16.6904 | 6.436378 | 1.064202 | -0.1129 | 0.16637 |
| 10 | 57669800 | 57671599 | 9.464227 | 16.80864 | 40.96816 | 18.08422 | -0.88382 | 0.46478 | 0.166387 |
| 18 | 35251000 | 35252799 | 9.963575 | 17.56822 | 43.55038 | 19.27354 | -0.91302 | 0.543941 | 0.166392 |

| | | | | | | | | | |
|---|---|---|---|---|---|---|---|---|---|
| 15 | 9403400 | 9404599 | 14.32231 | 27.15249 | 16.65794 | 6.727286 | 1.185726 | 0.049786 | 0.166397 |
| 7 | 7623800 | 7625799 | 49.67997 | 80.31839 | 57.64608 | 24.63172 | 0.83875 | 1.706412 | 0.166398 |
| 17 | 18357600 | 18361599 | 73.24355 | 116.6282 | 91.82311 | 38.40025 | 0.680748 | 2.210242 | 0.166401 |
| 10 | 129733200 | 129735399 | 25.96623 | 45.33426 | 29.32662 | 12.91534 | 1.012755 | 0.746088 | 0.166404 |
| 12 | 107991800 | 107992399 | 12.19925 | 23.15865 | 16.3097 | 6.567957 | 0.992873 | 0.019872 | 0.166409 |
| 1 | 48817800 | 48820199 | 20.59652 | 36.77576 | 24.04311 | 10.56545 | 1.007321 | 0.467976 | 0.166411 |
| 15 | 76052600 | 76086399 | 712.6092 | 909.6639 | 1809.898 | 759.5432 | -0.95612 | 6.031599 | 0.166444 |
| 7 | 8330200 | 8341399 | 61.98613 | 99.35423 | 70.80102 | 29.88788 | 0.83457 | 1.990104 | 0.166452 |
| 2 | 170198000 | 170202199 | 39.99369 | 63.98986 | 123.3272 | 51.92383 | -0.60104 | 2.03708 | 0.166453 |
| 14 | 106132200 | 106136999 | 26.47648 | 42.56641 | 116.1595 | 49.25867 | -1.09313 | 1.866516 | 0.166472 |
| 6 | 82860800 | 82868799 | 42.27329 | 67.48797 | 136.3354 | 57.22132 | -0.6724 | 2.109562 | 0.166478 |
| 3 | 90290200 | 90299399 | 38.0506 | 60.23313 | 164.7118 | 68.47764 | -1.12081 | 2.368338 | 0.166479 |
| 17 | 78571800 | 78574599 | 12.92805 | 22.63398 | 50.48323 | 22.28089 | -0.76407 | 0.670329 | 0.166486 |
| 4 | 119619200 | 119620599 | 8.86594 | 16.20387 | 37.71294 | 16.14738 | -0.78823 | 0.278708 | 0.166492 |
| 5 | 108512400 | 108523199 | 75.96988 | 116.8576 | 234.5912 | 97.42293 | -0.70817 | 2.94412 | 0.166505 |
| 1 | 132058800 | 132060399 | 43.36597 | 68.94493 | 140.8786 | 58.7939 | -0.69526 | 2.246706 | 0.166508 |
| 11 | 114741600 | 114744599 | 14.14556 | 23.89138 | 64.13687 | 28.05785 | -1.03771 | 1.069007 | 0.166511 |
| 2 | 179704800 | 179709199 | 34.20381 | 54.70828 | 123.3476 | 51.90853 | -0.8273 | 2.041771 | 0.166513 |
| X | 120274800 | 120277399 | 35.9865 | 60.39187 | 40.93933 | 17.86907 | 0.939841 | 1.174307 | 0.166536 |
| X | 5764000 | 5765599 | 16.1628 | 29.96135 | 20.96619 | 8.806225 | 0.953422 | 0.217846 | 0.166536 |
| X | 113319400 | 113322799 | 59.5077 | 95.64519 | 71.70552 | 30.39563 | 0.766226 | 1.903977 | 0.16654 |
| 6 | 52179800 | 52184599 | 26.49602 | 42.81347 | 98.49514 | 42.05698 | -0.83846 | 1.705605 | 0.166548 |
| 3 | 142410600 | 142411999 | 24.23954 | 42.91156 | 22.58082 | 9.985321 | 1.311878 | 0.547148 | 0.166553 |
| 17 | 4877600 | 4878399 | 11.21953 | 21.87681 | 12.30052 | 4.426722 | 1.459714 | -0.37092 | 0.166558 |
| 2 | 143932400 | 143934399 | 9.685579 | 17.4784 | 40.1043 | 17.44632 | -0.78305 | 0.36351 | 0.16656 |
| 12 | 3991800 | 3993599 | 9.118334 | 16.32218 | 39.47774 | 17.31117 | -0.86601 | 0.409389 | 0.166564 |
| 5 | 143176800 | 143182999 | 35.30449 | 56.52594 | 124.4946 | 52.46686 | -0.79204 | 2.009798 | 0.166567 |
| 9 | 20635600 | 20642599 | 39.2546 | 62.58555 | 137.1366 | 57.45074 | -0.79148 | 2.149061 | 0.166575 |
| Y | 311200 | 311599 | 6.330076 | 12.36012 | 4.762804 | 1.506063 | 2.147327 | -0.92514 | 0.166596 |
| 10 | 118143600 | 118146999 | 17.05509 | 28.47605 | 70.17446 | 30.53121 | -0.91546 | 1.19745 | 0.1666 |
| 18 | 11972000 | 11980799 | 45.2687 | 71.6036 | 156.5589 | 65.06256 | -0.79958 | 2.391881 | 0.16663 |
| 5 | 117751800 | 117773599 | 150.2637 | 220.5981 | 423.7003 | 174.506 | -0.70338 | 3.845371 | 0.166633 |
| 7 | 98391600 | 98397799 | 36.91983 | 58.72518 | 143.0476 | 59.73281 | -0.94753 | 2.225852 | 0.166638 |
| 4 | 135527200 | 135544399 | 132.2231 | 196.4954 | 361.9995 | 149.1327 | -0.62676 | 3.638959 | 0.166653 |
| 1 | 94163200 | 94164799 | 14.7036 | 27.77213 | 19.06435 | 7.759761 | 1.016533 | 0.086321 | 0.166656 |
| 7 | 27088600 | 27091199 | 17.87285 | 30.19256 | 63.91156 | 27.96094 | -0.69664 | 1.079759 | 0.166657 |
| 17 | 67668400 | 67670599 | 9.801471 | 17.32227 | 46.69233 | 20.65452 | -1.03229 | 0.519889 | 0.166658 |
| 16 | 20620800 | 20623599 | 14.87101 | 25.6407 | 55.92083 | 24.53126 | -0.73527 | 0.87328 | 0.166662 |
| 7 | 127834800 | 127837999 | 17.8266 | 29.77118 | 69.13386 | 30.08292 | -0.83007 | 1.196747 | 0.166666 |
| 5 | 84250600 | 84251999 | 17.74793 | 32.66234 | 20.67813 | 8.771239 | 1.086631 | 0.268846 | 0.166674 |
| 2 | 93840600 | 93841599 | 5.556282 | 10.65314 | 28.9952 | 11.40623 | -0.90902 | -0.08174 | 0.166707 |
| 18 | 29553600 | 29555399 | 23.14412 | 41.35563 | 21.84415 | 9.593066 | 1.313837 | 0.477628 | 0.166712 |
| 9 | 44678400 | 44679199 | 4.429992 | 8.538741 | 27.9925 | 10.8458 | -1.15336 | -0.14793 | 0.166713 |
| 11 | 53360400 | 53362399 | 8.877128 | 16.1548 | 38.15373 | 16.42408 | -0.81442 | 0.303818 | 0.166715 |
| 4 | 21241600 | 21244399 | 33.60214 | 56.99075 | 36.40062 | 15.94433 | 1.025502 | 1.068897 | 0.166724 |
| 7 | 8240200 | 8241199 | 21.45289 | 38.4316 | 21.6716 | 9.496828 | 1.221596 | 0.448403 | 0.166726 |
| 2 | 68301200 | 68302799 | 7.784634 | 13.91018 | 40.02175 | 17.57591 | -1.11578 | 0.413327 | 0.166726 |

| | | | | | | | | | |
|---|---|---|---|---|---|---|---|---|---|
| 8 | 85524200 | 85526999 | 20.38192 | 33.47289 | 80.20842 | 34.55161 | -0.87921 | 1.410788 | 0.16673 |
| 15 | 101307800 | 101312199 | 19.82536 | 32.41892 | 84.82443 | 36.47572 | -1.00868 | 1.468665 | 0.166746 |
| 12 | 17119800 | 17125399 | 44.25709 | 70.51436 | 139.0378 | 58.19362 | -0.64063 | 2.175796 | 0.166762 |
| 2 | 171907200 | 171908799 | 22.32755 | 39.07541 | 30.12189 | 13.29009 | 0.761096 | 0.691791 | 0.166775 |
| 16 | 18780200 | 18785399 | 73.10526 | 116.3762 | 92.02223 | 38.469 | 0.67408 | 2.217547 | 0.166776 |
| 4 | 9770400 | 9770999 | 3.818838 | 7.451083 | 23.95901 | 8.547343 | -1.01356 | -0.46043 | 0.166784 |
| 12 | 12986600 | 12988199 | 7.181755 | 13.24012 | 34.64452 | 14.66046 | -0.94283 | 0.210624 | 0.166787 |
| 9 | 51874000 | 51874999 | 4.920484 | 9.452002 | 30.13151 | 11.78896 | -1.12607 | -0.10017 | 0.166795 |
| 11 | 56294400 | 56295399 | 14.47401 | 27.15329 | 20.10061 | 8.281424 | 0.894274 | 0.129152 | 0.166799 |
| 10 | 13376200 | 13378399 | 9.913207 | 17.62457 | 40.95799 | 18.06371 | -0.81476 | 0.460616 | 0.166809 |
| 15 | 17419600 | 17420999 | 16.55129 | 30.3679 | 21.86804 | 9.314314 | 0.897948 | 0.278229 | 0.166812 |
| 2 | 45032200 | 45032999 | 11.663 | 22.29074 | 17.84336 | 7.083591 | 0.827629 | -0.02616 | 0.166815 |
| 16 | 30774800 | 30782399 | 43.52634 | 68.76592 | 161.1476 | 66.92929 | -0.9008 | 2.419936 | 0.16683 |
| 5 | 137464600 | 137467399 | 15.3772 | 26.5185 | 59.77036 | 26.22011 | -0.78254 | 0.879798 | 0.166849 |
| 16 | 35423000 | 35425399 | 9.708755 | 17.02576 | 46.72419 | 20.65054 | -1.06082 | 0.62085 | 0.166851 |
| Y | 9011000 | 9014199 | 18.22143 | 33.11522 | 11.15386 | 4.809497 | 1.97162 | 0.315169 | 0.166863 |
| 7 | 21157600 | 21162199 | 61.83519 | 99.2347 | 70.02153 | 29.61436 | 0.850842 | 1.951884 | 0.16687 |
| 16 | 20335400 | 20337199 | 9.055192 | 15.96188 | 43.58182 | 19.28858 | -1.05129 | 0.543324 | 0.166878 |
| 14 | 86213200 | 86214999 | 9.524316 | 17.24705 | 41.83272 | 18.11953 | -0.85863 | 0.345505 | 0.166885 |
| 9 | 21954000 | 21969799 | 101.3047 | 153.0198 | 292.8509 | 121.0698 | -0.66051 | 3.322747 | 0.166886 |
| X | 25730000 | 25731999 | 47.14538 | 76.84817 | 54.4271 | 23.37728 | 0.867951 | 1.553761 | 0.166886 |
| 14 | 102879800 | 102881799 | 14.08962 | 24.47434 | 58.16152 | 25.53248 | -0.85629 | 0.80335 | 0.166889 |
| 3 | 3513400 | 3515999 | 27.07739 | 47.15301 | 29.84715 | 13.12338 | 1.043696 | 0.787269 | 0.166915 |
| 17 | 89058000 | 89063199 | 35.41423 | 56.84519 | 114.0775 | 48.26191 | -0.65371 | 1.928257 | 0.16692 |
| 3 | 158561800 | 158562399 | 11.43804 | 22.12291 | 15.95205 | 6.095974 | 1.028962 | -0.15393 | 0.166921 |
| 7 | 61055200 | 61064399 | 249.1365 | 364.0382 | 292.4888 | 120.5003 | 0.546847 | 3.925066 | 0.166932 |
| 5 | 23571200 | 23573599 | 57.50568 | 92.22209 | 77.45847 | 32.72487 | 0.598827 | 1.969548 | 0.166934 |
| 17 | 40989800 | 40995399 | 81.10247 | 128.6226 | 97.5885 | 40.627 | 0.728257 | 2.336137 | 0.166946 |
| X | 156924200 | 156925399 | 16.46346 | 30.13509 | 22.47514 | 9.603892 | 0.844543 | 0.29246 | 0.166949 |
| X | 14079400 | 14080999 | 18.91623 | 35.28397 | 17.44044 | 7.255568 | 1.461187 | 0.196935 | 0.166959 |
| 5 | 147725200 | 147727399 | 14.26009 | 24.58281 | 54.79128 | 24.03561 | -0.76637 | 0.871975 | 0.166975 |
| 18 | 86852000 | 86852599 | 12.178 | 23.6865 | 12.61819 | 4.665097 | 1.501763 | -0.2672 | 0.166995 |
| 1 | 87420000 | 87423399 | 18.35196 | 30.98304 | 65.05811 | 28.45729 | -0.68515 | 1.087608 | 0.167008 |
| 7 | 19428400 | 19441799 | 82.09053 | 125.2919 | 254.7574 | 105.6454 | -0.73535 | 3.10399 | 0.167014 |
| 16 | 53141200 | 53142599 | 22.87642 | 39.99812 | 30.68426 | 13.52739 | 0.768163 | 0.711139 | 0.167017 |
| 2 | 47053000 | 47053799 | 12.12902 | 23.37678 | 17.31842 | 6.711499 | 0.970777 | -0.0981 | 0.167021 |
| 14 | 124062600 | 124064799 | 20.94748 | 37.33197 | 23.35086 | 10.27757 | 1.068955 | 0.479397 | 0.167033 |
| 11 | 47574800 | 47576199 | 20.59445 | 36.25026 | 28.00345 | 12.39742 | 0.758577 | 0.595284 | 0.167047 |
| 5 | 74264400 | 74265399 | 19.81035 | 35.76274 | 22.57118 | 9.814407 | 1.067783 | 0.394999 | 0.167051 |
| 14 | 76348000 | 76348799 | 2.547556 | 4.970332 | 22.43741 | 8.012906 | -1.49204 | -0.4751 | 0.16706 |
| 19 | 3383000 | 3393399 | 62.81969 | 97.04809 | 217.063 | 90.1633 | -0.85906 | 2.856757 | 0.167082 |
| 10 | 81362600 | 81390199 | 176.9949 | 253.5833 | 551.4306 | 227.7499 | -0.91681 | 4.21646 | 0.167094 |
| 9 | 111220200 | 111221999 | 10.04432 | 17.50971 | 49.92147 | 21.96427 | -1.11601 | 0.72248 | 0.167095 |
| 6 | 89481800 | 89483799 | 11.56998 | 20.15303 | 49.95861 | 21.96946 | -0.91588 | 0.740842 | 0.167111 |
| 12 | 3095000 | 3103599 | 99.89762 | 156.953 | 113.0659 | 46.89919 | 0.79231 | 2.562235 | 0.167128 |
| X | 126547000 | 126549999 | 47.75929 | 78.95235 | 39.96743 | 17.2731 | 1.359426 | 1.37021 | 0.167132 |
| 7 | 117968000 | 117971599 | 22.1891 | 36.71441 | 76.04069 | 32.88358 | -0.66655 | 1.320038 | 0.167133 |

| | | | | | | | | | |
|---|---|---|---|---|---|---|---|---|---|
| 2 | 30674400 | 30689599 | 71.98749 | 109.8371 | 263.1598 | 109.1217 | -0.97219 | 3.108222 | 0.167137 |
| X | 21981400 | 21983399 | 20.3433 | 36.47085 | 23.61148 | 10.34045 | 1.024731 | 0.44151 | 0.167144 |
| 13 | 49974000 | 49979799 | 36.9552 | 59.05379 | 126.4429 | 53.14067 | -0.75452 | 2.073345 | 0.167147 |
| 2 | 112353600 | 112354399 | 4.545433 | 8.785488 | 27.00262 | 10.35223 | -1.04656 | -0.1849 | 0.16715 |
| 2 | 34261000 | 34263599 | 14.99378 | 25.50796 | 60.56921 | 26.5472 | -0.86083 | 1.003021 | 0.167166 |
| 9 | 12997000 | 13000799 | 39.13927 | 65.65555 | 38.19236 | 16.66572 | 1.159841 | 1.187932 | 0.167173 |
| 4 | 62580600 | 62582399 | 7.520926 | 13.75586 | 36.64027 | 15.67325 | -0.98023 | 0.261361 | 0.167177 |
| 2 | 51290400 | 51291399 | 5.120294 | 9.901892 | 28.33928 | 10.83751 | -0.94267 | -0.18575 | 0.167182 |
| 6 | 53151400 | 53154799 | 31.0398 | 53.67138 | 29.72343 | 13.04016 | 1.234201 | 0.883803 | 0.167182 |
| 15 | 102352400 | 102355799 | 14.85702 | 25.23373 | 61.61578 | 27.0007 | -0.90132 | 1.018659 | 0.167184 |
| X | 131673000 | 131674999 | 39.57284 | 65.43131 | 48.29522 | 20.87489 | 0.817326 | 1.351832 | 0.16719 |
| 9 | 64644000 | 64644999 | 5.286438 | 10.12257 | 28.92988 | 11.42319 | -0.98394 | -0.07319 | 0.167194 |
| 8 | 52376400 | 52378399 | 25.58081 | 44.47854 | 29.75335 | 13.06953 | 0.964317 | 0.801252 | 0.167195 |
| 1 | 190833000 | 190834399 | 10.36604 | 18.44543 | 41.46954 | 18.27613 | -0.76674 | 0.458595 | 0.167195 |
| 2 | 94356400 | 94358199 | 10.02109 | 17.93426 | 41.54443 | 18.22076 | -0.8049 | 0.418676 | 0.167198 |
| 6 | 83293000 | 83294799 | 6.20241 | 11.70011 | 31.30431 | 12.77634 | -0.93295 | 0.052112 | 0.167224 |
| 2 | 160430400 | 160435399 | 29.32697 | 47.1215 | 114.1468 | 48.36401 | -0.92277 | 1.888558 | 0.167232 |
| 11 | 82873200 | 82874399 | 5.159567 | 9.863362 | 29.42237 | 11.67278 | -1.05168 | -0.05797 | 0.167234 |
| 2 | 23662200 | 23663799 | 18.49398 | 33.76131 | 21.02593 | 9.015688 | 1.099237 | 0.318991 | 0.167261 |
| 4 | 115530800 | 115532799 | 18.47364 | 33.80705 | 20.48103 | 8.754426 | 1.141671 | 0.30404 | 0.16727 |
| 16 | 94003800 | 94009999 | 39.29484 | 62.56128 | 137.1048 | 57.32528 | -0.79386 | 2.196485 | 0.167274 |
| 13 | 64298800 | 64301999 | 20.21539 | 33.64275 | 72.1471 | 31.32467 | -0.71584 | 1.243433 | 0.167274 |
| 14 | 43745000 | 43746799 | 9.032332 | 16.43227 | 38.14682 | 16.42733 | -0.7901 | 0.307107 | 0.167289 |
| 11 | 88121600 | 88126799 | 20.07164 | 32.98384 | 80.89591 | 34.85693 | -0.91234 | 1.401543 | 0.16729 |
| 16 | 4368800 | 4371799 | 12.70917 | 21.54339 | 62.43098 | 27.34164 | -1.14696 | 1.031581 | 0.167293 |
| 8 | 13672600 | 13678999 | 47.14303 | 74.74984 | 148.9608 | 62.00086 | -0.66302 | 2.329581 | 0.167293 |
| 19 | 21803400 | 21805999 | 11.97468 | 20.98198 | 48.85437 | 21.57687 | -0.82572 | 0.648118 | 0.167329 |
| 13 | 13214200 | 13217999 | 72.44467 | 115.3019 | 93.69715 | 39.15851 | 0.634464 | 2.22508 | 0.167353 |
| 8 | 21904200 | 21906799 | 32.87543 | 56.49373 | 30.79667 | 13.50845 | 1.255255 | 0.946731 | 0.167356 |
| 4 | 154455000 | 154461399 | 34.33547 | 54.9993 | 119.7931 | 50.52021 | -0.77519 | 1.994076 | 0.167367 |
| 11 | 119045400 | 119054399 | 40.78101 | 64.83033 | 141.9214 | 59.22456 | -0.79459 | 2.246508 | 0.167374 |
| 1 | 167084400 | 167093399 | 57.37371 | 89.53893 | 191.1391 | 79.31417 | -0.78041 | 2.671204 | 0.167396 |
| 10 | 80667400 | 80673199 | 58.61553 | 91.12148 | 198.7999 | 82.5385 | -0.81617 | 2.741571 | 0.167411 |
| 4 | 112389600 | 112391799 | 36.38261 | 60.49151 | 46.57259 | 20.20765 | 0.757165 | 1.27859 | 0.167414 |
| 9 | 22844200 | 22845199 | 11.90037 | 23.1301 | 13.88929 | 5.164536 | 1.324549 | -0.2472 | 0.167424 |
| 11 | 63484600 | 63487799 | 31.51102 | 53.44379 | 39.31417 | 17.2205 | 0.822701 | 1.066284 | 0.167427 |
| 3 | 137592800 | 137593399 | 3.935707 | 7.681824 | 23.45031 | 8.315073 | -0.9313 | -0.48138 | 0.167429 |
| 3 | 38754600 | 38755599 | 12.09446 | 23.46674 | 15.05557 | 5.668861 | 1.214374 | -0.19893 | 0.167434 |
| 19 | 13651800 | 13652599 | 12.09446 | 23.46674 | 15.05557 | 5.668861 | 1.214374 | -0.19893 | 0.167434 |
| 2 | 150580000 | 150581799 | 7.606844 | 14.07659 | 35.27879 | 14.85274 | -0.87637 | 0.191784 | 0.167435 |
| 14 | 78706400 | 78707799 | 7.606844 | 14.07659 | 35.27879 | 14.85274 | -0.87637 | 0.191784 | 0.167435 |
| 5 | 3091800 | 3092999 | 21.4992 | 37.61016 | 28.39438 | 12.52351 | 0.791096 | 0.691457 | 0.167449 |
| 14 | 30436600 | 30438999 | 13.38242 | 22.63024 | 65.2214 | 28.54292 | -1.13962 | 1.055735 | 0.16746 |
| 16 | 4680800 | 4685199 | 27.13143 | 43.82716 | 98.42757 | 42.01201 | -0.8043 | 1.716 | 0.167462 |
| 13 | 76203600 | 76205399 | 14.34785 | 24.48593 | 58.956 | 25.84668 | -0.87995 | 0.970076 | 0.167465 |
| 11 | 5109800 | 5112199 | 24.27706 | 39.59747 | 87.86718 | 37.74908 | -0.7746 | 1.515466 | 0.167466 |
| X | 5460800 | 5462199 | 19.02722 | 34.87432 | 19.91668 | 8.495203 | 1.228387 | 0.299017 | 0.167473 |

| | | | | | | | | | |
|---|---|---|---|---|---|---|---|---|---|
| 15 | 31580400 | 31584199 | 18.16988 | 30.70789 | 65.11098 | 28.4895 | -0.69912 | 1.076777 | 0.167479 |
| 12 | 19497400 | 19499799 | 29.31423 | 49.91962 | 33.70341 | 14.77322 | 0.946354 | 1.015118 | 0.16748 |
| 7 | 38461400 | 38468999 | 86.77827 | 136.8702 | 102.9837 | 42.73868 | 0.73384 | 2.475165 | 0.167501 |
| 6 | 108634400 | 108638799 | 20.65527 | 33.69861 | 87.62109 | 37.64683 | -1.00212 | 1.505158 | 0.167505 |
| 6 | 53161000 | 53163799 | 29.30263 | 50.0429 | 37.68328 | 16.52152 | 0.79017 | 0.994995 | 0.167507 |
| 6 | 30312800 | 30314399 | 26.6824 | 46.32089 | 32.1357 | 14.10203 | 0.911776 | 0.830834 | 0.167511 |
| 14 | 30798000 | 30801799 | 55.31683 | 88.82105 | 73.45925 | 31.0934 | 0.623108 | 1.926988 | 0.16752 |
| 15 | 103410800 | 103412599 | 9.310291 | 16.27458 | 51.49043 | 22.69531 | -1.26519 | 0.677931 | 0.167523 |
| 16 | 20095000 | 20098599 | 19.32748 | 32.50293 | 69.59608 | 30.40627 | -0.71364 | 1.128125 | 0.16753 |
| 12 | 11108400 | 11109399 | 8.395495 | 15.40093 | 36.17044 | 15.41014 | -0.79583 | 0.251003 | 0.167538 |
| 13 | 53547600 | 53551799 | 21.62778 | 35.56765 | 79.51371 | 34.27221 | -0.77872 | 1.394893 | 0.167539 |
| 11 | 50652000 | 50656999 | 27.57728 | 44.64021 | 95.89236 | 41.01956 | -0.73715 | 1.661023 | 0.167558 |
| 6 | 108958200 | 108958999 | 18.11305 | 33.39915 | 20.12469 | 8.513932 | 1.160133 | 0.260194 | 0.167561 |
| 14 | 57369800 | 57371399 | 6.400868 | 11.6235 | 38.1099 | 16.44925 | -1.28557 | 0.299447 | 0.167561 |
| 8 | 117600400 | 117602799 | 12.14458 | 21.24278 | 48.34738 | 21.32167 | -0.79294 | 0.677513 | 0.167581 |
| 1 | 38245800 | 38250399 | 26.38746 | 42.90386 | 91.39199 | 39.21818 | -0.71914 | 1.567503 | 0.167601 |
| X | 33396400 | 33397999 | 18.5776 | 34.31681 | 19.65842 | 8.296542 | 1.235192 | 0.253935 | 0.167616 |
| 10 | 58577800 | 58583199 | 30.68844 | 49.45423 | 105.8586 | 45.04381 | -0.73943 | 1.793627 | 0.167617 |
| 4 | 60304800 | 60309199 | 27.23581 | 46.80376 | 36.06495 | 15.81931 | 0.758335 | 0.928693 | 0.167619 |
| 3 | 6725400 | 6726999 | 29.78014 | 50.88756 | 37.09687 | 16.26517 | 0.836851 | 0.990971 | 0.167636 |
| 9 | 20138200 | 20140799 | 56.53158 | 90.98607 | 70.79182 | 30.06785 | 0.715159 | 1.855292 | 0.167667 |
| 7 | 20992800 | 20995199 | 41.5572 | 67.99165 | 54.88604 | 23.60637 | 0.683372 | 1.487874 | 0.167679 |
| 8 | 46381400 | 46381599 | 6.961226 | 13.60592 | 9.317914 | 2.996251 | 1.319551 | -0.82629 | 0.167689 |
| X | 56612400 | 56615799 | 30.6566 | 53.44136 | 25.88486 | 11.38865 | 1.427862 | 0.787024 | 0.167716 |
| 7 | 40701200 | 40702999 | 27.00748 | 46.33107 | 36.72558 | 16.10747 | 0.71735 | 0.944926 | 0.167719 |
| 8 | 70606200 | 70607599 | 17.41388 | 31.75536 | 22.49614 | 9.659091 | 0.913962 | 0.321665 | 0.16772 |
| 19 | 47371800 | 47376599 | 21.24608 | 34.66568 | 85.81137 | 36.8705 | -0.93124 | 1.505854 | 0.167726 |
| 4 | 22684200 | 22685999 | 22.72628 | 39.86267 | 28.92228 | 12.78505 | 0.847958 | 0.657196 | 0.167742 |
| 9 | 53637200 | 53637599 | 11.58433 | 22.16141 | 15.37787 | 6.088201 | 1.035605 | -0.04313 | 0.167744 |
| 6 | 42431200 | 42433799 | 14.96309 | 28.38859 | 17.89561 | 7.215661 | 1.149886 | 0.055643 | 0.167756 |
| 4 | 57320600 | 57324999 | 26.88952 | 43.65538 | 92.32601 | 39.57991 | -0.7107 | 1.597727 | 0.167761 |
| 9 | 79624000 | 79630399 | 38.44823 | 61.31557 | 134.3497 | 56.30311 | -0.791 | 2.139717 | 0.167762 |
| 3 | 82050200 | 82053199 | 14.05974 | 24.35635 | 53.7736 | 23.5927 | -0.75128 | 0.824459 | 0.167768 |
| 6 | 132495600 | 132498599 | 79.83074 | 126.7991 | 95.71667 | 39.90866 | 0.73772 | 2.292552 | 0.167774 |
| 4 | 115513200 | 115514999 | 11.50662 | 20.19173 | 48.29342 | 21.35583 | -0.86418 | 0.620427 | 0.16778 |
| 2 | 50424000 | 50425599 | 17.08937 | 31.11014 | 22.8803 | 9.845149 | 0.858026 | 0.328369 | 0.167786 |
| 2 | 122675400 | 122677599 | 12.12045 | 21.29528 | 49.34978 | 21.84586 | -0.81894 | 0.605892 | 0.167803 |
| 13 | 120495400 | 120498399 | 13.7231 | 23.89858 | 51.61521 | 22.69507 | -0.7185 | 0.755156 | 0.167815 |
| 7 | 15223400 | 15228599 | 91.92584 | 145.4053 | 98.81119 | 41.06411 | 0.883822 | 2.412209 | 0.167823 |
| X | 26583200 | 26585399 | 26.45665 | 46.61408 | 23.23236 | 10.28322 | 1.387708 | 0.613008 | 0.167846 |
| 14 | 96803200 | 96804399 | 14.24637 | 26.94052 | 17.81825 | 7.234353 | 1.072007 | 0.069242 | 0.167876 |
| 5 | 39799200 | 39802999 | 22.85411 | 37.30145 | 90.73892 | 38.99232 | -0.90612 | 1.503593 | 0.167885 |
| X | 134767200 | 134767999 | 13.27535 | 25.50709 | 17.16542 | 6.711585 | 1.09628 | -0.05866 | 0.167888 |
| 2 | 18253800 | 18254399 | 9.367596 | 18.20079 | 14.43714 | 5.383734 | 0.924501 | -0.26679 | 0.16789 |
| 7 | 78041800 | 78043999 | 22.34132 | 39.05182 | 30.74559 | 13.55129 | 0.730909 | 0.71239 | 0.167891 |
| 9 | 36462200 | 36463599 | 22.34132 | 39.05182 | 30.74559 | 13.55129 | 0.730909 | 0.71239 | 0.167891 |
| 18 | 23117400 | 23118999 | 7.298894 | 13.55153 | 34.25003 | 14.35479 | -0.88342 | 0.168654 | 0.167919 |

| | | | | | | | | | |
|---|---|---|---|---|---|---|---|---|---|
| 2 | 30174800 | 30177199 | 9.087769 | 15.97303 | 45.13281 | 19.98554 | -1.1023 | 0.577993 | 0.167929 |
| 2 | 78698600 | 78700399 | 8.965901 | 16.05106 | 39.48871 | 17.3144 | -0.89016 | 0.40854 | 0.16793 |
| 11 | 117898400 | 117902199 | 16.16357 | 27.25864 | 63.7195 | 27.86253 | -0.83917 | 1.088982 | 0.167932 |
| 18 | 86365600 | 86366799 | 14.54698 | 27.10525 | 20.55299 | 8.565876 | 0.846572 | 0.172503 | 0.167933 |
| 18 | 33919000 | 33922799 | 15.66462 | 26.50914 | 64.9432 | 28.42756 | -0.90649 | 1.055644 | 0.167935 |
| 14 | 52221200 | 52223599 | 12.42434 | 21.39023 | 55.21207 | 24.21924 | -0.97717 | 0.878142 | 0.167938 |
| 11 | 16176800 | 16179399 | 44.14629 | 72.35045 | 48.38675 | 20.8254 | 0.955174 | 1.467976 | 0.167938 |
| 14 | 28243200 | 28244799 | 6.718179 | 12.4839 | 34.8019 | 14.56931 | -1.02251 | 0.159634 | 0.16794 |
| 2 | 26421200 | 26426599 | 27.46981 | 44.33528 | 100.6052 | 42.89346 | -0.82069 | 1.742635 | 0.167948 |
| 6 | 120013800 | 120016399 | 15.27803 | 26.05084 | 60.11013 | 26.35082 | -0.81919 | 0.981824 | 0.167959 |
| 3 | 111489400 | 111490799 | 20.71836 | 38.2676 | 17.14365 | 7.236158 | 1.586369 | 0.265756 | 0.167962 |
| 6 | 125257200 | 125259999 | 25.75288 | 41.8718 | 88.66972 | 38.04991 | -0.71057 | 1.566522 | 0.167967 |
| 3 | 52022000 | 52024799 | 12.33953 | 21.46237 | 53.75174 | 23.616 | -0.93132 | 0.767725 | 0.167974 |
| 1 | 177890800 | 177892799 | 16.46034 | 29.91473 | 20.28605 | 8.747234 | 0.971851 | 0.325236 | 0.16799 |
| 11 | 116302600 | 116304399 | 5.559545 | 10.39362 | 34.92059 | 14.47198 | -1.27721 | 0.110729 | 0.167991 |
| 12 | 106157800 | 106167399 | 71.33671 | 110.1365 | 216.0654 | 89.74923 | -0.67086 | 2.87128 | 0.167996 |
| 7 | 103038800 | 103039799 | 13.91948 | 26.91147 | 14.99928 | 5.742762 | 1.393771 | -0.12371 | 0.168007 |
| 7 | 33334000 | 33338599 | 101.3119 | 158.7597 | 119.2781 | 49.4742 | 0.728859 | 2.612459 | 0.168009 |
| 7 | 40910200 | 40910999 | 17.27589 | 31.17051 | 23.945 | 10.41882 | 0.785467 | 0.382291 | 0.168027 |
| 7 | 31301000 | 31302999 | 40.86079 | 67.24596 | 51.08818 | 22.02695 | 0.77412 | 1.411378 | 0.168029 |
| 17 | 49298000 | 49300599 | 14.4032 | 24.87249 | 54.27748 | 23.81148 | -0.73549 | 0.856254 | 0.168044 |
| 9 | 12523200 | 12526399 | 31.99071 | 54.28821 | 38.73146 | 16.96818 | 0.866694 | 1.062499 | 0.168051 |
| 8 | 104439800 | 104441599 | 15.96379 | 29.05856 | 22.62774 | 9.737537 | 0.776169 | 0.320108 | 0.168067 |
| 17 | 57947600 | 57948599 | 15.96379 | 29.05856 | 22.62774 | 9.737537 | 0.776169 | 0.320108 | 0.168067 |
| 8 | 21683600 | 21687399 | 24.63353 | 43.50067 | 23.73217 | 10.49967 | 1.259056 | 0.578105 | 0.168072 |
| 7 | 101134600 | 101139399 | 26.58665 | 43.11814 | 92.27805 | 39.52857 | -0.72911 | 1.620976 | 0.168072 |
| 7 | 129193600 | 129193999 | 3.817963 | 7.451839 | 24.75135 | 8.77983 | -1.05269 | -0.47905 | 0.168073 |
| 3 | 123673200 | 123674199 | 16.80207 | 31.27407 | 19.79161 | 8.263327 | 1.10358 | 0.197168 | 0.168074 |
| 18 | 49964800 | 49966399 | 5.304589 | 10.17082 | 30.77827 | 12.1068 | -1.06149 | -0.08374 | 0.168074 |
| 12 | 60747200 | 60749199 | 17.69743 | 31.99879 | 18.54619 | 8.048376 | 1.191328 | 0.360605 | 0.168093 |
| 13 | 52264600 | 52278999 | 84.05533 | 128.3724 | 254.6269 | 105.6067 | -0.69889 | 3.09126 | 0.16811 |
| 5 | 124987800 | 124998199 | 70.45157 | 108.5758 | 222.8866 | 92.58348 | -0.7382 | 2.904574 | 0.168133 |
| 15 | 27590000 | 27591399 | 6.639605 | 12.13665 | 36.10128 | 15.45454 | -1.13825 | 0.258867 | 0.168142 |
| 2 | 157214800 | 157217399 | 13.89814 | 24.15498 | 52.15482 | 22.90349 | -0.71839 | 0.786006 | 0.16815 |
| 11 | 83337200 | 83341399 | 16.39188 | 27.45248 | 67.38389 | 29.37368 | -0.90961 | 1.160801 | 0.168151 |
| 12 | 19920600 | 19929399 | 302.4568 | 435.8417 | 333.8027 | 137.7448 | 0.596359 | 4.137363 | 0.168153 |
| 11 | 78061200 | 78063999 | 13.15592 | 22.76522 | 53.16144 | 23.32333 | -0.83188 | 0.832108 | 0.168155 |
| 10 | 67928800 | 67931199 | 15.8339 | 27.2532 | 59.7931 | 26.22854 | -0.74454 | 0.897825 | 0.168158 |
| 16 | 89873400 | 89877599 | 28.19686 | 45.69201 | 93.8653 | 40.18414 | -0.67144 | 1.639688 | 0.168161 |
| 1 | 58543200 | 58544599 | 13.10248 | 22.8444 | 49.98155 | 21.99286 | -0.73672 | 0.734096 | 0.168189 |
| 4 | 41034000 | 41038999 | 29.84569 | 48.21312 | 99.3331 | 42.40017 | -0.6803 | 1.718635 | 0.16819 |
| 2 | 40659400 | 40660999 | 19.18134 | 33.91079 | 25.96076 | 11.48306 | 0.773687 | 0.535033 | 0.1682 |
| 1 | 156300600 | 156302799 | 8.794546 | 15.81057 | 38.9907 | 17.0356 | -0.89028 | 0.377679 | 0.168209 |
| 9 | 94021600 | 94022999 | 8.896265 | 16.15702 | 37.57676 | 16.21655 | -0.79448 | 0.316207 | 0.168213 |
| 2 | 153516000 | 153516999 | 4.019189 | 7.761679 | 28.04737 | 10.79605 | -1.28294 | -0.17481 | 0.168217 |
| 7 | 8093600 | 8096599 | 37.97757 | 62.48208 | 51.56278 | 22.22873 | 0.654934 | 1.407153 | 0.168232 |
| 4 | 134759400 | 134763799 | 33.49503 | 53.75567 | 113.7117 | 48.09504 | -0.72975 | 1.932121 | 0.168237 |

| | | | | | | | | | |
|---|---|---|---|---|---|---|---|---|---|
| 5 | 93896200 | 93901799 | 122.3106 | 188.1362 | 155.2326 | 64.4661 | 0.573301 | 2.954033 | 0.168239 |
| 1 | 20207600 | 20208599 | 11.7244 | 22.75466 | 15.06711 | 5.666286 | 1.170714 | -0.2075 | 0.168243 |
| 17 | 86904600 | 86907199 | 25.65138 | 44.0586 | 33.69031 | 14.77748 | 0.769165 | 0.922769 | 0.168275 |
| 13 | 61695400 | 61697199 | 41.18066 | 68.08037 | 47.21641 | 20.40665 | 0.906848 | 1.356328 | 0.168287 |
| 8 | 91588200 | 91588999 | 3.431958 | 6.662184 | 25.99817 | 9.765039 | -1.35673 | -0.27112 | 0.168287 |
| 9 | 107558600 | 107563399 | 16.05257 | 26.51732 | 77.05998 | 33.28669 | -1.15312 | 1.326682 | 0.16829 |
| 12 | 45359000 | 45360199 | 8.374207 | 15.43131 | 36.28342 | 15.36373 | -0.79115 | 0.224882 | 0.168302 |
| X | 115561200 | 115563999 | 28.37348 | 48.3068 | 36.91621 | 16.18119 | 0.768447 | 1.019016 | 0.168311 |
| 16 | 22615400 | 22619599 | 20.27779 | 33.5768 | 75.29142 | 32.57622 | -0.78062 | 1.304903 | 0.168314 |
| 12 | 22421800 | 22423199 | 14.7444 | 28.11505 | 17.48999 | 6.977316 | 1.182362 | 0.019702 | 0.168323 |
| 14 | 41079400 | 41082599 | 38.28307 | 64.0264 | 39.80438 | 17.33621 | 1.064279 | 1.214296 | 0.168331 |
| 19 | 9554200 | 9556599 | 22.25175 | 39.20627 | 26.02319 | 11.51846 | 0.976533 | 0.589825 | 0.168344 |
| 8 | 118508800 | 118512399 | 18.833 | 31.28248 | 77.09699 | 33.42979 | -0.91611 | 1.264311 | 0.168351 |
| 6 | 113955400 | 113973399 | 126.7899 | 189.1202 | 346.7491 | 142.9128 | -0.61494 | 3.571826 | 0.168356 |
| X | 117850800 | 117852399 | 23.77061 | 41.6194 | 29.49833 | 13.01985 | 0.881734 | 0.692696 | 0.168362 |
| 7 | 3114000 | 3115399 | 25.03587 | 43.30649 | 34.13125 | 14.97411 | 0.727856 | 0.86014 | 0.168364 |
| 1 | 34897600 | 34904199 | 30.06718 | 48.13467 | 127.521 | 53.73253 | -1.05806 | 2.008996 | 0.168377 |
| 4 | 90146000 | 90149199 | 62.91966 | 101.1876 | 67.76474 | 28.74923 | 0.92968 | 1.888826 | 0.168391 |
| 5 | 74182000 | 74182999 | 15.23271 | 28.14786 | 21.22234 | 8.952331 | 0.842416 | 0.228732 | 0.168404 |
| 18 | 71398600 | 71399199 | 4.23529 | 8.242112 | 24.65981 | 9.078064 | -0.95234 | -0.33828 | 0.168411 |
| 5 | 66622400 | 66623999 | 10.02987 | 17.95467 | 40.496 | 17.75701 | -0.76618 | 0.4165 | 0.168424 |
| 4 | 6988800 | 6991599 | 16.56614 | 27.70524 | 68.4756 | 29.82303 | -0.91958 | 1.177274 | 0.168437 |
| 10 | 88894600 | 88896999 | 33.2234 | 56.59629 | 35.30314 | 15.47486 | 1.060145 | 1.029447 | 0.168441 |
| 4 | 123222400 | 123230399 | 48.81531 | 77.24028 | 154.8114 | 64.34764 | -0.6739 | 2.385119 | 0.168442 |
| 3 | 126011400 | 126012399 | 6.770011 | 12.84014 | 32.12883 | 12.96358 | -0.82343 | 0.019096 | 0.168473 |
| 10 | 30050800 | 30051399 | 14.86254 | 27.48169 | 21.22989 | 8.94761 | 0.808416 | 0.221815 | 0.168486 |
| 13 | 12934200 | 12934599 | 9.519167 | 18.55086 | 13.65547 | 4.950653 | 1.066476 | -0.36523 | 0.168491 |
| 2 | 155118000 | 155121999 | 17.08949 | 28.10617 | 80.91696 | 34.87804 | -1.14204 | 1.38533 | 0.168492 |
| 15 | 36878000 | 36879999 | 9.953872 | 17.58083 | 44.57534 | 19.72048 | -0.94465 | 0.525038 | 0.168494 |
| 4 | 40721200 | 40723399 | 10.16837 | 17.80581 | 47.27042 | 20.86626 | -1.01335 | 0.648402 | 0.168502 |
| 9 | 101961200 | 101966199 | 30.37157 | 48.94044 | 104.3826 | 44.41161 | -0.73433 | 1.795273 | 0.168526 |
| 4 | 133586600 | 133594799 | 37.42734 | 59.48108 | 146.2779 | 61.02426 | -0.96254 | 2.253064 | 0.168553 |
| 12 | 14553600 | 14554999 | 13.67968 | 26.1757 | 17.57475 | 6.953549 | 1.083577 | -0.01679 | 0.168561 |
| 15 | 84430800 | 84443399 | 119.3501 | 177.5835 | 359.0031 | 147.9168 | -0.75902 | 3.615539 | 0.168576 |
| 17 | 30095200 | 30100599 | 54.798 | 85.82945 | 180.3312 | 74.80149 | -0.75339 | 2.604788 | 0.168586 |
| 13 | 32073000 | 32075999 | 32.25608 | 54.49489 | 40.38442 | 17.66842 | 0.811899 | 1.10193 | 0.168595 |
| 16 | 57447200 | 57449199 | 14.94571 | 25.14252 | 65.16574 | 28.47355 | -0.98754 | 1.108897 | 0.168605 |
| 16 | 97789400 | 97790999 | 18.5134 | 33.26804 | 24.36986 | 10.65153 | 0.849559 | 0.416148 | 0.168607 |
| 13 | 100922200 | 100923399 | 8.937512 | 16.09399 | 38.49389 | 16.79062 | -0.84462 | 0.369095 | 0.168628 |
| 5 | 150153200 | 150153599 | 7.087619 | 13.86412 | 10.55802 | 3.485654 | 1.136756 | -0.73061 | 0.168636 |
| 7 | 62274400 | 62275599 | 11.13335 | 21.72117 | 10.98917 | 3.925207 | 1.617678 | -0.40234 | 0.168639 |
| 6 | 83253400 | 83260399 | 43.84027 | 69.72301 | 148.8928 | 62.16138 | -0.75835 | 2.237907 | 0.168643 |
| 7 | 55766600 | 55768199 | 18.54058 | 33.52615 | 23.3391 | 10.12724 | 0.929312 | 0.378636 | 0.168648 |
| 15 | 70109000 | 70110799 | 7.111725 | 13.33713 | 42.27599 | 17.42759 | -1.18925 | 0.119536 | 0.168657 |
| 6 | 60005600 | 60007199 | 18.65055 | 33.72078 | 23.25645 | 10.09295 | 0.942449 | 0.379518 | 0.168667 |
| 4 | 128608400 | 128612199 | 16.88737 | 28.21873 | 69.04475 | 30.05496 | -0.90512 | 1.187706 | 0.168667 |
| 1 | 82387000 | 82387999 | 4.950025 | 9.615062 | 26.95071 | 10.0721 | -0.88084 | -0.27029 | 0.168667 |

| | | | | | | | | | |
|---|---|---|---|---|---|---|---|---|---|
| 5 | 130310800 | 130320799 | 78.53831 | 120.8431 | 226.8761 | 94.22267 | -0.61132 | 2.939225 | 0.168668 |
| 3 | 9069000 | 9070999 | 9.583634 | 17.10972 | 40.47336 | 17.78742 | -0.83674 | 0.429972 | 0.168669 |
| 8 | 79680400 | 79680999 | 7.292166 | 13.56503 | 34.27715 | 14.32564 | -0.87952 | 0.159908 | 0.168691 |
| 16 | 75388800 | 75390399 | 9.224848 | 16.63641 | 39.02704 | 16.99118 | -0.81546 | 0.361264 | 0.16871 |
| 11 | 107359000 | 107361799 | 15.17081 | 25.3912 | 68.90256 | 30.02355 | -1.05374 | 1.164979 | 0.168722 |
| 5 | 24218800 | 24219399 | 3.129085 | 6.111084 | 22.21485 | 7.796324 | -1.16463 | -0.5336 | 0.168724 |
| 15 | 31297000 | 31299599 | 14.96784 | 25.53028 | 59.542 | 26.10245 | -0.8344 | 0.977383 | 0.168725 |
| 5 | 104753800 | 104758599 | 127.1263 | 195.5138 | 153.4398 | 63.72025 | 0.645317 | 2.958901 | 0.168728 |
| 7 | 31283600 | 31293799 | 170.7756 | 258.5757 | 183.2476 | 75.8153 | 0.774567 | 3.283291 | 0.168734 |
| 13 | 101004600 | 101009399 | 25.83505 | 42.21806 | 87.28087 | 37.52293 | -0.6709 | 1.491173 | 0.168735 |
| 17 | 84240400 | 84247599 | 33.80518 | 53.92809 | 133.9481 | 56.15939 | -0.97095 | 2.125529 | 0.168741 |
| 9 | 67737800 | 67739999 | 15.44199 | 26.29884 | 61.75838 | 27.07076 | -0.84481 | 0.994654 | 0.168742 |
| 15 | 46422800 | 46423399 | 12.94134 | 24.79271 | 16.38958 | 6.462272 | 1.109435 | -0.03983 | 0.168747 |
| 5 | 64728400 | 64728999 | 10.61568 | 20.62695 | 15.23053 | 5.676946 | 1.026862 | -0.25061 | 0.168752 |
| 2 | 12337600 | 12343399 | 67.50871 | 107.6091 | 90.16942 | 37.7634 | 0.592583 | 2.169879 | 0.168769 |
| 1 | 9305800 | 9308999 | 16.98989 | 28.51647 | 65.83513 | 28.73829 | -0.82156 | 1.138087 | 0.168776 |
| 1 | 158072600 | 158075999 | 23.25773 | 38.10876 | 82.34397 | 35.43733 | -0.73199 | 1.446174 | 0.16881 |
| 9 | 26252600 | 26253999 | 29.24915 | 49.46382 | 40.00663 | 17.5084 | 0.686217 | 1.082103 | 0.16882 |
| 2 | 71319200 | 71320399 | 5.876693 | 11.2206 | 30.01601 | 11.94109 | -0.90031 | -0.0411 | 0.168821 |
| 17 | 46169000 | 46198599 | 171.1304 | 246.0975 | 561.1108 | 231.6126 | -0.97974 | 4.161749 | 0.168822 |
| 1 | 136709200 | 136710599 | 6.177026 | 11.75763 | 31.08263 | 12.44653 | -0.89214 | -0.01458 | 0.168823 |
| 10 | 105341800 | 105343399 | 8.562228 | 15.6633 | 36.64814 | 15.67268 | -0.79437 | 0.269624 | 0.168837 |
| 7 | 88325400 | 88328799 | 33.03224 | 56.56532 | 31.30909 | 13.72987 | 1.232549 | 0.977104 | 0.168848 |
| 8 | 54483400 | 54484199 | 5.097049 | 9.858284 | 27.10774 | 10.35995 | -0.88372 | -0.189 | 0.168849 |
| 18 | 12037600 | 12038799 | 5.097049 | 9.858284 | 27.10774 | 10.35995 | -0.88372 | -0.189 | 0.168849 |
| 10 | 103628600 | 103628999 | 8.347747 | 16.28076 | 13.3732 | 4.798794 | 0.923904 | -0.41735 | 0.168891 |
| 12 | 52312400 | 52313399 | 5.948987 | 11.19488 | 33.48089 | 13.72204 | -1.098 | 0.067675 | 0.168908 |
| X | 77254800 | 77255999 | 24.79267 | 43.72669 | 24.48893 | 10.83721 | 1.221113 | 0.594709 | 0.168912 |
| 4 | 77669400 | 77670799 | 14.52073 | 27.82256 | 15.67131 | 6.176452 | 1.338937 | -0.02618 | 0.16892 |
| 16 | 93459400 | 93461399 | 25.78425 | 44.52587 | 35.04897 | 15.37595 | 0.729086 | 0.878606 | 0.168924 |
| 5 | 41914200 | 41922199 | 66.96372 | 103.7291 | 211.0597 | 87.66846 | -0.71976 | 2.80709 | 0.168935 |
| 10 | 117628200 | 117631399 | 16.45128 | 27.79554 | 66.10152 | 28.92044 | -0.864 | 1.073361 | 0.168945 |
| X | 122932200 | 122933799 | 21.03253 | 38.72518 | 17.8198 | 7.554006 | 1.544181 | 0.287123 | 0.168964 |
| 12 | 49547000 | 49548399 | 14.28787 | 26.81201 | 20.10599 | 8.279501 | 0.8763 | 0.125512 | 0.168966 |
| 18 | 5709000 | 5711199 | 24.87293 | 43.18499 | 33.4927 | 14.69867 | 0.751899 | 0.824004 | 0.168977 |
| 5 | 11267600 | 11269199 | 18.66597 | 34.05443 | 21.68752 | 9.306493 | 1.066889 | 0.32615 | 0.168993 |
| X | 4088400 | 4089599 | 23.88149 | 41.94705 | 27.30146 | 12.08271 | 1.003901 | 0.64074 | 0.169004 |
| 5 | 73860000 | 73862399 | 11.6793 | 20.43879 | 51.01899 | 22.50905 | -0.9257 | 0.669505 | 0.169006 |
| 5 | 146730600 | 146732199 | 11.7506 | 20.39931 | 52.08459 | 22.85627 | -0.95892 | 0.792745 | 0.169008 |
| 2 | 180608800 | 180609599 | 4.417586 | 8.548887 | 27.09167 | 10.32687 | -1.08252 | -0.20699 | 0.169021 |
| 12 | 111961200 | 111967999 | 50.2795 | 79.53019 | 155.8669 | 64.77531 | -0.64202 | 2.394189 | 0.169037 |
| 7 | 49227000 | 49230199 | 21.7463 | 35.86534 | 77.51126 | 33.4497 | -0.72839 | 1.357688 | 0.169044 |
| 10 | 93589800 | 93590599 | 4.758494 | 9.248076 | 28.2023 | 10.49346 | -0.99432 | -0.28146 | 0.169065 |
| 10 | 61563400 | 61567199 | 23.99073 | 39.04967 | 87.92816 | 37.74683 | -0.79769 | 1.546263 | 0.169066 |
| 3 | 40902400 | 40905599 | 19.50716 | 32.26302 | 75.00416 | 32.42575 | -0.83254 | 1.317618 | 0.169067 |
| 6 | 62604600 | 62606199 | 15.46083 | 28.82854 | 20.49188 | 8.530117 | 0.940648 | 0.176419 | 0.169068 |
| 13 | 64514200 | 64516799 | 13.40043 | 23.31941 | 54.3463 | 23.88482 | -0.82806 | 0.766056 | 0.169085 |

| | | | | | | | | | |
|---|---|---|---|---|---|---|---|---|---|
| 5 | 31293200 | 31297399 | 28.80091 | 46.35268 | 107.4212 | 45.62337 | -0.85598 | 1.833699 | 0.169113 |
| 14 | 58236200 | 58236999 | 8.470282 | 15.30004 | 38.01817 | 16.51888 | -0.89569 | 0.345307 | 0.169116 |
| 16 | 38655800 | 38656999 | 24.405 | 42.55574 | 31.08955 | 13.68106 | 0.83802 | 0.757085 | 0.16912 |
| X | 25286600 | 25287799 | 13.14668 | 25.42781 | 15.54405 | 5.939905 | 1.264016 | -0.13732 | 0.169127 |
| 4 | 146268800 | 146270599 | 31.22721 | 52.50412 | 42.11594 | 18.39918 | 0.697859 | 1.140149 | 0.169139 |
| 10 | 66932000 | 66933399 | 9.015833 | 16.02164 | 41.4889 | 18.30508 | -0.97028 | 0.457865 | 0.169142 |
| 3 | 16655400 | 16657199 | 20.30132 | 35.95365 | 26.28874 | 11.62407 | 0.840643 | 0.525168 | 0.169144 |
| 1 | 138890600 | 138891999 | 7.223912 | 13.2217 | 36.11956 | 15.43607 | -1.0152 | 0.254656 | 0.169147 |
| 10 | 99122800 | 99125399 | 13.27196 | 23.0492 | 52.14454 | 22.88806 | -0.7855 | 0.793212 | 0.169153 |
| 7 | 11530000 | 11532999 | 74.14141 | 117.733 | 91.65782 | 38.20679 | 0.692937 | 2.295129 | 0.169153 |
| 12 | 22596600 | 22600399 | 64.63339 | 103.5362 | 73.4221 | 30.96653 | 0.84061 | 2.012256 | 0.169158 |
| 4 | 32885800 | 32897799 | 131.1288 | 195.3417 | 350.859 | 144.5788 | -0.58696 | 3.595996 | 0.169159 |
| 6 | 48024200 | 48026199 | 12.62088 | 21.62528 | 57.29446 | 25.12636 | -1.01604 | 0.925272 | 0.169165 |
| 4 | 153374200 | 153425999 | 521.0388 | 691.4992 | 1172.657 | 490.1148 | -0.66949 | 5.399638 | 0.16917 |
| 6 | 117793600 | 117794199 | 4.360319 | 8.484817 | 24.65463 | 9.081406 | -0.91129 | -0.33384 | 0.169171 |
| 16 | 8828600 | 8830799 | 11.16422 | 19.71687 | 44.55962 | 19.71387 | -0.77989 | 0.53363 | 0.169178 |
| 5 | 58407400 | 58408999 | 16.01189 | 29.62196 | 20.85509 | 8.783567 | 0.941611 | 0.226263 | 0.169179 |
| 2 | 165356400 | 165370999 | 110.5137 | 165.9404 | 318.0527 | 131.2684 | -0.67022 | 3.44308 | 0.169187 |
| 3 | 31271600 | 31273799 | 10.75368 | 18.8707 | 46.70524 | 20.65373 | -0.913 | 0.614521 | 0.16919 |
| 12 | 20180600 | 20182199 | 17.94771 | 32.61468 | 19.57364 | 8.441071 | 1.147549 | 0.338254 | 0.1692 |
| 18 | 71770400 | 71773599 | 48.84986 | 79.13973 | 60.66112 | 25.96452 | 0.746943 | 1.661891 | 0.169218 |
| 13 | 73895200 | 73899399 | 18.70947 | 31.37902 | 69.48824 | 30.31852 | -0.76204 | 1.15187 | 0.169228 |
| 16 | 16799600 | 16802199 | 12.84983 | 22.21916 | 54.78452 | 24.03488 | -0.91037 | 0.837062 | 0.169237 |
| X | 5546200 | 5547399 | 25.95489 | 45.09592 | 32.39075 | 14.22149 | 0.862261 | 0.81844 | 0.16924 |
| 15 | 31243400 | 31249599 | 33.52653 | 53.72911 | 120.6912 | 50.93585 | -0.81877 | 1.977688 | 0.169247 |
| 6 | 3363400 | 3366599 | 49.33713 | 81.4794 | 38.24949 | 16.51926 | 1.467674 | 1.383504 | 0.169251 |
| 3 | 126933600 | 126935199 | 7.544313 | 13.73728 | 37.09639 | 15.95735 | -1.00514 | 0.289031 | 0.169251 |
| 4 | 82996400 | 82999399 | 12.16861 | 21.21336 | 51.04436 | 22.45864 | -0.87336 | 0.731118 | 0.169256 |
| 16 | 91443400 | 91445399 | 7.612664 | 13.42722 | 46.68454 | 20.65929 | -1.39741 | 0.53096 | 0.169273 |
| 12 | 23628400 | 23629999 | 24.95709 | 43.4826 | 26.89111 | 11.8274 | 1.077268 | 0.766285 | 0.16928 |
| 9 | 67165600 | 67167599 | 7.987323 | 14.53338 | 37.08298 | 15.96609 | -0.92494 | 0.296542 | 0.169294 |
| 7 | 44813200 | 44836599 | 242.3092 | 340.3475 | 655.4436 | 271.5915 | -0.77044 | 4.511304 | 0.169294 |
| 1 | 74881000 | 74884599 | 16.9371 | 28.56836 | 64.38757 | 28.15622 | -0.78686 | 1.089348 | 0.169297 |
| 7 | 89886800 | 89887999 | 5.831831 | 11.25037 | 31.88254 | 12.32274 | -0.94397 | -0.13229 | 0.169308 |
| 2 | 178606600 | 178607199 | 3.940043 | 7.677011 | 23.6128 | 8.579667 | -0.97362 | -0.39169 | 0.169321 |
| 13 | 8819800 | 8820599 | 3.940043 | 7.677011 | 23.6128 | 8.579667 | -0.97362 | -0.39169 | 0.169321 |
| 14 | 43225000 | 43226399 | 26.31992 | 45.16011 | 35.63754 | 15.6305 | 0.723677 | 0.937078 | 0.169321 |
| 9 | 20347800 | 20350799 | 56.8569 | 91.96087 | 56.49151 | 24.15064 | 1.063325 | 1.707863 | 0.169327 |
| 5 | 42010400 | 42011799 | 8.697387 | 15.93966 | 37.24037 | 15.8864 | -0.78995 | 0.260177 | 0.169338 |
| 13 | 100474800 | 100475999 | 10.35993 | 18.49134 | 42.03784 | 18.47861 | -0.78006 | 0.440651 | 0.169359 |
| X | 146989200 | 146992399 | 30.01318 | 52.34117 | 25.79296 | 11.35114 | 1.402996 | 0.777262 | 0.169362 |
| 3 | 88316200 | 88317599 | 6.064945 | 11.4673 | 31.95689 | 12.97899 | -0.98561 | 0.033422 | 0.169364 |
| 18 | 77063800 | 77066599 | 14.0796 | 24.31507 | 55.9193 | 24.53179 | -0.81097 | 0.85468 | 0.169367 |
| 4 | 135647800 | 135648999 | 6.419464 | 11.91008 | 33.67 | 14.12563 | -1.04392 | 0.166275 | 0.169382 |
| 8 | 74250800 | 74251799 | 26.30471 | 45.10003 | 34.80961 | 15.26645 | 0.755528 | 0.944574 | 0.169397 |
| 5 | 148878200 | 148881999 | 29.00347 | 46.97538 | 95.15158 | 40.71966 | -0.65188 | 1.651821 | 0.169399 |
| 10 | 99850600 | 99853399 | 13.12153 | 22.77517 | 52.67838 | 23.11423 | -0.8174 | 0.801093 | 0.169405 |

| | | | | | | | | | |
|---|---|---|---|---|---|---|---|---|---|
| 16 | 8838600 | 8840799 | 13.55126 | 23.63588 | 54.35162 | 23.92152 | -0.80847 | 0.734755 | 0.169406 |
| X | 54589600 | 54591399 | 44.65051 | 73.90047 | 42.78458 | 18.50571 | 1.166642 | 1.351275 | 0.169406 |
| 14 | 49168200 | 49169799 | 9.758431 | 17.34957 | 41.98793 | 18.51778 | -0.87316 | 0.459695 | 0.169406 |
| 11 | 49193400 | 49193799 | 3.454567 | 6.752729 | 21.29814 | 7.332274 | -0.93662 | -0.59746 | 0.169411 |
| 4 | 36904800 | 36906399 | 17.3873 | 31.93736 | 21.22797 | 9.027985 | 1.014217 | 0.277419 | 0.169414 |
| 2 | 166856600 | 166858199 | 6.132195 | 11.4099 | 33.71926 | 14.08691 | -1.10296 | 0.145443 | 0.16942 |
| 5 | 84695800 | 84696999 | 15.85464 | 29.91175 | 17.66824 | 7.209032 | 1.22787 | 0.10015 | 0.169431 |
| 17 | 88180800 | 88187999 | 40.02561 | 63.90207 | 129.4687 | 54.34222 | -0.67646 | 2.105074 | 0.169439 |
| 5 | 142094800 | 142111999 | 147.0653 | 217.7949 | 379.284 | 156.2014 | -0.55027 | 3.702222 | 0.169445 |
| 2 | 118526600 | 118528799 | 12.13016 | 20.86996 | 55.73081 | 24.4463 | -1.02617 | 0.883111 | 0.169447 |
| 14 | 118139200 | 118143799 | 21.80415 | 35.46406 | 90.30024 | 38.75459 | -0.97523 | 1.554026 | 0.169447 |
| 2 | 110192000 | 110193599 | 8.488241 | 15.29054 | 40.0689 | 17.46966 | -0.97545 | 0.36408 | 0.169454 |
| 14 | 98404200 | 98407599 | 18.77469 | 30.96176 | 78.72545 | 33.97246 | -0.96205 | 1.352298 | 0.169461 |
| 13 | 56514600 | 56518399 | 17.57534 | 29.22517 | 73.44581 | 31.8705 | -0.94381 | 1.246579 | 0.169473 |
| 2 | 19694600 | 19700199 | 34.09735 | 54.81747 | 110.346 | 46.79206 | -0.65532 | 1.875582 | 0.169482 |
| 1 | 126790800 | 126792199 | 17.64248 | 31.57461 | 24.56372 | 10.77306 | 0.759959 | 0.434973 | 0.169496 |
| 9 | 92423800 | 92425199 | 8.535318 | 15.23614 | 39.99392 | 17.57867 | -0.98552 | 0.424261 | 0.169505 |
| X | 116733600 | 116735599 | 21.63287 | 39.70028 | 17.64281 | 7.51238 | 1.58932 | 0.308611 | 0.169507 |
| 8 | 70241800 | 70244399 | 8.289024 | 14.69395 | 42.01656 | 18.57487 | -1.11412 | 0.471309 | 0.169514 |
| 7 | 10565000 | 10567199 | 35.599 | 59.33132 | 45.44008 | 19.7475 | 0.764703 | 1.247894 | 0.169515 |
| X | 5550800 | 5553599 | 44.66678 | 74.57337 | 35.35927 | 15.38337 | 1.453764 | 1.251676 | 0.169516 |
| 14 | 73790000 | 73790799 | 6.051523 | 11.44516 | 30.8264 | 12.51238 | -0.93548 | 0.030617 | 0.16953 |
| 10 | 10837800 | 10839599 | 43.26377 | 71.06906 | 47.28513 | 20.37129 | 0.963846 | 1.436471 | 0.169533 |
| 5 | 112691600 | 112697999 | 35.7461 | 57.33727 | 116.2969 | 49.14792 | -0.67028 | 1.951256 | 0.169539 |
| 16 | 46735000 | 46736399 | 14.15342 | 27.07804 | 16.25382 | 6.433756 | 1.242347 | -0.01506 | 0.169545 |
| 6 | 22355600 | 22356399 | 6.795785 | 12.80591 | 31.87032 | 13.03172 | -0.83229 | 0.064394 | 0.169562 |
| 8 | 86567000 | 86567999 | 6.795785 | 12.80591 | 31.87032 | 13.03172 | -0.83229 | 0.064394 | 0.169562 |
| 6 | 144747600 | 144750199 | 26.48666 | 43.24792 | 89.0739 | 38.2825 | -0.66634 | 1.505344 | 0.16957 |
| 2 | 36502000 | 36503999 | 26.29502 | 46.11693 | 25.26474 | 11.16713 | 1.250814 | 0.672607 | 0.169572 |
| 17 | 52231400 | 52233399 | 6.61791 | 12.56881 | 35.18611 | 14.15622 | -0.9809 | 0.013681 | 0.169577 |
| 11 | 69724000 | 69729599 | 25.22616 | 40.7963 | 99.29444 | 42.44489 | -0.91812 | 1.677745 | 0.169584 |
| 7 | 28251600 | 28265199 | 86.54601 | 131.203 | 280.4658 | 116.0826 | -0.81543 | 3.245465 | 0.169584 |
| 5 | 142535200 | 142549799 | 102.22 | 154.4987 | 288.9413 | 119.477 | -0.62651 | 3.310235 | 0.169591 |
| 12 | 110167400 | 110171599 | 20.20585 | 33.25199 | 79.32773 | 34.20195 | -0.8717 | 1.382836 | 0.169593 |
| 15 | 100392400 | 100397999 | 33.1178 | 53.29568 | 107.5038 | 45.65263 | -0.65646 | 1.841919 | 0.169594 |
| 7 | 8120600 | 8122599 | 21.53924 | 38.66958 | 20.51564 | 8.97394 | 1.310454 | 0.433377 | 0.169598 |
| 14 | 66187400 | 66195999 | 47.83978 | 75.68739 | 158.69 | 65.9557 | -0.73896 | 2.387179 | 0.169623 |
| 10 | 100950800 | 100952999 | 27.79139 | 47.6879 | 36.61534 | 16.05944 | 0.76315 | 0.945323 | 0.169629 |
| 10 | 79533600 | 79538399 | 20.95452 | 34.2231 | 86.71464 | 37.27177 | -0.96389 | 1.489359 | 0.169632 |
| 7 | 111935600 | 111938599 | 15.48739 | 26.26532 | 62.74766 | 27.48432 | -0.87012 | 1.034578 | 0.169653 |
| 13 | 39694400 | 39696199 | 14.99024 | 27.64222 | 21.67981 | 9.171244 | 0.782814 | 0.238529 | 0.169664 |
| 7 | 7483800 | 7486399 | 45.61818 | 74.75464 | 50.65249 | 21.79957 | 0.936307 | 1.473672 | 0.169672 |
| 6 | 90665600 | 90667799 | 9.140947 | 16.30169 | 40.47848 | 17.80595 | -0.90705 | 0.43427 | 0.169675 |
| 7 | 109302000 | 109303599 | 9.140947 | 16.30169 | 40.47848 | 17.80595 | -0.90705 | 0.43427 | 0.169675 |
| 2 | 170165600 | 170170199 | 35.24986 | 56.5982 | 112.9239 | 47.79551 | -0.64482 | 1.918536 | 0.169682 |
| 4 | 117000400 | 117007799 | 35.91891 | 57.21523 | 137.7321 | 57.62762 | -0.92823 | 2.176845 | 0.169696 |
| 14 | 41816600 | 41820599 | 57.12096 | 91.99809 | 62.66495 | 26.64269 | 0.907825 | 1.833148 | 0.169699 |

| | | | | | | | | | |
|---|---|---|---|---|---|---|---|---|---|
| 12 | 21868200 | 21871999 | 82.85414 | 130.7527 | 111.6542 | 46.34704 | 0.552139 | 2.461882 | 0.169702 |
| 4 | 40290600 | 40293199 | 57.35152 | 91.99777 | 77.51125 | 32.75957 | 0.59476 | 1.961815 | 0.169731 |
| 3 | 99125200 | 99127399 | 13.39292 | 23.35751 | 51.61143 | 22.71416 | -0.75109 | 0.731932 | 0.169732 |
| 1 | 117094600 | 117095599 | 22.35968 | 39.0206 | 29.49299 | 12.98505 | 0.789441 | 0.734342 | 0.169742 |
| 12 | 111406400 | 111418199 | 65.05405 | 99.87648 | 237.755 | 98.71104 | -0.95579 | 2.979387 | 0.169742 |
| 18 | 36587200 | 36587599 | 2.93588 | 5.742076 | 19.59394 | 6.625958 | -1.02359 | -0.67595 | 0.169745 |
| X | 91010600 | 91012199 | 22.3414 | 40.03644 | 21.98123 | 9.629845 | 1.260807 | 0.455429 | 0.169773 |
| 2 | 125700400 | 125700999 | 3.526763 | 6.887155 | 22.26286 | 7.824773 | -1.0003 | -0.52234 | 0.169773 |
| 18 | 88206000 | 88207399 | 15.3338 | 29.14789 | 17.33319 | 6.960534 | 1.238373 | 0.045183 | 0.169778 |
| X | 4512800 | 4515799 | 47.18703 | 77.65199 | 45.06279 | 19.42851 | 1.161448 | 1.424282 | 0.169778 |
| 4 | 93074600 | 93075599 | 15.93927 | 29.63439 | 20.58936 | 8.61298 | 0.968079 | 0.199779 | 0.169785 |
| 17 | 68671600 | 68672599 | 11.40902 | 22.22328 | 13.50948 | 4.928103 | 1.332319 | -0.31823 | 0.169798 |
| 5 | 122334200 | 122352799 | 160.9544 | 234.548 | 450.0929 | 185.4821 | -0.71237 | 3.96204 | 0.169798 |
| X | 126077600 | 126077999 | 8.145986 | 15.93442 | 10.35494 | 3.458926 | 1.346768 | -0.67349 | 0.169827 |
| 4 | 38305400 | 38308199 | 21.06061 | 37.95452 | 21.96079 | 9.569534 | 1.190443 | 0.412693 | 0.16984 |
| 9 | 36743800 | 36745599 | 9.715448 | 17.4049 | 39.98177 | 17.52042 | -0.7918 | 0.408053 | 0.169856 |
| 13 | 82793000 | 82793399 | 8.101239 | 15.83996 | 11.56985 | 3.955547 | 1.151683 | -0.59251 | 0.169858 |
| 6 | 131063600 | 131064199 | 10.68312 | 20.78722 | 14.70242 | 5.418154 | 1.103223 | -0.28961 | 0.169859 |
| 13 | 12944600 | 12944999 | 10.68312 | 20.78722 | 14.70242 | 5.418154 | 1.103223 | -0.28961 | 0.169859 |
| 5 | 43628800 | 43630999 | 12.75996 | 22.29277 | 54.89069 | 24.18406 | -0.90639 | 0.707022 | 0.169865 |
| 12 | 110124200 | 110127199 | 17.70532 | 29.96229 | 63.37221 | 27.7398 | -0.69532 | 1.063558 | 0.169879 |
| 15 | 78293000 | 78295199 | 11.37888 | 19.9253 | 46.74877 | 20.63578 | -0.83623 | 0.654703 | 0.169893 |
| 18 | 68366400 | 68369399 | 16.25738 | 27.58092 | 61.70009 | 27.02837 | -0.77569 | 1.033625 | 0.169896 |
| 8 | 12339400 | 12341199 | 10.72232 | 18.8864 | 46.14526 | 20.42618 | -0.89337 | 0.562243 | 0.169908 |
| 3 | 111472000 | 111474799 | 45.88011 | 74.96503 | 52.89306 | 22.73822 | 0.875663 | 1.51335 | 0.16991 |
| 18 | 23443000 | 23444399 | 11.86281 | 20.71186 | 48.88877 | 21.53088 | -0.84509 | 0.707966 | 0.16991 |
| 16 | 89291600 | 89292999 | 7.984157 | 14.52851 | 36.56104 | 15.74017 | -0.90475 | 0.296551 | 0.169919 |
| 8 | 129965800 | 129966999 | 13.05044 | 25.24898 | 14.18296 | 5.41246 | 1.385065 | -0.14793 | 0.169929 |
| 1 | 136793800 | 136795399 | 7.724211 | 14.38728 | 35.48172 | 14.79672 | -0.8431 | 0.154215 | 0.16993 |
| 1 | 83321400 | 83321999 | 10.27361 | 19.93361 | 15.83328 | 5.961909 | 0.909379 | -0.21846 | 0.169933 |
| 11 | 102982600 | 102986599 | 27.55929 | 44.65676 | 93.71352 | 40.11778 | -0.70214 | 1.639372 | 0.169935 |
| 10 | 12954200 | 12956599 | 10.53699 | 18.65585 | 44.03759 | 19.47315 | -0.84085 | 0.500663 | 0.169952 |
| 5 | 37978200 | 37982199 | 34.48961 | 55.25965 | 119.4712 | 50.40308 | -0.7641 | 1.986197 | 0.169954 |
| 18 | 76665600 | 76667799 | 15.64941 | 29.17166 | 20.48618 | 8.531957 | 0.957474 | 0.179975 | 0.169954 |
| Y | 16618800 | 16620799 | 13.0787 | 25.03579 | 8.574212 | 3.388525 | 2.042498 | -0.05674 | 0.169956 |
| 4 | 154781000 | 154797999 | 89.07051 | 134.4419 | 303.1506 | 125.299 | -0.8978 | 3.335501 | 0.16996 |
| 17 | 90212200 | 90212799 | 8.337201 | 16.29205 | 11.47561 | 3.98134 | 1.184 | -0.54075 | 0.169961 |
| 4 | 70177000 | 70179199 | 10.57283 | 18.63316 | 43.54143 | 19.27151 | -0.8287 | 0.553376 | 0.169964 |
| 3 | 51183400 | 51185199 | 9.466402 | 16.51682 | 50.9635 | 22.43517 | -1.22938 | 0.706538 | 0.169966 |
| Y | 8262400 | 8263399 | 9.335094 | 18.2008 | 8.17277 | 2.944532 | 1.777112 | -0.41208 | 0.169978 |
| 10 | 49910200 | 49912599 | 27.63312 | 48.10476 | 29.84885 | 13.12139 | 1.072318 | 0.795005 | 0.169981 |
| 15 | 100249400 | 100253399 | 28.75068 | 46.37324 | 101.6626 | 43.30895 | -0.77214 | 1.76484 | 0.169987 |
| 13 | 66001400 | 66017799 | 318.075 | 455.0857 | 355.6972 | 146.9404 | 0.556794 | 4.242307 | 0.169988 |
| 13 | 104909200 | 104911999 | 23.59822 | 41.61303 | 25.45426 | 11.26478 | 1.094314 | 0.58961 | 0.169996 |
| 1 | 109753400 | 109753999 | 10.31747 | 20.1124 | 12.93748 | 4.672617 | 1.263144 | -0.36962 | 0.169997 |
| 12 | 77832200 | 77834199 | 9.390582 | 16.89698 | 38.9865 | 17.02112 | -0.79427 | 0.376283 | 0.17003 |
| 1 | 37351200 | 37353999 | 10.41983 | 18.03772 | 54.12726 | 23.74735 | -1.19133 | 0.817042 | 0.170033 |

| | | | | | | | | | |
|---|---|---|---|---|---|---|---|---|---|
| 10 | 108215600 | 108220799 | 37.04054 | 59.42983 | 115.4451 | 48.80823 | -0.60762 | 1.944745 | 0.17004 |
| 18 | 67073200 | 67073999 | 6.199847 | 11.66179 | 32.31889 | 13.25573 | -0.98876 | 0.0749 | 0.17005 |
| 7 | 27172600 | 27174399 | 10.60998 | 18.60663 | 47.7697 | 21.11304 | -0.96562 | 0.625493 | 0.170051 |
| 19 | 56248400 | 56252399 | 21.09746 | 34.84289 | 76.96454 | 33.24012 | -0.75936 | 1.338872 | 0.170056 |
| 10 | 120163400 | 120164799 | 8.934076 | 16.09061 | 38.48871 | 16.78469 | -0.8447 | 0.366793 | 0.170061 |
| 7 | 11146400 | 11146799 | 16.89476 | 30.65649 | 21.74364 | 9.394003 | 0.906684 | 0.344064 | 0.170076 |
| 5 | 74354800 | 74356799 | 9.24131 | 16.22077 | 45.67853 | 20.20391 | -1.09746 | 0.602808 | 0.170078 |
| 11 | 52605400 | 52610799 | 39.80617 | 63.73849 | 123.3959 | 52.00994 | -0.60629 | 2.012665 | 0.170081 |
| 10 | 75932200 | 75932999 | 5.60572 | 10.69958 | 32.33704 | 12.8737 | -1.07627 | -0.03755 | 0.170094 |
| 2 | 63432800 | 63434399 | 19.20107 | 35.27146 | 19.3607 | 8.233063 | 1.288157 | 0.287195 | 0.170105 |
| X | 148368400 | 148370199 | 14.60387 | 27.58712 | 19.1415 | 7.789161 | 1.001702 | 0.085231 | 0.170126 |
| 11 | 57241400 | 57242199 | 12.46148 | 24.21537 | 13.87297 | 5.16761 | 1.389703 | -0.23409 | 0.170137 |
| 7 | 17381000 | 17393399 | 133.1745 | 204.9189 | 138.5976 | 57.56059 | 0.860185 | 2.949208 | 0.170149 |
| 3 | 4225800 | 4228999 | 60.77859 | 97.26505 | 82.07309 | 34.56458 | 0.588649 | 2.039599 | 0.170159 |
| 12 | 82521600 | 82523599 | 13.07754 | 22.88176 | 50.49594 | 22.27503 | -0.74886 | 0.68051 | 0.170164 |
| 2 | 30007200 | 30009599 | 6.627826 | 12.19453 | 36.75014 | 15.59381 | -1.1483 | 0.220005 | 0.170164 |
| 16 | 4476200 | 4479799 | 14.38721 | 24.8927 | 53.76187 | 23.58666 | -0.72004 | 0.837859 | 0.170165 |
| 11 | 12592400 | 12594599 | 23.46619 | 41.9262 | 21.03998 | 9.240823 | 1.386733 | 0.477922 | 0.170166 |
| 4 | 44065800 | 44067799 | 7.929915 | 14.59798 | 35.16175 | 14.90906 | -0.8268 | 0.225315 | 0.170175 |
| 15 | 56666000 | 56668999 | 39.60648 | 65.16849 | 51.63227 | 22.25962 | 0.713593 | 1.40981 | 0.17019 |
| 5 | 62704600 | 62706399 | 14.78742 | 27.9945 | 18.44611 | 7.471831 | 1.080871 | 0.069748 | 0.17019 |
| 15 | 92695400 | 92695999 | 5.609945 | 10.64217 | 30.33678 | 12.23507 | -1.00812 | 0.003903 | 0.170195 |
| 15 | 80733800 | 80736999 | 14.55368 | 25.09864 | 56.46628 | 24.77069 | -0.77987 | 0.870125 | 0.170196 |
| 12 | 83441000 | 83442999 | 16.54941 | 28.14172 | 60.66401 | 26.58778 | -0.72213 | 1.013007 | 0.170217 |
| 1 | 24040000 | 24041799 | 10.71021 | 18.90871 | 43.52177 | 19.256 | -0.80614 | 0.533351 | 0.170229 |
| 10 | 60567600 | 60575799 | 45.40012 | 71.89082 | 153.3515 | 63.76526 | -0.76272 | 2.364522 | 0.170233 |
| 9 | 107531200 | 107541199 | 41.04484 | 65.00637 | 154.8406 | 64.39053 | -0.92135 | 2.357406 | 0.170239 |
| 7 | 55624000 | 55624399 | 9.293576 | 18.1365 | 13.16124 | 4.679892 | 1.111667 | -0.43575 | 0.170243 |
| 7 | 61188400 | 61188999 | 9.293576 | 18.1365 | 13.16124 | 4.679892 | 1.111667 | -0.43575 | 0.170243 |
| 15 | 101071400 | 101074199 | 12.20084 | 21.19937 | 51.55393 | 22.63578 | -0.88876 | 0.783116 | 0.170259 |
| 9 | 94729000 | 94730599 | 19.63709 | 34.79553 | 26.36691 | 11.65742 | 0.789692 | 0.515449 | 0.170264 |
| 9 | 96593600 | 96596399 | 14.14456 | 24.26134 | 59.63823 | 26.15642 | -0.90873 | 0.921216 | 0.170278 |
| 12 | 22603600 | 22607199 | 62.69848 | 100.5193 | 70.32047 | 29.69548 | 0.861557 | 1.984254 | 0.170285 |
| 1 | 53372000 | 53375199 | 40.33959 | 66.37339 | 51.63223 | 22.25941 | 0.739964 | 1.412804 | 0.170285 |
| 11 | 10631200 | 10632999 | 23.78243 | 41.34679 | 32.93362 | 14.46545 | 0.71366 | 0.798375 | 0.170298 |
| 1 | 85290600 | 85290799 | 6.497161 | 12.70897 | 10.36725 | 3.467116 | 1.024508 | -0.69841 | 0.1703 |
| 2 | 125563400 | 125566999 | 15.58934 | 26.19131 | 66.33829 | 28.9745 | -0.95471 | 1.122289 | 0.170303 |
| 1 | 103853200 | 103855399 | 21.01572 | 38.04708 | 21.07711 | 9.132121 | 1.258144 | 0.385926 | 0.170311 |
| 4 | 11155800 | 11158399 | 10.56359 | 18.63335 | 44.04786 | 19.49221 | -0.84495 | 0.542948 | 0.170316 |
| 10 | 94447000 | 94448599 | 7.744422 | 14.36383 | 34.78731 | 14.60055 | -0.82406 | 0.1792 | 0.170336 |
| 10 | 31246200 | 31246399 | 7.657648 | 14.9691 | 12.28653 | 4.226245 | 0.978687 | -0.576 | 0.170341 |
| 17 | 29226000 | 29228599 | 11.1451 | 19.3744 | 53.68998 | 23.58085 | -1.07642 | 0.770534 | 0.170341 |
| 7 | 20795600 | 20804599 | 119.182 | 185.4949 | 121.2591 | 50.33594 | 0.921816 | 2.738803 | 0.170346 |
| 9 | 82582000 | 82585999 | 41.78978 | 66.74768 | 129.198 | 54.26205 | -0.61005 | 2.093322 | 0.170361 |
| 11 | 115880400 | 115885999 | 26.36011 | 42.55915 | 101.1861 | 43.16057 | -0.88722 | 1.730173 | 0.170372 |
| 3 | 68620000 | 68620799 | 10.6994 | 20.75077 | 16.41838 | 6.19475 | 0.912784 | -0.20384 | 0.170378 |
| 3 | 53064200 | 53064599 | 4.058147 | 7.917692 | 23.3173 | 8.324146 | -0.88866 | -0.45454 | 0.170387 |

| | | | | | | | | | |
|---|---|---|---|---|---|---|---|---|---|
| 2 | 52314200 | 52315599 | 7.841749 | 14.2694 | 37.61388 | 16.19339 | -0.97158 | 0.295613 | 0.170393 |
| 2 | 152919800 | 152926799 | 38.40993 | 61.34332 | 129.2144 | 54.26146 | -0.73192 | 2.092769 | 0.170397 |
| X | 117794400 | 117796799 | 30.30938 | 52.57637 | 29.93395 | 13.1338 | 1.195568 | 0.855074 | 0.170409 |
| X | 148834400 | 148837799 | 59.47958 | 96.08645 | 53.57671 | 22.86984 | 1.200796 | 1.746189 | 0.170423 |
| 3 | 120847400 | 120849999 | 31.54507 | 54.56533 | 29.79201 | 13.07039 | 1.254848 | 0.884149 | 0.170423 |
| 10 | 121758600 | 121760199 | 9.627445 | 17.06637 | 41.46131 | 18.32945 | -0.88087 | 0.478682 | 0.170424 |
| 6 | 108685200 | 108687199 | 7.514372 | 13.45813 | 44.76337 | 19.6202 | -1.32247 | 0.401912 | 0.170431 |
| X | 157614800 | 157617399 | 27.61038 | 47.38501 | 36.61523 | 16.05961 | 0.754021 | 0.942966 | 0.170442 |
| 5 | 114059200 | 114069399 | 79.47521 | 122.3422 | 223.5036 | 92.82804 | -0.57144 | 2.930463 | 0.170444 |
| 3 | 140087400 | 140087799 | 9.70268 | 18.86635 | 12.48451 | 4.626735 | 1.1919 | -0.29027 | 0.170444 |
| 18 | 5591000 | 5593599 | 14.31676 | 24.09444 | 65.09968 | 28.44826 | -1.04711 | 1.103638 | 0.170449 |
| 8 | 117585800 | 117587799 | 23.69281 | 41.126 | 32.50125 | 14.26137 | 0.724722 | 0.818698 | 0.17045 |
| 3 | 130014400 | 130014999 | 9.302654 | 18.12029 | 14.93155 | 5.447125 | 0.898677 | -0.34087 | 0.170457 |
| 15 | 85501800 | 85503399 | 15.03751 | 28.18544 | 20.08448 | 8.2872 | 0.947281 | 0.140047 | 0.170459 |
| 11 | 102494400 | 102503799 | 41.98262 | 66.57143 | 153.8135 | 64.02441 | -0.87597 | 2.324012 | 0.170479 |
| 8 | 105690600 | 105691799 | 6.337294 | 12.00665 | 30.90605 | 12.4983 | -0.86633 | 0.020255 | 0.170479 |
| 4 | 145436400 | 145438799 | 65.67749 | 104.9571 | 78.24315 | 32.88601 | 0.764865 | 2.086393 | 0.170486 |
| 11 | 98232400 | 98233799 | 5.540322 | 10.40522 | 34.51206 | 14.19219 | -1.24978 | 0.079254 | 0.170498 |
| 17 | 57747000 | 57747999 | 13.16022 | 25.39112 | 15.97335 | 6.171461 | 1.207792 | -0.10111 | 0.170505 |
| 4 | 118152000 | 118154199 | 7.640676 | 14.04022 | 35.12551 | 14.93024 | -0.88343 | 0.234047 | 0.170513 |
| 11 | 13382600 | 13386199 | 38.4071 | 63.93944 | 44.36272 | 19.26561 | 0.906864 | 1.268925 | 0.170513 |
| 4 | 148243000 | 148252999 | 66.485 | 102.069 | 248.5559 | 103.1948 | -0.98862 | 2.980469 | 0.170521 |
| 14 | 45720600 | 45721599 | 5.385556 | 10.43147 | 31.08464 | 11.79929 | -0.99058 | -0.20113 | 0.170529 |
| 1 | 181749200 | 181750199 | 8.46074 | 15.30815 | 38.05113 | 16.49897 | -0.89385 | 0.337902 | 0.170532 |
| 16 | 70480800 | 70481399 | 12.59604 | 23.77438 | 18.29507 | 7.457898 | 0.850964 | 0.068625 | 0.170532 |
| 5 | 60481400 | 60482199 | 6.309144 | 12.03359 | 30.5629 | 12.18581 | -0.82887 | -0.02664 | 0.170534 |
| 14 | 114530400 | 114532199 | 27.22641 | 47.57937 | 26.95074 | 11.87628 | 1.203263 | 0.735616 | 0.170539 |
| 10 | 25952600 | 25955199 | 14.45957 | 24.78206 | 57.98121 | 25.42806 | -0.83784 | 0.928832 | 0.170544 |
| 1 | 18251600 | 18254199 | 17.96329 | 33.38334 | 19.26185 | 8.062553 | 1.233331 | 0.215772 | 0.170549 |
| 7 | 40979200 | 40983199 | 67.0106 | 107.3261 | 77.35156 | 32.61539 | 0.817041 | 2.02193 | 0.170565 |
| 8 | 112370000 | 112371399 | 6.962908 | 13.05474 | 32.85522 | 13.54984 | -0.85858 | 0.100042 | 0.170566 |
| 7 | 5495200 | 5496999 | 21.39104 | 38.88849 | 19.76917 | 8.520848 | 1.38556 | 0.362184 | 0.170585 |
| 2 | 136924600 | 136927799 | 13.97943 | 24.0113 | 57.45197 | 25.2 | -0.8695 | 0.906064 | 0.170593 |
| 3 | 41517800 | 41521999 | 26.38681 | 42.90789 | 89.54062 | 38.42539 | -0.6894 | 1.565411 | 0.170593 |
| 12 | 23259800 | 23265399 | 105.7094 | 164.7774 | 130.2009 | 54.03545 | 0.650333 | 2.707306 | 0.170597 |
| 4 | 48595800 | 48603799 | 72.35935 | 112.4535 | 210.652 | 87.46322 | -0.59697 | 2.752624 | 0.17061 |
| 14 | 57101600 | 57103999 | 14.00144 | 23.99894 | 57.34812 | 25.15058 | -0.86784 | 0.928977 | 0.170615 |
| 12 | 80509800 | 80511599 | 6.249268 | 11.39588 | 38.70216 | 16.61755 | -1.33124 | 0.273532 | 0.170637 |
| 11 | 52053000 | 52060599 | 60.2344 | 94.15259 | 184.5889 | 76.58364 | -0.656 | 2.643801 | 0.170638 |
| 16 | 41993800 | 41996399 | 13.90199 | 24.11285 | 52.69236 | 23.11919 | -0.73608 | 0.811798 | 0.170642 |
| 19 | 59484400 | 59486399 | 9.400817 | 16.93487 | 40.06957 | 17.46939 | -0.82881 | 0.373626 | 0.170671 |
| 6 | 145587400 | 145588399 | 5.847152 | 11.22698 | 29.10593 | 11.40258 | -0.83435 | -0.09316 | 0.170672 |
| 18 | 62147400 | 62149999 | 14.0718 | 24.46112 | 53.77487 | 23.61754 | -0.74422 | 0.787126 | 0.170678 |
| 7 | 18003600 | 18013999 | 242.5648 | 356.1598 | 280.3375 | 115.4592 | 0.583142 | 3.848859 | 0.170696 |
| 1 | 92256800 | 92258399 | 9.087184 | 16.36154 | 38.48342 | 16.78857 | -0.8209 | 0.36997 | 0.170712 |
| 8 | 63345800 | 63349199 | 36.2344 | 60.33546 | 45.44872 | 19.73994 | 0.788582 | 1.259017 | 0.170717 |
| 7 | 32837200 | 32840399 | 68.17771 | 108.9763 | 77.59153 | 32.6222 | 0.83136 | 2.082629 | 0.170722 |

| | | | | | | | | | |
|---|---|---|---|---|---|---|---|---|---|
| 2 | 169892600 | 169912999 | 201.2956 | 290.1111 | 502.2388 | 207.2465 | -0.5795 | 4.130802 | 0.170744 |
| 2 | 95930400 | 95932399 | 19.37539 | 35.69251 | 17.58869 | 7.450787 | 1.445807 | 0.268855 | 0.170757 |
| 13 | 91721000 | 91721999 | 5.41325 | 10.44762 | 32.54093 | 12.56052 | -1.07752 | -0.14156 | 0.170775 |
| 14 | 88272000 | 88273999 | 33.00706 | 55.73229 | 37.20816 | 16.27601 | 0.961818 | 1.105766 | 0.17078 |
| 4 | 154446200 | 154448599 | 10.37203 | 18.08436 | 52.59759 | 23.14441 | -1.14515 | 0.722058 | 0.170782 |
| 15 | 96627800 | 96630399 | 12.13469 | 21.27781 | 46.71598 | 20.64511 | -0.74119 | 0.63542 | 0.1708 |
| X | 102364800 | 102366599 | 9.538849 | 17.21976 | 39.57989 | 17.21051 | -0.78494 | 0.360337 | 0.170809 |
| 12 | 83340800 | 83342599 | 11.1957 | 19.73257 | 44.57401 | 19.72819 | -0.7801 | 0.558696 | 0.170848 |
| 8 | 13725400 | 13733999 | 43.85432 | 69.65628 | 148.0845 | 61.74349 | -0.75371 | 2.274289 | 0.170864 |
| 17 | 38979000 | 38983399 | 82.77867 | 131.3381 | 97.12954 | 40.45355 | 0.765765 | 2.326424 | 0.170868 |
| 16 | 34973600 | 34982999 | 54.07376 | 84.5186 | 201.4551 | 83.58162 | -0.93757 | 2.64338 | 0.170875 |
| 5 | 31205000 | 31206399 | 7.212959 | 13.20251 | 35.59433 | 15.21027 | -0.99585 | 0.254653 | 0.170897 |
| 8 | 86426600 | 86428199 | 11.1327 | 19.37758 | 53.16008 | 23.36687 | -1.06175 | 0.751043 | 0.170909 |
| 4 | 57854200 | 57857999 | 46.10121 | 72.8557 | 158.6238 | 65.89107 | -0.79452 | 2.414091 | 0.170918 |
| 5 | 137013600 | 137014799 | 5.268231 | 10.14405 | 28.03417 | 10.89392 | -0.91426 | -0.12922 | 0.170922 |
| 9 | 71028200 | 71029199 | 5.268231 | 10.14405 | 28.03417 | 10.89392 | -0.91426 | -0.12922 | 0.170922 |
| 14 | 36856800 | 36857599 | 5.268231 | 10.14405 | 28.03417 | 10.89392 | -0.91426 | -0.12922 | 0.170922 |
| 5 | 136224200 | 136224999 | 10.4561 | 20.25917 | 16.38133 | 6.20896 | 0.875239 | -0.19271 | 0.170932 |
| 3 | 128911400 | 128912599 | 11.28014 | 21.81597 | 16.84196 | 6.438166 | 0.930038 | -0.15359 | 0.170935 |
| 15 | 14871600 | 14872199 | 11.85226 | 22.88823 | 16.70702 | 6.431331 | 1.000372 | -0.12522 | 0.170935 |
| 10 | 45878000 | 45880199 | 18.80405 | 33.3136 | 26.04265 | 11.51446 | 0.744935 | 0.511364 | 0.170943 |
| 16 | 35012800 | 35014799 | 8.622738 | 15.20455 | 44.09315 | 19.51361 | -1.13759 | 0.5367 | 0.170973 |
| 9 | 44114400 | 44117999 | 28.14514 | 45.76447 | 97.24004 | 41.72862 | -0.71556 | 1.573129 | 0.170997 |
| 14 | 79029000 | 79029999 | 10.43471 | 20.29191 | 14.66161 | 5.428889 | 1.066126 | -0.27749 | 0.170999 |
| 8 | 20091400 | 20093399 | 17.01771 | 32.12826 | 17.27383 | 7.037795 | 1.365038 | 0.106558 | 0.171013 |
| 19 | 44941400 | 44942599 | 16.74249 | 30.56345 | 22.58694 | 9.685215 | 0.854582 | 0.310803 | 0.171031 |
| 4 | 144913400 | 144916399 | 13.60736 | 23.53271 | 55.37332 | 24.29331 | -0.84336 | 0.839085 | 0.171051 |
| 8 | 72885200 | 72892599 | 40.89639 | 65.11276 | 140.2154 | 58.6279 | -0.76863 | 2.19646 | 0.171062 |
| 3 | 101945000 | 101946999 | 23.86292 | 39.22335 | 80.92137 | 34.8714 | -0.66362 | 1.40856 | 0.171062 |
| 18 | 77010000 | 77010399 | 2.836769 | 5.545803 | 20.63019 | 7.079321 | -1.16615 | -0.62292 | 0.171068 |
| 18 | 15892000 | 15893399 | 12.35745 | 21.45945 | 51.57101 | 22.63531 | -0.87198 | 0.790552 | 0.171073 |
| 6 | 26053400 | 26053999 | 11.80202 | 22.59 | 17.94422 | 7.093281 | 0.84485 | -0.03677 | 0.171075 |
| 16 | 11352400 | 11353399 | 11.80202 | 22.59 | 17.94422 | 7.093281 | 0.84485 | -0.03677 | 0.171075 |
| 3 | 124050200 | 124050999 | 12.31509 | 23.73031 | 17.31298 | 6.713528 | 0.991942 | -0.09403 | 0.171081 |
| 3 | 153618200 | 153618999 | 3.53074 | 6.883547 | 23.06965 | 8.318014 | -1.08503 | -0.42681 | 0.171083 |
| 5 | 100405000 | 100411399 | 39.01885 | 62.5251 | 121.3447 | 51.20427 | -0.60849 | 1.988995 | 0.171084 |
| 9 | 90201600 | 90202399 | 17.06267 | 32.53172 | 14.9075 | 5.948574 | 1.618186 | 0.03604 | 0.171087 |
| 10 | 39795800 | 39796599 | 3.417088 | 6.672634 | 24.19084 | 8.509977 | -1.16578 | -0.51636 | 0.1711 |
| 18 | 34102000 | 34105399 | 54.88853 | 88.25222 | 73.75109 | 31.27881 | 0.610508 | 1.885404 | 0.171101 |
| 8 | 11807800 | 11809999 | 8.816309 | 15.79002 | 39.48338 | 17.30581 | -0.91331 | 0.403165 | 0.171104 |
| 12 | 54687200 | 54690199 | 26.97788 | 46.20106 | 37.27031 | 16.34412 | 0.691794 | 0.960356 | 0.171104 |
| 15 | 47240200 | 47243399 | 59.47828 | 95.42711 | 77.43805 | 32.73792 | 0.649127 | 1.958593 | 0.171125 |
| 7 | 110618000 | 110619199 | 6.607208 | 12.59141 | 31.73432 | 12.67559 | -0.82062 | -0.01599 | 0.171128 |
| 4 | 154736800 | 154737599 | 3.623783 | 6.998104 | 29.75925 | 11.45486 | -1.51507 | -0.17525 | 0.171137 |
| X | 33340400 | 33342399 | 23.95347 | 42.37133 | 22.82616 | 10.09542 | 1.278538 | 0.556804 | 0.171144 |
| 2 | 103894400 | 103898799 | 33.24376 | 53.58063 | 109.2076 | 46.47875 | -0.66873 | 1.793601 | 0.171149 |
| 12 | 17592600 | 17596399 | 19.10772 | 31.82359 | 71.47408 | 31.04765 | -0.78221 | 1.231274 | 0.171149 |

| | | | | | | | | | |
|---|---|---|---|---|---|---|---|---|---|
| 12 | 82148200 | 82155599 | 61.24905 | 96.07477 | 178.6606 | 74.09734 | -0.57592 | 2.581169 | 0.171151 |
| 3 | 51057400 | 51062999 | 34.24709 | 55.07833 | 110.7015 | 46.96769 | -0.65248 | 1.863937 | 0.171157 |
| 11 | 68683800 | 68686199 | 15.45047 | 26.29256 | 61.73497 | 27.05878 | -0.84477 | 1.002103 | 0.17116 |
| 10 | 60477000 | 60478199 | 6.395063 | 11.973 | 33.34604 | 13.7855 | -1.00626 | 0.105125 | 0.171163 |
| 15 | 99669400 | 99670799 | 6.395063 | 11.973 | 33.34604 | 13.7855 | -1.00626 | 0.105125 | 0.171163 |
| 4 | 121767000 | 121767399 | 8.179372 | 15.97263 | 12.15747 | 4.274214 | 1.057538 | -0.49551 | 0.171166 |
| 8 | 121683000 | 121704599 | 167.5463 | 242.7711 | 482.7342 | 199.0565 | -0.77161 | 4.048563 | 0.171167 |
| 3 | 102152600 | 102153799 | 6.382537 | 11.95308 | 32.24296 | 13.32281 | -0.95937 | 0.102452 | 0.171171 |
| 7 | 10678800 | 10683999 | 82.85151 | 131.217 | 101.3015 | 42.1259 | 0.701517 | 2.378305 | 0.171173 |
| 7 | 13340800 | 13343599 | 58.53448 | 94.32369 | 65.79488 | 27.99449 | 0.874539 | 1.821263 | 0.171182 |
| 17 | 41190800 | 41191999 | 7.02328 | 13.00763 | 35.83486 | 15.0696 | -1.01059 | 0.18286 | 0.171184 |
| 5 | 115104000 | 115111599 | 41.12989 | 65.32317 | 146.5607 | 61.10273 | -0.83138 | 2.275047 | 0.1712 |
| X | 75596000 | 75596399 | 14.71418 | 27.37429 | 18.86148 | 7.881584 | 0.981332 | 0.181132 | 0.171206 |
| 17 | 28984600 | 29001199 | 101.8618 | 153.2669 | 313.9127 | 129.6318 | -0.76319 | 3.400491 | 0.171207 |
| X | 80094200 | 80096799 | 39.95873 | 65.91418 | 49.40077 | 21.32033 | 0.79498 | 1.381152 | 0.171211 |
| 2 | 178780400 | 178791599 | 87.53748 | 133.9271 | 247.0075 | 102.4885 | -0.59204 | 3.055973 | 0.171227 |
| 18 | 73812200 | 73813999 | 23.41017 | 40.96945 | 30.05993 | 13.26257 | 0.83208 | 0.698617 | 0.171246 |
| 16 | 46881200 | 46884399 | 33.22905 | 55.64064 | 43.7518 | 19.06991 | 0.72659 | 1.189656 | 0.171248 |
| 9 | 42367800 | 42369599 | 7.793166 | 14.30687 | 37.27421 | 15.86398 | -0.94359 | 0.242338 | 0.171255 |
| X | 147623200 | 147629399 | 111.902 | 174.396 | 125.3155 | 52.00946 | 0.78698 | 2.713741 | 0.171292 |
| 17 | 75766600 | 75767999 | 11.20772 | 19.66284 | 46.70787 | 20.65079 | -0.85416 | 0.620823 | 0.171299 |
| 17 | 18932800 | 18934999 | 44.48121 | 72.81655 | 52.83934 | 22.73085 | 0.837172 | 1.484964 | 0.171317 |
| 7 | 54328000 | 54329799 | 23.0256 | 40.1506 | 31.86293 | 14.02143 | 0.719331 | 0.752169 | 0.171326 |
| 14 | 118839600 | 118841999 | 15.58298 | 26.5901 | 59.03957 | 25.88303 | -0.76374 | 0.975639 | 0.171363 |
| 9 | 44351800 | 44352799 | 13.78479 | 23.83713 | 53.71056 | 23.56375 | -0.78087 | 0.841895 | 0.171378 |
| 5 | 109989200 | 109990199 | 15.94712 | 29.35327 | 21.99776 | 9.328814 | 0.845359 | 0.256606 | 0.171389 |
| 13 | 77247200 | 77248599 | 15.94712 | 29.35327 | 21.99776 | 9.328814 | 0.845359 | 0.256606 | 0.171389 |
| X | 18244000 | 18245599 | 22.75278 | 41.39557 | 17.97611 | 7.740642 | 1.611689 | 0.364398 | 0.171389 |
| 7 | 22290400 | 22290799 | 9.359796 | 18.24827 | 14.40472 | 5.192412 | 0.975477 | -0.38261 | 0.171396 |
| 18 | 77272600 | 77276999 | 34.23986 | 54.71858 | 129.5146 | 54.43704 | -0.89873 | 2.068614 | 0.171421 |
| 6 | 128905600 | 128909199 | 50.95504 | 82.33702 | 65.04433 | 27.77431 | 0.699902 | 1.728008 | 0.171438 |
| 13 | 75354000 | 75354599 | 3.341287 | 6.531581 | 20.67185 | 7.10872 | -0.93997 | -0.6062 | 0.171442 |
| 3 | 17181800 | 17183199 | 14.97805 | 28.34747 | 18.43946 | 7.473553 | 1.098639 | 0.073506 | 0.171452 |
| 11 | 59774600 | 59777799 | 16.58329 | 28.1021 | 61.68372 | 27.01195 | -0.74846 | 1.045277 | 0.171464 |
| 14 | 47350600 | 47357799 | 48.05751 | 76.3597 | 143.9388 | 60.0364 | -0.57999 | 2.267114 | 0.171471 |
| 7 | 62301200 | 62304199 | 69.07935 | 110.188 | 88.88315 | 37.25993 | 0.648438 | 2.150316 | 0.171475 |
| 18 | 50577600 | 50578399 | 11.98422 | 22.93396 | 17.94006 | 7.095885 | 0.866065 | -0.03284 | 0.171483 |
| 1 | 102607200 | 102609199 | 20.88703 | 37.44907 | 21.68422 | 9.495689 | 1.184216 | 0.43907 | 0.171494 |
| 7 | 85129000 | 85130599 | 25.44233 | 44.60745 | 26.42887 | 11.67858 | 1.138528 | 0.67439 | 0.171495 |
| 4 | 124324400 | 124339599 | 98.14081 | 148.2421 | 299.6122 | 123.8654 | -0.73914 | 3.321735 | 0.171498 |
| 7 | 77143000 | 77144399 | 16.05049 | 30.26452 | 18.3522 | 7.496152 | 1.189603 | 0.108087 | 0.1715 |
| 3 | 143907400 | 143908399 | 4.686885 | 9.014079 | 27.91001 | 10.88225 | -1.08007 | -0.12272 | 0.171511 |
| 7 | 43996800 | 43998199 | 7.058894 | 12.98101 | 36.2087 | 15.37476 | -1.03842 | 0.225336 | 0.171513 |
| 15 | 63292200 | 63293599 | 17.95682 | 32.95151 | 21.20994 | 9.032221 | 1.058992 | 0.287558 | 0.171516 |
| 13 | 40889600 | 40899799 | 94.18697 | 143.1919 | 270.5976 | 112.0934 | -0.63443 | 3.187856 | 0.171517 |
| 8 | 15214600 | 15217999 | 17.57271 | 29.64534 | 63.8346 | 27.9158 | -0.72122 | 1.091096 | 0.17152 |
| X | 96829800 | 96831599 | 29.04869 | 49.61481 | 37.7498 | 16.55083 | 0.775369 | 0.993872 | 0.171521 |

| | | | | | | | | | |
|---|---|---|---|---|---|---|---|---|---|
| 18 | 25385600 | 25388599 | 29.35439 | 50.41449 | 34.82774 | 15.27634 | 0.915031 | 0.941156 | 0.171542 |
| 5 | 39074000 | 39075599 | 16.75773 | 30.65985 | 22.39167 | 9.574112 | 0.873904 | 0.296742 | 0.171562 |
| 1 | 50441200 | 50441999 | 23.6823 | 41.30636 | 31.75403 | 13.97577 | 0.765006 | 0.751683 | 0.17157 |
| 2 | 158750600 | 158753799 | 20.03735 | 33.41366 | 70.49274 | 30.64555 | -0.69228 | 1.219949 | 0.171582 |
| 8 | 69713800 | 69715399 | 15.08597 | 28.14597 | 20.50329 | 8.526439 | 0.906568 | 0.169294 | 0.171585 |
| 10 | 26804000 | 26804399 | 7.660247 | 14.96549 | 12.91841 | 4.507309 | 0.88927 | -0.52467 | 0.171589 |
| 17 | 6378200 | 6380599 | 9.566733 | 17.14398 | 39.47211 | 17.29241 | -0.79461 | 0.404972 | 0.171597 |
| 2 | 119307400 | 119309399 | 8.310184 | 14.72414 | 44.59373 | 19.71616 | -1.19725 | 0.479785 | 0.171605 |
| 16 | 18630000 | 18631399 | 7.577892 | 14.12611 | 33.81242 | 14.08137 | -0.79831 | 0.145488 | 0.171612 |
| 5 | 117988400 | 117992399 | 17.50702 | 29.26169 | 69.15964 | 30.10987 | -0.85544 | 1.186232 | 0.171612 |
| 7 | 43681400 | 43682599 | 17.08002 | 30.98319 | 21.74101 | 9.396454 | 0.921735 | 0.347317 | 0.17162 |
| 2 | 178095000 | 178096199 | 15.65397 | 28.8269 | 22.07761 | 9.356832 | 0.814965 | 0.252231 | 0.171625 |
| 18 | 77400400 | 77401599 | 7.252088 | 13.61912 | 32.91111 | 13.53545 | -0.7972 | 0.092842 | 0.171685 |
| 4 | 66627200 | 66628799 | 21.22107 | 38.51119 | 19.62646 | 8.478106 | 1.379643 | 0.3696 | 0.171699 |
| 8 | 48126400 | 48129599 | 19.04098 | 31.13788 | 90.58282 | 38.95264 | -1.16112 | 1.469286 | 0.171702 |
| 16 | 34738000 | 34750999 | 78.42969 | 119.9241 | 252.3079 | 104.6739 | -0.78244 | 3.065143 | 0.171721 |
| 9 | 122687800 | 122691799 | 127.1728 | 195.3637 | 157.6014 | 65.43829 | 0.604402 | 2.981007 | 0.171723 |
| 7 | 24332400 | 24338599 | 40.68077 | 64.53984 | 150.321 | 62.60245 | -0.88677 | 2.307784 | 0.171732 |
| 1 | 87007000 | 87018799 | 83.08728 | 126.6768 | 256.3572 | 106.2823 | -0.72975 | 3.126614 | 0.171736 |
| 13 | 80197000 | 80197999 | 13.41489 | 23.36922 | 50.52783 | 22.22136 | -0.71993 | 0.748228 | 0.171748 |
| 11 | 68484800 | 68486199 | 11.34612 | 19.96234 | 44.56907 | 19.73299 | -0.76451 | 0.577802 | 0.17175 |
| 2 | 157366000 | 157366799 | 5.688131 | 10.94201 | 29.16941 | 11.36651 | -0.86659 | -0.1112 | 0.171765 |
| 6 | 53845600 | 53846799 | 15.37507 | 28.26846 | 21.71719 | 9.223958 | 0.808474 | 0.258437 | 0.1718 |
| 7 | 13792400 | 13796599 | 94.4362 | 148.5217 | 115.1277 | 47.76068 | 0.687827 | 2.539615 | 0.171814 |
| 4 | 39388400 | 39389799 | 39.64865 | 64.99432 | 47.2567 | 20.34102 | 0.835076 | 1.453475 | 0.171824 |
| 4 | 135022000 | 135042999 | 144.1689 | 211.392 | 451.9224 | 186.1438 | -0.85951 | 3.864829 | 0.171856 |
| 12 | 100191600 | 100193599 | 9.48436 | 16.79726 | 41.98487 | 18.56432 | -0.92221 | 0.48867 | 0.171863 |
| 1 | 168989400 | 168990999 | 16.4501 | 30.9286 | 18.20565 | 7.472734 | 1.226318 | 0.12847 | 0.171883 |
| 4 | 4249400 | 4250999 | 29.58899 | 50.60587 | 34.04808 | 14.92949 | 0.95225 | 0.978997 | 0.171922 |
| 4 | 4975400 | 4982399 | 47.86411 | 76.15467 | 144.0577 | 60.178 | -0.5831 | 2.225877 | 0.171926 |
| 11 | 95749200 | 95750799 | 6.630771 | 11.88311 | 41.59566 | 18.22207 | -1.394 | 0.388158 | 0.171927 |
| 3 | 47262000 | 47262599 | 9.537461 | 18.52733 | 15.43923 | 5.766208 | 0.851968 | -0.25604 | 0.171927 |
| 4 | 43451200 | 43457599 | 33.03063 | 52.97261 | 119.1972 | 50.36056 | -0.82 | 1.95373 | 0.171932 |
| 17 | 19658200 | 19665999 | 205.3544 | 305.5918 | 241.5579 | 99.52827 | 0.595032 | 3.620631 | 0.171936 |
| 14 | 57967000 | 57969599 | 10.23321 | 18.12825 | 42.47322 | 18.77927 | -0.82958 | 0.49053 | 0.17197 |
| 10 | 69932400 | 69936799 | 28.42073 | 45.88621 | 99.89503 | 42.61007 | -0.76047 | 1.732879 | 0.172001 |
| 3 | 78824000 | 78825399 | 10.70386 | 18.93547 | 42.4787 | 18.78696 | -0.76802 | 0.510179 | 0.172007 |
| 7 | 31417400 | 31420399 | 31.51884 | 54.69655 | 28.07011 | 12.31617 | 1.344488 | 0.853871 | 0.172008 |
| 2 | 74443800 | 74444999 | 21.54044 | 37.74577 | 28.51405 | 12.59419 | 0.789912 | 0.66358 | 0.172017 |
| 17 | 72030000 | 72031999 | 9.779174 | 17.33113 | 41.47301 | 18.33551 | -0.85918 | 0.483865 | 0.172026 |
| 11 | 64997600 | 64999399 | 9.370584 | 16.51405 | 44.07958 | 19.50977 | -1.0193 | 0.546911 | 0.17204 |
| 7 | 28722000 | 28738399 | 106.3171 | 159.8155 | 313.9418 | 129.6101 | -0.70426 | 3.420514 | 0.172069 |
| 17 | 17369000 | 17370799 | 33.01525 | 55.80329 | 35.29934 | 15.44584 | 1.039489 | 1.097164 | 0.172079 |
| 7 | 10611600 | 10615799 | 92.92366 | 146.0409 | 118.1689 | 49.01784 | 0.625222 | 2.552524 | 0.172132 |
| 15 | 78925200 | 78929599 | 21.95042 | 35.73922 | 89.98574 | 38.6343 | -0.95805 | 1.538258 | 0.172132 |
| 10 | 92928800 | 92930799 | 12.44579 | 21.84428 | 50.99273 | 22.55342 | -0.82947 | 0.627363 | 0.172136 |
| 14 | 41978000 | 41981399 | 36.07853 | 60.1539 | 44.8802 | 19.50897 | 0.8023 | 1.24466 | 0.172136 |

| | | | | | | | | | |
|---|---|---|---|---|---|---|---|---|---|
| 11 | 55138800 | 55144799 | 43.91714 | 70.02103 | 137.6612 | 57.67252 | -0.63533 | 2.152789 | 0.17214 |
| 6 | 61436600 | 61437799 | 20.71115 | 36.69139 | 23.75049 | 10.50105 | 1.015467 | 0.519161 | 0.172141 |
| 14 | 20044800 | 20046399 | 8.586527 | 15.6076 | 37.06249 | 15.97781 | -0.82331 | 0.307312 | 0.172167 |
| 3 | 136660600 | 136661799 | 26.47903 | 45.40785 | 36.76213 | 16.12302 | 0.686909 | 0.945691 | 0.172179 |
| 14 | 41990800 | 41994399 | 66.2187 | 105.6434 | 85.45015 | 35.82947 | 0.644787 | 2.142249 | 0.17218 |
| 12 | 22569000 | 22570799 | 32.87206 | 55.20118 | 38.96885 | 17.01599 | 0.88148 | 1.154305 | 0.172181 |
| 3 | 94638800 | 94643599 | 41.64069 | 66.50419 | 127.605 | 53.5869 | -0.59755 | 2.095794 | 0.172183 |
| 17 | 23950400 | 23980199 | 204.0265 | 289.9276 | 593.177 | 245.3216 | -0.8408 | 4.33803 | 0.172186 |
| 9 | 6466600 | 6468599 | 11.0821 | 19.38385 | 52.07261 | 22.96404 | -1.03086 | 0.677973 | 0.17222 |
| 7 | 11631200 | 11633399 | 58.16333 | 93.41399 | 74.21617 | 31.42522 | 0.681383 | 1.926095 | 0.172221 |
| 7 | 45542200 | 45547199 | 22.26203 | 36.25511 | 90.12516 | 38.69747 | -0.93946 | 1.53524 | 0.172223 |
| 15 | 61163400 | 61164799 | 10.83373 | 19.21598 | 44.04275 | 19.46267 | -0.79745 | 0.484996 | 0.172223 |
| 7 | 59744400 | 59746199 | 31.20427 | 52.67813 | 40.96151 | 17.9168 | 0.742711 | 1.10447 | 0.172224 |
| 1 | 23761400 | 23763799 | 13.39744 | 23.35285 | 51.07085 | 22.46866 | -0.7363 | 0.739133 | 0.172234 |
| 5 | 36369000 | 36375399 | 72.70261 | 112.4557 | 209.0817 | 86.84748 | -0.5914 | 2.837123 | 0.172237 |
| 6 | 51196600 | 51199199 | 11.05588 | 19.42235 | 45.12215 | 19.9722 | -0.82188 | 0.598457 | 0.17224 |
| 7 | 20666600 | 20671999 | 73.36922 | 116.5017 | 99.41519 | 41.43853 | 0.5607 | 2.296126 | 0.172243 |
| 6 | 76845800 | 76848199 | 21.69138 | 38.45601 | 25.08567 | 11.08978 | 1.004922 | 0.521133 | 0.172258 |
| 10 | 42044000 | 42047399 | 26.97642 | 43.57948 | 98.50629 | 42.04901 | -0.81352 | 1.713361 | 0.172259 |
| X | 74989000 | 74991799 | 35.976 | 60.66265 | 35.56177 | 15.54941 | 1.148679 | 1.127494 | 0.172263 |
| 17 | 34822200 | 34823799 | 6.323019 | 11.67111 | 35.72768 | 15.0939 | -1.16576 | 0.197398 | 0.172263 |
| 3 | 38357800 | 38359799 | 20.20938 | 35.68611 | 27.05568 | 11.9702 | 0.787282 | 0.558115 | 0.172264 |
| 14 | 112260200 | 112261199 | 10.50402 | 20.44023 | 14.70756 | 5.416838 | 1.079352 | -0.29406 | 0.172277 |
| 17 | 31015200 | 31024199 | 53.0618 | 83.406 | 175.1132 | 72.64051 | -0.74837 | 2.534663 | 0.172334 |
| 8 | 91784400 | 91786399 | 7.329444 | 13.50812 | 34.6397 | 14.66473 | -0.91436 | 0.214056 | 0.172341 |
| 8 | 47813400 | 47815799 | 17.44284 | 29.33358 | 64.84077 | 28.3287 | -0.75907 | 1.119533 | 0.172344 |
| 7 | 20736800 | 20739799 | 66.71 | 106.524 | 76.74701 | 32.22157 | 0.812802 | 2.111272 | 0.172348 |
| 14 | 91148600 | 91148799 | 19.78672 | 36.80772 | 15.39331 | 6.433396 | 1.693708 | 0.215102 | 0.172351 |
| 3 | 115164600 | 115165799 | 26.34899 | 46.37831 | 23.90352 | 10.58239 | 1.339246 | 0.624349 | 0.172357 |
| 4 | 19154000 | 19157199 | 28.01161 | 47.81752 | 37.71757 | 16.53598 | 0.72333 | 0.992343 | 0.17236 |
| 11 | 88491600 | 88495199 | 20.0927 | 33.35017 | 72.4923 | 31.41708 | -0.73551 | 1.275176 | 0.17236 |
| 19 | 56985000 | 56993799 | 45.37571 | 71.8982 | 156.5856 | 65.13112 | -0.79193 | 2.348542 | 0.172373 |
| 17 | 39782400 | 39786199 | 64.819 | 104.1501 | 68.51446 | 29.03012 | 0.953821 | 1.918002 | 0.172392 |
| 14 | 118471400 | 118475599 | 33.8774 | 54.20465 | 128.7365 | 54.2025 | -0.90169 | 2.03186 | 0.172409 |
| 12 | 22284000 | 22285999 | 47.05224 | 76.3723 | 49.99168 | 21.42769 | 0.976557 | 1.616213 | 0.172414 |
| 12 | 81071000 | 81075599 | 33.14093 | 53.25756 | 113.0567 | 47.91247 | -0.73258 | 1.887891 | 0.172423 |
| 2 | 15712600 | 15713799 | 17.9444 | 32.62218 | 22.7789 | 9.81824 | 0.930881 | 0.342399 | 0.172428 |
| 17 | 64170000 | 64171799 | 12.79465 | 22.28508 | 49.98408 | 21.97996 | -0.77252 | 0.746883 | 0.172447 |
| 12 | 69215000 | 69215999 | 6.077211 | 11.41355 | 31.74785 | 13.05547 | -0.99737 | 0.080631 | 0.172448 |
| 15 | 20795000 | 20795599 | 9.540796 | 18.57365 | 14.51644 | 5.328352 | 0.96727 | -0.31645 | 0.172477 |
| X | 32005400 | 32007399 | 18.75157 | 34.20434 | 21.7701 | 9.344063 | 1.067691 | 0.32854 | 0.172478 |
| X | 32239800 | 32243999 | 57.10473 | 92.88208 | 46.83491 | 20.10054 | 1.354501 | 1.601759 | 0.172504 |
| 4 | 139650800 | 139657599 | 43.02076 | 68.0225 | 166.5905 | 69.21839 | -0.96339 | 2.400747 | 0.172514 |
| 8 | 81639800 | 81640799 | 5.552917 | 10.70643 | 28.73955 | 11.12549 | -0.8681 | -0.13812 | 0.17252 |
| 17 | 88222200 | 88231999 | 77.23859 | 119.1989 | 217.1601 | 90.20413 | -0.56467 | 2.881288 | 0.172524 |
| 12 | 73040400 | 73042999 | 26.15982 | 44.91414 | 36.25693 | 15.90298 | 0.691336 | 0.933254 | 0.172541 |
| 2 | 125436200 | 125438599 | 19.36163 | 34.22028 | 26.43466 | 11.69332 | 0.760642 | 0.540055 | 0.172545 |

| | | | | | | | | | |
|---|---|---|---|---|---|---|---|---|---|
| 7 | 126073400 | 126074999 | 5.978368 | 11.17396 | 33.82967 | 14.02552 | -1.12881 | 0.113761 | 0.172551 |
| 13 | 41153800 | 41154999 | 5.978368 | 11.17396 | 33.82967 | 14.02552 | -1.12881 | 0.113761 | 0.172551 |
| 8 | 121710600 | 121713199 | 11.99958 | 20.96591 | 48.89208 | 21.54219 | -0.82787 | 0.695445 | 0.172551 |
| 19 | 6107400 | 6109599 | 11.99958 | 20.96591 | 48.89208 | 21.54219 | -0.82787 | 0.695445 | 0.172551 |
| 5 | 142699600 | 142708599 | 44.28037 | 69.73315 | 170.9936 | 70.94889 | -0.96979 | 2.492083 | 0.172557 |
| 14 | 68287400 | 68289199 | 9.049013 | 16.38611 | 39.65302 | 17.17572 | -0.85534 | 0.336312 | 0.172568 |
| 4 | 116414200 | 116415399 | 6.679858 | 12.10121 | 37.56759 | 16.26366 | -1.21002 | 0.315922 | 0.172568 |
| 11 | 119112400 | 119115399 | 15.42814 | 25.94399 | 66.36466 | 28.99398 | -0.96887 | 1.113366 | 0.172575 |
| 7 | 71961200 | 71962199 | 5.387517 | 10.42878 | 27.83921 | 10.5981 | -0.8367 | -0.19844 | 0.172581 |
| 13 | 95050600 | 95051799 | 5.678088 | 10.96084 | 29.39027 | 11.33436 | -0.86098 | -0.14554 | 0.172582 |
| 10 | 42453800 | 42463199 | 85.99864 | 130.7733 | 273.5413 | 113.3205 | -0.7805 | 3.181178 | 0.172588 |
| 1 | 90635600 | 90636399 | 13.73462 | 26.03422 | 19.21342 | 7.761912 | 0.922366 | 0.054081 | 0.17261 |
| 1 | 140961400 | 140965399 | 48.92236 | 79.09105 | 63.57908 | 27.16574 | 0.675634 | 1.711438 | 0.172634 |
| 12 | 106253600 | 106256199 | 23.88795 | 39.14527 | 83.11667 | 35.77137 | -0.70682 | 1.446272 | 0.172645 |
| 10 | 40424800 | 40427199 | 17.69469 | 29.89541 | 65.08261 | 28.47437 | -0.73705 | 1.077329 | 0.172645 |
| 3 | 104614000 | 104616999 | 20.11437 | 33.32733 | 73.46836 | 31.80136 | -0.75591 | 1.297445 | 0.172652 |
| 12 | 18397600 | 18423599 | 819.4565 | 1083.115 | 777.6703 | 325.1928 | 0.564357 | 5.467395 | 0.172664 |
| 3 | 39297800 | 39298199 | 3.231465 | 6.318769 | 20.79544 | 7.096852 | -0.98596 | -0.63601 | 0.172678 |
| 2 | 18092000 | 18094199 | 24.93121 | 43.28679 | 33.04206 | 14.50103 | 0.774794 | 0.824116 | 0.172684 |
| 5 | 108861600 | 108862599 | 5.215392 | 9.808726 | 31.74207 | 13.02213 | -1.21032 | 0.063818 | 0.172696 |
| 4 | 61269400 | 61271399 | 8.530562 | 15.67384 | 37.31341 | 15.86241 | -0.81329 | 0.243861 | 0.172696 |
| 12 | 38044200 | 38045599 | 8.530562 | 15.67384 | 37.31341 | 15.86241 | -0.81329 | 0.243861 | 0.172696 |
| 6 | 31888000 | 31889799 | 22.88272 | 40.2497 | 27.14678 | 12.01986 | 0.952887 | 0.614632 | 0.172715 |
| 14 | 22128200 | 22132399 | 41.49908 | 66.23133 | 128.7537 | 54.01534 | -0.61749 | 2.118087 | 0.172727 |
| 11 | 43106800 | 43108399 | 7.927119 | 14.58819 | 36.24481 | 15.37496 | -0.87202 | 0.227786 | 0.172731 |
| X | 108750200 | 108754399 | 34.80147 | 59.32418 | 33.60164 | 14.72981 | 1.198856 | 1.028519 | 0.172731 |
| 5 | 103219000 | 103219399 | 5.531417 | 10.69293 | 28.31696 | 10.84799 | -0.8339 | -0.17329 | 0.172741 |
| 19 | 4084200 | 4088199 | 14.71687 | 24.95659 | 61.55225 | 26.96006 | -0.91572 | 1.032701 | 0.172742 |
| 6 | 29205200 | 29207399 | 7.188392 | 13.25145 | 35.18226 | 14.8894 | -0.96412 | 0.210641 | 0.172759 |
| 17 | 55040800 | 55043599 | 34.11748 | 57.45412 | 37.80416 | 16.52338 | 0.982694 | 1.133843 | 0.172778 |
| 10 | 47537800 | 47538999 | 19.07556 | 33.87865 | 25.83342 | 11.40339 | 0.782331 | 0.493503 | 0.172789 |
| 17 | 65306800 | 65307999 | 4.653437 | 9.055708 | 30.61314 | 11.27108 | -1.12887 | -0.31829 | 0.172794 |
| 7 | 43004000 | 43006199 | 14.84637 | 25.66674 | 55.97009 | 24.55487 | -0.73431 | 0.848516 | 0.172802 |
| 11 | 114873800 | 114875799 | 14.19517 | 24.70202 | 54.91054 | 24.13255 | -0.76019 | 0.771918 | 0.172808 |
| 2 | 118619600 | 118624999 | 36.95793 | 59.05165 | 126.3397 | 53.08898 | -0.75357 | 2.076777 | 0.172816 |
| 18 | 75587600 | 75593799 | 30.38371 | 48.93375 | 109.8898 | 46.71843 | -0.80957 | 1.813121 | 0.172819 |
| 10 | 42149600 | 42154999 | 36.20037 | 57.75309 | 133.8528 | 56.12486 | -0.87121 | 2.124921 | 0.172821 |
| 2 | 88627600 | 88628999 | 15.71657 | 29.4365 | 19.32753 | 7.984843 | 1.06261 | 0.147808 | 0.172825 |
| 5 | 8516200 | 8521599 | 91.33585 | 143.8554 | 113.6505 | 47.15682 | 0.662032 | 2.508001 | 0.172841 |
| 6 | 133469400 | 133476199 | 214.3269 | 316.9922 | 258.3846 | 106.4096 | 0.542578 | 3.727861 | 0.172845 |
| 17 | 87088600 | 87093599 | 23.23707 | 38.12906 | 81.3456 | 35.02873 | -0.71285 | 1.427474 | 0.172847 |
| X | 122861800 | 122862999 | 19.05735 | 33.96352 | 25.4703 | 11.21038 | 0.809261 | 0.472953 | 0.17288 |
| 16 | 55335400 | 55336799 | 7.755595 | 14.35092 | 34.73055 | 14.62323 | -0.82629 | 0.192753 | 0.172884 |
| 4 | 137221800 | 137223999 | 9.213484 | 16.23998 | 43.57376 | 19.28523 | -1.02645 | 0.544093 | 0.172888 |
| 2 | 75230600 | 75231199 | 3.454054 | 6.753562 | 21.43073 | 7.326389 | -0.93624 | -0.62211 | 0.172896 |
| 2 | 156194400 | 156225799 | 286.1774 | 397.7146 | 719.3791 | 298.5772 | -0.6955 | 4.672592 | 0.172896 |
| 12 | 22266200 | 22266799 | 18.32251 | 33.68688 | 20.66005 | 8.775211 | 1.130896 | 0.279046 | 0.172904 |

| | | | | | | | | | |
|---|---|---|---|---|---|---|---|---|---|
| 2 | 36867000 | 36867999 | 19.58853 | 35.9598 | 19.34805 | 8.234971 | 1.315957 | 0.293955 | 0.172907 |
| 11 | 117004000 | 117006799 | 15.13113 | 25.78082 | 60.63544 | 26.57948 | -0.84684 | 0.988658 | 0.172911 |
| 7 | 19583800 | 19596399 | 91.22214 | 138.7523 | 270.7617 | 112.1777 | -0.68012 | 3.176983 | 0.172916 |
| 3 | 27868600 | 27870399 | 8.672969 | 15.5078 | 39.48673 | 17.33126 | -0.94046 | 0.413778 | 0.172933 |
| 5 | 82812200 | 82813999 | 14.69554 | 27.50035 | 19.97967 | 8.266359 | 0.916037 | 0.145495 | 0.172942 |
| 12 | 20186800 | 20197799 | 345.2712 | 489.9783 | 382.1992 | 158.1029 | 0.547871 | 4.362211 | 0.172945 |
| 14 | 42108400 | 42110999 | 30.8282 | 52.35804 | 39.30591 | 17.2241 | 0.793546 | 1.051904 | 0.172955 |
| 17 | 84486600 | 84491199 | 19.77081 | 32.47706 | 81.99462 | 35.32464 | -0.95424 | 1.405983 | 0.17296 |
| 2 | 104952600 | 104954999 | 13.71976 | 23.86342 | 52.16416 | 22.91873 | -0.73605 | 0.772777 | 0.172966 |
| 13 | 66097200 | 66103199 | 165.8592 | 251.4796 | 181.7139 | 75.21343 | 0.748254 | 3.254576 | 0.172967 |
| 9 | 63668600 | 63669199 | 13.9692 | 26.05021 | 20.71214 | 8.620115 | 0.780478 | 0.163347 | 0.172976 |
| 19 | 25323000 | 25330199 | 30.83148 | 49.71949 | 106.3555 | 45.29646 | -0.73732 | 1.772785 | 0.172981 |
| 1 | 139971800 | 139972199 | 8.608937 | 16.83856 | 8.148786 | 2.74218 | 1.749863 | -0.65193 | 0.172986 |
| 9 | 9559000 | 9560799 | 12.82891 | 24.36303 | 18.08094 | 7.275522 | 0.918555 | 0.028055 | 0.172991 |
| 8 | 106229600 | 106230799 | 6.346739 | 12.01324 | 33.77439 | 13.68546 | -0.99588 | 0.03129 | 0.173034 |
| 2 | 66578800 | 66580199 | 17.28619 | 32.78214 | 16.04731 | 6.476218 | 1.510431 | 0.076108 | 0.173052 |
| 5 | 111767800 | 111789199 | 159.8667 | 231.4353 | 492.1815 | 202.9763 | -0.86979 | 4.061456 | 0.173062 |
| 13 | 25627200 | 25628599 | 16.54253 | 30.64508 | 20.95551 | 8.80962 | 0.985644 | 0.224814 | 0.173064 |
| 18 | 90233200 | 90234399 | 17.19057 | 31.9873 | 19.10079 | 7.979077 | 1.18581 | 0.20017 | 0.173068 |
| 9 | 61218000 | 61227999 | 46.59661 | 73.62059 | 161.8846 | 67.23728 | -0.80911 | 2.419598 | 0.173074 |
| 7 | 5843600 | 5848599 | 116.403 | 180.0292 | 144.0611 | 59.83954 | 0.623438 | 2.848767 | 0.173082 |
| 15 | 97403600 | 97406399 | 26.52433 | 43.19156 | 88.6217 | 38.05033 | -0.66366 | 1.54485 | 0.17309 |
| 12 | 119201400 | 119203199 | 11.71271 | 20.10042 | 57.82412 | 25.36182 | -1.13386 | 0.906661 | 0.173094 |
| 3 | 119561800 | 119562999 | 19.45561 | 34.47929 | 26.36748 | 11.65731 | 0.776621 | 0.512468 | 0.173102 |
| 17 | 20330400 | 20331599 | 35.40215 | 58.73239 | 47.68201 | 20.65772 | 0.680679 | 1.301595 | 0.173107 |
| 12 | 23665200 | 23666799 | 15.00457 | 29.26649 | 7.818169 | 2.801934 | 2.519145 | -0.34887 | 0.173109 |
| 12 | 19595400 | 19597199 | 35.3662 | 58.86198 | 41.76493 | 18.13448 | 0.874559 | 1.258329 | 0.17313 |
| 12 | 6922800 | 6924199 | 21.83257 | 38.26547 | 29.25823 | 12.92511 | 0.772888 | 0.666077 | 0.173145 |
| X | 26633400 | 26635999 | 50.43953 | 81.4705 | 63.53096 | 27.11371 | 0.717929 | 1.740235 | 0.173148 |
| 12 | 22612200 | 22613799 | 44.36617 | 71.97741 | 58.5425 | 25.08379 | 0.663499 | 1.626105 | 0.173155 |
| 15 | 47401000 | 47404799 | 76.07791 | 120.6124 | 102.5305 | 42.6697 | 0.564024 | 2.345232 | 0.173158 |
| 2 | 75662000 | 75663199 | 8.245214 | 14.72305 | 40.51051 | 17.80087 | -1.05283 | 0.4203 | 0.17316 |
| 1 | 155629000 | 155630399 | 11.36082 | 19.93099 | 46.18336 | 20.41851 | -0.81848 | 0.621634 | 0.173168 |
| 5 | 97739800 | 97740799 | 7.497071 | 14.6599 | 11.69298 | 3.988923 | 1.029515 | -0.60943 | 0.173169 |
| 12 | 79957600 | 79958599 | 4.898316 | 9.378939 | 31.23187 | 12.33384 | -1.20332 | -0.0735 | 0.173181 |
| 2 | 36876400 | 36877999 | 42.86085 | 69.67183 | 56.92644 | 24.42621 | 0.6595 | 1.583184 | 0.173182 |
| 9 | 46238400 | 46244599 | 38.90402 | 62.09754 | 131.5459 | 55.1966 | -0.74099 | 2.111076 | 0.173197 |
| 1 | 31764400 | 31766799 | 34.49113 | 57.81782 | 39.01628 | 17.01772 | 0.946457 | 1.179609 | 0.173205 |
| 2 | 21038200 | 21039799 | 6.364551 | 11.99819 | 33.0055 | 13.48658 | -0.9751 | 0.058658 | 0.173211 |
| X | 89049400 | 89050999 | 40.37149 | 66.69529 | 46.6746 | 20.1626 | 0.893914 | 1.364073 | 0.173216 |
| 6 | 140369400 | 140371999 | 18.47857 | 30.79152 | 74.92598 | 32.55764 | -0.89752 | 1.225296 | 0.173223 |
| 4 | 141893400 | 141918399 | 162.6429 | 235.6652 | 490.1618 | 202.1194 | -0.83647 | 4.04822 | 0.173223 |
| 13 | 58287800 | 58288999 | 28.60202 | 48.62802 | 39.38722 | 17.26235 | 0.684487 | 1.033613 | 0.173269 |
| 17 | 72247800 | 72249399 | 7.460122 | 13.80599 | 35.83064 | 15.08373 | -0.9264 | 0.190814 | 0.173271 |
| 16 | 31751600 | 31757999 | 42.82619 | 68.16243 | 145.4627 | 60.79377 | -0.7562 | 2.209613 | 0.173284 |
| 10 | 18399600 | 18403399 | 26.84849 | 43.2871 | 102.8741 | 43.80154 | -0.88895 | 1.774159 | 0.17329 |
| 11 | 68878400 | 68885399 | 26.36261 | 42.3142 | 120.5672 | 51.00421 | -1.15818 | 1.920122 | 0.17331 |

| | | | | | | | | | |
|---|---|---|---|---|---|---|---|---|---|
| 11 | 61127400 | 61128999 | 6.922772 | 12.3384 | 41.05623 | 18.06931 | -1.32588 | 0.426384 | 0.173333 |
| 5 | 32144400 | 32146199 | 7.487713 | 13.75158 | 36.17495 | 15.38781 | -0.95573 | 0.240407 | 0.173341 |
| 9 | 39467000 | 39468599 | 15.83509 | 29.11483 | 20.25902 | 8.604392 | 0.949503 | 0.255899 | 0.173343 |
| 16 | 3895200 | 3895799 | 3.034268 | 5.910272 | 25.84786 | 9.430825 | -1.47984 | -0.38316 | 0.173348 |
| 11 | 62347400 | 62349599 | 12.28457 | 20.88457 | 66.96406 | 29.3484 | -1.29236 | 1.004709 | 0.17335 |
| 18 | 48805400 | 48807999 | 26.7754 | 46.83756 | 25.77906 | 11.36947 | 1.24421 | 0.715246 | 0.173353 |
| 1 | 100489200 | 100493999 | 48.44514 | 79.95041 | 39.95476 | 17.24741 | 1.37772 | 1.389565 | 0.173369 |
| 7 | 60304800 | 60310999 | 106.9802 | 166.2306 | 137.2218 | 56.97619 | 0.583778 | 2.762382 | 0.173373 |
| 16 | 4607200 | 4608799 | 11.27293 | 19.6363 | 51.55453 | 22.67177 | -0.9985 | 0.739206 | 0.173388 |
| 15 | 49488000 | 49488399 | 15.00577 | 27.6142 | 21.2517 | 9.015252 | 0.807167 | 0.24918 | 0.173393 |
| 4 | 61567200 | 61567999 | 16.69569 | 30.69146 | 21.97662 | 9.336242 | 0.908968 | 0.270217 | 0.173395 |
| 18 | 23400600 | 23402199 | 18.41487 | 34.45042 | 17.49888 | 7.243548 | 1.427856 | 0.174848 | 0.173399 |
| 18 | 38732400 | 38738599 | 36.46611 | 58.29159 | 125.2827 | 52.67783 | -0.75941 | 2.062184 | 0.173404 |
| 7 | 10367200 | 10375999 | 158.2313 | 240.3973 | 177.6795 | 73.58786 | 0.718012 | 3.212635 | 0.173411 |
| 14 | 34534200 | 34546999 | 112.8161 | 169.0281 | 332.7023 | 137.2421 | -0.71139 | 3.485713 | 0.173415 |
| 11 | 59354000 | 59358399 | 85.58373 | 135.1501 | 110.0894 | 45.7129 | 0.620897 | 2.448778 | 0.173422 |
| 15 | 83620400 | 83623199 | 14.39675 | 24.44171 | 61.00244 | 26.72852 | -0.93264 | 1.021108 | 0.17343 |
| 15 | 90825800 | 90828199 | 13.58228 | 23.59896 | 51.61199 | 22.66084 | -0.73682 | 0.787827 | 0.173432 |
| 17 | 6903400 | 6905199 | 15.80451 | 29.23881 | 21.50873 | 9.058646 | 0.879056 | 0.226792 | 0.173432 |
| X | 148897600 | 148903199 | 115.4474 | 180.4876 | 111.1486 | 46.11336 | 1.012789 | 2.660959 | 0.173459 |
| 5 | 95504800 | 95508199 | 31.59236 | 53.77339 | 37.58326 | 16.47324 | 0.896606 | 1.033093 | 0.173477 |
| 14 | 66279600 | 66282599 | 15.22198 | 26.12064 | 56.9471 | 24.977 | -0.73617 | 0.91981 | 0.173498 |
| 1 | 3116600 | 3119199 | 26.84765 | 46.95742 | 26.36315 | 11.62571 | 1.215858 | 0.719139 | 0.173509 |
| 12 | 67807400 | 67809199 | 21.68628 | 37.90647 | 29.41495 | 12.9647 | 0.751736 | 0.707285 | 0.173516 |
| 1 | 96689800 | 96690399 | 4.057298 | 7.918876 | 23.444 | 8.317762 | -0.88822 | -0.47663 | 0.173523 |
| 3 | 46753800 | 46756599 | 27.82921 | 48.8101 | 23.49865 | 10.38695 | 1.436011 | 0.677865 | 0.173541 |
| 18 | 24244200 | 24247599 | 14.74661 | 25.38376 | 55.31859 | 24.26584 | -0.73453 | 0.88905 | 0.173542 |
| 4 | 120504400 | 120510399 | 49.55026 | 78.16651 | 163.1961 | 67.72908 | -0.73623 | 2.459314 | 0.17356 |
| 7 | 31049800 | 31064799 | 299.4118 | 431.7027 | 335.0753 | 138.2594 | 0.577929 | 4.128084 | 0.173563 |
| 11 | 118350600 | 118352399 | 18.77608 | 33.3593 | 25.9144 | 11.43552 | 0.75604 | 0.48968 | 0.173568 |
| 17 | 85158200 | 85162399 | 24.08192 | 39.37568 | 83.28372 | 35.81197 | -0.70299 | 1.478508 | 0.17357 |
| 9 | 20619000 | 20626999 | 53.07399 | 83.8305 | 159.667 | 66.30655 | -0.60254 | 2.43068 | 0.173596 |
| 16 | 61140600 | 61142599 | 17.99272 | 32.23463 | 24.97129 | 10.94231 | 0.766869 | 0.432003 | 0.173603 |
| 7 | 44218200 | 44222199 | 21.76055 | 35.85409 | 77.35623 | 33.36911 | -0.72649 | 1.370533 | 0.173606 |
| 1 | 38732400 | 38735999 | 18.66578 | 31.02403 | 72.41794 | 31.41442 | -0.83778 | 1.254553 | 0.173609 |
| 13 | 18298400 | 18300599 | 9.586422 | 17.09955 | 43.08838 | 18.95083 | -0.92847 | 0.438696 | 0.173611 |
| 17 | 19428200 | 19432799 | 86.51061 | 137.1453 | 97.27313 | 40.4797 | 0.824823 | 2.354046 | 0.173626 |
| 18 | 69872400 | 69872999 | 5.090121 | 9.868212 | 26.71423 | 10.09727 | -0.8462 | -0.22846 | 0.173637 |
| 1 | 96799400 | 96800599 | 9.835519 | 17.30554 | 46.15078 | 20.43306 | -1.01933 | 0.569708 | 0.173647 |
| 4 | 148320400 | 148331399 | 101.7436 | 153.7786 | 288.7733 | 119.4075 | -0.6324 | 3.310065 | 0.173655 |
| 3 | 17935600 | 17936399 | 13.51587 | 25.42825 | 17.41504 | 7.140401 | 1.011506 | 0.096515 | 0.173668 |
| 4 | 107849400 | 107857199 | 45.88171 | 72.64998 | 155.9437 | 64.84159 | -0.77182 | 2.365564 | 0.173672 |
| 6 | 49132800 | 49135199 | 11.64647 | 20.538 | 45.0859 | 19.95263 | -0.73903 | 0.554841 | 0.173688 |
| 13 | 11398600 | 11400399 | 22.71204 | 40.2022 | 24.32871 | 10.75853 | 1.111506 | 0.541779 | 0.173703 |
| 4 | 106490400 | 106494399 | 16.66134 | 27.65819 | 73.7055 | 31.94561 | -1.02817 | 1.265386 | 0.173712 |
| 7 | 13388000 | 13391799 | 94.06135 | 147.0504 | 114.8063 | 47.63094 | 0.670865 | 2.647011 | 0.173712 |
| 7 | 110995800 | 110997199 | 8.363397 | 15.44685 | 36.34966 | 15.34371 | -0.7891 | 0.211616 | 0.173723 |

| | | | | | | | | | |
|---|---|---|---|---|---|---|---|---|---|
| 12 | 32323600 | 32323999 | 3.261101 | 6.377827 | 22.8445 | 7.759372 | -1.09932 | -0.64135 | 0.173724 |
| X | 139911400 | 139919199 | 30.84229 | 49.77753 | 104.8213 | 44.69008 | -0.71327 | 1.751997 | 0.173736 |
| 5 | 110945600 | 110946599 | 4.002493 | 7.772702 | 26.7622 | 10.02087 | -1.17608 | -0.27559 | 0.173747 |
| 9 | 55443200 | 55454399 | 91.03216 | 137.0537 | 332.4869 | 137.3239 | -1.00646 | 3.387905 | 0.173748 |
| 10 | 67337000 | 67340999 | 18.62127 | 31.06391 | 69.15255 | 30.07004 | -0.76936 | 1.21167 | 0.173752 |
| 7 | 62186000 | 62192199 | 113.0862 | 176.7372 | 115.1228 | 47.76381 | 0.931536 | 2.664269 | 0.173761 |
| 5 | 50215400 | 50217399 | 10.65469 | 18.59241 | 49.93775 | 21.98531 | -1.03063 | 0.708895 | 0.17377 |
| 11 | 10204400 | 10207199 | 26.92252 | 46.75756 | 32.13138 | 14.1039 | 0.92554 | 0.826689 | 0.173794 |
| 7 | 7129800 | 7134399 | 64.91811 | 103.5896 | 88.18615 | 36.98724 | 0.571417 | 2.133707 | 0.173806 |
| 7 | 14031000 | 14034799 | 60.54053 | 96.93762 | 81.44143 | 34.32599 | 0.595815 | 2.023349 | 0.17381 |
| 1 | 149771000 | 149773599 | 18.26482 | 32.73065 | 23.69391 | 10.38028 | 0.863897 | 0.427834 | 0.173842 |
| 6 | 114715200 | 114721199 | 35.34257 | 56.49378 | 124.5177 | 52.35378 | -0.79569 | 2.062937 | 0.173852 |
| 10 | 85343200 | 85346599 | 17.9909 | 30.45877 | 65.15643 | 28.52455 | -0.71175 | 1.060401 | 0.173855 |
| 7 | 27165400 | 27167999 | 10.0816 | 17.86027 | 42.99533 | 19.00999 | -0.86849 | 0.48967 | 0.173859 |
| 7 | 138661200 | 138663399 | 10.0816 | 17.86027 | 42.99533 | 19.00999 | -0.86849 | 0.48967 | 0.173859 |
| 11 | 51373400 | 51379999 | 34.59749 | 55.5727 | 111.7648 | 47.33338 | -0.6556 | 1.904788 | 0.17392 |
| 14 | 8132200 | 8132599 | 7.372478 | 14.41911 | 11.73732 | 3.973608 | 1.010679 | -0.63638 | 0.173952 |
| 19 | 39681200 | 39683799 | 22.53562 | 39.85265 | 24.90055 | 11.01346 | 1.065209 | 0.551482 | 0.173959 |
| 8 | 41087400 | 41088399 | 7.86873 | 14.19456 | 38.02712 | 16.55027 | -1.0044 | 0.351428 | 0.173968 |
| 6 | 124433400 | 124437599 | 20.97447 | 34.55552 | 78.45135 | 33.84011 | -0.79983 | 1.369627 | 0.173978 |
| 16 | 37682600 | 37683199 | 9.481314 | 18.44335 | 15.51124 | 5.729761 | 0.853725 | -0.29309 | 0.17399 |
| 5 | 64055400 | 64056399 | 6.11143 | 11.37198 | 33.17756 | 13.85912 | -1.08392 | 0.145404 | 0.174004 |
| 7 | 126843200 | 126844599 | 6.11143 | 11.37198 | 33.17756 | 13.85912 | -1.08392 | 0.145404 | 0.174004 |
| 8 | 20568800 | 20572799 | 141.7154 | 215.4387 | 180.0188 | 74.57055 | 0.541861 | 3.194675 | 0.174013 |
| 17 | 27396800 | 27450399 | 357.3283 | 481.1263 | 1099.781 | 458.7229 | -1.07926 | 5.167765 | 0.174023 |
| 2 | 42722200 | 42724399 | 26.12864 | 45.91397 | 24.76046 | 10.96216 | 1.273387 | 0.641987 | 0.174029 |
| X | 3152600 | 3154599 | 28.17927 | 49.10931 | 28.69737 | 12.6239 | 1.158546 | 0.781443 | 0.174036 |
| 1 | 172110600 | 172114999 | 21.75276 | 35.51199 | 88.68177 | 38.11149 | -0.94351 | 1.497392 | 0.174037 |
| 10 | 126645400 | 126658599 | 126.0803 | 188.5509 | 336.6152 | 138.8061 | -0.57312 | 3.522866 | 0.174065 |
| 8 | 127469000 | 127471999 | 14.57941 | 25.07733 | 56.41726 | 24.74721 | -0.7806 | 0.89447 | 0.174075 |
| 14 | 15527800 | 15530199 | 13.87438 | 24.14434 | 52.16703 | 22.92715 | -0.71926 | 0.766525 | 0.174082 |
| 13 | 56130200 | 56132999 | 14.16776 | 24.24703 | 57.92728 | 25.40146 | -0.8681 | 0.941503 | 0.174102 |
| 7 | 8464600 | 8465199 | 15.70238 | 28.72737 | 21.73815 | 9.295402 | 0.823277 | 0.288427 | 0.174105 |
| 1 | 59951600 | 59954599 | 18.52558 | 31.17716 | 65.58054 | 28.65824 | -0.6879 | 1.115698 | 0.174127 |
| 7 | 55711400 | 55714399 | 25.83717 | 45.17042 | 28.74688 | 12.67224 | 1.036269 | 0.725182 | 0.17413 |
| 14 | 84111400 | 84113999 | 25.83717 | 45.17042 | 28.74688 | 12.67224 | 1.036269 | 0.725182 | 0.17413 |
| 4 | 136160000 | 136164399 | 21.42111 | 35.35238 | 77.5571 | 33.47968 | -0.74963 | 1.348238 | 0.174133 |
| 6 | 132491200 | 132494399 | 69.90008 | 111.2237 | 94.92317 | 39.65694 | 0.563475 | 2.230451 | 0.174143 |
| 3 | 30358800 | 30359599 | 6.026129 | 11.4769 | 29.9286 | 11.97008 | -0.8705 | -0.02133 | 0.174157 |
| 1 | 79835800 | 79837399 | 6.58594 | 12.18582 | 34.16548 | 14.38656 | -1.03629 | 0.183395 | 0.174169 |
| 6 | 81107200 | 81109399 | 19.37646 | 35.67412 | 18.80014 | 7.970156 | 1.349509 | 0.275252 | 0.174186 |
| 2 | 167254800 | 167262199 | 57.16538 | 90.18161 | 167.0569 | 69.33218 | -0.56399 | 2.463591 | 0.174209 |
| 5 | 35652200 | 35654999 | 13.14016 | 22.40451 | 60.49314 | 26.51862 | -1.04465 | 0.975173 | 0.174223 |
| 13 | 4229200 | 4232199 | 57.15604 | 92.02581 | 69.59214 | 29.57445 | 0.756809 | 1.845607 | 0.174234 |
| 2 | 150601800 | 150607999 | 40.69341 | 64.90229 | 132.0071 | 55.32868 | -0.68367 | 2.138363 | 0.174235 |
| 11 | 5905000 | 5906399 | 6.310817 | 12.03863 | 30.59764 | 12.1957 | -0.82989 | -0.02935 | 0.174237 |
| 2 | 181327800 | 181329799 | 7.915162 | 14.59935 | 36.30591 | 15.35329 | -0.87015 | 0.214584 | 0.17424 |

| | | | | | | | | | |
|---|---|---|---|---|---|---|---|---|---|
| 13 | 29134600 | 29139399 | 27.46873 | 44.36077 | 102.368 | 43.67962 | -0.84391 | 1.723691 | 0.174243 |
| 15 | 89357000 | 89362599 | 36.04463 | 57.49342 | 133.5168 | 55.96888 | -0.87437 | 2.131285 | 0.174268 |
| 11 | 83331200 | 83336399 | 32.48905 | 52.24615 | 109.4307 | 46.42234 | -0.71196 | 1.864339 | 0.174279 |
| 19 | 6736600 | 6739999 | 30.71512 | 49.43483 | 106.8515 | 45.38324 | -0.75568 | 1.834804 | 0.174279 |
| 7 | 10524600 | 10527199 | 54.47859 | 88.18584 | 58.60773 | 25.07036 | 0.951206 | 1.689667 | 0.17428 |
| 7 | 8525400 | 8525799 | 7.979754 | 15.61016 | 9.704765 | 3.212437 | 1.419956 | -0.70954 | 0.174295 |
| 1 | 30647800 | 30649999 | 23.36611 | 40.81688 | 31.18856 | 13.74001 | 0.773848 | 0.728292 | 0.1743 |
| 14 | 53598000 | 53600199 | 13.72719 | 23.82766 | 54.34267 | 23.84527 | -0.79739 | 0.804141 | 0.174308 |
| X | 104163800 | 104165599 | 10.71732 | 18.96706 | 44.05295 | 19.4816 | -0.81749 | 0.510146 | 0.174308 |
| 2 | 37673200 | 37695799 | 146.2045 | 212.5643 | 476.9793 | 196.609 | -0.94115 | 3.99588 | 0.17431 |
| 11 | 36119000 | 36120199 | 6.716992 | 12.45641 | 33.66943 | 14.13416 | -0.98065 | 0.170782 | 0.174312 |
| 13 | 35222200 | 35224199 | 13.75438 | 23.89189 | 52.14194 | 22.88985 | -0.73377 | 0.794054 | 0.174318 |
| 8 | 89175200 | 89189399 | 124.6515 | 186.6229 | 332.0147 | 136.9389 | -0.56663 | 3.501637 | 0.174319 |
| 2 | 113912400 | 113913399 | 22.47819 | 39.23732 | 31.30658 | 13.78572 | 0.711642 | 0.733352 | 0.174329 |
| 17 | 81167800 | 81172199 | 35.2172 | 56.67193 | 109.9041 | 46.66926 | -0.59995 | 1.847189 | 0.17433 |
| 3 | 16594400 | 16595799 | 20.44394 | 36.36944 | 25.13268 | 11.07851 | 0.92491 | 0.49142 | 0.174331 |
| 6 | 100189600 | 100201599 | 55.9788 | 86.91305 | 207.9421 | 86.35041 | -0.95052 | 2.761905 | 0.174339 |
| 12 | 108050600 | 108051399 | 12.16586 | 23.28455 | 16.96555 | 6.707958 | 0.968356 | -0.03464 | 0.174345 |
| 11 | 71389800 | 71390999 | 7.261126 | 13.60601 | 32.84189 | 13.55758 | -0.79986 | 0.107303 | 0.174362 |
| 13 | 72627600 | 72628799 | 4.921046 | 9.362962 | 31.52292 | 12.63419 | -1.23814 | -0.02253 | 0.174387 |
| 18 | 67201000 | 67204199 | 16.84743 | 28.26108 | 72.1533 | 31.48484 | -0.96654 | 1.145605 | 0.174399 |
| 11 | 118968600 | 118983799 | 70.28285 | 107.1986 | 271.0339 | 112.3777 | -1.05013 | 3.115072 | 0.174402 |
| 18 | 15490600 | 15494199 | 17.59108 | 29.58047 | 66.0456 | 28.85438 | -0.77364 | 1.119769 | 0.174404 |
| 15 | 13506800 | 13508399 | 15.46894 | 28.49537 | 22.08252 | 9.354804 | 0.798483 | 0.248813 | 0.17443 |
| 18 | 44849800 | 44850199 | 4.357302 | 8.488789 | 24.25788 | 8.830824 | -0.87201 | -0.37776 | 0.174436 |
| 4 | 75422200 | 75423399 | 15.5272 | 29.09041 | 19.33359 | 7.983351 | 1.045747 | 0.14419 | 0.174438 |
| 1 | 100882800 | 100883999 | 12.6576 | 23.92767 | 19.29836 | 7.841106 | 0.787933 | 0.061719 | 0.174447 |
| 7 | 4582200 | 4586599 | 61.16323 | 97.82315 | 82.93246 | 34.89746 | 0.580984 | 2.058689 | 0.174471 |
| 1 | 77489400 | 77495799 | 28.56807 | 45.86544 | 119.7327 | 50.67018 | -1.03186 | 1.91425 | 0.174473 |
| 17 | 41748200 | 41749599 | 13.05932 | 24.90225 | 18.09049 | 7.216663 | 0.960045 | 0.003439 | 0.174479 |
| 3 | 106553600 | 106555599 | 21.25249 | 38.28861 | 21.95503 | 9.570681 | 1.203064 | 0.415864 | 0.174481 |
| 10 | 42082200 | 42086799 | 36.2534 | 58.09393 | 117.5078 | 49.58556 | -0.668 | 1.98363 | 0.174486 |
| 6 | 92717600 | 92721799 | 23.59331 | 38.60659 | 87.12266 | 37.47327 | -0.79552 | 1.465897 | 0.174491 |
| 12 | 51394600 | 51395599 | 6.35512 | 12.00607 | 33.09663 | 13.46531 | -0.97274 | 0.043698 | 0.174492 |
| 9 | 20886200 | 20890599 | 20.54267 | 33.75268 | 80.38186 | 34.63318 | -0.87014 | 1.404291 | 0.174496 |
| 14 | 20213400 | 20221199 | 39.13068 | 62.31002 | 137.4538 | 57.48409 | -0.80307 | 2.190483 | 0.174496 |
| 15 | 98927000 | 98928799 | 6.72929 | 12.47516 | 34.74827 | 14.5935 | -1.02459 | 0.173336 | 0.174498 |
| 17 | 48942800 | 48943999 | 8.523796 | 15.72606 | 37.99768 | 16.06303 | -0.82864 | 0.222507 | 0.174501 |
| 16 | 6854600 | 6856999 | 11.52225 | 20.23525 | 45.64269 | 20.19723 | -0.77964 | 0.606819 | 0.174508 |
| 3 | 34942400 | 34950999 | 84.82214 | 129.647 | 250.1888 | 103.7852 | -0.65839 | 3.074969 | 0.174508 |
| 19 | 39992800 | 39993399 | 3.706162 | 7.227601 | 25.74652 | 9.248746 | -1.16924 | -0.43295 | 0.174515 |
| 9 | 98773800 | 98780599 | 96.61301 | 151.3673 | 123.7619 | 51.33502 | 0.606765 | 2.613067 | 0.174524 |
| 9 | 120972400 | 120977599 | 43.80068 | 69.74494 | 135.9722 | 56.86289 | -0.62522 | 2.195078 | 0.17453 |
| 10 | 57407400 | 57408799 | 8.855365 | 16.1796 | 38.80737 | 16.62234 | -0.83202 | 0.2811 | 0.174555 |
| 2 | 32247600 | 32249599 | 7.120206 | 13.31676 | 32.78617 | 13.57797 | -0.83189 | 0.118 | 0.174559 |
| 9 | 114565400 | 114567399 | 9.546261 | 17.17242 | 38.98279 | 17.02341 | -0.77117 | 0.379449 | 0.174575 |
| 2 | 104240000 | 104241399 | 9.871003 | 17.29704 | 47.77997 | 21.1035 | -1.07039 | 0.635219 | 0.174586 |

| | | | | | | | | | |
|---|---|---|---|---|---|---|---|---|---|
| 19 | 8964600 | 8968599 | 21.17663 | 34.78286 | 81.08509 | 34.93131 | -0.83968 | 1.409904 | 0.174616 |
| 11 | 46876800 | 46878399 | 15.68161 | 29.16985 | 20.36599 | 8.512361 | 0.961435 | 0.192224 | 0.174635 |
| 14 | 108506800 | 108507599 | 11.59565 | 22.5858 | 13.50454 | 4.929221 | 1.355292 | -0.3137 | 0.174647 |
| 16 | 30003000 | 30010199 | 39.13386 | 62.31045 | 142.2647 | 59.48992 | -0.85277 | 2.193107 | 0.174661 |
| 7 | 42750400 | 42751999 | 32.61853 | 54.90159 | 42.05438 | 18.37819 | 0.764225 | 1.136625 | 0.174663 |
| 7 | 20976200 | 20977999 | 53.82482 | 86.59066 | 71.20869 | 30.22521 | 0.63466 | 1.86667 | 0.174664 |
| 18 | 77020800 | 77023199 | 12.93259 | 22.55218 | 49.98461 | 21.99666 | -0.75535 | 0.729519 | 0.174665 |
| X | 122443600 | 122445399 | 20.1208 | 36.08005 | 23.7848 | 10.41438 | 0.999171 | 0.439818 | 0.17468 |
| 5 | 111026400 | 111028199 | 10.75871 | 18.86847 | 46.71745 | 20.6489 | -0.91353 | 0.624339 | 0.174685 |
| 10 | 121461600 | 121462999 | 7.508146 | 13.76604 | 36.15361 | 15.41381 | -0.95653 | 0.246095 | 0.174702 |
| 14 | 53370400 | 53373799 | 22.88181 | 37.66062 | 78.63907 | 33.90691 | -0.68015 | 1.387221 | 0.174705 |
| 9 | 40854000 | 40856799 | 11.05679 | 19.40174 | 45.66081 | 20.19111 | -0.84065 | 0.616147 | 0.174737 |
| 11 | 3281800 | 3283599 | 12.48117 | 21.78492 | 48.9003 | 21.53187 | -0.77315 | 0.714637 | 0.174741 |
| 16 | 29764400 | 29768799 | 27.39046 | 44.40585 | 94.52344 | 40.47973 | -0.72197 | 1.628253 | 0.174748 |
| 17 | 87806600 | 87808199 | 10.25334 | 18.11066 | 43.01176 | 19.02855 | -0.85068 | 0.52465 | 0.174749 |
| 5 | 120020600 | 120022799 | 17.42975 | 29.34033 | 64.34349 | 28.1198 | -0.74765 | 1.1096 | 0.174756 |
| 10 | 7879200 | 7880399 | 14.99904 | 27.9203 | 21.10805 | 8.811753 | 0.849225 | 0.184676 | 0.174765 |
| 17 | 92089000 | 92089799 | 18.97473 | 33.88141 | 25.33897 | 11.13451 | 0.814631 | 0.459566 | 0.174767 |
| 16 | 38062600 | 38064199 | 6.486405 | 12.28232 | 32.04932 | 12.97592 | -0.88783 | 0.026852 | 0.174771 |
| 15 | 24643000 | 24644599 | 19.68495 | 35.81243 | 19.67348 | 8.472031 | 1.274648 | 0.343496 | 0.174772 |
| 5 | 66645200 | 66645999 | 4.824141 | 9.354927 | 26.72703 | 10.09016 | -0.92173 | -0.23696 | 0.174782 |
| 9 | 120871400 | 120875799 | 25.14494 | 40.82427 | 91.28941 | 39.13639 | -0.7908 | 1.593721 | 0.174784 |
| 7 | 98131800 | 98139599 | 35.58671 | 56.70124 | 135.8289 | 56.85238 | -0.92075 | 2.167708 | 0.174797 |
| 3 | 32880600 | 32882599 | 12.75483 | 22.29593 | 51.04992 | 22.50135 | -0.80184 | 0.6938 | 0.174808 |
| 8 | 47217400 | 47220199 | 18.36431 | 30.91058 | 65.57909 | 28.65897 | -0.70021 | 1.113579 | 0.174823 |
| 6 | 9626200 | 9627999 | 17.72412 | 31.83188 | 24.36403 | 10.65402 | 0.785978 | 0.412873 | 0.174858 |
| 19 | 8813800 | 8816999 | 14.39471 | 24.88321 | 53.75014 | 23.58087 | -0.72053 | 0.846223 | 0.174861 |
| 6 | 21120600 | 21121599 | 7.40568 | 13.90029 | 33.4609 | 13.77375 | -0.79273 | 0.099137 | 0.174863 |
| 10 | 45749800 | 45752199 | 14.46177 | 24.77347 | 60.7415 | 26.63753 | -0.90543 | 0.935283 | 0.174864 |
| 7 | 60283000 | 60283999 | 14.61343 | 27.60611 | 18.99178 | 7.727679 | 1.013564 | 0.083682 | 0.174879 |
| 17 | 32837600 | 32841199 | 58.45178 | 93.78607 | 76.97032 | 32.54328 | 0.633004 | 1.956673 | 0.174885 |
| 11 | 76900000 | 76904199 | 20.73917 | 34.37246 | 73.65962 | 31.8905 | -0.7153 | 1.294677 | 0.174885 |
| 12 | 23747200 | 23749399 | 37.23071 | 61.63987 | 42.92795 | 18.57185 | 0.900936 | 1.324788 | 0.174893 |
| X | 32414000 | 32416999 | 34.49014 | 59.19382 | 30.217 | 13.25334 | 1.349326 | 0.964533 | 0.174902 |
| 16 | 18226200 | 18230999 | 18.92832 | 30.9432 | 88.83 | 38.19418 | -1.1424 | 1.471935 | 0.174904 |
| 11 | 34261200 | 34261599 | 2.835618 | 5.546462 | 20.35581 | 6.832346 | -1.11808 | -0.70328 | 0.174906 |
| 9 | 95393200 | 95394399 | 7.300653 | 13.20475 | 39.62352 | 17.18797 | -1.1643 | 0.33113 | 0.174919 |
| 1 | 79533600 | 79534199 | 3.639564 | 7.110076 | 21.74811 | 7.595039 | -0.91296 | -0.54431 | 0.174929 |
| 5 | 6231200 | 6232599 | 15.04465 | 27.6801 | 21.57528 | 9.155056 | 0.788323 | 0.250147 | 0.174931 |
| 12 | 72347400 | 72352599 | 40.47256 | 64.7035 | 127.6783 | 53.694 | -0.63633 | 2.062696 | 0.174936 |
| 3 | 98220000 | 98224799 | 21.17884 | 34.77167 | 80.39213 | 34.62654 | -0.828 | 1.41556 | 0.17494 |
| 4 | 81360400 | 81361599 | 7.945294 | 14.58715 | 35.6435 | 15.16854 | -0.85094 | 0.243275 | 0.17495 |
| 10 | 42501200 | 42513799 | 97.84838 | 148.1331 | 282.3044 | 116.7928 | -0.65163 | 3.275796 | 0.174959 |
| 18 | 60792600 | 60795799 | 13.90665 | 23.69787 | 60.5413 | 26.53852 | -0.96544 | 0.983128 | 0.174963 |
| 5 | 135630200 | 135632399 | 43.56095 | 71.70744 | 48.92617 | 21.09699 | 0.928852 | 1.414285 | 0.174994 |
| 11 | 68481400 | 68482799 | 3.342522 | 6.449024 | 29.04552 | 11.22195 | -1.60081 | -0.1661 | 0.175001 |
| 3 | 27424000 | 27426999 | 21.48288 | 35.2994 | 79.98408 | 34.46258 | -0.79858 | 1.405472 | 0.175022 |

| | | | | | | | | | |
|---|---|---|---|---|---|---|---|---|---|
| 8 | 93178400 | 93190199 | 87.40797 | 133.3886 | 260.2951 | 107.9382 | -0.67628 | 3.108131 | 0.175027 |
| 11 | 115520800 | 115524199 | 15.16665 | 25.75403 | 61.64243 | 27.0106 | -0.87268 | 1.021432 | 0.175034 |
| 13 | 58400800 | 58403199 | 10.04891 | 17.926 | 40.46272 | 17.79477 | -0.77018 | 0.439191 | 0.175048 |
| 7 | 13324400 | 13325599 | 24.25693 | 42.18652 | 32.8326 | 14.42425 | 0.746954 | 0.795414 | 0.175055 |
| 5 | 107551200 | 107551999 | 6.374507 | 11.98995 | 32.92951 | 13.5069 | -0.97716 | 0.073444 | 0.175063 |
| 11 | 114801200 | 114804799 | 12.13919 | 21.24697 | 47.80715 | 21.09484 | -0.77651 | 0.666132 | 0.175076 |
| 1 | 93606600 | 93611999 | 37.00493 | 59.43781 | 117.7062 | 49.84722 | -0.63352 | 1.910082 | 0.175108 |
| 3 | 97188600 | 97191799 | 19.33521 | 32.46102 | 67.85028 | 29.62259 | -0.679 | 1.142449 | 0.175116 |
| 11 | 119564000 | 119564799 | 10.66095 | 20.71402 | 15.44999 | 5.760719 | 1.013387 | -0.24601 | 0.175126 |
| 1 | 132666400 | 132675999 | 45.72925 | 72.04299 | 171.0865 | 70.99101 | -0.92323 | 2.484469 | 0.175162 |
| 7 | 137495200 | 137497799 | 10.87275 | 19.26006 | 47.78325 | 21.12732 | -0.91185 | 0.506454 | 0.175173 |
| 7 | 30603200 | 30606799 | 21.41713 | 35.3663 | 75.76477 | 32.71906 | -0.71529 | 1.341443 | 0.17518 |
| 7 | 17584200 | 17586399 | 47.32992 | 76.49188 | 62.33923 | 26.62488 | 0.655328 | 1.715976 | 0.175191 |
| 4 | 47351400 | 47353999 | 15.7991 | 26.79227 | 61.0829 | 26.75462 | -0.80283 | 1.036044 | 0.175215 |
| 12 | 86993200 | 86995199 | 10.30195 | 18.07329 | 45.67637 | 20.19431 | -0.94268 | 0.616169 | 0.17523 |
| 18 | 63829400 | 63830399 | 16.97918 | 30.6482 | 24.02718 | 10.44902 | 0.756879 | 0.37822 | 0.175238 |
| 4 | 23071200 | 23073799 | 31.06672 | 53.04102 | 36.46682 | 15.98783 | 0.920806 | 1.002511 | 0.175243 |
| 5 | 7943200 | 7944399 | 17.05694 | 31.9061 | 17.38735 | 7.198875 | 1.326176 | 0.16218 | 0.175267 |
| 2 | 172037600 | 172046199 | 65.49063 | 101.8245 | 198.7644 | 82.52191 | -0.65578 | 2.740591 | 0.175278 |
| 16 | 90966600 | 90976399 | 52.50348 | 82.69894 | 171.0498 | 70.97399 | -0.72451 | 2.492011 | 0.175289 |
| 16 | 53953800 | 53961999 | 131.9235 | 202.7497 | 153.3487 | 63.67629 | 0.697785 | 2.973356 | 0.175293 |
| 7 | 133445200 | 133448799 | 24.66439 | 40.46986 | 83.41602 | 35.91982 | -0.66342 | 1.431996 | 0.175302 |
| 13 | 66442200 | 66444399 | 34.91599 | 58.42466 | 43.77313 | 19.07082 | 0.796213 | 1.204598 | 0.175303 |
| 13 | 37741000 | 37747199 | 32.70527 | 52.46261 | 116.6118 | 49.2851 | -0.80193 | 1.945929 | 0.175305 |
| 12 | 91815200 | 91816199 | 6.512733 | 11.88047 | 36.59986 | 15.70802 | -1.19088 | 0.270633 | 0.175305 |
| 9 | 30013400 | 30015799 | 15.67186 | 27.02681 | 59.2612 | 25.99676 | -0.74305 | 0.879307 | 0.175308 |
| 8 | 12541600 | 12543599 | 12.02494 | 20.95467 | 49.43733 | 21.7462 | -0.84484 | 0.73589 | 0.175314 |
| 9 | 120582400 | 120586999 | 40.2983 | 64.54884 | 123.688 | 52.16052 | -0.59091 | 2.00293 | 0.175319 |
| 8 | 48263000 | 48264599 | 9.594238 | 17.09916 | 40.46064 | 17.80768 | -0.8386 | 0.441226 | 0.175337 |
| 7 | 31604000 | 31604599 | 9.177053 | 17.93075 | 11.99991 | 4.178209 | 1.253143 | -0.51246 | 0.175342 |
| 15 | 86098200 | 86099999 | 8.616084 | 15.20836 | 43.57433 | 19.28006 | -1.11966 | 0.524186 | 0.175378 |
| 3 | 99265400 | 99268599 | 24.54044 | 42.96145 | 28.16611 | 12.43042 | 0.993352 | 0.695083 | 0.17538 |
| X | 5339800 | 5341799 | 36.71213 | 61.37582 | 41.67685 | 18.1474 | 0.937711 | 1.219225 | 0.175386 |
| 16 | 18030800 | 18032799 | 7.402312 | 13.47957 | 37.62625 | 16.18399 | -1.0526 | 0.288101 | 0.175389 |
| 6 | 11969200 | 11970799 | 38.2326 | 62.9675 | 51.55197 | 22.23391 | 0.666989 | 1.396315 | 0.17539 |
| 8 | 25923000 | 25925399 | 18.69742 | 31.45823 | 65.55037 | 28.64282 | -0.67432 | 1.119424 | 0.17541 |
| 5 | 125657600 | 125661399 | 27.8718 | 45.18246 | 92.66519 | 39.68151 | -0.66852 | 1.630937 | 0.175412 |
| 7 | 19102400 | 19109199 | 33.83536 | 54.24566 | 117.1641 | 49.47729 | -0.76158 | 1.964952 | 0.175417 |
| 4 | 144019200 | 144019999 | 23.97879 | 41.62596 | 30.66914 | 13.45817 | 0.825566 | 0.817897 | 0.175418 |
| 4 | 58435400 | 58437599 | 11.00924 | 19.46932 | 42.98901 | 19.01389 | -0.74559 | 0.516021 | 0.175437 |
| 12 | 3073400 | 3077999 | 85.40474 | 135.4769 | 95.97777 | 39.96566 | 0.827371 | 2.335357 | 0.175446 |
| 7 | 20960000 | 20961599 | 23.14557 | 40.22626 | 30.48454 | 13.38695 | 0.785041 | 0.79535 | 0.175447 |
| 6 | 75277800 | 75278799 | 5.829978 | 11.22694 | 31.05753 | 12.06493 | -0.91568 | -0.11529 | 0.175451 |
| 17 | 34517800 | 34520399 | 13.45106 | 23.29719 | 52.66029 | 23.10411 | -0.78497 | 0.823176 | 0.175455 |
| 3 | 20749000 | 20750999 | 16.15436 | 27.71953 | 61.44119 | 26.94802 | -0.76021 | 0.927908 | 0.175457 |
| 17 | 32332000 | 32338999 | 35.22152 | 56.22005 | 129.8282 | 54.47644 | -0.86512 | 2.107847 | 0.175459 |
| 4 | 111922400 | 111926999 | 67.03127 | 107.4025 | 75.37433 | 31.80121 | 0.855996 | 2.007659 | 0.175461 |

| | | | | | | | | | |
|---|---|---|---|---|---|---|---|---|---|
| 1 | 81589800 | 81591199 | 23.086 | 40.85147 | 24.32671 | 10.75831 | 1.134491 | 0.547683 | 0.175467 |
| 4 | 4528000 | 4529599 | 9.575979 | 17.107 | 43.65252 | 19.17426 | -0.94528 | 0.429606 | 0.175474 |
| 10 | 42059000 | 42062199 | 13.90024 | 23.69011 | 59.43634 | 26.05443 | -0.93931 | 0.981663 | 0.175474 |
| X | 42450400 | 42452399 | 29.09849 | 50.62439 | 28.63623 | 12.58343 | 1.20499 | 0.808745 | 0.17548 |
| 6 | 72919800 | 72920799 | 12.05303 | 22.91362 | 18.40032 | 7.387615 | 0.809265 | 0.015904 | 0.175483 |
| 13 | 98784200 | 98791199 | 42.96425 | 68.4438 | 140.481 | 58.78498 | -0.69866 | 2.179615 | 0.175484 |
| 7 | 10257800 | 10259799 | 28.23105 | 48.61152 | 34.87388 | 15.2978 | 0.861557 | 0.913958 | 0.175494 |
| 1 | 38665400 | 38668799 | 16.47312 | 27.45296 | 74.0169 | 32.16497 | -1.04474 | 1.218335 | 0.175495 |
| 1 | 132210200 | 132212199 | 7.513187 | 13.74598 | 35.58682 | 15.21619 | -0.93864 | 0.258952 | 0.175496 |
| 7 | 28243600 | 28249199 | 31.50258 | 50.40705 | 126.3127 | 53.18521 | -0.97878 | 2.026184 | 0.175509 |
| 8 | 53305000 | 53306199 | 13.7839 | 26.31414 | 18.18877 | 7.238247 | 1.03491 | 0.004725 | 0.175515 |
| 10 | 56531400 | 56532199 | 12.99189 | 24.65383 | 19.23675 | 7.75303 | 0.845375 | 0.038931 | 0.175572 |
| 10 | 95424000 | 95426399 | 16.503 | 28.16427 | 63.02193 | 27.62923 | -0.77513 | 0.980262 | 0.175573 |
| X | 38226400 | 38228799 | 35.25498 | 59.31774 | 40.40887 | 17.65716 | 0.932858 | 1.148654 | 0.175575 |
| 9 | 42515000 | 42515799 | 6.499327 | 12.25527 | 31.31191 | 12.78923 | -0.86823 | 0.056915 | 0.175577 |
| 7 | 15186000 | 15187399 | 71.58667 | 114.9868 | 61.9586 | 26.23991 | 1.240491 | 1.936172 | 0.175581 |
| 15 | 98321400 | 98322999 | 8.782482 | 15.82225 | 38.494 | 16.7808 | -0.86867 | 0.363613 | 0.1756 |
| 4 | 151552800 | 151563399 | 64.23432 | 99.40657 | 212.3186 | 88.19178 | -0.7907 | 2.823999 | 0.175607 |
| 9 | 95070000 | 95073399 | 18.66293 | 31.47624 | 65.65785 | 28.71084 | -0.67577 | 1.097119 | 0.175607 |
| 6 | 129803000 | 129803999 | 14.99753 | 28.25146 | 19.1647 | 7.84099 | 1.027672 | 0.107405 | 0.17563 |
| 6 | 132691000 | 132693599 | 30.19185 | 51.51743 | 35.90876 | 15.74243 | 0.900829 | 1.00165 | 0.175645 |
| 6 | 90836600 | 90838799 | 21.90019 | 38.35645 | 30.17372 | 13.32145 | 0.731994 | 0.678195 | 0.175648 |
| 19 | 6907600 | 6911999 | 22.86958 | 37.28167 | 88.0457 | 37.81834 | -0.86469 | 1.520805 | 0.175649 |
| 13 | 61134000 | 61134999 | 13.55144 | 26.08154 | 16.52385 | 6.429513 | 1.188657 | -0.07306 | 0.175654 |
| 7 | 26094400 | 26095599 | 23.24558 | 40.60805 | 31.24413 | 13.76486 | 0.763979 | 0.727609 | 0.175658 |
| 4 | 121253400 | 121254999 | 31.9555 | 54.03476 | 39.95697 | 17.48609 | 0.81516 | 1.095139 | 0.175662 |
| 8 | 122265800 | 122270799 | 58.48513 | 91.28265 | 187.5811 | 77.83694 | -0.72542 | 2.670659 | 0.175668 |
| 3 | 7114800 | 7119999 | 85.93667 | 135.6931 | 110.1135 | 45.72047 | 0.626225 | 2.450247 | 0.175679 |
| 17 | 71744800 | 71746399 | 8.479295 | 15.28326 | 37.99937 | 16.55539 | -0.89884 | 0.359576 | 0.175686 |
| 10 | 6728000 | 6728599 | 6.31948 | 12.02814 | 30.50655 | 12.21723 | -0.83283 | -0.01354 | 0.175694 |
| 1 | 194219000 | 194219799 | 14.40053 | 26.89293 | 20.73477 | 8.608706 | 0.827494 | 0.159067 | 0.175708 |
| 1 | 142176400 | 142179199 | 37.05045 | 61.36024 | 48.27171 | 20.88813 | 0.725701 | 1.324116 | 0.175709 |
| 5 | 143297400 | 143301599 | 19.17272 | 31.76936 | 73.98439 | 32.02629 | -0.8348 | 1.29334 | 0.175711 |
| 2 | 17770800 | 17771599 | 4.05921 | 7.916209 | 23.20393 | 8.333886 | -0.88973 | -0.43283 | 0.17573 |
| 13 | 47855000 | 47855599 | 4.05921 | 7.916209 | 23.20393 | 8.333886 | -0.88973 | -0.43283 | 0.17573 |
| 8 | 116791400 | 116792999 | 9.886466 | 17.64007 | 40.97886 | 18.01844 | -0.81142 | 0.436076 | 0.175737 |
| 19 | 8365200 | 8369399 | 75.56834 | 120.4742 | 85.19901 | 35.68916 | 0.837147 | 2.174628 | 0.175739 |
| 5 | 21559200 | 21561199 | 11.58809 | 20.15468 | 52.63526 | 23.12497 | -0.99123 | 0.763904 | 0.175741 |
| 16 | 15454400 | 15456599 | 10.01122 | 17.53259 | 46.75058 | 20.63873 | -1.0195 | 0.645754 | 0.175745 |
| 13 | 56564600 | 56568599 | 21.52187 | 35.2774 | 82.66715 | 35.58193 | -0.84839 | 1.437606 | 0.175748 |
| 19 | 24454000 | 24455999 | 11.2127 | 19.66007 | 46.7202 | 20.64604 | -0.85473 | 0.630604 | 0.175755 |
| 10 | 109852200 | 109852599 | 3.455067 | 6.751916 | 21.18055 | 7.341802 | -0.93773 | -0.5733 | 0.175758 |
| 2 | 108876600 | 108877799 | 12.6186 | 24.32118 | 15.31493 | 5.934514 | 1.201603 | -0.10083 | 0.175771 |
| 6 | 83483400 | 83484199 | 3.534294 | 6.880281 | 23.99278 | 8.808196 | -1.16531 | -0.35782 | 0.175776 |
| 15 | 64247600 | 64249399 | 10.85851 | 19.18768 | 42.99516 | 19.0196 | -0.76713 | 0.523676 | 0.175787 |
| 10 | 61770600 | 61773599 | 16.09412 | 27.33436 | 60.59842 | 26.55473 | -0.76257 | 1.022688 | 0.175787 |
| 14 | 51073400 | 51077399 | 16.07673 | 26.96341 | 68.02501 | 29.68005 | -0.94916 | 1.143228 | 0.175794 |

| | | | | | | | | | |
|---|---|---|---|---|---|---|---|---|---|
| 5 | 111401000 | 111429599 | 169.6866 | 243.8633 | 531.4607 | 219.396 | -0.91552 | 4.170653 | 0.175811 |
| 9 | 62764400 | 62768799 | 19.48178 | 31.94702 | 82.86136 | 35.67195 | -0.99399 | 1.428209 | 0.175819 |
| 11 | 63886200 | 63886999 | 4.06141 | 7.913142 | 23.60819 | 8.58292 | -0.93076 | -0.38714 | 0.175821 |
| 17 | 22612800 | 22613999 | 46.5066 | 75.21968 | 62.38994 | 26.67214 | 0.631262 | 1.692821 | 0.175823 |
| 7 | 10617800 | 10620199 | 42.1793 | 69.30707 | 51.67565 | 22.26545 | 0.800228 | 1.433839 | 0.175829 |
| 7 | 85676000 | 85676999 | 12.65902 | 24.55251 | 15.03826 | 5.672733 | 1.278438 | -0.18611 | 0.175831 |
| 10 | 82651800 | 82653199 | 13.781 | 26.00352 | 19.62716 | 8.005747 | 0.877856 | 0.085246 | 0.175839 |
| 11 | 117193200 | 117195599 | 12.49257 | 21.74891 | 49.97848 | 21.97114 | -0.8073 | 0.752559 | 0.17584 |
| 6 | 120005800 | 120006599 | 10.35796 | 20.15999 | 13.44355 | 4.943411 | 1.188681 | -0.30672 | 0.175853 |
| 1 | 133327800 | 133328999 | 5.6343 | 10.61524 | 31.25396 | 12.78374 | -1.07206 | 0.054802 | 0.175853 |
| 1 | 4289800 | 4291199 | 17.69591 | 32.32545 | 22.36559 | 9.582309 | 0.949645 | 0.313413 | 0.175858 |
| 3 | 6620600 | 6625999 | 95.84495 | 149.9402 | 126.8057 | 52.60559 | 0.556448 | 2.639157 | 0.175868 |
| X | 25818600 | 25820599 | 22.57061 | 40.13011 | 23.22224 | 10.24126 | 1.179299 | 0.507341 | 0.175884 |
| 14 | 54741400 | 54753199 | 94.15614 | 143.535 | 260.2739 | 107.9031 | -0.57171 | 3.131721 | 0.1759 |
| 14 | 45555400 | 45556399 | 6.073678 | 11.45961 | 31.88028 | 13.00579 | -0.98875 | 0.048472 | 0.175904 |
| 10 | 92820600 | 92823599 | 40.28388 | 66.22651 | 52.1981 | 22.49543 | 0.720614 | 1.422378 | 0.175908 |
| 7 | 49024000 | 49027599 | 18.18691 | 30.71657 | 63.94298 | 27.97288 | -0.67266 | 1.082442 | 0.175908 |
| 15 | 90526000 | 90528399 | 28.69302 | 46.45498 | 94.25627 | 40.32526 | -0.65478 | 1.659362 | 0.17591 |
| 12 | 80801800 | 80808399 | 41.03631 | 65.04264 | 155.4872 | 64.69296 | -0.92562 | 2.336595 | 0.175917 |
| 14 | 47866400 | 47866999 | 8.459786 | 16.50669 | 12.7023 | 4.52926 | 1.024189 | -0.44096 | 0.175923 |
| 6 | 133050600 | 133051799 | 17.94964 | 32.71927 | 21.5856 | 9.27277 | 1.014964 | 0.32389 | 0.175927 |
| 11 | 100309600 | 100315399 | 22.97185 | 37.21994 | 94.93178 | 40.63659 | -0.98275 | 1.634129 | 0.175929 |
| 5 | 106601000 | 106603599 | 22.70779 | 37.41408 | 78.16014 | 33.71561 | -0.6802 | 1.372837 | 0.175936 |
| 10 | 86052800 | 86054799 | 9.220817 | 16.64108 | 46.72469 | 20.32424 | -1.07295 | 0.369462 | 0.175942 |
| 8 | 74527200 | 74528399 | 15.32819 | 28.71812 | 20.00273 | 8.259383 | 0.978992 | 0.144716 | 0.175965 |
| 1 | 91042600 | 91043999 | 6.122394 | 11.41817 | 33.23098 | 13.82839 | -1.07644 | 0.128898 | 0.176011 |
| 14 | 50442000 | 50446599 | 102.865 | 159.9504 | 133.1711 | 55.28841 | 0.572254 | 2.748427 | 0.176014 |
| 9 | 62454800 | 62458599 | 19.91238 | 33.1114 | 71.46098 | 31.00927 | -0.72503 | 1.252095 | 0.176015 |
| 14 | 72033200 | 72035599 | 10.1777 | 18.2109 | 45.87174 | 20.12203 | -0.92553 | 0.428084 | 0.17603 |
| 2 | 143984400 | 143987399 | 24.75301 | 40.36863 | 87.34964 | 37.52499 | -0.73849 | 1.518036 | 0.176037 |
| 2 | 152893000 | 152897799 | 22.43308 | 36.39487 | 91.96765 | 39.40598 | -0.96745 | 1.606418 | 0.176048 |
| 4 | 95069400 | 95070999 | 10.27291 | 18.09199 | 43.55696 | 19.28107 | -0.87175 | 0.560023 | 0.17606 |
| 9 | 63534800 | 63535799 | 5.429744 | 10.43201 | 29.12718 | 11.39054 | -0.93815 | -0.10515 | 0.176062 |
| 1 | 5774200 | 5776199 | 28.90071 | 49.72923 | 34.96102 | 15.33569 | 0.890558 | 0.924747 | 0.176071 |
| 1 | 167627600 | 167631599 | 22.79533 | 37.32512 | 83.39228 | 35.87791 | -0.78048 | 1.454099 | 0.176074 |
| 4 | 148298000 | 148306199 | 53.57977 | 84.14329 | 177.82 | 73.75458 | -0.75882 | 2.553637 | 0.176075 |
| 18 | 74106000 | 74119999 | 94.56374 | 143.1748 | 295.2926 | 122.1701 | -0.76544 | 3.272882 | 0.176094 |
| 14 | 54680400 | 54682199 | 7.823921 | 14.27445 | 36.07199 | 15.47858 | -0.90764 | 0.278332 | 0.1761 |
| 17 | 29785200 | 29791999 | 31.59392 | 50.65309 | 116.8597 | 49.35018 | -0.85637 | 1.962445 | 0.176141 |
| 1 | 147573600 | 147574399 | 11.58988 | 22.60842 | 11.66567 | 4.175279 | 1.588668 | -0.38629 | 0.176143 |
| 8 | 89098600 | 89104199 | 42.51762 | 67.68365 | 136.4196 | 57.02872 | -0.67356 | 2.202968 | 0.176167 |
| 13 | 18289800 | 18290399 | 6.194216 | 11.71187 | 30.82049 | 12.51713 | -0.90294 | 0.034366 | 0.176194 |
| 1 | 116384800 | 116385399 | 7.587084 | 14.11341 | 33.7512 | 14.10623 | -0.80112 | 0.15939 | 0.176196 |
| 2 | 67628600 | 67629199 | 10.86735 | 21.12809 | 15.27086 | 5.66563 | 1.063831 | -0.26243 | 0.176222 |
| 12 | 47502000 | 47503599 | 21.24601 | 37.69232 | 25.01036 | 11.05023 | 0.980859 | 0.512959 | 0.176229 |
| 10 | 81106400 | 81132799 | 165.5385 | 238.5823 | 518.1914 | 213.8286 | -0.90653 | 4.128084 | 0.176238 |
| 2 | 179380200 | 179384799 | 28.87977 | 46.67741 | 96.74678 | 41.32433 | -0.68796 | 1.701345 | 0.176263 |

| | | | | | | | | | |
|---|---|---|---|---|---|---|---|---|---|
| 1 | 103338400 | 103339399 | 11.16464 | 21.64616 | 16.34029 | 6.174491 | 0.97779 | -0.194 | 0.176263 |
| 16 | 83157200 | 83157999 | 11.16464 | 21.64616 | 16.34029 | 6.174491 | 0.97779 | -0.194 | 0.176263 |
| 1 | 112670000 | 112670999 | 12.60178 | 24.27516 | 17.23583 | 6.689611 | 1.029385 | -0.08874 | 0.176308 |
| 17 | 79749400 | 79750399 | 12.60178 | 24.27516 | 17.23583 | 6.689611 | 1.029385 | -0.08874 | 0.176308 |
| 15 | 58826200 | 58828999 | 10.59959 | 18.60415 | 45.14766 | 19.96843 | -0.88441 | 0.609913 | 0.176331 |
| 3 | 109191600 | 109192399 | 14.76317 | 27.68405 | 20.01997 | 8.254109 | 0.92681 | 0.133838 | 0.176334 |
| 12 | 17630800 | 17631799 | 11.73842 | 22.42635 | 18.4085 | 7.313863 | 0.790932 | -0.02059 | 0.176365 |
| 17 | 81108000 | 81110399 | 11.08286 | 19.38555 | 48.34021 | 21.3184 | -0.92387 | 0.672037 | 0.176366 |
| 1 | 89866800 | 89870199 | 24.35318 | 39.52672 | 90.93231 | 38.97452 | -0.83198 | 1.599132 | 0.176376 |
| 7 | 24176200 | 24188199 | 81.55088 | 124.9262 | 239.4046 | 99.36303 | -0.64578 | 3.028191 | 0.176377 |
| 7 | 37783400 | 37786399 | 18.98761 | 31.53538 | 72.99897 | 31.65015 | -0.82589 | 1.264461 | 0.176389 |
| 1 | 101417200 | 101419399 | 30.33176 | 52.05929 | 34.21857 | 15.00851 | 0.986326 | 0.95011 | 0.176395 |
| 1 | 84647200 | 84648399 | 14.02329 | 26.57071 | 19.1353 | 7.736928 | 0.956186 | 0.058968 | 0.176403 |
| 12 | 23310200 | 23316599 | 161.0238 | 242.8416 | 193.5101 | 79.96073 | 0.600638 | 3.361315 | 0.176404 |
| 2 | 134460200 | 134461199 | 12.99584 | 24.97828 | 17.09853 | 6.679102 | 1.072732 | -0.06547 | 0.176408 |
| 11 | 59262200 | 59266599 | 22.43511 | 36.84451 | 81.33256 | 35.03574 | -0.76141 | 1.414499 | 0.176433 |
| 5 | 93736000 | 93741399 | 81.07463 | 128.9186 | 90.06207 | 37.59296 | 0.850826 | 2.259823 | 0.176434 |
| 12 | 22842600 | 22867399 | 576.8488 | 784.3016 | 588.4586 | 245.0822 | 0.541722 | 5.016363 | 0.176435 |
| 2 | 38533400 | 38534199 | 4.966226 | 9.595873 | 27.02631 | 10.3761 | -0.92424 | -0.17534 | 0.176464 |
| 14 | 51597000 | 51597799 | 4.966226 | 9.595873 | 27.02631 | 10.3761 | -0.92424 | -0.17534 | 0.176464 |
| 4 | 124522800 | 124525999 | 14.56863 | 25.1091 | 54.8061 | 24.04195 | -0.73646 | 0.875008 | 0.176465 |
| 18 | 24498000 | 24500199 | 25.73803 | 44.90296 | 30.52391 | 13.43644 | 0.941648 | 0.757517 | 0.176466 |
| 4 | 31612800 | 31614599 | 8.053507 | 14.88781 | 35.81719 | 15.10056 | -0.81949 | 0.20221 | 0.176467 |
| 6 | 107716600 | 107717999 | 8.053507 | 14.88781 | 35.81719 | 15.10056 | -0.81949 | 0.20221 | 0.176467 |
| 12 | 81676600 | 81682399 | 33.26381 | 53.55072 | 108.5522 | 46.12292 | -0.66292 | 1.829851 | 0.176489 |
| 7 | 117728200 | 117729399 | 6.752326 | 12.45712 | 35.18953 | 14.87696 | -1.05163 | 0.202787 | 0.176492 |
| 7 | 126773800 | 126778799 | 32.46024 | 52.29544 | 104.1467 | 44.29816 | -0.63604 | 1.805137 | 0.176501 |
| 2 | 171961800 | 171963399 | 16.48132 | 30.83238 | 19.30331 | 7.990819 | 1.128659 | 0.162237 | 0.176507 |
| Y | 6747400 | 6749999 | 15.45039 | 28.83476 | 10.13711 | 4.213389 | 1.946386 | 0.140152 | 0.176515 |
| 19 | 25586000 | 25593199 | 47.7728 | 75.75362 | 150.3778 | 62.59494 | -0.65737 | 2.327425 | 0.176516 |
| 13 | 65636600 | 65648199 | 204.108 | 305.8945 | 205.5899 | 84.81205 | 0.83816 | 3.490992 | 0.176518 |
| 16 | 13222400 | 13224199 | 9.167769 | 16.26093 | 40.97916 | 18.10896 | -0.93263 | 0.471925 | 0.176521 |
| 8 | 106387000 | 106387399 | 2.936757 | 5.741254 | 20.01327 | 6.868303 | -1.07325 | -0.62167 | 0.176521 |
| 7 | 113555800 | 113558399 | 14.54199 | 25.22283 | 53.78615 | 23.5991 | -0.701 | 0.816832 | 0.176524 |
| 1 | 191726600 | 191729999 | 18.36028 | 30.4625 | 76.70213 | 33.23239 | -0.94684 | 1.279612 | 0.176536 |
| 3 | 32583400 | 32583999 | 3.854075 | 7.419902 | 27.86912 | 10.83467 | -1.35052 | -0.14133 | 0.176541 |
| 3 | 9763800 | 9769599 | 31.21659 | 50.19326 | 112.2089 | 47.59181 | -0.806 | 1.867173 | 0.176542 |
| 12 | 63801600 | 63803199 | 15.35749 | 29.07087 | 16.51574 | 6.691299 | 1.291671 | 0.06974 | 0.17655 |
| 13 | 108070600 | 108072399 | 10.85344 | 19.17152 | 45.08986 | 19.94772 | -0.83705 | 0.529859 | 0.176555 |
| 2 | 23686600 | 23687999 | 28.1895 | 48.43998 | 36.06459 | 15.81912 | 0.807785 | 0.934438 | 0.176557 |
| 3 | 135096400 | 135097199 | 6.0102 | 11.47057 | 29.97748 | 11.93662 | -0.86761 | -0.0345 | 0.176576 |
| 13 | 53332200 | 53339999 | 54.22312 | 85.58862 | 162.2899 | 67.37316 | -0.5969 | 2.447445 | 0.176589 |
| 8 | 84755000 | 84759599 | 20.66154 | 33.72044 | 88.3377 | 37.95948 | -1.01259 | 1.499855 | 0.17659 |
| 3 | 40682800 | 40684599 | 11.12202 | 19.35268 | 50.47815 | 22.18347 | -0.98896 | 0.754674 | 0.176593 |
| X | 31995600 | 31997999 | 36.05551 | 60.95281 | 33.98237 | 14.87037 | 1.220916 | 1.10546 | 0.176599 |
| 9 | 3824200 | 3830199 | 95.25257 | 149.5271 | 120.1015 | 49.81201 | 0.634563 | 2.576967 | 0.17661 |
| 5 | 138883200 | 138887399 | 27.46584 | 44.32891 | 99.92416 | 42.60315 | -0.8111 | 1.742774 | 0.17661 |

| | | | | | | | | | |
|---|---|---|---|---|---|---|---|---|---|
| 9 | 45962600 | 45964599 | 9.596766 | 17.09217 | 42.53489 | 18.73122 | -0.91175 | 0.447714 | 0.176628 |
| 8 | 70301200 | 70312599 | 80.81168 | 124.0325 | 235.5313 | 97.7839 | -0.63049 | 2.989281 | 0.176632 |
| 2 | 87377000 | 87377799 | 5.280286 | 10.12983 | 28.44216 | 11.1701 | -0.95124 | -0.09257 | 0.176645 |
| 2 | 28025600 | 28038999 | 85.24194 | 130.0759 | 255.0243 | 105.7511 | -0.68309 | 3.108693 | 0.176654 |
| 3 | 52009200 | 52012999 | 18.72057 | 30.6898 | 85.05209 | 36.61054 | -1.08943 | 1.431069 | 0.176655 |
| 6 | 91732600 | 91737999 | 33.32911 | 53.50116 | 112.6704 | 47.67056 | -0.72296 | 1.924206 | 0.176658 |
| 6 | 116633200 | 116635199 | 9.527378 | 17.16682 | 40.06621 | 17.46289 | -0.80939 | 0.36989 | 0.176688 |
| 1 | 181279600 | 181287399 | 54.74664 | 86.28738 | 168.6642 | 69.98995 | -0.64238 | 2.480243 | 0.17669 |
| 3 | 58674000 | 58675799 | 15.47769 | 29.68254 | 15.06202 | 5.921174 | 1.491581 | -0.02553 | 0.17672 |
| 12 | 21714800 | 21717999 | 73.58229 | 116.7733 | 99.47465 | 41.43964 | 0.562406 | 2.313318 | 0.176741 |
| 6 | 88817800 | 88821399 | 18.62866 | 30.70793 | 79.24696 | 34.19138 | -0.98363 | 1.357473 | 0.176743 |
| 3 | 76750200 | 76750999 | 10.6198 | 20.61987 | 15.79749 | 5.923132 | 0.966636 | -0.22852 | 0.176774 |
| X | 42843000 | 42844399 | 18.85659 | 34.90851 | 19.00575 | 7.997231 | 1.3108 | 0.242644 | 0.176803 |
| 2 | 143129000 | 143131599 | 38.18331 | 63.93479 | 39.7858 | 17.341 | 1.062873 | 1.202404 | 0.176804 |
| 5 | 22765600 | 22766999 | 23.97958 | 42.22825 | 26.01576 | 11.51639 | 1.083452 | 0.60755 | 0.176812 |
| 13 | 103317600 | 103318799 | 13.71771 | 23.90589 | 51.07412 | 22.46702 | -0.70281 | 0.74436 | 0.176818 |
| 2 | 86414000 | 86416399 | 60.10567 | 96.15516 | 78.74957 | 33.14878 | 0.63111 | 2.048218 | 0.176822 |
| 3 | 106776200 | 106778999 | 33.86301 | 57.56125 | 35.82522 | 15.70098 | 1.062912 | 1.05125 | 0.176831 |
| 16 | 93774400 | 93784399 | 64.01469 | 98.88217 | 220.9048 | 91.75917 | -0.85753 | 2.859103 | 0.176833 |
| 16 | 22478400 | 22480199 | 10.57629 | 18.63757 | 43.54976 | 19.27555 | -0.82856 | 0.555494 | 0.176834 |
| 7 | 23771800 | 23774399 | 17.19289 | 28.744 | 68.55805 | 29.85277 | -0.86849 | 1.18231 | 0.176847 |
| 18 | 82491400 | 82495799 | 19.0701 | 31.18915 | 87.14285 | 37.47497 | -1.10294 | 1.464815 | 0.17685 |
| 2 | 33400000 | 33409399 | 69.90093 | 108.6513 | 197.4533 | 81.98098 | -0.55306 | 2.747781 | 0.176868 |
| 6 | 69063800 | 69065799 | 24.92901 | 43.54599 | 29.88994 | 13.16772 | 0.927697 | 0.734033 | 0.176869 |
| 18 | 77554000 | 77557399 | 21.40026 | 35.00251 | 84.69971 | 36.42523 | -0.89619 | 1.467436 | 0.176876 |
| 4 | 79562800 | 79564199 | 15.88934 | 29.46967 | 20.89815 | 8.771466 | 0.934862 | 0.211857 | 0.176896 |
| 4 | 133672800 | 133674399 | 18.82421 | 33.49707 | 25.93169 | 11.4281 | 0.762334 | 0.480914 | 0.176904 |
| 18 | 67120800 | 67121999 | 7.974905 | 14.5446 | 36.5917 | 15.70877 | -0.90198 | 0.281646 | 0.176909 |
| 15 | 52093200 | 52093799 | 10.41431 | 20.20169 | 16.48777 | 6.215419 | 0.869875 | -0.20944 | 0.176911 |
| 5 | 29707800 | 29709399 | 19.89282 | 34.98089 | 27.19265 | 12.04084 | 0.749062 | 0.600043 | 0.176916 |
| 2 | 153764000 | 153769199 | 24.62288 | 39.7329 | 102.1528 | 43.54261 | -1.00059 | 1.744621 | 0.176918 |
| 17 | 86469200 | 86471399 | 9.246776 | 16.61272 | 38.97421 | 17.03778 | -0.81941 | 0.384854 | 0.176954 |
| 2 | 3687200 | 3688399 | 8.240358 | 15.12487 | 35.6433 | 15.17391 | -0.79988 | 0.245189 | 0.176986 |
| 11 | 24077200 | 24080399 | 14.31371 | 24.51268 | 58.50695 | 25.65699 | -0.8667 | 0.935942 | 0.17699 |
| 4 | 153075200 | 153096799 | 220.1057 | 315.0225 | 535.4087 | 221.1685 | -0.56295 | 4.234156 | 0.177002 |
| 14 | 117958000 | 117959799 | 9.606583 | 17.08928 | 40.4612 | 17.83593 | -0.84083 | 0.454599 | 0.177013 |
| 1 | 27110400 | 27114399 | 56.9365 | 91.88246 | 63.7027 | 27.15573 | 0.885656 | 1.774977 | 0.177028 |
| X | 126853200 | 126854799 | 20.63032 | 37.36336 | 21.564 | 9.338698 | 1.199793 | 0.381782 | 0.177043 |
| 15 | 76832400 | 76835199 | 10.01144 | 17.56911 | 49.34737 | 21.82306 | -1.09459 | 0.612513 | 0.17705 |
| 11 | 3050000 | 3062999 | 692.0834 | 938.9543 | 498.9538 | 207.8743 | 1.036147 | 5.052543 | 0.177052 |
| 9 | 45593800 | 45598199 | 18.43963 | 30.19686 | 88.42884 | 38.04681 | -1.16938 | 1.445612 | 0.177054 |
| 8 | 33573400 | 33574799 | 8.04307 | 14.89945 | 36.43915 | 15.3194 | -0.84018 | 0.191247 | 0.177057 |
| 16 | 60736600 | 60738999 | 26.46575 | 45.85199 | 33.86031 | 14.85596 | 0.821693 | 0.851776 | 0.177058 |
| 6 | 88891400 | 88896199 | 26.95701 | 43.61406 | 100.6219 | 43.02665 | -0.84064 | 1.671309 | 0.177065 |
| 9 | 14012400 | 14013399 | 10.4868 | 18.3274 | 48.34534 | 21.30538 | -1.00433 | 0.684044 | 0.177071 |
| 9 | 108009600 | 108015199 | 29.02519 | 46.93793 | 100.371 | 42.90244 | -0.73207 | 1.685264 | 0.177076 |
| 14 | 116158000 | 116159399 | 7.248241 | 13.60464 | 34.62331 | 14.25248 | -0.87288 | 0.098187 | 0.177078 |

| | | | | | | | | | |
|---|---|---|---|---|---|---|---|---|---|
| 9 | 71736400 | 71740399 | 14.72624 | 24.64941 | 70.56778 | 30.75095 | -1.13072 | 1.163106 | 0.177089 |
| 17 | 71800000 | 71802999 | 23.68225 | 38.92651 | 84.10285 | 36.24244 | -0.73014 | 1.407435 | 0.177095 |
| 14 | 3675600 | 3676799 | 15.88679 | 29.38551 | 21.57808 | 9.090179 | 0.881527 | 0.229274 | 0.177103 |
| 4 | 101128400 | 101130199 | 17.10887 | 31.07222 | 23.40911 | 10.10246 | 0.82087 | 0.343002 | 0.177185 |
| 7 | 80673200 | 80675199 | 9.616434 | 17.07718 | 40.95511 | 18.08006 | -0.86087 | 0.465595 | 0.177204 |
| 5 | 126407400 | 126410599 | 30.17975 | 50.8575 | 41.63781 | 18.20274 | 0.668632 | 1.118259 | 0.177207 |
| 15 | 81899400 | 81900999 | 9.653722 | 17.02579 | 43.05474 | 19.05332 | -0.94113 | 0.540341 | 0.177235 |
| 6 | 58573200 | 58573999 | 36.43815 | 61.41263 | 36.92512 | 16.1432 | 1.112122 | 1.132353 | 0.177239 |
| 14 | 112964600 | 112965799 | 18.7601 | 33.91461 | 23.17231 | 10.05802 | 0.955613 | 0.380401 | 0.17725 |
| 6 | 132478800 | 132480599 | 33.21294 | 55.41386 | 44.90775 | 19.52935 | 0.683354 | 1.228048 | 0.177252 |
| 4 | 95385400 | 95385999 | 9.29566 | 18.13277 | 13.75197 | 4.923046 | 1.040483 | -0.41051 | 0.177256 |
| 1 | 13005800 | 13006399 | 10.3265 | 20.09626 | 14.71271 | 5.415523 | 1.055284 | -0.29852 | 0.177258 |
| 16 | 78049400 | 78049999 | 2.837908 | 5.545006 | 21.04315 | 7.323251 | -1.21299 | -0.57078 | 0.177275 |
| 12 | 22978200 | 22993399 | 353.5712 | 501.517 | 392.5972 | 162.4172 | 0.542038 | 4.369806 | 0.177279 |
| 3 | 152052000 | 152052799 | 3.835988 | 7.435998 | 25.95318 | 9.814718 | -1.20763 | -0.24254 | 0.177286 |
| 13 | 15112800 | 15113399 | 3.756568 | 7.340567 | 22.51254 | 7.818469 | -0.90968 | -0.55933 | 0.177288 |
| 5 | 122783400 | 122789199 | 34.5133 | 55.24101 | 121.0682 | 50.99857 | -0.78543 | 2.019462 | 0.17729 |
| 15 | 98065400 | 98068999 | 20.9721 | 34.2356 | 88.47331 | 38.02112 | -0.99278 | 1.496746 | 0.177295 |
| 9 | 38947800 | 38948999 | 14.04736 | 26.92494 | 16.31114 | 6.422198 | 1.236419 | -0.03239 | 0.177301 |
| 7 | 42757000 | 42766999 | 255.7811 | 372.9856 | 301.3095 | 124.1601 | 0.536175 | 3.956684 | 0.177303 |
| 18 | 17406600 | 17408199 | 15.3386 | 28.74555 | 19.33964 | 7.981859 | 1.028742 | 0.140569 | 0.177304 |
| 12 | 19384400 | 19386599 | 32.51874 | 54.25884 | 40.75975 | 17.72614 | 0.792405 | 1.221683 | 0.177306 |
| 7 | 4990400 | 4999799 | 124.4057 | 185.7988 | 337.8224 | 139.2679 | -0.60122 | 3.548234 | 0.177306 |
| 7 | 31962000 | 31965799 | 55.135 | 89.47426 | 53.0493 | 22.73499 | 1.118714 | 1.638642 | 0.177308 |
| 12 | 73249600 | 73250199 | 3.421111 | 6.669864 | 23.67577 | 8.535702 | -1.1672 | -0.42799 | 0.177313 |
| 5 | 93805400 | 93808999 | 70.33571 | 112.021 | 92.77785 | 38.8106 | 0.608103 | 2.200191 | 0.177325 |
| 5 | 70722600 | 70727799 | 74.62606 | 118.5744 | 97.96191 | 40.85932 | 0.608206 | 2.277564 | 0.17735 |
| 17 | 38062800 | 38069199 | 83.75178 | 132.9534 | 94.7767 | 39.49483 | 0.819448 | 2.31251 | 0.177357 |
| 12 | 19043000 | 19062199 | 533.458 | 729.123 | 549.2466 | 228.5652 | 0.543594 | 4.933224 | 0.177359 |
| 2 | 152251200 | 152254599 | 13.64767 | 23.50096 | 59.15033 | 25.94695 | -0.94154 | 0.882208 | 0.177364 |
| 8 | 109576200 | 109579799 | 22.51448 | 36.78183 | 84.71529 | 36.41637 | -0.82637 | 1.48728 | 0.177376 |
| 5 | 27132400 | 27135999 | 24.65734 | 40.4965 | 82.28905 | 35.44869 | -0.64233 | 1.41996 | 0.177383 |
| 8 | 57771000 | 57776999 | 45.97756 | 72.94312 | 152.4086 | 63.4718 | -0.73045 | 2.311254 | 0.177402 |
| 2 | 4721400 | 4724199 | 14.20628 | 24.22262 | 60.58651 | 26.55959 | -0.93498 | 0.978434 | 0.177403 |
| 19 | 23291600 | 23293799 | 11.6544 | 20.4659 | 46.68455 | 20.65703 | -0.79612 | 0.607395 | 0.177417 |
| 16 | 89993200 | 89997599 | 23.65692 | 38.55639 | 86.94679 | 37.34322 | -0.7985 | 1.525544 | 0.177429 |
| 10 | 22213200 | 22215799 | 13.87578 | 24.18965 | 52.15996 | 22.94961 | -0.71608 | 0.741651 | 0.177433 |
| 17 | 24908200 | 24909599 | 7.141681 | 13.29318 | 34.31535 | 14.32685 | -0.90952 | 0.151332 | 0.177435 |
| 1 | 63669000 | 63670399 | 74.59851 | 114.6512 | 227.5887 | 94.50394 | -0.69286 | 2.961203 | 0.177446 |
| 11 | 119397600 | 119401999 | 20.06956 | 32.67497 | 90.07334 | 38.67109 | -1.0883 | 1.536072 | 0.177461 |
| 3 | 94734600 | 94736799 | 21.63581 | 37.99268 | 29.04337 | 12.85079 | 0.77301 | 0.634121 | 0.177463 |
| 2 | 154013200 | 154016199 | 17.03076 | 28.49321 | 68.58497 | 29.8779 | -0.88167 | 1.173743 | 0.177463 |
| 14 | 89127800 | 89133599 | 105.5728 | 164.3587 | 133.8448 | 55.55811 | 0.605501 | 2.728651 | 0.177466 |
| 7 | 15269400 | 15273999 | 71.26981 | 114.4628 | 63.06883 | 26.70476 | 1.20815 | 1.940201 | 0.17748 |
| 7 | 55344400 | 55345399 | 13.37235 | 25.69098 | 17.08612 | 6.682437 | 1.112516 | -0.05746 | 0.177506 |
| 4 | 134836000 | 134852599 | 120.7903 | 180.1518 | 345.2116 | 142.2761 | -0.67871 | 3.573067 | 0.177526 |
| 6 | 41155000 | 41155999 | 6.302432 | 12.04882 | 30.6984 | 12.17704 | -0.82726 | -0.04537 | 0.177527 |

| | | | | | | | | | |
|---|---|---|---|---|---|---|---|---|---|
| 8 | 79853800 | 79854599 | 4.371425 | 8.470249 | 25.97699 | 9.843705 | -1.0265 | -0.22851 | 0.177532 |
| 6 | 25092800 | 25094999 | 26.79604 | 46.17921 | 30.31912 | 13.30014 | 0.988853 | 0.892599 | 0.177539 |
| 19 | 24810200 | 24812199 | 11.01471 | 19.49564 | 42.99215 | 19.01204 | -0.74318 | 0.507516 | 0.177546 |
| 3 | 35492400 | 35495199 | 20.2767 | 33.57771 | 74.71792 | 32.32989 | -0.76956 | 1.30359 | 0.177554 |
| 2 | 35605200 | 35607599 | 10.41237 | 18.37191 | 43.01847 | 19.03558 | -0.83094 | 0.538049 | 0.177558 |
| 7 | 58485800 | 58488199 | 15.81502 | 26.77264 | 62.16069 | 27.21234 | -0.82919 | 1.051855 | 0.177593 |
| 19 | 5049200 | 5052399 | 15.81502 | 26.77264 | 62.16069 | 27.21234 | -0.82919 | 1.051855 | 0.177593 |
| 2 | 169826000 | 169829999 | 24.1978 | 39.6589 | 88.69563 | 38.1748 | -0.78157 | 1.447123 | 0.177593 |
| 8 | 110848000 | 110855999 | 53.295 | 84.00112 | 165.7366 | 68.77528 | -0.65579 | 2.478922 | 0.177613 |
| 5 | 102752800 | 102753999 | 9.940341 | 17.58954 | 43.03325 | 19.03119 | -0.89175 | 0.506156 | 0.177615 |
| 9 | 99234200 | 99242999 | 56.1377 | 87.85811 | 187.875 | 77.93702 | -0.77972 | 2.619188 | 0.177623 |
| 13 | 63720000 | 63722199 | 12.8819 | 22.19376 | 54.7207 | 24.00422 | -0.91131 | 0.871937 | 0.177633 |
| 12 | 106477000 | 106489599 | 87.95841 | 134.0331 | 260.2743 | 107.8918 | -0.67088 | 3.138231 | 0.177633 |
| 9 | 107173400 | 107176199 | 10.30753 | 18.08143 | 46.7146 | 20.65173 | -0.97448 | 0.618058 | 0.177636 |
| 8 | 14936400 | 14939599 | 16.38042 | 27.88379 | 60.13702 | 26.35905 | -0.72258 | 1.001773 | 0.177643 |
| 15 | 68081000 | 68084599 | 15.24423 | 26.06461 | 62.99029 | 27.61951 | -0.88525 | 0.957685 | 0.177648 |
| 5 | 37882600 | 37883999 | 6.524748 | 12.24464 | 32.30696 | 13.30271 | -0.92405 | 0.089063 | 0.177651 |
| X | 33261400 | 33273199 | 154.2309 | 235.9848 | 148.5863 | 61.65278 | 0.95716 | 3.068508 | 0.177657 |
| 6 | 126410000 | 126410599 | 4.487133 | 8.737054 | 24.75341 | 9.071813 | -0.86846 | -0.34946 | 0.17766 |
| 13 | 118356600 | 118358999 | 29.70061 | 50.79165 | 37.13181 | 16.28152 | 0.833001 | 0.984541 | 0.177662 |
| 2 | 73216000 | 73217399 | 8.137511 | 14.45184 | 42.02395 | 18.55084 | -1.13672 | 0.457329 | 0.177667 |
| 10 | 69101200 | 69102999 | 13.34437 | 22.98174 | 54.73631 | 24.01079 | -0.8618 | 0.877379 | 0.177679 |
| 4 | 3834400 | 3836199 | 13.4005 | 23.3897 | 49.98273 | 22.00859 | -0.70282 | 0.71919 | 0.177687 |
| 14 | 25755600 | 25760799 | 24.76977 | 40.04502 | 102.1384 | 43.64205 | -0.98608 | 1.687471 | 0.177687 |
| 10 | 61735400 | 61739599 | 24.96912 | 40.58287 | 90.37035 | 38.77341 | -0.78331 | 1.570372 | 0.177688 |
| 8 | 79277800 | 79279399 | 8.073692 | 14.91962 | 35.7875 | 15.09562 | -0.8153 | 0.207159 | 0.17769 |
| 11 | 119141400 | 119146799 | 21.29636 | 34.67201 | 87.88596 | 37.73187 | -0.96763 | 1.538405 | 0.177704 |
| 16 | 70436000 | 70436799 | 10.93536 | 21.10525 | 17.41723 | 6.723047 | 0.821584 | -0.12383 | 0.177719 |
| 7 | 17926800 | 17928199 | 29.16578 | 49.85333 | 37.68235 | 16.52229 | 0.784947 | 0.986852 | 0.177741 |
| 12 | 22579200 | 22581599 | 41.70461 | 68.00235 | 57.00266 | 24.48651 | 0.62577 | 1.536856 | 0.177743 |
| 2 | 121621000 | 121621399 | 2.83864 | 5.55281 | 19.24112 | 6.359131 | -1.01324 | -0.75643 | 0.177753 |
| 8 | 77625400 | 77629399 | 24.90436 | 40.03715 | 116.1805 | 49.26446 | -1.18154 | 1.866132 | 0.177768 |
| 8 | 86939600 | 86943199 | 26.84183 | 43.7191 | 88.09734 | 37.82851 | -0.6374 | 1.541981 | 0.177773 |
| 10 | 70761000 | 70761799 | 6.621516 | 12.56471 | 31.56683 | 12.72351 | -0.82784 | 0.012546 | 0.177777 |
| 1 | 139050000 | 139052199 | 15.9164 | 26.66279 | 70.67786 | 30.8144 | -1.02017 | 1.159737 | 0.177782 |
| 17 | 39960000 | 39971999 | 33.81798 | 57.68017 | 31.55785 | 13.83451 | 1.248604 | 1.016583 | 0.17779 |
| 14 | 13183400 | 13184599 | 10.59162 | 18.61262 | 44.6061 | 19.74904 | -0.86632 | 0.590065 | 0.177805 |
| 15 | 17804800 | 17805799 | 12.52964 | 24.00441 | 17.89532 | 7.055617 | 0.939258 | -0.03724 | 0.177805 |
| 5 | 10181200 | 10181599 | 8.146647 | 15.93271 | 11.61508 | 3.943311 | 1.16392 | -0.6141 | 0.177821 |
| 11 | 95113200 | 95121399 | 37.54435 | 59.76093 | 141.0611 | 58.96119 | -0.90113 | 2.202187 | 0.177823 |
| 1 | 166419800 | 166422999 | 77.78436 | 123.5559 | 97.30686 | 40.57404 | 0.67674 | 2.288909 | 0.17783 |
| 10 | 19580800 | 19581999 | 5.958218 | 11.06088 | 33.66486 | 14.11211 | -1.14794 | 0.16075 | 0.177832 |
| 2 | 30145200 | 30152399 | 37.34356 | 59.51667 | 134.3606 | 56.25897 | -0.83493 | 2.159684 | 0.177835 |
| 18 | 46759400 | 46760399 | 11.77197 | 22.65276 | 17.8705 | 6.95698 | 0.874792 | -0.0863 | 0.177844 |
| 1 | 75496600 | 75498599 | 10.71445 | 18.89183 | 45.61795 | 20.18864 | -0.87587 | 0.54989 | 0.177848 |
| 9 | 57795400 | 57797799 | 11.2058 | 19.67441 | 45.12594 | 19.96419 | -0.80369 | 0.607079 | 0.17786 |
| 3 | 146060600 | 146063199 | 27.50105 | 44.76809 | 89.8106 | 38.55642 | -0.63157 | 1.553233 | 0.177862 |

| | | | | | | | | | |
|---|---|---|---|---|---|---|---|---|---|
| 2 | 181240200 | 181243399 | 13.52747 | 23.22412 | 57.416 | 25.18364 | -0.91653 | 0.908673 | 0.177864 |
| 6 | 132512800 | 132515999 | 60.51158 | 97.30841 | 70.52464 | 29.91721 | 0.815813 | 1.88751 | 0.177867 |
| 11 | 101865200 | 101869599 | 23.66114 | 38.53266 | 86.68513 | 37.22006 | -0.79581 | 1.538233 | 0.177871 |
| 3 | 130640000 | 130641199 | 8.988871 | 15.99642 | 41.01035 | 18.07229 | -0.95415 | 0.451257 | 0.177874 |
| 5 | 114635600 | 114640799 | 44.53867 | 70.67464 | 145.9723 | 60.80368 | -0.71345 | 2.304609 | 0.177886 |
| 4 | 45182600 | 45186799 | 19.30682 | 32.05003 | 75.9619 | 32.92378 | -0.86 | 1.272075 | 0.1779 |
| 4 | 151099000 | 151100799 | 8.126609 | 14.80878 | 38.18086 | 16.40829 | -0.93846 | 0.289705 | 0.177903 |
| 9 | 75633800 | 75635999 | 11.96278 | 21.01748 | 46.14398 | 20.42608 | -0.74121 | 0.601067 | 0.17793 |
| 1 | 179352200 | 179353799 | 23.43798 | 40.90206 | 30.02645 | 13.21985 | 0.831003 | 0.738292 | 0.177931 |
| 5 | 114237800 | 114239199 | 6.581047 | 12.0809 | 35.62911 | 15.16508 | -1.11932 | 0.235501 | 0.177943 |
| 7 | 20511200 | 20513599 | 58.02897 | 93.80205 | 57.09592 | 24.39185 | 1.075631 | 1.726005 | 0.177947 |
| 9 | 78355200 | 78358599 | 11.4682 | 19.88946 | 54.76197 | 24.02717 | -1.06772 | 0.803292 | 0.177947 |
| 18 | 7537600 | 7538399 | 13.71582 | 25.82614 | 19.58677 | 8.019406 | 0.866452 | 0.097357 | 0.17797 |
| 3 | 76700000 | 76702999 | 67.80263 | 108.0574 | 91.58461 | 38.34618 | 0.575875 | 2.177729 | 0.177989 |
| 4 | 151983000 | 151995999 | 86.71801 | 132.6432 | 244.5715 | 101.4724 | -0.59189 | 3.060346 | 0.178012 |
| 19 | 13721200 | 13721999 | 10.53334 | 20.38185 | 16.9881 | 6.478859 | 0.82401 | -0.16869 | 0.178023 |
| 9 | 101093400 | 101094199 | 5.855986 | 11.24333 | 29.72828 | 11.64821 | -0.86331 | -0.09302 | 0.178025 |
| 16 | 95615400 | 95625199 | 70.26961 | 108.3282 | 220.3981 | 91.54964 | -0.72497 | 2.898733 | 0.178034 |
| 8 | 32232000 | 32234199 | 28.15726 | 48.52554 | 34.91807 | 15.31758 | 0.857495 | 0.90712 | 0.178035 |
| 4 | 25429600 | 25430599 | 11.36471 | 21.77919 | 15.95658 | 6.285786 | 0.964917 | -0.0589 | 0.178047 |
| 2 | 156571800 | 156573199 | 6.658343 | 12.52238 | 31.81338 | 13.05164 | -0.8656 | 0.075466 | 0.178049 |
| 1 | 92926000 | 92937399 | 94.06305 | 143.5351 | 254.6641 | 105.6016 | -0.53917 | 3.111586 | 0.178064 |
| 18 | 59705400 | 59707399 | 27.1457 | 47.26465 | 30.41594 | 13.37217 | 1.020049 | 0.791012 | 0.178076 |
| 3 | 93439000 | 93440599 | 25.11862 | 43.40085 | 35.09055 | 15.39445 | 0.690778 | 0.870654 | 0.178077 |
| X | 32925000 | 32927199 | 27.39519 | 47.88813 | 27.75685 | 12.23449 | 1.170614 | 0.735541 | 0.178082 |
| 7 | 80799200 | 80809799 | 86.3136 | 131.8811 | 250.3563 | 103.8464 | -0.63513 | 3.082445 | 0.178084 |
| 17 | 86540600 | 86545199 | 33.16623 | 53.24013 | 116.1362 | 49.13755 | -0.77367 | 1.923253 | 0.178089 |
| 7 | 40923800 | 40924399 | 21.72272 | 38.3491 | 26.83415 | 11.87276 | 0.902198 | 0.569638 | 0.178091 |
| 7 | 44853600 | 44862399 | 51.98076 | 81.96881 | 167.1721 | 69.37524 | -0.70297 | 2.468349 | 0.178091 |
| 1 | 36140200 | 36146199 | 46.28551 | 73.5237 | 142.8242 | 59.55492 | -0.62366 | 2.273839 | 0.178109 |
| 16 | 86844800 | 86846599 | 9.854044 | 17.67141 | 41.57862 | 18.20433 | -0.82558 | 0.40407 | 0.178118 |
| 4 | 112575000 | 112576199 | 19.16331 | 35.38588 | 18.24731 | 7.704815 | 1.384486 | 0.25631 | 0.178129 |
| 13 | 61316600 | 61318199 | 28.3537 | 49.8134 | 22.40895 | 9.919311 | 1.533194 | 0.654448 | 0.178138 |
| X | 147784400 | 147785799 | 17.79578 | 32.9057 | 20.34789 | 8.575779 | 1.127274 | 0.244113 | 0.178143 |
| X | 24440600 | 24444599 | 55.75384 | 90.60301 | 50.21281 | 21.5412 | 1.217603 | 1.612868 | 0.178145 |
| 12 | 24321400 | 24326199 | 92.55083 | 145.3194 | 120.9131 | 50.15011 | 0.583981 | 2.570462 | 0.178151 |
| 6 | 35468000 | 35474199 | 51.33045 | 81.42689 | 152.3135 | 63.42554 | -0.57125 | 2.319022 | 0.178154 |
| 1 | 72592200 | 72593199 | 5.369712 | 10.4033 | 29.12762 | 11.04389 | -0.89902 | -0.20753 | 0.178173 |
| 3 | 88857000 | 88858599 | 10.62555 | 18.59156 | 47.28481 | 20.85857 | -0.95192 | 0.664096 | 0.178181 |
| 1 | 32304200 | 32307999 | 34.05589 | 54.89401 | 110.0033 | 46.82229 | -0.64419 | 1.793243 | 0.178207 |
| 3 | 3236800 | 3238999 | 43.92806 | 71.88243 | 53.87186 | 23.17157 | 0.79044 | 1.491058 | 0.178214 |
| 10 | 96452200 | 96454399 | 15.62173 | 26.18968 | 68.51282 | 29.88526 | -1.00123 | 1.14671 | 0.178214 |
| X | 3965000 | 3969599 | 49.03401 | 80.16817 | 50.13502 | 21.55746 | 1.049886 | 1.512054 | 0.178215 |
| 3 | 57435200 | 57437999 | 20.69743 | 34.41788 | 72.16483 | 31.31536 | -0.68355 | 1.254035 | 0.178222 |
| 6 | 52419400 | 52423199 | 20.69743 | 34.41788 | 72.16483 | 31.31536 | -0.68355 | 1.254035 | 0.178222 |
| 5 | 62263200 | 62265999 | 28.90134 | 49.00534 | 39.94141 | 17.49391 | 0.675268 | 1.058011 | 0.17824 |
| 17 | 47998400 | 48000599 | 11.61447 | 20.14491 | 53.68772 | 23.5559 | -1.02087 | 0.802268 | 0.178241 |

| | | | | | | | | | |
|---|---|---|---|---|---|---|---|---|---|
| 2 | 129271200 | 129272199 | 6.96292 | 13.05564 | 32.28949 | 13.315 | -0.83309 | 0.099986 | 0.178249 |
| 6 | 89619800 | 89621999 | 11.08098 | 19.38398 | 47.28928 | 20.8565 | -0.89226 | 0.670217 | 0.178259 |
| 15 | 74802400 | 74808999 | 37.8818 | 60.65702 | 119.3179 | 50.29096 | -0.62918 | 2.01024 | 0.17826 |
| 13 | 62811400 | 62818999 | 228.303 | 336.2767 | 271.3077 | 111.7268 | 0.551815 | 3.797548 | 0.178271 |
| 11 | 69436800 | 69438599 | 4.323818 | 8.005568 | 36.23776 | 15.2454 | -1.71826 | 0.166223 | 0.178296 |
| 8 | 67546000 | 67546999 | 14.58684 | 27.23292 | 20.73014 | 8.611036 | 0.845287 | 0.162645 | 0.178302 |
| 7 | 32244000 | 32246399 | 40.31192 | 66.43274 | 46.70489 | 20.14863 | 0.886362 | 1.390341 | 0.178335 |
| 11 | 79228600 | 79231599 | 13.73539 | 23.84803 | 52.15922 | 22.89089 | -0.73707 | 0.798918 | 0.178345 |
| 8 | 79235000 | 79238399 | 20.09716 | 33.34024 | 73.07037 | 31.65625 | -0.74746 | 1.281118 | 0.178356 |
| 2 | 106988800 | 106989999 | 22.2816 | 39.20414 | 27.78741 | 12.30277 | 0.881899 | 0.609476 | 0.178365 |
| 15 | 34702200 | 34703799 | 15.43689 | 29.28158 | 16.56657 | 6.68263 | 1.30323 | 0.057278 | 0.178384 |
| 3 | 109028800 | 109031199 | 10.70302 | 18.91574 | 42.99561 | 19.01923 | -0.78755 | 0.52074 | 0.178394 |
| 14 | 34936800 | 34938399 | 15.29003 | 28.4502 | 21.02914 | 8.784966 | 0.880616 | 0.189222 | 0.178398 |
| 5 | 115271800 | 115274399 | 12.76976 | 22.28456 | 49.97967 | 22.00462 | -0.77226 | 0.717768 | 0.178415 |
| 12 | 22184800 | 22189599 | 107.9545 | 167.2175 | 140.9974 | 58.5665 | 0.549698 | 2.817767 | 0.178431 |
| X | 63434600 | 63435399 | 14.41684 | 27.67555 | 15.7288 | 6.165699 | 1.333455 | -0.0437 | 0.178452 |
| X | 43203200 | 43204999 | 22.13935 | 39.75884 | 21.90747 | 9.580341 | 1.257273 | 0.440898 | 0.178459 |
| 5 | 123641000 | 123643399 | 10.52701 | 18.33324 | 51.52856 | 22.65743 | -1.09615 | 0.739041 | 0.178474 |
| 12 | 88993800 | 88994199 | 4.056634 | 7.919803 | 25.57587 | 9.030329 | -1.00654 | -0.48841 | 0.178499 |
| 9 | 53577600 | 53578999 | 4.565312 | 8.566324 | 34.94998 | 14.39427 | -1.54636 | 0.078672 | 0.178511 |
| 17 | 34235000 | 34236399 | 26.36126 | 45.36628 | 36.18018 | 15.87055 | 0.70943 | 0.91359 | 0.178515 |
| 3 | 117653000 | 117655799 | 36.44578 | 60.30535 | 49.35885 | 21.3455 | 0.668675 | 1.333901 | 0.178542 |
| 15 | 15277200 | 15278999 | 18.16475 | 33.23826 | 21.73851 | 9.293265 | 1.032285 | 0.306038 | 0.178542 |
| 6 | 142477600 | 142478199 | 5.539126 | 10.68044 | 28.11113 | 10.88045 | -0.83909 | -0.13828 | 0.178548 |
| 8 | 58616800 | 58617799 | 5.539126 | 10.68044 | 28.11113 | 10.88045 | -0.83909 | -0.13828 | 0.178548 |
| 17 | 48396000 | 48396799 | 5.539126 | 10.68044 | 28.11113 | 10.88045 | -0.83909 | -0.13828 | 0.178548 |
| 17 | 34196400 | 34198199 | 12.21512 | 20.87586 | 60.48646 | 26.52067 | -1.14534 | 0.949605 | 0.178556 |
| 11 | 55226200 | 55227599 | 8.334418 | 14.99152 | 38.50518 | 16.81576 | -0.94749 | 0.372808 | 0.17856 |
| 13 | 57575600 | 57578399 | 17.90303 | 29.73247 | 73.98911 | 32.07825 | -0.92985 | 1.262685 | 0.178567 |
| 18 | 7076000 | 7076999 | 11.25246 | 21.58929 | 17.92738 | 7.038483 | 0.790173 | -0.06497 | 0.178584 |
| 9 | 22378600 | 22379799 | 4.548743 | 8.575129 | 34.53942 | 14.11329 | -1.51884 | 0.04653 | 0.178589 |
| X | 125214200 | 125214999 | 19.58887 | 34.68361 | 25.65146 | 11.34283 | 0.824353 | 0.523317 | 0.178592 |
| X | 122415400 | 122417399 | 25.97564 | 45.40513 | 25.61816 | 11.29148 | 1.209282 | 0.725196 | 0.178593 |
| X | 26383200 | 26384999 | 17.62783 | 33.87923 | 11.97596 | 4.678835 | 2.014332 | -0.03055 | 0.178605 |
| 4 | 124047200 | 124048799 | 9.220291 | 16.66636 | 38.01845 | 16.50467 | -0.77243 | 0.347024 | 0.178622 |
| 15 | 76015000 | 76017999 | 12.33294 | 21.473 | 50.5136 | 22.20768 | -0.84102 | 0.749864 | 0.178636 |
| X | 25434800 | 25436599 | 18.18623 | 34.36872 | 15.96835 | 6.492153 | 1.576014 | 0.107188 | 0.178647 |
| 3 | 10448400 | 10452599 | 21.01665 | 34.91494 | 82.19848 | 35.64558 | -0.85049 | 1.271733 | 0.178658 |
| 6 | 109023000 | 109024399 | 11.48544 | 20.30708 | 46.15833 | 20.41655 | -0.78752 | 0.526816 | 0.178666 |
| 5 | 131535600 | 131536999 | 24.01968 | 39.00821 | 89.74612 | 38.48268 | -0.83125 | 1.58654 | 0.178666 |
| 8 | 14822000 | 14825399 | 17.64704 | 30.01501 | 62.97425 | 27.60081 | -0.68334 | 1.016841 | 0.178669 |
| 5 | 98313800 | 98318399 | 31.06452 | 49.93566 | 113.2584 | 48.01872 | -0.82728 | 1.875602 | 0.178677 |
| 2 | 58453200 | 58457799 | 26.3462 | 42.58871 | 102.5688 | 43.81943 | -0.90378 | 1.693306 | 0.178694 |
| 12 | 18760400 | 18797399 | 805.2148 | 1067.34 | 748.8882 | 313.0467 | 0.601706 | 5.421634 | 0.178725 |
| 9 | 118740600 | 118745799 | 32.16455 | 51.73411 | 109.5122 | 46.46966 | -0.72688 | 1.857625 | 0.17875 |
| 7 | 29345600 | 29349599 | 83.02356 | 131.1949 | 104.8186 | 43.53901 | 0.649554 | 2.431357 | 0.178767 |
| 1 | 63964800 | 63966999 | 19.98674 | 35.41285 | 25.65028 | 11.34072 | 0.854191 | 0.515721 | 0.178768 |

| | | | | | | | | | |
|---|---|---|---|---|---|---|---|---|---|
| 15 | 14705400 | 14705799 | 9.196078 | 17.88093 | 14.38428 | 5.331667 | 0.911985 | -0.29048 | 0.178781 |
| 10 | 53505400 | 53506799 | 10.28367 | 18.08979 | 45.6235 | 20.20056 | -0.93947 | 0.574141 | 0.178783 |
| 13 | 19495400 | 19499999 | 110.3053 | 170.6519 | 144.8613 | 60.17184 | 0.538705 | 2.84103 | 0.178798 |
| 4 | 57799800 | 57802399 | 17.66568 | 29.95325 | 62.91311 | 27.55504 | -0.68527 | 1.044354 | 0.178807 |
| 11 | 119864600 | 119865999 | 14.07487 | 24.45387 | 53.22765 | 23.36964 | -0.73 | 0.794037 | 0.178815 |
| 9 | 90021400 | 90024799 | 25.43165 | 41.35843 | 88.07955 | 37.79963 | -0.71848 | 1.562153 | 0.178819 |
| 2 | 39425600 | 39426199 | 6.919957 | 13.12093 | 32.11989 | 12.96749 | -0.79277 | 0.022884 | 0.178822 |
| 9 | 99318000 | 99321399 | 18.23217 | 30.23883 | 75.10242 | 32.53103 | -0.92725 | 1.279944 | 0.178823 |
| 7 | 14396800 | 14397599 | 24.74536 | 42.81483 | 34.54462 | 15.15558 | 0.69412 | 0.855705 | 0.178823 |
| 14 | 51181400 | 51182399 | 5.919307 | 11.11839 | 31.73738 | 13.0484 | -1.03372 | 0.079561 | 0.178828 |
| 11 | 50682600 | 50684999 | 13.48943 | 23.25044 | 54.76246 | 24.02285 | -0.84563 | 0.869834 | 0.178831 |
| 14 | 3258400 | 3263199 | 67.59155 | 107.8584 | 88.77177 | 37.23425 | 0.62009 | 2.136151 | 0.178839 |
| 8 | 112546600 | 112548399 | 8.321307 | 15.0354 | 38.02315 | 16.51492 | -0.92055 | 0.342094 | 0.178842 |
| 7 | 142120200 | 142137399 | 132.5128 | 196.263 | 377.9688 | 155.6599 | -0.6952 | 3.698477 | 0.178851 |
| 18 | 47011800 | 47013199 | 7.046225 | 12.99022 | 35.72257 | 15.11672 | -1.01446 | 0.209681 | 0.178859 |
| 8 | 55913000 | 55918199 | 92.64659 | 144.9726 | 124.1737 | 51.5124 | 0.538392 | 2.632206 | 0.17887 |
| X | 54460200 | 54460799 | 10.53435 | 20.59138 | 10.00005 | 3.44432 | 1.721707 | -0.53453 | 0.178877 |
| 2 | 166357800 | 166366999 | 58.12155 | 91.26404 | 174.95 | 72.56296 | -0.6185 | 2.558652 | 0.178894 |
| 2 | 80882800 | 80883799 | 24.92107 | 43.32459 | 29.40049 | 12.91325 | 0.943721 | 0.799857 | 0.178906 |
| 7 | 34573600 | 34578399 | 33.47773 | 53.76913 | 111.2868 | 47.12436 | -0.69705 | 1.906409 | 0.178908 |
| 7 | 19807800 | 19813199 | 100.484 | 156.5605 | 131.3632 | 54.52153 | 0.563308 | 2.713128 | 0.178913 |
| 1 | 181700800 | 181701599 | 4.789685 | 9.305002 | 25.68719 | 9.588334 | -0.85655 | -0.27806 | 0.178923 |
| 16 | 46214600 | 46215399 | 4.789685 | 9.305002 | 25.68719 | 9.588334 | -0.85655 | -0.27806 | 0.178923 |
| 9 | 65296200 | 65298999 | 17.2996 | 28.66144 | 74.20362 | 32.11798 | -0.9869 | 1.291116 | 0.178927 |
| 5 | 94786000 | 94786399 | 7.216721 | 14.10915 | 10.42367 | 3.572555 | 1.134529 | -0.6028 | 0.178935 |
| 11 | 33046800 | 33052399 | 39.56285 | 63.11642 | 130.4958 | 54.71599 | -0.70682 | 2.128212 | 0.178941 |
| 11 | 95019000 | 95022599 | 17.41258 | 28.94558 | 71.62325 | 31.07301 | -0.92141 | 1.24975 | 0.178949 |
| 1 | 93853400 | 93855799 | 10.90036 | 19.13428 | 48.28539 | 21.35833 | -0.94106 | 0.611319 | 0.178954 |
| 5 | 135603200 | 135611199 | 58.1184 | 90.85439 | 184.38 | 76.49615 | -0.70562 | 2.640838 | 0.178965 |
| 12 | 17917800 | 17919999 | 44.78672 | 72.69844 | 60.40949 | 25.89392 | 0.633342 | 1.616081 | 0.178999 |
| 12 | 23737400 | 23738999 | 13.67644 | 25.52662 | 19.82891 | 8.240298 | 0.815681 | 0.1536 | 0.179021 |
| 8 | 108122800 | 108125399 | 15.58189 | 26.63225 | 57.95007 | 25.40877 | -0.73422 | 0.960646 | 0.179031 |
| 1 | 39771200 | 39775799 | 41.67959 | 66.41533 | 134.7665 | 56.41494 | -0.6817 | 2.169033 | 0.179033 |
| 4 | 137648000 | 137652799 | 18.21003 | 30.24062 | 72.8586 | 31.58689 | -0.88333 | 1.263874 | 0.179037 |
| 4 | 146117600 | 146122399 | 65.88193 | 105.5036 | 72.13486 | 30.40929 | 0.892674 | 2.02382 | 0.179041 |
| X | 122863800 | 122864199 | 9.731859 | 19.03649 | 8.471658 | 2.796603 | 1.899127 | -0.69124 | 0.179043 |
| 3 | 42636200 | 42638199 | 20.37282 | 36.37653 | 24.69621 | 10.85239 | 0.953421 | 0.468506 | 0.179055 |
| 1 | 171762800 | 171765999 | 25.16771 | 41.20715 | 85.68014 | 36.86127 | -0.67726 | 1.46095 | 0.179082 |
| 2 | 36821200 | 36821999 | 15.13145 | 28.43672 | 18.79919 | 7.72216 | 1.059067 | 0.119941 | 0.179089 |
| 5 | 131803200 | 131806599 | 16.49027 | 28.25308 | 59.17825 | 25.95233 | -0.67914 | 0.941689 | 0.179096 |
| 14 | 12270400 | 12271399 | 17.56011 | 31.54605 | 24.12554 | 10.54727 | 0.787685 | 0.411198 | 0.179122 |
| 15 | 81953200 | 81958199 | 43.82986 | 69.73652 | 137.4846 | 57.43281 | -0.64254 | 2.221649 | 0.179125 |
| 12 | 68487400 | 68488199 | 7.452234 | 13.4002 | 38.52024 | 16.82708 | -1.1089 | 0.367892 | 0.179136 |
| 3 | 63494400 | 63495799 | 8.103929 | 14.84349 | 35.60309 | 15.19984 | -0.82788 | 0.257146 | 0.179138 |
| 14 | 102578200 | 102580599 | 34.66527 | 57.82433 | 41.18282 | 17.90699 | 0.869057 | 1.231733 | 0.179141 |
| 10 | 78535600 | 78537399 | 10.49394 | 18.72759 | 42.55772 | 18.70961 | -0.7799 | 0.441438 | 0.179144 |
| 5 | 32271200 | 32272999 | 9.020001 | 16.4483 | 37.66157 | 16.17095 | -0.76759 | 0.292258 | 0.179148 |

| | | | | | | | | | |
|---|---|---|---|---|---|---|---|---|---|
| 7 | 123182600 | 123188399 | 37.70295 | 60.3591 | 126.0045 | 53.09394 | -0.71518 | 2.016146 | 0.179157 |
| 7 | 6058600 | 6063599 | 99.91569 | 155.8034 | 132.2289 | 54.87409 | 0.547772 | 2.698441 | 0.179163 |
| 19 | 41704200 | 41706999 | 13.30821 | 23.00568 | 54.26107 | 23.80504 | -0.84675 | 0.840886 | 0.179164 |
| 5 | 138237200 | 138238199 | 6.504562 | 12.26429 | 31.88293 | 13.024 | -0.89363 | 0.056987 | 0.17917 |
| 19 | 58448200 | 58449599 | 6.504562 | 12.26429 | 31.88293 | 13.024 | -0.89363 | 0.056987 | 0.17917 |
| 14 | 26017400 | 26018999 | 8.854256 | 16.22204 | 37.23255 | 15.88987 | -0.76492 | 0.263542 | 0.17918 |
| 8 | 110594600 | 110596199 | 8.662499 | 15.5195 | 38.9815 | 17.08124 | -0.91925 | 0.400085 | 0.179181 |
| 11 | 115847800 | 115850199 | 7.29053 | 13.11087 | 38.53016 | 16.82986 | -1.14018 | 0.36702 | 0.179185 |
| 4 | 151419400 | 151420199 | 4.413767 | 8.551991 | 26.5985 | 10.08088 | -1.04781 | -0.22831 | 0.179195 |
| 14 | 119880000 | 119881799 | 12.12657 | 21.33101 | 46.6662 | 20.66235 | -0.73566 | 0.587899 | 0.17921 |
| 10 | 59163800 | 59166399 | 11.1521 | 19.76847 | 44.56488 | 19.70162 | -0.77418 | 0.493124 | 0.179212 |
| 4 | 136440600 | 136448999 | 48.89015 | 77.16001 | 161.2432 | 66.93388 | -0.73702 | 2.448047 | 0.17922 |
| 7 | 45261800 | 45269199 | 39.22873 | 62.61607 | 128.9164 | 54.09355 | -0.69991 | 2.111156 | 0.179224 |
| X | 147445400 | 147447199 | 24.07779 | 41.85001 | 33.03178 | 14.50632 | 0.726853 | 0.804729 | 0.179237 |
| 9 | 40302400 | 40305199 | 13.7565 | 23.4275 | 59.95605 | 26.28066 | -0.96801 | 0.988484 | 0.179251 |
| 9 | 15502600 | 15506399 | 22.75942 | 37.3566 | 80.05927 | 34.47922 | -0.71913 | 1.422461 | 0.179258 |
| 4 | 153068200 | 153074399 | 50.29622 | 79.48811 | 159.2408 | 66.14857 | -0.67464 | 2.414822 | 0.179264 |
| 3 | 142155400 | 142156199 | 4.652085 | 9.057387 | 25.42292 | 9.324249 | -0.85707 | -0.3447 | 0.179267 |
| 15 | 63440400 | 63441199 | 10.94951 | 21.18654 | 15.64745 | 5.968343 | 0.995424 | -0.16965 | 0.17928 |
| 4 | 93586200 | 93587799 | 12.1005 | 23.19518 | 17.69986 | 6.96741 | 0.907025 | -0.04948 | 0.179294 |
| 10 | 38908000 | 38909199 | 14.62784 | 27.24512 | 21.11874 | 8.80757 | 0.814468 | 0.177593 | 0.179309 |
| 16 | 30475200 | 30478999 | 19.24149 | 32.10637 | 69.87378 | 30.38832 | -0.73695 | 1.210095 | 0.179316 |
| X | 62209200 | 62213199 | 77.88244 | 123.9364 | 90.18631 | 37.68274 | 0.793245 | 2.235719 | 0.179325 |
| 8 | 122393800 | 122397199 | 14.18318 | 24.23146 | 57.8752 | 25.37509 | -0.86804 | 0.958663 | 0.17933 |
| 13 | 68742600 | 68744399 | 17.78324 | 32.07032 | 23.14999 | 10.07902 | 0.874463 | 0.388737 | 0.179333 |
| 3 | 51936800 | 51938399 | 7.868289 | 14.67939 | 36.12431 | 15.02345 | -0.83691 | 0.146387 | 0.179353 |
| 10 | 24471200 | 24473999 | 13.4054 | 22.93688 | 57.87087 | 25.37624 | -0.94641 | 0.941114 | 0.179359 |
| 8 | 115545600 | 115546999 | 7.260027 | 13.62862 | 33.44392 | 13.76382 | -0.81975 | 0.098074 | 0.179367 |
| 4 | 20504200 | 20504999 | 5.58921 | 10.60123 | 30.31437 | 12.22996 | -1.01245 | 0.006568 | 0.179372 |
| X | 136012400 | 136014199 | 5.837541 | 11.16599 | 35.39905 | 14.02021 | -1.13691 | -0.03949 | 0.179398 |
| 16 | 48703800 | 48705199 | 22.68161 | 40.0985 | 25.55636 | 11.30422 | 1.036801 | 0.557877 | 0.17941 |
| 7 | 27527400 | 27529599 | 20.73937 | 36.63585 | 24.01661 | 10.62483 | 0.995955 | 0.550925 | 0.179412 |
| 8 | 12170800 | 12171999 | 18.38585 | 33.25835 | 23.1835 | 10.05519 | 0.927545 | 0.373973 | 0.179412 |
| 4 | 106318400 | 106322599 | 26.00317 | 42.06936 | 96.54553 | 41.28248 | -0.83295 | 1.669808 | 0.179413 |
| 17 | 19366600 | 19368199 | 24.52274 | 43.84652 | 19.85426 | 8.714583 | 1.534471 | 0.47857 | 0.179434 |
| X | 5469600 | 5473199 | 89.18229 | 141.2652 | 91.56332 | 38.07926 | 0.95354 | 2.376817 | 0.179443 |
| 2 | 48838800 | 48840399 | 11.93764 | 20.65795 | 54.75172 | 24.02106 | -1.01374 | 0.825909 | 0.179451 |
| 6 | 149269800 | 149270199 | 4.056498 | 7.919993 | 24.2397 | 8.550056 | -0.92832 | -0.49555 | 0.179453 |
| X | 25674800 | 25678799 | 60.00531 | 96.75204 | 63.1972 | 26.91054 | 0.970079 | 1.804911 | 0.179496 |
| 4 | 112555800 | 112557199 | 28.2174 | 48.624 | 34.96139 | 15.33653 | 0.858642 | 0.908932 | 0.179509 |
| 10 | 67410200 | 67414399 | 27.02045 | 43.94139 | 90.34741 | 38.77262 | -0.66803 | 1.56556 | 0.17951 |
| 10 | 125801000 | 125803199 | 12.66107 | 21.98322 | 52.12602 | 22.87662 | -0.85294 | 0.79362 | 0.179529 |
| 12 | 72130800 | 72133199 | 13.02841 | 22.10246 | 61.96416 | 27.14388 | -1.09948 | 1.023848 | 0.179531 |
| 5 | 93666400 | 93667199 | 14.09091 | 26.82804 | 17.97832 | 7.195283 | 1.071539 | 0.023852 | 0.179536 |
| 4 | 152078200 | 152081399 | 13.53605 | 22.90411 | 65.82919 | 28.81429 | -1.13573 | 1.050653 | 0.179543 |
| 15 | 50273800 | 50275799 | 23.37013 | 41.21086 | 26.10798 | 11.55408 | 1.044164 | 0.590135 | 0.179544 |
| 10 | 31659000 | 31660399 | 14.73153 | 25.44794 | 54.81727 | 24.04853 | -0.71672 | 0.856438 | 0.17955 |

| | | | | | | | | | |
|---|---|---|---|---|---|---|---|---|---|
| 10 | 46680400 | 46680999 | 9.585093 | 18.66921 | 14.85518 | 5.423949 | 0.947161 | -0.33489 | 0.179559 |
| X | 16192400 | 16194399 | 27.52548 | 48.41216 | 23.21826 | 10.28058 | 1.442032 | 0.637998 | 0.179561 |
| 3 | 39727000 | 39729599 | 13.07468 | 22.84245 | 49.43838 | 21.78175 | -0.72095 | 0.702782 | 0.179567 |
| 8 | 101616800 | 101619599 | 30.32051 | 51.66562 | 37.58303 | 16.4746 | 0.83931 | 1.017316 | 0.179568 |
| 3 | 97557200 | 97559799 | 10.18437 | 18.17343 | 44.16746 | 19.41883 | -0.87644 | 0.441868 | 0.179576 |
| 12 | 18918600 | 18926199 | 111.6557 | 176.1304 | 85.47714 | 35.47399 | 1.365642 | 2.48904 | 0.179593 |
| 8 | 59871200 | 59872999 | 14.70259 | 25.45301 | 53.77453 | 23.59263 | -0.68824 | 0.834415 | 0.179596 |
| 6 | 46257000 | 46257599 | 5.083105 | 9.87829 | 28.3377 | 10.54958 | -0.90906 | -0.2787 | 0.179609 |
| 16 | 4265600 | 4267599 | 9.378321 | 16.91211 | 38.49813 | 16.76223 | -0.77235 | 0.362171 | 0.179632 |
| 3 | 15623600 | 15624999 | 17.77609 | 32.25019 | 23.31514 | 10.0751 | 0.878768 | 0.353671 | 0.179652 |
| 4 | 48042600 | 48049999 | 52.16493 | 81.83647 | 180.6867 | 74.93661 | -0.82306 | 2.573834 | 0.17966 |
| 8 | 89886000 | 89890199 | 26.06114 | 42.39424 | 91.45754 | 39.25338 | -0.73686 | 1.559293 | 0.179667 |
| X | 89568400 | 89569999 | 22.06695 | 39.5536 | 22.36439 | 9.795833 | 1.218659 | 0.452269 | 0.179684 |
| 6 | 30171400 | 30180199 | 70.49118 | 108.6024 | 225.5315 | 93.68091 | -0.75521 | 2.910477 | 0.179695 |
| 8 | 4623400 | 4625399 | 21.62295 | 38.02556 | 28.41418 | 12.58158 | 0.80584 | 0.612262 | 0.179711 |
| 14 | 57108000 | 57109599 | 27.16976 | 46.47763 | 37.81338 | 16.58078 | 0.679376 | 0.972003 | 0.179711 |
| 7 | 11383400 | 11385999 | 19.10231 | 35.08578 | 19.95683 | 8.487966 | 1.237197 | 0.288677 | 0.179715 |
| X | 54235800 | 54237399 | 19.27868 | 36.20941 | 15.72048 | 6.463935 | 1.659419 | 0.154225 | 0.179721 |
| 7 | 29610600 | 29612599 | 11.84097 | 20.72965 | 46.75167 | 20.63302 | -0.77962 | 0.660868 | 0.179723 |
| 14 | 112803000 | 112804399 | 21.63048 | 37.98954 | 28.35138 | 12.54634 | 0.807283 | 0.628733 | 0.179729 |
| 16 | 20958400 | 20959799 | 17.80266 | 32.51623 | 22.28424 | 9.549011 | 0.963032 | 0.314339 | 0.17974 |
| 18 | 36328800 | 36330799 | 11.62329 | 20.12402 | 53.12142 | 23.30616 | -1.00752 | 0.819411 | 0.179752 |
| 18 | 31892000 | 31894399 | 12.96588 | 22.16668 | 60.01246 | 26.31359 | -1.04793 | 0.948378 | 0.179759 |
| 6 | 29119000 | 29120399 | 6.821237 | 12.76076 | 32.7669 | 13.5643 | -0.89101 | 0.115907 | 0.179764 |
| 6 | 124185000 | 124186799 | 28.36683 | 48.81888 | 33.70604 | 14.78527 | 0.916286 | 0.916195 | 0.179782 |
| 2 | 156919200 | 156923999 | 23.80264 | 38.47273 | 98.50606 | 42.07765 | -0.99183 | 1.692692 | 0.179792 |
| 8 | 71878200 | 71878999 | 10.7529 | 20.95802 | 13.57595 | 4.912971 | 1.251709 | -0.35389 | 0.1798 |
| 11 | 43700600 | 43703599 | 29.52865 | 50.23912 | 39.35853 | 17.25076 | 0.732579 | 1.031824 | 0.179804 |
| 3 | 22645000 | 22646199 | 15.55244 | 29.41748 | 17.73307 | 7.196072 | 1.20565 | 0.080487 | 0.179806 |
| 11 | 22808200 | 22810599 | 16.96811 | 28.18004 | 73.23101 | 31.74976 | -0.992 | 1.260301 | 0.179806 |
| 19 | 40208000 | 40209799 | 9.830764 | 17.73408 | 40.6328 | 17.68411 | -0.78159 | 0.366677 | 0.179809 |
| 4 | 27639400 | 27640399 | 12.16007 | 23.10908 | 18.82255 | 7.56269 | 0.787779 | 0.019814 | 0.179814 |
| 10 | 8700600 | 8702399 | 6.384338 | 11.98184 | 32.86106 | 13.52954 | -0.97946 | 0.088058 | 0.179831 |
| 10 | 85763800 | 85764799 | 6.384338 | 11.98184 | 32.86106 | 13.52954 | -0.97946 | 0.088058 | 0.179831 |
| 8 | 87478600 | 87480599 | 26.56191 | 45.62422 | 36.2566 | 15.90314 | 0.714047 | 0.931881 | 0.179833 |
| 16 | 29967800 | 29970399 | 15.83077 | 27.25807 | 62.05306 | 27.22019 | -0.79755 | 0.896509 | 0.179859 |
| X | 126173400 | 126175399 | 24.95622 | 44.24406 | 22.10824 | 9.773529 | 1.386991 | 0.537013 | 0.179859 |
| X | 43860600 | 43861399 | 16.76982 | 31.35753 | 19.37841 | 8.028212 | 1.146671 | 0.168396 | 0.179873 |
| 8 | 69610200 | 69611799 | 7.415987 | 13.86108 | 33.33694 | 13.82103 | -0.79994 | 0.125244 | 0.17988 |
| 12 | 54380400 | 54381599 | 7.415987 | 13.86108 | 33.33694 | 13.82103 | -0.79994 | 0.125244 | 0.17988 |
| 8 | 4393600 | 4396799 | 13.41797 | 23.33177 | 51.07405 | 22.43571 | -0.738 | 0.774736 | 0.17989 |
| 12 | 14896800 | 14898599 | 24.15223 | 42.19918 | 30.69133 | 13.52279 | 0.844869 | 0.72984 | 0.179891 |
| 7 | 83540800 | 83541999 | 8.32325 | 14.71957 | 44.59709 | 19.72506 | -1.19883 | 0.501319 | 0.179894 |
| 13 | 24427000 | 24429599 | 10.31829 | 18.07798 | 47.26072 | 20.87139 | -0.99143 | 0.63949 | 0.179899 |
| 10 | 11398800 | 11400999 | 29.08461 | 50.03735 | 34.96093 | 15.33557 | 0.899401 | 0.927136 | 0.179923 |
| X | 3162200 | 3164599 | 53.77746 | 86.98475 | 60.66195 | 25.93455 | 0.880942 | 1.705789 | 0.179923 |
| 6 | 3223800 | 3225799 | 28.12091 | 48.32593 | 35.63612 | 15.63121 | 0.821667 | 0.933975 | 0.179934 |

| | | | | | | | | | |
|---|---|---|---|---|---|---|---|---|---|
| 5 | 93002000 | 93006399 | 22.85582 | 37.70837 | 77.09748 | 33.27604 | -0.64845 | 1.356378 | 0.179946 |
| 6 | 66314200 | 66314799 | 9.131942 | 17.78856 | 14.93575 | 5.445477 | 0.872528 | -0.3454 | 0.179953 |
| 7 | 63581600 | 63590399 | 46.62382 | 73.59957 | 162.7649 | 67.57318 | -0.81834 | 2.440935 | 0.179989 |
| 9 | 64928200 | 64929599 | 7.918177 | 14.6133 | 34.67404 | 14.65175 | -0.80167 | 0.209609 | 0.18001 |
| 18 | 34065600 | 34068399 | 12.12136 | 21.29395 | 46.1458 | 20.42516 | -0.7225 | 0.603885 | 0.180018 |
| 1 | 180760600 | 180765599 | 29.48542 | 47.38252 | 113.7228 | 48.19382 | -0.90923 | 1.884011 | 0.180027 |
| 7 | 10059000 | 10061399 | 71.36252 | 113.3769 | 97.43247 | 40.63419 | 0.551304 | 2.278036 | 0.180027 |
| 1 | 65965600 | 65968199 | 22.61607 | 37.08476 | 82.97867 | 35.72164 | -0.7818 | 1.434973 | 0.180027 |
| 9 | 8898400 | 8902399 | 30.36822 | 48.94149 | 105.8541 | 45.04617 | -0.75425 | 1.790796 | 0.180068 |
| 17 | 36403600 | 36406399 | 15.21288 | 26.13032 | 56.41936 | 24.74707 | -0.72202 | 0.910223 | 0.180077 |
| 5 | 70064400 | 70066799 | 38.78868 | 64.6541 | 43.20883 | 18.77906 | 0.960614 | 1.255112 | 0.180091 |
| 19 | 40540600 | 40546799 | 40.86153 | 65.15552 | 131.7919 | 55.22101 | -0.67607 | 2.146049 | 0.180106 |
| 7 | 106379800 | 106381399 | 19.12572 | 34.42227 | 23.003 | 10.04017 | 0.982131 | 0.406611 | 0.180107 |
| 15 | 84936400 | 84937999 | 7.300274 | 13.54909 | 34.22573 | 14.35181 | -0.88261 | 0.173659 | 0.180108 |
| 6 | 83137800 | 83139999 | 13.05451 | 22.44557 | 55.78058 | 24.46764 | -0.92337 | 0.892103 | 0.180134 |
| 5 | 125328400 | 125340599 | 78.87904 | 120.4619 | 249.9868 | 103.6813 | -0.76407 | 3.0912 | 0.180139 |
| 5 | 25701600 | 25705399 | 22.06588 | 36.01117 | 86.86037 | 37.3242 | -0.89389 | 1.502453 | 0.180139 |
| 12 | 99358000 | 99360799 | 12.7255 | 21.94386 | 55.82075 | 24.48744 | -0.95603 | 0.862918 | 0.180141 |
| 15 | 85933400 | 85936399 | 11.83922 | 20.71451 | 47.28582 | 20.85849 | -0.79708 | 0.670282 | 0.180152 |
| 5 | 43991000 | 43997199 | 58.60021 | 92.04623 | 173.4092 | 71.927 | -0.59309 | 2.552177 | 0.180161 |
| 8 | 5195600 | 5196799 | 15.67655 | 30.17214 | 14.64175 | 5.696878 | 1.5695 | -0.05584 | 0.180161 |
| 2 | 41786200 | 41786399 | 7.815975 | 15.28956 | 9.708572 | 3.211668 | 1.390468 | -0.71488 | 0.180179 |
| 7 | 38536600 | 38543999 | 182.7317 | 274.5535 | 212.9134 | 87.8898 | 0.6351 | 3.438554 | 0.180193 |
| 7 | 23855800 | 23860199 | 58.54224 | 94.04547 | 75.23005 | 31.86611 | 0.671943 | 1.918059 | 0.180201 |
| 9 | 114333200 | 114347599 | 250.0515 | 352.4419 | 618.0211 | 255.9429 | -0.63026 | 4.460249 | 0.180203 |
| 7 | 89168400 | 89170199 | 18.89391 | 33.56505 | 25.83668 | 11.40225 | 0.769004 | 0.49049 | 0.180233 |
| 9 | 93719000 | 93720999 | 22.23403 | 40.00786 | 21.2474 | 9.276216 | 1.311449 | 0.427977 | 0.180253 |
| 12 | 22239200 | 22240599 | 28.27643 | 48.51393 | 32.12975 | 14.09192 | 0.975557 | 0.943605 | 0.180271 |
| 19 | 13496400 | 13497599 | 14.03463 | 26.95335 | 15.74162 | 6.163161 | 1.295965 | -0.05168 | 0.180286 |
| 4 | 11384400 | 11389799 | 36.30886 | 58.03423 | 124.3385 | 52.27319 | -0.75503 | 2.065521 | 0.180301 |
| 14 | 18189400 | 18191799 | 24.94423 | 43.80131 | 25.83808 | 11.43233 | 1.144632 | 0.646753 | 0.180305 |
| 2 | 131018600 | 131022599 | 19.88143 | 32.74165 | 78.17039 | 33.71322 | -0.87234 | 1.372446 | 0.180309 |
| 17 | 18390600 | 18395599 | 74.46488 | 118.7495 | 86.04143 | 36.05654 | 0.80264 | 2.1636 | 0.180312 |
| 15 | 78140800 | 78142999 | 7.54456 | 13.71018 | 36.55157 | 15.76251 | -0.98877 | 0.301489 | 0.180317 |
| 17 | 48278800 | 48290399 | 68.12192 | 105.4737 | 209.5911 | 87.05865 | -0.68618 | 2.816203 | 0.180326 |
| 8 | 93174000 | 93177599 | 20.02186 | 33.01861 | 77.80562 | 33.57557 | -0.85274 | 1.354473 | 0.180328 |
| 13 | 54949600 | 54956399 | 53.49004 | 84.22158 | 168.1675 | 69.77466 | -0.67415 | 2.500101 | 0.18034 |
| X | 27549000 | 27549399 | 10.18306 | 19.83076 | 14.91053 | 5.455387 | 1.026286 | -0.31828 | 0.180344 |
| 5 | 139068400 | 139075999 | 41.3986 | 65.86928 | 139.5335 | 58.29005 | -0.74598 | 2.218477 | 0.180372 |
| 12 | 22556800 | 22557599 | 28.39893 | 48.44535 | 38.35794 | 16.8158 | 0.717898 | 1.00312 | 0.180394 |
| Y | 2209600 | 2210799 | 9.168711 | 17.88608 | 7.606862 | 2.721289 | 1.862248 | -0.44167 | 0.180405 |
| 14 | 89174800 | 89175999 | 15.36081 | 29.06313 | 17.74007 | 7.194661 | 1.188422 | 0.076738 | 0.180414 |
| 10 | 7161000 | 7162799 | 12.82235 | 22.24391 | 52.12211 | 22.8691 | -0.83609 | 0.804751 | 0.180416 |
| 4 | 80802400 | 80802799 | 3.040906 | 5.94472 | 20.00959 | 6.871206 | -1.02402 | -0.61666 | 0.180456 |
| 17 | 41641000 | 41641599 | 9.533576 | 18.5323 | 15.406 | 5.722485 | 0.862374 | -0.27522 | 0.180497 |
| 4 | 86306600 | 86307399 | 7.081659 | 13.36549 | 32.51738 | 13.2572 | -0.79664 | 0.056998 | 0.180503 |
| 14 | 108605400 | 108605999 | 7.081659 | 13.36549 | 32.51738 | 13.2572 | -0.79664 | 0.056998 | 0.180503 |

| | | | | | | | | | |
|---|---|---|---|---|---|---|---|---|---|
| 6 | 125630800 | 125635999 | 24.57481 | 40.13084 | 84.46676 | 36.30395 | -0.6973 | 1.497701 | 0.180529 |
| 3 | 37234000 | 37235599 | 9.736164 | 17.38102 | 39.95309 | 17.55963 | -0.79563 | 0.430853 | 0.18053 |
| 9 | 14347400 | 14349199 | 22.89732 | 40.21041 | 28.28015 | 12.51375 | 0.892544 | 0.638479 | 0.180544 |
| 16 | 23054400 | 23055999 | 13.17457 | 24.71582 | 19.8192 | 8.165331 | 0.779951 | 0.116946 | 0.18055 |
| 14 | 98778200 | 98779199 | 14.38655 | 26.92605 | 20.70293 | 8.563484 | 0.835651 | 0.144978 | 0.180557 |
| 3 | 34072000 | 34075199 | 19.3976 | 32.38494 | 68.7392 | 29.90543 | -0.70095 | 1.20444 | 0.180567 |
| 7 | 60712400 | 60713999 | 22.57083 | 40.23159 | 21.48927 | 9.456917 | 1.295151 | 0.485613 | 0.18057 |
| 17 | 39641000 | 39643799 | 40.45629 | 67.72166 | 38.72 | 16.87347 | 1.184744 | 1.214121 | 0.180573 |
| 3 | 131798600 | 131799599 | 15.10351 | 28.11128 | 21.03485 | 8.783041 | 0.863578 | 0.185688 | 0.180574 |
| 10 | 117429600 | 117431799 | 19.17128 | 31.74406 | 73.79927 | 31.93025 | -0.83233 | 1.303701 | 0.180575 |
| 3 | 11583600 | 11585599 | 25.55102 | 44.50964 | 29.84742 | 13.12622 | 0.960717 | 0.773492 | 0.180581 |
| 10 | 78355200 | 78356399 | 9.576266 | 17.13083 | 39.46038 | 17.31298 | -0.79673 | 0.416446 | 0.18059 |
| 17 | 5891400 | 5892399 | 7.17609 | 13.26239 | 34.69414 | 14.6322 | -0.9393 | 0.194813 | 0.1806 |
| 14 | 48934200 | 48934999 | 4.65599 | 9.052494 | 25.19479 | 9.344106 | -0.85962 | -0.305 | 0.18061 |
| 2 | 88601400 | 88602999 | 22.99083 | 40.0146 | 32.47626 | 14.27443 | 0.686863 | 0.778812 | 0.180634 |
| 10 | 19724600 | 19732199 | 47.97721 | 75.94286 | 163.0017 | 67.76491 | -0.77218 | 2.375328 | 0.180653 |
| 12 | 21713800 | 21717999 | 81.25536 | 128.3848 | 109.8372 | 45.62074 | 0.550834 | 2.432742 | 0.180661 |
| 8 | 74683200 | 74684399 | 14.95094 | 28.42881 | 18.02884 | 7.235199 | 1.147596 | 0.041882 | 0.180673 |
| 12 | 22026600 | 22038399 | 323.8184 | 462.6818 | 362.1262 | 149.6277 | 0.552905 | 4.262387 | 0.180678 |
| 9 | 21796800 | 21804799 | 55.04019 | 86.84566 | 164.3957 | 68.23436 | -0.59492 | 2.457043 | 0.180679 |
| 15 | 96691000 | 96691599 | 6.639221 | 12.54417 | 31.39602 | 12.77044 | -0.83387 | 0.043033 | 0.180682 |
| 6 | 8201600 | 8204999 | 55.69984 | 89.54426 | 74.47088 | 31.57775 | 0.617204 | 1.890924 | 0.180685 |
| X | 4097400 | 4099999 | 27.69051 | 48.26621 | 29.2686 | 12.87663 | 1.105446 | 0.777412 | 0.18069 |
| 8 | 17010400 | 17011399 | 12.8496 | 24.91888 | 15.03249 | 5.674026 | 1.299412 | -0.18185 | 0.180699 |
| 17 | 40857400 | 40858799 | 22.50309 | 39.74617 | 26.11922 | 11.55532 | 0.992425 | 0.567464 | 0.180703 |
| 1 | 110107800 | 110109399 | 33.21411 | 55.35013 | 43.92086 | 19.08589 | 0.713798 | 1.24202 | 0.180703 |
| X | 147669400 | 147678199 | 125.9819 | 195.2805 | 128.0272 | 53.17779 | 0.913121 | 2.811713 | 0.180715 |
| 3 | 126181800 | 126183399 | 19.22825 | 34.31912 | 25.41008 | 11.16986 | 0.828797 | 0.464874 | 0.180722 |
| 11 | 10349200 | 10350399 | 15.61585 | 28.62392 | 22.57095 | 9.626349 | 0.766655 | 0.27809 | 0.180744 |
| 13 | 9925200 | 9929599 | 31.12652 | 50.30168 | 98.52164 | 42.0748 | -0.60683 | 1.709529 | 0.180761 |
| 15 | 79019000 | 79020599 | 7.446374 | 13.82444 | 33.70508 | 14.12765 | -0.8319 | 0.169631 | 0.1808 |
| 5 | 143360800 | 143365799 | 43.1525 | 68.7268 | 134.5694 | 56.29211 | -0.63113 | 2.187746 | 0.180807 |
| 2 | 164362600 | 164372199 | 108.7566 | 163.851 | 301.3964 | 124.5058 | -0.60663 | 3.37645 | 0.180825 |
| 13 | 54490800 | 54503999 | 91.0787 | 138.0173 | 280.6139 | 116.1287 | -0.74373 | 3.254563 | 0.180833 |
| 9 | 54929000 | 54935399 | 36.64511 | 58.53357 | 127.2752 | 53.45849 | -0.77737 | 2.086268 | 0.180845 |
| 6 | 80283200 | 80283799 | 9.821737 | 19.14673 | 14.32003 | 5.169501 | 1.050336 | -0.37187 | 0.180846 |
| 10 | 41120800 | 41121799 | 5.716866 | 10.94127 | 28.94249 | 11.44464 | -0.87569 | -0.06413 | 0.18086 |
| 10 | 67368400 | 67377799 | 72.10718 | 110.7695 | 234.2548 | 97.26562 | -0.78464 | 2.969174 | 0.18086 |
| 8 | 110820200 | 110829799 | 59.39886 | 92.90602 | 184.5148 | 76.54785 | -0.67391 | 2.631512 | 0.180866 |
| 4 | 99944400 | 99945399 | 6.13645 | 11.79571 | 30.41853 | 11.87839 | -0.82301 | -0.09885 | 0.18087 |
| 19 | 55132200 | 55135399 | 13.00128 | 22.48298 | 53.70948 | 23.5633 | -0.86488 | 0.83606 | 0.18087 |
| 10 | 61740600 | 61745999 | 47.91053 | 76.09589 | 143.3343 | 59.76046 | -0.57946 | 2.27884 | 0.180879 |
| 14 | 94765200 | 94766999 | 19.74743 | 35.57435 | 23.02032 | 10.03603 | 1.029488 | 0.406723 | 0.180891 |
| 9 | 48214000 | 48215199 | 7.074652 | 13.40171 | 33.84015 | 13.68905 | -0.83985 | 0.034899 | 0.1809 |
| 1 | 116674200 | 116675799 | 15.85416 | 29.47424 | 21.01201 | 8.79075 | 0.930883 | 0.1998 | 0.18093 |
| 13 | 110775400 | 110776799 | 8.85137 | 15.73501 | 40.49029 | 17.8591 | -0.96048 | 0.454937 | 0.180933 |
| 10 | 84433600 | 84434199 | 3.818761 | 7.451143 | 23.32875 | 8.318447 | -0.9747 | -0.46393 | 0.180974 |

| | | | | | | | | | |
|---|---|---|---|---|---|---|---|---|---|
| 3 | 108747200 | 108747999 | 5.343177 | 10.1394 | 30.89945 | 12.45344 | -1.10295 | -0.00093 | 0.181 |
| 12 | 56624200 | 56626799 | 24.27791 | 42.38584 | 31.1927 | 13.73699 | 0.827829 | 0.741546 | 0.181004 |
| 9 | 32890800 | 32891799 | 5.578112 | 10.6779 | 28.94852 | 11.44029 | -0.91009 | -0.06806 | 0.181009 |
| 11 | 4798800 | 4799799 | 5.578112 | 10.6779 | 28.94852 | 11.44029 | -0.91009 | -0.06806 | 0.181009 |
| 16 | 27426400 | 27428799 | 25.08718 | 40.86896 | 94.18751 | 40.44641 | -0.83035 | 1.540239 | 0.181014 |
| 2 | 89995800 | 89997199 | 15.35717 | 28.70959 | 19.8817 | 8.23986 | 0.982725 | 0.157327 | 0.181016 |
| 16 | 84158800 | 84160399 | 15.35717 | 28.70959 | 19.8817 | 8.23986 | 0.982725 | 0.157327 | 0.181016 |
| 10 | 94082800 | 94083999 | 16.00578 | 29.13653 | 22.47211 | 9.669928 | 0.789702 | 0.318649 | 0.18103 |
| 3 | 56278600 | 56279799 | 8.145443 | 14.39797 | 42.56385 | 18.82764 | -1.16326 | 0.504954 | 0.181051 |
| 6 | 128026400 | 128028999 | 13.44475 | 23.28576 | 53.2082 | 23.34448 | -0.80064 | 0.823064 | 0.181066 |
| 2 | 35228600 | 35231199 | 13.89028 | 24.22226 | 53.79135 | 23.6719 | -0.75834 | 0.740108 | 0.181067 |
| 13 | 66459600 | 66462399 | 78.4843 | 124.6057 | 98.33493 | 40.98231 | 0.673123 | 2.305811 | 0.181072 |
| 17 | 32582800 | 32583799 | 5.865245 | 11.19952 | 29.4224 | 11.7032 | -0.87367 | -0.04122 | 0.181076 |
| 2 | 166945800 | 166950399 | 55.25148 | 86.168 | 190.4849 | 79.04834 | -0.83149 | 2.683531 | 0.181095 |
| 10 | 116772800 | 116777599 | 43.27549 | 69.07241 | 129.5465 | 54.35459 | -0.56573 | 2.117159 | 0.181097 |
| 6 | 60249800 | 60250399 | 8.965519 | 17.45717 | 14.96909 | 5.485749 | 0.836009 | -0.32963 | 0.181101 |
| 7 | 16272800 | 16274199 | 18.22103 | 33.18948 | 22.1543 | 9.525856 | 0.997513 | 0.331931 | 0.181128 |
| 12 | 107614600 | 107620599 | 43.12191 | 68.73079 | 134.3093 | 56.24298 | -0.62709 | 2.162065 | 0.181129 |
| 7 | 11051000 | 11055399 | 82.98822 | 131.555 | 99.50714 | 41.40741 | 0.732083 | 2.351321 | 0.18114 |
| 2 | 100634400 | 100636199 | 18.59897 | 33.85683 | 22.14293 | 9.528372 | 1.02614 | 0.338509 | 0.181151 |
| 14 | 99891400 | 99893199 | 24.41737 | 42.77272 | 28.15978 | 12.43419 | 0.987348 | 0.684906 | 0.181156 |
| 13 | 53751400 | 53753199 | 11.51553 | 20.18677 | 47.24956 | 20.87623 | -0.83308 | 0.636046 | 0.181162 |
| 13 | 58124600 | 58126199 | 16.07527 | 29.72442 | 21.57311 | 9.092131 | 0.897871 | 0.232736 | 0.181165 |
| 12 | 92601400 | 92601799 | 3.148213 | 6.158304 | 20.48254 | 6.835741 | -0.97005 | -0.71417 | 0.181172 |
| 18 | 76276200 | 76281199 | 22.19039 | 35.96894 | 95.51045 | 40.897 | -1.0401 | 1.623906 | 0.181195 |
| 5 | 117398600 | 117418599 | 136.8514 | 202.082 | 391.6531 | 161.2904 | -0.70842 | 3.748868 | 0.181209 |
| 6 | 84294600 | 84297999 | 14.13223 | 24.27579 | 56.3693 | 24.72517 | -0.82636 | 0.905107 | 0.181209 |
| 7 | 141290400 | 141303999 | 132.7409 | 196.0598 | 389.9645 | 160.5954 | -0.74548 | 3.7445 | 0.18121 |
| 16 | 75563200 | 75564599 | 8.272748 | 15.07773 | 37.11083 | 15.94494 | -0.8716 | 0.288263 | 0.181224 |
| 18 | 17596200 | 17599999 | 59.91043 | 96.19471 | 75.79322 | 32.0802 | 0.692889 | 1.936103 | 0.181265 |
| 11 | 88187800 | 88189399 | 11.69358 | 20.44895 | 47.29562 | 20.85355 | -0.81579 | 0.679018 | 0.181322 |
| 7 | 7769200 | 7770399 | 20.88991 | 37.02067 | 25.56889 | 11.30428 | 0.92263 | 0.51988 | 0.181331 |
| 16 | 69688400 | 69690599 | 19.9519 | 33.07361 | 73.41065 | 31.78664 | -0.76566 | 1.292452 | 0.181337 |
| 5 | 126868200 | 126868799 | 10.06043 | 19.52028 | 16.49677 | 6.211306 | 0.821514 | -0.21799 | 0.181351 |
| 3 | 66884400 | 66889999 | 45.44509 | 72.21673 | 147.405 | 61.48891 | -0.69446 | 2.261368 | 0.181358 |
| 13 | 102870600 | 102875399 | 21.75238 | 35.19976 | 98.46661 | 42.11059 | -1.11732 | 1.659508 | 0.181379 |
| 18 | 5873800 | 5875999 | 35.30861 | 58.93025 | 41.19496 | 17.91855 | 0.895931 | 1.229602 | 0.181387 |
| 13 | 93279200 | 93281399 | 10.18597 | 18.20014 | 40.46758 | 17.77247 | -0.74737 | 0.428736 | 0.181393 |
| 5 | 141328400 | 141329199 | 10.93186 | 21.32743 | 12.35742 | 4.415751 | 1.425993 | -0.39842 | 0.181396 |
| 6 | 114858400 | 114863999 | 25.58362 | 41.29048 | 102.6583 | 43.76803 | -0.95215 | 1.740117 | 0.181413 |
| 9 | 116839600 | 116840799 | 6.142005 | 11.78758 | 29.71207 | 11.65575 | -0.79643 | -0.08505 | 0.181418 |
| 14 | 92969000 | 92970199 | 8.574421 | 15.62183 | 36.57182 | 15.72056 | -0.80017 | 0.292502 | 0.18142 |
| 4 | 113949000 | 113953999 | 110.2459 | 171.85 | 125.9208 | 52.25908 | 0.759058 | 2.708063 | 0.181426 |
| 10 | 67815000 | 67817599 | 11.05157 | 19.40554 | 45.12404 | 19.96502 | -0.82338 | 0.604274 | 0.181433 |
| 7 | 7652600 | 7661399 | 118.2708 | 184.3856 | 117.1518 | 48.61876 | 0.964678 | 2.709428 | 0.181436 |
| 16 | 3246200 | 3251199 | 78.89765 | 124.748 | 107.199 | 44.54093 | 0.545265 | 2.413657 | 0.181439 |
| 13 | 50231600 | 50232999 | 15.58508 | 29.33414 | 18.8306 | 7.715023 | 1.104363 | 0.115375 | 0.181443 |

| | | | | | | | | | |
|---|---|---|---|---|---|---|---|---|---|
| 4 | 148457600 | 148484999 | 222.7277 | 316.2134 | 586.2111 | 242.4779 | -0.69994 | 4.352245 | 0.181447 |
| 8 | 71478600 | 71479599 | 6.027965 | 11.50029 | 31.77059 | 12.66035 | -0.94915 | -0.03146 | 0.181448 |
| 16 | 3322000 | 3325999 | 56.62285 | 91.87785 | 47.40865 | 20.31583 | 1.318421 | 1.642868 | 0.181473 |
| 13 | 38009000 | 38011999 | 16.64488 | 27.69231 | 73.34105 | 31.83644 | -1.01907 | 1.237812 | 0.18149 |
| 14 | 9452800 | 9454799 | 20.89585 | 37.29586 | 23.9498 | 10.52786 | 1.032655 | 0.47187 | 0.181503 |
| 15 | 38077000 | 38079399 | 12.05259 | 20.93248 | 52.64494 | 23.10836 | -0.9374 | 0.787158 | 0.181524 |
| X | 26298800 | 26300399 | 30.2415 | 51.46702 | 38.8132 | 17.01215 | 0.787417 | 1.031877 | 0.181532 |
| 10 | 107974800 | 107975799 | 11.92901 | 23.06068 | 16.7546 | 6.418023 | 1.013926 | -0.13983 | 0.181558 |
| 2 | 24472400 | 24477799 | 88.60769 | 139.2288 | 119.8549 | 49.71588 | 0.535754 | 2.55401 | 0.18156 |
| 1 | 30842800 | 30845399 | 15.91483 | 27.07613 | 61.2111 | 26.82863 | -0.79041 | 1.005586 | 0.181563 |
| 13 | 91447000 | 91448799 | 20.38407 | 36.63902 | 21.72046 | 9.491473 | 1.152489 | 0.420458 | 0.181593 |
| 17 | 46545400 | 46547999 | 8.968322 | 16.05715 | 39.9898 | 17.53249 | -0.90801 | 0.40624 | 0.181609 |
| 8 | 81584600 | 81586599 | 9.388133 | 16.92684 | 38.5016 | 16.76742 | -0.77127 | 0.364482 | 0.181622 |
| 11 | 8268400 | 8268799 | 8.564098 | 16.6927 | 14.52377 | 5.252764 | 0.832549 | -0.38153 | 0.181635 |
| 4 | 105955800 | 105961799 | 40.52003 | 64.6954 | 126.8401 | 53.24354 | -0.62904 | 2.10388 | 0.181639 |
| 11 | 75709200 | 75711399 | 9.849958 | 17.70813 | 39.49319 | 17.255 | -0.74645 | 0.385738 | 0.181645 |
| 9 | 63284200 | 63285399 | 5.09553 | 9.562771 | 32.23801 | 13.27369 | -1.27312 | 0.077465 | 0.181651 |
| 17 | 42334200 | 42336799 | 18.96402 | 31.55347 | 70.79578 | 30.73343 | -0.78073 | 1.241974 | 0.181654 |
| X | 33832200 | 33834399 | 35.13311 | 60.1001 | 31.30711 | 13.72785 | 1.319669 | 0.996669 | 0.181675 |
| 6 | 120631200 | 120632599 | 5.987602 | 11.16742 | 33.7736 | 14.05283 | -1.13141 | 0.12791 | 0.181692 |
| 6 | 134891600 | 134892799 | 5.987602 | 11.16742 | 33.7736 | 14.05283 | -1.13141 | 0.12791 | 0.181692 |
| 11 | 120958400 | 120963199 | 38.88532 | 62.11588 | 127.6539 | 53.62247 | -0.69607 | 2.086171 | 0.181721 |
| 5 | 85856600 | 85857999 | 13.59648 | 25.6624 | 19.6334 | 8.003632 | 0.859172 | 0.08153 | 0.181725 |
| 15 | 38837800 | 38839599 | 12.97057 | 22.51765 | 51.59472 | 22.64159 | -0.80373 | 0.796899 | 0.181738 |
| 5 | 73808000 | 73809599 | 8.73302 | 15.47725 | 41.49563 | 18.34547 | -1.02204 | 0.475975 | 0.181747 |
| X | 32802000 | 32802799 | 18.3555 | 33.57651 | 21.73253 | 9.294771 | 1.046777 | 0.309379 | 0.181753 |
| 1 | 150268200 | 150269599 | 9.95977 | 17.57709 | 44.05593 | 19.49382 | -0.92854 | 0.533457 | 0.181771 |
| 1 | 38578800 | 38583399 | 27.66729 | 44.98167 | 89.75844 | 38.51623 | -0.62521 | 1.570937 | 0.181784 |
| 19 | 28642800 | 28643599 | 4.531737 | 8.799346 | 25.60239 | 9.596538 | -0.93725 | -0.26765 | 0.181794 |
| 16 | 12913600 | 12917799 | 37.81892 | 62.52192 | 49.93127 | 21.57981 | 0.703945 | 1.348724 | 0.181794 |
| 11 | 27118200 | 27119999 | 21.81703 | 38.5227 | 26.0384 | 11.52026 | 0.951632 | 0.564837 | 0.181794 |
| 7 | 17130200 | 17130399 | 6.653675 | 12.99672 | 8.653343 | 2.753312 | 1.370761 | -0.8719 | 0.181803 |
| 4 | 154298400 | 154303199 | 36.94173 | 59.07677 | 122.3636 | 51.48113 | -0.70551 | 2.049677 | 0.181805 |
| 1 | 57620600 | 57623199 | 15.76081 | 26.82176 | 60.09244 | 26.33893 | -0.77733 | 1.001885 | 0.181805 |
| 14 | 38182200 | 38182599 | 7.8165 | 15.28867 | 11.00023 | 3.700146 | 1.193438 | -0.65384 | 0.18181 |
| 2 | 168560400 | 168574799 | 110.6331 | 165.305 | 356.2296 | 146.8693 | -0.84559 | 3.537927 | 0.18183 |
| 17 | 92601000 | 92602799 | 18.15724 | 33.14958 | 21.94954 | 9.410654 | 1.012224 | 0.320072 | 0.181844 |
| 15 | 91759000 | 91759799 | 6.685891 | 12.5095 | 33.90367 | 14.03121 | -0.96886 | 0.112401 | 0.18185 |
| 4 | 128195000 | 128201199 | 38.99326 | 62.41497 | 124.9823 | 52.65147 | -0.65548 | 2.022376 | 0.181852 |
| 5 | 94830400 | 94831799 | 20.48456 | 36.72991 | 22.85712 | 10.00875 | 1.081209 | 0.438708 | 0.181862 |
| 14 | 38332400 | 38333399 | 16.88763 | 31.11575 | 21.28496 | 9.014165 | 0.97733 | 0.256687 | 0.181863 |
| 11 | 49304800 | 49305799 | 13.32228 | 25.80539 | 12.91355 | 4.900886 | 1.556356 | -0.16647 | 0.181864 |
| 8 | 120753800 | 120761599 | 39.14628 | 62.30144 | 139.5383 | 58.31254 | -0.82579 | 2.208294 | 0.181864 |
| 17 | 15822800 | 15825399 | 39.51565 | 65.24766 | 45.07793 | 19.46592 | 0.912281 | 1.364472 | 0.181864 |
| 1 | 92837600 | 92848999 | 56.70274 | 87.88428 | 211.6892 | 87.92926 | -0.9622 | 2.792812 | 0.18187 |
| 1 | 63150800 | 63153999 | 15.28584 | 26.03693 | 61.75799 | 27.07092 | -0.85915 | 0.992387 | 0.181874 |
| 10 | 75037000 | 75037999 | 4.735845 | 9.074685 | 28.92053 | 11.39639 | -1.13654 | -0.08621 | 0.181877 |

| | | | | | | | | | |
|---|---|---|---|---|---|---|---|---|---|
| 14 | 103926000 | 103927399 | 4.735845 | 9.074685 | 28.92053 | 11.39639 | -1.13654 | -0.08621 | 0.181877 |
| 5 | 134807400 | 134809199 | 6.328024 | 12.01783 | 30.42474 | 12.24152 | -0.83609 | 0.002082 | 0.181882 |
| 12 | 106473600 | 106476199 | 21.16169 | 34.79362 | 80.03484 | 34.49367 | -0.81989 | 1.396331 | 0.181886 |
| 11 | 109517200 | 109519199 | 10.89356 | 19.14143 | 45.10613 | 19.96886 | -0.8425 | 0.591477 | 0.181902 |
| 7 | 139550200 | 139552999 | 11.45718 | 19.87838 | 52.60768 | 23.08224 | -1.01056 | 0.798088 | 0.181907 |
| X | 115460800 | 115462799 | 30.13903 | 51.43408 | 37.23995 | 16.3262 | 0.846546 | 1.005995 | 0.181914 |
| 8 | 92502200 | 92502799 | 4.233374 | 8.244158 | 24.15846 | 8.83942 | -0.91451 | -0.36177 | 0.181915 |
| 16 | 30418000 | 30422599 | 25.80684 | 41.81695 | 92.59162 | 39.6374 | -0.77925 | 1.63664 | 0.181922 |
| 11 | 54775600 | 54776999 | 7.732081 | 14.37814 | 34.30259 | 14.34511 | -0.79869 | 0.162954 | 0.181929 |
| 4 | 83859400 | 83860399 | 4.827819 | 9.35135 | 26.62699 | 10.10629 | -0.92419 | -0.21855 | 0.181931 |
| 7 | 137956800 | 137957599 | 4.827819 | 9.35135 | 26.62699 | 10.10629 | -0.92419 | -0.21855 | 0.181931 |
| 3 | 23829600 | 23830599 | 10.71007 | 20.60884 | 16.41288 | 6.389992 | 0.860971 | -0.10247 | 0.181939 |
| 18 | 60577800 | 60581799 | 61.94575 | 99.33821 | 77.73797 | 32.84072 | 0.700588 | 1.976469 | 0.181951 |
| 14 | 65313000 | 65315799 | 16.76799 | 27.86142 | 71.9631 | 31.21068 | -0.98298 | 1.255456 | 0.181952 |
| 4 | 140498800 | 140534799 | 262.1369 | 363.1152 | 791.8551 | 328.8264 | -0.96996 | 4.722094 | 0.181954 |
| 1 | 157239200 | 157240799 | 13.31062 | 23.01322 | 53.17541 | 23.32885 | -0.81709 | 0.839265 | 0.181969 |
| 9 | 12185400 | 12186399 | 13.04728 | 24.6516 | 19.14559 | 7.787531 | 0.840278 | 0.067651 | 0.181974 |
| 3 | 15532600 | 15534399 | 19.84427 | 36.15797 | 20.30107 | 8.725455 | 1.245517 | 0.33829 | 0.181993 |
| 2 | 156311000 | 156313199 | 9.677689 | 17.01992 | 44.09381 | 19.52417 | -0.97761 | 0.573119 | 0.182004 |
| 5 | 124804000 | 124821999 | 131.324 | 194.7625 | 369.8291 | 152.3267 | -0.67309 | 3.674777 | 0.182031 |
| 8 | 26403800 | 26406199 | 24.3137 | 40.01354 | 81.2684 | 35.03919 | -0.64045 | 1.393054 | 0.182031 |
| 16 | 49187000 | 49191399 | 32.42096 | 52.36428 | 102.9621 | 43.93951 | -0.61344 | 1.729473 | 0.182031 |
| 11 | 40645400 | 40647199 | 8.225624 | 15.11933 | 37.87764 | 16.09391 | -0.88596 | 0.239511 | 0.182044 |
| 13 | 70865000 | 70866199 | 5.093465 | 9.863417 | 26.61534 | 10.11375 | -0.84886 | -0.21012 | 0.182048 |
| 4 | 148162400 | 148170599 | 42.17314 | 66.81279 | 156.1521 | 64.94546 | -0.89373 | 2.350394 | 0.182049 |
| 11 | 117219800 | 117226199 | 28.40633 | 45.60374 | 117.6894 | 49.8021 | -1.01536 | 1.915128 | 0.182053 |
| 4 | 53667400 | 53668799 | 14.40156 | 27.27222 | 19.12273 | 7.740978 | 0.99306 | 0.066506 | 0.182054 |
| 4 | 59910400 | 59916999 | 50.40891 | 79.78787 | 155.617 | 64.69321 | -0.63429 | 2.377743 | 0.182061 |
| 8 | 12544400 | 12545999 | 9.384954 | 16.88235 | 40.03232 | 17.48164 | -0.83393 | 0.378516 | 0.182063 |
| 8 | 16950400 | 16950799 | 10.25036 | 19.99982 | 13.22967 | 4.719904 | 1.240953 | -0.40894 | 0.182071 |
| 9 | 64160000 | 64161799 | 10.45042 | 18.34385 | 45.65868 | 20.19587 | -0.92094 | 0.607036 | 0.182075 |
| 2 | 70885600 | 70886999 | 7.022018 | 13.0086 | 35.2907 | 14.83563 | -0.98814 | 0.180346 | 0.182085 |
| 7 | 108151800 | 108155999 | 103.5117 | 160.7395 | 138.1641 | 57.37288 | 0.524801 | 2.771293 | 0.182092 |
| 10 | 17193800 | 17194399 | 14.56835 | 26.83825 | 20.71225 | 8.773495 | 0.804447 | 0.23744 | 0.182101 |
| 18 | 18845600 | 18847399 | 30.98317 | 52.16853 | 42.6487 | 18.64003 | 0.67068 | 1.128602 | 0.182104 |
| 4 | 128589200 | 128592599 | 16.78798 | 28.29935 | 64.36227 | 28.13989 | -0.79992 | 1.094165 | 0.18211 |
| 11 | 68812200 | 68814399 | 6.891383 | 12.68182 | 34.61475 | 14.68399 | -1.00554 | 0.219443 | 0.182118 |
| 7 | 9776000 | 9777599 | 25.51029 | 44.44908 | 31.26594 | 13.75208 | 0.892576 | 0.775947 | 0.182139 |
| 1 | 74107200 | 74109399 | 10.97533 | 19.11071 | 51.01079 | 22.42734 | -1.02213 | 0.743071 | 0.182141 |
| 10 | 59389000 | 59390399 | 7.888785 | 14.68826 | 40.67414 | 16.97295 | -1.01011 | 0.169461 | 0.182142 |
| 9 | 65494000 | 65497399 | 15.95328 | 27.05223 | 61.10757 | 26.76498 | -0.78959 | 1.036581 | 0.182166 |
| 2 | 59020000 | 59022599 | 14.83084 | 25.68326 | 56.55664 | 24.81348 | -0.74789 | 0.83349 | 0.18217 |
| 13 | 97827400 | 97829199 | 12.76834 | 22.31405 | 48.36424 | 21.31313 | -0.72286 | 0.695644 | 0.182171 |
| 12 | 21884800 | 21887799 | 69.74853 | 110.9198 | 95.01795 | 39.66559 | 0.557236 | 2.248508 | 0.182179 |
| 15 | 83814200 | 83817999 | 21.31354 | 34.71352 | 88.67057 | 38.07415 | -0.97836 | 1.531943 | 0.182181 |
| 7 | 9920400 | 9929199 | 152.1791 | 232.2613 | 163.8108 | 67.93042 | 0.790814 | 3.118276 | 0.182184 |
| 3 | 138580800 | 138585399 | 38.22625 | 61.09466 | 129.3197 | 54.362 | -0.7378 | 2.069335 | 0.182192 |

| | | | | | | | | | |
|---|---|---|---|---|---|---|---|---|---|
| 6 | 10203800 | 10204999 | 10.75683 | 20.86911 | 14.5147 | 5.467935 | 1.097393 | -0.21646 | 0.182194 |
| 7 | 8662200 | 8662599 | 9.652396 | 18.78799 | 15.48327 | 5.690372 | 0.889325 | -0.30841 | 0.182208 |
| 6 | 54303000 | 54307399 | 22.94874 | 37.27277 | 95.19519 | 40.82225 | -0.9815 | 1.58275 | 0.182227 |
| 9 | 124246200 | 124249399 | 40.39881 | 66.69248 | 49.42271 | 21.33915 | 0.811449 | 1.372384 | 0.18223 |
| 11 | 75421400 | 75424999 | 16.95406 | 28.5529 | 64.33949 | 28.12204 | -0.78663 | 1.103222 | 0.182236 |
| 9 | 19059600 | 19060399 | 19.05321 | 33.71439 | 26.43808 | 11.6921 | 0.739718 | 0.52575 | 0.182253 |
| 8 | 12431000 | 12442399 | 76.24506 | 116.593 | 263.3699 | 109.2652 | -0.8848 | 3.064021 | 0.182254 |
| 2 | 172468800 | 172481799 | 105.7598 | 158.9584 | 316.6622 | 130.729 | -0.72461 | 3.422629 | 0.182263 |
| 17 | 88448400 | 88449799 | 10.72279 | 18.89636 | 43.53285 | 19.26794 | -0.80844 | 0.554141 | 0.182282 |
| 6 | 78657600 | 78659599 | 15.24591 | 28.23304 | 21.52469 | 9.053287 | 0.829085 | 0.216376 | 0.182284 |
| X | 53202400 | 53204199 | 36.26748 | 61.13564 | 38.70977 | 16.92432 | 1.038033 | 1.134876 | 0.182285 |
| 5 | 52628200 | 52632599 | 26.18946 | 42.68492 | 88.15723 | 37.86341 | -0.67215 | 1.530304 | 0.182294 |
| 17 | 88102200 | 88107799 | 50.87433 | 80.17221 | 166.4447 | 69.06756 | -0.72931 | 2.480966 | 0.182296 |
| 1 | 172152000 | 172157599 | 41.35193 | 65.9189 | 132.327 | 55.42361 | -0.66557 | 2.155827 | 0.182304 |
| 5 | 83696600 | 83697999 | 24.02644 | 41.97984 | 27.72779 | 12.21712 | 0.983073 | 0.723816 | 0.182319 |
| 4 | 120807800 | 120809199 | 10.52338 | 18.6917 | 41.44213 | 18.29182 | -0.74837 | 0.470455 | 0.18234 |
| 2 | 28902400 | 28907799 | 36.57735 | 58.60398 | 117.5192 | 49.5787 | -0.65582 | 1.988812 | 0.182342 |
| 12 | 105947400 | 105951199 | 36.04501 | 57.86836 | 114.5613 | 48.4802 | -0.63383 | 1.922753 | 0.182366 |
| 15 | 3118000 | 3130799 | 132.7331 | 203.9808 | 152.7048 | 63.40835 | 0.712501 | 2.974391 | 0.182368 |
| X | 92118400 | 92119999 | 23.07488 | 40.65601 | 26.74692 | 11.83883 | 0.989671 | 0.598738 | 0.18237 |
| 12 | 80005000 | 80009999 | 20.85189 | 34.27371 | 79.91988 | 34.43964 | -0.83962 | 1.399699 | 0.182384 |
| 11 | 41583800 | 41585199 | 9.683166 | 17.48225 | 39.04702 | 16.97641 | -0.74379 | 0.358929 | 0.182389 |
| 2 | 81623200 | 81626399 | 52.84145 | 85.08179 | 60.04087 | 25.51669 | 0.85638 | 1.832675 | 0.182392 |
| 10 | 18193600 | 18195599 | 11.45916 | 19.89247 | 54.23455 | 23.79906 | -1.05334 | 0.792919 | 0.182407 |
| 12 | 113733200 | 113737399 | 76.97024 | 122.0456 | 103.3396 | 43.01294 | 0.569919 | 2.340769 | 0.182412 |
| 3 | 34690400 | 34691799 | 16.42041 | 29.70873 | 21.26602 | 9.221322 | 0.889941 | 0.354804 | 0.182432 |
| 7 | 127825600 | 127826399 | 12.11386 | 23.41503 | 16.74877 | 6.419624 | 1.035496 | -0.13569 | 0.18244 |
| X | 30890600 | 30893399 | 26.36997 | 47.14224 | 18.16617 | 7.974646 | 1.765308 | 0.489343 | 0.182444 |
| 11 | 120523000 | 120528599 | 30.5157 | 49.21193 | 101.758 | 43.34032 | -0.68835 | 1.770674 | 0.182448 |
| 13 | 64721800 | 64724399 | 20.00914 | 33.42452 | 71.1955 | 30.98494 | -0.70616 | 1.200614 | 0.18248 |
| 12 | 16678600 | 16679199 | 5.262466 | 10.07553 | 28.9 | 11.41509 | -0.98895 | -0.07044 | 0.182488 |
| 12 | 4831000 | 4834399 | 17.77991 | 29.80967 | 67.65747 | 29.51329 | -0.79728 | 1.151604 | 0.182506 |
| 5 | 141106400 | 141107799 | 11.16731 | 19.74589 | 42.98848 | 19.01423 | -0.72542 | 0.518968 | 0.182508 |
| 1 | 184875200 | 184878399 | 13.49892 | 23.24035 | 56.38128 | 24.73213 | -0.88856 | 0.881251 | 0.182508 |
| 2 | 130498400 | 130501199 | 8.449122 | 14.93167 | 42.55447 | 18.82427 | -1.11108 | 0.508684 | 0.182511 |
| 12 | 18082600 | 18085599 | 139.0036 | 211.8887 | 166.0607 | 68.84362 | 0.637414 | 3.139521 | 0.182517 |
| 10 | 81340800 | 81354399 | 103.8937 | 156.4525 | 305.4451 | 126.1627 | -0.69306 | 3.384588 | 0.182523 |
| 10 | 82582800 | 82583799 | 4.997896 | 9.643306 | 28.72262 | 11.09579 | -1.01365 | -0.15161 | 0.182528 |
| 13 | 118258200 | 118258599 | 8.389723 | 16.38158 | 13.31706 | 4.690878 | 0.962108 | -0.48029 | 0.182541 |
| 17 | 69799200 | 69800399 | 7.104656 | 13.32662 | 36.92802 | 15.21765 | -0.99628 | 0.105094 | 0.182546 |
| 13 | 53911000 | 53912399 | 6.823357 | 12.76974 | 32.24106 | 13.33776 | -0.86634 | 0.110735 | 0.182555 |
| X | 67960200 | 67962799 | 25.82198 | 45.42023 | 24.76421 | 10.9649 | 1.258023 | 0.628505 | 0.18256 |
| 1 | 31859000 | 31860199 | 9.339435 | 17.00813 | 39.2627 | 16.88654 | -0.78097 | 0.305982 | 0.18256 |
| 13 | 60459600 | 60461399 | 9.339435 | 17.00813 | 39.2627 | 16.88654 | -0.78097 | 0.305982 | 0.18256 |
| 14 | 15653000 | 15654399 | 21.64702 | 37.96364 | 29.67103 | 13.11429 | 0.741331 | 0.655623 | 0.182561 |
| 2 | 33821000 | 33825599 | 31.45032 | 50.84812 | 99.9481 | 42.70909 | -0.61122 | 1.697795 | 0.182569 |
| 4 | 129396000 | 129398399 | 9.490533 | 16.79234 | 41.9941 | 18.57191 | -0.92346 | 0.499429 | 0.182585 |

| | | | | | | | | | |
|---|---|---|---|---|---|---|---|---|---|
| 6 | 48791600 | 48792799 | 9.548555 | 17.1447 | 39.4816 | 17.26824 | -0.79344 | 0.391114 | 0.182594 |
| 5 | 103906200 | 103908999 | 11.0498 | 19.39915 | 46.69335 | 20.6564 | -0.87312 | 0.608168 | 0.182599 |
| 8 | 13262600 | 13264399 | 8.344813 | 15.01505 | 38.49007 | 16.80459 | -0.94483 | 0.368217 | 0.182609 |
| 15 | 88406200 | 88409599 | 23.84318 | 39.23679 | 79.33833 | 34.20614 | -0.63404 | 1.392293 | 0.182611 |
| Y | 9429800 | 9430399 | 9.883452 | 19.23864 | 8.710035 | 3.199863 | 1.74132 | -0.33475 | 0.182615 |
| 15 | 27961800 | 27964599 | 11.66186 | 20.48249 | 45.10839 | 19.96248 | -0.74548 | 0.603493 | 0.182637 |
| 16 | 43427400 | 43431399 | 35.21335 | 56.63959 | 109.75 | 46.57263 | -0.59959 | 1.861158 | 0.18265 |
| 1 | 178603000 | 178605599 | 26.21982 | 45.76467 | 29.89424 | 13.16308 | 0.998865 | 0.75255 | 0.182659 |
| 1 | 36159800 | 36165199 | 38.5665 | 61.61019 | 126.1735 | 53.00498 | -0.69092 | 2.084068 | 0.18267 |
| 12 | 23344800 | 23347199 | 90.68065 | 142.3147 | 111.3937 | 46.19893 | 0.671498 | 2.578594 | 0.182677 |
| 5 | 115233000 | 115235599 | 13.07987 | 22.42694 | 56.81042 | 24.91545 | -0.95157 | 0.919289 | 0.182678 |
| 9 | 62442000 | 62444399 | 10.99304 | 19.10416 | 52.06882 | 22.86441 | -1.05241 | 0.773244 | 0.182705 |
| 7 | 15996400 | 16010999 | 92.1263 | 140.0932 | 263.5915 | 109.1991 | -0.62785 | 3.182461 | 0.182708 |
| 17 | 11830400 | 11833199 | 12.09486 | 20.91045 | 54.20381 | 23.78 | -0.9821 | 0.835043 | 0.182712 |
| 18 | 37666000 | 37666999 | 31.67256 | 53.99135 | 36.6589 | 16.07003 | 0.938656 | 1.021423 | 0.182733 |
| 11 | 114087400 | 114097999 | 70.04837 | 107.406 | 254.758 | 105.7517 | -0.95208 | 3.007263 | 0.182744 |
| 11 | 79695200 | 79698199 | 24.309 | 39.56591 | 87.41286 | 37.52499 | -0.7704 | 1.547143 | 0.182751 |
| 2 | 161223800 | 161225399 | 9.691544 | 17.51721 | 40.68424 | 17.66318 | -0.79849 | 0.356165 | 0.182753 |
| 4 | 130362600 | 130366599 | 19.21022 | 31.707 | 75.86132 | 32.75005 | -0.87405 | 1.342738 | 0.182766 |
| 12 | 44115600 | 44118199 | 20.56545 | 36.30408 | 27.31977 | 12.08776 | 0.79762 | 0.562496 | 0.182775 |
| 12 | 81140000 | 81142199 | 11.16776 | 19.68845 | 48.83338 | 21.61224 | -0.91471 | 0.558413 | 0.182794 |
| 3 | 74153800 | 74154799 | 14.29601 | 27.14308 | 18.51573 | 7.454001 | 1.038961 | 0.045891 | 0.182813 |
| 17 | 46608000 | 46619599 | 64.718 | 99.46938 | 234.1606 | 97.23332 | -0.93852 | 2.956973 | 0.182824 |
| 17 | 95024800 | 95025599 | 16.66286 | 30.78224 | 21.40984 | 9.034756 | 0.957368 | 0.241601 | 0.182836 |
| 1 | 114061200 | 114062799 | 30.62555 | 51.85539 | 33.81769 | 14.80542 | 0.995586 | 1.069777 | 0.182839 |
| 5 | 97255600 | 97257399 | 8.394314 | 15.40261 | 35.63715 | 15.17779 | -0.77399 | 0.248572 | 0.182843 |
| 7 | 42723400 | 42726199 | 52.08613 | 84.13906 | 64.92579 | 27.7128 | 0.733229 | 1.74034 | 0.182845 |
| 14 | 112488400 | 112489399 | 13.69757 | 26.16026 | 17.44195 | 6.934926 | 1.086712 | -0.0024 | 0.182845 |
| 1 | 105917400 | 105919799 | 13.78121 | 23.40926 | 60.97479 | 26.72076 | -0.99389 | 1.013873 | 0.182861 |
| 13 | 40853800 | 40860999 | 51.74185 | 81.79736 | 158.2302 | 65.73878 | -0.62388 | 2.408407 | 0.182866 |
| 14 | 40443600 | 40444999 | 22.78884 | 40.049 | 27.52085 | 12.18524 | 0.926045 | 0.627298 | 0.182874 |
| 17 | 12088800 | 12091199 | 16.50402 | 28.23317 | 58.59058 | 25.69143 | -0.66612 | 0.955555 | 0.182882 |
| 8 | 15033600 | 15035799 | 11.50923 | 20.22892 | 44.57506 | 19.73843 | -0.74618 | 0.590744 | 0.182883 |
| 11 | 121348800 | 121349999 | 15.94788 | 29.08443 | 23.05777 | 9.899206 | 0.752553 | 0.310038 | 0.182884 |
| 1 | 154608400 | 154609599 | 5.198321 | 9.906376 | 32.20919 | 12.86557 | -1.1846 | -0.03053 | 0.182888 |
| 4 | 26297000 | 26297599 | 12.52914 | 23.8569 | 18.13517 | 7.256546 | 0.89128 | 0.00757 | 0.182893 |
| 7 | 60141800 | 60142599 | 6.817089 | 12.78503 | 32.86189 | 13.54597 | -0.88822 | 0.096406 | 0.182895 |
| 14 | 87464400 | 87465599 | 14.05096 | 26.56563 | 19.01176 | 7.722215 | 0.959098 | 0.072474 | 0.182914 |
| 2 | 173288200 | 173300999 | 121.6693 | 182.2556 | 328.0533 | 135.3191 | -0.58275 | 3.490945 | 0.182919 |
| 2 | 3568600 | 3571999 | 19.90929 | 33.10597 | 72.07341 | 31.27484 | -0.73757 | 1.251955 | 0.18292 |
| 17 | 13524800 | 13542799 | 1207.207 | 1486.134 | 2697.986 | 1136.742 | -0.87692 | 6.616317 | 0.182921 |
| 16 | 85876200 | 85876999 | 12.49165 | 23.8338 | 18.76589 | 7.476785 | 0.846723 | -0.00379 | 0.182954 |
| 8 | 105328000 | 105329999 | 10.66015 | 18.59042 | 49.94049 | 21.97796 | -1.03094 | 0.718039 | 0.182964 |
| 11 | 7085600 | 7090799 | 30.38256 | 48.93422 | 108.6653 | 46.20102 | -0.79333 | 1.811208 | 0.182968 |
| 6 | 40958600 | 40959599 | 15.15536 | 28.33285 | 20.54313 | 8.513712 | 0.917315 | 0.157854 | 0.182969 |
| 5 | 150756400 | 150758999 | 10.43235 | 18.35411 | 48.80414 | 21.60819 | -1.01559 | 0.576878 | 0.182998 |
| 3 | 31202400 | 31208999 | 35.59595 | 57.06197 | 117.7885 | 49.73259 | -0.6966 | 1.970217 | 0.183012 |

| | | | | | | | | | |
|---|---|---|---|---|---|---|---|---|---|
| 4 | 148043200 | 148052599 | 60.86976 | 95.1818 | 185.7472 | 77.06118 | -0.64894 | 2.636901 | 0.183015 |
| 8 | 19617400 | 19619199 | 36.96031 | 61.07877 | 47.72662 | 20.62146 | 0.735342 | 1.351207 | 0.183026 |
| 18 | 82505800 | 82509999 | 20.29724 | 33.55676 | 74.49488 | 32.19698 | -0.76628 | 1.32426 | 0.183052 |
| 17 | 24979600 | 24981599 | 11.02107 | 19.45397 | 43.00106 | 19.0263 | -0.74814 | 0.537109 | 0.183065 |
| 17 | 86016600 | 86018399 | 17.23446 | 29.11655 | 62.77871 | 27.46602 | -0.72305 | 1.075128 | 0.183093 |
| 1 | 74914600 | 74925599 | 63.55054 | 98.47488 | 214.1192 | 88.93912 | -0.81502 | 2.799485 | 0.183103 |
| 7 | 11788600 | 11801199 | 239.4489 | 353.8681 | 251.1312 | 103.4216 | 0.741246 | 3.744019 | 0.183142 |
| X | 63540600 | 63541999 | 20.47251 | 37.00472 | 21.9224 | 9.517286 | 1.159863 | 0.39134 | 0.183155 |
| 12 | 82720400 | 82722799 | 11.9017 | 20.66111 | 51.55599 | 22.62396 | -0.92594 | 0.793493 | 0.183156 |
| 12 | 20725200 | 20739199 | 479.1036 | 661.3492 | 508.6734 | 211.3495 | 0.527991 | 4.779398 | 0.183175 |
| 13 | 66407000 | 66418999 | 130.8939 | 200.6758 | 139.9269 | 58.08356 | 0.812286 | 3.019173 | 0.183198 |
| 4 | 37621000 | 37625999 | 71.30305 | 113.9463 | 81.53735 | 34.27027 | 0.823839 | 2.095125 | 0.183198 |
| 6 | 85527200 | 85528999 | 20.7765 | 36.561 | 28.48582 | 12.61164 | 0.74619 | 0.600155 | 0.183199 |
| 18 | 67857000 | 67858799 | 10.92206 | 19.12779 | 48.32184 | 21.33245 | -0.94257 | 0.650282 | 0.183201 |
| 2 | 6469200 | 6489199 | 157.4827 | 231.227 | 422.6739 | 174.0818 | -0.63186 | 3.84387 | 0.183206 |
| 4 | 3676800 | 3679999 | 17.79539 | 29.80014 | 67.01224 | 29.20989 | -0.78388 | 1.165003 | 0.183213 |
| 7 | 16732000 | 16740999 | 39.42076 | 62.84108 | 133.6319 | 55.97094 | -0.74853 | 2.153742 | 0.18322 |
| 16 | 30868400 | 30871999 | 15.23024 | 25.70613 | 63.68364 | 27.85391 | -0.92253 | 1.077709 | 0.183225 |
| 7 | 19445600 | 19449999 | 22.00451 | 36.03034 | 81.97733 | 35.2692 | -0.80655 | 1.452405 | 0.183227 |
| 19 | 8022000 | 8024599 | 38.02407 | 62.7503 | 51.03299 | 22.02913 | 0.677649 | 1.370492 | 0.183245 |
| 7 | 24824800 | 24842399 | 108.7038 | 163.2081 | 316.6672 | 130.6935 | -0.68797 | 3.444508 | 0.183266 |
| 10 | 68605400 | 68606599 | 8.676264 | 15.96857 | 37.92755 | 16.08639 | -0.80765 | 0.236656 | 0.183266 |
| 3 | 3219200 | 3220399 | 25.94579 | 44.50439 | 34.31943 | 15.05201 | 0.756875 | 0.938225 | 0.183275 |
| 4 | 151496000 | 151498399 | 12.04482 | 20.93503 | 51.03725 | 22.41346 | -0.89242 | 0.774964 | 0.183276 |
| 7 | 26727600 | 26729199 | 25.68401 | 44.16461 | 35.63622 | 15.63154 | 0.692265 | 0.915333 | 0.183277 |
| 7 | 26999200 | 27002599 | 17.75709 | 29.8359 | 65.46439 | 28.59356 | -0.7486 | 1.127212 | 0.183279 |
| 11 | 101046600 | 101048599 | 7.04652 | 12.93114 | 35.10417 | 14.9483 | -1.00175 | 0.238367 | 0.183279 |
| X | 126628000 | 126630999 | 45.18594 | 74.52707 | 45.597 | 19.67762 | 1.086677 | 1.392558 | 0.183284 |
| 5 | 148504200 | 148508399 | 37.33888 | 59.9999 | 116.6263 | 49.4207 | -0.60592 | 1.897445 | 0.183285 |
| 5 | 122639200 | 122641199 | 8.348892 | 14.66124 | 45.67676 | 20.21114 | -1.24203 | 0.590112 | 0.183287 |
| 3 | 52472600 | 52474199 | 9.513074 | 17.21908 | 38.56866 | 16.71496 | -0.74487 | 0.341379 | 0.183299 |
| 17 | 29951600 | 29954199 | 21.18478 | 34.76648 | 80.33763 | 34.59678 | -0.82749 | 1.421051 | 0.183302 |
| 4 | 73747400 | 73748599 | 14.48575 | 27.49562 | 18.50903 | 7.455733 | 1.057263 | 0.049691 | 0.183318 |
| 10 | 94723200 | 94724999 | 27.37471 | 46.99309 | 36.57795 | 16.0436 | 0.743554 | 0.93883 | 0.183337 |
| X | 26568800 | 26571199 | 68.03224 | 109.2047 | 65.69868 | 27.79568 | 1.080844 | 1.955349 | 0.183346 |
| 6 | 18493600 | 18494599 | 4.5235 | 8.807675 | 25.42973 | 9.32131 | -0.8967 | -0.34919 | 0.183354 |
| 10 | 24165800 | 24166399 | 3.822349 | 7.448014 | 23.61742 | 8.576417 | -1.01647 | -0.39625 | 0.183362 |
| 3 | 154698600 | 154699599 | 13.75313 | 26.41125 | 17.07371 | 6.685775 | 1.151606 | -0.04946 | 0.183367 |
| 17 | 14334800 | 14336999 | 11.63267 | 20.51552 | 45.08187 | 19.95049 | -0.74052 | 0.552715 | 0.183367 |
| 16 | 61217600 | 61219599 | 20.42825 | 36.13695 | 26.76845 | 11.8389 | 0.821352 | 0.538687 | 0.183373 |
| 3 | 52120200 | 52126399 | 34.83763 | 55.75584 | 129.3155 | 54.46673 | -0.86718 | 2.022583 | 0.183382 |
| 4 | 124070600 | 124077199 | 39.47916 | 63.19839 | 123.2026 | 51.90831 | -0.61672 | 2.01998 | 0.183393 |
| 3 | 111082000 | 111083199 | 13.37632 | 25.30775 | 19.75368 | 8.011752 | 0.83717 | 0.064399 | 0.183394 |
| 19 | 10078000 | 10079399 | 8.468297 | 14.96344 | 43.56783 | 19.27308 | -1.14214 | 0.510712 | 0.1834 |
| 1 | 88903000 | 88905999 | 34.25322 | 55.08385 | 108.68 | 46.10477 | -0.62592 | 1.866214 | 0.183414 |
| 16 | 96144200 | 96146799 | 13.24325 | 23.07429 | 50.53217 | 22.22573 | -0.73834 | 0.743684 | 0.183419 |
| 15 | 96640200 | 96643799 | 19.686 | 32.96155 | 67.75645 | 29.53168 | -0.65493 | 1.173191 | 0.183438 |

| | | | | | | | | | |
|---|---|---|---|---|---|---|---|---|---|
| 7 | 82647000 | 82648599 | 11.17023 | 19.68622 | 46.14361 | 20.4232 | -0.83359 | 0.558481 | 0.183454 |
| 7 | 94833800 | 94835399 | 23.45156 | 40.85587 | 31.97577 | 14.06297 | 0.739263 | 0.768288 | 0.183457 |
| 12 | 23932000 | 23936199 | 61.62117 | 98.5826 | 83.89953 | 35.31811 | 0.575847 | 2.05073 | 0.183466 |
| 8 | 95465600 | 95468799 | 21.4922 | 35.71598 | 73.99125 | 32.09332 | -0.66641 | 1.264854 | 0.183468 |
| 8 | 53420000 | 53420999 | 12.69081 | 24.4438 | 17.30212 | 6.717591 | 1.033703 | -0.08592 | 0.18347 |
| 14 | 7285400 | 7287599 | 26.80558 | 46.87551 | 27.52296 | 12.13534 | 1.151775 | 0.724229 | 0.183473 |
| 1 | 119837600 | 119839199 | 12.68377 | 21.97725 | 52.61396 | 23.08406 | -0.86702 | 0.81816 | 0.183479 |
| 9 | 18769400 | 18769999 | 10.21546 | 19.90347 | 14.18226 | 5.156268 | 1.109754 | -0.34359 | 0.183486 |
| 14 | 41326200 | 41329199 | 44.71163 | 72.72678 | 59.33512 | 25.46566 | 0.662267 | 1.579589 | 0.183486 |
| 14 | 55633600 | 55635599 | 19.15307 | 34.91671 | 21.58679 | 9.272024 | 1.108473 | 0.333678 | 0.183489 |
| 15 | 82167000 | 82171199 | 24.84832 | 40.30473 | 92.78906 | 39.7517 | -0.83395 | 1.613047 | 0.183493 |
| 9 | 46767400 | 46770799 | 37.80012 | 62.42802 | 50.49097 | 21.80657 | 0.685647 | 1.360192 | 0.183501 |
| 12 | 52792800 | 52794999 | 14.6953 | 25.41999 | 53.7754 | 23.59243 | -0.69042 | 0.840957 | 0.183501 |
| 9 | 28682400 | 28686399 | 53.76456 | 87.09303 | 58.07443 | 24.85549 | 0.947348 | 1.6733 | 0.183514 |
| 10 | 129886800 | 129890999 | 56.35755 | 90.81356 | 68.14728 | 28.99438 | 0.769305 | 1.817428 | 0.183514 |
| 1 | 173495000 | 173496599 | 24.24084 | 42.21401 | 32.21585 | 14.165 | 0.775231 | 0.776007 | 0.183515 |
| 1 | 75161200 | 75163599 | 9.882941 | 17.26673 | 48.34179 | 21.29899 | -1.08952 | 0.686634 | 0.183519 |
| 12 | 20187800 | 20197599 | 334.1944 | 475.4219 | 374.7978 | 154.9774 | 0.53613 | 4.326004 | 0.183542 |
| 15 | 3103200 | 3106799 | 45.38866 | 73.88908 | 58.2835 | 25.02186 | 0.711644 | 1.569002 | 0.183564 |
| 18 | 24153400 | 24156599 | 16.69995 | 28.39618 | 60.68762 | 26.59801 | -0.70981 | 1.013555 | 0.183576 |
| X | 33275200 | 33276199 | 10.71946 | 20.9527 | 9.995824 | 3.445034 | 1.74644 | -0.52952 | 0.183577 |
| 11 | 114286000 | 114287599 | 10.33502 | 18.45282 | 42.03378 | 18.47181 | -0.78291 | 0.436171 | 0.183596 |
| 4 | 112758400 | 112759599 | 21.56415 | 38.09005 | 25.46955 | 11.26776 | 0.967173 | 0.557713 | 0.183597 |
| 8 | 13009600 | 13023199 | 138.5853 | 205.5715 | 365.7488 | 150.6491 | -0.57897 | 3.672439 | 0.183607 |
| 17 | 94160600 | 94161799 | 15.96876 | 30.03831 | 18.81839 | 7.717843 | 1.138201 | 0.122725 | 0.183613 |
| 9 | 105085800 | 105086599 | 5.249936 | 10.16478 | 27.24714 | 10.36163 | -0.8411 | -0.20572 | 0.183629 |
| 14 | 13161000 | 13162599 | 19.90717 | 35.3745 | 25.68365 | 11.33218 | 0.853065 | 0.495452 | 0.183632 |
| 8 | 118954200 | 118956799 | 29.93088 | 51.04569 | 38.16965 | 16.73291 | 0.799954 | 1.009786 | 0.183637 |
| 3 | 20861400 | 20863999 | 40.00945 | 65.53519 | 54.87406 | 23.61211 | 0.631584 | 1.468387 | 0.183639 |
| 9 | 27037800 | 27039599 | 19.50977 | 32.71305 | 67.24138 | 29.33354 | -0.65487 | 1.156465 | 0.183646 |
| 16 | 21547800 | 21550599 | 15.2405 | 26.15022 | 56.37223 | 24.72469 | -0.71986 | 0.92171 | 0.18365 |
| 9 | 67043600 | 67045999 | 10.08074 | 17.86085 | 42.47369 | 18.77891 | -0.85083 | 0.487545 | 0.183662 |
| 17 | 19512000 | 19514599 | 32.80424 | 56.46922 | 30.79829 | 13.50987 | 1.255081 | 0.932093 | 0.183668 |
| 6 | 22739600 | 22740999 | 17.93572 | 32.47756 | 22.62708 | 9.801218 | 0.929278 | 0.362885 | 0.183674 |
| X | 107962400 | 107963399 | 21.8559 | 38.60917 | 23.74495 | 10.50459 | 1.087947 | 0.559447 | 0.183676 |
| 6 | 117352000 | 117353399 | 15.90951 | 29.42899 | 21.43282 | 9.028377 | 0.893079 | 0.227774 | 0.183688 |
| 5 | 33431000 | 33433599 | 12.18554 | 21.20407 | 51.04611 | 22.43362 | -0.87428 | 0.757998 | 0.183693 |
| X | 31745200 | 31746399 | 16.2452 | 29.7435 | 22.62645 | 9.665038 | 0.816785 | 0.290585 | 0.183699 |
| 5 | 119693400 | 119700199 | 40.36498 | 64.3976 | 130.2104 | 54.59806 | -0.67471 | 2.128053 | 0.183702 |
| 13 | 94824200 | 94826399 | 17.67422 | 29.50347 | 68.50442 | 29.80059 | -0.82983 | 1.201306 | 0.183705 |
| 11 | 96055800 | 96067199 | 85.59654 | 130.7202 | 265.2709 | 110.0202 | -0.73187 | 3.092708 | 0.183709 |
| 12 | 44246200 | 44247599 | 17.54748 | 31.57448 | 24.53649 | 10.71229 | 0.766021 | 0.401811 | 0.183716 |
| 8 | 33710000 | 33713999 | 33.09391 | 53.33417 | 103.9624 | 44.24112 | -0.6046 | 1.795054 | 0.18372 |
| 18 | 23951400 | 23952599 | 5.84942 | 11.25058 | 29.8409 | 11.63319 | -0.86099 | -0.10982 | 0.183722 |
| 4 | 134698000 | 134707999 | 79.52909 | 122.3591 | 226.0104 | 93.86223 | -0.58792 | 2.940754 | 0.183727 |
| 15 | 43208200 | 43209799 | 22.1071 | 38.86309 | 28.34258 | 12.55077 | 0.840518 | 0.618664 | 0.183735 |
| 18 | 81040000 | 81042599 | 11.86676 | 20.71559 | 49.41306 | 21.75969 | -0.8601 | 0.71181 | 0.183744 |

| | | | | | | | | | |
|---|---|---|---|---|---|---|---|---|---|
| 3 | 70552200 | 70552999 | 12.42607 | 23.89385 | 17.78704 | 6.939136 | 0.954909 | -0.073 | 0.183746 |
| 17 | 55082000 | 55082999 | 12.42607 | 23.89385 | 17.78704 | 6.939136 | 0.954909 | -0.073 | 0.183746 |
| 6 | 127221000 | 127221599 | 11.30521 | 21.76594 | 17.94443 | 6.975291 | 0.813884 | -0.09561 | 0.183751 |
| 11 | 84069800 | 84071199 | 4.556671 | 8.570915 | 34.46624 | 14.13674 | -1.52084 | 0.061365 | 0.183753 |
| 1 | 49426000 | 49426399 | 8.562009 | 16.69626 | 14.4921 | 5.210046 | 0.843387 | -0.40265 | 0.183779 |
| 17 | 8089000 | 8097999 | 46.19454 | 73.15777 | 158.5444 | 65.92862 | -0.78544 | 2.361602 | 0.183788 |
| 17 | 33542600 | 33543999 | 25.47275 | 44.23921 | 32.99141 | 14.48131 | 0.808307 | 0.823117 | 0.183792 |
| 17 | 43935200 | 43943999 | 114.4978 | 172.1409 | 309.0876 | 127.6116 | -0.57449 | 3.418734 | 0.183806 |
| 6 | 118168200 | 118175599 | 32.38175 | 51.69048 | 138.8914 | 58.28453 | -1.08316 | 2.10397 | 0.183816 |
| 10 | 81933200 | 81940399 | 252.7256 | 370.79 | 279.3303 | 115.0499 | 0.645313 | 3.862288 | 0.183818 |
| 18 | 37594600 | 37596799 | 38.19391 | 63.19211 | 49.37473 | 21.35101 | 0.735494 | 1.340903 | 0.183838 |
| 7 | 8284800 | 8287799 | 62.11789 | 99.94878 | 66.80353 | 28.36223 | 0.933362 | 1.870404 | 0.183853 |
| 4 | 151684400 | 151708399 | 171.5762 | 249.7858 | 451.0893 | 185.9113 | -0.62615 | 3.977367 | 0.183857 |
| 1 | 116205600 | 116206199 | 10.29949 | 20.01343 | 15.98131 | 5.954722 | 0.916568 | -0.25152 | 0.183873 |
| 13 | 117160800 | 117164399 | 18.29129 | 30.55291 | 70.98216 | 30.89081 | -0.8309 | 1.195936 | 0.183889 |
| 7 | 32999000 | 33001999 | 74.10487 | 117.6468 | 100.472 | 41.87083 | 0.559295 | 2.303319 | 0.183897 |
| 7 | 31296000 | 31297399 | 39.65442 | 65.19784 | 48.29575 | 20.8145 | 0.810012 | 1.419995 | 0.183897 |
| 13 | 53162600 | 53165799 | 12.93024 | 22.53005 | 51.60696 | 22.69918 | -0.80288 | 0.742146 | 0.183928 |
| 10 | 69895800 | 69896999 | 7.567592 | 14.14038 | 33.32988 | 13.82482 | -0.7716 | 0.128836 | 0.183931 |
| 4 | 145576600 | 145581399 | 201.8846 | 299.5787 | 246.1556 | 101.3934 | 0.535436 | 3.669341 | 0.183953 |
| 7 | 85077200 | 85079399 | 28.11395 | 48.64674 | 33.20216 | 14.56665 | 0.934492 | 0.869231 | 0.18397 |
| 4 | 153288800 | 153296799 | 62.58856 | 97.26508 | 200.9348 | 83.42906 | -0.73798 | 2.747833 | 0.183971 |
| 11 | 55101800 | 55103599 | 12.46432 | 21.75849 | 48.90498 | 21.53579 | -0.77517 | 0.710701 | 0.183975 |
| 10 | 72939600 | 72942999 | 23.74966 | 41.39131 | 32.3631 | 14.2368 | 0.740661 | 0.764458 | 0.183978 |
| X | 93240000 | 93242199 | 17.75625 | 33.08132 | 19.2193 | 8.011559 | 1.227992 | 0.198596 | 0.183979 |
| 10 | 59289000 | 59291599 | 14.10222 | 24.35872 | 53.71328 | 23.56414 | -0.75024 | 0.853301 | 0.183997 |
| 10 | 42182600 | 42186199 | 20.01446 | 33.02379 | 77.28079 | 33.35652 | -0.84247 | 1.347435 | 0.183998 |
| 7 | 109350800 | 109352199 | 5.511233 | 10.42328 | 33.64668 | 13.65724 | -1.19519 | 0.029237 | 0.184005 |
| 19 | 54862600 | 54863399 | 10.97861 | 21.15381 | 16.78926 | 6.511491 | 0.870571 | -0.11122 | 0.184006 |
| 8 | 110458000 | 110460599 | 12.50839 | 21.30076 | 59.82414 | 26.22272 | -1.10115 | 0.98639 | 0.184008 |
| 17 | 23036400 | 23037799 | 7.38893 | 13.9282 | 34.20553 | 13.97881 | -0.81314 | 0.072496 | 0.184013 |
| 17 | 46750000 | 46752399 | 20.72914 | 34.42418 | 72.52925 | 31.44422 | -0.6906 | 1.272168 | 0.184016 |
| 5 | 102979600 | 102979999 | 2.6373 | 5.158191 | 18.93818 | 6.395011 | -1.12357 | -0.68674 | 0.184031 |
| 1 | 34339800 | 34342799 | 19.30159 | 32.05215 | 73.04969 | 31.66893 | -0.80357 | 1.266852 | 0.184033 |
| 12 | 96259200 | 96260799 | 7.722051 | 14.3898 | 34.37213 | 14.32295 | -0.79632 | 0.149084 | 0.184056 |
| 2 | 73044200 | 73045199 | 6.354182 | 12.0069 | 32.53106 | 13.22914 | -0.94731 | 0.040986 | 0.184062 |
| 2 | 161302400 | 161304599 | 24.65783 | 42.95451 | 32.21227 | 14.16639 | 0.800466 | 0.774244 | 0.18407 |
| X | 66762200 | 66762599 | 11.14421 | 21.58564 | 15.49397 | 5.881229 | 1.044744 | -0.18042 | 0.184084 |
| 2 | 20980400 | 20981999 | 17.34867 | 31.43405 | 23.29327 | 10.08195 | 0.841737 | 0.356949 | 0.184099 |
| 2 | 94194400 | 94196799 | 17.34867 | 31.43405 | 23.29327 | 10.08195 | 0.841737 | 0.356949 | 0.184099 |
| 12 | 12890600 | 12894199 | 19.09241 | 31.8434 | 69.2373 | 30.10305 | -0.73556 | 1.214613 | 0.184111 |
| 4 | 150327600 | 150333799 | 34.17097 | 54.74613 | 117.9294 | 49.74852 | -0.75891 | 1.987608 | 0.184113 |
| 3 | 90827800 | 90828399 | 11.03593 | 19.45834 | 43.53443 | 19.26662 | -0.76604 | 0.551706 | 0.18412 |
| 19 | 20174600 | 20175199 | 8.969219 | 17.45205 | 15.00381 | 5.529916 | 0.825032 | -0.30967 | 0.184145 |
| 5 | 134782200 | 134786399 | 22.35329 | 36.50668 | 83.94351 | 36.08035 | -0.82427 | 1.492254 | 0.184151 |
| 14 | 59686200 | 59687799 | 21.12192 | 37.23032 | 27.8695 | 12.3347 | 0.804478 | 0.58353 | 0.184152 |
| 17 | 41582800 | 41586599 | 36.79997 | 61.6265 | 40.90635 | 17.83086 | 0.970078 | 1.196991 | 0.184162 |

| | | | | | | | | | |
|---|---|---|---|---|---|---|---|---|---|
| 6 | 45810200 | 45811599 | 12.15467 | 21.32113 | 47.23795 | 20.88264 | -0.75409 | 0.631721 | 0.184169 |
| 10 | 72531000 | 72534199 | 22.88916 | 40.17825 | 27.64703 | 12.22901 | 0.923714 | 0.642919 | 0.184181 |
| 8 | 36568400 | 36573199 | 24.83415 | 40.30376 | 91.08048 | 39.03545 | -0.80647 | 1.601429 | 0.184194 |
| 5 | 47915200 | 47915999 | 9.187851 | 16.71413 | 37.60747 | 16.19636 | -0.74532 | 0.308036 | 0.184216 |
| 11 | 23202600 | 23207799 | 30.22047 | 48.67827 | 105.4376 | 44.8355 | -0.75724 | 1.807211 | 0.184216 |
| Y | 10611200 | 10614199 | 15.54499 | 29.01436 | 9.587136 | 3.984085 | 2.034502 | 0.138602 | 0.184234 |
| 7 | 24391200 | 24394999 | 16.51025 | 27.74618 | 64.75186 | 28.28396 | -0.83704 | 1.122117 | 0.184238 |
| 5 | 118171400 | 118181199 | 76.95202 | 118.6267 | 217.3297 | 90.27511 | -0.57393 | 2.901701 | 0.184243 |
| 12 | 23795000 | 23811399 | 568.7981 | 773.0674 | 573.1286 | 238.7113 | 0.558441 | 5.021988 | 0.184249 |
| 14 | 65750000 | 65751399 | 12.17544 | 23.08706 | 18.85376 | 7.610829 | 0.778271 | 0.035201 | 0.184251 |
| 18 | 77152600 | 77153599 | 5.100277 | 9.853667 | 27.02106 | 10.38011 | -0.88681 | -0.17121 | 0.184259 |
| 17 | 65919400 | 65921199 | 10.55619 | 18.32516 | 53.10879 | 23.30523 | -1.14065 | 0.788488 | 0.184263 |
| 1 | 130990400 | 130993199 | 12.50173 | 21.72163 | 52.12647 | 22.8868 | -0.87005 | 0.782405 | 0.184268 |
| 2 | 43182000 | 43182799 | 6.01984 | 11.50757 | 32.47946 | 12.88673 | -0.9744 | -0.04464 | 0.184272 |
| 1 | 132037600 | 132040399 | 12.21814 | 21.18547 | 52.63356 | 23.09316 | -0.9202 | 0.806905 | 0.184277 |
| 11 | 67388800 | 67389999 | 7.078725 | 13.39465 | 36.78387 | 14.91458 | -0.96332 | 0.048562 | 0.184279 |
| 10 | 30249200 | 30252399 | 22.47988 | 37.30448 | 79.24459 | 34.33836 | -0.70273 | 1.288514 | 0.18428 |
| 4 | 8754000 | 8755199 | 9.410685 | 16.87246 | 38.95838 | 17.06041 | -0.79827 | 0.399617 | 0.18429 |
| 5 | 61796200 | 61798199 | 21.95289 | 38.47519 | 29.48925 | 13.02683 | 0.76919 | 0.66538 | 0.184291 |
| 1 | 97798000 | 97799599 | 18.07494 | 32.9972 | 22.35551 | 9.585482 | 0.979123 | 0.320056 | 0.184291 |
| 15 | 34082400 | 34084599 | 11.52064 | 20.18405 | 47.26067 | 20.87093 | -0.83359 | 0.645721 | 0.184295 |
| 17 | 16045600 | 16047599 | 17.25755 | 29.08027 | 64.39591 | 28.15147 | -0.76161 | 1.098954 | 0.184295 |
| 4 | 147408200 | 147409399 | 11.37494 | 22.02551 | 16.21428 | 6.163849 | 1.00503 | -0.17323 | 0.184306 |
| 11 | 17102600 | 17102999 | 29.83717 | 50.75624 | 32.46625 | 14.22972 | 1.023948 | 1.027397 | 0.184307 |
| 6 | 83078200 | 83080799 | 12.70135 | 21.59168 | 60.90079 | 26.6909 | -1.10787 | 1.003274 | 0.18432 |
| 8 | 85638200 | 85643599 | 46.1482 | 73.20174 | 149.4532 | 62.23074 | -0.69736 | 2.316111 | 0.18437 |
| 12 | 69012600 | 69013799 | 14.99744 | 28.3231 | 18.30239 | 7.451866 | 1.102005 | 0.086888 | 0.184385 |
| 6 | 141092000 | 141092999 | 15.60187 | 29.29149 | 19.3767 | 7.972875 | 1.056594 | 0.132483 | 0.184393 |
| 6 | 42595200 | 42595999 | 11.49984 | 22.44353 | 10.32312 | 3.666538 | 1.759591 | -0.41813 | 0.1844 |
| 11 | 3091200 | 3105999 | 663.0089 | 894.9473 | 575.5282 | 239.9636 | 0.753609 | 5.131625 | 0.184405 |
| X | 3072800 | 3078399 | 57.47483 | 92.98083 | 56.92257 | 24.33888 | 1.068639 | 1.707772 | 0.184436 |
| 6 | 139985400 | 139990999 | 40.44421 | 64.74669 | 128.152 | 53.99768 | -0.63854 | 2.020571 | 0.18444 |
| 4 | 63398400 | 63401399 | 13.32861 | 22.98812 | 56.9573 | 24.98616 | -0.91835 | 0.864215 | 0.184444 |
| 4 | 21674000 | 21674999 | 8.21693 | 15.15396 | 35.20735 | 14.89138 | -0.77274 | 0.216507 | 0.184444 |
| 17 | 47747200 | 47752999 | 26.066 | 42.01187 | 103.7367 | 44.15032 | -0.94452 | 1.782809 | 0.18445 |
| 10 | 17509200 | 17509999 | 5.123744 | 9.898668 | 27.64186 | 10.62027 | -0.91375 | -0.17104 | 0.184451 |
| 2 | 103952000 | 103953799 | 9.604742 | 17.08872 | 40.45097 | 17.82903 | -0.84054 | 0.452383 | 0.184473 |
| 2 | 71730200 | 71731399 | 6.295409 | 12.05655 | 30.18189 | 11.91193 | -0.79479 | -0.06172 | 0.184486 |
| 3 | 124183400 | 124184599 | 6.295409 | 12.05655 | 30.18189 | 11.91193 | -0.79479 | -0.06172 | 0.184486 |
| 6 | 110428200 | 110429399 | 12.85572 | 24.90149 | 14.88487 | 5.649221 | 1.304396 | -0.16532 | 0.184489 |
| 5 | 71944400 | 71946399 | 24.24071 | 42.51452 | 28.21992 | 12.47345 | 0.975668 | 0.66635 | 0.184499 |
| 4 | 29799200 | 29801999 | 32.58467 | 55.9769 | 32.57137 | 14.28676 | 1.161738 | 0.951732 | 0.18452 |
| 12 | 107586200 | 107588599 | 12.92124 | 22.6466 | 51.01597 | 22.53665 | -0.77839 | 0.653103 | 0.18452 |
| 7 | 7268200 | 7269199 | 13.27809 | 25.18682 | 19.16041 | 7.728825 | 0.880472 | 0.043842 | 0.184529 |
| 14 | 104681800 | 104682999 | 13.27809 | 25.18682 | 19.16041 | 7.728825 | 0.880472 | 0.043842 | 0.184529 |
| 10 | 94350000 | 94351599 | 7.213021 | 12.9369 | 42.12299 | 18.44066 | -1.29026 | 0.386769 | 0.184555 |
| X | 103163400 | 103164599 | 6.442887 | 12.33373 | 31.39012 | 12.40398 | -0.82063 | -0.05479 | 0.184561 |

| | | | | | | | | | |
|---|---|---|---|---|---|---|---|---|---|
| 14 | 98505400 | 98506199 | 11.11666 | 21.68695 | 12.35253 | 4.416675 | 1.449759 | -0.39372 | 0.184572 |
| 1 | 15087600 | 15090999 | 24.58095 | 40.12164 | 86.88624 | 37.33733 | -0.73881 | 1.50492 | 0.184578 |
| 12 | 112663600 | 112668399 | 27.97645 | 45.10011 | 103.3153 | 43.98239 | -0.83635 | 1.778569 | 0.184583 |
| 5 | 134573000 | 134581799 | 43.64058 | 69.07992 | 155.7812 | 64.76384 | -0.84322 | 2.371724 | 0.18459 |
| 5 | 141225400 | 141229799 | 27.12521 | 43.8338 | 96.78887 | 41.33286 | -0.77927 | 1.704033 | 0.18459 |
| 11 | 102709200 | 102710799 | 8.75036 | 15.86794 | 37.02586 | 16.00916 | -0.8009 | 0.322885 | 0.184603 |
| 10 | 22259400 | 22259599 | 8.491812 | 16.57462 | 13.01078 | 4.613072 | 1.005594 | -0.45503 | 0.184616 |
| 4 | 106705200 | 106706799 | 8.561105 | 15.66513 | 36.11726 | 15.44045 | -0.77289 | 0.267217 | 0.184621 |
| 13 | 25333000 | 25334799 | 8.561105 | 15.66513 | 36.11726 | 15.44045 | -0.77289 | 0.267217 | 0.184621 |
| 11 | 20413200 | 20416599 | 16.78428 | 27.96703 | 74.06745 | 32.18368 | -1.01909 | 1.220791 | 0.184636 |
| 9 | 65185000 | 65187399 | 18.28943 | 30.5588 | 68.08812 | 29.63686 | -0.77064 | 1.192 | 0.184638 |
| 4 | 54626600 | 54628199 | 14.94393 | 28.09266 | 18.80523 | 7.720759 | 1.041698 | 0.11628 | 0.184641 |
| 14 | 88882400 | 88883799 | 13.02701 | 24.83911 | 18.37648 | 7.331698 | 0.934847 | 0.006571 | 0.184651 |
| X | 13145200 | 13147999 | 12.64741 | 22.13625 | 52.63076 | 23.22304 | -0.85566 | 0.678619 | 0.184667 |
| 10 | 94750800 | 94752399 | 10.72051 | 18.88754 | 45.09113 | 19.95866 | -0.85993 | 0.558161 | 0.184671 |
| 17 | 71317600 | 71319599 | 10.72051 | 18.88754 | 45.09113 | 19.95866 | -0.85993 | 0.558161 | 0.184671 |
| 18 | 56609000 | 56610799 | 10.39166 | 18.40882 | 41.44828 | 18.32613 | -0.7723 | 0.491447 | 0.184672 |
| 17 | 38695800 | 38696599 | 28.05164 | 47.81881 | 35.92492 | 15.74828 | 0.793241 | 1.005602 | 0.184696 |
| 18 | 7273200 | 7274999 | 11.46924 | 20.28508 | 45.08909 | 19.9423 | -0.7553 | 0.520468 | 0.184696 |
| 13 | 41418200 | 41419999 | 29.84367 | 51.23497 | 34.78231 | 15.25609 | 0.939995 | 0.946464 | 0.1847 |
| 10 | 76397000 | 76400199 | 19.6465 | 32.54592 | 74.65913 | 32.31254 | -0.81324 | 1.29745 | 0.184708 |
| 7 | 113659800 | 113661199 | 5.561496 | 10.6449 | 28.91873 | 11.43222 | -0.91295 | -0.06532 | 0.184711 |
| 16 | 89699000 | 89701999 | 23.26283 | 38.10373 | 81.70516 | 35.15719 | -0.72116 | 1.450352 | 0.184714 |
| 4 | 151611400 | 151616399 | 47.1376 | 74.74564 | 147.8036 | 61.52285 | -0.65178 | 2.327404 | 0.184728 |
| 2 | 131333400 | 131334999 | 6.88087 | 12.72576 | 34.1637 | 14.39467 | -0.97507 | 0.187859 | 0.184735 |
| 16 | 31917800 | 31918799 | 6.88087 | 12.72576 | 34.1637 | 14.39467 | -0.97507 | 0.187859 | 0.184735 |
| 6 | 142325000 | 142327799 | 12.9738 | 22.50238 | 52.66842 | 23.10867 | -0.83463 | 0.807156 | 0.184742 |
| 2 | 129486800 | 129489199 | 17.00787 | 31.17186 | 22.42182 | 9.56427 | 0.898277 | 0.289808 | 0.184752 |
| 8 | 23046600 | 23060399 | 91.76988 | 139.4534 | 271.1723 | 112.316 | -0.67616 | 3.196628 | 0.184752 |
| 7 | 12009800 | 12013399 | 57.76742 | 93.2747 | 61.54011 | 26.25248 | 0.957891 | 1.759841 | 0.184768 |
| 1 | 82326400 | 82328799 | 12.77086 | 22.26365 | 57.11193 | 25.12856 | -0.96569 | 0.73994 | 0.184768 |
| 13 | 40876200 | 40877199 | 7.915782 | 14.63994 | 34.71357 | 14.62611 | -0.79748 | 0.198704 | 0.184773 |
| 8 | 119935800 | 119940999 | 44.75847 | 71.26974 | 137.2109 | 57.38057 | -0.60717 | 2.195794 | 0.184779 |
| 9 | 5279200 | 5285199 | 87.15507 | 137.0853 | 116.5578 | 48.35461 | 0.55471 | 2.533172 | 0.184783 |
| 11 | 117640200 | 117642799 | 14.61348 | 25.0479 | 56.89756 | 24.95298 | -0.79535 | 0.927889 | 0.184795 |
| 12 | 23073400 | 23078999 | 157.9048 | 238.2905 | 191.9636 | 79.33775 | 0.585713 | 3.347175 | 0.184795 |
| 7 | 15231800 | 15246999 | 262.0411 | 383.0989 | 288.8302 | 118.9891 | 0.639325 | 3.918714 | 0.184801 |
| 4 | 148896200 | 148908599 | 90.0711 | 136.8201 | 268.7319 | 111.2926 | -0.69104 | 3.204354 | 0.184804 |
| 12 | 18493200 | 18510599 | 501.9934 | 690.527 | 521.8466 | 216.9435 | 0.548291 | 4.833792 | 0.184809 |
| 9 | 61453400 | 61457199 | 30.34911 | 48.96766 | 100.013 | 42.62448 | -0.66976 | 1.75565 | 0.184825 |
| 7 | 115373600 | 115374199 | 4.923087 | 9.569318 | 26.39776 | 9.810786 | -0.84988 | -0.28975 | 0.184833 |
| 7 | 103867800 | 103869399 | 8.422077 | 15.34892 | 36.57725 | 15.71694 | -0.8253 | 0.289196 | 0.184856 |
| X | 66665200 | 66668199 | 32.16115 | 54.51468 | 38.85768 | 17.01754 | 0.868069 | 1.074678 | 0.184862 |
| X | 109548200 | 109551799 | 59.82858 | 96.15247 | 72.95049 | 30.91943 | 0.748906 | 1.907411 | 0.184876 |
| 1 | 87122600 | 87123599 | 5.840837 | 11.2366 | 28.62271 | 11.15109 | -0.80161 | -0.11291 | 0.184879 |
| 8 | 13539600 | 13546399 | 36.48437 | 58.27853 | 127.2731 | 53.45984 | -0.78359 | 2.085089 | 0.184881 |
| 3 | 145608600 | 145611599 | 21.75139 | 35.86019 | 76.88496 | 33.17412 | -0.71713 | 1.362154 | 0.184882 |

| | | | | | | | | | |
|---|---|---|---|---|---|---|---|---|---|
| 9 | 107182800 | 107187399 | 18.91948 | 31.21843 | 77.05812 | 33.26103 | -0.91903 | 1.344547 | 0.184883 |
| 18 | 77636200 | 77646999 | 73.70655 | 113.2433 | 230.2362 | 95.599 | -0.72771 | 2.96671 | 0.184899 |
| 17 | 38591600 | 38592999 | 17.19032 | 32.52511 | 16.50463 | 6.693407 | 1.452761 | 0.091194 | 0.184899 |
| 6 | 67917800 | 67919599 | 40.08237 | 65.6179 | 54.81444 | 23.58087 | 0.634681 | 1.476384 | 0.1849 |
| 12 | 19948400 | 19950399 | 44.96331 | 73.25601 | 53.27882 | 22.88005 | 0.829206 | 1.553005 | 0.184914 |
| 9 | 47614400 | 47615399 | 5.677802 | 10.96139 | 28.79375 | 11.09906 | -0.83083 | -0.14854 | 0.184917 |
| X | 131117800 | 131121399 | 37.82395 | 63.58058 | 38.72632 | 16.91714 | 1.093707 | 1.1642 | 0.184917 |
| 8 | 30351200 | 30353199 | 21.28754 | 37.89906 | 24.48822 | 10.7875 | 1.021839 | 0.491215 | 0.184921 |
| 8 | 104373200 | 104376599 | 29.11691 | 49.39209 | 40.45629 | 17.72173 | 0.668204 | 1.056226 | 0.184928 |
| 1 | 71700200 | 71705599 | 64.95612 | 104.0382 | 79.56295 | 33.54915 | 0.731691 | 2.018613 | 0.184937 |
| 19 | 47478800 | 47482199 | 17.05294 | 28.8562 | 63.95543 | 27.99448 | -0.76264 | 1.06016 | 0.184945 |
| 9 | 84006400 | 84008199 | 16.0747 | 29.37556 | 22.67436 | 9.709806 | 0.793453 | 0.299898 | 0.18495 |
| 12 | 90725200 | 90726599 | 14.67638 | 27.84961 | 18.50231 | 7.45746 | 1.075405 | 0.053486 | 0.18496 |
| 7 | 44564200 | 44567399 | 12.52358 | 21.71064 | 52.10263 | 22.86007 | -0.87052 | 0.810157 | 0.184981 |
| 4 | 142938200 | 142939999 | 9.768255 | 17.34141 | 41.45589 | 18.30592 | -0.85672 | 0.468558 | 0.184985 |
| 7 | 13417800 | 13422199 | 61.2128 | 98.84784 | 57.66177 | 24.60123 | 1.135252 | 1.762361 | 0.184985 |
| 7 | 11995200 | 11996399 | 20.63179 | 37.45631 | 20.8274 | 8.992514 | 1.255512 | 0.36596 | 0.184986 |
| 11 | 67257800 | 67258799 | 5.378191 | 10.38591 | 28.17022 | 10.84437 | -0.87421 | -0.15693 | 0.184992 |
| 5 | 36064800 | 36068799 | 16.886 | 28.63545 | 61.18056 | 26.7974 | -0.70966 | 1.039258 | 0.185005 |
| 7 | 84408400 | 84409999 | 17.96123 | 32.06225 | 25.48723 | 11.20244 | 0.72686 | 0.454592 | 0.185012 |
| 19 | 46575800 | 46581799 | 31.67143 | 50.97408 | 107.9195 | 45.83698 | -0.72591 | 1.835621 | 0.185032 |
| 1 | 60749400 | 60749999 | 8.173766 | 15.97768 | 10.85685 | 3.736397 | 1.244597 | -0.57341 | 0.185037 |
| 14 | 11647000 | 11648799 | 11.15578 | 19.76299 | 42.97894 | 19.0028 | -0.72282 | 0.497563 | 0.185051 |
| 5 | 104193600 | 104195799 | 10.61446 | 18.60509 | 47.2525 | 20.87554 | -0.95007 | 0.633277 | 0.185054 |
| 8 | 78508200 | 78509599 | 7.962159 | 14.55654 | 35.57378 | 15.22563 | -0.85724 | 0.266588 | 0.185054 |
| 3 | 136421000 | 136422399 | 8.539542 | 15.66311 | 36.17106 | 15.41317 | -0.77204 | 0.251956 | 0.185055 |
| 7 | 61132200 | 61133999 | 20.6731 | 36.8985 | 24.6086 | 10.8174 | 0.978576 | 0.4724 | 0.185059 |
| 6 | 145691800 | 145693399 | 13.86116 | 23.72693 | 58.4678 | 25.63888 | -0.91257 | 0.938471 | 0.185066 |
| 14 | 79352200 | 79354199 | 8.915961 | 16.12682 | 37.51224 | 16.27158 | -0.79942 | 0.340605 | 0.185071 |
| 15 | 59186400 | 59187999 | 7.003237 | 12.67803 | 38.58812 | 16.72032 | -1.18329 | 0.322371 | 0.185076 |
| 6 | 136976400 | 136981999 | 52.23772 | 82.65963 | 154.0679 | 64.04022 | -0.56925 | 2.386765 | 0.185091 |
| 7 | 31003200 | 31007599 | 76.86262 | 121.8804 | 103.7598 | 43.18975 | 0.56219 | 2.339996 | 0.1851 |
| 17 | 17205600 | 17213999 | 220.9817 | 328.5402 | 235.1285 | 96.86998 | 0.737092 | 3.639049 | 0.185103 |
| 3 | 3114400 | 3116999 | 43.30008 | 71.47831 | 46.69684 | 20.16148 | 0.992644 | 1.377757 | 0.185104 |
| 7 | 37408400 | 37412399 | 20.75739 | 34.34788 | 74.73409 | 32.3201 | -0.73737 | 1.31376 | 0.185109 |
| 4 | 103474000 | 103480199 | 42.28876 | 67.46089 | 133.6747 | 56.04393 | -0.64579 | 2.132899 | 0.18511 |
| 6 | 88021400 | 88022799 | 7.388375 | 13.89464 | 34.71226 | 14.24215 | -0.84255 | 0.087283 | 0.185125 |
| 18 | 33051600 | 33052599 | 7.388375 | 13.89464 | 34.71226 | 14.24215 | -0.84255 | 0.087283 | 0.185125 |
| 10 | 56828600 | 56830799 | 22.03172 | 38.84525 | 27.3118 | 12.08717 | 0.894642 | 0.585649 | 0.185127 |
| 10 | 67240600 | 67242599 | 12.14277 | 21.3527 | 47.7307 | 21.13503 | -0.76671 | 0.594022 | 0.185128 |
| X | 33188800 | 33191199 | 25.86642 | 45.15356 | 30.08358 | 13.24751 | 0.970668 | 0.750528 | 0.185145 |
| 2 | 165033000 | 165040999 | 73.52725 | 113.3849 | 218.5558 | 90.78424 | -0.6468 | 2.89391 | 0.185153 |
| 14 | 48878800 | 48884399 | 57.0358 | 91.61821 | 74.1066 | 31.38664 | 0.655779 | 1.915996 | 0.185162 |
| 6 | 52236200 | 52243599 | 55.33615 | 86.9532 | 175.3131 | 72.72039 | -0.69039 | 2.542215 | 0.185162 |
| 17 | 50849200 | 50851399 | 25.36914 | 44.35545 | 29.51247 | 13.01054 | 0.972627 | 0.72432 | 0.185165 |
| 7 | 12085600 | 12087799 | 34.75028 | 58.45293 | 40.8941 | 17.86773 | 0.894488 | 1.147878 | 0.185187 |
| 5 | 38056800 | 38068599 | 95.47399 | 145.5131 | 259.0989 | 107.4107 | -0.5457 | 3.136377 | 0.185188 |

| | | | | | | | | | |
|---|---|---|---|---|---|---|---|---|---|
| 1 | 135006600 | 135007199 | 3.936538 | 7.680904 | 22.70265 | 8.096011 | -0.89238 | -0.46273 | 0.185204 |
| 6 | 149332600 | 149332999 | 3.936538 | 7.680904 | 22.70265 | 8.096011 | -0.89238 | -0.46273 | 0.185204 |
| 2 | 132319800 | 132320799 | 10.77442 | 20.89483 | 16.47876 | 6.219539 | 0.917406 | -0.20091 | 0.185218 |
| X | 75280200 | 75281199 | 10.77442 | 20.89483 | 16.47876 | 6.219539 | 0.917406 | -0.20091 | 0.185218 |
| 1 | 57597000 | 57597599 | 3.57358 | 6.978217 | 22.19596 | 7.80826 | -0.97687 | -0.51432 | 0.185227 |
| 13 | 14899400 | 14901399 | 29.15762 | 50.66199 | 29.15184 | 12.79964 | 1.180256 | 0.82658 | 0.185232 |
| 3 | 58394600 | 58395199 | 10.65271 | 20.74282 | 14.82736 | 5.432871 | 1.09637 | -0.30785 | 0.185232 |
| 13 | 38516600 | 38518799 | 10.01101 | 18.03373 | 42.76096 | 18.64324 | -0.83243 | 0.381286 | 0.185239 |
| 10 | 33651400 | 33652799 | 12.32811 | 23.8049 | 15.37545 | 5.921376 | 1.1736 | -0.12335 | 0.185239 |
| 4 | 64689000 | 64689999 | 8.844852 | 16.19163 | 39.42874 | 16.84578 | -0.85128 | 0.270783 | 0.185247 |
| 16 | 10879200 | 10886399 | 32.99281 | 52.99534 | 111.9884 | 47.42829 | -0.72681 | 1.902312 | 0.185249 |
| 16 | 97543600 | 97545399 | 8.448912 | 15.3187 | 39.14346 | 16.92834 | -0.93172 | 0.323358 | 0.185262 |
| 14 | 72626400 | 72627599 | 7.431355 | 13.8578 | 33.25947 | 13.84288 | -0.80116 | 0.141885 | 0.18527 |
| 18 | 13645200 | 13646599 | 18.01338 | 33.21132 | 20.71112 | 8.763475 | 1.11091 | 0.261691 | 0.185292 |
| 6 | 22173200 | 22174599 | 28.11771 | 47.97731 | 38.81043 | 17.0145 | 0.687061 | 0.998989 | 0.185302 |
| 1 | 110921400 | 110923399 | 23.5038 | 41.92038 | 20.90025 | 9.192928 | 1.394452 | 0.487626 | 0.185307 |
| 17 | 3125600 | 3135799 | 845.7912 | 1112.589 | 773.8925 | 323.7929 | 0.603234 | 5.544252 | 0.18532 |
| X | 31706200 | 31712599 | 147.2175 | 224.6537 | 153.1157 | 63.49301 | 0.839938 | 3.119285 | 0.18533 |
| 1 | 132368600 | 132368999 | 11.92441 | 22.99094 | 16.94662 | 6.56044 | 0.98003 | -0.10355 | 0.185349 |
| 18 | 56091000 | 56092999 | 17.27033 | 32.20035 | 19.14447 | 7.97016 | 1.196051 | 0.18896 | 0.185349 |
| 7 | 28388600 | 28400999 | 103.6257 | 156.6944 | 290.2641 | 120.0412 | -0.6122 | 3.30083 | 0.185354 |
| 6 | 47097800 | 47100199 | 16.86848 | 28.2506 | 70.92949 | 30.91921 | -0.94241 | 1.162676 | 0.185368 |
| 4 | 142640400 | 142641799 | 8.033799 | 14.90981 | 37.64353 | 15.78598 | -0.88329 | 0.182706 | 0.18537 |
| 9 | 60195600 | 60200199 | 25.76126 | 41.86372 | 88.60963 | 38.01354 | -0.71049 | 1.575119 | 0.185379 |
| 8 | 95399800 | 95400799 | 3.532933 | 6.881538 | 23.48917 | 8.571129 | -1.12671 | -0.38154 | 0.18538 |
| 16 | 35676400 | 35677599 | 8.619649 | 15.56886 | 37.49661 | 16.29355 | -0.85093 | 0.346218 | 0.185388 |
| 16 | 34134800 | 34135999 | 20.67123 | 36.3193 | 28.68676 | 12.69904 | 0.726109 | 0.619437 | 0.185393 |
| 4 | 51099600 | 51101199 | 17.40988 | 31.58484 | 20.55904 | 8.884452 | 1.029219 | 0.345358 | 0.185406 |
| 9 | 16514800 | 16515999 | 13.71507 | 26.12722 | 17.99148 | 7.191951 | 1.033983 | 0.016159 | 0.185412 |
| 3 | 107501800 | 107505399 | 26.83516 | 43.3248 | 100.0763 | 42.6855 | -0.84575 | 1.731418 | 0.185419 |
| 16 | 53546800 | 53548599 | 16.48349 | 30.84496 | 18.61924 | 7.704031 | 1.180955 | 0.158107 | 0.18544 |
| 14 | 42282400 | 42289399 | 65.22152 | 104.8429 | 67.40065 | 28.57637 | 0.987736 | 1.905348 | 0.185442 |
| 10 | 95473600 | 95475599 | 17.94271 | 32.33268 | 24.33259 | 10.60424 | 0.813466 | 0.394994 | 0.185444 |
| X | 47682800 | 47684399 | 20.10832 | 36.53839 | 20.84798 | 8.991416 | 1.220239 | 0.359135 | 0.185449 |
| 12 | 86928200 | 86931999 | 17.90878 | 29.36125 | 82.42576 | 35.4812 | -1.10771 | 1.428298 | 0.18545 |
| 17 | 84982600 | 84986399 | 18.34884 | 30.9042 | 66.75856 | 29.18004 | -0.72616 | 1.108046 | 0.185489 |
| 7 | 29168800 | 29177199 | 46.50205 | 73.68794 | 152.3332 | 63.36529 | -0.71676 | 2.347177 | 0.18551 |
| 5 | 136617000 | 136618999 | 9.651046 | 17.05544 | 43.02156 | 19.03096 | -0.93673 | 0.515069 | 0.185515 |
| 7 | 135554400 | 135555599 | 7.109955 | 13.32966 | 32.29988 | 13.32211 | -0.80434 | 0.101045 | 0.185536 |
| 4 | 61554200 | 61556599 | 20.60856 | 36.74587 | 19.67505 | 8.656152 | 1.292665 | 0.468463 | 0.185553 |
| 9 | 61200600 | 61206999 | 37.61372 | 60.06614 | 126.6681 | 53.18516 | -0.73366 | 2.094987 | 0.185556 |
| 5 | 65675200 | 65677399 | 13.7218 | 23.9003 | 50.53708 | 22.22335 | -0.68801 | 0.751494 | 0.185592 |
| 14 | 109902400 | 109903799 | 7.705881 | 14.44766 | 35.16427 | 14.50224 | -0.81114 | 0.111743 | 0.185597 |
| 9 | 89697400 | 89698799 | 11.49717 | 20.24557 | 44.04284 | 19.49663 | -0.72674 | 0.568445 | 0.185602 |
| 14 | 33640400 | 33642199 | 7.640871 | 14.04314 | 35.1301 | 14.92849 | -0.88332 | 0.231611 | 0.185603 |
| 14 | 103586200 | 103587799 | 7.640871 | 14.04314 | 35.1301 | 14.92849 | -0.88332 | 0.231611 | 0.185603 |
| 13 | 17233800 | 17234799 | 7.225965 | 13.65721 | 33.75202 | 13.71932 | -0.81543 | 0.051148 | 0.185609 |

| | | | | | | | | | |
|---|---|---|---|---|---|---|---|---|---|
| 8 | 112395800 | 112399399 | 28.70806 | 46.4266 | 96.51974 | 41.25798 | -0.6914 | 1.682913 | 0.185612 |
| 19 | 40500400 | 40503599 | 19.93234 | 33.09037 | 71.92225 | 31.17418 | -0.73537 | 1.271907 | 0.185615 |
| 11 | 88931200 | 88931999 | 4.666218 | 9.039789 | 26.00764 | 9.863796 | -0.93713 | -0.22297 | 0.18562 |
| X | 31673600 | 31676199 | 39.21233 | 66.53515 | 31.24711 | 13.6888 | 1.467199 | 1.083003 | 0.185621 |
| 9 | 3291400 | 3294999 | 84.97764 | 133.7174 | 106.7345 | 44.28182 | 0.646161 | 2.520249 | 0.185648 |
| 5 | 113714400 | 113718999 | 27.63041 | 44.58754 | 99.93126 | 42.59757 | -0.80311 | 1.747435 | 0.18565 |
| 18 | 84209400 | 84214599 | 37.83641 | 60.67827 | 117.5463 | 49.66113 | -0.60451 | 1.96 | 0.185678 |
| 17 | 73946800 | 73948999 | 11.40101 | 19.90265 | 48.36509 | 21.29828 | -0.88691 | 0.70537 | 0.185679 |
| 8 | 28594000 | 28598999 | 52.4532 | 84.55425 | 69.95961 | 29.78063 | 0.629374 | 1.801554 | 0.185682 |
| 18 | 15476800 | 15479799 | 22.18441 | 36.66942 | 77.85551 | 33.64482 | -0.70243 | 1.331386 | 0.185706 |
| 17 | 20765800 | 20769799 | 62.1797 | 100.116 | 65.48386 | 27.82926 | 0.965717 | 1.851517 | 0.185713 |
| 2 | 76153200 | 76154399 | 9.672793 | 17.4988 | 39.0913 | 16.95282 | -0.74173 | 0.346919 | 0.185718 |
| 1 | 174612000 | 174615799 | 33.42292 | 55.71393 | 44.43868 | 19.31436 | 0.706375 | 1.240599 | 0.185735 |
| 5 | 130474000 | 130485199 | 82.58474 | 126.4737 | 242.6998 | 100.724 | -0.64808 | 3.034156 | 0.185751 |
| 18 | 82964400 | 82976599 | 72.74676 | 111.6448 | 233.0754 | 96.76316 | -0.76707 | 2.988898 | 0.185761 |
| 19 | 39776400 | 39778399 | 25.92197 | 45.16204 | 30.97834 | 13.62475 | 0.928434 | 0.776654 | 0.185765 |
| X | 24883600 | 24883999 | 7.609489 | 14.88002 | 9.955742 | 3.238766 | 1.34007 | -0.7709 | 0.185771 |
| 6 | 31333000 | 31334399 | 7.617007 | 14.06642 | 34.68589 | 14.64407 | -0.85594 | 0.202712 | 0.185782 |
| 13 | 54452200 | 54453799 | 7.617007 | 14.06642 | 34.68589 | 14.64407 | -0.85594 | 0.202712 | 0.185782 |
| 6 | 47412600 | 47414799 | 23.2399 | 40.62258 | 31.18196 | 13.74364 | 0.767347 | 0.718385 | 0.18579 |
| 15 | 102884200 | 102886999 | 14.51754 | 24.73229 | 58.91967 | 25.82711 | -0.86505 | 0.987452 | 0.185794 |
| 8 | 84595000 | 84605599 | 70.77699 | 109.8559 | 200.6513 | 83.32658 | -0.56194 | 2.772782 | 0.185812 |
| 11 | 106918600 | 106920799 | 9.819753 | 17.30154 | 43.55722 | 19.27949 | -0.9356 | 0.551395 | 0.185816 |
| 2 | 167999200 | 168004599 | 23.95644 | 38.72092 | 97.91462 | 41.82446 | -0.97392 | 1.693163 | 0.18582 |
| 2 | 55888000 | 55888599 | 14.60606 | 27.01429 | 21.64824 | 9.120781 | 0.755996 | 0.218366 | 0.185825 |
| 1 | 94534200 | 94535199 | 14.25375 | 26.80775 | 20.2146 | 8.292823 | 0.87298 | 0.112593 | 0.18585 |
| 1 | 140339200 | 140340799 | 14.25375 | 26.80775 | 20.2146 | 8.292823 | 0.87298 | 0.112593 | 0.18585 |
| 6 | 113845600 | 113853399 | 48.40953 | 76.38851 | 161.4872 | 67.02943 | -0.75387 | 2.451876 | 0.185853 |
| 11 | 42309400 | 42312199 | 10.62551 | 18.60174 | 47.79838 | 21.09493 | -0.96681 | 0.654485 | 0.185859 |
| 10 | 99243200 | 99244999 | 25.2056 | 43.89417 | 32.21471 | 14.16451 | 0.831438 | 0.782013 | 0.185866 |
| 9 | 114197000 | 114197799 | 3.832687 | 7.438922 | 25.45551 | 9.570271 | -1.1713 | -0.26435 | 0.185871 |
| 8 | 74078800 | 74081599 | 17.30327 | 28.68402 | 74.29324 | 32.16955 | -0.98748 | 1.283627 | 0.185874 |
| 4 | 61642800 | 61643599 | 11.94739 | 22.81984 | 18.269 | 7.262311 | 0.82558 | -0.01826 | 0.185889 |
| 13 | 64440800 | 64441799 | 13.05976 | 24.83437 | 19.2806 | 7.7365 | 0.858245 | 0.02621 | 0.1859 |
| 15 | 53400200 | 53401799 | 9.357367 | 16.97806 | 38.08767 | 16.45689 | -0.74411 | 0.32598 | 0.1859 |
| 4 | 56864400 | 56866199 | 9.559484 | 17.13374 | 39.46505 | 17.28697 | -0.79523 | 0.402704 | 0.185922 |
| 18 | 25759200 | 25761199 | 9.559484 | 17.13374 | 39.46505 | 17.28697 | -0.79523 | 0.402704 | 0.185922 |
| 3 | 143954400 | 143957399 | 18.80975 | 31.76864 | 65.70892 | 28.74546 | -0.66353 | 1.085298 | 0.185923 |
| 9 | 38465000 | 38466799 | 23.22231 | 40.66844 | 29.99048 | 13.23956 | 0.824739 | 0.686458 | 0.185953 |
| 7 | 45787400 | 45790799 | 14.82933 | 25.25566 | 58.92068 | 25.82692 | -0.83506 | 0.991989 | 0.18596 |
| 18 | 22933400 | 22933799 | 2.455498 | 4.796231 | 22.11176 | 7.741788 | -1.49652 | -0.55135 | 0.185971 |
| 3 | 97861200 | 97863999 | 29.91341 | 50.42719 | 41.59684 | 18.18685 | 0.657783 | 1.112889 | 0.185972 |
| 11 | 43475200 | 43477799 | 12.01047 | 20.95853 | 48.90277 | 21.52985 | -0.82885 | 0.713903 | 0.185982 |
| X | 5250600 | 5254999 | 75.84181 | 120.6258 | 94.45557 | 39.43793 | 0.68683 | 2.250972 | 0.185985 |
| 19 | 56873200 | 56875599 | 10.47437 | 18.31746 | 47.29386 | 20.85333 | -0.97338 | 0.672851 | 0.185988 |
| 19 | 41137200 | 41139199 | 32.34499 | 54.91714 | 38.69763 | 16.95596 | 0.884529 | 1.059707 | 0.185996 |
| 16 | 21192000 | 21196399 | 25.26658 | 41.10931 | 88.21248 | 37.86316 | -0.72888 | 1.554287 | 0.186007 |

| | | | | | | | | | |
|---|---|---|---|---|---|---|---|---|---|
| 6 | 51246600 | 51250199 | 24.08386 | 39.36921 | 83.35112 | 35.83775 | -0.70467 | 1.481124 | 0.186008 |
| 8 | 65392600 | 65393799 | 16.54766 | 29.96115 | 23.1932 | 10.04767 | 0.778698 | 0.356667 | 0.186008 |
| 8 | 10026000 | 10028799 | 14.01817 | 23.98579 | 60.12424 | 26.36438 | -0.93734 | 0.943944 | 0.18604 |
| 12 | 118258000 | 118259399 | 14.323 | 27.60301 | 15.6034 | 6.048337 | 1.357582 | -0.07803 | 0.18604 |
| X | 78210400 | 78211399 | 11.44196 | 22.20839 | 15.69992 | 5.901927 | 1.078087 | -0.20997 | 0.186043 |
| 10 | 29409200 | 29412399 | 22.36381 | 36.51089 | 84.5274 | 36.32661 | -0.83447 | 1.497307 | 0.186045 |
| 11 | 29953800 | 29956199 | 11.09034 | 19.37707 | 47.83029 | 21.07422 | -0.90912 | 0.690878 | 0.186046 |
| 18 | 83845200 | 83847599 | 14.45032 | 24.78044 | 59.66104 | 26.16597 | -0.87897 | 0.924159 | 0.186069 |
| 4 | 132937400 | 132941199 | 17.77457 | 29.42623 | 74.21619 | 32.10676 | -0.94941 | 1.301347 | 0.186081 |
| 8 | 85806600 | 85808799 | 9.858999 | 17.27886 | 46.21919 | 20.41699 | -1.024 | 0.631404 | 0.186084 |
| 12 | 98241200 | 98242199 | 5.987262 | 11.52611 | 32.39726 | 12.59705 | -0.94081 | -0.11094 | 0.186084 |
| 7 | 29092800 | 29103399 | 68.68676 | 106.1009 | 212.3797 | 88.21804 | -0.69926 | 2.86054 | 0.186101 |
| 3 | 96309400 | 96319199 | 97.75701 | 148.736 | 270.133 | 111.9272 | -0.57629 | 3.172558 | 0.186111 |
| 5 | 8580200 | 8581399 | 15.37551 | 28.67422 | 20.41969 | 8.49767 | 0.938129 | 0.17388 | 0.186117 |
| 2 | 89531000 | 89533399 | 37.66468 | 63.05917 | 41.48769 | 18.08148 | 0.982952 | 1.204723 | 0.186124 |
| X | 88896200 | 88897199 | 20.48283 | 36.3677 | 25.43169 | 11.22874 | 0.906615 | 0.505941 | 0.186126 |
| 12 | 113766600 | 113767799 | 16.10271 | 30.58279 | 16.59756 | 6.677443 | 1.366422 | 0.056097 | 0.186131 |
| 8 | 48366600 | 48369199 | 15.39217 | 25.95529 | 63.66148 | 27.83636 | -0.90821 | 1.086854 | 0.186136 |
| 8 | 5465400 | 5467199 | 16.65791 | 30.55941 | 21.78646 | 9.281264 | 0.911995 | 0.279175 | 0.186137 |
| 14 | 115096200 | 115098599 | 15.99022 | 27.54185 | 59.81647 | 26.23936 | -0.72975 | 0.892159 | 0.186152 |
| 13 | 51977600 | 51982599 | 24.95415 | 40.59785 | 91.23616 | 39.15991 | -0.79552 | 1.555374 | 0.186187 |
| 7 | 33503600 | 33513199 | 201.1485 | 301.4625 | 209.8927 | 86.58727 | 0.787274 | 3.490321 | 0.186192 |
| 3 | 86888400 | 86888999 | 12.16857 | 23.27795 | 18.43455 | 7.299377 | 0.847024 | -0.02607 | 0.186192 |
| 17 | 20207200 | 20207999 | 13.41251 | 26.00912 | 12.96322 | 4.894216 | 1.569382 | -0.18169 | 0.186198 |
| 12 | 7574400 | 7574799 | 3.525782 | 6.888036 | 21.75333 | 7.592309 | -0.9579 | -0.54918 | 0.186202 |
| 12 | 49307400 | 49307799 | 10.1868 | 19.69811 | 16.02042 | 6.128934 | 0.854964 | -0.16533 | 0.186203 |
| 3 | 33729200 | 33730399 | 11.36473 | 21.95111 | 17.44751 | 6.710156 | 0.880631 | -0.13011 | 0.186213 |
| 1 | 91051800 | 91054799 | 18.76797 | 31.32801 | 69.26326 | 30.12994 | -0.75958 | 1.204214 | 0.186215 |
| 19 | 41632400 | 41635799 | 21.72168 | 35.88113 | 77.12564 | 33.3144 | -0.72017 | 1.337625 | 0.186216 |
| 6 | 131677400 | 131681199 | 75.0022 | 119.066 | 101.0718 | 42.11892 | 0.567922 | 2.304593 | 0.18622 |
| 1 | 145471200 | 145471799 | 10.00755 | 19.37386 | 16.48531 | 6.272841 | 0.797342 | -0.18434 | 0.186225 |
| 7 | 43874600 | 43883399 | 53.14641 | 83.72089 | 168.7195 | 70.00805 | -0.68693 | 2.48991 | 0.186226 |
| 11 | 101873800 | 101875999 | 8.409801 | 14.96185 | 40.49645 | 17.84992 | -1.03229 | 0.44583 | 0.186238 |
| 15 | 20763800 | 20764799 | 15.98152 | 29.49321 | 21.20723 | 8.961788 | 0.908674 | 0.24252 | 0.186253 |
| 17 | 72409400 | 72411399 | 18.99037 | 33.79807 | 25.22037 | 11.11314 | 0.81493 | 0.478171 | 0.186267 |
| 13 | 62535400 | 62539799 | 96.76035 | 151.5118 | 125.5676 | 52.08705 | 0.586632 | 2.622977 | 0.186271 |
| 5 | 134107000 | 134110399 | 25.94673 | 42.11052 | 95.16265 | 40.7858 | -0.80657 | 1.603296 | 0.186278 |
| 19 | 13653600 | 13654999 | 16.85036 | 31.59123 | 18.05054 | 7.444817 | 1.263159 | 0.148561 | 0.186285 |
| 5 | 144790400 | 144798999 | 97.9857 | 147.9866 | 291.003 | 120.3007 | -0.69976 | 3.324888 | 0.1863 |
| 4 | 32821200 | 32825399 | 21.2968 | 35.06632 | 78.45941 | 33.83533 | -0.77922 | 1.377539 | 0.186302 |
| 17 | 72199800 | 72201399 | 11.06938 | 19.39737 | 46.21859 | 20.41423 | -0.85817 | 0.641713 | 0.186322 |
| 17 | 40994000 | 40994999 | 24.0249 | 41.60697 | 33.54288 | 14.71652 | 0.695498 | 0.842313 | 0.186322 |
| 10 | 8974200 | 8975199 | 6.152745 | 11.78317 | 31.40949 | 12.39692 | -0.88533 | -0.06266 | 0.186323 |
| 18 | 80237200 | 80238199 | 12.78459 | 24.26644 | 19.37152 | 7.803279 | 0.813757 | 0.036688 | 0.186333 |
| 7 | 17190400 | 17194199 | 64.61487 | 103.4181 | 81.23024 | 34.21608 | 0.692083 | 2.041946 | 0.186337 |
| 1 | 190820800 | 190824999 | 21.84446 | 35.7964 | 84.54757 | 36.38709 | -0.85997 | 1.442024 | 0.186337 |
| 7 | 10786200 | 10788799 | 47.69363 | 77.18756 | 64.28469 | 27.49475 | 0.626204 | 1.68379 | 0.186338 |

| | | | | | | | | | |
|---|---|---|---|---|---|---|---|---|---|
| 4 | 58372600 | 58374399 | 8.843584 | 16.2387 | 37.29679 | 15.86901 | -0.76278 | 0.250644 | 0.186345 |
| 5 | 123858200 | 123859199 | 3.65255 | 7.094813 | 24.91784 | 9.312354 | -1.1992 | -0.28789 | 0.186346 |
| 4 | 138307800 | 138309199 | 5.911198 | 11.13047 | 31.24549 | 12.79282 | -1.00495 | 0.06219 | 0.186349 |
| 1 | 113135000 | 113136399 | 16.31114 | 30.45664 | 19.17634 | 7.963592 | 1.116543 | 0.171134 | 0.18635 |
| 12 | 69166800 | 69167999 | 8.030767 | 14.95314 | 34.90883 | 14.56627 | -0.76509 | 0.158748 | 0.186357 |
| 8 | 18307600 | 18308999 | 16.7586 | 30.51972 | 23.11196 | 9.940052 | 0.816886 | 0.325627 | 0.186365 |
| 9 | 19329000 | 19330599 | 31.92222 | 53.70159 | 42.67037 | 18.64444 | 0.711612 | 1.140768 | 0.186369 |
| 7 | 48704000 | 48706599 | 15.0119 | 25.49534 | 61.64114 | 27.01136 | -0.88714 | 1.019203 | 0.186371 |
| 12 | 83343600 | 83350799 | 45.78645 | 72.75483 | 141.7774 | 59.13787 | -0.62774 | 2.263268 | 0.18638 |
| 11 | 113645800 | 113649199 | 18.57003 | 31.13995 | 65.98444 | 28.78673 | -0.69846 | 1.152551 | 0.186387 |
| 7 | 38119200 | 38121599 | 38.64375 | 63.35701 | 53.18283 | 22.89337 | 0.628495 | 1.452208 | 0.186393 |
| 19 | 46568200 | 46571599 | 15.78579 | 26.81696 | 60.00806 | 26.2979 | -0.77583 | 1.020042 | 0.186396 |
| 8 | 92522800 | 92534599 | 85.97894 | 130.7196 | 269.3774 | 111.5895 | -0.75919 | 3.18487 | 0.186404 |
| 10 | 38841200 | 38842999 | 10.16984 | 18.26541 | 40.53783 | 17.72597 | -0.74033 | 0.396545 | 0.186426 |
| 14 | 24214000 | 24218199 | 21.77834 | 35.83291 | 77.77999 | 33.53193 | -0.736 | 1.388746 | 0.186441 |
| 5 | 75812200 | 75816999 | 34.11688 | 54.80591 | 110.3415 | 46.73706 | -0.65696 | 1.901091 | 0.186467 |
| 2 | 167857800 | 167872399 | 99.5587 | 150.6571 | 283.3259 | 117.1994 | -0.63303 | 3.284828 | 0.186479 |
| 7 | 10871600 | 10875999 | 139.6068 | 212.4993 | 174.7486 | 72.41464 | 0.566361 | 3.169118 | 0.186483 |
| 16 | 75852000 | 75853999 | 35.65282 | 58.87882 | 48.80568 | 21.07799 | 0.650289 | 1.353915 | 0.186486 |
| 12 | 45381600 | 45382999 | 14.16427 | 26.23937 | 20.13698 | 8.465513 | 0.819998 | 0.201423 | 0.186487 |
| 6 | 132590600 | 132591799 | 26.00167 | 44.97097 | 35.00792 | 15.35866 | 0.745518 | 0.866382 | 0.18649 |
| 1 | 12632800 | 12634599 | 17.83782 | 32.81313 | 21.2546 | 9.021237 | 1.05341 | 0.273738 | 0.186493 |
| 15 | 59054800 | 59063399 | 67.1192 | 104.7976 | 188.2126 | 78.09599 | -0.53097 | 2.667271 | 0.186504 |
| 10 | 85931200 | 85932599 | 8.551003 | 15.6495 | 36.12146 | 15.43841 | -0.77443 | 0.264808 | 0.186506 |
| 3 | 57431000 | 57431399 | 6.786948 | 12.81342 | 33.10177 | 13.48621 | -0.8818 | 0.052255 | 0.18651 |
| 8 | 48556000 | 48559399 | 71.58452 | 113.7752 | 98.22513 | 40.98121 | 0.545249 | 2.267335 | 0.18651 |
| 5 | 136758200 | 136759199 | 19.682 | 34.70576 | 27.19577 | 12.03558 | 0.739151 | 0.570122 | 0.186513 |
| 14 | 42431800 | 42435399 | 57.44336 | 92.4988 | 69.74676 | 29.64498 | 0.761182 | 1.842121 | 0.186522 |
| 13 | 74508400 | 74509599 | 7.105298 | 13.34824 | 32.34406 | 13.29416 | -0.80002 | 0.089125 | 0.186526 |
| 13 | 67000400 | 67010599 | 47.82036 | 77.79412 | 53.34849 | 22.89745 | 0.912322 | 1.580722 | 0.186532 |
| 10 | 31645400 | 31647399 | 29.0737 | 49.16906 | 40.52107 | 17.73375 | 0.659253 | 1.081445 | 0.186536 |
| 6 | 82981000 | 82985999 | 20.52885 | 34.16218 | 71.62547 | 31.09678 | -0.68339 | 1.244496 | 0.186539 |
| 3 | 121381600 | 121385399 | 30.3079 | 49.00553 | 97.45021 | 41.64659 | -0.62774 | 1.690985 | 0.186542 |
| 8 | 21356200 | 21356399 | 8.07281 | 15.78515 | 10.43348 | 3.562532 | 1.299283 | -0.59895 | 0.186564 |
| X | 24347200 | 24352999 | 50.15066 | 81.28896 | 59.93922 | 25.66527 | 0.803505 | 1.657657 | 0.186591 |
| 12 | 19914600 | 19916799 | 60.13385 | 97.51557 | 49.14908 | 21.05221 | 1.351425 | 1.662184 | 0.186601 |
| 3 | 107667000 | 107668399 | 5.888364 | 10.90217 | 35.77825 | 15.05239 | -1.26076 | 0.176014 | 0.186608 |
| 1 | 140963400 | 140965399 | 31.89167 | 53.5951 | 37.63504 | 16.43917 | 0.889245 | 1.144857 | 0.186625 |
| 6 | 37898600 | 37899799 | 14.99722 | 25.95331 | 54.89132 | 24.08232 | -0.69007 | 0.839152 | 0.186638 |
| 2 | 88257600 | 88261399 | 42.79885 | 70.51868 | 48.93088 | 21.10736 | 0.905143 | 1.400061 | 0.186639 |
| 11 | 51526400 | 51529199 | 12.33831 | 21.46937 | 50.51497 | 22.19985 | -0.84138 | 0.75876 | 0.186656 |
| 11 | 96575600 | 96587999 | 101.6413 | 154.1083 | 284.5333 | 117.7709 | -0.60413 | 3.245415 | 0.186666 |
| X | 64582000 | 64584399 | 17.80937 | 32.84427 | 20.88316 | 8.83295 | 1.083815 | 0.259761 | 0.186671 |
| 15 | 83446800 | 83448799 | 11.71153 | 20.42722 | 48.90197 | 21.52311 | -0.86566 | 0.719726 | 0.186674 |
| 12 | 80992000 | 80993399 | 7.650495 | 14.02705 | 35.09117 | 14.96083 | -0.88649 | 0.247036 | 0.186675 |
| X | 3772000 | 3776599 | 36.27894 | 61.42686 | 36.44836 | 15.95706 | 1.131472 | 1.095737 | 0.186681 |
| 7 | 141141400 | 141143199 | 10.62468 | 18.57852 | 47.29588 | 20.85253 | -0.95313 | 0.675547 | 0.186682 |

| | | | | | | | | | |
|---|---|---|---|---|---|---|---|---|---|
| 8 | 118024400 | 118029199 | 22.92234 | 37.22895 | 90.18549 | 38.67325 | -0.90531 | 1.582535 | 0.186687 |
| 13 | 62181200 | 62187999 | 181.4642 | 272.3578 | 219.3647 | 90.52697 | 0.579131 | 3.459471 | 0.186693 |
| 1 | 155119800 | 155122599 | 21.20825 | 35.16162 | 73.18071 | 31.69044 | -0.67326 | 1.291598 | 0.186697 |
| 8 | 27830600 | 27833199 | 21.43841 | 37.90633 | 25.47462 | 11.26772 | 0.960518 | 0.546462 | 0.186706 |
| 14 | 16275600 | 16277599 | 22.89726 | 40.16509 | 28.22093 | 12.47568 | 0.893836 | 0.654632 | 0.186708 |
| 10 | 119186400 | 119188599 | 22.39888 | 39.4082 | 27.71411 | 12.27046 | 0.893045 | 0.610222 | 0.186719 |
| 15 | 98052800 | 98055199 | 10.49299 | 18.77286 | 44.77271 | 19.64188 | -0.84717 | 0.429858 | 0.186729 |
| 1 | 153381400 | 153384999 | 17.63485 | 29.54811 | 66.37895 | 28.94424 | -0.78233 | 1.159605 | 0.186734 |
| 8 | 83773000 | 83774399 | 8.224218 | 15.12054 | 37.33003 | 15.85582 | -0.86457 | 0.237064 | 0.186754 |
| 5 | 94368000 | 94369399 | 16.65739 | 31.23875 | 18.05689 | 7.443684 | 1.247114 | 0.144932 | 0.186784 |
| 9 | 58461400 | 58462799 | 7.864408 | 14.25203 | 37.54894 | 16.24861 | -0.97558 | 0.320293 | 0.186786 |
| 5 | 67109400 | 67110599 | 16.33345 | 29.6988 | 23.57401 | 10.1587 | 0.747457 | 0.331381 | 0.186789 |
| 14 | 99919200 | 99920999 | 16.3502 | 29.66696 | 23.61811 | 10.20444 | 0.74088 | 0.343624 | 0.18679 |
| 1 | 147617800 | 147619399 | 15.53332 | 29.0224 | 20.53162 | 8.517407 | 0.951513 | 0.16503 | 0.186795 |
| 15 | 29551000 | 29552799 | 8.219837 | 15.23267 | 39.26879 | 16.5005 | -0.91494 | 0.19876 | 0.186801 |
| 14 | 21448400 | 21450799 | 21.90691 | 38.34299 | 30.73016 | 13.55958 | 0.705306 | 0.689595 | 0.186809 |
| 13 | 73305000 | 73308799 | 20.90762 | 34.6097 | 75.40215 | 32.61763 | -0.73925 | 1.309569 | 0.186809 |
| 15 | 47403000 | 47404799 | 43.74275 | 71.22654 | 58.72773 | 25.21476 | 0.648152 | 1.562552 | 0.18681 |
| 4 | 127491000 | 127498199 | 48.74846 | 77.28515 | 149.8773 | 62.37404 | -0.62403 | 2.334064 | 0.186848 |
| 1 | 58733800 | 58734799 | 5.253648 | 10.16058 | 27.14516 | 10.37705 | -0.84346 | -0.1877 | 0.186848 |
| 3 | 95529400 | 95530199 | 5.253648 | 10.16058 | 27.14516 | 10.37705 | -0.84346 | -0.1877 | 0.186848 |
| 12 | 49420800 | 49425799 | 25.684 | 41.92689 | 92.66128 | 39.82068 | -0.7681 | 1.507749 | 0.186859 |
| 3 | 154043200 | 154045799 | 16.72611 | 28.36886 | 61.17925 | 26.79813 | -0.72307 | 1.037047 | 0.186865 |
| 7 | 115803600 | 115804199 | 2.946156 | 5.762045 | 20.78752 | 7.043254 | -1.10354 | -0.65976 | 0.186871 |
| 13 | 66895000 | 66897599 | 40.54272 | 67.47879 | 39.41602 | 17.11479 | 1.15414 | 1.272122 | 0.186872 |
| 15 | 89420400 | 89435199 | 133.5106 | 197.8645 | 374.8271 | 154.3741 | -0.67062 | 3.687532 | 0.186898 |
| 16 | 87549200 | 87552399 | 27.53692 | 44.68971 | 90.41308 | 38.74914 | -0.64734 | 1.606862 | 0.186905 |
| 12 | 80481600 | 80483799 | 12.01428 | 20.96551 | 48.37337 | 21.29699 | -0.81262 | 0.714 | 0.186907 |
| 13 | 40649400 | 40652399 | 20.34517 | 33.96422 | 69.42431 | 30.20196 | -0.64683 | 1.20841 | 0.186909 |
| X | 126420600 | 126421799 | 18.02973 | 33.74061 | 18.20951 | 7.533656 | 1.342132 | 0.171781 | 0.186927 |
| 15 | 24320600 | 24322199 | 21.0199 | 36.98039 | 28.5518 | 12.64146 | 0.759207 | 0.605096 | 0.186931 |
| 1 | 171023000 | 171023999 | 6.510621 | 12.21414 | 32.27306 | 13.29609 | -0.92601 | 0.094225 | 0.186943 |
| X | 26413400 | 26415999 | 36.42712 | 62.95152 | 23.90648 | 10.48798 | 1.775069 | 0.906846 | 0.186944 |
| 11 | 53368800 | 53371599 | 25.09426 | 40.86027 | 86.37545 | 37.08502 | -0.7065 | 1.543066 | 0.186946 |
| 7 | 25936400 | 25953999 | 50.25166 | 81.24533 | 59.89884 | 25.5954 | 0.800178 | 1.708379 | 0.186947 |
| 4 | 33309000 | 33310799 | 9.573008 | 16.73824 | 47.81069 | 21.07811 | -1.11808 | 0.671991 | 0.186979 |
| 11 | 49548400 | 49551199 | 16.77792 | 28.30764 | 63.83333 | 27.91618 | -0.78758 | 1.085702 | 0.187026 |
| 1 | 52538800 | 52540199 | 11.23829 | 19.64601 | 47.8238 | 21.08032 | -0.88924 | 0.682233 | 0.187037 |
| 3 | 38725000 | 38726999 | 9.25706 | 16.59363 | 40.51649 | 17.74579 | -0.87887 | 0.400921 | 0.187048 |
| 19 | 58384200 | 58384999 | 5.004352 | 9.638332 | 28.62299 | 11.11482 | -1.01637 | -0.13435 | 0.187051 |
| 2 | 89098800 | 89102799 | 76.62583 | 121.5063 | 103.3367 | 43.01411 | 0.563656 | 2.33875 | 0.187051 |
| X | 136725000 | 136730799 | 85.64069 | 135.4498 | 105.8417 | 43.98359 | 0.682544 | 2.412558 | 0.187054 |
| 10 | 40664400 | 40665599 | 13.7984 | 25.97112 | 20.16074 | 8.260069 | 0.832552 | 0.102634 | 0.187054 |
| 8 | 57893000 | 57893799 | 3.816912 | 7.452736 | 24.2548 | 8.54517 | -1.01464 | -0.50515 | 0.187055 |
| 1 | 148290800 | 148291999 | 13.74911 | 26.4183 | 15.11864 | 5.910711 | 1.326312 | -0.06139 | 0.18706 |
| 9 | 124162000 | 124164599 | 71.47251 | 114.1125 | 84.81081 | 35.59984 | 0.767839 | 2.123787 | 0.187071 |
| 7 | 116170000 | 116170999 | 16.74024 | 30.4214 | 23.41906 | 10.099 | 0.790554 | 0.33647 | 0.187087 |

| | | | | | | | | | |
|---|---|---|---|---|---|---|---|---|---|
| 18 | 21086200 | 21087399 | 20.30719 | 35.70392 | 26.34086 | 11.66403 | 0.824156 | 0.603783 | 0.187088 |
| 1 | 97588400 | 97589999 | 13.18209 | 22.72529 | 55.29902 | 24.25837 | -0.89241 | 0.866683 | 0.187098 |
| 10 | 60109200 | 60113999 | 25.80979 | 41.82333 | 92.66558 | 39.67018 | -0.78015 | 1.63543 | 0.187151 |
| 10 | 99187800 | 99188399 | 10.168 | 19.7278 | 16.43323 | 6.189127 | 0.841424 | -0.21665 | 0.187162 |
| 2 | 178745200 | 178747399 | 9.885103 | 17.64133 | 40.45883 | 17.78665 | -0.79279 | 0.433884 | 0.187168 |
| 15 | 53382000 | 53384199 | 17.98095 | 32.53617 | 23.83832 | 10.33512 | 0.856677 | 0.371285 | 0.187179 |
| 17 | 80513600 | 80514999 | 8.874498 | 16.15775 | 37.08962 | 15.95646 | -0.77296 | 0.299093 | 0.187206 |
| 9 | 41287400 | 41287999 | 3.642148 | 7.107056 | 22.05092 | 7.852904 | -0.95861 | -0.4725 | 0.187214 |
| 11 | 68601600 | 68602399 | 3.642148 | 7.107056 | 22.05092 | 7.852904 | -0.95861 | -0.4725 | 0.187214 |
| 17 | 56865400 | 56866599 | 5.997042 | 11.5137 | 29.13747 | 11.41802 | -0.80062 | -0.09195 | 0.187218 |
| 7 | 29467600 | 29468799 | 5.387192 | 10.42922 | 27.24025 | 10.3651 | -0.80477 | -0.20152 | 0.187223 |
| 4 | 121257800 | 121264399 | 35.15876 | 59.70024 | 32.63402 | 14.29951 | 1.249153 | 1.061918 | 0.187226 |
| 2 | 130235200 | 130236999 | 8.122765 | 14.81333 | 36.06248 | 15.4828 | -0.85487 | 0.28258 | 0.187232 |
| 17 | 69008000 | 69009599 | 15.95703 | 29.66142 | 20.93584 | 8.760611 | 0.944796 | 0.200804 | 0.187254 |
| 9 | 45323400 | 45325799 | 10.51417 | 18.3363 | 50.46713 | 22.21008 | -1.06578 | 0.717265 | 0.187271 |
| 7 | 135502000 | 135504799 | 13.6991 | 23.88389 | 52.71413 | 23.19481 | -0.7496 | 0.738931 | 0.187278 |
| 12 | 13044400 | 13047399 | 13.58875 | 23.19756 | 61.12903 | 26.80001 | -1.00961 | 0.965023 | 0.187283 |
| X | 63501000 | 63503199 | 46.49293 | 75.70739 | 56.58664 | 24.2958 | 0.789443 | 1.568022 | 0.187283 |
| 15 | 90304800 | 90305399 | 3.760581 | 7.334584 | 22.55488 | 8.089859 | -0.95517 | -0.44246 | 0.1873 |
| 5 | 115457600 | 115461999 | 18.83887 | 31.69779 | 65.61512 | 28.67154 | -0.6649 | 1.118323 | 0.187307 |
| 12 | 74039800 | 74040599 | 5.40883 | 10.45157 | 32.08325 | 12.31491 | -1.049 | -0.16207 | 0.18731 |
| 16 | 8701600 | 8703399 | 16.84828 | 28.25007 | 66.40221 | 28.96725 | -0.84745 | 1.147692 | 0.187315 |
| 5 | 143571200 | 143573999 | 10.76851 | 18.86871 | 49.37732 | 21.808 | -0.99323 | 0.641793 | 0.187317 |
| 5 | 110954800 | 110959599 | 22.95042 | 37.57601 | 81.46293 | 35.04665 | -0.73713 | 1.456263 | 0.187319 |
| 16 | 91454400 | 91457799 | 34.57502 | 55.64131 | 107.1709 | 45.51195 | -0.58999 | 1.844151 | 0.187323 |
| 15 | 95836200 | 95848399 | 98.38846 | 149.2788 | 272.7318 | 112.9094 | -0.58813 | 3.23048 | 0.187324 |
| 17 | 19382000 | 19385799 | 52.20204 | 84.3253 | 65.59397 | 27.99757 | 0.721642 | 1.740521 | 0.187329 |
| 8 | 98557000 | 98558399 | 16.51163 | 30.44275 | 21.29712 | 9.011346 | 0.946007 | 0.249839 | 0.187338 |
| 7 | 141781600 | 141783999 | 12.80575 | 22.24482 | 52.67807 | 23.12569 | -0.85112 | 0.787385 | 0.187339 |
| 5 | 129937200 | 129939199 | 10.01991 | 17.5267 | 47.28917 | 20.85529 | -1.03642 | 0.666746 | 0.187346 |
| 7 | 53629400 | 53631799 | 22.62115 | 39.60678 | 29.99393 | 13.23858 | 0.786516 | 0.685758 | 0.187352 |
| 14 | 121959000 | 121962599 | 26.96367 | 43.59768 | 96.56696 | 41.26853 | -0.78282 | 1.685655 | 0.187355 |
| X | 61841200 | 61843599 | 24.41833 | 43.65518 | 20.45233 | 8.977894 | 1.485792 | 0.479933 | 0.187365 |
| 2 | 68691200 | 68692799 | 11.2241 | 19.65765 | 46.22486 | 20.40749 | -0.83944 | 0.650034 | 0.187367 |
| 7 | 23198200 | 23205199 | 106.3204 | 166.6146 | 115.6371 | 47.96378 | 0.843152 | 2.614441 | 0.187404 |
| 6 | 39529200 | 39536999 | 37.13598 | 59.2924 | 138.5991 | 58.17971 | -0.88235 | 2.101186 | 0.187407 |
| 16 | 47099800 | 47100599 | 14.15378 | 26.3807 | 21.1996 | 8.83077 | 0.76423 | 0.169475 | 0.187409 |
| 17 | 40182800 | 40183799 | 14.15378 | 26.3807 | 21.1996 | 8.83077 | 0.76423 | 0.169475 | 0.187409 |
| X | 148808200 | 148809599 | 20.40903 | 36.64985 | 23.63248 | 10.33508 | 1.031909 | 0.432328 | 0.187411 |
| 6 | 87698600 | 87699999 | 6.961895 | 13.05585 | 32.30611 | 13.31817 | -0.8338 | 0.09741 | 0.187413 |
| 17 | 75112600 | 75114199 | 16.35926 | 29.9056 | 21.23028 | 9.087595 | 0.912979 | 0.294036 | 0.187413 |
| 13 | 52206800 | 52216799 | 61.34832 | 95.56423 | 194.0377 | 80.53683 | -0.71038 | 2.705562 | 0.187424 |
| 6 | 49599200 | 49599599 | 23.44915 | 41.54698 | 21.85394 | 9.662282 | 1.312999 | 0.532463 | 0.187446 |
| 8 | 11604800 | 11608599 | 20.54901 | 33.39323 | 90.32628 | 38.74977 | -1.06304 | 1.566983 | 0.187469 |
| 8 | 67708000 | 67708799 | 19.20605 | 34.74344 | 20.43251 | 8.861569 | 1.171566 | 0.375367 | 0.187482 |
| 19 | 40050200 | 40051999 | 24.09402 | 42.42727 | 25.93097 | 11.47899 | 1.094775 | 0.6083 | 0.187486 |
| 12 | 52910200 | 52911399 | 18.47061 | 33.19666 | 24.0513 | 10.51083 | 0.865699 | 0.415331 | 0.187489 |

| | | | | | | | | | |
|---|---|---|---|---|---|---|---|---|---|
| 2 | 74291200 | 74293799 | 13.85965 | 24.16116 | 52.71578 | 23.19394 | -0.73308 | 0.741549 | 0.187495 |
| 7 | 17035000 | 17038399 | 59.33393 | 95.28263 | 75.91324 | 32.13796 | 0.677119 | 1.930128 | 0.187519 |
| 17 | 56483600 | 56491599 | 46.38816 | 73.40194 | 154.5271 | 64.23042 | -0.74462 | 2.382502 | 0.187538 |
| 9 | 20775600 | 20797599 | 176.9535 | 257.7579 | 445.7056 | 183.6819 | -0.56276 | 3.969452 | 0.187539 |
| 6 | 52208800 | 52213199 | 22.44369 | 36.50884 | 91.17331 | 39.13108 | -0.94726 | 1.552904 | 0.187542 |
| 5 | 142969000 | 142974599 | 47.1478 | 74.74442 | 148.0719 | 61.62014 | -0.65474 | 2.334617 | 0.187546 |
| 5 | 67068400 | 67069399 | 13.80363 | 25.69369 | 20.75502 | 8.665172 | 0.754206 | 0.173658 | 0.187558 |
| 16 | 16516600 | 16518399 | 18.21979 | 32.89248 | 23.71991 | 10.31346 | 0.87664 | 0.38497 | 0.187558 |
| 7 | 40834600 | 40836599 | 34.44332 | 57.58232 | 44.84625 | 19.52736 | 0.740476 | 1.211651 | 0.187559 |
| 6 | 109377000 | 109377399 | 9.188538 | 17.90636 | 14.97551 | 5.429875 | 0.88557 | -0.36288 | 0.187562 |
| 12 | 19533800 | 19545599 | 288.5308 | 416.404 | 316.7831 | 130.6954 | 0.608176 | 4.114084 | 0.187565 |
| 2 | 177524400 | 177525599 | 12.23163 | 23.61753 | 16.00292 | 6.164322 | 1.105272 | -0.1216 | 0.187575 |
| 4 | 137321000 | 137322399 | 5.957579 | 11.18876 | 32.85841 | 13.51239 | -1.07571 | 0.079688 | 0.187578 |
| 14 | 122709400 | 122714999 | 45.86275 | 72.66265 | 153.2323 | 63.7331 | -0.74562 | 2.350978 | 0.187583 |
| 1 | 110832200 | 110834399 | 24.36251 | 42.41645 | 32.82215 | 14.42957 | 0.755414 | 0.781682 | 0.187586 |
| 2 | 17068600 | 17072599 | 83.74595 | 131.9719 | 114.5776 | 47.54479 | 0.526697 | 2.492821 | 0.187587 |
| 4 | 78653400 | 78656599 | 27.4122 | 47.30858 | 35.00577 | 15.35671 | 0.817864 | 0.892353 | 0.187607 |
| 6 | 88161000 | 88169199 | 43.83673 | 69.66078 | 147.4092 | 61.49056 | -0.74638 | 2.260049 | 0.187635 |
| 10 | 69481200 | 69482999 | 14.03067 | 23.97422 | 58.43261 | 25.61976 | -0.89719 | 0.95623 | 0.187639 |
| 7 | 85021400 | 85025199 | 59.43211 | 95.26811 | 79.79608 | 33.68913 | 0.60206 | 1.987956 | 0.187639 |
| 1 | 24496200 | 24496799 | 3.379087 | 6.602843 | 23.66865 | 8.223322 | -1.1301 | -0.55528 | 0.187648 |
| 5 | 102227000 | 102231799 | 45.25836 | 73.51008 | 59.96037 | 25.71329 | 0.660877 | 1.604067 | 0.187649 |
| 6 | 37034200 | 37035399 | 5.691959 | 10.97408 | 28.14546 | 10.89674 | -0.80292 | -0.13703 | 0.187651 |
| 7 | 70006800 | 70011999 | 27.4994 | 44.33469 | 103.2237 | 43.94803 | -0.85955 | 1.775405 | 0.187688 |
| 2 | 165294600 | 165295999 | 7.072666 | 13.37696 | 32.61443 | 13.23703 | -0.79401 | 0.041967 | 0.187696 |
| 7 | 28992400 | 29007799 | 89.83396 | 136.1635 | 283.794 | 117.4536 | -0.77918 | 3.249216 | 0.187704 |
| 2 | 173152800 | 173158399 | 28.62232 | 46.12811 | 105.3149 | 44.81715 | -0.83204 | 1.788061 | 0.187715 |
| 8 | 127148800 | 127150399 | 9.796678 | 17.31267 | 42.00389 | 18.58081 | -0.88069 | 0.515981 | 0.187715 |
| 17 | 5989400 | 5992399 | 14.7783 | 25.30543 | 56.88282 | 24.94453 | -0.78049 | 0.938044 | 0.187716 |
| 1 | 36326400 | 36327999 | 12.96486 | 22.5228 | 51.06204 | 22.41725 | -0.78838 | 0.786455 | 0.187718 |
| 6 | 127742000 | 127749199 | 54.73002 | 85.90581 | 175.5823 | 72.82159 | -0.71127 | 2.564585 | 0.187724 |
| 1 | 108341600 | 108341999 | 9.711111 | 18.85411 | 14.02066 | 5.2601 | 1.009337 | -0.24236 | 0.18773 |
| 9 | 122363200 | 122364999 | 7.507605 | 13.73328 | 35.58607 | 15.21962 | -0.93992 | 0.261359 | 0.187746 |
| 19 | 39367000 | 39370599 | 61.89168 | 99.1669 | 79.93029 | 33.72311 | 0.656617 | 2.002635 | 0.187752 |
| 1 | 154066000 | 154069399 | 27.03897 | 43.92351 | 90.08425 | 38.63541 | -0.66571 | 1.585215 | 0.187763 |
| 18 | 32054400 | 32056199 | 11.50444 | 20.19255 | 46.69578 | 20.65335 | -0.81575 | 0.614497 | 0.187764 |
| 9 | 43042000 | 43043999 | 8.190442 | 14.74233 | 37.9892 | 16.57946 | -0.95171 | 0.365074 | 0.187788 |
| 16 | 44566000 | 44566799 | 4.368164 | 8.474525 | 25.4798 | 9.599031 | -0.99008 | -0.25018 | 0.187792 |
| 11 | 75622200 | 75624799 | 10.06884 | 17.56271 | 50.98495 | 22.44042 | -1.14219 | 0.713178 | 0.187799 |
| 4 | 134315200 | 134333999 | 106.9356 | 159.5459 | 353.9979 | 145.9076 | -0.88957 | 3.564184 | 0.187804 |
| 7 | 40934800 | 40936199 | 30.16143 | 50.74697 | 42.18251 | 18.43209 | 0.646744 | 1.131818 | 0.18781 |
| 2 | 15302000 | 15302399 | 3.34014 | 6.532984 | 23.14532 | 7.80173 | -1.07556 | -0.67034 | 0.187835 |
| 17 | 86952800 | 86959199 | 38.11672 | 60.80989 | 134.7902 | 56.51731 | -0.80706 | 2.126646 | 0.187858 |
| 10 | 20702600 | 20703199 | 10.03886 | 19.56027 | 14.18729 | 5.155056 | 1.085055 | -0.34814 | 0.187865 |
| 13 | 49008600 | 49012999 | 17.74078 | 29.45404 | 72.15944 | 31.27842 | -0.90729 | 1.265817 | 0.187884 |
| 4 | 9498800 | 9500999 | 18.66033 | 33.39227 | 25.3649 | 11.12556 | 0.794092 | 0.444373 | 0.187888 |
| 5 | 127354200 | 127368999 | 113.1309 | 170.4355 | 306.0109 | 126.4109 | -0.5718 | 3.377792 | 0.187915 |

| | | | | | | | | | |
|---|---|---|---|---|---|---|---|---|---|
| 11 | 95922000 | 95922799 | 9.253813 | 17.99038 | 15.48054 | 5.745245 | 0.814498 | -0.28099 | 0.187918 |
| 13 | 48980600 | 48982799 | 10.39652 | 18.37741 | 43.51821 | 19.2497 | -0.84625 | 0.516952 | 0.187931 |
| 13 | 64579600 | 64580999 | 11.37025 | 19.92432 | 46.20988 | 20.41014 | -0.81975 | 0.641115 | 0.187938 |
| 6 | 38915000 | 38915599 | 3.340267 | 6.532832 | 20.27406 | 6.87003 | -0.89276 | -0.66008 | 0.187943 |
| 1 | 180109000 | 180110399 | 12.3566 | 23.41889 | 19.34453 | 7.817183 | 0.760614 | 0.041729 | 0.187953 |
| 7 | 7255800 | 7258599 | 53.6412 | 86.33847 | 72.57851 | 30.82899 | 0.603907 | 1.851049 | 0.187956 |
| 1 | 54390200 | 54390999 | 5.8296 | 11.25381 | 28.83599 | 11.1183 | -0.79611 | -0.14728 | 0.187963 |
| 14 | 42383400 | 42385799 | 27.04145 | 46.19585 | 37.7827 | 16.56553 | 0.671441 | 0.982044 | 0.187978 |
| 11 | 35167000 | 35172199 | 26.40079 | 42.89118 | 91.85931 | 39.39965 | -0.7279 | 1.582102 | 0.187984 |
| 5 | 23971400 | 23972599 | 5.547284 | 10.71256 | 28.25589 | 10.87683 | -0.83529 | -0.15851 | 0.188014 |
| 1 | 112483600 | 112484999 | 39.13879 | 64.66007 | 38.31025 | 16.54957 | 1.133123 | 1.350423 | 0.18802 |
| 7 | 25748400 | 25767999 | 54.32871 | 87.49293 | 65.40718 | 27.80489 | 0.773712 | 1.831013 | 0.18804 |
| 17 | 52119200 | 52120999 | 13.64384 | 23.51566 | 54.22712 | 23.78798 | -0.81522 | 0.870557 | 0.188043 |
| 16 | 83133800 | 83137599 | 62.95219 | 100.6896 | 84.67371 | 35.63277 | 0.59273 | 2.059319 | 0.188044 |
| 16 | 91680400 | 91682599 | 11.54844 | 20.17219 | 47.83566 | 21.07313 | -0.8517 | 0.696909 | 0.188046 |
| 7 | 23331600 | 23336599 | 79.87571 | 126.8886 | 96.33624 | 40.1727 | 0.729645 | 2.28899 | 0.188048 |
| 5 | 111790400 | 111792799 | 17.63157 | 29.5441 | 68.18986 | 29.73482 | -0.82141 | 1.157624 | 0.188066 |
| 18 | 38531000 | 38536199 | 41.51667 | 67.84648 | 56.55191 | 24.31256 | 0.636104 | 1.504473 | 0.188067 |
| 2 | 127867200 | 127870199 | 12.19334 | 21.19769 | 53.74545 | 23.60596 | -0.94904 | 0.773952 | 0.188068 |
| 5 | 41689400 | 41690599 | 16.14653 | 30.00373 | 20.93027 | 8.762225 | 0.961196 | 0.20432 | 0.188085 |
| 11 | 68237200 | 68238199 | 4.955227 | 9.608905 | 26.12765 | 9.86494 | -0.85099 | -0.23584 | 0.188092 |
| 11 | 86574400 | 86575199 | 4.955227 | 9.608905 | 26.12765 | 9.86494 | -0.85099 | -0.23584 | 0.188092 |
| 13 | 30843200 | 30844599 | 7.710287 | 13.67616 | 42.54125 | 18.79882 | -1.23421 | 0.46363 | 0.188098 |
| 11 | 59117600 | 59119999 | 10.4013 | 18.07933 | 53.11605 | 23.32711 | -1.16007 | 0.768528 | 0.188107 |
| 19 | 6046000 | 6048599 | 15.03697 | 25.90632 | 54.82114 | 24.04827 | -0.69212 | 0.880455 | 0.188119 |
| 4 | 114515200 | 114517999 | 14.70893 | 25.37172 | 54.83377 | 24.05469 | -0.72206 | 0.867433 | 0.188121 |
| 9 | 37920800 | 37922599 | 18.52213 | 34.56115 | 17.94943 | 7.462815 | 1.391006 | 0.19238 | 0.188129 |
| 4 | 105075400 | 105079599 | 17.83089 | 29.77504 | 70.90254 | 30.85038 | -0.86631 | 1.197871 | 0.188138 |
| 14 | 83646200 | 83647399 | 19.57236 | 35.04677 | 24.81144 | 10.87693 | 0.895704 | 0.446314 | 0.188142 |
| 8 | 117095800 | 117099999 | 20.03009 | 33.40425 | 70.53387 | 30.665 | -0.69355 | 1.218425 | 0.188144 |
| 14 | 83124800 | 83126199 | 16.30275 | 30.146 | 20.76599 | 8.751579 | 0.97211 | 0.230707 | 0.188165 |
| 7 | 15172600 | 15175999 | 35.97478 | 61.27577 | 32.97551 | 14.45442 | 1.272183 | 1.039317 | 0.188171 |
| 16 | 93032800 | 93033799 | 19.00107 | 33.59699 | 25.72719 | 11.37942 | 0.773644 | 0.533534 | 0.188184 |
| 1 | 185574000 | 185576399 | 25.10167 | 43.25404 | 35.67529 | 15.64969 | 0.661407 | 0.895066 | 0.188191 |
| 5 | 112007600 | 112029999 | 292.8642 | 407.0026 | 707.0049 | 293.4418 | -0.6372 | 4.673054 | 0.188194 |
| 14 | 67952600 | 67954399 | 11.69457 | 20.43899 | 47.82949 | 21.07876 | -0.83271 | 0.688295 | 0.188201 |
| 3 | 69494000 | 69499599 | 42.27314 | 67.58685 | 126.1187 | 53.0434 | -0.55573 | 2.063029 | 0.188203 |
| 9 | 20298600 | 20306199 | 108.8038 | 168.8124 | 140.4392 | 58.32507 | 0.570927 | 2.788195 | 0.188216 |
| 8 | 48352600 | 48355599 | 19.62352 | 32.56258 | 73.63378 | 31.90581 | -0.79242 | 1.276681 | 0.188226 |
| X | 33346600 | 33348999 | 27.41615 | 48.17818 | 23.89576 | 10.57966 | 1.393702 | 0.649155 | 0.188229 |
| 9 | 3292200 | 3294999 | 79.77878 | 126.0415 | 98.43508 | 40.8832 | 0.682071 | 2.430386 | 0.188246 |
| 7 | 39824800 | 39827199 | 19.05827 | 33.90626 | 25.77757 | 11.36224 | 0.787971 | 0.482547 | 0.18825 |
| 1 | 113640400 | 113643599 | 25.60177 | 45.40659 | 18.99761 | 8.395702 | 1.640677 | 0.531179 | 0.18825 |
| 7 | 17819200 | 17823999 | 111.3801 | 172.844 | 138.6251 | 57.56995 | 0.623907 | 2.787101 | 0.18825 |
| 9 | 58136400 | 58138199 | 8.821067 | 15.79442 | 40.51267 | 17.76086 | -0.95029 | 0.405351 | 0.188254 |
| X | 54499200 | 54502999 | 64.08145 | 103.1637 | 63.81617 | 27.11505 | 1.045776 | 1.858698 | 0.18826 |
| 12 | 23277600 | 23287599 | 120.113 | 185.5017 | 137.8544 | 57.26184 | 0.728485 | 2.87593 | 0.188276 |

| | | | | | | | | | |
|---|---|---|---|---|---|---|---|---|---|
| 9 | 90175000 | 90176399 | 11.66453 | 22.28866 | 18.34469 | 7.286114 | 0.787207 | -0.02332 | 0.18828 |
| 4 | 86812600 | 86813399 | 5.531112 | 10.69343 | 27.72188 | 10.61497 | -0.80271 | -0.17634 | 0.188282 |
| 2 | 122544000 | 122544799 | 6.83685 | 12.73256 | 33.2062 | 13.85127 | -0.92207 | 0.146689 | 0.188286 |
| 8 | 121196400 | 121201199 | 20.1473 | 32.93211 | 84.25153 | 36.22405 | -0.97677 | 1.474861 | 0.188321 |
| 13 | 67666600 | 67668199 | 10.66053 | 18.99487 | 41.47393 | 18.25743 | -0.7238 | 0.451339 | 0.18833 |
| 14 | 78799200 | 78800999 | 8.885396 | 16.12382 | 40.25555 | 17.39074 | -0.89771 | 0.323195 | 0.188346 |
| 7 | 109267000 | 109268199 | 7.75076 | 14.40438 | 34.81668 | 14.57314 | -0.81793 | 0.170524 | 0.188351 |
| 16 | 14988200 | 14989199 | 17.92636 | 32.96978 | 21.33286 | 9.056779 | 1.054882 | 0.276183 | 0.188356 |
| 5 | 122529600 | 122530599 | 4.962517 | 9.60026 | 26.53242 | 10.12933 | -0.88944 | -0.19623 | 0.188364 |
| 4 | 7743600 | 7744999 | 23.63767 | 41.03307 | 32.45052 | 14.23972 | 0.723668 | 0.816815 | 0.188366 |
| 9 | 114027400 | 114028999 | 24.19625 | 42.40682 | 28.92928 | 12.77943 | 0.936601 | 0.67923 | 0.188384 |
| 7 | 120977200 | 120980399 | 13.73294 | 23.8231 | 53.24402 | 23.36133 | -0.76844 | 0.809176 | 0.188393 |
| 9 | 51808800 | 51810399 | 8.19796 | 15.22241 | 35.37902 | 14.82582 | -0.76348 | 0.178595 | 0.188396 |
| 7 | 16576600 | 16580199 | 17.38534 | 28.63508 | 78.85276 | 34.00958 | -1.07724 | 1.365603 | 0.188408 |
| X | 26844000 | 26848199 | 56.74198 | 92.21461 | 49.10507 | 21.06696 | 1.275111 | 1.612937 | 0.188419 |
| X | 61011600 | 61015199 | 43.77131 | 71.40276 | 57.10509 | 24.53433 | 0.69385 | 1.536489 | 0.188428 |
| 15 | 66894400 | 66896599 | 33.08364 | 55.0722 | 45.49468 | 19.7561 | 0.655895 | 1.252607 | 0.188438 |
| 4 | 147952000 | 147968999 | 139.47 | 206.778 | 374.2254 | 154.1325 | -0.60426 | 3.681223 | 0.188457 |
| 17 | 39746800 | 39749399 | 32.10406 | 55.40273 | 30.22461 | 13.25924 | 1.255314 | 0.906499 | 0.188472 |
| 12 | 23288400 | 23309399 | 413.8776 | 577.9482 | 451.6784 | 187.3289 | 0.521762 | 4.602572 | 0.188474 |
| 5 | 42333600 | 42336199 | 42.94429 | 70.63134 | 50.56381 | 21.79518 | 0.859301 | 1.42471 | 0.188494 |
| 1 | 48048200 | 48049599 | 13.77058 | 25.74326 | 20.68298 | 8.572097 | 0.770174 | 0.145753 | 0.188499 |
| 13 | 38113400 | 38116599 | 21.15959 | 34.44032 | 88.47012 | 37.97905 | -0.9871 | 1.541641 | 0.188508 |
| 17 | 19440600 | 19441799 | 21.30832 | 37.73305 | 23.56207 | 10.41864 | 1.066682 | 0.52669 | 0.188538 |
| 9 | 44709400 | 44714199 | 23.78797 | 38.83004 | 84.21013 | 36.18826 | -0.74069 | 1.502236 | 0.188547 |
| 16 | 9760400 | 9761799 | 8.247557 | 15.09986 | 36.66813 | 15.664 | -0.84674 | 0.260509 | 0.18855 |
| 3 | 33805800 | 33808999 | 27.7444 | 47.78626 | 35.43111 | 15.54236 | 0.814275 | 0.91024 | 0.188573 |
| 7 | 47085600 | 47087399 | 27.24579 | 46.45721 | 34.90475 | 15.30139 | 0.793084 | 0.997386 | 0.188573 |
| 17 | 33997200 | 34001199 | 18.25711 | 30.58243 | 67.67382 | 29.5038 | -0.76085 | 1.162863 | 0.188585 |
| 7 | 27117400 | 27119199 | 14.38441 | 24.93706 | 52.69336 | 23.11927 | -0.68798 | 0.819321 | 0.188587 |
| 13 | 116182000 | 116184199 | 17.86548 | 32.48459 | 22.70158 | 9.782698 | 0.929756 | 0.340033 | 0.188592 |
| 13 | 15010800 | 15011999 | 18.98894 | 35.67207 | 16.34956 | 6.720418 | 1.582612 | 0.152161 | 0.188596 |
| 13 | 47555000 | 47555599 | 10.94911 | 21.27038 | 15.90156 | 5.93704 | 1.007999 | -0.23709 | 0.188603 |
| 6 | 3367600 | 3376199 | 110.9615 | 174.0271 | 107.1303 | 44.43167 | 1.016848 | 2.603463 | 0.18861 |
| 2 | 52533800 | 52534199 | 1.870313 | 3.657518 | 17.99298 | 5.867795 | -1.48804 | -0.82878 | 0.188615 |
| 17 | 25179200 | 25180199 | 3.607852 | 7.006329 | 28.54221 | 10.6873 | -1.41593 | -0.27603 | 0.188618 |
| 3 | 29490600 | 29492399 | 21.44891 | 37.69134 | 28.96994 | 12.82603 | 0.765134 | 0.621418 | 0.188618 |
| X | 23428200 | 23430599 | 25.32053 | 45.0411 | 20.97611 | 9.246935 | 1.490488 | 0.517166 | 0.188628 |
| 1 | 139826600 | 139827799 | 12.9605 | 24.71128 | 18.68453 | 7.455241 | 0.902757 | 0.0051 | 0.188629 |
| 17 | 26310800 | 26316599 | 27.35549 | 44.0883 | 105.5586 | 44.92325 | -0.90034 | 1.785228 | 0.188634 |
| 4 | 51183000 | 51183599 | 11.12553 | 21.42456 | 17.94949 | 6.973 | 0.791613 | -0.09966 | 0.188645 |
| 17 | 31711600 | 31713599 | 16.66534 | 30.15604 | 24.06182 | 10.43183 | 0.734457 | 0.362178 | 0.18865 |
| X | 17768400 | 17773799 | 92.66712 | 146.093 | 110.0372 | 45.66326 | 0.731893 | 2.490978 | 0.188661 |
| 10 | 116587000 | 116588399 | 6.003232 | 11.48273 | 29.49414 | 11.68206 | -0.83588 | -0.05343 | 0.188665 |
| 12 | 111049400 | 111050999 | 6.003232 | 11.48273 | 29.49414 | 11.68206 | -0.83588 | -0.05343 | 0.188665 |
| 8 | 111099400 | 111102399 | 18.31224 | 31.04721 | 64.05462 | 28.05437 | -0.6596 | 1.049043 | 0.188674 |
| 17 | 62373600 | 62373999 | 11.6856 | 22.41595 | 17.00109 | 6.67855 | 0.919646 | -0.06005 | 0.18868 |

| | | | | | | | | | |
|---|---|---|---|---|---|---|---|---|---|
| 1 | 136872600 | 136880999 | 65.26833 | 102.0481 | 186.3177 | 77.2988 | -0.55315 | 2.641369 | 0.188688 |
| 2 | 119067200 | 119071399 | 24.90559 | 40.30095 | 98.84253 | 42.27033 | -0.92834 | 1.664507 | 0.188693 |
| 18 | 82153800 | 82157199 | 20.86583 | 34.65767 | 72.71395 | 31.52837 | -0.68462 | 1.268171 | 0.188707 |
| 6 | 18885600 | 18887199 | 17.6806 | 32.46123 | 21.13421 | 8.99372 | 1.043129 | 0.28175 | 0.188709 |
| 3 | 96261600 | 96262599 | 10.99025 | 21.28366 | 16.89794 | 6.419453 | 0.898494 | -0.17695 | 0.188718 |
| 4 | 136106000 | 136107199 | 7.981937 | 14.5219 | 36.56251 | 15.74427 | -0.90543 | 0.298922 | 0.18873 |
| 5 | 115378000 | 115380999 | 11.91793 | 20.66655 | 53.70229 | 23.56213 | -0.98471 | 0.805417 | 0.188741 |
| 12 | 40284000 | 40284599 | 8.832382 | 17.26821 | 10.74526 | 3.680918 | 1.375689 | -0.57771 | 0.188743 |
| 11 | 70206200 | 70208599 | 9.540314 | 16.77847 | 45.11611 | 19.97679 | -1.03121 | 0.572218 | 0.188744 |
| 19 | 25508400 | 25529999 | 200.9938 | 289.5352 | 510.3487 | 210.6065 | -0.60616 | 4.137765 | 0.188756 |
| 18 | 15096600 | 15098399 | 25.13169 | 43.37864 | 35.63569 | 15.63312 | 0.667621 | 0.880594 | 0.188762 |
| 4 | 8874800 | 8878799 | 23.51058 | 38.70258 | 78.81097 | 33.98365 | -0.64398 | 1.386038 | 0.188769 |
| 7 | 9891200 | 9893599 | 35.27631 | 59.62228 | 38.07184 | 16.65876 | 1.025878 | 1.107216 | 0.188772 |
| 15 | 91662400 | 91664399 | 12.43528 | 21.8147 | 46.7036 | 20.64749 | -0.70514 | 0.629156 | 0.188814 |
| 2 | 8414400 | 8414999 | 10.52735 | 20.39368 | 16.97075 | 6.437582 | 0.833437 | -0.1867 | 0.18885 |
| 2 | 88076600 | 88077999 | 11.92034 | 22.76232 | 18.90801 | 7.52042 | 0.77257 | -0.01387 | 0.188859 |
| 4 | 151184400 | 151189199 | 53.71971 | 84.82482 | 161.2487 | 66.9532 | -0.60012 | 2.438053 | 0.188859 |
| 2 | 11802800 | 11805599 | 15.49531 | 26.77089 | 55.97001 | 24.55434 | -0.67399 | 0.858355 | 0.188867 |
| 7 | 84639200 | 84641599 | 27.79854 | 48.20126 | 32.79795 | 14.39022 | 0.939577 | 0.850744 | 0.188877 |
| 7 | 138101600 | 138104799 | 18.46778 | 31.24688 | 66.31816 | 29.02772 | -0.70053 | 1.068346 | 0.18888 |
| 7 | 25705000 | 25709599 | 99.66845 | 155.728 | 129.3326 | 53.66075 | 0.581207 | 2.66305 | 0.1889 |
| 5 | 118687000 | 118689999 | 15.74267 | 26.90283 | 57.95052 | 25.40864 | -0.71974 | 0.962956 | 0.188906 |
| 4 | 139970000 | 139973599 | 26.16429 | 42.29572 | 96.1011 | 41.05763 | -0.81978 | 1.691864 | 0.188915 |
| 18 | 71014600 | 71016199 | 10.86535 | 19.16168 | 44.04028 | 19.48966 | -0.8048 | 0.546617 | 0.188929 |
| 7 | 27252400 | 27252999 | 3.419877 | 6.670722 | 23.17129 | 8.300452 | -1.12786 | -0.45297 | 0.188947 |
| 18 | 35659400 | 35663799 | 32.77339 | 52.80713 | 103.7982 | 44.15862 | -0.61697 | 1.800611 | 0.188958 |
| X | 116869400 | 116874599 | 84.91771 | 134.8326 | 92.1698 | 38.41657 | 0.880142 | 2.308298 | 0.188967 |
| 2 | 172763600 | 172771199 | 42.45825 | 67.33444 | 150.0898 | 62.48264 | -0.82406 | 2.320051 | 0.188978 |
| 7 | 50649600 | 50652799 | 38.07265 | 63.01075 | 48.83866 | 21.12365 | 0.747033 | 1.335278 | 0.188981 |
| 4 | 57854400 | 57859399 | 54.52469 | 85.7067 | 169.6245 | 70.3638 | -0.66325 | 2.535734 | 0.188981 |
| 3 | 11185600 | 11187199 | 16.3553 | 30.747 | 18.80619 | 7.720667 | 1.171462 | 0.130058 | 0.188989 |
| 5 | 117935400 | 117941599 | 34.61192 | 55.53427 | 117.6124 | 49.72596 | -0.73198 | 1.940233 | 0.189002 |
| 7 | 141849400 | 141850799 | 7.222673 | 13.22268 | 35.59584 | 15.20716 | -0.99376 | 0.252243 | 0.189004 |
| 7 | 13638000 | 13645599 | 51.96712 | 83.96121 | 60.45701 | 25.81106 | 0.833057 | 1.732044 | 0.189013 |
| 8 | 119696000 | 119702199 | 56.98819 | 89.95456 | 163.5249 | 67.88637 | -0.53615 | 2.44912 | 0.189025 |
| 15 | 30302400 | 30305399 | 29.76057 | 51.17172 | 34.91674 | 15.31593 | 0.933267 | 0.934876 | 0.18903 |
| 10 | 77494400 | 77500399 | 61.71332 | 96.86911 | 176.2344 | 73.09225 | -0.54359 | 2.568097 | 0.18903 |
| 8 | 60529800 | 60531599 | 17.76932 | 33.0251 | 19.76198 | 8.27113 | 1.181228 | 0.214789 | 0.189045 |
| 13 | 46905200 | 46906199 | 6.3018 | 12.04961 | 30.11974 | 11.94183 | -0.79921 | -0.04822 | 0.189049 |
| 4 | 128457800 | 128489999 | 335.6906 | 461.7885 | 791.018 | 328.8416 | -0.63187 | 4.830064 | 0.18905 |
| 1 | 146326400 | 146327999 | 22.5278 | 40.21555 | 22.10434 | 9.715222 | 1.255585 | 0.478104 | 0.189057 |
| 1 | 68364000 | 68364799 | 11.26325 | 21.57225 | 17.9567 | 7.085482 | 0.780103 | -0.04858 | 0.189079 |
| 13 | 21651600 | 21652799 | 7.548618 | 14.16679 | 33.47769 | 13.7836 | -0.76676 | 0.1002 | 0.189083 |
| 14 | 83798600 | 83799199 | 9.399658 | 18.312 | 12.46065 | 4.536776 | 1.172109 | -0.35752 | 0.189096 |
| 10 | 90800200 | 90806599 | 38.6918 | 61.52821 | 143.0498 | 59.71406 | -0.88083 | 2.234907 | 0.189113 |
| 17 | 14119200 | 14119799 | 2.837324 | 5.545412 | 20.53106 | 7.093819 | -1.16823 | -0.59845 | 0.189118 |
| 1 | 9299000 | 9299599 | 10.42538 | 20.17623 | 16.54119 | 6.303616 | 0.849034 | -0.17305 | 0.189119 |

| | | | | | | | | | |
|---|---|---|---|---|---|---|---|---|---|
| 4 | 123155000 | 123155799 | 10.42538 | 20.17623 | 16.54119 | 6.303616 | 0.849034 | -0.17305 | 0.189119 |
| 3 | 152493400 | 152494399 | 5.466011 | 10.39691 | 29.85625 | 11.9733 | -1.01145 | -0.01826 | 0.189122 |
| 6 | 82707800 | 82709399 | 8.366235 | 14.99677 | 38.98616 | 17.07554 | -0.9681 | 0.393839 | 0.189124 |
| 18 | 54831800 | 54834599 | 27.25578 | 44.12813 | 94.02466 | 40.22623 | -0.72516 | 1.655987 | 0.18914 |
| 19 | 23422800 | 23427199 | 22.39423 | 36.87154 | 82.89591 | 35.74783 | -0.78634 | 1.382123 | 0.189158 |
| 7 | 61212200 | 61213599 | 20.59884 | 37.57675 | 19.72279 | 8.464531 | 1.343403 | 0.336752 | 0.189164 |
| 1 | 74457400 | 74460799 | 18.03739 | 29.97564 | 71.11301 | 30.84554 | -0.86093 | 1.254707 | 0.189165 |
| X | 24539000 | 24541199 | 62.82578 | 100.631 | 79.31344 | 33.4464 | 0.688379 | 2.016451 | 0.189173 |
| 11 | 101704600 | 101707799 | 11.56515 | 20.15997 | 49.43201 | 21.7483 | -0.90014 | 0.72804 | 0.18918 |
| 8 | 79942200 | 79945199 | 28.30677 | 48.09247 | 39.95577 | 17.51055 | 0.647856 | 1.039705 | 0.189197 |
| 1 | 110460000 | 110462599 | 39.29969 | 65.23759 | 44.92895 | 19.47306 | 0.91717 | 1.301084 | 0.189212 |
| 4 | 79180200 | 79182199 | 18.44672 | 33.43182 | 23.21772 | 10.04616 | 0.935145 | 0.364298 | 0.189212 |
| 11 | 75238600 | 75239999 | 5.875863 | 11.22152 | 29.45107 | 11.71081 | -0.87232 | -0.04394 | 0.189213 |
| 2 | 118203200 | 118206599 | 19.72902 | 32.47032 | 83.31436 | 35.92219 | -0.9764 | 1.383312 | 0.189217 |
| 5 | 24063400 | 24065399 | 10.85773 | 19.18853 | 42.47575 | 18.7894 | -0.74953 | 0.521589 | 0.189222 |
| 7 | 31046800 | 31048599 | 25.44282 | 45.91357 | 17.2213 | 7.491678 | 1.812229 | 0.427333 | 0.189233 |
| 6 | 115771800 | 115773199 | 6.373482 | 11.9908 | 32.37548 | 13.27443 | -0.95223 | 0.070778 | 0.18925 |
| 7 | 25274400 | 25285599 | 92.86817 | 141.4408 | 257.4022 | 106.6846 | -0.57797 | 3.150359 | 0.189255 |
| 2 | 40994800 | 40995799 | 14.24459 | 27.29135 | 17.16757 | 6.767674 | 1.182354 | -0.0224 | 0.189262 |
| 6 | 44887200 | 44888999 | 23.55972 | 41.38803 | 27.63734 | 12.2329 | 0.966907 | 0.635777 | 0.189264 |
| 5 | 61770000 | 61771199 | 12.86908 | 24.21306 | 19.37115 | 7.941516 | 0.788972 | 0.095063 | 0.189273 |
| 7 | 16586800 | 16592599 | 26.47195 | 42.49923 | 114.8616 | 48.60713 | -1.08163 | 1.912644 | 0.189292 |
| 10 | 21099600 | 21104799 | 32.08906 | 51.8284 | 101.7466 | 43.42095 | -0.61114 | 1.727759 | 0.189297 |
| 7 | 11288800 | 11311999 | 542.3761 | 741.4984 | 561.2319 | 233.5436 | 0.537103 | 4.92986 | 0.189305 |
| 19 | 8653600 | 8659999 | 34.18829 | 54.73162 | 119.8208 | 50.48727 | -0.78352 | 2.012713 | 0.189319 |
| 6 | 47720800 | 47721799 | 4.007986 | 7.812036 | 24.26069 | 8.776115 | -0.98063 | -0.40158 | 0.189321 |
| 10 | 67744800 | 67745799 | 11.36521 | 19.94053 | 45.1436 | 19.96036 | -0.78495 | 0.619746 | 0.189329 |
| 6 | 130755000 | 130759799 | 71.3894 | 113.9896 | 84.43003 | 35.44434 | 0.77289 | 2.12024 | 0.189335 |
| 6 | 10048800 | 10053999 | 61.5827 | 98.81679 | 76.1579 | 32.20333 | 0.723602 | 1.956349 | 0.189337 |
| 8 | 4542600 | 4549799 | 40.31916 | 64.42301 | 128.535 | 54.01236 | -0.65308 | 2.079416 | 0.189337 |
| 7 | 9810200 | 9814599 | 89.42407 | 140.3534 | 122.0627 | 50.62428 | 0.519802 | 2.576708 | 0.18934 |
| 13 | 3537400 | 3539999 | 42.13715 | 69.9922 | 42.68374 | 18.50805 | 1.09213 | 1.302301 | 0.189343 |
| 12 | 62044600 | 62045799 | 20.59183 | 37.55261 | 20.41722 | 8.76587 | 1.292963 | 0.340592 | 0.189346 |
| 5 | 31018600 | 31022799 | 23.77115 | 38.84401 | 83.09936 | 35.7248 | -0.71994 | 1.486914 | 0.189348 |
| 6 | 10526200 | 10527599 | 14.63782 | 27.97957 | 17.54384 | 6.962355 | 1.177718 | 0.002766 | 0.189367 |
| 1 | 61179600 | 61181399 | 8.235253 | 15.11073 | 36.72322 | 15.64037 | -0.84476 | 0.247628 | 0.189404 |
| 15 | 66338800 | 66340399 | 16.93795 | 31.35851 | 20.7877 | 8.746316 | 1.029026 | 0.230261 | 0.189418 |
| 19 | 42299800 | 42308599 | 40.30677 | 64.06236 | 143.0783 | 59.69424 | -0.82376 | 2.252164 | 0.189419 |
| 16 | 3231000 | 3236999 | 69.89878 | 112.4491 | 58.55227 | 24.85288 | 1.292389 | 1.885371 | 0.189423 |
| 5 | 147334000 | 147350799 | 128.279 | 191.4244 | 351.3257 | 144.8113 | -0.61582 | 3.563626 | 0.189429 |
| 10 | 79687000 | 79691999 | 21.07375 | 34.47651 | 82.82941 | 35.62393 | -0.88552 | 1.463092 | 0.189434 |
| 1 | 89481600 | 89484199 | 15.03426 | 25.94788 | 54.29223 | 23.81755 | -0.6753 | 0.864207 | 0.189461 |
| 11 | 50685800 | 50690199 | 30.6995 | 49.44735 | 104.2317 | 44.31945 | -0.7182 | 1.809496 | 0.189493 |
| 7 | 60912000 | 60917599 | 104.3039 | 162.8586 | 127.9638 | 53.09659 | 0.660287 | 2.677176 | 0.189496 |
| 15 | 27930400 | 27932199 | 7.754863 | 14.33761 | 36.93305 | 15.56823 | -0.91736 | 0.200276 | 0.1895 |
| 8 | 117196000 | 117197399 | 5.97667 | 11.52933 | 31.2536 | 12.08103 | -0.88046 | -0.1312 | 0.189504 |
| 18 | 35694600 | 35696199 | 6.534162 | 12.23553 | 32.25181 | 13.32926 | -0.92692 | 0.103506 | 0.189508 |

| | | | | | | | | | |
|---|---|---|---|---|---|---|---|---|---|
| 12 | 17516800 | 17525399 | 57.21259 | 89.67149 | 179.4976 | 74.45059 | -0.6829 | 2.592994 | 0.189511 |
| X | 31977000 | 31980599 | 35.24732 | 60.55798 | 29.77918 | 13.06207 | 1.403414 | 0.959657 | 0.189516 |
| 5 | 137580200 | 137586599 | 43.34341 | 68.95492 | 137.2502 | 57.32613 | -0.65648 | 2.224583 | 0.189523 |
| 10 | 46466800 | 46467399 | 9.141465 | 17.77362 | 15.04136 | 5.578056 | 0.839769 | -0.28563 | 0.189541 |
| 4 | 135446600 | 135455399 | 86.96297 | 133.1545 | 238.4737 | 98.96593 | -0.54878 | 3.039507 | 0.189558 |
| 4 | 47330200 | 47331199 | 5.086455 | 9.873476 | 26.22232 | 9.852837 | -0.81071 | -0.25022 | 0.189574 |
| 10 | 77417400 | 77417999 | 5.086455 | 9.873476 | 26.22232 | 9.852837 | -0.81071 | -0.25022 | 0.189574 |
| 5 | 105606200 | 105607999 | 7.684807 | 14.0173 | 37.65987 | 16.16476 | -0.99613 | 0.279776 | 0.189576 |
| 1 | 35326200 | 35327199 | 19.2263 | 34.81224 | 22.53893 | 9.763829 | 1.034477 | 0.373461 | 0.18958 |
| 2 | 127111800 | 127112999 | 6.696422 | 12.47941 | 32.68831 | 13.61975 | -0.92729 | 0.13804 | 0.189584 |
| 8 | 77635200 | 77636399 | 10.24219 | 18.12954 | 41.45884 | 18.33372 | -0.79503 | 0.499247 | 0.189595 |
| 7 | 65294000 | 65295999 | 9.826584 | 17.3092 | 45.10418 | 19.96521 | -0.98543 | 0.555412 | 0.189611 |
| 9 | 33899200 | 33900999 | 15.64959 | 28.6945 | 21.93539 | 9.351414 | 0.81127 | 0.274194 | 0.189614 |
| X | 97637800 | 97639999 | 23.72057 | 42.52609 | 20.49233 | 8.974344 | 1.447642 | 0.458554 | 0.189617 |
| 4 | 59161200 | 59163199 | 10.26067 | 18.10231 | 44.57137 | 19.72316 | -0.90333 | 0.539205 | 0.189625 |
| 10 | 89984200 | 89986199 | 18.29884 | 34.40303 | 16.98408 | 6.971828 | 1.478015 | 0.142781 | 0.189626 |
| 14 | 47200000 | 47200999 | 8.936388 | 16.0957 | 37.97941 | 16.5614 | -0.82485 | 0.366821 | 0.189676 |
| 12 | 111130800 | 111133599 | 12.66627 | 21.64406 | 59.42862 | 26.05658 | -1.06816 | 0.951563 | 0.189676 |
| X | 12180000 | 12182399 | 24.81428 | 43.39493 | 30.0046 | 13.22997 | 0.917396 | 0.718171 | 0.18971 |
| 1 | 14606600 | 14609599 | 24.25738 | 42.8445 | 24.21962 | 10.71493 | 1.208165 | 0.574351 | 0.189715 |
| 14 | 83969800 | 83971599 | 24.25738 | 42.8445 | 24.21962 | 10.71493 | 1.208165 | 0.574351 | 0.189715 |
| 17 | 8564800 | 8566999 | 17.10577 | 28.82986 | 63.23433 | 27.64523 | -0.74775 | 1.097085 | 0.189743 |
| 6 | 18030000 | 18032999 | 20.65989 | 34.04401 | 78.44848 | 33.84172 | -0.82111 | 1.365977 | 0.189747 |
| 17 | 38756600 | 38758999 | 38.28791 | 64.0869 | 40.91286 | 17.82819 | 1.026225 | 1.208343 | 0.189751 |
| 17 | 76981400 | 76982799 | 13.92986 | 26.80352 | 15.80049 | 6.15222 | 1.29013 | -0.06953 | 0.189751 |
| 4 | 10905000 | 10906599 | 10.85483 | 19.22009 | 41.95218 | 18.55071 | -0.72819 | 0.500322 | 0.189752 |
| 17 | 22582800 | 22583399 | 19.70155 | 34.9061 | 22.85892 | 10.10667 | 0.99922 | 0.511426 | 0.189773 |
| 7 | 10611400 | 10620199 | 159.1997 | 241.4825 | 187.33 | 77.54762 | 0.64639 | 3.245005 | 0.189795 |
| 4 | 74257000 | 74258799 | 10.56306 | 18.6255 | 46.14823 | 20.42319 | -0.91285 | 0.549052 | 0.189796 |
| 6 | 62939400 | 62940999 | 20.71932 | 37.2799 | 22.322 | 9.745892 | 1.139314 | 0.418844 | 0.189839 |
| 16 | 87396400 | 87398199 | 10.35381 | 18.463 | 40.44745 | 17.79367 | -0.72774 | 0.443089 | 0.189848 |
| 5 | 17759000 | 17760199 | 7.928741 | 14.59949 | 34.63 | 14.67858 | -0.8044 | 0.222854 | 0.189881 |
| 3 | 63020000 | 63021399 | 16.5579 | 30.67507 | 20.80003 | 8.743671 | 0.997441 | 0.223309 | 0.189908 |
| 17 | 33836200 | 33837799 | 29.35266 | 50.39155 | 32.5082 | 14.25862 | 1.013433 | 0.943718 | 0.18991 |
| 3 | 122965200 | 122967999 | 13.89007 | 24.09473 | 52.70389 | 23.12432 | -0.7375 | 0.809921 | 0.189925 |
| 18 | 37770000 | 37774599 | 127.4834 | 195.3095 | 161.154 | 66.88731 | 0.568899 | 3.034557 | 0.189938 |
| 7 | 24606400 | 24613199 | 30.59593 | 49.15584 | 111.2983 | 47.15072 | -0.82572 | 1.892568 | 0.189943 |
| 1 | 39737400 | 39738399 | 4.548671 | 8.782209 | 26.92864 | 10.37636 | -1.05007 | -0.16724 | 0.189948 |
| 16 | 53473200 | 53475599 | 13.70745 | 23.91983 | 50.53008 | 22.2463 | -0.6864 | 0.724379 | 0.189968 |
| 8 | 85197600 | 85202999 | 31.67724 | 50.97699 | 105.7396 | 44.90289 | -0.69663 | 1.8401 | 0.189974 |
| 17 | 32675600 | 32677999 | 12.95129 | 22.63813 | 49.42073 | 21.78226 | -0.73309 | 0.696766 | 0.189984 |
| 15 | 89958200 | 89959399 | 13.08773 | 24.77163 | 19.82561 | 8.034145 | 0.802501 | 0.059539 | 0.189984 |
| 7 | 24981200 | 24985599 | 16.77901 | 27.66593 | 80.6712 | 34.8096 | -1.15895 | 1.35064 | 0.189994 |
| 6 | 64013200 | 64014199 | 12.24315 | 23.59899 | 16.56696 | 6.418535 | 1.047119 | -0.10142 | 0.190021 |
| 10 | 47590200 | 47590999 | 12.24315 | 23.59899 | 16.56696 | 6.418535 | 1.047119 | -0.10142 | 0.190021 |
| 17 | 83348600 | 83354199 | 30.72557 | 49.43165 | 108.6052 | 46.10106 | -0.77996 | 1.847632 | 0.190038 |
| 7 | 142848200 | 142850999 | 48.84622 | 77.21821 | 155.5785 | 64.63646 | -0.68249 | 2.407759 | 0.190058 |

| | | | | | | | | | |
|---|---|---|---|---|---|---|---|---|---|
| 10 | 116710000 | 116711199 | 6.02706 | 11.50113 | 31.19006 | 12.42314 | -0.92198 | -0.03428 | 0.190066 |
| 19 | 37462000 | 37465799 | 44.30771 | 72.16855 | 58.30767 | 25.03719 | 0.677658 | 1.557893 | 0.190067 |
| 4 | 140814000 | 140823799 | 61.39394 | 95.91026 | 183.7872 | 76.25551 | -0.62372 | 2.654551 | 0.19009 |
| 8 | 5493400 | 5495599 | 29.69697 | 51.08204 | 34.21985 | 15.01042 | 0.959634 | 0.928419 | 0.190092 |
| 17 | 89754200 | 89756999 | 35.91298 | 59.35833 | 49.36251 | 21.33085 | 0.64568 | 1.344434 | 0.190094 |
| X | 67076000 | 67076999 | 12.55941 | 24.01205 | 18.81115 | 7.460583 | 0.859928 | -0.01697 | 0.190101 |
| 7 | 32062600 | 32065999 | 51.79625 | 83.9146 | 60.22225 | 25.77702 | 0.841764 | 1.668481 | 0.190114 |
| 11 | 77772800 | 77777199 | 19.25573 | 32.09317 | 69.81156 | 30.33923 | -0.73622 | 1.222817 | 0.190127 |
| 13 | 3504600 | 3505799 | 26.9024 | 46.15473 | 37.1645 | 16.30009 | 0.694735 | 0.943029 | 0.19013 |
| 1 | 18885200 | 18886199 | 18.41401 | 32.73359 | 25.92045 | 11.43301 | 0.728933 | 0.483641 | 0.190149 |
| 5 | 57094400 | 57096199 | 29.36218 | 49.84982 | 39.85637 | 17.46336 | 0.702909 | 1.048527 | 0.190156 |
| 9 | 13282800 | 13283799 | 6.00331 | 11.50572 | 29.04374 | 11.4382 | -0.80369 | -0.07538 | 0.190185 |
| 7 | 64397800 | 64410199 | 69.77993 | 107.6476 | 221.6109 | 92.05308 | -0.7412 | 2.885302 | 0.190196 |
| 13 | 47575600 | 47577599 | 2.626039 | 5.131542 | 25.50381 | 8.85291 | -1.59045 | -0.56231 | 0.190199 |
| 2 | 35980400 | 35982199 | 12.51487 | 21.71824 | 51.57896 | 22.63103 | -0.85518 | 0.799823 | 0.190203 |
| 16 | 35320600 | 35321799 | 16.83186 | 30.51109 | 24.01512 | 10.38865 | 0.756033 | 0.353344 | 0.19021 |
| 9 | 50341000 | 50343199 | 18.24588 | 30.23105 | 75.02068 | 32.473 | -0.92616 | 1.292016 | 0.19021 |
| 8 | 54080400 | 54081199 | 14.51791 | 27.86149 | 16.3501 | 6.414591 | 1.28701 | -0.03801 | 0.190213 |
| 6 | 72346000 | 72348599 | 14.51718 | 25.16514 | 53.25763 | 23.36679 | -0.69037 | 0.819842 | 0.190229 |
| 1 | 86367600 | 86372799 | 20.09921 | 32.98392 | 81.94619 | 35.28902 | -0.93171 | 1.419181 | 0.190231 |
| 3 | 122187000 | 122187999 | 8.705915 | 15.92802 | 36.11494 | 15.4422 | -0.74932 | 0.268159 | 0.190255 |
| 17 | 60611200 | 60613999 | 38.38136 | 62.95822 | 52.59511 | 22.64515 | 0.635708 | 1.446146 | 0.190257 |
| X | 108419600 | 108421399 | 35.8077 | 59.68942 | 45.45994 | 19.75828 | 0.772825 | 1.249187 | 0.190271 |
| 7 | 25729400 | 25730999 | 14.69332 | 27.42189 | 21.15579 | 8.793173 | 0.825218 | 0.166251 | 0.190271 |
| 5 | 45333600 | 45334599 | 5.383588 | 10.43414 | 27.3529 | 10.35265 | -0.80275 | -0.21977 | 0.190275 |
| 5 | 103459000 | 103459599 | 5.383588 | 10.43414 | 27.3529 | 10.35265 | -0.80275 | -0.21977 | 0.190275 |
| 14 | 20335000 | 20335599 | 9.085453 | 17.66538 | 14.96998 | 5.549883 | 0.83773 | -0.28863 | 0.190279 |
| 4 | 19724200 | 19725799 | 13.41295 | 25.32297 | 19.63964 | 8.001518 | 0.840329 | 0.07781 | 0.190297 |
| 4 | 112206200 | 112210799 | 103.3496 | 160.5944 | 138.4215 | 57.47406 | 0.521561 | 2.760324 | 0.190328 |
| 2 | 181359800 | 181361199 | 11.67528 | 20.46823 | 45.67727 | 20.18272 | -0.76444 | 0.634983 | 0.190336 |
| 10 | 83062800 | 83066599 | 21.4124 | 35.35494 | 78.23687 | 33.7843 | -0.76206 | 1.3423 | 0.190342 |
| 2 | 166658800 | 166665199 | 44.00607 | 69.55512 | 158.8599 | 65.99418 | -0.86326 | 2.408432 | 0.190343 |
| 19 | 5789800 | 5794599 | 20.87882 | 34.2539 | 80.85723 | 34.81603 | -0.8583 | 1.42421 | 0.190372 |
| 17 | 14458400 | 14462399 | 26.22422 | 42.65182 | 88.35792 | 37.92054 | -0.67866 | 1.561545 | 0.190374 |
| 13 | 110981400 | 110982599 | 7.559689 | 14.15135 | 39.81285 | 16.46306 | -1.02222 | 0.12908 | 0.19038 |
| X | 37909000 | 37911199 | 19.71279 | 36.72618 | 17.41713 | 7.260432 | 1.518363 | 0.211136 | 0.190396 |
| 3 | 78556200 | 78557599 | 13.11633 | 24.56179 | 19.31486 | 7.982115 | 0.804287 | 0.126078 | 0.190412 |
| 6 | 128339400 | 128340399 | 6.49456 | 12.27417 | 31.40333 | 12.76611 | -0.86466 | 0.039296 | 0.190426 |
| 11 | 66915600 | 66916399 | 4.409881 | 8.555152 | 26.10427 | 9.835939 | -1.0124 | -0.24996 | 0.190426 |
| 11 | 63104000 | 63106599 | 16.35605 | 28.02522 | 58.58695 | 25.69317 | -0.67622 | 0.941234 | 0.190438 |
| 18 | 69321400 | 69323399 | 21.1706 | 37.38708 | 27.32095 | 12.08733 | 0.840064 | 0.562974 | 0.190467 |
| 4 | 4103600 | 4104399 | 6.487851 | 12.25777 | 31.35696 | 12.75545 | -0.86452 | 0.044566 | 0.190473 |
| 7 | 139493000 | 139495999 | 15.24939 | 25.71953 | 68.17921 | 29.81353 | -1.02009 | 1.086443 | 0.190482 |
| 10 | 43331400 | 43335799 | 36.06505 | 57.84099 | 116.4093 | 49.18521 | -0.65932 | 1.955549 | 0.190482 |
| 10 | 88566400 | 88568399 | 11.29606 | 19.62793 | 52.08181 | 22.86802 | -1.01407 | 0.776444 | 0.190483 |
| 1 | 176281600 | 176284599 | 29.58151 | 50.41868 | 38.72216 | 16.97427 | 0.76141 | 1.014296 | 0.190485 |
| 7 | 59070200 | 59071799 | 29.58151 | 50.41868 | 38.72216 | 16.97427 | 0.76141 | 1.014296 | 0.190485 |

| | | | | | | | | | |
|---|---|---|---|---|---|---|---|---|---|
| 12 | 9623000 | 9633599 | 83.33714 | 127.1808 | 261.0408 | 108.2483 | -0.74904 | 3.106211 | 0.190502 |
| 6 | 19306200 | 19307199 | 12.34434 | 23.65036 | 18.40716 | 7.256584 | 0.877698 | -0.03831 | 0.190503 |
| 5 | 146359200 | 146359999 | 3.825092 | 7.445599 | 24.03106 | 8.829765 | -1.05709 | -0.35192 | 0.190504 |
| 18 | 21262200 | 21262999 | 3.825092 | 7.445599 | 24.03106 | 8.829765 | -1.05709 | -0.35192 | 0.190504 |
| 2 | 144513000 | 144514399 | 19.4853 | 34.42437 | 27.53474 | 12.18096 | 0.710616 | 0.548514 | 0.190508 |
| X | 103874000 | 103874999 | 17.87736 | 33.95928 | 15.45923 | 6.217143 | 1.618527 | 0.069564 | 0.190508 |
| 5 | 111812800 | 111831999 | 123.2718 | 183.2901 | 361.9988 | 149.1404 | -0.72634 | 3.628097 | 0.190521 |
| 5 | 15657400 | 15658399 | 24.53097 | 42.52322 | 34.58673 | 15.17492 | 0.683164 | 0.839523 | 0.190521 |
| 15 | 19675600 | 19676199 | 9.972616 | 19.42302 | 14.14063 | 5.166476 | 1.072338 | -0.33082 | 0.190563 |
| 12 | 101179400 | 101180199 | 12.31776 | 23.77975 | 16.61769 | 6.406279 | 1.06053 | -0.11575 | 0.190592 |
| 16 | 16646600 | 16648199 | 8.259609 | 15.08924 | 36.6214 | 15.68843 | -0.84871 | 0.273262 | 0.190611 |
| 7 | 9848400 | 9849599 | 36.80393 | 61.06667 | 47.6838 | 20.66053 | 0.73672 | 1.304855 | 0.190614 |
| 13 | 13219800 | 13222599 | 82.86958 | 131.064 | 108.6877 | 45.16573 | 0.596864 | 2.410479 | 0.190619 |
| 2 | 113811200 | 113811599 | 2.936183 | 5.741812 | 19.49698 | 6.640791 | -1.02587 | -0.65044 | 0.190628 |
| 4 | 42734600 | 42734999 | 2.936183 | 5.741812 | 19.49698 | 6.640791 | -1.02587 | -0.65044 | 0.190628 |
| 10 | 23588800 | 23589399 | 2.936183 | 5.741812 | 19.49698 | 6.640791 | -1.02587 | -0.65044 | 0.190628 |
| 8 | 11245200 | 11245999 | 3.71083 | 7.174158 | 27.4315 | 10.5043 | -1.35279 | -0.19115 | 0.190637 |
| 7 | 27020200 | 27030799 | 63.96167 | 99.22924 | 203.6782 | 84.58974 | -0.73064 | 2.778387 | 0.190654 |
| 2 | 141383000 | 141384799 | 12.91019 | 22.57206 | 48.89361 | 21.55458 | -0.7221 | 0.688897 | 0.190671 |
| 9 | 48678800 | 48685399 | 53.54713 | 84.5871 | 159.5559 | 66.2641 | -0.58848 | 2.427945 | 0.190676 |
| 17 | 94192400 | 94193999 | 18.8087 | 34.23821 | 21.46977 | 9.238841 | 1.08615 | 0.337977 | 0.190686 |
| 9 | 22238400 | 22239799 | 10.58325 | 18.61766 | 44.06349 | 19.50966 | -0.84814 | 0.573775 | 0.190696 |
| X | 3205200 | 3212999 | 83.02266 | 132.0269 | 89.16462 | 37.22137 | 0.89971 | 2.261355 | 0.190715 |
| X | 127635800 | 127636999 | 19.27622 | 35.47419 | 20.08846 | 8.52135 | 1.246572 | 0.280683 | 0.190738 |
| X | 120458400 | 120461799 | 49.65255 | 80.51086 | 59.93121 | 25.66872 | 0.790335 | 1.649515 | 0.190763 |
| 7 | 61666000 | 61666599 | 11.23593 | 21.82527 | 15.82699 | 5.914284 | 1.0501 | -0.23139 | 0.190781 |
| X | 25942200 | 25944399 | 31.31627 | 53.68467 | 34.91718 | 15.31345 | 1.001459 | 0.968213 | 0.190784 |
| 8 | 101632000 | 101633199 | 13.40524 | 25.24514 | 20.30188 | 8.310071 | 0.783332 | 0.096802 | 0.190796 |
| 2 | 87425000 | 87427599 | 47.65567 | 76.99028 | 65.76196 | 28.07352 | 0.587147 | 1.728334 | 0.190796 |
| 6 | 49050600 | 49051399 | 4.844714 | 9.334914 | 27.44632 | 10.64464 | -0.99956 | -0.14154 | 0.190811 |
| 5 | 103752800 | 103754999 | 7.917296 | 14.24222 | 41.10467 | 17.94993 | -1.11528 | 0.370046 | 0.190817 |
| 16 | 56836200 | 56837999 | 26.07918 | 45.2796 | 33.25568 | 14.59471 | 0.830002 | 0.82807 | 0.190818 |
| 12 | 69010400 | 69011799 | 19.18521 | 34.31811 | 24.63961 | 10.811 | 0.874372 | 0.447777 | 0.190818 |
| X | 126813800 | 126815799 | 32.33482 | 55.18828 | 36.02931 | 15.79561 | 0.995271 | 1.012274 | 0.190832 |
| 7 | 61510400 | 61511399 | 14.52862 | 27.82754 | 16.9118 | 6.672338 | 1.229699 | -0.01886 | 0.190837 |
| 17 | 38106000 | 38107399 | 23.71776 | 41.22977 | 33.52694 | 14.72502 | 0.684024 | 0.801427 | 0.190839 |
| 7 | 13483400 | 13483999 | 13.73945 | 26.44283 | 15.80696 | 6.150944 | 1.270919 | -0.07356 | 0.190907 |
| 3 | 39063000 | 39065399 | 15.11417 | 25.42715 | 65.22858 | 28.50143 | -0.97286 | 1.107796 | 0.190911 |
| 11 | 107436600 | 107439399 | 15.11417 | 25.42715 | 65.22858 | 28.50143 | -0.97286 | 1.107796 | 0.190911 |
| 1 | 188386400 | 188387199 | 12.27309 | 23.47175 | 18.88031 | 7.481513 | 0.82343 | -0.02204 | 0.19092 |
| 14 | 6999800 | 7004399 | 61.38211 | 98.12247 | 83.63811 | 35.16863 | 0.572359 | 2.073574 | 0.19092 |
| 3 | 131075000 | 131085799 | 83.74939 | 127.8745 | 252.659 | 104.7848 | -0.69358 | 3.095877 | 0.190927 |
| 14 | 8341800 | 8342799 | 12.88773 | 24.81824 | 17.15548 | 6.664403 | 1.066292 | -0.08346 | 0.190933 |
| 7 | 81102200 | 81105799 | 27.42476 | 44.0628 | 122.2469 | 51.7921 | -1.11815 | 1.890539 | 0.190958 |
| X | 109907400 | 109910599 | 23.10892 | 40.44468 | 30.61393 | 13.50769 | 0.787386 | 0.697123 | 0.190988 |
| 5 | 62922800 | 62923999 | 6.728091 | 12.47611 | 34.21138 | 14.36305 | -1.00176 | 0.170813 | 0.190997 |
| 8 | 47852000 | 47855999 | 36.17916 | 58.19376 | 112.5975 | 47.78181 | -0.59751 | 1.862882 | 0.190998 |

| | | | | | | | | | |
|---|---|---|---|---|---|---|---|---|---|
| 15 | 87608800 | 87612799 | 27.36252 | 44.4335 | 90.6868 | 38.88376 | -0.65919 | 1.593141 | 0.191003 |
| 3 | 12902000 | 12905799 | 78.84261 | 124.65 | 108.6694 | 45.1499 | 0.524402 | 2.415368 | 0.19101 |
| 8 | 55079800 | 55081599 | 22.30689 | 39.11094 | 29.42007 | 13.00058 | 0.796075 | 0.661254 | 0.19102 |
| 7 | 99465000 | 99467799 | 12.63677 | 21.99876 | 50.52302 | 22.20976 | -0.80666 | 0.753118 | 0.191022 |
| 15 | 25698600 | 25701999 | 25.18117 | 41.18283 | 86.15397 | 37.04939 | -0.68709 | 1.476734 | 0.191022 |
| 7 | 19509400 | 19524599 | 114.4624 | 171.3623 | 329.2688 | 135.8054 | -0.67767 | 3.500466 | 0.191029 |
| 8 | 72082800 | 72089199 | 68.96042 | 106.2516 | 219.9029 | 91.34383 | -0.75021 | 2.908905 | 0.191029 |
| 10 | 10448600 | 10449199 | 7.187176 | 13.25254 | 34.65075 | 14.65945 | -0.94177 | 0.208167 | 0.191037 |
| 15 | 27680800 | 27681999 | 7.078367 | 12.93231 | 35.57333 | 15.2379 | -1.02699 | 0.264085 | 0.191039 |
| 15 | 25632800 | 25643199 | 64.61081 | 100.6254 | 198.5809 | 82.42505 | -0.66964 | 2.710638 | 0.19105 |
| 7 | 42239600 | 42242999 | 104.0808 | 161.5314 | 136.6467 | 56.74756 | 0.547169 | 2.781793 | 0.191091 |
| 11 | 70104600 | 70107399 | 16.01085 | 26.63611 | 71.47622 | 31.02584 | -1.03775 | 1.236614 | 0.191106 |
| 2 | 39812000 | 39812799 | 12.16719 | 23.69673 | 12.72013 | 4.64878 | 1.506396 | -0.30511 | 0.191116 |
| 16 | 17094000 | 17096599 | 19.09495 | 31.47393 | 78.15898 | 33.72077 | -0.92838 | 1.359026 | 0.191118 |
| X | 124937000 | 124939399 | 19.87929 | 36.46539 | 19.93138 | 8.492168 | 1.292626 | 0.302175 | 0.191119 |
| 17 | 19914000 | 19914999 | 12.83248 | 24.98946 | 12.02543 | 4.401819 | 1.659683 | -0.29348 | 0.191148 |
| 5 | 31245200 | 31258999 | 88.62081 | 134.5944 | 269.7667 | 111.7155 | -0.72042 | 3.207449 | 0.191151 |
| 2 | 10199400 | 10202199 | 21.23997 | 37.21263 | 30.22965 | 13.34994 | 0.68586 | 0.669287 | 0.191154 |
| 2 | 126391400 | 126395999 | 26.90573 | 43.64088 | 92.72355 | 39.72294 | -0.71863 | 1.616783 | 0.191174 |
| 9 | 101268200 | 101270199 | 25.79184 | 44.32967 | 36.72406 | 16.10849 | 0.654376 | 0.922258 | 0.191176 |
| X | 80802400 | 80804199 | 34.13628 | 57.77377 | 38.74694 | 16.96114 | 0.955533 | 1.094251 | 0.191176 |
| 1 | 80216400 | 80218199 | 11.81413 | 20.73518 | 45.64408 | 20.18742 | -0.74493 | 0.616125 | 0.191192 |
| 2 | 17312000 | 17313599 | 16.50077 | 30.05755 | 23.5289 | 10.11626 | 0.769482 | 0.322315 | 0.191226 |
| 5 | 116155000 | 116159999 | 25.7346 | 41.89417 | 86.51417 | 37.14176 | -0.6731 | 1.547094 | 0.19123 |
| 5 | 135901400 | 135904199 | 13.99866 | 23.66512 | 64.73702 | 28.32833 | -1.0648 | 1.059626 | 0.191238 |
| 6 | 16279800 | 16280199 | 7.781544 | 15.20219 | 12.83137 | 4.478296 | 0.920221 | -0.52303 | 0.191245 |
| 3 | 104959800 | 104960999 | 6.947497 | 12.66736 | 36.07252 | 15.49117 | -1.07903 | 0.275816 | 0.191245 |
| 1 | 17881600 | 17882399 | 11.63209 | 22.54526 | 16.25781 | 6.152501 | 1.040991 | -0.18419 | 0.191249 |
| 10 | 42783600 | 42784399 | 11.63209 | 22.54526 | 16.25781 | 6.152501 | 1.040991 | -0.18419 | 0.191249 |
| 2 | 154289400 | 154290199 | 5.855368 | 11.24403 | 29.14503 | 11.41418 | -0.83413 | -0.09594 | 0.191253 |
| 10 | 130385400 | 130388599 | 45.53596 | 74.07383 | 57.83946 | 24.82451 | 0.725433 | 1.577158 | 0.191256 |
| 19 | 36626000 | 36626599 | 4.229641 | 8.248099 | 23.87895 | 8.582892 | -0.87414 | -0.42785 | 0.191264 |
| 5 | 137458600 | 137460399 | 9.382002 | 16.97692 | 39.09974 | 16.95126 | -0.78485 | 0.345134 | 0.191273 |
| 12 | 73536000 | 73537399 | 24.69782 | 42.65289 | 34.715 | 15.22947 | 0.681204 | 0.873506 | 0.191285 |
| 2 | 49022800 | 49023599 | 12.26475 | 23.30514 | 18.75125 | 7.53601 | 0.804763 | 0.020945 | 0.19129 |
| 9 | 121094600 | 121098399 | 13.76447 | 23.40215 | 59.89471 | 26.25026 | -0.96837 | 1.005066 | 0.191292 |
| 4 | 136406400 | 136408799 | 8.857469 | 15.69259 | 41.51938 | 18.35719 | -1.00285 | 0.483336 | 0.191293 |
| 2 | 106852000 | 106854399 | 17.06526 | 28.86147 | 62.24771 | 27.24252 | -0.72342 | 1.064365 | 0.191297 |
| 8 | 3090400 | 3092399 | 17.27798 | 31.16688 | 24.42513 | 10.63089 | 0.756554 | 0.384573 | 0.191322 |
| 1 | 37278000 | 37281399 | 25.33834 | 41.45916 | 83.79134 | 36.03948 | -0.63694 | 1.469749 | 0.191326 |
| 13 | 114031200 | 114035399 | 21.84378 | 36.23625 | 73.26537 | 31.73979 | -0.63158 | 1.287174 | 0.191335 |
| 12 | 23812200 | 23820399 | 333.2071 | 474.0105 | 369.7863 | 152.9055 | 0.551233 | 4.326282 | 0.191336 |
| 15 | 80170200 | 80173999 | 29.1183 | 46.88387 | 106.7347 | 45.36039 | -0.8296 | 1.819034 | 0.191347 |
| 7 | 94500000 | 94501399 | 12.92941 | 22.53048 | 51.06704 | 22.46288 | -0.78771 | 0.740306 | 0.191369 |
| X | 130566400 | 130569599 | 48.66378 | 78.92809 | 59.85952 | 25.64293 | 0.763831 | 1.642575 | 0.191392 |
| 6 | 6279600 | 6282599 | 27.19758 | 44.19111 | 88.89666 | 38.1324 | -0.63755 | 1.580974 | 0.191399 |
| 1 | 53790200 | 53790799 | 9.419831 | 18.3817 | 13.87271 | 4.938386 | 1.055732 | -0.42587 | 0.191407 |

| | | | | | | | | | |
|---|---|---|---|---|---|---|---|---|---|
| 14 | 123689000 | 123691199 | 20.1888 | 36.20583 | 23.44075 | 10.26282 | 1.024688 | 0.436968 | 0.191409 |
| 7 | 16605000 | 16611199 | 33.44299 | 53.80436 | 106.8208 | 45.34472 | -0.6341 | 1.850739 | 0.191421 |
| 11 | 117318800 | 117329999 | 71.54726 | 110.2772 | 229.3005 | 95.24765 | -0.75656 | 2.903929 | 0.191426 |
| 10 | 29409000 | 29412399 | 23.83709 | 38.79182 | 87.27732 | 37.46469 | -0.79672 | 1.548498 | 0.191438 |
| 6 | 122836000 | 122839999 | 62.0757 | 99.20172 | 85.38821 | 35.88941 | 0.557834 | 2.084201 | 0.191445 |
| 15 | 79332800 | 79357599 | 181.0312 | 260.1116 | 548.295 | 226.3531 | -0.8678 | 4.174903 | 0.191447 |
| 6 | 6878000 | 6882799 | 39.76348 | 63.33664 | 135.5042 | 56.69226 | -0.75851 | 2.180636 | 0.191448 |
| 12 | 19598200 | 19598399 | 8.198379 | 16.01483 | 12.98963 | 4.541847 | 0.976563 | -0.50678 | 0.191457 |
| 12 | 83968000 | 83969199 | 7.001795 | 13.01255 | 33.17642 | 13.88647 | -0.89418 | 0.159048 | 0.191478 |
| 17 | 83821200 | 83822199 | 7.001795 | 13.01255 | 33.17642 | 13.88647 | -0.89418 | 0.159048 | 0.191478 |
| 3 | 112763400 | 112763799 | 10.60419 | 20.51966 | 16.53762 | 6.306012 | 0.872743 | -0.16887 | 0.191482 |
| 2 | 76235800 | 76237799 | 10.90875 | 19.13264 | 45.67324 | 20.18829 | -0.86107 | 0.623184 | 0.191498 |
| 14 | 51975200 | 51979199 | 33.01793 | 52.97822 | 115.0518 | 48.64543 | -0.76801 | 1.937308 | 0.191501 |
| 10 | 117440200 | 117445999 | 50.25226 | 79.52414 | 155.3575 | 64.57883 | -0.63688 | 2.382293 | 0.191513 |
| 1 | 76327800 | 76328399 | 7.48223 | 13.78693 | 35.17922 | 14.89722 | -0.90811 | 0.215047 | 0.191526 |
| 11 | 97671800 | 97673399 | 7.48223 | 13.78693 | 35.17922 | 14.89722 | -0.90811 | 0.215047 | 0.191526 |
| 15 | 85616000 | 85617399 | 7.48223 | 13.78693 | 35.17922 | 14.89722 | -0.90811 | 0.215047 | 0.191526 |
| 17 | 22294200 | 22296599 | 27.24894 | 47.5105 | 29.19835 | 12.8481 | 1.086165 | 0.770256 | 0.191543 |
| X | 24409400 | 24410999 | 34.15343 | 57.81188 | 38.65455 | 16.9215 | 0.959825 | 1.091473 | 0.191557 |
| 3 | 88592800 | 88594199 | 7.300497 | 13.15906 | 38.02528 | 16.56592 | -1.11379 | 0.350592 | 0.191563 |
| 3 | 22812000 | 22815199 | 37.8957 | 62.72484 | 48.83664 | 21.1244 | 0.740557 | 1.333398 | 0.191575 |
| 6 | 107544200 | 107547599 | 22.81565 | 37.79432 | 76.22504 | 32.98935 | -0.62825 | 1.308321 | 0.191595 |
| 12 | 106102400 | 106108199 | 38.47374 | 61.29781 | 133.7539 | 55.97957 | -0.78647 | 2.171427 | 0.191596 |
| 4 | 50170200 | 50171999 | 26.2829 | 45.76046 | 30.35561 | 13.3453 | 0.976288 | 0.78566 | 0.191601 |
| 7 | 141614000 | 141615399 | 10.72903 | 18.89033 | 43.53943 | 19.27432 | -0.8096 | 0.564412 | 0.19161 |
| 9 | 72018000 | 72019599 | 9.141188 | 15.99555 | 48.31667 | 21.31508 | -1.19836 | 0.656325 | 0.191614 |
| 15 | 15493800 | 15496799 | 42.06547 | 69.35321 | 49.45661 | 21.33972 | 0.866189 | 1.392179 | 0.191615 |
| 11 | 67079400 | 67080599 | 5.893461 | 11.20248 | 29.86304 | 11.99296 | -0.90717 | -0.00946 | 0.191628 |
| 4 | 91666000 | 91667199 | 12.93031 | 24.71017 | 18.79937 | 7.464868 | 0.90037 | -0.00916 | 0.191632 |
| 7 | 24649400 | 24658399 | 65.55893 | 101.6725 | 200.9705 | 83.46945 | -0.67667 | 2.785984 | 0.191641 |
| 13 | 96521200 | 96523399 | 30.09262 | 50.7152 | 42.11961 | 18.41387 | 0.648011 | 1.116505 | 0.191664 |
| X | 121267400 | 121270599 | 50.25512 | 81.35275 | 62.75647 | 26.84591 | 0.736812 | 1.684015 | 0.191669 |
| 5 | 40701600 | 40703199 | 15.26621 | 28.88121 | 18.34703 | 7.442416 | 1.131293 | 0.078358 | 0.19167 |
| 12 | 23903400 | 23903799 | 8.152293 | 15.92039 | 11.5073 | 4.042834 | 1.134942 | -0.49788 | 0.191673 |
| 3 | 117415400 | 117417199 | 9.503247 | 17.19215 | 40.17075 | 17.42085 | -0.80637 | 0.345981 | 0.191674 |
| 7 | 9726800 | 9738999 | 169.9037 | 257.8608 | 174.1667 | 72.11041 | 0.846656 | 3.236278 | 0.191679 |
| 13 | 106210000 | 106210599 | 8.734298 | 17.01608 | 14.55858 | 5.299349 | 0.84862 | -0.35621 | 0.191692 |
| 14 | 20332200 | 20333799 | 13.90999 | 26.05271 | 20.78111 | 8.585447 | 0.784473 | 0.136709 | 0.191698 |
| 9 | 72969200 | 72972199 | 12.9142 | 22.60669 | 47.80606 | 21.09691 | -0.68751 | 0.668206 | 0.191704 |
| 10 | 117680800 | 117682399 | 11.40833 | 19.90106 | 49.42637 | 21.75574 | -0.91844 | 0.716315 | 0.191711 |
| 6 | 100310000 | 100316399 | 38.66704 | 61.8966 | 124.9769 | 52.6538 | -0.66733 | 2.019916 | 0.191738 |
| 9 | 31942800 | 31944399 | 9.812769 | 17.30685 | 43.03604 | 19.04492 | -0.9173 | 0.538985 | 0.191744 |
| 11 | 95010600 | 95017799 | 39.03744 | 62.02501 | 145.5224 | 60.69446 | -0.89515 | 2.261021 | 0.191767 |
| 2 | 18775000 | 18776599 | 20.74066 | 36.4259 | 27.93664 | 12.36257 | 0.768044 | 0.622614 | 0.191771 |
| 11 | 100848000 | 100851199 | 13.8694 | 23.72162 | 58.4499 | 25.62935 | -0.91263 | 0.946218 | 0.191773 |
| 17 | 60927400 | 60927999 | 8.914911 | 17.34807 | 14.42072 | 5.310822 | 0.873689 | -0.31598 | 0.191795 |
| X | 3206000 | 3207999 | 26.3408 | 45.41168 | 36.1042 | 15.83807 | 0.714357 | 0.897472 | 0.191801 |

| | | | | | | | | | |
|---|---|---|---|---|---|---|---|---|---|
| 17 | 58815200 | 58816199 | 11.4488 | 22.19287 | 16.26369 | 6.151002 | 1.01869 | -0.18843 | 0.191801 |
| 11 | 116688400 | 116689399 | 6.768266 | 12.84249 | 30.99182 | 12.49316 | -0.77026 | 0.013582 | 0.191817 |
| 16 | 94943200 | 94945799 | 10.49456 | 18.77037 | 41.00317 | 17.99245 | -0.72118 | 0.425764 | 0.191828 |
| 1 | 35776600 | 35776999 | 9.182403 | 17.91933 | 13.80226 | 4.908832 | 1.026984 | -0.43378 | 0.191831 |
| 9 | 78259400 | 78261599 | 9.105085 | 16.33336 | 38.96455 | 17.05535 | -0.84463 | 0.393356 | 0.191844 |
| 1 | 85688000 | 85689999 | 13.22116 | 23.09347 | 50.52265 | 22.2565 | -0.73655 | 0.707234 | 0.19187 |
| 16 | 95344200 | 95355999 | 81.84863 | 125.5465 | 238.8688 | 99.16111 | -0.63397 | 3.001739 | 0.191876 |
| 5 | 103355000 | 103358199 | 17.42483 | 29.3171 | 68.34839 | 29.86543 | -0.83583 | 1.115101 | 0.191899 |
| 7 | 7668600 | 7672199 | 27.5436 | 48.21048 | 26.35403 | 11.62899 | 1.254258 | 0.712556 | 0.191923 |
| 9 | 114591000 | 114607199 | 105.4286 | 158.2964 | 324.3036 | 133.8464 | -0.76633 | 3.442988 | 0.191925 |
| 3 | 121254200 | 121255399 | 8.539827 | 15.6998 | 35.67918 | 15.15594 | -0.74558 | 0.238964 | 0.191944 |
| 6 | 8020000 | 8020799 | 14.38306 | 26.74411 | 21.08762 | 8.81971 | 0.786568 | 0.185292 | 0.191947 |
| 7 | 19257000 | 19258999 | 19.29967 | 32.05211 | 71.2595 | 30.89493 | -0.76778 | 1.26579 | 0.191963 |
| 2 | 124647000 | 124648799 | 22.77499 | 39.74207 | 31.90875 | 14.04792 | 0.702732 | 0.741778 | 0.191964 |
| 13 | 98419800 | 98421199 | 8.024616 | 14.47738 | 37.50666 | 16.32148 | -0.95642 | 0.350022 | 0.191967 |
| X | 38158400 | 38159199 | 17.0908 | 31.7294 | 19.6009 | 8.215903 | 1.134617 | 0.215794 | 0.191969 |
| 5 | 88182800 | 88184999 | 26.02685 | 45.63833 | 27.08148 | 11.9683 | 1.136588 | 0.678348 | 0.191978 |
| 13 | 55337800 | 55343799 | 28.65682 | 46.50343 | 91.30093 | 39.12732 | -0.60432 | 1.610548 | 0.192004 |
| 17 | 80786800 | 80788199 | 7.834146 | 14.26322 | 36.04638 | 15.50813 | -0.91017 | 0.290892 | 0.192011 |
| 10 | 117251400 | 117255599 | 19.07409 | 31.84326 | 71.64687 | 31.16947 | -0.78489 | 1.204332 | 0.192043 |
| 19 | 40727200 | 40728199 | 4.926414 | 9.56313 | 26.1783 | 9.833683 | -0.85294 | -0.25158 | 0.192044 |
| 7 | 31339800 | 31342199 | 44.38379 | 72.79121 | 47.83404 | 20.59369 | 0.980852 | 1.460466 | 0.192065 |
| 5 | 131288000 | 131289799 | 16.63309 | 29.98099 | 24.08346 | 10.49206 | 0.720324 | 0.383683 | 0.192068 |
| 3 | 124632000 | 124633199 | 15.23488 | 28.96441 | 17.2454 | 6.922973 | 1.236677 | 0.042491 | 0.192082 |
| 4 | 121257800 | 121258799 | 24.40555 | 42.31394 | 34.1294 | 14.97435 | 0.695291 | 0.83714 | 0.192085 |
| 1 | 118919400 | 118919999 | 8.684181 | 16.91144 | 14.53119 | 5.318141 | 0.83521 | -0.33889 | 0.192088 |
| 15 | 57835600 | 57836999 | 4.697152 | 9.00102 | 28.33701 | 11.16475 | -1.11797 | -0.08635 | 0.192092 |
| 12 | 19639200 | 19643799 | 102.8305 | 160.6277 | 128.2486 | 53.21238 | 0.637694 | 2.669139 | 0.192106 |
| 4 | 148831000 | 148832199 | 6.152093 | 11.78388 | 30.81832 | 12.15777 | -0.85733 | -0.06554 | 0.192111 |
| 15 | 84484600 | 84495599 | 94.54247 | 143.5385 | 268.8757 | 111.335 | -0.62333 | 3.215935 | 0.192121 |
| 1 | 164365000 | 164367999 | 23.05034 | 37.90331 | 78.59425 | 33.87447 | -0.67063 | 1.39884 | 0.19213 |
| 1 | 189637800 | 189648199 | 52.85955 | 82.81973 | 182.4322 | 75.6705 | -0.82115 | 2.597313 | 0.192136 |
| 3 | 16346000 | 16347799 | 24.22843 | 42.24121 | 32.25701 | 14.19341 | 0.774571 | 0.761382 | 0.192138 |
| 11 | 82084600 | 82088799 | 17.33788 | 29.52958 | 60.73099 | 26.62055 | -0.65421 | 1.002023 | 0.192138 |
| 4 | 16678400 | 16679599 | 14.78333 | 27.97955 | 19.11016 | 7.745033 | 1.029283 | 0.074028 | 0.192146 |
| 3 | 43907600 | 43909599 | 22.31499 | 39.0971 | 26.68436 | 11.78454 | 0.935764 | 0.668821 | 0.192147 |
| 17 | 71666400 | 71668399 | 27.64483 | 47.52157 | 36.5776 | 16.04433 | 0.760062 | 0.928975 | 0.19216 |
| 19 | 46044800 | 46045799 | 4.992509 | 9.564168 | 28.35505 | 11.18138 | -1.03392 | -0.0812 | 0.192165 |
| 10 | 57979200 | 58012799 | 2168.224 | 2572.894 | 3646.936 | 1544.346 | -0.57391 | 7.197647 | 0.192185 |
| 2 | 82869600 | 82870599 | 12.34165 | 23.44237 | 19.32124 | 7.771311 | 0.769544 | 0.026398 | 0.192187 |
| 3 | 9903200 | 9904399 | 12.34165 | 23.44237 | 19.32124 | 7.771311 | 0.769544 | 0.026398 | 0.192187 |
| 9 | 96567000 | 96571199 | 33.13928 | 53.25856 | 111.2116 | 47.13523 | -0.70873 | 1.885154 | 0.192191 |
| 8 | 106026600 | 106028599 | 7.180676 | 13.24146 | 34.11808 | 14.43275 | -0.92033 | 0.208149 | 0.192198 |
| 13 | 17057000 | 17057599 | 13.09432 | 24.84515 | 19.16662 | 7.726779 | 0.861137 | 0.040051 | 0.192207 |
| 15 | 100511400 | 100514599 | 16.46231 | 27.4142 | 70.47831 | 30.61263 | -0.97614 | 1.225739 | 0.192214 |
| 10 | 23356200 | 23357799 | 15.53223 | 26.67351 | 55.87778 | 24.50937 | -0.67864 | 0.91331 | 0.192216 |
| 1 | 30976800 | 30977799 | 6.632068 | 12.55089 | 31.98122 | 12.96416 | -0.85461 | 0.035915 | 0.192225 |

| | | | | | | | | | |
|---|---|---|---|---|---|---|---|---|---|
| 18 | 13983600 | 13984199 | 3.455613 | 6.751031 | 21.07685 | 7.354748 | -0.93952 | -0.54959 | 0.192259 |
| X | 120069600 | 120071999 | 24.22169 | 42.77491 | 24.88897 | 11.01139 | 1.166869 | 0.577787 | 0.19228 |
| 16 | 3400200 | 3405199 | 73.21118 | 117.0355 | 78.82696 | 33.14708 | 0.911573 | 2.08702 | 0.192286 |
| 11 | 9139400 | 9141999 | 17.81574 | 30.28055 | 61.83949 | 27.10192 | -0.64466 | 1.023311 | 0.192291 |
| 5 | 97033000 | 97033999 | 5.846603 | 11.22781 | 28.53133 | 11.17208 | -0.80497 | -0.09608 | 0.192292 |
| 9 | 64292800 | 64296999 | 36.3294 | 58.01881 | 126.9523 | 53.30923 | -0.78671 | 2.091637 | 0.1923 |
| 7 | 116979400 | 116983399 | 22.8495 | 37.28369 | 85.1811 | 36.60035 | -0.81602 | 1.506085 | 0.192347 |
| 7 | 31551200 | 31552599 | 22.26283 | 39.59196 | 23.80441 | 10.49597 | 1.124137 | 0.502796 | 0.192354 |
| 4 | 47091600 | 47094199 | 16.11202 | 26.92194 | 68.37294 | 29.7617 | -0.95857 | 1.185745 | 0.192384 |
| 12 | 114970200 | 114970599 | 18.63893 | 33.44027 | 19.91396 | 8.715662 | 1.145342 | 0.417712 | 0.192386 |
| X | 128002400 | 128003399 | 14.74723 | 28.11959 | 17.65169 | 7.042286 | 1.169909 | 0.021275 | 0.192389 |
| 9 | 77447800 | 77449599 | 16.88238 | 30.3636 | 23.97905 | 10.47302 | 0.742252 | 0.397166 | 0.192412 |
| 3 | 113423200 | 113424399 | 8.516236 | 15.69092 | 36.28937 | 15.36906 | -0.76786 | 0.225854 | 0.192416 |
| 17 | 45536400 | 45542199 | 29.49487 | 47.36777 | 113.1766 | 47.9178 | -0.90384 | 1.904551 | 0.192429 |
| 6 | 52252600 | 52254799 | 11.48276 | 20.20534 | 48.27453 | 21.37303 | -0.8621 | 0.580469 | 0.192453 |
| 11 | 49644200 | 49647799 | 20.16628 | 33.29436 | 80.38551 | 34.70992 | -0.88738 | 1.341256 | 0.192455 |
| 12 | 29255800 | 29258399 | 13.58572 | 23.54696 | 53.7729 | 23.59275 | -0.79966 | 0.816965 | 0.192471 |
| 2 | 108039200 | 108042999 | 38.5217 | 63.74753 | 48.29891 | 20.88875 | 0.779702 | 1.336727 | 0.192473 |
| 7 | 47733600 | 47735999 | 23.79924 | 41.31949 | 34.03892 | 14.93808 | 0.665173 | 0.819216 | 0.192475 |
| 12 | 105479800 | 105491799 | 91.71414 | 140.1932 | 255.8387 | 106.1311 | -0.57779 | 3.075366 | 0.192476 |
| 19 | 40811600 | 40816399 | 21.48619 | 35.70055 | 79.31763 | 34.39999 | -0.76714 | 1.269342 | 0.192478 |
| 9 | 15920200 | 15920799 | 11.81674 | 22.90016 | 16.25192 | 6.154002 | 1.063098 | -0.17995 | 0.192493 |
| 15 | 102967000 | 102969999 | 11.87501 | 20.68267 | 49.9717 | 21.9741 | -0.87922 | 0.742175 | 0.192499 |
| 8 | 122580200 | 122586999 | 42.47399 | 67.79423 | 127.8296 | 53.63548 | -0.57337 | 2.115874 | 0.192501 |
| 16 | 36599200 | 36599999 | 4.619383 | 8.993379 | 25.36016 | 9.304065 | -0.86325 | -0.34167 | 0.192504 |
| 17 | 61108000 | 61108799 | 10.8109 | 20.93903 | 16.90337 | 6.417662 | 0.875428 | -0.18117 | 0.19251 |
| 12 | 105883200 | 105892399 | 76.75378 | 118.4456 | 214.873 | 89.25414 | -0.55859 | 2.881608 | 0.192514 |
| 19 | 46344400 | 46345799 | 10.74428 | 18.88161 | 44.08333 | 19.51823 | -0.8289 | 0.588805 | 0.192528 |
| X | 32248000 | 32252799 | 99.9316 | 157.679 | 100.0281 | 41.51797 | 0.980133 | 2.475496 | 0.192545 |
| X | 38781400 | 38784399 | 43.78143 | 72.56647 | 43.29314 | 18.74431 | 1.12357 | 1.332811 | 0.192564 |
| 9 | 45574000 | 45578199 | 18.61288 | 30.71649 | 77.6273 | 33.50898 | -0.95288 | 1.342128 | 0.192569 |
| 3 | 43855000 | 43858999 | 84.56975 | 133.1301 | 116.1455 | 48.18883 | 0.518544 | 2.514097 | 0.192572 |
| 17 | 33741400 | 33745399 | 17.77951 | 29.81491 | 65.94989 | 28.77201 | -0.7602 | 1.148857 | 0.192572 |
| 15 | 84231800 | 84238799 | 61.40476 | 96.34186 | 175.5252 | 72.79519 | -0.54613 | 2.576714 | 0.192574 |
| 1 | 89071000 | 89077799 | 46.76367 | 73.86892 | 159.55 | 66.26053 | -0.78357 | 2.424737 | 0.192577 |
| 7 | 80480000 | 80482999 | 15.901 | 27.14984 | 57.95265 | 25.4078 | -0.70689 | 0.971156 | 0.192579 |
| 7 | 48007000 | 48008399 | 23.96035 | 42.5987 | 22.73256 | 10.02576 | 1.295081 | 0.514765 | 0.192582 |
| 9 | 14951200 | 14961199 | 57.58971 | 90.14044 | 186.7779 | 77.48103 | -0.73422 | 2.617693 | 0.192625 |
| 17 | 84364200 | 84369799 | 37.03836 | 59.41609 | 114.1236 | 48.2369 | -0.59167 | 1.949507 | 0.192625 |
| 13 | 53132400 | 53136399 | 20.67281 | 34.04239 | 80.77136 | 34.83387 | -0.86359 | 1.375297 | 0.192631 |
| 8 | 61679000 | 61679399 | 2.8356 | 5.54646 | 19.71003 | 6.61216 | -1.07121 | -0.7072 | 0.192646 |
| 2 | 85521400 | 85523199 | 16.92689 | 31.02922 | 22.34997 | 9.531198 | 0.896407 | 0.287387 | 0.192653 |
| 17 | 51460600 | 51462199 | 16.92689 | 31.02922 | 22.34997 | 9.531198 | 0.896407 | 0.287387 | 0.192653 |
| 9 | 5904600 | 5905599 | 12.62686 | 23.97449 | 19.24837 | 7.748577 | 0.805874 | 0.031331 | 0.192661 |
| 15 | 27421200 | 27422999 | 12.97707 | 22.51143 | 51.59001 | 22.63561 | -0.80417 | 0.805496 | 0.192662 |
| 10 | 67932000 | 67934599 | 23.86067 | 38.7188 | 90.57538 | 38.81522 | -0.85625 | 1.604116 | 0.192663 |
| 7 | 40847000 | 40848999 | 42.99389 | 70.29101 | 54.87076 | 23.59354 | 0.730937 | 1.502863 | 0.192668 |

| | | | | | | | | | |
|---|---|---|---|---|---|---|---|---|---|
| 3 | 142100600 | 142101799 | 8.277031 | 14.70179 | 41.0029 | 18.09787 | -1.07664 | 0.456042 | 0.192672 |
| 16 | 11378600 | 11379599 | 4.580238 | 8.860434 | 28.91491 | 11.04057 | -1.12787 | -0.19649 | 0.192686 |
| 16 | 13488600 | 13490799 | 10.09417 | 17.85086 | 42.48747 | 18.79211 | -0.85313 | 0.509028 | 0.192694 |
| 2 | 30438800 | 30452799 | 83.24431 | 126.922 | 253.6914 | 105.1782 | -0.71184 | 3.125597 | 0.192699 |
| 12 | 72991800 | 72993999 | 13.89693 | 24.15685 | 51.07903 | 22.43337 | -0.68828 | 0.782412 | 0.192707 |
| 7 | 47954800 | 47955999 | 18.60533 | 33.70709 | 23.36936 | 10.11635 | 0.937426 | 0.368986 | 0.192713 |
| 4 | 66650400 | 66650799 | 9.58824 | 18.66383 | 15.42214 | 5.666149 | 0.885392 | -0.31145 | 0.192735 |
| 4 | 86834000 | 86834599 | 9.58824 | 18.66383 | 15.42214 | 5.666149 | 0.885392 | -0.31145 | 0.192735 |
| 4 | 19651000 | 19651599 | 5.241243 | 10.11089 | 27.5082 | 10.63253 | -0.88397 | -0.1466 | 0.192741 |
| 2 | 94386600 | 94388199 | 11.97957 | 20.61671 | 55.19779 | 24.2127 | -1.02978 | 0.879017 | 0.192753 |
| 7 | 31431600 | 31434799 | 36.38776 | 61.46553 | 36.9109 | 16.14797 | 1.114051 | 1.114378 | 0.192759 |
| 5 | 22624000 | 22625599 | 18.9579 | 34.66637 | 20.36839 | 8.715418 | 1.184388 | 0.311649 | 0.192761 |
| 1 | 165541400 | 165542799 | 7.405175 | 13.8742 | 32.8538 | 13.56536 | -0.77293 | 0.108358 | 0.192774 |
| 2 | 5905200 | 5907599 | 10.71263 | 18.89297 | 44.56456 | 19.72155 | -0.84207 | 0.545778 | 0.192777 |
| 15 | 27937000 | 27939399 | 9.664742 | 17.03134 | 43.0539 | 19.05605 | -0.94123 | 0.548583 | 0.192794 |
| 19 | 58376800 | 58378399 | 10.50078 | 18.71915 | 40.956 | 18.02219 | -0.72632 | 0.446103 | 0.1928 |
| 14 | 3188800 | 3193599 | 100.3052 | 156.1216 | 135.4723 | 56.23562 | 0.513731 | 2.730323 | 0.19283 |
| X | 107248400 | 107249999 | 33.51557 | 55.75551 | 47.10623 | 20.44702 | 0.623576 | 1.263609 | 0.192833 |
| 15 | 66470800 | 66471799 | 14.98419 | 27.95308 | 21.08009 | 8.7679 | 0.856922 | 0.170844 | 0.192836 |
| 9 | 55038600 | 55040599 | 9.081934 | 16.35495 | 38.47825 | 16.78265 | -0.82125 | 0.367668 | 0.192838 |
| 2 | 178416000 | 178416599 | 5.755202 | 10.89133 | 30.27296 | 12.27199 | -0.9784 | 0.023023 | 0.192841 |
| 1 | 31879400 | 31880199 | 10.15486 | 19.74863 | 15.29155 | 5.659972 | 0.968123 | -0.28004 | 0.192845 |
| 9 | 71816000 | 71817399 | 16.52752 | 30.69137 | 20.91914 | 8.765455 | 0.993582 | 0.211341 | 0.192846 |
| 6 | 84166800 | 84172799 | 35.62689 | 57.0396 | 119.9338 | 50.54137 | -0.72531 | 2.010941 | 0.192851 |
| 7 | 59817400 | 59819599 | 41.91785 | 68.79701 | 52.81 | 22.74296 | 0.757685 | 1.446807 | 0.192859 |
| 8 | 44218200 | 44220599 | 20.20694 | 36.16479 | 23.98949 | 10.52108 | 0.988297 | 0.450563 | 0.192863 |
| 13 | 72317600 | 72318799 | 5.380249 | 10.4387 | 28.09554 | 10.58133 | -0.83381 | -0.23514 | 0.192866 |
| 11 | 5099000 | 5103999 | 18.9843 | 31.20062 | 80.713 | 34.77982 | -0.98887 | 1.397338 | 0.192888 |
| 3 | 125427400 | 125429199 | 14.09438 | 26.21811 | 21.16555 | 8.84615 | 0.753726 | 0.180755 | 0.192917 |
| 6 | 75059200 | 75060999 | 22.24667 | 39.64132 | 23.25182 | 10.23529 | 1.161128 | 0.489661 | 0.192923 |
| 7 | 14665000 | 14674399 | 161.8669 | 245.4371 | 187.1216 | 77.45274 | 0.670976 | 3.253265 | 0.19294 |
| 7 | 30461200 | 30469199 | 46.80605 | 74.22551 | 147.9277 | 61.57915 | -0.66294 | 2.324569 | 0.19294 |
| 7 | 21097600 | 21106999 | 127.1314 | 197.2386 | 124.4553 | 51.68729 | 0.969263 | 2.796949 | 0.192942 |
| 13 | 60573000 | 60576199 | 24.39227 | 39.90023 | 82.30528 | 35.39677 | -0.66654 | 1.471641 | 0.192952 |
| 13 | 113531000 | 113531999 | 4.33228 | 8.290111 | 29.3926 | 11.6277 | -1.29369 | -0.07599 | 0.192952 |
| 14 | 10642600 | 10646999 | 20.18973 | 33.66945 | 69.96852 | 30.41862 | -0.6706 | 1.219035 | 0.192962 |
| 3 | 106685600 | 106687399 | 16.85504 | 32.07149 | 14.73206 | 5.903549 | 1.608748 | 0.046129 | 0.192966 |
| 1 | 149420400 | 149421599 | 19.17177 | 33.92214 | 26.36239 | 11.65878 | 0.75254 | 0.526538 | 0.192967 |
| 16 | 94512400 | 94521799 | 69.69944 | 108.4415 | 197.6218 | 82.03928 | -0.55601 | 2.730582 | 0.192968 |
| 7 | 105391800 | 105393999 | 12.39541 | 21.42839 | 53.13924 | 23.3129 | -0.91845 | 0.83651 | 0.192974 |
| 17 | 47561800 | 47565599 | 19.07399 | 31.87433 | 67.60428 | 29.42896 | -0.69983 | 1.192695 | 0.192983 |
| 5 | 59160400 | 59164199 | 91.06889 | 142.9288 | 115.7816 | 48.01898 | 0.622082 | 2.579266 | 0.192996 |
| 3 | 32762400 | 32763599 | 8.856358 | 15.77228 | 40.9884 | 18.05134 | -0.97354 | 0.441548 | 0.193009 |
| 6 | 136398000 | 136403399 | 30.97641 | 49.98828 | 103.0174 | 43.91452 | -0.68247 | 1.751112 | 0.193027 |
| 11 | 53190400 | 53191599 | 6.830614 | 12.40081 | 37.06653 | 16.00389 | -1.15354 | 0.302206 | 0.193039 |
| X | 148351800 | 148354199 | 44.40708 | 72.89904 | 46.74231 | 20.13181 | 1.016718 | 1.448574 | 0.193064 |
| 10 | 93933200 | 93934199 | 5.534415 | 10.68807 | 27.62299 | 10.63152 | -0.80531 | -0.15868 | 0.193078 |

| | | | | | | | | | |
|---|---|---|---|---|---|---|---|---|---|
| 12 | 61923600 | 61925199 | 14.09536 | 26.39215 | 20.77644 | 8.587783 | 0.802812 | 0.140325 | 0.193079 |
| 19 | 8171400 | 8173199 | 22.50731 | 39.19306 | 31.06198 | 13.65719 | 0.721105 | 0.767529 | 0.193088 |
| 15 | 101187800 | 101189599 | 6.902981 | 12.70547 | 34.61845 | 14.68192 | -1.00309 | 0.216998 | 0.193091 |
| 16 | 28298800 | 28299999 | 14.88481 | 28.1754 | 18.36121 | 7.439375 | 1.096138 | 0.07085 | 0.193094 |
| 4 | 135360400 | 135363399 | 17.16074 | 28.7692 | 66.43949 | 28.98047 | -0.82212 | 1.150258 | 0.193118 |
| 4 | 125609200 | 125611199 | 11.98514 | 20.99485 | 46.21327 | 20.40593 | -0.74482 | 0.650069 | 0.193123 |
| 6 | 16352400 | 16353999 | 20.71509 | 36.45625 | 28.41876 | 12.58173 | 0.745425 | 0.598056 | 0.193136 |
| 1 | 36703400 | 36705399 | 8.832478 | 15.78723 | 40.49704 | 17.78019 | -0.95184 | 0.416792 | 0.193141 |
| 11 | 89190600 | 89194599 | 21.40485 | 34.99963 | 84.05746 | 36.14511 | -0.88564 | 1.47154 | 0.193145 |
| 7 | 107279600 | 107281399 | 12.59926 | 22.08721 | 46.71836 | 20.64206 | -0.68775 | 0.641668 | 0.193148 |
| 18 | 80749200 | 80753799 | 24.54497 | 39.79556 | 93.83161 | 40.18559 | -0.86891 | 1.621577 | 0.193157 |
| 14 | 53647000 | 53648399 | 16.17079 | 29.60529 | 23.04365 | 9.844144 | 0.783669 | 0.290577 | 0.193159 |
| X | 62056000 | 62057999 | 29.04755 | 50.87395 | 24.68461 | 10.89796 | 1.425264 | 0.709704 | 0.19316 |
| 16 | 35920800 | 35923999 | 46.35086 | 75.2859 | 60.53053 | 25.95814 | 0.681667 | 1.60755 | 0.1932 |
| 13 | 14787400 | 14788399 | 3.927718 | 7.529983 | 30.57193 | 12.02966 | -1.4797 | -0.09576 | 0.1932 |
| 7 | 113272600 | 113273799 | 8.433344 | 15.33743 | 36.54415 | 15.74373 | -0.82748 | 0.301676 | 0.193203 |
| 18 | 46222200 | 46222999 | 11.96595 | 22.9757 | 18.40169 | 7.208915 | 0.844898 | -0.06266 | 0.193207 |
| 9 | 11211800 | 11214599 | 29.02673 | 49.54302 | 38.16931 | 16.73383 | 0.757141 | 0.998393 | 0.193257 |
| 15 | 88950800 | 88953399 | 20.3058 | 33.17675 | 84.19212 | 36.19276 | -0.96555 | 1.481818 | 0.193271 |
| 7 | 31753200 | 31761599 | 163.9927 | 247.2144 | 207.7158 | 85.82017 | 0.523721 | 3.371163 | 0.193286 |
| 8 | 122936600 | 122946799 | 67.50503 | 104.4428 | 210.373 | 87.38373 | -0.7065 | 2.82948 | 0.19329 |
| 5 | 95537600 | 95541199 | 34.56209 | 58.03214 | 42.60351 | 18.60165 | 0.825298 | 1.166284 | 0.193297 |
| 7 | 66855400 | 66856599 | 6.629534 | 12.55541 | 30.91666 | 12.51297 | -0.80422 | 0.025145 | 0.193319 |
| 2 | 91540400 | 91541999 | 5.856324 | 10.92073 | 34.86485 | 14.51102 | -1.20943 | 0.129501 | 0.193327 |
| 2 | 76815600 | 76816399 | 5.543384 | 10.71678 | 28.36901 | 10.86451 | -0.83345 | -0.17616 | 0.193329 |
| 7 | 134036800 | 134037599 | 5.543384 | 10.71678 | 28.36901 | 10.86451 | -0.83345 | -0.17616 | 0.193329 |
| 6 | 74727200 | 74728999 | 21.57598 | 39.34067 | 17.2653 | 7.414216 | 1.598173 | 0.340857 | 0.193347 |
| 5 | 111612400 | 111633199 | 192.3973 | 276.7827 | 508.7965 | 210.0064 | -0.66863 | 4.156913 | 0.193347 |
| 12 | 86356000 | 86357999 | 12.90534 | 22.5763 | 48.88583 | 21.56141 | -0.72162 | 0.679502 | 0.193358 |
| 17 | 68185200 | 68186599 | 6.927702 | 13.13071 | 32.65669 | 13.19462 | -0.81617 | 0.028296 | 0.193369 |
| 12 | 109329600 | 109330399 | 12.73461 | 24.06423 | 19.85839 | 8.075052 | 0.754337 | 0.067072 | 0.193374 |
| 7 | 63917000 | 63918799 | 18.70216 | 31.42527 | 64.97411 | 28.37981 | -0.66321 | 1.129973 | 0.193385 |
| 7 | 62216400 | 62223399 | 190.1179 | 284.6279 | 226.0562 | 93.23592 | 0.596087 | 3.508762 | 0.193395 |
| 10 | 82656000 | 82658199 | 10.75628 | 18.86364 | 45.68781 | 20.1933 | -0.8815 | 0.626553 | 0.193399 |
| 7 | 75563800 | 75566599 | 13.13451 | 22.7531 | 53.74609 | 23.58062 | -0.84828 | 0.81983 | 0.193399 |
| 2 | 168795600 | 168815199 | 160.1047 | 235.4841 | 409.9061 | 168.805 | -0.55894 | 3.816904 | 0.193408 |
| 11 | 44196800 | 44197599 | 8.008669 | 14.47487 | 37.52955 | 16.29439 | -0.95521 | 0.340334 | 0.193411 |
| 4 | 118911000 | 118915399 | 25.75045 | 41.87499 | 87.56324 | 37.57743 | -0.69222 | 1.563025 | 0.193427 |
| 12 | 113861800 | 113863599 | 33.77838 | 56.63816 | 43.70634 | 19.0666 | 0.753602 | 1.177654 | 0.193435 |
| 7 | 32981800 | 32990999 | 196.3768 | 293.4625 | 230.0261 | 94.8397 | 0.612681 | 3.543749 | 0.193439 |
| 17 | 58046400 | 58046999 | 10.10284 | 19.71269 | 13.04321 | 4.650502 | 1.239781 | -0.41529 | 0.193442 |
| 2 | 12952000 | 12955399 | 53.91989 | 87.15934 | 62.53836 | 26.7167 | 0.838944 | 1.72164 | 0.193448 |
| 18 | 52660600 | 52664399 | 26.69399 | 43.42176 | 93.47108 | 40.11693 | -0.73384 | 1.561684 | 0.193457 |
| 13 | 38262800 | 38265599 | 15.11067 | 25.79622 | 59.0383 | 25.88344 | -0.80716 | 0.968758 | 0.19346 |
| 13 | 107345800 | 107347799 | 9.658624 | 17.47213 | 39.62734 | 17.18656 | -0.7637 | 0.346896 | 0.193506 |
| 5 | 143290400 | 143296599 | 29.99068 | 48.12138 | 115.1765 | 48.69888 | -0.90787 | 1.93399 | 0.193509 |
| 9 | 122634200 | 122634999 | 5.442187 | 10.41984 | 28.95454 | 11.43595 | -0.94463 | -0.072 | 0.193513 |

| | | | | | | | | | |
|---|---|---|---|---|---|---|---|---|---|
| 19 | 42030000 | 42035999 | 36.14979 | 57.77655 | 124.3423 | 52.27036 | -0.76158 | 2.066838 | 0.193529 |
| 6 | 79829600 | 79831399 | 16.42539 | 30.50603 | 20.99626 | 8.796082 | 0.979978 | 0.210343 | 0.193533 |
| 4 | 122359600 | 122360199 | 10.94014 | 21.30016 | 14.74426 | 5.407196 | 1.140741 | -0.30192 | 0.193535 |
| 17 | 18546000 | 18546599 | 10.94014 | 21.30016 | 14.74426 | 5.407196 | 1.140741 | -0.30192 | 0.193535 |
| X | 146945200 | 146945999 | 10.94014 | 21.30016 | 14.74426 | 5.407196 | 1.140741 | -0.30192 | 0.193535 |
| 9 | 87051000 | 87052799 | 9.170243 | 16.69602 | 40.92207 | 17.60548 | -0.86745 | 0.307402 | 0.193539 |
| 11 | 19282800 | 19284799 | 21.21402 | 37.57762 | 26.21179 | 11.58958 | 0.908411 | 0.529335 | 0.193544 |
| 7 | 9234400 | 9238399 | 29.29333 | 50.29472 | 35.94512 | 15.76624 | 0.866259 | 0.945661 | 0.193561 |
| X | 15522400 | 15526199 | 19.46282 | 34.7134 | 22.51816 | 9.904483 | 1.017806 | 0.464597 | 0.193581 |
| 2 | 170904600 | 170905799 | 14.97452 | 27.67066 | 22.12976 | 9.335167 | 0.757672 | 0.227776 | 0.19359 |
| 5 | 56207600 | 56208199 | 9.245537 | 16.61403 | 38.45635 | 16.80837 | -0.7999 | 0.382601 | 0.1936 |
| 7 | 128400200 | 128401599 | 9.245537 | 16.61403 | 38.45635 | 16.80837 | -0.7999 | 0.382601 | 0.1936 |
| 19 | 21035400 | 21036199 | 9.245537 | 16.61403 | 38.45635 | 16.80837 | -0.7999 | 0.382601 | 0.1936 |
| 4 | 149927600 | 149928399 | 4.109334 | 8.009941 | 24.3869 | 8.816485 | -0.95374 | -0.40776 | 0.193601 |
| 7 | 12036400 | 12037399 | 23.13253 | 40.38669 | 31.6919 | 13.95695 | 0.73553 | 0.734699 | 0.193605 |
| X | 33853200 | 33853999 | 15.21355 | 28.57151 | 20.04939 | 8.24523 | 0.973027 | 0.129361 | 0.193609 |
| 13 | 68525200 | 68528999 | 68.54433 | 109.1641 | 94.09022 | 39.35765 | 0.550612 | 2.20011 | 0.193616 |
| 8 | 81343600 | 81343999 | 7.375746 | 14.41438 | 12.96012 | 4.487209 | 0.841223 | -0.55484 | 0.193617 |
| 4 | 138217800 | 138229799 | 84.27202 | 128.8998 | 243.3116 | 100.9494 | -0.62571 | 3.05976 | 0.193625 |
| 6 | 47790000 | 47790799 | 5.558693 | 10.70009 | 28.0584 | 10.91284 | -0.84082 | -0.12394 | 0.193627 |
| 14 | 42372400 | 42386999 | 141.7441 | 216.9397 | 162.458 | 67.41292 | 0.707542 | 3.058245 | 0.193644 |
| 14 | 67521400 | 67522599 | 18.59999 | 33.34987 | 24.9796 | 10.9384 | 0.815988 | 0.432181 | 0.193646 |
| 2 | 19001600 | 19003999 | 15.63939 | 26.52845 | 61.10507 | 26.76648 | -0.81761 | 1.032155 | 0.19365 |
| 7 | 136255000 | 136257199 | 14.53343 | 25.14734 | 53.23473 | 23.35544 | -0.69129 | 0.836742 | 0.193696 |
| X | 48793600 | 48794599 | 23.34107 | 41.03943 | 26.96904 | 11.94212 | 0.990347 | 0.623816 | 0.193723 |
| 10 | 61008400 | 61012399 | 17.99745 | 30.02055 | 69.11327 | 30.05125 | -0.81765 | 1.210049 | 0.19375 |
| 19 | 21674600 | 21677999 | 48.26751 | 78.23246 | 62.1159 | 26.59627 | 0.696934 | 1.655408 | 0.193751 |
| 6 | 72012800 | 72014799 | 23.04694 | 37.9432 | 77.99994 | 33.63538 | -0.65745 | 1.384932 | 0.193767 |
| 13 | 10734400 | 10738199 | 34.91884 | 58.06321 | 47.64998 | 20.67646 | 0.665385 | 1.274398 | 0.193768 |
| 7 | 11681400 | 11696199 | 380.0666 | 537.3117 | 403.6351 | 167.0754 | 0.596663 | 4.419694 | 0.193784 |
| 8 | 26274800 | 26276799 | 7.343367 | 13.50575 | 34.6087 | 14.693 | -0.91652 | 0.224792 | 0.193787 |
| 15 | 76395200 | 76396599 | 7.343367 | 13.50575 | 34.6087 | 14.693 | -0.91652 | 0.224792 | 0.193787 |
| 13 | 58629600 | 58633199 | 20.43317 | 33.83861 | 73.57685 | 31.83732 | -0.73627 | 1.302859 | 0.193795 |
| 10 | 84258800 | 84267999 | 52.04587 | 81.91467 | 174.2725 | 72.30389 | -0.766 | 2.507477 | 0.193795 |
| 8 | 61670200 | 61673199 | 16.6387 | 28.502 | 63.68815 | 27.9297 | -0.7723 | 0.947593 | 0.193797 |
| 5 | 109798200 | 109800599 | 33.97654 | 56.47707 | 47.65549 | 20.67398 | 0.6253 | 1.27399 | 0.193801 |
| 2 | 123337200 | 123339399 | 12.80387 | 22.29659 | 54.83565 | 24.11029 | -0.90498 | 0.756356 | 0.193819 |
| 12 | 3116800 | 3119599 | 80.82195 | 127.6975 | 106.7956 | 44.35697 | 0.583539 | 2.433019 | 0.19382 |
| 6 | 10265200 | 10268799 | 46.91584 | 77.48466 | 42.77528 | 18.47452 | 1.234821 | 1.380308 | 0.193824 |
| 2 | 3066600 | 3067199 | 15.3322 | 28.082 | 20.8218 | 8.88981 | 0.854004 | 0.276908 | 0.19383 |
| 6 | 68150600 | 68154999 | 58.78616 | 94.71461 | 66.13602 | 28.13334 | 0.872722 | 1.823885 | 0.193831 |
| 7 | 43016200 | 43018199 | 19.55394 | 34.51432 | 27.4622 | 12.15107 | 0.717544 | 0.557576 | 0.19384 |
| 13 | 41512000 | 41512799 | 4.008344 | 7.768858 | 26.56821 | 10.05485 | -1.18073 | -0.23809 | 0.193848 |
| 14 | 73185400 | 73186599 | 12.23637 | 23.24527 | 19.38656 | 7.795548 | 0.753153 | 0.02527 | 0.193855 |
| 5 | 112183200 | 112186599 | 15.22667 | 25.71975 | 64.28513 | 28.1235 | -0.93531 | 1.070541 | 0.193876 |
| 6 | 128297400 | 128302399 | 35.2956 | 56.53314 | 119.5991 | 50.4328 | -0.7334 | 1.996781 | 0.193886 |
| 3 | 107237600 | 107241599 | 23.74496 | 38.50773 | 92.4205 | 39.58826 | -0.89403 | 1.615231 | 0.19389 |

| | | | | | | | | | |
|---|---|---|---|---|---|---|---|---|---|
| 8 | 66770800 | 66772199 | 15.28153 | 28.84068 | 18.88953 | 7.701596 | 1.081348 | 0.095937 | 0.193892 |
| 13 | 66484400 | 66486799 | 54.49888 | 88.27982 | 58.07466 | 24.85502 | 0.966799 | 1.676025 | 0.193919 |
| 5 | 151606400 | 151607799 | 5.861838 | 11.2369 | 29.05075 | 11.43419 | -0.8371 | -0.07934 | 0.19392 |
| 17 | 45816800 | 45817999 | 5.861838 | 11.2369 | 29.05075 | 11.43419 | -0.8371 | -0.07934 | 0.19392 |
| 1 | 32557800 | 32561599 | 38.95434 | 64.63961 | 45.4913 | 19.71159 | 0.886008 | 1.304615 | 0.193922 |
| 3 | 67441600 | 67442599 | 15.09094 | 28.48889 | 18.89663 | 7.699971 | 1.063922 | 0.092219 | 0.19393 |
| 14 | 56060400 | 56064799 | 23.78011 | 38.48602 | 95.39469 | 40.80074 | -0.94319 | 1.659113 | 0.193945 |
| 1 | 47714800 | 47715799 | 12.2067 | 23.6577 | 14.26369 | 5.399205 | 1.294708 | -0.18403 | 0.193951 |
| 14 | 41762600 | 41774799 | 173.6721 | 262.4903 | 192.2861 | 79.5098 | 0.724638 | 3.320764 | 0.193952 |
| 5 | 141794400 | 141797399 | 15.37371 | 25.97056 | 62.61431 | 27.39334 | -0.88341 | 1.069817 | 0.193961 |
| 10 | 18173800 | 18175799 | 23.84337 | 41.66612 | 31.07125 | 13.69171 | 0.808819 | 0.725096 | 0.193961 |
| 8 | 22166200 | 22171999 | 77.54072 | 123.818 | 77.63748 | 32.59517 | 1.012829 | 2.123706 | 0.193967 |
| 13 | 98305800 | 98308999 | 17.41951 | 29.35248 | 63.26467 | 27.65673 | -0.72267 | 1.099735 | 0.19398 |
| 18 | 83417400 | 83418799 | 9.127811 | 16.31322 | 39.46016 | 17.32664 | -0.86764 | 0.41856 | 0.193993 |
| 4 | 79660600 | 79661399 | 12.22982 | 23.34976 | 18.19047 | 7.237329 | 0.863343 | -0.01321 | 0.193996 |
| 6 | 116741000 | 116742399 | 5.557641 | 10.39478 | 33.82711 | 14.00802 | -1.23045 | 0.105517 | 0.194007 |
| 10 | 17069800 | 17070999 | 13.16164 | 25.02498 | 19.21366 | 7.711672 | 0.873632 | 0.027343 | 0.194011 |
| 14 | 53854000 | 53855399 | 6.751549 | 12.86489 | 31.76588 | 12.69152 | -0.79195 | -0.01484 | 0.194029 |
| 12 | 22129600 | 22138399 | 215.7233 | 318.5436 | 263.8272 | 108.6492 | 0.517387 | 3.754565 | 0.194035 |
| 8 | 109161800 | 109165399 | 21.26775 | 35.08925 | 76.92987 | 33.20407 | -0.7488 | 1.350891 | 0.194039 |
| 17 | 28977000 | 28978999 | 11.35137 | 19.95611 | 44.05539 | 19.50857 | -0.74871 | 0.5859 | 0.194047 |
| 12 | 102364400 | 102373399 | 58.93256 | 92.55502 | 171.9628 | 71.32574 | -0.57324 | 2.555192 | 0.194049 |
| 12 | 8673200 | 8680199 | 95.62458 | 149.5715 | 128.3312 | 53.23934 | 0.535486 | 2.641398 | 0.194051 |
| 15 | 94163600 | 94166999 | 29.02632 | 46.94935 | 94.72447 | 40.4968 | -0.64799 | 1.679454 | 0.194055 |
| 7 | 48895200 | 48900799 | 33.59711 | 53.70004 | 126.7857 | 53.30219 | -0.89482 | 2.062476 | 0.194059 |
| X | 118585000 | 118587599 | 37.61607 | 62.79933 | 43.77144 | 19.04267 | 0.900186 | 1.2371 | 0.194065 |
| 5 | 49879200 | 49879599 | 5.88652 | 11.16746 | 46.99425 | 18.95092 | -1.56762 | 0.03977 | 0.194071 |
| 8 | 46739000 | 46742799 | 24.13139 | 39.77393 | 80.83902 | 34.87649 | -0.6406 | 1.373019 | 0.194088 |
| 7 | 31715400 | 31724799 | 162.5988 | 244.9791 | 207.0056 | 85.51422 | 0.514865 | 3.381154 | 0.194121 |
| 12 | 112582000 | 112584999 | 26.5147 | 42.82435 | 99.62669 | 42.51544 | -0.85532 | 1.718658 | 0.19413 |
| 5 | 3396600 | 3397399 | 10.28034 | 20.06633 | 12.42594 | 4.402706 | 1.341849 | -0.436 | 0.194149 |
| 9 | 4252400 | 4254599 | 35.96596 | 60.97822 | 35.79423 | 15.67703 | 1.146891 | 1.081618 | 0.194152 |
| 15 | 77440000 | 77444399 | 32.08669 | 51.86904 | 102.03 | 43.5901 | -0.61258 | 1.703375 | 0.194161 |
| 3 | 87256800 | 87259199 | 12.76799 | 22.35325 | 47.27397 | 20.86408 | -0.68753 | 0.665574 | 0.194168 |
| 7 | 34637400 | 34639799 | 9.362818 | 16.5191 | 43.04544 | 19.04774 | -0.9841 | 0.532405 | 0.19418 |
| 7 | 121609200 | 121611799 | 9.362818 | 16.5191 | 43.04544 | 19.04774 | -0.9841 | 0.532405 | 0.19418 |
| 17 | 75788800 | 75789599 | 13.23608 | 24.87715 | 20.33646 | 8.352254 | 0.756054 | 0.107719 | 0.194185 |
| 10 | 96469600 | 96471399 | 7.949436 | 14.56772 | 35.61175 | 15.1933 | -0.85464 | 0.251364 | 0.194187 |
| 11 | 74816400 | 74817399 | 7.949436 | 14.56772 | 35.61175 | 15.1933 | -0.85464 | 0.251364 | 0.194187 |
| 4 | 32356600 | 32356999 | 3.638824 | 7.110932 | 21.23625 | 7.363331 | -0.86937 | -0.57163 | 0.194196 |
| 7 | 115811000 | 115811599 | 3.638824 | 7.110932 | 21.23625 | 7.363331 | -0.86937 | -0.57163 | 0.194196 |
| 4 | 152242600 | 152246199 | 33.66792 | 53.99518 | 114.7842 | 48.49682 | -0.7381 | 1.953745 | 0.1942 |
| 2 | 60205600 | 60207599 | 11.97571 | 21.00432 | 45.66085 | 20.1831 | -0.72692 | 0.628727 | 0.194201 |
| 6 | 133946400 | 133950599 | 22.50797 | 37.17767 | 78.45562 | 33.88742 | -0.69379 | 1.340813 | 0.194218 |
| 7 | 17521800 | 17528599 | 159.2394 | 241.8991 | 180.2522 | 74.65062 | 0.706123 | 3.215368 | 0.194238 |
| 2 | 104646000 | 104647799 | 11.52904 | 20.189 | 46.22462 | 20.40507 | -0.80125 | 0.653481 | 0.194257 |
| 9 | 54702200 | 54703999 | 21.05756 | 37.1203 | 27.79978 | 12.30362 | 0.803791 | 0.581411 | 0.194257 |

| | | | | | | | | | |
|---|---|---|---|---|---|---|---|---|---|
| 4 | 138405000 | 138405399 | 3.231419 | 6.318806 | 20.16067 | 6.876696 | -0.94086 | -0.63982 | 0.194272 |
| 9 | 16744800 | 16745399 | 4.485932 | 8.738999 | 24.25225 | 8.833857 | -0.83087 | -0.37322 | 0.194282 |
| 15 | 65935400 | 65937799 | 21.92856 | 38.82882 | 25.48109 | 11.26734 | 0.995571 | 0.535532 | 0.194283 |
| X | 25727600 | 25728799 | 27.56302 | 47.76014 | 32.45181 | 14.23805 | 0.941522 | 0.856157 | 0.194285 |
| 1 | 37757400 | 37759799 | 10.30134 | 18.0736 | 45.15893 | 19.96682 | -0.92627 | 0.614196 | 0.194295 |
| X | 39722400 | 39724399 | 19.18988 | 36.03865 | 16.34372 | 6.721458 | 1.597226 | 0.155808 | 0.194296 |
| 3 | 104326800 | 104333399 | 52.81068 | 83.65767 | 156.9454 | 65.27237 | -0.5774 | 2.360074 | 0.194306 |
| 14 | 41173000 | 41174799 | 15.65366 | 28.56033 | 23.13595 | 9.927345 | 0.722212 | 0.305802 | 0.194315 |
| 7 | 31358800 | 31376799 | 268.2475 | 395.5195 | 231.7918 | 95.45483 | 1.014137 | 3.784459 | 0.194318 |
| 4 | 151957600 | 151968399 | 97.04237 | 147.2723 | 270.2147 | 111.8755 | -0.59398 | 3.225131 | 0.194342 |
| 9 | 21712200 | 21714599 | 10.50834 | 18.31667 | 49.41213 | 21.73873 | -1.03688 | 0.724599 | 0.194352 |
| X | 25956600 | 25960399 | 37.01857 | 63.06668 | 31.82174 | 13.94891 | 1.364719 | 1.040606 | 0.194357 |
| 1 | 147732600 | 147735199 | 32.2536 | 55.19121 | 33.55052 | 14.71192 | 1.0979 | 0.983162 | 0.194363 |
| 7 | 4331600 | 4334199 | 31.60532 | 54.3288 | 32.84335 | 14.40633 | 1.107218 | 0.944072 | 0.194373 |
| X | 32235000 | 32238799 | 39.47128 | 67.46272 | 26.68981 | 11.70239 | 1.714232 | 1.017459 | 0.194382 |
| 6 | 6859400 | 6866199 | 53.61261 | 84.50025 | 166.8028 | 69.21753 | -0.65653 | 2.479182 | 0.194387 |
| 12 | 108035800 | 108038599 | 12.22625 | 21.18023 | 52.62461 | 23.08859 | -0.92057 | 0.815426 | 0.194426 |
| 5 | 42012800 | 42014399 | 20.17377 | 35.52365 | 27.93581 | 12.36651 | 0.733076 | 0.587466 | 0.194428 |
| 15 | 40305600 | 40306799 | 16.3874 | 30.66184 | 19.22111 | 7.954526 | 1.126907 | 0.159672 | 0.194447 |
| 5 | 21821000 | 21823199 | 9.570086 | 17.11086 | 41.53257 | 18.23015 | -0.8727 | 0.420787 | 0.194453 |
| 4 | 37902400 | 37903599 | 15.44575 | 28.86359 | 20.46208 | 8.486157 | 0.948628 | 0.162475 | 0.194455 |
| 1 | 139952600 | 139954799 | 34.26853 | 58.34987 | 34.07153 | 14.93444 | 1.154779 | 1.035247 | 0.194455 |
| 18 | 21134200 | 21135999 | 11.43271 | 19.60509 | 58.8799 | 25.82332 | -1.19588 | 0.913158 | 0.194456 |
| 2 | 167536000 | 167538399 | 21.94266 | 36.08604 | 78.41857 | 33.8004 | -0.73796 | 1.39462 | 0.194468 |
| 4 | 150244200 | 150249599 | 56.94581 | 89.51488 | 171.5823 | 71.17619 | -0.61716 | 2.535585 | 0.194474 |
| 7 | 98443000 | 98447999 | 18.81889 | 31.28047 | 70.68233 | 30.66286 | -0.79098 | 1.253938 | 0.194476 |
| 4 | 91044800 | 91045799 | 5.723725 | 10.97258 | 29.63363 | 11.66229 | -0.89957 | -0.0804 | 0.194477 |
| 18 | 76106000 | 76107799 | 11.16841 | 19.68754 | 45.08425 | 19.9534 | -0.79998 | 0.554385 | 0.194515 |
| 3 | 61615200 | 61615599 | 8.397878 | 16.36945 | 14.52713 | 5.250894 | 0.804935 | -0.38613 | 0.194529 |
| 19 | 57689200 | 57690999 | 11.97796 | 21.03348 | 45.11209 | 19.96076 | -0.70749 | 0.609111 | 0.194531 |
| X | 121239600 | 121241199 | 31.14182 | 53.0065 | 38.267 | 16.77293 | 0.850202 | 1.034177 | 0.194546 |
| 3 | 27484600 | 27486399 | 8.04978 | 14.9244 | 34.23112 | 14.37469 | -0.74658 | 0.183672 | 0.194549 |
| 1 | 45777400 | 45781799 | 55.45651 | 89.48278 | 64.88353 | 27.6546 | 0.82089 | 1.7749 | 0.194555 |
| X | 30139600 | 30140399 | 10.95799 | 21.06822 | 17.97299 | 7.014262 | 0.759658 | -0.08682 | 0.194579 |
| 6 | 132448200 | 132452399 | 54.16389 | 87.48435 | 63.70464 | 27.18712 | 0.8164 | 1.744123 | 0.194592 |
| 1 | 180441600 | 180442399 | 19.32016 | 34.08594 | 26.26304 | 11.62156 | 0.763636 | 0.560227 | 0.194594 |
| 8 | 46830600 | 46831999 | 10.14202 | 17.81825 | 44.60883 | 19.74625 | -0.9291 | 0.589541 | 0.194596 |
| 14 | 121892200 | 121896199 | 21.36688 | 35.39936 | 74.96411 | 32.44676 | -0.69836 | 1.300552 | 0.194605 |
| 8 | 85292200 | 85301799 | 69.18458 | 106.8977 | 211.9609 | 88.04393 | -0.68535 | 2.855474 | 0.194608 |
| 17 | 7521000 | 7523399 | 12.60338 | 22.08161 | 46.73099 | 20.6375 | -0.68849 | 0.65133 | 0.19461 |
| 14 | 116531000 | 116531799 | 6.613645 | 12.5811 | 30.49168 | 12.22552 | -0.76922 | -0.00587 | 0.194613 |
| 12 | 57277000 | 57277999 | 7.628295 | 14.05515 | 34.64085 | 14.67054 | -0.85843 | 0.216006 | 0.194627 |
| 11 | 109903800 | 109904399 | 11.03058 | 21.41725 | 14.49725 | 5.436055 | 1.145199 | -0.22484 | 0.194635 |
| 12 | 21507800 | 21510799 | 28.72433 | 49.20823 | 37.05915 | 16.25196 | 0.790557 | 0.96351 | 0.194636 |
| 8 | 127733000 | 127733999 | 6.318628 | 12.02919 | 29.94265 | 11.98577 | -0.80533 | -0.01634 | 0.194639 |
| 9 | 88468400 | 88468999 | 6.318628 | 12.02919 | 29.94265 | 11.98577 | -0.80533 | -0.01634 | 0.194639 |
| 11 | 63495800 | 63496999 | 6.318628 | 12.02919 | 29.94265 | 11.98577 | -0.80533 | -0.01634 | 0.194639 |

| | | | | | | | | | |
|---|---|---|---|---|---|---|---|---|---|
| 17 | 82460800 | 82461999 | 6.318628 | 12.02919 | 29.94265 | 11.98577 | -0.80533 | -0.01634 | 0.194639 |
| 2 | 132419400 | 132421999 | 17.47342 | 29.803 | 61.35315 | 26.89637 | -0.65537 | 0.989181 | 0.194656 |
| 4 | 21931000 | 21933799 | 14.06554 | 24.3504 | 52.68395 | 23.11416 | -0.72237 | 0.829409 | 0.194667 |
| 12 | 55060200 | 55061599 | 6.776884 | 12.83098 | 30.90933 | 12.5172 | -0.7735 | 0.028912 | 0.194667 |
| 13 | 114954400 | 114955199 | 6.776884 | 12.83098 | 30.90933 | 12.5172 | -0.7735 | 0.028912 | 0.194667 |
| 1 | 142331400 | 142332199 | 17.48006 | 31.2303 | 25.09629 | 11.02257 | 0.712071 | 0.445414 | 0.194672 |
| X | 114205800 | 114206999 | 15.45755 | 28.75237 | 21.1647 | 8.846566 | 0.886278 | 0.194265 | 0.194697 |
| 9 | 57168000 | 57172599 | 23.67862 | 38.54989 | 87.99444 | 37.77823 | -0.8171 | 1.541702 | 0.194699 |
| 7 | 31152400 | 31162399 | 297.4348 | 428.0789 | 345.5989 | 142.6365 | 0.518614 | 4.154958 | 0.194714 |
| 17 | 17235400 | 17235799 | 8.888444 | 17.38252 | 10.79363 | 3.671754 | 1.38812 | -0.59911 | 0.194722 |
| X | 61222600 | 61224599 | 28.51947 | 49.47831 | 32.07975 | 14.07534 | 1.008956 | 0.847422 | 0.194738 |
| 17 | 28293800 | 28297799 | 24.29435 | 39.55968 | 91.52274 | 39.29568 | -0.83629 | 1.543837 | 0.194739 |
| 10 | 60546800 | 60552399 | 100.6088 | 152.1982 | 281.6624 | 116.4998 | -0.61025 | 3.291915 | 0.194761 |
| 13 | 3497200 | 3498199 | 12.25213 | 23.51797 | 18.33332 | 7.188471 | 0.882242 | -0.05747 | 0.194804 |
| 15 | 86023800 | 86027599 | 17.8748 | 29.72662 | 70.58573 | 30.6319 | -0.86213 | 1.245201 | 0.194816 |
| 10 | 11663000 | 11665999 | 28.03777 | 48.57175 | 32.58936 | 14.2983 | 0.959273 | 0.857294 | 0.194825 |
| X | 119047800 | 119050199 | 25.7036 | 45.25654 | 24.77201 | 10.96619 | 1.252942 | 0.617742 | 0.194831 |
| 3 | 3461800 | 3462399 | 15.66514 | 28.54114 | 21.77912 | 9.361585 | 0.806383 | 0.31117 | 0.194848 |
| 10 | 112647400 | 112648999 | 22.4545 | 39.26422 | 31.28891 | 13.79509 | 0.71363 | 0.708968 | 0.194868 |
| 12 | 20606800 | 20619999 | 225.3749 | 331.8233 | 276.0683 | 113.6867 | 0.506939 | 3.803817 | 0.194871 |
| 6 | 48531200 | 48534399 | 17.06066 | 28.84979 | 63.37849 | 27.73627 | -0.74995 | 1.065751 | 0.194875 |
| 17 | 38743000 | 38747799 | 79.16494 | 125.5199 | 102.5096 | 42.66553 | 0.621928 | 2.345587 | 0.194888 |
| 14 | 101743400 | 101745199 | 7.720972 | 14.39107 | 33.82169 | 14.08815 | -0.77257 | 0.146514 | 0.194891 |
| 5 | 147366400 | 147368399 | 11.60515 | 20.14865 | 52.6257 | 23.09505 | -0.99162 | 0.790108 | 0.19491 |
| 5 | 144409200 | 144414999 | 49.86781 | 79.13404 | 145.9584 | 60.80024 | -0.55048 | 2.306814 | 0.194911 |
| 16 | 30125000 | 30128799 | 26.47344 | 42.84718 | 96.25569 | 41.17058 | -0.80176 | 1.66177 | 0.19492 |
| 10 | 57730400 | 57732199 | 12.92572 | 22.63849 | 47.79634 | 21.10178 | -0.68514 | 0.660683 | 0.194937 |
| 9 | 36324000 | 36330399 | 123.0105 | 188.9842 | 160.843 | 66.78497 | 0.527288 | 2.977374 | 0.194943 |
| 1 | 192050200 | 192051599 | 6.013583 | 11.51327 | 33.21804 | 13.12181 | -1.00003 | -0.05799 | 0.194974 |
| 4 | 72004600 | 72006799 | 12.11909 | 21.34409 | 49.36581 | 21.85229 | -0.81496 | 0.577717 | 0.194978 |
| 9 | 116492600 | 116494199 | 20.55225 | 36.38465 | 26.84531 | 11.87105 | 0.8276 | 0.532414 | 0.194983 |
| 19 | 6433400 | 6443999 | 50.74575 | 79.51099 | 180.3523 | 74.8075 | -0.86329 | 2.595851 | 0.194983 |
| 5 | 3154800 | 3156799 | 51.64094 | 83.17187 | 64.90602 | 27.59542 | 0.71207 | 1.825401 | 0.194986 |
| 4 | 17248600 | 17249599 | 14.77764 | 27.71881 | 19.35781 | 7.977389 | 0.976873 | 0.129681 | 0.194995 |
| 7 | 16453800 | 16460599 | 33.16797 | 53.2401 | 113.4819 | 48.01064 | -0.74044 | 1.924539 | 0.195 |
| 9 | 62528200 | 62542999 | 105.405 | 159.613 | 279.54 | 115.6583 | -0.52956 | 3.272209 | 0.195001 |
| 19 | 28839800 | 28842199 | 12.35029 | 21.49561 | 51.01918 | 22.42497 | -0.85354 | 0.759398 | 0.195006 |
| 11 | 98083600 | 98084599 | 4.429465 | 8.539168 | 27.42679 | 10.62146 | -1.12332 | -0.15091 | 0.195009 |
| 16 | 53414000 | 53414999 | 12.32901 | 23.75759 | 17.17427 | 6.659381 | 1.004508 | -0.09563 | 0.195019 |
| 8 | 117646800 | 117650599 | 19.78914 | 32.81415 | 74.79043 | 32.39114 | -0.80389 | 1.285857 | 0.195019 |
| 17 | 36113000 | 36114799 | 8.173986 | 14.77359 | 38.02812 | 16.51096 | -0.94558 | 0.338878 | 0.195023 |
| 11 | 10300200 | 10303599 | 33.70564 | 56.86362 | 40.91995 | 17.89507 | 0.854134 | 1.120593 | 0.195049 |
| 6 | 45442000 | 45442799 | 11.32686 | 22.05125 | 14.01008 | 5.141428 | 1.26131 | -0.2979 | 0.195052 |
| 13 | 98152600 | 98153999 | 7.639573 | 14.04169 | 34.59962 | 14.70172 | -0.86125 | 0.231601 | 0.195052 |
| 10 | 43385000 | 43393799 | 65.28974 | 100.8327 | 214.6777 | 89.17265 | -0.78837 | 2.863041 | 0.195052 |
| 18 | 81279200 | 81287799 | 63.2079 | 99.11184 | 180.8364 | 75.00249 | -0.54894 | 2.588526 | 0.195054 |
| 8 | 57772800 | 57776999 | 36.10719 | 57.80461 | 120.7701 | 50.88855 | -0.71627 | 2.013541 | 0.195055 |

| | | | | | | | | | |
|---|---|---|---|---|---|---|---|---|---|
| 4 | 130739000 | 130751999 | 122.4962 | 183.802 | 320.4199 | 132.2203 | -0.53446 | 3.458968 | 0.195065 |
| 3 | 69523000 | 69525599 | 27.09706 | 46.49958 | 36.02642 | 15.80123 | 0.750058 | 0.939119 | 0.195075 |
| 6 | 26842400 | 26843199 | 9.863959 | 19.22034 | 14.19232 | 5.153846 | 1.060158 | -0.35271 | 0.195089 |
| 5 | 125584600 | 125587199 | 24.69685 | 40.02152 | 92.36492 | 39.54266 | -0.83872 | 1.632869 | 0.1951 |
| 4 | 143413800 | 143415799 | 45.07697 | 73.47499 | 54.55789 | 23.43326 | 0.799743 | 1.544575 | 0.195117 |
| 3 | 100868400 | 100870399 | 6.894483 | 12.74039 | 36.32456 | 15.32237 | -1.06301 | 0.195376 | 0.195166 |
| 6 | 132881400 | 132882999 | 20.25542 | 35.60829 | 27.44723 | 12.15426 | 0.760848 | 0.604286 | 0.195175 |
| 2 | 5228600 | 5231599 | 19.9984 | 33.45874 | 68.37171 | 29.7853 | -0.64646 | 1.182137 | 0.195176 |
| 1 | 174364200 | 174366799 | 11.6223 | 20.52804 | 46.68848 | 20.65527 | -0.78917 | 0.538067 | 0.195176 |
| 1 | 58328200 | 58331999 | 17.90171 | 30.10007 | 65.52338 | 28.62543 | -0.73723 | 1.120915 | 0.195178 |
| X | 113308800 | 113309599 | 15.71088 | 29.99203 | 16.90906 | 6.729375 | 1.326285 | 0.020543 | 0.195182 |
| 11 | 100697800 | 100703199 | 31.10936 | 50.30881 | 97.68474 | 41.75472 | -0.5932 | 1.688294 | 0.195187 |
| 16 | 31018600 | 31020399 | 9.658697 | 17.01916 | 43.06354 | 19.06066 | -0.94248 | 0.5507 | 0.195187 |
| 1 | 143652400 | 143653799 | 9.743238 | 17.04326 | 48.8515 | 21.54485 | -1.12345 | 0.665072 | 0.195194 |
| 8 | 60725400 | 60726599 | 14.49597 | 27.93135 | 15.20384 | 5.896484 | 1.409255 | -0.07717 | 0.195195 |
| 7 | 16835000 | 16838599 | 58.27531 | 93.62568 | 75.69371 | 32.06698 | 0.656839 | 1.915561 | 0.195196 |
| 10 | 78096000 | 78101599 | 33.7404 | 54.35572 | 104.5963 | 44.48684 | -0.58673 | 1.807811 | 0.195199 |
| 10 | 62561400 | 62563799 | 8.879787 | 15.75765 | 41.48299 | 18.32424 | -0.99507 | 0.465856 | 0.1952 |
| 5 | 59957000 | 59957399 | 8.839007 | 17.25733 | 13.22105 | 4.664099 | 1.044407 | -0.46909 | 0.195208 |
| 10 | 61410800 | 61412999 | 10.68183 | 18.9365 | 41.93424 | 18.53832 | -0.74837 | 0.48434 | 0.195224 |
| 13 | 47300600 | 47301999 | 5.108273 | 9.738955 | 29.8527 | 11.91347 | -1.09687 | -0.03367 | 0.195229 |
| 4 | 118408600 | 118415799 | 55.24571 | 87.10867 | 161.8727 | 67.17517 | -0.56905 | 2.472702 | 0.195234 |
| 8 | 62839200 | 62839999 | 12.37136 | 23.39582 | 19.37361 | 7.865099 | 0.751365 | 0.056874 | 0.195238 |
| 6 | 18785400 | 18787599 | 24.83001 | 43.65484 | 22.90168 | 10.14118 | 1.312292 | 0.621428 | 0.195242 |
| 15 | 97140400 | 97144399 | 30.55834 | 49.17897 | 106.8557 | 45.38032 | -0.76334 | 1.836418 | 0.195245 |
| 9 | 40563000 | 40564399 | 8.619378 | 15.56746 | 38.00942 | 16.51759 | -0.87084 | 0.346195 | 0.195255 |
| 16 | 67429200 | 67429399 | 10.46284 | 20.41369 | 13.03318 | 4.65255 | 1.28944 | -0.40587 | 0.195263 |
| 16 | 33545600 | 33546199 | 3.040377 | 5.945425 | 19.49332 | 6.643604 | -0.97661 | -0.64537 | 0.195272 |
| 3 | 157352600 | 157353599 | 19.79536 | 35.5932 | 22.89777 | 10.00302 | 1.035807 | 0.416941 | 0.195275 |
| 4 | 83806400 | 83808399 | 11.20521 | 19.67489 | 44.60594 | 19.73541 | -0.78699 | 0.605091 | 0.195276 |
| 4 | 149689000 | 149690399 | 7.286309 | 13.56466 | 33.22342 | 13.86701 | -0.83376 | 0.149757 | 0.195277 |
| 7 | 31052400 | 31055199 | 47.59476 | 76.92149 | 65.35017 | 27.91159 | 0.595511 | 1.716115 | 0.195293 |
| 7 | 37009600 | 37014599 | 43.13501 | 68.76112 | 130.9017 | 54.82643 | -0.5892 | 2.157676 | 0.195299 |
| 11 | 120048600 | 120052599 | 18.34199 | 30.51387 | 70.71085 | 30.69334 | -0.82714 | 1.241857 | 0.195301 |
| 16 | 82460200 | 82461999 | 16.01788 | 29.34365 | 22.41408 | 9.567068 | 0.811221 | 0.283338 | 0.195305 |
| 1 | 139543200 | 139545799 | 23.54559 | 41.34467 | 26.99425 | 11.94373 | 0.998998 | 0.640237 | 0.195321 |
| 7 | 10056400 | 10058199 | 79.3836 | 125.4734 | 99.45789 | 41.31738 | 0.661197 | 2.420717 | 0.195321 |
| 18 | 13787200 | 13789199 | 9.990578 | 17.57139 | 47.73103 | 21.13439 | -1.04622 | 0.576528 | 0.195325 |
| 4 | 38241600 | 38245599 | 57.85139 | 93.25794 | 66.06068 | 28.12164 | 0.85296 | 1.807342 | 0.195337 |
| 14 | 111011600 | 111013599 | 22.15158 | 38.74274 | 30.85257 | 13.60514 | 0.714625 | 0.70369 | 0.195349 |
| 1 | 168754000 | 168755799 | 20.61851 | 37.42099 | 21.50662 | 9.28926 | 1.208162 | 0.369746 | 0.195352 |
| 15 | 88584600 | 88586399 | 7.069036 | 12.97396 | 35.63508 | 15.17199 | -1.01903 | 0.235949 | 0.195354 |
| X | 4530000 | 4536399 | 101.3083 | 159.2846 | 111.054 | 46.06902 | 0.840224 | 2.549368 | 0.195357 |
| 12 | 22909800 | 22912799 | 98.81922 | 154.1465 | 120.9796 | 50.20355 | 0.660868 | 2.691522 | 0.195367 |
| 11 | 50662600 | 50673399 | 73.16356 | 112.5373 | 243.0223 | 100.9238 | -0.81339 | 2.944892 | 0.195369 |
| 14 | 69408800 | 69413799 | 37.67901 | 60.39789 | 118.518 | 50.03641 | -0.62383 | 1.97431 | 0.19537 |
| 1 | 55882000 | 55882599 | 5.082592 | 9.879026 | 26.45224 | 9.833705 | -0.80831 | -0.28837 | 0.195395 |

| | | | | | | | | | |
|---|---|---|---|---|---|---|---|---|---|
| 7 | 17413000 | 17420399 | 209.2807 | 310.0129 | 258.3776 | 106.4089 | 0.512625 | 3.701653 | 0.195416 |
| 9 | 70283400 | 70286999 | 23.66558 | 38.97412 | 78.2822 | 33.76165 | -0.62399 | 1.380952 | 0.195418 |
| 11 | 51890400 | 51892399 | 9.181183 | 16.2832 | 41.98083 | 18.55306 | -0.9658 | 0.471818 | 0.195424 |
| 8 | 118297000 | 118313999 | 129.2016 | 192.2267 | 358.2938 | 147.6234 | -0.64206 | 3.617743 | 0.195429 |
| 11 | 88693000 | 88695999 | 15.80437 | 26.77879 | 62.20627 | 27.23975 | -0.82992 | 1.043009 | 0.195431 |
| 8 | 60933400 | 60935999 | 20.65943 | 34.45971 | 70.61609 | 30.70406 | -0.65053 | 1.218524 | 0.195439 |
| 8 | 106323000 | 106324599 | 8.716636 | 15.91345 | 36.07525 | 15.46748 | -0.75164 | 0.28086 | 0.195454 |
| 8 | 96533000 | 96534799 | 9.639366 | 17.05511 | 41.48259 | 18.34581 | -0.88359 | 0.500283 | 0.195482 |
| 1 | 120148600 | 120149399 | 5.010599 | 9.633235 | 28.53349 | 11.13672 | -1.01948 | -0.11733 | 0.195497 |
| 10 | 9799400 | 9800799 | 18.15633 | 33.74307 | 18.36923 | 7.688618 | 1.315776 | 0.213725 | 0.195501 |
| 11 | 120946400 | 120950599 | 22.73527 | 37.39028 | 77.90026 | 33.57728 | -0.67734 | 1.396775 | 0.195501 |
| Y | 2475000 | 2478199 | 16.66981 | 30.61723 | 11.72146 | 4.985483 | 1.8018 | 0.245602 | 0.19552 |
| 13 | 87717400 | 87718199 | 11.0798 | 19.38221 | 46.77005 | 20.62775 | -0.87641 | 0.670307 | 0.195523 |
| 10 | 82668000 | 82671399 | 18.86368 | 31.64855 | 66.66995 | 29.09519 | -0.69016 | 1.146722 | 0.195528 |
| 10 | 53541200 | 53542599 | 6.130212 | 11.80486 | 30.54931 | 11.86822 | -0.82113 | -0.11577 | 0.19554 |
| 4 | 142701800 | 142706999 | 36.07384 | 57.86325 | 113.7311 | 48.06393 | -0.62498 | 1.95133 | 0.195546 |
| 12 | 112959200 | 112960999 | 9.475906 | 16.80397 | 40.95139 | 18.09987 | -0.88504 | 0.473546 | 0.195549 |
| X | 133441800 | 133442799 | 15.95432 | 30.09303 | 18.11779 | 7.395984 | 1.20128 | 0.105877 | 0.195553 |
| 15 | 98923400 | 98925399 | 9.861288 | 17.66424 | 39.97217 | 17.51782 | -0.77047 | 0.408904 | 0.195584 |
| 16 | 37085800 | 37086599 | 11.51142 | 22.4089 | 14.0045 | 5.142457 | 1.284177 | -0.29342 | 0.195589 |
| 3 | 96328200 | 96330599 | 14.03379 | 24.4777 | 52.16194 | 22.95791 | -0.69907 | 0.735225 | 0.195598 |
| 13 | 62495000 | 62498599 | 72.93947 | 116.1778 | 92.51262 | 38.7042 | 0.66485 | 2.201089 | 0.19561 |
| 2 | 130900000 | 130906999 | 36.71042 | 58.49435 | 134.5327 | 56.3134 | -0.86206 | 2.166507 | 0.195622 |
| 17 | 53362000 | 53363399 | 16.16453 | 29.95672 | 21.46508 | 9.018978 | 0.919144 | 0.220356 | 0.195626 |
| 6 | 73213200 | 73215999 | 39.57069 | 65.11798 | 52.69461 | 22.71885 | 0.683308 | 1.410459 | 0.195648 |
| 16 | 86085400 | 86086599 | 19.07606 | 34.39227 | 23.61309 | 10.28421 | 0.945525 | 0.398596 | 0.195655 |
| 8 | 22742800 | 22744999 | 29.05443 | 50.5377 | 29.78681 | 13.08739 | 1.146114 | 0.813605 | 0.195672 |
| 5 | 52735200 | 52741999 | 43.33113 | 68.92255 | 140.8091 | 58.79746 | -0.69435 | 2.230129 | 0.195675 |
| 6 | 137625000 | 137626799 | 14.26544 | 24.54524 | 56.41668 | 24.7472 | -0.8113 | 0.889671 | 0.195675 |
| 1 | 85862600 | 85864999 | 15.32245 | 26.50729 | 54.89134 | 24.08209 | -0.65983 | 0.844103 | 0.19569 |
| 1 | 110469600 | 110471599 | 23.42116 | 41.2588 | 26.50838 | 11.73367 | 1.023338 | 0.600999 | 0.195708 |
| 9 | 47440600 | 47445399 | 27.25975 | 44.12468 | 93.95407 | 40.1894 | -0.72449 | 1.660611 | 0.195714 |
| 15 | 18642400 | 18643599 | 14.82712 | 27.28031 | 22.2074 | 9.423082 | 0.726352 | 0.250674 | 0.195714 |
| 18 | 35200800 | 35202999 | 11.84467 | 20.69904 | 48.35985 | 21.31051 | -0.83051 | 0.690881 | 0.195722 |
| 14 | 115845400 | 115846199 | 13.54202 | 25.37839 | 20.79044 | 8.58078 | 0.747315 | 0.129466 | 0.195728 |
| 4 | 113520000 | 113521799 | 37.54948 | 62.86957 | 39.76185 | 17.32988 | 1.039516 | 1.199686 | 0.195731 |
| 5 | 106211600 | 106214799 | 45.86721 | 74.28793 | 63.61325 | 27.22278 | 0.587209 | 1.664878 | 0.195732 |
| 17 | 38297400 | 38299999 | 35.67648 | 60.1125 | 39.76965 | 17.38535 | 0.9749 | 1.136694 | 0.195733 |
| 12 | 88544600 | 88545399 | 6.592382 | 12.61519 | 31.97395 | 12.64824 | -0.81621 | -0.048 | 0.195736 |
| 14 | 119948600 | 119949199 | 7.224184 | 13.65982 | 32.60504 | 13.24084 | -0.76434 | 0.045715 | 0.195739 |
| 18 | 76374000 | 76375599 | 12.49212 | 21.73647 | 49.98572 | 21.96691 | -0.80848 | 0.759521 | 0.195747 |
| 15 | 41878200 | 41879599 | 12.57726 | 24.05077 | 18.15697 | 7.1972 | 0.91309 | -0.02138 | 0.195759 |
| 4 | 152222800 | 152228399 | 27.89998 | 45.15103 | 95.97912 | 41.04894 | -0.72245 | 1.666308 | 0.195768 |
| 2 | 69727400 | 69728999 | 9.135702 | 15.99615 | 47.78808 | 21.09325 | -1.1825 | 0.644842 | 0.195772 |
| 17 | 30014000 | 30025999 | 99.73306 | 151.2509 | 273.2548 | 113.1107 | -0.57251 | 3.240039 | 0.195795 |
| 2 | 23818000 | 23819399 | 24.9953 | 43.18488 | 35.6356 | 15.63354 | 0.661483 | 0.871761 | 0.1958 |
| 10 | 122514200 | 122516799 | 16.85926 | 28.24857 | 67.52198 | 29.44264 | -0.87162 | 1.15565 | 0.195805 |

| | | | | | | | | | |
|---|---|---|---|---|---|---|---|---|---|
| 4 | 113183600 | 113186599 | 65.32006 | 104.4488 | 84.92556 | 35.72513 | 0.640828 | 2.070777 | 0.195806 |
| 3 | 12026200 | 12028799 | 23.98188 | 42.09624 | 27.56893 | 12.19444 | 0.994781 | 0.649852 | 0.195811 |
| 14 | 62333200 | 62335999 | 15.68959 | 26.91619 | 56.42181 | 24.7461 | -0.67984 | 0.923466 | 0.195817 |
| 8 | 123939200 | 123940399 | 6.054325 | 11.47678 | 30.9198 | 12.48949 | -0.93018 | 0.012697 | 0.195818 |
| 2 | 80086800 | 80087999 | 15.97103 | 29.69593 | 20.27653 | 8.480907 | 0.992422 | 0.196755 | 0.195818 |
| 2 | 180300600 | 180304199 | 16.83232 | 28.26289 | 65.34353 | 28.53063 | -0.8237 | 1.131362 | 0.195821 |
| 5 | 43389800 | 43392999 | 26.32757 | 42.54407 | 96.54989 | 41.22986 | -0.81868 | 1.703372 | 0.195822 |
| 4 | 112379000 | 112384399 | 93.33927 | 147.059 | 112.5144 | 46.6874 | 0.708602 | 2.502903 | 0.195828 |
| 12 | 20606800 | 20613799 | 103.0172 | 160.0659 | 138.939 | 57.68956 | 0.51135 | 2.761421 | 0.19583 |
| 1 | 136552000 | 136553799 | 10.02091 | 17.56786 | 48.29997 | 21.34154 | -1.06381 | 0.628182 | 0.195843 |
| X | 34588800 | 34589999 | 22.01922 | 38.89199 | 23.92641 | 10.58518 | 1.087135 | 0.560921 | 0.195846 |
| 18 | 37028200 | 37032399 | 98.7943 | 154.9143 | 121.6219 | 50.44303 | 0.666208 | 2.59997 | 0.195858 |
| 17 | 23382400 | 23384999 | 24.90888 | 43.89998 | 23.51501 | 10.40771 | 1.284011 | 0.598783 | 0.195889 |
| 8 | 22113600 | 22115799 | 18.88658 | 33.9124 | 24.84471 | 10.86616 | 0.848815 | 0.42467 | 0.195894 |
| 10 | 39163600 | 39168199 | 39.33899 | 63.07404 | 120.5744 | 50.92337 | -0.58576 | 1.971384 | 0.195904 |
| 1 | 190038800 | 190040599 | 15.01709 | 25.89426 | 55.39228 | 24.29944 | -0.70751 | 0.872182 | 0.195912 |
| 14 | 112286000 | 112286799 | 10.00175 | 19.48721 | 14.8894 | 5.412926 | 1.011199 | -0.34314 | 0.195913 |
| 11 | 62133400 | 62142399 | 63.86206 | 99.69767 | 185.0131 | 76.76952 | -0.57829 | 2.667977 | 0.195917 |
| 10 | 83950400 | 83952399 | 9.692234 | 17.46783 | 38.48611 | 16.76944 | -0.72622 | 0.368548 | 0.195921 |
| 18 | 83822800 | 83824999 | 15.65749 | 26.91616 | 59.23828 | 25.98418 | -0.74946 | 0.906197 | 0.195923 |
| 8 | 109206600 | 109208799 | 15.02801 | 25.91685 | 54.29327 | 23.81744 | -0.67733 | 0.870592 | 0.195932 |
| 7 | 80981600 | 80983599 | 8.456812 | 15.31362 | 37.00887 | 16.03057 | -0.85288 | 0.328596 | 0.195948 |
| 1 | 12390800 | 12391999 | 6.141419 | 11.78844 | 29.12991 | 11.42186 | -0.76728 | -0.08796 | 0.195968 |
| 4 | 107182400 | 107185799 | 16.0257 | 27.3825 | 59.12042 | 25.92129 | -0.72333 | 0.963843 | 0.195975 |
| 17 | 6845600 | 6846799 | 12.49308 | 23.57255 | 19.33643 | 7.887711 | 0.758782 | 0.072926 | 0.195986 |
| 11 | 21550200 | 21551399 | 17.51595 | 31.64526 | 24.45015 | 10.62112 | 0.778818 | 0.37822 | 0.196016 |
| 8 | 119362000 | 119366599 | 40.63606 | 65.01685 | 121.6697 | 51.22507 | -0.55865 | 2.037542 | 0.196018 |
| 13 | 58187600 | 58202199 | 126.6259 | 189.0176 | 340.2935 | 140.2726 | -0.58764 | 3.557416 | 0.196029 |
| 8 | 87812600 | 87814399 | 7.905327 | 14.65859 | 34.22606 | 14.36989 | -0.77169 | 0.182671 | 0.196044 |
| 11 | 104122000 | 104128399 | 36.85384 | 58.7619 | 132.7981 | 55.63845 | -0.83577 | 2.144246 | 0.196048 |
| 13 | 118453600 | 118455999 | 25.47635 | 44.05666 | 34.41387 | 15.09798 | 0.740408 | 0.862493 | 0.196056 |
| 2 | 125347400 | 125349399 | 12.07545 | 20.59123 | 59.86336 | 26.24276 | -1.15053 | 0.970315 | 0.19606 |
| 17 | 43676200 | 43680799 | 16.25144 | 27.57372 | 61.72453 | 27.03993 | -0.77667 | 1.031948 | 0.196062 |
| 2 | 130058600 | 130060999 | 15.26197 | 26.07813 | 57.41008 | 25.17144 | -0.75106 | 0.960329 | 0.196067 |
| 1 | 93099800 | 93113999 | 70.03307 | 107.2517 | 249.0453 | 103.3575 | -0.92282 | 3.030771 | 0.196084 |
| 7 | 126833600 | 126841999 | 81.63725 | 125.5906 | 223.0022 | 92.61172 | -0.53114 | 2.943608 | 0.196091 |
| 16 | 4793800 | 4798999 | 21.15772 | 34.80215 | 80.72766 | 34.79863 | -0.83173 | 1.390597 | 0.196091 |
| 18 | 80429200 | 80444799 | 130.432 | 193.7886 | 361.6076 | 148.9677 | -0.64551 | 3.642499 | 0.196099 |
| 1 | 116896200 | 116897999 | 30.18326 | 50.85132 | 42.67768 | 18.65599 | 0.632984 | 1.121221 | 0.196101 |
| 18 | 38623200 | 38625999 | 10.86387 | 19.18158 | 42.48365 | 18.79638 | -0.75082 | 0.532121 | 0.196123 |
| 3 | 130930200 | 130934799 | 19.17084 | 32.17206 | 67.85877 | 29.61874 | -0.69205 | 1.145494 | 0.196128 |
| 17 | 8741000 | 8749399 | 56.91356 | 89.14289 | 179.9435 | 74.64069 | -0.69575 | 2.604955 | 0.196146 |
| 8 | 122190600 | 122199999 | 70.09287 | 108.4271 | 206.7239 | 85.86807 | -0.62753 | 2.834223 | 0.196175 |
| 8 | 126216000 | 126219799 | 17.47313 | 29.28833 | 65.889 | 28.73753 | -0.78444 | 1.152957 | 0.196177 |
| 1 | 18978200 | 18979399 | 14.7239 | 27.35225 | 21.70943 | 9.091999 | 0.77612 | 0.196374 | 0.196207 |
| 18 | 61530600 | 61539199 | 60.46004 | 94.76652 | 176.8566 | 73.35478 | -0.58187 | 2.5937 | 0.196217 |
| 4 | 39817200 | 39819799 | 19.98217 | 35.38332 | 24.5142 | 10.83974 | 0.917391 | 0.518647 | 0.196218 |

| | | | | | | | | | |
|---|---|---|---|---|---|---|---|---|---|
| 8 | 66712800 | 66713599 | 12.67423 | 23.90249 | 19.82254 | 8.094322 | 0.741873 | 0.079328 | 0.196228 |
| 14 | 71003000 | 71004999 | 10.89288 | 19.14173 | 44.58502 | 19.73944 | -0.82577 | 0.589473 | 0.196232 |
| 7 | 11264200 | 11272999 | 133.5018 | 204.9164 | 158.575 | 65.83507 | 0.663432 | 2.999343 | 0.196246 |
| 4 | 49366000 | 49369599 | 93.83113 | 147.4023 | 119.7444 | 49.66854 | 0.619034 | 2.562554 | 0.196267 |
| 2 | 91801400 | 91810599 | 44.64112 | 70.93333 | 147.0275 | 61.32641 | -0.71669 | 2.262937 | 0.196281 |
| 4 | 55529600 | 55534199 | 28.7965 | 46.38074 | 106.8968 | 45.45056 | -0.84674 | 1.807978 | 0.196304 |
| 12 | 23932200 | 23936199 | 60.80178 | 97.30482 | 83.8794 | 35.32666 | 0.557937 | 2.039517 | 0.196312 |
| 2 | 173175800 | 173189599 | 96.20274 | 145.8563 | 278.1773 | 115.1387 | -0.65092 | 3.244875 | 0.19632 |
| 2 | 87381800 | 87382399 | 5.686601 | 10.98107 | 34.61122 | 13.32623 | -1.09129 | -0.14174 | 0.196333 |
| 2 | 89801000 | 89803799 | 22.30074 | 39.47558 | 25.47916 | 11.26719 | 1.019261 | 0.541451 | 0.196335 |
| 5 | 54334800 | 54338999 | 24.82399 | 40.70402 | 85.8547 | 36.95929 | -0.69703 | 1.441328 | 0.196365 |
| 11 | 120956200 | 120957599 | 5.519041 | 10.41827 | 32.99261 | 13.44556 | -1.17273 | 0.041616 | 0.196381 |
| 15 | 80160200 | 80163999 | 20.12404 | 33.31482 | 73.48468 | 31.78963 | -0.75691 | 1.306488 | 0.196386 |
| 9 | 95194400 | 95195999 | 13.88289 | 26.19262 | 19.55305 | 7.977551 | 0.893136 | 0.086331 | 0.196389 |
| 12 | 19930400 | 19944199 | 467.2989 | 646.0013 | 502.1663 | 208.5971 | 0.514831 | 4.756825 | 0.196397 |
| 19 | 32572200 | 32573799 | 12.76257 | 22.28982 | 48.9052 | 21.54237 | -0.74056 | 0.704824 | 0.196414 |
| 18 | 46859800 | 46861999 | 11.55864 | 20.16307 | 48.37301 | 21.29196 | -0.86858 | 0.717188 | 0.196415 |
| 6 | 120448800 | 120452799 | 18.76898 | 31.33286 | 68.1026 | 29.62693 | -0.735 | 1.202957 | 0.196441 |
| 8 | 104788200 | 104789999 | 12.78006 | 22.27669 | 49.45276 | 21.75677 | -0.75767 | 0.734105 | 0.196454 |
| 5 | 113608800 | 113610399 | 6.979574 | 13.02224 | 33.249 | 13.8261 | -0.88814 | 0.136254 | 0.196458 |
| 12 | 3158200 | 3158999 | 21.94183 | 38.52739 | 28.21808 | 12.4849 | 0.834628 | 0.63709 | 0.196466 |
| 10 | 74145000 | 74145799 | 11.43846 | 21.91433 | 16.50609 | 6.507535 | 0.924522 | -0.05326 | 0.196471 |
| 18 | 38942000 | 38943199 | 11.78558 | 22.46552 | 18.30205 | 7.306474 | 0.79533 | -0.00628 | 0.196483 |
| 8 | 13784400 | 13788999 | 18.75771 | 30.97206 | 76.53054 | 33.04775 | -0.92043 | 1.335609 | 0.196491 |
| 5 | 137058400 | 137061999 | 26.48574 | 42.79809 | 96.55153 | 41.2286 | -0.8102 | 1.704864 | 0.196499 |
| 4 | 16302400 | 16304199 | 14.73221 | 28.15734 | 17.46028 | 6.931074 | 1.192978 | 0.003926 | 0.196525 |
| 9 | 21688000 | 21694599 | 40.14566 | 63.81342 | 145.3573 | 60.65184 | -0.852 | 2.248551 | 0.196573 |
| 10 | 65344000 | 65344999 | 14.40201 | 26.88944 | 21.22921 | 8.817462 | 0.79308 | 0.16165 | 0.19658 |
| 18 | 55935400 | 55939199 | 22.06069 | 36.38443 | 76.41503 | 32.97685 | -0.68723 | 1.357408 | 0.196589 |
| 11 | 46327600 | 46328599 | 6.383319 | 11.98268 | 32.31279 | 13.29861 | -0.95474 | 0.085415 | 0.196601 |
| 17 | 13010600 | 13014799 | 19.91464 | 33.09461 | 73.22496 | 31.76111 | -0.76098 | 1.259351 | 0.196606 |
| 1 | 10937800 | 10939599 | 13.41507 | 23.41551 | 49.44282 | 21.77107 | -0.68554 | 0.719288 | 0.196637 |
| 9 | 36382400 | 36383199 | 12.20623 | 23.56312 | 17.36287 | 6.695294 | 0.985349 | -0.1124 | 0.196643 |
| 7 | 33894800 | 33896799 | 11.36058 | 19.94571 | 44.60783 | 19.7346 | -0.76743 | 0.607895 | 0.196647 |
| 4 | 132675800 | 132678399 | 10.4317 | 18.35255 | 43.56603 | 19.28909 | -0.8521 | 0.573095 | 0.196659 |
| 12 | 42613000 | 42613799 | 10.4317 | 18.35255 | 43.56603 | 19.28909 | -0.8521 | 0.573095 | 0.196659 |
| 16 | 16120600 | 16121599 | 5.045527 | 9.603518 | 29.84753 | 11.95292 | -1.12235 | -0.02708 | 0.196666 |
| 15 | 95787600 | 95788599 | 19.14766 | 33.95072 | 25.73488 | 11.37569 | 0.789375 | 0.502978 | 0.196719 |
| 5 | 28132800 | 28133399 | 4.782734 | 9.315576 | 26.18372 | 9.559144 | -0.85315 | -0.35718 | 0.196726 |
| 8 | 109658400 | 109661399 | 11.57395 | 20.15933 | 51.03176 | 22.44095 | -0.9462 | 0.74064 | 0.196733 |
| 7 | 38254000 | 38258599 | 76.50893 | 121.5924 | 91.3997 | 38.13054 | 0.74458 | 2.272309 | 0.196743 |
| X | 78092600 | 78094999 | 40.72991 | 68.29962 | 37.60354 | 16.40511 | 1.238952 | 1.196874 | 0.196748 |
| 19 | 8215800 | 8217599 | 30.67134 | 51.81873 | 42.06774 | 18.40501 | 0.680872 | 1.098419 | 0.196762 |
| 10 | 110594800 | 110603199 | 79.52396 | 122.3157 | 223.1088 | 92.65349 | -0.57012 | 2.946596 | 0.196775 |
| 12 | 24185600 | 24187399 | 50.61847 | 81.58749 | 70.28234 | 29.91367 | 0.571144 | 1.801764 | 0.196775 |
| 18 | 63582000 | 63583199 | 8.560483 | 15.17278 | 41.01326 | 18.1319 | -1.03328 | 0.475222 | 0.196776 |
| 15 | 95975800 | 95978799 | 15.0084 | 25.90218 | 55.40734 | 24.3067 | -0.7072 | 0.863975 | 0.196788 |

| | | | | | | | | | |
|---|---|---|---|---|---|---|---|---|---|
| 4 | 74791200 | 74792599 | 16.35272 | 30.75071 | 18.11516 | 7.433464 | 1.225228 | 0.125867 | 0.196806 |
| 4 | 24977400 | 24977999 | 2.163318 | 4.23085 | 20.01898 | 6.801251 | -1.4907 | -0.67043 | 0.196812 |
| 7 | 11014000 | 11025799 | 252.0433 | 367.6082 | 302.3024 | 124.5667 | 0.510768 | 3.953422 | 0.196837 |
| 9 | 14318000 | 14319599 | 20.80318 | 36.82419 | 26.43874 | 11.69156 | 0.866773 | 0.535442 | 0.196842 |
| 10 | 111134200 | 111135599 | 9.849897 | 17.67526 | 39.99274 | 17.50056 | -0.76885 | 0.397357 | 0.196873 |
| 1 | 7565400 | 7566999 | 16.00557 | 29.9651 | 19.23398 | 7.95189 | 1.094078 | 0.152461 | 0.196877 |
| X | 76214800 | 76216799 | 18.47634 | 34.9028 | 15.83878 | 6.445009 | 1.608366 | 0.112 | 0.196899 |
| 17 | 19019800 | 19021199 | 21.28475 | 35.07113 | 77.13257 | 33.27303 | -0.75396 | 1.370676 | 0.196919 |
| 2 | 164232800 | 164247999 | 153.4615 | 223.3136 | 481.6637 | 198.5215 | -0.88287 | 3.983384 | 0.196923 |
| 12 | 57362200 | 57362999 | 12.57239 | 24.35473 | 15.52827 | 5.891093 | 1.213131 | -0.16775 | 0.196925 |
| 11 | 16701000 | 16703999 | 19.26279 | 32.08664 | 69.77976 | 30.3145 | -0.73585 | 1.229132 | 0.196942 |
| 17 | 39144200 | 39145999 | 37.34244 | 62.39094 | 43.23694 | 18.81936 | 0.908466 | 1.228395 | 0.19695 |
| 17 | 80020000 | 80026199 | 39.42164 | 62.83506 | 135.6533 | 56.80833 | -0.77049 | 2.157498 | 0.196951 |
| 10 | 60514600 | 60523799 | 61.59056 | 95.7493 | 197.2161 | 81.88056 | -0.73326 | 2.740925 | 0.196961 |
| 14 | 67850600 | 67851799 | 7.749716 | 14.37835 | 33.69388 | 14.13533 | -0.77603 | 0.176646 | 0.196962 |
| 1 | 144833000 | 144834199 | 16.56354 | 31.05072 | 19.34614 | 7.98038 | 1.13974 | 0.150664 | 0.196964 |
| 13 | 48882200 | 48886999 | 33.71848 | 53.96969 | 122.3044 | 51.52046 | -0.83356 | 2.017592 | 0.196977 |
| 18 | 21721600 | 21724199 | 24.74707 | 42.81229 | 35.04996 | 15.37723 | 0.673136 | 0.85744 | 0.196978 |
| 15 | 96357800 | 96359799 | 12.42046 | 21.40961 | 54.1729 | 23.76428 | -0.94826 | 0.864777 | 0.19698 |
| 18 | 75137600 | 75139399 | 6.904724 | 13.14429 | 33.51976 | 13.41915 | -0.84085 | -0.00259 | 0.197033 |
| 13 | 32890600 | 32891199 | 3.346617 | 6.524956 | 22.43654 | 8.083616 | -1.11903 | -0.43175 | 0.197034 |
| 6 | 86709600 | 86710799 | 7.939426 | 14.23118 | 41.06078 | 17.98476 | -1.1178 | 0.393398 | 0.197047 |
| 14 | 64789600 | 64791799 | 7.939426 | 14.23118 | 41.06078 | 17.98476 | -1.1178 | 0.393398 | 0.197047 |
| 16 | 89662000 | 89674599 | 110.6707 | 166.8004 | 296.8191 | 122.6278 | -0.55835 | 3.369073 | 0.197049 |
| 6 | 15638600 | 15638999 | 3.340087 | 6.533037 | 20.40783 | 6.864678 | -0.89257 | -0.68593 | 0.197051 |
| 9 | 102731800 | 102739599 | 69.16598 | 107.7876 | 196.324 | 81.48059 | -0.55338 | 2.703945 | 0.197059 |
| 19 | 39476000 | 39478599 | 57.10284 | 91.80864 | 74.58801 | 31.63184 | 0.651083 | 1.890878 | 0.197065 |
| 3 | 101270600 | 101271199 | 4.232088 | 8.245512 | 23.6547 | 8.602207 | -0.87613 | -0.38567 | 0.197067 |
| 9 | 118225400 | 118231399 | 33.79339 | 54.27638 | 111.5768 | 47.24747 | -0.6873 | 1.905939 | 0.197074 |
| 18 | 54455000 | 54457999 | 38.56674 | 63.72028 | 50.48309 | 21.80934 | 0.715424 | 1.358876 | 0.197083 |
| 17 | 57051600 | 57070399 | 172.1295 | 248.9082 | 485.0391 | 200.0605 | -0.74536 | 4.078888 | 0.197087 |
| 10 | 50994200 | 50996199 | 22.49528 | 39.65785 | 26.51619 | 11.73534 | 0.966424 | 0.586645 | 0.197095 |
| 12 | 79439600 | 79440199 | 6.828696 | 12.74727 | 32.71736 | 13.59321 | -0.89454 | 0.130094 | 0.197108 |
| 15 | 81138000 | 81139999 | 11.33193 | 20.01524 | 42.9938 | 19.02071 | -0.70674 | 0.532464 | 0.197111 |
| 5 | 135302200 | 135308399 | 28.31572 | 45.61091 | 105.5012 | 44.86366 | -0.8517 | 1.80405 | 0.197114 |
| 7 | 25899800 | 25904999 | 74.33624 | 118.6284 | 83.63467 | 35.08259 | 0.843089 | 2.139732 | 0.197142 |
| 13 | 100971800 | 100974399 | 16.5569 | 28.11264 | 60.64727 | 26.57394 | -0.72349 | 1.026001 | 0.197148 |
| 1 | 50537600 | 50539399 | 23.78584 | 41.93252 | 25.94013 | 11.48042 | 1.078052 | 0.59453 | 0.197149 |
| 4 | 153132200 | 153180999 | 466.4917 | 625.9953 | 1025.91 | 428.1247 | -0.60421 | 5.222519 | 0.197164 |
| 6 | 115711600 | 115713799 | 22.02646 | 36.02347 | 83.0241 | 35.70345 | -0.82618 | 1.469426 | 0.19717 |
| X | 32999000 | 33003999 | 119.5907 | 184.7004 | 136.5129 | 56.70463 | 0.736385 | 2.877225 | 0.197172 |
| X | 89516000 | 89519399 | 69.11643 | 110.455 | 83.21248 | 34.97567 | 0.749806 | 2.093608 | 0.197192 |
| X | 4240400 | 4243999 | 25.2351 | 44.26347 | 27.72489 | 12.2558 | 1.059056 | 0.670595 | 0.197195 |
| 18 | 3029000 | 3031399 | 56.38792 | 90.71609 | 72.07334 | 30.59321 | 0.684346 | 1.870903 | 0.197196 |
| 4 | 90303800 | 90307599 | 53.08533 | 85.40807 | 73.65777 | 31.26848 | 0.566309 | 1.862811 | 0.19721 |
| 4 | 95941600 | 95941999 | 9.250016 | 17.99541 | 15.45273 | 5.703044 | 0.824567 | -0.30054 | 0.197221 |
| 13 | 79140600 | 79141999 | 15.72212 | 28.68705 | 22.73014 | 9.752443 | 0.754158 | 0.305814 | 0.197231 |

| | | | | | | | | | |
|---|---|---|---|---|---|---|---|---|---|
| 4 | 147347200 | 147349599 | 38.81078 | 64.58631 | 43.76953 | 19.00312 | 0.940546 | 1.271094 | 0.197234 |
| 6 | 4614200 | 4615199 | 12.65915 | 23.92532 | 19.79677 | 8.04737 | 0.750672 | 0.064432 | 0.197253 |
| 8 | 96297200 | 96306999 | 66.99319 | 104.073 | 198.6075 | 82.46714 | -0.62412 | 2.755722 | 0.197254 |
| 10 | 111308600 | 111310199 | 9.703176 | 17.0471 | 46.68549 | 20.66532 | -1.05802 | 0.58908 | 0.197259 |
| 2 | 180210800 | 180213399 | 12.14231 | 21.2266 | 48.88943 | 21.55024 | -0.81014 | 0.686812 | 0.197261 |
| 17 | 70133200 | 70136199 | 34.45681 | 58.24877 | 39.27668 | 17.18699 | 0.94774 | 1.105473 | 0.19727 |
| 10 | 37012000 | 37018399 | 39.19419 | 62.2978 | 152.4043 | 63.589 | -0.9549 | 2.250109 | 0.197283 |
| 10 | 76121400 | 76122999 | 5.574419 | 10.62449 | 29.33954 | 11.71608 | -0.94949 | -0.0303 | 0.197289 |
| 6 | 55411000 | 55413399 | 12.65881 | 21.97721 | 53.21268 | 23.35213 | -0.8831 | 0.795293 | 0.197311 |
| 6 | 124390200 | 124392999 | 16.63981 | 28.03237 | 64.33683 | 28.12353 | -0.813 | 1.098961 | 0.197312 |
| 17 | 88070000 | 88085999 | 132.0631 | 195.997 | 367.0414 | 151.1894 | -0.65214 | 3.662115 | 0.197355 |
| 12 | 23553800 | 23554799 | 14.75383 | 27.28443 | 21.30402 | 8.977172 | 0.793426 | 0.220832 | 0.197355 |
| 5 | 11213800 | 11215199 | 17.52045 | 32.41501 | 20.09777 | 8.463811 | 1.123646 | 0.236667 | 0.197366 |
| X | 148405200 | 148406599 | 22.718 | 40.76361 | 21.89634 | 9.582282 | 1.293186 | 0.450277 | 0.197371 |
| 7 | 142815000 | 142819999 | 30.20401 | 48.688 | 104.7166 | 44.57531 | -0.74578 | 1.783217 | 0.197375 |
| 18 | 75151600 | 75152399 | 4.235129 | 8.242281 | 24.06411 | 8.854562 | -0.9166 | -0.34157 | 0.197379 |
| 7 | 11166600 | 11170999 | 68.99156 | 109.9159 | 93.44958 | 39.10914 | 0.570902 | 2.190119 | 0.197422 |
| 13 | 49595200 | 49596799 | 7.568697 | 14.17089 | 33.37929 | 13.79829 | -0.76646 | 0.117166 | 0.197424 |
| 10 | 93735800 | 93740399 | 34.88824 | 56.1158 | 108.4307 | 46.01179 | -0.59557 | 1.86059 | 0.197432 |
| 5 | 135674400 | 135677999 | 22.75703 | 37.35495 | 80.75881 | 34.78139 | -0.73172 | 1.420953 | 0.197435 |
| 3 | 148224600 | 148225599 | 12.74436 | 24.77952 | 13.91584 | 5.159249 | 1.424712 | -0.24583 | 0.197447 |
| 4 | 123094400 | 123098199 | 16.62738 | 27.70091 | 71.72065 | 31.16063 | -0.98643 | 1.221133 | 0.197448 |
| 6 | 82108200 | 82109999 | 9.458437 | 16.82163 | 42.51702 | 18.74603 | -0.93523 | 0.455792 | 0.197469 |
| 11 | 50758400 | 50759599 | 21.79676 | 38.1329 | 30.67805 | 13.5314 | 0.699767 | 0.694996 | 0.197478 |
| 9 | 64437600 | 64440399 | 16.20227 | 27.6612 | 57.99083 | 25.42432 | -0.68115 | 0.972285 | 0.197508 |
| 11 | 73066400 | 73069799 | 17.7176 | 29.49685 | 72.43145 | 31.45577 | -0.91043 | 1.231824 | 0.197509 |
| 7 | 7681600 | 7683599 | 15.9553 | 30.08173 | 18.94616 | 7.740202 | 1.135346 | 0.10969 | 0.19752 |
| 2 | 93524800 | 93527599 | 7.630523 | 13.71276 | 40.04723 | 17.50276 | -1.13273 | 0.373252 | 0.197522 |
| 5 | 84600600 | 84601199 | 5.524332 | 10.70445 | 27.95316 | 10.59124 | -0.79875 | -0.21242 | 0.197523 |
| 14 | 37180400 | 37182399 | 31.13821 | 52.50346 | 42.60669 | 18.62962 | 0.681345 | 1.116988 | 0.197525 |
| 12 | 86170600 | 86172799 | 8.198315 | 14.75567 | 38.49402 | 16.80097 | -0.96967 | 0.365046 | 0.197532 |
| X | 4576600 | 4579599 | 64.45209 | 102.9939 | 84.63047 | 35.56826 | 0.624648 | 2.090729 | 0.197541 |
| 4 | 10853000 | 10855399 | 32.70878 | 54.52925 | 44.93296 | 19.53049 | 0.659433 | 1.23474 | 0.197544 |
| 16 | 29863400 | 29864799 | 7.557165 | 14.15488 | 32.84672 | 13.56907 | -0.74448 | 0.11198 | 0.19756 |
| 15 | 52751400 | 52752799 | 22.91555 | 40.16042 | 29.98251 | 13.24418 | 0.80705 | 0.673483 | 0.197564 |
| 6 | 102793000 | 102797399 | 69.58963 | 111.0269 | 89.12356 | 37.37233 | 0.655817 | 2.143899 | 0.197571 |
| 14 | 30074000 | 30075999 | 12.30438 | 21.51135 | 47.30034 | 20.85095 | -0.7434 | 0.685795 | 0.197605 |
| 8 | 79094000 | 79095799 | 21.63646 | 37.99084 | 29.53215 | 13.06619 | 0.748986 | 0.636101 | 0.197622 |
| 3 | 51647800 | 51651199 | 18.54831 | 31.14996 | 64.9492 | 28.36178 | -0.67531 | 1.134644 | 0.197627 |
| 16 | 62079800 | 62081799 | 20.61503 | 36.72799 | 25.07126 | 11.03769 | 0.943675 | 0.483444 | 0.197629 |
| 1 | 56419200 | 56419999 | 10.45991 | 18.32963 | 48.83369 | 21.56954 | -1.01857 | 0.640444 | 0.19763 |
| 13 | 64491200 | 64492399 | 5.730282 | 10.96637 | 29.54084 | 11.68301 | -0.90248 | -0.06401 | 0.197635 |
| 16 | 45727000 | 45728199 | 7.099575 | 13.34277 | 31.81292 | 13.06707 | -0.77636 | 0.083879 | 0.19765 |
| 18 | 56903200 | 56906399 | 22.67853 | 37.46486 | 75.44912 | 32.59211 | -0.62641 | 1.339096 | 0.197667 |
| 6 | 5518600 | 5520999 | 21.14129 | 37.3845 | 26.04645 | 11.52043 | 0.9089 | 0.54483 | 0.197668 |
| 4 | 7801000 | 7801799 | 11.05597 | 21.46159 | 16.39232 | 6.158118 | 0.969059 | -0.21379 | 0.19767 |
| 12 | 23454600 | 23456799 | 29.66397 | 50.65095 | 33.69964 | 14.77476 | 0.968065 | 0.99432 | 0.197673 |

| | | | | | | | | | |
|---|---|---|---|---|---|---|---|---|---|
| 2 | 141809400 | 141810999 | 8.666155 | 15.98247 | 38.56881 | 16.31311 | -0.82763 | 0.225957 | 0.19768 |
| 15 | 96662200 | 96672599 | 68.83413 | 106.8118 | 200.9315 | 83.44627 | -0.60474 | 2.776361 | 0.197695 |
| 2 | 22045000 | 22046199 | 13.68065 | 25.87984 | 19.02508 | 7.718579 | 0.922009 | 0.064982 | 0.197698 |
| 11 | 54682000 | 54682799 | 4.662669 | 9.044207 | 25.51096 | 9.619103 | -0.9008 | -0.24457 | 0.197701 |
| 2 | 23795600 | 23798999 | 41.40668 | 68.03871 | 52.2511 | 22.51356 | 0.757697 | 1.430717 | 0.197707 |
| 4 | 109830400 | 109846999 | 160.4909 | 234.248 | 452.1236 | 186.2879 | -0.71846 | 3.937162 | 0.197726 |
| 3 | 14493200 | 14496199 | 47.72488 | 77.4256 | 60.66238 | 25.99189 | 0.71726 | 1.632352 | 0.197728 |
| 17 | 51846600 | 51847399 | 4.523323 | 8.807852 | 24.81095 | 9.090135 | -0.8607 | -0.35251 | 0.197728 |
| 14 | 18102800 | 18104399 | 22.01508 | 38.57554 | 30.22802 | 13.35065 | 0.737679 | 0.672701 | 0.197732 |
| 7 | 109660200 | 109661599 | 5.819361 | 10.88552 | 32.20378 | 13.33324 | -1.0935 | 0.104898 | 0.197751 |
| 2 | 32851800 | 32852999 | 11.06317 | 19.40186 | 45.16201 | 19.96091 | -0.82472 | 0.626041 | 0.197753 |
| 10 | 84207400 | 84209199 | 9.29154 | 16.56849 | 40.4672 | 17.80259 | -0.88361 | 0.435103 | 0.197761 |
| 3 | 73968800 | 73971199 | 21.18877 | 37.54216 | 25.48496 | 11.26766 | 0.947243 | 0.523654 | 0.197771 |
| 14 | 56379200 | 56382599 | 25.02096 | 40.93184 | 83.72905 | 36.01012 | -0.65448 | 1.471608 | 0.197784 |
| 9 | 65037600 | 65039399 | 8.387614 | 14.97958 | 39.4867 | 17.3488 | -0.99113 | 0.418981 | 0.197791 |
| 7 | 6536400 | 6538999 | 28.90894 | 49.50496 | 34.54718 | 15.14987 | 0.899978 | 0.965411 | 0.197811 |
| 3 | 10362600 | 10365199 | 12.15121 | 21.25787 | 46.7642 | 20.62632 | -0.74395 | 0.673783 | 0.19782 |
| 7 | 16393400 | 16395599 | 12.15121 | 21.25787 | 46.7642 | 20.62632 | -0.74395 | 0.673783 | 0.19782 |
| 5 | 106128400 | 106129999 | 10.23155 | 18.12959 | 41.4365 | 18.32003 | -0.79382 | 0.486278 | 0.197825 |
| 19 | 23730800 | 23732599 | 12.28044 | 21.5342 | 46.18577 | 20.41146 | -0.70767 | 0.634184 | 0.197833 |
| 12 | 91103600 | 91109999 | 51.02109 | 80.88238 | 149.3743 | 62.16027 | -0.55377 | 2.337899 | 0.197841 |
| 17 | 60853200 | 60856799 | 59.65212 | 95.46129 | 82.97618 | 34.94436 | 0.545923 | 2.039145 | 0.197859 |
| 2 | 59290600 | 59291999 | 14.08627 | 26.50165 | 20.0842 | 8.23465 | 0.866006 | 0.107383 | 0.197884 |
| 17 | 40445400 | 40446399 | 14.08627 | 26.50165 | 20.0842 | 8.23465 | 0.866006 | 0.107383 | 0.197884 |
| 11 | 103950000 | 103953799 | 19.12113 | 31.81078 | 72.01717 | 31.26087 | -0.79375 | 1.243658 | 0.197898 |
| 10 | 41554600 | 41558799 | 21.7134 | 35.87907 | 78.96968 | 34.11856 | -0.75426 | 1.334223 | 0.1979 |
| 7 | 116898000 | 116898799 | 5.023167 | 9.622391 | 28.94103 | 11.41418 | -1.05547 | -0.08107 | 0.197912 |
| 7 | 60331400 | 60338399 | 91.76365 | 144.6739 | 112.0611 | 46.50337 | 0.691608 | 2.489353 | 0.197914 |
| 4 | 38845600 | 38846399 | 7.670719 | 14.00005 | 35.55764 | 15.25091 | -0.91415 | 0.275013 | 0.197943 |
| 8 | 86218800 | 86221999 | 14.83019 | 24.93943 | 69.13883 | 30.20634 | -1.08456 | 1.112273 | 0.197952 |
| 5 | 129023000 | 129034999 | 118.4069 | 177.3317 | 326.2831 | 134.5834 | -0.61471 | 3.494189 | 0.197961 |
| 7 | 32151000 | 32153199 | 28.43518 | 49.82878 | 23.39612 | 10.33389 | 1.471874 | 0.694341 | 0.197963 |
| 1 | 116834000 | 116834999 | 12.37815 | 24.01258 | 14.95619 | 5.636487 | 1.255117 | -0.19333 | 0.19797 |
| 7 | 10856200 | 10860599 | 75.74445 | 120.5683 | 86.47516 | 36.14627 | 0.814488 | 2.222743 | 0.197979 |
| 5 | 70428200 | 70431199 | 54.8633 | 88.29466 | 72.9341 | 30.97452 | 0.628887 | 1.855569 | 0.197982 |
| 15 | 83291600 | 83293599 | 9.315368 | 16.55124 | 40.96112 | 18.07469 | -0.90552 | 0.459552 | 0.197985 |
| 1 | 109540400 | 109541599 | 15.89182 | 29.04323 | 23.15757 | 9.915923 | 0.746943 | 0.298957 | 0.197997 |
| 17 | 87837200 | 87839999 | 12.03298 | 20.94085 | 51.04441 | 22.43439 | -0.89215 | 0.755416 | 0.198001 |
| X | 3621600 | 3626199 | 101.5403 | 158.1824 | 123.8924 | 51.42199 | 0.662364 | 2.716103 | 0.198004 |
| 19 | 8979000 | 8980199 | 6.050734 | 11.44615 | 30.28071 | 12.28524 | -0.90912 | 0.027892 | 0.198014 |
| 4 | 30239600 | 30242199 | 28.78148 | 49.52835 | 35.38825 | 15.52314 | 0.867231 | 0.922826 | 0.198017 |
| 7 | 8539000 | 8542199 | 50.22045 | 81.31837 | 62.82827 | 26.88193 | 0.734901 | 1.678233 | 0.198027 |
| 17 | 85915000 | 85916999 | 16.88477 | 28.63683 | 60.63271 | 26.55842 | -0.69664 | 1.037714 | 0.198032 |
| 9 | 37343400 | 37347399 | 27.06885 | 43.89468 | 92.06288 | 39.43277 | -0.70031 | 1.622031 | 0.198049 |
| 9 | 47473600 | 47474599 | 13.87706 | 25.82613 | 20.82379 | 8.696305 | 0.756705 | 0.176181 | 0.198051 |
| 7 | 8437000 | 8448399 | 242.4219 | 355.8082 | 286.3071 | 117.9207 | 0.55079 | 3.855213 | 0.198057 |
| 10 | 23561400 | 23563399 | 19.67537 | 32.94954 | 67.22925 | 29.30514 | -0.64425 | 1.1702 | 0.19807 |

| | | | | | | | | | |
|---|---|---|---|---|---|---|---|---|---|
| 4 | 92169800 | 92171599 | 39.27805 | 64.67112 | 46.17511 | 19.91255 | 0.863523 | 1.395601 | 0.19807 |
| 2 | 70721800 | 70722999 | 18.59327 | 34.24681 | 20.69287 | 8.767463 | 1.154911 | 0.271949 | 0.198073 |
| 1 | 168919000 | 168919799 | 6.903869 | 13.1456 | 32.92843 | 13.17618 | -0.81456 | -0.00539 | 0.198084 |
| 17 | 34225600 | 34227399 | 24.02844 | 42.00894 | 30.6844 | 13.52652 | 0.838719 | 0.719923 | 0.19809 |
| 7 | 100722400 | 100723999 | 8.19686 | 15.22429 | 34.82887 | 14.59016 | -0.74042 | 0.176061 | 0.19809 |
| 10 | 80846200 | 80873399 | 156.4837 | 226.2351 | 523.2186 | 215.8142 | -0.99265 | 4.081666 | 0.198091 |
| 8 | 71383400 | 71385999 | 15.36441 | 25.97832 | 62.09048 | 27.17161 | -0.87084 | 1.061221 | 0.198092 |
| 11 | 20492400 | 20493999 | 6.730422 | 11.84921 | 45.66288 | 20.21464 | -1.54497 | 0.546961 | 0.198127 |
| 10 | 44466000 | 44468599 | 13.85749 | 24.16349 | 51.08147 | 22.47844 | -0.68762 | 0.736005 | 0.198132 |
| 1 | 107225600 | 107227599 | 25.41335 | 44.13175 | 33.94782 | 14.9002 | 0.763702 | 0.824779 | 0.198133 |
| 2 | 173200400 | 173222599 | 198.7282 | 285.3257 | 518.9911 | 214.2806 | -0.65625 | 4.186275 | 0.198138 |
| 6 | 3248600 | 3249999 | 27.20313 | 46.41421 | 38.41844 | 16.84258 | 0.654106 | 0.996395 | 0.198149 |
| 11 | 36030000 | 36033399 | 26.33468 | 42.97331 | 85.32847 | 36.66454 | -0.61427 | 1.512184 | 0.198163 |
| 12 | 18045800 | 18050799 | 120.2003 | 185.19 | 153.9504 | 63.94784 | 0.564271 | 2.920716 | 0.198166 |
| 1 | 136873600 | 136883799 | 77.05946 | 119.0139 | 219.4943 | 91.17337 | -0.58148 | 2.866482 | 0.198167 |
| 5 | 88203400 | 88205399 | 23.56889 | 41.30135 | 28.72645 | 12.68829 | 0.908483 | 0.675192 | 0.198205 |
| 7 | 15356800 | 15368199 | 114.7849 | 178.0036 | 130.818 | 54.33339 | 0.749038 | 2.798576 | 0.198213 |
| 7 | 18155000 | 18168399 | 297.6419 | 428.3715 | 347.0182 | 143.2226 | 0.513671 | 4.154794 | 0.19822 |
| 5 | 109437400 | 109438999 | 29.43541 | 51.24929 | 26.80352 | 11.78369 | 1.317301 | 0.796523 | 0.198234 |
| 7 | 23651800 | 23652599 | 10.96605 | 21.33544 | 14.61133 | 5.389299 | 1.147745 | -0.28376 | 0.198236 |
| 1 | 134468400 | 134470199 | 10.55183 | 18.70372 | 41.9661 | 18.55286 | -0.76691 | 0.487913 | 0.198242 |
| 10 | 39887000 | 39889599 | 26.37041 | 42.95712 | 86.6417 | 37.20532 | -0.63851 | 1.539874 | 0.198251 |
| 14 | 20768600 | 20770199 | 6.686602 | 12.49045 | 32.19854 | 13.36238 | -0.89976 | 0.121374 | 0.198253 |
| 10 | 95399000 | 95401599 | 13.38474 | 23.33034 | 51.62009 | 22.71007 | -0.75314 | 0.739032 | 0.19826 |
| 16 | 24689000 | 24690799 | 27.07464 | 47.26802 | 29.2569 | 12.8844 | 1.076204 | 0.752841 | 0.19826 |
| 8 | 105994200 | 105999599 | 37.69225 | 60.01886 | 137.6049 | 57.54641 | -0.8586 | 2.189367 | 0.198271 |
| 4 | 3108400 | 3109799 | 10.96201 | 21.34413 | 14.02113 | 5.139334 | 1.214991 | -0.30687 | 0.198274 |
| 5 | 84772400 | 84773999 | 18.14466 | 32.374 | 25.96149 | 11.41555 | 0.713916 | 0.459846 | 0.198285 |
| 18 | 87983600 | 87985199 | 36.05257 | 59.71246 | 49.87275 | 21.58214 | 0.639708 | 1.323519 | 0.198285 |
| 9 | 3213600 | 3221599 | 137.2057 | 209.166 | 179.4698 | 74.40409 | 0.506939 | 3.137177 | 0.198317 |
| X | 38533400 | 38534199 | 12.06923 | 23.47274 | 14.22112 | 5.261698 | 1.320374 | -0.25882 | 0.198322 |
| 10 | 36115200 | 36116599 | 17.87078 | 32.70692 | 22.31983 | 9.539167 | 0.971745 | 0.30416 | 0.198329 |
| 5 | 71682800 | 71683799 | 14.56962 | 26.95252 | 21.30748 | 8.974665 | 0.776071 | 0.217366 | 0.198329 |
| 18 | 74347800 | 74354799 | 61.94478 | 95.91387 | 210.5549 | 87.45895 | -0.82962 | 2.813017 | 0.198344 |
| 17 | 15592200 | 15597399 | 30.91865 | 49.6663 | 111.308 | 47.1438 | -0.81128 | 1.898078 | 0.198373 |
| 1 | 167458600 | 167466999 | 60.54587 | 94.67503 | 181.8944 | 75.4628 | -0.62642 | 2.636735 | 0.198377 |
| 7 | 61945400 | 61947399 | 28.83375 | 50.20851 | 27.9752 | 12.29984 | 1.226609 | 0.793688 | 0.198384 |
| 2 | 40956400 | 40959199 | 32.26619 | 54.01722 | 44.85162 | 19.54674 | 0.648347 | 1.189892 | 0.198391 |
| 10 | 16789800 | 16790599 | 10.64524 | 20.76884 | 13.02817 | 4.653576 | 1.313948 | -0.40116 | 0.198404 |
| 10 | 82839200 | 82851399 | 108.109 | 163.0958 | 292.3873 | 120.831 | -0.56784 | 3.349336 | 0.198422 |
| 6 | 71683000 | 71688999 | 35.92514 | 57.55059 | 124.6908 | 52.59253 | -0.76755 | 1.992895 | 0.198426 |
| 7 | 20027200 | 20027799 | 13.1882 | 25.34748 | 16.39577 | 6.404963 | 1.152993 | -0.06597 | 0.198435 |
| 12 | 25230000 | 25231799 | 23.73053 | 41.20242 | 33.62888 | 14.75864 | 0.678606 | 0.818691 | 0.198442 |
| 2 | 58531800 | 58533799 | 10.15304 | 18.2421 | 40.00736 | 17.48846 | -0.72305 | 0.39201 | 0.198471 |
| 2 | 71599800 | 71601599 | 8.977659 | 16.05616 | 43.13827 | 18.93043 | -1.01811 | 0.417286 | 0.198475 |
| X | 88178600 | 88179999 | 28.47578 | 49.34702 | 32.74553 | 14.36697 | 0.975454 | 0.858839 | 0.198475 |
| 7 | 140423600 | 140441399 | 88.92686 | 134.2433 | 299.2693 | 123.7001 | -0.88118 | 3.332435 | 0.198477 |

| | | | | | | | | | |
|---|---|---|---|---|---|---|---|---|---|
| 7 | 32897400 | 32900599 | 49.69825 | 80.29429 | 58.66541 | 25.04569 | 0.812019 | 1.726298 | 0.198487 |
| 5 | 21629200 | 21630599 | 5.457134 | 10.4054 | 29.36954 | 11.71746 | -0.98018 | -0.03687 | 0.198487 |
| 10 | 103881600 | 103882399 | 5.457134 | 10.4054 | 29.36954 | 11.71746 | -0.98018 | -0.03687 | 0.198487 |
| 3 | 47494400 | 47495999 | 18.74567 | 34.37584 | 19.82618 | 8.450721 | 1.214741 | 0.293232 | 0.198503 |
| 13 | 85302400 | 85304399 | 19.95221 | 35.46394 | 25.03859 | 11.0451 | 0.893194 | 0.491908 | 0.198509 |
| 1 | 16106200 | 16109799 | 78.80658 | 124.5849 | 108.7784 | 45.1937 | 0.522125 | 2.416541 | 0.198511 |
| 6 | 136303000 | 136307799 | 78.73787 | 121.4407 | 215.6851 | 89.59174 | -0.52865 | 2.892726 | 0.198511 |
| 12 | 70044200 | 70045799 | 16.61207 | 30.92225 | 20.30126 | 8.475263 | 1.051044 | 0.196182 | 0.198514 |
| 18 | 9176000 | 9178399 | 25.83778 | 45.16805 | 29.24762 | 12.89214 | 1.011418 | 0.727068 | 0.198526 |
| 19 | 60877600 | 60879399 | 11.05132 | 19.43081 | 44.07223 | 19.51559 | -0.78726 | 0.584402 | 0.198533 |
| X | 93748800 | 93750599 | 17.3293 | 32.07192 | 20.10429 | 8.462769 | 1.108352 | 0.2332 | 0.19858 |
| 15 | 97602800 | 97603799 | 5.325035 | 10.15406 | 29.93152 | 11.94032 | -1.04228 | -0.03792 | 0.198585 |
| 3 | 142412400 | 142414199 | 9.470393 | 16.81455 | 42.49781 | 18.76418 | -0.93657 | 0.466825 | 0.198591 |
| 4 | 58030200 | 58030999 | 14.74777 | 27.64754 | 20.66513 | 8.524082 | 0.879338 | 0.137993 | 0.198595 |
| 4 | 153967600 | 153976599 | 76.50588 | 117.8934 | 219.1615 | 91.0362 | -0.5954 | 2.908308 | 0.198596 |
| 9 | 122397600 | 122403799 | 40.55961 | 64.26537 | 163.3511 | 67.94434 | -1.01448 | 2.347647 | 0.198598 |
| 8 | 72231600 | 72234599 | 25.6555 | 41.56616 | 92.59493 | 39.6349 | -0.78815 | 1.638568 | 0.198605 |
| 15 | 99359000 | 99362199 | 20.32183 | 34.02927 | 68.39253 | 29.81063 | -0.62258 | 1.174678 | 0.19861 |
| 5 | 60498800 | 60499399 | 9.239223 | 18.05156 | 12.65915 | 4.411386 | 1.186419 | -0.50552 | 0.198621 |
| 15 | 94803200 | 94803599 | 7.534416 | 14.72808 | 12.91859 | 4.444572 | 0.884437 | -0.57394 | 0.198621 |
| 11 | 69112000 | 69117399 | 47.89935 | 76.05844 | 143.9913 | 60.01881 | -0.58713 | 2.285957 | 0.198627 |
| 18 | 64965600 | 64967599 | 25.29247 | 43.92784 | 33.85675 | 14.86151 | 0.760822 | 0.821004 | 0.198632 |
| 13 | 61860200 | 61864799 | 104.4768 | 163.1178 | 128.6104 | 53.36526 | 0.655243 | 2.678537 | 0.198632 |
| 1 | 145713400 | 145713999 | 8.955552 | 17.47093 | 13.76117 | 4.920462 | 0.987734 | -0.41988 | 0.198661 |
| 4 | 24266600 | 24267199 | 8.955552 | 17.47093 | 13.76117 | 4.920462 | 0.987734 | -0.41988 | 0.198661 |
| 8 | 88466200 | 88475599 | 53.2253 | 83.65836 | 173.1571 | 71.82855 | -0.72789 | 2.535519 | 0.19868 |
| 7 | 31277800 | 31282599 | 92.31577 | 145.5779 | 110.9191 | 46.03083 | 0.715582 | 2.484183 | 0.198682 |
| 3 | 19364400 | 19366199 | 9.231169 | 16.25616 | 44.08544 | 19.51529 | -1.04248 | 0.554336 | 0.198682 |
| 8 | 127183800 | 127184799 | 5.267669 | 10.14468 | 27.46384 | 10.6671 | -0.88401 | -0.13218 | 0.198682 |
| 18 | 76722800 | 76724999 | 17.80193 | 29.79305 | 67.5786 | 29.4455 | -0.79635 | 1.171447 | 0.198683 |
| 7 | 42572600 | 42573599 | 32.7789 | 54.55289 | 44.45682 | 19.3026 | 0.675469 | 1.253685 | 0.198693 |
| X | 3956000 | 3956999 | 20.63836 | 36.43347 | 26.65571 | 11.7939 | 0.838382 | 0.564009 | 0.198695 |
| 8 | 17128200 | 17130399 | 18.34463 | 34.10142 | 17.13664 | 7.170306 | 1.430043 | 0.210661 | 0.198698 |
| 2 | 143939600 | 143944999 | 40.49938 | 64.70598 | 124.3289 | 52.23919 | -0.59894 | 2.081992 | 0.198699 |
| X | 143058200 | 143059799 | 11.32794 | 20.07286 | 45.09971 | 19.93951 | -0.7695 | 0.500427 | 0.198704 |
| X | 31262400 | 31263199 | 18.21382 | 33.08767 | 23.33733 | 10.06817 | 0.915806 | 0.350371 | 0.19871 |
| 5 | 55107800 | 55108399 | 10.12824 | 19.65049 | 16.55402 | 6.234867 | 0.826044 | -0.21505 | 0.198712 |
| 17 | 33308200 | 33308999 | 5.217046 | 10.07428 | 28.71601 | 11.05676 | -0.94498 | -0.15518 | 0.198726 |
| 12 | 94973200 | 94975199 | 17.55936 | 32.31748 | 21.18178 | 8.984053 | 1.037013 | 0.267887 | 0.198741 |
| 2 | 14059800 | 14061199 | 13.30951 | 23.0142 | 52.64195 | 23.0949 | -0.80249 | 0.837529 | 0.198754 |
| X | 89479800 | 89481599 | 20.72226 | 37.12153 | 24.08689 | 10.55553 | 1.020936 | 0.449948 | 0.198778 |
| 9 | 110330800 | 110332799 | 8.277066 | 15.08748 | 36.05889 | 15.49056 | -0.82878 | 0.288277 | 0.1988 |
| 6 | 105725000 | 105725599 | 10.45961 | 20.42387 | 11.79587 | 4.152675 | 1.448817 | -0.45709 | 0.198815 |
| X | 32367200 | 32370599 | 40.33515 | 67.86804 | 35.83591 | 15.65914 | 1.2989 | 1.161764 | 0.198828 |
| 2 | 145533400 | 145536399 | 16.13928 | 27.27701 | 66.09856 | 28.92191 | -0.89098 | 1.069023 | 0.198832 |
| 4 | 133004400 | 133005399 | 4.997283 | 9.643766 | 28.13733 | 10.86437 | -0.98337 | -0.15461 | 0.198835 |
| 14 | 78111800 | 78112399 | 4.997283 | 9.643766 | 28.13733 | 10.86437 | -0.98337 | -0.15461 | 0.198835 |

| | | | | | | | | | |
|---|---|---|---|---|---|---|---|---|---|
| 17 | 35183400 | 35190199 | 38.95414 | 62.06666 | 132.6117 | 55.5058 | -0.75608 | 2.170057 | 0.198845 |
| 4 | 105475000 | 105475799 | 15.42213 | 28.47394 | 22.18527 | 9.369242 | 0.79416 | 0.237147 | 0.198847 |
| 1 | 109771200 | 109771999 | 8.784033 | 17.1508 | 12.57419 | 4.432302 | 1.107286 | -0.47588 | 0.198851 |
| 17 | 7150200 | 7151199 | 4.353192 | 8.274532 | 30.30932 | 12.17267 | -1.35977 | -0.02235 | 0.198857 |
| 13 | 12664000 | 12665999 | 6.516746 | 11.93497 | 37.72584 | 16.10861 | -1.22327 | 0.24752 | 0.19886 |
| Y | 4897200 | 4898599 | 11.46672 | 22.13208 | 10.30024 | 3.9748 | 1.638528 | -0.14378 | 0.198874 |
| 6 | 137463200 | 137464599 | 7.869983 | 14.72209 | 34.49923 | 14.28241 | -0.76115 | 0.127046 | 0.198878 |
| 9 | 116013200 | 116014799 | 7.869983 | 14.72209 | 34.49923 | 14.28241 | -0.76115 | 0.127046 | 0.198878 |
| 16 | 93988800 | 93995399 | 40.69776 | 64.89869 | 130.5277 | 54.69634 | -0.66776 | 2.143366 | 0.198883 |
| 1 | 155404200 | 155410399 | 51.38762 | 81.36053 | 151.3493 | 62.92825 | -0.56577 | 2.371435 | 0.19889 |
| 7 | 142329400 | 142330799 | 13.5764 | 25.61797 | 20.27293 | 8.268798 | 0.810515 | 0.085766 | 0.198901 |
| 4 | 139531400 | 139547399 | 93.55254 | 141.4869 | 301.767 | 124.805 | -0.81517 | 3.295278 | 0.198912 |
| 5 | 49748200 | 49749799 | 8.663626 | 15.98594 | 37.45493 | 15.83102 | -0.7845 | 0.221007 | 0.198922 |
| 5 | 59277200 | 59278799 | 14.87724 | 27.56831 | 21.53508 | 9.049612 | 0.795085 | 0.209413 | 0.198934 |
| 4 | 77649600 | 77651199 | 21.09719 | 37.73055 | 22.72284 | 9.971694 | 1.125993 | 0.457939 | 0.198944 |
| 5 | 100993000 | 100994999 | 12.73281 | 21.94002 | 55.26924 | 24.24495 | -0.94221 | 0.869399 | 0.198945 |
| X | 26712200 | 26716999 | 55.35911 | 89.77155 | 55.29916 | 23.68628 | 1.062799 | 1.65517 | 0.19896 |
| 9 | 64575400 | 64576399 | 4.365711 | 8.477737 | 24.98035 | 9.35655 | -0.95338 | -0.27219 | 0.198961 |
| 1 | 148365600 | 148365999 | 9.657817 | 18.77572 | 14.06046 | 5.218413 | 1.013959 | -0.27893 | 0.198989 |
| 17 | 4802600 | 4805599 | 27.09144 | 46.1314 | 37.91663 | 16.61991 | 0.663741 | 1.012087 | 0.198989 |
| 10 | 37803600 | 37804799 | 11.93599 | 23.10561 | 15.49466 | 5.897765 | 1.13601 | -0.16531 | 0.19899 |
| 3 | 75502400 | 75503199 | 14.09984 | 26.77124 | 19.18227 | 7.721964 | 0.969088 | 0.046409 | 0.199 |
| 3 | 92546400 | 92547999 | 12.00181 | 20.99727 | 46.76511 | 20.62737 | -0.7616 | 0.673081 | 0.199013 |
| 4 | 144351600 | 144355799 | 75.93742 | 120.4281 | 103.3308 | 43.01646 | 0.551049 | 2.334704 | 0.199017 |
| 12 | 76415000 | 76418399 | 16.68485 | 27.99908 | 65.87098 | 28.75044 | -0.84874 | 1.137467 | 0.199033 |
| 14 | 44772000 | 44773599 | 12.11497 | 21.35163 | 46.1411 | 20.422 | -0.71695 | 0.565481 | 0.199034 |
| 10 | 119398200 | 119400599 | 24.49157 | 40.21919 | 81.12013 | 34.94328 | -0.6317 | 1.419867 | 0.199035 |
| 3 | 145544600 | 145547399 | 34.26476 | 55.10809 | 107.2667 | 45.51203 | -0.60623 | 1.863557 | 0.199049 |
| 3 | 84176200 | 84182199 | 33.31055 | 53.51249 | 110.2431 | 46.69779 | -0.68998 | 1.898359 | 0.199062 |
| X | 126168000 | 126169399 | 20.57949 | 38.58678 | 15.056 | 6.208608 | 1.808764 | 0.174455 | 0.199065 |
| 8 | 125446800 | 125449199 | 8.947967 | 15.75489 | 46.15656 | 20.43277 | -1.15352 | 0.556748 | 0.199071 |
| 11 | 108793600 | 108794999 | 6.951143 | 13.06761 | 31.81917 | 13.06322 | -0.80594 | 0.080212 | 0.199075 |
| 17 | 27204000 | 27206799 | 27.84175 | 45.23835 | 88.96739 | 38.16311 | -0.60498 | 1.582522 | 0.199099 |
| 2 | 128358200 | 128362199 | 26.22656 | 42.65589 | 87.0441 | 37.35677 | -0.65692 | 1.561831 | 0.199111 |
| 12 | 19834000 | 19852799 | 688.563 | 921.2441 | 696.4248 | 290.7551 | 0.507541 | 5.27184 | 0.199114 |
| 9 | 115565600 | 115567399 | 9.661097 | 17.51752 | 38.60756 | 16.69537 | -0.71966 | 0.332468 | 0.199117 |
| 4 | 83229000 | 83232599 | 20.86872 | 34.64145 | 73.87103 | 32.01666 | -0.70811 | 1.275487 | 0.199117 |
| 9 | 92615200 | 92619799 | 54.48475 | 87.5664 | 74.52441 | 31.58766 | 0.583544 | 1.899221 | 0.199119 |
| 7 | 93400000 | 93402599 | 29.11865 | 49.55629 | 39.85015 | 17.46666 | 0.69508 | 1.027204 | 0.199167 |
| 11 | 86298000 | 86299799 | 18.49418 | 33.66216 | 22.71969 | 9.777985 | 0.981013 | 0.339952 | 0.199188 |
| 12 | 111858200 | 111859599 | 7.770357 | 14.34604 | 34.14022 | 14.41858 | -0.80505 | 0.206144 | 0.199223 |
| 7 | 14509800 | 14513199 | 82.91046 | 132.1633 | 78.9924 | 33.08371 | 1.079207 | 2.183148 | 0.199235 |
| 10 | 22164600 | 22165999 | 14.13806 | 26.4087 | 21.17197 | 8.78688 | 0.771756 | 0.155521 | 0.199245 |
| 17 | 41691800 | 41693399 | 13.05369 | 24.54157 | 20.34076 | 8.349852 | 0.736824 | 0.104054 | 0.199258 |
| 2 | 74814600 | 74815799 | 8.708329 | 15.96475 | 36.15731 | 15.41775 | -0.74466 | 0.257753 | 0.199259 |
| 1 | 52152600 | 52153199 | 4.53445 | 8.796611 | 25.51576 | 9.615228 | -0.93999 | -0.24883 | 0.199268 |
| 4 | 78458400 | 78460999 | 23.64496 | 41.36004 | 29.31288 | 12.9276 | 0.881574 | 0.704171 | 0.199268 |

| | | | | | | | | | |
|---|---|---|---|---|---|---|---|---|---|
| 7 | 144489800 | 144497999 | 34.71356 | 55.19484 | 145.7512 | 60.83387 | -1.06431 | 2.236138 | 0.199274 |
| 4 | 24024000 | 24027599 | 26.00114 | 45.5698 | 26.92571 | 11.89429 | 1.142531 | 0.684566 | 0.199275 |
| 12 | 77921000 | 77922599 | 14.29674 | 26.51729 | 21.68767 | 9.102202 | 0.730768 | 0.200442 | 0.199276 |
| 7 | 27662200 | 27665399 | 59.86422 | 96.31251 | 70.55352 | 29.9498 | 0.801239 | 1.872283 | 0.199276 |
| 9 | 3115000 | 3117999 | 60.71527 | 97.3356 | 75.57564 | 31.91258 | 0.711421 | 1.986395 | 0.199279 |
| 8 | 12306800 | 12316999 | 64.07516 | 100.3573 | 179.0183 | 74.25711 | -0.51765 | 2.609483 | 0.19928 |
| 4 | 45141800 | 45143999 | 10.22486 | 18.1239 | 44.569 | 19.70364 | -0.89898 | 0.486018 | 0.199282 |
| 2 | 163295200 | 163296999 | 8.173497 | 14.74163 | 37.50307 | 16.32593 | -0.9307 | 0.353219 | 0.199297 |
| 8 | 3542800 | 3543999 | 5.51712 | 10.34913 | 31.70749 | 13.0656 | -1.13761 | 0.083179 | 0.199298 |
| 13 | 97850000 | 97852799 | 15.00613 | 26.05189 | 55.46777 | 24.34144 | -0.69867 | 0.810687 | 0.199305 |
| 1 | 108879200 | 108879999 | 12.38211 | 23.72774 | 17.61801 | 6.941257 | 0.944765 | -0.04434 | 0.199307 |
| 10 | 113005800 | 113007399 | 12.38211 | 23.72774 | 17.61801 | 6.941257 | 0.944765 | -0.04434 | 0.199307 |
| 5 | 104287400 | 104289199 | 7.86155 | 14.68838 | 37.94894 | 15.74306 | -0.90394 | 0.140086 | 0.199312 |
| 7 | 13317200 | 13320799 | 113.096 | 174.44 | 147.8332 | 61.40726 | 0.537751 | 2.89801 | 0.199314 |
| 7 | 47541200 | 47541799 | 20.21644 | 35.47028 | 28.29354 | 12.51008 | 0.71194 | 0.637299 | 0.199321 |
| 6 | 14037200 | 14038599 | 22.11585 | 39.986 | 20.04303 | 8.69842 | 1.399285 | 0.397676 | 0.199325 |
| 2 | 167222600 | 167224599 | 7.174869 | 13.26349 | 34.16095 | 14.40238 | -0.91657 | 0.192317 | 0.19934 |
| 14 | 63313200 | 63317999 | 25.27622 | 41.09752 | 87.42699 | 37.51687 | -0.71707 | 1.56436 | 0.199345 |
| 5 | 55872800 | 55875199 | 26.19301 | 45.87058 | 27.50987 | 12.1444 | 1.12129 | 0.698722 | 0.19935 |
| 2 | 12423000 | 12424399 | 7.108575 | 13.32311 | 33.42261 | 13.79213 | -0.855 | 0.103729 | 0.199363 |
| 3 | 115522400 | 115525199 | 16.25265 | 27.56198 | 64.52082 | 28.26063 | -0.84111 | 1.038002 | 0.199364 |
| 11 | 48670200 | 48672199 | 12.76819 | 22.28558 | 48.91072 | 21.53627 | -0.74107 | 0.714033 | 0.199366 |
| 3 | 22097600 | 22099999 | 31.66243 | 53.8952 | 37.51335 | 16.44266 | 0.902477 | 1.03194 | 0.199384 |
| 14 | 77393600 | 77395599 | 18.6172 | 31.06215 | 69.23074 | 30.10712 | -0.77114 | 1.208612 | 0.199393 |
| 10 | 104354600 | 104357199 | 13.69962 | 23.84706 | 55.47406 | 24.38496 | -0.82549 | 0.764072 | 0.1994 |
| 4 | 67766200 | 67767999 | 17.34549 | 31.70679 | 22.21853 | 9.511246 | 0.931829 | 0.305283 | 0.1994 |
| 5 | 30050600 | 30052199 | 8.210231 | 15.13292 | 36.8529 | 15.59957 | -0.84137 | 0.221466 | 0.199414 |
| 3 | 158793800 | 158795999 | 17.90554 | 32.40554 | 23.76509 | 10.30105 | 0.855396 | 0.36895 | 0.199427 |
| 2 | 28485800 | 28492599 | 37.41493 | 59.83461 | 122.9024 | 51.70921 | -0.69349 | 2.048536 | 0.199429 |
| 8 | 95615200 | 95617399 | 13.0177 | 22.47133 | 53.68834 | 23.55275 | -0.86557 | 0.852705 | 0.199439 |
| 15 | 103859200 | 103863199 | 37.56637 | 61.77479 | 52.60329 | 22.67252 | 0.609579 | 1.413963 | 0.19944 |
| 12 | 23564200 | 23565399 | 33.66264 | 56.08959 | 40.32703 | 17.5226 | 0.855622 | 1.241602 | 0.199442 |
| 5 | 46432400 | 46434799 | 27.62539 | 47.59743 | 34.83079 | 15.27918 | 0.833133 | 0.904934 | 0.199443 |
| 9 | 38692800 | 38694999 | 16.0827 | 27.36476 | 58.98483 | 25.85309 | -0.72175 | 1.004946 | 0.199444 |
| 5 | 111652000 | 111658799 | 47.77257 | 75.36859 | 163.4302 | 67.82914 | -0.7907 | 2.455665 | 0.19945 |
| 17 | 29290600 | 29293999 | 12.5943 | 21.66517 | 55.22934 | 24.22621 | -0.95968 | 0.885027 | 0.199451 |
| 18 | 90205600 | 90206599 | 10.33264 | 18.49357 | 39.95987 | 17.5244 | -0.70488 | 0.418235 | 0.199455 |
| 12 | 85262800 | 85266999 | 22.83674 | 37.29339 | 84.13906 | 36.1634 | -0.7971 | 1.493405 | 0.199471 |
| X | 48017400 | 48019399 | 17.12903 | 32.17197 | 18.08952 | 7.438019 | 1.289837 | 0.140498 | 0.199489 |
| 3 | 88442200 | 88444199 | 7.997065 | 14.52453 | 36.53323 | 15.76539 | -0.90646 | 0.306591 | 0.199504 |
| 18 | 5142400 | 5145799 | 30.15703 | 48.76356 | 95.41192 | 40.77764 | -0.60432 | 1.689901 | 0.199531 |
| 15 | 63610000 | 63611799 | 12.43024 | 21.82047 | 46.15932 | 20.41842 | -0.68793 | 0.617352 | 0.19955 |
| 16 | 20120000 | 20121399 | 5.595768 | 10.70759 | 30.69918 | 12.14551 | -0.9924 | -0.06221 | 0.199563 |
| 13 | 38194000 | 38195199 | 4.625285 | 8.829176 | 29.90068 | 11.90107 | -1.23689 | -0.05187 | 0.199619 |
| 4 | 129325200 | 129327399 | 13.75895 | 23.80296 | 53.21033 | 23.34461 | -0.76953 | 0.834604 | 0.199623 |
| 2 | 72078000 | 72078799 | 3.035319 | 5.90969 | 25.13604 | 9.218173 | -1.44655 | -0.36605 | 0.199633 |
| 4 | 80159400 | 80161399 | 19.46853 | 35.00211 | 24.13703 | 10.54526 | 0.936478 | 0.418863 | 0.199646 |

| | | | | | | | | | |
|---|---|---|---|---|---|---|---|---|---|
| 19 | 44241800 | 44243799 | 10.25976 | 18.10279 | 44.04872 | 19.49144 | -0.88626 | 0.537141 | 0.19966 |
| 7 | 3174600 | 3176199 | 24.81544 | 42.88132 | 35.63563 | 15.63363 | 0.651383 | 0.86931 | 0.199667 |
| 7 | 59245600 | 59251399 | 61.71373 | 98.82181 | 75.3677 | 31.76881 | 0.735199 | 2.019772 | 0.199671 |
| 13 | 18847400 | 18847999 | 8.669063 | 16.93308 | 12.62516 | 4.419758 | 1.09234 | -0.50028 | 0.199673 |
| 3 | 95017600 | 95018999 | 7.822745 | 14.27572 | 35.55163 | 15.25073 | -0.88636 | 0.275946 | 0.199673 |
| 8 | 123951000 | 123952799 | 7.822745 | 14.27572 | 35.55163 | 15.25073 | -0.88636 | 0.275946 | 0.199673 |
| X | 146536600 | 146538399 | 48.3367 | 79.31803 | 47.86985 | 20.61394 | 1.103615 | 1.464284 | 0.19968 |
| 2 | 95758600 | 95759799 | 14.41517 | 27.31204 | 17.77508 | 7.187618 | 1.100054 | 0.057931 | 0.199682 |
| 6 | 125307800 | 125311199 | 15.6263 | 26.55321 | 59.46622 | 26.06132 | -0.77693 | 1.016246 | 0.199694 |
| 4 | 130331000 | 130336199 | 40.17441 | 64.19765 | 123.6916 | 51.9839 | -0.60262 | 2.076324 | 0.199696 |
| 14 | 61793600 | 61795199 | 10.93784 | 19.12304 | 48.34605 | 21.31264 | -0.9436 | 0.678755 | 0.199723 |
| 14 | 90289000 | 90289999 | 11.944 | 23.0923 | 16.06562 | 6.150821 | 1.075856 | -0.14448 | 0.199731 |
| 4 | 140965600 | 140991399 | 176.2835 | 254.8545 | 491.6672 | 202.8007 | -0.73123 | 4.082548 | 0.199732 |
| 13 | 66481200 | 66487199 | 97.2002 | 153.8854 | 91.45317 | 38.02342 | 1.077946 | 2.39521 | 0.199733 |
| 17 | 88358800 | 88359399 | 4.228377 | 8.249436 | 24.01058 | 8.57828 | -0.874 | -0.4495 | 0.199737 |
| 10 | 47617800 | 47618599 | 10.31741 | 19.99737 | 15.18744 | 5.74713 | 0.966127 | -0.20477 | 0.199756 |
| 17 | 18717600 | 18724799 | 147.9455 | 224.487 | 191.6531 | 79.34521 | 0.508691 | 3.23544 | 0.199766 |
| 1 | 67575800 | 67577799 | 17.3695 | 31.97843 | 21.18825 | 8.982823 | 1.021878 | 0.264479 | 0.199768 |
| 11 | 95835000 | 95837399 | 13.83239 | 23.74533 | 56.8936 | 24.95431 | -0.87159 | 0.909972 | 0.199773 |
| 2 | 25706000 | 25714999 | 70.79864 | 110.2897 | 194.3437 | 80.66334 | -0.50615 | 2.709716 | 0.199785 |
| 11 | 102175000 | 102176399 | 6.277008 | 11.70245 | 33.7781 | 14.06434 | -1.06573 | 0.132507 | 0.199805 |
| 5 | 142771800 | 142775999 | 57.91023 | 90.632 | 179.8187 | 74.59502 | -0.67175 | 2.621245 | 0.199807 |
| 2 | 73267200 | 73270399 | 19.50107 | 32.72455 | 66.14183 | 28.86839 | -0.63066 | 1.146897 | 0.199811 |
| 5 | 138871200 | 138878599 | 45.21715 | 71.66143 | 148.7194 | 61.87035 | -0.72211 | 2.342822 | 0.199815 |
| 12 | 43938200 | 43939399 | 12.76714 | 24.6975 | 16.09236 | 6.14504 | 1.173635 | -0.1426 | 0.199826 |
| 5 | 24721000 | 24722999 | 9.77252 | 17.03931 | 50.44759 | 22.19855 | -1.17028 | 0.717131 | 0.19983 |
| 7 | 38658000 | 38658599 | 10.63304 | 20.59719 | 16.9088 | 6.415871 | 0.852167 | -0.18539 | 0.19985 |
| 8 | 59076000 | 59076399 | 10.63304 | 20.59719 | 16.9088 | 6.415871 | 0.852167 | -0.18539 | 0.19985 |
| 13 | 45172600 | 45173599 | 10.63304 | 20.59719 | 16.9088 | 6.415871 | 0.852167 | -0.18539 | 0.19985 |
| 5 | 126218800 | 126222799 | 36.8251 | 61.24309 | 47.10762 | 20.44346 | 0.758491 | 1.276646 | 0.199862 |
| 1 | 181297400 | 181300599 | 55.185 | 88.9955 | 67.95668 | 28.94379 | 0.745585 | 1.793986 | 0.199864 |
| 1 | 190681200 | 190682799 | 8.573169 | 15.62329 | 36.04691 | 15.49029 | -0.77898 | 0.290129 | 0.199869 |
| 5 | 78397000 | 78397799 | 11.19979 | 21.56507 | 18.00552 | 6.997092 | 0.796386 | -0.09684 | 0.199876 |
| 2 | 92487000 | 92495599 | 56.19074 | 88.19442 | 173.3994 | 71.91582 | -0.65542 | 2.56625 | 0.199896 |
| 1 | 75840000 | 75841799 | 25.25441 | 43.43272 | 36.21828 | 15.88694 | 0.645076 | 0.913433 | 0.199897 |
| 10 | 126814000 | 126816999 | 14.00197 | 23.63695 | 63.56655 | 27.80411 | -1.04026 | 1.072382 | 0.199902 |
| 13 | 65653600 | 65659399 | 145.3926 | 223.6803 | 136.4121 | 56.65157 | 1.009175 | 2.958549 | 0.199927 |
| 10 | 43096800 | 43098199 | 7.412241 | 13.47294 | 37.0714 | 15.98191 | -1.03407 | 0.298201 | 0.199938 |
| 5 | 107208600 | 107209599 | 14.35428 | 26.68314 | 21.71859 | 9.087706 | 0.740867 | 0.18934 | 0.199942 |
| 18 | 35064200 | 35065799 | 9.355785 | 16.52371 | 42.5249 | 18.81352 | -0.96565 | 0.519837 | 0.199945 |
| 13 | 65688000 | 65693799 | 125.1076 | 194.5523 | 119.4745 | 49.60143 | 1.010941 | 2.756537 | 0.199947 |
| 11 | 52321600 | 52329599 | 62.10609 | 96.86317 | 187.204 | 77.68632 | -0.63788 | 2.684468 | 0.199965 |
| 19 | 25383800 | 25384599 | 4.360166 | 8.485018 | 24.05946 | 8.857966 | -0.87557 | -0.33712 | 0.199968 |
| 5 | 135541400 | 135557199 | 121.7257 | 181.9821 | 334.1238 | 137.7705 | -0.61393 | 3.527719 | 0.199978 |
| 10 | 76239200 | 76240599 | 6.332356 | 11.66482 | 35.15617 | 14.89323 | -1.14619 | 0.208225 | 0.199979 |
| 5 | 23988000 | 23988799 | 5.609178 | 10.64304 | 29.79037 | 12.00915 | -0.98131 | 0.001143 | 0.199983 |
| 14 | 31749200 | 31753599 | 24.90022 | 40.28814 | 96.30587 | 41.18242 | -0.89148 | 1.6661 | 0.199983 |

| | | | | | | | | | |
|---|---|---|---|---|---|---|---|---|---|
| 7 | 142894200 | 142895799 | 9.440852 | 16.83376 | 39.94949 | 17.57569 | -0.84237 | 0.435981 | 0.199989 |
| 3 | 129134600 | 129135199 | 10.26848 | 19.94059 | 15.80776 | 5.920119 | 0.919167 | -0.23715 | 0.199993 |
| X | 153218400 | 153220399 | 39.6159 | 65.4869 | 49.39038 | 21.34475 | 0.786297 | 1.35584 | 0.200008 |
| 6 | 20830200 | 20831399 | 11.48744 | 20.30343 | 43.51029 | 19.24664 | -0.70303 | 0.526942 | 0.200012 |
| 1 | 54995200 | 54996799 | 9.197166 | 16.70036 | 37.03961 | 15.9885 | -0.72676 | 0.318041 | 0.200029 |
| 2 | 126527400 | 126530799 | 37.72991 | 62.33102 | 51.01376 | 22.03698 | 0.668666 | 1.358108 | 0.200034 |
| 15 | 41037000 | 41038599 | 13.58934 | 23.66893 | 50.52359 | 22.21703 | -0.70146 | 0.752765 | 0.200035 |
| 7 | 47291000 | 47294399 | 37.58214 | 63.60394 | 34.61649 | 15.15294 | 1.255395 | 1.10085 | 0.200044 |
| 19 | 43532600 | 43535999 | 17.57815 | 29.59159 | 64.40419 | 28.14675 | -0.73682 | 1.108496 | 0.20005 |
| 4 | 20211800 | 20214199 | 19.57224 | 35.07121 | 23.42985 | 10.26518 | 0.978868 | 0.436777 | 0.200053 |
| 12 | 65011600 | 65013399 | 11.28337 | 19.63237 | 51.02149 | 22.42149 | -0.98398 | 0.75523 | 0.200055 |
| 13 | 3250800 | 3252599 | 26.17373 | 45.30268 | 34.96677 | 15.34096 | 0.75797 | 0.860529 | 0.200067 |
| 6 | 80628600 | 80629199 | 9.797253 | 19.09152 | 13.55384 | 4.918091 | 1.116551 | -0.35918 | 0.200071 |
| 13 | 65589200 | 65591999 | 19.17603 | 34.46374 | 21.93066 | 9.584246 | 1.051541 | 0.415212 | 0.200123 |
| 12 | 61395400 | 61395999 | 9.747581 | 19.02845 | 12.44082 | 4.399936 | 1.266318 | -0.45032 | 0.200141 |
| 5 | 109219400 | 109222199 | 54.09482 | 87.31775 | 66.39713 | 28.31317 | 0.753082 | 1.764156 | 0.200155 |
| 10 | 19439400 | 19440599 | 12.14519 | 23.40839 | 17.18054 | 6.657708 | 0.983556 | -0.09969 | 0.20016 |
| 11 | 85775200 | 85777599 | 16.35118 | 27.48347 | 64.17241 | 28.03229 | -0.83772 | 1.120147 | 0.200162 |
| 8 | 3185200 | 3187599 | 26.3565 | 45.90179 | 31.54459 | 13.87051 | 0.925812 | 0.785589 | 0.20019 |
| 8 | 94489600 | 94490199 | 3.933764 | 7.683965 | 23.08895 | 8.078626 | -0.89144 | -0.53053 | 0.200197 |
| 3 | 103697800 | 103698999 | 5.642279 | 10.60659 | 31.21075 | 12.81436 | -1.07557 | 0.069577 | 0.200198 |
| 12 | 70870600 | 70873399 | 11.48415 | 19.88347 | 54.19907 | 23.77897 | -1.05377 | 0.818562 | 0.200203 |
| 2 | 163543200 | 163544799 | 7.91701 | 14.61484 | 34.14061 | 14.42138 | -0.77889 | 0.207126 | 0.20022 |
| 2 | 176850000 | 176851999 | 26.16329 | 45.14822 | 30.0892 | 13.19981 | 0.967932 | 0.879722 | 0.200235 |
| 12 | 115664000 | 115667799 | 58.36716 | 94.25586 | 60.51343 | 25.81902 | 0.997358 | 1.755905 | 0.200238 |
| 10 | 85335800 | 85342399 | 35.74076 | 57.32445 | 116.4044 | 49.18777 | -0.67207 | 1.952987 | 0.20024 |
| 19 | 25363400 | 25368799 | 37.19812 | 59.62556 | 117.6917 | 49.68567 | -0.63231 | 1.974338 | 0.200244 |
| 17 | 13134400 | 13135999 | 10.77355 | 18.86189 | 46.75235 | 20.6342 | -0.91505 | 0.653354 | 0.20025 |
| 7 | 24551600 | 24554599 | 17.45988 | 29.30467 | 64.79951 | 28.28582 | -0.75964 | 1.138267 | 0.200252 |
| 15 | 93233800 | 93235599 | 9.929569 | 17.59567 | 42.47415 | 18.77855 | -0.87223 | 0.484557 | 0.200261 |
| X | 5127200 | 5127999 | 11.77521 | 22.98839 | 10.36447 | 3.660739 | 1.795718 | -0.43221 | 0.200268 |
| X | 14272800 | 14274599 | 21.28598 | 39.7792 | 14.99379 | 6.217808 | 1.851917 | 0.20028 | 0.200279 |
| 7 | 9806600 | 9814599 | 140.6538 | 214.4807 | 178.6897 | 74.08532 | 0.549695 | 3.13276 | 0.200282 |
| 7 | 126101400 | 126102999 | 8.454593 | 14.9276 | 42.56436 | 18.8321 | -1.1123 | 0.519255 | 0.200284 |
| 11 | 87988000 | 87996599 | 41.7252 | 66.37541 | 138.7921 | 57.964 | -0.7276 | 2.225232 | 0.200286 |
| 7 | 9277800 | 9280199 | 34.89582 | 58.9774 | 35.63367 | 15.59177 | 1.105432 | 1.105137 | 0.200298 |
| 7 | 12281800 | 12285799 | 33.56512 | 56.58872 | 38.2812 | 16.7377 | 0.942865 | 1.122092 | 0.200312 |
| X | 8009400 | 8012199 | 47.98528 | 78.56574 | 48.3956 | 20.82066 | 1.07305 | 1.492342 | 0.200321 |
| 12 | 66976400 | 66977799 | 10.47743 | 18.79854 | 42.12397 | 18.43548 | -0.7552 | 0.407269 | 0.200324 |
| 3 | 11713000 | 11713599 | 11.18093 | 21.45453 | 17.86436 | 7.011424 | 0.786374 | -0.06775 | 0.200343 |
| 8 | 66963800 | 66966199 | 32.04238 | 51.44862 | 111.2395 | 47.08319 | -0.76041 | 1.914009 | 0.200346 |
| 4 | 121145000 | 121147399 | 35.90771 | 60.48533 | 38.02688 | 16.6221 | 1.047992 | 1.135786 | 0.200363 |
| 4 | 146496400 | 146496799 | 16.20756 | 30.77005 | 17.21027 | 6.9296 | 1.322687 | 0.061539 | 0.200364 |
| 2 | 48683000 | 48685199 | 29.5145 | 50.35672 | 38.13732 | 16.7193 | 0.781672 | 1.003514 | 0.200366 |
| 16 | 87228600 | 87229799 | 6.475367 | 12.29346 | 31.00758 | 12.48504 | -0.83213 | 0.005982 | 0.200391 |
| 10 | 79870800 | 79875799 | 18.67035 | 30.5845 | 81.80203 | 35.20038 | -1.03856 | 1.441253 | 0.200394 |
| 15 | 15524400 | 15526599 | 26.10173 | 45.23086 | 34.37135 | 15.08027 | 0.780621 | 0.848549 | 0.200399 |

| | | | | | | | | | |
|---|---|---|---|---|---|---|---|---|---|
| 1 | 13966200 | 13967799 | 14.20887 | 24.7365 | 51.62016 | 22.69261 | -0.66919 | 0.762929 | 0.200415 |
| 17 | 6320400 | 6322599 | 11.03268 | 19.43895 | 43.01562 | 19.03969 | -0.75073 | 0.557858 | 0.20043 |
| 2 | 27741600 | 27743999 | 10.60501 | 18.60385 | 45.15615 | 19.9633 | -0.88481 | 0.61771 | 0.200433 |
| 5 | 96647800 | 96648999 | 5.730066 | 10.86544 | 30.30932 | 12.23495 | -0.97771 | 0.010358 | 0.200435 |
| 1 | 102761800 | 102762999 | 15.32722 | 28.37092 | 22.08912 | 9.296345 | 0.798521 | 0.221366 | 0.200437 |
| 4 | 109346800 | 109348199 | 15.32722 | 28.37092 | 22.08912 | 9.296345 | 0.798521 | 0.221366 | 0.200437 |
| 11 | 59351600 | 59358399 | 104.4416 | 162.7753 | 133.2118 | 55.28633 | 0.599603 | 2.710188 | 0.200441 |
| 12 | 54977200 | 54978799 | 7.566598 | 14.14177 | 32.78223 | 13.59238 | -0.74722 | 0.126239 | 0.200446 |
| 4 | 23402400 | 23403199 | 9.069325 | 17.68424 | 13.71217 | 4.934405 | 1.00174 | -0.39693 | 0.200451 |
| 7 | 16550000 | 16551199 | 7.830134 | 14.25341 | 36.04932 | 15.51282 | -0.91123 | 0.293266 | 0.200463 |
| 4 | 135136000 | 135139399 | 24.66686 | 42.52762 | 34.75913 | 15.24804 | 0.674668 | 0.888047 | 0.200464 |
| 7 | 73323600 | 73325199 | 21.03791 | 36.93073 | 29.24033 | 12.93438 | 0.722733 | 0.636904 | 0.200465 |
| X | 165411800 | 165413999 | 25.00332 | 43.6928 | 30.57075 | 13.47107 | 0.90046 | 0.731976 | 0.200469 |
| 2 | 89307800 | 89309799 | 25.41939 | 44.44098 | 29.88021 | 13.17207 | 0.957499 | 0.724907 | 0.20047 |
| X | 31776800 | 31778999 | 24.59936 | 43.42846 | 24.88589 | 11.01071 | 1.188683 | 0.583597 | 0.20047 |
| 6 | 94602800 | 94603799 | 3.321202 | 6.460419 | 26.28424 | 9.710311 | -1.39457 | -0.33354 | 0.200475 |
| 13 | 93172200 | 93173999 | 9.366497 | 16.53649 | 44.58391 | 19.72591 | -1.03288 | 0.526007 | 0.200491 |
| 15 | 18706400 | 18708999 | 30.92059 | 51.98726 | 43.18416 | 18.86571 | 0.647642 | 1.140471 | 0.200496 |
| 19 | 16380600 | 16381799 | 15.83436 | 29.58056 | 19.78595 | 8.210088 | 1.030975 | 0.165532 | 0.200532 |
| 10 | 129859000 | 129860599 | 21.00482 | 37.22202 | 25.48593 | 11.26774 | 0.93496 | 0.520676 | 0.200545 |
| 7 | 34305000 | 34310799 | 41.54217 | 66.17465 | 132.5848 | 55.47646 | -0.66389 | 2.180415 | 0.200545 |
| 1 | 126795000 | 126796999 | 11.79646 | 20.78713 | 44.03018 | 19.48831 | -0.68816 | 0.561824 | 0.200547 |
| 11 | 69439400 | 69440999 | 5.182752 | 9.840451 | 29.78471 | 11.99003 | -1.09097 | -0.00759 | 0.200557 |
| 12 | 14047200 | 14048599 | 9.533326 | 17.19055 | 37.97208 | 16.5345 | -0.72923 | 0.363079 | 0.200561 |
| X | 9836200 | 9838399 | 20.30812 | 37.6257 | 17.07365 | 7.176445 | 1.571984 | 0.245621 | 0.200563 |
| 12 | 116266200 | 116268399 | 22.65614 | 39.76608 | 28.77967 | 12.72916 | 0.851431 | 0.645409 | 0.200582 |
| 9 | 18776000 | 18777599 | 26.95994 | 46.3687 | 36.21823 | 15.88695 | 0.739226 | 0.922562 | 0.200584 |
| X | 125176400 | 125177599 | 32.86989 | 55.22995 | 40.64804 | 17.75438 | 0.821783 | 1.153776 | 0.200588 |
| 1 | 62744200 | 62749799 | 28.09949 | 45.37333 | 98.01364 | 41.81441 | -0.74904 | 1.728068 | 0.200591 |
| X | 118079800 | 118083599 | 58.78155 | 94.86499 | 62.53119 | 26.65816 | 0.958475 | 1.777944 | 0.200612 |
| 13 | 22962800 | 22964999 | 23.23701 | 40.4364 | 32.87662 | 14.44891 | 0.68422 | 0.781935 | 0.200616 |
| 1 | 73865400 | 73870599 | 27.15031 | 43.84031 | 99.04163 | 42.25255 | -0.81352 | 1.726804 | 0.200623 |
| 11 | 32279400 | 32281999 | 11.8482 | 20.70704 | 47.30606 | 20.84769 | -0.79818 | 0.689082 | 0.200628 |
| 5 | 101420000 | 101421599 | 8.54526 | 15.63022 | 37.73747 | 16.1414 | -0.84006 | 0.269622 | 0.200629 |
| 4 | 13627800 | 13629199 | 23.16225 | 40.58618 | 30.04575 | 13.2703 | 0.819257 | 0.678253 | 0.200629 |
| 1 | 151675800 | 151677599 | 6.278348 | 11.65496 | 33.14698 | 13.89536 | -1.05163 | 0.160235 | 0.200641 |
| 4 | 85123400 | 85125199 | 12.1754 | 21.21777 | 48.91465 | 21.51687 | -0.81172 | 0.734717 | 0.200644 |
| 10 | 126547800 | 126550199 | 22.16075 | 36.75667 | 73.3175 | 31.75916 | -0.61214 | 1.28954 | 0.200644 |
| 3 | 9701000 | 9701999 | 6.991375 | 13.02381 | 32.68756 | 13.62878 | -0.86719 | 0.142622 | 0.200648 |
| 5 | 5685200 | 5686799 | 17.86707 | 33.21248 | 18.98307 | 7.943275 | 1.246518 | 0.211742 | 0.200648 |
| 9 | 7184200 | 7185399 | 7.764474 | 14.32958 | 35.76905 | 15.11452 | -0.87469 | 0.208664 | 0.20066 |
| 8 | 30292400 | 30294199 | 25.80897 | 45.26834 | 26.33947 | 11.643 | 1.164466 | 0.670274 | 0.200662 |
| 2 | 87415400 | 87417999 | 33.67444 | 58.19145 | 26.7099 | 11.71785 | 1.503267 | 0.896978 | 0.200663 |
| 6 | 132045400 | 132047999 | 37.70856 | 63.53268 | 36.88801 | 16.12535 | 1.162377 | 1.140359 | 0.200666 |
| 3 | 69865000 | 69866199 | 7.730169 | 14.40914 | 33.26044 | 13.85285 | -0.74636 | 0.146475 | 0.200667 |
| 12 | 84411000 | 84412799 | 10.136 | 17.82132 | 44.07865 | 19.51846 | -0.91152 | 0.577465 | 0.20067 |
| 10 | 72797400 | 72798799 | 18.91321 | 34.33221 | 22.58732 | 9.753831 | 1.014419 | 0.357255 | 0.200689 |

| | | | | | | | | | |
|---|---|---|---|---|---|---|---|---|---|
| 18 | 64039800 | 64040799 | 5.710313 | 10.98532 | 29.84958 | 11.6296 | -0.89478 | -0.11385 | 0.200692 |
| 1 | 71019200 | 71020599 | 16.33503 | 30.7963 | 17.55376 | 7.171888 | 1.277138 | 0.108638 | 0.200695 |
| 2 | 109357600 | 109358799 | 16.33503 | 30.7963 | 17.55376 | 7.171888 | 1.277138 | 0.108638 | 0.200695 |
| 7 | 97862200 | 97862799 | 5.245814 | 10.16945 | 26.75748 | 10.11641 | -0.80649 | -0.22713 | 0.200697 |
| 2 | 107120200 | 107121999 | 15.96027 | 27.03495 | 63.33553 | 27.73176 | -0.84219 | 1.046613 | 0.200708 |
| 10 | 22826000 | 22828399 | 24.94042 | 43.09466 | 35.59677 | 15.61655 | 0.660029 | 0.869923 | 0.200712 |
| 12 | 106135000 | 106148599 | 106.0894 | 160.1929 | 297.4348 | 122.9517 | -0.6173 | 3.330133 | 0.200715 |
| 17 | 87441000 | 87443999 | 19.28573 | 32.06748 | 70.20269 | 30.46201 | -0.74546 | 1.250755 | 0.200719 |
| 17 | 79524000 | 79525399 | 21.36258 | 37.68041 | 27.87255 | 12.33435 | 0.822003 | 0.578278 | 0.200719 |
| X | 3607800 | 3610599 | 21.36258 | 37.68041 | 27.87255 | 12.33435 | 0.822003 | 0.578278 | 0.200719 |
| 12 | 86146600 | 86148399 | 9.01746 | 16.45159 | 36.59463 | 15.70337 | -0.72546 | 0.287498 | 0.200731 |
| 7 | 25453200 | 25455799 | 14.89923 | 25.20109 | 62.06158 | 27.16201 | -0.91375 | 1.056334 | 0.200732 |
| 10 | 13000400 | 13004999 | 30.98977 | 49.99798 | 99.58485 | 42.4383 | -0.63374 | 1.758185 | 0.200741 |
| 6 | 119151400 | 119152799 | 9.784975 | 17.32805 | 41.45937 | 18.3323 | -0.85967 | 0.490328 | 0.200743 |
| 1 | 20798200 | 20798999 | 13.07455 | 24.80128 | 19.80901 | 7.990081 | 0.811171 | 0.044358 | 0.200776 |
| 7 | 15229600 | 15230399 | 11.79144 | 22.91632 | 15.245 | 5.672734 | 1.178933 | -0.24051 | 0.200786 |
| 19 | 24890000 | 24890399 | 7.863169 | 15.36066 | 13.52005 | 4.723947 | 0.860492 | -0.51289 | 0.200787 |
| 7 | 13842600 | 13845199 | 36.65231 | 61.77413 | 38.61857 | 16.8836 | 1.056273 | 1.136399 | 0.200789 |
| 7 | 9652600 | 9655199 | 19.65611 | 35.32949 | 24.13305 | 10.54601 | 0.949864 | 0.42201 | 0.200803 |
| 5 | 104897600 | 104902799 | 173.0852 | 260.5711 | 207.8607 | 85.84919 | 0.59687 | 3.399663 | 0.200811 |
| 4 | 128621000 | 128621999 | 7.469313 | 13.79806 | 34.69181 | 14.64023 | -0.88335 | 0.199258 | 0.200812 |
| 13 | 49564000 | 49565599 | 17.72621 | 31.82874 | 24.84539 | 10.86649 | 0.7576 | 0.415115 | 0.200824 |
| 1 | 64289400 | 64290399 | 10.90052 | 21.16271 | 15.71685 | 5.897917 | 1.009666 | -0.22286 | 0.200827 |
| 18 | 72258000 | 72258799 | 11.94515 | 23.02333 | 17.85283 | 6.917803 | 0.905581 | -0.09919 | 0.20083 |
| 8 | 36032800 | 36034799 | 15.53934 | 28.36229 | 22.73348 | 9.750109 | 0.737949 | 0.302496 | 0.200848 |
| 17 | 27627600 | 27659199 | 320.2215 | 440.5748 | 791.6882 | 329.1124 | -0.7007 | 4.827299 | 0.200854 |
| 8 | 80343600 | 80344799 | 8.688902 | 16.00101 | 36.26841 | 15.37069 | -0.73949 | 0.231729 | 0.200859 |
| 4 | 63935000 | 63936399 | 14.01852 | 23.9766 | 56.80069 | 24.90619 | -0.85596 | 0.947161 | 0.200859 |
| 13 | 107781400 | 107782999 | 10.53259 | 18.68052 | 40.91436 | 18.0813 | -0.73202 | 0.47926 | 0.200881 |
| 16 | 93149000 | 93150599 | 10.53259 | 18.68052 | 40.91436 | 18.0813 | -0.73202 | 0.47926 | 0.200881 |
| 1 | 27858800 | 27859399 | 10.34217 | 19.97539 | 15.8327 | 6.089994 | 0.883563 | -0.14801 | 0.200884 |
| 9 | 98400200 | 98401799 | 11.0119 | 19.43954 | 42.99192 | 19.02192 | -0.74896 | 0.534971 | 0.200885 |
| 14 | 8410000 | 8411399 | 13.19037 | 24.69063 | 19.86362 | 8.214427 | 0.771072 | 0.131259 | 0.200889 |
| 2 | 114020000 | 114022599 | 15.3149 | 26.51839 | 54.90662 | 24.08946 | -0.65936 | 0.835707 | 0.200925 |
| 1 | 171537200 | 171538799 | 17.38062 | 31.21869 | 25.07093 | 10.96204 | 0.717419 | 0.412545 | 0.200932 |
| 12 | 59906200 | 59909199 | 33.47187 | 57.0132 | 35.21389 | 15.4356 | 1.074237 | 1.030544 | 0.200936 |
| 10 | 46709200 | 46710399 | 14.72941 | 27.75377 | 19.45146 | 7.954737 | 0.980741 | 0.102224 | 0.200951 |
| 1 | 154811400 | 154812399 | 13.49843 | 25.19002 | 20.79246 | 8.643246 | 0.728262 | 0.155188 | 0.200961 |
| 16 | 97064400 | 97065799 | 5.834426 | 11.24639 | 28.13838 | 10.90058 | -0.76827 | -0.13296 | 0.20097 |
| 13 | 9143200 | 9144799 | 10.19892 | 17.81582 | 48.87165 | 21.53019 | -1.06057 | 0.689904 | 0.200979 |
| 15 | 102817200 | 102819999 | 14.54033 | 25.29124 | 56.00368 | 24.606 | -0.7547 | 0.782597 | 0.201008 |
| 4 | 62653600 | 62655399 | 10.09443 | 17.85386 | 41.46987 | 18.3415 | -0.81783 | 0.50699 | 0.201034 |
| 11 | 116006400 | 116008799 | 10.09443 | 17.85386 | 41.46987 | 18.3415 | -0.81783 | 0.50699 | 0.201034 |
| 11 | 22671600 | 22672199 | 9.7657 | 18.99338 | 15.97742 | 5.909923 | 0.851285 | -0.284 | 0.201035 |
| 3 | 4201400 | 4207399 | 76.0799 | 121.2617 | 87.48307 | 36.63419 | 0.808063 | 2.181614 | 0.201044 |
| 19 | 39367200 | 39370599 | 60.92252 | 97.66023 | 79.89896 | 33.73416 | 0.635937 | 1.987095 | 0.201053 |
| 15 | 56667200 | 56669399 | 33.13168 | 55.35919 | 45.95617 | 20.00373 | 0.648742 | 1.214263 | 0.201094 |

| | | | | | | | | | |
|---|---|---|---|---|---|---|---|---|---|
| 17 | 69662600 | 69666399 | 24.95958 | 41.01127 | 85.48939 | 36.83432 | -0.67888 | 1.415886 | 0.201097 |
| 14 | 42000400 | 42002999 | 17.33929 | 32.47531 | 18.63866 | 7.700439 | 1.255261 | 0.160992 | 0.201104 |
| 9 | 55138800 | 55140799 | 10.3765 | 18.40853 | 40.91784 | 18.07883 | -0.75297 | 0.476256 | 0.201105 |
| 6 | 70530000 | 70531999 | 26.21474 | 45.25041 | 35.47616 | 15.56316 | 0.734698 | 0.883976 | 0.201119 |
| 1 | 103657200 | 103658199 | 11.88621 | 22.82525 | 18.34515 | 7.184375 | 0.840037 | -0.06544 | 0.201119 |
| 15 | 3795600 | 3796999 | 11.88621 | 22.82525 | 18.34515 | 7.184375 | 0.840037 | -0.06544 | 0.201119 |
| 7 | 44397200 | 44398399 | 5.630443 | 10.68034 | 30.9136 | 12.46947 | -1.03072 | 0.003997 | 0.201125 |
| 8 | 80437800 | 80438599 | 5.630443 | 10.68034 | 30.9136 | 12.46947 | -1.03072 | 0.003997 | 0.201125 |
| 14 | 69969400 | 69970399 | 5.630443 | 10.68034 | 30.9136 | 12.46947 | -1.03072 | 0.003997 | 0.201125 |
| 11 | 117869800 | 117871199 | 7.788495 | 14.31352 | 34.59724 | 14.70336 | -0.83411 | 0.232564 | 0.201125 |
| 7 | 79370000 | 79373199 | 15.27828 | 26.06089 | 57.91717 | 25.39023 | -0.76503 | 0.977158 | 0.201154 |
| 1 | 12352000 | 12354799 | 27.30008 | 47.3573 | 33.20769 | 14.57019 | 0.896365 | 0.843761 | 0.201154 |
| 3 | 4123600 | 4124799 | 17.81409 | 31.69761 | 25.60671 | 11.28547 | 0.70117 | 0.473795 | 0.201155 |
| 7 | 49581800 | 49582799 | 6.175271 | 11.75951 | 29.94957 | 11.98156 | -0.83741 | -0.02018 | 0.201162 |
| 16 | 10951800 | 10952999 | 6.175271 | 11.75951 | 29.94957 | 11.98156 | -0.83741 | -0.02018 | 0.201162 |
| 3 | 47812000 | 47813799 | 24.83707 | 43.03826 | 35.05029 | 15.37814 | 0.681205 | 0.844395 | 0.201186 |
| 13 | 113454200 | 113457399 | 22.10785 | 36.33801 | 79.00261 | 34.04433 | -0.73896 | 1.401845 | 0.201194 |
| 13 | 113708400 | 113711399 | 20.19186 | 33.65029 | 71.11577 | 30.90422 | -0.69483 | 1.226615 | 0.201207 |
| 11 | 120965200 | 120966799 | 8.606504 | 15.60256 | 37.55513 | 16.24387 | -0.84541 | 0.32666 | 0.201225 |
| 10 | 43539600 | 43543999 | 28.46015 | 46.28589 | 90.63678 | 38.89779 | -0.5978 | 1.56777 | 0.201227 |
| 12 | 86204200 | 86206199 | 10.46023 | 18.34722 | 47.2397 | 20.88187 | -0.96973 | 0.620754 | 0.201243 |
| 5 | 138354200 | 138356799 | 39.86052 | 65.31654 | 55.42767 | 23.85423 | 0.612558 | 1.463841 | 0.201275 |
| 1 | 161231600 | 161232599 | 14.51557 | 27.35694 | 19.74453 | 8.071529 | 0.939934 | 0.101614 | 0.20128 |
| 7 | 121563600 | 121566599 | 14.08289 | 24.29565 | 57.01332 | 25.01094 | -0.84035 | 0.86453 | 0.201291 |
| 7 | 7571600 | 7575399 | 75.9224 | 120.9409 | 83.86322 | 35.09036 | 0.864222 | 2.199022 | 0.2013 |
| 6 | 69717000 | 69718599 | 18.90837 | 34.02348 | 24.14898 | 10.54304 | 0.895702 | 0.4094 | 0.201323 |
| 6 | 131821200 | 131825399 | 93.93599 | 147.9199 | 113.878 | 47.24993 | 0.699077 | 2.514715 | 0.201345 |
| 11 | 85578600 | 85579599 | 6.445327 | 12.33905 | 30.75616 | 12.1516 | -0.79013 | -0.05491 | 0.201346 |
| 14 | 99909600 | 99910599 | 6.445327 | 12.33905 | 30.75616 | 12.1516 | -0.79013 | -0.05491 | 0.201346 |
| 8 | 112956000 | 112956599 | 3.759614 | 7.336022 | 22.04677 | 7.856037 | -0.91377 | -0.46779 | 0.201363 |
| 12 | 106196600 | 106204399 | 51.01755 | 80.88617 | 148.8514 | 61.94987 | -0.54848 | 2.334361 | 0.201365 |
| 15 | 97861400 | 97862999 | 9.702233 | 17.45197 | 38.45899 | 16.7955 | -0.72862 | 0.380351 | 0.201368 |
| 5 | 53205400 | 53207599 | 12.35492 | 21.45824 | 51.04774 | 22.40771 | -0.85745 | 0.786874 | 0.201379 |
| 10 | 50310000 | 50310599 | 6.289024 | 12.06723 | 29.70397 | 11.65952 | -0.76327 | -0.08107 | 0.201413 |
| 4 | 130889400 | 130897999 | 57.15772 | 89.73449 | 172.0056 | 71.34006 | -0.61858 | 2.561735 | 0.201414 |
| 19 | 13492400 | 13492999 | 9.30995 | 18.08947 | 14.35192 | 5.350322 | 0.924052 | -0.26977 | 0.201416 |
| 4 | 95964400 | 95966199 | 12.12452 | 21.29003 | 45.10676 | 19.95775 | -0.69001 | 0.609848 | 0.201428 |
| 15 | 101909200 | 101914999 | 25.44457 | 41.05342 | 104.3863 | 44.48709 | -0.98501 | 1.749192 | 0.20143 |
| 9 | 119236200 | 119244799 | 104.6811 | 158.6348 | 275.6199 | 114.0632 | -0.51714 | 3.256898 | 0.201432 |
| 5 | 9659800 | 9660399 | 9.357838 | 18.25294 | 14.37921 | 5.151719 | 0.985885 | -0.40388 | 0.201469 |
| 3 | 15656000 | 15656999 | 20.51426 | 36.17507 | 24.55953 | 10.8688 | 0.94477 | 0.568706 | 0.201471 |
| 7 | 5422600 | 5431199 | 128.0347 | 198.113 | 133.793 | 55.57427 | 0.868469 | 2.844847 | 0.201473 |
| 6 | 33956000 | 33961199 | 33.07594 | 53.31132 | 106.7535 | 45.43559 | -0.64321 | 1.791402 | 0.201483 |
| 13 | 72098000 | 72112799 | 109.6124 | 164.7779 | 316.3756 | 130.6168 | -0.67125 | 3.4196 | 0.201485 |
| 12 | 20294000 | 20303599 | 295.2804 | 427.3193 | 320.1916 | 132.0536 | 0.633966 | 4.074474 | 0.201489 |
| 16 | 30783400 | 30785999 | 19.1273 | 31.80759 | 70.20062 | 30.46346 | -0.7571 | 1.248799 | 0.201492 |
| X | 33115000 | 33117799 | 38.9043 | 65.51182 | 37.04 | 16.18913 | 1.200752 | 1.152166 | 0.201497 |

| | | | | | | | | | |
|---|---|---|---|---|---|---|---|---|---|
| 17 | 91626800 | 91627399 | 9.021597 | 17.56463 | 15.5583 | 5.706208 | 0.789037 | -0.32327 | 0.201511 |
| 8 | 123622000 | 123624199 | 10.88964 | 19.15324 | 43.548 | 19.28211 | -0.79062 | 0.57745 | 0.201518 |
| 7 | 138202000 | 138205399 | 22.17659 | 36.75845 | 73.80638 | 31.95623 | -0.62168 | 1.298359 | 0.201518 |
| 1 | 10109600 | 10110399 | 4.838804 | 9.340696 | 26.95375 | 10.39476 | -0.96508 | -0.16187 | 0.201525 |
| 7 | 11953400 | 11956199 | 33.21159 | 56.60029 | 36.42762 | 15.96832 | 1.015394 | 1.028063 | 0.201526 |
| 2 | 177234400 | 177236199 | 20.22575 | 35.8582 | 26.13847 | 11.55516 | 0.845483 | 0.51225 | 0.201537 |
| 1 | 191251400 | 191254199 | 16.83257 | 28.66526 | 60.75503 | 26.63081 | -0.69757 | 0.999271 | 0.201537 |
| X | 4343400 | 4345199 | 32.33514 | 54.94589 | 38.66272 | 16.9441 | 0.886633 | 1.052463 | 0.201543 |
| 12 | 19791600 | 19806799 | 499.0613 | 685.6993 | 530.206 | 220.459 | 0.513563 | 4.85146 | 0.201553 |
| 1 | 58942200 | 58943399 | 6.643569 | 12.5434 | 33.68172 | 13.72094 | -0.93712 | 0.051233 | 0.201558 |
| 10 | 110844800 | 110848999 | 17.1971 | 29.11479 | 65.12534 | 28.51092 | -0.77592 | 1.056602 | 0.201561 |
| 15 | 8575600 | 8577999 | 30.46767 | 51.57234 | 41.49153 | 18.16336 | 0.693897 | 1.080653 | 0.201561 |
| 4 | 135319000 | 135322799 | 34.26948 | 55.10061 | 107.7073 | 45.68083 | -0.61281 | 1.872451 | 0.201581 |
| 18 | 84747200 | 84749199 | 14.83624 | 25.24967 | 59.48303 | 26.07194 | -0.84933 | 0.997614 | 0.201581 |
| 17 | 23764600 | 23766399 | 13.07843 | 22.83838 | 48.37574 | 21.30733 | -0.69001 | 0.708267 | 0.201585 |
| 12 | 114625600 | 114627199 | 29.88221 | 50.47507 | 42.12393 | 18.42844 | 0.641208 | 1.097909 | 0.201602 |
| 7 | 21036400 | 21041399 | 88.87049 | 140.3419 | 107.9877 | 44.84045 | 0.702996 | 2.451657 | 0.201606 |
| 7 | 44352400 | 44365799 | 112.1176 | 169.3309 | 299.1324 | 123.6617 | -0.54528 | 3.32735 | 0.201606 |
| 7 | 59128600 | 59129799 | 19.99442 | 35.20463 | 28.62069 | 12.66992 | 0.685518 | 0.590093 | 0.201608 |
| 14 | 48159200 | 48163599 | 31.12673 | 50.25524 | 101.4282 | 43.26414 | -0.65161 | 1.736511 | 0.201615 |
| 12 | 106492400 | 106495799 | 25.52175 | 41.69596 | 83.55286 | 35.91705 | -0.62618 | 1.492502 | 0.201632 |
| 18 | 65364200 | 65364999 | 5.677517 | 10.96195 | 28.20248 | 10.866 | -0.80033 | -0.15155 | 0.201635 |
| 14 | 106055000 | 106055999 | 7.570016 | 14.11373 | 33.82913 | 14.08444 | -0.80022 | 0.142954 | 0.201663 |
| 18 | 75238800 | 75241399 | 23.64009 | 39.00551 | 77.90129 | 33.62415 | -0.61475 | 1.356855 | 0.201687 |
| 7 | 114429800 | 114431399 | 7.68343 | 14.01815 | 37.12963 | 15.9324 | -0.97539 | 0.277393 | 0.2017 |
| 7 | 140870800 | 140880199 | 49.93933 | 78.61057 | 175.9802 | 73.01375 | -0.83917 | 2.504913 | 0.201709 |
| X | 55024800 | 55025599 | 12.58525 | 24.31853 | 16.6619 | 6.395733 | 1.094897 | -0.12613 | 0.201719 |
| 1 | 74826600 | 74834599 | 46.54587 | 73.64949 | 158.4168 | 65.84629 | -0.77564 | 2.382899 | 0.201739 |
| 9 | 25513600 | 25515199 | 13.23503 | 22.64982 | 59.98772 | 26.30489 | -1.01594 | 0.942822 | 0.20174 |
| 2 | 28797000 | 28803199 | 48.91558 | 77.13746 | 164.2199 | 68.15035 | -0.76474 | 2.465795 | 0.201769 |
| 13 | 19496400 | 19499599 | 99.82075 | 155.3375 | 135.7399 | 56.34833 | 0.503416 | 2.733452 | 0.201773 |
| 13 | 22410800 | 22412999 | 24.99688 | 43.39129 | 34.45886 | 15.12073 | 0.717832 | 0.828179 | 0.201787 |
| 6 | 90778200 | 90779999 | 9.756489 | 17.35771 | 39.93675 | 17.60325 | -0.79979 | 0.453255 | 0.201794 |
| X | 148377600 | 148379399 | 15.76372 | 28.88425 | 22.70821 | 9.689782 | 0.770724 | 0.28293 | 0.201797 |
| 18 | 72996600 | 72999599 | 27.20299 | 46.81653 | 36.61463 | 16.06114 | 0.737497 | 0.917439 | 0.201809 |
| 6 | 77972400 | 77973399 | 14.48565 | 27.96415 | 14.61694 | 5.635687 | 1.474622 | -0.0972 | 0.201817 |
| 6 | 149184000 | 149187599 | 73.20162 | 116.5342 | 94.3249 | 39.43375 | 0.640512 | 2.217738 | 0.20186 |
| 17 | 47757000 | 47759199 | 7.51832 | 13.75786 | 35.58801 | 15.21295 | -0.93746 | 0.256545 | 0.201865 |
| 10 | 103161400 | 103161999 | 12.12995 | 23.37451 | 17.84689 | 6.919762 | 0.926957 | -0.09513 | 0.20187 |
| 4 | 43957400 | 43958999 | 4.68065 | 8.790286 | 32.23288 | 13.25594 | -1.39146 | 0.06909 | 0.201883 |
| 15 | 96003600 | 96004199 | 4.922942 | 9.569596 | 25.78706 | 9.579292 | -0.81563 | -0.29299 | 0.20192 |
| 17 | 88001600 | 88007799 | 35.1052 | 56.31207 | 113.5286 | 47.98165 | -0.66144 | 1.948388 | 0.201927 |
| 2 | 143892000 | 143896399 | 21.03483 | 34.89829 | 73.26264 | 31.74112 | -0.68564 | 1.282162 | 0.201948 |
| 2 | 173621600 | 173625199 | 21.03483 | 34.89829 | 73.26264 | 31.74112 | -0.68564 | 1.282162 | 0.201948 |
| 7 | 13317200 | 13325599 | 179.7289 | 269.665 | 221.0209 | 91.20241 | 0.553546 | 3.466061 | 0.201949 |
| 5 | 70223400 | 70224399 | 13.35947 | 25.04361 | 20.79511 | 8.578449 | 0.728496 | 0.125838 | 0.201967 |
| 2 | 57425600 | 57427999 | 22.54297 | 39.5249 | 28.79168 | 12.7229 | 0.842061 | 0.660738 | 0.201976 |

| | | | | | | | | | |
|---|---|---|---|---|---|---|---|---|---|
| 18 | 33843200 | 33845199 | 22.24274 | 39.46862 | 25.01223 | 11.04916 | 1.047156 | 0.519331 | 0.201986 |
| 8 | 92669600 | 92671199 | 20.16639 | 35.53466 | 28.5576 | 12.64006 | 0.702477 | 0.582775 | 0.202002 |
| 12 | 22247800 | 22251999 | 78.08561 | 123.4587 | 104.9103 | 43.58906 | 0.561339 | 2.413841 | 0.202013 |
| 12 | 22248000 | 22251999 | 78.08561 | 123.4587 | 104.9103 | 43.58906 | 0.561339 | 2.413841 | 0.202013 |
| 5 | 110117600 | 110122599 | 136.365 | 208.4566 | 169.0529 | 70.13003 | 0.591343 | 3.082322 | 0.202017 |
| 12 | 114685200 | 114687999 | 48.62586 | 79.10765 | 56.83269 | 24.39324 | 0.845431 | 1.583151 | 0.202022 |
| 14 | 124733800 | 124734199 | 10.94327 | 21.28976 | 15.31746 | 5.652673 | 1.077636 | -0.27886 | 0.202054 |
| 14 | 20855400 | 20857199 | 7.331494 | 13.51777 | 34.11741 | 14.43509 | -0.89118 | 0.209127 | 0.202057 |
| 5 | 150368400 | 150380399 | 97.73421 | 148.9948 | 257.0254 | 106.5572 | -0.49975 | 3.13206 | 0.202062 |
| 5 | 66345400 | 66348599 | 14.06524 | 24.30537 | 57.60192 | 25.27043 | -0.85407 | 0.849718 | 0.202066 |
| X | 77004800 | 77005799 | 24.9959 | 43.36825 | 32.6516 | 14.32628 | 0.794506 | 0.832761 | 0.202066 |
| 2 | 174120800 | 174135199 | 204.5204 | 292.3985 | 537.2682 | 221.9744 | -0.67698 | 4.249031 | 0.202078 |
| 10 | 84661600 | 84667399 | 36.50695 | 58.66006 | 113.431 | 48.06312 | -0.59852 | 1.895888 | 0.202087 |
| 10 | 61707000 | 61710999 | 18.10665 | 30.31604 | 67.64531 | 29.48256 | -0.77281 | 1.167407 | 0.202102 |
| 8 | 69446600 | 69447799 | 9.667606 | 17.46133 | 38.52008 | 16.73871 | -0.72566 | 0.354313 | 0.20211 |
| 8 | 15702200 | 15702799 | 5.782136 | 10.91846 | 30.77753 | 12.53424 | -1.00487 | 0.038247 | 0.202113 |
| 17 | 20659400 | 20668999 | 133.9603 | 205.0565 | 156.9782 | 65.14122 | 0.675954 | 3.051967 | 0.202133 |
| 7 | 9417600 | 9419799 | 21.94841 | 39.11649 | 23.83694 | 10.49135 | 1.106609 | 0.487966 | 0.202142 |
| X | 43132000 | 43133199 | 12.36373 | 23.59737 | 18.45858 | 7.349317 | 0.857691 | -0.0062 | 0.20217 |
| 1 | 88495800 | 88497399 | 8.682503 | 15.95998 | 35.68041 | 15.1592 | -0.72245 | 0.239926 | 0.202206 |
| 5 | 88712400 | 88714399 | 11.97692 | 20.98014 | 46.73585 | 20.63642 | -0.76209 | 0.649981 | 0.202218 |
| Y | 324400 | 325199 | 10.0622 | 19.55598 | 9.816067 | 3.652018 | 1.579113 | -0.28446 | 0.202227 |
| 18 | 15195000 | 15199399 | 33.68912 | 53.98025 | 117.3063 | 49.49415 | -0.77136 | 1.982986 | 0.202233 |
| 15 | 83376600 | 83382399 | 40.18889 | 64.18148 | 125.0581 | 52.51265 | -0.61977 | 2.095776 | 0.202241 |
| 11 | 83363800 | 83364599 | 3.823491 | 7.447015 | 23.52736 | 8.592301 | -1.0185 | -0.37557 | 0.202247 |
| 17 | 10618400 | 10623399 | 25.42986 | 41.36031 | 86.90668 | 37.29786 | -0.69899 | 1.559912 | 0.202261 |
| 4 | 74850400 | 74852599 | 18.29654 | 33.01723 | 24.28184 | 10.56331 | 0.848146 | 0.38957 | 0.202273 |
| 5 | 133849400 | 133851199 | 7.580384 | 14.10323 | 33.7645 | 14.10791 | -0.80266 | 0.156872 | 0.202279 |
| 2 | 55433000 | 55434599 | 16.75974 | 30.65578 | 22.88417 | 9.788055 | 0.842178 | 0.29914 | 0.202282 |
| 4 | 8300600 | 8302999 | 30.42837 | 51.33216 | 42.14945 | 18.43248 | 0.664519 | 1.111313 | 0.202284 |
| 10 | 14680400 | 14681199 | 11.27386 | 21.82879 | 16.82693 | 6.3981 | 0.939239 | -0.17144 | 0.202332 |
| 18 | 77570600 | 77573799 | 14.11644 | 24.29974 | 54.23069 | 23.78864 | -0.76863 | 0.884127 | 0.202334 |
| 2 | 158626200 | 158627999 | 10.71172 | 18.89354 | 44.04079 | 19.48931 | -0.82496 | 0.543719 | 0.20235 |
| 15 | 19725600 | 19726599 | 13.66942 | 26.20212 | 17.5524 | 6.909728 | 1.093433 | -0.03296 | 0.202357 |
| 5 | 113910600 | 113915199 | 18.41996 | 30.47064 | 74.8969 | 32.3836 | -0.9126 | 1.311807 | 0.202373 |
| 15 | 46225800 | 46228799 | 33.37479 | 55.60669 | 46.00273 | 19.98802 | 0.653758 | 1.2439 | 0.202373 |
| 9 | 32571400 | 32572999 | 17.55738 | 31.85375 | 23.95638 | 10.35206 | 0.822352 | 0.354202 | 0.202374 |
| 14 | 59862200 | 59865799 | 21.82257 | 35.78895 | 79.7712 | 34.34301 | -0.77602 | 1.430986 | 0.202389 |
| 7 | 24313200 | 24331599 | 141.0292 | 208.4004 | 385.9006 | 158.9226 | -0.64173 | 3.737195 | 0.202405 |
| 5 | 146779400 | 146788599 | 57.2164 | 90.05069 | 174.0606 | 72.21577 | -0.62788 | 2.51085 | 0.202413 |
| 7 | 15097400 | 15099999 | 100.8306 | 158.8991 | 102.5064 | 42.53592 | 0.954601 | 2.500835 | 0.202424 |
| 9 | 31357400 | 31360199 | 10.98154 | 19.46857 | 45.0977 | 19.93667 | -0.81324 | 0.490072 | 0.202426 |
| 1 | 113905200 | 113906999 | 20.76991 | 37.52602 | 21.81826 | 9.47721 | 1.186148 | 0.395421 | 0.202429 |
| 12 | 43551600 | 43554599 | 31.92175 | 54.73533 | 32.97754 | 14.46306 | 1.111102 | 0.963227 | 0.202431 |
| 11 | 87856000 | 87858599 | 12.13391 | 21.318 | 45.09739 | 19.96269 | -0.68775 | 0.601968 | 0.202439 |
| 12 | 61288800 | 61290199 | 18.64385 | 33.17808 | 26.41376 | 11.63997 | 0.722204 | 0.480041 | 0.20244 |
| 9 | 107527000 | 107529399 | 8.239691 | 14.46736 | 47.74084 | 21.12186 | -1.32489 | 0.593202 | 0.202454 |

| | | | | | | | | | |
|---|---|---|---|---|---|---|---|---|---|
| 5 | 52781400 | 52783799 | 10.11469 | 17.83568 | 43.02641 | 19.04144 | -0.87404 | 0.542654 | 0.202458 |
| 10 | 46661800 | 46663599 | 28.81662 | 49.21947 | 38.75004 | 16.9894 | 0.726251 | 0.993054 | 0.202468 |
| 3 | 79535600 | 79537399 | 14.86282 | 25.22916 | 60.50924 | 26.51121 | -0.87552 | 1.022864 | 0.202471 |
| 1 | 176434400 | 176435399 | 16.57729 | 30.99938 | 19.88775 | 8.238393 | 1.093144 | 0.167396 | 0.202472 |
| 7 | 120151600 | 120152599 | 8.102818 | 14.84513 | 35.07813 | 14.97068 | -0.80603 | 0.254727 | 0.20248 |
| 3 | 19139000 | 19141799 | 38.87698 | 64.45904 | 47.16493 | 20.42536 | 0.830012 | 1.315609 | 0.202482 |
| 12 | 38968800 | 38969399 | 6.211608 | 11.72176 | 30.77815 | 12.55186 | -0.90539 | 0.046793 | 0.202511 |
| 16 | 92078000 | 92082399 | 20.99942 | 34.5354 | 78.24539 | 33.72599 | -0.79788 | 1.392195 | 0.202513 |
| 5 | 73412000 | 73415599 | 13.19916 | 22.71523 | 55.27294 | 24.24565 | -0.89287 | 0.882957 | 0.202514 |
| 12 | 4557200 | 4557799 | 3.028145 | 5.913473 | 24.78039 | 8.708752 | -1.37013 | -0.524 | 0.202527 |
| 7 | 20351600 | 20352399 | 11.63014 | 22.23118 | 17.5015 | 6.944975 | 0.852637 | -0.02743 | 0.202535 |
| 14 | 30138600 | 30139399 | 4.18439 | 8.159971 | 23.19878 | 8.336896 | -0.84681 | -0.42818 | 0.202537 |
| 15 | 63893800 | 63896199 | 11.525 | 20.18163 | 46.74499 | 20.63451 | -0.81797 | 0.653393 | 0.202541 |
| 9 | 87137200 | 87137799 | 5.249594 | 10.16517 | 26.65362 | 10.13106 | -0.80879 | -0.20881 | 0.202548 |
| 10 | 56582800 | 56584399 | 9.810962 | 17.31704 | 44.05629 | 19.4934 | -0.94986 | 0.530545 | 0.202551 |
| 10 | 117977400 | 117978799 | 7.424621 | 13.85064 | 32.73101 | 13.61393 | -0.7784 | 0.13677 | 0.202562 |
| 19 | 22893200 | 22894599 | 8.420776 | 15.35022 | 36.05197 | 15.48659 | -0.80412 | 0.286821 | 0.202565 |
| 1 | 90702600 | 90703999 | 5.88431 | 11.21237 | 29.37731 | 11.73724 | -0.87587 | -0.02799 | 0.202568 |
| 3 | 146226400 | 146227399 | 5.88431 | 11.21237 | 29.37731 | 11.73724 | -0.87587 | -0.02799 | 0.202568 |
| 1 | 104195400 | 104197199 | 20.13242 | 37.71514 | 13.09221 | 5.406902 | 1.971916 | 0.170414 | 0.202582 |
| 2 | 111244800 | 111247599 | 26.53905 | 45.91941 | 34.96614 | 15.34053 | 0.777349 | 0.865469 | 0.20259 |
| 7 | 73267800 | 73271799 | 16.65516 | 28.44309 | 59.1428 | 25.93 | -0.66942 | 0.97132 | 0.202616 |
| 7 | 48767600 | 48772799 | 23.66685 | 38.54845 | 86.2354 | 37.02924 | -0.78759 | 1.534537 | 0.202624 |
| 15 | 47015200 | 47017599 | 23.71709 | 41.45367 | 30.45009 | 13.42024 | 0.830329 | 0.719248 | 0.202641 |
| 15 | 72661400 | 72663599 | 11.06071 | 19.39873 | 45.15523 | 19.95894 | -0.82484 | 0.623993 | 0.202646 |
| 5 | 26019400 | 26025599 | 48.51888 | 76.70068 | 154.9354 | 64.36922 | -0.68621 | 2.407126 | 0.202649 |
| 7 | 4917400 | 4933199 | 94.30053 | 142.2016 | 302.3414 | 124.9273 | -0.81424 | 3.355479 | 0.202656 |
| X | 142735200 | 142736799 | 18.95582 | 35.87998 | 14.60876 | 5.919498 | 1.767948 | 0.100326 | 0.202659 |
| 12 | 24188800 | 24208599 | 406.8397 | 570.0425 | 446.7735 | 185.1972 | 0.522543 | 4.552147 | 0.202687 |
| 7 | 60280800 | 60281399 | 10.99681 | 21.26976 | 17.44493 | 6.666091 | 0.844206 | -0.15598 | 0.202704 |
| 6 | 39701400 | 39702599 | 6.498456 | 12.25647 | 30.76658 | 12.56141 | -0.84238 | 0.054223 | 0.202725 |
| 7 | 66068600 | 66070199 | 6.498456 | 12.25647 | 30.76658 | 12.56141 | -0.84238 | 0.054223 | 0.202725 |
| 17 | 18155400 | 18159999 | 48.26115 | 78.71128 | 54.06597 | 23.22694 | 0.912473 | 1.545487 | 0.202737 |
| 10 | 120918400 | 120919999 | 12.73805 | 22.30794 | 49.94883 | 22.05054 | -0.76975 | 0.661636 | 0.202739 |
| 13 | 31806800 | 31817599 | 81.23439 | 124.5793 | 244.8762 | 101.6521 | -0.68114 | 3.005584 | 0.202743 |
| 17 | 6129000 | 6130799 | 8.69365 | 15.94473 | 35.62965 | 15.18382 | -0.72492 | 0.252904 | 0.202745 |
| X | 31338400 | 31341799 | 92.59418 | 145.1551 | 120.6052 | 50.02264 | 0.584264 | 2.601542 | 0.202756 |
| 11 | 49473600 | 49475399 | 10.47626 | 18.80048 | 41.58962 | 18.19834 | -0.73653 | 0.40503 | 0.202765 |
| 13 | 12042800 | 12046399 | 22.12637 | 36.80818 | 74.68722 | 32.41851 | -0.63646 | 1.255471 | 0.202781 |
| 9 | 19237000 | 19238799 | 36.69142 | 60.71684 | 50.44471 | 21.81633 | 0.647202 | 1.335377 | 0.202801 |
| 18 | 77011800 | 77015799 | 32.43414 | 52.31386 | 101.1456 | 43.0881 | -0.59147 | 1.768627 | 0.202804 |
| 2 | 120458400 | 120459999 | 6.484567 | 12.28421 | 30.924 | 12.50874 | -0.83512 | 0.021373 | 0.20281 |
| 5 | 30359600 | 30360799 | 6.484567 | 12.28421 | 30.924 | 12.50874 | -0.83512 | 0.021373 | 0.20281 |
| 1 | 7788200 | 7790799 | 34.25868 | 58.34203 | 35.86576 | 15.72109 | 1.081145 | 1.038306 | 0.202819 |
| 5 | 95638800 | 95641199 | 52.39638 | 84.77383 | 61.96062 | 26.49506 | 0.813705 | 1.695922 | 0.202827 |
| 7 | 59400600 | 59401799 | 11.74655 | 22.70647 | 12.51761 | 4.796465 | 1.40879 | -0.15372 | 0.202827 |
| 16 | 33348200 | 33350799 | 27.31991 | 44.47405 | 89.43182 | 38.39407 | -0.635 | 1.550377 | 0.202832 |

| | | | | | | | | | |
|---|---|---|---|---|---|---|---|---|---|
| 12 | 72983600 | 72988799 | 36.7271 | 58.88983 | 114.272 | 48.26602 | -0.60712 | 1.963784 | 0.202837 |
| 5 | 18930600 | 18933399 | 22.61981 | 39.62117 | 30.60471 | 13.51299 | 0.75824 | 0.68152 | 0.202843 |
| 12 | 83131000 | 83133599 | 14.56715 | 25.08725 | 54.80731 | 24.04192 | -0.73802 | 0.88133 | 0.202845 |
| 9 | 121106000 | 121108999 | 16.39501 | 27.86813 | 59.55094 | 26.09911 | -0.70961 | 1.015003 | 0.202848 |
| 9 | 82369600 | 82370199 | 12.93449 | 25.16669 | 13.31022 | 4.907061 | 1.517319 | -0.26406 | 0.202848 |
| 12 | 18456400 | 18458399 | 57.9112 | 93.82619 | 53.14382 | 22.74809 | 1.182327 | 1.675365 | 0.202851 |
| 11 | 25461600 | 25463599 | 19.21747 | 34.70563 | 23.72572 | 10.31152 | 0.953847 | 0.391424 | 0.202873 |
| 1 | 6284400 | 6285799 | 12.56413 | 21.6909 | 55.2954 | 24.25784 | -0.95874 | 0.850519 | 0.202875 |
| 17 | 74175200 | 74178999 | 35.20059 | 56.62034 | 110.5535 | 46.91513 | -0.61056 | 1.859955 | 0.202878 |
| 9 | 62652400 | 62663199 | 81.46881 | 124.1743 | 254.9257 | 105.682 | -0.75075 | 3.131818 | 0.202886 |
| 7 | 117404800 | 117405999 | 10.00551 | 18.04497 | 40.08928 | 17.4528 | -0.73745 | 0.369899 | 0.202897 |
| 1 | 169374200 | 169377199 | 27.19144 | 46.38577 | 33.40954 | 14.64647 | 0.853844 | 0.990884 | 0.202904 |
| 16 | 76787000 | 76789199 | 35.42916 | 60.31143 | 34.07044 | 14.93367 | 1.202374 | 1.043067 | 0.202905 |
| 17 | 20573000 | 20574199 | 22.61155 | 40.1378 | 23.0943 | 10.19876 | 1.185426 | 0.513945 | 0.202919 |
| 14 | 11012600 | 11015799 | 44.30043 | 72.69078 | 51.75 | 22.28431 | 0.865938 | 1.455414 | 0.202921 |
| 2 | 27914600 | 27917199 | 10.82917 | 18.83587 | 50.47543 | 22.17692 | -1.02778 | 0.758379 | 0.202924 |
| 11 | 116664800 | 116676799 | 81.38982 | 125.1173 | 229.4971 | 95.29935 | -0.57879 | 2.957772 | 0.202926 |
| 16 | 89833200 | 89843999 | 89.59302 | 136.9364 | 249.5502 | 103.5198 | -0.5759 | 3.076478 | 0.202966 |
| 3 | 103127400 | 103127999 | 15.03459 | 27.84221 | 22.16274 | 9.321426 | 0.767538 | 0.216904 | 0.202971 |
| 9 | 93584400 | 93585999 | 15.22051 | 28.17729 | 22.15779 | 9.323486 | 0.784583 | 0.22038 | 0.202985 |
| 19 | 9853800 | 9856399 | 12.93619 | 22.16571 | 56.7951 | 24.90738 | -0.96815 | 0.92479 | 0.203005 |
| 19 | 5334600 | 5338599 | 15.816 | 26.77515 | 61.02213 | 26.71416 | -0.80235 | 1.052117 | 0.203011 |
| 1 | 25798200 | 25800399 | 21.66385 | 38.20375 | 27.80092 | 12.30318 | 0.845334 | 0.581928 | 0.203021 |
| 4 | 57258000 | 57258999 | 4.249935 | 8.226597 | 25.90895 | 9.864 | -1.07086 | -0.21445 | 0.203032 |
| 3 | 148503000 | 148506799 | 48.23529 | 76.60432 | 142.0607 | 59.22375 | -0.55714 | 2.281652 | 0.203038 |
| 10 | 122821000 | 122822599 | 8.283063 | 15.06866 | 36.54919 | 15.73997 | -0.85268 | 0.298389 | 0.203048 |
| 3 | 48109800 | 48112399 | 22.70847 | 39.90737 | 28.20172 | 12.4866 | 0.885585 | 0.625786 | 0.20306 |
| 3 | 88288200 | 88292399 | 17.5076 | 29.24884 | 67.38341 | 29.32781 | -0.81853 | 1.191581 | 0.203099 |
| 5 | 42155800 | 42157599 | 22.28256 | 39.20126 | 28.27679 | 12.51974 | 0.856679 | 0.611489 | 0.203114 |
| 13 | 65423200 | 65424599 | 31.63622 | 53.3158 | 42.1375 | 18.4216 | 0.719411 | 1.124624 | 0.203121 |
| 7 | 48072200 | 48075199 | 41.41259 | 68.39992 | 48.88223 | 21.113 | 0.86367 | 1.369247 | 0.203121 |
| 12 | 35806800 | 35807399 | 4.182022 | 8.164079 | 26.24693 | 9.277426 | -1.00165 | -0.48006 | 0.203134 |
| 7 | 15405200 | 15410599 | 133.5413 | 204.7273 | 163.299 | 67.785 | 0.618456 | 3.022464 | 0.203141 |
| 8 | 92571800 | 92575399 | 17.71515 | 29.91803 | 62.23241 | 27.22408 | -0.67146 | 1.080109 | 0.203149 |
| 17 | 34036000 | 34038999 | 14.77717 | 25.30652 | 56.34418 | 24.70856 | -0.76672 | 0.936406 | 0.203152 |
| 10 | 53613000 | 53615399 | 17.41053 | 29.33649 | 64.42694 | 28.16458 | -0.74973 | 1.099467 | 0.203158 |
| 15 | 102905800 | 102909999 | 22.41763 | 36.85855 | 79.11564 | 34.09748 | -0.72041 | 1.39848 | 0.203168 |
| 9 | 95737000 | 95738599 | 17.02851 | 30.56068 | 24.19538 | 10.58666 | 0.737183 | 0.413354 | 0.203179 |
| 8 | 8885200 | 8886599 | 7.939231 | 14.58595 | 34.59231 | 14.70755 | -0.80733 | 0.235962 | 0.203182 |
| 17 | 29433000 | 29436199 | 13.07021 | 22.43447 | 55.75545 | 24.45489 | -0.92389 | 0.908071 | 0.203198 |
| 2 | 150209600 | 150210399 | 13.85947 | 26.56077 | 17.54573 | 6.911309 | 1.112679 | -0.029 | 0.203202 |
| 14 | 111564600 | 111566799 | 13.85947 | 24.16166 | 56.59218 | 24.90001 | -0.83511 | 0.745447 | 0.203208 |
| 1 | 124755400 | 124756599 | 14.00979 | 26.60443 | 19.11806 | 7.693569 | 0.965093 | 0.043726 | 0.203228 |
| 1 | 133162200 | 133169399 | 36.81801 | 58.77921 | 134.4997 | 56.4514 | -0.85171 | 2.104702 | 0.203234 |
| 8 | 53748800 | 53749599 | 12.97922 | 24.40562 | 20.27775 | 8.321643 | 0.733417 | 0.101454 | 0.203241 |
| 11 | 74474600 | 74482999 | 62.74039 | 97.52221 | 197.801 | 82.12522 | -0.71127 | 2.743866 | 0.203245 |
| 5 | 41901200 | 41904599 | 20.01929 | 33.4345 | 68.2897 | 29.7164 | -0.64572 | 1.201591 | 0.20325 |

| | | | | | | | | | |
|---|---|---|---|---|---|---|---|---|---|
| 17 | 31181000 | 31185999 | 28.5504 | 46.16752 | 95.84789 | 40.96609 | -0.68954 | 1.684906 | 0.203283 |
| 4 | 112507600 | 112510399 | 33.61638 | 57.06465 | 38.14417 | 16.71247 | 0.960398 | 1.066429 | 0.203303 |
| X | 167911800 | 167913799 | 24.65024 | 44.51441 | 18.45112 | 8.019665 | 1.670375 | 0.420593 | 0.203319 |
| Y | 5576600 | 5579999 | 19.71181 | 35.04992 | 14.70096 | 6.48286 | 1.638633 | 0.463656 | 0.203319 |
| 16 | 89366000 | 89368799 | 17.41369 | 29.34568 | 63.2925 | 27.66976 | -0.72367 | 1.098113 | 0.203323 |
| 5 | 30244800 | 30246999 | 10.01581 | 17.93998 | 39.44671 | 17.28839 | -0.72928 | 0.409794 | 0.203332 |
| 5 | 110128200 | 110129799 | 27.17627 | 46.2981 | 38.45094 | 16.85476 | 0.64897 | 1.009648 | 0.203342 |
| 7 | 10065000 | 10066799 | 43.81047 | 72.16685 | 48.91842 | 21.09967 | 0.938668 | 1.407675 | 0.203346 |
| 17 | 23771600 | 23781999 | 57.28167 | 89.6205 | 183.9401 | 76.30715 | -0.72095 | 2.624976 | 0.203351 |
| 1 | 117656800 | 117659599 | 62.47769 | 100.2446 | 76.76552 | 32.45619 | 0.732691 | 1.960914 | 0.203366 |
| 15 | 36578200 | 36579199 | 5.404397 | 10.40583 | 27.45842 | 10.67138 | -0.84815 | -0.12813 | 0.203366 |
| 1 | 178877600 | 178878199 | 3.576494 | 6.972282 | 23.01563 | 8.305813 | -1.06299 | -0.4191 | 0.203385 |
| 8 | 21249400 | 21250599 | 9.54755 | 18.51438 | 15.52168 | 5.86005 | 0.82935 | -0.21856 | 0.203393 |
| 17 | 22606200 | 22612199 | 85.17852 | 134.5865 | 110.7307 | 45.99239 | 0.607149 | 2.434738 | 0.203399 |
| 7 | 12143400 | 12146199 | 29.39741 | 50.82147 | 32.52845 | 14.27068 | 1.026433 | 0.881645 | 0.203403 |
| 2 | 7878600 | 7879199 | 4.782637 | 9.315725 | 25.54739 | 9.322403 | -0.81723 | -0.36052 | 0.20341 |
| 18 | 16283400 | 16283999 | 4.486988 | 8.737289 | 24.14828 | 8.845798 | -0.83228 | -0.35279 | 0.203413 |
| 18 | 18962000 | 18964599 | 56.6424 | 90.79319 | 75.97627 | 32.07478 | 0.60333 | 1.982674 | 0.20344 |
| X | 64718600 | 64720199 | 20.6728 | 39.05389 | 13.41956 | 5.460043 | 2.005142 | 0.125905 | 0.203446 |
| 3 | 4690000 | 4691599 | 21.8066 | 38.33834 | 28.90143 | 12.798 | 0.793072 | 0.617099 | 0.203461 |
| 9 | 102742200 | 102743999 | 6.423479 | 11.94975 | 33.18739 | 13.86888 | -1.01461 | 0.147445 | 0.203466 |
| X | 88559200 | 88561999 | 39.62711 | 66.17135 | 42.08342 | 18.31085 | 1.031893 | 1.239637 | 0.203478 |
| 11 | 82019000 | 82020199 | 16.35212 | 29.6633 | 24.09697 | 10.41447 | 0.711635 | 0.345954 | 0.203492 |
| 18 | 13390800 | 13392199 | 8.173256 | 15.20967 | 36.03782 | 15.05207 | -0.78778 | 0.164878 | 0.203492 |
| 12 | 20155000 | 20159199 | 136.6348 | 208.0401 | 177.2166 | 73.44416 | 0.515936 | 3.161674 | 0.203493 |
| 17 | 75380600 | 75383199 | 13.10859 | 22.76757 | 52.69225 | 23.12981 | -0.81824 | 0.790569 | 0.203502 |
| 11 | 5470800 | 5473399 | 13.25322 | 23.06543 | 50.00145 | 21.97661 | -0.72385 | 0.759713 | 0.203502 |
| 10 | 88491000 | 88492999 | 16.97023 | 31.34519 | 20.66119 | 8.720677 | 1.033656 | 0.242045 | 0.203515 |
| 10 | 63431200 | 63431799 | 12.68207 | 24.47253 | 17.28414 | 6.674798 | 1.043866 | -0.10296 | 0.203518 |
| 7 | 13465000 | 13467199 | 34.45465 | 58.95716 | 32.40062 | 14.20769 | 1.24288 | 0.992407 | 0.203531 |
| 2 | 36723200 | 36725599 | 40.93303 | 67.79277 | 46.63314 | 20.1796 | 0.918755 | 1.334124 | 0.203555 |
| 14 | 65272600 | 65275399 | 11.90352 | 20.32006 | 58.79093 | 25.77488 | -1.14323 | 0.958943 | 0.203559 |
| 7 | 42037000 | 42040599 | 74.54356 | 118.8235 | 88.5482 | 37.08477 | 0.761512 | 2.177148 | 0.203561 |
| 5 | 29798600 | 29799599 | 21.54649 | 37.99903 | 27.87158 | 12.3343 | 0.834076 | 0.581166 | 0.203565 |
| 15 | 84177000 | 84179999 | 17.69397 | 29.48296 | 68.96776 | 29.96787 | -0.84049 | 1.221815 | 0.203584 |
| 14 | 124072200 | 124074199 | 19.44246 | 34.90609 | 23.45761 | 10.26059 | 0.972021 | 0.424463 | 0.203594 |
| 15 | 82113600 | 82121399 | 35.7064 | 56.98771 | 128.7008 | 53.99492 | -0.83322 | 2.111261 | 0.203601 |
| 5 | 134483200 | 134486199 | 15.11396 | 25.44575 | 65.83682 | 28.77369 | -0.98519 | 1.100783 | 0.203604 |
| 1 | 191872000 | 191874399 | 14.24921 | 24.59492 | 53.1917 | 23.33481 | -0.72268 | 0.858631 | 0.203611 |
| 17 | 39631200 | 39639799 | 202.7184 | 302.2928 | 222.3526 | 91.63786 | 0.701547 | 3.583476 | 0.203615 |
| 18 | 20716000 | 20717799 | 31.46052 | 53.8319 | 35.86464 | 15.72662 | 0.966448 | 0.984484 | 0.203616 |
| 9 | 62314800 | 62321399 | 73.35039 | 113.978 | 199.2897 | 82.74727 | -0.49779 | 2.755042 | 0.203641 |
| 10 | 38722600 | 38723199 | 7.26911 | 13.59447 | 32.23733 | 13.35198 | -0.77818 | 0.118996 | 0.203655 |
| 6 | 142696000 | 142699799 | 24.6614 | 40.04085 | 94.68866 | 40.5949 | -0.87123 | 1.594393 | 0.203659 |
| 8 | 126051400 | 126062999 | 90.92772 | 138.8289 | 247.8296 | 102.7809 | -0.54735 | 3.098715 | 0.203673 |
| 9 | 69052000 | 69057199 | 48.34685 | 78.17721 | 65.99376 | 28.20518 | 0.60558 | 1.704216 | 0.203677 |
| 10 | 81557800 | 81558399 | 17.46458 | 32.54395 | 19.81501 | 8.257285 | 1.161332 | 0.196594 | 0.203705 |

| | | | | | | | | | |
|---|---|---|---|---|---|---|---|---|---|
| 13 | 58213200 | 58216999 | 23.59262 | 38.60829 | 81.65582 | 35.1228 | -0.70203 | 1.463172 | 0.203735 |
| 17 | 19330000 | 19332399 | 31.54133 | 53.86104 | 37.02403 | 16.23236 | 0.921189 | 1.004608 | 0.203739 |
| X | 3734200 | 3740199 | 60.28819 | 97.65181 | 51.9955 | 22.25227 | 1.270963 | 1.686385 | 0.203743 |
| 19 | 26050600 | 26052599 | 20.85685 | 34.66648 | 71.60665 | 31.06293 | -0.66209 | 1.259324 | 0.203747 |
| 5 | 120546600 | 120558799 | 112.1311 | 168.729 | 302.6139 | 124.9679 | -0.57172 | 3.401037 | 0.203747 |
| 11 | 70008000 | 70012199 | 61.80966 | 99.29247 | 73.75091 | 31.23868 | 0.778684 | 1.923911 | 0.203775 |
| X | 33915200 | 33917199 | 32.27374 | 54.11827 | 44.31832 | 19.33425 | 0.668267 | 1.174511 | 0.203775 |
| 7 | 129444000 | 129445399 | 7.234637 | 13.61906 | 33.01432 | 13.52271 | -0.7973 | 0.075607 | 0.203778 |
| 12 | 104717400 | 104719399 | 10.88589 | 19.14618 | 44.05549 | 19.50691 | -0.80793 | 0.577375 | 0.20381 |
| 6 | 58567800 | 58568399 | 20.21738 | 35.46918 | 28.76868 | 12.71934 | 0.687986 | 0.639265 | 0.20381 |
| 14 | 18626600 | 18630399 | 24.84879 | 42.82339 | 35.75304 | 15.68385 | 0.644005 | 0.893863 | 0.203813 |
| 7 | 20953000 | 20961599 | 79.91911 | 127.2247 | 88.81725 | 37.12979 | 0.853542 | 2.224507 | 0.203814 |
| 5 | 137230600 | 137233999 | 19.35992 | 32.40993 | 67.19928 | 29.28562 | -0.66733 | 1.172678 | 0.203817 |
| 19 | 56224000 | 56228999 | 31.59004 | 51.07101 | 99.597 | 42.55585 | -0.59999 | 1.698086 | 0.203832 |
| 4 | 3447800 | 3450799 | 29.99347 | 51.3882 | 35.86496 | 15.7284 | 0.899859 | 0.9659 | 0.203846 |
| 8 | 78191800 | 78201599 | 70.76664 | 109.112 | 237.8517 | 98.79949 | -0.82428 | 2.897051 | 0.203846 |
| 9 | 70437800 | 70439799 | 12.89977 | 22.5811 | 48.34072 | 21.33259 | -0.70521 | 0.668115 | 0.203851 |
| 14 | 66682000 | 66685599 | 23.35901 | 38.44284 | 77.54207 | 33.43273 | -0.63043 | 1.388752 | 0.203858 |
| 3 | 120797800 | 120799199 | 15.50415 | 29.10485 | 19.96804 | 8.217707 | 1.004381 | 0.134064 | 0.203861 |
| 10 | 8626400 | 8627199 | 3.939952 | 7.67711 | 23.0177 | 8.359474 | -0.93637 | -0.39507 | 0.203868 |
| 6 | 107645800 | 107647999 | 18.74325 | 33.95452 | 23.12874 | 10.01309 | 0.962316 | 0.368424 | 0.203879 |
| 13 | 104389200 | 104393599 | 49.18338 | 79.74305 | 61.75708 | 26.44902 | 0.733202 | 1.649455 | 0.203884 |
| 3 | 13211000 | 13211999 | 15.18738 | 28.25617 | 21.60234 | 9.026059 | 0.832218 | 0.190759 | 0.203888 |
| 7 | 46031600 | 46032999 | 6.465391 | 12.30059 | 30.46941 | 12.21342 | -0.79976 | -0.00701 | 0.203905 |
| 2 | 27523000 | 27527999 | 23.48922 | 38.317 | 86.55357 | 37.18634 | -0.80005 | 1.511213 | 0.203914 |
| 4 | 8423800 | 8429399 | 29.93955 | 48.54375 | 97.02603 | 41.5548 | -0.63131 | 1.631513 | 0.203914 |
| 19 | 39799600 | 39802199 | 42.59196 | 69.35615 | 59.1989 | 25.41751 | 0.598416 | 1.55974 | 0.203925 |
| 10 | 79795600 | 79796599 | 4.004603 | 7.771336 | 26.07327 | 9.809961 | -1.14531 | -0.25985 | 0.203926 |
| 2 | 172155800 | 172164799 | 90.65724 | 138.7211 | 246.388 | 102.235 | -0.53756 | 3.053632 | 0.203929 |
| 1 | 100801800 | 100803199 | 13.14022 | 25.11631 | 17.46159 | 6.930231 | 1.028911 | -0.01408 | 0.203938 |
| 4 | 21041400 | 21042799 | 16.82036 | 31.22042 | 20.83438 | 8.735554 | 1.023114 | 0.215841 | 0.203938 |
| 17 | 29271400 | 29273599 | 9.942267 | 17.58737 | 41.46997 | 18.341 | -0.83932 | 0.504035 | 0.203952 |
| 12 | 63445600 | 63447399 | 15.21555 | 29.12128 | 16.97519 | 6.71187 | 1.28702 | -0.0043 | 0.203959 |
| 12 | 3089600 | 3091799 | 57.59157 | 92.31593 | 75.26318 | 31.77442 | 0.640906 | 1.984745 | 0.203969 |
| 17 | 19490400 | 19492999 | 31.46626 | 54.48598 | 29.65713 | 13.01163 | 1.259455 | 0.874012 | 0.203974 |
| X | 25599800 | 25604599 | 54.06563 | 87.81286 | 54.13596 | 23.21512 | 1.06372 | 1.621554 | 0.203976 |
| 5 | 3230600 | 3234199 | 48.24643 | 78.07598 | 58.71362 | 25.11051 | 0.772989 | 1.682648 | 0.203983 |
| 4 | 103242000 | 103242799 | 5.546922 | 10.71294 | 27.66537 | 10.64449 | -0.80428 | -0.16152 | 0.203989 |
| 3 | 145403200 | 145405799 | 13.85677 | 24.16426 | 50.54014 | 22.2414 | -0.67224 | 0.734154 | 0.204002 |
| 2 | 171524200 | 171527199 | 22.8004 | 37.75864 | 75.70712 | 32.75897 | -0.61986 | 1.310063 | 0.204009 |
| 2 | 6071600 | 6081599 | 84.32745 | 129.2656 | 236.8804 | 98.327 | -0.58056 | 3.015204 | 0.204017 |
| Y | 21307400 | 21307799 | 12.30623 | 23.70503 | 8.043346 | 3.1221 | 2.078452 | -0.12408 | 0.204018 |
| 13 | 99147000 | 99148799 | 7.675209 | 14.01073 | 35.55564 | 15.24667 | -0.91303 | 0.272622 | 0.20402 |
| 13 | 44630400 | 44633799 | 29.66814 | 47.96357 | 97.29732 | 41.57284 | -0.65666 | 1.693999 | 0.204026 |
| 3 | 16090000 | 16091799 | 28.94348 | 49.4343 | 35.18273 | 15.42532 | 0.87126 | 0.989959 | 0.204052 |
| 3 | 51456400 | 51458399 | 7.62854 | 14.05656 | 34.10782 | 14.44346 | -0.83565 | 0.215991 | 0.204055 |
| 19 | 3983800 | 3985599 | 7.62854 | 14.05656 | 34.10782 | 14.44346 | -0.83565 | 0.215991 | 0.204055 |

| | | | | | | | | | |
|---|---|---|---|---|---|---|---|---|---|
| 4 | 125248000 | 125251199 | 21.26667 | 35.09007 | 76.35402 | 32.95656 | -0.7379 | 1.349615 | 0.204075 |
| 15 | 59578000 | 59579999 | 10.57562 | 18.61827 | 45.09319 | 19.96256 | -0.88101 | 0.565542 | 0.204087 |
| 7 | 25836200 | 25838199 | 25.28415 | 44.32038 | 28.15071 | 12.43713 | 1.038959 | 0.680827 | 0.204098 |
| 13 | 66133200 | 66141599 | 144.6607 | 220.0769 | 165.0126 | 68.36937 | 0.698874 | 3.179098 | 0.204099 |
| 8 | 16578000 | 16581199 | 33.66367 | 56.55561 | 43.12902 | 18.83721 | 0.770575 | 1.157312 | 0.204102 |
| 18 | 70415200 | 70415999 | 15.00062 | 27.9168 | 21.60811 | 9.024038 | 0.815045 | 0.187227 | 0.204106 |
| 10 | 75686000 | 75689799 | 24.54123 | 40.17501 | 81.25722 | 34.95609 | -0.63747 | 1.460356 | 0.204109 |
| 4 | 146036600 | 146041799 | 90.25568 | 142.4044 | 110.9099 | 46.0334 | 0.684528 | 2.471221 | 0.204115 |
| 7 | 30835000 | 30843399 | 42.30329 | 67.07756 | 149.8405 | 62.36848 | -0.8274 | 2.324788 | 0.204127 |
| 1 | 59965200 | 59966599 | 5.151886 | 9.780366 | 29.76468 | 11.98581 | -1.09855 | -0.00492 | 0.204128 |
| 9 | 43016400 | 43017999 | 9.962033 | 17.56953 | 42.52369 | 18.81831 | -0.87849 | 0.539824 | 0.204141 |
| 12 | 104755200 | 104772999 | 136.2992 | 202.0681 | 366.1526 | 150.8069 | -0.60608 | 3.681693 | 0.204145 |
| 14 | 65595400 | 65596799 | 6.65267 | 12.53608 | 33.01848 | 13.50146 | -0.91398 | 0.063447 | 0.204156 |
| 4 | 10926000 | 10927599 | 10.34212 | 18.4451 | 40.43953 | 17.78778 | -0.72884 | 0.440856 | 0.204161 |
| 6 | 72517200 | 72519199 | 10.34212 | 18.4451 | 40.43953 | 17.78778 | -0.72884 | 0.440856 | 0.204161 |
| 2 | 158603400 | 158606199 | 14.34668 | 24.48927 | 58.4353 | 25.61885 | -0.867 | 0.96674 | 0.204188 |
| 7 | 126852400 | 126853799 | 6.15153 | 11.39369 | 33.64428 | 14.14738 | -1.10868 | 0.172926 | 0.204189 |
| 18 | 11484000 | 11485799 | 8.727194 | 15.89915 | 36.04185 | 15.49398 | -0.75404 | 0.293434 | 0.204194 |
| 1 | 134160000 | 134165799 | 29.08587 | 46.62831 | 121.9829 | 51.50742 | -1.03747 | 1.962045 | 0.204195 |
| 1 | 34537000 | 34542799 | 35.61713 | 57.06026 | 116.3949 | 49.09754 | -0.68054 | 1.990699 | 0.204205 |
| 15 | 18635600 | 18636599 | 13.24323 | 24.73642 | 19.76169 | 8.202034 | 0.776639 | 0.143895 | 0.204206 |
| 19 | 33868800 | 33869799 | 14.34196 | 27.03238 | 20.12034 | 8.223811 | 0.895568 | 0.09905 | 0.204206 |
| X | 89224000 | 89225599 | 25.39104 | 44.80534 | 24.20895 | 10.71219 | 1.272644 | 0.591759 | 0.20422 |
| 14 | 26254400 | 26256799 | 15.01898 | 25.92747 | 53.76508 | 23.58645 | -0.66238 | 0.860659 | 0.204223 |
| 13 | 60242800 | 60245399 | 14.12019 | 24.27537 | 56.94363 | 24.97772 | -0.84079 | 0.896995 | 0.204228 |
| 12 | 21844000 | 21847199 | 63.44013 | 101.3988 | 86.91339 | 36.53896 | 0.56415 | 2.081598 | 0.204236 |
| 14 | 42292600 | 42292999 | 8.893741 | 17.3705 | 13.26855 | 4.651632 | 1.057019 | -0.48856 | 0.204249 |
| 8 | 80081400 | 80084399 | 10.59638 | 18.61039 | 45.64047 | 20.19728 | -0.8998 | 0.597838 | 0.204259 |
| 6 | 21770400 | 21771199 | 4.688216 | 9.113362 | 27.72837 | 10.30005 | -0.99034 | -0.29503 | 0.204261 |
| 19 | 39329200 | 39333999 | 98.28539 | 154.0384 | 123.8384 | 51.36498 | 0.631452 | 2.608513 | 0.204264 |
| 2 | 173172200 | 173174399 | 8.794563 | 15.47717 | 44.61093 | 19.75049 | -1.13014 | 0.560123 | 0.204271 |
| 16 | 32582600 | 32591999 | 66.1144 | 102.8542 | 192.6404 | 79.97242 | -0.59552 | 2.730481 | 0.204294 |
| 12 | 22085000 | 22099999 | 364.0344 | 514.0368 | 410.9005 | 170.116 | 0.505348 | 4.436329 | 0.204306 |
| 9 | 109945600 | 109946999 | 5.214595 | 9.809487 | 31.20838 | 12.79819 | -1.1854 | 0.061151 | 0.204306 |
| 1 | 85275000 | 85280199 | 85.31331 | 134.2624 | 117.0732 | 48.57193 | 0.518978 | 2.521074 | 0.204328 |
| 7 | 5053400 | 5065599 | 77.50744 | 118.7413 | 241.319 | 100.1593 | -0.73034 | 3.02507 | 0.20433 |
| 3 | 117419400 | 117420199 | 5.686506 | 10.98116 | 27.65857 | 10.64806 | -0.76932 | -0.15741 | 0.204342 |
| 2 | 77565200 | 77565799 | 9.41629 | 18.33046 | 15.42689 | 5.664545 | 0.859878 | -0.31592 | 0.204352 |
| 13 | 15510400 | 15510799 | 9.41629 | 18.33046 | 15.42689 | 5.664545 | 0.859878 | -0.31592 | 0.204352 |
| 10 | 39508000 | 39510999 | 13.88206 | 24.07414 | 54.34474 | 23.84405 | -0.78298 | 0.813344 | 0.204382 |
| X | 122589000 | 122594199 | 61.32081 | 98.81822 | 64.44233 | 27.42085 | 0.971505 | 1.823184 | 0.204391 |
| 17 | 43415800 | 43416599 | 6.61109 | 12.59687 | 31.69412 | 12.66366 | -0.81823 | -0.01329 | 0.204395 |
| 17 | 37938200 | 37942799 | 68.98332 | 110.1275 | 87.93298 | 36.9053 | 0.664438 | 2.124804 | 0.204406 |
| 4 | 125703800 | 125706799 | 20.89533 | 34.65097 | 71.94408 | 31.15999 | -0.66969 | 1.288052 | 0.204424 |
| 14 | 51551400 | 51552399 | 8.893975 | 16.16054 | 36.52903 | 15.75502 | -0.75298 | 0.311514 | 0.204425 |
| 16 | 64671000 | 64673199 | 17.32506 | 29.02331 | 67.00592 | 29.21399 | -0.82172 | 1.158831 | 0.204435 |
| 9 | 76986800 | 76991799 | 44.9227 | 71.56402 | 132.9662 | 55.64073 | -0.55524 | 2.180708 | 0.20445 |

| | | | | | | | | | |
|---|---|---|---|---|---|---|---|---|---|
| 16 | 60425000 | 60426999 | 16.56194 | 27.72193 | 67.42848 | 29.38285 | -0.8966 | 1.16632 | 0.204464 |
| 7 | 62389800 | 62393199 | 21.33073 | 37.69123 | 24.68176 | 10.91846 | 0.997484 | 0.552014 | 0.204467 |
| X | 115631000 | 115634799 | 53.54645 | 86.57863 | 62.98783 | 26.91792 | 0.819482 | 1.715066 | 0.204471 |
| 15 | 100704200 | 100712399 | 51.12139 | 80.74861 | 158.3429 | 65.75763 | -0.64446 | 2.428163 | 0.204472 |
| 4 | 60546600 | 60549799 | 28.73642 | 48.56585 | 38.04035 | 16.64467 | 0.732305 | 1.083654 | 0.204496 |
| 15 | 102871200 | 102877599 | 31.04582 | 49.93972 | 106.6096 | 45.24457 | -0.73877 | 1.852661 | 0.204505 |
| 5 | 139216400 | 139217199 | 6.301169 | 12.05039 | 29.54549 | 11.70858 | -0.77084 | -0.05107 | 0.204509 |
| 7 | 134446200 | 134447399 | 6.301169 | 12.05039 | 29.54549 | 11.70858 | -0.77084 | -0.05107 | 0.204509 |
| 5 | 93770000 | 93775799 | 99.71305 | 156.4876 | 118.9364 | 49.32705 | 0.713875 | 2.584548 | 0.204523 |
| 12 | 28707000 | 28710399 | 16.19346 | 27.67186 | 57.46428 | 25.19535 | -0.66727 | 0.963045 | 0.204527 |
| 18 | 36954800 | 36959199 | 107.9449 | 167.0394 | 146.0246 | 60.65496 | 0.496597 | 2.835196 | 0.20453 |
| 12 | 34012000 | 34014599 | 19.83552 | 33.19341 | 67.80467 | 29.54232 | -0.64592 | 1.178678 | 0.204543 |
| 17 | 39168800 | 39172799 | 61.37003 | 99.04455 | 58.92229 | 25.12031 | 1.106023 | 1.777725 | 0.204543 |
| 8 | 80666200 | 80668399 | 29.89482 | 51.06058 | 38.2 | 16.74824 | 0.79961 | 0.997646 | 0.204545 |
| 12 | 16860000 | 16861799 | 9.818952 | 17.3021 | 43.04429 | 19.05202 | -0.91846 | 0.549349 | 0.204563 |
| 3 | 3369600 | 3374999 | 97.25561 | 151.5767 | 133.042 | 55.21582 | 0.498722 | 2.705574 | 0.204565 |
| 9 | 96405000 | 96405999 | 11.31627 | 21.74414 | 18.4776 | 7.223493 | 0.763033 | -0.07586 | 0.204568 |
| 1 | 34823400 | 34827799 | 21.20897 | 35.14758 | 73.19006 | 31.68502 | -0.67409 | 1.296152 | 0.204581 |
| 3 | 130971000 | 130974999 | 29.35694 | 47.12169 | 115.353 | 48.80053 | -0.93967 | 1.921179 | 0.204609 |
| 5 | 149812000 | 149813399 | 24.36241 | 42.19349 | 35.13065 | 15.41315 | 0.649319 | 0.847209 | 0.204611 |
| 10 | 77987000 | 77991199 | 24.07568 | 39.37453 | 82.89019 | 35.64574 | -0.69603 | 1.473351 | 0.204632 |
| 4 | 110569800 | 110572599 | 28.0299 | 48.6424 | 32.645 | 14.32347 | 0.959499 | 0.843544 | 0.204642 |
| 5 | 25108400 | 25110999 | 9.798598 | 17.31763 | 41.99248 | 18.57543 | -0.87999 | 0.513828 | 0.204645 |
| 9 | 51421200 | 51421999 | 5.704067 | 10.99118 | 30.57898 | 11.85824 | -0.92238 | -0.12794 | 0.204651 |
| 7 | 65432000 | 65435399 | 15.86642 | 26.7413 | 65.40569 | 28.59459 | -0.90441 | 1.092881 | 0.20466 |
| 16 | 91139600 | 91145999 | 45.93625 | 73.02807 | 139.0568 | 58.03106 | -0.59375 | 2.248651 | 0.20467 |
| 14 | 116148400 | 116150199 | 23.2656 | 41.20312 | 24.23806 | 10.71734 | 1.15242 | 0.540043 | 0.204674 |
| 8 | 4374000 | 4375799 | 10.32254 | 18.07645 | 46.74915 | 20.63681 | -0.97588 | 0.64718 | 0.204675 |
| 2 | 22095400 | 22097999 | 26.91784 | 46.19952 | 36.02635 | 15.8014 | 0.740781 | 0.93676 | 0.204687 |
| 3 | 3191600 | 3192999 | 28.33323 | 48.47133 | 37.61461 | 16.49372 | 0.74719 | 0.97413 | 0.204691 |
| 12 | 11693000 | 11694599 | 20.0753 | 35.72544 | 25.62508 | 11.29241 | 0.871678 | 0.487517 | 0.204692 |
| 1 | 95763800 | 95764799 | 11.81951 | 22.65961 | 18.2984 | 7.200231 | 0.826714 | -0.05175 | 0.204703 |
| 7 | 15194400 | 15197399 | 49.3202 | 80.11786 | 54.48798 | 23.36876 | 0.92263 | 1.608339 | 0.204707 |
| 7 | 26623200 | 26625199 | 30.59192 | 51.37538 | 43.22228 | 18.87401 | 0.629329 | 1.149775 | 0.204713 |
| 5 | 111659800 | 111668399 | 60.73593 | 94.90232 | 183.2781 | 76.04262 | -0.63471 | 2.650046 | 0.204723 |
| 16 | 21023000 | 21025599 | 18.76431 | 30.96798 | 76.48504 | 33.01708 | -0.91981 | 1.341418 | 0.204727 |
| 7 | 10716200 | 10720999 | 90.47628 | 142.4608 | 116.4657 | 48.32311 | 0.612441 | 2.514556 | 0.204747 |
| 8 | 96044400 | 96049599 | 27.61483 | 44.60253 | 103.4801 | 44.16156 | -0.8514 | 1.721817 | 0.204758 |
| 9 | 37839800 | 37841799 | 15.40712 | 28.51345 | 22.15279 | 9.325526 | 0.801482 | 0.223852 | 0.204761 |
| 7 | 73417200 | 73419999 | 27.86221 | 47.54097 | 39.38243 | 17.26525 | 0.652955 | 0.999468 | 0.204764 |
| 13 | 47434200 | 47440199 | 42.77684 | 68.25749 | 128.71 | 53.98231 | -0.57393 | 2.124388 | 0.20477 |
| 9 | 20060600 | 20066399 | 82.02229 | 130.1015 | 100.9661 | 42.03965 | 0.69596 | 2.33516 | 0.204776 |
| 10 | 80662200 | 80665399 | 15.70285 | 26.47674 | 63.14511 | 27.6098 | -0.86796 | 1.088021 | 0.20478 |
| 18 | 75499400 | 75504799 | 29.78835 | 47.89161 | 109.7186 | 46.5283 | -0.84138 | 1.868424 | 0.204796 |
| 10 | 82534400 | 82536399 | 12.52527 | 21.7071 | 54.27057 | 23.81104 | -0.92964 | 0.81346 | 0.204797 |
| 11 | 115330400 | 115336599 | 41.69633 | 66.40941 | 133.4613 | 55.83064 | -0.66856 | 2.185122 | 0.204799 |
| 8 | 26669400 | 26672799 | 34.28963 | 57.03902 | 47.10773 | 20.44663 | 0.656309 | 1.268234 | 0.204802 |

| | | | | | | | | | |
|---|---|---|---|---|---|---|---|---|---|
| 6 | 113666000 | 113668399 | 10.83737 | 19.21159 | 41.41357 | 18.30827 | -0.70973 | 0.485202 | 0.204817 |
| 8 | 26647800 | 26649199 | 6.924435 | 12.64056 | 37.66139 | 16.14912 | -1.14231 | 0.272725 | 0.204835 |
| 11 | 94317800 | 94321799 | 14.64809 | 24.63367 | 64.59118 | 28.21966 | -1.00418 | 1.110044 | 0.204842 |
| 5 | 140605800 | 140628599 | 131.2914 | 191.9573 | 467.1041 | 192.4317 | -1.05015 | 3.909205 | 0.204844 |
| 12 | 22024400 | 22039799 | 356.4787 | 505.2861 | 396.2441 | 163.9452 | 0.538484 | 4.380718 | 0.204846 |
| 14 | 61564200 | 61568999 | 24.04706 | 39.04477 | 91.72971 | 39.32736 | -0.86172 | 1.589728 | 0.204852 |
| 7 | 17116800 | 17121799 | 103.2708 | 160.7715 | 136.8492 | 56.80537 | 0.541691 | 2.728313 | 0.204866 |
| 2 | 103833200 | 103834199 | 5.564877 | 10.69302 | 27.97401 | 10.93553 | -0.84428 | -0.107 | 0.20488 |
| 2 | 84773400 | 84775399 | 12.61728 | 22.10461 | 46.7341 | 20.63782 | -0.687 | 0.653359 | 0.204886 |
| 3 | 125947800 | 125948799 | 7.062648 | 13.38987 | 32.14508 | 12.98098 | -0.76546 | 0.024005 | 0.204894 |
| 19 | 32187000 | 32188399 | 7.062648 | 13.38987 | 32.14508 | 12.98098 | -0.76546 | 0.024005 | 0.204894 |
| 15 | 6565400 | 6566199 | 7.05632 | 12.95469 | 34.5855 | 14.71883 | -0.97746 | 0.233502 | 0.204896 |
| X | 121306400 | 121307399 | 18.88484 | 34.13413 | 23.01499 | 9.991916 | 0.974938 | 0.384208 | 0.204911 |
| 7 | 13914800 | 13917799 | 81.26294 | 128.2415 | 112.7766 | 46.81296 | 0.509795 | 2.461526 | 0.204912 |
| 17 | 20465600 | 20468999 | 61.30186 | 98.54051 | 71.79925 | 30.44056 | 0.807448 | 1.900764 | 0.204923 |
| 16 | 89816800 | 89832199 | 171.0953 | 249.5651 | 442.7011 | 182.4178 | -0.59767 | 3.947883 | 0.204939 |
| 17 | 28306600 | 28308599 | 8.792092 | 15.81289 | 37.95921 | 16.57897 | -0.85119 | 0.373156 | 0.204942 |
| 2 | 91885600 | 91886799 | 5.435422 | 10.42648 | 28.46776 | 11.18314 | -0.91212 | -0.09137 | 0.204947 |
| 11 | 60242200 | 60243799 | 6.327176 | 12.01886 | 29.86786 | 12.01187 | -0.80885 | -0.00069 | 0.204953 |
| 16 | 92751000 | 92752199 | 6.327176 | 12.01886 | 29.86786 | 12.01187 | -0.80885 | -0.00069 | 0.204953 |
| 8 | 48364600 | 48368799 | 28.46377 | 46.32074 | 90.03136 | 38.64677 | -0.58641 | 1.559248 | 0.204973 |
| 7 | 138284000 | 138286599 | 14.04016 | 24.37563 | 51.63314 | 22.65909 | -0.69114 | 0.800308 | 0.205004 |
| 8 | 25857400 | 25863399 | 32.62764 | 52.51568 | 106.3666 | 45.18068 | -0.66207 | 1.834829 | 0.205014 |
| 18 | 80352600 | 80355599 | 29.55635 | 47.67133 | 104.2539 | 44.41117 | -0.76881 | 1.764867 | 0.205016 |
| 14 | 55113200 | 55115199 | 10.85695 | 19.18939 | 41.9585 | 18.56015 | -0.73179 | 0.5195 | 0.205017 |
| 4 | 88671200 | 88683599 | 87.19341 | 133.2862 | 241.6801 | 100.2581 | -0.56849 | 3.073628 | 0.205024 |
| 5 | 75313400 | 75317799 | 24.1236 | 39.0274 | 97.92983 | 41.87355 | -0.96142 | 1.667391 | 0.205032 |
| 1 | 166419600 | 166422999 | 79.12225 | 125.5689 | 100.8351 | 42.00765 | 0.647485 | 2.316549 | 0.205042 |
| 8 | 45701600 | 45703599 | 10.50024 | 18.81626 | 44.81171 | 19.63216 | -0.84365 | 0.420696 | 0.205053 |
| 2 | 74521400 | 74525399 | 16.93711 | 28.19926 | 70.56702 | 30.6473 | -0.93745 | 1.228648 | 0.205069 |
| 15 | 58175800 | 58179599 | 27.35395 | 44.46722 | 87.77668 | 37.66429 | -0.60969 | 1.571798 | 0.205073 |
| 18 | 86192200 | 86192799 | 9.680618 | 18.89862 | 11.76609 | 4.157918 | 1.335952 | -0.45709 | 0.205077 |
| 13 | 58275000 | 58277399 | 12.98469 | 22.504 | 51.5831 | 22.63194 | -0.80472 | 0.814037 | 0.205081 |
| X | 109982800 | 109983999 | 14.05052 | 26.92119 | 17.53905 | 6.912891 | 1.131757 | -0.02505 | 0.205104 |
| 8 | 52154000 | 52155399 | 17.09727 | 31.81899 | 19.00969 | 7.938778 | 1.185172 | 0.197646 | 0.205108 |
| 1 | 61566200 | 61568599 | 16.34352 | 27.92342 | 58.02726 | 25.44184 | -0.66838 | 0.965228 | 0.205118 |
| 18 | 8011200 | 8012799 | 19.7114 | 34.88174 | 27.4703 | 12.14841 | 0.733847 | 0.532525 | 0.205127 |
| 11 | 3991200 | 3994999 | 19.0418 | 31.50586 | 75.04796 | 32.45358 | -0.86751 | 1.310525 | 0.205183 |
| 17 | 65352600 | 65353999 | 12.59367 | 24.02543 | 18.69703 | 7.451332 | 0.862887 | -0.00264 | 0.205186 |
| 13 | 87551200 | 87552199 | 14.55024 | 27.75359 | 17.32653 | 6.907497 | 1.177438 | 0.014263 | 0.205203 |
| 18 | 37362600 | 37366199 | 47.19202 | 77.03802 | 53.50705 | 22.99449 | 0.897389 | 1.529225 | 0.20521 |
| 11 | 27558800 | 27559199 | 10.96856 | 21.051 | 17.99761 | 7.059879 | 0.749804 | -0.07015 | 0.205216 |
| X | 8390200 | 8391199 | 27.97687 | 48.54817 | 32.74772 | 14.36852 | 0.952286 | 0.84479 | 0.205217 |
| 9 | 17561400 | 17563199 | 7.230648 | 13.62325 | 36.58092 | 14.96181 | -0.9425 | 0.076907 | 0.20523 |
| 1 | 100470800 | 100471999 | 13.81841 | 26.01594 | 19.51023 | 7.989818 | 0.882102 | 0.098431 | 0.205244 |
| 1 | 108508800 | 108509999 | 13.81841 | 26.01594 | 19.51023 | 7.989818 | 0.882102 | 0.098431 | 0.205244 |
| 6 | 76024800 | 76025999 | 16.48418 | 30.09193 | 23.49192 | 10.07325 | 0.77579 | 0.309852 | 0.205268 |

| | | | | | | | | | |
|---|---|---|---|---|---|---|---|---|---|
| 7 | 38090200 | 38092599 | 31.87114 | 53.94916 | 40.32332 | 17.65208 | 0.799647 | 1.083525 | 0.205289 |
| 6 | 144284800 | 144287199 | 27.48681 | 44.72959 | 88.77308 | 38.10625 | -0.61647 | 1.555884 | 0.205309 |
| 10 | 59520600 | 59524199 | 20.53292 | 34.17497 | 69.87916 | 30.3423 | -0.64728 | 1.241907 | 0.205323 |
| 8 | 84058600 | 84059399 | 4.971591 | 9.483079 | 29.30959 | 11.68444 | -1.10725 | -0.04036 | 0.205344 |
| 9 | 74547400 | 74549999 | 35.89134 | 60.30136 | 41.45566 | 18.10664 | 0.919669 | 1.163496 | 0.205354 |
| 11 | 110823200 | 110823799 | 4.818156 | 9.360663 | 25.73839 | 9.604538 | -0.85121 | -0.28102 | 0.205356 |
| 19 | 24177800 | 24182399 | 25.15565 | 41.21943 | 82.78795 | 35.62871 | -0.62694 | 1.449594 | 0.205373 |
| 5 | 120405600 | 120412599 | 47.69693 | 75.87327 | 139.2136 | 58.13812 | -0.53946 | 2.231138 | 0.205377 |
| 2 | 165247200 | 165249799 | 27.74676 | 44.88578 | 93.81713 | 40.11289 | -0.69859 | 1.674378 | 0.205383 |
| Y | 84296800 | 84298399 | 11.65882 | 22.50482 | 9.184484 | 3.542774 | 1.825191 | -0.14726 | 0.205387 |
| 15 | 27556200 | 27558999 | 13.03506 | 22.4591 | 54.20109 | 23.77633 | -0.88057 | 0.870843 | 0.205395 |
| 17 | 39493600 | 39506399 | 198.1856 | 296.8393 | 218.702 | 90.2106 | 0.70492 | 3.501623 | 0.205398 |
| 1 | 4857800 | 4860399 | 41.71741 | 68.18545 | 57.11263 | 24.55506 | 0.629252 | 1.503447 | 0.205398 |
| 11 | 26508600 | 26512999 | 57.80935 | 93.12123 | 68.32002 | 29.05353 | 0.801178 | 1.829781 | 0.205425 |
| 2 | 36291600 | 36293799 | 25.10535 | 43.62933 | 30.85454 | 13.54885 | 0.884437 | 0.807776 | 0.205427 |
| 14 | 50970400 | 50971999 | 15.15642 | 28.7408 | 18.40687 | 7.429771 | 1.125957 | 0.062178 | 0.205434 |
| 17 | 81975000 | 81979599 | 26.44851 | 42.86183 | 94.22742 | 40.34122 | -0.76888 | 1.634249 | 0.205454 |
| 18 | 13134600 | 13138599 | 22.00252 | 36.44215 | 74.44763 | 32.21692 | -0.64671 | 1.305378 | 0.205458 |
| 12 | 79104000 | 79109599 | 44.9567 | 71.53933 | 135.5424 | 56.63235 | -0.58508 | 2.217519 | 0.205466 |
| 2 | 60040000 | 60040999 | 4.413445 | 8.552243 | 26.01722 | 9.855852 | -1.0154 | -0.23142 | 0.205474 |
| 6 | 87288800 | 87289999 | 4.413445 | 8.552243 | 26.01722 | 9.855852 | -1.0154 | -0.23142 | 0.205474 |
| 19 | 42133800 | 42137399 | 19.44245 | 32.32688 | 70.24664 | 30.48031 | -0.73481 | 1.251214 | 0.205486 |
| 7 | 126443200 | 126444999 | 8.671232 | 15.97544 | 35.73757 | 15.13671 | -0.72017 | 0.226813 | 0.205488 |
| 16 | 34071600 | 34072799 | 5.85228 | 11.21918 | 28.44964 | 11.19588 | -0.80867 | -0.07947 | 0.205488 |
| 17 | 92613000 | 92614199 | 13.6987 | 25.85223 | 19.55972 | 7.97563 | 0.874595 | 0.082617 | 0.20549 |
| 7 | 10791800 | 10793999 | 44.96124 | 73.33175 | 57.16039 | 24.55661 | 0.730656 | 1.541768 | 0.205505 |
| 3 | 16961600 | 16962999 | 11.32035 | 20.03473 | 42.46027 | 18.77718 | -0.68631 | 0.509165 | 0.205514 |
| 19 | 39158200 | 39158799 | 15.01847 | 27.87747 | 17.94956 | 7.525423 | 1.074912 | 0.194257 | 0.205519 |
| 14 | 42310400 | 42311999 | 14.06907 | 26.84668 | 18.25115 | 7.269869 | 1.05786 | 0.011329 | 0.205519 |
| 16 | 11585800 | 11590999 | 75.4329 | 119.4919 | 103.6074 | 43.08919 | 0.534451 | 2.365465 | 0.205533 |
| 15 | 3366800 | 3368399 | 20.28415 | 36.28913 | 24.55885 | 10.77425 | 0.959402 | 0.455025 | 0.205533 |
| 9 | 100180200 | 100182599 | 14.66365 | 25.41422 | 53.81431 | 23.6109 | -0.69129 | 0.822232 | 0.205539 |
| 7 | 31529000 | 31533999 | 65.61978 | 105.3887 | 65.28101 | 27.6415 | 1.039661 | 1.933443 | 0.205544 |
| 13 | 113122000 | 113127799 | 36.13278 | 57.78781 | 122.4655 | 51.52995 | -0.73835 | 2.04523 | 0.205563 |
| 13 | 66187400 | 66192199 | 62.30215 | 100.6516 | 55.98616 | 23.90001 | 1.204432 | 1.751821 | 0.205572 |
| 17 | 24381600 | 24385399 | 15.31252 | 25.69318 | 68.53617 | 29.91152 | -1.02924 | 1.135856 | 0.205576 |
| 16 | 21750200 | 21753799 | 16.20086 | 27.63605 | 58.5367 | 25.66194 | -0.69615 | 0.979739 | 0.205585 |
| X | 39116400 | 39116999 | 16.50646 | 30.22186 | 21.74664 | 9.289219 | 0.896719 | 0.291538 | 0.205595 |
| 2 | 105053000 | 105055199 | 11.01847 | 19.43331 | 42.99857 | 19.02835 | -0.75014 | 0.545398 | 0.205609 |
| 5 | 24552200 | 24559999 | 42.89293 | 68.13099 | 142.8501 | 59.56251 | -0.73368 | 2.272312 | 0.205617 |
| 17 | 15627400 | 15632599 | 27.99848 | 45.48317 | 89.04484 | 38.19023 | -0.59878 | 1.586517 | 0.20563 |
| 6 | 101353000 | 101355599 | 13.61671 | 23.54012 | 52.65355 | 23.09983 | -0.77049 | 0.840619 | 0.205636 |
| 14 | 121086400 | 121087999 | 10.17696 | 18.25318 | 39.46612 | 17.27727 | -0.70391 | 0.40366 | 0.205649 |
| 16 | 66065400 | 66069599 | 29.81092 | 51.08371 | 35.86501 | 15.72862 | 0.89134 | 0.963566 | 0.205662 |
| 3 | 83243400 | 83244599 | 6.913042 | 13.10983 | 32.15451 | 12.97726 | -0.79544 | 0.02022 | 0.205677 |
| 17 | 19025000 | 19028599 | 34.1782 | 58.32106 | 34.68971 | 15.20803 | 1.128828 | 1.018602 | 0.205683 |
| 6 | 148845800 | 148848399 | 15.27734 | 25.40619 | 74.23725 | 32.21591 | -1.16029 | 1.239847 | 0.205702 |

| | | | | | | | | | |
|---|---|---|---|---|---|---|---|---|---|
| 17 | 21089400 | 21089999 | 11.89089 | 23.10881 | 15.16434 | 5.644292 | 1.197665 | -0.23914 | 0.205707 |
| 9 | 102909000 | 102912999 | 35.8942 | 57.60972 | 112.339 | 47.51903 | -0.61254 | 1.931261 | 0.205721 |
| 1 | 45100400 | 45102399 | 27.90188 | 47.74068 | 38.19897 | 16.75016 | 0.703293 | 0.972296 | 0.205723 |
| 17 | 23614400 | 23618199 | 31.53112 | 53.16361 | 42.02497 | 18.37502 | 0.719004 | 1.117037 | 0.205731 |
| 7 | 23607200 | 23608999 | 50.59711 | 81.67175 | 67.47512 | 28.77278 | 0.63372 | 1.760652 | 0.205743 |
| 17 | 29345600 | 29348999 | 19.14577 | 32.19569 | 67.93602 | 29.68053 | -0.69261 | 1.125217 | 0.205744 |
| 1 | 164279000 | 164281199 | 9.444857 | 16.82959 | 42.01558 | 18.49515 | -0.91592 | 0.442484 | 0.205747 |
| 7 | 31912200 | 31912599 | 16.88303 | 30.4098 | 24.62358 | 10.73695 | 0.708197 | 0.391439 | 0.205747 |
| X | 73257000 | 73257799 | 14.11593 | 27.28486 | 15.51604 | 5.947131 | 1.363663 | -0.11653 | 0.205756 |
| X | 48919400 | 48921399 | 16.07685 | 29.71961 | 22.06756 | 9.303807 | 0.864797 | 0.235225 | 0.205769 |
| 17 | 27287800 | 27293199 | 29.0841 | 46.88555 | 101.0045 | 42.99848 | -0.74791 | 1.780405 | 0.205773 |
| 3 | 32190800 | 32192799 | 19.66696 | 34.73591 | 28.01566 | 12.39438 | 0.698568 | 0.553509 | 0.205773 |
| 7 | 8604000 | 8605799 | 36.39617 | 60.51105 | 38.22352 | 16.58446 | 1.041613 | 1.271443 | 0.205775 |
| 2 | 59441600 | 59444199 | 14.2675 | 24.19993 | 63.21272 | 27.68922 | -0.9983 | 1.029021 | 0.205801 |
| 3 | 152186000 | 152193599 | 46.06981 | 73.30281 | 137.5481 | 57.46191 | -0.57136 | 2.221429 | 0.20581 |
| 7 | 30164600 | 30168199 | 21.19452 | 34.76989 | 80.34228 | 34.59348 | -0.82766 | 1.425159 | 0.205816 |
| 1 | 169053600 | 169056199 | 30.96638 | 52.05273 | 43.72721 | 19.10167 | 0.631502 | 1.14353 | 0.205817 |
| 8 | 36715000 | 36716999 | 12.02118 | 20.95127 | 48.91289 | 21.51756 | -0.8298 | 0.732102 | 0.205821 |
| 2 | 18809000 | 18811199 | 19.09811 | 31.83751 | 68.64508 | 29.83645 | -0.72344 | 1.219585 | 0.205826 |
| 14 | 58018000 | 58019199 | 14.52374 | 27.04834 | 21.68695 | 9.046176 | 0.765973 | 0.179135 | 0.205848 |
| 10 | 82830200 | 82834599 | 27.47991 | 44.35499 | 100.8545 | 43.00403 | -0.82353 | 1.739216 | 0.205877 |
| 17 | 36966000 | 36966599 | 4.521255 | 8.809959 | 28.34546 | 10.2919 | -1.03966 | -0.37675 | 0.205882 |
| 16 | 91280800 | 91284999 | 21.87684 | 36.13619 | 77.18197 | 33.33793 | -0.71106 | 1.338056 | 0.205888 |
| 18 | 10244200 | 10245999 | 18.34074 | 33.62566 | 21.69259 | 9.249111 | 1.054598 | 0.296713 | 0.205902 |
| 8 | 47128000 | 47129199 | 15.08767 | 28.1426 | 21.00819 | 8.739848 | 0.8711 | 0.171867 | 0.20591 |
| 8 | 105350400 | 105352199 | 7.285325 | 13.21293 | 38.08847 | 16.46598 | -1.10347 | 0.311966 | 0.205919 |
| 1 | 5540400 | 5542199 | 25.96818 | 44.82454 | 36.06715 | 15.82242 | 0.697444 | 0.884469 | 0.205923 |
| 8 | 93166000 | 93166999 | 5.103544 | 9.849002 | 26.94421 | 10.40316 | -0.89031 | -0.15368 | 0.205945 |
| 8 | 96735000 | 96735399 | 5.103544 | 9.849002 | 26.94421 | 10.40316 | -0.89031 | -0.15368 | 0.205945 |
| 17 | 94991800 | 94995199 | 51.14152 | 82.52644 | 67.34365 | 28.70592 | 0.65096 | 1.766261 | 0.205949 |
| 7 | 41232000 | 41234799 | 44.18167 | 71.81506 | 58.30005 | 25.01509 | 0.66867 | 1.585001 | 0.20596 |
| 9 | 119168600 | 119170999 | 13.83942 | 23.7408 | 55.78765 | 24.46785 | -0.8437 | 0.916348 | 0.205962 |
| 5 | 91908800 | 91909799 | 6.14984 | 11.78611 | 35.213 | 13.86934 | -1.04626 | -0.06175 | 0.205965 |
| 5 | 129390400 | 129395799 | 47.45384 | 75.28693 | 144.4438 | 60.15617 | -0.60751 | 2.311677 | 0.205972 |
| 11 | 15832600 | 15834799 | 20.74854 | 36.6174 | 27.80662 | 12.30383 | 0.784483 | 0.567461 | 0.20598 |
| 8 | 110084400 | 110085999 | 15.73512 | 28.58906 | 23.08182 | 9.956092 | 0.722065 | 0.329145 | 0.205981 |
| 16 | 32995800 | 33000599 | 28.99935 | 46.96356 | 94.63547 | 40.49551 | -0.64458 | 1.65301 | 0.205981 |
| 10 | 81602800 | 81604799 | 29.42742 | 51.27208 | 26.20882 | 11.52743 | 1.35013 | 0.785876 | 0.205991 |
| 14 | 42276200 | 42277799 | 16.80943 | 31.19861 | 20.98581 | 8.799629 | 1.012011 | 0.217352 | 0.205997 |
| 1 | 157993400 | 157994799 | 23.6626 | 41.336 | 30.45555 | 13.41734 | 0.825929 | 0.726488 | 0.206005 |
| 18 | 24176000 | 24179799 | 17.73273 | 29.46065 | 71.07305 | 30.82193 | -0.88506 | 1.256984 | 0.206013 |
| 13 | 109981400 | 109982599 | 8.661106 | 15.98942 | 36.35004 | 15.35339 | -0.74046 | 0.216047 | 0.206037 |
| 7 | 84571000 | 84572599 | 16.99201 | 31.21385 | 22.38869 | 9.520559 | 0.905445 | 0.276996 | 0.206037 |
| 7 | 73021400 | 73028199 | 38.62792 | 61.55871 | 132.9496 | 55.66203 | -0.77129 | 2.162826 | 0.206042 |
| 2 | 143494400 | 143495999 | 8.500377 | 15.76446 | 37.05255 | 15.55326 | -0.78166 | 0.190684 | 0.206052 |
| 13 | 24418800 | 24420399 | 8.500377 | 15.76446 | 37.05255 | 15.55326 | -0.78166 | 0.190684 | 0.206052 |
| X | 114366200 | 114367399 | 14.55359 | 27.75189 | 18.16552 | 7.246435 | 1.110089 | 0.020219 | 0.206064 |

| | | | | | | | | | |
|---|---|---|---|---|---|---|---|---|---|
| 1 | 135667000 | 135669199 | 8.506773 | 15.24537 | 38.48428 | 16.85882 | -0.92588 | 0.397 | 0.206068 |
| X | 5271600 | 5273999 | 41.54457 | 68.8489 | 46.06784 | 19.94364 | 0.958623 | 1.329535 | 0.206069 |
| 7 | 139640600 | 139645599 | 31.37559 | 50.44685 | 108.3631 | 45.95795 | -0.74857 | 1.868024 | 0.206077 |
| 8 | 118581200 | 118583599 | 10.66945 | 18.94915 | 41.42981 | 18.288 | -0.72852 | 0.471269 | 0.206096 |
| 2 | 36894800 | 36895599 | 12.69035 | 23.87818 | 19.85848 | 8.140493 | 0.733477 | 0.094048 | 0.206105 |
| 5 | 100370200 | 100374999 | 59.8378 | 93.73658 | 176.0871 | 73.03998 | -0.59197 | 2.604429 | 0.206106 |
| 13 | 52350400 | 52352399 | 6.666997 | 12.47377 | 32.75692 | 13.55732 | -0.92254 | 0.114859 | 0.206107 |
| 3 | 48969200 | 48971399 | 30.06662 | 50.75125 | 42.07363 | 18.40255 | 0.65065 | 1.102653 | 0.206113 |
| 12 | 71021000 | 71022599 | 7.296502 | 13.55403 | 33.17451 | 13.89434 | -0.83661 | 0.163581 | 0.206126 |
| 5 | 30120600 | 30123199 | 30.2022 | 51.63765 | 36.98733 | 16.21787 | 0.862176 | 0.986803 | 0.206129 |
| 17 | 27916400 | 27933799 | 151.2256 | 221.1271 | 427.8656 | 176.2655 | -0.71951 | 3.908594 | 0.20613 |
| 12 | 49394600 | 49397199 | 18.69303 | 31.59071 | 69.01333 | 30.1966 | -0.74206 | 1.084685 | 0.206135 |
| 19 | 8532600 | 8534999 | 14.02031 | 23.98167 | 57.37101 | 25.15644 | -0.87014 | 0.945287 | 0.206157 |
| 10 | 127167600 | 127170399 | 19.23528 | 32.12374 | 67.61486 | 29.42195 | -0.68886 | 1.19963 | 0.206162 |
| 1 | 62609400 | 62610999 | 15.69531 | 29.45495 | 19.96186 | 8.219302 | 1.02142 | 0.137707 | 0.206166 |
| 1 | 8962200 | 8964599 | 36.1329 | 59.91427 | 49.33511 | 21.36641 | 0.660219 | 1.309254 | 0.206186 |
| 9 | 9544600 | 9545399 | 6.654549 | 12.5161 | 31.2526 | 12.82009 | -0.84029 | 0.0754 | 0.206187 |
| 5 | 85624200 | 85626799 | 31.61151 | 53.04142 | 44.27733 | 19.32409 | 0.640584 | 1.163572 | 0.2062 |
| 6 | 66678600 | 66680999 | 31.61151 | 53.04142 | 44.27733 | 19.32409 | 0.640584 | 1.163572 | 0.2062 |
| 8 | 21155400 | 21156599 | 12.53761 | 24.05856 | 18.26218 | 7.1669 | 0.918958 | -0.05229 | 0.206229 |
| 17 | 21867600 | 21870399 | 16.41499 | 27.82045 | 62.83468 | 27.51703 | -0.78962 | 1.042813 | 0.206237 |
| 6 | 68182400 | 68184199 | 19.91035 | 35.64006 | 24.5672 | 10.77289 | 0.933428 | 0.448835 | 0.206238 |
| 12 | 83168400 | 83170599 | 11.66561 | 20.47844 | 44.6033 | 19.7303 | -0.72955 | 0.611431 | 0.206245 |
| 7 | 83226400 | 83229399 | 26.11674 | 44.99181 | 36.61532 | 16.06192 | 0.680566 | 0.903027 | 0.206266 |
| 17 | 92682200 | 92684399 | 29.72207 | 50.60843 | 39.24544 | 17.2023 | 0.747306 | 1.023845 | 0.206288 |
| 3 | 84489200 | 84490599 | 9.138364 | 16.30386 | 39.45376 | 17.3491 | -0.86965 | 0.42989 | 0.206295 |
| 2 | 9956600 | 9958199 | 8.719396 | 15.88396 | 36.56687 | 15.7209 | -0.77645 | 0.293426 | 0.206303 |
| 7 | 42270000 | 42277599 | 196.0916 | 293.4943 | 224.4298 | 92.56044 | 0.650384 | 3.514584 | 0.206329 |
| 4 | 134069600 | 134071999 | 17.43189 | 29.32797 | 63.79717 | 27.87645 | -0.73605 | 1.113375 | 0.206355 |
| 18 | 74456200 | 74460199 | 46.1272 | 74.99594 | 59.49859 | 25.53168 | 0.702049 | 1.587876 | 0.20636 |
| 2 | 28915000 | 28920799 | 34.89938 | 56.08097 | 111.0887 | 47.0755 | -0.63304 | 1.890808 | 0.206368 |
| X | 33037600 | 33040399 | 28.14238 | 49.26845 | 26.52052 | 11.70444 | 1.276637 | 0.714617 | 0.206371 |
| 11 | 22909400 | 22912399 | 12.11149 | 21.35765 | 47.75493 | 21.13388 | -0.76566 | 0.561323 | 0.206384 |
| 18 | 75087400 | 75090199 | 13.89733 | 24.08823 | 52.15605 | 22.8834 | -0.72309 | 0.81669 | 0.206407 |
| 12 | 36065600 | 36067199 | 7.899262 | 14.61477 | 34.73535 | 14.62795 | -0.8008 | 0.193745 | 0.206418 |
| 12 | 100052400 | 100060199 | 47.50153 | 75.18512 | 155.0658 | 64.46367 | -0.71475 | 2.375724 | 0.206427 |
| 13 | 107736800 | 107737999 | 6.372457 | 11.99166 | 31.82507 | 13.04362 | -0.92705 | 0.06811 | 0.20644 |
| 5 | 60936200 | 60937999 | 30.04813 | 50.64551 | 42.10101 | 18.40632 | 0.646646 | 1.114886 | 0.206443 |
| 6 | 131148800 | 131149799 | 14.66204 | 27.88452 | 18.62824 | 7.472885 | 1.073774 | 0.039706 | 0.206471 |
| 7 | 128660600 | 128661399 | 4.815994 | 9.362721 | 25.85027 | 9.594339 | -0.84998 | -0.30036 | 0.206477 |
| 15 | 6737000 | 6738999 | 13.06659 | 22.44677 | 56.31891 | 24.70312 | -0.93744 | 0.899992 | 0.206478 |
| 15 | 16939800 | 16940999 | 14.33703 | 27.09568 | 18.92505 | 7.693481 | 0.992711 | 0.077309 | 0.206485 |
| 15 | 44926200 | 44927599 | 14.33703 | 27.09568 | 18.92505 | 7.693481 | 0.992711 | 0.077309 | 0.206485 |
| X | 62364400 | 62367199 | 36.58171 | 62.38346 | 32.39933 | 14.20321 | 1.323495 | 1.031433 | 0.206487 |
| 7 | 12377000 | 12378599 | 32.82223 | 54.955 | 43.83498 | 19.10519 | 0.706182 | 1.192397 | 0.206494 |
| 2 | 38176400 | 38179399 | 18.68685 | 31.39867 | 66.15332 | 28.89463 | -0.69038 | 1.129702 | 0.206511 |
| 7 | 143102200 | 143103199 | 7.055144 | 12.9185 | 35.07945 | 14.98258 | -1.00523 | 0.251291 | 0.20652 |

| | | | | | | | | | |
|---|---|---|---|---|---|---|---|---|---|
| 7 | 91881200 | 91882199 | 24.8028 | 43.44891 | 26.44925 | 11.67912 | 1.10005 | 0.685991 | 0.20652 |
| 7 | 10809600 | 10810999 | 36.25262 | 59.82201 | 50.96068 | 21.99686 | 0.610953 | 1.3674 | 0.206524 |
| 7 | 15215600 | 15216799 | 32.56237 | 54.52046 | 45.39779 | 19.78646 | 0.644342 | 1.191495 | 0.206535 |
| 8 | 85702200 | 85706999 | 20.58894 | 33.73709 | 81.84241 | 35.22203 | -0.89829 | 1.439422 | 0.206541 |
| 7 | 82974800 | 82977599 | 37.45062 | 63.01495 | 39.21911 | 17.13752 | 1.062704 | 1.154728 | 0.206543 |
| 7 | 46367600 | 46371199 | 22.6587 | 37.45 | 76.75388 | 33.16668 | -0.65165 | 1.332969 | 0.206552 |
| 1 | 50043200 | 50046999 | 61.71062 | 98.84721 | 82.46264 | 34.7757 | 0.60658 | 2.013904 | 0.206586 |
| 10 | 82210000 | 82212599 | 79.07629 | 126.1578 | 80.80061 | 33.88055 | 0.980953 | 2.154508 | 0.206588 |
| 3 | 34840800 | 34843199 | 19.5043 | 32.7898 | 66.14835 | 28.89716 | -0.62787 | 1.129837 | 0.206614 |
| 7 | 75803600 | 75805399 | 9.432663 | 16.83681 | 42.03847 | 18.47839 | -0.91466 | 0.431242 | 0.206614 |
| 12 | 77902600 | 77904399 | 8.827267 | 16.21171 | 37.31076 | 15.87099 | -0.76563 | 0.248212 | 0.206639 |
| X | 118108000 | 118109399 | 17.27579 | 32.83715 | 15.98842 | 6.421434 | 1.523861 | 0.060896 | 0.206641 |
| 5 | 94573800 | 94576599 | 22.18642 | 39.15834 | 26.52525 | 11.73665 | 0.948365 | 0.572709 | 0.206642 |
| X | 127107600 | 127108399 | 12.86073 | 24.85086 | 15.84624 | 6.082609 | 1.198881 | -0.12358 | 0.206645 |
| 14 | 29518000 | 29519799 | 12.2755 | 21.5799 | 45.07716 | 19.96119 | -0.66961 | 0.592689 | 0.206651 |
| 3 | 50744400 | 50746199 | 10.22162 | 18.13854 | 40.92133 | 18.07635 | -0.77407 | 0.473248 | 0.206677 |
| 8 | 22173000 | 22175199 | 38.08708 | 63.69382 | 42.61547 | 18.55966 | 0.958897 | 1.219621 | 0.20669 |
| 18 | 81012600 | 81019799 | 85.74483 | 129.7849 | 297.3453 | 123.0081 | -0.91728 | 3.278282 | 0.206695 |
| 1 | 92703600 | 92706199 | 14.40748 | 24.8416 | 53.72945 | 23.56995 | -0.72318 | 0.869173 | 0.206698 |
| 7 | 138967800 | 138970799 | 14.69115 | 24.98414 | 60.00444 | 26.2989 | -0.87723 | 1.004358 | 0.206707 |
| 12 | 104387200 | 104390199 | 14.22621 | 24.56988 | 55.39087 | 24.30001 | -0.78237 | 0.853573 | 0.206707 |
| 10 | 14319800 | 14320599 | 10.07299 | 19.4972 | 17.05799 | 6.495978 | 0.756825 | -0.17836 | 0.20674 |
| 17 | 19838400 | 19851199 | 256.532 | 375.5954 | 290.5827 | 119.7001 | 0.604092 | 3.895059 | 0.206755 |
| 12 | 74849800 | 74851999 | 147.1142 | 227.8504 | 93.99315 | 39.04222 | 1.579625 | 2.832461 | 0.206768 |
| 8 | 46744200 | 46750199 | 44.17058 | 70.1618 | 146.4127 | 61.04919 | -0.72688 | 2.272588 | 0.206771 |
| 6 | 28333000 | 28334199 | 9.207586 | 16.68501 | 37.00706 | 16.0155 | -0.72919 | 0.330285 | 0.206792 |
| 17 | 66035000 | 66035999 | 9.207586 | 16.68501 | 37.00706 | 16.0155 | -0.72919 | 0.330285 | 0.206792 |
| 1 | 80190200 | 80191799 | 11.86934 | 20.6834 | 51.0418 | 22.4519 | -0.90967 | 0.73492 | 0.206797 |
| 8 | 103779400 | 103780799 | 19.66958 | 36.19694 | 18.6808 | 7.924577 | 1.378651 | 0.279613 | 0.206817 |
| 11 | 6224200 | 6226999 | 11.54655 | 20.17028 | 47.83567 | 21.07403 | -0.85192 | 0.69493 | 0.206833 |
| 4 | 19208200 | 19210399 | 18.06483 | 32.51419 | 22.40611 | 9.779157 | 0.93837 | 0.398529 | 0.20684 |
| 9 | 80543800 | 80544599 | 12.15779 | 23.38568 | 17.73046 | 6.910525 | 0.929569 | -0.07983 | 0.206848 |
| 9 | 73061000 | 73062999 | 10.52151 | 18.35404 | 53.15152 | 23.39541 | -1.13903 | 0.715479 | 0.206848 |
| 14 | 62821800 | 62825599 | 24.55957 | 40.16603 | 82.18187 | 35.34118 | -0.65499 | 1.476012 | 0.206858 |
| 13 | 58143400 | 58147599 | 22.75592 | 37.3558 | 80.17759 | 34.53212 | -0.72125 | 1.41973 | 0.206861 |
| X | 81580400 | 81582199 | 16.24629 | 30.61707 | 18.85642 | 7.709198 | 1.166404 | 0.114462 | 0.206878 |
| 17 | 20358200 | 20361599 | 52.76189 | 85.55081 | 58.08121 | 24.87598 | 0.922729 | 1.652448 | 0.206879 |
| 11 | 66058400 | 66059799 | 10.71492 | 18.90401 | 42.49241 | 18.80315 | -0.77245 | 0.539664 | 0.206883 |
| 6 | 83446000 | 83447399 | 15.94994 | 29.08109 | 23.54295 | 10.11059 | 0.722247 | 0.312411 | 0.206894 |
| 9 | 27132200 | 27133399 | 17.54263 | 31.33701 | 25.17019 | 11.05654 | 0.712719 | 0.447659 | 0.206897 |
| 7 | 14605800 | 14615199 | 240.9683 | 353.5839 | 284.5583 | 117.2023 | 0.550229 | 3.859605 | 0.206905 |
| 2 | 57957800 | 57959599 | 16.45716 | 30.48472 | 21.3834 | 8.99039 | 0.948884 | 0.224827 | 0.206912 |
| 9 | 21629400 | 21633399 | 17.72295 | 29.85314 | 65.01573 | 28.41915 | -0.73778 | 1.103598 | 0.206913 |
| 13 | 98640600 | 98642999 | 17.9732 | 30.04054 | 67.52564 | 29.39827 | -0.7833 | 1.186541 | 0.206951 |
| 12 | 79066400 | 79068599 | 11.63023 | 20.51836 | 43.50303 | 19.25045 | -0.68888 | 0.546539 | 0.206966 |
| 3 | 156149400 | 156150199 | 8.370633 | 15.43642 | 34.66337 | 14.66261 | -0.72399 | 0.217469 | 0.206966 |
| 15 | 65658600 | 65659799 | 8.370633 | 15.43642 | 34.66337 | 14.66261 | -0.72399 | 0.217469 | 0.206966 |

| | | | | | | | | | |
|---|---|---|---|---|---|---|---|---|---|
| 12 | 105698200 | 105702399 | 23.622 | 38.58043 | 83.10564 | 35.72081 | -0.7304 | 1.493026 | 0.206978 |
| 6 | 84287600 | 84288999 | 6.111522 | 11.42737 | 32.20661 | 13.34362 | -1.02527 | 0.109549 | 0.206981 |
| 1 | 39790000 | 39792399 | 12.26555 | 21.51546 | 48.3037 | 21.35364 | -0.77349 | 0.630227 | 0.206986 |
| 2 | 173415000 | 173421599 | 47.54098 | 75.55019 | 144.3077 | 60.19921 | -0.59882 | 2.261434 | 0.207 |
| X | 95988200 | 95991199 | 51.78341 | 83.8753 | 61.20436 | 26.19314 | 0.817569 | 1.675048 | 0.207017 |
| 8 | 110291800 | 110292799 | 6.393034 | 11.97469 | 32.25719 | 13.32502 | -0.95748 | 0.099885 | 0.207018 |
| 7 | 97823600 | 97824999 | 4.903182 | 9.375513 | 29.99526 | 11.89178 | -1.15107 | -0.06294 | 0.207027 |
| 11 | 33463000 | 33465599 | 24.32374 | 42.51173 | 28.54039 | 12.57801 | 0.959605 | 0.721705 | 0.207034 |
| 7 | 60230600 | 60243999 | 25.19464 | 43.68938 | 31.41282 | 13.78252 | 0.860237 | 0.834001 | 0.207038 |
| 17 | 19072400 | 19074199 | 21.45803 | 38.55517 | 22.97663 | 10.04352 | 1.145359 | 0.435114 | 0.207072 |
| 13 | 17994600 | 17996399 | 30.86556 | 49.70213 | 103.8032 | 44.1209 | -0.70528 | 1.819401 | 0.207073 |
| 5 | 7846000 | 7846799 | 8.704676 | 15.9297 | 35.58506 | 15.2105 | -0.72758 | 0.265751 | 0.207081 |
| 11 | 102791400 | 102793599 | 9.97638 | 18.03612 | 39.61151 | 17.19259 | -0.71825 | 0.353337 | 0.207086 |
| 5 | 66988200 | 66989599 | 5.681934 | 10.9535 | 28.10458 | 10.88448 | -0.80344 | -0.13421 | 0.207114 |
| 1 | 29152000 | 29153799 | 14.66434 | 25.514 | 54.37849 | 23.89611 | -0.69982 | 0.775908 | 0.207137 |
| Y | 21513600 | 21513999 | 7.005409 | 13.70344 | 7.131068 | 2.380379 | 1.662618 | -0.71481 | 0.207142 |
| 19 | 49929800 | 49931599 | 19.24158 | 35.06298 | 22.16118 | 9.523857 | 1.07651 | 0.338424 | 0.207175 |
| 6 | 38989600 | 38996999 | 52.51103 | 83.12611 | 153.3065 | 63.73694 | -0.55355 | 2.375743 | 0.207179 |
| 1 | 4855400 | 4856199 | 5.848178 | 11.252 | 28.66297 | 11.16312 | -0.8017 | -0.11571 | 0.207197 |
| 19 | 44010000 | 44012199 | 19.27588 | 34.39863 | 25.74155 | 11.3171 | 0.813209 | 0.465649 | 0.207215 |
| 8 | 9056000 | 9056999 | 13.63219 | 25.60147 | 20.69775 | 8.511978 | 0.77011 | 0.11615 | 0.207229 |
| X | 88723000 | 88726399 | 33.55826 | 56.46987 | 42.58551 | 18.60896 | 0.786719 | 1.143446 | 0.207255 |
| 4 | 4890200 | 4890399 | 11.62308 | 22.24463 | 18.95704 | 7.496409 | 0.743587 | -0.03498 | 0.207265 |
| 10 | 61253000 | 61255199 | 17.50063 | 29.25697 | 66.85782 | 29.11147 | -0.80687 | 1.183686 | 0.207268 |
| 10 | 58867800 | 58869199 | 11.81476 | 22.56354 | 18.97056 | 7.543288 | 0.755837 | -0.01503 | 0.207274 |
| 17 | 86509600 | 86512999 | 14.02427 | 23.98233 | 59.01295 | 25.87587 | -0.91082 | 0.948416 | 0.207275 |
| 10 | 88689000 | 88690199 | 16.57782 | 30.06517 | 23.95053 | 10.35443 | 0.739181 | 0.347751 | 0.207293 |
| 15 | 97626400 | 97645199 | 144.8866 | 213.1533 | 405.7027 | 167.0714 | -0.68738 | 3.811313 | 0.207295 |
| 9 | 39026200 | 39029799 | 73.22177 | 116.4114 | 98.98667 | 41.31103 | 0.567525 | 2.260329 | 0.207323 |
| 2 | 166446200 | 166455599 | 49.95817 | 78.59593 | 167.9987 | 69.69717 | -0.77316 | 2.51513 | 0.207328 |
| 9 | 83645400 | 83646799 | 8.028752 | 14.9562 | 33.80682 | 14.09558 | -0.71784 | 0.153624 | 0.207343 |
| 7 | 28714400 | 28721199 | 48.08235 | 76.27303 | 147.0122 | 61.21696 | -0.61439 | 2.315869 | 0.207346 |
| 8 | 11728600 | 11738399 | 72.17053 | 112.2234 | 202.4758 | 84.06134 | -0.54223 | 2.741684 | 0.207357 |
| 11 | 27371800 | 27372999 | 15.51537 | 29.55236 | 17.29162 | 6.914385 | 1.266726 | 0.033551 | 0.207361 |
| X | 118166000 | 118167799 | 15.51537 | 29.55236 | 17.29162 | 6.914385 | 1.266726 | 0.033551 | 0.207361 |
| 4 | 46347600 | 46354199 | 38.329 | 61.03683 | 135.9483 | 56.85884 | -0.81684 | 2.187496 | 0.207362 |
| 3 | 27642600 | 27643599 | 7.108975 | 13.3309 | 31.75475 | 13.09215 | -0.77931 | 0.098413 | 0.207368 |
| 8 | 8070800 | 8071799 | 7.108975 | 13.3309 | 31.75475 | 13.09215 | -0.77931 | 0.098413 | 0.207368 |
| 3 | 138201600 | 138204799 | 20.1259 | 33.7696 | 68.50094 | 29.89611 | -0.63596 | 1.151143 | 0.20737 |
| 8 | 127460200 | 127464199 | 18.09796 | 30.32243 | 67.11141 | 29.26262 | -0.76112 | 1.159371 | 0.207379 |
| 17 | 30026800 | 30028399 | 6.636521 | 12.18937 | 35.64396 | 15.16016 | -1.10742 | 0.22821 | 0.207382 |
| 7 | 26404200 | 26406799 | 35.52388 | 58.63437 | 49.33592 | 21.29909 | 0.628729 | 1.360544 | 0.207398 |
| 1 | 18492200 | 18493399 | 13.87034 | 26.53043 | 18.09448 | 7.167014 | 1.059752 | -0.01002 | 0.207402 |
| 14 | 88269600 | 88270399 | 10.90732 | 21.15146 | 16.28135 | 6.146513 | 0.950637 | -0.20119 | 0.207402 |
| 6 | 22124800 | 22125399 | 4.655824 | 9.0527 | 24.59756 | 9.118307 | -0.82455 | -0.30824 | 0.207415 |
| 8 | 109292600 | 109293199 | 4.655824 | 9.0527 | 24.59756 | 9.118307 | -0.82455 | -0.30824 | 0.207415 |
| 18 | 12116200 | 12116999 | 4.655824 | 9.0527 | 24.59756 | 9.118307 | -0.82455 | -0.30824 | 0.207415 |

| | | | | | | | | | |
|---|---|---|---|---|---|---|---|---|---|
| 13 | 59881000 | 59882599 | 12.27044 | 21.54377 | 45.62845 | 20.18661 | -0.68962 | 0.61257 | 0.207432 |
| 18 | 61971200 | 61973199 | 10.54031 | 18.65283 | 41.95037 | 18.55196 | -0.77153 | 0.502948 | 0.207448 |
| 9 | 124090200 | 124091799 | 61.7338 | 98.79529 | 84.40745 | 35.54849 | 0.57079 | 2.042262 | 0.207463 |
| 16 | 73345000 | 73345599 | 13.33347 | 25.81564 | 15.50317 | 5.895878 | 1.295752 | -0.1509 | 0.207465 |
| 17 | 27388000 | 27395599 | 41.98758 | 66.56516 | 150.1014 | 62.46598 | -0.84119 | 2.329986 | 0.207483 |
| 6 | 130693600 | 130703999 | 18.63448 | 33.44967 | 22.47332 | 9.831498 | 0.972647 | 0.419598 | 0.207488 |
| 3 | 149035600 | 149037999 | 13.06662 | 22.81545 | 49.44665 | 21.77765 | -0.72303 | 0.710046 | 0.207496 |
| 13 | 24984000 | 24986599 | 10.98743 | 19.08502 | 51.00254 | 22.38925 | -1.02404 | 0.780281 | 0.207521 |
| 2 | 28181600 | 28188199 | 40.443 | 64.33217 | 138.8197 | 57.97111 | -0.77298 | 2.226059 | 0.207548 |
| 19 | 54419800 | 54424799 | 65.05473 | 104.2682 | 77.57815 | 32.74659 | 0.77239 | 1.996345 | 0.207551 |
| 13 | 69957800 | 69960599 | 11.98079 | 20.63875 | 55.22389 | 24.22486 | -1.02874 | 0.86908 | 0.207558 |
| 15 | 25489800 | 25490399 | 3.705849 | 7.22779 | 23.83815 | 8.550839 | -1.0569 | -0.44332 | 0.2076 |
| 10 | 35143200 | 35145999 | 32.88147 | 55.18682 | 44.262 | 19.32061 | 0.698136 | 1.16635 | 0.207617 |
| 8 | 92134400 | 92143599 | 75.93678 | 117.1789 | 212.9504 | 88.45556 | -0.56118 | 2.882128 | 0.207618 |
| 5 | 120348600 | 120349599 | 5.549961 | 10.66315 | 27.9424 | 10.92674 | -0.84656 | -0.10422 | 0.20762 |
| 9 | 64381000 | 64385799 | 26.15794 | 42.30182 | 94.46854 | 40.38028 | -0.79426 | 1.679699 | 0.207621 |
| 18 | 77484600 | 77487999 | 22.29512 | 36.57355 | 80.58529 | 34.69736 | -0.75937 | 1.427958 | 0.207648 |
| 11 | 94631800 | 94634199 | 11.58163 | 20.15033 | 49.96779 | 21.95775 | -0.91658 | 0.756753 | 0.207654 |
| 4 | 134658000 | 134658799 | 5.422165 | 10.43946 | 28.06493 | 10.90884 | -0.87566 | -0.12798 | 0.207679 |
| 5 | 148653000 | 148655999 | 48.16605 | 77.74442 | 63.87265 | 27.23516 | 0.641663 | 1.754847 | 0.207685 |
| 4 | 34785000 | 34786999 | 12.30886 | 21.50704 | 47.3103 | 20.84569 | -0.74398 | 0.695127 | 0.207686 |
| 3 | 33319400 | 33320599 | 12.39568 | 23.70849 | 18.16318 | 7.195349 | 0.892825 | -0.02529 | 0.207687 |
| 19 | 39364800 | 39365799 | 13.32262 | 25.86119 | 14.33768 | 5.386863 | 1.425599 | -0.19349 | 0.207718 |
| 16 | 15326200 | 15330999 | 47.03131 | 76.57152 | 57.85819 | 24.84026 | 0.773595 | 1.57047 | 0.207721 |
| 16 | 21094400 | 21106799 | 79.82341 | 122.2003 | 241.178 | 100.084 | -0.68895 | 3.040698 | 0.207735 |
| 8 | 49090200 | 49091799 | 9.506966 | 17.28003 | 38.12612 | 16.4371 | -0.71824 | 0.316952 | 0.207738 |
| 4 | 85663200 | 85664599 | 14.10433 | 26.46914 | 20.61853 | 8.489714 | 0.821877 | 0.124503 | 0.207746 |
| 5 | 139855800 | 139857399 | 6.825975 | 12.77671 | 32.25465 | 13.33971 | -0.86617 | 0.108173 | 0.207751 |
| 3 | 126526000 | 126527199 | 6.75865 | 12.85537 | 30.51381 | 12.23756 | -0.74005 | -0.00472 | 0.207753 |
| 6 | 36121800 | 36123199 | 30.86493 | 51.91961 | 43.73555 | 19.10948 | 0.627614 | 1.137852 | 0.207756 |
| 17 | 75763200 | 75764199 | 7.070831 | 13.37934 | 31.48002 | 12.7648 | -0.74176 | 0.036517 | 0.207759 |
| 6 | 54957000 | 54958199 | 17.16549 | 30.73454 | 25.11507 | 11.01077 | 0.689828 | 0.430137 | 0.207762 |
| 10 | 84752600 | 84754199 | 9.917971 | 17.6098 | 40.42679 | 17.84927 | -0.79843 | 0.467277 | 0.207766 |
| 4 | 71216800 | 71218599 | 51.26024 | 82.52282 | 71.81303 | 30.51275 | 0.554439 | 1.840039 | 0.207766 |
| 3 | 126917200 | 126918599 | 7.300568 | 13.5502 | 35.3552 | 14.82459 | -0.92992 | 0.171218 | 0.207766 |
| 7 | 23024600 | 23025399 | 18.87947 | 35.64421 | 15.82524 | 6.447192 | 1.638288 | 0.119441 | 0.207767 |
| 6 | 89896400 | 89898199 | 24.6587 | 42.95237 | 32.71812 | 14.38809 | 0.77801 | 0.77607 | 0.207792 |
| 9 | 14397400 | 14397799 | 3.128089 | 6.111779 | 21.08939 | 7.345914 | -1.08023 | -0.56424 | 0.207798 |
| 8 | 107593600 | 107595599 | 10.15804 | 18.28573 | 39.51003 | 17.24103 | -0.69983 | 0.380324 | 0.207818 |
| 13 | 8935400 | 8936399 | 5.976404 | 11.17537 | 32.74719 | 13.56628 | -1.08102 | 0.108555 | 0.207827 |
| 13 | 66344200 | 66345999 | 18.16645 | 33.23292 | 22.2324 | 9.507692 | 0.999512 | 0.308435 | 0.207829 |
| 17 | 38506200 | 38507399 | 18.16645 | 33.23292 | 22.2324 | 9.507692 | 0.999512 | 0.308435 | 0.207829 |
| 7 | 18271800 | 18276399 | 106.2472 | 164.704 | 144.0851 | 59.84695 | 0.497531 | 2.802693 | 0.207836 |
| 5 | 94470400 | 94475799 | 95.45542 | 150.6105 | 106.4541 | 44.18431 | 0.824335 | 2.474296 | 0.207842 |
| 5 | 145819600 | 145823199 | 90.39822 | 141.8193 | 124.2199 | 51.51791 | 0.509038 | 2.585692 | 0.207865 |
| 7 | 9593800 | 9593999 | 6.497577 | 12.70859 | 10.93841 | 3.687746 | 0.938031 | -0.66829 | 0.207866 |
| 11 | 108832600 | 108842399 | 59.24157 | 91.93916 | 203.7293 | 84.60648 | -0.84052 | 2.770258 | 0.20789 |

| | | | | | | | | | |
|---|---|---|---|---|---|---|---|---|---|
| X | 54585000 | 54593199 | 149.0347 | 228.4793 | 147.107 | 61.0637 | 0.927176 | 3.02887 | 0.207898 |
| 9 | 12459200 | 12461199 | 17.82055 | 32.057 | 20.58223 | 8.988003 | 1.03921 | 0.395729 | 0.207907 |
| 1 | 182829600 | 182830599 | 5.016647 | 9.628024 | 28.45334 | 11.16129 | -1.02296 | -0.10053 | 0.207915 |
| 7 | 8291000 | 8298399 | 145.0762 | 220.7909 | 184.2011 | 76.32818 | 0.545429 | 3.174075 | 0.207924 |
| 8 | 14027000 | 14044399 | 116.1704 | 174.2825 | 320.1704 | 132.1117 | -0.6102 | 3.460734 | 0.207926 |
| 7 | 13345200 | 13348199 | 52.32151 | 84.50209 | 66.81423 | 28.51137 | 0.697805 | 1.746098 | 0.207948 |
| 15 | 85215000 | 85216799 | 10.35035 | 18.41191 | 43.03212 | 18.96964 | -0.82302 | 0.460658 | 0.20795 |
| 3 | 90340200 | 90341199 | 6.704902 | 12.49441 | 33.23515 | 13.8491 | -0.94986 | 0.138082 | 0.207981 |
| 14 | 18082000 | 18084399 | 17.22075 | 31.55354 | 22.26448 | 9.500626 | 0.92519 | 0.291689 | 0.207981 |
| 13 | 14237600 | 14239599 | 12.79796 | 21.92405 | 58.96631 | 25.85911 | -1.038 | 0.927255 | 0.208 |
| 11 | 113792800 | 113796199 | 18.31813 | 30.53302 | 69.12523 | 30.04253 | -0.79358 | 1.218877 | 0.208 |
| 4 | 152958800 | 152977999 | 221.8898 | 317.0909 | 538.3905 | 222.4535 | -0.56372 | 4.256405 | 0.208016 |
| 13 | 64093800 | 64094999 | 8.467552 | 15.29957 | 36.99381 | 16.06794 | -0.85593 | 0.343028 | 0.208019 |
| 8 | 11497600 | 11504599 | 37.41681 | 59.8389 | 121.5499 | 51.14168 | -0.6774 | 2.047844 | 0.208019 |
| 10 | 45516200 | 45517199 | 13.02737 | 25.41748 | 10.28658 | 3.671508 | 1.936881 | -0.38027 | 0.208022 |
| 8 | 89856400 | 89857799 | 9.914977 | 17.65209 | 40.98564 | 18.05485 | -0.81207 | 0.453957 | 0.208047 |
| 5 | 63281800 | 63282799 | 13.43785 | 25.04445 | 19.34114 | 8.057632 | 0.821056 | 0.158882 | 0.208049 |
| 5 | 82342000 | 82344199 | 21.62961 | 38.30312 | 26.05288 | 11.51996 | 0.944295 | 0.533885 | 0.20807 |
| 10 | 25212200 | 25214399 | 10.65801 | 18.96092 | 41.44767 | 18.26977 | -0.72685 | 0.460238 | 0.208077 |
| 11 | 55068600 | 55072999 | 21.68145 | 35.91531 | 75.0192 | 32.46735 | -0.67864 | 1.302896 | 0.208088 |
| 7 | 18797800 | 18802199 | 39.68851 | 63.40454 | 124.7398 | 52.40495 | -0.63309 | 2.083715 | 0.208089 |
| 18 | 12490200 | 12491199 | 5.983589 | 11.53084 | 28.76151 | 11.15051 | -0.76543 | -0.12873 | 0.208102 |
| 12 | 88401600 | 88403199 | 11.18974 | 19.71277 | 43.01539 | 19.04015 | -0.73073 | 0.560736 | 0.208104 |
| 7 | 140301200 | 140302399 | 6.930788 | 13.08996 | 31.40425 | 12.78431 | -0.77459 | 0.047867 | 0.20812 |
| 13 | 115664400 | 115667599 | 20.98007 | 35.03024 | 69.50261 | 30.23936 | -0.60405 | 1.208501 | 0.208132 |
| 14 | 65010000 | 65010799 | 3.031915 | 5.911513 | 24.23772 | 8.729941 | -1.3706 | -0.4353 | 0.208135 |
| 17 | 50914600 | 50917199 | 19.15074 | 32.29135 | 64.54375 | 28.22164 | -0.6144 | 1.100559 | 0.208153 |
| 11 | 77497400 | 77499799 | 10.62659 | 18.60977 | 49.91688 | 22.03603 | -1.02869 | 0.650617 | 0.208157 |
| 12 | 96635800 | 96636399 | 10.53388 | 20.38077 | 17.51346 | 6.682462 | 0.779677 | -0.16562 | 0.208173 |
| 8 | 12161600 | 12164799 | 18.18958 | 30.68347 | 62.81915 | 27.47104 | -0.64879 | 1.091801 | 0.208173 |
| 12 | 109149200 | 109150999 | 17.40313 | 32.28012 | 20.14719 | 8.455254 | 1.117768 | 0.222303 | 0.208175 |
| 3 | 98692600 | 98694599 | 12.89396 | 22.58665 | 51.58159 | 22.77618 | -0.79819 | 0.660599 | 0.208182 |
| 13 | 53595800 | 53596999 | 6.960881 | 13.05696 | 31.76054 | 13.08815 | -0.80877 | 0.094775 | 0.208189 |
| 7 | 140532400 | 140537599 | 23.75801 | 38.85904 | 83.47212 | 35.89735 | -0.72495 | 1.471101 | 0.208192 |
| 2 | 166985800 | 166995799 | 83.00485 | 127.2241 | 234.3076 | 97.25757 | -0.58787 | 3.017046 | 0.2082 |
| 4 | 54975000 | 54976999 | 10.4923 | 18.6953 | 42.52705 | 18.72088 | -0.78276 | 0.450414 | 0.208204 |
| 17 | 31283200 | 31287399 | 23.49554 | 38.28581 | 85.00025 | 36.50457 | -0.77618 | 1.528105 | 0.208218 |
| 4 | 46660600 | 46662199 | 18.74496 | 33.20785 | 26.19309 | 11.58105 | 0.73226 | 0.512748 | 0.208258 |
| 3 | 111871200 | 111873599 | 25.82666 | 44.8799 | 32.63759 | 14.33123 | 0.844215 | 0.813283 | 0.208263 |
| 8 | 45907800 | 45909999 | 24.69076 | 42.56989 | 35.1321 | 15.41167 | 0.660636 | 0.886714 | 0.208267 |
| 9 | 50278200 | 50279399 | 6.342967 | 12.0166 | 31.4926 | 12.7373 | -0.8927 | 0.020342 | 0.208273 |
| 18 | 39362000 | 39363999 | 9.544367 | 17.129 | 41.05784 | 17.96487 | -0.85156 | 0.395567 | 0.208282 |
| 15 | 82134200 | 82138199 | 23.06915 | 37.89602 | 78.37638 | 33.76616 | -0.66744 | 1.412697 | 0.208284 |
| 12 | 70033200 | 70034199 | 4.559088 | 8.771579 | 27.35738 | 10.65531 | -1.08892 | -0.1297 | 0.208285 |
| 3 | 107514600 | 107515799 | 5.279746 | 10.13049 | 27.88218 | 10.94494 | -0.92195 | -0.09548 | 0.208303 |
| 4 | 156338000 | 156342399 | 29.56486 | 47.6658 | 102.7174 | 43.73387 | -0.74825 | 1.776818 | 0.208304 |
| 1 | 102392600 | 102394799 | 30.50109 | 52.12313 | 37.5803 | 16.47723 | 0.852763 | 0.994188 | 0.208315 |

| | | | | | | | | | |
|---|---|---|---|---|---|---|---|---|---|
| 8 | 121212600 | 121217399 | 25.82539 | 41.82953 | 93.85942 | 40.16776 | -0.79901 | 1.64451 | 0.208322 |
| 15 | 31530200 | 31532199 | 10.6074 | 18.5961 | 45.17783 | 19.96204 | -0.88595 | 0.629536 | 0.20834 |
| 6 | 48907000 | 48910799 | 25.20447 | 41.16975 | 83.49063 | 35.88779 | -0.64357 | 1.49432 | 0.208359 |
| 8 | 118814600 | 118820399 | 49.43178 | 78.24954 | 154.0542 | 64.04694 | -0.64766 | 2.374071 | 0.208361 |
| 4 | 139774000 | 139801599 | 275.8509 | 385.473 | 668.5657 | 277.2505 | -0.62695 | 4.589162 | 0.208363 |
| 2 | 20971600 | 20975199 | 22.47863 | 36.8037 | 80.88503 | 34.79808 | -0.75676 | 1.453876 | 0.208363 |
| 5 | 59160200 | 59163999 | 91.59157 | 143.6372 | 119.2251 | 49.44832 | 0.586117 | 2.594243 | 0.208365 |
| 12 | 85469600 | 85473199 | 21.4945 | 35.30279 | 81.20827 | 34.98346 | -0.82061 | 1.410719 | 0.208381 |
| 2 | 158536400 | 158538399 | 11.89871 | 20.6562 | 51.02032 | 22.38909 | -0.91116 | 0.793601 | 0.208382 |
| 5 | 112632600 | 112633399 | 3.709612 | 7.225269 | 24.12568 | 8.809929 | -1.09765 | -0.37673 | 0.20839 |
| 19 | 34504400 | 34504999 | 3.709612 | 7.225269 | 24.12568 | 8.809929 | -1.09765 | -0.37673 | 0.20839 |
| 14 | 42222200 | 42224199 | 33.31922 | 55.79162 | 40.57586 | 17.68562 | 0.838779 | 1.186598 | 0.208396 |
| 10 | 72354800 | 72355199 | 2.937277 | 5.740742 | 19.92832 | 6.886245 | -1.07616 | -0.59721 | 0.208398 |
| 14 | 73663600 | 73664599 | 13.44154 | 25.03983 | 20.28274 | 8.456471 | 0.75175 | 0.164012 | 0.208415 |
| 5 | 109513800 | 109515599 | 32.44002 | 54.78353 | 41.44383 | 18.1297 | 0.782269 | 1.107083 | 0.208425 |
| 16 | 80932000 | 80933999 | 23.87245 | 41.87384 | 28.71518 | 12.6937 | 0.928866 | 0.662778 | 0.208445 |
| 7 | 37265400 | 37269399 | 28.12651 | 45.3504 | 100.9097 | 42.96859 | -0.79413 | 1.772956 | 0.208475 |
| 7 | 61125000 | 61128399 | 41.81737 | 68.61961 | 53.90015 | 23.21065 | 0.724504 | 1.450189 | 0.208476 |
| 4 | 15265200 | 15266999 | 14.10754 | 24.30854 | 53.70641 | 23.55937 | -0.75385 | 0.87431 | 0.208477 |
| 14 | 40526600 | 40529199 | 24.19666 | 42.40425 | 29.42249 | 12.99645 | 0.912253 | 0.681167 | 0.208481 |
| 1 | 174874400 | 174879199 | 56.41962 | 90.50567 | 79.0202 | 33.39887 | 0.543519 | 1.960782 | 0.208484 |
| 2 | 178728000 | 178731399 | 19.07054 | 31.51473 | 81.46466 | 35.19859 | -0.98542 | 1.327831 | 0.208487 |
| 2 | 56399000 | 56399999 | 13.31117 | 25.1283 | 20.21931 | 8.239433 | 0.787537 | 0.079417 | 0.208494 |
| X | 65922000 | 65924199 | 22.07312 | 39.07273 | 26.12992 | 11.55496 | 0.968529 | 0.541978 | 0.208498 |
| 15 | 36561400 | 36562199 | 4.523545 | 8.589798 | 32.54915 | 13.09972 | -1.4121 | -0.0101 | 0.208506 |
| 6 | 90521000 | 90522199 | 5.315974 | 10.16143 | 29.44649 | 11.6854 | -1.01115 | -0.0568 | 0.208516 |
| 11 | 69648800 | 69652599 | 15.17261 | 25.74945 | 60.53398 | 26.52031 | -0.84685 | 1.025628 | 0.208538 |
| 5 | 137572200 | 137573599 | 8.964201 | 16.05735 | 38.45607 | 16.85336 | -0.85121 | 0.401828 | 0.208561 |
| 1 | 53213400 | 53215999 | 33.47688 | 55.81762 | 47.06985 | 20.46113 | 0.626253 | 1.238611 | 0.208577 |
| 18 | 19198600 | 19199199 | 10.8801 | 21.09626 | 16.94974 | 6.402267 | 0.889419 | -0.19646 | 0.208579 |
| 4 | 104222600 | 104226199 | 22.55725 | 37.14198 | 76.90241 | 33.16398 | -0.66771 | 1.379844 | 0.208583 |
| 4 | 148083000 | 148086599 | 26.08928 | 42.37334 | 89.87286 | 38.53976 | -0.71425 | 1.586581 | 0.208592 |
| 17 | 47502600 | 47507999 | 48.638 | 76.97335 | 153.2695 | 63.71246 | -0.66427 | 2.379817 | 0.208614 |
| 2 | 171660400 | 171661199 | 4.281481 | 8.324666 | 27.05197 | 10.02979 | -1.08122 | -0.30844 | 0.208621 |
| 6 | 3250800 | 3260999 | 107.0229 | 166.2605 | 140.0741 | 58.16242 | 0.554151 | 2.766819 | 0.208623 |
| 14 | 68320000 | 68322799 | 18.53488 | 30.75957 | 73.48515 | 31.84387 | -0.87133 | 1.272236 | 0.20863 |
| X | 132292400 | 132294599 | 23.10154 | 41.60747 | 20.56327 | 8.967324 | 1.415445 | 0.427611 | 0.208634 |
| 14 | 48397000 | 48403599 | 46.86673 | 74.16923 | 152.9909 | 63.59625 | -0.71509 | 2.37913 | 0.208641 |
| 7 | 20845800 | 20846999 | 37.23021 | 61.55199 | 51.00017 | 22.0426 | 0.651031 | 1.348146 | 0.208663 |
| 16 | 58490200 | 58491399 | 6.748565 | 12.12507 | 39.04093 | 17.0662 | -1.27187 | 0.368408 | 0.208669 |
| 14 | 122727600 | 122728799 | 13.2269 | 23.05773 | 51.61846 | 22.71091 | -0.76996 | 0.736415 | 0.208671 |
| 6 | 97536400 | 97537399 | 4.988938 | 9.650058 | 27.74936 | 10.60068 | -0.94792 | -0.19298 | 0.208672 |
| 6 | 137550200 | 137552599 | 10.9977 | 19.45297 | 42.45818 | 18.77833 | -0.72893 | 0.51176 | 0.208685 |
| 13 | 46776000 | 46777199 | 11.76279 | 22.74486 | 16.07184 | 6.149531 | 1.054353 | -0.14863 | 0.208692 |
| 10 | 56261600 | 56265599 | 35.59378 | 57.07024 | 114.6888 | 48.43901 | -0.65761 | 1.963174 | 0.208692 |
| 5 | 143404000 | 143405599 | 5.448272 | 10.41391 | 28.8821 | 11.46272 | -0.94836 | -0.05574 | 0.208694 |
| 5 | 145125600 | 145128399 | 11.06473 | 19.39275 | 46.20606 | 20.41182 | -0.85845 | 0.635609 | 0.208695 |

| | | | | | | | | | |
|---|---|---|---|---|---|---|---|---|---|
| 4 | 96236400 | 96237799 | 11.81654 | 22.42886 | 17.89345 | 7.207961 | 0.814307 | 0.022105 | 0.2087 |
| 9 | 35566400 | 35570599 | 74.37803 | 118.3367 | 96.05375 | 40.12353 | 0.635557 | 2.238216 | 0.20872 |
| 16 | 94090600 | 94093399 | 14.22383 | 24.62346 | 52.14824 | 22.8791 | -0.69166 | 0.830147 | 0.208724 |
| 6 | 57050200 | 57055799 | 105.5819 | 163.6577 | 143.8284 | 59.74113 | 0.490813 | 2.803811 | 0.208727 |
| 16 | 32617400 | 32625599 | 52.52129 | 83.13258 | 152.3994 | 63.35604 | -0.54502 | 2.378632 | 0.208734 |
| 4 | 114762400 | 114771199 | 60.23984 | 94.1504 | 184.2215 | 76.4321 | -0.6533 | 2.645179 | 0.208746 |
| 17 | 84407200 | 84413199 | 32.96941 | 53.01178 | 108.3139 | 45.93999 | -0.67649 | 1.86841 | 0.208747 |
| 16 | 17751200 | 17752599 | 7.012236 | 13.00131 | 33.13811 | 13.91493 | -0.89706 | 0.17276 | 0.208748 |
| 1 | 24138600 | 24140399 | 11.75319 | 20.40681 | 51.56268 | 22.62934 | -0.94398 | 0.789075 | 0.208774 |
| 6 | 79607600 | 79609199 | 32.72319 | 57.42106 | 19.05807 | 8.427501 | 1.968931 | 0.693507 | 0.20878 |
| 14 | 63201800 | 63203799 | 11.67833 | 20.44967 | 46.22244 | 20.40346 | -0.78286 | 0.654192 | 0.208794 |
| 6 | 72574200 | 72575999 | 6.761883 | 12.44968 | 34.62288 | 14.67752 | -1.03198 | 0.21358 | 0.208804 |
| 12 | 99017600 | 99019399 | 7.719893 | 14.39233 | 33.27432 | 13.85481 | -0.74856 | 0.143942 | 0.208807 |
| 1 | 102354600 | 102357199 | 27.57285 | 47.96522 | 30.86884 | 13.56361 | 1.019808 | 0.805455 | 0.208809 |
| 7 | 50096200 | 50096799 | 11.70517 | 22.48235 | 18.35106 | 7.182329 | 0.818664 | -0.06944 | 0.208829 |
| 16 | 91341200 | 91344399 | 15.39975 | 26.32993 | 58.55125 | 25.67307 | -0.76555 | 0.954719 | 0.208831 |
| 11 | 58636800 | 58640599 | 36.6232 | 60.71149 | 49.34636 | 21.36745 | 0.678873 | 1.313506 | 0.208848 |
| 19 | 47483000 | 47484799 | 7.753168 | 14.35373 | 33.6564 | 14.16116 | -0.78017 | 0.187739 | 0.208848 |
| 5 | 120237400 | 120244999 | 57.67728 | 90.06427 | 185.8363 | 77.10919 | -0.73079 | 2.660462 | 0.208855 |
| 12 | 20475200 | 20477199 | 77.15709 | 122.1051 | 95.53296 | 39.7117 | 0.681396 | 2.388751 | 0.208856 |
| 7 | 39258200 | 39258799 | 10.69993 | 20.74959 | 16.95523 | 6.400469 | 0.866006 | -0.20071 | 0.208863 |
| 19 | 13682000 | 13683199 | 11.62384 | 20.52603 | 43.49867 | 19.24497 | -0.68769 | 0.536098 | 0.208866 |
| 5 | 144601000 | 144602199 | 7.911235 | 14.60317 | 34.68222 | 14.65154 | -0.80298 | 0.207144 | 0.208877 |
| 6 | 124713400 | 124718199 | 22.34924 | 36.53438 | 83.03 | 35.69994 | -0.80652 | 1.476805 | 0.20889 |
| 19 | 45130200 | 45137399 | 65.02264 | 101.4528 | 185.8088 | 77.10416 | -0.55996 | 2.67835 | 0.208896 |
| 7 | 130307600 | 130308799 | 6.9215 | 13.10035 | 31.48834 | 12.76082 | -0.77161 | 0.032758 | 0.208901 |
| X | 20157600 | 20158799 | 19.58451 | 35.22028 | 23.0385 | 10.063 | 1.012883 | 0.41813 | 0.208904 |
| 8 | 124238800 | 124244399 | 26.74786 | 43.07012 | 107.3004 | 45.60527 | -0.95912 | 1.813755 | 0.208904 |
| 17 | 90201200 | 90202999 | 21.07509 | 37.33851 | 26.05573 | 11.52015 | 0.907723 | 0.524975 | 0.208924 |
| 13 | 101370400 | 101371999 | 8.868304 | 16.13775 | 38.69696 | 16.65921 | -0.8365 | 0.303807 | 0.208928 |
| 17 | 94299800 | 94300599 | 14.86607 | 27.79717 | 21.12613 | 8.752576 | 0.850358 | 0.155821 | 0.208938 |
| 14 | 54781000 | 54792399 | 70.77495 | 109.1401 | 219.4777 | 91.16719 | -0.70785 | 2.89276 | 0.208956 |
| 14 | 24910400 | 24912599 | 10.38457 | 18.40012 | 40.92121 | 18.09233 | -0.7546 | 0.487139 | 0.208957 |
| 11 | 85243400 | 85244399 | 6.171116 | 11.71947 | 30.37989 | 12.22429 | -0.86922 | 0.003624 | 0.208964 |
| Y | 10553200 | 10555599 | 14.27447 | 26.77509 | 12.33514 | 5.083305 | 1.571353 | 0.105018 | 0.208965 |
| 11 | 27083800 | 27085399 | 35.9679 | 59.29245 | 47.74583 | 20.60041 | 0.691282 | 1.375196 | 0.208975 |
| 12 | 36410000 | 36413599 | 33.50912 | 56.39308 | 42.56798 | 18.60189 | 0.785332 | 1.141841 | 0.208988 |
| 5 | 33020800 | 33021599 | 5.174539 | 9.848554 | 29.29558 | 11.7332 | -1.0596 | -0.02603 | 0.208993 |
| 10 | 95301800 | 95303199 | 6.333502 | 12.02477 | 31.58364 | 12.71555 | -0.89009 | 0.004945 | 0.209005 |
| 7 | 15202200 | 15203799 | 14.61843 | 27.5406 | 20.17815 | 8.253971 | 0.9175 | 0.105365 | 0.209024 |
| 5 | 29675800 | 29685199 | 76.01031 | 117.1434 | 217.6883 | 90.42418 | -0.59474 | 2.905838 | 0.209027 |
| 1 | 100044800 | 100046999 | 19.25695 | 34.58263 | 23.46183 | 10.26004 | 0.958609 | 0.421328 | 0.209032 |
| 9 | 121883200 | 121884799 | 9.344969 | 16.91887 | 39.10926 | 16.94415 | -0.78998 | 0.338126 | 0.209042 |
| 5 | 73965800 | 73968799 | 27.94404 | 47.84802 | 38.19815 | 16.7508 | 0.706726 | 0.966357 | 0.209044 |
| 14 | 17995800 | 17997599 | 11.31864 | 19.99591 | 42.46427 | 18.78566 | -0.6903 | 0.528237 | 0.209046 |
| 4 | 139808200 | 139814199 | 27.70776 | 44.61169 | 110.7996 | 47.08628 | -0.95496 | 1.818385 | 0.209055 |
| 15 | 96529000 | 96537799 | 96.76772 | 146.5797 | 291.0471 | 120.4312 | -0.71011 | 3.264831 | 0.209058 |

| | | | | | | | | | |
|---|---|---|---|---|---|---|---|---|---|
| 10 | 40316800 | 40317399 | 3.415904 | 6.673416 | 22.39258 | 7.814637 | -1.04456 | -0.5504 | 0.209063 |
| 11 | 74820400 | 74821999 | 6.513227 | 12.25585 | 31.27299 | 12.8186 | -0.8708 | 0.069122 | 0.209068 |
| 2 | 98052600 | 98053999 | 23.2405 | 41.28537 | 23.10515 | 10.19707 | 1.225939 | 0.514322 | 0.209086 |
| 3 | 54749200 | 54750399 | 12.78589 | 24.26347 | 19.87696 | 8.010678 | 0.776083 | 0.039446 | 0.209094 |
| 7 | 8398000 | 8405599 | 118.0993 | 183.4818 | 131.2292 | 54.48913 | 0.790263 | 2.769618 | 0.209096 |
| 4 | 60107400 | 60107999 | 9.129691 | 17.7921 | 14.91281 | 5.405492 | 0.882232 | -0.36598 | 0.209115 |
| 16 | 11499400 | 11500199 | 9.129691 | 17.7921 | 14.91281 | 5.405492 | 0.882232 | -0.36598 | 0.209115 |
| 7 | 19730000 | 19734999 | 65.02399 | 103.9631 | 79.55778 | 33.46151 | 0.727924 | 2.071365 | 0.209136 |
| 1 | 34258600 | 34261399 | 13.10762 | 22.78526 | 50.53686 | 22.19554 | -0.75687 | 0.776583 | 0.209156 |
| 1 | 31559600 | 31562999 | 42.25336 | 68.96311 | 57.18134 | 24.57184 | 0.642367 | 1.523043 | 0.20916 |
| 15 | 34798000 | 34799199 | 7.693409 | 14.4233 | 35.76968 | 14.75362 | -0.83845 | 0.111794 | 0.209165 |
| X | 89419400 | 89421599 | 36.88826 | 62.03267 | 39.86026 | 17.41462 | 1.01687 | 1.159337 | 0.209177 |
| 3 | 14291800 | 14302399 | 225.5713 | 331.9019 | 279.2928 | 115.0139 | 0.48968 | 3.814409 | 0.209184 |
| 11 | 70350000 | 70356999 | 40.95886 | 65.07424 | 142.2143 | 59.31141 | -0.793 | 2.26387 | 0.209186 |
| 19 | 42713400 | 42722599 | 49.19563 | 77.32813 | 175.3726 | 72.74799 | -0.85948 | 2.532512 | 0.209189 |
| 19 | 34115000 | 34116399 | 19.00556 | 33.58903 | 27.60452 | 12.21094 | 0.671953 | 0.54188 | 0.209198 |
| 14 | 32692600 | 32694599 | 8.312906 | 14.67961 | 42.56409 | 18.83143 | -1.13623 | 0.516332 | 0.209201 |
| 6 | 84379600 | 84380599 | 8.554035 | 15.71909 | 36.20694 | 15.38922 | -0.76532 | 0.243848 | 0.209219 |
| 10 | 124764400 | 124767199 | 30.63716 | 51.7189 | 42.59929 | 18.63297 | 0.659997 | 1.105263 | 0.209239 |
| 14 | 41986600 | 41989999 | 35.50269 | 60.3669 | 35.21284 | 15.43297 | 1.156174 | 1.055188 | 0.209261 |
| 10 | 121575400 | 121576599 | 8.54369 | 15.22812 | 39.48312 | 17.37622 | -0.96904 | 0.433347 | 0.209267 |
| 1 | 74049400 | 74050599 | 15.34207 | 28.06485 | 23.17012 | 9.909319 | 0.698103 | 0.288941 | 0.209274 |
| 5 | 140870600 | 140879999 | 74.67103 | 115.1412 | 215.5568 | 89.53853 | -0.6048 | 2.895336 | 0.209278 |
| 17 | 82831400 | 82833399 | 22.17935 | 39.30163 | 25.48437 | 11.26715 | 1.013217 | 0.530011 | 0.209284 |
| 4 | 41676000 | 41678199 | 9.62546 | 17.0682 | 40.44044 | 17.8749 | -0.84466 | 0.474409 | 0.2093 |
| 11 | 43419200 | 43421199 | 9.62546 | 17.0682 | 40.44044 | 17.8749 | -0.84466 | 0.474409 | 0.2093 |
| 8 | 27532000 | 27533199 | 5.691378 | 10.97483 | 27.5644 | 10.66655 | -0.77222 | -0.14001 | 0.2093 |
| 6 | 109910200 | 109910799 | 11.20432 | 21.42494 | 17.92484 | 7.105595 | 0.766478 | -0.03514 | 0.209307 |
| 16 | 15297600 | 15298799 | 9.310472 | 16.54673 | 39.95324 | 17.62619 | -0.86973 | 0.457446 | 0.209308 |
| 5 | 77428800 | 77430799 | 9.033592 | 16.00846 | 40.97421 | 18.11529 | -0.95569 | 0.477643 | 0.209316 |
| 13 | 3362400 | 3364799 | 41.47908 | 67.71165 | 57.66149 | 24.7798 | 0.604187 | 1.519706 | 0.209325 |
| 3 | 35246200 | 35247999 | 17.38632 | 31.62919 | 23.28664 | 10.02852 | 0.855655 | 0.334918 | 0.209343 |
| 14 | 68706000 | 68707799 | 20.11393 | 35.92231 | 25.1065 | 11.03061 | 0.911816 | 0.465383 | 0.209367 |
| 16 | 92034200 | 92035399 | 9.039502 | 16.42293 | 36.52125 | 15.75412 | -0.72991 | 0.312426 | 0.209367 |
| 1 | 171910000 | 171910999 | 5.577539 | 10.67857 | 28.38824 | 11.21355 | -0.8813 | -0.07093 | 0.209372 |
| X | 6689400 | 6692199 | 38.59335 | 63.92454 | 45.04894 | 19.49548 | 0.884143 | 1.329244 | 0.209372 |
| 17 | 39429400 | 39430799 | 27.11847 | 47.25962 | 29.13317 | 12.81543 | 1.081603 | 0.775301 | 0.209406 |
| 7 | 88320600 | 88321599 | 8.348147 | 15.42545 | 35.81702 | 15.10986 | -0.76947 | 0.206656 | 0.209414 |
| 8 | 23778400 | 23779999 | 6.775427 | 12.82456 | 32.06313 | 12.99228 | -0.82793 | 0.031729 | 0.209416 |
| 10 | 115588600 | 115588999 | 10.00938 | 19.46515 | 16.02865 | 5.934194 | 0.881156 | -0.27657 | 0.209423 |
| 19 | 5167000 | 5169799 | 14.42475 | 24.82499 | 54.24426 | 23.79443 | -0.7384 | 0.887134 | 0.209426 |
| 1 | 44417600 | 44420999 | 41.98143 | 68.39412 | 58.72265 | 25.21735 | 0.590368 | 1.549845 | 0.209426 |
| 16 | 87084000 | 87084999 | 12.19215 | 23.06325 | 19.37813 | 7.862364 | 0.731208 | 0.053124 | 0.20943 |
| 5 | 108410400 | 108411599 | 5.901674 | 11.19365 | 29.79954 | 12.0228 | -0.91105 | 0.006065 | 0.209432 |
| 8 | 34089400 | 34090199 | 5.901674 | 11.19365 | 29.79954 | 12.0228 | -0.91105 | 0.006065 | 0.209432 |
| 9 | 107749400 | 107750399 | 5.901674 | 11.19365 | 29.79954 | 12.0228 | -0.91105 | 0.006065 | 0.209432 |
| 16 | 49284800 | 49287799 | 12.48889 | 21.772 | 52.64699 | 23.16397 | -0.88034 | 0.736753 | 0.209433 |

| | | | | | | | | | |
|---|---|---|---|---|---|---|---|---|---|
| 17 | 89789800 | 89791599 | 27.94335 | 48.599 | 30.86993 | 13.56221 | 1.038612 | 0.810566 | 0.209461 |
| 15 | 36893800 | 36897599 | 20.63166 | 34.55988 | 69.02761 | 30.09393 | -0.61368 | 1.172062 | 0.209476 |
| 16 | 8580800 | 8582999 | 14.20916 | 24.6813 | 51.08994 | 22.42772 | -0.6579 | 0.794307 | 0.209481 |
| 11 | 120120200 | 120122599 | 9.568997 | 17.12418 | 38.93624 | 17.07768 | -0.77792 | 0.411974 | 0.209485 |
| 12 | 105660400 | 105663399 | 19.98194 | 33.45079 | 69.00203 | 30.071 | -0.66005 | 1.175451 | 0.209485 |
| 10 | 50407800 | 50408999 | 24.93709 | 42.93417 | 36.21775 | 15.88724 | 0.628826 | 0.902422 | 0.209488 |
| 4 | 137205800 | 137218399 | 61.12668 | 94.26257 | 220.6238 | 91.64324 | -0.92646 | 2.889928 | 0.209508 |
| 15 | 18151600 | 18152799 | 16.80366 | 31.68071 | 15.66547 | 6.40009 | 1.479413 | 0.107533 | 0.209508 |
| 2 | 36628200 | 36634399 | 104.2088 | 162.547 | 132.1493 | 54.83895 | 0.610039 | 2.695439 | 0.209508 |
| 7 | 47016000 | 47022799 | 174.2592 | 261.8717 | 219.873 | 90.76713 | 0.520784 | 3.433779 | 0.209535 |
| 19 | 55274800 | 55276399 | 7.079366 | 13.39345 | 31.45335 | 12.7581 | -0.73913 | 0.039163 | 0.209535 |
| 4 | 99544000 | 99546599 | 22.59071 | 37.10733 | 78.37712 | 33.76541 | -0.69776 | 1.411508 | 0.209536 |
| 5 | 80997400 | 80998399 | 15.84942 | 27.18934 | 55.8801 | 24.50843 | -0.65146 | 0.924173 | 0.209541 |
| 7 | 43417600 | 43428399 | 79.57662 | 122.3288 | 225.4591 | 93.62241 | -0.58569 | 2.95731 | 0.209546 |
| 2 | 167336800 | 167339599 | 13.40563 | 23.34427 | 49.46693 | 21.74963 | -0.69106 | 0.751576 | 0.209555 |
| 19 | 25641400 | 25652399 | 79.28804 | 121.5511 | 241.661 | 100.3126 | -0.69794 | 3.011917 | 0.209564 |
| 3 | 154052200 | 154055799 | 75.06981 | 119.0947 | 104.1564 | 43.37565 | 0.524057 | 2.32494 | 0.209573 |
| 11 | 77279800 | 77282199 | 33.34106 | 56.40194 | 39.87708 | 17.45341 | 0.879693 | 1.096764 | 0.209581 |
| 1 | 121965000 | 121965799 | 6.705188 | 12.46959 | 32.64813 | 13.65136 | -0.93073 | 0.151959 | 0.209592 |
| 7 | 24918200 | 24921199 | 17.01206 | 28.50635 | 66.41217 | 28.95984 | -0.83471 | 1.154843 | 0.209592 |
| X | 33808200 | 33811799 | 34.97253 | 59.30839 | 37.5593 | 16.4513 | 1.037987 | 1.079172 | 0.209605 |
| 2 | 115217400 | 115219999 | 12.89746 | 22.62831 | 48.3007 | 21.35581 | -0.701 | 0.631556 | 0.209606 |
| 14 | 71093400 | 71094199 | 4.527421 | 8.80366 | 24.60269 | 9.11497 | -0.86401 | -0.31264 | 0.209629 |
| 17 | 48405800 | 48410399 | 30.5108 | 48.88865 | 119.4486 | 50.39969 | -0.93995 | 1.979059 | 0.209645 |
| 4 | 84695600 | 84696199 | 3.93369 | 7.684044 | 22.43915 | 7.847337 | -0.84987 | -0.53416 | 0.209649 |
| 7 | 60043800 | 60046399 | 24.38341 | 42.15237 | 35.2114 | 15.44771 | 0.644136 | 0.8639 | 0.209658 |
| 4 | 71009600 | 71010799 | 12.83065 | 24.36089 | 18.59044 | 7.48408 | 0.878061 | 0.030818 | 0.209658 |
| 12 | 3385800 | 3388799 | 47.44989 | 76.7166 | 63.98094 | 27.34022 | 0.622829 | 1.706974 | 0.20967 |
| 9 | 76096200 | 76096999 | 7.41253 | 13.88891 | 32.28743 | 13.32999 | -0.74603 | 0.108305 | 0.209684 |
| 7 | 47382000 | 47384199 | 53.85918 | 86.71628 | 72.61998 | 30.86051 | 0.60993 | 1.842382 | 0.209707 |
| 3 | 131195400 | 131196799 | 8.305525 | 15.04504 | 36.50345 | 15.80458 | -0.85786 | 0.325374 | 0.209711 |
| 6 | 131675600 | 131676599 | 20.59676 | 36.24591 | 28.97331 | 12.82739 | 0.709411 | 0.599326 | 0.209724 |
| 5 | 25702800 | 25705199 | 17.29781 | 28.70849 | 73.85505 | 32.00561 | -0.97765 | 1.268675 | 0.209727 |
| 8 | 47839600 | 47846199 | 41.39717 | 65.8633 | 139.66 | 58.33839 | -0.74751 | 2.219855 | 0.20973 |
| 12 | 115577200 | 115582199 | 101.1681 | 158.1726 | 128.2567 | 53.21083 | 0.616408 | 2.65283 | 0.209761 |
| 7 | 93038800 | 93039599 | 15.43336 | 28.56342 | 22.01899 | 9.268581 | 0.812441 | 0.222352 | 0.209786 |
| 3 | 85530200 | 85532399 | 26.1988 | 44.93837 | 37.19962 | 16.31522 | 0.654934 | 0.941445 | 0.209795 |
| 8 | 85386600 | 85391999 | 21.81211 | 35.50473 | 90.5761 | 38.88404 | -0.97722 | 1.541495 | 0.209797 |
| 2 | 30600600 | 30602399 | 8.331165 | 15.02681 | 37.48424 | 16.31664 | -0.90288 | 0.35179 | 0.20981 |
| 9 | 96994400 | 96996599 | 8.331165 | 15.02681 | 37.48424 | 16.31664 | -0.90288 | 0.35179 | 0.20981 |
| 10 | 32939800 | 32940199 | 10.72306 | 20.5935 | 16.94608 | 6.634746 | 0.806759 | -0.08275 | 0.209812 |
| 2 | 29666000 | 29674799 | 46.46754 | 73.75038 | 143.271 | 59.69064 | -0.62492 | 2.299103 | 0.209813 |
| 7 | 24849000 | 24866399 | 98.97031 | 149.486 | 292.8533 | 121.0688 | -0.69427 | 3.322819 | 0.209823 |
| X | 147535800 | 147545599 | 156.0384 | 237.2807 | 180.1343 | 74.62483 | 0.680485 | 3.193498 | 0.209836 |
| 8 | 29163600 | 29166799 | 17.12628 | 29.28162 | 59.18274 | 25.94994 | -0.62839 | 0.962955 | 0.209844 |
| 7 | 47110400 | 47113999 | 36.26008 | 60.43795 | 45.97127 | 19.97936 | 0.774559 | 1.24939 | 0.209861 |
| 10 | 71042600 | 71044399 | 10.15367 | 18.2018 | 41.56857 | 18.20578 | -0.78325 | 0.408049 | 0.209873 |

| | | | | | | | | | |
|---|---|---|---|---|---|---|---|---|---|
| 13 | 66736200 | 66740399 | 125.9247 | 193.7548 | 156.7143 | 65.08418 | 0.60236 | 2.949333 | 0.209873 |
| 11 | 45537600 | 45538999 | 15.66693 | 29.12787 | 21.52016 | 9.002444 | 0.879784 | 0.198817 | 0.209877 |
| 13 | 36174600 | 36176199 | 14.18889 | 24.66241 | 51.09692 | 22.44011 | -0.65917 | 0.783709 | 0.209879 |
| 7 | 31195200 | 31210799 | 267.7268 | 392.1713 | 278.1501 | 114.5753 | 0.730066 | 3.888421 | 0.209882 |
| 11 | 100417800 | 100420199 | 9.725401 | 17.04484 | 47.25435 | 20.87439 | -1.07556 | 0.630515 | 0.209894 |
| 2 | 124076600 | 124077799 | 7.692311 | 14.42459 | 35.19653 | 14.5115 | -0.81467 | 0.109165 | 0.2099 |
| X | 25426000 | 25431599 | 97.32871 | 153.1918 | 112.41 | 46.63802 | 0.767727 | 2.526235 | 0.209902 |
| 4 | 119928400 | 119931599 | 17.5291 | 29.63445 | 62.8562 | 27.50559 | -0.69953 | 1.06911 | 0.209903 |
| 12 | 7946200 | 7947799 | 12.15526 | 21.253 | 46.77554 | 20.62153 | -0.74461 | 0.683199 | 0.209921 |
| 6 | 69909800 | 69912199 | 35.84713 | 59.79571 | 42.24466 | 18.36868 | 0.880759 | 1.238353 | 0.209952 |
| X | 19537400 | 19538799 | 17.03847 | 31.38029 | 21.58917 | 9.148312 | 0.969184 | 0.263089 | 0.209959 |
| 5 | 64852200 | 64856399 | 28.61727 | 46.53757 | 91.30782 | 39.18459 | -0.60071 | 1.569252 | 0.209994 |
| 10 | 15795000 | 15796999 | 24.6064 | 43.17319 | 28.14229 | 12.44335 | 1.00167 | 0.662204 | 0.210007 |
| Y | 5545400 | 5547399 | 14.65891 | 27.50789 | 10.15688 | 4.182633 | 1.886665 | 0.098717 | 0.210022 |
| 2 | 174122400 | 174143599 | 479.1855 | 636.5349 | 1134.392 | 474.0686 | -0.73978 | 5.384545 | 0.210062 |
| 13 | 101270200 | 101271399 | 17.37055 | 31.88861 | 22.37652 | 9.523754 | 0.936045 | 0.283762 | 0.210097 |
| 14 | 101006600 | 101007799 | 12.03569 | 23.2371 | 17.23819 | 6.642943 | 0.97587 | -0.1181 | 0.2101 |
| 13 | 51353000 | 51360399 | 58.67365 | 91.96379 | 191.0348 | 79.2357 | -0.7363 | 2.594975 | 0.210114 |
| 5 | 105625600 | 105630799 | 50.61136 | 80.02093 | 154.3249 | 64.12061 | -0.61932 | 2.404182 | 0.210114 |
| 13 | 68145200 | 68156599 | 224.7093 | 333.0154 | 251.8022 | 103.7056 | 0.653584 | 3.695268 | 0.210129 |
| 7 | 32303000 | 32306399 | 77.81277 | 123.1284 | 106.4488 | 44.24663 | 0.537452 | 2.395257 | 0.210133 |
| 8 | 118388400 | 118393199 | 38.54938 | 61.65371 | 120.5193 | 50.71223 | -0.62203 | 2.047655 | 0.210153 |
| 7 | 5141600 | 5142799 | 14.48165 | 27.48073 | 19.16957 | 7.726031 | 1.006098 | 0.054006 | 0.210157 |
| 12 | 20120800 | 20126799 | 199.8833 | 296.7686 | 235.0759 | 96.83492 | 0.588961 | 3.659932 | 0.210185 |
| 18 | 30610200 | 30611599 | 11.35407 | 19.9355 | 44.59956 | 19.73196 | -0.768 | 0.605828 | 0.210207 |
| 1 | 163230800 | 163231599 | 5.556689 | 10.39537 | 33.2853 | 13.77835 | -1.20673 | 0.102907 | 0.210221 |
| 9 | 108673000 | 108674999 | 7.505527 | 13.76806 | 35.09777 | 14.95358 | -0.91303 | 0.241237 | 0.210228 |
| 18 | 32551800 | 32553399 | 7.505527 | 13.76806 | 35.09777 | 14.95358 | -0.91303 | 0.241237 | 0.210228 |
| 1 | 137453200 | 137455399 | 33.80891 | 56.57786 | 41.16381 | 17.93515 | 0.838339 | 1.196771 | 0.210232 |
| 7 | 100566600 | 100570999 | 21.809 | 35.80244 | 78.72704 | 33.90781 | -0.75594 | 1.417612 | 0.210237 |
| 12 | 24619800 | 24623599 | 21.809 | 35.80244 | 78.72704 | 33.90781 | -0.75594 | 1.417612 | 0.210237 |
| 5 | 142295600 | 142307199 | 86.95934 | 133.159 | 235.8109 | 97.86258 | -0.53245 | 3.037787 | 0.210245 |
| 3 | 43332800 | 43334599 | 18.35426 | 33.66084 | 21.02416 | 8.960347 | 1.101036 | 0.292815 | 0.210247 |
| 10 | 48622200 | 48622999 | 11.02928 | 21.20282 | 18.03085 | 7.039175 | 0.764045 | -0.08402 | 0.210252 |
| 7 | 88414200 | 88415799 | 9.491865 | 17.3087 | 40.41344 | 17.3526 | -0.79582 | 0.301542 | 0.21026 |
| 10 | 119301400 | 119304399 | 34.91942 | 56.08446 | 109.9351 | 46.55172 | -0.6188 | 1.907562 | 0.210285 |
| 11 | 30919600 | 30922599 | 22.72011 | 39.84384 | 26.83703 | 11.86125 | 0.954489 | 0.655691 | 0.210287 |
| 9 | 38719200 | 38721399 | 22.39038 | 39.4294 | 27.64688 | 12.23845 | 0.897755 | 0.600135 | 0.21029 |
| 12 | 50925800 | 50926799 | 18.56478 | 32.8937 | 26.19323 | 11.58075 | 0.718676 | 0.509768 | 0.210301 |
| 7 | 11091200 | 11093999 | 24.75392 | 43.65867 | 24.69535 | 10.92854 | 1.206546 | 0.593265 | 0.210316 |
| 1 | 60597400 | 60598799 | 16.19923 | 30.90255 | 16.8017 | 6.696601 | 1.376154 | 0.02944 | 0.210329 |
| 14 | 14524200 | 14528399 | 21.2454 | 34.76653 | 84.82423 | 36.48539 | -0.90755 | 1.459069 | 0.210344 |
| 8 | 90688000 | 90689199 | 14.70368 | 27.00854 | 21.62994 | 9.198705 | 0.747207 | 0.256347 | 0.210351 |
| 13 | 88948600 | 88949399 | 6.596684 | 12.60828 | 30.10375 | 11.94955 | -0.73512 | -0.04041 | 0.210358 |
| 7 | 55393000 | 55395199 | 14.36319 | 24.85322 | 54.33134 | 23.83656 | -0.73757 | 0.835893 | 0.21037 |
| 6 | 7692200 | 7693999 | 9.451751 | 16.82445 | 39.94231 | 17.59785 | -0.84431 | 0.447171 | 0.210378 |
| 1 | 112344200 | 112345199 | 8.56093 | 15.63772 | 35.55425 | 15.2337 | -0.75517 | 0.275139 | 0.210389 |

| | | | | | | | | | |
|---|---|---|---|---|---|---|---|---|---|
| 2 | 94267600 | 94268999 | 9.105676 | 15.96565 | 45.70316 | 20.19658 | -1.12095 | 0.61971 | 0.210403 |
| 8 | 118220400 | 118223999 | 19.81559 | 32.80024 | 77.02122 | 33.31678 | -0.84703 | 1.307662 | 0.210409 |
| 6 | 79660000 | 79662399 | 19.73094 | 35.61725 | 22.97393 | 9.990277 | 1.036384 | 0.394962 | 0.210412 |
| 6 | 89257600 | 89260799 | 27.25436 | 44.12932 | 92.83784 | 39.72079 | -0.70672 | 1.653875 | 0.210429 |
| 1 | 178689200 | 178690599 | 18.87692 | 33.6368 | 26.42927 | 11.63396 | 0.742216 | 0.474249 | 0.210432 |
| 17 | 17137400 | 17142599 | 25.54853 | 44.16511 | 29.01102 | 12.72751 | 0.989074 | 0.858862 | 0.21045 |
| 2 | 19630000 | 19631399 | 22.944 | 40.38602 | 27.64249 | 12.23762 | 0.932204 | 0.608694 | 0.210462 |
| 14 | 117940800 | 117941999 | 6.766712 | 12.83302 | 32.74096 | 13.21245 | -0.85183 | 0.019184 | 0.210468 |
| X | 12096000 | 12100599 | 107.2792 | 166.4409 | 140.4777 | 58.34141 | 0.550112 | 2.789795 | 0.21047 |
| 4 | 17202800 | 17203999 | 9.492983 | 17.30655 | 40.96349 | 17.59411 | -0.8157 | 0.303916 | 0.210475 |
| 2 | 174982800 | 174984199 | 8.892089 | 16.11819 | 38.06673 | 16.46784 | -0.81915 | 0.326101 | 0.210475 |
| 15 | 59180200 | 59181999 | 8.892089 | 16.11819 | 38.06673 | 16.46784 | -0.81915 | 0.326101 | 0.210475 |
| 2 | 16587200 | 16591999 | 80.29541 | 126.8462 | 111.188 | 46.17777 | 0.515661 | 2.436582 | 0.210479 |
| 12 | 105442600 | 105444599 | 16.4769 | 28.18914 | 59.18145 | 25.95069 | -0.68287 | 0.953681 | 0.21048 |
| 3 | 89047400 | 89053199 | 31.54737 | 50.69411 | 109.2363 | 46.29 | -0.75411 | 1.886999 | 0.210489 |
| 17 | 28534000 | 28537999 | 21.38376 | 34.98217 | 81.67753 | 35.12796 | -0.84447 | 1.464161 | 0.210497 |
| 13 | 54181000 | 54183199 | 18.86282 | 31.694 | 64.42054 | 28.13769 | -0.63866 | 1.130611 | 0.2105 |
| 1 | 85001000 | 85002399 | 26.63965 | 45.74281 | 37.71113 | 16.54096 | 0.661093 | 0.935661 | 0.210506 |
| 4 | 153977600 | 154003399 | 236.9684 | 336.2923 | 578.3068 | 239.225 | -0.59219 | 4.358868 | 0.21052 |
| 9 | 98025000 | 98027199 | 21.29857 | 37.31013 | 30.77715 | 13.59014 | 0.663915 | 0.673248 | 0.210526 |
| 7 | 3122800 | 3126199 | 65.06515 | 103.9845 | 86.83912 | 36.50191 | 0.601488 | 2.08824 | 0.210531 |
| 6 | 32178600 | 32180599 | 8.755751 | 15.84827 | 36.99445 | 16.03276 | -0.80379 | 0.332729 | 0.210536 |
| 14 | 5286200 | 5288399 | 19.93686 | 36.84204 | 18.86966 | 7.959429 | 1.395815 | 0.260911 | 0.210538 |
| 4 | 34777800 | 34778599 | 4.109227 | 8.010025 | 23.76624 | 8.587994 | -0.91613 | -0.41117 | 0.210539 |
| 7 | 34420200 | 34422399 | 11.10343 | 19.38384 | 49.94979 | 21.9966 | -0.9713 | 0.703714 | 0.210552 |
| 12 | 19590600 | 19591399 | 22.05501 | 38.72567 | 24.74409 | 10.94774 | 1.030327 | 0.630404 | 0.210564 |
| 5 | 71490000 | 71490799 | 11.50309 | 21.93592 | 18.37839 | 7.330677 | 0.756465 | -0.0113 | 0.210565 |
| 13 | 98595000 | 98595599 | 12.75407 | 24.7325 | 15.65887 | 5.91172 | 1.230493 | -0.18004 | 0.210569 |
| 3 | 45071800 | 45072999 | 30.00144 | 50.57811 | 41.55725 | 18.16982 | 0.663398 | 1.11178 | 0.210574 |
| 12 | 113216400 | 113218199 | 9.683198 | 17.41217 | 40.02126 | 17.48247 | -0.78967 | 0.382547 | 0.210577 |
| 19 | 28698000 | 28700399 | 10.89724 | 19.13468 | 46.16405 | 20.42528 | -0.87643 | 0.603377 | 0.210589 |
| 19 | 53459800 | 53466999 | 51.03401 | 80.85186 | 150.0482 | 62.40724 | -0.56163 | 2.355456 | 0.21059 |
| 3 | 110761600 | 110764399 | 35.78594 | 59.75894 | 45.41578 | 19.76116 | 0.775812 | 1.230208 | 0.210605 |
| 18 | 60733000 | 60745199 | 75.0056 | 114.7751 | 251.4657 | 104.3428 | -0.84 | 3.048351 | 0.210606 |
| 13 | 29256600 | 29257999 | 13.11488 | 22.82592 | 49.45998 | 21.73937 | -0.72302 | 0.759158 | 0.210615 |
| 15 | 86285400 | 86286999 | 5.882157 | 10.61587 | 40.6353 | 17.67513 | -1.51448 | 0.337009 | 0.210615 |
| 15 | 58932800 | 58943799 | 87.35663 | 132.9355 | 259.7655 | 107.6372 | -0.68151 | 3.166129 | 0.21063 |
| 13 | 81931000 | 81931999 | 6.980999 | 13.03506 | 32.19855 | 13.37161 | -0.83973 | 0.125998 | 0.210631 |
| 13 | 62188800 | 62190999 | 54.38493 | 87.9524 | 61.93342 | 26.47432 | 0.866766 | 1.710353 | 0.210636 |
| 5 | 95526800 | 95528599 | 36.73879 | 60.75507 | 50.98411 | 22.03955 | 0.632767 | 1.343981 | 0.210639 |
| 14 | 21556200 | 21556999 | 4.82028 | 9.358649 | 25.63789 | 9.617847 | -0.8529 | -0.26198 | 0.210649 |
| 3 | 45899000 | 45902799 | 28.34242 | 48.84395 | 34.78562 | 15.25947 | 0.872228 | 0.906507 | 0.210661 |
| 5 | 9310400 | 9311999 | 8.037939 | 14.9052 | 34.24265 | 14.37592 | -0.74888 | 0.181177 | 0.210665 |
| 1 | 27223800 | 27226599 | 34.22333 | 57.35716 | 39.40136 | 17.18346 | 0.920675 | 1.178219 | 0.210711 |
| 7 | 102227600 | 102228599 | 8.364995 | 14.99771 | 38.47563 | 16.84894 | -0.94889 | 0.391604 | 0.210724 |
| 17 | 30949800 | 30957199 | 48.17969 | 76.62802 | 140.6337 | 58.71716 | -0.5401 | 2.237191 | 0.210738 |
| 14 | 114207200 | 114209599 | 34.97249 | 58.78086 | 40.85425 | 17.8461 | 0.903838 | 1.154032 | 0.210757 |

| | | | | | | | | | |
|---|---|---|---|---|---|---|---|---|---|
| 4 | 138160200 | 138162399 | 9.730017 | 17.37283 | 39.9444 | 17.55345 | -0.796 | 0.428612 | 0.210761 |
| 19 | 3241200 | 3243399 | 17.69655 | 29.91684 | 61.72837 | 27.01186 | -0.6598 | 1.0699 | 0.210766 |
| 15 | 52652600 | 52653799 | 15.51066 | 29.12514 | 19.29919 | 7.938721 | 1.054254 | 0.129892 | 0.210766 |
| 12 | 23063200 | 23065599 | 40.74675 | 66.48128 | 57.50011 | 24.70572 | 0.581347 | 1.526778 | 0.210771 |
| 9 | 39956600 | 39958199 | 20.10983 | 36.27903 | 22.96074 | 9.99217 | 1.062962 | 0.40134 | 0.210772 |
| 7 | 40312200 | 40314199 | 19.82713 | 34.89195 | 28.00308 | 12.39781 | 0.70354 | 0.591896 | 0.210787 |
| 5 | 120562200 | 120579199 | 136.1529 | 200.2753 | 416.0668 | 171.341 | -0.8138 | 3.812738 | 0.210792 |
| 11 | 76754400 | 76756199 | 10.99031 | 19.5386 | 41.44593 | 18.27965 | -0.68419 | 0.468516 | 0.21081 |
| 16 | 33420600 | 33420999 | 7.65641 | 14.97082 | 12.23763 | 4.179502 | 0.993195 | -0.60056 | 0.210817 |
| 1 | 16069600 | 16070599 | 8.826724 | 16.26511 | 39.67987 | 16.80199 | -0.84446 | 0.234325 | 0.210831 |
| 3 | 143848800 | 143849999 | 8.826724 | 16.26511 | 39.67987 | 16.80199 | -0.84446 | 0.234325 | 0.210831 |
| 15 | 36792600 | 36794799 | 13.72024 | 23.45427 | 57.33672 | 25.14018 | -0.90128 | 0.950107 | 0.210834 |
| 8 | 57784200 | 57787799 | 25.37422 | 41.05151 | 106.2407 | 45.42786 | -1.00602 | 1.672463 | 0.210837 |
| 8 | 4784200 | 4788599 | 23.88227 | 41.59047 | 29.80321 | 13.10436 | 0.865636 | 0.770585 | 0.21085 |
| 14 | 121185600 | 121200199 | 132.563 | 197.2769 | 358.6034 | 147.7547 | -0.60559 | 3.613655 | 0.21085 |
| 2 | 59004600 | 59005399 | 10.80417 | 20.82212 | 16.79415 | 6.509418 | 0.848286 | -0.11529 | 0.210854 |
| 13 | 77825000 | 77826199 | 10.80417 | 20.82212 | 16.79415 | 6.509418 | 0.848286 | -0.11529 | 0.210854 |
| 19 | 13304800 | 13305799 | 14.17033 | 27.04432 | 17.34049 | 6.904744 | 1.14062 | 0.006518 | 0.210877 |
| 11 | 106084000 | 106084799 | 5.723097 | 10.9732 | 29.05777 | 11.43019 | -0.87067 | -0.0833 | 0.210894 |
| 2 | 113843200 | 113845599 | 19.09025 | 31.84558 | 68.1149 | 29.61844 | -0.71194 | 1.211837 | 0.210904 |
| 14 | 55172000 | 55174599 | 13.5134 | 23.23144 | 54.72777 | 24.00574 | -0.84664 | 0.894128 | 0.210905 |
| 19 | 5785600 | 5788999 | 13.5134 | 23.23144 | 54.72777 | 24.00574 | -0.84664 | 0.894128 | 0.210905 |
| 15 | 91182000 | 91183799 | 7.319566 | 13.53009 | 33.62603 | 14.17811 | -0.86547 | 0.193283 | 0.210909 |
| 15 | 96422400 | 96422999 | 5.253309 | 10.16096 | 26.5607 | 10.14879 | -0.81152 | -0.19076 | 0.210913 |
| 14 | 9215800 | 9217599 | 21.25024 | 37.3903 | 27.71685 | 12.27148 | 0.817342 | 0.601131 | 0.210929 |
| 12 | 81495000 | 81499399 | 78.29261 | 124.7511 | 87.05252 | 36.43878 | 0.855578 | 2.193593 | 0.210934 |
| 12 | 87526600 | 87532999 | 49.65776 | 78.46442 | 156.6371 | 65.06083 | -0.66976 | 2.418654 | 0.210942 |
| 6 | 23038400 | 23039199 | 11.63382 | 22.22425 | 18.97521 | 7.540639 | 0.73449 | -0.01892 | 0.210948 |
| 13 | 15495400 | 15497199 | 8.844842 | 16.23671 | 42.96189 | 18.28598 | -0.9665 | 0.262045 | 0.210967 |
| 5 | 57149000 | 57153599 | 56.43528 | 90.85888 | 71.95858 | 30.57697 | 0.690168 | 1.846854 | 0.210992 |
| 7 | 122367600 | 122369599 | 13.07262 | 22.84447 | 47.8369 | 21.0797 | -0.67344 | 0.697121 | 0.211016 |
| 3 | 96614400 | 96615999 | 7.354032 | 13.49527 | 34.57803 | 14.72455 | -0.91948 | 0.237847 | 0.211026 |
| 12 | 98594000 | 98594999 | 6.157543 | 11.77814 | 29.55296 | 11.70465 | -0.80315 | -0.05499 | 0.211032 |
| 13 | 97636800 | 97638199 | 6.157543 | 11.77814 | 29.55296 | 11.70465 | -0.80315 | -0.05499 | 0.211032 |
| 17 | 57536800 | 57537799 | 6.157543 | 11.77814 | 29.55296 | 11.70465 | -0.80315 | -0.05499 | 0.211032 |
| 7 | 6431400 | 6433599 | 34.23931 | 57.34062 | 44.8282 | 19.54087 | 0.735003 | 1.19432 | 0.211042 |
| 16 | 53391800 | 53392199 | 7.499819 | 14.65668 | 12.87809 | 4.460171 | 0.872958 | -0.55317 | 0.211044 |
| 2 | 136519800 | 136522199 | 12.94592 | 22.52003 | 50.53392 | 22.20408 | -0.77351 | 0.765226 | 0.21105 |
| 1 | 118090800 | 118092799 | 36.09895 | 60.65717 | 40.85514 | 17.84498 | 0.949029 | 1.161448 | 0.21105 |
| 7 | 53741600 | 53743399 | 29.43746 | 50.4912 | 33.29632 | 14.60323 | 0.981476 | 0.950685 | 0.211063 |
| 15 | 55255600 | 55259799 | 27.61707 | 45.02532 | 90.36284 | 38.81499 | -0.63067 | 1.529587 | 0.211063 |
| 10 | 31631400 | 31633399 | 34.62097 | 59.34843 | 31.24382 | 13.70241 | 1.304871 | 0.976138 | 0.211066 |
| 13 | 57565600 | 57567999 | 17.29837 | 29.49873 | 59.69585 | 26.16942 | -0.63079 | 0.987924 | 0.211083 |
| 6 | 40267400 | 40267999 | 9.089314 | 17.66092 | 15.01498 | 5.597114 | 0.826116 | -0.26931 | 0.211093 |
| X | 33886600 | 33887199 | 20.01293 | 35.45463 | 25.65244 | 11.34197 | 0.856245 | 0.520738 | 0.211095 |
| X | 63977200 | 63981599 | 46.37992 | 75.36557 | 58.10127 | 24.92428 | 0.742762 | 1.600553 | 0.211097 |
| 14 | 114743000 | 114743599 | 13.72773 | 25.80803 | 18.16019 | 7.455113 | 0.970844 | 0.103972 | 0.211097 |

| | | | | | | | | | |
|---|---|---|---|---|---|---|---|---|---|
| 13 | 98739200 | 98742799 | 40.17088 | 64.15416 | 127.3826 | 53.49043 | -0.64691 | 2.094463 | 0.211099 |
| 3 | 14815600 | 14816399 | 12.97081 | 24.60752 | 19.87148 | 8.013192 | 0.795948 | 0.04324 | 0.211112 |
| 12 | 46003200 | 46003999 | 11.7941 | 22.90449 | 15.8114 | 5.918848 | 1.118439 | -0.21839 | 0.211117 |
| 19 | 15909000 | 15909599 | 11.04018 | 21.50597 | 14.07169 | 5.129623 | 1.227987 | -0.32394 | 0.211134 |
| 16 | 19871200 | 19873399 | 11.93608 | 21.04626 | 45.07623 | 19.94881 | -0.70388 | 0.556376 | 0.211136 |
| 4 | 19122200 | 19123399 | 16.18234 | 29.70248 | 22.37412 | 9.524845 | 0.833877 | 0.273901 | 0.211147 |
| 4 | 35065000 | 35066399 | 7.548897 | 14.13519 | 33.98041 | 14.04448 | -0.79605 | 0.11465 | 0.211167 |
| 5 | 99063200 | 99064599 | 8.019305 | 14.97055 | 33.87724 | 14.07393 | -0.71529 | 0.139632 | 0.21118 |
| 3 | 137328000 | 137330199 | 14.23124 | 24.58833 | 53.21284 | 23.34479 | -0.72341 | 0.848513 | 0.211204 |
| 18 | 64464200 | 64466999 | 25.04364 | 40.91038 | 83.48947 | 35.88837 | -0.65256 | 1.492623 | 0.211209 |
| 10 | 62206400 | 62212999 | 52.68567 | 83.41128 | 151.2315 | 62.87756 | -0.52886 | 2.373517 | 0.211221 |
| 11 | 79750000 | 79753799 | 18.9227 | 31.57893 | 71.04348 | 30.89981 | -0.78469 | 1.207339 | 0.211228 |
| 18 | 43045200 | 43047599 | 25.90536 | 44.62384 | 34.96322 | 15.33717 | 0.734907 | 0.89879 | 0.211233 |
| X | 54594400 | 54597999 | 52.63617 | 85.8635 | 50.17586 | 21.55727 | 1.145239 | 1.552387 | 0.211239 |
| 16 | 37663200 | 37664199 | 4.62572 | 8.980869 | 25.55235 | 9.5861 | -0.90554 | -0.26039 | 0.211243 |
| 5 | 94887000 | 94888199 | 18.18394 | 33.27514 | 21.5707 | 9.22086 | 1.044748 | 0.304561 | 0.211244 |
| 2 | 27028600 | 27032799 | 22.8358 | 37.29408 | 83.55781 | 35.91436 | -0.78703 | 1.492237 | 0.211253 |
| 5 | 103171800 | 103173399 | 23.04205 | 40.11897 | 33.46608 | 14.713 | 0.647559 | 0.775532 | 0.21126 |
| 9 | 121242000 | 121257399 | 103.3336 | 155.9009 | 296.4416 | 122.5088 | -0.65265 | 3.346683 | 0.211281 |
| 6 | 114362200 | 114363399 | 5.386685 | 10.36879 | 27.9923 | 10.88772 | -0.8814 | -0.12237 | 0.211294 |
| 7 | 85001800 | 85005999 | 61.89919 | 99.30112 | 78.24722 | 33.07489 | 0.691308 | 1.966192 | 0.211304 |
| 7 | 44065800 | 44067599 | 8.525345 | 15.67999 | 35.15047 | 14.92253 | -0.72553 | 0.234073 | 0.211316 |
| 9 | 102898400 | 102899999 | 11.81422 | 20.76371 | 44.04885 | 19.5044 | -0.69175 | 0.592314 | 0.211316 |
| 7 | 78810400 | 78812799 | 28.07664 | 48.69986 | 31.39842 | 13.77631 | 1.016713 | 0.844134 | 0.211336 |
| 8 | 80551000 | 80552399 | 7.872854 | 14.67331 | 33.26733 | 13.85859 | -0.7211 | 0.147502 | 0.211339 |
| 7 | 31411800 | 31414199 | 42.68532 | 69.48612 | 59.8299 | 25.68546 | 0.585508 | 1.563684 | 0.211344 |
| 1 | 103739400 | 103740799 | 14.80411 | 27.95234 | 19.50038 | 7.94302 | 0.99231 | 0.090107 | 0.211347 |
| 12 | 104412600 | 104415599 | 23.06642 | 37.87995 | 79.07404 | 34.06231 | -0.68104 | 1.415328 | 0.211351 |
| 1 | 106408600 | 106409599 | 6.278829 | 12.08438 | 31.7583 | 12.35749 | -0.84546 | -0.10582 | 0.211356 |
| 12 | 86916400 | 86919599 | 19.5369 | 32.64087 | 68.2827 | 29.72025 | -0.68012 | 1.195537 | 0.21136 |
| 2 | 21901200 | 21902999 | 13.11795 | 25.11243 | 18.1213 | 7.160298 | 0.982015 | -0.02571 | 0.211366 |
| 10 | 87236000 | 87237199 | 13.11795 | 25.11243 | 18.1213 | 7.160298 | 0.982015 | -0.02571 | 0.211366 |
| 17 | 82476800 | 82477399 | 6.230512 | 11.7027 | 31.21941 | 12.83587 | -0.93764 | 0.079051 | 0.211367 |
| 12 | 23530400 | 23533599 | 58.93358 | 94.40375 | 75.16737 | 31.70161 | 0.673817 | 2.007396 | 0.21138 |
| 11 | 32889000 | 32891799 | 14.24933 | 24.57293 | 53.72898 | 23.56986 | -0.73874 | 0.866741 | 0.211387 |
| 6 | 107932400 | 107932999 | 8.852106 | 17.2359 | 15.5622 | 5.704269 | 0.762362 | -0.32775 | 0.21139 |
| 17 | 4958000 | 4959599 | 7.107474 | 13.32412 | 32.86796 | 13.55796 | -0.83041 | 0.101101 | 0.211393 |
| 5 | 4368000 | 4370799 | 60.77785 | 97.42352 | 74.91256 | 31.62647 | 0.725142 | 1.987296 | 0.211448 |
| 7 | 10141200 | 10150599 | 201.9794 | 300.3091 | 247.9486 | 102.1524 | 0.531094 | 3.636218 | 0.21145 |
| 4 | 125193800 | 125194999 | 7.905167 | 14.63045 | 33.65085 | 14.16499 | -0.75301 | 0.191217 | 0.211453 |
| 8 | 3680600 | 3681599 | 7.905167 | 14.63045 | 33.65085 | 14.16499 | -0.75301 | 0.191217 | 0.211453 |
| 11 | 89925600 | 89926599 | 7.905167 | 14.63045 | 33.65085 | 14.16499 | -0.75301 | 0.191217 | 0.211453 |
| 6 | 99764000 | 99766599 | 18.9621 | 32.05773 | 64.05251 | 28.02927 | -0.61372 | 1.075891 | 0.211454 |
| 7 | 10824400 | 10834399 | 159.3559 | 241.4333 | 193.4439 | 80.04936 | 0.598378 | 3.268281 | 0.211462 |
| 6 | 125549200 | 125550199 | 6.441032 | 12.33635 | 29.63282 | 11.69275 | -0.73569 | -0.06341 | 0.211466 |
| 7 | 16446000 | 16452799 | 47.70986 | 75.87947 | 138.2229 | 57.70922 | -0.52933 | 2.2374 | 0.211471 |
| 12 | 65908600 | 65910599 | 21.19007 | 37.10301 | 30.72578 | 13.56218 | 0.658233 | 0.678729 | 0.211473 |

| | | | | | | | | | |
|---|---|---|---|---|---|---|---|---|---|
| 14 | 99750200 | 99751599 | 14.06791 | 26.4598 | 20.72579 | 8.501681 | 0.818557 | 0.111578 | 0.211477 |
| 2 | 129198200 | 129198599 | 3.340254 | 6.532846 | 19.63722 | 6.650805 | -0.84636 | -0.66393 | 0.21148 |
| 14 | 74460800 | 74461199 | 3.340254 | 6.532846 | 19.63722 | 6.650805 | -0.84636 | -0.66393 | 0.21148 |
| 7 | 10195200 | 10196999 | 17.96554 | 33.38028 | 19.57956 | 8.197961 | 1.209073 | 0.216786 | 0.211483 |
| 10 | 7241800 | 7243599 | 11.82817 | 20.72348 | 45.68792 | 20.17691 | -0.74706 | 0.645384 | 0.211483 |
| 10 | 115498000 | 115501599 | 20.84466 | 34.66115 | 72.82984 | 31.60294 | -0.68668 | 1.254353 | 0.211497 |
| 7 | 52505400 | 52506799 | 14.34493 | 24.94589 | 51.09796 | 22.4475 | -0.64272 | 0.777513 | 0.211508 |
| 2 | 36571400 | 36575999 | 74.58443 | 118.5381 | 99.62962 | 41.56541 | 0.583816 | 2.271693 | 0.211515 |
| 2 | 33776400 | 33779199 | 16.84897 | 28.67767 | 59.06669 | 25.88963 | -0.65647 | 1.003684 | 0.211526 |
| 6 | 35109800 | 35111599 | 10.03715 | 17.56621 | 48.32442 | 21.32144 | -1.06463 | 0.657076 | 0.211527 |
| 8 | 43236600 | 43237599 | 8.22 | 15.12417 | 35.70144 | 15.14856 | -0.79909 | 0.229709 | 0.211535 |
| 8 | 48588200 | 48594599 | 47.25603 | 75.03775 | 142.3328 | 59.32819 | -0.58987 | 2.284991 | 0.211551 |
| 6 | 87913000 | 87914799 | 8.813817 | 15.79221 | 38.4584 | 16.85057 | -0.87495 | 0.398709 | 0.211563 |
| 5 | 142442400 | 142446799 | 18.19241 | 29.89908 | 79.33153 | 34.18451 | -1.02517 | 1.395617 | 0.211563 |
| 3 | 53354000 | 53354599 | 4.107929 | 8.01111 | 23.88516 | 8.58012 | -0.91547 | -0.43249 | 0.211569 |
| 11 | 21443400 | 21443999 | 4.107929 | 8.01111 | 23.88516 | 8.58012 | -0.91547 | -0.43249 | 0.211569 |
| 10 | 100138600 | 100142799 | 88.64903 | 134.4884 | 271.3597 | 112.338 | -0.73135 | 3.230523 | 0.211578 |
| 16 | 70594800 | 70596799 | 25.89342 | 44.42429 | 37.23381 | 16.3305 | 0.637147 | 0.938508 | 0.211582 |
| 14 | 3600000 | 3605599 | 97.21187 | 151.8917 | 131.5732 | 54.59033 | 0.520532 | 2.661996 | 0.211591 |
| 16 | 37943200 | 37944199 | 5.26112 | 10.15212 | 26.97252 | 10.41697 | -0.84943 | -0.15242 | 0.211605 |
| 8 | 47414800 | 47417799 | 27.84816 | 45.21831 | 88.84445 | 38.09146 | -0.60454 | 1.595879 | 0.211617 |
| 4 | 127991800 | 127993799 | 16.83648 | 28.25063 | 64.70834 | 28.24115 | -0.81017 | 1.139822 | 0.211618 |
| 8 | 21486000 | 21487199 | 14.9521 | 28.42454 | 18.54604 | 7.446348 | 1.106307 | 0.044657 | 0.211623 |
| 3 | 145802800 | 145812199 | 111.289 | 168.0962 | 287.599 | 118.8975 | -0.49901 | 3.32541 | 0.211633 |
| 4 | 26091400 | 26092399 | 17.02858 | 31.51209 | 21.3648 | 8.994645 | 0.996365 | 0.235245 | 0.211635 |
| 1 | 117801000 | 117802199 | 21.63111 | 38.00465 | 29.46181 | 13.0418 | 0.752876 | 0.626226 | 0.211646 |
| 14 | 33548200 | 33549799 | 8.333879 | 15.48965 | 38.29052 | 16.02497 | -0.85099 | 0.178593 | 0.211647 |
| 1 | 113220200 | 113221199 | 18.69576 | 33.08394 | 27.13146 | 11.9986 | 0.675723 | 0.525485 | 0.211653 |
| 14 | 37076800 | 37078999 | 16.62947 | 28.56629 | 58.13291 | 25.49876 | -0.63707 | 0.917148 | 0.211677 |
| 3 | 101443400 | 101444599 | 4.794196 | 9.298153 | 25.50617 | 9.622982 | -0.86172 | -0.24031 | 0.21168 |
| 5 | 113271800 | 113282799 | 62.62725 | 97.19828 | 199.0124 | 82.64698 | -0.72656 | 2.770807 | 0.211684 |
| X | 26084000 | 26084999 | 14.39944 | 27.86265 | 14.83547 | 5.657932 | 1.463818 | -0.13171 | 0.211694 |
| X | 31784600 | 31787399 | 40.49224 | 66.44999 | 48.33022 | 20.81276 | 0.835231 | 1.443003 | 0.211702 |
| 11 | 22578600 | 22580399 | 8.879525 | 16.12849 | 38.11053 | 16.44452 | -0.81751 | 0.313833 | 0.211721 |
| 13 | 87039600 | 87040799 | 8.879525 | 16.12849 | 38.11053 | 16.44452 | -0.81751 | 0.313833 | 0.211721 |
| 11 | 9920400 | 9922599 | 20.22521 | 35.63949 | 28.42538 | 12.58145 | 0.713243 | 0.581418 | 0.211743 |
| 8 | 41827000 | 41828399 | 9.567139 | 17.11279 | 40.48387 | 17.76353 | -0.83544 | 0.416364 | 0.211748 |
| 14 | 60689200 | 60690599 | 10.19752 | 18.16082 | 40.42883 | 17.80486 | -0.75195 | 0.44899 | 0.211771 |
| 18 | 80205800 | 80207199 | 10.19752 | 18.16082 | 40.42883 | 17.80486 | -0.75195 | 0.44899 | 0.211771 |
| 1 | 14901600 | 14902999 | 20.67796 | 36.30336 | 29.72041 | 13.14811 | 0.674716 | 0.633296 | 0.21179 |
| 7 | 110443200 | 110444599 | 7.86304 | 14.68641 | 33.33271 | 13.83561 | -0.71848 | 0.133462 | 0.211791 |
| 16 | 79898000 | 79898799 | 10.97766 | 21.3104 | 16.33186 | 6.133511 | 0.964198 | -0.21674 | 0.211794 |
| X | 24521200 | 24521999 | 25.90256 | 44.9608 | 31.1493 | 13.66798 | 0.914121 | 0.831506 | 0.211795 |
| 3 | 152108600 | 152109599 | 12.095 | 23.46528 | 15.60082 | 5.877095 | 1.162808 | -0.19579 | 0.211804 |
| 6 | 24941600 | 24943599 | 21.20601 | 37.60341 | 26.14028 | 11.55411 | 0.913892 | 0.518607 | 0.211814 |
| 6 | 83751800 | 83752999 | 5.738923 | 10.91245 | 29.29712 | 11.75719 | -0.91587 | -0.01339 | 0.211831 |
| 8 | 111216000 | 111222199 | 44.75179 | 71.31051 | 131.9144 | 55.22068 | -0.5485 | 2.171679 | 0.211843 |

| | | | | | | | | | |
|---|---|---|---|---|---|---|---|---|---|
| 10 | 34082400 | 34083599 | 7.886607 | 14.23507 | 37.50347 | 16.31051 | -0.98018 | 0.344506 | 0.211853 |
| 2 | 128705600 | 128708599 | 16.2422 | 27.61306 | 58.98695 | 25.85228 | -0.70904 | 1.012891 | 0.211871 |
| 5 | 136431400 | 136433399 | 10.87798 | 19.15149 | 43.53057 | 19.27061 | -0.78975 | 0.565171 | 0.211877 |
| 11 | 78380000 | 78382199 | 10.87798 | 19.15149 | 43.53057 | 19.27061 | -0.78975 | 0.565171 | 0.211877 |
| 13 | 109396800 | 109397599 | 4.233274 | 8.244259 | 23.56045 | 8.616561 | -0.87793 | -0.3651 | 0.21188 |
| 9 | 67539800 | 67541799 | 11.77155 | 20.38247 | 52.06477 | 22.84263 | -0.96027 | 0.818399 | 0.211883 |
| 14 | 62028800 | 62032199 | 37.49231 | 61.68434 | 53.19974 | 22.93453 | 0.5915 | 1.406871 | 0.211887 |
| 5 | 9916600 | 9918399 | 15.57886 | 28.68661 | 22.50379 | 9.538355 | 0.780307 | 0.252235 | 0.211899 |
| 7 | 139778800 | 139783999 | 32.05624 | 51.44481 | 112.9049 | 47.75129 | -0.7828 | 1.929935 | 0.211899 |
| 19 | 5059800 | 5061199 | 7.693677 | 14.46711 | 33.55469 | 13.76992 | -0.73621 | 0.089307 | 0.211903 |
| 2 | 163912200 | 163914799 | 12.09121 | 20.92245 | 54.76721 | 24.02781 | -0.99599 | 0.826535 | 0.211909 |
| 15 | 101352400 | 101362199 | 67.64473 | 105.0555 | 195.2458 | 81.07469 | -0.58629 | 2.761306 | 0.211925 |
| 12 | 69229600 | 69231999 | 10.30027 | 18.08346 | 45.13697 | 19.96806 | -0.92493 | 0.602251 | 0.211929 |
| X | 165991800 | 165993999 | 20.02355 | 36.36422 | 22.18671 | 9.575417 | 1.123401 | 0.363775 | 0.211935 |
| 12 | 115945600 | 115947399 | 23.90707 | 41.77121 | 31.62986 | 13.93639 | 0.786908 | 0.728955 | 0.211962 |
| 16 | 52996600 | 52997799 | 12.96256 | 24.43005 | 20.25002 | 8.278179 | 0.741147 | 0.086798 | 0.211967 |
| 7 | 80775600 | 80779399 | 42.69252 | 67.94715 | 134.6923 | 56.28956 | -0.64993 | 2.210563 | 0.211973 |
| 2 | 120807400 | 120808799 | 7.221435 | 13.22367 | 35.07484 | 14.97957 | -0.97212 | 0.249827 | 0.211977 |
| 12 | 100494200 | 100497599 | 17.0857 | 28.87068 | 61.62888 | 26.96451 | -0.70854 | 1.073226 | 0.211977 |
| 1 | 179460800 | 179464199 | 22.75564 | 37.35823 | 78.96057 | 34.00907 | -0.69907 | 1.41865 | 0.211995 |
| 13 | 61490400 | 61495799 | 141.1707 | 215.7536 | 170.4428 | 70.70288 | 0.628872 | 3.087068 | 0.212012 |
| 7 | 5318600 | 5321999 | 45.10436 | 73.42121 | 59.35841 | 25.4827 | 0.676156 | 1.570518 | 0.212017 |
| 7 | 129522800 | 129526399 | 23.30339 | 38.54037 | 75.63509 | 32.69778 | -0.58915 | 1.327561 | 0.212036 |
| 6 | 57118000 | 57119199 | 15.83969 | 29.94878 | 17.62885 | 7.159868 | 1.238433 | 0.085325 | 0.212037 |
| 18 | 82931400 | 82933399 | 6.600701 | 12.21228 | 34.70591 | 14.61551 | -1.05618 | 0.183421 | 0.212052 |
| 8 | 31548600 | 31550399 | 29.10658 | 49.40929 | 40.94951 | 17.94161 | 0.651305 | 1.050545 | 0.212057 |
| 2 | 147035000 | 147038399 | 31.21377 | 50.1945 | 109.1487 | 46.3016 | -0.76593 | 1.862551 | 0.212072 |
| 4 | 111210600 | 111211599 | 7.691214 | 14.42588 | 34.62617 | 14.27073 | -0.79063 | 0.106533 | 0.212083 |
| 14 | 78024000 | 78025799 | 8.727164 | 15.92984 | 36.07449 | 15.4706 | -0.75015 | 0.283252 | 0.212086 |
| 13 | 49883800 | 49889599 | 27.94922 | 45.56505 | 88.5045 | 38.0162 | -0.58363 | 1.530766 | 0.212092 |
| 2 | 55695400 | 55697399 | 28.32099 | 49.21057 | 31.44736 | 13.80792 | 1.029925 | 0.826083 | 0.212105 |
| X | 12152200 | 12155399 | 14.54158 | 25.28859 | 53.25092 | 23.39373 | -0.68237 | 0.782343 | 0.212108 |
| 5 | 29158400 | 29158799 | 3.453566 | 6.754373 | 20.26881 | 6.872467 | -0.84554 | -0.65496 | 0.212109 |
| 15 | 62960800 | 62963599 | 32.60642 | 55.85637 | 34.12043 | 14.96316 | 1.091263 | 0.977016 | 0.212122 |
| 16 | 34391200 | 34394799 | 17.37984 | 28.98541 | 69.03495 | 30.01397 | -0.86641 | 1.209957 | 0.21213 |
| 2 | 156614400 | 156625199 | 58.59788 | 91.26333 | 193.0867 | 80.14424 | -0.76983 | 2.707383 | 0.21213 |
| 8 | 20044200 | 20044599 | 9.580321 | 18.67739 | 14.25049 | 5.139768 | 1.022238 | -0.37961 | 0.212138 |
| 13 | 92729400 | 92732399 | 14.18863 | 24.59445 | 59.3611 | 26.04465 | -0.87947 | 0.823819 | 0.212142 |
| 6 | 86477000 | 86480799 | 21.64262 | 35.97061 | 72.88899 | 31.61805 | -0.63459 | 1.262612 | 0.212146 |
| X | 5110800 | 5111599 | 12.277 | 23.83954 | 15.01514 | 5.626016 | 1.247034 | -0.2132 | 0.21215 |
| 12 | 54429600 | 54430599 | 6.287972 | 12.06658 | 29.74629 | 11.67232 | -0.76535 | -0.08385 | 0.212161 |
| 7 | 15374000 | 15377199 | 71.81233 | 114.7228 | 82.48078 | 34.6508 | 0.816529 | 2.104322 | 0.212162 |
| 17 | 31172800 | 31179799 | 46.21143 | 73.15868 | 151.6962 | 63.06962 | -0.72218 | 2.369886 | 0.212164 |
| 7 | 44812800 | 44829799 | 200.0414 | 286.8152 | 521.4236 | 215.3315 | -0.65748 | 4.206597 | 0.212172 |
| 2 | 15720800 | 15722999 | 23.85263 | 42.08598 | 25.94616 | 11.48114 | 1.083462 | 0.586415 | 0.212174 |
| 5 | 119323800 | 119328399 | 27.78012 | 45.30882 | 88.56817 | 38.04951 | -0.59235 | 1.524003 | 0.212177 |
| 4 | 147158600 | 147165999 | 128.7231 | 196.9182 | 171.2415 | 71.04992 | 0.49154 | 3.063831 | 0.212178 |

| | | | | | | | | | |
|---|---|---|---|---|---|---|---|---|---|
| 1 | 174115800 | 174122199 | 82.93655 | 130.6235 | 112.544 | 46.69757 | 0.537086 | 2.503065 | 0.212179 |
| 10 | 47261000 | 47263599 | 19.66183 | 34.99931 | 26.27086 | 11.5743 | 0.806879 | 0.485012 | 0.212183 |
| 7 | 132460800 | 132461599 | 4.531562 | 8.799511 | 25.0167 | 9.372657 | -0.9034 | -0.27082 | 0.212214 |
| 5 | 24312000 | 24313399 | 9.368466 | 16.89731 | 38.49503 | 16.75721 | -0.77344 | 0.35986 | 0.212222 |
| 4 | 151959000 | 151968399 | 90.85741 | 138.457 | 252.3857 | 104.6232 | -0.57965 | 3.137966 | 0.212227 |
| 2 | 180471800 | 180473799 | 12.01508 | 20.95531 | 48.38077 | 21.29283 | -0.8137 | 0.721178 | 0.212232 |
| 4 | 133559200 | 133559599 | 3.713145 | 7.222822 | 24.5372 | 9.064944 | -1.13763 | -0.33309 | 0.21224 |
| X | 53749200 | 53750999 | 40.09422 | 66.80992 | 39.4997 | 17.16374 | 1.136854 | 1.260859 | 0.212243 |
| 1 | 16831800 | 16834199 | 26.29986 | 45.04318 | 36.69141 | 16.09035 | 0.677696 | 0.954194 | 0.212253 |
| 10 | 60316600 | 60318199 | 16.14382 | 27.7804 | 55.93873 | 24.53764 | -0.62125 | 0.897086 | 0.212257 |
| 8 | 28119000 | 28120399 | 16.80949 | 31.28919 | 19.61921 | 8.190768 | 1.116472 | 0.195652 | 0.212262 |
| 16 | 91982000 | 91983599 | 8.205858 | 15.20887 | 34.22476 | 14.37832 | -0.71971 | 0.187165 | 0.212276 |
| X | 24194600 | 24197199 | 30.32587 | 52.47805 | 32.07182 | 14.07077 | 1.093243 | 0.878756 | 0.212281 |
| 7 | 85444800 | 85446199 | 22.60093 | 39.9983 | 25.39331 | 11.22971 | 1.042802 | 0.54519 | 0.212281 |
| 11 | 73776800 | 73777399 | 8.382023 | 16.39303 | 10.80687 | 3.669328 | 1.304784 | -0.61445 | 0.212281 |
| 6 | 71065600 | 71066799 | 5.20518 | 9.901837 | 30.98074 | 12.42147 | -1.13414 | -0.02034 | 0.212288 |
| 9 | 93507400 | 93509399 | 17.87171 | 32.33478 | 24.402 | 10.58103 | 0.814227 | 0.372703 | 0.212291 |
| 17 | 49868600 | 49873799 | 29.22846 | 47.16457 | 100.6076 | 42.88378 | -0.73211 | 1.749684 | 0.212303 |
| 5 | 130046400 | 130050799 | 18.78114 | 31.31357 | 69.80486 | 30.34327 | -0.77145 | 1.216844 | 0.212305 |
| 1 | 57813200 | 57814999 | 10.17017 | 17.82156 | 46.73594 | 20.64265 | -0.99588 | 0.634794 | 0.212332 |
| 2 | 57127600 | 57130999 | 19.19547 | 32.1323 | 68.92482 | 30.03624 | -0.71622 | 1.173341 | 0.21234 |
| 7 | 5570600 | 5575999 | 103.7953 | 162.5468 | 121.7956 | 50.52174 | 0.732021 | 2.62514 | 0.212347 |
| 19 | 58594000 | 58595199 | 8.27005 | 15.08003 | 36.05703 | 15.4829 | -0.82943 | 0.283509 | 0.212376 |
| 6 | 70181400 | 70182999 | 17.34803 | 31.04626 | 25.58563 | 11.22167 | 0.677292 | 0.435455 | 0.212379 |
| 12 | 23288600 | 23290599 | 46.71909 | 75.51132 | 65.08584 | 27.8011 | 0.57486 | 1.71415 | 0.212385 |
| 7 | 75410400 | 75412799 | 19.27419 | 32.08516 | 69.10538 | 30.00945 | -0.72186 | 1.237066 | 0.212391 |
| 10 | 7407200 | 7414199 | 121.4155 | 187.6257 | 145.7453 | 60.53937 | 0.665344 | 2.866297 | 0.212396 |
| 4 | 29301400 | 29302599 | 8.203251 | 15.17135 | 34.18232 | 14.40262 | -0.72471 | 0.198183 | 0.212401 |
| 13 | 60194400 | 60198799 | 19.47677 | 32.29291 | 71.76012 | 31.07656 | -0.7673 | 1.285839 | 0.212402 |
| 1 | 165931000 | 165935799 | 27.90007 | 45.1511 | 96.65619 | 41.33846 | -0.73258 | 1.666166 | 0.212425 |
| 7 | 111944600 | 111946399 | 9.341513 | 16.53307 | 41.48265 | 18.34474 | -0.92802 | 0.494355 | 0.212428 |
| 6 | 64138200 | 64139599 | 16.44745 | 29.89506 | 23.98639 | 10.34118 | 0.731503 | 0.334636 | 0.212435 |
| 17 | 72009600 | 72020399 | 78.89609 | 121.0275 | 234.495 | 97.3465 | -0.66006 | 2.999002 | 0.212447 |
| 4 | 45466600 | 45474999 | 44.25054 | 70.11851 | 153.4758 | 63.83908 | -0.79906 | 2.346079 | 0.212449 |
| 8 | 127602200 | 127603999 | 7.185961 | 13.25362 | 34.1222 | 14.43091 | -0.91922 | 0.20569 | 0.212458 |
| 8 | 15801000 | 15801799 | 17.82983 | 31.88074 | 25.98551 | 11.40542 | 0.692341 | 0.444691 | 0.212467 |
| 11 | 102554400 | 102557999 | 55.89207 | 87.66117 | 173.4354 | 71.93116 | -0.66527 | 2.580507 | 0.212469 |
| 16 | 22991000 | 22992399 | 18.74594 | 33.20562 | 26.66113 | 11.78827 | 0.706677 | 0.514865 | 0.21247 |
| 18 | 43523000 | 43527199 | 21.5435 | 35.65833 | 73.1178 | 31.62701 | -0.65205 | 1.311893 | 0.212482 |
| 14 | 18199400 | 18201999 | 36.53549 | 60.27573 | 52.08343 | 22.47943 | 0.590328 | 1.370789 | 0.212489 |
| 14 | 46037200 | 46038799 | 23.56293 | 41.09284 | 32.80702 | 14.43804 | 0.710744 | 0.754709 | 0.212497 |
| 8 | 52738000 | 52739199 | 15.47745 | 28.8607 | 20.33996 | 8.466286 | 0.952607 | 0.174901 | 0.212512 |
| 10 | 42176000 | 42177199 | 6.016353 | 11.48921 | 28.88482 | 11.48695 | -0.81082 | -0.04288 | 0.212528 |
| 3 | 111982600 | 111983999 | 17.90228 | 33.66967 | 17.50261 | 7.180276 | 1.404777 | 0.138101 | 0.212534 |
| 16 | 95576000 | 95579599 | 23.19073 | 38.22141 | 76.4251 | 32.97142 | -0.61698 | 1.370238 | 0.212537 |
| 12 | 42652200 | 42654599 | 29.2831 | 50.32109 | 35.90505 | 15.74916 | 0.868835 | 0.937788 | 0.212554 |
| 19 | 50414600 | 50415199 | 5.96366 | 11.12851 | 32.16251 | 13.37078 | -1.06474 | 0.122699 | 0.212556 |

| | | | | | | | | | |
|---|---|---|---|---|---|---|---|---|---|
| 8 | 116224400 | 116225599 | 13.69616 | 25.77589 | 20.73677 | 8.497633 | 0.781351 | 0.104227 | 0.212568 |
| 13 | 75430600 | 75431599 | 13.69616 | 25.77589 | 20.73677 | 8.497633 | 0.781351 | 0.104227 | 0.212568 |
| 6 | 135030600 | 135033599 | 11.68008 | 20.43776 | 47.81107 | 21.09166 | -0.83237 | 0.667481 | 0.212582 |
| 17 | 22299800 | 22300599 | 6.576821 | 12.22806 | 34.26603 | 14.33345 | -1.02903 | 0.153564 | 0.212588 |
| 8 | 106736400 | 106738199 | 11.21626 | 19.65788 | 45.68576 | 20.1816 | -0.82268 | 0.636414 | 0.212593 |
| 15 | 91486400 | 91487199 | 5.481833 | 10.44251 | 32.23461 | 12.88655 | -1.11181 | -0.02552 | 0.212604 |
| 13 | 3064800 | 3070199 | 47.84121 | 77.51574 | 57.0132 | 24.40545 | 0.806573 | 1.657788 | 0.212606 |
| 3 | 88069600 | 88070399 | 3.423318 | 6.668317 | 23.49283 | 8.567583 | -1.17122 | -0.38606 | 0.212614 |
| 2 | 72359400 | 72360999 | 8.091772 | 14.86152 | 34.58738 | 14.71176 | -0.78074 | 0.239356 | 0.212615 |
| 2 | 72171200 | 72174199 | 37.26082 | 61.97726 | 47.10498 | 20.44448 | 0.775769 | 1.277372 | 0.21262 |
| 1 | 120399200 | 120402799 | 21.84191 | 36.23911 | 72.12156 | 31.24782 | -0.6088 | 1.284513 | 0.212621 |
| 18 | 5048000 | 5049599 | 7.096899 | 13.33374 | 32.9413 | 13.53744 | -0.82827 | 0.086594 | 0.212622 |
| 18 | 46657600 | 46659799 | 14.14726 | 24.26366 | 55.80349 | 24.47437 | -0.81292 | 0.919295 | 0.212624 |
| 13 | 53376400 | 53384799 | 102.1876 | 154.6196 | 279.612 | 115.6594 | -0.57671 | 3.287316 | 0.212645 |
| 1 | 106345200 | 106346799 | 21.55408 | 37.97263 | 28.42277 | 12.58086 | 0.804445 | 0.593426 | 0.212648 |
| 12 | 20802600 | 20817999 | 426.6308 | 596.5965 | 452.6512 | 187.6912 | 0.566505 | 4.58251 | 0.212652 |
| 11 | 35647000 | 35648799 | 8.026064 | 14.91851 | 34.30303 | 14.35489 | -0.7468 | 0.167493 | 0.21266 |
| 2 | 121303000 | 121304999 | 10.33143 | 18.49532 | 39.43902 | 17.29289 | -0.68571 | 0.416017 | 0.212688 |
| 3 | 43639800 | 43642799 | 38.0097 | 62.92109 | 49.9238 | 21.59644 | 0.713449 | 1.33444 | 0.212691 |
| 14 | 56090400 | 56092999 | 12.04851 | 20.93572 | 53.73894 | 23.59588 | -0.96679 | 0.780144 | 0.212697 |
| 16 | 90897600 | 90899599 | 17.49816 | 31.41948 | 25.47748 | 11.14288 | 0.703181 | 0.415643 | 0.212701 |
| 14 | 6965000 | 6967199 | 30.1408 | 50.77903 | 43.18602 | 18.87812 | 0.613888 | 1.121085 | 0.212705 |
| 7 | 13511200 | 13519199 | 131.6053 | 202.2384 | 157.0168 | 65.19845 | 0.659961 | 2.975797 | 0.21271 |
| 5 | 29778800 | 29781399 | 22.06319 | 38.70676 | 29.97177 | 13.25105 | 0.754549 | 0.652086 | 0.212722 |
| 16 | 14397800 | 14398399 | 9.693449 | 18.87986 | 14.77992 | 5.397982 | 0.969599 | -0.33344 | 0.212725 |
| 10 | 123207600 | 123208999 | 8.180419 | 15.20049 | 34.29587 | 14.35821 | -0.72011 | 0.171015 | 0.212739 |
| 1 | 154673800 | 154678599 | 50.18964 | 79.6451 | 144.8006 | 60.31822 | -0.52973 | 2.306659 | 0.212743 |
| 7 | 62191000 | 62192199 | 32.37723 | 54.272 | 44.27009 | 19.30877 | 0.673797 | 1.175814 | 0.212745 |
| 4 | 150129400 | 150154999 | 148.5494 | 215.5153 | 500.96 | 206.5393 | -0.99498 | 4.028983 | 0.212757 |
| 1 | 118143200 | 118144799 | 42.37442 | 69.05936 | 59.28367 | 25.46163 | 0.591 | 1.546769 | 0.212769 |
| 1 | 79934000 | 79939399 | 28.7219 | 46.4142 | 101.2874 | 43.25413 | -0.7625 | 1.705675 | 0.212771 |
| 18 | 19103200 | 19104399 | 17.63213 | 32.43527 | 21.91251 | 9.300281 | 0.993425 | 0.274271 | 0.212773 |
| 14 | 69366800 | 69374999 | 83.9377 | 128.7977 | 231.9666 | 96.3006 | -0.55446 | 2.994992 | 0.212795 |
| 1 | 142655200 | 142656999 | 22.45957 | 40.3201 | 21.59577 | 9.446752 | 1.297247 | 0.443445 | 0.212814 |
| 5 | 132691000 | 132693799 | 16.02698 | 27.44045 | 56.39373 | 24.72983 | -0.65183 | 0.943805 | 0.212821 |
| 10 | 82458400 | 82459399 | 9.356043 | 16.90868 | 38.5336 | 16.7321 | -0.7716 | 0.347891 | 0.212845 |
| 13 | 91648600 | 91649999 | 21.08517 | 37.31552 | 26.61906 | 11.77179 | 0.875621 | 0.537735 | 0.212881 |
| 10 | 46781800 | 46783799 | 18.41166 | 33.51806 | 22.64254 | 9.742564 | 0.979798 | 0.337577 | 0.2129 |
| 12 | 100919200 | 100927399 | 39.96069 | 63.56879 | 139.4485 | 58.23853 | -0.79658 | 2.223552 | 0.212908 |
| 5 | 121815600 | 121818599 | 18.15266 | 30.69665 | 62.9096 | 27.53114 | -0.65016 | 1.069073 | 0.21291 |
| 8 | 45034800 | 45037399 | 34.7704 | 58.22561 | 44.25229 | 19.2889 | 0.775546 | 1.194836 | 0.212915 |
| 3 | 4987000 | 4988799 | 26.04138 | 45.11834 | 34.9677 | 15.34183 | 0.752372 | 0.851535 | 0.212917 |
| X | 135390400 | 135391999 | 26.70249 | 46.28656 | 34.37095 | 15.08024 | 0.813936 | 0.849466 | 0.212927 |
| 3 | 154052400 | 154055799 | 73.42939 | 116.6159 | 101.9014 | 42.48165 | 0.526761 | 2.29257 | 0.212957 |
| 18 | 50768200 | 50770199 | 18.51081 | 33.00236 | 26.43591 | 11.63156 | 0.714938 | 0.468146 | 0.212981 |
| 4 | 129903000 | 129904199 | 6.595831 | 11.8988 | 39.06908 | 17.00296 | -1.29574 | 0.341507 | 0.212986 |
| 14 | 3070200 | 3073199 | 82.74035 | 130.3427 | 115.395 | 47.88188 | 0.498178 | 2.498798 | 0.212988 |

| | | | | | | | | | |
|---|---|---|---|---|---|---|---|---|---|
| 8 | 72193200 | 72195199 | 12.63061 | 22.02076 | 48.38891 | 21.29151 | -0.74287 | 0.729736 | 0.212989 |
| 17 | 37628200 | 37631799 | 52.55275 | 84.68957 | 71.20266 | 30.29798 | 0.605817 | 1.80996 | 0.213 |
| 8 | 121205600 | 121206999 | 8.122423 | 14.81178 | 36.5881 | 15.70969 | -0.87611 | 0.282575 | 0.213011 |
| 14 | 31379600 | 31381599 | 8.122423 | 14.81178 | 36.5881 | 15.70969 | -0.87611 | 0.282575 | 0.213011 |
| 7 | 9955000 | 9957799 | 38.48815 | 65.20763 | 33.45682 | 14.6501 | 1.340136 | 1.094578 | 0.213013 |
| 3 | 13213000 | 13217399 | 68.41117 | 109.1369 | 90.72298 | 38.03827 | 0.605333 | 2.145703 | 0.213021 |
| X | 125518600 | 125520199 | 32.51928 | 55.14243 | 39.3431 | 17.23209 | 0.866703 | 1.07345 | 0.213025 |
| 7 | 15877000 | 15881999 | 27.95569 | 45.12577 | 101.7068 | 43.37091 | -0.81078 | 1.737956 | 0.213028 |
| 4 | 102699600 | 102700799 | 13.74146 | 25.68181 | 21.30977 | 8.835952 | 0.723787 | 0.149868 | 0.213031 |
| 12 | 47795000 | 47797399 | 30.36359 | 51.59027 | 39.77212 | 17.43015 | 0.755321 | 1.045645 | 0.213033 |
| 12 | 80564800 | 80569199 | 23.86363 | 38.78313 | 88.25675 | 37.86445 | -0.81453 | 1.569321 | 0.213037 |
| 14 | 123366800 | 123369199 | 16.2973 | 28.07504 | 58.17306 | 25.51857 | -0.66213 | 0.890096 | 0.213047 |
| 3 | 66853600 | 66854799 | 8.497206 | 15.25849 | 37.97899 | 16.60964 | -0.90401 | 0.383143 | 0.213052 |
| 4 | 140150400 | 140175399 | 200.2357 | 287.2965 | 511.6446 | 211.2697 | -0.62674 | 4.196545 | 0.213088 |
| X | 54463000 | 54467999 | 84.92079 | 134.6591 | 100.0208 | 41.63368 | 0.758706 | 2.346511 | 0.213094 |
| 17 | 5585600 | 5588399 | 15.22026 | 26.14991 | 54.78478 | 24.0303 | -0.67851 | 0.907007 | 0.213099 |
| 12 | 98224800 | 98225999 | 6.20227 | 11.73142 | 30.29323 | 12.29341 | -0.8752 | 0.029002 | 0.213102 |
| X | 65272600 | 65275599 | 38.38071 | 64.8795 | 35.85204 | 15.68822 | 1.233724 | 1.11692 | 0.213106 |
| 7 | 15157400 | 15164199 | 214.1779 | 319.7456 | 221.8012 | 91.43086 | 0.787548 | 3.564764 | 0.213123 |
| 1 | 78277600 | 78279399 | 12.59898 | 22.04993 | 46.75243 | 20.62802 | -0.69126 | 0.668429 | 0.213124 |
| 9 | 27386800 | 27390399 | 30.13628 | 48.75526 | 95.18797 | 40.70146 | -0.60046 | 1.676221 | 0.213126 |
| 12 | 24008200 | 24008799 | 13.35081 | 25.74873 | 16.50395 | 6.383648 | 1.179679 | -0.09405 | 0.213143 |
| X | 75577400 | 75578799 | 21.08642 | 38.33063 | 21.04554 | 9.072189 | 1.275588 | 0.364158 | 0.213149 |
| 13 | 23906000 | 23906999 | 6.741677 | 12.87819 | 31.29628 | 12.43805 | -0.76244 | -0.03356 | 0.21315 |
| 14 | 123868000 | 123869999 | 38.25146 | 62.81128 | 54.24989 | 23.36805 | 0.588382 | 1.434413 | 0.21316 |
| 6 | 22719800 | 22720399 | 3.233215 | 6.317154 | 20.48068 | 7.134596 | -0.99111 | -0.56301 | 0.213163 |
| 11 | 116259800 | 116260399 | 3.233215 | 6.317154 | 20.48068 | 7.134596 | -0.99111 | -0.56301 | 0.213163 |
| 2 | 53762200 | 53763199 | 4.520702 | 8.810494 | 25.06591 | 9.079357 | -0.86018 | -0.3937 | 0.213165 |
| 4 | 152048400 | 152050999 | 21.34273 | 34.98052 | 84.73665 | 36.47126 | -0.89609 | 1.442029 | 0.213184 |
| 15 | 94047800 | 94049799 | 30.0663 | 51.4514 | 36.98673 | 16.21878 | 0.857203 | 0.9786 | 0.213208 |
| 6 | 42496600 | 42497399 | 6.964111 | 13.05311 | 33.9885 | 14.02428 | -0.90835 | 0.102727 | 0.21321 |
| X | 98201400 | 98203599 | 9.38413 | 17.02732 | 41.88289 | 18.09579 | -0.87609 | 0.334791 | 0.213215 |
| 2 | 87778000 | 87779199 | 24.20507 | 42.3059 | 28.60913 | 12.6087 | 0.949239 | 0.721019 | 0.213218 |
| 18 | 66780400 | 66782199 | 10.51313 | 18.67646 | 41.43404 | 18.2858 | -0.74925 | 0.46825 | 0.213232 |
| 16 | 20490800 | 20494599 | 21.69528 | 35.51842 | 81.33966 | 34.99253 | -0.81598 | 1.453094 | 0.213241 |
| 3 | 67878200 | 67879399 | 18.55607 | 33.33919 | 24.69976 | 10.79723 | 0.832995 | 0.416964 | 0.213255 |
| 13 | 63420000 | 63422199 | 8.746408 | 15.89439 | 36.5365 | 15.75579 | -0.77675 | 0.310601 | 0.213265 |
| 9 | 111515600 | 111519599 | 20.4943 | 33.79467 | 77.34496 | 33.37551 | -0.81089 | 1.355888 | 0.213266 |
| 7 | 117935400 | 117936599 | 22.8901 | 40.17673 | 28.14256 | 12.44737 | 0.898177 | 0.644891 | 0.213288 |
| 7 | 31271800 | 31274999 | 66.34217 | 105.8445 | 91.25287 | 38.26461 | 0.55296 | 2.142008 | 0.213293 |
| 2 | 172386000 | 172389799 | 31.34992 | 50.46345 | 105.3898 | 44.77137 | -0.70598 | 1.830168 | 0.2133 |
| 15 | 37542600 | 37544399 | 8.592864 | 15.58976 | 37.0285 | 16.00077 | -0.826 | 0.31728 | 0.213313 |
| 10 | 21704400 | 21706399 | 10.54701 | 18.64716 | 41.95914 | 18.55932 | -0.77278 | 0.513616 | 0.213332 |
| 1 | 164853000 | 164854399 | 13.66878 | 23.49519 | 57.95544 | 25.42141 | -0.91285 | 0.900358 | 0.213336 |
| 9 | 67642600 | 67644199 | 8.820673 | 16.27464 | 36.89248 | 15.59469 | -0.7372 | 0.22197 | 0.213339 |
| 12 | 112748200 | 112752999 | 37.42872 | 59.82071 | 124.3773 | 52.28522 | -0.71196 | 2.067411 | 0.213344 |
| 10 | 101018000 | 101018799 | 10.34709 | 18.51645 | 39.96639 | 17.52964 | -0.70334 | 0.420494 | 0.21335 |

| | | | | | | | | | |
|---|---|---|---|---|---|---|---|---|---|
| 18 | 37683800 | 37684799 | 18.08884 | 32.57663 | 24.95749 | 10.88467 | 0.78743 | 0.401871 | 0.213366 |
| 1 | 135433600 | 135438999 | 25.01394 | 40.57261 | 93.53172 | 40.06931 | -0.83596 | 1.613256 | 0.213367 |
| 8 | 50310800 | 50312599 | 20.66089 | 36.61045 | 26.70069 | 11.80494 | 0.844606 | 0.522526 | 0.213367 |
| 4 | 89190600 | 89191999 | 12.43625 | 21.77969 | 46.73848 | 20.63352 | -0.70855 | 0.656168 | 0.21337 |
| 14 | 30547400 | 30548199 | 4.246558 | 8.230179 | 25.41081 | 9.618898 | -1.03454 | -0.23586 | 0.21337 |
| 10 | 81922400 | 81925399 | 89.09363 | 140.75 | 108.8419 | 45.20418 | 0.696274 | 2.441822 | 0.213372 |
| 19 | 25773000 | 25774399 | 13.9027 | 24.19636 | 50.53467 | 22.21131 | -0.6703 | 0.764916 | 0.213372 |
| 17 | 27092400 | 27094399 | 10.27213 | 18.09105 | 43.05409 | 19.05837 | -0.85498 | 0.560095 | 0.213373 |
| X | 28405400 | 28405599 | 7.571889 | 14.80143 | 9.135248 | 2.944104 | 1.463297 | -0.80758 | 0.213373 |
| 5 | 105955200 | 105962399 | 111.3579 | 167.0613 | 314.6382 | 129.8287 | -0.64607 | 3.460446 | 0.213379 |
| 5 | 3895000 | 3895799 | 5.974817 | 11.53271 | 28.82806 | 11.12171 | -0.76147 | -0.14318 | 0.213381 |
| 16 | 96162400 | 96166399 | 21.59637 | 35.60622 | 75.5755 | 32.60973 | -0.70253 | 1.362016 | 0.213425 |
| 2 | 96147800 | 96148599 | 5.678975 | 10.99094 | 27.87708 | 10.62166 | -0.7652 | -0.19295 | 0.213438 |
| 11 | 110296200 | 110298599 | 17.84053 | 30.20711 | 61.21783 | 26.80107 | -0.6339 | 1.056325 | 0.213461 |
| 7 | 22120800 | 22121799 | 11.47529 | 22.42133 | 10.63379 | 3.702026 | 1.743835 | -0.48241 | 0.213462 |
| 7 | 19767000 | 19769799 | 37.27094 | 61.6987 | 49.92401 | 21.59667 | 0.685195 | 1.331367 | 0.213463 |
| 14 | 4330800 | 4332399 | 16.01183 | 29.52777 | 22.52961 | 9.529982 | 0.822333 | 0.248438 | 0.213465 |
| 1 | 118588600 | 118589999 | 7.233557 | 13.62021 | 32.45169 | 13.28699 | -0.77205 | 0.072934 | 0.213472 |
| 12 | 62251400 | 62252999 | 16.82184 | 30.83296 | 22.92252 | 9.77696 | 0.851013 | 0.28886 | 0.213484 |
| 1 | 75544600 | 75545999 | 7.061725 | 13.39107 | 31.57274 | 12.74391 | -0.73896 | 0.021255 | 0.21349 |
| 17 | 28832800 | 28834999 | 12.28792 | 21.56198 | 45.12495 | 19.9541 | -0.67235 | 0.622518 | 0.213493 |
| 5 | 71252600 | 71253999 | 11.03908 | 19.41497 | 43.53858 | 19.27816 | -0.77097 | 0.578202 | 0.213501 |
| 3 | 138797000 | 138797599 | 11.49771 | 22.38077 | 14.72866 | 5.411187 | 1.210881 | -0.28848 | 0.213501 |
| 16 | 35588800 | 35591799 | 14.86249 | 24.92794 | 69.02323 | 30.11637 | -1.08283 | 1.139356 | 0.213503 |
| 10 | 94777600 | 94781999 | 26.05291 | 42.06843 | 100.666 | 42.94567 | -0.89629 | 1.725079 | 0.213506 |
| 7 | 60547800 | 60549599 | 23.93852 | 41.68276 | 33.41902 | 14.69296 | 0.704553 | 0.779604 | 0.213507 |
| 17 | 19251400 | 19259199 | 181.3621 | 272.4229 | 218.7495 | 90.29246 | 0.584546 | 3.444053 | 0.213523 |
| 11 | 88871800 | 88872999 | 5.299171 | 10.10768 | 28.80123 | 11.48236 | -0.99221 | -0.04085 | 0.213525 |
| 6 | 111661600 | 111662199 | 11.01063 | 21.50648 | 11.1414 | 3.901342 | 1.610083 | -0.46883 | 0.213551 |
| 18 | 64630000 | 64632199 | 24.84812 | 43.24698 | 33.28177 | 14.62845 | 0.763238 | 0.789306 | 0.213573 |
| 10 | 117706000 | 117708799 | 14.80866 | 25.70687 | 57.72641 | 25.33127 | -0.77535 | 0.811493 | 0.213581 |
| 3 | 67272000 | 67273199 | 12.416 | 23.91824 | 17.77577 | 6.898373 | 0.964161 | -0.08983 | 0.213601 |
| X | 62297800 | 62300599 | 53.30871 | 86.46694 | 51.72978 | 22.16126 | 1.105172 | 1.646365 | 0.213613 |
| 8 | 57830800 | 57831399 | 5.115352 | 9.90647 | 26.65995 | 10.12751 | -0.84524 | -0.21302 | 0.213636 |
| 15 | 57221600 | 57222199 | 5.115352 | 9.90647 | 26.65995 | 10.12751 | -0.84524 | -0.21302 | 0.213636 |
| 8 | 3053800 | 3054599 | 10.85055 | 21.04323 | 16.02648 | 6.047797 | 0.968014 | -0.2011 | 0.213638 |
| 6 | 141191000 | 141192399 | 7.699439 | 14.41624 | 33.4134 | 13.81148 | -0.74377 | 0.11565 | 0.213641 |
| 8 | 75944800 | 75946199 | 8.43205 | 15.33872 | 36.02291 | 15.51456 | -0.80644 | 0.299318 | 0.213645 |
| 12 | 24449600 | 24461999 | 215.277 | 319.149 | 257.3145 | 105.9796 | 0.561504 | 3.688947 | 0.213652 |
| 3 | 152315600 | 152315999 | 1.946575 | 3.807342 | 17.30987 | 5.690713 | -1.38559 | -0.79904 | 0.213654 |
| 9 | 10150200 | 10151399 | 10.14771 | 18.30349 | 39.55417 | 17.21741 | -0.69774 | 0.368494 | 0.213666 |
| 17 | 34308800 | 34317799 | 64.55443 | 99.80899 | 219.5308 | 91.18799 | -0.83405 | 2.842819 | 0.213672 |
| 9 | 70129800 | 70130599 | 4.505198 | 8.708017 | 26.39861 | 10.11906 | -1.02568 | -0.18508 | 0.213674 |
| 6 | 131708800 | 131713599 | 68.67769 | 109.8051 | 83.64683 | 35.1815 | 0.734522 | 2.078112 | 0.213684 |
| 10 | 13493000 | 13493999 | 11.95611 | 22.99823 | 18.39289 | 7.168279 | 0.85379 | -0.07933 | 0.213686 |
| 14 | 27632000 | 27633999 | 28.50679 | 49.52798 | 31.44795 | 13.80723 | 1.039133 | 0.828613 | 0.213688 |
| 9 | 41987800 | 41988999 | 6.134682 | 11.79831 | 28.6478 | 11.1706 | -0.73465 | -0.10766 | 0.213705 |

| | | | | | | | | | |
|---|---|---|---|---|---|---|---|---|---|
| 18 | 40718600 | 40719399 | 10.67712 | 20.80063 | 14.08267 | 5.12756 | 1.180553 | -0.33302 | 0.213716 |
| 7 | 41123600 | 41124599 | 29.62929 | 51.31017 | 31.96229 | 14.02303 | 1.065879 | 0.867644 | 0.213717 |
| 1 | 14253400 | 14254799 | 9.184475 | 16.68027 | 37.03871 | 15.98458 | -0.72842 | 0.315683 | 0.213723 |
| 16 | 76287000 | 76288599 | 9.64928 | 17.53651 | 38.12343 | 16.43852 | -0.69731 | 0.317863 | 0.213753 |
| 17 | 34863000 | 34868999 | 31.65509 | 50.98909 | 103.3754 | 43.95857 | -0.66207 | 1.808542 | 0.213763 |
| 1 | 43888400 | 43889399 | 13.65018 | 25.87238 | 19.6654 | 7.945651 | 0.879711 | 0.054156 | 0.213766 |
| 10 | 22415600 | 22422799 | 45.57906 | 74.55436 | 50.07744 | 21.53637 | 0.94753 | 1.496293 | 0.213768 |
| 5 | 149102800 | 149109399 | 38.49732 | 61.68267 | 117.1047 | 49.40692 | -0.57699 | 1.98892 | 0.213777 |
| 1 | 154598800 | 154604199 | 39.09598 | 62.32679 | 135.126 | 56.60105 | -0.77631 | 2.151228 | 0.213781 |
| 10 | 53318000 | 53319199 | 12.52432 | 23.77741 | 19.82667 | 7.983079 | 0.751593 | 0.032969 | 0.213783 |
| 14 | 13872600 | 13873999 | 15.67306 | 28.5279 | 23.65901 | 10.18197 | 0.685843 | 0.320621 | 0.213814 |
| 3 | 81980200 | 81982799 | 10.78352 | 18.8656 | 48.33631 | 21.32016 | -0.96278 | 0.666627 | 0.213814 |
| 13 | 58298200 | 58303599 | 38.79855 | 62.25378 | 115.6306 | 48.89062 | -0.54305 | 1.944422 | 0.213822 |
| 11 | 6158800 | 6162199 | 46.77473 | 75.67651 | 64.61302 | 27.62875 | 0.590023 | 1.689579 | 0.213826 |
| 7 | 99004200 | 99007799 | 14.39817 | 24.85054 | 53.2033 | 23.33986 | -0.70822 | 0.859244 | 0.213827 |
| 5 | 141761600 | 141768999 | 54.32703 | 85.91041 | 155.6709 | 64.68546 | -0.52937 | 2.402499 | 0.213828 |
| 10 | 112024800 | 112025399 | 8.973163 | 17.44662 | 15.55164 | 5.766691 | 0.765512 | -0.28678 | 0.213831 |
| 10 | 42719600 | 42722599 | 43.82019 | 71.03703 | 59.6084 | 25.52603 | 0.617573 | 1.640051 | 0.213849 |
| 7 | 44739000 | 44741599 | 11.41095 | 19.8953 | 48.37719 | 21.28698 | -0.88796 | 0.723665 | 0.213857 |
| 5 | 3146400 | 3150999 | 126.1629 | 193.2814 | 157.7611 | 65.47871 | 0.584491 | 3.034476 | 0.213861 |
| 11 | 59014800 | 59019599 | 42.4458 | 67.7739 | 125.8516 | 52.83275 | -0.55077 | 2.103239 | 0.213862 |
| 8 | 119049000 | 119051199 | 11.78347 | 20.80479 | 43.49801 | 19.24525 | -0.66839 | 0.539016 | 0.213868 |
| 5 | 117253000 | 117254599 | 9.824785 | 17.30998 | 44.06528 | 19.50446 | -0.95173 | 0.551325 | 0.213882 |
| 2 | 112068600 | 112075199 | 33.22802 | 53.21605 | 121.4502 | 51.20542 | -0.84273 | 1.997804 | 0.213883 |
| X | 102550000 | 102552599 | 9.34068 | 16.95885 | 37.03261 | 15.98797 | -0.70478 | 0.318949 | 0.213888 |
| 5 | 128196400 | 128198799 | 26.81108 | 45.72175 | 38.92688 | 17.06477 | 0.61346 | 0.999222 | 0.213892 |
| X | 24283200 | 24284199 | 14.39031 | 27.91721 | 13.65065 | 5.14343 | 1.600951 | -0.17388 | 0.213894 |
| 10 | 49814200 | 49817399 | 52.26962 | 84.29816 | 70.09719 | 29.8574 | 0.622948 | 1.787459 | 0.213913 |
| 6 | 103808200 | 103810399 | 16.14789 | 27.68815 | 57.54842 | 25.23917 | -0.66786 | 0.930294 | 0.213929 |
| 7 | 74394400 | 74398799 | 28.31619 | 46.09725 | 92.33621 | 39.64138 | -0.62937 | 1.555948 | 0.213937 |
| 10 | 118898200 | 118901599 | 20.65003 | 34.05058 | 76.77142 | 33.12765 | -0.78934 | 1.356455 | 0.213958 |
| 7 | 27092000 | 27104399 | 100.1266 | 152.107 | 265.8819 | 110.115 | -0.52293 | 3.204675 | 0.213958 |
| 1 | 90923000 | 90928599 | 38.30603 | 61.04901 | 132.2013 | 55.35284 | -0.77504 | 2.160704 | 0.213964 |
| 2 | 98285200 | 98289199 | 96.89606 | 151.7837 | 126.0543 | 52.2868 | 0.584079 | 2.616117 | 0.213974 |
| 12 | 55031400 | 55033199 | 8.853523 | 15.51041 | 49.34809 | 21.79173 | -1.27296 | 0.635549 | 0.213975 |
| 3 | 128990200 | 128991199 | 13.57157 | 25.32234 | 20.85817 | 8.672983 | 0.731121 | 0.15773 | 0.213981 |
| 18 | 65257800 | 65258999 | 9.329202 | 16.9732 | 37.07247 | 15.96369 | -0.70271 | 0.306595 | 0.213983 |
| 8 | 103717600 | 103718599 | 14.03992 | 26.06108 | 21.13396 | 8.860485 | 0.743827 | 0.191916 | 0.213986 |
| 18 | 3216600 | 3222599 | 127.7606 | 195.3507 | 164.4765 | 68.23536 | 0.537567 | 3.073593 | 0.213996 |
| 5 | 66711800 | 66713199 | 8.207442 | 15.13543 | 35.76 | 15.12655 | -0.79718 | 0.216522 | 0.213997 |
| 7 | 14783000 | 14792999 | 232.293 | 343.6972 | 255.4121 | 105.1859 | 0.676413 | 3.724105 | 0.21401 |
| 14 | 77467400 | 77469799 | 17.79043 | 30.2714 | 60.78068 | 26.64028 | -0.62004 | 1.011107 | 0.214012 |
| X | 33565600 | 33572799 | 146.3782 | 222.6538 | 169.8238 | 70.35938 | 0.674056 | 3.185718 | 0.214016 |
| 13 | 22995600 | 22998399 | 61.92741 | 99.37345 | 76.82448 | 32.48695 | 0.719207 | 1.956041 | 0.214035 |
| 5 | 3390400 | 3395399 | 32.52017 | 52.22192 | 111.0086 | 46.99387 | -0.73577 | 1.910103 | 0.21405 |
| 17 | 56423000 | 56434199 | 119.8575 | 179.0149 | 333.3235 | 137.4145 | -0.63544 | 3.545909 | 0.214053 |
| 17 | 20187600 | 20189199 | 34.9074 | 57.94236 | 49.29211 | 21.36311 | 0.613489 | 1.295254 | 0.214065 |

| | | | | | | | | | |
|---|---|---|---|---|---|---|---|---|---|
| 1 | 81175400 | 81176599 | 9.899932 | 17.64193 | 39.42884 | 17.35526 | -0.75674 | 0.44298 | 0.214082 |
| 9 | 114804600 | 114812399 | 45.34401 | 71.91088 | 151.7797 | 63.18608 | -0.74554 | 2.321629 | 0.214106 |
| 13 | 63289800 | 63293999 | 48.07572 | 76.30593 | 143.5124 | 59.79367 | -0.57824 | 2.298766 | 0.214144 |
| 1 | 131080800 | 131081599 | 4.650487 | 9.059377 | 24.30381 | 8.855085 | -0.78362 | -0.37176 | 0.214145 |
| 8 | 42728400 | 42728799 | 4.650487 | 9.059377 | 24.30381 | 8.855085 | -0.78362 | -0.37176 | 0.214145 |
| 13 | 59977000 | 59977599 | 4.650487 | 9.059377 | 24.30381 | 8.855085 | -0.78362 | -0.37176 | 0.214145 |
| 15 | 38812600 | 38815199 | 9.615718 | 17.07861 | 40.94507 | 18.07321 | -0.86042 | 0.463391 | 0.214167 |
| 7 | 80707200 | 80710999 | 25.39272 | 41.39633 | 85.0083 | 36.51416 | -0.6636 | 1.523927 | 0.214175 |
| 4 | 145622400 | 145623199 | 18.50673 | 34.10448 | 19.92596 | 8.436782 | 1.203145 | 0.265543 | 0.214179 |
| 16 | 31197200 | 31198799 | 35.19262 | 58.25632 | 50.40944 | 21.80616 | 0.588739 | 1.327238 | 0.214179 |
| 4 | 88533600 | 88534999 | 12.19249 | 23.68069 | 13.05393 | 4.882444 | 1.437493 | -0.22733 | 0.214188 |
| 18 | 84494800 | 84495199 | 2.45244 | 4.797474 | 20.43473 | 6.796425 | -1.31646 | -0.74228 | 0.214192 |
| 1 | 73152400 | 73152999 | 4.113602 | 8.006343 | 24.06877 | 8.851161 | -0.95767 | -0.34603 | 0.214193 |
| 7 | 126622200 | 126625599 | 13.74554 | 23.43553 | 58.35984 | 25.58334 | -0.92844 | 0.976195 | 0.214205 |
| 9 | 25759400 | 25762599 | 38.22271 | 62.79701 | 52.69632 | 22.70389 | 0.630062 | 1.424309 | 0.214217 |
| 1 | 173815600 | 173816199 | 4.230776 | 8.246895 | 23.14923 | 8.365875 | -0.83684 | -0.40999 | 0.214219 |
| 17 | 18394600 | 18396999 | 41.06569 | 67.46076 | 53.8477 | 23.19893 | 0.702048 | 1.435913 | 0.21422 |
| 7 | 47367200 | 47369199 | 29.61369 | 50.551 | 38.69043 | 16.96246 | 0.766678 | 1.001041 | 0.214223 |
| 5 | 135572200 | 135574799 | 14.54719 | 24.72105 | 60.52557 | 26.52572 | -0.90504 | 1.011066 | 0.214224 |
| 15 | 79697400 | 79703999 | 41.68946 | 66.40752 | 133.0566 | 55.67186 | -0.66402 | 2.180309 | 0.214229 |
| 13 | 66391600 | 66393399 | 41.202 | 67.69737 | 48.87913 | 21.06001 | 0.846428 | 1.430789 | 0.214232 |
| 5 | 3852200 | 3853999 | 11.98691 | 20.97372 | 46.75952 | 20.62699 | -0.76322 | 0.66907 | 0.214236 |
| 3 | 135190200 | 135191599 | 6.995949 | 13.02855 | 33.77745 | 14.09012 | -0.91465 | 0.145227 | 0.214237 |
| 6 | 123280600 | 123281999 | 13.98859 | 26.01856 | 21.7285 | 9.083068 | 0.705092 | 0.18229 | 0.214256 |
| X | 25388000 | 25390999 | 57.30689 | 92.42495 | 65.70941 | 27.98959 | 0.848574 | 1.795612 | 0.214259 |
| 18 | 45985200 | 45988999 | 49.84616 | 80.59089 | 66.13499 | 28.2607 | 0.646055 | 1.710887 | 0.214267 |
| 11 | 104565400 | 104566199 | 7.231506 | 13.18039 | 35.5518 | 15.27451 | -1.00167 | 0.280061 | 0.214277 |
| 12 | 85335000 | 85337799 | 15.2354 | 26.10629 | 55.83659 | 24.4871 | -0.70893 | 0.932226 | 0.214278 |
| 14 | 42668800 | 42671599 | 35.21725 | 59.12496 | 41.98255 | 18.32911 | 0.873127 | 1.167311 | 0.214288 |
| 13 | 70686600 | 70687999 | 7.38702 | 13.48959 | 36.08508 | 15.46662 | -0.98818 | 0.268356 | 0.214308 |
| 5 | 110244800 | 110256399 | 161.8323 | 235.1881 | 459.0909 | 189.2436 | -0.74078 | 4.003105 | 0.214313 |
| 4 | 19575000 | 19576799 | 9.380765 | 16.88608 | 38.46151 | 16.78406 | -0.77543 | 0.371717 | 0.214319 |
| 7 | 66223200 | 66226599 | 19.50669 | 32.7171 | 65.55999 | 28.60513 | -0.61825 | 1.15214 | 0.214323 |
| 5 | 133865600 | 133874199 | 56.48052 | 88.73627 | 186.4059 | 77.32002 | -0.74977 | 2.548082 | 0.214332 |
| X | 8381200 | 8383199 | 36.72036 | 61.65596 | 40.97497 | 17.89059 | 0.968465 | 1.17355 | 0.214336 |
| 8 | 36690000 | 36692199 | 9.265835 | 16.58707 | 39.45837 | 17.30192 | -0.84251 | 0.407979 | 0.214365 |
| 13 | 66383600 | 66389999 | 23.51427 | 41.21425 | 27.39949 | 12.10431 | 0.973449 | 0.670695 | 0.214371 |
| 12 | 104654600 | 104658199 | 14.78489 | 25.2989 | 56.33013 | 24.70076 | -0.76696 | 0.944173 | 0.21438 |
| 7 | 49310000 | 49314399 | 24.12946 | 39.73153 | 82.62121 | 35.63051 | -0.67412 | 1.38533 | 0.214391 |
| 7 | 19478400 | 19494799 | 100.2278 | 151.2475 | 298.122 | 123.2116 | -0.70421 | 3.341668 | 0.214399 |
| 13 | 41847000 | 41853199 | 53.6389 | 84.93758 | 153.4432 | 63.80366 | -0.52344 | 2.369279 | 0.214409 |
| 1 | 157388200 | 157388999 | 5.246274 | 10.10233 | 27.43152 | 10.65802 | -0.88814 | -0.12937 | 0.214409 |
| 6 | 117822000 | 117824199 | 11.6395 | 20.47619 | 44.55503 | 19.72722 | -0.72784 | 0.579307 | 0.214412 |
| 17 | 33048800 | 33049999 | 7.72026 | 14.42482 | 32.77562 | 13.59625 | -0.71909 | 0.12983 | 0.214415 |
| 2 | 59566800 | 59569799 | 16.22094 | 27.59759 | 60.13824 | 26.35845 | -0.7376 | 1.00524 | 0.214422 |
| X | 78793000 | 78794399 | 21.71677 | 38.37045 | 27.25025 | 12.05487 | 0.881325 | 0.561308 | 0.214428 |
| 14 | 20979000 | 20980599 | 7.920303 | 14.20959 | 38.49225 | 16.84705 | -1.02616 | 0.38448 | 0.214437 |

| | | | | | | | | | |
|---|---|---|---|---|---|---|---|---|---|
| 7 | 15862200 | 15866799 | 29.92404 | 48.15106 | 105.8947 | 44.96259 | -0.78097 | 1.840258 | 0.21444 |
| 6 | 142090000 | 142090999 | 12.60272 | 24.27274 | 17.7694 | 6.900153 | 0.984964 | -0.08578 | 0.21444 |
| 18 | 8072800 | 8073599 | 12.60272 | 24.27274 | 17.7694 | 6.900153 | 0.984964 | -0.08578 | 0.21444 |
| 4 | 124608200 | 124609199 | 4.935975 | 9.352703 | 30.82609 | 12.46318 | -1.21858 | 0.005804 | 0.214441 |
| 1 | 28928400 | 28930999 | 41.74781 | 67.93158 | 56.86444 | 24.40865 | 0.625707 | 1.566987 | 0.214457 |
| 1 | 183411400 | 183412999 | 8.5209 | 15.65111 | 38.42778 | 16.34322 | -0.85829 | 0.246308 | 0.214458 |
| 9 | 49337400 | 49338599 | 5.58854 | 10.60227 | 29.77096 | 12.00517 | -0.98575 | 0.003809 | 0.214463 |
| 13 | 29868600 | 29869999 | 25.14 | 43.36466 | 31.32967 | 13.74382 | 0.852053 | 0.881588 | 0.21447 |
| 9 | 113088000 | 113089199 | 18.83416 | 33.88643 | 24.45996 | 10.67975 | 0.871905 | 0.412268 | 0.214472 |
| 2 | 42642400 | 42643399 | 18.96749 | 33.68305 | 26.1341 | 11.53718 | 0.757678 | 0.496292 | 0.214473 |
| 15 | 12984600 | 12985399 | 12.23048 | 23.56591 | 17.78214 | 6.896593 | 0.943174 | -0.09389 | 0.214485 |
| 11 | 69855200 | 69858799 | 23.33324 | 38.05482 | 85.25368 | 36.62515 | -0.78832 | 1.513659 | 0.214486 |
| 11 | 8004200 | 8005599 | 7.556165 | 14.15627 | 32.29678 | 13.33668 | -0.71964 | 0.109359 | 0.214489 |
| 7 | 89778800 | 89780399 | 7.554864 | 13.75238 | 38.15368 | 16.41829 | -1.04455 | 0.29139 | 0.214508 |
| X | 54521000 | 54523599 | 25.55069 | 44.59651 | 28.55078 | 12.57149 | 1.027778 | 0.747467 | 0.214514 |
| 4 | 85487200 | 85488999 | 17.78509 | 30.32817 | 61.37851 | 26.90703 | -0.63094 | 0.992019 | 0.214535 |
| 15 | 31351400 | 31358199 | 51.33047 | 81.35144 | 152.8889 | 63.60607 | -0.57933 | 2.345802 | 0.214542 |
| 7 | 40893400 | 40900999 | 163.4372 | 246.5004 | 208.6885 | 86.23474 | 0.513437 | 3.360199 | 0.214547 |
| 10 | 72010600 | 72011399 | 12.14854 | 21.29163 | 45.68296 | 20.18021 | -0.708 | 0.643257 | 0.214547 |
| 11 | 33019200 | 33022599 | 17.58469 | 29.57618 | 65.52058 | 28.6269 | -0.76239 | 1.116693 | 0.214563 |
| 4 | 38236000 | 38237399 | 13.53279 | 26.13571 | 15.92723 | 6.128774 | 1.258633 | -0.11013 | 0.214569 |
| 13 | 24948000 | 24949599 | 9.990885 | 18.07449 | 42.36147 | 18.3707 | -0.81075 | 0.355167 | 0.214575 |
| 7 | 11024400 | 11025799 | 30.55804 | 51.86274 | 40.33107 | 17.66957 | 0.742793 | 1.056658 | 0.214577 |
| 8 | 5549800 | 5551599 | 18.14512 | 33.78734 | 19.02231 | 7.936785 | 1.27158 | 0.204156 | 0.214579 |
| 14 | 50526800 | 50527599 | 11.71104 | 22.72367 | 16.30834 | 6.139444 | 1.055146 | -0.19959 | 0.214583 |
| 3 | 30704200 | 30705199 | 6.934766 | 13.09833 | 30.82886 | 12.54835 | -0.74663 | 0.047791 | 0.214586 |
| 7 | 84836400 | 84840199 | 87.7918 | 138.0014 | 121.7655 | 50.51101 | 0.500646 | 2.546122 | 0.214589 |
| 2 | 142496600 | 142497999 | 16.24394 | 27.59396 | 59.5315 | 26.08868 | -0.72346 | 1.020115 | 0.21459 |
| 7 | 141907600 | 141909799 | 11.08642 | 19.38383 | 47.29771 | 20.85141 | -0.89262 | 0.677642 | 0.214605 |
| 14 | 25041400 | 25043399 | 7.336888 | 13.52339 | 35.18512 | 14.89331 | -0.93558 | 0.211618 | 0.214614 |
| 10 | 71469000 | 71470799 | 23.29197 | 40.75789 | 31.11246 | 13.72561 | 0.775294 | 0.701737 | 0.214621 |
| 1 | 184550600 | 184553799 | 19.46811 | 32.76838 | 65.11802 | 28.45745 | -0.60621 | 1.116859 | 0.214631 |
| 6 | 123855000 | 123856999 | 43.13406 | 70.08077 | 60.35573 | 25.88856 | 0.583076 | 1.594185 | 0.214636 |
| 15 | 90140400 | 90141199 | 14.62823 | 27.12719 | 20.856 | 8.755321 | 0.819564 | 0.203897 | 0.214662 |
| 13 | 56446600 | 56464999 | 148.2592 | 218.9431 | 382.5863 | 157.5566 | -0.55902 | 3.748777 | 0.214666 |
| 2 | 38240000 | 38248199 | 65.14534 | 100.6336 | 214.6195 | 89.14837 | -0.79057 | 2.858141 | 0.214667 |
| 11 | 94966400 | 94968999 | 8.397539 | 14.97159 | 39.48494 | 17.37314 | -0.99325 | 0.430284 | 0.214674 |
| 19 | 35465000 | 35466999 | 22.40435 | 39.15999 | 31.12736 | 13.71896 | 0.71705 | 0.712981 | 0.214688 |
| 2 | 26502800 | 26504199 | 7.603087 | 14.08038 | 33.66196 | 14.15734 | -0.80751 | 0.184258 | 0.214704 |
| 5 | 70349800 | 70350999 | 19.26644 | 33.97839 | 28.07831 | 12.42576 | 0.662871 | 0.566319 | 0.214705 |
| 10 | 78810200 | 78811599 | 11.45268 | 20.26241 | 44.02204 | 19.46909 | -0.7224 | 0.514105 | 0.214709 |
| 16 | 44953000 | 44954599 | 22.17793 | 38.95346 | 29.46458 | 13.03966 | 0.788204 | 0.634649 | 0.214715 |
| 4 | 69231800 | 69233199 | 17.07844 | 31.44899 | 21.77925 | 9.230815 | 0.958936 | 0.262559 | 0.214725 |
| X | 50930600 | 50931799 | 14.91087 | 28.65472 | 16.48067 | 6.440525 | 1.32168 | -0.0449 | 0.214761 |
| 16 | 66228400 | 66230199 | 12.76358 | 22.41972 | 48.27695 | 21.3665 | -0.71347 | 0.613162 | 0.214779 |
| 11 | 75604000 | 75605399 | 22.09423 | 38.63893 | 31.78156 | 14.01373 | 0.66811 | 0.705492 | 0.214791 |
| 3 | 121750400 | 121750999 | 8.554492 | 16.70909 | 13.2779 | 4.649222 | 1.001926 | -0.49827 | 0.214798 |

| | | | | | | | | | |
|---|---|---|---|---|---|---|---|---|---|
| 4 | 10555800 | 10556399 | 8.554492 | 16.70909 | 13.2779 | 4.649222 | 1.001926 | -0.49827 | 0.214798 |
| 11 | 116313000 | 116315199 | 14.25683 | 24.58635 | 53.18243 | 23.33012 | -0.72318 | 0.866857 | 0.21481 |
| 14 | 108852400 | 108855199 | 27.62767 | 47.2551 | 37.64882 | 16.50846 | 0.709303 | 0.972532 | 0.21481 |
| 14 | 104702200 | 104705799 | 31.52252 | 50.71151 | 106.2732 | 45.10824 | -0.71199 | 1.849656 | 0.214815 |
| 3 | 98516400 | 98517399 | 8.983031 | 16.4967 | 36.75694 | 15.63681 | -0.71925 | 0.249164 | 0.214828 |
| 4 | 58991400 | 58992799 | 8.983031 | 16.4967 | 36.75694 | 15.63681 | -0.71925 | 0.249164 | 0.214828 |
| 7 | 40975800 | 40980399 | 79.74274 | 126.0459 | 111.7197 | 46.41217 | 0.500325 | 2.422242 | 0.214834 |
| 7 | 16624800 | 16629599 | 21.25141 | 34.76387 | 84.76332 | 36.45442 | -0.90699 | 1.464386 | 0.214867 |
| 9 | 36352400 | 36355199 | 42.72312 | 69.51582 | 60.33513 | 25.89804 | 0.573578 | 1.571045 | 0.214872 |
| 4 | 3633000 | 3637199 | 107.503 | 166.2605 | 145.5146 | 60.44328 | 0.494278 | 2.845852 | 0.214876 |
| 13 | 66442000 | 66456199 | 270.9264 | 396.4225 | 282.2384 | 116.2674 | 0.723068 | 3.906349 | 0.214877 |
| 11 | 94338800 | 94343599 | 19.25327 | 31.72223 | 76.96562 | 33.2003 | -0.89515 | 1.364039 | 0.214883 |
| 1 | 49849800 | 49851799 | 20.38511 | 35.98694 | 27.80869 | 12.304 | 0.759575 | 0.561652 | 0.21489 |
| 1 | 101011800 | 101012399 | 9.944486 | 19.32525 | 16.52687 | 6.153759 | 0.819266 | -0.25703 | 0.214892 |
| 16 | 22503600 | 22504799 | 7.738337 | 14.35152 | 35.35048 | 14.83836 | -0.8487 | 0.179228 | 0.214898 |
| X | 90903600 | 90905399 | 21.40128 | 38.30169 | 23.98776 | 10.52047 | 1.070947 | 0.460085 | 0.214898 |
| 12 | 98181800 | 98185199 | 20.79101 | 34.76258 | 69.59548 | 30.30618 | -0.61706 | 1.192033 | 0.214901 |
| 4 | 156214600 | 156225399 | 120.3565 | 179.4871 | 344.8864 | 142.1394 | -0.68281 | 3.574381 | 0.214904 |
| 2 | 79904600 | 79906399 | 11.33116 | 19.93279 | 46.13611 | 20.42745 | -0.81661 | 0.579673 | 0.214907 |
| 19 | 3413600 | 3432599 | 149.3696 | 220.0712 | 392.7988 | 161.7583 | -0.59273 | 3.787513 | 0.214926 |
| 12 | 87806200 | 87808399 | 15.25317 | 28.09836 | 22.71253 | 9.621483 | 0.738315 | 0.249341 | 0.214927 |
| 6 | 8626600 | 8631199 | 34.57222 | 55.57504 | 109.3192 | 46.34957 | -0.62231 | 1.87857 | 0.21496 |
| 1 | 171849400 | 171850399 | 5.146417 | 9.876695 | 27.88728 | 10.94048 | -0.95774 | -0.09946 | 0.214963 |
| 6 | 29347600 | 29348599 | 5.146417 | 9.876695 | 27.88728 | 10.94048 | -0.95774 | -0.09946 | 0.214963 |
| 2 | 162999000 | 163012199 | 106.4427 | 160.4827 | 306.6478 | 126.7 | -0.66055 | 3.361072 | 0.214995 |
| 17 | 18555000 | 18567599 | 331.7132 | 472.2333 | 386.3394 | 159.7304 | 0.483736 | 4.315644 | 0.215004 |
| 1 | 116670800 | 116671599 | 11.85289 | 22.88708 | 17.24424 | 6.641332 | 0.954392 | -0.12221 | 0.215005 |
| 12 | 55349000 | 55350399 | 10.0035 | 17.95366 | 38.94188 | 17.03849 | -0.70803 | 0.396011 | 0.215008 |
| 4 | 153309000 | 153340599 | 285.7514 | 399.2745 | 670.2926 | 277.9703 | -0.58006 | 4.590685 | 0.21503 |
| 7 | 26427200 | 26428399 | 24.27312 | 42.40123 | 31.63132 | 13.93506 | 0.808342 | 0.734273 | 0.215043 |
| 13 | 67958200 | 67960399 | 17.39397 | 29.36489 | 62.23511 | 27.22225 | -0.6984 | 1.081176 | 0.215055 |
| 4 | 102783400 | 102786799 | 19.96059 | 33.50278 | 67.36467 | 29.40605 | -0.6233 | 1.146165 | 0.215065 |
| 12 | 74274600 | 74276399 | 12.6243 | 22.05333 | 47.31371 | 20.84431 | -0.7081 | 0.700469 | 0.215067 |
| 2 | 47854000 | 47855399 | 10.96419 | 19.52863 | 41.45914 | 18.25668 | -0.68394 | 0.455216 | 0.215068 |
| 1 | 134686800 | 134688999 | 11.82963 | 20.77541 | 44.5963 | 19.73447 | -0.70858 | 0.606384 | 0.215071 |
| 14 | 47588800 | 47592999 | 29.8071 | 48.25143 | 94.0517 | 40.23327 | -0.59731 | 1.663075 | 0.215076 |
| 15 | 82782800 | 82784599 | 9.100806 | 15.96758 | 45.17725 | 19.97525 | -1.10415 | 0.607922 | 0.215094 |
| 1 | 97196200 | 97196799 | 10.16313 | 19.73552 | 16.41841 | 6.150058 | 0.850315 | -0.23532 | 0.215105 |
| 7 | 15399600 | 15403999 | 96.72744 | 153.1557 | 92.62593 | 38.51341 | 1.052892 | 2.392397 | 0.215106 |
| 9 | 122126200 | 122126799 | 3.041386 | 5.944079 | 19.92513 | 6.88929 | -1.02699 | -0.59226 | 0.215107 |
| 8 | 92406800 | 92408399 | 7.568902 | 14.11486 | 33.28131 | 13.85103 | -0.77621 | 0.140379 | 0.215145 |
| 9 | 7294000 | 7295799 | 16.09878 | 29.76163 | 21.92881 | 9.244684 | 0.875628 | 0.233726 | 0.215164 |
| 12 | 73237600 | 73239999 | 24.34241 | 42.04243 | 35.71273 | 15.66721 | 0.619779 | 0.872048 | 0.215166 |
| 17 | 38564400 | 38565999 | 27.06554 | 47.61093 | 24.08024 | 10.66191 | 1.365769 | 0.634256 | 0.215166 |
| 8 | 21606600 | 21610599 | 69.38722 | 110.9145 | 83.36019 | 35.04975 | 0.753574 | 2.0845 | 0.215169 |
| 12 | 105718000 | 105724599 | 51.64968 | 81.87967 | 151.9684 | 63.24036 | -0.56092 | 2.337922 | 0.215174 |
| 10 | 60181800 | 60186199 | 21.77649 | 35.83509 | 76.63232 | 33.03919 | -0.71442 | 1.386251 | 0.215182 |

| | | | | | | | | | |
|---|---|---|---|---|---|---|---|---|---|
| 15 | 96133800 | 96136199 | 17.00722 | 28.98902 | 58.53871 | 25.66121 | -0.62769 | 0.99114 | 0.215183 |
| X | 148164800 | 148166599 | 23.76019 | 41.38304 | 29.38435 | 12.9212 | 0.878848 | 0.7681 | 0.215184 |
| 18 | 37947800 | 37955199 | 85.90807 | 130.8953 | 253.1996 | 104.956 | -0.6654 | 3.140021 | 0.215197 |
| 7 | 34346600 | 34349399 | 25.41166 | 41.37987 | 85.34142 | 36.64197 | -0.67094 | 1.541539 | 0.215219 |
| 16 | 22946200 | 22946999 | 4.651747 | 9.057806 | 24.19434 | 8.865134 | -0.7847 | -0.35134 | 0.215233 |
| 16 | 64782000 | 64783199 | 12.92052 | 24.73496 | 18.78874 | 7.423621 | 0.909145 | -0.0252 | 0.215237 |
| 6 | 68749000 | 68749999 | 22.29575 | 38.90132 | 30.5652 | 13.45544 | 0.733847 | 0.739346 | 0.215251 |
| 4 | 9363000 | 9364199 | 15.49029 | 28.45474 | 23.09874 | 9.822738 | 0.728076 | 0.266711 | 0.215267 |
| 7 | 134417800 | 134419599 | 15.49029 | 28.45474 | 23.09874 | 9.822738 | 0.728076 | 0.266711 | 0.215267 |
| 17 | 7402800 | 7406999 | 27.52139 | 44.69448 | 88.91118 | 38.12431 | -0.6222 | 1.591442 | 0.215293 |
| 9 | 55418800 | 55419599 | 6.101482 | 11.4359 | 31.71558 | 13.08711 | -0.99748 | 0.092575 | 0.215301 |
| 2 | 80145600 | 80146599 | 4.69229 | 9.110271 | 26.24066 | 9.84296 | -0.92462 | -0.26311 | 0.215317 |
| 6 | 123851400 | 123852999 | 19.50751 | 35.7065 | 20.98344 | 8.966632 | 1.185875 | 0.312946 | 0.215321 |
| 15 | 92015800 | 92016799 | 21.23256 | 38.99324 | 17.40113 | 7.402337 | 1.583672 | 0.300373 | 0.215325 |
| 14 | 95297600 | 95298799 | 11.72499 | 22.75337 | 15.61353 | 5.874707 | 1.119089 | -0.20435 | 0.21534 |
| 3 | 15492600 | 15493199 | 8.432429 | 16.49298 | 10.85676 | 3.659981 | 1.316594 | -0.63649 | 0.215342 |
| 4 | 123684200 | 123685199 | 6.277678 | 12.08629 | 30.53944 | 11.87156 | -0.78774 | -0.11173 | 0.215344 |
| 14 | 62788600 | 62789599 | 8.882812 | 16.17775 | 36.03679 | 15.49768 | -0.72928 | 0.296736 | 0.215357 |
| 17 | 42125800 | 42128599 | 23.63862 | 41.46367 | 29.34781 | 12.97334 | 0.883578 | 0.663296 | 0.215362 |
| 1 | 25513400 | 25513999 | 9.307429 | 18.11182 | 13.06089 | 4.810187 | 1.077788 | -0.31392 | 0.215364 |
| 7 | 5834600 | 5835199 | 12.67967 | 24.07026 | 19.4363 | 7.827332 | 0.797877 | 0.035565 | 0.215395 |
| 7 | 9174200 | 9176799 | 28.70778 | 50.34549 | 24.77093 | 10.94831 | 1.405344 | 0.688585 | 0.215397 |
| 4 | 67958800 | 67959799 | 9.995986 | 19.50396 | 13.69902 | 4.885334 | 1.155037 | -0.41333 | 0.215401 |
| 7 | 11922400 | 11923799 | 20.18521 | 37.76559 | 16.14428 | 6.681849 | 1.674329 | 0.186492 | 0.215402 |
| 17 | 6837200 | 6843399 | 29.64277 | 47.98415 | 95.92615 | 41.03431 | -0.63377 | 1.663504 | 0.215405 |
| 10 | 126734000 | 126737599 | 21.88293 | 36.13147 | 76.55902 | 33.05952 | -0.69963 | 1.342589 | 0.215435 |
| 1 | 103360600 | 103362599 | 26.15352 | 44.92541 | 37.7437 | 16.55546 | 0.634061 | 0.930338 | 0.215442 |
| 7 | 23059200 | 23062799 | 75.66821 | 120.7621 | 82.70852 | 34.69736 | 0.884898 | 2.138938 | 0.215453 |
| 10 | 43451800 | 43459399 | 51.31958 | 81.3781 | 150.7845 | 62.76451 | -0.55813 | 2.329617 | 0.215459 |
| 6 | 130959600 | 130962999 | 92.21036 | 144.3337 | 127.7655 | 53.00078 | 0.491284 | 2.627813 | 0.215471 |
| X | 61950800 | 61954799 | 50.39887 | 83.14221 | 40.52703 | 17.49119 | 1.413518 | 1.401963 | 0.215483 |
| 8 | 105984200 | 105988799 | 34.93467 | 56.07775 | 110.7137 | 46.84178 | -0.63007 | 1.925766 | 0.215498 |
| X | 63587400 | 63590199 | 34.18091 | 58.81625 | 30.79652 | 13.50894 | 1.313668 | 0.942651 | 0.215503 |
| 16 | 97872000 | 97873999 | 50.34864 | 81.88172 | 56.31447 | 24.16773 | 0.907871 | 1.592058 | 0.21555 |
| 9 | 72040000 | 72041199 | 5.861225 | 11.2376 | 28.4803 | 11.2043 | -0.8079 | -0.08223 | 0.215553 |
| 11 | 96661400 | 96664199 | 10.30073 | 18.0739 | 44.64365 | 19.74026 | -0.90974 | 0.612222 | 0.215559 |
| 13 | 62482800 | 62490599 | 197.0703 | 293.6083 | 245.824 | 101.2981 | 0.51367 | 3.599216 | 0.215571 |
| 5 | 74271200 | 74272999 | 29.19991 | 49.25467 | 40.64869 | 17.77562 | 0.65722 | 1.105722 | 0.215575 |
| 9 | 88168400 | 88170399 | 12.42111 | 21.79118 | 46.70036 | 20.64713 | -0.70667 | 0.627104 | 0.215584 |
| 8 | 5659400 | 5660599 | 14.33482 | 24.95853 | 51.09614 | 22.46381 | -0.64179 | 0.759837 | 0.215585 |
| 15 | 89182000 | 89185199 | 14.62161 | 24.65396 | 63.03186 | 27.5604 | -0.9677 | 1.084862 | 0.21561 |
| 19 | 9240400 | 9242999 | 40.27713 | 65.97276 | 55.9433 | 24.07206 | 0.613428 | 1.471189 | 0.215618 |
| 5 | 149173600 | 149186199 | 93.00883 | 141.8694 | 255.5346 | 105.9527 | -0.56137 | 3.118853 | 0.215623 |
| 2 | 57519400 | 57520599 | 62.84436 | 101.3227 | 58.48077 | 24.89999 | 1.147904 | 1.80647 | 0.215624 |
| X | 122228400 | 122230399 | 25.48664 | 44.78685 | 26.57239 | 11.76458 | 1.136732 | 0.642211 | 0.215625 |
| 5 | 108776400 | 108777999 | 10.57486 | 18.62375 | 43.0243 | 19.04528 | -0.81274 | 0.559507 | 0.215632 |
| 19 | 10327800 | 10335599 | 42.94917 | 68.50931 | 128.3937 | 53.82427 | -0.56561 | 2.13545 | 0.215642 |

| | | | | | | | | | |
|---|---|---|---|---|---|---|---|---|---|
| 11 | 72204800 | 72209399 | 22.48373 | 36.8117 | 82.16647 | 35.34923 | -0.77913 | 1.45359 | 0.215643 |
| 9 | 104019800 | 104022199 | 13.07772 | 22.83907 | 47.84494 | 21.07466 | -0.67409 | 0.706389 | 0.215654 |
| 11 | 83446000 | 83450399 | 20.43439 | 33.48705 | 81.84104 | 35.22287 | -0.90891 | 1.437675 | 0.215655 |
| 6 | 76594800 | 76595599 | 7.536927 | 14.18314 | 32.4365 | 13.29392 | -0.71447 | 0.080304 | 0.215656 |
| 15 | 32840200 | 32842199 | 17.19799 | 31.50217 | 22.91039 | 9.78033 | 0.881719 | 0.295578 | 0.215659 |
| 2 | 83982800 | 83983399 | 4.617137 | 8.997831 | 24.98885 | 9.059419 | -0.82596 | -0.38603 | 0.215677 |
| 7 | 99593400 | 99597799 | 57.44314 | 90.33284 | 166.74 | 69.17111 | -0.56235 | 2.528376 | 0.21568 |
| 12 | 112378000 | 112386399 | 52.31328 | 82.91779 | 150.5542 | 62.64139 | -0.52954 | 2.343428 | 0.215689 |
| 3 | 15670200 | 15672199 | 29.41542 | 49.92254 | 40.92831 | 17.93112 | 0.666879 | 1.053295 | 0.215705 |
| 1 | 119677400 | 119678599 | 15.94202 | 29.69779 | 20.90857 | 8.717457 | 0.952476 | 0.187102 | 0.215721 |
| 19 | 3226800 | 3228799 | 18.71495 | 33.24908 | 26.32002 | 11.61498 | 0.728869 | 0.489622 | 0.215735 |
| 5 | 18523800 | 18525999 | 27.874 | 47.51668 | 39.92555 | 17.50204 | 0.632354 | 1.008551 | 0.215754 |
| 14 | 104101200 | 104101999 | 13.72433 | 26.50223 | 15.92092 | 6.130017 | 1.278377 | -0.10601 | 0.215755 |
| 11 | 116788000 | 116789599 | 8.244774 | 15.10224 | 35.60336 | 15.1989 | -0.80348 | 0.255679 | 0.215761 |
| 17 | 18328800 | 18333799 | 64.52232 | 104.0315 | 59.51648 | 25.34205 | 1.160765 | 1.811821 | 0.215763 |
| 9 | 120479400 | 120480599 | 14.77252 | 27.37096 | 22.10496 | 9.29062 | 0.747327 | 0.210932 | 0.215771 |
| 17 | 94528600 | 94529799 | 14.77252 | 27.37096 | 22.10496 | 9.29062 | 0.747327 | 0.210932 | 0.215771 |
| 6 | 132512600 | 132515999 | 62.66358 | 100.6061 | 75.1754 | 31.8143 | 0.769067 | 1.941678 | 0.215776 |
| 18 | 63665800 | 63667399 | 10.10032 | 17.84763 | 41.48133 | 18.34999 | -0.81923 | 0.517614 | 0.215779 |
| 4 | 153342000 | 153356199 | 117.9901 | 176.522 | 327.5913 | 135.0954 | -0.62845 | 3.513826 | 0.21578 |
| 3 | 65864800 | 65866199 | 6.258748 | 11.67443 | 32.16262 | 13.37999 | -0.99737 | 0.127306 | 0.215783 |
| 8 | 102856200 | 102857999 | 19.73618 | 34.82229 | 28.14599 | 12.4546 | 0.695032 | 0.566037 | 0.215801 |
| 3 | 49031000 | 49031799 | 7.682053 | 14.01899 | 36.60185 | 15.70123 | -0.95445 | 0.275007 | 0.215802 |
| 3 | 2999800 | 3002599 | 188.9175 | 288.0906 | 119.2944 | 49.46035 | 1.557309 | 3.143551 | 0.21581 |
| 2 | 17342200 | 17344399 | 11.94641 | 21.08182 | 49.3945 | 21.8563 | -0.83245 | 0.554241 | 0.215827 |
| 7 | 130616400 | 130620199 | 39.56105 | 65.8911 | 44.31931 | 19.2524 | 0.951469 | 1.267238 | 0.215854 |
| 4 | 130065800 | 130066399 | 3.419782 | 6.670782 | 22.56732 | 8.080204 | -1.08935 | -0.45645 | 0.215865 |
| 11 | 98650800 | 98652799 | 10.73295 | 18.87994 | 44.05552 | 19.50635 | -0.82795 | 0.574528 | 0.215869 |
| 14 | 87804800 | 87806199 | 10.69411 | 18.92437 | 41.43317 | 18.32336 | -0.73298 | 0.503802 | 0.21588 |
| 6 | 13467800 | 13469199 | 19.95806 | 35.27597 | 27.88282 | 12.3338 | 0.727846 | 0.546741 | 0.215892 |
| 7 | 33320600 | 33333199 | 316.2065 | 454.6608 | 347.4081 | 143.4047 | 0.596588 | 4.170078 | 0.215902 |
| 4 | 148211600 | 148219999 | 65.28899 | 102.0255 | 182.9585 | 75.90932 | -0.52784 | 2.650597 | 0.215904 |
| 10 | 117734600 | 117736599 | 7.627055 | 14.0564 | 34.11131 | 14.44135 | -0.83582 | 0.213537 | 0.215913 |
| 1 | 146920800 | 146921999 | 12.98494 | 25.00298 | 17.07847 | 6.63635 | 1.082654 | -0.08223 | 0.215918 |
| 18 | 38492600 | 38494599 | 15.0576 | 26.00261 | 54.81791 | 24.04857 | -0.68587 | 0.861342 | 0.215931 |
| 15 | 86414200 | 86415399 | 6.590582 | 12.61807 | 30.20568 | 11.93157 | -0.73228 | -0.0566 | 0.215939 |
| 5 | 81530600 | 81532999 | 19.41063 | 34.35686 | 27.54 | 12.17872 | 0.708615 | 0.528779 | 0.215952 |
| 8 | 18947200 | 18950599 | 14.71678 | 25.34032 | 55.36297 | 24.28553 | -0.73822 | 0.883625 | 0.21596 |
| 9 | 114559000 | 114560799 | 8.085564 | 14.85283 | 34.58922 | 14.70888 | -0.78162 | 0.236923 | 0.215967 |
| 2 | 70301400 | 70307799 | 43.41577 | 68.86459 | 148.9346 | 62.01219 | -0.78035 | 2.31508 | 0.215974 |
| 17 | 84169000 | 84173199 | 25.39319 | 41.38948 | 88.08636 | 37.83376 | -0.71527 | 1.527078 | 0.215975 |
| X | 146630600 | 146633399 | 24.3091 | 43.23475 | 20.73729 | 9.142933 | 1.447539 | 0.509347 | 0.21598 |
| 7 | 8386400 | 8397199 | 203.1973 | 302.777 | 240.6951 | 99.18597 | 0.588584 | 3.598511 | 0.215989 |
| 10 | 64675000 | 64675799 | 12.45022 | 24.2572 | 12.76415 | 4.641681 | 1.541618 | -0.31772 | 0.215993 |
| 11 | 63197400 | 63198199 | 16.04025 | 29.73688 | 21.18933 | 8.898844 | 0.928071 | 0.218361 | 0.215999 |
| 3 | 54461200 | 54464399 | 30.0787 | 51.42317 | 37.54583 | 16.46268 | 0.834681 | 0.98785 | 0.216007 |
| 10 | 71242800 | 71243399 | 13.19369 | 24.68535 | 20.83055 | 8.621079 | 0.701722 | 0.136476 | 0.216009 |

| | | | | | | | | | |
|---|---|---|---|---|---|---|---|---|---|
| 7 | 10946600 | 10957599 | 256.8381 | 375.7416 | 294.1305 | 121.1676 | 0.586054 | 3.907968 | 0.216014 |
| 16 | 31354800 | 31355999 | 13.37771 | 25.30475 | 20.26563 | 8.223257 | 0.79976 | 0.067119 | 0.216031 |
| 10 | 16020800 | 16022199 | 7.259237 | 13.60875 | 31.74895 | 13.09616 | -0.75006 | 0.102048 | 0.216036 |
| 9 | 3191400 | 3197399 | 91.65239 | 145.2721 | 92.6868 | 38.56461 | 0.977053 | 2.360899 | 0.21604 |
| 18 | 61687800 | 61689199 | 8.123151 | 14.45948 | 40.50055 | 17.84381 | -1.08098 | 0.439743 | 0.216043 |
| 4 | 154396200 | 154400399 | 27.94385 | 45.10918 | 97.78134 | 41.70001 | -0.75448 | 1.736512 | 0.21606 |
| 1 | 187590400 | 187593799 | 31.67686 | 53.0713 | 45.38902 | 19.79051 | 0.605833 | 1.181254 | 0.216072 |
| 4 | 56742800 | 56744999 | 23.747 | 38.87476 | 80.49318 | 34.62738 | -0.67115 | 1.45852 | 0.216084 |
| 8 | 125234400 | 125237999 | 23.747 | 38.87476 | 80.49318 | 34.62738 | -0.67115 | 1.45852 | 0.216084 |
| 1 | 145763000 | 145763399 | 8.730509 | 17.02343 | 15.02922 | 5.408995 | 0.817787 | -0.39457 | 0.216085 |
| 18 | 72591200 | 72593999 | 43.69252 | 70.85276 | 61.28298 | 26.24933 | 0.5744 | 1.634705 | 0.216102 |
| 13 | 86920400 | 86920799 | 9.46475 | 18.47676 | 13.11364 | 4.636026 | 1.150442 | -0.45312 | 0.216104 |
| 7 | 63465000 | 63468999 | 23.10133 | 37.84799 | 81.18327 | 34.93605 | -0.72151 | 1.446065 | 0.216105 |
| 4 | 95334400 | 95338199 | 26.3052 | 42.991 | 85.05126 | 36.56584 | -0.60746 | 1.488953 | 0.216112 |
| 6 | 16277200 | 16278399 | 14.32688 | 27.00293 | 20.2511 | 8.277785 | 0.885054 | 0.100593 | 0.216118 |
| 7 | 23364600 | 23367399 | 47.66463 | 77.35029 | 61.73525 | 26.45724 | 0.691049 | 1.628325 | 0.216129 |
| 16 | 94646800 | 94648799 | 18.59004 | 33.08053 | 26.55573 | 11.70307 | 0.710415 | 0.481441 | 0.216146 |
| 14 | 68170000 | 68171799 | 10.46485 | 18.33712 | 45.70048 | 20.18454 | -0.92273 | 0.636402 | 0.216174 |
| 3 | 32783200 | 32786599 | 15.08408 | 25.44668 | 63.11275 | 27.60002 | -0.92416 | 1.081061 | 0.216182 |
| 11 | 6629600 | 6631399 | 8.237892 | 15.12789 | 34.5839 | 14.71292 | -0.75554 | 0.240315 | 0.216196 |
| 4 | 143935200 | 143937199 | 31.75681 | 54.37969 | 35.25722 | 15.4612 | 1.005639 | 0.978087 | 0.216216 |
| 4 | 60366000 | 60366599 | 20.78988 | 36.78297 | 24.85831 | 10.99373 | 0.953122 | 0.536203 | 0.216235 |
| 19 | 30535800 | 30539199 | 42.25062 | 69.43546 | 52.77659 | 22.74137 | 0.772686 | 1.433536 | 0.216243 |
| 7 | 20864800 | 20865399 | 14.60302 | 27.19148 | 21.74777 | 9.074091 | 0.76938 | 0.181663 | 0.216248 |
| 17 | 69213800 | 69215599 | 11.14531 | 19.77858 | 41.4171 | 18.29463 | -0.66727 | 0.480307 | 0.216256 |
| X | 25861800 | 25864999 | 36.72117 | 62.49958 | 33.49528 | 14.68136 | 1.278036 | 1.04986 | 0.216264 |
| 17 | 30142600 | 30146399 | 31.75665 | 51.36768 | 97.23421 | 41.57101 | -0.55623 | 1.684221 | 0.216264 |
| 8 | 15076200 | 15079599 | 22.05525 | 36.00251 | 83.88448 | 36.04862 | -0.84377 | 1.49795 | 0.216298 |
| 15 | 5097800 | 5099199 | 8.693977 | 15.90908 | 36.11921 | 15.44019 | -0.75112 | 0.265751 | 0.216307 |
| 7 | 17958200 | 17965999 | 159.3363 | 240.8651 | 203.8188 | 84.28325 | 0.516937 | 3.31394 | 0.216308 |
| 17 | 20135600 | 20144199 | 176.3515 | 265.4815 | 212.1723 | 87.62932 | 0.594155 | 3.400279 | 0.216308 |
| 2 | 16434400 | 16435199 | 4.947422 | 9.618106 | 25.23464 | 9.366813 | -0.77702 | -0.29921 | 0.216318 |
| 5 | 21732200 | 21734199 | 9.869743 | 17.63628 | 42.04905 | 18.46598 | -0.8477 | 0.426925 | 0.216323 |
| 14 | 66598000 | 66599999 | 10.81453 | 19.23826 | 40.92249 | 18.03916 | -0.68774 | 0.46111 | 0.216332 |
| 2 | 111341400 | 111346199 | 40.82148 | 67.34032 | 50.5651 | 21.82324 | 0.79223 | 1.381917 | 0.21636 |
| 5 | 146371400 | 146372599 | 7.057581 | 13.43155 | 33.5085 | 13.42252 | -0.81014 | 0.001251 | 0.216362 |
| 7 | 61467400 | 61468199 | 11.30618 | 21.76405 | 18.46814 | 7.182422 | 0.771909 | -0.09265 | 0.216362 |
| 6 | 99964600 | 99966799 | 9.765467 | 17.34358 | 40.42979 | 17.84664 | -0.82017 | 0.464259 | 0.216371 |
| X | 26222200 | 26225599 | 39.64345 | 66.8736 | 34.58542 | 15.12434 | 1.328627 | 1.135648 | 0.216376 |
| 18 | 69314000 | 69315399 | 17.13016 | 31.03098 | 24.42328 | 10.57409 | 0.75525 | 0.359795 | 0.216387 |
| 19 | 50817800 | 50818799 | 12.93255 | 24.76272 | 18.128 | 7.158621 | 0.962163 | -0.02964 | 0.216411 |
| 7 | 59784400 | 59790199 | 28.4443 | 48.64022 | 35.18111 | 15.42611 | 0.848296 | 0.977183 | 0.216415 |
| 10 | 107231000 | 107232199 | 9.229422 | 16.6147 | 38.98828 | 17.01259 | -0.81815 | 0.370814 | 0.216417 |
| 12 | 28715000 | 28718599 | 18.83576 | 31.67533 | 65.06251 | 28.42538 | -0.6538 | 1.122076 | 0.216431 |
| 14 | 41472400 | 41473599 | 15.94224 | 29.62469 | 21.57988 | 9.033942 | 0.899507 | 0.20485 | 0.216438 |
| 5 | 75085200 | 75087199 | 9.98472 | 18.02334 | 38.50658 | 16.74533 | -0.6804 | 0.360722 | 0.216444 |
| 3 | 152568400 | 152570999 | 31.48044 | 50.74747 | 101.264 | 43.10769 | -0.63781 | 1.783414 | 0.216447 |

| | | | | | | | | | |
|---|---|---|---|---|---|---|---|---|---|
| 10 | 67283200 | 67285199 | 22.46402 | 39.25892 | 31.68332 | 13.96238 | 0.695323 | 0.716851 | 0.216464 |
| 4 | 101365200 | 101366999 | 9.905769 | 17.62143 | 39.92204 | 17.59833 | -0.77786 | 0.454077 | 0.216472 |
| 9 | 113849200 | 113853399 | 28.86066 | 46.6837 | 96.75133 | 41.36482 | -0.68662 | 1.677512 | 0.216483 |
| 12 | 33315600 | 33316799 | 6.474443 | 12.29441 | 30.44391 | 12.25238 | -0.80509 | 0.003215 | 0.216486 |
| 6 | 30355400 | 30357399 | 8.41103 | 14.99181 | 40.49196 | 17.82006 | -1.02793 | 0.432426 | 0.216493 |
| 1 | 102088600 | 102090599 | 23.24716 | 40.60378 | 32.2412 | 14.20259 | 0.718699 | 0.731358 | 0.216516 |
| 9 | 114860800 | 114870999 | 57.66191 | 90.46116 | 176.0664 | 73.02031 | -0.6414 | 2.575786 | 0.216521 |
| 1 | 25053000 | 25057999 | 46.52355 | 74.12786 | 135.292 | 56.62837 | -0.52927 | 2.176181 | 0.21653 |
| 11 | 113307400 | 113316999 | 68.6077 | 106.6195 | 196.4983 | 81.58696 | -0.5735 | 2.749865 | 0.216531 |
| 4 | 130617200 | 130619599 | 8.953128 | 16.06934 | 37.95013 | 16.60386 | -0.82947 | 0.388012 | 0.216536 |
| 8 | 126864400 | 126866799 | 16.95354 | 28.55911 | 62.66272 | 27.3912 | -0.74826 | 1.100388 | 0.21654 |
| 10 | 75427400 | 75429999 | 11.04758 | 19.39972 | 45.11711 | 19.96298 | -0.8237 | 0.602205 | 0.216547 |
| 14 | 76024000 | 76025799 | 10.6785 | 18.97004 | 40.40669 | 17.83256 | -0.69067 | 0.469171 | 0.216548 |
| 3 | 17239600 | 17240199 | 21.98247 | 38.40309 | 31.46182 | 13.86113 | 0.673939 | 0.721458 | 0.216553 |
| 1 | 168680600 | 168682599 | 18.37837 | 32.78386 | 25.81183 | 11.35183 | 0.739919 | 0.461502 | 0.216571 |
| 9 | 30810400 | 30813999 | 31.32489 | 50.49005 | 101.0989 | 43.03009 | -0.64299 | 1.787328 | 0.216585 |
| 5 | 131317200 | 131318399 | 15.22045 | 27.79885 | 22.59343 | 9.681943 | 0.718482 | 0.294383 | 0.21659 |
| 7 | 27730200 | 27731599 | 15.15692 | 28.32912 | 21.05104 | 8.727671 | 0.881689 | 0.160446 | 0.216595 |
| 7 | 105420600 | 105422599 | 10.93341 | 19.11841 | 46.77621 | 20.62343 | -0.89636 | 0.67504 | 0.216599 |
| 13 | 96903600 | 96906799 | 20.82764 | 34.29148 | 77.19752 | 33.29019 | -0.78802 | 1.37664 | 0.216608 |
| 12 | 3087600 | 3088799 | 30.40074 | 51.06991 | 41.22107 | 18.00347 | 0.689003 | 1.146144 | 0.216622 |
| X | 33935800 | 33937199 | 19.09675 | 35.05565 | 21.13012 | 8.99393 | 1.153725 | 0.295016 | 0.216624 |
| 10 | 100138000 | 100142599 | 92.65693 | 140.3807 | 274.9193 | 113.7649 | -0.68995 | 3.259068 | 0.216627 |
| 4 | 108074200 | 108077399 | 14.92847 | 25.56419 | 56.36049 | 24.7156 | -0.75265 | 0.936984 | 0.216637 |
| 4 | 114856800 | 114857799 | 5.745958 | 10.95176 | 29.38328 | 11.7328 | -0.90912 | -0.03185 | 0.216638 |
| 7 | 60736800 | 60738799 | 27.11486 | 46.79484 | 32.94852 | 14.45421 | 0.889086 | 0.887608 | 0.216642 |
| 14 | 111131600 | 111132999 | 19.52261 | 34.7702 | 25.6373 | 11.29004 | 0.832708 | 0.478407 | 0.216653 |
| 5 | 18183000 | 18185599 | 35.89593 | 59.3895 | 50.96454 | 22.03833 | 0.600663 | 1.336059 | 0.216657 |
| 17 | 56723000 | 56736599 | 74.33072 | 112.9415 | 272.7708 | 112.9747 | -0.9886 | 3.195135 | 0.216666 |
| 6 | 14300800 | 14305999 | 127.0905 | 194.3844 | 167.2806 | 69.4035 | 0.506389 | 3.066352 | 0.216707 |
| 11 | 67812200 | 67813799 | 9.299273 | 16.55821 | 39.448 | 17.37548 | -0.84887 | 0.444172 | 0.216716 |
| 1 | 87770400 | 87774999 | 28.78434 | 46.83754 | 90.10526 | 38.67702 | -0.57174 | 1.561249 | 0.216721 |
| 9 | 22704800 | 22707399 | 11.22669 | 19.65217 | 46.23331 | 20.40244 | -0.84019 | 0.657586 | 0.216722 |
| 6 | 120531000 | 120532199 | 9.741477 | 17.36333 | 39.93321 | 17.57405 | -0.79781 | 0.439874 | 0.216737 |
| 4 | 108767200 | 108768599 | 12.73003 | 24.16741 | 19.17906 | 7.72269 | 0.821977 | 0.032456 | 0.216772 |
| X | 41503200 | 41504999 | 34.12739 | 57.26772 | 43.71998 | 19.08187 | 0.769109 | 1.172769 | 0.21679 |
| 5 | 135717400 | 135718999 | 10.56484 | 18.64247 | 41.99055 | 18.58043 | -0.77515 | 0.536825 | 0.216797 |
| 13 | 66088800 | 66089999 | 12.96481 | 25.04842 | 15.94617 | 6.125046 | 1.198381 | -0.1225 | 0.216802 |
| 17 | 84373200 | 84376599 | 29.6235 | 47.64212 | 108.6735 | 46.10689 | -0.83453 | 1.857244 | 0.216811 |
| 17 | 20538600 | 20544799 | 64.28167 | 102.8039 | 79.32841 | 33.37778 | 0.716369 | 2.063159 | 0.216816 |
| 5 | 36737200 | 36740199 | 13.15032 | 22.37198 | 59.33535 | 26.00642 | -1.01932 | 0.996193 | 0.216855 |
| 19 | 3366800 | 3379999 | 80.50536 | 123.5256 | 231.4393 | 96.08114 | -0.61145 | 2.994716 | 0.216861 |
| 4 | 129905000 | 129911399 | 31.19604 | 49.89246 | 124.4501 | 52.36513 | -0.97289 | 2.040699 | 0.216874 |
| 10 | 97021800 | 97022599 | 4.521904 | 8.809278 | 24.30994 | 8.852222 | -0.8233 | -0.37629 | 0.216886 |
| 7 | 4782800 | 4784999 | 10.09995 | 17.84652 | 41.98076 | 18.57101 | -0.83667 | 0.517551 | 0.216888 |
| 1 | 116419200 | 116420799 | 20.33733 | 36.32109 | 24.44978 | 10.74213 | 0.96547 | 0.464853 | 0.216892 |
| 10 | 69984800 | 69998599 | 104.8501 | 158.2645 | 289.3277 | 119.5869 | -0.59535 | 3.337338 | 0.216898 |

| | | | | | | | | | |
|---|---|---|---|---|---|---|---|---|---|
| 9 | 64485400 | 64488199 | 10.50072 | 18.35459 | 52.07612 | 22.95254 | -1.10937 | 0.683987 | 0.216905 |
| 8 | 27224000 | 27225199 | 6.12327 | 11.81505 | 30.08012 | 11.61917 | -0.78997 | -0.13593 | 0.216922 |
| 15 | 93101800 | 93102999 | 6.12327 | 11.81505 | 30.08012 | 11.61917 | -0.78997 | -0.13593 | 0.216922 |
| 12 | 23727400 | 23731399 | 52.76457 | 84.986 | 72.77317 | 30.94255 | 0.578523 | 1.827779 | 0.216928 |
| 8 | 14382400 | 14385799 | 21.89206 | 36.1321 | 75.30955 | 32.511 | -0.67593 | 1.347247 | 0.21693 |
| 19 | 12933600 | 12938599 | 71.68211 | 114.5352 | 83.30989 | 35.00795 | 0.800076 | 2.100829 | 0.216937 |
| 5 | 82565600 | 82566799 | 15.92267 | 28.97515 | 23.24823 | 10.00825 | 0.733158 | 0.323925 | 0.216968 |
| 12 | 25147800 | 25152199 | 21.27966 | 35.07977 | 76.25676 | 32.90199 | -0.737 | 1.361113 | 0.21698 |
| 18 | 46684000 | 46690399 | 35.68654 | 57.37486 | 111.7271 | 47.38068 | -0.60757 | 1.875699 | 0.21699 |
| 8 | 47955600 | 47957799 | 9.613864 | 17.07963 | 39.93553 | 17.62385 | -0.82409 | 0.461289 | 0.216991 |
| 17 | 5171200 | 5172799 | 7.7993 | 14.30155 | 34.56497 | 14.73414 | -0.83709 | 0.245562 | 0.216992 |
| 2 | 173668400 | 173676999 | 81.72124 | 125.9749 | 218.8845 | 90.92095 | -0.49765 | 2.903613 | 0.217012 |
| 6 | 95863400 | 95864799 | 15.76765 | 28.7576 | 23.54764 | 10.1087 | 0.706247 | 0.309102 | 0.217032 |
| 9 | 6531600 | 6532799 | 7.222406 | 13.66243 | 31.4717 | 12.76877 | -0.7121 | 0.040271 | 0.217038 |
| 10 | 21604000 | 21604999 | 11.27492 | 21.55387 | 18.48376 | 7.333074 | 0.730289 | -0.02955 | 0.217064 |
| 7 | 64306800 | 64309399 | 16.14874 | 27.8312 | 56.50148 | 24.78564 | -0.63254 | 0.884405 | 0.217066 |
| 19 | 25491400 | 25493999 | 10.84206 | 19.18055 | 42.45872 | 18.778 | -0.74911 | 0.508806 | 0.217067 |
| 10 | 42624800 | 42625599 | 11.54907 | 22.10553 | 18.90221 | 7.472263 | 0.738912 | -0.03772 | 0.217073 |
| 6 | 132861200 | 132864999 | 52.29173 | 84.57344 | 63.59897 | 27.18827 | 0.772277 | 1.704405 | 0.217074 |
| 17 | 93185800 | 93187999 | 25.76331 | 44.56527 | 35.47877 | 15.5653 | 0.713291 | 0.863801 | 0.217075 |
| 6 | 50315800 | 50318599 | 16.68173 | 28.4464 | 57.9937 | 25.42318 | -0.64129 | 0.984992 | 0.217077 |
| 3 | 26825600 | 26826199 | 6.138955 | 11.79788 | 31.68084 | 12.3793 | -0.88239 | -0.09578 | 0.217093 |
| 4 | 13124600 | 13125199 | 5.398137 | 10.41433 | 26.96708 | 10.42114 | -0.81344 | -0.14834 | 0.217096 |
| 12 | 20377400 | 20397399 | 634.054 | 853.3525 | 646.5834 | 269.7098 | 0.512794 | 5.177106 | 0.217099 |
| 3 | 11993400 | 11995599 | 20.15929 | 37.04562 | 19.96554 | 8.485916 | 1.315435 | 0.295211 | 0.217104 |
| 2 | 109965200 | 109965799 | 10.43126 | 20.16274 | 17.07582 | 6.544943 | 0.794836 | -0.15221 | 0.217107 |
| 7 | 40851600 | 40855599 | 48.69654 | 78.93898 | 62.83999 | 26.90911 | 0.69336 | 1.652826 | 0.217107 |
| 13 | 64685000 | 64689399 | 45.68089 | 74.22656 | 61.0939 | 26.20725 | 0.648455 | 1.59864 | 0.21711 |
| 7 | 38572000 | 38572799 | 13.74367 | 26.44121 | 17.05166 | 6.642311 | 1.161883 | -0.06608 | 0.21711 |
| 14 | 97354000 | 97354399 | 3.132727 | 6.108471 | 22.44294 | 8.076597 | -1.21224 | -0.44099 | 0.21713 |
| 4 | 62639200 | 62646999 | 59.42739 | 93.29571 | 171.9666 | 71.32369 | -0.56222 | 2.563027 | 0.21714 |
| X | 126937000 | 126938199 | 15.10113 | 29.07278 | 15.91216 | 6.186204 | 1.399333 | -0.06046 | 0.21714 |
| 9 | 89936000 | 89937199 | 8.383151 | 15.38877 | 35.11006 | 14.94452 | -0.75346 | 0.243711 | 0.217148 |
| 7 | 120483000 | 120484199 | 14.6637 | 26.92692 | 22.25395 | 9.467993 | 0.701956 | 0.260261 | 0.217148 |
| 1 | 186500800 | 186503599 | 26.48467 | 46.03827 | 32.01569 | 14.06102 | 0.908445 | 0.811114 | 0.217155 |
| 7 | 29979000 | 29984399 | 26.28655 | 42.60517 | 92.36922 | 39.54966 | -0.74864 | 1.630691 | 0.217156 |
| 3 | 43855400 | 43858999 | 79.54205 | 125.586 | 111.6137 | 46.34149 | 0.495124 | 2.448254 | 0.217164 |
| 16 | 64876600 | 64876999 | 8.48807 | 16.58529 | 13.40471 | 4.662095 | 0.987482 | -0.51988 | 0.21717 |
| 5 | 137431200 | 137437599 | 42.75511 | 68.32052 | 124.6851 | 52.40035 | -0.52457 | 2.079403 | 0.217189 |
| 12 | 95657800 | 95663199 | 30.86087 | 49.70029 | 105.6302 | 44.90497 | -0.73033 | 1.813666 | 0.217199 |
| 5 | 76451600 | 76453199 | 11.6989 | 20.4359 | 47.31328 | 20.84357 | -0.81735 | 0.695714 | 0.217213 |
| 7 | 27911800 | 27914599 | 16.61844 | 28.06946 | 61.58047 | 26.93993 | -0.74783 | 1.075424 | 0.217213 |
| 14 | 122047400 | 122048399 | 28.023 | 47.5364 | 40.55046 | 17.76298 | 0.609869 | 1.053002 | 0.217216 |
| 7 | 27869000 | 27869999 | 4.314686 | 8.303397 | 27.92353 | 10.86814 | -1.1959 | -0.13481 | 0.217234 |
| 9 | 14411400 | 14412399 | 4.314686 | 8.303397 | 27.92353 | 10.86814 | -1.1959 | -0.13481 | 0.217234 |
| 5 | 75679000 | 75680999 | 10.34075 | 18.44652 | 39.92079 | 17.55652 | -0.71 | 0.438668 | 0.217245 |
| 11 | 110921000 | 110922599 | 10.16116 | 18.23167 | 38.93801 | 17.0407 | -0.68607 | 0.399146 | 0.217259 |

| | | | | | | | | | |
|---|---|---|---|---|---|---|---|---|---|
| 9 | 107431000 | 107432599 | 5.685786 | 10.63869 | 32.20796 | 13.32855 | -1.12599 | 0.101291 | 0.21727 |
| 9 | 38094800 | 38097599 | 47.87439 | 77.44912 | 65.97918 | 28.21117 | 0.593103 | 1.692992 | 0.217281 |
| 9 | 50570200 | 50571399 | 6.671331 | 12.48963 | 31.6973 | 13.10498 | -0.87264 | 0.107069 | 0.217301 |
| 7 | 40717000 | 40719399 | 50.90272 | 82.10584 | 69.72158 | 29.70309 | 0.593023 | 1.782415 | 0.217311 |
| 9 | 69846400 | 69847599 | 6.89522 | 13.15891 | 33.06178 | 13.16629 | -0.81287 | -0.02119 | 0.217312 |
| 10 | 127401800 | 127403399 | 8.042539 | 14.4837 | 37.48408 | 16.3483 | -0.95723 | 0.359624 | 0.217322 |
| 11 | 18976000 | 18978599 | 12.46868 | 21.73866 | 49.4494 | 21.75827 | -0.79265 | 0.728864 | 0.217325 |
| 3 | 137929600 | 137932799 | 25.26367 | 41.11173 | 86.4511 | 37.10949 | -0.69958 | 1.550909 | 0.21736 |
| 5 | 109794800 | 109795399 | 21.22401 | 37.2284 | 29.79184 | 13.16966 | 0.707504 | 0.651416 | 0.217361 |
| 5 | 139502000 | 139505599 | 20.80624 | 34.30727 | 76.19899 | 32.88028 | -0.76778 | 1.357021 | 0.217377 |
| 17 | 73036400 | 73037399 | 13.10792 | 25.08843 | 18.78238 | 7.425592 | 0.929187 | -0.02127 | 0.217384 |
| 5 | 92510200 | 92514399 | 19.978 | 33.51501 | 66.18168 | 28.88042 | -0.59723 | 1.151552 | 0.217418 |
| 7 | 102977600 | 102979399 | 19.78332 | 35.48102 | 24.59497 | 10.76756 | 0.927008 | 0.436605 | 0.217419 |
| 14 | 60993800 | 60999999 | 32.53999 | 52.21117 | 113.6269 | 48.04269 | -0.77104 | 1.936464 | 0.217429 |
| 11 | 50591200 | 50597199 | 40.59937 | 65.03995 | 120.2381 | 50.71617 | -0.53902 | 1.997375 | 0.21743 |
| 14 | 26929400 | 26932999 | 19.34317 | 32.42688 | 66.12641 | 28.84331 | -0.64346 | 1.156595 | 0.217433 |
| 10 | 49904600 | 49906999 | 30.66519 | 52.48392 | 33.95146 | 14.88798 | 1.008856 | 0.977576 | 0.217433 |
| 7 | 31621600 | 31626599 | 83.14016 | 131.579 | 105.6152 | 43.90367 | 0.644557 | 2.392851 | 0.217437 |
| 7 | 40787000 | 40789399 | 42.95453 | 71.27641 | 43.23942 | 18.73566 | 1.099583 | 1.31631 | 0.217438 |
| 1 | 138792800 | 138794799 | 24.09768 | 42.29785 | 27.49247 | 12.16022 | 1.005559 | 0.650577 | 0.21746 |
| 16 | 94276600 | 94278199 | 9.315138 | 17.04919 | 37.76048 | 16.13735 | -0.71544 | 0.273615 | 0.217473 |
| 12 | 113841000 | 113843399 | 40.4117 | 66.21958 | 55.98725 | 24.09503 | 0.61788 | 1.466239 | 0.217488 |
| 4 | 128240400 | 128241399 | 4.493924 | 8.726099 | 24.97624 | 9.360313 | -0.9126 | -0.26788 | 0.217495 |
| 14 | 64546000 | 64549399 | 22.48658 | 37.21575 | 74.46567 | 32.20792 | -0.61695 | 1.315552 | 0.217505 |
| 6 | 131017400 | 131020399 | 43.34161 | 71.77198 | 44.9619 | 19.45456 | 1.053254 | 1.341885 | 0.217512 |
| 9 | 30424000 | 30424999 | 18.93665 | 33.50219 | 27.46972 | 12.14878 | 0.675701 | 0.529161 | 0.217524 |
| 14 | 32905400 | 32908399 | 23.29301 | 38.52603 | 75.68646 | 32.72165 | -0.59068 | 1.32612 | 0.217526 |
| 1 | 180884000 | 180885799 | 20.57715 | 36.28227 | 28.36109 | 12.55143 | 0.742487 | 0.576931 | 0.217542 |
| 2 | 35348800 | 35352599 | 21.80446 | 36.29669 | 71.18391 | 30.91179 | -0.58735 | 1.244875 | 0.217558 |
| 9 | 114301400 | 114303399 | 9.35769 | 16.54003 | 43.53987 | 19.25954 | -0.99787 | 0.511281 | 0.217574 |
| 5 | 148923400 | 148930799 | 82.55711 | 127.2008 | 219.0922 | 91.00708 | -0.4856 | 2.913283 | 0.217584 |
| 8 | 27653200 | 27654799 | 6.071903 | 11.4612 | 30.784 | 12.54711 | -0.93715 | 0.043072 | 0.217587 |
| 4 | 107225000 | 107225599 | 4.409556 | 8.555405 | 25.52056 | 9.611357 | -0.97924 | -0.2531 | 0.217599 |
| 8 | 35248800 | 35256999 | 57.79399 | 91.31167 | 161.3195 | 67.00244 | -0.49394 | 2.427491 | 0.217607 |
| 11 | 113574000 | 113576399 | 13.33228 | 22.98901 | 53.69157 | 23.55337 | -0.83334 | 0.863993 | 0.21761 |
| 8 | 25722000 | 25723399 | 13.7773 | 25.92426 | 20.79502 | 8.523921 | 0.785438 | 0.106835 | 0.217618 |
| 4 | 64334400 | 64335399 | 10.49792 | 20.45243 | 14.08815 | 5.12653 | 1.156537 | -0.33757 | 0.217636 |
| 8 | 65838400 | 65840599 | 37.1119 | 61.56954 | 47.15372 | 20.429 | 0.764405 | 1.302527 | 0.21764 |
| 11 | 49668800 | 49670799 | 17.8387 | 30.17942 | 61.77413 | 27.03794 | -0.64831 | 1.063328 | 0.217652 |
| 7 | 27941000 | 27949199 | 96.61049 | 146.7361 | 263.4641 | 109.1071 | -0.5618 | 3.208308 | 0.217667 |
| 3 | 46967200 | 46973399 | 85.45126 | 134.5909 | 118.5045 | 49.17104 | 0.505904 | 2.504909 | 0.217672 |
| 6 | 20164200 | 20166199 | 23.9878 | 42.64987 | 22.52731 | 9.934779 | 1.309593 | 0.513363 | 0.217681 |
| 15 | 54434200 | 54436399 | 16.72822 | 27.97999 | 67.99427 | 29.61587 | -0.89533 | 1.174787 | 0.217682 |
| 7 | 18869200 | 18871599 | 16.47693 | 27.79136 | 63.66073 | 27.83813 | -0.80989 | 1.09119 | 0.217685 |
| X | 24594600 | 24596599 | 29.04655 | 49.8709 | 33.05211 | 14.4973 | 0.974723 | 0.942427 | 0.217687 |
| X | 53189600 | 53190599 | 23.34141 | 40.92033 | 29.97806 | 13.24593 | 0.834234 | 0.671415 | 0.217689 |
| 4 | 138327200 | 138328199 | 13.16193 | 24.73606 | 20.77391 | 8.532922 | 0.717057 | 0.107773 | 0.217699 |

| | | | | | | | | | |
|---|---|---|---|---|---|---|---|---|---|
| 1 | 45602000 | 45603599 | 14.96916 | 27.9864 | 21.05723 | 8.725901 | 0.864357 | 0.156855 | 0.217713 |
| 17 | 57634400 | 57642399 | 46.06604 | 73.27467 | 139.3651 | 58.19667 | -0.5913 | 2.231171 | 0.217714 |
| 17 | 72907600 | 72913599 | 53.14443 | 84.17487 | 152.1379 | 63.26859 | -0.52384 | 2.362961 | 0.217723 |
| 11 | 41933400 | 41935799 | 21.91177 | 39.60778 | 21.23432 | 9.218114 | 1.303042 | 0.400501 | 0.217732 |
| 17 | 27294000 | 27300999 | 38.61034 | 61.5734 | 128.9784 | 54.05127 | -0.72617 | 2.139737 | 0.217742 |
| 7 | 17629000 | 17634399 | 102.5374 | 161.5885 | 103.5859 | 42.98383 | 0.963725 | 2.504868 | 0.217754 |
| 5 | 70817800 | 70821799 | 105.1176 | 162.8222 | 143.5791 | 59.63887 | 0.485153 | 2.816393 | 0.217759 |
| 13 | 54552400 | 54556199 | 33.47423 | 56.48145 | 41.42814 | 18.11811 | 0.826671 | 1.118319 | 0.217764 |
| 10 | 78069200 | 78069999 | 4.52319 | 8.807983 | 24.19996 | 8.862083 | -0.8243 | -0.35583 | 0.217767 |
| 7 | 27781800 | 27807999 | 239.9247 | 339.0871 | 598.5708 | 247.7768 | -0.63591 | 4.419595 | 0.217779 |
| 18 | 37497000 | 37500399 | 59.65181 | 95.52785 | 78.63124 | 33.14855 | 0.625536 | 2.018413 | 0.217801 |
| 1 | 65817800 | 65819999 | 9.970198 | 17.99186 | 39.56333 | 17.21118 | -0.7223 | 0.362985 | 0.217802 |
| 7 | 34314600 | 34321999 | 54.99545 | 86.90488 | 157.242 | 65.31452 | -0.5281 | 2.420295 | 0.217803 |
| 3 | 89459400 | 89461999 | 37.86503 | 60.63808 | 117.7623 | 49.64514 | -0.61052 | 2.004775 | 0.217815 |
| 3 | 16692000 | 16692799 | 11.78137 | 22.7169 | 17.19306 | 6.654366 | 0.94112 | -0.10783 | 0.21783 |
| 15 | 83137800 | 83141399 | 25.82134 | 42.20552 | 86.23068 | 37.0746 | -0.65369 | 1.486189 | 0.217841 |
| 2 | 160076800 | 160078399 | 20.06056 | 35.28989 | 28.27436 | 12.51882 | 0.706104 | 0.600284 | 0.217844 |
| 11 | 120447200 | 120450199 | 40.50041 | 66.55658 | 53.86323 | 23.20922 | 0.6824 | 1.42922 | 0.217844 |
| 10 | 4322000 | 4325399 | 13.9174 | 24.05099 | 53.75025 | 23.5807 | -0.76954 | 0.845321 | 0.217881 |
| 14 | 20211200 | 20212399 | 5.465218 | 10.39769 | 29.30717 | 11.74764 | -0.9841 | -0.02105 | 0.217883 |
| 14 | 4048600 | 4050799 | 33.76001 | 56.37585 | 40.71201 | 17.71448 | 0.849131 | 1.218091 | 0.217887 |
| 4 | 136636400 | 136636999 | 4.182519 | 8.163229 | 22.80596 | 8.092204 | -0.80534 | -0.47542 | 0.217888 |
| 12 | 73620000 | 73620599 | 2.453679 | 4.797153 | 20.58282 | 7.052363 | -1.36552 | -0.63466 | 0.217908 |
| 2 | 113715000 | 113715599 | 6.666603 | 12.51302 | 31.21725 | 12.85024 | -0.84337 | 0.087426 | 0.217909 |
| 17 | 35927200 | 35934599 | 58.44047 | 91.36855 | 181.4024 | 75.25988 | -0.67389 | 2.638371 | 0.217911 |
| 16 | 52270600 | 52274799 | 26.25669 | 42.33066 | 106.7485 | 45.448 | -0.97449 | 1.774095 | 0.217917 |
| 4 | 142608200 | 142611199 | 15.04381 | 25.84676 | 56.43561 | 24.7544 | -0.73808 | 0.906055 | 0.21792 |
| 2 | 160737000 | 160754999 | 122.6872 | 182.0024 | 369.0961 | 152.029 | -0.76756 | 3.668705 | 0.217927 |
| 1 | 9150800 | 9154599 | 47.58472 | 77.09682 | 63.39181 | 27.13506 | 0.646075 | 1.659648 | 0.217933 |
| 5 | 97412200 | 97413599 | 11.51788 | 20.24446 | 44.06479 | 19.51274 | -0.72842 | 0.590822 | 0.217934 |
| 7 | 63403000 | 63409599 | 54.0935 | 84.88643 | 177.4001 | 73.57322 | -0.74358 | 2.568742 | 0.217935 |
| 18 | 27365800 | 27366999 | 7.053421 | 13.40178 | 32.25451 | 12.9648 | -0.76321 | 0.00858 | 0.217948 |
| 14 | 42399600 | 42402999 | 34.02637 | 56.40363 | 48.75671 | 21.11135 | 0.590335 | 1.306998 | 0.217965 |
| 15 | 83802200 | 83810199 | 47.64182 | 75.46981 | 149.7734 | 62.28968 | -0.65831 | 2.35551 | 0.21797 |
| 2 | 73722200 | 73724599 | 12.53025 | 21.7047 | 52.10469 | 22.86042 | -0.87128 | 0.816797 | 0.217975 |
| 12 | 90405200 | 90405999 | 10.3531 | 20.2241 | 11.85146 | 4.143131 | 1.437284 | -0.48144 | 0.217977 |
| 10 | 46883800 | 46886999 | 36.9604 | 62.53852 | 36.93741 | 16.16786 | 1.138076 | 1.104287 | 0.217991 |
| 5 | 127860400 | 127865599 | 47.18167 | 75.18172 | 136.1636 | 56.99867 | -0.51808 | 2.175005 | 0.218005 |
| 1 | 59961000 | 59962399 | 15.53449 | 28.83941 | 20.24809 | 8.487445 | 0.950042 | 0.200832 | 0.218015 |
| 2 | 98054800 | 98056999 | 32.73582 | 55.76644 | 36.91353 | 16.18078 | 0.974867 | 1.029112 | 0.218016 |
| 4 | 96260800 | 96264199 | 38.14153 | 61.24731 | 113.3855 | 47.99824 | -0.53693 | 1.918653 | 0.218028 |
| 3 | 52002800 | 52005599 | 18.3056 | 30.54295 | 73.13889 | 31.80716 | -0.87436 | 1.211013 | 0.218032 |
| 19 | 25561400 | 25567799 | 45.56517 | 72.56837 | 133.7403 | 55.94425 | -0.54395 | 2.189619 | 0.218038 |
| 8 | 80111200 | 80118399 | 45.08126 | 71.79507 | 133.5056 | 55.84324 | -0.5569 | 2.190343 | 0.218051 |
| 7 | 122737800 | 122739599 | 14.51232 | 27.1591 | 20.49274 | 8.47779 | 0.861909 | 0.144494 | 0.218059 |
| 3 | 141525200 | 141526399 | 5.28915 | 10.18376 | 28.65108 | 11.13542 | -0.9407 | -0.12906 | 0.218075 |
| 19 | 30022200 | 30025599 | 33.18711 | 55.90691 | 42.56132 | 18.60461 | 0.773154 | 1.132403 | 0.218077 |

| | | | | | | | | | |
|---|---|---|---|---|---|---|---|---|---|
| 18 | 56262000 | 56262799 | 13.85031 | 26.59016 | 17.67371 | 6.92649 | 1.111102 | -0.04376 | 0.218082 |
| 6 | 54571800 | 54572799 | 5.166216 | 9.856792 | 28.80584 | 11.47752 | -1.02769 | -0.04473 | 0.21809 |
| 7 | 122248600 | 122248999 | 2.835583 | 5.546459 | 19.07798 | 6.39681 | -1.02384 | -0.71113 | 0.2181 |
| 7 | 43069600 | 43074999 | 23.66348 | 38.95876 | 77.70718 | 33.50932 | -0.61414 | 1.383952 | 0.218101 |
| X | 39398800 | 39400999 | 34.77091 | 59.08237 | 36.95139 | 16.19405 | 1.056005 | 1.061039 | 0.218101 |
| 13 | 56270200 | 56281599 | 78.60054 | 120.3712 | 250.3329 | 103.8917 | -0.76399 | 3.034206 | 0.218102 |
| X | 61505200 | 61507799 | 26.37863 | 46.4699 | 24.09481 | 10.66526 | 1.330829 | 0.61495 | 0.218124 |
| X | 62306000 | 62308599 | 27.10809 | 47.6116 | 24.56904 | 10.87288 | 1.336542 | 0.654658 | 0.21814 |
| 11 | 118924200 | 118930399 | 40.22888 | 64.11413 | 131.7597 | 55.16872 | -0.6997 | 2.161767 | 0.218145 |
| 12 | 31744600 | 31748999 | 54.29468 | 87.32771 | 75.44328 | 32.0126 | 0.563314 | 1.8751 | 0.218154 |
| 8 | 118322800 | 118345799 | 248.6757 | 351.5894 | 592.7095 | 245.335 | -0.56895 | 4.41439 | 0.218162 |
| 14 | 42038000 | 42042399 | 67.48738 | 108.3074 | 66.69475 | 28.20811 | 1.046881 | 1.957838 | 0.218169 |
| 1 | 9868200 | 9870399 | 8.681531 | 15.92145 | 36.17137 | 15.41607 | -0.74898 | 0.252908 | 0.218177 |
| 9 | 88363000 | 88365399 | 15.5352 | 26.2219 | 62.04957 | 27.13953 | -0.85657 | 1.077511 | 0.21818 |
| 2 | 98684400 | 98685599 | 11.10073 | 21.4695 | 17.38436 | 6.644776 | 0.861875 | -0.1547 | 0.218182 |
| 17 | 19246400 | 19249799 | 76.40443 | 121.5405 | 95.72447 | 39.97679 | 0.678732 | 2.245625 | 0.218182 |
| 13 | 86525200 | 86526999 | 16.0612 | 29.28019 | 23.47028 | 10.081 | 0.736206 | 0.313332 | 0.218196 |
| 5 | 114010400 | 114014799 | 42.51387 | 67.69714 | 132.7281 | 55.50764 | -0.63329 | 2.193065 | 0.218198 |
| 4 | 131195600 | 131198399 | 15.30925 | 26.52677 | 59.94451 | 26.30031 | -0.785 | 0.83458 | 0.218201 |
| 8 | 45467800 | 45469599 | 10.16652 | 18.2712 | 38.96158 | 17.02749 | -0.68235 | 0.389793 | 0.218203 |
| X | 108111600 | 108113799 | 19.33085 | 34.55242 | 25.76334 | 11.31128 | 0.819778 | 0.456681 | 0.218216 |
| 5 | 17494400 | 17495399 | 8.819463 | 16.27655 | 36.34171 | 15.35656 | -0.71506 | 0.219492 | 0.21822 |
| 10 | 40049400 | 40050599 | 7.385362 | 13.93415 | 31.93857 | 13.03016 | -0.71201 | 0.061737 | 0.218235 |
| 2 | 160714200 | 160715199 | 7.45585 | 13.8131 | 33.1293 | 13.92592 | -0.81126 | 0.180746 | 0.218249 |
| 8 | 117943000 | 117949399 | 38.39713 | 61.38186 | 121.3623 | 51.03355 | -0.63916 | 2.062027 | 0.218254 |
| X | 8459600 | 8462199 | 61.53464 | 99.12313 | 66.09673 | 28.1085 | 0.938774 | 1.836816 | 0.218267 |
| 14 | 48470600 | 48471799 | 6.45565 | 12.31339 | 30.62477 | 12.20971 | -0.79954 | -0.02819 | 0.218268 |
| 18 | 37845200 | 37845999 | 26.52316 | 46.28293 | 30.33581 | 13.35549 | 0.993927 | 0.75771 | 0.218271 |
| 7 | 79676200 | 79679399 | 13.39975 | 23.35026 | 48.93037 | 21.52312 | -0.67492 | 0.740774 | 0.218273 |
| 12 | 115554000 | 115555799 | 52.3289 | 84.1862 | 73.36537 | 31.14175 | 0.551206 | 1.864289 | 0.218274 |
| 3 | 44237400 | 44238599 | 14.42985 | 27.64159 | 16.82792 | 6.637018 | 1.227287 | -0.02162 | 0.218302 |
| 13 | 87521800 | 87526199 | 78.85113 | 124.9681 | 106.5192 | 44.31512 | 0.559697 | 2.357255 | 0.21831 |
| 13 | 87522000 | 87526199 | 78.85113 | 124.9681 | 106.5192 | 44.31512 | 0.559697 | 2.357255 | 0.21831 |
| 18 | 47146000 | 47147999 | 34.20528 | 58.57998 | 32.97548 | 14.46075 | 1.208641 | 0.985246 | 0.218315 |
| 1 | 132973400 | 132975599 | 12.17754 | 21.20821 | 49.44991 | 21.74377 | -0.82819 | 0.743607 | 0.218325 |
| 12 | 13659200 | 13660399 | 8.214472 | 15.15709 | 34.13483 | 14.42783 | -0.72729 | 0.211552 | 0.218327 |
| 6 | 83188600 | 83190399 | 7.468029 | 13.79918 | 34.15722 | 14.40968 | -0.86057 | 0.196766 | 0.218336 |
| X | 5114600 | 5116599 | 38.91752 | 64.69572 | 42.75134 | 18.54755 | 0.976782 | 1.28369 | 0.218337 |
| 12 | 17200200 | 17210799 | 79.25921 | 122.2072 | 216.6491 | 89.99229 | -0.52636 | 2.898244 | 0.218341 |
| 4 | 69138400 | 69140999 | 25.41496 | 44.3791 | 30.33247 | 13.35969 | 0.933754 | 0.741884 | 0.218353 |
| 14 | 42075600 | 42085399 | 88.66497 | 140.3329 | 103.2699 | 42.93031 | 0.769801 | 2.396802 | 0.218387 |
| 9 | 83027600 | 83028799 | 10.2944 | 18.08527 | 44.60265 | 19.74287 | -0.90769 | 0.590285 | 0.218419 |
| 10 | 12672000 | 12672999 | 5.410229 | 10.45024 | 27.67216 | 10.64092 | -0.8394 | -0.16564 | 0.218435 |
| 12 | 12763200 | 12763999 | 5.410229 | 10.45024 | 27.67216 | 10.64092 | -0.8394 | -0.16564 | 0.218435 |
| 16 | 8310000 | 8311399 | 6.317775 | 12.03023 | 29.38314 | 11.75629 | -0.77753 | -0.01913 | 0.218443 |
| 9 | 19757000 | 19757999 | 12.84504 | 24.93714 | 14.99702 | 5.629129 | 1.31104 | -0.20027 | 0.218447 |
| 5 | 144293800 | 144297199 | 19.25943 | 32.08363 | 69.82034 | 30.33367 | -0.73685 | 1.227618 | 0.218459 |

| | | | | | | | | | |
|---|---|---|---|---|---|---|---|---|---|
| 2 | 36572600 | 36575199 | 58.70626 | 94.06564 | 82.67383 | 34.88211 | 0.532046 | 1.999867 | 0.218462 |
| 7 | 43400800 | 43402599 | 10.96544 | 19.48442 | 43.53736 | 19.21035 | -0.75958 | 0.472719 | 0.21847 |
| 13 | 57651400 | 57656599 | 35.84092 | 57.64286 | 111.9897 | 47.51571 | -0.60363 | 1.865111 | 0.218495 |
| 1 | 45889000 | 45889799 | 11.34764 | 21.99579 | 16.87708 | 6.384344 | 0.953069 | -0.18664 | 0.218501 |
| 1 | 26378800 | 26380999 | 24.0673 | 41.87585 | 33.92955 | 14.91045 | 0.689305 | 0.790882 | 0.218508 |
| X | 149005400 | 149009799 | 84.29224 | 133.7114 | 99.36769 | 41.37701 | 0.75848 | 2.335607 | 0.218512 |
| 1 | 188524400 | 188526599 | 12.32837 | 21.46649 | 50.51969 | 22.2113 | -0.8417 | 0.747934 | 0.218515 |
| 9 | 55233800 | 55235799 | 15.41082 | 26.35606 | 56.34775 | 24.70744 | -0.70877 | 0.95172 | 0.218524 |
| 16 | 48093600 | 48095799 | 24.96134 | 42.90091 | 36.29335 | 15.91955 | 0.624255 | 0.918427 | 0.218525 |
| 8 | 100979000 | 100982799 | 46.24345 | 75.34731 | 52.72544 | 22.64303 | 0.884843 | 1.555916 | 0.218528 |
| 5 | 85919400 | 85920399 | 10.66394 | 20.70347 | 16.52188 | 6.199814 | 0.908475 | -0.22078 | 0.218539 |
| 9 | 26299000 | 26301399 | 26.64304 | 46.61182 | 28.66658 | 12.64429 | 1.085418 | 0.71934 | 0.21855 |
| 7 | 92523200 | 92524799 | 10.98535 | 19.10679 | 50.48324 | 22.1798 | -1.00762 | 0.759035 | 0.218557 |
| 10 | 100016000 | 100017599 | 25.54934 | 44.06895 | 36.68762 | 16.09425 | 0.648375 | 0.890696 | 0.218572 |
| 7 | 53416400 | 53418799 | 39.38804 | 65.82559 | 42.03623 | 18.2997 | 1.025899 | 1.229725 | 0.218572 |
| 9 | 122147200 | 122152199 | 32.42426 | 52.32493 | 99.55951 | 42.44066 | -0.56754 | 1.752748 | 0.218573 |
| 7 | 19419200 | 19427599 | 47.65976 | 75.43324 | 156.6246 | 65.11067 | -0.72447 | 2.376473 | 0.218583 |
| 3 | 6620800 | 6626199 | 94.86272 | 148.3858 | 130.294 | 54.0534 | 0.502196 | 2.641174 | 0.218586 |
| X | 31685800 | 31689199 | 74.22336 | 118.8855 | 65.7178 | 27.71258 | 1.199035 | 2.026124 | 0.218592 |
| 7 | 41100000 | 41107599 | 92.83718 | 145.2477 | 129.2863 | 53.63418 | 0.482813 | 2.635902 | 0.218598 |
| 6 | 3233600 | 3238399 | 62.53572 | 100.3286 | 77.84409 | 32.90764 | 0.713667 | 1.964686 | 0.218637 |
| 4 | 133074600 | 133076999 | 17.90908 | 31.96065 | 26.10688 | 11.47737 | 0.687804 | 0.458188 | 0.218638 |
| 6 | 116527400 | 116529399 | 42.9487 | 68.08952 | 147.6776 | 61.45788 | -0.78512 | 2.330449 | 0.218641 |
| 7 | 5128600 | 5132399 | 36.54644 | 58.6364 | 112.2656 | 47.46319 | -0.58676 | 1.943005 | 0.218642 |
| 2 | 32550800 | 32553799 | 14.38253 | 24.94019 | 51.62024 | 22.64861 | -0.65813 | 0.815797 | 0.218658 |
| 17 | 39249400 | 39256999 | 69.9427 | 111.769 | 78.81208 | 33.12338 | 0.844948 | 2.093617 | 0.218658 |
| 5 | 140323600 | 140328599 | 35.2787 | 56.5727 | 113.1526 | 47.80003 | -0.65054 | 1.959173 | 0.21866 |
| 2 | 137880400 | 137882999 | 20.80996 | 36.51483 | 29.54063 | 13.06245 | 0.691561 | 0.640437 | 0.218664 |
| 5 | 139682800 | 139699799 | 128.4369 | 190.5086 | 365.819 | 150.6773 | -0.68901 | 3.671568 | 0.218668 |
| 1 | 193756600 | 193757399 | 12.75888 | 24.70993 | 16.2225 | 6.161527 | 1.170711 | -0.15882 | 0.218696 |
| 9 | 105371600 | 105372999 | 8.877309 | 15.7343 | 40.48137 | 17.89676 | -0.96279 | 0.474704 | 0.218717 |
| X | 164486800 | 164490399 | 35.59441 | 61.20563 | 28.97915 | 12.71147 | 1.457788 | 0.951893 | 0.21873 |
| 7 | 30730200 | 30744799 | 97.87425 | 147.8825 | 286.774 | 118.5687 | -0.67913 | 3.315518 | 0.218733 |
| 7 | 53032800 | 53038199 | 53.087 | 85.79097 | 64.76343 | 27.65809 | 0.765441 | 1.72834 | 0.218765 |
| 18 | 31177200 | 31177999 | 11.08746 | 21.23639 | 18.46221 | 7.284113 | 0.717965 | -0.04965 | 0.218776 |
| X | 98544000 | 98547799 | 13.78193 | 23.99119 | 56.48608 | 24.83037 | -0.84245 | 0.770399 | 0.218799 |
| 2 | 11124400 | 11126799 | 15.90123 | 27.08693 | 59.04372 | 25.88129 | -0.7377 | 0.991819 | 0.218801 |
| 9 | 61278000 | 61282599 | 23.78018 | 38.83794 | 82.52624 | 35.47149 | -0.71078 | 1.493553 | 0.218802 |
| 8 | 115575400 | 115577399 | 9.091786 | 16.34566 | 37.94387 | 16.5776 | -0.80357 | 0.377187 | 0.218822 |
| 8 | 4662200 | 4663799 | 10.43374 | 18.35274 | 45.61927 | 20.19892 | -0.91877 | 0.574894 | 0.218827 |
| 12 | 84873800 | 84878999 | 29.64301 | 47.63384 | 110.5832 | 46.8606 | -0.86135 | 1.884713 | 0.21883 |
| 9 | 3917600 | 3922199 | 48.43058 | 78.62756 | 60.17073 | 25.79505 | 0.752109 | 1.619719 | 0.21883 |
| 4 | 32888800 | 32898599 | 117.0078 | 175.9101 | 305.0304 | 125.9357 | -0.52425 | 3.419381 | 0.218844 |
| 7 | 62410600 | 62414999 | 96.70325 | 151.3556 | 128.5799 | 53.33898 | 0.550506 | 2.631899 | 0.218846 |
| 7 | 9429400 | 9432799 | 29.47164 | 51.50468 | 26.27666 | 11.57955 | 1.353491 | 0.747321 | 0.218859 |
| 4 | 137422800 | 137423599 | 3.030747 | 5.912132 | 23.73557 | 8.494394 | -1.33214 | -0.46037 | 0.218871 |
| 12 | 16001600 | 16003399 | 17.41965 | 31.40375 | 25.04492 | 10.90876 | 0.73102 | 0.391473 | 0.218872 |

| | | | | | | | | | |
|---|---|---|---|---|---|---|---|---|---|
| 3 | 86131000 | 86132799 | 9.435656 | 16.52038 | 47.27156 | 20.86423 | -1.12076 | 0.646352 | 0.21888 |
| 8 | 110251800 | 110252199 | 3.817653 | 7.452083 | 22.19945 | 7.859159 | -0.89411 | -0.49322 | 0.218883 |
| 18 | 38220400 | 38222799 | 10.90822 | 19.13141 | 46.19107 | 20.41691 | -0.87748 | 0.623101 | 0.218898 |
| 6 | 5252400 | 5253799 | 5.558103 | 10.70076 | 27.48526 | 10.68473 | -0.81045 | -0.1269 | 0.218908 |
| 12 | 90037000 | 90037799 | 5.558103 | 10.70076 | 27.48526 | 10.68473 | -0.81045 | -0.1269 | 0.218908 |
| 14 | 123846400 | 123847199 | 5.558103 | 10.70076 | 27.48526 | 10.68473 | -0.81045 | -0.1269 | 0.218908 |
| 9 | 118539000 | 118546399 | 51.56266 | 81.57836 | 152.1373 | 63.23262 | -0.57024 | 2.388777 | 0.218915 |
| 1 | 80922400 | 80922799 | 3.2609 | 6.378611 | 21.83617 | 7.299718 | -1.01371 | -0.70102 | 0.218918 |
| 7 | 16205600 | 16215599 | 76.31436 | 116.7963 | 251.239 | 104.2525 | -0.81336 | 3.045359 | 0.21892 |
| 15 | 14606800 | 14607799 | 14.31709 | 26.48524 | 22.21273 | 9.356009 | 0.691131 | 0.216411 | 0.218921 |
| 4 | 48446200 | 48447799 | 14.08048 | 26.01318 | 21.21971 | 8.953813 | 0.728945 | 0.218837 | 0.218957 |
| X | 61936200 | 61940199 | 22.65367 | 39.82915 | 25.9458 | 11.48911 | 1.002681 | 0.617132 | 0.218967 |
| 14 | 27331600 | 27334199 | 25.40992 | 44.47946 | 29.23369 | 12.90056 | 0.990014 | 0.704355 | 0.218972 |
| 11 | 99837200 | 99837999 | 11.36523 | 21.9498 | 17.97621 | 6.916824 | 0.837159 | -0.12709 | 0.218984 |
| 1 | 116644200 | 116646199 | 26.34432 | 45.94258 | 30.90966 | 13.60287 | 0.956377 | 0.765942 | 0.218986 |
| 19 | 52918200 | 52918599 | 9.466604 | 18.47299 | 13.71439 | 4.881963 | 1.077885 | -0.4275 | 0.218995 |
| 2 | 84899600 | 84903599 | 21.81011 | 35.80664 | 78.7844 | 33.93347 | -0.75681 | 1.416245 | 0.219003 |
| 9 | 113463800 | 113474999 | 101.5385 | 153.7458 | 278.3886 | 115.179 | -0.57769 | 3.27253 | 0.219011 |
| 4 | 112873000 | 112873399 | 15.70971 | 28.71228 | 23.64649 | 10.12445 | 0.700593 | 0.298016 | 0.219023 |
| 10 | 68793400 | 68796199 | 34.70481 | 57.70849 | 48.73411 | 21.14719 | 0.624197 | 1.273659 | 0.219029 |
| 7 | 11070800 | 11076199 | 77.6253 | 123.4372 | 88.56203 | 36.96787 | 0.812496 | 2.257319 | 0.219034 |
| 13 | 66843600 | 66848599 | 81.55613 | 129.2537 | 97.40912 | 40.5246 | 0.737042 | 2.358675 | 0.219066 |
| 16 | 31965000 | 31968399 | 24.08096 | 39.39834 | 84.89407 | 36.5126 | -0.72903 | 1.472844 | 0.219073 |
| 13 | 59239200 | 59245999 | 26.50516 | 42.83919 | 98.21248 | 41.95012 | -0.83308 | 1.696604 | 0.219076 |
| 13 | 13173600 | 13173999 | 6.802427 | 13.30441 | 12.46991 | 4.215899 | 0.812761 | -0.64633 | 0.219105 |
| 12 | 106185400 | 106195799 | 71.8607 | 111.2886 | 204.2767 | 84.85109 | -0.57158 | 2.812828 | 0.219113 |
| 8 | 125913000 | 125915999 | 16.8741 | 28.64897 | 59.55715 | 26.09565 | -0.67023 | 1.027344 | 0.219121 |
| 11 | 65697800 | 65698399 | 4.996781 | 9.644138 | 27.55742 | 10.63535 | -0.95277 | -0.15761 | 0.219122 |
| 12 | 11315400 | 11316599 | 10.63491 | 18.59864 | 47.29314 | 20.85342 | -0.95173 | 0.671552 | 0.219132 |
| X | 146720800 | 146725199 | 69.26486 | 110.3198 | 92.31615 | 38.61951 | 0.593337 | 2.200323 | 0.219141 |
| X | 33703200 | 33707999 | 50.85997 | 82.80608 | 54.56595 | 23.42692 | 0.970566 | 1.578526 | 0.219156 |
| X | 26416800 | 26418999 | 19.83334 | 37.69948 | 12.21836 | 4.906983 | 2.103109 | 0.075457 | 0.219158 |
| 3 | 75298400 | 75299999 | 16.69633 | 28.3999 | 59.04744 | 25.87988 | -0.67012 | 1.008846 | 0.219158 |
| 16 | 37340200 | 37341399 | 13.89579 | 25.91163 | 21.14013 | 8.799217 | 0.743199 | 0.163381 | 0.219179 |
| 19 | 14020000 | 14020399 | 2.452497 | 4.797301 | 20.58929 | 6.794986 | -1.31726 | -0.76967 | 0.219183 |
| 4 | 130827400 | 130830399 | 19.31182 | 32.04181 | 71.231 | 30.86232 | -0.76779 | 1.276526 | 0.219186 |
| 8 | 119231600 | 119232799 | 8.824527 | 16.21482 | 36.22205 | 15.39737 | -0.72211 | 0.243331 | 0.219196 |
| 7 | 20962800 | 20970199 | 155.6851 | 236.3942 | 176.4591 | 73.06994 | 0.703062 | 3.222165 | 0.219199 |
| 11 | 105236000 | 105243199 | 32.57824 | 52.19593 | 118.2325 | 49.87864 | -0.83121 | 1.984216 | 0.2192 |
| 11 | 77787800 | 77789399 | 7.492593 | 13.77838 | 34.6071 | 14.69496 | -0.88826 | 0.225758 | 0.219204 |
| 2 | 90746400 | 90748599 | 9.392185 | 16.8757 | 38.44474 | 16.80486 | -0.7773 | 0.383466 | 0.219208 |
| 1 | 156833400 | 156834999 | 15.31034 | 26.47189 | 53.80213 | 23.60384 | -0.63313 | 0.847138 | 0.219223 |
| 10 | 50447400 | 50451399 | 44.06481 | 72.12198 | 55.03813 | 23.6751 | 0.764623 | 1.488538 | 0.219234 |
| 5 | 127472000 | 127485799 | 127.147 | 190.5153 | 327.4806 | 135.0903 | -0.51597 | 3.486344 | 0.219248 |
| 19 | 39885600 | 39887799 | 18.54397 | 33.51615 | 24.31027 | 10.55499 | 0.869877 | 0.383227 | 0.219248 |
| 18 | 34236200 | 34238599 | 17.21568 | 29.11774 | 61.72447 | 27.01405 | -0.6986 | 1.063372 | 0.21927 |
| 7 | 19146600 | 19148799 | 24.11977 | 39.31756 | 84.48192 | 36.28686 | -0.72874 | 1.524729 | 0.219272 |

| | | | | | | | | | |
|---|---|---|---|---|---|---|---|---|---|
| 13 | 89982800 | 89983599 | 5.822104 | 11.26537 | 28.47647 | 10.86165 | -0.76187 | -0.18569 | 0.219277 |
| 12 | 98315600 | 98317199 | 12.78473 | 22.273 | 48.92631 | 21.51798 | -0.74258 | 0.741275 | 0.219277 |
| 6 | 116155200 | 116158399 | 25.77485 | 44.49148 | 36.17957 | 15.87174 | 0.682372 | 0.885082 | 0.219329 |
| 10 | 81394200 | 81397399 | 21.12921 | 34.81701 | 76.78057 | 33.12207 | -0.75791 | 1.366262 | 0.219337 |
| 8 | 82900400 | 82901399 | 6.910204 | 13.11289 | 36.41648 | 14.67735 | -0.9721 | 0.024137 | 0.219346 |
| 2 | 69596200 | 69596999 | 7.585116 | 14.11613 | 32.67434 | 13.64571 | -0.75338 | 0.154282 | 0.219356 |
| 13 | 81907000 | 81907999 | 7.585116 | 14.11613 | 32.67434 | 13.64571 | -0.75338 | 0.154282 | 0.219356 |
| 18 | 76328200 | 76329999 | 14.12215 | 26.43716 | 21.1494 | 8.744824 | 0.779022 | 0.14141 | 0.219367 |
| 10 | 67382600 | 67385599 | 15.44872 | 26.32391 | 57.88272 | 25.37298 | -0.74992 | 0.98869 | 0.219393 |
| 11 | 93846000 | 93849599 | 17.58954 | 29.59207 | 63.24828 | 27.63629 | -0.71073 | 1.114017 | 0.219402 |
| 7 | 44929200 | 44932799 | 20.8944 | 34.65226 | 71.37704 | 30.91614 | -0.65824 | 1.286725 | 0.219415 |
| 4 | 84784400 | 84785999 | 13.3881 | 23.32766 | 50.00233 | 21.99328 | -0.70756 | 0.742513 | 0.219418 |
| 1 | 179947600 | 179949199 | 7.128234 | 13.30532 | 32.74332 | 13.60591 | -0.8354 | 0.129623 | 0.219438 |
| 7 | 7643200 | 7647599 | 44.07113 | 72.66401 | 48.37707 | 20.87442 | 0.965024 | 1.395979 | 0.219449 |
| 4 | 50745000 | 50745599 | 10.22879 | 19.87564 | 16.46436 | 6.135752 | 0.863423 | -0.2513 | 0.219453 |
| 5 | 151510800 | 151513799 | 26.29294 | 45.74187 | 31.4409 | 13.81557 | 0.925226 | 0.798063 | 0.219454 |
| 9 | 123959000 | 123960999 | 62.5024 | 99.8361 | 88.65019 | 37.23534 | 0.512299 | 2.100988 | 0.219472 |
| 4 | 146632800 | 146636799 | 55.93367 | 89.98405 | 66.89574 | 28.39488 | 0.779944 | 1.867572 | 0.219484 |
| 11 | 54944600 | 54947399 | 13.88177 | 24.18035 | 49.46785 | 21.75395 | -0.64052 | 0.750429 | 0.219484 |
| 12 | 38246400 | 38247799 | 16.9052 | 30.63117 | 24.56642 | 10.63273 | 0.728753 | 0.358009 | 0.219516 |
| X | 120219800 | 120220799 | 16.9052 | 30.63117 | 24.56642 | 10.63273 | 0.728753 | 0.358009 | 0.219516 |
| 1 | 182510400 | 182512399 | 11.65165 | 20.46737 | 44.5776 | 19.72989 | -0.72962 | 0.599424 | 0.219518 |
| 5 | 145154600 | 145161199 | 34.62463 | 55.53978 | 112.7191 | 47.63769 | -0.67105 | 1.948424 | 0.219543 |
| 1 | 102194800 | 102195799 | 14.23868 | 27.28118 | 16.83492 | 6.635736 | 1.208663 | -0.02556 | 0.219575 |
| 1 | 100484600 | 100485999 | 12.29556 | 23.51522 | 18.75821 | 7.432672 | 0.834934 | -0.02361 | 0.219586 |
| 17 | 46348600 | 46353599 | 24.37107 | 39.56469 | 93.65724 | 40.14913 | -0.87289 | 1.593733 | 0.219603 |
| 12 | 113628600 | 113635199 | 97.29634 | 152.7623 | 120.6886 | 50.05663 | 0.65858 | 2.5753 | 0.219604 |
| 1 | 156349600 | 156351799 | 16.46195 | 28.24481 | 57.01769 | 25.0077 | -0.62579 | 0.925495 | 0.219616 |
| 15 | 35295600 | 35297399 | 10.07483 | 17.58739 | 54.22139 | 23.87704 | -1.22873 | 0.704068 | 0.219632 |
| X | 159777000 | 159778799 | 23.29378 | 40.79074 | 29.74991 | 13.13258 | 0.840873 | 0.689885 | 0.219634 |
| 4 | 110484600 | 110487999 | 46.53076 | 76.49402 | 48.42188 | 20.86775 | 1.035934 | 1.436964 | 0.219645 |
| 10 | 37861600 | 37862599 | 11.4053 | 19.8994 | 47.84842 | 21.0636 | -0.87177 | 0.712686 | 0.21965 |
| 15 | 79944600 | 79951999 | 51.50553 | 81.20539 | 161.8059 | 67.14681 | -0.66961 | 2.471188 | 0.219655 |
| 1 | 48610800 | 48613399 | 33.34278 | 55.72835 | 46.48926 | 20.23954 | 0.641727 | 1.21279 | 0.219663 |
| 8 | 83869400 | 83872599 | 38.21041 | 62.88809 | 53.7688 | 23.18332 | 0.604218 | 1.405524 | 0.219668 |
| 1 | 125289200 | 125292799 | 32.92812 | 53.05147 | 103.2866 | 43.93502 | -0.60339 | 1.803028 | 0.219684 |
| 19 | 25337000 | 25339399 | 10.25804 | 18.10646 | 41.99076 | 18.57942 | -0.81685 | 0.531002 | 0.219687 |
| X | 91284800 | 91286799 | 40.14527 | 65.84915 | 54.83663 | 23.60988 | 0.640272 | 1.453628 | 0.219692 |
| 4 | 80385000 | 80385999 | 11.83221 | 22.64064 | 18.83468 | 7.452032 | 0.776938 | -0.03262 | 0.219703 |
| 13 | 65605600 | 65610999 | 79.51418 | 126.1035 | 104.0516 | 43.31758 | 0.607421 | 2.337968 | 0.219709 |
| 7 | 37080000 | 37082999 | 15.80751 | 27.24114 | 54.86381 | 24.06718 | -0.62108 | 0.880787 | 0.219718 |
| 14 | 47168400 | 47170199 | 8.134473 | 14.8028 | 36.55422 | 15.73621 | -0.87806 | 0.2951 | 0.219723 |
| 11 | 5443200 | 5447199 | 24.03834 | 39.43601 | 79.02973 | 34.02861 | -0.62291 | 1.43142 | 0.219724 |
| 5 | 95543400 | 95544399 | 20.67793 | 36.51773 | 25.32412 | 11.20385 | 0.915019 | 0.554999 | 0.219728 |
| 6 | 145976800 | 145990799 | 131.4645 | 196.8108 | 334.2044 | 137.8404 | -0.49956 | 3.503854 | 0.219736 |
| 15 | 100507800 | 100508999 | 5.234103 | 10.12313 | 26.52188 | 10.13714 | -0.81457 | -0.18789 | 0.219737 |
| 3 | 152668600 | 152675999 | 62.91742 | 98.56656 | 176.91 | 73.38087 | -0.52626 | 2.604186 | 0.219749 |

| | | | | | | | | | |
|---|---|---|---|---|---|---|---|---|---|
| 13 | 60499200 | 60499799 | 5.846054 | 11.22864 | 27.9619 | 10.94383 | -0.77528 | -0.099 | 0.219755 |
| 8 | 106099200 | 106110799 | 75.05215 | 114.3793 | 270.3832 | 112.0861 | -0.95417 | 3.133931 | 0.219819 |
| 2 | 54707200 | 54707799 | 10.97978 | 21.15243 | 17.30674 | 6.715514 | 0.826374 | -0.10826 | 0.219823 |
| 16 | 54863200 | 54865799 | 31.43558 | 53.64064 | 37.51161 | 16.44515 | 0.896289 | 1.010264 | 0.219825 |
| 3 | 145453000 | 145453999 | 4.729222 | 9.080617 | 27.87367 | 10.91931 | -1.07474 | -0.10864 | 0.219842 |
| 1 | 185094800 | 185095799 | 5.592087 | 10.66216 | 28.81324 | 11.49681 | -0.91796 | -0.03582 | 0.219872 |
| 16 | 80875400 | 80877199 | 19.10633 | 33.82971 | 27.40275 | 12.1172 | 0.693582 | 0.521473 | 0.219882 |
| 2 | 46209200 | 46210199 | 12.36514 | 23.68668 | 18.80784 | 7.417716 | 0.847961 | -0.03701 | 0.219904 |
| 17 | 39400000 | 39402599 | 27.29721 | 47.93879 | 24.56925 | 10.87202 | 1.346384 | 0.657451 | 0.219906 |
| 17 | 51485400 | 51486799 | 8.002758 | 14.53056 | 37.57425 | 16.22064 | -0.94681 | 0.308923 | 0.219915 |
| 7 | 16963600 | 16969799 | 154.2551 | 233.3856 | 200.777 | 83.04802 | 0.494196 | 3.295821 | 0.219931 |
| 10 | 20274600 | 20277599 | 19.09474 | 31.83446 | 68.68376 | 29.85479 | -0.72442 | 1.218063 | 0.219933 |
| 7 | 19070800 | 19072999 | 9.635949 | 17.04951 | 40.9873 | 18.12667 | -0.86662 | 0.500343 | 0.219944 |
| 12 | 75715400 | 75716999 | 8.589958 | 15.6185 | 36.02214 | 15.52351 | -0.78085 | 0.30497 | 0.219955 |
| 19 | 47394200 | 47404999 | 72.51985 | 111.2157 | 235.1099 | 97.5981 | -0.78596 | 3.002792 | 0.219958 |
| X | 80416800 | 80417999 | 28.71595 | 49.8466 | 32.18727 | 14.12386 | 1.015079 | 0.844575 | 0.219958 |
| 10 | 128875600 | 128879399 | 54.74224 | 87.85926 | 75.97749 | 32.1375 | 0.558173 | 1.941754 | 0.21996 |
| 7 | 38535000 | 38544599 | 202.6721 | 302.3182 | 238.0296 | 98.10476 | 0.603857 | 3.578235 | 0.219968 |
| 18 | 41595600 | 41596599 | 4.687817 | 9.11365 | 26.47409 | 9.824857 | -0.92265 | -0.3015 | 0.219984 |
| 7 | 59286400 | 59289799 | 45.15688 | 74.2753 | 47.86537 | 20.63261 | 1.010431 | 1.424646 | 0.219991 |
| X | 150052600 | 150053599 | 13.93596 | 26.80538 | 17.04495 | 6.643803 | 1.181255 | -0.06205 | 0.219992 |
| 8 | 55809000 | 55816999 | 76.18039 | 117.3191 | 216.8459 | 90.068 | -0.5878 | 2.920615 | 0.219995 |
| 6 | 41402000 | 41403999 | 32.30065 | 54.44288 | 42.62539 | 18.63566 | 0.732932 | 1.126123 | 0.219998 |
| 16 | 33757000 | 33758599 | 7.949608 | 14.5726 | 34.56046 | 14.73849 | -0.81043 | 0.248936 | 0.220001 |
| X | 117205200 | 117206599 | 20.67998 | 36.64409 | 26.44174 | 11.69046 | 0.860161 | 0.523979 | 0.220005 |
| X | 33711400 | 33721999 | 142.4909 | 218.7114 | 151.2836 | 62.80783 | 0.82517 | 3.003069 | 0.220009 |
| 1 | 40693200 | 40694999 | 7.692703 | 14.46867 | 32.99013 | 13.53287 | -0.71123 | 0.086646 | 0.220017 |
| 4 | 133799600 | 133806399 | 49.78429 | 78.34657 | 166.058 | 68.89478 | -0.76064 | 2.508811 | 0.220022 |
| 6 | 117502000 | 117504999 | 18.62822 | 31.62051 | 64.66132 | 28.3303 | -0.64681 | 1.040539 | 0.220024 |
| 10 | 52044800 | 52047199 | 9.709602 | 17.53245 | 47.29173 | 20.55732 | -1.01456 | 0.374082 | 0.220029 |
| 7 | 108379400 | 108381999 | 28.9535 | 50.16227 | 32.53456 | 14.2743 | 1.008034 | 0.861329 | 0.220035 |
| 1 | 114575600 | 114576999 | 21.87117 | 38.41503 | 26.97623 | 11.93852 | 0.894722 | 0.627651 | 0.220041 |
| 14 | 62043600 | 62047199 | 27.93726 | 45.55853 | 89.83368 | 38.59115 | -0.60503 | 1.526677 | 0.220043 |
| 1 | 139612000 | 139615399 | 38.32282 | 63.16644 | 52.70981 | 22.74095 | 0.639987 | 1.386541 | 0.220047 |
| 7 | 54485200 | 54487199 | 12.57069 | 22.07596 | 47.21038 | 20.89051 | -0.70368 | 0.612158 | 0.22005 |
| 18 | 38128800 | 38132799 | 17.24216 | 28.7054 | 69.45364 | 30.1626 | -0.88898 | 1.229857 | 0.220057 |
| 7 | 4483400 | 4493199 | 47.6076 | 75.12581 | 167.4874 | 69.52124 | -0.83043 | 2.449665 | 0.220064 |
| 1 | 152922400 | 152924599 | 12.63385 | 22.04042 | 47.85527 | 21.06512 | -0.72548 | 0.720739 | 0.220075 |
| 9 | 35930800 | 35937199 | 83.26608 | 132.0275 | 102.0473 | 42.47474 | 0.701303 | 2.347268 | 0.220082 |
| 6 | 112085000 | 112086799 | 5.070941 | 9.666476 | 38.33311 | 15.30215 | -1.4689 | -0.02213 | 0.220086 |
| 1 | 26748400 | 26749199 | 16.88491 | 30.40602 | 25.09864 | 10.94733 | 0.68033 | 0.393707 | 0.220095 |
| 17 | 55395000 | 55395999 | 5.220573 | 10.1465 | 26.44495 | 9.83666 | -0.77045 | -0.28399 | 0.220121 |
| 5 | 25211800 | 25213399 | 6.110334 | 11.164 | 39.82011 | 17.05644 | -1.3994 | 0.262312 | 0.220124 |
| 7 | 86692800 | 86695399 | 42.51936 | 69.50039 | 57.19292 | 24.59012 | 0.654764 | 1.5041 | 0.220124 |
| 6 | 39396400 | 39401199 | 28.36125 | 45.96846 | 91.33678 | 39.10799 | -0.62311 | 1.635903 | 0.220126 |
| 16 | 3330400 | 3335799 | 87.64292 | 137.6971 | 122.5753 | 50.84354 | 0.487354 | 2.556263 | 0.220128 |
| 2 | 5711800 | 5720399 | 54.35689 | 85.89503 | 155.4091 | 64.5529 | -0.52804 | 2.420151 | 0.220128 |

| | | | | | | | | | |
|---|---|---|---|---|---|---|---|---|---|
| 6 | 39282600 | 39289399 | 41.91526 | 67.00949 | 126.3485 | 53.13423 | -0.57084 | 2.063783 | 0.220132 |
| 1 | 45964400 | 45965599 | 9.239402 | 16.63842 | 37.44959 | 16.33411 | -0.7575 | 0.368601 | 0.220137 |
| 7 | 14855000 | 14859799 | 81.94017 | 130.3644 | 90.11243 | 37.64021 | 0.866906 | 2.245661 | 0.220141 |
| 4 | 19868800 | 19870599 | 9.689352 | 17.43536 | 37.93225 | 16.56022 | -0.71 | 0.37579 | 0.220156 |
| 7 | 31384600 | 31397799 | 342.561 | 486.1598 | 398.1045 | 164.6808 | 0.477872 | 4.361438 | 0.22016 |
| 2 | 169009000 | 169016599 | 56.55423 | 88.66565 | 185.2552 | 76.83637 | -0.74432 | 2.588671 | 0.220164 |
| 11 | 100383200 | 100385199 | 10.2668 | 18.10005 | 42.53452 | 18.82456 | -0.83625 | 0.54774 | 0.220173 |
| 11 | 83740200 | 83744599 | 27.73255 | 44.59323 | 110.4499 | 46.84876 | -0.95332 | 1.860929 | 0.220186 |
| 17 | 7786000 | 7786799 | 5.245469 | 10.16984 | 26.16086 | 9.886053 | -0.77342 | -0.23025 | 0.220192 |
| 7 | 15509800 | 15510799 | 23.36834 | 40.81326 | 31.8483 | 14.02904 | 0.743808 | 0.733432 | 0.220192 |
| 6 | 140677600 | 140685199 | 47.09314 | 74.84027 | 161.0179 | 67.1716 | -0.77027 | 2.264732 | 0.220197 |
| 12 | 30049200 | 30054799 | 30.87312 | 49.69331 | 104.5184 | 44.39773 | -0.71623 | 1.831329 | 0.220208 |
| 7 | 34121800 | 34122599 | 6.584991 | 12.62706 | 30.91114 | 12.15562 | -0.75836 | -0.0701 | 0.220212 |
| 6 | 52548800 | 52549599 | 11.64056 | 22.32121 | 18.30425 | 7.198195 | 0.805466 | -0.05571 | 0.220216 |
| 10 | 19588000 | 19588799 | 4.237001 | 8.240277 | 23.98046 | 8.87275 | -0.91925 | -0.3217 | 0.220222 |
| 11 | 82881600 | 82882799 | 7.240069 | 13.63661 | 31.31897 | 12.81663 | -0.71842 | 0.070191 | 0.220236 |
| 19 | 50018000 | 50019399 | 12.74806 | 22.33596 | 46.20061 | 20.40393 | -0.65571 | 0.650103 | 0.220239 |
| 12 | 60751800 | 60755799 | 59.70881 | 95.51946 | 84.49192 | 35.56553 | 0.520212 | 2.050596 | 0.22025 |
| 10 | 4175800 | 4183799 | 44.39012 | 70.82328 | 129.926 | 54.48312 | -0.53455 | 2.129729 | 0.22025 |
| 14 | 75361400 | 75363599 | 11.92945 | 21.05394 | 45.07684 | 19.94542 | -0.70288 | 0.545998 | 0.220254 |
| X | 90448200 | 90450799 | 31.22572 | 54.12032 | 30.23902 | 13.26731 | 1.222237 | 0.866166 | 0.22027 |
| 2 | 37312400 | 37313199 | 5.127135 | 9.895503 | 26.97792 | 10.41283 | -0.88557 | -0.15651 | 0.220272 |
| 7 | 4828400 | 4842199 | 103.1658 | 156.2344 | 277.5232 | 114.8267 | -0.54986 | 3.269685 | 0.220276 |
| 17 | 44415800 | 44416999 | 5.954611 | 11.13788 | 31.66971 | 13.1134 | -1.03676 | 0.105894 | 0.220279 |
| 19 | 5414200 | 5418199 | 17.93077 | 29.73281 | 74.48755 | 32.26011 | -0.9397 | 1.280837 | 0.220284 |
| 13 | 103403200 | 103404599 | 9.029766 | 16.40424 | 37.04481 | 15.98119 | -0.75224 | 0.312413 | 0.220293 |
| 10 | 62485600 | 62487599 | 10.55928 | 18.62743 | 44.04129 | 19.48895 | -0.84526 | 0.540818 | 0.220299 |
| X | 5095000 | 5103999 | 99.87791 | 156.955 | 115.1774 | 47.77635 | 0.765916 | 2.559556 | 0.2203 |
| 14 | 118594600 | 118596399 | 14.66581 | 25.51167 | 51.64172 | 22.69073 | -0.6257 | 0.775663 | 0.220313 |
| 6 | 137228400 | 137230399 | 13.64822 | 23.50519 | 54.22889 | 23.7882 | -0.81619 | 0.87689 | 0.220315 |
| 4 | 37695400 | 37696999 | 31.22522 | 52.64553 | 36.45933 | 15.94128 | 0.909162 | 1.111854 | 0.22032 |
| 8 | 8502000 | 8503999 | 9.642631 | 17.4913 | 38.07831 | 16.45796 | -0.70165 | 0.327788 | 0.220326 |
| 4 | 143551200 | 143552199 | 38.09287 | 62.50242 | 54.21752 | 23.34645 | 0.58172 | 1.443555 | 0.22033 |
| 9 | 36243400 | 36245599 | 22.38212 | 39.56804 | 26.61262 | 11.77148 | 0.959645 | 0.558331 | 0.220332 |
| 12 | 103817200 | 103823799 | 69.18866 | 107.7058 | 190.8922 | 79.2391 | -0.51527 | 2.720489 | 0.220339 |
| 4 | 65509200 | 65510399 | 6.335607 | 12.00871 | 29.80174 | 12.04058 | -0.81267 | 0.014763 | 0.220339 |
| 5 | 90907000 | 90907799 | 6.335607 | 12.00871 | 29.80174 | 12.04058 | -0.81267 | 0.014763 | 0.220339 |
| 4 | 131112200 | 131116799 | 49.8328 | 79.15281 | 143.1833 | 59.70037 | -0.52117 | 2.278213 | 0.22034 |
| 3 | 86699000 | 86700599 | 8.742857 | 15.86088 | 36.50228 | 15.77463 | -0.78078 | 0.318202 | 0.220345 |
| 14 | 78935600 | 78936799 | 19.15388 | 33.75468 | 27.73562 | 12.27593 | 0.670716 | 0.573793 | 0.220372 |
| 4 | 148956400 | 148967799 | 67.26251 | 104.2257 | 201.6997 | 83.78042 | -0.64713 | 2.800702 | 0.220375 |
| 7 | 17342600 | 17348599 | 116.4893 | 180.5793 | 140.2585 | 58.2576 | 0.668926 | 2.806962 | 0.220378 |
| 8 | 110210600 | 110211999 | 10.17172 | 18.21819 | 38.92391 | 17.0601 | -0.68807 | 0.410684 | 0.220379 |
| 9 | 44248800 | 44251799 | 14.48706 | 25.15348 | 53.83598 | 23.62697 | -0.70622 | 0.802611 | 0.220395 |
| 1 | 187729800 | 187730599 | 4.695579 | 9.107714 | 26.13932 | 9.857684 | -0.9267 | -0.24435 | 0.2204 |
| 9 | 61943800 | 61944999 | 4.695579 | 9.107714 | 26.13932 | 9.857684 | -0.9267 | -0.24435 | 0.2204 |
| 11 | 97851400 | 97853799 | 13.94324 | 24.11324 | 52.64264 | 23.0953 | -0.73541 | 0.840894 | 0.220418 |

| | | | | | | | | | |
|---|---|---|---|---|---|---|---|---|---|
| 2 | 60871600 | 60872799 | 14.78221 | 27.64498 | 21.06343 | 8.724133 | 0.846877 | 0.15326 | 0.220444 |
| 14 | 111188000 | 111191199 | 34.10712 | 57.47887 | 41.41524 | 18.10574 | 0.852216 | 1.130489 | 0.220446 |
| 12 | 116448400 | 116452999 | 32.79498 | 52.76314 | 106.8189 | 45.34558 | -0.66217 | 1.848235 | 0.220475 |
| 11 | 58818800 | 58824999 | 26.81944 | 43.35162 | 98.37021 | 42.02309 | -0.81812 | 1.693936 | 0.220481 |
| 3 | 53030600 | 53033799 | 22.62892 | 37.47849 | 76.33654 | 33.03364 | -0.6425 | 1.308026 | 0.220484 |
| 12 | 103389400 | 103393399 | 24.3718 | 39.92391 | 80.14827 | 34.48863 | -0.62633 | 1.45087 | 0.220494 |
| 11 | 63687400 | 63688599 | 13.24713 | 25.17791 | 19.79241 | 7.950383 | 0.839104 | 0.032809 | 0.220506 |
| 9 | 118322600 | 118324999 | 19.45616 | 34.27228 | 28.62201 | 12.66937 | 0.647114 | 0.581502 | 0.220509 |
| 18 | 54370800 | 54371799 | 16.1731 | 29.82167 | 22.16696 | 9.378083 | 0.859945 | 0.250895 | 0.220511 |
| 3 | 19848800 | 19850799 | 21.43999 | 37.51612 | 30.67599 | 13.53274 | 0.67641 | 0.689592 | 0.220523 |
| 12 | 8818400 | 8819399 | 10.95901 | 21.06656 | 18.48779 | 7.218716 | 0.718442 | -0.08388 | 0.220527 |
| 2 | 155839800 | 155841399 | 7.740764 | 14.36807 | 33.16606 | 13.90556 | -0.754 | 0.171615 | 0.220528 |
| 8 | 34445800 | 34447399 | 7.740764 | 14.36807 | 33.16606 | 13.90556 | -0.754 | 0.171615 | 0.220528 |
| 15 | 57621200 | 57627599 | 40.64045 | 64.93226 | 126.0244 | 52.95265 | -0.61342 | 2.081048 | 0.220528 |
| 19 | 19893600 | 19894399 | 11.82643 | 22.53921 | 18.50413 | 7.390047 | 0.784594 | -0.00201 | 0.220536 |
| 16 | 12171600 | 12173999 | 42.05882 | 68.36784 | 59.65224 | 25.59577 | 0.565012 | 1.58317 | 0.220555 |
| 7 | 9463200 | 9463799 | 9.195324 | 17.89215 | 14.08645 | 5.191441 | 0.951681 | -0.30877 | 0.220576 |
| 2 | 5968000 | 5968599 | 3.932908 | 7.684906 | 23.24779 | 8.080333 | -0.89248 | -0.55399 | 0.220577 |
| X | 32481800 | 32484199 | 28.12117 | 49.10142 | 28.61024 | 12.60063 | 1.162911 | 0.755775 | 0.220599 |
| 4 | 26582600 | 26584199 | 22.05882 | 38.74578 | 29.52785 | 13.06809 | 0.777504 | 0.633916 | 0.220603 |
| X | 118248800 | 118249999 | 24.33628 | 42.59669 | 29.20909 | 12.88883 | 0.929458 | 0.70224 | 0.220651 |
| 6 | 83738800 | 83740999 | 11.48332 | 20.26505 | 42.47168 | 18.79298 | -0.67195 | 0.541638 | 0.22067 |
| 5 | 20813000 | 20813399 | 1.949275 | 3.804394 | 15.26553 | 4.802748 | -1.15057 | -0.96579 | 0.220674 |
| 1 | 103621800 | 103623399 | 18.2345 | 33.52275 | 21.07436 | 8.952029 | 1.095183 | 0.279042 | 0.220681 |
| 5 | 112755000 | 112767599 | 83.95811 | 128.4966 | 243.3813 | 100.9922 | -0.62996 | 3.047007 | 0.22069 |
| 1 | 33519200 | 33520399 | 5.987316 | 11.48855 | 30.2167 | 11.86012 | -0.85722 | -0.08058 | 0.2207 |
| 7 | 31898000 | 31902999 | 81.83663 | 129.5199 | 101.2178 | 42.07634 | 0.683144 | 2.392061 | 0.220701 |
| 7 | 62718000 | 62718999 | 11.33411 | 21.822 | 17.82231 | 6.927444 | 0.826804 | -0.09719 | 0.220705 |
| 16 | 90310600 | 90313199 | 12.27452 | 21.53978 | 45.11786 | 19.952 | -0.67369 | 0.620461 | 0.220709 |
| 2 | 23642000 | 23645599 | 52.34339 | 84.38587 | 70.16871 | 29.87359 | 0.622349 | 1.797114 | 0.220717 |
| 4 | 145384800 | 145388199 | 42.97358 | 70.6495 | 51.72759 | 22.29282 | 0.826656 | 1.428978 | 0.220725 |
| 17 | 86260800 | 86263399 | 20.07961 | 32.95916 | 78.95106 | 34.00244 | -0.87899 | 1.4158 | 0.220735 |
| 2 | 3109400 | 3112999 | 41.04552 | 67.27132 | 54.45838 | 23.439 | 0.680415 | 1.46193 | 0.220738 |
| 10 | 50452600 | 50457999 | 85.04883 | 134.3064 | 113.3797 | 47.07933 | 0.569454 | 2.448388 | 0.220754 |
| 9 | 70759400 | 70761599 | 13.4921 | 22.92233 | 62.08203 | 27.2073 | -1.05004 | 1.010836 | 0.220757 |
| 9 | 60458600 | 60460199 | 9.722809 | 17.45092 | 38.97147 | 17.0506 | -0.74937 | 0.39657 | 0.22076 |
| 6 | 137791400 | 137794599 | 17.78382 | 29.80661 | 65.95831 | 28.76623 | -0.76081 | 1.153943 | 0.220762 |
| 8 | 12193000 | 12194799 | 14.96534 | 25.99094 | 53.84534 | 23.6351 | -0.65939 | 0.801575 | 0.220777 |
| 4 | 20076400 | 20079599 | 44.0391 | 71.48205 | 62.54151 | 26.807 | 0.559121 | 1.617441 | 0.220784 |
| 5 | 74413600 | 74414799 | 4.711199 | 9.095491 | 26.95848 | 10.3906 | -1.00263 | -0.16597 | 0.220804 |
| 16 | 70464000 | 70464799 | 7.897947 | 14.61606 | 34.19493 | 14.39539 | -0.77779 | 0.191241 | 0.220808 |
| 4 | 141263600 | 141266399 | 14.50987 | 24.74378 | 58.97849 | 25.85497 | -0.86572 | 0.976487 | 0.220812 |
| 8 | 11035800 | 11037799 | 14.50987 | 24.74378 | 58.97849 | 25.85497 | -0.86572 | 0.976487 | 0.220812 |
| 1 | 103029400 | 103031599 | 32.19496 | 54.43458 | 40.86117 | 17.88129 | 0.793432 | 1.094788 | 0.220822 |
| 6 | 38653200 | 38659599 | 39.5663 | 63.13222 | 126.4997 | 53.05251 | -0.66138 | 2.122935 | 0.220823 |
| 7 | 129251000 | 129253999 | 18.49845 | 30.77557 | 70.19237 | 30.46926 | -0.8043 | 1.240956 | 0.220824 |
| 6 | 87787600 | 87789599 | 12.4107 | 21.41683 | 53.12205 | 23.3041 | -0.91927 | 0.853131 | 0.22083 |

| | | | | | | | | | |
|---|---|---|---|---|---|---|---|---|---|
| 9 | 9388400 | 9391399 | 56.64659 | 91.00384 | 70.27608 | 29.76943 | 0.722728 | 1.911814 | 0.220838 |
| 7 | 128569200 | 128570999 | 16.90932 | 31.06779 | 22.31848 | 9.488285 | 0.903324 | 0.27456 | 0.220844 |
| X | 120314000 | 120314799 | 15.0702 | 28.51005 | 19.17245 | 7.777446 | 1.050659 | 0.080436 | 0.220859 |
| 10 | 121200200 | 121201399 | 6.297015 | 11.68878 | 33.68292 | 14.12139 | -1.07101 | 0.160269 | 0.22086 |
| 16 | 93680400 | 93681599 | 7.091462 | 12.95852 | 35.5723 | 15.2341 | -1.02414 | 0.261687 | 0.220861 |
| 6 | 47754400 | 47755599 | 10.6187 | 18.5848 | 48.85019 | 21.55422 | -0.99918 | 0.662364 | 0.220886 |
| 8 | 37049600 | 37052399 | 33.64001 | 53.60542 | 140.016 | 58.61389 | -1.04509 | 2.164236 | 0.22089 |
| 3 | 37400800 | 37403799 | 13.89613 | 24.08922 | 51.62031 | 22.64843 | -0.70814 | 0.814929 | 0.220904 |
| 3 | 156436600 | 156437799 | 13.0949 | 22.4154 | 60.57632 | 26.56351 | -1.04503 | 0.938611 | 0.220915 |
| 11 | 101969200 | 101976999 | 40.16153 | 63.79195 | 142.1151 | 59.25559 | -0.82098 | 2.270803 | 0.220919 |
| 7 | 31946600 | 31950999 | 61.59908 | 99.22231 | 64.64983 | 27.49133 | 0.971922 | 1.835155 | 0.22092 |
| 12 | 115458800 | 115459999 | 31.37825 | 52.67778 | 44.82105 | 19.5679 | 0.613181 | 1.158934 | 0.220921 |
| 5 | 146749000 | 146750999 | 31.03344 | 52.07987 | 44.83562 | 19.56975 | 0.596309 | 1.161458 | 0.220938 |
| 3 | 90602200 | 90603199 | 12.14604 | 23.12923 | 19.30991 | 7.725331 | 0.757677 | 0.007031 | 0.220955 |
| X | 57889200 | 57891999 | 48.35576 | 78.22927 | 61.22358 | 26.17813 | 0.715263 | 1.693615 | 0.220956 |
| 9 | 5055200 | 5059199 | 42.58092 | 69.761 | 54.50516 | 23.45533 | 0.731267 | 1.470869 | 0.220982 |
| 16 | 34591600 | 34592599 | 17.8328 | 31.66997 | 26.13434 | 11.53669 | 0.669111 | 0.486933 | 0.220987 |
| 7 | 141487000 | 141489399 | 13.15262 | 22.74953 | 52.10954 | 22.86153 | -0.80439 | 0.833318 | 0.221016 |
| 10 | 56152000 | 56154199 | 16.97667 | 28.98993 | 58.04772 | 25.44973 | -0.61518 | 0.972701 | 0.221017 |
| 4 | 3574600 | 3575199 | 5.544058 | 10.67251 | 27.45291 | 10.67564 | -0.81245 | -0.12409 | 0.22103 |
| X | 5029400 | 5034599 | 176.4992 | 266.5825 | 195.3579 | 80.76273 | 0.723267 | 3.335394 | 0.221034 |
| 10 | 20664400 | 20665199 | 11.73913 | 22.72247 | 16.7462 | 6.376843 | 1.001303 | -0.16175 | 0.221048 |
| 12 | 26991200 | 26991799 | 10.83462 | 21.10568 | 14.79677 | 5.39351 | 1.130594 | -0.32349 | 0.221054 |
| 4 | 126048200 | 126059599 | 66.1523 | 102.8451 | 198.0364 | 82.22065 | -0.63622 | 2.741676 | 0.221063 |
| 6 | 143303400 | 143304399 | 12.58871 | 23.94963 | 19.87116 | 7.965721 | 0.764413 | 0.0202 | 0.221064 |
| 3 | 138330000 | 138331599 | 14.57505 | 26.83011 | 22.14364 | 9.389559 | 0.707132 | 0.244816 | 0.221084 |
| 5 | 12711000 | 12712399 | 14.38985 | 26.61486 | 22.27526 | 9.384735 | 0.694004 | 0.218892 | 0.221085 |
| 3 | 13284400 | 13286599 | 20.7545 | 37.18126 | 23.91684 | 10.48054 | 1.033222 | 0.448526 | 0.22109 |
| 6 | 3057600 | 3060599 | 78.78265 | 124.6471 | 110.0173 | 45.72699 | 0.507463 | 2.399324 | 0.221098 |
| 9 | 85756800 | 85757999 | 5.119228 | 9.902855 | 26.56647 | 10.14506 | -0.84787 | -0.19493 | 0.221106 |
| 13 | 113247200 | 113250799 | 43.46407 | 71.81308 | 46.68008 | 20.16666 | 0.999955 | 1.368092 | 0.221108 |
| 8 | 72210800 | 72213399 | 12.30501 | 21.57631 | 45.67029 | 20.18287 | -0.68856 | 0.636314 | 0.221116 |
| 7 | 42702800 | 42709199 | 147.8828 | 224.0679 | 194.3124 | 80.41068 | 0.484408 | 3.264867 | 0.221117 |
| 5 | 114028600 | 114031199 | 14.35982 | 24.92734 | 51.09912 | 22.42289 | -0.64405 | 0.803582 | 0.221124 |
| 8 | 11180000 | 11184199 | 24.08068 | 39.36845 | 83.47851 | 35.894 | -0.70682 | 1.478481 | 0.221127 |
| 17 | 22944600 | 22956599 | 272.8485 | 396.7912 | 316.0602 | 130.2753 | 0.552677 | 3.99947 | 0.221131 |
| 12 | 55478600 | 55480999 | 13.76627 | 23.79723 | 52.66451 | 23.1046 | -0.75527 | 0.841239 | 0.221147 |
| 17 | 13105200 | 13111199 | 32.85585 | 52.72336 | 112.4963 | 47.57205 | -0.74241 | 1.933496 | 0.22117 |
| 18 | 42146200 | 42150199 | 21.59662 | 35.59212 | 78.09902 | 33.69279 | -0.75073 | 1.365755 | 0.221178 |
| 4 | 112961400 | 112962399 | 25.91624 | 44.45762 | 36.86803 | 16.1699 | 0.652538 | 0.941597 | 0.22119 |
| 7 | 43318400 | 43328599 | 111.2509 | 167.6777 | 295.6366 | 122.1398 | -0.54501 | 3.368458 | 0.221195 |
| 14 | 92664000 | 92665199 | 15.17743 | 29.11637 | 14.95479 | 5.87343 | 1.474926 | -0.03228 | 0.221196 |
| 2 | 118001000 | 118001399 | 11.46858 | 22.14811 | 17.91874 | 6.896585 | 0.853964 | -0.12583 | 0.221207 |
| 18 | 10229600 | 10231799 | 16.42692 | 30.50083 | 21.4975 | 9.009298 | 0.945566 | 0.212869 | 0.22121 |
| 1 | 183315800 | 183317199 | 8.658595 | 15.99292 | 35.25506 | 14.8806 | -0.6955 | 0.211077 | 0.221214 |
| 19 | 41074200 | 41078599 | 30.21837 | 48.6795 | 103.0166 | 43.81206 | -0.72357 | 1.803371 | 0.221221 |
| 9 | 77949800 | 77951199 | 14.16903 | 24.64025 | 52.18836 | 22.92667 | -0.69086 | 0.776649 | 0.221246 |

| | | | | | | | | | |
|---|---|---|---|---|---|---|---|---|---|
| 2 | 135724600 | 135726199 | 10.03114 | 17.90003 | 40.44761 | 17.78333 | -0.77173 | 0.434719 | 0.221247 |
| 1 | 75416600 | 75436199 | 101.5237 | 151.6476 | 351.2435 | 144.805 | -0.94996 | 3.541243 | 0.221254 |
| 9 | 99733800 | 99734399 | 4.649239 | 9.060932 | 24.42565 | 8.848336 | -0.7831 | -0.39252 | 0.221262 |
| 5 | 48139600 | 48143999 | 26.09717 | 42.03431 | 104.7018 | 44.51914 | -0.95828 | 1.80348 | 0.221283 |
| 12 | 63809600 | 63810399 | 4.943384 | 9.622933 | 26.10286 | 9.586481 | -0.81076 | -0.33548 | 0.221305 |
| 18 | 73490600 | 73493799 | 14.18721 | 24.59454 | 54.90701 | 24.09049 | -0.76723 | 0.818246 | 0.221307 |
| 5 | 42000000 | 42002799 | 12.89642 | 22.19868 | 55.80877 | 24.48084 | -0.93973 | 0.879831 | 0.221317 |
| 6 | 133834000 | 133836399 | 16.50593 | 30.8198 | 18.47447 | 7.672358 | 1.186403 | 0.170606 | 0.221332 |
| 6 | 82611400 | 82613199 | 18.88813 | 33.9078 | 25.33013 | 11.0805 | 0.82069 | 0.426907 | 0.221359 |
| 18 | 60174000 | 60175599 | 18.88813 | 33.9078 | 25.33013 | 11.0805 | 0.82069 | 0.426907 | 0.221359 |
| 1 | 189060600 | 189064199 | 22.61201 | 37.55317 | 74.10597 | 32.12907 | -0.59662 | 1.275459 | 0.22137 |
| X | 30991400 | 30994399 | 47.10498 | 78.2675 | 38.17486 | 16.55638 | 1.413291 | 1.312294 | 0.221381 |
| 11 | 53597400 | 53598399 | 5.629582 | 10.68104 | 30.35988 | 12.24046 | -1.00408 | 0.001238 | 0.221382 |
| 2 | 72515000 | 72518799 | 48.34556 | 78.95342 | 53.50176 | 22.99565 | 0.933325 | 1.527798 | 0.221389 |
| 15 | 20059400 | 20060599 | 16.32092 | 29.48333 | 24.59174 | 10.68396 | 0.668919 | 0.370025 | 0.221389 |
| 4 | 41754400 | 41761399 | 32.26526 | 51.68943 | 119.3664 | 50.34949 | -0.85916 | 1.987071 | 0.221389 |
| X | 3979000 | 3984599 | 57.06111 | 92.273 | 59.82729 | 25.57139 | 0.985435 | 1.716583 | 0.221406 |
| 1 | 185805800 | 185808199 | 52.06197 | 83.99078 | 64.31657 | 27.40676 | 0.74209 | 1.775084 | 0.22142 |
| 7 | 29445200 | 29446399 | 11.86168 | 22.59688 | 19.37848 | 7.746131 | 0.720493 | 0.002037 | 0.221439 |
| 9 | 42609800 | 42610599 | 11.86168 | 22.59688 | 19.37848 | 7.746131 | 0.720493 | 0.002037 | 0.221439 |
| 9 | 20334000 | 20338199 | 82.71013 | 130.2822 | 116.1412 | 48.19085 | 0.488113 | 2.500508 | 0.221442 |
| 5 | 76622800 | 76623799 | 7.118257 | 13.31921 | 31.70358 | 13.11944 | -0.7825 | 0.112782 | 0.221465 |
| 7 | 22423000 | 22423399 | 9.246155 | 18.00058 | 15.43163 | 5.662942 | 0.834165 | -0.3204 | 0.221466 |
| 8 | 40835600 | 40836999 | 16.53715 | 29.98259 | 24.57574 | 10.62864 | 0.698168 | 0.351527 | 0.221468 |
| 12 | 24712400 | 24715599 | 13.40488 | 23.29255 | 52.70927 | 23.14375 | -0.78598 | 0.785067 | 0.221472 |
| 7 | 14254800 | 14259599 | 98.85414 | 155.5484 | 110.7662 | 45.95433 | 0.810604 | 2.530193 | 0.221484 |
| X | 54585400 | 54588399 | 57.1427 | 92.08672 | 67.83908 | 28.86689 | 0.796088 | 1.819066 | 0.221488 |
| 6 | 73251600 | 73252799 | 6.970498 | 13.04648 | 31.70892 | 13.11529 | -0.81183 | 0.109171 | 0.221498 |
| 7 | 7706000 | 7707799 | 17.19153 | 32.52003 | 17.01788 | 6.904861 | 1.408185 | 0.093908 | 0.221502 |
| 5 | 63874600 | 63876399 | 9.974356 | 18.03922 | 43.47122 | 18.85897 | -0.85119 | 0.357447 | 0.221515 |
| 6 | 116346800 | 116348999 | 12.48611 | 21.75633 | 48.39205 | 21.2852 | -0.7603 | 0.738099 | 0.221542 |
| 14 | 34031400 | 34033599 | 13.72205 | 23.86094 | 50.01387 | 21.97024 | -0.67569 | 0.774341 | 0.221543 |
| 2 | 104541800 | 104542999 | 5.790596 | 10.91086 | 30.72697 | 12.56594 | -1.00858 | 0.053207 | 0.221549 |
| 3 | 46026200 | 46026999 | 5.675463 | 10.99551 | 28.62124 | 10.85262 | -0.79579 | -0.208 | 0.221566 |
| X | 147799400 | 147801399 | 25.4448 | 44.96788 | 21.75102 | 9.621684 | 1.431443 | 0.570431 | 0.221583 |
| 17 | 83005000 | 83006799 | 19.50833 | 34.5775 | 22.13034 | 9.783758 | 1.031972 | 0.502747 | 0.221598 |
| 17 | 27977800 | 27980599 | 11.58677 | 20.14725 | 49.9697 | 21.95036 | -0.91693 | 0.765586 | 0.221615 |
| 1 | 131922600 | 131923799 | 13.064 | 24.52298 | 18.94664 | 7.797974 | 0.834618 | 0.110614 | 0.221618 |
| 8 | 69214600 | 69215199 | 11.56958 | 22.07422 | 17.00609 | 6.775822 | 0.878354 | -0.01704 | 0.22163 |
| 1 | 131838000 | 131839199 | 8.953145 | 16.06854 | 38.45224 | 16.82427 | -0.84865 | 0.387978 | 0.22164 |
| 10 | 8144800 | 8147999 | 19.80565 | 33.34047 | 66.23638 | 28.94722 | -0.60581 | 1.118806 | 0.22165 |
| 1 | 8222400 | 8230199 | 39.27931 | 64.56595 | 47.25008 | 20.36182 | 0.827258 | 1.418553 | 0.221655 |
| 3 | 102300200 | 102301199 | 7.981913 | 14.53134 | 35.52639 | 15.28563 | -0.86267 | 0.291822 | 0.221673 |
| 15 | 84969200 | 84969999 | 7.981913 | 14.53134 | 35.52639 | 15.28563 | -0.86267 | 0.291822 | 0.221673 |
| X | 27112000 | 27112399 | 8.435244 | 16.48702 | 12.73173 | 4.393877 | 1.061178 | -0.55059 | 0.221685 |
| 10 | 15326000 | 15327199 | 12.49314 | 23.83122 | 19.28699 | 7.688103 | 0.806751 | -0.00097 | 0.221696 |
| 2 | 152179200 | 152183999 | 24.9842 | 40.5722 | 88.93202 | 38.13193 | -0.76174 | 1.587727 | 0.221702 |

| | | | | | | | | | |
|---|---|---|---|---|---|---|---|---|---|
| 6 | 3579000 | 3581599 | 19.60694 | 35.47106 | 23.01855 | 9.982281 | 1.030267 | 0.382207 | 0.221735 |
| 2 | 132219400 | 132221799 | 13.91407 | 24.07342 | 52.1391 | 22.8747 | -0.72402 | 0.833599 | 0.221735 |
| 5 | 10701800 | 10704199 | 32.95405 | 55.97322 | 38.6017 | 16.91594 | 0.91544 | 1.057979 | 0.221741 |
| 14 | 68360200 | 68363399 | 25.24294 | 40.78849 | 97.657 | 41.69885 | -0.89586 | 1.702755 | 0.22176 |
| 13 | 86669600 | 86670399 | 10.00148 | 19.38241 | 16.98566 | 6.431659 | 0.761658 | -0.19938 | 0.22176 |
| 11 | 121728400 | 121729799 | 9.223179 | 16.26038 | 43.05375 | 19.05431 | -1.00744 | 0.539917 | 0.221771 |
| 4 | 90138000 | 90138799 | 13.20903 | 24.77562 | 20.16838 | 8.31194 | 0.757605 | 0.118052 | 0.221786 |
| 9 | 43201000 | 43202599 | 7.729217 | 14.41064 | 32.71871 | 13.62197 | -0.72218 | 0.143903 | 0.221787 |
| 1 | 72573800 | 72576599 | 19.37788 | 32.00762 | 75.54523 | 32.63041 | -0.85451 | 1.331786 | 0.221794 |
| 1 | 21582400 | 21583599 | 12.51109 | 23.54781 | 19.86238 | 8.138035 | 0.713764 | 0.090368 | 0.221796 |
| 17 | 19908600 | 19912999 | 64.69599 | 104.001 | 69.1932 | 29.33753 | 0.938395 | 1.904079 | 0.221803 |
| 17 | 46980000 | 46986199 | 27.13594 | 43.8414 | 96.20535 | 41.07191 | -0.7707 | 1.709614 | 0.221819 |
| 10 | 57306400 | 57307999 | 20.17072 | 35.96191 | 25.00039 | 10.9999 | 0.917917 | 0.475121 | 0.221821 |
| 1 | 25442200 | 25443399 | 10.31177 | 18.58514 | 40.60808 | 17.69314 | -0.71452 | 0.376243 | 0.221839 |
| 6 | 47812400 | 47813599 | 6.611878 | 12.20493 | 34.66378 | 14.6434 | -1.05851 | 0.196859 | 0.221842 |
| 9 | 105807600 | 105809399 | 11.13135 | 19.75288 | 41.92904 | 18.52149 | -0.68694 | 0.480251 | 0.221854 |
| 9 | 43935800 | 43936599 | 14.78793 | 27.11091 | 20.82964 | 8.88166 | 0.804247 | 0.266816 | 0.221866 |
| 18 | 81475800 | 81476799 | 14.86799 | 27.4705 | 22.69191 | 9.573871 | 0.711264 | 0.229283 | 0.221867 |
| 4 | 125290000 | 125301599 | 76.73748 | 117.3697 | 242.3982 | 100.5693 | -0.75537 | 3.058611 | 0.221894 |
| 4 | 111908200 | 111909999 | 25.19368 | 43.92868 | 32.09749 | 14.12414 | 0.8379 | 0.764194 | 0.2219 |
| 17 | 67106400 | 67106999 | 15.7307 | 29.81236 | 17.69166 | 7.149922 | 1.233093 | 0.069213 | 0.221912 |
| X | 31691000 | 31692399 | 47.07001 | 76.73641 | 55.50148 | 23.84 | 0.837911 | 1.553189 | 0.221928 |
| 4 | 155753000 | 155753999 | 6.016555 | 11.45946 | 29.34747 | 11.73594 | -0.84389 | -0.02144 | 0.221931 |
| 15 | 74670400 | 74672799 | 25.96627 | 42.07963 | 90.89841 | 38.91552 | -0.74354 | 1.633779 | 0.221936 |
| 18 | 76931400 | 76935799 | 33.36743 | 53.46922 | 115.9706 | 48.93674 | -0.76841 | 1.979745 | 0.221944 |
| 5 | 136868800 | 136870999 | 22.38223 | 36.89594 | 75.84749 | 32.72486 | -0.65673 | 1.364129 | 0.221953 |
| 13 | 61912200 | 61914599 | 55.37289 | 88.88864 | 78.65814 | 33.28118 | 0.52534 | 1.937052 | 0.221961 |
| 4 | 22812600 | 22814399 | 15.98591 | 29.9875 | 19.87746 | 8.190468 | 1.052189 | 0.142396 | 0.221965 |
| 9 | 80115800 | 80118599 | 14.35716 | 24.9826 | 50.55156 | 22.2161 | -0.62501 | 0.76881 | 0.221969 |
| 12 | 22274000 | 22275599 | 34.87491 | 57.64132 | 48.78126 | 21.07936 | 0.620562 | 1.343096 | 0.221977 |
| 13 | 55875000 | 55875999 | 5.611177 | 10.69544 | 29.94798 | 11.95694 | -0.97024 | -0.0329 | 0.221992 |
| 9 | 122564800 | 122569599 | 39.30981 | 63.04416 | 115.2653 | 48.70093 | -0.52113 | 1.960551 | 0.222005 |
| 8 | 121462600 | 121469399 | 44.75815 | 70.86517 | 158.8825 | 66.06082 | -0.83462 | 2.368166 | 0.22201 |
| 3 | 90467400 | 90468199 | 9.821792 | 19.06102 | 16.44313 | 6.185387 | 0.792871 | -0.22521 | 0.222024 |
| 2 | 43395400 | 43397799 | 31.06519 | 52.82872 | 36.62405 | 16.05164 | 0.908101 | 1.037925 | 0.22203 |
| 11 | 74213400 | 74214599 | 15.19662 | 27.928 | 23.17154 | 9.847589 | 0.697501 | 0.262366 | 0.222038 |
| 9 | 4980400 | 4981999 | 16.7303 | 31.05917 | 20.75968 | 8.701983 | 1.020907 | 0.213335 | 0.222048 |
| X | 135382800 | 135384199 | 16.62884 | 32.14611 | 11.45935 | 4.389687 | 2.027493 | -0.10309 | 0.222062 |
| 1 | 148578000 | 148578599 | 9.289793 | 18.14329 | 12.5065 | 4.387584 | 1.201154 | -0.4841 | 0.222066 |
| 11 | 61298800 | 61299599 | 5.697868 | 10.92418 | 28.4254 | 11.18335 | -0.84472 | -0.08067 | 0.222071 |
| 10 | 115915400 | 115917199 | 14.95047 | 27.57541 | 21.46031 | 9.075469 | 0.793775 | 0.234578 | 0.222072 |
| 6 | 143212000 | 143216399 | 33.39696 | 53.80911 | 108.6205 | 46.20672 | -0.65546 | 1.803302 | 0.222081 |
| 5 | 45968600 | 45969199 | 9.989815 | 19.52227 | 11.22145 | 3.889148 | 1.474606 | -0.51823 | 0.222089 |
| 2 | 7841200 | 7841599 | 7.53558 | 14.7261 | 13.49788 | 4.678658 | 0.812339 | -0.54619 | 0.222098 |
| 5 | 6611400 | 6618799 | 65.5021 | 104.8523 | 76.57988 | 32.26341 | 0.796958 | 2.034978 | 0.222111 |
| 8 | 56158000 | 56158999 | 13.38432 | 25.65911 | 18.16785 | 7.149214 | 1.014907 | -0.03522 | 0.222122 |
| 17 | 87410600 | 87412399 | 12.64705 | 21.99199 | 49.99458 | 21.96199 | -0.79212 | 0.769006 | 0.222125 |

| | | | | | | | | | |
|---|---|---|---|---|---|---|---|---|---|
| Y | 1951800 | 1955399 | 29.31828 | 50.25518 | 19.23883 | 8.437416 | 1.761095 | 0.932738 | 0.222149 |
| 7 | 42560800 | 42564199 | 104.1086 | 161.4241 | 142.5509 | 59.20776 | 0.484214 | 2.798114 | 0.222157 |
| 1 | 14405800 | 14407599 | 32.30477 | 55.32977 | 35.34393 | 15.49949 | 1.027175 | 0.980312 | 0.222164 |
| 1 | 71122800 | 71125799 | 13.21298 | 23.1001 | 48.36325 | 21.31971 | -0.67328 | 0.690381 | 0.222165 |
| 11 | 107212200 | 107214599 | 24.514 | 40.22457 | 79.14508 | 34.08123 | -0.59643 | 1.429892 | 0.222167 |
| X | 58883600 | 58885999 | 35.53605 | 60.62119 | 34.16901 | 14.97942 | 1.206167 | 1.027177 | 0.22217 |
| 5 | 125703400 | 125726799 | 253.7711 | 358.6543 | 603.4922 | 249.8142 | -0.56682 | 4.420235 | 0.222203 |
| 3 | 120617000 | 120620599 | 41.46078 | 66.32773 | 121.0556 | 50.95714 | -0.52282 | 2.042794 | 0.222257 |
| 16 | 55634200 | 55636799 | 22.51758 | 37.16906 | 76.04358 | 32.83067 | -0.64923 | 1.347293 | 0.22226 |
| 6 | 52146800 | 52149999 | 14.01783 | 23.9807 | 56.2825 | 24.67941 | -0.84254 | 0.943782 | 0.222264 |
| 6 | 88495200 | 88496599 | 7.374099 | 13.49815 | 35.5929 | 15.2086 | -0.96454 | 0.25319 | 0.222271 |
| 14 | 81919600 | 81921799 | 24.96043 | 43.49437 | 32.76407 | 14.41172 | 0.794152 | 0.773642 | 0.222278 |
| 17 | 58266600 | 58268399 | 26.19568 | 45.03229 | 37.74349 | 16.55575 | 0.637726 | 0.9242 | 0.222295 |
| 18 | 38273200 | 38282199 | 52.34033 | 82.45158 | 164.1293 | 68.10347 | -0.66917 | 2.488938 | 0.222321 |
| 8 | 93154400 | 93156399 | 13.88453 | 23.71186 | 57.87594 | 25.3746 | -0.89934 | 0.959963 | 0.222322 |
| 1 | 3244200 | 3247999 | 56.04449 | 90.29215 | 70.41933 | 29.95292 | 0.713486 | 1.823709 | 0.22233 |
| 11 | 98436200 | 98438799 | 17.66023 | 30.09325 | 60.75359 | 26.6315 | -0.62747 | 0.999088 | 0.22233 |
| 13 | 77920000 | 77922999 | 30.40729 | 51.7679 | 38.07351 | 16.68842 | 0.823294 | 1.025145 | 0.222332 |
| 4 | 114407600 | 114408399 | 4.82347 | 9.355561 | 25.54659 | 9.635193 | -0.85549 | -0.24323 | 0.222339 |
| 6 | 96093600 | 96094399 | 4.82347 | 9.355561 | 25.54659 | 9.635193 | -0.85549 | -0.24323 | 0.222339 |
| 19 | 20269200 | 20271199 | 19.88857 | 35.52004 | 25.7517 | 11.31402 | 0.859443 | 0.465912 | 0.222351 |
| 1 | 150923400 | 150924199 | 7.52193 | 13.73594 | 35.04489 | 15.02079 | -0.91994 | 0.269295 | 0.222367 |
| 7 | 8010400 | 8012799 | 17.30491 | 32.09402 | 20.74035 | 8.705641 | 1.067938 | 0.223826 | 0.222373 |
| 9 | 116747200 | 116748999 | 8.281721 | 15.0698 | 36.02759 | 15.51073 | -0.83164 | 0.296029 | 0.22239 |
| 4 | 61133800 | 61137199 | 20.28629 | 34.04399 | 74.30078 | 32.43052 | -0.74131 | 1.154307 | 0.222401 |
| 3 | 85463600 | 85464799 | 13.86528 | 25.8477 | 21.76793 | 9.064698 | 0.697527 | 0.167464 | 0.222417 |
| 14 | 113413800 | 113415599 | 19.43522 | 34.93273 | 24.78081 | 10.82901 | 0.895878 | 0.422541 | 0.222424 |
| X | 21477600 | 21478799 | 19.43522 | 34.93273 | 24.78081 | 10.82901 | 0.895878 | 0.422541 | 0.222424 |
| 7 | 27958800 | 27960599 | 11.52462 | 20.23048 | 44.61805 | 19.73645 | -0.74731 | 0.61276 | 0.222425 |
| 18 | 12466600 | 12468999 | 15.05003 | 25.86427 | 54.79749 | 24.03607 | -0.6946 | 0.902832 | 0.222443 |
| 7 | 31702800 | 31707199 | 51.72845 | 84.27685 | 52.99042 | 22.75608 | 1.038657 | 1.568656 | 0.222445 |
| 9 | 19252800 | 19253999 | 16.80113 | 30.40541 | 23.26488 | 10.0848 | 0.794982 | 0.362233 | 0.222446 |
| 5 | 62402800 | 62404199 | 17.85245 | 32.84173 | 21.08809 | 8.949941 | 1.065669 | 0.272248 | 0.222447 |
| 11 | 11914800 | 11917799 | 13.8804 | 24.10277 | 51.09418 | 22.42522 | -0.69219 | 0.795998 | 0.22245 |
| 5 | 111297400 | 111299599 | 10.53945 | 18.6535 | 41.4335 | 18.32296 | -0.75359 | 0.500838 | 0.222468 |
| 9 | 60952400 | 60953199 | 3.71696 | 7.220192 | 24.95879 | 9.324862 | -1.17742 | -0.29085 | 0.222472 |
| 4 | 122009000 | 122009599 | 11.21903 | 21.64235 | 17.87659 | 6.909979 | 0.818201 | -0.1155 | 0.222476 |
| 1 | 133985400 | 133986399 | 6.917424 | 13.12481 | 30.42183 | 12.26478 | -0.7126 | 0.014619 | 0.2225 |
| 17 | 50388800 | 50389999 | 6.917424 | 13.12481 | 30.42183 | 12.26478 | -0.7126 | 0.014619 | 0.2225 |
| 18 | 49911800 | 49913199 | 8.670711 | 15.51026 | 38.46998 | 16.87903 | -0.90238 | 0.409351 | 0.2225 |
| 9 | 79700000 | 79700999 | 5.982967 | 11.53161 | 28.17221 | 10.91669 | -0.73497 | -0.13171 | 0.222502 |
| 3 | 76450400 | 76452199 | 17.87224 | 32.79407 | 21.62728 | 9.210805 | 1.023949 | 0.287551 | 0.222506 |
| 10 | 21092400 | 21093399 | 14.97647 | 27.66712 | 22.62585 | 9.547787 | 0.725347 | 0.230261 | 0.222511 |
| 2 | 89399200 | 89400599 | 19.97068 | 36.19122 | 22.46797 | 9.722609 | 1.095731 | 0.37437 | 0.222523 |
| 2 | 30091200 | 30094199 | 17.33835 | 29.45011 | 59.59705 | 26.11661 | -0.63158 | 1.025032 | 0.222534 |
| 6 | 99145200 | 99146999 | 11.30694 | 19.97474 | 42.97596 | 19.01213 | -0.70916 | 0.528172 | 0.222579 |
| 7 | 114013800 | 114015999 | 13.17867 | 22.72831 | 53.14752 | 23.31484 | -0.83494 | 0.861675 | 0.222596 |

| | | | | | | | | | |
|---|---|---|---|---|---|---|---|---|---|
| 4 | 10978800 | 10983999 | 33.54408 | 54.13115 | 103.0219 | 43.9157 | -0.56799 | 1.755823 | 0.222601 |
| X | 112145400 | 112146999 | 19.65484 | 36.24373 | 19.33243 | 8.178257 | 1.334598 | 0.270575 | 0.222612 |
| 19 | 26043000 | 26044199 | 7.995791 | 14.52565 | 36.0167 | 15.53789 | -0.88561 | 0.304243 | 0.222613 |
| 14 | 30723000 | 30724599 | 8.522142 | 15.24857 | 38.47181 | 16.87604 | -0.92663 | 0.406247 | 0.222631 |
| 16 | 34654800 | 34659199 | 24.0082 | 39.44579 | 78.72317 | 33.91817 | -0.61614 | 1.410991 | 0.222633 |
| 1 | 37197000 | 37204799 | 99.56677 | 155.2148 | 136.0932 | 56.48016 | 0.500489 | 2.703216 | 0.222634 |
| 17 | 34848800 | 34852799 | 19.59091 | 32.24083 | 78.6947 | 33.92939 | -0.90455 | 1.379986 | 0.222637 |
| 9 | 60776600 | 60780199 | 19.8119 | 32.78431 | 72.73078 | 31.45437 | -0.7651 | 1.311694 | 0.222644 |
| 17 | 40073200 | 40074999 | 27.67068 | 47.94595 | 33.10527 | 14.52475 | 0.918129 | 0.85407 | 0.222683 |
| 6 | 87557400 | 87564399 | 33.96781 | 54.52817 | 110.5173 | 46.7619 | -0.66777 | 1.922786 | 0.222697 |
| 19 | 54407400 | 54408599 | 14.50896 | 27.07942 | 21.66542 | 9.0041 | 0.773175 | 0.165258 | 0.222713 |
| 8 | 3877400 | 3883399 | 123.1833 | 189.2287 | 163.4604 | 67.87072 | 0.505766 | 2.979836 | 0.222728 |
| 5 | 124867800 | 124893399 | 225.6386 | 320.5269 | 567.4328 | 234.6883 | -0.63263 | 4.344 | 0.222739 |
| 17 | 49314000 | 49317599 | 26.5919 | 43.11944 | 96.67147 | 41.40545 | -0.79632 | 1.624863 | 0.222747 |
| 3 | 96371400 | 96371799 | 2.936718 | 5.741289 | 19.41317 | 6.658976 | -1.02893 | -0.62543 | 0.222766 |
| 3 | 49687400 | 49689599 | 23.71949 | 41.22592 | 34.53109 | 15.16445 | 0.641476 | 0.805011 | 0.222775 |
| X | 153797800 | 153800999 | 61.68929 | 99.12244 | 73.73704 | 31.24346 | 0.776909 | 1.916929 | 0.222827 |
| 1 | 163607000 | 163607599 | 4.974965 | 9.585508 | 26.87961 | 10.42603 | -0.93184 | -0.14045 | 0.222839 |
| 18 | 3037600 | 3048199 | 263.4447 | 386.0126 | 263.9475 | 108.723 | 0.78321 | 3.883403 | 0.222852 |
| 8 | 122424000 | 122437399 | 93.75316 | 141.8411 | 280.3523 | 115.9614 | -0.70498 | 3.288159 | 0.222852 |
| 18 | 28299200 | 28301199 | 24.85613 | 43.23014 | 30.73099 | 13.50126 | 0.876984 | 0.794626 | 0.222866 |
| 3 | 10544400 | 10548599 | 50.68945 | 81.92601 | 66.73284 | 28.50346 | 0.656193 | 1.722686 | 0.222878 |
| 18 | 75078000 | 75079999 | 12.7118 | 21.95047 | 53.68701 | 23.5524 | -0.89912 | 0.847805 | 0.222891 |
| 6 | 32025800 | 32027399 | 9.565665 | 17.11375 | 39.96255 | 17.53167 | -0.81655 | 0.41415 | 0.222896 |
| 12 | 32750600 | 32751999 | 14.98337 | 27.76113 | 21.46104 | 9.020202 | 0.80967 | 0.210405 | 0.222901 |
| 14 | 26482600 | 26487199 | 95.27701 | 148.8172 | 130.1609 | 54.00624 | 0.506219 | 2.665433 | 0.222909 |
| 14 | 119009800 | 119011799 | 27.5838 | 46.96178 | 39.49341 | 17.31082 | 0.63089 | 1.019028 | 0.222917 |
| 12 | 114540400 | 114544599 | 64.73884 | 104.0991 | 67.04207 | 28.43706 | 0.985671 | 1.893624 | 0.222949 |
| 2 | 147200600 | 147212599 | 93.81867 | 142.2964 | 289.9703 | 120.0337 | -0.74595 | 3.237328 | 0.222954 |
| 1 | 188116800 | 188118399 | 14.71593 | 25.4799 | 52.68425 | 23.11439 | -0.65716 | 0.832781 | 0.222958 |
| 18 | 37385000 | 37389999 | 79.10607 | 125.5988 | 99.94862 | 41.6568 | 0.661159 | 2.303467 | 0.222962 |
| 8 | 87617800 | 87619599 | 11.32515 | 20.02666 | 41.94839 | 18.55388 | -0.6699 | 0.517724 | 0.222987 |
| 13 | 70576800 | 70578399 | 17.43256 | 31.69816 | 23.99253 | 10.3387 | 0.815791 | 0.341096 | 0.222993 |
| 16 | 9921400 | 9923599 | 19.39064 | 34.59963 | 25.66416 | 11.28412 | 0.825714 | 0.466477 | 0.223006 |
| 3 | 64609400 | 64611999 | 23.23631 | 40.93456 | 27.00224 | 11.95219 | 0.985539 | 0.600165 | 0.223008 |
| 10 | 12230400 | 12231599 | 11.84374 | 20.74825 | 45.6989 | 20.1797 | -0.7455 | 0.649442 | 0.223016 |
| 10 | 41097000 | 41100999 | 27.86471 | 45.19046 | 89.76979 | 38.45387 | -0.62198 | 1.620817 | 0.223024 |
| 17 | 57168200 | 57184799 | 156.9353 | 230.668 | 401.1501 | 165.2061 | -0.55851 | 3.82698 | 0.223035 |
| 6 | 137541200 | 137545399 | 26.44833 | 43.35014 | 84.74483 | 36.47534 | -0.58792 | 1.451 | 0.223035 |
| 6 | 41581400 | 41584199 | 18.93825 | 31.57769 | 68.08311 | 29.59614 | -0.7234 | 1.216208 | 0.223038 |
| 10 | 59940000 | 59947799 | 42.77565 | 67.84744 | 149.3184 | 62.17031 | -0.80552 | 2.315393 | 0.223042 |
| 5 | 79653800 | 79656999 | 16.11932 | 27.71653 | 60.39679 | 26.49254 | -0.7352 | 0.906922 | 0.223044 |
| 3 | 132285200 | 132289199 | 45.49035 | 73.70248 | 64.25912 | 27.50603 | 0.561716 | 1.657006 | 0.223053 |
| 14 | 106138800 | 106142999 | 20.94504 | 35.00896 | 71.35433 | 31.06585 | -0.64285 | 1.196738 | 0.223053 |
| 7 | 81816000 | 81817799 | 48.39923 | 78.26779 | 66.52327 | 28.43424 | 0.595882 | 1.703085 | 0.223059 |
| 9 | 41184600 | 41186799 | 10.95335 | 19.54308 | 41.48292 | 18.24055 | -0.68228 | 0.444031 | 0.223062 |
| 13 | 8840000 | 8841399 | 16.50339 | 30.79935 | 19.67977 | 8.179968 | 1.094514 | 0.177283 | 0.223078 |

| | | | | | | | | | |
|---|---|---|---|---|---|---|---|---|---|
| 7 | 119622800 | 119623799 | 20.00816 | 35.47236 | 26.51743 | 11.72269 | 0.80972 | 0.51423 | 0.2231 |
| 7 | 20542600 | 20542999 | 7.901968 | 15.45836 | 11.09547 | 3.676631 | 1.217251 | -0.69989 | 0.223101 |
| 4 | 144563200 | 144563999 | 7.069913 | 13.38054 | 30.91836 | 12.53125 | -0.7152 | 0.033787 | 0.223106 |
| 2 | 6490200 | 6498999 | 66.62978 | 103.9992 | 190.3337 | 78.97879 | -0.55852 | 2.672229 | 0.22311 |
| 9 | 117616000 | 117622399 | 69.4166 | 107.8664 | 211.4002 | 87.77553 | -0.66222 | 2.754871 | 0.223112 |
| 4 | 65949000 | 65950399 | 20.25538 | 35.59249 | 29.58733 | 13.0989 | 0.6526 | 0.615716 | 0.223144 |
| 8 | 124549600 | 124551999 | 13.95841 | 24.03458 | 53.69607 | 23.55433 | -0.77005 | 0.880025 | 0.223155 |
| 5 | 140315400 | 140318199 | 12.52142 | 21.71241 | 51.04349 | 22.39555 | -0.84078 | 0.806628 | 0.223179 |
| X | 78959000 | 78960399 | 34.47678 | 58.16739 | 40.91915 | 17.89498 | 0.8868 | 1.123858 | 0.223192 |
| 5 | 64787200 | 64790999 | 21.65214 | 36.01188 | 70.54653 | 30.61687 | -0.58581 | 1.254003 | 0.223196 |
| 9 | 48020400 | 48023399 | 16.16928 | 27.85671 | 57.58144 | 25.25911 | -0.65852 | 0.889613 | 0.223196 |
| 10 | 8761000 | 8763399 | 11.98729 | 20.95917 | 48.3548 | 21.31765 | -0.81253 | 0.682224 | 0.223216 |
| 3 | 151783000 | 151783799 | 4.400266 | 8.562872 | 24.70669 | 9.097824 | -0.90171 | -0.33685 | 0.223219 |
| 13 | 67419600 | 67420799 | 19.11053 | 34.044 | 22.60366 | 9.952199 | 0.983469 | 0.469614 | 0.223225 |
| 2 | 30010600 | 30012199 | 6.793692 | 12.80707 | 31.33778 | 12.80533 | -0.80751 | 0.059094 | 0.22325 |
| 8 | 12554400 | 12559799 | 32.58542 | 52.53521 | 106.2138 | 45.22198 | -0.65656 | 1.781939 | 0.223269 |
| 10 | 95498600 | 95499799 | 13.00863 | 24.94944 | 18.18139 | 7.145841 | 0.975209 | -0.04314 | 0.223282 |
| 19 | 14424000 | 14424799 | 13.00863 | 24.94944 | 18.18139 | 7.145841 | 0.975209 | -0.04314 | 0.223282 |
| X | 64664800 | 64667399 | 30.3006 | 52.63134 | 30.30341 | 13.30262 | 1.179642 | 0.839867 | 0.223282 |
| 7 | 99591800 | 99597799 | 66.88229 | 104.1762 | 187.7695 | 77.93611 | -0.53884 | 2.709796 | 0.223286 |
| 17 | 14819200 | 14820599 | 8.668948 | 15.93399 | 36.2299 | 15.39407 | -0.74703 | 0.239934 | 0.223286 |
| 4 | 73761800 | 73764999 | 36.15221 | 60.32657 | 45.95199 | 19.98528 | 0.772493 | 1.237831 | 0.223288 |
| 17 | 59221600 | 59221799 | 2.839277 | 5.551914 | 18.19013 | 5.909651 | -0.9109 | -0.8211 | 0.223296 |
| 4 | 70503400 | 70505599 | 28.74565 | 50.34436 | 25.76204 | 11.37347 | 1.349068 | 0.710123 | 0.223306 |
| 2 | 156704000 | 156705599 | 10.05602 | 17.91533 | 39.42604 | 17.35792 | -0.7348 | 0.446035 | 0.223315 |
| 3 | 130968400 | 130973599 | 45.67727 | 72.42691 | 147.4202 | 61.3606 | -0.69348 | 2.326893 | 0.223321 |
| 15 | 89475600 | 89476799 | 7.126117 | 12.93488 | 36.54321 | 15.78161 | -1.07319 | 0.30404 | 0.223331 |
| 12 | 83734800 | 83735399 | 4.665926 | 9.040149 | 25.43409 | 9.641648 | -0.90441 | -0.22607 | 0.223345 |
| 13 | 88522200 | 88522999 | 13.07025 | 24.51173 | 20.86497 | 8.601152 | 0.69387 | 0.121153 | 0.223366 |
| 3 | 48006000 | 48007599 | 14.05255 | 26.56273 | 19.52912 | 7.936091 | 0.919923 | 0.075186 | 0.223366 |
| 2 | 113639200 | 113643399 | 37.62147 | 62.22873 | 48.8561 | 21.12292 | 0.72845 | 1.33896 | 0.223369 |
| 10 | 5536600 | 5538799 | 17.88853 | 32.29106 | 24.9257 | 10.83917 | 0.779242 | 0.386908 | 0.223376 |
| 2 | 131077400 | 131078999 | 7.708644 | 14.40547 | 32.79045 | 13.59864 | -0.7216 | 0.12728 | 0.223377 |
| 10 | 39968200 | 39969399 | 7.708644 | 14.40547 | 32.79045 | 13.59864 | -0.7216 | 0.12728 | 0.223377 |
| 13 | 56626000 | 56627399 | 4.204898 | 8.074214 | 30.1202 | 11.79789 | -1.35325 | -0.11016 | 0.223381 |
| 10 | 21867200 | 21867799 | 4.815805 | 9.362901 | 25.24079 | 9.363712 | -0.8151 | -0.3036 | 0.223382 |
| 16 | 13835600 | 13836599 | 4.815805 | 9.362901 | 25.24079 | 9.363712 | -0.8151 | -0.3036 | 0.223382 |
| 19 | 60744000 | 60745799 | 9.831102 | 17.276 | 43.59776 | 19.30245 | -0.9402 | 0.584043 | 0.223383 |
| 6 | 43888200 | 43889399 | 10.35141 | 18.46644 | 39.41511 | 17.33341 | -0.68996 | 0.438714 | 0.223394 |
| 9 | 76591800 | 76592999 | 13.32821 | 25.09832 | 20.74775 | 8.49359 | 0.743496 | 0.096859 | 0.223433 |
| X | 30755800 | 30756599 | 20.26175 | 38.00904 | 14.92009 | 6.147629 | 1.800519 | 0.168244 | 0.223445 |
| 11 | 64411000 | 64412599 | 9.207906 | 16.65471 | 36.9737 | 16.03817 | -0.73281 | 0.340081 | 0.223447 |
| 18 | 80542200 | 80544199 | 7.935006 | 14.58085 | 34.59454 | 14.70485 | -0.8079 | 0.233527 | 0.223449 |
| X | 33938200 | 33944799 | 84.98433 | 135.5049 | 77.19102 | 32.34106 | 1.148806 | 2.178356 | 0.223463 |
| X | 125307800 | 125310999 | 35.97277 | 61.83543 | 28.97827 | 12.71096 | 1.472468 | 0.956633 | 0.223467 |
| 10 | 125975800 | 125978799 | 27.78271 | 45.30533 | 86.72157 | 37.25388 | -0.56229 | 1.525661 | 0.223468 |
| 12 | 36254200 | 36255799 | 22.3336 | 39.05131 | 31.83593 | 14.03581 | 0.680981 | 0.710171 | 0.223482 |

| | | | | | | | | | |
|---|---|---|---|---|---|---|---|---|---|
| 15 | 75370400 | 75373599 | 15.40719 | 26.32436 | 57.98875 | 25.42501 | -0.75211 | 0.960773 | 0.2235 |
| 5 | 62360800 | 62361599 | 10.68717 | 20.77807 | 15.82737 | 5.871477 | 0.988865 | -0.26359 | 0.223507 |
| X | 142818600 | 142819799 | 7.393187 | 13.88879 | 31.81727 | 13.07808 | -0.72018 | 0.088611 | 0.22351 |
| 13 | 86077800 | 86078799 | 12.74829 | 24.41504 | 18.1347 | 7.156945 | 0.942144 | -0.03358 | 0.223518 |
| 17 | 44889000 | 44890399 | 6.342024 | 12.01744 | 30.93134 | 12.50451 | -0.8662 | 0.017599 | 0.223526 |
| 11 | 97961200 | 97969399 | 83.45786 | 128.39 | 222.2345 | 92.29274 | -0.49442 | 2.944031 | 0.223535 |
| 4 | 140787000 | 140788399 | 7.552671 | 13.73058 | 36.0329 | 15.53083 | -0.96588 | 0.296773 | 0.22354 |
| 14 | 121284000 | 121285599 | 9.888927 | 17.6564 | 38.92331 | 17.10485 | -0.73542 | 0.429538 | 0.223544 |
| X | 60833400 | 60834399 | 12.06149 | 23.54673 | 9.633104 | 3.403807 | 1.931832 | -0.42941 | 0.223545 |
| 7 | 11106000 | 11106999 | 13.57377 | 26.01679 | 18.16108 | 7.150902 | 1.034496 | -0.03126 | 0.223553 |
| 6 | 69681600 | 69687999 | 149.3357 | 226.8883 | 173.8706 | 72.01092 | 0.665897 | 3.208776 | 0.223574 |
| 12 | 18586400 | 18591799 | 133.7809 | 204.3061 | 177.6562 | 73.68233 | 0.489695 | 3.100813 | 0.223593 |
| 8 | 83529600 | 83529999 | 4.181709 | 8.164645 | 22.9332 | 8.086013 | -0.8049 | -0.49791 | 0.223595 |
| 13 | 31981400 | 31983199 | 12.15867 | 21.21805 | 48.37753 | 21.29918 | -0.79573 | 0.712717 | 0.223598 |
| 7 | 98156800 | 98157799 | 6.920556 | 13.10143 | 30.92616 | 12.52719 | -0.74504 | 0.030026 | 0.223617 |
| X | 32999000 | 33006599 | 141.3514 | 216.0123 | 162.9349 | 67.58722 | 0.695449 | 3.090575 | 0.223622 |
| 3 | 121174000 | 121176199 | 11.93694 | 21.00673 | 46.13205 | 20.42404 | -0.74103 | 0.578833 | 0.223627 |
| 4 | 53614600 | 53615399 | 5.971451 | 11.53883 | 28.94818 | 11.11109 | -0.75968 | -0.16069 | 0.22363 |
| 7 | 59047800 | 59050199 | 19.62447 | 35.42096 | 23.55284 | 10.2435 | 0.992772 | 0.396369 | 0.223637 |
| 8 | 97723600 | 97724599 | 12.24188 | 23.54064 | 18.32607 | 7.148595 | 0.890973 | -0.07408 | 0.22365 |
| 6 | 50478200 | 50480999 | 18.13343 | 30.79668 | 60.73971 | 26.61632 | -0.5947 | 1.03029 | 0.223656 |
| 14 | 124428600 | 124431599 | 24.12533 | 39.78169 | 77.3301 | 33.36893 | -0.57618 | 1.365427 | 0.223691 |
| 14 | 19260400 | 19261799 | 8.887889 | 15.75263 | 40.97417 | 18.11472 | -0.97871 | 0.474649 | 0.223714 |
| 14 | 47567800 | 47569399 | 7.89319 | 14.64629 | 33.16048 | 13.90931 | -0.72673 | 0.175122 | 0.223726 |
| 10 | 30353200 | 30354799 | 19.98731 | 36.13698 | 23.00591 | 9.984471 | 1.057076 | 0.388632 | 0.22373 |
| 14 | 55822000 | 55823199 | 6.016488 | 11.14711 | 33.64805 | 14.14285 | -1.13974 | 0.169438 | 0.223738 |
| 5 | 72720200 | 72741399 | 241.8489 | 344.3307 | 562.5867 | 232.5957 | -0.51368 | 4.311865 | 0.223753 |
| 14 | 36523200 | 36524999 | 15.67007 | 27.03027 | 54.30664 | 23.82347 | -0.61719 | 0.872131 | 0.223755 |
| 12 | 13148800 | 13155799 | 52.5896 | 83.01121 | 161.2299 | 66.93349 | -0.63149 | 2.445328 | 0.223778 |
| 1 | 169024600 | 169025399 | 6.286465 | 12.07607 | 31.55398 | 12.34525 | -0.84413 | -0.0862 | 0.223782 |
| 7 | 67363600 | 67365599 | 8.184169 | 14.76603 | 37.4885 | 16.31247 | -0.92781 | 0.348593 | 0.223787 |
| 18 | 30167800 | 30168999 | 6.28122 | 12.0749 | 29.86233 | 11.6587 | -0.76314 | -0.10056 | 0.223789 |
| 5 | 109344800 | 109347799 | 41.53443 | 68.4107 | 50.60664 | 21.8269 | 0.812829 | 1.400998 | 0.223794 |
| 8 | 66525800 | 66528399 | 39.89358 | 65.9914 | 45.56503 | 19.70095 | 0.913219 | 1.343796 | 0.223802 |
| 2 | 17656800 | 17657599 | 3.711107 | 7.224241 | 24.03524 | 8.82621 | -1.09976 | -0.35638 | 0.22381 |
| 17 | 71183800 | 71184799 | 7.40294 | 13.90484 | 31.80042 | 13.07479 | -0.71781 | 0.091201 | 0.223813 |
| 4 | 127261000 | 127263199 | 9.862744 | 17.27682 | 45.71615 | 20.18628 | -1.00841 | 0.639135 | 0.223831 |
| 5 | 52532400 | 52535599 | 16.84847 | 28.25534 | 65.88097 | 28.74305 | -0.83588 | 1.144669 | 0.223837 |
| 5 | 74108200 | 74109999 | 9.354575 | 16.90999 | 38.00771 | 16.4989 | -0.75147 | 0.345587 | 0.223845 |
| 8 | 13755800 | 13759199 | 20.21993 | 33.62593 | 70.41725 | 30.55831 | -0.68172 | 1.250299 | 0.223846 |
| 9 | 16581400 | 16582599 | 7.206503 | 13.68583 | 32.82699 | 13.20882 | -0.75975 | 0.015162 | 0.223848 |
| 13 | 61178800 | 61181199 | 33.53155 | 56.31162 | 43.1044 | 18.82242 | 0.765117 | 1.159251 | 0.223872 |
| 11 | 36535000 | 36535599 | 4.482536 | 8.744507 | 23.99694 | 8.583501 | -0.79134 | -0.44013 | 0.223875 |
| 8 | 41966200 | 41968999 | 28.3679 | 45.95642 | 91.79238 | 39.28869 | -0.63131 | 1.646347 | 0.223882 |
| 6 | 136792800 | 136794199 | 5.709947 | 10.9504 | 27.89351 | 10.96527 | -0.8136 | -0.08633 | 0.223888 |
| 16 | 8948000 | 8949399 | 7.855355 | 14.65766 | 36.15388 | 15.03158 | -0.84012 | 0.143847 | 0.223901 |
| 12 | 88421400 | 88426599 | 35.06934 | 56.31523 | 113.3085 | 47.96723 | -0.65676 | 1.9131 | 0.22392 |

| | | | | | | | | | |
|---|---|---|---|---|---|---|---|---|---|
| X | 46772400 | 46774999 | 31.70831 | 53.9404 | 39.21669 | 17.18841 | 0.839894 | 1.041182 | 0.223937 |
| X | 49085800 | 49087399 | 22.56162 | 39.5183 | 30.72106 | 13.56409 | 0.749176 | 0.682836 | 0.223941 |
| 5 | 29379600 | 29380399 | 7.052492 | 13.40299 | 31.67502 | 12.7259 | -0.73646 | 0.005805 | 0.223943 |
| 1 | 135586800 | 135588399 | 7.606193 | 14.08712 | 33.12448 | 13.92995 | -0.78335 | 0.184232 | 0.223951 |
| 2 | 31009200 | 31010399 | 7.606193 | 14.08712 | 33.12448 | 13.92995 | -0.78335 | 0.184232 | 0.223951 |
| 14 | 47979600 | 47981399 | 7.943298 | 14.59076 | 34.59054 | 14.71045 | -0.8068 | 0.238396 | 0.223985 |
| 18 | 9182200 | 9184999 | 20.41263 | 36.11817 | 26.0502 | 11.52068 | 0.859485 | 0.533049 | 0.223998 |
| 19 | 57470000 | 57473999 | 29.68214 | 47.94857 | 98.21496 | 41.92248 | -0.67194 | 1.717243 | 0.224008 |
| X | 44695800 | 44696399 | 24.82453 | 43.34979 | 31.63366 | 13.93313 | 0.84001 | 0.742228 | 0.224015 |
| 2 | 85472800 | 85474599 | 19.24707 | 34.76101 | 23.56597 | 10.24138 | 0.965648 | 0.38998 | 0.224016 |
| 18 | 10725400 | 10726799 | 11.69325 | 20.43971 | 46.78081 | 20.61917 | -0.80078 | 0.684504 | 0.224018 |
| 6 | 143330600 | 143332799 | 16.27978 | 27.55904 | 60.55202 | 26.508 | -0.7499 | 1.052181 | 0.224032 |
| 7 | 32277400 | 32279999 | 32.31434 | 55.72593 | 31.94794 | 14.01494 | 1.184073 | 0.917045 | 0.22404 |
| X | 24673400 | 24675399 | 17.80683 | 33.65432 | 17.19911 | 6.991397 | 1.440558 | 0.106687 | 0.224047 |
| 1 | 98697200 | 98699799 | 31.46017 | 52.59666 | 45.43191 | 19.78274 | 0.591607 | 1.202192 | 0.224069 |
| 2 | 4819400 | 4827799 | 52.49663 | 83.14037 | 151.0992 | 62.8334 | -0.53189 | 2.364712 | 0.224073 |
| 2 | 8592000 | 8592599 | 11.71671 | 22.70528 | 16.86513 | 6.387562 | 0.997984 | -0.17816 | 0.224076 |
| 16 | 33607000 | 33611799 | 40.14415 | 64.27087 | 118.0963 | 49.76589 | -0.5313 | 2.016554 | 0.224109 |
| 10 | 70234000 | 70235999 | 43.49025 | 70.6991 | 60.98055 | 26.16632 | 0.581613 | 1.585098 | 0.224151 |
| 5 | 93190000 | 93194399 | 36.8003 | 58.78993 | 128.4512 | 53.96344 | -0.78413 | 2.081113 | 0.224165 |
| 7 | 32274000 | 32276199 | 22.71161 | 39.97697 | 28.27917 | 12.51948 | 0.885046 | 0.609217 | 0.224166 |
| 4 | 133483800 | 133484999 | 6.522769 | 12.24658 | 31.22223 | 12.84602 | -0.87387 | 0.083776 | 0.224174 |
| 15 | 98766200 | 98770599 | 20.41776 | 33.47885 | 80.10821 | 34.48539 | -0.8781 | 1.429975 | 0.224178 |
| 3 | 134090400 | 134092599 | 22.65188 | 39.74113 | 28.14781 | 12.44507 | 0.882219 | 0.649812 | 0.224214 |
| 9 | 121542400 | 121543399 | 10.82443 | 20.9133 | 17.9926 | 6.910832 | 0.768869 | -0.13947 | 0.22422 |
| 1 | 13440800 | 13444999 | 26.95191 | 43.61161 | 94.47916 | 40.40392 | -0.74981 | 1.666619 | 0.224226 |
| 10 | 44951600 | 44952999 | 7.865342 | 14.22917 | 36.50564 | 15.83124 | -0.93914 | 0.330163 | 0.22425 |
| 7 | 140906400 | 140908199 | 6.737889 | 12.46843 | 33.63948 | 14.1628 | -0.98132 | 0.181883 | 0.224277 |
| 15 | 96687800 | 96689999 | 11.064 | 19.39293 | 45.68384 | 20.18242 | -0.8421 | 0.633656 | 0.224277 |
| 7 | 105081400 | 105085399 | 39.86233 | 63.64435 | 125.5893 | 52.71738 | -0.63838 | 2.102508 | 0.224284 |
| 10 | 82464800 | 82466199 | 8.147252 | 14.48679 | 43.58334 | 19.21946 | -1.18484 | 0.450537 | 0.224298 |
| 18 | 5795400 | 5797199 | 21.38778 | 37.58331 | 29.39634 | 13.01463 | 0.739967 | 0.621334 | 0.224301 |
| 12 | 86288000 | 86289199 | 8.218595 | 15.12538 | 35.16364 | 14.91527 | -0.77681 | 0.227253 | 0.224302 |
| 19 | 23429400 | 23431399 | 27.59126 | 47.98441 | 32.06357 | 14.08621 | 0.966012 | 0.812221 | 0.224304 |
| 9 | 24478000 | 24479399 | 13.09063 | 22.91364 | 47.2777 | 20.86228 | -0.65209 | 0.671016 | 0.224306 |
| 2 | 89534400 | 89535799 | 16.52917 | 30.68635 | 21.42412 | 8.980272 | 0.958813 | 0.213866 | 0.224307 |
| 4 | 118159400 | 118164399 | 29.64823 | 47.99884 | 92.7804 | 39.69223 | -0.58512 | 1.660629 | 0.22432 |
| 3 | 98231400 | 98233999 | 11.77521 | 20.77051 | 44.02002 | 19.4799 | -0.68852 | 0.54937 | 0.224321 |
| 3 | 127048000 | 127049599 | 14.83595 | 27.87231 | 20.57414 | 8.455924 | 0.901141 | 0.124696 | 0.224322 |
| 4 | 59412000 | 59413999 | 16.56984 | 29.86526 | 24.48648 | 10.66842 | 0.69078 | 0.38365 | 0.22435 |
| 12 | 72771400 | 72774399 | 18.48693 | 31.1665 | 65.69453 | 28.72327 | -0.69087 | 1.098672 | 0.224352 |
| 10 | 82796400 | 82799399 | 17.47087 | 29.68625 | 61.86569 | 27.1128 | -0.67389 | 1.022797 | 0.224355 |
| 18 | 87948400 | 87952999 | 52.63374 | 84.9167 | 69.59711 | 29.65849 | 0.644388 | 1.777681 | 0.224356 |
| 7 | 85581400 | 85585999 | 108.0489 | 167.5838 | 144.5997 | 60.05601 | 0.517954 | 2.794802 | 0.224365 |
| 12 | 114690200 | 114692399 | 40.15462 | 66.58921 | 47.16118 | 20.42624 | 0.876947 | 1.318432 | 0.224378 |
| 13 | 65974800 | 65978399 | 44.91663 | 73.23092 | 53.88687 | 23.14704 | 0.813061 | 1.542769 | 0.224389 |
| 4 | 136873600 | 136879999 | 55.6848 | 87.90997 | 157.6317 | 65.44467 | -0.5162 | 2.444038 | 0.224413 |

| | | | | | | | | | |
|---|---|---|---|---|---|---|---|---|---|
| 7 | 100557400 | 100559799 | 9.039755 | 16.00422 | 40.98685 | 18.1243 | -0.95702 | 0.488472 | 0.224425 |
| 7 | 7511400 | 7518999 | 115.6969 | 181.2567 | 106.5029 | 44.17644 | 1.082724 | 2.622833 | 0.224429 |
| 6 | 67587200 | 67589199 | 19.91599 | 35.13449 | 28.43114 | 12.58115 | 0.69309 | 0.567497 | 0.22444 |
| 14 | 42659400 | 42661399 | 19.91599 | 35.13449 | 28.43114 | 12.58115 | 0.69309 | 0.567497 | 0.22444 |
| 7 | 108837600 | 108838999 | 8.20607 | 15.13672 | 35.21741 | 14.89192 | -0.77474 | 0.214047 | 0.224454 |
| 15 | 39791200 | 39793199 | 10.4556 | 18.77449 | 40.52498 | 17.72366 | -0.70088 | 0.398266 | 0.224458 |
| 13 | 66801000 | 66802799 | 39.34168 | 65.03293 | 49.92348 | 21.57498 | 0.760876 | 1.356268 | 0.224461 |
| 12 | 88106200 | 88109799 | 50.40554 | 81.77501 | 59.71063 | 25.58421 | 0.818682 | 1.637429 | 0.224463 |
| 7 | 102133400 | 102134599 | 7.668613 | 14.02761 | 36.11072 | 15.44429 | -0.93135 | 0.259897 | 0.224469 |
| 10 | 68646800 | 68647199 | 11.28745 | 21.80109 | 17.92466 | 6.894777 | 0.831636 | -0.12995 | 0.224496 |
| 10 | 116983800 | 116984999 | 5.423913 | 10.16023 | 36.11652 | 14.91731 | -1.35425 | 0.098022 | 0.2245 |
| 4 | 101149200 | 101150999 | 8.030713 | 14.49427 | 36.9893 | 16.08816 | -0.93454 | 0.345415 | 0.224507 |
| X | 5392600 | 5393399 | 15.1934 | 29.1363 | 16.2325 | 6.381971 | 1.358349 | -0.02491 | 0.224535 |
| 17 | 89564000 | 89565399 | 8.874834 | 16.29808 | 41.20581 | 17.52929 | -0.90022 | 0.259609 | 0.224545 |
| 2 | 44998600 | 45001399 | 12.34705 | 21.45864 | 51.05528 | 22.42027 | -0.85764 | 0.776264 | 0.22455 |
| 17 | 83104600 | 83107399 | 18.02713 | 32.75067 | 23.83466 | 10.28189 | 0.870945 | 0.349444 | 0.224558 |
| 10 | 50395000 | 50398199 | 37.81369 | 62.40443 | 52.15417 | 22.51343 | 0.638424 | 1.369781 | 0.224562 |
| 2 | 35387800 | 35388599 | 11.92956 | 23.05934 | 17.29609 | 6.62867 | 0.967702 | -0.1368 | 0.224572 |
| 3 | 92696200 | 92696799 | 4.653722 | 9.055331 | 24.09563 | 8.878857 | -0.78666 | -0.33125 | 0.224587 |
| 2 | 173276400 | 173290799 | 108.6413 | 163.5957 | 304.8044 | 125.8968 | -0.62573 | 3.3857 | 0.22459 |
| 9 | 110253400 | 110254399 | 6.300538 | 12.05118 | 28.97585 | 11.47738 | -0.74216 | -0.05393 | 0.224596 |
| 17 | 75749200 | 75750399 | 6.300538 | 12.05118 | 28.97585 | 11.47738 | -0.74216 | -0.05393 | 0.224596 |
| 3 | 146054800 | 146056799 | 16.3848 | 27.87903 | 58.48341 | 25.63334 | -0.6828 | 1.004473 | 0.224596 |
| 8 | 46996600 | 46999999 | 14.46281 | 24.77255 | 57.44008 | 25.18951 | -0.82509 | 0.933242 | 0.224624 |
| 15 | 5371200 | 5373599 | 22.28464 | 38.94925 | 31.68232 | 13.9631 | 0.683978 | 0.71418 | 0.224638 |
| 2 | 27286400 | 27290999 | 46.91868 | 74.61235 | 136.8621 | 57.13235 | -0.5395 | 2.240828 | 0.224646 |
| 12 | 3095000 | 3098799 | 41.52758 | 67.82748 | 58.22056 | 25.02499 | 0.593322 | 1.513159 | 0.224654 |
| 1 | 12092200 | 12093199 | 20.21925 | 35.46699 | 29.73038 | 13.14281 | 0.64069 | 0.643192 | 0.224659 |
| 1 | 135357600 | 135358599 | 6.192813 | 11.74124 | 29.80762 | 12.03604 | -0.8445 | 0.010979 | 0.224661 |
| 13 | 56669800 | 56670599 | 6.192813 | 11.74124 | 29.80762 | 12.03604 | -0.8445 | 0.010979 | 0.224661 |
| 12 | 114450600 | 114455599 | 67.53626 | 108.5592 | 66.90711 | 28.36538 | 1.048413 | 1.911074 | 0.224671 |
| 15 | 11944400 | 11944999 | 8.960781 | 17.46371 | 14.91749 | 5.40401 | 0.855821 | -0.37056 | 0.224688 |
| 5 | 135013000 | 135016999 | 25.35845 | 41.05991 | 93.63891 | 40.06381 | -0.82276 | 1.650391 | 0.224691 |
| 8 | 37224600 | 37225399 | 12.34316 | 23.43999 | 19.83255 | 7.980748 | 0.731391 | 0.029165 | 0.224698 |
| 12 | 32460000 | 32461399 | 12.34316 | 23.43999 | 19.83255 | 7.980748 | 0.731391 | 0.029165 | 0.224698 |
| 7 | 108340000 | 108341999 | 22.33817 | 39.28296 | 26.25764 | 11.6267 | 0.965633 | 0.61213 | 0.224738 |
| 18 | 38637000 | 38638399 | 8.334522 | 15.44017 | 35.3361 | 14.85459 | -0.74505 | 0.190703 | 0.224746 |
| 15 | 43740600 | 43745199 | 54.4637 | 86.20127 | 153.9776 | 64.00993 | -0.50775 | 2.381629 | 0.224751 |
| 1 | 116945600 | 116946399 | 9.849693 | 19.24396 | 10.46342 | 3.64733 | 1.54431 | -0.50506 | 0.224756 |
| 11 | 6565400 | 6572199 | 28.08439 | 45.37976 | 95.17337 | 40.6383 | -0.70533 | 1.707586 | 0.224758 |
| 7 | 3135800 | 3137999 | 33.74858 | 56.26601 | 47.06873 | 20.45958 | 0.637765 | 1.239544 | 0.224762 |
| 15 | 45418800 | 45419999 | 14.16893 | 26.23446 | 21.0993 | 8.87629 | 0.752101 | 0.206439 | 0.224763 |
| 7 | 114797800 | 114798799 | 15.24155 | 28.96296 | 18.46069 | 7.418491 | 1.13851 | 0.049598 | 0.224768 |
| 2 | 17413400 | 17414399 | 12.40556 | 23.60781 | 19.8771 | 7.96338 | 0.744095 | 0.016366 | 0.224782 |
| 6 | 55428600 | 55431799 | 19.91328 | 32.75036 | 79.27704 | 34.1712 | -0.89301 | 1.389072 | 0.22479 |
| 8 | 127738400 | 127739599 | 7.859334 | 14.74136 | 33.46163 | 13.79055 | -0.71023 | 0.107485 | 0.224794 |
| 18 | 38236400 | 38243199 | 46.13732 | 73.24777 | 141.468 | 58.95674 | -0.6161 | 2.289875 | 0.224802 |

| | | | | | | | | | |
|---|---|---|---|---|---|---|---|---|---|
| 6 | 92332600 | 92333999 | 5.517296 | 10.41947 | 31.88291 | 12.98149 | -1.12228 | 0.036205 | 0.224806 |
| 1 | 154078200 | 154082799 | 28.07739 | 45.39388 | 95.02574 | 40.60458 | -0.70176 | 1.68958 | 0.224817 |
| 5 | 9210800 | 9212599 | 11.00356 | 19.44736 | 41.9466 | 18.55511 | -0.71236 | 0.520285 | 0.224824 |
| 16 | 25923200 | 25925199 | 11.00356 | 19.44736 | 41.9466 | 18.55511 | -0.71236 | 0.520285 | 0.224824 |
| 16 | 97282800 | 97283999 | 11.64588 | 22.20133 | 18.50792 | 7.387186 | 0.763367 | -0.00587 | 0.224825 |
| 18 | 24227600 | 24229599 | 11.04503 | 19.41015 | 43.55113 | 19.28259 | -0.77205 | 0.58829 | 0.224833 |
| 7 | 40609400 | 40610399 | 11.20069 | 21.56308 | 18.52374 | 7.202043 | 0.754953 | -0.09388 | 0.224834 |
| 16 | 27945200 | 27945999 | 9.84633 | 17.713 | 37.92878 | 16.56251 | -0.68743 | 0.378957 | 0.224835 |
| 13 | 13611600 | 13613599 | 12.5422 | 21.69983 | 53.70784 | 23.56294 | -0.91561 | 0.828634 | 0.224836 |
| 18 | 55869400 | 55870199 | 10.65294 | 20.74201 | 15.37581 | 5.6366 | 1.043772 | -0.30453 | 0.224838 |
| 2 | 75638600 | 75641199 | 17.79266 | 29.7977 | 67.64081 | 29.4852 | -0.79751 | 1.1633 | 0.224852 |
| X | 122522000 | 122525599 | 42.06304 | 69.8057 | 44.92572 | 19.46798 | 1.014683 | 1.314681 | 0.224853 |
| 12 | 71607200 | 71610599 | 20.06876 | 33.35808 | 70.94831 | 30.77829 | -0.70409 | 1.255894 | 0.224857 |
| 8 | 64936600 | 64937999 | 7.698348 | 14.41752 | 32.85799 | 13.5766 | -0.71912 | 0.113032 | 0.224869 |
| 7 | 51779800 | 51780199 | 3.933616 | 7.684122 | 21.79968 | 7.61989 | -0.80777 | -0.5378 | 0.224903 |
| 13 | 120134600 | 120136199 | 27.55676 | 46.84587 | 39.52245 | 17.32147 | 0.625965 | 1.032059 | 0.224906 |
| 2 | 130914400 | 130918599 | 25.97874 | 42.06798 | 91.87016 | 39.3096 | -0.76027 | 1.649934 | 0.224906 |
| 19 | 8614600 | 8615399 | 5.53501 | 10.72587 | 28.01241 | 10.61032 | -0.79903 | -0.21533 | 0.224941 |
| 4 | 34661000 | 34662399 | 8.942051 | 16.08166 | 37.44455 | 16.35499 | -0.80744 | 0.374057 | 0.224942 |
| 9 | 65121600 | 65123199 | 8.942051 | 16.08166 | 37.44455 | 16.35499 | -0.80744 | 0.374057 | 0.224942 |
| 19 | 25577800 | 25580799 | 15.81934 | 27.15303 | 58.11438 | 25.48952 | -0.70976 | 0.919351 | 0.22496 |
| 12 | 24401600 | 24415599 | 267.5278 | 388.0526 | 325.8866 | 134.3616 | 0.472817 | 4.037471 | 0.224963 |
| 10 | 94778000 | 94781999 | 25.0566 | 40.55267 | 95.27022 | 40.74807 | -0.86612 | 1.659738 | 0.224964 |
| 11 | 34642400 | 34644199 | 19.00452 | 34.89293 | 21.04693 | 8.956219 | 1.152789 | 0.292584 | 0.224965 |
| 7 | 13403400 | 13410599 | 113.9993 | 176.9286 | 139.37 | 57.87836 | 0.64997 | 2.786124 | 0.224968 |
| 16 | 65918400 | 65920799 | 17.36758 | 31.24582 | 25.49926 | 11.13411 | 0.695644 | 0.403167 | 0.22497 |
| 19 | 32288000 | 32289999 | 17.36758 | 31.24582 | 25.49926 | 11.13411 | 0.695644 | 0.403167 | 0.22497 |
| 7 | 27331400 | 27334599 | 17.15755 | 28.76952 | 64.74961 | 28.24687 | -0.78499 | 1.147511 | 0.224984 |
| 10 | 75811200 | 75815999 | 45.23949 | 72.06196 | 132.4584 | 55.4211 | -0.53989 | 2.183599 | 0.224984 |
| 1 | 93270200 | 93271799 | 6.988601 | 12.68607 | 37.06068 | 16.00382 | -1.12117 | 0.303118 | 0.224985 |
| 2 | 88271200 | 88272799 | 21.4079 | 37.73563 | 27.08804 | 11.98876 | 0.864362 | 0.581649 | 0.225023 |
| 3 | 83935400 | 83939199 | 24.49326 | 42.21935 | 36.25401 | 15.90371 | 0.603636 | 0.890722 | 0.225026 |
| 7 | 126970200 | 126975399 | 55.34821 | 86.959 | 168.4697 | 69.88035 | -0.63306 | 2.545297 | 0.225033 |
| 15 | 17254200 | 17256999 | 23.68739 | 41.08605 | 35.13024 | 15.41368 | 0.611536 | 0.830649 | 0.225033 |
| 14 | 64750600 | 64751799 | 11.21005 | 21.54211 | 18.53526 | 7.244043 | 0.745825 | -0.07708 | 0.225036 |
| 9 | 120742600 | 120748999 | 42.65487 | 67.94628 | 132.1471 | 55.29235 | -0.62114 | 2.180523 | 0.225045 |
| 4 | 133299000 | 133302999 | 28.36753 | 45.93162 | 92.54865 | 39.597 | -0.64467 | 1.655358 | 0.225048 |
| 14 | 49845000 | 49845399 | 9.542002 | 18.52154 | 16.51027 | 6.205151 | 0.74746 | -0.23084 | 0.225049 |
| 7 | 11697000 | 11698199 | 27.48196 | 46.98444 | 39.32538 | 17.24297 | 0.638478 | 0.979733 | 0.225075 |
| 8 | 86418400 | 86419599 | 14.64767 | 27.52676 | 20.58039 | 8.454237 | 0.883368 | 0.121038 | 0.225087 |
| 5 | 45577000 | 45578199 | 6.984343 | 13.03749 | 33.29337 | 13.83312 | -0.88837 | 0.128632 | 0.225089 |
| 1 | 126530600 | 126532599 | 9.178325 | 16.28479 | 40.95689 | 18.09431 | -0.92982 | 0.467534 | 0.225089 |
| 7 | 8408800 | 8410599 | 22.1416 | 39.74532 | 22.87061 | 10.00525 | 1.194778 | 0.445361 | 0.225115 |
| 7 | 135850200 | 135853399 | 14.84296 | 25.24859 | 62.81082 | 27.52944 | -0.92794 | 1.003661 | 0.225118 |
| 14 | 66707200 | 66710999 | 29.98139 | 48.49852 | 93.84193 | 40.12215 | -0.58771 | 1.678367 | 0.225121 |
| 2 | 95526000 | 95532199 | 116.7226 | 180.5775 | 146.5329 | 60.86619 | 0.603633 | 2.843712 | 0.225139 |
| 6 | 34924600 | 34927199 | 19.75269 | 32.858 | 69.15921 | 30.01812 | -0.68887 | 1.246201 | 0.22516 |

| | | | | | | | | | |
|---|---|---|---|---|---|---|---|---|---|
| 18 | 32295200 | 32296799 | 9.602159 | 17.09119 | 39.43025 | 17.37309 | -0.80325 | 0.448044 | 0.225182 |
| 15 | 46842000 | 46843199 | 18.81531 | 33.78265 | 25.25888 | 11.0478 | 0.81945 | 0.424626 | 0.225194 |
| 1 | 181302800 | 181306799 | 22.30974 | 36.98935 | 72.8031 | 31.52783 | -0.59301 | 1.293229 | 0.225199 |
| 3 | 103554200 | 103560799 | 57.77761 | 91.26164 | 161.3304 | 66.998 | -0.49513 | 2.433235 | 0.22521 |
| 7 | 12989200 | 12991599 | 33.77971 | 57.13776 | 35.75293 | 15.64783 | 1.055174 | 1.094925 | 0.225249 |
| 9 | 37466200 | 37468199 | 10.28795 | 18.08748 | 44.07061 | 19.51512 | -0.89015 | 0.578215 | 0.22525 |
| 6 | 80813800 | 80815999 | 22.55208 | 39.5283 | 30.47904 | 13.46492 | 0.760657 | 0.669933 | 0.225252 |
| 17 | 38508600 | 38515199 | 72.51371 | 115.7772 | 85.58596 | 35.92623 | 0.775634 | 2.124859 | 0.225266 |
| 16 | 4595200 | 4596399 | 7.74899 | 14.40815 | 33.73228 | 14.10898 | -0.77131 | 0.165436 | 0.225267 |
| 9 | 62463600 | 62465999 | 13.32487 | 22.99978 | 52.62587 | 23.0866 | -0.80346 | 0.854162 | 0.225289 |
| 4 | 114125200 | 114127999 | 33.5678 | 56.67266 | 41.44488 | 18.12867 | 0.831021 | 1.114646 | 0.225299 |
| 11 | 108798800 | 108814999 | 112.668 | 168.4485 | 349.4881 | 144.107 | -0.79009 | 3.526329 | 0.225337 |
| 1 | 13231800 | 13233399 | 7.060802 | 13.39227 | 31.00413 | 12.50857 | -0.71215 | 0.018501 | 0.225349 |
| X | 33461200 | 33468199 | 106.9356 | 168.3254 | 102.6304 | 42.58096 | 1.034663 | 2.530417 | 0.225359 |
| 5 | 74282600 | 74289799 | 60.09836 | 94.28938 | 172.9954 | 71.74624 | -0.55626 | 2.575957 | 0.225371 |
| 11 | 55295600 | 55299599 | 29.12722 | 47.26631 | 90.9284 | 38.96738 | -0.57507 | 1.610517 | 0.225385 |
| 9 | 82933200 | 82935399 | 16.21963 | 27.59858 | 59.59148 | 26.11905 | -0.72438 | 1.003666 | 0.225393 |
| 4 | 69286200 | 69288399 | 21.77186 | 38.9006 | 22.60852 | 9.929385 | 1.176219 | 0.468014 | 0.225407 |
| 7 | 58715600 | 58717799 | 26.77037 | 46.64684 | 30.85607 | 13.5722 | 0.980563 | 0.778808 | 0.22541 |
| 15 | 96935000 | 96937199 | 10.07047 | 17.87953 | 39.91432 | 17.62386 | -0.75833 | 0.468143 | 0.225413 |
| X | 66116600 | 66118399 | 20.98328 | 38.23287 | 20.895 | 8.980863 | 1.284637 | 0.349546 | 0.225413 |
| 7 | 23436400 | 23446999 | 55.73776 | 89.65608 | 68.63504 | 29.12713 | 0.737482 | 1.871933 | 0.225422 |
| 7 | 14534200 | 14539199 | 100.8836 | 158.3603 | 110.8175 | 45.96124 | 0.833122 | 2.579908 | 0.225429 |
| 11 | 58396800 | 58398199 | 18.09974 | 33.03605 | 22.69257 | 9.732376 | 0.958871 | 0.320962 | 0.22547 |
| 4 | 88502200 | 88503799 | 20.25557 | 33.60785 | 70.75932 | 30.65807 | -0.68965 | 1.279184 | 0.22547 |
| 8 | 30505200 | 30508799 | 46.55101 | 74.19357 | 132.602 | 55.52515 | -0.49859 | 2.16705 | 0.225474 |
| 11 | 38799800 | 38800599 | 8.276041 | 15.08922 | 35.53961 | 15.26291 | -0.80749 | 0.2859 | 0.225501 |
| 10 | 77716200 | 77727799 | 89.23838 | 136.0651 | 251.6752 | 104.343 | -0.60007 | 3.125777 | 0.225523 |
| 13 | 24684400 | 24685799 | 9.44184 | 16.8312 | 40.97366 | 18.03017 | -0.87934 | 0.43812 | 0.225527 |
| 2 | 143851600 | 143853799 | 12.34817 | 21.46264 | 49.98823 | 21.95301 | -0.82689 | 0.774571 | 0.225531 |
| 13 | 68389000 | 68396199 | 134.0185 | 205.5103 | 163.3253 | 67.79821 | 0.623999 | 3.017634 | 0.225551 |
| 6 | 31196800 | 31198199 | 10.32008 | 18.51179 | 38.93414 | 17.04291 | -0.66427 | 0.402279 | 0.225557 |
| 1 | 131056400 | 131058399 | 25.20903 | 41.16512 | 82.84685 | 35.60726 | -0.63288 | 1.498352 | 0.225596 |
| 3 | 65299200 | 65303599 | 28.9314 | 46.62897 | 100.9295 | 42.95417 | -0.75511 | 1.786321 | 0.22561 |
| 4 | 13749400 | 13751999 | 17.90266 | 30.11259 | 63.27888 | 27.64791 | -0.68641 | 1.116639 | 0.225622 |
| 9 | 42963600 | 42968399 | 51.20134 | 81.35903 | 152.0613 | 63.4274 | -0.56769 | 2.271019 | 0.225626 |
| 5 | 146603400 | 146604999 | 58.58861 | 94.27192 | 71.20032 | 30.22979 | 0.757468 | 1.86556 | 0.225637 |
| 5 | 148231200 | 148254399 | 239.4166 | 339.3698 | 578.0248 | 239.1551 | -0.58004 | 4.376081 | 0.225655 |
| 9 | 89619400 | 89620799 | 11.45096 | 20.26415 | 42.96554 | 19.00095 | -0.68723 | 0.509896 | 0.225664 |
| 8 | 81605400 | 81608199 | 16.56465 | 27.69034 | 70.67255 | 30.77116 | -0.96576 | 1.18448 | 0.225675 |
| 16 | 57051000 | 57052599 | 9.530773 | 16.78226 | 43.56453 | 19.28505 | -0.97989 | 0.55593 | 0.22568 |
| 12 | 22450600 | 22451799 | 20.0129 | 35.40346 | 27.88615 | 12.33314 | 0.733366 | 0.538359 | 0.225692 |
| 4 | 141449000 | 141451199 | 10.86918 | 19.17554 | 41.97819 | 18.57572 | -0.7345 | 0.540505 | 0.225717 |
| 11 | 102196000 | 102198799 | 11.3996 | 19.90353 | 47.31937 | 20.83998 | -0.85539 | 0.701619 | 0.225719 |
| 3 | 31481600 | 31482799 | 11.97223 | 21.09182 | 44.03686 | 19.49219 | -0.66753 | 0.566817 | 0.225724 |
| 16 | 70067000 | 70067799 | 17.50058 | 31.19896 | 26.09683 | 11.48308 | 0.652786 | 0.461003 | 0.225735 |
| 13 | 79166000 | 79166599 | 5.105111 | 9.915975 | 27.0052 | 10.09518 | -0.84063 | -0.26892 | 0.225739 |

| | | | | | | | | | |
|---|---|---|---|---|---|---|---|---|---|
| 7 | 35017200 | 35019399 | 13.40492 | 23.34496 | 48.93538 | 21.51704 | -0.67547 | 0.749746 | 0.225764 |
| 15 | 92826200 | 92826999 | 13.48594 | 25.80109 | 18.76964 | 7.429537 | 0.968742 | -0.01342 | 0.225772 |
| 9 | 82130200 | 82130999 | 4.657727 | 9.050332 | 24.51074 | 9.13542 | -0.82699 | -0.28882 | 0.225777 |
| 15 | 89491800 | 89493799 | 15.81623 | 27.28077 | 54.32016 | 23.82935 | -0.60436 | 0.872739 | 0.225783 |
| 7 | 127375400 | 127377599 | 9.888034 | 17.31542 | 48.84232 | 21.56066 | -1.10061 | 0.646716 | 0.225796 |
| 4 | 3262400 | 3263799 | 29.22313 | 50.22336 | 35.86462 | 15.73148 | 0.867646 | 0.936001 | 0.225809 |
| 13 | 95499600 | 95501599 | 10.68676 | 18.93156 | 40.90703 | 18.08682 | -0.71348 | 0.490926 | 0.225811 |
| 5 | 30226400 | 30229199 | 21.8564 | 38.22911 | 31.72691 | 13.99152 | 0.655187 | 0.700804 | 0.225826 |
| 9 | 61190800 | 61196799 | 48.50897 | 77.152 | 138.7126 | 57.91536 | -0.51046 | 2.237152 | 0.225861 |
| 8 | 110709000 | 110714599 | 38.97246 | 62.46415 | 116.2502 | 49.07052 | -0.54777 | 1.978986 | 0.225861 |
| 10 | 88159200 | 88159999 | 4.988568 | 9.650324 | 27.15778 | 10.36975 | -0.91631 | -0.19604 | 0.225884 |
| 11 | 78766200 | 78769199 | 19.45006 | 32.79251 | 64.60654 | 28.24817 | -0.59373 | 1.101587 | 0.22589 |
| 8 | 64704000 | 64706399 | 32.38184 | 55.37871 | 35.21302 | 15.44022 | 1.033311 | 0.991495 | 0.225892 |
| 3 | 4482600 | 4485999 | 30.73022 | 52.37187 | 38.63028 | 16.93388 | 0.819409 | 1.01928 | 0.225911 |
| 5 | 123988200 | 123999599 | 79.11217 | 121.6034 | 226.7858 | 94.17211 | -0.60282 | 2.958783 | 0.225911 |
| 3 | 74173800 | 74176199 | 49.91987 | 80.89035 | 62.84936 | 26.90535 | 0.72779 | 1.663674 | 0.225921 |
| 11 | 27247000 | 27248599 | 15.0614 | 28.5577 | 19.00939 | 7.674628 | 1.07068 | 0.06381 | 0.225922 |
| 4 | 108435600 | 108436999 | 6.265186 | 11.71059 | 32.75103 | 13.57723 | -1.01535 | 0.113198 | 0.225932 |
| 13 | 84370200 | 84370999 | 6.265186 | 11.71059 | 32.75103 | 13.57723 | -1.01535 | 0.113198 | 0.225932 |
| 9 | 124154400 | 124155399 | 15.17302 | 28.2905 | 21.58101 | 8.984222 | 0.839491 | 0.176975 | 0.22594 |
| 18 | 74100800 | 74104599 | 27.70948 | 44.91855 | 91.55477 | 39.2055 | -0.65964 | 1.629518 | 0.225942 |
| 3 | 74674400 | 74675799 | 16.38995 | 30.67507 | 18.52821 | 7.66419 | 1.18011 | 0.155541 | 0.225946 |
| 16 | 38142000 | 38145799 | 30.81142 | 49.74779 | 97.56063 | 41.61619 | -0.61014 | 1.734218 | 0.225948 |
| 10 | 123757400 | 123759799 | 25.8445 | 45.14511 | 30.32849 | 13.36091 | 0.958624 | 0.740018 | 0.22595 |
| 1 | 100774800 | 100776399 | 27.41487 | 46.82879 | 39.40802 | 17.27799 | 0.630586 | 0.98851 | 0.22595 |
| X | 130568000 | 130569799 | 31.69184 | 53.3621 | 42.17836 | 18.43449 | 0.719265 | 1.133087 | 0.225952 |
| 9 | 89321200 | 89321999 | 17.41145 | 31.15467 | 25.65552 | 11.25388 | 0.678351 | 0.437712 | 0.225956 |
| 7 | 60968400 | 60968999 | 18.18279 | 32.30796 | 23.84239 | 10.52042 | 0.829927 | 0.482227 | 0.225963 |
| 7 | 38554800 | 38555199 | 15.25743 | 28.44315 | 21.64214 | 9.012211 | 0.843008 | 0.179513 | 0.225973 |
| 5 | 112149400 | 112151399 | 12.94676 | 22.53913 | 48.9327 | 21.5111 | -0.7258 | 0.752821 | 0.225989 |
| X | 153281600 | 153284599 | 58.27355 | 93.9219 | 67.49523 | 28.72579 | 0.832025 | 1.816312 | 0.225993 |
| 10 | 115379800 | 115382199 | 37.75239 | 62.65039 | 44.97531 | 19.48914 | 0.857537 | 1.301342 | 0.225999 |
| 7 | 27838600 | 27840599 | 11.23319 | 19.65011 | 46.25343 | 20.39951 | -0.84073 | 0.6711 | 0.226008 |
| 8 | 116257800 | 116258999 | 4.831102 | 9.34815 | 25.96235 | 9.901202 | -0.89491 | -0.20347 | 0.226022 |
| 17 | 9049800 | 9050799 | 4.831102 | 9.34815 | 25.96235 | 9.901202 | -0.89491 | -0.20347 | 0.226022 |
| 14 | 3177400 | 3182399 | 74.41517 | 118.5434 | 86.06496 | 36.01306 | 0.798109 | 2.196987 | 0.226025 |
| 4 | 32129600 | 32130999 | 7.399301 | 13.85907 | 33.98883 | 14.04109 | -0.82412 | 0.111032 | 0.226035 |
| 3 | 132653200 | 132656599 | 20.96827 | 34.56546 | 78.61569 | 33.91937 | -0.80214 | 1.361108 | 0.226036 |
| 12 | 94404200 | 94404799 | 8.518943 | 16.58979 | 15.0452 | 5.505695 | 0.758053 | -0.34006 | 0.226042 |
| 1 | 4865400 | 4866399 | 12.87734 | 24.55657 | 18.6195 | 7.426837 | 0.898879 | 0.002386 | 0.226048 |
| 5 | 137438800 | 137445999 | 53.7147 | 84.81725 | 158.6511 | 65.87475 | -0.57684 | 2.437397 | 0.22605 |
| 8 | 4889200 | 4891199 | 9.308011 | 17.06126 | 39.48314 | 16.83977 | -0.77644 | 0.268062 | 0.226063 |
| 10 | 45822600 | 45823999 | 13.26087 | 23.12384 | 48.38943 | 21.2942 | -0.67269 | 0.731167 | 0.226071 |
| 17 | 45883200 | 45885399 | 14.8269 | 25.25616 | 59.52362 | 26.09159 | -0.84981 | 0.988438 | 0.226081 |
| 4 | 99035000 | 99035799 | 10.19476 | 19.68558 | 17.55549 | 6.760841 | 0.714138 | -0.13865 | 0.226081 |
| 9 | 64867800 | 64869799 | 7.553451 | 13.75308 | 37.62472 | 16.18586 | -1.02414 | 0.289024 | 0.226088 |
| 1 | 43972400 | 43973799 | 9.479266 | 16.80915 | 41.45201 | 18.32185 | -0.90225 | 0.473492 | 0.226132 |

| | | | | | | | | | |
|---|---|---|---|---|---|---|---|---|---|
| 17 | 78943200 | 78946799 | 20.87898 | 34.62278 | 74.41898 | 32.23121 | -0.71965 | 1.287488 | 0.226151 |
| 2 | 19393400 | 19394599 | 7.232477 | 13.62136 | 31.89255 | 13.05289 | -0.74651 | 0.070258 | 0.226159 |
| 6 | 117812000 | 117813999 | 13.25262 | 23.09674 | 48.39686 | 21.2901 | -0.67474 | 0.738263 | 0.226168 |
| 4 | 117054000 | 117054399 | 9.886496 | 19.18324 | 17.03346 | 6.412672 | 0.750864 | -0.21912 | 0.226177 |
| 1 | 112192000 | 112194199 | 36.67941 | 60.30122 | 52.04377 | 22.429 | 0.590067 | 1.413503 | 0.226192 |
| 15 | 79953400 | 79954799 | 5.457859 | 10.4585 | 32.60035 | 12.84537 | -1.10691 | -0.07388 | 0.226199 |
| 1 | 175424400 | 175425199 | 4.980446 | 9.656464 | 27.37541 | 10.34278 | -0.91241 | -0.2325 | 0.226204 |
| 12 | 110409000 | 110411799 | 14.3954 | 24.81052 | 56.46676 | 24.77044 | -0.7967 | 0.874052 | 0.226218 |
| 12 | 110260400 | 110269399 | 78.12945 | 120.406 | 215.1401 | 89.36573 | -0.53822 | 2.907067 | 0.22623 |
| 7 | 16997800 | 16999999 | 22.81739 | 40.6161 | 23.7246 | 10.45254 | 1.166358 | 0.500932 | 0.226269 |
| 3 | 42226800 | 42229199 | 20.35384 | 36.86279 | 22.45529 | 9.724638 | 1.122253 | 0.38083 | 0.226271 |
| 6 | 42347600 | 42350399 | 11.50952 | 19.87412 | 54.16539 | 23.76225 | -1.05432 | 0.843712 | 0.226272 |
| 4 | 113017000 | 113018599 | 49.31398 | 79.68581 | 67.76618 | 28.93682 | 0.593807 | 1.726167 | 0.22628 |
| 1 | 111341200 | 111343599 | 28.77724 | 49.7801 | 32.47714 | 14.24843 | 0.999038 | 0.872431 | 0.226286 |
| 6 | 37080400 | 37081999 | 8.84733 | 16.18864 | 35.58087 | 15.21142 | -0.70466 | 0.266694 | 0.226294 |
| 7 | 17085200 | 17098399 | 366.5823 | 517.406 | 422.4267 | 174.9007 | 0.474233 | 4.442834 | 0.226302 |
| 14 | 29948000 | 29950199 | 23.87859 | 41.96121 | 28.19985 | 12.48515 | 0.957871 | 0.63484 | 0.226325 |
| X | 54499800 | 54503199 | 57.13856 | 92.32531 | 61.97295 | 26.46151 | 0.934294 | 1.740791 | 0.226333 |
| 13 | 28961400 | 28962199 | 6.008889 | 11.49865 | 28.39736 | 11.23332 | -0.77822 | -0.06189 | 0.226336 |
| 17 | 15511800 | 15513199 | 7.106372 | 13.32513 | 32.31668 | 13.32535 | -0.80556 | 0.09847 | 0.226338 |
| 6 | 27636000 | 27636799 | 6.929844 | 13.09102 | 30.8488 | 12.55243 | -0.74827 | 0.045158 | 0.226338 |
| 10 | 9893400 | 9894199 | 9.325045 | 16.54416 | 40.44547 | 17.86923 | -0.88913 | 0.468399 | 0.22637 |
| 12 | 12961600 | 12962399 | 6.134073 | 11.79379 | 29.19007 | 11.39132 | -0.76278 | -0.10192 | 0.226374 |
| 2 | 35629400 | 35632399 | 16.17642 | 27.24476 | 62.581 | 27.35388 | -0.81401 | 1.099881 | 0.226385 |
| 11 | 96678000 | 96680999 | 12.74987 | 22.29891 | 47.2925 | 20.85438 | -0.69171 | 0.680495 | 0.2264 |
| 10 | 82782600 | 82791999 | 89.86592 | 137.3625 | 240.7465 | 99.86937 | -0.51957 | 3.075214 | 0.226417 |
| X | 32271200 | 32271799 | 10.7215 | 20.94224 | 11.84151 | 4.144792 | 1.486926 | -0.47168 | 0.22645 |
| 2 | 163239000 | 163241799 | 16.34996 | 27.88886 | 58.55684 | 25.67065 | -0.68362 | 0.980299 | 0.226458 |
| 15 | 13510400 | 13511199 | 11.26533 | 21.56896 | 18.96718 | 7.491447 | 0.700159 | -0.04284 | 0.226468 |
| 7 | 13701400 | 13712599 | 332.8681 | 473.954 | 388.9395 | 160.801 | 0.479543 | 4.313642 | 0.226472 |
| X | 32175600 | 32177399 | 25.14858 | 44.75007 | 20.74344 | 9.141811 | 1.497092 | 0.51274 | 0.226473 |
| 5 | 45265400 | 45265999 | 9.232849 | 18.01832 | 13.06826 | 4.645402 | 1.112015 | -0.43893 | 0.226475 |
| 6 | 78396800 | 78397799 | 23.29576 | 40.36011 | 34.17287 | 14.99309 | 0.625263 | 0.837208 | 0.226479 |
| 2 | 48572400 | 48573799 | 19.59624 | 34.64703 | 27.88422 | 12.33366 | 0.70208 | 0.540871 | 0.226481 |
| 10 | 89941600 | 89942599 | 12.74526 | 24.35788 | 19.32979 | 7.674598 | 0.84007 | -0.01022 | 0.226514 |
| X | 18636800 | 18637799 | 12.74526 | 24.35788 | 19.32979 | 7.674598 | 0.84007 | -0.01022 | 0.226514 |
| 2 | 84740000 | 84744999 | 30.08766 | 48.82266 | 98.52625 | 42.22204 | -0.6441 | 1.616225 | 0.226518 |
| 7 | 46211000 | 46220599 | 69.05636 | 107.8573 | 191.295 | 79.37233 | -0.51256 | 2.662286 | 0.226521 |
| X | 90658200 | 90658799 | 11.38226 | 21.90797 | 18.51859 | 7.204442 | 0.777299 | -0.08984 | 0.226524 |
| 5 | 135697600 | 135702199 | 22.04052 | 36.39356 | 75.40643 | 32.56298 | -0.66739 | 1.341877 | 0.226542 |
| 3 | 3196800 | 3201399 | 74.30654 | 118.3325 | 91.45944 | 38.2534 | 0.707538 | 2.210143 | 0.226545 |
| 2 | 57004200 | 57006999 | 23.92097 | 39.11308 | 82.24732 | 35.3665 | -0.69462 | 1.475732 | 0.226558 |
| 2 | 32526400 | 32528399 | 8.558862 | 15.66878 | 35.06322 | 14.97973 | -0.7293 | 0.262396 | 0.226561 |
| 13 | 85825800 | 85825999 | 9.124892 | 17.83324 | 12.71298 | 4.398328 | 1.172574 | -0.53069 | 0.226566 |
| 17 | 45784000 | 45786199 | 10.28801 | 18.07463 | 43.5931 | 19.29653 | -0.87535 | 0.590439 | 0.226567 |
| 6 | 105006600 | 105010399 | 63.73265 | 101.6674 | 90.30388 | 37.85954 | 0.509798 | 2.141986 | 0.226572 |
| 2 | 152872400 | 152875199 | 20.79084 | 34.31864 | 75.14726 | 32.4436 | -0.74681 | 1.342932 | 0.22658 |

| | | | | | | | | | |
|---|---|---|---|---|---|---|---|---|---|
| 3 | 38798800 | 38801399 | 38.99945 | 64.226 | 53.27798 | 22.97765 | 0.648017 | 1.399305 | 0.226583 |
| 8 | 62171000 | 62171999 | 14.32391 | 26.74184 | 21.67125 | 9.002075 | 0.755342 | 0.161685 | 0.226585 |
| 14 | 54385800 | 54387999 | 14.0351 | 24.34793 | 52.17601 | 22.89275 | -0.70805 | 0.808789 | 0.226587 |
| 18 | 80347600 | 80350799 | 28.50014 | 46.22502 | 89.98496 | 38.5485 | -0.59276 | 1.619904 | 0.226602 |
| 3 | 26399800 | 26403199 | 27.20562 | 44.17628 | 88.18416 | 37.80918 | -0.6273 | 1.593239 | 0.226607 |
| 5 | 127759000 | 127773599 | 119.4831 | 179.5019 | 311.188 | 128.4532 | -0.52485 | 3.433807 | 0.226609 |
| 19 | 6062400 | 6066599 | 17.89977 | 29.68475 | 71.47936 | 30.95968 | -0.88252 | 1.280069 | 0.226612 |
| 1 | 16540000 | 16541799 | 16.66792 | 28.61934 | 59.19502 | 25.96421 | -0.6605 | 0.924043 | 0.226626 |
| 16 | 94324600 | 94327999 | 16.62829 | 28.03742 | 64.39044 | 28.15441 | -0.81394 | 1.090398 | 0.226636 |
| 5 | 10548600 | 10550599 | 41.94318 | 68.73731 | 55.49554 | 23.88463 | 0.684337 | 1.468628 | 0.226636 |
| Y | 41344000 | 41345599 | 11.67749 | 22.49833 | 9.733002 | 3.775831 | 1.735031 | -0.12701 | 0.226638 |
| 13 | 65882600 | 65885799 | 28.83461 | 50.20572 | 28.48289 | 12.52239 | 1.200724 | 0.795503 | 0.22665 |
| 19 | 5802800 | 5812199 | 49.39371 | 77.55727 | 175.5533 | 72.81294 | -0.85788 | 2.554256 | 0.226652 |
| 18 | 81742200 | 81743399 | 10.18219 | 18.20487 | 38.91301 | 17.08065 | -0.69017 | 0.422118 | 0.226654 |
| 2 | 78778200 | 78779999 | 18.52776 | 32.96809 | 26.96726 | 11.88492 | 0.683326 | 0.481385 | 0.226661 |
| 3 | 30653600 | 30655199 | 5.686161 | 10.9816 | 27.07426 | 10.41818 | -0.73797 | -0.16042 | 0.226663 |
| 11 | 20780200 | 20781799 | 11.67765 | 20.45003 | 45.70005 | 20.17415 | -0.76649 | 0.652259 | 0.226679 |
| X | 113480400 | 113481799 | 28.28854 | 49.14797 | 31.73131 | 13.93149 | 1.015443 | 0.830485 | 0.226697 |
| 9 | 102365800 | 102372799 | 38.94771 | 62.06862 | 132.5545 | 55.49698 | -0.75513 | 2.162531 | 0.226697 |
| 11 | 27267600 | 27268999 | 22.63575 | 39.39859 | 33.46937 | 14.71141 | 0.621294 | 0.777287 | 0.226704 |
| 11 | 15557200 | 15559799 | 30.36274 | 51.69987 | 39.79645 | 17.44383 | 0.758021 | 1.028192 | 0.226708 |
| 5 | 122249000 | 122255399 | 40.65198 | 64.55398 | 142.7611 | 59.50928 | -0.81079 | 2.278939 | 0.226716 |
| 7 | 47137200 | 47138999 | 30.34192 | 51.75603 | 35.64422 | 15.62603 | 0.918263 | 1.008108 | 0.226726 |
| 4 | 62448200 | 62449999 | 8.495729 | 15.7721 | 34.83028 | 14.59993 | -0.69067 | 0.180579 | 0.226729 |
| 17 | 23155800 | 23158199 | 26.59039 | 45.77712 | 37.19767 | 16.31705 | 0.682671 | 0.913136 | 0.226754 |
| 15 | 93016400 | 93017999 | 7.787276 | 14.31485 | 34.0726 | 14.47548 | -0.81167 | 0.230116 | 0.226762 |
| 1 | 134133000 | 134138999 | 34.49008 | 55.26006 | 115.57 | 48.75677 | -0.71627 | 1.985708 | 0.226767 |
| 5 | 112832400 | 112837199 | 29.73519 | 47.9122 | 101.714 | 43.27252 | -0.72767 | 1.795246 | 0.226819 |
| X | 64984000 | 64985399 | 23.76219 | 41.75832 | 28.27114 | 12.51713 | 0.947349 | 0.634105 | 0.226822 |
| 4 | 56669000 | 56671799 | 42.5459 | 69.56509 | 57.18968 | 24.59147 | 0.6565 | 1.500197 | 0.226835 |
| 7 | 60145800 | 60146799 | 18.02626 | 31.95809 | 25.72952 | 11.37442 | 0.703159 | 0.49868 | 0.226835 |
| 5 | 119818000 | 119822199 | 20.26422 | 33.58456 | 73.09052 | 31.64363 | -0.73751 | 1.292184 | 0.226853 |
| 3 | 119771200 | 119773199 | 17.86919 | 33.19716 | 20.17748 | 8.449725 | 1.158147 | 0.218317 | 0.226856 |
| 13 | 118993000 | 118996199 | 26.80294 | 43.74961 | 84.93652 | 36.49984 | -0.58163 | 1.508313 | 0.226856 |
| 10 | 63407400 | 63411199 | 33.23692 | 53.60315 | 100.719 | 42.90066 | -0.55053 | 1.772579 | 0.226868 |
| 8 | 5636400 | 5637199 | 14.31285 | 27.10238 | 19.57503 | 7.925213 | 0.950218 | 0.066576 | 0.226883 |
| 13 | 42130000 | 42131199 | 14.31285 | 27.10238 | 19.57503 | 7.925213 | 0.950218 | 0.066576 | 0.226883 |
| 8 | 13596200 | 13606199 | 56.94104 | 89.12768 | 181.6139 | 75.33862 | -0.71005 | 2.616371 | 0.226899 |
| 14 | 37399400 | 37400199 | 6.584387 | 12.62803 | 30.31792 | 11.91658 | -0.72978 | -0.073 | 0.226904 |
| 3 | 110906400 | 110908999 | 37.49663 | 62.33095 | 47.66424 | 20.67838 | 0.76688 | 1.284279 | 0.226905 |
| 3 | 89798800 | 89801399 | 15.02015 | 26.02451 | 52.70576 | 23.12431 | -0.62723 | 0.827451 | 0.226926 |
| X | 24187800 | 24188999 | 22.83818 | 40.14754 | 25.94537 | 11.48927 | 1.014072 | 0.619959 | 0.22693 |
| 12 | 101872200 | 101872799 | 3.654028 | 7.093091 | 24.85343 | 9.336974 | -1.20273 | -0.26887 | 0.226931 |
| 7 | 80752800 | 80755799 | 14.86955 | 25.6775 | 52.6782 | 23.10987 | -0.64678 | 0.856607 | 0.226932 |
| 10 | 127022600 | 127027599 | 32.20649 | 51.7077 | 111.4379 | 47.16105 | -0.75592 | 1.916075 | 0.226933 |
| 2 | 36544400 | 36547799 | 71.64187 | 113.9846 | 98.21782 | 41.02809 | 0.549519 | 2.236535 | 0.226956 |
| 1 | 48132800 | 48134999 | 26.86042 | 45.79579 | 39.46439 | 17.3001 | 0.596063 | 1.00253 | 0.226957 |

| | | | | | | | | | |
|---|---|---|---|---|---|---|---|---|---|
| 4 | 41634000 | 41640599 | 43.05208 | 68.3928 | 140.3021 | 58.50783 | -0.70203 | 2.26836 | 0.226981 |
| 2 | 122125200 | 122129599 | 38.41389 | 61.33612 | 126.4993 | 53.10469 | -0.70184 | 2.098798 | 0.226985 |
| 11 | 109363000 | 109368999 | 28.41415 | 45.89647 | 97.80953 | 41.74499 | -0.72903 | 1.717727 | 0.226992 |
| 11 | 118646800 | 118658799 | 104.5255 | 158.0737 | 279.7962 | 115.7164 | -0.54646 | 3.29851 | 0.227028 |
| 1 | 57109400 | 57110199 | 8.725992 | 15.90077 | 35.51989 | 15.26503 | -0.73265 | 0.291062 | 0.227073 |
| 5 | 42470800 | 42472199 | 13.68437 | 26.18525 | 17.41632 | 6.890438 | 1.096513 | -0.01837 | 0.227078 |
| 6 | 113828600 | 113829999 | 12.92351 | 24.25025 | 20.24482 | 8.337515 | 0.722425 | 0.113355 | 0.227083 |
| 7 | 17948600 | 17950799 | 36.34153 | 60.27512 | 49.87976 | 21.60599 | 0.653107 | 1.307945 | 0.227087 |
| 14 | 48910400 | 48912999 | 17.94022 | 30.06781 | 65.40536 | 28.52451 | -0.73615 | 1.154572 | 0.22709 |
| 9 | 35861800 | 35867399 | 81.46338 | 129.3741 | 97.46686 | 40.63266 | 0.740385 | 2.298109 | 0.227091 |
| 7 | 22022000 | 22022399 | 10.43941 | 20.28471 | 15.78274 | 5.882155 | 0.952111 | -0.25176 | 0.227092 |
| 14 | 5457800 | 5458799 | 10.43941 | 20.28471 | 15.78274 | 5.882155 | 0.952111 | -0.25176 | 0.227092 |
| 17 | 43020800 | 43021999 | 9.215237 | 16.62587 | 38.50139 | 16.75376 | -0.79657 | 0.356664 | 0.227101 |
| 17 | 19549600 | 19551199 | 37.46916 | 61.83328 | 52.0802 | 22.48085 | 0.627301 | 1.37134 | 0.227102 |
| 15 | 79222600 | 79228399 | 36.70966 | 58.91192 | 111.1199 | 46.99536 | -0.56484 | 1.93598 | 0.227103 |
| 12 | 30422600 | 30423799 | 7.246526 | 13.23387 | 36.6139 | 15.69204 | -1.03645 | 0.267419 | 0.227106 |
| 18 | 29874400 | 29876199 | 10.12775 | 18.27501 | 39.02565 | 16.97862 | -0.6803 | 0.363894 | 0.227108 |
| 14 | 52713200 | 52716999 | 62.72774 | 100.8145 | 72.4089 | 30.69106 | 0.827828 | 1.909465 | 0.227111 |
| 13 | 98164400 | 98165599 | 8.282805 | 15.07269 | 35.51368 | 15.28831 | -0.81044 | 0.296038 | 0.227113 |
| 10 | 11552000 | 11553199 | 13.92077 | 26.43951 | 19.05274 | 7.664744 | 0.961234 | 0.041043 | 0.227129 |
| 14 | 3093400 | 3099999 | 101.3641 | 159.1019 | 118.173 | 49.01138 | 0.747198 | 2.583866 | 0.227133 |
| 14 | 7001400 | 7004199 | 51.30226 | 82.54717 | 68.3634 | 29.02497 | 0.624777 | 1.853656 | 0.227138 |
| 10 | 130385200 | 130388199 | 41.2467 | 67.42245 | 56.62722 | 24.3472 | 0.625382 | 1.498829 | 0.22715 |
| 6 | 75203000 | 75203799 | 15.5047 | 28.54792 | 18.68525 | 7.920644 | 1.039693 | 0.243211 | 0.227153 |
| 1 | 18989800 | 18990999 | 12.58499 | 22.10658 | 45.08821 | 19.95564 | -0.63549 | 0.606265 | 0.22716 |
| 19 | 13517800 | 13519399 | 15.59584 | 29.83707 | 16.78589 | 6.644719 | 1.335706 | 0.001956 | 0.227165 |
| 14 | 4275600 | 4277599 | 28.64407 | 49.75543 | 32.07283 | 14.07962 | 1.017561 | 0.83417 | 0.227168 |
| 6 | 83473000 | 83475599 | 13.87518 | 23.71782 | 57.35471 | 25.14795 | -0.88576 | 0.950678 | 0.227174 |
| 6 | 59892400 | 59893599 | 29.2787 | 49.30053 | 42.17787 | 18.43447 | 0.605329 | 1.12124 | 0.227175 |
| 13 | 36917600 | 36922399 | 36.87236 | 59.08934 | 120.8212 | 50.98842 | -0.68354 | 1.981492 | 0.227177 |
| 8 | 12172800 | 12173399 | 5.08225 | 9.879517 | 25.22849 | 9.369919 | -0.73905 | -0.29483 | 0.22719 |
| 9 | 17696600 | 17697399 | 5.08225 | 9.879517 | 25.22849 | 9.369919 | -0.73905 | -0.29483 | 0.22719 |
| 5 | 72368200 | 72369999 | 10.48786 | 18.69911 | 40.94957 | 18.01693 | -0.72762 | 0.443872 | 0.227193 |
| 8 | 115681200 | 115687599 | 45.53378 | 72.57989 | 133.4643 | 55.89431 | -0.53944 | 2.16162 | 0.227222 |
| 17 | 68497000 | 68500399 | 41.99687 | 68.45905 | 59.31279 | 25.47617 | 0.577933 | 1.542174 | 0.227227 |
| 17 | 32253000 | 32256399 | 17.53779 | 29.24766 | 69.07595 | 30.03103 | -0.85436 | 1.210429 | 0.22723 |
| X | 5309400 | 5315399 | 79.62435 | 127.0787 | 79.76939 | 33.46542 | 1.01049 | 2.144137 | 0.227235 |
| 11 | 50486400 | 50496399 | 83.97874 | 128.385 | 241.5581 | 100.2099 | -0.62164 | 3.070009 | 0.22724 |
| 12 | 3199200 | 3209199 | 152.8157 | 231.593 | 197.8195 | 81.86766 | 0.506513 | 3.261904 | 0.227266 |
| 8 | 22630200 | 22634999 | 35.18831 | 56.66802 | 105.5029 | 44.84916 | -0.53986 | 1.821117 | 0.227268 |
| 13 | 11412600 | 11414399 | 16.68187 | 30.51901 | 22.8139 | 9.755531 | 0.840284 | 0.296732 | 0.227275 |
| 17 | 19122200 | 19123199 | 15.60797 | 26.59116 | 57.88322 | 25.37288 | -0.73545 | 0.990961 | 0.227279 |
| 2 | 3612600 | 3614599 | 22.05054 | 38.73456 | 29.33441 | 12.98342 | 0.786317 | 0.630505 | 0.227284 |
| 10 | 19050000 | 19051599 | 22.05054 | 38.73456 | 29.33441 | 12.98342 | 0.786317 | 0.630505 | 0.227284 |
| 12 | 22649800 | 22655199 | 105.5994 | 164.0593 | 129.7832 | 53.88885 | 0.644957 | 2.762881 | 0.227318 |
| 8 | 9721400 | 9725199 | 46.7277 | 75.78546 | 62.28959 | 26.68329 | 0.647965 | 1.63687 | 0.227321 |
| 14 | 12583400 | 12584199 | 8.231057 | 15.11433 | 35.11619 | 14.94073 | -0.77908 | 0.240323 | 0.227324 |

| | | | | | | | | | |
|---|---|---|---|---|---|---|---|---|---|
| 1 | 128447800 | 128449399 | 8.072789 | 14.86648 | 34.09885 | 14.45112 | -0.75628 | 0.221362 | 0.227331 |
| 9 | 31776600 | 31777199 | 8.072789 | 14.86648 | 34.09885 | 14.45112 | -0.75628 | 0.221362 | 0.227331 |
| X | 53892400 | 53895199 | 32.53042 | 55.20909 | 39.7509 | 17.41528 | 0.853555 | 1.065001 | 0.227332 |
| 1 | 35899000 | 35899999 | 10.75937 | 20.76608 | 17.94821 | 6.92724 | 0.755501 | -0.12492 | 0.227334 |
| 4 | 119939400 | 119940199 | 4.817965 | 9.360834 | 25.139 | 9.376485 | -0.81676 | -0.28423 | 0.227353 |
| 5 | 105958200 | 105962599 | 98.47972 | 149.1419 | 274.2126 | 113.4579 | -0.59948 | 3.268348 | 0.227379 |
| 19 | 46704800 | 46705799 | 22.10057 | 38.62512 | 32.33737 | 14.25104 | 0.642746 | 0.716675 | 0.227399 |
| 14 | 115946800 | 115947599 | 9.948618 | 19.24722 | 14.51506 | 5.539989 | 0.965869 | -0.18155 | 0.227406 |
| 1 | 77820800 | 77824599 | 20.9465 | 35.0814 | 69.07049 | 30.10944 | -0.59306 | 1.174603 | 0.227416 |
| 9 | 95286600 | 95290199 | 25.55748 | 41.30209 | 97.21929 | 41.51888 | -0.87118 | 1.699076 | 0.227426 |
| 19 | 59021800 | 59023599 | 19.128 | 34.01043 | 26.81739 | 11.8264 | 0.735261 | 0.48907 | 0.227427 |
| 9 | 60918200 | 60928599 | 77.79935 | 119.8867 | 214.6889 | 89.17832 | -0.54151 | 2.908698 | 0.227451 |
| 17 | 45087000 | 45088599 | 10.83738 | 19.2472 | 40.40319 | 17.83499 | -0.66995 | 0.47219 | 0.227465 |
| 12 | 24214200 | 24218599 | 63.57148 | 102.5048 | 60.14081 | 25.60994 | 1.124546 | 1.807685 | 0.227469 |
| 12 | 81212600 | 81215999 | 20.8496 | 34.65702 | 71.65066 | 31.08362 | -0.66339 | 1.257832 | 0.22749 |
| 7 | 91600000 | 91603399 | 25.66649 | 44.80625 | 30.90209 | 13.60827 | 0.920857 | 0.748357 | 0.2275 |
| 1 | 24690600 | 24692199 | 14.4107 | 26.80276 | 20.44245 | 8.545957 | 0.83439 | 0.181241 | 0.227511 |
| 4 | 127955400 | 127956199 | 6.943052 | 13.08573 | 30.7663 | 12.57654 | -0.75035 | 0.062727 | 0.227515 |
| 7 | 34598400 | 34601999 | 15.71575 | 26.84269 | 58.55367 | 25.67201 | -0.73816 | 0.96526 | 0.22752 |
| 7 | 106908200 | 106910799 | 21.54212 | 37.73103 | 31.15916 | 13.75705 | 0.662214 | 0.676595 | 0.227522 |
| 5 | 138912000 | 138918999 | 37.06932 | 59.34184 | 118.5057 | 49.93076 | -0.65131 | 2.015635 | 0.227525 |
| 6 | 147591000 | 147592199 | 8.443206 | 15.32737 | 35.99882 | 15.54423 | -0.80893 | 0.311694 | 0.227543 |
| 18 | 37050000 | 37050399 | 21.53289 | 37.73482 | 27.5146 | 12.15337 | 0.841155 | 0.664457 | 0.227571 |
| 12 | 64360600 | 64361999 | 17.73772 | 31.61487 | 25.94999 | 11.42055 | 0.679545 | 0.462654 | 0.227574 |
| 7 | 143004800 | 143021799 | 234.1819 | 332.3563 | 565.0713 | 233.7478 | -0.57569 | 4.358073 | 0.227577 |
| 18 | 89939200 | 89940399 | 13.0093 | 24.46889 | 19.64067 | 8.056456 | 0.782979 | 0.097264 | 0.227583 |
| 8 | 72270800 | 72273599 | 82.8614 | 131.0788 | 108.2757 | 44.99935 | 0.602724 | 2.403971 | 0.227616 |
| 7 | 136694600 | 136695599 | 6.591518 | 12.59983 | 30.13976 | 11.95995 | -0.73771 | -0.04314 | 0.227637 |
| 5 | 33479600 | 33480599 | 12.17852 | 23.08264 | 19.85282 | 8.021663 | 0.702837 | 0.040694 | 0.227639 |
| 12 | 3198800 | 3206199 | 126.0941 | 193.4074 | 167.3431 | 69.46848 | 0.501763 | 3.011291 | 0.227644 |
| 1 | 8019000 | 8020599 | 20.78622 | 36.54196 | 25.56171 | 11.31927 | 0.900205 | 0.604753 | 0.227671 |
| 7 | 21067600 | 21075399 | 122.626 | 189.2545 | 140.8149 | 58.4918 | 0.725989 | 2.889541 | 0.227687 |
| 15 | 94013400 | 94014799 | 8.193547 | 15.14848 | 35.2775 | 14.87063 | -0.77284 | 0.200702 | 0.227722 |
| 17 | 37054200 | 37055799 | 29.05162 | 49.05765 | 39.63205 | 17.33674 | 0.687932 | 1.095248 | 0.22773 |
| X | 146905400 | 146905599 | 6.669283 | 13.04421 | 11.52466 | 3.891022 | 0.899913 | -0.65587 | 0.227734 |
| 16 | 97089600 | 97090799 | 6.755913 | 12.84358 | 32.27455 | 12.95774 | -0.8236 | 0.000942 | 0.227738 |
| 19 | 43869000 | 43870399 | 14.00052 | 26.16619 | 20.39607 | 8.456975 | 0.813638 | 0.150827 | 0.22775 |
| 19 | 43662400 | 43664399 | 9.215949 | 16.26424 | 42.5342 | 18.82053 | -0.98908 | 0.527427 | 0.22775 |
| 17 | 38677400 | 38680599 | 74.61597 | 118.2457 | 105.4736 | 43.87629 | 0.494136 | 2.356843 | 0.22776 |
| 14 | 16041800 | 16043799 | 15.41098 | 25.90124 | 67.37014 | 29.42848 | -0.99277 | 1.121114 | 0.227794 |
| 7 | 94491400 | 94491999 | 10.46405 | 20.24705 | 17.46127 | 6.660415 | 0.774594 | -0.16851 | 0.227795 |
| 2 | 7142600 | 7143399 | 11.80299 | 22.93886 | 15.08643 | 5.613545 | 1.194335 | -0.24214 | 0.227803 |
| 1 | 34760600 | 34764199 | 24.68686 | 40.02999 | 90.15228 | 38.61294 | -0.80267 | 1.619118 | 0.227807 |
| 6 | 144611400 | 144615799 | 40.84477 | 65.18051 | 126.0266 | 52.86374 | -0.60983 | 2.11939 | 0.227827 |
| 7 | 14398400 | 14401799 | 71.20498 | 113.6367 | 87.92684 | 36.88546 | 0.709187 | 2.136206 | 0.227846 |
| 2 | 84930600 | 84933999 | 27.77076 | 45.29879 | 88.04209 | 37.82509 | -0.58398 | 1.521558 | 0.227863 |
| 12 | 109049200 | 109064599 | 135.981 | 202.2106 | 352.4051 | 145.1896 | -0.54584 | 3.627482 | 0.227866 |

| | | | | | | | | | |
|---|---|---|---|---|---|---|---|---|---|
| 8 | 79754600 | 79757799 | 16.78327 | 27.94895 | 70.006 | 30.40766 | -0.93878 | 1.225298 | 0.227871 |
| 11 | 111684000 | 111685199 | 6.596088 | 12.60923 | 29.53056 | 11.71646 | -0.70677 | -0.04326 | 0.227897 |
| 4 | 142076600 | 142079999 | 19.62651 | 33.00798 | 65.12575 | 28.45376 | -0.59598 | 1.124202 | 0.227904 |
| 6 | 26839200 | 26841599 | 24.30996 | 42.62663 | 29.2842 | 12.94148 | 0.926413 | 0.672291 | 0.227912 |
| 15 | 58369600 | 58370199 | 4.060192 | 7.91484 | 22.5065 | 8.127439 | -0.85321 | -0.41491 | 0.227914 |
| 19 | 6955800 | 6958199 | 13.04072 | 22.44982 | 53.64867 | 23.53339 | -0.86659 | 0.879093 | 0.227925 |
| 12 | 18308800 | 18323999 | 404.6512 | 568.2727 | 438.5238 | 181.713 | 0.548249 | 4.518738 | 0.227926 |
| 16 | 68204600 | 68206599 | 20.1464 | 35.35949 | 29.78454 | 13.17314 | 0.633794 | 0.634747 | 0.227928 |
| 12 | 23317400 | 23327399 | 220.8713 | 325.0638 | 264.3466 | 108.8592 | 0.539282 | 3.80899 | 0.227928 |
| 4 | 137219200 | 137220999 | 8.166313 | 14.76748 | 36.49057 | 15.83298 | -0.88603 | 0.334284 | 0.227949 |
| 1 | 138763600 | 138779599 | 115.3979 | 173.1859 | 320.3479 | 132.1977 | -0.61964 | 3.453315 | 0.22795 |
| 11 | 62441600 | 62444199 | 10.9099 | 19.13525 | 44.65158 | 19.73721 | -0.82811 | 0.62335 | 0.227964 |
| 5 | 24770400 | 24774599 | 18.05829 | 29.99663 | 72.33593 | 31.36606 | -0.88462 | 1.259103 | 0.227974 |
| 17 | 37336600 | 37338199 | 17.03403 | 30.55085 | 25.60644 | 11.21018 | 0.654856 | 0.420058 | 0.227979 |
| 1 | 23798400 | 23803199 | 35.4088 | 56.83042 | 111.6274 | 47.21775 | -0.623 | 1.930281 | 0.227982 |
| 16 | 33500600 | 33502599 | 10.22556 | 18.15011 | 39.91156 | 17.62655 | -0.7369 | 0.471157 | 0.227995 |
| 18 | 74181600 | 74184599 | 20.80688 | 34.77535 | 67.82929 | 29.5297 | -0.5795 | 1.195837 | 0.22802 |
| 5 | 108430400 | 108432599 | 26.6623 | 46.41409 | 29.54903 | 12.98948 | 1.035508 | 0.79038 | 0.228031 |
| 9 | 35304800 | 35306599 | 13.48055 | 25.49617 | 20.20179 | 8.199466 | 0.814564 | 0.068219 | 0.228036 |
| 1 | 182694000 | 182697199 | 21.46771 | 34.98385 | 86.47268 | 37.13784 | -0.93052 | 1.522682 | 0.228044 |
| 10 | 68531000 | 68532599 | 10.14608 | 17.81615 | 44.11119 | 19.51742 | -0.91312 | 0.597535 | 0.228061 |
| 8 | 58023800 | 58026599 | 15.98587 | 27.41802 | 57.00085 | 24.99958 | -0.66813 | 0.926288 | 0.228062 |
| 3 | 41567600 | 41569399 | 8.225604 | 15.1431 | 34.0935 | 14.45508 | -0.73007 | 0.224781 | 0.228082 |
| 9 | 45935000 | 45936799 | 6.42248 | 11.95057 | 32.65715 | 13.64211 | -0.99094 | 0.14489 | 0.228089 |
| 3 | 88638000 | 88641599 | 32.58688 | 52.58683 | 98.9573 | 42.1834 | -0.55164 | 1.753205 | 0.228105 |
| 6 | 140436800 | 140438999 | 8.524041 | 15.68153 | 34.61622 | 14.6907 | -0.70304 | 0.231621 | 0.228113 |
| 3 | 96508200 | 96509599 | 6.511912 | 12.27155 | 33.59175 | 13.73453 | -0.96916 | 0.062431 | 0.228122 |
| 7 | 22160000 | 22160799 | 11.96978 | 23.33453 | 12.83105 | 4.631072 | 1.488394 | -0.34897 | 0.228126 |
| 12 | 108899200 | 108901199 | 7.799815 | 14.30155 | 35.08971 | 14.95917 | -0.85907 | 0.245568 | 0.228134 |
| 5 | 10524000 | 10525799 | 23.60755 | 41.47302 | 27.48519 | 12.16579 | 0.977619 | 0.634437 | 0.228148 |
| 6 | 94258200 | 94261999 | 21.66268 | 35.55003 | 79.94142 | 34.42585 | -0.7884 | 1.421043 | 0.22815 |
| 19 | 31060000 | 31062399 | 48.76478 | 77.30787 | 145.5122 | 60.55496 | -0.58106 | 2.335223 | 0.228172 |
| 13 | 104240200 | 104241599 | 24.37824 | 42.74369 | 29.35099 | 12.97031 | 0.927155 | 0.67433 | 0.228177 |
| 1 | 98456800 | 98458199 | 17.14925 | 30.99418 | 24.94691 | 10.83192 | 0.720526 | 0.3741 | 0.228223 |
| 5 | 12007200 | 12008999 | 22.25437 | 39.2017 | 27.00442 | 11.95333 | 0.923157 | 0.593891 | 0.228228 |
| 2 | 39199000 | 39200599 | 15.42589 | 28.7217 | 19.11418 | 7.976182 | 1.031243 | 0.179559 | 0.228229 |
| 10 | 116311600 | 116313799 | 26.17221 | 45.13541 | 35.98961 | 15.78796 | 0.710113 | 0.892016 | 0.228234 |
| 10 | 111233800 | 111237399 | 26.47718 | 43.21441 | 87.96266 | 37.79916 | -0.64984 | 1.509442 | 0.228268 |
| 15 | 12057400 | 12057599 | 10.2936 | 20.02974 | 15.95216 | 5.877642 | 0.934646 | -0.29032 | 0.228275 |
| 3 | 123441400 | 123442999 | 24.76258 | 42.78907 | 36.10601 | 15.83998 | 0.629398 | 0.869192 | 0.228275 |
| X | 12096400 | 12100599 | 103.0698 | 160.2945 | 137.6611 | 57.15116 | 0.52775 | 2.746801 | 0.228282 |
| 5 | 56573600 | 56576599 | 28.1311 | 45.87864 | 96.36597 | 41.39863 | -0.6959 | 1.530844 | 0.228285 |
| 1 | 60723600 | 60725599 | 20.50944 | 35.96799 | 29.85414 | 13.19503 | 0.65503 | 0.650055 | 0.228304 |
| 7 | 23595200 | 23598799 | 73.81458 | 117.5234 | 94.84661 | 39.65812 | 0.644931 | 2.215858 | 0.228308 |
| 4 | 112336000 | 112338399 | 60.90602 | 97.44312 | 85.92634 | 36.17375 | 0.524824 | 2.050198 | 0.228342 |
| 9 | 67490800 | 67492999 | 9.680391 | 17.41459 | 38.96789 | 17.01618 | -0.75079 | 0.378024 | 0.22836 |
| 7 | 11607600 | 11609199 | 30.70416 | 52.13091 | 40.90242 | 17.92199 | 0.730089 | 1.056007 | 0.228364 |

| | | | | | | | | | |
|---|---|---|---|---|---|---|---|---|---|
| 1 | 48929600 | 48930999 | 14.6063 | 27.57941 | 20.1609 | 8.21171 | 0.925816 | 0.09063 | 0.228374 |
| 15 | 96513000 | 96523799 | 124.5726 | 187.2621 | 312.0766 | 128.841 | -0.46718 | 3.423335 | 0.228383 |
| 5 | 143096400 | 143099399 | 17.36804 | 29.37033 | 62.30207 | 27.2654 | -0.69964 | 1.065418 | 0.228397 |
| X | 5490000 | 5490799 | 25.33286 | 44.15559 | 29.50624 | 12.98098 | 0.966454 | 0.769231 | 0.228401 |
| 16 | 89782600 | 89786199 | 33.61503 | 54.05347 | 105.8784 | 44.91136 | -0.6156 | 1.8672 | 0.228402 |
| 7 | 92945200 | 92945599 | 10.30925 | 19.88514 | 16.04447 | 6.203732 | 0.851944 | -0.12906 | 0.228406 |
| 19 | 45990600 | 46000799 | 96.4319 | 146.8234 | 263.3538 | 109.1413 | -0.55747 | 3.157142 | 0.228411 |
| 15 | 50281600 | 50282399 | 10.60181 | 20.51899 | 15.65786 | 5.965209 | 0.950139 | -0.17802 | 0.228424 |
| 17 | 37914800 | 37917999 | 40.56708 | 67.70682 | 42.04026 | 18.28626 | 1.066329 | 1.24703 | 0.228433 |
| 18 | 59695800 | 59696799 | 6.61882 | 12.5679 | 29.87755 | 12.0258 | -0.74675 | 0.004253 | 0.228438 |
| 7 | 59597000 | 59597799 | 17.53532 | 31.35062 | 23.33878 | 10.2447 | 0.822457 | 0.438858 | 0.228443 |
| 1 | 154424600 | 154426199 | 18.5657 | 34.37888 | 19.41847 | 8.168021 | 1.258248 | 0.239181 | 0.228445 |
| 18 | 27194800 | 27196199 | 18.5657 | 34.37888 | 19.41847 | 8.168021 | 1.258248 | 0.239181 | 0.228445 |
| 12 | 6858200 | 6862199 | 36.26734 | 60.21303 | 49.30598 | 21.37239 | 0.668401 | 1.297042 | 0.228447 |
| 1 | 136185200 | 136192799 | 45.00466 | 71.40731 | 150.9815 | 62.88425 | -0.74739 | 2.306621 | 0.228448 |
| 18 | 53117400 | 53119399 | 11.76708 | 20.82722 | 45.0867 | 19.9383 | -0.71707 | 0.513559 | 0.228453 |
| 9 | 48599400 | 48600799 | 8.755369 | 15.87798 | 36.51164 | 15.78563 | -0.77963 | 0.322902 | 0.228456 |
| 1 | 37878400 | 37879999 | 9.316268 | 16.98941 | 36.58432 | 15.70741 | -0.67959 | 0.291742 | 0.22846 |
| 15 | 9394400 | 9394999 | 10.06475 | 17.92429 | 39.94827 | 17.59329 | -0.75316 | 0.450436 | 0.228461 |
| 16 | 51851600 | 51853599 | 10.31681 | 18.47212 | 39.4312 | 17.28712 | -0.68722 | 0.413756 | 0.228472 |
| 1 | 86403800 | 86405999 | 10.38378 | 18.40089 | 40.40854 | 17.86524 | -0.73635 | 0.485011 | 0.228495 |
| 13 | 13357800 | 13358599 | 13.53532 | 25.71524 | 19.72304 | 7.929309 | 0.873146 | 0.037751 | 0.228507 |
| 19 | 23157200 | 23158199 | 9.496115 | 17.19954 | 37.51353 | 16.24449 | -0.70605 | 0.334406 | 0.228507 |
| 12 | 5563400 | 5564199 | 12.4154 | 23.96685 | 16.5367 | 6.376791 | 1.078013 | -0.1145 | 0.228509 |
| 10 | 61232200 | 61234399 | 9.137072 | 16.30495 | 38.94472 | 17.12226 | -0.85072 | 0.427697 | 0.22852 |
| 12 | 55882800 | 55883999 | 6.213458 | 11.74671 | 31.97684 | 12.99036 | -0.95273 | 0.034529 | 0.228523 |
| 12 | 79853200 | 79856599 | 19.46273 | 32.30728 | 71.92409 | 31.17265 | -0.76988 | 1.270741 | 0.228531 |
| 17 | 19310600 | 19315799 | 68.76081 | 110.4396 | 68.20479 | 28.88199 | 1.044084 | 1.936555 | 0.228535 |
| 5 | 53890800 | 53892399 | 9.632685 | 17.56326 | 40.42091 | 17.35857 | -0.77545 | 0.302466 | 0.228543 |
| 5 | 148052200 | 148053799 | 6.533474 | 12.20868 | 31.66219 | 13.132 | -0.90727 | 0.117736 | 0.228553 |
| 5 | 95418400 | 95419999 | 35.541 | 59.1013 | 48.74114 | 21.15082 | 0.658292 | 1.275748 | 0.228554 |
| 8 | 123559600 | 123562199 | 12.14699 | 20.90322 | 55.76108 | 24.45973 | -1.02477 | 0.879502 | 0.228555 |
| 8 | 121872000 | 121875599 | 23.45753 | 38.33294 | 82.64562 | 35.53022 | -0.73112 | 1.483618 | 0.22856 |
| 10 | 24929200 | 24930999 | 11.17997 | 19.69958 | 42.48976 | 18.80672 | -0.71401 | 0.556577 | 0.228566 |
| 2 | 119126200 | 119128599 | 10.82629 | 18.85633 | 49.95216 | 21.96566 | -1.01117 | 0.736805 | 0.228566 |
| 17 | 18600400 | 18601999 | 41.98379 | 68.38054 | 58.54619 | 25.13924 | 0.594222 | 1.555862 | 0.228576 |
| 16 | 90314000 | 90325999 | 82.42448 | 126.0099 | 251.36 | 104.2764 | -0.70589 | 3.069309 | 0.228602 |
| 13 | 52268200 | 52270199 | 19.17367 | 32.1693 | 65.58716 | 28.62301 | -0.64316 | 1.146404 | 0.228618 |
| X | 48050000 | 48052199 | 30.68384 | 52.71688 | 35.30056 | 15.48391 | 0.960013 | 0.945417 | 0.228623 |
| 6 | 113303200 | 113306399 | 16.78846 | 28.74717 | 57.55134 | 25.23767 | -0.61433 | 0.945709 | 0.228624 |
| 10 | 56799800 | 56801199 | 12.95017 | 25.08293 | 14.78173 | 5.611657 | 1.323841 | -0.16402 | 0.228641 |
| 5 | 3207000 | 3212799 | 64.49243 | 103.0924 | 87.67992 | 36.86712 | 0.575546 | 2.080174 | 0.228671 |
| 15 | 64829800 | 64831599 | 20.91287 | 37.44442 | 24.56975 | 10.77163 | 1.004523 | 0.455263 | 0.228671 |
| 17 | 29708600 | 29711999 | 14.80809 | 24.90432 | 64.07703 | 27.99557 | -0.97703 | 1.109078 | 0.228674 |
| 17 | 39857400 | 39862399 | 74.48103 | 119.2889 | 71.36246 | 30.08931 | 1.085173 | 2.029664 | 0.228675 |
| 17 | 45895400 | 45901599 | 37.34926 | 59.50837 | 139.82 | 58.52128 | -0.89297 | 2.170649 | 0.228679 |
| 1 | 32101000 | 32103199 | 25.0727 | 43.69619 | 32.04884 | 14.09837 | 0.832306 | 0.769257 | 0.228685 |

| | | | | | | | | | |
|---|---|---|---|---|---|---|---|---|---|
| 11 | 109913200 | 109913999 | 11.96689 | 22.97358 | 18.92835 | 7.418884 | 0.803707 | -0.05975 | 0.228691 |
| 3 | 123013600 | 123014799 | 5.736395 | 10.96065 | 28.89677 | 11.47825 | -0.8774 | -0.05067 | 0.228692 |
| 11 | 105259800 | 105260799 | 5.736395 | 10.96065 | 28.89677 | 11.47825 | -0.8774 | -0.05067 | 0.228692 |
| 8 | 60005200 | 60006599 | 21.12696 | 36.98432 | 28.51979 | 12.58612 | 0.760788 | 0.679917 | 0.228693 |
| 1 | 151303600 | 151304999 | 6.244462 | 11.39795 | 36.60651 | 15.69905 | -1.24986 | 0.263987 | 0.228693 |
| 5 | 103839600 | 103841599 | 16.74485 | 28.33532 | 61.14781 | 26.76566 | -0.72418 | 1.056956 | 0.228704 |
| 9 | 20571600 | 20576199 | 23.29351 | 38.08157 | 82.70348 | 35.56076 | -0.74115 | 1.476658 | 0.228736 |
| 17 | 26894200 | 26895999 | 7.479675 | 13.78914 | 34.11645 | 14.43731 | -0.86311 | 0.210105 | 0.22874 |
| 13 | 37452400 | 37453399 | 9.937701 | 19.34006 | 15.9628 | 5.874595 | 0.884994 | -0.29923 | 0.228763 |
| 2 | 152783000 | 152786399 | 20.15906 | 33.28783 | 74.46569 | 32.15737 | -0.77746 | 1.337081 | 0.228763 |
| 1 | 87254200 | 87255599 | 7.018232 | 13.01151 | 33.67582 | 14.14208 | -0.91946 | 0.172789 | 0.228774 |
| 7 | 10384200 | 10386599 | 48.33807 | 78.61374 | 58.35194 | 25.04234 | 0.798208 | 1.59044 | 0.228781 |
| 7 | 8789600 | 8790999 | 11.61801 | 22.58101 | 15.09255 | 5.612508 | 1.17197 | -0.24651 | 0.228786 |
| 9 | 11556400 | 11557399 | 17.89354 | 31.77956 | 25.98617 | 11.47076 | 0.682093 | 0.485541 | 0.228789 |
| 12 | 52776200 | 52777399 | 10.0237 | 19.48281 | 14.62684 | 5.438122 | 1.005562 | -0.26991 | 0.228791 |
| 18 | 58792200 | 58793199 | 7.454963 | 13.81048 | 33.66751 | 14.15351 | -0.83505 | 0.180773 | 0.228794 |
| 1 | 29628200 | 29630399 | 17.00168 | 30.86651 | 24.46153 | 10.56137 | 0.747979 | 0.34677 | 0.228798 |
| 6 | 11255000 | 11257999 | 36.96233 | 61.49121 | 45.99692 | 19.9659 | 0.798547 | 1.270372 | 0.228803 |
| 8 | 126534600 | 126535799 | 6.285612 | 11.69657 | 33.19231 | 13.86433 | -1.04497 | 0.143908 | 0.228816 |
| 16 | 45664000 | 45665199 | 6.285612 | 11.69657 | 33.19231 | 13.86433 | -1.04497 | 0.143908 | 0.228816 |
| 7 | 114344000 | 114345999 | 10.87705 | 19.15203 | 43.01396 | 19.04148 | -0.77248 | 0.563137 | 0.228819 |
| 12 | 57621800 | 57623599 | 10.87705 | 19.15203 | 43.01396 | 19.04148 | -0.77248 | 0.563137 | 0.228819 |
| 4 | 138518600 | 138525399 | 44.63318 | 70.96545 | 138.1679 | 57.66737 | -0.62559 | 2.243277 | 0.228823 |
| 5 | 93189600 | 93194199 | 37.9383 | 60.56268 | 130.1638 | 54.62513 | -0.76163 | 2.105868 | 0.228826 |
| 14 | 84054800 | 84056599 | 22.7007 | 39.92624 | 28.13237 | 12.45646 | 0.890081 | 0.615814 | 0.228831 |
| 12 | 24316400 | 24320399 | 68.39182 | 109.0086 | 93.96349 | 39.34831 | 0.55173 | 2.174523 | 0.228855 |
| 14 | 6219800 | 6220599 | 4.111741 | 8.007923 | 23.56502 | 8.613316 | -0.91909 | -0.3696 | 0.228858 |
| 12 | 19628600 | 19637999 | 177.7221 | 267.6186 | 197.9784 | 81.77312 | 0.705875 | 3.394194 | 0.228862 |
| 3 | 16354200 | 16357999 | 60.37753 | 96.72303 | 82.07309 | 34.61749 | 0.58228 | 2.006678 | 0.22887 |
| 2 | 150038800 | 150041199 | 74.62372 | 118.3014 | 105.7677 | 44.0148 | 0.491314 | 2.346313 | 0.228875 |
| 5 | 86694400 | 86696399 | 20.15439 | 33.76571 | 66.1381 | 28.83704 | -0.58553 | 1.16691 | 0.228879 |
| 9 | 108746800 | 108748599 | 7.15972 | 12.91207 | 37.5207 | 16.33538 | -1.12114 | 0.345113 | 0.228891 |
| 5 | 76366200 | 76367399 | 11.99268 | 23.28855 | 15.6606 | 5.865627 | 1.154379 | -0.21569 | 0.228891 |
| 3 | 83692600 | 83697399 | 30.91907 | 50.07937 | 95.0996 | 40.69824 | -0.55903 | 1.65397 | 0.228895 |
| 13 | 13108000 | 13109199 | 13.62674 | 25.96303 | 17.91557 | 7.159231 | 1.031105 | 0.013451 | 0.228897 |
| 8 | 83590000 | 83591199 | 7.444061 | 13.82723 | 32.63681 | 13.67035 | -0.78463 | 0.164559 | 0.228897 |
| 19 | 4875400 | 4876799 | 7.444061 | 13.82723 | 32.63681 | 13.67035 | -0.78463 | 0.164559 | 0.228897 |
| 13 | 78708400 | 78711799 | 34.19747 | 57.58866 | 41.39947 | 18.0948 | 0.855436 | 1.136223 | 0.228899 |
| 2 | 28850800 | 28861799 | 68.57388 | 105.9262 | 213.1279 | 88.52882 | -0.70671 | 2.859712 | 0.228914 |
| 15 | 50482600 | 50483799 | 7.403105 | 13.87672 | 31.75652 | 13.10205 | -0.72299 | 0.103106 | 0.228916 |
| 10 | 82401400 | 82411399 | 198.4953 | 295.8793 | 244.9294 | 100.9347 | 0.530698 | 3.590707 | 0.228919 |
| 19 | 15961800 | 15962599 | 5.754462 | 10.8923 | 29.7335 | 12.04777 | -0.95187 | 0.02029 | 0.228928 |
| 14 | 122687000 | 122692599 | 42.17235 | 67.15962 | 134.4571 | 56.23802 | -0.66329 | 2.188857 | 0.22893 |
| X | 157934400 | 157935399 | 13.86864 | 26.04479 | 19.90271 | 8.18541 | 0.851422 | 0.118643 | 0.228933 |
| 15 | 88938600 | 88940399 | 9.655895 | 17.0522 | 42.00278 | 18.58286 | -0.90292 | 0.521464 | 0.228943 |
| 16 | 33928800 | 33934199 | 26.37272 | 42.9116 | 87.97242 | 37.76095 | -0.66323 | 1.55455 | 0.228951 |
| 3 | 85535400 | 85537599 | 34.53503 | 57.16829 | 46.11863 | 19.95016 | 0.689592 | 1.322541 | 0.228952 |

| | | | | | | | | | |
|---|---|---|---|---|---|---|---|---|---|
| 7 | 41465200 | 41472199 | 133.9279 | 205.2386 | 165.4561 | 68.67664 | 0.602712 | 3.031911 | 0.228954 |
| 7 | 5600600 | 5607399 | 103.7343 | 162.8668 | 115.1584 | 47.7628 | 0.818418 | 2.578603 | 0.228958 |
| 8 | 100591800 | 100592999 | 11.62173 | 22.57148 | 15.67288 | 5.863348 | 1.109925 | -0.22434 | 0.228964 |
| 15 | 26894400 | 26896199 | 10.92289 | 19.12437 | 45.71459 | 20.17708 | -0.86294 | 0.652222 | 0.228968 |
| 7 | 38380200 | 38381199 | 13.33026 | 25.8736 | 15.0706 | 5.665253 | 1.355456 | -0.18871 | 0.228978 |
| 8 | 127799800 | 127800399 | 4.107821 | 8.011194 | 23.26151 | 8.35206 | -0.8769 | -0.43595 | 0.228984 |
| 7 | 9164800 | 9173399 | 106.9447 | 167.8532 | 112.353 | 46.59731 | 0.897061 | 2.588086 | 0.22899 |
| 14 | 33666600 | 33670599 | 25.79917 | 41.85095 | 92.56968 | 39.66046 | -0.77632 | 1.611746 | 0.228994 |
| 16 | 91074400 | 91078399 | 25.77395 | 41.8578 | 88.55583 | 37.97271 | -0.71069 | 1.587283 | 0.228996 |
| 2 | 31249000 | 31252999 | 25.02004 | 40.54714 | 91.33751 | 39.10416 | -0.80381 | 1.631421 | 0.228997 |
| 15 | 75712200 | 75713999 | 7.642214 | 14.04466 | 35.6679 | 15.15841 | -0.90533 | 0.231622 | 0.228998 |
| 10 | 75509800 | 75512399 | 9.90965 | 17.62918 | 39.42518 | 17.3786 | -0.75905 | 0.454127 | 0.22901 |
| 4 | 5661800 | 5664599 | 17.61667 | 30.0112 | 59.69811 | 26.16828 | -0.60634 | 0.998253 | 0.229013 |
| 18 | 64471600 | 64479799 | 74.08651 | 114.4442 | 207.8278 | 86.32726 | -0.55894 | 2.861586 | 0.229039 |
| 1 | 136610200 | 136611599 | 8.096922 | 14.84071 | 34.55493 | 14.73876 | -0.78448 | 0.249887 | 0.229041 |
| 19 | 39861400 | 39864399 | 31.4434 | 53.29827 | 39.73319 | 17.40099 | 0.80338 | 1.068848 | 0.229047 |
| 14 | 118461200 | 118469599 | 58.26154 | 91.13031 | 194.2333 | 80.57939 | -0.77566 | 2.631492 | 0.229047 |
| 18 | 38289200 | 38291599 | 11.17981 | 19.67888 | 43.5214 | 19.26734 | -0.75068 | 0.568792 | 0.229058 |
| 12 | 115335600 | 115336999 | 27.66818 | 46.94977 | 40.54865 | 17.76394 | 0.592084 | 1.048581 | 0.229081 |
| 4 | 27296600 | 27297599 | 18.5098 | 31.26318 | 62.28549 | 27.24812 | -0.60951 | 1.082244 | 0.229084 |
| X | 15332800 | 15333599 | 16.53774 | 30.66028 | 19.49777 | 8.188042 | 1.090752 | 0.217286 | 0.229089 |
| 2 | 155614200 | 155618199 | 25.73486 | 41.8873 | 86.65282 | 37.19892 | -0.67587 | 1.548284 | 0.229112 |
| 1 | 186890800 | 186893399 | 29.01654 | 49.39256 | 39.79748 | 17.44386 | 0.692199 | 1.023802 | 0.229119 |
| 1 | 116375800 | 116377599 | 20.4414 | 35.89108 | 30.14545 | 13.3371 | 0.63784 | 0.630415 | 0.229131 |
| 13 | 11696800 | 11697799 | 8.121027 | 14.81278 | 36.06209 | 15.47921 | -0.85492 | 0.280194 | 0.229143 |
| 10 | 106051000 | 106052199 | 10.32815 | 18.45996 | 39.41609 | 17.30612 | -0.68906 | 0.425168 | 0.229167 |
| 13 | 54856200 | 54857399 | 10.32815 | 18.45996 | 39.41609 | 17.30612 | -0.68906 | 0.425168 | 0.229167 |
| 16 | 3330200 | 3335599 | 88.81985 | 139.4225 | 124.9009 | 51.80222 | 0.47724 | 2.574806 | 0.229171 |
| 3 | 84249000 | 84253199 | 20.76757 | 34.33598 | 74.12933 | 32.04005 | -0.72628 | 1.322828 | 0.22919 |
| 4 | 35464400 | 35466199 | 17.93742 | 31.90024 | 26.47837 | 11.67482 | 0.661765 | 0.47893 | 0.229202 |
| X | 73547600 | 73550599 | 31.53329 | 53.5052 | 39.91128 | 17.48439 | 0.802814 | 1.065115 | 0.229206 |
| 1 | 109429800 | 109430399 | 7.557283 | 14.12666 | 32.79747 | 13.59494 | -0.74936 | 0.123686 | 0.22923 |
| 3 | 143927400 | 143929799 | 12.59344 | 22.09506 | 45.1215 | 19.95026 | -0.63726 | 0.626031 | 0.229259 |
| 10 | 64229000 | 64230999 | 16.32674 | 30.10388 | 21.80532 | 9.225538 | 0.896241 | 0.248877 | 0.229265 |
| 7 | 56048800 | 56051599 | 28.9058 | 49.44384 | 38.74835 | 16.99066 | 0.733195 | 0.981325 | 0.229273 |
| 10 | 22021200 | 22027999 | 53.454 | 84.24071 | 163.926 | 68.02277 | -0.63601 | 2.480926 | 0.229283 |
| 14 | 7988400 | 7990199 | 30.88674 | 52.17848 | 38.90573 | 17.02119 | 0.803063 | 1.095799 | 0.229291 |
| 7 | 12771000 | 12772999 | 15.74359 | 26.82255 | 59.0433 | 25.88143 | -0.75175 | 0.989549 | 0.229315 |
| 17 | 38663200 | 38673399 | 324.7323 | 463.6455 | 380.9739 | 157.4348 | 0.481668 | 4.273049 | 0.229321 |
| 6 | 52201000 | 52204199 | 13.13553 | 22.40707 | 58.33862 | 25.5751 | -0.9922 | 0.968775 | 0.229323 |
| 1 | 62681000 | 62683199 | 22.87058 | 39.79693 | 34.01699 | 14.94966 | 0.612465 | 0.783596 | 0.229328 |
| 15 | 43080400 | 43081599 | 9.090143 | 16.3716 | 37.46295 | 16.33748 | -0.78088 | 0.367726 | 0.229329 |
| 5 | 14996600 | 14998399 | 27.08552 | 46.44995 | 38.19688 | 16.75233 | 0.664454 | 0.948674 | 0.229337 |
| X | 151749600 | 151751399 | 19.21 | 34.54334 | 24.45412 | 10.68222 | 0.899378 | 0.418578 | 0.229339 |
| 5 | 145236800 | 145239999 | 31.95628 | 53.915 | 41.42999 | 18.11839 | 0.759654 | 1.111796 | 0.229346 |
| 3 | 133054600 | 133056799 | 14.49428 | 25.18952 | 51.10754 | 22.44271 | -0.62917 | 0.786908 | 0.229351 |
| 3 | 99676400 | 99677399 | 12.68229 | 24.5046 | 16.5841 | 6.367618 | 1.111776 | -0.12487 | 0.229353 |

| | | | | | | | | | |
|---|---|---|---|---|---|---|---|---|---|
| 13 | 12937000 | 12937399 | 10.87626 | 20.95142 | 17.89782 | 6.946377 | 0.764644 | -0.10653 | 0.229354 |
| 5 | 94467200 | 94469199 | 26.92728 | 47.05222 | 28.53468 | 12.5733 | 1.105408 | 0.737903 | 0.229354 |
| 7 | 32193800 | 32195599 | 26.43467 | 45.68817 | 32.94871 | 14.45494 | 0.855089 | 0.87145 | 0.229363 |
| 17 | 84389000 | 84394199 | 25.24405 | 40.80364 | 97.88462 | 41.81369 | -0.89815 | 1.692615 | 0.229364 |
| 18 | 80219000 | 80219999 | 9.544667 | 17.14867 | 37.92461 | 16.57831 | -0.73476 | 0.384364 | 0.229369 |
| 17 | 85264400 | 85266199 | 10.4374 | 18.34962 | 43.55649 | 19.28759 | -0.85265 | 0.57904 | 0.229371 |
| 1 | 18031600 | 18036799 | 80.16321 | 127.0273 | 106.8472 | 44.44446 | 0.578591 | 2.362854 | 0.229375 |
| 7 | 86585600 | 86586799 | 15.38322 | 28.1416 | 23.03119 | 9.8492 | 0.710453 | 0.28747 | 0.229376 |
| 19 | 59332600 | 59334799 | 9.624324 | 16.79447 | 49.91798 | 21.97665 | -1.17577 | 0.703484 | 0.229399 |
| X | 32753800 | 32755999 | 22.7197 | 41.82502 | 17.08464 | 7.243914 | 1.714419 | 0.299967 | 0.22941 |
| 7 | 33417800 | 33419199 | 35.06255 | 58.60991 | 45.93365 | 19.99919 | 0.731527 | 1.21716 | 0.22941 |
| 2 | 91924000 | 91924399 | 61.42278 | 99.23479 | 43.58653 | 18.60764 | 1.543706 | 1.736006 | 0.229417 |
| 3 | 130693000 | 130695399 | 19.06453 | 31.88609 | 65.95813 | 28.72804 | -0.66384 | 1.181963 | 0.229424 |
| 7 | 42687800 | 42689999 | 28.09751 | 47.69858 | 40.994 | 17.96135 | 0.599093 | 1.046173 | 0.229424 |
| 7 | 41867800 | 41873199 | 27.6092 | 47.86224 | 31.18393 | 13.68197 | 1.001694 | 0.848315 | 0.229426 |
| 3 | 25360200 | 25361799 | 20.49347 | 37.04465 | 21.73228 | 9.434097 | 1.173662 | 0.389892 | 0.229434 |
| 9 | 21913000 | 21928799 | 80.50743 | 122.7977 | 251.3698 | 104.2252 | -0.74575 | 3.116206 | 0.229437 |
| 9 | 120029600 | 120039399 | 77.08842 | 118.9458 | 211.4209 | 87.82025 | -0.52932 | 2.883948 | 0.229452 |
| 17 | 63603400 | 63605199 | 20.31315 | 35.92692 | 27.81672 | 12.30296 | 0.757846 | 0.542088 | 0.229465 |
| 7 | 30852600 | 30856599 | 23.96877 | 39.06988 | 84.03262 | 36.09336 | -0.73024 | 1.523144 | 0.229465 |
| 7 | 136010200 | 136012199 | 9.329332 | 16.93319 | 38.09272 | 16.45178 | -0.74801 | 0.321273 | 0.229467 |
| 7 | 106309400 | 106310799 | 15.53894 | 28.75483 | 21.94737 | 9.240176 | 0.826352 | 0.223337 | 0.229507 |
| 7 | 60095000 | 60096799 | 22.30148 | 39.35907 | 26.44541 | 11.70146 | 0.959921 | 0.573476 | 0.229507 |
| 14 | 118784400 | 118786199 | 17.34662 | 29.44008 | 59.58172 | 26.10342 | -0.63174 | 1.03233 | 0.229508 |
| 5 | 118831000 | 118833399 | 13.77876 | 23.92906 | 52.65934 | 23.11384 | -0.74563 | 0.799624 | 0.229516 |
| 5 | 145510600 | 145515799 | 147.3781 | 223.7179 | 192.0068 | 79.50142 | 0.501569 | 3.227597 | 0.229518 |
| 7 | 28290600 | 28297599 | 32.00181 | 51.14508 | 124.6731 | 52.40663 | -0.9409 | 2.063986 | 0.229522 |
| 17 | 5386200 | 5393399 | 51.60009 | 81.58839 | 152.3919 | 63.31911 | -0.57315 | 2.404114 | 0.229546 |
| 1 | 58885000 | 58885999 | 6.734437 | 12.88799 | 31.4181 | 12.42571 | -0.76045 | -0.04968 | 0.229566 |
| 2 | 170316000 | 170316799 | 7.28634 | 13.56079 | 34.3248 | 14.33374 | -0.88186 | 0.152352 | 0.229566 |
| 14 | 40812400 | 40813799 | 7.710228 | 14.44074 | 32.29014 | 13.34046 | -0.6914 | 0.112981 | 0.229567 |
| 3 | 17236400 | 17236999 | 14.51886 | 26.7802 | 22.73081 | 9.61233 | 0.669962 | 0.235631 | 0.229571 |
| 8 | 15871600 | 15872999 | 14.51886 | 26.7802 | 22.73081 | 9.61233 | 0.669962 | 0.235631 | 0.229571 |
| 15 | 102279000 | 102281199 | 8.315547 | 15.03288 | 36.48698 | 15.8373 | -0.8607 | 0.337506 | 0.229589 |
| X | 155304600 | 155307599 | 32.3593 | 54.87522 | 40.31067 | 17.65628 | 0.824713 | 1.071515 | 0.229616 |
| X | 129487400 | 129491399 | 47.28688 | 78.02241 | 44.46391 | 19.19177 | 1.188996 | 1.3935 | 0.229648 |
| 8 | 103097200 | 103099599 | 16.2428 | 29.96245 | 21.22713 | 8.975474 | 0.928424 | 0.243947 | 0.22966 |
| 7 | 113309800 | 113310999 | 7.249239 | 13.62326 | 31.2599 | 12.84118 | -0.72148 | 0.084888 | 0.229661 |
| 10 | 31319600 | 31322599 | 19.49128 | 32.73749 | 64.48733 | 28.16222 | -0.59365 | 1.135808 | 0.229661 |
| 19 | 43848800 | 43849999 | 14.06971 | 26.53161 | 20.06233 | 8.193619 | 0.873549 | 0.092726 | 0.229664 |
| 17 | 38408000 | 38412999 | 65.80758 | 105.4647 | 78.7308 | 33.2283 | 0.767441 | 2.001403 | 0.229666 |
| 7 | 31576600 | 31581199 | 86.84154 | 136.827 | 117.5726 | 48.78652 | 0.541398 | 2.499683 | 0.229669 |
| 7 | 45359000 | 45362999 | 18.35384 | 30.50435 | 70.0803 | 30.39953 | -0.8148 | 1.252918 | 0.229693 |
| 5 | 142185400 | 142201599 | 112.6357 | 168.9596 | 321.6618 | 132.7229 | -0.66176 | 3.462309 | 0.229729 |
| 16 | 66267000 | 66272199 | 134.4297 | 204.8623 | 181.1896 | 75.11042 | 0.462821 | 3.143406 | 0.229731 |
| 17 | 40750200 | 40752399 | 26.12891 | 45.38292 | 31.45583 | 13.80807 | 0.912942 | 0.818361 | 0.22974 |
| 8 | 82447800 | 82448599 | 7.673985 | 14.01185 | 35.03767 | 15.02007 | -0.89155 | 0.270231 | 0.229749 |

| | | | | | | | | | |
|---|---|---|---|---|---|---|---|---|---|
| 13 | 58050000 | 58065799 | 136.8341 | 203.4172 | 349.9628 | 144.1789 | -0.52764 | 3.633195 | 0.229752 |
| 12 | 107089400 | 107094399 | 33.1168 | 53.27996 | 105.6046 | 44.82892 | -0.6317 | 1.8491 | 0.229753 |
| 6 | 45903600 | 45905199 | 13.02019 | 24.92341 | 18.72431 | 7.400171 | 0.924327 | -0.02402 | 0.229757 |
| 13 | 36913600 | 36916399 | 20.64122 | 34.05649 | 75.67296 | 32.66195 | -0.768 | 1.348149 | 0.229778 |
| 16 | 61041600 | 61044199 | 77.80339 | 123.0085 | 108.4417 | 45.05554 | 0.508351 | 2.415416 | 0.22978 |
| 6 | 125386800 | 125388999 | 13.38434 | 23.41268 | 47.28899 | 20.85594 | -0.62162 | 0.681879 | 0.229781 |
| 19 | 53890400 | 53892799 | 10.97579 | 19.12442 | 51.01764 | 22.43987 | -1.02132 | 0.732151 | 0.229785 |
| X | 3620000 | 3623799 | 88.45411 | 139.2137 | 112.141 | 46.526 | 0.633249 | 2.522709 | 0.229803 |
| 11 | 107251400 | 107252399 | 5.204348 | 9.902433 | 30.42182 | 12.1917 | -1.10732 | -0.02313 | 0.229818 |
| 7 | 88209800 | 88210399 | 12.80424 | 24.32086 | 17.74475 | 7.13734 | 0.943996 | 0.026842 | 0.229825 |
| 7 | 15378600 | 15383799 | 117.4517 | 181.5319 | 149.5323 | 62.11235 | 0.580982 | 2.862045 | 0.229835 |
| 4 | 106495800 | 106502799 | 54.93846 | 86.61211 | 160.5455 | 66.62318 | -0.56557 | 2.475765 | 0.229836 |
| 12 | 44758600 | 44760999 | 17.09247 | 30.98484 | 23.22271 | 10.04532 | 0.825833 | 0.351364 | 0.229846 |
| 4 | 123904000 | 123907199 | 19.93996 | 33.56823 | 66.30333 | 28.97692 | -0.59753 | 1.117703 | 0.22985 |
| 5 | 144424800 | 144427999 | 16.43813 | 28.27301 | 57.62005 | 25.27477 | -0.63883 | 0.903321 | 0.229851 |
| 6 | 140568200 | 140570799 | 18.81646 | 31.27928 | 69.51941 | 30.16026 | -0.76712 | 1.252725 | 0.229859 |
| 9 | 96787000 | 96787799 | 11.71098 | 22.22628 | 18.44845 | 7.433101 | 0.757501 | 0.02374 | 0.229864 |
| 1 | 14201800 | 14203799 | 29.806 | 50.30477 | 43.15988 | 18.87692 | 0.60134 | 1.104613 | 0.229868 |
| 13 | 25568800 | 25570199 | 21.33915 | 37.75111 | 27.18348 | 12.02228 | 0.862092 | 0.544946 | 0.229885 |
| 10 | 45210600 | 45212399 | 12.26442 | 21.54945 | 44.55854 | 19.72584 | -0.65515 | 0.598631 | 0.229897 |
| 13 | 4014400 | 4014999 | 4.063869 | 7.909717 | 23.44028 | 8.618552 | -0.93579 | -0.3463 | 0.229897 |
| 12 | 110393000 | 110399799 | 52.48324 | 82.73493 | 160.65 | 66.66589 | -0.63256 | 2.474741 | 0.229898 |
| 19 | 59507200 | 59509599 | 9.600341 | 17.08706 | 44.64995 | 19.67377 | -0.98264 | 0.454181 | 0.229905 |
| 7 | 121887200 | 121890199 | 20.06747 | 33.01499 | 79.96634 | 34.47319 | -0.89374 | 1.385122 | 0.229907 |
| 11 | 115039800 | 115046599 | 48.51141 | 76.6894 | 154.6893 | 64.26724 | -0.68413 | 2.406118 | 0.229913 |
| X | 4576400 | 4584199 | 126.7255 | 195.4306 | 148.4856 | 61.67831 | 0.695 | 2.907265 | 0.229919 |
| 1 | 37191000 | 37192999 | 9.155322 | 16.82993 | 38.45166 | 16.33551 | -0.75375 | 0.249324 | 0.229921 |
| 7 | 85169400 | 85174999 | 197.6585 | 293.5689 | 250.899 | 103.3566 | 0.479882 | 3.654001 | 0.229924 |
| 13 | 51253800 | 51261199 | 59.828 | 94.20138 | 164.9377 | 68.43459 | -0.4849 | 2.503507 | 0.229928 |
| X | 24039000 | 24041199 | 50.81154 | 82.23007 | 63.87502 | 27.31867 | 0.726465 | 1.691857 | 0.229951 |
| 9 | 63767000 | 63770999 | 17.68821 | 29.49788 | 67.9246 | 29.53009 | -0.81788 | 1.211178 | 0.229953 |
| 4 | 145550000 | 145550999 | 13.33223 | 25.3357 | 19.84739 | 7.975101 | 0.843907 | 0.035498 | 0.229954 |
| 7 | 11970400 | 11973199 | 61.57173 | 98.69745 | 81.49853 | 34.40766 | 0.622576 | 1.991078 | 0.229965 |
| 2 | 167263200 | 167264199 | 6.392019 | 11.97553 | 31.71811 | 13.09726 | -0.93273 | 0.097261 | 0.229967 |
| 3 | 134637400 | 134638199 | 6.392019 | 11.97553 | 31.71811 | 13.09726 | -0.93273 | 0.097261 | 0.229967 |
| 9 | 62413600 | 62414999 | 6.392019 | 11.97553 | 31.71811 | 13.09726 | -0.93273 | 0.097261 | 0.229967 |
| 4 | 85649600 | 85651999 | 27.02901 | 46.74238 | 35.43691 | 15.54682 | 0.783497 | 0.872741 | 0.229983 |
| 9 | 120465800 | 120466799 | 11.05294 | 21.47009 | 16.38148 | 6.120838 | 0.977483 | -0.23254 | 0.229983 |
| 7 | 72527400 | 72528999 | 23.80127 | 41.31621 | 35.04974 | 15.38009 | 0.623 | 0.822759 | 0.230005 |
| 14 | 3426200 | 3431199 | 45.76697 | 72.78888 | 136.0975 | 56.82126 | -0.56689 | 2.235649 | 0.230026 |
| X | 20444600 | 20447199 | 27.18202 | 47.76819 | 24.56407 | 10.87582 | 1.341562 | 0.6469 | 0.230036 |
| 7 | 44090800 | 44099399 | 53.14553 | 83.34232 | 175.1894 | 72.65962 | -0.7527 | 2.580865 | 0.230044 |
| 5 | 9502200 | 9503399 | 13.9944 | 26.0091 | 18.94032 | 7.927294 | 0.900195 | 0.180752 | 0.230053 |
| 5 | 73979200 | 73981199 | 14.19179 | 24.2315 | 57.89535 | 25.38268 | -0.86878 | 0.962832 | 0.230073 |
| 19 | 5587800 | 5592199 | 22.04253 | 36.03659 | 84.25453 | 36.22775 | -0.84725 | 1.47442 | 0.230083 |
| 5 | 52769800 | 52777999 | 65.32893 | 101.9784 | 183.9105 | 76.31275 | -0.53719 | 2.669745 | 0.230085 |
| 16 | 93131600 | 93138799 | 78.3656 | 120.9877 | 209.9784 | 87.2208 | -0.49427 | 2.873541 | 0.230119 |

| | | | | | | | | | |
|---|---|---|---|---|---|---|---|---|---|
| 11 | 55409800 | 55410199 | 4.228214 | 8.249599 | 22.74915 | 8.119691 | -0.79534 | -0.45648 | 0.230121 |
| 3 | 11499800 | 11501399 | 18.88776 | 33.387 | 27.46701 | 12.14966 | 0.670387 | 0.537575 | 0.230131 |
| 18 | 5170200 | 5173199 | 33.69117 | 56.68368 | 43.10375 | 18.83443 | 0.774739 | 1.145414 | 0.230138 |
| 2 | 74469600 | 74470399 | 8.479221 | 15.73626 | 35.95703 | 15.079 | -0.74033 | 0.183135 | 0.23014 |
| 3 | 151237200 | 151237999 | 13.95489 | 26.74107 | 17.46524 | 6.881494 | 1.128284 | -0.02782 | 0.230145 |
| 13 | 15219200 | 15220399 | 13.95489 | 26.74107 | 17.46524 | 6.881494 | 1.128284 | -0.02782 | 0.230145 |
| 2 | 134407200 | 134411599 | 59.385 | 95.2747 | 80.30567 | 33.94875 | 0.594438 | 1.96097 | 0.23015 |
| 1 | 25515600 | 25516799 | 13.57773 | 25.61483 | 20.7829 | 8.480759 | 0.774159 | 0.088456 | 0.230152 |
| 1 | 110137800 | 110139399 | 29.3227 | 49.36366 | 42.71416 | 18.6676 | 0.589022 | 1.124327 | 0.230155 |
| 7 | 5167600 | 5168799 | 12.79986 | 24.23103 | 20.39951 | 8.263627 | 0.730619 | 0.057401 | 0.230163 |
| 13 | 93155400 | 93159799 | 34.19869 | 55.12209 | 105.1893 | 44.77103 | -0.57385 | 1.791987 | 0.230166 |
| 5 | 91862400 | 91863399 | 6.156896 | 11.77885 | 28.98279 | 11.47338 | -0.77446 | -0.05784 | 0.23017 |
| 9 | 65734000 | 65735199 | 8.528024 | 15.64484 | 36.17875 | 15.41261 | -0.77396 | 0.249531 | 0.230171 |
| 7 | 134781400 | 134784199 | 14.27135 | 24.53081 | 57.50691 | 25.22466 | -0.84003 | 0.899266 | 0.230183 |
| 2 | 121892800 | 121893399 | 3.642083 | 7.10713 | 21.45613 | 7.637369 | -0.91875 | -0.47601 | 0.23019 |
| 1 | 19202400 | 19203199 | 12.30065 | 23.41108 | 19.93477 | 7.984381 | 0.728491 | 0.015227 | 0.230193 |
| 3 | 66948200 | 66949399 | 17.07541 | 30.99478 | 24.52639 | 10.59176 | 0.750046 | 0.349126 | 0.230196 |
| 4 | 60046400 | 60048399 | 33.59719 | 56.17084 | 46.48811 | 20.24297 | 0.653167 | 1.211454 | 0.230211 |
| 12 | 32254000 | 32256399 | 25.65063 | 41.54901 | 91.20228 | 39.03238 | -0.76719 | 1.64353 | 0.230215 |
| 4 | 114757000 | 114760599 | 27.19233 | 44.18981 | 87.19652 | 37.40763 | -0.60958 | 1.576395 | 0.230218 |
| 6 | 70933000 | 70934799 | 13.29058 | 23.03184 | 51.06249 | 22.40346 | -0.75693 | 0.815412 | 0.230222 |
| 12 | 91554800 | 91558799 | 20.3978 | 33.51997 | 81.07793 | 34.93543 | -0.89246 | 1.400985 | 0.230245 |
| 18 | 54161600 | 54162799 | 13.3487 | 25.36779 | 19.72983 | 7.927331 | 0.853868 | 0.033937 | 0.230246 |
| 9 | 68156200 | 68157799 | 10.54616 | 18.64783 | 41.44395 | 18.33102 | -0.7549 | 0.51152 | 0.230251 |
| 16 | 44703000 | 44704999 | 24.12996 | 42.2867 | 29.85551 | 13.18755 | 0.887229 | 0.681073 | 0.230257 |
| 13 | 23174200 | 23174999 | 10.5403 | 20.36811 | 18.05188 | 6.927609 | 0.727757 | -0.14487 | 0.230333 |
| 8 | 125811200 | 125818599 | 48.81191 | 77.22112 | 152.2427 | 63.27427 | -0.65036 | 2.388399 | 0.230344 |
| 7 | 90075600 | 90076799 | 9.542677 | 17.17742 | 37.43096 | 16.33447 | -0.71171 | 0.37265 | 0.230345 |
| 1 | 112459200 | 112460399 | 19.43423 | 34.51885 | 24.41371 | 10.77587 | 0.890401 | 0.493095 | 0.230362 |
| 5 | 150910400 | 150913799 | 24.23168 | 39.64581 | 80.50047 | 34.62321 | -0.64358 | 1.46769 | 0.230377 |
| 16 | 37721600 | 37722199 | 4.519363 | 8.811844 | 25.21356 | 9.079225 | -0.86077 | -0.41482 | 0.230385 |
| 12 | 68492000 | 68492799 | 6.585265 | 12.60722 | 30.83536 | 12.18217 | -0.76369 | -0.05648 | 0.230389 |
| 7 | 8977400 | 8979399 | 16.148 | 31.18092 | 13.30284 | 5.123146 | 1.765745 | -0.08627 | 0.230392 |
| 3 | 122788000 | 122792399 | 58.81096 | 94.53623 | 75.31907 | 31.93273 | 0.678855 | 1.900191 | 0.23042 |
| X | 147433200 | 147434599 | 14.52957 | 27.82459 | 17.4439 | 6.885665 | 1.184611 | -0.01599 | 0.230435 |
| 3 | 104418600 | 104419799 | 8.28577 | 14.69587 | 40.49778 | 17.89691 | -1.0606 | 0.464891 | 0.230447 |
| 18 | 70871000 | 70873599 | 25.30544 | 44.15365 | 31.47333 | 13.85543 | 0.873435 | 0.75401 | 0.230457 |
| 7 | 41898600 | 41906399 | 249.4241 | 364.3655 | 304.072 | 125.2752 | 0.49179 | 3.929787 | 0.230519 |
| 17 | 28931800 | 28933799 | 12.29062 | 21.60699 | 44.56015 | 19.73215 | -0.65112 | 0.592767 | 0.230536 |
| X | 33343400 | 33345799 | 29.02256 | 50.698 | 26.78189 | 11.7987 | 1.30342 | 0.751046 | 0.230553 |
| 13 | 37884000 | 37887399 | 34.2508 | 55.08632 | 105.6259 | 44.81666 | -0.58459 | 1.861583 | 0.230557 |
| 19 | 36381400 | 36383799 | 19.962 | 35.06053 | 29.71576 | 13.15054 | 0.624852 | 0.622095 | 0.230565 |
| 10 | 79078600 | 79079999 | 23.62875 | 41.42508 | 29.22097 | 12.91167 | 0.888308 | 0.669575 | 0.230565 |
| 17 | 85815800 | 85822599 | 39.43436 | 63.23909 | 120.1483 | 50.7582 | -0.57661 | 1.962902 | 0.230607 |
| 12 | 39048400 | 39048999 | 3.524971 | 6.888739 | 20.61648 | 7.14052 | -0.87087 | -0.58026 | 0.230614 |
| 13 | 43263000 | 43268599 | 50.70152 | 80.36931 | 146.4163 | 60.92441 | -0.53422 | 2.339655 | 0.230624 |
| 10 | 5790200 | 5792999 | 12.89774 | 22.58265 | 46.74101 | 20.63059 | -0.65672 | 0.662338 | 0.230626 |

| | | | | | | | | | |
|---|---|---|---|---|---|---|---|---|---|
| 1 | 136342800 | 136343399 | 3.601427 | 7.009586 | 26.45719 | 9.728445 | -1.28242 | -0.34763 | 0.230647 |
| 4 | 17717400 | 17717999 | 5.107183 | 9.914083 | 26.27324 | 9.868987 | -0.80786 | -0.25316 | 0.230658 |
| 19 | 56390800 | 56399399 | 87.57915 | 133.5107 | 269.1186 | 111.5702 | -0.72421 | 3.130337 | 0.230661 |
| 13 | 81823400 | 81825999 | 21.84244 | 38.25178 | 31.10468 | 13.7314 | 0.68438 | 0.680029 | 0.230668 |
| 13 | 11403000 | 11404599 | 15.70237 | 29.0388 | 22.5791 | 9.514372 | 0.799219 | 0.230796 | 0.230673 |
| 11 | 69870800 | 69872399 | 7.331165 | 13.18817 | 38.51437 | 16.81842 | -1.13146 | 0.362433 | 0.230677 |
| 6 | 86501000 | 86504399 | 15.16135 | 25.75794 | 58.90693 | 25.81579 | -0.80706 | 1.015325 | 0.230688 |
| 7 | 85801600 | 85802599 | 21.75486 | 37.98143 | 32.4472 | 14.28957 | 0.613662 | 0.729757 | 0.230688 |
| 2 | 8203000 | 8204799 | 10.82422 | 19.22687 | 40.39171 | 17.82753 | -0.67105 | 0.469986 | 0.230709 |
| 11 | 104514600 | 104517199 | 13.8546 | 24.16657 | 48.92687 | 21.53486 | -0.62541 | 0.728591 | 0.23071 |
| 7 | 67600000 | 67603999 | 21.99754 | 35.73894 | 90.17693 | 38.67914 | -0.96347 | 1.572886 | 0.230725 |
| 8 | 14779000 | 14783199 | 20.36719 | 33.53332 | 80.13425 | 34.55658 | -0.87396 | 1.376071 | 0.230747 |
| 6 | 91828400 | 91829799 | 8.859038 | 16.17527 | 35.54296 | 15.23708 | -0.70694 | 0.279403 | 0.23076 |
| 15 | 99534200 | 99535199 | 4.406711 | 8.557671 | 25.02127 | 9.369062 | -0.9428 | -0.27514 | 0.230781 |
| 14 | 6103400 | 6105799 | 16.70245 | 27.9945 | 66.40815 | 28.96262 | -0.86067 | 1.15071 | 0.230784 |
| 10 | 84817400 | 84818199 | 5.555263 | 10.65474 | 27.87198 | 10.95388 | -0.85079 | -0.08753 | 0.230792 |
| 18 | 81632800 | 81633999 | 7.295307 | 13.55528 | 32.64164 | 13.6664 | -0.81285 | 0.16104 | 0.230793 |
| X | 25479400 | 25484199 | 55.77541 | 90.74952 | 50.23216 | 21.56113 | 1.220666 | 1.594596 | 0.230794 |
| 8 | 21688800 | 21691799 | 28.46635 | 49.03459 | 35.90672 | 15.75104 | 0.832227 | 0.913503 | 0.230797 |
| 14 | 17909400 | 17911799 | 13.68309 | 25.51544 | 21.77297 | 9.062353 | 0.67918 | 0.163904 | 0.230797 |
| X | 33142000 | 33143799 | 15.66462 | 30.08899 | 14.99596 | 5.868269 | 1.523162 | -0.03792 | 0.2308 |
| 2 | 113328600 | 113329999 | 8.323445 | 15.45217 | 35.96584 | 15.07604 | -0.76635 | 0.179628 | 0.23082 |
| 3 | 4646400 | 4650599 | 60.60118 | 96.90119 | 83.01613 | 34.92091 | 0.565544 | 2.066404 | 0.230821 |
| 4 | 26787800 | 26789799 | 22.51695 | 39.60899 | 28.13395 | 12.45678 | 0.878611 | 0.612978 | 0.230823 |
| 14 | 3439200 | 3440799 | 16.6932 | 31.67008 | 17.09567 | 6.893549 | 1.371495 | 0.07024 | 0.230824 |
| 18 | 63840400 | 63842199 | 11.93296 | 21.10257 | 44.01937 | 19.46968 | -0.66425 | 0.522959 | 0.230844 |
| 1 | 181669200 | 181670799 | 11.21557 | 19.65816 | 45.16565 | 19.95312 | -0.80619 | 0.634462 | 0.230847 |
| X | 42965600 | 42968999 | 26.90848 | 47.00519 | 29.88614 | 13.1663 | 1.037871 | 0.746215 | 0.23085 |
| 15 | 60728600 | 60729599 | 13.25636 | 25.56462 | 17.12727 | 6.626209 | 1.11647 | -0.0923 | 0.23085 |
| 7 | 33928200 | 33932199 | 23.43823 | 38.34713 | 81.65854 | 35.12113 | -0.71209 | 1.465431 | 0.230877 |
| 17 | 92441800 | 92442599 | 11.10773 | 21.45668 | 17.93059 | 6.892971 | 0.809119 | -0.13407 | 0.230884 |
| 17 | 33629400 | 33631399 | 26.29039 | 46.13317 | 25.6623 | 11.34792 | 1.228854 | 0.664896 | 0.230908 |
| X | 24112400 | 24114199 | 38.05186 | 63.0824 | 48.23803 | 20.88812 | 0.766758 | 1.319531 | 0.230913 |
| 7 | 93658000 | 93659799 | 10.96101 | 19.48868 | 41.95537 | 18.5026 | -0.70555 | 0.466252 | 0.230915 |
| 18 | 37000800 | 37001799 | 16.94984 | 30.52472 | 24.6936 | 10.76987 | 0.709445 | 0.393743 | 0.23092 |
| 3 | 110616400 | 110617399 | 7.213462 | 13.64163 | 32.63784 | 13.24989 | -0.76758 | 0.043073 | 0.230928 |
| 3 | 119439400 | 119440399 | 15.51536 | 28.70233 | 22.58492 | 9.512593 | 0.782568 | 0.227333 | 0.230933 |
| 12 | 19311000 | 19324799 | 370.0801 | 521.7296 | 425.5059 | 176.2098 | 0.474056 | 4.460392 | 0.230936 |
| 16 | 26276200 | 26276999 | 6.428071 | 12.35462 | 29.85314 | 11.66211 | -0.7307 | -0.09655 | 0.230944 |
| 17 | 80090400 | 80091399 | 6.428071 | 12.35462 | 29.85314 | 11.66211 | -0.7307 | -0.09655 | 0.230944 |
| 3 | 4781000 | 4790599 | 113.2916 | 174.6594 | 153.2462 | 63.65589 | 0.48722 | 2.907353 | 0.230952 |
| 2 | 156925400 | 156927599 | 11.97552 | 20.98085 | 45.68434 | 20.1747 | -0.72932 | 0.6461 | 0.230967 |
| 7 | 142203200 | 142215399 | 72.32461 | 110.763 | 243.5069 | 101.0594 | -0.84397 | 3.030598 | 0.231009 |
| 7 | 130536800 | 130541599 | 30.08527 | 48.40395 | 105.2089 | 44.66264 | -0.76431 | 1.844741 | 0.231014 |
| 4 | 7524400 | 7525599 | 13.89655 | 25.79061 | 21.82803 | 9.153578 | 0.682786 | 0.194944 | 0.231019 |
| X | 107538600 | 107540199 | 17.78776 | 33.61943 | 16.9956 | 6.908169 | 1.455572 | 0.105126 | 0.231068 |
| 12 | 59321000 | 59323999 | 29.34919 | 49.75436 | 42.04639 | 18.41471 | 0.623305 | 1.064708 | 0.231071 |

| | | | | | | | | | |
|---|---|---|---|---|---|---|---|---|---|
| 3 | 137660400 | 137661799 | 5.573995 | 10.72531 | 29.85827 | 11.62601 | -0.92871 | -0.11788 | 0.231081 |
| 14 | 3758000 | 3760199 | 31.96928 | 54.56202 | 37.58056 | 16.47564 | 0.918283 | 1.012493 | 0.231082 |
| 15 | 19621400 | 19623399 | 19.15631 | 34.13633 | 25.64546 | 11.28847 | 0.806253 | 0.472317 | 0.231083 |
| 14 | 110888800 | 110889599 | 11.81098 | 22.9162 | 16.23806 | 6.11473 | 1.072626 | -0.19826 | 0.231085 |
| 14 | 26389400 | 26391399 | 10.51258 | 18.31441 | 48.89428 | 21.50378 | -1.02196 | 0.731792 | 0.231086 |
| 17 | 43186400 | 43187999 | 12.73828 | 22.34661 | 45.64343 | 20.17908 | -0.63762 | 0.628727 | 0.231091 |
| 8 | 89099000 | 89104199 | 41.84153 | 66.69376 | 129.0715 | 54.05378 | -0.61298 | 2.158616 | 0.231112 |
| 6 | 48672200 | 48672799 | 12.1315 | 23.04374 | 19.95267 | 8.024899 | 0.699372 | 0.02691 | 0.231135 |
| X | 65470800 | 65471399 | 15.87561 | 29.29422 | 21.75974 | 9.196779 | 0.862057 | 0.244022 | 0.23114 |
| 2 | 171814000 | 171833199 | 144.5984 | 214.5942 | 369.8757 | 152.3564 | -0.53259 | 3.664114 | 0.23115 |
| 10 | 81929400 | 81932399 | 108.8814 | 168.8688 | 144.8571 | 60.16328 | 0.526358 | 2.795893 | 0.231156 |
| 17 | 23958400 | 23965999 | 66.78389 | 103.1617 | 213.5454 | 88.70224 | -0.74762 | 2.859021 | 0.231157 |
| 9 | 14186400 | 14188399 | 11.68788 | 20.44324 | 46.24747 | 20.39499 | -0.78406 | 0.673204 | 0.231162 |
| 8 | 116858800 | 116860799 | 11.11113 | 19.77584 | 43.03289 | 18.96141 | -0.72016 | 0.46255 | 0.231169 |
| 11 | 54298200 | 54298799 | 6.43398 | 12.34629 | 29.15291 | 11.44078 | -0.70376 | -0.08277 | 0.23117 |
| 16 | 71824600 | 71826999 | 12.65618 | 21.97798 | 51.59988 | 22.64782 | -0.83875 | 0.789947 | 0.23117 |
| 12 | 91367000 | 91368199 | 7.983483 | 14.53676 | 35.52164 | 15.28021 | -0.86197 | 0.289447 | 0.23117 |
| 9 | 21854400 | 21861799 | 42.86713 | 68.14946 | 138.5872 | 57.83556 | -0.68841 | 2.245499 | 0.231188 |
| 9 | 20107400 | 20110799 | 52.74042 | 85.92611 | 50.79462 | 21.81317 | 1.127343 | 1.567986 | 0.231189 |
| 5 | 78191800 | 78193399 | 22.58993 | 39.44197 | 32.84538 | 14.46553 | 0.650329 | 0.73375 | 0.2312 |
| 6 | 46857200 | 46859399 | 27.03645 | 46.53288 | 33.99608 | 14.91253 | 0.835266 | 0.911647 | 0.231201 |
| 8 | 94915800 | 94922599 | 36.89242 | 59.07936 | 120.8685 | 50.95311 | -0.68556 | 2.004189 | 0.231208 |
| 8 | 3524800 | 3526399 | 6.554244 | 12.24311 | 33.29658 | 13.81824 | -0.97695 | 0.12043 | 0.231209 |
| 17 | 31101800 | 31113599 | 100.6173 | 152.8258 | 263.5463 | 109.142 | -0.50362 | 3.208329 | 0.231219 |
| 4 | 137225000 | 137231999 | 25.63823 | 41.30889 | 106.1516 | 45.15744 | -1.00256 | 1.791735 | 0.231228 |
| 8 | 39304600 | 39306999 | 29.80248 | 51.3375 | 34.73881 | 15.23873 | 0.945645 | 0.917716 | 0.231231 |
| X | 29359600 | 29365199 | 70.77105 | 113.9518 | 56.83204 | 24.15326 | 1.355944 | 1.861236 | 0.231232 |
| 5 | 123272600 | 123278199 | 21.46109 | 34.98709 | 85.95443 | 36.92066 | -0.92133 | 1.516439 | 0.231269 |
| 14 | 42676600 | 42679799 | 38.32842 | 63.76115 | 46.54878 | 20.20485 | 0.833617 | 1.277532 | 0.23128 |
| 6 | 106776400 | 106777199 | 4.288339 | 8.320412 | 26.1096 | 9.832141 | -1.05172 | -0.25423 | 0.231285 |
| X | 24479800 | 24483999 | 88.2299 | 139.4175 | 108.1856 | 44.93684 | 0.691555 | 2.436846 | 0.231289 |
| 2 | 33300800 | 33302799 | 10.26492 | 18.09955 | 43.01678 | 19.0374 | -0.85281 | 0.543424 | 0.23129 |
| 6 | 89500600 | 89503399 | 13.41018 | 23.2899 | 52.16562 | 22.89771 | -0.77129 | 0.791963 | 0.23129 |
| 4 | 62905600 | 62909199 | 31.90299 | 53.27343 | 46.52245 | 20.24237 | 0.575985 | 1.218537 | 0.231307 |
| 14 | 65386800 | 65387799 | 6.210748 | 11.72266 | 30.23644 | 12.32532 | -0.8792 | 0.044092 | 0.231328 |
| 15 | 102880600 | 102881999 | 6.210748 | 11.72266 | 30.23644 | 12.32532 | -0.8792 | 0.044092 | 0.231328 |
| 15 | 61919800 | 61923199 | 24.27834 | 40.00892 | 86.35788 | 37.25465 | -0.72735 | 1.377698 | 0.231329 |
| 5 | 10428200 | 10428999 | 9.295843 | 18.13244 | 14.31025 | 5.125438 | 0.983023 | -0.40703 | 0.231344 |
| 16 | 15303200 | 15304199 | 21.80576 | 38.12113 | 31.73324 | 13.98829 | 0.650821 | 0.708137 | 0.231382 |
| 14 | 30620400 | 30624399 | 38.98665 | 62.4869 | 114.8078 | 48.46165 | -0.52924 | 1.979252 | 0.231394 |
| 13 | 65584000 | 65585399 | 17.5348 | 32.03492 | 22.71096 | 9.728847 | 0.914573 | 0.311001 | 0.231396 |
| 4 | 149644000 | 149652999 | 56.49098 | 88.7308 | 168.8887 | 70.05175 | -0.60789 | 2.551098 | 0.231409 |
| 3 | 31627600 | 31631199 | 41.44759 | 68.94101 | 43.19386 | 18.74693 | 1.053403 | 1.28518 | 0.231414 |
| 7 | 28522000 | 28527399 | 20.36958 | 33.91319 | 68.74831 | 29.85694 | -0.63485 | 1.237202 | 0.23142 |
| 4 | 73415400 | 73416999 | 9.955579 | 18.06813 | 40.27012 | 17.39426 | -0.73494 | 0.331404 | 0.231436 |
| 16 | 93472800 | 93482999 | 81.17547 | 124.8751 | 221.1678 | 91.84938 | -0.52752 | 2.943568 | 0.231443 |
| 3 | 95074000 | 95074999 | 7.116822 | 13.31565 | 32.25645 | 13.34973 | -0.80814 | 0.112833 | 0.231456 |

| | | | | | | | | | |
|---|---|---|---|---|---|---|---|---|---|
| 14 | 31216000 | 31217399 | 7.116822 | 13.31565 | 32.25645 | 13.34973 | -0.80814 | 0.112833 | 0.231456 |
| 12 | 19405000 | 19409599 | 92.54182 | 145.1517 | 125.9026 | 52.21692 | 0.522897 | 2.592315 | 0.231467 |
| 3 | 9202000 | 9202999 | 5.421538 | 10.44002 | 27.4912 | 10.68064 | -0.84528 | -0.13094 | 0.231472 |
| 6 | 100636600 | 100637599 | 5.421538 | 10.44002 | 27.4912 | 10.68064 | -0.84528 | -0.13094 | 0.231472 |
| 11 | 106548400 | 106549399 | 5.421538 | 10.44002 | 27.4912 | 10.68064 | -0.84528 | -0.13094 | 0.231472 |
| 2 | 80081200 | 80081999 | 9.38615 | 16.52814 | 44.07826 | 19.51233 | -1.01862 | 0.555098 | 0.231482 |
| 2 | 156329200 | 156331199 | 11.35988 | 19.9204 | 45.66798 | 20.183 | -0.80319 | 0.627408 | 0.231488 |
| 2 | 136955400 | 136959599 | 26.5803 | 42.79178 | 105.1469 | 44.67707 | -0.93957 | 1.819926 | 0.231489 |
| 8 | 71579400 | 71580199 | 4.689871 | 9.112077 | 25.74452 | 9.601344 | -0.88933 | -0.28537 | 0.23149 |
| 14 | 36696400 | 36697199 | 4.689871 | 9.112077 | 25.74452 | 9.601344 | -0.88933 | -0.28537 | 0.23149 |
| 14 | 45626000 | 45626799 | 4.689871 | 9.112077 | 25.74452 | 9.601344 | -0.88933 | -0.28537 | 0.23149 |
| 9 | 53922000 | 53925799 | 45.53755 | 73.6345 | 63.93495 | 27.32524 | 0.565035 | 1.698278 | 0.231498 |
| X | 88283600 | 88285999 | 29.98867 | 52.16622 | 29.71715 | 13.05733 | 1.195065 | 0.817689 | 0.231537 |
| X | 14282600 | 14284199 | 38.0266 | 62.76918 | 50.98008 | 22.00985 | 0.67967 | 1.369991 | 0.231538 |
| 2 | 81624000 | 81626399 | 49.40561 | 79.67878 | 59.98839 | 25.55145 | 0.765961 | 1.778487 | 0.231546 |
| 15 | 52931400 | 52932599 | 8.202016 | 15.17292 | 33.64568 | 14.1716 | -0.70147 | 0.195683 | 0.23155 |
| 5 | 135029200 | 135031399 | 15.17813 | 25.72732 | 60.42695 | 26.4636 | -0.84538 | 1.03959 | 0.231558 |
| 7 | 7252600 | 7258599 | 84.60771 | 133.9115 | 107.5156 | 44.69532 | 0.644297 | 2.395149 | 0.231567 |
| 9 | 22109000 | 22115799 | 134.4944 | 205.7822 | 172.404 | 71.53509 | 0.545456 | 3.063258 | 0.231592 |
| 4 | 129270000 | 129271999 | 8.443136 | 15.3227 | 37.03386 | 15.99709 | -0.85064 | 0.31402 | 0.231595 |
| 14 | 120388400 | 120392399 | 21.44312 | 35.7716 | 70.74237 | 30.76151 | -0.59943 | 1.219095 | 0.231605 |
| 16 | 96116400 | 96119599 | 27.79187 | 45.25741 | 87.78854 | 37.69316 | -0.58285 | 1.545624 | 0.231608 |
| 12 | 81417600 | 81420599 | 30.05601 | 48.41732 | 107.5844 | 45.75383 | -0.79381 | 1.806822 | 0.231624 |
| 3 | 20142800 | 20144199 | 20.13082 | 35.60991 | 27.81751 | 12.30293 | 0.745148 | 0.539143 | 0.231632 |
| 15 | 31384200 | 31384799 | 11.13324 | 21.62524 | 16.43796 | 6.143848 | 0.982891 | -0.22956 | 0.231646 |
| 19 | 3117800 | 3120399 | 21.21527 | 37.25505 | 30.14342 | 13.33778 | 0.691585 | 0.633822 | 0.231652 |
| 3 | 116601400 | 116602199 | 6.464183 | 12.30478 | 29.963 | 11.99703 | -0.77456 | -0.01519 | 0.231656 |
| 8 | 125671000 | 125672199 | 6.464183 | 12.30478 | 29.963 | 11.99703 | -0.77456 | -0.01519 | 0.231656 |
| 9 | 35165400 | 35166199 | 10.47397 | 20.37922 | 15.94684 | 5.879166 | 0.959152 | -0.28587 | 0.23167 |
| 1 | 3771200 | 3772399 | 13.67651 | 26.16016 | 18.76327 | 7.431511 | 0.988255 | -0.00951 | 0.23167 |
| 2 | 123446200 | 123448799 | 35.74183 | 59.85912 | 43.6739 | 19.03815 | 0.834217 | 1.196025 | 0.231674 |
| 7 | 108382800 | 108383799 | 11.53736 | 22.33223 | 17.42207 | 6.633769 | 0.920684 | -0.16126 | 0.231675 |
| 15 | 91858600 | 91861199 | 15.96644 | 27.47578 | 64.93823 | 28.4853 | -0.85177 | 0.903726 | 0.231681 |
| 10 | 14984400 | 14989999 | 74.61505 | 118.4729 | 103.2666 | 43.04135 | 0.530049 | 2.300422 | 0.231699 |
| 16 | 44377600 | 44379599 | 10.05974 | 17.5636 | 48.87469 | 21.52452 | -1.08119 | 0.69651 | 0.231703 |
| 17 | 4582000 | 4582999 | 12.22357 | 23.268 | 19.88304 | 7.961041 | 0.723599 | 0.012527 | 0.231704 |
| 3 | 65881000 | 65882399 | 6.766522 | 12.84484 | 29.8707 | 12.03001 | -0.71592 | 0.008057 | 0.231705 |
| 13 | 98427400 | 98428799 | 6.766522 | 12.84484 | 29.8707 | 12.03001 | -0.71592 | 0.008057 | 0.231705 |
| 14 | 89128400 | 89133799 | 102.906 | 160.38 | 136.3229 | 56.57762 | 0.544956 | 2.709465 | 0.231711 |
| 6 | 144771200 | 144772999 | 18.13141 | 30.30188 | 66.3465 | 28.88305 | -0.74542 | 1.188844 | 0.231713 |
| X | 124891800 | 124892599 | 15.22534 | 28.45776 | 20.74846 | 8.601828 | 0.909523 | 0.161937 | 0.231718 |
| 17 | 38620800 | 38626999 | 83.90807 | 132.9453 | 104.9465 | 43.65079 | 0.669913 | 2.369221 | 0.23173 |
| 1 | 115404000 | 115404799 | 11.63515 | 22.22174 | 19.48328 | 7.746269 | 0.695865 | -0.01608 | 0.231769 |
| 9 | 36187800 | 36191999 | 48.93525 | 79.3921 | 61.30951 | 26.26987 | 0.73815 | 1.634938 | 0.231777 |
| 12 | 70356400 | 70356999 | 4.106545 | 8.012258 | 23.3815 | 8.34455 | -0.87629 | -0.45768 | 0.231788 |
| 7 | 34560800 | 34561999 | 7.253126 | 13.59952 | 31.76271 | 13.09821 | -0.75162 | 0.099473 | 0.23179 |
| X | 66209600 | 66211399 | 29.6345 | 50.397 | 40.44579 | 17.72668 | 0.697898 | 1.038341 | 0.231826 |

| | | | | | | | | | |
|---|---|---|---|---|---|---|---|---|---|
| 7 | 62127200 | 62132599 | 77.53216 | 123.3613 | 94.39153 | 39.43198 | 0.720674 | 2.239202 | 0.231846 |
| 2 | 80668200 | 80668799 | 3.234901 | 6.315552 | 20.91347 | 7.383611 | -1.03869 | -0.51306 | 0.231864 |
| 3 | 9079800 | 9080799 | 15.89007 | 29.3763 | 22.57328 | 9.516152 | 0.815728 | 0.234256 | 0.231871 |
| 5 | 94494600 | 94496999 | 41.84365 | 69.41239 | 45.48645 | 19.70525 | 0.988669 | 1.318629 | 0.231873 |
| 1 | 30958800 | 30961999 | 74.72975 | 118.8045 | 97.98622 | 40.892 | 0.6114 | 2.261203 | 0.231879 |
| 2 | 158123000 | 158128199 | 24.86468 | 40.30667 | 92.06619 | 39.42428 | -0.82345 | 1.624999 | 0.23189 |
| 7 | 105202800 | 105209399 | 35.15939 | 56.2628 | 120.782 | 50.86226 | -0.75601 | 2.025819 | 0.231908 |
| 8 | 111321600 | 111323799 | 11.1577 | 19.69579 | 42.98232 | 19.01789 | -0.72998 | 0.535748 | 0.231944 |
| 2 | 169815000 | 169817199 | 16.93632 | 29.04071 | 58.13603 | 25.49693 | -0.61398 | 0.934095 | 0.231947 |
| 7 | 11029600 | 11032199 | 46.98872 | 77.012 | 47.32037 | 20.36679 | 1.077046 | 1.473691 | 0.231947 |
| X | 147825000 | 147825799 | 12.83511 | 25.01723 | 12.12817 | 4.387047 | 1.664799 | -0.33245 | 0.231953 |
| 7 | 66601600 | 66602999 | 5.620008 | 10.68853 | 29.87493 | 11.98387 | -0.97356 | -0.01711 | 0.231957 |
| 9 | 37880200 | 37881199 | 14.85581 | 25.65645 | 52.68842 | 23.11442 | -0.64826 | 0.85477 | 0.231994 |
| X | 153141800 | 153143999 | 48.68918 | 78.99701 | 61.02248 | 26.14789 | 0.737868 | 1.636482 | 0.231996 |
| 6 | 146789600 | 146791199 | 7.503599 | 13.46149 | 40.55318 | 17.74962 | -1.17871 | 0.384041 | 0.23203 |
| 11 | 67397000 | 67399199 | 17.10145 | 29.31563 | 58.68844 | 25.73862 | -0.6141 | 0.938091 | 0.232031 |
| 7 | 13881600 | 13887999 | 93.99114 | 148.1319 | 112.928 | 46.86092 | 0.714144 | 2.498592 | 0.232032 |
| 17 | 70990400 | 70991199 | 5.084202 | 9.87671 | 25.12783 | 9.383028 | -0.74085 | -0.27554 | 0.232037 |
| 11 | 30721400 | 30722599 | 9.710446 | 17.43898 | 37.93182 | 16.58751 | -0.71116 | 0.389797 | 0.232043 |
| 7 | 9110400 | 9112399 | 46.20793 | 74.68008 | 65.61399 | 28.02424 | 0.547179 | 1.715163 | 0.232043 |
| 7 | 27054000 | 27076199 | 171.3049 | 249.4255 | 441.2407 | 181.8497 | -0.5962 | 3.975002 | 0.232059 |
| 16 | 17822200 | 17825799 | 20.85493 | 34.63898 | 73.95517 | 32.06619 | -0.70981 | 1.26789 | 0.232068 |
| 5 | 53215200 | 53217199 | 12.90265 | 22.57836 | 46.75323 | 20.62604 | -0.65735 | 0.671847 | 0.232072 |
| 11 | 45913600 | 45915199 | 8.056388 | 14.86159 | 35.76745 | 15.12315 | -0.82321 | 0.213085 | 0.232074 |
| 3 | 109960200 | 109962199 | 28.08807 | 48.16128 | 38.22783 | 16.76568 | 0.715357 | 0.956148 | 0.232079 |
| 17 | 8043200 | 8048799 | 29.89422 | 48.1731 | 103.8643 | 44.19353 | -0.74985 | 1.792049 | 0.232079 |
| 7 | 21952200 | 21952799 | 12.12081 | 23.39743 | 17.84097 | 6.88076 | 0.935823 | -0.11225 | 0.232092 |
| 1 | 15936400 | 15938999 | 26.09827 | 44.75168 | 37.87388 | 16.61051 | 0.623251 | 0.94724 | 0.23211 |
| 8 | 43022800 | 43023399 | 4.618061 | 8.996003 | 24.24661 | 8.836893 | -0.78984 | -0.36869 | 0.232121 |
| 5 | 93732400 | 93734599 | 30.02169 | 51.33102 | 37.5112 | 16.44763 | 0.833412 | 0.98611 | 0.23213 |
| 9 | 72199800 | 72200999 | 8.01877 | 14.50497 | 36.49417 | 15.82867 | -0.91155 | 0.331059 | 0.232139 |
| 19 | 13489800 | 13491399 | 25.11883 | 43.22914 | 37.2321 | 16.33198 | 0.598866 | 0.908189 | 0.23214 |
| 18 | 84938000 | 84938799 | 4.575262 | 8.863473 | 27.23037 | 10.32812 | -1.03211 | -0.22388 | 0.23214 |
| 1 | 173400000 | 173408799 | 254.5767 | 370.6908 | 313.4945 | 129.2006 | 0.468086 | 3.978699 | 0.232159 |
| 11 | 79227200 | 79227799 | 12.36607 | 23.59108 | 19.46285 | 7.756632 | 0.780192 | -0.00056 | 0.232174 |
| 19 | 39768600 | 39770999 | 30.31747 | 52.41595 | 32.47203 | 14.24596 | 1.073358 | 0.885666 | 0.232176 |
| 10 | 20040600 | 20041799 | 16.62191 | 30.39843 | 23.58874 | 10.09178 | 0.786745 | 0.302078 | 0.232182 |
| 1 | 119932800 | 119936799 | 23.97374 | 39.07771 | 85.32762 | 36.64949 | -0.75202 | 1.522858 | 0.232197 |
| 1 | 95785400 | 95787399 | 20.44761 | 36.36053 | 26.10507 | 11.51126 | 0.869771 | 0.495726 | 0.232207 |
| 14 | 109989600 | 109989999 | 8.314207 | 16.26325 | 10.913 | 3.649613 | 1.299895 | -0.66428 | 0.232209 |
| 17 | 33839800 | 33842999 | 33.58787 | 55.75422 | 47.13616 | 20.42926 | 0.622466 | 1.289616 | 0.23222 |
| 8 | 19461800 | 19464799 | 33.79396 | 56.60144 | 45.37814 | 19.78205 | 0.698841 | 1.192727 | 0.232228 |
| Y | 84075400 | 84076399 | 8.090408 | 15.80313 | 8.223795 | 2.877918 | 1.60403 | -0.53087 | 0.232229 |
| 4 | 146322400 | 146325199 | 45.66067 | 74.97839 | 50.66471 | 21.82383 | 0.941408 | 1.449413 | 0.232248 |
| 1 | 95241200 | 95242199 | 5.434832 | 10.42709 | 27.90459 | 10.95667 | -0.88271 | -0.09427 | 0.232251 |
| 3 | 5640800 | 5641799 | 9.811158 | 17.30798 | 42.01361 | 18.59161 | -0.88251 | 0.534861 | 0.232259 |
| 19 | 27987800 | 27989799 | 11.74297 | 20.4046 | 49.97316 | 21.94225 | -0.8989 | 0.776911 | 0.232265 |

| | | | | | | | | | |
|---|---|---|---|---|---|---|---|---|---|
| 8 | 84923800 | 84932399 | 38.86436 | 61.77931 | 141.4296 | 59.01544 | -0.85909 | 2.245867 | 0.232276 |
| 3 | 14290800 | 14301599 | 229.7494 | 337.5403 | 288.6471 | 118.8772 | 0.464401 | 3.838544 | 0.232283 |
| 2 | 156656600 | 156664399 | 55.65733 | 87.92372 | 156.4699 | 64.98367 | -0.50457 | 2.428067 | 0.232284 |
| 18 | 35980800 | 35984399 | 18.01284 | 30.0143 | 69.08953 | 30.02079 | -0.81746 | 1.221234 | 0.232286 |
| 17 | 19151400 | 19153399 | 44.1104 | 71.92851 | 58.38471 | 25.08062 | 0.672084 | 1.541116 | 0.232312 |
| 7 | 107597600 | 107598999 | 25.61639 | 44.02436 | 37.74369 | 16.5557 | 0.605059 | 0.923213 | 0.232314 |
| 1 | 113368600 | 113369599 | 11.8049 | 22.58409 | 19.4685 | 7.707006 | 0.725734 | -0.02819 | 0.232322 |
| 11 | 23952800 | 23953799 | 11.8049 | 22.58409 | 19.4685 | 7.707006 | 0.725734 | -0.02819 | 0.232322 |
| 16 | 4586200 | 4586999 | 4.357112 | 8.489036 | 23.04996 | 8.383136 | -0.79748 | -0.3845 | 0.232333 |
| 17 | 8556800 | 8562799 | 33.50199 | 53.74189 | 113.8412 | 48.11585 | -0.7326 | 1.944937 | 0.232334 |
| 4 | 124373200 | 124374799 | 9.689833 | 17.47165 | 37.44897 | 16.30801 | -0.6861 | 0.363986 | 0.232346 |
| 4 | 130296000 | 130298999 | 17.50959 | 29.25958 | 66.90004 | 29.12209 | -0.80772 | 1.187086 | 0.232371 |
| 1 | 21003600 | 21006199 | 14.51109 | 25.21911 | 50.56267 | 22.20364 | -0.61201 | 0.787015 | 0.232387 |
| 2 | 176799000 | 176802999 | 45.20834 | 73.67694 | 53.91085 | 23.15396 | 0.820742 | 1.548568 | 0.232389 |
| 16 | 37516200 | 37518199 | 10.01764 | 17.91081 | 39.94373 | 17.53295 | -0.75124 | 0.421191 | 0.232392 |
| 1 | 8110600 | 8118999 | 195.3814 | 291.9155 | 235.9873 | 97.29363 | 0.56791 | 3.54762 | 0.232405 |
| 17 | 40958800 | 40959999 | 14.96041 | 27.44547 | 23.14463 | 9.859774 | 0.671591 | 0.269453 | 0.232409 |
| 17 | 85461000 | 85462399 | 7.504217 | 13.76906 | 34.57398 | 14.72545 | -0.89098 | 0.238805 | 0.232411 |
| 1 | 85037600 | 85042399 | 86.01247 | 135.6757 | 113.7516 | 47.20802 | 0.578052 | 2.473335 | 0.232416 |
| 2 | 8109000 | 8110399 | 15.47429 | 29.18987 | 19.40085 | 7.918442 | 1.059088 | 0.102336 | 0.232476 |
| 1 | 153725000 | 153727199 | 14.96841 | 25.98728 | 52.20338 | 22.90939 | -0.61509 | 0.804858 | 0.23248 |
| 7 | 61763200 | 61768799 | 108.605 | 168.0633 | 149.0033 | 61.89223 | 0.47632 | 2.835494 | 0.23249 |
| 14 | 7030200 | 7032599 | 17.8069 | 29.78923 | 66.43787 | 28.94102 | -0.77202 | 1.17516 | 0.232491 |
| 5 | 34734400 | 34735599 | 5.281615 | 10.18935 | 28.16669 | 10.88526 | -0.90782 | -0.14926 | 0.232493 |
| 1 | 146271200 | 146272999 | 25.20713 | 43.88939 | 33.22957 | 14.60918 | 0.786724 | 0.78565 | 0.232494 |
| 7 | 23491400 | 23495399 | 79.43444 | 126.5385 | 87.86508 | 36.76606 | 0.862303 | 2.202644 | 0.232502 |
| 17 | 63140800 | 63142199 | 17.92445 | 33.04502 | 21.13237 | 8.942579 | 1.074598 | 0.261692 | 0.232513 |
| 14 | 71215800 | 71218799 | 12.32285 | 21.46073 | 53.77308 | 23.64434 | -0.93192 | 0.748044 | 0.232529 |
| 17 | 5966200 | 5969599 | 15.97616 | 27.02937 | 60.50876 | 26.48439 | -0.77674 | 1.056609 | 0.232531 |
| 3 | 72444000 | 72445799 | 24.81789 | 43.52516 | 26.8412 | 11.86367 | 1.080911 | 0.666031 | 0.232539 |
| 4 | 97639200 | 97641199 | 7.751955 | 14.35513 | 33.12398 | 13.93232 | -0.75676 | 0.185229 | 0.232548 |
| 1 | 49290000 | 49291799 | 19.27686 | 35.04503 | 22.53242 | 9.711825 | 1.049086 | 0.351596 | 0.232554 |
| 6 | 3100400 | 3102599 | 15.28295 | 28.83695 | 19.40848 | 7.91691 | 1.041792 | 0.098626 | 0.232557 |
| 6 | 144882600 | 144890799 | 69.43605 | 107.8685 | 194.2295 | 80.64923 | -0.54038 | 2.756409 | 0.232561 |
| 13 | 66678200 | 66682599 | 98.21722 | 155.5046 | 91.73323 | 38.141 | 1.08869 | 2.394787 | 0.232575 |
| 9 | 22322600 | 22323199 | 5.096452 | 9.859137 | 25.95286 | 9.909143 | -0.81981 | -0.19512 | 0.232578 |
| 17 | 6085600 | 6088399 | 11.88501 | 20.67683 | 49.44934 | 21.72904 | -0.86465 | 0.758172 | 0.232602 |
| 9 | 67921600 | 67923799 | 10.10615 | 17.84146 | 41.49337 | 18.3587 | -0.82064 | 0.528151 | 0.232603 |
| 7 | 16930600 | 16935999 | 118.7 | 182.3734 | 158.4605 | 65.79625 | 0.497537 | 2.973652 | 0.232606 |
| 5 | 21887800 | 21895599 | 95.09327 | 145.4356 | 246.8438 | 102.4122 | -0.47268 | 3.065492 | 0.232607 |
| 17 | 85235800 | 85240199 | 47.18702 | 75.20206 | 133.5985 | 55.9371 | -0.49005 | 2.169638 | 0.232611 |
| 16 | 83009600 | 83010999 | 17.69921 | 32.31551 | 23.35253 | 10.01134 | 0.886744 | 0.318178 | 0.232611 |
| 4 | 103085400 | 103087199 | 18.13646 | 32.54911 | 24.68516 | 10.80097 | 0.798413 | 0.419958 | 0.232621 |
| 12 | 115811400 | 115812799 | 21.63472 | 37.80444 | 31.79189 | 14.00861 | 0.636217 | 0.714735 | 0.232633 |
| 1 | 90016600 | 90024599 | 53.30679 | 84.48356 | 148.6226 | 61.83115 | -0.48413 | 2.347407 | 0.232638 |
| 13 | 43185600 | 43188599 | 49.03666 | 79.10035 | 70.14253 | 29.88459 | 0.530757 | 1.777267 | 0.232662 |
| 3 | 55929200 | 55930599 | 15.78403 | 30.24951 | 16.2116 | 6.385523 | 1.411561 | -0.01304 | 0.232667 |

| | | | | | | | | | |
|---|---|---|---|---|---|---|---|---|---|
| 11 | 74237400 | 74238399 | 6.492732 | 12.27604 | 30.2992 | 12.30595 | -0.81189 | 0.03386 | 0.232671 |
| 17 | 37462000 | 37464399 | 29.27184 | 49.43287 | 42.63054 | 18.64879 | 0.59395 | 1.096665 | 0.232701 |
| 4 | 152742800 | 152764399 | 244.3366 | 345.7985 | 583.6349 | 241.5392 | -0.5692 | 4.399494 | 0.232702 |
| 17 | 86121200 | 86127199 | 64.98806 | 101.4826 | 183.5453 | 76.15842 | -0.54095 | 2.663047 | 0.23271 |
| 1 | 161296800 | 161297199 | 7.051564 | 13.4042 | 31.09927 | 12.48872 | -0.70941 | 0.003026 | 0.23272 |
| 7 | 116102600 | 116103999 | 6.542481 | 12.2275 | 31.67025 | 13.13174 | -0.90548 | 0.115186 | 0.23272 |
| 12 | 27505000 | 27508399 | 31.85123 | 54.37182 | 37.51097 | 16.4454 | 0.915894 | 1.009046 | 0.232723 |
| 13 | 84125000 | 84126199 | 5.059812 | 9.841071 | 26.51794 | 9.807098 | -0.80984 | -0.30488 | 0.23273 |
| 17 | 43726600 | 43728599 | 10.2299 | 18.13094 | 40.40873 | 17.86479 | -0.75748 | 0.482019 | 0.232733 |
| 17 | 50416200 | 50417999 | 8.667527 | 15.93537 | 35.68875 | 15.15896 | -0.72493 | 0.237487 | 0.232747 |
| 5 | 93239000 | 93242799 | 22.00111 | 36.04972 | 80.9967 | 34.85751 | -0.7881 | 1.441938 | 0.232763 |
| 7 | 83799200 | 83801999 | 13.62852 | 23.51818 | 53.18009 | 23.32955 | -0.78673 | 0.857105 | 0.232775 |
| 7 | 41543800 | 41544599 | 20.09983 | 35.44902 | 28.62534 | 12.66778 | 0.696202 | 0.573879 | 0.232779 |
| 12 | 56505000 | 56506999 | 10.45983 | 18.82761 | 43.29338 | 18.89369 | -0.78958 | 0.388469 | 0.232825 |
| 12 | 115106600 | 115107799 | 24.58863 | 42.88317 | 32.70653 | 14.39414 | 0.776403 | 0.759471 | 0.232826 |
| 12 | 22644400 | 22646199 | 26.07131 | 45.07532 | 32.94883 | 14.45514 | 0.835755 | 0.866537 | 0.232838 |
| 3 | 132773400 | 132775399 | 16.96675 | 31.71018 | 19.71385 | 8.17398 | 1.136804 | 0.173047 | 0.232839 |
| 14 | 96537800 | 96539599 | 19.22484 | 34.03748 | 27.33219 | 12.08615 | 0.705966 | 0.522265 | 0.232842 |
| 16 | 45172400 | 45175399 | 23.10763 | 38.25848 | 76.98057 | 33.30461 | -0.62502 | 1.313564 | 0.232849 |
| 3 | 38432000 | 38432599 | 6.742586 | 12.90427 | 30.78183 | 12.17221 | -0.72872 | -0.0498 | 0.232889 |
| 10 | 85908800 | 85910399 | 14.13974 | 26.4057 | 21.67707 | 9.000051 | 0.737358 | 0.158107 | 0.232914 |
| 7 | 10963600 | 10984799 | 596.8947 | 809.274 | 625.8865 | 260.7835 | 0.493899 | 5.060997 | 0.232921 |
| 11 | 35869800 | 35871599 | 13.10058 | 22.79037 | 49.47092 | 21.73779 | -0.72578 | 0.764173 | 0.232929 |
| 2 | 168403400 | 168409599 | 37.29076 | 59.55244 | 124.7965 | 52.40187 | -0.72453 | 2.092704 | 0.232938 |
| 10 | 37696400 | 37697999 | 32.09894 | 54.79513 | 36.91273 | 16.18313 | 0.950076 | 1.008566 | 0.23294 |
| 15 | 23769800 | 23773199 | 32.09894 | 54.79513 | 36.91273 | 16.18313 | 0.950076 | 1.008566 | 0.23294 |
| 1 | 189239000 | 189250399 | 115.4915 | 174.5277 | 298.5995 | 123.464 | -0.49843 | 3.316495 | 0.232944 |
| X | 75541800 | 75543799 | 25.52982 | 44.35528 | 34.36371 | 15.08729 | 0.753471 | 0.818606 | 0.232944 |
| 7 | 9789000 | 9792399 | 60.63848 | 97.61928 | 69.48937 | 29.52484 | 0.843636 | 1.85492 | 0.232948 |
| 15 | 20338800 | 20339999 | 13.51228 | 25.80778 | 17.97713 | 7.146885 | 1.024168 | -0.00346 | 0.232956 |
| 12 | 107087000 | 107088599 | 10.03615 | 17.91734 | 38.90828 | 17.0998 | -0.71405 | 0.430375 | 0.232957 |
| 4 | 48407600 | 48409599 | 8.329849 | 15.02792 | 36.97302 | 16.08944 | -0.88277 | 0.349501 | 0.23296 |
| 11 | 116938600 | 116955999 | 132.641 | 197.1278 | 351.8956 | 144.9711 | -0.58123 | 3.637531 | 0.232966 |
| X | 156406200 | 156407799 | 17.21315 | 29.26946 | 60.64093 | 26.57759 | -0.66505 | 1.018 | 0.232967 |
| 2 | 129166200 | 129167599 | 9.834532 | 17.8475 | 40.25729 | 17.38987 | -0.75187 | 0.332854 | 0.232988 |
| 3 | 76384200 | 76385399 | 15.67504 | 28.52459 | 24.14135 | 10.3927 | 0.65645 | 0.322978 | 0.232996 |
| 18 | 3404800 | 3406799 | 43.47744 | 70.70843 | 60.83839 | 26.10938 | 0.585642 | 1.581186 | 0.233009 |
| 1 | 142303200 | 142304999 | 23.35493 | 40.84236 | 31.56553 | 13.91809 | 0.75745 | 0.711681 | 0.233021 |
| 18 | 40991200 | 40993999 | 24.55018 | 42.90147 | 29.18705 | 12.86148 | 0.940239 | 0.723563 | 0.233029 |
| 7 | 35145400 | 35147999 | 13.55701 | 23.5926 | 49.47606 | 21.7448 | -0.67628 | 0.761135 | 0.233033 |
| 2 | 97277600 | 97278999 | 11.15286 | 19.72908 | 41.41944 | 18.31861 | -0.67305 | 0.510512 | 0.233034 |
| 14 | 79063800 | 79066199 | 11.15286 | 19.72908 | 41.41944 | 18.31861 | -0.67305 | 0.510512 | 0.233034 |
| 19 | 18726800 | 18728199 | 6.089962 | 11.48187 | 34.15195 | 13.95231 | -1.08695 | 0.056624 | 0.233043 |
| 4 | 28740600 | 28743199 | 31.47131 | 54.09775 | 34.12145 | 14.96695 | 1.046034 | 0.942496 | 0.233044 |
| 12 | 105245400 | 105249199 | 24.77379 | 40.35392 | 85.98828 | 36.92395 | -0.71763 | 1.534795 | 0.233046 |
| 13 | 64916800 | 64918399 | 15.68629 | 28.88049 | 22.43137 | 9.509417 | 0.794265 | 0.253198 | 0.233051 |
| 12 | 109836000 | 109841999 | 45.7142 | 72.86825 | 130.6904 | 54.73344 | -0.50345 | 2.161467 | 0.23307 |

| | | | | | | | | | |
|---|---|---|---|---|---|---|---|---|---|
| 4 | 83715000 | 83717599 | 15.01671 | 25.93007 | 52.68881 | 23.11448 | -0.63308 | 0.857221 | 0.233074 |
| 18 | 76879400 | 76882199 | 26.71247 | 45.86997 | 37.64525 | 16.5121 | 0.667537 | 0.934465 | 0.233077 |
| 4 | 147330800 | 147334599 | 89.80152 | 141.6981 | 106.0124 | 44.00232 | 0.742508 | 2.46886 | 0.233088 |
| 7 | 23989000 | 24000599 | 121.8138 | 182.7521 | 312.2152 | 128.8298 | -0.50549 | 3.461426 | 0.233091 |
| 3 | 79540600 | 79541799 | 21.27902 | 37.34789 | 29.30546 | 12.9614 | 0.735759 | 0.641753 | 0.233094 |
| 6 | 10990600 | 10991999 | 12.45298 | 21.73068 | 52.14531 | 22.95738 | -0.86949 | 0.71798 | 0.233128 |
| 6 | 119082400 | 119086999 | 25.31971 | 41.08183 | 90.64436 | 38.84159 | -0.77242 | 1.606204 | 0.233137 |
| 5 | 85667200 | 85671599 | 44.87562 | 73.42651 | 55.08639 | 23.6932 | 0.788839 | 1.494462 | 0.233148 |
| 7 | 39041400 | 39043399 | 22.13917 | 39.43716 | 24.32057 | 10.70863 | 1.089158 | 0.493174 | 0.233155 |
| 6 | 59330000 | 59332399 | 15.32542 | 26.01091 | 59.45178 | 26.04684 | -0.80637 | 1.024747 | 0.233161 |
| 2 | 3513200 | 3514399 | 11.03819 | 19.41561 | 43.02376 | 19.04977 | -0.75376 | 0.576183 | 0.233163 |
| 8 | 20084200 | 20086199 | 17.26533 | 31.56602 | 22.14112 | 9.475949 | 0.930498 | 0.302879 | 0.23317 |
| 16 | 89415000 | 89416799 | 10.63392 | 19.09217 | 44.32155 | 19.38624 | -0.8055 | 0.40779 | 0.233171 |
| 10 | 104394000 | 104394999 | 15.06051 | 28.56341 | 18.33009 | 7.396324 | 1.123196 | 0.059505 | 0.233177 |
| 1 | 158807000 | 158808599 | 9.956526 | 18.00766 | 39.08168 | 16.95387 | -0.70083 | 0.348702 | 0.233182 |
| 6 | 89927000 | 89929199 | 28.04849 | 48.97798 | 28.53687 | 12.56888 | 1.162912 | 0.753808 | 0.233185 |
| 8 | 56560000 | 56561799 | 17.16552 | 31.59525 | 22.35378 | 9.480071 | 0.927829 | 0.26746 | 0.233194 |
| 9 | 45923600 | 45924799 | 5.666123 | 10.65315 | 31.22398 | 12.81626 | -1.07009 | 0.066984 | 0.233198 |
| 4 | 120873200 | 120875399 | 15.09936 | 25.83197 | 55.79514 | 24.46578 | -0.72332 | 0.947198 | 0.233204 |
| 4 | 47536400 | 47538199 | 8.247488 | 15.08753 | 38.84103 | 16.61047 | -0.93197 | 0.267738 | 0.23323 |
| 10 | 108737200 | 108738599 | 13.31982 | 24.85816 | 21.2868 | 8.84786 | 0.67565 | 0.154195 | 0.233238 |
| 9 | 112924600 | 112928199 | 43.98133 | 71.46981 | 61.38091 | 26.33168 | 0.587378 | 1.595684 | 0.233259 |
| 9 | 62320400 | 62321399 | 12.35938 | 21.45516 | 50.51855 | 22.1684 | -0.84273 | 0.793745 | 0.233262 |
| 17 | 75926000 | 75927599 | 15.09106 | 27.61954 | 23.10842 | 9.876188 | 0.67952 | 0.283186 | 0.233264 |
| 9 | 32187600 | 32192799 | 43.07198 | 68.79399 | 125.8709 | 52.8629 | -0.52903 | 2.094212 | 0.233269 |
| 12 | 69176400 | 69177799 | 11.49085 | 20.22274 | 43.00352 | 19.03875 | -0.69444 | 0.572481 | 0.233286 |
| 13 | 97722000 | 97724199 | 18.13646 | 32.18379 | 27.0053 | 11.92912 | 0.644451 | 0.495033 | 0.233296 |
| 12 | 63468000 | 63469999 | 21.77276 | 38.42958 | 27.66408 | 12.24047 | 0.861243 | 0.572391 | 0.233304 |
| 19 | 46362800 | 46363599 | 3.941038 | 7.675902 | 22.9387 | 8.378162 | -0.93897 | -0.3744 | 0.233309 |
| 8 | 63878000 | 63879799 | 20.56325 | 36.6907 | 25.66244 | 11.28369 | 0.910166 | 0.475938 | 0.233311 |
| 10 | 116793000 | 116800999 | 46.70718 | 74.44337 | 132.084 | 55.30916 | -0.48809 | 2.166521 | 0.233312 |
| 7 | 139523400 | 139524999 | 15.76525 | 26.81763 | 58.44465 | 25.6158 | -0.73763 | 1.004581 | 0.233313 |
| 1 | 168434800 | 168440999 | 65.36359 | 105.0787 | 68.69843 | 29.12958 | 0.963641 | 1.903539 | 0.233317 |
| 7 | 28009200 | 28012799 | 16.22208 | 27.23272 | 65.86493 | 28.7546 | -0.8885 | 1.131203 | 0.233321 |
| 1 | 159406000 | 159407599 | 9.415777 | 16.84657 | 40.48794 | 17.7613 | -0.85785 | 0.413263 | 0.233323 |
| 8 | 21581400 | 21582199 | 16.50429 | 29.7985 | 25.06796 | 10.8982 | 0.656059 | 0.375519 | 0.233329 |
| 10 | 81762200 | 81764199 | 21.55598 | 38.50679 | 23.78274 | 10.44694 | 1.08913 | 0.470735 | 0.233333 |
| 6 | 57679600 | 57680999 | 5.887721 | 11.24086 | 30.62159 | 12.18382 | -0.92704 | -0.04099 | 0.233342 |
| 13 | 23205000 | 23206199 | 12.50325 | 24.13228 | 17.15423 | 6.620241 | 1.034773 | -0.10865 | 0.233344 |
| 9 | 99298600 | 99302799 | 19.22525 | 32.15093 | 65.96016 | 28.72669 | -0.65199 | 1.183999 | 0.233354 |
| 19 | 10917200 | 10924399 | 41.5384 | 66.14168 | 133.2422 | 55.72011 | -0.67239 | 2.192636 | 0.233359 |
| 11 | 44322000 | 44322599 | 6.308244 | 12.04179 | 28.89755 | 11.50175 | -0.74566 | -0.03783 | 0.233377 |
| 9 | 34485200 | 34486999 | 7.72886 | 14.38189 | 32.67678 | 13.64954 | -0.72725 | 0.155301 | 0.233396 |
| 7 | 78842400 | 78845199 | 13.64737 | 23.50036 | 55.31685 | 24.26535 | -0.84517 | 0.878468 | 0.233398 |
| 1 | 57356400 | 57357999 | 28.24663 | 49.24614 | 29.70279 | 13.06942 | 1.113214 | 0.77814 | 0.2334 |
| 6 | 44941600 | 44943399 | 18.59356 | 33.74843 | 23.8175 | 10.28641 | 0.91403 | 0.359219 | 0.233468 |
| 7 | 103084200 | 103084999 | 14.40093 | 26.59354 | 20.89128 | 8.820489 | 0.782722 | 0.224746 | 0.233477 |

| | | | | | | | | | |
|---|---|---|---|---|---|---|---|---|---|
| 13 | 75151200 | 75153399 | 14.33341 | 24.96023 | 50.01686 | 21.99162 | -0.61096 | 0.756184 | 0.233478 |
| 16 | 42532800 | 42534199 | 13.30925 | 25.26451 | 17.30013 | 6.967577 | 1.031888 | 0.03247 | 0.233485 |
| 8 | 6908000 | 6909199 | 14.92848 | 27.2775 | 22.67279 | 9.710646 | 0.6869 | 0.290122 | 0.233497 |
| 6 | 5069200 | 5071199 | 24.41623 | 42.87003 | 28.06193 | 12.41529 | 0.995569 | 0.64968 | 0.233501 |
| 9 | 38542400 | 38544799 | 31.846 | 54.12182 | 39.74404 | 17.41818 | 0.825314 | 1.050579 | 0.233511 |
| X | 18154600 | 18156599 | 22.08224 | 39.48307 | 23.88559 | 10.48401 | 1.119844 | 0.470181 | 0.233512 |
| 7 | 5222600 | 5224799 | 25.3619 | 44.01967 | 34.33063 | 15.06334 | 0.743713 | 0.830079 | 0.233529 |
| 1 | 160733600 | 160734799 | 8.409653 | 14.99246 | 39.9815 | 17.59235 | -1.00946 | 0.43024 | 0.233537 |
| 4 | 93195600 | 93197799 | 16.29943 | 28.07158 | 55.40855 | 24.30569 | -0.59233 | 0.889732 | 0.233547 |
| X | 32730600 | 32734199 | 35.63221 | 61.12182 | 31.36874 | 13.75792 | 1.341792 | 0.977078 | 0.233553 |
| 17 | 38587800 | 38590799 | 27.90742 | 48.93259 | 25.07418 | 11.08056 | 1.346745 | 0.685742 | 0.233564 |
| 12 | 103651000 | 103658999 | 52.50352 | 83.08557 | 155.2632 | 64.53917 | -0.57294 | 2.383695 | 0.233578 |
| X | 6879000 | 6882199 | 58.75547 | 95.00183 | 58.76702 | 25.11442 | 1.052918 | 1.722689 | 0.233597 |
| 2 | 36530000 | 36531199 | 18.29846 | 33.01281 | 24.77209 | 10.7798 | 0.819019 | 0.39185 | 0.233601 |
| 14 | 113237000 | 113241199 | 51.253 | 82.76788 | 68.27921 | 29.13265 | 0.636595 | 1.74895 | 0.233607 |
| 14 | 47118800 | 47123799 | 24.98348 | 40.57268 | 88.34546 | 37.88157 | -0.75214 | 1.586627 | 0.233611 |
| 12 | 61675000 | 61675999 | 13.93716 | 24.22839 | 53.20624 | 23.3686 | -0.74252 | 0.786585 | 0.233633 |
| 15 | 96768800 | 96769999 | 15.09241 | 28.48532 | 19.41612 | 7.915378 | 1.024347 | 0.094912 | 0.233634 |
| 7 | 139660200 | 139663999 | 26.67318 | 43.44546 | 86.39659 | 37.09855 | -0.61837 | 1.539404 | 0.233644 |
| 15 | 80975000 | 80977199 | 12.11086 | 21.26898 | 44.57462 | 19.72461 | -0.67446 | 0.605789 | 0.233648 |
| 11 | 27376800 | 27381199 | 61.95607 | 99.06754 | 87.41581 | 36.77159 | 0.522994 | 2.067586 | 0.233657 |
| 12 | 45505400 | 45506399 | 5.53464 | 10.72627 | 27.39999 | 10.37334 | -0.76661 | -0.21844 | 0.233665 |
| 10 | 91006600 | 91008599 | 9.963869 | 17.57435 | 42.51297 | 18.81323 | -0.87782 | 0.537696 | 0.233667 |
| 13 | 60336200 | 60338199 | 12.2459 | 21.56844 | 45.06981 | 19.95043 | -0.6692 | 0.570302 | 0.233677 |
| 2 | 93777800 | 93780799 | 28.65615 | 49.13833 | 36.98624 | 16.22123 | 0.791482 | 0.953988 | 0.23368 |
| 9 | 77298600 | 77298999 | 4.353511 | 8.493774 | 22.74348 | 8.122445 | -0.75402 | -0.45179 | 0.233694 |
| 7 | 44089000 | 44089999 | 6.127838 | 11.80836 | 28.1646 | 10.92033 | -0.70143 | -0.12765 | 0.233701 |
| 1 | 96282600 | 96282999 | 7.185637 | 14.0516 | 11.70023 | 3.986434 | 0.969482 | -0.61958 | 0.233714 |
| 7 | 10785400 | 10788799 | 55.16557 | 88.79815 | 73.98923 | 31.43136 | 0.616764 | 1.852051 | 0.233719 |
| 15 | 32886600 | 32887999 | 15.7882 | 28.72687 | 24.072 | 10.36449 | 0.670458 | 0.323891 | 0.233726 |
| 13 | 44630000 | 44633599 | 30.63696 | 49.49134 | 97.86051 | 41.77158 | -0.62117 | 1.718371 | 0.233733 |
| 14 | 37599000 | 37603199 | 77.89542 | 123.2213 | 111.1761 | 46.20441 | 0.475584 | 2.402443 | 0.233734 |
| 1 | 132041200 | 132047199 | 28.77807 | 46.36982 | 101.3689 | 43.12552 | -0.76984 | 1.794281 | 0.233735 |
| 7 | 37176400 | 37180199 | 25.6466 | 41.55352 | 90.68363 | 38.81802 | -0.75846 | 1.637785 | 0.233743 |
| 19 | 53128800 | 53132199 | 22.37227 | 36.50355 | 83.24813 | 35.76928 | -0.81338 | 1.505325 | 0.233745 |
| 3 | 3747400 | 3750999 | 44.53369 | 72.62194 | 57.32727 | 24.62676 | 0.712203 | 1.539949 | 0.233747 |
| 3 | 32086600 | 32090199 | 18.3263 | 30.90056 | 65.69301 | 28.72397 | -0.70315 | 1.096532 | 0.23376 |
| 13 | 62376000 | 62380599 | 93.78202 | 147.6461 | 115.7357 | 48.01937 | 0.672851 | 2.517738 | 0.233762 |
| 3 | 158441400 | 158441799 | 8.235762 | 16.04757 | 14.06552 | 5.108483 | 0.816783 | -0.37324 | 0.233762 |
| X | 13185200 | 13188599 | 43.67248 | 71.51112 | 54.96565 | 23.64815 | 0.75461 | 1.483407 | 0.233774 |
| 17 | 33903600 | 33904599 | 5.601351 | 10.70319 | 29.46403 | 11.70226 | -0.93922 | -0.05172 | 0.233775 |
| 18 | 65849800 | 65854599 | 28.09164 | 45.37129 | 101.6759 | 43.38994 | -0.80153 | 1.723211 | 0.233776 |
| 3 | 8208800 | 8215799 | 58.21575 | 93.55153 | 71.07011 | 30.11829 | 0.746826 | 1.906243 | 0.233789 |
| 10 | 35494200 | 35495399 | 25.03233 | 43.11629 | 37.26556 | 16.34708 | 0.594114 | 0.901352 | 0.233791 |
| 2 | 36987400 | 36989399 | 20.89764 | 36.75683 | 23.2086 | 10.27621 | 1.047404 | 0.595447 | 0.233794 |
| 1 | 190956000 | 190957799 | 10.2643 | 18.10099 | 42.00152 | 18.58757 | -0.81815 | 0.541435 | 0.233795 |
| 1 | 194561600 | 194562599 | 5.386541 | 10.4301 | 26.06059 | 9.906724 | -0.73982 | -0.2077 | 0.233804 |

| | | | | | | | | | |
|---|---|---|---|---|---|---|---|---|---|
| 6 | 13124400 | 13125799 | 6.939233 | 13.08068 | 30.78176 | 12.57935 | -0.75162 | 0.060112 | 0.233817 |
| 13 | 30138600 | 30142399 | 20.55743 | 34.15949 | 69.73991 | 30.24745 | -0.6452 | 1.262017 | 0.233823 |
| 9 | 97209200 | 97210999 | 21.19279 | 37.64473 | 26.08779 | 11.51418 | 0.919719 | 0.507788 | 0.233834 |
| 7 | 64198400 | 64204799 | 50.94148 | 80.51414 | 154.204 | 64.05883 | -0.60976 | 2.412701 | 0.233838 |
| 16 | 31481200 | 31483599 | 12.13656 | 21.24874 | 45.70035 | 20.17033 | -0.71159 | 0.65843 | 0.233853 |
| 3 | 142804600 | 142806999 | 14.78 | 25.2931 | 56.34589 | 24.70787 | -0.76773 | 0.942419 | 0.233854 |
| 1 | 183487600 | 183490799 | 19.15716 | 32.18473 | 65.07133 | 28.42091 | -0.63114 | 1.13148 | 0.233863 |
| 8 | 38098400 | 38101399 | 25.86419 | 45.08083 | 32.04623 | 14.09825 | 0.87733 | 0.772685 | 0.233898 |
| 12 | 59256800 | 59260599 | 63.16518 | 101.1559 | 83.38467 | 35.1533 | 0.623456 | 2.021994 | 0.233908 |
| 2 | 156324200 | 156328399 | 37.3727 | 59.88659 | 114.8328 | 48.45562 | -0.59112 | 1.984185 | 0.233913 |
| 13 | 31898000 | 31901399 | 23.16186 | 38.22224 | 74.8987 | 32.33037 | -0.58724 | 1.352128 | 0.233931 |
| 4 | 112454200 | 112457599 | 61.20884 | 98.42243 | 72.37889 | 30.70053 | 0.794723 | 1.89169 | 0.233936 |
| 6 | 97315200 | 97317999 | 16.69512 | 28.40114 | 58.50368 | 25.64179 | -0.65672 | 1.007274 | 0.233961 |
| 7 | 18290400 | 18291599 | 20.77392 | 36.35965 | 30.89948 | 13.63419 | 0.621206 | 0.68408 | 0.233995 |
| 6 | 136304000 | 136307799 | 74.65281 | 115.5474 | 202.4494 | 84.09232 | -0.50514 | 2.824738 | 0.23401 |
| 19 | 8872200 | 8875999 | 16.96746 | 28.53819 | 63.76953 | 27.86188 | -0.77467 | 1.113875 | 0.234018 |
| 18 | 20191200 | 20192799 | 9.312594 | 16.55312 | 39.94058 | 17.61825 | -0.86875 | 0.455237 | 0.234023 |
| 6 | 49821000 | 49822399 | 18.91024 | 34.01939 | 24.64015 | 10.75947 | 0.866483 | 0.411656 | 0.234023 |
| 4 | 145592400 | 145595599 | 140.8473 | 214.1597 | 188.9467 | 78.2733 | 0.463733 | 3.191597 | 0.234023 |
| 12 | 9239800 | 9240599 | 11.57683 | 22.06315 | 19.44457 | 7.765899 | 0.682623 | -0.00297 | 0.234026 |
| 2 | 55961000 | 55961799 | 10.2599 | 19.95223 | 15.21507 | 5.633232 | 0.98913 | -0.27864 | 0.234031 |
| 5 | 147204400 | 147210199 | 45.42481 | 72.20816 | 145.3196 | 60.63889 | -0.67364 | 2.250769 | 0.234038 |
| 14 | 48174400 | 48175799 | 9.789849 | 17.78975 | 38.12045 | 16.43981 | -0.67692 | 0.318774 | 0.234043 |
| 13 | 65531400 | 65538999 | 49.82624 | 80.52958 | 58.03538 | 24.78803 | 0.832388 | 1.717746 | 0.234043 |
| 6 | 31193000 | 31195799 | 19.39979 | 32.00789 | 77.22555 | 33.33193 | -0.8864 | 1.345829 | 0.23405 |
| 3 | 9026400 | 9030199 | 16.8684 | 28.6116 | 61.22793 | 26.81985 | -0.71205 | 1.035909 | 0.234063 |
| 7 | 10751200 | 10755999 | 50.362 | 81.84086 | 57.99677 | 24.88047 | 0.863972 | 1.605145 | 0.234069 |
| 5 | 41817800 | 41819599 | 20.01796 | 35.67629 | 26.21366 | 11.53735 | 0.838485 | 0.480051 | 0.234073 |
| 3 | 19643200 | 19645199 | 26.29956 | 45.82624 | 32.10186 | 14.12012 | 0.898444 | 0.779823 | 0.2341 |
| 11 | 15811400 | 15812799 | 34.13182 | 56.70849 | 48.72704 | 21.13198 | 0.599234 | 1.282624 | 0.234107 |
| 5 | 54603400 | 54603999 | 4.281101 | 8.324898 | 25.81807 | 9.563285 | -1.01296 | -0.31493 | 0.234116 |
| 18 | 36892800 | 36894399 | 9.014919 | 16.45488 | 35.53752 | 15.24056 | -0.6825 | 0.282729 | 0.23412 |
| 10 | 45502200 | 45504199 | 7.706861 | 14.37812 | 37.30516 | 15.51474 | -0.91308 | 0.145436 | 0.234136 |
| 15 | 46367200 | 46369599 | 25.13109 | 43.57721 | 31.52304 | 13.83086 | 0.851604 | 0.835243 | 0.234139 |
| 4 | 155682600 | 155683399 | 7.088074 | 13.35735 | 30.77593 | 12.58319 | -0.72193 | 0.063814 | 0.234149 |
| 7 | 125829200 | 125830599 | 7.088074 | 13.35735 | 30.77593 | 12.58319 | -0.72193 | 0.063814 | 0.234149 |
| 18 | 20814800 | 20816199 | 12.76263 | 22.3618 | 45.67915 | 20.1739 | -0.63765 | 0.65019 | 0.23415 |
| 12 | 5292200 | 5297999 | 30.52013 | 49.20039 | 102.3821 | 43.58478 | -0.69815 | 1.781052 | 0.234168 |
| 4 | 123864000 | 123869999 | 36.18387 | 58.16834 | 107.8576 | 45.73281 | -0.5372 | 1.879234 | 0.234171 |
| 10 | 4417200 | 4419399 | 16.03054 | 27.40428 | 56.38112 | 24.72123 | -0.65377 | 0.957498 | 0.234174 |
| 5 | 134499800 | 134501399 | 14.08272 | 26.01065 | 21.70051 | 9.159885 | 0.696323 | 0.221323 | 0.234181 |
| 1 | 7734200 | 7735199 | 8.406767 | 15.3645 | 35.03518 | 15.00106 | -0.75868 | 0.269407 | 0.234184 |
| 17 | 38907200 | 38914999 | 74.92928 | 119.2653 | 88.57894 | 37.02331 | 0.764134 | 2.222507 | 0.234184 |
| 3 | 4955200 | 4958999 | 63.43946 | 101.4807 | 86.12206 | 36.24822 | 0.579757 | 2.056642 | 0.234184 |
| 8 | 71340200 | 71351799 | 78.33122 | 119.61 | 248.1542 | 102.9182 | -0.76384 | 3.092854 | 0.234185 |
| 14 | 89028600 | 89029199 | 10.12456 | 19.65939 | 17.07109 | 6.397805 | 0.789127 | -0.23054 | 0.234186 |
| 11 | 109374800 | 109377599 | 21.30128 | 35.48124 | 70.07736 | 30.44325 | -0.59751 | 1.234707 | 0.234187 |

| | | | | | | | | | |
|---|---|---|---|---|---|---|---|---|---|
| 7 | 43029200 | 43029999 | 10.62643 | 20.60849 | 16.90097 | 6.377926 | 0.86086 | -0.20365 | 0.234193 |
| 4 | 112064600 | 112066599 | 42.33429 | 69.73168 | 51.10333 | 22.0416 | 0.82644 | 1.406194 | 0.234204 |
| 12 | 99606200 | 99607999 | 9.115716 | 16.32272 | 39.46168 | 17.29946 | -0.86545 | 0.404868 | 0.234204 |
| X | 32764200 | 32765199 | 13.97049 | 26.66197 | 18.32308 | 7.296556 | 1.04291 | 0.010178 | 0.234208 |
| 12 | 20432000 | 20435399 | 78.42819 | 123.874 | 110.0223 | 45.68946 | 0.496472 | 2.438829 | 0.234221 |
| 3 | 100512000 | 100512999 | 8.565798 | 15.61262 | 36.04586 | 15.4864 | -0.77991 | 0.287748 | 0.234241 |
| X | 24505200 | 24514999 | 131.2088 | 203.4421 | 124.9244 | 51.88676 | 1.007809 | 2.80964 | 0.234252 |
| 9 | 11597400 | 11599199 | 21.71364 | 38.54454 | 25.74775 | 11.3704 | 0.97213 | 0.516061 | 0.234255 |
| 7 | 24582400 | 24585999 | 19.87506 | 32.76965 | 77.23461 | 33.32621 | -0.85316 | 1.355696 | 0.234258 |
| 9 | 22920800 | 22921799 | 5.123039 | 9.899306 | 26.48336 | 10.16556 | -0.85091 | -0.1771 | 0.234261 |
| 14 | 54491000 | 54491799 | 5.123039 | 9.899306 | 26.48336 | 10.16556 | -0.85091 | -0.1771 | 0.234261 |
| 8 | 47764200 | 47765999 | 13.23168 | 23.17471 | 46.73523 | 20.63358 | -0.61932 | 0.66026 | 0.234266 |
| 17 | 19234400 | 19235399 | 31.77014 | 53.18136 | 45.95106 | 20.02459 | 0.591198 | 1.192352 | 0.234271 |
| X | 23805800 | 23807599 | 20.08554 | 37.49199 | 16.59952 | 6.897329 | 1.619604 | 0.200407 | 0.234274 |
| 7 | 5280400 | 5286599 | 139.4475 | 212.7896 | 179.6211 | 74.48351 | 0.531582 | 3.118757 | 0.234277 |
| 10 | 25853600 | 25854799 | 24.04864 | 41.69306 | 35.63543 | 15.63511 | 0.61198 | 0.83741 | 0.23428 |
| 1 | 175747600 | 175749199 | 8.963008 | 16.05859 | 37.94734 | 16.62746 | -0.83178 | 0.3996 | 0.234289 |
| 13 | 95740200 | 95748799 | 127.043 | 189.7134 | 331.2446 | 136.5528 | -0.54298 | 3.550054 | 0.234331 |
| 17 | 94962000 | 94964199 | 23.01353 | 40.17371 | 32.74926 | 14.42143 | 0.68084 | 0.737751 | 0.234334 |
| 7 | 49426200 | 49430399 | 20.75935 | 34.34211 | 73.59846 | 31.82321 | -0.71562 | 1.315627 | 0.234346 |
| 19 | 17410600 | 17411199 | 5.831202 | 11.27 | 29.01322 | 11.12602 | -0.79589 | -0.16769 | 0.234355 |
| 2 | 165765400 | 165771999 | 63.6771 | 98.65941 | 201.609 | 83.74277 | -0.72566 | 2.801435 | 0.234372 |
| 7 | 111932600 | 111934599 | 11.36754 | 19.98099 | 44.0727 | 19.51688 | -0.74737 | 0.590075 | 0.234376 |
| 15 | 87445800 | 87456599 | 73.69276 | 113.6868 | 215.4989 | 89.51426 | -0.62221 | 2.885645 | 0.234386 |
| 5 | 42476000 | 42477199 | 15.84028 | 28.64948 | 23.72648 | 10.29256 | 0.680592 | 0.358114 | 0.234389 |
| 5 | 112166600 | 112167999 | 15.84028 | 28.64948 | 23.72648 | 10.29256 | 0.680592 | 0.358114 | 0.234389 |
| 14 | 102121400 | 102125799 | 40.28706 | 64.54034 | 116.807 | 49.26975 | -0.50839 | 1.996222 | 0.234391 |
| 10 | 62077400 | 62080799 | 24.83186 | 40.72614 | 79.2116 | 34.10279 | -0.58013 | 1.436142 | 0.234411 |
| 1 | 120268000 | 120274399 | 54.15758 | 85.20338 | 170.129 | 70.58103 | -0.67508 | 2.51636 | 0.234412 |
| 14 | 100382200 | 100383399 | 14.85127 | 27.50455 | 22.66686 | 9.532583 | 0.717868 | 0.21593 | 0.234424 |
| 2 | 89453800 | 89455599 | 20.64573 | 36.98249 | 24.49466 | 10.7347 | 0.991327 | 0.449899 | 0.234439 |
| 6 | 10146800 | 10147999 | 46.12023 | 75.16379 | 54.83474 | 23.55084 | 0.825207 | 1.556057 | 0.234463 |
| 10 | 126558600 | 126559199 | 5.285365 | 10.12386 | 27.81959 | 10.97436 | -0.92628 | -0.07894 | 0.234465 |
| 15 | 59519200 | 59523599 | 26.39843 | 42.58254 | 103.3708 | 44.04115 | -0.91873 | 1.75681 | 0.234468 |
| 12 | 22751400 | 22776799 | 781.5793 | 1034.359 | 795.9489 | 332.7843 | 0.466227 | 5.446727 | 0.234479 |
| 7 | 31476600 | 31481999 | 49.76949 | 80.88291 | 53.99991 | 23.16687 | 0.949798 | 1.599844 | 0.234489 |
| 17 | 18783600 | 18787399 | 96.10166 | 150.3476 | 129.652 | 53.7862 | 0.528424 | 2.639873 | 0.234496 |
| 18 | 50813200 | 50814399 | 11.40445 | 22.05759 | 16.08429 | 6.146955 | 1.010827 | -0.15696 | 0.2345 |
| 15 | 31348000 | 31350399 | 9.429306 | 16.8501 | 38.43413 | 16.87777 | -0.78342 | 0.420328 | 0.234517 |
| 2 | 151821400 | 151822599 | 8.921686 | 15.73386 | 42.55445 | 18.83213 | -1.03754 | 0.534191 | 0.234529 |
| 4 | 48279000 | 48280199 | 8.921686 | 15.73386 | 42.55445 | 18.83213 | -1.03754 | 0.534191 | 0.234529 |
| 2 | 177069400 | 177071399 | 20.15903 | 35.55042 | 28.98354 | 12.82651 | 0.682267 | 0.574567 | 0.234532 |
| 15 | 82480600 | 82483199 | 32.80946 | 52.80041 | 112.1108 | 47.61024 | -0.73028 | 1.843954 | 0.234547 |
| 8 | 13028800 | 13036799 | 62.98137 | 98.50714 | 177.6665 | 73.70695 | -0.53509 | 2.633495 | 0.23455 |
| 9 | 81160600 | 81163799 | 34.91228 | 58.56233 | 41.97857 | 18.31739 | 0.859479 | 1.1738 | 0.234554 |
| 10 | 12846800 | 12847399 | 10.2453 | 19.97219 | 12.82816 | 4.631712 | 1.26479 | -0.37274 | 0.234567 |
| X | 89752400 | 89755399 | 38.80956 | 65.23173 | 39.16746 | 17.1094 | 1.114253 | 1.168295 | 0.234583 |

| | | | | | | | | | |
|---|---|---|---|---|---|---|---|---|---|
| 14 | 42459800 | 42462999 | 39.90045 | 65.31373 | 55.48674 | 23.8691 | 0.610327 | 1.474882 | 0.234583 |
| 9 | 99336400 | 99337199 | 4.234967 | 8.242451 | 23.47661 | 8.634282 | -0.88048 | -0.34487 | 0.234597 |
| 15 | 11035200 | 11038199 | 25.18112 | 40.79671 | 91.27073 | 39.06747 | -0.7943 | 1.637673 | 0.234602 |
| 4 | 117162800 | 117164799 | 9.428113 | 16.83925 | 40.47022 | 17.77964 | -0.85929 | 0.424649 | 0.234619 |
| 10 | 67598600 | 67600999 | 24.7705 | 42.98549 | 33.81853 | 14.83837 | 0.73103 | 0.827422 | 0.234636 |
| 18 | 4839200 | 4841799 | 12.28304 | 21.48388 | 48.88614 | 21.55833 | -0.79284 | 0.678121 | 0.234651 |
| 11 | 24558000 | 24558999 | 8.535649 | 15.66768 | 34.57437 | 14.71781 | -0.70571 | 0.244661 | 0.234656 |
| X | 31753200 | 31754599 | 28.69574 | 49.21832 | 31.70503 | 13.9053 | 1.015399 | 0.948435 | 0.234684 |
| 12 | 5423400 | 5427599 | 27.45765 | 44.3602 | 96.36598 | 41.13932 | -0.75624 | 1.710267 | 0.234697 |
| 6 | 28279600 | 28280199 | 4.483573 | 8.742827 | 23.24456 | 8.360592 | -0.75309 | -0.42203 | 0.234702 |
| 2 | 91022200 | 91024199 | 7.933264 | 14.60881 | 34.09852 | 14.45196 | -0.78093 | 0.222841 | 0.234707 |
| 2 | 30988400 | 30992799 | 33.00901 | 52.98381 | 110.8433 | 46.89279 | -0.71352 | 1.92452 | 0.234712 |
| 13 | 77770400 | 77772999 | 38.32983 | 63.57832 | 48.80589 | 21.14208 | 0.761142 | 1.311804 | 0.234713 |
| 5 | 29885000 | 29887999 | 35.52728 | 58.85233 | 50.96487 | 22.05731 | 0.587687 | 1.320463 | 0.234717 |
| 7 | 31966600 | 31968599 | 33.43485 | 56.82163 | 38.00573 | 16.65634 | 0.959468 | 1.054157 | 0.234719 |
| 6 | 135628600 | 135630199 | 7.398143 | 13.8601 | 33.42931 | 13.80456 | -0.79973 | 0.10841 | 0.234722 |
| 3 | 136764600 | 136765399 | 4.753106 | 9.261039 | 27.56324 | 10.01776 | -0.92858 | -0.36793 | 0.234731 |
| 6 | 7281400 | 7289599 | 162.3209 | 244.383 | 213.6289 | 88.23317 | 0.465026 | 3.395521 | 0.234735 |
| 6 | 51687200 | 51690999 | 24.36393 | 39.91343 | 80.85014 | 34.79154 | -0.63926 | 1.449383 | 0.234748 |
| 7 | 139702800 | 139707599 | 21.48739 | 35.30147 | 78.25877 | 33.71704 | -0.76721 | 1.405941 | 0.234749 |
| 1 | 180082200 | 180084199 | 9.192546 | 16.25705 | 41.00026 | 18.13417 | -0.9356 | 0.502166 | 0.23475 |
| 11 | 70140800 | 70147199 | 35.97689 | 57.52855 | 122.369 | 51.47674 | -0.74396 | 2.050067 | 0.234754 |
| 12 | 32944400 | 32947999 | 16.4459 | 27.81377 | 60.53507 | 26.49292 | -0.73632 | 1.0615 | 0.234756 |
| 15 | 35975000 | 35975599 | 9.81287 | 19.07206 | 15.86857 | 5.902179 | 0.859041 | -0.26623 | 0.234761 |
| 12 | 32870000 | 32870999 | 6.232761 | 11.40344 | 36.11192 | 15.44051 | -1.22678 | 0.248825 | 0.234789 |
| 17 | 13900000 | 13911999 | 73.12052 | 112.9629 | 209.2934 | 86.93598 | -0.58774 | 2.859053 | 0.234795 |
| 18 | 36250400 | 36252799 | 13.06403 | 22.44785 | 55.24779 | 24.23348 | -0.90967 | 0.896648 | 0.234801 |
| 19 | 6111200 | 6114599 | 13.06403 | 22.44785 | 55.24779 | 24.23348 | -0.90967 | 0.896648 | 0.234801 |
| 1 | 31651800 | 31654399 | 26.18667 | 45.11436 | 36.54224 | 16.02989 | 0.687282 | 0.901816 | 0.234802 |
| 17 | 19069000 | 19070199 | 12.25311 | 23.51583 | 18.86495 | 7.400561 | 0.84054 | -0.05456 | 0.234817 |
| 13 | 41911400 | 41914399 | 17.84743 | 30.19829 | 60.65106 | 26.54748 | -0.62097 | 1.061938 | 0.234831 |
| 8 | 3869800 | 3871799 | 19.62607 | 35.82992 | 21.43459 | 9.190288 | 1.157091 | 0.328791 | 0.234833 |
| 12 | 28622000 | 28624199 | 21.96804 | 36.53184 | 70.59387 | 30.63426 | -0.5662 | 1.256417 | 0.234836 |
| 2 | 164354600 | 164360999 | 39.26862 | 62.58798 | 136.9595 | 57.3513 | -0.79003 | 2.160785 | 0.234858 |
| 18 | 52838800 | 52841799 | 24.448 | 42.47686 | 30.97118 | 13.59804 | 0.840576 | 0.808164 | 0.23486 |
| X | 89633600 | 89634199 | 8.381116 | 16.39433 | 9.473384 | 3.164824 | 1.510263 | -0.67381 | 0.234872 |
| 14 | 83746800 | 83751599 | 46.41762 | 75.63251 | 58.39186 | 25.07665 | 0.743442 | 1.558859 | 0.234882 |
| 3 | 30937600 | 30942399 | 32.48418 | 52.2884 | 104.0034 | 44.18134 | -0.63577 | 1.834108 | 0.234897 |
| 1 | 8016600 | 8017799 | 16.42283 | 30.5718 | 20.81796 | 8.690101 | 0.998804 | 0.195195 | 0.234915 |
| 2 | 163673600 | 163680399 | 55.14703 | 86.7638 | 164.2393 | 68.13797 | -0.59809 | 2.516893 | 0.234939 |
| 7 | 45406400 | 45410999 | 25.04649 | 40.93006 | 81.61187 | 35.08596 | -0.61869 | 1.48783 | 0.234947 |
| 2 | 122206000 | 122207199 | 6.961846 | 13.05488 | 32.87504 | 13.55426 | -0.85942 | 0.097467 | 0.234987 |
| 7 | 14793800 | 14795599 | 36.32085 | 59.93086 | 50.39736 | 21.75313 | 0.629556 | 1.368959 | 0.234988 |
| 3 | 84187600 | 84190199 | 10.56556 | 18.62369 | 43.52492 | 19.26351 | -0.82902 | 0.549137 | 0.235003 |
| 1 | 115372600 | 115375999 | 53.31758 | 85.83744 | 74.39212 | 31.60994 | 0.560406 | 1.844069 | 0.235006 |
| 7 | 65531000 | 65535799 | 27.81939 | 45.25636 | 86.27815 | 37.03098 | -0.55889 | 1.5609 | 0.235007 |
| 12 | 18353400 | 18356799 | 79.64416 | 126.6931 | 93.78105 | 39.16471 | 0.768075 | 2.249169 | 0.235017 |

| | | | | | | | | | |
|---|---|---|---|---|---|---|---|---|---|
| 19 | 13643200 | 13645599 | 31.31289 | 54.14215 | 31.95476 | 14.01907 | 1.143071 | 0.889811 | 0.235041 |
| 4 | 152745600 | 152776599 | 367.6052 | 502.4429 | 824.9469 | 343.2955 | -0.58016 | 4.930892 | 0.235045 |
| 8 | 57965200 | 57967199 | 43.72166 | 70.8886 | 62.94296 | 26.95715 | 0.53646 | 1.639143 | 0.235063 |
| 9 | 71631800 | 71635599 | 22.59468 | 37.10005 | 79.01698 | 34.03572 | -0.71 | 1.415509 | 0.235065 |
| 3 | 90155000 | 90156799 | 8.466453 | 15.30383 | 36.47452 | 15.83429 | -0.83495 | 0.338398 | 0.235077 |
| 13 | 67457800 | 67458599 | 11.98236 | 23.32137 | 13.89188 | 5.108678 | 1.350375 | -0.28305 | 0.235077 |
| 1 | 78473600 | 78482199 | 96.12611 | 146.2113 | 259.2913 | 107.4255 | -0.54225 | 3.177697 | 0.235093 |
| 2 | 44262200 | 44263799 | 14.8969 | 27.72082 | 22.17847 | 9.263911 | 0.767544 | 0.188764 | 0.235094 |
| 4 | 22526600 | 22527799 | 14.8969 | 27.72082 | 22.17847 | 9.263911 | 0.767544 | 0.188764 | 0.235094 |
| 5 | 149409200 | 149410199 | 14.8969 | 27.72082 | 22.17847 | 9.263911 | 0.767544 | 0.188764 | 0.235094 |
| 9 | 11821600 | 11824999 | 111.7782 | 172.2502 | 148.6043 | 61.72785 | 0.511035 | 2.913133 | 0.235094 |
| 7 | 122140200 | 122140799 | 2.838802 | 5.552222 | 18.93186 | 6.400703 | -1.01961 | -0.67647 | 0.2351 |
| 3 | 10340600 | 10342599 | 11.33926 | 20.04743 | 42.47715 | 18.78853 | -0.68643 | 0.521976 | 0.235102 |
| 18 | 44670200 | 44671399 | 6.512232 | 12.25683 | 30.73065 | 12.59135 | -0.84511 | 0.066452 | 0.235114 |
| 3 | 101575000 | 101575799 | 4.969253 | 9.592288 | 26.38484 | 10.17646 | -0.89656 | -0.16078 | 0.235117 |
| 9 | 87611400 | 87613799 | 19.03188 | 31.52849 | 78.15468 | 33.82597 | -0.92481 | 1.296289 | 0.235138 |
| 18 | 86247200 | 86248199 | 11.75284 | 22.69286 | 17.85326 | 6.877333 | 0.892573 | -0.12046 | 0.23514 |
| 9 | 11012200 | 11012599 | 10.97996 | 21.03247 | 18.52451 | 7.306728 | 0.699928 | -0.05083 | 0.235144 |
| 9 | 36542000 | 36544999 | 37.78852 | 63.29664 | 41.98149 | 18.30219 | 0.971304 | 1.200107 | 0.235148 |
| 5 | 23542000 | 23544599 | 15.9694 | 27.51904 | 54.32107 | 23.82907 | -0.59221 | 0.88151 | 0.235165 |
| 10 | 120849800 | 120850999 | 37.19648 | 61.62636 | 50.45154 | 21.83733 | 0.668509 | 1.323286 | 0.235176 |
| Y | 4046400 | 4048399 | 12.56018 | 24.00456 | 10.23722 | 4.06439 | 1.725234 | -0.04219 | 0.235191 |
| 12 | 22309000 | 22312599 | 48.87046 | 78.81515 | 65.98882 | 28.10701 | 0.613077 | 1.78175 | 0.235214 |
| 14 | 52157600 | 52162199 | 27.0708 | 43.89303 | 90.2198 | 38.64022 | -0.67139 | 1.623542 | 0.235233 |
| 12 | 14946000 | 14947799 | 29.66161 | 50.10366 | 43.15445 | 18.87941 | 0.59581 | 1.09712 | 0.235239 |
| 18 | 85942800 | 85945399 | 32.14794 | 55.20381 | 32.92234 | 14.44052 | 1.126059 | 0.95175 | 0.235271 |
| X | 26022600 | 26026399 | 51.1259 | 83.42054 | 51.77511 | 22.24637 | 1.058906 | 1.548218 | 0.235271 |
| 7 | 10887800 | 10892199 | 85.54763 | 135.3189 | 108.9142 | 45.26317 | 0.640061 | 2.409375 | 0.235272 |
| 16 | 42835000 | 42836799 | 26.16206 | 45.71777 | 30.38637 | 13.38985 | 0.974144 | 0.737462 | 0.235282 |
| 1 | 100600800 | 100603399 | 43.30404 | 71.28554 | 50.62403 | 21.82924 | 0.871278 | 1.413439 | 0.235282 |
| 7 | 98772800 | 98774599 | 22.8636 | 39.80928 | 33.9674 | 14.93364 | 0.615061 | 0.774768 | 0.235286 |
| 6 | 10429800 | 10431599 | 17.88773 | 33.15233 | 20.04329 | 8.420851 | 1.162158 | 0.230267 | 0.235302 |
| X | 17692400 | 17693399 | 19.50905 | 34.66902 | 26.54171 | 11.71021 | 0.777462 | 0.496567 | 0.235315 |
| 4 | 5183600 | 5186799 | 48.16384 | 77.82964 | 68.20096 | 29.12359 | 0.550726 | 1.723549 | 0.23534 |
| 11 | 50267600 | 50270399 | 11.15918 | 19.38714 | 54.23723 | 23.81215 | -1.09025 | 0.779173 | 0.235353 |
| 6 | 65820800 | 65821799 | 11.46797 | 21.86335 | 19.00358 | 7.584155 | 0.703616 | -0.00693 | 0.235366 |
| 15 | 57870000 | 57873599 | 15.6551 | 26.51608 | 59.96099 | 26.25362 | -0.7911 | 1.045196 | 0.235371 |
| 15 | 11948200 | 11951799 | 26.23241 | 42.64397 | 86.46526 | 37.10155 | -0.64825 | 1.568087 | 0.235396 |
| 4 | 100335000 | 100336999 | 8.806745 | 16.23543 | 39.14481 | 16.56551 | -0.82714 | 0.229401 | 0.235397 |
| 19 | 20952200 | 20953799 | 22.57389 | 39.26582 | 33.61086 | 14.76766 | 0.610424 | 0.787349 | 0.235407 |
| 7 | 90125000 | 90126199 | 8.069321 | 14.89503 | 33.60505 | 14.19642 | -0.72905 | 0.20809 | 0.235414 |
| 7 | 85510600 | 85511199 | 9.312805 | 18.10223 | 16.59515 | 6.166687 | 0.722679 | -0.27153 | 0.235416 |
| 9 | 5033200 | 5034199 | 13.68127 | 25.80704 | 20.72079 | 8.457573 | 0.788663 | 0.08954 | 0.235418 |
| 14 | 83510200 | 83511599 | 13.68127 | 25.80704 | 20.72079 | 8.457573 | 0.788663 | 0.08954 | 0.235418 |
| 14 | 46988400 | 46988799 | 9.884784 | 19.30235 | 13.15523 | 4.627468 | 1.215337 | -0.46274 | 0.235422 |
| 4 | 113669800 | 113672799 | 37.5871 | 63.05768 | 40.83194 | 17.81858 | 1.005672 | 1.182052 | 0.235425 |
| 17 | 48077400 | 48084799 | 78.79494 | 120.9919 | 225.0114 | 93.42221 | -0.59988 | 2.97936 | 0.235437 |

| | | | | | | | | | |
|---|---|---|---|---|---|---|---|---|---|
| 9 | 103621000 | 103623399 | 15.34901 | 26.46838 | 53.21692 | 23.3452 | -0.61829 | 0.872042 | 0.235489 |
| 16 | 84570600 | 84572199 | 9.259304 | 16.27363 | 45.6397 | 20.20907 | -1.09193 | 0.578667 | 0.235491 |
| 4 | 46986800 | 46997799 | 69.44331 | 107.1132 | 231.3977 | 96.11844 | -0.81082 | 2.888859 | 0.235496 |
| 15 | 68149200 | 68149799 | 4.227227 | 8.250632 | 22.86955 | 8.112319 | -0.79474 | -0.4786 | 0.235497 |
| 10 | 95626400 | 95626799 | 3.932807 | 7.685009 | 21.92999 | 7.614606 | -0.80754 | -0.56135 | 0.235498 |
| 18 | 23884600 | 23887799 | 23.67466 | 38.96293 | 76.44514 | 32.96 | -0.5906 | 1.388477 | 0.23552 |
| 2 | 36477400 | 36477999 | 13.84303 | 25.63502 | 21.80186 | 9.169911 | 0.672664 | 0.205973 | 0.235529 |
| 14 | 42245400 | 42248799 | 50.16346 | 81.29535 | 57.34402 | 24.55084 | 0.866969 | 1.656168 | 0.235533 |
| 10 | 29485400 | 29486799 | 17.96036 | 32.94114 | 22.20621 | 9.463073 | 0.992074 | 0.292398 | 0.235545 |
| 12 | 22288200 | 22290799 | 25.77417 | 44.77186 | 34.40833 | 15.1052 | 0.765076 | 0.823029 | 0.235547 |
| 5 | 10592600 | 10593799 | 16.28495 | 29.53966 | 24.51542 | 10.59613 | 0.680754 | 0.34593 | 0.235547 |
| 1 | 96852800 | 96853799 | 12.17034 | 23.68548 | 13.88581 | 5.109508 | 1.372441 | -0.27859 | 0.235552 |
| 10 | 9549000 | 9553399 | 25.81479 | 42.24095 | 81.51262 | 35.06005 | -0.57033 | 1.468834 | 0.235572 |
| 1 | 28420400 | 28424599 | 50.59021 | 81.71967 | 67.92889 | 28.99331 | 0.626053 | 1.737796 | 0.235575 |
| 9 | 92917200 | 92918199 | 4.226458 | 8.251448 | 24.9879 | 8.820944 | -0.9153 | -0.49039 | 0.235591 |
| 6 | 130538000 | 130551199 | 190.4096 | 287.592 | 181.7817 | 75.15016 | 0.93645 | 3.338008 | 0.2356 |
| 10 | 77369000 | 77370999 | 10.91756 | 19.12739 | 45.18333 | 19.95349 | -0.84592 | 0.640671 | 0.235624 |
| 7 | 31220200 | 31223399 | 54.31016 | 87.74203 | 65.36388 | 27.90402 | 0.784067 | 1.739715 | 0.235626 |
| 10 | 52257400 | 52258799 | 8.960129 | 15.752 | 45.12573 | 19.9826 | -1.12214 | 0.573068 | 0.235628 |
| 2 | 169711000 | 169716199 | 30.16253 | 48.72312 | 96.41995 | 41.15469 | -0.62236 | 1.718641 | 0.235647 |
| 7 | 119143200 | 119144999 | 29.28357 | 49.77026 | 40.98866 | 17.96374 | 0.660617 | 1.042743 | 0.235663 |
| 7 | 38496400 | 38499199 | 62.21007 | 99.8346 | 77.89271 | 32.94247 | 0.706142 | 1.955613 | 0.235677 |
| 5 | 30637600 | 30644199 | 34.66568 | 55.50317 | 116.8408 | 49.24129 | -0.72687 | 2.008618 | 0.235678 |
| 13 | 106134600 | 106136799 | 30.98601 | 52.99699 | 36.34816 | 15.93808 | 0.924579 | 0.985623 | 0.23568 |
| 19 | 6814200 | 6818999 | 26.70129 | 43.08494 | 101.1291 | 43.10397 | -0.86974 | 1.747107 | 0.235693 |
| 3 | 39973600 | 39974199 | 3.880933 | 7.572489 | 22.04262 | 7.859173 | -0.86893 | -0.46309 | 0.235721 |
| 4 | 145545200 | 145545999 | 17.90051 | 32.16069 | 22.94449 | 10.03011 | 0.887146 | 0.409283 | 0.235725 |
| 9 | 122751800 | 122753199 | 5.405786 | 10.45421 | 27.18271 | 10.3942 | -0.80558 | -0.18641 | 0.235729 |
| 11 | 101192600 | 101193399 | 5.405786 | 10.45421 | 27.18271 | 10.3942 | -0.80558 | -0.18641 | 0.235729 |
| 17 | 18696400 | 18699199 | 41.7922 | 69.58735 | 42.04646 | 18.26178 | 1.105549 | 1.274043 | 0.235756 |
| 16 | 18694000 | 18697799 | 19.41558 | 32.34567 | 68.69628 | 29.84644 | -0.70179 | 1.226848 | 0.235759 |
| 12 | 24066000 | 24068199 | 19.25151 | 34.20349 | 24.41625 | 10.77455 | 0.877282 | 0.490081 | 0.235765 |
| 6 | 28479600 | 28480999 | 10.2437 | 18.11891 | 40.94544 | 18.10935 | -0.77849 | 0.505679 | 0.235768 |
| 8 | 110267800 | 110269199 | 5.871886 | 11.18948 | 28.80316 | 11.50617 | -0.84981 | -0.0281 | 0.235772 |
| 5 | 133765000 | 133766399 | 10.83517 | 19.21407 | 40.38267 | 17.84877 | -0.67307 | 0.480932 | 0.235775 |
| 12 | 112059800 | 112061999 | 19.36762 | 34.21041 | 28.56598 | 12.63761 | 0.648863 | 0.551888 | 0.235813 |
| 1 | 89826200 | 89829399 | 20.57918 | 34.10464 | 71.95647 | 31.15145 | -0.69291 | 1.293356 | 0.235813 |
| 2 | 93790800 | 93796599 | 32.06818 | 51.45049 | 115.4701 | 48.81676 | -0.81548 | 1.938 | 0.235814 |
| 7 | 9255600 | 9256599 | 26.71608 | 46.04393 | 29.68485 | 13.02193 | 1.015304 | 0.893448 | 0.235834 |
| 16 | 87515600 | 87519599 | 27.24692 | 44.13603 | 100.3085 | 42.93058 | -0.8174 | 1.647982 | 0.235848 |
| 10 | 26614400 | 26615399 | 15.58241 | 28.54672 | 23.6819 | 10.10782 | 0.693367 | 0.284396 | 0.235851 |
| 1 | 74644400 | 74650799 | 37.57123 | 60.09603 | 119.8915 | 50.45602 | -0.65117 | 2.04212 | 0.235852 |
| 1 | 78791400 | 78793399 | 9.629254 | 17.50741 | 37.59316 | 16.20142 | -0.67923 | 0.313149 | 0.235862 |
| 11 | 15209800 | 15218399 | 125.7849 | 193.4934 | 159.453 | 66.2194 | 0.575186 | 2.954433 | 0.235862 |
| 15 | 27183000 | 27185399 | 16.63522 | 28.61916 | 62.64105 | 27.47742 | -0.7414 | 0.908392 | 0.235864 |
| 13 | 116842200 | 116846399 | 29.46599 | 47.78856 | 90.13256 | 38.60996 | -0.54735 | 1.623359 | 0.235874 |
| 9 | 69446000 | 69448999 | 18.87435 | 31.65648 | 64.38595 | 28.09285 | -0.63959 | 1.149211 | 0.235885 |

| | | | | | | | | | |
|---|---|---|---|---|---|---|---|---|---|
| 1 | 137298000 | 137298599 | 4.054594 | 7.922651 | 22.57673 | 7.847878 | -0.80707 | -0.55275 | 0.235896 |
| 16 | 31718400 | 31719199 | 15.59779 | 28.51143 | 23.7143 | 10.15184 | 0.686678 | 0.297084 | 0.235898 |
| X | 29404600 | 29405999 | 18.85406 | 36.26227 | 11.30913 | 4.409108 | 2.195437 | -0.02589 | 0.235898 |
| X | 20910000 | 20910999 | 29.20058 | 50.74064 | 31.50298 | 13.83263 | 1.071515 | 0.831406 | 0.235933 |
| 7 | 34686000 | 34691599 | 26.43863 | 42.86426 | 90.66509 | 38.82842 | -0.71271 | 1.624308 | 0.235942 |
| 5 | 37113800 | 37116799 | 15.12306 | 25.80359 | 56.82427 | 24.9117 | -0.75176 | 0.973394 | 0.235947 |
| 7 | 3338800 | 3342799 | 23.18965 | 38.17606 | 76.44087 | 32.96231 | -0.61969 | 1.38305 | 0.235947 |
| 6 | 11978600 | 11978999 | 3.571355 | 6.982889 | 20.77958 | 7.104494 | -0.84446 | -0.62086 | 0.23596 |
| 7 | 113623200 | 113625199 | 26.02796 | 44.70679 | 38.28815 | 16.79423 | 0.606482 | 0.931412 | 0.235961 |
| 5 | 10235400 | 10238199 | 37.98892 | 63.13254 | 47.67043 | 20.67728 | 0.785041 | 1.288607 | 0.235978 |
| 15 | 86510000 | 86516799 | 41.90431 | 67.01895 | 124.7564 | 52.49462 | -0.55174 | 2.05088 | 0.235978 |
| 7 | 61545600 | 61547599 | 22.60952 | 39.65013 | 30.47263 | 13.46816 | 0.765396 | 0.662341 | 0.235985 |
| 12 | 32053800 | 32056599 | 22.60952 | 39.65013 | 30.47263 | 13.46816 | 0.765396 | 0.662341 | 0.235985 |
| 4 | 88489400 | 88489999 | 4.647965 | 9.062522 | 24.56072 | 8.845036 | -0.78315 | -0.41363 | 0.235993 |
| 17 | 38610800 | 38611599 | 23.15025 | 40.35599 | 33.31557 | 14.65794 | 0.662554 | 0.758366 | 0.236 |
| 7 | 29033800 | 29044399 | 51.82617 | 81.3283 | 173.1032 | 71.79115 | -0.7699 | 2.567743 | 0.236035 |
| 10 | 62935200 | 62936399 | 9.648501 | 17.05688 | 41.48142 | 18.34839 | -0.884 | 0.508793 | 0.236037 |
| X | 136015600 | 136016799 | 5.664721 | 10.98667 | 28.70067 | 10.82743 | -0.79345 | -0.22346 | 0.23605 |
| 1 | 172453600 | 172456599 | 31.24565 | 52.76539 | 42.54574 | 18.61162 | 0.690582 | 1.103628 | 0.236058 |
| 11 | 7824200 | 7825199 | 10.75743 | 20.94471 | 15.30668 | 5.61286 | 1.063284 | -0.3031 | 0.236059 |
| 6 | 94929000 | 94932399 | 29.63613 | 50.14719 | 42.58435 | 18.63977 | 0.616232 | 1.081537 | 0.236062 |
| 11 | 45946600 | 45948199 | 8.641228 | 15.54909 | 37.4497 | 16.34976 | -0.85549 | 0.367724 | 0.236065 |
| 16 | 58764000 | 58764399 | 7.363673 | 13.92495 | 32.05715 | 13.01562 | -0.71248 | 0.0441 | 0.236069 |
| 5 | 73350200 | 73352399 | 14.11611 | 24.27729 | 55.31957 | 24.26562 | -0.79894 | 0.891958 | 0.236072 |
| X | 23239600 | 23240999 | 19.60072 | 35.86118 | 21.56585 | 9.221083 | 1.15242 | 0.317753 | 0.236072 |
| 14 | 79378800 | 79381599 | 19.33359 | 32.43658 | 65.03097 | 28.37972 | -0.61901 | 1.147037 | 0.236079 |
| 8 | 3808800 | 3819599 | 174.396 | 261.3697 | 228.9552 | 94.45126 | 0.456204 | 3.48602 | 0.236082 |
| 4 | 126909400 | 126911399 | 7.735619 | 14.35382 | 34.25611 | 14.36899 | -0.80258 | 0.174175 | 0.236101 |
| 5 | 134836000 | 134839399 | 26.17077 | 42.33537 | 95.43984 | 40.80604 | -0.80748 | 1.671448 | 0.236103 |
| 7 | 59284200 | 59285199 | 17.35506 | 32.41672 | 19.70007 | 8.176454 | 1.168313 | 0.18021 | 0.236176 |
| 12 | 105009200 | 105019199 | 73.93057 | 114.6152 | 198.6609 | 82.51456 | -0.48778 | 2.794455 | 0.236176 |
| 12 | 23530800 | 23533599 | 55.79398 | 89.54455 | 71.27525 | 30.14712 | 0.677477 | 1.943474 | 0.23618 |
| 12 | 84741000 | 84745999 | 29.15517 | 47.21887 | 93.51678 | 40.01867 | -0.61958 | 1.651233 | 0.236186 |
| 3 | 156266200 | 156268999 | 16.28571 | 27.54679 | 61.10139 | 26.74193 | -0.76362 | 1.059189 | 0.236187 |
| 3 | 4328000 | 4330599 | 32.14656 | 54.05954 | 44.21849 | 19.31876 | 0.66985 | 1.144563 | 0.236196 |
| 9 | 46330400 | 46333799 | 24.09409 | 41.52822 | 34.75913 | 15.24791 | 0.64036 | 0.887097 | 0.236198 |
| 7 | 18851600 | 18854399 | 26.33877 | 42.56025 | 95.44809 | 40.757 | -0.8017 | 1.70347 | 0.236199 |
| 17 | 27248400 | 27250799 | 15.15127 | 26.3002 | 52.19528 | 22.90334 | -0.59764 | 0.809267 | 0.2362 |
| 2 | 168682800 | 168688599 | 49.67699 | 78.90272 | 141.0103 | 58.79239 | -0.50373 | 2.278842 | 0.236207 |
| 7 | 52733000 | 52734399 | 27.57555 | 47.17402 | 39.35185 | 17.25545 | 0.643486 | 0.975577 | 0.236209 |
| 1 | 181651800 | 181652799 | 13.25366 | 24.84259 | 21.35673 | 8.811733 | 0.678702 | 0.127411 | 0.236227 |
| 16 | 12604800 | 12605999 | 14.77956 | 27.71587 | 19.87034 | 8.191876 | 0.938862 | 0.132308 | 0.236229 |
| 1 | 88833200 | 88836999 | 18.2329 | 30.61498 | 63.8165 | 27.86411 | -0.67489 | 1.134296 | 0.236229 |
| 2 | 108017200 | 108017799 | 13.79196 | 25.88676 | 21.32071 | 8.776636 | 0.742794 | 0.12397 | 0.236251 |
| 10 | 122793400 | 122805599 | 101.4296 | 153.8477 | 270.8587 | 112.1241 | -0.53509 | 3.236745 | 0.236255 |
| 3 | 98632800 | 98634399 | 22.10877 | 38.85834 | 29.32807 | 12.98668 | 0.791227 | 0.622664 | 0.23627 |
| 4 | 155922000 | 155928599 | 30.91187 | 49.67917 | 109.9309 | 46.59284 | -0.79213 | 1.88152 | 0.236279 |

| | | | | | | | | | |
|---|---|---|---|---|---|---|---|---|---|
| 12 | 35834800 | 35836399 | 18.15112 | 33.28026 | 22.19933 | 9.464306 | 1.00678 | 0.295764 | 0.23628 |
| 8 | 36406400 | 36408199 | 9.350726 | 16.94631 | 36.47676 | 15.78377 | -0.6862 | 0.32886 | 0.236284 |
| 13 | 44373600 | 44379799 | 54.66764 | 86.55482 | 151.7558 | 63.09767 | -0.48048 | 2.374217 | 0.236289 |
| 12 | 4512400 | 4513999 | 16.61914 | 30.00216 | 24.99725 | 10.86914 | 0.669646 | 0.376398 | 0.236291 |
| 3 | 124024200 | 124024999 | 5.104906 | 9.916151 | 26.38809 | 9.859872 | -0.80683 | -0.27211 | 0.236292 |
| 1 | 117284600 | 117286599 | 26.48149 | 45.33601 | 37.71465 | 16.53854 | 0.647411 | 0.959787 | 0.236296 |
| 13 | 33812600 | 33819399 | 84.99483 | 134.4992 | 107.952 | 44.87249 | 0.644549 | 2.39891 | 0.236309 |
| 9 | 113449600 | 113456599 | 56.79644 | 89.74738 | 156.4828 | 64.99788 | -0.47481 | 2.422614 | 0.236319 |
| 14 | 91755000 | 91756599 | 18.9335 | 34.28034 | 23.61689 | 10.23144 | 0.945477 | 0.373963 | 0.236327 |
| 17 | 18380800 | 18381599 | 12.86624 | 24.85399 | 17.26478 | 6.636495 | 1.073782 | -0.11611 | 0.236328 |
| 7 | 144576600 | 144581199 | 23.23634 | 38.12494 | 79.02764 | 34.02948 | -0.67145 | 1.426705 | 0.236331 |
| 3 | 77256400 | 77256999 | 3.420922 | 6.669983 | 22.4811 | 8.097261 | -1.09167 | -0.43486 | 0.236351 |
| 12 | 96697600 | 96698599 | 12.31097 | 23.79323 | 16.5976 | 6.365185 | 1.069953 | -0.13314 | 0.236358 |
| 7 | 9084200 | 9086999 | 19.45048 | 34.4889 | 24.4835 | 10.8222 | 0.883717 | 0.503963 | 0.23636 |
| 9 | 54408400 | 54410399 | 9.715604 | 17.38465 | 38.92297 | 17.07342 | -0.75601 | 0.412821 | 0.236364 |
| 9 | 52414600 | 52416399 | 31.92626 | 54.0235 | 41.40981 | 18.12577 | 0.763435 | 1.088211 | 0.236373 |
| 13 | 42985000 | 42986799 | 10.19424 | 18.22349 | 38.92396 | 17.08777 | -0.68911 | 0.424369 | 0.236375 |
| 11 | 34098800 | 34100999 | 11.78191 | 20.80689 | 42.45238 | 18.78175 | -0.63314 | 0.534868 | 0.236392 |
| 17 | 38448800 | 38451199 | 26.08364 | 45.64334 | 29.16763 | 12.86772 | 1.030353 | 0.713159 | 0.236395 |
| 18 | 21851200 | 21853399 | 14.48521 | 25.25596 | 53.28425 | 23.44655 | -0.6849 | 0.742895 | 0.236402 |
| 3 | 92612400 | 92613999 | 22.3343 | 38.87974 | 33.5134 | 14.73215 | 0.600412 | 0.77405 | 0.236403 |
| 12 | 52573000 | 52573799 | 3.709511 | 7.22533 | 23.53146 | 8.588905 | -1.06125 | -0.38008 | 0.23641 |
| 18 | 76946200 | 76948999 | 27.93129 | 45.12956 | 95.92733 | 40.95693 | -0.72479 | 1.708094 | 0.236436 |
| 7 | 11175200 | 11209799 | 872.891 | 1146.041 | 867.5971 | 363.0761 | 0.478355 | 5.575365 | 0.236437 |
| 5 | 147762000 | 147767599 | 33.60104 | 54.04018 | 106.8434 | 45.33186 | -0.62874 | 1.860946 | 0.236438 |
| 11 | 73457800 | 73459399 | 20.28588 | 36.28438 | 25.0492 | 10.99137 | 0.930695 | 0.457228 | 0.236441 |
| 7 | 10680000 | 10683999 | 70.45038 | 112.1956 | 93.72862 | 39.2044 | 0.595586 | 2.205512 | 0.236448 |
| 13 | 66837400 | 66838999 | 16.93548 | 31.81034 | 18.61119 | 7.651743 | 1.233063 | 0.139486 | 0.236464 |
| 16 | 93986400 | 93987799 | 5.239076 | 9.879196 | 31.26675 | 12.77017 | -1.17387 | 0.043693 | 0.236481 |
| Y | 87089000 | 87089799 | 8.905463 | 17.36443 | 8.747761 | 3.148642 | 1.615108 | -0.4194 | 0.236484 |
| 5 | 129055600 | 129070599 | 107.6285 | 161.6874 | 310.2216 | 128.0531 | -0.67107 | 3.432329 | 0.236484 |
| 1 | 75290000 | 75294799 | 28.23592 | 45.64975 | 93.76256 | 40.06435 | -0.6744 | 1.690193 | 0.23649 |
| 4 | 52187000 | 52191399 | 55.94709 | 89.76972 | 80.19554 | 33.90831 | 0.510923 | 1.953193 | 0.2365 |
| 1 | 20856600 | 20859799 | 22.59981 | 37.53572 | 74.1536 | 32.15138 | -0.59822 | 1.273976 | 0.236508 |
| 1 | 101453200 | 101454799 | 17.2501 | 30.98771 | 25.4116 | 11.10995 | 0.687271 | 0.41022 | 0.236509 |
| 18 | 36061600 | 36071999 | 48.52639 | 76.67509 | 156.2373 | 64.89407 | -0.69932 | 2.418103 | 0.236514 |
| 6 | 92155200 | 92159199 | 44.85439 | 72.7674 | 63.73575 | 27.30837 | 0.556852 | 1.629818 | 0.236517 |
| 4 | 11044000 | 11045599 | 8.48552 | 15.78893 | 34.90334 | 14.5811 | -0.68848 | 0.166846 | 0.236523 |
| 1 | 162769000 | 162769599 | 5.363552 | 10.41609 | 26.92893 | 10.08125 | -0.76745 | -0.25731 | 0.236528 |
| 4 | 60319800 | 60321199 | 18.51805 | 33.7074 | 22.55808 | 9.707783 | 0.992961 | 0.338488 | 0.236528 |
| 16 | 67229600 | 67232399 | 35.67858 | 59.66114 | 45.36702 | 19.75698 | 0.774893 | 1.214312 | 0.236528 |
| 18 | 24203600 | 24205199 | 10.64509 | 19.01621 | 39.40845 | 17.3106 | -0.64664 | 0.431341 | 0.236529 |
| 7 | 108036000 | 108038199 | 22.22326 | 40.06053 | 21.66934 | 9.439367 | 1.287133 | 0.418599 | 0.236533 |
| 9 | 3613800 | 3614999 | 21.01986 | 36.78455 | 31.2799 | 13.80029 | 0.620222 | 0.687398 | 0.236535 |
| 17 | 19982200 | 19983199 | 21.01986 | 36.78455 | 31.2799 | 13.80029 | 0.620222 | 0.687398 | 0.236535 |
| 12 | 33808000 | 33809599 | 9.471413 | 17.22565 | 37.6004 | 16.1984 | -0.70243 | 0.309867 | 0.236556 |
| X | 116999600 | 117000799 | 16.91017 | 32.23328 | 16.36655 | 6.534082 | 1.472444 | 0.040945 | 0.236565 |

| | | | | | | | | | |
|---|---|---|---|---|---|---|---|---|---|
| 8 | 9665000 | 9665999 | 11.63263 | 22.5439 | 16.80667 | 6.363355 | 0.992778 | -0.18108 | 0.236566 |
| 3 | 72657200 | 72657799 | 17.08367 | 30.69057 | 25.62226 | 11.20148 | 0.661938 | 0.410788 | 0.236581 |
| 7 | 7017200 | 7020999 | 69.47983 | 110.5405 | 98.88598 | 41.30503 | 0.495501 | 2.235771 | 0.236581 |
| 15 | 73055600 | 73062799 | 37.29665 | 59.54774 | 125.9463 | 52.86693 | -0.73824 | 2.100129 | 0.236582 |
| 5 | 9980200 | 9982999 | 46.2109 | 75.04573 | 56.49323 | 24.22371 | 0.776177 | 1.609184 | 0.236593 |
| 8 | 117959400 | 117966799 | 52.92699 | 83.54376 | 156.9393 | 65.15229 | -0.5834 | 2.44564 | 0.236599 |
| 5 | 146070200 | 146072999 | 78.22579 | 123.5422 | 108.7738 | 45.16874 | 0.508935 | 2.44041 | 0.236601 |
| 6 | 60396600 | 60398399 | 14.94486 | 27.72441 | 19.20869 | 8.059809 | 0.968767 | 0.199328 | 0.236603 |
| 13 | 13407000 | 13411399 | 42.06633 | 68.53248 | 60.35526 | 25.91845 | 0.553848 | 1.552057 | 0.236612 |
| 6 | 145939400 | 145946199 | 54.92613 | 86.52993 | 162.7681 | 67.53915 | -0.58757 | 2.489037 | 0.236616 |
| 7 | 129677400 | 129679599 | 10.99628 | 19.42948 | 43.50133 | 19.24384 | -0.76625 | 0.524394 | 0.236628 |
| 10 | 8735600 | 8736799 | 6.281968 | 12.07905 | 28.64022 | 11.17435 | -0.70141 | -0.10364 | 0.236634 |
| 6 | 26160800 | 26163799 | 40.26396 | 66.28708 | 53.28127 | 22.97593 | 0.693226 | 1.407727 | 0.236636 |
| 1 | 128030000 | 128031799 | 6.131243 | 11.45828 | 33.84287 | 14.03391 | -1.09401 | 0.114805 | 0.236665 |
| 14 | 24540200 | 24541399 | 5.545688 | 10.40202 | 32.25687 | 13.29349 | -1.15563 | 0.083241 | 0.236667 |
| 7 | 12058400 | 12061799 | 67.44747 | 107.8029 | 86.50037 | 36.36481 | 0.659198 | 2.086649 | 0.236671 |
| 11 | 40539200 | 40544599 | 63.8564 | 100.1411 | 175.0636 | 72.60728 | -0.48715 | 2.586055 | 0.236699 |
| 19 | 7661000 | 7661999 | 6.002536 | 11.52362 | 29.76348 | 11.66515 | -0.83054 | -0.09181 | 0.236708 |
| 12 | 77087800 | 77093399 | 44.40466 | 70.77241 | 131.6158 | 55.1119 | -0.55582 | 2.16368 | 0.236714 |
| 4 | 154956000 | 154957399 | 15.42466 | 28.16347 | 22.34029 | 9.577014 | 0.753437 | 0.299185 | 0.236718 |
| 2 | 10290800 | 10291599 | 4.012151 | 7.802496 | 24.4798 | 9.062045 | -1.02535 | -0.31685 | 0.236739 |
| 7 | 20514400 | 20519799 | 146.8286 | 222.3701 | 194.5385 | 80.49276 | 0.471209 | 3.273057 | 0.236771 |
| 1 | 134222800 | 134226999 | 28.90276 | 46.65195 | 103.518 | 44.14015 | -0.78842 | 1.743282 | 0.236772 |
| 10 | 19350000 | 19351399 | 7.257565 | 13.6335 | 31.79001 | 13.06771 | -0.74503 | 0.090133 | 0.236781 |
| 16 | 18467000 | 18468599 | 18.29051 | 32.34879 | 27.13023 | 11.99939 | 0.643149 | 0.528036 | 0.236786 |
| 8 | 22107400 | 22108999 | 21.74194 | 38.66407 | 25.53331 | 11.25867 | 0.989813 | 0.503795 | 0.236791 |
| 17 | 55261000 | 55263999 | 25.72218 | 44.96076 | 30.32221 | 13.36451 | 0.953094 | 0.73015 | 0.236795 |
| 11 | 109583600 | 109584599 | 7.858404 | 14.74304 | 32.90347 | 13.55518 | -0.68546 | 0.10485 | 0.236817 |
| 11 | 114460800 | 114480799 | 218.5947 | 313.6773 | 514.6245 | 212.496 | -0.50826 | 4.195421 | 0.236821 |
| 6 | 83388200 | 83389599 | 6.23309 | 11.7328 | 31.83091 | 13.03961 | -0.95802 | 0.064426 | 0.236822 |
| 7 | 12174800 | 12175999 | 15.18892 | 28.25251 | 22.10778 | 9.240858 | 0.798442 | 0.193303 | 0.236825 |
| 12 | 25133400 | 25134199 | 3.830549 | 7.440745 | 24.38434 | 9.111174 | -1.10108 | -0.28971 | 0.236829 |
| 13 | 34119400 | 34120199 | 4.240813 | 8.236248 | 24.41014 | 9.133564 | -0.96022 | -0.27969 | 0.236831 |
| 1 | 185849000 | 185850599 | 6.763786 | 12.83595 | 31.02084 | 12.50082 | -0.77226 | 0.010913 | 0.236838 |
| 2 | 10374200 | 10380799 | 48.39026 | 76.77992 | 145.5828 | 60.63632 | -0.59044 | 2.308405 | 0.236839 |
| 5 | 52834600 | 52836199 | 8.582766 | 15.58775 | 39.73225 | 17.14665 | -0.92643 | 0.314328 | 0.236848 |
| 9 | 99756800 | 99764199 | 71.59669 | 111.1237 | 196.3367 | 81.53529 | -0.51412 | 2.77326 | 0.236848 |
| 15 | 55682200 | 55683999 | 19.60257 | 34.47789 | 27.52164 | 12.18599 | 0.710936 | 0.595033 | 0.236859 |
| 7 | 61293800 | 61298199 | 42.74619 | 70.18814 | 49.48369 | 21.31453 | 0.880299 | 1.441516 | 0.236868 |
| 11 | 44206400 | 44207399 | 6.27501 | 12.08254 | 30.59617 | 11.8897 | -0.79081 | -0.11454 | 0.236872 |
| 11 | 69711200 | 69713199 | 16.50709 | 28.22871 | 56.39663 | 24.72872 | -0.61151 | 0.95676 | 0.236877 |
| 16 | 94563000 | 94563599 | 4.649135 | 9.06106 | 23.80094 | 8.617902 | -0.74531 | -0.39592 | 0.23689 |
| X | 148468800 | 148473199 | 42.27807 | 70.22961 | 44.34925 | 19.23086 | 1.042053 | 1.304672 | 0.236895 |
| 2 | 130892200 | 130899199 | 39.47102 | 62.81014 | 137.0182 | 57.24917 | -0.78803 | 2.212015 | 0.236897 |
| 1 | 5634400 | 5638199 | 34.06853 | 56.76586 | 42.82394 | 18.60753 | 0.786402 | 1.243578 | 0.236908 |
| 1 | 45286000 | 45287199 | 12.5151 | 24.10681 | 17.70845 | 6.874179 | 0.980165 | -0.0886 | 0.236913 |
| 19 | 27338400 | 27339399 | 15.99564 | 29.56738 | 22.50533 | 9.489296 | 0.829021 | 0.235235 | 0.236915 |

| | | | | | | | | | |
|---|---|---|---|---|---|---|---|---|---|
| 7 | 5564200 | 5567799 | 61.58344 | 98.77988 | 79.9283 | 33.77783 | 0.652878 | 1.97099 | 0.236916 |
| 6 | 133652000 | 133652799 | 10.34426 | 18.41637 | 40.93528 | 18.03267 | -0.75018 | 0.451989 | 0.236939 |
| 15 | 96956000 | 96959799 | 31.14708 | 50.23923 | 99.68241 | 42.46619 | -0.62866 | 1.764838 | 0.236943 |
| 7 | 114410200 | 114412199 | 31.71091 | 53.15135 | 45.91561 | 20.02558 | 0.591415 | 1.177663 | 0.236949 |
| 7 | 128636800 | 128639999 | 31.71091 | 53.15135 | 45.91561 | 20.02558 | 0.591415 | 1.177663 | 0.236949 |
| 11 | 71161000 | 71161399 | 13.18124 | 24.88518 | 20.4056 | 8.313824 | 0.76128 | 0.079872 | 0.236966 |
| X | 125025800 | 125026399 | 25.51741 | 44.08624 | 33.2897 | 14.60437 | 0.789166 | 0.873575 | 0.236969 |
| 4 | 148941800 | 148944999 | 18.02847 | 30.00397 | 69.58418 | 30.21241 | -0.8282 | 1.235194 | 0.236986 |
| 13 | 62809200 | 62809799 | 11.62407 | 22.24277 | 19.47405 | 7.704587 | 0.704274 | -0.03212 | 0.237002 |
| 17 | 19624200 | 19633199 | 90.59691 | 142.6221 | 110.0028 | 45.6404 | 0.696094 | 2.517286 | 0.237017 |
| 7 | 30804400 | 30808799 | 22.60154 | 37.09554 | 78.32969 | 33.7336 | -0.69782 | 1.421145 | 0.237026 |
| 1 | 16817600 | 16820799 | 21.83776 | 36.18203 | 72.7216 | 31.47766 | -0.6232 | 1.299529 | 0.237067 |
| 1 | 169389400 | 169392399 | 23.58914 | 41.29624 | 30.99245 | 13.67781 | 0.799606 | 0.695734 | 0.237083 |
| 9 | 22372200 | 22373799 | 15.24267 | 27.77325 | 23.11861 | 9.936531 | 0.681534 | 0.30929 | 0.237086 |
| 1 | 143577800 | 143578999 | 9.064626 | 16.01423 | 42.52493 | 18.81242 | -1.01041 | 0.513977 | 0.237099 |
| 1 | 34881600 | 34882599 | 4.579162 | 8.861057 | 27.13096 | 10.34464 | -1.03442 | -0.20569 | 0.237116 |
| 10 | 42420800 | 42422799 | 12.78396 | 22.27352 | 48.39792 | 21.2867 | -0.72692 | 0.739436 | 0.237117 |
| 2 | 170504200 | 170506399 | 18.12131 | 30.87075 | 60.80183 | 26.65208 | -0.59208 | 1.002464 | 0.237121 |
| 2 | 169544000 | 169547799 | 32.08828 | 51.81359 | 97.96756 | 41.79414 | -0.55748 | 1.733753 | 0.237124 |
| 2 | 51844000 | 51844999 | 11.95071 | 22.70664 | 19.95775 | 8.022218 | 0.678595 | 0.023098 | 0.237126 |
| 12 | 22680600 | 22681999 | 11.95071 | 22.70664 | 19.95775 | 8.022218 | 0.678595 | 0.023098 | 0.237126 |
| 8 | 46941000 | 46942399 | 7.601835 | 14.08164 | 33.12915 | 13.92843 | -0.78411 | 0.181744 | 0.237128 |
| 7 | 16482600 | 16485199 | 13.89493 | 24.09021 | 51.08633 | 22.41423 | -0.69309 | 0.813167 | 0.237131 |
| 16 | 91393800 | 91396199 | 13.89493 | 24.09021 | 51.08633 | 22.41423 | -0.69309 | 0.813167 | 0.237131 |
| 1 | 189899800 | 189902999 | 22.147 | 36.30594 | 79.261 | 34.11737 | -0.74644 | 1.436007 | 0.237148 |
| 18 | 88963800 | 88965599 | 11.96148 | 21.11401 | 43.50516 | 19.24934 | -0.64749 | 0.544069 | 0.237158 |
| 9 | 70892200 | 70897199 | 35.84749 | 57.65421 | 107.493 | 45.60873 | -0.54428 | 1.863685 | 0.237162 |
| 2 | 130951800 | 130952999 | 6.152855 | 11.77174 | 28.39139 | 11.23756 | -0.7451 | -0.05797 | 0.237175 |
| 3 | 58543800 | 58544799 | 6.152855 | 11.77174 | 28.39139 | 11.23756 | -0.7451 | -0.05797 | 0.237175 |
| 17 | 18170400 | 18175999 | 78.29533 | 124.304 | 101.5459 | 42.31994 | 0.623311 | 2.304977 | 0.237191 |
| 4 | 145251800 | 145254799 | 53.47217 | 86.36319 | 61.29985 | 26.1593 | 0.853125 | 1.742989 | 0.237191 |
| 10 | 60587200 | 60594799 | 59.63784 | 93.97274 | 163.6629 | 67.91234 | -0.47639 | 2.487999 | 0.237222 |
| 19 | 57398200 | 57401199 | 16.9108 | 28.22964 | 69.06028 | 30.04277 | -0.90494 | 1.197538 | 0.237228 |
| 4 | 26236200 | 26236799 | 7.775161 | 14.32842 | 33.58004 | 14.21769 | -0.78588 | 0.214496 | 0.237232 |
| 7 | 143568800 | 143570799 | 9.985139 | 17.57178 | 45.10173 | 19.96793 | -0.9644 | 0.56641 | 0.237241 |
| 11 | 5230800 | 5232399 | 6.891061 | 12.35851 | 38.53301 | 16.87011 | -1.22722 | 0.380992 | 0.237247 |
| 7 | 22886400 | 22886999 | 10.91031 | 21.29272 | 13.073 | 4.644252 | 1.352023 | -0.41497 | 0.237247 |
| 11 | 116742800 | 116745199 | 11.54206 | 20.17764 | 46.26148 | 20.39217 | -0.80315 | 0.68191 | 0.237268 |
| 17 | 19117000 | 19121399 | 81.66152 | 125.9765 | 214.7563 | 89.20583 | -0.46935 | 2.890502 | 0.237281 |
| 9 | 58159800 | 58161399 | 8.430755 | 15.34001 | 35.50428 | 15.28664 | -0.78521 | 0.296958 | 0.237306 |
| 2 | 72576800 | 72578599 | 10.17228 | 18.15858 | 43.63572 | 19.17905 | -0.86002 | 0.437438 | 0.237321 |
| 6 | 39499400 | 39503199 | 23.29657 | 38.47434 | 76.82916 | 33.18139 | -0.61438 | 1.343389 | 0.237329 |
| 7 | 18283000 | 18283599 | 17.13161 | 30.58998 | 23.78345 | 10.45107 | 0.758984 | 0.443939 | 0.237335 |
| 1 | 17353200 | 17354799 | 15.4152 | 28.59813 | 21.9938 | 9.227893 | 0.81909 | 0.208908 | 0.237335 |
| 7 | 58860000 | 58861399 | 28.4446 | 48.84829 | 34.49894 | 15.13209 | 0.883383 | 0.938522 | 0.23734 |
| 3 | 23928600 | 23930599 | 25.68245 | 44.82582 | 31.58176 | 13.90596 | 0.890404 | 0.753467 | 0.23734 |
| 7 | 43563200 | 43570799 | 54.27844 | 85.50681 | 163.1433 | 67.69457 | -0.60806 | 2.489288 | 0.237341 |

| | | | | | | | | | |
|---|---|---|---|---|---|---|---|---|---|
| 16 | 87372600 | 87374199 | 7.602943 | 14.07196 | 35.29286 | 14.85494 | -0.8774 | 0.189302 | 0.237343 |
| 13 | 73682800 | 73687199 | 27.70772 | 44.57294 | 114.3617 | 48.53901 | -1.00331 | 1.848584 | 0.237343 |
| 2 | 180716600 | 180720399 | 24.01867 | 39.46342 | 77.45168 | 33.3705 | -0.59201 | 1.411277 | 0.237344 |
| 7 | 31815400 | 31818599 | 66.55978 | 106.0198 | 92.47837 | 38.69166 | 0.533662 | 2.194624 | 0.237347 |
| 8 | 33830800 | 33832799 | 20.39044 | 36.3203 | 26.20546 | 11.53905 | 0.864199 | 0.48614 | 0.237375 |
| 8 | 111797600 | 111802799 | 30.85333 | 49.71243 | 100.6889 | 42.83466 | -0.66006 | 1.7979 | 0.237384 |
| 4 | 122396000 | 122397999 | 23.84645 | 42.11513 | 25.87018 | 11.44534 | 1.08912 | 0.576097 | 0.237395 |
| 3 | 7899200 | 7902399 | 64.31752 | 102.513 | 90.18852 | 37.76664 | 0.52228 | 2.170715 | 0.2374 |
| 17 | 13775000 | 13777999 | 15.40972 | 26.35732 | 55.81079 | 24.47223 | -0.69489 | 0.950095 | 0.237415 |
| 10 | 77070200 | 77078199 | 71.79645 | 111.3405 | 198.224 | 82.32933 | -0.52604 | 2.787949 | 0.237416 |
| 15 | 52189000 | 52191399 | 18.53782 | 33.66337 | 23.09537 | 9.968555 | 0.954709 | 0.353074 | 0.237426 |
| 3 | 3184200 | 3194399 | 91.99889 | 145.2247 | 102.9368 | 42.73458 | 0.820802 | 2.460758 | 0.237435 |
| X | 92993600 | 92996199 | 10.80152 | 19.36946 | 42.11252 | 18.43895 | -0.71234 | 0.413526 | 0.23747 |
| 14 | 110610000 | 110611399 | 18.46271 | 33.10027 | 25.805 | 11.30083 | 0.758278 | 0.432091 | 0.23747 |
| 4 | 134223400 | 134228199 | 21.18878 | 34.79242 | 81.15036 | 34.955 | -0.84066 | 1.41268 | 0.237478 |
| 5 | 5743600 | 5744599 | 7.766434 | 13.96507 | 37.98969 | 16.59542 | -1.03029 | 0.367371 | 0.237487 |
| 19 | 8037200 | 8038399 | 28.48131 | 48.24338 | 42.08892 | 18.42874 | 0.577469 | 1.068045 | 0.237494 |
| 1 | 41009000 | 41009799 | 13.71507 | 25.58204 | 21.14548 | 8.797132 | 0.725014 | 0.159818 | 0.237494 |
| 5 | 116067000 | 116071599 | 48.43538 | 76.79137 | 144.8683 | 60.29147 | -0.58422 | 2.332521 | 0.237498 |
| 16 | 77638400 | 77639799 | 18.74521 | 33.94985 | 23.62325 | 10.23027 | 0.931523 | 0.370735 | 0.2375 |
| 11 | 103002200 | 103004199 | 8.772343 | 15.48684 | 42.54428 | 18.82366 | -1.05932 | 0.520816 | 0.237502 |
| 10 | 61459000 | 61464199 | 33.8501 | 54.23147 | 115.1684 | 48.58298 | -0.73838 | 1.987308 | 0.237508 |
| 7 | 31934800 | 31937799 | 53.42841 | 86.04798 | 65.7482 | 27.95218 | 0.742381 | 1.829068 | 0.237508 |
| 11 | 62071400 | 62075199 | 45.97974 | 75.15098 | 55.14318 | 23.70819 | 0.819647 | 1.511572 | 0.237515 |
| 13 | 65437000 | 65448399 | 359.328 | 511.7485 | 376.4888 | 155.6473 | 0.637452 | 4.311898 | 0.237519 |
| 8 | 75466800 | 75467799 | 17.35159 | 31.27923 | 25.46323 | 11.09177 | 0.701379 | 0.391438 | 0.237524 |
| 14 | 122705800 | 122707199 | 9.473246 | 16.80612 | 39.93664 | 17.64528 | -0.84844 | 0.469264 | 0.23755 |
| 7 | 18135000 | 18138199 | 106.0073 | 164.1468 | 143.5866 | 59.64206 | 0.496402 | 2.823565 | 0.237566 |
| 8 | 34143400 | 34145999 | 14.54717 | 25.13223 | 52.14266 | 22.87524 | -0.66278 | 0.849975 | 0.237569 |
| 14 | 87583800 | 87585799 | 14.54717 | 25.13223 | 52.14266 | 22.87524 | -0.66278 | 0.849975 | 0.237569 |
| 2 | 162759600 | 162763399 | 33.10376 | 53.28987 | 104.109 | 44.23119 | -0.60985 | 1.830028 | 0.237578 |
| 12 | 19456000 | 19460599 | 63.30586 | 102.1605 | 59.4412 | 25.3389 | 1.137928 | 1.785916 | 0.23758 |
| 12 | 23609600 | 23610799 | 19.38931 | 34.16532 | 27.33611 | 12.09941 | 0.708937 | 0.576745 | 0.237591 |
| 12 | 24284200 | 24293399 | 280.2107 | 404.811 | 339.6957 | 140.1295 | 0.468214 | 4.098477 | 0.237594 |
| 17 | 15875800 | 15878399 | 11.77828 | 20.8701 | 44.02444 | 19.46425 | -0.67914 | 0.500764 | 0.237596 |
| 7 | 137138600 | 137140399 | 11.96943 | 21.0105 | 44.07824 | 19.49705 | -0.6758 | 0.61298 | 0.237603 |
| 10 | 37012000 | 37015199 | 26.04465 | 42.08346 | 99.85382 | 42.63627 | -0.88301 | 1.704157 | 0.237616 |
| 5 | 141188200 | 141190199 | 12.56438 | 22.08163 | 46.1267 | 20.42407 | -0.66995 | 0.598194 | 0.237616 |
| 18 | 37537400 | 37540799 | 56.08302 | 90.63541 | 63.13195 | 26.9624 | 0.881311 | 1.733937 | 0.237628 |
| 9 | 100348200 | 100351599 | 20.02147 | 33.02753 | 76.13873 | 32.86028 | -0.82103 | 1.349237 | 0.237654 |
| 17 | 19151800 | 19162799 | 211.7639 | 313.5427 | 267.0491 | 109.9792 | 0.480718 | 3.710124 | 0.237656 |
| 7 | 31757200 | 31761399 | 99.6869 | 154.9595 | 139.4751 | 57.90836 | 0.459566 | 2.752254 | 0.237665 |
| 4 | 43380400 | 43382399 | 9.378462 | 16.5313 | 43.55747 | 19.2779 | -1.00071 | 0.54274 | 0.237677 |
| 7 | 63697000 | 63700199 | 14.64482 | 25.01263 | 59.03702 | 25.88385 | -0.85135 | 0.961856 | 0.237679 |
| 17 | 79920400 | 79923199 | 14.36982 | 24.87785 | 51.62298 | 22.6487 | -0.66238 | 0.829027 | 0.23768 |
| 17 | 49512200 | 49515799 | 21.28504 | 35.07548 | 75.63512 | 32.62859 | -0.7256 | 1.365562 | 0.237687 |
| 2 | 167231000 | 167239599 | 88.36036 | 135.5013 | 229.9353 | 95.44933 | -0.46933 | 3.008821 | 0.237698 |

| | | | | | | | | | |
|---|---|---|---|---|---|---|---|---|---|
| 13 | 78417800 | 78419799 | 24.83732 | 43.25119 | 30.92568 | 13.59742 | 0.868527 | 0.775655 | 0.237712 |
| 9 | 73146400 | 73147799 | 10.13482 | 18.32573 | 38.54155 | 16.72459 | -0.65586 | 0.351947 | 0.237719 |
| 1 | 63265000 | 63265799 | 6.584593 | 12.60803 | 30.24657 | 11.94383 | -0.73528 | -0.05935 | 0.237724 |
| 11 | 73955200 | 73956599 | 25.70538 | 44.04375 | 35.91503 | 15.75043 | 0.676477 | 0.949802 | 0.237743 |
| 14 | 8506800 | 8509799 | 16.03896 | 27.37057 | 57.44934 | 25.18604 | -0.68296 | 0.974248 | 0.237744 |
| 9 | 20588600 | 20590799 | 15.12108 | 26.22414 | 53.29573 | 23.38383 | -0.63216 | 0.817602 | 0.237749 |
| 14 | 34223800 | 34225399 | 10.75589 | 18.86926 | 44.63358 | 19.73291 | -0.84786 | 0.616468 | 0.237757 |
| 10 | 44334400 | 44334999 | 3.605272 | 7.007546 | 26.22666 | 9.750023 | -1.28476 | -0.3081 | 0.237765 |
| 18 | 35660800 | 35663199 | 24.31865 | 40.00671 | 77.08349 | 33.22959 | -0.56479 | 1.395548 | 0.237774 |
| 14 | 62569400 | 62570199 | 4.413122 | 8.552495 | 25.44263 | 9.633573 | -0.98265 | -0.23453 | 0.237791 |
| 3 | 25124800 | 25126799 | 11.12665 | 19.87001 | 42.56723 | 18.70324 | -0.69471 | 0.442474 | 0.237804 |
| 15 | 74378400 | 74395999 | 116.9703 | 174.4664 | 345.0038 | 142.1924 | -0.72398 | 3.570589 | 0.237805 |
| 14 | 60488400 | 60490199 | 12.00499 | 20.95216 | 48.37583 | 21.2999 | -0.81377 | 0.71007 | 0.237811 |
| 18 | 60320800 | 60322399 | 17.87314 | 32.20543 | 24.748 | 10.78607 | 0.783348 | 0.394958 | 0.23782 |
| 7 | 118012400 | 118015999 | 26.10455 | 42.81316 | 81.85191 | 35.23942 | -0.5555 | 1.44011 | 0.237824 |
| 1 | 40345400 | 40348599 | 37.03461 | 59.4382 | 110.5986 | 46.78064 | -0.54512 | 1.933891 | 0.237839 |
| X | 122080400 | 122081399 | 13.49691 | 25.76504 | 19.30417 | 7.682892 | 0.919569 | 0.005334 | 0.237841 |
| 10 | 81133600 | 81139399 | 24.09335 | 38.9579 | 94.44412 | 40.36298 | -0.91249 | 1.680603 | 0.237842 |
| 7 | 28834000 | 28835199 | 5.906533 | 11.22656 | 30.44543 | 12.22861 | -0.93245 | -0.00936 | 0.237868 |
| 9 | 53616600 | 53618199 | 11.44222 | 19.8961 | 51.03037 | 22.41643 | -0.96525 | 0.764724 | 0.237876 |
| 9 | 20050000 | 20050799 | 17.92498 | 31.92638 | 26.63455 | 11.72872 | 0.655975 | 0.471505 | 0.237876 |
| X | 157833800 | 157835399 | 17.59821 | 33.34389 | 17.16091 | 6.938821 | 1.436902 | 0.088027 | 0.237888 |
| 13 | 30132400 | 30137799 | 60.51512 | 95.20427 | 165.5583 | 68.6847 | -0.47597 | 2.520932 | 0.237896 |
| 7 | 46950800 | 46951199 | 10.13857 | 19.55334 | 17.02026 | 6.583529 | 0.742706 | -0.12716 | 0.237897 |
| X | 45290200 | 45293199 | 33.29481 | 57.72934 | 28.56774 | 12.53419 | 1.396705 | 0.871524 | 0.237901 |
| 7 | 31740600 | 31741399 | 13.50728 | 26.27702 | 12.49652 | 4.612976 | 1.665904 | -0.25565 | 0.237908 |
| 14 | 124918800 | 124922399 | 35.99936 | 59.55804 | 51.53899 | 22.28607 | 0.588584 | 1.33635 | 0.237909 |
| 3 | 24285600 | 24289799 | 73.96752 | 117.4651 | 103.1556 | 43.00256 | 0.519534 | 2.294974 | 0.237949 |
| 14 | 47205200 | 47206799 | 9.667947 | 17.50656 | 36.98655 | 16.02111 | -0.66047 | 0.337664 | 0.237958 |
| 10 | 33513000 | 33514999 | 21.78182 | 38.40497 | 28.22245 | 12.49017 | 0.831123 | 0.584734 | 0.237962 |
| 17 | 71691200 | 71693599 | 21.78182 | 38.40497 | 28.22245 | 12.49017 | 0.831123 | 0.584734 | 0.237962 |
| 12 | 18087000 | 18094399 | 288.0009 | 415.0282 | 345.8862 | 142.7303 | 0.474574 | 4.135891 | 0.237966 |
| 2 | 179444600 | 179449999 | 45.46188 | 72.27258 | 136.6632 | 57.03109 | -0.58372 | 2.248857 | 0.237975 |
| 15 | 99328000 | 99332799 | 19.80909 | 32.42987 | 80.91044 | 34.80761 | -0.93905 | 1.449688 | 0.237977 |
| 5 | 30877800 | 30880799 | 14.64659 | 25.02285 | 56.28639 | 24.67845 | -0.78188 | 0.959002 | 0.238008 |
| 6 | 90407000 | 90416199 | 86.82585 | 132.5582 | 243.9677 | 101.1778 | -0.59139 | 3.098631 | 0.238017 |
| 16 | 91100000 | 91103999 | 23.72667 | 38.88187 | 80.78376 | 34.76767 | -0.67554 | 1.443754 | 0.238051 |
| 1 | 90885800 | 90892599 | 36.20715 | 58.12901 | 110.1562 | 46.61646 | -0.57087 | 1.921645 | 0.238057 |
| 13 | 61885200 | 61889799 | 93.67786 | 147.7809 | 110.8296 | 45.99647 | 0.738798 | 2.478008 | 0.238063 |
| 5 | 113600600 | 113607799 | 54.35877 | 85.43798 | 168.0949 | 69.72845 | -0.65484 | 2.536225 | 0.238081 |
| 14 | 36024600 | 36026799 | 27.17983 | 47.51321 | 28.52934 | 12.57533 | 1.119712 | 0.733352 | 0.238087 |
| 9 | 86576000 | 86578199 | 15.68615 | 26.92007 | 54.80034 | 24.03564 | -0.6376 | 0.918497 | 0.238099 |
| 5 | 34426800 | 34434399 | 43.60972 | 69.45892 | 134.646 | 56.32854 | -0.61663 | 2.185052 | 0.238105 |
| 15 | 35583400 | 35584399 | 3.040314 | 5.945493 | 22.50636 | 7.337785 | -1.12479 | -0.79429 | 0.238118 |
| 15 | 43733000 | 43734199 | 5.756663 | 10.98142 | 33.55637 | 13.37526 | -1.09413 | -0.03074 | 0.238125 |
| 9 | 98269000 | 98270999 | 11.61617 | 20.4954 | 43.49507 | 19.24658 | -0.69027 | 0.544407 | 0.238128 |
| 17 | 39226000 | 39230199 | 86.45811 | 136.6413 | 110.8333 | 46.04063 | 0.627932 | 2.431057 | 0.238128 |

| | | | | | | | | | |
|---|---|---|---|---|---|---|---|---|---|
| 12 | 23962200 | 23965999 | 104.0855 | 161.2064 | 134.3815 | 55.81845 | 0.566059 | 2.816222 | 0.238145 |
| 7 | 13250600 | 13253399 | 27.71129 | 48.18186 | 29.55808 | 12.98358 | 1.088664 | 0.812575 | 0.238146 |
| 17 | 20437000 | 20439799 | 38.25239 | 63.76622 | 44.84433 | 19.49103 | 0.887308 | 1.253295 | 0.238158 |
| 17 | 76244600 | 76245399 | 6.819646 | 12.76364 | 31.69273 | 13.10945 | -0.84191 | 0.110683 | 0.23816 |
| 4 | 72423800 | 72424199 | 8.102668 | 15.83775 | 12.74109 | 4.39166 | 1.004138 | -0.56057 | 0.238167 |
| 7 | 130088800 | 130090199 | 30.00654 | 50.71056 | 43.13612 | 18.87413 | 0.613755 | 1.094432 | 0.238168 |
| 1 | 10457600 | 10459199 | 22.76009 | 40.06366 | 28.13832 | 12.45704 | 0.895196 | 0.607864 | 0.238191 |
| 6 | 8509000 | 8510599 | 11.04968 | 19.40669 | 43.57052 | 19.28143 | -0.77293 | 0.5983 | 0.238197 |
| 8 | 12475000 | 12485799 | 70.25805 | 108.771 | 206.829 | 85.91136 | -0.62281 | 2.818076 | 0.238199 |
| 15 | 3065200 | 3070999 | 121.8079 | 188.1734 | 148.359 | 61.62516 | 0.643593 | 2.873033 | 0.238199 |
| 9 | 40796800 | 40799399 | 29.41727 | 49.91951 | 41.4547 | 18.16106 | 0.648419 | 1.054838 | 0.238212 |
| 8 | 74478600 | 74480599 | 21.16085 | 37.89845 | 23.2375 | 10.18529 | 1.101504 | 0.451098 | 0.238216 |
| 17 | 18245600 | 18250999 | 78.9964 | 125.7583 | 90.78223 | 37.95227 | 0.805352 | 2.222659 | 0.238217 |
| Y | 40947200 | 40948399 | 10.09127 | 19.62935 | 8.12866 | 3.003299 | 1.859838 | -0.31531 | 0.23822 |
| 10 | 32052000 | 32053599 | 16.82208 | 31.21539 | 21.34282 | 8.951942 | 0.987987 | 0.218361 | 0.238239 |
| 1 | 59788400 | 59790199 | 16.59278 | 30.14104 | 24.6048 | 10.61578 | 0.706309 | 0.341703 | 0.238251 |
| 12 | 82157800 | 82158599 | 10.94038 | 21.29934 | 15.30095 | 5.614116 | 1.087117 | -0.29861 | 0.238265 |
| 7 | 42243800 | 42250599 | 194.9014 | 290.1962 | 250.4641 | 103.2036 | 0.46903 | 3.610113 | 0.238265 |
| 6 | 143045000 | 143046199 | 10.3738 | 18.41131 | 39.8949 | 17.62075 | -0.71602 | 0.47197 | 0.238267 |
| 3 | 87432600 | 87433599 | 5.33254 | 10.14804 | 29.3122 | 11.74287 | -1.01843 | -0.02489 | 0.238274 |
| 15 | 94420800 | 94421599 | 5.33254 | 10.14804 | 29.3122 | 11.74287 | -1.01843 | -0.02489 | 0.238274 |
| 7 | 20971000 | 20974599 | 104.1284 | 161.3173 | 135.9211 | 56.45788 | 0.550943 | 2.812304 | 0.238275 |
| 7 | 20971200 | 20974599 | 104.1284 | 161.3173 | 135.9211 | 56.45788 | 0.550943 | 2.812304 | 0.238275 |
| 7 | 92837000 | 92839399 | 27.12831 | 46.79537 | 35.35087 | 15.50787 | 0.787751 | 0.893088 | 0.238275 |
| 17 | 18554600 | 18573599 | 422.1608 | 589.3894 | 471.2671 | 195.4595 | 0.488498 | 4.60644 | 0.238276 |
| 11 | 68164200 | 68169799 | 30.98851 | 49.98529 | 98.99565 | 42.17562 | -0.62593 | 1.763538 | 0.238304 |
| 2 | 20430200 | 20431599 | 15.95238 | 29.20585 | 23.67206 | 10.1124 | 0.725855 | 0.291107 | 0.238326 |
| 4 | 19528800 | 19529199 | 2.835351 | 5.546527 | 18.55499 | 6.171186 | -0.97349 | -0.74184 | 0.238333 |
| 8 | 11578800 | 11586399 | 46.29749 | 73.51024 | 140.3139 | 58.48229 | -0.59899 | 2.286554 | 0.238333 |
| 7 | 104508400 | 104512399 | 52.7014 | 84.69958 | 75.1115 | 31.83613 | 0.524279 | 1.89541 | 0.238344 |
| 4 | 6508000 | 6510399 | 14.19088 | 24.77297 | 49.99169 | 22.0146 | -0.62062 | 0.721296 | 0.238348 |
| 2 | 153690600 | 153704799 | 124.258 | 185.4946 | 332.1943 | 136.9357 | -0.57995 | 3.555734 | 0.238359 |
| 2 | 111694200 | 111696799 | 24.61628 | 43.05334 | 30.31799 | 13.36937 | 0.890928 | 0.714084 | 0.238363 |
| 1 | 121973400 | 121975199 | 18.41966 | 33.36481 | 24.35049 | 10.54341 | 0.863591 | 0.370378 | 0.238364 |
| 18 | 63676800 | 63678199 | 11.1587 | 19.75863 | 40.90601 | 18.08801 | -0.65234 | 0.499861 | 0.238364 |
| 7 | 62093000 | 62095999 | 36.37447 | 60.54791 | 43.83744 | 19.03598 | 0.845533 | 1.261611 | 0.238366 |
| 9 | 96376200 | 96378999 | 14.82515 | 25.80171 | 51.64331 | 22.69854 | -0.60944 | 0.769413 | 0.238402 |
| 11 | 110297400 | 110303199 | 55.65816 | 87.90545 | 156.9924 | 65.19333 | -0.50993 | 2.433292 | 0.23844 |
| 7 | 120441800 | 120444799 | 13.52903 | 23.21894 | 54.70672 | 23.99465 | -0.8473 | 0.910077 | 0.238447 |
| 18 | 19256000 | 19259399 | 62.47381 | 99.88413 | 88.0332 | 37.02538 | 0.524509 | 2.071996 | 0.238454 |
| 7 | 29347200 | 29349399 | 70.72957 | 112.4236 | 95.61608 | 39.89527 | 0.566919 | 2.265552 | 0.238461 |
| 7 | 136688800 | 136689999 | 6.692792 | 12.50404 | 32.20921 | 13.36313 | -0.8987 | 0.118827 | 0.23847 |
| X | 136304800 | 136307399 | 38.6164 | 63.75032 | 52.13236 | 22.50917 | 0.669807 | 1.369906 | 0.238472 |
| 17 | 36912800 | 36915399 | 24.76387 | 43.1919 | 32.04217 | 14.1024 | 0.816032 | 0.757011 | 0.238476 |
| 1 | 31852600 | 31853999 | 19.32268 | 35.85021 | 17.60524 | 7.385769 | 1.46067 | 0.230706 | 0.238478 |
| 13 | 57669800 | 57673799 | 37.6803 | 60.38414 | 116.471 | 49.15262 | -0.5995 | 1.981778 | 0.238483 |
| 9 | 3074800 | 3077799 | 98.53375 | 153.2839 | 132.0781 | 54.83097 | 0.523276 | 2.738374 | 0.238485 |

| | | | | | | | | | |
|---|---|---|---|---|---|---|---|---|---|
| 9 | 108469200 | 108471399 | 12.95188 | 22.53443 | 48.93724 | 21.50493 | -0.72631 | 0.761702 | 0.238498 |
| 1 | 142223800 | 142225599 | 18.4632 | 33.24829 | 24.08465 | 10.505 | 0.867164 | 0.400772 | 0.238507 |
| 18 | 3171400 | 3172999 | 26.07334 | 45.03152 | 36.61671 | 16.06378 | 0.682568 | 0.881644 | 0.238509 |
| 1 | 37903400 | 37904799 | 12.33602 | 21.47067 | 48.92795 | 21.50573 | -0.79534 | 0.753299 | 0.23851 |
| 5 | 132569200 | 132573399 | 26.21022 | 42.31801 | 98.7984 | 42.14324 | -0.86098 | 1.727921 | 0.238511 |
| 4 | 77624000 | 77626599 | 31.80742 | 53.4501 | 43.12844 | 18.83835 | 0.689306 | 1.14667 | 0.238533 |
| 2 | 168605600 | 168609199 | 28.74791 | 46.39667 | 98.03761 | 41.79712 | -0.71817 | 1.743608 | 0.238537 |
| X | 54302800 | 54304399 | 16.38825 | 31.57667 | 14.67274 | 5.690917 | 1.636165 | -0.05763 | 0.238548 |
| 7 | 14065800 | 14070399 | 98.04129 | 153.0176 | 135.1881 | 56.09655 | 0.490878 | 2.681904 | 0.238556 |
| 3 | 40599800 | 40601199 | 10.85502 | 19.1699 | 41.95623 | 18.56226 | -0.73374 | 0.527902 | 0.238558 |
| X | 116544600 | 116545999 | 26.40858 | 45.73301 | 35.47915 | 15.56601 | 0.750844 | 0.858243 | 0.238562 |
| 4 | 105273800 | 105274799 | 12.78719 | 24.2605 | 20.38977 | 8.221243 | 0.738818 | 0.042201 | 0.238567 |
| 1 | 173399600 | 173411799 | 283.0769 | 408.6849 | 343.285 | 141.6224 | 0.465865 | 4.10828 | 0.238569 |
| 14 | 61122600 | 61129599 | 67.19074 | 105.2044 | 180.1642 | 74.73482 | -0.45925 | 2.616788 | 0.23858 |
| 11 | 72120600 | 72121599 | 5.409852 | 10.45057 | 27.08675 | 10.41089 | -0.80802 | -0.16866 | 0.238588 |
| 17 | 14929000 | 14930399 | 10.17046 | 18.21978 | 38.40718 | 16.83062 | -0.66857 | 0.408459 | 0.238589 |
| 3 | 93614200 | 93615799 | 14.71095 | 25.32351 | 57.01459 | 25.00983 | -0.7816 | 0.886888 | 0.238603 |
| 5 | 139555400 | 139558799 | 25.20552 | 41.17657 | 81.67842 | 35.11042 | -0.61159 | 1.492124 | 0.238607 |
| X | 114458200 | 114466399 | 111.6754 | 173.1996 | 137.7586 | 57.21518 | 0.635253 | 2.798569 | 0.238616 |
| 7 | 89246400 | 89247799 | 13.80732 | 26.28687 | 19.11441 | 7.650733 | 0.954759 | 0.024455 | 0.238624 |
| 1 | 188924400 | 188928599 | 31.24684 | 50.58379 | 95.03933 | 40.65677 | -0.54441 | 1.665112 | 0.238631 |
| 7 | 28508000 | 28511199 | 14.18886 | 24.22975 | 56.80476 | 24.90482 | -0.84143 | 0.961346 | 0.238643 |
| 18 | 37560800 | 37564199 | 68.88599 | 110.3904 | 76.64416 | 32.34412 | 0.871733 | 2.00763 | 0.238645 |
| 16 | 82191600 | 82193599 | 23.2713 | 40.82443 | 29.8453 | 13.19465 | 0.837109 | 0.659719 | 0.238649 |
| 17 | 66825600 | 66827199 | 9.630051 | 16.7941 | 49.92047 | 21.96923 | -1.17595 | 0.712643 | 0.238679 |
| 17 | 78231200 | 78234599 | 43.76087 | 71.27518 | 54.90651 | 23.57642 | 0.746151 | 1.554919 | 0.238679 |
| X | 112073000 | 112074799 | 21.48134 | 38.16473 | 24.50075 | 10.81194 | 1.030003 | 0.508038 | 0.238684 |
| 3 | 158265400 | 158268199 | 20.78929 | 34.31903 | 73.95445 | 31.93051 | -0.72373 | 1.341792 | 0.238696 |
| 7 | 41668200 | 41675199 | 101.6278 | 159.2888 | 122.9788 | 51.00806 | 0.689871 | 2.609266 | 0.238697 |
| 13 | 78171800 | 78174199 | 21.09798 | 34.85281 | 73.3751 | 31.69557 | -0.69009 | 1.335436 | 0.238723 |
| 5 | 123266600 | 123271799 | 22.92226 | 37.26642 | 88.18036 | 37.83878 | -0.87024 | 1.560852 | 0.238749 |
| 4 | 108316400 | 108317999 | 9.677756 | 17.05344 | 44.0655 | 19.50399 | -0.97308 | 0.548444 | 0.238757 |
| 7 | 18319000 | 18320999 | 43.35575 | 70.66763 | 58.65748 | 25.19385 | 0.639587 | 1.548938 | 0.238808 |
| 7 | 110680600 | 110682799 | 13.33104 | 22.98641 | 54.23884 | 23.79343 | -0.84817 | 0.863878 | 0.238814 |
| 3 | 13213400 | 13217399 | 64.298 | 102.7732 | 88.29819 | 37.12275 | 0.560828 | 2.082292 | 0.238822 |
| 12 | 93543000 | 93543999 | 100.386 | 158.3336 | 97.30921 | 40.38378 | 1.025313 | 2.481234 | 0.238849 |
| 1 | 85818800 | 85823799 | 34.79084 | 55.77707 | 115.8261 | 48.91167 | -0.70501 | 1.964976 | 0.238853 |
| 15 | 36881600 | 36886399 | 28.82354 | 46.73539 | 89.99369 | 38.54363 | -0.57752 | 1.626571 | 0.23886 |
| 12 | 18169600 | 18170799 | 28.5651 | 48.39362 | 41.52729 | 18.18377 | 0.601204 | 1.065385 | 0.238865 |
| 5 | 122201400 | 122202799 | 7.811859 | 14.29223 | 35.05582 | 14.98942 | -0.8617 | 0.258431 | 0.238882 |
| 18 | 35947200 | 35949199 | 7.811859 | 14.29223 | 35.05582 | 14.98942 | -0.8617 | 0.258431 | 0.238882 |
| 6 | 131545400 | 131558199 | 66.84322 | 106.8014 | 81.03287 | 34.05017 | 0.739266 | 2.095358 | 0.23889 |
| 4 | 14587200 | 14589199 | 17.78841 | 32.55398 | 22.74497 | 9.721754 | 0.937742 | 0.304149 | 0.238893 |
| 18 | 39141000 | 39144199 | 15.99337 | 27.41126 | 56.44043 | 24.75237 | -0.65442 | 0.932485 | 0.238894 |
| 6 | 118022600 | 118027399 | 37.53516 | 60.16039 | 113.6864 | 47.98821 | -0.56971 | 1.977371 | 0.238927 |
| 7 | 98062400 | 98065199 | 13.93217 | 24.10271 | 51.58925 | 22.6334 | -0.70692 | 0.835565 | 0.238934 |
| 17 | 27762200 | 27766599 | 23.15218 | 37.79377 | 82.27962 | 35.35917 | -0.74612 | 1.499788 | 0.238943 |

| | | | | | | | | | |
|---|---|---|---|---|---|---|---|---|---|
| 14 | 31362400 | 31363399 | 6.075515 | 11.41563 | 30.67887 | 12.60603 | -0.94701 | 0.075326 | 0.238949 |
| 8 | 27325000 | 27327799 | 16.25636 | 27.56975 | 60.06479 | 26.30928 | -0.73767 | 1.034568 | 0.238968 |
| X | 67175000 | 67178399 | 68.4293 | 109.197 | 90.35679 | 37.89991 | 0.612416 | 2.138635 | 0.238971 |
| 12 | 72709400 | 72711999 | 14.40683 | 24.82133 | 53.7309 | 23.57003 | -0.72467 | 0.87552 | 0.238977 |
| 14 | 47656600 | 47660799 | 18.96061 | 31.55514 | 70.31536 | 30.52978 | -0.77089 | 1.2376 | 0.23898 |
| 7 | 139909800 | 139910799 | 10.90066 | 20.91954 | 18.44753 | 7.23751 | 0.704944 | -0.07002 | 0.238999 |
| 12 | 7434000 | 7434599 | 10.90066 | 20.91954 | 18.44753 | 7.23751 | 0.704944 | -0.07002 | 0.238999 |
| 5 | 66850600 | 66853799 | 18.83655 | 31.65221 | 65.62737 | 28.66546 | -0.66736 | 1.128742 | 0.23901 |
| 6 | 97412000 | 97414399 | 12.77693 | 22.25907 | 49.46038 | 21.75266 | -0.75917 | 0.741174 | 0.239019 |
| 12 | 38147000 | 38147399 | 3.814826 | 7.454495 | 22.59066 | 7.843219 | -0.8935 | -0.56258 | 0.239028 |
| 13 | 101036200 | 101037599 | 7.363492 | 13.97079 | 34.58202 | 13.94276 | -0.80751 | 0.02725 | 0.239029 |
| 18 | 53536800 | 53538999 | 24.042 | 41.448 | 34.22262 | 15.01266 | 0.659951 | 0.88358 | 0.239042 |
| 4 | 27276200 | 27277199 | 12.1944 | 23.32002 | 19.34896 | 7.6684 | 0.778554 | -0.02194 | 0.239047 |
| 17 | 19532600 | 19534199 | 35.04768 | 57.92594 | 51.49253 | 22.25071 | 0.549838 | 1.345572 | 0.239048 |
| 16 | 91335200 | 91339799 | 19.71377 | 32.50956 | 76.03417 | 32.81072 | -0.84193 | 1.352729 | 0.239062 |
| 9 | 119350000 | 119354199 | 38.53042 | 61.63011 | 120.3175 | 50.63514 | -0.62001 | 2.043011 | 0.239071 |
| 11 | 45733600 | 45734599 | 13.75553 | 25.76594 | 21.16104 | 8.740954 | 0.742441 | 0.134181 | 0.239091 |
| 1 | 183668400 | 183671399 | 28.35147 | 48.56001 | 38.71755 | 16.97897 | 0.708649 | 0.966684 | 0.239108 |
| 5 | 140293200 | 140294399 | 6.627747 | 12.55752 | 29.80827 | 12.05321 | -0.75039 | 0.019662 | 0.239116 |
| 13 | 33661000 | 33663799 | 33.36993 | 56.25624 | 40.27549 | 17.60934 | 0.861289 | 1.121546 | 0.23913 |
| X | 9462000 | 9466799 | 77.35957 | 122.9885 | 97.69263 | 40.77052 | 0.665648 | 2.265521 | 0.239156 |
| 7 | 9067200 | 9069999 | 48.96112 | 79.14546 | 61.42764 | 26.24295 | 0.725959 | 1.711955 | 0.239158 |
| 19 | 40249800 | 40250799 | 9.65879 | 17.58745 | 38.16922 | 16.41887 | -0.6925 | 0.307845 | 0.23917 |
| 7 | 38817000 | 38818999 | 48.63571 | 78.56138 | 68.80088 | 29.36493 | 0.550943 | 1.735537 | 0.239175 |
| 19 | 38807200 | 38808199 | 6.671367 | 12.52112 | 31.77212 | 13.08014 | -0.86822 | 0.087487 | 0.239189 |
| 18 | 74581000 | 74590399 | 85.70364 | 131.9676 | 223.3393 | 92.76341 | -0.46094 | 2.926753 | 0.239196 |
| 6 | 131629600 | 131630799 | 25.23449 | 43.62696 | 36.03141 | 15.8075 | 0.660347 | 0.866444 | 0.23921 |
| 10 | 95433000 | 95434399 | 8.804029 | 15.80518 | 38.45524 | 16.82175 | -0.87225 | 0.384837 | 0.239211 |
| 11 | 75541200 | 75543199 | 10.88403 | 19.14724 | 43.02357 | 19.04913 | -0.77365 | 0.573333 | 0.239212 |
| 4 | 65702000 | 65703399 | 14.11688 | 26.18544 | 22.18731 | 9.311053 | 0.680126 | 0.199466 | 0.239218 |
| 18 | 15860800 | 15864399 | 35.25409 | 56.60453 | 109.059 | 46.15911 | -0.5945 | 1.91804 | 0.239223 |
| 1 | 79526600 | 79529599 | 23.87908 | 41.85385 | 29.79045 | 13.16062 | 0.875495 | 0.6763 | 0.239225 |
| 6 | 51569200 | 51571199 | 7.316852 | 13.52601 | 33.0973 | 13.95386 | -0.84275 | 0.19326 | 0.239237 |
| 6 | 134608800 | 134610999 | 10.82615 | 19.19346 | 41.40164 | 18.30266 | -0.71074 | 0.483012 | 0.239239 |
| 5 | 144703600 | 144706599 | 15.14272 | 25.77191 | 58.42354 | 25.6076 | -0.7941 | 0.997267 | 0.239249 |
| 2 | 152324600 | 152329799 | 43.23942 | 69.12346 | 123.2078 | 51.81033 | -0.48992 | 2.066768 | 0.239251 |
| 8 | 105320200 | 105321599 | 10.25703 | 18.10423 | 42.49313 | 18.80179 | -0.83427 | 0.530936 | 0.239254 |
| 15 | 77195800 | 77198799 | 18.76337 | 33.38013 | 26.82498 | 11.82436 | 0.708414 | 0.483017 | 0.239264 |
| 10 | 55832200 | 55833199 | 12.55009 | 24.42091 | 13.87367 | 5.111173 | 1.41599 | -0.26968 | 0.239276 |
| 12 | 91679000 | 91682399 | 18.66252 | 31.4202 | 63.97105 | 27.9583 | -0.64102 | 1.110163 | 0.239276 |
| 16 | 43132000 | 43133599 | 14.02594 | 24.4919 | 48.91461 | 21.54409 | -0.60564 | 0.7149 | 0.239286 |
| 9 | 108268800 | 108271799 | 22.00082 | 36.42666 | 73.88186 | 31.96441 | -0.63639 | 1.308574 | 0.239289 |
| 16 | 65928600 | 65931599 | 24.29148 | 42.34095 | 33.31539 | 14.65689 | 0.731623 | 0.767004 | 0.239298 |
| 8 | 14017200 | 14018199 | 11.90901 | 22.64176 | 18.76191 | 7.531429 | 0.763966 | 0.013307 | 0.239303 |
| 13 | 117513000 | 117514599 | 14.92037 | 27.84626 | 20.35932 | 8.461807 | 0.901544 | 0.164207 | 0.239303 |
| 13 | 66887600 | 66888999 | 13.61393 | 26.38754 | 14.81812 | 5.606589 | 1.397838 | -0.16646 | 0.23933 |
| 18 | 61902600 | 61906399 | 39.37918 | 64.61733 | 55.99537 | 24.11269 | 0.583195 | 1.446321 | 0.239333 |

| | | | | | | | | | |
|---|---|---|---|---|---|---|---|---|---|
| 4 | 22476400 | 22480799 | 28.65622 | 46.46273 | 92.51729 | 39.62362 | -0.62583 | 1.626872 | 0.239338 |
| 11 | 100972000 | 100974399 | 11.15933 | 19.72201 | 41.42987 | 18.32665 | -0.67444 | 0.521134 | 0.239342 |
| 7 | 62159600 | 62166599 | 140.1004 | 213.8441 | 177.8862 | 73.76874 | 0.553001 | 3.112576 | 0.239363 |
| 14 | 70573600 | 70574399 | 5.447688 | 10.41451 | 28.32864 | 11.23776 | -0.91986 | -0.05859 | 0.239368 |
| 13 | 8352400 | 8353599 | 20.22646 | 35.63754 | 28.9151 | 12.79851 | 0.688586 | 0.583456 | 0.239372 |
| 16 | 59336200 | 59338599 | 26.5052 | 45.20436 | 39.46387 | 17.3003 | 0.577449 | 0.997979 | 0.239398 |
| 7 | 72656200 | 72657799 | 7.697258 | 14.41881 | 32.30589 | 13.34326 | -0.69421 | 0.110412 | 0.239401 |
| 2 | 90770400 | 90771599 | 9.889855 | 17.62227 | 40.43637 | 17.8002 | -0.79497 | 0.442885 | 0.239408 |
| 6 | 63035600 | 63038599 | 34.2759 | 57.24206 | 47.04326 | 20.47038 | 0.663308 | 1.225785 | 0.23943 |
| 5 | 38044200 | 38054199 | 69.14588 | 107.355 | 203.0794 | 84.3313 | -0.61214 | 2.766345 | 0.239432 |
| 13 | 32285200 | 32287599 | 18.71106 | 34.36114 | 21.64345 | 9.207501 | 1.091148 | 0.290685 | 0.239437 |
| 3 | 109900600 | 109901599 | 9.051855 | 16.36485 | 39.07586 | 16.95989 | -0.83816 | 0.343761 | 0.239454 |
| 18 | 15074200 | 15075199 | 8.060015 | 14.88058 | 33.6085 | 14.19433 | -0.73056 | 0.205624 | 0.239478 |
| 16 | 47888000 | 47889599 | 17.92731 | 32.36284 | 24.78409 | 10.77684 | 0.790451 | 0.385467 | 0.239493 |
| 17 | 75470400 | 75473399 | 13.40308 | 23.29321 | 51.62738 | 22.67104 | -0.75608 | 0.781478 | 0.239502 |
| 3 | 94294000 | 94299199 | 32.02928 | 51.46086 | 108.0317 | 45.76968 | -0.7168 | 1.892046 | 0.239503 |
| 4 | 15499600 | 15500999 | 18.87464 | 34.83093 | 20.55508 | 8.686682 | 1.191139 | 0.263288 | 0.239504 |
| 8 | 114756400 | 114757999 | 18.20461 | 30.66494 | 62.22601 | 27.19986 | -0.63607 | 1.104186 | 0.239507 |
| 10 | 26549400 | 26551399 | 13.2234 | 23.05935 | 49.46082 | 21.76618 | -0.70841 | 0.729015 | 0.239509 |
| 17 | 40090600 | 40093599 | 45.01833 | 73.25951 | 60.56876 | 25.9995 | 0.643614 | 1.573927 | 0.239524 |
| 1 | 48530400 | 48532599 | 20.14111 | 35.36877 | 30.20308 | 13.36418 | 0.614085 | 0.626884 | 0.239524 |
| 7 | 10771600 | 10777799 | 162.578 | 244.9433 | 215.1749 | 88.88853 | 0.458835 | 3.381989 | 0.239531 |
| 14 | 50250000 | 50251999 | 21.13955 | 38.24989 | 21.75225 | 9.43123 | 1.219538 | 0.389958 | 0.239544 |
| 1 | 3107200 | 3108599 | 28.55944 | 48.25977 | 41.66069 | 18.22796 | 0.592616 | 1.089421 | 0.239554 |
| 7 | 26557800 | 26566199 | 228.1601 | 335.8021 | 285.2836 | 117.4812 | 0.476293 | 3.810645 | 0.239561 |
| 2 | 173474800 | 173479799 | 34.5506 | 55.60526 | 105.0038 | 44.59541 | -0.56142 | 1.840143 | 0.239565 |
| 4 | 120906600 | 120915399 | 59.60219 | 93.1325 | 184.5867 | 76.58541 | -0.67207 | 2.648637 | 0.23959 |
| 7 | 60476400 | 60477599 | 16.01289 | 30.47077 | 17.88501 | 7.166187 | 1.260192 | 0.046724 | 0.239608 |
| 8 | 121910400 | 121920599 | 58.84345 | 91.80089 | 196.1696 | 81.40449 | -0.78231 | 2.678784 | 0.239611 |
| 19 | 15997200 | 15997599 | 9.609545 | 18.62747 | 17.10474 | 6.469262 | 0.696984 | -0.20626 | 0.239619 |
| 7 | 18124800 | 18131799 | 103.9683 | 162.1767 | 133.2964 | 55.31478 | 0.594328 | 2.694178 | 0.239628 |
| 14 | 6507200 | 6507799 | 17.7844 | 31.74171 | 26.4492 | 11.62676 | 0.65914 | 0.455899 | 0.239632 |
| 2 | 150207400 | 150208799 | 14.38952 | 26.91901 | 21.71201 | 8.988009 | 0.76617 | 0.150187 | 0.239641 |
| 3 | 28644400 | 28645799 | 13.23914 | 24.87115 | 21.33609 | 8.769813 | 0.686017 | 0.113017 | 0.239652 |
| 7 | 24775600 | 24777199 | 10.87521 | 19.16873 | 41.98893 | 18.58386 | -0.73588 | 0.55088 | 0.239664 |
| 17 | 71781600 | 71782999 | 10.87521 | 19.16873 | 41.98893 | 18.58386 | -0.73588 | 0.55088 | 0.239664 |
| 19 | 36347400 | 36353199 | 35.39681 | 56.85098 | 108.9043 | 46.11497 | -0.58571 | 1.907646 | 0.239675 |
| 3 | 40912800 | 40913599 | 7.382095 | 13.90234 | 31.33037 | 12.82406 | -0.69182 | 0.071274 | 0.239676 |
| 3 | 87523600 | 87525199 | 7.382095 | 13.90234 | 31.33037 | 12.82406 | -0.69182 | 0.071274 | 0.239676 |
| 14 | 52122200 | 52123799 | 7.382095 | 13.90234 | 31.33037 | 12.82406 | -0.69182 | 0.071274 | 0.239676 |
| X | 153214200 | 153217999 | 37.81813 | 63.21008 | 40.48856 | 17.62738 | 1.021498 | 1.22177 | 0.239687 |
| 1 | 149274000 | 149277199 | 27.62281 | 47.17784 | 39.84638 | 17.47 | 0.625246 | 0.989506 | 0.239702 |
| 1 | 165549800 | 165551599 | 10.53859 | 18.65417 | 40.91888 | 18.09495 | -0.73551 | 0.498727 | 0.23971 |
| 3 | 104683400 | 104684999 | 10.53859 | 18.65417 | 40.91888 | 18.09495 | -0.73551 | 0.498727 | 0.23971 |
| 3 | 150715200 | 150715599 | 3.025754 | 5.914609 | 23.96507 | 8.247125 | -1.29462 | -0.60244 | 0.239727 |
| 7 | 26616000 | 26626599 | 135.8845 | 207.6721 | 175.7287 | 72.89539 | 0.529807 | 3.085878 | 0.239732 |
| 7 | 3412800 | 3417999 | 28.34646 | 45.95709 | 89.85109 | 38.47964 | -0.59952 | 1.627824 | 0.239751 |

| | | | | | | | | | |
|---|---|---|---|---|---|---|---|---|---|
| 4 | 61551200 | 61552999 | 23.6494 | 41.37866 | 30.9412 | 13.64875 | 0.80478 | 0.703827 | 0.239779 |
| 12 | 20128600 | 20134799 | 104.8669 | 165.4443 | 98.51976 | 40.88916 | 1.071149 | 2.483611 | 0.239781 |
| 11 | 101697000 | 101700599 | 19.62595 | 32.56066 | 71.32357 | 30.90088 | -0.74663 | 1.277426 | 0.239781 |
| 10 | 57202600 | 57203599 | 12.02933 | 23.17884 | 18.44166 | 7.151937 | 0.867984 | -0.09339 | 0.239786 |
| X | 33075400 | 33077199 | 15.09345 | 28.99991 | 16.29597 | 6.37192 | 1.353492 | -0.04249 | 0.239807 |
| 12 | 57581200 | 57583999 | 22.37647 | 36.88324 | 77.68936 | 33.51832 | -0.69186 | 1.36511 | 0.239831 |
| 1 | 145975400 | 145976599 | 12.70662 | 23.96448 | 19.68644 | 8.038243 | 0.755098 | 0.077791 | 0.239837 |
| 5 | 98450000 | 98451599 | 9.134488 | 16.73523 | 36.69542 | 15.66094 | -0.69975 | 0.263018 | 0.239841 |
| 13 | 113798400 | 113800399 | 15.29964 | 26.4841 | 52.72924 | 23.13396 | -0.60315 | 0.835283 | 0.239843 |
| 2 | 144171600 | 144175599 | 30.38708 | 48.93062 | 103.4672 | 43.97676 | -0.72321 | 1.817065 | 0.239848 |
| 12 | 99161800 | 99168399 | 50.0887 | 79.32677 | 147.6044 | 61.38012 | -0.56578 | 2.363327 | 0.239852 |
| 16 | 86263600 | 86265399 | 21.97077 | 38.90482 | 25.31188 | 11.19245 | 1.00776 | 0.534147 | 0.239854 |
| X | 63801800 | 63804599 | 33.36011 | 57.04769 | 35.21207 | 15.43971 | 1.076077 | 0.997156 | 0.239869 |
| 2 | 98977000 | 98978399 | 14.01309 | 27.08512 | 15.96832 | 6.1215 | 1.311436 | -0.11637 | 0.239882 |
| 5 | 147669600 | 147670999 | 17.42579 | 31.12508 | 25.71194 | 11.2955 | 0.672421 | 0.44902 | 0.239903 |
| 18 | 52386800 | 52390199 | 28.77255 | 46.83191 | 87.79594 | 37.68974 | -0.53425 | 1.555478 | 0.239934 |
| 8 | 119728200 | 119736799 | 66.50064 | 103.2846 | 205.1104 | 85.17017 | -0.68176 | 2.760696 | 0.239944 |
| 2 | 154418800 | 154419999 | 6.052544 | 11.47842 | 29.81351 | 12.0315 | -0.87649 | 0.007192 | 0.239951 |
| 8 | 83500200 | 83501399 | 6.052544 | 11.47842 | 29.81351 | 12.0315 | -0.87649 | 0.007192 | 0.239951 |
| X | 32812000 | 32822399 | 130.0867 | 201.7595 | 125.7508 | 52.22777 | 0.98666 | 2.8049 | 0.239958 |
| 17 | 39333800 | 39337799 | 99.6943 | 155.1081 | 139.2183 | 57.79412 | 0.464593 | 2.737714 | 0.239977 |
| 4 | 3633600 | 3637399 | 105.4909 | 163.2795 | 146.1582 | 60.71037 | 0.462725 | 2.830484 | 0.239993 |
| 18 | 40793200 | 40794599 | 15.74338 | 28.6438 | 24.20548 | 10.42265 | 0.658548 | 0.32535 | 0.240001 |
| 2 | 52713400 | 52714399 | 12.97855 | 24.67821 | 19.74341 | 7.923377 | 0.814806 | 0.026297 | 0.240015 |
| 8 | 56346400 | 56348399 | 22.2096 | 40.12864 | 21.12208 | 9.174391 | 1.328752 | 0.404577 | 0.240021 |
| 3 | 129295400 | 129297599 | 13.51984 | 23.67227 | 48.33926 | 21.33632 | -0.63748 | 0.667471 | 0.240025 |
| 1 | 6991600 | 6994799 | 22.0711 | 39.12795 | 24.1522 | 10.67718 | 1.083653 | 0.520012 | 0.240029 |
| 8 | 102811400 | 102812999 | 7.844214 | 14.76885 | 35.36787 | 14.48295 | -0.77973 | 0.086322 | 0.240029 |
| 19 | 7183200 | 7184799 | 7.347504 | 13.51582 | 34.61203 | 14.69077 | -0.9156 | 0.22235 | 0.240065 |
| 8 | 125516400 | 125527599 | 61.45454 | 95.49137 | 211.1826 | 87.68558 | -0.83633 | 2.750486 | 0.240067 |
| 16 | 94592200 | 94600199 | 55.00595 | 86.86327 | 158.0259 | 65.61789 | -0.53673 | 2.436107 | 0.240078 |
| 7 | 74040000 | 74043999 | 29.30988 | 47.50509 | 90.59708 | 38.79071 | -0.56418 | 1.635837 | 0.240078 |
| 3 | 110718000 | 110718599 | 9.647761 | 18.79849 | 15.45542 | 5.614806 | 0.907181 | -0.34889 | 0.240091 |
| 5 | 77505200 | 77506799 | 7.44175 | 13.82195 | 33.1772 | 13.89806 | -0.80913 | 0.164591 | 0.240094 |
| 5 | 100259400 | 100260599 | 7.44175 | 13.82195 | 33.1772 | 13.89806 | -0.80913 | 0.164591 | 0.240094 |
| 4 | 139735600 | 139742399 | 46.52021 | 73.67046 | 150.4153 | 62.54691 | -0.69957 | 2.361915 | 0.240101 |
| X | 147739600 | 147742599 | 48.71209 | 79.31223 | 57.39739 | 24.6429 | 0.83576 | 1.573208 | 0.240104 |
| 7 | 38606400 | 38606599 | 7.060784 | 13.80734 | 12.34209 | 4.205995 | 0.868907 | -0.61751 | 0.240118 |
| 18 | 6283800 | 6285199 | 19.16432 | 33.73835 | 28.27437 | 12.51696 | 0.641881 | 0.586032 | 0.24012 |
| 13 | 14696400 | 14697799 | 11.31409 | 19.6204 | 51.54148 | 22.6141 | -1 | 0.800614 | 0.24013 |
| 9 | 30573600 | 30578199 | 76.85179 | 121.8276 | 107.4526 | 44.71378 | 0.510783 | 2.348984 | 0.240132 |
| 2 | 55786800 | 55789999 | 33.1014 | 55.72381 | 43.08474 | 18.82944 | 0.750796 | 1.137483 | 0.240136 |
| 4 | 150938400 | 150939999 | 7.029792 | 13.00271 | 33.63564 | 14.1699 | -0.92197 | 0.18635 | 0.240138 |
| 7 | 30737200 | 30742399 | 41.02501 | 65.42204 | 127.2749 | 53.33499 | -0.6198 | 2.142673 | 0.240148 |
| 7 | 67278400 | 67279999 | 7.094685 | 13.33577 | 31.83058 | 13.0707 | -0.77789 | 0.081285 | 0.240165 |
| 9 | 61578200 | 61579199 | 16.7778 | 30.46777 | 24.60011 | 10.61784 | 0.721717 | 0.344966 | 0.240173 |
| 7 | 10759400 | 10760799 | 29.42827 | 49.50565 | 42.70255 | 18.65844 | 0.593472 | 1.130052 | 0.240201 |

| | | | | | | | | | |
|---|---|---|---|---|---|---|---|---|---|
| 6 | 72763600 | 72766799 | 17.7517 | 29.47894 | 71.68584 | 31.07806 | -0.89664 | 1.261508 | 0.240203 |
| 1 | 33874800 | 33879199 | 52.6127 | 83.00204 | 159.786 | 66.31564 | -0.61931 | 2.458574 | 0.240204 |
| 6 | 103487600 | 103491399 | 27.12111 | 44.26423 | 85.00482 | 36.52777 | -0.56608 | 1.510376 | 0.24022 |
| 1 | 103073600 | 103075599 | 21.69718 | 38.08719 | 27.53306 | 12.17895 | 0.853142 | 0.636684 | 0.240223 |
| 10 | 80234400 | 80236199 | 9.090503 | 16.34684 | 37.43058 | 16.35041 | -0.78372 | 0.374927 | 0.240229 |
| 5 | 126446000 | 126447399 | 12.75431 | 22.37508 | 45.12331 | 19.94939 | -0.61923 | 0.628811 | 0.240249 |
| 13 | 73719600 | 73721999 | 12.75431 | 22.37508 | 45.12331 | 19.94939 | -0.61923 | 0.628811 | 0.240249 |
| 12 | 28259600 | 28260799 | 15.87103 | 29.8429 | 20.0735 | 8.237998 | 1.036299 | 0.12851 | 0.240258 |
| 7 | 7576200 | 7581199 | 85.32264 | 135.374 | 98.35287 | 40.96294 | 0.791342 | 2.329239 | 0.24026 |
| 8 | 24780200 | 24781999 | 38.63211 | 63.87773 | 51.02066 | 22.05337 | 0.703839 | 1.351395 | 0.240263 |
| 10 | 76833600 | 76834199 | 4.39671 | 8.56576 | 24.31608 | 8.849362 | -0.86304 | -0.38083 | 0.240263 |
| 10 | 73378400 | 73379799 | 6.1208 | 11.82111 | 31.399 | 12.08188 | -0.84512 | -0.14447 | 0.240267 |
| 2 | 27533000 | 27534999 | 8.006705 | 14.5158 | 35.99868 | 15.56984 | -0.88823 | 0.316553 | 0.240269 |
| 7 | 88987600 | 88988199 | 4.947242 | 9.618318 | 24.63307 | 9.139203 | -0.74176 | -0.30245 | 0.240277 |
| 17 | 38341200 | 38343599 | 36.63069 | 60.76573 | 49.86085 | 21.6005 | 0.665383 | 1.308242 | 0.240295 |
| 13 | 41054000 | 41055599 | 6.939933 | 13.07191 | 33.03044 | 13.51595 | -0.85564 | 0.068228 | 0.2403 |
| 12 | 3949200 | 3952999 | 21.23869 | 35.12674 | 72.41813 | 31.30772 | -0.65986 | 1.320108 | 0.240309 |
| 4 | 46775800 | 46787999 | 101.4601 | 154.1246 | 263.8481 | 109.271 | -0.49293 | 3.205871 | 0.240316 |
| X | 24024600 | 24028399 | 78.71411 | 125.1515 | 96.67342 | 40.34822 | 0.705955 | 2.263153 | 0.240322 |
| 7 | 18901000 | 18907799 | 42.55043 | 67.62507 | 141.657 | 59.09836 | -0.73155 | 2.254381 | 0.240326 |
| 2 | 7585400 | 7587399 | 15.89435 | 29.83614 | 18.59659 | 7.653822 | 1.140855 | 0.133012 | 0.24033 |
| 7 | 111754800 | 111758399 | 16.80127 | 28.28825 | 62.66626 | 27.38853 | -0.7621 | 1.103545 | 0.240338 |
| 17 | 25556000 | 25563199 | 50.32647 | 79.44935 | 159.6272 | 66.27257 | -0.68009 | 2.440553 | 0.240356 |
| 4 | 83907000 | 83908999 | 10.37378 | 18.38022 | 46.1715 | 20.41379 | -0.92996 | 0.493205 | 0.24036 |
| 1 | 189465800 | 189467399 | 16.94329 | 28.57054 | 61.59226 | 26.93149 | -0.72284 | 1.090469 | 0.240361 |
| 2 | 173549400 | 173551999 | 18.47149 | 31.20818 | 62.36381 | 27.28496 | -0.61394 | 1.077321 | 0.240362 |
| 6 | 113354200 | 113355799 | 10.42957 | 18.35461 | 42.53349 | 18.83074 | -0.8174 | 0.566943 | 0.240362 |
| 13 | 64105200 | 64108999 | 56.60234 | 90.84484 | 80.45978 | 34.03356 | 0.523768 | 1.946017 | 0.240372 |
| 18 | 87439000 | 87441199 | 14.79968 | 27.68525 | 20.41105 | 8.450001 | 0.894176 | 0.149137 | 0.240375 |
| 9 | 34943000 | 34945199 | 10.9934 | 19.48985 | 40.37966 | 17.85089 | -0.65271 | 0.483931 | 0.240376 |
| 6 | 118456000 | 118458599 | 14.27958 | 24.52701 | 56.94498 | 24.97762 | -0.82633 | 0.905579 | 0.240388 |
| 3 | 102131800 | 102132599 | 6.121879 | 11.81709 | 28.27209 | 10.90497 | -0.69881 | -0.14492 | 0.240389 |
| 7 | 104879000 | 104879999 | 6.121879 | 11.81709 | 28.27209 | 10.90497 | -0.69881 | -0.14492 | 0.240389 |
| 17 | 90900000 | 90901599 | 8.769521 | 15.84225 | 36.46666 | 15.84263 | -0.78575 | 0.344829 | 0.240389 |
| 7 | 31174000 | 31177399 | 67.33647 | 107.7377 | 82.29747 | 34.64902 | 0.731649 | 2.052978 | 0.240396 |
| 1 | 171282400 | 171284799 | 12.31811 | 21.48239 | 47.85914 | 21.06239 | -0.76254 | 0.722629 | 0.240401 |
| 9 | 94901400 | 94903399 | 14.48681 | 25.25363 | 50.55247 | 22.24166 | -0.60937 | 0.742689 | 0.240404 |
| X | 5042000 | 5043799 | 58.78715 | 94.83291 | 65.38966 | 27.86119 | 0.892963 | 1.791412 | 0.240415 |
| 17 | 3277600 | 3279199 | 18.0225 | 31.96408 | 24.81641 | 10.96727 | 0.755614 | 0.494415 | 0.24044 |
| 3 | 41984800 | 41985399 | 6.140249 | 11.79016 | 27.98043 | 10.96056 | -0.70798 | -0.09379 | 0.240444 |
| 2 | 36678400 | 36680599 | 34.84301 | 57.93123 | 49.28635 | 21.38502 | 0.613507 | 1.275463 | 0.240488 |
| 7 | 117890800 | 117895399 | 90.01704 | 141.3311 | 125.2587 | 51.95203 | 0.49294 | 2.571905 | 0.240492 |
| 7 | 23305400 | 23310199 | 74.93043 | 119.7622 | 79.3484 | 33.35779 | 0.934949 | 2.093516 | 0.240503 |
| 7 | 120422600 | 120423999 | 6.286693 | 12.06819 | 28.58044 | 11.20402 | -0.70648 | -0.08966 | 0.240509 |
| 7 | 97990000 | 97994399 | 18.76993 | 31.32138 | 68.14832 | 29.63903 | -0.73654 | 1.206301 | 0.240518 |
| 6 | 137173600 | 137174999 | 8.023718 | 14.92115 | 33.21276 | 13.88825 | -0.69932 | 0.162402 | 0.240524 |
| 3 | 67422000 | 67424999 | 19.75285 | 32.84328 | 69.77059 | 30.27395 | -0.70229 | 1.250775 | 0.24054 |

| | | | | | | | | | |
|---|---|---|---|---|---|---|---|---|---|
| 4 | 151141800 | 151142799 | 5.449747 | 10.34403 | 29.23619 | 11.77731 | -0.9937 | -0.00275 | 0.240548 |
| 8 | 30001000 | 30006999 | 63.36214 | 101.2846 | 88.57419 | 37.24261 | 0.535443 | 2.079721 | 0.240562 |
| 7 | 81583000 | 81586599 | 18.8106 | 31.29284 | 69.67573 | 30.24363 | -0.76974 | 1.242019 | 0.240572 |
| 7 | 15720200 | 15723599 | 22.11296 | 36.3312 | 77.05646 | 33.1998 | -0.70349 | 1.407822 | 0.240599 |
| 7 | 118204200 | 118206799 | 27.86522 | 47.53471 | 40.41805 | 17.71893 | 0.61541 | 1.002654 | 0.240601 |
| X | 23243400 | 23245799 | 22.86536 | 41.46217 | 20.10036 | 8.691465 | 1.450764 | 0.388162 | 0.240604 |
| 10 | 126204200 | 126208399 | 30.25224 | 49.08405 | 92.35 | 39.57192 | -0.54334 | 1.616512 | 0.240632 |
| 2 | 98824200 | 98825999 | 17.71551 | 32.26741 | 23.88179 | 10.26914 | 0.849773 | 0.332976 | 0.240642 |
| 8 | 95327800 | 95329599 | 12.69061 | 21.96174 | 51.56895 | 22.62472 | -0.83977 | 0.825993 | 0.240646 |
| 7 | 116032400 | 116036399 | 64.99986 | 100.8289 | 205.0371 | 85.15546 | -0.71728 | 2.783817 | 0.240646 |
| X | 15301400 | 15302799 | 16.71915 | 31.63282 | 18.34958 | 7.44202 | 1.262128 | 0.092308 | 0.240654 |
| 13 | 8091800 | 8094799 | 31.98574 | 54.37734 | 38.56904 | 16.90368 | 0.87515 | 1.045176 | 0.24066 |
| 11 | 73168000 | 73172999 | 33.76607 | 54.30513 | 105.5362 | 44.77668 | -0.60398 | 1.861513 | 0.240671 |
| 16 | 60745400 | 60748999 | 41.77245 | 68.96787 | 50.00187 | 21.59088 | 0.842971 | 1.375486 | 0.240686 |
| 5 | 137135800 | 137149399 | 109.6765 | 163.4524 | 367.3108 | 151.3645 | -0.9094 | 3.586063 | 0.240693 |
| X | 26722400 | 26723799 | 15.80382 | 30.52037 | 14.33594 | 5.51759 | 1.631451 | -0.08826 | 0.240702 |
| 14 | 7898800 | 7900199 | 8.499243 | 15.7116 | 34.17762 | 14.40957 | -0.67519 | 0.202637 | 0.240705 |
| 16 | 91250600 | 91257399 | 53.69196 | 84.88037 | 152.9414 | 63.54152 | -0.5216 | 2.409083 | 0.240734 |
| 15 | 102061400 | 102063799 | 12.62409 | 22.02674 | 47.3298 | 20.83581 | -0.71055 | 0.716915 | 0.24074 |
| 4 | 35922800 | 35923599 | 6.910193 | 13.11309 | 30.44726 | 12.27107 | -0.715 | 0.011949 | 0.240741 |
| 15 | 74718600 | 74720199 | 11.76945 | 20.82391 | 42.43892 | 18.76875 | -0.63049 | 0.51371 | 0.240749 |
| 5 | 146706400 | 146707999 | 7.625815 | 14.05764 | 33.58481 | 14.21359 | -0.81299 | 0.211065 | 0.240769 |
| 1 | 57979200 | 57982599 | 16.77105 | 28.30564 | 62.75867 | 27.44902 | -0.76327 | 1.081068 | 0.24077 |
| 3 | 37365200 | 37367399 | 14.50083 | 25.23366 | 50.02152 | 21.98375 | -0.59554 | 0.767651 | 0.240773 |
| X | 21746800 | 21749199 | 25.34357 | 44.50634 | 27.48389 | 12.16142 | 1.079237 | 0.65187 | 0.240775 |
| 10 | 81452600 | 81471199 | 150.1464 | 219.6055 | 427.28 | 176.0214 | -0.72714 | 3.903901 | 0.240785 |
| 7 | 98331000 | 98332399 | 8.090656 | 14.86318 | 34.06355 | 14.48403 | -0.7583 | 0.236914 | 0.240793 |
| 10 | 108192000 | 108196199 | 52.05367 | 82.3907 | 149.1664 | 62.01134 | -0.52705 | 2.378923 | 0.240802 |
| 8 | 121897400 | 121905599 | 39.79788 | 63.31495 | 136.7933 | 57.13607 | -0.77451 | 2.220389 | 0.240804 |
| 4 | 139589000 | 139605999 | 117.5625 | 176.4344 | 310.8174 | 128.2647 | -0.5493 | 3.453762 | 0.240808 |
| 19 | 24600000 | 24601999 | 11.80345 | 20.71929 | 45.11241 | 19.9546 | -0.72928 | 0.612084 | 0.24083 |
| X | 81249400 | 81252399 | 34.35526 | 57.60608 | 41.07925 | 17.92067 | 0.86712 | 1.178246 | 0.240837 |
| 11 | 88804600 | 88805199 | 10.0019 | 19.48677 | 15.44497 | 5.617634 | 0.958198 | -0.33977 | 0.240843 |
| 4 | 32979600 | 32985599 | 33.0701 | 53.32969 | 100.724 | 42.89817 | -0.5581 | 1.774289 | 0.240852 |
| 12 | 59177600 | 59178599 | 6.748763 | 12.45993 | 33.60697 | 14.19337 | -0.98413 | 0.195335 | 0.240853 |
| X | 121111400 | 121114799 | 41.69061 | 70.73644 | 29.41286 | 12.88503 | 1.641902 | 1.09155 | 0.24087 |
| 3 | 101321800 | 101324399 | 11.33578 | 19.93003 | 44.55991 | 19.73172 | -0.76712 | 0.583734 | 0.240884 |
| 12 | 23962000 | 23965999 | 104.429 | 161.6921 | 134.895 | 56.03185 | 0.564582 | 2.821667 | 0.240898 |
| 16 | 8330800 | 8335399 | 24.20333 | 39.65415 | 80.84865 | 34.79231 | -0.64855 | 1.447641 | 0.2409 |
| 10 | 78829400 | 78830999 | 19.39817 | 35.60356 | 20.3485 | 8.663025 | 1.229243 | 0.295425 | 0.240907 |
| 7 | 135534400 | 135536799 | 17.53314 | 29.65235 | 60.63065 | 26.5347 | -0.64673 | 1.06469 | 0.240908 |
| 3 | 80943600 | 80945199 | 11.25106 | 19.63273 | 47.32704 | 20.83571 | -0.87525 | 0.710139 | 0.240908 |
| 1 | 171078200 | 171081599 | 14.13224 | 23.92304 | 61.48858 | 26.9178 | -0.97507 | 1.045528 | 0.240914 |
| 5 | 88701600 | 88703799 | 12.03047 | 20.94165 | 49.45164 | 21.73772 | -0.84646 | 0.749948 | 0.240916 |
| 15 | 31363000 | 31369799 | 43.02641 | 68.04894 | 165.9714 | 68.97012 | -0.95718 | 2.393623 | 0.240924 |
| 2 | 158827400 | 158828599 | 8.013176 | 14.93301 | 33.27396 | 13.86423 | -0.69671 | 0.148528 | 0.240933 |
| 6 | 104209600 | 104211799 | 17.16093 | 31.37148 | 22.72322 | 9.726499 | 0.884448 | 0.304341 | 0.240944 |

| | | | | | | | | | |
|---|---|---|---|---|---|---|---|---|---|
| 5 | 66407600 | 66410399 | 23.1542 | 38.23126 | 74.36442 | 32.1123 | -0.57653 | 1.34506 | 0.240967 |
| 2 | 36768000 | 36770399 | 23.74734 | 42.38204 | 22.58709 | 9.92949 | 1.299913 | 0.489774 | 0.240976 |
| 5 | 113033600 | 113041199 | 59.91165 | 93.29954 | 193.305 | 80.23615 | -0.7398 | 2.709377 | 0.240987 |
| 5 | 23637800 | 23640599 | 17.37759 | 28.98689 | 67.92091 | 29.53299 | -0.84292 | 1.207179 | 0.240988 |
| 18 | 80348000 | 80350599 | 26.84166 | 43.71925 | 84.50229 | 36.28495 | -0.57739 | 1.540204 | 0.240989 |
| 18 | 63235200 | 63236199 | 13.35613 | 25.26308 | 20.38297 | 8.271455 | 0.789444 | 0.068668 | 0.24099 |
| 12 | 18966200 | 19014599 | 1188.318 | 1520.77 | 1129.198 | 473.7633 | 0.469385 | 5.991958 | 0.241001 |
| 15 | 21775000 | 21777999 | 48.15126 | 77.69038 | 69.60536 | 29.66472 | 0.516302 | 1.767956 | 0.241009 |
| 5 | 146634000 | 146641599 | 48.87978 | 77.54416 | 145.5931 | 60.63087 | -0.5765 | 2.313568 | 0.24105 |
| 7 | 23150800 | 23153999 | 63.60733 | 102.0538 | 77.03477 | 32.56843 | 0.753347 | 1.963123 | 0.241056 |
| 7 | 54181800 | 54182599 | 8.235421 | 15.1584 | 34.09122 | 14.45772 | -0.72857 | 0.227229 | 0.241066 |
| 10 | 26782400 | 26783599 | 8.007704 | 14.98823 | 32.843 | 13.58368 | -0.66395 | 0.120263 | 0.241071 |
| 3 | 101944600 | 101950799 | 53.79402 | 85.23263 | 149.2844 | 62.0926 | -0.47818 | 2.355143 | 0.241085 |
| 8 | 104210800 | 104212599 | 8.821788 | 16.21792 | 35.14316 | 14.92855 | -0.67769 | 0.238441 | 0.241086 |
| 13 | 89954600 | 89955599 | 11.82026 | 20.71088 | 45.68247 | 20.17554 | -0.74786 | 0.643353 | 0.241094 |
| X | 139697400 | 139701399 | 22.86325 | 37.82752 | 78.87815 | 34.10917 | -0.67629 | 1.326594 | 0.241095 |
| 7 | 32121600 | 32131999 | 289.8463 | 416.927 | 349.6755 | 144.3315 | 0.462841 | 4.162355 | 0.241098 |
| 14 | 18162400 | 18165399 | 21.63971 | 38.2746 | 27.10868 | 11.98971 | 0.885644 | 0.548672 | 0.241105 |
| 10 | 95548800 | 95550399 | 7.263257 | 13.58885 | 31.7086 | 13.12428 | -0.75454 | 0.11383 | 0.241112 |
| 4 | 130994200 | 130995999 | 7.148609 | 13.28697 | 32.64648 | 13.66244 | -0.84127 | 0.157518 | 0.241118 |
| 7 | 20809800 | 20824999 | 293.1248 | 424.271 | 323.4123 | 133.3789 | 0.609449 | 4.07222 | 0.241124 |
| 18 | 37488000 | 37490599 | 93.42094 | 145.9591 | 120.127 | 49.84156 | 0.59419 | 2.65987 | 0.241127 |
| 6 | 131576400 | 131579599 | 46.69452 | 76.26132 | 55.69838 | 23.94026 | 0.8256 | 1.524665 | 0.241143 |
| 15 | 77452600 | 77457399 | 57.86987 | 92.81697 | 81.77406 | 34.55426 | 0.530223 | 1.968025 | 0.241151 |
| 18 | 6168000 | 6176999 | 110.9057 | 171.8189 | 147.3186 | 61.19211 | 0.525813 | 2.816193 | 0.241156 |
| 6 | 83831200 | 83833599 | 11.44809 | 20.22995 | 45.61581 | 20.1788 | -0.77658 | 0.528831 | 0.241166 |
| 17 | 39744600 | 39745999 | 19.59198 | 35.94759 | 20.34128 | 8.663788 | 1.243115 | 0.298796 | 0.24117 |
| 13 | 76144200 | 76145399 | 20.39081 | 38.42276 | 15.14945 | 6.19499 | 1.803536 | 0.143749 | 0.241193 |
| 2 | 70531400 | 70535999 | 19.5541 | 32.60326 | 71.1056 | 30.90926 | -0.74009 | 1.218658 | 0.241201 |
| 7 | 45654200 | 45659799 | 29.74197 | 47.91056 | 102.2418 | 43.48059 | -0.73565 | 1.803461 | 0.241227 |
| 3 | 93340400 | 93341399 | 10.39005 | 18.39448 | 40.422 | 17.87459 | -0.73784 | 0.495816 | 0.241232 |
| 11 | 73561800 | 73573799 | 367.7916 | 518.544 | 424.7579 | 175.8978 | 0.467874 | 4.459207 | 0.241239 |
| 1 | 192352000 | 192354399 | 17.35101 | 28.99843 | 66.31002 | 28.86766 | -0.80782 | 1.185197 | 0.241246 |
| 14 | 5960400 | 5964199 | 33.99724 | 57.01741 | 44.77772 | 19.53663 | 0.7284 | 1.177546 | 0.24126 |
| 1 | 109621600 | 109622199 | 13.36403 | 25.33452 | 20.25919 | 8.18249 | 0.807648 | 0.052006 | 0.241263 |
| 2 | 38225200 | 38227599 | 19.31017 | 32.43359 | 66.22613 | 28.92173 | -0.64541 | 1.1349 | 0.241279 |
| 5 | 3142000 | 3151799 | 170.1673 | 255.6869 | 220.4119 | 90.98621 | 0.48262 | 3.436387 | 0.241283 |
| 3 | 83945600 | 83948199 | 16.07489 | 27.33868 | 58.46483 | 25.62421 | -0.71057 | 1.00738 | 0.241285 |
| 11 | 117422600 | 117428799 | 29.62957 | 47.64703 | 108.6794 | 46.1026 | -0.83465 | 1.86018 | 0.241292 |
| 11 | 59414600 | 59416999 | 16.76325 | 28.77635 | 57.60207 | 25.26504 | -0.61361 | 0.922212 | 0.241297 |
| 5 | 85943200 | 85944999 | 17.42134 | 31.39935 | 25.5278 | 11.12238 | 0.703149 | 0.393747 | 0.2413 |
| 13 | 34868200 | 34870599 | 12.43487 | 21.78056 | 45.68442 | 20.17076 | -0.6757 | 0.652297 | 0.241324 |
| 6 | 57359600 | 57361599 | 23.78956 | 41.54122 | 32.08631 | 14.13223 | 0.758242 | 0.73605 | 0.241326 |
| 11 | 3274200 | 3275199 | 10.67478 | 20.52648 | 18.54936 | 7.236379 | 0.677881 | -0.08912 | 0.241344 |
| X | 89504000 | 89504999 | 13.43292 | 26.00361 | 16.12611 | 6.137831 | 1.249202 | -0.14491 | 0.241348 |
| 17 | 65832600 | 65834799 | 24.31397 | 42.26811 | 34.52873 | 15.16544 | 0.677529 | 0.805876 | 0.241352 |
| 17 | 45921400 | 45924799 | 17.28373 | 29.43884 | 62.49506 | 27.39392 | -0.69998 | 0.999928 | 0.241388 |

| | | | | | | | | | |
|---|---|---|---|---|---|---|---|---|---|
| X | 26704400 | 26704999 | 10.24574 | 20.02128 | 11.90877 | 4.13338 | 1.4255 | -0.50629 | 0.241403 |
| X | 29307800 | 29310199 | 22.55459 | 39.94922 | 25.97259 | 11.48399 | 1.009268 | 0.538065 | 0.241416 |
| 11 | 68791600 | 68793999 | 9.845356 | 17.30182 | 44.08959 | 19.52095 | -0.95448 | 0.581884 | 0.241434 |
| 19 | 31172200 | 31173599 | 9.845356 | 17.30182 | 44.08959 | 19.52095 | -0.95448 | 0.581884 | 0.241434 |
| 5 | 146912800 | 146919199 | 53.49307 | 84.67582 | 150.2085 | 62.44713 | -0.4976 | 2.37753 | 0.241438 |
| 13 | 12915000 | 12915399 | 9.348522 | 18.27561 | 11.29401 | 3.878206 | 1.383091 | -0.55882 | 0.24144 |
| 11 | 99187400 | 99188199 | 11.6069 | 22.54125 | 16.36424 | 6.125148 | 1.046523 | -0.21954 | 0.241442 |
| 13 | 110101600 | 110102199 | 6.59085 | 12.60064 | 29.5631 | 11.72555 | -0.70925 | -0.04599 | 0.241454 |
| 16 | 35257800 | 35263199 | 53.269 | 84.01729 | 159.7897 | 66.31336 | -0.6021 | 2.464162 | 0.24147 |
| 10 | 11430400 | 11431599 | 18.0265 | 30.44461 | 61.16297 | 26.75645 | -0.62148 | 1.079778 | 0.241473 |
| 1 | 5088000 | 5091799 | 38.89415 | 62.12292 | 122.8482 | 51.59543 | -0.64078 | 2.089224 | 0.241478 |
| 9 | 4587400 | 4589399 | 20.76852 | 36.36845 | 31.32625 | 13.82837 | 0.60181 | 0.676519 | 0.24148 |
| 9 | 91524800 | 91526999 | 20.83227 | 37.27319 | 22.63208 | 9.928178 | 1.114424 | 0.452597 | 0.241486 |
| 10 | 65743400 | 65744799 | 20.03345 | 35.63389 | 26.75052 | 11.79313 | 0.80613 | 0.493231 | 0.24149 |
| 11 | 99227800 | 99231599 | 23.2898 | 38.53006 | 73.95137 | 31.97686 | -0.55712 | 1.322219 | 0.241493 |
| 18 | 19115000 | 19115999 | 18.66361 | 33.14217 | 27.42815 | 12.10884 | 0.664855 | 0.495305 | 0.241493 |
| 3 | 48896600 | 48898999 | 66.7157 | 106.1717 | 94.97594 | 39.68672 | 0.495928 | 2.222449 | 0.241495 |
| 7 | 38218400 | 38224199 | 206.1803 | 305.1234 | 263.8243 | 108.6505 | 0.458312 | 3.717222 | 0.241496 |
| 7 | 16899400 | 16902999 | 50.61035 | 81.52228 | 73.0684 | 31.0714 | 0.51292 | 1.822132 | 0.241508 |
| 10 | 81039200 | 81042399 | 18.70715 | 31.01286 | 73.45694 | 31.80855 | -0.85913 | 1.284823 | 0.241521 |
| 5 | 129890600 | 129896999 | 37.38617 | 59.8848 | 115.0699 | 48.52687 | -0.59479 | 1.996968 | 0.241521 |
| 1 | 112984800 | 112986399 | 23.21406 | 40.69054 | 27.95518 | 12.35044 | 0.926319 | 0.669177 | 0.241523 |
| 17 | 88872600 | 88873999 | 8.577997 | 15.60227 | 36.01469 | 15.51344 | -0.78203 | 0.300236 | 0.241524 |
| 2 | 130398600 | 130399599 | 5.538316 | 10.72226 | 26.68521 | 10.1518 | -0.73583 | -0.2033 | 0.241542 |
| 5 | 113468600 | 113483999 | 195.1257 | 281.7259 | 475.3166 | 196.0904 | -0.53978 | 4.103378 | 0.241546 |
| 8 | 87537800 | 87539799 | 14.83333 | 25.78689 | 51.10284 | 22.4451 | -0.59528 | 0.78519 | 0.241546 |
| 7 | 128005400 | 128007199 | 8.207505 | 14.76807 | 39.52383 | 17.25585 | -1.00691 | 0.367287 | 0.241555 |
| 12 | 19579800 | 19582799 | 44.20746 | 71.96255 | 53.72332 | 23.06369 | 0.790797 | 1.564757 | 0.241559 |
| 8 | 70663200 | 70665599 | 10.79358 | 18.86356 | 47.82647 | 21.07674 | -0.94765 | 0.683538 | 0.241571 |
| 1 | 113174600 | 113175799 | 18.02968 | 31.95266 | 27.13445 | 11.99636 | 0.626409 | 0.505063 | 0.241578 |
| 4 | 63411000 | 63414599 | 17.46958 | 29.29026 | 64.81491 | 28.27386 | -0.76073 | 1.1485 | 0.241583 |
| 11 | 67010800 | 67013199 | 29.5976 | 50.38729 | 40.32263 | 17.6742 | 0.702089 | 1.026441 | 0.241588 |
| 18 | 87944400 | 87945199 | 11.4907 | 22.41372 | 12.89983 | 4.620275 | 1.433102 | -0.38086 | 0.241595 |
| 12 | 100462400 | 100464599 | 13.37157 | 23.37858 | 47.29831 | 20.85118 | -0.62406 | 0.689271 | 0.241603 |
| 14 | 38741400 | 38744399 | 27.83647 | 48.46655 | 31.48755 | 13.84315 | 1.006549 | 0.797421 | 0.24161 |
| 17 | 47385400 | 47390599 | 147.4483 | 226.7604 | 124.6603 | 51.7685 | 1.158299 | 2.9661 | 0.241623 |
| X | 87659000 | 87660999 | 21.21275 | 38.36387 | 22.51497 | 9.767806 | 1.174314 | 0.396038 | 0.241625 |
| 8 | 67581600 | 67583399 | 33.54296 | 55.71169 | 43.99351 | 19.07478 | 0.72064 | 1.280116 | 0.241627 |
| 6 | 133255000 | 133259199 | 18.17949 | 30.76521 | 80.76231 | 35.3521 | -1.00637 | 1.07855 | 0.24163 |
| 2 | 92598000 | 92599799 | 6.610658 | 12.20574 | 34.13586 | 14.41546 | -1.03601 | 0.194372 | 0.241641 |
| 9 | 26083400 | 26085199 | 24.85756 | 43.88571 | 24.02125 | 10.62808 | 1.254028 | 0.5817 | 0.241643 |
| 6 | 21214000 | 21217399 | 36.2648 | 58.06929 | 116.4278 | 49.07466 | -0.65657 | 2.006257 | 0.241644 |
| 16 | 3237800 | 3242399 | 27.41521 | 46.9263 | 34.72578 | 15.22765 | 0.81565 | 0.963403 | 0.241651 |
| 6 | 60281800 | 60282599 | 11.19586 | 21.67928 | 16.76536 | 6.372635 | 0.934673 | -0.17435 | 0.24166 |
| 6 | 28795600 | 28796599 | 6.002114 | 11.50727 | 27.90895 | 10.98077 | -0.745 | -0.08116 | 0.241661 |
| 14 | 106347800 | 106349999 | 18.02493 | 32.34556 | 25.25213 | 11.05002 | 0.756857 | 0.42134 | 0.241667 |
| 10 | 100467000 | 100469399 | 27.29137 | 46.70501 | 39.29682 | 17.23183 | 0.631133 | 0.969763 | 0.241669 |

| | | | | | | | | | |
|---|---|---|---|---|---|---|---|---|---|
| X | 92069800 | 92070799 | 14.01866 | 27.06241 | 16.5391 | 6.377035 | 1.25247 | -0.09605 | 0.241672 |
| 13 | 110752200 | 110752999 | 6.983222 | 13.03836 | 32.74947 | 13.6019 | -0.86417 | 0.126044 | 0.241672 |
| 17 | 33184200 | 33185399 | 6.983222 | 13.03836 | 32.74947 | 13.6019 | -0.86417 | 0.126044 | 0.241672 |
| 5 | 140108400 | 140111199 | 12.50159 | 21.72876 | 49.46299 | 21.722 | -0.79401 | 0.775326 | 0.241688 |
| 2 | 121636000 | 121638999 | 18.66791 | 31.39384 | 66.2126 | 28.92807 | -0.69188 | 1.1213 | 0.241696 |
| 6 | 137782800 | 137787599 | 44.03468 | 70.32541 | 128.8561 | 54.10987 | -0.53125 | 2.098282 | 0.241698 |
| 5 | 121644400 | 121648599 | 24.62288 | 40.64408 | 80.34335 | 34.68695 | -0.59988 | 1.357148 | 0.241704 |
| 1 | 32964400 | 32964999 | 16.69856 | 30.54065 | 23.16752 | 9.910674 | 0.819508 | 0.302089 | 0.241748 |
| 9 | 86173000 | 86174599 | 24.99124 | 43.40391 | 34.92278 | 15.32928 | 0.699085 | 0.821392 | 0.241748 |
| 11 | 24833800 | 24835199 | 16.22372 | 29.91031 | 22.38654 | 9.472999 | 0.849274 | 0.250374 | 0.241781 |
| 4 | 43681000 | 43682599 | 7.563442 | 13.72204 | 36.01615 | 15.5634 | -0.96847 | 0.309135 | 0.241791 |
| 17 | 29378400 | 29382599 | 61.43555 | 95.84948 | 183.224 | 76.03207 | -0.62168 | 2.678577 | 0.241801 |
| 4 | 141807000 | 141815399 | 42.77756 | 67.84653 | 147.9142 | 61.58209 | -0.792 | 2.3169 | 0.241822 |
| 9 | 3879000 | 3879999 | 19.24495 | 34.2147 | 26.74423 | 11.79521 | 0.74761 | 0.489883 | 0.241853 |
| 13 | 44423200 | 44424199 | 11.7292 | 22.44185 | 19.4187 | 7.684753 | 0.720528 | -0.03094 | 0.24186 |
| 2 | 25150600 | 25154399 | 18.2946 | 30.55551 | 66.35536 | 28.87552 | -0.73365 | 1.195791 | 0.241862 |
| 9 | 102393600 | 102395999 | 17.35398 | 29.40446 | 60.11733 | 26.32502 | -0.64647 | 1.046673 | 0.241863 |
| 9 | 98861000 | 98862199 | 5.982345 | 11.53238 | 27.5882 | 10.68515 | -0.70415 | -0.13469 | 0.241877 |
| 4 | 154086400 | 154091999 | 28.46997 | 45.85629 | 102.3247 | 43.50861 | -0.80004 | 1.80591 | 0.241908 |
| 10 | 25987800 | 25989199 | 9.945607 | 18.02036 | 39.67377 | 17.17197 | -0.71934 | 0.338918 | 0.24193 |
| 11 | 121150400 | 121151799 | 7.491277 | 13.7794 | 34.08165 | 14.46694 | -0.86582 | 0.223305 | 0.241946 |
| 6 | 121523800 | 121525599 | 22.75906 | 39.55962 | 33.66229 | 14.78362 | 0.618881 | 0.798634 | 0.241963 |
| 7 | 10409200 | 10415599 | 141.7218 | 215.8017 | 185.811 | 77.00767 | 0.50057 | 3.162751 | 0.241977 |
| 4 | 53905600 | 53907199 | 10.07729 | 17.86317 | 40.40892 | 17.86433 | -0.77875 | 0.479025 | 0.241979 |
| 1 | 118036400 | 118038799 | 35.25886 | 59.22127 | 42.51386 | 18.56631 | 0.857394 | 1.163661 | 0.241981 |
| 15 | 99011400 | 99014599 | 18.93451 | 32.09999 | 62.44061 | 27.34689 | -0.57501 | 1.049906 | 0.242018 |
| 5 | 145219200 | 145225799 | 33.48418 | 53.74914 | 113.1625 | 47.87751 | -0.72266 | 1.923182 | 0.242022 |
| 5 | 143268800 | 143269799 | 13.21092 | 24.65799 | 21.35258 | 8.873334 | 0.660036 | 0.153166 | 0.242029 |
| 10 | 70369200 | 70376399 | 56.5436 | 89.07978 | 162.9406 | 67.61089 | -0.54725 | 2.489524 | 0.242035 |
| 5 | 94521600 | 94528399 | 137.9292 | 211.1408 | 168.6027 | 69.96574 | 0.615196 | 3.053639 | 0.242037 |
| 11 | 3401600 | 3402999 | 6.70425 | 12.47065 | 32.12214 | 13.42632 | -0.90684 | 0.149407 | 0.242043 |
| 10 | 121871600 | 121874799 | 22.35036 | 36.94251 | 73.14667 | 31.6092 | -0.60193 | 1.330204 | 0.24205 |
| 8 | 115784600 | 115785599 | 8.026739 | 14.95925 | 32.71704 | 13.63069 | -0.6696 | 0.148489 | 0.242097 |
| 16 | 95882400 | 95884799 | 10.04697 | 17.56505 | 47.81658 | 21.07878 | -1.04949 | 0.674087 | 0.242106 |
| 2 | 157045600 | 157046999 | 4.611396 | 8.839487 | 28.36903 | 11.16721 | -1.14506 | -0.09305 | 0.242107 |
| 4 | 98121400 | 98124599 | 26.05878 | 42.3958 | 86.65901 | 37.19559 | -0.65907 | 1.555282 | 0.242125 |
| 16 | 45153400 | 45155599 | 19.04465 | 31.87695 | 66.05006 | 28.78183 | -0.66634 | 1.172304 | 0.242128 |
| 9 | 64511600 | 64513599 | 9.300025 | 16.56217 | 39.43532 | 17.36758 | -0.8481 | 0.441949 | 0.242129 |
| 13 | 92462200 | 92464199 | 9.314935 | 16.99107 | 36.05333 | 15.47477 | -0.65817 | 0.289363 | 0.242143 |
| 8 | 80605800 | 80607999 | 26.53624 | 45.68297 | 37.67825 | 16.52785 | 0.661221 | 0.913014 | 0.242156 |
| 15 | 101911200 | 101913999 | 15.29014 | 25.70208 | 65.81632 | 28.75108 | -0.97046 | 1.115012 | 0.242171 |
| 10 | 33746600 | 33748999 | 37.28986 | 61.42106 | 53.70358 | 23.16287 | 0.572478 | 1.394639 | 0.242185 |
| 7 | 49281800 | 49286799 | 52.93972 | 83.51451 | 159.1403 | 66.04628 | -0.60466 | 2.459807 | 0.242185 |
| 7 | 33489000 | 33499199 | 249.9652 | 365.3447 | 306.7055 | 126.3553 | 0.483938 | 3.921648 | 0.242189 |
| 13 | 24174400 | 24175799 | 7.686827 | 14.43106 | 32.37509 | 13.32199 | -0.69177 | 0.09598 | 0.242192 |
| X | 153766600 | 153768399 | 19.97693 | 36.61867 | 21.01271 | 8.961485 | 1.22221 | 0.309424 | 0.242224 |
| 6 | 69315800 | 69320199 | 41.45516 | 68.70538 | 46.61754 | 20.18256 | 0.938535 | 1.327937 | 0.242259 |

| | | | | | | | | | |
|---|---|---|---|---|---|---|---|---|---|
| 12 | 20411400 | 20414199 | 77.41627 | 122.3635 | 103.8867 | 43.15714 | 0.562282 | 2.420156 | 0.242271 |
| 8 | 47258600 | 47260999 | 34.21469 | 57.21845 | 46.47233 | 20.23949 | 0.68019 | 1.210002 | 0.24229 |
| 6 | 48997800 | 48998799 | 5.664427 | 10.98725 | 28.07805 | 10.58738 | -0.76124 | -0.2266 | 0.242291 |
| 12 | 80048200 | 80052799 | 23.53745 | 38.64431 | 80.41756 | 34.63759 | -0.67675 | 1.418997 | 0.242293 |
| 5 | 66179200 | 66185999 | 95.87962 | 145.2307 | 273.3831 | 113.1212 | -0.63319 | 3.263833 | 0.242294 |
| 9 | 72292200 | 72293599 | 7.231396 | 13.62251 | 31.33705 | 12.82046 | -0.7207 | 0.06758 | 0.242307 |
| 4 | 115000800 | 115001399 | 6.919611 | 13.1025 | 30.36788 | 12.29537 | -0.71818 | 0.027291 | 0.242324 |
| 11 | 35101000 | 35106999 | 40.80996 | 65.22971 | 120.9907 | 50.86699 | -0.54746 | 2.06827 | 0.242328 |
| X | 55117200 | 55119799 | 46.68898 | 75.91482 | 59.9064 | 25.70736 | 0.709892 | 1.59119 | 0.242344 |
| 4 | 17996200 | 17999199 | 28.51153 | 48.69712 | 39.82011 | 17.45853 | 0.671784 | 0.993406 | 0.242347 |
| 17 | 32659600 | 32662799 | 15.00761 | 25.9408 | 52.16117 | 22.88371 | -0.61771 | 0.847195 | 0.242361 |
| 3 | 51999800 | 52001399 | 8.505425 | 15.27815 | 37.95782 | 16.59589 | -0.9015 | 0.378577 | 0.242369 |
| 9 | 39026000 | 39029599 | 71.73387 | 114.0867 | 100.1189 | 41.7998 | 0.522594 | 2.248642 | 0.242379 |
| 7 | 4575600 | 4579999 | 51.82039 | 83.35108 | 74.42449 | 31.58197 | 0.515855 | 1.870075 | 0.242381 |
| 10 | 96509400 | 96510399 | 6.813954 | 12.78799 | 31.22402 | 12.85575 | -0.81312 | 0.088492 | 0.242383 |
| 3 | 14537200 | 14538199 | 27.23131 | 46.36096 | 37.51514 | 16.44368 | 0.686249 | 1.015627 | 0.242393 |
| 11 | 83315000 | 83318999 | 20.72269 | 34.37917 | 70.89668 | 30.71529 | -0.65991 | 1.280532 | 0.242417 |
| 13 | 48414200 | 48416599 | 16.10324 | 26.96411 | 67.42259 | 29.38713 | -0.93632 | 1.160163 | 0.242424 |
| 5 | 4358800 | 4360599 | 9.169821 | 16.69627 | 36.02215 | 15.49625 | -0.6841 | 0.298579 | 0.24248 |
| 14 | 70401800 | 70405799 | 17.39952 | 28.98491 | 69.56908 | 30.22427 | -0.87754 | 1.222511 | 0.242481 |
| 4 | 12851000 | 12851999 | 9.494688 | 17.20102 | 36.98855 | 16.01243 | -0.6854 | 0.332085 | 0.242483 |
| 8 | 105985400 | 105988599 | 29.84747 | 48.21657 | 94.9861 | 40.54528 | -0.61571 | 1.717115 | 0.242491 |
| 17 | 60851800 | 60852199 | 8.611971 | 16.8005 | 15.07565 | 5.391094 | 0.803004 | -0.41751 | 0.242496 |
| 18 | 27816600 | 27819999 | 26.43704 | 45.48166 | 37.64552 | 16.51316 | 0.655868 | 0.917417 | 0.242514 |
| 9 | 35571400 | 35575599 | 58.51704 | 93.99849 | 77.69481 | 32.90684 | 0.624687 | 1.920622 | 0.242518 |
| 7 | 48798200 | 48806999 | 60.0958 | 94.855 | 168.1935 | 69.82269 | -0.50037 | 2.453211 | 0.242518 |
| 6 | 34680000 | 34683999 | 76.02949 | 120.962 | 90.11272 | 37.64089 | 0.759083 | 2.239114 | 0.242521 |
| 5 | 78928000 | 78930399 | 59.71428 | 95.67217 | 84.03645 | 35.45217 | 0.532743 | 2.00445 | 0.242534 |
| 13 | 62483400 | 62493799 | 229.7351 | 338.7802 | 277.8157 | 114.4082 | 0.529904 | 3.77866 | 0.242534 |
| X | 102177600 | 102179199 | 19.66106 | 37.08763 | 15.0329 | 6.135652 | 1.76553 | 0.130037 | 0.242543 |
| X | 64236400 | 64240399 | 82.83078 | 131.1747 | 107.1784 | 44.56814 | 0.619737 | 2.380748 | 0.242546 |
| 1 | 143305000 | 143306199 | 9.482395 | 17.214 | 37.02529 | 15.9873 | -0.68337 | 0.319857 | 0.242549 |
| 16 | 96266000 | 96267799 | 9.482395 | 17.214 | 37.02529 | 15.9873 | -0.68337 | 0.319857 | 0.242549 |
| 3 | 151819200 | 151820799 | 6.612047 | 12.58365 | 29.37031 | 11.76486 | -0.71395 | -0.01145 | 0.242549 |
| 15 | 36833000 | 36834399 | 6.612047 | 12.58365 | 29.37031 | 11.76486 | -0.71395 | -0.01145 | 0.242549 |
| 4 | 37762000 | 37763999 | 18.72633 | 34.30162 | 22.17874 | 9.468013 | 1.050185 | 0.305838 | 0.242551 |
| 1 | 115493400 | 115493999 | 4.943018 | 9.623365 | 24.84977 | 9.117513 | -0.73888 | -0.3421 | 0.242552 |
| 15 | 48736800 | 48737999 | 5.815647 | 11.25906 | 31.68841 | 12.05577 | -0.91201 | -0.18541 | 0.242557 |
| 11 | 94271200 | 94272399 | 5.860611 | 11.2383 | 27.91494 | 10.97663 | -0.77839 | -0.08512 | 0.242576 |
| X | 147539400 | 147542199 | 49.41784 | 80.1246 | 62.87349 | 26.92751 | 0.714335 | 1.65088 | 0.242586 |
| 12 | 18018600 | 18023199 | 76.77847 | 122.1816 | 91.88788 | 38.39455 | 0.745761 | 2.230739 | 0.242602 |
| 8 | 17091600 | 17092599 | 12.25322 | 23.21547 | 20.41604 | 8.255816 | 0.670207 | 0.046098 | 0.242603 |
| 9 | 55420400 | 55421599 | 7.530635 | 14.24337 | 36.20289 | 14.69298 | -0.85393 | 0.054449 | 0.242606 |
| 5 | 99372000 | 99373599 | 8.511015 | 15.69715 | 34.12805 | 14.43385 | -0.67766 | 0.21597 | 0.242608 |
| 6 | 140041200 | 140044799 | 20.93783 | 35.09467 | 67.95867 | 29.63959 | -0.56918 | 1.165138 | 0.242622 |
| 5 | 31115200 | 31117999 | 14.51708 | 24.73931 | 58.41988 | 25.60869 | -0.85243 | 0.98243 | 0.242625 |
| 7 | 60940400 | 60941799 | 26.90339 | 46.54522 | 30.56204 | 13.4083 | 0.990185 | 0.863131 | 0.242633 |

| | | | | | | | | | |
|---|---|---|---|---|---|---|---|---|---|
| 17 | 82990200 | 82990799 | 19.91986 | 34.94665 | 29.78945 | 13.17046 | 0.616655 | 0.639694 | 0.242653 |
| 3 | 18880600 | 18881799 | 9.452431 | 16.82485 | 40.44423 | 17.81971 | -0.86242 | 0.447123 | 0.242665 |
| 13 | 112018400 | 112021199 | 28.38408 | 45.91561 | 93.53261 | 39.9905 | -0.66166 | 1.674677 | 0.242666 |
| 7 | 108269000 | 108271399 | 25.76488 | 44.56226 | 35.99359 | 15.7911 | 0.692463 | 0.865531 | 0.242666 |
| 6 | 100113000 | 100113799 | 3.032882 | 5.910988 | 23.53872 | 8.52355 | -1.33562 | -0.41748 | 0.242676 |
| 15 | 89909000 | 89911399 | 20.69668 | 36.6527 | 25.39936 | 11.23089 | 0.917204 | 0.524051 | 0.242695 |
| 14 | 18240000 | 18241999 | 15.70819 | 28.84723 | 22.95897 | 9.766881 | 0.7558 | 0.268616 | 0.24274 |
| 3 | 106841000 | 106843399 | 49.10158 | 79.80932 | 54.54093 | 23.40154 | 0.916426 | 1.595766 | 0.242746 |
| 10 | 30656600 | 30658199 | 13.36256 | 23.34544 | 48.91069 | 21.54928 | -0.67438 | 0.702274 | 0.242755 |
| 19 | 57302800 | 57304799 | 9.989834 | 17.96916 | 37.91607 | 16.55851 | -0.66659 | 0.379827 | 0.242759 |
| 6 | 54476000 | 54477399 | 13.50184 | 25.18476 | 21.77801 | 9.06001 | 0.660679 | 0.160341 | 0.242762 |
| 10 | 100232000 | 100232999 | 7.068996 | 13.38173 | 30.36059 | 12.29949 | -0.68835 | 0.031055 | 0.242787 |
| 18 | 76770000 | 76771599 | 9.129108 | 16.8018 | 36.27002 | 15.38365 | -0.67073 | 0.23708 | 0.242812 |
| 10 | 4134400 | 4136399 | 11.28497 | 19.99487 | 42.43553 | 18.76197 | -0.68792 | 0.494058 | 0.242818 |
| 14 | 33523800 | 33525199 | 18.70674 | 33.06401 | 28.15009 | 12.45204 | 0.621347 | 0.540251 | 0.242823 |
| 6 | 30901800 | 30902799 | 11.77058 | 22.48335 | 18.78924 | 7.468258 | 0.764198 | -0.01932 | 0.242825 |
| 16 | 79473000 | 79473999 | 11.77058 | 22.48335 | 18.78924 | 7.468258 | 0.764198 | -0.01932 | 0.242825 |
| 12 | 21718800 | 21721199 | 21.74643 | 39.13102 | 22.81407 | 9.960077 | 1.177845 | 0.427564 | 0.242825 |
| 17 | 13760200 | 13773599 | 807.1441 | 1031.954 | 1751.258 | 734.8224 | -0.7244 | 6.006638 | 0.242825 |
| 8 | 35080400 | 35082599 | 13.37536 | 23.42549 | 46.73391 | 20.63235 | -0.6039 | 0.660971 | 0.242833 |
| 7 | 75105000 | 75105999 | 6.201407 | 11.73233 | 29.74929 | 12.06713 | -0.84849 | 0.026276 | 0.242847 |
| 10 | 60742600 | 60745199 | 32.18746 | 54.85185 | 37.44127 | 16.41258 | 0.930756 | 1.023916 | 0.242857 |
| 6 | 117307800 | 117308799 | 6.578094 | 12.63817 | 30.44098 | 11.90469 | -0.72765 | -0.08961 | 0.242872 |
| 1 | 83036600 | 83039199 | 22.20179 | 36.69555 | 73.04778 | 31.56564 | -0.60959 | 1.331206 | 0.242873 |
| 19 | 38042600 | 38043799 | 8.951951 | 16.07063 | 37.44114 | 16.37824 | -0.80977 | 0.385766 | 0.242875 |
| 19 | 12981600 | 12984799 | 51.24543 | 82.62177 | 71.71339 | 30.53408 | 0.560725 | 1.794533 | 0.242883 |
| 12 | 103943000 | 103947999 | 28.41208 | 45.89562 | 95.93835 | 40.94925 | -0.70113 | 1.715837 | 0.242884 |
| 13 | 83994400 | 83996999 | 19.76752 | 32.82795 | 71.44167 | 30.97316 | -0.73719 | 1.265667 | 0.242898 |
| 13 | 108445000 | 108448799 | 33.22283 | 53.5698 | 103.9424 | 44.2623 | -0.59715 | 1.77838 | 0.242902 |
| 2 | 68671600 | 68673599 | 14.01576 | 24.51012 | 51.06359 | 22.51517 | -0.66634 | 0.694608 | 0.242929 |
| X | 113845000 | 113846799 | 24.48757 | 43.22334 | 25.94735 | 11.48076 | 1.121983 | 0.586945 | 0.242934 |
| 14 | 32555000 | 32558999 | 27.53537 | 44.68577 | 88.05905 | 37.73937 | -0.60953 | 1.606075 | 0.242938 |
| 1 | 10899000 | 10900399 | 15.54493 | 28.35176 | 24.17327 | 10.37711 | 0.64849 | 0.309631 | 0.242955 |
| 4 | 5887400 | 5889199 | 19.70672 | 35.55446 | 24.12211 | 10.4966 | 0.963582 | 0.400964 | 0.242975 |
| 1 | 176807200 | 176809399 | 12.24969 | 21.61769 | 44.01513 | 19.47589 | -0.63072 | 0.547683 | 0.242979 |
| 15 | 49992600 | 49994799 | 20.55172 | 34.18597 | 69.20188 | 30.03591 | -0.63281 | 1.249783 | 0.242983 |
| 12 | 20372400 | 20374999 | 58.16228 | 94.02378 | 59.94669 | 25.61191 | 1.009083 | 1.726855 | 0.242987 |
| 16 | 37203800 | 37205799 | 12.51069 | 21.71279 | 52.13135 | 22.87346 | -0.871 | 0.797832 | 0.24299 |
| 5 | 134830800 | 134835199 | 41.48081 | 66.22318 | 124.0535 | 52.06689 | -0.56359 | 2.106586 | 0.24299 |
| 7 | 47052200 | 47054799 | 36.77418 | 61.54634 | 43.67216 | 19.02925 | 0.87428 | 1.206699 | 0.243003 |
| 11 | 24081200 | 24083999 | 16.4752 | 28.27495 | 55.36792 | 24.28463 | -0.58192 | 0.922305 | 0.243006 |
| 2 | 14305400 | 14307199 | 14.88498 | 27.83366 | 20.99331 | 8.696954 | 0.860993 | 0.154296 | 0.243011 |
| 4 | 25866200 | 25869399 | 24.44009 | 40.33247 | 76.8488 | 33.17262 | -0.54698 | 1.356342 | 0.243014 |
| 12 | 86816800 | 86824199 | 47.43662 | 75.24523 | 144.7085 | 60.2549 | -0.61123 | 2.316308 | 0.243015 |
| X | 33066400 | 33067599 | 16.29997 | 31.16062 | 16.85265 | 6.685643 | 1.389783 | 0.016429 | 0.243023 |
| 5 | 135932400 | 135933799 | 6.212498 | 11.74745 | 31.41796 | 12.75746 | -0.92674 | 0.031809 | 0.243025 |
| 10 | 127368000 | 127371399 | 27.9093 | 45.14156 | 92.63913 | 39.60459 | -0.67244 | 1.677195 | 0.243026 |

| | | | | | | | | | |
|---|---|---|---|---|---|---|---|---|---|
| 17 | 40164600 | 40166599 | 14.85048 | 25.82624 | 52.17694 | 22.92217 | -0.6229 | 0.781948 | 0.243064 |
| 12 | 50627600 | 50629399 | 19.47929 | 34.67644 | 26.76273 | 11.79042 | 0.766998 | 0.484138 | 0.243083 |
| 12 | 17622600 | 17629599 | 44.24404 | 70.4947 | 133.307 | 55.7963 | -0.58036 | 2.173599 | 0.243085 |
| 3 | 74382800 | 74385599 | 29.34618 | 50.43185 | 35.82362 | 15.71347 | 0.875191 | 0.936588 | 0.243103 |
| 10 | 119636600 | 119638799 | 12.89707 | 22.58317 | 46.21168 | 20.39825 | -0.64031 | 0.660409 | 0.243116 |
| 1 | 101388000 | 101389599 | 25.40029 | 44.51876 | 28.57872 | 12.62498 | 1.023667 | 0.683476 | 0.24312 |
| 7 | 25856200 | 25857199 | 23.58049 | 40.83779 | 34.25913 | 15.03076 | 0.638547 | 0.843421 | 0.243122 |
| 12 | 13484800 | 13487799 | 58.85783 | 92.72417 | 161.9146 | 67.18502 | -0.4804 | 2.492307 | 0.243135 |
| 2 | 25461200 | 25475999 | 82.15808 | 124.0671 | 316.4633 | 130.821 | -1.07491 | 3.325057 | 0.24316 |
| 6 | 59968400 | 59971799 | 32.20893 | 54.59259 | 40.26421 | 17.63322 | 0.818844 | 1.073671 | 0.243177 |
| 19 | 45782600 | 45785799 | 12.63548 | 21.65986 | 56.26022 | 24.67368 | -0.9876 | 0.918439 | 0.243189 |
| 7 | 26578800 | 26595999 | 438.3023 | 608.7859 | 495.565 | 205.7015 | 0.455216 | 4.684216 | 0.24319 |
| 7 | 11139200 | 11140399 | 16.7519 | 30.92481 | 22.48048 | 9.495703 | 0.893251 | 0.249028 | 0.243192 |
| 13 | 59703000 | 59705599 | 12.72903 | 21.94165 | 53.66735 | 23.54252 | -0.89968 | 0.864288 | 0.243193 |
| 5 | 103039800 | 103040999 | 10.05338 | 17.88213 | 39.90706 | 17.59329 | -0.75648 | 0.454899 | 0.243207 |
| 6 | 47998400 | 48006799 | 115.8391 | 170.8903 | 413.4621 | 170.27 | -1.02954 | 3.763867 | 0.243211 |
| 7 | 11515800 | 11526999 | 38.21438 | 63.14534 | 44.55346 | 19.24919 | 0.882099 | 1.352646 | 0.24322 |
| 13 | 71768800 | 71770799 | 16.22862 | 29.2162 | 24.06756 | 10.50118 | 0.682677 | 0.386807 | 0.243234 |
| 7 | 7063600 | 7071799 | 156.9572 | 237.4115 | 204.98 | 84.77949 | 0.488666 | 3.301541 | 0.243247 |
| 18 | 73274200 | 73277799 | 22.40097 | 39.40268 | 28.70105 | 12.70802 | 0.842513 | 0.614244 | 0.243258 |
| 7 | 11166400 | 11168999 | 47.03307 | 75.94913 | 67.84171 | 28.944 | 0.521984 | 1.741981 | 0.243266 |
| 8 | 122099000 | 122101599 | 18.99664 | 31.52838 | 70.66397 | 30.62302 | -0.77948 | 1.271336 | 0.243267 |
| 13 | 65820000 | 65832399 | 172.3543 | 261.0583 | 187.1704 | 77.44692 | 0.758249 | 3.27753 | 0.243276 |
| 10 | 113495200 | 113496599 | 9.170536 | 16.7993 | 36.14318 | 15.42771 | -0.67231 | 0.265434 | 0.243283 |
| 5 | 56055800 | 56058799 | 60.06556 | 96.40578 | 79.35269 | 33.56665 | 0.629308 | 1.949303 | 0.243283 |
| 3 | 91388600 | 91390999 | 27.06553 | 46.90704 | 34.23669 | 15.02125 | 0.838521 | 0.851341 | 0.243294 |
| 13 | 120576000 | 120577799 | 23.32912 | 40.90229 | 29.78832 | 13.16304 | 0.842517 | 0.668054 | 0.243299 |
| 4 | 3402200 | 3406799 | 46.23691 | 75.28471 | 58.49022 | 25.11241 | 0.733621 | 1.566221 | 0.243306 |
| 13 | 52089000 | 52090999 | 12.21336 | 21.18574 | 50.50833 | 22.16098 | -0.86069 | 0.80171 | 0.243329 |
| 12 | 49157000 | 49157399 | 4.68323 | 9.117286 | 26.75725 | 9.820064 | -0.92263 | -0.34112 | 0.243331 |
| 11 | 106585800 | 106591799 | 30.85872 | 49.70692 | 101.1321 | 43.0054 | -0.66706 | 1.807201 | 0.24334 |
| 10 | 77884000 | 77888799 | 27.91548 | 45.14187 | 95.17197 | 40.67843 | -0.71164 | 1.682363 | 0.24334 |
| 7 | 32280800 | 32292399 | 197.2041 | 293.7818 | 245.0224 | 100.9669 | 0.519116 | 3.601729 | 0.243342 |
| 9 | 104282200 | 104284199 | 12.26115 | 21.6008 | 43.49743 | 19.25342 | -0.61577 | 0.566298 | 0.243352 |
| 4 | 147364200 | 147366999 | 32.18878 | 54.14074 | 39.55021 | 17.28021 | 0.832441 | 1.139769 | 0.243372 |
| 5 | 84977400 | 84978399 | 14.86757 | 27.79346 | 21.6367 | 8.967664 | 0.8155 | 0.158416 | 0.243397 |
| 16 | 78112200 | 78114199 | 11.14244 | 19.70781 | 42.44777 | 18.77395 | -0.71004 | 0.51255 | 0.243404 |
| 8 | 21658600 | 21660999 | 32.693 | 55.57433 | 39.15867 | 17.16196 | 0.884744 | 1.051194 | 0.243415 |
| 1 | 128700000 | 128701599 | 14.72383 | 25.38508 | 52.66099 | 23.10119 | -0.66322 | 0.870621 | 0.243429 |
| 8 | 72140000 | 72148599 | 90.42622 | 137.5107 | 261.4209 | 108.3005 | -0.64273 | 3.180852 | 0.243431 |
| 1 | 92834400 | 92835399 | 5.542291 | 10.71795 | 26.58808 | 10.16793 | -0.73831 | -0.1853 | 0.243435 |
| 2 | 63808600 | 63811399 | 25.49022 | 41.29696 | 96.06939 | 41.1211 | -0.85057 | 1.641142 | 0.243438 |
| 5 | 93998400 | 94000999 | 26.94602 | 46.2589 | 34.18286 | 14.99313 | 0.818309 | 0.934892 | 0.243445 |
| 8 | 112962400 | 112964199 | 15.24205 | 26.09919 | 55.28705 | 24.24472 | -0.69524 | 0.938385 | 0.243448 |
| 2 | 34683600 | 34685399 | 12.73766 | 22.34723 | 45.11618 | 19.94727 | -0.62088 | 0.626758 | 0.24345 |
| 4 | 112252200 | 112253799 | 44.76783 | 72.51835 | 64.58618 | 27.64223 | 0.530885 | 1.661009 | 0.243458 |
| 3 | 3197600 | 3200799 | 61.02132 | 97.76583 | 82.0784 | 34.6259 | 0.59788 | 2.007309 | 0.243465 |

| | | | | | | | | | |
|---|---|---|---|---|---|---|---|---|---|
| 10 | 81746800 | 81749399 | 45.46383 | 74.51652 | 47.88006 | 20.60812 | 1.012706 | 1.467663 | 0.243479 |
| 19 | 42478600 | 42482399 | 22.67441 | 37.45138 | 73.73271 | 31.84675 | -0.59384 | 1.339633 | 0.24348 |
| 8 | 72117400 | 72121599 | 22.99828 | 37.56059 | 83.68643 | 35.97004 | -0.77901 | 1.49131 | 0.243483 |
| 7 | 74516000 | 74517599 | 16.61463 | 31.44003 | 18.2691 | 7.406908 | 1.259796 | 0.089653 | 0.243494 |
| 5 | 120342200 | 120347799 | 31.58108 | 50.68206 | 112.197 | 47.45165 | -0.79525 | 1.928956 | 0.243508 |
| 4 | 156343600 | 156344599 | 5.306502 | 10.09918 | 28.7493 | 11.5157 | -0.99674 | -0.02499 | 0.243516 |
| 6 | 8113400 | 8117799 | 47.73941 | 77.84032 | 56.80859 | 24.40289 | 0.82508 | 1.550993 | 0.243517 |
| 17 | 38262200 | 38266199 | 37.13374 | 61.22635 | 53.15884 | 22.93782 | 0.58315 | 1.381636 | 0.243529 |
| 12 | 59477400 | 59478999 | 21.03074 | 37.24973 | 27.18938 | 12.02204 | 0.84324 | 0.530587 | 0.243533 |
| 9 | 117494600 | 117499599 | 52.6837 | 83.45508 | 147.6063 | 61.38778 | -0.49248 | 2.359897 | 0.243534 |
| X | 131060600 | 131062999 | 34.57218 | 59.34124 | 32.4569 | 14.2359 | 1.250485 | 0.96806 | 0.243537 |
| 5 | 117263600 | 117265799 | 13.55033 | 23.6874 | 47.30174 | 20.84992 | -0.60524 | 0.693948 | 0.243541 |
| 1 | 6540600 | 6541799 | 13.91295 | 26.04604 | 21.7844 | 9.007091 | 0.715619 | 0.141934 | 0.243549 |
| 1 | 121618400 | 121618999 | 5.576966 | 10.67924 | 27.83309 | 10.98904 | -0.85222 | -0.07381 | 0.243555 |
| 8 | 79291600 | 79292399 | 5.576966 | 10.67924 | 27.83309 | 10.98904 | -0.85222 | -0.07381 | 0.243555 |
| 11 | 46297400 | 46298399 | 5.576966 | 10.67924 | 27.83309 | 10.98904 | -0.85222 | -0.07381 | 0.243555 |
| 2 | 145664000 | 145665399 | 7.850993 | 14.70254 | 32.29884 | 13.34687 | -0.66653 | 0.114033 | 0.243564 |
| 8 | 8982800 | 8984199 | 7.850993 | 14.70254 | 32.29884 | 13.34687 | -0.66653 | 0.114033 | 0.243564 |
| 5 | 100460600 | 100468199 | 50.17555 | 79.6213 | 143.6449 | 59.83571 | -0.51864 | 2.306547 | 0.243573 |
| X | 26560400 | 26564199 | 87.81269 | 138.7871 | 109.8862 | 45.64801 | 0.662788 | 2.431274 | 0.243581 |
| 13 | 108254200 | 108254999 | 4.81343 | 9.365224 | 24.74165 | 9.123753 | -0.7782 | -0.32655 | 0.243587 |
| 18 | 69556600 | 69556999 | 5.218539 | 10.15004 | 25.95225 | 9.595958 | -0.73509 | -0.30673 | 0.243595 |
| 3 | 44568400 | 44571199 | 31.03521 | 52.69611 | 40.27778 | 17.64861 | 0.767606 | 1.054393 | 0.243596 |
| 5 | 129044600 | 129053999 | 76.94063 | 118.2786 | 236.3291 | 98.13646 | -0.70206 | 2.957188 | 0.243612 |
| 11 | 95282600 | 95285199 | 9.735086 | 17.04364 | 46.7547 | 20.63508 | -1.06034 | 0.647862 | 0.243623 |
| 7 | 32867200 | 32875599 | 99.61451 | 156.8733 | 102.533 | 42.5431 | 0.934987 | 2.514395 | 0.243625 |
| 6 | 6289000 | 6291199 | 12.88712 | 22.59188 | 46.18419 | 20.40605 | -0.63894 | 0.641155 | 0.243643 |
| 17 | 94449800 | 94451399 | 15.22924 | 28.26242 | 21.99977 | 9.226273 | 0.8022 | 0.205409 | 0.24365 |
| 17 | 74166000 | 74166599 | 11.63906 | 22.52774 | 17.36235 | 6.612871 | 0.937377 | -0.15997 | 0.243679 |
| 4 | 152868000 | 152901799 | 418.1996 | 566.381 | 904.6064 | 376.9786 | -0.55369 | 5.075935 | 0.243689 |
| 17 | 65045600 | 65046799 | 6.88537 | 13.17411 | 31.4079 | 12.42905 | -0.72932 | -0.04576 | 0.243704 |
| 12 | 50183000 | 50184999 | 18.05816 | 32.90478 | 21.99059 | 9.451168 | 0.995883 | 0.327171 | 0.243721 |
| 3 | 111169000 | 111169799 | 12.65681 | 24.13039 | 19.91653 | 7.948123 | 0.777704 | 0.007273 | 0.243736 |
| 11 | 66250600 | 66252199 | 8.972352 | 16.57915 | 38.08059 | 16.06436 | -0.75406 | 0.21709 | 0.243742 |
| 7 | 32176600 | 32177199 | 12.90303 | 24.15722 | 20.41738 | 8.440797 | 0.700845 | 0.129212 | 0.243745 |
| 10 | 17595000 | 17600599 | 33.09444 | 52.97301 | 130.2239 | 54.85971 | -0.95075 | 2.016248 | 0.243747 |
| 3 | 151915400 | 151916799 | 8.918016 | 16.10856 | 36.4532 | 15.83919 | -0.76164 | 0.345718 | 0.243753 |
| 17 | 38825000 | 38834399 | 149.5965 | 228.8036 | 161.981 | 67.20451 | 0.787778 | 3.075585 | 0.243757 |
| 2 | 168337200 | 168339599 | 10.2528 | 18.11648 | 40.9727 | 18.12511 | -0.78006 | 0.518462 | 0.243757 |
| 7 | 18892800 | 18894599 | 7.616324 | 14.07325 | 33.09061 | 13.95964 | -0.78654 | 0.19771 | 0.243765 |
| 5 | 52752600 | 52753999 | 12.93016 | 22.58613 | 46.79016 | 20.61279 | -0.65792 | 0.70194 | 0.243767 |
| 1 | 133979000 | 133981399 | 9.418244 | 16.86332 | 37.92778 | 16.62794 | -0.76162 | 0.406684 | 0.243771 |
| 5 | 30437200 | 30438399 | 9.418244 | 16.86332 | 37.92778 | 16.62794 | -0.76162 | 0.406684 | 0.243771 |
| 2 | 39777600 | 39778599 | 12.57941 | 24.33492 | 16.64722 | 6.356208 | 1.104134 | -0.14372 | 0.243772 |
| 1 | 116970000 | 116971999 | 26.94691 | 46.49022 | 35.3515 | 15.50823 | 0.778361 | 0.89066 | 0.243782 |
| 14 | 111459200 | 111461599 | 24.80196 | 43.44954 | 29.79404 | 13.15661 | 0.929145 | 0.689983 | 0.243786 |
| 7 | 31728000 | 31730599 | 32.49504 | 55.89408 | 31.76673 | 13.93436 | 1.195631 | 0.937406 | 0.243805 |

| | | | | | | | | | |
|---|---|---|---|---|---|---|---|---|---|
| 7 | 6737200 | 6739199 | 15.06255 | 27.51257 | 22.81345 | 9.77529 | 0.690215 | 0.294919 | 0.243812 |
| 12 | 20796600 | 20800799 | 106.3545 | 164.6151 | 144.6165 | 60.06999 | 0.489745 | 2.8315 | 0.243819 |
| 7 | 23407400 | 23409199 | 39.56171 | 65.24313 | 51.677 | 22.30037 | 0.715239 | 1.38163 | 0.243821 |
| 13 | 73661000 | 73663199 | 10.56674 | 18.6305 | 41.98914 | 18.58285 | -0.77659 | 0.5451 | 0.243821 |
| 5 | 148303400 | 148317799 | 131.8178 | 196.3928 | 341.6533 | 140.7872 | -0.54072 | 3.593174 | 0.243838 |
| 18 | 46437400 | 46440199 | 17.92064 | 30.53981 | 60.84912 | 26.67358 | -0.60869 | 0.99677 | 0.243844 |
| 14 | 106485600 | 106486599 | 16.0059 | 28.94701 | 23.49882 | 10.19469 | 0.708817 | 0.35756 | 0.243846 |
| 1 | 90893600 | 90895999 | 12.12074 | 21.26113 | 44.60942 | 19.72026 | -0.67607 | 0.625523 | 0.243847 |
| 1 | 13447000 | 13449199 | 15.71153 | 28.70855 | 24.1362 | 10.3373 | 0.670726 | 0.300407 | 0.24385 |
| 8 | 33949600 | 33951399 | 15.71153 | 28.70855 | 24.1362 | 10.3373 | 0.670726 | 0.300407 | 0.24385 |
| 12 | 96927400 | 96928999 | 18.54981 | 33.90417 | 22.71871 | 9.726817 | 0.996135 | 0.317495 | 0.243858 |
| 13 | 83666000 | 83666799 | 5.308525 | 10.16769 | 28.39931 | 11.20475 | -0.95049 | -0.07881 | 0.243861 |
| 8 | 22211800 | 22217399 | 67.93153 | 108.4856 | 87.25945 | 36.6404 | 0.654378 | 2.110963 | 0.243877 |
| 8 | 106552600 | 106554599 | 10.65375 | 18.96517 | 39.89088 | 17.57431 | -0.67099 | 0.453736 | 0.243882 |
| 7 | 26603000 | 26606599 | 67.16534 | 106.8794 | 95.22509 | 39.78676 | 0.501627 | 2.226561 | 0.243904 |
| 10 | 43241600 | 43245799 | 27.65585 | 44.9606 | 88.71725 | 38.06868 | -0.60917 | 1.569865 | 0.243907 |
| 7 | 17944600 | 17946999 | 32.22985 | 53.89695 | 47.02766 | 20.48082 | 0.577128 | 1.204672 | 0.243923 |
| 8 | 85442400 | 85445799 | 13.30235 | 22.6698 | 58.87724 | 25.80922 | -0.98895 | 0.97849 | 0.243924 |
| 5 | 77497800 | 77498599 | 6.456723 | 12.31717 | 29.42517 | 11.72797 | -0.7406 | -0.02841 | 0.243933 |
| 6 | 66649000 | 66651199 | 25.3398 | 43.63057 | 37.74335 | 16.55649 | 0.592724 | 0.906043 | 0.243943 |
| 7 | 41231200 | 41235399 | 52.2645 | 84.25519 | 69.68844 | 29.66753 | 0.629916 | 1.797022 | 0.243946 |
| 7 | 13572600 | 13575199 | 30.97312 | 53.85489 | 29.07018 | 12.76951 | 1.27229 | 0.827375 | 0.243947 |
| 7 | 45529800 | 45531199 | 12.0987 | 21.27897 | 44.02284 | 19.49251 | -0.65558 | 0.583732 | 0.243951 |
| 5 | 73856000 | 73856599 | 4.651607 | 9.05798 | 23.59207 | 8.640345 | -0.7479 | -0.35466 | 0.243956 |
| 14 | 56820200 | 56820799 | 4.651607 | 9.05798 | 23.59207 | 8.640345 | -0.7479 | -0.35466 | 0.243956 |
| 2 | 91172400 | 91173599 | 18.0757 | 32.07421 | 27.14354 | 11.99073 | 0.632348 | 0.496424 | 0.243958 |
| 13 | 75662800 | 75666599 | 27.97583 | 45.47158 | 89.9793 | 38.60679 | -0.61348 | 1.574075 | 0.243963 |
| 3 | 68655000 | 68656799 | 18.6639 | 33.3843 | 26.33739 | 11.55631 | 0.739429 | 0.448786 | 0.243976 |
| 14 | 3078200 | 3080599 | 67.00703 | 106.6571 | 96.00703 | 40.12888 | 0.487275 | 2.21776 | 0.243997 |
| 13 | 78357800 | 78360799 | 21.56469 | 35.2309 | 87.43044 | 37.58382 | -0.93367 | 1.486884 | 0.244012 |
| 11 | 23074400 | 23075999 | 6.792636 | 12.80807 | 30.78758 | 12.57552 | -0.78151 | 0.056406 | 0.244019 |
| 2 | 176896000 | 176898599 | 13.23553 | 23.05212 | 49.46902 | 21.75295 | -0.70925 | 0.747045 | 0.24402 |
| 5 | 114296600 | 114298799 | 8.85274 | 15.78876 | 42.03557 | 18.48985 | -1.00728 | 0.432468 | 0.244022 |
| 9 | 64481800 | 64484199 | 12.71672 | 21.94278 | 52.60551 | 23.07743 | -0.87048 | 0.854472 | 0.244044 |
| 13 | 98486400 | 98489199 | 18.25723 | 31.07897 | 60.82744 | 26.6617 | -0.58326 | 1.008751 | 0.244058 |
| 15 | 89613800 | 89615199 | 13.72925 | 23.89013 | 48.94347 | 21.50947 | -0.64266 | 0.763818 | 0.24408 |
| 4 | 146205000 | 146205599 | 14.68177 | 28.43995 | 14.12547 | 5.359781 | 1.569191 | -0.1471 | 0.244081 |
| 12 | 53881800 | 53883199 | 5.518496 | 10.66832 | 27.67532 | 10.60048 | -0.80353 | -0.17347 | 0.244105 |
| 19 | 56385400 | 56389999 | 25.52557 | 41.32814 | 94.9778 | 40.63494 | -0.83394 | 1.651648 | 0.244117 |
| 8 | 126507600 | 126509599 | 11.19454 | 19.70575 | 42.51157 | 18.8201 | -0.71473 | 0.568939 | 0.244121 |
| 7 | 31939000 | 31942399 | 30.39763 | 52.94748 | 27.89041 | 12.26486 | 1.30738 | 0.798645 | 0.244138 |
| 7 | 13512200 | 13520399 | 137.0728 | 209.9538 | 167.4071 | 69.47907 | 0.617994 | 3.041776 | 0.244158 |
| X | 39270400 | 39271599 | 17.47662 | 31.56493 | 25.07312 | 10.89574 | 0.738955 | 0.381954 | 0.244179 |
| 17 | 82490400 | 82490999 | 8.470725 | 16.49687 | 14.46119 | 5.28787 | 0.807104 | -0.34645 | 0.244187 |
| 1 | 99657400 | 99659599 | 25.3715 | 44.262 | 32.03882 | 14.10299 | 0.851434 | 0.757817 | 0.24419 |
| 2 | 105298200 | 105298999 | 8.454248 | 15.31616 | 35.97954 | 15.57556 | -0.81156 | 0.323951 | 0.244191 |
| 6 | 47940200 | 47941599 | 8.454248 | 15.31616 | 35.97954 | 15.57556 | -0.81156 | 0.323951 | 0.244191 |

| | | | | | | | | | |
|---|---|---|---|---|---|---|---|---|---|
| 2 | 168535000 | 168545999 | 86.56459 | 132.071 | 247.0265 | 102.4306 | -0.61541 | 3.112133 | 0.244211 |
| 17 | 87312800 | 87314999 | 21.03969 | 36.92519 | 30.69837 | 13.57665 | 0.652658 | 0.642822 | 0.244251 |
| 13 | 95250400 | 95250799 | 2.946045 | 5.762991 | 19.2364 | 6.361511 | -0.96069 | -0.75111 | 0.244254 |
| X | 33025200 | 33029399 | 44.53467 | 73.93773 | 42.67735 | 18.49891 | 1.17124 | 1.316659 | 0.244263 |
| 7 | 40823200 | 40823599 | 5.848044 | 11.18877 | 28.90758 | 11.44127 | -0.84267 | -0.05747 | 0.244302 |
| 5 | 137046000 | 137048399 | 13.1933 | 23.11667 | 48.86603 | 21.58157 | -0.68719 | 0.65436 | 0.244305 |
| 1 | 22330000 | 22330799 | 8.468191 | 16.49915 | 14.42042 | 5.242827 | 0.818423 | -0.36692 | 0.244307 |
| 4 | 149391000 | 149392799 | 9.822986 | 17.31076 | 43.03467 | 19.04742 | -0.91751 | 0.547231 | 0.244343 |
| 14 | 54887200 | 54888599 | 6.071022 | 11.46201 | 30.24182 | 12.32052 | -0.91095 | 0.040368 | 0.244349 |
| 1 | 74751400 | 74752999 | 9.206548 | 16.65617 | 36.45604 | 15.80898 | -0.71215 | 0.33777 | 0.244352 |
| 7 | 84965800 | 84969399 | 47.21712 | 77.32289 | 52.37054 | 22.53258 | 0.936592 | 1.487493 | 0.244352 |
| 7 | 7102200 | 7105799 | 39.65163 | 65.09352 | 55.49899 | 23.90346 | 0.606795 | 1.440468 | 0.244356 |
| 3 | 99688000 | 99688999 | 7.081952 | 13.42236 | 32.07766 | 12.96215 | -0.75917 | 0.02933 | 0.244372 |
| 1 | 100278400 | 100280799 | 25.94876 | 45.38652 | 29.10423 | 12.83479 | 1.025281 | 0.718487 | 0.244374 |
| 17 | 47695000 | 47708399 | 111.4868 | 167.4569 | 311.1123 | 128.4153 | -0.62487 | 3.435975 | 0.24438 |
| 2 | 36749400 | 36750199 | 12.95115 | 25.07926 | 16.065 | 6.105148 | 1.204363 | -0.15623 | 0.244407 |
| 6 | 3653800 | 3654799 | 12.95115 | 25.07926 | 16.065 | 6.105148 | 1.204363 | -0.15623 | 0.244407 |
| 2 | 167241000 | 167251599 | 73.94206 | 113.8993 | 219.7842 | 91.29147 | -0.64957 | 2.914335 | 0.244414 |
| 11 | 5345400 | 5346799 | 7.355052 | 13.93565 | 32.738 | 13.23764 | -0.73638 | 0.031649 | 0.244425 |
| 14 | 55101000 | 55101799 | 4.285411 | 8.322217 | 25.61256 | 9.589483 | -1.01612 | -0.27627 | 0.244426 |
| 1 | 71931800 | 71933799 | 16.95249 | 31.07777 | 22.19704 | 9.46531 | 0.90827 | 0.285907 | 0.244432 |
| 5 | 6321600 | 6325799 | 97.26623 | 151.7493 | 136.2447 | 56.53723 | 0.467238 | 2.688794 | 0.244439 |
| 19 | 20948000 | 20948799 | 16.82305 | 30.08426 | 25.17857 | 11.0501 | 0.654513 | 0.435315 | 0.24444 |
| 8 | 85572400 | 85573799 | 6.604719 | 12.168 | 33.58329 | 14.22459 | -1.01974 | 0.207672 | 0.244443 |
| 4 | 96171600 | 96176799 | 87.09425 | 137.8887 | 105.2843 | 43.77443 | 0.717017 | 2.389608 | 0.244454 |
| 13 | 66570000 | 66572199 | 38.46563 | 63.80123 | 48.81166 | 21.14404 | 0.765989 | 1.312248 | 0.244455 |
| 1 | 66621600 | 66623999 | 20.64059 | 36.19688 | 30.09963 | 13.30319 | 0.65213 | 0.649363 | 0.244471 |
| 2 | 149431200 | 149433999 | 26.69199 | 46.66789 | 29.1064 | 12.83142 | 1.065231 | 0.729208 | 0.244482 |
| 13 | 65562600 | 65563199 | 29.25994 | 49.44756 | 38.62874 | 16.9018 | 0.736235 | 1.089027 | 0.244488 |
| 7 | 59019800 | 59021799 | 19.44618 | 34.49665 | 27.27535 | 12.05173 | 0.729433 | 0.505926 | 0.244495 |
| 10 | 130089200 | 130089999 | 9.261838 | 17.97978 | 16.57409 | 6.223859 | 0.700878 | -0.23648 | 0.244504 |
| 10 | 32069600 | 32071599 | 22.19331 | 38.90093 | 31.14941 | 13.76151 | 0.706602 | 0.669737 | 0.244531 |
| 9 | 31481800 | 31483599 | 14.21528 | 24.72381 | 50.01133 | 21.97203 | -0.62451 | 0.77516 | 0.244541 |
| 14 | 27431400 | 27433599 | 7.702024 | 14.38226 | 35.03325 | 14.5473 | -0.82064 | 0.135103 | 0.244552 |
| 11 | 74162600 | 74163399 | 10.23642 | 18.12523 | 40.42212 | 17.87411 | -0.75892 | 0.492843 | 0.244558 |
| X | 110063200 | 110064599 | 31.78391 | 53.4943 | 44.24796 | 19.33653 | 0.653889 | 1.137869 | 0.244568 |
| 17 | 23631200 | 23634999 | 68.63995 | 109.4743 | 92.94944 | 38.95822 | 0.574488 | 2.15502 | 0.244575 |
| 8 | 118595600 | 118596999 | 8.990791 | 16.48645 | 35.08799 | 14.957 | -0.65547 | 0.254804 | 0.244591 |
| 4 | 78398600 | 78405799 | 156.6801 | 236.3821 | 208.6581 | 86.23059 | 0.4536 | 3.353414 | 0.244614 |
| 17 | 6842400 | 6843399 | 11.87824 | 22.57244 | 19.90436 | 7.998134 | 0.674109 | 0.020413 | 0.244628 |
| 14 | 23257400 | 23259199 | 13.7098 | 23.97554 | 47.82491 | 21.08749 | -0.60371 | 0.689179 | 0.244638 |
| 6 | 133924600 | 133926599 | 13.70003 | 23.99415 | 49.41058 | 21.81409 | -0.64955 | 0.666536 | 0.244639 |
| 12 | 73876400 | 73877599 | 7.239114 | 13.638 | 30.7701 | 12.58703 | -0.69243 | 0.067512 | 0.244648 |
| 4 | 92302400 | 92304399 | 28.7691 | 49.84482 | 33.05187 | 14.50128 | 0.976267 | 0.860589 | 0.244682 |
| 10 | 83752800 | 83757799 | 35.49795 | 57.19284 | 106.5359 | 45.31833 | -0.53982 | 1.807194 | 0.244694 |
| 2 | 88301000 | 88302399 | 20.19394 | 35.6886 | 27.74408 | 12.27272 | 0.751497 | 0.548298 | 0.244703 |
| 14 | 25456400 | 25462199 | 32.49961 | 52.23551 | 107.2642 | 45.46845 | -0.6844 | 1.880354 | 0.244708 |

| | | | | | | | | | |
|---|---|---|---|---|---|---|---|---|---|
| 1 | 8261800 | 8264399 | 23.7455 | 41.70351 | 28.63202 | 12.67037 | 0.927111 | 0.642546 | 0.24476 |
| 16 | 93769600 | 93772999 | 29.32879 | 47.51752 | 90.90505 | 38.91158 | -0.56917 | 1.645224 | 0.244766 |
| 8 | 3368000 | 3369799 | 19.75123 | 35.56422 | 23.99957 | 10.46919 | 0.96883 | 0.411248 | 0.244788 |
| 3 | 124114400 | 124116199 | 10.04683 | 17.90552 | 38.90324 | 17.12244 | -0.71623 | 0.441628 | 0.244797 |
| 11 | 30209600 | 30210399 | 10.15519 | 19.7481 | 15.84455 | 5.867489 | 0.916696 | -0.27679 | 0.2448 |
| 19 | 47799000 | 47800599 | 14.35516 | 26.42575 | 22.7685 | 9.654311 | 0.645737 | 0.245343 | 0.2448 |
| 3 | 134584600 | 134588199 | 36.41125 | 58.39391 | 112.2195 | 47.41209 | -0.59286 | 1.957905 | 0.244807 |
| 8 | 4002600 | 4013799 | 30.94865 | 52.22975 | 39.47945 | 17.26685 | 0.783438 | 1.105908 | 0.244826 |
| 4 | 144209200 | 144210999 | 28.15082 | 48.09639 | 39.81991 | 17.45889 | 0.654 | 0.988815 | 0.244829 |
| 4 | 6356200 | 6357399 | 16.79066 | 30.12754 | 25.54061 | 11.17724 | 0.63876 | 0.414655 | 0.24484 |
| 18 | 5842000 | 5845399 | 37.6865 | 62.0399 | 54.32333 | 23.42467 | 0.570039 | 1.401361 | 0.244842 |
| 7 | 63504400 | 63509599 | 24.88523 | 40.30928 | 93.70474 | 40.10342 | -0.84979 | 1.641099 | 0.244853 |
| 1 | 100752400 | 100753199 | 7.050636 | 13.40541 | 30.52742 | 12.25332 | -0.68205 | 0.000245 | 0.244879 |
| 2 | 105343400 | 105344999 | 17.94857 | 32.08267 | 26.39822 | 11.58973 | 0.678447 | 0.44773 | 0.244884 |
| 13 | 79117400 | 79118999 | 16.27609 | 30.51882 | 19.79032 | 8.160451 | 1.082668 | 0.147075 | 0.244885 |
| 16 | 82004000 | 82005599 | 11.11702 | 19.76901 | 40.90522 | 18.0372 | -0.64849 | 0.464963 | 0.244894 |
| 15 | 102068200 | 102071799 | 18.68268 | 31.0247 | 72.43513 | 31.40271 | -0.83824 | 1.263892 | 0.244899 |
| 2 | 166001200 | 166004599 | 25.6995 | 41.92874 | 82.79468 | 35.57331 | -0.60644 | 1.512431 | 0.244902 |
| 10 | 32596200 | 32597799 | 18.9653 | 34.56813 | 22.58502 | 9.702986 | 1.029093 | 0.335106 | 0.244908 |
| 6 | 67243200 | 67245199 | 15.13324 | 25.77904 | 57.90169 | 25.38072 | -0.7806 | 0.988216 | 0.244919 |
| 5 | 25229200 | 25234399 | 31.48229 | 50.74585 | 99.98001 | 42.55612 | -0.61963 | 1.785692 | 0.244924 |
| 15 | 98950800 | 98955599 | 31.18256 | 50.21336 | 102.4373 | 43.54131 | -0.6715 | 1.816908 | 0.244925 |
| 1 | 132123000 | 132129999 | 37.4535 | 59.80538 | 124.804 | 52.39638 | -0.7187 | 2.096333 | 0.244928 |
| 8 | 79224200 | 79225199 | 12.39236 | 23.97617 | 16.6541 | 6.355005 | 1.083046 | -0.1479 | 0.244931 |
| 15 | 20768000 | 20769399 | 12.39236 | 23.97617 | 16.6541 | 6.355005 | 1.083046 | -0.1479 | 0.244931 |
| 5 | 41622400 | 41623199 | 9.75436 | 17.40151 | 40.49287 | 17.80721 | -0.81335 | 0.43976 | 0.244938 |
| 8 | 35551200 | 35552199 | 6.899789 | 13.18917 | 34.35954 | 13.63422 | -0.85997 | -0.02877 | 0.244946 |
| 15 | 99755000 | 99756999 | 12.10518 | 21.27343 | 44.03274 | 19.496 | -0.65659 | 0.593804 | 0.244946 |
| 6 | 32383000 | 32385199 | 13.75525 | 23.80558 | 51.60532 | 22.64071 | -0.72521 | 0.829376 | 0.244953 |
| 7 | 116323200 | 116325799 | 20.6986 | 34.43447 | 68.72156 | 29.82994 | -0.61241 | 1.247383 | 0.244954 |
| 6 | 124445800 | 124452399 | 39.96456 | 63.92396 | 122.9432 | 51.74006 | -0.59853 | 2.04459 | 0.244963 |
| 13 | 112358200 | 112364199 | 77.77564 | 120.3517 | 204.5901 | 84.98162 | -0.46187 | 2.830441 | 0.244978 |
| 5 | 25977200 | 25981199 | 24.79172 | 40.77536 | 77.77967 | 33.52636 | -0.55053 | 1.399414 | 0.244978 |
| 4 | 131539800 | 131543399 | 19.57346 | 32.60769 | 68.13723 | 29.60297 | -0.67843 | 1.227454 | 0.244979 |
| 16 | 50729600 | 50731199 | 17.20191 | 30.89284 | 26.03454 | 11.38487 | 0.648104 | 0.41389 | 0.244981 |
| 4 | 86666400 | 86667999 | 13.63673 | 23.53356 | 55.85304 | 24.50219 | -0.85619 | 0.861099 | 0.244995 |
| 10 | 76843400 | 76846199 | 13.24561 | 23.07194 | 48.40414 | 21.28601 | -0.67664 | 0.745313 | 0.244997 |
| 2 | 60115400 | 60116799 | 18.31629 | 32.66967 | 22.51801 | 9.904109 | 0.930591 | 0.455166 | 0.245008 |
| 5 | 53212600 | 53214199 | 8.826168 | 16.17247 | 37.79976 | 16.12937 | -0.79122 | 0.261126 | 0.245039 |
| 4 | 40702800 | 40703399 | 3.417723 | 6.672175 | 21.54934 | 7.614048 | -1.00575 | -0.5079 | 0.245049 |
| 11 | 66597000 | 66597399 | 3.417723 | 6.672175 | 21.54934 | 7.614048 | -1.00575 | -0.5079 | 0.245049 |
| 15 | 11889000 | 11889599 | 3.417723 | 6.672175 | 21.54934 | 7.614048 | -1.00575 | -0.5079 | 0.245049 |
| X | 80023000 | 80026799 | 74.94308 | 118.9969 | 99.54797 | 41.49255 | 0.589287 | 2.300767 | 0.245057 |
| 5 | 38403600 | 38406599 | 19.02771 | 31.93212 | 63.8557 | 27.87164 | -0.61524 | 1.143147 | 0.245088 |
| 7 | 138528600 | 138531999 | 21.25958 | 35.6046 | 72.00788 | 31.38928 | -0.63174 | 1.177695 | 0.24511 |
| 7 | 6289000 | 6292999 | 25.44528 | 41.35177 | 86.85127 | 37.26022 | -0.69929 | 1.572214 | 0.245113 |
| 6 | 57925600 | 57927399 | 17.18203 | 31.33424 | 23.25759 | 9.985361 | 0.846731 | 0.319246 | 0.245116 |

| | | | | | | | | | |
|---|---|---|---|---|---|---|---|---|---|
| 14 | 14226600 | 14228999 | 16.15271 | 27.6833 | 55.89769 | 24.51427 | -0.62634 | 0.933188 | 0.245119 |
| 5 | 98669800 | 98671399 | 10.28671 | 18.56067 | 38.49071 | 16.74722 | -0.6383 | 0.364803 | 0.245122 |
| 15 | 78830400 | 78833799 | 12.04279 | 20.93631 | 49.98483 | 21.95445 | -0.86242 | 0.769455 | 0.245125 |
| 5 | 92385800 | 92389599 | 26.63847 | 43.48991 | 83.75797 | 35.99602 | -0.56987 | 1.504306 | 0.245127 |
| 12 | 23188400 | 23197199 | 238.9637 | 351.5896 | 283.1169 | 116.589 | 0.5531 | 3.817268 | 0.245127 |
| 19 | 47922400 | 47923599 | 9.663496 | 17.4662 | 36.95125 | 16.04285 | -0.66469 | 0.347413 | 0.245133 |
| 1 | 26456400 | 26458199 | 18.58449 | 33.89522 | 22.59797 | 9.700962 | 1.000742 | 0.328503 | 0.245142 |
| 1 | 84448000 | 84449599 | 16.31239 | 29.86212 | 22.82671 | 9.75254 | 0.809111 | 0.29004 | 0.24515 |
| 11 | 46440200 | 46441999 | 14.04996 | 26.17836 | 22.26342 | 9.279207 | 0.682505 | 0.173583 | 0.245154 |
| 15 | 76858600 | 76861199 | 13.55195 | 23.56733 | 50.5529 | 22.20832 | -0.70894 | 0.771672 | 0.245171 |
| 11 | 115326000 | 115327599 | 8.108158 | 14.82875 | 34.52634 | 14.77058 | -0.78753 | 0.262722 | 0.245196 |
| 11 | 103185000 | 103186999 | 10.11097 | 18.29706 | 41.83398 | 18.13534 | -0.77511 | 0.351415 | 0.245201 |
| 11 | 5133800 | 5135799 | 8.697694 | 15.50418 | 38.96361 | 17.14613 | -0.92435 | 0.431995 | 0.245218 |
| 13 | 31834000 | 31838399 | 28.01776 | 45.45211 | 88.72732 | 38.01559 | -0.59646 | 1.614238 | 0.245223 |
| 1 | 154683800 | 154684799 | 12.44764 | 23.6355 | 19.77008 | 7.957681 | 0.747277 | 0.03029 | 0.245233 |
| 1 | 81960000 | 81964399 | 57.09166 | 91.63474 | 80.59727 | 34.0943 | 0.533858 | 1.945096 | 0.245236 |
| 9 | 48839800 | 48840999 | 5.874202 | 11.22336 | 28.33537 | 11.25652 | -0.81544 | -0.04961 | 0.245255 |
| 11 | 35173000 | 35175599 | 12.96272 | 22.52458 | 49.47189 | 21.72247 | -0.74273 | 0.781078 | 0.245274 |
| 6 | 137761400 | 137764799 | 18.98323 | 31.90633 | 67.94138 | 29.67835 | -0.70576 | 1.128311 | 0.245276 |
| 6 | 68394400 | 68397399 | 40.5146 | 67.16542 | 47.72159 | 20.66455 | 0.872336 | 1.323552 | 0.245293 |
| 9 | 108473600 | 108474999 | 8.375385 | 14.96159 | 38.48798 | 16.90633 | -0.95561 | 0.416819 | 0.245316 |
| 7 | 41790400 | 41794599 | 52.29621 | 84.6884 | 57.90152 | 24.77587 | 0.910954 | 1.67692 | 0.245321 |
| 13 | 118997600 | 119002199 | 73.08415 | 116.2361 | 99.78648 | 41.66195 | 0.554299 | 2.249531 | 0.245327 |
| 14 | 34348800 | 34350199 | 8.809493 | 16.23209 | 35.19819 | 14.90586 | -0.6755 | 0.225319 | 0.245332 |
| 4 | 83606200 | 83608199 | 9.794205 | 17.85228 | 40.4112 | 17.36106 | -0.75205 | 0.305773 | 0.245333 |
| 1 | 12898400 | 12901999 | 43.1402 | 70.23748 | 60.49616 | 25.97296 | 0.58519 | 1.562195 | 0.245334 |
| 7 | 30822000 | 30833999 | 76.08889 | 116.8116 | 230.1992 | 95.5733 | -0.68364 | 2.982397 | 0.245334 |
| 12 | 55923200 | 55925199 | 31.93264 | 53.30251 | 46.51865 | 20.23582 | 0.576824 | 1.220801 | 0.245346 |
| 4 | 120991200 | 120995199 | 57.49655 | 92.76106 | 66.61121 | 28.38565 | 0.834679 | 1.785353 | 0.245351 |
| 1 | 133172400 | 133175599 | 13.3007 | 23.00089 | 53.1986 | 23.33917 | -0.81857 | 0.835567 | 0.245358 |
| 3 | 22573000 | 22576199 | 43.64488 | 71.12677 | 59.85024 | 25.70456 | 0.619753 | 1.551956 | 0.245364 |
| 8 | 128025800 | 128027799 | 15.18447 | 26.3048 | 52.70602 | 23.12431 | -0.6119 | 0.829945 | 0.245392 |
| X | 149161800 | 149163599 | 38.2268 | 63.6534 | 45.99839 | 19.97874 | 0.848423 | 1.268886 | 0.245393 |
| 18 | 33598600 | 33599999 | 9.049566 | 16.44221 | 36.0042 | 15.5299 | -0.70749 | 0.31244 | 0.245395 |
| Y | 10371800 | 10373599 | 10.91981 | 21.1616 | 8.654658 | 3.27747 | 1.846076 | -0.21898 | 0.245398 |
| 4 | 83335600 | 83336599 | 8.229659 | 15.11553 | 34.5858 | 14.71007 | -0.75675 | 0.237883 | 0.24542 |
| 9 | 3832400 | 3832999 | 9.97283 | 19.42272 | 14.69585 | 5.37192 | 1.016731 | -0.32748 | 0.245422 |
| 4 | 36884000 | 36886199 | 16.20938 | 29.7612 | 21.72238 | 9.243546 | 0.87919 | 0.269991 | 0.245424 |
| 4 | 35631600 | 35633399 | 17.79034 | 32.9678 | 20.64252 | 8.674052 | 1.111996 | 0.231787 | 0.245438 |
| 5 | 128634400 | 128640199 | 52.41305 | 84.36413 | 74.93421 | 31.83095 | 0.524655 | 1.848166 | 0.245458 |
| 2 | 79185000 | 79186199 | 9.627864 | 17.50907 | 37.06046 | 15.96707 | -0.65831 | 0.310796 | 0.245468 |
| 1 | 156494200 | 156497799 | 17.54129 | 28.94024 | 76.83909 | 33.1645 | -1.02388 | 1.34233 | 0.245486 |
| 7 | 30179000 | 30182199 | 19.42883 | 32.34709 | 68.0281 | 29.54423 | -0.68758 | 1.234792 | 0.245488 |
| 7 | 62284400 | 62285399 | 30.10621 | 51.04092 | 42.04579 | 18.4148 | 0.660076 | 1.068 | 0.24549 |
| 10 | 125189600 | 125190799 | 7.831783 | 14.72838 | 33.00741 | 13.54288 | -0.68698 | 0.08772 | 0.245491 |
| 12 | 113037200 | 113040799 | 19.17008 | 31.77763 | 71.76358 | 31.07349 | -0.79054 | 1.286584 | 0.245493 |
| 7 | 15333200 | 15333999 | 11.70095 | 22.75718 | 15.14717 | 5.602965 | 1.185143 | -0.26277 | 0.245523 |

| | | | | | | | | | |
|---|---|---|---|---|---|---|---|---|---|
| 11 | 102163000 | 102164599 | 6.183342 | 11.76884 | 31.69313 | 12.69284 | -0.91927 | -0.01448 | 0.245526 |
| 9 | 48197400 | 48198599 | 14.47717 | 25.20746 | 50.56328 | 22.23014 | -0.61251 | 0.758673 | 0.24554 |
| 6 | 53792600 | 53799599 | 38.65517 | 61.54374 | 145.0861 | 60.66355 | -0.89899 | 2.196161 | 0.24554 |
| 9 | 5248000 | 5249799 | 17.40493 | 32.27495 | 20.65661 | 8.672155 | 1.081429 | 0.224816 | 0.245545 |
| 14 | 115537600 | 115539799 | 29.6096 | 48.07897 | 89.27308 | 38.27553 | -0.52324 | 1.598793 | 0.245595 |
| 17 | 50712000 | 50713999 | 17.56323 | 31.29897 | 26.63969 | 11.72564 | 0.627624 | 0.465408 | 0.245635 |
| 7 | 10902600 | 10903999 | 48.22298 | 77.81154 | 69.49943 | 29.62224 | 0.520872 | 1.767044 | 0.245648 |
| 12 | 49907400 | 49909999 | 20.45626 | 34.26855 | 66.2093 | 28.86655 | -0.56588 | 1.167889 | 0.24565 |
| 16 | 34572800 | 34573599 | 6.142626 | 11.79408 | 28.58795 | 11.20029 | -0.73897 | -0.09365 | 0.245668 |
| 2 | 67464800 | 67468399 | 38.16866 | 64.08978 | 40.25287 | 17.57565 | 1.049527 | 1.17513 | 0.245699 |
| 2 | 168633200 | 168633799 | 8.976418 | 17.44215 | 15.0902 | 5.623624 | 0.801914 | -0.27069 | 0.245707 |
| 15 | 32899800 | 32901799 | 25.9132 | 44.93522 | 32.34888 | 14.19322 | 0.858478 | 0.839045 | 0.24572 |
| 8 | 97353000 | 97354799 | 33.53109 | 55.85558 | 48.70499 | 21.15939 | 0.578182 | 1.250528 | 0.24576 |
| 6 | 35245400 | 35249599 | 58.03871 | 90.94 | 173.2912 | 71.87846 | -0.61226 | 2.599118 | 0.245762 |
| 2 | 130226400 | 130228599 | 13.63501 | 23.59585 | 58.04733 | 25.46695 | -0.90684 | 0.84028 | 0.245767 |
| 13 | 32521800 | 32522999 | 7.396985 | 13.86112 | 32.87298 | 13.56952 | -0.77507 | 0.105787 | 0.245776 |
| 16 | 41021000 | 41023599 | 28.17733 | 48.24119 | 34.23279 | 15.01173 | 0.876014 | 0.965123 | 0.245783 |
| 4 | 5350600 | 5353399 | 24.15443 | 42.91355 | 23.69221 | 10.4552 | 1.245844 | 0.522057 | 0.245811 |
| 7 | 138764600 | 138773399 | 85.84144 | 131.14 | 242.903 | 100.7513 | -0.59985 | 3.084821 | 0.245813 |
| 3 | 127433000 | 127435399 | 14.33657 | 24.19032 | 65.85144 | 28.79937 | -1.05804 | 1.077803 | 0.245816 |
| 4 | 21840200 | 21840999 | 11.45606 | 22.17639 | 17.36882 | 6.611358 | 0.915105 | -0.16416 | 0.245818 |
| 7 | 24054600 | 24077599 | 190.9848 | 275.9374 | 475.7055 | 196.233 | -0.56991 | 4.092634 | 0.245838 |
| 15 | 15062600 | 15065599 | 46.35927 | 76.40621 | 46.72032 | 20.15614 | 1.087098 | 1.405006 | 0.245843 |
| 6 | 46788400 | 46788999 | 6.293719 | 12.05935 | 28.46081 | 11.21654 | -0.70824 | -0.07036 | 0.245848 |
| 4 | 54155200 | 54155999 | 8.285022 | 15.07407 | 35.5191 | 15.29402 | -0.81052 | 0.298409 | 0.24585 |
| 7 | 122512000 | 122515599 | 20.88421 | 36.59099 | 31.21083 | 13.78377 | 0.615774 | 0.667682 | 0.245851 |
| 18 | 9211600 | 9215399 | 29.86535 | 48.19163 | 97.61403 | 41.60707 | -0.65763 | 1.750518 | 0.245859 |
| 4 | 143485600 | 143486199 | 17.882 | 32.01914 | 26.48783 | 11.61208 | 0.672304 | 0.437811 | 0.245871 |
| 4 | 72052000 | 72052999 | 7.19657 | 13.7005 | 32.36352 | 12.95499 | -0.73126 | -0.00319 | 0.245881 |
| 4 | 9296800 | 9297999 | 8.393715 | 15.37787 | 34.54377 | 14.7429 | -0.73388 | 0.254205 | 0.245886 |
| 16 | 65819800 | 65820599 | 9.364637 | 18.20413 | 15.94429 | 5.921812 | 0.787766 | -0.27637 | 0.245906 |
| 16 | 15311600 | 15312199 | 6.890921 | 13.16554 | 30.12278 | 11.96733 | -0.67551 | -0.03534 | 0.245951 |
| 10 | 70142800 | 70154999 | 89.26021 | 136.178 | 253.4174 | 105.0796 | -0.60816 | 3.11349 | 0.245965 |
| 1 | 95434000 | 95435799 | 21.79867 | 38.13046 | 31.67298 | 13.96845 | 0.653855 | 0.698806 | 0.245969 |
| 13 | 8817400 | 8818399 | 19.97456 | 35.04046 | 29.85033 | 13.1967 | 0.617646 | 0.641745 | 0.245974 |
| 4 | 129384400 | 129385999 | 8.975311 | 16.05167 | 38.44108 | 16.86942 | -0.85264 | 0.411044 | 0.245977 |
| 14 | 18519200 | 18521199 | 8.975311 | 16.05167 | 38.44108 | 16.86942 | -0.85264 | 0.411044 | 0.245977 |
| 7 | 31814400 | 31819199 | 75.53613 | 119.7729 | 105.2355 | 43.80254 | 0.51669 | 2.341674 | 0.24599 |
| 4 | 67420400 | 67421799 | 35.6905 | 59.29852 | 43.93914 | 19.05563 | 0.812152 | 1.282489 | 0.245992 |
| 14 | 11024400 | 11025599 | 5.682077 | 10.9869 | 26.58181 | 10.17147 | -0.70328 | -0.18115 | 0.245992 |
| 4 | 123133000 | 123134199 | 5.599919 | 10.6534 | 28.75822 | 11.52889 | -0.92229 | -0.02001 | 0.246002 |
| 6 | 5859000 | 5860599 | 8.189419 | 15.18899 | 33.15589 | 13.91607 | -0.67525 | 0.179628 | 0.246013 |
| X | 54224400 | 54228199 | 30.99555 | 54.07891 | 26.86939 | 11.82823 | 1.391548 | 0.779346 | 0.246014 |
| 14 | 34107400 | 34108199 | 14.79869 | 27.09662 | 23.18357 | 9.90225 | 0.648174 | 0.278893 | 0.246034 |
| 3 | 108092000 | 108096199 | 21.0168 | 34.52138 | 77.51731 | 33.39474 | -0.7857 | 1.407645 | 0.246036 |
| 2 | 11114600 | 11115399 | 4.815617 | 9.363081 | 24.63868 | 9.136031 | -0.77982 | -0.30684 | 0.246048 |
| 9 | 81132400 | 81134199 | 19.95622 | 35.84697 | 24.53787 | 10.72845 | 0.946305 | 0.428249 | 0.246055 |

| | | | | | | | | | |
|---|---|---|---|---|---|---|---|---|---|
| 14 | 87407800 | 87409799 | 13.52114 | 23.22734 | 54.72981 | 24.00608 | -0.84723 | 0.900338 | 0.246056 |
| 2 | 16988600 | 16989199 | 9.889426 | 19.17844 | 14.09333 | 5.319353 | 1.01855 | -0.21944 | 0.246056 |
| 15 | 92789800 | 92791799 | 13.67235 | 23.90935 | 48.89919 | 21.56065 | -0.63976 | 0.688991 | 0.24606 |
| 13 | 90109400 | 90110199 | 13.72859 | 25.70844 | 21.78991 | 9.004894 | 0.697094 | 0.13832 | 0.246068 |
| 5 | 128871600 | 128894599 | 201.0652 | 289.8337 | 493.1728 | 203.5003 | -0.55435 | 4.127429 | 0.24609 |
| 5 | 117689600 | 117694799 | 54.4742 | 85.7401 | 163.4888 | 67.83022 | -0.60815 | 2.507399 | 0.246101 |
| 9 | 91258600 | 91261599 | 23.88717 | 39.12963 | 84.42594 | 36.3292 | -0.73016 | 1.450023 | 0.246102 |
| 3 | 35512400 | 35516199 | 42.86896 | 69.94662 | 59.39687 | 25.52106 | 0.607788 | 1.531447 | 0.246105 |
| X | 104965000 | 104968399 | 33.28156 | 56.33949 | 40.87509 | 17.8939 | 0.842562 | 1.091091 | 0.24611 |
| 19 | 45164800 | 45169999 | 31.78888 | 51.2697 | 99.17063 | 42.24313 | -0.59219 | 1.768481 | 0.246119 |
| 7 | 132271600 | 132272999 | 7.564609 | 14.14453 | 31.69712 | 13.13227 | -0.6977 | 0.121037 | 0.246121 |
| 6 | 41274000 | 41276799 | 24.10439 | 39.87966 | 76.94606 | 33.27291 | -0.56474 | 1.326322 | 0.246124 |
| 2 | 132711600 | 132712799 | 6.91658 | 13.1261 | 29.86385 | 12.03422 | -0.68529 | 0.011858 | 0.246125 |
| 18 | 49925600 | 49927999 | 33.1522 | 55.46347 | 41.72164 | 18.17501 | 0.790407 | 1.198102 | 0.246128 |
| 16 | 36053200 | 36056799 | 67.20402 | 107.3132 | 90.03926 | 37.80238 | 0.593453 | 2.11643 | 0.24613 |
| 1 | 102689800 | 102694199 | 40.94185 | 66.89912 | 57.70805 | 24.80853 | 0.586604 | 1.504298 | 0.246138 |
| 1 | 156884200 | 156884599 | 3.148454 | 6.157783 | 18.97084 | 6.419808 | -0.87836 | -0.66956 | 0.246145 |
| 14 | 52602600 | 52606199 | 73.13768 | 116.0013 | 104.4869 | 43.50274 | 0.481233 | 2.3293 | 0.246152 |
| 6 | 73679000 | 73679799 | 5.103157 | 9.849554 | 26.38062 | 10.18069 | -0.85924 | -0.15668 | 0.246156 |
| 3 | 88318600 | 88321399 | 9.42337 | 16.52145 | 45.6969 | 20.19254 | -1.07198 | 0.621161 | 0.246158 |
| 13 | 61094200 | 61097799 | 41.57279 | 67.68455 | 58.16598 | 24.97241 | 0.588386 | 1.556642 | 0.246161 |
| 3 | 93744200 | 93745799 | 57.25813 | 91.74982 | 82.90077 | 34.9805 | 0.492295 | 1.99704 | 0.246175 |
| 2 | 122727800 | 122730199 | 39.95942 | 66.54638 | 44.29403 | 19.23992 | 0.966472 | 1.268617 | 0.246196 |
| 7 | 5092800 | 5106599 | 91.27884 | 138.378 | 272.8668 | 112.9375 | -0.6991 | 3.245481 | 0.246205 |
| 4 | 140599400 | 140604999 | 70.81617 | 109.8058 | 196.5414 | 81.63213 | -0.53389 | 2.790088 | 0.246216 |
| 7 | 78866000 | 78869799 | 33.71314 | 54.36807 | 100.8945 | 42.97221 | -0.53275 | 1.774637 | 0.246242 |
| 14 | 31296000 | 31299999 | 23.40901 | 38.36596 | 81.36349 | 35.01897 | -0.70498 | 1.441466 | 0.246245 |
| 14 | 65722200 | 65723799 | 17.79553 | 32.40726 | 23.94819 | 10.30003 | 0.851916 | 0.335355 | 0.246253 |
| 9 | 5717200 | 5718799 | 21.81935 | 38.30207 | 30.52137 | 13.50091 | 0.713429 | 0.642845 | 0.246268 |
| 13 | 28059800 | 28061599 | 22.95954 | 40.33553 | 29.26579 | 12.95419 | 0.847869 | 0.634658 | 0.246269 |
| 18 | 24131800 | 24138199 | 129.1539 | 199.0954 | 139.8812 | 58.10429 | 0.807349 | 2.913715 | 0.246286 |
| 1 | 156408800 | 156409399 | 4.054548 | 7.922715 | 21.92364 | 7.617011 | -0.76435 | -0.55643 | 0.246293 |
| 13 | 89363800 | 89364199 | 4.230675 | 8.246999 | 22.54692 | 8.144312 | -0.7984 | -0.41341 | 0.246316 |
| 2 | 61085600 | 61086599 | 7.069764 | 13.41017 | 30.41447 | 12.26892 | -0.68217 | 0.018413 | 0.246329 |
| 18 | 74261800 | 74262799 | 16.04718 | 29.80763 | 22.01655 | 9.221824 | 0.878922 | 0.208381 | 0.246329 |
| 11 | 20477200 | 20479599 | 13.36531 | 23.43997 | 49.39025 | 21.83033 | -0.68254 | 0.641756 | 0.246342 |
| 9 | 114836400 | 114839799 | 41.76354 | 67.99529 | 60.82698 | 26.11488 | 0.530684 | 1.561194 | 0.246344 |
| 13 | 23805600 | 23806599 | 5.279207 | 10.13114 | 27.32761 | 10.72212 | -0.89237 | -0.09839 | 0.246349 |
| 7 | 26975600 | 26976799 | 12.88406 | 24.09058 | 18.82634 | 7.800078 | 0.81017 | 0.131523 | 0.246358 |
| 5 | 144899000 | 144903399 | 22.30601 | 36.5655 | 79.3206 | 34.14259 | -0.73731 | 1.436407 | 0.246384 |
| 19 | 5046800 | 5048199 | 7.908618 | 14.60571 | 33.61401 | 14.19052 | -0.75708 | 0.202173 | 0.246395 |
| 8 | 122129600 | 122132199 | 27.39523 | 44.40409 | 89.49572 | 38.32057 | -0.64357 | 1.63154 | 0.246395 |
| 2 | 101601200 | 101602999 | 9.868462 | 17.65715 | 38.40534 | 16.84719 | -0.7145 | 0.413708 | 0.246404 |
| 7 | 139624000 | 139630999 | 47.69283 | 75.78787 | 143.1527 | 59.71449 | -0.58279 | 2.262387 | 0.246411 |
| 13 | 101089800 | 101093599 | 30.25509 | 49.10228 | 91.14617 | 39.06406 | -0.52353 | 1.610779 | 0.246422 |
| 4 | 9834400 | 9835199 | 6.008918 | 11.51778 | 29.07845 | 11.44815 | -0.80391 | -0.07814 | 0.246438 |
| 1 | 131034600 | 131035999 | 11.84332 | 20.74909 | 45.18008 | 19.95187 | -0.72902 | 0.6475 | 0.246444 |

| | | | | | | | | | |
|---|---|---|---|---|---|---|---|---|---|
| 10 | 34372800 | 34373399 | 10.97057 | 21.04772 | 19.01727 | 7.467135 | 0.669327 | -0.06435 | 0.246448 |
| 10 | 69063800 | 69065399 | 11.25293 | 19.63733 | 47.32585 | 20.83608 | -0.87498 | 0.708175 | 0.246467 |
| 6 | 29745200 | 29750199 | 259.9133 | 362.5031 | 674.7487 | 279.8984 | -0.73162 | 4.618188 | 0.246481 |
| 19 | 36210400 | 36212999 | 14.3179 | 24.97963 | 50.00646 | 22.01276 | -0.60931 | 0.729148 | 0.246505 |
| 16 | 23917400 | 23918199 | 12.16119 | 23.29645 | 19.45358 | 7.671968 | 0.776379 | -0.0365 | 0.246506 |
| 12 | 11702200 | 11704199 | 16.06725 | 30.21239 | 19.24904 | 7.899413 | 1.113183 | 0.126441 | 0.24651 |
| 6 | 139491000 | 139492399 | 15.04834 | 28.15613 | 19.14361 | 7.922568 | 1.011215 | 0.149021 | 0.246531 |
| 14 | 62799400 | 62802399 | 16.47816 | 28.32548 | 55.38034 | 24.2919 | -0.57921 | 0.908211 | 0.246534 |
| 15 | 84487200 | 84493799 | 73.6156 | 114.0914 | 198.0706 | 82.2729 | -0.49044 | 2.799736 | 0.246537 |
| X | 26792800 | 26793399 | 21.38085 | 37.67724 | 27.66782 | 12.24606 | 0.832221 | 0.589958 | 0.246554 |
| 8 | 103874800 | 103875999 | 5.469989 | 10.4504 | 30.60795 | 12.16096 | -1.02861 | -0.04994 | 0.246564 |
| 3 | 94633200 | 94634799 | 17.01139 | 31.16245 | 23.41598 | 9.995254 | 0.834951 | 0.294631 | 0.246597 |
| 12 | 14610200 | 14611799 | 29.29412 | 49.74157 | 41.53355 | 18.19905 | 0.640708 | 1.051564 | 0.246603 |
| 17 | 81661400 | 81665199 | 80.91832 | 127.9774 | 112.4758 | 46.73981 | 0.513175 | 2.410031 | 0.246612 |
| 10 | 120007400 | 120008799 | 14.63121 | 27.23881 | 22.12638 | 9.235025 | 0.74645 | 0.182704 | 0.246616 |
| 11 | 41461800 | 41462999 | 14.63121 | 27.23881 | 22.12638 | 9.235025 | 0.74645 | 0.182704 | 0.246616 |
| 13 | 40116000 | 40118599 | 39.97792 | 66.49132 | 45.43544 | 19.71876 | 0.928678 | 1.283679 | 0.246622 |
| 17 | 83334600 | 83341399 | 55.92394 | 88.59986 | 162.2203 | 67.4702 | -0.54223 | 2.360972 | 0.246644 |
| 5 | 60930000 | 60933399 | 32.17372 | 54.30404 | 41.96086 | 18.35283 | 0.751735 | 1.110123 | 0.246649 |
| 16 | 33827600 | 33828999 | 8.193563 | 15.22995 | 33.19621 | 13.89159 | -0.66984 | 0.168442 | 0.246654 |
| 4 | 37837000 | 37838399 | 14.93518 | 28.46507 | 17.84336 | 7.125573 | 1.169661 | 0.024794 | 0.246659 |
| 15 | 28034600 | 28037999 | 14.33945 | 24.87022 | 52.18701 | 22.89742 | -0.67799 | 0.811936 | 0.24666 |
| 11 | 115148600 | 115150199 | 14.66984 | 24.99871 | 57.88192 | 25.37279 | -0.82414 | 0.983114 | 0.246675 |
| 1 | 138922200 | 138923199 | 13.84019 | 25.75691 | 21.10352 | 8.813473 | 0.733146 | 0.174702 | 0.246682 |
| 13 | 113966600 | 113972999 | 56.09253 | 88.85978 | 160.1205 | 66.59416 | -0.51933 | 2.362753 | 0.246701 |
| 2 | 135911200 | 135913599 | 13.04962 | 22.92394 | 47.20271 | 20.89243 | -0.64934 | 0.610653 | 0.246704 |
| 3 | 9412000 | 9412999 | 5.593855 | 10.70925 | 28.97913 | 11.44869 | -0.90748 | -0.07081 | 0.246731 |
| 12 | 19368000 | 19370999 | 57.23211 | 91.93667 | 62.31448 | 26.39359 | 0.910324 | 1.911946 | 0.246737 |
| 12 | 61527400 | 61528599 | 11.33442 | 22.03087 | 15.73932 | 5.850776 | 1.077677 | -0.24904 | 0.246739 |
| 16 | 61130400 | 61132199 | 18.63921 | 32.97907 | 28.02147 | 12.39256 | 0.624672 | 0.52752 | 0.246748 |
| 19 | 18460200 | 18461999 | 18.63921 | 32.97907 | 28.02147 | 12.39256 | 0.624672 | 0.52752 | 0.246748 |
| 5 | 137771800 | 137773199 | 7.418778 | 13.84199 | 32.74324 | 13.61527 | -0.77979 | 0.134232 | 0.246748 |
| 9 | 50236200 | 50237999 | 21.98537 | 38.42847 | 32.23479 | 14.2073 | 0.639855 | 0.712705 | 0.246751 |
| 1 | 136172400 | 136173399 | 13.59297 | 25.57893 | 21.30972 | 8.734021 | 0.731265 | 0.105819 | 0.246753 |
| 3 | 132768600 | 132769799 | 13.59297 | 25.57893 | 21.30972 | 8.734021 | 0.731265 | 0.105819 | 0.246753 |
| 14 | 45095000 | 45097199 | 16.57435 | 28.06984 | 62.28956 | 27.27221 | -0.76433 | 1.049006 | 0.246758 |
| 18 | 12959800 | 12962799 | 38.34937 | 62.90991 | 55.95044 | 24.09049 | 0.545652 | 1.446635 | 0.246762 |
| 1 | 153527800 | 153532999 | 25.75379 | 41.87327 | 87.70417 | 37.63557 | -0.69481 | 1.564192 | 0.246777 |
| 14 | 55315000 | 55318399 | 16.01077 | 27.39568 | 56.41137 | 24.73632 | -0.65485 | 0.948043 | 0.246784 |
| X | 33293000 | 33296599 | 36.30794 | 60.96672 | 42.55145 | 18.57954 | 0.898 | 1.172144 | 0.246805 |
| 13 | 56094800 | 56106799 | 81.10436 | 124.2937 | 236.5003 | 98.16584 | -0.63504 | 3.018769 | 0.246812 |
| 18 | 63214600 | 63216199 | 9.892095 | 17.63445 | 38.90604 | 17.11983 | -0.738 | 0.438568 | 0.246825 |
| X | 149602000 | 149604399 | 28.10744 | 48.3697 | 37.09313 | 16.271 | 0.765753 | 0.923091 | 0.246851 |
| 8 | 69740200 | 69745799 | 113.3842 | 175.2017 | 151.4619 | 62.91395 | 0.511083 | 2.865974 | 0.246854 |
| 13 | 66215800 | 66219399 | 72.58139 | 115.6577 | 93.46589 | 39.12651 | 0.644283 | 2.186923 | 0.246865 |
| 1 | 69144400 | 69148399 | 28.15279 | 45.35209 | 110.437 | 46.968 | -0.9256 | 1.803104 | 0.246865 |
| 13 | 51894600 | 51900599 | 31.74365 | 50.92807 | 111.9219 | 47.31421 | -0.78523 | 1.938852 | 0.246873 |

| | | | | | | | | | |
|---|---|---|---|---|---|---|---|---|---|
| 8 | 4873200 | 4875799 | 27.28157 | 46.84695 | 38.25734 | 16.78057 | 0.674805 | 0.940099 | 0.246885 |
| 18 | 81253000 | 81259399 | 46.72871 | 74.25028 | 142.6546 | 59.50232 | -0.60739 | 2.263737 | 0.246891 |
| 15 | 85494000 | 85497999 | 28.26457 | 45.62835 | 96.75389 | 41.26351 | -0.72277 | 1.735147 | 0.246903 |
| 2 | 102439000 | 102440399 | 15.13521 | 25.78282 | 57.3466 | 25.13743 | -0.76647 | 0.988342 | 0.246922 |
| 14 | 99548800 | 99551399 | 25.34473 | 41.48011 | 80.16136 | 34.48137 | -0.57216 | 1.465279 | 0.246932 |
| 8 | 96220000 | 96221399 | 8.81583 | 16.28226 | 34.70452 | 14.64953 | -0.64727 | 0.212042 | 0.246932 |
| 5 | 142146200 | 142149199 | 35.84978 | 57.652 | 106.7955 | 45.30608 | -0.53514 | 1.866752 | 0.246935 |
| 4 | 55008400 | 55014999 | 44.26715 | 70.10631 | 157.7174 | 65.58515 | -0.83921 | 2.361065 | 0.246937 |
| X | 4564600 | 4569799 | 89.69451 | 142.2354 | 96.43435 | 40.14268 | 0.890211 | 2.34801 | 0.24696 |
| 10 | 42971600 | 42973799 | 11.29117 | 19.98911 | 41.91779 | 18.53616 | -0.67114 | 0.502712 | 0.246966 |
| 3 | 106942800 | 106946199 | 26.00863 | 42.44323 | 83.56848 | 35.90891 | -0.60208 | 1.5092 | 0.246989 |
| 1 | 46564400 | 46566399 | 12.598 | 22.14317 | 44.55262 | 19.72894 | -0.61587 | 0.596346 | 0.246991 |
| 12 | 60171600 | 60175599 | 57.51287 | 92.55449 | 73.64791 | 31.27558 | 0.682655 | 1.863036 | 0.246995 |
| 17 | 42520000 | 42520999 | 8.315311 | 15.51676 | 33.88015 | 14.08493 | -0.66507 | 0.144239 | 0.246996 |
| 11 | 121710600 | 121713199 | 38.93254 | 64.4357 | 50.48084 | 21.83435 | 0.731721 | 1.340692 | 0.246997 |
| 1 | 100568400 | 100570599 | 16.18202 | 30.87067 | 16.60932 | 6.619492 | 1.390532 | 0.02786 | 0.247006 |
| 1 | 50913600 | 50915399 | 11.43954 | 20.32752 | 41.41733 | 18.2744 | -0.62701 | 0.473173 | 0.247015 |
| 7 | 41504000 | 41508799 | 78.96892 | 125.6542 | 93.85233 | 39.20971 | 0.755548 | 2.239584 | 0.247015 |
| 7 | 110226800 | 110228599 | 13.80109 | 23.77046 | 53.16306 | 23.32072 | -0.77149 | 0.875915 | 0.247023 |
| 6 | 121861000 | 121862599 | 10.27392 | 18.51912 | 41.15348 | 17.92812 | -0.73892 | 0.373904 | 0.247029 |
| 4 | 147101400 | 147103999 | 27.00724 | 46.40044 | 34.54426 | 15.15217 | 0.807689 | 0.927459 | 0.247035 |
| 14 | 41854400 | 41856999 | 19.58947 | 34.99181 | 26.31798 | 11.56116 | 0.806943 | 0.464199 | 0.247038 |
| 9 | 31593400 | 31595399 | 9.70005 | 17.04685 | 44.60296 | 19.74862 | -0.99243 | 0.581061 | 0.247048 |
| 17 | 5933000 | 5933599 | 6.501707 | 12.2671 | 30.23949 | 12.33664 | -0.81572 | 0.048908 | 0.247054 |
| 8 | 117327800 | 117335599 | 86.54861 | 133.2009 | 223.7061 | 92.90927 | -0.45046 | 2.937039 | 0.247056 |
| X | 54652400 | 54653199 | 13.14807 | 25.42489 | 16.76791 | 6.414184 | 1.154905 | -0.12961 | 0.247056 |
| 9 | 36466200 | 36471799 | 64.67987 | 104.1758 | 64.10204 | 27.25501 | 1.054197 | 1.844673 | 0.247059 |
| Y | 2061400 | 2062999 | 10.55564 | 20.45663 | 9.781205 | 3.703435 | 1.624889 | -0.21963 | 0.247068 |
| 7 | 83181400 | 83182199 | 5.378928 | 10.44049 | 25.6604 | 9.645366 | -0.70084 | -0.24774 | 0.247077 |
| 2 | 99878800 | 99879999 | 13.45763 | 25.89949 | 16.98454 | 6.609855 | 1.138732 | -0.07292 | 0.247085 |
| 15 | 36954000 | 36957199 | 25.35281 | 41.43317 | 81.86156 | 35.1941 | -0.60552 | 1.485967 | 0.247094 |
| 12 | 33521200 | 33521999 | 8.63131 | 16.8156 | 14.94835 | 5.440544 | 0.792898 | -0.35901 | 0.24711 |
| 5 | 74044800 | 74048599 | 25.19492 | 43.22232 | 36.76159 | 16.12328 | 0.616033 | 0.935279 | 0.247115 |
| 7 | 21123200 | 21126599 | 30.45739 | 52.54432 | 33.55578 | 14.72045 | 1.029148 | 0.906034 | 0.247121 |
| 5 | 114165000 | 114171199 | 42.54377 | 67.66363 | 143.136 | 59.76113 | -0.74464 | 2.234375 | 0.247125 |
| 7 | 38922800 | 38923199 | 9.145788 | 17.76685 | 16.61208 | 6.203155 | 0.688152 | -0.25672 | 0.247134 |
| 12 | 81148200 | 81153399 | 30.30109 | 48.98337 | 97.54485 | 41.6748 | -0.63019 | 1.696494 | 0.247163 |
| 1 | 23664200 | 23665199 | 18.03077 | 32.16063 | 26.99611 | 11.87364 | 0.648174 | 0.463384 | 0.247173 |
| 17 | 13017000 | 13030599 | 130.3866 | 199.4255 | 173.0249 | 71.78677 | 0.49463 | 3.067687 | 0.247182 |
| 17 | 63652200 | 63656799 | 41.78931 | 66.80688 | 120.6782 | 50.74832 | -0.50906 | 2.064254 | 0.247189 |
| 2 | 165220800 | 165228799 | 119.2316 | 178.3346 | 322.1106 | 132.8288 | -0.58972 | 3.519178 | 0.247216 |
| 1 | 4561800 | 4563999 | 21.71109 | 35.51997 | 81.28629 | 34.96053 | -0.81561 | 1.462425 | 0.24722 |
| 9 | 94957200 | 94960399 | 14.26848 | 24.53188 | 56.41769 | 24.74707 | -0.81238 | 0.895914 | 0.247223 |
| 10 | 31124000 | 31127599 | 35.37812 | 56.86799 | 106.9721 | 45.3515 | -0.55802 | 1.880559 | 0.247242 |
| 7 | 16186600 | 16200399 | 96.8399 | 146.5954 | 278.9413 | 115.3988 | -0.64949 | 3.276293 | 0.247247 |
| 3 | 107704000 | 107705399 | 11.77113 | 22.37166 | 19.96283 | 8.019539 | 0.657638 | 0.019281 | 0.24726 |
| 18 | 37116400 | 37116999 | 11.77113 | 22.37166 | 19.96283 | 8.019539 | 0.657638 | 0.019281 | 0.24726 |

| | | | | | | | | | |
|---|---|---|---|---|---|---|---|---|---|
| 18 | 20727000 | 20728999 | 13.73959 | 23.79652 | 52.69428 | 23.11901 | -0.75564 | 0.82257 | 0.247262 |
| 12 | 24449400 | 24455999 | 130.3495 | 199.4861 | 174.1399 | 72.2592 | 0.486485 | 3.057588 | 0.247297 |
| 9 | 3109800 | 3113399 | 58.59376 | 94.13859 | 77.61683 | 32.88235 | 0.628607 | 1.915997 | 0.247299 |
| 2 | 166686800 | 166689999 | 14.70369 | 25.34933 | 53.75433 | 23.58078 | -0.69493 | 0.870389 | 0.2473 |
| 8 | 24398800 | 24399399 | 3.230961 | 6.319175 | 19.01788 | 6.433788 | -0.8465 | -0.67291 | 0.247306 |
| 8 | 55941600 | 55941999 | 3.230961 | 6.319175 | 19.01788 | 6.433788 | -0.8465 | -0.67291 | 0.247306 |
| 5 | 115939400 | 115943399 | 35.87685 | 57.62619 | 108.4784 | 45.93973 | -0.56045 | 1.905794 | 0.247306 |
| 7 | 139407600 | 139409599 | 10.40319 | 18.38112 | 40.95829 | 18.11894 | -0.75891 | 0.519246 | 0.247316 |
| 10 | 52364400 | 52365999 | 12.73821 | 22.30718 | 46.20987 | 20.39913 | -0.65791 | 0.657678 | 0.247319 |
| 12 | 62201400 | 62202399 | 12.72411 | 24.40708 | 18.79205 | 7.382964 | 0.897205 | -0.04541 | 0.247322 |
| 2 | 37091200 | 37092199 | 16.10328 | 29.21903 | 24.52042 | 10.59424 | 0.665134 | 0.342678 | 0.247341 |
| 15 | 58066000 | 58066399 | 3.040341 | 5.945473 | 18.89276 | 6.435627 | -0.93108 | -0.64918 | 0.247341 |
| 13 | 65468200 | 65477999 | 232.9333 | 342.084 | 293.1022 | 120.7149 | 0.46106 | 3.845101 | 0.247343 |
| 8 | 81773000 | 81774799 | 23.28648 | 40.50562 | 34.56467 | 15.18681 | 0.614904 | 0.792452 | 0.247353 |
| 13 | 94925400 | 94926199 | 7.097605 | 13.34527 | 30.72231 | 12.60919 | -0.72509 | 0.078563 | 0.247365 |
| 18 | 12849200 | 12850799 | 7.097605 | 13.34527 | 30.72231 | 12.60919 | -0.72509 | 0.078563 | 0.247365 |
| 6 | 149466600 | 149468199 | 12.88209 | 22.5963 | 46.16997 | 20.40988 | -0.63823 | 0.631419 | 0.247368 |
| 3 | 87518800 | 87519599 | 4.988198 | 9.650591 | 26.57223 | 10.14134 | -0.88434 | -0.1991 | 0.24738 |
| 3 | 83764800 | 83766799 | 12.75647 | 22.32674 | 45.70238 | 20.16562 | -0.64068 | 0.667304 | 0.247386 |
| 1 | 101293000 | 101294199 | 24.01843 | 41.47087 | 35.1317 | 15.41218 | 0.623468 | 0.87062 | 0.247391 |
| 16 | 30455400 | 30469199 | 112.7733 | 170.6901 | 295.1104 | 122.0823 | -0.51158 | 3.283589 | 0.247404 |
| 7 | 9980400 | 9983399 | 75.86974 | 120.6541 | 98.11012 | 40.95635 | 0.632294 | 2.256171 | 0.247414 |
| 10 | 76083800 | 76085799 | 8.968565 | 16.06016 | 39.98368 | 17.52744 | -0.90756 | 0.403979 | 0.247432 |
| 7 | 98916000 | 98918999 | 19.79046 | 32.81358 | 70.68114 | 30.60966 | -0.72252 | 1.285547 | 0.247441 |
| X | 95877600 | 95877999 | 7.247537 | 13.69379 | 33.67768 | 13.69797 | -0.80865 | 0.056424 | 0.247441 |
| 5 | 97884000 | 97884999 | 6.589383 | 12.61998 | 29.04851 | 11.46349 | -0.6747 | -0.06234 | 0.247448 |
| 4 | 152777400 | 152787999 | 85.18104 | 130.2483 | 245.4401 | 101.8249 | -0.62364 | 3.066009 | 0.247451 |
| 6 | 3741200 | 3741799 | 4.917835 | 9.578993 | 24.9119 | 9.09143 | -0.74137 | -0.35935 | 0.24746 |
| 6 | 33345800 | 33346399 | 4.917835 | 9.578993 | 24.9119 | 9.09143 | -0.74137 | -0.35935 | 0.24746 |
| 6 | 6836200 | 6837599 | 7.840694 | 14.67212 | 34.55092 | 14.29586 | -0.76801 | 0.121924 | 0.247472 |
| 9 | 12846400 | 12847999 | 22.40944 | 39.38121 | 29.2636 | 12.95658 | 0.813539 | 0.626221 | 0.247493 |
| 7 | 40969800 | 40974599 | 81.19519 | 128.4999 | 110.684 | 46.01055 | 0.542929 | 2.394123 | 0.247499 |
| 17 | 31849400 | 31852399 | 14.14492 | 24.2655 | 54.73437 | 24.00594 | -0.78493 | 0.915985 | 0.247504 |
| 14 | 40322400 | 40324599 | 22.70276 | 40.37396 | 22.42791 | 9.889112 | 1.237091 | 0.501737 | 0.247515 |
| 12 | 86511200 | 86518199 | 42.00394 | 66.93229 | 129.6049 | 54.25435 | -0.61426 | 2.168396 | 0.247518 |
| 14 | 9325000 | 9331999 | 86.71938 | 131.5957 | 268.3521 | 111.1017 | -0.7463 | 3.224209 | 0.247519 |
| 3 | 59694000 | 59695599 | 16.03725 | 29.57614 | 22.3931 | 9.471589 | 0.833169 | 0.246945 | 0.247539 |
| 15 | 45563600 | 45565399 | 22.86312 | 37.68294 | 74.70362 | 32.22926 | -0.60459 | 1.367088 | 0.247543 |
| 19 | 60455400 | 60457799 | 22.08952 | 36.353 | 76.22437 | 32.86438 | -0.68612 | 1.384442 | 0.247552 |
| 4 | 40899800 | 40901399 | 23.53187 | 40.7422 | 35.71096 | 15.6676 | 0.575415 | 0.846795 | 0.247554 |
| X | 80871200 | 80872599 | 20.44173 | 37.00621 | 23.03158 | 9.979519 | 1.091172 | 0.385474 | 0.247595 |
| 19 | 55087000 | 55088599 | 7.597572 | 13.69519 | 36.99719 | 16.11705 | -1.01737 | 0.350095 | 0.247601 |
| 13 | 79815800 | 79817799 | 19.30377 | 34.11266 | 26.8017 | 11.85605 | 0.736661 | 0.544949 | 0.24762 |
| 19 | 13288600 | 13289599 | 15.05427 | 27.79846 | 23.1847 | 9.789542 | 0.696691 | 0.235215 | 0.247631 |
| 5 | 33786200 | 33787199 | 4.299586 | 8.313211 | 26.44135 | 10.12054 | -1.09327 | -0.19624 | 0.24765 |
| 12 | 107487600 | 107491199 | 22.75588 | 37.8504 | 73.63507 | 31.9589 | -0.57594 | 1.257588 | 0.247655 |
| 4 | 59396000 | 59397599 | 8.691164 | 15.91181 | 35.05809 | 14.97649 | -0.70734 | 0.26093 | 0.247655 |

| | | | | | | | | | |
|---|---|---|---|---|---|---|---|---|---|
| 5 | 18003600 | 18004999 | 10.32901 | 18.49882 | 38.40384 | 16.83299 | -0.64686 | 0.411574 | 0.247673 |
| 11 | 95109400 | 95110599 | 6.488914 | 12.26963 | 29.7385 | 12.07655 | -0.78526 | 0.033778 | 0.247674 |
| 15 | 47289600 | 47292199 | 27.90193 | 47.97115 | 35.86598 | 15.73226 | 0.801675 | 0.925494 | 0.247685 |
| 17 | 10816000 | 10817199 | 9.798742 | 17.77757 | 37.01238 | 15.99378 | -0.63736 | 0.326377 | 0.247711 |
| 14 | 110149000 | 110150199 | 20.30716 | 35.71577 | 28.90553 | 12.79885 | 0.691172 | 0.602365 | 0.247717 |
| 7 | 11739600 | 11742999 | 56.36938 | 91.09613 | 63.21526 | 26.99719 | 0.886662 | 1.734587 | 0.247722 |
| 14 | 24606400 | 24608799 | 11.43536 | 20.28068 | 42.43547 | 18.7543 | -0.66693 | 0.486202 | 0.247736 |
| 7 | 89640800 | 89641599 | 12.14628 | 23.40642 | 17.72206 | 6.870991 | 0.938395 | -0.09672 | 0.247742 |
| 2 | 71865200 | 71867999 | 11.52843 | 20.20289 | 44.65954 | 19.72652 | -0.75065 | 0.640058 | 0.247748 |
| 15 | 98670200 | 98672399 | 11.41791 | 19.89657 | 48.3843 | 21.28281 | -0.88825 | 0.730774 | 0.247754 |
| 4 | 82422400 | 82425599 | 21.08965 | 34.48395 | 81.621 | 35.09732 | -0.86462 | 1.469988 | 0.247757 |
| 1 | 165287000 | 165289599 | 15.04937 | 25.46011 | 60.43829 | 26.45315 | -0.8609 | 1.053947 | 0.247762 |
| 6 | 61453600 | 61455599 | 20.84682 | 37.41906 | 23.27346 | 10.18196 | 1.082855 | 0.435731 | 0.247763 |
| 2 | 163637600 | 163644599 | 55.56832 | 87.56023 | 161.2298 | 66.90271 | -0.5566 | 2.484986 | 0.247764 |
| 5 | 73092600 | 73093599 | 11.6073 | 22.60853 | 13.29242 | 4.851675 | 1.377925 | -0.31533 | 0.247766 |
| 10 | 61025000 | 61025599 | 11.78366 | 22.35082 | 19.4852 | 7.860765 | 0.685805 | 0.031925 | 0.247773 |
| 9 | 58670400 | 58675199 | 34.87054 | 56.13355 | 103.9513 | 44.16366 | -0.53288 | 1.832864 | 0.247773 |
| 4 | 63482200 | 63506999 | 201.1088 | 287.4092 | 574.5594 | 237.3959 | -0.79914 | 4.254856 | 0.247774 |
| X | 79522800 | 79526399 | 38.21236 | 63.93345 | 43.11896 | 18.78452 | 0.947369 | 1.215054 | 0.247775 |
| 13 | 73990000 | 73993199 | 18.44612 | 31.27312 | 60.7547 | 26.60881 | -0.57314 | 1.045884 | 0.247777 |
| 9 | 110359800 | 110362399 | 11.66446 | 20.47956 | 43.56721 | 19.27449 | -0.69568 | 0.607462 | 0.247786 |
| 15 | 95625600 | 95627399 | 8.837596 | 16.24808 | 34.6039 | 14.69813 | -0.65253 | 0.238433 | 0.247788 |
| 2 | 17788200 | 17789999 | 11.01001 | 19.42015 | 42.98916 | 19.02384 | -0.75086 | 0.54327 | 0.247789 |
| 14 | 119174800 | 119176199 | 8.418174 | 15.35283 | 35.00928 | 15.02969 | -0.76115 | 0.282063 | 0.247798 |
| 6 | 145610800 | 145613399 | 22.92279 | 38.04368 | 74.75154 | 32.3939 | -0.59055 | 1.283793 | 0.247798 |
| X | 4519200 | 4521999 | 58.493 | 94.15907 | 66.14558 | 28.10141 | 0.86255 | 1.852886 | 0.247798 |
| 7 | 105072800 | 105076399 | 25.36935 | 41.41113 | 83.49977 | 35.8829 | -0.63603 | 1.503771 | 0.247803 |
| 10 | 68194000 | 68195599 | 20.72756 | 36.42973 | 30.13374 | 13.34313 | 0.659846 | 0.617556 | 0.247804 |
| 3 | 27970600 | 27977399 | 68.88371 | 107.6314 | 183.8881 | 76.29587 | -0.45821 | 2.654416 | 0.247808 |
| X | 125056200 | 125059999 | 27.55403 | 47.46481 | 31.64347 | 13.88091 | 0.967171 | 0.905408 | 0.247818 |
| 4 | 92886800 | 92887999 | 11.09943 | 21.21925 | 18.98913 | 7.532709 | 0.669336 | -0.03061 | 0.247825 |
| 3 | 55065000 | 55065999 | 7.454699 | 13.81448 | 32.60167 | 13.69938 | -0.78769 | 0.178228 | 0.247845 |
| 19 | 8336000 | 8338199 | 29.04906 | 49.86257 | 37.57724 | 16.48178 | 0.790079 | 0.949452 | 0.247849 |
| 1 | 8387600 | 8388799 | 15.68237 | 29.94637 | 15.99275 | 6.360766 | 1.403058 | 0.014564 | 0.247852 |
| 2 | 57279600 | 57281399 | 22.03438 | 38.76951 | 28.70413 | 12.70862 | 0.81923 | 0.608566 | 0.24786 |
| 18 | 12699400 | 12700199 | 3.93986 | 7.677209 | 22.43165 | 8.142778 | -0.89874 | -0.39845 | 0.247861 |
| X | 81652200 | 81653999 | 21.35836 | 37.69522 | 25.4867 | 11.27722 | 0.951295 | 0.568003 | 0.247862 |
| 6 | 42042200 | 42043199 | 7.697173 | 14.52 | 36.63136 | 14.95007 | -0.85032 | 0.075836 | 0.247864 |
| 10 | 48643400 | 48644799 | 18.58176 | 33.06596 | 26.82878 | 11.82334 | 0.694842 | 0.479985 | 0.247875 |
| 5 | 22915200 | 22920399 | 116.2197 | 178.6415 | 159.846 | 66.37559 | 0.455678 | 2.965111 | 0.24788 |
| 14 | 10031400 | 10033999 | 16.06537 | 27.3471 | 57.94124 | 25.39657 | -0.69699 | 0.998383 | 0.247883 |
| 14 | 106130400 | 106131399 | 5.954605 | 11.19091 | 31.22857 | 12.82712 | -1.00101 | 0.071727 | 0.247888 |
| 5 | 131517600 | 131529799 | 109.3637 | 165.4525 | 282.0973 | 116.6818 | -0.49205 | 3.291596 | 0.247896 |
| 8 | 10876200 | 10878799 | 11.25329 | 19.6215 | 47.85367 | 21.05736 | -0.89198 | 0.723088 | 0.247903 |
| 5 | 26574400 | 26575999 | 22.48991 | 39.21952 | 33.35913 | 14.68054 | 0.619699 | 0.74985 | 0.247903 |
| 13 | 52686600 | 52687599 | 15.03264 | 27.56591 | 23.69666 | 10.10096 | 0.643599 | 0.274302 | 0.247912 |
| 16 | 64061200 | 64062199 | 6.221968 | 11.74055 | 31.34418 | 12.78444 | -0.92983 | 0.046869 | 0.247957 |

| | | | | | | | | | |
|---|---|---|---|---|---|---|---|---|---|
| 1 | 65152000 | 65154399 | 8.467637 | 15.30508 | 36.97984 | 16.05484 | -0.85487 | 0.338378 | 0.247958 |
| 8 | 109555200 | 109556999 | 8.467637 | 15.30508 | 36.97984 | 16.05484 | -0.85487 | 0.338378 | 0.247958 |
| 11 | 94771800 | 94774399 | 13.65548 | 23.50025 | 53.68534 | 23.54923 | -0.80221 | 0.883305 | 0.247971 |
| 3 | 24617000 | 24618799 | 47.9788 | 78.1163 | 57.97354 | 24.89004 | 0.799482 | 1.571498 | 0.247976 |
| 1 | 5567000 | 5567799 | 17.93904 | 31.89625 | 26.68784 | 11.76932 | 0.650368 | 0.482539 | 0.247977 |
| 13 | 25672200 | 25672599 | 3.341205 | 6.531679 | 19.44545 | 6.680186 | -0.85096 | -0.61371 | 0.247986 |
| 13 | 52250000 | 52253199 | 15.91696 | 27.07425 | 58.46438 | 25.62433 | -0.72449 | 1.005131 | 0.247993 |
| 8 | 3084800 | 3089599 | 37.21117 | 62.10843 | 44.82262 | 19.49703 | 0.850059 | 1.236585 | 0.247997 |
| 4 | 49365000 | 49369599 | 104.9688 | 163.6861 | 134.2393 | 55.70826 | 0.597183 | 2.701811 | 0.248016 |
| 7 | 60718400 | 60719999 | 20.52571 | 36.63959 | 24.9368 | 10.96062 | 0.949347 | 0.470108 | 0.24805 |
| X | 91986800 | 91988199 | 7.841391 | 14.77398 | 33.63882 | 13.75842 | -0.70588 | 0.078297 | 0.248056 |
| 7 | 59380600 | 59381199 | 14.60308 | 26.8822 | 21.8756 | 9.275671 | 0.727863 | 0.246295 | 0.24806 |
| 15 | 53258000 | 53259199 | 15.1441 | 27.76659 | 23.63107 | 10.07483 | 0.657674 | 0.275248 | 0.248062 |
| 2 | 77231800 | 77233599 | 9.014603 | 16.41863 | 36.02791 | 15.49286 | -0.70803 | 0.295281 | 0.248063 |
| 10 | 117464800 | 117465999 | 8.791265 | 15.81395 | 37.94989 | 16.57265 | -0.85079 | 0.370863 | 0.248074 |
| 6 | 22480600 | 22483399 | 32.97312 | 55.69478 | 41.95493 | 18.35438 | 0.788389 | 1.108066 | 0.248075 |
| 17 | 34740600 | 34742399 | 16.43718 | 30.96867 | 18.68164 | 7.641149 | 1.195133 | 0.116607 | 0.248085 |
| 11 | 50351600 | 50353199 | 9.588013 | 17.0979 | 39.42295 | 17.34256 | -0.80103 | 0.434627 | 0.24809 |
| 4 | 49711200 | 49712999 | 18.43489 | 33.31517 | 24.87548 | 10.80195 | 0.828254 | 0.38464 | 0.248097 |
| 4 | 32091000 | 32094399 | 21.77463 | 35.83727 | 75.48934 | 32.54842 | -0.69262 | 1.383753 | 0.248102 |
| 18 | 9449200 | 9451399 | 19.95305 | 33.47874 | 66.2471 | 28.91201 | -0.60027 | 1.148386 | 0.248112 |
| 2 | 55553400 | 55554199 | 12.71341 | 24.19007 | 19.16497 | 7.680479 | 0.830203 | 0.016991 | 0.248114 |
| X | 125461600 | 125464599 | 44.86947 | 73.44365 | 55.10544 | 23.7046 | 0.789044 | 1.489191 | 0.248124 |
| 3 | 3464600 | 3465999 | 28.26074 | 47.87593 | 41.6273 | 18.22733 | 0.582413 | 1.067118 | 0.24817 |
| 8 | 111623800 | 111624799 | 6.58392 | 12.60885 | 29.66213 | 11.70743 | -0.70654 | -0.06222 | 0.24817 |
| 17 | 19853200 | 19862999 | 243.8687 | 358.1785 | 268.404 | 110.5423 | 0.654412 | 3.844393 | 0.248192 |
| 3 | 23431400 | 23432599 | 6.433379 | 12.33616 | 32.17803 | 12.65434 | -0.84982 | -0.07104 | 0.248198 |
| 7 | 63501600 | 63503399 | 10.20283 | 18.17872 | 38.90045 | 17.12505 | -0.69462 | 0.444685 | 0.248211 |
| 11 | 100509600 | 100511199 | 10.20283 | 18.17872 | 38.90045 | 17.12505 | -0.69462 | 0.444685 | 0.248211 |
| 14 | 42319200 | 42324399 | 45.35743 | 73.53353 | 64.86058 | 27.77641 | 0.545851 | 1.644899 | 0.248228 |
| 4 | 89573600 | 89577199 | 24.97803 | 40.9435 | 85.33554 | 36.73083 | -0.68007 | 1.444578 | 0.248231 |
| 12 | 25015600 | 25021999 | 69.3793 | 110.5666 | 94.77784 | 39.68138 | 0.559519 | 2.180231 | 0.248237 |
| 13 | 66403800 | 66405799 | 30.54232 | 52.86109 | 31.26603 | 13.71731 | 1.140002 | 0.875176 | 0.248241 |
| 6 | 27497200 | 27499199 | 19.738 | 34.81732 | 29.11008 | 12.88182 | 0.646352 | 0.570147 | 0.248261 |
| 2 | 151420400 | 151424999 | 25.8482 | 42.21504 | 81.74474 | 35.13546 | -0.57722 | 1.497594 | 0.248267 |
| 7 | 28741800 | 28748599 | 48.10636 | 76.2754 | 144.225 | 60.02819 | -0.58741 | 2.329814 | 0.248273 |
| 4 | 132832400 | 132834399 | 12.45032 | 21.76894 | 46.24909 | 20.38946 | -0.69407 | 0.682691 | 0.248284 |
| 14 | 107730600 | 107731199 | 11.82064 | 22.65791 | 18.82694 | 7.411821 | 0.785198 | -0.04886 | 0.248286 |
| 12 | 92557600 | 92558399 | 7.21819 | 13.712 | 31.6303 | 12.71948 | -0.70262 | 0.012302 | 0.248288 |
| 16 | 66266200 | 66272199 | 140.4031 | 213.3833 | 191.0152 | 79.11896 | 0.442246 | 3.200914 | 0.248303 |
| 1 | 54182800 | 54184199 | 13.0475 | 24.86196 | 19.79537 | 7.9087 | 0.827438 | 0.013425 | 0.248304 |
| 18 | 70701000 | 70702199 | 9.612828 | 17.0805 | 39.92261 | 17.61579 | -0.82357 | 0.459083 | 0.248329 |
| 15 | 69822400 | 69824399 | 18.20561 | 33.22496 | 22.61094 | 9.698942 | 0.97194 | 0.321884 | 0.248336 |
| 11 | 83670800 | 83673199 | 19.27227 | 32.08772 | 67.98798 | 29.52755 | -0.69828 | 1.234327 | 0.248355 |
| 1 | 3156800 | 3159399 | 36.73006 | 60.98656 | 48.78679 | 21.1486 | 0.701899 | 1.294552 | 0.248375 |
| 3 | 7897800 | 7903599 | 93.03739 | 145.8048 | 128.8674 | 53.45123 | 0.494878 | 2.607592 | 0.248376 |
| 13 | 16232400 | 16233999 | 16.63171 | 31.32572 | 18.67469 | 7.642208 | 1.211547 | 0.120295 | 0.248384 |

| | | | | | | | | | |
|---|---|---|---|---|---|---|---|---|---|
| 11 | 13301200 | 13302599 | 21.27369 | 37.15437 | 32.00329 | 14.09718 | 0.601893 | 0.7219 | 0.248395 |
| 10 | 30709800 | 30710399 | 4.965601 | 9.596604 | 25.88759 | 9.929541 | -0.86105 | -0.18141 | 0.248395 |
| 13 | 33130000 | 33133999 | 19.52673 | 34.55199 | 24.94291 | 11.03089 | 0.858425 | 0.526146 | 0.248396 |
| 7 | 78039800 | 78040799 | 7.860488 | 14.65265 | 33.28136 | 13.86061 | -0.72365 | 0.14497 | 0.248399 |
| 8 | 118794800 | 118796399 | 7.860488 | 14.65265 | 33.28136 | 13.86061 | -0.72365 | 0.14497 | 0.248399 |
| 1 | 52529000 | 52529799 | 6.453783 | 12.31534 | 29.4812 | 11.74268 | -0.74349 | -0.03384 | 0.2484 |
| 19 | 9239600 | 9242999 | 43.80809 | 71.30802 | 60.93176 | 26.15773 | 0.596455 | 1.569595 | 0.248409 |
| 7 | 44048200 | 44049999 | 9.887731 | 17.65796 | 38.41343 | 16.87781 | -0.71617 | 0.427339 | 0.248423 |
| 1 | 170895400 | 170898999 | 58.5257 | 91.68981 | 174.9254 | 72.55739 | -0.61414 | 2.60085 | 0.248429 |
| 1 | 111460000 | 111462799 | 43.70566 | 71.57248 | 55.62239 | 23.93169 | 0.738765 | 1.480985 | 0.248442 |
| 4 | 34249400 | 34250799 | 10.79526 | 19.31314 | 39.41891 | 17.29353 | -0.62356 | 0.423071 | 0.248467 |
| 3 | 36952800 | 36955599 | 32.53185 | 55.20465 | 40.28086 | 17.64679 | 0.834403 | 1.06654 | 0.248483 |
| 11 | 81337200 | 81339199 | 10.61869 | 19.00141 | 39.93841 | 17.5182 | -0.66567 | 0.419928 | 0.248496 |
| 3 | 95307200 | 95309199 | 15.29332 | 28.44318 | 22.0415 | 9.215088 | 0.812004 | 0.194311 | 0.2485 |
| 3 | 75740200 | 75741199 | 13.82675 | 25.6592 | 22.25716 | 9.334453 | 0.647275 | 0.194967 | 0.248506 |
| 9 | 95440200 | 95443199 | 21.72995 | 35.50857 | 82.26655 | 35.36679 | -0.83453 | 1.480626 | 0.24851 |
| 6 | 116388800 | 116393399 | 28.27295 | 45.62726 | 97.87599 | 41.72483 | -0.73995 | 1.744722 | 0.248542 |
| 5 | 48583200 | 48585199 | 19.33257 | 34.54808 | 26.25324 | 11.52846 | 0.792403 | 0.458877 | 0.248545 |
| 9 | 36072800 | 36077999 | 152.295 | 230.6043 | 201.9087 | 83.53718 | 0.469787 | 3.278535 | 0.248551 |
| 3 | 145587400 | 145590199 | 16.15192 | 27.65182 | 57.53327 | 25.22935 | -0.66974 | 0.944134 | 0.248551 |
| X | 54441800 | 54444399 | 48.5628 | 79.28726 | 54.626 | 23.4766 | 0.909241 | 1.53309 | 0.248556 |
| 1 | 170891000 | 170899199 | 172.2745 | 249.135 | 475.3819 | 196.0754 | -0.71495 | 4.077633 | 0.248564 |
| 13 | 86953400 | 86954799 | 15.10517 | 27.82897 | 23.08372 | 9.777167 | 0.701118 | 0.246892 | 0.248578 |
| 17 | 23029800 | 23031199 | 15.10517 | 27.82897 | 23.08372 | 9.777167 | 0.701118 | 0.246892 | 0.248578 |
| 10 | 21989400 | 21990399 | 14.57006 | 27.06958 | 22.08514 | 9.247899 | 0.736448 | 0.193886 | 0.248587 |
| 8 | 12103200 | 12106399 | 17.04169 | 28.90883 | 58.99952 | 25.84463 | -0.64376 | 1.035292 | 0.248597 |
| 17 | 4792800 | 4793199 | 7.825265 | 15.27444 | 14.58713 | 5.167293 | 0.726101 | -0.46344 | 0.248607 |
| 6 | 66323800 | 66324599 | 6.015865 | 11.5114 | 28.98974 | 11.46938 | -0.80691 | -0.06176 | 0.248613 |
| 10 | 12711800 | 12712399 | 3.882749 | 7.569099 | 22.4665 | 8.110727 | -0.91294 | -0.41641 | 0.248618 |
| 4 | 10202800 | 10204799 | 20.45141 | 36.12594 | 28.50136 | 12.60894 | 0.730402 | 0.558964 | 0.24862 |
| X | 145693400 | 145695199 | 19.82064 | 36.07537 | 22.63756 | 9.74302 | 1.085578 | 0.350644 | 0.248647 |
| 15 | 97545800 | 97547999 | 10.4849 | 18.70165 | 39.90757 | 17.55229 | -0.69003 | 0.439501 | 0.248655 |
| 17 | 82708600 | 82711399 | 18.76092 | 33.62187 | 25.72738 | 11.27052 | 0.784652 | 0.436053 | 0.248661 |
| 1 | 45989400 | 45991199 | 13.84948 | 24.23084 | 48.35245 | 21.32829 | -0.60436 | 0.682381 | 0.248668 |
| 12 | 51524000 | 51526199 | 13.84948 | 24.23084 | 48.35245 | 21.32829 | -0.60436 | 0.682381 | 0.248668 |
| 14 | 94693600 | 94695199 | 16.63653 | 30.49534 | 23.42897 | 9.992063 | 0.803968 | 0.287911 | 0.248685 |
| 11 | 33871400 | 33873599 | 16.37452 | 27.89001 | 57.41824 | 25.16855 | -0.65557 | 0.993865 | 0.248716 |
| 15 | 3332200 | 3333199 | 14.34527 | 26.44114 | 20.41376 | 8.640885 | 0.80493 | 0.233035 | 0.248727 |
| 7 | 22273400 | 22274399 | 11.15164 | 21.67717 | 15.74554 | 5.849635 | 1.054666 | -0.25342 | 0.24874 |
| 13 | 51320800 | 51328799 | 68.49667 | 106.3315 | 197.4314 | 81.98767 | -0.58545 | 2.768445 | 0.248755 |
| 18 | 75910000 | 75913199 | 29.64074 | 49.75071 | 42.767 | 18.66874 | 0.598406 | 1.15241 | 0.248757 |
| 11 | 118039800 | 118048399 | 77.40523 | 119.4657 | 209.2315 | 86.9107 | -0.50774 | 2.879007 | 0.248768 |
| 7 | 42962000 | 42962999 | 9.425779 | 16.84649 | 38.42112 | 16.86978 | -0.78327 | 0.41808 | 0.248773 |
| 3 | 138731000 | 138736199 | 56.60517 | 89.49497 | 155.8334 | 64.7464 | -0.47223 | 2.408796 | 0.248776 |
| 12 | 20484800 | 20487399 | 50.43615 | 82.44308 | 51.23849 | 22.03023 | 1.058702 | 1.521127 | 0.248783 |
| 1 | 180035000 | 180036199 | 6.002184 | 11.48453 | 28.37165 | 11.22678 | -0.77866 | -0.05915 | 0.248786 |
| 15 | 12406400 | 12407799 | 10.94939 | 19.54833 | 39.91069 | 17.53989 | -0.62584 | 0.437456 | 0.248788 |

| | | | | | | | | | |
|---|---|---|---|---|---|---|---|---|---|
| 19 | 33689200 | 33690799 | 10.94939 | 19.54833 | 39.91069 | 17.53989 | -0.62584 | 0.437456 | 0.248788 |
| 1 | 8799000 | 8803599 | 71.61702 | 114.2888 | 88.57432 | 37.15457 | 0.70682 | 2.140496 | 0.248792 |
| Y | 4386200 | 4388399 | 15.16974 | 28.10412 | 12.82568 | 5.391315 | 1.563006 | 0.188702 | 0.2488 |
| 3 | 12583800 | 12587999 | 47.92713 | 78.48712 | 49.0555 | 21.10644 | 1.052015 | 1.484005 | 0.248801 |
| 1 | 114138000 | 114144599 | 82.48154 | 130.263 | 114.9546 | 47.73526 | 0.505627 | 2.444097 | 0.24881 |
| 2 | 163751800 | 163756399 | 29.70505 | 47.9347 | 97.61224 | 41.60825 | -0.66523 | 1.749066 | 0.248812 |
| 5 | 105281800 | 105283799 | 11.3067 | 20.1205 | 41.9789 | 18.49964 | -0.65965 | 0.465693 | 0.248817 |
| 19 | 20146600 | 20148199 | 16.67005 | 30.29197 | 23.83938 | 10.28106 | 0.758729 | 0.336449 | 0.248817 |
| 6 | 103430200 | 103431999 | 21.39903 | 38.4576 | 22.70159 | 9.921463 | 1.15877 | 0.431395 | 0.248819 |
| 19 | 28905600 | 28909199 | 25.45513 | 41.77355 | 79.44095 | 34.21027 | -0.54745 | 1.42957 | 0.248828 |
| 12 | 9023800 | 9025599 | 10.36154 | 18.42431 | 39.3895 | 17.36984 | -0.69512 | 0.458805 | 0.248831 |
| 18 | 33868800 | 33869199 | 2.735727 | 5.351235 | 17.36931 | 5.727989 | -0.91655 | -0.77387 | 0.248831 |
| 5 | 135916400 | 135920199 | 21.13806 | 34.79866 | 75.37877 | 32.50433 | -0.7326 | 1.379298 | 0.248832 |
| 13 | 82173800 | 82175399 | 32.52246 | 54.99297 | 41.96696 | 18.36566 | 0.769817 | 1.097876 | 0.248833 |
| 17 | 87711200 | 87713799 | 20.43476 | 34.29501 | 66.27666 | 28.92918 | -0.56628 | 1.14793 | 0.24884 |
| 4 | 63548400 | 63549599 | 5.323401 | 10.15545 | 28.82332 | 11.48746 | -0.98673 | -0.04356 | 0.248844 |
| 9 | 19223200 | 19223799 | 10.43223 | 20.16047 | 18.10396 | 6.945706 | 0.709618 | -0.14614 | 0.24885 |
| 8 | 48253200 | 48255999 | 44.67368 | 72.27381 | 62.83324 | 26.86677 | 0.56391 | 1.684107 | 0.248854 |
| 2 | 29640400 | 29643599 | 25.56436 | 41.64675 | 83.65478 | 35.92353 | -0.63251 | 1.534737 | 0.248856 |
| 5 | 148915800 | 148922399 | 73.00977 | 113.4725 | 193.7967 | 80.46399 | -0.46368 | 2.749972 | 0.248858 |
| X | 119175200 | 119177199 | 24.38994 | 42.67848 | 31.05417 | 13.69971 | 0.844167 | 0.708566 | 0.248869 |
| 7 | 7484000 | 7489799 | 82.63584 | 131.6324 | 85.33685 | 35.70595 | 0.961122 | 2.206905 | 0.248882 |
| 18 | 69478800 | 69479999 | 6.928905 | 13.09209 | 30.29726 | 12.32235 | -0.72167 | 0.042447 | 0.248891 |
| 8 | 93419800 | 93421199 | 15.96004 | 27.48171 | 54.3218 | 23.82883 | -0.59445 | 0.887812 | 0.248896 |
| 10 | 71427400 | 71429599 | 27.8614 | 47.36905 | 41.52947 | 18.20125 | 0.570718 | 1.033754 | 0.248903 |
| 12 | 80767200 | 80770799 | 25.44193 | 41.34318 | 86.06646 | 36.92146 | -0.68657 | 1.574864 | 0.248919 |
| 6 | 117579000 | 117580199 | 18.56506 | 33.84961 | 23.25284 | 9.985829 | 0.957799 | 0.33234 | 0.248938 |
| 10 | 38988800 | 38993399 | 31.7614 | 51.30769 | 96.2226 | 41.07234 | -0.545 | 1.720105 | 0.24895 |
| 17 | 58579200 | 58579799 | 5.216948 | 10.15278 | 27.36596 | 10.07179 | -0.80463 | -0.31996 | 0.248956 |
| 15 | 19786000 | 19793199 | 81.96944 | 129.811 | 108.3396 | 45.05109 | 0.589159 | 2.379936 | 0.248969 |
| 15 | 78468800 | 78474399 | 46.30407 | 73.51479 | 148.7558 | 61.99589 | -0.68321 | 2.290637 | 0.248974 |
| 8 | 48555200 | 48559399 | 76.08775 | 120.5241 | 109.5712 | 45.56848 | 0.46619 | 2.369479 | 0.248991 |
| 6 | 146941800 | 146944799 | 17.88566 | 30.11353 | 62.77846 | 27.43743 | -0.67493 | 1.106691 | 0.249004 |
| X | 8399200 | 8401799 | 50.11559 | 81.6108 | 56.30003 | 24.17321 | 0.904821 | 1.574093 | 0.249014 |
| X | 149624800 | 149627999 | 13.90794 | 24.25056 | 49.98655 | 21.9961 | -0.6511 | 0.738158 | 0.249019 |
| 4 | 120535200 | 120543799 | 51.9888 | 81.97223 | 171.601 | 71.21217 | -0.74072 | 2.471612 | 0.249029 |
| 12 | 64392400 | 64394199 | 25.47071 | 44.2473 | 31.84327 | 13.97965 | 0.859323 | 0.816418 | 0.249037 |
| 4 | 21521800 | 21523999 | 26.39468 | 45.37384 | 34.13591 | 14.97332 | 0.792838 | 0.919858 | 0.249044 |
| 8 | 31438000 | 31439799 | 15.85887 | 28.84327 | 24.6236 | 10.60759 | 0.643406 | 0.328613 | 0.249048 |
| 13 | 65578000 | 65579999 | 20.63512 | 37.44399 | 21.81276 | 9.423738 | 1.188289 | 0.37061 | 0.249056 |
| 8 | 21849200 | 21855599 | 67.08055 | 107.0129 | 92.39976 | 38.7407 | 0.550656 | 2.146662 | 0.249059 |
| 3 | 121394400 | 121396199 | 7.589012 | 14.09454 | 32.63717 | 13.67306 | -0.75765 | 0.165569 | 0.249069 |
| 9 | 74882000 | 74883599 | 7.589012 | 14.09454 | 32.63717 | 13.67306 | -0.75765 | 0.165569 | 0.249069 |
| 11 | 107598600 | 107601799 | 20.40359 | 33.87514 | 69.71203 | 30.21766 | -0.65676 | 1.270865 | 0.249072 |
| X | 125288200 | 125290399 | 46.19998 | 75.10361 | 60.41112 | 25.92192 | 0.682149 | 1.59361 | 0.249073 |
| 9 | 90826200 | 90827999 | 17.06103 | 31.49061 | 21.75044 | 9.189276 | 0.965918 | 0.249467 | 0.249073 |
| 10 | 67804000 | 67806399 | 25.23563 | 44.11735 | 30.88894 | 13.61613 | 0.899338 | 0.726007 | 0.24908 |

| | | | | | | | | | |
|---|---|---|---|---|---|---|---|---|---|
| 3 | 11602000 | 11603799 | 12.09027 | 21.25154 | 47.19222 | 20.89833 | -0.75779 | 0.593717 | 0.249086 |
| 6 | 123079600 | 123082599 | 32.60241 | 55.58104 | 37.44062 | 16.41282 | 0.949883 | 1.022726 | 0.249096 |
| 17 | 46249200 | 46251999 | 14.39581 | 24.8527 | 52.13468 | 22.87124 | -0.67883 | 0.855809 | 0.249099 |
| 5 | 23074600 | 23077199 | 14.81278 | 25.75503 | 50.57042 | 22.21335 | -0.58203 | 0.781482 | 0.249104 |
| 13 | 77960400 | 77963799 | 47.60783 | 77.14383 | 64.91457 | 27.78942 | 0.613012 | 1.660401 | 0.249124 |
| 3 | 123162800 | 123163399 | 3.343152 | 6.529268 | 20.39841 | 7.156729 | -0.94729 | -0.53477 | 0.249128 |
| 8 | 69173600 | 69174199 | 3.343152 | 6.529268 | 20.39841 | 7.156729 | -0.94729 | -0.53477 | 0.249128 |
| 6 | 47724800 | 47725599 | 10.04655 | 17.96142 | 38.93796 | 17.09235 | -0.71017 | 0.423527 | 0.249134 |
| 7 | 13729200 | 13731799 | 45.03452 | 73.22704 | 61.65544 | 26.45822 | 0.61659 | 1.586028 | 0.249135 |
| 13 | 53709400 | 53713599 | 31.32391 | 50.47577 | 102.4057 | 43.57142 | -0.66235 | 1.794354 | 0.249142 |
| 12 | 99798800 | 99800999 | 17.63268 | 30.15125 | 59.7341 | 26.19196 | -0.59961 | 0.965591 | 0.249147 |
| 2 | 22076800 | 22077199 | 7.219662 | 14.10556 | 13.53359 | 4.717173 | 0.74003 | -0.53244 | 0.249151 |
| 5 | 52092000 | 52093799 | 13.85717 | 24.21876 | 47.83624 | 21.08182 | -0.58961 | 0.699214 | 0.249162 |
| 8 | 91348000 | 91348399 | 4.226216 | 8.251689 | 23.00402 | 8.108527 | -0.79479 | -0.50111 | 0.249166 |
| 9 | 12069400 | 12072799 | 26.51863 | 46.21657 | 30.79456 | 13.54669 | 0.97006 | 0.774271 | 0.249171 |
| 5 | 148783000 | 148792799 | 81.34869 | 125.5121 | 212.2 | 88.1439 | -0.45726 | 2.886598 | 0.249171 |
| 15 | 27103400 | 27105399 | 17.45963 | 32.12176 | 22.27101 | 9.450908 | 0.956093 | 0.271822 | 0.249181 |
| 2 | 122097000 | 122101399 | 32.09068 | 51.81117 | 97.29094 | 41.49897 | -0.54776 | 1.737129 | 0.249183 |
| 11 | 115872800 | 115875999 | 19.76476 | 32.84384 | 69.09368 | 29.96811 | -0.68824 | 1.258592 | 0.249195 |
| 2 | 121207200 | 121210199 | 16.69671 | 27.99037 | 64.72048 | 28.23008 | -0.82379 | 1.147966 | 0.24924 |
| 14 | 91788200 | 91789799 | 13.37802 | 25.01132 | 21.8182 | 9.041409 | 0.652683 | 0.145279 | 0.249248 |
| 7 | 33581200 | 33585799 | 88.2041 | 138.9226 | 119.7833 | 49.70146 | 0.536044 | 2.506929 | 0.249251 |
| 16 | 58033600 | 58035599 | 17.28153 | 31.64122 | 23.47401 | 10.02696 | 0.852723 | 0.3004 | 0.249262 |
| 11 | 13938800 | 13941399 | 22.17265 | 38.72475 | 32.79455 | 14.44581 | 0.626148 | 0.726476 | 0.249272 |
| 1 | 173696600 | 173698799 | 34.36365 | 57.18298 | 43.36402 | 18.82644 | 0.778907 | 1.258357 | 0.249286 |
| 7 | 11218200 | 11225599 | 244.9875 | 357.3513 | 303.9447 | 125.2421 | 0.462281 | 3.951141 | 0.249314 |
| 1 | 91232800 | 91234199 | 8.145017 | 14.79492 | 36.00743 | 15.5363 | -0.85927 | 0.305157 | 0.249315 |
| 7 | 28465600 | 28467599 | 8.563505 | 15.23997 | 39.47104 | 17.39346 | -0.96895 | 0.442316 | 0.249318 |
| 16 | 7722600 | 7723999 | 17.03686 | 31.98767 | 19.21426 | 7.905165 | 1.194884 | 0.144716 | 0.249337 |
| 10 | 82761800 | 82766399 | 51.13101 | 82.41981 | 72.73194 | 30.95928 | 0.536565 | 1.801488 | 0.249359 |
| 1 | 6054400 | 6056199 | 49.78958 | 80.17528 | 71.05439 | 30.20136 | 0.528626 | 1.829885 | 0.249359 |
| 8 | 7747200 | 7748999 | 20.15994 | 36.12855 | 25.07858 | 10.98643 | 0.924452 | 0.445058 | 0.249367 |
| 4 | 104703800 | 104704999 | 10.55752 | 18.65103 | 40.95824 | 18.11947 | -0.73808 | 0.522178 | 0.249384 |
| 7 | 29824800 | 29828799 | 25.29075 | 41.62439 | 79.57315 | 34.30929 | -0.55329 | 1.394527 | 0.249393 |
| 3 | 89738000 | 89741799 | 28.04088 | 45.4215 | 91.42751 | 39.12925 | -0.64263 | 1.645502 | 0.249413 |
| 1 | 174347400 | 174350999 | 23.6408 | 41.16985 | 30.40838 | 13.37043 | 0.822272 | 0.77109 | 0.249417 |
| 13 | 67037600 | 67043399 | 103.2359 | 160.6778 | 139.6916 | 57.98736 | 0.51095 | 2.732242 | 0.249433 |
| 6 | 34936200 | 34937399 | 7.094436 | 13.36714 | 30.76088 | 12.58054 | -0.72018 | 0.066427 | 0.249437 |
| 1 | 119271600 | 119279999 | 60.63094 | 95.00861 | 173.3064 | 71.88424 | -0.54927 | 2.598109 | 0.249437 |
| 17 | 22501000 | 22503799 | 65.90741 | 105.5539 | 81.92735 | 34.54215 | 0.710246 | 2.023914 | 0.249453 |
| 10 | 23846000 | 23847799 | 10.13456 | 18.21965 | 38.94947 | 17.01755 | -0.68595 | 0.385214 | 0.24947 |
| 17 | 26768200 | 26769999 | 10.13456 | 18.21965 | 38.94947 | 17.01755 | -0.68595 | 0.385214 | 0.24947 |
| 12 | 106443600 | 106444999 | 9.338039 | 16.96212 | 35.98467 | 15.52631 | -0.6627 | 0.314263 | 0.249473 |
| 8 | 48685600 | 48706199 | 237.1225 | 337.2493 | 551.2021 | 227.9288 | -0.51567 | 4.326771 | 0.249476 |
| 3 | 15537200 | 15538599 | 16.90389 | 31.91233 | 17.48912 | 7.126003 | 1.336694 | 0.105059 | 0.249482 |
| 10 | 103573800 | 103575399 | 17.09991 | 32.27336 | 17.48121 | 7.126997 | 1.352758 | 0.108774 | 0.249488 |
| 10 | 58333400 | 58334599 | 9.076544 | 16.34809 | 38.47344 | 16.77688 | -0.82164 | 0.365366 | 0.249492 |

| | | | | | | | | | |
|---|---|---|---|---|---|---|---|---|---|
| 10 | 14923400 | 14924999 | 16.05019 | 30.25779 | 18.69555 | 7.639034 | 1.16188 | 0.109218 | 0.249505 |
| 2 | 8891800 | 8893599 | 19.36366 | 35.18778 | 23.10903 | 9.965686 | 1.018368 | 0.356299 | 0.249519 |
| 1 | 40140600 | 40142599 | 10.5453 | 18.6485 | 40.93101 | 18.10372 | -0.73688 | 0.509422 | 0.249522 |
| 11 | 68341200 | 68343799 | 11.4285 | 19.89934 | 49.96891 | 21.96895 | -0.9346 | 0.743336 | 0.24953 |
| 17 | 39320800 | 39331199 | 225.366 | 331.7893 | 284.7554 | 117.264 | 0.462035 | 3.80554 | 0.249538 |
| 3 | 9315000 | 9317199 | 13.56473 | 22.89183 | 63.53946 | 27.78968 | -1.08567 | 1.072937 | 0.249541 |
| X | 125255200 | 125259599 | 101.9116 | 158.5775 | 137.5411 | 57.09681 | 0.514082 | 2.737402 | 0.249549 |
| X | 29430200 | 29433999 | 26.25908 | 45.62045 | 28.52772 | 12.5247 | 1.061247 | 0.815974 | 0.249552 |
| 3 | 43514400 | 43514999 | 11.21095 | 21.54014 | 19.05286 | 7.449996 | 0.705581 | -0.07415 | 0.249562 |
| 8 | 108314400 | 108318399 | 18.1292 | 30.72084 | 60.73486 | 26.59619 | -0.5983 | 1.048683 | 0.249572 |
| 1 | 39487600 | 39488999 | 6.665646 | 12.51411 | 30.68105 | 12.62459 | -0.81791 | 0.084782 | 0.249581 |
| 14 | 42134200 | 42138599 | 74.93745 | 118.8247 | 107.4318 | 44.71706 | 0.475488 | 2.340343 | 0.249587 |
| 7 | 8369600 | 8376199 | 84.01319 | 133.6041 | 91.40086 | 38.15546 | 0.881164 | 2.26216 | 0.249589 |
| 1 | 7927000 | 7929399 | 28.73466 | 49.95464 | 30.74883 | 13.50551 | 1.083661 | 0.819498 | 0.249594 |
| 14 | 121689800 | 121696599 | 49.07891 | 77.80015 | 158.5636 | 65.98143 | -0.69581 | 2.339254 | 0.249597 |
| 15 | 16877800 | 16879799 | 34.0454 | 57.818 | 38.57038 | 16.90143 | 0.963245 | 1.064042 | 0.249606 |
| 4 | 86550200 | 86551599 | 7.588734 | 14.0822 | 34.809 | 14.5996 | -0.85288 | 0.1732 | 0.249609 |
| X | 155620000 | 155621399 | 9.798676 | 17.84421 | 37.63158 | 16.18529 | -0.65135 | 0.3064 | 0.249617 |
| 7 | 112570400 | 112572599 | 21.13929 | 35.19212 | 71.22686 | 30.93193 | -0.63271 | 1.240237 | 0.249626 |
| 14 | 37589000 | 37590799 | 20.51623 | 36.78865 | 22.66764 | 9.925334 | 1.095215 | 0.437284 | 0.249637 |
| 2 | 11869800 | 11870399 | 3.878018 | 7.578007 | 21.2258 | 7.368611 | -0.77931 | -0.56177 | 0.249645 |
| 18 | 38458000 | 38459599 | 17.43413 | 31.69329 | 24.48413 | 10.55375 | 0.786205 | 0.34344 | 0.249649 |
| 11 | 30216800 | 30219799 | 18.16966 | 30.27439 | 69.13103 | 30.03806 | -0.80595 | 1.221702 | 0.249655 |
| 14 | 116831800 | 116834599 | 30.43275 | 51.64027 | 41.41243 | 18.14228 | 0.698604 | 1.059207 | 0.24966 |
| 3 | 121516600 | 121518599 | 9.600868 | 17.09243 | 38.92319 | 17.14669 | -0.78437 | 0.445872 | 0.249677 |
| 3 | 152617000 | 152618799 | 15.00211 | 25.90758 | 52.69911 | 23.11907 | -0.63463 | 0.855385 | 0.249679 |
| 8 | 11683800 | 11686999 | 16.06996 | 26.96716 | 64.70625 | 28.24202 | -0.87698 | 1.134535 | 0.249681 |
| 8 | 114525800 | 114528399 | 13.78867 | 23.76931 | 53.71705 | 23.56435 | -0.78629 | 0.867641 | 0.249695 |
| 8 | 64051600 | 64051999 | 5.118873 | 9.903176 | 25.98771 | 9.919332 | -0.81557 | -0.19799 | 0.249701 |
| 14 | 51595600 | 51596199 | 5.118873 | 9.903176 | 25.98771 | 9.919332 | -0.81557 | -0.19799 | 0.249701 |
| X | 108349400 | 108351199 | 32.86101 | 55.26112 | 43.72398 | 19.10273 | 0.717736 | 1.149036 | 0.249706 |
| 2 | 12959200 | 12961399 | 9.790278 | 17.78924 | 43.11018 | 18.59336 | -0.8538 | 0.327574 | 0.249714 |
| 9 | 53949800 | 53952199 | 15.27075 | 26.05678 | 56.85767 | 24.92643 | -0.73867 | 0.972229 | 0.249719 |
| 5 | 9013800 | 9014999 | 7.334263 | 13.52534 | 34.12159 | 14.43328 | -0.89057 | 0.206669 | 0.249754 |
| 3 | 91453600 | 91455199 | 28.4448 | 49.09218 | 35.30759 | 15.48913 | 0.858588 | 0.89562 | 0.249764 |
| X | 148181200 | 148182999 | 21.96223 | 39.34448 | 23.91492 | 10.47979 | 1.114659 | 0.458073 | 0.249779 |
| 7 | 140593800 | 140596399 | 18.95003 | 32.03057 | 61.83398 | 27.05672 | -0.56425 | 1.075257 | 0.249801 |
| 9 | 120467800 | 120473399 | 49.55925 | 78.59026 | 143.6997 | 59.81813 | -0.5389 | 2.326536 | 0.249805 |
| X | 3592200 | 3593199 | 10.61073 | 20.50282 | 18.0994 | 6.948068 | 0.733328 | -0.142 | 0.249808 |
| 18 | 83941200 | 83944599 | 18.75806 | 31.34432 | 65.90236 | 28.68691 | -0.68723 | 1.190902 | 0.24981 |
| 4 | 154272800 | 154278999 | 42.3185 | 67.50248 | 125.9451 | 52.79691 | -0.55919 | 2.135449 | 0.249817 |
| 14 | 57372600 | 57373799 | 7.614159 | 13.72114 | 38.00485 | 16.56197 | -1.05386 | 0.352354 | 0.249826 |
| 6 | 26148600 | 26149599 | 14.73703 | 28.10919 | 16.60366 | 6.619655 | 1.255869 | 0.013748 | 0.249828 |
| 5 | 66155600 | 66159399 | 19.4144 | 32.34662 | 68.13516 | 29.60433 | -0.68995 | 1.225471 | 0.24983 |
| 7 | 15572200 | 15578199 | 82.51982 | 130.0942 | 118.0906 | 49.00521 | 0.462997 | 2.483629 | 0.249841 |
| 1 | 170497400 | 170499199 | 24.6819 | 42.91029 | 33.75668 | 14.8269 | 0.731417 | 0.802856 | 0.24985 |
| 1 | 4029400 | 4030999 | 17.73163 | 32.22004 | 24.40859 | 10.52694 | 0.813701 | 0.347614 | 0.249875 |

| | | | | | | | | | |
|---|---|---|---|---|---|---|---|---|---|
| 13 | 116429800 | 116431599 | 16.82013 | 28.71067 | 56.94224 | 24.96422 | -0.60151 | 0.974791 | 0.249875 |
| 18 | 85291400 | 85292399 | 8.500381 | 15.66854 | 35.75698 | 15.13459 | -0.74826 | 0.220931 | 0.249902 |
| 6 | 41195000 | 41197199 | 27.44445 | 47.20736 | 37.67791 | 16.52729 | 0.708214 | 0.924979 | 0.249909 |
| 12 | 59058200 | 59060599 | 12.69049 | 21.96106 | 52.10744 | 22.86095 | -0.85485 | 0.825882 | 0.24991 |
| 13 | 68343600 | 68352999 | 257.97 | 375.9309 | 316.7299 | 130.5233 | 0.474633 | 3.966794 | 0.249911 |
| 5 | 110324600 | 110325199 | 4.834657 | 9.3447 | 25.89182 | 9.925408 | -0.89857 | -0.18555 | 0.249928 |
| 8 | 118895600 | 118896999 | 8.913941 | 16.10026 | 36.95852 | 16.05992 | -0.78241 | 0.345695 | 0.249934 |
| 2 | 178834800 | 178836399 | 11.17916 | 19.70043 | 41.97616 | 18.57897 | -0.6964 | 0.554531 | 0.249946 |
| 15 | 18203200 | 18204599 | 15.9269 | 29.73452 | 20.88669 | 8.676171 | 0.959893 | 0.173247 | 0.249947 |
| 18 | 89643800 | 89645399 | 15.9269 | 29.73452 | 20.88669 | 8.676171 | 0.959893 | 0.173247 | 0.249947 |
| 11 | 88968400 | 88975399 | 64.9747 | 101.5282 | 178.8249 | 74.19476 | -0.50201 | 2.651751 | 0.249949 |
| X | 32256200 | 32257799 | 15.98376 | 30.80533 | 15.21215 | 5.895717 | 1.55013 | -0.06279 | 0.249954 |
| 8 | 44053800 | 44055199 | 15.63426 | 30.20204 | 12.64143 | 4.858544 | 1.794569 | -0.10038 | 0.249965 |
| 17 | 56503800 | 56506199 | 28.73857 | 46.39173 | 96.59469 | 41.19354 | -0.69655 | 1.738595 | 0.249965 |
| 5 | 93813400 | 93819399 | 70.91828 | 113.258 | 86.88633 | 36.48491 | 0.72261 | 2.115046 | 0.24997 |
| 11 | 76280800 | 76284599 | 32.93325 | 53.07058 | 100.1657 | 42.6201 | -0.5583 | 1.796227 | 0.249976 |
| 15 | 86419600 | 86422999 | 26.83126 | 43.76306 | 83.44601 | 35.84953 | -0.55641 | 1.520177 | 0.250006 |
| 11 | 56253800 | 56254999 | 8.319389 | 15.45658 | 34.30309 | 14.36394 | -0.69687 | 0.172024 | 0.250009 |
| X | 66390200 | 66392599 | 21.06789 | 38.46426 | 20.93266 | 8.974466 | 1.293093 | 0.339573 | 0.250009 |
| 5 | 142920000 | 142922999 | 17.28906 | 29.46917 | 58.59875 | 25.68751 | -0.60568 | 0.990537 | 0.250015 |
| 15 | 14085000 | 14087399 | 16.74314 | 30.78186 | 21.66838 | 9.204008 | 0.932959 | 0.267245 | 0.250018 |
| 9 | 113940200 | 113946199 | 49.39445 | 78.33108 | 143.698 | 59.81909 | -0.5436 | 2.325521 | 0.250036 |
| 14 | 3922600 | 3922999 | 3.453944 | 6.75375 | 19.53035 | 6.666438 | -0.80114 | -0.63351 | 0.250036 |
| 16 | 45083000 | 45083399 | 3.453944 | 6.75375 | 19.53035 | 6.666438 | -0.80114 | -0.63351 | 0.250036 |
| 4 | 63099600 | 63101399 | 7.684616 | 14.00186 | 35.01972 | 15.0518 | -0.89425 | 0.282849 | 0.250054 |
| 9 | 113213000 | 113213799 | 7.045955 | 13.45192 | 31.27707 | 12.44511 | -0.7006 | -0.02579 | 0.250065 |
| 17 | 40000400 | 40000799 | 8.509374 | 16.60173 | 14.40271 | 5.144759 | 0.851418 | -0.42731 | 0.250075 |
| 8 | 13058800 | 13065799 | 52.47989 | 82.74456 | 158.7119 | 65.86318 | -0.6147 | 2.471017 | 0.250079 |
| 14 | 51201000 | 51203199 | 21.1081 | 37.02103 | 31.14319 | 13.76576 | 0.635657 | 0.653196 | 0.25009 |
| 2 | 139609600 | 139610399 | 12.53863 | 24.0564 | 18.79892 | 7.381172 | 0.876724 | -0.04939 | 0.25009 |
| 4 | 149137200 | 149137999 | 4.238838 | 8.238312 | 23.90707 | 8.893721 | -0.92234 | -0.30214 | 0.250092 |
| 12 | 41866200 | 41866799 | 4.238838 | 8.238312 | 23.90707 | 8.893721 | -0.92234 | -0.30214 | 0.250092 |
| 17 | 27066000 | 27071199 | 31.26959 | 50.54317 | 95.15133 | 40.64616 | -0.54949 | 1.701377 | 0.250095 |
| 7 | 57206000 | 57207199 | 11.94467 | 23.09119 | 16.6112 | 6.362793 | 1.027418 | -0.14143 | 0.250125 |
| 8 | 67791600 | 67794799 | 32.09787 | 51.95519 | 104.1321 | 44.5426 | -0.63763 | 1.67487 | 0.250128 |
| 7 | 59100000 | 59107199 | 119.278 | 184.0905 | 143.4175 | 59.57287 | 0.65974 | 2.88744 | 0.250135 |
| 10 | 100157000 | 100157799 | 10.90423 | 21.03758 | 18.57606 | 7.17788 | 0.723965 | -0.11625 | 0.25014 |
| 17 | 38831600 | 38833399 | 42.04322 | 68.59406 | 57.47328 | 24.69387 | 0.627048 | 1.532141 | 0.250155 |
| 6 | 88958400 | 88968199 | 53.64279 | 84.48114 | 162.4623 | 67.40946 | -0.61972 | 2.494754 | 0.250165 |
| 16 | 13595400 | 13598999 | 46.36383 | 74.90225 | 67.29799 | 28.72875 | 0.514325 | 1.727429 | 0.250165 |
| 1 | 117673000 | 117682399 | 193.6185 | 289.0939 | 242.3148 | 99.88975 | 0.51491 | 3.559042 | 0.250187 |
| 3 | 23061600 | 23064199 | 27.003 | 46.34021 | 38.7455 | 16.99388 | 0.640757 | 0.94369 | 0.250198 |
| 9 | 13555600 | 13559599 | 30.73878 | 52.0414 | 42.56322 | 18.6336 | 0.670321 | 1.08209 | 0.250208 |
| 14 | 95190800 | 95192399 | 20.65694 | 36.62116 | 24.77896 | 10.95424 | 0.952432 | 0.514039 | 0.250213 |
| 19 | 57598400 | 57600399 | 12.54507 | 21.69171 | 51.55437 | 22.61774 | -0.85729 | 0.833773 | 0.250215 |
| 2 | 47963200 | 47964799 | 16.27437 | 30.50714 | 20.4576 | 8.439572 | 1.034516 | 0.151231 | 0.250226 |
| 14 | 37080600 | 37082199 | 16.27437 | 30.50714 | 20.4576 | 8.439572 | 1.034516 | 0.151231 | 0.250226 |

| | | | | | | | | | |
|---|---|---|---|---|---|---|---|---|---|
| 1 | 5375600 | 5380199 | 56.08202 | 90.31774 | 72.65076 | 30.88477 | 0.668321 | 1.836857 | 0.250237 |
| 1 | 91019200 | 91020999 | 8.368259 | 15.43984 | 33.59748 | 14.20197 | -0.6781 | 0.212515 | 0.250237 |
| 2 | 85858800 | 85860599 | 25.31658 | 43.34906 | 36.90581 | 16.18407 | 0.614339 | 0.954398 | 0.250245 |
| 18 | 17459600 | 17460399 | 23.50112 | 40.62417 | 35.25069 | 15.46492 | 0.589425 | 0.859899 | 0.250245 |
| 2 | 105657800 | 105659599 | 20.99773 | 37.03938 | 28.85578 | 12.76929 | 0.747624 | 0.576598 | 0.250256 |
| 3 | 11551200 | 11555599 | 64.68076 | 103.5302 | 84.64054 | 35.65582 | 0.634512 | 2.039932 | 0.250278 |
| 7 | 38780200 | 38783599 | 49.2138 | 81.30537 | 41.6072 | 17.97392 | 1.343863 | 1.383076 | 0.250288 |
| 4 | 58068200 | 58073999 | 75.44427 | 116.8525 | 200.4852 | 83.27615 | -0.47415 | 2.81199 | 0.250289 |
| 7 | 60064400 | 60068599 | 48.70366 | 79.11891 | 59.98835 | 25.72837 | 0.766325 | 1.611587 | 0.250296 |
| 8 | 47004600 | 47005999 | 6.96949 | 13.04758 | 31.17265 | 12.88848 | -0.78676 | 0.106556 | 0.250298 |
| 14 | 79086400 | 79087199 | 5.203516 | 9.903029 | 29.86726 | 11.96389 | -1.08023 | -0.02593 | 0.250299 |
| 7 | 5974000 | 5975799 | 42.09696 | 68.57445 | 60.88961 | 26.14682 | 0.54194 | 1.554468 | 0.25031 |
| X | 32772200 | 32773999 | 21.06492 | 37.93493 | 23.41803 | 10.21374 | 1.096819 | 0.419481 | 0.250321 |
| 15 | 59636000 | 59646199 | 144.5383 | 215.0475 | 349.902 | 144.166 | -0.44614 | 3.619747 | 0.250322 |
| 15 | 87458200 | 87465799 | 54.18075 | 85.72682 | 150.3446 | 62.48986 | -0.48163 | 2.389188 | 0.250342 |
| 18 | 49309800 | 49311199 | 14.55175 | 27.75009 | 17.85841 | 7.122998 | 1.133435 | 0.017073 | 0.250358 |
| 15 | 49702200 | 49703799 | 16.69181 | 31.12595 | 20.85882 | 8.681858 | 1.025214 | 0.187507 | 0.250368 |
| 19 | 30152200 | 30153199 | 8.304701 | 15.04972 | 35.98348 | 15.57146 | -0.83653 | 0.320706 | 0.250374 |
| 10 | 81815800 | 81821799 | 95.59177 | 149.4891 | 119.7871 | 49.69582 | 0.633726 | 2.643631 | 0.250379 |
| 3 | 89150800 | 89154199 | 13.25803 | 22.67784 | 56.25062 | 24.66516 | -0.92192 | 0.941641 | 0.250384 |
| X | 100835800 | 100840799 | 40.66937 | 65.10937 | 120.3759 | 50.72367 | -0.54022 | 2.020338 | 0.250397 |
| 10 | 91251400 | 91252799 | 12.8209 | 24.54238 | 19.37985 | 7.658832 | 0.85355 | -0.02351 | 0.2504 |
| 13 | 6963200 | 6964199 | 9.503647 | 17.2864 | 36.53798 | 15.73828 | -0.65574 | 0.309894 | 0.250425 |
| 8 | 60603000 | 60604599 | 15.89333 | 29.46166 | 21.91086 | 9.203073 | 0.86596 | 0.216885 | 0.250439 |
| 9 | 81604000 | 81605199 | 14.02601 | 26.56394 | 20.17545 | 8.16523 | 0.878851 | 0.064436 | 0.250439 |
| X | 122984400 | 122989799 | 50.22546 | 81.95765 | 53.54758 | 23.00861 | 0.985038 | 1.54362 | 0.25045 |
| 8 | 130025000 | 130027799 | 45.97505 | 76.19593 | 33.10929 | 14.33425 | 1.578472 | 1.327107 | 0.250456 |
| 7 | 113862400 | 113863399 | 26.15692 | 45.04372 | 37.26551 | 16.34695 | 0.656978 | 0.909541 | 0.250457 |
| 16 | 17398600 | 17401999 | 32.08177 | 53.55541 | 47.59226 | 20.70381 | 0.550896 | 1.223327 | 0.250479 |
| 3 | 122086000 | 122087999 | 11.24268 | 19.64543 | 47.308 | 20.84555 | -0.87378 | 0.687696 | 0.250485 |
| 5 | 23788000 | 23789599 | 9.609094 | 17.53187 | 39.33756 | 16.87932 | -0.73836 | 0.295324 | 0.250507 |
| 4 | 41135000 | 41136799 | 10.51818 | 18.6979 | 39.38648 | 17.37238 | -0.67408 | 0.461835 | 0.250519 |
| 2 | 148879600 | 148882399 | 46.03374 | 75.31813 | 54.58364 | 23.47658 | 0.838575 | 1.497361 | 0.25052 |
| 4 | 119025600 | 119029399 | 23.11285 | 37.83088 | 79.67762 | 34.27447 | -0.6957 | 1.459753 | 0.250522 |
| 9 | 112470200 | 112471799 | 9.164029 | 16.29333 | 39.9431 | 17.61546 | -0.89131 | 0.452206 | 0.250525 |
| 10 | 81212800 | 81214999 | 9.164029 | 16.29333 | 39.9431 | 17.61546 | -0.89131 | 0.452206 | 0.250525 |
| 11 | 11982800 | 11983199 | 2.94601 | 5.762861 | 19.11893 | 6.371919 | -0.962 | -0.72394 | 0.250536 |
| 4 | 111882000 | 111883199 | 12.59284 | 24.29892 | 17.76496 | 6.861634 | 0.99386 | -0.10281 | 0.250537 |
| 4 | 138105000 | 138108999 | 21.33739 | 35.42488 | 71.06329 | 30.80303 | -0.62017 | 1.273107 | 0.25054 |
| 10 | 62904600 | 62913599 | 91.79713 | 140.0625 | 245.0515 | 101.613 | -0.51911 | 3.112315 | 0.250542 |
| 4 | 46342200 | 46346199 | 36.56949 | 58.59359 | 114.5359 | 48.32289 | -0.61898 | 1.986214 | 0.250555 |
| 6 | 38610600 | 38614599 | 21.17928 | 34.79685 | 80.04507 | 34.48731 | -0.8204 | 1.404695 | 0.250557 |
| 4 | 141008800 | 141011399 | 10.70101 | 18.90074 | 41.95655 | 18.56184 | -0.75396 | 0.524977 | 0.250565 |
| 3 | 71258400 | 71260599 | 17.2029 | 31.29756 | 23.78914 | 10.24409 | 0.809986 | 0.333988 | 0.250578 |
| 1 | 20026800 | 20029199 | 30.9021 | 52.15413 | 43.09896 | 18.85021 | 0.655231 | 1.107789 | 0.25058 |
| 12 | 22333600 | 22333999 | 7.991876 | 15.58715 | 14.11957 | 5.064737 | 0.786185 | -0.41846 | 0.25059 |
| 3 | 48830800 | 48831999 | 18.83789 | 33.25858 | 28.62837 | 12.66644 | 0.604948 | 0.553574 | 0.250594 |

| | | | | | | | | | |
|---|---|---|---|---|---|---|---|---|---|
| 11 | 43380000 | 43390999 | 71.51803 | 110.4522 | 211.0325 | 87.65849 | -0.6324 | 2.864213 | 0.250598 |
| 11 | 79692400 | 79696599 | 7.48342 | 13.37136 | 41.05333 | 18.0296 | -1.20877 | 0.413807 | 0.250607 |
| 5 | 68262600 | 68264999 | 11.81303 | 20.7652 | 43.01323 | 19.04755 | -0.65741 | 0.588297 | 0.250612 |
| 3 | 137130600 | 137132399 | 17.79872 | 32.20077 | 24.82643 | 10.76588 | 0.783354 | 0.372672 | 0.250623 |
| 18 | 29448000 | 29448999 | 7.681319 | 14.48689 | 31.94601 | 13.04391 | -0.65791 | 0.06654 | 0.25065 |
| 17 | 95128600 | 95134399 | 132.5003 | 203.4004 | 163.1733 | 67.7451 | 0.611645 | 2.997226 | 0.250653 |
| 1 | 38692400 | 38694599 | 21.44699 | 35.76713 | 69.00209 | 29.99899 | -0.56375 | 1.221296 | 0.250655 |
| 6 | 10502600 | 10504799 | 23.47084 | 41.72905 | 24.28813 | 10.71168 | 1.170703 | 0.514362 | 0.250688 |
| 8 | 43989000 | 43991999 | 35.66021 | 59.7086 | 44.78527 | 19.51976 | 0.794528 | 1.199937 | 0.250704 |
| 12 | 49176000 | 49179199 | 35.66021 | 59.7086 | 44.78527 | 19.51976 | 0.794528 | 1.199937 | 0.250704 |
| 9 | 85630800 | 85631999 | 5.89627 | 11.23439 | 29.96341 | 11.97316 | -0.90206 | -0.02788 | 0.250707 |
| 17 | 64451800 | 64452999 | 5.89627 | 11.23439 | 29.96341 | 11.97316 | -0.90206 | -0.02788 | 0.250707 |
| 6 | 14089600 | 14090799 | 16.04596 | 29.30239 | 20.1636 | 8.642236 | 0.956985 | 0.296682 | 0.250759 |
| 16 | 95681400 | 95682599 | 18.20923 | 33.544 | 21.70857 | 9.196115 | 1.056687 | 0.270027 | 0.250776 |
| 12 | 3830400 | 3834999 | 25.72529 | 41.90556 | 83.65587 | 35.92295 | -0.62369 | 1.536391 | 0.250797 |
| 1 | 178187600 | 178189199 | 12.74585 | 22.38864 | 44.56389 | 19.7232 | -0.60047 | 0.607085 | 0.250801 |
| 14 | 86756200 | 86760199 | 47.89416 | 77.31908 | 69.5818 | 29.67622 | 0.510836 | 1.7523 | 0.250803 |
| 4 | 126968800 | 126972599 | 8.91802 | 16.1437 | 44.56558 | 19.31138 | -1.04442 | 0.348826 | 0.250813 |
| 16 | 61308600 | 61309599 | 11.58042 | 22.32603 | 18.40426 | 7.12626 | 0.818898 | -0.10436 | 0.250819 |
| 8 | 26210000 | 26211399 | 6.350432 | 12.01024 | 30.30551 | 12.30149 | -0.84283 | 0.030112 | 0.250822 |
| X | 113905000 | 113907599 | 26.0833 | 45.9002 | 25.7472 | 11.39913 | 1.217608 | 0.627458 | 0.250825 |
| 7 | 46130800 | 46132999 | 11.60944 | 20.15744 | 52.63643 | 23.10073 | -0.99137 | 0.788219 | 0.250836 |
| 15 | 46528800 | 46529599 | 11.42337 | 22.25127 | 13.29826 | 4.850938 | 1.355203 | -0.31986 | 0.250851 |
| 5 | 106262200 | 106264199 | 19.36372 | 34.05388 | 28.61745 | 12.67148 | 0.637194 | 0.597565 | 0.250859 |
| 3 | 88049200 | 88050999 | 9.286729 | 16.57105 | 38.43591 | 16.89938 | -0.80865 | 0.428577 | 0.250867 |
| 1 | 74275200 | 74276599 | 6.735352 | 12.43602 | 33.08528 | 13.97173 | -0.96397 | 0.195313 | 0.250868 |
| 5 | 100186000 | 100188599 | 17.86494 | 29.75456 | 69.05552 | 29.99755 | -0.82923 | 1.225566 | 0.250884 |
| 1 | 146030000 | 146030399 | 10.7246 | 20.69545 | 18.58126 | 7.175476 | 0.700865 | -0.12034 | 0.250887 |
| 4 | 38144600 | 38145999 | 15.18637 | 28.65181 | 20.00721 | 8.16288 | 0.989014 | 0.100232 | 0.250892 |
| 9 | 40963800 | 40966199 | 20.83148 | 34.67192 | 70.00738 | 30.39798 | -0.62924 | 1.241327 | 0.250902 |
| 3 | 111005200 | 111006599 | 19.05248 | 34.28847 | 23.40711 | 10.21729 | 0.951386 | 0.406572 | 0.250908 |
| 17 | 39004800 | 39006999 | 33.40667 | 57.19261 | 34.02054 | 14.91862 | 1.129276 | 0.985123 | 0.25093 |
| 9 | 13912600 | 13913999 | 16.93601 | 30.49254 | 25.44188 | 11.09954 | 0.664509 | 0.39454 | 0.250931 |
| 13 | 29439600 | 29440199 | 9.404564 | 16.89969 | 37.43683 | 16.36285 | -0.73667 | 0.383532 | 0.250937 |
| 7 | 61405400 | 61407999 | 29.20296 | 50.04545 | 37.51015 | 16.4502 | 0.797469 | 0.962648 | 0.25095 |
| 6 | 130701800 | 130704199 | 74.19082 | 117.7755 | 104.6378 | 43.60402 | 0.502253 | 2.305272 | 0.250959 |
| X | 33353200 | 33355799 | 32.10124 | 54.45262 | 36.27971 | 15.89203 | 0.965079 | 1.065344 | 0.250974 |
| 3 | 101127000 | 101128799 | 38.24779 | 62.93596 | 54.86564 | 23.65408 | 0.576156 | 1.409395 | 0.250985 |
| 8 | 23574600 | 23579199 | 36.75807 | 58.81888 | 118.7327 | 49.99665 | -0.66747 | 2.028274 | 0.250997 |
| 13 | 4029000 | 4029399 | 10.80407 | 20.69987 | 18.52874 | 7.304027 | 0.677529 | -0.05478 | 0.251003 |
| 7 | 60651800 | 60653399 | 20.15056 | 35.56872 | 28.92488 | 12.79816 | 0.686386 | 0.564239 | 0.251007 |
| 7 | 116546000 | 116546599 | 4.409231 | 8.555659 | 24.94384 | 9.389627 | -0.94574 | -0.25625 | 0.251015 |
| X | 148782200 | 148785599 | 29.21412 | 49.96112 | 35.18102 | 15.42618 | 0.886884 | 0.980528 | 0.251016 |
| 15 | 84040000 | 84046599 | 38.6778 | 61.8891 | 119.4734 | 50.30665 | -0.60325 | 2.031028 | 0.251029 |
| 15 | 90562800 | 90564599 | 12.61569 | 21.99407 | 49.45477 | 21.76007 | -0.77612 | 0.729537 | 0.251048 |
| 10 | 77961200 | 77962399 | 15.17424 | 27.66023 | 22.56824 | 9.694865 | 0.710577 | 0.304539 | 0.251061 |
| 11 | 35865400 | 35866199 | 5.1047 | 9.916326 | 25.77764 | 9.627268 | -0.77261 | -0.27531 | 0.251063 |

| | | | | | | | | | |
|---|---|---|---|---|---|---|---|---|---|
| 8 | 121807600 | 121814399 | 59.03361 | 92.43523 | 175.1507 | 72.65491 | -0.60494 | 2.61103 | 0.251064 |
| 17 | 88249600 | 88254599 | 67.06768 | 106.6997 | 96.65228 | 40.37034 | 0.477414 | 2.233 | 0.251084 |
| 2 | 48096400 | 48099799 | 39.73737 | 65.02427 | 56.0279 | 24.09884 | 0.589633 | 1.479647 | 0.251091 |
| 17 | 93095800 | 93097599 | 21.40877 | 37.54317 | 31.08886 | 13.74014 | 0.658243 | 0.656679 | 0.251092 |
| 7 | 61420000 | 61430199 | 126.195 | 193.7497 | 166.5034 | 69.12906 | 0.512621 | 2.991973 | 0.251108 |
| 12 | 55554400 | 55557399 | 14.34284 | 24.94841 | 49.48792 | 21.74359 | -0.5965 | 0.772091 | 0.25111 |
| 12 | 107838200 | 107848399 | 85.82271 | 130.8086 | 251.1355 | 104.1103 | -0.65417 | 3.132399 | 0.251125 |
| 9 | 46806400 | 46808199 | 10.33802 | 18.44935 | 38.88968 | 17.09702 | -0.67183 | 0.434287 | 0.251126 |
| 8 | 52396000 | 52398999 | 35.9243 | 59.22003 | 53.12639 | 22.92182 | 0.535967 | 1.380233 | 0.251142 |
| 2 | 148522600 | 148523799 | 6.173854 | 11.43424 | 34.17893 | 14.37357 | -1.12674 | 0.172956 | 0.251149 |
| 6 | 116626000 | 116629199 | 20.17256 | 33.29089 | 76.24152 | 32.91113 | -0.81137 | 1.343885 | 0.251165 |
| 9 | 98130800 | 98133399 | 13.23498 | 23.16894 | 46.21962 | 20.39734 | -0.60371 | 0.667876 | 0.251189 |
| 7 | 28372000 | 28373599 | 14.97211 | 26.04279 | 51.11089 | 22.45656 | -0.58131 | 0.777076 | 0.251193 |
| 12 | 19911000 | 19913799 | 72.99982 | 116.7388 | 80.32203 | 33.79263 | 0.881407 | 2.076161 | 0.251193 |
| 3 | 107125200 | 107126999 | 10.18519 | 17.83473 | 48.31949 | 21.3288 | -1.04285 | 0.648228 | 0.251194 |
| 2 | 99104600 | 99106199 | 18.01693 | 32.89089 | 22.61743 | 9.697933 | 0.957367 | 0.318569 | 0.251196 |
| X | 24581400 | 24583399 | 39.75892 | 65.37238 | 54.91762 | 23.66881 | 0.629014 | 1.422752 | 0.251198 |
| 14 | 42672800 | 42675799 | 41.2936 | 67.54492 | 52.69054 | 22.6602 | 0.73235 | 1.487993 | 0.251206 |
| 7 | 106124200 | 106126399 | 26.29508 | 45.11598 | 38.77509 | 17.00658 | 0.601119 | 0.942103 | 0.251221 |
| 4 | 45717200 | 45720999 | 22.61268 | 37.47025 | 77.01135 | 33.33712 | -0.65545 | 1.301905 | 0.251223 |
| 14 | 47514000 | 47514799 | 5.28096 | 10.18983 | 27.58391 | 10.65482 | -0.87709 | -0.15226 | 0.251223 |
| 7 | 10141400 | 10142399 | 39.97639 | 65.31057 | 53.59512 | 23.03898 | 0.658328 | 1.505151 | 0.251225 |
| 12 | 60889000 | 60889599 | 10.65877 | 20.72153 | 16.50076 | 6.124443 | 0.925594 | -0.25873 | 0.251227 |
| 15 | 73425600 | 73427199 | 7.242654 | 13.20618 | 35.03489 | 15.04463 | -0.97763 | 0.275299 | 0.251233 |
| 12 | 87973800 | 87978799 | 52.90973 | 85.51858 | 65.38637 | 27.93007 | 0.747353 | 1.723427 | 0.251236 |
| 2 | 48904600 | 48906399 | 20.81502 | 36.72267 | 28.85701 | 12.76945 | 0.735301 | 0.573706 | 0.251253 |
| 4 | 61225200 | 61226799 | 13.54517 | 25.3724 | 21.79541 | 9.002697 | 0.67841 | 0.134703 | 0.251258 |
| 7 | 131534400 | 131536799 | 22.685 | 39.707 | 31.55543 | 13.92436 | 0.717369 | 0.693524 | 0.251258 |
| 11 | 89092800 | 89094999 | 11.42936 | 20.2874 | 44.58247 | 19.68895 | -0.73669 | 0.483856 | 0.251283 |
| 12 | 18076000 | 18077799 | 59.6304 | 96.95464 | 47.41493 | 20.34676 | 1.398148 | 1.612257 | 0.251292 |
| 10 | 91160400 | 91168999 | 64.61404 | 101.5386 | 176.8226 | 73.34716 | -0.4793 | 2.546851 | 0.251299 |
| Y | 3835200 | 3837599 | 20.47587 | 36.24498 | 14.33044 | 6.337103 | 1.720072 | 0.502589 | 0.251312 |
| 17 | 26600000 | 26608799 | 54.11688 | 84.85217 | 172.4537 | 71.52614 | -0.70432 | 2.584389 | 0.251318 |
| 10 | 11270400 | 11271599 | 14.37844 | 27.35807 | 18.4177 | 7.381058 | 1.063139 | 0.031612 | 0.251326 |
| 5 | 112968400 | 112969599 | 6.659916 | 12.53029 | 31.28313 | 12.8255 | -0.84013 | 0.070201 | 0.251329 |
| 14 | 60919000 | 60920599 | 6.659916 | 12.53029 | 31.28313 | 12.8255 | -0.84013 | 0.070201 | 0.251329 |
| 13 | 111625200 | 111627999 | 16.75498 | 27.70461 | 78.10471 | 33.76067 | -1.10975 | 1.313647 | 0.251341 |
| 5 | 67764400 | 67767399 | 15.93717 | 26.68603 | 70.7262 | 30.82706 | -1.01998 | 1.163211 | 0.251344 |
| 16 | 3385400 | 3391199 | 71.27405 | 114.2234 | 74.53502 | 31.45662 | 0.961148 | 2.008851 | 0.251349 |
| 2 | 59213400 | 59215399 | 13.28947 | 23.03283 | 50.53365 | 22.17152 | -0.74186 | 0.813652 | 0.251357 |
| 2 | 56768800 | 56769199 | 3.523609 | 6.889916 | 20.21004 | 6.89989 | -0.82348 | -0.63302 | 0.251357 |
| 18 | 62125200 | 62125599 | 3.523609 | 6.889916 | 20.21004 | 6.89989 | -0.82348 | -0.63302 | 0.251357 |
| 6 | 138420400 | 138420999 | 9.190825 | 17.90153 | 16.08484 | 5.871861 | 0.7745 | -0.33562 | 0.25136 |
| 6 | 3333400 | 3334799 | 16.81982 | 30.3554 | 24.71941 | 10.75233 | 0.702479 | 0.381066 | 0.251368 |
| 14 | 35915600 | 35916599 | 13.86689 | 25.84474 | 22.26887 | 9.27693 | 0.6643 | 0.17003 | 0.251381 |
| 1 | 170522000 | 170526999 | 41.8203 | 66.72983 | 124.896 | 52.38118 | -0.5632 | 2.123833 | 0.251382 |
| 19 | 33417600 | 33419799 | 26.11527 | 44.99473 | 32.66204 | 14.32781 | 0.844861 | 0.897579 | 0.251387 |

| | | | | | | | | | |
|---|---|---|---|---|---|---|---|---|---|
| 13 | 105779200 | 105779999 | 9.6032 | 17.09387 | 38.93783 | 17.15543 | -0.78477 | 0.448093 | 0.251394 |
| 3 | 38194600 | 38196399 | 10.28057 | 18.51162 | 38.97295 | 17.00054 | -0.66248 | 0.376627 | 0.251408 |
| 7 | 22848400 | 22857399 | 137.5651 | 210.1539 | 165.2105 | 68.5257 | 0.635421 | 3.09367 | 0.251412 |
| 1 | 183677600 | 183679199 | 7.308835 | 13.5441 | 33.67307 | 14.14969 | -0.86276 | 0.177284 | 0.251414 |
| 7 | 34408200 | 34411999 | 23.68332 | 38.92527 | 77.56084 | 33.4222 | -0.61364 | 1.404954 | 0.251428 |
| 1 | 46153200 | 46154399 | 14.40409 | 27.26478 | 20.16077 | 8.168929 | 0.9158 | 0.071961 | 0.251435 |
| 6 | 29746800 | 29749199 | 246.0399 | 344.2665 | 654.8301 | 271.499 | -0.75814 | 4.569114 | 0.251446 |
| 9 | 63924000 | 63931599 | 36.37562 | 58.00288 | 130.1738 | 54.54349 | -0.82564 | 2.141011 | 0.251457 |
| 7 | 48856600 | 48862799 | 34.60557 | 55.54327 | 110.8177 | 46.87781 | -0.64542 | 1.927722 | 0.251488 |
| 15 | 56745000 | 56749599 | 40.45913 | 64.91449 | 121.5624 | 51.39415 | -0.55496 | 1.951169 | 0.251488 |
| 10 | 70022000 | 70025999 | 24.65496 | 40.06601 | 86.64692 | 37.1716 | -0.74135 | 1.571559 | 0.251489 |
| 19 | 18340200 | 18343799 | 28.48951 | 48.74656 | 39.24102 | 17.20711 | 0.694638 | 0.976569 | 0.251493 |
| 13 | 64670400 | 64671799 | 15.94132 | 28.87405 | 24.56097 | 10.63526 | 0.643718 | 0.351572 | 0.251508 |
| 3 | 43108200 | 43109399 | 13.35397 | 25.46027 | 17.31449 | 6.909293 | 1.053268 | 0.004078 | 0.251516 |
| 16 | 64858200 | 64858999 | 13.35397 | 25.46027 | 17.31449 | 6.909293 | 1.053268 | 0.004078 | 0.251516 |
| 5 | 136070000 | 136079199 | 58.19376 | 90.81318 | 184.8936 | 76.72306 | -0.71225 | 2.671679 | 0.251521 |
| 10 | 121885400 | 121887999 | 16.48913 | 28.14734 | 58.59538 | 25.6891 | -0.67119 | 0.973287 | 0.251544 |
| 19 | 56219400 | 56222799 | 28.65207 | 46.48521 | 88.93923 | 38.10885 | -0.56747 | 1.613319 | 0.251548 |
| 1 | 162659800 | 162661999 | 11.92541 | 21.11425 | 42.95915 | 18.99623 | -0.62776 | 0.508085 | 0.25155 |
| 8 | 21996200 | 21996999 | 11.70739 | 22.73563 | 16.29863 | 6.102272 | 1.063808 | -0.21819 | 0.251558 |
| 9 | 70449400 | 70450799 | 9.960489 | 17.57084 | 41.50588 | 18.36705 | -0.84342 | 0.535701 | 0.251571 |
| 18 | 13377200 | 13380199 | 24.68887 | 43.14631 | 31.46005 | 13.86376 | 0.841079 | 0.729109 | 0.251585 |
| 7 | 20975400 | 20996799 | 380.9791 | 539.5861 | 400.1802 | 165.5954 | 0.617802 | 4.393383 | 0.251602 |
| 1 | 32245200 | 32246799 | 30.89847 | 52.31914 | 38.83462 | 17.002 | 0.809649 | 1.076282 | 0.251617 |
| 3 | 41696600 | 41697399 | 5.441013 | 10.42104 | 27.83814 | 10.98458 | -0.88673 | -0.07775 | 0.251627 |
| 12 | 111455000 | 111456199 | 5.441013 | 10.42104 | 27.83814 | 10.98458 | -0.88673 | -0.07775 | 0.251627 |
| 4 | 100454600 | 100459399 | 39.45702 | 63.33733 | 113.4155 | 47.98516 | -0.48961 | 1.931911 | 0.251635 |
| 10 | 12698200 | 12702999 | 21.75052 | 35.85275 | 75.79369 | 32.70095 | -0.69698 | 1.362524 | 0.251637 |
| 7 | 73300200 | 73304599 | 33.57514 | 54.15263 | 99.42212 | 42.35229 | -0.51704 | 1.771458 | 0.251652 |
| 19 | 44423800 | 44427199 | 20.33023 | 33.95759 | 66.64452 | 29.00253 | -0.58833 | 1.199883 | 0.251653 |
| 11 | 88890000 | 88895799 | 32.6972 | 52.46767 | 120.7018 | 51.04034 | -0.85088 | 1.93611 | 0.251654 |
| 7 | 44325400 | 44342599 | 142.5439 | 211.593 | 359.4782 | 148.0769 | -0.51147 | 3.659664 | 0.251699 |
| 16 | 44502200 | 44503199 | 14.48625 | 27.96239 | 15.15713 | 5.846809 | 1.422113 | -0.0942 | 0.251707 |
| X | 60672800 | 60674999 | 22.98863 | 41.48485 | 21.09394 | 9.176354 | 1.377016 | 0.41734 | 0.251715 |
| 5 | 128512800 | 128515399 | 18.2548 | 30.59312 | 64.30347 | 28.04285 | -0.6868 | 1.15583 | 0.251717 |
| 5 | 79306800 | 79307999 | 16.74447 | 30.41436 | 24.40185 | 10.52918 | 0.730725 | 0.34115 | 0.25173 |
| 3 | 41975400 | 41976399 | 12.08096 | 23.14543 | 19.40567 | 7.650532 | 0.770771 | -0.03927 | 0.251731 |
| 19 | 16446800 | 16447599 | 12.01195 | 22.87087 | 19.99605 | 8.00207 | 0.691887 | 0.010244 | 0.251738 |
| 11 | 13952200 | 13953599 | 16.87162 | 31.15118 | 21.75743 | 9.18814 | 0.950346 | 0.246027 | 0.251739 |
| 7 | 102679800 | 102681199 | 26.54768 | 45.65663 | 38.22663 | 16.76789 | 0.639359 | 0.922683 | 0.251747 |
| 6 | 54626400 | 54627199 | 12.52579 | 23.77488 | 20.34487 | 8.195612 | 0.71389 | 0.035729 | 0.251751 |
| 8 | 95153800 | 95156399 | 11.38123 | 19.92255 | 45.20383 | 19.94801 | -0.78809 | 0.658472 | 0.251755 |
| 19 | 15285600 | 15286199 | 5.529635 | 10.64173 | 27.98665 | 10.89204 | -0.84475 | -0.11833 | 0.251788 |
| 5 | 84865600 | 84867199 | 17.06459 | 31.88014 | 20.30588 | 8.425225 | 1.101553 | 0.178188 | 0.251814 |
| 11 | 10461200 | 10463799 | 24.08238 | 42.0769 | 31.46384 | 13.86312 | 0.804776 | 0.728344 | 0.251816 |
| 8 | 122148000 | 122154599 | 47.65165 | 75.46907 | 158.1637 | 65.77194 | -0.73712 | 2.36224 | 0.251817 |
| 12 | 108237800 | 108248199 | 51.84178 | 81.33472 | 174.5416 | 72.38756 | -0.78197 | 2.571663 | 0.25182 |

| | | | | | | | | | |
|---|---|---|---|---|---|---|---|---|---|
| 1 | 74403600 | 74404399 | 4.659597 | 9.048019 | 24.43429 | 9.155394 | -0.82988 | -0.26969 | 0.25182 |
| 13 | 86946200 | 86947399 | 13.60978 | 25.38294 | 21.71396 | 9.035267 | 0.675744 | 0.161366 | 0.25182 |
| 7 | 38891000 | 38892799 | 24.31183 | 42.29434 | 34.35216 | 15.0946 | 0.685712 | 0.79399 | 0.25182 |
| 6 | 68896200 | 68899399 | 44.39047 | 72.46574 | 58.37224 | 25.08543 | 0.684249 | 1.527554 | 0.251856 |
| 6 | 125325200 | 125327599 | 8.562517 | 15.21024 | 39.48841 | 17.42808 | -0.9737 | 0.455623 | 0.251858 |
| 4 | 136641400 | 136645199 | 23.73836 | 38.88519 | 78.81455 | 33.91432 | -0.64006 | 1.449569 | 0.251858 |
| 6 | 98197600 | 98199399 | 9.302059 | 16.95835 | 37.65975 | 16.17528 | -0.72435 | 0.294111 | 0.251859 |
| X | 81875800 | 81877399 | 20.16835 | 36.35852 | 24.18385 | 10.5339 | 0.991255 | 0.409638 | 0.251877 |
| 5 | 113808400 | 113811399 | 18.02435 | 30.42031 | 61.16815 | 26.7532 | -0.62281 | 1.085156 | 0.251886 |
| 8 | 129470000 | 129472999 | 71.4801 | 113.689 | 100.4447 | 41.9388 | 0.512972 | 2.247574 | 0.251899 |
| 18 | 73835800 | 73837399 | 11.04348 | 19.42537 | 42.52329 | 18.82809 | -0.73609 | 0.576257 | 0.251901 |
| 2 | 92266600 | 92267599 | 12.19522 | 23.05888 | 20.37575 | 8.274852 | 0.657824 | 0.058563 | 0.251908 |
| 15 | 47252400 | 47253199 | 10.77533 | 20.89216 | 17.54883 | 6.629963 | 0.825849 | -0.19465 | 0.25192 |
| 14 | 48895600 | 48900399 | 29.61402 | 47.99569 | 95.75727 | 40.99865 | -0.62928 | 1.63796 | 0.251924 |
| 17 | 30473000 | 30475199 | 8.940868 | 16.08296 | 36.93522 | 16.12963 | -0.78747 | 0.371793 | 0.25193 |
| 7 | 27991600 | 27998599 | 38.15242 | 60.79513 | 133.5846 | 55.9205 | -0.79629 | 2.164884 | 0.251932 |
| 1 | 156357000 | 156360199 | 29.09763 | 47.33999 | 87.80282 | 37.68659 | -0.51929 | 1.56246 | 0.251935 |
| 4 | 125124400 | 125126199 | 21.02776 | 36.77456 | 31.8341 | 14.03711 | 0.59466 | 0.698696 | 0.251941 |
| 11 | 113020200 | 113027799 | 64.15051 | 100.24 | 179.6208 | 74.52501 | -0.52681 | 2.650774 | 0.252018 |
| 1 | 24009000 | 24011799 | 29.58342 | 49.94233 | 43.14326 | 18.87116 | 0.591475 | 1.100833 | 0.252019 |
| 1 | 152927200 | 152928799 | 7.478398 | 13.79025 | 33.58957 | 14.20949 | -0.84028 | 0.20763 | 0.252031 |
| 5 | 98357600 | 98360999 | 20.43679 | 34.38442 | 66.87257 | 29.2205 | -0.57532 | 1.125525 | 0.252047 |
| X | 16245600 | 16248399 | 25.64718 | 45.38303 | 23.54734 | 10.41366 | 1.332145 | 0.563601 | 0.252054 |
| 6 | 14898000 | 14901999 | 32.55837 | 52.20854 | 115.2052 | 48.66448 | -0.79205 | 1.955869 | 0.252057 |
| 8 | 70980000 | 70980999 | 4.541223 | 8.789733 | 25.37079 | 9.662758 | -0.9474 | -0.21204 | 0.252064 |
| 7 | 14419000 | 14426599 | 154.1004 | 234.787 | 176.3972 | 73.11891 | 0.69774 | 3.153686 | 0.252076 |
| 13 | 52302000 | 52309399 | 47.53055 | 75.63491 | 134.2058 | 56.07374 | -0.49066 | 2.218856 | 0.252078 |
| 10 | 115220000 | 115221999 | 13.53403 | 23.22385 | 56.33866 | 24.71011 | -0.88946 | 0.911497 | 0.25209 |
| 15 | 54607400 | 54611599 | 20.09891 | 33.32812 | 74.31905 | 32.1872 | -0.7725 | 1.285424 | 0.252112 |
| 9 | 116805600 | 116807399 | 13.69028 | 23.85262 | 49.47357 | 21.75982 | -0.66038 | 0.74395 | 0.252124 |
| 6 | 125132200 | 125136199 | 15.50016 | 26.25698 | 61.09039 | 26.74915 | -0.83215 | 1.042737 | 0.252128 |
| 1 | 147331600 | 147332399 | 12.23921 | 23.24025 | 20.40765 | 8.213923 | 0.678112 | 0.030803 | 0.252151 |
| 17 | 56068200 | 56072999 | 34.02172 | 54.89216 | 102.5122 | 43.6884 | -0.54106 | 1.761226 | 0.252156 |
| 4 | 33516200 | 33518799 | 31.67161 | 52.83534 | 47.06743 | 20.46685 | 0.54736 | 1.227611 | 0.252163 |
| 9 | 57503600 | 57506999 | 13.53904 | 23.21631 | 55.24021 | 24.22697 | -0.86168 | 0.917859 | 0.252183 |
| 13 | 22743600 | 22744799 | 20.46292 | 35.85464 | 28.50855 | 12.59088 | 0.716757 | 0.661477 | 0.252184 |
| 13 | 115429400 | 115430199 | 5.979144 | 11.52489 | 27.55138 | 10.67439 | -0.70315 | -0.13187 | 0.252197 |
| 1 | 24382400 | 24384399 | 21.47647 | 38.08496 | 25.32361 | 11.19265 | 0.977365 | 0.516688 | 0.252204 |
| 13 | 40870200 | 40873599 | 20.15524 | 33.73261 | 65.56435 | 28.56843 | -0.5744 | 1.177082 | 0.252209 |
| 17 | 39910600 | 39917999 | 95.21044 | 149.914 | 115.8495 | 48.06741 | 0.693547 | 2.517033 | 0.252209 |
| 3 | 94190000 | 94192199 | 18.60668 | 34.28393 | 19.82619 | 8.396114 | 1.217479 | 0.266505 | 0.252217 |
| 7 | 38509800 | 38514199 | 86.46114 | 136.7061 | 109.8939 | 45.66056 | 0.641365 | 2.419344 | 0.252224 |
| 2 | 171092600 | 171093999 | 5.887042 | 11.16664 | 29.23934 | 11.79512 | -0.88661 | 0.005972 | 0.252225 |
| 7 | 22239200 | 22239599 | 9.589966 | 18.75983 | 8.743766 | 2.909557 | 1.82024 | -0.66732 | 0.252226 |
| 14 | 41984600 | 41985799 | 21.22395 | 37.54508 | 27.74763 | 12.27193 | 0.824805 | 0.546255 | 0.252246 |
| 6 | 90288000 | 90297999 | 112.0039 | 169.0038 | 288.1492 | 119.0867 | -0.49526 | 3.345564 | 0.25226 |
| 7 | 20843200 | 20844999 | 56.11706 | 89.96549 | 76.81596 | 32.43715 | 0.574615 | 1.978937 | 0.252261 |

| | | | | | | | | | |
|---|---|---|---|---|---|---|---|---|---|
| 7 | 78221000 | 78230199 | 66.27296 | 103.1307 | 193.8114 | 80.45508 | -0.60007 | 2.724703 | 0.252277 |
| X | 116662400 | 116664599 | 18.86036 | 34.88615 | 20.6874 | 8.714803 | 1.18774 | 0.251626 | 0.252279 |
| 5 | 125806400 | 125818199 | 132.9857 | 199.2557 | 328.2464 | 135.4052 | -0.45473 | 3.488407 | 0.252281 |
| 17 | 9167600 | 9170599 | 24.7762 | 40.34672 | 84.04501 | 36.08588 | -0.68521 | 1.538988 | 0.252285 |
| 5 | 3950000 | 3953799 | 56.03127 | 90.12017 | 70.49522 | 29.91268 | 0.706254 | 1.875293 | 0.252295 |
| 10 | 77044400 | 77045399 | 5.506852 | 10.42629 | 30.85006 | 12.4934 | -1.06725 | 0.015586 | 0.252317 |
| 2 | 109553000 | 109554999 | 19.4245 | 32.78229 | 64.09922 | 28.03489 | -0.58281 | 1.091505 | 0.25232 |
| 4 | 69964600 | 69965599 | 5.830424 | 11.27079 | 27.8027 | 10.65165 | -0.73334 | -0.17377 | 0.252325 |
| 3 | 83919000 | 83924199 | 41.13077 | 65.72542 | 121.6326 | 51.11806 | -0.5446 | 2.076466 | 0.252326 |
| 7 | 41427600 | 41431599 | 71.28225 | 113.5627 | 95.42394 | 39.93411 | 0.587764 | 2.193241 | 0.252331 |
| 3 | 48008400 | 48008999 | 10.88061 | 21.09501 | 17.49701 | 6.61225 | 0.843208 | -0.19333 | 0.252351 |
| 5 | 34442800 | 34447399 | 33.74393 | 54.32414 | 103.0712 | 43.79463 | -0.56764 | 1.828493 | 0.252361 |
| 6 | 90979800 | 90981199 | 7.857475 | 14.74472 | 32.34865 | 13.32137 | -0.66042 | 0.102213 | 0.252363 |
| 2 | 34912400 | 34913799 | 7.823578 | 14.28319 | 35.03014 | 15.0192 | -0.86413 | 0.271166 | 0.252375 |
| 18 | 21094200 | 21095999 | 7.823578 | 14.28319 | 35.03014 | 15.0192 | -0.86413 | 0.271166 | 0.252375 |
| 7 | 34616200 | 34626199 | 64.3168 | 100.0955 | 191.3329 | 79.42515 | -0.62446 | 2.722331 | 0.252383 |
| 16 | 68013200 | 68013599 | 9.826006 | 19.13574 | 16.01672 | 5.858906 | 0.873088 | -0.32063 | 0.252395 |
| 19 | 8717800 | 8719799 | 9.814147 | 17.27702 | 42.56653 | 18.84417 | -0.90473 | 0.561807 | 0.252396 |
| 18 | 18490600 | 18492599 | 28.45621 | 49.06116 | 35.86752 | 15.73427 | 0.83481 | 0.905439 | 0.252413 |
| 17 | 35904800 | 35909199 | 35.21852 | 56.5815 | 109.8772 | 46.54745 | -0.60485 | 1.897379 | 0.252417 |
| 13 | 66577200 | 66579999 | 81.76529 | 130.8285 | 68.05825 | 28.6555 | 1.284844 | 2.068587 | 0.252418 |
| 8 | 12721600 | 12722599 | 7.137343 | 13.29711 | 32.15674 | 13.40509 | -0.81401 | 0.141081 | 0.252422 |
| 7 | 5201200 | 5202999 | 25.65407 | 44.84449 | 30.25911 | 13.33739 | 0.952295 | 0.728166 | 0.252428 |
| X | 114728400 | 114730399 | 22.5782 | 40.85261 | 21.25325 | 9.214987 | 1.347425 | 0.400597 | 0.252443 |
| 17 | 73497000 | 73499399 | 13.39084 | 23.46477 | 46.72428 | 20.63713 | -0.60114 | 0.653398 | 0.252455 |
| 18 | 74071200 | 74072199 | 4.68967 | 9.112214 | 25.14459 | 9.373218 | -0.85486 | -0.28857 | 0.252495 |
| 8 | 37014400 | 37016799 | 13.402 | 23.38749 | 47.32827 | 20.83706 | -0.6245 | 0.719014 | 0.25251 |
| 2 | 109503800 | 109508799 | 50.72314 | 80.33066 | 147.48 | 61.32802 | -0.54648 | 2.363466 | 0.252535 |
| 6 | 113207200 | 113208799 | 22.95427 | 40.07306 | 32.69926 | 14.40004 | 0.679382 | 0.735785 | 0.252559 |
| 15 | 20435800 | 20436799 | 7.525655 | 14.2528 | 32.69108 | 13.23063 | -0.70294 | 0.038078 | 0.252562 |
| 11 | 53363600 | 53365999 | 15.60008 | 27.01579 | 53.84164 | 23.62231 | -0.60455 | 0.833464 | 0.252563 |
| 4 | 46363000 | 46371399 | 51.54318 | 81.54424 | 153.1801 | 63.66472 | -0.58074 | 2.389081 | 0.252566 |
| 13 | 15558200 | 15561199 | 27.77997 | 47.75832 | 36.42699 | 15.97831 | 0.773008 | 0.926573 | 0.252568 |
| 7 | 85280000 | 85281599 | 35.32314 | 58.61616 | 50.4125 | 21.84449 | 0.597727 | 1.299257 | 0.252573 |
| 3 | 82528400 | 82529799 | 16.52352 | 29.76607 | 25.58709 | 11.15529 | 0.622536 | 0.389628 | 0.252592 |
| 1 | 78464800 | 78466399 | 7.849477 | 14.66347 | 33.35023 | 13.83905 | -0.72139 | 0.130914 | 0.252605 |
| 7 | 144712200 | 144715599 | 17.382 | 29.3438 | 63.39296 | 27.72845 | -0.72604 | 1.08091 | 0.252609 |
| 7 | 105334800 | 105337599 | 15.26821 | 26.03933 | 60.15507 | 26.37095 | -0.82096 | 0.978379 | 0.252629 |
| 16 | 58432800 | 58433399 | 3.936394 | 7.681056 | 21.49131 | 7.656577 | -0.81247 | -0.46974 | 0.25263 |
| 17 | 30970000 | 30972799 | 17.10278 | 28.8153 | 62.1698 | 27.17711 | -0.72414 | 1.097794 | 0.252647 |
| 7 | 127531800 | 127533799 | 10.20611 | 18.15274 | 39.39252 | 17.3673 | -0.71632 | 0.455771 | 0.252655 |
| 18 | 83353200 | 83354399 | 6.042675 | 11.48725 | 29.32718 | 11.7753 | -0.84539 | -0.0111 | 0.252658 |
| 17 | 32384600 | 32386599 | 10.28999 | 18.08061 | 43.07154 | 19.06788 | -0.8577 | 0.586357 | 0.252659 |
| 15 | 98901600 | 98904799 | 26.03005 | 42.43156 | 83.32403 | 35.78954 | -0.59935 | 1.525856 | 0.252659 |
| 1 | 124568400 | 124572999 | 73.12974 | 115.9255 | 104.6293 | 43.53826 | 0.477589 | 2.346711 | 0.252669 |
| 14 | 22227400 | 22228399 | 11.85415 | 22.80038 | 18.98273 | 7.399775 | 0.796192 | -0.0776 | 0.252669 |
| 5 | 60979000 | 60979799 | 8.047128 | 14.89494 | 33.11766 | 13.9383 | -0.70445 | 0.189704 | 0.25267 |

| | | | | | | | | | |
|---|---|---|---|---|---|---|---|---|---|
| 6 | 5866600 | 5868199 | 8.047128 | 14.89494 | 33.11766 | 13.9383 | -0.70445 | 0.189704 | 0.25267 |
| 1 | 190713400 | 190722599 | 74.26576 | 114.6617 | 220.8867 | 91.75189 | -0.64463 | 2.872515 | 0.252679 |
| 10 | 16962400 | 16964399 | 12.40301 | 21.80478 | 45.06494 | 19.95565 | -0.65449 | 0.591348 | 0.252693 |
| 12 | 44808200 | 44814999 | 70.43031 | 111.9953 | 98.87949 | 41.27438 | 0.513632 | 2.250695 | 0.252693 |
| 1 | 3872400 | 3875999 | 53.56073 | 86.30595 | 72.50293 | 30.84712 | 0.606826 | 1.815057 | 0.252696 |
| 4 | 62936600 | 62944999 | 54.48537 | 85.71095 | 167.3867 | 69.44247 | -0.64321 | 2.518236 | 0.252698 |
| 13 | 3241600 | 3243599 | 36.00723 | 59.33591 | 53.16488 | 22.93503 | 0.537503 | 1.383524 | 0.252704 |
| X | 4944000 | 4946199 | 23.48206 | 41.66727 | 24.83757 | 10.97211 | 1.134966 | 0.527317 | 0.252706 |
| 6 | 87444400 | 87448999 | 33.37778 | 53.94185 | 99.78786 | 42.61765 | -0.52464 | 1.712214 | 0.252707 |
| 15 | 99076200 | 99078999 | 14.4953 | 25.24151 | 49.48155 | 21.7559 | -0.5794 | 0.75695 | 0.252734 |
| 6 | 32745600 | 32748799 | 18.64132 | 31.46915 | 61.79223 | 27.02812 | -0.58875 | 1.085002 | 0.252739 |
| 4 | 25848600 | 25849599 | 14.78737 | 27.91882 | 19.77661 | 8.056703 | 0.971114 | 0.093206 | 0.252744 |
| 6 | 56987800 | 56989799 | 24.13293 | 42.21471 | 30.88443 | 13.621 | 0.836129 | 0.709911 | 0.252756 |
| 13 | 85326000 | 85327999 | 23.91677 | 41.53985 | 34.96404 | 15.34786 | 0.634284 | 0.814819 | 0.252757 |
| 7 | 28195000 | 28198599 | 24.35533 | 39.95664 | 77.93396 | 33.55702 | -0.58395 | 1.431152 | 0.252774 |
| 7 | 85349800 | 85352199 | 27.0923 | 47.18786 | 31.99606 | 14.0699 | 0.945139 | 0.788913 | 0.25278 |
| 7 | 106167000 | 106169399 | 39.06604 | 63.8108 | 56.90673 | 24.46079 | 0.538399 | 1.506255 | 0.252789 |
| 2 | 25261200 | 25263999 | 12.53426 | 21.35647 | 58.26452 | 25.54022 | -1.05943 | 0.978197 | 0.252792 |
| X | 9818800 | 9820199 | 16.52418 | 31.3491 | 17.97894 | 7.249868 | 1.285953 | 0.072333 | 0.252792 |
| X | 38270600 | 38271599 | 6.726977 | 12.89812 | 30.9505 | 12.17438 | -0.73047 | -0.0689 | 0.252794 |
| 1 | 33708400 | 33709799 | 10.08406 | 17.85814 | 40.42223 | 17.87362 | -0.78014 | 0.489867 | 0.252796 |
| 3 | 33838800 | 33839599 | 10.76398 | 20.92533 | 16.44397 | 6.105062 | 0.943781 | -0.25734 | 0.252799 |
| 5 | 89475200 | 89478199 | 47.52478 | 75.64164 | 133.7927 | 55.91741 | -0.48576 | 2.211895 | 0.252801 |
| 1 | 34404400 | 34405799 | 20.19635 | 35.71823 | 28.37566 | 12.55032 | 0.72097 | 0.543395 | 0.252812 |
| 1 | 181987200 | 181987799 | 9.37963 | 18.18727 | 16.53536 | 6.245121 | 0.712679 | -0.21656 | 0.25282 |
| 4 | 62118400 | 62120399 | 50.29945 | 81.00172 | 73.49218 | 31.24056 | 0.494947 | 1.829635 | 0.252833 |
| 14 | 99205000 | 99206799 | 24.10345 | 42.04454 | 33.26113 | 14.63988 | 0.723929 | 0.755406 | 0.252838 |
| 8 | 102553400 | 102554399 | 17.38315 | 30.98641 | 26.64225 | 11.7241 | 0.613286 | 0.462355 | 0.252842 |
| 11 | 21725200 | 21730199 | 86.4677 | 136.1506 | 121.1783 | 50.27851 | 0.490026 | 2.512075 | 0.252848 |
| 13 | 94846000 | 94847599 | 14.7128 | 25.34288 | 53.74358 | 23.57544 | -0.69526 | 0.878543 | 0.252868 |
| 8 | 108707200 | 108709599 | 10.49553 | 18.72544 | 38.88638 | 17.09942 | -0.65061 | 0.437358 | 0.252879 |
| 2 | 113355400 | 113356199 | 5.817756 | 11.27209 | 27.3839 | 10.37955 | -0.69631 | -0.20998 | 0.252885 |
| 5 | 107996000 | 107999199 | 19.11871 | 32.4011 | 70.38374 | 30.82249 | -0.73391 | 1.060896 | 0.252889 |
| 11 | 114142600 | 114146599 | 24.79155 | 40.34176 | 85.79969 | 36.82931 | -0.71592 | 1.549864 | 0.252889 |
| 6 | 63966200 | 63967199 | 15.51241 | 29.12149 | 19.81818 | 8.155493 | 1.015643 | 0.132528 | 0.252903 |
| 17 | 29634800 | 29636399 | 7.896247 | 14.2277 | 36.98069 | 16.11819 | -0.96276 | 0.354164 | 0.252903 |
| 7 | 53017000 | 53017999 | 14.60654 | 26.8749 | 22.82725 | 9.685891 | 0.665678 | 0.251201 | 0.252913 |
| 15 | 89510800 | 89511399 | 14.60654 | 26.8749 | 22.82725 | 9.685891 | 0.665678 | 0.251201 | 0.252913 |
| 8 | 34852000 | 34853799 | 9.173345 | 16.65904 | 37.56794 | 16.21421 | -0.75087 | 0.315668 | 0.252921 |
| 7 | 17014400 | 17017999 | 39.62842 | 65.69242 | 44.98174 | 19.47734 | 0.925433 | 1.319409 | 0.252921 |
| 10 | 28806200 | 28806799 | 9.309455 | 18.1082 | 16.5896 | 6.130412 | 0.730773 | -0.29098 | 0.252925 |
| 6 | 26734200 | 26734999 | 14.41176 | 27.28632 | 19.50316 | 7.898171 | 0.964639 | 0.06768 | 0.252929 |
| 1 | 69319000 | 69320399 | 8.012351 | 14.98114 | 41.00486 | 16.98966 | -0.98546 | 0.142537 | 0.25295 |
| 7 | 8548200 | 8549799 | 17.26521 | 32.89317 | 15.93456 | 6.368521 | 1.53698 | 0.045491 | 0.25295 |
| 5 | 5133800 | 5135199 | 14.98116 | 27.41515 | 23.66513 | 10.11575 | 0.634681 | 0.284652 | 0.252956 |
| 13 | 61245400 | 61247199 | 22.29377 | 39.49713 | 25.8931 | 11.44827 | 0.997345 | 0.532917 | 0.252965 |
| 12 | 18705000 | 18708399 | 129.2324 | 197.6398 | 176.0277 | 73.03088 | 0.456757 | 3.071188 | 0.252972 |

| | | | | | | | | | |
|---|---|---|---|---|---|---|---|---|---|
| 7 | 37539800 | 37542399 | 15.1714 | 26.17238 | 52.69043 | 23.11452 | -0.62008 | 0.866074 | 0.252972 |
| 8 | 95600800 | 95603199 | 12.1942 | 21.199 | 49.45958 | 21.72341 | -0.82932 | 0.770212 | 0.252975 |
| 4 | 46450400 | 46454199 | 15.65364 | 26.51594 | 60.54724 | 26.51101 | -0.8052 | 1.043401 | 0.252977 |
| X | 8526200 | 8532199 | 125.1804 | 193.5763 | 141.6954 | 58.85692 | 0.75172 | 2.856912 | 0.25298 |
| 10 | 53278400 | 53278799 | 7.816919 | 15.28798 | 12.19052 | 4.135668 | 1.03677 | -0.6205 | 0.252981 |
| Y | 5419800 | 5420999 | 10.27736 | 19.94418 | 9.793703 | 3.684895 | 1.595364 | -0.24374 | 0.253022 |
| X | 21234400 | 21236599 | 26.42491 | 46.53443 | 25.34966 | 11.22153 | 1.260155 | 0.622458 | 0.253029 |
| 4 | 37603200 | 37604799 | 30.95452 | 53.42112 | 33.60597 | 14.74262 | 1.05098 | 0.906412 | 0.253035 |
| 17 | 46900200 | 46901999 | 5.247853 | 9.873654 | 31.22221 | 12.80033 | -1.17697 | 0.058553 | 0.253066 |
| 19 | 12771000 | 12774199 | 18.56128 | 31.09512 | 66.62698 | 29.04928 | -0.71458 | 1.160979 | 0.253074 |
| 15 | 52335000 | 52337399 | 27.14795 | 47.2565 | 31.94657 | 14.04285 | 0.9493 | 0.796451 | 0.253074 |
| 17 | 13793400 | 13798399 | 36.34054 | 58.37123 | 112.029 | 47.44367 | -0.58837 | 1.906238 | 0.253075 |
| 5 | 115413000 | 115415199 | 14.16479 | 24.68858 | 48.93798 | 21.5293 | -0.59521 | 0.740942 | 0.253079 |
| 6 | 121604400 | 121604799 | 3.524273 | 6.889339 | 20.09968 | 6.910554 | -0.82484 | -0.60825 | 0.253081 |
| 11 | 53327600 | 53331999 | 24.50808 | 40.18838 | 79.21397 | 34.1014 | -0.5994 | 1.436707 | 0.253083 |
| 4 | 73185800 | 73189199 | 51.29238 | 82.80979 | 69.61552 | 29.69278 | 0.608966 | 1.75542 | 0.253111 |
| 11 | 73562800 | 73573799 | 356.2282 | 503.4276 | 417.359 | 172.7665 | 0.453973 | 4.424162 | 0.253112 |
| 2 | 28530200 | 28532199 | 7.980791 | 14.53275 | 35.01313 | 15.06031 | -0.84134 | 0.289455 | 0.253125 |
| 13 | 112382600 | 112386399 | 50.20274 | 81.23941 | 65.70296 | 28.09908 | 0.668296 | 1.691432 | 0.253137 |
| 2 | 169832000 | 169833599 | 11.48998 | 20.2235 | 42.48806 | 18.81012 | -0.67698 | 0.570452 | 0.253158 |
| 14 | 90386000 | 90387999 | 26.06826 | 45.33003 | 30.79077 | 13.52597 | 0.942474 | 0.80506 | 0.253163 |
| 10 | 54565600 | 54569399 | 46.67314 | 76.16664 | 56.82679 | 24.41829 | 0.794172 | 1.53593 | 0.253179 |
| 9 | 23170600 | 23171999 | 22.42935 | 39.32995 | 28.09383 | 12.41566 | 0.870448 | 0.659484 | 0.253201 |
| 17 | 65296000 | 65297599 | 16.63949 | 29.97131 | 25.51774 | 11.12676 | 0.636058 | 0.390499 | 0.25321 |
| 5 | 131147800 | 131149599 | 19.8674 | 35.00697 | 29.59355 | 13.09865 | 0.629878 | 0.583219 | 0.253211 |
| 10 | 61555200 | 61556999 | 11.20839 | 19.67182 | 43.59155 | 19.27946 | -0.75412 | 0.611029 | 0.25323 |
| 14 | 37930600 | 37931999 | 14.61635 | 27.59934 | 20.03038 | 8.157956 | 0.935826 | 0.089081 | 0.253231 |
| 7 | 32323800 | 32328799 | 69.40695 | 111.2437 | 76.15749 | 32.14689 | 0.892274 | 2.003248 | 0.253244 |
| 7 | 111088400 | 111091799 | 21.12325 | 35.25985 | 67.85389 | 29.51704 | -0.56014 | 1.209666 | 0.253247 |
| 11 | 21956600 | 21958599 | 16.78984 | 28.7051 | 57.53624 | 25.22791 | -0.61642 | 0.959398 | 0.253258 |
| 3 | 130069600 | 130070999 | 15.62426 | 29.17374 | 21.61169 | 8.97513 | 0.884481 | 0.17275 | 0.253263 |
| 7 | 5411200 | 5414599 | 64.35392 | 103.0375 | 84.08637 | 35.43756 | 0.637573 | 2.030495 | 0.253286 |
| 1 | 140760400 | 140761999 | 17.26119 | 30.78547 | 26.02955 | 11.44931 | 0.637578 | 0.455712 | 0.253293 |
| 13 | 62427200 | 62433599 | 157.4112 | 238.1881 | 204.3961 | 84.54807 | 0.497948 | 3.294172 | 0.253305 |
| 13 | 65449200 | 65455799 | 180.7908 | 272.0094 | 214.6861 | 88.65401 | 0.611473 | 3.412717 | 0.253313 |
| 4 | 112045600 | 112046799 | 21.89854 | 38.60847 | 28.15236 | 12.45929 | 0.842181 | 0.585494 | 0.253316 |
| 7 | 56432200 | 56433399 | 14.1867 | 24.66467 | 49.48998 | 21.73776 | -0.61305 | 0.778311 | 0.25333 |
| 17 | 12377600 | 12385799 | 51.52125 | 81.57381 | 150.5833 | 62.61043 | -0.55467 | 2.369945 | 0.253339 |
| 9 | 67701800 | 67704799 | 12.91239 | 22.66358 | 45.10861 | 19.95121 | -0.60039 | 0.621775 | 0.25335 |
| 5 | 125004600 | 125007199 | 22.85118 | 37.2823 | 82.79945 | 35.57557 | -0.77535 | 1.506606 | 0.253352 |
| 3 | 82345000 | 82345799 | 10.40336 | 20.22723 | 16.4556 | 6.10219 | 0.895647 | -0.26613 | 0.253353 |
| 15 | 99428000 | 99430999 | 16.39991 | 27.88335 | 57.90722 | 25.37923 | -0.66845 | 1.012 | 0.253356 |
| 12 | 107497400 | 107498399 | 16.98477 | 31.11205 | 23.54938 | 10.05292 | 0.824664 | 0.29611 | 0.253364 |
| 16 | 25016400 | 25016999 | 5.382605 | 10.43547 | 25.56499 | 9.661926 | -0.7036 | -0.22913 | 0.253412 |
| 17 | 71551800 | 71552599 | 5.382605 | 10.43547 | 25.56499 | 9.661926 | -0.7036 | -0.22913 | 0.253412 |
| 17 | 34482200 | 34485399 | 16.71137 | 28.38416 | 58.46938 | 25.62218 | -0.65703 | 1.022004 | 0.253422 |
| 5 | 38464000 | 38468599 | 26.5225 | 43.17138 | 87.51634 | 37.57028 | -0.6467 | 1.548795 | 0.253424 |

| | | | | | | | | | |
|---|---|---|---|---|---|---|---|---|---|
| 7 | 11094800 | 11098399 | 55.02275 | 88.44346 | 77.59363 | 32.89525 | 0.540042 | 1.89707 | 0.253435 |
| 14 | 13761800 | 13762399 | 9.357994 | 18.25256 | 14.94254 | 5.356195 | 0.930341 | -0.40041 | 0.253441 |
| 2 | 87305600 | 87306599 | 5.96939 | 11.54849 | 29.14193 | 11.12385 | -0.76096 | -0.18133 | 0.253447 |
| 5 | 14713400 | 14717199 | 41.2081 | 67.68259 | 54.99612 | 23.6919 | 0.676306 | 1.438761 | 0.253457 |
| X | 146567400 | 146568799 | 29.06651 | 49.36235 | 42.03511 | 18.41964 | 0.612421 | 1.049214 | 0.25348 |
| 14 | 106329600 | 106333599 | 40.26929 | 64.59204 | 115.1402 | 48.6585 | -0.48441 | 1.956859 | 0.253486 |
| 4 | 78981200 | 78982399 | 15.8795 | 29.07047 | 24.10439 | 10.29803 | 0.693009 | 0.291099 | 0.253491 |
| 13 | 70619600 | 70620999 | 15.8795 | 29.07047 | 24.10439 | 10.29803 | 0.693009 | 0.291099 | 0.253491 |
| 3 | 48541600 | 48542999 | 9.850571 | 17.30067 | 44.10723 | 19.51892 | -0.95513 | 0.591912 | 0.253503 |
| 18 | 52454800 | 52456399 | 12.58388 | 22.10805 | 44.03808 | 19.49341 | -0.60151 | 0.602273 | 0.253548 |
| 14 | 63427600 | 63431999 | 20.60193 | 34.08861 | 71.79657 | 31.04845 | -0.69052 | 1.312722 | 0.25355 |
| 2 | 114980800 | 114981999 | 12.31477 | 23.38011 | 20.45705 | 8.236508 | 0.683061 | 0.033483 | 0.253562 |
| 16 | 90840400 | 90844199 | 20.58409 | 34.09535 | 72.53376 | 31.3903 | -0.70488 | 1.299214 | 0.253576 |
| 9 | 87308000 | 87309799 | 10.91689 | 19.12649 | 44.6756 | 19.72952 | -0.82962 | 0.640757 | 0.253579 |
| 3 | 28859400 | 28859999 | 6.146309 | 11.78129 | 27.90296 | 10.98491 | -0.71178 | -0.0772 | 0.25359 |
| 5 | 58573200 | 58573799 | 6.146309 | 11.78129 | 27.90296 | 10.98491 | -0.71178 | -0.0772 | 0.25359 |
| 13 | 61616400 | 61620199 | 49.26791 | 80.44562 | 54.07378 | 23.24003 | 0.944822 | 1.533518 | 0.253595 |
| 19 | 42158800 | 42159599 | 6.771226 | 12.8518 | 29.84868 | 12.02502 | -0.71415 | 0.01073 | 0.253602 |
| 14 | 14769200 | 14770799 | 23.54293 | 40.72399 | 35.74906 | 15.68388 | 0.572997 | 0.855158 | 0.253616 |
| X | 147386000 | 147393199 | 60.58789 | 97.52717 | 70.46693 | 29.93544 | 0.822002 | 1.859734 | 0.253617 |
| X | 32392800 | 32395799 | 23.21664 | 40.88855 | 27.86289 | 12.33383 | 0.939146 | 0.607015 | 0.253617 |
| 7 | 119319400 | 119320199 | 3.894453 | 7.548291 | 27.15841 | 10.27711 | -1.25335 | -0.23921 | 0.253627 |
| 9 | 19114600 | 19123199 | 126.5919 | 194.7646 | 159.6086 | 66.2854 | 0.583386 | 2.951482 | 0.253638 |
| 16 | 4994000 | 4995199 | 8.353017 | 15.5173 | 33.73097 | 14.12655 | -0.66683 | 0.174503 | 0.253651 |
| 9 | 6392600 | 6395599 | 53.01884 | 85.49174 | 62.98351 | 26.82142 | 0.796953 | 1.792155 | 0.253653 |
| 8 | 115842600 | 115844799 | 16.99555 | 28.93235 | 57.99774 | 25.42147 | -0.61749 | 1.00108 | 0.253662 |
| 12 | 78320000 | 78321999 | 19.80153 | 35.43389 | 25.63031 | 11.24212 | 0.86419 | 0.452412 | 0.253666 |
| 2 | 162767600 | 162768999 | 14.17268 | 26.64707 | 21.17693 | 8.69214 | 0.796937 | 0.115299 | 0.253671 |
| 1 | 147206200 | 147210999 | 62.27994 | 99.81872 | 81.87529 | 34.55937 | 0.631947 | 1.995089 | 0.253672 |
| 3 | 159007400 | 159009199 | 40.07877 | 65.33606 | 58.53721 | 25.14356 | 0.529771 | 1.537687 | 0.253672 |
| 9 | 3506000 | 3507799 | 23.05186 | 40.61099 | 27.50005 | 12.17245 | 0.947947 | 0.599331 | 0.25368 |
| 12 | 25805800 | 25807399 | 13.69626 | 23.84845 | 49.47763 | 21.75323 | -0.66083 | 0.752902 | 0.253687 |
| 7 | 139868000 | 139869199 | 12.2204 | 23.58829 | 17.7784 | 6.858494 | 0.951817 | -0.111 | 0.253709 |
| 14 | 52434200 | 52435199 | 4.849892 | 9.329836 | 26.82329 | 10.45128 | -0.97358 | -0.12738 | 0.253746 |
| 15 | 30171800 | 30173999 | 15.20698 | 26.13619 | 53.72495 | 23.56567 | -0.65111 | 0.901872 | 0.25375 |
| 3 | 73285400 | 73287399 | 20.66807 | 34.45007 | 67.72551 | 29.43427 | -0.59075 | 1.2243 | 0.25376 |
| 5 | 93285800 | 93287599 | 24.05007 | 41.69012 | 36.14347 | 15.85795 | 0.591504 | 0.839165 | 0.253765 |
| 16 | 16211400 | 16211799 | 9.23933 | 18.05123 | 13.22943 | 4.612487 | 1.122922 | -0.50186 | 0.253765 |
| 4 | 122668800 | 122671399 | 29.59224 | 50.59833 | 38.62827 | 16.93748 | 0.770647 | 0.985898 | 0.253766 |
| 3 | 41176600 | 41179799 | 49.88485 | 80.40497 | 71.50482 | 30.43395 | 0.525283 | 1.800023 | 0.253769 |
| 15 | 103271600 | 103275599 | 16.25946 | 27.57878 | 58.95873 | 25.82588 | -0.71041 | 1.033152 | 0.253776 |
| 16 | 11752600 | 11754399 | 8.724791 | 15.90239 | 35.0006 | 15.03736 | -0.71107 | 0.288688 | 0.253785 |
| 6 | 39496400 | 39497799 | 24.44718 | 42.75163 | 31.50845 | 13.8927 | 0.825719 | 0.718485 | 0.253787 |
| 1 | 4560800 | 4563999 | 27.43993 | 44.37114 | 93.7538 | 40.07025 | -0.71483 | 1.682651 | 0.253796 |
| 10 | 82274200 | 82280399 | 157.9264 | 239.509 | 194.157 | 80.36952 | 0.582756 | 3.248035 | 0.253796 |
| 2 | 157869800 | 157872199 | 13.4714 | 23.27534 | 51.57531 | 22.62639 | -0.75706 | 0.844876 | 0.253806 |
| 6 | 29609000 | 29609999 | 6.52178 | 12.24755 | 30.68561 | 12.6203 | -0.84841 | 0.081128 | 0.25382 |

| | | | | | | | | | |
|---|---|---|---|---|---|---|---|---|---|
| 12 | 72244800 | 72249999 | 34.11445 | 54.78138 | 109.7559 | 46.46266 | -0.65061 | 1.911536 | 0.253832 |
| 2 | 121391200 | 121393799 | 14.97432 | 25.93292 | 52.18851 | 22.89705 | -0.61841 | 0.828564 | 0.253843 |
| 7 | 53293600 | 53295599 | 22.65902 | 40.17919 | 25.32063 | 11.19199 | 1.05447 | 0.526046 | 0.253844 |
| 9 | 91657400 | 91663199 | 61.77585 | 96.87745 | 188.7654 | 78.29014 | -0.64349 | 2.588288 | 0.253854 |
| 2 | 127850200 | 127851399 | 7.523834 | 14.16084 | 32.45997 | 13.29983 | -0.7177 | 0.077699 | 0.253855 |
| 9 | 51014600 | 51015799 | 7.523834 | 14.16084 | 32.45997 | 13.29983 | -0.7177 | 0.077699 | 0.253855 |
| 11 | 120536400 | 120539399 | 12.99432 | 22.15202 | 58.32088 | 25.56601 | -1.00834 | 0.974057 | 0.253855 |
| 14 | 61834800 | 61837199 | 13.8901 | 24.16743 | 48.94511 | 21.50873 | -0.62615 | 0.766395 | 0.253856 |
| 13 | 49572800 | 49575399 | 16.34806 | 27.91845 | 56.38362 | 24.72028 | -0.62739 | 0.968025 | 0.253861 |
| 17 | 35276600 | 35280799 | 18.21882 | 30.23418 | 70.53205 | 30.56356 | -0.83699 | 1.270747 | 0.253863 |
| 2 | 21372200 | 21374599 | 20.16461 | 33.28349 | 73.85404 | 31.88431 | -0.76581 | 1.341608 | 0.253879 |
| 7 | 8466000 | 8466999 | 12.15887 | 23.38373 | 18.27013 | 7.124396 | 0.8859 | -0.07689 | 0.253879 |
| 1 | 49911200 | 49913199 | 12.75093 | 22.44813 | 48.82216 | 21.61332 | -0.72695 | 0.58503 | 0.253882 |
| 5 | 147829600 | 147832799 | 20.1124 | 33.3225 | 72.48049 | 31.37274 | -0.73674 | 1.294966 | 0.25389 |
| 7 | 20904400 | 20908799 | 34.47617 | 58.89888 | 35.21144 | 15.43823 | 1.121855 | 1.010929 | 0.25389 |
| 7 | 27659800 | 27661199 | 28.06573 | 47.98863 | 39.84484 | 17.47092 | 0.650087 | 0.982383 | 0.253891 |
| 3 | 38508000 | 38512599 | 82.43893 | 125.8685 | 255.9241 | 106.1379 | -0.73491 | 3.097825 | 0.253893 |
| 7 | 67834200 | 67835599 | 10.22907 | 18.13162 | 39.89834 | 17.63872 | -0.73909 | 0.479887 | 0.253897 |
| 5 | 135694400 | 135696399 | 23.37431 | 40.80188 | 32.09027 | 14.13088 | 0.73217 | 0.737919 | 0.253911 |
| 17 | 6371000 | 6371799 | 3.530562 | 6.883702 | 21.89482 | 7.886882 | -1.00877 | -0.43367 | 0.253921 |
| 12 | 4818800 | 4819599 | 6.184379 | 11.72604 | 29.24553 | 11.8074 | -0.81842 | 0.010888 | 0.253931 |
| 12 | 33423400 | 33424399 | 6.184379 | 11.72604 | 29.24553 | 11.8074 | -0.81842 | 0.010888 | 0.253931 |
| 6 | 90160400 | 90162399 | 23.94854 | 42.07812 | 28.76569 | 12.73413 | 0.933441 | 0.638907 | 0.253936 |
| 13 | 65903400 | 65911999 | 135.6112 | 209.9195 | 127.4021 | 52.91956 | 1.022861 | 2.841533 | 0.253966 |
| 16 | 26736000 | 26739199 | 29.2506 | 49.76632 | 40.87446 | 17.9152 | 0.664601 | 1.030885 | 0.253969 |
| 19 | 38111200 | 38113799 | 16.64059 | 28.45831 | 56.42898 | 24.74299 | -0.60077 | 0.955584 | 0.253971 |
| 1 | 49644000 | 49645799 | 21.38419 | 37.76898 | 26.4532 | 11.70313 | 0.90062 | 0.558938 | 0.253973 |
| 11 | 63556000 | 63557999 | 15.40546 | 28.3402 | 20.61458 | 8.748981 | 0.887287 | 0.252061 | 0.253976 |
| 10 | 85567400 | 85568199 | 4.980436 | 9.578984 | 26.8192 | 10.4559 | -0.93644 | -0.12336 | 0.254 |
| 11 | 13157600 | 13159199 | 19.01562 | 34.41447 | 24.18682 | 10.4839 | 0.916544 | 0.378605 | 0.254008 |
| 8 | 66991400 | 66993999 | 20.01396 | 36.41584 | 22.63146 | 9.744207 | 1.099103 | 0.353923 | 0.254013 |
| 14 | 85240000 | 85242999 | 36.68821 | 60.48425 | 53.75003 | 23.1917 | 0.54958 | 1.380718 | 0.254021 |
| 6 | 115056000 | 115056999 | 7.37532 | 13.95095 | 30.90276 | 12.53936 | -0.65611 | 0.041299 | 0.254024 |
| 1 | 103487800 | 103489799 | 25.76811 | 44.58613 | 34.8409 | 15.28559 | 0.73997 | 0.858989 | 0.254036 |
| 13 | 70155400 | 70156799 | 8.224366 | 15.14462 | 33.56628 | 14.22668 | -0.70719 | 0.22232 | 0.254051 |
| 16 | 92294400 | 92296599 | 18.4225 | 30.82999 | 65.98332 | 28.74945 | -0.71285 | 1.172244 | 0.254057 |
| 7 | 86238400 | 86239199 | 12.20915 | 23.29369 | 19.87944 | 7.921236 | 0.731403 | -0.00318 | 0.254065 |
| X | 96137600 | 96139399 | 21.61772 | 38.51503 | 25.56168 | 11.25344 | 0.984243 | 0.492012 | 0.254066 |
| 8 | 69856400 | 69858199 | 20.12756 | 35.39259 | 30.07502 | 13.31722 | 0.621271 | 0.606985 | 0.25407 |
| 6 | 73264400 | 73269599 | 74.66542 | 115.0839 | 211.7244 | 87.94684 | -0.58012 | 2.903365 | 0.254073 |
| 4 | 112660200 | 112662799 | 47.37202 | 77.1628 | 58.48494 | 25.11368 | 0.769639 | 1.564899 | 0.254074 |
| 14 | 122640000 | 122646199 | 49.48121 | 78.63076 | 141.0866 | 58.85343 | -0.5088 | 2.263451 | 0.254075 |
| 8 | 14688000 | 14690199 | 10.20255 | 18.13805 | 40.92362 | 18.04934 | -0.7728 | 0.460044 | 0.254113 |
| 1 | 155292400 | 155293399 | 7.041224 | 12.99404 | 33.60115 | 14.19959 | -0.92467 | 0.199767 | 0.254117 |
| 11 | 3487000 | 3495999 | 59.46071 | 93.23864 | 174.5278 | 72.38578 | -0.58573 | 2.584872 | 0.254126 |
| 7 | 24141200 | 24147599 | 45.94909 | 72.99056 | 138.9994 | 57.96106 | -0.59501 | 2.271771 | 0.254132 |
| 13 | 36008800 | 36009799 | 7.220628 | 13.66505 | 30.3533 | 12.30362 | -0.65871 | 0.034817 | 0.254136 |

| | | | | | | | | | |
|---|---|---|---|---|---|---|---|---|---|
| 8 | 124261000 | 124263999 | 11.27448 | 19.63519 | 48.91084 | 21.50654 | -0.92288 | 0.739006 | 0.254145 |
| 18 | 55123600 | 55126199 | 10.4606 | 18.72358 | 39.94302 | 17.51624 | -0.68675 | 0.416823 | 0.254169 |
| 3 | 145674400 | 145677599 | 13.56401 | 23.56029 | 50.55544 | 22.19285 | -0.7096 | 0.789142 | 0.254215 |
| 1 | 13442200 | 13445199 | 23.34091 | 38.05852 | 84.67448 | 36.37326 | -0.77862 | 1.516333 | 0.254215 |
| 15 | 83131800 | 83136999 | 42.79642 | 68.18864 | 132.0839 | 55.28488 | -0.61495 | 2.171843 | 0.254215 |
| 17 | 88563400 | 88564999 | 8.241992 | 15.10462 | 34.54905 | 14.73888 | -0.75937 | 0.250838 | 0.254221 |
| 9 | 26054400 | 26055999 | 20.58017 | 36.52145 | 26.56742 | 11.73536 | 0.849218 | 0.509561 | 0.254225 |
| 14 | 92275000 | 92278599 | 39.14084 | 64.31107 | 55.39659 | 23.86817 | 0.592601 | 1.430635 | 0.254236 |
| X | 26751600 | 26766799 | 56.29221 | 90.96454 | 57.27534 | 24.45806 | 1.026468 | 1.735274 | 0.25425 |
| 12 | 88000200 | 88003399 | 62.64821 | 100.3763 | 75.9237 | 32.03137 | 0.748052 | 2.002618 | 0.254256 |
| 3 | 110698800 | 110699999 | 7.991702 | 14.95728 | 33.4182 | 13.82314 | -0.69221 | 0.120312 | 0.254256 |
| 12 | 23658000 | 23659999 | 17.14729 | 31.13612 | 23.75181 | 10.25278 | 0.802521 | 0.343882 | 0.254261 |
| 4 | 38066400 | 38069799 | 38.48073 | 61.8476 | 112.5653 | 47.71423 | -0.51084 | 1.891522 | 0.254264 |
| 3 | 121352400 | 121357199 | 35.1813 | 56.62011 | 107.2035 | 45.52943 | -0.56528 | 1.841168 | 0.254266 |
| 11 | 121433000 | 121436599 | 16.00328 | 27.00761 | 62.13021 | 27.17347 | -0.81616 | 1.079072 | 0.254278 |
| 9 | 121156600 | 121159599 | 28.00522 | 45.45658 | 88.33633 | 37.86331 | -0.58925 | 1.602325 | 0.254292 |
| X | 36832200 | 36833799 | 15.16635 | 28.20935 | 22.23069 | 9.292894 | 0.788529 | 0.194816 | 0.254316 |
| 9 | 60185000 | 60187199 | 8.279458 | 14.73616 | 40.48847 | 17.84007 | -1.05366 | 0.44057 | 0.25432 |
| 11 | 120112600 | 120118999 | 27.52339 | 44.35671 | 103.8432 | 44.19037 | -0.86811 | 1.785729 | 0.254349 |
| 10 | 82602600 | 82605399 | 14.2573 | 24.53679 | 55.89009 | 24.51638 | -0.79829 | 0.886181 | 0.254352 |
| 14 | 52494600 | 52495599 | 5.891819 | 11.20429 | 28.76607 | 11.54167 | -0.85202 | -0.01503 | 0.254366 |
| 3 | 59888000 | 59893399 | 69.9678 | 111.8178 | 85.68086 | 36.01417 | 0.725384 | 2.093165 | 0.254377 |
| 12 | 56727800 | 56731199 | 23.27911 | 38.51519 | 74.00054 | 31.99974 | -0.55864 | 1.320775 | 0.254393 |
| 4 | 17495000 | 17496199 | 12.01482 | 23.22013 | 17.89922 | 6.865372 | 0.927887 | -0.13089 | 0.254396 |
| 9 | 99875800 | 99878799 | 17.67654 | 29.88921 | 61.77849 | 27.0355 | -0.66236 | 1.066602 | 0.254399 |
| 8 | 73372400 | 73377399 | 34.70177 | 55.89535 | 102.3551 | 43.5233 | -0.51564 | 1.813246 | 0.254407 |
| 6 | 123283200 | 123283799 | 10.70045 | 20.74841 | 17.50306 | 6.610537 | 0.819768 | -0.19759 | 0.254447 |
| 4 | 118965600 | 118966799 | 8.131049 | 14.80452 | 35.00288 | 15.05826 | -0.81473 | 0.290378 | 0.254453 |
| 2 | 49457400 | 49463199 | 37.33084 | 59.8811 | 115.2503 | 48.70794 | -0.59475 | 1.950645 | 0.254464 |
| 9 | 87621000 | 87625199 | 17.92196 | 29.70419 | 76.67085 | 33.19487 | -0.98266 | 1.289075 | 0.254494 |
| 1 | 65181400 | 65183599 | 34.04114 | 57.20538 | 38.76049 | 16.93059 | 0.940325 | 1.149617 | 0.254499 |
| 14 | 106500400 | 106501399 | 6.617922 | 12.56896 | 29.32286 | 11.79704 | -0.71914 | 0.001483 | 0.254504 |
| 10 | 14726600 | 14729199 | 20.77271 | 37.13052 | 24.95797 | 10.95747 | 0.968412 | 0.46423 | 0.254513 |
| 7 | 8906400 | 8907799 | 10.04518 | 19.54855 | 15.90214 | 5.853919 | 0.904979 | -0.29781 | 0.254531 |
| 10 | 119075000 | 119076999 | 17.70294 | 29.88726 | 61.16557 | 26.75463 | -0.64814 | 1.080845 | 0.254531 |
| 7 | 20730000 | 20732799 | 56.2944 | 90.54591 | 70.07177 | 29.73421 | 0.721745 | 1.873889 | 0.254531 |
| 10 | 22695000 | 22696199 | 7.467057 | 13.79929 | 33.09321 | 13.95818 | -0.81436 | 0.196723 | 0.254536 |
| 6 | 147084200 | 147085599 | 9.400254 | 16.85561 | 39.46209 | 17.2794 | -0.81839 | 0.397309 | 0.254536 |
| 19 | 17021800 | 17025399 | 44.17344 | 72.79061 | 49.51415 | 21.35983 | 0.933779 | 1.403153 | 0.254537 |
| 11 | 106678400 | 106680199 | 10.29294 | 18.0855 | 43.57167 | 19.28902 | -0.87404 | 0.58628 | 0.254538 |
| 1 | 40403800 | 40406399 | 14.38643 | 24.836 | 52.67825 | 23.10968 | -0.69478 | 0.855694 | 0.254543 |
| 11 | 53683400 | 53687999 | 21.68849 | 35.92374 | 71.49466 | 30.94006 | -0.60902 | 1.30244 | 0.254557 |
| 5 | 115295200 | 115297599 | 11.25541 | 19.64135 | 47.84668 | 21.06518 | -0.89053 | 0.708079 | 0.254562 |
| 10 | 93425000 | 93426599 | 7.273265 | 13.57834 | 31.66134 | 13.15253 | -0.75769 | 0.128027 | 0.25457 |
| 7 | 13695800 | 13700399 | 107.6593 | 166.6019 | 150.6831 | 62.58996 | 0.447596 | 2.835772 | 0.254573 |
| 6 | 123850200 | 123850599 | 8.02496 | 15.69908 | 10.97587 | 3.638189 | 1.253033 | -0.69803 | 0.254581 |
| 7 | 60684800 | 60685399 | 12.47865 | 23.85637 | 19.28267 | 7.648762 | 0.814691 | -0.01686 | 0.254583 |

| | | | | | | | | | |
|---|---|---|---|---|---|---|---|---|---|
| 19 | 58018000 | 58019399 | 22.57627 | 39.85742 | 27.66384 | 12.23977 | 0.913867 | 0.575797 | 0.25459 |
| 6 | 128243600 | 128248399 | 33.56499 | 54.10782 | 99.97019 | 42.5569 | -0.52704 | 1.786081 | 0.254593 |
| 2 | 132539400 | 132541999 | 11.68928 | 20.43398 | 46.77823 | 20.61908 | -0.8012 | 0.682514 | 0.254596 |
| 3 | 58858800 | 58859399 | 4.615944 | 9.000213 | 23.86029 | 8.591227 | -0.75047 | -0.41395 | 0.254622 |
| 8 | 56587200 | 56587799 | 4.615944 | 9.000213 | 23.86029 | 8.591227 | -0.75047 | -0.41395 | 0.254622 |
| 19 | 15176200 | 15176799 | 4.615944 | 9.000213 | 23.86029 | 8.591227 | -0.75047 | -0.41395 | 0.254622 |
| 15 | 81864600 | 81866799 | 19.59376 | 32.57941 | 70.94136 | 30.78237 | -0.73787 | 1.250036 | 0.254622 |
| 3 | 48563000 | 48563799 | 7.437916 | 13.84699 | 32.14015 | 13.41472 | -0.75633 | 0.150703 | 0.254627 |
| 6 | 130893000 | 130895999 | 46.38407 | 76.09101 | 51.79651 | 22.30202 | 0.930301 | 1.464272 | 0.254641 |
| 1 | 28747000 | 28748799 | 15.43816 | 26.79335 | 52.76283 | 23.15053 | -0.58665 | 0.812313 | 0.254642 |
| 13 | 14739200 | 14740199 | 22.16681 | 38.56131 | 31.18014 | 13.70048 | 0.692219 | 0.780047 | 0.254663 |
| 5 | 144047600 | 144048399 | 5.397597 | 10.41507 | 26.40051 | 10.1972 | -0.78221 | -0.15133 | 0.254711 |
| 6 | 64696600 | 64697399 | 6.420806 | 12.36489 | 29.37818 | 11.41209 | -0.69892 | -0.11641 | 0.254721 |
| X | 120148400 | 120151199 | 45.33017 | 74.41614 | 52.29178 | 22.52218 | 0.885042 | 1.454574 | 0.25473 |
| 4 | 17065600 | 17066799 | 13.04853 | 22.9256 | 46.13196 | 20.42119 | -0.61625 | 0.606662 | 0.254749 |
| 2 | 37028000 | 37030399 | 28.44305 | 48.85154 | 37.54353 | 16.46759 | 0.762004 | 0.940589 | 0.254753 |
| 10 | 101466400 | 101467199 | 9.186315 | 16.28004 | 40.4523 | 17.88433 | -0.91326 | 0.476318 | 0.254767 |
| 10 | 107026400 | 107031799 | 92.20411 | 146.0462 | 98.84145 | 41.1053 | 0.891294 | 2.38142 | 0.254778 |
| 4 | 82453400 | 82458599 | 27.128 | 43.85428 | 94.12597 | 40.21459 | -0.73785 | 1.690898 | 0.254783 |
| 7 | 16571400 | 16575799 | 26.9474 | 43.60952 | 92.03512 | 39.36219 | -0.71195 | 1.663615 | 0.254798 |
| 2 | 111389200 | 111389599 | 8.839147 | 17.2571 | 13.78659 | 4.86606 | 0.983992 | -0.46551 | 0.254799 |
| 18 | 10324000 | 10325999 | 12.64557 | 22.00654 | 48.40627 | 21.27466 | -0.74455 | 0.756646 | 0.254811 |
| 15 | 35164600 | 35167799 | 48.56084 | 78.81345 | 62.34672 | 26.72149 | 0.704002 | 1.628988 | 0.254827 |
| 13 | 53249400 | 53250199 | 6.636555 | 12.5473 | 29.74741 | 12.08321 | -0.75432 | 0.034885 | 0.254836 |
| X | 115314600 | 115316999 | 31.97907 | 54.52051 | 38.6598 | 16.94733 | 0.876306 | 1.023131 | 0.25484 |
| 14 | 41367600 | 41372199 | 43.36913 | 70.71998 | 59.97404 | 25.76339 | 0.609092 | 1.541392 | 0.254842 |
| 6 | 50585200 | 50585599 | 9.607404 | 18.63112 | 15.11247 | 5.704587 | 0.877302 | -0.21877 | 0.254858 |
| 9 | 98601200 | 98602399 | 14.40498 | 26.88246 | 22.23631 | 9.243038 | 0.725376 | 0.16681 | 0.254885 |
| 10 | 14839800 | 14841199 | 14.40498 | 26.88246 | 22.23631 | 9.243038 | 0.725376 | 0.16681 | 0.254885 |
| 14 | 119247000 | 119251199 | 31.62835 | 51.02882 | 97.35298 | 41.48779 | -0.57172 | 1.757747 | 0.254885 |
| 19 | 6230000 | 6235199 | 35.71659 | 57.36354 | 108.3899 | 45.89587 | -0.566 | 1.908347 | 0.254903 |
| 16 | 89780600 | 89796799 | 184.1417 | 267.6924 | 445.2954 | 183.551 | -0.5097 | 3.999947 | 0.254911 |
| 5 | 95572600 | 95576799 | 48.19751 | 78.0603 | 64.06327 | 27.41488 | 0.648276 | 1.667909 | 0.254922 |
| 7 | 116032600 | 116037399 | 69.8548 | 108.0033 | 212.8872 | 88.4284 | -0.67607 | 2.846416 | 0.254924 |
| 6 | 105197400 | 105199799 | 24.91869 | 43.65235 | 29.72953 | 13.12769 | 0.938889 | 0.690689 | 0.254931 |
| 17 | 53419000 | 53424999 | 89.97785 | 141.2909 | 126.2612 | 52.36883 | 0.48121 | 2.568675 | 0.254939 |
| 13 | 85290800 | 85291999 | 16.29737 | 28.07502 | 54.32143 | 23.82899 | -0.56358 | 0.886345 | 0.254959 |
| 7 | 8354000 | 8355799 | 33.97806 | 57.06188 | 44.1944 | 19.29849 | 0.748306 | 1.162796 | 0.254971 |
| 6 | 89799000 | 89801199 | 40.74451 | 66.48511 | 59.77283 | 25.68329 | 0.525761 | 1.526269 | 0.254977 |
| Y | 4206800 | 4207199 | 7.394776 | 14.46238 | 7.106088 | 2.410651 | 1.723188 | -0.65389 | 0.254994 |
| 19 | 47514800 | 47516399 | 7.252057 | 13.60065 | 31.21901 | 12.86897 | -0.72626 | 0.09684 | 0.255009 |
| 12 | 11707000 | 11708599 | 11.19851 | 19.66733 | 43.56157 | 19.27684 | -0.75359 | 0.59904 | 0.255019 |
| 8 | 61144800 | 61146799 | 26.09134 | 44.76693 | 38.86115 | 17.04434 | 0.586868 | 0.942714 | 0.255021 |
| 12 | 46653400 | 46655599 | 10.84224 | 19.16018 | 43.50199 | 19.24356 | -0.78623 | 0.52146 | 0.255029 |
| 5 | 143220800 | 143221799 | 7.281457 | 13.58612 | 32.17877 | 13.37999 | -0.78065 | 0.135695 | 0.255032 |
| 16 | 89819800 | 89832199 | 142.4869 | 211.2729 | 362.1037 | 149.1415 | -0.52563 | 3.679336 | 0.255043 |
| 10 | 6573400 | 6575599 | 24.85732 | 43.22441 | 31.39559 | 13.79167 | 0.846217 | 0.799594 | 0.255046 |

| | | | | | | | | | |
|---|---|---|---|---|---|---|---|---|---|
| 4 | 126426600 | 126427799 | 8.627616 | 15.55955 | 36.4571 | 15.87203 | -0.81314 | 0.353605 | 0.255061 |
| 7 | 43429200 | 43434599 | 38.2383 | 61.09856 | 121.7983 | 51.18201 | -0.65176 | 2.07606 | 0.25507 |
| 19 | 8917200 | 8920199 | 16.63533 | 28.03838 | 61.56947 | 26.91777 | -0.74935 | 1.093141 | 0.255073 |
| 7 | 7614200 | 7619599 | 91.50529 | 143.1888 | 130.9688 | 54.33119 | 0.443804 | 2.632248 | 0.255076 |
| 3 | 95152600 | 95153199 | 5.110686 | 9.910804 | 25.57658 | 9.65487 | -0.77649 | -0.23763 | 0.255087 |
| 6 | 40447800 | 40448599 | 5.110686 | 9.910804 | 25.57658 | 9.65487 | -0.77649 | -0.23763 | 0.255087 |
| 17 | 27263800 | 27264399 | 5.110686 | 9.910804 | 25.57658 | 9.65487 | -0.77649 | -0.23763 | 0.255087 |
| 5 | 59087600 | 59089599 | 29.43475 | 50.71457 | 35.26044 | 15.46765 | 0.906738 | 0.914592 | 0.255095 |
| 2 | 7511200 | 7513199 | 19.95499 | 33.50923 | 64.55389 | 28.18751 | -0.56169 | 1.138902 | 0.255113 |
| 9 | 77238400 | 77242599 | 71.47227 | 113.4026 | 100.4537 | 41.83929 | 0.50584 | 2.317771 | 0.255119 |
| 13 | 3077400 | 3081199 | 43.77165 | 71.70179 | 55.59459 | 23.92273 | 0.742269 | 1.478341 | 0.255126 |
| 7 | 17967000 | 17969799 | 45.13746 | 73.53671 | 59.54041 | 25.56864 | 0.674807 | 1.55708 | 0.255141 |
| 8 | 57939600 | 57943999 | 23.26588 | 38.10019 | 81.83325 | 35.20897 | -0.72373 | 1.451692 | 0.255152 |
| 6 | 92111200 | 92111999 | 12.75926 | 24.70805 | 16.7679 | 6.371834 | 1.122668 | -0.15574 | 0.255159 |
| 7 | 138430200 | 138433799 | 25.67591 | 41.9974 | 80.68791 | 34.70042 | -0.56435 | 1.475213 | 0.25516 |
| 2 | 39668600 | 39669999 | 20.60257 | 36.04262 | 30.96031 | 13.65555 | 0.605836 | 0.690801 | 0.255164 |
| 8 | 58922200 | 58924799 | 15.01099 | 26.12342 | 54.92443 | 24.13967 | -0.6802 | 0.775971 | 0.255168 |
| X | 28475600 | 28476199 | 11.014 | 21.23381 | 15.10656 | 5.849539 | 1.030485 | -0.11871 | 0.255179 |
| 4 | 117751800 | 117752799 | 3.999798 | 7.774377 | 24.49127 | 9.104934 | -1.03904 | -0.30765 | 0.25519 |
| 18 | 60981400 | 60982199 | 3.999798 | 7.774377 | 24.49127 | 9.104934 | -1.03904 | -0.30765 | 0.25519 |
| 16 | 68738400 | 68739599 | 13.45124 | 25.26312 | 21.2141 | 8.723458 | 0.715818 | 0.11514 | 0.255193 |
| 16 | 19361600 | 19364599 | 36.29355 | 60.91716 | 42.50747 | 18.55545 | 0.898243 | 1.174548 | 0.255202 |
| 4 | 3443400 | 3443799 | 14.60135 | 26.88586 | 21.40813 | 9.074312 | 0.759401 | 0.243838 | 0.255212 |
| 8 | 35387200 | 35388399 | 13.35072 | 24.81475 | 20.8785 | 8.737563 | 0.693273 | 0.17934 | 0.255229 |
| 7 | 108199200 | 108201399 | 33.34977 | 55.65718 | 47.06202 | 20.46968 | 0.622425 | 1.231103 | 0.255241 |
| 13 | 101228600 | 101230599 | 16.66167 | 28.51542 | 55.87035 | 24.4995 | -0.58333 | 0.949742 | 0.255245 |
| 10 | 88073000 | 88076799 | 45.89483 | 74.42411 | 64.89626 | 27.79701 | 0.562743 | 1.641378 | 0.255249 |
| 17 | 34694200 | 34697199 | 14.93083 | 25.54979 | 56.36218 | 24.71491 | -0.7537 | 0.942994 | 0.255268 |
| 4 | 9710400 | 9713199 | 31.12133 | 50.28123 | 94.85838 | 40.49785 | -0.55325 | 1.714454 | 0.25527 |
| 2 | 23176000 | 23177999 | 24.18017 | 42.37569 | 29.72691 | 13.13101 | 0.896317 | 0.679753 | 0.255292 |
| 18 | 86128800 | 86130599 | 26.03671 | 45.43837 | 31.4132 | 13.83064 | 0.917423 | 0.755485 | 0.255294 |
| 2 | 67395400 | 67397399 | 14.36863 | 24.87894 | 51.08928 | 22.41463 | -0.64733 | 0.827278 | 0.255304 |
| 17 | 42174200 | 42176999 | 32.20278 | 54.64978 | 40.38017 | 17.69065 | 0.816448 | 1.066613 | 0.255304 |
| 5 | 43372800 | 43373799 | 11.04065 | 21.17973 | 19.06637 | 7.488888 | 0.674429 | -0.06157 | 0.255307 |
| 5 | 114309800 | 114314599 | 24.98685 | 40.57028 | 87.69603 | 37.59771 | -0.74186 | 1.590384 | 0.255315 |
| 1 | 85587600 | 85589399 | 21.09536 | 37.04619 | 30.5176 | 13.50387 | 0.665663 | 0.631668 | 0.255329 |
| 1 | 90165400 | 90169199 | 24.49344 | 40.20172 | 78.73848 | 33.91003 | -0.58973 | 1.424586 | 0.255331 |
| 12 | 103575400 | 103575999 | 10.19529 | 19.81372 | 15.1654 | 5.644338 | 0.976703 | -0.26232 | 0.255335 |
| 8 | 64252600 | 64253399 | 10.55621 | 20.33729 | 18.59192 | 7.213316 | 0.668835 | -0.10727 | 0.255351 |
| 1 | 53852200 | 53853199 | 6.191945 | 11.74217 | 29.26144 | 11.81003 | -0.81725 | 0.008226 | 0.255366 |
| 4 | 96887800 | 96888599 | 11.24528 | 21.79638 | 17.4849 | 6.61568 | 0.889476 | -0.18484 | 0.255366 |
| X | 4313800 | 4315799 | 26.48034 | 46.16538 | 32.04292 | 14.09883 | 0.911723 | 0.773513 | 0.255366 |
| X | 43762600 | 43765799 | 21.53401 | 39.50585 | 18.50059 | 7.881154 | 1.513688 | 0.311013 | 0.255372 |
| 13 | 86799400 | 86800799 | 20.58502 | 36.80519 | 24.96267 | 10.95698 | 0.955719 | 0.461148 | 0.255381 |
| 8 | 113559000 | 113560199 | 12.56736 | 22.07866 | 44.55348 | 19.72129 | -0.62023 | 0.602193 | 0.255385 |
| 10 | 14960600 | 14961399 | 18.57722 | 32.86833 | 28.1525 | 12.45059 | 0.613283 | 0.528897 | 0.255406 |
| 9 | 110073800 | 110076999 | 15.48461 | 25.90877 | 66.15387 | 28.82048 | -0.96618 | 1.166969 | 0.25542 |

| | | | | | | | | | |
|---|---|---|---|---|---|---|---|---|---|
| 8 | 63504800 | 63506599 | 17.93601 | 33.48141 | 18.98716 | 7.888717 | 1.265843 | 0.186825 | 0.255422 |
| 7 | 10699400 | 10701999 | 17.95473 | 33.02536 | 20.99895 | 8.913765 | 1.079384 | 0.273203 | 0.255435 |
| 17 | 8909600 | 8916199 | 88.58228 | 139.0158 | 126.4253 | 52.4328 | 0.455276 | 2.578883 | 0.255447 |
| 19 | 42169800 | 42174399 | 20.01804 | 33.03681 | 76.81421 | 33.15898 | -0.83316 | 1.343315 | 0.255453 |
| 11 | 77690200 | 77696199 | 29.37439 | 47.43385 | 95.28087 | 40.65066 | -0.64436 | 1.728101 | 0.255458 |
| 3 | 63649200 | 63650399 | 13.65163 | 25.86933 | 20.19013 | 8.161535 | 0.841248 | 0.056896 | 0.255462 |
| 8 | 122088000 | 122091999 | 35.57099 | 57.09973 | 109.17 | 46.18932 | -0.58385 | 1.925237 | 0.255462 |
| 14 | 76463000 | 76465599 | 15.80476 | 27.16444 | 55.93961 | 24.53713 | -0.65393 | 0.90474 | 0.255467 |
| 16 | 12154800 | 12155599 | 10.54661 | 20.35571 | 18.58637 | 7.173111 | 0.677538 | -0.12443 | 0.255473 |
| 3 | 20111600 | 20112399 | 3.822154 | 7.448174 | 22.43526 | 8.139417 | -0.94155 | -0.40301 | 0.255483 |
| 4 | 150998600 | 150999799 | 3.822154 | 7.448174 | 22.43526 | 8.139417 | -0.94155 | -0.40301 | 0.255483 |
| 9 | 103078800 | 103079399 | 3.822154 | 7.448174 | 22.43526 | 8.139417 | -0.94155 | -0.40301 | 0.255483 |
| 11 | 16541400 | 16541999 | 3.822154 | 7.448174 | 22.43526 | 8.139417 | -0.94155 | -0.40301 | 0.255483 |
| 2 | 94716200 | 94716799 | 14.47929 | 28.0881 | 13.52461 | 5.09741 | 1.621885 | -0.17256 | 0.25549 |
| X | 26079200 | 26081599 | 29.42392 | 51.58471 | 24.47683 | 10.81536 | 1.457145 | 0.696207 | 0.255521 |
| 14 | 90663600 | 90664399 | 11.81734 | 22.55821 | 19.99372 | 7.957781 | 0.679028 | -0.00937 | 0.255534 |
| 9 | 42963800 | 42968999 | 53.02676 | 84.17551 | 156.3013 | 65.13082 | -0.55971 | 2.302734 | 0.25554 |
| 4 | 26299000 | 26301199 | 39.52898 | 64.94682 | 48.84949 | 21.04701 | 0.787624 | 1.426767 | 0.255541 |
| 1 | 11975200 | 11976399 | 8.042659 | 15.05032 | 37.9775 | 15.71318 | -0.86661 | 0.134645 | 0.255544 |
| 5 | 71286200 | 71288599 | 26.61942 | 46.27823 | 33.75359 | 14.82544 | 0.840271 | 0.814787 | 0.255555 |
| 8 | 11759000 | 11762599 | 17.85628 | 29.75969 | 68.52862 | 29.78102 | -0.81798 | 1.217866 | 0.255564 |
| 17 | 36730400 | 36733599 | 43.96192 | 71.70922 | 58.95346 | 25.32788 | 0.654068 | 1.536806 | 0.255575 |
| 18 | 57382400 | 57383399 | 5.526828 | 10.7004 | 26.05481 | 9.910356 | -0.70368 | -0.20349 | 0.255582 |
| 12 | 29191000 | 29195999 | 25.27936 | 41.09825 | 85.67725 | 36.76459 | -0.68806 | 1.564693 | 0.255583 |
| 11 | 96002200 | 96004799 | 12.25878 | 21.49228 | 48.31428 | 21.34883 | -0.77542 | 0.637928 | 0.255592 |
| 11 | 10731000 | 10734799 | 30.16446 | 51.68988 | 37.50939 | 16.44978 | 0.844017 | 0.968345 | 0.255598 |
| 15 | 36141800 | 36143599 | 11.42877 | 20.34306 | 41.43856 | 18.25748 | -0.62524 | 0.46213 | 0.255613 |
| 17 | 20527000 | 20528999 | 30.53662 | 51.79903 | 41.98478 | 18.39114 | 0.683334 | 1.06409 | 0.255634 |
| 7 | 44070200 | 44083199 | 88.49711 | 134.801 | 252.9628 | 104.8487 | -0.62203 | 3.145864 | 0.255636 |
| 8 | 72128600 | 72134799 | 56.47386 | 88.34851 | 178.515 | 74.05724 | -0.69849 | 2.625814 | 0.25564 |
| 11 | 47912800 | 47914399 | 11.44194 | 20.38981 | 42.52375 | 18.71755 | -0.65805 | 0.457579 | 0.255645 |
| X | 5197800 | 5201999 | 42.45349 | 70.55063 | 44.33251 | 19.22918 | 1.049191 | 1.300529 | 0.255654 |
| 9 | 83117800 | 83119799 | 11.78728 | 20.79956 | 41.94208 | 18.55898 | -0.6167 | 0.543253 | 0.255666 |
| 5 | 84563200 | 84565599 | 17.39184 | 29.33756 | 63.37077 | 27.7122 | -0.72588 | 1.087907 | 0.255667 |
| 7 | 36173800 | 36174999 | 9.908465 | 17.63071 | 38.9186 | 17.15231 | -0.74016 | 0.45196 | 0.255675 |
| 2 | 103754600 | 103756399 | 20.04305 | 35.15871 | 30.21562 | 13.35758 | 0.604893 | 0.642382 | 0.255682 |
| 17 | 76863400 | 76864599 | 14.44979 | 26.7772 | 22.79679 | 9.579347 | 0.672326 | 0.210467 | 0.255689 |
| 10 | 128247000 | 128248399 | 7.158506 | 12.91286 | 37.01531 | 16.1107 | -1.10129 | 0.342824 | 0.255699 |
| 8 | 3810400 | 3819399 | 164.9536 | 248.0304 | 221.0618 | 91.27199 | 0.435453 | 3.420865 | 0.255732 |
| 12 | 103992000 | 104000199 | 62.31192 | 97.48664 | 177.441 | 73.61133 | -0.54796 | 2.627588 | 0.25574 |
| 17 | 39092600 | 39094199 | 25.46689 | 44.65952 | 28.57268 | 12.62806 | 1.028518 | 0.675926 | 0.255741 |
| 1 | 117924800 | 117925599 | 20.58874 | 36.2607 | 29.40594 | 13.01685 | 0.689122 | 0.591223 | 0.255759 |
| 7 | 12065200 | 12066399 | 26.84548 | 46.11687 | 37.80799 | 16.58368 | 0.669377 | 0.934903 | 0.255762 |
| X | 62682600 | 62683599 | 13.62602 | 26.31422 | 15.83412 | 6.094622 | 1.275977 | -0.10887 | 0.255764 |
| 1 | 40609600 | 40611199 | 13.8668 | 24.11366 | 49.48674 | 21.73926 | -0.64538 | 0.773189 | 0.255764 |
| 4 | 146154800 | 146155199 | 7.952056 | 15.50273 | 14.09475 | 5.084736 | 0.773262 | -0.40026 | 0.255767 |
| 12 | 83921400 | 83922799 | 5.600493 | 10.7039 | 28.90275 | 11.47391 | -0.91091 | -0.05456 | 0.255772 |

| | | | | | | | | | |
|---|---|---|---|---|---|---|---|---|---|
| 4 | 148949200 | 148955399 | 24.08008 | 39.04401 | 91.98559 | 39.39662 | -0.86764 | 1.618384 | 0.255772 |
| 7 | 35380000 | 35384599 | 26.27758 | 42.99994 | 84.11714 | 36.18083 | -0.59 | 1.469453 | 0.255778 |
| 4 | 40675400 | 40678399 | 15.48103 | 26.7942 | 52.17918 | 22.89253 | -0.57134 | 0.837878 | 0.255783 |
| 6 | 126432000 | 126433799 | 11.31425 | 20.10379 | 41.4374 | 18.28409 | -0.64342 | 0.474565 | 0.255784 |
| 1 | 48454000 | 48457999 | 34.03275 | 57.26188 | 42.50081 | 18.57131 | 0.809492 | 1.143854 | 0.255796 |
| 10 | 63582000 | 63583799 | 17.84108 | 32.41477 | 24.33722 | 10.49783 | 0.826295 | 0.348517 | 0.255805 |
| 7 | 6037400 | 6054799 | 319.9982 | 457.0937 | 384.2153 | 158.7622 | 0.449585 | 4.266154 | 0.255808 |
| 1 | 183226200 | 183232799 | 59.70907 | 93.87182 | 165.3862 | 68.60717 | -0.49554 | 2.533808 | 0.25581 |
| 11 | 84476000 | 84478599 | 13.25097 | 23.06722 | 48.41001 | 21.28045 | -0.67718 | 0.754247 | 0.255824 |
| 5 | 23950000 | 23951199 | 6.201944 | 11.7552 | 30.93868 | 12.50029 | -0.89743 | 0.01382 | 0.255825 |
| 6 | 113581200 | 113583399 | 11.64846 | 20.44925 | 45.11059 | 19.95547 | -0.74805 | 0.609286 | 0.255835 |
| 3 | 71215400 | 71217399 | 27.53354 | 47.15422 | 35.63966 | 15.62908 | 0.785345 | 0.959515 | 0.255836 |
| 5 | 139475000 | 139477599 | 15.5682 | 26.6052 | 55.81296 | 24.47146 | -0.68174 | 0.958358 | 0.255836 |
| 15 | 60693200 | 60696999 | 23.8079 | 38.82847 | 84.22126 | 36.18146 | -0.74188 | 1.513782 | 0.255842 |
| 12 | 87743400 | 87754199 | 145.98 | 216.2382 | 364.1438 | 149.9667 | -0.50156 | 3.697393 | 0.255843 |
| 5 | 37764200 | 37766999 | 36.00301 | 60.84536 | 39.10386 | 17.1099 | 1.016658 | 1.11186 | 0.255845 |
| 12 | 34956200 | 34958599 | 14.43068 | 24.79164 | 55.32182 | 24.26581 | -0.76925 | 0.902952 | 0.255863 |
| 7 | 117714800 | 117715799 | 6.0614 | 11.47055 | 29.75468 | 12.06243 | -0.88035 | 0.022519 | 0.255871 |
| 2 | 36518800 | 36520399 | 23.01043 | 39.98342 | 35.07936 | 15.40278 | 0.574828 | 0.805293 | 0.255879 |
| 12 | 60753000 | 60755799 | 52.93591 | 85.02597 | 75.35398 | 31.91131 | 0.524149 | 1.915287 | 0.255881 |
| 5 | 135738400 | 135739999 | 10.30259 | 18.48734 | 38.40528 | 16.8061 | -0.64634 | 0.397764 | 0.255887 |
| 6 | 88985200 | 88991999 | 40.76062 | 64.84137 | 137.2818 | 57.33374 | -0.74549 | 2.22345 | 0.255891 |
| 5 | 150183000 | 150183799 | 4.817774 | 9.361006 | 24.54693 | 9.15137 | -0.78192 | -0.28743 | 0.255894 |
| 6 | 32470400 | 32471199 | 4.817774 | 9.361006 | 24.54693 | 9.15137 | -0.78192 | -0.28743 | 0.255894 |
| 11 | 3723200 | 3728999 | 39.83925 | 63.67402 | 120.672 | 50.72837 | -0.57851 | 2.068953 | 0.255898 |
| 17 | 79805600 | 79807799 | 22.37547 | 38.99376 | 33.95897 | 14.93822 | 0.585791 | 0.760011 | 0.255946 |
| X | 16816800 | 16817399 | 10.24967 | 19.83252 | 18.10379 | 6.905045 | 0.693955 | -0.16803 | 0.255962 |
| 1 | 79142400 | 79144999 | 40.72774 | 67.35031 | 49.97871 | 21.60465 | 0.809632 | 1.354224 | 0.255978 |
| 1 | 125289000 | 125290799 | 23.34229 | 38.46379 | 74.75972 | 32.25031 | -0.57632 | 1.371163 | 0.25598 |
| 13 | 74498000 | 74498999 | 7.072512 | 13.35605 | 31.41957 | 12.79376 | -0.74707 | 0.048966 | 0.255982 |
| X | 57251400 | 57252999 | 16.86344 | 30.97235 | 23.09587 | 9.823579 | 0.849884 | 0.279964 | 0.255983 |
| 17 | 19209200 | 19210599 | 24.07952 | 42.19401 | 30.47971 | 13.46211 | 0.854503 | 0.684328 | 0.255986 |
| X | 11922800 | 11929599 | 32.26944 | 52.20525 | 100.855 | 43.12395 | -0.58538 | 1.684895 | 0.255987 |
| 5 | 73599200 | 73603599 | 23.58111 | 39.0786 | 75.31946 | 32.60681 | -0.56301 | 1.303376 | 0.255991 |
| 11 | 102482200 | 102484199 | 10.07639 | 17.85688 | 41.94367 | 18.5438 | -0.83312 | 0.483233 | 0.255995 |
| 1 | 163821800 | 163824199 | 13.84304 | 24.13304 | 48.93482 | 21.53086 | -0.62777 | 0.735708 | 0.255997 |
| X | 42915400 | 42919199 | 68.2326 | 109.2491 | 80.04719 | 33.73598 | 0.792806 | 2.035787 | 0.256022 |
| 2 | 106523200 | 106526399 | 18.81273 | 30.96213 | 77.38074 | 33.36376 | -0.93839 | 1.376862 | 0.256034 |
| 2 | 180818200 | 180819599 | 9.144205 | 16.72443 | 35.56945 | 15.21572 | -0.6583 | 0.270984 | 0.256057 |
| 1 | 88729600 | 88735599 | 50.10982 | 79.66158 | 146.5792 | 61.16912 | -0.54452 | 2.252935 | 0.256068 |
| 16 | 80839400 | 80840799 | 19.41928 | 35.13385 | 23.52126 | 10.19946 | 0.985659 | 0.381291 | 0.256072 |
| 2 | 152526800 | 152534799 | 67.71925 | 105.4204 | 188.342 | 78.18028 | -0.52676 | 2.718959 | 0.25608 |
| 11 | 33144800 | 33149599 | 71.99 | 110.773 | 218.4504 | 90.72403 | -0.68207 | 2.935561 | 0.256089 |
| 1 | 144612400 | 144613599 | 15.75469 | 29.35056 | 21.42861 | 8.933383 | 0.900517 | 0.186113 | 0.256112 |
| 10 | 71526000 | 71527599 | 16.60912 | 30.10042 | 25.12511 | 10.87195 | 0.671716 | 0.356195 | 0.25612 |
| 19 | 37184000 | 37185999 | 10.2162 | 17.83454 | 49.40472 | 21.75625 | -1.07488 | 0.699072 | 0.256125 |
| 7 | 14878800 | 14883999 | 64.36233 | 103.5401 | 68.16288 | 28.93124 | 0.954868 | 1.88085 | 0.256127 |

| | | | | | | | | | |
|---|---|---|---|---|---|---|---|---|---|
| 15 | 63520000 | 63523399 | 17.76929 | 30.21419 | 61.3601 | 26.89272 | -0.63664 | 1.016666 | 0.256135 |
| X | 54352400 | 54355999 | 47.5541 | 77.84055 | 52.87379 | 22.74489 | 0.932251 | 1.495721 | 0.256144 |
| 5 | 138617600 | 138618199 | 4.485755 | 8.739289 | 23.04539 | 8.386295 | -0.75639 | -0.37995 | 0.256144 |
| 10 | 85626800 | 85627999 | 4.485755 | 8.739289 | 23.04539 | 8.386295 | -0.75639 | -0.37995 | 0.256144 |
| 9 | 73009400 | 73012599 | 23.74056 | 38.87478 | 79.45477 | 34.1846 | -0.65234 | 1.45345 | 0.256177 |
| 6 | 127643800 | 127648199 | 26.8059 | 43.34346 | 93.52556 | 39.96982 | -0.7451 | 1.683501 | 0.256179 |
| 6 | 145581000 | 145586599 | 50.26733 | 79.5025 | 152.1341 | 63.22062 | -0.60772 | 2.39536 | 0.256189 |
| 4 | 148122200 | 148126399 | 27.09319 | 43.88213 | 94.32123 | 40.3648 | -0.73725 | 1.646198 | 0.256193 |
| 13 | 34027000 | 34029999 | 16.42748 | 28.23671 | 57.06896 | 25.03256 | -0.62708 | 0.90785 | 0.256202 |
| 18 | 64621600 | 64624599 | 15.88674 | 27.11869 | 56.3525 | 24.70576 | -0.6684 | 0.970475 | 0.256203 |
| 3 | 14114200 | 14114999 | 9.513295 | 17.26809 | 36.5051 | 15.76498 | -0.65836 | 0.322229 | 0.256218 |
| 16 | 43958800 | 43961199 | 15.75243 | 26.84477 | 56.84436 | 24.91806 | -0.69579 | 0.986625 | 0.25622 |
| 15 | 27461800 | 27464399 | 16.95035 | 29.07591 | 55.92685 | 24.5288 | -0.55638 | 0.929268 | 0.256221 |
| 19 | 27384600 | 27386799 | 19.00278 | 34.10234 | 25.74702 | 11.266 | 0.80513 | 0.429993 | 0.256222 |
| 1 | 128171600 | 128172799 | 8.516258 | 15.27047 | 37.95365 | 16.61893 | -0.90352 | 0.390225 | 0.256231 |
| 5 | 110079000 | 110079999 | 22.0862 | 38.45097 | 33.60228 | 14.77198 | 0.580773 | 0.772735 | 0.256245 |
| 3 | 142936200 | 142938199 | 18.26307 | 33.09314 | 24.21239 | 10.47887 | 0.860185 | 0.365693 | 0.256262 |
| 18 | 19256400 | 19259199 | 58.5146 | 93.74632 | 84.49181 | 35.64209 | 0.495604 | 2.003824 | 0.256266 |
| 18 | 6116600 | 6117999 | 15.20138 | 28.6074 | 20.54516 | 8.419759 | 0.943735 | 0.117544 | 0.256275 |
| 8 | 25588000 | 25590199 | 9.925979 | 17.59761 | 40.40911 | 17.86387 | -0.80017 | 0.476027 | 0.256275 |
| 14 | 66125400 | 66128199 | 24.23729 | 39.63097 | 80.50785 | 34.61874 | -0.6449 | 1.475615 | 0.256288 |
| 5 | 75764200 | 75765999 | 7.786058 | 14.31618 | 33.55093 | 14.24902 | -0.78902 | 0.227666 | 0.256301 |
| 2 | 9885800 | 9892799 | 52.68294 | 82.93538 | 165.9429 | 68.84996 | -0.67733 | 2.502426 | 0.256304 |
| X | 120367400 | 120370199 | 39.96826 | 66.29853 | 48.2553 | 20.90412 | 0.837767 | 1.317123 | 0.256327 |
| 14 | 50304600 | 50307599 | 35.16407 | 58.08699 | 52.0631 | 22.48888 | 0.537903 | 1.351676 | 0.25635 |
| 12 | 8139600 | 8141999 | 13.8674 | 24.27275 | 48.88037 | 21.56909 | -0.61744 | 0.67685 | 0.256362 |
| 4 | 120515600 | 120516799 | 14.12726 | 26.68702 | 20.75692 | 8.446361 | 0.83827 | 0.084788 | 0.25637 |
| 17 | 18702400 | 18703599 | 26.18883 | 45.88693 | 27.9286 | 12.33462 | 1.100069 | 0.691168 | 0.256372 |
| 17 | 19751400 | 19753799 | 29.24959 | 50.16297 | 37.50954 | 16.45093 | 0.80104 | 0.956648 | 0.256393 |
| 13 | 61228600 | 61230199 | 19.9556 | 36.24051 | 22.42455 | 9.678168 | 1.102807 | 0.362286 | 0.256418 |
| 10 | 82219800 | 82224199 | 89.94954 | 141.3443 | 124.9281 | 51.82061 | 0.497808 | 2.554696 | 0.256428 |
| 12 | 63617000 | 63617999 | 8.820419 | 16.21947 | 34.60765 | 14.69605 | -0.65514 | 0.235993 | 0.256432 |
| 13 | 63290600 | 63294599 | 48.0497 | 76.33184 | 138.5863 | 57.79218 | -0.52619 | 2.271704 | 0.256442 |
| 11 | 80712000 | 80713799 | 18.6117 | 32.84043 | 28.14604 | 12.45446 | 0.610804 | 0.556862 | 0.256447 |
| 2 | 89384800 | 89386199 | 17.84931 | 32.51664 | 23.16063 | 9.95693 | 0.904588 | 0.330065 | 0.256452 |
| 11 | 101237800 | 101238599 | 4.945696 | 9.528035 | 26.85116 | 10.41838 | -0.93866 | -0.13764 | 0.256455 |
| 10 | 29841200 | 29842999 | 15.33455 | 27.99682 | 23.00087 | 9.861135 | 0.702482 | 0.297699 | 0.256489 |
| 12 | 111540400 | 111541999 | 7.384392 | 13.49125 | 35.04498 | 15.01182 | -0.94542 | 0.263565 | 0.256493 |
| 10 | 35096200 | 35102199 | 85.82292 | 135.6922 | 110.8214 | 46.0451 | 0.618508 | 2.421443 | 0.256505 |
| 18 | 37755600 | 37763399 | 90.65167 | 142.9919 | 114.5456 | 47.53745 | 0.643706 | 2.478318 | 0.256508 |
| 4 | 32614800 | 32615799 | 5.619144 | 10.68924 | 29.32317 | 11.75707 | -0.94611 | -0.0199 | 0.256508 |
| 12 | 113602200 | 113604999 | 36.49082 | 60.35074 | 52.11239 | 22.52877 | 0.591652 | 1.341825 | 0.256518 |
| 17 | 28476000 | 28482399 | 51.80432 | 81.32557 | 179.235 | 74.33795 | -0.81968 | 2.56127 | 0.256546 |
| 7 | 32929800 | 32934799 | 95.1357 | 150.189 | 107.4014 | 44.58999 | 0.808168 | 2.461476 | 0.256548 |
| 5 | 35730800 | 35732599 | 9.638481 | 17.49624 | 36.49255 | 15.75852 | -0.6393 | 0.320781 | 0.256556 |
| 12 | 60462200 | 60463199 | 14.60222 | 28.21285 | 15.94915 | 6.125259 | 1.369243 | -0.10396 | 0.256581 |
| 7 | 20852800 | 20858599 | 131.0369 | 200.4442 | 173.8209 | 72.11901 | 0.495505 | 3.067714 | 0.256588 |

| | | | | | | | | | |
|---|---|---|---|---|---|---|---|---|---|
| 10 | 94085600 | 94090199 | 28.85169 | 46.71172 | 90.70257 | 38.80481 | -0.59151 | 1.658001 | 0.256605 |
| 4 | 112324000 | 112328399 | 91.28119 | 143.4241 | 124.6457 | 51.70285 | 0.52202 | 2.55728 | 0.256611 |
| 2 | 73359200 | 73360199 | 6.391006 | 11.97637 | 31.18277 | 12.87121 | -0.90773 | 0.094635 | 0.256646 |
| 13 | 72511000 | 72516799 | 38.627 | 61.95772 | 113.5984 | 48.00684 | -0.52495 | 1.953526 | 0.256646 |
| 5 | 131136000 | 131137399 | 19.34875 | 34.50827 | 26.78968 | 11.78377 | 0.760143 | 0.472245 | 0.25665 |
| 9 | 32451400 | 32458999 | 45.05532 | 71.38544 | 151.1717 | 62.87696 | -0.75161 | 2.348809 | 0.256654 |
| 7 | 79163600 | 79167599 | 38.38083 | 61.3556 | 121.7933 | 51.21456 | -0.64491 | 2.060766 | 0.256665 |
| 13 | 13488800 | 13491199 | 35.31578 | 58.91493 | 43.82688 | 19.05779 | 0.806537 | 1.236525 | 0.256713 |
| 17 | 20513000 | 20517799 | 81.77498 | 129.9286 | 97.43962 | 40.63999 | 0.747602 | 2.286348 | 0.256743 |
| 12 | 18127000 | 18168599 | 1173.54 | 1503.407 | 1127.599 | 473.0449 | 0.45636 | 5.975154 | 0.256754 |
| 15 | 67046000 | 67050399 | 30.02538 | 48.44938 | 97.01809 | 41.35253 | -0.64115 | 1.750975 | 0.256776 |
| 13 | 57659200 | 57664399 | 29.04296 | 46.92296 | 98.20671 | 41.92694 | -0.70269 | 1.711308 | 0.25678 |
| 9 | 106392200 | 106393999 | 9.480962 | 17.21549 | 36.49864 | 15.7552 | -0.66239 | 0.317517 | 0.256789 |
| 1 | 130461600 | 130462199 | 10.12918 | 19.64817 | 17.61415 | 6.640706 | 0.735674 | -0.20876 | 0.256799 |
| 15 | 14572400 | 14575199 | 30.75693 | 52.31404 | 40.27294 | 17.65114 | 0.757607 | 1.03875 | 0.256816 |
| 9 | 19220600 | 19222399 | 26.60606 | 45.80529 | 37.61254 | 16.49905 | 0.667514 | 0.911806 | 0.256818 |
| 5 | 150924800 | 150927999 | 17.62447 | 29.54871 | 64.84776 | 28.28743 | -0.74887 | 1.148994 | 0.256818 |
| 2 | 111422000 | 111422799 | 12.93084 | 24.4768 | 20.22225 | 8.193361 | 0.756739 | 0.056958 | 0.256829 |
| 6 | 42189200 | 42189799 | 6.159537 | 11.76201 | 28.3244 | 11.26535 | -0.74923 | -0.04183 | 0.256835 |
| 6 | 41344200 | 41345999 | 23.29998 | 40.7224 | 32.07866 | 14.13734 | 0.730062 | 0.720858 | 0.256836 |
| 3 | 71621200 | 71621599 | 4.18531 | 8.158382 | 22.50239 | 8.130661 | -0.81037 | -0.41031 | 0.25684 |
| 2 | 4806800 | 4809199 | 13.24978 | 23.205 | 46.73839 | 20.63392 | -0.61747 | 0.662283 | 0.256841 |
| 5 | 126596200 | 126597599 | 14.68611 | 27.13625 | 23.19553 | 9.785462 | 0.662305 | 0.228315 | 0.256866 |
| 5 | 121823000 | 121823799 | 4.363692 | 8.480372 | 23.90341 | 8.89743 | -0.88145 | -0.29777 | 0.256882 |
| 9 | 69428000 | 69429599 | 9.495507 | 16.77229 | 41.01462 | 18.14496 | -0.89202 | 0.518664 | 0.256884 |
| 16 | 7775200 | 7777799 | 27.5412 | 47.76327 | 33.62555 | 14.75341 | 0.89052 | 0.846774 | 0.256887 |
| 8 | 58163200 | 58164999 | 15.31463 | 26.01994 | 58.38927 | 25.58906 | -0.77985 | 1.014309 | 0.256898 |
| 14 | 98010600 | 98011799 | 12.13902 | 22.91199 | 19.83724 | 8.086408 | 0.682202 | 0.068186 | 0.256908 |
| 12 | 22361800 | 22362599 | 23.72751 | 41.44962 | 29.71427 | 13.09081 | 0.865499 | 0.727982 | 0.256912 |
| 11 | 83535800 | 83537799 | 22.07709 | 38.67616 | 31.60012 | 13.95421 | 0.677575 | 0.677187 | 0.25692 |
| 4 | 46580400 | 46582599 | 81.56802 | 128.983 | 102.1995 | 42.46579 | 0.662275 | 2.412766 | 0.256939 |
| 7 | 47083800 | 47088999 | 70.00403 | 111.3381 | 100.5783 | 41.99289 | 0.480917 | 2.247314 | 0.25695 |
| 7 | 133734800 | 133737399 | 19.9598 | 33.54039 | 63.98807 | 27.9496 | -0.54765 | 1.132278 | 0.256954 |
| 10 | 74570800 | 74571599 | 10.93738 | 21.10193 | 18.47786 | 7.139561 | 0.735413 | -0.11784 | 0.256967 |
| 8 | 13099800 | 13101399 | 18.12185 | 32.20586 | 27.15839 | 11.98167 | 0.638768 | 0.487714 | 0.256972 |
| 9 | 74610000 | 74611999 | 8.630592 | 15.97315 | 36.44501 | 15.341 | -0.74233 | 0.200193 | 0.257002 |
| 3 | 127439600 | 127441199 | 8.632945 | 16.02876 | 34.28801 | 14.37028 | -0.64503 | 0.179052 | 0.257012 |
| 19 | 10356600 | 10357999 | 14.26574 | 26.44477 | 22.80142 | 9.577036 | 0.654553 | 0.206979 | 0.257014 |
| 7 | 5608400 | 5615199 | 105.5386 | 165.7251 | 114.0186 | 47.29102 | 0.857983 | 2.576365 | 0.257029 |
| 3 | 6939200 | 6940999 | 23.28775 | 40.7711 | 31.5516 | 13.92529 | 0.755591 | 0.694236 | 0.257034 |
| 2 | 169139800 | 169150799 | 102.8651 | 156.2193 | 271.228 | 112.3188 | -0.51354 | 3.211281 | 0.257039 |
| 4 | 16330000 | 16330999 | 14.28649 | 26.30421 | 22.70539 | 9.625119 | 0.643211 | 0.242893 | 0.257039 |
| 17 | 88261600 | 88262599 | 13.0313 | 24.32978 | 21.35687 | 8.870769 | 0.641071 | 0.14959 | 0.257057 |
| 18 | 82827400 | 82828599 | 12.12274 | 22.93043 | 19.81295 | 8.039922 | 0.690715 | 0.053199 | 0.257073 |
| 15 | 82485800 | 82507199 | 224.9908 | 322.7894 | 516.1029 | 213.1133 | -0.4714 | 4.19882 | 0.257078 |
| 18 | 24475000 | 24478599 | 18.91547 | 32.07498 | 61.9092 | 27.11602 | -0.56386 | 1.046696 | 0.25708 |
| 6 | 82350200 | 82354399 | 36.00084 | 59.55561 | 52.09342 | 22.52455 | 0.573127 | 1.337647 | 0.257082 |

| | | | | | | | | | |
|---|---|---|---|---|---|---|---|---|---|
| 4 | 135220800 | 135223799 | 12.76527 | 21.92332 | 54.68986 | 23.98838 | -0.92909 | 0.899988 | 0.257107 |
| 15 | 94705600 | 94708399 | 20.61271 | 34.58968 | 65.68414 | 28.66847 | -0.54103 | 1.150148 | 0.257112 |
| X | 164452400 | 164453399 | 12.02485 | 23.30582 | 14.81509 | 5.6073 | 1.219252 | -0.18516 | 0.257122 |
| 18 | 37548400 | 37550999 | 25.30159 | 44.42803 | 28.13221 | 12.44721 | 1.043413 | 0.655182 | 0.257126 |
| 2 | 155322800 | 155325199 | 24.32305 | 42.07478 | 36.65364 | 16.08075 | 0.583929 | 0.858934 | 0.257129 |
| 14 | 52865200 | 52867599 | 23.57556 | 41.69219 | 25.21494 | 11.15249 | 1.111948 | 0.558527 | 0.257139 |
| 3 | 137369600 | 137375599 | 39.25965 | 64.59614 | 49.96517 | 21.54067 | 0.748091 | 1.408071 | 0.257147 |
| 12 | 26117400 | 26119799 | 11.37583 | 19.914 | 45.70997 | 20.17213 | -0.80492 | 0.656368 | 0.257147 |
| 11 | 103765400 | 103766199 | 8.782729 | 15.82818 | 36.97136 | 16.11005 | -0.80935 | 0.361388 | 0.257159 |
| 17 | 84290200 | 84292599 | 14.65745 | 25.41975 | 51.11107 | 22.42529 | -0.61666 | 0.813426 | 0.257162 |
| 6 | 131584000 | 131587599 | 59.74966 | 95.88654 | 80.42831 | 34.01504 | 0.601935 | 1.953276 | 0.257202 |
| 3 | 67769400 | 67771599 | 19.43741 | 35.08563 | 24.05432 | 10.46074 | 0.948984 | 0.395458 | 0.257209 |
| 4 | 134019600 | 134024199 | 23.83131 | 39.25496 | 75.33821 | 32.49473 | -0.55835 | 1.377888 | 0.257227 |
| 13 | 29272800 | 29274599 | 21.56008 | 38.17896 | 26.46748 | 11.70335 | 0.916883 | 0.533817 | 0.257237 |
| 1 | 67791800 | 67792599 | 11.86319 | 22.59465 | 19.89493 | 7.956366 | 0.682078 | 0.004838 | 0.257256 |
| 15 | 85763000 | 85763999 | 5.905617 | 11.22729 | 29.88706 | 11.99874 | -0.9052 | -0.01214 | 0.257259 |
| 9 | 108139600 | 108141799 | 9.879654 | 17.64636 | 38.3999 | 16.86959 | -0.71661 | 0.425096 | 0.257261 |
| 16 | 10592400 | 10596999 | 18.9093 | 31.59543 | 66.52895 | 28.95873 | -0.68944 | 1.191243 | 0.257274 |
| 1 | 192505000 | 192509199 | 29.60134 | 48.06941 | 88.74937 | 38.05325 | -0.51496 | 1.596456 | 0.257279 |
| 11 | 86705400 | 86706799 | 5.38274 | 10.1808 | 34.22565 | 13.89068 | -1.25328 | 0.028212 | 0.257291 |
| 15 | 96611400 | 96614199 | 14.62965 | 25.49945 | 51.10377 | 22.48246 | -0.61113 | 0.745202 | 0.257309 |
| 1 | 115026200 | 115028999 | 36.04766 | 59.59383 | 52.62998 | 22.74621 | 0.559267 | 1.346232 | 0.257318 |
| 3 | 62898800 | 62900799 | 25.60315 | 43.95151 | 38.37906 | 16.83358 | 0.578508 | 0.935759 | 0.257326 |
| 9 | 63943000 | 63944799 | 16.37214 | 29.40385 | 24.74215 | 10.81934 | 0.650374 | 0.405325 | 0.257331 |
| 18 | 37559200 | 37564999 | 94.51923 | 149.7037 | 97.02876 | 40.35015 | 0.953526 | 2.385462 | 0.257335 |
| 10 | 81949000 | 81950199 | 29.01857 | 49.95446 | 36.42663 | 15.9788 | 0.838139 | 0.922385 | 0.257341 |
| 4 | 152770400 | 152776599 | 64.66056 | 100.5665 | 189.6594 | 78.73793 | -0.60574 | 2.733895 | 0.257387 |
| 6 | 18433800 | 18436199 | 13.62239 | 23.53482 | 51.58392 | 22.63013 | -0.74147 | 0.845494 | 0.25739 |
| 9 | 75456800 | 75459599 | 18.4209 | 31.3564 | 60.27398 | 26.41903 | -0.55734 | 1.00943 | 0.257402 |
| 5 | 64880400 | 64884399 | 53.47123 | 86.26468 | 70.29429 | 29.95417 | 0.652656 | 1.780456 | 0.257406 |
| 7 | 10337800 | 10344199 | 64.7252 | 104.4198 | 59.45344 | 25.33436 | 1.168648 | 1.7974 | 0.257412 |
| 13 | 90680600 | 90681599 | 19.1054 | 34.7545 | 21.86663 | 9.417677 | 1.080492 | 0.344516 | 0.257415 |
| 13 | 54207000 | 54210399 | 21.66391 | 35.54211 | 78.10346 | 33.63037 | -0.75536 | 1.424219 | 0.257427 |
| 2 | 84948400 | 84950599 | 25.01458 | 43.64107 | 32.70268 | 14.39557 | 0.801835 | 0.757656 | 0.257442 |
| 9 | 88430200 | 88430799 | 4.865104 | 9.401329 | 27.65443 | 10.61278 | -0.98664 | -0.1793 | 0.25746 |
| 2 | 147030600 | 147033399 | 27.20937 | 44.16136 | 90.03771 | 38.59036 | -0.65842 | 1.602474 | 0.257476 |
| 7 | 43082800 | 43097999 | 107.5603 | 162.5045 | 288.4343 | 119.2521 | -0.55166 | 3.318555 | 0.257482 |
| 7 | 32293800 | 32301999 | 84.77286 | 134.5817 | 97.09539 | 40.4633 | 0.802269 | 2.313684 | 0.257492 |
| 7 | 9988200 | 9990799 | 30.15257 | 51.25626 | 37.78882 | 16.56161 | 0.819677 | 1.040029 | 0.257497 |
| 14 | 103568000 | 103569199 | 12.67781 | 24.17115 | 19.80908 | 7.904732 | 0.787494 | 0.005707 | 0.257503 |
| 17 | 83483600 | 83487599 | 18.17315 | 30.63061 | 64.54745 | 28.21981 | -0.69046 | 1.098352 | 0.257505 |
| 2 | 14006200 | 14008199 | 29.22895 | 49.99323 | 38.62827 | 16.93791 | 0.753404 | 0.981282 | 0.257506 |
| 2 | 169044400 | 169067999 | 234.8312 | 335.414 | 534.5537 | 220.8865 | -0.47314 | 4.264457 | 0.257511 |
| 5 | 92582000 | 92583399 | 8.454012 | 15.31483 | 36.48673 | 15.7963 | -0.83207 | 0.323934 | 0.257532 |
| 6 | 114791800 | 114792799 | 13.91507 | 25.75698 | 22.34886 | 9.405668 | 0.643477 | 0.210966 | 0.25754 |
| 3 | 31262600 | 31264599 | 9.537067 | 17.13413 | 38.4329 | 16.80124 | -0.7553 | 0.384331 | 0.257564 |
| 1 | 26536000 | 26537199 | 15.11538 | 28.3773 | 21.14537 | 8.701055 | 0.886527 | 0.13358 | 0.257571 |

| | | | | | | | | | |
|---|---|---|---|---|---|---|---|---|---|
| 7 | 30287400 | 30291199 | 23.25052 | 37.73733 | 88.31066 | 37.85107 | -0.85644 | 1.595745 | 0.257574 |
| 5 | 81983600 | 81984399 | 16.30265 | 31.00935 | 17.79042 | 7.134506 | 1.291558 | 0.051682 | 0.257596 |
| 18 | 75952200 | 75955399 | 22.75861 | 37.35198 | 79.02341 | 34.03184 | -0.70066 | 1.421407 | 0.257597 |
| 4 | 146300800 | 146309399 | 180.718 | 272.4488 | 205.0031 | 84.70117 | 0.682898 | 3.372816 | 0.257603 |
| 2 | 69523000 | 69524599 | 9.310329 | 17.05739 | 35.60713 | 15.19646 | -0.62899 | 0.263964 | 0.257609 |
| 16 | 65856000 | 65859799 | 62.04519 | 99.1481 | 89.92555 | 37.79386 | 0.482441 | 2.088241 | 0.257611 |
| 13 | 115721800 | 115722999 | 13.31876 | 25.43296 | 19.19191 | 7.633129 | 0.909638 | -0.00012 | 0.25762 |
| 2 | 35623200 | 35625599 | 24.73829 | 42.82548 | 36.03315 | 15.8089 | 0.634006 | 0.852562 | 0.257624 |
| 2 | 89510200 | 89512799 | 26.93346 | 46.33632 | 36.46597 | 15.99575 | 0.728231 | 0.916438 | 0.257629 |
| 5 | 131346200 | 131346599 | 8.439677 | 16.47786 | 14.5026 | 5.115199 | 0.847742 | -0.46991 | 0.257638 |
| 17 | 49248400 | 49250599 | 10.97277 | 19.4453 | 42.96419 | 18.995 | -0.74562 | 0.490211 | 0.257651 |
| X | 148221600 | 148223599 | 15.08772 | 27.72455 | 23.72869 | 10.08601 | 0.652894 | 0.263849 | 0.257675 |
| 15 | 92633000 | 92633999 | 13.43797 | 25.17478 | 21.85397 | 9.024948 | 0.663756 | 0.133656 | 0.257689 |
| 4 | 18471600 | 18478399 | 87.15594 | 137.0263 | 122.4727 | 50.80554 | 0.48227 | 2.540982 | 0.257711 |
| 13 | 67673600 | 67674399 | 7.696168 | 14.42009 | 31.75723 | 13.11152 | -0.66904 | 0.10779 | 0.257712 |
| 2 | 46271600 | 46273199 | 23.37665 | 40.79344 | 29.84773 | 13.14084 | 0.836058 | 0.740966 | 0.257713 |
| 13 | 100400200 | 100403199 | 17.25039 | 29.61625 | 58.72684 | 25.75864 | -0.60006 | 0.924731 | 0.25772 |
| 15 | 72667400 | 72683199 | 194.9129 | 280.8332 | 482.7433 | 199.2091 | -0.56967 | 4.134279 | 0.257725 |
| X | 33094200 | 33095199 | 15.91696 | 29.70778 | 21.53849 | 8.9511 | 0.914411 | 0.177347 | 0.257734 |
| 15 | 87330000 | 87330599 | 12.52079 | 23.6659 | 19.98749 | 8.123398 | 0.722156 | 0.064877 | 0.257745 |
| 10 | 81551600 | 81553199 | 14.36788 | 26.97039 | 19.1897 | 7.898097 | 0.952377 | 0.122828 | 0.257751 |
| 12 | 64192600 | 64193599 | 14.36788 | 26.97039 | 19.1897 | 7.898097 | 0.952377 | 0.122828 | 0.257751 |
| 3 | 132288000 | 132289199 | 22.30458 | 38.75997 | 33.62003 | 14.76274 | 0.591213 | 0.798426 | 0.257756 |
| 8 | 83267200 | 83268599 | 16.03135 | 29.48014 | 23.55618 | 10.0027 | 0.752248 | 0.266394 | 0.257762 |
| 18 | 27310600 | 27313799 | 21.8883 | 38.74222 | 27.11208 | 11.98949 | 0.903347 | 0.543222 | 0.257775 |
| 9 | 57588800 | 57590199 | 8.899172 | 15.77035 | 41.98554 | 18.56238 | -1.0124 | 0.476727 | 0.257777 |
| 13 | 51251000 | 51261399 | 85.89647 | 132.1989 | 220.7405 | 91.67777 | -0.44209 | 2.936194 | 0.25778 |
| 3 | 16599200 | 16600199 | 24.81517 | 42.71074 | 37.77544 | 16.57038 | 0.560761 | 0.906876 | 0.257788 |
| 10 | 7769800 | 7771199 | 15.39258 | 28.95854 | 20.53846 | 8.42128 | 0.961141 | 0.121219 | 0.257791 |
| 4 | 78524600 | 78524999 | 2.637119 | 5.15834 | 17.8247 | 5.968518 | -1.02574 | -0.7208 | 0.257796 |
| 5 | 95642200 | 95645599 | 41.35973 | 68.57451 | 47.16556 | 20.42543 | 0.919058 | 1.324312 | 0.257797 |
| 1 | 5639000 | 5640599 | 16.89615 | 30.55149 | 24.88636 | 10.79911 | 0.703788 | 0.36856 | 0.2578 |
| 19 | 28901000 | 28903399 | 15.27385 | 26.51371 | 52.21643 | 22.91302 | -0.58671 | 0.808021 | 0.257826 |
| 17 | 38663400 | 38673999 | 326.3509 | 465.5864 | 389.3097 | 160.901 | 0.455327 | 4.284778 | 0.257827 |
| 19 | 18422600 | 18423399 | 8.819726 | 15.77591 | 37.96357 | 16.65765 | -0.85903 | 0.410237 | 0.257827 |
| 17 | 19477800 | 19480199 | 56.35399 | 90.4553 | 80.66485 | 34.1255 | 0.514069 | 1.941662 | 0.25783 |
| 11 | 102199800 | 102202199 | 10.11326 | 17.55466 | 51.50606 | 22.60063 | -1.1581 | 0.786092 | 0.257833 |
| 12 | 23956400 | 23957999 | 37.32265 | 61.39705 | 45.14252 | 19.45995 | 0.820964 | 1.403199 | 0.257845 |
| 6 | 115685800 | 115687999 | 13.35636 | 23.39363 | 46.73081 | 20.63204 | -0.60583 | 0.658948 | 0.257872 |
| 17 | 53624000 | 53626799 | 32.29538 | 55.06283 | 38.00448 | 16.66013 | 0.914987 | 1.021491 | 0.257876 |
| 13 | 13238600 | 13240199 | 34.25974 | 56.7036 | 50.91397 | 22.02214 | 0.535574 | 1.327491 | 0.257877 |
| 1 | 175863000 | 175863599 | 11.43684 | 22.01765 | 17.75566 | 6.903454 | 0.844284 | -0.09596 | 0.257883 |
| 2 | 116893200 | 116894999 | 12.14601 | 21.24125 | 45.72781 | 20.16318 | -0.71292 | 0.677395 | 0.257893 |
| X | 14160600 | 14161399 | 13.28704 | 25.42536 | 18.8955 | 7.475294 | 0.939352 | -0.01577 | 0.257906 |
| 7 | 122477400 | 122480399 | 18.07505 | 30.82653 | 60.31614 | 26.44157 | -0.58244 | 0.989949 | 0.257924 |
| 5 | 122020400 | 122029799 | 94.19907 | 142.5416 | 278.3166 | 115.126 | -0.68716 | 3.283909 | 0.257927 |
| 5 | 112874400 | 112876599 | 10.59317 | 18.61076 | 43.06342 | 19.06207 | -0.81608 | 0.589925 | 0.257927 |

| | | | | | | | | | |
|---|---|---|---|---|---|---|---|---|---|
| 7 | 41169200 | 41176399 | 96.12383 | 150.6412 | 128.4006 | 53.25763 | 0.547121 | 2.608816 | 0.257937 |
| 6 | 109297400 | 109299999 | 14.66729 | 25.58272 | 53.82775 | 23.69111 | -0.68093 | 0.740276 | 0.257941 |
| 12 | 7981800 | 7983999 | 25.91332 | 44.67053 | 37.71132 | 16.54303 | 0.628165 | 0.898938 | 0.257948 |
| 9 | 26940600 | 26942399 | 9.197432 | 16.0326 | 50.43542 | 22.18972 | -1.25721 | 0.717672 | 0.257966 |
| 6 | 14371400 | 14373599 | 25.77482 | 44.76927 | 34.92077 | 15.32935 | 0.743749 | 0.824806 | 0.257969 |
| 15 | 100789400 | 100794799 | 57.865 | 91.15297 | 160.3438 | 66.53256 | -0.49105 | 2.492533 | 0.257972 |
| 5 | 107088200 | 107089599 | 16.89175 | 30.66338 | 25.02327 | 10.80546 | 0.705913 | 0.348202 | 0.25798 |
| 4 | 33602800 | 33605199 | 23.19511 | 40.48269 | 33.25691 | 14.64316 | 0.669744 | 0.742253 | 0.257981 |
| 5 | 95118000 | 95120999 | 64.94101 | 104.4426 | 67.68339 | 28.71642 | 0.977028 | 1.889036 | 0.257982 |
| 7 | 84847200 | 84848399 | 24.58934 | 42.88073 | 33.21535 | 14.61729 | 0.754145 | 0.761315 | 0.257983 |
| 7 | 108213200 | 108214799 | 18.3403 | 32.45711 | 28.08798 | 12.42145 | 0.598536 | 0.523775 | 0.257993 |
| 2 | 39470200 | 39471999 | 15.93403 | 30.12344 | 17.52868 | 7.121035 | 1.254308 | 0.086418 | 0.257997 |
| 3 | 131246400 | 131250599 | 51.94968 | 82.50164 | 143.1838 | 59.69074 | -0.46173 | 2.284798 | 0.258003 |
| 2 | 64781800 | 64782599 | 9.59244 | 18.65664 | 16.53282 | 6.11472 | 0.776903 | -0.28516 | 0.258004 |
| 6 | 117564000 | 117564599 | 15.41616 | 28.18124 | 24.20567 | 10.36135 | 0.640605 | 0.296142 | 0.25801 |
| 14 | 30671200 | 30673799 | 15.26913 | 26.51901 | 52.76421 | 23.1608 | -0.60137 | 0.801185 | 0.258011 |
| 17 | 66678200 | 66680599 | 12.88506 | 22.6442 | 45.06716 | 19.95416 | -0.60041 | 0.599839 | 0.258018 |
| 2 | 87625400 | 87628599 | 36.29807 | 60.52053 | 47.06003 | 20.45665 | 0.742874 | 1.247369 | 0.258027 |
| 4 | 88671200 | 88671999 | 11.7115 | 22.72214 | 16.86207 | 6.351069 | 1.00665 | -0.19644 | 0.258035 |
| 17 | 9085800 | 9088799 | 17.39765 | 29.3785 | 60.56979 | 26.49585 | -0.65865 | 1.078298 | 0.258036 |
| 14 | 15986200 | 15988399 | 42.90409 | 69.68953 | 63.05705 | 27.04221 | 0.511712 | 1.601465 | 0.258042 |
| 2 | 147238400 | 147241399 | 29.01817 | 49.11692 | 43.14145 | 18.88541 | 0.567747 | 1.077648 | 0.258043 |
| 9 | 18937400 | 18942399 | 45.1625 | 73.86512 | 51.71066 | 22.23789 | 0.887939 | 1.498665 | 0.258058 |
| 2 | 177598200 | 177599399 | 7.708309 | 14.01995 | 39.75769 | 17.12646 | -1.0773 | 0.299394 | 0.258073 |
| 16 | 97751000 | 97753999 | 19.37055 | 32.41776 | 64.90912 | 28.28207 | -0.61704 | 1.175111 | 0.258081 |
| Y | 43619400 | 43621999 | 17.12258 | 31.06991 | 13.83568 | 5.977483 | 1.571759 | 0.322477 | 0.258089 |
| X | 18421200 | 18423999 | 28.02421 | 49.15207 | 26.33308 | 11.63983 | 1.283251 | 0.685449 | 0.25809 |
| 11 | 70191800 | 70192999 | 19.39468 | 34.39617 | 27.89947 | 12.33026 | 0.692724 | 0.509448 | 0.258101 |
| 7 | 4144000 | 4155399 | 78.25453 | 119.6647 | 240.3764 | 99.72555 | -0.71563 | 3.062956 | 0.25811 |
| 4 | 49058400 | 49061399 | 17.20633 | 28.74309 | 66.8682 | 29.10201 | -0.83262 | 1.189516 | 0.258131 |
| 14 | 3572200 | 3574399 | 25.01066 | 43.36091 | 36.03419 | 15.81006 | 0.652301 | 0.841954 | 0.258136 |
| 7 | 72725000 | 72728199 | 36.97082 | 61.84288 | 44.80982 | 19.51848 | 0.844342 | 1.215716 | 0.258143 |
| 9 | 19303600 | 19306599 | 36.97082 | 61.84288 | 44.80982 | 19.51848 | 0.844342 | 1.215716 | 0.258143 |
| 11 | 100115200 | 100116999 | 19.83079 | 36.73022 | 19.42198 | 8.166862 | 1.353447 | 0.249198 | 0.258157 |
| X | 122180600 | 122180999 | 9.255434 | 18.0052 | 15.63796 | 5.773352 | 0.809252 | -0.2959 | 0.25816 |
| 4 | 151576600 | 151584599 | 64.21807 | 100.1711 | 182.3284 | 75.66196 | -0.55131 | 2.682365 | 0.258178 |
| 1 | 40879600 | 40881799 | 10.86154 | 19.18413 | 40.94351 | 18.1137 | -0.69736 | 0.525888 | 0.258193 |
| X | 117838000 | 117840399 | 33.72699 | 57.33648 | 38.56755 | 16.903 | 0.951483 | 1.054038 | 0.258197 |
| 7 | 12044600 | 12049399 | 95.78614 | 150.4947 | 122.0917 | 50.64312 | 0.621135 | 2.560983 | 0.258198 |
| 11 | 45587400 | 45588999 | 11.45814 | 20.22193 | 42.97202 | 19.01412 | -0.69201 | 0.539435 | 0.25822 |
| 11 | 69087600 | 69089199 | 10.31405 | 18.475 | 38.39284 | 16.82588 | -0.6483 | 0.40931 | 0.258223 |
| 3 | 23708000 | 23709399 | 14.10504 | 25.98506 | 22.22394 | 9.413663 | 0.657215 | 0.237014 | 0.258227 |
| 12 | 29156000 | 29157799 | 8.497946 | 15.7132 | 33.63951 | 14.17779 | -0.65189 | 0.20014 | 0.25823 |
| 9 | 72950800 | 72953399 | 11.68516 | 20.47554 | 44.66148 | 19.72575 | -0.73147 | 0.642809 | 0.258251 |
| 19 | 35632600 | 35634599 | 18.04904 | 29.94722 | 74.37698 | 32.22581 | -0.92703 | 1.276508 | 0.258252 |
| 7 | 19294400 | 19300199 | 36.08967 | 57.82213 | 114.3368 | 48.23552 | -0.63563 | 1.987093 | 0.258265 |
| 1 | 88255800 | 88256999 | 8.534351 | 15.66921 | 34.04761 | 14.48866 | -0.68317 | 0.242224 | 0.258268 |

| | | | | | | | | | |
|---|---|---|---|---|---|---|---|---|---|
| 8 | 109611600 | 109615399 | 20.29432 | 33.56439 | 74.12257 | 32.04428 | -0.7588 | 1.317198 | 0.258269 |
| X | 52401200 | 52401599 | 9.772092 | 19.1021 | 11.97812 | 4.121743 | 1.361311 | -0.54169 | 0.258276 |
| 13 | 68584200 | 68586399 | 29.06519 | 49.03354 | 43.74364 | 19.12976 | 0.545266 | 1.10722 | 0.258305 |
| 15 | 45770800 | 45774199 | 71.41984 | 113.2951 | 103.0379 | 42.90642 | 0.467579 | 2.32389 | 0.258312 |
| 15 | 13951000 | 13952199 | 13.146 | 24.98316 | 20.3738 | 8.185678 | 0.786443 | 0.034453 | 0.258318 |
| 12 | 106735200 | 106739999 | 32.73947 | 52.82752 | 98.44456 | 41.95914 | -0.53777 | 1.755718 | 0.258322 |
| 1 | 148073200 | 148074199 | 14.35797 | 26.67604 | 22.71877 | 9.512979 | 0.675215 | 0.194412 | 0.258328 |
| 1 | 20547800 | 20548599 | 6.275376 | 12.09012 | 28.157 | 10.92398 | -0.66809 | -0.12359 | 0.258338 |
| 2 | 127291200 | 127292999 | 8.106032 | 14.8239 | 35.04466 | 14.99364 | -0.80972 | 0.262722 | 0.25834 |
| 8 | 13573000 | 13574999 | 8.106032 | 14.8239 | 35.04466 | 14.99364 | -0.80972 | 0.262722 | 0.25834 |
| 1 | 141780200 | 141781999 | 34.68706 | 58.07026 | 42.16606 | 18.37639 | 0.841187 | 1.196764 | 0.258349 |
| 7 | 99375600 | 99377599 | 24.27594 | 43.04053 | 24.14949 | 10.67555 | 1.220968 | 0.535749 | 0.258351 |
| X | 4022600 | 4028999 | 104.6813 | 163.2525 | 133.866 | 55.55256 | 0.597477 | 2.700564 | 0.258356 |
| 17 | 67935400 | 67941799 | 39.9654 | 63.98628 | 116.3375 | 49.02657 | -0.51604 | 2.014437 | 0.25837 |
| 6 | 107927800 | 107928799 | 10.72684 | 18.89266 | 46.1471 | 20.42736 | -0.89288 | 0.564354 | 0.258396 |
| X | 49275000 | 49275999 | 9.952479 | 19.3079 | 17.61925 | 6.638618 | 0.711024 | -0.21304 | 0.258396 |
| 12 | 88438800 | 88439799 | 8.001365 | 14.94635 | 32.79099 | 13.60832 | -0.66982 | 0.131907 | 0.258399 |
| 2 | 89476000 | 89477199 | 19.58969 | 34.46614 | 29.58794 | 13.10007 | 0.606887 | 0.596453 | 0.258411 |
| 16 | 17317600 | 17321799 | 23.88924 | 39.15047 | 78.35008 | 33.72106 | -0.62153 | 1.443376 | 0.258413 |
| 13 | 70029600 | 70032199 | 42.96592 | 68.52343 | 126.0695 | 52.83651 | -0.53927 | 2.141495 | 0.258415 |
| 18 | 10609400 | 10610599 | 9.900523 | 17.61489 | 39.91032 | 17.5908 | -0.77798 | 0.451862 | 0.258418 |
| 19 | 55572400 | 55573999 | 9.900523 | 17.61489 | 39.91032 | 17.5908 | -0.77798 | 0.451862 | 0.258418 |
| 2 | 152403200 | 152404399 | 14.02768 | 25.95515 | 22.77507 | 9.590202 | 0.626688 | 0.214455 | 0.258419 |
| 11 | 68446600 | 68449799 | 20.35159 | 33.50338 | 75.27173 | 32.45795 | -0.78508 | 1.37728 | 0.258427 |
| 18 | 45114200 | 45115599 | 15.97174 | 30.02951 | 19.85417 | 8.149256 | 1.060188 | 0.128029 | 0.258438 |
| 5 | 59371400 | 59374599 | 29.5617 | 50.11426 | 41.98951 | 18.38975 | 0.635538 | 1.064105 | 0.258439 |
| 1 | 34657400 | 34659999 | 26.29093 | 43.04751 | 80.46038 | 34.61607 | -0.52389 | 1.460983 | 0.258446 |
| 5 | 55712800 | 55714599 | 19.98089 | 35.47726 | 27.21769 | 12.01709 | 0.773458 | 0.504083 | 0.258446 |
| 3 | 151445400 | 151446799 | 16.02665 | 28.95717 | 25.14611 | 10.92224 | 0.611394 | 0.368229 | 0.25847 |
| 4 | 26194200 | 26196599 | 32.02822 | 53.80918 | 45.31693 | 19.79294 | 0.627885 | 1.153725 | 0.258475 |
| 5 | 19615200 | 19616999 | 18.656 | 33.70633 | 24.72796 | 10.74125 | 0.853052 | 0.386399 | 0.258505 |
| 8 | 79417600 | 79419599 | 9.413286 | 16.85714 | 37.91468 | 16.61995 | -0.76168 | 0.404422 | 0.258522 |
| 1 | 175975600 | 175977199 | 24.72875 | 43.03372 | 34.34829 | 15.09601 | 0.710864 | 0.792274 | 0.258548 |
| 8 | 27974600 | 27976799 | 24.53054 | 43.25443 | 26.35023 | 11.66143 | 1.100247 | 0.597875 | 0.258582 |
| 2 | 167900400 | 167909399 | 54.81077 | 86.22096 | 166.0875 | 68.9033 | -0.62345 | 2.518817 | 0.258583 |
| 13 | 28124600 | 28126799 | 26.08258 | 45.05436 | 36.50914 | 16.01666 | 0.687448 | 0.878586 | 0.258586 |
| X | 17762800 | 17774399 | 156.0495 | 237.8839 | 170.9288 | 70.86255 | 0.762645 | 3.144563 | 0.258594 |
| 4 | 34501400 | 34502599 | 23.69211 | 40.90534 | 35.78867 | 15.70036 | 0.577258 | 0.872334 | 0.258602 |
| 8 | 95079400 | 95084199 | 38.18846 | 61.13542 | 117.0642 | 49.32572 | -0.59078 | 2.015247 | 0.258609 |
| 19 | 36811200 | 36812399 | 8.380454 | 15.39149 | 34.05291 | 14.48471 | -0.70859 | 0.238834 | 0.25862 |
| 7 | 47468400 | 47469999 | 18.71921 | 33.53809 | 26.35962 | 11.55011 | 0.746215 | 0.439694 | 0.258648 |
| 13 | 46970600 | 46974599 | 18.56334 | 31.1047 | 64.92218 | 28.30951 | -0.6768 | 1.158249 | 0.258651 |
| 1 | 86452600 | 86457999 | 30.87297 | 49.42336 | 123.5072 | 52.03935 | -0.97403 | 2.008977 | 0.258661 |
| 19 | 44919400 | 44920199 | 5.995237 | 11.51604 | 27.41953 | 10.72927 | -0.71115 | -0.10071 | 0.258663 |
| 3 | 91277200 | 91278599 | 13.17925 | 25.05925 | 19.08127 | 7.658023 | 0.885084 | 0.025783 | 0.258684 |
| 1 | 95183400 | 95184399 | 8.537978 | 15.63625 | 35.06417 | 14.97271 | -0.7322 | 0.257571 | 0.258686 |
| 11 | 98840600 | 98843799 | 15.51662 | 26.24358 | 60.487 | 26.47232 | -0.81863 | 1.057188 | 0.258689 |

| | | | | | | | | | |
|---|---|---|---|---|---|---|---|---|---|
| 6 | 105232200 | 105235799 | 41.2144 | 67.41712 | 58.26329 | 25.05707 | 0.585044 | 1.491533 | 0.258701 |
| 17 | 61688800 | 61690799 | 21.36435 | 37.88223 | 25.9007 | 11.44871 | 0.937399 | 0.518015 | 0.258703 |
| 1 | 85299200 | 85301799 | 41.98981 | 68.26512 | 60.11369 | 25.79507 | 0.551935 | 1.582774 | 0.258707 |
| 2 | 10681200 | 10682799 | 14.28978 | 26.87757 | 19.9366 | 8.177716 | 0.89587 | 0.109519 | 0.258713 |
| 13 | 66488400 | 66491399 | 45.22847 | 75.39451 | 38.63389 | 16.79163 | 1.342409 | 1.272141 | 0.25873 |
| 8 | 89674200 | 89685199 | 139.2731 | 207.6291 | 338.9427 | 139.6809 | -0.4482 | 3.583296 | 0.258751 |
| 15 | 31197400 | 31201599 | 39.29283 | 63.07095 | 112.0837 | 47.41845 | -0.47873 | 1.932644 | 0.258772 |
| 5 | 24976800 | 24991199 | 205.3955 | 296.0112 | 485.1093 | 200.1788 | -0.50047 | 4.131032 | 0.258773 |
| 7 | 132048800 | 132050399 | 9.6387 | 17.45152 | 38.03384 | 16.4772 | -0.70554 | 0.337622 | 0.258777 |
| 4 | 14114600 | 14116999 | 32.54553 | 54.93013 | 43.08214 | 18.8431 | 0.73042 | 1.114829 | 0.258796 |
| 13 | 8920600 | 8921599 | 7.329137 | 13.52021 | 33.07037 | 13.98254 | -0.84543 | 0.204172 | 0.258801 |
| 1 | 160039400 | 160040999 | 8.342793 | 15.43125 | 33.64578 | 14.17426 | -0.67767 | 0.196673 | 0.258805 |
| 9 | 111598000 | 111598999 | 5.853517 | 11.24614 | 27.42553 | 10.72523 | -0.74463 | -0.10471 | 0.258805 |
| X | 157917200 | 157921799 | 45.76509 | 75.2353 | 50.65055 | 21.82822 | 0.947396 | 1.437099 | 0.258806 |
| 8 | 73888400 | 73889399 | 8.165285 | 15.17519 | 34.31064 | 14.36077 | -0.72308 | 0.168505 | 0.258809 |
| 5 | 104886200 | 104892399 | 177.7488 | 266.454 | 232.7516 | 96.02153 | 0.460565 | 3.483012 | 0.258823 |
| 1 | 189723200 | 189726199 | 21.80037 | 36.30299 | 68.90862 | 29.93055 | -0.54035 | 1.239403 | 0.258823 |
| 2 | 50076200 | 50077399 | 5.565424 | 10.73177 | 28.7861 | 11.13986 | -0.8671 | -0.14094 | 0.258826 |
| 14 | 89572400 | 89573999 | 18.60286 | 34.25735 | 21.02369 | 8.910156 | 1.132014 | 0.272906 | 0.258826 |
| 19 | 3395600 | 3399399 | 24.4441 | 40.32658 | 75.64033 | 32.64637 | -0.52453 | 1.359552 | 0.258847 |
| 8 | 114413800 | 114415199 | 8.166661 | 15.21816 | 32.72123 | 13.63526 | -0.64565 | 0.149515 | 0.258855 |
| 9 | 104783800 | 104784999 | 14.57622 | 27.64959 | 19.74765 | 7.965473 | 0.971557 | 0.059434 | 0.258878 |
| 19 | 17333400 | 17334999 | 8.526879 | 15.27908 | 39.99976 | 17.52495 | -0.97894 | 0.396892 | 0.258882 |
| 19 | 44627600 | 44630399 | 14.20918 | 24.60694 | 51.08879 | 22.41454 | -0.66306 | 0.824789 | 0.25891 |
| 10 | 116268600 | 116281399 | 135.5678 | 201.9625 | 344.3298 | 141.8889 | -0.51177 | 3.595074 | 0.258914 |
| X | 4335600 | 4339999 | 49.38082 | 80.77644 | 47.34154 | 20.36008 | 1.143707 | 1.504375 | 0.258932 |
| 7 | 7602800 | 7622999 | 235.0958 | 347.0522 | 270.3057 | 111.3168 | 0.605632 | 3.762276 | 0.258932 |
| 10 | 47213200 | 47214999 | 21.6693 | 37.76256 | 33.00669 | 14.51918 | 0.580535 | 0.754871 | 0.258937 |
| X | 33629000 | 33632399 | 41.29263 | 68.64701 | 44.91889 | 19.48852 | 0.991091 | 1.295297 | 0.258945 |
| 4 | 120954000 | 120958799 | 25.41411 | 41.79424 | 84.54913 | 36.44253 | -0.63529 | 1.40493 | 0.258951 |
| 3 | 104911200 | 104913399 | 12.97476 | 22.49153 | 51.60082 | 22.63986 | -0.8061 | 0.810295 | 0.258963 |
| 10 | 60458600 | 60461399 | 12.97476 | 22.49153 | 51.60082 | 22.63986 | -0.8061 | 0.810295 | 0.258963 |
| 18 | 54115200 | 54118999 | 37.16198 | 59.78315 | 107.1417 | 45.47602 | -0.48692 | 1.858665 | 0.258968 |
| 13 | 95408400 | 95412199 | 19.4231 | 32.78383 | 63.53978 | 27.7912 | -0.57012 | 1.090006 | 0.258968 |
| 16 | 95281000 | 95282599 | 7.672762 | 14.01297 | 34.52255 | 14.79481 | -0.86987 | 0.267838 | 0.258976 |
| 1 | 52856200 | 52857599 | 9.100816 | 16.33179 | 38.43986 | 16.82041 | -0.82514 | 0.388839 | 0.258981 |
| 14 | 65047800 | 65049799 | 9.100816 | 16.33179 | 38.43986 | 16.82041 | -0.82514 | 0.388839 | 0.258981 |
| 9 | 79871800 | 79873399 | 9.313602 | 16.99273 | 35.52485 | 15.24335 | -0.63653 | 0.286981 | 0.258982 |
| 16 | 74352400 | 74353399 | 9.313602 | 16.99273 | 35.52485 | 15.24335 | -0.63653 | 0.286981 | 0.258982 |
| 18 | 88634400 | 88635199 | 9.313602 | 16.99273 | 35.52485 | 15.24335 | -0.63653 | 0.286981 | 0.258982 |
| 7 | 13744400 | 13751799 | 153.8838 | 233.4425 | 196.4127 | 81.30997 | 0.529447 | 3.242363 | 0.258999 |
| 3 | 45112800 | 45116199 | 43.26009 | 69.07122 | 123.4586 | 51.82513 | -0.49588 | 2.104977 | 0.259017 |
| 1 | 143245800 | 143247999 | 26.32421 | 42.97146 | 82.4601 | 35.43662 | -0.56475 | 1.505087 | 0.259054 |
| 13 | 49364000 | 49371599 | 35.88916 | 57.25497 | 131.343 | 55.07032 | -0.85646 | 2.124294 | 0.259058 |
| X | 4557600 | 4561999 | 79.30318 | 126.2526 | 84.75561 | 35.43511 | 0.910006 | 2.221408 | 0.259059 |
| 1 | 30793000 | 30793999 | 6.128568 | 11.8091 | 28.79882 | 11.16805 | -0.73394 | -0.12749 | 0.259062 |
| 15 | 88617200 | 88618999 | 10.04434 | 17.93286 | 38.41087 | 16.88049 | -0.69413 | 0.430419 | 0.259067 |

| | | | | | | | | | |
|---|---|---|---|---|---|---|---|---|---|
| 6 | 52198200 | 52199799 | 6.901069 | 12.73622 | 34.13178 | 14.42256 | -0.97581 | 0.198805 | 0.259071 |
| 5 | 118712400 | 118714399 | 8.558415 | 15.64066 | 34.50945 | 14.77714 | -0.71149 | 0.270347 | 0.259075 |
| 15 | 97210400 | 97212599 | 14.15387 | 24.65192 | 50.02017 | 21.99551 | -0.62895 | 0.751655 | 0.259097 |
| 13 | 58286200 | 58289399 | 41.13039 | 67.07394 | 59.30868 | 25.47819 | 0.549014 | 1.533868 | 0.2591 |
| 7 | 15685800 | 15702599 | 129.5423 | 193.4547 | 333.8949 | 137.6466 | -0.52626 | 3.549201 | 0.259101 |
| 2 | 131642800 | 131643599 | 5.24891 | 10.16593 | 25.48564 | 9.677821 | -0.74307 | -0.215 | 0.259103 |
| 9 | 20688800 | 20692599 | 28.15373 | 45.75316 | 86.15917 | 36.9632 | -0.54231 | 1.577699 | 0.259114 |
| 2 | 173042000 | 173054399 | 101.9887 | 154.5316 | 270.938 | 112.1203 | -0.5304 | 3.258575 | 0.259121 |
| 4 | 139578000 | 139585199 | 53.6348 | 84.49431 | 160.6555 | 66.66242 | -0.60304 | 2.488525 | 0.259124 |
| 8 | 25244800 | 25247799 | 37.53774 | 63.0926 | 40.80581 | 17.82495 | 1.007405 | 1.164738 | 0.259132 |
| 12 | 17371600 | 17375999 | 28.13172 | 45.76488 | 85.82602 | 36.83704 | -0.53526 | 1.560509 | 0.25914 |
| 9 | 115783400 | 115784399 | 9.876616 | 17.67262 | 37.90696 | 16.62786 | -0.69429 | 0.413749 | 0.259148 |
| 4 | 74572800 | 74574199 | 15.27291 | 28.55015 | 20.31986 | 8.422414 | 0.94319 | 0.157363 | 0.25916 |
| 11 | 88422600 | 88424399 | 7.988182 | 15.01811 | 32.99892 | 13.5461 | -0.65926 | 0.09139 | 0.259163 |
| 13 | 62451600 | 62460599 | 344.9186 | 488.3343 | 407.317 | 168.5588 | 0.447841 | 4.39666 | 0.259178 |
| 3 | 103494400 | 103495999 | 11.15851 | 19.72287 | 40.91578 | 18.09882 | -0.65636 | 0.519044 | 0.25919 |
| 15 | 100615800 | 100618999 | 26.89114 | 43.65168 | 88.12221 | 37.77372 | -0.64391 | 1.598487 | 0.259194 |
| 11 | 64407000 | 64407799 | 5.21662 | 10.15335 | 26.087 | 9.591855 | -0.73464 | -0.32655 | 0.259209 |
| 12 | 23474600 | 23474999 | 8.755087 | 17.02548 | 14.97666 | 5.545913 | 0.785693 | -0.29743 | 0.259224 |
| 12 | 70629000 | 70630999 | 23.58409 | 41.31166 | 30.92748 | 13.6559 | 0.803104 | 0.686265 | 0.259225 |
| 5 | 114910600 | 114913799 | 10.84943 | 18.85008 | 50.48261 | 22.16453 | -1.02715 | 0.774045 | 0.259241 |
| 10 | 78845600 | 78847199 | 26.17823 | 45.29203 | 32.99833 | 14.47717 | 0.840767 | 0.861924 | 0.259247 |
| 7 | 15037200 | 15052399 | 579.2046 | 790.3383 | 577.2398 | 240.2762 | 0.58577 | 4.960504 | 0.259249 |
| 1 | 182019000 | 182022199 | 18.91706 | 32.12684 | 61.92045 | 27.13354 | -0.56169 | 1.033863 | 0.259253 |
| 17 | 8644600 | 8646599 | 12.56109 | 22.1397 | 44.538 | 19.71215 | -0.61421 | 0.563752 | 0.259273 |
| 2 | 18804400 | 18807999 | 29.0681 | 46.91117 | 99.48526 | 42.41165 | -0.72353 | 1.745103 | 0.259276 |
| 1 | 108734400 | 108735399 | 19.65043 | 34.76875 | 28.378 | 12.5502 | 0.682375 | 0.534556 | 0.259295 |
| 13 | 3086000 | 3088399 | 98.84506 | 154.8611 | 127.4705 | 52.8734 | 0.59709 | 2.615854 | 0.259302 |
| 9 | 88513200 | 88514399 | 19.03201 | 34.36467 | 24.71508 | 10.74386 | 0.880891 | 0.392797 | 0.259304 |
| 16 | 37688000 | 37689799 | 7.476754 | 13.78816 | 33.06632 | 13.98684 | -0.81757 | 0.207612 | 0.259314 |
| 4 | 154566200 | 154570599 | 22.11805 | 36.33085 | 77.80181 | 33.51754 | -0.71759 | 1.409091 | 0.259323 |
| 4 | 108857400 | 108858599 | 5.272957 | 10.13854 | 26.83481 | 10.47103 | -0.85781 | -0.11811 | 0.259325 |
| 11 | 6575600 | 6576799 | 5.272957 | 10.13854 | 26.83481 | 10.47103 | -0.85781 | -0.11811 | 0.259325 |
| 12 | 106232400 | 106233199 | 5.272957 | 10.13854 | 26.83481 | 10.47103 | -0.85781 | -0.11811 | 0.259325 |
| X | 29426000 | 29427199 | 21.76454 | 38.18014 | 27.68472 | 12.23917 | 0.848904 | 0.648511 | 0.25934 |
| 12 | 70257600 | 70259999 | 8.848069 | 16.15423 | 36.07183 | 15.46503 | -0.73028 | 0.279399 | 0.259346 |
| 1 | 136726200 | 136728399 | 11.15587 | 19.69712 | 41.94366 | 18.55747 | -0.69459 | 0.531602 | 0.25935 |
| 16 | 11223400 | 11224799 | 7.442904 | 13.82862 | 32.10774 | 13.44406 | -0.76064 | 0.162019 | 0.259365 |
| 2 | 64560800 | 64561799 | 13.65161 | 25.56854 | 21.73551 | 8.979885 | 0.69299 | 0.13575 | 0.25937 |
| 7 | 22427400 | 22427799 | 9.905619 | 19.29298 | 14.05375 | 5.133169 | 1.071255 | -0.33388 | 0.259385 |
| 13 | 117758400 | 117760999 | 21.9493 | 38.77201 | 27.59775 | 12.20918 | 0.877826 | 0.564927 | 0.259406 |
| 10 | 81760000 | 81761199 | 14.82743 | 28.32942 | 17.90968 | 7.114883 | 1.164139 | 0.007917 | 0.259411 |
| 16 | 11002600 | 11004999 | 14.27808 | 24.53689 | 54.23477 | 23.78922 | -0.75539 | 0.899015 | 0.259421 |
| 14 | 5521800 | 5524599 | 15.47282 | 26.88282 | 56.61511 | 24.85699 | -0.68297 | 0.802707 | 0.259422 |
| 13 | 44296000 | 44301799 | 76.56214 | 121.2664 | 110.4217 | 45.92046 | 0.463808 | 2.371544 | 0.259422 |
| 10 | 74894000 | 74901799 | 42.03284 | 66.87404 | 139.5152 | 58.27712 | -0.72411 | 2.220017 | 0.259428 |
| 7 | 11533800 | 11544399 | 367.661 | 517.8743 | 433.215 | 179.4278 | 0.436218 | 4.472613 | 0.259429 |

| | | | | | | | | | |
|---|---|---|---|---|---|---|---|---|---|
| 11 | 16025800 | 16027399 | 20.3142 | 35.92449 | 28.30929 | 12.52112 | 0.732498 | 0.544178 | 0.259439 |
| 12 | 107923400 | 107928399 | 24.52497 | 39.79436 | 88.5982 | 37.96622 | -0.78523 | 1.603512 | 0.259452 |
| 13 | 92287800 | 92288399 | 4.061239 | 7.913381 | 22.42804 | 8.146143 | -0.85592 | -0.3939 | 0.259454 |
| 9 | 116423400 | 116426799 | 33.90695 | 56.52883 | 48.69467 | 21.16605 | 0.595708 | 1.243444 | 0.259458 |
| 17 | 20146600 | 20158399 | 157.4558 | 239.1483 | 176.3652 | 73.03708 | 0.720851 | 3.217625 | 0.259459 |
| 5 | 102409600 | 102411199 | 13.20792 | 23.15144 | 45.65827 | 20.17162 | -0.58734 | 0.644719 | 0.259466 |
| 18 | 24784600 | 24788199 | 19.80645 | 32.80656 | 76.5376 | 33.12253 | -0.83764 | 1.298928 | 0.259469 |
| 9 | 19175400 | 19176399 | 19.69609 | 34.91835 | 27.82737 | 12.30183 | 0.717729 | 0.51324 | 0.259476 |
| 10 | 115708000 | 115709599 | 8.664686 | 15.93814 | 34.61421 | 14.69251 | -0.68006 | 0.232586 | 0.259481 |
| 17 | 91832200 | 91833799 | 16.11993 | 29.71538 | 22.96453 | 9.719176 | 0.803333 | 0.251871 | 0.259491 |
| 14 | 95384200 | 95390799 | 89.43582 | 141.1264 | 114.0042 | 47.32034 | 0.632028 | 2.468592 | 0.259496 |
| 6 | 58992600 | 58994799 | 12.41731 | 21.89261 | 43.49177 | 19.24798 | -0.5959 | 0.55889 | 0.259496 |
| 18 | 81150800 | 81153399 | 17.65593 | 29.90555 | 61.26447 | 26.82449 | -0.64947 | 1.050808 | 0.259498 |
| 3 | 48268600 | 48271199 | 18.63178 | 33.45513 | 25.75545 | 11.26442 | 0.77758 | 0.423729 | 0.259499 |
| 6 | 18981600 | 18983999 | 18.63178 | 33.45513 | 25.75545 | 11.26442 | 0.77758 | 0.423729 | 0.259499 |
| 17 | 21403200 | 21405599 | 18.63178 | 33.45513 | 25.75545 | 11.26442 | 0.77758 | 0.423729 | 0.259499 |
| 1 | 135074200 | 135076199 | 9.668779 | 17.05695 | 43.02689 | 19.03997 | -0.93788 | 0.533954 | 0.259519 |
| 8 | 115797800 | 115799199 | 8.498907 | 15.66978 | 35.21311 | 14.89927 | -0.72577 | 0.218459 | 0.259535 |
| 12 | 95323200 | 95324399 | 8.498907 | 15.66978 | 35.21311 | 14.89927 | -0.72577 | 0.218459 | 0.259535 |
| 17 | 31863200 | 31867999 | 59.55845 | 93.16225 | 176.1108 | 73.06084 | -0.60267 | 2.629908 | 0.259538 |
| 10 | 113108200 | 113109399 | 20.28509 | 35.54659 | 28.50715 | 12.59132 | 0.704419 | 0.658733 | 0.25954 |
| X | 33121800 | 33123599 | 19.69979 | 36.69484 | 17.69234 | 7.377583 | 1.49381 | 0.212161 | 0.259543 |
| 5 | 43251400 | 43252399 | 7.395828 | 13.86215 | 32.31997 | 13.33604 | -0.75015 | 0.10316 | 0.259543 |
| 7 | 35227800 | 35228399 | 14.32442 | 26.61939 | 22.33696 | 9.350386 | 0.697066 | 0.193385 | 0.259546 |
| 4 | 32826800 | 32832799 | 33.42973 | 53.80676 | 102.8184 | 43.67365 | -0.57825 | 1.834985 | 0.259553 |
| 4 | 146717600 | 146719199 | 19.55477 | 35.61983 | 21.8947 | 9.413916 | 1.115599 | 0.341161 | 0.259556 |
| 15 | 56761800 | 56766199 | 44.52003 | 72.379 | 62.19542 | 26.6884 | 0.587115 | 1.587199 | 0.259561 |
| 4 | 153198400 | 153205199 | 49.05123 | 77.84289 | 141.0903 | 58.77591 | -0.5252 | 2.303442 | 0.259563 |
| 10 | 92824800 | 92826199 | 6.590181 | 12.60145 | 28.99098 | 11.49316 | -0.68047 | -0.04884 | 0.259573 |
| 13 | 105826400 | 105827799 | 14.85639 | 28.2406 | 19.00597 | 7.632149 | 1.061504 | 0.044736 | 0.259581 |
| 17 | 81721600 | 81722199 | 6.133504 | 11.80004 | 27.49249 | 10.70969 | -0.67406 | -0.11355 | 0.2596 |
| 5 | 128618200 | 128622599 | 27.71175 | 44.91181 | 92.82362 | 39.74205 | -0.68001 | 1.633948 | 0.259617 |
| 14 | 63780800 | 63783799 | 11.82072 | 20.69639 | 47.27212 | 20.86457 | -0.79812 | 0.656745 | 0.259617 |
| 13 | 44021800 | 44023999 | 11.159 | 19.86992 | 49.59154 | 21.84308 | -0.91658 | 0.473057 | 0.259618 |
| 3 | 35233200 | 35234599 | 5.866992 | 11.23128 | 27.84559 | 11.00334 | -0.78232 | -0.06868 | 0.259618 |
| 8 | 9037200 | 9039199 | 21.31563 | 35.49165 | 68.3065 | 29.66684 | -0.56022 | 1.238383 | 0.259621 |
| 1 | 92276000 | 92278799 | 18.7421 | 31.81209 | 61.92583 | 27.13134 | -0.57603 | 1.037239 | 0.259627 |
| 6 | 122263800 | 122266399 | 12.60835 | 21.99674 | 48.91693 | 21.53295 | -0.76012 | 0.718583 | 0.259666 |
| 6 | 132916600 | 132918399 | 20.61948 | 37.18212 | 22.17718 | 9.656485 | 1.147129 | 0.40486 | 0.259667 |
| 18 | 36960400 | 36973199 | 298.5242 | 428.4961 | 365.528 | 150.9251 | 0.435311 | 4.194133 | 0.259676 |
| 13 | 67382000 | 67383199 | 28.39842 | 48.25387 | 42.05161 | 18.42922 | 0.57926 | 1.042005 | 0.259676 |
| 7 | 66769600 | 66770799 | 8.02259 | 14.92242 | 32.67243 | 13.65708 | -0.6752 | 0.159853 | 0.259676 |
| 1 | 131570000 | 131572199 | 28.72067 | 49.02726 | 31.82451 | 13.95235 | 1.003405 | 0.989848 | 0.25968 |
| 9 | 42078000 | 42079399 | 19.81445 | 35.18359 | 27.42679 | 12.10887 | 0.750822 | 0.504673 | 0.259693 |
| 12 | 24416400 | 24423199 | 149.3741 | 227.3066 | 186.4775 | 77.26892 | 0.569522 | 3.177857 | 0.259711 |
| 12 | 17214400 | 17227599 | 123.2838 | 185.0447 | 307.9787 | 127.0977 | -0.46722 | 3.452234 | 0.259732 |
| 1 | 127495600 | 127498399 | 33.94041 | 54.56213 | 105.1972 | 44.60516 | -0.5933 | 1.87586 | 0.259739 |

| | | | | | | | | | |
|---|---|---|---|---|---|---|---|---|---|
| 11 | 110684600 | 110686999 | 32.4573 | 54.22854 | 47.57969 | 20.70949 | 0.569229 | 1.216082 | 0.259745 |
| 4 | 86460000 | 86461799 | 11.28403 | 20.09596 | 40.4061 | 17.79472 | -0.60597 | 0.456967 | 0.259757 |
| 16 | 23003800 | 23004799 | 13.2263 | 24.63365 | 21.38604 | 8.919671 | 0.652269 | 0.167095 | 0.259772 |
| 3 | 73769400 | 73771599 | 29.33331 | 49.74425 | 34.8764 | 15.27609 | 0.891645 | 1.053102 | 0.259775 |
| 8 | 30512000 | 30514999 | 11.02222 | 21.3302 | 14.71727 | 5.609899 | 1.094093 | -0.1729 | 0.25978 |
| 16 | 60741400 | 60742999 | 19.02201 | 34.51691 | 22.99817 | 9.935858 | 0.995745 | 0.36048 | 0.259793 |
| 8 | 53787000 | 53790399 | 62.56267 | 100.0581 | 80.38046 | 33.82202 | 0.658454 | 2.059727 | 0.259794 |
| 13 | 114023400 | 114028399 | 31.61326 | 51.022 | 97.66441 | 41.63896 | -0.57596 | 1.746827 | 0.259803 |
| 17 | 4001600 | 4003199 | 17.26302 | 31.31811 | 25.01248 | 10.80911 | 0.736176 | 0.354708 | 0.259809 |
| 17 | 4888400 | 4889799 | 17.26302 | 31.31811 | 25.01248 | 10.80911 | 0.736176 | 0.354708 | 0.259809 |
| 7 | 52955800 | 52957399 | 21.52506 | 38.06956 | 27.67622 | 12.24088 | 0.848249 | 0.549979 | 0.259816 |
| 10 | 73994200 | 73996599 | 24.29588 | 39.97147 | 78.41395 | 33.80392 | -0.59071 | 1.393874 | 0.259826 |
| 8 | 12444400 | 12448599 | 24.25478 | 39.24884 | 99.11334 | 42.38694 | -0.97024 | 1.664205 | 0.259835 |
| 3 | 81404000 | 81406599 | 20.71171 | 34.40219 | 69.22823 | 30.01729 | -0.62452 | 1.265786 | 0.259839 |
| 4 | 35654800 | 35657599 | 26.77476 | 46.35848 | 35.35844 | 15.51296 | 0.775097 | 0.860106 | 0.25984 |
| 17 | 19307200 | 19308599 | 18.93231 | 35.2034 | 19.50385 | 8.157179 | 1.292069 | 0.220806 | 0.259875 |
| 2 | 158251000 | 158252599 | 10.8252 | 19.26619 | 39.38549 | 17.35484 | -0.63013 | 0.456829 | 0.259888 |
| 4 | 93574800 | 93576199 | 9.725293 | 17.36353 | 40.4557 | 17.77888 | -0.81524 | 0.428569 | 0.259899 |
| 2 | 31025400 | 31028399 | 11.17207 | 19.70811 | 41.45256 | 18.34346 | -0.67724 | 0.542118 | 0.2599 |
| 4 | 151969200 | 151971199 | 16.28158 | 27.56582 | 59.44688 | 26.02526 | -0.72299 | 1.050782 | 0.2599 |
| 5 | 117451400 | 117455199 | 22.39775 | 36.87449 | 75.75582 | 32.66769 | -0.65657 | 1.379778 | 0.259908 |
| 4 | 115391000 | 115395599 | 64.09113 | 102.3975 | 90.00276 | 37.81588 | 0.527329 | 2.095331 | 0.259914 |
| 4 | 48645800 | 48646399 | 5.373097 | 10.44845 | 27.91751 | 10.34024 | -0.80116 | -0.29556 | 0.259921 |
| 5 | 148841000 | 148851399 | 68.93354 | 107.1244 | 191.754 | 79.61729 | -0.53148 | 2.749894 | 0.259941 |
| 13 | 61915600 | 61922399 | 126.4109 | 193.8912 | 169.6274 | 70.41661 | 0.485808 | 3.011038 | 0.259941 |
| 16 | 45228800 | 45229999 | 6.909243 | 13.11418 | 29.88612 | 12.0393 | -0.68759 | 0.009187 | 0.259959 |
| Y | 16627200 | 16630399 | 22.89893 | 40.17395 | 14.72352 | 6.509665 | 1.825315 | 0.623837 | 0.259962 |
| 2 | 174689600 | 174691399 | 8.291935 | 15.06083 | 35.4881 | 15.31344 | -0.81293 | 0.306081 | 0.259965 |
| 18 | 11138400 | 11138999 | 14.23939 | 26.22951 | 21.88169 | 9.270157 | 0.693065 | 0.239459 | 0.25997 |
| 7 | 6023400 | 6027999 | 96.23166 | 150.3811 | 134.5521 | 55.82547 | 0.474026 | 2.659537 | 0.259974 |
| 10 | 66291400 | 66293199 | 19.63212 | 35.06987 | 26.17526 | 11.49794 | 0.817805 | 0.462786 | 0.259981 |
| 7 | 8172600 | 8174999 | 19.40921 | 36.16035 | 18.32111 | 7.637354 | 1.423443 | 0.210163 | 0.259981 |
| 1 | 115385400 | 115386399 | 12.92094 | 24.78415 | 18.11836 | 7.118691 | 0.970786 | -0.04592 | 0.259987 |
| 11 | 88062000 | 88064999 | 13.32023 | 22.99705 | 52.10536 | 22.85841 | -0.78933 | 0.850608 | 0.259989 |
| 11 | 87248800 | 87250199 | 9.775212 | 17.33629 | 39.42842 | 17.42542 | -0.78574 | 0.473134 | 0.259989 |
| 1 | 16566200 | 16571199 | 42.79457 | 68.2397 | 126.7066 | 53.09227 | -0.55275 | 2.145872 | 0.259995 |
| X | 42932200 | 42933799 | 20.02597 | 36.35075 | 23.16622 | 10.0043 | 1.060395 | 0.368432 | 0.260004 |
| 5 | 37329200 | 37333999 | 27.99024 | 45.10193 | 101.2807 | 43.09009 | -0.80842 | 1.791412 | 0.260006 |
| 1 | 129860600 | 129861999 | 20.8241 | 36.70015 | 29.4095 | 13.01666 | 0.706669 | 0.586015 | 0.260007 |
| 10 | 78424400 | 78428799 | 26.78997 | 43.36489 | 94.03032 | 40.22245 | -0.75065 | 1.658241 | 0.260031 |
| 1 | 66080600 | 66082799 | 20.74048 | 36.63337 | 28.23598 | 12.49176 | 0.76353 | 0.559141 | 0.260037 |
| 7 | 4053000 | 4061599 | 100.0279 | 156.4339 | 131.2499 | 54.45108 | 0.567556 | 2.647892 | 0.260039 |
| 3 | 3295200 | 3298599 | 51.99107 | 84.14368 | 64.20668 | 27.46166 | 0.752156 | 1.69116 | 0.260044 |
| 8 | 92290000 | 92291399 | 11.04312 | 19.47376 | 42.51369 | 18.81436 | -0.7306 | 0.549721 | 0.260048 |
| 10 | 125185600 | 125187599 | 19.27815 | 34.15771 | 27.82522 | 12.3023 | 0.685833 | 0.515848 | 0.260052 |
| 1 | 98022400 | 98025399 | 19.38618 | 32.38331 | 65.97808 | 28.71394 | -0.64205 | 1.195903 | 0.260059 |
| 11 | 77927800 | 77929599 | 6.740206 | 12.49452 | 35.30344 | 14.82809 | -1.04556 | 0.173365 | 0.260066 |

| | | | | | | | | | |
|---|---|---|---|---|---|---|---|---|---|
| 6 | 140834400 | 140836199 | 10.12361 | 18.2804 | 37.44846 | 16.27811 | -0.61979 | 0.357021 | 0.260067 |
| 19 | 13042400 | 13043599 | 21.44378 | 37.703 | 29.89067 | 13.23588 | 0.720781 | 0.615449 | 0.26007 |
| 6 | 76740200 | 76741199 | 7.361737 | 13.92738 | 30.92624 | 12.54484 | -0.6595 | 0.038654 | 0.260082 |
| 14 | 122341800 | 122342999 | 7.361737 | 13.92738 | 30.92624 | 12.54484 | -0.6595 | 0.038654 | 0.260082 |
| 7 | 49336200 | 49339599 | 15.99232 | 27.38292 | 57.53292 | 25.22952 | -0.68374 | 0.9418 | 0.260085 |
| 16 | 70981800 | 70982799 | 12.4706 | 24.18505 | 16.14417 | 6.092265 | 1.154946 | -0.18412 | 0.260096 |
| 12 | 21791000 | 21804599 | 453.1312 | 626.3999 | 499.2558 | 207.3889 | 0.478735 | 4.756322 | 0.260119 |
| 3 | 32216400 | 32217599 | 20.3241 | 36.97492 | 21.86752 | 9.416952 | 1.169599 | 0.354292 | 0.260129 |
| 2 | 16587600 | 16591799 | 73.64647 | 116.7282 | 106.5063 | 44.31305 | 0.461721 | 2.351759 | 0.260133 |
| 10 | 90559000 | 90560599 | 7.665786 | 14.02934 | 35.06069 | 14.98516 | -0.88808 | 0.255079 | 0.260134 |
| 12 | 45647200 | 45649799 | 32.7013 | 54.73525 | 47.0061 | 20.48322 | 0.599932 | 1.196232 | 0.260134 |
| 9 | 3222600 | 3225999 | 41.54789 | 68.83716 | 47.73566 | 20.6621 | 0.907226 | 1.333463 | 0.260151 |
| X | 94970000 | 94971399 | 22.75469 | 39.56843 | 31.23866 | 13.7228 | 0.726607 | 0.789649 | 0.260155 |
| 7 | 115637800 | 115639999 | 13.42018 | 23.32932 | 48.9492 | 21.49867 | -0.67706 | 0.776295 | 0.26016 |
| 2 | 10286800 | 10288599 | 10.34915 | 18.43746 | 38.88368 | 17.1192 | -0.67395 | 0.445513 | 0.260169 |
| 4 | 142626800 | 142630599 | 31.56109 | 51.12571 | 94.05011 | 40.25286 | -0.51313 | 1.65217 | 0.260187 |
| 4 | 122900200 | 122902199 | 12.32577 | 21.46728 | 48.92605 | 21.51399 | -0.79551 | 0.742439 | 0.260191 |
| 3 | 78951800 | 78952799 | 15.25496 | 27.96611 | 23.56423 | 10.04816 | 0.6717 | 0.276193 | 0.260203 |
| 5 | 55612400 | 55614399 | 15.84198 | 28.87387 | 24.5933 | 10.56641 | 0.649091 | 0.31623 | 0.260209 |
| X | 61126000 | 61128799 | 43.87186 | 71.25892 | 60.68953 | 26.0262 | 0.598836 | 1.605185 | 0.260213 |
| 1 | 95592000 | 95595799 | 20.55124 | 34.13944 | 69.22612 | 30.01867 | -0.63548 | 1.263846 | 0.260222 |
| X | 3295600 | 3301999 | 64.56003 | 103.5986 | 76.68954 | 32.43018 | 0.781756 | 1.962356 | 0.260225 |
| 12 | 49398200 | 49400999 | 33.00983 | 55.60379 | 43.63179 | 19.07207 | 0.729624 | 1.133235 | 0.260228 |
| 14 | 100056400 | 100057399 | 13.89879 | 26.45083 | 19.70616 | 7.894008 | 0.919274 | 0.029946 | 0.26023 |
| 8 | 120591800 | 120594199 | 8.639881 | 15.55019 | 36.94085 | 16.12337 | -0.83548 | 0.365455 | 0.260235 |
| 2 | 164562400 | 164564199 | 20.43412 | 35.87805 | 29.8406 | 13.20215 | 0.652109 | 0.631835 | 0.260239 |
| 13 | 74711000 | 74714599 | 19.75811 | 33.24079 | 67.47368 | 29.48268 | -0.63689 | 1.122159 | 0.260251 |
| 16 | 92134400 | 92139199 | 25.43401 | 41.36237 | 86.45966 | 37.10448 | -0.69165 | 1.559963 | 0.260261 |
| 10 | 102268600 | 102269199 | 5.314781 | 10.16249 | 28.33368 | 11.2332 | -0.95442 | -0.06249 | 0.260274 |
| 12 | 57097400 | 57101399 | 31.07437 | 53.25788 | 36.94807 | 16.20383 | 0.908686 | 0.970653 | 0.260276 |
| 1 | 92196000 | 92201799 | 34.72828 | 55.79964 | 113.7565 | 48.19805 | -0.67469 | 1.893891 | 0.260289 |
| 16 | 61828000 | 61828599 | 9.201327 | 17.87587 | 15.96919 | 5.960068 | 0.753225 | -0.26153 | 0.260294 |
| 17 | 46744000 | 46746399 | 16.39159 | 27.89428 | 57.38509 | 25.15222 | -0.65466 | 1.003025 | 0.260303 |
| 7 | 14304800 | 14305799 | 32.59219 | 54.19102 | 46.58783 | 20.21469 | 0.598556 | 1.267385 | 0.260309 |
| 12 | 112043800 | 112048199 | 24.3942 | 39.88393 | 81.79273 | 35.17009 | -0.65867 | 1.477041 | 0.260317 |
| 12 | 24267600 | 24283199 | 421.934 | 589.5432 | 470.5764 | 195.1488 | 0.492149 | 4.594666 | 0.260326 |
| 2 | 74575600 | 74578199 | 19.27169 | 32.07075 | 69.79677 | 30.30325 | -0.73696 | 1.238665 | 0.260331 |
| 5 | 116525400 | 116526799 | 6.491819 | 12.27698 | 29.75324 | 12.07867 | -0.78509 | 0.031138 | 0.260345 |
| 18 | 56907200 | 56907999 | 6.491819 | 12.27698 | 29.75324 | 12.07867 | -0.78509 | 0.031138 | 0.260345 |
| 7 | 15336000 | 15344999 | 299.7011 | 430.2888 | 363.8342 | 150.2201 | 0.448377 | 4.190994 | 0.260352 |
| 8 | 86610400 | 86613399 | 26.10401 | 45.2102 | 36.03206 | 15.80864 | 0.712164 | 0.856883 | 0.260353 |
| 9 | 66619400 | 66623999 | 24.01434 | 39.42711 | 78.09655 | 33.63781 | -0.60574 | 1.419419 | 0.260364 |
| 13 | 68279000 | 68291399 | 255.2816 | 373.6927 | 299.1659 | 123.2373 | 0.554531 | 3.897695 | 0.26037 |
| 8 | 125347400 | 125349399 | 10.37721 | 18.39092 | 40.39358 | 17.8583 | -0.73667 | 0.482825 | 0.260384 |
| 19 | 58746800 | 58751999 | 31.62672 | 51.00261 | 99.77811 | 42.49593 | -0.60869 | 1.771178 | 0.260386 |
| 8 | 73799200 | 73800399 | 26.70498 | 46.27878 | 35.40026 | 15.53173 | 0.771241 | 0.852964 | 0.260397 |
| 5 | 142379400 | 142380999 | 14.53691 | 25.23243 | 50.55908 | 22.18464 | -0.61125 | 0.806214 | 0.260407 |

| | | | | | | | | | |
|---|---|---|---|---|---|---|---|---|---|
| 6 | 42630200 | 42631399 | 11.44078 | 20.39203 | 41.99481 | 18.48145 | -0.63973 | 0.455401 | 0.260446 |
| 2 | 21241600 | 21243199 | 9.958995 | 18.06286 | 37.00593 | 15.99702 | -0.61463 | 0.329632 | 0.260446 |
| 7 | 5053200 | 5060999 | 51.2548 | 81.01183 | 153.9561 | 63.95683 | -0.59854 | 2.411678 | 0.260462 |
| 7 | 19676800 | 19678999 | 19.75075 | 32.86025 | 68.03931 | 29.53543 | -0.66526 | 1.243478 | 0.260464 |
| 9 | 103479800 | 103481199 | 7.750742 | 14.35654 | 32.5948 | 13.705 | -0.73313 | 0.182716 | 0.260466 |
| 7 | 4059800 | 4061599 | 33.26718 | 55.38344 | 49.24365 | 21.38549 | 0.55021 | 1.257212 | 0.260495 |
| 1 | 133115000 | 133118799 | 27.76644 | 45.35363 | 83.94727 | 36.08381 | -0.51226 | 1.500582 | 0.260498 |
| 5 | 147796200 | 147798199 | 12.31732 | 21.48281 | 47.33482 | 20.83273 | -0.74666 | 0.720771 | 0.260503 |
| 2 | 81043200 | 81044199 | 16.638 | 30.0292 | 25.19805 | 10.95965 | 0.659514 | 0.380156 | 0.260504 |
| 17 | 18151400 | 18152399 | 13.06249 | 24.82833 | 20.32436 | 8.163114 | 0.7812 | 0.031764 | 0.260505 |
| 7 | 42648400 | 42648799 | 19.73735 | 34.64602 | 26.96234 | 11.92623 | 0.747397 | 0.625081 | 0.260516 |
| 9 | 47540600 | 47541399 | 11.41429 | 21.71039 | 19.46867 | 7.808128 | 0.652566 | 0.008852 | 0.26054 |
| 7 | 61318200 | 61319999 | 25.49707 | 44.44706 | 32.55699 | 14.32419 | 0.834284 | 0.769309 | 0.260574 |
| X | 63719800 | 63720799 | 15.38511 | 29.88799 | 12.38746 | 4.625053 | 1.848108 | -0.19567 | 0.260576 |
| 6 | 62323400 | 62325199 | 18.98025 | 33.86954 | 26.7979 | 11.78196 | 0.733302 | 0.466125 | 0.260599 |
| 17 | 47364000 | 47365799 | 9.793996 | 17.73157 | 37.49843 | 16.24654 | -0.66241 | 0.338539 | 0.260612 |
| 18 | 5093000 | 5094599 | 10.65233 | 18.96658 | 39.37604 | 17.34445 | -0.65204 | 0.451565 | 0.260621 |
| 7 | 107521000 | 107522199 | 29.30807 | 49.51176 | 39.65564 | 17.3493 | 0.700397 | 1.093679 | 0.260624 |
| 14 | 24548800 | 24551199 | 10.52049 | 18.6697 | 39.87798 | 17.61483 | -0.69562 | 0.472781 | 0.260625 |
| 14 | 66658600 | 66660199 | 19.8268 | 34.87092 | 30.1353 | 13.34337 | 0.597229 | 0.603404 | 0.26063 |
| 2 | 57013400 | 57014999 | 10.13774 | 17.82799 | 43.55584 | 19.28618 | -0.89385 | 0.573356 | 0.260637 |
| 5 | 107414800 | 107417399 | 7.759341 | 13.99894 | 39.57467 | 17.22116 | -1.0822 | 0.348159 | 0.260654 |
| 18 | 15392800 | 15394799 | 26.77749 | 45.65227 | 39.51747 | 17.3233 | 0.589671 | 1.003692 | 0.26067 |
| 7 | 31772000 | 31776999 | 81.50227 | 129.7072 | 84.11946 | 35.15175 | 0.959173 | 2.233605 | 0.260677 |
| 16 | 6211000 | 6211799 | 12.84752 | 24.01021 | 19.91843 | 8.259558 | 0.723972 | 0.138235 | 0.260681 |
| 13 | 72784400 | 72790399 | 51.71397 | 81.83606 | 149.1729 | 62.00756 | -0.53708 | 2.382915 | 0.260689 |
| 14 | 44515000 | 44516999 | 19.03762 | 34.76933 | 22.62543 | 9.696095 | 1.037225 | 0.325068 | 0.260715 |
| 12 | 108144400 | 108146599 | 14.03176 | 24.38336 | 49.49183 | 21.73189 | -0.62964 | 0.784496 | 0.260726 |
| 9 | 122906000 | 122910399 | 19.71924 | 32.50636 | 75.41571 | 32.53867 | -0.83053 | 1.3572 | 0.26073 |
| 13 | 113226000 | 113227799 | 10.81043 | 19.24297 | 39.37252 | 17.34679 | -0.63137 | 0.45461 | 0.260768 |
| 12 | 69322200 | 69324199 | 13.73545 | 23.82074 | 50.55358 | 22.18069 | -0.69408 | 0.808894 | 0.260774 |
| 17 | 39104000 | 39108599 | 58.91832 | 95.02414 | 66.2262 | 28.20982 | 0.877174 | 1.795906 | 0.260778 |
| 7 | 8084400 | 8085399 | 13.68475 | 25.51265 | 22.27433 | 9.274655 | 0.64594 | 0.166474 | 0.260793 |
| 7 | 33150400 | 33157799 | 184.8366 | 277.3234 | 226.3855 | 93.41562 | 0.559322 | 3.466925 | 0.260796 |
| 7 | 37194800 | 37199599 | 28.10249 | 45.87323 | 85.06922 | 36.55756 | -0.51563 | 1.51146 | 0.260846 |
| 16 | 64632200 | 64632599 | 7.654979 | 14.97276 | 11.57062 | 3.89162 | 1.091686 | -0.65513 | 0.260858 |
| 3 | 22706600 | 22708799 | 24.83664 | 43.26175 | 32.56584 | 14.32101 | 0.794871 | 0.775457 | 0.260865 |
| 18 | 7174800 | 7176399 | 17.89377 | 32.74241 | 22.66601 | 9.689775 | 0.95069 | 0.305082 | 0.260873 |
| 17 | 20521000 | 20521999 | 18.44718 | 32.93812 | 27.00861 | 11.86857 | 0.682751 | 0.460571 | 0.260879 |
| 9 | 24691400 | 24693399 | 10.73822 | 18.8768 | 43.03388 | 19.0565 | -0.79481 | 0.580615 | 0.260894 |
| 12 | 18856600 | 18888999 | 1088.22 | 1401.724 | 1066.878 | 447.3357 | 0.443077 | 5.886241 | 0.260905 |
| 5 | 119395000 | 119396999 | 10.81032 | 18.84494 | 47.8549 | 21.05496 | -0.95011 | 0.722345 | 0.260919 |
| 9 | 3204800 | 3206599 | 31.65358 | 54.07336 | 38.10241 | 16.70567 | 0.885735 | 1.003729 | 0.260922 |
| 17 | 18540400 | 18543399 | 32.30684 | 55.31962 | 36.38564 | 15.956 | 0.985141 | 0.98356 | 0.26093 |
| 16 | 55736600 | 55737799 | 3.601088 | 7.00975 | 25.23285 | 9.269445 | -1.21316 | -0.35424 | 0.260935 |
| 3 | 143783400 | 143784799 | 8.835002 | 16.16503 | 36.11665 | 15.44184 | -0.7285 | 0.266694 | 0.26094 |
| 15 | 42265000 | 42266799 | 18.91168 | 35.20104 | 17.0439 | 7.116008 | 1.48565 | 0.207612 | 0.260942 |

| | | | | | | | | | |
|---|---|---|---|---|---|---|---|---|---|
| 8 | 93082400 | 93087399 | 43.54737 | 69.14205 | 140.7166 | 58.6488 | -0.69137 | 2.284581 | 0.260942 |
| 12 | 23413400 | 23415399 | 36.99505 | 61.12207 | 47.2046 | 20.39266 | 0.752092 | 1.351877 | 0.260945 |
| 12 | 57792200 | 57794199 | 25.80356 | 44.70029 | 32.94888 | 14.45603 | 0.824338 | 0.848605 | 0.26095 |
| 7 | 61249000 | 61250199 | 16.73221 | 31.05452 | 21.27249 | 8.920138 | 0.985388 | 0.215857 | 0.260956 |
| 2 | 98519600 | 98520599 | 12.16523 | 23.70358 | 12.6498 | 4.59592 | 1.52164 | -0.32538 | 0.260963 |
| 17 | 17460200 | 17472999 | 377.3328 | 533.8576 | 411.2345 | 170.1994 | 0.56158 | 4.40886 | 0.260968 |
| 7 | 25458000 | 25466399 | 42.03555 | 66.88203 | 136.0958 | 56.85312 | -0.68809 | 2.218055 | 0.260977 |
| 5 | 75234600 | 75237999 | 27.71115 | 44.91451 | 90.96715 | 38.94956 | -0.65071 | 1.631949 | 0.260981 |
| 9 | 59959600 | 59960799 | 13.197 | 24.68008 | 21.82329 | 9.039059 | 0.633772 | 0.141682 | 0.261001 |
| 2 | 87214800 | 87215799 | 12.18006 | 23.08043 | 20.36345 | 8.231858 | 0.665725 | 0.043436 | 0.261011 |
| 9 | 91603800 | 91604399 | 12.18006 | 23.08043 | 20.36345 | 8.231858 | 0.665725 | 0.043436 | 0.261011 |
| 10 | 89273200 | 89275799 | 13.5496 | 23.76118 | 48.84778 | 21.58903 | -0.64705 | 0.64461 | 0.261013 |
| 2 | 160272000 | 160273199 | 14.57663 | 26.93767 | 23.25678 | 9.809469 | 0.648309 | 0.227336 | 0.261033 |
| 7 | 38403000 | 38405399 | 26.74535 | 46.87951 | 27.34135 | 12.07907 | 1.161259 | 0.687916 | 0.261041 |
| 10 | 82374200 | 82376799 | 26.74535 | 46.87951 | 27.34135 | 12.07907 | 1.161259 | 0.687916 | 0.261041 |
| 7 | 89174600 | 89176799 | 12.40403 | 21.85126 | 43.48817 | 19.24955 | -0.59903 | 0.567059 | 0.26105 |
| 1 | 15729600 | 15730399 | 6.755124 | 12.88124 | 29.43891 | 11.7434 | -0.6786 | -0.02338 | 0.261053 |
| 1 | 32949200 | 32950799 | 16.94152 | 31.34872 | 21.8031 | 9.180071 | 0.959552 | 0.235255 | 0.261056 |
| X | 152868200 | 152871999 | 91.38627 | 143.9206 | 117.8589 | 48.90236 | 0.610213 | 2.512355 | 0.261076 |
| 7 | 23630800 | 23631999 | 24.61421 | 42.81394 | 34.31005 | 15.07468 | 0.704969 | 0.797201 | 0.261115 |
| 11 | 21583200 | 21585199 | 15.81212 | 27.23538 | 53.23001 | 23.35012 | -0.57804 | 0.883825 | 0.261117 |
| 11 | 73818400 | 73819999 | 14.75133 | 27.01527 | 22.84096 | 9.753272 | 0.665765 | 0.277956 | 0.261126 |
| 13 | 47316800 | 47317599 | 14.75133 | 27.01527 | 22.84096 | 9.753272 | 0.665765 | 0.277956 | 0.261126 |
| 4 | 35225000 | 35226599 | 12.49763 | 21.72334 | 49.46683 | 21.72461 | -0.79455 | 0.773421 | 0.261128 |
| 17 | 87492200 | 87494799 | 12.49763 | 21.72334 | 49.46683 | 21.72461 | -0.79455 | 0.773421 | 0.261128 |
| 6 | 33176600 | 33178399 | 11.60719 | 20.50327 | 42.44253 | 18.77649 | -0.65387 | 0.529799 | 0.261131 |
| 19 | 46529000 | 46530199 | 8.977765 | 16.50358 | 34.60109 | 14.6996 | -0.63041 | 0.239395 | 0.261134 |
| X | 22706600 | 22713399 | 78.25117 | 124.5505 | 93.55054 | 39.10031 | 0.747961 | 2.230056 | 0.261137 |
| X | 6196800 | 6199199 | 26.02422 | 45.65417 | 27.9919 | 12.37534 | 1.089595 | 0.672618 | 0.261144 |
| 15 | 6373400 | 6374999 | 21.22794 | 37.05022 | 32.49905 | 14.30968 | 0.575666 | 0.73096 | 0.261147 |
| 4 | 141529000 | 141534999 | 50.36542 | 79.95972 | 140.4146 | 58.51732 | -0.4791 | 2.293536 | 0.26116 |
| 6 | 42686000 | 42687199 | 9.039123 | 16.39528 | 35.96797 | 15.54736 | -0.71246 | 0.320029 | 0.261172 |
| 17 | 29537600 | 29543999 | 36.93608 | 59.07794 | 118.1309 | 49.71252 | -0.65449 | 2.042959 | 0.26119 |
| 19 | 40492000 | 40497599 | 33.3766 | 53.87284 | 98.76008 | 42.11103 | -0.51366 | 1.747397 | 0.261199 |
| 7 | 100250000 | 100250799 | 12.8786 | 24.3763 | 20.4527 | 8.287891 | 0.735259 | 0.060056 | 0.261205 |
| 5 | 131013000 | 131014599 | 11.93597 | 21.09793 | 41.92344 | 18.5444 | -0.59469 | 0.525182 | 0.261256 |
| 11 | 18963200 | 18966399 | 16.41487 | 27.48982 | 69.76434 | 30.41054 | -0.9578 | 1.160254 | 0.261276 |
| 10 | 42347600 | 42355199 | 53.81303 | 85.11611 | 153.9846 | 63.99182 | -0.52681 | 2.396082 | 0.261278 |
| 2 | 36360600 | 36361199 | 12.93507 | 24.36574 | 21.38322 | 8.749025 | 0.658736 | 0.093689 | 0.261288 |
| 5 | 39241000 | 39241999 | 7.039925 | 13.46254 | 32.59179 | 12.91759 | -0.7535 | -0.03613 | 0.261312 |
| 2 | 89375400 | 89375999 | 7.058957 | 13.39467 | 29.87881 | 12.04333 | -0.65764 | 0.012986 | 0.261316 |
| 13 | 38358600 | 38359599 | 7.058957 | 13.39467 | 29.87881 | 12.04333 | -0.65764 | 0.012986 | 0.261316 |
| 1 | 146281400 | 146282599 | 22.18113 | 38.71147 | 33.35178 | 14.68457 | 0.601392 | 0.737485 | 0.261316 |
| 12 | 110854600 | 110857799 | 11.84177 | 20.70009 | 46.25766 | 20.38924 | -0.76653 | 0.683302 | 0.261326 |
| 6 | 130066600 | 130067799 | 12.47659 | 24.16549 | 16.71196 | 6.344655 | 1.09647 | -0.16286 | 0.261328 |
| 13 | 44371600 | 44372799 | 6.480071 | 12.28187 | 29.25062 | 11.81925 | -0.75379 | 0.015795 | 0.261339 |
| 9 | 5115800 | 5116799 | 9.408675 | 18.30569 | 14.15466 | 5.220243 | 0.973826 | -0.30729 | 0.261359 |

| | | | | | | | | | |
|---|---|---|---|---|---|---|---|---|---|
| 1 | 167135600 | 167136599 | 5.238205 | 9.879802 | 30.7291 | 12.54547 | -1.14838 | 0.040999 | 0.261365 |
| 7 | 26642200 | 26646399 | 51.13907 | 82.49512 | 71.74248 | 30.56788 | 0.558838 | 1.779477 | 0.261375 |
| 4 | 146245600 | 146245999 | 13.94438 | 26.09089 | 18.28173 | 7.565718 | 0.967986 | 0.136346 | 0.261375 |
| X | 32101600 | 32103199 | 51.71826 | 84.80985 | 41.74907 | 17.97306 | 1.396095 | 1.475558 | 0.261396 |
| 6 | 3222400 | 3224799 | 28.98466 | 49.02489 | 42.16998 | 18.45611 | 0.597851 | 1.08447 | 0.261415 |
| 3 | 39755800 | 39756999 | 18.0305 | 33.05337 | 22.9248 | 9.775899 | 0.951049 | 0.298721 | 0.26142 |
| 6 | 73196600 | 73200399 | 25.71885 | 41.89955 | 84.43303 | 36.25808 | -0.63715 | 1.533684 | 0.261426 |
| 9 | 121157000 | 121159799 | 25.71885 | 41.89955 | 84.43303 | 36.25808 | -0.63715 | 1.533684 | 0.261426 |
| 7 | 42736000 | 42740199 | 73.19651 | 116.8395 | 86.63448 | 36.35408 | 0.770865 | 2.13169 | 0.261428 |
| 5 | 65592400 | 65593799 | 9.24182 | 16.61797 | 36.91724 | 16.12697 | -0.74033 | 0.375827 | 0.261428 |
| 19 | 6170200 | 6174399 | 19.10942 | 31.82582 | 67.46688 | 29.3066 | -0.69902 | 1.229468 | 0.261444 |
| X | 125440600 | 125441599 | 15.19212 | 29.23991 | 16.35374 | 6.362904 | 1.367026 | -0.05629 | 0.261449 |
| 7 | 137196600 | 137201999 | 27.70134 | 44.921 | 90.50848 | 38.76742 | -0.64255 | 1.621413 | 0.261452 |
| 4 | 121630800 | 121632399 | 18.96463 | 33.83265 | 23.729 | 10.43503 | 0.905708 | 0.460551 | 0.261455 |
| 5 | 64029200 | 64032599 | 27.45285 | 44.88666 | 84.52129 | 36.34391 | -0.53591 | 1.486628 | 0.261458 |
| 9 | 107107600 | 107111599 | 28.11528 | 45.80335 | 84.83129 | 36.42902 | -0.51581 | 1.539489 | 0.26146 |
| 14 | 16266200 | 16271999 | 45.64623 | 73.96595 | 65.23387 | 27.92631 | 0.545428 | 1.657037 | 0.261463 |
| 2 | 19427000 | 19428399 | 7.554167 | 14.15906 | 31.20749 | 12.8768 | -0.66919 | 0.104108 | 0.261476 |
| 5 | 3977800 | 3978999 | 7.554167 | 14.15906 | 31.20749 | 12.8768 | -0.66919 | 0.104108 | 0.261476 |
| 16 | 61002200 | 61002999 | 11.34793 | 21.68202 | 19.54389 | 7.763669 | 0.657455 | -0.02114 | 0.261486 |
| 3 | 15983000 | 15989999 | 99.32531 | 155.8996 | 122.3393 | 50.73922 | 0.667963 | 2.58308 | 0.261493 |
| 14 | 29912000 | 29912599 | 6.924764 | 13.13672 | 30.39715 | 12.25879 | -0.71021 | 0.017288 | 0.261494 |
| 12 | 106904400 | 106908199 | 22.52488 | 37.19049 | 73.06846 | 31.5517 | -0.59088 | 1.343733 | 0.261504 |
| 19 | 41707800 | 41709999 | 11.62059 | 20.52984 | 41.41563 | 18.32167 | -0.61663 | 0.527783 | 0.261523 |
| 2 | 22279400 | 22281599 | 24.25493 | 42.0055 | 36.07071 | 15.82557 | 0.60486 | 0.846979 | 0.261534 |
| 19 | 17886400 | 17889599 | 47.76648 | 77.23371 | 68.12546 | 29.1138 | 0.542263 | 1.707016 | 0.261549 |
| X | 67487600 | 67489599 | 22.69117 | 40.95594 | 21.69588 | 9.435941 | 1.318662 | 0.415561 | 0.261573 |
| 6 | 39254600 | 39255999 | 10.59218 | 19.09037 | 40.06485 | 17.45665 | -0.65695 | 0.375717 | 0.261575 |
| 4 | 44652400 | 44656399 | 18.72982 | 31.36099 | 64.90101 | 28.28738 | -0.66452 | 1.166917 | 0.261584 |
| 14 | 35794600 | 35798599 | 89.05803 | 139.6174 | 127.6542 | 52.94608 | 0.446415 | 2.599013 | 0.261585 |
| 12 | 115955600 | 115958799 | 72.09407 | 114.325 | 104.818 | 43.63256 | 0.455462 | 2.334958 | 0.261598 |
| 7 | 12259200 | 12260999 | 42.40668 | 68.83407 | 58.41389 | 25.03824 | 0.603102 | 1.612058 | 0.261603 |
| 17 | 12160200 | 12161799 | 15.44844 | 28.39181 | 22.39831 | 9.518512 | 0.769291 | 0.260355 | 0.261605 |
| 12 | 104662800 | 104663399 | 5.709386 | 10.95115 | 27.33668 | 10.74123 | -0.78391 | -0.08923 | 0.26161 |
| 13 | 55931400 | 55932399 | 9.394404 | 16.91648 | 36.93453 | 16.11073 | -0.71391 | 0.369505 | 0.261613 |
| 4 | 31963200 | 31964599 | 10.38069 | 18.42441 | 39.40196 | 17.40234 | -0.69696 | 0.472023 | 0.261614 |
| 14 | 82595800 | 82596999 | 13.63774 | 25.67731 | 20.02987 | 8.201803 | 0.825769 | 0.097414 | 0.261638 |
| 12 | 107700200 | 107703999 | 25.15796 | 41.20277 | 80.39509 | 34.59165 | -0.58559 | 1.454192 | 0.261644 |
| 7 | 31859800 | 31862599 | 35.24401 | 60.5831 | 31.24405 | 13.70482 | 1.335208 | 0.956776 | 0.261649 |
| 2 | 153528200 | 153529199 | 5.821107 | 11.26691 | 26.67151 | 10.15855 | -0.66575 | -0.19491 | 0.26165 |
| 10 | 63404600 | 63405399 | 12.1476 | 23.12698 | 19.83058 | 7.937422 | 0.718837 | 0.009827 | 0.261666 |
| X | 78185600 | 78195599 | 141.1179 | 215.9522 | 168.5712 | 69.94745 | 0.647585 | 3.062631 | 0.261668 |
| 7 | 4641200 | 4643399 | 11.65529 | 20.46472 | 43.55628 | 19.27064 | -0.69646 | 0.605395 | 0.261674 |
| 2 | 6326400 | 6328399 | 10.42192 | 18.35665 | 43.5313 | 19.26924 | -0.85037 | 0.556563 | 0.261684 |
| 4 | 41093200 | 41099999 | 44.1211 | 70.19768 | 136.3101 | 56.93968 | -0.62076 | 2.220925 | 0.261687 |
| 15 | 99961000 | 99962399 | 8.928942 | 16.09629 | 36.43817 | 15.87203 | -0.76443 | 0.357678 | 0.261687 |
| 5 | 89034200 | 89035999 | 12.83258 | 22.22898 | 51.05487 | 22.39984 | -0.80779 | 0.816413 | 0.26169 |

| | | | | | | | | | |
|---|---|---|---|---|---|---|---|---|---|
| 13 | 76531800 | 76533799 | 17.86006 | 29.74938 | 67.90232 | 29.49844 | -0.80521 | 1.22433 | 0.261693 |
| 7 | 57239600 | 57242199 | 25.79054 | 41.84559 | 88.29762 | 37.83589 | -0.70815 | 1.606601 | 0.261695 |
| 16 | 91977400 | 91981399 | 32.78276 | 52.77104 | 103.4164 | 43.9344 | -0.61439 | 1.830407 | 0.261699 |
| 1 | 36533800 | 36535399 | 6.655455 | 12.52567 | 30.18778 | 12.37012 | -0.78854 | 0.067464 | 0.261708 |
| 2 | 88335800 | 88339599 | 51.82735 | 83.7171 | 68.56567 | 29.26466 | 0.647385 | 1.741471 | 0.261709 |
| 2 | 124603400 | 124605799 | 23.35567 | 40.84055 | 32.07209 | 14.14053 | 0.734533 | 0.713576 | 0.261712 |
| 7 | 96730400 | 96732999 | 13.7437 | 23.87056 | 49.48408 | 21.72149 | -0.65987 | 0.791895 | 0.261712 |
| 11 | 106602800 | 106604799 | 11.00907 | 19.42072 | 42.47034 | 18.79381 | -0.7333 | 0.541209 | 0.261714 |
| 8 | 49906800 | 49908199 | 16.55226 | 31.09302 | 19.83308 | 8.152996 | 1.109955 | 0.139003 | 0.261716 |
| 8 | 112468800 | 112473999 | 24.20787 | 39.29331 | 88.66048 | 38.00253 | -0.80401 | 1.595521 | 0.261718 |
| 2 | 168575600 | 168578799 | 26.22729 | 42.25853 | 107.2654 | 45.63208 | -0.98484 | 1.794245 | 0.261754 |
| 7 | 41372600 | 41380999 | 141.1203 | 215.2863 | 169.2331 | 70.17171 | 0.633965 | 3.12358 | 0.261777 |
| 10 | 121547800 | 121549599 | 14.02777 | 26.16761 | 19.91795 | 8.2852 | 0.842783 | 0.157747 | 0.261826 |
| 6 | 87428800 | 87435199 | 36.354 | 58.44268 | 105.9314 | 44.91702 | -0.50444 | 1.878878 | 0.26183 |
| 3 | 97168800 | 97171199 | 19.43644 | 32.81073 | 62.39852 | 27.29315 | -0.54283 | 1.088639 | 0.261869 |
| 2 | 76532400 | 76533399 | 8.176861 | 15.20506 | 32.66593 | 13.66093 | -0.64857 | 0.163384 | 0.261869 |
| 1 | 58916800 | 58920199 | 53.63026 | 86.18223 | 78.59795 | 33.30818 | 0.483753 | 1.902838 | 0.261871 |
| 1 | 66983800 | 66984599 | 6.498503 | 12.28082 | 31.98832 | 13.00572 | -0.89092 | 0.039375 | 0.261885 |
| X | 126862200 | 126866599 | 83.50323 | 132.5549 | 92.36697 | 38.48951 | 0.852164 | 2.312072 | 0.26189 |
| 2 | 88591800 | 88593599 | 21.13994 | 37.17333 | 26.89413 | 11.90869 | 0.852002 | 0.606414 | 0.261897 |
| 10 | 41798200 | 41803599 | 49.374 | 78.35236 | 141.2175 | 58.82686 | -0.51716 | 2.304683 | 0.26191 |
| 15 | 46738200 | 46738799 | 8.669211 | 16.93287 | 13.19124 | 4.620278 | 1.029148 | -0.49665 | 0.261926 |
| 13 | 49444800 | 49445599 | 11.0038 | 21.25497 | 18.52659 | 7.123582 | 0.748799 | -0.13232 | 0.261942 |
| 4 | 48248000 | 48250199 | 21.91779 | 38.56213 | 29.26909 | 12.95891 | 0.783591 | 0.609833 | 0.261943 |
| 9 | 26855000 | 26857799 | 39.11617 | 64.91141 | 48.79646 | 21.14446 | 0.791353 | 1.310361 | 0.261948 |
| X | 109948200 | 109949799 | 18.29615 | 33.80732 | 19.88771 | 8.389794 | 1.196948 | 0.24899 | 0.261949 |
| 6 | 41845000 | 41847599 | 27.57829 | 47.67537 | 35.31285 | 15.49222 | 0.816736 | 0.877039 | 0.261955 |
| 13 | 93634600 | 93636599 | 12.48928 | 21.35575 | 56.18842 | 24.63722 | -1.00643 | 0.943003 | 0.261966 |
| 7 | 51044000 | 51045199 | 23.87619 | 41.56954 | 34.01137 | 14.95233 | 0.675488 | 0.782692 | 0.261976 |
| 7 | 13484800 | 13493399 | 189.8008 | 283.6169 | 223.2297 | 92.03637 | 0.606628 | 3.542989 | 0.261985 |
| 19 | 39340000 | 39344199 | 138.7475 | 213.0616 | 156.0313 | 64.78291 | 0.743398 | 2.9918 | 0.261986 |
| 13 | 44601200 | 44602199 | 12.45908 | 23.81444 | 19.92279 | 7.906999 | 0.765165 | -0.01245 | 0.261987 |
| 10 | 93517200 | 93523399 | 42.37116 | 67.80733 | 125.4728 | 52.84211 | -0.54219 | 2.032434 | 0.262003 |
| 1 | 168552400 | 168554799 | 27.93134 | 48.08199 | 36.98692 | 16.22457 | 0.761267 | 0.917689 | 0.262004 |
| 11 | 53055200 | 53059399 | 24.11868 | 39.76426 | 75.59562 | 32.61824 | -0.54426 | 1.365936 | 0.262015 |
| 16 | 69496800 | 69498399 | 16.37257 | 30.68726 | 20.38301 | 8.410637 | 1.047758 | 0.152275 | 0.26204 |
| 4 | 8579000 | 8580599 | 19.13869 | 33.77856 | 29.05034 | 12.85398 | 0.605989 | 0.559319 | 0.262082 |
| 10 | 118694400 | 118704399 | 166.0066 | 242.6455 | 417.5721 | 172.0291 | -0.55104 | 3.915613 | 0.262085 |
| 10 | 81791800 | 81794199 | 19.94827 | 35.08253 | 30.07583 | 13.31719 | 0.608664 | 0.60415 | 0.26209 |
| 4 | 124753000 | 124758399 | 38.67045 | 61.93369 | 114.8706 | 48.43298 | -0.54411 | 2.00216 | 0.26209 |
| 8 | 112666600 | 112668199 | 6.925681 | 12.72113 | 34.5904 | 14.71103 | -1.00335 | 0.22768 | 0.262107 |
| 14 | 32422400 | 32423999 | 9.822085 | 17.31115 | 42.5226 | 18.82034 | -0.90021 | 0.545181 | 0.262129 |
| 7 | 13210600 | 13211799 | 13.91749 | 26.86897 | 16.46237 | 6.345386 | 1.248876 | -0.0989 | 0.262161 |
| 17 | 76404800 | 76405799 | 15.51142 | 28.41475 | 24.1158 | 10.29396 | 0.660454 | 0.28439 | 0.262174 |
| 9 | 31296000 | 31298399 | 14.43989 | 24.78648 | 55.30921 | 24.25924 | -0.76944 | 0.910906 | 0.26218 |
| 17 | 93439600 | 93440999 | 16.69622 | 31.14278 | 20.18871 | 8.39902 | 1.072885 | 0.183418 | 0.262181 |
| 2 | 97498400 | 97499399 | 11.59281 | 22.03888 | 19.96791 | 8.016862 | 0.636501 | 0.01546 | 0.262187 |

| | | | | | | | | | |
|---|---|---|---|---|---|---|---|---|---|
| 13 | 63387600 | 63388599 | 7.577061 | 14.10661 | 32.14699 | 13.41665 | -0.73048 | 0.149202 | 0.262195 |
| 11 | 3811800 | 3814199 | 14.76598 | 25.29249 | 56.37599 | 24.72291 | -0.76839 | 0.932883 | 0.262202 |
| 6 | 26139800 | 26140599 | 10.27489 | 19.93187 | 16.91291 | 6.374726 | 0.813591 | -0.21218 | 0.262209 |
| 2 | 152210600 | 152211999 | 7.021436 | 12.99144 | 32.58372 | 13.72074 | -0.87681 | 0.18382 | 0.262219 |
| 7 | 7635400 | 7636399 | 19.14309 | 34.02271 | 27.50587 | 12.13439 | 0.699309 | 0.494827 | 0.262221 |
| 7 | 8543000 | 8547399 | 112.0767 | 173.4717 | 139.6817 | 58.02022 | 0.615293 | 2.832594 | 0.262244 |
| 13 | 115360800 | 115362599 | 9.110397 | 16.76199 | 37.38349 | 15.86284 | -0.71818 | 0.239543 | 0.262245 |
| 17 | 19250200 | 19259999 | 204.0226 | 304.1907 | 243.6844 | 100.4263 | 0.578303 | 3.587793 | 0.262259 |
| 1 | 119281800 | 119284999 | 24.94561 | 41.08245 | 76.71363 | 33.08525 | -0.51911 | 1.38326 | 0.26226 |
| 9 | 21210400 | 21214199 | 36.31432 | 58.42419 | 107.9359 | 45.81843 | -0.53054 | 1.853593 | 0.262266 |
| 9 | 113936000 | 113938399 | 26.16 | 45.10715 | 32.66099 | 14.32769 | 0.848741 | 0.891284 | 0.262276 |
| 11 | 99920400 | 99920799 | 2.453051 | 4.797427 | 19.46155 | 6.614558 | -1.27467 | -0.66741 | 0.262278 |
| 4 | 5184400 | 5186599 | 40.71705 | 66.35059 | 60.28649 | 25.89124 | 0.509168 | 1.545166 | 0.262286 |
| 12 | 70982400 | 70985199 | 16.08199 | 27.44904 | 57.91615 | 25.39107 | -0.69022 | 0.976914 | 0.262295 |
| 9 | 36495200 | 36499399 | 24.94037 | 43.56913 | 29.09804 | 12.81946 | 0.967203 | 0.733555 | 0.2623 |
| 7 | 12358800 | 12360799 | 39.87953 | 66.16139 | 48.25938 | 20.90808 | 0.834644 | 1.314193 | 0.262309 |
| 6 | 136365200 | 136376999 | 129.7527 | 194.2667 | 318.5831 | 131.3999 | -0.44917 | 3.501407 | 0.262309 |
| 14 | 118492600 | 118494999 | 20.26659 | 34.07072 | 64.6027 | 28.22179 | -0.53887 | 1.129561 | 0.262317 |
| X | 62660600 | 62662999 | 25.34099 | 44.89547 | 23.55497 | 10.41452 | 1.316833 | 0.549299 | 0.262326 |
| 7 | 121494200 | 121496399 | 10.50625 | 18.71237 | 38.88065 | 17.12171 | -0.65282 | 0.448565 | 0.262328 |
| 12 | 62583200 | 62583799 | 14.76719 | 26.93988 | 22.25621 | 9.551222 | 0.693846 | 0.297003 | 0.262335 |
| 16 | 49886800 | 49887799 | 10.27398 | 18.57945 | 38.00526 | 16.48954 | -0.616 | 0.350532 | 0.262335 |
| 2 | 98624800 | 98630399 | 27.35883 | 47.21666 | 35.87165 | 15.73657 | 0.779804 | 0.890894 | 0.262352 |
| 12 | 3057600 | 3072599 | 860.645 | 1128.25 | 847.5595 | 354.7508 | 0.487308 | 5.59973 | 0.262353 |
| 9 | 21662400 | 21664199 | 10.9056 | 19.14185 | 43.60913 | 19.28847 | -0.79367 | 0.609561 | 0.262365 |
| 12 | 19659800 | 19677999 | 568.9082 | 773.065 | 607.6331 | 253.0903 | 0.473832 | 5.02562 | 0.262369 |
| 16 | 92631800 | 92633799 | 14.33709 | 24.95556 | 48.95007 | 21.51656 | -0.58028 | 0.761433 | 0.26238 |
| 6 | 108669000 | 108670199 | 7.230317 | 13.62367 | 30.78533 | 12.58975 | -0.69462 | 0.0649 | 0.262384 |
| 6 | 118318600 | 118319399 | 7.230317 | 13.62367 | 30.78533 | 12.58975 | -0.69462 | 0.0649 | 0.262384 |
| 13 | 22174000 | 22175999 | 25.71754 | 44.97858 | 30.25285 | 13.34044 | 0.956895 | 0.720895 | 0.262385 |
| 11 | 113037600 | 113039999 | 11.62142 | 20.49065 | 42.45777 | 18.79014 | -0.65627 | 0.55067 | 0.262403 |
| 12 | 6449600 | 6451199 | 19.17262 | 34.94558 | 21.90831 | 9.412404 | 1.087964 | 0.334573 | 0.262408 |
| 1 | 31913200 | 31913799 | 9.747723 | 19.02817 | 13.00955 | 4.603402 | 1.20199 | -0.44677 | 0.262411 |
| 13 | 39546000 | 39547999 | 9.836855 | 17.65846 | 39.46277 | 17.2632 | -0.75087 | 0.392836 | 0.262425 |
| 7 | 11356600 | 11372399 | 266.3563 | 391.0476 | 272.8709 | 112.3848 | 0.756664 | 3.853781 | 0.262431 |
| 17 | 74174800 | 74178999 | 37.62962 | 60.43796 | 111.7106 | 47.3043 | -0.5343 | 1.909947 | 0.262432 |
| 1 | 91055600 | 91058799 | 25.13729 | 40.81494 | 92.39421 | 39.61224 | -0.80832 | 1.593552 | 0.262432 |
| 5 | 117288400 | 117294999 | 46.10736 | 73.28007 | 146.7517 | 61.22554 | -0.66679 | 2.258366 | 0.262442 |
| 13 | 102141600 | 102143999 | 13.03287 | 22.88571 | 46.14232 | 20.41644 | -0.61913 | 0.614529 | 0.262452 |
| 12 | 19680400 | 19682199 | 18.71125 | 35.05105 | 18.54929 | 7.661244 | 1.371963 | 0.172171 | 0.262458 |
| 1 | 173306600 | 173309599 | 15.97351 | 27.39633 | 57.56654 | 25.24765 | -0.68353 | 0.926166 | 0.262462 |
| 12 | 77998000 | 77999599 | 9.116341 | 16.82106 | 35.79096 | 15.12798 | -0.64641 | 0.221433 | 0.262463 |
| 16 | 95502600 | 95503999 | 8.630646 | 15.23893 | 42.54416 | 18.82305 | -1.08238 | 0.517895 | 0.262468 |
| 8 | 35568600 | 35572799 | 29.8214 | 48.25698 | 91.38142 | 39.08047 | -0.55613 | 1.670463 | 0.262473 |
| 13 | 65740200 | 65770999 | 646.3268 | 871.0793 | 669.4369 | 279.1846 | 0.494365 | 5.156015 | 0.262473 |
| 18 | 81983400 | 81985199 | 11.64382 | 20.47257 | 43.00197 | 19.04167 | -0.67737 | 0.583371 | 0.262476 |
| 7 | 4576000 | 4579999 | 49.05176 | 79.07247 | 71.69535 | 30.52017 | 0.49765 | 1.797159 | 0.262477 |

| | | | | | | | | | |
|---|---|---|---|---|---|---|---|---|---|
| 12 | 77072200 | 77074599 | 13.04454 | 22.93167 | 46.65463 | 20.65977 | -0.63201 | 0.598658 | 0.26251 |
| 9 | 54050600 | 54052399 | 18.69904 | 33.07793 | 28.57066 | 12.63598 | 0.601079 | 0.531782 | 0.262517 |
| 13 | 14035400 | 14035799 | 24.41754 | 43.21081 | 19.72821 | 8.724528 | 1.514405 | 0.545258 | 0.262518 |
| 9 | 115110800 | 115112799 | 8.576592 | 15.60342 | 35.49455 | 15.28484 | -0.76074 | 0.297877 | 0.262523 |
| 5 | 61848800 | 61849799 | 16.03066 | 29.77549 | 22.14482 | 9.276161 | 0.869288 | 0.209892 | 0.262524 |
| 10 | 33637400 | 33637999 | 9.149147 | 17.7616 | 16.63153 | 6.24375 | 0.679169 | -0.23783 | 0.262526 |
| 11 | 119664200 | 119668399 | 56.8175 | 90.9979 | 82.25804 | 34.68271 | 0.490782 | 2.010548 | 0.262527 |
| X | 55003400 | 55005999 | 53.93855 | 86.98805 | 70.63155 | 30.08426 | 0.657243 | 1.793037 | 0.262545 |
| 12 | 62397600 | 62398799 | 17.44949 | 31.64667 | 25.00709 | 10.81093 | 0.751125 | 0.357956 | 0.262545 |
| 4 | 141110200 | 141112799 | 17.96102 | 30.04857 | 65.91853 | 28.71571 | -0.7483 | 1.174226 | 0.262545 |
| 18 | 64494200 | 64495999 | 11.78619 | 20.75927 | 42.46597 | 18.79773 | -0.63843 | 0.563854 | 0.262548 |
| 12 | 85449000 | 85451399 | 23.26631 | 40.84043 | 29.77487 | 13.17025 | 0.841007 | 0.649987 | 0.262548 |
| 9 | 115236200 | 115239399 | 17.65422 | 29.93447 | 59.61066 | 26.10878 | -0.60862 | 1.04066 | 0.262564 |
| 9 | 98661400 | 98662599 | 5.729865 | 10.92427 | 28.26276 | 11.27843 | -0.85562 | -0.03477 | 0.262582 |
| 15 | 43086000 | 43086599 | 9.666233 | 18.71592 | 17.01048 | 6.46812 | 0.703833 | -0.18965 | 0.262589 |
| 4 | 151409000 | 151413399 | 26.70858 | 43.41504 | 86.01691 | 36.90824 | -0.61508 | 1.569319 | 0.26259 |
| 14 | 120453000 | 120454199 | 14.85965 | 27.59468 | 22.01172 | 9.223036 | 0.767997 | 0.1984 | 0.262592 |
| 10 | 116644000 | 116646999 | 19.3313 | 32.43885 | 63.91816 | 27.8974 | -0.59407 | 1.144141 | 0.262608 |
| 4 | 105708600 | 105709999 | 15.55125 | 28.97207 | 22.08024 | 9.204766 | 0.839331 | 0.186642 | 0.262613 |
| 4 | 93952000 | 93954199 | 43.68598 | 71.47839 | 56.74708 | 24.40728 | 0.707395 | 1.492599 | 0.262622 |
| 17 | 36139000 | 36140999 | 10.48342 | 18.70293 | 39.38964 | 17.32144 | -0.67099 | 0.437312 | 0.262629 |
| 11 | 109629400 | 109631999 | 16.76488 | 30.38052 | 24.92664 | 10.78703 | 0.695985 | 0.355639 | 0.262652 |
| 13 | 116204000 | 116206199 | 21.19112 | 37.53724 | 26.46973 | 11.70362 | 0.892559 | 0.527882 | 0.262657 |
| 4 | 101744400 | 101747199 | 36.4597 | 60.5358 | 46.05623 | 19.96451 | 0.774081 | 1.293054 | 0.262666 |
| 2 | 90507400 | 90510199 | 15.80839 | 26.77339 | 61.11651 | 26.75885 | -0.80484 | 1.045469 | 0.262667 |
| 13 | 25491600 | 25491999 | 4.647754 | 9.06278 | 23.2961 | 8.381505 | -0.70608 | -0.42051 | 0.262683 |
| 3 | 14723800 | 14725199 | 6.598777 | 12.59134 | 28.90922 | 11.51616 | -0.68375 | -0.03279 | 0.262696 |
| 19 | 8043400 | 8047799 | 51.19034 | 82.58571 | 71.6822 | 30.54597 | 0.561796 | 1.777452 | 0.262697 |
| 1 | 34055400 | 34057399 | 11.94192 | 21.08875 | 41.93225 | 18.55175 | -0.59612 | 0.535713 | 0.262709 |
| 1 | 119042600 | 119043999 | 6.691727 | 12.50489 | 31.67389 | 13.13598 | -0.87408 | 0.116229 | 0.262743 |
| 8 | 114377200 | 114384599 | 41.97748 | 66.91272 | 131.6526 | 55.13731 | -0.63677 | 2.156688 | 0.262744 |
| 2 | 172555400 | 172561599 | 44.74761 | 71.29669 | 129.2795 | 54.11019 | -0.51987 | 2.174165 | 0.262753 |
| 7 | 16018000 | 16020799 | 12.17381 | 21.20316 | 49.45368 | 21.74635 | -0.82872 | 0.741673 | 0.262757 |
| 12 | 4803200 | 4805799 | 32.3132 | 54.19782 | 45.9055 | 20.02979 | 0.619793 | 1.173769 | 0.262757 |
| 14 | 9183400 | 9185599 | 15.54394 | 26.65837 | 54.24052 | 23.78883 | -0.63695 | 0.924142 | 0.262758 |
| 12 | 106082200 | 106097799 | 126.7584 | 189.3915 | 327.1796 | 134.8913 | -0.52691 | 3.539491 | 0.262765 |
| 7 | 86690600 | 86695399 | 62.43372 | 100.2227 | 77.94016 | 32.97715 | 0.711352 | 1.9476 | 0.262789 |
| 6 | 85458200 | 85462399 | 30.85586 | 49.70582 | 100.7006 | 42.82629 | -0.66082 | 1.804045 | 0.262792 |
| 7 | 32909000 | 32913599 | 63.05941 | 101.5967 | 64.7274 | 27.53302 | 1.004646 | 1.832005 | 0.262806 |
| 5 | 148526600 | 148528399 | 7.465463 | 13.8014 | 33.09727 | 13.95293 | -0.81434 | 0.191775 | 0.262807 |
| 10 | 95057600 | 95059599 | 7.465463 | 13.8014 | 33.09727 | 13.95293 | -0.81434 | 0.191775 | 0.262807 |
| 7 | 117665800 | 117668799 | 32.10712 | 51.77307 | 99.86777 | 42.5296 | -0.58855 | 1.774355 | 0.262814 |
| 3 | 43763000 | 43764599 | 17.91281 | 32.69699 | 23.20092 | 9.949582 | 0.912381 | 0.320024 | 0.26283 |
| 7 | 142324200 | 142325599 | 21.27872 | 37.14579 | 31.09323 | 13.69229 | 0.64297 | 0.727329 | 0.262838 |
| 4 | 67050400 | 67051799 | 11.59261 | 20.62487 | 41.95905 | 18.49423 | -0.62383 | 0.467333 | 0.262839 |
| 1 | 17922800 | 17923599 | 12.4136 | 23.68319 | 19.23 | 7.662853 | 0.802206 | -0.00369 | 0.262858 |
| 14 | 79898600 | 79901199 | 13.03154 | 22.46135 | 52.60972 | 23.07851 | -0.83744 | 0.865735 | 0.262863 |

| | | | | | | | | | |
|---|---|---|---|---|---|---|---|---|---|
| 12 | 31792800 | 31795599 | 36.75836 | 60.58703 | 54.26908 | 23.41355 | 0.538108 | 1.384788 | 0.262865 |
| 11 | 27377400 | 27381399 | 56.831 | 91.11139 | 82.71314 | 34.92813 | 0.486337 | 1.979675 | 0.262871 |
| 12 | 22617800 | 22621799 | 122.8432 | 188.7695 | 145.1604 | 60.27509 | 0.673745 | 2.972161 | 0.262874 |
| 12 | 80026200 | 80027599 | 7.737117 | 14.39818 | 32.1356 | 13.42203 | -0.70126 | 0.155265 | 0.262895 |
| 1 | 189471800 | 189475599 | 19.89922 | 33.10547 | 69.28231 | 30.07389 | -0.68072 | 1.24366 | 0.262912 |
| 14 | 42494800 | 42495799 | 17.60361 | 31.63955 | 23.03353 | 10.06574 | 0.858555 | 0.405275 | 0.262914 |
| 17 | 13699200 | 13705799 | 65.51092 | 104.3471 | 94.2422 | 39.42854 | 0.483482 | 2.192204 | 0.26293 |
| 4 | 80212200 | 80214199 | 29.97934 | 48.51177 | 91.45666 | 39.11217 | -0.54977 | 1.6708 | 0.262937 |
| 2 | 104847000 | 104847799 | 3.905056 | 7.542953 | 27.48009 | 10.56667 | -1.2931 | -0.18194 | 0.26296 |
| 12 | 73591400 | 73593399 | 18.95823 | 31.55654 | 69.18896 | 30.04415 | -0.74756 | 1.234866 | 0.262962 |
| 18 | 37946000 | 37946999 | 4.961887 | 9.600997 | 25.38935 | 9.683722 | -0.82482 | -0.20236 | 0.262972 |
| 12 | 111419400 | 111422799 | 14.02013 | 24.39433 | 48.95179 | 21.51205 | -0.61305 | 0.765133 | 0.262976 |
| 5 | 31397800 | 31399199 | 8.156826 | 14.78615 | 35.98742 | 15.56736 | -0.86168 | 0.317458 | 0.262979 |
| 6 | 7010200 | 7013999 | 92.31523 | 144.8822 | 127.2742 | 52.78525 | 0.505262 | 2.581063 | 0.26298 |
| 17 | 31651400 | 31653799 | 11.71449 | 20.42488 | 47.33833 | 20.83009 | -0.81912 | 0.72326 | 0.262985 |
| 16 | 20588800 | 20591599 | 32.56054 | 55.21608 | 40.23503 | 17.62311 | 0.836234 | 1.07091 | 0.262988 |
| 4 | 73716600 | 73717999 | 15.17457 | 28.28681 | 22.09352 | 9.201205 | 0.805226 | 0.179541 | 0.262996 |
| 6 | 96140400 | 96143199 | 21.69522 | 35.91688 | 71.45765 | 30.91334 | -0.6086 | 1.308405 | 0.263 |
| 15 | 28859400 | 28861599 | 16.51582 | 30.43478 | 22.30579 | 9.444763 | 0.878627 | 0.254764 | 0.263005 |
| 1 | 8934000 | 8934399 | 8.073993 | 15.73248 | 14.57033 | 5.29129 | 0.738004 | -0.37442 | 0.263013 |
| 19 | 31750400 | 31751599 | 8.361243 | 15.382 | 35.70359 | 15.15221 | -0.77532 | 0.230676 | 0.263014 |
| 6 | 149240400 | 149242599 | 34.69757 | 58.04396 | 46.45897 | 20.23763 | 0.70118 | 1.208068 | 0.263015 |
| 15 | 93236600 | 93238599 | 11.79338 | 20.79127 | 41.95205 | 18.56679 | -0.61811 | 0.553631 | 0.263032 |
| 8 | 78523400 | 78525599 | 15.65961 | 28.81515 | 23.56852 | 9.999149 | 0.71963 | 0.259588 | 0.263046 |
| X | 5107800 | 5108599 | 15.2239 | 28.16894 | 23.15974 | 9.751975 | 0.720034 | 0.225378 | 0.263049 |
| 12 | 76146000 | 76147999 | 14.2995 | 24.52974 | 54.2029 | 23.77277 | -0.75534 | 0.916692 | 0.263049 |
| 9 | 18202000 | 18204199 | 15.60629 | 27.00877 | 52.74059 | 23.13878 | -0.5754 | 0.838389 | 0.263061 |
| 4 | 136758600 | 136761399 | 16.12155 | 27.75873 | 53.78085 | 23.59151 | -0.56565 | 0.888531 | 0.263069 |
| 3 | 130586200 | 130590399 | 24.04889 | 39.39107 | 80.21019 | 34.51384 | -0.64686 | 1.450076 | 0.263089 |
| 16 | 16727800 | 16730799 | 14.45893 | 24.7748 | 55.82107 | 24.48022 | -0.78374 | 0.928315 | 0.263089 |
| 19 | 44386400 | 44390399 | 22.53973 | 37.14019 | 75.90759 | 32.74506 | -0.64866 | 1.368851 | 0.263094 |
| 11 | 101335600 | 101336399 | 3.125213 | 6.113417 | 19.90042 | 6.631838 | -0.93929 | -0.72679 | 0.263094 |
| X | 64173800 | 64180399 | 100.9726 | 158.9319 | 111.045 | 46.07186 | 0.838372 | 2.528943 | 0.263107 |
| 14 | 3400400 | 3402199 | 26.43967 | 45.91552 | 34.23517 | 15.02663 | 0.808383 | 0.828102 | 0.263129 |
| 3 | 84721000 | 84723199 | 13.20388 | 22.71218 | 53.66295 | 23.53907 | -0.85067 | 0.885974 | 0.263134 |
| 6 | 40418800 | 40421199 | 14.79846 | 25.7165 | 50.57831 | 22.20941 | -0.58453 | 0.788309 | 0.263137 |
| 4 | 144211800 | 144213199 | 15.38028 | 29.01707 | 19.32885 | 7.886453 | 1.056066 | 0.099708 | 0.263164 |
| X | 5185600 | 5187199 | 15.38028 | 29.01707 | 19.32885 | 7.886453 | 1.056066 | 0.099708 | 0.263164 |
| 11 | 96577800 | 96585599 | 76.40837 | 118.3999 | 203.4335 | 84.50058 | -0.47573 | 2.803651 | 0.263164 |
| 2 | 71338400 | 71340199 | 13.22164 | 23.06015 | 48.39296 | 21.29849 | -0.67695 | 0.725305 | 0.263169 |
| 3 | 131060200 | 131061599 | 8.79576 | 16.2482 | 34.71438 | 14.65002 | -0.65061 | 0.209572 | 0.263171 |
| 2 | 50054800 | 50058599 | 19.41393 | 32.79414 | 63.56147 | 27.80732 | -0.57013 | 1.082997 | 0.263175 |
| 17 | 52931800 | 52933399 | 14.83747 | 27.86853 | 21.09376 | 8.67298 | 0.864768 | 0.127338 | 0.263183 |
| 6 | 79475000 | 79476199 | 9.797411 | 17.77938 | 36.48647 | 15.76183 | -0.61638 | 0.324042 | 0.263184 |
| 12 | 11043400 | 11044599 | 15.21638 | 28.56323 | 21.08035 | 8.676199 | 0.899884 | 0.134635 | 0.263192 |
| 8 | 31374600 | 31375999 | 16.01787 | 30.34428 | 17.58741 | 7.113886 | 1.26557 | 0.074089 | 0.263195 |
| X | 31736000 | 31737599 | 17.65038 | 32.94525 | 20.285 | 8.429229 | 1.14851 | 0.18891 | 0.2632 |

| | | | | | | | | | |
|---|---|---|---|---|---|---|---|---|---|
| 10 | 15040600 | 15043999 | 41.48071 | 68.29455 | 52.82317 | 22.779 | 0.748555 | 1.408358 | 0.2632 |
| X | 30930200 | 30933399 | 18.00004 | 32.49186 | 20.70879 | 9.005572 | 1.053898 | 0.379078 | 0.263206 |
| 9 | 96549000 | 96552999 | 32.60596 | 52.54096 | 100.1681 | 42.61851 | -0.57285 | 1.796489 | 0.263207 |
| 2 | 172624600 | 172637599 | 138.5293 | 205.7453 | 350.4504 | 144.3661 | -0.51448 | 3.649626 | 0.263208 |
| 4 | 60242200 | 60243799 | 17.98538 | 32.40369 | 24.67057 | 10.75398 | 0.796408 | 0.395828 | 0.263216 |
| 17 | 37356400 | 37359799 | 18.61985 | 31.05993 | 66.96364 | 29.11728 | -0.72331 | 1.209438 | 0.263235 |
| 15 | 65302800 | 65303999 | 8.681393 | 15.53107 | 38.956 | 17.0907 | -0.91879 | 0.407057 | 0.263247 |
| 9 | 63661600 | 63669199 | 47.44886 | 75.22676 | 145.508 | 60.55725 | -0.62023 | 2.331258 | 0.263249 |
| 9 | 124252600 | 124256599 | 816.3868 | 1094.06 | 550.0846 | 229.6026 | 1.098141 | 5.246907 | 0.263251 |
| 7 | 30169600 | 30176799 | 47.53125 | 75.57487 | 136.8441 | 57.11608 | -0.52108 | 2.244541 | 0.263263 |
| 14 | 27646400 | 27647799 | 21.98594 | 38.66579 | 26.97023 | 11.94209 | 0.904721 | 0.611906 | 0.263276 |
| 9 | 102946200 | 102952399 | 50.67558 | 80.37816 | 143.2934 | 59.66209 | -0.50212 | 2.320404 | 0.263288 |
| 8 | 42107800 | 42111999 | 40.98387 | 66.76657 | 59.78993 | 25.67534 | 0.529755 | 1.54853 | 0.263292 |
| 2 | 125253000 | 125254799 | 18.65762 | 31.02613 | 68.44211 | 29.70009 | -0.75631 | 1.246561 | 0.263299 |
| 7 | 41896000 | 41906599 | 280.4052 | 405.7237 | 339.5216 | 140.0288 | 0.474413 | 4.075862 | 0.263301 |
| 14 | 46780600 | 46782799 | 25.77116 | 44.20083 | 38.80456 | 17.019 | 0.570488 | 0.942872 | 0.26331 |
| 18 | 70449400 | 70451199 | 16.00283 | 29.93756 | 20.93557 | 8.666358 | 0.970379 | 0.161799 | 0.263311 |
| 1 | 110670400 | 110671199 | 9.953909 | 19.45704 | 11.97301 | 4.122578 | 1.387499 | -0.53667 | 0.263314 |
| 14 | 32318800 | 32323799 | 40.69232 | 64.92171 | 133.4773 | 55.97016 | -0.69869 | 2.128554 | 0.263342 |
| 1 | 12903800 | 12906599 | 16.32021 | 27.88593 | 58.61349 | 25.69904 | -0.68484 | 0.963364 | 0.263362 |
| 8 | 15577800 | 15580599 | 22.5257 | 39.36767 | 31.57472 | 13.91481 | 0.704114 | 0.715555 | 0.263379 |
| 12 | 24380000 | 24384199 | 58.85391 | 94.57095 | 77.27072 | 32.74294 | 0.641894 | 1.910738 | 0.26339 |
| 3 | 13356000 | 13358799 | 31.027 | 52.90951 | 38.56759 | 16.90722 | 0.836513 | 1.014652 | 0.263391 |
| 14 | 106878600 | 106880399 | 15.81185 | 29.58903 | 20.94261 | 8.664931 | 0.953652 | 0.158198 | 0.263408 |
| 1 | 104509200 | 104511599 | 37.04327 | 60.74992 | 53.09015 | 22.85617 | 0.570735 | 1.445518 | 0.26341 |
| 3 | 19123800 | 19125599 | 14.40176 | 24.84766 | 58.66378 | 25.73504 | -0.84905 | 0.86815 | 0.263429 |
| 8 | 88556200 | 88562799 | 28.21071 | 45.67362 | 90.6288 | 38.77304 | -0.62269 | 1.656484 | 0.263439 |
| 9 | 119394200 | 119397599 | 29.81416 | 47.8793 | 109.5367 | 46.37484 | -0.84134 | 1.904621 | 0.263441 |
| 11 | 70979400 | 70980599 | 8.509725 | 15.69874 | 33.59477 | 14.20342 | -0.65454 | 0.213494 | 0.263442 |
| 13 | 52100000 | 52101999 | 10.71958 | 18.89936 | 41.47886 | 18.35714 | -0.73847 | 0.545938 | 0.263453 |
| 5 | 133567400 | 133570399 | 24.08854 | 41.85883 | 34.91835 | 15.33256 | 0.647204 | 0.808729 | 0.263458 |
| 11 | 102035800 | 102040199 | 32.52416 | 52.21917 | 109.003 | 46.13255 | -0.70988 | 1.915118 | 0.26348 |
| X | 145232600 | 145233199 | 14.24688 | 26.72669 | 20.86954 | 8.5985 | 0.81846 | 0.131207 | 0.263486 |
| 18 | 11306200 | 11307399 | 16.27932 | 31.12182 | 16.66734 | 6.61171 | 1.403201 | 0.014843 | 0.263489 |
| 14 | 28264200 | 28266599 | 13.06087 | 22.85644 | 45.70061 | 20.16255 | -0.60703 | 0.670724 | 0.263504 |
| 4 | 86546800 | 86548599 | 16.88867 | 30.39845 | 26.06104 | 11.37381 | 0.625442 | 0.398236 | 0.263522 |
| 4 | 65530200 | 65533999 | 52.09071 | 82.37114 | 150.5926 | 62.5742 | -0.54219 | 2.40233 | 0.263525 |
| 4 | 126628400 | 126630999 | 21.05475 | 34.51929 | 79.71491 | 34.31439 | -0.82714 | 1.431722 | 0.263532 |
| X | 88451000 | 88453399 | 37.77226 | 62.72325 | 49.32563 | 21.38365 | 0.726585 | 1.299283 | 0.263542 |
| 4 | 103516000 | 103517199 | 13.71857 | 23.90462 | 47.87425 | 21.0579 | -0.60984 | 0.742314 | 0.263557 |
| 7 | 38517800 | 38527199 | 84.93256 | 134.4616 | 100.6411 | 41.84317 | 0.745668 | 2.386658 | 0.263574 |
| 17 | 22692800 | 22701399 | 133.9674 | 206.3252 | 148.7582 | 61.78801 | 0.769083 | 2.933024 | 0.263587 |
| 8 | 117169000 | 117169599 | 8.516864 | 16.59238 | 15.55301 | 5.661752 | 0.717305 | -0.35711 | 0.263589 |
| 2 | 177515200 | 177516999 | 20.38789 | 36.0128 | 28.92774 | 12.79764 | 0.704495 | 0.558892 | 0.263596 |
| 19 | 39446000 | 39447599 | 26.87304 | 46.60075 | 33.05865 | 14.50464 | 0.87939 | 0.842236 | 0.263612 |
| X | 16461600 | 16464399 | 23.58452 | 42.42428 | 21.62367 | 9.443046 | 1.370249 | 0.44072 | 0.263632 |
| 12 | 109144400 | 109146999 | 49.40479 | 80.16356 | 62.39066 | 26.7355 | 0.727114 | 1.635639 | 0.263648 |

| | | | | | | | | | |
|---|---|---|---|---|---|---|---|---|---|
| 8 | 27573000 | 27579199 | 34.11638 | 54.78574 | 108.4401 | 45.90727 | -0.63307 | 1.910773 | 0.263655 |
| 5 | 113381400 | 113384599 | 20.56222 | 34.1526 | 69.14435 | 29.98014 | -0.63319 | 1.266817 | 0.263663 |
| 8 | 45473400 | 45498399 | 494.0262 | 660.379 | 1016.915 | 424.6109 | -0.51999 | 5.284594 | 0.26367 |
| 5 | 76969000 | 76970999 | 10.19328 | 18.16459 | 38.88671 | 17.1167 | -0.69523 | 0.442458 | 0.263673 |
| 18 | 14335200 | 14338599 | 30.75535 | 49.88249 | 91.0188 | 38.99122 | -0.49968 | 1.625062 | 0.263683 |
| 2 | 169869600 | 169877399 | 72.12073 | 111.8467 | 195.5967 | 81.23776 | -0.50024 | 2.787426 | 0.263691 |
| 5 | 17570400 | 17571999 | 21.77467 | 39.64887 | 19.46388 | 8.372607 | 1.436959 | 0.355046 | 0.263693 |
| 3 | 93780600 | 93781399 | 5.533901 | 10.72705 | 26.19322 | 9.906876 | -0.70056 | -0.22468 | 0.263703 |
| 8 | 96380200 | 96381199 | 5.824729 | 11.26132 | 26.57555 | 10.17503 | -0.66842 | -0.17699 | 0.263714 |
| 1 | 84309600 | 84310199 | 5.211942 | 9.897373 | 29.80542 | 11.99453 | -1.0839 | -0.01026 | 0.263715 |
| 7 | 16954200 | 16973999 | 390.9573 | 550.002 | 441.8197 | 183.029 | 0.492852 | 4.492299 | 0.263729 |
| 5 | 75974800 | 75975999 | 10.32541 | 18.46281 | 38.38359 | 16.84681 | -0.65035 | 0.420752 | 0.263732 |
| 3 | 49281200 | 49282199 | 6.879445 | 13.18328 | 32.14378 | 12.66354 | -0.75554 | -0.05917 | 0.263734 |
| 4 | 114068000 | 114070399 | 23.47192 | 41.46148 | 26.36903 | 11.6656 | 1.039347 | 0.57234 | 0.263743 |
| 8 | 25560200 | 25561999 | 8.378979 | 15.42445 | 33.56134 | 14.23072 | -0.68117 | 0.225739 | 0.263745 |
| Y | 6488400 | 6491599 | 17.96632 | 32.23206 | 15.25106 | 6.67513 | 1.47324 | 0.397939 | 0.263756 |
| 5 | 118162400 | 118170599 | 58.19464 | 90.4335 | 207.2573 | 86.05467 | -0.88757 | 2.747565 | 0.263758 |
| 15 | 9070800 | 9072399 | 13.60725 | 23.53214 | 51.59809 | 22.63703 | -0.74181 | 0.835299 | 0.263761 |
| 7 | 25661600 | 25668799 | 91.67286 | 143.5891 | 130.9479 | 54.31716 | 0.449132 | 2.615872 | 0.263773 |
| 10 | 47027000 | 47031199 | 47.69127 | 77.29414 | 59.84226 | 25.6217 | 0.733132 | 1.653428 | 0.263782 |
| 3 | 96503000 | 96506799 | 25.2956 | 41.07841 | 86.52726 | 37.10118 | -0.70439 | 1.587769 | 0.263808 |
| 5 | 6976200 | 6978399 | 21.77917 | 38.73241 | 25.37106 | 11.18661 | 1.001344 | 0.502393 | 0.26382 |
| 19 | 23664400 | 23666199 | 8.950504 | 16.07061 | 37.42857 | 16.37053 | -0.80934 | 0.383485 | 0.263827 |
| 1 | 113684600 | 113688199 | 33.64407 | 57.29242 | 38.03763 | 16.67302 | 0.970633 | 1.038995 | 0.263828 |
| 17 | 37944400 | 37950999 | 178.1132 | 266.9066 | 234.2979 | 96.65288 | 0.453006 | 3.489618 | 0.263852 |
| 17 | 18265600 | 18267999 | 23.58121 | 41.92069 | 24.19509 | 10.67164 | 1.182594 | 0.515168 | 0.263859 |
| 17 | 17848600 | 17849799 | 15.25654 | 28.09016 | 23.70026 | 10.0469 | 0.676137 | 0.254256 | 0.263873 |
| 19 | 33378400 | 33380599 | 27.74912 | 47.77625 | 36.98715 | 16.22475 | 0.752129 | 0.915291 | 0.2639 |
| 7 | 41948800 | 41951399 | 45.14424 | 74.4163 | 48.94583 | 21.11779 | 0.982389 | 1.402495 | 0.263903 |
| 5 | 51936800 | 51938199 | 8.641195 | 16.08978 | 37.80913 | 15.77642 | -0.77492 | 0.169339 | 0.263914 |
| 10 | 75181800 | 75182599 | 14.06215 | 26.26516 | 21.6182 | 8.973091 | 0.733739 | 0.155557 | 0.26392 |
| 8 | 21808400 | 21811599 | 32.21563 | 55.30035 | 34.02127 | 14.92214 | 1.081413 | 0.956826 | 0.263922 |
| X | 145809000 | 145810599 | 18.88361 | 34.74555 | 21.76246 | 9.233657 | 1.102214 | 0.282702 | 0.263935 |
| 16 | 19461200 | 19463199 | 23.37728 | 41.65206 | 23.64494 | 10.40916 | 1.208085 | 0.499058 | 0.263938 |
| 5 | 125974800 | 125976999 | 17.15384 | 29.17388 | 58.54391 | 25.65889 | -0.61927 | 1.010621 | 0.26394 |
| 4 | 98098200 | 98099199 | 6.115631 | 11.82627 | 31.00961 | 11.86084 | -0.81882 | -0.16799 | 0.263946 |
| 1 | 48987400 | 48988999 | 17.90669 | 32.24302 | 25.93545 | 11.31363 | 0.717307 | 0.40323 | 0.263949 |
| 6 | 105442800 | 105445599 | 34.97589 | 59.29191 | 39.14295 | 17.14301 | 0.97824 | 1.083759 | 0.26396 |
| 2 | 172938200 | 172940199 | 14.64109 | 25.01849 | 56.31742 | 24.69246 | -0.78299 | 0.955521 | 0.263961 |
| 8 | 104525400 | 104527599 | 14.24569 | 24.57583 | 52.12701 | 22.86737 | -0.69492 | 0.861611 | 0.263962 |
| 19 | 28815800 | 28818399 | 14.24569 | 24.57583 | 52.12701 | 22.86737 | -0.69492 | 0.861611 | 0.263962 |
| 2 | 36234600 | 36241399 | 121.1191 | 185.8133 | 168.853 | 70.09718 | 0.431293 | 3.005142 | 0.26398 |
| 9 | 96328800 | 96338599 | 89.4557 | 137.3629 | 227.3737 | 94.40313 | -0.43206 | 2.982694 | 0.263987 |
| 7 | 20475600 | 20478599 | 41.92725 | 68.56213 | 53.28239 | 22.91239 | 0.737493 | 1.493366 | 0.264014 |
| 6 | 60805000 | 60806599 | 9.298631 | 16.56311 | 38.92779 | 17.14106 | -0.82922 | 0.439772 | 0.26402 |
| 3 | 71584400 | 71585199 | 11.95208 | 22.70415 | 20.46815 | 8.231277 | 0.641655 | 0.025868 | 0.264029 |
| 5 | 151456600 | 151458999 | 73.62695 | 116.837 | 106.3426 | 44.29799 | 0.466941 | 2.316782 | 0.264031 |

| | | | | | | | | | |
|---|---|---|---|---|---|---|---|---|---|
| 4 | 72368800 | 72371399 | 27.21003 | 47.39094 | 31.94127 | 14.04543 | 0.953717 | 0.789569 | 0.264035 |
| 10 | 7830600 | 7832799 | 12.23083 | 21.177 | 51.03184 | 22.38942 | -0.87673 | 0.818644 | 0.264036 |
| 16 | 60456800 | 60458199 | 18.97931 | 33.44038 | 29.16854 | 12.91067 | 0.584843 | 0.577038 | 0.264044 |
| 13 | 73870400 | 73872399 | 11.18542 | 19.69371 | 41.98751 | 18.58734 | -0.69777 | 0.5648 | 0.264046 |
| 14 | 59438200 | 59439199 | 6.080063 | 11.45417 | 30.19504 | 12.35225 | -0.91468 | 0.05531 | 0.264051 |
| 16 | 26158200 | 26158799 | 4.830772 | 9.348462 | 25.39358 | 9.679687 | -0.86242 | -0.20654 | 0.264057 |
| 15 | 47383800 | 47387399 | 43.27882 | 70.66889 | 58.39997 | 25.09966 | 0.647651 | 1.520011 | 0.26406 |
| 14 | 57613600 | 57614999 | 8.539433 | 15.25543 | 38.45949 | 16.89308 | -0.92702 | 0.415416 | 0.264062 |
| 17 | 88466400 | 88474399 | 65.54044 | 102.6373 | 174.9977 | 72.59045 | -0.45272 | 2.612066 | 0.264069 |
| 13 | 74687400 | 74689999 | 20.03548 | 35.97548 | 25.10838 | 10.98145 | 0.91829 | 0.432769 | 0.26408 |
| 5 | 93890800 | 93892599 | 39.77738 | 66.30939 | 44.8425 | 19.49043 | 0.943681 | 1.259607 | 0.264083 |
| X | 24046800 | 24048599 | 33.70774 | 56.69008 | 43.69661 | 19.09129 | 0.755344 | 1.151177 | 0.264101 |
| 6 | 133318600 | 133320399 | 28.73255 | 49.37346 | 34.04324 | 14.93249 | 0.918108 | 0.934909 | 0.264102 |
| 8 | 18650600 | 18650999 | 11.83556 | 22.84867 | 18.45086 | 7.113253 | 0.854507 | -0.11461 | 0.264119 |
| 12 | 49594200 | 49596799 | 23.20669 | 40.78821 | 29.26013 | 12.95664 | 0.864208 | 0.629643 | 0.264119 |
| 6 | 138947400 | 138948799 | 8.46938 | 15.81565 | 36.78816 | 15.27522 | -0.75457 | 0.146693 | 0.264121 |
| X | 41063600 | 41064999 | 26.54528 | 45.86906 | 36.14214 | 15.85582 | 0.728017 | 0.880267 | 0.264135 |
| 7 | 10762400 | 10769799 | 197.4141 | 292.9959 | 259.0411 | 106.703 | 0.430069 | 3.666364 | 0.264142 |
| 13 | 3082400 | 3087799 | 147.5768 | 225.0933 | 178.2473 | 73.90614 | 0.623171 | 3.130326 | 0.264149 |
| 1 | 35744400 | 35747799 | 24.07617 | 39.8453 | 75.88382 | 32.82059 | -0.54597 | 1.320826 | 0.264153 |
| 13 | 111855800 | 111857999 | 10.84887 | 19.15634 | 42.98336 | 19.01722 | -0.76971 | 0.529913 | 0.264155 |
| X | 128049000 | 128049399 | 9.773946 | 19.09367 | 13.21378 | 4.615588 | 1.202715 | -0.48717 | 0.264165 |
| 6 | 101888000 | 101890199 | 20.97732 | 37.19047 | 27.26556 | 12.05343 | 0.83757 | 0.521261 | 0.264169 |
| 9 | 77332200 | 77335199 | 16.41163 | 27.83534 | 58.97973 | 25.83465 | -0.69766 | 1.033691 | 0.264171 |
| 8 | 72674800 | 72677399 | 17.43213 | 29.31212 | 63.83129 | 27.88666 | -0.73769 | 1.117001 | 0.264174 |
| 14 | 84403400 | 84405799 | 26.62718 | 46.52697 | 29.62169 | 13.05212 | 1.035697 | 0.738313 | 0.264176 |
| 13 | 65177600 | 65179799 | 24.64831 | 42.74729 | 36.0349 | 15.81054 | 0.631871 | 0.836956 | 0.264181 |
| 7 | 61167800 | 61169999 | 39.88131 | 65.38844 | 57.13143 | 24.59349 | 0.570915 | 1.458048 | 0.26419 |
| 7 | 73601600 | 73602599 | 11.85352 | 22.88593 | 17.79153 | 6.855419 | 0.90899 | -0.11921 | 0.264205 |
| 8 | 21819000 | 21821599 | 54.36043 | 87.37511 | 78.0583 | 33.09019 | 0.513873 | 1.897612 | 0.264212 |
| 1 | 110676400 | 110678399 | 27.94986 | 47.36619 | 42.086 | 18.43017 | 0.551189 | 1.061465 | 0.264231 |
| 7 | 130086000 | 130090199 | 54.31951 | 87.37059 | 77.22202 | 32.76897 | 0.530535 | 1.874952 | 0.264257 |
| 16 | 32315200 | 32318799 | 15.84896 | 26.39551 | 68.73142 | 29.85579 | -0.99438 | 1.221513 | 0.264264 |
| 3 | 44042800 | 44043199 | 3.75584 | 7.341664 | 20.09037 | 6.915762 | -0.73484 | -0.59826 | 0.26427 |
| 4 | 104800400 | 104800799 | 3.75584 | 7.341664 | 20.09037 | 6.915762 | -0.73484 | -0.59826 | 0.26427 |
| 7 | 10759400 | 10769799 | 241.5974 | 352.9046 | 307.2504 | 126.5857 | 0.430599 | 3.929677 | 0.264275 |
| 3 | 157446800 | 157447999 | 15.66635 | 28.9109 | 22.40623 | 9.468771 | 0.800557 | 0.240076 | 0.26428 |
| 18 | 38886000 | 38889599 | 18.51667 | 31.18773 | 61.70557 | 26.97161 | -0.59972 | 1.105343 | 0.264285 |
| 7 | 29950000 | 29953999 | 18.52874 | 30.74425 | 69.97642 | 30.31991 | -0.80157 | 1.273266 | 0.264305 |
| 15 | 27950400 | 27951599 | 5.745223 | 10.98893 | 31.90681 | 12.63862 | -1.01273 | -0.05531 | 0.264316 |
| 4 | 146342800 | 146343399 | 10.72344 | 20.81917 | 16.27598 | 6.107093 | 0.93638 | -0.224 | 0.264327 |
| 6 | 78299800 | 78300599 | 10.72344 | 20.81917 | 16.27598 | 6.107093 | 0.93638 | -0.224 | 0.264327 |
| 11 | 82725600 | 82727399 | 12.78391 | 24.09477 | 20.76421 | 8.487535 | 0.685496 | 0.085779 | 0.26433 |
| 14 | 43864200 | 43867799 | 35.93037 | 60.26634 | 41.94154 | 18.30051 | 0.90199 | 1.177997 | 0.264335 |
| 6 | 133513000 | 133515799 | 35.73297 | 59.52171 | 43.87754 | 19.0618 | 0.819717 | 1.254581 | 0.264358 |
| 12 | 24248200 | 24255199 | 236.9783 | 347.0106 | 294.6692 | 121.3825 | 0.469986 | 3.89152 | 0.264376 |
| 5 | 96495600 | 96496799 | 6.749102 | 12.9267 | 33.85906 | 13.35945 | -0.85968 | -0.04897 | 0.264388 |

| | | | | | | | | | |
|---|---|---|---|---|---|---|---|---|---|
| X | 125086800 | 125090399 | 56.06626 | 90.90787 | 57.06448 | 24.44035 | 1.035559 | 1.659911 | 0.26439 |
| 7 | 61783200 | 61787999 | 84.70788 | 133.2886 | 121.9931 | 50.61207 | 0.448999 | 2.523342 | 0.264393 |
| 15 | 31129200 | 31131599 | 13.21442 | 23.03754 | 51.62687 | 22.71559 | -0.77152 | 0.734467 | 0.264397 |
| 1 | 122677000 | 122679999 | 31.75967 | 54.37119 | 36.30717 | 15.92131 | 0.963223 | 0.981334 | 0.26441 |
| 7 | 10882800 | 10886199 | 77.87148 | 124.042 | 84.43099 | 35.32647 | 0.890919 | 2.201586 | 0.264415 |
| 14 | 111718200 | 111718799 | 5.23474 | 10.18194 | 26.01532 | 9.616347 | -0.7343 | -0.30974 | 0.26442 |
| 13 | 100743200 | 100745399 | 14.35153 | 25.06657 | 49.98931 | 22.02082 | -0.60356 | 0.714758 | 0.26442 |
| 17 | 38576600 | 38579999 | 43.80145 | 71.90966 | 49.49409 | 21.3176 | 0.914788 | 1.446647 | 0.264425 |
| 7 | 102289400 | 102290399 | 6.340141 | 12.01913 | 29.82094 | 12.04447 | -0.81234 | 0.012103 | 0.264434 |
| 1 | 68905800 | 68906999 | 9.785834 | 17.79522 | 36.5236 | 15.73702 | -0.61417 | 0.311724 | 0.264445 |
| 19 | 39581800 | 39583999 | 51.54658 | 83.48928 | 63.60091 | 27.21818 | 0.755603 | 1.675334 | 0.264451 |
| 5 | 15888400 | 15889399 | 23.72608 | 41.45246 | 32.64101 | 14.38137 | 0.730705 | 0.730049 | 0.264467 |
| 5 | 70427200 | 70431999 | 67.93787 | 108.4949 | 89.92916 | 37.76101 | 0.611117 | 2.113499 | 0.264497 |
| 7 | 31820000 | 31825599 | 74.29515 | 118.9363 | 75.83225 | 31.95167 | 0.992727 | 2.045878 | 0.26451 |
| 13 | 85386800 | 85387399 | 5.084033 | 9.876953 | 24.53682 | 9.158052 | -0.70605 | -0.27874 | 0.264513 |
| 2 | 108843600 | 108845199 | 16.88926 | 31.01344 | 23.46409 | 9.982973 | 0.828567 | 0.280908 | 0.264516 |
| 2 | 78490800 | 78492599 | 16.37751 | 29.88638 | 24.55074 | 10.53204 | 0.702438 | 0.313685 | 0.26452 |
| 10 | 71970200 | 71970799 | 9.87037 | 19.20977 | 15.90789 | 5.852572 | 0.880161 | -0.30227 | 0.26452 |
| 2 | 8406800 | 8409599 | 67.80812 | 107.9683 | 98.49792 | 41.18906 | 0.468489 | 2.208094 | 0.264526 |
| 4 | 89924800 | 89926799 | 20.44944 | 36.35608 | 26.59693 | 11.73025 | 0.842651 | 0.497876 | 0.264526 |
| 7 | 127067000 | 127068799 | 26.74232 | 45.67271 | 39.87234 | 17.48134 | 0.577674 | 0.98561 | 0.264545 |
| 5 | 66422000 | 66423399 | 6.738532 | 12.88247 | 28.98356 | 11.49699 | -0.64929 | -0.04493 | 0.26455 |
| 18 | 74753600 | 74760399 | 39.32211 | 62.93232 | 117.1882 | 49.35802 | -0.55105 | 2.025486 | 0.264571 |
| 3 | 139730800 | 139731599 | 10.04909 | 19.54128 | 16.46723 | 6.099323 | 0.846694 | -0.27493 | 0.264572 |
| 6 | 11365000 | 11366799 | 19.79801 | 34.73154 | 30.33128 | 13.40958 | 0.581962 | 0.640934 | 0.26458 |
| X | 75524400 | 75525799 | 24.21134 | 42.45168 | 29.54594 | 13.05662 | 0.90799 | 0.674942 | 0.264586 |
| 9 | 60347800 | 60348999 | 6.519362 | 12.26656 | 31.83791 | 13.05168 | -0.89575 | 0.06919 | 0.264589 |
| 17 | 18469800 | 18470599 | 11.04034 | 21.50549 | 14.63905 | 5.33905 | 1.170954 | -0.32059 | 0.264591 |
| 7 | 31564000 | 31565199 | 35.96516 | 59.25634 | 53.09723 | 22.90417 | 0.537365 | 1.387309 | 0.264608 |
| 7 | 63909800 | 63913599 | 53.14683 | 85.46724 | 77.56833 | 32.90636 | 0.492025 | 1.879411 | 0.264641 |
| 8 | 76510400 | 76513799 | 49.88825 | 80.40426 | 72.92527 | 31.03544 | 0.49687 | 1.805065 | 0.26465 |
| 19 | 8045000 | 8046999 | 23.85077 | 41.23648 | 36.72252 | 16.11072 | 0.552198 | 0.861702 | 0.264679 |
| 9 | 97186000 | 97186599 | 6.117718 | 11.8232 | 27.78687 | 10.65806 | -0.66559 | -0.16548 | 0.264681 |
| 9 | 43867400 | 43869799 | 15.96735 | 27.47382 | 53.76978 | 23.58614 | -0.58038 | 0.894197 | 0.264682 |
| 14 | 73987800 | 73988999 | 8.784667 | 16.26139 | 35.32751 | 14.86676 | -0.67168 | 0.198659 | 0.26469 |
| 1 | 84911400 | 84915399 | 26.27413 | 42.61935 | 92.12811 | 39.46986 | -0.74335 | 1.612942 | 0.264693 |
| 4 | 153834000 | 153836599 | 18.19401 | 30.67796 | 61.15022 | 26.73778 | -0.61031 | 1.09428 | 0.264702 |
| 10 | 31548400 | 31551199 | 17.64702 | 29.97518 | 58.52813 | 25.64839 | -0.58017 | 1.02469 | 0.26472 |
| 7 | 59341400 | 59344999 | 57.013 | 91.51957 | 72.94758 | 30.86401 | 0.675813 | 1.938114 | 0.264722 |
| 19 | 36604400 | 36605799 | 9.947864 | 18.08005 | 37.04673 | 15.97326 | -0.61245 | 0.317353 | 0.264726 |
| 15 | 88563600 | 88571199 | 55.42151 | 86.87844 | 181.6227 | 75.33075 | -0.74525 | 2.59025 | 0.264738 |
| 1 | 119408200 | 119412199 | 28.30025 | 46.01989 | 85.76422 | 36.80494 | -0.52665 | 1.567094 | 0.264754 |
| 7 | 18734400 | 18735599 | 5.96409 | 11.18415 | 31.18617 | 12.85743 | -1.00419 | 0.086309 | 0.264759 |
| 12 | 74960400 | 74961799 | 9.765014 | 17.82384 | 37.69326 | 16.16527 | -0.65283 | 0.291496 | 0.264777 |
| 19 | 28967000 | 28968799 | 11.48341 | 20.18413 | 45.60384 | 20.19448 | -0.78246 | 0.588504 | 0.264792 |
| 10 | 66571400 | 66574399 | 42.01489 | 68.20272 | 60.14079 | 25.77973 | 0.547976 | 1.609688 | 0.264795 |
| 7 | 69944600 | 69946399 | 9.457926 | 17.37394 | 39.00669 | 16.58721 | -0.72988 | 0.256118 | 0.264795 |

| | | | | | | | | | |
|---|---|---|---|---|---|---|---|---|---|
| 3 | 123678400 | 123679199 | 16.84004 | 30.06167 | 25.23934 | 11.09322 | 0.648577 | 0.44661 | 0.264795 |
| X | 22363200 | 22368199 | 42.04822 | 70.8174 | 35.21393 | 15.39176 | 1.385275 | 1.160558 | 0.264814 |
| 8 | 35535200 | 35538199 | 21.91078 | 36.5343 | 72.56161 | 31.54309 | -0.60567 | 1.226183 | 0.264814 |
| 4 | 40471800 | 40474799 | 17.1234 | 28.8031 | 62.10097 | 27.13361 | -0.72319 | 1.113178 | 0.26482 |
| 6 | 132326200 | 132328399 | 23.42702 | 41.42068 | 26.45634 | 11.70205 | 1.033814 | 0.563292 | 0.264822 |
| 3 | 93985600 | 93987399 | 13.70835 | 23.91849 | 47.33022 | 20.83595 | -0.59241 | 0.722334 | 0.264837 |
| 8 | 46837200 | 46841399 | 27.42765 | 44.81285 | 83.41995 | 35.86082 | -0.52017 | 1.49471 | 0.264842 |
| 2 | 108114200 | 108117799 | 28.75037 | 49.60817 | 35.26413 | 15.46996 | 0.875291 | 0.898653 | 0.264851 |
| 7 | 105229800 | 105231399 | 8.550632 | 15.62545 | 35.02808 | 15.0004 | -0.73459 | 0.270345 | 0.264856 |
| 12 | 10752400 | 10755399 | 24.0544 | 41.913 | 33.17626 | 14.59246 | 0.722802 | 0.768176 | 0.264895 |
| 5 | 109350000 | 109353599 | 80.54477 | 127.5747 | 109.715 | 45.62661 | 0.546119 | 2.37459 | 0.26491 |
| 13 | 66701800 | 66719799 | 415.8811 | 584.536 | 428.9566 | 177.7253 | 0.62198 | 4.507037 | 0.264918 |
| 7 | 60402600 | 60404399 | 22.79708 | 39.68901 | 34.55604 | 15.19136 | 0.586112 | 0.777857 | 0.264941 |
| 15 | 27970800 | 27973599 | 13.26576 | 23.03393 | 50.01541 | 21.95081 | -0.72658 | 0.79284 | 0.264944 |
| 14 | 69978000 | 69979399 | 16.04383 | 28.92966 | 25.18562 | 10.9671 | 0.605492 | 0.380119 | 0.264953 |
| 11 | 61892800 | 61896399 | 20.26144 | 34.03584 | 65.16925 | 28.46254 | -0.55297 | 1.136202 | 0.264959 |
| X | 31648400 | 31652199 | 152.061 | 231.1791 | 169.6003 | 70.25601 | 0.729513 | 3.197016 | 0.264961 |
| 11 | 63436400 | 63437399 | 5.236795 | 10.1796 | 25.27385 | 9.389318 | -0.69979 | -0.29343 | 0.264971 |
| 9 | 76985800 | 76991599 | 50.65911 | 80.488 | 138.9284 | 57.94262 | -0.45321 | 2.270358 | 0.264974 |
| X | 3865600 | 3868799 | 31.95847 | 55.54364 | 28.41192 | 12.4769 | 1.349367 | 0.837018 | 0.264995 |
| 19 | 12426400 | 12427199 | 42.63604 | 69.48016 | 60.91949 | 26.16343 | 0.560528 | 1.550637 | 0.26501 |
| 2 | 112379600 | 112381399 | 33.8583 | 56.01251 | 49.84072 | 21.55084 | 0.548502 | 1.330037 | 0.265033 |
| 1 | 145581200 | 145582799 | 21.33963 | 37.88515 | 26.72208 | 11.79949 | 0.894146 | 0.514328 | 0.265034 |
| 14 | 110835800 | 110836199 | 11.48158 | 21.8399 | 19.03302 | 7.632288 | 0.693966 | 0.008759 | 0.265086 |
| 4 | 72257000 | 72259399 | 31.65472 | 52.90988 | 44.42558 | 19.34177 | 0.632481 | 1.207105 | 0.265091 |
| 7 | 35365200 | 35371799 | 41.11099 | 65.71122 | 122.0472 | 51.31157 | -0.54941 | 2.0677 | 0.265096 |
| 10 | 14945200 | 14945599 | 6.779488 | 13.2596 | 12.39158 | 4.188136 | 0.816337 | -0.64971 | 0.265098 |
| 8 | 117243000 | 117245199 | 54.23753 | 87.18042 | 76.74411 | 32.53462 | 0.535174 | 1.89655 | 0.265099 |
| 1 | 98338400 | 98340199 | 16.09636 | 27.78882 | 53.81585 | 23.60876 | -0.56427 | 0.86403 | 0.265103 |
| 12 | 7874200 | 7878599 | 46.89822 | 76.06601 | 58.10868 | 24.89326 | 0.753389 | 1.636651 | 0.265103 |
| 7 | 130918400 | 130920399 | 12.24952 | 21.52459 | 44.54997 | 19.72307 | -0.65662 | 0.596551 | 0.265118 |
| 10 | 61618200 | 61620999 | 17.60857 | 29.57974 | 62.62769 | 27.35268 | -0.69717 | 1.127779 | 0.26512 |
| 7 | 61609400 | 61618599 | 113.3481 | 175.954 | 141.8518 | 58.90718 | 0.616807 | 2.782787 | 0.265121 |
| 1 | 77730800 | 77731399 | 13.38529 | 25.65658 | 18.70822 | 7.365483 | 0.972183 | -0.03233 | 0.265126 |
| 8 | 45051800 | 45052799 | 13.38529 | 25.65658 | 18.70822 | 7.365483 | 0.972183 | -0.03233 | 0.265126 |
| 7 | 4660000 | 4662599 | 15.07289 | 25.80958 | 56.93453 | 24.96719 | -0.75362 | 0.93744 | 0.265126 |
| 7 | 80332400 | 80335599 | 19.29819 | 32.50654 | 62.33531 | 27.24751 | -0.55486 | 1.107338 | 0.265128 |
| 7 | 8480600 | 8482599 | 28.69976 | 49.81504 | 33.05599 | 14.50442 | 0.975775 | 0.844974 | 0.265141 |
| 1 | 24118800 | 24120599 | 14.64923 | 27.52315 | 21.10047 | 8.671373 | 0.846981 | 0.123684 | 0.265142 |
| 7 | 37614400 | 37620399 | 45.93854 | 72.96662 | 141.1635 | 58.85767 | -0.61789 | 2.27439 | 0.265149 |
| Y | 39882000 | 39883399 | 13.87382 | 26.18053 | 10.1776 | 4.15073 | 1.824068 | 0.056124 | 0.265166 |
| 2 | 178357000 | 178360999 | 24.42138 | 39.86456 | 83.9533 | 36.07807 | -0.69854 | 1.501294 | 0.265172 |
| 12 | 78954800 | 78958999 | 40.41337 | 64.73469 | 119.7291 | 50.49283 | -0.53989 | 1.99959 | 0.26518 |
| 10 | 97578800 | 97580599 | 10.88011 | 19.13989 | 43.52914 | 19.27307 | -0.79112 | 0.573259 | 0.265187 |
| 17 | 76572200 | 76574199 | 24.42687 | 42.35835 | 34.37439 | 15.08193 | 0.686295 | 0.832673 | 0.265202 |
| 1 | 8799800 | 8803599 | 62.84065 | 100.7147 | 82.04115 | 34.62795 | 0.641892 | 1.999062 | 0.265217 |
| 15 | 37106200 | 37107399 | 10.1467 | 18.25026 | 37.39438 | 16.33149 | -0.62446 | 0.380728 | 0.265229 |

| | | | | | | | | | |
|---|---|---|---|---|---|---|---|---|---|
| 18 | 35665400 | 35666399 | 10.1467 | 18.25026 | 37.39438 | 16.33149 | -0.62446 | 0.380728 | 0.265229 |
| 12 | 13209400 | 13210999 | 9.505437 | 17.18968 | 36.4382 | 15.80956 | -0.66687 | 0.341889 | 0.265244 |
| 15 | 19536200 | 19538399 | 16.20476 | 27.65833 | 62.96642 | 27.60514 | -0.7998 | 0.979138 | 0.26525 |
| 9 | 23756200 | 23758199 | 24.05794 | 42.18833 | 29.71994 | 13.13499 | 0.890288 | 0.669443 | 0.265251 |
| 7 | 81819000 | 81829999 | 108.7792 | 164.9882 | 270.2758 | 111.8813 | -0.43127 | 3.239655 | 0.265257 |
| 5 | 127627000 | 127634399 | 59.6573 | 93.88482 | 165.4569 | 68.64548 | -0.49485 | 2.512962 | 0.265262 |
| 17 | 69247200 | 69248799 | 18.46607 | 32.85671 | 27.38698 | 12.07039 | 0.656177 | 0.481328 | 0.265271 |
| 11 | 105051800 | 105063199 | 178.5764 | 259.988 | 432.7969 | 178.3716 | -0.50867 | 3.977337 | 0.265279 |
| 7 | 29144000 | 29148599 | 24.63826 | 40.06434 | 84.89405 | 36.43129 | -0.71123 | 1.560857 | 0.26528 |
| 17 | 69349600 | 69353199 | 36.66079 | 60.92368 | 49.3172 | 21.39117 | 0.684989 | 1.286221 | 0.265292 |
| 4 | 143528200 | 143530599 | 54.13627 | 88.10588 | 51.95405 | 22.30237 | 1.129889 | 1.587005 | 0.265294 |
| 17 | 24888400 | 24889199 | 3.706889 | 7.227077 | 22.51881 | 8.117795 | -0.98178 | -0.42873 | 0.265298 |
| 16 | 5702400 | 5704999 | 18.55622 | 30.72048 | 71.50487 | 30.93396 | -0.83402 | 1.301351 | 0.265298 |
| 6 | 37444400 | 37445599 | 14.18906 | 26.61115 | 21.70603 | 8.947172 | 0.754874 | 0.132349 | 0.265321 |
| 10 | 77598400 | 77599399 | 14.18906 | 26.61115 | 21.70603 | 8.947172 | 0.754874 | 0.132349 | 0.265321 |
| 11 | 113514000 | 113516999 | 28.71999 | 46.4154 | 95.12263 | 40.62636 | -0.67185 | 1.701074 | 0.265322 |
| 15 | 19843200 | 19845399 | 31.89672 | 53.76551 | 44.20203 | 19.32533 | 0.662683 | 1.124694 | 0.265329 |
| 4 | 148416200 | 148427999 | 84.21182 | 129.3719 | 224.5003 | 93.21617 | -0.49958 | 2.970944 | 0.265336 |
| 1 | 149877400 | 149879199 | 16.41167 | 30.63331 | 19.59797 | 8.14366 | 1.09282 | 0.174737 | 0.265343 |
| 14 | 42194800 | 42196999 | 24.45122 | 42.49101 | 33.76131 | 14.82493 | 0.717019 | 0.807083 | 0.265345 |
| 1 | 7644400 | 7644999 | 7.107016 | 13.33338 | 30.67566 | 12.63738 | -0.72849 | 0.093142 | 0.265352 |
| 12 | 22349600 | 22351999 | 35.76288 | 59.64847 | 43.80851 | 19.04732 | 0.824911 | 1.239452 | 0.265353 |
| X | 54567600 | 54568599 | 16.81895 | 30.53675 | 24.96179 | 10.77649 | 0.703592 | 0.345845 | 0.265381 |
| 7 | 79234200 | 79243799 | 72.92037 | 112.6812 | 217.2167 | 90.22703 | -0.64463 | 2.855788 | 0.265388 |
| 10 | 112346400 | 112349999 | 38.33538 | 63.14413 | 54.37744 | 23.45367 | 0.594299 | 1.39596 | 0.265388 |
| 5 | 143402000 | 143402999 | 13.60816 | 25.54329 | 21.83444 | 8.987212 | 0.689608 | 0.122961 | 0.265391 |
| 17 | 23373600 | 23374199 | 9.873012 | 19.14008 | 15.80149 | 5.973594 | 0.848367 | -0.21145 | 0.265392 |
| 3 | 55829800 | 55830199 | 8.398231 | 16.36892 | 15.59802 | 5.643445 | 0.701905 | -0.37931 | 0.265394 |
| 1 | 111380600 | 111383199 | 28.56455 | 49.92469 | 28.5275 | 12.57257 | 1.191046 | 0.744801 | 0.265417 |
| 1 | 105813600 | 105815799 | 12.43675 | 21.81279 | 44.09247 | 19.48974 | -0.62253 | 0.629108 | 0.265443 |
| 1 | 72543800 | 72546999 | 19.53515 | 32.6275 | 67.1675 | 29.22896 | -0.65703 | 1.197644 | 0.265459 |
| 19 | 3438200 | 3443199 | 44.09375 | 70.26943 | 128.6342 | 53.85552 | -0.53323 | 2.166602 | 0.265469 |
| 11 | 73859000 | 73861799 | 46.58129 | 75.44422 | 66.52862 | 28.47032 | 0.544956 | 1.66825 | 0.265478 |
| 16 | 14240000 | 14242199 | 12.60037 | 22.21494 | 46.67622 | 20.65739 | -0.67644 | 0.565888 | 0.265486 |
| 15 | 76053400 | 76068399 | 611.7835 | 794.3495 | 1425.974 | 597.437 | -0.78312 | 5.754207 | 0.265491 |
| 7 | 11646800 | 11653999 | 83.24668 | 132.4634 | 90.00935 | 37.60516 | 0.891792 | 2.241773 | 0.265492 |
| 7 | 49159800 | 49161999 | 12.12703 | 21.23333 | 45.69498 | 20.16899 | -0.71255 | 0.656412 | 0.265502 |
| 9 | 19654400 | 19659999 | 86.66869 | 137.4901 | 99.3304 | 41.36418 | 0.799262 | 2.336025 | 0.265505 |
| 19 | 17813000 | 17815399 | 15.05767 | 25.83623 | 54.78722 | 24.02949 | -0.69634 | 0.916937 | 0.265509 |
| 9 | 18956800 | 18961799 | 98.17337 | 152.8015 | 140.5106 | 58.32717 | 0.430121 | 2.730192 | 0.265528 |
| X | 102704000 | 102706799 | 29.00152 | 50.01363 | 35.94663 | 15.7691 | 0.859476 | 0.907632 | 0.265558 |
| 15 | 101117400 | 101124999 | 36.49715 | 58.27346 | 125.0992 | 52.51417 | -0.75959 | 2.097784 | 0.265568 |
| 14 | 33073000 | 33077599 | 27.46615 | 44.36182 | 96.29819 | 41.09592 | -0.75564 | 1.717986 | 0.265584 |
| 1 | 67076400 | 67078199 | 7.902822 | 14.63355 | 32.59001 | 13.70895 | -0.70597 | 0.186199 | 0.265588 |
| 16 | 89848600 | 89853599 | 57.45261 | 89.86919 | 179.0819 | 74.29334 | -0.67861 | 2.628153 | 0.265589 |
| 11 | 36274600 | 36276199 | 9.274817 | 17.04185 | 39.59898 | 16.83333 | -0.77901 | 0.252729 | 0.265598 |
| 4 | 31133800 | 31134799 | 8.786378 | 17.148 | 13.73487 | 4.877486 | 0.972112 | -0.44644 | 0.265602 |

| | | | | | | | | | |
|---|---|---|---|---|---|---|---|---|---|
| 1 | 135579400 | 135583199 | 36.01706 | 60.27241 | 44.22018 | 19.26667 | 0.826245 | 1.204916 | 0.265603 |
| 6 | 73091400 | 73092399 | 6.725659 | 12.89993 | 29.76233 | 11.6957 | -0.67279 | -0.07469 | 0.265608 |
| X | 54893800 | 54897799 | 76.76621 | 122.0439 | 98.5568 | 41.13074 | 0.64185 | 2.266085 | 0.265627 |
| 13 | 31726200 | 31727199 | 5.66384 | 10.98841 | 26.85031 | 10.11455 | -0.69561 | -0.23288 | 0.265632 |
| 8 | 44034200 | 44035999 | 16.67863 | 31.18617 | 19.64138 | 8.137535 | 1.118824 | 0.166899 | 0.265637 |
| 6 | 131576000 | 131578199 | 40.50992 | 66.82019 | 52.23605 | 22.54352 | 0.734277 | 1.383461 | 0.265642 |
| 19 | 26951400 | 26952999 | 9.269258 | 17.04884 | 37.37383 | 15.86561 | -0.69392 | 0.24295 | 0.265642 |
| 6 | 26881200 | 26882399 | 9.323926 | 17.10874 | 36.72963 | 15.64258 | -0.66693 | 0.258363 | 0.265652 |
| 3 | 30884800 | 30886199 | 14.24309 | 26.22299 | 22.83485 | 9.680664 | 0.630835 | 0.244378 | 0.265663 |
| 6 | 3498000 | 3498999 | 11.04531 | 21.07736 | 18.45619 | 7.35107 | 0.695188 | -0.0202 | 0.265676 |
| 14 | 83025200 | 83026399 | 23.63097 | 40.83574 | 36.21602 | 15.88827 | 0.557941 | 0.863965 | 0.26568 |
| 15 | 76523600 | 76524399 | 4.529239 | 8.801802 | 23.93512 | 8.911627 | -0.83137 | -0.29639 | 0.265692 |
| 8 | 3879200 | 3883399 | 109.1458 | 168.7212 | 152.8932 | 63.50836 | 0.443644 | 2.855311 | 0.265695 |
| 8 | 75510400 | 75513599 | 101.6038 | 157.5291 | 142.5592 | 59.21185 | 0.448847 | 2.797671 | 0.265707 |
| 2 | 39537400 | 39538199 | 18.45314 | 32.66243 | 26.80286 | 11.85271 | 0.675085 | 0.521801 | 0.265714 |
| 3 | 21737200 | 21740599 | 31.97677 | 54.22237 | 41.43756 | 18.14944 | 0.767951 | 1.072988 | 0.265717 |
| 8 | 80845200 | 80847399 | 11.94684 | 20.9989 | 43.50906 | 19.26702 | -0.65804 | 0.588967 | 0.265724 |
| 12 | 102865600 | 102867999 | 18.60111 | 31.47535 | 62.43138 | 27.32721 | -0.6032 | 1.063681 | 0.265734 |
| 13 | 66027000 | 66030399 | 33.74189 | 57.61928 | 36.30643 | 15.91776 | 1.046205 | 1.012497 | 0.265736 |
| X | 147323400 | 147323999 | 9.428205 | 18.31248 | 17.09047 | 6.390247 | 0.688756 | -0.24783 | 0.265743 |
| 2 | 87691400 | 87693799 | 25.90923 | 45.19322 | 31.36064 | 13.80314 | 0.911902 | 0.7606 | 0.265744 |
| 7 | 140372000 | 140373399 | 10.42099 | 18.35965 | 42.00036 | 18.59063 | -0.7984 | 0.552548 | 0.265757 |
| X | 98889000 | 98891999 | 14.96617 | 25.98975 | 50.58141 | 22.201 | -0.56952 | 0.79952 | 0.265759 |
| 8 | 93082200 | 93087399 | 44.37095 | 70.40379 | 142.281 | 59.26214 | -0.68217 | 2.304881 | 0.265771 |
| 2 | 41940400 | 41943199 | 25.12801 | 41.26882 | 77.68651 | 33.46576 | -0.53233 | 1.418244 | 0.265772 |
| 16 | 96240400 | 96245399 | 27.22966 | 44.15181 | 88.38482 | 37.85507 | -0.63329 | 1.623098 | 0.265772 |
| 18 | 11923600 | 11926199 | 34.63726 | 57.69624 | 48.71873 | 21.1649 | 0.624377 | 1.253551 | 0.265775 |
| 8 | 77558000 | 77558399 | 3.339861 | 6.533287 | 18.59024 | 6.196434 | -0.74706 | -0.72422 | 0.265782 |
| 10 | 47907400 | 47911999 | 85.12522 | 134.4698 | 106.8213 | 44.36316 | 0.657397 | 2.439284 | 0.265785 |
| 11 | 18825000 | 18827999 | 18.51756 | 30.76197 | 69.52954 | 30.15082 | -0.79143 | 1.258194 | 0.265806 |
| 8 | 25607200 | 25612999 | 42.73333 | 68.26149 | 124.7144 | 52.38632 | -0.52673 | 2.089327 | 0.265814 |
| 18 | 47972400 | 47973999 | 14.83308 | 27.27046 | 23.67608 | 10.05671 | 0.632936 | 0.258014 | 0.265816 |
| 12 | 91504000 | 91505399 | 14.37697 | 24.87085 | 51.08344 | 22.41146 | -0.64789 | 0.835695 | 0.265817 |
| 3 | 67758000 | 67759399 | 6.364016 | 12.01687 | 34.85429 | 14.1954 | -1.04745 | 0.049189 | 0.265818 |
| X | 79751400 | 79753399 | 27.17409 | 47.22697 | 33.75596 | 14.82341 | 0.869318 | 0.822411 | 0.26582 |
| 9 | 63156200 | 63159799 | 14.34289 | 24.90366 | 50.03514 | 21.95917 | -0.61518 | 0.798139 | 0.265837 |
| 4 | 89488000 | 89488999 | 15.20782 | 28.62747 | 19.88231 | 8.144277 | 0.991809 | 0.11334 | 0.265874 |
| 2 | 122110000 | 122118399 | 55.02103 | 86.44151 | 167.3116 | 69.39961 | -0.63177 | 2.545148 | 0.265883 |
| 13 | 65108800 | 65109999 | 21.59635 | 37.64073 | 32.12355 | 14.13195 | 0.614974 | 0.752469 | 0.265893 |
| X | 26393200 | 26394799 | 20.79335 | 38.859 | 16.62576 | 6.894446 | 1.671165 | 0.199839 | 0.265895 |
| 12 | 4859000 | 4860799 | 27.7105 | 47.02313 | 40.99642 | 17.96028 | 0.578454 | 1.04731 | 0.265905 |
| X | 33914200 | 33917999 | 44.84956 | 73.16 | 58.82965 | 25.27505 | 0.686027 | 1.54104 | 0.265905 |
| 4 | 115070400 | 115072799 | 23.84714 | 42.11129 | 26.36512 | 11.66459 | 1.061753 | 0.578161 | 0.265906 |
| 8 | 19308400 | 19310199 | 16.99045 | 29.06544 | 56.41215 | 24.73622 | -0.56986 | 0.95602 | 0.265911 |
| 18 | 38944800 | 38945599 | 6.141972 | 11.7948 | 28.01192 | 10.96932 | -0.70902 | -0.09657 | 0.265914 |
| 7 | 7865400 | 7865999 | 10.00208 | 19.38156 | 17.52383 | 6.638683 | 0.716297 | -0.19627 | 0.265932 |
| 15 | 34041600 | 34042799 | 26.77055 | 45.50549 | 40.0055 | 17.53306 | 0.566672 | 1.028626 | 0.265943 |

| | | | | | | | | | |
|---|---|---|---|---|---|---|---|---|---|
| 1 | 51477600 | 51478999 | 19.12479 | 34.4473 | 25.30746 | 11.03529 | 0.847703 | 0.409146 | 0.265945 |
| 12 | 114044200 | 114045799 | 22.75216 | 40.08674 | 28.07426 | 12.42639 | 0.899752 | 0.597744 | 0.26595 |
| 8 | 74405200 | 74406199 | 8.65483 | 15.99817 | 33.63323 | 14.18132 | -0.62631 | 0.203604 | 0.26596 |
| 9 | 36337600 | 36339799 | 27.74552 | 48.32046 | 30.73615 | 13.51496 | 1.036671 | 0.790483 | 0.265962 |
| 3 | 30561200 | 30566999 | 31.65539 | 50.9799 | 102.7829 | 43.69462 | -0.6544 | 1.813684 | 0.26598 |
| 1 | 103667400 | 103668799 | 29.03069 | 49.91839 | 33.94844 | 14.89121 | 0.938021 | 0.929952 | 0.265982 |
| 6 | 11081000 | 11082199 | 7.981336 | 15.02863 | 34.81542 | 14.25413 | -0.73216 | 0.084728 | 0.26599 |
| 7 | 123773800 | 123777399 | 37.00261 | 61.37542 | 50.44778 | 21.85297 | 0.662844 | 1.309432 | 0.265993 |
| 19 | 23691600 | 23692599 | 22.31705 | 38.85246 | 32.66887 | 14.36155 | 0.636231 | 0.773605 | 0.26601 |
| 14 | 52770600 | 52772799 | 33.74609 | 56.07038 | 50.35548 | 21.83894 | 0.535655 | 1.281294 | 0.266017 |
| 7 | 5442200 | 5444999 | 68.97151 | 109.5755 | 98.94024 | 41.262 | 0.480038 | 2.277774 | 0.266038 |
| 17 | 3471400 | 3472199 | 4.917745 | 9.579165 | 24.29154 | 8.860821 | -0.70457 | -0.3627 | 0.26605 |
| 15 | 63923200 | 63927399 | 22.85499 | 37.29058 | 82.86225 | 35.60327 | -0.77609 | 1.505306 | 0.266051 |
| 3 | 20055200 | 20055599 | 3.342321 | 6.530303 | 19.88475 | 6.927688 | -0.90147 | -0.56179 | 0.266054 |
| 7 | 79887200 | 79889799 | 14.66891 | 24.99795 | 56.7774 | 24.88869 | -0.79634 | 0.983366 | 0.266055 |
| 18 | 60419800 | 60423999 | 52.16067 | 84.19372 | 70.21515 | 29.94033 | 0.620142 | 1.763692 | 0.266058 |
| 8 | 108411200 | 108415199 | 24.40596 | 40.31183 | 78.80146 | 34.03631 | -0.58378 | 1.345759 | 0.266069 |
| 1 | 87109800 | 87121199 | 60.89375 | 93.78118 | 238.1722 | 98.92655 | -1.04561 | 2.915821 | 0.266078 |
| 18 | 63054800 | 63058799 | 21.63588 | 35.97737 | 69.46131 | 30.14156 | -0.56494 | 1.254495 | 0.266094 |
| 3 | 54730000 | 54734999 | 77.2611 | 119.6177 | 200.7801 | 83.39881 | -0.44301 | 2.820384 | 0.2661 |
| 15 | 96565200 | 96566599 | 8.254187 | 15.09387 | 34.51815 | 14.76961 | -0.76217 | 0.263663 | 0.266108 |
| 12 | 105180800 | 105192599 | 99.20992 | 150.6987 | 262.8726 | 108.8653 | -0.52005 | 3.206029 | 0.266108 |
| 1 | 51321200 | 51323399 | 13.88076 | 24.24352 | 47.84514 | 21.07585 | -0.5884 | 0.710462 | 0.266117 |
| 5 | 10324200 | 10327799 | 38.89454 | 65.16017 | 40.80949 | 17.79312 | 1.05365 | 1.200363 | 0.26612 |
| 1 | 135117800 | 135120199 | 7.612849 | 14.06487 | 34.15841 | 14.41282 | -0.83372 | 0.197753 | 0.266136 |
| 9 | 18348000 | 18348999 | 10.16346 | 19.735 | 16.9694 | 6.359462 | 0.802456 | -0.23215 | 0.266188 |
| 3 | 83933600 | 83935199 | 9.013177 | 16.41996 | 35.50287 | 15.26251 | -0.68655 | 0.292911 | 0.266194 |
| 3 | 130424800 | 130425599 | 9.013177 | 16.41996 | 35.50287 | 15.26251 | -0.68655 | 0.292911 | 0.266194 |
| 7 | 107553200 | 107554399 | 15.51713 | 28.69814 | 23.09138 | 9.729349 | 0.75021 | 0.229826 | 0.266221 |
| 1 | 144792400 | 144795199 | 20.06106 | 35.75504 | 26.06529 | 11.47152 | 0.849654 | 0.478645 | 0.266229 |
| 5 | 130036600 | 130041999 | 65.96445 | 103.0262 | 179.1064 | 74.31518 | -0.4837 | 2.660216 | 0.266239 |
| 8 | 125737800 | 125742999 | 34.51655 | 55.70378 | 100.7573 | 42.95365 | -0.49466 | 1.755094 | 0.26624 |
| X | 106803000 | 106805599 | 30.56123 | 53.06558 | 31.39035 | 13.77675 | 1.140904 | 0.847804 | 0.266244 |
| 7 | 9698400 | 9701599 | 37.94457 | 64.2071 | 36.82116 | 16.11723 | 1.180409 | 1.106974 | 0.266266 |
| 7 | 32329800 | 32330199 | 9.30301 | 18.11965 | 16.03247 | 5.854383 | 0.795696 | -0.33417 | 0.266289 |
| 3 | 113717400 | 113718999 | 11.61516 | 20.65737 | 42.48995 | 18.73406 | -0.63985 | 0.471709 | 0.266294 |
| 1 | 74838400 | 74848999 | 65.16924 | 100.9325 | 214.4785 | 89.0886 | -0.78247 | 2.810081 | 0.266306 |
| 8 | 122076600 | 122078999 | 10.62305 | 18.59274 | 45.21467 | 19.9504 | -0.88745 | 0.656412 | 0.266315 |
| 7 | 38820000 | 38825999 | 147.0277 | 222.5679 | 199.9109 | 82.70351 | 0.432652 | 3.282785 | 0.26633 |
| 18 | 84737400 | 84738999 | 12.58826 | 22.00409 | 48.36443 | 21.31711 | -0.74312 | 0.688989 | 0.26633 |
| X | 25645200 | 25645999 | 23.44069 | 40.6516 | 32.79612 | 14.38949 | 0.695102 | 0.82908 | 0.266338 |
| 13 | 92047600 | 92050799 | 28.99542 | 49.27145 | 40.87598 | 17.91409 | 0.649879 | 1.039913 | 0.266338 |
| 13 | 58307600 | 58318599 | 103.968 | 157.112 | 279.5022 | 115.5673 | -0.55467 | 3.31348 | 0.266341 |
| 6 | 118028200 | 118031399 | 26.89543 | 43.64845 | 88.05801 | 37.7397 | -0.64328 | 1.603305 | 0.266352 |
| 3 | 9539800 | 9541999 | 15.42195 | 26.31333 | 56.87471 | 24.93379 | -0.72508 | 0.972791 | 0.266365 |
| 8 | 126437600 | 126440399 | 22.87018 | 37.63493 | 78.81638 | 33.98086 | -0.68455 | 1.387279 | 0.266386 |
| 5 | 100148000 | 100149599 | 9.082245 | 16.36782 | 36.43427 | 15.87619 | -0.74063 | 0.360866 | 0.266394 |

| | | | | | | | | | |
|---|---|---|---|---|---|---|---|---|---|
| 6 | 82377400 | 82381199 | 28.92774 | 46.64891 | 100.901 | 42.97272 | -0.75322 | 1.768155 | 0.266422 |
| 16 | 18868200 | 18869999 | 20.64734 | 36.97733 | 24.9889 | 10.95331 | 0.962313 | 0.452112 | 0.266444 |
| 12 | 106821800 | 106824599 | 18.60272 | 31.5114 | 60.76213 | 26.60509 | -0.56247 | 1.053612 | 0.266446 |
| 1 | 18031200 | 18036999 | 84.68257 | 133.8125 | 113.9223 | 47.31976 | 0.557988 | 2.433187 | 0.266447 |
| 1 | 101740400 | 101741599 | 16.62128 | 29.99891 | 25.4844 | 11.08425 | 0.641532 | 0.378685 | 0.266465 |
| 4 | 135610000 | 135613799 | 17.94031 | 30.07388 | 63.71969 | 27.79281 | -0.69825 | 1.151831 | 0.266486 |
| 13 | 45541800 | 45544599 | 14.20251 | 24.22997 | 58.43653 | 25.61826 | -0.88249 | 0.970313 | 0.266486 |
| 7 | 85081400 | 85088199 | 99.29451 | 156.3589 | 111.6303 | 46.31736 | 0.807735 | 2.51681 | 0.266491 |
| 4 | 116350000 | 116353399 | 34.11492 | 56.95302 | 47.58891 | 20.70318 | 0.639381 | 1.227426 | 0.266507 |
| 5 | 67046600 | 67048399 | 16.28027 | 27.55873 | 64.41742 | 28.19981 | -0.83898 | 1.055723 | 0.266521 |
| 1 | 52470400 | 52472399 | 11.30302 | 20.01534 | 40.37655 | 17.85369 | -0.61516 | 0.498543 | 0.266525 |
| 11 | 11314400 | 11315199 | 10.61215 | 20.63301 | 15.76418 | 5.846222 | 0.984467 | -0.26659 | 0.266561 |
| 17 | 47846600 | 47847799 | 6.163673 | 11.77171 | 28.34608 | 11.27033 | -0.74916 | -0.04455 | 0.266563 |
| 6 | 88521400 | 88525799 | 24.0357 | 39.40861 | 78.51896 | 33.80032 | -0.61495 | 1.437013 | 0.266568 |
| 10 | 22848000 | 22849199 | 23.02387 | 37.9469 | 74.13269 | 31.9826 | -0.58352 | 1.367652 | 0.266583 |
| 8 | 86872000 | 86874599 | 14.48479 | 25.25648 | 48.93556 | 21.53368 | -0.56242 | 0.737147 | 0.266588 |
| X | 22747000 | 22749799 | 49.30528 | 79.54802 | 69.45331 | 29.59784 | 0.553419 | 1.773664 | 0.266596 |
| 13 | 4415600 | 4418199 | 11.00406 | 19.63873 | 48.5508 | 21.34364 | -0.90088 | 0.456746 | 0.266601 |
| 7 | 117321200 | 117323999 | 35.98061 | 59.716 | 50.42727 | 21.85312 | 0.624053 | 1.299415 | 0.266606 |
| 4 | 90072400 | 90073999 | 16.96786 | 30.66832 | 25.44362 | 11.0467 | 0.67714 | 0.373186 | 0.26662 |
| 7 | 59167400 | 59170799 | 24.73537 | 43.01516 | 34.90978 | 15.33604 | 0.68689 | 0.802819 | 0.266647 |
| X | 146958000 | 146961399 | 45.2174 | 74.20788 | 52.89623 | 22.77952 | 0.864268 | 1.457135 | 0.266657 |
| 8 | 114519600 | 114524799 | 36.66008 | 59.02978 | 105.6958 | 44.92187 | -0.48397 | 1.828428 | 0.266674 |
| 2 | 3398000 | 3398399 | 3.040055 | 5.945853 | 18.37305 | 6.210514 | -0.88101 | -0.67856 | 0.266679 |
| 15 | 96606400 | 96608199 | 8.228261 | 15.11673 | 34.05821 | 14.48077 | -0.7342 | 0.235441 | 0.266684 |
| 7 | 7416600 | 7419999 | 44.13931 | 72.70047 | 50.08517 | 21.6019 | 0.915239 | 1.408628 | 0.266692 |
| 7 | 31732600 | 31738599 | 75.49706 | 119.9686 | 102.0483 | 42.56742 | 0.566325 | 2.278496 | 0.266699 |
| 7 | 47412600 | 47421399 | 213.9478 | 317.0502 | 266.8707 | 109.911 | 0.498878 | 3.695535 | 0.266707 |
| 4 | 146294600 | 146296799 | 86.62522 | 136.8391 | 113.2127 | 47.01751 | 0.59885 | 2.441424 | 0.26673 |
| 2 | 55325000 | 55328199 | 26.87335 | 43.66954 | 85.95672 | 36.87457 | -0.60605 | 1.575855 | 0.266741 |
| 16 | 91779800 | 91791599 | 105.8662 | 159.187 | 324.1603 | 133.8399 | -0.75573 | 3.414566 | 0.266755 |
| 1 | 13120400 | 13123799 | 14.95547 | 26.00209 | 50.57821 | 22.21621 | -0.5686 | 0.782133 | 0.266756 |
| 4 | 11086600 | 11087199 | 4.752806 | 9.261811 | 24.98207 | 9.062224 | -0.78498 | -0.38146 | 0.266758 |
| 15 | 55962000 | 55962799 | 5.114636 | 9.907117 | 25.49087 | 9.674124 | -0.77949 | -0.21921 | 0.266763 |
| 9 | 70180000 | 70182799 | 19.63506 | 33.0344 | 62.86973 | 27.47114 | -0.54405 | 1.119925 | 0.266767 |
| 1 | 34373400 | 34374599 | 17.62415 | 31.40244 | 26.70632 | 11.75657 | 0.628737 | 0.467635 | 0.266767 |
| 1 | 8361600 | 8366799 | 59.99172 | 96.46824 | 74.12072 | 31.44073 | 0.731662 | 1.888988 | 0.266778 |
| Y | 39320800 | 39322599 | 12.81227 | 24.29971 | 11.84411 | 4.779073 | 1.515481 | 0.017199 | 0.266792 |
| 7 | 106035000 | 106035599 | 9.720967 | 17.45506 | 37.9425 | 16.59444 | -0.71022 | 0.39208 | 0.266793 |
| 10 | 22236600 | 22239399 | 11.62844 | 20.48443 | 42.46632 | 18.79734 | -0.65749 | 0.56098 | 0.266794 |
| 13 | 64605200 | 64611999 | 57.10244 | 89.76987 | 165.7199 | 68.7452 | -0.56285 | 2.532843 | 0.266795 |
| 7 | 12614200 | 12615599 | 13.71456 | 23.86815 | 48.41625 | 21.27978 | -0.62845 | 0.760078 | 0.266797 |
| 12 | 28683400 | 28686999 | 19.56391 | 32.61629 | 67.04786 | 29.1435 | -0.65487 | 1.218367 | 0.266805 |
| 9 | 18337800 | 18340599 | 23.04063 | 37.90368 | 75.76614 | 32.66149 | -0.61763 | 1.391243 | 0.266829 |
| 3 | 79337200 | 79339599 | 13.55233 | 23.6341 | 47.33736 | 20.83235 | -0.60987 | 0.728808 | 0.266854 |
| 11 | 12699200 | 12701999 | 19.37055 | 34.40408 | 24.79932 | 10.94652 | 0.862993 | 0.493036 | 0.266862 |
| 9 | 92578800 | 92580199 | 21.68326 | 38.14582 | 26.57525 | 11.76644 | 0.906686 | 0.606312 | 0.266891 |

| | | | | | | | | | |
|---|---|---|---|---|---|---|---|---|---|
| 7 | 27214400 | 27221199 | 54.36192 | 85.02974 | 177.4685 | 73.62442 | -0.74535 | 2.62849 | 0.266895 |
| 4 | 147495200 | 147500599 | 20.2263 | 35.63773 | 25.18583 | 11.14781 | 0.886822 | 0.577374 | 0.266896 |
| 12 | 99574000 | 99578199 | 25.56621 | 41.63497 | 86.19471 | 37.00945 | -0.6764 | 1.539211 | 0.266898 |
| 1 | 182236200 | 182237199 | 6.444302 | 12.32503 | 28.99839 | 11.48934 | -0.71182 | -0.05275 | 0.26691 |
| 11 | 54328400 | 54329599 | 6.444302 | 12.32503 | 28.99839 | 11.48934 | -0.71182 | -0.05275 | 0.26691 |
| 13 | 105288600 | 105289199 | 4.919009 | 9.57683 | 24.18297 | 8.871309 | -0.70585 | -0.34238 | 0.266929 |
| 10 | 83891200 | 83892999 | 9.158577 | 16.67078 | 37.08107 | 15.95779 | -0.72844 | 0.300933 | 0.266945 |
| 14 | 6744400 | 6745999 | 23.27189 | 40.82228 | 30.34785 | 13.41594 | 0.813072 | 0.661677 | 0.266983 |
| X | 30925800 | 30928999 | 35.92114 | 61.14455 | 35.69393 | 15.64518 | 1.155324 | 1.048307 | 0.26699 |
| 8 | 88055000 | 88056399 | 10.61071 | 19.06414 | 38.4127 | 16.79158 | -0.6015 | 0.392419 | 0.266994 |
| 5 | 65525800 | 65527199 | 7.820379 | 14.74379 | 32.52825 | 13.28985 | -0.65961 | 0.070316 | 0.267006 |
| 13 | 57854600 | 57856999 | 13.13755 | 22.79233 | 49.99814 | 21.93723 | -0.74117 | 0.804581 | 0.267019 |
| X | 22800600 | 22802399 | 17.72363 | 32.66684 | 22.3089 | 9.443576 | 0.980445 | 0.264686 | 0.267026 |
| 8 | 119343400 | 119346999 | 34.53151 | 55.64021 | 101.2942 | 43.09364 | -0.5066 | 1.801595 | 0.267034 |
| 14 | 32721200 | 32722799 | 8.034094 | 14.90952 | 32.62617 | 13.68297 | -0.67793 | 0.173598 | 0.267034 |
| 12 | 9733000 | 9734199 | 10.96674 | 19.58027 | 39.40082 | 17.31509 | -0.60486 | 0.437502 | 0.267036 |
| 1 | 127434400 | 127437199 | 16.46539 | 28.24065 | 54.8286 | 24.04689 | -0.56975 | 0.926754 | 0.267039 |
| 17 | 39112400 | 39114599 | 33.64753 | 57.31385 | 37.40426 | 16.39575 | 0.995243 | 1.034565 | 0.267055 |
| 1 | 192105400 | 192106199 | 5.981723 | 11.53315 | 27.00971 | 10.45597 | -0.67299 | -0.13767 | 0.267056 |
| 3 | 64081000 | 64083199 | 23.40511 | 41.19282 | 26.83594 | 11.88083 | 1.00273 | 0.609667 | 0.267057 |
| 2 | 79297400 | 79298599 | 7.980301 | 15.03023 | 34.23687 | 14.01167 | -0.70749 | 0.082055 | 0.267084 |
| 19 | 24524600 | 24526399 | 36.50286 | 60.02592 | 54.25473 | 23.3843 | 0.523761 | 1.411106 | 0.267094 |
| 7 | 59901800 | 59904199 | 31.05278 | 52.85478 | 39.68891 | 17.39617 | 0.793495 | 1.032628 | 0.2671 |
| 13 | 57787600 | 57791199 | 18.79873 | 31.78419 | 60.70522 | 26.56517 | -0.54879 | 1.071704 | 0.267113 |
| 9 | 78049000 | 78049999 | 18.9004 | 33.55547 | 27.2874 | 12.04887 | 0.689704 | 0.49692 | 0.267127 |
| 2 | 166657200 | 166657999 | 20.16708 | 36.36915 | 19.61856 | 8.54157 | 1.292112 | 0.398664 | 0.267136 |
| 4 | 14985800 | 14986399 | 5.914839 | 11.22032 | 29.8194 | 12.02696 | -0.90864 | 0.003402 | 0.26714 |
| 17 | 93268400 | 93271599 | 31.53226 | 53.27108 | 40.8445 | 17.86961 | 0.762834 | 1.097407 | 0.267149 |
| 7 | 37804800 | 37808999 | 24.84652 | 40.6836 | 79.6958 | 34.28752 | -0.59136 | 1.45622 | 0.267159 |
| 4 | 96166800 | 96170799 | 76.26985 | 121.4908 | 92.79016 | 38.82401 | 0.725051 | 2.20226 | 0.267165 |
| 7 | 31484400 | 31485399 | 17.98269 | 32.25409 | 26.37531 | 11.54623 | 0.690173 | 0.427227 | 0.26717 |
| 7 | 31180200 | 31181199 | 45.67112 | 73.81624 | 61.06924 | 26.08485 | 0.633821 | 1.710273 | 0.267174 |
| 13 | 61313600 | 61314999 | 27.10186 | 46.13765 | 38.04341 | 16.67515 | 0.659224 | 1.016653 | 0.267187 |
| 17 | 22183200 | 22183799 | 10.02235 | 19.35312 | 18.07565 | 6.960396 | 0.64803 | -0.13975 | 0.267198 |
| 4 | 131266400 | 131269799 | 21.58833 | 35.60832 | 73.3995 | 31.6766 | -0.66018 | 1.354108 | 0.267203 |
| 7 | 25994000 | 25995999 | 37.9775 | 62.48221 | 54.91074 | 23.67208 | 0.564442 | 1.409021 | 0.267207 |
| 14 | 47433800 | 47434799 | 12.22595 | 23.01544 | 20.42268 | 8.364954 | 0.641628 | 0.088277 | 0.26723 |
| 2 | 64857400 | 64859999 | 15.66363 | 26.88041 | 55.91024 | 24.52237 | -0.66868 | 0.918255 | 0.267235 |
| 3 | 3125200 | 3140199 | 230.4519 | 341.5163 | 237.4601 | 97.79847 | 0.774441 | 3.696567 | 0.267244 |
| 6 | 23252400 | 23255799 | 26.97574 | 43.97165 | 84.82308 | 36.43297 | -0.57382 | 1.527885 | 0.267264 |
| 1 | 115310800 | 115311399 | 9.7208 | 18.84004 | 17.58089 | 6.654456 | 0.672542 | -0.20183 | 0.267264 |
| 13 | 43545400 | 43546199 | 9.035485 | 17.55596 | 15.45056 | 5.760449 | 0.775938 | -0.26908 | 0.267279 |
| 12 | 49428600 | 49432799 | 34.40545 | 54.94526 | 123.4798 | 51.84981 | -0.82509 | 2.090083 | 0.267287 |
| 15 | 12370000 | 12377199 | 49.18519 | 77.68537 | 154.6975 | 64.24169 | -0.66665 | 2.427904 | 0.267311 |
| 13 | 66678200 | 66684999 | 120.7103 | 188.7993 | 109.6166 | 45.47567 | 1.098145 | 2.655521 | 0.267316 |
| 16 | 87697800 | 87698999 | 6.325481 | 12.02094 | 28.76742 | 11.55853 | -0.75353 | -0.00624 | 0.267322 |
| 8 | 10280200 | 10282399 | 10.16867 | 18.18743 | 38.3869 | 16.84445 | -0.67181 | 0.417655 | 0.267324 |

| | | | | | | | | | |
|---|---|---|---|---|---|---|---|---|---|
| 10 | 8203200 | 8204999 | 28.14129 | 47.63839 | 42.63229 | 18.6632 | 0.540856 | 1.072321 | 0.267333 |
| X | 61121200 | 61123399 | 27.61876 | 47.83506 | 34.7536 | 15.24774 | 0.845002 | 0.860538 | 0.267336 |
| 3 | 82204600 | 82206199 | 11.88369 | 20.67654 | 48.40029 | 21.26936 | -0.83372 | 0.756458 | 0.267341 |
| 2 | 21359600 | 21364399 | 74.55187 | 118.3851 | 103.5821 | 43.17744 | 0.524744 | 2.298321 | 0.267364 |
| 16 | 65001200 | 65001799 | 24.87681 | 43.53826 | 24.8674 | 10.97205 | 1.191748 | 0.699243 | 0.267365 |
| 7 | 57532600 | 57533599 | 20.41031 | 35.94403 | 26.57693 | 11.76472 | 0.821693 | 0.586346 | 0.267368 |
| 3 | 134583400 | 134588199 | 41.65444 | 66.54181 | 121.735 | 51.13888 | -0.5285 | 2.08798 | 0.267368 |
| X | 23677800 | 23680799 | 26.42441 | 46.0051 | 33.13103 | 14.56511 | 0.858615 | 0.792966 | 0.26739 |
| 6 | 109687400 | 109688199 | 21.90835 | 38.67112 | 23.20779 | 10.26859 | 1.122524 | 0.566409 | 0.267395 |
| 16 | 61040200 | 61044999 | 96.82387 | 151.0086 | 137.4225 | 57.02798 | 0.447414 | 2.695037 | 0.267407 |
| 1 | 49801600 | 49802199 | 16.65056 | 29.99841 | 24.76953 | 10.79728 | 0.680648 | 0.389698 | 0.267413 |
| 4 | 46998800 | 47001399 | 12.88576 | 22.59293 | 45.12724 | 19.94162 | -0.60559 | 0.637245 | 0.26744 |
| 3 | 155465600 | 155467199 | 18.27797 | 32.89168 | 25.92386 | 11.31712 | 0.745871 | 0.409559 | 0.267453 |
| 2 | 148235600 | 148239999 | 22.44317 | 36.83878 | 79.59972 | 34.28181 | -0.73096 | 1.41909 | 0.267468 |
| 10 | 117265400 | 117268999 | 27.28687 | 44.5235 | 83.17951 | 35.74011 | -0.52668 | 1.513731 | 0.267508 |
| 8 | 34514600 | 34519199 | 47.16381 | 75.12794 | 132.7695 | 55.55198 | -0.48327 | 2.184534 | 0.267522 |
| 18 | 3130400 | 3135199 | 29.73197 | 51.2542 | 32.12247 | 14.09135 | 1.0559 | 0.907463 | 0.267533 |
| 5 | 57823400 | 57825399 | 21.55777 | 38.50167 | 24.27611 | 10.6656 | 1.05934 | 0.472923 | 0.267539 |
| 8 | 75125600 | 75127599 | 17.47581 | 31.856 | 23.17356 | 9.954751 | 0.875009 | 0.323469 | 0.267541 |
| 17 | 78778000 | 78786399 | 12.06652 | 23.17453 | 17.30192 | 6.773275 | 0.946139 | -0.06728 | 0.267571 |
| 3 | 95180400 | 95182199 | 7.428661 | 13.86224 | 31.64932 | 13.1575 | -0.72809 | 0.134145 | 0.267576 |
| 17 | 16509200 | 16512199 | 27.32046 | 47.04573 | 37.57893 | 16.48442 | 0.707282 | 0.913382 | 0.267579 |
| 1 | 110426600 | 110427999 | 12.10627 | 22.94925 | 19.79469 | 7.995563 | 0.6989 | 0.03803 | 0.267586 |
| 5 | 103449800 | 103451399 | 19.11889 | 33.80718 | 28.63259 | 12.66437 | 0.629172 | 0.539831 | 0.267608 |
| 11 | 66413200 | 66419399 | 50.22624 | 79.53325 | 148.7689 | 61.85987 | -0.57354 | 2.365853 | 0.267619 |
| 4 | 56594200 | 56595999 | 10.42152 | 18.81727 | 41.20948 | 17.91422 | -0.71598 | 0.365211 | 0.26763 |
| Y | 9718800 | 9721599 | 15.20176 | 28.09071 | 13.35348 | 5.632964 | 1.501275 | 0.204662 | 0.26766 |
| 7 | 27472600 | 27476599 | 91.39887 | 142.9138 | 132.0284 | 54.77504 | 0.428578 | 2.645974 | 0.267662 |
| 14 | 118373200 | 118374999 | 11.97411 | 20.98156 | 44.64117 | 19.71654 | -0.69606 | 0.642212 | 0.267676 |
| X | 154844200 | 154847999 | 50.43129 | 81.65634 | 62.59625 | 26.78183 | 0.746198 | 1.681982 | 0.267678 |
| 1 | 78385000 | 78386399 | 8.189413 | 15.1524 | 33.65206 | 14.17073 | -0.70364 | 0.193202 | 0.267691 |
| 14 | 72878000 | 72879399 | 9.024747 | 16.44206 | 34.98602 | 15.03915 | -0.6633 | 0.292919 | 0.2677 |
| 3 | 47289200 | 47290199 | 14.11011 | 26.78555 | 19.57637 | 7.883845 | 0.939768 | 0.047607 | 0.267705 |
| 16 | 65270200 | 65270999 | 8.132665 | 14.79871 | 35.52818 | 15.29554 | -0.83693 | 0.297489 | 0.267706 |
| 9 | 61434600 | 61436199 | 8.763476 | 15.83318 | 36.96347 | 16.05612 | -0.80625 | 0.342484 | 0.267719 |
| 5 | 95133000 | 95133799 | 12.25353 | 23.91036 | 10.84625 | 3.863744 | 1.776467 | -0.39565 | 0.267721 |
| 10 | 23786800 | 23788399 | 12.75298 | 24.02986 | 21.38839 | 8.746747 | 0.639038 | 0.089995 | 0.267727 |
| 3 | 15170800 | 15172399 | 10.8215 | 19.22999 | 39.36601 | 17.36877 | -0.6335 | 0.46568 | 0.267734 |
| 17 | 3375400 | 3377599 | 13.9705 | 24.01649 | 53.68875 | 23.54996 | -0.77147 | 0.894356 | 0.267739 |
| 2 | 152031000 | 152032599 | 8.440247 | 15.3247 | 35.9932 | 15.53838 | -0.80895 | 0.309338 | 0.267743 |
| 2 | 172390800 | 172393799 | 25.07595 | 40.55865 | 95.1307 | 40.66967 | -0.86465 | 1.67232 | 0.267743 |
| 9 | 50029800 | 50031199 | 8.304164 | 15.53311 | 33.40144 | 13.82973 | -0.63846 | 0.127525 | 0.267748 |
| 7 | 14542400 | 14550999 | 208.6655 | 308.5649 | 272.3192 | 112.148 | 0.427758 | 3.73049 | 0.267761 |
| 9 | 66900000 | 66900999 | 13.12023 | 25.05449 | 19.84838 | 7.893813 | 0.840526 | 0.000392 | 0.267768 |
| 5 | 37885600 | 37887199 | 9.277261 | 16.58365 | 37.94404 | 16.6581 | -0.78741 | 0.417277 | 0.267769 |
| 9 | 64014400 | 64015999 | 14.96708 | 27.23192 | 22.7747 | 9.806338 | 0.673301 | 0.315139 | 0.267769 |
| 2 | 167393400 | 167395199 | 8.213662 | 15.19553 | 33.10847 | 13.94399 | -0.67589 | 0.19566 | 0.267777 |

| | | | | | | | | | |
|---|---|---|---|---|---|---|---|---|---|
| 7 | 41630400 | 41632399 | 22.94582 | 40.37976 | 28.63448 | 12.67738 | 0.881274 | 0.612727 | 0.267791 |
| 19 | 51531600 | 51533999 | 17.77896 | 32.11422 | 24.14818 | 10.49249 | 0.817092 | 0.378538 | 0.267792 |
| 12 | 48941800 | 48942799 | 11.6645 | 22.55143 | 17.24299 | 6.602246 | 0.941029 | -0.14382 | 0.267794 |
| 6 | 51216800 | 51219399 | 49.49009 | 79.7097 | 72.78573 | 30.94665 | 0.485961 | 1.823904 | 0.267811 |
| 17 | 19096400 | 19097599 | 35.45359 | 58.62442 | 52.58007 | 22.72804 | 0.537093 | 1.343071 | 0.267822 |
| 4 | 139293200 | 139299199 | 33.71879 | 53.9649 | 117.6781 | 49.56474 | -0.77828 | 2.019838 | 0.267822 |
| 12 | 33499000 | 33502199 | 20.88564 | 34.26249 | 78.56072 | 33.8261 | -0.81652 | 1.423443 | 0.267826 |
| 4 | 9480600 | 9482599 | 19.86217 | 35.60105 | 25.65534 | 11.23747 | 0.870953 | 0.443322 | 0.267831 |
| X | 78005400 | 78005999 | 12.4126 | 23.85923 | 18.96454 | 7.406272 | 0.860243 | -0.06555 | 0.267836 |
| 6 | 64312800 | 64316399 | 36.84244 | 60.59475 | 54.82841 | 23.63329 | 0.522298 | 1.410366 | 0.267854 |
| 1 | 113996200 | 113997799 | 19.66061 | 35.71171 | 23.01919 | 9.932398 | 1.044553 | 0.360461 | 0.267857 |
| 10 | 121203000 | 121207799 | 29.66409 | 47.95482 | 96.18157 | 41.09367 | -0.64041 | 1.694103 | 0.267882 |
| 2 | 152384800 | 152385399 | 9.709387 | 18.85664 | 17.03423 | 6.374639 | 0.733963 | -0.24214 | 0.26789 |
| 16 | 26399200 | 26401399 | 14.64831 | 25.42822 | 50.57948 | 22.19492 | -0.60089 | 0.803084 | 0.267897 |
| 18 | 53620200 | 53623999 | 27.30473 | 44.50355 | 83.58563 | 35.90017 | -0.53545 | 1.530207 | 0.267907 |
| 16 | 39328400 | 39333599 | 68.66609 | 109.5094 | 94.1034 | 39.43859 | 0.557091 | 2.157463 | 0.26792 |
| X | 143268000 | 143270399 | 21.02499 | 37.63277 | 24.97942 | 10.95428 | 0.987656 | 0.458308 | 0.267921 |
| 7 | 32189200 | 32192999 | 75.929 | 120.1691 | 111.4662 | 46.3364 | 0.436252 | 2.389809 | 0.267924 |
| 18 | 68035000 | 68035799 | 10.86432 | 21.13569 | 15.81378 | 5.83686 | 1.020961 | -0.27862 | 0.267926 |
| 18 | 58526800 | 58529399 | 17.54394 | 29.64097 | 69.01672 | 30.19567 | -0.83377 | 1.080327 | 0.267933 |
| 8 | 22204800 | 22208199 | 46.93335 | 76.99367 | 50.71298 | 21.83567 | 0.977632 | 1.46345 | 0.267935 |
| X | 4808600 | 4813399 | 12.51843 | 23.78753 | 17.83044 | 7.165718 | 0.9066 | 0.0219 | 0.267957 |
| 8 | 126970800 | 126972799 | 7.489961 | 13.78041 | 33.55918 | 14.24033 | -0.84317 | 0.220849 | 0.267965 |
| 3 | 105996400 | 106001399 | 52.67367 | 84.73421 | 76.95761 | 32.66285 | 0.49159 | 1.868577 | 0.267968 |
| 5 | 44563200 | 44564599 | 12.08496 | 21.40385 | 45.08757 | 19.93608 | -0.67754 | 0.508782 | 0.267975 |
| 17 | 41070200 | 41073799 | 33.8423 | 57.58265 | 37.9707 | 16.64258 | 0.980181 | 1.045726 | 0.267982 |
| 9 | 54834600 | 54836999 | 19.29603 | 34.98843 | 23.56886 | 10.19177 | 0.979408 | 0.368398 | 0.268003 |
| 13 | 18875600 | 18877999 | 19.29603 | 34.98843 | 23.56886 | 10.19177 | 0.979408 | 0.368398 | 0.268003 |
| 3 | 74776200 | 74778999 | 25.92195 | 45.35401 | 30.37225 | 13.39756 | 0.963331 | 0.717579 | 0.268019 |
| 10 | 111341800 | 111344799 | 16.70686 | 28.349 | 60.12869 | 26.34156 | -0.69921 | 1.030598 | 0.268023 |
| 5 | 109372200 | 109375399 | 39.03565 | 65.69757 | 39.12908 | 17.09942 | 1.125955 | 1.158527 | 0.268029 |
| 17 | 20180200 | 20182799 | 54.1955 | 87.5641 | 66.48955 | 28.38763 | 0.756722 | 1.73816 | 0.268033 |
| 15 | 89111200 | 89112599 | 7.262195 | 13.58997 | 31.17056 | 12.89658 | -0.7294 | 0.111219 | 0.268035 |
| 1 | 14160400 | 14163199 | 33.7136 | 56.59916 | 44.18844 | 19.29189 | 0.736757 | 1.164228 | 0.268037 |
| 3 | 7291600 | 7292199 | 4.80848 | 9.370048 | 26.42259 | 9.588193 | -0.85073 | -0.38086 | 0.268059 |
| 17 | 38470400 | 38473399 | 34.79581 | 59.83924 | 31.18141 | 13.67753 | 1.320379 | 0.950248 | 0.26806 |
| 3 | 91170800 | 91174199 | 40.04207 | 65.48223 | 58.74389 | 25.26133 | 0.531057 | 1.49257 | 0.268076 |
| 7 | 124382400 | 124387199 | 57.26596 | 92.1104 | 76.00647 | 32.25345 | 0.629443 | 1.878214 | 0.26808 |
| 7 | 138792800 | 138793599 | 5.122686 | 9.899624 | 25.91332 | 9.942061 | -0.81899 | -0.18013 | 0.268092 |
| 11 | 70404400 | 70405399 | 5.122686 | 9.899624 | 25.91332 | 9.942061 | -0.81899 | -0.18013 | 0.268092 |
| 7 | 5176600 | 5179799 | 37.55683 | 61.79147 | 55.40394 | 23.88492 | 0.53563 | 1.409027 | 0.268093 |
| 12 | 119870200 | 119873399 | 42.38643 | 69.03841 | 61.52956 | 26.42057 | 0.536474 | 1.556339 | 0.268124 |
| 1 | 138945000 | 138946999 | 31.1311 | 52.10792 | 46.47534 | 20.25215 | 0.545488 | 1.189505 | 0.268125 |
| 11 | 52654200 | 52657599 | 21.16449 | 34.79793 | 77.15903 | 33.25412 | -0.76705 | 1.394329 | 0.268138 |
| 3 | 65687800 | 65688399 | 8.977349 | 17.44089 | 16.61599 | 6.201031 | 0.662018 | -0.26107 | 0.268152 |
| 11 | 113050800 | 113054999 | 24.98407 | 40.58637 | 88.54817 | 37.97727 | -0.75457 | 1.579258 | 0.268159 |
| 2 | 169821400 | 169821999 | 4.231887 | 8.245717 | 22.46335 | 8.161225 | -0.80075 | -0.39242 | 0.268161 |

| | | | | | | | | | |
|---|---|---|---|---|---|---|---|---|---|
| 9 | 44345000 | 44347399 | 15.72835 | 26.89707 | 55.27648 | 24.23756 | -0.65197 | 0.953112 | 0.268162 |
| 6 | 39337000 | 39339599 | 12.22511 | 21.59129 | 45.07272 | 19.94072 | -0.66626 | 0.539238 | 0.268169 |
| 15 | 103630800 | 103631199 | 3.81652 | 7.453043 | 21.07315 | 7.405875 | -0.80983 | -0.52311 | 0.268186 |
| 7 | 59907400 | 59910799 | 47.49688 | 76.95508 | 65.62682 | 28.09192 | 0.593596 | 1.661302 | 0.268193 |
| 19 | 23962000 | 23963999 | 8.689139 | 15.48968 | 38.47554 | 16.93066 | -0.90734 | 0.432039 | 0.2682 |
| Y | 4296000 | 4299799 | 17.68562 | 31.85992 | 14.87874 | 6.48663 | 1.495822 | 0.370537 | 0.268202 |
| 7 | 9822200 | 9824799 | 52.23474 | 84.50088 | 65.25952 | 27.90292 | 0.734287 | 1.701663 | 0.268206 |
| 8 | 117112200 | 117113599 | 8.78468 | 16.33148 | 36.02292 | 15.07081 | -0.68727 | 0.176482 | 0.268209 |
| X | 54631400 | 54633999 | 26.84769 | 47.14178 | 26.90467 | 11.90384 | 1.192469 | 0.662441 | 0.268214 |
| 19 | 10432400 | 10436399 | 21.51037 | 35.29343 | 78.73344 | 33.90294 | -0.777 | 1.422312 | 0.268227 |
| 5 | 100647600 | 100648799 | 4.466979 | 8.622666 | 29.29108 | 11.29779 | -1.19919 | -0.16215 | 0.268229 |
| 7 | 85563200 | 85565199 | 26.09915 | 45.57361 | 31.40785 | 13.8333 | 0.922038 | 0.748373 | 0.268239 |
| 17 | 35005400 | 35036599 | 367.6719 | 504.9229 | 769.7448 | 320.0839 | -0.46624 | 4.857268 | 0.26826 |
| 4 | 112034800 | 112037799 | 21.00761 | 37.01412 | 26.10995 | 11.55686 | 0.88961 | 0.584835 | 0.268278 |
| 15 | 78253800 | 78256599 | 17.92465 | 30.58619 | 58.08784 | 25.46581 | -0.53937 | 0.983 | 0.268279 |
| 14 | 103913000 | 103917399 | 34.89753 | 56.06249 | 109.7855 | 46.50462 | -0.61702 | 1.898623 | 0.268283 |
| 4 | 131903600 | 131906799 | 18.05573 | 30.34111 | 65.00516 | 28.39395 | -0.71443 | 1.125157 | 0.268283 |
| 14 | 70693200 | 70697199 | 31.8147 | 51.26964 | 99.13885 | 42.18502 | -0.59305 | 1.793985 | 0.268336 |
| 17 | 23848800 | 23853799 | 30.50496 | 49.23361 | 95.8989 | 40.8866 | -0.60097 | 1.745808 | 0.26835 |
| 4 | 74516600 | 74519599 | 31.89354 | 54.23403 | 40.27236 | 17.65054 | 0.809455 | 1.046536 | 0.268354 |
| 2 | 30536200 | 30541799 | 23.44468 | 38.00343 | 90.45374 | 38.73278 | -0.88262 | 1.623098 | 0.268362 |
| 7 | 13713800 | 13718599 | 57.53521 | 92.96042 | 63.30853 | 27.02961 | 0.913353 | 1.742631 | 0.268381 |
| X | 33239200 | 33240999 | 18.27765 | 34.52156 | 16.86983 | 6.867038 | 1.502078 | 0.113665 | 0.268388 |
| 13 | 30971400 | 30976599 | 41.74882 | 66.79002 | 118.986 | 50.08753 | -0.48796 | 2.040644 | 0.268399 |
| 17 | 38284400 | 38285399 | 17.35348 | 31.27521 | 25.95286 | 11.30841 | 0.673601 | 0.393711 | 0.268406 |
| 19 | 7837200 | 7838799 | 29.20064 | 49.25345 | 44.25753 | 19.35347 | 0.53488 | 1.109225 | 0.268409 |
| 3 | 70015800 | 70017799 | 12.41191 | 21.76524 | 46.17882 | 20.40878 | -0.6923 | 0.632863 | 0.268411 |
| 4 | 62041400 | 62047799 | 118.1243 | 181.3038 | 164.6251 | 68.34639 | 0.432943 | 2.994804 | 0.268412 |
| 7 | 56531000 | 56533199 | 15.32866 | 28.53104 | 20.21821 | 8.441709 | 0.940872 | 0.18365 | 0.268426 |
| 15 | 56322000 | 56325599 | 39.70843 | 65.22085 | 50.47416 | 21.74407 | 0.746445 | 1.432216 | 0.268434 |
| 13 | 3279000 | 3282199 | 62.37768 | 99.8068 | 87.61686 | 36.89024 | 0.531424 | 2.049355 | 0.26844 |
| 14 | 33687000 | 33689199 | 14.34214 | 24.94925 | 48.95501 | 21.51055 | -0.58087 | 0.77028 | 0.268441 |
| 3 | 88270800 | 88280799 | 129.0085 | 192.4919 | 329.1704 | 135.6763 | -0.51402 | 3.565679 | 0.268448 |
| 5 | 111531200 | 111532599 | 6.712656 | 12.4883 | 32.12651 | 13.42448 | -0.90503 | 0.146892 | 0.268466 |
| 4 | 75158600 | 75158999 | 4.180753 | 8.166322 | 21.78819 | 7.625034 | -0.72154 | -0.52808 | 0.268477 |
| 7 | 112177400 | 112180199 | 15.9628 | 27.0375 | 59.46547 | 26.03673 | -0.75127 | 1.044729 | 0.26849 |
| 11 | 40804800 | 40806999 | 30.39604 | 52.47722 | 34.12777 | 14.97171 | 1.003319 | 0.900828 | 0.268491 |
| 7 | 14514000 | 14523799 | 287.1362 | 414.4604 | 350.3992 | 144.5613 | 0.456233 | 4.111683 | 0.268497 |
| 17 | 14360800 | 14362799 | 11.80295 | 20.82532 | 41.9543 | 18.56472 | -0.61529 | 0.545386 | 0.268504 |
| 13 | 114261000 | 114264999 | 35.19931 | 56.69859 | 101.7466 | 43.26662 | -0.48645 | 1.813437 | 0.268508 |
| X | 32181200 | 32182599 | 24.13232 | 42.27889 | 27.95974 | 12.34722 | 0.981164 | 0.68288 | 0.268509 |
| 3 | 32790400 | 32791799 | 11.5888 | 20.15377 | 49.44889 | 21.7236 | -0.90147 | 0.761865 | 0.26852 |
| 5 | 122785600 | 122787199 | 14.92488 | 25.56721 | 54.7514 | 24.01067 | -0.71072 | 0.932065 | 0.268525 |
| 2 | 48968000 | 48969999 | 29.92208 | 51.28825 | 37.47475 | 16.43494 | 0.834141 | 0.964249 | 0.268532 |
| 12 | 59270200 | 59270999 | 5.908965 | 11.18584 | 29.20667 | 11.83186 | -0.88833 | 0.018671 | 0.268535 |
| 15 | 45185000 | 45185799 | 17.61985 | 31.40655 | 22.4394 | 9.875078 | 0.878833 | 0.455556 | 0.268545 |
| 7 | 41737200 | 41740399 | 72.94086 | 116.0781 | 98.87748 | 41.31682 | 0.566444 | 2.230274 | 0.268545 |

| | | | | | | | | | |
|---|---|---|---|---|---|---|---|---|---|
| 2 | 33744800 | 33747199 | 9.971189 | 18.11489 | 37.57655 | 16.21016 | -0.63043 | 0.322062 | 0.268546 |
| 14 | 57014200 | 57015999 | 9.645123 | 17.0664 | 41.97199 | 18.56201 | -0.89976 | 0.498057 | 0.268551 |
| 4 | 146372000 | 146375799 | 35.92028 | 60.28414 | 38.71537 | 16.89897 | 1.017398 | 1.169364 | 0.268556 |
| 1 | 71141400 | 71142399 | 9.19298 | 16.27643 | 40.46638 | 17.89395 | -0.91459 | 0.487154 | 0.26856 |
| 11 | 27363600 | 27368599 | 62.71355 | 100.3774 | 86.6145 | 36.48531 | 0.55672 | 2.039538 | 0.268567 |
| 5 | 139725600 | 139728599 | 12.94531 | 22.54033 | 47.88006 | 21.05054 | -0.69442 | 0.749167 | 0.268567 |
| 8 | 89032400 | 89033999 | 12.91192 | 22.6645 | 44.58362 | 19.72029 | -0.5835 | 0.619797 | 0.268577 |
| 7 | 33187600 | 33192199 | 118.5353 | 183.5484 | 145.2214 | 60.3211 | 0.64111 | 2.828272 | 0.268599 |
| 1 | 53335800 | 53336599 | 5.933827 | 11.20599 | 30.24721 | 12.31571 | -0.94286 | 0.036641 | 0.268605 |
| 8 | 84481800 | 84484599 | 571.734 | 790.5191 | 370.7074 | 153.9853 | 1.244006 | 4.755648 | 0.26862 |
| 1 | 118360400 | 118362999 | 21.32867 | 37.29295 | 32.2819 | 14.23204 | 0.594675 | 0.703975 | 0.268648 |
| X | 33194000 | 33196799 | 33.48839 | 57.24099 | 36.34616 | 15.9364 | 1.035407 | 1.003844 | 0.268668 |
| 7 | 37052600 | 37059199 | 40.52483 | 64.63363 | 127.6053 | 53.484 | -0.64078 | 2.136545 | 0.268687 |
| 18 | 7713200 | 7717399 | 112.1577 | 172.7737 | 157.6319 | 65.47571 | 0.430032 | 2.921644 | 0.268711 |
| 6 | 130294400 | 130296999 | 46.09734 | 75.05042 | 59.61227 | 25.59327 | 0.701712 | 1.568106 | 0.268714 |
| 4 | 147944400 | 147951199 | 45.7122 | 72.39287 | 151.6977 | 63.08105 | -0.73697 | 2.360314 | 0.268722 |
| 13 | 120209400 | 120211999 | 21.00001 | 34.91154 | 70.55622 | 30.61156 | -0.63072 | 1.255583 | 0.268725 |
| 15 | 35953800 | 35956799 | 48.34947 | 78.08257 | 70.40686 | 30.04222 | 0.508597 | 1.74231 | 0.268728 |
| 7 | 11226600 | 11240599 | 374.2251 | 526.8795 | 441.7187 | 182.9636 | 0.432385 | 4.479763 | 0.268739 |
| 18 | 37685600 | 37690999 | 158.7651 | 239.8456 | 207.5972 | 85.82806 | 0.483549 | 3.328083 | 0.268751 |
| X | 148854600 | 148858199 | 42.62162 | 70.61097 | 47.18091 | 20.42095 | 0.960641 | 1.337568 | 0.268751 |
| 5 | 109586000 | 109588599 | 35.42084 | 59.78573 | 40.21452 | 17.58911 | 0.951087 | 1.12067 | 0.268773 |
| 13 | 30660400 | 30661199 | 11.59405 | 22.59201 | 14.02111 | 5.090153 | 1.308675 | -0.33069 | 0.268775 |
| 1 | 110781000 | 110782799 | 16.37277 | 30.32621 | 21.82429 | 9.176665 | 0.911901 | 0.224839 | 0.268776 |
| 4 | 39347600 | 39348799 | 13.72428 | 26.06102 | 20.245 | 8.147885 | 0.853582 | 0.044323 | 0.268781 |
| 13 | 100664600 | 100667399 | 14.1934 | 24.76802 | 48.38815 | 21.30341 | -0.57407 | 0.724848 | 0.268793 |
| 17 | 23630200 | 23635199 | 78.28837 | 124.2933 | 103.5633 | 43.16081 | 0.594873 | 2.30663 | 0.268803 |
| 16 | 17883200 | 17884799 | 7.875937 | 14.77007 | 33.43809 | 13.78469 | -0.7065 | 0.110065 | 0.268815 |
| 2 | 170220000 | 170221799 | 12.57164 | 22.07465 | 44.04728 | 19.48861 | -0.60405 | 0.61015 | 0.268831 |
| 17 | 16044400 | 16047799 | 24.63279 | 40.50602 | 76.56835 | 33.00149 | -0.53757 | 1.400717 | 0.268832 |
| 11 | 17100200 | 17104999 | 52.34775 | 84.23705 | 76.41206 | 32.44645 | 0.49393 | 1.857954 | 0.268834 |
| 12 | 6386800 | 6388999 | 29.81781 | 50.68881 | 41.48799 | 18.18288 | 0.669555 | 1.043692 | 0.268837 |
| 7 | 41236400 | 41237599 | 22.99283 | 40.21401 | 32.0703 | 14.14211 | 0.712376 | 0.708204 | 0.268861 |
| 1 | 57654400 | 57655799 | 15.0759 | 28.509 | 20.07164 | 8.149365 | 0.983454 | 0.084311 | 0.268874 |
| 13 | 112515200 | 112517199 | 13.97416 | 24.0389 | 53.13579 | 23.30788 | -0.75484 | 0.890033 | 0.268883 |
| 6 | 133528000 | 133528799 | 11.78392 | 22.94975 | 14.61476 | 5.342834 | 1.263525 | -0.3025 | 0.268884 |
| 6 | 23078800 | 23079999 | 7.067361 | 13.45712 | 31.70713 | 12.68706 | -0.72639 | -0.00444 | 0.268895 |
| 4 | 43032600 | 43034799 | 9.843857 | 17.71632 | 36.89855 | 16.10582 | -0.64719 | 0.374426 | 0.26891 |
| 10 | 11885200 | 11885799 | 9.242406 | 18.04227 | 14.41907 | 5.099633 | 0.981864 | -0.44915 | 0.268916 |
| 15 | 73511800 | 73513999 | 9.237073 | 16.60857 | 37.41635 | 16.34572 | -0.76065 | 0.375796 | 0.268927 |
| 17 | 8501400 | 8503599 | 11.4072 | 19.90376 | 48.37377 | 21.29557 | -0.88735 | 0.710627 | 0.268929 |
| 12 | 102246600 | 102250199 | 18.78113 | 31.72836 | 61.83854 | 27.05438 | -0.57804 | 1.078499 | 0.268929 |
| 7 | 21017200 | 21020399 | 49.23943 | 79.60687 | 68.32453 | 29.19487 | 0.581446 | 1.712511 | 0.268937 |
| 17 | 88970800 | 88972599 | 11.16476 | 19.66958 | 44.02926 | 19.48877 | -0.76767 | 0.558478 | 0.268939 |
| 9 | 66882200 | 66882799 | 3.526551 | 6.887332 | 20.43521 | 7.171748 | -0.87553 | -0.53315 | 0.268946 |
| 18 | 73938400 | 73939999 | 9.622808 | 17.57929 | 36.08252 | 15.45795 | -0.60877 | 0.283398 | 0.26898 |
| X | 116459600 | 116461399 | 20.77164 | 37.9309 | 21.52798 | 9.227413 | 1.233253 | 0.337715 | 0.268986 |

| | | | | | | | | | |
|---|---|---|---|---|---|---|---|---|---|
| 16 | 65852600 | 65854999 | 32.11105 | 54.12932 | 40.46972 | 17.6938 | 0.799126 | 1.120599 | 0.268989 |
| 12 | 104575800 | 104577999 | 14.67483 | 25.40335 | 51.09772 | 22.41818 | -0.61773 | 0.830391 | 0.268996 |
| 13 | 29445600 | 29446799 | 8.411122 | 15.40415 | 34.54693 | 14.75156 | -0.73166 | 0.259036 | 0.269001 |
| 4 | 60267200 | 60270199 | 51.05772 | 82.1512 | 74.65003 | 31.69268 | 0.491175 | 1.857421 | 0.269006 |
| 13 | 3679800 | 3682999 | 25.50878 | 44.15979 | 36.54836 | 16.03494 | 0.657766 | 0.857698 | 0.269065 |
| 18 | 4332200 | 4335199 | 14.6648 | 25.45719 | 50.03838 | 21.95767 | -0.58372 | 0.803193 | 0.269081 |
| 2 | 167261200 | 167262599 | 16.69169 | 29.89302 | 26.14044 | 11.4581 | 0.592805 | 0.427512 | 0.269082 |
| 13 | 97350200 | 97352599 | 9.762957 | 17.34187 | 40.4183 | 17.83918 | -0.81992 | 0.462055 | 0.269089 |
| 1 | 102344000 | 102346399 | 31.21514 | 52.51414 | 45.31619 | 19.80074 | 0.59295 | 1.139672 | 0.269115 |
| 2 | 179434400 | 179443799 | 51.21687 | 80.66186 | 163.5837 | 67.87489 | -0.69625 | 2.492768 | 0.269116 |
| 7 | 25334400 | 25339799 | 26.00726 | 42.02668 | 94.03442 | 40.15592 | -0.79824 | 1.701931 | 0.269128 |
| 9 | 105651200 | 105651999 | 4.791139 | 9.302793 | 24.43022 | 9.159058 | -0.79066 | -0.26538 | 0.269133 |
| 7 | 127738600 | 127742599 | 28.11908 | 45.76232 | 85.95608 | 36.89816 | -0.53713 | 1.554261 | 0.269135 |
| 4 | 115285800 | 115288399 | 22.34286 | 39.54665 | 25.80514 | 11.41154 | 1.003386 | 0.542167 | 0.269138 |
| 19 | 25099200 | 25100999 | 8.662339 | 15.5315 | 37.44472 | 16.39689 | -0.85986 | 0.391118 | 0.269145 |
| 5 | 145350000 | 145354799 | 120.8829 | 185.7034 | 167.4626 | 69.53291 | 0.443851 | 2.978919 | 0.269153 |
| 4 | 121388200 | 121390799 | 70.95232 | 112.9549 | 96.74436 | 40.44597 | 0.559148 | 2.217926 | 0.269155 |
| 8 | 119490600 | 119492199 | 16.24051 | 29.21769 | 25.22637 | 11.01509 | 0.614905 | 0.395061 | 0.269164 |
| 11 | 51319200 | 51330999 | 157.3162 | 230.9497 | 397.6532 | 163.7783 | -0.54578 | 3.845896 | 0.269176 |
| 19 | 46128400 | 46131999 | 21.3642 | 35.41538 | 70.34746 | 30.46067 | -0.60612 | 1.291449 | 0.269178 |
| X | 169273600 | 169273999 | 3.534011 | 6.880538 | 22.84379 | 8.378427 | -1.09357 | -0.36445 | 0.269186 |
| 10 | 86096000 | 86098199 | 28.44765 | 48.63337 | 39.76692 | 17.43651 | 0.671968 | 0.98408 | 0.269189 |
| X | 3279600 | 3287199 | 77.3466 | 123.4605 | 83.616 | 35.08606 | 0.901309 | 2.138964 | 0.26924 |
| X | 3313600 | 3317199 | 23.10401 | 41.59181 | 21.53501 | 9.396763 | 1.348252 | 0.432121 | 0.269286 |
| 8 | 58003200 | 58005199 | 11.44067 | 20.27486 | 40.86662 | 18.06757 | -0.6137 | 0.490666 | 0.269312 |
| 2 | 22958000 | 22959599 | 11.7074 | 20.42653 | 47.32896 | 20.83495 | -0.81873 | 0.712181 | 0.26933 |
| 10 | 84324200 | 84326399 | 11.7074 | 20.42653 | 47.32896 | 20.83495 | -0.81873 | 0.712181 | 0.26933 |
| 4 | 25751200 | 25752799 | 20.85079 | 37.25588 | 25.52944 | 11.21196 | 0.94059 | 0.468669 | 0.269345 |
| 1 | 102229000 | 102231599 | 46.83271 | 75.98817 | 63.97423 | 27.41357 | 0.614004 | 1.630763 | 0.269346 |
| 8 | 65145400 | 65146399 | 7.624575 | 14.05888 | 33.06144 | 13.98732 | -0.78994 | 0.20859 | 0.269355 |
| 9 | 122682200 | 122686999 | 85.83902 | 135.3665 | 118.7215 | 49.26889 | 0.512883 | 2.480427 | 0.26936 |
| 5 | 43991000 | 43994999 | 52.19618 | 82.62547 | 147.6243 | 61.37373 | -0.50785 | 2.375564 | 0.269381 |
| 7 | 30844200 | 30851799 | 54.9587 | 86.10548 | 173.7474 | 72.06843 | -0.69485 | 2.598291 | 0.269391 |
| 8 | 21639200 | 21639599 | 10.09562 | 19.60517 | 16.36025 | 6.126388 | 0.845862 | -0.23829 | 0.269399 |
| 13 | 120557000 | 120559199 | 14.26357 | 24.56065 | 52.64387 | 23.09279 | -0.71053 | 0.879654 | 0.269407 |
| 15 | 3109200 | 3112799 | 60.05332 | 96.0631 | 88.20093 | 37.12267 | 0.466398 | 2.054457 | 0.269411 |
| 4 | 43964000 | 43966999 | 14.08818 | 24.30807 | 52.12639 | 22.86721 | -0.71059 | 0.859172 | 0.269429 |
| 2 | 8378400 | 8379799 | 22.76917 | 39.5984 | 34.48383 | 15.14952 | 0.585549 | 0.790345 | 0.269435 |
| 13 | 30123600 | 30127199 | 33.75923 | 54.34756 | 101.6 | 43.16813 | -0.54633 | 1.829853 | 0.269452 |
| X | 117786600 | 117787399 | 13.53702 | 26.17025 | 16.75637 | 6.417517 | 1.195713 | -0.12129 | 0.269457 |
| 10 | 41993000 | 41995199 | 14.8253 | 25.26056 | 56.77796 | 24.88862 | -0.78137 | 0.985641 | 0.269458 |
| 5 | 121716400 | 121721199 | 38.5059 | 61.7466 | 111.6907 | 47.18106 | -0.5059 | 1.963006 | 0.269483 |
| 19 | 12909400 | 12910199 | 10.18278 | 19.70444 | 18.06012 | 6.886183 | 0.688225 | -0.17092 | 0.269492 |
| 5 | 78710600 | 78712799 | 30.00627 | 50.88429 | 38.40427 | 16.8212 | 0.785829 | 1.061097 | 0.269512 |
| 13 | 18114600 | 18115799 | 5.814007 | 11.27789 | 27.51258 | 10.37198 | -0.6949 | -0.2284 | 0.269524 |
| 15 | 43032800 | 43034599 | 16.03645 | 27.37275 | 56.36863 | 24.71273 | -0.65547 | 0.971039 | 0.269528 |
| 12 | 91599800 | 91601399 | 22.35149 | 38.8558 | 34.61172 | 15.20172 | 0.553072 | 0.79522 | 0.269537 |

| | | | | | | | | | |
|---|---|---|---|---|---|---|---|---|---|
| 12 | 93794600 | 93798199 | 24.66516 | 42.53037 | 37.74383 | 16.55743 | 0.556405 | 0.890112 | 0.269554 |
| 13 | 115225000 | 115227399 | 17.11922 | 28.81351 | 65.97129 | 28.82995 | -0.80962 | 1.111402 | 0.269563 |
| 12 | 81654400 | 81655799 | 15.82737 | 28.63278 | 23.50304 | 10.19267 | 0.693225 | 0.35434 | 0.269577 |
| 4 | 41313200 | 41314999 | 8.810255 | 15.80551 | 40.5288 | 17.73907 | -0.94844 | 0.391534 | 0.269579 |
| 5 | 137280600 | 137283599 | 25.77259 | 42.34469 | 78.37657 | 33.76884 | -0.50781 | 1.415556 | 0.269579 |
| 5 | 136087200 | 136090599 | 13.6648 | 23.49656 | 53.13646 | 23.30787 | -0.78782 | 0.891485 | 0.269592 |
| 3 | 4995800 | 4999999 | 50.44594 | 81.31341 | 73.26487 | 31.18508 | 0.506551 | 1.800993 | 0.269596 |
| 2 | 83642600 | 83644199 | 8.953229 | 16.06787 | 38.96011 | 17.04722 | -0.86779 | 0.387944 | 0.269596 |
| 5 | 95613400 | 95614199 | 9.186692 | 17.91025 | 15.5211 | 5.596997 | 0.841513 | -0.38037 | 0.269599 |
| 5 | 120487800 | 120492799 | 27.30161 | 44.11205 | 95.84765 | 40.92277 | -0.75641 | 1.706599 | 0.26961 |
| 1 | 93063000 | 93066799 | 18.14883 | 30.28309 | 66.33247 | 28.84816 | -0.74603 | 1.205097 | 0.269618 |
| 11 | 10727200 | 10728799 | 17.38502 | 30.98321 | 27.11969 | 11.93632 | 0.587472 | 0.464531 | 0.269626 |
| 2 | 89151600 | 89152199 | 8.9001 | 17.36315 | 13.67957 | 4.889788 | 0.987036 | -0.42303 | 0.269645 |
| 3 | 94072200 | 94073999 | 14.87096 | 25.64041 | 52.13802 | 22.87182 | -0.63452 | 0.869524 | 0.26967 |
| 8 | 27750000 | 27752199 | 14.87096 | 25.64041 | 52.13802 | 22.87182 | -0.63452 | 0.869524 | 0.26967 |
| 18 | 26695600 | 26696999 | 18.73444 | 33.49779 | 26.89383 | 11.80423 | 0.714092 | 0.453241 | 0.269681 |
| 7 | 100757000 | 100759399 | 15.96394 | 27.02886 | 62.23853 | 27.24768 | -0.81743 | 1.049065 | 0.269704 |
| 5 | 46423600 | 46426399 | 27.57096 | 47.41619 | 38.16421 | 16.7405 | 0.696049 | 0.927241 | 0.269705 |
| 2 | 97812400 | 97815399 | 33.8628 | 57.51024 | 39.10148 | 17.13437 | 0.93592 | 1.063372 | 0.269709 |
| 10 | 41762600 | 41763799 | 6.626854 | 12.55857 | 29.26044 | 11.82622 | -0.72305 | 0.016917 | 0.269713 |
| 3 | 59487400 | 59488399 | 14.1249 | 25.96234 | 22.26518 | 9.460457 | 0.650177 | 0.250085 | 0.269719 |
| 5 | 31336600 | 31337999 | 8.188178 | 15.19058 | 32.62062 | 13.68664 | -0.65136 | 0.177102 | 0.269739 |
| 1 | 39166600 | 39168599 | 12.57639 | 22.0081 | 48.34702 | 21.32971 | -0.74237 | 0.670166 | 0.269749 |
| 9 | 6483200 | 6487399 | 55.99243 | 90.46078 | 64.95319 | 27.72906 | 0.837082 | 1.742345 | 0.269752 |
| 8 | 30893400 | 30895199 | 16.94998 | 30.97861 | 23.87792 | 10.22375 | 0.795308 | 0.307263 | 0.269752 |
| 1 | 51905800 | 51910799 | 59.82786 | 94.20155 | 161.5431 | 67.02618 | -0.45492 | 2.502979 | 0.269753 |
| 7 | 143633800 | 143638799 | 50.41256 | 79.78464 | 148.7794 | 61.84713 | -0.56972 | 2.379402 | 0.269759 |
| 15 | 55797800 | 55799599 | 10.80518 | 19.36205 | 39.44342 | 17.2808 | -0.61938 | 0.413885 | 0.269766 |
| 13 | 66053400 | 66057199 | 90.39289 | 143.8597 | 81.51561 | 34.06478 | 1.153677 | 2.243416 | 0.269785 |
| 9 | 59842200 | 59844399 | 13.19255 | 23.17053 | 48.28276 | 21.36696 | -0.66657 | 0.615373 | 0.269788 |
| 3 | 64825600 | 64828199 | 30.50142 | 51.63122 | 41.96105 | 18.36908 | 0.679285 | 1.079391 | 0.269799 |
| 9 | 19231800 | 19238799 | 90.44869 | 142.8987 | 110.5574 | 45.918 | 0.695641 | 2.441945 | 0.269802 |
| 3 | 91261400 | 91264399 | 51.70538 | 83.37065 | 71.87708 | 30.60723 | 0.570568 | 1.792351 | 0.269804 |
| 13 | 72110000 | 72116599 | 54.84349 | 86.6173 | 154.6475 | 64.21904 | -0.50953 | 2.432711 | 0.269805 |
| 13 | 65540600 | 65542999 | 52.52912 | 85.45803 | 55.78487 | 23.94354 | 0.983379 | 1.593234 | 0.269808 |
| 17 | 16896400 | 16897199 | 9.895687 | 17.66935 | 38.42717 | 16.88612 | -0.71574 | 0.429582 | 0.269827 |
| 18 | 5681200 | 5682999 | 10.04552 | 17.90692 | 38.39489 | 16.89574 | -0.69704 | 0.439445 | 0.269862 |
| 4 | 25158600 | 25159799 | 8.64444 | 16.0837 | 34.34159 | 14.34783 | -0.63877 | 0.167829 | 0.269864 |
| X | 25227600 | 25230399 | 34.98159 | 60.44632 | 28.98588 | 12.71632 | 1.44096 | 0.908842 | 0.269892 |
| X | 94323800 | 94326799 | 30.29295 | 52.68299 | 30.16285 | 13.25119 | 1.187796 | 0.822677 | 0.26992 |
| 2 | 8015600 | 8016799 | 9.107512 | 16.76515 | 36.28291 | 15.38525 | -0.67428 | 0.234633 | 0.269949 |
| 9 | 36506400 | 36507799 | 26.28484 | 45.89268 | 31.40845 | 13.83257 | 0.932031 | 0.751015 | 0.26995 |
| 6 | 20920200 | 20923199 | 37.16587 | 62.0504 | 45.92759 | 19.98124 | 0.813752 | 1.235594 | 0.269967 |
| 2 | 30333400 | 30344799 | 76.07957 | 117.0775 | 216.1016 | 89.74981 | -0.58663 | 2.934607 | 0.269969 |
| 4 | 139300400 | 139304799 | 20.32999 | 33.21421 | 82.52955 | 35.47725 | -0.93532 | 1.4809 | 0.26997 |
| 10 | 71989200 | 71991599 | 22.582 | 39.30445 | 33.92738 | 14.91259 | 0.598279 | 0.777181 | 0.269973 |
| 18 | 3720200 | 3725399 | 79.23476 | 125.0059 | 115.3826 | 47.88791 | 0.439741 | 2.466562 | 0.269984 |

| | | | | | | | | | |
|---|---|---|---|---|---|---|---|---|---|
| 17 | 31404000 | 31406799 | 24.86333 | 40.67768 | 79.52636 | 34.20377 | -0.58919 | 1.468211 | 0.26999 |
| 6 | 144051000 | 144052799 | 9.443276 | 17.25551 | 36.62878 | 15.6872 | -0.65689 | 0.28006 | 0.269996 |
| 10 | 81909800 | 81914199 | 92.89136 | 145.3867 | 132.5039 | 54.96691 | 0.449191 | 2.630474 | 0.269998 |
| 17 | 24946400 | 24947999 | 7.204373 | 13.2567 | 34.61696 | 14.68658 | -0.94312 | 0.218939 | 0.270011 |
| 10 | 46240400 | 46241399 | 12.08957 | 21.2868 | 42.97329 | 19.02352 | -0.61974 | 0.569518 | 0.270017 |
| 5 | 88364600 | 88365799 | 16.23515 | 29.76958 | 24.07177 | 10.26152 | 0.731293 | 0.285077 | 0.270036 |
| 7 | 85298200 | 85302399 | 49.72333 | 80.67444 | 62.43065 | 26.75118 | 0.735233 | 1.63756 | 0.270042 |
| 10 | 47510800 | 47514999 | 81.7645 | 128.8329 | 119.7179 | 49.677 | 0.428643 | 2.49472 | 0.270048 |
| 14 | 82814400 | 82821199 | 94.59621 | 148.2924 | 128.9279 | 53.47466 | 0.518909 | 2.602407 | 0.270055 |
| 4 | 13742600 | 13745799 | 20.95968 | 34.56759 | 80.92879 | 34.92434 | -0.84349 | 1.357539 | 0.270058 |
| 12 | 3079400 | 3093999 | 36.60445 | 60.74481 | 39.31794 | 17.03752 | 1.006447 | 1.294369 | 0.270071 |
| 4 | 139354400 | 139357799 | 22.83673 | 37.27198 | 81.58129 | 35.05672 | -0.75401 | 1.501733 | 0.270084 |
| 2 | 181437400 | 181439799 | 8.413746 | 15.3438 | 35.52668 | 15.253 | -0.78335 | 0.282058 | 0.270114 |
| 14 | 41421200 | 41425599 | 47.76616 | 77.27573 | 61.47653 | 26.28637 | 0.6916 | 1.689047 | 0.270138 |
| 1 | 189750400 | 189754199 | 23.87508 | 39.16618 | 77.30201 | 33.28398 | -0.601 | 1.430105 | 0.270142 |
| 15 | 69019200 | 69022799 | 32.22438 | 54.5605 | 41.36299 | 18.10856 | 0.779279 | 1.083836 | 0.270169 |
| 1 | 157455600 | 157457199 | 12.59628 | 22.02334 | 46.24701 | 20.38792 | -0.67741 | 0.683387 | 0.270182 |
| 16 | 21075600 | 21078799 | 16.45862 | 28.17084 | 57.55224 | 25.23724 | -0.64358 | 0.947042 | 0.270183 |
| 14 | 50237600 | 50240799 | 53.74019 | 86.25561 | 78.85342 | 33.35669 | 0.478242 | 1.939417 | 0.270193 |
| 12 | 106177600 | 106180599 | 23.73784 | 39.35727 | 74.77861 | 32.38342 | -0.54237 | 1.297957 | 0.270194 |
| 8 | 48577600 | 48583199 | 37.6769 | 60.39944 | 112.8419 | 47.64599 | -0.55295 | 1.970229 | 0.270199 |
| 16 | 32098800 | 32100399 | 28.20943 | 47.97829 | 42.03138 | 18.42176 | 0.571899 | 1.032458 | 0.27021 |
| X | 70211200 | 70212799 | 8.494054 | 15.8518 | 37.90177 | 15.75432 | -0.79516 | 0.154405 | 0.270238 |
| 7 | 62498200 | 62500599 | 40.70668 | 66.77619 | 56.63763 | 24.38626 | 0.613906 | 1.453441 | 0.270243 |
| 17 | 17192800 | 17204799 | 346.6868 | 495.008 | 374.888 | 154.919 | 0.599336 | 4.274358 | 0.270297 |
| 6 | 117972000 | 117980199 | 53.12218 | 83.78601 | 156.5342 | 64.96233 | -0.57639 | 2.464424 | 0.270299 |
| 2 | 131670400 | 131670999 | 5.376606 | 10.44365 | 25.16195 | 9.403668 | -0.66464 | -0.26983 | 0.270306 |
| 19 | 6189200 | 6192999 | 19.53962 | 32.63829 | 66.01421 | 28.72893 | -0.6316 | 1.196381 | 0.270314 |
| 6 | 24249400 | 24250999 | 12.58333 | 22.10879 | 43.51619 | 19.26366 | -0.58433 | 0.600274 | 0.270331 |
| 1 | 11483800 | 11485399 | 14.16486 | 24.64315 | 49.49011 | 21.74694 | -0.61428 | 0.767603 | 0.270332 |
| 3 | 33330400 | 33332599 | 33.7916 | 55.98573 | 48.74301 | 21.09794 | 0.580047 | 1.313552 | 0.270343 |
| 2 | 103059400 | 103060399 | 5.673538 | 10.96965 | 26.54341 | 10.15998 | -0.70336 | -0.17828 | 0.270364 |
| 16 | 78411200 | 78412399 | 11.68648 | 20.45657 | 48.33147 | 21.32793 | -0.84644 | 0.663922 | 0.270367 |
| 17 | 56198400 | 56199599 | 8.510601 | 15.65967 | 34.62078 | 14.68897 | -0.70516 | 0.229177 | 0.270367 |
| 1 | 104113000 | 104114399 | 18.17957 | 33.67731 | 19.94295 | 8.384078 | 1.191057 | 0.234732 | 0.270368 |
| X | 87642400 | 87643599 | 17.35029 | 31.40524 | 25.10639 | 10.88046 | 0.732339 | 0.369125 | 0.270391 |
| 15 | 96623200 | 96626799 | 20.0074 | 33.40672 | 67.21021 | 29.24228 | -0.62403 | 1.202151 | 0.270423 |
| 8 | 20622400 | 20623799 | 15.45126 | 29.92148 | 12.7551 | 4.846635 | 1.784158 | -0.13804 | 0.27043 |
| 2 | 83058200 | 83058599 | 3.414998 | 6.673979 | 20.62113 | 7.137827 | -0.91572 | -0.58519 | 0.270431 |
| 4 | 153297800 | 153307999 | 86.95702 | 132.8586 | 238.5619 | 98.95322 | -0.55496 | 3.082886 | 0.270453 |
| 13 | 48278600 | 48279599 | 4.472998 | 8.619696 | 29.18644 | 11.3157 | -1.2015 | -0.14479 | 0.270454 |
| 11 | 20403200 | 20406799 | 20.92119 | 34.59429 | 73.03845 | 31.57127 | -0.69417 | 1.320667 | 0.270455 |
| 5 | 139630600 | 139635399 | 37.14475 | 59.28652 | 124.0589 | 52.05145 | -0.72329 | 2.110212 | 0.270457 |
| 10 | 95774200 | 95777399 | 21.99324 | 36.41679 | 73.93058 | 31.98711 | -0.63775 | 1.307122 | 0.270458 |
| 6 | 109101000 | 109103399 | 25.88777 | 44.29108 | 38.83646 | 17.02959 | 0.571701 | 0.964952 | 0.270486 |
| X | 62252400 | 62254399 | 22.47701 | 40.07718 | 24.91578 | 10.96188 | 1.078573 | 0.491808 | 0.27049 |
| 1 | 84958200 | 84961799 | 55.98624 | 90.196 | 72.62904 | 30.89168 | 0.667458 | 1.826381 | 0.270491 |

| | | | | | | | | | |
|---|---|---|---|---|---|---|---|---|---|
| 17 | 23479000 | 23481599 | 26.84939 | 43.72082 | 93.45271 | 40.12454 | -0.72256 | 1.548436 | 0.270502 |
| 3 | 14125000 | 14126199 | 17.05548 | 30.75121 | 26.02547 | 11.33402 | 0.645907 | 0.389658 | 0.270504 |
| 11 | 23908400 | 23916999 | 111.0752 | 171.8401 | 151.3541 | 62.86878 | 0.485497 | 2.841331 | 0.270507 |
| 6 | 128773200 | 128776999 | 16.04728 | 26.99674 | 67.08637 | 29.31312 | -0.92744 | 1.110023 | 0.270509 |
| 19 | 39103800 | 39105599 | 19.66036 | 34.74893 | 28.93077 | 12.79741 | 0.653324 | 0.547194 | 0.270514 |
| 9 | 97312200 | 97313199 | 6.414687 | 12.37353 | 30.11812 | 11.64417 | -0.72723 | -0.13059 | 0.270536 |
| 12 | 67552400 | 67553399 | 6.908322 | 13.09711 | 33.35228 | 13.46886 | -0.85042 | 0.023057 | 0.270541 |
| 8 | 111172600 | 111174799 | 23.08785 | 40.09505 | 35.04577 | 15.38251 | 0.58007 | 0.812669 | 0.270542 |
| 7 | 8434400 | 8453799 | 387.1098 | 544.5954 | 444.6478 | 184.2002 | 0.469419 | 4.491948 | 0.270545 |
| 16 | 13277800 | 13282999 | 28.01904 | 45.43293 | 90.58394 | 38.79769 | -0.62756 | 1.623331 | 0.270547 |
| 5 | 98399600 | 98404199 | 35.56075 | 57.07766 | 110.592 | 46.78397 | -0.60324 | 1.927761 | 0.27055 |
| 8 | 71986600 | 71990399 | 19.04229 | 31.8789 | 64.9367 | 28.29998 | -0.64179 | 1.169456 | 0.270555 |
| 7 | 52727400 | 52730999 | 59.15727 | 94.68544 | 86.66525 | 36.50673 | 0.471796 | 2.034349 | 0.270563 |
| 14 | 110642000 | 110646799 | 42.29043 | 69.67198 | 50.08397 | 21.60358 | 0.854132 | 1.398987 | 0.270564 |
| 6 | 9709600 | 9711599 | 17.95008 | 32.60865 | 24.26457 | 10.46816 | 0.838886 | 0.34942 | 0.270567 |
| 13 | 65294200 | 65296999 | 30.97748 | 52.94054 | 38.65817 | 16.94991 | 0.834588 | 0.99789 | 0.270568 |
| 7 | 108387800 | 108389199 | 9.637116 | 17.05711 | 39.96234 | 17.67224 | -0.82955 | 0.493953 | 0.270575 |
| 8 | 87786800 | 87787799 | 12.48837 | 24.12771 | 17.82716 | 6.847805 | 0.986355 | -0.12135 | 0.270576 |
| 1 | 34840600 | 34852199 | 72.55726 | 111.2939 | 232.4204 | 96.48412 | -0.76815 | 2.998479 | 0.270607 |
| 13 | 84620200 | 84620999 | 11.89456 | 22.54841 | 20.42704 | 8.250605 | 0.629056 | 0.038543 | 0.270614 |
| 7 | 140947200 | 140948399 | 7.660677 | 14.46191 | 35.54962 | 14.48575 | -0.81153 | 0.065192 | 0.270614 |
| 16 | 19671200 | 19672599 | 13.48203 | 25.49331 | 20.725 | 8.415782 | 0.777198 | 0.070935 | 0.270633 |
| 18 | 22346600 | 22347999 | 13.48203 | 25.49331 | 20.725 | 8.415782 | 0.777198 | 0.070935 | 0.270633 |
| 5 | 120085400 | 120086599 | 10.68613 | 18.95953 | 39.38107 | 17.39851 | -0.65555 | 0.475844 | 0.270653 |
| 12 | 71479400 | 71481799 | 14.02558 | 24.38916 | 48.95652 | 21.506 | -0.61357 | 0.773946 | 0.270672 |
| 9 | 115598400 | 115605399 | 54.41538 | 86.30778 | 148.8346 | 61.96767 | -0.45433 | 2.324988 | 0.270677 |
| 5 | 36292600 | 36294399 | 9.001071 | 16.47296 | 34.52016 | 14.75429 | -0.63596 | 0.265228 | 0.270705 |
| 18 | 35800600 | 35802199 | 9.001071 | 16.47296 | 34.52016 | 14.75429 | -0.63596 | 0.265228 | 0.270705 |
| 4 | 112544600 | 112547399 | 52.93502 | 85.70242 | 62.64312 | 26.80007 | 0.814512 | 1.684455 | 0.270719 |
| 8 | 94738200 | 94739599 | 8.799983 | 15.79701 | 37.4687 | 16.39101 | -0.83512 | 0.387184 | 0.27073 |
| 9 | 88908400 | 88910799 | 21.0659 | 37.35925 | 26.47988 | 11.70308 | 0.885934 | 0.516352 | 0.270743 |
| 8 | 75216200 | 75218799 | 26.45963 | 45.86641 | 35.36119 | 15.51472 | 0.759989 | 0.848609 | 0.270757 |
| 1 | 114002000 | 114004999 | 34.64853 | 57.54827 | 50.39581 | 21.85079 | 0.57184 | 1.286968 | 0.270765 |
| 12 | 28945200 | 28954199 | 68.55042 | 106.6729 | 187.5381 | 77.85112 | -0.50401 | 2.72607 | 0.270773 |
| 11 | 60110800 | 60114199 | 13.82787 | 23.74575 | 55.8306 | 24.48865 | -0.84441 | 0.904858 | 0.270775 |
| 7 | 31428800 | 31429199 | 15.43999 | 28.13127 | 22.39751 | 9.627246 | 0.745731 | 0.311698 | 0.270776 |
| 5 | 90242400 | 90244799 | 19.57012 | 33.12835 | 63.5985 | 27.84148 | -0.55626 | 1.065587 | 0.270778 |
| 18 | 3442800 | 3444799 | 9.115066 | 16.82298 | 35.24393 | 14.89161 | -0.62376 | 0.218954 | 0.270779 |
| 13 | 56420800 | 56423599 | 22.65057 | 37.4576 | 72.71429 | 31.43329 | -0.57347 | 1.3239 | 0.270783 |
| 12 | 84911800 | 84915599 | 19.77598 | 32.47384 | 77.86326 | 33.53966 | -0.88014 | 1.408431 | 0.270801 |
| 12 | 47382000 | 47382999 | 7.35044 | 13.99268 | 31.76717 | 12.71773 | -0.67447 | -0.00215 | 0.27082 |
| 3 | 82414600 | 82416199 | 19.89798 | 35.14829 | 27.17773 | 12.02337 | 0.758769 | 0.549473 | 0.27082 |
| 14 | 83362000 | 83362399 | 9.412027 | 18.40822 | 10.69139 | 3.616824 | 1.489949 | -0.60888 | 0.270831 |
| 11 | 58495800 | 58497799 | 11.15122 | 19.7308 | 40.39013 | 17.86254 | -0.63662 | 0.506304 | 0.270832 |
| 19 | 55792200 | 55796399 | 23.03642 | 37.93079 | 74.61079 | 32.17566 | -0.59393 | 1.38027 | 0.270835 |
| 17 | 84381600 | 84388199 | 38.52138 | 61.67303 | 115.1037 | 48.50504 | -0.5537 | 2.013552 | 0.270844 |
| 7 | 16687200 | 16692399 | 34.36949 | 55.33103 | 105.9561 | 45.02019 | -0.581 | 1.829385 | 0.270851 |

| | | | | | | | | | |
|---|---|---|---|---|---|---|---|---|---|
| 13 | 114593000 | 114595999 | 21.00146 | 34.53365 | 75.90613 | 32.71566 | -0.75433 | 1.392819 | 0.270871 |
| 12 | 55472200 | 55474399 | 11.95211 | 20.99551 | 43.52918 | 19.26613 | -0.65892 | 0.598989 | 0.270879 |
| 15 | 69109000 | 69111799 | 25.99679 | 45.00512 | 35.3596 | 15.5134 | 0.73237 | 0.856704 | 0.270908 |
| 3 | 115641800 | 115642599 | 6.426193 | 12.35729 | 28.08877 | 10.95685 | -0.641 | -0.10535 | 0.270921 |
| 7 | 7744000 | 7747799 | 68.44113 | 109.0802 | 88.08863 | 36.88486 | 0.645564 | 2.175869 | 0.270935 |
| 5 | 38691400 | 38694399 | 17.45563 | 29.77983 | 56.96068 | 24.97125 | -0.54958 | 0.982224 | 0.270946 |
| 10 | 3315600 | 3317799 | 16.53425 | 28.1126 | 58.50486 | 25.64122 | -0.67158 | 1.010723 | 0.27095 |
| 10 | 74329800 | 74331599 | 18.45723 | 33.52312 | 23.01842 | 9.93309 | 0.953567 | 0.350715 | 0.270954 |
| 7 | 21152800 | 21156799 | 67.97729 | 108.6051 | 82.54544 | 34.682 | 0.73663 | 2.09902 | 0.270963 |
| 11 | 77794200 | 77798599 | 18.32578 | 30.5333 | 69.17267 | 30.0554 | -0.79461 | 1.222171 | 0.27098 |
| 13 | 45487200 | 45489399 | 12.72634 | 22.31561 | 45.12546 | 19.94249 | -0.62327 | 0.63448 | 0.270984 |
| 17 | 37992400 | 37998399 | 22.36128 | 39.27227 | 25.73569 | 11.38863 | 0.994189 | 0.631444 | 0.271016 |
| 4 | 112425400 | 112430799 | 127.2951 | 195.0771 | 173.1233 | 71.85928 | 0.464243 | 3.028974 | 0.27102 |
| X | 25542400 | 25547199 | 57.88708 | 93.43148 | 65.27338 | 27.83106 | 0.874997 | 1.775574 | 0.271047 |
| 18 | 11711000 | 11713799 | 48.6312 | 79.22423 | 57.93724 | 24.87969 | 0.821272 | 1.566539 | 0.27107 |
| 4 | 55529400 | 55532199 | 20.44922 | 33.82235 | 72.98396 | 31.55318 | -0.7255 | 1.316475 | 0.271092 |
| 2 | 154222400 | 154223199 | 4.800315 | 9.288898 | 25.36326 | 9.671163 | -0.86954 | -0.20366 | 0.271094 |
| Y | 2293000 | 2294199 | 10.5667 | 20.45266 | 10.32742 | 3.932999 | 1.539716 | -0.19833 | 0.271102 |
| 5 | 151156400 | 151157599 | 7.307511 | 13.54509 | 33.13949 | 13.92066 | -0.83935 | 0.174764 | 0.271145 |
| 11 | 51827000 | 51828599 | 9.676002 | 17.41837 | 37.40743 | 16.32308 | -0.69107 | 0.371224 | 0.271147 |
| 12 | 54113600 | 54116199 | 21.01824 | 34.50905 | 76.84032 | 33.09713 | -0.7738 | 1.413306 | 0.271153 |
| 10 | 116545400 | 116549799 | 33.24383 | 53.59536 | 98.67521 | 42.01095 | -0.52179 | 1.782397 | 0.271168 |
| 12 | 22678200 | 22679799 | 18.21999 | 32.43971 | 26.83638 | 11.82131 | 0.667399 | 0.473912 | 0.271182 |
| 14 | 120456800 | 120458999 | 18.95258 | 33.45884 | 28.27749 | 12.51136 | 0.631395 | 0.555784 | 0.271189 |
| 16 | 28646000 | 28654599 | 61.89943 | 96.65717 | 181.457 | 75.29208 | -0.59466 | 2.661555 | 0.271222 |
| 17 | 21452000 | 21453599 | 33.71214 | 55.98835 | 48.75588 | 21.13853 | 0.579834 | 1.282527 | 0.27123 |
| 13 | 87243800 | 87245399 | 16.84728 | 30.40651 | 24.85227 | 10.80943 | 0.696718 | 0.378156 | 0.27123 |
| X | 75418800 | 75420599 | 18.47834 | 34.89076 | 16.86195 | 6.867926 | 1.517279 | 0.117385 | 0.271232 |
| 9 | 38476200 | 38479399 | 46.17539 | 75.90815 | 50.11817 | 21.59859 | 0.975348 | 1.437089 | 0.271232 |
| 7 | 124621000 | 124622399 | 19.51174 | 34.65916 | 27.49929 | 12.13702 | 0.725843 | 0.500857 | 0.271238 |
| 5 | 150127800 | 150130599 | 27.08284 | 44.30449 | 83.59689 | 35.95317 | -0.53838 | 1.47554 | 0.271259 |
| 1 | 143189800 | 143191999 | 18.67161 | 31.02996 | 70.77357 | 30.69853 | -0.80451 | 1.253693 | 0.271267 |
| 8 | 22004000 | 22004799 | 12.56142 | 24.00781 | 19.86916 | 7.887832 | 0.780038 | -0.01128 | 0.27127 |
| 6 | 98521400 | 98523399 | 17.8579 | 31.81892 | 26.35757 | 11.60305 | 0.666075 | 0.465653 | 0.271273 |
| 7 | 27597400 | 27599199 | 16.7736 | 31.34238 | 20.90745 | 8.672072 | 1.035874 | 0.176163 | 0.271285 |
| 19 | 47056000 | 47056999 | 6.049157 | 11.44814 | 29.20217 | 11.83679 | -0.85563 | 0.022433 | 0.271286 |
| 17 | 39352200 | 39353199 | 18.15283 | 33.27487 | 22.70464 | 9.683108 | 0.973949 | 0.298177 | 0.271289 |
| 4 | 128305200 | 128309199 | 28.02994 | 45.42985 | 89.78685 | 38.44334 | -0.61552 | 1.632925 | 0.271293 |
| 15 | 13845800 | 13848399 | 27.00907 | 46.60555 | 34.74649 | 15.24289 | 0.806676 | 0.887738 | 0.271316 |
| 7 | 15419400 | 15439599 | 428.5431 | 598.5738 | 471.7198 | 195.6334 | 0.510081 | 4.599997 | 0.27132 |
| 3 | 133405600 | 133406999 | 11.75857 | 20.90582 | 43.00967 | 18.96826 | -0.64053 | 0.474683 | 0.271322 |
| X | 26709800 | 26711199 | 18.36565 | 34.69134 | 16.23268 | 6.606245 | 1.56402 | 0.112053 | 0.271324 |
| 5 | 82965800 | 82967999 | 12.46217 | 21.86592 | 44.61265 | 19.72617 | -0.63575 | 0.625386 | 0.271333 |
| X | 38877200 | 38880399 | 38.37782 | 65.46453 | 31.69598 | 13.89519 | 1.424305 | 1.035939 | 0.271335 |
| 7 | 20634800 | 20635999 | 15.17702 | 29.31543 | 15.19344 | 5.842469 | 1.49094 | -0.09621 | 0.271341 |
| 9 | 19154000 | 19157799 | 42.57176 | 70.11254 | 52.2698 | 22.54332 | 0.801698 | 1.408125 | 0.271345 |
| 2 | 69023400 | 69024199 | 13.18124 | 24.70518 | 21.80641 | 8.998308 | 0.640561 | 0.127457 | 0.271353 |

| | | | | | | | | | |
|---|---|---|---|---|---|---|---|---|---|
| 16 | 54935200 | 54936999 | 18.48741 | 34.1299 | 21.07849 | 8.903063 | 1.126489 | 0.258886 | 0.271358 |
| 6 | 34167800 | 34170399 | 26.13791 | 42.81655 | 79.81039 | 34.34308 | -0.51967 | 1.455379 | 0.271358 |
| 8 | 52037200 | 52038599 | 18.60285 | 33.93887 | 21.92875 | 9.410144 | 1.0457 | 0.324662 | 0.271363 |
| 11 | 121811800 | 121816599 | 29.44241 | 47.7689 | 89.81906 | 38.4867 | -0.54244 | 1.613836 | 0.271378 |
| 7 | 47784200 | 47785799 | 20.28158 | 35.99758 | 27.14026 | 11.98672 | 0.798077 | 0.507892 | 0.271379 |
| 11 | 107206400 | 107206799 | 3.637457 | 7.112501 | 19.57552 | 6.683924 | -0.73244 | -0.63177 | 0.27138 |
| 14 | 47885600 | 47887999 | 13.84803 | 24.12887 | 48.40704 | 21.2915 | -0.61249 | 0.74288 | 0.271381 |
| 13 | 62320800 | 62336199 | 413.304 | 582.5488 | 411.0677 | 170.2363 | 0.682578 | 4.465334 | 0.271385 |
| 14 | 11492200 | 11492799 | 9.449367 | 17.30984 | 35.60392 | 15.19781 | -0.60814 | 0.26491 | 0.271386 |
| 5 | 140732200 | 140746799 | 91.42507 | 137.4609 | 325.6485 | 134.4513 | -0.97393 | 3.414123 | 0.271388 |
| 3 | 34439200 | 34441199 | 10.29535 | 18.54795 | 37.41112 | 16.30856 | -0.60016 | 0.372112 | 0.271393 |
| 16 | 30423400 | 30425199 | 10.29535 | 18.54795 | 37.41112 | 16.30856 | -0.60016 | 0.372112 | 0.271393 |
| X | 120998800 | 121001199 | 34.70817 | 58.95112 | 38.60053 | 16.91529 | 0.990104 | 1.066821 | 0.2714 |
| 7 | 8350000 | 8368799 | 330.0687 | 472.5104 | 372.0262 | 153.6725 | 0.547065 | 4.235303 | 0.271441 |
| 12 | 114681000 | 114683799 | 63.29784 | 101.1354 | 89.85022 | 37.75459 | 0.51202 | 2.094971 | 0.271448 |
| 7 | 85092000 | 85092799 | 29.93663 | 50.22358 | 43.33053 | 18.90909 | 0.593285 | 1.160681 | 0.271454 |
| 7 | 10172800 | 10174399 | 21.40402 | 38.28991 | 24.96995 | 10.95526 | 1.012626 | 0.46449 | 0.271474 |
| 11 | 82918400 | 82919199 | 8.392373 | 15.37922 | 34.0212 | 14.51498 | -0.71151 | 0.251785 | 0.271475 |
| 9 | 10961600 | 10965199 | 44.45939 | 71.98811 | 65.03811 | 27.82693 | 0.509732 | 1.669445 | 0.271475 |
| 5 | 118876600 | 118879599 | 32.29955 | 52.03334 | 99.75671 | 42.42708 | -0.58134 | 1.805334 | 0.271491 |
| 1 | 124880600 | 124881399 | 12.69534 | 23.87066 | 21.35147 | 8.763004 | 0.627754 | 0.102028 | 0.271491 |
| 11 | 39958400 | 39959999 | 29.40079 | 49.37392 | 43.3061 | 18.91035 | 0.569504 | 1.147742 | 0.271492 |
| 7 | 17516800 | 17519599 | 46.49771 | 75.88202 | 57.4012 | 24.66532 | 0.774336 | 1.535409 | 0.271495 |
| X | 31300800 | 31303999 | 25.79486 | 45.25178 | 28.63885 | 12.66135 | 1.044231 | 0.673138 | 0.271498 |
| 14 | 29862800 | 29865799 | 15.39328 | 26.33453 | 55.84038 | 24.4856 | -0.69696 | 0.946594 | 0.271525 |
| 15 | 48748600 | 48753599 | 49.95593 | 79.5178 | 138.188 | 57.76109 | -0.46025 | 2.212054 | 0.271532 |
| 14 | 41972200 | 41976599 | 36.37291 | 60.90825 | 44.77364 | 19.51597 | 0.823518 | 1.20158 | 0.271544 |
| X | 5959800 | 5960999 | 16.17066 | 30.32264 | 20.16403 | 8.313851 | 1.047706 | 0.149101 | 0.271549 |
| Y | 88976800 | 88977799 | 11.32938 | 21.78613 | 11.38037 | 4.439115 | 1.45995 | -0.10765 | 0.27155 |
| 6 | 134186400 | 134189999 | 24.21289 | 41.89855 | 36.07024 | 15.82497 | 0.600957 | 0.853498 | 0.27156 |
| 2 | 144448000 | 144449599 | 18.30502 | 32.51582 | 27.63222 | 12.19563 | 0.627311 | 0.492879 | 0.271561 |
| 5 | 149229400 | 149232199 | 21.2446 | 35.54132 | 68.56718 | 29.86918 | -0.56384 | 1.185697 | 0.271563 |
| 9 | 20230800 | 20233599 | 61.54016 | 98.23938 | 90.444 | 37.9561 | 0.459285 | 2.119186 | 0.271576 |
| 17 | 88985000 | 88985799 | 6.607964 | 12.58059 | 28.83657 | 11.5419 | -0.68733 | -0.01694 | 0.271576 |
| 1 | 173270600 | 173272799 | 11.4301 | 20.24358 | 44.01639 | 19.46253 | -0.72308 | 0.501221 | 0.271588 |
| 14 | 122344200 | 122345199 | 5.655173 | 10.66125 | 30.19596 | 12.33788 | -1.01567 | 0.046818 | 0.27161 |
| 5 | 87577000 | 87578999 | 23.51283 | 41.21702 | 30.29407 | 13.38451 | 0.829386 | 0.672749 | 0.271623 |
| 1 | 169733400 | 169737599 | 22.13834 | 36.86825 | 70.03872 | 30.42269 | -0.54153 | 1.241331 | 0.271644 |
| 1 | 33206200 | 33208399 | 21.77024 | 38.16642 | 31.59156 | 13.95903 | 0.658894 | 0.664144 | 0.271648 |
| 11 | 80273600 | 80274799 | 5.690216 | 10.97634 | 26.41926 | 10.21341 | -0.7098 | -0.14599 | 0.271654 |
| 7 | 43213200 | 43215399 | 10.49841 | 18.76411 | 38.39306 | 16.85706 | -0.6277 | 0.426095 | 0.271669 |
| 2 | 151372400 | 151374399 | 11.60677 | 20.54645 | 40.87683 | 18.07609 | -0.59556 | 0.504425 | 0.27167 |
| 6 | 132477800 | 132481399 | 48.65373 | 78.70885 | 67.32538 | 28.78649 | 0.587224 | 1.694555 | 0.271725 |
| 4 | 146074200 | 146075599 | 31.3765 | 52.30034 | 46.02707 | 20.00416 | 0.564922 | 1.235047 | 0.271729 |
| 8 | 53887600 | 53889199 | 13.55206 | 25.1078 | 21.87339 | 9.194215 | 0.638932 | 0.201498 | 0.27174 |
| 1 | 111791800 | 111792999 | 8.346484 | 15.02748 | 37.45785 | 16.34174 | -0.90408 | 0.361378 | 0.271776 |
| 17 | 37888600 | 37889999 | 19.59616 | 36.5876 | 17.74856 | 7.371663 | 1.489691 | 0.197397 | 0.271789 |

| | | | | | | | | | |
|---|---|---|---|---|---|---|---|---|---|
| 3 | 9891000 | 9892999 | 12.27853 | 21.63196 | 42.98569 | 19.02682 | -0.59658 | 0.566371 | 0.271791 |
| 1 | 151085800 | 151086399 | 3.885983 | 7.552378 | 26.2733 | 9.775551 | -1.1824 | -0.30234 | 0.271794 |
| 16 | 4936200 | 4940399 | 27.8619 | 45.18262 | 88.66586 | 37.97894 | -0.60453 | 1.620967 | 0.271798 |
| 4 | 132967600 | 132968999 | 7.677921 | 14.02151 | 35.03413 | 15.01523 | -0.89046 | 0.267837 | 0.271803 |
| 3 | 156256400 | 156258199 | 24.40624 | 43.12519 | 25.78861 | 11.40812 | 1.127935 | 0.574462 | 0.271828 |
| X | 52885000 | 52886799 | 27.89582 | 47.88666 | 34.18312 | 14.99317 | 0.868054 | 0.940618 | 0.271838 |
| 2 | 94389600 | 94392799 | 37.73512 | 62.19211 | 53.28729 | 22.98921 | 0.601863 | 1.385202 | 0.271856 |
| Y | 90830600 | 90831999 | 23.763 | 42.92904 | 20.51421 | 8.911997 | 1.467232 | 0.416072 | 0.271869 |
| 1 | 162837200 | 162841799 | 42.10994 | 69.05016 | 55.63028 | 23.94842 | 0.687641 | 1.460014 | 0.271882 |
| 7 | 65868000 | 65869799 | 9.238605 | 16.59632 | 40.01025 | 17.49222 | -0.85894 | 0.384788 | 0.271887 |
| 16 | 15088200 | 15088999 | 8.545839 | 15.65555 | 34.01632 | 14.51908 | -0.68621 | 0.255151 | 0.271902 |
| 5 | 128176400 | 128177599 | 24.82187 | 42.55368 | 36.90362 | 16.1847 | 0.588061 | 0.941365 | 0.27191 |
| 17 | 23923800 | 23926399 | 14.23605 | 24.584 | 51.60237 | 22.63791 | -0.6796 | 0.851638 | 0.27191 |
| 5 | 93805200 | 93812199 | 94.06503 | 147.7025 | 125.3759 | 52.00136 | 0.555242 | 2.571433 | 0.271911 |
| 9 | 80109200 | 80111599 | 16.95572 | 28.9366 | 58.06829 | 25.45751 | -0.61854 | 0.976832 | 0.271928 |
| 17 | 17476200 | 17479199 | 94.78499 | 150.3128 | 91.75979 | 38.1981 | 1.041389 | 2.350994 | 0.27195 |
| 5 | 12251400 | 12253799 | 39.63057 | 64.83551 | 57.66757 | 24.80215 | 0.543757 | 1.484138 | 0.271956 |
| X | 38801600 | 38805399 | 33.78836 | 57.84463 | 35.2995 | 15.47944 | 1.092823 | 0.991208 | 0.271959 |
| 5 | 51110600 | 51111599 | 13.78803 | 25.58613 | 22.37919 | 9.38625 | 0.635584 | 0.196469 | 0.271966 |
| 6 | 72875600 | 72879399 | 18.52083 | 30.76715 | 70.13151 | 30.41169 | -0.80363 | 1.258054 | 0.271967 |
| 13 | 8627600 | 8628599 | 22.44566 | 38.95723 | 33.6763 | 14.77603 | 0.595954 | 0.816303 | 0.271979 |
| 16 | 38504400 | 38505399 | 11.06619 | 21.26692 | 18.95835 | 7.408128 | 0.694726 | -0.07974 | 0.271983 |
| 3 | 110797200 | 110797999 | 9.906156 | 19.15108 | 18.11765 | 6.938633 | 0.637268 | -0.15858 | 0.271991 |
| 16 | 81936000 | 81936599 | 9.906156 | 19.15108 | 18.11765 | 6.938633 | 0.637268 | -0.15858 | 0.271991 |
| 1 | 86079000 | 86082799 | 19.32227 | 32.42333 | 64.50395 | 28.15287 | -0.60792 | 1.144005 | 0.271993 |
| 7 | 133384400 | 133391599 | 46.09382 | 73.27943 | 135.2451 | 56.44103 | -0.54874 | 2.247568 | 0.272004 |
| 15 | 27678000 | 27679799 | 7.071483 | 13.35701 | 30.86171 | 12.56084 | -0.72067 | 0.046259 | 0.272028 |
| 15 | 82063600 | 82067799 | 44.75247 | 71.27515 | 130.2223 | 54.47383 | -0.53141 | 2.187539 | 0.27203 |
| 7 | 102012400 | 102013999 | 13.54538 | 23.69563 | 46.2323 | 20.39081 | -0.57188 | 0.680762 | 0.272037 |
| 6 | 122008800 | 122010199 | 16.18997 | 29.25718 | 24.45459 | 10.61964 | 0.666059 | 0.365376 | 0.272038 |
| 4 | 117044200 | 117044999 | 11.77947 | 22.35778 | 18.04579 | 7.269816 | 0.79804 | 0.023653 | 0.272059 |
| 17 | 23351000 | 23353799 | 33.39989 | 56.45257 | 41.35283 | 18.09456 | 0.828571 | 1.102772 | 0.272073 |
| 1 | 183167600 | 183169399 | 24.70673 | 42.63574 | 37.74389 | 16.55786 | 0.560209 | 0.883778 | 0.272075 |
| 14 | 105282600 | 105284399 | 22.03746 | 38.53616 | 32.73194 | 14.43168 | 0.621916 | 0.707291 | 0.272075 |
| 9 | 90334400 | 90335999 | 19.79883 | 34.92165 | 29.47705 | 13.04443 | 0.632483 | 0.570838 | 0.272076 |
| 5 | 62013200 | 62017599 | 90.02091 | 140.8851 | 130.4793 | 54.12737 | 0.425979 | 2.629047 | 0.272079 |
| 4 | 3524000 | 3527399 | 25.9398 | 45.11783 | 33.65708 | 14.7874 | 0.80782 | 0.803322 | 0.272088 |
| 10 | 98619200 | 98619799 | 5.963911 | 11.5526 | 27.98647 | 10.61938 | -0.69381 | -0.20266 | 0.272095 |
| 16 | 50543600 | 50545599 | 19.7694 | 34.77315 | 30.07665 | 13.31715 | 0.595973 | 0.601311 | 0.272134 |
| 17 | 50269200 | 50269799 | 8.916726 | 16.10999 | 35.94288 | 15.61289 | -0.74098 | 0.343418 | 0.272134 |
| 6 | 6278400 | 6282799 | 34.87227 | 56.13177 | 102.6532 | 43.60703 | -0.51497 | 1.835058 | 0.272154 |
| 5 | 132685600 | 132689199 | 44.45345 | 72.13071 | 63.81173 | 27.34477 | 0.54272 | 1.624219 | 0.272158 |
| 17 | 12487400 | 12492599 | 33.32336 | 53.49752 | 109.294 | 46.24939 | -0.67908 | 1.918578 | 0.272187 |
| 7 | 20709400 | 20715999 | 133.7846 | 204.3392 | 163.943 | 67.99694 | 0.605857 | 3.096245 | 0.272192 |
| 4 | 143417000 | 143417599 | 9.541757 | 18.57104 | 16.62916 | 6.115734 | 0.770089 | -0.30323 | 0.272196 |
| 12 | 63261800 | 63262999 | 14.69788 | 27.74183 | 20.73626 | 8.452825 | 0.893138 | 0.095959 | 0.272203 |
| 4 | 27169000 | 27170399 | 10.25622 | 18.60578 | 40.26237 | 17.40023 | -0.69323 | 0.335563 | 0.272217 |

| | | | | | | | | | |
|---|---|---|---|---|---|---|---|---|---|
| 7 | 42727200 | 42729399 | 43.9602 | 71.40909 | 63.6654 | 27.30426 | 0.53307 | 1.603706 | 0.272217 |
| 13 | 65797000 | 65801199 | 76.21369 | 121.654 | 84.09618 | 35.28826 | 0.871818 | 2.134934 | 0.272217 |
| 2 | 4405600 | 4407399 | 8.643318 | 15.55204 | 37.9529 | 16.57018 | -0.87463 | 0.367695 | 0.272218 |
| 2 | 71744000 | 71744999 | 9.229521 | 16.63124 | 36.41822 | 15.87159 | -0.71742 | 0.361744 | 0.272218 |
| 13 | 35937800 | 35939599 | 11.28938 | 19.99066 | 40.87781 | 18.07543 | -0.63479 | 0.498483 | 0.272228 |
| 12 | 61236800 | 61238799 | 24.85012 | 43.53495 | 29.66371 | 13.09932 | 0.938129 | 0.688666 | 0.27223 |
| 2 | 157398000 | 157405599 | 166.1626 | 241.9978 | 432.7199 | 178.3335 | -0.61132 | 3.970903 | 0.272271 |
| X | 38136200 | 38138399 | 33.2062 | 55.45269 | 47.05339 | 20.4741 | 0.617384 | 1.224262 | 0.272287 |
| 2 | 9637400 | 9639399 | 10.96702 | 19.4825 | 39.86133 | 17.59194 | -0.63313 | 0.468649 | 0.272328 |
| 3 | 63097800 | 63098799 | 11.93011 | 23.058 | 17.84772 | 6.843408 | 0.922069 | -0.13376 | 0.272328 |
| 15 | 94684400 | 94685199 | 13.24066 | 24.86815 | 21.84554 | 8.982805 | 0.651566 | 0.115662 | 0.272339 |
| 4 | 8850600 | 8855799 | 27.68513 | 44.94396 | 87.08601 | 37.33164 | -0.58482 | 1.598192 | 0.27234 |
| 10 | 103309200 | 103311399 | 10.76419 | 19.24746 | 41.51359 | 18.22054 | -0.70353 | 0.427406 | 0.272342 |
| 17 | 85792600 | 85794799 | 18.18459 | 30.68956 | 60.62249 | 26.51424 | -0.59725 | 1.085808 | 0.272374 |
| 3 | 91098000 | 91099799 | 30.08751 | 51.91708 | 34.07909 | 14.95011 | 0.989714 | 0.902886 | 0.272384 |
| 7 | 10664200 | 10677799 | 68.73842 | 109.5503 | 88.86951 | 37.21006 | 0.638989 | 2.177097 | 0.272395 |
| 7 | 60290800 | 60301999 | 145.1591 | 222.5985 | 157.2357 | 65.27054 | 0.793926 | 3.021917 | 0.272398 |
| 9 | 98222200 | 98237199 | 124.0085 | 185.8001 | 314.4415 | 129.7098 | -0.4936 | 3.486639 | 0.272403 |
| 17 | 65863400 | 65869599 | 48.40341 | 76.78268 | 142.528 | 59.34999 | -0.56015 | 2.315149 | 0.272416 |
| 7 | 60171000 | 60179799 | 128.0251 | 196.4918 | 156.8255 | 65.10792 | 0.618801 | 2.998367 | 0.272428 |
| 1 | 10604800 | 10605799 | 29.81412 | 50.11508 | 43.29947 | 18.91473 | 0.591209 | 1.141628 | 0.272432 |
| 16 | 59277800 | 59280999 | 33.84306 | 54.24103 | 113.9123 | 48.08122 | -0.7217 | 1.97343 | 0.272443 |
| 5 | 97756000 | 97757799 | 16.37417 | 30.23943 | 23.00093 | 9.710329 | 0.828812 | 0.244618 | 0.272446 |
| 12 | 73594600 | 73596599 | 11.47901 | 20.31965 | 40.90533 | 18.0888 | -0.6123 | 0.505802 | 0.272446 |
| 8 | 121797400 | 121806399 | 76.04857 | 117.123 | 214.0022 | 88.88757 | -0.57112 | 2.918591 | 0.272468 |
| 13 | 39251600 | 39252199 | 3.595577 | 7.012514 | 25.10891 | 9.010105 | -1.17515 | -0.44105 | 0.272471 |
| 16 | 19916000 | 19917399 | 10.27759 | 18.5149 | 37.9244 | 16.5335 | -0.6225 | 0.372082 | 0.27248 |
| 2 | 35402200 | 35411599 | 111.8668 | 168.3426 | 295.1606 | 121.8921 | -0.53898 | 3.399133 | 0.272486 |
| 1 | 140553200 | 140555399 | 22.10508 | 38.83128 | 29.20263 | 12.92535 | 0.796225 | 0.629167 | 0.2725 |
| 17 | 38655800 | 38662199 | 190.152 | 283.9343 | 214.9788 | 88.62199 | 0.661581 | 3.556473 | 0.272504 |
| 4 | 127575000 | 127577799 | 14.53363 | 25.11462 | 51.61787 | 22.64517 | -0.64922 | 0.846407 | 0.272504 |
| 2 | 130508800 | 130510399 | 10.03865 | 17.8938 | 38.88975 | 17.1142 | -0.71668 | 0.439399 | 0.272513 |
| 7 | 10733800 | 10736399 | 35.54056 | 59.10272 | 41.85346 | 18.16242 | 0.877419 | 1.270509 | 0.27252 |
| 10 | 65921600 | 65922399 | 10.35751 | 20.20735 | 14.26012 | 5.094145 | 1.146388 | -0.40041 | 0.272529 |
| 9 | 45061600 | 45063999 | 16.10685 | 27.30184 | 59.50372 | 26.05937 | -0.73823 | 1.038054 | 0.272535 |
| 1 | 10857000 | 10858799 | 18.43568 | 33.56358 | 22.47875 | 9.671442 | 0.991931 | 0.336113 | 0.272536 |
| 13 | 12475400 | 12476799 | 8.143635 | 14.7959 | 35.49206 | 15.3094 | -0.83819 | 0.302808 | 0.272543 |
| 5 | 13938400 | 13941399 | 39.10186 | 64.96174 | 48.2281 | 20.91465 | 0.809412 | 1.29693 | 0.272559 |
| X | 145354000 | 145355199 | 16.00271 | 30.52227 | 17.86402 | 7.122663 | 1.270281 | 0.031191 | 0.272578 |
| 7 | 83600600 | 83602199 | 10.12583 | 17.8273 | 42.02465 | 18.60395 | -0.8417 | 0.559186 | 0.27258 |
| 17 | 64081400 | 64082599 | 14.11894 | 26.18245 | 22.68803 | 9.524549 | 0.647608 | 0.201985 | 0.272593 |
| 14 | 25707800 | 25712599 | 20.54365 | 33.73742 | 76.782 | 33.07515 | -0.80507 | 1.409345 | 0.2726 |
| 1 | 28248400 | 28250199 | 15.37789 | 28.58912 | 22.61215 | 9.460022 | 0.782115 | 0.199367 | 0.272612 |
| 16 | 29809400 | 29815399 | 32.61754 | 52.5228 | 101.6718 | 43.2144 | -0.59606 | 1.819049 | 0.272614 |
| 4 | 123155600 | 123158599 | 18.55085 | 31.14688 | 62.7195 | 27.38405 | -0.62517 | 1.135568 | 0.272615 |
| 14 | 101670600 | 101671999 | 7.671596 | 14.50251 | 31.47224 | 12.78696 | -0.62867 | 0.048872 | 0.272621 |
| 3 | 107361600 | 107368199 | 44.04339 | 69.88804 | 143.8015 | 59.89889 | -0.70801 | 2.302586 | 0.272627 |

| | | | | | | | | | |
|---|---|---|---|---|---|---|---|---|---|
| 3 | 153282400 | 153284199 | 24.23754 | 42.21616 | 34.34004 | 15.10076 | 0.683759 | 0.777634 | 0.272672 |
| 18 | 52661200 | 52664199 | 23.14144 | 38.18779 | 76.7942 | 33.14815 | -0.62458 | 1.351701 | 0.272679 |
| 15 | 59748600 | 59749999 | 8.776192 | 16.27149 | 37.0835 | 15.57209 | -0.73824 | 0.192681 | 0.272679 |
| 3 | 158558000 | 158560799 | 33.03441 | 54.94639 | 49.80077 | 21.61426 | 0.522543 | 1.266018 | 0.272679 |
| 5 | 60885600 | 60887199 | 8.792316 | 16.31937 | 34.28047 | 14.37345 | -0.6194 | 0.182561 | 0.27268 |
| 18 | 82927800 | 82930599 | 12.15067 | 21.23756 | 45.74113 | 20.15955 | -0.71357 | 0.686774 | 0.272682 |
| 5 | 112649600 | 112651999 | 15.39132 | 26.37879 | 54.22994 | 23.7829 | -0.65196 | 0.929639 | 0.272691 |
| 9 | 44643800 | 44647599 | 18.12874 | 29.9393 | 73.54057 | 31.76083 | -0.91172 | 1.330426 | 0.272691 |
| 18 | 23186000 | 23188199 | 23.73266 | 41.74551 | 27.98488 | 12.3909 | 0.961626 | 0.620721 | 0.272695 |
| 8 | 8196400 | 8198999 | 26.53082 | 46.25468 | 31.93356 | 14.05096 | 0.919305 | 0.772233 | 0.272715 |
| 6 | 72593000 | 72594199 | 5.256542 | 9.868234 | 31.18426 | 12.83257 | -1.18029 | 0.073243 | 0.272725 |
| 5 | 95501000 | 95503799 | 36.44745 | 61.4483 | 40.80802 | 17.84252 | 0.969521 | 1.136062 | 0.272726 |
| 19 | 53034800 | 53035999 | 9.625085 | 17.51239 | 36.00211 | 15.50181 | -0.61586 | 0.306083 | 0.272735 |
| 12 | 24188800 | 24213199 | 511.4691 | 701.4822 | 570.1299 | 237.1267 | 0.439053 | 4.879711 | 0.272735 |
| 5 | 76427800 | 76428999 | 10.25519 | 18.54 | 39.06816 | 16.95697 | -0.65916 | 0.352812 | 0.272736 |
| X | 30968000 | 30972999 | 64.67529 | 104.1192 | 66.77012 | 28.36996 | 0.993942 | 1.861567 | 0.272758 |
| 1 | 102841000 | 102841799 | 11.03331 | 21.31438 | 17.87854 | 6.870761 | 0.803838 | -0.13693 | 0.272771 |
| 1 | 176069200 | 176070199 | 11.03331 | 21.31438 | 17.87854 | 6.870761 | 0.803838 | -0.13693 | 0.272771 |
| 2 | 40021000 | 40023199 | 29.68581 | 50.83332 | 38.62692 | 16.9389 | 0.777704 | 0.974084 | 0.272774 |
| 19 | 54123200 | 54125199 | 16.21763 | 27.70376 | 61.82779 | 27.10781 | -0.77091 | 0.97179 | 0.272778 |
| 6 | 57063000 | 57066799 | 86.11816 | 135.2953 | 125.5343 | 52.07064 | 0.427576 | 2.556221 | 0.2728 |
| 15 | 58645200 | 58648199 | 18.59029 | 31.57162 | 59.68543 | 26.15409 | -0.53385 | 1.030491 | 0.272831 |
| X | 80731400 | 80733399 | 36.27055 | 60.61036 | 45.4116 | 19.76802 | 0.796327 | 1.226052 | 0.272841 |
| 16 | 10843400 | 10844399 | 5.564728 | 10.73226 | 28.19503 | 10.90578 | -0.83661 | -0.14393 | 0.272847 |
| 4 | 46729400 | 46730999 | 8.764502 | 15.83966 | 35.9468 | 15.6088 | -0.76506 | 0.340203 | 0.27285 |
| X | 33565600 | 33569999 | 129.0245 | 197.4804 | 158.7767 | 65.88597 | 0.605167 | 3.055107 | 0.272852 |
| 17 | 40764400 | 40771399 | 66.85821 | 106.6804 | 92.56576 | 38.82155 | 0.543879 | 2.137728 | 0.272865 |
| 5 | 118478400 | 118480399 | 11.57409 | 20.58662 | 43.01362 | 18.96316 | -0.66248 | 0.469442 | 0.272868 |
| 10 | 13690800 | 13691799 | 11.4116 | 21.96107 | 18.94924 | 7.37523 | 0.746708 | -0.08847 | 0.272868 |
| 8 | 122238400 | 122240999 | 9.722349 | 17.04462 | 45.18046 | 19.96333 | -1.01104 | 0.622693 | 0.272883 |
| 19 | 7932800 | 7938599 | 76.73798 | 121.8335 | 104.6208 | 43.60214 | 0.551458 | 2.304594 | 0.272897 |
| 11 | 63350600 | 63353399 | 17.27484 | 29.52739 | 56.44655 | 24.74964 | -0.54837 | 0.963099 | 0.272899 |
| 17 | 22354200 | 22357199 | 92.60759 | 144.7145 | 133.702 | 55.47288 | 0.427786 | 2.657998 | 0.272909 |
| 3 | 15500600 | 15503199 | 44.77983 | 72.49597 | 66.22479 | 28.33149 | 0.493619 | 1.673418 | 0.272911 |
| X | 5629400 | 5632599 | 64.20589 | 103.0623 | 76.8012 | 32.49276 | 0.772707 | 1.952234 | 0.272911 |
| 2 | 51036200 | 51039399 | 21.15654 | 34.80261 | 76.63624 | 33.03594 | -0.75678 | 1.38749 | 0.272928 |
| 13 | 36670600 | 36671799 | 14.70754 | 27.76102 | 20.61712 | 8.403649 | 0.902085 | 0.09441 | 0.27294 |
| 6 | 130982600 | 130989199 | 99.82649 | 157.4983 | 106.354 | 44.14151 | 0.890072 | 2.480145 | 0.272941 |
| 8 | 113529600 | 113532799 | 26.74042 | 43.84479 | 81.98229 | 35.29173 | -0.52371 | 1.444416 | 0.272952 |
| 19 | 42622400 | 42623999 | 9.130205 | 16.73989 | 35.08396 | 14.95797 | -0.63379 | 0.255755 | 0.27297 |
| 5 | 118004600 | 118007399 | 29.38263 | 47.4249 | 95.65544 | 40.78578 | -0.65106 | 1.742232 | 0.272994 |
| 18 | 56477400 | 56479599 | 40.26908 | 65.67148 | 60.24549 | 25.88094 | 0.496139 | 1.533912 | 0.27301 |
| 5 | 77552800 | 77558399 | 66.24102 | 103.6139 | 175.9601 | 72.9984 | -0.44828 | 2.633434 | 0.273038 |
| 15 | 66819000 | 66820399 | 17.88376 | 31.97202 | 26.33089 | 11.55858 | 0.677185 | 0.445481 | 0.273064 |
| 17 | 66550800 | 66553799 | 25.22185 | 41.14753 | 82.15026 | 35.29583 | -0.62229 | 1.511383 | 0.273066 |
| 14 | 19308200 | 19310799 | 22.00445 | 38.83387 | 27.51783 | 12.17605 | 0.883669 | 0.573925 | 0.273076 |
| 13 | 28244600 | 28245999 | 7.044193 | 13.45512 | 36.25489 | 14.4109 | -0.91097 | -0.01909 | 0.273083 |

| | | | | | | | | | |
|---|---|---|---|---|---|---|---|---|---|
| 1 | 171764400 | 171766399 | 20.92043 | 35.03555 | 66.84825 | 29.13957 | -0.54792 | 1.172349 | 0.273123 |
| 17 | 59653000 | 59653799 | 5.967085 | 11.54681 | 27.25992 | 10.39377 | -0.66339 | -0.18765 | 0.273131 |
| 4 | 56759800 | 56761999 | 15.47248 | 28.36448 | 22.92632 | 9.77634 | 0.731064 | 0.275665 | 0.273131 |
| 11 | 92330400 | 92333199 | 31.59445 | 53.37351 | 43.06077 | 18.83896 | 0.689781 | 1.102455 | 0.273135 |
| 3 | 35986000 | 35987799 | 12.28222 | 21.50458 | 45.174 | 19.93867 | -0.67784 | 0.657211 | 0.273147 |
| 17 | 88812800 | 88814199 | 12.12495 | 21.2577 | 44.10689 | 19.48973 | -0.65999 | 0.633322 | 0.273148 |
| 17 | 61696000 | 61697799 | 17.54385 | 32.35613 | 21.65629 | 9.159367 | 1.009817 | 0.257303 | 0.273162 |
| 4 | 113153400 | 113154399 | 19.82643 | 35.14285 | 27.96073 | 12.36125 | 0.720136 | 0.517616 | 0.27317 |
| 7 | 28986400 | 28990999 | 25.7014 | 41.54503 | 93.66599 | 40.01427 | -0.80869 | 1.691081 | 0.27317 |
| 1 | 135284800 | 135288599 | 18.39741 | 30.89057 | 62.69058 | 27.37216 | -0.63635 | 1.135068 | 0.273182 |
| 16 | 89581400 | 89583799 | 13.43192 | 23.27781 | 51.07851 | 22.41052 | -0.74226 | 0.814174 | 0.273187 |
| 7 | 32844000 | 32846199 | 25.69256 | 44.77461 | 33.22016 | 14.61326 | 0.816008 | 0.776965 | 0.27319 |
| 5 | 25876400 | 25878799 | 11.77427 | 20.73539 | 43.49593 | 19.25419 | -0.6747 | 0.565819 | 0.273197 |
| 6 | 141791600 | 141795399 | 39.33269 | 64.48435 | 57.13255 | 24.59324 | 0.550795 | 1.456848 | 0.273198 |
| 2 | 134627600 | 134629399 | 12.13242 | 21.23072 | 45.70893 | 20.16547 | -0.71315 | 0.665939 | 0.273214 |
| 11 | 80615800 | 80616399 | 3.935417 | 7.682132 | 20.97887 | 7.424142 | -0.76904 | -0.49557 | 0.273215 |
| 3 | 52814200 | 52818399 | 49.19865 | 79.59219 | 67.32449 | 28.78659 | 0.60335 | 1.695735 | 0.27323 |
| X | 8344000 | 8347799 | 46.39267 | 76.06059 | 52.92597 | 22.78269 | 0.898384 | 1.472051 | 0.273241 |
| X | 25887600 | 25888599 | 28.36872 | 48.40879 | 39.94941 | 17.51392 | 0.65853 | 1.003279 | 0.273243 |
| X | 91089200 | 91091599 | 20.20759 | 37.60314 | 18.29175 | 7.640445 | 1.479654 | 0.224296 | 0.273248 |
| 12 | 44666400 | 44668799 | 12.79134 | 22.28856 | 47.34834 | 20.82635 | -0.69432 | 0.737696 | 0.273248 |
| 1 | 133350200 | 133353799 | 19.57644 | 32.59678 | 68.14521 | 29.59734 | -0.67909 | 1.232236 | 0.273257 |
| 5 | 19373800 | 19375999 | 21.17767 | 37.32106 | 28.71777 | 12.71065 | 0.764697 | 0.586251 | 0.273262 |
| 14 | 25069000 | 25070599 | 7.716656 | 14.39612 | 31.65186 | 13.16402 | -0.67506 | 0.136211 | 0.273274 |
| 7 | 92390200 | 92391399 | 7.895248 | 14.25315 | 37.49701 | 16.30415 | -0.97814 | 0.342188 | 0.273295 |
| 12 | 3057600 | 3078599 | 968.1897 | 1257.108 | 959.6577 | 402.0657 | 0.450312 | 5.755978 | 0.273304 |
| 8 | 25385800 | 25386199 | 14.34896 | 27.42706 | 18.54334 | 7.359292 | 1.069511 | 0.0018 | 0.273307 |
| 3 | 123960000 | 123960399 | 5.828289 | 11.25583 | 26.49012 | 10.19451 | -0.6715 | -0.15933 | 0.273317 |
| 18 | 81438000 | 81447199 | 77.65053 | 120.0641 | 203.6998 | 84.61214 | -0.4599 | 2.844553 | 0.273327 |
| X | 63697200 | 63698999 | 35.19927 | 58.59393 | 48.71193 | 21.15318 | 0.64691 | 1.26501 | 0.273346 |
| 8 | 72863200 | 72864999 | 13.0553 | 22.91531 | 44.57682 | 19.71664 | -0.56756 | 0.620529 | 0.273346 |
| 5 | 109598400 | 109599399 | 11.78507 | 22.62544 | 19.47008 | 7.630331 | 0.741549 | -0.06083 | 0.273348 |
| 12 | 23732400 | 23732999 | 11.78507 | 22.62544 | 19.47008 | 7.630331 | 0.741549 | -0.06083 | 0.273348 |
| 4 | 151061000 | 151062999 | 17.33229 | 29.426 | 58.512 | 25.63747 | -0.60638 | 1.027544 | 0.273351 |
| 18 | 68175600 | 68177199 | 9.805188 | 17.72014 | 36.9391 | 16.04002 | -0.6438 | 0.348303 | 0.273361 |
| 7 | 14441000 | 14473399 | 589.4055 | 802.5718 | 595.3724 | 247.9043 | 0.560263 | 4.993677 | 0.273367 |
| 16 | 60985000 | 60988999 | 37.83648 | 62.62581 | 51.59295 | 22.3164 | 0.65937 | 1.336464 | 0.273368 |
| 5 | 138278000 | 138280599 | 38.66879 | 61.96951 | 112.6657 | 47.54819 | -0.51429 | 1.982495 | 0.27337 |
| 16 | 89804000 | 89809599 | 64.8979 | 100.3419 | 216.5362 | 89.9441 | -0.80656 | 2.842066 | 0.273371 |
| 9 | 117550400 | 117554399 | 41.30749 | 65.94546 | 123.5403 | 51.85037 | -0.5637 | 2.106145 | 0.273374 |
| 5 | 87133000 | 87134199 | 13.97647 | 25.80471 | 22.74453 | 9.605479 | 0.617162 | 0.22531 | 0.273375 |
| 9 | 35857800 | 35860799 | 25.37489 | 44.32078 | 32.08123 | 14.13225 | 0.851695 | 0.742912 | 0.273396 |
| 3 | 142230600 | 142231599 | 14.54124 | 27.78773 | 18.53564 | 7.360695 | 1.088076 | 0.005697 | 0.273423 |
| 1 | 44326400 | 44334999 | 181.9815 | 272.0273 | 242.3705 | 99.93477 | 0.428327 | 3.535963 | 0.273437 |
| 7 | 24957600 | 24960599 | 15.92099 | 27.05957 | 59.00788 | 25.86089 | -0.7388 | 1.01239 | 0.273445 |
| 16 | 29227400 | 29229399 | 10.81411 | 19.28353 | 38.87979 | 17.10423 | -0.60868 | 0.443491 | 0.273462 |
| 1 | 78324000 | 78326199 | 14.51468 | 25.1303 | 51.0973 | 22.41811 | -0.63319 | 0.827905 | 0.273473 |

| | | | | | | | | | |
|---|---|---|---|---|---|---|---|---|---|
| 9 | 5552600 | 5555999 | 82.21517 | 129.4636 | 119.7225 | 49.67494 | 0.434943 | 2.506548 | 0.273489 |
| 6 | 136212000 | 136213799 | 11.63564 | 20.47872 | 42.47609 | 18.80504 | -0.65871 | 0.571209 | 0.2735 |
| 7 | 60604400 | 60608999 | 76.20931 | 122.0334 | 72.10929 | 30.39529 | 1.102628 | 2.03828 | 0.273503 |
| 17 | 88397400 | 88400399 | 20.94845 | 36.6743 | 32.31645 | 14.26244 | 0.569166 | 0.683087 | 0.273505 |
| 13 | 33425800 | 33427799 | 27.80898 | 48.45818 | 30.73106 | 13.51749 | 1.041095 | 0.783563 | 0.273521 |
| 19 | 10138800 | 10140199 | 18.89105 | 34.76103 | 21.60458 | 9.166026 | 1.113046 | 0.281208 | 0.273545 |
| 17 | 20228600 | 20229399 | 15.19046 | 28.24885 | 22.61879 | 9.458153 | 0.765062 | 0.195845 | 0.273545 |
| 19 | 40923600 | 40926399 | 15.34693 | 26.4714 | 52.14667 | 22.87588 | -0.58883 | 0.868626 | 0.273547 |
| 9 | 52699200 | 52700799 | 16.53047 | 30.2973 | 24.00047 | 10.23747 | 0.760023 | 0.289384 | 0.273566 |
| 16 | 37997400 | 37999799 | 13.56134 | 23.73382 | 46.75137 | 20.62729 | -0.58557 | 0.675233 | 0.273569 |
| 2 | 67149600 | 67150199 | 13.98497 | 25.79356 | 20.41834 | 8.635289 | 0.769894 | 0.22617 | 0.273573 |
| 12 | 97629600 | 97630799 | 11.37244 | 21.76832 | 19.43063 | 7.680273 | 0.677518 | -0.03879 | 0.273581 |
| 9 | 90756600 | 90758199 | 22.20869 | 39.10618 | 28.64136 | 12.67889 | 0.835179 | 0.601328 | 0.273582 |
| 9 | 49031600 | 49035599 | 32.88942 | 53.16731 | 95.46279 | 40.78503 | -0.48118 | 1.701717 | 0.273587 |
| 1 | 182698000 | 182704399 | 109.2008 | 165.2142 | 277.8734 | 114.9365 | -0.47232 | 3.29033 | 0.273587 |
| 5 | 83287200 | 83290599 | 33.7287 | 56.17583 | 43.36407 | 18.83719 | 0.753368 | 1.247267 | 0.273589 |
| 11 | 75597800 | 75600199 | 18.1172 | 30.87829 | 58.59897 | 25.68745 | -0.53853 | 0.996121 | 0.273598 |
| 13 | 108752400 | 108754799 | 19.40504 | 34.66444 | 26.81286 | 11.77796 | 0.766761 | 0.463316 | 0.273602 |
| 17 | 9199800 | 9200799 | 5.266544 | 10.14595 | 26.34033 | 10.22079 | -0.82256 | -0.13812 | 0.273603 |
| 12 | 113709200 | 113710799 | 30.96127 | 51.87799 | 46.47305 | 20.26455 | 0.539254 | 1.180106 | 0.273623 |
| 5 | 11401000 | 11402199 | 10.74845 | 20.56364 | 18.00284 | 7.126147 | 0.703603 | -0.04414 | 0.273629 |
| 3 | 134898800 | 134899199 | 9.641433 | 18.81294 | 14.28297 | 5.089978 | 1.044679 | -0.41926 | 0.273632 |
| 14 | 25317000 | 25320799 | 18.93494 | 31.58075 | 66.4118 | 28.87456 | -0.68749 | 1.212049 | 0.273644 |
| X | 5168200 | 5170199 | 38.85556 | 64.89669 | 44.24202 | 19.25265 | 0.931975 | 1.235657 | 0.27365 |
| 4 | 140198600 | 140211399 | 103.7062 | 156.0147 | 299.6966 | 123.7553 | -0.67098 | 3.403635 | 0.273653 |
| 9 | 67513200 | 67514999 | 9.136684 | 16.7904 | 34.59453 | 14.70314 | -0.60588 | 0.242794 | 0.273663 |
| 5 | 121073800 | 121081599 | 61.2161 | 95.67061 | 178.4839 | 74.05205 | -0.58468 | 2.646086 | 0.273668 |
| 5 | 126856400 | 126858999 | 24.20143 | 42.08233 | 34.31399 | 15.07334 | 0.67995 | 0.798902 | 0.273669 |
| 3 | 50423000 | 50425799 | 27.87819 | 47.93376 | 38.13263 | 16.72654 | 0.712756 | 0.930213 | 0.273675 |
| 1 | 31325000 | 31326999 | 16.79655 | 28.90994 | 55.95455 | 24.54474 | -0.56433 | 0.898642 | 0.273675 |
| 3 | 59952400 | 59953599 | 19.33899 | 31.99907 | 71.51355 | 30.92616 | -0.77563 | 1.310803 | 0.273678 |
| 5 | 148095000 | 148097799 | 11.65332 | 20.44726 | 44.60391 | 19.72276 | -0.73194 | 0.617178 | 0.273681 |
| 7 | 40108600 | 40109799 | 16.56729 | 29.69256 | 24.80159 | 10.86435 | 0.659311 | 0.419996 | 0.273684 |
| 8 | 43107400 | 43108799 | 19.87698 | 35.55535 | 26.19558 | 11.49375 | 0.837608 | 0.456871 | 0.27369 |
| 16 | 49675200 | 49676799 | 12.98579 | 22.49349 | 50.52795 | 22.16863 | -0.77589 | 0.817154 | 0.273701 |
| 3 | 53147200 | 53148999 | 15.72128 | 28.42021 | 21.34978 | 9.267263 | 0.819722 | 0.353952 | 0.273702 |
| 5 | 115100000 | 115100999 | 4.987828 | 9.650857 | 25.99293 | 9.915539 | -0.85202 | -0.20217 | 0.273722 |
| X | 147546600 | 147549999 | 28.11312 | 49.14858 | 28.45193 | 12.54204 | 1.172234 | 0.737486 | 0.273734 |
| 3 | 77617200 | 77618999 | 21.27006 | 38.08977 | 23.14183 | 10.14431 | 1.114435 | 0.451938 | 0.273743 |
| 9 | 55587800 | 55589199 | 6.77632 | 12.47542 | 35.1555 | 14.90468 | -1.05137 | 0.213596 | 0.273747 |
| 13 | 97466800 | 97468799 | 13.30081 | 23.01026 | 51.05924 | 22.40087 | -0.75875 | 0.830461 | 0.273749 |
| X | 44688800 | 44693199 | 50.94398 | 82.66827 | 61.33891 | 26.2865 | 0.795981 | 1.636911 | 0.273752 |
| 18 | 17991400 | 17996399 | 78.80375 | 124.6772 | 113.1812 | 47.04138 | 0.466938 | 2.40065 | 0.273755 |
| 7 | 126972400 | 126974999 | 35.46482 | 56.78917 | 114.7898 | 48.38535 | -0.66821 | 2.004904 | 0.273759 |
| 4 | 122742000 | 122745199 | 18.7232 | 31.34344 | 67.19948 | 29.28536 | -0.71547 | 1.169495 | 0.273766 |
| 6 | 40674200 | 40675399 | 7.202005 | 13.69248 | 29.95179 | 12.02364 | -0.6245 | 0.001288 | 0.273789 |
| 17 | 53707200 | 53708399 | 7.202005 | 13.69248 | 29.95179 | 12.02364 | -0.6245 | 0.001288 | 0.273789 |

| | | | | | | | | | |
|---|---|---|---|---|---|---|---|---|---|
| 19 | 35866200 | 35867199 | 12.14383 | 21.2253 | 46.25764 | 20.38695 | -0.73064 | 0.68668 | 0.273802 |
| 4 | 53016200 | 53018799 | 36.82545 | 61.61527 | 44.77877 | 19.50806 | 0.839957 | 1.211071 | 0.273808 |
| X | 34412600 | 34413799 | 17.75538 | 32.41044 | 23.56007 | 10.10329 | 0.878376 | 0.322045 | 0.27386 |
| 8 | 25506400 | 25508799 | 9.846387 | 17.3148 | 45.1055 | 19.97255 | -0.98607 | 0.573723 | 0.273864 |
| 9 | 115446400 | 115451399 | 38.78456 | 62.32651 | 110.204 | 46.70312 | -0.46948 | 1.895244 | 0.273866 |
| 9 | 31136800 | 31138599 | 11.27434 | 20.05178 | 39.86383 | 17.57578 | -0.59092 | 0.463674 | 0.273868 |
| 15 | 83414000 | 83422399 | 49.03238 | 77.75854 | 149.6089 | 62.281 | -0.61225 | 2.324154 | 0.273876 |
| 10 | 47367200 | 47368799 | 30.39057 | 51.24302 | 44.76725 | 19.57434 | 0.575344 | 1.117129 | 0.273899 |
| 17 | 58324600 | 58327199 | 25.1609 | 43.74256 | 34.95292 | 15.35221 | 0.709241 | 0.809826 | 0.273907 |
| 1 | 108590200 | 108590799 | 10.57733 | 20.43988 | 17.94837 | 6.887563 | 0.740451 | -0.14646 | 0.273907 |
| 11 | 118931200 | 118933599 | 18.43705 | 30.81484 | 67.08092 | 29.20407 | -0.7373 | 1.186574 | 0.273915 |
| 14 | 9850400 | 9851199 | 7.210934 | 13.67931 | 29.8715 | 12.04736 | -0.62789 | 0.016781 | 0.273927 |
| 1 | 180259400 | 180262799 | 13.57871 | 23.57423 | 49.4893 | 21.71813 | -0.67808 | 0.79612 | 0.273938 |
| 10 | 28615000 | 28616599 | 18.53094 | 33.55402 | 24.7731 | 10.73133 | 0.846514 | 0.373577 | 0.273955 |
| X | 88981000 | 88982599 | 22.72675 | 40.03558 | 28.42109 | 12.5803 | 0.880576 | 0.60257 | 0.273985 |
| 1 | 119291200 | 119291599 | 5.230429 | 10.12945 | 25.45081 | 9.667642 | -0.74606 | -0.2121 | 0.273994 |
| 4 | 94617200 | 94618999 | 25.91222 | 44.92777 | 35.99632 | 15.79349 | 0.704872 | 0.845988 | 0.273996 |
| 8 | 53194800 | 53196999 | 20.45528 | 35.89688 | 30.03146 | 13.28087 | 0.643363 | 0.636844 | 0.274001 |
| 3 | 121591200 | 121592999 | 14.00406 | 24.46162 | 47.31396 | 20.84334 | -0.5596 | 0.707292 | 0.27401 |
| 3 | 69795000 | 69795599 | 4.106377 | 8.012383 | 22.13772 | 7.892971 | -0.79664 | -0.46468 | 0.274016 |
| 12 | 44086800 | 44087199 | 4.106377 | 8.012383 | 22.13772 | 7.892971 | -0.79664 | -0.46468 | 0.274016 |
| 7 | 35696600 | 35697799 | 10.63932 | 18.97996 | 38.87032 | 17.0939 | -0.63086 | 0.438192 | 0.274023 |
| 13 | 61823400 | 61829599 | 136.6652 | 209.6676 | 163.6241 | 67.92664 | 0.650754 | 3.010236 | 0.274034 |
| 8 | 11153000 | 11155399 | 13.40939 | 23.31213 | 48.95184 | 21.50074 | -0.67826 | 0.774388 | 0.274051 |
| 10 | 119301800 | 119307999 | 65.89959 | 103.094 | 176.2448 | 73.11546 | -0.45771 | 2.630429 | 0.274087 |
| 4 | 65522800 | 65523999 | 5.280306 | 10.1903 | 27.00676 | 10.42678 | -0.84602 | -0.15525 | 0.274099 |
| 1 | 85675600 | 85686799 | 78.84221 | 121.1224 | 226.842 | 94.18844 | -0.60949 | 2.969075 | 0.274132 |
| 17 | 37697000 | 37698999 | 21.27141 | 37.63026 | 27.60698 | 12.20965 | 0.835212 | 0.544876 | 0.274142 |
| 6 | 7543200 | 7543999 | 12.58706 | 23.6701 | 21.40929 | 8.784825 | 0.612201 | 0.100959 | 0.274151 |
| 3 | 48336800 | 48339199 | 30.66344 | 52.23682 | 40.27149 | 17.65274 | 0.755891 | 1.025173 | 0.274169 |
| 7 | 107472600 | 107474399 | 20.73898 | 36.69632 | 28.31152 | 12.5207 | 0.763282 | 0.541671 | 0.274174 |
| 14 | 6231400 | 6232999 | 15.14536 | 28.02888 | 23.10389 | 9.725971 | 0.716456 | 0.222894 | 0.274178 |
| 3 | 128996200 | 128999799 | 29.51419 | 47.38374 | 123.2385 | 52.15265 | -1.02662 | 1.916839 | 0.274191 |
| 3 | 129878800 | 129879399 | 12.92124 | 24.39404 | 21.37534 | 8.709142 | 0.665903 | 0.078912 | 0.274195 |
| 12 | 18210600 | 18211199 | 8.774017 | 17.15825 | 12.06525 | 4.107365 | 1.211367 | -0.59319 | 0.274198 |
| 8 | 99600200 | 99601999 | 15.84325 | 29.50376 | 22.00775 | 9.181029 | 0.869191 | 0.191199 | 0.274207 |
| 9 | 22867000 | 22868599 | 15.84325 | 29.50376 | 22.00775 | 9.181029 | 0.869191 | 0.191199 | 0.274207 |
| 7 | 11634200 | 11635199 | 20.25708 | 35.58988 | 30.57151 | 13.53281 | 0.605524 | 0.619706 | 0.274219 |
| 2 | 45451000 | 45451799 | 15.21637 | 27.65212 | 24.25681 | 10.46027 | 0.603584 | 0.327838 | 0.27422 |
| 17 | 43311600 | 43313199 | 10.13173 | 18.22236 | 37.9031 | 16.5544 | -0.64625 | 0.380696 | 0.274227 |
| 17 | 91803000 | 91803999 | 10.13173 | 18.22236 | 37.9031 | 16.5544 | -0.64625 | 0.380696 | 0.274227 |
| 4 | 57266200 | 57266999 | 4.924334 | 9.567019 | 24.50158 | 9.142439 | -0.74854 | -0.28012 | 0.274242 |
| 11 | 109833800 | 109834399 | 4.924334 | 9.567019 | 24.50158 | 9.142439 | -0.74854 | -0.28012 | 0.274242 |
| 2 | 168875800 | 168876599 | 11.81226 | 20.76619 | 42.4997 | 18.81972 | -0.64001 | 0.586286 | 0.274245 |
| 2 | 11504800 | 11508999 | 22.80049 | 37.32089 | 79.22139 | 34.07864 | -0.70697 | 1.456426 | 0.274253 |
| 13 | 12580400 | 12581599 | 9.962123 | 17.5753 | 41.49245 | 18.36078 | -0.84268 | 0.53357 | 0.274259 |
| 16 | 76543800 | 76545199 | 17.97979 | 32.88529 | 23.24224 | 9.94208 | 0.920499 | 0.309886 | 0.274268 |

| | | | | | | | | | |
|---|---|---|---|---|---|---|---|---|---|
| 13 | 45284800 | 45286799 | 14.20933 | 26.65649 | 21.06633 | 8.679445 | 0.800337 | 0.12819 | 0.274286 |
| 7 | 16924000 | 16927399 | 39.28951 | 64.92 | 51.65498 | 22.31693 | 0.709212 | 1.359213 | 0.274298 |
| 12 | 69556000 | 69556999 | 11.84123 | 20.71254 | 45.21114 | 19.93886 | -0.73268 | 0.672085 | 0.274304 |
| 1 | 8254800 | 8256399 | 18.2849 | 32.54313 | 27.39078 | 12.06932 | 0.642418 | 0.478295 | 0.274314 |
| 7 | 5998200 | 6001799 | 53.00316 | 85.35627 | 71.10654 | 30.22672 | 0.6179 | 1.835684 | 0.274319 |
| 10 | 75311000 | 75313399 | 10.89002 | 19.14304 | 42.52305 | 18.83042 | -0.75755 | 0.581439 | 0.274333 |
| 12 | 113592200 | 113593399 | 21.98867 | 38.25875 | 30.70283 | 13.49237 | 0.703078 | 0.775666 | 0.274334 |
| 13 | 65787000 | 65803199 | 308.7562 | 447.3002 | 314.8212 | 129.8203 | 0.725781 | 4.059642 | 0.274363 |
| 17 | 23649800 | 23653399 | 72.3277 | 114.9931 | 102.2451 | 42.67205 | 0.503263 | 2.260315 | 0.274369 |
| 19 | 46036000 | 46037199 | 6.742011 | 12.85731 | 30.06663 | 11.99168 | -0.71201 | -0.02599 | 0.274381 |
| 17 | 39876000 | 39876199 | 7.608498 | 14.88033 | 13.00338 | 4.412308 | 0.908558 | -0.61679 | 0.274412 |
| 6 | 115718800 | 115720999 | 13.85242 | 24.16888 | 47.3305 | 20.83553 | -0.57753 | 0.723013 | 0.274419 |
| 11 | 111471000 | 111472999 | 12.72334 | 22.42465 | 48.29597 | 21.37559 | -0.71233 | 0.57067 | 0.274422 |
| 8 | 3510200 | 3514999 | 21.72126 | 35.90102 | 71.86453 | 31.05415 | -0.61756 | 1.328875 | 0.274423 |
| 14 | 4658400 | 4659599 | 8.845638 | 16.29681 | 34.64472 | 14.67378 | -0.64681 | 0.227761 | 0.274443 |
| 8 | 64716200 | 64716999 | 6.414047 | 12.37442 | 29.50793 | 11.40262 | -0.69709 | -0.13358 | 0.274445 |
| X | 153155000 | 153155999 | 25.44411 | 43.61128 | 38.44079 | 16.85857 | 0.56473 | 0.949063 | 0.274456 |
| 1 | 102969600 | 102972199 | 29.04545 | 50.13282 | 33.5102 | 14.70067 | 0.963578 | 0.893532 | 0.274487 |
| 12 | 104436200 | 104443599 | 47.76253 | 75.37377 | 167.7206 | 69.61964 | -0.82762 | 2.449711 | 0.274488 |
| 6 | 50261400 | 50262399 | 14.44963 | 26.65064 | 23.1655 | 9.797079 | 0.635595 | 0.235641 | 0.274504 |
| 5 | 89264400 | 89266199 | 17.15495 | 31.26786 | 24.39976 | 10.48286 | 0.774594 | 0.325449 | 0.274515 |
| 6 | 128032200 | 128035999 | 17.4267 | 29.3329 | 61.60527 | 26.92249 | -0.68546 | 1.107469 | 0.274516 |
| 14 | 14733400 | 14734399 | 7.185585 | 13.71678 | 31.3092 | 12.46119 | -0.67478 | -0.02462 | 0.27452 |
| 3 | 38611600 | 38614999 | 19.77245 | 32.82467 | 70.27508 | 30.46005 | -0.71366 | 1.269107 | 0.274523 |
| 9 | 47229600 | 47235599 | 41.93175 | 67.03233 | 119.6492 | 50.31318 | -0.49193 | 2.06445 | 0.274535 |
| 18 | 65106600 | 65110599 | 33.37227 | 53.84065 | 100.5202 | 42.83567 | -0.54075 | 1.761086 | 0.274535 |
| 5 | 26272400 | 26273999 | 9.31458 | 17.13021 | 36.79437 | 15.6221 | -0.66448 | 0.245386 | 0.274538 |
| 4 | 43560200 | 43563199 | 10.58662 | 18.61301 | 43.03357 | 19.05581 | -0.81491 | 0.577776 | 0.274553 |
| 9 | 61188200 | 61194999 | 52.54711 | 83.04081 | 154.4931 | 64.17459 | -0.56812 | 2.416293 | 0.274563 |
| 10 | 7845200 | 7847399 | 16.57912 | 28.57856 | 54.90688 | 24.08628 | -0.55336 | 0.88112 | 0.274576 |
| 12 | 18653600 | 18655999 | 38.78786 | 63.41679 | 58.00913 | 24.94331 | 0.502854 | 1.493505 | 0.274606 |
| 7 | 108526000 | 108528599 | 17.07747 | 29.29174 | 56.50522 | 24.78231 | -0.56065 | 0.929729 | 0.274634 |
| 2 | 58947400 | 58949599 | 18.45667 | 30.80985 | 65.81117 | 28.62913 | -0.70989 | 1.201305 | 0.274634 |
| 6 | 140232600 | 140234799 | 15.4156 | 26.75922 | 56.09528 | 24.61478 | -0.67659 | 0.81249 | 0.274638 |
| 1 | 66855400 | 66857399 | 9.35403 | 16.54124 | 41.48148 | 18.34732 | -0.92789 | 0.502898 | 0.274642 |
| 19 | 3091400 | 3093999 | 23.11271 | 40.95901 | 25.89344 | 11.44781 | 1.049757 | 0.536314 | 0.274644 |
| 18 | 62121600 | 62124399 | 27.87907 | 45.17414 | 88.91689 | 38.06391 | -0.60992 | 1.638633 | 0.274655 |
| 7 | 80036200 | 80038199 | 10.25521 | 18.10519 | 41.46688 | 18.34686 | -0.79892 | 0.52679 | 0.274657 |
| 12 | 11530000 | 11530399 | 5.67011 | 11.00247 | 26.296 | 9.89776 | -0.66328 | -0.23896 | 0.274683 |
| 13 | 111359400 | 111360599 | 15.26077 | 28.51293 | 21.49815 | 8.918489 | 0.860001 | 0.164067 | 0.274683 |
| 4 | 6453400 | 6454399 | 7.428338 | 13.83364 | 32.15266 | 13.41258 | -0.75819 | 0.145656 | 0.274689 |
| 5 | 141926000 | 141931199 | 38.48458 | 61.6947 | 112.4252 | 47.4706 | -0.51706 | 1.971007 | 0.274693 |
| 7 | 40877600 | 40885199 | 133.7863 | 204.3165 | 181.6702 | 75.34732 | 0.457574 | 3.100395 | 0.274699 |
| 3 | 131001600 | 131002999 | 8.965532 | 16.06149 | 38.94851 | 17.0676 | -0.86939 | 0.399526 | 0.274702 |
| 7 | 81946600 | 81952399 | 36.04866 | 57.86497 | 108.7525 | 46.01083 | -0.55923 | 1.926209 | 0.274713 |
| Y | 88673400 | 88674399 | 9.554602 | 18.6012 | 8.712158 | 3.196912 | 1.694131 | -0.34369 | 0.274731 |
| 2 | 146686200 | 146687799 | 11.77853 | 20.76697 | 41.93956 | 18.56093 | -0.61941 | 0.551508 | 0.274746 |

| | | | | | | | | | |
|---|---|---|---|---|---|---|---|---|---|
| 10 | 70057600 | 70059399 | 11.59905 | 20.51044 | 41.91306 | 18.53835 | -0.63475 | 0.51714 | 0.274757 |
| 5 | 30864200 | 30865599 | 7.97823 | 15.03341 | 33.08889 | 13.53104 | -0.6573 | 0.0767 | 0.274759 |
| Y | 20706600 | 20706799 | 6.19286 | 12.11199 | 7.751076 | 2.541268 | 1.392976 | -0.78721 | 0.274782 |
| 13 | 67766400 | 67768999 | 33.81616 | 58.28984 | 31.88717 | 13.98809 | 1.251368 | 0.92828 | 0.274782 |
| 14 | 41801200 | 41809199 | 103.8094 | 162.1947 | 131.5852 | 54.59549 | 0.614938 | 2.665884 | 0.27479 |
| 2 | 87142800 | 87147399 | 63.21404 | 101.2487 | 84.59399 | 35.67094 | 0.604295 | 2.018442 | 0.274798 |
| 2 | 6082400 | 6084599 | 27.49659 | 44.73103 | 84.45286 | 36.2475 | -0.5449 | 1.559239 | 0.274798 |
| 9 | 46693000 | 46694799 | 11.18729 | 19.74952 | 41.46766 | 18.34744 | -0.67413 | 0.535949 | 0.274815 |
| 5 | 56793400 | 56796399 | 29.95872 | 50.80087 | 42.5179 | 18.62268 | 0.637171 | 1.064532 | 0.274818 |
| 11 | 95538200 | 95539999 | 6.481812 | 12.28704 | 29.26628 | 11.82178 | -0.75393 | 0.013136 | 0.274819 |
| 18 | 23936800 | 23937599 | 6.481812 | 12.28704 | 29.26628 | 11.82178 | -0.75393 | 0.013136 | 0.274819 |
| 2 | 82293000 | 82298999 | 87.67127 | 138.5112 | 112.7907 | 46.84561 | 0.621896 | 2.439237 | 0.274842 |
| Y | 5013600 | 5014799 | 11.79213 | 22.67632 | 9.721337 | 3.791821 | 1.740669 | -0.10877 | 0.27485 |
| 19 | 54872600 | 54874799 | 22.91284 | 40.22152 | 30.46049 | 13.47397 | 0.786567 | 0.649781 | 0.274861 |
| 14 | 42902600 | 42905199 | 30.05176 | 51.44329 | 38.62692 | 16.93846 | 0.794798 | 0.978729 | 0.274874 |
| 7 | 26828000 | 26832999 | 39.69035 | 63.39337 | 122.0419 | 51.25539 | -0.6022 | 2.089788 | 0.274879 |
| 7 | 43323400 | 43328599 | 81.86661 | 125.7879 | 219.2089 | 91.02096 | -0.50551 | 2.967613 | 0.274887 |
| 7 | 48201400 | 48203199 | 18.31856 | 32.96657 | 25.78873 | 11.25734 | 0.756503 | 0.408125 | 0.274891 |
| 17 | 39452800 | 39453199 | 20.43754 | 35.86801 | 28.95848 | 12.80705 | 0.694848 | 0.637705 | 0.274894 |
| 12 | 29024400 | 29026599 | 10.39761 | 18.3682 | 42.47438 | 18.79025 | -0.81252 | 0.521203 | 0.274905 |
| 7 | 41257400 | 41259399 | 21.05055 | 36.89448 | 31.80734 | 14.05108 | 0.600602 | 0.666121 | 0.274906 |
| 1 | 108343400 | 108344799 | 34.23172 | 57.07252 | 38.7241 | 16.83257 | 0.938823 | 1.234555 | 0.274909 |
| 18 | 83662000 | 83663999 | 12.23568 | 21.63844 | 42.95314 | 18.99799 | -0.59309 | 0.522502 | 0.274917 |
| 14 | 14082800 | 14086399 | 54.75835 | 87.83529 | 81.12242 | 34.28417 | 0.462484 | 1.960964 | 0.274919 |
| 6 | 134694800 | 134696199 | 6.645256 | 12.53727 | 29.69469 | 12.11574 | -0.75854 | 0.049927 | 0.274923 |
| 7 | 66678600 | 66679799 | 6.645256 | 12.53727 | 29.69469 | 12.11574 | -0.75854 | 0.049927 | 0.274923 |
| 1 | 156582000 | 156584199 | 11.47514 | 20.23856 | 41.43841 | 18.33796 | -0.63885 | 0.545789 | 0.274927 |
| 11 | 53659600 | 53662399 | 11.29328 | 19.64081 | 50.49898 | 22.18562 | -0.96871 | 0.760348 | 0.27494 |
| 14 | 59361400 | 59362799 | 18.71941 | 33.88487 | 24.76668 | 10.73267 | 0.860631 | 0.376802 | 0.274961 |
| 3 | 31022600 | 31023799 | 8.216303 | 14.76365 | 38.47551 | 16.81759 | -0.96983 | 0.374528 | 0.274968 |
| 10 | 47738200 | 47739199 | 17.225 | 32.34025 | 18.50873 | 7.615162 | 1.263606 | 0.144187 | 0.274972 |
| 5 | 14762400 | 14763199 | 6.880178 | 13.14704 | 30.15871 | 11.97769 | -0.67915 | -0.03807 | 0.274981 |
| X | 26830400 | 26831399 | 26.68823 | 46.45938 | 32.32092 | 14.208 | 0.90848 | 0.797031 | 0.27499 |
| 2 | 127089000 | 127090799 | 9.241129 | 16.277 | 43.56456 | 19.28393 | -1.02359 | 0.550185 | 0.275022 |
| 7 | 125274800 | 125278199 | 14.43833 | 24.8118 | 53.16 | 23.31777 | -0.71053 | 0.899847 | 0.275042 |
| 1 | 86182600 | 86187599 | 23.14547 | 37.8203 | 81.4253 | 35.00468 | -0.72914 | 1.483101 | 0.275048 |
| 14 | 60127000 | 60129999 | 14.99081 | 25.91776 | 51.63438 | 22.65276 | -0.60437 | 0.843618 | 0.275053 |
| 8 | 92402200 | 92404799 | 12.0088 | 20.94369 | 49.44435 | 21.7605 | -0.84596 | 0.720981 | 0.275054 |
| 9 | 58199600 | 58201999 | 12.0088 | 20.94369 | 49.44435 | 21.7605 | -0.84596 | 0.720981 | 0.275054 |
| 7 | 93419600 | 93420999 | 14.32946 | 27.06239 | 20.63273 | 8.400146 | 0.865877 | 0.086967 | 0.275068 |
| 7 | 32057000 | 32063799 | 131.9928 | 201.4849 | 182.9631 | 75.8755 | 0.426717 | 3.109633 | 0.275069 |
| 7 | 29286800 | 29288399 | 48.33128 | 78.02966 | 70.95231 | 30.26319 | 0.496014 | 1.750905 | 0.275073 |
| 3 | 23312400 | 23313999 | 21.63842 | 37.98533 | 31.01972 | 13.72162 | 0.67824 | 0.64203 | 0.275096 |
| 2 | 79837600 | 79839199 | 20.21409 | 35.67333 | 29.47928 | 13.04401 | 0.663315 | 0.568434 | 0.2751 |
| 18 | 66875200 | 66876999 | 20.21409 | 35.67333 | 29.47928 | 13.04401 | 0.663315 | 0.568434 | 0.2751 |
| 6 | 142740800 | 142743399 | 13.36607 | 23.34269 | 47.31677 | 20.84204 | -0.62695 | 0.705882 | 0.275104 |
| 4 | 88171200 | 88175999 | 46.20228 | 73.61679 | 129.2256 | 54.09001 | -0.47311 | 2.174403 | 0.275106 |

| | | | | | | | | | |
|---|---|---|---|---|---|---|---|---|---|
| 2 | 158015400 | 158019399 | 26.2532 | 42.62582 | 86.80555 | 37.21556 | -0.65612 | 1.592395 | 0.275111 |
| 19 | 22701000 | 22703599 | 29.14686 | 48.96194 | 43.27507 | 18.90031 | 0.558418 | 1.142467 | 0.275122 |
| 1 | 92448000 | 92450199 | 13.44712 | 23.28347 | 50.5365 | 22.17212 | -0.72675 | 0.822768 | 0.275137 |
| 8 | 45199800 | 45201999 | 31.72946 | 51.40949 | 92.84386 | 39.74289 | -0.48628 | 1.648131 | 0.275139 |
| 7 | 43910000 | 43929799 | 124.7662 | 186.2549 | 331.2057 | 136.5284 | -0.56974 | 3.555196 | 0.275148 |
| X | 61441800 | 61443999 | 40.78198 | 67.0139 | 55.50969 | 23.91785 | 0.648936 | 1.433289 | 0.275196 |
| 17 | 89455000 | 89459799 | 34.54365 | 58.96034 | 35.69432 | 15.64879 | 1.103554 | 1.018857 | 0.275211 |
| 4 | 76722200 | 76724199 | 27.85404 | 47.76097 | 39.23918 | 17.20926 | 0.6658 | 0.955319 | 0.275213 |
| 2 | 144928200 | 144930999 | 17.90175 | 30.61838 | 58.69297 | 25.73593 | -0.55228 | 0.961743 | 0.275218 |
| 10 | 46075800 | 46077599 | 20.77878 | 36.74596 | 27.03458 | 11.95732 | 0.83077 | 0.543128 | 0.275234 |
| 5 | 49894200 | 49895799 | 13.55822 | 23.67441 | 46.79422 | 20.60976 | -0.59059 | 0.710611 | 0.275246 |
| 10 | 44614000 | 44614599 | 15.00138 | 27.38573 | 24.18641 | 10.37074 | 0.598982 | 0.299682 | 0.275261 |
| 2 | 162533600 | 162535999 | 15.64871 | 26.96166 | 52.69281 | 23.11467 | -0.57788 | 0.879712 | 0.275262 |
| 19 | 33369000 | 33369799 | 7.282827 | 13.56344 | 32.69507 | 13.63752 | -0.81039 | 0.144672 | 0.275268 |
| 4 | 7294400 | 7296599 | 19.34186 | 34.26328 | 28.38324 | 12.54955 | 0.661741 | 0.520145 | 0.27528 |
| 14 | 82167600 | 82168399 | 11.35428 | 21.97837 | 17.97937 | 6.845212 | 0.852882 | -0.16238 | 0.275285 |
| 10 | 98611400 | 98613199 | 8.484728 | 15.72948 | 33.15029 | 13.92241 | -0.62573 | 0.184126 | 0.275291 |
| 5 | 149165600 | 149169999 | 36.1449 | 58.18741 | 105.7127 | 44.91475 | -0.5053 | 1.833306 | 0.275326 |
| 11 | 96075800 | 96081599 | 42.58026 | 67.94874 | 131.1753 | 55.02195 | -0.60771 | 2.121027 | 0.275336 |
| 3 | 115341200 | 115341999 | 5.537585 | 10.68294 | 26.3956 | 10.20145 | -0.7464 | -0.14724 | 0.275339 |
| 8 | 64371800 | 64372999 | 13.0673 | 25.20103 | 17.68458 | 6.841025 | 1.0503 | -0.0934 | 0.275378 |
| 3 | 65770600 | 65775999 | 41.48596 | 66.20433 | 123.7499 | 51.90938 | -0.56112 | 2.119042 | 0.275381 |
| 19 | 56902000 | 56906799 | 41.48596 | 66.20433 | 123.7499 | 51.90938 | -0.56112 | 2.119042 | 0.275381 |
| 1 | 54233200 | 54234399 | 11.82626 | 20.72483 | 44.13028 | 19.49275 | -0.69709 | 0.639547 | 0.275382 |
| 4 | 88908400 | 88909599 | 5.859998 | 11.23899 | 27.35486 | 10.75125 | -0.74856 | -0.08802 | 0.275391 |
| 14 | 61021800 | 61022399 | 5.378675 | 10.44084 | 25.06792 | 9.418181 | -0.66664 | -0.2509 | 0.275394 |
| 6 | 75338200 | 75339399 | 14.61545 | 27.26841 | 22.11345 | 9.195869 | 0.752975 | 0.168859 | 0.275396 |
| 19 | 27645000 | 27647999 | 15.93567 | 27.63637 | 57.74858 | 25.33754 | -0.67224 | 0.827205 | 0.275398 |
| 7 | 61459000 | 61459999 | 14.57765 | 27.25656 | 22.21929 | 9.206229 | 0.749946 | 0.156383 | 0.275411 |
| 1 | 37035400 | 37037799 | 13.33974 | 23.36063 | 48.33814 | 21.33869 | -0.65655 | 0.662724 | 0.275412 |
| 3 | 49221200 | 49222199 | 10.13122 | 19.56222 | 18.02091 | 6.941186 | 0.667104 | -0.13849 | 0.275418 |
| 15 | 12117200 | 12119199 | 13.47425 | 23.26204 | 51.57759 | 22.62685 | -0.75824 | 0.851337 | 0.275419 |
| 4 | 17825200 | 17827599 | 33.13385 | 55.35565 | 43.31475 | 18.85244 | 0.733775 | 1.213691 | 0.275423 |
| 16 | 97803200 | 97809399 | 132.2211 | 203.487 | 155.3266 | 64.50978 | 0.686059 | 2.947658 | 0.275426 |
| 15 | 24246800 | 24248799 | 23.36953 | 40.80875 | 33.20413 | 14.62479 | 0.683602 | 0.735779 | 0.275432 |
| 2 | 75327600 | 75328399 | 9.604064 | 18.63679 | 17.62945 | 6.634447 | 0.661094 | -0.22159 | 0.275452 |
| 5 | 5066600 | 5068799 | 20.54429 | 35.99286 | 29.621 | 13.08086 | 0.666982 | 0.662632 | 0.275467 |
| 1 | 8465600 | 8466399 | 12.72691 | 23.82333 | 21.39544 | 8.847745 | 0.613435 | 0.130824 | 0.275468 |
| X | 125457800 | 125460399 | 43.72174 | 72.2812 | 47.78667 | 20.65309 | 0.975704 | 1.363786 | 0.27548 |
| 2 | 50923600 | 50924599 | 4.815428 | 9.363261 | 24.04422 | 8.911403 | -0.74414 | -0.31008 | 0.27548 |
| 18 | 76012600 | 76018599 | 45.21426 | 72.16131 | 125.0798 | 52.47471 | -0.45232 | 2.120176 | 0.275481 |
| 7 | 130848000 | 130848799 | 10.98196 | 21.02922 | 19.54331 | 7.716037 | 0.621697 | -0.04509 | 0.275481 |
| 9 | 109602800 | 109604599 | 11.78651 | 20.8006 | 41.42626 | 18.33031 | -0.59878 | 0.541187 | 0.275481 |
| 7 | 121445600 | 121450199 | 30.09784 | 48.8771 | 88.7485 | 38.05369 | -0.49092 | 1.597643 | 0.275482 |
| 3 | 52039800 | 52041599 | 13.54835 | 23.76446 | 47.23383 | 20.87979 | -0.59855 | 0.63873 | 0.275493 |
| 7 | 96745000 | 96747199 | 25.37878 | 44.23891 | 33.1212 | 14.57202 | 0.802912 | 0.770481 | 0.275498 |
| 12 | 60947800 | 60948999 | 9.431759 | 17.26784 | 37.23028 | 15.90425 | -0.67674 | 0.269736 | 0.275501 |

| | | | | | | | | | |
|---|---|---|---|---|---|---|---|---|---|
| 8 | 21756400 | 21756999 | 11.53783 | 22.33081 | 17.97282 | 6.846835 | 0.87541 | -0.15818 | 0.275519 |
| 1 | 64571400 | 64572999 | 10.27336 | 18.09558 | 44.05039 | 19.49889 | -0.88795 | 0.55573 | 0.275529 |
| 7 | 42605200 | 42611999 | 164.8474 | 249.4588 | 201.374 | 83.29991 | 0.586165 | 3.294244 | 0.275531 |
| 2 | 129456600 | 129462199 | 58.15341 | 91.68824 | 158.1775 | 65.64123 | -0.46204 | 2.474374 | 0.275542 |
| 4 | 148066000 | 148081999 | 139.103 | 206.8627 | 350.1134 | 144.2461 | -0.50358 | 3.626732 | 0.27555 |
| 12 | 64672800 | 64674999 | 23.61588 | 41.01991 | 35.5147 | 15.59022 | 0.593776 | 0.81343 | 0.275564 |
| 17 | 38115600 | 38116999 | 22.99984 | 40.20024 | 32.63762 | 14.38441 | 0.686756 | 0.719385 | 0.275604 |
| 8 | 18938400 | 18940799 | 13.98129 | 24.38413 | 50.01183 | 22.01045 | -0.64423 | 0.731022 | 0.27563 |
| 9 | 70771800 | 70775599 | 19.23149 | 32.64568 | 63.06733 | 27.63222 | -0.56512 | 1.039727 | 0.275643 |
| 3 | 149364800 | 149365999 | 21.68507 | 37.74342 | 33.11173 | 14.5541 | 0.575136 | 0.772678 | 0.275657 |
| X | 24019600 | 24022399 | 64.1976 | 102.922 | 80.56596 | 34.02167 | 0.699773 | 1.990907 | 0.275669 |
| 9 | 38791400 | 38794599 | 56.25501 | 90.21934 | 83.14475 | 35.12897 | 0.465279 | 1.96934 | 0.275669 |
| 3 | 12663800 | 12665599 | 24.8089 | 42.89264 | 37.13323 | 16.2909 | 0.59275 | 0.866206 | 0.275675 |
| 12 | 71557800 | 71558599 | 5.68173 | 10.98735 | 26.00071 | 9.944379 | -0.67086 | -0.1842 | 0.275679 |
| 2 | 10157400 | 10158999 | 8.616117 | 15.56793 | 37.48605 | 16.28183 | -0.85059 | 0.341572 | 0.275684 |
| 7 | 97240600 | 97245799 | 42.08761 | 67.38208 | 117.4812 | 49.50635 | -0.45573 | 2.01862 | 0.275688 |
| 7 | 30747000 | 30763199 | 98.99081 | 149.4289 | 285.628 | 118.0599 | -0.65952 | 3.334569 | 0.27569 |
| 10 | 126587400 | 126589599 | 11.93761 | 21.04432 | 41.93917 | 18.5613 | -0.60043 | 0.554398 | 0.275692 |
| 4 | 90205800 | 90208199 | 40.79742 | 66.58507 | 59.33153 | 25.49557 | 0.538748 | 1.521599 | 0.275706 |
| 1 | 32778400 | 32780399 | 19.56447 | 34.74742 | 27.85309 | 12.29479 | 0.710864 | 0.501603 | 0.275708 |
| 12 | 82582600 | 82584399 | 24.13172 | 42.42433 | 28.69302 | 12.70583 | 0.948893 | 0.631815 | 0.275725 |
| 14 | 77863000 | 77864399 | 9.611717 | 17.59731 | 36.13247 | 15.43528 | -0.60648 | 0.270683 | 0.27573 |
| 15 | 68115200 | 68119599 | 20.83779 | 34.29762 | 77.25741 | 33.31167 | -0.7891 | 1.379499 | 0.275743 |
| 2 | 8137400 | 8138999 | 25.58614 | 44.7285 | 30.19545 | 13.31003 | 0.951527 | 0.72618 | 0.275754 |
| 2 | 30592600 | 30599799 | 43.40222 | 69.27666 | 124.8473 | 52.38505 | -0.50822 | 2.114105 | 0.275759 |
| X | 32424800 | 32425999 | 22.66659 | 39.785 | 24.83549 | 10.98479 | 1.063941 | 0.640844 | 0.275787 |
| 18 | 56768000 | 56768799 | 16.12594 | 29.57331 | 24.13531 | 10.28672 | 0.718324 | 0.284133 | 0.275788 |
| 8 | 16492000 | 16493799 | 23.35232 | 41.13165 | 27.42184 | 12.13834 | 0.970155 | 0.602946 | 0.2758 |
| 2 | 131713400 | 131718199 | 26.68048 | 43.4689 | 95.99687 | 41.22488 | -0.76902 | 1.541262 | 0.275816 |
| 19 | 8432400 | 8435399 | 22.23619 | 39.04211 | 27.93815 | 12.36047 | 0.867732 | 0.637215 | 0.27582 |
| 15 | 75109400 | 75111199 | 10.14358 | 18.21083 | 37.89008 | 16.57388 | -0.64815 | 0.392383 | 0.275822 |
| 18 | 24584600 | 24584999 | 9.968336 | 19.27395 | 17.14375 | 6.561425 | 0.726806 | -0.1618 | 0.275823 |
| 14 | 27286000 | 27287999 | 28.18472 | 47.85837 | 42.57897 | 18.65719 | 0.549431 | 1.046923 | 0.275828 |
| 11 | 52763400 | 52767999 | 45.71894 | 72.82113 | 130.0509 | 54.40905 | -0.49847 | 2.185334 | 0.275853 |
| 6 | 81528200 | 81529999 | 9.288374 | 17.02453 | 35.07703 | 14.96142 | -0.60976 | 0.259126 | 0.275906 |
| 14 | 42234000 | 42237199 | 32.45486 | 55.17729 | 39.66123 | 17.38241 | 0.856104 | 1.048833 | 0.275914 |
| 2 | 80565200 | 80568199 | 24.94262 | 41.00694 | 77.84343 | 33.5481 | -0.54382 | 1.404016 | 0.275915 |
| 17 | 39972800 | 39974799 | 24.66891 | 43.32973 | 28.55763 | 12.63855 | 0.985805 | 0.646709 | 0.275921 |
| 3 | 54525200 | 54526799 | 16.75218 | 28.84678 | 54.89272 | 24.0793 | -0.53987 | 0.891653 | 0.27594 |
| 7 | 116600800 | 116603599 | 30.08754 | 48.39655 | 103.7501 | 44.02878 | -0.74481 | 1.852007 | 0.275943 |
| 2 | 36881200 | 36884199 | 26.75873 | 46.4106 | 35.40148 | 15.53266 | 0.775542 | 0.846403 | 0.275955 |
| 15 | 82148000 | 82151199 | 16.03829 | 27.35025 | 57.45096 | 25.18533 | -0.68426 | 0.980087 | 0.27596 |
| 9 | 99800200 | 99801599 | 19.91477 | 34.95319 | 30.6981 | 13.57727 | 0.573735 | 0.633847 | 0.275971 |
| 14 | 53309400 | 53310999 | 9.298346 | 17.07779 | 35.1222 | 14.93894 | -0.60408 | 0.24863 | 0.275974 |
| 8 | 105034600 | 105038799 | 26.01248 | 42.43974 | 82.33987 | 35.37801 | -0.5811 | 1.512033 | 0.275977 |
| 2 | 161094000 | 161094999 | 7.563615 | 14.14591 | 31.15993 | 12.90471 | -0.67257 | 0.118432 | 0.275979 |
| 7 | 11704600 | 11707399 | 67.69715 | 107.6596 | 96.63206 | 40.34038 | 0.490044 | 2.247343 | 0.275983 |

| | | | | | | | | | |
|---|---|---|---|---|---|---|---|---|---|
| 11 | 27078200 | 27085399 | 164.9204 | 248.8969 | 197.6237 | 81.67903 | 0.606705 | 3.347938 | 0.275989 |
| X | 120362200 | 120362999 | 16.16316 | 29.85801 | 23.13048 | 9.761343 | 0.803194 | 0.24267 | 0.27599 |
| 4 | 21369600 | 21371199 | 13.36133 | 23.44561 | 45.62905 | 20.17707 | -0.56832 | 0.627998 | 0.275999 |
| X | 151267000 | 151267199 | 8.441328 | 16.47447 | 15.07565 | 5.354231 | 0.783482 | -0.44407 | 0.276 |
| 12 | 69134400 | 69134799 | 4.184229 | 8.160257 | 21.99364 | 7.896184 | -0.769 | -0.43509 | 0.276008 |
| 18 | 90282600 | 90284599 | 21.72334 | 38.03882 | 32.16129 | 14.19646 | 0.628453 | 0.683173 | 0.276017 |
| 4 | 121531600 | 121533399 | 33.68922 | 56.1006 | 48.69644 | 21.1513 | 0.584818 | 1.25656 | 0.276022 |
| 8 | 129843400 | 129846399 | 36.79655 | 61.14699 | 49.32365 | 21.39341 | 0.690046 | 1.286672 | 0.276035 |
| X | 90315600 | 90316799 | 18.18906 | 34.35071 | 17.49947 | 7.124886 | 1.442645 | 0.115277 | 0.276041 |
| 14 | 105413800 | 105415199 | 10.12125 | 17.83477 | 42.5115 | 18.8162 | -0.85732 | 0.548833 | 0.276043 |
| 7 | 139553800 | 139560399 | 40.36925 | 64.37306 | 128.0178 | 53.64254 | -0.65155 | 2.142734 | 0.276044 |
| 10 | 41735600 | 41737599 | 12.42262 | 21.73787 | 49.4214 | 21.80273 | -0.79172 | 0.671289 | 0.276061 |
| 4 | 137234200 | 137240199 | 24.06782 | 39.01848 | 89.42892 | 38.29753 | -0.82814 | 1.621257 | 0.276077 |
| 18 | 32120600 | 32123399 | 13.13184 | 22.76632 | 50.00588 | 21.94016 | -0.74339 | 0.809385 | 0.276136 |
| 4 | 71701000 | 71704199 | 51.98661 | 83.71987 | 75.49804 | 32.09386 | 0.50373 | 1.832965 | 0.27614 |
| 9 | 89439200 | 89439799 | 14.55046 | 26.66215 | 22.63787 | 9.659281 | 0.660028 | 0.270705 | 0.276157 |
| 4 | 6637200 | 6639199 | 25.70181 | 44.43968 | 37.09899 | 16.27591 | 0.645042 | 0.870216 | 0.276158 |
| 7 | 42908000 | 42909799 | 20.17379 | 36.95794 | 21.50683 | 9.179502 | 1.201368 | 0.315185 | 0.276161 |
| X | 148974400 | 148976399 | 31.78545 | 55.10696 | 30.68807 | 13.46445 | 1.227124 | 0.866991 | 0.276166 |
| 7 | 44966200 | 44969999 | 18.61284 | 31.06928 | 65.84701 | 28.64398 | -0.69866 | 1.201781 | 0.276171 |
| 19 | 51807200 | 51808599 | 15.6784 | 28.7764 | 24.09019 | 10.25592 | 0.682806 | 0.274957 | 0.276172 |
| 6 | 88614200 | 88615399 | 20.50635 | 36.51498 | 26.62222 | 11.72575 | 0.848899 | 0.489121 | 0.276174 |
| 9 | 34469200 | 34470799 | 8.856397 | 16.17824 | 34.49567 | 14.77921 | -0.66315 | 0.274619 | 0.276178 |
| 14 | 64314000 | 64315599 | 8.856397 | 16.17824 | 34.49567 | 14.77921 | -0.66315 | 0.274619 | 0.276178 |
| 11 | 116654600 | 116659199 | 41.01109 | 65.40634 | 126.356 | 52.95719 | -0.60956 | 2.138418 | 0.276186 |
| 10 | 68527000 | 68527799 | 5.39667 | 10.46237 | 26.20009 | 9.903609 | -0.73591 | -0.22892 | 0.276188 |
| 11 | 11607800 | 11608599 | 5.39667 | 10.46237 | 26.20009 | 9.903609 | -0.73591 | -0.22892 | 0.276188 |
| 3 | 62455600 | 62457799 | 21.26025 | 37.97679 | 24.86435 | 10.92247 | 1.005544 | 0.471212 | 0.27619 |
| 1 | 6187000 | 6188199 | 7.814572 | 14.75164 | 34.94928 | 14.24275 | -0.75879 | 0.066215 | 0.276204 |
| 1 | 156891400 | 156896799 | 49.64095 | 78.89683 | 138.6381 | 57.84098 | -0.47849 | 2.258626 | 0.276221 |
| 7 | 108003400 | 108008199 | 86.93595 | 136.4337 | 126.4629 | 52.44847 | 0.427845 | 2.577706 | 0.276226 |
| X | 121358200 | 121361399 | 61.82027 | 99.25387 | 78.39778 | 33.17886 | 0.689215 | 1.942555 | 0.276229 |
| 15 | 82417000 | 82418199 | 11.27282 | 20.11496 | 39.90205 | 17.54404 | -0.58497 | 0.443602 | 0.276231 |
| 17 | 79142400 | 79143199 | 7.140654 | 13.29642 | 33.7854 | 14.09633 | -0.88634 | 0.14625 | 0.276232 |
| 4 | 151225800 | 151229999 | 53.51553 | 83.77626 | 175.022 | 72.6032 | -0.74578 | 2.613072 | 0.276239 |
| 4 | 117105400 | 117106399 | 9.155014 | 16.7125 | 35.0044 | 15.00978 | -0.63855 | 0.281274 | 0.276244 |
| 11 | 57432600 | 57433799 | 9.155014 | 16.7125 | 35.0044 | 15.00978 | -0.63855 | 0.281274 | 0.276244 |
| 11 | 39954800 | 39957599 | 40.46825 | 66.012 | 59.27463 | 25.46607 | 0.527147 | 1.528491 | 0.276265 |
| 7 | 41308800 | 41310999 | 48.01295 | 77.51946 | 70.89652 | 30.24127 | 0.487785 | 1.749125 | 0.276268 |
| 17 | 17327400 | 17328199 | 10.17139 | 19.86926 | 12.43047 | 4.345095 | 1.344168 | -0.48268 | 0.276269 |
| 4 | 121115400 | 121116999 | 21.53866 | 37.71597 | 32.31335 | 14.26375 | 0.609593 | 0.683781 | 0.276274 |
| 19 | 17129400 | 17130599 | 12.96222 | 24.20479 | 21.29607 | 8.843048 | 0.637903 | 0.147043 | 0.276282 |
| 2 | 73099800 | 73105199 | 40.14962 | 63.80404 | 147.4188 | 61.49692 | -0.87283 | 2.25698 | 0.276283 |
| 17 | 48848200 | 48859599 | 100.1743 | 152.2407 | 261.796 | 108.4359 | -0.49886 | 3.195698 | 0.276289 |
| 7 | 108078600 | 108083399 | 126.3601 | 193.4602 | 175.6801 | 72.90521 | 0.43004 | 3.047572 | 0.276297 |
| 14 | 4095600 | 4097799 | 13.82728 | 24.1971 | 47.2797 | 20.85883 | -0.57417 | 0.676547 | 0.276302 |
| 8 | 70895200 | 70898999 | 24.08636 | 39.83204 | 74.06324 | 32.01645 | -0.51157 | 1.328722 | 0.276308 |

| | | | | | | | | | |
|---|---|---|---|---|---|---|---|---|---|
| 19 | 9220800 | 9221999 | 17.96842 | 32.56545 | 24.79237 | 10.72731 | 0.80349 | 0.363881 | 0.276312 |
| 11 | 35795200 | 35795999 | 6.607662 | 12.57357 | 30.55257 | 12.24199 | -0.77283 | -0.01125 | 0.276323 |
| 7 | 7120400 | 7123199 | 27.48304 | 46.81516 | 41.48593 | 18.18496 | 0.555679 | 1.014505 | 0.276326 |
| 5 | 91950600 | 91951199 | 8.56235 | 16.69567 | 15.58949 | 5.61006 | 0.737724 | -0.39576 | 0.27633 |
| 14 | 26482200 | 26486999 | 95.85289 | 149.4879 | 136.2417 | 56.53827 | 0.445203 | 2.693414 | 0.276332 |
| 2 | 46066400 | 46067999 | 15.30062 | 28.7904 | 20.47582 | 8.393197 | 0.957282 | 0.118611 | 0.276354 |
| 7 | 7009800 | 7016199 | 85.8628 | 135.2668 | 121.239 | 50.307 | 0.480474 | 2.500764 | 0.27636 |
| 16 | 21035400 | 21043799 | 41.47727 | 65.82169 | 143.3016 | 59.69569 | -0.78916 | 2.298381 | 0.276376 |
| 10 | 108191600 | 108196199 | 54.35507 | 85.93303 | 150.3608 | 62.47098 | -0.47926 | 2.408384 | 0.276385 |
| 6 | 143126400 | 143129599 | 14.32755 | 25.03284 | 48.37667 | 21.31479 | -0.55836 | 0.707137 | 0.276399 |
| 2 | 58732800 | 58736399 | 20.86965 | 34.66941 | 82.08179 | 35.59404 | -0.85855 | 1.272765 | 0.2764 |
| 5 | 130953200 | 130956999 | 35.72637 | 58.97801 | 53.71351 | 23.19152 | 0.514873 | 1.36249 | 0.276403 |
| 18 | 77920600 | 77923399 | 18.07135 | 30.83116 | 58.65735 | 25.71512 | -0.54202 | 0.98516 | 0.27641 |
| 15 | 100753200 | 100757999 | 34.38036 | 55.33694 | 103.8706 | 44.12069 | -0.55258 | 1.835803 | 0.276411 |
| 10 | 75658000 | 75659199 | 16.75243 | 29.99693 | 26.20631 | 11.48855 | 0.594142 | 0.429775 | 0.276417 |
| 15 | 97916000 | 97919199 | 17.18399 | 29.12412 | 60.13806 | 26.33621 | -0.66076 | 1.042803 | 0.27642 |
| 12 | 69698400 | 69701199 | 106.7304 | 166.4545 | 132.5914 | 55.02334 | 0.639277 | 2.698195 | 0.27643 |
| 17 | 14463200 | 14466399 | 22.04819 | 36.37123 | 76.57737 | 33.04975 | -0.69076 | 1.353163 | 0.276432 |
| 1 | 113273800 | 113275599 | 18.40619 | 33.0503 | 26.39006 | 11.5419 | 0.725372 | 0.42426 | 0.27644 |
| 9 | 13823000 | 13824399 | 7.674058 | 14.44612 | 31.33446 | 12.83485 | -0.6381 | 0.076043 | 0.27644 |
| 7 | 59920000 | 59933599 | 179.0617 | 270.1563 | 204.3479 | 84.45004 | 0.676268 | 3.358249 | 0.276445 |
| 5 | 149866400 | 149867799 | 8.010907 | 14.93558 | 32.18569 | 13.40045 | -0.64782 | 0.143385 | 0.276453 |
| 19 | 23838000 | 23839599 | 8.010907 | 14.93558 | 32.18569 | 13.40045 | -0.64782 | 0.143385 | 0.276453 |
| X | 96264800 | 96266599 | 22.86054 | 40.91458 | 23.12914 | 10.14364 | 1.217638 | 0.467667 | 0.276464 |
| 5 | 109193400 | 109195599 | 42.01747 | 68.42868 | 61.53112 | 26.42 | 0.523529 | 1.556786 | 0.276477 |
| 2 | 156695000 | 156698199 | 13.24619 | 23.14932 | 46.25847 | 20.38529 | -0.60608 | 0.696112 | 0.276478 |
| 16 | 75530800 | 75531999 | 9.808748 | 19.07717 | 15.85601 | 5.864836 | 0.867617 | -0.28561 | 0.276479 |
| 3 | 54539800 | 54548599 | 124.2037 | 186.4024 | 305.5969 | 126.1106 | -0.44564 | 3.454965 | 0.276481 |
| 6 | 68211800 | 68214399 | 43.711 | 71.18233 | 61.63858 | 26.46554 | 0.577743 | 1.561927 | 0.276487 |
| 14 | 16240600 | 16241999 | 14.99461 | 25.50674 | 57.86814 | 25.36312 | -0.79532 | 1.000868 | 0.276502 |
| 4 | 15496200 | 15496999 | 11.09874 | 19.84855 | 39.40188 | 17.29159 | -0.58414 | 0.427012 | 0.276516 |
| 14 | 13165600 | 13166999 | 15.38409 | 28.24909 | 24.15832 | 10.27855 | 0.652958 | 0.270626 | 0.276527 |
| 9 | 96003600 | 96010399 | 48.15753 | 76.56995 | 140.0158 | 58.43933 | -0.53529 | 2.245279 | 0.276548 |
| 9 | 54455400 | 54457199 | 8.982543 | 16.03758 | 37.95093 | 16.67798 | -0.83667 | 0.422466 | 0.276562 |
| 11 | 80319200 | 80320399 | 7.702989 | 14.00543 | 35.51003 | 15.30332 | -0.91658 | 0.295335 | 0.276591 |
| 17 | 39770600 | 39781599 | 211.6793 | 313.5761 | 244.4793 | 100.6854 | 0.608861 | 3.700864 | 0.276609 |
| 12 | 16059200 | 16063599 | 27.44355 | 44.7939 | 82.65352 | 35.51837 | -0.5085 | 1.509104 | 0.276615 |
| 7 | 10990400 | 10992999 | 84.33022 | 132.4968 | 116.8719 | 48.47801 | 0.500569 | 2.55274 | 0.276623 |
| 8 | 27664000 | 27668599 | 31.77334 | 51.27231 | 97.67292 | 41.63414 | -0.56931 | 1.75148 | 0.276648 |
| 17 | 18633800 | 18634399 | 20.75676 | 36.38769 | 31.70007 | 14.00634 | 0.585444 | 0.6593 | 0.276658 |
| 7 | 17005000 | 17012599 | 108.8738 | 169.1105 | 143.8305 | 59.72351 | 0.540346 | 2.770888 | 0.27668 |
| 5 | 103707600 | 103709599 | 13.99201 | 26.01299 | 22.72955 | 9.508912 | 0.639329 | 0.187376 | 0.276682 |
| 5 | 33570800 | 33572199 | 8.569414 | 15.62766 | 34.48807 | 14.80711 | -0.71423 | 0.282998 | 0.276693 |
| 10 | 48499000 | 48501199 | 19.5046 | 35.0611 | 24.46992 | 10.69221 | 0.918904 | 0.419678 | 0.276694 |
| 3 | 12509600 | 12510399 | 9.759155 | 19.00599 | 15.9665 | 5.838751 | 0.867739 | -0.3237 | 0.276695 |
| 1 | 30157400 | 30158599 | 16.16266 | 29.13349 | 25.59629 | 11.15093 | 0.591852 | 0.383271 | 0.276704 |
| 13 | 12716600 | 12718399 | 11.37508 | 19.91418 | 45.1908 | 19.94428 | -0.78848 | 0.65444 | 0.276708 |

| | | | | | | | | | |
|---|---|---|---|---|---|---|---|---|---|
| 4 | 135577800 | 135580199 | 10.61596 | 18.604 | 44.66383 | 19.72857 | -0.86911 | 0.633223 | 0.276713 |
| 4 | 65316000 | 65317199 | 14.4839 | 27.47256 | 20.21539 | 8.155336 | 0.928421 | 0.059507 | 0.276716 |
| 3 | 77078800 | 77079999 | 13.10308 | 24.3715 | 20.33788 | 8.501761 | 0.706088 | 0.170755 | 0.276725 |
| 13 | 103541800 | 103543799 | 18.90834 | 34.21635 | 24.76026 | 10.73401 | 0.874637 | 0.380023 | 0.276733 |
| 3 | 16147200 | 16148999 | 18.89622 | 33.59138 | 27.93193 | 12.32005 | 0.659016 | 0.491749 | 0.276741 |
| 14 | 42066200 | 42067399 | 18.09459 | 32.92933 | 24.36896 | 10.48999 | 0.849 | 0.341906 | 0.276754 |
| 17 | 55964600 | 55966399 | 13.35752 | 23.4511 | 46.14581 | 20.41454 | -0.58419 | 0.620131 | 0.276766 |
| 7 | 62241800 | 62253999 | 200.6836 | 300.1949 | 230.6863 | 95.13013 | 0.642545 | 3.526523 | 0.276778 |
| 15 | 97364800 | 97371399 | 54.97569 | 86.88986 | 153.6127 | 63.81303 | -0.49448 | 2.414881 | 0.276784 |
| 2 | 33566600 | 33571399 | 21.80698 | 35.43592 | 86.48119 | 37.09902 | -0.91514 | 1.568368 | 0.276793 |
| X | 33472200 | 33478199 | 97.58683 | 154.2831 | 100.9164 | 41.92846 | 0.93829 | 2.429425 | 0.276808 |
| 12 | 52867400 | 52868799 | 22.77186 | 39.53454 | 33.76176 | 14.81616 | 0.613625 | 0.815904 | 0.276809 |
| X | 154615000 | 154616599 | 17.747 | 33.12137 | 20.20293 | 8.396935 | 1.161532 | 0.18993 | 0.276814 |
| 12 | 114987400 | 114990599 | 57.50825 | 92.88563 | 64.46406 | 27.51104 | 0.885616 | 1.752276 | 0.276818 |
| 3 | 41755000 | 41758399 | 49.43879 | 80.27528 | 60.85 | 26.08886 | 0.765986 | 1.619494 | 0.276819 |
| 7 | 108418200 | 108422999 | 49.7944 | 80.95758 | 58.62675 | 25.15749 | 0.833354 | 1.5953 | 0.276828 |
| 1 | 81869600 | 81872199 | 28.04775 | 47.76737 | 40.89665 | 17.92637 | 0.605076 | 1.016838 | 0.276842 |
| 2 | 169847400 | 169852199 | 47.69137 | 75.95314 | 131.0449 | 54.81527 | -0.449 | 2.192157 | 0.276844 |
| 5 | 45780200 | 45782799 | 58.34319 | 93.82281 | 75.67368 | 32.10217 | 0.661884 | 1.885415 | 0.276866 |
| 8 | 118704800 | 118705599 | 10.73657 | 18.8915 | 42.02646 | 18.60693 | -0.75889 | 0.570629 | 0.276868 |
| 12 | 65831600 | 65834199 | 51.02002 | 82.12015 | 75.68488 | 32.14883 | 0.471476 | 1.848198 | 0.276881 |
| 12 | 83538000 | 83539199 | 12.60475 | 23.91566 | 20.91612 | 8.432498 | 0.6819 | 0.041161 | 0.276885 |
| 13 | 85874200 | 85878399 | 28.26248 | 48.77948 | 35.83021 | 15.71842 | 0.828305 | 0.894894 | 0.276885 |
| 9 | 12821600 | 12823399 | 17.31989 | 31.0945 | 26.45461 | 11.57177 | 0.634123 | 0.416986 | 0.276888 |
| 17 | 80790200 | 80790999 | 5.132732 | 9.890218 | 26.34471 | 10.21638 | -0.85855 | -0.14218 | 0.27691 |
| 17 | 81251000 | 81253199 | 13.51175 | 23.59013 | 48.92051 | 21.54445 | -0.65985 | 0.712194 | 0.276913 |
| 7 | 90229800 | 90230399 | 8.566644 | 16.68836 | 16.14822 | 5.883036 | 0.671342 | -0.35067 | 0.276916 |
| 2 | 11896000 | 11896599 | 5.523092 | 10.70648 | 25.5592 | 9.665458 | -0.66738 | -0.22488 | 0.276917 |
| 9 | 41837000 | 41839199 | 9.779696 | 17.74616 | 37.01085 | 15.98957 | -0.63978 | 0.324025 | 0.276926 |
| 10 | 63055600 | 63056399 | 8.47287 | 15.80986 | 33.3187 | 13.85288 | -0.61436 | 0.145231 | 0.276932 |
| X | 34819000 | 34823599 | 88.80093 | 140.4675 | 108.2874 | 45.00307 | 0.702204 | 2.412059 | 0.276956 |
| 16 | 34296800 | 34298399 | 10.70501 | 18.88558 | 43.51111 | 19.25412 | -0.80817 | 0.539528 | 0.276958 |
| 9 | 51959200 | 51960599 | 9.599576 | 17.09366 | 38.41843 | 16.92137 | -0.76533 | 0.443698 | 0.276986 |
| 1 | 15874600 | 15876399 | 14.84402 | 25.66876 | 51.09219 | 22.41503 | -0.60295 | 0.841263 | 0.276996 |
| 11 | 35327200 | 35329399 | 14.63601 | 25.439 | 50.03867 | 21.97505 | -0.58466 | 0.783965 | 0.276999 |
| X | 88448600 | 88453399 | 70.64244 | 112.968 | 83.86637 | 35.28338 | 0.77196 | 2.075875 | 0.277001 |
| 4 | 62204400 | 62206599 | 10.33969 | 18.41971 | 39.38308 | 17.33978 | -0.69384 | 0.445465 | 0.277011 |
| 2 | 164056800 | 164057599 | 4.864722 | 9.401542 | 27.07111 | 10.38403 | -0.95537 | -0.18233 | 0.277014 |
| 3 | 55163200 | 55164599 | 5.989788 | 11.44647 | 31.70496 | 12.58983 | -0.94666 | -0.03657 | 0.277026 |
| 2 | 94179000 | 94179999 | 5.413623 | 10.44718 | 26.43006 | 10.20528 | -0.77953 | -0.15416 | 0.277043 |
| 16 | 90681200 | 90682599 | 5.413623 | 10.44718 | 26.43006 | 10.20528 | -0.77953 | -0.15416 | 0.277043 |
| 7 | 62179000 | 62182199 | 43.88672 | 72.63206 | 46.6446 | 20.17448 | 1.017535 | 1.351412 | 0.277052 |
| 7 | 54584800 | 54588999 | 61.10476 | 97.76379 | 87.62163 | 36.88889 | 0.501443 | 2.048534 | 0.277083 |
| 16 | 52340800 | 52341399 | 10.87037 | 20.82801 | 18.0933 | 7.131273 | 0.720954 | -0.0549 | 0.277112 |
| 7 | 31407200 | 31410599 | 63.6434 | 102.0577 | 80.6934 | 34.08778 | 0.685744 | 1.982422 | 0.277127 |
| X | 136284400 | 136287199 | 23.011 | 41.33161 | 22.68975 | 9.920934 | 1.262707 | 0.447275 | 0.277145 |
| 17 | 54138400 | 54140799 | 35.3546 | 58.68966 | 50.95691 | 22.08592 | 0.58413 | 1.296009 | 0.277149 |

| | | | | | | | | | |
|---|---|---|---|---|---|---|---|---|---|
| 12 | 20743400 | 20752199 | 290.9416 | 417.9081 | 354.81 | 146.4834 | 0.443164 | 4.183245 | 0.277165 |
| 4 | 26184800 | 26188199 | 50.88651 | 81.8784 | 74.64836 | 31.69347 | 0.486474 | 1.856055 | 0.277188 |
| 9 | 118771200 | 118772199 | 6.420838 | 12.3861 | 29.45378 | 11.38545 | -0.69316 | -0.13075 | 0.277198 |
| 4 | 79350000 | 79350599 | 11.08959 | 21.23332 | 19.49071 | 7.697341 | 0.638803 | -0.0439 | 0.277213 |
| 1 | 166234800 | 166240399 | 45.50274 | 72.68868 | 127.5008 | 53.5603 | -0.46802 | 2.091967 | 0.27723 |
| 16 | 19229400 | 19231999 | 37.1331 | 61.10019 | 55.33533 | 23.85631 | 0.521312 | 1.407792 | 0.277241 |
| 6 | 87357400 | 87361599 | 23.02376 | 37.93223 | 74.1375 | 31.97951 | -0.5844 | 1.371944 | 0.277245 |
| 2 | 70480400 | 70481399 | 14.10942 | 26.0605 | 21.43455 | 9.047364 | 0.717268 | 0.222801 | 0.277256 |
| 5 | 69582200 | 69586999 | 131.4598 | 200.3345 | 181.0694 | 75.06037 | 0.43155 | 3.142802 | 0.277271 |
| 17 | 36205400 | 36208999 | 13.33745 | 22.9831 | 53.16096 | 23.32015 | -0.81967 | 0.868653 | 0.277273 |
| 8 | 4215600 | 4217199 | 10.29669 | 18.0904 | 44.59454 | 19.7403 | -0.90714 | 0.588207 | 0.277282 |
| 7 | 8103600 | 8107599 | 35.82079 | 60.53985 | 39.6402 | 17.34478 | 0.989846 | 1.111206 | 0.277293 |
| 16 | 25850800 | 25852799 | 28.83655 | 50.03668 | 33.67174 | 14.77392 | 0.955681 | 0.850484 | 0.277299 |
| 6 | 145959400 | 145961399 | 12.28886 | 21.68285 | 44.0283 | 19.48243 | -0.62675 | 0.551949 | 0.277302 |
| 1 | 110345200 | 110346199 | 14.78168 | 27.11939 | 23.64167 | 10.07061 | 0.624048 | 0.268503 | 0.277304 |
| 19 | 12903200 | 12907399 | 38.91006 | 64.32218 | 52.16372 | 22.54378 | 0.6819 | 1.354833 | 0.277305 |
| 10 | 119914400 | 119925999 | 117.7483 | 177.084 | 299.0943 | 123.4962 | -0.48589 | 3.412209 | 0.277313 |
| 5 | 6053400 | 6054999 | 21.07195 | 37.65081 | 24.86932 | 10.92211 | 0.993093 | 0.468138 | 0.277336 |
| 3 | 148415400 | 148416199 | 6.75602 | 12.85893 | 28.83021 | 11.54608 | -0.6564 | -0.0131 | 0.277346 |
| 6 | 55243800 | 55244799 | 6.75602 | 12.85893 | 28.83021 | 11.54608 | -0.6564 | -0.0131 | 0.277346 |
| 13 | 120507800 | 120508599 | 6.75602 | 12.85893 | 28.83021 | 11.54608 | -0.6564 | -0.0131 | 0.277346 |
| 1 | 174050800 | 174052599 | 20.1779 | 35.51856 | 29.40358 | 13.01717 | 0.65919 | 0.593545 | 0.277352 |
| 13 | 27883400 | 27886399 | 53.49855 | 86.00162 | 78.64454 | 33.34535 | 0.480523 | 1.891841 | 0.277356 |
| 18 | 37601600 | 37603399 | 36.32067 | 59.93108 | 53.68446 | 23.17204 | 0.538667 | 1.370849 | 0.277358 |
| 7 | 8282200 | 8304199 | 369.8234 | 526.0271 | 382.3579 | 158.1075 | 0.652991 | 4.331077 | 0.277363 |
| 19 | 8050800 | 8055199 | 52.75274 | 85.20403 | 68.18102 | 29.09703 | 0.680657 | 1.744693 | 0.27737 |
| 17 | 38851600 | 38858199 | 99.12806 | 154.9087 | 133.7591 | 55.49641 | 0.525307 | 2.661679 | 0.277394 |
| 12 | 61446400 | 61446999 | 9.313091 | 18.10172 | 17.13831 | 6.37157 | 0.675921 | -0.2683 | 0.277395 |
| 10 | 92406400 | 92406799 | 8.673273 | 16.92706 | 14.38216 | 5.108395 | 0.888234 | -0.44447 | 0.277396 |
| 7 | 85678800 | 85682799 | 36.15962 | 59.61652 | 53.65635 | 23.15189 | 0.531397 | 1.380443 | 0.277397 |
| 12 | 84931400 | 84939599 | 53.72237 | 84.41285 | 175.9974 | 73.00376 | -0.73876 | 2.543282 | 0.277405 |
| 12 | 109453000 | 109454199 | 9.453296 | 17.38299 | 36.79954 | 15.62714 | -0.64404 | 0.246347 | 0.277414 |
| 8 | 71708000 | 71711799 | 16.02669 | 27.38142 | 55.8438 | 24.48446 | -0.64136 | 0.961804 | 0.277425 |
| 14 | 3786600 | 3788999 | 26.3301 | 46.15269 | 28.50278 | 12.59242 | 1.079274 | 0.68763 | 0.277426 |
| 11 | 102555400 | 102557999 | 51.0112 | 80.46988 | 156.2416 | 64.84232 | -0.63166 | 2.460023 | 0.277431 |
| 9 | 38480400 | 38481999 | 23.14727 | 40.58438 | 30.29267 | 13.38617 | 0.807203 | 0.667247 | 0.277431 |
| X | 54605400 | 54609399 | 46.58184 | 76.41747 | 52.2994 | 22.5188 | 0.92265 | 1.466844 | 0.277436 |
| 12 | 58314200 | 58316999 | 17.95982 | 30.04933 | 65.36444 | 28.47599 | -0.73612 | 1.172806 | 0.277446 |
| 1 | 119980800 | 119985199 | 25.67934 | 41.95008 | 80.64246 | 34.66494 | -0.56648 | 1.49225 | 0.277461 |
| X | 114458200 | 114461799 | 69.98962 | 111.4141 | 95.66554 | 39.99089 | 0.555409 | 2.219466 | 0.277461 |
| Y | 2275600 | 2277199 | 12.21828 | 23.29468 | 11.8715 | 4.737424 | 1.465605 | -0.02396 | 0.277462 |
| 3 | 9592600 | 9593799 | 9.103154 | 16.34112 | 36.93256 | 16.15656 | -0.76623 | 0.386662 | 0.277496 |
| 10 | 116262000 | 116265999 | 59.08587 | 92.7962 | 166.8901 | 69.22175 | -0.52633 | 2.554117 | 0.277498 |
| 2 | 172436400 | 172438599 | 13.42139 | 23.30356 | 49.48764 | 21.71884 | -0.69461 | 0.79359 | 0.277505 |
| 7 | 73723000 | 73726199 | 30.3901 | 49.35542 | 92.09271 | 39.48981 | -0.53011 | 1.596519 | 0.277513 |
| 3 | 134854000 | 134855399 | 14.61226 | 27.00486 | 23.13883 | 9.759059 | 0.658961 | 0.22584 | 0.277514 |
| 2 | 82440800 | 82443799 | 33.48343 | 56.09195 | 46.43564 | 20.24547 | 0.652638 | 1.188838 | 0.277518 |

| | | | | | | | | | |
|---|---|---|---|---|---|---|---|---|---|
| 5 | 4993200 | 4993999 | 7.727312 | 14.41365 | 31.64544 | 13.16498 | -0.67308 | 0.138749 | 0.277523 |
| 7 | 26492200 | 26493199 | 20.97566 | 36.83453 | 31.0727 | 13.74905 | 0.631611 | 0.632919 | 0.277543 |
| 1 | 4510600 | 4511799 | 14.14212 | 24.71313 | 48.37608 | 21.31708 | -0.57685 | 0.702484 | 0.277548 |
| 9 | 51181800 | 51182799 | 8.005668 | 14.99133 | 31.74403 | 13.1189 | -0.61388 | 0.115033 | 0.277549 |
| 11 | 68310000 | 68311799 | 8.005668 | 14.99133 | 31.74403 | 13.1189 | -0.61388 | 0.115033 | 0.277549 |
| X | 146952400 | 146955199 | 21.95353 | 38.89209 | 27.11654 | 11.98934 | 0.909176 | 0.534766 | 0.277555 |
| 1 | 52932600 | 52933599 | 8.880642 | 16.15069 | 34.96845 | 15.06516 | -0.69027 | 0.302092 | 0.277562 |
| 12 | 78953200 | 78953999 | 5.988255 | 11.52496 | 26.92903 | 10.47881 | -0.67666 | -0.12053 | 0.277573 |
| 17 | 56381000 | 56382199 | 6.200545 | 11.73325 | 29.20969 | 11.84282 | -0.82151 | 0.023546 | 0.277578 |
| 17 | 69724000 | 69724999 | 6.200545 | 11.73325 | 29.20969 | 11.84282 | -0.82151 | 0.023546 | 0.277578 |
| 19 | 8480000 | 8483199 | 27.35743 | 47.02172 | 35.00499 | 15.35446 | 0.807885 | 0.92595 | 0.277586 |
| 5 | 56929800 | 56930599 | 10.62046 | 20.61873 | 16.90062 | 6.342761 | 0.868832 | -0.22218 | 0.277595 |
| 12 | 66254000 | 66256799 | 36.29486 | 60.91346 | 43.052 | 18.792 | 0.879867 | 1.175988 | 0.277596 |
| 6 | 123031600 | 123033199 | 15.62439 | 29.67459 | 18.28822 | 7.357495 | 1.18484 | 0.05565 | 0.277618 |
| 3 | 112445800 | 112447399 | 20.11338 | 36.26801 | 23.94951 | 10.42874 | 1.001561 | 0.405887 | 0.277623 |
| 4 | 112376400 | 112377799 | 30.25441 | 51.33973 | 41.94348 | 18.37515 | 0.671896 | 1.058539 | 0.277633 |
| 11 | 28222400 | 28223999 | 18.22443 | 32.26185 | 27.47188 | 12.14825 | 0.621696 | 0.51733 | 0.277641 |
| 3 | 26838200 | 26838799 | 6.362637 | 11.64601 | 34.56259 | 14.76952 | -1.13299 | 0.244942 | 0.277645 |
| 1 | 133153400 | 133155999 | 11.98411 | 20.95984 | 46.24345 | 20.39196 | -0.74829 | 0.674605 | 0.27765 |
| 19 | 36386600 | 36387799 | 10.22825 | 18.13229 | 39.39035 | 17.41374 | -0.72055 | 0.477752 | 0.277655 |
| 13 | 22787200 | 22790999 | 67.48321 | 107.8217 | 89.35988 | 37.54807 | 0.612069 | 2.098438 | 0.277656 |
| 7 | 11172400 | 11174399 | 53.48251 | 86.13106 | 65.658 | 27.91193 | 0.745679 | 1.831247 | 0.277672 |
| 15 | 33601600 | 33604399 | 31.44343 | 53.0578 | 44.20685 | 19.33367 | 0.643546 | 1.114635 | 0.277672 |
| 15 | 100600800 | 100603199 | 12.35788 | 21.45601 | 49.46683 | 21.70909 | -0.81239 | 0.790184 | 0.277679 |
| 7 | 42221400 | 42229799 | 192.5168 | 288.92 | 219.8292 | 90.71639 | 0.661082 | 3.464289 | 0.277706 |
| 11 | 39376400 | 39378199 | 27.24419 | 47.5244 | 30.14607 | 13.26902 | 1.040774 | 0.764968 | 0.277728 |
| 7 | 116780800 | 116781999 | 7.030818 | 13.43114 | 30.14972 | 11.9812 | -0.64883 | -0.03418 | 0.277729 |
| 9 | 72437600 | 72440399 | 14.84857 | 25.24003 | 57.83327 | 25.34635 | -0.80965 | 1.007719 | 0.277743 |
| 15 | 93280200 | 93282999 | 29.58053 | 49.65523 | 44.86913 | 19.58776 | 0.526706 | 1.15418 | 0.277754 |
| X | 148906600 | 148909999 | 49.72121 | 80.5443 | 65.17944 | 27.89667 | 0.668828 | 1.669805 | 0.277766 |
| 1 | 43318400 | 43320999 | 33.72891 | 56.29399 | 47.59364 | 20.70185 | 0.622484 | 1.228262 | 0.277779 |
| 13 | 83877400 | 83881999 | 36.3926 | 58.32932 | 114.3012 | 48.24798 | -0.62199 | 1.974767 | 0.277789 |
| 10 | 121168000 | 121173799 | 58.08112 | 91.87708 | 156.1023 | 64.87546 | -0.43627 | 2.397778 | 0.277798 |
| 11 | 17754800 | 17755599 | 7.360769 | 13.92859 | 30.36646 | 12.31208 | -0.63257 | 0.035927 | 0.27786 |
| 9 | 45761000 | 45762599 | 4.203372 | 8.075314 | 28.41903 | 11.1156 | -1.26765 | -0.11892 | 0.277864 |
| 2 | 163510200 | 163512999 | 14.37451 | 25.10069 | 51.06705 | 22.49219 | -0.63226 | 0.720483 | 0.277868 |
| 2 | 57996200 | 57999999 | 17.78373 | 30.20149 | 59.66362 | 26.14444 | -0.5971 | 1.02666 | 0.277879 |
| 11 | 90207600 | 90210999 | 18.32548 | 30.9252 | 61.21946 | 26.77477 | -0.60043 | 1.086194 | 0.27788 |
| 17 | 44815600 | 44817999 | 19.9812 | 33.45132 | 65.03615 | 28.34363 | -0.57483 | 1.172052 | 0.277881 |
| 7 | 38003400 | 38005599 | 19.77726 | 35.30997 | 26.80458 | 11.77978 | 0.793269 | 0.469459 | 0.277901 |
| 12 | 8638200 | 8640399 | 17.69155 | 29.92266 | 59.54151 | 26.05868 | -0.60759 | 1.063829 | 0.27791 |
| 7 | 55600600 | 55602599 | 36.70397 | 60.98563 | 49.86702 | 21.62725 | 0.67051 | 1.289045 | 0.277997 |
| 8 | 57777800 | 57783399 | 47.16867 | 75.08182 | 134.7807 | 56.33755 | -0.50693 | 2.207854 | 0.278003 |
| 5 | 109147200 | 109151999 | 58.31066 | 94.05949 | 67.30939 | 28.67728 | 0.839419 | 1.791259 | 0.278005 |
| 18 | 13175000 | 13177599 | 11.94919 | 21.1393 | 41.92661 | 18.54264 | -0.59143 | 0.516729 | 0.278008 |
| 4 | 132059400 | 132063599 | 16.84815 | 28.24368 | 63.5977 | 27.73999 | -0.78561 | 1.148733 | 0.27801 |
| 9 | 10366000 | 10367599 | 11.84415 | 20.69018 | 47.31268 | 20.84366 | -0.79976 | 0.694424 | 0.278023 |

| | | | | | | | | | |
|---|---|---|---|---|---|---|---|---|---|
| 2 | 164009400 | 164010999 | 20.73551 | 34.39843 | 69.63976 | 30.16953 | -0.63328 | 1.282287 | 0.278032 |
| 7 | 15441600 | 15457999 | 306.8529 | 443.3748 | 334.3768 | 137.9323 | 0.622436 | 4.097985 | 0.278035 |
| 7 | 15601400 | 15602399 | 21.36194 | 37.2514 | 30.15221 | 13.26902 | 0.691009 | 0.740524 | 0.278049 |
| 6 | 54968800 | 54970399 | 8.536522 | 15.63745 | 34.53717 | 14.74271 | -0.70999 | 0.255154 | 0.278059 |
| 3 | 57289800 | 57291999 | 32.6557 | 54.30867 | 49.26071 | 21.37773 | 0.521398 | 1.265444 | 0.278069 |
| 12 | 106253800 | 106263199 | 74.03697 | 114.547 | 203.7663 | 84.63948 | -0.52747 | 2.831335 | 0.278079 |
| 12 | 106057600 | 106073799 | 155.6255 | 228.5805 | 396.3263 | 163.2268 | -0.55515 | 3.837749 | 0.278107 |
| 9 | 114767000 | 114778199 | 108.1655 | 163.6258 | 274.359 | 113.4781 | -0.46806 | 3.292982 | 0.27811 |
| X | 65335000 | 65336599 | 17.09948 | 32.48783 | 17.75965 | 7.139408 | 1.357893 | 0.066951 | 0.278147 |
| 5 | 95532800 | 95536599 | 41.97638 | 69.87241 | 43.15353 | 18.73895 | 1.074101 | 1.278714 | 0.278161 |
| 5 | 137097000 | 137098799 | 10.93577 | 19.56651 | 38.88538 | 17.05799 | -0.58531 | 0.421702 | 0.278184 |
| 12 | 51968600 | 51970599 | 16.52553 | 29.76267 | 26.07107 | 11.36964 | 0.595216 | 0.391897 | 0.278201 |
| 12 | 23528400 | 23529599 | 23.3327 | 40.66129 | 30.88746 | 13.58696 | 0.782058 | 0.760746 | 0.278215 |
| 11 | 19286800 | 19288799 | 14.48037 | 25.20394 | 48.95418 | 21.51783 | -0.56624 | 0.762092 | 0.278232 |
| 5 | 123226400 | 123229399 | 27.27426 | 44.58489 | 81.5891 | 35.08027 | -0.49502 | 1.486891 | 0.278245 |
| 1 | 47119400 | 47120199 | 12.51127 | 23.68889 | 20.97079 | 8.491881 | 0.659229 | 0.055261 | 0.278248 |
| 7 | 24691800 | 24694399 | 16.62964 | 28.46982 | 55.35896 | 24.27589 | -0.57242 | 0.944599 | 0.278273 |
| 1 | 44323400 | 44324999 | 11.42999 | 20.41262 | 41.49395 | 18.23098 | -0.61942 | 0.441996 | 0.278288 |
| 9 | 43990000 | 43992399 | 12.18619 | 21.19912 | 49.46039 | 21.73281 | -0.82934 | 0.759487 | 0.278294 |
| 2 | 159249800 | 159251199 | 18.41393 | 32.5265 | 27.80541 | 12.30108 | 0.615262 | 0.544962 | 0.278314 |
| X | 61337600 | 61339999 | 22.49956 | 39.95915 | 26.00535 | 11.47943 | 1.009835 | 0.517986 | 0.278329 |
| 19 | 49472000 | 49473199 | 13.16455 | 25.01921 | 20.26694 | 8.142197 | 0.795697 | 0.032892 | 0.278339 |
| 13 | 107804000 | 107805399 | 7.543593 | 14.17382 | 30.71712 | 12.62215 | -0.6402 | 0.086968 | 0.278341 |
| 6 | 67576400 | 67577999 | 20.47745 | 36.9868 | 23.40059 | 10.16824 | 1.064858 | 0.398204 | 0.278355 |
| 18 | 83518200 | 83520799 | 23.82805 | 41.27297 | 36.6543 | 16.08169 | 0.556631 | 0.844991 | 0.278359 |
| 5 | 29582000 | 29585399 | 17.86314 | 30.13261 | 61.17202 | 26.75069 | -0.63662 | 1.088362 | 0.278359 |
| 5 | 101668400 | 101669599 | 8.941911 | 16.55137 | 34.76939 | 14.6332 | -0.62356 | 0.199629 | 0.278376 |
| 4 | 150420000 | 150420999 | 6.740846 | 12.9074 | 29.04102 | 11.46733 | -0.64287 | -0.0584 | 0.27838 |
| 2 | 74036200 | 74038199 | 14.49727 | 25.30828 | 48.92157 | 21.5436 | -0.55886 | 0.720888 | 0.27838 |
| 13 | 65274200 | 65281999 | 101.5076 | 158.3522 | 136.9668 | 56.8374 | 0.520895 | 2.691999 | 0.278397 |
| X | 8368800 | 8374799 | 125.5335 | 194.0903 | 143.5228 | 59.61603 | 0.736897 | 2.860402 | 0.278403 |
| 4 | 112532800 | 112533199 | 10.14016 | 19.62178 | 18.15135 | 6.92128 | 0.675572 | -0.16932 | 0.278414 |
| 7 | 140259600 | 140260799 | 26.18013 | 44.81892 | 38.77777 | 17.0047 | 0.590995 | 0.960144 | 0.278427 |
| 10 | 17390400 | 17392399 | 18.4844 | 32.82586 | 27.91831 | 12.32462 | 0.625704 | 0.494437 | 0.278451 |
| 2 | 173916400 | 173919399 | 16.31232 | 27.52744 | 60.47814 | 26.45042 | -0.74994 | 1.080509 | 0.278455 |
| 4 | 55552400 | 55553599 | 14.11462 | 26.05007 | 22.86413 | 9.660879 | 0.623003 | 0.230245 | 0.278457 |
| 2 | 58975600 | 58976199 | 10.77026 | 20.90674 | 17.55524 | 6.595639 | 0.833712 | -0.21311 | 0.278467 |
| 5 | 21893600 | 21894999 | 18.76092 | 31.74833 | 61.32252 | 26.84251 | -0.565 | 1.062818 | 0.278467 |
| 9 | 3202000 | 3206599 | 69.27605 | 110.9519 | 80.0681 | 33.75965 | 0.81517 | 2.027016 | 0.278479 |
| X | 67978400 | 67979999 | 19.26659 | 35.5215 | 21.04884 | 8.906498 | 1.184084 | 0.272607 | 0.278488 |
| 8 | 126324400 | 126324999 | 9.268481 | 17.97095 | 17.11291 | 6.464609 | 0.64646 | -0.21476 | 0.278509 |
| 3 | 154552000 | 154553999 | 13.34578 | 23.40412 | 46.17072 | 20.40614 | -0.58789 | 0.637687 | 0.278513 |
| 4 | 90892600 | 90893799 | 16.20997 | 30.76058 | 18.26367 | 7.360844 | 1.236125 | 0.067045 | 0.278516 |
| 1 | 115372800 | 115375999 | 51.18859 | 82.49037 | 74.33867 | 31.63194 | 0.505906 | 1.810785 | 0.278521 |
| 1 | 53823400 | 53824599 | 6.500788 | 12.26801 | 29.69995 | 12.11101 | -0.78918 | 0.046209 | 0.278523 |
| 2 | 165235800 | 165238199 | 28.0117 | 45.45841 | 87.02584 | 37.29634 | -0.56791 | 1.606187 | 0.278523 |
| 5 | 115509000 | 115511199 | 11.99623 | 21.00571 | 44.67346 | 19.71922 | -0.69524 | 0.655879 | 0.278523 |

| | | | | | | | | | |
|---|---|---|---|---|---|---|---|---|---|
| 18 | 84606800 | 84608199 | 8.749276 | 15.84305 | 36.47093 | 15.79776 | -0.78349 | 0.328074 | 0.278523 |
| 17 | 66905000 | 66906999 | 9.758201 | 17.76829 | 38.7255 | 16.65838 | -0.69907 | 0.306376 | 0.278534 |
| 3 | 8279000 | 8282599 | 34.20597 | 55.15326 | 99.00574 | 42.15833 | -0.48518 | 1.780244 | 0.278535 |
| 7 | 11882800 | 11883199 | 12.94483 | 24.22516 | 21.26948 | 8.799403 | 0.645062 | 0.132913 | 0.278539 |
| 15 | 80294600 | 80296799 | 14.03515 | 24.55179 | 48.35361 | 21.32596 | -0.58543 | 0.687086 | 0.278548 |
| 16 | 37874400 | 37875999 | 16.25935 | 29.41658 | 25.65303 | 11.12397 | 0.606961 | 0.364097 | 0.278551 |
| 1 | 156293400 | 156295399 | 17.124 | 30.86925 | 26.08625 | 11.36303 | 0.647958 | 0.391968 | 0.278576 |
| 13 | 7458400 | 7461399 | 30.37672 | 51.95854 | 38.00408 | 16.66427 | 0.832206 | 0.984716 | 0.278577 |
| 17 | 12803000 | 12803999 | 7.152151 | 13.2892 | 33.72724 | 14.12104 | -0.88856 | 0.160096 | 0.278595 |
| 2 | 73326000 | 73328199 | 11.59176 | 20.51692 | 41.9054 | 18.53146 | -0.63352 | 0.506474 | 0.278597 |
| 17 | 16027400 | 16029399 | 11.59176 | 20.51692 | 41.9054 | 18.53146 | -0.63352 | 0.506474 | 0.278597 |
| 13 | 67025600 | 67028399 | 62.31166 | 99.75872 | 86.22601 | 36.33439 | 0.554664 | 2.030763 | 0.278599 |
| 5 | 71377400 | 71378799 | 14.37757 | 27.34309 | 19.62714 | 7.873862 | 0.970592 | 0.038739 | 0.278614 |
| 17 | 39154600 | 39165599 | 227.3622 | 334.5208 | 288.4975 | 118.8047 | 0.454196 | 3.816293 | 0.278617 |
| 1 | 151407800 | 151411799 | 37.63764 | 60.43136 | 110.2119 | 46.64681 | -0.51554 | 1.919715 | 0.278623 |
| 3 | 130720000 | 130726999 | 54.52437 | 85.69787 | 174.6028 | 72.428 | -0.70521 | 2.537803 | 0.278627 |
| 16 | 19552600 | 19553399 | 8.426341 | 15.33469 | 35.49921 | 15.28109 | -0.7855 | 0.294587 | 0.278629 |
| 19 | 42544400 | 42546399 | 17.38551 | 31.28496 | 24.75681 | 10.80792 | 0.740329 | 0.402378 | 0.278636 |
| 7 | 11035800 | 11040399 | 62.0136 | 99.61548 | 76.75312 | 32.50815 | 0.725846 | 1.926896 | 0.278665 |
| 4 | 101296600 | 101297399 | 4.97329 | 9.661734 | 25.78427 | 9.624248 | -0.80947 | -0.27965 | 0.278666 |
| 8 | 86025400 | 86027199 | 8.942618 | 15.7562 | 43.06239 | 19.06044 | -1.05324 | 0.544496 | 0.278667 |
| 4 | 70922000 | 70923599 | 20.77535 | 36.57503 | 30.07968 | 13.31613 | 0.669002 | 0.599567 | 0.278677 |
| 4 | 125997000 | 125998799 | 7.477121 | 13.79135 | 33.06584 | 13.98316 | -0.81723 | 0.205153 | 0.278684 |
| 16 | 9883000 | 9885999 | 18.95501 | 31.92628 | 65.17563 | 28.48845 | -0.64494 | 1.10726 | 0.2787 |
| 1 | 167192400 | 167195399 | 15.98953 | 27.4146 | 54.8159 | 24.04062 | -0.61215 | 0.927544 | 0.27871 |
| 13 | 29213800 | 29215199 | 8.727117 | 15.87627 | 34.97276 | 15.06134 | -0.71468 | 0.298806 | 0.278726 |
| 13 | 17508200 | 17509799 | 22.06461 | 38.97333 | 27.52473 | 12.17702 | 0.88896 | 0.565699 | 0.278738 |
| 8 | 45514000 | 45522599 | 58.15944 | 91.64843 | 159.8587 | 66.33425 | -0.4785 | 2.485083 | 0.278738 |
| 2 | 129822400 | 129822799 | 3.523559 | 6.889954 | 19.58016 | 6.681416 | -0.77744 | -0.63682 | 0.27874 |
| 7 | 14467600 | 14470799 | 72.86941 | 115.9427 | 99.65674 | 41.6317 | 0.553181 | 2.237178 | 0.278745 |
| 2 | 142743000 | 142745399 | 10.49688 | 18.33972 | 46.79091 | 20.61593 | -0.95649 | 0.685715 | 0.278748 |
| 13 | 4230000 | 4232399 | 47.74995 | 77.27171 | 67.71025 | 28.95794 | 0.552949 | 1.687856 | 0.278749 |
| 8 | 15575400 | 15576599 | 13.10276 | 24.45351 | 21.90637 | 9.105108 | 0.611331 | 0.154718 | 0.278756 |
| 4 | 38116400 | 38119599 | 32.99926 | 55.62988 | 43.61219 | 19.06807 | 0.730969 | 1.126378 | 0.278767 |
| 10 | 98002400 | 98002999 | 7.20948 | 13.64603 | 30.37422 | 12.30813 | -0.66158 | 0.032168 | 0.27878 |
| 7 | 25879000 | 25884199 | 91.57049 | 143.2579 | 133.0443 | 55.19352 | 0.421591 | 2.637175 | 0.278781 |
| 1 | 142954600 | 142955199 | 8.303091 | 15.53468 | 32.84529 | 13.59415 | -0.61375 | 0.124919 | 0.278789 |
| 5 | 96308400 | 96311199 | 13.49218 | 22.94247 | 62.13094 | 27.23065 | -1.04979 | 1.001761 | 0.278792 |
| 4 | 114522400 | 114526999 | 24.58488 | 40.11925 | 81.44209 | 34.99503 | -0.64593 | 1.506109 | 0.278804 |
| 10 | 66789800 | 66791799 | 11.05365 | 19.40365 | 43.07737 | 19.05487 | -0.75676 | 0.60625 | 0.278812 |
| 14 | 113358400 | 113360199 | 22.95768 | 40.28337 | 29.71604 | 13.14002 | 0.824014 | 0.652846 | 0.278815 |
| 1 | 64190600 | 64192999 | 34.98603 | 57.91081 | 52.06865 | 22.52296 | 0.533575 | 1.327819 | 0.278822 |
| 12 | 115508200 | 115512399 | 88.53479 | 139.6956 | 116.5377 | 48.37038 | 0.585615 | 2.470608 | 0.278829 |
| 18 | 65472800 | 65476599 | 21.57438 | 36.04105 | 69.1278 | 30.08603 | -0.55547 | 1.202578 | 0.278833 |
| X | 116384400 | 116390999 | 52.33056 | 84.60033 | 61.72182 | 26.37192 | 0.81518 | 1.715977 | 0.278835 |
| X | 152930400 | 152931799 | 17.86115 | 34.44274 | 12.57722 | 4.866742 | 1.981373 | -0.05529 | 0.278849 |
| 12 | 95764200 | 95765199 | 11.98957 | 23.29838 | 15.64541 | 5.827795 | 1.163319 | -0.23456 | 0.278849 |

| | | | | | | | | | |
|---|---|---|---|---|---|---|---|---|---|
| 4 | 66093000 | 66095199 | 13.06001 | 22.98 | 46.12678 | 20.42227 | -0.61205 | 0.588656 | 0.278853 |
| 13 | 32744600 | 32746399 | 30.57178 | 51.71236 | 43.64528 | 19.10213 | 0.625079 | 1.090018 | 0.27889 |
| 6 | 36020400 | 36021799 | 9.629067 | 17.65185 | 37.81273 | 16.12472 | -0.66537 | 0.267538 | 0.278917 |
| 17 | 55232200 | 55233799 | 10.25217 | 18.61179 | 38.63852 | 16.68348 | -0.63278 | 0.32856 | 0.278921 |
| 4 | 112474400 | 112476399 | 29.55259 | 50.94607 | 35.82554 | 15.71581 | 0.890676 | 0.911848 | 0.278922 |
| 12 | 17972000 | 17981199 | 325.4459 | 463.5871 | 396.9193 | 164.0881 | 0.41899 | 4.306965 | 0.278925 |
| 2 | 76110200 | 76110599 | 4.121516 | 7.999685 | 24.34989 | 9.153693 | -1.00454 | -0.26503 | 0.278927 |
| 12 | 86847800 | 86848999 | 4.121516 | 7.999685 | 24.34989 | 9.153693 | -1.00454 | -0.26503 | 0.278927 |
| 13 | 3683800 | 3684599 | 5.151665 | 9.871418 | 27.27657 | 10.74872 | -0.9327 | -0.08577 | 0.27895 |
| 7 | 136927800 | 136929199 | 19.78584 | 34.7492 | 30.69055 | 13.58129 | 0.565687 | 0.623264 | 0.278951 |
| 7 | 85232000 | 85234199 | 32.73791 | 56.0421 | 36.30617 | 15.92079 | 1.006818 | 0.987015 | 0.278978 |
| 7 | 35151800 | 35157999 | 31.80082 | 51.25226 | 99.4716 | 42.33223 | -0.59824 | 1.789554 | 0.278982 |
| 1 | 166367400 | 166370399 | 39.16637 | 65.67758 | 40.79247 | 17.79631 | 1.065626 | 1.192433 | 0.279004 |
| 1 | 102287800 | 102293599 | 94.9444 | 148.5879 | 132.1066 | 54.80254 | 0.484817 | 2.633657 | 0.279004 |
| 1 | 110103800 | 110106999 | 41.60791 | 69.60617 | 40.8753 | 17.80573 | 1.146458 | 1.227002 | 0.279009 |
| 6 | 7111400 | 7111999 | 9.466737 | 18.47271 | 14.28868 | 5.08894 | 1.018713 | -0.42398 | 0.27901 |
| 6 | 142907400 | 142911599 | 27.07368 | 43.89101 | 90.29672 | 38.66881 | -0.67291 | 1.625842 | 0.279018 |
| 11 | 3751800 | 3755399 | 29.10798 | 47.30477 | 86.43225 | 37.08084 | -0.49889 | 1.578717 | 0.279018 |
| 12 | 13133400 | 13136799 | 27.25932 | 44.57902 | 81.70908 | 35.13719 | -0.49692 | 1.480312 | 0.279023 |
| 5 | 111736800 | 111738799 | 9.81284 | 17.90038 | 38.21712 | 16.40225 | -0.66692 | 0.298659 | 0.279043 |
| X | 66119200 | 66120199 | 14.18704 | 26.98837 | 19.63489 | 7.872257 | 0.952029 | 0.034912 | 0.279047 |
| 1 | 128939200 | 128939799 | 10.38981 | 19.98464 | 18.6085 | 7.253059 | 0.636406 | -0.09437 | 0.279058 |
| 10 | 120416600 | 120418599 | 14.15239 | 24.70194 | 47.85587 | 21.072 | -0.56209 | 0.719054 | 0.279072 |
| 7 | 18176800 | 18180199 | 64.31114 | 102.9823 | 84.45525 | 35.60004 | 0.630736 | 2.027809 | 0.279091 |
| 3 | 6706400 | 6707799 | 14.31907 | 27.48633 | 16.16806 | 6.338909 | 1.283663 | -0.04358 | 0.279112 |
| 9 | 11380200 | 11380799 | 9.146085 | 17.76639 | 17.144 | 6.404813 | 0.642371 | -0.25353 | 0.279126 |
| 7 | 60884400 | 60892199 | 120.0334 | 184.4991 | 166.9389 | 69.32067 | 0.439586 | 2.968164 | 0.279128 |
| 18 | 20580000 | 20580599 | 8.14537 | 15.87987 | 13.34526 | 4.813299 | 0.884265 | -0.40379 | 0.27913 |
| 4 | 147770000 | 147775399 | 91.60873 | 144.7449 | 109.7125 | 45.56922 | 0.725293 | 2.44046 | 0.279134 |
| 10 | 52128800 | 52129799 | 6.874072 | 13.15399 | 31.45575 | 12.44391 | -0.73358 | -0.0485 | 0.279142 |
| 9 | 26596800 | 26598799 | 12.70406 | 22.3852 | 45.5963 | 20.18171 | -0.63238 | 0.568603 | 0.279144 |
| 11 | 58334800 | 58339799 | 47.04104 | 74.88767 | 131.1212 | 54.81706 | -0.47079 | 2.204249 | 0.279173 |
| 7 | 102918600 | 102919199 | 19.3614 | 34.01517 | 29.8329 | 13.20551 | 0.575817 | 0.615001 | 0.279183 |
| 4 | 117334200 | 117335799 | 9.491836 | 17.20398 | 35.94568 | 15.55178 | -0.64351 | 0.327435 | 0.279193 |
| 18 | 62681200 | 62682799 | 9.491836 | 17.20398 | 35.94568 | 15.55178 | -0.64351 | 0.327435 | 0.279193 |
| 10 | 22060400 | 22063799 | 14.64178 | 25.43391 | 50.04119 | 21.96796 | -0.58509 | 0.792665 | 0.279193 |
| 15 | 20095200 | 20096599 | 13.88009 | 25.93266 | 21.62441 | 8.971239 | 0.715598 | 0.151974 | 0.279204 |
| 3 | 74601600 | 74602999 | 13.59755 | 25.9479 | 19.77711 | 7.877062 | 0.893893 | 0.009308 | 0.279215 |
| 8 | 47595200 | 47597599 | 24.81863 | 43.07585 | 35.999 | 15.79517 | 0.644515 | 0.830966 | 0.27925 |
| 17 | 20192000 | 20194799 | 48.15773 | 78.13794 | 58.87639 | 25.22906 | 0.773641 | 1.631175 | 0.279253 |
| 8 | 47542200 | 47544599 | 15.41646 | 26.75804 | 52.7744 | 23.15611 | -0.58889 | 0.810431 | 0.279256 |
| 2 | 43390200 | 43391599 | 21.05493 | 37.69657 | 24.31854 | 10.66202 | 1.028529 | 0.45468 | 0.279262 |
| 17 | 49305000 | 49307199 | 12.4338 | 21.86492 | 42.99181 | 19.03658 | -0.58251 | 0.587478 | 0.279265 |
| 5 | 100700200 | 100701199 | 6.329715 | 12.02817 | 29.33556 | 11.7885 | -0.78132 | -0.00614 | 0.27927 |
| 1 | 125383800 | 125385399 | 11.76709 | 20.8271 | 40.87634 | 18.07642 | -0.57615 | 0.507391 | 0.279272 |
| 19 | 20116400 | 20119999 | 44.82369 | 73.13671 | 59.0117 | 25.35565 | 0.68132 | 1.534953 | 0.279283 |
| 18 | 82720400 | 82726199 | 44.36922 | 70.76547 | 130.5893 | 54.73509 | -0.54361 | 2.140058 | 0.279283 |

| | | | | | | | | | |
|---|---|---|---|---|---|---|---|---|---|
| 7 | 19361400 | 19363799 | 15.14899 | 25.76498 | 58.42522 | 25.60693 | -0.79472 | 1.002991 | 0.279284 |
| 2 | 99130800 | 99132199 | 22.32175 | 39.57718 | 26.55122 | 11.73692 | 0.964913 | 0.527902 | 0.279289 |
| X | 107614800 | 107616599 | 22.32175 | 39.57718 | 26.55122 | 11.73692 | 0.964913 | 0.527902 | 0.279289 |
| 2 | 145265600 | 145266999 | 20.55072 | 35.94866 | 31.88296 | 14.06143 | 0.55985 | 0.692612 | 0.279292 |
| 14 | 79157200 | 79159799 | 16.62945 | 28.56635 | 54.2991 | 23.81714 | -0.53891 | 0.91343 | 0.279294 |
| 12 | 105488000 | 105494199 | 51.61151 | 81.98413 | 140.7214 | 58.67823 | -0.44546 | 2.276901 | 0.279299 |
| 4 | 147725800 | 147727399 | 24.79492 | 43.7046 | 26.92393 | 11.9162 | 1.084136 | 0.604905 | 0.279303 |
| 19 | 52318200 | 52318599 | 10.8146 | 20.68438 | 18.55571 | 7.350921 | 0.667851 | -0.03854 | 0.279303 |
| 4 | 50465200 | 50465599 | 8.383473 | 16.39087 | 12.01895 | 4.115102 | 1.143491 | -0.58195 | 0.279315 |
| 18 | 34035800 | 34037199 | 17.31858 | 31.34766 | 25.40536 | 11.0101 | 0.7124 | 0.367647 | 0.279316 |
| 3 | 72470600 | 72471999 | 12.83543 | 24.57073 | 19.2709 | 7.615313 | 0.862855 | -0.02509 | 0.279318 |
| 14 | 6785800 | 6786799 | 12.83543 | 24.57073 | 19.2709 | 7.615313 | 0.862855 | -0.02509 | 0.279318 |
| 12 | 24161800 | 24182999 | 509.7518 | 698.6448 | 571.4528 | 237.7028 | 0.428859 | 4.890049 | 0.279324 |
| 12 | 23406600 | 23410799 | 78.21881 | 123.9319 | 94.4252 | 39.27474 | 0.720944 | 2.362413 | 0.279336 |
| 6 | 6955800 | 6956599 | 6.369385 | 11.99423 | 30.19691 | 12.36172 | -0.84996 | 0.06009 | 0.279351 |
| 9 | 102848400 | 102852999 | 25.71079 | 41.90573 | 83.32555 | 35.78851 | -0.61746 | 1.526329 | 0.279351 |
| 1 | 93570800 | 93573999 | 33.82631 | 56.54325 | 47.567 | 20.7103 | 0.629635 | 1.213671 | 0.27937 |
| 2 | 160105200 | 160110399 | 32.64587 | 52.49439 | 105.8787 | 44.91064 | -0.65776 | 1.864935 | 0.279386 |
| 8 | 22307000 | 22310599 | 39.53178 | 65.29318 | 51.12784 | 22.08285 | 0.732132 | 1.363149 | 0.279388 |
| 7 | 10864400 | 10881999 | 390.958 | 550.1921 | 437.8533 | 181.3759 | 0.506855 | 4.487615 | 0.279432 |
| 3 | 22395200 | 22397599 | 27.84458 | 47.87761 | 35.82545 | 15.71453 | 0.800479 | 0.923702 | 0.279442 |
| 15 | 12696600 | 12698799 | 10.91232 | 19.53742 | 42.07546 | 18.44715 | -0.70033 | 0.421319 | 0.279449 |
| 9 | 95678000 | 95678799 | 12.36908 | 24.01456 | 16.2103 | 6.081559 | 1.146998 | -0.2039 | 0.279473 |
| 5 | 113500600 | 113502599 | 12.94934 | 22.56143 | 47.35001 | 20.82567 | -0.67691 | 0.740306 | 0.279475 |
| 13 | 120150200 | 120152999 | 16.67554 | 28.3702 | 59.08843 | 25.89826 | -0.67267 | 1.005445 | 0.279484 |
| 5 | 79923800 | 79924199 | 3.756403 | 7.340819 | 19.99511 | 6.930183 | -0.73705 | -0.57408 | 0.279485 |
| 3 | 20263000 | 20264999 | 23.23658 | 40.63073 | 32.63185 | 14.38686 | 0.702341 | 0.714779 | 0.279493 |
| 14 | 51361400 | 51362199 | 5.25263 | 10.16171 | 25.41074 | 9.700154 | -0.74659 | -0.19689 | 0.279504 |
| X | 94913400 | 94914599 | 16.72485 | 30.44178 | 24.21081 | 10.41969 | 0.746207 | 0.331806 | 0.279508 |
| 11 | 57362600 | 57375199 | 117.5345 | 176.7627 | 297.2246 | 122.7243 | -0.47947 | 3.412255 | 0.279511 |
| 18 | 80897200 | 80898199 | 7.532486 | 14.15198 | 31.27626 | 12.85339 | -0.66871 | 0.087029 | 0.279513 |
| X | 36842200 | 36845799 | 32.45173 | 55.10519 | 40.83116 | 17.89147 | 0.812383 | 1.062016 | 0.279515 |
| 12 | 35567200 | 35568799 | 8.205122 | 14.75057 | 36.95865 | 16.14809 | -0.91269 | 0.370041 | 0.279518 |
| 6 | 89017200 | 89022599 | 37.0403 | 59.36279 | 113.0765 | 47.7253 | -0.58132 | 1.978097 | 0.279524 |
| 1 | 101854400 | 101858399 | 36.60184 | 61.6732 | 40.20007 | 17.57364 | 0.996221 | 1.139213 | 0.279525 |
| 4 | 57255800 | 57257199 | 5.427371 | 10.43446 | 26.85453 | 10.48156 | -0.81861 | -0.11688 | 0.27953 |
| 13 | 38134000 | 38135199 | 5.427371 | 10.43446 | 26.85453 | 10.48156 | -0.81861 | -0.11688 | 0.27953 |
| 6 | 30985800 | 30991399 | 37.70334 | 60.48884 | 126.9939 | 53.68491 | -0.71968 | 1.951668 | 0.279533 |
| 3 | 73496200 | 73497599 | 16.6911 | 30.37264 | 25.00327 | 10.76416 | 0.696114 | 0.332678 | 0.279542 |
| 4 | 106141000 | 106142399 | 16.6911 | 30.37264 | 25.00327 | 10.76416 | 0.696114 | 0.332678 | 0.279542 |
| 6 | 59054200 | 59056999 | 70.31731 | 111.6408 | 103.8225 | 43.26989 | 0.436651 | 2.298995 | 0.279559 |
| 6 | 129866400 | 129869399 | 38.11582 | 64.27817 | 37.91505 | 16.57896 | 1.13976 | 1.135485 | 0.27956 |
| 3 | 156584000 | 156584799 | 11.10832 | 21.4556 | 18.47613 | 7.106116 | 0.765508 | -0.13105 | 0.279561 |
| 7 | 30517400 | 30524599 | 45.4457 | 72.21611 | 138.0774 | 57.59672 | -0.6003 | 2.260354 | 0.279563 |
| 19 | 39533400 | 39534199 | 22.85972 | 39.63767 | 35.70985 | 15.66783 | 0.536385 | 0.830256 | 0.279566 |
| 18 | 45298200 | 45299799 | 16.44388 | 30.96512 | 19.89084 | 8.142927 | 1.104703 | 0.123489 | 0.279568 |
| 14 | 39662400 | 39663599 | 6.113843 | 11.8289 | 27.90553 | 10.64794 | -0.6639 | -0.18328 | 0.27957 |

| | | | | | | | | | |
|---|---|---|---|---|---|---|---|---|---|
| 2 | 16521600 | 16522799 | 8.496649 | 15.7148 | 33.10444 | 13.94745 | -0.62836 | 0.197641 | 0.279572 |
| 13 | 50457400 | 50458999 | 8.496649 | 15.7148 | 33.10444 | 13.94745 | -0.62836 | 0.197641 | 0.279572 |
| 13 | 38969000 | 38971599 | 18.92688 | 31.58837 | 65.88487 | 28.65752 | -0.67569 | 1.204265 | 0.279594 |
| 12 | 69618800 | 69620799 | 10.75332 | 19.37955 | 39.52439 | 17.22253 | -0.61594 | 0.376623 | 0.279621 |
| 11 | 96141200 | 96143599 | 11.03642 | 19.4169 | 42.00044 | 18.59583 | -0.71893 | 0.572138 | 0.279622 |
| 9 | 21748200 | 21754599 | 28.29795 | 45.62566 | 99.97017 | 42.56709 | -0.77204 | 1.772209 | 0.279632 |
| X | 159073400 | 159075999 | 21.38794 | 37.59919 | 30.50715 | 13.50927 | 0.687563 | 0.618797 | 0.279639 |
| 10 | 66877800 | 66882999 | 33.89547 | 54.66959 | 98.30155 | 41.87525 | -0.48704 | 1.77124 | 0.279661 |
| 3 | 19171800 | 19174799 | 106.895 | 165.1261 | 151.4396 | 62.90479 | 0.425601 | 2.867513 | 0.279685 |
| 6 | 19874600 | 19875199 | 4.781067 | 9.318122 | 23.17924 | 8.393644 | -0.66747 | -0.39473 | 0.279685 |
| 11 | 7214400 | 7217799 | 21.25757 | 35.52747 | 67.38036 | 29.33366 | -0.53925 | 1.195868 | 0.279687 |
| 2 | 15973200 | 15974199 | 6.44315 | 12.34313 | 28.45432 | 11.22062 | -0.67541 | -0.06641 | 0.279687 |
| 1 | 89267000 | 89269599 | 14.17652 | 24.63395 | 49.49733 | 21.73377 | -0.61518 | 0.785143 | 0.27969 |
| 8 | 3269400 | 3271599 | 13.06357 | 22.90279 | 44.61202 | 19.71255 | -0.56944 | 0.640094 | 0.279705 |
| 11 | 91944600 | 91944999 | 3.572777 | 6.979886 | 20.4697 | 7.143598 | -0.85012 | -0.54837 | 0.279713 |
| 2 | 11987000 | 11988799 | 19.41372 | 34.71171 | 24.96485 | 10.95773 | 0.871393 | 0.451653 | 0.279716 |
| 14 | 63476800 | 63479399 | 23.8846 | 39.14822 | 77.83084 | 33.49952 | -0.61198 | 1.440821 | 0.279724 |
| 17 | 39489200 | 39492199 | 30.00188 | 52.47125 | 26.76743 | 11.80341 | 1.353728 | 0.740121 | 0.279727 |
| 5 | 73088800 | 73091399 | 29.05876 | 46.91271 | 101.0989 | 43.11926 | -0.74601 | 1.737517 | 0.279728 |
| 3 | 27756000 | 27759799 | 20.52196 | 34.15241 | 68.23522 | 29.6259 | -0.61402 | 1.24105 | 0.279738 |
| 2 | 106750400 | 106751199 | 10.46692 | 18.32368 | 47.79702 | 21.09392 | -0.98814 | 0.655792 | 0.279746 |
| 6 | 56357000 | 56357999 | 12.04699 | 22.69056 | 19.9384 | 8.159269 | 0.656653 | 0.081916 | 0.279754 |
| 5 | 109396000 | 109400199 | 60.42738 | 97.30831 | 70.2326 | 29.85347 | 0.824242 | 1.845258 | 0.279764 |
| 3 | 99312200 | 99314799 | 24.12009 | 42.24474 | 30.24073 | 13.35112 | 0.867269 | 0.68918 | 0.279764 |
| 7 | 31270600 | 31282799 | 210.884 | 313.7465 | 251.2311 | 103.5114 | 0.576086 | 3.626564 | 0.279799 |
| 6 | 101495000 | 101499199 | 19.7701 | 33.2297 | 64.60453 | 28.22095 | -0.57488 | 1.128442 | 0.279809 |
| 8 | 33371200 | 33375199 | 29.22279 | 47.16844 | 99.04423 | 42.23246 | -0.709 | 1.740074 | 0.279831 |
| 2 | 59980800 | 59981199 | 3.933468 | 7.684279 | 20.55344 | 7.177099 | -0.72209 | -0.54508 | 0.279843 |
| 4 | 87147800 | 87148999 | 9.388164 | 16.87926 | 36.90135 | 16.12152 | -0.71759 | 0.376696 | 0.279849 |
| 13 | 58792600 | 58804599 | 87.38558 | 133.5999 | 251.614 | 104.3827 | -0.623 | 3.070914 | 0.279858 |
| 10 | 94098600 | 94099999 | 5.877483 | 11.24891 | 28.99668 | 11.46538 | -0.83952 | -0.06569 | 0.27986 |
| 15 | 56035200 | 56036799 | 15.16651 | 26.34473 | 51.10671 | 22.43536 | -0.56475 | 0.798977 | 0.279875 |
| 7 | 130428000 | 130432199 | 26.75393 | 43.37764 | 88.3154 | 37.82355 | -0.65769 | 1.623136 | 0.279878 |
| 18 | 32524200 | 32530999 | 42.14535 | 67.19834 | 126.4862 | 52.99324 | -0.57252 | 2.147854 | 0.279886 |
| 7 | 63723000 | 63740599 | 228.4866 | 324.8564 | 544.7244 | 225.263 | -0.55311 | 4.331372 | 0.279889 |
| 4 | 31618000 | 31620199 | 10.28945 | 18.45889 | 39.44369 | 17.26435 | -0.68723 | 0.399967 | 0.279891 |
| 18 | 61795400 | 61799799 | 22.39566 | 36.89062 | 73.98358 | 31.91063 | -0.62152 | 1.373143 | 0.279911 |
| 4 | 103236800 | 103237399 | 6.560755 | 12.20991 | 31.60343 | 13.1982 | -0.91273 | 0.143314 | 0.27992 |
| 13 | 98452800 | 98453799 | 6.820599 | 12.77335 | 30.6389 | 12.66023 | -0.79138 | 0.102884 | 0.279933 |
| 2 | 71549600 | 71551999 | 11.29589 | 20.02425 | 39.84925 | 17.61674 | -0.59505 | 0.4856 | 0.279946 |
| X | 14281200 | 14284799 | 57.0252 | 91.91512 | 69.58774 | 29.61853 | 0.757102 | 1.814412 | 0.279949 |
| 13 | 66466600 | 66473799 | 229.1219 | 335.7723 | 290.5787 | 119.6916 | 0.443755 | 3.87752 | 0.279983 |
| 8 | 123384000 | 123388199 | 21.33081 | 35.05249 | 77.73895 | 33.49642 | -0.76768 | 1.401546 | 0.279988 |
| 16 | 89868800 | 89872599 | 20.98893 | 34.9391 | 68.31089 | 29.66415 | -0.58289 | 1.239196 | 0.279989 |
| 7 | 48022400 | 48027199 | 60.96663 | 98.15376 | 70.51176 | 29.96199 | 0.830618 | 1.855454 | 0.280011 |
| 6 | 107986800 | 107988799 | 24.26395 | 42.43218 | 32.01615 | 14.11682 | 0.791986 | 0.717696 | 0.280026 |
| 3 | 34702000 | 34705999 | 37.59085 | 60.07244 | 121.1687 | 50.92753 | -0.66845 | 2.070498 | 0.280026 |

| | | | | | | | | | |
|---|---|---|---|---|---|---|---|---|---|
| 10 | 93870800 | 93872599 | 9.883042 | 17.29383 | 45.73674 | 20.17709 | -1.00775 | 0.65628 | 0.280036 |
| 2 | 100742200 | 100742999 | 13.00007 | 24.92397 | 19.37722 | 7.623795 | 0.881742 | -0.03578 | 0.28004 |
| 16 | 29974200 | 29981999 | 59.80551 | 94.12916 | 166.5255 | 69.08991 | -0.50026 | 2.510514 | 0.280041 |
| 12 | 102123600 | 102125399 | 9.223419 | 16.61929 | 36.91818 | 16.09063 | -0.73827 | 0.361717 | 0.280044 |
| 3 | 9200400 | 9201199 | 4.785425 | 9.311483 | 23.49939 | 8.660042 | -0.71094 | -0.3301 | 0.280051 |
| 15 | 37668600 | 37669399 | 7.248019 | 13.65253 | 30.75545 | 12.58454 | -0.69024 | 0.070123 | 0.280051 |
| 7 | 12057000 | 12061599 | 75.94777 | 120.8704 | 97.13356 | 40.59144 | 0.65044 | 2.231033 | 0.280058 |
| 1 | 59630400 | 59631999 | 9.285489 | 16.57229 | 37.93178 | 16.6748 | -0.78939 | 0.426382 | 0.280068 |
| 17 | 55450000 | 55450999 | 10.65091 | 18.968 | 38.86335 | 17.11563 | -0.63292 | 0.449392 | 0.280071 |
| 10 | 118889200 | 118890799 | 8.948608 | 16.48204 | 35.74449 | 15.14518 | -0.67643 | 0.228754 | 0.280071 |
| 17 | 44566800 | 44568999 | 12.90241 | 22.52047 | 48.91731 | 21.53954 | -0.72636 | 0.712748 | 0.280073 |
| 13 | 62295800 | 62296199 | 15.14651 | 27.89138 | 23.76123 | 10.07089 | 0.662575 | 0.253293 | 0.280088 |
| 3 | 131789800 | 131793399 | 27.56514 | 44.64996 | 90.10049 | 38.56071 | -0.64624 | 1.644607 | 0.280093 |
| 1 | 24624200 | 24626599 | 19.74104 | 32.88352 | 66.94786 | 29.08437 | -0.64091 | 1.229714 | 0.280099 |
| 3 | 56089800 | 56091599 | 11.24779 | 19.64361 | 46.79541 | 20.61261 | -0.85831 | 0.695105 | 0.2801 |
| 3 | 54599600 | 54600599 | 8.214379 | 15.129 | 33.56686 | 14.22339 | -0.70865 | 0.21987 | 0.280112 |
| 3 | 119133800 | 119135399 | 17.1285 | 29.23432 | 56.41736 | 24.73392 | -0.56227 | 0.976044 | 0.280112 |
| 6 | 133186400 | 133189199 | 34.72674 | 57.97211 | 48.13521 | 20.94031 | 0.648526 | 1.231511 | 0.280114 |
| 8 | 77828600 | 77830399 | 15.43337 | 28.14501 | 24.72596 | 10.6159 | 0.605537 | 0.311083 | 0.28012 |
| 1 | 119444400 | 119446199 | 18.00204 | 31.81521 | 27.26817 | 12.06217 | 0.611693 | 0.534878 | 0.280123 |
| 4 | 41502000 | 41507999 | 44.42073 | 70.75888 | 129.6956 | 54.2663 | -0.53577 | 2.181389 | 0.280129 |
| 1 | 148108800 | 148110199 | 14.51915 | 27.34402 | 21.26931 | 8.705459 | 0.831286 | 0.109634 | 0.280148 |
| 17 | 17307600 | 17309599 | 28.75497 | 50.77512 | 21.8471 | 9.665813 | 1.599552 | 0.60273 | 0.280156 |
| 4 | 10133600 | 10135599 | 22.54542 | 39.34568 | 32.70312 | 14.3986 | 0.6528 | 0.737653 | 0.280156 |
| 2 | 36868800 | 36878199 | 129.2779 | 198.8023 | 161.2021 | 66.94282 | 0.598124 | 2.962146 | 0.280167 |
| 4 | 131118800 | 131133199 | 100.0759 | 151.4958 | 272.0332 | 112.5413 | -0.56593 | 3.276657 | 0.280168 |
| 12 | 3917800 | 3925199 | 34.38541 | 54.98548 | 121.9358 | 51.29689 | -0.80387 | 2.046897 | 0.280176 |
| 7 | 10325800 | 10329999 | 50.84854 | 82.8649 | 55.76756 | 23.95097 | 0.941179 | 1.565821 | 0.28018 |
| 14 | 71682400 | 71684599 | 33.77189 | 56.3008 | 49.2359 | 21.40062 | 0.574016 | 1.244354 | 0.280189 |
| 1 | 35792600 | 35794199 | 6.074667 | 11.41667 | 30.15047 | 12.38407 | -0.92148 | 0.072669 | 0.280193 |
| 6 | 48774400 | 48776199 | 9.809316 | 17.67879 | 38.99521 | 16.99133 | -0.72832 | 0.367089 | 0.280206 |
| 10 | 17930800 | 17932199 | 9.646803 | 17.05784 | 40.46606 | 17.89844 | -0.84817 | 0.5046 | 0.280234 |
| 4 | 73590600 | 73591199 | 7.942612 | 15.5154 | 13.93593 | 4.871707 | 0.82948 | -0.51087 | 0.280251 |
| 6 | 110622000 | 110626599 | 34.41959 | 55.29318 | 108.9846 | 46.16354 | -0.6264 | 1.898676 | 0.280256 |
| 5 | 55138000 | 55141399 | 38.00953 | 63.26358 | 47.64167 | 20.68667 | 0.789039 | 1.271655 | 0.280259 |
| X | 148451400 | 148454399 | 46.10229 | 75.11324 | 60.08115 | 25.80144 | 0.692428 | 1.560242 | 0.280265 |
| 15 | 91948400 | 91950199 | 11.87102 | 20.68225 | 48.39228 | 21.28381 | -0.8331 | 0.734725 | 0.280267 |
| X | 30902200 | 30902999 | 12.16614 | 23.67433 | 15.19068 | 5.595338 | 1.24349 | -0.27038 | 0.28027 |
| 19 | 30526000 | 30527999 | 14.8408 | 25.77342 | 50.0327 | 21.9609 | -0.56572 | 0.798987 | 0.280273 |
| 10 | 113742200 | 113744199 | 21.17022 | 37.13085 | 31.03111 | 13.71649 | 0.644968 | 0.65187 | 0.280274 |
| 13 | 107595400 | 107596399 | 10.75834 | 20.86765 | 15.59105 | 5.879214 | 0.993802 | -0.21019 | 0.280277 |
| 6 | 137774000 | 137777799 | 52.55011 | 83.11677 | 148.7991 | 61.8362 | -0.51169 | 2.396028 | 0.280295 |
| 12 | 113197400 | 113198799 | 9.698485 | 17.04672 | 43.5745 | 19.29576 | -0.9589 | 0.577041 | 0.280299 |
| 1 | 183562000 | 183563799 | 22.35496 | 39.38794 | 28.91212 | 12.79718 | 0.832648 | 0.598871 | 0.28031 |
| 18 | 37586600 | 37590999 | 54.9818 | 89.32936 | 55.32433 | 23.7303 | 1.057564 | 1.618489 | 0.280315 |
| 7 | 38382400 | 38389999 | 116.3352 | 179.7106 | 155.0059 | 64.38616 | 0.514056 | 2.872029 | 0.280358 |
| 18 | 67597000 | 67598399 | 9.85441 | 17.67061 | 37.38633 | 16.36996 | -0.67296 | 0.397752 | 0.28036 |

| | | | | | | | | | |
|---|---|---|---|---|---|---|---|---|---|
| 19 | 59137000 | 59138999 | 14.9493 | 26.00918 | 50.03616 | 21.98759 | -0.55262 | 0.771539 | 0.280366 |
| 19 | 8779400 | 8781999 | 13.44938 | 23.25662 | 53.21082 | 23.34449 | -0.8031 | 0.836189 | 0.28037 |
| 4 | 135332800 | 135335799 | 18.38355 | 30.87244 | 62.74459 | 27.39861 | -0.63851 | 1.131883 | 0.280374 |
| 13 | 78058400 | 78060799 | 26.78844 | 46.10583 | 38.1336 | 16.72747 | 0.657072 | 0.915884 | 0.280389 |
| 1 | 29647200 | 29648999 | 34.61556 | 58.24712 | 43.04012 | 18.80743 | 0.815939 | 1.146457 | 0.280403 |
| 6 | 24203400 | 24204599 | 14.81794 | 27.57208 | 22.63206 | 9.454418 | 0.730522 | 0.188789 | 0.280404 |
| 8 | 89674600 | 89685399 | 139.3981 | 207.902 | 334.3365 | 137.7938 | -0.42604 | 3.575409 | 0.280405 |
| 9 | 14017400 | 14021199 | 20.63455 | 34.43636 | 68.42232 | 29.7601 | -0.60614 | 1.209909 | 0.280405 |
| 2 | 110408600 | 110410199 | 18.84572 | 34.95471 | 19.96918 | 8.380935 | 1.244469 | 0.234298 | 0.280417 |
| 5 | 30866400 | 30871199 | 27.08733 | 43.88151 | 90.02423 | 38.53164 | -0.66982 | 1.641167 | 0.280441 |
| 7 | 66487400 | 66492199 | 20.22299 | 33.27705 | 78.29506 | 33.75537 | -0.85188 | 1.381793 | 0.280454 |
| 6 | 21213200 | 21217399 | 40.16898 | 64.17731 | 120.482 | 50.62196 | -0.56549 | 2.080695 | 0.280476 |
| 6 | 134542800 | 134545199 | 11.8109 | 20.73748 | 43.04208 | 19.04279 | -0.66038 | 0.606212 | 0.280484 |
| 9 | 98868000 | 98868999 | 6.37034 | 11.96741 | 30.15018 | 12.39263 | -0.85521 | 0.077406 | 0.280484 |
| 7 | 11002200 | 11003399 | 13.61653 | 26.44509 | 14.93983 | 5.59023 | 1.404504 | -0.20163 | 0.280508 |
| 2 | 107420400 | 107426199 | 46.40214 | 74.93195 | 66.26131 | 28.27054 | 0.536498 | 1.737593 | 0.28052 |
| 17 | 55905600 | 55909599 | 46.53423 | 75.93265 | 57.94936 | 24.89979 | 0.761517 | 1.537854 | 0.280542 |
| 9 | 3165600 | 3170399 | 44.37719 | 73.16307 | 50.04176 | 21.59202 | 0.926233 | 1.40012 | 0.280558 |
| 16 | 4564800 | 4565399 | 11.87636 | 22.84353 | 18.88205 | 7.360116 | 0.806041 | -0.07919 | 0.28058 |
| 8 | 105950000 | 105952199 | 10.47655 | 18.67803 | 41.46885 | 18.24616 | -0.74751 | 0.443825 | 0.28062 |
| 17 | 83883600 | 83884999 | 9.276106 | 17.04008 | 35.12733 | 14.93739 | -0.60737 | 0.246196 | 0.28063 |
| 13 | 66612800 | 66624799 | 295.1076 | 423.6355 | 365.4459 | 150.8888 | 0.41933 | 4.19253 | 0.280645 |
| 8 | 37551200 | 37552399 | 14.25769 | 26.14398 | 22.23696 | 9.479181 | 0.65859 | 0.263971 | 0.280647 |
| 13 | 63578200 | 63580399 | 16.96639 | 28.96215 | 56.41696 | 24.73406 | -0.57567 | 0.97375 | 0.280651 |
| 13 | 40850000 | 40852399 | 12.88166 | 22.55756 | 46.20746 | 20.39743 | -0.64189 | 0.658389 | 0.280656 |
| 13 | 36910400 | 36912599 | 18.30726 | 30.91449 | 62.36836 | 27.28261 | -0.62769 | 1.080554 | 0.280664 |
| 7 | 25735800 | 25747399 | 147.6488 | 225.5823 | 172.0054 | 71.34584 | 0.67958 | 3.096324 | 0.280668 |
| X | 43511000 | 43511799 | 11.33523 | 21.91623 | 18.07341 | 6.918184 | 0.834798 | -0.14301 | 0.280675 |
| 12 | 72871800 | 72873199 | 7.405645 | 13.85353 | 31.71336 | 13.129 | -0.72763 | 0.114878 | 0.280683 |
| 4 | 56559600 | 56560199 | 4.234805 | 8.242621 | 22.89759 | 8.41733 | -0.844 | -0.34817 | 0.280684 |
| 6 | 13361400 | 13364599 | 28.65235 | 49.63264 | 34.14073 | 14.97892 | 0.923588 | 0.863452 | 0.28069 |
| 7 | 13723400 | 13724599 | 56.69457 | 90.91936 | 73.89609 | 31.2188 | 0.646136 | 1.970301 | 0.280722 |
| 4 | 7939000 | 7939399 | 4.486813 | 8.737576 | 22.9629 | 8.403665 | -0.75882 | -0.35944 | 0.280724 |
| 1 | 101347600 | 101348599 | 15.3336 | 27.85494 | 24.67086 | 10.6437 | 0.589258 | 0.331089 | 0.280728 |
| 8 | 22998400 | 22999799 | 8.070235 | 14.86924 | 33.04676 | 13.99569 | -0.71029 | 0.21643 | 0.280747 |
| 5 | 17615200 | 17616599 | 18.35994 | 33.17772 | 25.30494 | 10.98926 | 0.797596 | 0.384606 | 0.280748 |
| 9 | 62655800 | 62661799 | 61.79671 | 96.34201 | 182.5687 | 75.76719 | -0.60996 | 2.690722 | 0.280757 |
| 2 | 71564800 | 71568399 | 22.47486 | 37.25227 | 70.48519 | 30.51792 | -0.53628 | 1.295867 | 0.280771 |
| 16 | 21016800 | 21019199 | 12.57333 | 22.03664 | 45.13975 | 19.94065 | -0.64177 | 0.641428 | 0.280774 |
| 13 | 20330000 | 20332799 | 17.00513 | 28.87236 | 60.16937 | 26.36416 | -0.6739 | 1.026072 | 0.280776 |
| 2 | 174136000 | 174144599 | 264.0285 | 369.1684 | 645.424 | 267.6294 | -0.63761 | 4.580021 | 0.280794 |
| 14 | 113915400 | 113918199 | 59.23712 | 94.70515 | 87.66483 | 36.86818 | 0.453816 | 2.070274 | 0.280794 |
| 10 | 111808200 | 111809199 | 5.751932 | 10.98465 | 30.62958 | 12.17986 | -0.95971 | -0.04488 | 0.280795 |
| 16 | 47968000 | 47970799 | 23.15425 | 40.56937 | 30.86701 | 13.63223 | 0.779771 | 0.678776 | 0.280797 |
| 12 | 97009800 | 97010199 | 8.604741 | 16.81756 | 13.30583 | 4.597245 | 1.025291 | -0.54189 | 0.280797 |
| 14 | 52855800 | 52857999 | 27.87545 | 47.34369 | 42.59485 | 18.6654 | 0.533377 | 1.044163 | 0.280807 |
| 2 | 94479800 | 94481399 | 19.85896 | 35.02508 | 26.03309 | 11.52055 | 0.81489 | 0.565424 | 0.28082 |

| | | | | | | | | | |
|---|---|---|---|---|---|---|---|---|---|
| X | 153235800 | 153238199 | 59.60776 | 96.10529 | 67.77729 | 28.8586 | 0.859664 | 1.808657 | 0.280833 |
| 18 | 70102600 | 70103599 | 14.11547 | 26.7232 | 20.75713 | 8.407406 | 0.845931 | 0.069824 | 0.280836 |
| 4 | 15957200 | 15960199 | 17.41268 | 29.90104 | 57.06915 | 25.0319 | -0.54507 | 0.922119 | 0.28084 |
| 15 | 66194200 | 66196599 | 17.41268 | 29.90104 | 57.06915 | 25.0319 | -0.54507 | 0.922119 | 0.28084 |
| 7 | 84648200 | 84649199 | 16.46073 | 30.47538 | 22.40415 | 9.426263 | 0.880919 | 0.229831 | 0.280845 |
| 3 | 137816600 | 137817399 | 4.404277 | 8.559564 | 23.93921 | 8.908072 | -0.8708 | -0.30075 | 0.280848 |
| 13 | 54322200 | 54330999 | 67.15499 | 104.7393 | 181.7963 | 75.44212 | -0.483 | 2.68562 | 0.28085 |
| 18 | 66634800 | 66636999 | 15.10388 | 27.41351 | 23.21945 | 10.02855 | 0.65221 | 0.330707 | 0.280858 |
| 16 | 50348800 | 50350999 | 13.1572 | 22.43804 | 60.53888 | 26.5391 | -1.04369 | 0.971727 | 0.28088 |
| 6 | 63232800 | 63235199 | 23.4828 | 38.30712 | 81.7115 | 35.10622 | -0.71747 | 1.509707 | 0.280883 |
| 4 | 113857800 | 113860199 | 29.09062 | 50.01608 | 37.54346 | 16.46808 | 0.796235 | 0.935482 | 0.280891 |
| 5 | 117607000 | 117616399 | 66.50941 | 103.3224 | 192.4098 | 79.8936 | -0.58884 | 2.754487 | 0.280896 |
| 14 | 19821400 | 19824799 | 15.82049 | 27.15186 | 54.82705 | 24.04744 | -0.62607 | 0.917304 | 0.280904 |
| X | 43825800 | 43827599 | 17.72041 | 33.23681 | 19.10415 | 7.873585 | 1.255909 | 0.156677 | 0.280908 |
| 5 | 122159200 | 122165599 | 75.82042 | 117.29 | 200.1825 | 83.15094 | -0.46795 | 2.835261 | 0.28091 |
| 2 | 89924800 | 89927799 | 35.38553 | 59.26619 | 44.76185 | 19.51319 | 0.784522 | 1.194777 | 0.280912 |
| 9 | 114813400 | 114822399 | 89.71184 | 137.1996 | 235.8398 | 97.84586 | -0.49099 | 3.063675 | 0.280912 |
| 1 | 6052200 | 6056199 | 76.69042 | 121.2957 | 112.8577 | 46.89952 | 0.431078 | 2.405042 | 0.280916 |
| 7 | 126830200 | 126832799 | 12.1453 | 21.21125 | 47.84271 | 21.07404 | -0.78019 | 0.699661 | 0.280929 |
| 13 | 65518000 | 65530599 | 241.8529 | 352.9997 | 305.635 | 125.9305 | 0.43749 | 3.941843 | 0.280943 |
| 12 | 86259400 | 86264199 | 31.30993 | 50.48945 | 98.41055 | 41.91103 | -0.60343 | 1.771734 | 0.28095 |
| 6 | 90408400 | 90415999 | 81.48782 | 124.9886 | 224.8715 | 93.34999 | -0.55337 | 3.001829 | 0.280959 |
| 4 | 53462600 | 53463799 | 8.344081 | 15.47482 | 32.61508 | 13.69031 | -0.62499 | 0.180603 | 0.280963 |
| 16 | 95928000 | 95929599 | 12.44 | 21.75219 | 46.24515 | 20.38876 | -0.69514 | 0.680696 | 0.280976 |
| 2 | 88874600 | 88875399 | 13.37963 | 25.00852 | 22.32328 | 9.254355 | 0.619265 | 0.147875 | 0.280984 |
| 7 | 93455200 | 93456399 | 13.37963 | 25.00852 | 22.32328 | 9.254355 | 0.619265 | 0.147875 | 0.280984 |
| 17 | 57188000 | 57199599 | 150.9618 | 222.7523 | 374.6194 | 154.2743 | -0.50491 | 3.762414 | 0.280999 |
| 9 | 18962600 | 18964799 | 34.61147 | 58.06369 | 44.74331 | 19.52505 | 0.75561 | 1.176036 | 0.280999 |
| 2 | 153114000 | 153115799 | 7.69719 | 14.38641 | 32.80585 | 13.60127 | -0.7241 | 0.124728 | 0.281002 |
| 16 | 3546000 | 3547599 | 8.831223 | 16.20703 | 34.03569 | 14.49239 | -0.63509 | 0.246567 | 0.281006 |
| 12 | 63057400 | 63058799 | 19.02784 | 33.54977 | 29.65606 | 13.12458 | 0.566166 | 0.570941 | 0.281014 |
| 7 | 7553200 | 7555999 | 21.32335 | 37.79693 | 27.04673 | 11.95713 | 0.872033 | 0.523652 | 0.281025 |
| 7 | 19597200 | 19603399 | 40.38108 | 64.37772 | 126.9522 | 53.17982 | -0.63973 | 2.150281 | 0.281026 |
| 13 | 95385000 | 95386999 | 17.78331 | 30.28051 | 57.49087 | 25.19978 | -0.53939 | 1.001648 | 0.281035 |
| 12 | 69408000 | 69408599 | 3.415676 | 6.673549 | 20.52364 | 7.151691 | -0.91767 | -0.56136 | 0.281048 |
| 16 | 49985600 | 49988199 | 13.05625 | 22.82232 | 46.24858 | 20.38466 | -0.62635 | 0.689456 | 0.281051 |
| 12 | 8723600 | 8724999 | 10.51904 | 18.67095 | 39.36844 | 17.38675 | -0.67687 | 0.470636 | 0.281068 |
| 15 | 82458800 | 82459399 | 12.28893 | 23.43691 | 20.45536 | 8.15734 | 0.69851 | 0.002353 | 0.281073 |
| 19 | 50342400 | 50343399 | 12.28893 | 23.43691 | 20.45536 | 8.15734 | 0.69851 | 0.002353 | 0.281073 |
| 11 | 121035800 | 121038599 | 19.10749 | 31.83843 | 66.86784 | 29.0555 | -0.68557 | 1.224814 | 0.281086 |
| 9 | 28314200 | 28316199 | 13.29099 | 23.01694 | 50.53577 | 22.17191 | -0.74321 | 0.820277 | 0.281089 |
| 4 | 91258200 | 91263799 | 48.22299 | 76.56653 | 138.7601 | 57.83395 | -0.52432 | 2.287477 | 0.281095 |
| 16 | 34965200 | 34968799 | 31.52696 | 53.20141 | 44.18224 | 19.3232 | 0.64819 | 1.113521 | 0.281096 |
| 6 | 146330200 | 146332199 | 10.80907 | 19.24454 | 38.86008 | 17.11802 | -0.61225 | 0.452439 | 0.2811 |
| 1 | 156416000 | 156416999 | 10.97158 | 21.04608 | 19.54006 | 7.67615 | 0.629716 | -0.06145 | 0.2811 |
| 7 | 63306200 | 63306799 | 6.299276 | 12.05276 | 27.85112 | 11.02139 | -0.68386 | -0.05964 | 0.281117 |
| 1 | 84943400 | 84944599 | 37.17081 | 61.2889 | 54.34506 | 23.4499 | 0.55283 | 1.380363 | 0.281135 |

| | | | | | | | | | |
|---|---|---|---|---|---|---|---|---|---|
| 18 | 81232600 | 81234399 | 10.03353 | 17.92017 | 37.88823 | 16.64562 | -0.67529 | 0.425978 | 0.281179 |
| 7 | 8359800 | 8365399 | 109.0189 | 168.6075 | 153.7972 | 63.88367 | 0.434723 | 2.847484 | 0.281184 |
| 4 | 60643800 | 60645599 | 21.6086 | 37.81449 | 32.77147 | 14.45824 | 0.593062 | 0.693839 | 0.281193 |
| 11 | 44368000 | 44370599 | 21.6086 | 37.81449 | 32.77147 | 14.45824 | 0.593062 | 0.693839 | 0.281193 |
| 4 | 12252400 | 12252999 | 13.64975 | 25.44636 | 22.38756 | 9.322449 | 0.635176 | 0.167988 | 0.281196 |
| 10 | 47510400 | 47514999 | 84.12793 | 132.3302 | 123.7249 | 51.32827 | 0.417916 | 2.530754 | 0.281207 |
| 4 | 140652800 | 140668399 | 145.8653 | 215.0813 | 379.8739 | 156.437 | -0.57646 | 3.773422 | 0.281215 |
| 1 | 90817200 | 90821599 | 55.92722 | 90.50937 | 60.53065 | 25.88841 | 0.941861 | 1.697173 | 0.281241 |
| Y | 88735200 | 88736799 | 12.26842 | 23.27126 | 13.33436 | 5.378917 | 1.28498 | 0.015635 | 0.281254 |
| 5 | 122258000 | 122261399 | 22.21771 | 36.63923 | 73.555 | 31.74107 | -0.62236 | 1.357221 | 0.281267 |
| X | 146566400 | 146569599 | 44.28408 | 72.28101 | 59.4557 | 25.54962 | 0.653983 | 1.530783 | 0.28127 |
| 17 | 47320400 | 47321599 | 7.092472 | 13.3378 | 30.73445 | 12.61067 | -0.72644 | 0.075966 | 0.281271 |
| 6 | 73036200 | 73038599 | 24.61007 | 42.3583 | 37.86991 | 16.61182 | 0.545371 | 0.907487 | 0.281277 |
| 4 | 127049600 | 127051799 | 21.27604 | 35.58009 | 66.20739 | 28.83415 | -0.51181 | 1.189658 | 0.281283 |
| 8 | 48646000 | 48676399 | 354.7402 | 487.3785 | 780.1988 | 324.4084 | -0.53606 | 4.849797 | 0.281288 |
| 1 | 26654600 | 26656599 | 32.23699 | 54.05979 | 42.16133 | 18.39392 | 0.738331 | 1.169586 | 0.281292 |
| 1 | 192530800 | 192531999 | 12.60196 | 23.64483 | 21.43031 | 8.827045 | 0.604964 | 0.115452 | 0.281293 |
| 12 | 23496200 | 23497999 | 25.83397 | 45.18051 | 30.18977 | 13.3122 | 0.966271 | 0.721585 | 0.281309 |
| 6 | 122399200 | 122401399 | 31.65988 | 53.60938 | 42.49182 | 18.60324 | 0.715364 | 1.083119 | 0.28131 |
| 3 | 46970400 | 46973399 | 59.07848 | 94.48445 | 87.80685 | 36.94614 | 0.44867 | 2.059602 | 0.281331 |
| 5 | 59501200 | 59502799 | 20.69641 | 37.00023 | 24.87927 | 10.9214 | 0.967917 | 0.461981 | 0.281339 |
| X | 4913600 | 4914599 | 40.87988 | 68.84387 | 36.81023 | 16.08909 | 1.280916 | 1.15748 | 0.281375 |
| 13 | 97059600 | 97063599 | 26.30937 | 42.59735 | 91.50854 | 39.15078 | -0.73684 | 1.651805 | 0.28138 |
| 13 | 83877000 | 83880999 | 33.7465 | 54.30047 | 104.1377 | 44.21495 | -0.58402 | 1.844507 | 0.281385 |
| 10 | 19637800 | 19638199 | 4.651505 | 9.058105 | 22.99824 | 8.418898 | -0.7107 | -0.35799 | 0.28139 |
| 11 | 99812200 | 99812799 | 4.651505 | 9.058105 | 22.99824 | 8.418898 | -0.7107 | -0.35799 | 0.28139 |
| 9 | 34380000 | 34381199 | 12.97704 | 24.28046 | 21.4484 | 8.876506 | 0.636468 | 0.136997 | 0.281392 |
| 9 | 11721400 | 11723199 | 20.68108 | 36.15482 | 31.8904 | 14.05752 | 0.567725 | 0.702657 | 0.281392 |
| 2 | 79083600 | 79086599 | 16.27604 | 27.56261 | 58.92801 | 25.79986 | -0.71057 | 1.047589 | 0.281398 |
| 6 | 66018400 | 66019799 | 15.81062 | 28.69347 | 25.08329 | 10.83486 | 0.606842 | 0.340854 | 0.281399 |
| 8 | 21988000 | 21989799 | 23.428 | 40.91552 | 32.59486 | 14.35747 | 0.713858 | 0.732618 | 0.281409 |
| X | 38263600 | 38266399 | 47.83155 | 77.69259 | 62.2511 | 26.69641 | 0.686694 | 1.612827 | 0.28141 |
| 12 | 85317000 | 85318799 | 9.277641 | 17.1132 | 35.78256 | 15.13103 | -0.62183 | 0.224872 | 0.28141 |
| 1 | 147626400 | 147626999 | 9.362172 | 18.24256 | 16.08189 | 5.840139 | 0.808346 | -0.35154 | 0.28142 |
| 4 | 106032200 | 106034999 | 23.95139 | 39.59088 | 73.34659 | 31.69673 | -0.50632 | 1.334299 | 0.281435 |
| 13 | 88377200 | 88381399 | 50.1444 | 81.05597 | 66.42149 | 28.37539 | 0.647718 | 1.714996 | 0.281442 |
| 18 | 49282400 | 49283199 | 10.06812 | 19.50613 | 18.11242 | 6.86723 | 0.677456 | -0.19025 | 0.281456 |
| 5 | 146068200 | 146072999 | 104.805 | 162.4154 | 149.5184 | 62.10572 | 0.423664 | 2.808705 | 0.281457 |
| 4 | 4729800 | 4732999 | 20.96591 | 34.93705 | 69.5438 | 30.21935 | -0.60877 | 1.227854 | 0.281469 |
| 9 | 60366000 | 60367399 | 15.91053 | 27.11318 | 56.30934 | 24.68346 | -0.6679 | 0.987308 | 0.281469 |
| 9 | 14898800 | 14900599 | 10.53688 | 18.65551 | 39.89656 | 17.64204 | -0.69891 | 0.494499 | 0.281489 |
| 1 | 138917800 | 138919999 | 12.74104 | 22.34344 | 44.08647 | 19.48484 | -0.58791 | 0.632605 | 0.281499 |
| 6 | 65000200 | 65004199 | 40.54288 | 66.16129 | 60.37089 | 25.941 | 0.504474 | 1.52445 | 0.281502 |
| 12 | 80268800 | 80271999 | 21.48148 | 35.72789 | 68.2875 | 29.63638 | -0.55037 | 1.251323 | 0.281508 |
| 12 | 56392200 | 56394799 | 11.74367 | 20.80248 | 41.89762 | 18.52505 | -0.61298 | 0.4987 | 0.281511 |
| 2 | 99056000 | 99060199 | 83.71983 | 132.0762 | 119.4176 | 49.56153 | 0.469443 | 2.471482 | 0.28152 |
| 6 | 134967400 | 134969799 | 13.0701 | 22.84686 | 45.72857 | 20.15571 | -0.60848 | 0.68955 | 0.28153 |

| | | | | | | | | | |
|---|---|---|---|---|---|---|---|---|---|
| 3 | 112995200 | 112995999 | 13.02677 | 24.5903 | 21.3201 | 8.68885 | 0.680603 | 0.080001 | 0.281544 |
| X | 4935200 | 4938399 | 124.6197 | 193.331 | 133.812 | 55.57363 | 0.835932 | 2.79906 | 0.281558 |
| 2 | 153233400 | 153235999 | 17.26765 | 29.49055 | 56.99271 | 24.98744 | -0.56423 | 0.970614 | 0.281565 |
| 6 | 76605600 | 76607799 | 17.26765 | 29.49055 | 56.99271 | 24.98744 | -0.56423 | 0.970614 | 0.281565 |
| 15 | 44829600 | 44830199 | 12.08896 | 22.85568 | 20.94122 | 8.506291 | 0.60583 | 0.060179 | 0.281569 |
| 10 | 8632600 | 8633599 | 6.022538 | 11.50535 | 28.35201 | 11.26609 | -0.78151 | -0.04844 | 0.281578 |
| 6 | 89989000 | 89991999 | 40.88551 | 67.18644 | 55.55577 | 23.93807 | 0.65144 | 1.432324 | 0.281582 |
| 3 | 32450400 | 32452199 | 11.34593 | 19.92548 | 43.55257 | 19.27224 | -0.73468 | 0.59978 | 0.281582 |
| 8 | 127332400 | 127333599 | 5.873372 | 11.22428 | 27.78497 | 11.03266 | -0.78657 | -0.05245 | 0.281589 |
| 15 | 48978400 | 48978799 | 4.226055 | 8.251849 | 21.72923 | 7.651591 | -0.71178 | -0.50828 | 0.281605 |
| 2 | 167812200 | 167814799 | 11.54576 | 20.17433 | 45.75642 | 20.16144 | -0.78759 | 0.68936 | 0.281607 |
| 4 | 153437400 | 153440799 | 27.27025 | 44.5649 | 81.59326 | 35.07821 | -0.49605 | 1.49081 | 0.28162 |
| 10 | 21933800 | 21935199 | 11.5497 | 20.16818 | 46.28012 | 20.3839 | -0.80457 | 0.698549 | 0.281622 |
| 2 | 5955800 | 5957199 | 9.328163 | 16.5523 | 40.43302 | 17.86136 | -0.88801 | 0.466202 | 0.28163 |
| 17 | 48050200 | 48056799 | 52.0092 | 82.32496 | 149.9473 | 62.33777 | -0.53567 | 2.376223 | 0.281645 |
| 19 | 44487400 | 44492199 | 22.76628 | 37.74221 | 72.95152 | 31.58975 | -0.56704 | 1.294566 | 0.28166 |
| 7 | 10778600 | 10783199 | 91.33806 | 144.5511 | 104.0073 | 43.23494 | 0.802122 | 2.401056 | 0.281664 |
| 1 | 85297600 | 85301799 | 55.37527 | 88.976 | 79.26652 | 33.5858 | 0.517302 | 1.910266 | 0.281674 |
| 3 | 54687800 | 54690199 | 21.09789 | 35.28945 | 66.23899 | 28.85166 | -0.52432 | 1.186073 | 0.281694 |
| 11 | 35435200 | 35436599 | 7.39706 | 13.95713 | 31.35958 | 12.79219 | -0.68292 | 0.061666 | 0.281702 |
| 1 | 101778400 | 101781399 | 35.17124 | 59.10088 | 43.6067 | 19.04707 | 0.818035 | 1.160953 | 0.281708 |
| 6 | 45151000 | 45151799 | 6.167415 | 11.75057 | 28.26592 | 11.29575 | -0.75364 | -0.0259 | 0.281709 |
| 1 | 23479800 | 23480999 | 15.30036 | 29.32781 | 17.39714 | 6.84898 | 1.267357 | -0.01637 | 0.281717 |
| 1 | 176100600 | 176103399 | 18.38107 | 30.51492 | 72.39372 | 31.37843 | -0.86127 | 1.266148 | 0.281722 |
| 14 | 61884800 | 61886799 | 14.93304 | 25.56012 | 54.74039 | 24.00425 | -0.711 | 0.939852 | 0.281734 |
| 12 | 85157200 | 85158199 | 5.59986 | 10.70444 | 28.34633 | 11.24772 | -0.8823 | -0.05741 | 0.281738 |
| 13 | 105034800 | 105036199 | 16.20278 | 29.85479 | 23.53287 | 9.965753 | 0.774534 | 0.25679 | 0.281747 |
| 16 | 56607000 | 56608799 | 13.79245 | 24.03708 | 64.95939 | 28.57572 | -1.04057 | 0.764142 | 0.281756 |
| X | 80726600 | 80732599 | 89.95224 | 142.3223 | 105.6204 | 43.89996 | 0.757306 | 2.4082 | 0.281764 |
| 13 | 31745800 | 31746999 | 7.351157 | 13.94055 | 30.44854 | 12.28896 | -0.62945 | 0.02066 | 0.281764 |
| 5 | 48459600 | 48460599 | 11.66176 | 22.07812 | 19.39171 | 7.854181 | 0.669725 | 0.041852 | 0.281775 |
| 9 | 95330400 | 95333599 | 29.40707 | 47.84387 | 87.34956 | 37.49218 | -0.49662 | 1.562069 | 0.281779 |
| 7 | 38232000 | 38232999 | 22.36692 | 38.95741 | 33.15185 | 14.57813 | 0.61911 | 0.768482 | 0.281819 |
| 11 | 117448400 | 117451199 | 23.90136 | 39.58993 | 74.17529 | 32.10063 | -0.52235 | 1.309066 | 0.28182 |
| 1 | 74120600 | 74125999 | 24.47863 | 39.8102 | 84.31354 | 36.18397 | -0.71036 | 1.558109 | 0.281821 |
| 2 | 137348600 | 137349599 | 13.65014 | 26.25466 | 18.21411 | 7.100254 | 1.056794 | -0.06142 | 0.281824 |
| 10 | 82255400 | 82257199 | 34.88814 | 58.5048 | 45.31156 | 19.76811 | 0.748418 | 1.182684 | 0.281831 |
| 7 | 65610200 | 65612799 | 14.1633 | 24.69012 | 47.87153 | 21.06236 | -0.56341 | 0.737252 | 0.281832 |
| 13 | 93609600 | 93612599 | 28.2274 | 48.12579 | 41.46394 | 18.17662 | 0.596354 | 1.01038 | 0.281834 |
| 12 | 52963400 | 52966799 | 31.73621 | 51.39909 | 92.108 | 39.41583 | -0.47567 | 1.656388 | 0.281835 |
| X | 43821200 | 43821799 | 18.25626 | 33.21958 | 24.00383 | 10.33435 | 0.883347 | 0.344301 | 0.281851 |
| 13 | 116061600 | 116062999 | 14.81512 | 27.30014 | 23.65543 | 10.01655 | 0.638911 | 0.244965 | 0.281856 |
| 1 | 136646000 | 136647799 | 16.04584 | 28.92645 | 25.66391 | 11.17867 | 0.578067 | 0.382396 | 0.281861 |
| 11 | 10588800 | 10590799 | 29.19572 | 49.12932 | 44.82084 | 19.58581 | 0.512962 | 1.129711 | 0.281863 |
| 5 | 105851200 | 105852799 | 9.249809 | 16.59032 | 39.46593 | 17.27715 | -0.84107 | 0.394179 | 0.281873 |
| 14 | 20132200 | 20133599 | 6.04178 | 11.48807 | 28.77826 | 11.54952 | -0.81757 | -0.01389 | 0.281889 |
| 4 | 74314200 | 74315599 | 17.00883 | 30.7514 | 24.81398 | 10.76938 | 0.71708 | 0.369445 | 0.281926 |

| | | | | | | | | | |
|---|---|---|---|---|---|---|---|---|---|
| 8 | 13366800 | 13372599 | 36.55304 | 58.7153 | 107.6765 | 45.60364 | -0.52269 | 1.906571 | 0.281944 |
| 4 | 149610400 | 149612799 | 8.679942 | 15.53173 | 38.44571 | 16.86385 | -0.89961 | 0.404839 | 0.281948 |
| Y | 2184600 | 2186199 | 15.46495 | 28.66127 | 11.20568 | 4.708056 | 1.783559 | 0.18415 | 0.281962 |
| 9 | 111213800 | 111215399 | 7.838799 | 14.71893 | 31.26302 | 12.86059 | -0.61293 | 0.094343 | 0.281969 |
| 9 | 48000200 | 48004399 | 24.29358 | 40.01551 | 74.925 | 32.3162 | -0.52289 | 1.377886 | 0.281977 |
| 15 | 83346000 | 83349799 | 22.81145 | 37.7226 | 72.14438 | 31.18633 | -0.55201 | 1.324478 | 0.281978 |
| 4 | 151376600 | 151378799 | 9.622906 | 17.07334 | 39.41468 | 17.41319 | -0.80697 | 0.467932 | 0.28198 |
| 5 | 37742400 | 37745999 | 26.25954 | 42.62422 | 88.6403 | 37.99435 | -0.6867 | 1.598042 | 0.281983 |
| 14 | 110589800 | 110591399 | 24.91409 | 43.6703 | 29.6574 | 13.10244 | 0.942907 | 0.681157 | 0.281988 |
| 9 | 35871200 | 35871599 | 13.6614 | 25.41937 | 21.92298 | 9.158636 | 0.66007 | 0.179256 | 0.28199 |
| 14 | 52987800 | 52988599 | 7.986124 | 15.02127 | 31.87631 | 13.0749 | -0.60835 | 0.08607 | 0.282 |
| 17 | 19864200 | 19869399 | 83.97537 | 133.7715 | 85.04094 | 35.58408 | 0.989401 | 2.206926 | 0.282 |
| 13 | 94701000 | 94702599 | 8.199547 | 15.17607 | 32.58179 | 13.71399 | -0.6543 | 0.190674 | 0.282001 |
| 5 | 103276600 | 103276999 | 3.93743 | 7.679907 | 21.41295 | 7.67505 | -0.81524 | -0.44786 | 0.282006 |
| 5 | 146167600 | 146168799 | 8.861921 | 15.76848 | 38.94975 | 17.16609 | -0.90126 | 0.444014 | 0.282014 |
| 18 | 38256600 | 38257399 | 11.39303 | 21.8816 | 19.57343 | 7.661963 | 0.688062 | -0.06723 | 0.282015 |
| 2 | 120626200 | 120627599 | 40.90659 | 66.8883 | 59.34268 | 25.51776 | 0.546932 | 1.497039 | 0.282042 |
| 1 | 140152000 | 140153399 | 9.494541 | 17.2453 | 35.45945 | 15.30091 | -0.6179 | 0.315188 | 0.282043 |
| 6 | 117665200 | 117666199 | 12.00172 | 22.74876 | 20.36936 | 8.229431 | 0.645265 | 0.039659 | 0.282057 |
| 2 | 36106000 | 36108599 | 13.27933 | 23.04227 | 49.48353 | 21.71156 | -0.71073 | 0.801566 | 0.282062 |
| 8 | 69952400 | 69955399 | 48.83187 | 78.72075 | 72.97162 | 31.06486 | 0.465874 | 1.795461 | 0.282072 |
| 12 | 74017800 | 74020399 | 26.93941 | 44.15798 | 81.79842 | 35.20816 | -0.51036 | 1.450215 | 0.282073 |
| 4 | 14267600 | 14268599 | 7.497404 | 14.23893 | 31.68174 | 12.74675 | -0.65215 | 0.014564 | 0.282089 |
| 6 | 22147600 | 22148999 | 7.497404 | 14.23893 | 31.68174 | 12.74675 | -0.65215 | 0.014564 | 0.282089 |
| 12 | 114107400 | 114109399 | 29.20033 | 50.61013 | 33.57166 | 14.7295 | 0.975909 | 0.860783 | 0.282091 |
| 12 | 20802600 | 20822999 | 512.1366 | 705.0634 | 540.1495 | 224.523 | 0.529828 | 4.821546 | 0.282097 |
| 9 | 113585200 | 113588999 | 30.92949 | 50.08564 | 90.69732 | 38.80821 | -0.49087 | 1.657052 | 0.282098 |
| 9 | 109881000 | 109882799 | 8.939685 | 16.08426 | 36.42849 | 15.90548 | -0.76732 | 0.369527 | 0.282112 |
| X | 4053600 | 4058999 | 45.59706 | 75.98264 | 39.1837 | 17.02688 | 1.333311 | 1.277441 | 0.282143 |
| 11 | 116636400 | 116638199 | 14.48812 | 24.76182 | 55.22168 | 24.2124 | -0.76938 | 0.953456 | 0.282151 |
| 17 | 72047200 | 72053999 | 53.11474 | 84.14417 | 149.2896 | 62.0902 | -0.49692 | 2.357225 | 0.282157 |
| 9 | 118816800 | 118818399 | 8.077809 | 14.84499 | 34.06352 | 14.47683 | -0.75999 | 0.232044 | 0.282165 |
| 11 | 36834600 | 36836199 | 7.663489 | 14.45862 | 31.40845 | 12.81443 | -0.63562 | 0.061226 | 0.282168 |
| 17 | 65139400 | 65141399 | 9.987204 | 17.97214 | 36.88214 | 16.10108 | -0.62626 | 0.375297 | 0.282169 |
| 3 | 36743000 | 36746199 | 18.70411 | 31.38336 | 63.30744 | 27.62917 | -0.62777 | 1.142657 | 0.28218 |
| 10 | 85607800 | 85614199 | 60.75539 | 97.46104 | 80.81384 | 34.15705 | 0.617794 | 1.965447 | 0.282194 |
| 10 | 63282000 | 63283599 | 9.50855 | 17.15589 | 37.44101 | 16.29135 | -0.71151 | 0.356167 | 0.282197 |
| 9 | 20069800 | 20071399 | 19.31748 | 35.35201 | 21.99535 | 9.401453 | 1.10384 | 0.314398 | 0.282197 |
| 2 | 100194800 | 100196799 | 28.68366 | 49.01799 | 40.31906 | 17.67806 | 0.663439 | 0.989731 | 0.282222 |
| 5 | 109619600 | 109623999 | 96.33944 | 150.2191 | 138.3309 | 57.4068 | 0.430114 | 2.698788 | 0.282236 |
| 17 | 38098400 | 38100799 | 41.76502 | 68.26484 | 58.36007 | 25.09159 | 0.59982 | 1.502227 | 0.282238 |
| 15 | 42032200 | 42032999 | 5.690533 | 11.00287 | 27.21304 | 10.41469 | -0.7356 | -0.18096 | 0.282254 |
| 13 | 66866600 | 66872399 | 125.6745 | 193.6214 | 156.6131 | 65.05375 | 0.603719 | 2.923682 | 0.282256 |
| 4 | 82154200 | 82155399 | 31.31148 | 52.83806 | 44.21854 | 19.33911 | 0.63722 | 1.114121 | 0.28226 |
| 17 | 20394200 | 20401399 | 86.64804 | 137.7328 | 91.88502 | 38.34045 | 0.916777 | 2.277092 | 0.282261 |
| 10 | 122283000 | 122285999 | 18.21427 | 30.23772 | 75.13356 | 32.56578 | -0.92777 | 1.269242 | 0.28227 |
| 12 | 18018400 | 18026799 | 127.403 | 196.3706 | 150.7244 | 62.60843 | 0.679746 | 2.914659 | 0.282281 |

| | | | | | | | | | |
|---|---|---|---|---|---|---|---|---|---|
| 12 | 65877000 | 65882799 | 93.26231 | 146.392 | 127.359 | 52.82154 | 0.519415 | 2.578083 | 0.282283 |
| 4 | 148802600 | 148806199 | 23.32013 | 38.45266 | 74.33579 | 32.07639 | -0.56821 | 1.36137 | 0.282285 |
| 1 | 20959600 | 20962199 | 14.3772 | 24.81954 | 53.22588 | 23.34991 | -0.71068 | 0.855579 | 0.282303 |
| 8 | 84462800 | 84469799 | 44.46953 | 70.6813 | 138.6999 | 57.8698 | -0.63739 | 2.25254 | 0.282305 |
| 7 | 12334400 | 12338599 | 83.77785 | 131.9764 | 122.1018 | 50.66473 | 0.434732 | 2.500955 | 0.282326 |
| 2 | 36707600 | 36711399 | 77.09536 | 122.3497 | 106.2304 | 44.25441 | 0.534977 | 2.317505 | 0.282334 |
| 2 | 130681000 | 130682399 | 7.830635 | 14.26701 | 34.50566 | 14.83203 | -0.84619 | 0.283781 | 0.282334 |
| 1 | 160972600 | 160974599 | 24.94655 | 43.24216 | 36.61958 | 16.06682 | 0.625239 | 0.844947 | 0.282353 |
| 11 | 23131200 | 23132199 | 10.45984 | 20.16709 | 16.80392 | 6.505275 | 0.803203 | -0.12342 | 0.282358 |
| Y | 42464800 | 42466799 | 16.63124 | 29.78962 | 26.07407 | 11.42744 | 0.59151 | 0.425249 | 0.282406 |
| 12 | 22179600 | 22181199 | 17.29081 | 33.28316 | 14.52828 | 5.650728 | 1.721619 | -0.04022 | 0.282441 |
| 10 | 19653800 | 19655799 | 12.88507 | 22.59345 | 44.60186 | 19.71072 | -0.58872 | 0.635288 | 0.282446 |
| 6 | 38188600 | 38210599 | 202.9557 | 294.7104 | 461.8339 | 190.3905 | -0.42522 | 4.019016 | 0.282456 |
| 10 | 68035400 | 68038599 | 22.57196 | 37.56732 | 70.79523 | 30.73927 | -0.53008 | 1.247325 | 0.282458 |
| X | 78190800 | 78193799 | 41.73063 | 68.71572 | 53.36883 | 23.01568 | 0.742728 | 1.408589 | 0.282462 |
| 9 | 33836200 | 33837199 | 7.85815 | 14.65496 | 32.1922 | 13.39669 | -0.67475 | 0.139821 | 0.282464 |
| 17 | 85634200 | 85636599 | 11.77336 | 20.81852 | 40.88784 | 18.08492 | -0.57766 | 0.518055 | 0.28247 |
| 4 | 44493200 | 44499599 | 48.00845 | 76.41854 | 132.1188 | 55.22489 | -0.45277 | 2.208959 | 0.282483 |
| 17 | 94850800 | 94852999 | 22.94044 | 40.09242 | 33.24385 | 14.65036 | 0.656576 | 0.722567 | 0.282499 |
| 19 | 55750800 | 55752999 | 17.73609 | 29.8707 | 61.0832 | 26.69896 | -0.64708 | 1.10335 | 0.282508 |
| 8 | 106840600 | 106843599 | 17.08006 | 29.41285 | 57.10527 | 25.0499 | -0.56878 | 0.895149 | 0.282521 |
| Y | 4183200 | 4184399 | 11.14069 | 21.46992 | 10.84437 | 4.203706 | 1.515824 | -0.13167 | 0.282531 |
| 5 | 58462200 | 58464599 | 23.54895 | 40.89912 | 36.10139 | 15.84666 | 0.566072 | 0.81646 | 0.282532 |
| 17 | 52177400 | 52179799 | 23.54895 | 40.89912 | 36.10139 | 15.84666 | 0.566072 | 0.81646 | 0.282532 |
| 11 | 17105800 | 17107599 | 23.557 | 40.70088 | 36.7909 | 16.14038 | 0.530588 | 0.866913 | 0.282534 |
| 4 | 10954800 | 10956399 | 9.831435 | 17.7331 | 36.40551 | 15.84676 | -0.62394 | 0.360297 | 0.28254 |
| 16 | 59221600 | 59222999 | 10.35195 | 18.41019 | 39.37578 | 17.36146 | -0.69572 | 0.456592 | 0.282542 |
| 7 | 128346600 | 128348199 | 7.917366 | 14.58388 | 34.64088 | 14.67588 | -0.80633 | 0.217953 | 0.282547 |
| 17 | 45037200 | 45037999 | 5.405026 | 10.45487 | 26.01214 | 9.937136 | -0.74103 | -0.19253 | 0.282557 |
| 17 | 57548600 | 57557999 | 61.38917 | 95.54006 | 188.2912 | 78.1624 | -0.66844 | 2.72084 | 0.28256 |
| 13 | 57399600 | 57405999 | 66.729 | 104.3606 | 176.2426 | 73.11679 | -0.44043 | 2.636243 | 0.282561 |
| 12 | 73837200 | 73838999 | 10.05047 | 17.88434 | 38.8862 | 17.13726 | -0.71871 | 0.450566 | 0.282584 |
| 13 | 112320400 | 112321399 | 10.05047 | 17.88434 | 38.8862 | 17.13726 | -0.71871 | 0.450566 | 0.282584 |
| 8 | 87793800 | 87798199 | 34.19972 | 55.1847 | 98.09112 | 41.80945 | -0.46974 | 1.757997 | 0.282585 |
| 18 | 37002600 | 37012199 | 250.468 | 366.2907 | 309.6839 | 127.5778 | 0.474524 | 3.912693 | 0.282592 |
| 10 | 43064000 | 43065199 | 9.772791 | 17.81311 | 36.0343 | 15.48085 | -0.59065 | 0.296912 | 0.282599 |
| 14 | 45667000 | 45669799 | 40.70847 | 66.654 | 58.28183 | 25.07429 | 0.569051 | 1.477136 | 0.282601 |
| 12 | 27270400 | 27272199 | 14.88748 | 27.18128 | 24.2513 | 10.39689 | 0.584648 | 0.298754 | 0.282617 |
| 7 | 103183200 | 103185999 | 40.67059 | 67.2768 | 50.51081 | 21.83937 | 0.792903 | 1.352117 | 0.28262 |
| 4 | 77491400 | 77493599 | 29.13056 | 49.72198 | 37.20484 | 16.31104 | 0.799179 | 0.998651 | 0.282626 |
| 3 | 27211600 | 27213999 | 7.401036 | 13.87925 | 30.67376 | 12.64545 | -0.67201 | 0.097843 | 0.282658 |
| 2 | 3829600 | 3831799 | 23.35181 | 40.87758 | 32.11094 | 14.17049 | 0.734174 | 0.698164 | 0.282665 |
| 1 | 134954600 | 134960799 | 41.11334 | 65.92336 | 116.2087 | 49.08455 | -0.46893 | 1.969719 | 0.282684 |
| 19 | 13014200 | 13015599 | 28.847 | 48.67245 | 44.25553 | 19.35447 | 0.517886 | 1.104942 | 0.282685 |
| 1 | 124569400 | 124572999 | 66.51009 | 105.8155 | 94.86879 | 39.62689 | 0.492311 | 2.232353 | 0.282687 |
| 9 | 118435400 | 118438199 | 13.48988 | 23.24486 | 52.63093 | 23.08761 | -0.789 | 0.869375 | 0.282701 |
| 10 | 71576200 | 71578399 | 17.80912 | 32.90617 | 21.69685 | 9.153457 | 1.034009 | 0.250061 | 0.282704 |

| | | | | | | | | | |
|---|---|---|---|---|---|---|---|---|---|
| 15 | 19530800 | 19532799 | 23.01679 | 40.40907 | 29.70963 | 13.14322 | 0.828815 | 0.64514 | 0.282705 |
| 8 | 126622200 | 126625399 | 17.34292 | 29.44453 | 57.95114 | 25.39185 | -0.59157 | 1.027674 | 0.282709 |
| 8 | 49442000 | 49445799 | 46.79451 | 75.77436 | 60.88974 | 26.05336 | 0.678403 | 1.668661 | 0.282711 |
| 7 | 3074200 | 3077799 | 73.37908 | 116.3262 | 102.4627 | 42.63867 | 0.512776 | 2.346029 | 0.282712 |
| 6 | 11036600 | 11038999 | 25.36625 | 41.41348 | 81.74899 | 35.13282 | -0.60525 | 1.500284 | 0.282723 |
| 10 | 130232200 | 130232999 | 7.828584 | 14.7327 | 31.32736 | 12.83829 | -0.61019 | 0.079727 | 0.282728 |
| 13 | 21579800 | 21580999 | 7.828584 | 14.7327 | 31.32736 | 12.83829 | -0.61019 | 0.079727 | 0.282728 |
| 14 | 62868000 | 62870199 | 27.28381 | 47.4783 | 33.08449 | 14.54022 | 0.90586 | 0.803902 | 0.28273 |
| 6 | 26544400 | 26545799 | 13.03367 | 24.89311 | 19.79923 | 7.871646 | 0.834983 | -0.00234 | 0.282734 |
| 10 | 15448200 | 15451399 | 41.59194 | 66.58985 | 116.1323 | 48.93943 | -0.45609 | 2.016821 | 0.282737 |
| 5 | 114627400 | 114631599 | 41.10136 | 65.79531 | 115.9357 | 48.8467 | -0.47116 | 2.020284 | 0.282756 |
| 5 | 108134200 | 108135399 | 8.021461 | 14.9237 | 32.13537 | 13.42745 | -0.65083 | 0.157302 | 0.282786 |
| 8 | 125442400 | 125445599 | 11.83607 | 20.71675 | 44.67957 | 19.71506 | -0.7154 | 0.660684 | 0.282806 |
| 13 | 22825200 | 22826799 | 51.89633 | 83.45688 | 74.19383 | 31.47515 | 0.521824 | 1.877244 | 0.282813 |
| 3 | 96250000 | 96252799 | 28.16018 | 45.73277 | 86.04091 | 36.89411 | -0.54188 | 1.591098 | 0.28283 |
| 15 | 42751000 | 42751999 | 8.202579 | 15.21448 | 32.61643 | 13.68832 | -0.64896 | 0.1796 | 0.282836 |
| 11 | 90938200 | 90938399 | 7.655414 | 14.93709 | 8.534389 | 2.677518 | 1.605988 | -0.89608 | 0.282838 |
| 3 | 125669800 | 125671399 | 16.06331 | 29.83903 | 21.88388 | 9.165651 | 0.888841 | 0.20687 | 0.282843 |
| 5 | 66175600 | 66185599 | 126.7025 | 189.331 | 322.9087 | 133.1336 | -0.50826 | 3.537171 | 0.282864 |
| 15 | 102835400 | 102840599 | 35.14797 | 56.66567 | 106.9438 | 45.53179 | -0.55755 | 1.786462 | 0.282887 |
| Y | 88474800 | 88476999 | 12.61201 | 23.6664 | 16.84857 | 6.938498 | 0.950303 | 0.100533 | 0.282908 |
| 11 | 45904800 | 45905999 | 8.673502 | 16.02972 | 33.62794 | 14.18265 | -0.62332 | 0.206079 | 0.282921 |
| 12 | 44729600 | 44730599 | 9.636009 | 18.82545 | 12.44722 | 4.342879 | 1.267246 | -0.49735 | 0.282927 |
| 10 | 116899200 | 116899999 | 4.527071 | 8.804008 | 23.4307 | 8.672586 | -0.79268 | -0.31914 | 0.282932 |
| 14 | 64559400 | 64560199 | 4.527071 | 8.804008 | 23.4307 | 8.672586 | -0.79268 | -0.31914 | 0.282932 |
| 12 | 113588600 | 113589799 | 19.67479 | 34.51669 | 30.38826 | 13.4352 | 0.570411 | 0.64021 | 0.28294 |
| 5 | 32790200 | 32793799 | 29.13612 | 50.58848 | 33.05186 | 14.5055 | 0.998179 | 0.84338 | 0.282945 |
| X | 107573800 | 107575599 | 22.42211 | 41.16901 | 18.51698 | 7.879308 | 1.572897 | 0.314214 | 0.282949 |
| 4 | 74445400 | 74449599 | 130.2557 | 198.6942 | 180.1419 | 74.69343 | 0.428214 | 3.121994 | 0.282956 |
| 7 | 26090000 | 26095599 | 85.57906 | 135.6321 | 105.0334 | 43.69735 | 0.697922 | 2.362863 | 0.28299 |
| X | 88719800 | 88721199 | 16.22196 | 30.7753 | 18.95146 | 7.642262 | 1.183774 | 0.071359 | 0.283001 |
| 2 | 119600000 | 119602399 | 11.11209 | 19.38134 | 47.85193 | 21.06072 | -0.90989 | 0.714594 | 0.283008 |
| 3 | 51018400 | 51024199 | 46.67411 | 74.49421 | 128.2804 | 53.82055 | -0.44285 | 2.121 | 0.283043 |
| 9 | 32868200 | 32869799 | 15.4581 | 28.3797 | 24.21377 | 10.30475 | 0.65616 | 0.273059 | 0.283048 |
| 17 | 46680600 | 46682999 | 33.17517 | 55.06224 | 50.36532 | 21.82692 | 0.509138 | 1.290817 | 0.283058 |
| 11 | 97970400 | 97974999 | 35.37739 | 56.90049 | 103.4689 | 43.90395 | -0.50817 | 1.861242 | 0.283062 |
| 19 | 61268000 | 61269799 | 12.74483 | 22.3903 | 43.51798 | 19.26282 | -0.56622 | 0.603094 | 0.283091 |
| 3 | 158407400 | 158408999 | 24.09217 | 42.3107 | 27.91183 | 12.35126 | 0.984217 | 0.642756 | 0.28312 |
| 11 | 6615000 | 6617399 | 26.15844 | 42.78109 | 80.17079 | 34.47639 | -0.52865 | 1.47785 | 0.283129 |
| 2 | 159629800 | 159631199 | 7.382809 | 13.93842 | 30.27705 | 12.33529 | -0.63312 | 0.05364 | 0.283129 |
| 1 | 191670400 | 191681199 | 102.1423 | 154.1324 | 285.2964 | 117.9098 | -0.61361 | 3.342266 | 0.28313 |
| 2 | 177507800 | 177511199 | 27.92354 | 48.35163 | 34.70888 | 15.22801 | 0.862191 | 0.863636 | 0.283136 |
| 5 | 150871000 | 150873799 | 17.24869 | 29.55946 | 55.94443 | 24.53462 | -0.53341 | 0.938266 | 0.283141 |
| 7 | 59733000 | 59736799 | 61.09878 | 98.53831 | 64.67382 | 27.54767 | 0.96354 | 1.799705 | 0.28316 |
| 4 | 41679800 | 41688399 | 43.17234 | 68.67032 | 134.1012 | 55.99683 | -0.62928 | 2.22874 | 0.28317 |
| 1 | 118529200 | 118531199 | 9.159076 | 16.75688 | 34.5145 | 14.75815 | -0.61166 | 0.26858 | 0.283174 |
| 3 | 15585600 | 15589199 | 49.77728 | 81.15191 | 55.28469 | 23.74708 | 0.923958 | 1.555062 | 0.283185 |

| | | | | | | | | | |
|---|---|---|---|---|---|---|---|---|---|
| 7 | 106946000 | 106947399 | 13.3651 | 23.44017 | 45.11614 | 19.9416 | -0.55238 | 0.635812 | 0.283188 |
| 11 | 60265400 | 60266799 | 7.241453 | 13.20713 | 34.52463 | 14.82111 | -0.95616 | 0.272917 | 0.283192 |
| 7 | 3399600 | 3401399 | 10.82401 | 19.26806 | 38.87458 | 17.12671 | -0.61105 | 0.45466 | 0.283192 |
| 15 | 25883400 | 25885799 | 12.26137 | 21.49084 | 46.19087 | 20.40576 | -0.71088 | 0.639818 | 0.283206 |
| 5 | 142515200 | 142517999 | 33.35724 | 53.91734 | 97.52012 | 41.65781 | -0.49194 | 1.705787 | 0.283213 |
| 6 | 65938600 | 65939399 | 14.35134 | 27.42554 | 19.21217 | 7.628946 | 1.018546 | 0.006213 | 0.283228 |
| 6 | 42929000 | 42930999 | 18.41268 | 32.75302 | 28.00483 | 12.34546 | 0.619569 | 0.484886 | 0.283237 |
| 12 | 15054600 | 15056199 | 15.43921 | 28.28772 | 24.05718 | 10.26634 | 0.657641 | 0.281973 | 0.28325 |
| 2 | 154581600 | 154582599 | 8.513458 | 15.74298 | 33.10108 | 13.94953 | -0.6257 | 0.20012 | 0.283251 |
| 19 | 53378400 | 53379399 | 6.998652 | 13.01595 | 31.59354 | 13.20935 | -0.82235 | 0.151406 | 0.283262 |
| X | 33663200 | 33666799 | 34.16385 | 57.99206 | 39.69397 | 17.39131 | 0.926359 | 1.070477 | 0.283265 |
| 1 | 125559200 | 125560199 | 14.07081 | 26.45266 | 21.75751 | 8.932083 | 0.747649 | 0.116887 | 0.283271 |
| 8 | 128384400 | 128385199 | 13.32171 | 25.36754 | 20.36988 | 8.149722 | 0.813827 | 0.022781 | 0.283283 |
| 10 | 40444200 | 40448799 | 36.87385 | 59.08837 | 115.6518 | 48.80263 | -0.62066 | 1.981963 | 0.283291 |
| 3 | 15791800 | 15793199 | 14.66054 | 27.40696 | 22.27145 | 9.230452 | 0.754323 | 0.158942 | 0.283298 |
| 11 | 8270000 | 8271199 | 14.66054 | 27.40696 | 22.27145 | 9.230452 | 0.754323 | 0.158942 | 0.283298 |
| 13 | 3291600 | 3293199 | 19.39493 | 36.66386 | 16.37056 | 6.651488 | 1.633878 | 0.117504 | 0.283306 |
| X | 31243000 | 31243999 | 14.95439 | 28.41594 | 19.60376 | 7.878624 | 1.025303 | 0.050196 | 0.283311 |
| 2 | 179901000 | 179904399 | 15.22739 | 25.71909 | 60.98776 | 26.68038 | -0.85956 | 1.068536 | 0.283312 |
| 12 | 87328600 | 87329999 | 8.904388 | 16.12386 | 35.44718 | 15.35439 | -0.7172 | 0.329009 | 0.283313 |
| 12 | 115697400 | 115702799 | 120.9594 | 186.1677 | 163.6025 | 67.9475 | 0.483102 | 2.94273 | 0.283323 |
| 15 | 98392800 | 98397999 | 37.51546 | 60.16646 | 110.4542 | 46.66823 | -0.52696 | 1.956323 | 0.283341 |
| 1 | 147219000 | 147220399 | 19.72728 | 34.83536 | 28.86345 | 12.76997 | 0.659631 | 0.556282 | 0.28335 |
| 11 | 68321000 | 68323999 | 18.42009 | 30.83166 | 64.87334 | 28.26911 | -0.68836 | 1.169398 | 0.28336 |
| 14 | 42104600 | 42107599 | 46.90393 | 76.05255 | 65.56291 | 28.0804 | 0.579052 | 1.645892 | 0.283367 |
| 14 | 17855400 | 17856399 | 19.91713 | 35.13267 | 25.18579 | 11.14528 | 0.866954 | 0.563443 | 0.283431 |
| 6 | 131864600 | 131871799 | 148.5653 | 226.625 | 165.9704 | 68.8175 | 0.735919 | 3.127557 | 0.283451 |
| 1 | 178938800 | 178942199 | 34.50889 | 55.68762 | 99.54805 | 42.4344 | -0.47781 | 1.756297 | 0.283452 |
| 13 | 9837600 | 9839399 | 11.57876 | 20.5286 | 42.95047 | 18.98689 | -0.66719 | 0.489143 | 0.283455 |
| 8 | 110891800 | 110897199 | 35.40868 | 56.82166 | 109.7046 | 46.39378 | -0.59845 | 1.934205 | 0.283469 |
| 19 | 50960800 | 50962199 | 16.57464 | 29.90598 | 26.10122 | 11.35668 | 0.602635 | 0.382408 | 0.283506 |
| 17 | 29872400 | 29873799 | 14.43075 | 26.93166 | 22.12009 | 9.194092 | 0.735264 | 0.165291 | 0.283507 |
| 8 | 69135200 | 69137399 | 13.59282 | 23.72555 | 48.37862 | 21.30234 | -0.63529 | 0.720165 | 0.283516 |
| X | 109146800 | 109147799 | 16.8131 | 31.79944 | 18.78795 | 7.625461 | 1.23473 | 0.096115 | 0.283528 |
| X | 4048000 | 4049999 | 43.41382 | 72.07811 | 39.54828 | 17.14279 | 1.244107 | 1.311744 | 0.28353 |
| 5 | 110099800 | 110102199 | 28.60097 | 48.49807 | 43.14522 | 18.89738 | 0.549518 | 1.06178 | 0.283538 |
| 8 | 125673000 | 125677199 | 24.60986 | 40.09963 | 83.00713 | 35.64795 | -0.67557 | 1.528722 | 0.283548 |
| 6 | 107069200 | 107069999 | 7.543273 | 14.14096 | 31.2183 | 12.87769 | -0.67147 | 0.101534 | 0.28355 |
| 3 | 37986000 | 37987399 | 7.248215 | 13.5894 | 32.32702 | 13.33243 | -0.77839 | 0.099529 | 0.28355 |
| 18 | 53928200 | 53929399 | 13.27808 | 25.4991 | 18.7744 | 7.351634 | 0.965681 | -0.04992 | 0.28356 |
| 10 | 92995400 | 92998199 | 13.15494 | 22.7351 | 52.65354 | 23.09962 | -0.82062 | 0.839726 | 0.283567 |
| 18 | 20681200 | 20683799 | 22.96573 | 37.97334 | 72.76689 | 31.45291 | -0.55488 | 1.32621 | 0.283567 |
| 14 | 111643200 | 111644599 | 16.93438 | 30.30796 | 26.68077 | 11.70135 | 0.58281 | 0.435094 | 0.28359 |
| 14 | 11051000 | 11052799 | 10.23198 | 18.11902 | 40.40693 | 17.86706 | -0.75895 | 0.490667 | 0.283598 |
| 7 | 4887800 | 4895599 | 57.70799 | 90.93185 | 158.2624 | 65.67138 | -0.4754 | 2.486833 | 0.28361 |
| 1 | 48516200 | 48517399 | 14.3053 | 27.07502 | 20.74971 | 8.409367 | 0.864479 | 0.073583 | 0.283618 |
| X | 49209200 | 49210599 | 14.3053 | 27.07502 | 20.74971 | 8.409367 | 0.864479 | 0.073583 | 0.283618 |

| | | | | | | | | | |
|---|---|---|---|---|---|---|---|---|---|
| 14 | 118434400 | 118435399 | 6.278648 | 12.07813 | 27.51782 | 10.72248 | -0.6428 | -0.11232 | 0.283626 |
| 5 | 78588800 | 78590399 | 17.98545 | 33.30953 | 21.14927 | 8.894188 | 1.091267 | 0.237767 | 0.283636 |
| 10 | 119447800 | 119449799 | 11.4356 | 20.33318 | 39.85982 | 17.57795 | -0.571 | 0.466705 | 0.283661 |
| 18 | 17730800 | 17732799 | 13.66872 | 23.97247 | 48.31042 | 21.35334 | -0.61857 | 0.641328 | 0.283666 |
| 15 | 4365200 | 4366399 | 21.82073 | 38.29932 | 27.67902 | 12.24222 | 0.853693 | 0.640804 | 0.283684 |
| 12 | 100776600 | 100778599 | 9.766751 | 17.75942 | 37.05465 | 15.96707 | -0.63801 | 0.31171 | 0.283686 |
| 7 | 29374200 | 29383799 | 72.55617 | 115.5257 | 91.03142 | 38.06556 | 0.679431 | 2.211379 | 0.283689 |
| 7 | 65133000 | 65136599 | 22.28711 | 37.08294 | 68.9252 | 29.92156 | -0.51018 | 1.25005 | 0.283707 |
| 10 | 40058400 | 40068399 | 59.9656 | 94.42293 | 162.7422 | 67.52414 | -0.46215 | 2.501603 | 0.28371 |
| 10 | 122802400 | 122804199 | 24.49577 | 42.06493 | 37.87152 | 16.61136 | 0.534593 | 0.92601 | 0.283715 |
| 17 | 50672600 | 50673999 | 8.440629 | 15.32994 | 34.97217 | 15.092 | -0.76656 | 0.307006 | 0.28372 |
| 8 | 48463600 | 48465199 | 12.2773 | 21.5369 | 43.5722 | 19.26262 | -0.62374 | 0.624366 | 0.283729 |
| 6 | 111765000 | 111766999 | 18.12254 | 33.0794 | 22.53455 | 9.663968 | 0.970546 | 0.319491 | 0.283729 |
| 12 | 93694000 | 93695799 | 18.12254 | 33.0794 | 22.53455 | 9.663968 | 0.970546 | 0.319491 | 0.283729 |
| 7 | 23384600 | 23386999 | 63.08413 | 100.7725 | 91.69731 | 38.51907 | 0.477197 | 2.101131 | 0.283738 |
| 15 | 103666600 | 103667199 | 23.79468 | 41.54086 | 29.36453 | 12.93162 | 0.885684 | 0.737241 | 0.283767 |
| 6 | 125065600 | 125068999 | 14.03005 | 24.35137 | 50.03396 | 21.9535 | -0.64741 | 0.801702 | 0.283776 |
| 1 | 194214000 | 194215399 | 19.98388 | 36.65417 | 19.62221 | 8.361394 | 1.321714 | 0.301629 | 0.283781 |
| 12 | 18013800 | 18017799 | 65.64365 | 104.8569 | 91.33516 | 38.36475 | 0.540041 | 2.104127 | 0.283785 |
| 18 | 37438200 | 37450199 | 225.9451 | 333.8524 | 253.101 | 104.2326 | 0.645784 | 3.745473 | 0.283803 |
| 1 | 33909800 | 33911799 | 17.52291 | 31.16503 | 27.58028 | 12.16003 | 0.569974 | 0.478584 | 0.283804 |
| 10 | 62448400 | 62451199 | 48.27017 | 77.96274 | 69.97434 | 29.86174 | 0.515408 | 1.737859 | 0.283808 |
| 17 | 33165000 | 33166799 | 8.721951 | 15.86213 | 36.00614 | 15.51217 | -0.75833 | 0.301153 | 0.283816 |
| 12 | 49328000 | 49330199 | 22.6042 | 37.60174 | 69.53833 | 30.18286 | -0.50299 | 1.25383 | 0.283833 |
| 3 | 91370800 | 91372799 | 22.91691 | 40.20113 | 31.02624 | 13.71628 | 0.759475 | 0.661476 | 0.283839 |
| 2 | 36712200 | 36716399 | 105.8527 | 163.9416 | 148.8674 | 61.8355 | 0.442782 | 2.820476 | 0.283857 |
| 17 | 42976600 | 42978599 | 11.45624 | 20.22327 | 41.93083 | 18.55256 | -0.65653 | 0.535296 | 0.283861 |
| 6 | 83844000 | 83846799 | 15.93308 | 27.56866 | 52.19943 | 22.90121 | -0.53114 | 0.841603 | 0.283867 |
| 3 | 107511000 | 107513799 | 15.86758 | 27.11383 | 55.84336 | 24.48457 | -0.65543 | 0.959496 | 0.283869 |
| 3 | 101342400 | 101343199 | 3.99541 | 7.777318 | 23.4849 | 8.627366 | -0.96259 | -0.35379 | 0.283881 |
| 7 | 98098800 | 98100999 | 9.359596 | 16.521 | 41.01429 | 18.14728 | -0.91419 | 0.524125 | 0.283888 |
| 14 | 76695800 | 76696199 | 3.63808 | 7.111783 | 19.48061 | 6.698377 | -0.7347 | -0.607 | 0.283899 |
| 10 | 96450800 | 96455199 | 24.25861 | 39.6111 | 85.83964 | 36.89287 | -0.73942 | 1.496606 | 0.283907 |
| X | 90618400 | 90621399 | 45.83937 | 75.1519 | 53.98895 | 23.24006 | 0.852522 | 1.47287 | 0.283922 |
| 10 | 54897000 | 54897999 | 20.35027 | 35.84461 | 29.2298 | 12.9386 | 0.68156 | 0.591147 | 0.283929 |
| 14 | 58012800 | 58013999 | 13.36542 | 25.33149 | 20.78681 | 8.399574 | 0.770064 | 0.054748 | 0.283929 |
| 17 | 36714600 | 36716799 | 21.61511 | 38.10303 | 29.40952 | 13.01606 | 0.760754 | 0.589419 | 0.283946 |
| 1 | 156145200 | 156146399 | 12.00365 | 23.25445 | 17.35231 | 6.581175 | 0.989326 | -0.16951 | 0.283947 |
| 4 | 147105600 | 147107999 | 61.51129 | 99.05458 | 59.91102 | 25.46642 | 1.078536 | 1.84071 | 0.283961 |
| 2 | 28871800 | 28880999 | 80.0142 | 123.5087 | 206.4678 | 85.76267 | -0.44047 | 2.877446 | 0.283972 |
| X | 89999600 | 90000799 | 30.28292 | 51.59102 | 40.8724 | 17.91621 | 0.716807 | 1.025101 | 0.283984 |
| 1 | 20922600 | 20924999 | 35.09425 | 58.35459 | 50.37418 | 21.85848 | 0.592628 | 1.276939 | 0.283986 |
| 5 | 57172400 | 57176199 | 42.012 | 68.42603 | 61.35372 | 26.3447 | 0.527777 | 1.557783 | 0.28399 |
| 15 | 51875600 | 51878799 | 29.57904 | 48.18567 | 89.25409 | 38.32762 | -0.5161 | 1.545904 | 0.284026 |
| 1 | 17714800 | 17716599 | 11.30114 | 19.95014 | 42.4509 | 18.78254 | -0.69375 | 0.534453 | 0.284035 |
| 14 | 6582800 | 6583999 | 7.810595 | 14.75705 | 32.62983 | 13.27292 | -0.65733 | 0.055399 | 0.284048 |
| 17 | 73222400 | 73226599 | 27.74319 | 44.89218 | 91.00974 | 38.92034 | -0.65429 | 1.666776 | 0.284053 |

| | | | | | | | | | |
|---|---|---|---|---|---|---|---|---|---|
| 4 | 54944800 | 54949799 | 26.50486 | 42.83915 | 94.56916 | 40.39447 | -0.77861 | 1.694932 | 0.284069 |
| 17 | 19620600 | 19622399 | 44.54508 | 72.33374 | 64.36912 | 27.59892 | 0.535275 | 1.611058 | 0.284072 |
| 2 | 87872400 | 87875399 | 56.33349 | 90.75711 | 66.93091 | 28.46882 | 0.793984 | 1.822714 | 0.284094 |
| X | 33927800 | 33932399 | 75.99104 | 121.0157 | 93.7683 | 39.21939 | 0.703939 | 2.212208 | 0.284094 |
| 6 | 60001000 | 60002799 | 18.98054 | 33.43865 | 29.65338 | 13.12557 | 0.561038 | 0.579076 | 0.284101 |
| 17 | 94332600 | 94333199 | 9.79447 | 18.93822 | 17.65181 | 6.755135 | 0.659905 | -0.16291 | 0.284106 |
| 3 | 100378200 | 100380999 | 31.65785 | 51.01065 | 98.97694 | 42.10868 | -0.5982 | 1.797871 | 0.284116 |
| 4 | 24295200 | 24296199 | 11.42608 | 21.69421 | 18.52431 | 7.457381 | 0.718152 | 0.018811 | 0.284125 |
| 4 | 45341200 | 45342999 | 7.677381 | 14.03253 | 38.25327 | 16.37737 | -1.0149 | 0.267108 | 0.284127 |
| 1 | 58554000 | 58556399 | 33.11797 | 56.61925 | 36.87322 | 16.16776 | 0.999057 | 1.000855 | 0.284142 |
| 5 | 120985400 | 121001799 | 137.5363 | 204.9404 | 335.2704 | 138.1588 | -0.45193 | 3.591525 | 0.284144 |
| 3 | 11414000 | 11415199 | 19.19055 | 34.79314 | 24.14976 | 10.44448 | 0.936461 | 0.369824 | 0.284171 |
| 4 | 85843400 | 85844799 | 11.92304 | 22.85203 | 18.76509 | 7.38948 | 0.801512 | -0.04767 | 0.28418 |
| 7 | 38407400 | 38412999 | 128.0252 | 195.9649 | 177.1428 | 73.50835 | 0.436377 | 3.053066 | 0.284185 |
| 14 | 25550600 | 25554199 | 16.78691 | 27.94677 | 68.30444 | 29.66305 | -0.90346 | 1.227474 | 0.284213 |
| 13 | 10499800 | 10500599 | 5.666838 | 10.98261 | 26.14181 | 9.896425 | -0.66474 | -0.21753 | 0.284215 |
| 17 | 61020600 | 61021399 | 10.55713 | 20.33555 | 19.11533 | 7.420065 | 0.628278 | -0.1043 | 0.28423 |
| 11 | 102964200 | 102965999 | 10.86076 | 19.18498 | 40.43471 | 17.88819 | -0.67925 | 0.523807 | 0.284257 |
| 13 | 48267600 | 48268999 | 10.86076 | 19.18498 | 40.43471 | 17.88819 | -0.67925 | 0.523807 | 0.284257 |
| X | 4067000 | 4068399 | 15.50622 | 29.65826 | 17.9466 | 7.109255 | 1.230892 | 0.006485 | 0.28428 |
| 8 | 126746200 | 126748799 | 16.19476 | 27.64191 | 55.83116 | 24.47702 | -0.62763 | 0.971728 | 0.284289 |
| 17 | 26873400 | 26875199 | 8.223128 | 15.14615 | 33.04209 | 13.99973 | -0.68408 | 0.219855 | 0.284298 |
| 16 | 4983400 | 4986999 | 20.80024 | 34.3031 | 73.19878 | 31.5879 | -0.70995 | 1.354906 | 0.2843 |
| 14 | 92460200 | 92462399 | 27.88597 | 47.96049 | 37.51064 | 16.45385 | 0.737154 | 0.925548 | 0.284303 |
| 7 | 20482400 | 20483399 | 18.2066 | 32.45696 | 23.98077 | 10.55215 | 0.830605 | 0.46088 | 0.284315 |
| 12 | 58897000 | 58900599 | 35.46198 | 58.83947 | 50.97667 | 22.08713 | 0.587166 | 1.301031 | 0.284351 |
| 3 | 28777600 | 28781799 | 32.11266 | 51.83009 | 95.70433 | 40.80539 | -0.52406 | 1.747271 | 0.284369 |
| 3 | 113149800 | 113151599 | 23.35285 | 40.65451 | 33.79089 | 14.85452 | 0.652539 | 0.777397 | 0.284386 |
| 14 | 32854800 | 32855799 | 6.781131 | 12.81912 | 29.75605 | 12.08976 | -0.72469 | 0.035992 | 0.284393 |
| 18 | 77861200 | 77862399 | 6.781131 | 12.81912 | 29.75605 | 12.08976 | -0.72469 | 0.035992 | 0.284393 |
| 14 | 46903200 | 46907599 | 44.17337 | 71.82272 | 62.7698 | 26.93587 | 0.56288 | 1.585078 | 0.284394 |
| 3 | 48884800 | 48886599 | 26.11689 | 45.76374 | 27.85174 | 12.30133 | 1.100077 | 0.689141 | 0.284399 |
| 7 | 43300200 | 43312599 | 96.62799 | 147.645 | 240.4093 | 99.72112 | -0.41393 | 3.084378 | 0.284404 |
| 15 | 101767200 | 101769999 | 12.49446 | 21.71752 | 50.00003 | 21.95857 | -0.81044 | 0.773316 | 0.284406 |
| 7 | 41953600 | 41956599 | 85.04722 | 133.7037 | 123.7743 | 51.34513 | 0.431627 | 2.542898 | 0.284412 |
| 7 | 50101000 | 50101799 | 8.723589 | 15.904 | 34.48409 | 14.81101 | -0.68928 | 0.286311 | 0.284423 |
| 9 | 19082000 | 19084199 | 32.91364 | 55.84291 | 40.20547 | 17.61284 | 0.853541 | 1.06814 | 0.284438 |
| 3 | 67335000 | 67338399 | 23.39101 | 38.38873 | 86.81958 | 37.38883 | -0.79678 | 1.427912 | 0.28444 |
| 5 | 147529200 | 147532199 | 22.75228 | 37.78363 | 71.28567 | 30.9049 | -0.53201 | 1.273839 | 0.284446 |
| 9 | 104285200 | 104286599 | 9.621574 | 17.5813 | 35.55054 | 15.22543 | -0.58698 | 0.281004 | 0.28445 |
| 3 | 54731200 | 54735199 | 72.15971 | 112.2456 | 188.0338 | 78.06144 | -0.43489 | 2.73468 | 0.284453 |
| 9 | 112036800 | 112039799 | 16.79935 | 28.7802 | 54.81791 | 24.03996 | -0.54252 | 0.939283 | 0.284456 |
| 7 | 142717400 | 142720799 | 17.03922 | 28.88779 | 58.46261 | 25.60491 | -0.63171 | 1.039317 | 0.284458 |
| 17 | 18368600 | 18370199 | 36.44311 | 59.96199 | 55.31355 | 23.84644 | 0.494789 | 1.406183 | 0.284478 |
| 10 | 93507400 | 93509199 | 11.75148 | 20.75209 | 43.48269 | 19.23777 | -0.67162 | 0.534734 | 0.28449 |
| 8 | 5845400 | 5849799 | 79.77091 | 125.8777 | 118.1385 | 49.03681 | 0.41599 | 2.462914 | 0.284493 |
| 18 | 80142400 | 80145199 | 19.63502 | 32.96622 | 63.41587 | 27.68756 | -0.55958 | 1.138551 | 0.284507 |

| | | | | | | | | | |
|---|---|---|---|---|---|---|---|---|---|
| 11 | 57294400 | 57297599 | 30.23895 | 49.1526 | 89.07559 | 38.22035 | -0.48688 | 1.578483 | 0.284512 |
| 4 | 9140800 | 9150999 | 71.59655 | 111.131 | 194.001 | 80.5645 | -0.4967 | 2.771469 | 0.28452 |
| 17 | 78911600 | 78913999 | 13.20743 | 23.06817 | 47.31518 | 20.84278 | -0.64388 | 0.70322 | 0.284527 |
| 1 | 192983600 | 192986199 | 13.49162 | 23.24833 | 52.09371 | 22.85196 | -0.77395 | 0.869491 | 0.284531 |
| 17 | 49772800 | 49775599 | 14.82951 | 25.6132 | 53.24885 | 23.36035 | -0.66599 | 0.852818 | 0.284535 |
| 8 | 5846600 | 5849799 | 72.08688 | 114.2759 | 106.4979 | 44.31716 | 0.431479 | 2.344768 | 0.284542 |
| 1 | 157378000 | 157380599 | 23.0958 | 39.94112 | 35.21068 | 15.44757 | 0.56675 | 0.853128 | 0.284542 |
| 15 | 99834600 | 99847799 | 114.6443 | 171.5506 | 317.1373 | 130.7894 | -0.6226 | 3.509681 | 0.284548 |
| 1 | 84058400 | 84059799 | 17.53909 | 31.14049 | 27.62726 | 12.19832 | 0.565029 | 0.489545 | 0.284548 |
| 4 | 7775400 | 7776399 | 17.53909 | 31.14049 | 27.62726 | 12.19832 | 0.565029 | 0.489545 | 0.284548 |
| 9 | 20006200 | 20006799 | 15.64385 | 28.40653 | 24.31416 | 10.49536 | 0.638066 | 0.336734 | 0.284568 |
| 7 | 11669600 | 11698799 | 645.5515 | 871.5388 | 656.372 | 273.6635 | 0.526061 | 5.128867 | 0.284571 |
| 5 | 118982200 | 118984199 | 10.07833 | 18.34002 | 38.70785 | 16.66348 | -0.65369 | 0.312959 | 0.284586 |
| 8 | 107224600 | 107234999 | 66.14137 | 102.8403 | 205.446 | 85.29545 | -0.6891 | 2.74074 | 0.284592 |
| 12 | 18395000 | 18396799 | 82.78158 | 130.2334 | 110.4316 | 45.8144 | 0.558838 | 2.524714 | 0.2846 |
| 8 | 106441200 | 106444199 | 14.95498 | 25.55036 | 55.23674 | 24.21829 | -0.72487 | 0.958647 | 0.284612 |
| 16 | 97378400 | 97380799 | 25.83593 | 44.25905 | 39.89392 | 17.49509 | 0.532392 | 0.955706 | 0.284618 |
| 13 | 120296400 | 120302399 | 44.82111 | 71.17288 | 141.0686 | 58.80248 | -0.65313 | 2.281326 | 0.284619 |
| 10 | 45389000 | 45390399 | 15.52249 | 29.05298 | 21.54056 | 8.909507 | 0.887626 | 0.156159 | 0.284625 |
| 7 | 12375600 | 12385999 | 152.8535 | 232.5495 | 186.7027 | 77.35709 | 0.600396 | 3.183597 | 0.284626 |
| 6 | 131231200 | 131233999 | 49.41265 | 79.67684 | 72.96035 | 31.07004 | 0.483905 | 1.790398 | 0.284643 |
| 7 | 19469400 | 19477199 | 47.09191 | 74.76585 | 138.9056 | 57.89088 | -0.56019 | 2.288461 | 0.284645 |
| 19 | 17058000 | 17059399 | 9.03771 | 16.3966 | 35.45104 | 15.31939 | -0.69127 | 0.317694 | 0.284655 |
| 4 | 131072400 | 131073999 | 5.118518 | 9.903497 | 25.41544 | 9.696289 | -0.78291 | -0.20106 | 0.284666 |
| 19 | 6720200 | 6726199 | 32.02418 | 51.46476 | 106.4803 | 45.12777 | -0.69546 | 1.8833 | 0.284679 |
| 16 | 18252400 | 18255399 | 23.53603 | 38.67333 | 75.67237 | 32.60491 | -0.58759 | 1.407767 | 0.284689 |
| 3 | 13409200 | 13411999 | 29.85571 | 51.74449 | 31.79245 | 13.94886 | 1.085741 | 0.860523 | 0.284691 |
| 4 | 84070600 | 84072199 | 8.037844 | 14.87672 | 33.65833 | 14.16721 | -0.72979 | 0.189728 | 0.2847 |
| 12 | 91469000 | 91471199 | 14.0319 | 24.56011 | 48.34413 | 21.33206 | -0.58468 | 0.677646 | 0.28471 |
| 14 | 7381600 | 7383599 | 21.04301 | 36.93128 | 31.0236 | 13.72057 | 0.637578 | 0.641404 | 0.284727 |
| 2 | 36954000 | 36955599 | 20.63678 | 37.43783 | 22.3128 | 9.642724 | 1.155317 | 0.372932 | 0.284738 |
| 5 | 149559000 | 149560399 | 7.586626 | 13.73548 | 36.50496 | 15.81576 | -0.98909 | 0.321363 | 0.284739 |
| X | 3180000 | 3187599 | 160.8412 | 245.3003 | 167.2498 | 69.34468 | 0.838776 | 3.136383 | 0.284742 |
| Y | 5506800 | 5509199 | 26.7802 | 45.80188 | 28.45709 | 12.47794 | 1.06646 | 0.957937 | 0.284746 |
| 14 | 70526400 | 70527799 | 8.789445 | 15.8157 | 36.44509 | 15.91125 | -0.79184 | 0.368654 | 0.284759 |
| 10 | 75811000 | 75815799 | 45.54026 | 72.59425 | 127.5285 | 53.41253 | -0.47352 | 2.158851 | 0.284762 |
| X | 50686200 | 50687399 | 15.49761 | 29.69954 | 17.38949 | 6.850182 | 1.285246 | -0.01241 | 0.284766 |
| 19 | 55890000 | 55892599 | 16.00673 | 27.46813 | 53.72857 | 23.56541 | -0.58025 | 0.919913 | 0.284773 |
| 14 | 36122200 | 36122999 | 23.49332 | 40.74486 | 35.74655 | 15.68403 | 0.574655 | 0.833021 | 0.284791 |
| 3 | 44974800 | 44976799 | 18.41547 | 33.50855 | 23.64457 | 10.17996 | 0.916782 | 0.342281 | 0.284792 |
| 3 | 63871200 | 63872999 | 11.01537 | 19.4164 | 41.96387 | 18.57265 | -0.7168 | 0.549526 | 0.284803 |
| 2 | 51808600 | 51811599 | 31.68304 | 52.93367 | 48.11807 | 20.94351 | 0.518238 | 1.212768 | 0.284814 |
| 8 | 91508000 | 91510999 | 28.28624 | 46.13439 | 84.88149 | 36.46572 | -0.50512 | 1.5247 | 0.284817 |
| 9 | 19998400 | 19999999 | 23.8921 | 41.57612 | 34.30678 | 15.07739 | 0.662966 | 0.786939 | 0.28483 |
| 7 | 73717600 | 73719599 | 22.31437 | 37.00945 | 69.90671 | 30.28055 | -0.53373 | 1.288065 | 0.284836 |
| 15 | 26917800 | 26919199 | 21.54574 | 35.65575 | 70.80642 | 30.62351 | -0.60593 | 1.3126 | 0.28484 |
| 10 | 53109600 | 53111599 | 15.12663 | 26.34299 | 50.56732 | 22.23441 | -0.54931 | 0.760135 | 0.284854 |

| | | | | | | | | | |
|---|---|---|---|---|---|---|---|---|---|
| 8 | 113951800 | 113953999 | 18.31581 | 32.67062 | 26.22766 | 11.53519 | 0.711886 | 0.46145 | 0.284865 |
| 17 | 37958600 | 37969199 | 163.3884 | 247.5405 | 197.7596 | 81.83503 | 0.602581 | 3.270969 | 0.284869 |
| 7 | 61595000 | 61596599 | 15.713 | 29.40084 | 21.53346 | 8.91103 | 0.904619 | 0.159763 | 0.284871 |
| 2 | 32529400 | 32533999 | 17.44021 | 29.30698 | 63.25696 | 27.6304 | -0.72497 | 1.122347 | 0.284875 |
| 8 | 67156000 | 67158199 | 14.94384 | 26.0155 | 50.033 | 21.9946 | -0.5521 | 0.762687 | 0.284878 |
| 17 | 6300400 | 6301999 | 15.64346 | 28.71079 | 24.2084 | 10.30697 | 0.672687 | 0.276439 | 0.284883 |
| 17 | 75731000 | 75731999 | 8.379106 | 15.39285 | 33.52865 | 14.25687 | -0.68582 | 0.236393 | 0.284884 |
| 18 | 84178800 | 84187999 | 77.66989 | 119.2654 | 221.9149 | 92.13671 | -0.6006 | 2.978347 | 0.284901 |
| 11 | 30599000 | 30599599 | 4.282876 | 8.32373 | 24.52449 | 9.124927 | -0.94532 | -0.30189 | 0.284907 |
| 5 | 116195600 | 116196199 | 4.518917 | 8.812274 | 22.68272 | 8.152123 | -0.70656 | -0.42859 | 0.284916 |
| 4 | 38702600 | 38703399 | 12.67925 | 23.89491 | 21.33745 | 8.723448 | 0.634504 | 0.087378 | 0.284918 |
| 7 | 5893400 | 5899599 | 75.49443 | 120.6071 | 82.86676 | 34.81375 | 0.881893 | 2.109602 | 0.284921 |
| 6 | 103460600 | 103461999 | 7.978458 | 14.97232 | 32.37914 | 13.33306 | -0.64022 | 0.100691 | 0.28493 |
| 10 | 59576400 | 59577599 | 7.978458 | 14.97232 | 32.37914 | 13.33306 | -0.64022 | 0.100691 | 0.28493 |
| 7 | 37589600 | 37604399 | 134.4136 | 200.5775 | 334.2955 | 137.7915 | -0.47685 | 3.564045 | 0.284939 |
| 7 | 129892800 | 129894799 | 12.31792 | 21.47027 | 47.86641 | 21.05832 | -0.76372 | 0.729752 | 0.28494 |
| 15 | 19035000 | 19035999 | 6.170424 | 11.77851 | 30.06668 | 11.96751 | -0.83471 | -0.03877 | 0.284946 |
| 2 | 20262000 | 20264199 | 44.5975 | 72.43051 | 64.32408 | 27.58288 | 0.538436 | 1.608709 | 0.28495 |
| 1 | 119447000 | 119462599 | 140.8867 | 208.9012 | 354.8129 | 146.1387 | -0.51257 | 3.679038 | 0.284961 |
| 7 | 61815800 | 61818199 | 33.10866 | 56.11097 | 40.77436 | 17.856 | 0.84021 | 1.079023 | 0.284968 |
| 1 | 37821800 | 37823199 | 12.57652 | 23.9756 | 20.39765 | 8.141349 | 0.733869 | 0.007368 | 0.284974 |
| 9 | 11829400 | 11830599 | 12.57652 | 23.9756 | 20.39765 | 8.141349 | 0.733869 | 0.007368 | 0.284974 |
| 12 | 114795800 | 114799199 | 65.34981 | 104.5858 | 85.95538 | 36.20221 | 0.626698 | 2.045516 | 0.284986 |
| 10 | 129770200 | 129773399 | 30.41046 | 51.80542 | 40.84943 | 17.90606 | 0.723541 | 1.02565 | 0.284992 |
| 9 | 34892400 | 34893599 | 7.272208 | 13.57945 | 31.1288 | 12.92632 | -0.73277 | 0.125437 | 0.285005 |
| 15 | 21718200 | 21722399 | 61.57188 | 98.31256 | 90.41727 | 37.95721 | 0.46112 | 2.110692 | 0.285007 |
| 4 | 30591400 | 30591999 | 4.970869 | 9.663488 | 25.9015 | 9.615965 | -0.80859 | -0.29896 | 0.28505 |
| 11 | 114580400 | 114590799 | 83.90489 | 128.5361 | 232.4802 | 96.48976 | -0.56239 | 3.027101 | 0.285054 |
| 10 | 89146800 | 89147599 | 13.46834 | 25.85895 | 18.76701 | 7.353247 | 0.985538 | -0.04592 | 0.285054 |
| 7 | 78226200 | 78229999 | 30.51682 | 49.19754 | 102.5536 | 43.66064 | -0.70058 | 1.778823 | 0.285069 |
| 16 | 18006200 | 18008999 | 23.78589 | 39.32474 | 72.7697 | 31.45153 | -0.50463 | 1.331136 | 0.285079 |
| 4 | 63213000 | 63214399 | 8.475997 | 15.29417 | 35.95478 | 15.64302 | -0.81726 | 0.348118 | 0.285079 |
| 4 | 37895800 | 37896199 | 9.325356 | 18.07999 | 17.16258 | 6.485445 | 0.650924 | -0.21184 | 0.285098 |
| 10 | 93926200 | 93928599 | 26.97823 | 46.59161 | 37.09632 | 16.27416 | 0.712826 | 0.887361 | 0.285191 |
| 17 | 89334800 | 89336199 | 17.04798 | 31.44859 | 22.91295 | 9.687931 | 0.888918 | 0.255904 | 0.285199 |
| 6 | 131652600 | 131656399 | 52.65302 | 85.37488 | 60.95729 | 26.10793 | 0.85026 | 1.653968 | 0.285207 |
| 8 | 120062400 | 120089999 | 235.5763 | 334.417 | 556.4628 | 230.1773 | -0.54428 | 4.354175 | 0.285215 |
| 12 | 73081400 | 73082799 | 12.79394 | 22.26569 | 47.88508 | 21.04616 | -0.71224 | 0.755485 | 0.285216 |
| 7 | 112527800 | 112528999 | 7.679513 | 14.48979 | 30.83085 | 12.57704 | -0.60545 | 0.061154 | 0.285218 |
| 2 | 59216400 | 59218199 | 12.24831 | 21.56559 | 42.45346 | 18.79352 | -0.58359 | 0.57035 | 0.285228 |
| 11 | 10548000 | 10549399 | 8.629939 | 16.11082 | 36.77691 | 15.27803 | -0.72819 | 0.150271 | 0.285244 |
| 14 | 110541600 | 110543399 | 22.7866 | 40.1958 | 26.85903 | 11.88563 | 0.967565 | 0.582083 | 0.28525 |
| X | 153063000 | 153072199 | 246.5501 | 362.2108 | 288.2362 | 118.7082 | 0.56813 | 3.841868 | 0.285256 |
| 13 | 77856600 | 77857199 | 5.94301 | 11.19922 | 30.19922 | 12.3479 | -0.94657 | 0.051612 | 0.28526 |
| 14 | 87646600 | 87647399 | 18.00327 | 31.81364 | 27.73923 | 12.27084 | 0.586969 | 0.536961 | 0.285262 |
| 10 | 56771200 | 56772199 | 16.48575 | 30.81883 | 20.84802 | 8.640629 | 1.016471 | 0.17002 | 0.285263 |
| 9 | 96488800 | 96490399 | 32.41442 | 54.71453 | 39.45157 | 17.25572 | 0.851339 | 1.110802 | 0.285279 |

| | | | | | | | | | |
|---|---|---|---|---|---|---|---|---|---|
| 1 | 49788200 | 49789799 | 29.84335 | 50.20947 | 42.26884 | 18.46956 | 0.628612 | 1.133514 | 0.285282 |
| 8 | 33861600 | 33865999 | 60.36813 | 96.80954 | 75.80421 | 32.02442 | 0.699731 | 1.972546 | 0.285288 |
| 9 | 120616600 | 120629999 | 129.1061 | 193.257 | 319.9543 | 131.9599 | -0.46316 | 3.504911 | 0.28529 |
| 16 | 76984600 | 76984999 | 9.645742 | 18.80309 | 15.45329 | 5.580768 | 0.915104 | -0.36964 | 0.285293 |
| 10 | 77974800 | 77976999 | 9.761405 | 17.34267 | 39.9069 | 17.61042 | -0.80138 | 0.459901 | 0.285295 |
| 10 | 87065800 | 87066599 | 6.271755 | 12.08659 | 27.61095 | 10.70337 | -0.63971 | -0.12938 | 0.285299 |
| 5 | 80815200 | 80816199 | 13.02384 | 24.47297 | 21.45259 | 8.81387 | 0.655902 | 0.11089 | 0.285306 |
| 6 | 134500600 | 134502999 | 20.26001 | 35.79354 | 28.80126 | 12.74121 | 0.701959 | 0.554601 | 0.285307 |
| 3 | 104107400 | 104109399 | 40.37873 | 64.41925 | 124.0172 | 52.00091 | -0.60399 | 2.127961 | 0.28531 |
| 12 | 44374200 | 44377399 | 23.12047 | 37.83793 | 81.13683 | 34.90034 | -0.72182 | 1.459456 | 0.285316 |
| 15 | 83862200 | 83864599 | 20.16783 | 33.7168 | 64.95936 | 28.28531 | -0.5617 | 1.188716 | 0.285316 |
| 7 | 61159000 | 61163599 | 55.76492 | 90.31883 | 60.43671 | 25.86346 | 0.94216 | 1.68295 | 0.285317 |
| 7 | 45554400 | 45561999 | 45.91558 | 73.10533 | 130.0115 | 54.36273 | -0.49303 | 2.199296 | 0.285319 |
| 9 | 122676600 | 122681399 | 77.78584 | 123.6371 | 99.75126 | 41.61986 | 0.64284 | 2.270984 | 0.285335 |
| 7 | 56590200 | 56591199 | 13.88457 | 26.11029 | 21.76389 | 8.930208 | 0.729101 | 0.11322 | 0.285359 |
| 1 | 46506800 | 46508399 | 19.30089 | 35.41158 | 21.45499 | 9.138809 | 1.145332 | 0.299321 | 0.285361 |
| 4 | 86690200 | 86691199 | 12.15775 | 22.89085 | 20.36227 | 8.337867 | 0.638198 | 0.085668 | 0.285373 |
| 1 | 18322600 | 18325399 | 30.27346 | 50.69421 | 45.96263 | 20.03418 | 0.521957 | 1.183617 | 0.285376 |
| 2 | 177463000 | 177465199 | 33.2536 | 55.26144 | 49.28445 | 21.37694 | 0.545611 | 1.274912 | 0.285412 |
| 3 | 14221600 | 14222199 | 9.291918 | 18.13947 | 13.69449 | 4.840873 | 1.062382 | -0.45435 | 0.285414 |
| Y | 35933200 | 35933799 | 9.504387 | 18.38593 | 16.52907 | 6.311651 | 0.713847 | -0.17894 | 0.285425 |
| 14 | 93201000 | 93203399 | 24.51313 | 42.80943 | 32.54189 | 14.33388 | 0.781235 | 0.739375 | 0.285439 |
| 16 | 34437000 | 34437999 | 15.33273 | 28.7063 | 21.54766 | 8.907985 | 0.870487 | 0.152551 | 0.285445 |
| 17 | 10247800 | 10252199 | 30.12045 | 48.73708 | 94.81997 | 40.54867 | -0.59525 | 1.671617 | 0.285448 |
| 13 | 70526600 | 70528199 | 8.974802 | 16.57479 | 34.1576 | 14.41996 | -0.59902 | 0.213008 | 0.285451 |
| 13 | 97404000 | 97406199 | 44.4811 | 72.55951 | 59.55847 | 25.58825 | 0.656394 | 1.537861 | 0.285453 |
| 14 | 120200600 | 120204599 | 29.27078 | 47.50214 | 91.1481 | 39.06328 | -0.57133 | 1.608408 | 0.28546 |
| 12 | 60206400 | 60209199 | 38.68091 | 63.85366 | 53.2995 | 23.01232 | 0.640244 | 1.370043 | 0.285477 |
| 18 | 24338000 | 24339599 | 14.0682 | 24.32288 | 51.07684 | 22.40817 | -0.67987 | 0.839126 | 0.285502 |
| 11 | 50731600 | 50732399 | 5.216293 | 10.15393 | 24.83626 | 9.123201 | -0.66283 | -0.33316 | 0.285516 |
| 18 | 53309200 | 53310799 | 8.971635 | 16.04926 | 37.44427 | 16.42839 | -0.8147 | 0.408869 | 0.285528 |
| 5 | 41183000 | 41184199 | 15.33156 | 29.18953 | 19.0493 | 7.624248 | 1.109965 | 0.039654 | 0.285532 |
| 13 | 46597200 | 46598599 | 8.791093 | 16.32131 | 33.73315 | 14.13901 | -0.59576 | 0.18003 | 0.285546 |
| 6 | 22169000 | 22176199 | 66.20516 | 105.6749 | 93.33275 | 39.16242 | 0.51892 | 2.128482 | 0.28555 |
| 6 | 136339400 | 136341999 | 17.05099 | 28.85757 | 59.53671 | 26.0616 | -0.65954 | 1.055096 | 0.285566 |
| 5 | 79002200 | 79005799 | 57.95407 | 92.99122 | 81.57696 | 34.49309 | 0.537095 | 1.954169 | 0.28558 |
| 9 | 90256200 | 90257999 | 22.10824 | 39.16978 | 26.39133 | 11.66848 | 0.958073 | 0.532057 | 0.285593 |
| 4 | 6283800 | 6284999 | 12.7095 | 24.11144 | 20.86326 | 8.413379 | 0.696695 | 0.04228 | 0.285598 |
| 8 | 67312800 | 67313999 | 8.704663 | 15.51906 | 38.95018 | 17.13793 | -0.92253 | 0.429764 | 0.285603 |
| 10 | 94923400 | 94926199 | 11.92884 | 20.98041 | 45.06757 | 19.95506 | -0.70967 | 0.582829 | 0.285606 |
| X | 43350400 | 43351999 | 27.79494 | 48.21533 | 34.7994 | 15.26856 | 0.85503 | 0.849319 | 0.285615 |
| 6 | 115841800 | 115843199 | 6.770184 | 12.87336 | 30.45548 | 12.22209 | -0.73542 | 0.000654 | 0.285621 |
| X | 126241000 | 126241599 | 18.36387 | 32.62878 | 26.7515 | 11.80435 | 0.679087 | 0.490791 | 0.285621 |
| 8 | 65481800 | 65483599 | 12.06093 | 21.31451 | 42.9497 | 18.99912 | -0.61509 | 0.527987 | 0.285622 |
| 14 | 66331400 | 66333199 | 11.74988 | 20.79609 | 41.38237 | 18.30029 | -0.59609 | 0.50733 | 0.285639 |
| 11 | 10946000 | 10950999 | 60.66994 | 97.50202 | 76.72417 | 32.5176 | 0.696322 | 1.910655 | 0.285646 |
| 5 | 113454000 | 113466199 | 77.89318 | 119.4545 | 241.3819 | 100.2015 | -0.72096 | 3.005303 | 0.285659 |

| | | | | | | | | | |
|---|---|---|---|---|---|---|---|---|---|
| 7 | 20786200 | 20793199 | 64.82833 | 104.2678 | 70.13805 | 29.76053 | 0.923532 | 1.891463 | 0.285662 |
| 11 | 106514600 | 106526399 | 118.6247 | 178.5931 | 294.1562 | 121.4921 | -0.44829 | 3.391339 | 0.285665 |
| 18 | 38245800 | 38255799 | 69.38669 | 108.3808 | 180.3087 | 74.81334 | -0.42025 | 2.660275 | 0.285671 |
| 8 | 23023400 | 23032999 | 66.80147 | 103.8592 | 191.6135 | 79.55344 | -0.57451 | 2.740443 | 0.285681 |
| 13 | 34996000 | 34999199 | 51.371 | 82.74454 | 75.36453 | 32.04612 | 0.48976 | 1.826553 | 0.285684 |
| 8 | 46371400 | 46372199 | 7.913509 | 14.61945 | 32.5589 | 13.73907 | -0.70923 | 0.199663 | 0.285713 |
| 5 | 57174200 | 57176399 | 32.65825 | 54.3168 | 47.11096 | 20.44582 | 0.585893 | 1.265552 | 0.285722 |
| 7 | 131880400 | 131881399 | 12.10611 | 23.09476 | 20.46187 | 8.155066 | 0.677696 | -0.00152 | 0.285726 |
| 13 | 66604200 | 66608599 | 20.41736 | 35.93208 | 25.72537 | 11.3893 | 0.86764 | 0.592441 | 0.285726 |
| X | 111644000 | 111646199 | 23.13159 | 40.85793 | 27.59403 | 12.20768 | 0.953358 | 0.574164 | 0.285729 |
| 9 | 74859600 | 74860999 | 9.798459 | 17.32297 | 40.95844 | 18.11686 | -0.84366 | 0.507487 | 0.285751 |
| X | 76257600 | 76259599 | 23.01699 | 40.65797 | 27.66661 | 12.23964 | 0.942646 | 0.573394 | 0.285756 |
| 6 | 50804200 | 50806199 | 36.927 | 60.88212 | 54.84291 | 23.66394 | 0.530124 | 1.382191 | 0.285768 |
| 14 | 48290800 | 48292399 | 18.37769 | 32.59815 | 27.34488 | 12.08341 | 0.644386 | 0.498779 | 0.28577 |
| 5 | 73745200 | 73746399 | 10.93424 | 21.31954 | 14.10324 | 5.078521 | 1.227851 | -0.36675 | 0.285776 |
| 7 | 19241000 | 19242799 | 7.421549 | 13.85437 | 31.12392 | 12.93053 | -0.70437 | 0.129018 | 0.28578 |
| 8 | 40656000 | 40656599 | 7.421549 | 13.85437 | 31.12392 | 12.93053 | -0.70437 | 0.129018 | 0.28578 |
| 2 | 166658800 | 166662599 | 38.91818 | 62.09198 | 124.5248 | 52.21707 | -0.66279 | 2.124102 | 0.285785 |
| 12 | 105664800 | 105666999 | 7.689835 | 14.01384 | 35.01377 | 15.04593 | -0.89283 | 0.280469 | 0.28581 |
| 4 | 50062400 | 50063399 | 14.10047 | 26.07824 | 23.32611 | 9.828185 | 0.598721 | 0.219433 | 0.285815 |
| 9 | 101127800 | 101129799 | 12.13205 | 21.21465 | 47.29561 | 20.85267 | -0.76337 | 0.679116 | 0.285818 |
| 1 | 111346600 | 111347999 | 14.02539 | 26.31919 | 21.07298 | 8.677847 | 0.78217 | 0.124551 | 0.285821 |
| X | 25569200 | 25575399 | 93.28969 | 148.1471 | 88.68817 | 36.97079 | 1.071443 | 2.310755 | 0.285828 |
| 8 | 118270400 | 118270999 | 5.25875 | 10.20603 | 27.54734 | 10.36032 | -0.83648 | -0.2455 | 0.28584 |
| 8 | 45677200 | 45677999 | 12.1621 | 23.29423 | 19.98659 | 7.88602 | 0.736881 | -0.03363 | 0.285875 |
| 12 | 117267000 | 117269799 | 15.44078 | 26.73359 | 51.65929 | 22.66467 | -0.56019 | 0.832417 | 0.285887 |
| 2 | 38859000 | 38859599 | 11.03205 | 21.19726 | 19.58476 | 7.657097 | 0.643271 | -0.07523 | 0.285909 |
| 7 | 18170000 | 18173799 | 75.64013 | 120.2199 | 102.3401 | 42.69764 | 0.565501 | 2.272723 | 0.285936 |
| 7 | 17192000 | 17193799 | 45.02652 | 73.00641 | 64.60685 | 27.67036 | 0.541361 | 1.644012 | 0.285939 |
| 5 | 115119600 | 115123799 | 30.94825 | 50.03164 | 92.67897 | 39.59805 | -0.52594 | 1.695431 | 0.285954 |
| 16 | 80217000 | 80217799 | 20.55706 | 36.37111 | 25.49207 | 11.27401 | 0.901003 | 0.536728 | 0.285966 |
| 10 | 75169600 | 75172799 | 24.63075 | 40.48759 | 76.57369 | 32.9986 | -0.53855 | 1.404903 | 0.285969 |
| 7 | 49044800 | 49045799 | 18.23466 | 32.21849 | 27.33675 | 12.0931 | 0.62614 | 0.539959 | 0.285984 |
| 13 | 60545200 | 60547799 | 23.90312 | 41.78228 | 32.58451 | 14.36194 | 0.744571 | 0.723498 | 0.285991 |
| 10 | 124414000 | 124415599 | 13.93334 | 25.7238 | 22.86796 | 9.658258 | 0.605116 | 0.226801 | 0.285993 |
| 14 | 106719200 | 106720799 | 13.93334 | 25.7238 | 22.86796 | 9.658258 | 0.605116 | 0.226801 | 0.285993 |
| 15 | 61363600 | 61364599 | 11.29689 | 21.78243 | 19.00908 | 7.357533 | 0.738076 | -0.10669 | 0.285998 |
| 3 | 58911200 | 58914799 | 26.09643 | 42.37174 | 88.70335 | 38.0295 | -0.69588 | 1.591611 | 0.286005 |
| 3 | 142222400 | 142224799 | 28.78738 | 46.8326 | 85.23818 | 36.58505 | -0.49214 | 1.562208 | 0.286008 |
| 19 | 8289800 | 8291399 | 37.55137 | 61.69522 | 56.49244 | 24.33962 | 0.504601 | 1.426731 | 0.286008 |
| 14 | 111912000 | 111919799 | 59.56209 | 92.79228 | 202.2965 | 83.96325 | -0.81274 | 2.703501 | 0.286016 |
| X | 22349600 | 22351199 | 27.24198 | 47.50574 | 31.14323 | 13.70524 | 0.993936 | 0.774559 | 0.28602 |
| 7 | 6021200 | 6022399 | 26.49883 | 45.21407 | 40.98357 | 17.96717 | 0.523527 | 0.995243 | 0.286041 |
| X | 17896400 | 17899799 | 40.55291 | 67.23479 | 48.81353 | 21.13866 | 0.841383 | 1.325612 | 0.286043 |
| 18 | 84514400 | 84515599 | 15.28898 | 28.35172 | 23.19849 | 9.739063 | 0.730086 | 0.214454 | 0.286052 |
| 8 | 48188200 | 48189199 | 7.892056 | 14.68234 | 32.13032 | 13.42627 | -0.67357 | 0.158802 | 0.286053 |
| 4 | 153854000 | 153867199 | 117.1098 | 176.481 | 292.9408 | 121.0211 | -0.45825 | 3.372147 | 0.28606 |

| | | | | | | | | | |
|---|---|---|---|---|---|---|---|---|---|
| X | 164579000 | 164579999 | 12.42141 | 23.8292 | 19.49529 | 7.65671 | 0.811445 | -0.0461 | 0.286065 |
| 12 | 22310600 | 22312399 | 36.81415 | 60.47973 | 51.97538 | 22.39279 | 0.595881 | 1.424524 | 0.286067 |
| 4 | 9824400 | 9825999 | 12.27166 | 21.58543 | 42.47573 | 18.80708 | -0.58347 | 0.582617 | 0.28608 |
| 11 | 114459400 | 114478599 | 212.4535 | 306.3094 | 483.4885 | 199.4991 | -0.44574 | 4.125327 | 0.286083 |
| 8 | 23873800 | 23878599 | 27.51244 | 44.71236 | 84.85316 | 36.40373 | -0.55337 | 1.575182 | 0.286086 |
| 1 | 168981200 | 168982399 | 4.655493 | 9.053111 | 23.4266 | 8.676045 | -0.75326 | -0.31474 | 0.28609 |
| 13 | 78377800 | 78378399 | 4.655493 | 9.053111 | 23.4266 | 8.676045 | -0.75326 | -0.31474 | 0.28609 |
| 18 | 31923600 | 31925199 | 7.50149 | 14.18387 | 32.07753 | 13.0272 | -0.68757 | 0.045205 | 0.286133 |
| 11 | 58275200 | 58275799 | 4.782248 | 9.31632 | 23.08076 | 8.406324 | -0.66908 | -0.37394 | 0.28614 |
| 11 | 88781800 | 88787399 | 156.8374 | 229.1756 | 417.6394 | 172.06 | -0.63403 | 3.91948 | 0.286141 |
| 13 | 99095400 | 99096999 | 14.37129 | 25.11114 | 51.06495 | 22.50031 | -0.63152 | 0.711263 | 0.286142 |
| 4 | 156045600 | 156047799 | 7.763967 | 13.96656 | 36.97724 | 16.14737 | -0.99096 | 0.362841 | 0.286142 |
| 13 | 56020800 | 56022799 | 15.75773 | 27.37283 | 53.32026 | 23.40769 | -0.5709 | 0.80193 | 0.286151 |
| X | 88226600 | 88229999 | 60.67391 | 97.21803 | 84.5594 | 35.67745 | 0.547058 | 2.004585 | 0.286165 |
| 3 | 106261000 | 106266199 | 90.53254 | 143.1988 | 107.2256 | 44.56057 | 0.744085 | 2.414974 | 0.286173 |
| X | 21934200 | 21938599 | 64.60301 | 103.7911 | 74.05038 | 31.37047 | 0.836862 | 1.925688 | 0.286174 |
| X | 62510400 | 62512999 | 28.18419 | 49.29846 | 28.44598 | 12.54493 | 1.176921 | 0.730236 | 0.286199 |
| 1 | 106381600 | 106384199 | 35.76429 | 59.36812 | 50.43263 | 21.85828 | 0.615493 | 1.296022 | 0.286212 |
| 14 | 61676600 | 61677999 | 12.11738 | 21.29853 | 42.48755 | 18.81457 | -0.60346 | 0.589868 | 0.286216 |
| X | 3414800 | 3415999 | 46.82369 | 76.17785 | 51.02683 | 21.90152 | 0.946267 | 1.578647 | 0.286224 |
| 17 | 79718000 | 79719799 | 16.19664 | 27.66794 | 55.29324 | 24.2428 | -0.61213 | 0.964235 | 0.286237 |
| 14 | 104826600 | 104828199 | 12.91145 | 22.66542 | 44.06073 | 19.49027 | -0.56648 | 0.617817 | 0.286239 |
| 7 | 142896600 | 142899199 | 11.83729 | 20.69196 | 46.77759 | 20.61816 | -0.78325 | 0.683209 | 0.286247 |
| 1 | 91339400 | 91342999 | 16.55905 | 28.11045 | 57.91428 | 25.37481 | -0.65725 | 1.025533 | 0.286252 |
| 8 | 13795800 | 13799999 | 31.62178 | 51.00599 | 98.04708 | 41.77066 | -0.58302 | 1.763949 | 0.286255 |
| 14 | 44053400 | 44054399 | 7.510784 | 14.28115 | 34.11828 | 13.69291 | -0.75086 | 0.01284 | 0.28628 |
| 16 | 13327200 | 13334399 | 48.80632 | 77.77696 | 133.5623 | 55.91617 | -0.4413 | 2.173321 | 0.286296 |
| 5 | 134067800 | 134080199 | 106.2289 | 161.1411 | 269.9412 | 111.7471 | -0.46334 | 3.235804 | 0.286296 |
| 15 | 85940600 | 85951399 | 59.2544 | 92.634 | 176.1948 | 73.09995 | -0.61218 | 2.639677 | 0.286332 |
| 11 | 6757400 | 6758999 | 30.37978 | 51.26537 | 44.75144 | 19.57212 | 0.576506 | 1.110209 | 0.286334 |
| 11 | 43349800 | 43350999 | 14.85282 | 27.23798 | 24.1953 | 10.31258 | 0.596638 | 0.273352 | 0.286335 |
| 6 | 89864600 | 89865799 | 8.291512 | 15.48696 | 37.41227 | 15.5285 | -0.80861 | 0.143151 | 0.286336 |
| X | 18028200 | 18029999 | 12.42707 | 21.94838 | 43.49127 | 19.241 | -0.59119 | 0.54022 | 0.286368 |
| 10 | 115348400 | 115352199 | 33.65824 | 55.76305 | 50.93056 | 22.04433 | 0.511091 | 1.312983 | 0.286393 |
| 3 | 49757400 | 49757799 | 5.081908 | 9.880008 | 24.03406 | 8.917883 | -0.66816 | -0.3013 | 0.286406 |
| 19 | 47888200 | 47892199 | 21.42451 | 35.00488 | 81.62977 | 35.09137 | -0.84402 | 1.481284 | 0.286408 |
| X | 15629400 | 15630199 | 12.37502 | 23.93964 | 17.45219 | 6.663429 | 1.014549 | -0.14162 | 0.286431 |
| 1 | 97161400 | 97163799 | 30.20248 | 51.43666 | 40.24699 | 17.64159 | 0.734448 | 1.024652 | 0.286439 |
| 6 | 12472800 | 12474199 | 18.03422 | 33.13797 | 22.75867 | 9.674074 | 0.968089 | 0.284445 | 0.286442 |
| 7 | 9930000 | 9930999 | 18.03422 | 33.13797 | 22.75867 | 9.674074 | 0.968089 | 0.284445 | 0.286442 |
| 18 | 85635600 | 85637199 | 18.03422 | 33.13797 | 22.75867 | 9.674074 | 0.968089 | 0.284445 | 0.286442 |
| 11 | 21343000 | 21347599 | 68.40047 | 108.8513 | 100.4046 | 41.95431 | 0.451752 | 2.225945 | 0.286455 |
| 5 | 143335600 | 143352399 | 165.5731 | 243.2148 | 393.2393 | 161.9544 | -0.45427 | 3.837356 | 0.286456 |
| 7 | 61151000 | 61155599 | 70.97777 | 112.9271 | 101.2434 | 42.2915 | 0.492382 | 2.236737 | 0.286468 |
| 4 | 143547800 | 143550199 | 49.26864 | 80.06672 | 60.75723 | 26.06533 | 0.765601 | 1.604415 | 0.286477 |
| 11 | 38977400 | 38978799 | 16.67234 | 30.28881 | 24.33624 | 10.49851 | 0.72873 | 0.33879 | 0.28648 |
| 13 | 60829800 | 60833199 | 24.2533 | 39.61459 | 81.61749 | 35.08216 | -0.66632 | 1.491056 | 0.286492 |

| | | | | | | | | | |
|---|---|---|---|---|---|---|---|---|---|
| 17 | 84541600 | 84543199 | 9.904371 | 17.67832 | 38.94792 | 17.12029 | -0.73452 | 0.434019 | 0.286504 |
| 9 | 60947600 | 60951199 | 24.43759 | 39.85203 | 81.90314 | 35.1845 | -0.66435 | 1.515863 | 0.286519 |
| 6 | 48266200 | 48270399 | 103.8606 | 160.7818 | 145.263 | 60.33847 | 0.449542 | 2.827264 | 0.286522 |
| 2 | 76451800 | 76457999 | 97.7547 | 152.4477 | 139.6006 | 57.93252 | 0.438324 | 2.696432 | 0.286539 |
| X | 24384800 | 24385599 | 22.81554 | 39.64524 | 35.19568 | 15.45397 | 0.557995 | 0.805904 | 0.286544 |
| 1 | 164602800 | 164607399 | 48.90251 | 78.83995 | 72.85462 | 31.01776 | 0.470492 | 1.79458 | 0.286546 |
| 15 | 83616800 | 83619599 | 15.73836 | 26.82617 | 56.87729 | 24.9329 | -0.69768 | 0.983189 | 0.286573 |
| X | 41968600 | 41969599 | 18.64242 | 32.95316 | 27.80646 | 12.29983 | 0.634356 | 0.539539 | 0.286591 |
| 12 | 24380000 | 24400799 | 371.7027 | 529.0176 | 377.4898 | 156.0787 | 0.680516 | 4.322515 | 0.2866 |
| 10 | 129654000 | 129656399 | 54.30107 | 87.14627 | 81.16416 | 34.32845 | 0.451273 | 1.94509 | 0.2866 |
| 13 | 31958600 | 31960199 | 10.37111 | 18.41409 | 38.88049 | 17.16676 | -0.67844 | 0.467675 | 0.286601 |
| 10 | 41251400 | 41256799 | 38.65411 | 61.95008 | 112.2453 | 47.37554 | -0.50926 | 1.97965 | 0.286605 |
| 2 | 3770800 | 3771799 | 7.058035 | 13.39587 | 29.32242 | 11.81356 | -0.62994 | 0.010224 | 0.286614 |
| 14 | 122066600 | 122068799 | 16.39728 | 28.32014 | 54.93677 | 24.10081 | -0.56664 | 0.862274 | 0.286616 |
| 13 | 100144400 | 100145999 | 6.488102 | 12.27076 | 29.19983 | 11.85236 | -0.7583 | 0.031055 | 0.28663 |
| 4 | 16020800 | 16022199 | 23.42683 | 40.48876 | 35.82448 | 15.71656 | 0.561458 | 0.862852 | 0.286722 |
| 2 | 71482600 | 71483799 | 6.134342 | 11.803 | 27.52481 | 10.71878 | -0.67537 | -0.11635 | 0.286726 |
| 1 | 12881200 | 12882199 | 8.011796 | 14.89803 | 33.7694 | 14.12254 | -0.72576 | 0.162436 | 0.286726 |
| 2 | 79258800 | 79259599 | 4.380093 | 8.458904 | 25.78922 | 9.925389 | -1.03914 | -0.17454 | 0.286756 |
| 12 | 109803600 | 109806599 | 29.97783 | 48.50228 | 90.86901 | 38.8567 | -0.54097 | 1.67315 | 0.286756 |
| 11 | 99957200 | 99958999 | 10.2761 | 18.51654 | 37.40329 | 16.30156 | -0.60222 | 0.369806 | 0.28677 |
| 15 | 70004600 | 70005799 | 14.6329 | 27.23565 | 22.6387 | 9.452553 | 0.713034 | 0.185255 | 0.286772 |
| 12 | 76874200 | 76874799 | 4.057028 | 7.919252 | 20.97478 | 7.427084 | -0.72605 | -0.49081 | 0.286782 |
| 18 | 64969200 | 64969599 | 4.057028 | 7.919252 | 20.97478 | 7.427084 | -0.72605 | -0.49081 | 0.286782 |
| 14 | 83854000 | 83855599 | 14.63481 | 27.48845 | 21.73839 | 8.937716 | 0.802328 | 0.127863 | 0.286798 |
| 4 | 3197000 | 3198999 | 22.57848 | 39.46759 | 33.2418 | 14.65189 | 0.634073 | 0.717223 | 0.286799 |
| 9 | 53295200 | 53296999 | 8.549214 | 15.62662 | 34.50507 | 14.77198 | -0.71259 | 0.267946 | 0.286812 |
| 4 | 113430600 | 113432599 | 39.33431 | 64.70059 | 51.04261 | 22.00252 | 0.719603 | 1.413317 | 0.286814 |
| 2 | 10876600 | 10877399 | 5.721214 | 10.97508 | 27.36036 | 10.74705 | -0.78209 | -0.09199 | 0.286816 |
| 1 | 136272800 | 136273999 | 7.43916 | 13.82421 | 32.1109 | 13.44201 | -0.76125 | 0.159511 | 0.286816 |
| X | 64174200 | 64177199 | 53.31086 | 86.07848 | 69.02119 | 29.44366 | 0.677269 | 1.757638 | 0.286818 |
| 16 | 92304600 | 92305199 | 5.533531 | 10.72745 | 25.5994 | 9.677597 | -0.66695 | -0.2278 | 0.28682 |
| 13 | 28008200 | 28009999 | 15.35175 | 29.08985 | 19.45731 | 7.865967 | 1.061963 | 0.071434 | 0.286825 |
| 8 | 90974800 | 90976199 | 11.4677 | 20.24635 | 40.91322 | 18.1018 | -0.6194 | 0.53329 | 0.286828 |
| 15 | 85751800 | 85756399 | 19.51976 | 32.26966 | 72.18296 | 31.19662 | -0.77715 | 1.320031 | 0.286829 |
| 1 | 145327800 | 145329599 | 17.32662 | 31.94609 | 22.96769 | 9.717903 | 0.907461 | 0.261796 | 0.286838 |
| 1 | 107942600 | 107943999 | 17.05215 | 31.92833 | 20.28426 | 8.384391 | 1.109477 | 0.164164 | 0.286848 |
| 7 | 4618600 | 4620599 | 14.37578 | 24.87194 | 50.55281 | 22.17874 | -0.63277 | 0.833954 | 0.286868 |
| 13 | 61387200 | 61390999 | 42.90632 | 69.60463 | 64.63663 | 27.69366 | 0.472666 | 1.627506 | 0.286876 |
| 1 | 164324200 | 164325399 | 14.3924 | 26.36803 | 23.32406 | 9.95437 | 0.601449 | 0.27365 | 0.28688 |
| 3 | 50088600 | 50090399 | 18.44871 | 32.93382 | 27.4958 | 12.08484 | 0.656734 | 0.462759 | 0.286886 |
| 12 | 117339200 | 117341799 | 19.01901 | 33.56538 | 29.60013 | 13.0974 | 0.570017 | 0.560611 | 0.286891 |
| 9 | 117263400 | 117265599 | 25.53201 | 44.10751 | 37.64935 | 16.51693 | 0.612862 | 0.87774 | 0.286896 |
| 10 | 63693800 | 63694199 | 7.221487 | 14.1033 | 14.09727 | 4.948775 | 0.672622 | -0.50562 | 0.286901 |
| 4 | 151998800 | 152008399 | 96.6323 | 147.189 | 246.9695 | 102.3638 | -0.46079 | 3.148675 | 0.286925 |
| 4 | 11253600 | 11254399 | 5.420945 | 10.38315 | 27.26826 | 10.75808 | -0.86149 | -0.07788 | 0.286933 |
| 17 | 89393200 | 89395599 | 18.41174 | 33.20513 | 25.19279 | 10.96785 | 0.802697 | 0.395001 | 0.286936 |

| | | | | | | | | | |
|---|---|---|---|---|---|---|---|---|---|
| 5 | 51989400 | 51992399 | 21.6047 | 38.02841 | 28.64009 | 12.67928 | 0.794825 | 0.600774 | 0.286938 |
| 1 | 15827600 | 15833399 | 71.43788 | 111.2956 | 187.0168 | 77.62123 | -0.43758 | 2.707033 | 0.286945 |
| 2 | 166464000 | 166465999 | 9.220678 | 16.26154 | 41.5315 | 18.37935 | -0.95539 | 0.533742 | 0.286958 |
| 5 | 149447600 | 149450599 | 16.39815 | 27.86468 | 57.37116 | 25.14315 | -0.65619 | 1.016106 | 0.286967 |
| 7 | 8505800 | 8511599 | 49.17479 | 80.79643 | 48.44444 | 20.87265 | 1.113712 | 1.451228 | 0.286969 |
| 9 | 36206000 | 36208599 | 68.4714 | 109.1044 | 87.84957 | 36.77355 | 0.649466 | 2.18316 | 0.286974 |
| 1 | 114172600 | 114174799 | 25.45143 | 44.57945 | 29.02073 | 12.81322 | 1.003534 | 0.693603 | 0.286988 |
| 8 | 67622600 | 67624599 | 21.73031 | 38.02605 | 32.7232 | 14.43649 | 0.603146 | 0.694538 | 0.286998 |
| 2 | 93678200 | 93680399 | 10.83928 | 19.16163 | 41.93554 | 18.54932 | -0.73321 | 0.51519 | 0.287005 |
| 3 | 57482200 | 57484199 | 11.79954 | 20.3986 | 52.07009 | 22.84386 | -0.95992 | 0.831533 | 0.287005 |
| 7 | 25946600 | 25953799 | 143.0894 | 217.8083 | 190.8289 | 79.07906 | 0.47509 | 3.169854 | 0.287006 |
| 18 | 90421600 | 90422799 | 19.72328 | 34.62033 | 30.38204 | 13.43854 | 0.575032 | 0.632475 | 0.287016 |
| 12 | 23308400 | 23308999 | 20.42108 | 35.74213 | 28.58614 | 12.61346 | 0.708377 | 0.678529 | 0.287018 |
| 1 | 116173800 | 116174999 | 8.188036 | 15.1537 | 33.11621 | 13.94027 | -0.6801 | 0.190697 | 0.287053 |
| 2 | 164875200 | 164876999 | 8.188036 | 15.1537 | 33.11621 | 13.94027 | -0.6801 | 0.190697 | 0.287053 |
| 5 | 118903600 | 118905799 | 10.50494 | 18.71393 | 38.37148 | 16.89453 | -0.63358 | 0.446389 | 0.287054 |
| 10 | 4381800 | 4383999 | 9.699619 | 17.39774 | 37.38916 | 16.36748 | -0.69517 | 0.394621 | 0.287057 |
| 5 | 140285400 | 140286399 | 4.586413 | 8.856554 | 26.3975 | 10.16377 | -1.00913 | -0.17309 | 0.287098 |
| 1 | 184577600 | 184579199 | 8.094368 | 14.84338 | 33.51519 | 14.28565 | -0.73963 | 0.245035 | 0.287109 |
| 9 | 66257200 | 66258399 | 8.094368 | 14.84338 | 33.51519 | 14.28565 | -0.73963 | 0.245035 | 0.287109 |
| 4 | 101705600 | 101710799 | 104.3311 | 162.5362 | 138.505 | 57.4867 | 0.540869 | 2.716339 | 0.287126 |
| 2 | 106030400 | 106031599 | 15.16452 | 27.99099 | 23.62742 | 9.981841 | 0.678797 | 0.238667 | 0.287129 |
| 1 | 31508600 | 31513599 | 106.3041 | 164.39 | 150.4211 | 62.48137 | 0.430065 | 2.846659 | 0.287143 |
| X | 128711400 | 128712999 | 15.1591 | 28.73287 | 19.46558 | 7.864632 | 1.04437 | 0.06766 | 0.287158 |
| 19 | 42966200 | 42967599 | 11.74857 | 20.92397 | 43.04201 | 18.95504 | -0.63894 | 0.463626 | 0.287188 |
| 7 | 38617400 | 38617999 | 8.809523 | 17.12008 | 14.60864 | 5.438449 | 0.823022 | -0.27826 | 0.287205 |
| 1 | 99140200 | 99141399 | 15.66703 | 28.32449 | 25.15469 | 10.91773 | 0.579865 | 0.361805 | 0.287215 |
| 3 | 156022000 | 156022799 | 7.830024 | 14.79473 | 32.02907 | 13.02673 | -0.62658 | 0.055325 | 0.287216 |
| 4 | 110920800 | 110923599 | 23.31415 | 40.69971 | 32.03681 | 14.10796 | 0.730911 | 0.735954 | 0.287228 |
| 3 | 94685000 | 94687199 | 9.829651 | 17.294 | 42.05009 | 18.61777 | -0.88638 | 0.565782 | 0.287239 |
| 7 | 19496000 | 19501199 | 39.9924 | 63.92948 | 119.0118 | 50.04227 | -0.55253 | 2.063309 | 0.287252 |
| 17 | 77612400 | 77615399 | 39.29638 | 64.85667 | 51.14447 | 22.07882 | 0.721916 | 1.369983 | 0.287267 |
| 8 | 8986800 | 8990399 | 22.89508 | 38.03995 | 74.8722 | 32.4711 | -0.59291 | 1.270098 | 0.28728 |
| X | 11992400 | 11996599 | 19.16853 | 32.41723 | 63.99022 | 28.00476 | -0.59618 | 1.075935 | 0.287282 |
| 10 | 39468200 | 39471999 | 86.25188 | 135.383 | 125.8215 | 52.1836 | 0.424217 | 2.573653 | 0.287305 |
| 19 | 58452800 | 58456399 | 41.22574 | 66.08063 | 115.8222 | 48.89546 | -0.46131 | 1.978927 | 0.287316 |
| 13 | 66312200 | 66314399 | 49.93622 | 80.98617 | 63.5258 | 27.2121 | 0.715224 | 1.647224 | 0.287317 |
| 4 | 134579600 | 134581599 | 10.45552 | 18.34731 | 44.11408 | 19.50773 | -0.87129 | 0.608921 | 0.287317 |
| X | 23954800 | 23958999 | 86.7318 | 137.2367 | 108.7955 | 45.22126 | 0.662294 | 2.404219 | 0.287331 |
| 6 | 130777400 | 130789199 | 213.1493 | 316.4203 | 260.1797 | 107.1744 | 0.534945 | 3.665642 | 0.287335 |
| 9 | 94054400 | 94054799 | 8.432451 | 16.49292 | 11.43567 | 3.857188 | 1.242061 | -0.6326 | 0.287341 |
| 10 | 100989400 | 100989799 | 8.432451 | 16.49292 | 11.43567 | 3.857188 | 1.242061 | -0.6326 | 0.287341 |
| 3 | 37819200 | 37820199 | 12.84782 | 24.54513 | 19.8066 | 7.869843 | 0.814995 | -0.00623 | 0.287357 |
| 1 | 93947600 | 93955799 | 69.19081 | 107.3112 | 196.034 | 81.41845 | -0.56301 | 2.784369 | 0.28736 |
| 1 | 54048800 | 54050799 | 11.46357 | 20.21765 | 41.94035 | 18.56017 | -0.65775 | 0.54572 | 0.287366 |
| 1 | 23486200 | 23487399 | 11.29703 | 21.87387 | 16.6954 | 6.34781 | 0.952702 | -0.17305 | 0.28737 |
| 3 | 104724800 | 104725999 | 5.762186 | 10.93683 | 28.72053 | 11.57011 | -0.88944 | -0.00325 | 0.287372 |

| | | | | | | | | | |
|---|---|---|---|---|---|---|---|---|---|
| 3 | 144109600 | 144114999 | 36.80125 | 59.17477 | 107.791 | 45.72018 | -0.51122 | 1.871544 | 0.287373 |
| 3 | 36724400 | 36726799 | 18.38058 | 30.91288 | 61.63795 | 26.93001 | -0.6109 | 1.118564 | 0.287373 |
| 10 | 62701000 | 62702599 | 9.371811 | 16.9201 | 35.91918 | 15.61655 | -0.67067 | 0.350724 | 0.287374 |
| 5 | 77370400 | 77371199 | 6.089393 | 11.44621 | 30.15772 | 12.38364 | -0.91823 | 0.070077 | 0.287376 |
| 18 | 17280400 | 17282599 | 37.37741 | 61.99898 | 50.41057 | 21.83731 | 0.678534 | 1.312663 | 0.28738 |
| 2 | 168842400 | 168849799 | 64.52949 | 101.1565 | 170.709 | 70.80411 | -0.43676 | 2.593167 | 0.287421 |
| 11 | 21726400 | 21729999 | 73.43135 | 116.4677 | 108.5707 | 45.20325 | 0.431806 | 2.331351 | 0.287436 |
| 7 | 67821400 | 67821799 | 8.789856 | 17.11567 | 15.16673 | 5.556757 | 0.790754 | -0.32946 | 0.287439 |
| X | 54639600 | 54641599 | 39.89731 | 66.10541 | 49.94661 | 21.61976 | 0.783903 | 1.330078 | 0.287441 |
| 1 | 115255800 | 115256799 | 12.45467 | 23.89809 | 18.7448 | 7.357471 | 0.871541 | -0.05216 | 0.28746 |
| 9 | 73177600 | 73179599 | 10.14544 | 18.25189 | 36.87883 | 16.10342 | -0.60421 | 0.378465 | 0.287461 |
| 17 | 60854000 | 60858799 | 75.56779 | 119.6616 | 111.8897 | 46.52506 | 0.425306 | 2.376772 | 0.28751 |
| 5 | 106610800 | 106620399 | 67.84185 | 105.347 | 199.3391 | 82.77612 | -0.61237 | 2.762472 | 0.28751 |
| 3 | 92657600 | 92658399 | 4.163422 | 8.095221 | 27.6869 | 10.26414 | -1.15438 | -0.30959 | 0.287512 |
| 16 | 68370200 | 68371599 | 12.96602 | 24.42496 | 21.27406 | 8.703809 | 0.66925 | 0.092152 | 0.287517 |
| 7 | 38163000 | 38168199 | 94.96492 | 149.8202 | 111.4988 | 46.27559 | 0.749938 | 2.477272 | 0.287519 |
| 7 | 34910400 | 34915599 | 31.64149 | 50.97738 | 104.4128 | 44.41258 | -0.67646 | 1.800943 | 0.287527 |
| 11 | 30960800 | 30963799 | 54.19387 | 87.42812 | 69.97082 | 29.81528 | 0.67851 | 1.781037 | 0.287533 |
| 18 | 18083800 | 18087199 | 25.18481 | 44.30811 | 26.15889 | 11.58197 | 1.143847 | 0.623932 | 0.287535 |
| 7 | 12464000 | 12467599 | 40.27079 | 65.58265 | 60.72253 | 26.0726 | 0.481441 | 1.55397 | 0.28754 |
| 7 | 121529600 | 121531799 | 13.9303 | 24.10557 | 50.53272 | 22.17005 | -0.67691 | 0.832081 | 0.287543 |
| 10 | 25983400 | 25983799 | 4.352429 | 8.495192 | 21.61609 | 7.664694 | -0.67165 | -0.48093 | 0.287561 |
| 18 | 76494600 | 76497399 | 23.20392 | 38.20024 | 74.56025 | 32.1516 | -0.58285 | 1.383479 | 0.287568 |
| 3 | 79390400 | 79393999 | 40.38087 | 65.78668 | 60.82025 | 26.11815 | 0.483953 | 1.54804 | 0.287573 |
| 2 | 173226000 | 173237599 | 93.89276 | 143.4394 | 241.3143 | 100.0923 | -0.4612 | 3.086842 | 0.287576 |
| X | 125973000 | 125975599 | 34.06546 | 58.59112 | 32.92176 | 14.44098 | 1.212302 | 0.948729 | 0.287582 |
| X | 149067600 | 149069599 | 37.10647 | 61.81608 | 48.17369 | 20.92957 | 0.739797 | 1.261038 | 0.287585 |
| 9 | 92616400 | 92618799 | 40.76668 | 66.28016 | 58.96922 | 25.30447 | 0.537165 | 1.577802 | 0.287587 |
| 13 | 13186200 | 13189199 | 46.83819 | 76.3342 | 59.08641 | 25.37692 | 0.739908 | 1.556079 | 0.287603 |
| 10 | 81012400 | 81037999 | 261.7033 | 367.0329 | 621.4503 | 257.5666 | -0.58697 | 4.534586 | 0.287612 |
| 1 | 16457000 | 16457999 | 13.65964 | 26.22063 | 18.75961 | 7.354861 | 1.005217 | -0.04194 | 0.287619 |
| 16 | 17831200 | 17833599 | 70.76292 | 112.2411 | 102.7399 | 42.77779 | 0.458142 | 2.326394 | 0.287641 |
| 17 | 13950800 | 13952799 | 12.59639 | 22.05202 | 44.65703 | 19.70849 | -0.62519 | 0.660735 | 0.287654 |
| 14 | 94563200 | 94567999 | 60.48455 | 97.2633 | 75.13549 | 31.87239 | 0.723982 | 1.890886 | 0.287663 |
| 12 | 114564200 | 114567399 | 56.90939 | 91.88682 | 66.61847 | 28.41788 | 0.822205 | 1.762575 | 0.287663 |
| 17 | 37291600 | 37292799 | 7.481486 | 13.79154 | 34.65276 | 14.66563 | -0.88562 | 0.210122 | 0.28767 |
| 18 | 73005400 | 73012599 | 96.82444 | 151.0966 | 137.805 | 57.18329 | 0.444893 | 2.685189 | 0.28767 |
| 7 | 26679200 | 26686599 | 100.73 | 156.465 | 145.661 | 60.48336 | 0.41014 | 2.766301 | 0.287674 |
| 13 | 61971600 | 61976599 | 92.79567 | 146.7595 | 104.7757 | 43.53961 | 0.812596 | 2.417481 | 0.28768 |
| 7 | 28112200 | 28113799 | 7.514217 | 14.21509 | 30.35871 | 12.31603 | -0.60375 | 0.039682 | 0.287701 |
| 16 | 16768000 | 16768599 | 10.97263 | 21.19453 | 18.61757 | 7.158913 | 0.738228 | -0.13072 | 0.287732 |
| 18 | 23279400 | 23281999 | 24.1295 | 39.7312 | 75.50566 | 32.56177 | -0.54448 | 1.381587 | 0.287733 |
| 6 | 33067400 | 33070999 | 36.97081 | 61.1639 | 48.27202 | 20.87308 | 0.721026 | 1.336933 | 0.28774 |
| 5 | 34526600 | 34528199 | 10.71417 | 18.89153 | 41.46512 | 18.35073 | -0.73865 | 0.543814 | 0.287753 |
| 2 | 37173400 | 37177199 | 39.452 | 65.39758 | 49.39294 | 21.38687 | 0.784446 | 1.322065 | 0.287762 |
| 13 | 44295800 | 44299399 | 57.94287 | 92.84912 | 85.45698 | 36.06033 | 0.465757 | 1.998175 | 0.287768 |
| 4 | 79558000 | 79560199 | 29.86308 | 51.19249 | 37.43986 | 16.41978 | 0.832775 | 0.962486 | 0.287784 |

| | | | | | | | | | |
|---|---|---|---|---|---|---|---|---|---|
| 17 | 40498400 | 40498999 | 11.55316 | 21.87516 | 19.45874 | 7.879316 | 0.652077 | 0.040742 | 0.287785 |
| 17 | 39590800 | 39594599 | 39.21319 | 66.12554 | 37.34016 | 16.32734 | 1.202461 | 1.139906 | 0.287799 |
| 11 | 105671800 | 105673199 | 6.420298 | 12.38719 | 28.85241 | 11.14748 | -0.66277 | -0.13374 | 0.28782 |
| 7 | 78853800 | 78855599 | 21.28915 | 37.12788 | 33.09271 | 14.56388 | 0.552653 | 0.74386 | 0.287828 |
| 6 | 37686000 | 37688999 | 49.07995 | 79.51602 | 64.72157 | 27.7034 | 0.660615 | 1.66396 | 0.287841 |
| 15 | 19744000 | 19746599 | 32.07802 | 54.05598 | 44.73957 | 19.55869 | 0.653048 | 1.1283 | 0.287871 |
| 2 | 174167800 | 174172999 | 58.11799 | 90.45234 | 199.1135 | 82.65446 | -0.82778 | 2.720581 | 0.287871 |
| 16 | 89448000 | 89448999 | 16.12985 | 27.80139 | 52.70244 | 23.11917 | -0.53397 | 0.878833 | 0.287882 |
| 6 | 29083000 | 29084799 | 8.151277 | 15.18851 | 33.28032 | 13.86975 | -0.67312 | 0.149553 | 0.287884 |
| 9 | 81779400 | 81780199 | 8.504452 | 15.82878 | 33.78452 | 14.10673 | -0.6373 | 0.164259 | 0.287907 |
| 6 | 97880800 | 97884399 | 27.44329 | 46.58975 | 42.13079 | 18.45965 | 0.526079 | 1.046578 | 0.287927 |
| 7 | 107783800 | 107784999 | 20.77952 | 36.3908 | 31.16142 | 13.75659 | 0.610083 | 0.673187 | 0.287927 |
| 9 | 71499000 | 71501199 | 9.559769 | 17.11761 | 37.89898 | 16.61459 | -0.73933 | 0.405273 | 0.287934 |
| 14 | 4352800 | 4356999 | 47.26738 | 76.88926 | 60.2332 | 25.85176 | 0.720781 | 1.583055 | 0.287936 |
| X | 4303600 | 4304799 | 13.8196 | 26.77668 | 16.09547 | 6.099992 | 1.29953 | -0.15439 | 0.287938 |
| 17 | 79327600 | 79329199 | 9.331799 | 16.53926 | 39.4424 | 17.43755 | -0.85421 | 0.475046 | 0.287967 |
| 4 | 125255600 | 125261199 | 37.08623 | 59.31482 | 119.8972 | 50.4519 | -0.67024 | 2.043597 | 0.287975 |
| 9 | 19663400 | 19666399 | 42.03989 | 69.43355 | 50.58658 | 21.8486 | 0.835937 | 1.37186 | 0.287987 |
| 4 | 132459800 | 132461999 | 15.84931 | 27.15635 | 54.25498 | 23.79336 | -0.61104 | 0.933148 | 0.288002 |
| 3 | 109317800 | 109320599 | 20.18989 | 35.50338 | 29.95487 | 13.26393 | 0.631393 | 0.60564 | 0.288037 |
| 8 | 74246800 | 74252799 | 70.73701 | 112.4725 | 102.8684 | 42.93543 | 0.462678 | 2.25583 | 0.288053 |
| 2 | 93533400 | 93534399 | 4.856071 | 9.406707 | 26.67261 | 10.12042 | -0.91845 | -0.22144 | 0.288054 |
| 2 | 168207600 | 168227399 | 225.734 | 323.8358 | 505.1505 | 208.5927 | -0.43588 | 4.199658 | 0.288068 |
| 16 | 11801000 | 11806599 | 37.99467 | 60.53694 | 129.9911 | 54.40771 | -0.76328 | 2.168156 | 0.288072 |
| 17 | 95120200 | 95125399 | 126.9245 | 196.2563 | 143.8954 | 59.77078 | 0.749234 | 2.858379 | 0.288078 |
| X | 147635200 | 147638399 | 44.65932 | 72.63049 | 62.65306 | 26.88815 | 0.581947 | 1.584766 | 0.288086 |
| 8 | 57849800 | 57850599 | 5.262793 | 10.20306 | 26.80841 | 10.13245 | -0.80464 | -0.23004 | 0.288103 |
| 11 | 62770000 | 62771199 | 5.703927 | 10.69016 | 33.90522 | 13.99199 | -1.19012 | 0.092202 | 0.288134 |
| 1 | 135505600 | 135508999 | 15.56342 | 26.23091 | 63.17618 | 27.61996 | -0.88219 | 1.089586 | 0.288152 |
| 13 | 50078200 | 50079599 | 36.52879 | 60.08237 | 55.36228 | 23.86399 | 0.496181 | 1.409401 | 0.288153 |
| 7 | 59327600 | 59329599 | 38.30273 | 62.82284 | 49.32434 | 21.23599 | 0.724974 | 1.446612 | 0.288154 |
| 15 | 28096800 | 28097199 | 7.53238 | 14.73156 | 12.86424 | 4.362407 | 0.908514 | -0.62353 | 0.288168 |
| 15 | 82926200 | 82931399 | 49.16939 | 78.11262 | 137.9258 | 57.51824 | -0.48607 | 2.271309 | 0.288175 |
| 7 | 116347400 | 116347999 | 9.202914 | 17.8761 | 17.18698 | 6.422838 | 0.6476 | -0.25056 | 0.288187 |
| X | 26296800 | 26301799 | 63.99148 | 102.7799 | 75.87892 | 32.13192 | 0.78715 | 1.932885 | 0.288195 |
| 19 | 26879800 | 26882599 | 22.25904 | 37.1788 | 69.6592 | 30.31422 | -0.52175 | 1.208684 | 0.288198 |
| 17 | 39334200 | 39338599 | 102.2699 | 158.7556 | 146.6891 | 60.91608 | 0.420239 | 2.778405 | 0.288199 |
| 13 | 99038600 | 99042999 | 45.52941 | 72.74731 | 125.4716 | 52.72429 | -0.44338 | 2.085536 | 0.288258 |
| 10 | 49768400 | 49769599 | 11.72289 | 22.45245 | 16.41823 | 6.478516 | 0.965498 | -0.04813 | 0.288258 |
| 14 | 32128800 | 32132999 | 25.68129 | 42.00638 | 80.04763 | 34.42512 | -0.55256 | 1.475356 | 0.288278 |
| 11 | 6149200 | 6150399 | 7.869041 | 14.64427 | 32.14143 | 13.42353 | -0.67765 | 0.153765 | 0.28828 |
| 3 | 145268600 | 145270599 | 29.41393 | 50.80926 | 35.26535 | 15.47102 | 0.91001 | 0.893289 | 0.288292 |
| X | 24537600 | 24541599 | 80.09623 | 127.1194 | 103.2745 | 43.02551 | 0.630814 | 2.319377 | 0.288303 |
| 1 | 48311600 | 48312399 | 9.762554 | 18.99945 | 16.53158 | 6.083275 | 0.809607 | -0.30039 | 0.288303 |
| 4 | 76318600 | 76319399 | 11.24009 | 21.81223 | 17.49306 | 6.582098 | 0.897236 | -0.20319 | 0.288316 |
| 11 | 34656600 | 34657399 | 11.24009 | 21.81223 | 17.49306 | 6.582098 | 0.897236 | -0.20319 | 0.288316 |
| 8 | 46918400 | 46921399 | 28.5875 | 48.82327 | 40.29528 | 17.66675 | 0.658382 | 0.993838 | 0.288338 |

| | | | | | | | | | |
|---|---|---|---|---|---|---|---|---|---|
| 13 | 54446600 | 54447999 | 16.39661 | 29.37357 | 22.81283 | 9.996932 | 0.763097 | 0.415771 | 0.288341 |
| 9 | 117849600 | 117852799 | 49.10298 | 79.46856 | 67.26667 | 28.77018 | 0.602888 | 1.688661 | 0.288362 |
| 8 | 22533600 | 22534599 | 5.382277 | 10.43592 | 24.98228 | 9.437133 | -0.66979 | -0.23226 | 0.288369 |
| 17 | 4865400 | 4865999 | 5.382277 | 10.43592 | 24.98228 | 9.437133 | -0.66979 | -0.23226 | 0.288369 |
| 7 | 98634800 | 98636599 | 10.99986 | 19.4268 | 41.42791 | 18.32831 | -0.69649 | 0.526604 | 0.288377 |
| X | 33957600 | 33959999 | 46.86849 | 76.35931 | 59.47487 | 25.5408 | 0.730697 | 1.563233 | 0.2884 |
| 8 | 74122600 | 74124999 | 23.58471 | 41.30954 | 31.43473 | 13.87897 | 0.779682 | 0.688191 | 0.288407 |
| 13 | 60323400 | 60327799 | 34.37566 | 55.3547 | 101.5918 | 43.18134 | -0.51934 | 1.820806 | 0.288409 |
| 9 | 43452000 | 43452799 | 5.529408 | 10.73194 | 25.69983 | 9.66288 | -0.66456 | -0.2464 | 0.288433 |
| 3 | 58881400 | 58883999 | 25.48037 | 44.58968 | 30.81671 | 13.59575 | 0.917973 | 0.712087 | 0.288456 |
| 18 | 19949600 | 19951999 | 20.53772 | 36.35474 | 27.61129 | 12.21004 | 0.78575 | 0.533068 | 0.288457 |
| 16 | 60704200 | 60706199 | 32.68651 | 54.26985 | 49.80956 | 21.59119 | 0.504288 | 1.285182 | 0.288467 |
| 15 | 84270400 | 84276799 | 84.75257 | 130.408 | 215.4014 | 89.45745 | -0.4267 | 2.940175 | 0.28848 |
| 17 | 41804600 | 41806799 | 26.23452 | 45.70829 | 33.07491 | 14.5472 | 0.851804 | 0.781543 | 0.288489 |
| 18 | 82136600 | 82138199 | 16.29042 | 30.09045 | 22.9412 | 9.682628 | 0.825566 | 0.242146 | 0.288496 |
| X | 66858400 | 66863399 | 54.43299 | 88.31446 | 58.58113 | 25.10141 | 0.957052 | 1.647486 | 0.288499 |
| 15 | 13382800 | 13383999 | 12.60937 | 22.07341 | 44.66506 | 19.71103 | -0.62397 | 0.662751 | 0.288499 |
| 7 | 32315400 | 32317999 | 36.5778 | 60.91684 | 48.72776 | 21.16623 | 0.70228 | 1.262731 | 0.2885 |
| 13 | 81033600 | 81034599 | 9.639328 | 17.06257 | 39.9457 | 17.66456 | -0.82859 | 0.491782 | 0.288523 |
| 13 | 62308800 | 62317999 | 266.9371 | 389.9945 | 304.4661 | 125.436 | 0.588109 | 3.92994 | 0.288543 |
| 13 | 66520800 | 66524799 | 46.05239 | 75.10788 | 54.44334 | 23.38894 | 0.83508 | 1.542529 | 0.288545 |
| 1 | 104015000 | 104016799 | 23.44552 | 41.17598 | 29.70445 | 13.14476 | 0.856117 | 0.642979 | 0.288547 |
| 5 | 147910200 | 147915999 | 42.41703 | 67.74824 | 122.5112 | 51.45229 | -0.5121 | 2.092295 | 0.288552 |
| 17 | 39585800 | 39588799 | 31.87038 | 54.23021 | 40.32188 | 17.67318 | 0.807799 | 1.042065 | 0.288596 |
| 6 | 91890800 | 91892199 | 10.48206 | 18.74184 | 37.86766 | 16.62118 | -0.60942 | 0.421638 | 0.288609 |
| 5 | 3700200 | 3701999 | 19.61529 | 34.43235 | 30.75291 | 13.60311 | 0.549651 | 0.630328 | 0.288621 |
| 2 | 110325200 | 110328599 | 54.66747 | 87.68886 | 80.73961 | 34.12199 | 0.466861 | 1.9598 | 0.288688 |
| 13 | 120652400 | 120654599 | 19.95015 | 35.26641 | 27.67539 | 12.24174 | 0.737998 | 0.543214 | 0.288713 |
| 19 | 24327600 | 24330199 | 16.75252 | 28.91739 | 55.47018 | 24.3344 | -0.55067 | 0.870715 | 0.288714 |
| 2 | 80594600 | 80596599 | 9.420013 | 17.28044 | 37.85206 | 16.12645 | -0.69671 | 0.259318 | 0.288729 |
| 17 | 43468600 | 43470199 | 7.270643 | 13.57256 | 32.20763 | 13.38094 | -0.78328 | 0.128072 | 0.288731 |
| 10 | 82244400 | 82254599 | 210.4721 | 313.1056 | 252.4384 | 104.0079 | 0.566107 | 3.62833 | 0.288759 |
| 8 | 42490400 | 42491399 | 10.08633 | 18.32941 | 37.0402 | 15.97294 | -0.59284 | 0.318264 | 0.288768 |
| 4 | 71279600 | 71281199 | 11.24591 | 20.16071 | 41.56296 | 18.20262 | -0.63648 | 0.416123 | 0.288773 |
| 5 | 80648600 | 80648999 | 13.93976 | 26.76964 | 16.18287 | 6.336941 | 1.246058 | -0.05156 | 0.28878 |
| 2 | 173500000 | 173502199 | 13.77551 | 23.77635 | 52.12514 | 22.86689 | -0.74223 | 0.854285 | 0.288783 |
| 2 | 53372200 | 53373799 | 24.63298 | 42.50651 | 37.80607 | 16.58511 | 0.553542 | 0.884997 | 0.288795 |
| 12 | 34888000 | 34888999 | 5.667373 | 11.00602 | 26.41525 | 9.889625 | -0.66217 | -0.25777 | 0.288796 |
| 7 | 82841400 | 82842799 | 18.3766 | 33.13201 | 25.82745 | 11.24829 | 0.763721 | 0.398714 | 0.288812 |
| 2 | 58771200 | 58772399 | 13.8244 | 26.24788 | 20.18467 | 8.125738 | 0.867565 | 0.045445 | 0.288814 |
| 11 | 84770400 | 84771999 | 8.672947 | 15.52323 | 37.44746 | 16.42239 | -0.86217 | 0.402657 | 0.288822 |
| 6 | 39004400 | 39005399 | 6.309401 | 12.04585 | 28.923 | 11.50815 | -0.74642 | -0.04055 | 0.288831 |
| 12 | 19654800 | 19658599 | 176.18 | 263.6144 | 219.9805 | 90.72023 | 0.524059 | 3.516811 | 0.288845 |
| 4 | 124770400 | 124780799 | 80.77597 | 124.4524 | 212.3816 | 88.21918 | -0.47227 | 2.912258 | 0.288848 |
| X | 109579600 | 109581399 | 18.34827 | 34.13011 | 20.7229 | 8.661576 | 1.161939 | 0.216836 | 0.288852 |
| 8 | 66736400 | 66737999 | 17.06295 | 31.48596 | 22.26293 | 9.409119 | 0.931991 | 0.251939 | 0.288861 |
| 2 | 55718000 | 55720999 | 14.42106 | 24.85005 | 55.8723 | 24.50952 | -0.77907 | 0.877803 | 0.288864 |

| | | | | | | | | | |
|---|---|---|---|---|---|---|---|---|---|
| 13 | 120133200 | 120136399 | 39.71985 | 64.88776 | 59.79298 | 25.70272 | 0.491425 | 1.507166 | 0.288874 |
| 12 | 87593200 | 87594199 | 14.04178 | 26.94787 | 18.06964 | 7.089112 | 1.096602 | -0.03847 | 0.288889 |
| 6 | 54268800 | 54271199 | 9.481522 | 17.41846 | 38.43228 | 16.34144 | -0.70465 | 0.256115 | 0.288897 |
| 1 | 182574800 | 182577399 | 19.57836 | 32.6032 | 66.98975 | 29.09732 | -0.65417 | 1.231004 | 0.288907 |
| 1 | 79809600 | 79811199 | 10.74986 | 19.25996 | 41.01264 | 17.97062 | -0.68351 | 0.413764 | 0.288918 |
| 12 | 20825200 | 20836799 | 505.2449 | 691.663 | 561.1885 | 233.4761 | 0.438819 | 4.908174 | 0.288921 |
| 2 | 180828400 | 180831399 | 21.74876 | 35.85801 | 73.45528 | 31.6951 | -0.6515 | 1.358863 | 0.288925 |
| 7 | 137281200 | 137282599 | 8.940587 | 16.55316 | 34.22234 | 14.39795 | -0.60026 | 0.197123 | 0.288931 |
| 1 | 85268400 | 85269799 | 24.20561 | 42.04951 | 36.02654 | 15.81615 | 0.608914 | 0.816085 | 0.288945 |
| 5 | 53134200 | 53137799 | 23.38991 | 38.38969 | 86.22681 | 37.13468 | -0.78684 | 1.426693 | 0.288947 |
| 2 | 162999600 | 163006599 | 73.36886 | 113.5661 | 205.0187 | 85.15952 | -0.54819 | 2.822253 | 0.288958 |
| 8 | 86281400 | 86282799 | 8.83196 | 16.16747 | 35.05237 | 14.97671 | -0.68462 | 0.261875 | 0.288971 |
| 7 | 17931400 | 17935999 | 79.21161 | 125.5566 | 109.264 | 45.46328 | 0.530011 | 2.354282 | 0.288995 |
| 7 | 62355800 | 62371799 | 623.8947 | 839.7245 | 682.5963 | 284.7296 | 0.411474 | 5.176471 | 0.288995 |
| 9 | 80684000 | 80685199 | 24.8128 | 42.80968 | 37.80612 | 16.58505 | 0.563713 | 0.887429 | 0.288996 |
| 15 | 38247000 | 38250399 | 20.42314 | 34.40826 | 64.08544 | 28.01419 | -0.51299 | 1.111282 | 0.289003 |
| 3 | 123611000 | 123611599 | 9.453077 | 17.24501 | 35.51729 | 15.24252 | -0.6154 | 0.28791 | 0.289005 |
| 9 | 109592000 | 109593799 | 25.49306 | 44.1669 | 36.61886 | 16.06625 | 0.655554 | 0.852416 | 0.289012 |
| 2 | 111801800 | 111805599 | 40.82994 | 67.03991 | 56.13402 | 24.17881 | 0.632923 | 1.441765 | 0.289018 |
| 6 | 149181200 | 149190199 | 152.2394 | 231.027 | 198.3387 | 82.11571 | 0.500787 | 3.234614 | 0.289025 |
| 1 | 35692800 | 35696199 | 31.00563 | 52.12882 | 45.85666 | 20.03306 | 0.565265 | 1.144209 | 0.289026 |
| X | 133592200 | 133595599 | 19.776 | 33.46043 | 64.10408 | 28.05863 | -0.55328 | 1.072601 | 0.289033 |
| 13 | 101903600 | 101905399 | 13.77109 | 23.79327 | 51.06355 | 22.40188 | -0.71124 | 0.844384 | 0.289046 |
| 4 | 145607800 | 145610399 | 127.0054 | 194.0203 | 178.4285 | 74.00807 | 0.409337 | 3.091843 | 0.289057 |
| 7 | 31905400 | 31921599 | 198.6229 | 297.5631 | 225.6075 | 93.06318 | 0.663921 | 3.49784 | 0.289058 |
| 13 | 65540000 | 65550999 | 170.8219 | 259.3262 | 178.6142 | 73.95793 | 0.818817 | 3.232683 | 0.289068 |
| 6 | 23768000 | 23769599 | 26.78456 | 46.68454 | 32.50361 | 14.29938 | 0.907233 | 0.778614 | 0.289078 |
| 7 | 11205400 | 11208999 | 84.83643 | 133.3481 | 125.171 | 51.92331 | 0.411419 | 2.546026 | 0.289081 |
| 1 | 50092000 | 50092599 | 9.71428 | 18.84953 | 17.58226 | 6.618554 | 0.680463 | -0.22025 | 0.289088 |
| Y | 665600 | 667599 | 14.47368 | 26.97053 | 12.3082 | 5.129089 | 1.571911 | 0.14711 | 0.289098 |
| 16 | 35616200 | 35623599 | 95.01833 | 145.5163 | 236.4939 | 98.14865 | -0.4086 | 3.037658 | 0.289105 |
| 9 | 14080600 | 14082199 | 14.5754 | 27.40312 | 19.85301 | 8.149307 | 0.928653 | 0.114253 | 0.289115 |
| 19 | 55303800 | 55305199 | 8.502813 | 15.27993 | 36.94108 | 16.14558 | -0.86196 | 0.374067 | 0.289122 |
| 7 | 17552400 | 17556599 | 95.43436 | 150.2893 | 116.3301 | 48.26778 | 0.691368 | 2.51609 | 0.289143 |
| 15 | 5271200 | 5274799 | 28.65535 | 46.10283 | 110.9848 | 47.11379 | -0.91138 | 1.844999 | 0.289149 |
| Y | 88345400 | 88346799 | 13.60203 | 25.67206 | 10.72764 | 4.37346 | 1.721755 | 0.05401 | 0.289152 |
| 3 | 7619000 | 7619599 | 8.557157 | 15.64213 | 33.9913 | 14.55088 | -0.68934 | 0.267947 | 0.289154 |
| 14 | 68944600 | 68946199 | 8.557157 | 15.64213 | 33.9913 | 14.55088 | -0.68934 | 0.267947 | 0.289154 |
| 19 | 52091800 | 52092999 | 14.05242 | 26.91558 | 18.61972 | 7.346064 | 1.044648 | -0.01931 | 0.289156 |
| 10 | 101694200 | 101695199 | 14.54405 | 27.00583 | 23.22141 | 9.73126 | 0.660665 | 0.200455 | 0.289157 |
| 6 | 94750600 | 94752199 | 9.936933 | 18.03033 | 36.47665 | 15.76004 | -0.59618 | 0.324948 | 0.289159 |
| 6 | 133160400 | 133169399 | 166.6434 | 250.7437 | 223.0363 | 92.10418 | 0.439196 | 3.407377 | 0.289169 |
| 7 | 11738200 | 11743399 | 76.42646 | 122.1023 | 82.24192 | 34.55374 | 0.910627 | 2.109763 | 0.289171 |
| X | 24109200 | 24110799 | 16.79043 | 31.91487 | 17.68398 | 7.102301 | 1.339329 | 0.060446 | 0.289181 |
| 15 | 44221800 | 44223399 | 19.9795 | 33.51288 | 63.37239 | 27.65208 | -0.53489 | 1.151011 | 0.289198 |
| 12 | 115724200 | 115727199 | 46.21987 | 75.57432 | 56.28396 | 24.20226 | 0.798707 | 1.507829 | 0.289199 |
| 10 | 59102800 | 59105399 | 11.74098 | 20.76023 | 46.15504 | 20.41483 | -0.75609 | 0.524072 | 0.2892 |

| | | | | | | | | | |
|---|---|---|---|---|---|---|---|---|---|
| 9 | 22804000 | 22805599 | 17.66901 | 32.40504 | 23.30307 | 9.931219 | 0.899493 | 0.292923 | 0.289213 |
| 3 | 30547000 | 30548399 | 17.80106 | 32.88102 | 22.35451 | 9.434912 | 0.990017 | 0.254042 | 0.289261 |
| 7 | 128449200 | 128452599 | 45.21535 | 73.38782 | 64.95833 | 27.84195 | 0.542366 | 1.622062 | 0.289271 |
| 8 | 76164800 | 76166799 | 28.65038 | 49.09685 | 39.18327 | 17.1839 | 0.707342 | 0.963492 | 0.289278 |
| 1 | 65166600 | 65168199 | 12.1183 | 21.41253 | 42.45442 | 18.78062 | -0.59165 | 0.532355 | 0.289299 |
| 2 | 177100400 | 177102199 | 14.98218 | 28.3362 | 20.011 | 8.122053 | 0.979256 | 0.081661 | 0.289303 |
| X | 136327400 | 136328199 | 12.36841 | 23.5848 | 20.49867 | 8.177317 | 0.704312 | 0.00507 | 0.28931 |
| 5 | 115038800 | 115040799 | 9.660757 | 17.46945 | 35.91697 | 15.58464 | -0.62309 | 0.342803 | 0.28932 |
| 8 | 33875200 | 33876999 | 13.19292 | 22.71845 | 52.61302 | 23.0793 | -0.82153 | 0.874498 | 0.289323 |
| 7 | 41405000 | 41407599 | 47.6449 | 77.51313 | 60.23135 | 25.85215 | 0.732594 | 1.58263 | 0.289323 |
| 5 | 143148600 | 143155199 | 46.54764 | 74.01609 | 136.7179 | 57.06735 | -0.54967 | 2.241172 | 0.28933 |
| 2 | 130416800 | 130426599 | 83.69216 | 127.9679 | 238.0387 | 98.75362 | -0.60498 | 3.065902 | 0.289332 |
| 7 | 66171200 | 66172999 | 8.818522 | 16.17861 | 35.09782 | 14.95107 | -0.68238 | 0.249004 | 0.289344 |
| 10 | 11406400 | 11408799 | 27.76201 | 45.30887 | 82.71925 | 35.54523 | -0.49332 | 1.511166 | 0.289348 |
| 13 | 50965800 | 50968999 | 35.9908 | 57.52124 | 120.6258 | 50.70656 | -0.72428 | 2.06579 | 0.289353 |
| 2 | 146005200 | 146006199 | 7.34869 | 13.99561 | 30.60705 | 12.24169 | -0.61957 | -0.00774 | 0.289353 |
| 10 | 120913000 | 120916999 | 42.72141 | 68.27083 | 122.5005 | 51.48669 | -0.50007 | 2.075851 | 0.289357 |
| 1 | 110135600 | 110139599 | 46.20814 | 74.84802 | 67.15139 | 28.73913 | 0.520259 | 1.665029 | 0.289381 |
| 13 | 67302400 | 67303999 | 46.82159 | 75.6682 | 68.38472 | 29.20582 | 0.50652 | 1.720472 | 0.289385 |
| 5 | 104236800 | 104237199 | 5.975945 | 11.54033 | 26.51775 | 10.20787 | -0.63809 | -0.15806 | 0.289386 |
| 10 | 11517200 | 11518599 | 19.61704 | 35.10585 | 26.13112 | 11.46121 | 0.823124 | 0.451524 | 0.289392 |
| 12 | 15105000 | 15106599 | 19.61704 | 35.10585 | 26.13112 | 11.46121 | 0.823124 | 0.451524 | 0.289392 |
| X | 32348800 | 32350399 | 41.09369 | 67.80913 | 51.61311 | 22.2783 | 0.772973 | 1.38291 | 0.289394 |
| 2 | 36237200 | 36241399 | 97.339 | 151.3444 | 139.742 | 58.0172 | 0.423045 | 2.74682 | 0.289403 |
| 9 | 52588400 | 52591799 | 30.00247 | 48.45438 | 96.43077 | 41.14815 | -0.63085 | 1.723698 | 0.289404 |
| 7 | 92252600 | 92254399 | 31.19676 | 52.023 | 47.60664 | 20.69626 | 0.508722 | 1.230879 | 0.289415 |
| 6 | 133467600 | 133481399 | 297.8582 | 428.5933 | 363.2636 | 149.9323 | 0.448143 | 4.158711 | 0.289416 |
| 14 | 33374000 | 33380599 | 45.87098 | 73.14083 | 126.4765 | 52.99135 | -0.45037 | 2.150728 | 0.289467 |
| X | 126536400 | 126537399 | 14.9667 | 28.36883 | 20.13737 | 8.135671 | 0.978016 | 0.068182 | 0.289471 |
| 11 | 77919000 | 77920599 | 5.840793 | 10.92986 | 31.68295 | 13.10897 | -1.06395 | 0.099729 | 0.289479 |
| 1 | 154868800 | 154870399 | 9.999537 | 17.92509 | 37.8837 | 16.59223 | -0.67184 | 0.400841 | 0.289503 |
| 4 | 111965400 | 111966999 | 39.62827 | 64.75574 | 59.72616 | 25.67654 | 0.490405 | 1.50421 | 0.289507 |
| 15 | 75387000 | 75389199 | 21.88074 | 38.18979 | 33.90057 | 14.92683 | 0.558503 | 0.73603 | 0.289523 |
| 16 | 51563200 | 51565999 | 24.44238 | 42.50142 | 33.7094 | 14.80778 | 0.719632 | 0.798399 | 0.289527 |
| 1 | 95400200 | 95402199 | 21.03653 | 36.76352 | 32.88515 | 14.49179 | 0.547625 | 0.71179 | 0.289542 |
| 2 | 68182400 | 68183599 | 7.705391 | 14.40929 | 31.1617 | 12.90814 | -0.64671 | 0.119477 | 0.289542 |
| X | 5074800 | 5075399 | 11.75937 | 22.57565 | 19.5621 | 7.666837 | 0.732065 | -0.05924 | 0.289547 |
| 10 | 86050600 | 86051399 | 11.12503 | 21.65589 | 15.86469 | 5.827315 | 1.057837 | -0.29082 | 0.289549 |
| 7 | 9053000 | 9062399 | 123.1566 | 190.3147 | 149.5395 | 62.11539 | 0.648818 | 2.868629 | 0.289556 |
| 3 | 32288400 | 32291399 | 28.69509 | 48.991 | 40.87087 | 17.91864 | 0.642928 | 0.998897 | 0.289564 |
| 13 | 8004600 | 8008599 | 39.25838 | 64.34661 | 58.18183 | 25.04229 | 0.521431 | 1.46155 | 0.289574 |
| X | 60857000 | 60860399 | 46.36318 | 75.60836 | 58.92373 | 25.31294 | 0.730824 | 1.548349 | 0.289577 |
| 1 | 155595800 | 155601399 | 43.99222 | 69.94416 | 134.518 | 56.14227 | -0.60792 | 2.243164 | 0.289582 |
| 1 | 30756000 | 30760599 | 36.69224 | 58.91611 | 107.1866 | 45.36951 | -0.5119 | 1.917264 | 0.289596 |
| 5 | 80513800 | 80516199 | 17.35538 | 31.42352 | 24.77256 | 10.73191 | 0.752159 | 0.36389 | 0.289603 |
| 9 | 39571000 | 39577199 | 75.83807 | 120.4661 | 103.739 | 43.25667 | 0.548091 | 2.288315 | 0.289608 |
| 17 | 83607200 | 83609199 | 9.471009 | 17.27881 | 34.99356 | 15.01658 | -0.59101 | 0.287917 | 0.289627 |

| | | | | | | | | | |
|---|---|---|---|---|---|---|---|---|---|
| 5 | 81099200 | 81101599 | 20.17674 | 33.64608 | 67.75627 | 29.4559 | -0.6254 | 1.216803 | 0.289634 |
| 18 | 37481400 | 37482999 | 30.11438 | 50.8231 | 44.7715 | 19.58155 | 0.563448 | 1.10918 | 0.289642 |
| 4 | 98444400 | 98445999 | 10.11755 | 18.23625 | 37.40961 | 16.29827 | -0.624 | 0.366622 | 0.289655 |
| 15 | 10618200 | 10622399 | 41.05891 | 65.84638 | 115.4618 | 48.78066 | -0.46108 | 1.962965 | 0.289659 |
| 16 | 88158800 | 88160199 | 10.32578 | 18.43046 | 38.8774 | 17.08932 | -0.67284 | 0.432048 | 0.289663 |
| X | 121810400 | 121812199 | 17.29442 | 31.23042 | 26.05251 | 11.32381 | 0.668478 | 0.38333 | 0.289668 |
| 10 | 129536400 | 129537399 | 14.37684 | 26.3921 | 23.77468 | 10.1192 | 0.578047 | 0.263222 | 0.289702 |
| 4 | 31825600 | 31827599 | 19.09451 | 33.86944 | 28.45776 | 12.57524 | 0.642327 | 0.507781 | 0.28971 |
| 5 | 21346000 | 21346999 | 11.67472 | 22.14691 | 19.54811 | 7.884108 | 0.668584 | 0.030804 | 0.289711 |
| 12 | 63757600 | 63760199 | 32.40061 | 54.19812 | 48.10012 | 20.95169 | 0.552706 | 1.203042 | 0.289715 |
| 12 | 29398600 | 29401599 | 17.50106 | 29.65917 | 59.04825 | 25.85833 | -0.60828 | 1.044134 | 0.289716 |
| 7 | 7700200 | 7702199 | 21.54257 | 38.37921 | 25.97988 | 11.43804 | 0.955763 | 0.491953 | 0.289717 |
| 7 | 128907400 | 128909599 | 10.84691 | 19.15731 | 41.94445 | 18.55671 | -0.73434 | 0.525758 | 0.289724 |
| 4 | 147779200 | 147783599 | 54.37972 | 87.58547 | 74.31612 | 31.59756 | 0.591647 | 1.831008 | 0.28973 |
| 19 | 3493200 | 3497799 | 35.70527 | 57.42415 | 103.473 | 43.90161 | -0.49516 | 1.863956 | 0.289734 |
| 16 | 69733000 | 69733599 | 8.72678 | 17.03069 | 15.01578 | 5.337753 | 0.835025 | -0.43776 | 0.289765 |
| 8 | 92346000 | 92347599 | 7.784839 | 14.31751 | 33.03235 | 14.02402 | -0.76616 | 0.225214 | 0.289768 |
| 11 | 88781600 | 88787199 | 157.8559 | 230.5967 | 418.1401 | 172.2706 | -0.62725 | 3.924073 | 0.28979 |
| X | 59244400 | 59246599 | 28.87992 | 49.80059 | 36.50294 | 16.0131 | 0.831296 | 0.908726 | 0.289802 |
| 8 | 74876800 | 74877999 | 7.823211 | 14.80721 | 33.28605 | 13.48898 | -0.67606 | 0.045686 | 0.289818 |
| 8 | 75187800 | 75188799 | 12.44952 | 23.83745 | 19.93037 | 7.870727 | 0.772586 | -0.02851 | 0.289822 |
| 8 | 23361400 | 23362199 | 9.322312 | 16.51944 | 39.47771 | 17.4538 | -0.85698 | 0.479419 | 0.289829 |
| 5 | 135878400 | 135887599 | 58.44094 | 91.3808 | 179.3643 | 74.41336 | -0.65728 | 2.635108 | 0.289835 |
| 11 | 16562000 | 16564599 | 25.61167 | 44.20703 | 37.61643 | 16.50205 | 0.617055 | 0.884725 | 0.289839 |
| 13 | 97518200 | 97522399 | 28.76965 | 48.81637 | 43.12543 | 18.8925 | 0.559608 | 1.056787 | 0.289863 |
| 12 | 42828200 | 42829199 | 28.80578 | 48.5688 | 43.26798 | 18.91855 | 0.547321 | 1.111629 | 0.289871 |
| 7 | 41455600 | 41460999 | 78.58416 | 125.5093 | 79.77232 | 33.50095 | 0.993632 | 2.11966 | 0.289876 |
| 19 | 46408400 | 46409399 | 4.949123 | 9.616095 | 23.95792 | 8.932701 | -0.70873 | -0.2863 | 0.289883 |
| 2 | 173867400 | 173870199 | 19.0873 | 32.39597 | 60.24101 | 26.39268 | -0.51016 | 1.038709 | 0.289907 |
| 9 | 121202400 | 121205799 | 16.58616 | 27.73429 | 67.44366 | 29.37063 | -0.89633 | 1.176313 | 0.289917 |
| 1 | 82264400 | 82270799 | 51.41343 | 80.14541 | 198.818 | 82.51211 | -0.99809 | 2.68986 | 0.289922 |
| 15 | 28139800 | 28141199 | 8.61963 | 16.04746 | 33.25689 | 13.8801 | -0.5949 | 0.160198 | 0.289923 |
| 3 | 8946200 | 8950399 | 59.83177 | 94.23379 | 159.5917 | 66.2189 | -0.4366 | 2.497504 | 0.289929 |
| X | 75684800 | 75685799 | 12.93745 | 25.15004 | 15.02468 | 5.579131 | 1.334599 | -0.23496 | 0.289976 |
| 3 | 20208600 | 20209399 | 7.521569 | 13.76969 | 35.06557 | 14.9809 | -0.91463 | 0.251723 | 0.28998 |
| 1 | 30285200 | 30286799 | 8.285493 | 15.49358 | 34.54852 | 14.31187 | -0.69139 | 0.130174 | 0.289981 |
| 1 | 133345200 | 133349199 | 16.57218 | 28.08346 | 58.98569 | 25.83051 | -0.68513 | 1.041461 | 0.289987 |
| 4 | 126892400 | 126896999 | 27.63075 | 44.98096 | 85.36289 | 36.6475 | -0.55164 | 1.548213 | 0.289993 |
| 16 | 81713800 | 81714399 | 11.18244 | 21.32281 | 19.56533 | 7.805389 | 0.626657 | -0.00913 | 0.289995 |
| 5 | 135769200 | 135781399 | 108.4821 | 164.073 | 273.1949 | 112.9897 | -0.45823 | 3.297039 | 0.289996 |
| 6 | 82973400 | 82975799 | 13.06352 | 22.78876 | 47.86028 | 21.06661 | -0.67801 | 0.720813 | 0.290016 |
| 15 | 28119400 | 28121199 | 10.81289 | 19.43011 | 41.07756 | 17.95437 | -0.67026 | 0.399742 | 0.290044 |
| X | 78810800 | 78814199 | 51.15397 | 82.99754 | 61.88606 | 26.51828 | 0.788754 | 1.640785 | 0.290047 |
| 7 | 25581400 | 25581999 | 4.648926 | 9.061316 | 22.57767 | 8.167156 | -0.66838 | -0.40272 | 0.290052 |
| X | 79382400 | 79383599 | 17.27492 | 33.05386 | 16.05685 | 6.353659 | 1.546052 | 0.015672 | 0.290073 |
| 5 | 27856400 | 27859599 | 22.30249 | 36.97337 | 70.5266 | 30.53972 | -0.54793 | 1.292493 | 0.290077 |
| 8 | 16097600 | 16098199 | 14.10769 | 26.06398 | 20.96954 | 8.848003 | 0.749282 | 0.220314 | 0.290092 |

| | | | | | | | | | |
|---|---|---|---|---|---|---|---|---|---|
| 2 | 61640600 | 61643399 | 24.62428 | 40.51582 | 75.46193 | 32.5327 | -0.51594 | 1.392641 | 0.290095 |
| 8 | 55367600 | 55368599 | 12.82179 | 24.53991 | 19.9164 | 7.874723 | 0.813653 | -0.02065 | 0.290101 |
| 7 | 60539200 | 60544799 | 57.55519 | 93.07062 | 61.65736 | 26.3531 | 0.954446 | 1.718508 | 0.290101 |
| 16 | 45125000 | 45125799 | 11.44991 | 22.19032 | 17.37348 | 6.577232 | 0.92284 | -0.18221 | 0.290107 |
| 5 | 139012600 | 139016599 | 19.55227 | 32.64074 | 65.39858 | 28.45074 | -0.61796 | 1.202956 | 0.290118 |
| 17 | 84218000 | 84221599 | 20.63325 | 34.43733 | 67.85645 | 29.51567 | -0.59415 | 1.208516 | 0.290122 |
| 15 | 59634000 | 59639399 | 58.63888 | 92.42471 | 158.9651 | 65.96539 | -0.45803 | 2.48072 | 0.290125 |
| 11 | 50692800 | 50701399 | 98.184 | 149.8039 | 243.3865 | 100.922 | -0.4124 | 3.114623 | 0.290134 |
| 1 | 146632200 | 146635199 | 33.93739 | 57.88149 | 37.40322 | 16.39699 | 1.009761 | 1.025426 | 0.290137 |
| 5 | 60817600 | 60819799 | 24.19308 | 42.33861 | 31.38768 | 13.84811 | 0.817159 | 0.704459 | 0.290143 |
| 5 | 10450400 | 10451599 | 14.59257 | 27.21827 | 22.7422 | 9.46172 | 0.709943 | 0.17295 | 0.290149 |
| 4 | 112899000 | 112900999 | 32.33243 | 54.71834 | 42.47744 | 18.59387 | 0.745263 | 1.090177 | 0.290154 |
| 14 | 41883200 | 41885399 | 17.02369 | 31.58058 | 21.18685 | 8.889938 | 1.014463 | 0.220347 | 0.290166 |
| 14 | 78029400 | 78031199 | 9.581085 | 17.56595 | 38.39365 | 16.36983 | -0.69441 | 0.265116 | 0.290185 |
| 19 | 21576600 | 21578399 | 10.58662 | 19.09825 | 37.94746 | 16.51469 | -0.57725 | 0.366592 | 0.290197 |
| 10 | 83195800 | 83196799 | 8.600624 | 15.5831 | 35.45503 | 15.34634 | -0.76575 | 0.322527 | 0.290209 |
| 19 | 40576200 | 40578199 | 8.600624 | 15.5831 | 35.45503 | 15.34634 | -0.76575 | 0.322527 | 0.290209 |
| 7 | 94247600 | 94248399 | 9.701499 | 18.8681 | 16.47886 | 6.096462 | 0.796941 | -0.28376 | 0.290224 |
| 5 | 110093600 | 110095999 | 53.68291 | 86.44766 | 75.68575 | 32.17153 | 0.546243 | 1.837551 | 0.290237 |
| 15 | 4912600 | 4917399 | 60.10013 | 96.22491 | 87.33221 | 36.80446 | 0.484505 | 2.027385 | 0.290264 |
| 9 | 73810400 | 73814799 | 44.18138 | 71.80932 | 63.31292 | 27.16452 | 0.549823 | 1.591089 | 0.290285 |
| 10 | 46489200 | 46490199 | 12.58532 | 22.16789 | 42.97473 | 19.02275 | -0.56129 | 0.570005 | 0.290287 |
| 13 | 104953000 | 104954399 | 8.318819 | 15.04406 | 36.47024 | 15.82311 | -0.85899 | 0.332849 | 0.290292 |
| 17 | 64798400 | 64800199 | 8.333515 | 15.55532 | 32.76239 | 13.61345 | -0.61254 | 0.141616 | 0.290302 |
| 7 | 15104600 | 15105399 | 11.09816 | 21.65791 | 13.63773 | 4.8503 | 1.314695 | -0.4067 | 0.290306 |
| 16 | 44676800 | 44679199 | 20.64025 | 36.80621 | 26.71615 | 11.75289 | 0.856575 | 0.482524 | 0.290309 |
| 10 | 68982600 | 68985199 | 17.1189 | 29.24507 | 55.88972 | 24.50443 | -0.54803 | 0.966826 | 0.29031 |
| 9 | 22474400 | 22478599 | 54.34846 | 87.30641 | 80.26606 | 33.99818 | 0.47158 | 1.915774 | 0.290312 |
| 10 | 6059000 | 6059999 | 11.69502 | 22.49953 | 18.88899 | 7.358319 | 0.784556 | -0.08322 | 0.290314 |
| 16 | 45139800 | 45141599 | 18.45238 | 33.41782 | 24.70658 | 10.70009 | 0.844621 | 0.371237 | 0.290328 |
| 8 | 126790200 | 126792999 | 20.05926 | 35.30768 | 29.96057 | 13.2639 | 0.623733 | 0.594816 | 0.290334 |
| 12 | 50732600 | 50733199 | 4.810817 | 9.367777 | 23.63319 | 8.65778 | -0.70339 | -0.35322 | 0.290355 |
| 1 | 181057800 | 181060399 | 14.23214 | 24.69042 | 50.0176 | 21.94575 | -0.62702 | 0.809864 | 0.290362 |
| 1 | 15408200 | 15409599 | 11.57146 | 22.34318 | 18.41035 | 7.091144 | 0.82649 | -0.12156 | 0.290362 |
| 9 | 53035600 | 53038399 | 26.57785 | 45.66976 | 38.68657 | 16.9692 | 0.622254 | 0.929299 | 0.290374 |
| 14 | 48957000 | 48959399 | 14.52252 | 25.0949 | 52.16126 | 22.88355 | -0.66548 | 0.846293 | 0.290386 |
| 4 | 33695200 | 33696199 | 14.5487 | 27.33131 | 21.68646 | 8.913828 | 0.797675 | 0.125268 | 0.290392 |
| X | 4003800 | 4008799 | 54.32252 | 87.70755 | 68.79146 | 29.34376 | 0.708938 | 1.758869 | 0.290393 |
| 14 | 114764000 | 114764999 | 14.6406 | 26.75013 | 23.70859 | 10.15492 | 0.594863 | 0.290641 | 0.290402 |
| 6 | 135243400 | 135244599 | 6.597869 | 12.59243 | 28.34935 | 11.28779 | -0.65496 | -0.03561 | 0.290407 |
| 14 | 34800800 | 34803399 | 18.91226 | 32.13423 | 59.70392 | 26.16532 | -0.50886 | 1.027638 | 0.290407 |
| 4 | 150762200 | 150768999 | 69.12968 | 107.7677 | 183.0193 | 75.9549 | -0.45213 | 2.694804 | 0.290424 |
| 10 | 30678600 | 30680199 | 13.24574 | 23.04576 | 48.41708 | 21.27639 | -0.67887 | 0.761201 | 0.290426 |
| 16 | 59406600 | 59411199 | 34.58185 | 55.60106 | 112.3739 | 47.65782 | -0.66098 | 1.874452 | 0.290434 |
| 3 | 76437200 | 76438799 | 21.61663 | 38.01038 | 29.20013 | 12.92986 | 0.765842 | 0.612841 | 0.290456 |
| 8 | 26243800 | 26246199 | 10.3641 | 18.40077 | 39.37143 | 17.38421 | -0.69768 | 0.467623 | 0.29046 |
| 17 | 23377200 | 23380599 | 38.45869 | 63.99318 | 45.97707 | 19.96 | 0.856533 | 1.272605 | 0.290466 |

| | | | | | | | | | |
|---|---|---|---|---|---|---|---|---|---|
| 4 | 87388800 | 87393199 | 29.13132 | 47.35779 | 86.87658 | 37.27841 | -0.50424 | 1.575018 | 0.290471 |
| 8 | 117873000 | 117876399 | 26.49602 | 43.23598 | 81.17515 | 34.87974 | -0.53366 | 1.510911 | 0.290476 |
| 3 | 67009600 | 67012399 | 46.10821 | 74.52646 | 69.00362 | 29.47583 | 0.471897 | 1.714369 | 0.290479 |
| 1 | 141949600 | 141952399 | 30.63651 | 51.57051 | 45.30376 | 19.79878 | 0.567256 | 1.131735 | 0.290493 |
| 16 | 59278000 | 59281199 | 33.83539 | 54.24498 | 112.2589 | 47.40487 | -0.70002 | 1.963436 | 0.290498 |
| 7 | 111581600 | 111581999 | 5.057748 | 9.845645 | 24.786 | 9.102204 | -0.70299 | -0.33459 | 0.2905 |
| 12 | 75747800 | 75749199 | 6.528922 | 12.26013 | 31.23474 | 12.84762 | -0.8729 | 0.081191 | 0.290505 |
| 7 | 58103200 | 58103799 | 9.884883 | 19.30197 | 13.73519 | 4.833942 | 1.153166 | -0.45914 | 0.290534 |
| X | 146041600 | 146043799 | 21.18943 | 37.06607 | 32.92028 | 14.51896 | 0.55796 | 0.699134 | 0.290541 |
| 4 | 38964000 | 38965199 | 14.82449 | 27.83644 | 21.73201 | 8.939596 | 0.820235 | 0.131514 | 0.290548 |
| 7 | 11545200 | 11548799 | 51.65994 | 84.33078 | 51.92521 | 22.31462 | 1.070699 | 1.539825 | 0.29055 |
| 1 | 66464400 | 66465599 | 6.110981 | 11.83311 | 29.91254 | 11.36987 | -0.75775 | -0.19206 | 0.290553 |
| 4 | 156074600 | 156078199 | 23.37267 | 38.42568 | 75.09418 | 32.36319 | -0.58547 | 1.400567 | 0.290555 |
| 6 | 65019400 | 65020999 | 9.552901 | 17.14008 | 36.89625 | 16.14733 | -0.69709 | 0.391523 | 0.290597 |
| 17 | 19376800 | 19380599 | 42.67735 | 70.25164 | 52.85915 | 22.79272 | 0.788127 | 1.413987 | 0.290618 |
| 18 | 33071600 | 33072999 | 12.13999 | 21.24586 | 44.67553 | 19.71105 | -0.67915 | 0.664109 | 0.290618 |
| 3 | 32428600 | 32430399 | 18.49922 | 33.02575 | 27.36774 | 12.02801 | 0.667312 | 0.461349 | 0.290627 |
| 7 | 77490600 | 77491599 | 7.029879 | 13.43238 | 29.57122 | 11.74571 | -0.62029 | -0.03701 | 0.290629 |
| 9 | 101157400 | 101158999 | 10.84054 | 19.20786 | 39.37199 | 17.40741 | -0.6376 | 0.487536 | 0.290631 |
| 8 | 126533000 | 126533599 | 6.471674 | 12.29727 | 28.77851 | 11.56595 | -0.72223 | -0.0051 | 0.290647 |
| 16 | 70407000 | 70408199 | 17.08225 | 31.43979 | 22.79564 | 9.668378 | 0.89246 | 0.26746 | 0.290662 |
| 16 | 77467200 | 77467599 | 7.785685 | 15.19745 | 14.5312 | 5.145907 | 0.724105 | -0.46663 | 0.290664 |
| X | 25853800 | 25854799 | 34.3747 | 56.97796 | 49.27 | 21.3328 | 0.58993 | 1.311536 | 0.290679 |
| 1 | 141401600 | 141402399 | 14.43727 | 27.64098 | 18.60416 | 7.348885 | 1.08239 | -0.01144 | 0.290697 |
| 5 | 12839000 | 12842999 | 41.84708 | 69.40055 | 47.16092 | 20.42708 | 0.936434 | 1.322586 | 0.290702 |
| 3 | 36274800 | 36277199 | 15.55858 | 26.5946 | 55.29137 | 24.2432 | -0.6688 | 0.954996 | 0.290718 |
| 8 | 125866200 | 125867999 | 8.854061 | 15.75524 | 38.46458 | 16.95159 | -0.88423 | 0.444061 | 0.290725 |
| 11 | 106185600 | 106190199 | 64.08399 | 100.3458 | 171.9087 | 71.30989 | -0.45969 | 2.61317 | 0.29073 |
| 1 | 57613000 | 57613999 | 10.6959 | 20.49107 | 19.08986 | 7.52683 | 0.620211 | -0.05309 | 0.290732 |
| 2 | 133753000 | 133753799 | 10.63071 | 20.36852 | 19.03907 | 7.504338 | 0.615595 | -0.05585 | 0.290738 |
| 19 | 5874600 | 5875599 | 4.178425 | 8.08808 | 27.10228 | 10.31888 | -1.16082 | -0.21537 | 0.290754 |
| 13 | 100151600 | 100152599 | 6.339199 | 12.01997 | 29.27213 | 11.81735 | -0.78498 | 0.009351 | 0.29078 |
| 5 | 85519200 | 85520199 | 12.52799 | 24.07919 | 18.80397 | 7.345188 | 0.884471 | -0.06592 | 0.290796 |
| 4 | 146387000 | 146392799 | 17.21672 | 31.56219 | 21.27768 | 9.073264 | 0.991245 | 0.286862 | 0.290797 |
| 17 | 50420200 | 50424999 | 35.8437 | 57.64032 | 104.9673 | 44.52833 | -0.51063 | 1.866904 | 0.290809 |
| X | 114799200 | 114801399 | 23.65769 | 41.459 | 31.09413 | 13.73464 | 0.800699 | 0.682107 | 0.290812 |
| 17 | 27432000 | 27435799 | 39.64864 | 63.46982 | 114.8025 | 48.36931 | -0.5088 | 2.018428 | 0.29085 |
| 18 | 75983400 | 75984799 | 17.43558 | 31.29869 | 26.38709 | 11.54333 | 0.647005 | 0.417841 | 0.290877 |
| 4 | 79446000 | 79447599 | 20.42724 | 37.15775 | 21.77358 | 9.378017 | 1.182553 | 0.355196 | 0.290921 |
| 7 | 104480400 | 104483199 | 32.43644 | 53.99305 | 49.77539 | 21.61455 | 0.498202 | 1.256481 | 0.290926 |
| 19 | 39613200 | 39613999 | 22.76113 | 39.55578 | 31.29787 | 13.7436 | 0.723381 | 0.798362 | 0.290928 |
| 7 | 142942400 | 142944399 | 12.78032 | 22.30064 | 46.28689 | 20.37744 | -0.66076 | 0.715828 | 0.290938 |
| 8 | 15025000 | 15032799 | 41.34821 | 65.88292 | 130.9832 | 54.82616 | -0.65238 | 2.169201 | 0.29094 |
| 13 | 33857200 | 33858799 | 25.49019 | 43.90897 | 38.3174 | 16.80853 | 0.580402 | 0.905099 | 0.290941 |
| 7 | 60340600 | 60342599 | 21.00534 | 37.51152 | 26.169 | 11.49814 | 0.914678 | 0.475344 | 0.290945 |
| X | 33277200 | 33284599 | 56.64697 | 91.04008 | 69.51574 | 29.46405 | 0.739688 | 1.9013 | 0.290945 |
| X | 54266600 | 54267399 | 11.8808 | 23.08632 | 16.40906 | 6.113716 | 1.083138 | -0.23109 | 0.290955 |

| | | | | | | | | | |
|---|---|---|---|---|---|---|---|---|---|
| 16 | 84868200 | 84869799 | 12.11495 | 21.35185 | 41.95141 | 18.56761 | -0.58007 | 0.559404 | 0.290964 |
| 2 | 100632200 | 100633399 | 14.43613 | 26.80919 | 23.27843 | 9.753293 | 0.646996 | 0.199454 | 0.290966 |
| 16 | 20732000 | 20733599 | 30.65344 | 51.85534 | 39.95468 | 17.48737 | 0.756007 | 1.085354 | 0.290971 |
| 16 | 50418800 | 50420199 | 11.94156 | 20.97502 | 44.5644 | 19.72277 | -0.6942 | 0.600905 | 0.290978 |
| 19 | 7924200 | 7927199 | 35.09203 | 58.89459 | 44.16925 | 19.27964 | 0.794572 | 1.17259 | 0.290996 |
| 4 | 16889000 | 16891799 | 28.33406 | 48.86101 | 35.78767 | 15.69935 | 0.832108 | 0.901805 | 0.290997 |
| 8 | 12404200 | 12404999 | 5.275675 | 10.19367 | 26.51294 | 10.17963 | -0.81151 | -0.17582 | 0.291002 |
| 15 | 66157400 | 66159399 | 11.87707 | 20.68023 | 48.39781 | 21.27812 | -0.83348 | 0.743701 | 0.291007 |
| 17 | 19005600 | 19014799 | 189.0335 | 283.9033 | 220.7971 | 91.13394 | 0.63044 | 3.44908 | 0.291007 |
| 6 | 59793400 | 59797399 | 54.52734 | 88.18574 | 64.3531 | 27.50986 | 0.815482 | 1.710304 | 0.291013 |
| 2 | 121711200 | 121713599 | 22.87072 | 40.01855 | 32.7227 | 14.43532 | 0.676632 | 0.703365 | 0.291014 |
| 7 | 13659400 | 13660799 | 29.41957 | 50.19589 | 40.87118 | 17.91785 | 0.677739 | 1.008027 | 0.291018 |
| 9 | 102566800 | 102569399 | 10.89886 | 19.13352 | 43.577 | 19.27723 | -0.79358 | 0.60337 | 0.291021 |
| 14 | 91838000 | 91838199 | 9.444818 | 18.28747 | 17.64364 | 6.708429 | 0.619093 | -0.18918 | 0.291046 |
| 13 | 41758600 | 41759799 | 6.616778 | 12.56593 | 29.90074 | 12.03125 | -0.74803 | 0.001578 | 0.29107 |
| 7 | 20826000 | 20833999 | 92.3587 | 146.5288 | 93.00917 | 38.73505 | 0.985777 | 2.336139 | 0.291077 |
| 16 | 10113000 | 10117999 | 35.61681 | 57.03087 | 119.8313 | 50.50816 | -0.72411 | 2.005221 | 0.291082 |
| 18 | 88570600 | 88571599 | 9.425633 | 17.34408 | 35.71954 | 15.15429 | -0.60365 | 0.239002 | 0.291091 |
| 8 | 36152600 | 36154199 | 6.264182 | 12.09568 | 32.80191 | 12.63314 | -0.87701 | -0.14003 | 0.291113 |
| 1 | 43137400 | 43138199 | 5.685471 | 10.98249 | 25.92351 | 9.965954 | -0.67423 | -0.16647 | 0.291119 |
| 13 | 101803600 | 101805199 | 17.44315 | 32.64099 | 20.2694 | 8.386704 | 1.141081 | 0.171373 | 0.291126 |
| 5 | 100396000 | 100398599 | 14.04968 | 26.50495 | 19.18851 | 7.830291 | 0.937316 | 0.088897 | 0.291127 |
| 14 | 15519600 | 15520799 | 13.22596 | 24.89715 | 21.83764 | 8.945004 | 0.658103 | 0.101122 | 0.291128 |
| 15 | 20168400 | 20171199 | 34.64322 | 57.99311 | 46.43853 | 20.23755 | 0.7005 | 1.200024 | 0.291132 |
| 16 | 90505800 | 90515999 | 143.772 | 213.3215 | 351.1814 | 144.6528 | -0.46666 | 3.667627 | 0.291134 |
| X | 32435800 | 32438199 | 28.61311 | 49.93817 | 29.55783 | 13.01433 | 1.140354 | 0.765429 | 0.291141 |
| 8 | 63468600 | 63471999 | 74.48751 | 118.0062 | 109.6754 | 45.61513 | 0.434558 | 2.363785 | 0.291153 |
| 11 | 97003400 | 97006999 | 18.9477 | 31.56565 | 66.94558 | 29.08549 | -0.6997 | 1.224597 | 0.291164 |
| 13 | 84655400 | 84656999 | 16.70776 | 31.07347 | 21.91074 | 9.160591 | 0.947303 | 0.206334 | 0.291167 |
| 2 | 36772200 | 36775799 | 44.43148 | 72.24489 | 63.26799 | 27.15 | 0.560002 | 1.586663 | 0.291189 |
| 7 | 58862200 | 58866799 | 70.27159 | 112.05 | 94.57275 | 39.62558 | 0.582682 | 2.164308 | 0.291202 |
| 3 | 65149600 | 65151199 | 9.750269 | 17.84418 | 36.66999 | 15.67302 | -0.6084 | 0.274034 | 0.29123 |
| 12 | 98941600 | 98945599 | 34.55501 | 55.58799 | 103.5468 | 43.9487 | -0.54253 | 1.853372 | 0.29123 |
| 17 | 78506800 | 78510599 | 19.26009 | 32.07774 | 68.14109 | 29.60005 | -0.70209 | 1.228281 | 0.291238 |
| X | 148561000 | 148563599 | 48.2381 | 78.06948 | 67.8027 | 28.99962 | 0.565899 | 1.686666 | 0.291242 |
| 12 | 33479400 | 33480399 | 7.060029 | 13.42717 | 29.37324 | 11.7828 | -0.6233 | -0.00264 | 0.291257 |
| 14 | 80659000 | 80659799 | 16.76371 | 29.97655 | 25.32426 | 11.11453 | 0.641239 | 0.436727 | 0.291268 |
| 8 | 120924000 | 120940999 | 139.678 | 206.379 | 370.6682 | 152.6493 | -0.59787 | 3.738685 | 0.291293 |
| 4 | 83404000 | 83405199 | 8.226863 | 15.11793 | 33.53354 | 14.25285 | -0.71143 | 0.232996 | 0.291298 |
| 18 | 27434800 | 27437399 | 48.55879 | 78.29148 | 72.85143 | 31.0193 | 0.46062 | 1.791736 | 0.29131 |
| 1 | 58484200 | 58484999 | 6.141318 | 11.79553 | 27.44114 | 10.74062 | -0.67873 | -0.09949 | 0.29131 |
| 8 | 80173600 | 80174999 | 8.634268 | 16.1027 | 33.86347 | 14.0917 | -0.61235 | 0.151377 | 0.291314 |
| 14 | 44033000 | 44034799 | 16.48966 | 28.21188 | 54.79445 | 24.02709 | -0.57084 | 0.950135 | 0.291318 |
| 4 | 82100000 | 82100999 | 9.116053 | 16.75557 | 34.59794 | 14.70092 | -0.60891 | 0.240357 | 0.29132 |
| 17 | 21040600 | 21043599 | 51.97171 | 83.84517 | 66.53275 | 28.35113 | 0.690854 | 1.776683 | 0.291328 |
| 6 | 13793800 | 13794399 | 7.501402 | 14.65481 | 14.01105 | 4.890523 | 0.742678 | -0.52225 | 0.291336 |
| 6 | 131401400 | 131402599 | 10.4348 | 18.86908 | 37.47134 | 16.25918 | -0.57356 | 0.351446 | 0.29134 |

| | | | | | | | | | |
|---|---|---|---|---|---|---|---|---|---|
| X | 8517000 | 8522199 | 108.2159 | 168.4071 | 139.4223 | 57.87669 | 0.58131 | 2.738762 | 0.291345 |
| 7 | 17135000 | 17158799 | 65.39478 | 104.7352 | 77.28573 | 32.58625 | 0.783012 | 2.019622 | 0.291356 |
| 7 | 41741400 | 41746799 | 87.36582 | 137.8879 | 117.0561 | 48.59209 | 0.560759 | 2.463589 | 0.291365 |
| 10 | 126806800 | 126809599 | 13.29619 | 23.02644 | 50.00385 | 21.93855 | -0.72732 | 0.820388 | 0.291366 |
| 7 | 28578600 | 28583999 | 37.74352 | 60.35199 | 117.6366 | 49.48428 | -0.6182 | 2.052316 | 0.291377 |
| 13 | 52346600 | 52348399 | 15.55584 | 28.09639 | 23.63549 | 10.27072 | 0.65633 | 0.362233 | 0.291382 |
| 7 | 73713000 | 73713999 | 6.265445 | 12.10718 | 27.77895 | 10.66126 | -0.63199 | -0.16134 | 0.291389 |
| 3 | 154302200 | 154303599 | 11.33963 | 19.92787 | 43.0157 | 19.04553 | -0.71669 | 0.587776 | 0.291396 |
| 3 | 154738800 | 154739799 | 6.690923 | 12.52095 | 33.92579 | 14.03649 | -0.96849 | 0.109842 | 0.291396 |
| 12 | 110703800 | 110705199 | 10.27688 | 18.09221 | 42.012 | 18.59856 | -0.82026 | 0.559946 | 0.291401 |
| 5 | 71061000 | 71063999 | 30.42064 | 51.77461 | 41.4021 | 18.14696 | 0.703192 | 1.034601 | 0.291419 |
| 15 | 103354000 | 103355199 | 6.924057 | 13.11466 | 29.24875 | 11.83511 | -0.66186 | 0.024467 | 0.291419 |
| 17 | 40528600 | 40529199 | 6.924057 | 13.11466 | 29.24875 | 11.83511 | -0.66186 | 0.024467 | 0.291419 |
| 4 | 83050800 | 83052399 | 17.29 | 31.73045 | 23.31779 | 9.928755 | 0.869269 | 0.286181 | 0.291424 |
| 13 | 30668400 | 30670399 | 11.56092 | 20.60293 | 42.51463 | 18.71269 | -0.64298 | 0.456177 | 0.291439 |
| 17 | 50970800 | 50973399 | 70.6927 | 112.3399 | 103.8645 | 43.32715 | 0.446341 | 2.27319 | 0.291442 |
| 17 | 54157800 | 54160399 | 25.36511 | 44.01427 | 35.36515 | 15.51694 | 0.700832 | 0.833608 | 0.291444 |
| 16 | 15400600 | 15401199 | 5.670073 | 10.97635 | 26.04903 | 9.913992 | -0.66772 | -0.1993 | 0.291446 |
| 18 | 83440800 | 83441799 | 5.670073 | 10.97635 | 26.04903 | 9.913992 | -0.66772 | -0.1993 | 0.291446 |
| 4 | 47049200 | 47053599 | 26.77149 | 43.3632 | 89.22447 | 38.18502 | -0.67424 | 1.644089 | 0.29145 |
| 1 | 117644200 | 117648599 | 80.46665 | 128.2662 | 85.07425 | 35.63033 | 0.929285 | 2.181491 | 0.291457 |
| 12 | 114066200 | 114068399 | 33.3331 | 57.15497 | 34.54669 | 15.15124 | 1.106649 | 0.972542 | 0.291464 |
| 9 | 19307400 | 19308799 | 31.68115 | 53.22196 | 44.75229 | 19.54586 | 0.630033 | 1.150539 | 0.291503 |
| 18 | 79519200 | 79519999 | 11.30525 | 21.86246 | 17.25677 | 6.599641 | 0.896978 | -0.15214 | 0.291506 |
| 2 | 117797400 | 117798599 | 19.98271 | 35.47302 | 27.71084 | 12.237 | 0.747354 | 0.506221 | 0.291511 |
| 17 | 4381800 | 4382599 | 6.42922 | 12.34063 | 28.60822 | 11.21794 | -0.67664 | -0.08845 | 0.291522 |
| 8 | 12449600 | 12454999 | 37.84638 | 60.64101 | 113.1925 | 47.75793 | -0.55252 | 1.986077 | 0.291527 |
| 16 | 67414600 | 67418399 | 44.92496 | 71.52712 | 129.8955 | 54.31323 | -0.52319 | 2.198118 | 0.291539 |
| 17 | 13656400 | 13669599 | 310.538 | 432.4045 | 727.6666 | 301.9262 | -0.5886 | 4.644688 | 0.29154 |
| 3 | 145201800 | 145203399 | 13.88293 | 25.5763 | 22.84246 | 9.675443 | 0.595397 | 0.23754 | 0.291541 |
| 16 | 9548800 | 9553799 | 54.52916 | 87.72553 | 77.00138 | 32.68492 | 0.540824 | 1.867418 | 0.291546 |
| 5 | 36878400 | 36882199 | 21.43308 | 35.82005 | 66.2194 | 28.82787 | -0.50241 | 1.196624 | 0.291547 |
| 7 | 22579000 | 22592799 | 345.5362 | 493.9374 | 371.7812 | 153.6062 | 0.60989 | 4.257445 | 0.29155 |
| 4 | 57089800 | 57090399 | 11.29264 | 21.65976 | 18.60726 | 7.311816 | 0.741181 | -0.05772 | 0.291552 |
| 13 | 60381600 | 60382199 | 9.549961 | 18.5113 | 17.58593 | 6.65238 | 0.647693 | -0.20607 | 0.291562 |
| 13 | 37606400 | 37609799 | 30.73199 | 49.86619 | 89.43954 | 38.32692 | -0.47439 | 1.614596 | 0.291564 |
| 11 | 31554800 | 31560399 | 40.76603 | 65.25641 | 115.9379 | 48.84542 | -0.48308 | 2.020427 | 0.291568 |
| 15 | 5802400 | 5804599 | 13.74007 | 24.0728 | 54.32629 | 23.99149 | -0.78112 | 0.668864 | 0.291581 |
| 13 | 107793000 | 107796199 | 18.41503 | 31.31108 | 58.60434 | 25.68486 | -0.51927 | 1.017803 | 0.291582 |
| 3 | 73208000 | 73210199 | 20.56498 | 34.56466 | 65.22856 | 28.49327 | -0.53211 | 1.132031 | 0.291584 |
| 10 | 123332400 | 123334199 | 17.91433 | 30.60066 | 57.00966 | 24.99525 | -0.51152 | 0.9722 | 0.29159 |
| 9 | 52713200 | 52715599 | 15.95633 | 27.48555 | 52.70319 | 23.11907 | -0.55062 | 0.882733 | 0.291596 |
| 3 | 122831800 | 122833999 | 18.94113 | 32.00122 | 60.73554 | 26.57246 | -0.53983 | 1.077645 | 0.291605 |
| 17 | 37982600 | 37990399 | 113.7204 | 176.4575 | 142.2221 | 59.06469 | 0.616687 | 2.788535 | 0.291605 |
| X | 122089600 | 122092599 | 21.88734 | 39.73126 | 21.12228 | 9.121111 | 1.319733 | 0.376589 | 0.291639 |
| 3 | 48892800 | 48899199 | 105.1417 | 162.8543 | 151.3322 | 62.85929 | 0.409615 | 2.818066 | 0.29164 |
| 11 | 108849000 | 108856399 | 55.07125 | 87.23548 | 149.9483 | 62.3612 | -0.4514 | 2.361164 | 0.291649 |

| | | | | | | | | | |
|---|---|---|---|---|---|---|---|---|---|
| 4 | 142562000 | 142566999 | 33.85936 | 54.72826 | 98.1259 | 41.91588 | -0.47939 | 1.707985 | 0.29165 |
| 2 | 33846000 | 33848399 | 19.59025 | 33.09909 | 61.31111 | 26.82457 | -0.50491 | 1.080769 | 0.291651 |
| 1 | 111609000 | 111612799 | 43.7318 | 72.56571 | 45.47417 | 19.70389 | 1.053051 | 1.322659 | 0.291664 |
| 1 | 6943200 | 6945199 | 20.75039 | 36.39813 | 31.63982 | 13.98674 | 0.588542 | 0.649619 | 0.291691 |
| 5 | 106649000 | 106651199 | 14.05031 | 24.40153 | 48.96271 | 21.49176 | -0.61312 | 0.79329 | 0.291712 |
| 8 | 111757600 | 111758999 | 9.05796 | 16.42754 | 35.46833 | 15.33362 | -0.68932 | 0.322395 | 0.291723 |
| 13 | 25225600 | 25226399 | 12.76816 | 24.69428 | 17.19895 | 6.574055 | 1.077109 | -0.1365 | 0.291736 |
| 6 | 92068000 | 92069599 | 7.797129 | 13.94633 | 37.99144 | 16.67034 | -1.03663 | 0.40223 | 0.291744 |
| 19 | 23481400 | 23483399 | 10.40326 | 18.84035 | 39.11529 | 16.94061 | -0.63583 | 0.343972 | 0.291745 |
| 5 | 7578000 | 7580599 | 14.81626 | 27.03079 | 22.10101 | 9.484004 | 0.70849 | 0.295541 | 0.29175 |
| 18 | 81215200 | 81219799 | 26.14109 | 42.74719 | 79.71355 | 34.27815 | -0.52173 | 1.477968 | 0.291751 |
| 1 | 128673600 | 128676599 | 21.10092 | 35.25251 | 66.78749 | 29.06872 | -0.53769 | 1.198903 | 0.291756 |
| 13 | 65773600 | 65774399 | 20.91617 | 36.73234 | 31.01601 | 13.72468 | 0.630175 | 0.630851 | 0.29176 |
| 5 | 24105200 | 24106599 | 19.38723 | 34.16918 | 30.14342 | 13.34139 | 0.568831 | 0.57875 | 0.291768 |
| 1 | 72958400 | 72960399 | 24.93438 | 43.58999 | 31.96513 | 14.09033 | 0.832946 | 0.726419 | 0.291782 |
| 7 | 9059200 | 9061999 | 53.83278 | 87.04354 | 65.45561 | 27.97333 | 0.771913 | 1.715349 | 0.291784 |
| X | 61395800 | 61396999 | 22.90805 | 39.88343 | 33.64675 | 14.79191 | 0.631548 | 0.77807 | 0.29179 |
| 2 | 157497000 | 157503799 | 57.45182 | 90.75534 | 153.1173 | 63.5902 | -0.42774 | 2.42972 | 0.291801 |
| 18 | 19091000 | 19094199 | 31.27857 | 53.4883 | 38.59619 | 16.92358 | 0.851783 | 0.993167 | 0.291817 |
| 17 | 18239800 | 18240399 | 12.08801 | 23.01211 | 20.04561 | 8.024543 | 0.696984 | 0.012947 | 0.291823 |
| X | 157883200 | 157887999 | 59.85446 | 96.55401 | 66.26214 | 28.23269 | 0.899745 | 1.792562 | 0.291831 |
| 17 | 18407400 | 18411999 | 63.28007 | 101.7085 | 74.27272 | 31.48387 | 0.804075 | 1.909419 | 0.291836 |
| 8 | 30599400 | 30600199 | 10.0517 | 19.4575 | 16.87955 | 6.425688 | 0.768337 | -0.18285 | 0.291841 |
| 5 | 130947800 | 130948199 | 4.646344 | 9.064538 | 22.78913 | 8.145871 | -0.66586 | -0.44556 | 0.291849 |
| 4 | 77070200 | 77071199 | 5.673699 | 10.99779 | 25.5931 | 9.680947 | -0.63177 | -0.22356 | 0.291872 |
| 11 | 88348400 | 88351799 | 15.06511 | 25.83051 | 54.24081 | 23.78847 | -0.68239 | 0.923165 | 0.291881 |
| 14 | 63454200 | 63455799 | 16.84341 | 30.78023 | 24.45091 | 10.47061 | 0.752076 | 0.308723 | 0.291883 |
| 5 | 141062000 | 141063599 | 11.57857 | 20.58093 | 40.36201 | 17.80412 | -0.57198 | 0.46968 | 0.291884 |
| X | 89576000 | 89577799 | 31.05175 | 52.96105 | 39.79427 | 17.44521 | 0.793171 | 1.017838 | 0.291887 |
| 13 | 77747600 | 77749399 | 15.73115 | 28.66855 | 25.15157 | 10.80784 | 0.606629 | 0.317679 | 0.291894 |
| X | 147655200 | 147656799 | 31.3218 | 52.8072 | 44.76769 | 19.57385 | 0.618604 | 1.122517 | 0.291896 |
| 17 | 56774600 | 56776599 | 19.78695 | 32.81526 | 70.20488 | 30.40821 | -0.71274 | 1.28128 | 0.291896 |
| 4 | 35603000 | 35605199 | 23.82018 | 41.72201 | 30.81044 | 13.60335 | 0.822641 | 0.687676 | 0.291904 |
| 2 | 49584800 | 49586399 | 8.793041 | 16.25144 | 33.63231 | 14.18355 | -0.60413 | 0.20459 | 0.291906 |
| 14 | 124447600 | 124451199 | 80.46077 | 127.1982 | 115.7478 | 48.08887 | 0.462572 | 2.420712 | 0.29191 |
| 2 | 123710000 | 123712199 | 22.07521 | 38.67915 | 30.92756 | 13.65909 | 0.708384 | 0.671909 | 0.291913 |
| 1 | 105957600 | 105958999 | 6.241126 | 11.72725 | 30.69472 | 12.61174 | -0.90992 | 0.073809 | 0.291918 |
| 5 | 147158200 | 147159599 | 6.241126 | 11.72725 | 30.69472 | 12.61174 | -0.90992 | 0.073809 | 0.291918 |
| 5 | 62191400 | 62194999 | 36.63008 | 60.36464 | 54.33151 | 23.43868 | 0.531008 | 1.384849 | 0.291935 |
| 4 | 113802200 | 113804599 | 18.82045 | 33.50452 | 24.98614 | 11.00747 | 0.816294 | 0.47659 | 0.291947 |
| 17 | 35176000 | 35179399 | 12.63967 | 21.98424 | 48.9409 | 21.50224 | -0.76203 | 0.763475 | 0.291949 |
| 12 | 69100800 | 69103199 | 27.39658 | 47.36314 | 35.83591 | 15.72171 | 0.78618 | 0.876308 | 0.291962 |
| 19 | 58453400 | 58456399 | 38.11089 | 61.25872 | 107.8182 | 45.70805 | -0.4624 | 1.885203 | 0.291969 |
| 17 | 43834400 | 43839799 | 64.86883 | 103.5733 | 93.22229 | 39.13288 | 0.492167 | 2.117701 | 0.292 |
| 1 | 140127000 | 140128799 | 18.22613 | 32.88036 | 25.1998 | 10.96674 | 0.788524 | 0.391815 | 0.292016 |
| 10 | 80177800 | 80181199 | 24.78847 | 40.34378 | 84.0519 | 36.08136 | -0.68604 | 1.54648 | 0.292041 |
| 3 | 145642400 | 145645399 | 15.85751 | 27.12192 | 55.31821 | 24.25614 | -0.64121 | 0.950188 | 0.292043 |

| | | | | | | | | | |
|---|---|---|---|---|---|---|---|---|---|
| 2 | 59316800 | 59318399 | 8.929717 | 16.56784 | 34.83844 | 14.61528 | -0.62167 | 0.186091 | 0.29205 |
| 5 | 108460000 | 108460999 | 5.126435 | 9.896135 | 25.84845 | 9.967538 | -0.8228 | -0.16252 | 0.292051 |
| 6 | 12297600 | 12300599 | 42.89679 | 69.54322 | 64.56857 | 27.65103 | 0.472032 | 1.64048 | 0.292055 |
| 13 | 4673800 | 4674999 | 17.05785 | 31.17168 | 23.80737 | 10.1952 | 0.8082 | 0.308191 | 0.29206 |
| 6 | 131102000 | 131104399 | 25.58116 | 44.49198 | 34.82478 | 15.29999 | 0.73889 | 0.804978 | 0.292062 |
| 13 | 119330800 | 119333799 | 26.9731 | 46.20976 | 35.75959 | 15.68209 | 0.751601 | 0.955456 | 0.292063 |
| 7 | 17117200 | 17122399 | 100.2937 | 156.2555 | 141.5344 | 58.7435 | 0.452973 | 2.714706 | 0.292077 |
| 15 | 15970600 | 15970999 | 3.413556 | 6.674882 | 20.21525 | 6.897446 | -0.86841 | -0.63808 | 0.292083 |
| 9 | 64308600 | 64310799 | 8.775826 | 15.82861 | 35.93514 | 15.64261 | -0.76789 | 0.3522 | 0.29209 |
| X | 90943200 | 90944199 | 14.40003 | 26.76188 | 21.90748 | 9.169687 | 0.73283 | 0.193362 | 0.292093 |
| 10 | 121321800 | 121323599 | 12.29872 | 21.54647 | 44.13597 | 19.49043 | -0.64146 | 0.647792 | 0.292097 |
| 6 | 38169000 | 38173999 | 34.542 | 55.59792 | 102.6557 | 43.60556 | -0.52883 | 1.835296 | 0.292103 |
| 8 | 83322800 | 83324199 | 14.97875 | 25.97491 | 50.04253 | 21.95641 | -0.55525 | 0.814914 | 0.292127 |
| 10 | 14635600 | 14636399 | 7.553927 | 14.1301 | 31.1674 | 12.90424 | -0.67446 | 0.115874 | 0.292143 |
| 8 | 128109400 | 128110199 | 16.10259 | 29.75322 | 22.94827 | 9.681305 | 0.809392 | 0.238697 | 0.292148 |
| 6 | 33342000 | 33343199 | 17.47973 | 31.55546 | 26.04716 | 11.32579 | 0.683298 | 0.386528 | 0.292152 |
| 2 | 117538400 | 117539599 | 6.724341 | 12.90174 | 28.59224 | 11.22505 | -0.61373 | -0.08049 | 0.292154 |
| 9 | 115935200 | 115939999 | 45.53457 | 72.6481 | 124.9944 | 52.41397 | -0.44219 | 2.130612 | 0.292164 |
| 18 | 63100200 | 63102999 | 13.09643 | 22.77291 | 48.94855 | 21.50218 | -0.71175 | 0.769264 | 0.292165 |
| 7 | 32226200 | 32233599 | 37.03482 | 61.17336 | 48.27703 | 20.85256 | 0.720985 | 1.35587 | 0.292168 |
| 13 | 94720000 | 94725599 | 41.23644 | 66.0441 | 115.8959 | 48.86616 | -0.46441 | 2.004346 | 0.292168 |
| 19 | 6384600 | 6385799 | 5.021482 | 9.623935 | 27.28073 | 10.74405 | -0.96849 | -0.08972 | 0.292182 |
| Y | 1375000 | 1378199 | 24.76407 | 42.78602 | 24.7692 | 10.86639 | 1.170089 | 0.851644 | 0.292185 |
| 5 | 16658600 | 16661599 | 68.77309 | 109.4893 | 100.0433 | 41.8271 | 0.465998 | 2.213553 | 0.292189 |
| 12 | 72304000 | 72305399 | 10.78321 | 19.23056 | 39.37082 | 17.31862 | -0.6308 | 0.441212 | 0.292196 |
| 15 | 60493600 | 60495999 | 27.16465 | 46.9321 | 35.83492 | 15.72097 | 0.772865 | 0.880261 | 0.292205 |
| 2 | 148247800 | 148253999 | 52.29172 | 82.49908 | 155.5088 | 64.54185 | -0.58891 | 2.457028 | 0.292205 |
| 13 | 86631600 | 86632799 | 14.59568 | 27.61351 | 20.68234 | 8.389197 | 0.896144 | 0.078436 | 0.292208 |
| 8 | 72062800 | 72080599 | 175.7888 | 255.8175 | 435.1274 | 179.3402 | -0.54034 | 3.983145 | 0.292241 |
| 11 | 101239400 | 101244399 | 38.46368 | 61.75825 | 108.3688 | 45.8674 | -0.45996 | 1.920771 | 0.292258 |
| 8 | 4781000 | 4781999 | 10.3964 | 20.24354 | 15.89004 | 5.822678 | 0.961958 | -0.30876 | 0.292264 |
| 3 | 64879600 | 64880599 | 11.50218 | 22.17778 | 18.35385 | 7.104368 | 0.813403 | -0.10719 | 0.29229 |
| 13 | 50600600 | 50602399 | 16.25491 | 30.09487 | 23.05138 | 9.698339 | 0.822385 | 0.230362 | 0.292294 |
| 12 | 77382000 | 77385199 | 48.04306 | 77.95811 | 64.05338 | 27.44906 | 0.649178 | 1.63227 | 0.29232 |
| 3 | 4640000 | 4643599 | 34.03784 | 57.84571 | 39.0688 | 17.12222 | 0.945556 | 1.058353 | 0.292325 |
| 4 | 133337200 | 133342799 | 46.75724 | 74.29531 | 135.5436 | 56.53603 | -0.53282 | 2.263768 | 0.292348 |
| 5 | 139559800 | 139577799 | 159.8291 | 234.0625 | 431.3539 | 177.6857 | -0.64717 | 3.885518 | 0.292349 |
| 3 | 118355400 | 118356399 | 9.141364 | 16.72752 | 34.51203 | 14.75343 | -0.61399 | 0.266171 | 0.292368 |
| 1 | 53379200 | 53382599 | 87.18245 | 136.8661 | 128.2621 | 53.19698 | 0.412424 | 2.572417 | 0.29238 |
| 17 | 30912400 | 30916799 | 28.6062 | 46.51499 | 86.55868 | 37.14517 | -0.52461 | 1.567679 | 0.292386 |
| 16 | 19342600 | 19344199 | 17.26384 | 31.46227 | 24.33063 | 10.45484 | 0.7873 | 0.326366 | 0.292389 |
| 2 | 46907200 | 46907599 | 3.419592 | 6.670902 | 21.3888 | 7.650872 | -1.01114 | -0.4634 | 0.292401 |
| 5 | 98390600 | 98391599 | 7.105159 | 13.31888 | 33.44522 | 13.79764 | -0.85639 | 0.101156 | 0.292402 |
| 7 | 31070800 | 31112799 | 88.59137 | 139.6485 | 110.9545 | 46.04313 | 0.654774 | 2.490262 | 0.292407 |
| 11 | 46300800 | 46302199 | 13.8652 | 24.11483 | 48.42501 | 21.27504 | -0.61409 | 0.769574 | 0.292409 |
| 8 | 84334200 | 84335599 | 7.292921 | 13.55778 | 31.58629 | 13.21532 | -0.76466 | 0.155952 | 0.292423 |
| 12 | 18347400 | 18351999 | 120.3177 | 186.2788 | 145.2252 | 60.32274 | 0.662188 | 2.83338 | 0.292424 |

| | | | | | | | | | |
|---|---|---|---|---|---|---|---|---|---|
| 6 | 121103800 | 121105399 | 6.572488 | 12.23097 | 32.13099 | 13.41996 | -0.93452 | 0.143354 | 0.29243 |
| X | 149878600 | 149880599 | 18.70216 | 34.32529 | 22.80484 | 9.709083 | 1.014269 | 0.297021 | 0.292445 |
| 1 | 175452600 | 175453399 | 6.360719 | 11.97874 | 29.65495 | 12.13845 | -0.82529 | 0.060019 | 0.29245 |
| 7 | 144871200 | 144880199 | 58.77646 | 92.67668 | 160.2927 | 66.51958 | -0.46568 | 2.473045 | 0.292457 |
| 7 | 10261200 | 10262999 | 20.56344 | 37.1264 | 23.97798 | 10.42465 | 1.034982 | 0.402807 | 0.292459 |
| 11 | 56819200 | 56821199 | 19.17816 | 33.71675 | 30.25093 | 13.39562 | 0.543232 | 0.604192 | 0.292478 |
| 8 | 3820400 | 3823799 | 37.50045 | 61.8908 | 49.37848 | 21.31603 | 0.705213 | 1.366896 | 0.292481 |
| 7 | 141632600 | 141639399 | 36.58163 | 58.25553 | 134.2143 | 56.13533 | -0.86545 | 2.184204 | 0.29249 |
| 5 | 116124600 | 116134399 | 77.79741 | 119.5223 | 219.2211 | 91.02463 | -0.57933 | 2.968222 | 0.292491 |
| 7 | 47330000 | 47341399 | 152.4073 | 231.1946 | 183.9457 | 76.14812 | 0.609981 | 3.241935 | 0.292502 |
| 1 | 51989000 | 51990599 | 8.464777 | 15.30705 | 35.95462 | 15.60063 | -0.81374 | 0.333763 | 0.292502 |
| 2 | 134542200 | 134543199 | 16.90003 | 31.42187 | 21.90331 | 9.162021 | 0.963237 | 0.209855 | 0.292513 |
| 3 | 83110800 | 83112399 | 22.26498 | 38.97265 | 32.66968 | 14.41631 | 0.640817 | 0.693242 | 0.292522 |
| 1 | 55811800 | 55812799 | 6.760862 | 12.83888 | 29.33603 | 11.80536 | -0.69 | 0.002617 | 0.292524 |
| 13 | 52129000 | 52133399 | 37.99295 | 60.92928 | 110.653 | 46.74878 | -0.51151 | 1.958148 | 0.292548 |
| 15 | 87386200 | 87402199 | 136.4739 | 204.0074 | 328.9255 | 135.624 | -0.42677 | 3.53185 | 0.292554 |
| 7 | 34129800 | 34133399 | 26.74732 | 43.80928 | 80.70126 | 34.72457 | -0.50282 | 1.456817 | 0.292559 |
| 4 | 17732000 | 17732999 | 14.13981 | 26.15229 | 23.21174 | 9.779341 | 0.609623 | 0.217937 | 0.292589 |
| 7 | 130521000 | 130521999 | 14.43038 | 26.67738 | 23.14464 | 9.7572 | 0.641532 | 0.222383 | 0.292601 |
| 18 | 34240400 | 34242799 | 9.998223 | 17.95949 | 36.87619 | 16.12309 | -0.62853 | 0.387024 | 0.292603 |
| 10 | 43048200 | 43052199 | 27.91635 | 45.14128 | 92.72453 | 39.63007 | -0.67421 | 1.682755 | 0.292616 |
| 2 | 160994800 | 160996799 | 10.90356 | 19.60991 | 39.47313 | 17.24058 | -0.59943 | 0.389262 | 0.292635 |
| 4 | 54148400 | 54149399 | 10.90356 | 19.60991 | 39.47313 | 17.24058 | -0.59943 | 0.389262 | 0.292635 |
| 15 | 14960400 | 14963199 | 26.28593 | 45.80307 | 32.45495 | 14.27555 | 0.881857 | 0.77844 | 0.292636 |
| 14 | 42890600 | 42895199 | 52.86375 | 85.70482 | 61.50934 | 26.34171 | 0.842672 | 1.657802 | 0.292645 |
| 19 | 42767600 | 42771999 | 30.45045 | 49.27364 | 91.08874 | 38.95456 | -0.52158 | 1.670764 | 0.292658 |
| 4 | 128008600 | 128026799 | 147.8418 | 218.5945 | 368.9891 | 151.959 | -0.50747 | 3.727802 | 0.292672 |
| 4 | 116733800 | 116735399 | 10.39809 | 18.37591 | 40.43486 | 17.88665 | -0.74084 | 0.515007 | 0.292675 |
| 8 | 129410600 | 129412999 | 10.39809 | 18.37591 | 40.43486 | 17.88665 | -0.74084 | 0.515007 | 0.292675 |
| 13 | 41385200 | 41387399 | 12.90812 | 22.51825 | 48.38973 | 21.29999 | -0.71097 | 0.720037 | 0.292679 |
| 12 | 32781400 | 32782599 | 12.76914 | 23.99976 | 21.91108 | 8.998809 | 0.597815 | 0.107281 | 0.292684 |
| 8 | 16681000 | 16684399 | 32.39841 | 55.08696 | 39.63351 | 17.37041 | 0.854741 | 1.047145 | 0.292694 |
| 13 | 66366600 | 66377599 | 191.3483 | 288.9668 | 187.2381 | 77.40225 | 0.900519 | 3.342113 | 0.292696 |
| 4 | 14319200 | 14322399 | 31.78912 | 53.82517 | 42.47441 | 18.5953 | 0.721705 | 1.083644 | 0.292701 |
| 5 | 125799000 | 125816599 | 235.9689 | 337.6165 | 521.7379 | 215.5323 | -0.42623 | 4.2394 | 0.292704 |
| 18 | 84302600 | 84304599 | 20.95177 | 36.91401 | 30.02403 | 13.28945 | 0.685309 | 0.59228 | 0.292713 |
| 16 | 62830200 | 62830999 | 5.852901 | 11.24685 | 26.86314 | 10.50023 | -0.71415 | -0.10763 | 0.292719 |
| 12 | 18169600 | 18207999 | 1171.671 | 1499.002 | 1151.494 | 483.1325 | 0.419934 | 5.997014 | 0.292725 |
| 6 | 120538000 | 120539599 | 7.624016 | 14.05689 | 33.58654 | 14.21079 | -0.81313 | 0.208608 | 0.292746 |
| 3 | 121537000 | 121545199 | 109.3512 | 165.6488 | 272.087 | 112.5876 | -0.43665 | 3.266249 | 0.292749 |
| 12 | 28995600 | 28999799 | 23.91603 | 39.11682 | 79.34991 | 34.12437 | -0.64253 | 1.469817 | 0.292752 |
| 14 | 27424200 | 27424799 | 6.462021 | 12.30698 | 28.27257 | 11.30792 | -0.68919 | -0.02089 | 0.292753 |
| 3 | 6513400 | 6514199 | 15.64551 | 28.40271 | 24.79072 | 10.70429 | 0.609744 | 0.33907 | 0.292778 |
| 4 | 138731000 | 138744399 | 78.72906 | 121.1268 | 216.1411 | 89.76335 | -0.53815 | 2.939673 | 0.292795 |
| 16 | 59008600 | 59009199 | 9.432438 | 18.30611 | 17.63455 | 6.632365 | 0.63582 | -0.22587 | 0.292798 |
| 15 | 28159200 | 28161799 | 14.15163 | 24.70271 | 47.32333 | 20.83862 | -0.54595 | 0.717185 | 0.292841 |
| 17 | 22618000 | 22619799 | 29.86216 | 51.43614 | 35.17272 | 15.42903 | 0.930473 | 0.91819 | 0.292842 |

| | | | | | | | | | |
|---|---|---|---|---|---|---|---|---|---|
| 5 | 123821600 | 123823199 | 10.87444 | 19.154 | 41.47767 | 18.36014 | -0.71986 | 0.557022 | 0.292861 |
| 11 | 94218600 | 94219399 | 4.691693 | 9.110681 | 24.47035 | 9.166149 | -0.82249 | -0.27264 | 0.292875 |
| 9 | 35966000 | 35970999 | 61.06534 | 97.8993 | 76.95254 | 32.49525 | 0.693724 | 1.983055 | 0.29289 |
| 2 | 25702600 | 25704599 | 10.46835 | 18.71639 | 38.36769 | 16.84136 | -0.63039 | 0.421595 | 0.292898 |
| 3 | 72598400 | 72599399 | 10.77254 | 20.74648 | 19.0191 | 7.38682 | 0.6629 | -0.10183 | 0.29291 |
| 18 | 80155000 | 80155999 | 5.226977 | 10.1354 | 24.95233 | 9.423222 | -0.7089 | -0.23359 | 0.29291 |
| 5 | 110113000 | 110123799 | 216.301 | 320.7168 | 264.9816 | 109.1392 | 0.526445 | 3.68684 | 0.292924 |
| X | 71317800 | 71319599 | 15.70518 | 28.89212 | 23.74621 | 10.0778 | 0.71275 | 0.263522 | 0.292927 |
| X | 117662400 | 117663599 | 17.85318 | 32.64806 | 23.95482 | 10.24931 | 0.866952 | 0.312826 | 0.29293 |
| 17 | 93093200 | 93094199 | 13.56604 | 25.64247 | 21.30254 | 8.65722 | 0.745402 | 0.076292 | 0.292941 |
| 9 | 124250200 | 124256599 | 853.4897 | 1139.011 | 590.9322 | 246.7979 | 1.046875 | 5.313755 | 0.292987 |
| 12 | 83678600 | 83679799 | 7.58238 | 14.14505 | 31.6444 | 13.16182 | -0.69949 | 0.137718 | 0.292989 |
| 1 | 183126400 | 183127399 | 5.314185 | 10.16302 | 27.78485 | 11.01043 | -0.92562 | -0.06535 | 0.292991 |
| 1 | 149268200 | 149270799 | 48.04959 | 77.46057 | 71.58336 | 30.47434 | 0.470327 | 1.793966 | 0.292995 |
| 13 | 85752000 | 85755399 | 36.61623 | 61.17771 | 41.53966 | 18.08062 | 0.93759 | 1.220695 | 0.293009 |
| 5 | 10925200 | 10926599 | 11.96308 | 21.01675 | 42.50809 | 18.81567 | -0.62325 | 0.59707 | 0.293021 |
| 15 | 90506200 | 90507199 | 6.886086 | 13.21763 | 32.23455 | 12.63973 | -0.7492 | -0.07287 | 0.293026 |
| 7 | 46669400 | 46671199 | 10.1852 | 18.14933 | 39.38661 | 17.33745 | -0.71497 | 0.442411 | 0.293032 |
| 16 | 85894000 | 85895199 | 15.72161 | 29.4251 | 20.87776 | 8.635647 | 0.950252 | 0.155631 | 0.293049 |
| 14 | 6616200 | 6618399 | 30.60315 | 49.65017 | 89.74776 | 38.45502 | -0.48573 | 1.619012 | 0.293053 |
| 2 | 70264600 | 70269799 | 50.57066 | 82.18157 | 59.73212 | 25.62531 | 0.827574 | 1.606442 | 0.293091 |
| 12 | 110661600 | 110663799 | 11.81228 | 20.69814 | 45.67706 | 20.17418 | -0.74866 | 0.641322 | 0.29311 |
| 7 | 44297200 | 44300399 | 18.03092 | 30.35605 | 63.38754 | 27.70306 | -0.67736 | 1.107067 | 0.293112 |
| 1 | 122371400 | 122372599 | 7.653486 | 14.47048 | 31.49853 | 12.79351 | -0.63289 | 0.046231 | 0.293141 |
| 7 | 36394600 | 36398999 | 29.60205 | 47.6559 | 103.214 | 43.85499 | -0.75807 | 1.82246 | 0.29315 |
| 7 | 28251000 | 28253799 | 33.91061 | 54.60871 | 100.0348 | 42.52279 | -0.5165 | 1.819086 | 0.293168 |
| 16 | 93860600 | 93865599 | 45.01366 | 71.84012 | 126.0482 | 52.87993 | -0.46989 | 2.119479 | 0.293179 |
| 10 | 8331600 | 8335799 | 26.94039 | 44.01985 | 81.5951 | 35.07726 | -0.51394 | 1.491318 | 0.293182 |
| 8 | 23512600 | 23513799 | 6.602384 | 12.59911 | 28.3283 | 11.28307 | -0.65317 | -0.0329 | 0.293185 |
| 15 | 76470200 | 76474799 | 20.74379 | 33.98766 | 78.34367 | 33.71497 | -0.82479 | 1.439337 | 0.293201 |
| 14 | 96339000 | 96340999 | 23.77352 | 41.60847 | 32.00689 | 14.12248 | 0.764212 | 0.70227 | 0.293213 |
| 13 | 66964200 | 66965399 | 15.61092 | 29.30265 | 19.73716 | 8.123927 | 1.029946 | 0.133582 | 0.293233 |
| 1 | 192378600 | 192379599 | 6.66967 | 12.49224 | 30.63731 | 12.65691 | -0.82259 | 0.101829 | 0.293241 |
| 8 | 72203600 | 72208999 | 40.80909 | 65.16395 | 123.7453 | 51.95238 | -0.58292 | 2.098351 | 0.293245 |
| 15 | 90526200 | 90532399 | 56.42625 | 89.27252 | 150.3685 | 62.49722 | -0.42359 | 2.391782 | 0.293252 |
| 4 | 5412800 | 5413399 | 8.302018 | 15.53626 | 32.29238 | 13.36008 | -0.58878 | 0.12231 | 0.293258 |
| 10 | 7280200 | 7281399 | 14.93661 | 28.46124 | 18.38245 | 7.344423 | 1.126317 | 0.027594 | 0.293271 |
| 2 | 112040400 | 112042799 | 17.55748 | 30.14426 | 57.63931 | 25.28168 | -0.54781 | 0.924367 | 0.293276 |
| 13 | 105934400 | 105936199 | 15.93593 | 29.10392 | 24.50256 | 10.50083 | 0.667947 | 0.304652 | 0.293281 |
| 9 | 72187200 | 72187599 | 3.75522 | 7.342595 | 19.57087 | 6.686436 | -0.68738 | -0.62672 | 0.293306 |
| 15 | 10874800 | 10875199 | 3.75522 | 7.342595 | 19.57087 | 6.686436 | -0.68738 | -0.62672 | 0.293306 |
| 12 | 18576200 | 18582399 | 144.8447 | 222.1669 | 158.1453 | 65.65019 | 0.783068 | 3.01809 | 0.293314 |
| 7 | 33193400 | 33199399 | 184.727 | 275.9283 | 246.9802 | 101.8209 | 0.420735 | 3.550627 | 0.293318 |
| 9 | 38324200 | 38330999 | 166.4786 | 250.0889 | 227.8663 | 94.0578 | 0.402469 | 3.439859 | 0.293323 |
| 3 | 3005000 | 3026599 | 1018.401 | 1316.836 | 1012.322 | 424.3034 | 0.434229 | 5.823647 | 0.293351 |
| 18 | 34102800 | 34106399 | 54.59413 | 87.62371 | 81.97017 | 34.67363 | 0.445041 | 1.942738 | 0.293365 |
| 2 | 179173400 | 179177199 | 25.64134 | 42.00921 | 78.06919 | 33.59588 | -0.51522 | 1.450584 | 0.293384 |

| | | | | | | | | | |
|---|---|---|---|---|---|---|---|---|---|
| 8 | 12922600 | 12925799 | 16.39429 | 27.47628 | 64.14517 | 27.97457 | -0.83757 | 1.14905 | 0.29339 |
| 12 | 18360200 | 18363999 | 82.24951 | 130.6553 | 99.45028 | 41.46993 | 0.725928 | 2.292921 | 0.293399 |
| 5 | 71813200 | 71815999 | 24.19052 | 39.66406 | 77.99933 | 33.57721 | -0.59609 | 1.436203 | 0.293403 |
| 2 | 111568000 | 111568599 | 12.46215 | 23.39573 | 20.81844 | 8.566813 | 0.632002 | 0.107642 | 0.293425 |
| 1 | 73053800 | 73055199 | 17.83208 | 30.18631 | 59.03556 | 25.84399 | -0.5827 | 1.055705 | 0.293425 |
| X | 5019800 | 5020399 | 22.82523 | 39.86434 | 33.84924 | 14.91036 | 0.622471 | 0.732958 | 0.293441 |
| 18 | 87511400 | 87512799 | 15.70391 | 29.46523 | 20.33633 | 8.376309 | 0.994531 | 0.138801 | 0.293448 |
| 2 | 101155600 | 101156199 | 8.745599 | 15.87431 | 34.96496 | 15.1002 | -0.71693 | 0.31354 | 0.293451 |
| 5 | 53745800 | 53749199 | 19.86937 | 32.77235 | 74.37889 | 32.099 | -0.79859 | 1.349335 | 0.293453 |
| 13 | 48448600 | 48451199 | 28.12294 | 45.75844 | 84.71579 | 36.3628 | -0.51655 | 1.556995 | 0.293461 |
| 6 | 131754400 | 131759199 | 86.04827 | 136.4934 | 101.4484 | 42.2423 | 0.758256 | 2.33756 | 0.293467 |
| X | 15413400 | 15414999 | 23.87573 | 42.23798 | 26.45889 | 11.70186 | 1.062108 | 0.560838 | 0.293469 |
| 6 | 131625800 | 131627399 | 22.44933 | 39.71194 | 27.60524 | 12.209 | 0.912715 | 0.55416 | 0.293471 |
| 12 | 78272600 | 78274599 | 15.72856 | 26.46821 | 62.01671 | 27.10032 | -0.84261 | 1.104193 | 0.293474 |
| 2 | 39843600 | 39844199 | 7.717388 | 14.42937 | 31.15462 | 12.90878 | -0.64445 | 0.122033 | 0.293478 |
| 6 | 134100200 | 134102399 | 12.70396 | 22.33184 | 44.52954 | 19.71892 | -0.60315 | 0.592919 | 0.293487 |
| 6 | 128412400 | 128416399 | 19.0874 | 31.49403 | 74.27071 | 32.05724 | -0.85344 | 1.342947 | 0.293498 |
| 6 | 62981800 | 62985399 | 37.83553 | 63.20443 | 45.92182 | 19.98592 | 0.84048 | 1.233464 | 0.293509 |
| 14 | 121615600 | 121617599 | 11.35839 | 19.948 | 42.54982 | 18.82902 | -0.69936 | 0.599937 | 0.293517 |
| 13 | 59268600 | 59283799 | 179.3661 | 261.8178 | 420.8344 | 173.3944 | -0.4545 | 3.937385 | 0.293521 |
| X | 143604000 | 143605199 | 13.4376 | 23.371 | 52.67165 | 23.14068 | -0.77973 | 0.775192 | 0.293544 |
| 18 | 76204600 | 76208599 | 23.60541 | 38.59477 | 79.11654 | 34.01951 | -0.65793 | 1.475646 | 0.293561 |
| 13 | 23176400 | 23182199 | 90.32023 | 142.0106 | 125.2844 | 51.97238 | 0.501137 | 2.5438 | 0.293574 |
| 4 | 106590600 | 106596599 | 35.59943 | 57.0722 | 110.7246 | 46.75588 | -0.60709 | 1.966396 | 0.29359 |
| 13 | 10824400 | 10829799 | 92.71033 | 145.3027 | 131.4473 | 54.52087 | 0.461429 | 2.606505 | 0.293598 |
| 11 | 3504400 | 3506199 | 12.09158 | 21.39244 | 41.39391 | 18.30684 | -0.55603 | 0.515417 | 0.293599 |
| 8 | 97492200 | 97494399 | 24.32407 | 42.81724 | 26.75332 | 11.84386 | 1.062802 | 0.613836 | 0.293611 |
| 18 | 3101000 | 3102999 | 53.46643 | 85.89453 | 80.26282 | 33.99969 | 0.448256 | 1.912008 | 0.293634 |
| 4 | 152120800 | 152121399 | 10.85366 | 20.85891 | 19.59042 | 7.654667 | 0.620582 | -0.07924 | 0.293634 |
| 7 | 23479200 | 23484399 | 96.96746 | 152.2669 | 124.6855 | 51.71531 | 0.607111 | 2.572937 | 0.293637 |
| 8 | 34471800 | 34472399 | 8.74023 | 17.00462 | 16.69475 | 6.158607 | 0.634694 | -0.30266 | 0.293654 |
| 7 | 17570000 | 17579599 | 159.0158 | 242.5203 | 169.7244 | 70.37092 | 0.801197 | 3.135836 | 0.293658 |
| 14 | 82396600 | 82397799 | 19.01176 | 31.98216 | 61.68898 | 26.95428 | -0.56324 | 1.118576 | 0.293674 |
| 14 | 76821400 | 76822399 | 6.917724 | 13.10466 | 29.26371 | 11.83735 | -0.66361 | 0.021812 | 0.293692 |
| 2 | 13187600 | 13190399 | 26.84378 | 45.54005 | 41.10627 | 18.00693 | 0.528623 | 1.047972 | 0.293697 |
| 17 | 38686000 | 38687199 | 29.49479 | 49.61522 | 44.33081 | 19.36962 | 0.542986 | 1.135168 | 0.293698 |
| 11 | 68476600 | 68479199 | 12.75783 | 22.29288 | 45.73839 | 20.15386 | -0.64402 | 0.693516 | 0.293704 |
| 4 | 112051000 | 112051599 | 12.61513 | 23.62256 | 20.96934 | 8.66628 | 0.63101 | 0.127164 | 0.293705 |
| 1 | 38173800 | 38174599 | 12.7424 | 23.80023 | 21.42485 | 8.892772 | 0.6059 | 0.144976 | 0.293705 |
| 13 | 92980600 | 92981999 | 10.109 | 18.29953 | 36.43659 | 15.78901 | -0.57605 | 0.340373 | 0.293707 |
| 7 | 118167400 | 118169599 | 14.46823 | 25.16721 | 49.49713 | 21.74801 | -0.58434 | 0.77083 | 0.293711 |
| 7 | 10102200 | 10104799 | 36.78906 | 62.16949 | 39.63977 | 17.34402 | 1.028099 | 1.11664 | 0.293727 |
| 5 | 109577800 | 109580399 | 33.42917 | 56.77556 | 40.22318 | 17.62606 | 0.876847 | 1.063176 | 0.293728 |
| 4 | 145494200 | 145495799 | 21.58521 | 37.86347 | 28.63816 | 12.6601 | 0.788324 | 0.649661 | 0.293737 |
| 11 | 57899200 | 57901599 | 13.96984 | 24.00563 | 56.40153 | 24.73958 | -0.84328 | 0.899144 | 0.29374 |
| 11 | 32006600 | 32008799 | 14.06012 | 24.43136 | 48.95583 | 21.49582 | -0.61103 | 0.786503 | 0.293745 |
| 3 | 91250400 | 91252599 | 21.27213 | 37.3675 | 28.07486 | 12.42603 | 0.797827 | 0.624543 | 0.29376 |

| | | | | | | | | | |
|---|---|---|---|---|---|---|---|---|---|
| 8 | 114244400 | 114249199 | 40.45317 | 64.85706 | 112.817 | 47.64327 | -0.44993 | 1.971426 | 0.293761 |
| 14 | 122033000 | 122036199 | 19.69113 | 32.53052 | 73.9401 | 31.94082 | -0.80033 | 1.328781 | 0.293766 |
| 18 | 64397400 | 64399999 | 26.34021 | 42.9551 | 81.70103 | 35.09777 | -0.55298 | 1.519354 | 0.293769 |
| 3 | 4568800 | 4573199 | 81.83621 | 128.9792 | 119.7753 | 49.70245 | 0.429866 | 2.488802 | 0.293802 |
| 8 | 76366800 | 76368599 | 21.85843 | 38.48719 | 29.83776 | 13.20866 | 0.753697 | 0.603124 | 0.293808 |
| 8 | 126704800 | 126706799 | 12.43159 | 21.93853 | 42.97176 | 19.01425 | -0.575 | 0.548595 | 0.293824 |
| 12 | 20588000 | 20590399 | 51.50559 | 82.98057 | 63.45639 | 26.99167 | 0.74179 | 1.812915 | 0.293841 |
| 4 | 123811000 | 123811999 | 6.968687 | 13.04694 | 30.63477 | 12.66466 | -0.76138 | 0.106502 | 0.293846 |
| 7 | 105284600 | 105290399 | 25.02174 | 40.55792 | 89.06403 | 38.13638 | -0.76687 | 1.625919 | 0.293889 |
| 7 | 49177200 | 49184399 | 33.54115 | 54.08124 | 100.2995 | 42.70917 | -0.53215 | 1.778499 | 0.293893 |
| 13 | 51053400 | 51053999 | 4.055197 | 7.921808 | 20.54883 | 7.179797 | -0.67901 | -0.54021 | 0.293908 |
| 4 | 137284000 | 137285199 | 5.890998 | 11.20519 | 28.22468 | 11.31918 | -0.82404 | -0.01782 | 0.293914 |
| 17 | 31307000 | 31318999 | 116.0581 | 174.6221 | 294.8403 | 121.7547 | -0.48483 | 3.402786 | 0.293921 |
| 7 | 99272400 | 99276799 | 25.56845 | 41.3118 | 95.37244 | 40.71996 | -0.84355 | 1.70379 | 0.29394 |
| X | 77829200 | 77832999 | 56.67918 | 91.33122 | 71.26114 | 30.31894 | 0.713236 | 1.822816 | 0.293948 |
| 4 | 115049800 | 115052599 | 37.96295 | 62.73658 | 48.84058 | 21.10298 | 0.740594 | 1.352401 | 0.293958 |
| 14 | 6964200 | 6967399 | 33.51514 | 55.75619 | 50.35765 | 21.85871 | 0.5276 | 1.265554 | 0.293973 |
| 1 | 150218800 | 150219799 | 14.24692 | 26.59633 | 22.12673 | 9.192316 | 0.717406 | 0.161719 | 0.293978 |
| 10 | 16113000 | 16115799 | 33.01002 | 55.97163 | 41.39483 | 18.13045 | 0.815079 | 1.077349 | 0.294005 |
| 9 | 18284000 | 18284999 | 14.66379 | 28.30767 | 13.8273 | 5.331566 | 1.570086 | -0.0977 | 0.294006 |
| X | 80351600 | 80355399 | 39.86105 | 66.7182 | 41.93759 | 18.27441 | 1.048544 | 1.214939 | 0.294018 |
| 1 | 120519000 | 120540399 | 177.683 | 257.1709 | 501.4229 | 206.7935 | -0.7449 | 4.065993 | 0.294021 |
| 4 | 3856200 | 3857199 | 6.58716 | 12.13647 | 33.07045 | 14.00609 | -1.00089 | 0.207654 | 0.294036 |
| 3 | 109649600 | 109650799 | 9.302883 | 17.15684 | 40.8852 | 17.29273 | -0.80975 | 0.236377 | 0.294043 |
| 13 | 113780600 | 113782599 | 18.76217 | 31.83618 | 59.12269 | 25.90076 | -0.50826 | 1.03844 | 0.294048 |
| 8 | 93824000 | 93825999 | 20.78864 | 36.78538 | 28.17777 | 12.46146 | 0.773426 | 0.54029 | 0.294059 |
| 9 | 94105800 | 94107399 | 15.10859 | 28.10126 | 22.56312 | 9.432224 | 0.76124 | 0.193327 | 0.294063 |
| 1 | 189903800 | 189907999 | 26.81405 | 43.34675 | 102.6516 | 43.85709 | -0.87949 | 1.693306 | 0.29407 |
| 7 | 136217600 | 136219399 | 11.77766 | 20.76782 | 41.424 | 18.33234 | -0.6015 | 0.549454 | 0.294091 |
| 17 | 64618800 | 64619599 | 8.649142 | 15.95379 | 33.59175 | 14.20474 | -0.63168 | 0.214472 | 0.294093 |
| 10 | 41988600 | 41990799 | 27.56811 | 47.9786 | 31.83581 | 13.99246 | 0.975965 | 0.800324 | 0.294099 |
| 7 | 59737600 | 59738999 | 14.80694 | 27.94523 | 20.55163 | 8.37911 | 0.915613 | 0.095494 | 0.294106 |
| 7 | 19574400 | 19582999 | 45.76565 | 72.74849 | 135.131 | 56.38624 | -0.55818 | 2.250661 | 0.294113 |
| 9 | 88566800 | 88569199 | 26.2469 | 45.19623 | 38.13409 | 16.72794 | 0.628533 | 0.908686 | 0.294129 |
| 6 | 13488600 | 13489999 | 18.81581 | 33.47525 | 26.68294 | 11.76121 | 0.719985 | 0.481618 | 0.29414 |
| 11 | 101355600 | 101356799 | 6.200017 | 11.75669 | 29.82727 | 12.04011 | -0.84356 | 0.008318 | 0.294154 |
| 15 | 96552600 | 96554199 | 12.27029 | 21.51171 | 44.09925 | 19.48574 | -0.64278 | 0.634044 | 0.294163 |
| 14 | 50805200 | 50805999 | 19.29375 | 34.13979 | 29.0591 | 12.85126 | 0.622355 | 0.534294 | 0.294166 |
| 3 | 33477600 | 33479399 | 18.4653 | 33.74751 | 23.15216 | 9.913205 | 0.962195 | 0.31748 | 0.294175 |
| 17 | 37999400 | 38000399 | 13.15401 | 25.40248 | 17.74458 | 6.830223 | 1.063743 | -0.10772 | 0.294186 |
| 19 | 3128800 | 3130399 | 34.25608 | 56.60283 | 52.54657 | 22.70208 | 0.487445 | 1.347528 | 0.294189 |
| 3 | 134430600 | 134430999 | 5.221933 | 10.14414 | 24.53177 | 9.161398 | -0.66831 | -0.2744 | 0.294209 |
| 7 | 102443200 | 102444399 | 6.425567 | 12.35818 | 27.51083 | 10.72619 | -0.61041 | -0.10829 | 0.294219 |
| 4 | 62723800 | 62725399 | 7.441748 | 13.83001 | 31.58219 | 13.2194 | -0.73643 | 0.159476 | 0.294226 |
| 17 | 40019800 | 40022999 | 83.83408 | 131.88 | 124.1254 | 51.49503 | 0.408452 | 2.529217 | 0.294234 |
| 1 | 180102000 | 180107599 | 39.10438 | 62.69809 | 114.2503 | 48.2527 | -0.51679 | 1.966407 | 0.294241 |
| 3 | 3511400 | 3514199 | 24.15247 | 41.94445 | 31.08745 | 13.64521 | 0.817085 | 0.813612 | 0.294251 |

| | | | | | | | | | |
|---|---|---|---|---|---|---|---|---|---|
| 6 | 146143400 | 146146599 | 33.23232 | 55.45438 | 49.22447 | 21.40858 | 0.552513 | 1.232067 | 0.294253 |
| 10 | 68980200 | 68981199 | 9.478095 | 17.21847 | 35.45266 | 15.2946 | -0.61982 | 0.312828 | 0.294273 |
| 10 | 85247200 | 85248999 | 9.478095 | 17.21847 | 35.45266 | 15.2946 | -0.61982 | 0.312828 | 0.294273 |
| 3 | 35208400 | 35210399 | 12.71799 | 22.28597 | 46.19199 | 20.40222 | -0.65882 | 0.646041 | 0.294277 |
| X | 102149400 | 102150199 | 12.59333 | 24.24574 | 18.96314 | 7.371482 | 0.889501 | -0.07827 | 0.294278 |
| 5 | 89581000 | 89582599 | 15.81942 | 27.27581 | 52.15673 | 22.88003 | -0.54627 | 0.874098 | 0.294286 |
| 13 | 60429400 | 60430599 | 9.135422 | 16.7923 | 34.06262 | 14.47213 | -0.5831 | 0.240349 | 0.294303 |
| 13 | 74852600 | 74854599 | 9.135422 | 16.7923 | 34.06262 | 14.47213 | -0.5831 | 0.240349 | 0.294303 |
| 9 | 59917800 | 59920399 | 14.34347 | 25.08588 | 48.89625 | 21.56146 | -0.57063 | 0.692386 | 0.294306 |
| 1 | 153124800 | 153126399 | 24.47902 | 42.14581 | 36.89793 | 16.18561 | 0.575635 | 0.903522 | 0.294312 |
| 7 | 105266800 | 105268799 | 13.64322 | 23.50818 | 52.10277 | 22.8559 | -0.75832 | 0.870098 | 0.294323 |
| 6 | 131155400 | 131156399 | 13.48249 | 25.84041 | 18.64305 | 7.341837 | 0.986781 | -0.03115 | 0.29433 |
| 7 | 14578000 | 14603999 | 484.1263 | 672.6442 | 476.4094 | 197.7407 | 0.656396 | 4.681468 | 0.294335 |
| 7 | 37306000 | 37311799 | 34.70036 | 55.83411 | 105.1098 | 44.62632 | -0.55737 | 1.846396 | 0.294367 |
| 13 | 79669600 | 79670999 | 15.94524 | 29.68998 | 21.94043 | 9.154882 | 0.882212 | 0.19221 | 0.294372 |
| 17 | 71067000 | 71076399 | 71.61681 | 111.102 | 211.6756 | 87.91151 | -0.62337 | 2.782931 | 0.294375 |
| 19 | 7890200 | 7891999 | 9.246794 | 18.0297 | 13.44317 | 4.839665 | 1.059213 | -0.39098 | 0.294402 |
| 2 | 132925800 | 132927399 | 15.16003 | 28.32166 | 22.08491 | 9.16358 | 0.811727 | 0.165623 | 0.294418 |
| 7 | 130873200 | 130874799 | 8.68867 | 15.9515 | 33.51886 | 14.26492 | -0.63517 | 0.243175 | 0.294422 |
| 16 | 89845200 | 89847199 | 17.42766 | 29.94942 | 57.06875 | 25.0323 | -0.54248 | 0.915971 | 0.294432 |
| 3 | 24140800 | 24144999 | 57.32559 | 92.0692 | 80.76686 | 34.19935 | 0.538736 | 1.926574 | 0.294434 |
| 1 | 92351200 | 92354999 | 27.21563 | 44.17377 | 86.16229 | 36.93073 | -0.59452 | 1.602094 | 0.294448 |
| 12 | 27319800 | 27320999 | 9.304092 | 17.15445 | 36.31647 | 15.36598 | -0.6401 | 0.229805 | 0.294459 |
| 16 | 16964800 | 16966599 | 11.32372 | 19.93559 | 42.47627 | 18.80401 | -0.69714 | 0.565486 | 0.294501 |
| 2 | 30482600 | 30488799 | 34.62137 | 55.52878 | 110.9893 | 46.89924 | -0.64925 | 1.950273 | 0.294505 |
| 3 | 101420800 | 101425199 | 21.51726 | 35.67073 | 69.82714 | 30.23633 | -0.58512 | 1.290571 | 0.29451 |
| 14 | 77776200 | 77777799 | 9.765248 | 17.76094 | 36.52277 | 15.73317 | -0.61684 | 0.309356 | 0.294515 |
| 14 | 24052200 | 24055599 | 22.60969 | 37.08788 | 82.32719 | 35.44667 | -0.77009 | 1.431033 | 0.294528 |
| 9 | 70528000 | 70530999 | 22.29973 | 37.20515 | 71.83743 | 31.2401 | -0.56498 | 1.223712 | 0.29453 |
| 6 | 128819000 | 128822599 | 18.48994 | 31.14134 | 63.44257 | 27.73049 | -0.64191 | 1.104941 | 0.294542 |
| 2 | 92621800 | 92623199 | 23.8968 | 41.16292 | 37.36429 | 16.39042 | 0.523742 | 0.896179 | 0.294552 |
| 12 | 114586400 | 114587799 | 53.39305 | 85.76421 | 75.7487 | 32.08074 | 0.529181 | 1.91627 | 0.294558 |
| 17 | 85236400 | 85240599 | 48.80786 | 77.77471 | 132.1748 | 55.33067 | -0.42639 | 2.175008 | 0.294563 |
| 7 | 42403200 | 42408399 | 125.5101 | 191.7686 | 175.2399 | 72.68839 | 0.418693 | 3.087779 | 0.294577 |
| 3 | 151068800 | 151069399 | 10.29017 | 19.91107 | 18.00971 | 6.868853 | 0.706484 | -0.16962 | 0.29458 |
| 8 | 5406600 | 5409199 | 35.77411 | 59.26142 | 52.60649 | 22.76784 | 0.552085 | 1.32325 | 0.294582 |
| 7 | 39184200 | 39188199 | 87.5526 | 137.1594 | 127.4446 | 52.86304 | 0.42239 | 2.610168 | 0.294587 |
| 1 | 110659800 | 110661999 | 31.52854 | 53.02384 | 45.84716 | 20.03073 | 0.590064 | 1.143743 | 0.29459 |
| 2 | 171034200 | 171038399 | 43.048 | 68.89564 | 118.3426 | 49.84454 | -0.43484 | 2.030089 | 0.294595 |
| 11 | 105532000 | 105533999 | 17.81919 | 32.1563 | 25.84718 | 11.24407 | 0.720725 | 0.389157 | 0.294613 |
| 5 | 89169600 | 89171399 | 13.48949 | 23.64801 | 48.31969 | 21.34971 | -0.63847 | 0.646229 | 0.294629 |
| 7 | 41985800 | 41986599 | 15.84857 | 28.63726 | 25.63373 | 11.13315 | 0.567955 | 0.367316 | 0.294646 |
| 7 | 37607400 | 37611599 | 26.97781 | 43.98299 | 82.41331 | 35.40027 | -0.53176 | 1.52459 | 0.294646 |
| 14 | 110397000 | 110399799 | 30.31985 | 51.39011 | 43.05365 | 18.85476 | 0.635746 | 1.07046 | 0.294653 |
| 2 | 69574600 | 69575599 | 18.30554 | 32.80622 | 26.98966 | 11.82336 | 0.681038 | 0.437112 | 0.294664 |
| 7 | 13740200 | 13743199 | 36.6953 | 61.78553 | 41.93567 | 18.32831 | 0.938213 | 1.149559 | 0.294669 |
| 1 | 15827400 | 15832599 | 65.88483 | 103.0801 | 176.6717 | 73.29192 | -0.46129 | 2.628423 | 0.294674 |

| | | | | | | | | | |
|---|---|---|---|---|---|---|---|---|---|
| 18 | 37480800 | 37492199 | 196.8117 | 293.257 | 250.7773 | 103.3404 | 0.483338 | 3.59842 | 0.294698 |
| 9 | 6582600 | 6586199 | 67.26539 | 106.9352 | 99.81863 | 41.65571 | 0.43308 | 2.256749 | 0.294714 |
| 6 | 10357600 | 10361399 | 50.92729 | 82.19735 | 71.3463 | 30.42002 | 0.562466 | 1.76303 | 0.294718 |
| X | 121922000 | 121923599 | 35.12195 | 59.41572 | 40.78507 | 17.85092 | 0.922109 | 1.102981 | 0.294746 |
| 6 | 115657400 | 115659799 | 14.13925 | 24.66314 | 48.39835 | 21.30226 | -0.58071 | 0.728085 | 0.294763 |
| 9 | 13510400 | 13513599 | 14.13925 | 24.66314 | 48.39835 | 21.30226 | -0.58071 | 0.728085 | 0.294763 |
| 10 | 77567200 | 77567999 | 5.68207 | 11.0102 | 28.06384 | 10.63273 | -0.76507 | -0.21401 | 0.29477 |
| 5 | 128903200 | 128905199 | 12.45531 | 21.88337 | 43.53697 | 19.27249 | -0.59955 | 0.601611 | 0.29479 |
| 2 | 117035000 | 117038399 | 26.79188 | 43.72894 | 83.24001 | 35.77001 | -0.55334 | 1.507389 | 0.294798 |
| 15 | 25982200 | 25985599 | 19.89231 | 33.12071 | 67.03043 | 29.11204 | -0.63243 | 1.233445 | 0.29481 |
| 10 | 62525200 | 62527799 | 17.6709 | 29.91822 | 59.03434 | 25.84471 | -0.59549 | 1.053515 | 0.294821 |
| 5 | 119044600 | 119046599 | 12.1138 | 21.26639 | 43.03343 | 19.03675 | -0.62403 | 0.609749 | 0.294833 |
| 10 | 41067800 | 41069399 | 12.1138 | 21.26639 | 43.03343 | 19.03675 | -0.62403 | 0.609749 | 0.294833 |
| 14 | 57777400 | 57781799 | 51.09452 | 80.89189 | 146.5886 | 60.94709 | -0.52808 | 2.372947 | 0.294844 |
| 6 | 65393600 | 65396199 | 19.25091 | 34.75124 | 24.56074 | 10.68011 | 0.905438 | 0.394548 | 0.294849 |
| 1 | 117589600 | 117593999 | 65.9153 | 105.3713 | 88.87305 | 37.37098 | 0.587626 | 2.08012 | 0.294853 |
| 1 | 116980800 | 116982799 | 31.73046 | 53.43894 | 45.29078 | 19.79614 | 0.618907 | 1.132382 | 0.294866 |
| 4 | 53050200 | 53052199 | 25.34112 | 43.30484 | 35.88071 | 15.73202 | 0.652843 | 0.969547 | 0.294889 |
| 9 | 18457000 | 18458399 | 13.7759 | 25.73525 | 22.20059 | 9.211947 | 0.66691 | 0.153531 | 0.294894 |
| 14 | 121132000 | 121139799 | 61.79867 | 97.32821 | 161.7115 | 67.09818 | -0.40912 | 2.499836 | 0.294909 |
| 13 | 95002200 | 95005999 | 33.37377 | 53.85666 | 97.48323 | 41.55504 | -0.49571 | 1.752943 | 0.29492 |
| 5 | 65356800 | 65357599 | 5.409097 | 10.45122 | 25.93386 | 9.958369 | -0.74423 | -0.17472 | 0.294923 |
| 9 | 103879200 | 103880599 | 8.77837 | 15.83043 | 35.9483 | 15.65175 | -0.76827 | 0.354512 | 0.294925 |
| 7 | 11475600 | 11478399 | 70.63844 | 112.5495 | 97.19432 | 40.68123 | 0.548551 | 2.189641 | 0.294936 |
| 2 | 173006400 | 173011199 | 26.72473 | 43.39949 | 86.4222 | 37.0563 | -0.62365 | 1.588756 | 0.294937 |
| 5 | 39091800 | 39094199 | 16.07666 | 27.88203 | 53.32101 | 23.39494 | -0.54481 | 0.822251 | 0.294951 |
| 18 | 50105600 | 50106999 | 20.54739 | 36.39834 | 28.2508 | 12.49103 | 0.755215 | 0.528077 | 0.29496 |
| 17 | 66174600 | 66177599 | 14.33576 | 24.91004 | 48.96695 | 21.50055 | -0.58368 | 0.785897 | 0.295005 |
| 18 | 17602000 | 17603199 | 16.2574 | 29.18845 | 25.75281 | 11.26432 | 0.582195 | 0.408953 | 0.295012 |
| 13 | 112113600 | 112116599 | 18.06888 | 30.83425 | 57.55865 | 25.23394 | -0.51462 | 0.981961 | 0.295016 |
| 7 | 12304400 | 12310799 | 112.4655 | 174.7299 | 141.8554 | 58.90135 | 0.607637 | 2.767956 | 0.295028 |
| 2 | 178409400 | 178412199 | 15.98825 | 27.41571 | 54.27767 | 23.8048 | -0.59787 | 0.925894 | 0.295034 |
| 18 | 41968000 | 41970199 | 30.34446 | 51.96918 | 38.6566 | 16.95212 | 0.808478 | 0.976875 | 0.295041 |
| 7 | 14648600 | 14656599 | 242.1309 | 353.9098 | 310.3432 | 127.8536 | 0.421143 | 3.920646 | 0.295056 |
| 4 | 57623600 | 57624399 | 4.490241 | 8.732056 | 23.39566 | 8.662232 | -0.8018 | -0.31615 | 0.295075 |
| 14 | 120464200 | 120466199 | 21.53976 | 37.55546 | 33.99788 | 14.95987 | 0.530216 | 0.748897 | 0.295123 |
| 1 | 113553600 | 113557399 | 73.76989 | 117.3538 | 92.94855 | 38.81941 | 0.670657 | 2.240196 | 0.295126 |
| 12 | 22195200 | 22196999 | 62.57127 | 99.79043 | 91.80237 | 38.47482 | 0.45891 | 2.150636 | 0.29514 |
| 13 | 16048000 | 16049599 | 8.780053 | 16.33883 | 38.98384 | 16.28261 | -0.79817 | 0.175461 | 0.295145 |
| 3 | 143066800 | 143067799 | 19.41113 | 34.11859 | 30.30861 | 13.42141 | 0.557483 | 0.609108 | 0.295145 |
| 1 | 6631600 | 6635599 | 34.99615 | 58.41699 | 48.13886 | 20.9408 | 0.659384 | 1.232453 | 0.295148 |
| 19 | 53870400 | 53872399 | 9.274724 | 17.04181 | 34.59098 | 14.70429 | -0.58477 | 0.243754 | 0.295213 |
| 1 | 96606400 | 96609799 | 46.03055 | 74.49348 | 68.1948 | 29.16638 | 0.490038 | 1.682824 | 0.295214 |
| X | 127580600 | 127581199 | 12.32963 | 23.5886 | 17.58622 | 6.961328 | 0.934508 | -0.02628 | 0.295222 |
| 1 | 63070200 | 63072599 | 13.63117 | 23.50746 | 52.64884 | 23.09604 | -0.77319 | 0.861795 | 0.295229 |
| 13 | 36681800 | 36682199 | 9.071368 | 17.69403 | 15.57785 | 5.581728 | 0.827366 | -0.40314 | 0.295231 |
| 18 | 77800800 | 77802399 | 12.46891 | 21.77603 | 45.75722 | 20.15518 | -0.67819 | 0.701416 | 0.295239 |

| | | | | | | | | | |
|---|---|---|---|---|---|---|---|---|---|
| X | 37980200 | 37982599 | 26.64057 | 47.00303 | 24.50275 | 10.84253 | 1.323956 | 0.601655 | 0.295244 |
| 8 | 111504800 | 111514599 | 67.88982 | 105.6666 | 185.4329 | 76.97483 | -0.50116 | 2.721969 | 0.295258 |
| 5 | 62008400 | 62009799 | 20.93407 | 36.92614 | 29.2898 | 12.96142 | 0.721715 | 0.576522 | 0.295258 |
| 4 | 11965000 | 11966999 | 13.83464 | 24.1395 | 47.32921 | 20.83586 | -0.5793 | 0.721052 | 0.295265 |
| 13 | 51052000 | 51052399 | 10.24689 | 20.01579 | 13.11304 | 4.587491 | 1.278444 | -0.47578 | 0.29527 |
| 2 | 30071200 | 30085599 | 114.4844 | 172.2561 | 292.6321 | 120.8431 | -0.49364 | 3.402527 | 0.295276 |
| 11 | 109611800 | 109616599 | 36.49322 | 58.63827 | 107.891 | 45.71601 | -0.52692 | 1.893439 | 0.295288 |
| 10 | 13067600 | 13069599 | 11.41791 | 20.25295 | 42.4295 | 18.74915 | -0.66866 | 0.484008 | 0.295288 |
| 10 | 75949000 | 75951799 | 29.45647 | 49.54419 | 45.36833 | 19.82215 | 0.507628 | 1.136498 | 0.295295 |
| 9 | 90967800 | 90968799 | 8.891925 | 16.13791 | 34.95103 | 15.09652 | -0.69307 | 0.314449 | 0.295298 |
| 5 | 37382200 | 37384599 | 11.85679 | 20.68633 | 47.3296 | 20.8347 | -0.8007 | 0.712861 | 0.295309 |
| 2 | 21564200 | 21566799 | 15.63297 | 27.18348 | 54.40255 | 23.89794 | -0.60954 | 0.789384 | 0.295314 |
| 11 | 74614200 | 74615399 | 6.708207 | 12.20361 | 38.1371 | 16.4239 | -1.21539 | 0.28886 | 0.295314 |
| 1 | 110974800 | 110977599 | 40.71761 | 66.88369 | 51.10936 | 22.01845 | 0.764645 | 1.432155 | 0.295327 |
| 8 | 57442600 | 57443999 | 14.69794 | 27.41137 | 22.68464 | 9.439774 | 0.723329 | 0.173972 | 0.295351 |
| 14 | 41407800 | 41409799 | 24.01607 | 42.00545 | 31.96116 | 14.09449 | 0.779858 | 0.713016 | 0.295352 |
| 19 | 44411200 | 44413799 | 12.28467 | 21.48266 | 46.24329 | 20.3896 | -0.71299 | 0.678002 | 0.295352 |
| 4 | 112687800 | 112690799 | 75.4998 | 119.6139 | 111.1343 | 46.22267 | 0.435072 | 2.365872 | 0.295361 |
| 4 | 51327800 | 51329799 | 24.1899 | 42.43669 | 30.28313 | 13.38842 | 0.871869 | 0.665828 | 0.295374 |
| 2 | 180294200 | 180294799 | 4.965288 | 9.596969 | 25.32787 | 9.710268 | -0.82897 | -0.18445 | 0.295377 |
| 5 | 11513000 | 11515199 | 18.67261 | 34.03289 | 23.68015 | 10.17413 | 0.938905 | 0.33562 | 0.295405 |
| 15 | 63869200 | 63870599 | 10.5436 | 18.64983 | 39.91212 | 17.65224 | -0.70042 | 0.505221 | 0.295407 |
| 17 | 19173800 | 19195599 | 627.8267 | 845.7763 | 687.7516 | 286.8419 | 0.41228 | 5.162466 | 0.29542 |
| 4 | 110687800 | 110688999 | 12.52727 | 23.77235 | 20.87015 | 8.411199 | 0.676631 | 0.038487 | 0.295426 |
| 3 | 105866200 | 105867599 | 7.919076 | 14.241 | 37.46881 | 16.36647 | -0.98253 | 0.365979 | 0.295433 |
| 13 | 60224000 | 60226999 | 13.10225 | 22.81301 | 47.35847 | 20.82089 | -0.66141 | 0.749932 | 0.295433 |
| 3 | 37981000 | 37981799 | 15.25822 | 28.08606 | 24.20154 | 10.26298 | 0.645556 | 0.256709 | 0.29544 |
| 13 | 116095800 | 116096199 | 4.05383 | 7.923717 | 20.76612 | 7.158563 | -0.67628 | -0.58784 | 0.295442 |
| 12 | 83851200 | 83854599 | 42.42954 | 69.27598 | 60.4985 | 26.00075 | 0.568173 | 1.522369 | 0.295449 |
| 16 | 20462200 | 20464199 | 11.15046 | 19.77088 | 39.35644 | 17.39688 | -0.59517 | 0.482659 | 0.29548 |
| 6 | 132711400 | 132713199 | 22.54595 | 39.52094 | 30.29702 | 13.38547 | 0.768794 | 0.666578 | 0.29548 |
| 2 | 77416800 | 77418199 | 31.95676 | 53.80586 | 45.30135 | 19.79917 | 0.628429 | 1.136109 | 0.295485 |
| 1 | 133256800 | 133259799 | 16.79071 | 28.74447 | 54.81952 | 24.03926 | -0.54454 | 0.945298 | 0.295488 |
| 2 | 172381200 | 172383399 | 34.59711 | 57.07488 | 52.53894 | 22.67554 | 0.499308 | 1.367135 | 0.295506 |
| 9 | 26203000 | 26205199 | 27.35465 | 46.59349 | 42.02957 | 18.42317 | 0.530154 | 1.015532 | 0.295517 |
| 7 | 15431400 | 15439399 | 197.7089 | 293.2866 | 264.6515 | 109.0084 | 0.399942 | 3.67551 | 0.295526 |
| 5 | 140728600 | 140731199 | 17.79925 | 30.25998 | 57.45875 | 25.18251 | -0.5399 | 1.016461 | 0.295539 |
| 1 | 26393000 | 26394999 | 20.70235 | 36.24619 | 32.4145 | 14.30663 | 0.548045 | 0.681679 | 0.295545 |
| 9 | 19483400 | 19489399 | 49.32789 | 79.66075 | 64.10626 | 27.35294 | 0.672296 | 1.742141 | 0.295576 |
| 15 | 63336800 | 63340799 | 43.08412 | 70.14373 | 62.67896 | 26.90977 | 0.532264 | 1.56516 | 0.295581 |
| 7 | 62343800 | 62346799 | 69.52854 | 110.5212 | 103.1249 | 43.03086 | 0.433518 | 2.263747 | 0.295582 |
| 10 | 52479000 | 52481199 | 24.55132 | 42.9391 | 30.7665 | 13.56698 | 0.865973 | 0.713985 | 0.295585 |
| 10 | 71718200 | 71719199 | 21.64673 | 37.73676 | 34.08481 | 14.99682 | 0.533555 | 0.752791 | 0.295589 |
| 4 | 59826400 | 59830399 | 26.77758 | 43.75742 | 81.59395 | 35.07782 | -0.52244 | 1.489617 | 0.295592 |
| 14 | 63589600 | 63593199 | 20.74642 | 34.35163 | 70.79622 | 30.63058 | -0.65924 | 1.303145 | 0.295607 |
| 17 | 31391600 | 31394399 | 21.82489 | 36.1938 | 69.90859 | 30.27916 | -0.56585 | 1.286893 | 0.295633 |
| 9 | 3270400 | 3276199 | 107.7446 | 166.4012 | 155.3199 | 64.51667 | 0.399981 | 2.8726 | 0.295653 |

| | | | | | | | | | |
|---|---|---|---|---|---|---|---|---|---|
| 17 | 66050800 | 66064399 | 179.3359 | 262.6197 | 409.2995 | 168.6013 | -0.40556 | 3.887415 | 0.295659 |
| 3 | 101538400 | 101540399 | 11.18415 | 19.67569 | 41.98722 | 18.59019 | -0.69961 | 0.572892 | 0.295663 |
| X | 54509600 | 54518599 | 151.0169 | 229.7136 | 187.3089 | 77.60402 | 0.577824 | 3.187544 | 0.295666 |
| 14 | 52377400 | 52379399 | 11.35144 | 19.937 | 42.53626 | 18.82399 | -0.6998 | 0.597865 | 0.29568 |
| 14 | 79367600 | 79368599 | 12.93655 | 24.36273 | 21.89659 | 8.96268 | 0.624095 | 0.096362 | 0.295681 |
| X | 121180800 | 121182599 | 26.42526 | 46.52396 | 26.33103 | 11.65651 | 1.205245 | 0.626492 | 0.295684 |
| 18 | 3364200 | 3373599 | 119.7475 | 183.8887 | 169.9122 | 70.54633 | 0.408345 | 2.986089 | 0.295697 |
| 8 | 66450800 | 66452799 | 21.19461 | 37.83908 | 26.16458 | 11.49888 | 0.92719 | 0.478411 | 0.295698 |
| 1 | 128635400 | 128636799 | 12.89223 | 22.63482 | 43.52874 | 19.2572 | -0.55104 | 0.61377 | 0.295715 |
| 15 | 73317600 | 73320599 | 14.50186 | 25.10755 | 51.64093 | 22.6566 | -0.64978 | 0.827793 | 0.295716 |
| 17 | 72243000 | 72245199 | 11.91583 | 21.1291 | 40.85697 | 18.05934 | -0.55377 | 0.488793 | 0.295728 |
| 1 | 130155400 | 130155999 | 10.42243 | 20.0842 | 18.4959 | 7.171304 | 0.658577 | -0.11291 | 0.295741 |
| 11 | 74774000 | 74775599 | 9.946058 | 17.58379 | 39.9623 | 17.67627 | -0.78669 | 0.508398 | 0.295754 |
| 8 | 101217400 | 101219999 | 23.10218 | 40.24295 | 34.34054 | 15.102 | 0.614887 | 0.769032 | 0.295756 |
| 15 | 45770200 | 45774199 | 75.10651 | 118.8371 | 112.296 | 46.67534 | 0.40922 | 2.395278 | 0.295767 |
| 14 | 25405600 | 25407999 | 12.90377 | 22.54504 | 46.25902 | 20.38165 | -0.64424 | 0.696095 | 0.295782 |
| 9 | 114387800 | 114393799 | 39.92814 | 63.58117 | 134.3805 | 56.20042 | -0.74142 | 2.188542 | 0.295782 |
| 6 | 145546000 | 145548399 | 20.26266 | 34.07606 | 62.93038 | 27.49415 | -0.50088 | 1.125161 | 0.29579 |
| 3 | 94674400 | 94677999 | 23.31901 | 38.45366 | 73.76169 | 31.82966 | -0.55697 | 1.3601 | 0.295798 |
| 7 | 15667400 | 15678399 | 65.35568 | 101.5688 | 190.96 | 79.28706 | -0.60217 | 2.747158 | 0.295801 |
| X | 132748200 | 132749799 | 10.62263 | 19.20218 | 41.79121 | 18.14135 | -0.70551 | 0.363406 | 0.295801 |
| 10 | 84273000 | 84277999 | 27.63238 | 44.62316 | 96.98046 | 41.37854 | -0.75764 | 1.722586 | 0.295803 |
| 14 | 77200200 | 77204799 | 35.36305 | 57.02328 | 100.9867 | 43.00612 | -0.46559 | 1.781556 | 0.295828 |
| 14 | 8112000 | 8129399 | 484.4442 | 648.7229 | 986.8241 | 411.9418 | -0.49978 | 5.256135 | 0.295835 |
| 13 | 62297200 | 62306999 | 233.478 | 344.4168 | 279.3877 | 115.056 | 0.546023 | 3.773346 | 0.295837 |
| 5 | 149583600 | 149585199 | 19.88865 | 33.58756 | 63.03441 | 27.57463 | -0.52382 | 1.086331 | 0.29584 |
| 7 | 63936000 | 63937999 | 14.51387 | 25.21514 | 48.96592 | 21.49774 | -0.56612 | 0.79034 | 0.295845 |
| 4 | 73157400 | 73157799 | 7.201106 | 13.69381 | 29.38879 | 11.79209 | -0.59653 | -0.0015 | 0.295854 |
| 5 | 44432600 | 44433799 | 7.201106 | 13.69381 | 29.38879 | 11.79209 | -0.59653 | -0.0015 | 0.295854 |
| 4 | 34729800 | 34730999 | 8.327798 | 15.44748 | 32.61933 | 13.68865 | -0.62767 | 0.178106 | 0.295862 |
| 3 | 90767800 | 90768599 | 11.37167 | 19.93314 | 43.63093 | 19.27623 | -0.73644 | 0.633399 | 0.295863 |
| 2 | 176905600 | 176907399 | 20.934 | 37.85992 | 23.42939 | 10.16445 | 1.098154 | 0.395085 | 0.295893 |
| 2 | 111734800 | 111737199 | 33.6169 | 57.41784 | 36.22542 | 15.88251 | 1.04434 | 1.009043 | 0.295894 |
| X | 135393200 | 135398999 | 53.96507 | 87.3059 | 59.10189 | 25.27136 | 0.923958 | 1.699813 | 0.295905 |
| 5 | 5890200 | 5891799 | 15.62937 | 28.75068 | 24.18992 | 10.26708 | 0.67893 | 0.263523 | 0.295911 |
| 5 | 43157600 | 43159399 | 13.83974 | 26.21262 | 20.71388 | 8.38217 | 0.822175 | 0.063401 | 0.295916 |
| 17 | 39768000 | 39769599 | 39.65787 | 64.80041 | 59.69497 | 25.6626 | 0.492103 | 1.505523 | 0.295916 |
| 1 | 102247400 | 102248399 | 6.286153 | 12.0665 | 29.20053 | 11.4489 | -0.73809 | -0.08953 | 0.295917 |
| 10 | 51901000 | 51904199 | 26.80345 | 43.76635 | 80.77188 | 34.71476 | -0.50831 | 1.501485 | 0.295926 |
| 14 | 64185000 | 64186599 | 12.63354 | 22.01782 | 46.82553 | 20.59887 | -0.69587 | 0.73324 | 0.295935 |
| 13 | 106820800 | 106822399 | 15.68597 | 29.50604 | 19.79012 | 8.116526 | 1.040245 | 0.121758 | 0.295952 |
| 2 | 29972800 | 29975399 | 13.37124 | 23.33898 | 46.79748 | 20.60623 | -0.61137 | 0.713216 | 0.295952 |
| 14 | 41360000 | 41362999 | 37.00724 | 61.90571 | 45.33063 | 19.74572 | 0.829127 | 1.214506 | 0.295952 |
| 16 | 20683200 | 20683799 | 4.355774 | 8.490794 | 21.94995 | 7.929475 | -0.71837 | -0.41222 | 0.295954 |
| 3 | 116638000 | 116638799 | 6.568381 | 12.62737 | 28.70367 | 11.20521 | -0.6425 | -0.10119 | 0.295954 |
| 4 | 7472000 | 7473799 | 18.94816 | 33.73675 | 27.95336 | 12.31316 | 0.665461 | 0.482975 | 0.295962 |
| 3 | 158264800 | 158268199 | 24.70696 | 40.41307 | 78.89215 | 33.92831 | -0.58726 | 1.470603 | 0.295973 |

| | | | | | | | | | |
|---|---|---|---|---|---|---|---|---|---|
| X | 62619000 | 62622999 | 76.42306 | 121.4827 | 100.9723 | 42.13336 | 0.600156 | 2.269855 | 0.295976 |
| 16 | 10746800 | 10755199 | 81.20193 | 125.2579 | 208.5956 | 86.64684 | -0.43573 | 2.890104 | 0.295988 |
| 7 | 42234800 | 42238599 | 86.21171 | 136.9255 | 95.99691 | 40.02294 | 0.844108 | 2.300441 | 0.296002 |
| 6 | 140475200 | 140476799 | 15.20281 | 28.60322 | 21.07193 | 8.639143 | 0.906816 | 0.120202 | 0.296009 |
| 6 | 13766600 | 13768999 | 28.17314 | 47.47655 | 42.22523 | 18.4596 | 0.549591 | 1.111207 | 0.296018 |
| 8 | 92585800 | 92589599 | 27.62003 | 45.06265 | 81.94754 | 35.20949 | -0.48796 | 1.516073 | 0.296022 |
| 1 | 21184600 | 21185799 | 12.42294 | 23.31576 | 21.43467 | 8.824466 | 0.585121 | 0.111808 | 0.296028 |
| 13 | 111951400 | 111952599 | 13.90202 | 25.78066 | 23.31032 | 9.785546 | 0.586931 | 0.202511 | 0.296029 |
| Y | 42513000 | 42513399 | 7.127658 | 13.94003 | 7.695457 | 2.609516 | 1.559311 | -0.65692 | 0.296038 |
| 5 | 9072400 | 9073399 | 19.01289 | 33.80097 | 23.6735 | 10.4412 | 0.905323 | 0.484156 | 0.296038 |
| 4 | 73462400 | 73466199 | 50.48402 | 81.50898 | 71.27413 | 30.40227 | 0.552374 | 1.753403 | 0.296042 |
| 12 | 97697600 | 97698799 | 14.8561 | 27.0204 | 23.78803 | 10.24721 | 0.599368 | 0.318934 | 0.296043 |
| 1 | 74830800 | 74834599 | 25.74224 | 41.88063 | 84.83623 | 36.41316 | -0.64605 | 1.553622 | 0.296046 |
| 8 | 99029000 | 99030199 | 8.284703 | 15.56176 | 34.14581 | 14.03799 | -0.6593 | 0.101424 | 0.29605 |
| 17 | 29669000 | 29671799 | 20.83408 | 34.68893 | 67.14573 | 29.16221 | -0.56844 | 1.235995 | 0.296058 |
| 6 | 134587000 | 134589599 | 13.20053 | 23.07196 | 46.77507 | 20.6148 | -0.62708 | 0.692044 | 0.296073 |
| 17 | 19173600 | 19195399 | 627.5743 | 845.4589 | 687.7531 | 286.8413 | 0.411774 | 5.162045 | 0.296113 |
| 11 | 97479400 | 97481799 | 13.21451 | 23.09843 | 45.72513 | 20.15357 | -0.59272 | 0.690244 | 0.296134 |
| 9 | 8281400 | 8283199 | 19.02211 | 33.43391 | 28.68239 | 12.70099 | 0.607284 | 0.601939 | 0.296141 |
| 5 | 93925600 | 93928599 | 44.79119 | 73.03392 | 60.58026 | 26.02316 | 0.640913 | 1.546967 | 0.296143 |
| 5 | 93838200 | 93839999 | 42.12136 | 69.02895 | 56.75418 | 24.42627 | 0.658108 | 1.467786 | 0.296147 |
| 9 | 51399200 | 51401399 | 18.31652 | 32.70852 | 27.37186 | 12.02705 | 0.65345 | 0.458278 | 0.296158 |
| 3 | 86791800 | 86792999 | 19.72396 | 34.61884 | 30.86569 | 13.65157 | 0.55229 | 0.634449 | 0.296177 |
| 3 | 104126400 | 104128599 | 15.56186 | 26.57507 | 56.3673 | 24.71307 | -0.69784 | 0.964144 | 0.29618 |
| 7 | 17589000 | 17592599 | 66.55629 | 106.0272 | 98.27375 | 41.12332 | 0.446404 | 2.191885 | 0.296188 |
| 10 | 93621200 | 93623799 | 65.06546 | 103.8267 | 93.97219 | 39.41511 | 0.483218 | 2.134371 | 0.296192 |
| 13 | 33727000 | 33727999 | 17.72354 | 31.9153 | 26.43391 | 11.53014 | 0.675396 | 0.402326 | 0.296193 |
| 12 | 12242600 | 12244399 | 9.80185 | 17.92688 | 42.23551 | 18.06779 | -0.80541 | 0.290171 | 0.296197 |
| 3 | 119575000 | 119577399 | 11.96723 | 20.95055 | 47.80462 | 21.09761 | -0.79679 | 0.659382 | 0.296218 |
| 7 | 103091000 | 103092999 | 14.00775 | 24.52492 | 46.73796 | 20.62835 | -0.53822 | 0.669865 | 0.296226 |
| 11 | 89887200 | 89890599 | 30.62654 | 49.4899 | 95.008 | 40.57026 | -0.5781 | 1.707668 | 0.296252 |
| 2 | 43544400 | 43546599 | 21.16555 | 37.40034 | 28.73658 | 12.71212 | 0.768373 | 0.558766 | 0.296256 |
| 16 | 87863600 | 87865999 | 18.39505 | 32.97725 | 26.23115 | 11.48796 | 0.729297 | 0.432045 | 0.296275 |
| X | 125382000 | 125394799 | 150.9196 | 229.979 | 177.8718 | 73.73384 | 0.656118 | 3.151241 | 0.29628 |
| 13 | 45848200 | 45853999 | 40.23672 | 64.52541 | 114.019 | 48.16838 | -0.4722 | 1.962295 | 0.296284 |
| 7 | 30623400 | 30627199 | 25.22259 | 41.15134 | 80.92904 | 34.77193 | -0.60051 | 1.510371 | 0.296286 |
| 1 | 93608200 | 93610799 | 27.22306 | 44.62611 | 82.04946 | 35.31728 | -0.49962 | 1.448313 | 0.296301 |
| 13 | 13567600 | 13568999 | 9.923059 | 17.59945 | 39.39407 | 17.40945 | -0.76312 | 0.47175 | 0.296311 |
| 12 | 60505800 | 60506999 | 11.93649 | 22.83419 | 19.30313 | 7.643126 | 0.752719 | -0.02859 | 0.296311 |
| 14 | 12968000 | 12968799 | 9.956436 | 18.06681 | 35.95725 | 15.53445 | -0.57246 | 0.324965 | 0.296314 |
| 3 | 34407600 | 34412199 | 31.91574 | 51.53464 | 96.84722 | 41.31582 | -0.54871 | 1.732365 | 0.296353 |
| 11 | 111772200 | 111772999 | 5.374312 | 10.44678 | 24.6618 | 9.162889 | -0.62768 | -0.29229 | 0.29636 |
| 7 | 92799800 | 92802599 | 34.94989 | 58.14778 | 50.36342 | 21.86244 | 0.587894 | 1.270319 | 0.296361 |
| 5 | 121518800 | 121519999 | 14.82621 | 25.73675 | 49.50712 | 21.7242 | -0.55269 | 0.803941 | 0.296363 |
| 7 | 130263600 | 130269399 | 58.6905 | 92.39938 | 159.5649 | 66.20434 | -0.46512 | 2.50446 | 0.29637 |
| 9 | 115365800 | 115367199 | 11.29252 | 19.95572 | 41.92248 | 18.54488 | -0.67478 | 0.521854 | 0.296379 |
| 8 | 15786800 | 15792999 | 100.5806 | 156.4733 | 142.5571 | 59.18093 | 0.442963 | 2.738614 | 0.29639 |

| | | | | | | | | | |
|---|---|---|---|---|---|---|---|---|---|
| 7 | 84568000 | 84569999 | 29.91388 | 51.08225 | 39.68712 | 17.39994 | 0.745205 | 0.999549 | 0.29639 |
| 4 | 73062000 | 73063199 | 6.575957 | 12.6739 | 31.89403 | 12.36069 | -0.77798 | -0.11474 | 0.296394 |
| 3 | 51250200 | 51252999 | 23.60792 | 40.86522 | 36.65426 | 16.08136 | 0.542145 | 0.849044 | 0.296396 |
| 1 | 13136200 | 13138199 | 13.19865 | 23.11162 | 45.15591 | 19.93249 | -0.57394 | 0.659987 | 0.296401 |
| 18 | 26904800 | 26905999 | 13.88584 | 26.48714 | 19.71022 | 7.857257 | 0.926944 | 0.01433 | 0.296408 |
| 7 | 59425600 | 59427599 | 16.30128 | 29.32857 | 24.82626 | 10.8399 | 0.643441 | 0.395097 | 0.296412 |
| X | 122976200 | 122979399 | 37.47351 | 62.37025 | 48.73835 | 21.16258 | 0.735919 | 1.272537 | 0.296419 |
| X | 139330600 | 139336599 | 39.03309 | 62.61547 | 115.4597 | 48.80101 | -0.53293 | 1.95469 | 0.296436 |
| 6 | 141646200 | 141648399 | 22.17235 | 39.32515 | 26.40223 | 11.66784 | 0.963894 | 0.523521 | 0.296437 |
| 15 | 65826000 | 65827599 | 15.876 | 29.87002 | 18.55974 | 7.609093 | 1.149588 | 0.118576 | 0.296459 |
| 5 | 20853000 | 20855199 | 20.22184 | 34.08151 | 64.1427 | 28.04232 | -0.52803 | 1.105911 | 0.296465 |
| 7 | 23780800 | 23788599 | 99.9568 | 152.5986 | 245.4319 | 101.784 | -0.39716 | 3.103329 | 0.296465 |
| 4 | 132611200 | 132612399 | 6.782801 | 12.82312 | 29.20086 | 11.86254 | -0.69668 | 0.035911 | 0.296468 |
| 5 | 140440000 | 140455799 | 129.0082 | 191.948 | 361.8705 | 149.0977 | -0.65841 | 3.617345 | 0.296485 |
| 6 | 135473400 | 135474599 | 14.52644 | 26.54495 | 23.77551 | 10.18195 | 0.580039 | 0.289708 | 0.296496 |
| 13 | 86901400 | 86903799 | 21.98446 | 39.0025 | 26.40732 | 11.66742 | 0.951998 | 0.52053 | 0.296497 |
| 5 | 81597200 | 81599399 | 23.29022 | 40.53752 | 34.90264 | 15.34135 | 0.601976 | 0.782296 | 0.296501 |
| 1 | 76557400 | 76558199 | 13.38048 | 25.29874 | 21.30996 | 8.655161 | 0.726265 | 0.072551 | 0.296507 |
| 11 | 22600600 | 22602599 | 10.26376 | 18.09913 | 43.00736 | 19.03289 | -0.8526 | 0.541299 | 0.296569 |
| 8 | 5020000 | 5022199 | 31.36554 | 53.00093 | 43.04334 | 18.83285 | 0.680269 | 1.09866 | 0.29658 |
| 3 | 7834200 | 7835599 | 20.91637 | 37.04047 | 28.24886 | 12.49094 | 0.780283 | 0.534013 | 0.296581 |
| X | 62711000 | 62713799 | 36.93672 | 62.62885 | 37.9821 | 16.63564 | 1.100225 | 1.088899 | 0.296582 |
| 3 | 76073600 | 76074999 | 16.13889 | 29.39501 | 25.02065 | 10.75918 | 0.649182 | 0.322811 | 0.296592 |
| 5 | 11506200 | 11508999 | 28.59137 | 46.59187 | 83.78449 | 35.98313 | -0.47311 | 1.535837 | 0.296595 |
| 15 | 93915600 | 93918399 | 35.26329 | 58.86349 | 47.58941 | 20.70192 | 0.686809 | 1.232006 | 0.296609 |
| 4 | 134117800 | 134121399 | 28.0477 | 47.35954 | 40.18341 | 17.57761 | 0.617426 | 1.092452 | 0.296614 |
| 5 | 95585800 | 95589999 | 59.90669 | 95.98731 | 84.49092 | 35.64778 | 0.52981 | 2.001041 | 0.296615 |
| 4 | 132260200 | 132260999 | 5.104289 | 9.916677 | 24.57782 | 9.170579 | -0.70295 | -0.28171 | 0.29663 |
| 13 | 30871400 | 30872399 | 5.104289 | 9.916677 | 24.57782 | 9.170579 | -0.70295 | -0.28171 | 0.29663 |
| 7 | 20668600 | 20671599 | 58.00279 | 92.78922 | 86.74407 | 36.51296 | 0.440542 | 2.048799 | 0.296641 |
| 3 | 44168600 | 44169399 | 12.72856 | 23.82087 | 21.8922 | 9.05674 | 0.579927 | 0.133433 | 0.296647 |
| 4 | 35597600 | 35598999 | 15.49235 | 26.72397 | 51.62227 | 22.64588 | -0.56068 | 0.867526 | 0.29665 |
| 17 | 57013400 | 57015199 | 7.833795 | 14.27528 | 34.49748 | 14.82525 | -0.84505 | 0.281401 | 0.296664 |
| 10 | 4113600 | 4116799 | 16.74988 | 28.85 | 53.80468 | 23.60219 | -0.51084 | 0.88827 | 0.296667 |
| 15 | 4427600 | 4428599 | 13.92982 | 26.0081 | 22.81633 | 9.476496 | 0.642071 | 0.161216 | 0.296713 |
| 3 | 44708000 | 44709199 | 14.9545 | 27.24682 | 24.15873 | 10.38424 | 0.590997 | 0.309805 | 0.29672 |
| X | 52867800 | 52869799 | 22.7353 | 39.96772 | 29.51973 | 13.07202 | 0.822648 | 0.626691 | 0.296728 |
| 8 | 25145600 | 25147199 | 18.07816 | 32.76021 | 24.72014 | 10.69775 | 0.815983 | 0.364773 | 0.296781 |
| 8 | 47339600 | 47347399 | 45.83398 | 73.21813 | 125.6031 | 52.76341 | -0.43595 | 2.091918 | 0.296787 |
| X | 88226800 | 88229999 | 59.35201 | 95.17806 | 83.4467 | 35.25198 | 0.536981 | 1.977404 | 0.296795 |
| 7 | 60181200 | 60182599 | 20.02128 | 35.66697 | 27.19987 | 11.97572 | 0.784788 | 0.484388 | 0.296802 |
| 13 | 60421000 | 60423999 | 46.00202 | 74.73024 | 63.44345 | 27.21026 | 0.603561 | 1.605004 | 0.296839 |
| 12 | 87175800 | 87177399 | 7.532886 | 14.18881 | 30.22596 | 12.36832 | -0.61072 | 0.069612 | 0.296847 |
| 15 | 65066800 | 65069799 | 26.14208 | 42.76438 | 79.12924 | 34.0314 | -0.51017 | 1.472829 | 0.29685 |
| 18 | 77328000 | 77336999 | 58.94071 | 92.13044 | 179.2205 | 74.35632 | -0.64475 | 2.641534 | 0.296861 |
| 1 | 31509000 | 31513599 | 103.4179 | 160.175 | 146.9304 | 61.03093 | 0.428174 | 2.816637 | 0.296881 |
| 9 | 38066400 | 38070999 | 58.78733 | 94.1637 | 86.6163 | 36.52675 | 0.465908 | 2.012389 | 0.296885 |

| | | | | | | | | | |
|---|---|---|---|---|---|---|---|---|---|
| 10 | 43272400 | 43276599 | 27.92857 | 45.62298 | 82.18405 | 35.32719 | -0.47307 | 1.498774 | 0.296887 |
| 4 | 105374200 | 105376399 | 13.50992 | 23.59093 | 47.84966 | 21.0751 | -0.62795 | 0.708445 | 0.296903 |
| X | 114484400 | 114487999 | 30.24146 | 52.69609 | 29.56601 | 13.00491 | 1.217066 | 0.795899 | 0.296904 |
| 4 | 18275800 | 18280799 | 60.38938 | 96.94212 | 73.73388 | 31.19798 | 0.743278 | 1.940998 | 0.296907 |
| 11 | 67188000 | 67190999 | 21.14431 | 35.29893 | 66.09557 | 28.75417 | -0.52071 | 1.208549 | 0.296908 |
| 9 | 35812800 | 35813999 | 24.7998 | 42.63519 | 38.40737 | 16.84698 | 0.53417 | 0.919549 | 0.296915 |
| 7 | 18654800 | 18667599 | 89.18812 | 136.1347 | 241.2404 | 100.0417 | -0.53706 | 3.103122 | 0.296916 |
| 4 | 97562000 | 97562799 | 5.526905 | 10.73462 | 25.80904 | 9.652614 | -0.66326 | -0.26528 | 0.296918 |
| 3 | 79498400 | 79499199 | 5.838856 | 11.26198 | 26.44285 | 10.22535 | -0.67516 | -0.14473 | 0.296927 |
| 10 | 76210200 | 76211199 | 5.838856 | 11.26198 | 26.44285 | 10.22535 | -0.67516 | -0.14473 | 0.296927 |
| 17 | 22371600 | 22373999 | 20.2131 | 35.88186 | 27.07091 | 11.95445 | 0.797168 | 0.505694 | 0.296951 |
| 7 | 81640800 | 81643599 | 20.12092 | 32.94988 | 84.55276 | 36.37898 | -0.97945 | 1.448283 | 0.296962 |
| 11 | 108882200 | 108886199 | 38.00078 | 60.95633 | 109.4109 | 46.24074 | -0.49419 | 1.950927 | 0.296967 |
| 5 | 137682000 | 137686599 | 21.9756 | 36.06907 | 78.32901 | 33.73366 | -0.73817 | 1.418188 | 0.297012 |
| 16 | 44544400 | 44545399 | 5.184752 | 9.915873 | 28.33872 | 11.22865 | -0.98908 | -0.0664 | 0.297013 |
| 1 | 149135600 | 149136799 | 15.40038 | 27.96451 | 25.21706 | 10.88506 | 0.563113 | 0.335788 | 0.297016 |
| 2 | 107582800 | 107585599 | 34.40107 | 57.24867 | 49.28541 | 21.39809 | 0.596523 | 1.262392 | 0.297022 |
| 9 | 110609600 | 110612599 | 14.0104 | 24.36441 | 48.95916 | 21.50807 | -0.61516 | 0.772035 | 0.297043 |
| 7 | 4313200 | 4314199 | 12.59684 | 24.02054 | 19.75724 | 7.881437 | 0.782449 | 0.002993 | 0.297047 |
| 16 | 82016600 | 82018799 | 21.98955 | 38.25811 | 34.56271 | 15.18806 | 0.532899 | 0.781348 | 0.297047 |
| 12 | 70252400 | 70253999 | 7.698135 | 14.45999 | 30.71135 | 12.62598 | -0.61185 | 0.090628 | 0.29706 |
| 2 | 168857000 | 168858799 | 21.86501 | 38.04296 | 34.6031 | 15.20619 | 0.523166 | 0.780695 | 0.297087 |
| 7 | 10837800 | 10855199 | 346.2734 | 491.3129 | 416.5269 | 172.3082 | 0.427502 | 4.365586 | 0.297095 |
| 13 | 67167000 | 67174999 | 146.2055 | 222.7056 | 173.1692 | 71.77579 | 0.647921 | 3.154764 | 0.297101 |
| 14 | 108270200 | 108271399 | 12.71288 | 23.84433 | 21.8745 | 9.015518 | 0.586732 | 0.119141 | 0.297119 |
| 17 | 39459800 | 39460199 | 12.71288 | 23.84433 | 21.8745 | 9.015518 | 0.586732 | 0.119141 | 0.297119 |
| 9 | 12437800 | 12438999 | 12.99635 | 24.64572 | 20.80165 | 8.395665 | 0.731121 | 0.047186 | 0.297127 |
| 6 | 4902200 | 4904399 | 11.71899 | 20.42159 | 47.34528 | 20.82601 | -0.81973 | 0.730371 | 0.297133 |
| 7 | 92317800 | 92319799 | 11.71899 | 20.42159 | 47.34528 | 20.82601 | -0.81973 | 0.730371 | 0.297133 |
| 13 | 5076000 | 5083399 | 94.0503 | 147.9326 | 122.0311 | 50.62535 | 0.598334 | 2.537459 | 0.297143 |
| X | 123056600 | 123061399 | 60.40369 | 97.79527 | 56.57291 | 24.20328 | 1.151184 | 1.698335 | 0.29715 |
| X | 122293200 | 122293799 | 12.71811 | 23.96895 | 20.99367 | 8.582414 | 0.662864 | 0.088854 | 0.297152 |
| 8 | 59257400 | 59258399 | 8.483424 | 15.73109 | 32.61343 | 13.69217 | -0.60177 | 0.181605 | 0.297152 |
| 7 | 98199200 | 98200799 | 8.568354 | 15.68719 | 34.54242 | 14.75592 | -0.70582 | 0.262395 | 0.297171 |
| 19 | 27378000 | 27380599 | 24.3192 | 41.76851 | 37.87138 | 16.61139 | 0.524473 | 0.923639 | 0.297172 |
| 12 | 86708800 | 86711599 | 17.12727 | 29.23567 | 55.8764 | 24.49699 | -0.54832 | 0.974439 | 0.297203 |
| 16 | 48304600 | 48306399 | 19.33204 | 34.28345 | 28.32608 | 12.5193 | 0.666113 | 0.509385 | 0.297208 |
| 16 | 5925000 | 5926799 | 11.24505 | 20.03297 | 40.3885 | 17.78216 | -0.6098 | 0.452518 | 0.297211 |
| 14 | 102548200 | 102548999 | 7.239523 | 13.66952 | 30.26771 | 12.32839 | -0.65997 | 0.052543 | 0.297225 |
| 9 | 66852800 | 66853999 | 8.625548 | 15.56215 | 35.9387 | 15.63839 | -0.79204 | 0.349003 | 0.297225 |
| 7 | 13889200 | 13906799 | 180.0495 | 271.7862 | 186.0427 | 76.89758 | 0.820838 | 3.345668 | 0.297232 |
| 6 | 100707200 | 100720399 | 160.4377 | 236.677 | 378.5616 | 155.8974 | -0.43291 | 3.767365 | 0.297232 |
| 14 | 58490400 | 58491599 | 9.157542 | 16.82564 | 34.59159 | 14.70557 | -0.60283 | 0.245231 | 0.297235 |
| 2 | 83320200 | 83321199 | 9.818889 | 17.75011 | 35.91202 | 15.58831 | -0.60037 | 0.346023 | 0.297241 |
| 2 | 135899600 | 135901799 | 18.33408 | 30.94383 | 60.10078 | 26.28657 | -0.57299 | 1.084826 | 0.297257 |
| 2 | 152014200 | 152017399 | 26.43419 | 43.28974 | 79.41363 | 34.1683 | -0.4969 | 1.457647 | 0.297257 |
| 4 | 32452800 | 32454599 | 14.97884 | 28.32257 | 20.66657 | 8.392718 | 0.932156 | 0.085928 | 0.29726 |

| | | | | | | | | | |
|---|---|---|---|---|---|---|---|---|---|
| 14 | 110070400 | 110074199 | 45.97313 | 74.34505 | 68.16751 | 29.13734 | 0.48671 | 1.69747 | 0.297275 |
| 15 | 59549400 | 59550399 | 10.77115 | 19.35127 | 37.88746 | 16.5628 | -0.56032 | 0.393317 | 0.297278 |
| 7 | 122812800 | 122814799 | 10.10398 | 17.84335 | 40.46572 | 17.90287 | -0.7844 | 0.521856 | 0.297279 |
| 17 | 19218200 | 19225799 | 117.4611 | 182.1374 | 142.8822 | 59.34489 | 0.654929 | 2.804595 | 0.297283 |
| 16 | 91613000 | 91615799 | 21.53221 | 35.65666 | 70.32187 | 30.42767 | -0.59596 | 1.303888 | 0.2973 |
| 6 | 70701800 | 70702199 | 2.736355 | 5.350392 | 16.31053 | 5.287231 | -0.80459 | -0.83905 | 0.297316 |
| 15 | 100787600 | 100788399 | 4.81305 | 9.365587 | 23.53995 | 8.67236 | -0.70548 | -0.33312 | 0.297333 |
| 5 | 126241600 | 126243999 | 21.76032 | 38.18104 | 30.34829 | 13.41918 | 0.716835 | 0.647172 | 0.297337 |
| 4 | 156206800 | 156213399 | 45.22018 | 72.10377 | 125.6201 | 52.6332 | -0.46113 | 2.149574 | 0.29735 |
| 18 | 80660200 | 80664799 | 30.00744 | 48.47111 | 93.21053 | 39.78324 | -0.5811 | 1.718826 | 0.297353 |
| 4 | 75290600 | 75292599 | 20.34634 | 36.60515 | 23.8136 | 10.39801 | 1.020315 | 0.419292 | 0.297363 |
| 7 | 12099000 | 12099999 | 19.9206 | 35.42197 | 27.80425 | 12.26094 | 0.741955 | 0.496723 | 0.297366 |
| 5 | 114043600 | 114048599 | 24.14074 | 39.02171 | 104.2953 | 44.55423 | -1.05366 | 1.692441 | 0.297366 |
| 14 | 118612600 | 118615599 | 22.43103 | 37.34453 | 68.43016 | 29.71837 | -0.48969 | 1.242907 | 0.29738 |
| 14 | 35892200 | 35893199 | 4.268198 | 8.325746 | 25.13495 | 8.999473 | -0.92719 | -0.43156 | 0.297398 |
| 8 | 47862200 | 47863399 | 6.286755 | 12.07099 | 27.40753 | 10.73736 | -0.64473 | -0.09272 | 0.297404 |
| 1 | 13541400 | 13545199 | 53.80188 | 86.64046 | 75.80949 | 32.22478 | 0.547114 | 1.836764 | 0.297421 |
| 2 | 28366800 | 28370199 | 18.08573 | 30.34036 | 62.67432 | 27.34977 | -0.66182 | 1.142759 | 0.297428 |
| 13 | 100046600 | 100048999 | 14.52655 | 25.15458 | 50.02786 | 21.94901 | -0.60128 | 0.826268 | 0.297437 |
| 4 | 36275600 | 36276799 | 12.86352 | 24.51272 | 20.33761 | 8.1244 | 0.768588 | 0.012373 | 0.297444 |
| 4 | 71293200 | 71294599 | 12.86352 | 24.51272 | 20.33761 | 8.1244 | 0.768588 | 0.012373 | 0.297444 |
| 14 | 41061000 | 41063199 | 25.18235 | 43.96141 | 32.4924 | 14.30866 | 0.821508 | 0.747985 | 0.297493 |
| X | 80021400 | 80026399 | 18.2237 | 32.72158 | 23.11126 | 10.10816 | 0.901346 | 0.416992 | 0.297502 |
| 8 | 80058400 | 80062799 | 37.61522 | 60.55074 | 107.2216 | 45.55185 | -0.46852 | 1.838379 | 0.297515 |
| 6 | 100080200 | 100081999 | 7.707332 | 14.37779 | 32.19872 | 13.39294 | -0.70186 | 0.136254 | 0.297523 |
| 9 | 74753200 | 74756999 | 34.20913 | 55.09681 | 101.0619 | 42.96773 | -0.5182 | 1.814328 | 0.297524 |
| 11 | 49904200 | 49904799 | 4.93988 | 9.627084 | 23.84648 | 8.641822 | -0.66321 | -0.38985 | 0.297524 |
| 8 | 3109400 | 3111199 | 47.0542 | 75.91671 | 70.5022 | 30.0451 | 0.464568 | 1.768639 | 0.297531 |
| 11 | 73864400 | 73867999 | 42.13344 | 69.5316 | 51.6945 | 22.31461 | 0.806772 | 1.382974 | 0.297534 |
| 4 | 3192400 | 3195199 | 48.11456 | 77.74179 | 61.85962 | 26.41207 | 0.689377 | 1.723357 | 0.297544 |
| 4 | 71411600 | 71413999 | 27.25218 | 47.21011 | 35.27714 | 15.47754 | 0.804674 | 0.857319 | 0.297554 |
| 5 | 148605200 | 148628999 | 219.1447 | 313.5693 | 554.2986 | 228.9805 | -0.61998 | 4.237824 | 0.297558 |
| 14 | 106454200 | 106455799 | 17.70424 | 31.47247 | 28.06035 | 12.37668 | 0.55892 | 0.483765 | 0.297581 |
| 18 | 31330800 | 31332799 | 19.45515 | 32.74785 | 61.79004 | 27.0034 | -0.53165 | 1.113249 | 0.297589 |
| 11 | 88720000 | 88722599 | 18.09383 | 30.80282 | 57.5094 | 25.20747 | -0.51536 | 1.004459 | 0.297603 |
| 10 | 68216200 | 68217799 | 16.7572 | 30.17253 | 25.93589 | 11.31415 | 0.62184 | 0.393636 | 0.297621 |
| 17 | 21516200 | 21517599 | 21.44862 | 37.36146 | 33.01084 | 14.51698 | 0.564915 | 0.759276 | 0.297623 |
| 8 | 71347800 | 71349999 | 15.78331 | 26.80994 | 57.32012 | 25.11914 | -0.71031 | 1.01789 | 0.297624 |
| 16 | 44390600 | 44394799 | 60.56825 | 96.80366 | 90.88163 | 38.20447 | 0.433003 | 2.081989 | 0.297631 |
| 10 | 50367000 | 50373199 | 73.52815 | 116.8358 | 94.22495 | 39.30265 | 0.643005 | 2.273439 | 0.29764 |
| 9 | 75289800 | 75291399 | 10.6413 | 19.02144 | 37.86459 | 16.62359 | -0.58828 | 0.424731 | 0.297652 |
| 11 | 85813800 | 85816599 | 12.41995 | 21.73807 | 47.80986 | 21.09579 | -0.74404 | 0.665532 | 0.297663 |
| 7 | 93097800 | 93099799 | 13.69898 | 25.60174 | 21.63061 | 8.969388 | 0.697308 | 0.148387 | 0.29767 |
| 7 | 29413600 | 29421399 | 53.82642 | 85.13724 | 149.0672 | 61.94825 | -0.47968 | 2.395763 | 0.297681 |
| 16 | 97899400 | 97903599 | 51.60546 | 83.44879 | 67.43376 | 28.82168 | 0.668513 | 1.711146 | 0.297715 |
| 10 | 41527000 | 41529199 | 13.20081 | 23.22908 | 45.58985 | 20.18431 | -0.57978 | 0.587387 | 0.29772 |
| 14 | 42524600 | 42526599 | 31.30248 | 52.66659 | 41.60823 | 18.18114 | 0.719811 | 1.135099 | 0.297724 |

| | | | | | | | | | |
|---|---|---|---|---|---|---|---|---|---|
| 3 | 7736600 | 7738799 | 27.03708 | 46.47444 | 35.175 | 15.42907 | 0.784272 | 0.925972 | 0.297739 |
| 16 | 29398000 | 29398799 | 3.033828 | 5.910474 | 22.88058 | 8.328636 | -1.30177 | -0.39994 | 0.297739 |
| 14 | 121803200 | 121805799 | 14.37352 | 24.84515 | 51.08507 | 22.41163 | -0.64971 | 0.842213 | 0.297762 |
| 6 | 84150800 | 84152399 | 17.34404 | 30.8552 | 27.58332 | 12.15861 | 0.55568 | 0.475554 | 0.297769 |
| 14 | 54704800 | 54715599 | 64.55974 | 99.88068 | 214.6526 | 89.16141 | -0.79993 | 2.830049 | 0.297788 |
| 11 | 95035200 | 95037199 | 14.56494 | 24.71996 | 59.43003 | 26.03723 | -0.87898 | 1.020929 | 0.297794 |
| 5 | 55479800 | 55481399 | 12.61918 | 23.88498 | 21.4357 | 8.686464 | 0.638527 | 0.059097 | 0.297802 |
| 7 | 69929800 | 69933799 | 33.22846 | 53.56559 | 99.45164 | 42.33644 | -0.53406 | 1.784262 | 0.29781 |
| 12 | 31139800 | 31143199 | 21.3028 | 35.47953 | 67.21797 | 29.19887 | -0.53761 | 1.234131 | 0.297822 |
| 3 | 53227800 | 53228399 | 10.96798 | 21.20937 | 18.62136 | 7.122884 | 0.745971 | -0.14828 | 0.297826 |
| 7 | 65492000 | 65495399 | 49.88717 | 80.40241 | 73.2713 | 31.18266 | 0.489979 | 1.802926 | 0.297828 |
| 3 | 27428400 | 27429399 | 7.662468 | 14.45983 | 30.84978 | 12.58072 | -0.60916 | 0.05853 | 0.29784 |
| 7 | 82018800 | 82019799 | 7.662468 | 14.45983 | 30.84978 | 12.58072 | -0.60916 | 0.05853 | 0.29784 |
| 16 | 87292000 | 87293199 | 7.906001 | 14.60826 | 32.55821 | 13.73535 | -0.71028 | 0.197191 | 0.297855 |
| 3 | 44211400 | 44214599 | 44.47642 | 72.69147 | 57.93278 | 24.90724 | 0.700745 | 1.509974 | 0.297868 |
| 4 | 20391200 | 20392999 | 16.41101 | 28.30407 | 53.27336 | 23.37008 | -0.52363 | 0.873582 | 0.297877 |
| Y | 4284800 | 4286599 | 15.02871 | 27.6936 | 14.94404 | 6.32788 | 1.31661 | 0.223429 | 0.297883 |
| 6 | 34890400 | 34891399 | 4.402017 | 8.561418 | 23.43481 | 8.66913 | -0.83217 | -0.32355 | 0.297899 |
| 17 | 15718200 | 15718799 | 4.402017 | 8.561418 | 23.43481 | 8.66913 | -0.83217 | -0.32355 | 0.297899 |
| 8 | 48891000 | 48892999 | 12.09857 | 21.45792 | 43.49703 | 19.22268 | -0.62133 | 0.49379 | 0.297899 |
| 6 | 10239600 | 10241199 | 16.89937 | 28.66494 | 62.80065 | 27.50899 | -0.7457 | 1.039997 | 0.297906 |
| X | 4038400 | 4043399 | 62.22205 | 100.5876 | 56.70644 | 24.22782 | 1.186149 | 1.732858 | 0.29791 |
| 10 | 32088200 | 32090999 | 31.82372 | 54.46172 | 37.4035 | 16.40161 | 0.922857 | 0.986323 | 0.297933 |
| 16 | 35614400 | 35623599 | 105.2237 | 160.2081 | 258.1781 | 106.9961 | -0.40311 | 3.15999 | 0.297937 |
| 13 | 94736200 | 94736799 | 4.397827 | 8.564853 | 23.00728 | 8.412599 | -0.78993 | -0.36698 | 0.297977 |
| 3 | 34391400 | 34393199 | 15.05361 | 27.66844 | 23.54732 | 10.00569 | 0.660917 | 0.259928 | 0.29799 |
| 11 | 121046000 | 121049199 | 21.07175 | 34.86537 | 70.76308 | 30.60114 | -0.6373 | 1.312873 | 0.297995 |
| 15 | 25674600 | 25675999 | 15.40075 | 26.32894 | 55.29089 | 24.2433 | -0.68318 | 0.952681 | 0.297998 |
| 16 | 80995800 | 80998399 | 14.22606 | 24.56931 | 51.61031 | 22.64147 | -0.68071 | 0.849801 | 0.298002 |
| 9 | 115906000 | 115907599 | 12.55786 | 22.14425 | 42.43552 | 18.77987 | -0.54414 | 0.555552 | 0.298005 |
| 6 | 12154400 | 12156599 | 17.62573 | 31.39877 | 27.46934 | 12.09457 | 0.587706 | 0.468351 | 0.298009 |
| 12 | 84021800 | 84023799 | 17.62573 | 31.39877 | 27.46934 | 12.09457 | 0.587706 | 0.468351 | 0.298009 |
| 9 | 110884600 | 110887599 | 13.9445 | 24.0839 | 51.05105 | 22.39615 | -0.69343 | 0.850521 | 0.298021 |
| 4 | 111964200 | 111967799 | 58.38304 | 93.80048 | 79.14874 | 33.53157 | 0.595318 | 1.917377 | 0.298044 |
| 12 | 52357400 | 52360799 | 32.72804 | 55.46167 | 41.91247 | 18.35398 | 0.783983 | 1.080492 | 0.298047 |
| 6 | 6209800 | 6213199 | 27.24643 | 44.59569 | 80.65071 | 34.69277 | -0.47675 | 1.467351 | 0.29805 |
| 15 | 15418400 | 15419999 | 20.01488 | 35.85509 | 26.28786 | 11.51985 | 0.845982 | 0.450071 | 0.298055 |
| 17 | 61622600 | 61623399 | 14.2468 | 26.21647 | 23.80916 | 10.1007 | 0.569835 | 0.249287 | 0.298062 |
| 7 | 13222600 | 13226199 | 71.88324 | 114.1001 | 105.7049 | 44.04402 | 0.441837 | 2.308282 | 0.298069 |
| 13 | 51717000 | 51723199 | 48.35603 | 76.80502 | 138.8437 | 57.88933 | -0.52022 | 2.276434 | 0.298106 |
| 7 | 17877400 | 17881199 | 74.31551 | 118.2884 | 98.62272 | 41.22134 | 0.59763 | 2.225828 | 0.298108 |
| 14 | 63262400 | 63263799 | 13.08584 | 22.83033 | 46.28655 | 20.37537 | -0.62714 | 0.719148 | 0.298112 |
| 1 | 155544200 | 155545199 | 11.51546 | 22.39672 | 15.72548 | 5.816263 | 1.108906 | -0.26394 | 0.298113 |
| 2 | 121279400 | 121281599 | 11.87214 | 20.68047 | 49.98157 | 21.98031 | -0.87981 | 0.738302 | 0.298133 |
| 7 | 50015000 | 50015399 | 9.135312 | 17.78327 | 16.59745 | 6.094967 | 0.712188 | -0.31521 | 0.298141 |
| 9 | 38456200 | 38458199 | 28.78733 | 49.51339 | 37.47567 | 16.43854 | 0.784333 | 0.929518 | 0.298142 |
| 2 | 11376600 | 11378799 | 11.90037 | 21.03639 | 42.42435 | 18.76572 | -0.61611 | 0.522962 | 0.298147 |

| | | | | | | | | | |
|---|---|---|---|---|---|---|---|---|---|
| 4 | 147988800 | 147989399 | 6.014542 | 11.51268 | 27.86298 | 11.01312 | -0.74857 | -0.06749 | 0.298154 |
| 7 | 130730000 | 130732399 | 18.89667 | 32.15831 | 59.74441 | 26.18639 | -0.50843 | 1.012913 | 0.298158 |
| 2 | 154248600 | 154250599 | 11.4743 | 20.23941 | 40.92639 | 18.11095 | -0.62085 | 0.543729 | 0.298181 |
| 10 | 43354000 | 43356799 | 13.15355 | 22.73554 | 52.12079 | 22.86598 | -0.80592 | 0.837993 | 0.298189 |
| 12 | 22732800 | 22750399 | 547.761 | 744.6297 | 603.1152 | 251.1928 | 0.431128 | 5.018273 | 0.298198 |
| 3 | 31610600 | 31611199 | 9.599995 | 18.64372 | 17.63463 | 6.600658 | 0.668234 | -0.24032 | 0.298201 |
| 17 | 6404600 | 6406199 | 6.790612 | 12.81 | 29.69994 | 12.1209 | -0.72861 | 0.051021 | 0.298223 |
| 8 | 63467800 | 63472199 | 82.97323 | 130.7646 | 120.2422 | 49.89588 | 0.44401 | 2.490458 | 0.298231 |
| 1 | 48920600 | 48922199 | 40.40597 | 67.43001 | 38.95928 | 16.94487 | 1.169934 | 1.247172 | 0.298234 |
| 11 | 71952000 | 71956999 | 47.40881 | 75.30844 | 136.1931 | 56.79018 | -0.52062 | 2.273006 | 0.298266 |
| 17 | 39563600 | 39569199 | 138.8608 | 211.6531 | 170.0105 | 70.4797 | 0.601831 | 3.140784 | 0.298287 |
| 4 | 43495000 | 43499799 | 19.86133 | 32.77641 | 73.85717 | 31.8813 | -0.78799 | 1.34228 | 0.298295 |
| 1 | 84261800 | 84263599 | 13.38211 | 23.41581 | 45.18014 | 19.93092 | -0.55584 | 0.67423 | 0.298296 |
| 13 | 5482000 | 5482999 | 17.15256 | 31.11413 | 25.57201 | 11.0525 | 0.69503 | 0.356137 | 0.298308 |
| 15 | 17686800 | 17689399 | 21.59078 | 38.10515 | 28.66136 | 12.68195 | 0.798203 | 0.573607 | 0.298313 |
| 2 | 168268000 | 168273199 | 65.27045 | 101.6127 | 184.0037 | 76.37897 | -0.54611 | 2.717223 | 0.29832 |
| 2 | 56242200 | 56243399 | 14.4601 | 27.10053 | 22.27017 | 9.190578 | 0.743091 | 0.141182 | 0.298326 |
| 1 | 32356400 | 32357399 | 21.87799 | 38.02579 | 33.70627 | 14.80236 | 0.560157 | 0.794589 | 0.298333 |
| 13 | 60820600 | 60826999 | 42.20952 | 67.59029 | 117.3645 | 49.47169 | -0.44955 | 2.011527 | 0.298353 |
| 3 | 108152800 | 108154599 | 10.56689 | 19.12637 | 39.64482 | 17.18153 | -0.63415 | 0.349518 | 0.298395 |
| 6 | 140176200 | 140179599 | 23.71593 | 39.31064 | 77.80447 | 33.6877 | -0.60124 | 1.301639 | 0.298413 |
| 4 | 113451600 | 113455599 | 66.76794 | 106.5875 | 86.29922 | 36.218 | 0.644331 | 2.12242 | 0.298433 |
| 1 | 11715400 | 11716399 | 7.520425 | 14.16436 | 30.78786 | 12.59973 | -0.64004 | 0.06968 | 0.298452 |
| 3 | 53793600 | 53796799 | 46.47009 | 75.34985 | 66.01052 | 28.27232 | 0.555932 | 1.645926 | 0.298459 |
| 7 | 10147400 | 10149599 | 72.00875 | 114.1702 | 101.7395 | 42.3438 | 0.496225 | 2.34014 | 0.29846 |
| 14 | 38176800 | 38180799 | 59.18093 | 94.68744 | 85.89793 | 36.16411 | 0.484076 | 2.047893 | 0.298463 |
| 1 | 181619200 | 181622599 | 26.97256 | 43.97426 | 83.06158 | 35.67868 | -0.54334 | 1.524447 | 0.298466 |
| 6 | 132365800 | 132368199 | 35.71309 | 59.92205 | 43.60224 | 19.02917 | 0.837994 | 1.176276 | 0.298469 |
| 5 | 117198600 | 117199799 | 8.331922 | 15.49251 | 32.12324 | 13.43528 | -0.59777 | 0.164365 | 0.298481 |
| 7 | 86655200 | 86655999 | 5.810732 | 11.28297 | 27.02681 | 10.12913 | -0.66079 | -0.25027 | 0.298486 |
| 4 | 146363200 | 146367999 | 150.4521 | 228.6401 | 194.3655 | 80.50502 | 0.51666 | 3.205714 | 0.298492 |
| 14 | 92134400 | 92135399 | 10.96104 | 21.06325 | 19.544 | 7.638626 | 0.637337 | -0.07803 | 0.298495 |
| 1 | 44120600 | 44122999 | 10.10867 | 18.37115 | 36.51062 | 15.74337 | -0.56867 | 0.31828 | 0.298563 |
| 14 | 99570400 | 99572199 | 10.10867 | 18.37115 | 36.51062 | 15.74337 | -0.56867 | 0.31828 | 0.298563 |
| 16 | 47797600 | 47799599 | 25.62467 | 44.18408 | 38.16724 | 16.74316 | 0.595147 | 0.894581 | 0.298564 |
| X | 40416800 | 40419799 | 27.22121 | 47.90325 | 25.65456 | 11.35809 | 1.284146 | 0.634488 | 0.298605 |
| 10 | 46734400 | 46736599 | 21.96768 | 38.71765 | 29.21736 | 12.93148 | 0.792863 | 0.591692 | 0.298608 |
| 1 | 154997000 | 154998599 | 8.95133 | 16.53869 | 33.62567 | 14.18693 | -0.57917 | 0.208048 | 0.298611 |
| 17 | 30334600 | 30336399 | 17.49878 | 31.02236 | 27.61201 | 12.20666 | 0.559089 | 0.498222 | 0.298614 |
| 12 | 14589200 | 14590799 | 18.49932 | 33.64675 | 24.21895 | 10.43283 | 0.887932 | 0.346999 | 0.298614 |
| 3 | 72520000 | 72521199 | 21.14268 | 36.84953 | 30.63015 | 13.47486 | 0.652727 | 0.746826 | 0.298618 |
| 14 | 18324200 | 18325599 | 15.38372 | 27.99305 | 25.18738 | 10.8437 | 0.568644 | 0.323489 | 0.298622 |
| 15 | 47099600 | 47100599 | 15.38372 | 27.99305 | 25.18738 | 10.8437 | 0.568644 | 0.323489 | 0.298622 |
| 10 | 21171600 | 21172599 | 5.916965 | 11.17728 | 29.16548 | 11.86824 | -0.89303 | 0.03385 | 0.298625 |
| 17 | 76919800 | 76921999 | 26.20826 | 45.47333 | 35.92241 | 15.7614 | 0.725326 | 0.840655 | 0.298631 |
| 1 | 87014400 | 87018799 | 47.72344 | 75.40778 | 151.3345 | 62.85829 | -0.67734 | 2.417462 | 0.298636 |
| 5 | 30444600 | 30445599 | 11.95344 | 22.70166 | 20.98573 | 8.443419 | 0.605119 | 0.028636 | 0.298655 |

| | | | | | | | | | |
|---|---|---|---|---|---|---|---|---|---|
| X | 148491600 | 148493399 | 20.41452 | 36.62758 | 25.59655 | 11.2023 | 0.915864 | 0.441255 | 0.298656 |
| 9 | 48570600 | 48573399 | 27.62884 | 45.02988 | 82.41778 | 35.39803 | -0.49832 | 1.531241 | 0.298679 |
| 17 | 23206600 | 23208199 | 29.13978 | 49.91644 | 39.15488 | 17.17092 | 0.732089 | 0.968731 | 0.298681 |
| 18 | 50380800 | 50383599 | 24.09851 | 42.05921 | 33.71915 | 14.84664 | 0.704837 | 0.748212 | 0.298686 |
| 7 | 36456200 | 36458999 | 21.56195 | 36.09488 | 66.89319 | 29.15151 | -0.50606 | 1.178983 | 0.298687 |
| 15 | 36712800 | 36714799 | 13.22514 | 23.13031 | 45.19357 | 19.92896 | -0.57387 | 0.680963 | 0.298701 |
| 7 | 5408800 | 5414599 | 99.79542 | 156.7481 | 121.8735 | 50.5507 | 0.682057 | 2.569297 | 0.298705 |
| 3 | 55358200 | 55360199 | 9.771447 | 17.81495 | 35.505 | 15.24886 | -0.56896 | 0.294537 | 0.298707 |
| 11 | 115588200 | 115589199 | 5.981102 | 11.53392 | 26.43695 | 10.22923 | -0.64147 | -0.14065 | 0.29873 |
| 3 | 54539600 | 54544799 | 98.93463 | 150.2508 | 256.2457 | 106.1146 | -0.48775 | 3.210478 | 0.298735 |
| 12 | 18057400 | 18074999 | 514.0428 | 705.6939 | 569.1662 | 236.6898 | 0.451536 | 4.864919 | 0.298746 |
| X | 26802800 | 26816799 | 65.83633 | 105.4891 | 73.93881 | 31.19608 | 0.857867 | 2.008362 | 0.298748 |
| X | 3267600 | 3268599 | 24.98654 | 42.91727 | 37.93392 | 16.63885 | 0.561239 | 0.928195 | 0.298771 |
| 16 | 31766600 | 31767999 | 8.774566 | 15.82542 | 36.43092 | 15.85956 | -0.78813 | 0.352175 | 0.298772 |
| 5 | 18450600 | 18453399 | 30.66488 | 52.23301 | 40.80301 | 17.88555 | 0.73692 | 1.026749 | 0.298782 |
| 3 | 93461800 | 93463599 | 8.561563 | 15.61611 | 34.48124 | 14.8009 | -0.715 | 0.280612 | 0.298796 |
| 12 | 115797400 | 115798399 | 16.95763 | 30.46093 | 26.46194 | 11.5696 | 0.604599 | 0.410708 | 0.298801 |
| 1 | 7659200 | 7660199 | 15.45989 | 28.25571 | 24.57809 | 10.52332 | 0.622031 | 0.297065 | 0.298803 |
| 11 | 27640200 | 27641799 | 18.44885 | 33.12862 | 26.2565 | 11.48298 | 0.735905 | 0.422832 | 0.298809 |
| 7 | 45995400 | 46003399 | 50.25447 | 79.47715 | 153.8214 | 63.91975 | -0.62415 | 2.396317 | 0.298819 |
| 9 | 67263800 | 67265599 | 17.83552 | 31.87046 | 27.44562 | 12.0528 | 0.612915 | 0.451291 | 0.29885 |
| 3 | 19522800 | 19523999 | 12.48674 | 23.74786 | 20.97136 | 8.412579 | 0.674422 | 0.024632 | 0.298853 |
| 14 | 80004800 | 80011799 | 69.77589 | 107.991 | 199.7343 | 82.96457 | -0.5832 | 2.823291 | 0.298855 |
| 9 | 35576800 | 35577599 | 19.80405 | 35.21854 | 27.87162 | 12.28953 | 0.730374 | 0.495912 | 0.298872 |
| 14 | 112151000 | 112152399 | 16.70876 | 30.07654 | 26.5535 | 11.5878 | 0.583363 | 0.397332 | 0.298872 |
| 1 | 183301200 | 183304599 | 61.05662 | 97.44638 | 88.91039 | 37.30106 | 0.471907 | 2.125052 | 0.298874 |
| 16 | 39220000 | 39221199 | 7.406052 | 13.83752 | 35.05181 | 14.54087 | -0.87565 | 0.127933 | 0.298882 |
| X | 21895200 | 21897399 | 25.64922 | 44.57621 | 35.42955 | 15.5582 | 0.716852 | 0.817343 | 0.29889 |
| 5 | 83699000 | 83701599 | 18.40863 | 32.5363 | 29.17697 | 12.90634 | 0.54685 | 0.540706 | 0.298909 |
| Y | 6858400 | 6859999 | 18.36327 | 32.60485 | 18.41869 | 8.132223 | 1.211775 | 0.472753 | 0.298932 |
| 16 | 3525400 | 3529799 | 42.3059 | 69.12595 | 59.91932 | 25.75887 | 0.57966 | 1.511802 | 0.298946 |
| 11 | 97252800 | 97254599 | 9.458537 | 16.81848 | 38.4244 | 16.9444 | -0.79001 | 0.451784 | 0.29895 |
| 3 | 97516600 | 97518199 | 7.683539 | 14.43494 | 30.72242 | 12.62703 | -0.61479 | 0.088038 | 0.298952 |
| 8 | 105047600 | 105049799 | 13.36565 | 23.31037 | 48.39586 | 21.29922 | -0.66161 | 0.725981 | 0.298953 |
| 14 | 9646200 | 9648399 | 18.34496 | 30.51779 | 67.92096 | 29.48167 | -0.76905 | 1.239779 | 0.298971 |
| 10 | 48445200 | 48446599 | 18.67501 | 33.65564 | 25.75033 | 11.2207 | 0.789949 | 0.402762 | 0.298973 |
| 12 | 9274800 | 9275599 | 7.985095 | 15.02285 | 31.32027 | 12.84173 | -0.58248 | 0.083407 | 0.298979 |
| 6 | 136149000 | 136151999 | 16.95112 | 28.54527 | 62.17344 | 27.17451 | -0.73781 | 1.100958 | 0.299022 |
| 15 | 90911000 | 90911999 | 14.55326 | 27.7466 | 18.39863 | 7.342033 | 1.090052 | 0.019881 | 0.299038 |
| 19 | 21629600 | 21631199 | 11.04991 | 19.39826 | 43.06531 | 19.05051 | -0.75687 | 0.604186 | 0.299056 |
| 4 | 147372200 | 147374799 | 56.48458 | 91.63386 | 57.07839 | 24.4557 | 1.04736 | 1.651362 | 0.299067 |
| 12 | 9628800 | 9630199 | 12.60079 | 22.02059 | 45.73653 | 20.15466 | -0.66161 | 0.69084 | 0.299091 |
| 5 | 74863600 | 74865399 | 9.15363 | 16.71399 | 34.48086 | 14.78084 | -0.61649 | 0.278885 | 0.299096 |
| 5 | 142448000 | 142450399 | 17.45782 | 29.29921 | 62.63984 | 27.34192 | -0.71119 | 1.136011 | 0.299102 |
| 10 | 123236600 | 123237599 | 8.067071 | 14.8984 | 32.55449 | 13.74316 | -0.68239 | 0.203118 | 0.299115 |
| 8 | 15368600 | 15369199 | 4.104188 | 8.014168 | 21.73491 | 7.649027 | -0.7532 | -0.51309 | 0.29913 |
| 13 | 77684000 | 77686399 | 25.73004 | 44.0012 | 40.48538 | 17.75198 | 0.502532 | 0.972168 | 0.299134 |

| | | | | | | | | | |
|---|---|---|---|---|---|---|---|---|---|
| 12 | 114165200 | 114171599 | 94.22728 | 148.6029 | 114.5876 | 47.5537 | 0.698427 | 2.485268 | 0.299135 |
| 13 | 66302800 | 66307599 | 87.8138 | 139.3572 | 97.32167 | 40.54238 | 0.848601 | 2.324773 | 0.29914 |
| 8 | 101773400 | 101775199 | 20.82261 | 37.34908 | 24.93487 | 10.91535 | 0.981006 | 0.443843 | 0.299153 |
| 7 | 109444200 | 109446399 | 19.28283 | 32.07358 | 67.36969 | 29.23961 | -0.68582 | 1.245462 | 0.299174 |
| 8 | 19492000 | 19495199 | 33.78366 | 57.2235 | 40.75395 | 17.84409 | 0.869131 | 1.08611 | 0.299175 |
| 6 | 64711200 | 64713399 | 21.89129 | 36.51928 | 73.79667 | 32.08967 | -0.63059 | 1.221079 | 0.299186 |
| 12 | 29403000 | 29406599 | 20.95498 | 34.94523 | 68.43724 | 29.75268 | -0.58537 | 1.218757 | 0.299186 |
| X | 31999000 | 31999999 | 21.45904 | 38.06772 | 24.31496 | 10.74359 | 1.03579 | 0.51528 | 0.299198 |
| 13 | 68977000 | 68979199 | 26.98708 | 45.87652 | 42.06998 | 18.43794 | 0.506055 | 1.031466 | 0.299202 |
| 1 | 184526000 | 184528199 | 16.22257 | 28.05187 | 54.40292 | 23.86748 | -0.56592 | 0.849789 | 0.29921 |
| 4 | 112887200 | 112890799 | 59.22014 | 95.44035 | 69.14804 | 29.42634 | 0.820123 | 1.818171 | 0.299217 |
| 17 | 20617800 | 20626799 | 159.9596 | 242.197 | 204.0509 | 84.42207 | 0.525224 | 3.282654 | 0.299229 |
| 3 | 71305600 | 71306799 | 14.03682 | 26.20459 | 22.76114 | 9.455585 | 0.655949 | 0.162244 | 0.299243 |
| 12 | 116851200 | 116856799 | 37.34196 | 59.91551 | 108.8771 | 46.03383 | -0.51148 | 1.941128 | 0.299244 |
| X | 91052400 | 91053799 | 18.09069 | 34.08519 | 17.96781 | 7.348731 | 1.388193 | 0.129956 | 0.299255 |
| 9 | 63825600 | 63830799 | 26.78889 | 43.36038 | 90.92823 | 38.89367 | -0.70256 | 1.65879 | 0.29926 |
| 14 | 82705200 | 82706599 | 15.81541 | 28.75574 | 25.23575 | 10.87532 | 0.6037 | 0.332421 | 0.299268 |
| 1 | 28432800 | 28435999 | 49.42948 | 80.08969 | 65.71294 | 28.12966 | 0.649258 | 1.665544 | 0.299288 |
| 6 | 89816000 | 89819199 | 52.02937 | 83.72428 | 77.53242 | 32.92194 | 0.464177 | 1.858161 | 0.299289 |
| 14 | 30270000 | 30271999 | 9.697556 | 17.04674 | 43.06419 | 19.07008 | -0.94192 | 0.575028 | 0.299301 |
| 18 | 63084800 | 63086599 | 12.4328 | 21.78186 | 44.11901 | 19.48333 | -0.62548 | 0.646478 | 0.299308 |
| 9 | 115741000 | 115743799 | 11.06695 | 19.38955 | 45.17217 | 19.94895 | -0.82624 | 0.639358 | 0.299328 |
| 2 | 106013800 | 106014599 | 5.515244 | 10.71934 | 25.26052 | 9.395181 | -0.6266 | -0.28469 | 0.299329 |
| 4 | 122958400 | 122961199 | 13.53628 | 23.60971 | 46.80901 | 20.6018 | -0.59526 | 0.724998 | 0.299333 |
| 11 | 37126200 | 37127999 | 10.31129 | 18.47788 | 37.36355 | 16.36894 | -0.60867 | 0.404856 | 0.299342 |
| 13 | 45055800 | 45056999 | 7.261132 | 13.59109 | 30.63629 | 12.67061 | -0.704 | 0.108606 | 0.299343 |
| 9 | 57428400 | 57429199 | 4.32181 | 8.392654 | 24.37167 | 9.127076 | -0.93121 | -0.27242 | 0.299361 |
| 13 | 104436400 | 104437599 | 9.913733 | 18.1331 | 37.2031 | 15.91308 | -0.60689 | 0.279786 | 0.299363 |
| 13 | 3268600 | 3276199 | 124.3224 | 190.0664 | 172.0108 | 71.35874 | 0.433317 | 3.074838 | 0.299371 |
| 6 | 62735600 | 62737399 | 23.22532 | 40.42003 | 35.49477 | 15.60052 | 0.573716 | 0.785376 | 0.299399 |
| 1 | 44288400 | 44290399 | 28.60431 | 49.20677 | 37.47596 | 16.43874 | 0.775433 | 0.927133 | 0.299409 |
| 4 | 56739800 | 56741399 | 22.8214 | 37.29729 | 79.51554 | 34.18586 | -0.71476 | 1.48012 | 0.29941 |
| 1 | 110215600 | 110217599 | 24.57514 | 42.89123 | 32.49613 | 14.30817 | 0.785845 | 0.747192 | 0.299422 |
| 5 | 10768000 | 10770599 | 22.56835 | 39.76424 | 28.57699 | 12.64882 | 0.862697 | 0.596909 | 0.299468 |
| X | 54615600 | 54618399 | 24.25103 | 42.93584 | 25.88656 | 11.44692 | 1.117398 | 0.554104 | 0.29948 |
| 9 | 78661400 | 78664399 | 39.30109 | 64.89615 | 52.7808 | 22.79291 | 0.677616 | 1.367551 | 0.299485 |
| 17 | 60724000 | 60727999 | 53.2896 | 84.5054 | 142.6589 | 59.38458 | -0.42393 | 2.329173 | 0.299487 |
| X | 153841600 | 153843199 | 24.63855 | 43.82182 | 23.51642 | 10.36801 | 1.287337 | 0.518036 | 0.299527 |
| 6 | 89708000 | 89708999 | 29.01288 | 49.24857 | 41.96142 | 18.38478 | 0.611474 | 1.050281 | 0.299535 |
| 2 | 35991600 | 35995399 | 35.60244 | 57.05706 | 111.8684 | 47.21376 | -0.62289 | 1.977212 | 0.299536 |
| 1 | 46426400 | 46430399 | 43.95098 | 71.96941 | 56.26886 | 24.20915 | 0.730032 | 1.481623 | 0.299539 |
| 2 | 84710200 | 84721599 | 64.67024 | 100.2937 | 204.563 | 84.96159 | -0.72191 | 2.783661 | 0.299539 |
| 3 | 78390800 | 78396199 | 36.95392 | 59.42577 | 108.5872 | 46.06408 | -0.51557 | 1.868778 | 0.299573 |
| 5 | 109582600 | 109585199 | 37.40108 | 62.06702 | 49.91485 | 21.6295 | 0.694236 | 1.303423 | 0.299591 |
| 11 | 45316400 | 45319599 | 44.63125 | 72.48115 | 63.91123 | 27.40468 | 0.54859 | 1.607842 | 0.299594 |
| X | 15606200 | 15607399 | 21.59299 | 37.83093 | 32.30674 | 14.26715 | 0.614276 | 0.676284 | 0.299596 |
| 4 | 72735800 | 72737199 | 12.20712 | 23.61835 | 17.2214 | 6.570746 | 1.013765 | -0.14901 | 0.299625 |

| | | | | | | | | | |
|---|---|---|---|---|---|---|---|---|---|
| 16 | 15379200 | 15382599 | 36.90064 | 61.40425 | 49.29534 | 21.40142 | 0.697082 | 1.272711 | 0.299631 |
| 5 | 117321400 | 117331599 | 96.93326 | 148.2399 | 242.595 | 100.6479 | -0.4202 | 3.066678 | 0.299633 |
| 13 | 6697200 | 6699399 | 14.42704 | 24.44795 | 59.37572 | 26.0012 | -0.89357 | 1.03492 | 0.299636 |
| 11 | 50806400 | 50811599 | 43.88607 | 70.20547 | 120.0415 | 50.52802 | -0.42896 | 2.04574 | 0.299646 |
| 4 | 142967800 | 142981399 | 126.0186 | 189.3957 | 303.0611 | 125.1118 | -0.40885 | 3.426982 | 0.299651 |
| 16 | 54952400 | 54952999 | 6.432114 | 12.34893 | 27.42822 | 10.74837 | -0.61398 | -0.0915 | 0.29966 |
| 2 | 99055600 | 99060199 | 85.23692 | 134.279 | 122.8119 | 50.95961 | 0.45133 | 2.501149 | 0.299668 |
| 15 | 7868000 | 7869399 | 9.32192 | 16.93986 | 35.45801 | 15.29114 | -0.64308 | 0.309555 | 0.299671 |
| 12 | 3476400 | 3477799 | 11.82927 | 20.70609 | 44.67173 | 19.7126 | -0.71598 | 0.658671 | 0.2997 |
| X | 117478000 | 117483399 | 69.38193 | 110.8076 | 88.88313 | 37.32492 | 0.658472 | 2.115732 | 0.2997 |
| 3 | 55686600 | 55688399 | 14.35707 | 24.93065 | 48.96865 | 21.49241 | -0.58256 | 0.796466 | 0.299714 |
| X | 54541800 | 54544799 | 48.17455 | 78.83069 | 49.00488 | 21.07811 | 1.059312 | 1.497265 | 0.299716 |
| 8 | 126390800 | 126399199 | 119.7483 | 179.8338 | 299.7907 | 123.7363 | -0.46886 | 3.440963 | 0.299717 |
| 1 | 75206000 | 75209799 | 20.31149 | 33.97877 | 64.46066 | 28.07953 | -0.53952 | 1.181271 | 0.299725 |
| 16 | 56218400 | 56219799 | 13.2982 | 25.4386 | 19.84948 | 7.859897 | 0.867874 | -0.01164 | 0.299731 |
| 9 | 6582400 | 6586199 | 68.09691 | 108.195 | 101.5047 | 42.335 | 0.425025 | 2.274538 | 0.299734 |
| 6 | 131785600 | 131791599 | 109.5594 | 170.6567 | 137.0384 | 56.87919 | 0.626315 | 2.722183 | 0.299739 |
| 15 | 67842400 | 67842999 | 9.393205 | 18.17192 | 16.59594 | 6.335457 | 0.692016 | -0.18023 | 0.299742 |
| 7 | 46174400 | 46175599 | 7.247026 | 13.5903 | 31.77702 | 13.10048 | -0.75321 | 0.096897 | 0.299744 |
| 16 | 87078800 | 87080399 | 18.08148 | 33.47219 | 21.73811 | 9.147474 | 1.058482 | 0.242753 | 0.299765 |
| 1 | 104310200 | 104312599 | 30.67343 | 53.02148 | 32.30331 | 14.17186 | 1.097151 | 0.887499 | 0.299768 |
| 1 | 191632400 | 191633999 | 7.189507 | 13.26591 | 33.59911 | 14.20131 | -0.89537 | 0.20075 | 0.299779 |
| 12 | 47761200 | 47763399 | 22.00395 | 38.93982 | 27.53644 | 12.17759 | 0.888206 | 0.546104 | 0.299783 |
| 17 | 53344400 | 53346399 | 22.00395 | 38.93982 | 27.53644 | 12.17759 | 0.888206 | 0.546104 | 0.299783 |
| 6 | 79395800 | 79397799 | 21.42271 | 38.03684 | 26.4226 | 11.66616 | 0.915797 | 0.511536 | 0.299795 |
| 15 | 80138600 | 80141599 | 11.64803 | 20.16302 | 55.30524 | 24.26413 | -1.06304 | 0.820829 | 0.299799 |
| X | 123092600 | 123093199 | 18.63839 | 33.89383 | 23.99368 | 10.33802 | 0.912105 | 0.350855 | 0.299816 |
| 13 | 35642400 | 35643799 | 12.21059 | 23.29113 | 20.41153 | 8.13717 | 0.692818 | -0.00036 | 0.299863 |
| 19 | 3204400 | 3206999 | 42.90724 | 69.60327 | 65.20893 | 27.93801 | 0.459896 | 1.628586 | 0.299871 |
| X | 13247600 | 13251399 | 41.33919 | 68.67209 | 47.12522 | 20.43501 | 0.922563 | 1.304219 | 0.299878 |
| 5 | 106088800 | 106091199 | 19.12991 | 32.21043 | 61.78739 | 27.00479 | -0.55536 | 1.109003 | 0.29989 |
| X | 121224400 | 121226799 | 27.09765 | 47.94705 | 22.84397 | 10.10271 | 1.454378 | 0.572389 | 0.299905 |
| 18 | 73168200 | 73169799 | 26.88329 | 46.02057 | 39.79248 | 17.44961 | 0.591914 | 0.964992 | 0.299923 |
| 8 | 27754000 | 27761599 | 59.76704 | 94.32631 | 156.3941 | 64.92098 | -0.40372 | 2.453709 | 0.29994 |
| 13 | 18359800 | 18361799 | 12.22299 | 21.65732 | 42.94601 | 18.98758 | -0.59056 | 0.501102 | 0.299955 |
| 16 | 78279000 | 78281799 | 25.39647 | 41.87788 | 76.3919 | 32.96413 | -0.48479 | 1.367603 | 0.299968 |
| 10 | 60016800 | 60024799 | 61.55237 | 96.59589 | 167.1746 | 69.33285 | -0.47183 | 2.571003 | 0.299968 |
| 16 | 93860800 | 93867599 | 67.92763 | 106.0412 | 180.757 | 75.00375 | -0.45592 | 2.669203 | 0.299971 |
| 6 | 66983600 | 66991999 | 86.71328 | 132.5457 | 233.7557 | 96.96969 | -0.52851 | 3.07391 | 0.29999 |
| 7 | 23120200 | 23137799 | 509.1236 | 697.3936 | 580.9388 | 241.6699 | 0.401713 | 4.899119 | 0.299991 |
| X | 78811000 | 78814199 | 47.84388 | 78.01083 | 58.47294 | 25.11785 | 0.786721 | 1.553216 | 0.299997 |
| X | 26542600 | 26547599 | 66.68022 | 107.0262 | 76.05577 | 32.17379 | 0.840961 | 1.957224 | 0.3 |
| 2 | 69204000 | 69205199 | 10.18181 | 18.2449 | 37.39488 | 16.38339 | -0.62785 | 0.40627 | 0.300009 |
| 8 | 57446600 | 57448599 | 24.685 | 42.57643 | 37.1316 | 16.289 | 0.581421 | 0.885374 | 0.30001 |
| 13 | 58254000 | 58256999 | 23.75985 | 39.26774 | 73.45157 | 31.73845 | -0.52024 | 1.333982 | 0.300015 |
| 6 | 123214800 | 123216999 | 22.1902 | 39.26333 | 27.53527 | 12.17745 | 0.900077 | 0.549053 | 0.300021 |
| 2 | 86083000 | 86083999 | 19.3204 | 33.90437 | 29.90543 | 13.22483 | 0.567675 | 0.632681 | 0.300023 |

| | | | | | | | | | |
|---|---|---|---|---|---|---|---|---|---|
| 7 | 38046200 | 38049199 | 44.37887 | 72.49456 | 58.99875 | 25.36076 | 0.670117 | 1.518415 | 0.300043 |
| 2 | 177609000 | 177610799 | 22.71627 | 40.19025 | 26.30202 | 11.63227 | 0.999023 | 0.550162 | 0.300047 |
| 6 | 74091400 | 74093399 | 29.1444 | 49.33284 | 36.90135 | 16.15399 | 0.798509 | 1.06969 | 0.300058 |
| 7 | 62516000 | 62523599 | 89.01639 | 140.5838 | 114.2015 | 47.42259 | 0.624963 | 2.449234 | 0.300071 |
| 5 | 151445000 | 151446799 | 31.27425 | 53.06778 | 41.90376 | 18.35748 | 0.72085 | 1.062908 | 0.300077 |
| 12 | 21194400 | 21198799 | 29.50911 | 47.70405 | 92.37404 | 39.46666 | -0.58977 | 1.693267 | 0.300092 |
| 7 | 30529000 | 30537599 | 86.59946 | 133.1687 | 217.8493 | 90.46657 | -0.41348 | 2.952612 | 0.300111 |
| 17 | 94642400 | 94644399 | 30.45899 | 51.03522 | 46.46041 | 20.25875 | 0.515999 | 1.176314 | 0.300134 |
| 15 | 43869400 | 43871399 | 25.32846 | 44.58852 | 26.75015 | 11.84228 | 1.121332 | 0.620115 | 0.300162 |
| 11 | 114684800 | 114688399 | 19.98755 | 33.44513 | 64.45645 | 28.08208 | -0.5622 | 1.17719 | 0.300173 |
| 2 | 26725200 | 26739799 | 182.6228 | 266.836 | 420.3195 | 173.1649 | -0.42398 | 3.922605 | 0.300175 |
| 11 | 16389000 | 16393999 | 59.86095 | 94.145 | 161.7591 | 67.10573 | -0.45899 | 2.526112 | 0.300179 |
| 7 | 92926800 | 92927399 | 5.960041 | 11.55969 | 27.50389 | 10.3749 | -0.65993 | -0.22414 | 0.300188 |
| 10 | 60078000 | 60079799 | 23.58695 | 39.1078 | 71.23054 | 30.84729 | -0.48152 | 1.295591 | 0.300196 |
| 5 | 55931200 | 55932599 | 16.21857 | 29.41745 | 24.94373 | 10.78208 | 0.649775 | 0.345907 | 0.300197 |
| 13 | 93977400 | 93982199 | 55.27318 | 87.08272 | 157.5633 | 65.38006 | -0.53147 | 2.488356 | 0.300222 |
| 3 | 36574600 | 36577799 | 20.87646 | 34.61487 | 72.13519 | 31.23977 | -0.67515 | 1.286755 | 0.300222 |
| 1 | 52062200 | 52065199 | 23.7525 | 39.33773 | 71.78093 | 31.06136 | -0.4843 | 1.309338 | 0.300233 |
| 17 | 12575200 | 12577199 | 13.42477 | 23.39653 | 47.34687 | 20.82808 | -0.6246 | 0.739134 | 0.300233 |
| 1 | 186393200 | 186394399 | 7.862886 | 14.62198 | 34.3265 | 14.35412 | -0.77597 | 0.161455 | 0.300238 |
| 6 | 132132000 | 132134399 | 45.72454 | 74.62477 | 59.0394 | 25.36981 | 0.709893 | 1.53439 | 0.300239 |
| 1 | 42296200 | 42297599 | 7.359801 | 13.92981 | 29.81069 | 12.08115 | -0.60534 | 0.033197 | 0.300246 |
| 11 | 10305400 | 10309199 | 37.57855 | 62.29465 | 51.56083 | 22.3262 | 0.652816 | 1.319248 | 0.300249 |
| 11 | 59779000 | 59781599 | 16.90282 | 29.08271 | 54.3484 | 23.83994 | -0.5141 | 0.898628 | 0.300249 |
| 11 | 83551000 | 83554399 | 16.0842 | 27.80327 | 52.20013 | 22.90085 | -0.51931 | 0.850576 | 0.300253 |
| 6 | 133546200 | 133548599 | 30.69723 | 51.35195 | 47.56178 | 20.71801 | 0.491299 | 1.195849 | 0.300255 |
| 7 | 31225600 | 31227599 | 35.00658 | 58.39446 | 48.69931 | 21.17547 | 0.642215 | 1.240174 | 0.300257 |
| 4 | 152789000 | 152812599 | 216.0133 | 309.7575 | 506.7058 | 209.249 | -0.50503 | 4.205091 | 0.30027 |
| 12 | 62377200 | 62378799 | 19.18261 | 34.14676 | 27.82157 | 12.25735 | 0.689261 | 0.484609 | 0.300282 |
| 12 | 115544600 | 115547399 | 53.33842 | 85.59466 | 77.68243 | 32.85194 | 0.488376 | 1.943372 | 0.300282 |
| 2 | 179088000 | 179125399 | 672.6499 | 883.0685 | 1234.194 | 516.4959 | -0.40616 | 5.575747 | 0.300301 |
| X | 147754400 | 147759999 | 99.49139 | 156.2695 | 122.2832 | 50.72056 | 0.672809 | 2.569536 | 0.300306 |
| 17 | 13670400 | 13679999 | 255.6276 | 361.1107 | 618.0757 | 255.8706 | -0.59209 | 4.427664 | 0.300307 |
| 12 | 11149400 | 11150199 | 6.162792 | 11.77266 | 27.79315 | 11.04526 | -0.72017 | -0.04739 | 0.300315 |
| 8 | 81981200 | 81983999 | 30.43413 | 52.32044 | 36.91133 | 16.19075 | 0.885305 | 0.94134 | 0.300324 |
| 5 | 98327400 | 98329399 | 17.68657 | 29.86225 | 61.76444 | 27.0173 | -0.66344 | 1.079075 | 0.300347 |
| 19 | 16151200 | 16153399 | 20.35135 | 35.83846 | 30.51823 | 13.50953 | 0.619064 | 0.593773 | 0.300348 |
| 7 | 60532000 | 60534199 | 32.67118 | 54.64609 | 48.10229 | 20.95153 | 0.564449 | 1.204001 | 0.300354 |
| 9 | 123657200 | 123659399 | 12.14438 | 21.26594 | 44.15151 | 19.48945 | -0.66076 | 0.654658 | 0.300356 |
| 1 | 153137400 | 153139399 | 7.507297 | 13.77722 | 34.5745 | 14.72203 | -0.89015 | 0.236379 | 0.300363 |
| 11 | 65678200 | 65679399 | 6.007701 | 11.50019 | 27.28808 | 10.78426 | -0.71955 | -0.06762 | 0.300372 |
| 5 | 32402200 | 32404999 | 24.28105 | 40.09058 | 73.32131 | 31.66116 | -0.48792 | 1.348243 | 0.300378 |
| 9 | 32253400 | 32257999 | 31.5996 | 51.04602 | 93.75251 | 40.01769 | -0.51491 | 1.718994 | 0.300381 |
| 3 | 10705400 | 10706799 | 11.56908 | 20.66315 | 40.95602 | 17.99307 | -0.58318 | 0.440635 | 0.300431 |
| 7 | 24867800 | 24869399 | 6.873564 | 12.75386 | 32.61945 | 13.68303 | -0.90118 | 0.164236 | 0.300448 |
| 16 | 62558400 | 62559799 | 13.44156 | 25.1438 | 20.55464 | 8.509937 | 0.746077 | 0.137039 | 0.300451 |
| 15 | 64021000 | 64022999 | 7.427198 | 13.83465 | 31.6198 | 13.18525 | -0.73364 | 0.143092 | 0.30046 |

| | | | | | | | | | |
|---|---|---|---|---|---|---|---|---|---|
| 13 | 90735400 | 90737599 | 17.58862 | 31.4497 | 26.89503 | 11.80454 | 0.623336 | 0.443758 | 0.300472 |
| 14 | 3250600 | 3256999 | 81.5129 | 129.3558 | 103.8864 | 43.27646 | 0.64728 | 2.321245 | 0.300481 |
| 18 | 36811800 | 36812599 | 4.285023 | 8.32242 | 24.4465 | 9.144424 | -0.94799 | -0.28264 | 0.300495 |
| 11 | 33856600 | 33864999 | 66.14141 | 103.6662 | 175.2563 | 72.69139 | -0.43954 | 2.596754 | 0.300502 |
| 7 | 20572200 | 20575199 | 36.99224 | 62.19061 | 41.88839 | 18.29918 | 0.949069 | 1.161374 | 0.300509 |
| 13 | 16191800 | 16193599 | 11.90914 | 21.07726 | 40.86164 | 18.07013 | -0.55863 | 0.508185 | 0.30052 |
| 1 | 169901400 | 169905199 | 37.17386 | 59.69379 | 106.7601 | 45.19397 | -0.48716 | 1.915426 | 0.300523 |
| 7 | 26938000 | 26939599 | 8.535066 | 15.63866 | 34.01294 | 14.51404 | -0.68756 | 0.252734 | 0.300548 |
| 10 | 40921800 | 40923599 | 10.47676 | 18.79975 | 37.37959 | 16.35735 | -0.58327 | 0.398696 | 0.300553 |
| 15 | 22742400 | 22743599 | 13.65308 | 25.86628 | 20.72177 | 8.380415 | 0.803274 | 0.059632 | 0.300556 |
| 11 | 12413800 | 12415599 | 12.4644 | 21.74026 | 46.81187 | 20.60313 | -0.71365 | 0.71959 | 0.300564 |
| 10 | 61729000 | 61731999 | 20.93541 | 34.58684 | 71.7757 | 31.00629 | -0.66944 | 1.331253 | 0.300599 |
| 14 | 6333400 | 6336999 | 28.25445 | 46.05282 | 83.84698 | 36.01281 | -0.49072 | 1.531234 | 0.300608 |
| 1 | 34544400 | 34545199 | 4.888189 | 9.386542 | 27.34149 | 10.70857 | -1.00002 | -0.11034 | 0.300614 |
| 4 | 151736200 | 151741799 | 37.18175 | 59.6675 | 108.2559 | 45.78172 | -0.50895 | 1.936182 | 0.300618 |
| 3 | 16843400 | 16845799 | 24.57804 | 42.91604 | 33.10631 | 14.58106 | 0.760158 | 0.743211 | 0.300621 |
| 18 | 43141800 | 43145799 | 45.40009 | 73.6426 | 66.04534 | 28.29508 | 0.522664 | 1.635322 | 0.300625 |
| X | 75198000 | 75201399 | 55.4803 | 89.27339 | 75.41916 | 32.0225 | 0.596303 | 1.865635 | 0.300625 |
| 2 | 89178800 | 89180799 | 26.51258 | 46.3206 | 30.7119 | 13.53137 | 0.977494 | 0.74138 | 0.300629 |
| 9 | 40920400 | 40921399 | 7.21004 | 13.68063 | 29.31562 | 11.81766 | -0.60019 | 0.014023 | 0.300637 |
| 17 | 29019400 | 29020799 | 9.083902 | 16.03133 | 43.55129 | 19.27053 | -1.04391 | 0.52652 | 0.300647 |
| 6 | 124949400 | 124951199 | 12.74467 | 22.46308 | 43.49043 | 19.24854 | -0.5591 | 0.564661 | 0.300668 |
| 11 | 11063200 | 11066399 | 33.23863 | 53.5584 | 99.82478 | 42.4701 | -0.54039 | 1.797855 | 0.300672 |
| 7 | 78545600 | 78550799 | 51.95432 | 82.00252 | 153.8313 | 63.85971 | -0.58147 | 2.446124 | 0.300672 |
| 7 | 12055200 | 12055799 | 11.51716 | 22.29058 | 18.06774 | 6.883303 | 0.865862 | -0.15658 | 0.300677 |
| 9 | 98548600 | 98552399 | 37.54188 | 60.1183 | 113.8574 | 47.99618 | -0.5744 | 2.004437 | 0.300689 |
| 4 | 93046600 | 93047199 | 13.56955 | 25.08118 | 21.91481 | 9.241134 | 0.631477 | 0.214923 | 0.300708 |
| 7 | 41573600 | 41576999 | 63.21006 | 101.3738 | 81.51236 | 34.43941 | 0.661691 | 1.977745 | 0.300711 |
| 7 | 8455200 | 8465199 | 62.42236 | 100.37 | 67.2848 | 28.53827 | 0.927884 | 1.8918 | 0.300748 |
| 7 | 43055800 | 43067799 | 92.3217 | 141.2173 | 234.0677 | 97.11486 | -0.43831 | 3.060559 | 0.300758 |
| 6 | 39787400 | 39788999 | 11.42349 | 20.35066 | 39.35466 | 17.32712 | -0.54985 | 0.453444 | 0.30077 |
| 11 | 15679200 | 15679999 | 11.42349 | 20.35066 | 39.35466 | 17.32712 | -0.54985 | 0.453444 | 0.30077 |
| 1 | 75649600 | 75654799 | 41.07444 | 65.82764 | 113.667 | 47.97403 | -0.44004 | 1.983434 | 0.300778 |
| 6 | 46088800 | 46089399 | 5.51929 | 10.71269 | 25.06224 | 9.421545 | -0.63031 | -0.24661 | 0.300783 |
| 15 | 10651800 | 10653199 | 7.802813 | 14.76762 | 33.91079 | 13.7441 | -0.70702 | 0.045763 | 0.300788 |
| 4 | 61278200 | 61280999 | 24.06992 | 42.14935 | 31.38027 | 13.85213 | 0.811062 | 0.69434 | 0.300798 |
| 7 | 33594800 | 33604799 | 186.2481 | 278.7356 | 240.6885 | 99.26415 | 0.475042 | 3.514915 | 0.300812 |
| X | 4738200 | 4741999 | 50.00592 | 81.70772 | 54.10726 | 23.26034 | 0.967043 | 1.526119 | 0.300822 |
| 2 | 13549000 | 13549599 | 4.108905 | 8.010276 | 21.95806 | 7.923236 | -0.80075 | -0.42143 | 0.300826 |
| 11 | 30323600 | 30325599 | 42.2108 | 68.5927 | 63.65705 | 27.30812 | 0.475807 | 1.591289 | 0.300827 |
| 1 | 49539000 | 49540599 | 23.90917 | 41.51235 | 36.61725 | 16.07007 | 0.567053 | 0.823283 | 0.300839 |
| 8 | 94723200 | 94725799 | 13.22511 | 23.26706 | 46.12556 | 20.42233 | -0.59427 | 0.591504 | 0.300841 |
| 19 | 34874600 | 34875599 | 5.876808 | 11.24947 | 28.43029 | 11.23609 | -0.81049 | -0.06856 | 0.300857 |
| 11 | 111214000 | 111215599 | 11.22887 | 20.11036 | 39.93563 | 17.50444 | -0.58358 | 0.418658 | 0.300861 |
| 14 | 93259800 | 93261199 | 11.22887 | 20.11036 | 39.93563 | 17.50444 | -0.58358 | 0.418658 | 0.300861 |
| 4 | 121538400 | 121539799 | 30.86268 | 51.78013 | 46.97254 | 20.49917 | 0.521064 | 1.165154 | 0.300866 |
| 13 | 25232200 | 25241199 | 97.49292 | 148.5154 | 246.7439 | 102.2735 | -0.44642 | 3.146359 | 0.300877 |

| | | | | | | | | | |
|---|---|---|---|---|---|---|---|---|---|
| 12 | 21236200 | 21239599 | 18.25007 | 31.08866 | 57.52606 | 25.21608 | -0.50237 | 0.99927 | 0.300879 |
| 6 | 145359800 | 145364799 | 35.52353 | 57.08838 | 108.3986 | 45.94071 | -0.57198 | 1.887398 | 0.30088 |
| X | 125534000 | 125536199 | 53.52227 | 86.37073 | 70.01461 | 29.84555 | 0.66066 | 1.775178 | 0.300885 |
| 14 | 13980800 | 13983599 | 13.9809 | 24.49167 | 47.25771 | 20.86767 | -0.55615 | 0.660221 | 0.300898 |
| 18 | 4021600 | 4022799 | 11.05926 | 21.142 | 19.10231 | 7.578076 | 0.656017 | -0.02905 | 0.300906 |
| 1 | 93034600 | 93037399 | 37.36973 | 61.24663 | 56.97735 | 24.52312 | 0.480423 | 1.456104 | 0.300915 |
| X | 29802400 | 29803999 | 12.59141 | 23.94394 | 20.92227 | 8.395095 | 0.689044 | 0.025765 | 0.300922 |
| 10 | 52764800 | 52766599 | 17.08196 | 31.8361 | 20.69508 | 8.621679 | 1.067474 | 0.193091 | 0.300948 |
| 17 | 22596800 | 22604999 | 173.6397 | 263.1298 | 188.4964 | 78.00724 | 0.760049 | 3.268627 | 0.300951 |
| 1 | 55817200 | 55817999 | 10.03196 | 19.3396 | 18.60615 | 7.205647 | 0.598022 | -0.11946 | 0.300952 |
| 2 | 74564200 | 74567399 | 19.00028 | 31.89403 | 63.94292 | 27.91409 | -0.61902 | 1.138384 | 0.300961 |
| 18 | 56406200 | 56408399 | 22.36442 | 39.49446 | 27.44871 | 12.14391 | 0.911828 | 0.569382 | 0.300974 |
| 3 | 122024800 | 122025399 | 4.483393 | 8.743122 | 22.02832 | 7.915489 | -0.67474 | -0.42892 | 0.300982 |
| 17 | 38170400 | 38178599 | 115.9698 | 178.5759 | 164.3305 | 68.25445 | 0.417231 | 2.931141 | 0.300988 |
| 2 | 5340000 | 5340999 | 8.52013 | 15.68614 | 33.03066 | 14.00345 | -0.6342 | 0.224248 | 0.300997 |
| 3 | 71030400 | 71032599 | 19.52804 | 32.68897 | 70.01511 | 30.51134 | -0.71406 | 1.176838 | 0.300999 |
| 1 | 182175800 | 182179999 | 44.8026 | 72.54071 | 67.36894 | 28.82328 | 0.470003 | 1.672108 | 0.301015 |
| 2 | 13984400 | 13985799 | 21.53214 | 37.56584 | 33.94845 | 14.94403 | 0.53276 | 0.739853 | 0.301025 |
| 18 | 82202600 | 82212599 | 107.3144 | 161.6766 | 287.868 | 118.9169 | -0.55969 | 3.375673 | 0.301041 |
| 14 | 40768000 | 40770199 | 21.60921 | 38.12834 | 29.35293 | 12.98858 | 0.764947 | 0.579161 | 0.301042 |
| 3 | 15185400 | 15186399 | 6.888083 | 13.13811 | 28.9198 | 11.53014 | -0.62478 | -0.02775 | 0.301059 |
| 18 | 5038400 | 5040799 | 27.88093 | 47.0573 | 42.73093 | 18.68949 | 0.519814 | 1.097798 | 0.301064 |
| X | 24349800 | 24350599 | 18.57961 | 33.17083 | 26.65736 | 11.71534 | 0.711734 | 0.462864 | 0.301074 |
| 6 | 101127400 | 101132199 | 50.74727 | 80.29427 | 147.2462 | 61.20201 | -0.54591 | 2.385553 | 0.301078 |
| 2 | 181374600 | 181376199 | 16.66077 | 29.93931 | 26.53136 | 11.59615 | 0.576438 | 0.406704 | 0.301091 |
| 1 | 87434800 | 87441199 | 49.8171 | 79.07275 | 142.851 | 59.52026 | -0.52026 | 2.297711 | 0.301134 |
| 17 | 18682400 | 18685199 | 38.3459 | 64.14194 | 44.20414 | 19.25444 | 0.916319 | 1.218344 | 0.301144 |
| 10 | 82630600 | 82631999 | 9.585796 | 17.09311 | 39.92676 | 17.56491 | -0.81987 | 0.434582 | 0.301151 |
| 14 | 76979200 | 76980399 | 15.12783 | 27.79261 | 24.10864 | 10.25034 | 0.633092 | 0.264804 | 0.301152 |
| 15 | 17994800 | 17996199 | 15.12783 | 27.79261 | 24.10864 | 10.25034 | 0.633092 | 0.264804 | 0.301152 |
| 9 | 33187800 | 33190199 | 28.54139 | 49.09004 | 38.06882 | 16.69871 | 0.749493 | 0.930008 | 0.301158 |
| 15 | 11567000 | 11567999 | 7.176784 | 13.72987 | 30.84023 | 12.20905 | -0.64477 | -0.04352 | 0.301171 |
| 5 | 9083400 | 9084999 | 13.97858 | 25.80199 | 23.23872 | 9.817603 | 0.58583 | 0.227789 | 0.301175 |
| 12 | 113847200 | 113853799 | 88.80133 | 140.0673 | 118.5422 | 49.19585 | 0.564488 | 2.479602 | 0.301181 |
| 2 | 52835600 | 52839399 | 18.91198 | 31.95714 | 64.70847 | 28.31206 | -0.63304 | 1.077881 | 0.301192 |
| 17 | 18436200 | 18440599 | 99.89717 | 155.7525 | 140.3647 | 58.25162 | 0.461104 | 2.701424 | 0.301202 |
| 18 | 44389800 | 44399599 | 83.46749 | 131.6961 | 109.9831 | 45.6491 | 0.584003 | 2.465974 | 0.301204 |
| 15 | 25646200 | 25649399 | 29.27745 | 47.63185 | 85.89307 | 36.8667 | -0.47878 | 1.563255 | 0.301209 |
| 3 | 153357200 | 153360799 | 19.58546 | 32.59677 | 66.96007 | 29.07403 | -0.65381 | 1.237278 | 0.301245 |
| 2 | 150157400 | 150157999 | 22.28574 | 38.71713 | 34.7388 | 15.2515 | 0.542702 | 0.805152 | 0.301245 |
| 4 | 40849600 | 40850799 | 6.050764 | 11.48006 | 28.72472 | 11.5814 | -0.8217 | 0.001677 | 0.301262 |
| X | 31653000 | 31655399 | 50.4548 | 81.85547 | 63.01369 | 26.9996 | 0.742698 | 1.6414 | 0.301266 |
| 12 | 19446400 | 19447799 | 24.95474 | 43.26085 | 29.66993 | 13.01771 | 0.928114 | 0.834188 | 0.301266 |
| 19 | 17396200 | 17398399 | 11.94491 | 21.00009 | 42.47439 | 18.80839 | -0.62325 | 0.584945 | 0.301271 |
| 16 | 94795800 | 94799199 | 24.20087 | 39.66439 | 77.31413 | 33.27634 | -0.5836 | 1.441761 | 0.301296 |
| 12 | 11141400 | 11143399 | 15.79239 | 27.25959 | 51.63735 | 22.65295 | -0.53245 | 0.862352 | 0.301315 |
| 1 | 157559600 | 157560199 | 4.521576 | 8.809595 | 22.49447 | 8.179489 | -0.7101 | -0.38639 | 0.301319 |

| | | | | | | | | | |
|---|---|---|---|---|---|---|---|---|---|
| X | 125903600 | 125907599 | 77.44592 | 123.2019 | 97.94155 | 40.90482 | 0.665391 | 2.247633 | 0.30132 |
| 14 | 88092400 | 88094199 | 23.46143 | 41.11954 | 31.4272 | 13.88304 | 0.773405 | 0.677952 | 0.301326 |
| 2 | 174969000 | 174971199 | 15.66149 | 28.81123 | 24.07537 | 10.21775 | 0.688575 | 0.262036 | 0.301329 |
| 6 | 130477400 | 130478199 | 23.03914 | 40.07528 | 35.14669 | 15.44251 | 0.575647 | 0.792956 | 0.301339 |
| 5 | 109314400 | 109322799 | 150.3867 | 228.2144 | 200.1645 | 82.8715 | 0.469916 | 3.234861 | 0.30135 |
| 12 | 13099600 | 13100599 | 7.039837 | 13.38693 | 30.55764 | 12.26147 | -0.68534 | -0.00244 | 0.301356 |
| 4 | 55136800 | 55138199 | 13.24977 | 25.17155 | 20.84964 | 8.383069 | 0.763024 | 0.038349 | 0.301359 |
| 13 | 31409000 | 31410599 | 7.848117 | 14.83897 | 35.58461 | 14.45209 | -0.77139 | 0.059224 | 0.301363 |
| 4 | 103264000 | 103265199 | 14.71972 | 27.4595 | 22.04994 | 9.171407 | 0.766715 | 0.169883 | 0.301368 |
| 7 | 79873400 | 79875999 | 13.25162 | 23.0422 | 48.4226 | 21.27081 | -0.67933 | 0.770022 | 0.301372 |
| 5 | 70151600 | 70153599 | 23.525 | 40.90639 | 35.61626 | 15.64546 | 0.586008 | 0.800521 | 0.301373 |
| 14 | 120985600 | 120988199 | 11.01398 | 19.49679 | 46.70266 | 20.65239 | -0.86204 | 0.511576 | 0.30138 |
| 5 | 100435400 | 100437199 | 27.78372 | 48.26789 | 34.18365 | 15.00759 | 0.882458 | 0.830532 | 0.301393 |
| 10 | 19809200 | 19811399 | 15.63291 | 26.22005 | 66.91504 | 29.19647 | -0.9657 | 1.139217 | 0.301404 |
| 9 | 64718200 | 64719599 | 7.942345 | 14.22932 | 37.97311 | 16.63934 | -1.00602 | 0.39157 | 0.301408 |
| 15 | 98673400 | 98678999 | 45.31867 | 71.9596 | 137.5271 | 57.32357 | -0.60072 | 2.2834 | 0.301408 |
| 2 | 18036000 | 18038799 | 13.7191 | 23.86363 | 47.89533 | 21.04394 | -0.61323 | 0.767109 | 0.301409 |
| 5 | 145635400 | 145638999 | 30.97346 | 53.32949 | 35.78224 | 15.69622 | 0.957801 | 0.929269 | 0.301418 |
| 4 | 111029400 | 111032599 | 31.6557 | 54.0638 | 39.15539 | 17.167 | 0.846265 | 1.006928 | 0.301429 |
| 8 | 61445600 | 61446799 | 7.014448 | 13.01442 | 32.08968 | 13.462 | -0.84876 | 0.165211 | 0.301454 |
| X | 121956800 | 121957999 | 15.47232 | 30.03391 | 12.87132 | 4.836158 | 1.792141 | -0.17261 | 0.301471 |
| 10 | 47328600 | 47330599 | 11.87759 | 20.68177 | 47.35587 | 20.82042 | -0.80188 | 0.743903 | 0.301485 |
| 12 | 84727000 | 84729199 | 12.15067 | 21.28789 | 44.14347 | 19.49436 | -0.65894 | 0.647076 | 0.301502 |
| 14 | 121324000 | 121325199 | 17.4037 | 29.849 | 55.41296 | 24.30311 | -0.5055 | 0.93114 | 0.301512 |
| 2 | 11453200 | 11455599 | 17.78853 | 31.73559 | 27.42376 | 12.05949 | 0.606598 | 0.46027 | 0.301514 |
| 3 | 23344000 | 23345799 | 22.19054 | 38.69676 | 34.41532 | 15.14518 | 0.555683 | 0.750254 | 0.301517 |
| 13 | 42204600 | 42206999 | 15.69963 | 26.90464 | 53.70937 | 23.55442 | -0.60979 | 0.930924 | 0.301518 |
| 14 | 96387800 | 96388999 | 12.79783 | 23.94656 | 21.94529 | 9.081245 | 0.583861 | 0.135997 | 0.30152 |
| 17 | 17537600 | 17543199 | 134.8727 | 207.8882 | 148.621 | 61.73471 | 0.782191 | 2.919731 | 0.30152 |
| 5 | 99951200 | 99953199 | 12.15089 | 21.20948 | 47.32351 | 20.83822 | -0.76468 | 0.707012 | 0.301521 |
| 10 | 103520000 | 103520999 | 12.15089 | 21.20948 | 47.32351 | 20.83822 | -0.76468 | 0.707012 | 0.301521 |
| 2 | 80276600 | 80279599 | 18.75696 | 31.71588 | 61.88324 | 27.08128 | -0.57963 | 1.069791 | 0.301532 |
| 19 | 11776200 | 11778399 | 10.89448 | 19.62235 | 40.58461 | 17.69759 | -0.63691 | 0.382043 | 0.301533 |
| 2 | 172584200 | 172585199 | 8.410119 | 15.40613 | 34.02841 | 14.52525 | -0.7094 | 0.256618 | 0.301539 |
| 4 | 42999600 | 43000599 | 5.279825 | 10.19064 | 26.43534 | 10.20131 | -0.81463 | -0.15825 | 0.30156 |
| 19 | 56312200 | 56313399 | 6.479254 | 12.28301 | 28.70991 | 11.5954 | -0.72629 | 0.013046 | 0.301574 |
| 19 | 17345200 | 17346399 | 7.644415 | 14.48138 | 32.17419 | 13.01447 | -0.65711 | 0.033797 | 0.301589 |
| 7 | 42654800 | 42656599 | 21.46865 | 37.43044 | 34.08369 | 14.99736 | 0.521898 | 0.750176 | 0.301595 |
| 16 | 91352000 | 91359599 | 38.27054 | 61.06574 | 125.9859 | 52.83833 | -0.70342 | 2.119237 | 0.30161 |
| 11 | 117063200 | 117075599 | 115.2211 | 173.735 | 286.4838 | 118.3719 | -0.44798 | 3.360357 | 0.301629 |
| 17 | 17408600 | 17412599 | 85.91036 | 135.7205 | 114.419 | 47.52036 | 0.571868 | 2.441649 | 0.301636 |
| 3 | 11921400 | 11923199 | 20.85234 | 37.2506 | 26.02682 | 11.43249 | 0.912545 | 0.470859 | 0.301643 |
| 6 | 11838600 | 11841199 | 16.86087 | 30.73782 | 24.976 | 10.72782 | 0.716907 | 0.323597 | 0.301669 |
| 1 | 10713000 | 10714399 | 21.1516 | 38.06585 | 24.48646 | 10.68856 | 1.037191 | 0.426292 | 0.301673 |
| 5 | 73013000 | 73017399 | 43.84304 | 70.13975 | 121.7232 | 51.23912 | -0.4503 | 2.043563 | 0.301675 |
| 5 | 77122000 | 77122999 | 5.770277 | 10.973 | 29.89345 | 11.99438 | -0.93761 | -0.01597 | 0.301678 |
| 4 | 84840800 | 84842599 | 11.76301 | 20.78267 | 40.88522 | 18.08681 | -0.5806 | 0.526482 | 0.301681 |

| | | | | | | | | | |
|---|---|---|---|---|---|---|---|---|---|
| 3 | 91928000 | 91930599 | 31.1721 | 52.24728 | 47.00028 | 20.49841 | 0.533241 | 1.17857 | 0.301701 |
| 17 | 17584800 | 17589999 | 71.81143 | 114.5703 | 92.04157 | 38.5953 | 0.654661 | 2.147831 | 0.301706 |
| 12 | 18719200 | 18757799 | 1296.016 | 1643.52 | 1265.255 | 531.3259 | 0.403991 | 6.140185 | 0.30171 |
| 16 | 22137800 | 22142399 | 41.65862 | 68.18453 | 58.37463 | 25.11043 | 0.599163 | 1.483479 | 0.301732 |
| 17 | 40188800 | 40189799 | 11.95801 | 22.99395 | 19.47527 | 7.597651 | 0.770354 | -0.07345 | 0.301737 |
| 7 | 126417800 | 126419199 | 15.23532 | 27.62238 | 24.77494 | 10.71476 | 0.569089 | 0.342401 | 0.301742 |
| 12 | 21676600 | 21687399 | 357.8746 | 504.3714 | 424.6352 | 175.858 | 0.427774 | 4.463958 | 0.301744 |
| 19 | 52369600 | 52372599 | 21.21359 | 37.49095 | 28.102 | 12.43103 | 0.803818 | 0.555315 | 0.301754 |
| 3 | 130902600 | 130906599 | 21.22135 | 34.76469 | 78.46151 | 33.76345 | -0.79459 | 1.441878 | 0.301758 |
| 5 | 95482000 | 95483599 | 16.54886 | 30.62475 | 22.98043 | 9.674658 | 0.851076 | 0.234815 | 0.301764 |
| 1 | 27382400 | 27384199 | 19.48966 | 34.94514 | 26.16237 | 11.4563 | 0.816443 | 0.439314 | 0.301765 |
| 7 | 59002200 | 59003199 | 20.82634 | 36.45912 | 32.86316 | 14.50325 | 0.53667 | 0.684304 | 0.301782 |
| 7 | 35729400 | 35732199 | 24.97044 | 41.02021 | 75.89525 | 32.69755 | -0.50728 | 1.414154 | 0.301783 |
| 14 | 14084600 | 14086399 | 38.74678 | 63.22472 | 57.94315 | 24.89702 | 0.498132 | 1.52068 | 0.301793 |
| 2 | 71042000 | 71042799 | 4.238678 | 8.238479 | 23.33918 | 8.678404 | -0.88719 | -0.30536 | 0.301801 |
| 7 | 68044000 | 68046199 | 13.02571 | 22.80552 | 47.27077 | 20.86349 | -0.65897 | 0.662957 | 0.301815 |
| 5 | 54717200 | 54721799 | 54.58374 | 88.23904 | 65.01029 | 27.77541 | 0.801005 | 1.721608 | 0.301816 |
| 7 | 20563000 | 20570599 | 148.2561 | 224.8949 | 201.6936 | 83.495 | 0.437373 | 3.242298 | 0.301831 |
| 13 | 60211000 | 60217799 | 37.94998 | 60.9481 | 109.2515 | 46.2579 | -0.49026 | 1.911814 | 0.301854 |
| 2 | 121512200 | 121513399 | 6.89455 | 13.10845 | 31.7216 | 12.73305 | -0.76959 | -0.0007 | 0.301855 |
| 2 | 87155400 | 87156999 | 23.99183 | 41.42968 | 37.74432 | 16.55838 | 0.519113 | 0.874023 | 0.30192 |
| 15 | 4008600 | 4009799 | 7.882658 | 14.69922 | 31.64014 | 13.16914 | -0.64529 | 0.142316 | 0.301921 |
| 7 | 95972200 | 95975399 | 54.4881 | 86.19414 | 147.9063 | 61.46955 | -0.4505 | 2.393632 | 0.301924 |
| 4 | 63353200 | 63361799 | 72.60368 | 112.6175 | 192.7557 | 80.05547 | -0.46903 | 2.78386 | 0.301925 |
| 16 | 34576400 | 34579199 | 12.1255 | 21.23385 | 44.65146 | 19.7109 | -0.6793 | 0.652549 | 0.301929 |
| X | 54742200 | 54744399 | 41.80394 | 68.99054 | 51.06816 | 22.04484 | 0.813135 | 1.383266 | 0.301942 |
| 12 | 34005800 | 34008999 | 25.32902 | 41.08449 | 88.819 | 38.05092 | -0.74345 | 1.61129 | 0.301947 |
| 7 | 38641400 | 38641599 | 7.867553 | 15.35067 | 15.19376 | 5.42046 | 0.666668 | -0.43468 | 0.301948 |
| 7 | 31893600 | 31895399 | 21.00846 | 37.21495 | 26.96915 | 11.92436 | 0.853378 | 0.527023 | 0.301963 |
| 7 | 15904800 | 15913799 | 64.42767 | 100.813 | 172.9273 | 71.73762 | -0.46231 | 2.625643 | 0.301976 |
| 13 | 7810800 | 7811399 | 16.86299 | 30.03135 | 26.70171 | 11.75945 | 0.56438 | 0.464326 | 0.301994 |
| 12 | 119031800 | 119032599 | 6.148188 | 11.78819 | 27.36697 | 10.76525 | -0.68245 | -0.08285 | 0.302008 |
| 14 | 34489600 | 34492799 | 23.01664 | 37.92586 | 73.61685 | 31.75682 | -0.57444 | 1.369279 | 0.302016 |
| 2 | 19718200 | 19718599 | 4.524864 | 8.806262 | 22.92486 | 8.434544 | -0.7531 | -0.34228 | 0.302017 |
| 2 | 42542200 | 42543799 | 15.89877 | 27.07352 | 57.41853 | 25.16821 | -0.69836 | 0.992831 | 0.302026 |
| 4 | 144317600 | 144320599 | 57.43947 | 92.17868 | 83.01621 | 35.10927 | 0.499543 | 1.951214 | 0.302039 |
| 14 | 42266600 | 42275399 | 155.6922 | 235.3283 | 212.6741 | 87.94174 | 0.421956 | 3.316104 | 0.302043 |
| 6 | 118209600 | 118211199 | 24.9108 | 43.09357 | 37.68241 | 16.53212 | 0.578604 | 0.863325 | 0.302054 |
| 13 | 25089400 | 25090799 | 7.701625 | 14.00626 | 34.99816 | 15.07826 | -0.89535 | 0.292977 | 0.302055 |
| 11 | 28541000 | 28541399 | 3.936322 | 7.681133 | 20.90107 | 7.442672 | -0.77189 | -0.47325 | 0.30207 |
| 2 | 163043400 | 163046199 | 30.75983 | 49.8517 | 89.04208 | 38.12296 | -0.46998 | 1.639518 | 0.302082 |
| 17 | 78590800 | 78594599 | 19.11024 | 32.19782 | 62.94236 | 27.5149 | -0.58257 | 1.103455 | 0.302083 |
| 2 | 71173600 | 71174799 | 16.93609 | 30.30454 | 27.16152 | 11.91434 | 0.556842 | 0.437305 | 0.302084 |
| 18 | 74646600 | 74651599 | 35.31671 | 56.97416 | 114.1714 | 48.64913 | -0.64287 | 1.771116 | 0.302085 |
| 8 | 75156400 | 75157799 | 7.801816 | 14.76899 | 33.32527 | 13.50048 | -0.68129 | 0.043034 | 0.302103 |
| 18 | 89772800 | 89773399 | 10.15553 | 19.74757 | 16.41014 | 6.079868 | 0.865919 | -0.27354 | 0.30211 |
| 9 | 8171400 | 8174999 | 56.50958 | 90.78031 | 80.6815 | 34.17537 | 0.520372 | 1.918208 | 0.302135 |

| | | | | | | | | | |
|---|---|---|---|---|---|---|---|---|---|
| 5 | 60821400 | 60821999 | 5.359642 | 10.42435 | 25.32729 | 9.365847 | -0.66211 | -0.30558 | 0.302147 |
| 7 | 59872600 | 59874399 | 19.9679 | 35.27636 | 29.54413 | 13.06993 | 0.64476 | 0.556497 | 0.302148 |
| 8 | 8562400 | 8568199 | 36.64054 | 58.92974 | 105.3242 | 44.68859 | -0.48344 | 1.863567 | 0.302158 |
| 8 | 107453000 | 107454399 | 11.07608 | 19.88294 | 38.91587 | 17.02403 | -0.5607 | 0.401814 | 0.302161 |
| 11 | 12466200 | 12467399 | 6.160829 | 11.78575 | 29.58693 | 11.7143 | -0.80384 | -0.05772 | 0.302172 |
| 12 | 20371400 | 20425599 | 1397.441 | 1760.909 | 1346.958 | 566.0064 | 0.403989 | 6.243086 | 0.302173 |
| 12 | 109186800 | 109190799 | 44.89609 | 73.26575 | 58.52623 | 25.14843 | 0.695858 | 1.531589 | 0.302179 |
| 1 | 189122600 | 189123799 | 9.460242 | 17.36964 | 35.11522 | 14.94241 | -0.57995 | 0.252019 | 0.30219 |
| 2 | 98878600 | 98881399 | 37.86951 | 62.70263 | 52.14443 | 22.56043 | 0.645888 | 1.333313 | 0.30219 |
| 10 | 26406800 | 26408199 | 15.03663 | 28.64073 | 18.99383 | 7.594919 | 1.087808 | 0.033102 | 0.302196 |
| 7 | 12411600 | 12419199 | 78.52146 | 124.2646 | 113.8423 | 47.3227 | 0.454076 | 2.393056 | 0.302199 |
| X | 148693200 | 148695199 | 24.42384 | 43.25781 | 23.95071 | 10.59012 | 1.239227 | 0.545811 | 0.302203 |
| 6 | 59185000 | 59186199 | 8.997847 | 16.43455 | 34.48591 | 14.77737 | -0.64053 | 0.275555 | 0.302203 |
| 2 | 150135600 | 150136999 | 19.10007 | 34.33625 | 25.63011 | 11.19852 | 0.822748 | 0.41931 | 0.302205 |
| 2 | 98284200 | 98289799 | 106.9346 | 166.3575 | 142.4084 | 59.11925 | 0.532743 | 2.742676 | 0.302209 |
| 7 | 62044800 | 62046999 | 24.99376 | 44.2464 | 23.94646 | 10.58923 | 1.27172 | 0.554685 | 0.302212 |
| X | 24112200 | 24118999 | 115.7642 | 178.8965 | 150.2954 | 62.4294 | 0.552415 | 2.866154 | 0.302212 |
| 2 | 61000600 | 61001999 | 10.82017 | 19.19839 | 39.34675 | 17.38256 | -0.63653 | 0.474458 | 0.302216 |
| 16 | 56536400 | 56538199 | 10.82017 | 19.19839 | 39.34675 | 17.38256 | -0.63653 | 0.474458 | 0.302216 |
| 3 | 110224000 | 110224799 | 11.33823 | 22.02061 | 16.8795 | 6.315379 | 0.968845 | -0.22366 | 0.302236 |
| 5 | 124562800 | 124564199 | 7.931007 | 14.58441 | 33.02455 | 14.02323 | -0.73977 | 0.226181 | 0.302237 |
| 17 | 19693800 | 19703399 | 129.7427 | 199.7588 | 157.8025 | 65.54193 | 0.637089 | 2.939057 | 0.302256 |
| 4 | 4553200 | 4553799 | 17.29212 | 31.403 | 22.92078 | 9.892521 | 0.866234 | 0.343928 | 0.302258 |
| X | 108327200 | 108327999 | 11.03193 | 21.4109 | 17.59135 | 6.615307 | 0.864058 | -0.20586 | 0.302275 |
| 3 | 67544800 | 67546399 | 15.5682 | 29.00484 | 21.95528 | 9.152031 | 0.848841 | 0.185124 | 0.302284 |
| 19 | 29077600 | 29079399 | 12.57801 | 22.03378 | 44.6324 | 19.70816 | -0.62569 | 0.649134 | 0.302289 |
| 5 | 21305200 | 21305799 | 17.50392 | 31.19159 | 26.77457 | 11.78554 | 0.615572 | 0.466811 | 0.3023 |
| 6 | 131536200 | 131538399 | 16.93274 | 31.01628 | 23.85903 | 10.18531 | 0.801111 | 0.294622 | 0.302303 |
| 8 | 91898200 | 91901199 | 13.7247 | 23.94739 | 46.80593 | 20.60534 | -0.57458 | 0.722433 | 0.302319 |
| 12 | 114537600 | 114538999 | 22.57599 | 37.69852 | 70.2768 | 30.57803 | -0.51451 | 1.212568 | 0.302336 |
| 7 | 32829200 | 32833599 | 63.28141 | 101.5508 | 73.03473 | 30.88649 | 0.823351 | 1.957327 | 0.302344 |
| 4 | 27857600 | 27858199 | 5.817905 | 11.28387 | 28.83079 | 10.87609 | -0.76265 | -0.22506 | 0.302349 |
| 7 | 27586400 | 27587599 | 26.92572 | 45.83831 | 42.08641 | 18.44729 | 0.504661 | 1.019588 | 0.30235 |
| 2 | 144494000 | 144495999 | 10.60099 | 19.01961 | 37.8679 | 16.57329 | -0.58563 | 0.399517 | 0.302354 |
| 13 | 31978200 | 31980199 | 10.60099 | 19.01961 | 37.8679 | 16.57329 | -0.58563 | 0.399517 | 0.302354 |
| 6 | 89205000 | 89207999 | 32.50964 | 54.63344 | 45.83788 | 20.02219 | 0.63331 | 1.153134 | 0.302361 |
| 6 | 48303800 | 48305399 | 17.27086 | 30.99814 | 27.07147 | 11.84429 | 0.596557 | 0.420649 | 0.30237 |
| X | 72095000 | 72097999 | 24.51812 | 42.94789 | 31.04124 | 13.70617 | 0.853833 | 0.693745 | 0.302372 |
| 1 | 135159600 | 135159999 | 3.880113 | 7.574044 | 20.93753 | 7.411741 | -0.78561 | -0.49239 | 0.302373 |
| 17 | 94134400 | 94134999 | 10.76423 | 20.92456 | 17.01112 | 6.318681 | 0.89463 | -0.25411 | 0.302375 |
| 1 | 87184000 | 87187199 | 20.90269 | 35.14861 | 64.66314 | 28.25027 | -0.49545 | 1.129583 | 0.302378 |
| 19 | 30449200 | 30450799 | 9.49041 | 17.20546 | 35.42797 | 15.32327 | -0.62227 | 0.325107 | 0.302386 |
| 16 | 54252600 | 54252999 | 5.061011 | 9.838435 | 24.58219 | 9.128344 | -0.70687 | -0.29507 | 0.302389 |
| 11 | 116912400 | 116920599 | 72.67759 | 112.9714 | 188.2483 | 78.15873 | -0.42803 | 2.747006 | 0.302395 |
| 19 | 47379600 | 47383399 | 35.87185 | 57.68298 | 103.393 | 43.85765 | -0.48785 | 1.869299 | 0.302395 |
| 17 | 39480600 | 39483399 | 46.21549 | 75.42591 | 53.91676 | 23.16855 | 0.855825 | 1.532859 | 0.302398 |
| 17 | 80307000 | 80310199 | 49.83123 | 80.18755 | 74.28296 | 31.5424 | 0.46368 | 1.852916 | 0.302399 |

| | | | | | | | | | |
|---|---|---|---|---|---|---|---|---|---|
| 1 | 65853000 | 65856799 | 69.10353 | 109.7179 | 103.0979 | 42.96951 | 0.42176 | 2.296098 | 0.302403 |
| 12 | 107032800 | 107035799 | 17.38122 | 29.76639 | 59.2872 | 25.99918 | -0.6071 | 0.946915 | 0.302406 |
| 3 | 85992000 | 85992599 | 4.783405 | 9.314556 | 22.99371 | 8.422053 | -0.67121 | -0.3535 | 0.302412 |
| 5 | 27720600 | 27721999 | 14.32232 | 25.04187 | 47.29649 | 20.85109 | -0.52537 | 0.69405 | 0.302424 |
| 11 | 109861000 | 109865599 | 53.65913 | 86.30055 | 78.88428 | 33.46994 | 0.481971 | 1.877358 | 0.302429 |
| 8 | 20108800 | 20114399 | 142.1099 | 211.8895 | 335.3596 | 138.2082 | -0.4034 | 3.581492 | 0.302474 |
| X | 9174600 | 9176399 | 19.41965 | 35.34829 | 23.79758 | 10.24128 | 0.985048 | 0.350247 | 0.302483 |
| 13 | 115640200 | 115641199 | 14.84048 | 25.85108 | 49.47686 | 21.75833 | -0.54489 | 0.755149 | 0.302493 |
| 5 | 132954200 | 132955999 | 16.57539 | 28.65302 | 53.84118 | 23.62081 | -0.52057 | 0.854832 | 0.302496 |
| 5 | 120776000 | 120778999 | 18.50536 | 31.15118 | 61.73621 | 26.98041 | -0.60221 | 1.109 | 0.302531 |
| 4 | 35077800 | 35079599 | 21.21694 | 37.24921 | 31.62839 | 13.99206 | 0.622325 | 0.639758 | 0.302531 |
| 4 | 23354200 | 23355599 | 15.77389 | 29.30752 | 22.47824 | 9.411913 | 0.825093 | 0.204889 | 0.302538 |
| 6 | 63667200 | 63667599 | 3.692265 | 7.221593 | 22.41973 | 7.567291 | -0.88855 | -0.65874 | 0.302539 |
| X | 149482000 | 149483399 | 15.23075 | 29.0022 | 18.98564 | 7.596224 | 1.105678 | 0.036946 | 0.302543 |
| 3 | 24645200 | 24646999 | 17.35537 | 31.2712 | 26.44631 | 11.52684 | 0.646127 | 0.395981 | 0.30256 |
| X | 24364200 | 24367799 | 36.6945 | 61.45527 | 45.3323 | 19.7612 | 0.818612 | 1.203195 | 0.302597 |
| 2 | 176853000 | 176853399 | 8.033858 | 15.67461 | 15.19025 | 5.422443 | 0.696155 | -0.42998 | 0.302606 |
| 1 | 49370400 | 49370999 | 7.907113 | 15.42858 | 14.48415 | 5.163108 | 0.741577 | -0.44296 | 0.30263 |
| 2 | 53613400 | 53613999 | 8.638186 | 16.80836 | 16.06348 | 5.919241 | 0.673759 | -0.31199 | 0.302639 |
| X | 153288800 | 153290799 | 40.41252 | 67.05242 | 48.81816 | 21.15164 | 0.837352 | 1.315242 | 0.302658 |
| 17 | 8017800 | 8022999 | 34.50568 | 55.62584 | 101.0901 | 43.03526 | -0.50325 | 1.785799 | 0.302661 |
| 13 | 28950000 | 28952599 | 30.21857 | 50.6586 | 46.99459 | 20.49836 | 0.488943 | 1.171733 | 0.302662 |
| 10 | 122360800 | 122364599 | 30.25491 | 49.10251 | 87.51262 | 37.50707 | -0.46493 | 1.609044 | 0.302685 |
| X | 121856600 | 121858399 | 27.415 | 48.18701 | 26.22837 | 11.61461 | 1.260362 | 0.649235 | 0.30269 |
| 7 | 17500600 | 17508199 | 171.2475 | 257.4908 | 226.6554 | 93.58111 | 0.453384 | 3.423333 | 0.302693 |
| 4 | 112292400 | 112296199 | 69.90788 | 111.4479 | 95.81105 | 40.13334 | 0.555897 | 2.171816 | 0.302694 |
| X | 146728200 | 146730399 | 23.44501 | 41.31455 | 28.64393 | 12.6781 | 0.914509 | 0.602443 | 0.302695 |
| 13 | 33232000 | 33236599 | 27.37806 | 44.87232 | 86.93369 | 37.41686 | -0.57507 | 1.452957 | 0.302704 |
| 9 | 114966000 | 114968999 | 17.15995 | 29.16824 | 57.44115 | 25.17445 | -0.5923 | 1.014863 | 0.302704 |
| 3 | 108081200 | 108083199 | 9.355338 | 16.88446 | 37.44699 | 16.28791 | -0.73428 | 0.352968 | 0.302712 |
| 3 | 31316200 | 31317999 | 10.95629 | 19.53905 | 38.35039 | 16.86988 | -0.57012 | 0.442137 | 0.30273 |
| 3 | 129336400 | 129338399 | 28.76936 | 48.78845 | 43.08225 | 18.87037 | 0.560104 | 1.059508 | 0.302744 |
| 18 | 54438800 | 54442599 | 46.20275 | 75.09326 | 62.69875 | 26.90062 | 0.628237 | 1.598038 | 0.302762 |
| 13 | 12917800 | 12918199 | 9.201691 | 17.87553 | 16.50377 | 6.162501 | 0.705469 | -0.25833 | 0.30277 |
| 14 | 120438200 | 120439599 | 15.40837 | 28.90814 | 21.59807 | 8.897343 | 0.881338 | 0.140906 | 0.302773 |
| 18 | 76869600 | 76874799 | 38.31813 | 61.39264 | 114.7342 | 48.40332 | -0.55441 | 1.988588 | 0.302784 |
| 8 | 69331800 | 69334399 | 13.99347 | 24.47531 | 46.22924 | 20.38652 | -0.52543 | 0.68486 | 0.302796 |
| 19 | 10176800 | 10178999 | 11.97844 | 20.97868 | 44.14034 | 19.48712 | -0.68002 | 0.649902 | 0.302822 |
| 16 | 4924400 | 4928599 | 33.00551 | 52.98536 | 107.892 | 45.65365 | -0.67438 | 1.919039 | 0.302832 |
| 17 | 62318400 | 62318999 | 4.181521 | 8.164983 | 21.0593 | 7.414249 | -0.68073 | -0.50867 | 0.302856 |
| 16 | 12597000 | 12598799 | 31.66169 | 53.2449 | 45.84038 | 20.02744 | 0.596238 | 1.144245 | 0.302862 |
| 13 | 65574800 | 65576999 | 16.49082 | 31.43113 | 17.76239 | 7.092688 | 1.318415 | 0.040052 | 0.302866 |
| 14 | 36872400 | 36877399 | 82.39192 | 131.1193 | 92.69433 | 38.73347 | 0.834977 | 2.238796 | 0.302874 |
| 12 | 34006000 | 34008999 | 24.99158 | 40.58276 | 87.24789 | 37.40884 | -0.73391 | 1.5868 | 0.30288 |
| 7 | 60305800 | 60306999 | 30.48276 | 51.38154 | 45.32026 | 19.81416 | 0.561609 | 1.121854 | 0.302888 |
| 17 | 49570800 | 49571799 | 5.114279 | 9.90744 | 24.91639 | 9.451567 | -0.74608 | -0.22232 | 0.302889 |
| 1 | 42916400 | 42918399 | 10.89531 | 19.55438 | 40.5055 | 17.72252 | -0.64243 | 0.403122 | 0.30289 |

| | | | | | | | | | |
|---|---|---|---|---|---|---|---|---|---|
| 6 | 67049600 | 67051199 | 6.402944 | 11.9952 | 32.27197 | 13.32746 | -0.95573 | 0.097318 | 0.302898 |
| 6 | 28422800 | 28426799 | 22.17497 | 36.30795 | 80.42893 | 34.60243 | -0.76816 | 1.450545 | 0.302911 |
| 9 | 113873600 | 113881999 | 89.8232 | 137.4943 | 231.2356 | 95.95577 | -0.45845 | 3.045023 | 0.302958 |
| 4 | 7284000 | 7284999 | 10.14342 | 19.61404 | 16.16396 | 6.18413 | 0.836737 | -0.16374 | 0.302982 |
| 11 | 94993000 | 94994399 | 15.61288 | 26.95361 | 52.18213 | 22.89204 | -0.56378 | 0.859791 | 0.302984 |
| 7 | 31590000 | 31594799 | 50.08193 | 81.5459 | 58.0592 | 24.92742 | 0.859283 | 1.576198 | 0.30301 |
| 19 | 13646400 | 13650799 | 49.81192 | 80.46024 | 70.71505 | 30.1814 | 0.545806 | 1.738929 | 0.303028 |
| 1 | 73900800 | 73902399 | 10.83112 | 19.21871 | 38.85555 | 17.16488 | -0.6168 | 0.474512 | 0.30303 |
| 1 | 90538600 | 90540599 | 10.83112 | 19.21871 | 38.85555 | 17.16488 | -0.6168 | 0.474512 | 0.30303 |
| 7 | 4523200 | 4528599 | 33.83469 | 54.24505 | 112.3561 | 47.44753 | -0.70123 | 1.962435 | 0.303051 |
| 13 | 65650200 | 65669799 | 464.6894 | 645.4169 | 496.9549 | 206.2794 | 0.535407 | 4.686133 | 0.303067 |
| 16 | 31537200 | 31538999 | 14.25215 | 24.87572 | 53.77564 | 23.67841 | -0.71945 | 0.731114 | 0.303068 |
| 8 | 3731000 | 3733799 | 24.66031 | 40.51328 | 75.7845 | 32.6507 | -0.523 | 1.413338 | 0.303075 |
| 15 | 3092400 | 3095199 | 16.79046 | 30.3383 | 22.62411 | 9.826378 | 0.829751 | 0.368144 | 0.303083 |
| 16 | 58058000 | 58060799 | 23.416 | 38.81176 | 71.23797 | 30.84359 | -0.49265 | 1.298233 | 0.303089 |
| 12 | 19235200 | 19246599 | 341.0372 | 484.3842 | 412.9475 | 170.7978 | 0.420966 | 4.3504 | 0.303095 |
| 6 | 95391800 | 95393399 | 16.58329 | 30.41274 | 22.71682 | 9.681849 | 0.844577 | 0.281592 | 0.303099 |
| 2 | 135379000 | 135381799 | 37.45006 | 61.7453 | 54.3864 | 23.46709 | 0.56235 | 1.381203 | 0.303101 |
| 17 | 20159600 | 20164999 | 65.95778 | 106.0038 | 72.22139 | 30.61076 | 0.903804 | 1.915076 | 0.30312 |
| 8 | 102352600 | 102353199 | 11.30812 | 21.76027 | 19.54244 | 7.607765 | 0.689384 | -0.08675 | 0.303129 |
| 15 | 46427400 | 46430399 | 40.99097 | 66.83626 | 60.9828 | 26.1959 | 0.503706 | 1.536929 | 0.303144 |
| 6 | 62609200 | 62610999 | 16.92164 | 30.33303 | 27.11866 | 11.87757 | 0.561896 | 0.425893 | 0.303152 |
| 10 | 121704200 | 121705799 | 16.92164 | 30.33303 | 27.11866 | 11.87757 | 0.561896 | 0.425893 | 0.303152 |
| 7 | 23601000 | 23601999 | 30.07902 | 50.31851 | 45.9799 | 20.02912 | 0.51074 | 1.193082 | 0.303163 |
| 9 | 121232200 | 121235599 | 27.33026 | 44.45387 | 84.5209 | 36.27474 | -0.55517 | 1.560047 | 0.303166 |
| 9 | 70695200 | 70696999 | 18.32241 | 32.48475 | 28.45184 | 12.57756 | 0.582209 | 0.504431 | 0.303173 |
| 13 | 92709200 | 92712999 | 26.71376 | 43.40392 | 86.61436 | 37.15273 | -0.62603 | 1.577751 | 0.303173 |
| 7 | 104382800 | 104383999 | 12.60351 | 23.6422 | 21.93031 | 9.036578 | 0.571282 | 0.118085 | 0.303174 |
| 10 | 86820200 | 86821399 | 12.60351 | 23.6422 | 21.93031 | 9.036578 | 0.571282 | 0.118085 | 0.303174 |
| 2 | 148247800 | 148249799 | 19.74839 | 32.84677 | 67.52656 | 29.30696 | -0.65504 | 1.245343 | 0.303176 |
| 13 | 35677800 | 35680399 | 47.6779 | 76.95707 | 71.96941 | 30.6879 | 0.455225 | 1.756899 | 0.303177 |
| 5 | 67379800 | 67390199 | 99.32531 | 151.2472 | 248.4109 | 102.9523 | -0.43032 | 3.154987 | 0.30318 |
| 1 | 130485400 | 130487399 | 16.7617 | 31.39043 | 20.893 | 8.63325 | 1.043312 | 0.162122 | 0.303187 |
| 6 | 26887400 | 26888599 | 22.49196 | 39.21642 | 34.37445 | 15.12575 | 0.576503 | 0.753541 | 0.303195 |
| 19 | 56814000 | 56817999 | 27.8794 | 45.17727 | 94.30696 | 40.37302 | -0.69449 | 1.640066 | 0.303196 |
| 14 | 55918600 | 55919999 | 8.340118 | 15.43415 | 32.57805 | 13.71495 | -0.63037 | 0.191667 | 0.303202 |
| 11 | 97284200 | 97287599 | 15.61938 | 26.53796 | 57.88995 | 25.37025 | -0.73927 | 1.013837 | 0.303203 |
| 12 | 94521000 | 94522999 | 20.23948 | 36.40902 | 24.40029 | 10.65514 | 0.97775 | 0.420685 | 0.303216 |
| 14 | 79526400 | 79527999 | 7.660615 | 14.03935 | 37.23364 | 15.88387 | -0.97175 | 0.249476 | 0.303218 |
| 4 | 148445400 | 148456399 | 96.8495 | 147.2839 | 255.6014 | 105.8908 | -0.51127 | 3.181145 | 0.303242 |
| 18 | 61943200 | 61944599 | 6.179449 | 11.77187 | 29.41445 | 11.75817 | -0.80942 | -0.02568 | 0.303256 |
| 7 | 31099800 | 31111999 | 237.9003 | 348.9956 | 303.5414 | 125.0229 | 0.437938 | 3.863097 | 0.30326 |
| 6 | 58407200 | 58408999 | 12.39487 | 21.86263 | 41.91557 | 18.54879 | -0.54465 | 0.5506 | 0.303262 |
| 6 | 56338400 | 56339599 | 6.561245 | 12.23847 | 31.63915 | 13.16262 | -0.90696 | 0.126793 | 0.303266 |
| 4 | 78398800 | 78402999 | 133.7202 | 203.447 | 184.7704 | 76.55718 | 0.422693 | 3.176836 | 0.303273 |
| 5 | 81167400 | 81170999 | 25.06223 | 40.5182 | 98.98361 | 42.29452 | -0.92394 | 1.687666 | 0.303275 |
| 15 | 97712800 | 97715599 | 19.79392 | 32.81048 | 69.56507 | 30.12073 | -0.6998 | 1.287467 | 0.303282 |

| | | | | | | | | | |
|---|---|---|---|---|---|---|---|---|---|
| 1 | 36283000 | 36284199 | 4.174317 | 8.09007 | 26.60914 | 10.07309 | -1.12628 | -0.23676 | 0.303285 |
| 7 | 47121200 | 47126599 | 83.71314 | 132.1795 | 117.3554 | 48.7269 | 0.496665 | 2.452516 | 0.303293 |
| 2 | 128337600 | 128338999 | 8.195287 | 15.1196 | 35.22795 | 14.89265 | -0.77674 | 0.211586 | 0.303297 |
| 19 | 13620800 | 13622599 | 23.98108 | 41.83333 | 33.67847 | 14.82407 | 0.698688 | 0.753348 | 0.3033 |
| 6 | 135128800 | 135132399 | 19.04325 | 31.8642 | 64.94493 | 28.29442 | -0.64265 | 1.174461 | 0.30331 |
| 10 | 116807400 | 116810599 | 27.39965 | 44.3979 | 88.98785 | 38.09466 | -0.63601 | 1.636244 | 0.303314 |
| 2 | 4508600 | 4509999 | 6.517143 | 12.26808 | 30.74382 | 12.59326 | -0.84445 | 0.063848 | 0.30332 |
| 19 | 38277000 | 38281199 | 26.54477 | 43.65201 | 83.49216 | 35.98638 | -0.55465 | 1.408053 | 0.303322 |
| 1 | 112941800 | 112945199 | 34.072 | 56.44251 | 52.56282 | 22.74914 | 0.483163 | 1.317664 | 0.303325 |
| 10 | 72994200 | 72996399 | 26.3098 | 44.93539 | 40.96039 | 17.95843 | 0.515637 | 0.985399 | 0.303329 |
| 18 | 3400200 | 3403799 | 52.53168 | 84.80722 | 70.85024 | 30.21787 | 0.617925 | 1.760071 | 0.303336 |
| 9 | 104989400 | 104990399 | 12.98139 | 24.9826 | 18.8495 | 7.335951 | 0.938945 | -0.07178 | 0.303339 |
| 7 | 106614600 | 106616399 | 23.92797 | 41.47029 | 37.20009 | 16.32144 | 0.542485 | 0.842044 | 0.303341 |
| 1 | 19027200 | 19028399 | 11.90833 | 22.4527 | 18.3622 | 7.501046 | 0.761129 | 0.068642 | 0.303359 |
| 2 | 136094400 | 136094799 | 3.339847 | 6.5333 | 17.96306 | 5.984246 | -0.69722 | -0.7282 | 0.303365 |
| 10 | 78595400 | 78597599 | 17.37777 | 29.8219 | 57.08831 | 25.03906 | -0.54957 | 0.926457 | 0.303375 |
| 3 | 51546400 | 51548599 | 14.79995 | 25.77246 | 48.41776 | 21.28842 | -0.5183 | 0.75655 | 0.303381 |
| X | 5337600 | 5342399 | 63.64698 | 102.2199 | 76.09768 | 32.22135 | 0.775055 | 1.936028 | 0.303396 |
| 2 | 151793800 | 151794799 | 13.03469 | 24.32488 | 22.34922 | 9.290209 | 0.574767 | 0.154754 | 0.303404 |
| 12 | 18171800 | 18198399 | 908.3464 | 1185.046 | 935.3599 | 391.6976 | 0.409025 | 5.67828 | 0.303407 |
| 5 | 128288000 | 128288799 | 6.717413 | 12.91118 | 28.69511 | 11.2086 | -0.61103 | -0.09718 | 0.303412 |
| 17 | 6207400 | 6209799 | 11.65613 | 20.56985 | 41.45372 | 18.34258 | -0.61556 | 0.542548 | 0.303413 |
| 7 | 113363600 | 113367599 | 25.3829 | 41.39211 | 84.03262 | 36.09881 | -0.64688 | 1.517755 | 0.303417 |
| 9 | 16535000 | 16536999 | 19.83863 | 35.47717 | 26.828 | 11.77392 | 0.800186 | 0.46049 | 0.303419 |
| 14 | 17515800 | 17517999 | 17.2055 | 30.88589 | 27.01163 | 11.81647 | 0.594563 | 0.41837 | 0.30342 |
| 3 | 24034800 | 24035999 | 17.32725 | 30.87873 | 27.53888 | 12.12132 | 0.560477 | 0.464479 | 0.303456 |
| 1 | 24234000 | 24235599 | 12.08765 | 21.2883 | 41.9364 | 18.56365 | -0.58441 | 0.565447 | 0.303463 |
| 11 | 95695200 | 95703599 | 66.14189 | 103.2456 | 178.9605 | 74.25891 | -0.4801 | 2.669719 | 0.303471 |
| X | 26005600 | 26009999 | 55.95018 | 90.56094 | 62.13931 | 26.57954 | 0.905205 | 1.696775 | 0.303481 |
| 16 | 26055800 | 26057199 | 14.6658 | 27.87984 | 19.55268 | 7.850773 | 1.002625 | 0.043689 | 0.303484 |
| 8 | 22103200 | 22106599 | 30.30363 | 51.80427 | 39.73933 | 17.42436 | 0.763833 | 0.993645 | 0.303491 |
| 7 | 107842800 | 107845399 | 28.20444 | 48.66457 | 36.95363 | 16.2111 | 0.780566 | 0.89952 | 0.303492 |
| 5 | 118564800 | 118569799 | 36.36786 | 58.37762 | 107.6272 | 45.53509 | -0.53154 | 1.926012 | 0.303506 |
| 9 | 113476600 | 113478599 | 14.61678 | 25.45587 | 48.95305 | 21.52504 | -0.55191 | 0.753852 | 0.303508 |
| 7 | 10877200 | 10878599 | 29.38629 | 49.92286 | 42.07079 | 18.43723 | 0.627532 | 1.049221 | 0.30352 |
| 1 | 180045600 | 180046599 | 12.70651 | 22.38209 | 42.95378 | 19.01335 | -0.54693 | 0.568653 | 0.303531 |
| 9 | 48847200 | 48848999 | 12.70651 | 22.38209 | 42.95378 | 19.01335 | -0.54693 | 0.568653 | 0.303531 |
| 5 | 24507800 | 24509199 | 7.715577 | 14.39738 | 31.11801 | 12.93701 | -0.65006 | 0.133629 | 0.303535 |
| 6 | 90750800 | 90752399 | 7.715577 | 14.39738 | 31.11801 | 12.93701 | -0.65006 | 0.133629 | 0.303535 |
| 11 | 40590200 | 40591799 | 28.48734 | 48.42782 | 42.59197 | 18.66674 | 0.566226 | 1.039639 | 0.303538 |
| X | 61497400 | 61499199 | 18.45793 | 34.23843 | 21.18348 | 8.889516 | 1.130738 | 0.233855 | 0.303544 |
| 19 | 58610600 | 58612399 | 20.84579 | 34.78001 | 66.55378 | 28.94272 | -0.55188 | 1.215389 | 0.303548 |
| 13 | 86984600 | 86985999 | 15.06743 | 27.88991 | 23.04757 | 9.699782 | 0.712933 | 0.220408 | 0.303559 |
| 18 | 38479000 | 38479599 | 5.370164 | 10.45246 | 25.51392 | 9.379243 | -0.66158 | -0.32842 | 0.303577 |
| 5 | 81443600 | 81445199 | 32.31758 | 54.54937 | 40.43799 | 17.687 | 0.811435 | 1.11053 | 0.303602 |
| 8 | 89266800 | 89280199 | 88.53184 | 135.1196 | 240.6142 | 99.77972 | -0.54421 | 3.104975 | 0.303624 |
| 1 | 119825400 | 119831199 | 40.87965 | 65.14656 | 126.5241 | 52.9712 | -0.61838 | 2.164265 | 0.303628 |

| | | | | | | | | | |
|---|---|---|---|---|---|---|---|---|---|
| 6 | 116572200 | 116572599 | 3.309749 | 6.466252 | 22.39838 | 7.811979 | -1.08928 | -0.55528 | 0.303638 |
| 13 | 9813600 | 9815399 | 12.21201 | 21.19055 | 49.99542 | 21.93617 | -0.84575 | 0.796177 | 0.30367 |
| 19 | 18646600 | 18648999 | 12.21201 | 21.19055 | 49.99542 | 21.93617 | -0.84575 | 0.796177 | 0.30367 |
| 14 | 95656400 | 95658999 | 26.23408 | 42.6423 | 84.06834 | 36.0717 | -0.60792 | 1.56856 | 0.303685 |
| 13 | 93126800 | 93127599 | 5.992689 | 11.53285 | 27.5318 | 10.71508 | -0.70811 | -0.12038 | 0.303686 |
| 18 | 87275600 | 87277599 | 12.05909 | 21.31624 | 41.89953 | 18.5337 | -0.57926 | 0.523812 | 0.303689 |
| 11 | 101731200 | 101732199 | 14.16772 | 26.5638 | 22.33116 | 9.209144 | 0.711434 | 0.136506 | 0.303691 |
| 13 | 20720400 | 20723999 | 25.08976 | 41.24131 | 79.71404 | 34.35162 | -0.56995 | 1.406756 | 0.3037 |
| 2 | 144059200 | 144064799 | 7.783277 | 15.20019 | 12.36883 | 4.343306 | 0.965113 | -0.50356 | 0.303705 |
| 4 | 80177800 | 80180599 | 54.41329 | 87.29385 | 82.37924 | 34.82263 | 0.431673 | 1.956402 | 0.303712 |
| 13 | 60301400 | 60302399 | 22.13104 | 38.81098 | 31.69435 | 14.00811 | 0.678444 | 0.663376 | 0.303721 |
| 2 | 145516800 | 145519999 | 29.73092 | 48.3522 | 86.76288 | 37.23515 | -0.4721 | 1.567901 | 0.303727 |
| 13 | 3103200 | 3107799 | 69.01997 | 110.0365 | 96.431 | 40.39514 | 0.528302 | 2.169288 | 0.303734 |
| 5 | 17629800 | 17632199 | 18.22704 | 30.59438 | 63.30136 | 27.63306 | -0.66425 | 1.136403 | 0.303736 |
| 19 | 39054400 | 39055799 | 12.18978 | 23.40148 | 18.1472 | 7.112715 | 0.889353 | -0.06183 | 0.303742 |
| 18 | 46342600 | 46345999 | 26.70553 | 43.41758 | 84.26272 | 36.15782 | -0.58515 | 1.565972 | 0.303749 |
| 7 | 13946600 | 13948799 | 23.10524 | 41.58398 | 22.02794 | 9.614757 | 1.315305 | 0.434372 | 0.30376 |
| 2 | 89460600 | 89461199 | 9.12504 | 17.79942 | 14.88985 | 5.332482 | 0.899907 | -0.40823 | 0.303763 |
| 5 | 116291600 | 116297399 | 40.28728 | 64.43147 | 119.1012 | 50.11357 | -0.54162 | 2.048122 | 0.303766 |
| 15 | 63529000 | 63533199 | 29.08137 | 47.2949 | 86.69415 | 37.20539 | -0.5028 | 1.566347 | 0.303794 |
| X | 30767200 | 30768599 | 19.81486 | 35.96796 | 23.81837 | 10.28578 | 1.005644 | 0.368967 | 0.303815 |
| 19 | 29680600 | 29682999 | 21.84658 | 36.19045 | 69.79649 | 30.20795 | -0.56376 | 1.300391 | 0.303817 |
| X | 62204600 | 62212799 | 121.132 | 188.2024 | 133.5861 | 55.46739 | 0.801292 | 2.7725 | 0.303835 |
| 11 | 102360200 | 102361599 | 8.34327 | 15.3955 | 34.14009 | 14.42908 | -0.70549 | 0.210069 | 0.303837 |
| 14 | 33168400 | 33169599 | 8.34327 | 15.3955 | 34.14009 | 14.42908 | -0.70549 | 0.210069 | 0.303837 |
| 17 | 88521000 | 88523199 | 12.57683 | 22.11771 | 42.4629 | 18.80217 | -0.54826 | 0.586212 | 0.303841 |
| 19 | 48008600 | 48010199 | 9.269376 | 16.98858 | 35.61589 | 15.19263 | -0.63495 | 0.259128 | 0.303846 |
| 12 | 88149200 | 88151399 | 23.61693 | 41.01834 | 36.02943 | 15.81535 | 0.573041 | 0.815209 | 0.30385 |
| 10 | 121674600 | 121679399 | 37.28413 | 59.94821 | 107.2748 | 45.4979 | -0.48571 | 1.873611 | 0.303854 |
| 6 | 19676000 | 19676999 | 6.878547 | 13.14894 | 29.00504 | 11.50855 | -0.62171 | -0.04376 | 0.303871 |
| 8 | 119263600 | 119264799 | 12.96406 | 24.20263 | 21.79314 | 9.052988 | 0.604252 | 0.149628 | 0.303877 |
| 12 | 111351800 | 111356199 | 30.64072 | 49.52808 | 91.7877 | 39.21071 | -0.52677 | 1.699413 | 0.303888 |
| 6 | 60871800 | 60876599 | 60.60148 | 97.19384 | 83.35507 | 35.21994 | 0.568939 | 1.974286 | 0.303898 |
| 11 | 44506600 | 44508199 | 7.785356 | 14.3094 | 35.13459 | 14.92938 | -0.85698 | 0.230138 | 0.3039 |
| 3 | 91968400 | 91969199 | 14.17642 | 26.34051 | 23.23288 | 9.727364 | 0.625055 | 0.193432 | 0.303903 |
| 7 | 5351800 | 5356599 | 68.83847 | 109.7112 | 89.73857 | 37.57462 | 0.627099 | 2.176317 | 0.303903 |
| X | 33848200 | 33852199 | 46.10558 | 74.9636 | 57.08392 | 24.49585 | 0.761127 | 1.58809 | 0.303946 |
| 12 | 96838000 | 96839399 | 16.59213 | 29.87026 | 26.6176 | 11.614 | 0.570254 | 0.396465 | 0.303947 |
| 1 | 153207600 | 153209999 | 14.46556 | 24.44613 | 61.49931 | 26.90877 | -0.94444 | 1.060848 | 0.303968 |
| 1 | 177893800 | 177895799 | 24.10418 | 39.80892 | 72.79308 | 31.43871 | -0.48764 | 1.34368 | 0.303986 |
| 2 | 178999200 | 179002799 | 25.18943 | 41.15953 | 81.92661 | 35.21999 | -0.61639 | 1.486865 | 0.303995 |
| 4 | 140706800 | 140709599 | 32.40021 | 52.37204 | 92.91669 | 39.69291 | -0.46407 | 1.702095 | 0.303999 |
| 17 | 57607800 | 57610999 | 24.84698 | 40.70767 | 77.31955 | 33.27325 | -0.54657 | 1.448742 | 0.304001 |
| 19 | 18815600 | 18817599 | 15.35498 | 26.66092 | 52.72108 | 23.13794 | -0.59235 | 0.808743 | 0.304024 |
| 18 | 25546400 | 25547599 | 13.71106 | 26.095 | 20.25167 | 8.111588 | 0.860895 | 0.028896 | 0.304028 |
| 10 | 108247400 | 108248799 | 6.607061 | 12.58167 | 28.28399 | 11.31542 | -0.65884 | -0.01974 | 0.304032 |
| 14 | 14468000 | 14468799 | 6.607061 | 12.58167 | 28.28399 | 11.31542 | -0.65884 | -0.01974 | 0.304032 |

| | | | | | | | | | |
|---|---|---|---|---|---|---|---|---|---|
| 13 | 109600000 | 109602599 | 30.71806 | 52.19919 | 41.33896 | 18.11701 | 0.716694 | 1.047476 | 0.304033 |
| 6 | 25688200 | 25690399 | 18.13895 | 30.29207 | 65.28925 | 28.40953 | -0.72284 | 1.194341 | 0.304046 |
| 1 | 103663600 | 103664599 | 12.35539 | 23.7058 | 19.35152 | 7.597263 | 0.814354 | -0.05046 | 0.304068 |
| 14 | 118448800 | 118452199 | 17.90522 | 30.51018 | 58.09092 | 25.46428 | -0.5434 | 0.994606 | 0.304069 |
| 2 | 37941400 | 37944799 | 19.04089 | 31.93815 | 62.70556 | 27.36093 | -0.58876 | 1.148535 | 0.304097 |
| 5 | 76142800 | 76142999 | 6.214339 | 12.15474 | 11.04841 | 3.721345 | 0.862225 | -0.67525 | 0.304105 |
| 8 | 48602200 | 48613799 | 118.4442 | 178.5407 | 293.9534 | 121.4483 | -0.44615 | 3.366952 | 0.304124 |
| 7 | 9257600 | 9260199 | 27.11043 | 47.0592 | 34.71552 | 15.23317 | 0.823664 | 0.838053 | 0.304125 |
| 12 | 73821000 | 73823599 | 14.29521 | 24.89917 | 49.48741 | 21.76183 | -0.59927 | 0.750488 | 0.304127 |
| 15 | 36851800 | 36853999 | 11.12835 | 19.95318 | 41.03635 | 17.96954 | -0.63264 | 0.415312 | 0.304164 |
| 8 | 11473600 | 11477199 | 12.56351 | 21.37499 | 58.24957 | 25.53065 | -1.0582 | 0.991457 | 0.304174 |
| 19 | 15924000 | 15928399 | 38.50195 | 61.67815 | 112.4925 | 47.44705 | -0.51953 | 1.993884 | 0.304177 |
| 18 | 71254800 | 71256799 | 19.28998 | 34.1374 | 28.87377 | 12.76898 | 0.631282 | 0.530748 | 0.304184 |
| 7 | 116666400 | 116669799 | 31.70992 | 53.79871 | 41.32675 | 18.10385 | 0.760396 | 1.064642 | 0.304186 |
| 4 | 115932000 | 115935199 | 21.36724 | 35.38706 | 70.9305 | 30.69317 | -0.61929 | 1.301849 | 0.304191 |
| 13 | 69173600 | 69175399 | 15.93528 | 29.37696 | 23.48225 | 9.937383 | 0.755026 | 0.250909 | 0.304191 |
| 15 | 86670200 | 86675399 | 40.45184 | 64.80337 | 112.9927 | 47.6598 | -0.45473 | 1.995998 | 0.304191 |
| 5 | 131164200 | 131164799 | 4.777742 | 9.3232 | 24.86559 | 8.851713 | -0.74528 | -0.45225 | 0.304193 |
| 10 | 48014800 | 48016799 | 22.58676 | 39.71428 | 29.69989 | 13.15129 | 0.804546 | 0.621088 | 0.304225 |
| 3 | 153317400 | 153319399 | 10.76092 | 19.30129 | 37.86403 | 16.57539 | -0.56461 | 0.402649 | 0.304242 |
| 8 | 110976000 | 110977199 | 9.794748 | 17.783 | 35.44196 | 15.30151 | -0.57381 | 0.319365 | 0.304254 |
| 15 | 97252800 | 97253999 | 9.794748 | 17.783 | 35.44196 | 15.30151 | -0.57381 | 0.319365 | 0.304254 |
| 16 | 70749800 | 70751999 | 13.2128 | 23.06495 | 46.79586 | 20.60694 | -0.62827 | 0.710566 | 0.304264 |
| 14 | 120402200 | 120404199 | 11.59852 | 20.55635 | 39.83181 | 17.60977 | -0.557 | 0.489399 | 0.304274 |
| 2 | 40785000 | 40786999 | 19.23309 | 32.7092 | 61.40908 | 26.91459 | -0.5237 | 1.022116 | 0.304277 |
| 3 | 134212000 | 134212399 | 10.69795 | 20.90326 | 12.57666 | 4.374315 | 1.407888 | -0.48944 | 0.304295 |
| 13 | 85453000 | 85455799 | 25.43949 | 44.08561 | 36.47924 | 16.00512 | 0.658304 | 0.847451 | 0.304304 |
| 10 | 46682400 | 46684399 | 15.27243 | 28.1863 | 23.56603 | 9.957766 | 0.692184 | 0.239641 | 0.304313 |
| 4 | 49842800 | 49845999 | 26.03206 | 42.4102 | 83.98017 | 36.06373 | -0.612 | 1.53329 | 0.304319 |
| 15 | 43494800 | 43497399 | 36.51405 | 60.57077 | 51.54743 | 22.33078 | 0.612864 | 1.307814 | 0.304339 |
| X | 64233800 | 64240399 | 116.9677 | 181.356 | 143.9276 | 59.77988 | 0.638128 | 2.806015 | 0.304356 |
| 10 | 95010200 | 95013199 | 16.75454 | 28.31663 | 60.04398 | 26.27262 | -0.69901 | 1.066482 | 0.30436 |
| 13 | 26854200 | 26856799 | 25.48731 | 44.77368 | 28.55359 | 12.63831 | 1.033171 | 0.650152 | 0.304365 |
| 1 | 81536200 | 81537399 | 15.17604 | 27.81152 | 24.30287 | 10.36816 | 0.619487 | 0.281564 | 0.304368 |
| 5 | 10548800 | 10552399 | 62.4012 | 99.84937 | 88.83746 | 37.40678 | 0.512196 | 2.048448 | 0.304371 |
| X | 3614400 | 3616599 | 19.13938 | 34.39187 | 26.33308 | 11.50991 | 0.786203 | 0.425274 | 0.304376 |
| 1 | 104947200 | 104950199 | 42.39529 | 69.12257 | 54.7868 | 23.5339 | 0.706253 | 1.538927 | 0.304378 |
| X | 102524800 | 102525999 | 9.912599 | 17.61473 | 38.40113 | 16.94315 | -0.72345 | 0.458664 | 0.304388 |
| 17 | 72563000 | 72564999 | 12.57031 | 22.07572 | 43.0053 | 19.03007 | -0.56956 | 0.606173 | 0.304401 |
| 7 | 27520800 | 27527199 | 68.48088 | 109.9413 | 73.44676 | 31.08081 | 0.930273 | 1.951626 | 0.304422 |
| 15 | 98227000 | 98227399 | 14.34598 | 26.22967 | 23.29985 | 9.971352 | 0.592506 | 0.283976 | 0.304429 |
| X | 28665000 | 28665999 | 13.23639 | 25.68742 | 16.24832 | 6.112382 | 1.237083 | -0.18375 | 0.304434 |
| 8 | 74263600 | 74266199 | 25.10688 | 43.62569 | 35.35401 | 15.52325 | 0.688784 | 0.815207 | 0.304445 |
| 2 | 114843000 | 114844599 | 20.08542 | 35.99312 | 25.50218 | 11.1725 | 0.894898 | 0.445009 | 0.304454 |
| 16 | 30675000 | 30676599 | 11.22457 | 19.65376 | 48.32569 | 21.33064 | -0.90374 | 0.655747 | 0.304473 |
| 18 | 85120600 | 85125999 | 53.64319 | 86.54711 | 71.53144 | 30.48363 | 0.632345 | 1.779588 | 0.304494 |
| 9 | 37248800 | 37251799 | 40.52753 | 66.27991 | 53.17103 | 22.86535 | 0.69194 | 1.489845 | 0.304509 |

| | | | | | | | | | |
|---|---|---|---|---|---|---|---|---|---|
| 12 | 84408400 | 84409199 | 3.997449 | 7.775945 | 23.41028 | 8.647785 | -0.96558 | -0.33379 | 0.304536 |
| X | 33672800 | 33677599 | 62.18799 | 100.0766 | 71.96127 | 30.55865 | 0.828475 | 1.869918 | 0.304537 |
| 1 | 44913200 | 44914799 | 19.06072 | 33.69941 | 29.54741 | 13.06938 | 0.579285 | 0.54185 | 0.304544 |
| 11 | 57064200 | 57065799 | 7.672861 | 14.40721 | 32.39504 | 13.3265 | -0.69496 | 0.093394 | 0.304552 |
| 11 | 68409400 | 68411399 | 23.69071 | 41.26988 | 34.97387 | 15.38093 | 0.625047 | 0.774889 | 0.304558 |
| 15 | 87263400 | 87276599 | 118.1738 | 178.4777 | 284.5325 | 117.6261 | -0.3973 | 3.328109 | 0.304559 |
| 1 | 97339800 | 97340199 | 5.232056 | 10.18499 | 24.89427 | 9.141717 | -0.66203 | -0.33619 | 0.304559 |
| 10 | 81472000 | 81483599 | 88.24173 | 134.4133 | 252.7745 | 104.7709 | -0.6251 | 3.145084 | 0.30456 |
| 10 | 46789800 | 46790999 | 15.21727 | 29.05083 | 18.43734 | 7.336998 | 1.156612 | 0.018513 | 0.30459 |
| 13 | 33760000 | 33761199 | 26.18969 | 45.04451 | 39.26622 | 17.22394 | 0.58141 | 0.921255 | 0.304627 |
| 13 | 35058000 | 35060199 | 14.19044 | 24.6605 | 48.43386 | 21.26809 | -0.58232 | 0.783455 | 0.304646 |
| 11 | 15617200 | 15617799 | 13.1926 | 24.46399 | 21.40239 | 8.985148 | 0.634121 | 0.192001 | 0.304649 |
| 6 | 140062600 | 140063599 | 14.08304 | 26.78694 | 20.34211 | 8.158114 | 0.890929 | 0.038128 | 0.304649 |
| 2 | 44246600 | 44249599 | 18.55474 | 31.18514 | 67.16543 | 29.33911 | -0.7219 | 1.130136 | 0.304653 |
| 10 | 103049200 | 103051199 | 18.55474 | 31.18514 | 67.16543 | 29.33911 | -0.7219 | 1.130136 | 0.304653 |
| 13 | 57512400 | 57517599 | 31.54378 | 51.08034 | 94.13983 | 40.28119 | -0.51626 | 1.657862 | 0.304661 |
| 7 | 66280200 | 66286599 | 49.98472 | 79.539 | 133.6551 | 55.84247 | -0.41228 | 2.221785 | 0.304671 |
| 5 | 102627800 | 102629999 | 14.69093 | 25.33312 | 53.22767 | 23.34991 | -0.68167 | 0.86686 | 0.304671 |
| X | 63965200 | 63966999 | 23.36123 | 40.86993 | 31.82909 | 14.03911 | 0.746775 | 0.708955 | 0.304677 |
| 2 | 165636200 | 165638799 | 12.23768 | 21.17248 | 50.50433 | 22.15746 | -0.8623 | 0.82534 | 0.30468 |
| 9 | 16150400 | 16151399 | 7.038465 | 13.42122 | 28.91245 | 11.53397 | -0.5946 | -0.02388 | 0.304687 |
| 13 | 79022200 | 79023999 | 14.48618 | 25.15352 | 49.50783 | 21.72833 | -0.58566 | 0.797 | 0.304696 |
| 16 | 65583600 | 65586599 | 27.24649 | 46.33219 | 42.60885 | 18.67459 | 0.502089 | 1.031317 | 0.304702 |
| 9 | 88911600 | 88913399 | 14.79695 | 27.33026 | 23.63989 | 9.978291 | 0.644639 | 0.231777 | 0.304703 |
| 7 | 31887000 | 31890399 | 42.01109 | 69.00193 | 55.63217 | 23.96598 | 0.687532 | 1.440626 | 0.304703 |
| 6 | 57073800 | 57075799 | 28.23413 | 47.91009 | 42.54877 | 18.64016 | 0.551903 | 1.051346 | 0.304706 |
| 10 | 22033800 | 22036199 | 18.58464 | 31.08771 | 64.29895 | 28.00781 | -0.66365 | 1.176576 | 0.304707 |
| 5 | 137178000 | 137201599 | 511.2057 | 679.3563 | 1076.004 | 449.6424 | -0.56904 | 5.377992 | 0.304707 |
| 7 | 24496000 | 24499399 | 47.01383 | 75.83327 | 70.51189 | 30.03987 | 0.462379 | 1.774953 | 0.304709 |
| 2 | 148865000 | 148865799 | 10.8986 | 21.05081 | 19.1183 | 7.352641 | 0.689956 | -0.13056 | 0.304724 |
| 15 | 58202000 | 58203999 | 16.12028 | 27.71481 | 53.23286 | 23.35005 | -0.55367 | 0.901144 | 0.304726 |
| 17 | 25491600 | 25493399 | 8.083221 | 14.48088 | 37.46821 | 16.4189 | -0.96169 | 0.39246 | 0.304729 |
| 10 | 48897400 | 48899999 | 22.35222 | 39.25277 | 30.27609 | 13.39683 | 0.760026 | 0.638007 | 0.304734 |
| 17 | 38808800 | 38813599 | 48.06429 | 78.93657 | 50.15013 | 21.60347 | 1.030184 | 1.453825 | 0.304735 |
| 9 | 5378800 | 5382199 | 25.96446 | 44.64129 | 34.71999 | 15.22959 | 0.745181 | 0.917962 | 0.304738 |
| 3 | 31936400 | 31941399 | 38.5938 | 62.06633 | 108.8668 | 46.18642 | -0.45659 | 1.87035 | 0.304743 |
| 2 | 99395600 | 99398599 | 41.64567 | 67.91041 | 54.2563 | 23.30728 | 0.694909 | 1.533634 | 0.304757 |
| 8 | 30506000 | 30508799 | 42.59303 | 68.13021 | 117.5415 | 49.46879 | -0.44199 | 2.046972 | 0.304764 |
| 2 | 56789800 | 56793199 | 36.94732 | 60.96762 | 54.85394 | 23.67724 | 0.532338 | 1.37218 | 0.304765 |
| 7 | 73710000 | 73710999 | 6.581904 | 12.6113 | 27.93677 | 11.01061 | -0.61831 | -0.07085 | 0.304784 |
| 19 | 53513400 | 53515399 | 15.03538 | 25.87723 | 52.13369 | 22.86866 | -0.62176 | 0.886395 | 0.304805 |
| 2 | 150093400 | 150096399 | 75.97777 | 120.2602 | 112.1217 | 46.61159 | 0.429019 | 2.386113 | 0.304821 |
| 7 | 64041400 | 64042799 | 10.32613 | 18.0879 | 45.19006 | 19.95337 | -0.92632 | 0.639342 | 0.304823 |
| 8 | 3191600 | 3192799 | 23.54345 | 40.84864 | 33.80965 | 14.83439 | 0.658525 | 0.827848 | 0.304845 |
| 15 | 4943600 | 4944999 | 13.85487 | 26.17797 | 21.23902 | 8.638689 | 0.778103 | 0.081121 | 0.304873 |
| 17 | 33258400 | 33260999 | 43.20252 | 70.14631 | 64.82385 | 27.79174 | 0.480805 | 1.610297 | 0.304876 |
| X | 148957800 | 148960599 | 47.82191 | 77.57101 | 65.01551 | 27.85389 | 0.620107 | 1.640917 | 0.304877 |

| | | | | | | | | | |
|---|---|---|---|---|---|---|---|---|---|
| 18 | 62373600 | 62375399 | 17.16059 | 32.06429 | 19.53751 | 8.104363 | 1.164689 | 0.175102 | 0.304879 |
| 7 | 10737200 | 10739799 | 29.88545 | 52.40753 | 24.96925 | 11.03538 | 1.451385 | 0.696127 | 0.304893 |
| X | 107263800 | 107266199 | 25.76448 | 45.2024 | 28.42204 | 12.56642 | 1.053345 | 0.66976 | 0.304897 |
| 17 | 19445800 | 19450799 | 90.99076 | 143.3215 | 120.3142 | 49.9222 | 0.574664 | 2.507245 | 0.304897 |
| 14 | 90700600 | 90702199 | 30.60502 | 51.45749 | 46.41103 | 20.27551 | 0.529405 | 1.144166 | 0.304905 |
| 14 | 124684000 | 124686199 | 25.1535 | 43.76081 | 34.78091 | 15.28349 | 0.716858 | 0.797948 | 0.304907 |
| X | 34559400 | 34561199 | 13.74553 | 26.43555 | 18.14831 | 7.07659 | 1.071151 | -0.0602 | 0.304927 |
| 13 | 43713000 | 43713999 | 7.856982 | 14.65611 | 31.65267 | 13.16709 | -0.64992 | 0.137243 | 0.304942 |
| X | 119293400 | 119295799 | 44.75251 | 72.60777 | 62.28994 | 26.69043 | 0.586895 | 1.63077 | 0.304963 |
| 15 | 84129400 | 84134199 | 40.90422 | 65.50099 | 116.428 | 49.07794 | -0.48319 | 2.008951 | 0.30497 |
| 4 | 80845400 | 80849399 | 46.53828 | 74.15829 | 127.3271 | 53.30144 | -0.4411 | 2.17204 | 0.304974 |
| 8 | 73324400 | 73327599 | 19.71189 | 32.86433 | 67.12285 | 29.17684 | -0.64565 | 1.217315 | 0.304974 |
| 9 | 71504000 | 71507399 | 21.25754 | 35.09092 | 74.16123 | 32.0191 | -0.69586 | 1.341655 | 0.30499 |
| 1 | 60101200 | 60103599 | 27.19126 | 46.43661 | 41.4622 | 18.1783 | 0.545378 | 0.9909 | 0.305006 |
| 6 | 78932400 | 78934399 | 17.98775 | 32.01962 | 27.95093 | 12.31441 | 0.590286 | 0.476603 | 0.305016 |
| 13 | 98636600 | 98639799 | 22.72335 | 37.37513 | 75.76828 | 32.65998 | -0.63798 | 1.391857 | 0.305028 |
| 14 | 33998800 | 33999999 | 6.120822 | 11.81864 | 26.51155 | 10.21141 | -0.60442 | -0.15395 | 0.30504 |
| 4 | 60739600 | 60740799 | 14.12933 | 25.95736 | 23.23081 | 9.877537 | 0.588306 | 0.254967 | 0.305054 |
| 16 | 71954800 | 71956399 | 11.02997 | 19.40729 | 41.98694 | 18.58952 | -0.71924 | 0.57004 | 0.305062 |
| 19 | 58931400 | 58933799 | 11.02997 | 19.40729 | 41.98694 | 18.58952 | -0.71924 | 0.57004 | 0.305062 |
| 19 | 31060200 | 31063999 | 62.32075 | 98.02604 | 162.7169 | 67.50419 | -0.40916 | 2.52561 | 0.305064 |
| 10 | 100065000 | 100067599 | 24.70748 | 40.3991 | 80.11219 | 34.4484 | -0.61026 | 1.4756 | 0.305084 |
| 12 | 61424200 | 61425799 | 22.54112 | 39.55333 | 30.85076 | 13.64203 | 0.744067 | 0.652553 | 0.305087 |
| 5 | 33791800 | 33793599 | 18.0139 | 31.9628 | 27.27898 | 12.05112 | 0.620327 | 0.497814 | 0.305115 |
| 1 | 3122600 | 3126199 | 74.26792 | 117.6513 | 111.6591 | 46.43879 | 0.404337 | 2.366426 | 0.305118 |
| 1 | 31044000 | 31046799 | 56.51934 | 90.65196 | 83.59114 | 35.32273 | 0.464637 | 1.968057 | 0.305163 |
| 7 | 119718600 | 119719799 | 7.864435 | 14.7321 | 31.19598 | 12.88464 | -0.61293 | 0.111361 | 0.305165 |
| 18 | 5187600 | 5189999 | 17.18673 | 30.87224 | 25.83249 | 11.29446 | 0.6582 | 0.41018 | 0.305193 |
| 3 | 38642200 | 38644199 | 26.03573 | 45.13319 | 36.47874 | 16.00506 | 0.692214 | 0.848366 | 0.3052 |
| 10 | 19254800 | 19257599 | 56.80187 | 90.98122 | 85.54593 | 36.07402 | 0.434263 | 2.010405 | 0.305208 |
| 1 | 35797600 | 35802999 | 35.24003 | 56.53965 | 115.5477 | 48.84984 | -0.68074 | 1.940575 | 0.305234 |
| 19 | 41371800 | 41374199 | 12.2781 | 21.48484 | 45.70552 | 20.16334 | -0.69604 | 0.666643 | 0.305235 |
| 14 | 50727600 | 50729799 | 19.10813 | 34.65 | 24.07943 | 10.4117 | 0.934808 | 0.367476 | 0.305246 |
| 19 | 35217200 | 35219399 | 19.10813 | 34.65 | 24.07943 | 10.4117 | 0.934808 | 0.367476 | 0.305246 |
| 8 | 107249200 | 107257199 | 57.66528 | 90.87309 | 158.4247 | 65.73954 | -0.47772 | 2.48366 | 0.305254 |
| 8 | 4262800 | 4268199 | 39.04999 | 62.36433 | 121.1017 | 50.85329 | -0.61477 | 2.092001 | 0.30526 |
| 19 | 8259000 | 8263799 | 60.74705 | 97.8165 | 70.8928 | 30.13116 | 0.818154 | 1.847885 | 0.305264 |
| 17 | 80368600 | 80370399 | 11.5322 | 20.17739 | 44.69142 | 19.7141 | -0.75364 | 0.664757 | 0.305269 |
| 1 | 82538000 | 82539599 | 15.75993 | 27.36935 | 51.13522 | 22.4447 | -0.51093 | 0.803437 | 0.305278 |
| 5 | 93682200 | 93683399 | 15.41274 | 29.41555 | 18.42916 | 7.338201 | 1.1744 | 0.022391 | 0.30528 |
| 7 | 8796200 | 8798399 | 15.41274 | 29.41555 | 18.42916 | 7.338201 | 1.1744 | 0.022391 | 0.30528 |
| 9 | 57545000 | 57548599 | 12.37272 | 21.44624 | 49.99338 | 21.93455 | -0.82878 | 0.807261 | 0.305309 |
| 19 | 11820000 | 11821199 | 13.76665 | 25.95164 | 21.81639 | 8.91492 | 0.721723 | 0.097531 | 0.305318 |
| 13 | 106725800 | 106727399 | 15.56711 | 26.99328 | 52.77477 | 23.15514 | -0.57665 | 0.819612 | 0.305328 |
| 15 | 56844800 | 56846599 | 19.13039 | 33.79165 | 29.48421 | 13.04382 | 0.585679 | 0.550971 | 0.305356 |
| X | 153658200 | 153661799 | 70.46113 | 112.1602 | 99.04284 | 41.40041 | 0.514987 | 2.221597 | 0.305364 |
| 8 | 78696400 | 78699999 | 23.26535 | 38.47892 | 74.06032 | 32.01783 | -0.56123 | 1.32379 | 0.305368 |

| | | | | | | | | | |
|---|---|---|---|---|---|---|---|---|---|
| 12 | 84272800 | 84274399 | 10.23779 | 18.5595 | 37.51895 | 16.22865 | -0.59633 | 0.333765 | 0.305388 |
| 8 | 21961600 | 21970999 | 122.8644 | 189.7928 | 150.0718 | 62.33665 | 0.639319 | 2.873327 | 0.305393 |
| 11 | 78688000 | 78689399 | 11.44581 | 20.26936 | 39.84647 | 17.61909 | -0.57814 | 0.497223 | 0.305409 |
| 6 | 137121400 | 137125199 | 37.709 | 60.40855 | 120.3232 | 50.75075 | -0.64639 | 1.995372 | 0.305427 |
| 11 | 80780800 | 80783399 | 11.40347 | 19.90255 | 46.28887 | 20.38115 | -0.82394 | 0.705118 | 0.30543 |
| 8 | 61049800 | 61051799 | 16.77202 | 30.95549 | 22.85845 | 9.658107 | 0.870206 | 0.250004 | 0.305445 |
| 15 | 47501200 | 47502799 | 16.77202 | 30.95549 | 22.85845 | 9.658107 | 0.870206 | 0.250004 | 0.305445 |
| 12 | 17811600 | 17814399 | 16.55634 | 28.54586 | 54.3744 | 23.85285 | -0.54097 | 0.877599 | 0.305454 |
| 15 | 90351600 | 90352399 | 11.52651 | 22.36377 | 17.43398 | 6.565746 | 0.936323 | -0.19758 | 0.305488 |
| 19 | 42690400 | 42692999 | 22.50828 | 37.19287 | 71.44676 | 30.8685 | -0.55841 | 1.332616 | 0.305513 |
| 1 | 129015600 | 129016399 | 7.331533 | 13.96508 | 30.64097 | 12.25127 | -0.62417 | -0.01044 | 0.305514 |
| 18 | 82407000 | 82409999 | 20.92226 | 34.6021 | 70.6787 | 30.55494 | -0.6465 | 1.31835 | 0.305529 |
| 5 | 128668400 | 128671799 | 24.27764 | 39.59672 | 81.32409 | 34.93752 | -0.66323 | 1.513039 | 0.305529 |
| 16 | 56268800 | 56270999 | 26.72823 | 46.47007 | 33.56465 | 14.74298 | 0.854095 | 0.812269 | 0.305539 |
| 4 | 152422800 | 152424599 | 14.58142 | 25.05883 | 53.18036 | 23.32674 | -0.69696 | 0.898648 | 0.30554 |
| 5 | 9774400 | 9776199 | 14.58142 | 25.05883 | 53.18036 | 23.32674 | -0.69696 | 0.898648 | 0.30554 |
| 3 | 40512800 | 40515199 | 14.85731 | 25.62274 | 51.61414 | 22.64216 | -0.62097 | 0.865993 | 0.305545 |
| X | 26002600 | 26004599 | 17.38006 | 33.09132 | 17.09735 | 6.842274 | 1.44376 | 0.053781 | 0.30556 |
| 12 | 113146600 | 113160999 | 139.6393 | 205.9779 | 375.9196 | 154.8092 | -0.62322 | 3.766519 | 0.30556 |
| 6 | 125083600 | 125084999 | 20.03231 | 35.12875 | 31.85247 | 14.07742 | 0.528107 | 0.651629 | 0.305567 |
| 10 | 17718400 | 17719199 | 8.451683 | 15.31871 | 34.96064 | 15.12559 | -0.76949 | 0.319297 | 0.305594 |
| 5 | 54297400 | 54302399 | 80.66375 | 127.091 | 120.2938 | 49.91554 | 0.402058 | 2.495763 | 0.3056 |
| X | 21100600 | 21101999 | 23.26965 | 42.04873 | 21.62077 | 9.389665 | 1.36271 | 0.413374 | 0.305619 |
| 8 | 7037200 | 7038999 | 15.43511 | 28.14133 | 25.21729 | 10.83016 | 0.576842 | 0.313455 | 0.30562 |
| 8 | 74515600 | 74516599 | 15.43511 | 28.14133 | 25.21729 | 10.83016 | 0.576842 | 0.313455 | 0.30562 |
| 2 | 107359400 | 107362199 | 25.07153 | 43.45043 | 37.10207 | 16.27841 | 0.613249 | 0.847324 | 0.305622 |
| 16 | 45257000 | 45259199 | 16.18402 | 27.62636 | 55.84587 | 24.48367 | -0.62885 | 0.969997 | 0.305651 |
| 4 | 97665400 | 97667999 | 14.11988 | 24.27485 | 53.16842 | 23.32177 | -0.74213 | 0.893274 | 0.305655 |
| 12 | 4642600 | 4643799 | 7.571243 | 14.09488 | 32.69191 | 13.64555 | -0.7562 | 0.14924 | 0.305665 |
| 8 | 25744800 | 25753199 | 76.80175 | 118.8114 | 198.6524 | 82.51534 | -0.43828 | 2.834992 | 0.305671 |
| 7 | 39183200 | 39188599 | 97.21212 | 151.2872 | 142.4518 | 59.13404 | 0.395865 | 2.73228 | 0.305671 |
| 5 | 117811600 | 117812599 | 11.89605 | 22.54622 | 20.94098 | 8.462166 | 0.592719 | 0.041287 | 0.305683 |
| 12 | 115121600 | 115122399 | 23.89179 | 41.50995 | 35.66184 | 15.65737 | 0.605148 | 0.814277 | 0.305694 |
| 3 | 96400200 | 96404999 | 36.67973 | 59.02618 | 103.3664 | 43.89387 | -0.45301 | 1.847068 | 0.305699 |
| 7 | 26465600 | 26468199 | 32.05746 | 53.60032 | 48.64824 | 21.18452 | 0.520472 | 1.206867 | 0.305713 |
| 19 | 32344000 | 32345799 | 9.386824 | 16.88045 | 36.39191 | 15.8961 | -0.69733 | 0.374435 | 0.305719 |
| 2 | 161690800 | 161692399 | 17.46624 | 31.59596 | 26.02204 | 11.28583 | 0.688851 | 0.374621 | 0.305731 |
| 2 | 15202600 | 15204999 | 24.65539 | 42.54625 | 38.22897 | 16.77065 | 0.538794 | 0.88366 | 0.305743 |
| 1 | 117279200 | 117281799 | 23.72481 | 41.77013 | 27.91966 | 12.35975 | 0.966274 | 0.61068 | 0.305746 |
| 16 | 30015800 | 30026999 | 92.12175 | 141.0098 | 232.4777 | 96.47073 | -0.42982 | 3.046189 | 0.305747 |
| 5 | 142709200 | 142715599 | 69.45493 | 107.8487 | 189.5692 | 78.7195 | -0.50614 | 2.763871 | 0.305778 |
| 11 | 101765800 | 101767799 | 10.77244 | 19.28764 | 37.85385 | 16.59578 | -0.5667 | 0.414162 | 0.305782 |
| 5 | 148340400 | 148342799 | 19.57576 | 34.46065 | 30.6863 | 13.58487 | 0.554798 | 0.593834 | 0.305793 |
| 7 | 59883400 | 59884399 | 12.41794 | 23.91365 | 18.87191 | 7.33109 | 0.876959 | -0.08391 | 0.305795 |
| 10 | 75294200 | 75295199 | 12.41794 | 23.91365 | 18.87191 | 7.33109 | 0.876959 | -0.08391 | 0.305795 |
| 19 | 35318200 | 35319999 | 23.89217 | 41.81455 | 31.95362 | 14.09852 | 0.773653 | 0.702967 | 0.305796 |
| X | 60227200 | 60230799 | 63.95948 | 103.0686 | 65.81318 | 28.00344 | 1.001833 | 1.829542 | 0.305801 |

| | | | | | | | | | |
|---|---|---|---|---|---|---|---|---|---|
| 1 | 85181200 | 85196199 | 482.6159 | 663.4692 | 557.7446 | 231.8893 | 0.393801 | 4.842499 | 0.305803 |
| 13 | 96202000 | 96203199 | 12.78569 | 24.09239 | 21.27999 | 8.701865 | 0.649741 | 0.088458 | 0.305831 |
| 2 | 116342600 | 116345199 | 31.83385 | 54.44795 | 36.79848 | 16.13556 | 0.945705 | 0.989413 | 0.305847 |
| 11 | 48486000 | 48486799 | 8.01498 | 15.03941 | 31.79065 | 13.09287 | -0.60719 | 0.103219 | 0.305854 |
| X | 148735400 | 148735999 | 13.17054 | 24.58731 | 20.99684 | 8.722687 | 0.680731 | 0.152099 | 0.305856 |
| 10 | 114986800 | 114988199 | 9.776705 | 17.74916 | 35.95772 | 15.52536 | -0.59751 | 0.319349 | 0.305859 |
| 7 | 10501800 | 10515599 | 375.4327 | 527.9121 | 451.4007 | 187.0118 | 0.402106 | 4.498734 | 0.30586 |
| 10 | 82076400 | 82079199 | 23.70822 | 39.35478 | 74.31858 | 32.21055 | -0.53347 | 1.283055 | 0.305889 |
| 6 | 8595800 | 8597199 | 10.61304 | 18.96505 | 39.38936 | 17.29753 | -0.64978 | 0.426849 | 0.305892 |
| 6 | 100076800 | 100077799 | 5.541563 | 10.71872 | 25.43117 | 9.716246 | -0.67307 | -0.19142 | 0.305895 |
| 1 | 180640000 | 180641999 | 13.57181 | 23.57993 | 48.43124 | 21.26382 | -0.64655 | 0.78388 | 0.305899 |
| 7 | 20694800 | 20695599 | 21.24672 | 37.46978 | 25.18719 | 11.14639 | 0.959371 | 0.575502 | 0.305914 |
| 12 | 22158600 | 22168599 | 229.7678 | 336.5846 | 299.2443 | 123.264 | 0.404321 | 3.884257 | 0.305935 |
| 12 | 21617800 | 21619799 | 27.9232 | 47.28457 | 39.94914 | 17.49032 | 0.623453 | 1.063535 | 0.305938 |
| 19 | 31740400 | 31742599 | 12.64906 | 21.65267 | 55.17452 | 24.19479 | -0.96033 | 0.930806 | 0.305964 |
| 7 | 38249400 | 38252599 | 29.11724 | 50.24933 | 36.39098 | 15.96437 | 0.848739 | 0.901821 | 0.305994 |
| 6 | 75759600 | 75759999 | 6.181486 | 11.75309 | 28.22914 | 11.33042 | -0.75737 | -0.01285 | 0.305998 |
| 13 | 68644800 | 68651199 | 92.8354 | 145.624 | 128.6554 | 53.35815 | 0.49649 | 2.589508 | 0.306001 |
| 7 | 23368400 | 23372199 | 63.6443 | 102.8552 | 56.72063 | 24.22427 | 1.217395 | 1.744729 | 0.306002 |
| 3 | 65917600 | 65919799 | 18.12372 | 30.72642 | 58.54258 | 25.63993 | -0.54511 | 1.04253 | 0.306003 |
| 14 | 41177400 | 41179799 | 38.64573 | 64.26276 | 48.21773 | 20.92337 | 0.794195 | 1.28398 | 0.306003 |
| 5 | 95562200 | 95566999 | 63.86539 | 102.2404 | 80.1588 | 33.77499 | 0.695027 | 2.03345 | 0.306007 |
| 10 | 120129200 | 120130999 | 10.49246 | 18.69467 | 38.35592 | 16.8854 | -0.63447 | 0.444163 | 0.30601 |
| 3 | 137351400 | 137353999 | 25.11968 | 43.16631 | 37.16405 | 16.3014 | 0.598857 | 0.919363 | 0.306017 |
| 17 | 13246600 | 13250399 | 26.35793 | 42.93704 | 82.1062 | 35.25778 | -0.56181 | 1.535745 | 0.30602 |
| 12 | 82609400 | 82610799 | 22.76343 | 39.55082 | 36.16799 | 15.87992 | 0.515355 | 0.80758 | 0.30602 |
| 5 | 64385800 | 64388399 | 14.04511 | 24.29457 | 53.78853 | 23.59817 | -0.75595 | 0.835738 | 0.306025 |
| 1 | 3132800 | 3133999 | 12.98438 | 24.39806 | 21.80095 | 8.957464 | 0.627812 | 0.109411 | 0.306046 |
| 13 | 66879800 | 66886599 | 171.2364 | 256.3424 | 220.9249 | 91.11832 | 0.478185 | 3.505604 | 0.306047 |
| 2 | 145650600 | 145653399 | 20.23752 | 34.06334 | 62.97988 | 27.52302 | -0.50253 | 1.116719 | 0.30605 |
| 2 | 170824200 | 170826999 | 26.18327 | 44.87204 | 40.38894 | 17.71275 | 0.534525 | 0.954257 | 0.306059 |
| 15 | 16573200 | 16575199 | 22.65055 | 39.53486 | 34.40575 | 15.14962 | 0.587061 | 0.741309 | 0.306067 |
| 10 | 82819200 | 82823599 | 25.05972 | 40.89118 | 80.98895 | 34.7994 | -0.61058 | 1.507391 | 0.306087 |
| 12 | 31581800 | 31582399 | 4.653388 | 9.055746 | 22.92076 | 8.437909 | -0.71363 | -0.33783 | 0.306091 |
| 19 | 5985000 | 5986199 | 4.454003 | 8.628762 | 27.83519 | 10.55638 | -1.10193 | -0.225 | 0.306093 |
| 17 | 18413200 | 18415199 | 25.59329 | 44.45138 | 35.95361 | 15.78217 | 0.691628 | 0.825832 | 0.306101 |
| 5 | 43227800 | 43230999 | 29.9463 | 48.59192 | 87.30816 | 37.41391 | -0.47685 | 1.613134 | 0.306107 |
| 9 | 73731600 | 73732599 | 10.11602 | 18.23773 | 36.89036 | 16.06736 | -0.60353 | 0.364341 | 0.306111 |
| 13 | 56725400 | 56728199 | 13.8687 | 24.27029 | 46.22258 | 20.39296 | -0.53718 | 0.676731 | 0.306125 |
| 3 | 13073600 | 13074399 | 12.45982 | 24.20623 | 16.26903 | 6.072102 | 1.160225 | -0.21965 | 0.306125 |
| 14 | 118989600 | 118991399 | 11.96481 | 20.96653 | 44.63314 | 19.714 | -0.69692 | 0.640179 | 0.306128 |
| 7 | 116215400 | 116218199 | 18.56597 | 31.5073 | 60.26514 | 26.40227 | -0.55079 | 1.035962 | 0.306128 |
| 15 | 57791000 | 57792999 | 10.53602 | 18.65618 | 39.38898 | 17.41718 | -0.68039 | 0.492382 | 0.306132 |
| 14 | 124310800 | 124312399 | 24.49472 | 42.60068 | 35.38469 | 15.5456 | 0.653524 | 0.800153 | 0.306151 |
| 6 | 57801400 | 57802399 | 7.292581 | 13.55618 | 32.11571 | 13.43815 | -0.78909 | 0.155987 | 0.306163 |
| 5 | 145836800 | 145840999 | 76.77294 | 122.0313 | 102.2098 | 42.64706 | 0.589 | 2.270877 | 0.306164 |
| 7 | 140087200 | 140087799 | 5.256286 | 10.15757 | 25.34553 | 9.725268 | -0.75051 | -0.17903 | 0.306168 |

| | | | | | | | | | |
|---|---|---|---|---|---|---|---|---|---|
| 11 | 83308600 | 83314199 | 30.73555 | 49.43201 | 104.807 | 44.46832 | -0.72925 | 1.855992 | 0.306169 |
| 9 | 86897200 | 86898799 | 14.06693 | 24.32374 | 50.54738 | 22.17597 | -0.66479 | 0.83739 | 0.306172 |
| 5 | 133989800 | 133991799 | 17.88544 | 32.01154 | 27.46776 | 12.04604 | 0.619488 | 0.442237 | 0.306174 |
| 6 | 56808800 | 56809799 | 7.369294 | 13.91804 | 29.74429 | 12.10878 | -0.60903 | 0.048305 | 0.306183 |
| 7 | 31499200 | 31504399 | 74.45741 | 118.1713 | 108.0171 | 45.0018 | 0.460988 | 2.313763 | 0.306184 |
| 16 | 86522000 | 86522799 | 10.0092 | 17.94713 | 36.8736 | 16.14625 | -0.63089 | 0.398644 | 0.306202 |
| 4 | 23449800 | 23452399 | 31.14453 | 53.07982 | 40.22026 | 17.63091 | 0.780802 | 1.022826 | 0.306203 |
| 9 | 50637200 | 50640399 | 18.3362 | 30.97212 | 59.5511 | 26.05246 | -0.55838 | 1.077939 | 0.306204 |
| 9 | 85530200 | 85531399 | 14.44626 | 27.21237 | 21.10063 | 8.633398 | 0.835604 | 0.105475 | 0.306234 |
| 12 | 52625000 | 52628599 | 44.9181 | 73.04749 | 63.34391 | 27.18412 | 0.574329 | 1.585346 | 0.30625 |
| 6 | 87085400 | 87089399 | 18.41002 | 30.83887 | 63.79156 | 27.81142 | -0.66385 | 1.159945 | 0.306257 |
| 5 | 47304800 | 47305999 | 10.37992 | 20.1344 | 16.85192 | 6.353527 | 0.832484 | -0.21084 | 0.306285 |
| 2 | 115972400 | 115972799 | 3.755798 | 7.341728 | 19.47651 | 6.701039 | -0.6897 | -0.60201 | 0.3063 |
| 8 | 3939800 | 3943399 | 53.68224 | 86.69297 | 69.32061 | 29.57792 | 0.681528 | 1.751954 | 0.306304 |
| 7 | 34332000 | 34334999 | 17.60945 | 30.06896 | 55.89089 | 24.50402 | -0.50823 | 0.973729 | 0.306306 |
| 5 | 26710600 | 26712999 | 36.61134 | 60.65021 | 52.63499 | 22.78067 | 0.58463 | 1.325317 | 0.306323 |
| 4 | 71299800 | 71300399 | 6.777968 | 12.8395 | 29.23981 | 11.82849 | -0.69112 | 0.02335 | 0.306324 |
| 1 | 8110600 | 8114599 | 85.61081 | 134.7575 | 124.7934 | 51.77628 | 0.432493 | 2.51807 | 0.306336 |
| 1 | 181637600 | 181637999 | 3.636737 | 7.113279 | 19.16098 | 6.44459 | -0.6821 | -0.68676 | 0.306336 |
| 7 | 102237000 | 102238399 | 9.933269 | 17.9815 | 37.48942 | 16.24514 | -0.64226 | 0.339435 | 0.306362 |
| 1 | 178320400 | 178321999 | 10.44227 | 18.73993 | 37.87177 | 16.57119 | -0.60681 | 0.396382 | 0.306363 |
| 13 | 44685200 | 44690199 | 32.24453 | 52.03864 | 97.31337 | 41.48704 | -0.54248 | 1.74824 | 0.306366 |
| 11 | 20121200 | 20125599 | 32.06558 | 51.81913 | 94.12374 | 40.19654 | -0.49834 | 1.707782 | 0.30637 |
| 12 | 62235800 | 62236199 | 8.572206 | 16.67893 | 16.69875 | 6.156472 | 0.607395 | -0.3071 | 0.306379 |
| 1 | 49764600 | 49767199 | 56.92772 | 91.18203 | 86.30405 | 36.39325 | 0.424711 | 2.010536 | 0.306394 |
| 7 | 84338400 | 84340599 | 11.05302 | 19.40403 | 42.56741 | 18.83051 | -0.73961 | 0.604266 | 0.306397 |
| 8 | 113377800 | 113379999 | 19.86585 | 33.16105 | 72.26316 | 31.43518 | -0.73892 | 1.209359 | 0.306414 |
| 18 | 77678400 | 77689799 | 87.44631 | 134.4012 | 218.8782 | 90.88734 | -0.40756 | 2.963164 | 0.30642 |
| 1 | 62799400 | 62800199 | 5.106572 | 9.914606 | 24.49249 | 9.187286 | -0.70525 | -0.26266 | 0.306423 |
| 2 | 36632600 | 36633799 | 40.75529 | 66.2223 | 60.05689 | 25.76221 | 0.508891 | 1.588574 | 0.306426 |
| 4 | 32797600 | 32798799 | 9.413567 | 17.36157 | 35.78739 | 15.13585 | -0.60171 | 0.225847 | 0.30644 |
| 5 | 103090600 | 103095599 | 46.67718 | 74.45812 | 125.8634 | 52.76568 | -0.41701 | 2.137605 | 0.306452 |
| 7 | 17992600 | 17994999 | 41.07164 | 67.44572 | 56.67536 | 24.41335 | 0.6279 | 1.442017 | 0.306454 |
| 11 | 78384600 | 78385999 | 13.10817 | 24.57231 | 21.75315 | 8.973805 | 0.636467 | 0.124881 | 0.306455 |
| 13 | 48885400 | 48888399 | 23.27415 | 38.09356 | 78.13356 | 33.60983 | -0.65768 | 1.457893 | 0.30646 |
| 17 | 83029800 | 83030399 | 5.549882 | 10.70971 | 25.86051 | 9.98649 | -0.71287 | -0.15307 | 0.306463 |
| 14 | 69344200 | 69347799 | 16.72065 | 28.89137 | 55.50296 | 24.34953 | -0.55257 | 0.860628 | 0.306464 |
| 5 | 104478000 | 104482599 | 54.19862 | 85.18147 | 163.936 | 68.00276 | -0.62312 | 2.536692 | 0.306465 |
| 14 | 111990200 | 111991799 | 22.22492 | 38.8317 | 33.80168 | 14.89322 | 0.586753 | 0.723008 | 0.306469 |
| 16 | 51460200 | 51463399 | 41.08556 | 67.27774 | 58.84958 | 25.31955 | 0.568574 | 1.477801 | 0.30648 |
| 5 | 144268600 | 144271799 | 18.53183 | 31.58967 | 61.47972 | 26.95057 | -0.57495 | 0.996523 | 0.306511 |
| 9 | 17318200 | 17320399 | 18.38833 | 31.34382 | 58.10943 | 25.47308 | -0.50504 | 0.993928 | 0.306554 |
| 5 | 4774400 | 4776199 | 14.98898 | 27.94251 | 22.61871 | 9.418986 | 0.754096 | 0.17855 | 0.306554 |
| 6 | 74436800 | 74438199 | 14.98898 | 27.94251 | 22.61871 | 9.418986 | 0.754096 | 0.17855 | 0.306554 |
| 7 | 5479000 | 5484799 | 61.66635 | 99.60948 | 61.1984 | 26.12001 | 1.061167 | 1.757296 | 0.306556 |
| 6 | 22655800 | 22660799 | 82.55942 | 130.1453 | 121.8096 | 50.54797 | 0.418797 | 2.486253 | 0.306563 |
| 2 | 30113200 | 30114799 | 27.08158 | 46.37136 | 35.09433 | 15.3897 | 0.783314 | 0.957442 | 0.306564 |

| | | | | | | | | | |
|---|---|---|---|---|---|---|---|---|---|
| 1 | 59442400 | 59445199 | 20.88977 | 34.62458 | 80.80575 | 34.98612 | -0.83814 | 1.297494 | 0.306604 |
| 18 | 65316600 | 65318799 | 15.84432 | 26.74624 | 64.29025 | 28.12039 | -0.87922 | 1.079185 | 0.306611 |
| 2 | 129040200 | 129042199 | 9.459934 | 16.82016 | 38.91105 | 17.15969 | -0.80815 | 0.451735 | 0.306614 |
| X | 153620400 | 153621399 | 13.9477 | 26.76527 | 18.68988 | 7.334057 | 1.038588 | -0.03676 | 0.306627 |
| 7 | 32121600 | 32143399 | 481.1544 | 662.914 | 552.6883 | 229.7071 | 0.408929 | 4.808978 | 0.30663 |
| 18 | 29001800 | 29002999 | 12.10816 | 21.27079 | 42.49611 | 18.80976 | -0.60569 | 0.597812 | 0.30663 |
| 16 | 43519200 | 43523399 | 32.71434 | 52.85168 | 94.73934 | 40.44376 | -0.47983 | 1.718045 | 0.306631 |
| 4 | 25919800 | 25927399 | 90.39351 | 141.7793 | 117.4973 | 48.73098 | 0.588536 | 2.588769 | 0.306638 |
| 17 | 19391200 | 19394999 | 55.90644 | 90.57028 | 58.87399 | 25.19927 | 0.984117 | 1.675683 | 0.306649 |
| 10 | 73987600 | 73990199 | 19.63247 | 34.80461 | 28.25568 | 12.49125 | 0.691031 | 0.513176 | 0.306653 |
| 4 | 140219000 | 140220599 | 15.58271 | 26.5902 | 55.254 | 24.22347 | -0.66838 | 0.972002 | 0.306671 |
| 7 | 139915200 | 139917399 | 16.12878 | 27.80301 | 52.16618 | 22.88401 | -0.51913 | 0.877131 | 0.306675 |
| 7 | 120020200 | 120021399 | 7.360972 | 14.03915 | 32.43418 | 12.93313 | -0.69424 | -0.01239 | 0.306677 |
| 1 | 115677800 | 115682399 | 44.85761 | 72.53203 | 67.93259 | 29.03225 | 0.45595 | 1.699484 | 0.306698 |
| 8 | 20588600 | 20592799 | 66.0917 | 103.3674 | 174.0879 | 72.22263 | -0.43643 | 2.635788 | 0.306703 |
| 10 | 47924200 | 47926399 | 22.20979 | 39.10352 | 29.14301 | 12.90126 | 0.810108 | 0.603352 | 0.306718 |
| 10 | 39277200 | 39282599 | 49.1371 | 78.24435 | 131.6466 | 55.05598 | -0.41303 | 2.197719 | 0.306727 |
| 5 | 85688000 | 85689399 | 16.15805 | 29.17725 | 25.75097 | 11.19327 | 0.587693 | 0.375167 | 0.306743 |
| 17 | 27832600 | 27839599 | 48.38765 | 76.86116 | 135.8039 | 56.62629 | -0.48715 | 2.274772 | 0.306768 |
| 14 | 41351400 | 41353799 | 33.9577 | 56.39198 | 50.94021 | 22.08443 | 0.527165 | 1.286676 | 0.306768 |
| 4 | 112183200 | 112183799 | 13.52301 | 25.74439 | 20.25951 | 8.109877 | 0.841671 | 0.025058 | 0.306773 |
| 4 | 14304200 | 14304799 | 9.43491 | 17.33066 | 34.58401 | 14.70767 | -0.56082 | 0.247148 | 0.30678 |
| 17 | 18750400 | 18752199 | 57.77218 | 92.41777 | 85.75036 | 36.09328 | 0.451333 | 2.05033 | 0.306785 |
| 14 | 76625000 | 76628599 | 37.10667 | 61.28933 | 54.31638 | 23.46046 | 0.554188 | 1.361234 | 0.306789 |
| 4 | 108637000 | 108638199 | 8.897258 | 16.11378 | 35.4367 | 15.34648 | -0.71765 | 0.32667 | 0.306801 |
| 5 | 115464600 | 115466199 | 6.730743 | 12.39711 | 33.06775 | 14.01094 | -0.97078 | 0.211081 | 0.306818 |
| 18 | 83024200 | 83026799 | 18.86377 | 31.63072 | 63.31756 | 27.62228 | -0.61673 | 1.149843 | 0.306827 |
| 4 | 22626400 | 22629399 | 24.95141 | 43.52641 | 33.67645 | 14.82326 | 0.755858 | 0.759401 | 0.306845 |
| 8 | 61842200 | 61844999 | 17.22924 | 29.52915 | 55.95983 | 24.54158 | -0.5353 | 0.9365 | 0.306857 |
| 6 | 127662000 | 127664599 | 11.65392 | 20.46545 | 42.52643 | 18.81745 | -0.66201 | 0.601416 | 0.306868 |
| 18 | 32595200 | 32596799 | 11.65392 | 20.46545 | 42.52643 | 18.81745 | -0.66201 | 0.601416 | 0.306868 |
| 15 | 8707400 | 8708199 | 9.062217 | 17.69251 | 13.03176 | 4.600068 | 1.09829 | -0.46591 | 0.306876 |
| 14 | 21897200 | 21899199 | 8.995773 | 16.03622 | 37.94002 | 16.69508 | -0.83783 | 0.431519 | 0.306893 |
| 4 | 121353000 | 121358999 | 97.5618 | 152.448 | 136.9984 | 56.84089 | 0.467651 | 2.661906 | 0.306934 |
| 6 | 89829600 | 89830399 | 14.79363 | 26.9116 | 23.7212 | 10.21614 | 0.597687 | 0.316565 | 0.306949 |
| 12 | 18550600 | 18574999 | 706.0282 | 941.1246 | 764.9785 | 319.5376 | 0.397583 | 5.331163 | 0.306951 |
| 4 | 28734400 | 28734999 | 7.113537 | 13.31865 | 30.64119 | 12.66649 | -0.7327 | 0.104991 | 0.306954 |
| 1 | 43200800 | 43204599 | 26.43907 | 43.26342 | 79.94738 | 34.38709 | -0.50803 | 1.468166 | 0.306971 |
| 12 | 9762800 | 9764999 | 11.43539 | 20.40185 | 48.57366 | 21.35957 | -0.84745 | 0.459234 | 0.306971 |
| 7 | 99438600 | 99441799 | 19.65981 | 32.96797 | 62.79593 | 27.396 | -0.5453 | 1.152094 | 0.306975 |
| 7 | 59548800 | 59550399 | 13.06463 | 24.28169 | 21.90827 | 9.169192 | 0.593142 | 0.17989 | 0.306979 |
| 1 | 113288200 | 113289599 | 16.83594 | 30.64139 | 24.30336 | 10.46063 | 0.74932 | 0.329775 | 0.306981 |
| 3 | 38896000 | 38897399 | 16.83594 | 30.64139 | 24.30336 | 10.46063 | 0.74932 | 0.329775 | 0.306981 |
| 15 | 103429600 | 103432199 | 17.7513 | 30.38101 | 55.93172 | 24.52646 | -0.49396 | 0.953043 | 0.306983 |
| 13 | 46612400 | 46614799 | 20.77098 | 36.36442 | 32.81476 | 14.48264 | 0.53499 | 0.68229 | 0.306994 |
| 8 | 3633200 | 3636199 | 11.51564 | 20.18595 | 43.61167 | 19.26809 | -0.71792 | 0.632082 | 0.307005 |
| 7 | 35253400 | 35255599 | 10.05951 | 17.86574 | 39.8877 | 17.6074 | -0.75849 | 0.463739 | 0.307028 |

| | | | | | | | | | |
|---|---|---|---|---|---|---|---|---|---|
| 5 | 10276000 | 10277799 | 19.52065 | 35.21259 | 25.12425 | 10.93507 | 0.89115 | 0.40233 | 0.30703 |
| 1 | 3224400 | 3228199 | 30.38348 | 51.25304 | 40.51253 | 17.71628 | 0.719111 | 1.107779 | 0.307073 |
| 11 | 58058800 | 58059599 | 4.117314 | 8.003153 | 23.34236 | 8.674647 | -0.9281 | -0.30973 | 0.307074 |
| 6 | 120340400 | 120342199 | 11.22397 | 19.65328 | 44.16615 | 19.49514 | -0.77433 | 0.649868 | 0.307084 |
| 6 | 48642000 | 48643999 | 12.92769 | 22.51083 | 48.40812 | 21.2834 | -0.71223 | 0.747134 | 0.307087 |
| 2 | 172101000 | 172102399 | 9.438913 | 17.26008 | 35.02579 | 14.98705 | -0.59139 | 0.272865 | 0.30709 |
| 5 | 24158000 | 24160199 | 22.77703 | 39.52431 | 36.24869 | 15.90462 | 0.511032 | 0.824753 | 0.30709 |
| 5 | 128835200 | 128840399 | 51.37787 | 81.28743 | 149.1795 | 62.00381 | -0.54708 | 2.386893 | 0.307098 |
| 7 | 53785400 | 53786599 | 14.61174 | 26.86394 | 24.2938 | 10.31888 | 0.574712 | 0.258542 | 0.307101 |
| 9 | 73294200 | 73295199 | 12.82139 | 24.0448 | 21.81742 | 8.993924 | 0.602073 | 0.120196 | 0.307104 |
| 12 | 67779800 | 67781799 | 18.25051 | 32.97619 | 25.87302 | 11.23759 | 0.756975 | 0.386018 | 0.307116 |
| 11 | 76153400 | 76155999 | 10.3025 | 18.08223 | 43.09939 | 19.06208 | -0.85859 | 0.604228 | 0.307128 |
| 14 | 57250200 | 57252599 | 23.46665 | 41.10114 | 31.99839 | 14.12737 | 0.746957 | 0.689412 | 0.307138 |
| 4 | 23258400 | 23264199 | 115.9618 | 178.0536 | 164.5903 | 68.33558 | 0.407599 | 2.986781 | 0.307158 |
| 7 | 139765600 | 139767599 | 8.649307 | 15.54229 | 36.43468 | 15.92503 | -0.81775 | 0.374965 | 0.307166 |
| 15 | 25897600 | 25903799 | 43.86448 | 70.00993 | 127.1921 | 53.364 | -0.51999 | 2.116283 | 0.307166 |
| 5 | 6938000 | 6941799 | 85.07519 | 133.5961 | 124.108 | 51.47597 | 0.425338 | 2.56545 | 0.307176 |
| 2 | 166204200 | 166220399 | 112.5677 | 168.9818 | 324.0462 | 133.7331 | -0.67128 | 3.448461 | 0.307209 |
| 3 | 3228200 | 3238999 | 148.3377 | 226.5284 | 175.576 | 72.81894 | 0.65547 | 3.106279 | 0.307235 |
| 1 | 6230200 | 6239399 | 151.3143 | 228.6972 | 210.4556 | 87.02273 | 0.39579 | 3.315647 | 0.307235 |
| 6 | 59549600 | 59550399 | 11.34242 | 22.00947 | 17.44112 | 6.564429 | 0.913653 | -0.20186 | 0.307246 |
| 7 | 106443800 | 106444599 | 11.34242 | 22.00947 | 17.44112 | 6.564429 | 0.913653 | -0.20186 | 0.307246 |
| 7 | 40222600 | 40224599 | 19.89835 | 35.66778 | 25.50721 | 11.17209 | 0.881792 | 0.441898 | 0.307255 |
| 1 | 134470600 | 134473599 | 33.46404 | 55.50432 | 50.93216 | 22.06232 | 0.504454 | 1.299085 | 0.307256 |
| 4 | 51311200 | 51312799 | 16.14085 | 29.66987 | 23.9994 | 10.19634 | 0.734065 | 0.269789 | 0.307267 |
| 18 | 3145600 | 3146999 | 36.43138 | 60.70393 | 48.16157 | 20.92702 | 0.714114 | 1.258729 | 0.307284 |
| 1 | 22069600 | 22073399 | 65.53825 | 104.304 | 96.68365 | 40.40354 | 0.444805 | 2.221357 | 0.307284 |
| 9 | 57567200 | 57569399 | 14.14535 | 24.70948 | 46.78243 | 20.60974 | -0.52893 | 0.706097 | 0.307289 |
| 5 | 67988800 | 67989999 | 8.674463 | 15.92833 | 33.51581 | 14.26008 | -0.63706 | 0.240745 | 0.307293 |
| 8 | 79482000 | 79484199 | 19.93487 | 35.37869 | 28.33684 | 12.51624 | 0.711416 | 0.50975 | 0.307296 |
| 10 | 9041200 | 9044999 | 45.71925 | 73.89384 | 69.63423 | 29.74317 | 0.446427 | 1.714546 | 0.307296 |
| 11 | 11013800 | 11015199 | 15.74916 | 29.76122 | 19.30133 | 7.848269 | 1.098636 | 0.092354 | 0.307299 |
| 10 | 128639800 | 128643399 | 19.54629 | 32.26359 | 73.23697 | 31.61838 | -0.79848 | 1.338719 | 0.307301 |
| 3 | 77296000 | 77297199 | 11.26038 | 20.06965 | 38.8426 | 17.09481 | -0.55059 | 0.448211 | 0.307332 |
| 2 | 128529000 | 128530399 | 9.947894 | 18.01756 | 35.92166 | 15.55544 | -0.57726 | 0.33482 | 0.307335 |
| 11 | 83722200 | 83725199 | 15.12241 | 26.22242 | 50.04858 | 21.95902 | -0.54187 | 0.815548 | 0.307343 |
| 7 | 16027800 | 16034199 | 35.5859 | 57.069 | 109.8738 | 46.41946 | -0.59552 | 1.955204 | 0.307366 |
| 4 | 99717200 | 99718199 | 5.999859 | 11.52622 | 27.4476 | 10.73675 | -0.71136 | -0.10349 | 0.307392 |
| 10 | 85550000 | 85551599 | 26.87065 | 46.03816 | 36.18375 | 15.86821 | 0.729244 | 0.95357 | 0.307399 |
| 6 | 130453600 | 130456799 | 42.88357 | 71.07668 | 47.17384 | 20.42418 | 0.970317 | 1.332517 | 0.307408 |
| 15 | 36891000 | 36892599 | 6.711981 | 12.50963 | 33.24794 | 13.85064 | -0.94868 | 0.135553 | 0.307416 |
| X | 121526400 | 121527599 | 22.18093 | 38.94418 | 28.08014 | 12.42304 | 0.856992 | 0.63855 | 0.30742 |
| 13 | 34741000 | 34742999 | 10.90862 | 19.68372 | 40.6412 | 17.68185 | -0.63221 | 0.372518 | 0.30742 |
| 11 | 49025000 | 49025999 | 9.45841 | 17.29684 | 34.49991 | 14.76082 | -0.56636 | 0.272867 | 0.307425 |
| 7 | 32202800 | 32203599 | 10.28655 | 20.04908 | 15.37474 | 5.563913 | 1.011203 | -0.35435 | 0.307428 |
| 18 | 65863800 | 65873399 | 55.46381 | 87.71457 | 148.8194 | 61.84081 | -0.43449 | 2.400525 | 0.307434 |
| 5 | 138263600 | 138264599 | 6.300234 | 12.05396 | 28.43735 | 11.25508 | -0.71422 | -0.05952 | 0.307438 |

| | | | | | | | | | |
|---|---|---|---|---|---|---|---|---|---|
| 9 | 19373000 | 19376599 | 66.18344 | 105.4621 | 98.53984 | 41.25029 | 0.435226 | 2.182407 | 0.307448 |
| 8 | 39126600 | 39128599 | 14.49048 | 25.19325 | 48.43285 | 21.27366 | -0.55159 | 0.777917 | 0.307487 |
| 15 | 42096000 | 42098599 | 43.54643 | 71.75506 | 51.75645 | 22.32683 | 0.849479 | 1.403705 | 0.307495 |
| 5 | 14983800 | 14984399 | 7.499397 | 14.18611 | 30.94171 | 12.5543 | -0.63443 | 0.039762 | 0.307504 |
| 15 | 27200600 | 27203799 | 20.38376 | 34.27101 | 68.69029 | 30.00834 | -0.61881 | 1.130324 | 0.307514 |
| 7 | 62198200 | 62202199 | 75.67996 | 120.5573 | 95.01771 | 39.75884 | 0.679864 | 2.201184 | 0.307516 |
| 11 | 109058600 | 109063199 | 33.04359 | 53.33698 | 96.94931 | 41.34057 | -0.50135 | 1.744696 | 0.307517 |
| 8 | 25404800 | 25409399 | 57.09306 | 91.54492 | 85.0415 | 35.91935 | 0.453464 | 1.977304 | 0.307528 |
| 6 | 17965800 | 17967999 | 17.04845 | 29.32429 | 55.45429 | 24.32481 | -0.53132 | 0.902586 | 0.307532 |
| 8 | 89638400 | 89639799 | 14.22401 | 24.65935 | 49.4923 | 21.71481 | -0.6139 | 0.814764 | 0.307544 |
| 16 | 3339000 | 3340599 | 35.55041 | 58.50454 | 54.75802 | 23.60888 | 0.473993 | 1.400145 | 0.307553 |
| 7 | 93155200 | 93155399 | 7.412296 | 14.47476 | 14.68727 | 5.163176 | 0.650084 | -0.49396 | 0.307561 |
| 9 | 44328800 | 44331399 | 16.47312 | 28.15951 | 55.88815 | 24.50498 | -0.60222 | 0.957588 | 0.307566 |
| 4 | 34478000 | 34480599 | 16.54314 | 28.55952 | 56.60915 | 24.83415 | -0.59785 | 0.868039 | 0.30758 |
| 18 | 72760600 | 72761199 | 16.6685 | 29.95321 | 26.23773 | 11.46781 | 0.593366 | 0.408171 | 0.307583 |
| 4 | 4028000 | 4029399 | 20.56931 | 35.92639 | 30.61009 | 13.48401 | 0.617543 | 0.714843 | 0.307586 |
| X | 58768800 | 58769799 | 12.57907 | 24.29873 | 18.433 | 7.086265 | 0.947906 | -0.11546 | 0.30759 |
| 15 | 78472200 | 78475199 | 23.14942 | 38.19801 | 73.23231 | 31.60963 | -0.55584 | 1.351232 | 0.307609 |
| 12 | 88415400 | 88417599 | 14.46737 | 25.16779 | 48.96178 | 21.5139 | -0.56866 | 0.769019 | 0.307615 |
| X | 54666400 | 54668199 | 36.70145 | 61.42332 | 45.89566 | 19.99783 | 0.800141 | 1.211108 | 0.307619 |
| 1 | 175267800 | 175268999 | 12.10794 | 22.94731 | 20.31017 | 8.207637 | 0.661365 | 0.040774 | 0.307623 |
| 11 | 52246600 | 52249399 | 23.71565 | 41.23395 | 36.01788 | 15.82093 | 0.581225 | 0.801704 | 0.307631 |
| 5 | 71989000 | 71989599 | 10.39956 | 19.96774 | 18.62674 | 7.297463 | 0.627028 | -0.07763 | 0.307642 |
| 7 | 59920800 | 59924799 | 69.33768 | 110.3706 | 101.1085 | 42.26321 | 0.462055 | 2.220045 | 0.307671 |
| 12 | 21249600 | 21250399 | 4.358143 | 8.48768 | 22.38692 | 8.184349 | -0.76253 | -0.36727 | 0.307684 |
| 7 | 59716800 | 59720799 | 45.67355 | 74.93877 | 53.50695 | 23.04022 | 0.861711 | 1.460894 | 0.307688 |
| 7 | 40678600 | 40682199 | 44.10325 | 73.3623 | 42.58363 | 18.48279 | 1.163272 | 1.293105 | 0.307694 |
| 11 | 117839000 | 117840999 | 10.32425 | 18.43158 | 38.36358 | 16.86049 | -0.65346 | 0.429853 | 0.307711 |
| 11 | 109369800 | 109373799 | 23.17031 | 38.19526 | 73.09252 | 31.53629 | -0.55373 | 1.364136 | 0.307726 |
| 7 | 41148400 | 41149199 | 7.091366 | 13.33882 | 30.19221 | 12.38331 | -0.70032 | 0.073302 | 0.307727 |
| 6 | 132666200 | 132670399 | 76.91569 | 121.5999 | 115.5918 | 48.02526 | 0.399715 | 2.416235 | 0.307729 |
| 3 | 6807800 | 6814799 | 108.9349 | 169.1644 | 135.9013 | 56.43308 | 0.622238 | 2.772931 | 0.30773 |
| 8 | 21696600 | 21701399 | 63.03039 | 101.3763 | 72.17316 | 30.62415 | 0.841824 | 1.886576 | 0.307737 |
| 4 | 128340400 | 128348599 | 47.28688 | 75.03149 | 147.8938 | 61.60075 | -0.64624 | 2.308313 | 0.307742 |
| 17 | 19682000 | 19691599 | 124.8305 | 192.8673 | 149.935 | 62.27974 | 0.66402 | 2.872688 | 0.30775 |
| 1 | 178325800 | 178326999 | 18.50571 | 33.2866 | 26.28219 | 11.47795 | 0.742767 | 0.413538 | 0.307756 |
| 7 | 104126000 | 104128599 | 38.49753 | 61.71682 | 110.3839 | 46.61022 | -0.4901 | 1.970408 | 0.307804 |
| 13 | 91968000 | 91970999 | 29.54019 | 49.72003 | 45.87029 | 20.04575 | 0.496882 | 1.130057 | 0.307809 |
| 14 | 41708200 | 41709799 | 12.14916 | 23.12473 | 20.35872 | 8.152707 | 0.68046 | 0.012619 | 0.307812 |
| 19 | 5911000 | 5913999 | 35.93019 | 57.62224 | 110.0955 | 46.51451 | -0.58447 | 1.957199 | 0.307813 |
| 16 | 20496000 | 20499199 | 14.19678 | 24.61736 | 49.50084 | 21.7187 | -0.61666 | 0.811017 | 0.307823 |
| 1 | 114141800 | 114143199 | 17.95391 | 31.71448 | 25.87918 | 11.44888 | 0.681883 | 0.537001 | 0.30783 |
| 3 | 49402800 | 49404399 | 18.85158 | 33.69626 | 27.32077 | 11.99491 | 0.699633 | 0.456275 | 0.307852 |
| 3 | 107742800 | 107744399 | 16.37974 | 29.39217 | 26.64273 | 11.6589 | 0.542889 | 0.414287 | 0.30786 |
| 12 | 13605000 | 13605399 | 3.986054 | 7.78331 | 23.01774 | 8.103689 | -0.87773 | -0.51072 | 0.30786 |
| 2 | 87422000 | 87427999 | 85.09334 | 133.8068 | 126.008 | 52.2733 | 0.40721 | 2.537545 | 0.307906 |
| 19 | 48920800 | 48921599 | 7.431111 | 13.84282 | 31.089 | 12.96251 | -0.70808 | 0.143052 | 0.307907 |

| | | | | | | | | | |
|---|---|---|---|---|---|---|---|---|---|
| 4 | 58411600 | 58412399 | 14.42171 | 26.84304 | 23.26868 | 9.715309 | 0.653179 | 0.185782 | 0.307911 |
| 11 | 66783400 | 66784999 | 9.759889 | 17.34793 | 38.40318 | 16.9403 | -0.74521 | 0.455636 | 0.307914 |
| 19 | 34723000 | 34726399 | 25.61848 | 41.99511 | 78.23615 | 33.67566 | -0.5185 | 1.442496 | 0.30792 |
| 7 | 20871000 | 20884599 | 218.5693 | 324.1522 | 245.2094 | 100.9978 | 0.653906 | 3.682434 | 0.307928 |
| 11 | 27235800 | 27236999 | 31.81368 | 52.95771 | 48.1561 | 20.9117 | 0.517952 | 1.253769 | 0.307937 |
| 1 | 171248600 | 171249799 | 7.047854 | 13.40905 | 28.83702 | 11.55853 | -0.59811 | -0.00811 | 0.307948 |
| 15 | 97257600 | 97258999 | 13.19058 | 24.46656 | 20.9303 | 8.783874 | 0.666624 | 0.189479 | 0.307955 |
| 4 | 46901800 | 46908399 | 52.11352 | 82.79625 | 138.2693 | 57.66596 | -0.40568 | 2.272066 | 0.307957 |
| 1 | 101508600 | 101510199 | 13.2061 | 23.15356 | 44.08215 | 19.47906 | -0.5367 | 0.63886 | 0.307962 |
| 4 | 146224800 | 146226799 | 16.87259 | 31.53205 | 20.15768 | 8.358963 | 1.096523 | 0.173077 | 0.307973 |
| 4 | 89200200 | 89201399 | 6.4614 | 12.30767 | 28.28986 | 11.31118 | -0.68996 | -0.02359 | 0.307986 |
| 6 | 51989000 | 51989799 | 6.4614 | 12.30767 | 28.28986 | 11.31118 | -0.68996 | -0.02359 | 0.307986 |
| 7 | 132220000 | 132222799 | 15.71102 | 26.92141 | 53.69867 | 23.54953 | -0.60858 | 0.93269 | 0.308014 |
| 14 | 76109000 | 76112999 | 30.61367 | 49.54816 | 89.8989 | 38.45167 | -0.49436 | 1.66679 | 0.30802 |
| 6 | 67182200 | 67183199 | 8.495265 | 15.84905 | 33.30285 | 13.85013 | -0.61024 | 0.147771 | 0.308027 |
| 4 | 53011400 | 53012799 | 12.44469 | 21.74989 | 45.73466 | 20.15547 | -0.67931 | 0.688161 | 0.308062 |
| 2 | 98201000 | 98204599 | 53.85599 | 86.43629 | 80.35779 | 33.98997 | 0.453967 | 1.943376 | 0.308072 |
| 9 | 72723800 | 72724599 | 9.960389 | 19.23478 | 18.02586 | 6.939021 | 0.643287 | -0.1426 | 0.308076 |
| 3 | 93777200 | 93779399 | 24.15893 | 39.68904 | 76.46997 | 32.94583 | -0.56558 | 1.410633 | 0.308085 |
| 10 | 81855600 | 81859399 | 145.564 | 220.2891 | 202.0241 | 83.5702 | 0.402306 | 3.28834 | 0.308094 |
| X | 77183000 | 77184799 | 17.72354 | 32.57205 | 23.47995 | 9.978603 | 0.899249 | 0.284029 | 0.308096 |
| 9 | 68930800 | 68931999 | 11.8246 | 22.41924 | 20.37526 | 8.227006 | 0.624637 | 0.035877 | 0.308097 |
| 7 | 10097000 | 10100599 | 75.13225 | 119.4729 | 101.9833 | 42.57026 | 0.562252 | 2.257591 | 0.308098 |
| 17 | 39396600 | 39397199 | 10.28226 | 20.06091 | 14.18769 | 5.066698 | 1.142961 | -0.40359 | 0.308104 |
| 2 | 181311200 | 181313199 | 18.18138 | 30.66705 | 60.08197 | 26.27339 | -0.58548 | 1.089663 | 0.308109 |
| X | 26669200 | 26674199 | 45.62375 | 75.60949 | 44.3546 | 19.20202 | 1.147793 | 1.340019 | 0.308172 |
| 7 | 82499800 | 82500599 | 5.218052 | 10.15089 | 24.11994 | 8.905715 | -0.62807 | -0.31653 | 0.308182 |
| 1 | 34713600 | 34719599 | 36.96626 | 59.04026 | 127.2172 | 53.42607 | -0.76451 | 2.088111 | 0.3082 |
| 2 | 105970200 | 105972599 | 22.30908 | 36.94667 | 71.0698 | 30.75333 | -0.56014 | 1.304348 | 0.308216 |
| 14 | 44697200 | 44700399 | 54.27621 | 85.96399 | 145.942 | 60.69072 | -0.43368 | 2.363619 | 0.308238 |
| 5 | 135678800 | 135680999 | 9.667131 | 17.04486 | 41.01469 | 18.15149 | -0.87017 | 0.53825 | 0.308253 |
| 10 | 22378200 | 22379799 | 13.50184 | 23.69309 | 45.09381 | 19.94105 | -0.53635 | 0.626755 | 0.308266 |
| 3 | 38939200 | 38941999 | 23.24845 | 37.78899 | 86.50487 | 37.11693 | -0.82283 | 1.563173 | 0.30828 |
| 19 | 9235200 | 9237799 | 45.66993 | 74.14086 | 64.86882 | 27.80779 | 0.559352 | 1.620575 | 0.308292 |
| 4 | 54748200 | 54750399 | 23.74071 | 41.41393 | 34.28226 | 15.08981 | 0.65878 | 0.753857 | 0.308294 |
| 7 | 19482400 | 19484399 | 13.66039 | 23.4938 | 53.15625 | 23.3167 | -0.7886 | 0.887886 | 0.308311 |
| 13 | 118850000 | 118852599 | 18.73053 | 31.34899 | 64.35523 | 28.04206 | -0.65293 | 1.170508 | 0.308327 |
| X | 33749800 | 33753199 | 76.32059 | 121.8612 | 85.53178 | 35.90594 | 0.8504 | 2.128648 | 0.308344 |
| 4 | 129359000 | 129360599 | 9.793979 | 17.85284 | 35.50748 | 15.25333 | -0.56607 | 0.296921 | 0.308356 |
| 4 | 115775000 | 115783599 | 67.35058 | 104.9473 | 185.5733 | 77.01995 | -0.51082 | 2.701315 | 0.308363 |
| 6 | 17829800 | 17830799 | 13.92638 | 26.88025 | 17.71623 | 6.835696 | 1.143924 | -0.0913 | 0.308374 |
| 7 | 22443400 | 22443999 | 21.09963 | 37.2309 | 20.4171 | 9.034425 | 1.251308 | 0.557532 | 0.308377 |
| 7 | 55043800 | 55044599 | 8.29909 | 16.19278 | 13.77957 | 4.915308 | 0.879921 | -0.43866 | 0.308378 |
| 10 | 93514600 | 93515799 | 9.693204 | 17.38167 | 38.4072 | 16.81686 | -0.73547 | 0.396823 | 0.308384 |
| 17 | 60750400 | 60751399 | 16.53143 | 29.55515 | 26.67043 | 11.70737 | 0.546296 | 0.438 | 0.30839 |
| 19 | 41105000 | 41107599 | 16.26525 | 28.06137 | 52.18311 | 22.89201 | -0.50612 | 0.869556 | 0.308391 |
| X | 31641600 | 31643599 | 53.15269 | 86.49307 | 50.18766 | 21.54302 | 1.152927 | 1.586297 | 0.308393 |

| | | | | | | | | | |
|---|---|---|---|---|---|---|---|---|---|
| 4 | 8909200 | 8913999 | 22.95641 | 37.9607 | 72.81479 | 31.47521 | -0.55632 | 1.324771 | 0.308394 |
| 17 | 94816200 | 94816599 | 12.56122 | 23.47945 | 20.46651 | 8.484202 | 0.653503 | 0.136201 | 0.308396 |
| 15 | 4481600 | 4484199 | 29.45975 | 50.62754 | 38.06767 | 16.69778 | 0.793666 | 0.941895 | 0.308406 |
| 14 | 122771400 | 122772199 | 5.122333 | 9.899943 | 25.34972 | 9.721239 | -0.78674 | -0.18317 | 0.308414 |
| 2 | 123130800 | 123135399 | 39.60375 | 64.86866 | 57.73831 | 24.84426 | 0.543628 | 1.468218 | 0.308424 |
| 7 | 41476000 | 41480399 | 58.82684 | 95.09612 | 62.22353 | 26.58397 | 0.971781 | 1.731156 | 0.308434 |
| 8 | 108355200 | 108356999 | 13.08003 | 22.83641 | 45.75402 | 20.14995 | -0.61003 | 0.708106 | 0.308451 |
| 2 | 162748000 | 162750799 | 21.83332 | 36.22685 | 68.72877 | 29.77897 | -0.53994 | 1.281114 | 0.308454 |
| X | 78246600 | 78248199 | 24.21435 | 42.31369 | 33.19594 | 14.62775 | 0.736167 | 0.732013 | 0.308487 |
| 4 | 8567600 | 8569799 | 18.40775 | 33.04589 | 26.88542 | 11.76072 | 0.698325 | 0.426497 | 0.308492 |
| 13 | 10473600 | 10476599 | 41.8349 | 68.56718 | 57.72876 | 24.8469 | 0.624093 | 1.469017 | 0.308516 |
| 15 | 4184400 | 4186799 | 31.94305 | 54.41526 | 39.63257 | 17.37264 | 0.837544 | 1.029123 | 0.308516 |
| 6 | 69501600 | 69504199 | 26.10632 | 45.20245 | 37.06657 | 16.26239 | 0.67118 | 0.860364 | 0.308518 |
| 15 | 88759600 | 88760999 | 6.251143 | 11.72031 | 30.65544 | 12.64251 | -0.9132 | 0.088373 | 0.308525 |
| 11 | 84657400 | 84659599 | 12.62557 | 21.98981 | 47.8769 | 21.05289 | -0.72988 | 0.742038 | 0.308529 |
| 7 | 14166400 | 14179999 | 191.1229 | 287.1107 | 219.2 | 90.48061 | 0.657411 | 3.444986 | 0.308554 |
| 1 | 130389800 | 130391399 | 9.563756 | 17.12921 | 36.89756 | 16.17201 | -0.6995 | 0.403086 | 0.308554 |
| 5 | 8472200 | 8473199 | 9.563756 | 17.12921 | 36.89756 | 16.17201 | -0.6995 | 0.403086 | 0.308554 |
| 5 | 133813200 | 133815199 | 14.58631 | 25.08969 | 52.10978 | 22.85777 | -0.66556 | 0.89086 | 0.308563 |
| 14 | 42239000 | 42240399 | 15.07713 | 28.50492 | 20.60406 | 8.369508 | 0.945182 | 0.087022 | 0.308571 |
| 4 | 144247800 | 144251799 | 49.22178 | 79.90601 | 63.03637 | 27.02202 | 0.708314 | 1.625611 | 0.308574 |
| 5 | 112588600 | 112590999 | 14.56073 | 25.11737 | 51.06927 | 22.4033 | -0.63414 | 0.863063 | 0.308576 |
| 17 | 15523600 | 15527399 | 27.43934 | 44.36706 | 91.24801 | 38.99574 | -0.67605 | 1.68422 | 0.308621 |
| 3 | 131252400 | 131255999 | 40.91257 | 65.58707 | 112.5067 | 47.50507 | -0.43004 | 1.974073 | 0.308636 |
| 5 | 142202400 | 142203399 | 7.052921 | 12.95808 | 33.04059 | 14.04714 | -0.91037 | 0.226175 | 0.308668 |
| 8 | 71278000 | 71280199 | 18.54937 | 31.13089 | 62.17865 | 27.14047 | -0.61346 | 1.13926 | 0.308672 |
| 3 | 98810600 | 98813199 | 11.70185 | 20.43416 | 49.42168 | 21.76779 | -0.88023 | 0.70324 | 0.308677 |
| 6 | 96759400 | 96761999 | 22.35218 | 39.30181 | 30.32838 | 13.42959 | 0.759616 | 0.621609 | 0.308678 |
| 1 | 142291600 | 142293599 | 20.45705 | 35.86446 | 32.24906 | 14.24964 | 0.540024 | 0.657777 | 0.308682 |
| 4 | 38066800 | 38069999 | 37.64925 | 60.59635 | 105.6371 | 44.86886 | -0.44624 | 1.843805 | 0.308702 |
| 13 | 63297400 | 63300199 | 15.94717 | 27.41435 | 54.86578 | 24.06624 | -0.61288 | 0.901986 | 0.308703 |
| 1 | 151119800 | 151125399 | 33.66768 | 54.37084 | 101.1393 | 43.15437 | -0.53387 | 1.73242 | 0.308703 |
| 3 | 60676200 | 60677999 | 11.26029 | 20.13628 | 38.87719 | 17.06216 | -0.54427 | 0.427898 | 0.308722 |
| 9 | 45749800 | 45752599 | 11.14246 | 19.3072 | 50.46944 | 22.14319 | -0.99277 | 0.808741 | 0.308754 |
| 12 | 115575000 | 115576599 | 25.21585 | 44.38765 | 27.3327 | 12.10032 | 1.084 | 0.621385 | 0.30876 |
| 5 | 90663400 | 90664999 | 10.75974 | 19.45687 | 40.70809 | 17.6634 | -0.64859 | 0.35969 | 0.308763 |
| 9 | 81135200 | 81138999 | 81.22207 | 127.9766 | 121.1087 | 50.25401 | 0.402376 | 2.494314 | 0.308774 |
| 12 | 85846200 | 85847399 | 6.927027 | 13.09423 | 29.20652 | 11.86782 | -0.66763 | 0.037017 | 0.308775 |
| 18 | 56485400 | 56486199 | 6.927027 | 13.09423 | 29.20652 | 11.86782 | -0.66763 | 0.037017 | 0.308775 |
| 17 | 33921800 | 33922799 | 6.274237 | 12.09205 | 26.99679 | 10.46359 | -0.60615 | -0.12953 | 0.308786 |
| 13 | 42496600 | 42503599 | 51.44372 | 81.88072 | 143.1966 | 59.84931 | -0.46948 | 2.216845 | 0.308806 |
| 3 | 47827800 | 47831599 | 36.11097 | 60.85432 | 41.30697 | 18.05827 | 0.937993 | 1.137417 | 0.30881 |
| 5 | 65528800 | 65532399 | 23.80437 | 39.20652 | 74.92602 | 32.31553 | -0.55235 | 1.3767 | 0.308814 |
| 13 | 45166600 | 45167399 | 11.53049 | 22.0146 | 20.56978 | 8.18394 | 0.604029 | -0.01159 | 0.308822 |
| 9 | 122122800 | 122123999 | 9.876929 | 17.64887 | 37.38407 | 16.41753 | -0.67753 | 0.42069 | 0.308824 |
| 18 | 64455200 | 64455599 | 8.568972 | 16.68441 | 16.69752 | 6.121025 | 0.615306 | -0.32704 | 0.308826 |
| 10 | 48847400 | 48848799 | 15.33271 | 28.35898 | 23.6096 | 9.946789 | 0.701419 | 0.228856 | 0.308828 |

| | | | | | | | | | |
|---|---|---|---|---|---|---|---|---|---|
| 6 | 59530400 | 59531599 | 15.79307 | 29.26459 | 23.009 | 9.669356 | 0.78588 | 0.220915 | 0.308834 |
| X | 169304600 | 169304799 | 2.542772 | 4.973606 | 16.78071 | 5.52419 | -0.96939 | -0.78859 | 0.308835 |
| 7 | 9200800 | 9204399 | 37.75408 | 63.00139 | 42.70804 | 18.57477 | 0.940166 | 1.241139 | 0.308839 |
| 9 | 18771400 | 18772399 | 19.13844 | 33.60596 | 30.32005 | 13.41524 | 0.535142 | 0.622024 | 0.30884 |
| 14 | 77302800 | 77304199 | 16.90722 | 30.96004 | 24.49318 | 10.46039 | 0.760722 | 0.298507 | 0.308843 |
| 16 | 20833800 | 20837599 | 51.95803 | 83.68481 | 76.20848 | 32.40203 | 0.489868 | 1.828713 | 0.308846 |
| X | 3727800 | 3728599 | 15.03648 | 27.55902 | 24.68795 | 10.53074 | 0.58376 | 0.279139 | 0.308867 |
| 12 | 64847600 | 64852399 | 46.78148 | 76.97507 | 50.11655 | 21.60627 | 0.996013 | 1.428634 | 0.308874 |
| 9 | 5995400 | 5998399 | 68.89893 | 109.5552 | 103.0888 | 43.0295 | 0.421804 | 2.252591 | 0.308875 |
| 6 | 108611600 | 108613399 | 14.87235 | 27.20598 | 24.71258 | 10.56879 | 0.561172 | 0.288519 | 0.308881 |
| 13 | 68932200 | 68933599 | 16.03068 | 28.9507 | 26.11618 | 11.35033 | 0.55622 | 0.372815 | 0.308881 |
| 7 | 86316800 | 86319599 | 36.16497 | 60.4818 | 46.44662 | 20.22868 | 0.760714 | 1.215196 | 0.308904 |
| X | 151350200 | 151352199 | 28.09033 | 48.92209 | 32.50806 | 14.29429 | 0.974278 | 0.796754 | 0.308924 |
| 2 | 36549400 | 36552999 | 41.69197 | 68.82841 | 51.71475 | 22.32903 | 0.791582 | 1.37647 | 0.308937 |
| 17 | 78066600 | 78072599 | 41.22006 | 67.40625 | 54.29269 | 23.3469 | 0.686067 | 1.493787 | 0.308937 |
| 12 | 18357800 | 18359399 | 39.50119 | 65.63454 | 48.77904 | 21.15566 | 0.807936 | 1.297526 | 0.308948 |
| 19 | 61162200 | 61162999 | 14.69471 | 27.79745 | 20.62078 | 8.366404 | 0.909416 | 0.079533 | 0.308951 |
| 14 | 8113600 | 8128999 | 470.1033 | 630.7138 | 964.5063 | 402.5124 | -0.50465 | 5.225785 | 0.308964 |
| 12 | 117114200 | 117120199 | 48.47986 | 76.72002 | 157.7896 | 65.59012 | -0.71095 | 2.381497 | 0.308965 |
| 7 | 78547200 | 78557599 | 78.58782 | 120.8933 | 216.8022 | 90.03628 | -0.54549 | 2.942071 | 0.308969 |
| 7 | 10216600 | 10218599 | 16.71943 | 30.62599 | 24.50013 | 10.45878 | 0.74518 | 0.295154 | 0.308978 |
| 4 | 21107200 | 21109399 | 21.79756 | 38.88543 | 26.00203 | 11.4349 | 0.974516 | 0.486178 | 0.30899 |
| 5 | 150874400 | 150877399 | 22.58268 | 37.11572 | 80.80079 | 34.81812 | -0.74096 | 1.405361 | 0.308991 |
| 11 | 113514400 | 113517199 | 29.02519 | 46.93418 | 92.43753 | 39.50728 | -0.61373 | 1.684223 | 0.308992 |
| X | 33316200 | 33318399 | 36.40122 | 60.57776 | 49.26883 | 21.39102 | 0.678446 | 1.27238 | 0.30903 |
| 18 | 37968400 | 37979399 | 76.55385 | 117.9228 | 212.9235 | 88.44177 | -0.55383 | 2.914592 | 0.309032 |
| 13 | 50189200 | 50191799 | 15.99143 | 29.26504 | 20.27262 | 8.665152 | 0.950287 | 0.285502 | 0.309088 |
| 10 | 41903000 | 41905599 | 13.62347 | 23.5212 | 51.05826 | 22.39907 | -0.72782 | 0.850232 | 0.309091 |
| X | 89444400 | 89445599 | 18.72936 | 34.2896 | 23.19148 | 9.907249 | 0.985004 | 0.310644 | 0.309092 |
| 16 | 10455400 | 10458199 | 10.59357 | 18.61088 | 44.07103 | 19.50756 | -0.84957 | 0.589771 | 0.309106 |
| 12 | 23674200 | 23675999 | 48.09467 | 77.53943 | 68.60349 | 29.20876 | 0.533147 | 1.791326 | 0.309107 |
| 1 | 160861800 | 160862999 | 5.88402 | 11.24384 | 28.35794 | 11.26185 | -0.81402 | -0.05233 | 0.309117 |
| 9 | 115457800 | 115458999 | 5.88402 | 11.24384 | 28.35794 | 11.26185 | -0.81402 | -0.05233 | 0.309117 |
| 12 | 115654000 | 115655999 | 41.20917 | 67.22223 | 61.5486 | 26.44327 | 0.499173 | 1.532144 | 0.309118 |
| 15 | 57762000 | 57765799 | 47.47647 | 76.58223 | 71.83313 | 30.60416 | 0.44985 | 1.77471 | 0.30913 |
| 10 | 106116000 | 106117799 | 19.99539 | 35.00732 | 31.48797 | 13.89732 | 0.539732 | 0.676443 | 0.309139 |
| 17 | 12092800 | 12093799 | 5.13234 | 9.949351 | 28.03834 | 10.60129 | -0.90558 | -0.22804 | 0.309142 |
| 2 | 177275000 | 177275799 | 11.23098 | 21.39158 | 17.07106 | 6.826343 | 0.823417 | -0.00896 | 0.309152 |
| 6 | 105291400 | 105293399 | 29.98682 | 51.40162 | 37.40462 | 16.40425 | 0.839945 | 0.963062 | 0.309154 |
| 4 | 36411800 | 36412199 | 3.877428 | 7.579158 | 20.08571 | 6.91837 | -0.6898 | -0.59327 | 0.309164 |
| X | 148859800 | 148861199 | 14.98091 | 27.85964 | 23.24975 | 9.721713 | 0.706033 | 0.196374 | 0.309178 |
| 9 | 92120400 | 92129599 | 86.34905 | 135.9949 | 113.9876 | 47.29639 | 0.576753 | 2.505452 | 0.309203 |
| 15 | 89671600 | 89673399 | 16.45633 | 28.44594 | 53.81514 | 23.60936 | -0.53022 | 0.856059 | 0.309204 |
| 7 | 5191000 | 5192599 | 16.59479 | 30.93444 | 21.96947 | 9.14944 | 0.941572 | 0.191617 | 0.309238 |
| 7 | 108426800 | 108434199 | 23.79519 | 41.32608 | 32.05603 | 14.0708 | 0.751659 | 0.812112 | 0.309238 |
| 5 | 137271200 | 137273399 | 37.26102 | 61.24465 | 57.02236 | 24.57243 | 0.480882 | 1.421634 | 0.309255 |
| 2 | 80026200 | 80026999 | 6.740042 | 12.85931 | 28.92715 | 11.52631 | -0.65514 | -0.03162 | 0.309264 |

| | | | | | | | | | |
|---|---|---|---|---|---|---|---|---|---|
| 9 | 93531600 | 93533799 | 46.25 | 74.90383 | 55.28964 | 23.6598 | 0.800794 | 1.661555 | 0.309269 |
| 6 | 76906000 | 76906999 | 6.265108 | 12.10777 | 27.18368 | 10.42783 | -0.60019 | -0.16438 | 0.309278 |
| 17 | 69222000 | 69223999 | 20.91525 | 36.73963 | 29.83527 | 13.20491 | 0.685966 | 0.623471 | 0.30928 |
| X | 117306000 | 117307599 | 26.65934 | 45.82748 | 39.32011 | 17.24729 | 0.604141 | 0.929605 | 0.309282 |
| 7 | 130964000 | 130965199 | 5.923033 | 11.21417 | 29.21955 | 11.83329 | -0.88538 | 0.016023 | 0.309285 |
| 5 | 95513000 | 95515799 | 35.61547 | 59.62738 | 45.87223 | 19.99218 | 0.75808 | 1.201214 | 0.30929 |
| 12 | 11603600 | 11607399 | 35.00935 | 58.97741 | 43.01513 | 18.80305 | 0.834757 | 1.138679 | 0.309292 |
| 16 | 40433000 | 40435199 | 66.79654 | 107.0432 | 76.51044 | 32.28824 | 0.829857 | 2.000627 | 0.309294 |
| 7 | 15069200 | 15087199 | 531.8916 | 729.2552 | 571.824 | 237.8435 | 0.490306 | 4.885604 | 0.309298 |
| 14 | 47835400 | 47837999 | 10.57659 | 19.04559 | 37.90455 | 16.53348 | -0.58181 | 0.376133 | 0.309309 |
| 8 | 41043000 | 41045599 | 18.16437 | 30.71461 | 59.02253 | 25.82957 | -0.5575 | 1.067192 | 0.309319 |
| 9 | 88443400 | 88446999 | 63.2018 | 101.1732 | 87.45895 | 36.84838 | 0.554339 | 2.035979 | 0.309331 |
| 14 | 26294200 | 26295399 | 10.45537 | 18.83504 | 36.88834 | 16.08194 | -0.55798 | 0.373017 | 0.309332 |
| 1 | 156892000 | 156896999 | 48.96443 | 77.91006 | 133.661 | 55.83959 | -0.44226 | 2.221988 | 0.309335 |
| 10 | 12142200 | 12142799 | 8.049342 | 14.86736 | 33.08099 | 13.96325 | -0.70872 | 0.200669 | 0.309345 |
| 2 | 33927200 | 33929799 | 15.79285 | 27.21178 | 52.16663 | 22.8839 | -0.5502 | 0.878618 | 0.309354 |
| 2 | 52268200 | 52270199 | 17.22932 | 29.58327 | 54.8804 | 24.07173 | -0.50422 | 0.919236 | 0.309365 |
| 18 | 68422000 | 68423799 | 13.6581 | 23.92251 | 45.66727 | 20.16275 | -0.54069 | 0.658525 | 0.309376 |
| 16 | 18507600 | 18512399 | 32.06302 | 51.80571 | 95.33213 | 40.70319 | -0.51739 | 1.713123 | 0.309392 |
| 15 | 29291000 | 29291999 | 18.65071 | 32.93285 | 28.34175 | 12.53926 | 0.605554 | 0.552098 | 0.309412 |
| 14 | 124565800 | 124567599 | 15.61756 | 27.1254 | 50.58631 | 22.20545 | -0.5082 | 0.801013 | 0.309422 |
| 13 | 19449200 | 19451199 | 26.79793 | 43.8777 | 80.39924 | 34.58979 | -0.49529 | 1.463515 | 0.309429 |
| 2 | 17822800 | 17825799 | 13.17185 | 23.08752 | 46.71294 | 20.63703 | -0.62429 | 0.644545 | 0.309435 |
| 13 | 100460000 | 100465199 | 42.79346 | 69.80042 | 61.58461 | 26.45789 | 0.552432 | 1.538251 | 0.309442 |
| 10 | 96168200 | 96170399 | 11.38241 | 19.90628 | 49.41455 | 21.78921 | -0.91772 | 0.675256 | 0.309457 |
| 16 | 55869000 | 55869599 | 13.80232 | 23.77893 | 51.56948 | 22.62186 | -0.72705 | 0.872636 | 0.30946 |
| 12 | 33059600 | 33061399 | 11.27631 | 20.1888 | 39.94714 | 17.51426 | -0.57846 | 0.423182 | 0.309466 |
| 7 | 107656000 | 107659399 | 47.33747 | 76.56273 | 62.46673 | 26.70476 | 0.65495 | 1.694665 | 0.309472 |
| 4 | 122439600 | 122440599 | 14.91604 | 27.77682 | 20.54241 | 8.569868 | 0.881336 | 0.179596 | 0.309495 |
| 2 | 39547600 | 39550799 | 33.47255 | 55.96069 | 48.11095 | 20.94921 | 0.598376 | 1.211033 | 0.309502 |
| 3 | 145706800 | 145711999 | 46.35816 | 73.77318 | 134.3213 | 56.12608 | -0.52772 | 2.21487 | 0.309502 |
| 1 | 117696200 | 117714199 | 249.1782 | 368.6846 | 247.9376 | 102.1077 | 0.820449 | 3.725907 | 0.309512 |
| X | 85080600 | 85082599 | 23.98733 | 43.12663 | 21.40342 | 9.350929 | 1.40779 | 0.44557 | 0.309538 |
| 1 | 118017200 | 118022399 | 86.21394 | 136.2734 | 112.5826 | 46.77042 | 0.601621 | 2.426015 | 0.309549 |
| 3 | 20986000 | 20988199 | 26.06768 | 42.93839 | 78.10098 | 33.68626 | -0.48128 | 1.384374 | 0.309555 |
| 7 | 85341800 | 85342999 | 15.18429 | 29.27976 | 16.33093 | 6.319871 | 1.377924 | -0.07312 | 0.309583 |
| 4 | 11771200 | 11772999 | 16.09173 | 27.74148 | 52.72226 | 23.12801 | -0.53764 | 0.875194 | 0.309592 |
| 8 | 44423400 | 44427399 | 34.64568 | 58.38349 | 43.01307 | 18.80398 | 0.820274 | 1.13444 | 0.3096 |
| 14 | 51619400 | 51622199 | 76.33099 | 118.0169 | 199.5959 | 82.90753 | -0.4554 | 2.845438 | 0.309611 |
| 16 | 76345400 | 76347599 | 25.8726 | 44.20064 | 41.00357 | 17.97775 | 0.490527 | 0.981975 | 0.309616 |
| 6 | 130463400 | 130466999 | 52.72048 | 85.51683 | 59.91575 | 25.66905 | 0.877918 | 1.645033 | 0.309638 |
| 12 | 54365400 | 54368399 | 33.90858 | 56.04085 | 52.54301 | 22.70396 | 0.473203 | 1.343805 | 0.309684 |
| 5 | 93926400 | 93929199 | 44.13718 | 72.06455 | 60.02485 | 25.79724 | 0.63629 | 1.526431 | 0.309693 |
| 15 | 96137200 | 96142999 | 47.1618 | 75.17952 | 127.4704 | 53.38864 | -0.42247 | 2.160212 | 0.309696 |
| X | 20524400 | 20525999 | 22.43337 | 39.41647 | 29.64946 | 13.12551 | 0.796426 | 0.633062 | 0.309719 |
| 11 | 95681800 | 95683599 | 18.90133 | 32.04392 | 59.70029 | 26.14677 | -0.51301 | 1.046081 | 0.309729 |
| 7 | 111817600 | 111820599 | 25.09749 | 41.40982 | 75.11455 | 32.4212 | -0.47629 | 1.35989 | 0.309731 |

| | | | | | | | | | |
|---|---|---|---|---|---|---|---|---|---|
| 1 | 27889200 | 27890399 | 14.78147 | 27.45467 | 21.94881 | 9.19432 | 0.76489 | 0.195892 | 0.309731 |
| 14 | 109285600 | 109287199 | 15.85557 | 29.90788 | 17.99574 | 7.345582 | 1.200372 | 0.101287 | 0.309742 |
| 3 | 53370000 | 53371399 | 8.760493 | 15.83518 | 35.93616 | 15.60077 | -0.76503 | 0.337877 | 0.309758 |
| 10 | 71247000 | 71249999 | 17.57203 | 29.9722 | 58.65795 | 25.7149 | -0.58273 | 0.984129 | 0.309768 |
| 17 | 86248200 | 86253399 | 35.18438 | 56.60287 | 104.5128 | 44.36147 | -0.5298 | 1.852526 | 0.309775 |
| 8 | 12582400 | 12584199 | 10.08976 | 18.26363 | 36.9552 | 16.01493 | -0.59938 | 0.340352 | 0.309794 |
| 10 | 121297400 | 121300799 | 19.62167 | 32.56447 | 68.47301 | 29.67274 | -0.68769 | 1.272225 | 0.309832 |
| 8 | 48562200 | 48563799 | 10.18363 | 18.15029 | 38.87318 | 17.10845 | -0.69586 | 0.44023 | 0.309844 |
| 7 | 4992400 | 4998599 | 106.8153 | 161.6198 | 270.5145 | 111.8959 | -0.46521 | 3.288859 | 0.309845 |
| 6 | 127329600 | 127329999 | 4.229232 | 8.248516 | 21.44286 | 7.692425 | -0.71728 | -0.44168 | 0.309856 |
| 12 | 22394800 | 22395799 | 14.99992 | 27.62468 | 24.15476 | 10.23604 | 0.625126 | 0.250844 | 0.309858 |
| 16 | 33200800 | 33201999 | 7.866223 | 14.26786 | 35.9953 | 15.55917 | -0.91247 | 0.310951 | 0.309864 |
| 4 | 141427200 | 141430599 | 14.50046 | 25.10832 | 51.10498 | 22.42155 | -0.6347 | 0.826045 | 0.309874 |
| 15 | 68931200 | 68932999 | 10.39068 | 18.85636 | 39.17636 | 16.92341 | -0.63446 | 0.331801 | 0.309883 |
| 11 | 51646800 | 51647799 | 6.979861 | 13.04098 | 31.13857 | 12.91791 | -0.7901 | 0.118265 | 0.309893 |
| 15 | 69934400 | 69937599 | 36.4923 | 58.65526 | 105.5174 | 44.73071 | -0.49389 | 1.882987 | 0.309894 |
| 12 | 53298200 | 53300599 | 14.06905 | 24.30214 | 51.60976 | 22.64134 | -0.69635 | 0.847346 | 0.309902 |
| 1 | 35153600 | 35157799 | 109.2701 | 168.687 | 157.7816 | 65.53934 | 0.396527 | 2.88151 | 0.309913 |
| 7 | 135107800 | 135109399 | 21.54698 | 36.11337 | 66.93935 | 29.19383 | -0.50636 | 1.165804 | 0.309924 |
| 7 | 10184800 | 10185399 | 11.5429 | 21.98788 | 20.57044 | 8.223939 | 0.59624 | 0.004202 | 0.309928 |
| 2 | 101026600 | 101028799 | 27.89692 | 48.49636 | 33.56276 | 14.74109 | 0.915577 | 0.820785 | 0.309937 |
| 12 | 110328800 | 110337399 | 74.66793 | 115.5671 | 198.9453 | 82.63681 | -0.47976 | 2.824274 | 0.309954 |
| 16 | 89761200 | 89775399 | 121.6666 | 182.9691 | 295.3766 | 121.9598 | -0.42078 | 3.413756 | 0.309962 |
| 11 | 91081800 | 91084399 | 19.48296 | 34.66824 | 27.74971 | 12.22961 | 0.714359 | 0.488458 | 0.309984 |
| X | 126700200 | 126704399 | 87.51633 | 138.1948 | 116.1665 | 48.23506 | 0.575538 | 2.453156 | 0.309992 |
| 7 | 42876200 | 42880199 | 70.81506 | 112.9141 | 95.19858 | 39.88659 | 0.584221 | 2.16612 | 0.310024 |
| 8 | 100449400 | 100451799 | 15.5667 | 27.05239 | 51.68538 | 22.69598 | -0.54302 | 0.792127 | 0.310059 |
| 14 | 66688800 | 66691599 | 29.43496 | 50.05586 | 38.81436 | 17.0116 | 0.747181 | 1.034211 | 0.310063 |
| 9 | 18921400 | 18922599 | 16.77143 | 31.32963 | 21.55818 | 8.946186 | 0.991202 | 0.180276 | 0.310084 |
| 9 | 19065800 | 19069599 | 62.34651 | 99.7801 | 88.45888 | 37.25702 | 0.517624 | 2.042168 | 0.310102 |
| 5 | 89540400 | 89544599 | 59.3381 | 95.09646 | 78.68528 | 33.20902 | 0.619259 | 1.993273 | 0.310104 |
| X | 94393000 | 94393999 | 13.71231 | 25.61437 | 22.4204 | 9.304427 | 0.646521 | 0.15657 | 0.310104 |
| 7 | 41835600 | 41838399 | 56.62702 | 91.03734 | 77.71478 | 32.95457 | 0.57973 | 1.895488 | 0.310121 |
| 4 | 109430400 | 109433199 | 18.38444 | 31.41021 | 58.14329 | 25.4927 | -0.50231 | 0.972971 | 0.310123 |
| 1 | 163407200 | 163408199 | 15.26891 | 27.78767 | 25.24893 | 10.86847 | 0.554826 | 0.322576 | 0.310131 |
| 14 | 3195200 | 3202999 | 168.227 | 252.5809 | 232.2414 | 95.85009 | 0.388667 | 3.450481 | 0.310163 |
| 1 | 68390400 | 68391999 | 24.97378 | 43.44502 | 35.42016 | 15.56369 | 0.680412 | 0.800365 | 0.310183 |
| 6 | 32631800 | 32633399 | 10.07967 | 18.42186 | 38.28501 | 16.39265 | -0.62639 | 0.287923 | 0.310206 |
| X | 63604800 | 63605999 | 19.41717 | 34.63013 | 22.70314 | 9.986298 | 1.002809 | 0.466004 | 0.310211 |
| 12 | 106851600 | 106853999 | 19.14449 | 32.26073 | 60.65865 | 26.51789 | -0.52648 | 1.102371 | 0.310223 |
| 13 | 30362600 | 30363799 | 6.285445 | 12.06716 | 28.61621 | 11.21438 | -0.70835 | -0.09244 | 0.310231 |
| 12 | 18760400 | 18762199 | 41.80113 | 67.8445 | 61.09114 | 26.18391 | 0.517661 | 1.61486 | 0.310235 |
| 4 | 52954200 | 52955599 | 7.682374 | 14.55203 | 31.5372 | 12.75913 | -0.62114 | 0.036372 | 0.310238 |
| 8 | 58859200 | 58861399 | 21.62032 | 38.50311 | 26.5495 | 11.69256 | 0.929076 | 0.496338 | 0.310251 |
| 1 | 52738200 | 52739399 | 15.23258 | 28.5157 | 22.13483 | 9.152407 | 0.822373 | 0.154118 | 0.310261 |
| 10 | 129056800 | 129058999 | 36.20558 | 60.20036 | 50.4069 | 21.87214 | 0.636454 | 1.279937 | 0.310262 |
| 6 | 129801400 | 129801999 | 11.83611 | 22.84735 | 19.00375 | 7.329943 | 0.811523 | -0.11161 | 0.310263 |

| | | | | | | | | | |
|---|---|---|---|---|---|---|---|---|---|
| 9 | 116513400 | 116513999 | 11.83611 | 22.84735 | 19.00375 | 7.329943 | 0.811523 | -0.11161 | 0.310263 |
| 14 | 105749600 | 105752399 | 18.2247 | 31.19041 | 58.15936 | 25.50324 | -0.51241 | 0.95706 | 0.310306 |
| 18 | 22196800 | 22199199 | 52.87976 | 84.93808 | 78.67958 | 33.32103 | 0.460548 | 1.916815 | 0.310315 |
| 4 | 54373800 | 54375199 | 8.342882 | 15.47656 | 32.08352 | 13.46249 | -0.60087 | 0.178078 | 0.310346 |
| 1 | 40345800 | 40349599 | 40.44886 | 64.80751 | 112.4664 | 47.44775 | -0.44768 | 1.991501 | 0.310379 |
| 18 | 68048800 | 68051199 | 26.52487 | 45.95542 | 36.54797 | 16.03516 | 0.71538 | 0.857071 | 0.310379 |
| 7 | 98351800 | 98353799 | 8.00713 | 14.53762 | 38.10245 | 16.44991 | -0.96646 | 0.308911 | 0.310382 |
| 5 | 119890600 | 119892199 | 12.31507 | 21.52314 | 45.23129 | 19.93347 | -0.67819 | 0.689548 | 0.31039 |
| 18 | 60831400 | 60833799 | 8.926309 | 15.76136 | 41.51808 | 18.36898 | -0.99909 | 0.517442 | 0.310404 |
| 18 | 41647400 | 41649399 | 15.28716 | 28.84588 | 19.25561 | 7.853976 | 1.053176 | 0.097078 | 0.310422 |
| 11 | 8559200 | 8561599 | 10.71327 | 18.89209 | 40.95585 | 18.12493 | -0.72078 | 0.541757 | 0.310436 |
| 16 | 84465000 | 84467599 | 63.76955 | 101.8756 | 92.47915 | 38.85196 | 0.480779 | 2.099852 | 0.310445 |
| 15 | 44516000 | 44517799 | 18.17149 | 33.21709 | 23.74415 | 10.16418 | 0.903616 | 0.315611 | 0.310454 |
| 14 | 68514400 | 68516599 | 12.87355 | 22.6034 | 43.5175 | 19.25322 | -0.55274 | 0.611703 | 0.310454 |
| 7 | 32955400 | 32969799 | 439.3732 | 609.9265 | 514.6998 | 213.663 | 0.40245 | 4.693064 | 0.310457 |
| 11 | 76069600 | 76070999 | 8.264655 | 15.08463 | 33.98133 | 14.57413 | -0.74269 | 0.273974 | 0.310472 |
| 13 | 100787200 | 100788399 | 8.264655 | 15.08463 | 33.98133 | 14.57413 | -0.74269 | 0.273974 | 0.310472 |
| 3 | 53795000 | 53797199 | 37.98485 | 62.43299 | 57.02249 | 24.57225 | 0.508537 | 1.424614 | 0.31048 |
| 16 | 67540400 | 67540999 | 9.659631 | 18.77146 | 17.68208 | 6.584548 | 0.681166 | -0.25634 | 0.310487 |
| 10 | 43612600 | 43617799 | 28.6819 | 46.43406 | 92.18611 | 39.43777 | -0.62359 | 1.657221 | 0.31049 |
| 19 | 3719800 | 3728999 | 78.70405 | 121.4733 | 204.1142 | 84.78509 | -0.44801 | 2.878923 | 0.310503 |
| 13 | 79800000 | 79803599 | 33.54416 | 55.79483 | 49.28475 | 21.39045 | 0.559485 | 1.264997 | 0.310507 |
| 6 | 78638800 | 78639999 | 6.574336 | 12.67733 | 30.04997 | 11.62874 | -0.69013 | -0.12368 | 0.310515 |
| 7 | 46769200 | 46770199 | 6.574336 | 12.67733 | 30.04997 | 11.62874 | -0.69013 | -0.12368 | 0.310515 |
| 3 | 16640400 | 16640999 | 11.80682 | 22.58015 | 20.52538 | 8.133267 | 0.648463 | -0.02249 | 0.310522 |
| 17 | 24688800 | 24694399 | 23.72421 | 38.54815 | 86.51347 | 37.11058 | -0.79487 | 1.571696 | 0.310536 |
| 2 | 177057600 | 177058799 | 11.23699 | 21.70757 | 17.20044 | 6.610577 | 0.884579 | -0.13742 | 0.310552 |
| 5 | 110211000 | 110211999 | 20.74396 | 36.40869 | 31.57905 | 13.96695 | 0.591672 | 0.639863 | 0.310563 |
| X | 121316800 | 121318799 | 28.03036 | 49.02246 | 29.67397 | 13.08492 | 1.108427 | 0.734636 | 0.310592 |
| 6 | 90389000 | 90401999 | 87.74713 | 134.2149 | 233.0653 | 96.6981 | -0.50546 | 3.059785 | 0.310604 |
| 18 | 4532000 | 4534199 | 46.13679 | 74.70589 | 60.82033 | 26.02278 | 0.659548 | 1.668336 | 0.310607 |
| X | 125097400 | 125098399 | 13.30983 | 25.40349 | 20.37988 | 8.114014 | 0.821188 | 0.007098 | 0.310616 |
| 1 | 53999600 | 54001199 | 18.32035 | 32.96238 | 26.28673 | 11.47723 | 0.728684 | 0.410382 | 0.310654 |
| 1 | 171280400 | 171281599 | 9.157826 | 16.75874 | 33.9919 | 14.52979 | -0.58923 | 0.266172 | 0.310655 |
| 3 | 104114800 | 104115999 | 13.0124 | 24.61643 | 21.3248 | 8.651046 | 0.687488 | 0.065058 | 0.310659 |
| 15 | 27041600 | 27044199 | 21.50846 | 37.80497 | 30.94793 | 13.7045 | 0.674684 | 0.621538 | 0.310659 |
| 10 | 39245400 | 39246599 | 8.733424 | 15.85404 | 35.4603 | 15.31191 | -0.73922 | 0.311172 | 0.310668 |
| 19 | 19554000 | 19554999 | 19.68929 | 34.69025 | 30.5783 | 13.53474 | 0.569776 | 0.584421 | 0.310671 |
| 7 | 11530000 | 11544399 | 453.7386 | 626.8598 | 530.4573 | 220.3712 | 0.391482 | 4.766115 | 0.310688 |
| 18 | 19367200 | 19368599 | 17.17821 | 32.0026 | 21.28909 | 8.875188 | 1.033867 | 0.198182 | 0.310707 |
| 7 | 65758000 | 65760199 | 13.51545 | 23.58772 | 47.32605 | 20.83734 | -0.61239 | 0.715766 | 0.310707 |
| 2 | 75452200 | 75453999 | 10.96408 | 19.48533 | 38.83735 | 17.13405 | -0.59517 | 0.464341 | 0.310713 |
| 4 | 38219200 | 38220799 | 17.49949 | 31.8155 | 24.21106 | 10.43465 | 0.807468 | 0.34052 | 0.310724 |
| 17 | 3464400 | 3466599 | 14.30132 | 24.53628 | 53.13468 | 23.30452 | -0.72623 | 0.915155 | 0.310737 |
| 3 | 90432800 | 90436399 | 20.61384 | 34.07367 | 74.18524 | 32.05893 | -0.73847 | 1.323959 | 0.31074 |
| 1 | 101875800 | 101877599 | 18.71534 | 33.27181 | 28.4269 | 12.53774 | 0.620122 | 0.490878 | 0.310743 |
| 1 | 111636200 | 111638199 | 27.92667 | 47.53818 | 36.38288 | 15.94721 | 0.766199 | 1.016931 | 0.310758 |

| | | | | | | | | | |
|---|---|---|---|---|---|---|---|---|---|
| X | 42819400 | 42827599 | 116.4419 | 180.6318 | 143.6703 | 59.66855 | 0.6355 | 2.797068 | 0.31076 |
| 3 | 92709400 | 92710199 | 11.28609 | 21.67829 | 19.61973 | 7.677975 | 0.671906 | -0.06844 | 0.310764 |
| 4 | 53900400 | 53901799 | 13.94926 | 25.64118 | 22.74951 | 9.665381 | 0.601529 | 0.249131 | 0.310769 |
| 13 | 100469800 | 100470999 | 12.90915 | 24.80598 | 18.11613 | 7.081258 | 0.978954 | -0.06242 | 0.310781 |
| 1 | 79340600 | 79341799 | 8.281885 | 15.49755 | 32.86317 | 13.59777 | -0.61786 | 0.122357 | 0.310785 |
| 9 | 8169800 | 8170399 | 15.56756 | 28.09116 | 25.26173 | 10.99031 | 0.559852 | 0.372859 | 0.310798 |
| 8 | 52885600 | 52886799 | 11.15971 | 21.65776 | 17.44827 | 6.563113 | 0.890783 | -0.20614 | 0.310806 |
| 7 | 63857600 | 63860999 | 24.93919 | 40.98906 | 77.26747 | 33.29567 | -0.53394 | 1.406958 | 0.310809 |
| 17 | 50470200 | 50476399 | 83.5681 | 131.5046 | 124.1976 | 51.52717 | 0.403861 | 2.521636 | 0.310812 |
| 6 | 128599600 | 128600999 | 21.11784 | 36.81323 | 32.65533 | 14.36749 | 0.5595 | 0.748417 | 0.310828 |
| 13 | 20220200 | 20221599 | 10.78384 | 19.27416 | 37.84683 | 16.61731 | -0.56888 | 0.425572 | 0.310844 |
| 12 | 52190800 | 52192799 | 15.04333 | 28.16974 | 22.14199 | 9.150789 | 0.804954 | 0.150507 | 0.310854 |
| 17 | 38632400 | 38640599 | 129.4201 | 198.3966 | 161.7674 | 67.14865 | 0.586692 | 3.022482 | 0.310858 |
| 5 | 128506000 | 128511599 | 43.61961 | 69.46365 | 129.1349 | 54.01114 | -0.55655 | 2.189421 | 0.310868 |
| 7 | 22485400 | 22504399 | 790.1505 | 1046.229 | 820.8982 | 343.1942 | 0.438891 | 5.438984 | 0.310874 |
| 4 | 148158800 | 148161199 | 13.8712 | 24.08016 | 49.50052 | 21.72199 | -0.64805 | 0.797401 | 0.310881 |
| 2 | 99373000 | 99374399 | 12.80268 | 24.59477 | 19.3964 | 7.587551 | 0.86884 | -0.0562 | 0.310885 |
| 3 | 80534600 | 80535799 | 12.80268 | 24.59477 | 19.3964 | 7.587551 | 0.86884 | -0.0562 | 0.310885 |
| X | 25790000 | 25790599 | 12.80268 | 24.59477 | 19.3964 | 7.587551 | 0.86884 | -0.0562 | 0.310885 |
| 17 | 37809400 | 37811599 | 23.57388 | 41.34298 | 30.7958 | 13.61145 | 0.810214 | 0.667111 | 0.310929 |
| 13 | 65855000 | 65865399 | 83.93692 | 133.0006 | 100.291 | 41.7159 | 0.735936 | 2.365698 | 0.310929 |
| 16 | 19711400 | 19711999 | 5.079745 | 9.883125 | 23.52978 | 8.678651 | -0.6294 | -0.32424 | 0.31093 |
| 2 | 58456000 | 58458599 | 17.88266 | 30.08816 | 63.36343 | 27.68742 | -0.68959 | 1.111815 | 0.310945 |
| 1 | 58965000 | 58966199 | 21.19008 | 37.06439 | 33.41908 | 14.73801 | 0.536297 | 0.701037 | 0.310946 |
| 7 | 27616800 | 27620199 | 31.24543 | 50.68482 | 89.89772 | 38.51514 | -0.45867 | 1.622863 | 0.310947 |
| 2 | 95704600 | 95705399 | 5.513211 | 10.72267 | 25.37703 | 9.386824 | -0.6255 | -0.30417 | 0.310961 |
| 7 | 21083200 | 21085799 | 23.7148 | 42.0076 | 25.7241 | 11.37394 | 1.095086 | 0.543836 | 0.310963 |
| 19 | 35946600 | 35947399 | 7.730185 | 14.35844 | 32.10415 | 13.44758 | -0.70749 | 0.164042 | 0.310974 |
| 15 | 89586200 | 89586999 | 7.868681 | 14.67889 | 31.11275 | 12.94107 | -0.62262 | 0.137201 | 0.31099 |
| 7 | 30209000 | 30213199 | 27.47663 | 44.75461 | 82.29826 | 35.33887 | -0.50566 | 1.539702 | 0.31101 |
| 11 | 49039000 | 49039999 | 13.56204 | 23.66816 | 46.28063 | 20.37657 | -0.57517 | 0.717926 | 0.311015 |
| 7 | 142960800 | 142964599 | 27.80884 | 45.27038 | 82.82399 | 35.55736 | -0.49891 | 1.549154 | 0.311015 |
| 2 | 165074600 | 165078399 | 31.77849 | 51.26837 | 96.32296 | 41.04606 | -0.54977 | 1.75813 | 0.311027 |
| 3 | 12500200 | 12501199 | 15.20314 | 27.67295 | 25.19211 | 10.84157 | 0.552209 | 0.320202 | 0.311037 |
| 17 | 32769200 | 32770399 | 8.29059 | 15.06104 | 35.48098 | 15.30699 | -0.81262 | 0.303722 | 0.311053 |
| 19 | 34601000 | 34602799 | 14.05863 | 24.35699 | 49.49365 | 21.71498 | -0.63191 | 0.818899 | 0.311062 |
| 18 | 32958400 | 32960199 | 22.44071 | 39.29926 | 32.61351 | 14.3974 | 0.655247 | 0.686499 | 0.311074 |
| 11 | 85613400 | 85615199 | 8.470813 | 15.81327 | 32.22197 | 13.3865 | -0.56509 | 0.140064 | 0.311086 |
| 2 | 172790800 | 172803999 | 107.1431 | 162.2079 | 268.5911 | 111.1202 | -0.44899 | 3.276285 | 0.311093 |
| 4 | 156239200 | 156264999 | 357.135 | 491.0313 | 752.8794 | 313.0125 | -0.47281 | 4.838453 | 0.311093 |
| 4 | 134778600 | 134782199 | 37.85733 | 60.64456 | 112.2936 | 47.36206 | -0.54133 | 1.993508 | 0.311112 |
| 9 | 40769000 | 40770999 | 10.85348 | 19.15351 | 41.43944 | 18.33624 | -0.71765 | 0.534173 | 0.311126 |
| 6 | 102356600 | 102357799 | 10.82048 | 20.80505 | 18.38061 | 7.172119 | 0.709137 | -0.08948 | 0.31113 |
| 8 | 16297000 | 16299399 | 21.80163 | 38.35105 | 30.33312 | 13.43048 | 0.724429 | 0.613111 | 0.311133 |
| 9 | 52837000 | 52838199 | 12.19722 | 23.20663 | 21.02525 | 8.426975 | 0.638939 | 0.019666 | 0.31115 |
| X | 126148600 | 126150999 | 33.89793 | 57.64197 | 39.68742 | 17.39416 | 0.91814 | 1.054934 | 0.311161 |
| 11 | 108158600 | 108160799 | 11.12285 | 19.76226 | 38.83409 | 17.13644 | -0.57502 | 0.467365 | 0.31118 |

| | | | | | | | | | |
|---|---|---|---|---|---|---|---|---|---|
| 18 | 37952000 | 37961199 | 89.9532 | 137.0302 | 246.1108 | 102.0112 | -0.55871 | 3.144256 | 0.311198 |
| 10 | 77656000 | 77665799 | 86.64758 | 133.1254 | 218.634 | 90.78145 | -0.42009 | 2.969939 | 0.31123 |
| 2 | 79984200 | 79984999 | 10.2061 | 19.66774 | 19.12578 | 7.415069 | 0.581224 | -0.11242 | 0.311255 |
| 18 | 44802600 | 44803399 | 10.2061 | 19.66774 | 19.12578 | 7.415069 | 0.581224 | -0.11242 | 0.311255 |
| 14 | 106933600 | 106935799 | 12.37044 | 21.89365 | 43.47137 | 19.22423 | -0.59316 | 0.514747 | 0.311256 |
| 4 | 136025600 | 136027599 | 16.91245 | 30.76757 | 24.87084 | 10.71057 | 0.721767 | 0.334499 | 0.31126 |
| 11 | 44360600 | 44361799 | 7.201688 | 13.25836 | 33.56779 | 14.2318 | -0.89802 | 0.214018 | 0.311295 |
| 13 | 106905200 | 106905799 | 5.817109 | 11.28468 | 27.57354 | 10.39165 | -0.69726 | -0.23133 | 0.311308 |
| 10 | 46419800 | 46427999 | 229.2647 | 337.0715 | 298.0947 | 122.7626 | 0.416951 | 3.827543 | 0.311309 |
| 8 | 13592600 | 13593999 | 16.59006 | 29.66785 | 26.05732 | 11.43568 | 0.585164 | 0.434383 | 0.31132 |
| 8 | 90095600 | 90099799 | 25.5546 | 42.11816 | 76.39975 | 32.96074 | -0.477 | 1.373732 | 0.311324 |
| X | 78641400 | 78642599 | 16.23588 | 30.05816 | 23.16918 | 9.748558 | 0.813534 | 0.231866 | 0.311333 |
| 12 | 76268800 | 76269199 | 7.381905 | 14.40542 | 14.1554 | 5.036037 | 0.680481 | -0.45586 | 0.311335 |
| 9 | 44089600 | 44092399 | 24.13436 | 39.79653 | 73.19062 | 31.58267 | -0.49642 | 1.363686 | 0.311337 |
| 8 | 3728600 | 3729799 | 13.53547 | 23.21853 | 53.65044 | 23.53042 | -0.81945 | 0.912895 | 0.31134 |
| 12 | 22151200 | 22173999 | 500.5469 | 685.5263 | 570.4689 | 237.321 | 0.402936 | 4.901403 | 0.311348 |
| 5 | 103470800 | 103472199 | 9.01111 | 16.45981 | 33.9719 | 14.55597 | -0.6165 | 0.275558 | 0.31136 |
| 7 | 10070400 | 10073799 | 44.96348 | 73.88388 | 53.44942 | 23.02998 | 0.843734 | 1.442577 | 0.311373 |
| 7 | 32906600 | 32908199 | 16.00375 | 30.51719 | 18.40463 | 7.341811 | 1.226816 | 0.033997 | 0.311386 |
| 5 | 131686400 | 131692199 | 40.91414 | 65.53474 | 113.9244 | 48.04253 | -0.45066 | 2.000292 | 0.31139 |
| 9 | 50097000 | 50099399 | 13.4303 | 23.29746 | 54.31441 | 23.83096 | -0.82906 | 0.809563 | 0.3114 |
| 7 | 5456400 | 5458599 | 37.97408 | 62.85036 | 48.27818 | 20.88187 | 0.760002 | 1.334986 | 0.311409 |
| 7 | 17943000 | 17946399 | 49.15366 | 79.30307 | 73.27816 | 31.22764 | 0.471762 | 1.772823 | 0.31143 |
| 14 | 122578200 | 122579399 | 12.21325 | 23.03335 | 21.40391 | 8.739921 | 0.578922 | 0.078888 | 0.311437 |
| 5 | 31733800 | 31737199 | 21.97766 | 36.44585 | 69.95361 | 30.29791 | -0.55682 | 1.28734 | 0.311472 |
| 13 | 114606800 | 114608199 | 18.86849 | 33.20907 | 30.25367 | 13.39388 | 0.521934 | 0.590524 | 0.311483 |
| 19 | 39187000 | 39189199 | 31.97365 | 54.543 | 39.68624 | 17.3981 | 0.839356 | 1.018851 | 0.311489 |
| 8 | 34418800 | 34420599 | 10.89809 | 19.13362 | 43.06341 | 19.05126 | -0.77653 | 0.601384 | 0.31149 |
| 16 | 30206400 | 30212399 | 40.57664 | 64.94351 | 120.6262 | 50.80981 | -0.54737 | 2.024751 | 0.311496 |
| 15 | 43574800 | 43575599 | 14.40968 | 26.3414 | 23.36489 | 10.00143 | 0.594539 | 0.286379 | 0.311497 |
| 2 | 172912400 | 172916799 | 46.32423 | 73.9022 | 126.082 | 52.86436 | -0.43016 | 2.133502 | 0.311497 |
| 19 | 5657600 | 5660399 | 11.78824 | 20.4024 | 51.02449 | 22.38592 | -0.93014 | 0.819637 | 0.311513 |
| 15 | 36887200 | 36889799 | 13.37735 | 23.4226 | 44.64146 | 19.70403 | -0.5382 | 0.662805 | 0.311522 |
| 1 | 184887400 | 184888199 | 5.378493 | 10.44108 | 24.48249 | 9.193958 | -0.63209 | -0.25407 | 0.311524 |
| 3 | 106267600 | 106271999 | 59.40294 | 95.29755 | 81.88165 | 34.61171 | 0.566671 | 1.961693 | 0.311535 |
| 10 | 22344200 | 22348199 | 114.0204 | 175.4147 | 163.6297 | 67.95494 | 0.396648 | 2.947316 | 0.31157 |
| 3 | 130122000 | 130123199 | 13.53171 | 25.39804 | 22.30205 | 9.180759 | 0.650748 | 0.123009 | 0.311572 |
| 5 | 93826400 | 93836599 | 143.3431 | 218.397 | 189.1702 | 78.41347 | 0.492658 | 3.151543 | 0.311574 |
| 3 | 34947800 | 34951999 | 60.54004 | 95.12657 | 163.1663 | 67.6815 | -0.45754 | 2.544908 | 0.311578 |
| 8 | 48153600 | 48154799 | 12.41523 | 23.6809 | 19.76453 | 7.879645 | 0.762224 | -0.00087 | 0.311579 |
| 12 | 80940400 | 80943399 | 17.08405 | 28.8423 | 59.4654 | 26.01092 | -0.65865 | 1.077996 | 0.311597 |
| 3 | 15674000 | 15675599 | 17.76495 | 32.27445 | 24.77543 | 10.68769 | 0.794294 | 0.348493 | 0.311604 |
| 9 | 7396600 | 7398599 | 23.69345 | 41.07523 | 37.1684 | 16.30768 | 0.529982 | 0.837681 | 0.311606 |
| 11 | 58648400 | 58649399 | 6.368361 | 11.99509 | 29.66237 | 12.13812 | -0.82376 | 0.057411 | 0.311606 |
| 2 | 168816000 | 168823999 | 87.05447 | 133.9957 | 219.9656 | 91.35731 | -0.41745 | 2.933878 | 0.311628 |
| X | 131692400 | 131693199 | 7.514081 | 14.27475 | 30.50227 | 12.26467 | -0.59304 | 0.011703 | 0.311638 |
| 17 | 83261600 | 83264399 | 20.59507 | 34.08772 | 70.75711 | 30.60542 | -0.66958 | 1.30721 | 0.311662 |

| | | | | | | | | | |
|---|---|---|---|---|---|---|---|---|---|
| 7 | 102812000 | 102813399 | 13.94185 | 26.09506 | 22.1867 | 9.176448 | 0.691381 | 0.143017 | 0.31167 |
| 1 | 139381600 | 139383199 | 9.583634 | 17.10076 | 37.89579 | 16.66198 | -0.74349 | 0.428056 | 0.311681 |
| 7 | 104440000 | 104441199 | 17.81621 | 32.92953 | 21.02308 | 8.865443 | 1.080285 | 0.246078 | 0.311684 |
| 9 | 24121800 | 24123199 | 15.17535 | 28.35806 | 21.43713 | 8.890658 | 0.856397 | 0.161512 | 0.311696 |
| 1 | 113045000 | 113046199 | 5.808044 | 11.27667 | 27.58678 | 10.347 | -0.69153 | -0.24415 | 0.311714 |
| 13 | 9270200 | 9275199 | 45.87563 | 73.13499 | 125.6342 | 52.62527 | -0.44113 | 2.15646 | 0.311723 |
| 7 | 38110400 | 38117599 | 65.77475 | 105.2745 | 86.48343 | 36.42884 | 0.627472 | 2.044603 | 0.311726 |
| 14 | 29873600 | 29874599 | 12.58222 | 22.11027 | 42.47898 | 18.80698 | -0.54955 | 0.596271 | 0.311742 |
| 4 | 24792400 | 24793399 | 6.711043 | 12.91986 | 29.40826 | 11.43383 | -0.63904 | -0.11113 | 0.311744 |
| 19 | 45228600 | 45236199 | 51.75668 | 81.748 | 153.5815 | 63.77865 | -0.58281 | 2.428377 | 0.311757 |
| 18 | 46927000 | 46927999 | 17.97471 | 31.81967 | 27.80807 | 12.29692 | 0.584813 | 0.519297 | 0.311757 |
| 7 | 59317600 | 59323599 | 99.95054 | 155.869 | 140.761 | 58.41424 | 0.458362 | 2.698535 | 0.311765 |
| 1 | 131994800 | 131997599 | 15.00707 | 25.90286 | 51.08884 | 22.4123 | -0.5904 | 0.858492 | 0.311769 |
| 12 | 66974200 | 66975399 | 14.61978 | 25.51107 | 47.86039 | 21.07053 | -0.51603 | 0.72502 | 0.311773 |
| 13 | 47420600 | 47421799 | 12.88087 | 24.36976 | 21.47333 | 8.709669 | 0.663895 | 0.065505 | 0.311784 |
| 3 | 48850200 | 48853799 | 94.66689 | 147.9856 | 120.762 | 50.10226 | 0.607057 | 2.650539 | 0.311814 |
| Y | 2098400 | 2102399 | 27.96899 | 47.83674 | 22.79813 | 9.996613 | 1.446674 | 0.951746 | 0.311831 |
| 4 | 145092400 | 145094399 | 21.68831 | 37.73963 | 34.602 | 15.20675 | 0.511714 | 0.778122 | 0.311837 |
| 4 | 143619600 | 143626799 | 149.613 | 228.0154 | 183.7347 | 76.16696 | 0.597148 | 3.147117 | 0.311847 |
| X | 5388200 | 5391799 | 49.86297 | 81.24135 | 57.99308 | 24.90483 | 0.85621 | 1.568038 | 0.311849 |
| 7 | 56161200 | 56161799 | 13.43841 | 24.9071 | 21.46536 | 9.017845 | 0.655135 | 0.19799 | 0.31185 |
| 4 | 88608200 | 88609599 | 6.73055 | 12.86867 | 29.01293 | 11.5049 | -0.65219 | -0.04766 | 0.31185 |
| 10 | 3161200 | 3161999 | 6.73055 | 12.86867 | 29.01293 | 11.5049 | -0.65219 | -0.04766 | 0.31185 |
| 15 | 61856200 | 61860199 | 27.31471 | 44.11994 | 97.05079 | 41.42039 | -0.77464 | 1.715211 | 0.311888 |
| 10 | 78687200 | 78691599 | 63.29834 | 101.5214 | 81.11609 | 34.27506 | 0.670889 | 1.975143 | 0.311901 |
| 14 | 33653800 | 33656599 | 18.84669 | 31.27442 | 69.54706 | 30.1356 | -0.76822 | 1.271355 | 0.311918 |
| 12 | 23310200 | 23327399 | 395.0842 | 551.8731 | 456.914 | 189.4913 | 0.439001 | 4.597166 | 0.311921 |
| 12 | 76276000 | 76277799 | 8.547795 | 15.62779 | 33.98491 | 14.54491 | -0.69038 | 0.265545 | 0.311926 |
| 15 | 96715000 | 96716999 | 14.13517 | 24.72039 | 46.75892 | 20.61815 | -0.52756 | 0.687463 | 0.311939 |
| X | 118764400 | 118766799 | 34.88038 | 58.31922 | 47.56406 | 20.71216 | 0.674286 | 1.217064 | 0.311939 |
| 1 | 171122000 | 171124199 | 12.41196 | 21.98153 | 44.0153 | 19.46223 | -0.60476 | 0.510499 | 0.311942 |
| 4 | 138327400 | 138334399 | 42.60018 | 68.00918 | 121.3738 | 50.94108 | -0.49385 | 2.107095 | 0.311952 |
| 6 | 122671200 | 122681199 | 87.47406 | 134.3943 | 219.9048 | 91.30901 | -0.41479 | 2.970296 | 0.311952 |
| 15 | 32250200 | 32251999 | 13.36471 | 23.31067 | 47.86358 | 21.06607 | -0.64567 | 0.724124 | 0.31198 |
| 4 | 51571800 | 51572999 | 14.92413 | 27.35676 | 24.74679 | 10.55393 | 0.570075 | 0.278195 | 0.311985 |
| 16 | 8704400 | 8707999 | 38.1317 | 61.28602 | 105.7073 | 44.80663 | -0.43345 | 1.888496 | 0.312002 |
| 4 | 99629200 | 99629799 | 6.438561 | 12.33985 | 27.35509 | 10.77332 | -0.6179 | -0.07493 | 0.312007 |
| 7 | 94051800 | 94052399 | 6.438561 | 12.33985 | 27.35509 | 10.77332 | -0.6179 | -0.07493 | 0.312007 |
| 8 | 66375400 | 66384199 | 53.59947 | 86.23699 | 70.73305 | 30.02778 | 0.638582 | 1.862242 | 0.31201 |
| 3 | 91358000 | 91361599 | 42.58159 | 70.08095 | 53.96125 | 23.26611 | 0.754871 | 1.417444 | 0.312016 |
| 11 | 115914200 | 115915399 | 7.063337 | 13.00126 | 34.62207 | 14.68247 | -0.97078 | 0.215524 | 0.312024 |
| 3 | 90590800 | 90592599 | 20.49551 | 35.99724 | 31.62427 | 13.99504 | 0.57335 | 0.628567 | 0.312024 |
| 10 | 13544600 | 13545399 | 18.239 | 32.21099 | 29.235 | 12.93406 | 0.52916 | 0.548265 | 0.312036 |
| 19 | 34033600 | 34034399 | 11.865 | 22.77234 | 20.05172 | 7.864761 | 0.707834 | -0.05514 | 0.312039 |
| 4 | 126689200 | 126691799 | 26.42433 | 45.22954 | 40.95742 | 17.96028 | 0.525639 | 0.96771 | 0.312041 |
| 15 | 33571600 | 33572399 | 10.28863 | 20.04333 | 15.95412 | 5.810726 | 0.950016 | -0.3304 | 0.312054 |
| 15 | 40517600 | 40519799 | 15.8713 | 26.7308 | 60.95693 | 26.64399 | -0.80358 | 1.096459 | 0.312059 |

| | | | | | | | | | |
|---|---|---|---|---|---|---|---|---|---|
| 7 | 59130400 | 59131799 | 21.24822 | 37.46363 | 26.11101 | 11.55576 | 0.907308 | 0.579597 | 0.312063 |
| 15 | 77766000 | 77768399 | 11.35064 | 19.92347 | 43.05702 | 19.04464 | -0.71835 | 0.60772 | 0.312063 |
| 8 | 120596400 | 120600199 | 60.09895 | 94.28876 | 166.8548 | 69.19964 | -0.5042 | 2.575516 | 0.312084 |
| 1 | 25452400 | 25453599 | 23.849 | 41.65289 | 30.63624 | 13.49516 | 0.828577 | 0.732411 | 0.312092 |
| 1 | 5644400 | 5646399 | 18.07006 | 32.42825 | 25.10263 | 10.98411 | 0.768872 | 0.419916 | 0.312117 |
| 7 | 46778800 | 46784799 | 44.11327 | 70.23735 | 129.0312 | 53.95455 | -0.53972 | 2.195888 | 0.312117 |
| 13 | 43547600 | 43552399 | 58.13568 | 91.67276 | 156.5006 | 64.94651 | -0.44678 | 2.471024 | 0.312123 |
| 13 | 37883400 | 37887599 | 38.15705 | 61.20783 | 108.8945 | 46.02346 | -0.48143 | 1.950325 | 0.312126 |
| 7 | 26578800 | 26607199 | 619.7343 | 835.1326 | 689.1568 | 287.4144 | 0.391517 | 5.15745 | 0.312139 |
| 10 | 52493800 | 52496399 | 29.03302 | 50.18257 | 34.61201 | 15.18468 | 0.919053 | 0.883623 | 0.312148 |
| 17 | 13706600 | 13709999 | 45.54085 | 72.53658 | 129.5367 | 54.19303 | -0.49843 | 2.18489 | 0.312151 |
| 5 | 139471600 | 139472799 | 24.92115 | 42.86803 | 39.37362 | 17.27112 | 0.506404 | 0.915513 | 0.312166 |
| 19 | 9637400 | 9640399 | 46.2455 | 74.69779 | 70.66542 | 30.15935 | 0.439829 | 1.734331 | 0.312176 |
| 1 | 15873400 | 15876599 | 22.16423 | 36.67114 | 71.56429 | 30.94654 | -0.58099 | 1.315658 | 0.312198 |
| 6 | 23892600 | 23894999 | 34.19793 | 56.9415 | 45.49145 | 19.75785 | 0.704051 | 1.254948 | 0.312202 |
| 3 | 17721400 | 17725399 | 44.0625 | 71.98122 | 59.54191 | 25.59469 | 0.64676 | 1.517509 | 0.312206 |
| X | 91481400 | 91488999 | 76.66364 | 121.7928 | 95.09944 | 39.65918 | 0.689303 | 2.285291 | 0.312209 |
| 12 | 116251600 | 116252199 | 13.48193 | 25.08114 | 22.90916 | 9.573394 | 0.577401 | 0.18082 | 0.312216 |
| 1 | 136599600 | 136603999 | 26.73133 | 43.8266 | 79.63646 | 34.2808 | -0.48256 | 1.443642 | 0.312218 |
| 15 | 19963200 | 19965399 | 22.23929 | 39.07353 | 30.95144 | 13.70131 | 0.721984 | 0.632804 | 0.31222 |
| 6 | 48244200 | 48244799 | 13.74028 | 25.78899 | 21.66544 | 8.919 | 0.713783 | 0.122487 | 0.31222 |
| 19 | 46774800 | 46776399 | 6.947071 | 13.07209 | 29.65212 | 12.15441 | -0.70234 | 0.069557 | 0.312237 |
| 19 | 7657800 | 7658799 | 19.94871 | 34.91029 | 31.01048 | 13.68151 | 0.557671 | 0.681985 | 0.312245 |
| 13 | 35940600 | 35950199 | 88.01463 | 135.5137 | 219.9063 | 91.33636 | -0.4005 | 2.928719 | 0.312249 |
| 17 | 30182400 | 30186399 | 22.397 | 36.88158 | 73.41906 | 31.66305 | -0.61103 | 1.376157 | 0.312275 |
| 5 | 109250800 | 109251399 | 11.33561 | 21.91467 | 18.61017 | 7.127114 | 0.792132 | -0.13996 | 0.312286 |
| 18 | 76263800 | 76268599 | 51.97358 | 82.47215 | 139.5405 | 58.12434 | -0.42624 | 2.308426 | 0.3123 |
| 1 | 83015600 | 83016399 | 5.529035 | 10.73233 | 25.10294 | 9.433853 | -0.63014 | -0.24955 | 0.312317 |
| 6 | 18988000 | 18989599 | 25.08877 | 41.34772 | 74.50147 | 32.14089 | -0.46734 | 1.372972 | 0.312354 |
| 6 | 143231600 | 143235199 | 22.72515 | 37.94602 | 70.32007 | 30.59606 | -0.506 | 1.213042 | 0.312362 |
| 6 | 130640200 | 130643999 | 64.16254 | 103.1857 | 71.14453 | 30.18279 | 0.88775 | 1.890993 | 0.312382 |
| 7 | 28023000 | 28023999 | 6.442606 | 12.34414 | 27.89098 | 10.99321 | -0.64594 | -0.06929 | 0.312385 |
| 14 | 83896400 | 83897199 | 15.58301 | 28.28071 | 25.70086 | 11.10147 | 0.55137 | 0.341382 | 0.312391 |
| 15 | 97361400 | 97370799 | 85.80402 | 131.5565 | 225.696 | 93.68714 | -0.48528 | 3.010569 | 0.312397 |
| 13 | 66572200 | 66590799 | 399.7564 | 567.2712 | 373.9029 | 154.6722 | 0.790702 | 4.366995 | 0.312405 |
| 13 | 19619200 | 19619999 | 13.45475 | 25.13891 | 22.88144 | 9.492145 | 0.590716 | 0.153022 | 0.312422 |
| 10 | 129889400 | 129890399 | 19.90872 | 34.96092 | 31.12936 | 13.77412 | 0.553976 | 0.625983 | 0.312431 |
| 1 | 8362600 | 8366399 | 45.65786 | 74.27136 | 62.37477 | 26.77095 | 0.620681 | 1.581262 | 0.312432 |
| 12 | 71701200 | 71702399 | 6.730508 | 12.89337 | 27.92986 | 11.01441 | -0.58706 | -0.06691 | 0.312441 |
| 4 | 105231400 | 105234599 | 19.51912 | 32.26152 | 76.72846 | 33.1561 | -0.86543 | 1.325959 | 0.312448 |
| 1 | 90729000 | 90731999 | 25.19366 | 41.17173 | 79.48031 | 34.16993 | -0.57215 | 1.484803 | 0.312469 |
| 8 | 87720600 | 87721999 | 11.8047 | 20.77608 | 41.46406 | 18.35702 | -0.60325 | 0.572113 | 0.312483 |
| 3 | 111658800 | 111663399 | 54.5819 | 87.57017 | 83.047 | 35.10846 | 0.424712 | 1.954809 | 0.312498 |
| 15 | 35229600 | 35231199 | 8.868067 | 16.20058 | 33.98332 | 14.55855 | -0.63941 | 0.274622 | 0.312503 |
| 14 | 87305800 | 87306999 | 22.42799 | 38.90554 | 35.28411 | 15.48123 | 0.52702 | 0.824726 | 0.312514 |
| 9 | 121524000 | 121530399 | 46.67838 | 73.93895 | 154.5278 | 64.26494 | -0.73292 | 2.357306 | 0.312519 |
| 1 | 153520200 | 153523799 | 34.12311 | 56.69001 | 50.94217 | 22.09155 | 0.534732 | 1.282508 | 0.312538 |

| | | | | | | | | | |
|---|---|---|---|---|---|---|---|---|---|
| 17 | 69001600 | 69002799 | 18.62146 | 31.57732 | 58.54105 | 25.641 | -0.50577 | 1.0436 | 0.312539 |
| 4 | 114077000 | 114080599 | 54.8605 | 88.24847 | 78.47559 | 33.30514 | 0.521909 | 1.873952 | 0.31254 |
| X | 95736000 | 95736799 | 18.30817 | 32.50778 | 28.59584 | 12.6254 | 0.577407 | 0.497165 | 0.312541 |
| X | 21011800 | 21015199 | 69.11041 | 110.099 | 98.12605 | 41.06303 | 0.502897 | 2.194194 | 0.312566 |
| 7 | 87602000 | 87604999 | 30.01327 | 50.6885 | 43.62892 | 19.08594 | 0.596679 | 1.09854 | 0.312578 |
| 9 | 41860800 | 41863799 | 15.57766 | 26.55281 | 57.97535 | 25.41429 | -0.73995 | 0.982378 | 0.312589 |
| 7 | 60431400 | 60433799 | 21.44438 | 38.12507 | 27.09467 | 11.94973 | 0.884396 | 0.50641 | 0.31259 |
| 8 | 46681800 | 46683799 | 15.37267 | 26.40063 | 52.65239 | 23.09467 | -0.60786 | 0.908887 | 0.312595 |
| 14 | 49470800 | 49473599 | 9.672971 | 17.42099 | 36.38015 | 15.86562 | -0.6503 | 0.36668 | 0.312598 |
| 11 | 11552800 | 11559999 | 125.6559 | 192.5338 | 176.6037 | 73.30186 | 0.41646 | 3.032179 | 0.312604 |
| 19 | 29808000 | 29809199 | 9.802225 | 17.7231 | 35.90195 | 15.58051 | -0.6021 | 0.343694 | 0.312613 |
| 12 | 87747800 | 87754199 | 104.6464 | 157.984 | 278.0386 | 114.927 | -0.54031 | 3.333436 | 0.312617 |
| 1 | 49132200 | 49133199 | 13.49921 | 25.75761 | 20.37242 | 8.115928 | 0.840828 | 0.010974 | 0.312633 |
| 7 | 60166200 | 60168399 | 24.77385 | 43.19027 | 34.24301 | 15.06686 | 0.720699 | 0.767611 | 0.312633 |
| 1 | 106318600 | 106320799 | 8.830934 | 15.7969 | 38.94294 | 17.08652 | -0.89423 | 0.407905 | 0.312633 |
| 6 | 4365800 | 4368399 | 19.71534 | 34.87202 | 29.42959 | 13.01613 | 0.634314 | 0.540908 | 0.312641 |
| 18 | 25355200 | 25356999 | 23.7473 | 41.83837 | 28.5726 | 12.64707 | 0.936014 | 0.606061 | 0.312646 |
| 3 | 21278200 | 21279999 | 12.94753 | 22.51858 | 47.89348 | 21.04139 | -0.69643 | 0.764994 | 0.312656 |
| 18 | 12758200 | 12759999 | 10.66413 | 18.92651 | 39.34999 | 17.38013 | -0.6569 | 0.471449 | 0.312658 |
| X | 4451000 | 4459199 | 190.6717 | 285.3531 | 235.4504 | 97.10366 | 0.540585 | 3.517044 | 0.312693 |
| 5 | 45740000 | 45742399 | 17.06423 | 28.88087 | 58.41447 | 25.57059 | -0.63091 | 1.055358 | 0.312695 |
| 7 | 45737400 | 45739199 | 27.8795 | 47.35569 | 41.12141 | 18.02032 | 0.584471 | 1.044019 | 0.312699 |
| 7 | 33547200 | 33561599 | 355.9173 | 504.1585 | 425.2608 | 175.9699 | 0.43239 | 4.390282 | 0.312702 |
| 19 | 35148400 | 35150999 | 26.31596 | 45.25901 | 39.2389 | 17.21192 | 0.589192 | 0.921846 | 0.312708 |
| 11 | 51030400 | 51032199 | 14.85488 | 27.82506 | 22.14916 | 9.149173 | 0.787384 | 0.146893 | 0.312711 |
| 7 | 62342800 | 62354999 | 199.3038 | 296.7909 | 257.3957 | 106.0613 | 0.462249 | 3.608774 | 0.312739 |
| 17 | 66036800 | 66039199 | 19.58037 | 33.06371 | 60.76451 | 26.58087 | -0.49361 | 1.08465 | 0.31275 |
| X | 80263400 | 80264399 | 23.77935 | 41.10063 | 37.80545 | 16.58566 | 0.505684 | 0.866382 | 0.312774 |
| 15 | 18714400 | 18716399 | 33.71886 | 56.26373 | 44.40106 | 19.31025 | 0.721649 | 1.230408 | 0.312777 |
| 1 | 27259200 | 27260999 | 17.93141 | 32.35479 | 25.77831 | 11.21585 | 0.733149 | 0.390032 | 0.312785 |
| 13 | 95562600 | 95565199 | 11.35624 | 19.93361 | 42.55785 | 18.82411 | -0.70075 | 0.607801 | 0.312796 |
| Y | 3340200 | 3341399 | 10.52059 | 20.23215 | 13.87806 | 5.412383 | 1.071472 | -0.10869 | 0.312802 |
| 17 | 48339200 | 48346199 | 48.68927 | 77.28989 | 140.4211 | 58.51336 | -0.52825 | 2.293768 | 0.312812 |
| 2 | 131210400 | 131211199 | 8.608049 | 16.06379 | 32.77477 | 13.62429 | -0.56787 | 0.14368 | 0.312812 |
| 5 | 106565000 | 106567399 | 20.04345 | 33.87444 | 62.49734 | 27.34592 | -0.49924 | 1.079955 | 0.312816 |
| 6 | 134903800 | 134907399 | 15.84819 | 27.1055 | 55.87154 | 24.49874 | -0.65645 | 0.950064 | 0.312847 |
| 3 | 132389400 | 132390199 | 6.932264 | 13.10214 | 29.19551 | 11.86715 | -0.6663 | 0.039654 | 0.312859 |
| 8 | 21734200 | 21736999 | 31.08185 | 53.5348 | 34.55278 | 15.15707 | 1.013461 | 0.923491 | 0.312884 |
| 9 | 100720200 | 100722599 | 14.86098 | 25.6893 | 55.9528 | 24.5472 | -0.73258 | 0.850362 | 0.312897 |
| 2 | 21339600 | 21340199 | 6.838082 | 12.77682 | 31.67724 | 13.1401 | -0.84396 | 0.117272 | 0.312943 |
| 8 | 79415800 | 79416799 | 6.838082 | 12.77682 | 31.67724 | 13.1401 | -0.84396 | 0.117272 | 0.312943 |
| 7 | 48156000 | 48158199 | 25.29064 | 44.28812 | 30.74887 | 13.57371 | 0.91132 | 0.69998 | 0.31295 |
| 2 | 63635800 | 63636599 | 10.85998 | 19.18583 | 39.92829 | 17.66374 | -0.66101 | 0.521724 | 0.312955 |
| 8 | 104766000 | 104767399 | 10.85998 | 19.18583 | 39.92829 | 17.66374 | -0.66101 | 0.521724 | 0.312955 |
| 13 | 65899800 | 65901599 | 22.48429 | 39.52683 | 30.44651 | 13.48239 | 0.76227 | 0.625782 | 0.31296 |
| 10 | 75937400 | 75938799 | 14.59602 | 26.89683 | 24.27588 | 10.27905 | 0.580699 | 0.245493 | 0.312981 |
| 6 | 39357800 | 39358599 | 3.709308 | 7.225453 | 22.36939 | 8.157093 | -0.98735 | -0.38678 | 0.313004 |

| | | | | | | | | | |
|---|---|---|---|---|---|---|---|---|---|
| 7 | 19822800 | 19825799 | 85.89914 | 134.7193 | 125.542 | 52.0679 | 0.419444 | 2.589768 | 0.313004 |
| 1 | 24864200 | 24866799 | 21.21919 | 35.665 | 65.82354 | 28.75336 | -0.50012 | 1.135119 | 0.313037 |
| 6 | 130645000 | 130645599 | 22.268 | 38.75026 | 35.60875 | 15.64967 | 0.508543 | 0.782577 | 0.313041 |
| 15 | 63461200 | 63463799 | 20.5884 | 34.68532 | 63.52805 | 27.77021 | -0.48889 | 1.111928 | 0.313041 |
| 17 | 30083000 | 30090799 | 94.68401 | 144.2476 | 244.4888 | 101.3411 | -0.47511 | 3.144022 | 0.313048 |
| 17 | 88546400 | 88548199 | 19.77835 | 34.98021 | 29.48904 | 13.04267 | 0.63589 | 0.543069 | 0.313055 |
| 9 | 100819800 | 100821399 | 21.50985 | 38.11692 | 27.54508 | 12.17812 | 0.857817 | 0.52875 | 0.313056 |
| 11 | 69986600 | 69990199 | 34.39619 | 55.35093 | 101.1082 | 42.93439 | -0.51374 | 1.842328 | 0.31306 |
| 10 | 72363400 | 72364999 | 9.264002 | 17.13652 | 34.75361 | 14.63939 | -0.57399 | 0.206574 | 0.313071 |
| 12 | 45820400 | 45824399 | 34.86999 | 57.64864 | 53.70428 | 23.21099 | 0.482459 | 1.34303 | 0.313074 |
| 12 | 115000000 | 115006799 | 94.14421 | 148.3632 | 117.6632 | 48.82539 | 0.657144 | 2.500544 | 0.313077 |
| 19 | 46050600 | 46053999 | 17.68586 | 29.50701 | 65.7473 | 28.59148 | -0.77058 | 1.204091 | 0.313077 |
| 6 | 42772800 | 42773999 | 10.14417 | 18.25353 | 36.36581 | 15.87653 | -0.58377 | 0.3762 | 0.313078 |
| 9 | 109872600 | 109874399 | 17.79206 | 32.54413 | 23.75867 | 10.16206 | 0.874122 | 0.308942 | 0.313079 |
| 19 | 55010400 | 55013199 | 35.38178 | 59.30915 | 45.85596 | 20.00063 | 0.750893 | 1.188718 | 0.313087 |
| 9 | 40815600 | 40816399 | 4.89664 | 9.518667 | 25.13365 | 9.334992 | -0.78546 | -0.29323 | 0.313091 |
| 17 | 17413000 | 17449399 | 921.9417 | 1213.868 | 799.6837 | 334.5497 | 0.682932 | 5.530977 | 0.313104 |
| 4 | 53767000 | 53768599 | 9.806218 | 17.76734 | 35.41804 | 15.33046 | -0.57647 | 0.331599 | 0.313108 |
| 18 | 30493800 | 30495399 | 13.28864 | 24.68474 | 22.27092 | 9.327798 | 0.592158 | 0.184442 | 0.313127 |
| X | 16268400 | 16269599 | 36.61476 | 60.50523 | 47.71006 | 20.61372 | 0.722347 | 1.352777 | 0.313129 |
| 17 | 74622800 | 74623799 | 7.001871 | 13.02404 | 31.5952 | 13.20644 | -0.82153 | 0.148892 | 0.313143 |
| 3 | 153678400 | 153681199 | 19.91751 | 33.08672 | 69.22571 | 30.01852 | -0.68054 | 1.260741 | 0.313153 |
| 2 | 86443200 | 86446599 | 66.47475 | 105.948 | 98.28391 | 41.15483 | 0.445902 | 2.175704 | 0.313156 |
| X | 25831000 | 25836799 | 64.21723 | 103.7444 | 57.85641 | 24.69807 | 1.200716 | 1.756717 | 0.313156 |
| 5 | 19093200 | 19094799 | 25.40601 | 44.15711 | 35.50637 | 15.59273 | 0.700251 | 0.81606 | 0.313164 |
| 18 | 59055400 | 59056799 | 13.27137 | 24.92678 | 21.73762 | 8.937842 | 0.661762 | 0.11413 | 0.313242 |
| 7 | 27841600 | 27849199 | 64.59647 | 101.0729 | 172.1353 | 71.40936 | -0.45203 | 2.626585 | 0.313247 |
| 2 | 33759800 | 33761999 | 17.31892 | 29.40995 | 57.98705 | 25.4092 | -0.59422 | 1.024305 | 0.313251 |
| 15 | 15935000 | 15936999 | 17.62475 | 32.01429 | 25.47417 | 10.99145 | 0.742923 | 0.351386 | 0.313262 |
| X | 132782400 | 132784999 | 11.44427 | 20.4723 | 45.87754 | 20.12861 | -0.7581 | 0.43924 | 0.313276 |
| 9 | 46760000 | 46761399 | 8.263277 | 15.07756 | 35.00942 | 15.02074 | -0.78675 | 0.276358 | 0.313276 |
| 15 | 27934200 | 27936199 | 8.263277 | 15.07756 | 35.00942 | 15.02074 | -0.78675 | 0.276358 | 0.313276 |
| 14 | 58289200 | 58293199 | 51.70696 | 83.13291 | 78.3062 | 33.20848 | 0.438134 | 1.885244 | 0.313278 |
| X | 83856000 | 83858399 | 19.61268 | 35.87579 | 21.89976 | 9.366433 | 1.130419 | 0.318666 | 0.313284 |
| 1 | 139071800 | 139072999 | 8.068972 | 14.87064 | 32.52553 | 13.77024 | -0.68696 | 0.21396 | 0.313288 |
| 5 | 75345000 | 75346799 | 8.068972 | 14.87064 | 32.52553 | 13.77024 | -0.68696 | 0.21396 | 0.313288 |
| 19 | 60336000 | 60337199 | 14.47854 | 24.77147 | 54.17404 | 23.75507 | -0.74104 | 0.942493 | 0.313293 |
| 13 | 68530200 | 68546999 | 328.9253 | 470.3455 | 385.4964 | 159.2646 | 0.487535 | 4.251337 | 0.313297 |
| 14 | 112931600 | 112932199 | 9.672051 | 18.74342 | 18.20447 | 6.894145 | 0.615232 | -0.19711 | 0.3133 |
| 7 | 73243600 | 73245199 | 6.923899 | 13.08462 | 30.31492 | 12.32557 | -0.72324 | 0.039813 | 0.313304 |
| 10 | 8430000 | 8431399 | 33.05801 | 56.09374 | 41.38962 | 18.13216 | 0.818442 | 1.071856 | 0.313323 |
| 16 | 36788600 | 36791799 | 24.72707 | 40.38566 | 80.52206 | 34.61044 | -0.61922 | 1.492494 | 0.313323 |
| 2 | 122523200 | 122524399 | 180.2324 | 276.2807 | 111.927 | 46.45998 | 1.594661 | 3.039909 | 0.313325 |
| 14 | 41785200 | 41792199 | 135.9653 | 208.0994 | 176.3132 | 73.16252 | 0.529643 | 3.057623 | 0.313336 |
| 9 | 87894800 | 87896999 | 17.01695 | 31.5499 | 22.36831 | 9.393035 | 0.934893 | 0.226882 | 0.313351 |
| 8 | 27235000 | 27237599 | 16.48404 | 28.18203 | 54.79608 | 24.02641 | -0.5726 | 0.956092 | 0.31336 |
| 4 | 145624000 | 145626399 | 66.14834 | 106.3184 | 65.17676 | 27.62809 | 1.055887 | 1.909425 | 0.313365 |

| | | | | | | | | | |
|---|---|---|---|---|---|---|---|---|---|
| 10 | 89142400 | 89145999 | 42.63516 | 70.56221 | 48.33011 | 20.90379 | 0.924897 | 1.348048 | 0.313369 |
| 2 | 85910400 | 85911399 | 12.06492 | 22.66768 | 20.46031 | 8.408315 | 0.613403 | 0.099215 | 0.313378 |
| 18 | 38272800 | 38274799 | 20.88993 | 36.42243 | 32.60553 | 14.34719 | 0.546316 | 0.743852 | 0.313401 |
| 7 | 9400600 | 9403599 | 61.08645 | 98.12569 | 72.50312 | 30.70788 | 0.786128 | 1.923529 | 0.313419 |
| Y | 204200 | 204999 | 10.872 | 20.95495 | 11.39611 | 4.415865 | 1.411143 | -0.13401 | 0.313423 |
| 2 | 29172000 | 29174799 | 13.6557 | 23.87537 | 46.76585 | 20.61647 | -0.57772 | 0.688762 | 0.313425 |
| 2 | 51774200 | 51777599 | 69.43802 | 110.2076 | 104.1552 | 43.39387 | 0.412953 | 2.307538 | 0.313426 |
| 10 | 119919400 | 119925999 | 81.78098 | 125.5674 | 220.0826 | 91.37487 | -0.51455 | 2.980968 | 0.313429 |
| 10 | 118694200 | 118700599 | 148.9863 | 219.9667 | 367.3923 | 151.299 | -0.49416 | 3.752414 | 0.313431 |
| 11 | 78440000 | 78445999 | 45.83511 | 73.21655 | 123.6038 | 51.92015 | -0.41294 | 2.092913 | 0.313433 |
| 2 | 78162800 | 78164199 | 8.438765 | 15.78655 | 32.85483 | 13.60097 | -0.59156 | 0.125963 | 0.313449 |
| 10 | 32571600 | 32572599 | 6.267783 | 12.12064 | 29.67781 | 11.35084 | -0.72014 | -0.16705 | 0.313451 |
| 15 | 62490800 | 62492199 | 10.45794 | 20.25637 | 18.01216 | 6.837121 | 0.737294 | -0.18341 | 0.313453 |
| 4 | 112315600 | 112322799 | 174.3859 | 261.7442 | 234.3344 | 96.70561 | 0.426703 | 3.457846 | 0.313462 |
| 7 | 14690000 | 14697799 | 170.4483 | 255.9724 | 232.6924 | 96.04187 | 0.405382 | 3.44696 | 0.313462 |
| 1 | 117203400 | 117205999 | 29.34498 | 49.60122 | 44.74035 | 19.5771 | 0.52947 | 1.091946 | 0.31348 |
| 14 | 12427000 | 12431599 | 32.68548 | 52.96067 | 96.24167 | 41.19862 | -0.49523 | 1.649785 | 0.313492 |
| 1 | 102516800 | 102517599 | 12.09319 | 23.12082 | 20.4724 | 8.119321 | 0.684732 | -0.01741 | 0.313505 |
| 12 | 110153800 | 110156199 | 22.42005 | 36.86284 | 74.36441 | 32.0499 | -0.63103 | 1.396906 | 0.313513 |
| 6 | 130179000 | 130182199 | 35.56207 | 59.60375 | 45.85813 | 19.99996 | 0.757961 | 1.190773 | 0.313516 |
| 11 | 107090200 | 107092199 | 38.25456 | 63.05249 | 55.47246 | 23.93262 | 0.56393 | 1.389552 | 0.313529 |
| 1 | 69865800 | 69866799 | 7.400202 | 13.90939 | 30.17299 | 12.39103 | -0.64054 | 0.083242 | 0.313538 |
| 15 | 100192800 | 100193999 | 7.400202 | 13.90939 | 30.17299 | 12.39103 | -0.64054 | 0.083242 | 0.313538 |
| 10 | 72045000 | 72047799 | 38.16325 | 64.10889 | 41.30702 | 18.04091 | 1.012787 | 1.171352 | 0.313552 |
| 5 | 43471000 | 43472199 | 12.36723 | 23.68263 | 19.88665 | 7.850852 | 0.766582 | -0.03127 | 0.313563 |
| X | 74012000 | 74014799 | 21.15172 | 35.50029 | 71.90896 | 31.38876 | -0.63392 | 1.156982 | 0.313567 |
| 7 | 31871800 | 31875199 | 60.9246 | 97.8133 | 74.07483 | 31.34809 | 0.749764 | 1.94023 | 0.313577 |
| 7 | 61040800 | 61045599 | 97.04069 | 150.8047 | 140.3915 | 58.29124 | 0.410654 | 2.75546 | 0.313584 |
| 12 | 44912200 | 44914799 | 23.44447 | 40.87398 | 34.8853 | 15.34978 | 0.614863 | 0.761416 | 0.313597 |
| 18 | 40592600 | 40594399 | 17.5842 | 32.25556 | 23.23804 | 9.900971 | 0.896984 | 0.290497 | 0.3136 |
| 1 | 62742000 | 62742999 | 7.149421 | 13.29093 | 32.65274 | 13.66164 | -0.84115 | 0.155008 | 0.313609 |
| 4 | 7945600 | 7948199 | 24.59304 | 42.39336 | 38.83106 | 17.03422 | 0.510944 | 0.894723 | 0.313613 |
| 1 | 122393800 | 122395599 | 12.21612 | 21.66758 | 43.48533 | 19.21663 | -0.60713 | 0.492402 | 0.313626 |
| 18 | 53714000 | 53716199 | 11.54617 | 20.62121 | 41.4825 | 18.22514 | -0.6046 | 0.440588 | 0.313644 |
| 16 | 41739800 | 41740599 | 10.60084 | 20.52816 | 18.64569 | 7.085631 | 0.706067 | -0.1745 | 0.313648 |
| 9 | 13737600 | 13739799 | 12.19808 | 21.19666 | 48.41667 | 21.25961 | -0.79888 | 0.775371 | 0.313656 |
| 14 | 51556800 | 51562199 | 136.2672 | 207.1096 | 192.3979 | 79.69307 | 0.3889 | 3.198841 | 0.313663 |
| 1 | 115925600 | 115927199 | 18.27394 | 32.36997 | 28.44818 | 12.57839 | 0.576774 | 0.51302 | 0.313667 |
| X | 67917400 | 67918599 | 18.93614 | 34.57131 | 23.71628 | 10.16824 | 0.961267 | 0.328903 | 0.313674 |
| 17 | 12898000 | 12913599 | 112.0791 | 168.7559 | 290.7552 | 120.0908 | -0.51306 | 3.387775 | 0.313688 |
| 11 | 79148200 | 79149799 | 18.00194 | 32.21648 | 27.40355 | 12.01901 | 0.631837 | 0.443077 | 0.31369 |
| 4 | 107338400 | 107340999 | 21.76165 | 36.32062 | 66.20837 | 28.79879 | -0.4822 | 1.211979 | 0.313694 |
| 7 | 126689600 | 126692199 | 14.16865 | 24.24642 | 54.16148 | 23.74997 | -0.77137 | 0.939588 | 0.313734 |
| 17 | 39470600 | 39473799 | 52.75476 | 85.35884 | 60.04627 | 25.67761 | 0.868897 | 1.696863 | 0.313742 |
| 2 | 30567800 | 30571399 | 33.27429 | 55.51468 | 49.76826 | 21.64269 | 0.538301 | 1.234976 | 0.313747 |
| 3 | 71503200 | 71507199 | 45.84587 | 74.32404 | 66.68554 | 28.55756 | 0.52128 | 1.64687 | 0.313748 |
| 7 | 138139800 | 138142199 | 12.55827 | 22.08705 | 42.45035 | 18.79627 | -0.54986 | 0.584119 | 0.313761 |

| | | | | | | | | | |
|---|---|---|---|---|---|---|---|---|---|
| 9 | 118661400 | 118664599 | 21.39691 | 35.75191 | 69.16368 | 30.10555 | -0.56783 | 1.199018 | 0.313782 |
| 5 | 122334200 | 122335599 | 24.45227 | 39.82597 | 82.11813 | 35.26072 | -0.67028 | 1.536116 | 0.313782 |
| 6 | 62798400 | 62801199 | 31.60158 | 53.75065 | 40.75641 | 17.86304 | 0.779398 | 1.042479 | 0.313796 |
| 17 | 20062800 | 20066799 | 58.36935 | 94.3741 | 61.74108 | 26.38417 | 0.972281 | 1.723656 | 0.313798 |
| 9 | 18901200 | 18906799 | 130.2163 | 198.6428 | 185.7107 | 77.00324 | 0.384012 | 3.122593 | 0.313803 |
| 7 | 42474200 | 42483999 | 184.1253 | 276.6943 | 223.9992 | 92.47018 | 0.573347 | 3.436381 | 0.313805 |
| 1 | 108304400 | 108307999 | 40.5991 | 65.01771 | 113.6344 | 47.90631 | -0.45871 | 2.006021 | 0.313806 |
| 7 | 125789800 | 125790599 | 11.99975 | 23.18836 | 18.54599 | 7.117251 | 0.875014 | -0.12625 | 0.313814 |
| 9 | 37166200 | 37167199 | 5.145233 | 9.941855 | 27.09923 | 10.40361 | -0.87859 | -0.17692 | 0.313815 |
| 12 | 7909400 | 7911799 | 23.87035 | 39.19537 | 75.57229 | 32.5522 | -0.56678 | 1.418218 | 0.313828 |
| 7 | 124009000 | 124009999 | 12.42896 | 23.88702 | 19.41139 | 7.584114 | 0.827469 | -0.06421 | 0.313836 |
| 5 | 33720200 | 33722199 | 9.202781 | 16.69207 | 34.94315 | 15.10444 | -0.64503 | 0.320967 | 0.313847 |
| 1 | 170377000 | 170378599 | 20.619 | 36.01833 | 33.03034 | 14.55146 | 0.511953 | 0.716991 | 0.313851 |
| 18 | 3139000 | 3162799 | 401.9528 | 568.266 | 403.9002 | 167.1845 | 0.676533 | 4.421167 | 0.313864 |
| 5 | 96039600 | 96042199 | 37.38507 | 61.7448 | 54.35722 | 23.47695 | 0.563729 | 1.362088 | 0.313864 |
| 5 | 139089400 | 139090999 | 13.74283 | 23.84015 | 48.96467 | 21.48373 | -0.64682 | 0.805467 | 0.313878 |
| 3 | 84096800 | 84099199 | 14.86536 | 25.68316 | 50.55067 | 22.17686 | -0.58701 | 0.849712 | 0.313899 |
| 14 | 91176600 | 91177999 | 20.14767 | 36.16491 | 25.53035 | 11.16913 | 0.901572 | 0.435853 | 0.313902 |
| 3 | 147857800 | 147858399 | 8.986862 | 16.49158 | 33.50525 | 14.2678 | -0.58766 | 0.247517 | 0.313918 |
| 4 | 118434600 | 118438999 | 41.48336 | 66.27024 | 119.2486 | 50.09621 | -0.50462 | 2.086062 | 0.313925 |
| 7 | 18256800 | 18257799 | 19.45253 | 34.10943 | 30.87704 | 13.64544 | 0.530415 | 0.647082 | 0.313935 |
| X | 22439400 | 22441199 | 19.94536 | 36.81703 | 21.07439 | 8.902789 | 1.235557 | 0.272306 | 0.313944 |
| 14 | 89239600 | 89240799 | 18.9732 | 34.17841 | 25.67122 | 11.19221 | 0.815918 | 0.406802 | 0.313971 |
| 2 | 89557400 | 89558599 | 18.3778 | 33.50211 | 24.26852 | 10.42438 | 0.881496 | 0.33377 | 0.313972 |
| 3 | 8693800 | 8697999 | 29.15615 | 47.21036 | 90.47731 | 38.71188 | -0.57255 | 1.654034 | 0.314002 |
| 3 | 124363400 | 124365599 | 28.66635 | 48.85516 | 41.9521 | 18.39004 | 0.601033 | 1.015338 | 0.314003 |
| 3 | 33037600 | 33038999 | 10.99399 | 19.53226 | 38.86954 | 17.15286 | -0.59294 | 0.468755 | 0.314012 |
| 6 | 53319600 | 53321799 | 23.28893 | 40.76688 | 32.56727 | 14.37168 | 0.709962 | 0.698068 | 0.314036 |
| 18 | 56062600 | 56065399 | 46.19932 | 74.96223 | 65.63185 | 28.12436 | 0.557667 | 1.63016 | 0.314036 |
| 17 | 16138200 | 16139599 | 9.21539 | 16.64651 | 35.4146 | 15.38697 | -0.67311 | 0.345218 | 0.314041 |
| 10 | 75615200 | 75616999 | 13.05838 | 22.91064 | 43.55057 | 19.25558 | -0.53435 | 0.626412 | 0.314045 |
| 15 | 47393600 | 47394999 | 13.13475 | 24.66656 | 22.41259 | 9.217654 | 0.603114 | 0.117244 | 0.314047 |
| 14 | 44049400 | 44052199 | 44.08332 | 71.41091 | 67.45014 | 28.86839 | 0.446305 | 1.660133 | 0.314049 |
| 19 | 50455600 | 50457199 | 14.5791 | 27.25287 | 22.73815 | 9.424952 | 0.716219 | 0.158977 | 0.314056 |
| 14 | 52068800 | 52070399 | 18.52211 | 33.69826 | 23.56652 | 10.14799 | 0.929358 | 0.343189 | 0.314059 |
| 10 | 60831200 | 60833399 | 11.0748 | 19.8849 | 38.39141 | 16.79157 | -0.54089 | 0.399571 | 0.314065 |
| Y | 5319800 | 5321999 | 14.19781 | 26.46125 | 12.84801 | 5.352407 | 1.48385 | 0.145098 | 0.314068 |
| 2 | 101611200 | 101612399 | 7.867878 | 14.64542 | 31.60734 | 13.19543 | -0.65303 | 0.151207 | 0.314074 |
| 3 | 36704200 | 36705999 | 7.867878 | 14.64542 | 31.60734 | 13.19543 | -0.65303 | 0.151207 | 0.314074 |
| 6 | 124274400 | 124275199 | 7.867878 | 14.64542 | 31.60734 | 13.19543 | -0.65303 | 0.151207 | 0.314074 |
| 18 | 32872000 | 32873199 | 14.54425 | 26.52065 | 23.81612 | 10.22669 | 0.573865 | 0.302284 | 0.314077 |
| 9 | 12794400 | 12794999 | 16.05237 | 29.42956 | 20.74676 | 8.846095 | 0.927777 | 0.277572 | 0.314082 |
| 11 | 110149400 | 110152199 | 28.87732 | 49.2883 | 41.39986 | 18.15015 | 0.633092 | 1.002779 | 0.314105 |
| 7 | 10490000 | 10494999 | 103.5375 | 162.0819 | 129.0949 | 53.55164 | 0.64362 | 2.632953 | 0.314117 |
| 4 | 113884400 | 113887599 | 59.36311 | 95.32147 | 80.74263 | 34.17812 | 0.588777 | 1.934467 | 0.314133 |
| 6 | 47746800 | 47748999 | 17.18868 | 29.16715 | 57.39742 | 25.14518 | -0.59138 | 1.031187 | 0.314133 |
| 15 | 79863800 | 79871199 | 49.97499 | 78.95435 | 153.0441 | 63.56386 | -0.62753 | 2.420941 | 0.314141 |

| | | | | | | | | | |
|---|---|---|---|---|---|---|---|---|---|
| 18 | 50110000 | 50113399 | 24.16598 | 39.68341 | 76.42011 | 32.91831 | -0.56509 | 1.41615 | 0.314165 |
| 15 | 48907000 | 48907999 | 7.68328 | 14.3983 | 31.77088 | 13.11353 | -0.67175 | 0.105219 | 0.314169 |
| 16 | 4756800 | 4761599 | 22.53566 | 36.76517 | 80.95052 | 34.78034 | -0.76296 | 1.506226 | 0.314174 |
| 19 | 7359400 | 7361999 | 11.68674 | 20.15833 | 55.24973 | 24.23715 | -1.06295 | 0.854202 | 0.314178 |
| 9 | 15601000 | 15602199 | 8.443906 | 15.72021 | 34.38603 | 14.35151 | -0.67261 | 0.159248 | 0.314184 |
| 5 | 5542000 | 5542999 | 11.49904 | 21.80997 | 20.54279 | 8.288228 | 0.574981 | 0.03252 | 0.314193 |
| 2 | 70655200 | 70656399 | 6.098607 | 11.43836 | 30.12675 | 12.41705 | -0.92201 | 0.084675 | 0.314198 |
| 16 | 64731400 | 64732599 | 12.08157 | 21.25558 | 42.97051 | 19.02535 | -0.62229 | 0.577597 | 0.314202 |
| 12 | 37558800 | 37560799 | 20.90486 | 36.98878 | 28.10925 | 12.43131 | 0.784735 | 0.541075 | 0.314206 |
| 3 | 145272600 | 145273599 | 30.5102 | 51.20179 | 46.9687 | 20.50079 | 0.505011 | 1.161005 | 0.314209 |
| 12 | 115471000 | 115472599 | 32.30754 | 54.03514 | 48.64131 | 21.18563 | 0.532325 | 1.20552 | 0.314212 |
| 14 | 53685600 | 53687999 | 27.07062 | 46.92711 | 35.99457 | 15.79263 | 0.767532 | 0.854387 | 0.314221 |
| 1 | 112082800 | 112084199 | 15.68027 | 28.77252 | 24.5956 | 10.47471 | 0.652488 | 0.277383 | 0.314224 |
| 11 | 103120200 | 103122399 | 14.26866 | 24.57294 | 51.57907 | 22.6258 | -0.68031 | 0.878077 | 0.314245 |
| 5 | 96203400 | 96211999 | 35.39877 | 58.94131 | 45.46038 | 19.74443 | 0.754598 | 1.256921 | 0.314247 |
| 12 | 107112800 | 107115399 | 16.59626 | 28.61944 | 52.72209 | 23.12814 | -0.49274 | 0.876129 | 0.314252 |
| 16 | 11607600 | 11612599 | 45.66142 | 72.87783 | 125.008 | 52.44751 | -0.43722 | 2.118286 | 0.314257 |
| 6 | 47430400 | 47432199 | 12.6518 | 21.97926 | 48.94909 | 21.48982 | -0.76275 | 0.781008 | 0.314263 |
| 13 | 58981000 | 58982999 | 14.91543 | 27.29306 | 24.09659 | 10.3011 | 0.602137 | 0.284646 | 0.31427 |
| 11 | 109085800 | 109087599 | 13.22389 | 23.27025 | 45.06371 | 19.95127 | -0.56047 | 0.587464 | 0.31427 |
| 13 | 117357000 | 117357999 | 9.346784 | 16.95117 | 34.9282 | 15.10031 | -0.62263 | 0.321872 | 0.31428 |
| 7 | 133425600 | 133426599 | 18.27857 | 32.33913 | 29.1797 | 12.90494 | 0.538572 | 0.529355 | 0.314282 |
| 7 | 23231000 | 23237999 | 191.4707 | 285.0471 | 258.1325 | 106.3619 | 0.399546 | 3.612697 | 0.314285 |
| 10 | 67353800 | 67355799 | 30.53915 | 52.01551 | 41.35846 | 18.12894 | 0.711503 | 1.027774 | 0.314286 |
| 7 | 17008200 | 17012399 | 79.80094 | 126.0709 | 118.2712 | 49.12064 | 0.41791 | 2.436222 | 0.314287 |
| 8 | 84271000 | 84271799 | 12.52441 | 24.02038 | 19.98581 | 7.855384 | 0.786062 | -0.04201 | 0.314288 |
| 7 | 12611400 | 12613399 | 19.85784 | 35.0269 | 29.35446 | 12.99009 | 0.642666 | 0.569476 | 0.314294 |
| 18 | 43483400 | 43486199 | 28.4456 | 46.32633 | 83.01184 | 35.64344 | -0.46859 | 1.54442 | 0.314307 |
| 4 | 81581400 | 81583199 | 15.86221 | 29.88961 | 20.44409 | 8.36002 | 1.015962 | 0.115092 | 0.314308 |
| 13 | 68334600 | 68338599 | 91.85648 | 143.5435 | 135.4867 | 56.21327 | 0.396983 | 2.656422 | 0.314327 |
| X | 15421400 | 15425399 | 93.68533 | 147.4729 | 121.1391 | 50.26038 | 0.605283 | 2.523107 | 0.314358 |
| 3 | 57073800 | 57076399 | 36.05269 | 60.01864 | 49.83539 | 21.64198 | 0.6486 | 1.265779 | 0.314376 |
| 11 | 35378000 | 35380399 | 10.48074 | 18.74342 | 37.35739 | 16.3943 | -0.58963 | 0.419427 | 0.31438 |
| 13 | 33804200 | 33807199 | 35.71159 | 59.57599 | 48.69993 | 21.17697 | 0.671017 | 1.241813 | 0.31439 |
| 5 | 117365400 | 117369399 | 23.48254 | 38.32642 | 81.95522 | 35.21754 | -0.72057 | 1.500649 | 0.314401 |
| 4 | 14237400 | 14238199 | 11.9038 | 21.14779 | 40.353 | 17.80812 | -0.5331 | 0.475725 | 0.31441 |
| 4 | 115142000 | 115142799 | 5.98725 | 11.21655 | 33.91559 | 14.00599 | -1.12289 | 0.096886 | 0.314418 |
| 14 | 7904200 | 7908199 | 34.76149 | 58.50607 | 44.15022 | 19.29385 | 0.785795 | 1.147416 | 0.31443 |
| 16 | 13917600 | 13918799 | 9.322568 | 16.98144 | 34.45142 | 14.81286 | -0.59528 | 0.294775 | 0.314432 |
| 1 | 44556200 | 44556799 | 4.778529 | 9.321999 | 22.78296 | 8.148458 | -0.62616 | -0.44087 | 0.314453 |
| 6 | 147090200 | 147091599 | 13.61712 | 25.23666 | 20.99443 | 8.820797 | 0.705739 | 0.198964 | 0.314462 |
| 14 | 29261600 | 29265799 | 53.69106 | 86.12658 | 79.9322 | 33.78311 | 0.455528 | 1.956995 | 0.314474 |
| 16 | 93100400 | 93104199 | 26.88156 | 44.14179 | 79.79141 | 34.37021 | -0.47416 | 1.424963 | 0.314476 |
| 1 | 58785200 | 58789599 | 58.94349 | 94.22258 | 88.63493 | 37.26881 | 0.430399 | 2.074852 | 0.314476 |
| 10 | 30475200 | 30476999 | 12.39806 | 21.80847 | 42.4507 | 18.79588 | -0.56802 | 0.581274 | 0.314484 |
| 6 | 5495800 | 5496799 | 6.780168 | 12.82008 | 29.21231 | 11.86339 | -0.69752 | 0.033272 | 0.314546 |
| 16 | 20244200 | 20245199 | 6.780168 | 12.82008 | 29.21231 | 11.86339 | -0.69752 | 0.033272 | 0.314546 |

| | | | | | | | | | |
|---|---|---|---|---|---|---|---|---|---|
| X | 62482800 | 62485599 | 39.70962 | 65.5617 | 53.26049 | 22.99787 | 0.679359 | 1.373432 | 0.314566 |
| 8 | 48370000 | 48370799 | 5.404646 | 10.4552 | 25.43636 | 9.712554 | -0.70822 | -0.19559 | 0.314567 |
| 6 | 143596800 | 143601199 | 49.08261 | 79.81871 | 60.81569 | 26.09898 | 0.760496 | 1.59304 | 0.31457 |
| 4 | 128660000 | 128662999 | 20.95036 | 34.57473 | 73.49376 | 31.72515 | -0.70416 | 1.343891 | 0.31458 |
| 8 | 117264800 | 117266199 | 11.62537 | 20.52404 | 40.40161 | 17.87556 | -0.58177 | 0.534134 | 0.314581 |
| 17 | 83937200 | 83940399 | 23.79202 | 38.84098 | 82.07807 | 35.27346 | -0.70332 | 1.49754 | 0.314589 |
| 4 | 129273400 | 129277599 | 27.37546 | 44.91015 | 82.16228 | 35.37725 | -0.49204 | 1.439232 | 0.314596 |
| 7 | 78452800 | 78454799 | 12.78595 | 22.32482 | 45.76075 | 20.15396 | -0.6426 | 0.706736 | 0.314601 |
| 16 | 3224800 | 3225799 | 11.50319 | 22.35451 | 16.98555 | 6.324092 | 0.988499 | -0.2366 | 0.314616 |
| 4 | 132811000 | 132815799 | 20.17917 | 33.29378 | 75.10164 | 32.41521 | -0.78969 | 1.345693 | 0.314618 |
| 1 | 153615800 | 153617799 | 9.967931 | 17.57163 | 41.00008 | 18.14577 | -0.82624 | 0.541913 | 0.314626 |
| 1 | 113671200 | 113673599 | 33.20429 | 55.51168 | 48.65559 | 21.18624 | 0.570635 | 1.211482 | 0.314642 |
| 1 | 35908200 | 35910199 | 12.89436 | 22.58524 | 44.11506 | 19.47775 | -0.57342 | 0.652675 | 0.314649 |
| 18 | 81945200 | 81947199 | 12.89436 | 22.58524 | 44.11506 | 19.47775 | -0.57342 | 0.652675 | 0.314649 |
| 7 | 25859600 | 25874399 | 223.95 | 331.8497 | 264.9841 | 109.1332 | 0.5748 | 3.69788 | 0.314649 |
| 9 | 82271200 | 82272599 | 15.08848 | 27.85586 | 23.57267 | 9.956172 | 0.675296 | 0.236199 | 0.31465 |
| 10 | 31885800 | 31886399 | 6.454236 | 12.31259 | 30.66077 | 12.2201 | -0.80127 | -0.03091 | 0.314655 |
| 4 | 146310600 | 146312199 | 24.16311 | 42.56444 | 27.92311 | 12.35996 | 0.993503 | 0.608397 | 0.314657 |
| 18 | 37699600 | 37717799 | 299.8525 | 431.3001 | 368.6055 | 152.1461 | 0.435627 | 4.164347 | 0.31468 |
| 2 | 45309200 | 45311199 | 19.70483 | 32.89707 | 64.87942 | 28.22578 | -0.59527 | 1.202035 | 0.314693 |
| 10 | 92354400 | 92358199 | 55.77531 | 89.64349 | 72.77062 | 30.84577 | 0.651594 | 1.897109 | 0.314721 |
| 17 | 19971400 | 19972199 | 18.85048 | 33.23818 | 30.14543 | 13.34107 | 0.529275 | 0.570103 | 0.314722 |
| 13 | 120251600 | 120253599 | 12.93406 | 22.5083 | 48.41381 | 21.27782 | -0.71263 | 0.756042 | 0.31473 |
| 4 | 96076400 | 96077999 | 31.92032 | 53.85629 | 44.14642 | 19.30657 | 0.666881 | 1.115062 | 0.314749 |
| 15 | 68586000 | 68588999 | 24.22456 | 40.18141 | 73.08411 | 31.65895 | -0.47952 | 1.293691 | 0.314772 |
| 11 | 42442600 | 42443399 | 9.475723 | 18.45455 | 16.59695 | 6.066975 | 0.771131 | -0.32632 | 0.31478 |
| 18 | 73112600 | 73113199 | 9.475723 | 18.45455 | 16.59695 | 6.066975 | 0.771131 | -0.32632 | 0.31478 |
| 5 | 98800800 | 98801799 | 14.94577 | 27.46695 | 24.11479 | 10.24848 | 0.616246 | 0.261412 | 0.314806 |
| 12 | 115319400 | 115320999 | 20.88918 | 36.80281 | 30.46288 | 13.48377 | 0.660064 | 0.592201 | 0.314809 |
| 6 | 100428000 | 100450399 | 271.6266 | 379.0024 | 699.4896 | 290.142 | -0.71875 | 4.611735 | 0.314836 |
| 7 | 80642400 | 80644599 | 11.52911 | 20.17168 | 45.20087 | 19.93858 | -0.77044 | 0.664668 | 0.314837 |
| 17 | 87253600 | 87266799 | 107.332 | 162.0545 | 301.0012 | 124.4225 | -0.61794 | 3.332733 | 0.31484 |
| 3 | 44140400 | 44141599 | 17.35176 | 31.69476 | 23.72854 | 10.16707 | 0.836226 | 0.312445 | 0.314841 |
| 5 | 145123400 | 145124399 | 6.151701 | 11.77343 | 27.28308 | 10.78862 | -0.68645 | -0.06369 | 0.314854 |
| 15 | 49292600 | 49294599 | 17.52074 | 31.77933 | 24.7405 | 10.69423 | 0.772183 | 0.355048 | 0.314856 |
| 4 | 149940800 | 149941799 | 5.666547 | 10.98318 | 25.55341 | 9.668995 | -0.63134 | -0.22064 | 0.314859 |
| 13 | 76077600 | 76079599 | 12.23279 | 21.58246 | 41.39765 | 18.31862 | -0.54515 | 0.545639 | 0.314862 |
| 8 | 84267600 | 84268399 | 5.98765 | 11.52575 | 26.36482 | 10.25428 | -0.64551 | -0.12349 | 0.314876 |
| 19 | 39901600 | 39901999 | 8.517056 | 16.59215 | 16.10402 | 5.865154 | 0.666879 | -0.35374 | 0.314879 |
| 3 | 43453400 | 43455199 | 19.68354 | 34.94776 | 28.27926 | 12.48626 | 0.696999 | 0.504549 | 0.31488 |
| 17 | 37180200 | 37181199 | 7.249921 | 13.60292 | 30.14308 | 12.41576 | -0.67475 | 0.091566 | 0.314903 |
| 7 | 17105200 | 17105999 | 11.35653 | 21.79059 | 17.35322 | 6.811593 | 0.850094 | -0.06667 | 0.314904 |
| 17 | 79359400 | 79360999 | 9.061743 | 16.37382 | 35.4185 | 15.38296 | -0.69661 | 0.342005 | 0.314915 |
| 14 | 116926400 | 116927199 | 13.53807 | 25.70884 | 20.78862 | 8.365699 | 0.79626 | 0.043281 | 0.314939 |
| 18 | 83142200 | 83147799 | 44.90471 | 71.49907 | 130.2414 | 54.46227 | -0.52751 | 2.195502 | 0.314942 |
| 13 | 22165200 | 22168199 | 18.60895 | 30.69823 | 72.94409 | 31.48188 | -0.86431 | 1.344892 | 0.314948 |
| 15 | 88764200 | 88767199 | 24.46613 | 40.22679 | 75.41746 | 32.50368 | -0.52581 | 1.400637 | 0.314988 |

| | | | | | | | | | |
|---|---|---|---|---|---|---|---|---|---|
| 7 | 81112400 | 81118599 | 39.86004 | 63.62946 | 122.3727 | 51.35435 | -0.60162 | 2.106783 | 0.315001 |
| 8 | 107619000 | 107620799 | 10.24629 | 18.10568 | 41.9575 | 18.56061 | -0.81543 | 0.516185 | 0.315008 |
| 17 | 14319800 | 14322799 | 17.03286 | 28.91989 | 62.3208 | 27.30697 | -0.72198 | 1.03004 | 0.315032 |
| 1 | 126967200 | 126967999 | 11.29423 | 21.52357 | 20.53248 | 8.201209 | 0.569191 | -0.00236 | 0.315033 |
| 3 | 140084200 | 140084799 | 11.29423 | 21.52357 | 20.53248 | 8.201209 | 0.569191 | -0.00236 | 0.315033 |
| 18 | 4481200 | 4481999 | 4.819712 | 9.359159 | 23.89096 | 8.950038 | -0.74972 | -0.27151 | 0.315038 |
| 8 | 70972200 | 70975999 | 14.76548 | 25.30421 | 53.68669 | 23.5445 | -0.69724 | 0.926431 | 0.315041 |
| 18 | 37642200 | 37643799 | 49.29689 | 79.46388 | 73.32983 | 31.21406 | 0.47225 | 1.800609 | 0.315046 |
| 10 | 8110200 | 8115199 | 32.20569 | 52.14714 | 91.66967 | 39.22143 | -0.44834 | 1.660988 | 0.31505 |
| 1 | 195031400 | 195032799 | 18.63364 | 33.19658 | 27.83459 | 12.25467 | 0.648702 | 0.475488 | 0.315079 |
| 10 | 39802200 | 39803999 | 8.652321 | 16.00167 | 32.56705 | 13.72227 | -0.57901 | 0.19861 | 0.315084 |
| 4 | 132493600 | 132496799 | 15.60224 | 26.96271 | 51.65238 | 22.66035 | -0.54832 | 0.849783 | 0.315095 |
| 19 | 57613000 | 57616199 | 33.58721 | 54.11401 | 98.29082 | 41.8107 | -0.5034 | 1.803156 | 0.315123 |
| 9 | 19626600 | 19627599 | 11.76417 | 22.7428 | 17.17474 | 6.577923 | 0.958087 | -0.14254 | 0.315132 |
| 14 | 50819800 | 50822399 | 43.13046 | 70.3928 | 61.07506 | 26.24478 | 0.577157 | 1.530235 | 0.31514 |
| 17 | 38365400 | 38367199 | 26.23369 | 45.15522 | 39.26624 | 17.22436 | 0.585182 | 0.915063 | 0.315154 |
| 10 | 99444400 | 99445799 | 18.74302 | 31.8109 | 58.59431 | 25.67104 | -0.49651 | 1.035213 | 0.315162 |
| 13 | 66375200 | 66377599 | 47.28938 | 77.29649 | 56.18528 | 24.15679 | 0.833382 | 1.517687 | 0.315176 |
| 17 | 92483000 | 92483399 | 10.88271 | 20.80532 | 19.60655 | 7.777223 | 0.595789 | -0.03011 | 0.315197 |
| 4 | 101345200 | 101345799 | 7.415342 | 13.84503 | 31.12855 | 12.9289 | -0.70551 | 0.126475 | 0.315206 |
| 7 | 78650200 | 78651399 | 7.415342 | 13.84503 | 31.12855 | 12.9289 | -0.70551 | 0.126475 | 0.315206 |
| 7 | 106065400 | 106067399 | 26.41905 | 45.96777 | 33.5578 | 14.74708 | 0.838874 | 0.800362 | 0.315207 |
| 1 | 119816000 | 119817599 | 11.00624 | 19.42244 | 40.92696 | 18.10959 | -0.67976 | 0.535015 | 0.315207 |
| 15 | 83083800 | 83084199 | 3.955553 | 7.659716 | 24.77111 | 9.413822 | -1.10499 | -0.22614 | 0.315209 |
| 7 | 137037800 | 137040199 | 14.84708 | 25.63092 | 51.0883 | 22.41217 | -0.6055 | 0.856046 | 0.315213 |
| 1 | 19076600 | 19077999 | 25.11016 | 42.97528 | 39.0049 | 17.10392 | 0.522381 | 0.961523 | 0.315218 |
| 6 | 67352800 | 67353599 | 7.038805 | 13.38794 | 29.98659 | 12.02669 | -0.65756 | -0.00523 | 0.315233 |
| 7 | 9639000 | 9641199 | 27.36036 | 47.47676 | 35.28098 | 15.48001 | 0.813137 | 0.8442 | 0.315246 |
| 7 | 14366000 | 14368399 | 37.49722 | 61.7813 | 55.47755 | 23.93106 | 0.534255 | 1.390794 | 0.315247 |
| 14 | 87609200 | 87611599 | 18.53322 | 32.95694 | 28.43095 | 12.53668 | 0.606475 | 0.487855 | 0.315259 |
| 10 | 41231200 | 41232999 | 7.917865 | 14.59678 | 32.52991 | 13.76616 | -0.71336 | 0.210528 | 0.315268 |
| 13 | 96224600 | 96226199 | 10.2927 | 18.55184 | 36.37912 | 15.84943 | -0.55912 | 0.367557 | 0.315277 |
| 7 | 40888000 | 40903399 | 258.3941 | 377.1198 | 318.7651 | 131.3413 | 0.472067 | 3.944289 | 0.315288 |
| 19 | 3069000 | 3074199 | 67.15255 | 107.7317 | 77.85116 | 32.90883 | 0.81601 | 1.972946 | 0.315333 |
| 7 | 3694000 | 3698599 | 35.24198 | 56.56657 | 107.9563 | 45.66967 | -0.58142 | 1.926439 | 0.315361 |
| 15 | 95718200 | 95720199 | 16.65855 | 28.52034 | 54.25722 | 23.79282 | -0.54085 | 0.944846 | 0.315368 |
| 18 | 24080200 | 24081599 | 26.35154 | 45.19148 | 40.3644 | 17.70297 | 0.545732 | 0.948815 | 0.315375 |
| 4 | 24268600 | 24270999 | 27.49106 | 46.96762 | 35.3492 | 15.49868 | 0.791042 | 0.982073 | 0.315378 |
| 4 | 124092200 | 124094999 | 14.29131 | 24.52068 | 54.22653 | 23.78353 | -0.75657 | 0.913137 | 0.315388 |
| 3 | 40698600 | 40700599 | 13.37128 | 23.3077 | 47.87109 | 21.06118 | -0.64616 | 0.733197 | 0.315396 |
| 4 | 146096200 | 146101199 | 62.64968 | 100.4199 | 83.77617 | 35.36631 | 0.607698 | 1.993844 | 0.315429 |
| 12 | 7161400 | 7161799 | 7.99046 | 15.58977 | 15.69607 | 5.604063 | 0.640721 | -0.42964 | 0.31543 |
| 8 | 123859400 | 123863399 | 30.48648 | 49.25687 | 92.17088 | 39.35185 | -0.5414 | 1.712362 | 0.315466 |
| 3 | 22420200 | 22423799 | 40.86772 | 66.91619 | 58.72981 | 25.26725 | 0.563752 | 1.479687 | 0.315469 |
| 9 | 114680800 | 114687799 | 42.15287 | 67.15462 | 129.8517 | 54.34086 | -0.61259 | 2.174702 | 0.31547 |
| 2 | 76477600 | 76479399 | 12.17215 | 20.9155 | 55.2175 | 24.22019 | -1.01035 | 0.892125 | 0.315475 |
| 7 | 19453000 | 19463199 | 104.8401 | 158.9163 | 262.1629 | 108.5028 | -0.4421 | 3.251051 | 0.315487 |

| | | | | | | | | | |
|---|---|---|---|---|---|---|---|---|---|
| 3 | 103512800 | 103514799 | 13.69234 | 24.00934 | 45.11967 | 19.93982 | -0.51804 | 0.641348 | 0.315491 |
| 9 | 78479200 | 78481399 | 13.69234 | 24.00934 | 45.11967 | 19.93982 | -0.51804 | 0.641348 | 0.315491 |
| 10 | 118946800 | 118947999 | 9.93114 | 18.19742 | 37.80794 | 16.13254 | -0.62232 | 0.27185 | 0.315506 |
| 14 | 111302200 | 111304999 | 31.59936 | 53.21468 | 45.83715 | 20.03456 | 0.595635 | 1.13335 | 0.315513 |
| 16 | 16539400 | 16543199 | 32.34767 | 55.17837 | 39.06796 | 17.12699 | 0.878474 | 1.018925 | 0.315516 |
| 13 | 17712800 | 17713799 | 7.976154 | 14.97496 | 31.27 | 12.86621 | -0.58902 | 0.095408 | 0.315518 |
| 15 | 59865800 | 59866799 | 7.976154 | 14.97496 | 31.27 | 12.86621 | -0.58902 | 0.095408 | 0.315518 |
| 4 | 69991800 | 69993399 | 14.88365 | 27.55872 | 23.05409 | 9.698216 | 0.695826 | 0.216934 | 0.315526 |
| 8 | 38044000 | 38045599 | 14.88365 | 27.55872 | 23.05409 | 9.698216 | 0.695826 | 0.216934 | 0.315526 |
| 7 | 15093000 | 15100199 | 197.6327 | 296.6873 | 217.0064 | 89.55877 | 0.718344 | 3.458457 | 0.315534 |
| 17 | 57920800 | 57922599 | 22.35803 | 39.2806 | 30.89423 | 13.67553 | 0.73218 | 0.633538 | 0.315548 |
| 5 | 58733800 | 58738199 | 79.84466 | 126.2378 | 116.199 | 48.27897 | 0.446159 | 2.416707 | 0.315552 |
| 9 | 8236800 | 8237999 | 18.78348 | 33.12282 | 29.77348 | 13.17624 | 0.542288 | 0.569413 | 0.315553 |
| 15 | 38739000 | 38741599 | 18.58408 | 31.49073 | 59.68218 | 26.13497 | -0.53763 | 1.048879 | 0.315567 |
| X | 154802800 | 154804399 | 25.77705 | 44.76505 | 35.55297 | 15.6051 | 0.717949 | 0.82972 | 0.31557 |
| 16 | 20297000 | 20300999 | 40.36342 | 65.89326 | 60.96946 | 26.2018 | 0.484809 | 1.518792 | 0.3156 |
| 8 | 88360600 | 88361599 | 6.442933 | 12.32649 | 27.86746 | 11.03077 | -0.65327 | -0.05846 | 0.315613 |
| 13 | 111189200 | 111190999 | 22.32574 | 39.07362 | 33.22604 | 14.6601 | 0.620347 | 0.697164 | 0.315639 |
| 3 | 23633600 | 23634199 | 6.292592 | 12.06121 | 27.33838 | 10.76386 | -0.64897 | -0.07613 | 0.315655 |
| 7 | 39709400 | 39711199 | 17.73388 | 31.42388 | 28.63837 | 12.66163 | 0.525 | 0.507649 | 0.315671 |
| 9 | 26616000 | 26623199 | 99.90478 | 155.2294 | 146.8292 | 60.96571 | 0.387536 | 2.761118 | 0.315677 |
| 7 | 43098800 | 43104799 | 33.63109 | 54.0074 | 108.5333 | 45.94798 | -0.65489 | 1.90862 | 0.315695 |
| Y | 16600200 | 16602399 | 17.01011 | 30.7353 | 14.88698 | 6.465861 | 1.446884 | 0.348146 | 0.315711 |
| 15 | 11786000 | 11789399 | 24.11723 | 39.79205 | 72.7124 | 31.38425 | -0.48683 | 1.355255 | 0.315722 |
| 1 | 27718200 | 27719399 | 13.37679 | 24.77833 | 22.84989 | 9.60718 | 0.557059 | 0.203036 | 0.315722 |
| 9 | 99597600 | 99599599 | 14.82771 | 25.64649 | 50.56665 | 22.18462 | -0.58931 | 0.837676 | 0.315733 |
| 12 | 103809600 | 103837599 | 297.8775 | 417.7063 | 622.9606 | 258.1983 | -0.40417 | 4.537807 | 0.315741 |
| 3 | 9747600 | 9750999 | 43.41541 | 69.30199 | 121.492 | 50.98164 | -0.46834 | 2.111992 | 0.315741 |
| 8 | 62629000 | 62630599 | 26.19489 | 45.51405 | 35.31266 | 15.50463 | 0.751385 | 0.821799 | 0.315753 |
| 12 | 84067200 | 84068999 | 8.388413 | 15.41098 | 33.01222 | 14.036 | -0.66153 | 0.236384 | 0.315789 |
| 8 | 64980800 | 64982799 | 9.286809 | 17.19382 | 37.59127 | 15.81577 | -0.68053 | 0.208139 | 0.315792 |
| 18 | 75877000 | 75879599 | 39.63705 | 64.73892 | 60.82747 | 26.14541 | 0.463259 | 1.511731 | 0.315792 |
| X | 5246000 | 5248999 | 41.39004 | 69.09724 | 43.67634 | 19.00084 | 1.04074 | 1.249552 | 0.315816 |
| 13 | 56655400 | 56657399 | 21.48653 | 37.46525 | 33.95399 | 14.9412 | 0.528571 | 0.746948 | 0.315825 |
| 16 | 68895000 | 68897199 | 18.15245 | 32.3604 | 27.90833 | 12.28039 | 0.608863 | 0.468566 | 0.315836 |
| 3 | 122038800 | 122039999 | 6.496026 | 12.25981 | 29.15676 | 11.88788 | -0.76308 | 0.046133 | 0.315837 |
| 9 | 122508800 | 122510599 | 9.790572 | 17.69269 | 37.45682 | 16.26603 | -0.66624 | 0.348279 | 0.315838 |
| X | 125185600 | 125192199 | 112.1081 | 175.0662 | 128.6271 | 53.37112 | 0.757158 | 2.680691 | 0.315841 |
| 7 | 17217600 | 17220999 | 39.32971 | 64.48888 | 52.64934 | 22.66479 | 0.668546 | 1.452597 | 0.315841 |
| 16 | 59054400 | 59057399 | 36.28656 | 60.24122 | 50.99574 | 22.10389 | 0.620543 | 1.297063 | 0.315844 |
| 5 | 26239800 | 26241599 | 17.78274 | 31.55979 | 27.97563 | 12.35652 | 0.56575 | 0.493302 | 0.315856 |
| 3 | 154468000 | 154470799 | 21.41206 | 35.36575 | 71.21644 | 30.75955 | -0.62618 | 1.335551 | 0.315866 |
| 14 | 46655800 | 46657799 | 13.22537 | 23.18593 | 44.09189 | 19.48236 | -0.53494 | 0.640901 | 0.31587 |
| 12 | 22291600 | 22291799 | 8.031333 | 15.68123 | 15.18139 | 5.350869 | 0.713492 | -0.47432 | 0.315873 |
| 15 | 98122200 | 98123799 | 10.14074 | 18.21349 | 36.8591 | 16.11712 | -0.60795 | 0.387888 | 0.315881 |
| 13 | 90766600 | 90767199 | 8.391913 | 16.37832 | 15.02643 | 5.335112 | 0.779561 | -0.44725 | 0.315924 |
| 15 | 34039400 | 34042999 | 39.52194 | 64.48135 | 60.30693 | 25.91149 | 0.468763 | 1.52465 | 0.315927 |

| | | | | | | | | | |
|---|---|---|---|---|---|---|---|---|---|
| 10 | 118054200 | 118055199 | 12.14234 | 22.90822 | 20.84203 | 8.503871 | 0.609996 | 0.073575 | 0.315938 |
| X | 15346200 | 15346599 | 2.838691 | 5.552546 | 17.81942 | 5.974101 | -0.9218 | -0.7104 | 0.315943 |
| X | 154290400 | 154292599 | 23.89188 | 42.08526 | 28.70782 | 12.70739 | 0.937741 | 0.61028 | 0.315944 |
| 7 | 49181600 | 49182599 | 10.04246 | 19.32468 | 18.61713 | 7.24764 | 0.589187 | -0.10243 | 0.315963 |
| 14 | 114515800 | 114518599 | 25.25469 | 44.04867 | 33.63086 | 14.80176 | 0.774888 | 0.76269 | 0.315966 |
| 14 | 90537000 | 90539599 | 22.0065 | 38.23816 | 35.1574 | 15.43684 | 0.507519 | 0.800607 | 0.315976 |
| 2 | 99767400 | 99769199 | 16.69291 | 30.36854 | 25.50499 | 10.98351 | 0.667149 | 0.33503 | 0.315981 |
| 14 | 109274800 | 109276199 | 14.70807 | 26.82383 | 23.13052 | 9.930222 | 0.631887 | 0.299371 | 0.315992 |
| 4 | 44038800 | 44041999 | 14.15775 | 24.25258 | 54.19422 | 23.76628 | -0.77179 | 0.928281 | 0.315995 |
| 19 | 15493600 | 15494799 | 13.09095 | 24.38398 | 21.75349 | 9.071398 | 0.613064 | 0.1646 | 0.315998 |
| 4 | 151502400 | 151505799 | 16.41774 | 27.82998 | 57.88631 | 25.35135 | -0.67103 | 1.037846 | 0.316006 |
| 2 | 160657400 | 160662999 | 86.23691 | 136.1482 | 118.1009 | 49.03347 | 0.530019 | 2.456036 | 0.316018 |
| 9 | 55056200 | 55058799 | 17.12604 | 29.23703 | 55.33708 | 24.26078 | -0.53427 | 0.972834 | 0.316024 |
| 2 | 75156200 | 75156999 | 7.338971 | 14.01199 | 30.13175 | 11.98821 | -0.58879 | -0.0264 | 0.316034 |
| 15 | 41596200 | 41597599 | 7.338971 | 14.01199 | 30.13175 | 11.98821 | -0.58879 | -0.0264 | 0.316034 |
| 3 | 15825000 | 15826799 | 20.16317 | 36.11862 | 26.0745 | 11.42692 | 0.867866 | 0.44949 | 0.316049 |
| 4 | 122466000 | 122467599 | 20.16317 | 36.11862 | 26.0745 | 11.42692 | 0.867866 | 0.44949 | 0.316049 |
| 7 | 33457200 | 33462399 | 100.7506 | 156.5867 | 146.4441 | 60.80334 | 0.404176 | 2.757203 | 0.316057 |
| Y | 3846600 | 3848599 | 12.96032 | 24.46692 | 12.36269 | 5.037424 | 1.451991 | 0.051382 | 0.316084 |
| 4 | 110816600 | 110817399 | 13.00658 | 22.48743 | 53.68446 | 23.55217 | -0.86384 | 0.839757 | 0.316092 |
| 2 | 67948200 | 67949599 | 19.10148 | 33.61108 | 30.31019 | 13.41954 | 0.536485 | 0.595474 | 0.316132 |
| 12 | 85675000 | 85676999 | 13.11535 | 22.77918 | 48.42988 | 21.25872 | -0.6961 | 0.786843 | 0.316136 |
| X | 153028200 | 153030599 | 37.86628 | 62.6115 | 53.19688 | 22.99404 | 0.614988 | 1.351532 | 0.316138 |
| 4 | 148528200 | 148529399 | 8.383446 | 15.36565 | 34.0182 | 14.51011 | -0.7126 | 0.249366 | 0.316159 |
| 7 | 125416800 | 125418399 | 8.383446 | 15.36565 | 34.0182 | 14.51011 | -0.7126 | 0.249366 | 0.316159 |
| 13 | 43262800 | 43265999 | 39.17769 | 62.69725 | 114.3108 | 48.14181 | -0.52071 | 2.027376 | 0.316162 |
| 4 | 100892400 | 100892799 | 7.466919 | 14.58814 | 13.38559 | 4.668411 | 0.801638 | -0.52915 | 0.316173 |
| X | 33334000 | 33337599 | 34.08468 | 57.5336 | 43.08036 | 18.84289 | 0.797148 | 1.121365 | 0.316195 |
| 18 | 55478800 | 55481199 | 33.39491 | 55.93898 | 43.2555 | 18.85797 | 0.750758 | 1.18793 | 0.316198 |
| X | 126290400 | 126293799 | 42.07368 | 70.66358 | 37.90578 | 16.55332 | 1.276228 | 1.185253 | 0.316202 |
| 2 | 9019600 | 9020999 | 19.8363 | 35.07539 | 26.51087 | 11.72595 | 0.792362 | 0.539703 | 0.316208 |
| 2 | 13949000 | 13950799 | 16.42607 | 30.56361 | 21.85593 | 9.130575 | 0.927854 | 0.200287 | 0.316216 |
| 5 | 118929000 | 118933399 | 77.11965 | 118.911 | 206.2819 | 85.68593 | -0.49504 | 2.896524 | 0.316217 |
| 5 | 74915600 | 74918799 | 29.87116 | 49.9699 | 43.89935 | 19.12261 | 0.567315 | 1.190348 | 0.316235 |
| 14 | 92568000 | 92569799 | 25.49165 | 44.46208 | 33.01655 | 14.53141 | 0.814689 | 0.762181 | 0.316242 |
| 16 | 93937600 | 93940999 | 29.08023 | 47.39343 | 83.84865 | 36.01204 | -0.44956 | 1.535745 | 0.316242 |
| 3 | 33664400 | 33665599 | 11.63883 | 20.50794 | 40.94001 | 18.12081 | -0.60298 | 0.556966 | 0.316246 |
| 13 | 75443800 | 75444799 | 11.63883 | 20.50794 | 40.94001 | 18.12081 | -0.60298 | 0.556966 | 0.316246 |
| 9 | 4330200 | 4338399 | 134.4419 | 207.7567 | 139.9151 | 58.11766 | 0.871081 | 2.872183 | 0.31627 |
| 3 | 12282200 | 12284599 | 18.50215 | 33.74286 | 23.02891 | 9.886749 | 0.967001 | 0.328484 | 0.316271 |
| 13 | 94193400 | 94194999 | 9.02912 | 16.38003 | 35.96167 | 15.54121 | -0.71351 | 0.317679 | 0.31628 |
| 14 | 79881000 | 79882999 | 10.46801 | 18.88768 | 37.44185 | 16.29098 | -0.57342 | 0.36574 | 0.316299 |
| 2 | 16708800 | 16710599 | 19.21447 | 34.52108 | 26.39705 | 11.53948 | 0.788067 | 0.42756 | 0.316321 |
| 4 | 94326600 | 94328199 | 8.470114 | 15.74755 | 32.12186 | 13.43746 | -0.57475 | 0.165378 | 0.316325 |
| 14 | 68791400 | 68793799 | 16.75576 | 28.91176 | 53.27304 | 23.37026 | -0.49292 | 0.87208 | 0.316334 |
| 3 | 158401000 | 158403599 | 18.77136 | 32.02407 | 62.54464 | 27.4192 | -0.57973 | 0.993844 | 0.316334 |
| 5 | 95556200 | 95558599 | 25.54853 | 44.31916 | 35.88574 | 15.74521 | 0.689689 | 0.837875 | 0.316346 |

| | | | | | | | | | |
|---|---|---|---|---|---|---|---|---|---|
| 12 | 11485800 | 11488399 | 19.73172 | 32.86022 | 66.47121 | 28.87299 | -0.63172 | 1.230097 | 0.316354 |
| 2 | 98459400 | 98461599 | 34.4978 | 56.98304 | 53.0661 | 22.92121 | 0.482924 | 1.353525 | 0.316355 |
| 10 | 79151200 | 79152399 | 10.47893 | 20.22439 | 19.08113 | 7.365224 | 0.630124 | -0.12428 | 0.316359 |
| 14 | 37757000 | 37759199 | 31.3934 | 52.38689 | 49.20835 | 21.40185 | 0.471145 | 1.22586 | 0.31637 |
| 7 | 12458000 | 12463199 | 64.30606 | 103.1471 | 80.96283 | 34.21507 | 0.69667 | 1.975381 | 0.316383 |
| 19 | 37622600 | 37622999 | 3.523508 | 6.889993 | 18.96342 | 6.467644 | -0.73091 | -0.64063 | 0.316402 |
| 4 | 115491800 | 115494199 | 21.09846 | 35.25488 | 65.66245 | 28.58237 | -0.51314 | 1.196095 | 0.316407 |
| 5 | 98332400 | 98334799 | 22.65302 | 37.0581 | 78.53685 | 33.77706 | -0.70522 | 1.464158 | 0.316415 |
| 13 | 83671800 | 83673199 | 6.800737 | 12.80045 | 29.65666 | 12.15031 | -0.73209 | 0.065874 | 0.316422 |
| 10 | 43755200 | 43756399 | 30.88621 | 51.50885 | 48.13787 | 20.92816 | 0.478487 | 1.229801 | 0.316438 |
| 17 | 20312200 | 20313999 | 36.5548 | 60.97881 | 48.14702 | 20.9357 | 0.720982 | 1.246639 | 0.316448 |
| 7 | 18966200 | 18969199 | 15.73281 | 26.86692 | 54.73378 | 23.99688 | -0.63973 | 0.965084 | 0.316488 |
| 7 | 11658600 | 11664999 | 91.48739 | 145.2528 | 92.9638 | 38.74453 | 0.974904 | 2.315398 | 0.31649 |
| 5 | 151387600 | 151388999 | 15.81185 | 29.22202 | 23.53528 | 9.92617 | 0.747752 | 0.236752 | 0.316496 |
| 7 | 96096800 | 96097599 | 4.562088 | 8.871402 | 24.64989 | 9.129697 | -0.85619 | -0.31563 | 0.316516 |
| 3 | 106742600 | 106744199 | 29.48517 | 49.48841 | 45.41037 | 19.82234 | 0.504673 | 1.156717 | 0.31652 |
| 4 | 153618000 | 153620599 | 19.33039 | 32.46675 | 61.70228 | 26.94568 | -0.54205 | 1.133181 | 0.316531 |
| 12 | 72349400 | 72351999 | 29.12475 | 47.24255 | 87.90695 | 37.66093 | -0.5277 | 1.616979 | 0.316537 |
| 9 | 20145600 | 20147599 | 42.75178 | 69.30534 | 58.86626 | 25.2083 | 0.600131 | 1.638839 | 0.316548 |
| 1 | 10302000 | 10303599 | 10.67525 | 18.61927 | 49.95019 | 21.9841 | -1.02921 | 0.714128 | 0.316549 |
| 5 | 141922800 | 141925199 | 15.72629 | 26.89995 | 54.21106 | 23.77085 | -0.62376 | 0.949862 | 0.316555 |
| 7 | 60229800 | 60233799 | 64.45264 | 102.7056 | 97.73013 | 40.91214 | 0.408974 | 2.179193 | 0.316564 |
| X | 34849800 | 34853999 | 66.62567 | 107.2597 | 66.51794 | 28.26305 | 1.042162 | 1.863625 | 0.316583 |
| 4 | 142893200 | 142893999 | 4.061153 | 7.9135 | 21.85184 | 7.9331 | -0.81795 | -0.39728 | 0.316583 |
| 13 | 87011200 | 87011799 | 5.840763 | 11.27688 | 28.93885 | 11.15094 | -0.79835 | -0.15296 | 0.316598 |
| 4 | 153477600 | 153483599 | 64.76629 | 101.2521 | 180.5422 | 74.90353 | -0.51917 | 2.641358 | 0.316609 |
| 4 | 55418000 | 55420399 | 23.8193 | 39.24942 | 73.09784 | 31.53326 | -0.5149 | 1.371448 | 0.316619 |
| 3 | 142385800 | 142389399 | 38.94653 | 64.0745 | 56.57586 | 24.38959 | 0.557705 | 1.414725 | 0.316621 |
| 5 | 140306200 | 140308199 | 10.50435 | 18.684 | 38.3545 | 16.90904 | -0.63664 | 0.4553 | 0.316634 |
| X | 40183000 | 40184399 | 13.92323 | 26.30214 | 21.39011 | 8.703633 | 0.774797 | 0.08531 | 0.316646 |
| 8 | 115241600 | 115244199 | 23.00117 | 37.94019 | 72.56531 | 31.31712 | -0.55259 | 1.355295 | 0.31665 |
| 13 | 46388600 | 46390199 | 11.42535 | 20.42161 | 39.39318 | 17.29564 | -0.54342 | 0.433196 | 0.316675 |
| 5 | 121683200 | 121684399 | 6.219102 | 11.74273 | 29.71064 | 12.10161 | -0.85097 | 0.038763 | 0.316684 |
| 17 | 81463200 | 81464799 | 16.4107 | 29.36131 | 25.58315 | 11.223 | 0.59699 | 0.429078 | 0.316691 |
| 5 | 38328400 | 38334199 | 56.43708 | 89.17612 | 152.7259 | 63.43617 | -0.44904 | 2.421841 | 0.316697 |
| 13 | 60887400 | 60888399 | 12.14854 | 23.00971 | 21.49762 | 8.699119 | 0.582796 | 0.050461 | 0.316713 |
| 9 | 49689600 | 49690999 | 15.97536 | 28.65367 | 25.20203 | 11.03262 | 0.58568 | 0.410586 | 0.316729 |
| 12 | 49370800 | 49372399 | 18.95144 | 33.48064 | 29.48495 | 13.04375 | 0.572435 | 0.548049 | 0.316753 |
| 7 | 122033600 | 122036799 | 21.11025 | 35.27495 | 65.06937 | 28.32439 | -0.49924 | 1.196234 | 0.316758 |
| 4 | 107540200 | 107544199 | 43.77577 | 69.8278 | 123.2664 | 51.67482 | -0.47944 | 2.135852 | 0.316759 |
| 17 | 31865200 | 31868599 | 38.56032 | 61.62275 | 118.0768 | 49.6292 | -0.5945 | 2.071086 | 0.316763 |
| 4 | 126861000 | 126862199 | 8.126855 | 15.21184 | 32.87151 | 13.59457 | -0.64434 | 0.118748 | 0.316781 |
| 7 | 53007200 | 53009599 | 30.64363 | 52.28862 | 40.21971 | 17.63237 | 0.759478 | 1.010276 | 0.316797 |
| 7 | 3331800 | 3335199 | 17.88068 | 30.64807 | 56.52323 | 24.78882 | -0.49624 | 0.939718 | 0.31681 |
| 11 | 34103800 | 34105599 | 9.219187 | 16.60687 | 37.94235 | 16.54299 | -0.77917 | 0.363967 | 0.316821 |
| 18 | 56705800 | 56707199 | 8.439341 | 15.33122 | 34.46297 | 14.86786 | -0.74509 | 0.304658 | 0.31683 |
| 2 | 149198600 | 149200799 | 23.74756 | 41.39608 | 34.84573 | 15.3309 | 0.634679 | 0.764692 | 0.316854 |

| | | | | | | | | | |
|---|---|---|---|---|---|---|---|---|---|
| 17 | 49677400 | 49678799 | 15.93032 | 28.71494 | 26.21058 | 11.41841 | 0.537251 | 0.383799 | 0.31687 |
| 7 | 23252600 | 23253399 | 19.3511 | 34.22615 | 23.83823 | 10.54328 | 0.909308 | 0.526077 | 0.316881 |
| 14 | 6640400 | 6645399 | 64.73924 | 103.8565 | 80.01855 | 33.82323 | 0.723656 | 1.970728 | 0.316892 |
| 5 | 114632800 | 114634799 | 16.1681 | 27.74398 | 53.1953 | 23.33145 | -0.55155 | 0.92062 | 0.316893 |
| 17 | 57917000 | 57918799 | 22.80974 | 39.91125 | 33.18116 | 14.63738 | 0.652741 | 0.703268 | 0.316895 |
| 17 | 19573600 | 19581399 | 168.8497 | 254.3179 | 223.2485 | 92.21651 | 0.459511 | 3.38947 | 0.316898 |
| X | 67475200 | 67477399 | 30.2206 | 52.55638 | 30.61551 | 13.44992 | 1.162901 | 0.822563 | 0.316905 |
| 5 | 4930400 | 4932199 | 24.74262 | 42.56794 | 39.37359 | 17.27119 | 0.496349 | 0.913121 | 0.316906 |
| 7 | 138528400 | 138531599 | 21.54408 | 36.16843 | 67.54007 | 29.48665 | -0.51705 | 1.148825 | 0.316907 |
| 15 | 93803400 | 93807599 | 47.63312 | 75.86934 | 131.7419 | 55.11583 | -0.45807 | 2.186896 | 0.316934 |
| 3 | 99163600 | 99166799 | 25.67222 | 41.94586 | 79.54291 | 34.19449 | -0.54669 | 1.488594 | 0.316944 |
| 13 | 44822200 | 44824199 | 12.70561 | 22.38307 | 42.43159 | 18.78177 | -0.52922 | 0.566614 | 0.316955 |
| 4 | 124436400 | 124444599 | 67.49117 | 105.6698 | 174.6781 | 72.45947 | -0.40838 | 2.616328 | 0.316983 |
| 16 | 14157600 | 14159799 | 14.0531 | 26.17212 | 23.2813 | 9.711045 | 0.617181 | 0.178702 | 0.316983 |
| 14 | 123229200 | 123231999 | 16.71214 | 28.34529 | 58.47925 | 25.61549 | -0.65939 | 1.033221 | 0.317 |
| 10 | 78322000 | 78326199 | 24.85721 | 40.65745 | 80.22976 | 34.50314 | -0.60288 | 1.470707 | 0.317001 |
| 7 | 7447400 | 7450999 | 38.77915 | 65.38758 | 39.02699 | 17.06444 | 1.122914 | 1.141202 | 0.317002 |
| X | 77781200 | 77790799 | 52.99349 | 85.9371 | 55.62618 | 23.8267 | 0.991582 | 1.651963 | 0.317003 |
| 9 | 116320000 | 116321999 | 28.5188 | 49.16507 | 36.31318 | 15.92973 | 0.820015 | 0.905918 | 0.317012 |
| 15 | 34217000 | 34220799 | 28.1178 | 45.73434 | 85.96436 | 36.8941 | -0.53875 | 1.561681 | 0.317013 |
| 5 | 60630000 | 60632599 | 20.78329 | 36.54717 | 31.11907 | 13.77941 | 0.618543 | 0.613659 | 0.317055 |
| 8 | 44050600 | 44051799 | 16.15966 | 29.62916 | 24.52102 | 10.45399 | 0.697774 | 0.285071 | 0.31706 |
| 4 | 70167600 | 70168599 | 7.798829 | 14.7731 | 31.5881 | 12.77885 | -0.60227 | 0.034829 | 0.317076 |
| 19 | 22698000 | 22699799 | 16.21232 | 29.51622 | 25.57846 | 11.00488 | 0.623096 | 0.327548 | 0.317092 |
| 8 | 109984400 | 109987399 | 18.19323 | 32.10315 | 29.23304 | 12.93536 | 0.523939 | 0.556542 | 0.317098 |
| 4 | 39031400 | 39032199 | 12.22643 | 23.26294 | 20.93618 | 8.390718 | 0.648164 | 0.018106 | 0.317099 |
| 12 | 114701800 | 114705799 | 66.833 | 106.5453 | 97.83294 | 40.97997 | 0.461004 | 2.168414 | 0.317105 |
| X | 4791600 | 4793599 | 26.17321 | 45.6583 | 33.06292 | 14.55331 | 0.850953 | 0.764949 | 0.317109 |
| 8 | 84952400 | 84954599 | 9.835646 | 17.77727 | 35.92239 | 15.59625 | -0.59865 | 0.34835 | 0.317121 |
| 11 | 7941000 | 7944199 | 18.70053 | 31.86176 | 59.23272 | 25.96455 | -0.50915 | 0.999929 | 0.317136 |
| 7 | 67583600 | 67584599 | 7.021826 | 13.44293 | 29.09145 | 11.49337 | -0.58852 | -0.05603 | 0.317146 |
| 10 | 55157600 | 55158999 | 15.46227 | 29.22428 | 19.92317 | 8.098072 | 1.027839 | 0.090235 | 0.317159 |
| 10 | 114999600 | 115002199 | 22.26596 | 37.03999 | 67.84195 | 29.44704 | -0.48911 | 1.250547 | 0.317174 |
| 14 | 64774200 | 64775399 | 5.446522 | 10.4157 | 27.23722 | 10.79462 | -0.86201 | -0.06429 | 0.317199 |
| 13 | 108294600 | 108295799 | 8.507144 | 15.70191 | 32.53749 | 13.74696 | -0.6076 | 0.208533 | 0.317206 |
| 17 | 15045000 | 15045599 | 4.05788 | 7.918064 | 20.89747 | 7.44576 | -0.72895 | -0.46854 | 0.317228 |
| 3 | 127218200 | 127219799 | 13.17103 | 23.08785 | 46.17889 | 20.40241 | -0.60774 | 0.642594 | 0.317265 |
| 7 | 30928400 | 30935599 | 222.1217 | 328.3922 | 277.8166 | 114.4118 | 0.48845 | 3.736403 | 0.317277 |
| 3 | 98214600 | 98218199 | 27.09479 | 44.25245 | 82.83747 | 35.60557 | -0.52859 | 1.497938 | 0.317315 |
| 13 | 46880200 | 46881799 | 10.39721 | 18.37648 | 39.92823 | 17.66214 | -0.72261 | 0.512917 | 0.317321 |
| 1 | 125102400 | 125106399 | 46.18769 | 73.56897 | 129.1356 | 54.02737 | -0.47357 | 2.184518 | 0.317323 |
| 16 | 42839000 | 42840799 | 20.52487 | 36.17189 | 30.46543 | 13.48408 | 0.635251 | 0.58646 | 0.31736 |
| 15 | 33220800 | 33222799 | 17.22791 | 29.53046 | 55.41417 | 24.30251 | -0.52111 | 0.934856 | 0.317361 |
| 4 | 58255200 | 58256599 | 16.84247 | 31.15349 | 22.90727 | 9.650014 | 0.879435 | 0.239259 | 0.317361 |
| 13 | 46519400 | 46520999 | 20.61643 | 36.18318 | 31.79058 | 14.05995 | 0.573394 | 0.642526 | 0.317373 |
| 7 | 110721200 | 110722599 | 9.39419 | 16.85901 | 37.39497 | 16.36257 | -0.74008 | 0.388351 | 0.317382 |
| X | 80437600 | 80440199 | 84.78698 | 134.5093 | 102.9576 | 42.87602 | 0.716084 | 2.335115 | 0.317382 |

| | | | | | | | | | |
|---|---|---|---|---|---|---|---|---|---|
| 18 | 89637400 | 89640399 | 38.57958 | 63.68843 | 49.38233 | 21.32154 | 0.746342 | 1.367867 | 0.317391 |
| 17 | 19442600 | 19443799 | 15.80499 | 30.27328 | 17.99982 | 7.100909 | 1.261787 | -0.00283 | 0.317391 |
| 7 | 130588400 | 130590199 | 12.85538 | 22.62001 | 43.47307 | 19.24734 | -0.54969 | 0.581632 | 0.317395 |
| 5 | 148956800 | 148957999 | 7.057113 | 13.39708 | 28.77045 | 11.58578 | -0.60194 | 0.00746 | 0.317396 |
| 13 | 62908000 | 62908999 | 7.057113 | 13.39708 | 28.77045 | 11.58578 | -0.60194 | 0.00746 | 0.317396 |
| 1 | 102880000 | 102880999 | 16.27235 | 29.16265 | 25.80676 | 11.30509 | 0.576484 | 0.420481 | 0.317401 |
| 9 | 103690200 | 103691599 | 16.27235 | 29.16265 | 25.80676 | 11.30509 | 0.576484 | 0.420481 | 0.317401 |
| 12 | 18325000 | 18335999 | 383.1029 | 536.7525 | 458.0378 | 189.8683 | 0.399761 | 4.552468 | 0.31743 |
| 7 | 28235400 | 28239199 | 24.94511 | 40.60022 | 82.89292 | 35.58432 | -0.65689 | 1.545862 | 0.317435 |
| 11 | 66626200 | 66629599 | 39.58565 | 64.83748 | 59.32355 | 25.52612 | 0.504008 | 1.471428 | 0.31745 |
| 12 | 106056800 | 106066799 | 102.5027 | 155.2211 | 266.2843 | 110.1745 | -0.49971 | 3.270232 | 0.317452 |
| 4 | 96525800 | 96526999 | 16.22413 | 30.7045 | 19.34796 | 7.842475 | 1.144051 | 0.087585 | 0.317458 |
| 7 | 32002400 | 32017599 | 422.4915 | 587.5249 | 502.7552 | 208.6474 | 0.384903 | 4.665995 | 0.317475 |
| 9 | 5995200 | 5998399 | 69.40107 | 110.319 | 104.257 | 43.50337 | 0.415144 | 2.262436 | 0.31748 |
| 12 | 99840400 | 99841999 | 11.45949 | 20.43497 | 40.42693 | 17.78441 | -0.58153 | 0.451089 | 0.317489 |
| 9 | 90590600 | 90592199 | 9.901864 | 18.15167 | 37.27342 | 15.89504 | -0.60501 | 0.267012 | 0.317494 |
| 16 | 72523000 | 72525199 | 20.12112 | 35.22078 | 32.41721 | 14.30515 | 0.506644 | 0.68099 | 0.317517 |
| 17 | 18548800 | 18553799 | 55.87642 | 90.11167 | 70.69306 | 30.10975 | 0.706699 | 1.794091 | 0.317529 |
| 17 | 40278800 | 40283399 | 38.04393 | 63.85832 | 42.45281 | 18.53218 | 0.967905 | 1.180404 | 0.31753 |
| 3 | 74378000 | 74378799 | 9.497927 | 16.79904 | 39.9618 | 17.67446 | -0.85193 | 0.499546 | 0.317565 |
| 6 | 18848400 | 18848999 | 5.570656 | 10.72796 | 26.94686 | 10.46693 | -0.77784 | -0.13266 | 0.317594 |
| 2 | 162008000 | 162008599 | 9.586451 | 17.10662 | 37.40908 | 16.4473 | -0.72422 | 0.428099 | 0.317607 |
| 5 | 72875000 | 72876599 | 10.76519 | 19.44565 | 38.512 | 16.73316 | -0.57098 | 0.362474 | 0.317616 |
| 10 | 66262000 | 66263799 | 18.77475 | 33.52641 | 26.09776 | 11.46733 | 0.756936 | 0.457222 | 0.317623 |
| 10 | 24609600 | 24612399 | 23.94794 | 41.65588 | 36.01298 | 15.82309 | 0.59614 | 0.79745 | 0.317627 |
| 18 | 45284400 | 45285599 | 11.34709 | 21.89144 | 17.13871 | 6.62297 | 0.894249 | -0.11882 | 0.317633 |
| 9 | 19723800 | 19733999 | 57.40364 | 92.3625 | 70.75193 | 30.03811 | 0.73708 | 1.866157 | 0.317638 |
| X | 26237200 | 26239399 | 22.75537 | 40.71311 | 24.21721 | 10.62374 | 1.14454 | 0.471235 | 0.317642 |
| 12 | 114049600 | 114051399 | 19.03487 | 34.85265 | 21.92402 | 9.364441 | 1.088612 | 0.308641 | 0.317647 |
| 7 | 17584200 | 17592399 | 138.7442 | 211.2576 | 192.4361 | 79.75227 | 0.419337 | 3.163402 | 0.317654 |
| 18 | 88321600 | 88321999 | 4.647544 | 9.063038 | 22.06753 | 7.931825 | -0.62712 | -0.4274 | 0.317657 |
| 11 | 113195200 | 113198399 | 28.9861 | 46.99503 | 86.98108 | 37.25105 | -0.52062 | 1.627492 | 0.317658 |
| 3 | 151409400 | 151410599 | 7.378959 | 13.90619 | 29.68848 | 12.13757 | -0.61277 | 0.063233 | 0.317661 |
| 14 | 50238000 | 50240599 | 47.65011 | 76.78819 | 70.38976 | 29.95168 | 0.481352 | 1.802676 | 0.317683 |
| 6 | 52769600 | 52771799 | 10.03572 | 17.89601 | 37.87227 | 16.6606 | -0.67805 | 0.435029 | 0.317689 |
| X | 30886600 | 30887599 | 16.51058 | 31.4605 | 18.4585 | 7.374758 | 1.264671 | 0.044429 | 0.317689 |
| 7 | 83634800 | 83636399 | 8.089119 | 14.83645 | 33.51048 | 14.28011 | -0.74007 | 0.242613 | 0.31769 |
| 5 | 42656200 | 42657799 | 17.58367 | 30.0032 | 56.45039 | 24.74787 | -0.52594 | 0.979459 | 0.317719 |
| 16 | 72459800 | 72461599 | 11.1775 | 19.66352 | 42.48688 | 18.81151 | -0.71763 | 0.572818 | 0.317723 |
| X | 8630600 | 8636199 | 122.4652 | 187.8645 | 171.1534 | 71.05141 | 0.427542 | 3.008764 | 0.317728 |
| 19 | 23413000 | 23413399 | 4.351398 | 8.496546 | 21.10234 | 7.43159 | -0.62815 | -0.50706 | 0.317739 |
| 1 | 116733400 | 116735199 | 16.52873 | 30.14398 | 25.16334 | 10.80306 | 0.678947 | 0.320839 | 0.317741 |
| 13 | 8754200 | 8756799 | 13.22534 | 23.08916 | 45.75063 | 20.14793 | -0.59419 | 0.70879 | 0.317745 |
| 9 | 82495800 | 82496199 | 8.119205 | 15.81213 | 15.68768 | 5.737051 | 0.631205 | -0.34413 | 0.317745 |
| 3 | 27059800 | 27061599 | 18.60822 | 33.51514 | 23.33853 | 10.1774 | 0.923577 | 0.397941 | 0.317766 |
| 5 | 137455800 | 137457599 | 10.27208 | 18.47275 | 37.88989 | 16.55017 | -0.62639 | 0.381564 | 0.317773 |
| 14 | 48533000 | 48534599 | 10.27208 | 18.47275 | 37.88989 | 16.55017 | -0.62639 | 0.381564 | 0.317773 |

| | | | | | | | | | |
|---|---|---|---|---|---|---|---|---|---|
| 12 | 72582200 | 72583599 | 9.347236 | 16.91657 | 35.41274 | 15.3499 | -0.64794 | 0.334039 | 0.317776 |
| 7 | 68169600 | 68170799 | 8.44857 | 15.77432 | 32.23329 | 13.38749 | -0.56902 | 0.137519 | 0.317779 |
| 10 | 14722200 | 14724999 | 29.31968 | 50.48585 | 37.51001 | 16.45416 | 0.811394 | 0.923746 | 0.317798 |
| 7 | 112032400 | 112037599 | 56.85208 | 89.22633 | 176.2358 | 73.09151 | -0.66212 | 2.569567 | 0.317803 |
| X | 33688000 | 33688999 | 14.30159 | 27.7378 | 15.41675 | 5.816018 | 1.417121 | -0.16581 | 0.317804 |
| 13 | 45660600 | 45662399 | 11.03748 | 19.40334 | 42.00057 | 18.59884 | -0.72047 | 0.580175 | 0.317811 |
| 17 | 56019200 | 56019799 | 3.992211 | 7.779397 | 22.47148 | 8.154808 | -0.8831 | -0.40153 | 0.317832 |
| 5 | 69128000 | 69129599 | 15.14819 | 27.76007 | 24.62799 | 10.50708 | 0.597377 | 0.280083 | 0.317857 |
| 19 | 58061400 | 58063799 | 12.81144 | 22.24122 | 48.9546 | 21.48291 | -0.74598 | 0.792218 | 0.317859 |
| X | 118830800 | 118832599 | 21.40457 | 38.05219 | 27.23508 | 12.01222 | 0.87435 | 0.507811 | 0.317874 |
| 9 | 20251600 | 20255199 | 58.25561 | 93.56509 | 81.11898 | 34.34968 | 0.555707 | 1.926477 | 0.317881 |
| 7 | 14711800 | 14728199 | 359.6597 | 513.3886 | 372.9057 | 154.107 | 0.659161 | 4.279062 | 0.317899 |
| 3 | 4842800 | 4843999 | 12.24387 | 23.53632 | 19.41888 | 7.582396 | 0.806509 | -0.06822 | 0.3179 |
| 10 | 103701400 | 103704399 | 16.5674 | 28.12068 | 61.72149 | 27.04177 | -0.74832 | 1.03079 | 0.317903 |
| 12 | 63209800 | 63212799 | 29.40934 | 50.10321 | 41.35909 | 18.12956 | 0.657622 | 1.019935 | 0.317908 |
| 19 | 48037800 | 48042799 | 32.26157 | 52.02699 | 97.54135 | 41.54388 | -0.54741 | 1.770743 | 0.317909 |
| 6 | 9778200 | 9780399 | 28.92578 | 49.94149 | 35.74896 | 15.68294 | 0.865469 | 0.894639 | 0.317912 |
| 16 | 52358800 | 52361199 | 17.20308 | 31.48745 | 24.42159 | 10.43628 | 0.788426 | 0.302791 | 0.317915 |
| X | 88541600 | 88543199 | 16.44409 | 30.51811 | 22.39082 | 9.389785 | 0.887077 | 0.216404 | 0.317918 |
| 4 | 106446600 | 106449599 | 21.77677 | 36.23841 | 67.83496 | 29.45086 | -0.52038 | 1.24466 | 0.317927 |
| 13 | 36766200 | 36768799 | 15.6975 | 26.85566 | 54.78105 | 24.02114 | -0.64129 | 0.946237 | 0.317927 |
| 13 | 71237800 | 71238199 | 29.72616 | 50.14937 | 39.02584 | 17.06667 | 0.741579 | 1.103304 | 0.317931 |
| 17 | 16978200 | 16983599 | 81.66415 | 128.731 | 122.2645 | 50.73659 | 0.397665 | 2.487115 | 0.317941 |
| X | 63504000 | 63506399 | 43.82824 | 72.34367 | 50.60994 | 21.84961 | 0.894315 | 1.384532 | 0.317943 |
| 15 | 66749200 | 66750599 | 7.997577 | 14.90976 | 32.73987 | 13.63268 | -0.67517 | 0.143425 | 0.317958 |
| 16 | 7689800 | 7692199 | 29.97475 | 50.6119 | 40.55316 | 17.73919 | 0.699673 | 1.099275 | 0.317976 |
| 3 | 116850000 | 116852799 | 19.95457 | 33.71148 | 62.94618 | 27.53916 | -0.51638 | 1.085868 | 0.318017 |
| 4 | 17851600 | 17854799 | 19.94357 | 33.45778 | 64.01514 | 27.93527 | -0.55187 | 1.147711 | 0.318025 |
| 7 | 81355400 | 81360999 | 48.33489 | 76.9237 | 130.7481 | 54.63772 | -0.42854 | 2.215032 | 0.318029 |
| 2 | 82438400 | 82439999 | 21.73749 | 39.00537 | 25.01838 | 10.94829 | 1.03905 | 0.449455 | 0.318032 |
| 6 | 71623400 | 71624799 | 15.26244 | 28.50936 | 22.02195 | 9.139412 | 0.824893 | 0.166652 | 0.318035 |
| X | 113914200 | 113917399 | 22.71324 | 40.79489 | 23.78063 | 10.39832 | 1.176737 | 0.447761 | 0.318036 |
| 16 | 17326400 | 17327599 | 5.617416 | 10.69066 | 28.23373 | 11.30973 | -0.89038 | -0.02549 | 0.318038 |
| 4 | 135139000 | 135140799 | 7.582166 | 14.12021 | 31.08453 | 12.96687 | -0.68 | 0.146602 | 0.318066 |
| 13 | 73447400 | 73448999 | 9.373495 | 16.89256 | 35.89281 | 15.64042 | -0.67421 | 0.360343 | 0.318067 |
| 17 | 45454000 | 45459999 | 48.61965 | 77.35537 | 136.3407 | 56.95287 | -0.48128 | 2.224139 | 0.318076 |
| 1 | 131309600 | 131310399 | 5.219936 | 10.14763 | 24.02899 | 8.921129 | -0.63037 | -0.29692 | 0.318079 |
| 13 | 34221200 | 34221999 | 5.219936 | 10.14763 | 24.02899 | 8.921129 | -0.63037 | -0.29692 | 0.318079 |
| 7 | 39526000 | 39526599 | 4.644422 | 9.06693 | 24.37923 | 8.614633 | -0.74728 | -0.4828 | 0.318088 |
| X | 150515600 | 150517799 | 27.75481 | 48.71367 | 26.65606 | 11.78957 | 1.25272 | 0.672947 | 0.318088 |
| 8 | 48877600 | 48879399 | 16.94535 | 28.9837 | 54.8215 | 24.03843 | -0.53291 | 0.953604 | 0.318099 |
| 17 | 37506200 | 37508799 | 19.43437 | 32.77108 | 60.727 | 26.55314 | -0.50557 | 1.096465 | 0.318101 |
| 5 | 17143600 | 17146199 | 17.60054 | 30.08181 | 55.36287 | 24.27434 | -0.49376 | 0.964487 | 0.31812 |
| 7 | 10020400 | 10035799 | 264.0151 | 386.465 | 299.5582 | 123.3991 | 0.601063 | 3.900031 | 0.318123 |
| 9 | 40293000 | 40297599 | 44.04058 | 70.45513 | 118.8946 | 50.04779 | -0.40997 | 2.044343 | 0.31813 |
| 17 | 22449400 | 22459599 | 191.5123 | 278.8801 | 428.2477 | 176.4928 | -0.39199 | 3.974917 | 0.318166 |
| 16 | 19920800 | 19924599 | 72.42848 | 115.7639 | 85.99484 | 36.1522 | 0.770268 | 2.093594 | 0.318169 |

| | | | | | | | | | |
|---|---|---|---|---|---|---|---|---|---|
| 11 | 76968000 | 76970199 | 9.353722 | 16.88548 | 36.92768 | 16.05734 | -0.71386 | 0.350675 | 0.318186 |
| 7 | 107743600 | 107751999 | 119.5673 | 184.4295 | 159.7052 | 66.33865 | 0.506743 | 2.901197 | 0.3182 |
| 16 | 72498800 | 72499999 | 7.642365 | 14.48388 | 31.02535 | 12.53807 | -0.60349 | 0.028312 | 0.318212 |
| 5 | 117440400 | 117444999 | 28.39666 | 46.31561 | 84.03466 | 36.10223 | -0.48509 | 1.521418 | 0.318212 |
| 2 | 94235800 | 94237199 | 8.983647 | 16.44814 | 33.99301 | 14.52048 | -0.61559 | 0.260406 | 0.318216 |
| 1 | 92086000 | 92087199 | 9.985098 | 18.17491 | 37.61994 | 16.19007 | -0.62499 | 0.31207 | 0.318247 |
| 16 | 90523600 | 90530199 | 60.7176 | 95.80478 | 159.0342 | 66.00936 | -0.40571 | 2.459365 | 0.318263 |
| 7 | 38291600 | 38292799 | 31.66643 | 52.74499 | 48.14854 | 20.91653 | 0.51238 | 1.247061 | 0.318297 |
| 7 | 37280200 | 37282799 | 12.22073 | 21.50614 | 45.59181 | 20.18486 | -0.69053 | 0.578316 | 0.318301 |
| 11 | 117076400 | 117078599 | 10.56841 | 18.63819 | 40.48183 | 17.91486 | -0.72317 | 0.541054 | 0.318303 |
| 16 | 7848800 | 7853799 | 49.23053 | 79.52034 | 71.67707 | 30.5916 | 0.509445 | 1.739658 | 0.31832 |
| 16 | 91321800 | 91324799 | 26.94263 | 44.06075 | 79.25289 | 34.08086 | -0.46979 | 1.478756 | 0.318321 |
| 9 | 122032600 | 122035999 | 19.84775 | 33.12764 | 67.217 | 29.23827 | -0.63625 | 1.205025 | 0.318344 |
| X | 81532600 | 81534399 | 22.06702 | 40.11906 | 21.76004 | 9.378468 | 1.293088 | 0.37153 | 0.318348 |
| 4 | 63360000 | 63361999 | 30.44851 | 50.85875 | 46.53107 | 20.24966 | 0.509034 | 1.211461 | 0.31835 |
| 19 | 3438000 | 3441199 | 30.33398 | 48.96708 | 94.34646 | 40.23471 | -0.58499 | 1.737929 | 0.318355 |
| 10 | 30171800 | 30172999 | 16.25358 | 29.72131 | 25.09407 | 10.73746 | 0.665527 | 0.302587 | 0.31836 |
| 17 | 19928400 | 19930399 | 27.45744 | 48.12831 | 28.42584 | 12.55833 | 1.143389 | 0.694533 | 0.318396 |
| 18 | 19495000 | 19506599 | 236.1977 | 345.7543 | 311.4305 | 128.2896 | 0.384535 | 3.896366 | 0.31841 |
| 12 | 4073000 | 4074599 | 9.869936 | 17.6355 | 37.87588 | 16.63421 | -0.69759 | 0.420648 | 0.318431 |
| 1 | 88137800 | 88220799 | 9337.087 | 9867.762 | 11331.05 | 4890.692 | -0.44033 | 9.029664 | 0.318441 |
| 17 | 48899000 | 48902999 | 35.47407 | 57.19399 | 100.7343 | 42.89056 | -0.45796 | 1.783791 | 0.318445 |
| 18 | 19498600 | 19505399 | 192.0688 | 285.2381 | 259.7209 | 106.9897 | 0.388415 | 3.654842 | 0.318457 |
| 2 | 90409400 | 90411599 | 13.17602 | 22.73267 | 51.03266 | 22.38736 | -0.77606 | 0.856653 | 0.318474 |
| 1 | 17086200 | 17088999 | 27.63296 | 44.9926 | 82.95012 | 35.61398 | -0.50979 | 1.544962 | 0.318488 |
| 11 | 95588400 | 95592199 | 18.84821 | 31.25901 | 68.87133 | 29.83099 | -0.75487 | 1.27905 | 0.318489 |
| 10 | 78205600 | 78212199 | 65.85525 | 103.132 | 172.0087 | 71.35049 | -0.42095 | 2.611222 | 0.318491 |
| 11 | 92484600 | 92486399 | 11.97931 | 21.03141 | 42.54384 | 18.82103 | -0.62334 | 0.609095 | 0.318499 |
| 2 | 145481600 | 145483599 | 11.18379 | 19.69533 | 40.97014 | 18.13613 | -0.66226 | 0.560728 | 0.31851 |
| 15 | 65817400 | 65819599 | 19.50206 | 35.65819 | 23.16086 | 9.911404 | 1.041361 | 0.323998 | 0.318511 |
| 10 | 44974400 | 44977999 | 26.62134 | 43.47464 | 82.12451 | 35.29778 | -0.54175 | 1.498199 | 0.318527 |
| 10 | 54995400 | 54996799 | 15.28579 | 28.82947 | 20.46691 | 8.356519 | 0.96426 | 0.103996 | 0.318536 |
| 5 | 151379000 | 151381799 | 18.67616 | 33.65365 | 21.95625 | 9.568874 | 1.018344 | 0.396349 | 0.318541 |
| 17 | 20529800 | 20533399 | 37.98278 | 63.65475 | 44.18057 | 19.26715 | 0.906106 | 1.198965 | 0.318543 |
| X | 134501600 | 134503599 | 39.08582 | 65.0271 | 48.19703 | 20.92161 | 0.811974 | 1.282588 | 0.318562 |
| 10 | 43406000 | 43407799 | 9.258758 | 16.59186 | 36.90172 | 16.16652 | -0.74494 | 0.396848 | 0.318591 |
| 12 | 113139600 | 113141399 | 21.20549 | 36.9244 | 32.18181 | 14.14913 | 0.584623 | 0.763207 | 0.318611 |
| 11 | 84654600 | 84656399 | 10.99257 | 19.45757 | 39.3703 | 17.41018 | -0.61967 | 0.499122 | 0.318613 |
| 7 | 31762400 | 31770599 | 88.29475 | 139.9727 | 103.3 | 42.98744 | 0.767641 | 2.357613 | 0.318622 |
| 10 | 116416400 | 116418399 | 12.1638 | 21.20571 | 47.34025 | 20.82935 | -0.7656 | 0.725283 | 0.318634 |
| 2 | 165186800 | 165190599 | 22.0167 | 36.41321 | 70.91038 | 30.65572 | -0.57794 | 1.319829 | 0.318639 |
| 15 | 66797200 | 66798199 | 14.74928 | 28.09853 | 19.61579 | 7.840595 | 1.015013 | 0.030911 | 0.318645 |
| 16 | 55467800 | 55469199 | 14.74928 | 28.09853 | 19.61579 | 7.840595 | 1.015013 | 0.030911 | 0.318645 |
| X | 26629000 | 26630199 | 14.74928 | 28.09853 | 19.61579 | 7.840595 | 1.015013 | 0.030911 | 0.318645 |
| 1 | 181464800 | 181466199 | 7.706131 | 14.37882 | 31.65875 | 13.16327 | -0.67703 | 0.133673 | 0.318655 |
| 3 | 11636000 | 11637399 | 18.79009 | 33.10781 | 29.35122 | 12.99146 | 0.561742 | 0.577634 | 0.318662 |
| 3 | 6388000 | 6391199 | 33.33868 | 55.59636 | 49.78742 | 21.6447 | 0.53983 | 1.238725 | 0.318701 |

| | | | | | | | | | |
|---|---|---|---|---|---|---|---|---|---|
| 1 | 19699000 | 19700799 | 28.4249 | 48.03869 | 44.78323 | 19.59495 | 0.482116 | 1.090919 | 0.318704 |
| 8 | 100190600 | 100191599 | 11.09065 | 21.23178 | 20.01727 | 7.909115 | 0.599867 | -0.04104 | 0.318715 |
| X | 120309400 | 120311599 | 31.26668 | 53.22879 | 37.20752 | 16.30879 | 0.896573 | 1.031585 | 0.318717 |
| 17 | 83542600 | 83544599 | 10.87357 | 19.15466 | 40.9701 | 18.13505 | -0.70204 | 0.554981 | 0.318733 |
| 19 | 47943600 | 47945199 | 10.39296 | 18.78676 | 39.06223 | 16.957 | -0.64024 | 0.3537 | 0.318744 |
| 9 | 81512600 | 81516599 | 34.575 | 55.55416 | 106.0183 | 44.92032 | -0.57951 | 1.891566 | 0.318749 |
| 1 | 50964000 | 50968399 | 29.34415 | 47.45488 | 101.6628 | 43.45597 | -0.73469 | 1.685822 | 0.318754 |
| 7 | 32820800 | 32823199 | 27.19944 | 47.43271 | 31.81822 | 14.0025 | 0.960618 | 0.77178 | 0.318757 |
| 1 | 22572200 | 22573399 | 10.71797 | 18.90085 | 40.46545 | 17.90782 | -0.70267 | 0.541825 | 0.318776 |
| 18 | 18700000 | 18702399 | 33.02957 | 55.37499 | 43.23042 | 18.85735 | 0.73697 | 1.177238 | 0.318782 |
| 7 | 20896000 | 20899599 | 30.5101 | 52.80959 | 34.08712 | 14.95505 | 1.014855 | 0.87985 | 0.318784 |
| 2 | 10241600 | 10243799 | 8.062111 | 14.85698 | 33.04564 | 13.99173 | -0.71139 | 0.213976 | 0.318786 |
| 7 | 9908000 | 9914399 | 74.48022 | 119.2505 | 77.45211 | 32.6415 | 0.966374 | 2.04353 | 0.318824 |
| 12 | 114228600 | 114235399 | 109.0885 | 169.7803 | 140.8334 | 58.46168 | 0.578596 | 2.736553 | 0.318824 |
| 9 | 49739000 | 49740799 | 7.245838 | 13.59121 | 31.23062 | 12.8702 | -0.72775 | 0.094262 | 0.318825 |
| 17 | 93384000 | 93384799 | 10.79482 | 20.96977 | 16.89994 | 6.311935 | 0.899269 | -0.23666 | 0.318826 |
| 4 | 121317400 | 121320599 | 65.8939 | 105.4667 | 86.64349 | 36.49698 | 0.627451 | 2.043898 | 0.318831 |
| 7 | 60946000 | 60948999 | 28.88914 | 49.7665 | 37.47658 | 16.43972 | 0.792119 | 0.91714 | 0.318849 |
| 1 | 165063400 | 165065199 | 9.987574 | 17.57043 | 42.53246 | 18.83163 | -0.88034 | 0.566464 | 0.318864 |
| 2 | 89940600 | 89942599 | 25.71519 | 44.20132 | 39.84055 | 17.47554 | 0.533236 | 0.925521 | 0.318876 |
| 8 | 51385800 | 51387799 | 18.20426 | 33.11651 | 24.80399 | 10.68225 | 0.831237 | 0.345163 | 0.318889 |
| 7 | 94034600 | 94035599 | 17.75051 | 31.39672 | 28.2802 | 12.50778 | 0.540918 | 0.52692 | 0.318903 |
| 3 | 53164200 | 53166599 | 18.43459 | 31.39494 | 57.50859 | 25.2079 | -0.48786 | 1.003245 | 0.31891 |
| 9 | 26164400 | 26168599 | 51.72628 | 83.42213 | 73.10945 | 31.14057 | 0.54734 | 1.786945 | 0.318913 |
| X | 82408800 | 82410399 | 24.90819 | 44.36053 | 23.53924 | 10.36641 | 1.304646 | 0.512371 | 0.318922 |
| 17 | 62879800 | 62881199 | 9.503677 | 17.13412 | 39.5392 | 17.22061 | -0.79272 | 0.362996 | 0.318924 |
| X | 67408000 | 67411199 | 38.10827 | 63.6797 | 45.92146 | 19.98965 | 0.851296 | 1.232158 | 0.318934 |
| 7 | 44020800 | 44025199 | 25.57378 | 42.03673 | 77.37997 | 33.34572 | -0.49968 | 1.407179 | 0.318947 |
| 3 | 26649800 | 26651199 | 14.27802 | 26.16195 | 23.39019 | 9.980793 | 0.58644 | 0.272705 | 0.318952 |
| 13 | 56576200 | 56579199 | 16.78399 | 28.29693 | 60.53246 | 26.46667 | -0.71178 | 1.089047 | 0.318953 |
| 4 | 40196400 | 40199599 | 16.83951 | 28.65855 | 56.36014 | 24.70311 | -0.59 | 1.001526 | 0.318968 |
| 1 | 100917200 | 100919399 | 21.64306 | 38.92355 | 24.3931 | 10.65439 | 1.073988 | 0.433453 | 0.318976 |
| 19 | 43479800 | 43480399 | 4.657395 | 9.050735 | 23.3593 | 8.698128 | -0.75665 | -0.29525 | 0.318987 |
| 9 | 121389800 | 121393999 | 24.6778 | 40.41958 | 78.59242 | 33.81715 | -0.5805 | 1.450746 | 0.319007 |
| 10 | 81645800 | 81648399 | 26.70452 | 45.72244 | 36.22288 | 15.8845 | 0.717671 | 0.958981 | 0.319011 |
| 2 | 68376400 | 68383999 | 181.1596 | 263.4083 | 431.8525 | 178.0062 | -0.48848 | 3.996626 | 0.319015 |
| 5 | 75633200 | 75637199 | 28.12089 | 45.76044 | 83.54241 | 35.86067 | -0.49629 | 1.554745 | 0.319016 |
| X | 90150200 | 90152599 | 36.91197 | 61.57205 | 48.15296 | 20.93773 | 0.734667 | 1.246518 | 0.319027 |
| 16 | 63699600 | 63701199 | 24.98933 | 43.24274 | 33.44909 | 14.67501 | 0.75466 | 0.853473 | 0.319028 |
| 1 | 106999200 | 106999599 | 13.50735 | 24.95806 | 22.81721 | 9.625936 | 0.565922 | 0.217428 | 0.319032 |
| 6 | 116457200 | 116460199 | 12.63844 | 22.01356 | 50.51538 | 22.21395 | -0.80532 | 0.746116 | 0.319042 |
| 12 | 21122000 | 21124999 | 43.10731 | 70.0969 | 64.29902 | 27.59317 | 0.493377 | 1.583372 | 0.31906 |
| 12 | 76474400 | 76477199 | 13.37371 | 23.37621 | 45.20379 | 19.92341 | -0.55906 | 0.691031 | 0.319098 |
| 16 | 43458800 | 43460799 | 12.25288 | 21.4934 | 44.59473 | 19.71415 | -0.66017 | 0.624199 | 0.319107 |
| X | 24407800 | 24418399 | 142.0413 | 219.1582 | 137.7894 | 57.23516 | 0.968437 | 2.904079 | 0.319112 |
| 15 | 53274800 | 53277599 | 29.96617 | 51.09792 | 41.40025 | 18.14892 | 0.684755 | 1.016435 | 0.319117 |
| 2 | 28575800 | 28578199 | 8.474727 | 15.29543 | 35.45148 | 15.41958 | -0.79661 | 0.345825 | 0.319117 |

| | | | | | | | | | |
|---|---|---|---|---|---|---|---|---|---|
| 5 | 60502200 | 60504199 | 8.105039 | 14.84185 | 36.13071 | 15.4303 | -0.84952 | 0.259562 | 0.319119 |
| 18 | 7382800 | 7386799 | 51.4951 | 83.36052 | 65.96174 | 28.21719 | 0.700201 | 1.686343 | 0.319123 |
| 10 | 83068200 | 83071399 | 17.0285 | 28.87611 | 57.94045 | 25.37795 | -0.61942 | 1.036102 | 0.319135 |
| 6 | 22784200 | 22785999 | 21.33419 | 37.47618 | 31.17926 | 13.79994 | 0.651666 | 0.632064 | 0.319137 |
| 17 | 7502400 | 7506999 | 41.8211 | 66.70228 | 123.3427 | 51.69868 | -0.54644 | 2.136963 | 0.319143 |
| 19 | 9760000 | 9763399 | 41.08619 | 67.72981 | 53.92104 | 23.26476 | 0.707758 | 1.394124 | 0.319156 |
| 7 | 60636400 | 60638599 | 30.09859 | 51.37807 | 37.09776 | 16.26419 | 0.850385 | 1.001378 | 0.31919 |
| 1 | 110667400 | 110667999 | 9.445371 | 18.28664 | 18.16979 | 6.911834 | 0.576281 | -0.18609 | 0.319206 |
| 14 | 110732800 | 110733999 | 22.53206 | 39.15574 | 36.13119 | 15.86552 | 0.502359 | 0.803143 | 0.319217 |
| 13 | 15454000 | 15456799 | 38.7828 | 64.06621 | 53.82845 | 23.25181 | 0.630927 | 1.364054 | 0.31923 |
| 18 | 3365400 | 3373399 | 108.2398 | 167.1153 | 158.183 | 65.70602 | 0.379494 | 2.878272 | 0.31923 |
| 2 | 80899200 | 80900599 | 13.98767 | 26.37503 | 20.52558 | 8.383826 | 0.832024 | 0.092786 | 0.319261 |
| 3 | 121608200 | 121609999 | 10.79514 | 19.26084 | 37.84288 | 16.63995 | -0.57116 | 0.436881 | 0.319261 |
| 9 | 66174400 | 66176199 | 10.79514 | 19.26084 | 37.84288 | 16.63995 | -0.57116 | 0.436881 | 0.319261 |
| 2 | 168255400 | 168257399 | 13.71806 | 23.8331 | 48.43439 | 21.26468 | -0.63138 | 0.784527 | 0.31929 |
| 4 | 8030600 | 8033599 | 28.32374 | 47.92206 | 44.21388 | 19.35249 | 0.497006 | 1.079061 | 0.319294 |
| 3 | 38538400 | 38539599 | 9.272255 | 16.57917 | 38.41112 | 16.85933 | -0.80565 | 0.412736 | 0.3193 |
| 5 | 116958800 | 116964599 | 56.57888 | 89.55057 | 148.1602 | 61.59212 | -0.39732 | 2.381848 | 0.319308 |
| 5 | 67220000 | 67221999 | 17.44851 | 31.07818 | 28.14506 | 12.39373 | 0.538207 | 0.471102 | 0.319314 |
| 11 | 88956000 | 88956999 | 6.71675 | 12.91209 | 28.10906 | 10.97434 | -0.58067 | -0.10011 | 0.31932 |
| 17 | 69382600 | 69386199 | 32.42679 | 52.28404 | 97.46798 | 41.50303 | -0.53953 | 1.776434 | 0.31933 |
| 10 | 111401200 | 111404199 | 20.93573 | 34.98726 | 65.66016 | 28.58359 | -0.52405 | 1.194072 | 0.31933 |
| 17 | 19950400 | 19953799 | 56.28948 | 91.19554 | 60.49872 | 25.89567 | 0.954984 | 1.678331 | 0.319357 |
| 2 | 17087800 | 17091599 | 52.91664 | 85.25375 | 75.59656 | 32.1555 | 0.52876 | 1.822589 | 0.319359 |
| 1 | 56213400 | 56215199 | 10.74797 | 19.38809 | 37.41651 | 16.28492 | -0.53547 | 0.367501 | 0.319365 |
| 13 | 45590400 | 45591399 | 10.74797 | 19.38809 | 37.41651 | 16.28492 | -0.53547 | 0.367501 | 0.319365 |
| 17 | 38875600 | 38882799 | 93.0546 | 147.3541 | 102.1554 | 42.47773 | 0.85634 | 2.388893 | 0.319365 |
| 12 | 19130000 | 19159399 | 1216.279 | 1549.197 | 1207.655 | 506.9159 | 0.392329 | 6.068813 | 0.319374 |
| 17 | 17554600 | 17560799 | 78.30047 | 124.4734 | 101.3409 | 42.28933 | 0.63 | 2.269096 | 0.31938 |
| 7 | 31350400 | 31354799 | 84.54204 | 133.0098 | 125.5228 | 52.07556 | 0.404753 | 2.52794 | 0.319391 |
| 13 | 33453200 | 33453599 | 12.67368 | 23.76987 | 21.98048 | 9.059795 | 0.57559 | 0.120667 | 0.3194 |
| 4 | 117159000 | 117161999 | 19.34212 | 32.03444 | 71.21597 | 30.81753 | -0.76809 | 1.296269 | 0.319401 |
| 1 | 88136600 | 88240199 | 11261.85 | 11728.91 | 13185.91 | 5704.714 | -0.43118 | 9.265055 | 0.319401 |
| 5 | 124280000 | 124283999 | 30.94742 | 50.12227 | 89.95341 | 38.4942 | -0.47772 | 1.654951 | 0.319406 |
| 4 | 86062400 | 86064199 | 19.56701 | 35.22639 | 25.00295 | 10.90926 | 0.896213 | 0.412604 | 0.319412 |
| 6 | 71219200 | 71223999 | 24.56516 | 40.70982 | 74.21258 | 32.12787 | -0.48288 | 1.306168 | 0.319412 |
| 13 | 10199400 | 10201599 | 27.66458 | 47.10583 | 42.53802 | 18.64554 | 0.528504 | 1.019915 | 0.319413 |
| 11 | 113955600 | 113959599 | 30.87589 | 49.69318 | 102.2034 | 43.40856 | -0.68416 | 1.832471 | 0.319417 |
| 8 | 35600200 | 35602199 | 47.88769 | 77.54115 | 61.64092 | 26.37391 | 0.69385 | 1.671057 | 0.319421 |
| 16 | 84876000 | 84877599 | 8.883327 | 16.12515 | 34.94102 | 15.08888 | -0.69377 | 0.312095 | 0.319422 |
| 4 | 152835200 | 152865199 | 356.1777 | 491.6367 | 722.8973 | 300.3603 | -0.4067 | 4.778284 | 0.319423 |
| 5 | 109857800 | 109858799 | 23.62683 | 41.19488 | 34.24413 | 15.06834 | 0.652616 | 0.758962 | 0.319429 |
| 12 | 48492000 | 48494199 | 41.83288 | 67.95755 | 63.99628 | 27.44738 | 0.454403 | 1.59868 | 0.31944 |
| 4 | 21399600 | 21400599 | 16.29954 | 27.9708 | 53.21544 | 23.34036 | -0.54048 | 0.919428 | 0.319453 |
| 3 | 58165600 | 58168599 | 22.74228 | 39.82191 | 32.5646 | 14.37414 | 0.676347 | 0.689924 | 0.319475 |
| 4 | 39957800 | 39961999 | 42.51693 | 69.29335 | 62.20448 | 26.71633 | 0.526682 | 1.547786 | 0.319478 |
| 5 | 75140200 | 75159999 | 231.174 | 331.2929 | 514.9174 | 212.6607 | -0.43218 | 4.215138 | 0.319489 |

| | | | | | | | | | |
|---|---|---|---|---|---|---|---|---|---|
| 12 | 84193200 | 84195799 | 12.95963 | 22.5111 | 48.42821 | 21.25939 | -0.71304 | 0.784303 | 0.319517 |
| 13 | 3307000 | 3308199 | 19.84282 | 34.84167 | 31.73324 | 14.04012 | 0.521603 | 0.6293 | 0.319528 |
| 5 | 109798000 | 109804599 | 81.53913 | 129.2037 | 110.3587 | 45.90402 | 0.55645 | 2.365754 | 0.319529 |
| 18 | 3721400 | 3724599 | 66.78064 | 106.1496 | 99.53364 | 41.53091 | 0.426382 | 2.260451 | 0.319552 |
| 12 | 99094600 | 99096999 | 21.54577 | 36.07078 | 66.92786 | 29.16813 | -0.5078 | 1.177423 | 0.319555 |
| 6 | 38020200 | 38030199 | 81.55194 | 125.7733 | 210.3655 | 87.38211 | -0.44221 | 2.893481 | 0.319572 |
| 8 | 14145000 | 14149799 | 39.21853 | 62.61744 | 131.2381 | 55.08975 | -0.72516 | 2.102543 | 0.319573 |
| X | 44694000 | 44697399 | 55.92353 | 90.28705 | 68.845 | 29.36432 | 0.749469 | 1.764441 | 0.319573 |
| 16 | 28009800 | 28010399 | 5.062573 | 9.834894 | 24.497 | 9.145954 | -0.70953 | -0.27578 | 0.319575 |
| 13 | 114734000 | 114736199 | 15.56422 | 28.71155 | 23.49634 | 9.93456 | 0.722165 | 0.244049 | 0.319578 |
| 6 | 109525400 | 109527199 | 16.23 | 29.23588 | 26.62764 | 11.60966 | 0.539566 | 0.390122 | 0.319582 |
| 3 | 148129600 | 148132399 | 18.92606 | 32.11299 | 58.57645 | 25.66115 | -0.48247 | 1.039109 | 0.319631 |
| 10 | 89760600 | 89762599 | 34.23909 | 57.55067 | 45.26847 | 19.7746 | 0.726108 | 1.15755 | 0.319645 |
| 14 | 31234600 | 31237799 | 30.54985 | 51.98546 | 41.91351 | 18.37086 | 0.691332 | 1.036713 | 0.319646 |
| 13 | 62627600 | 62633199 | 93.49927 | 146.6464 | 131.1528 | 54.39466 | 0.478737 | 2.594003 | 0.319669 |
| 16 | 90470400 | 90472599 | 31.13506 | 50.28975 | 92.77588 | 39.59489 | -0.52146 | 1.722586 | 0.319673 |
| 6 | 19314400 | 19315199 | 8.994507 | 16.53963 | 33.54023 | 14.24274 | -0.58192 | 0.236938 | 0.319685 |
| 2 | 120649000 | 120652199 | 56.91667 | 91.45868 | 80.48229 | 34.10417 | 0.535106 | 1.909358 | 0.319698 |
| 17 | 66620200 | 66621799 | 9.412511 | 17.28835 | 35.12514 | 14.93916 | -0.58689 | 0.247154 | 0.319702 |
| 7 | 58543400 | 58546199 | 23.04457 | 40.63125 | 27.93592 | 12.36337 | 0.926506 | 0.591182 | 0.319703 |
| 15 | 43172600 | 43174799 | 18.33721 | 33.07666 | 25.12544 | 10.93607 | 0.801064 | 0.392689 | 0.319712 |
| 14 | 61926200 | 61929199 | 18.9076 | 32.08505 | 58.59563 | 25.67037 | -0.48422 | 1.037431 | 0.319723 |
| X | 3992600 | 3995399 | 31.54875 | 54.74146 | 31.79139 | 13.94888 | 1.167177 | 0.860839 | 0.319739 |
| 12 | 24260800 | 24262599 | 28.6839 | 49.88307 | 32.36013 | 14.21621 | 1.008425 | 0.817475 | 0.319739 |
| 7 | 110372000 | 110373599 | 10.47899 | 18.70674 | 37.84898 | 16.63514 | -0.61283 | 0.430734 | 0.31974 |
| 13 | 54562600 | 54575199 | 106.6181 | 161.8742 | 260.2816 | 107.7768 | -0.40322 | 3.217586 | 0.319741 |
| 4 | 57681400 | 57684199 | 18.05797 | 30.84799 | 56.48055 | 24.76339 | -0.48656 | 0.971155 | 0.319781 |
| 3 | 12636600 | 12639999 | 38.07902 | 62.76545 | 54.43053 | 23.48347 | 0.584594 | 1.385691 | 0.319783 |
| 6 | 119124600 | 119128199 | 31.99835 | 54.14304 | 43.61411 | 19.09037 | 0.692192 | 1.093286 | 0.319795 |
| 4 | 140670000 | 140680999 | 100.3054 | 152.0595 | 262.5411 | 108.6632 | -0.50762 | 3.247363 | 0.319827 |
| 6 | 19367200 | 19369599 | 18.2535 | 31.21442 | 57.57255 | 25.24428 | -0.49685 | 0.964881 | 0.319836 |
| 9 | 9206400 | 9207799 | 13.05998 | 24.53023 | 22.36709 | 9.196348 | 0.598224 | 0.114651 | 0.31985 |
| 1 | 99295800 | 99297999 | 26.07566 | 45.30393 | 32.33958 | 14.19866 | 0.871095 | 0.81908 | 0.319851 |
| 9 | 99245400 | 99246599 | 8.841908 | 16.19476 | 33.48187 | 14.29478 | -0.61537 | 0.257044 | 0.319858 |
| 13 | 60116600 | 60117199 | 3.705433 | 7.228041 | 21.42014 | 7.668704 | -0.90098 | -0.45719 | 0.319858 |
| 13 | 66053400 | 66060799 | 159.867 | 244.8534 | 149.7729 | 62.15953 | 1.00001 | 3.051142 | 0.319868 |
| 3 | 31311000 | 31312999 | 15.09486 | 26.19829 | 50.59683 | 22.20018 | -0.55866 | 0.806883 | 0.31987 |
| 4 | 25901600 | 25903199 | 17.60933 | 32.52906 | 22.87708 | 9.654579 | 0.941552 | 0.253074 | 0.319872 |
| 3 | 28988600 | 28989199 | 8.142553 | 15.24928 | 31.2034 | 12.89316 | -0.56478 | 0.113466 | 0.319892 |
| 7 | 109462400 | 109466999 | 22.15541 | 36.67779 | 71.03134 | 30.72836 | -0.56991 | 1.308415 | 0.319898 |
| 7 | 69929600 | 69933799 | 34.85035 | 56.11751 | 101.1631 | 42.99738 | -0.49362 | 1.821848 | 0.3199 |
| 11 | 32061400 | 32066799 | 53.73938 | 85.27027 | 142.4739 | 59.34354 | -0.40824 | 2.311088 | 0.319913 |
| 17 | 13706600 | 13713199 | 68.48873 | 106.7333 | 181.7884 | 75.44776 | -0.45667 | 2.699904 | 0.319924 |
| 4 | 153465800 | 153473599 | 91.4185 | 139.4727 | 238.4725 | 98.88297 | -0.48605 | 3.114071 | 0.319925 |
| 1 | 119850400 | 119853599 | 18.83065 | 31.63631 | 63.40542 | 27.69334 | -0.61855 | 1.128078 | 0.319945 |
| 13 | 13888000 | 13889199 | 20.08011 | 36.36718 | 23.38086 | 10.12542 | 1.044817 | 0.379878 | 0.319947 |
| 8 | 64093800 | 64094599 | 11.09081 | 19.94226 | 38.94378 | 17.01256 | -0.55599 | 0.392455 | 0.31995 |

| | | | | | | | | | |
|---|---|---|---|---|---|---|---|---|---|
| 16 | 72203400 | 72204199 | 17.67093 | 31.75838 | 26.90231 | 11.75709 | 0.641233 | 0.413939 | 0.319953 |
| 5 | 42888600 | 42891799 | 44.05431 | 72.31845 | 55.13605 | 23.74677 | 0.767668 | 1.4524 | 0.319962 |
| 7 | 23176000 | 23178199 | 34.41606 | 57.45232 | 49.22904 | 21.4159 | 0.603274 | 1.231817 | 0.319981 |
| 1 | 46210600 | 46212399 | 21.43547 | 37.52128 | 32.11481 | 14.17165 | 0.610757 | 0.685899 | 0.319998 |
| 9 | 3432200 | 3434599 | 16.14484 | 29.76281 | 22.8327 | 9.662715 | 0.813657 | 0.250379 | 0.320007 |
| 14 | 40180200 | 40181999 | 17.51411 | 31.81738 | 25.53775 | 11.01712 | 0.730736 | 0.350484 | 0.320022 |
| 3 | 40848200 | 40849199 | 8.62599 | 15.56205 | 36.43515 | 15.85555 | -0.81202 | 0.348979 | 0.320027 |
| 6 | 8209000 | 8209799 | 8.62599 | 15.56205 | 36.43515 | 15.85555 | -0.81202 | 0.348979 | 0.320027 |
| 7 | 98789000 | 98790999 | 9.351734 | 16.5261 | 39.99536 | 17.69238 | -0.87695 | 0.509407 | 0.320029 |
| 11 | 11984800 | 11988199 | 21.18216 | 35.14339 | 69.89965 | 30.28449 | -0.60797 | 1.279214 | 0.320035 |
| 7 | 43577000 | 43590999 | 109.4246 | 165.4296 | 274.2144 | 113.3965 | -0.45224 | 3.30527 | 0.320037 |
| 3 | 93766200 | 93768199 | 16.73799 | 30.19866 | 25.90868 | 11.27387 | 0.626865 | 0.381846 | 0.320043 |
| 17 | 20579400 | 20589199 | 140.5074 | 215.5556 | 163.7926 | 67.99632 | 0.689163 | 3.012927 | 0.320066 |
| 12 | 115061400 | 115065799 | 87.72533 | 138.0265 | 125.1997 | 51.94191 | 0.461881 | 2.529095 | 0.320087 |
| 3 | 91987800 | 91989399 | 13.08856 | 22.77661 | 47.88878 | 21.04674 | -0.67993 | 0.756804 | 0.320098 |
| 9 | 102502600 | 102504799 | 9.207946 | 16.28722 | 40.97246 | 18.12122 | -0.93219 | 0.497842 | 0.320104 |
| 3 | 3904600 | 3906599 | 23.42157 | 41.44633 | 26.88317 | 11.88882 | 1.012165 | 0.554878 | 0.320116 |
| 5 | 110076400 | 110076999 | 14.20677 | 26.57247 | 22.75168 | 9.421192 | 0.680201 | 0.151788 | 0.320128 |
| 6 | 66560800 | 66562799 | 23.31137 | 41.4503 | 24.60733 | 10.8514 | 1.14224 | 0.51209 | 0.320145 |
| 2 | 40064000 | 40066199 | 26.99547 | 46.35571 | 39.76338 | 17.44116 | 0.604091 | 0.939142 | 0.320147 |
| 7 | 61198400 | 61200599 | 29.31204 | 50.24619 | 39.12562 | 17.15905 | 0.742764 | 0.96333 | 0.32015 |
| 7 | 64081600 | 64083599 | 34.07736 | 56.23457 | 52.53585 | 22.67715 | 0.478214 | 1.361615 | 0.320151 |
| 7 | 58625000 | 58625599 | 12.78508 | 23.97015 | 22.43126 | 9.251251 | 0.557653 | 0.12435 | 0.320158 |
| 5 | 5806000 | 5806599 | 3.938565 | 7.678644 | 21.34541 | 7.696339 | -0.81857 | -0.42636 | 0.320161 |
| 8 | 126299200 | 126300399 | 13.76969 | 25.3711 | 22.90682 | 9.700919 | 0.580128 | 0.236571 | 0.320174 |
| 7 | 13300800 | 13302399 | 30.95491 | 52.24051 | 45.28513 | 19.80606 | 0.586593 | 1.112821 | 0.320181 |
| 7 | 98178800 | 98181399 | 14.77124 | 25.69052 | 49.50414 | 21.74908 | -0.55506 | 0.774052 | 0.320203 |
| 6 | 86605000 | 86606399 | 10.58544 | 18.61558 | 41.52114 | 18.38518 | -0.76302 | 0.573816 | 0.320218 |
| 1 | 89028600 | 89031599 | 23.29397 | 38.58877 | 70.4966 | 30.51129 | -0.48588 | 1.305385 | 0.320221 |
| 7 | 33260000 | 33284999 | 135.2427 | 207.26 | 158.935 | 65.97019 | 0.674823 | 3.030935 | 0.320223 |
| 5 | 75964800 | 75966599 | 27.58106 | 46.65639 | 43.74018 | 19.14402 | 0.474064 | 1.078697 | 0.320226 |
| 12 | 18901200 | 18905599 | 178.3602 | 266.3038 | 227.6299 | 93.83564 | 0.486608 | 3.556803 | 0.320227 |
| 7 | 39302400 | 39305399 | 26.39481 | 43.40018 | 77.43207 | 33.37269 | -0.45454 | 1.406393 | 0.320251 |
| 16 | 89661600 | 89665599 | 44.23311 | 70.57434 | 122.6644 | 51.43966 | -0.45672 | 2.127503 | 0.320258 |
| 7 | 14733800 | 14742999 | 162.9407 | 247.9729 | 178.8464 | 74.10884 | 0.755298 | 3.17966 | 0.32026 |
| 16 | 97122400 | 97123999 | 15.44714 | 26.72711 | 50.57944 | 22.19047 | -0.53034 | 0.837339 | 0.320267 |
| 2 | 168641000 | 168644199 | 30.60102 | 49.51836 | 91.2377 | 39.01889 | -0.5168 | 1.669913 | 0.320294 |
| 2 | 177227400 | 177227999 | 18.2935 | 32.30808 | 28.81632 | 12.74876 | 0.554285 | 0.548336 | 0.320324 |
| 17 | 57494000 | 57498199 | 31.42265 | 50.81394 | 91.1097 | 38.94276 | -0.47853 | 1.686631 | 0.320336 |
| 2 | 93652400 | 93653999 | 7.825265 | 14.28879 | 35.02662 | 15.01438 | -0.86346 | 0.268773 | 0.320337 |
| 2 | 88423800 | 88428799 | 63.76675 | 102.1014 | 86.90825 | 36.62904 | 0.57698 | 2.030074 | 0.320343 |
| 10 | 74260800 | 74262399 | 16.45546 | 28.44805 | 53.27185 | 23.3711 | -0.51548 | 0.85433 | 0.320346 |
| 3 | 145513400 | 145514799 | 12.26118 | 21.55225 | 41.96042 | 18.58146 | -0.56859 | 0.588606 | 0.32035 |
| 8 | 19086600 | 19088599 | 25.16237 | 43.02493 | 39.97136 | 17.52642 | 0.488578 | 0.969525 | 0.320357 |
| 14 | 37224400 | 37227999 | 28.23301 | 46.05243 | 84.03346 | 36.10278 | -0.49318 | 1.519746 | 0.320357 |
| 14 | 42294800 | 42305599 | 107.0524 | 166.769 | 139.6692 | 57.97024 | 0.565819 | 2.719252 | 0.32036 |
| 7 | 17845600 | 17854399 | 129.8852 | 200.0056 | 158.343 | 65.76767 | 0.634088 | 2.936212 | 0.320374 |

| | | | | | | | | | |
|---|---|---|---|---|---|---|---|---|---|
| 14 | 84604800 | 84606799 | 14.67996 | 25.65095 | 49.98826 | 22.02615 | -0.57039 | 0.710851 | 0.320393 |
| 8 | 114932200 | 114933599 | 9.551606 | 17.14139 | 36.38993 | 15.9229 | -0.67694 | 0.38928 | 0.320394 |
| 5 | 85070800 | 85073399 | 28.17486 | 48.76048 | 35.79739 | 15.70532 | 0.829853 | 0.87143 | 0.320397 |
| 5 | 6032600 | 6033999 | 16.34692 | 29.44288 | 26.5643 | 11.58381 | 0.55288 | 0.390992 | 0.320403 |
| 12 | 21962800 | 21983399 | 659.2023 | 883.0518 | 729.4695 | 304.4954 | 0.381389 | 5.252054 | 0.32041 |
| 9 | 36321800 | 36331199 | 141.9354 | 216.0458 | 193.7105 | 80.27254 | 0.441742 | 3.170247 | 0.320411 |
| 7 | 97841000 | 97846199 | 32.15168 | 51.74738 | 108.6216 | 46.13569 | -0.71357 | 1.836624 | 0.320414 |
| 14 | 44062200 | 44063199 | 4.970447 | 9.663796 | 24.6801 | 9.15388 | -0.73802 | -0.30544 | 0.320416 |
| 1 | 24804800 | 24806999 | 17.89569 | 30.6965 | 56.52171 | 24.78963 | -0.49374 | 0.933658 | 0.320422 |
| 10 | 20677800 | 20678799 | 14.08406 | 26.29346 | 22.41085 | 9.309655 | 0.683564 | 0.16373 | 0.320435 |
| 4 | 66693200 | 66694599 | 17.6784 | 29.48064 | 70.06988 | 30.46217 | -0.86337 | 1.214934 | 0.320438 |
| 6 | 61481000 | 61481999 | 10.70734 | 20.73317 | 18.60461 | 7.071988 | 0.722787 | -0.1732 | 0.320442 |
| 16 | 97882800 | 97884199 | 12.90621 | 22.51868 | 47.32956 | 20.83552 | -0.67907 | 0.716313 | 0.320445 |
| 7 | 102327600 | 102329599 | 10.66934 | 18.90191 | 41.39895 | 18.30408 | -0.73311 | 0.488687 | 0.320446 |
| 18 | 81836800 | 81840599 | 19.20905 | 32.12164 | 64.36883 | 28.03243 | -0.61824 | 1.181618 | 0.320469 |
| 5 | 29508800 | 29510599 | 29.56072 | 50.16364 | 43.6089 | 19.10488 | 0.582693 | 1.058814 | 0.320469 |
| 10 | 66363000 | 66364399 | 15.50857 | 29.09237 | 21.5353 | 8.873423 | 0.894204 | 0.141956 | 0.32047 |
| 14 | 42014600 | 42015999 | 25.32119 | 43.82441 | 37.68234 | 16.53233 | 0.602952 | 0.861786 | 0.320507 |
| 13 | 12367800 | 12369999 | 19.25026 | 34.00134 | 29.42356 | 13.01714 | 0.597422 | 0.551746 | 0.32051 |
| 9 | 78015200 | 78016399 | 7.505831 | 13.77794 | 34.05155 | 14.49447 | -0.86783 | 0.233942 | 0.320518 |
| 7 | 126545400 | 126555199 | 45.44348 | 71.85346 | 159.2775 | 66.17997 | -0.81981 | 2.400581 | 0.320521 |
| X | 138511000 | 138511999 | 12.46813 | 23.66922 | 20.0538 | 8.075627 | 0.729429 | 0.036079 | 0.320527 |
| 4 | 16695000 | 16695599 | 5.131924 | 9.949574 | 27.42411 | 10.36404 | -0.87311 | -0.23117 | 0.320549 |
| 2 | 177664800 | 177666399 | 11.13427 | 19.74936 | 38.83237 | 17.15995 | -0.57727 | 0.478328 | 0.32057 |
| 4 | 120297400 | 120299399 | 25.95775 | 45.07432 | 32.79617 | 14.39416 | 0.843511 | 0.825663 | 0.32057 |
| 1 | 162442800 | 162444599 | 23.16805 | 40.32539 | 31.4957 | 13.84395 | 0.742032 | 0.777759 | 0.32058 |
| 4 | 148114000 | 148115799 | 7.704306 | 14.41056 | 30.62603 | 12.68133 | -0.62124 | 0.116871 | 0.320593 |
| 6 | 112582200 | 112583399 | 7.704306 | 14.41056 | 30.62603 | 12.68133 | -0.62124 | 0.116871 | 0.320593 |
| 13 | 53707400 | 53708599 | 7.704306 | 14.41056 | 30.62603 | 12.68133 | -0.62124 | 0.116871 | 0.320593 |
| 8 | 91875400 | 91881599 | 68.893 | 107.6198 | 176.7466 | 73.33466 | -0.40154 | 2.657977 | 0.320604 |
| 6 | 49962200 | 49965199 | 13.20918 | 23.21457 | 43.49121 | 19.25692 | -0.51353 | 0.599481 | 0.320648 |
| 13 | 33594000 | 33599199 | 49.38394 | 79.95709 | 67.44332 | 28.85473 | 0.608481 | 1.680174 | 0.320649 |
| 13 | 40385800 | 40386399 | 4.618885 | 8.994378 | 22.95883 | 8.406978 | -0.71788 | -0.35495 | 0.320653 |
| 1 | 40166800 | 40167399 | 8.07662 | 15.72956 | 15.63523 | 5.716145 | 0.628384 | -0.34718 | 0.320664 |
| 1 | 61677200 | 61679199 | 13.88174 | 24.05042 | 51.09563 | 22.41784 | -0.69605 | 0.817937 | 0.320692 |
| 7 | 17264400 | 17269799 | 64.27048 | 102.5457 | 95.54556 | 40.06816 | 0.441261 | 2.139688 | 0.320702 |
| 12 | 80759800 | 80762399 | 18.98516 | 32.02256 | 60.10581 | 26.28375 | -0.52388 | 1.093424 | 0.320703 |
| 2 | 177295200 | 177297999 | 41.55758 | 67.77539 | 62.12337 | 26.68803 | 0.497371 | 1.536625 | 0.320703 |
| 12 | 117099200 | 117100399 | 9.634333 | 17.50119 | 34.92853 | 15.06968 | -0.57513 | 0.313753 | 0.320714 |
| 2 | 49509200 | 49515199 | 51.40395 | 81.79854 | 140.6416 | 58.76264 | -0.44535 | 2.222866 | 0.320727 |
| X | 146580000 | 146581599 | 22.39487 | 39.14418 | 33.88495 | 14.93458 | 0.594877 | 0.719721 | 0.320743 |
| 17 | 38324600 | 38325399 | 16.67925 | 29.91163 | 27.06178 | 11.84828 | 0.545076 | 0.420501 | 0.320743 |
| 7 | 37019600 | 37020599 | 5.101764 | 9.918982 | 24.07507 | 8.930391 | -0.66523 | -0.30429 | 0.320759 |
| 1 | 67794600 | 67797799 | 53.23521 | 85.42916 | 73.34892 | 31.01937 | 0.568257 | 1.942736 | 0.320765 |
| 13 | 76246200 | 76247399 | 9.297772 | 17.01262 | 33.98172 | 14.52802 | -0.5676 | 0.267114 | 0.32077 |
| 14 | 60323800 | 60325199 | 9.297772 | 17.01262 | 33.98172 | 14.52802 | -0.5676 | 0.267114 | 0.32077 |
| 19 | 58339200 | 58340999 | 9.297772 | 17.01262 | 33.98172 | 14.52802 | -0.5676 | 0.267114 | 0.32077 |

| | | | | | | | | | |
|---|---|---|---|---|---|---|---|---|---|
| 5 | 96088000 | 96089399 | 14.47076 | 26.42336 | 21.41866 | 9.181233 | 0.722272 | 0.287985 | 0.3208 |
| 18 | 3025000 | 3031399 | 100.8141 | 158.1178 | 125.4994 | 52.04998 | 0.650757 | 2.599296 | 0.320807 |
| 5 | 127199800 | 127201799 | 25.61962 | 44.11813 | 39.32017 | 17.24825 | 0.549987 | 0.909071 | 0.32082 |
| 12 | 68421800 | 68422599 | 11.37484 | 21.92636 | 19.59639 | 7.591183 | 0.703127 | -0.10088 | 0.320832 |
| 17 | 92759600 | 92762399 | 29.88372 | 50.4664 | 44.17793 | 19.32572 | 0.572407 | 1.099523 | 0.320851 |
| 1 | 34416600 | 34419199 | 20.97044 | 34.98122 | 65.55143 | 28.50334 | -0.52184 | 1.214875 | 0.320868 |
| 7 | 40546400 | 40553399 | 63.57593 | 99.15159 | 181.8173 | 75.45129 | -0.56221 | 2.684286 | 0.320877 |
| 15 | 61709200 | 61710799 | 12.15412 | 21.28172 | 44.1608 | 19.49256 | -0.65991 | 0.656678 | 0.32089 |
| 16 | 26811800 | 26812999 | 7.891147 | 14.68397 | 31.60117 | 13.2001 | -0.64912 | 0.156248 | 0.32091 |
| 5 | 99875200 | 99877799 | 13.12042 | 22.74842 | 50.54622 | 22.17729 | -0.7601 | 0.807402 | 0.320926 |
| 2 | 10314000 | 10317199 | 24.439 | 40.37005 | 72.78868 | 31.4412 | -0.46738 | 1.343006 | 0.320936 |
| 15 | 97973000 | 97974199 | 6.401818 | 11.99584 | 31.73032 | 13.09869 | -0.9309 | 0.094693 | 0.320939 |
| 5 | 123068600 | 123071399 | 17.17423 | 29.21629 | 56.34562 | 24.69594 | -0.56197 | 1.007735 | 0.320944 |
| 6 | 66873000 | 66874599 | 12.01253 | 20.94252 | 47.86297 | 21.05967 | -0.79933 | 0.72451 | 0.320949 |
| 18 | 54287000 | 54294399 | 87.96433 | 135.5506 | 217.8435 | 90.48869 | -0.38559 | 2.912638 | 0.320964 |
| 7 | 75915800 | 75919599 | 31.07605 | 53.01845 | 39.63235 | 17.37465 | 0.800465 | 1.012073 | 0.320971 |
| 17 | 71359600 | 71361599 | 7.462898 | 13.80364 | 32.05035 | 13.50229 | -0.76723 | 0.186775 | 0.320974 |
| 8 | 92452800 | 92484399 | 347.6758 | 480.5088 | 711.6025 | 295.6079 | -0.41517 | 4.758653 | 0.320975 |
| 10 | 61884800 | 61886599 | 18.81243 | 31.76479 | 59.57625 | 26.06054 | -0.52272 | 1.082781 | 0.320975 |
| 3 | 64492200 | 64496399 | 65.19397 | 103.9088 | 98.42929 | 41.21683 | 0.415815 | 2.17426 | 0.320978 |
| 8 | 67303600 | 67308399 | 54.74847 | 88.19265 | 74.96561 | 31.88066 | 0.58934 | 1.827595 | 0.320981 |
| 4 | 62442600 | 62443799 | 6.7221 | 12.48086 | 31.57596 | 13.2337 | -0.88426 | 0.15816 | 0.320996 |
| 19 | 9235400 | 9238399 | 50.14311 | 81.01643 | 70.54228 | 30.11785 | 0.559713 | 1.734098 | 0.320997 |
| 12 | 19539600 | 19545399 | 186.0242 | 277.3359 | 226.1804 | 93.21132 | 0.552412 | 3.586134 | 0.321006 |
| 3 | 86453800 | 86456199 | 23.51524 | 38.26548 | 88.30811 | 37.90624 | -0.83331 | 1.550018 | 0.321006 |
| 6 | 121106400 | 121108399 | 10.70969 | 18.88316 | 41.96436 | 18.57177 | -0.75663 | 0.543745 | 0.321011 |
| X | 146485600 | 146488399 | 44.13644 | 71.9222 | 62.20536 | 26.71527 | 0.580138 | 1.555138 | 0.321051 |
| 19 | 13306600 | 13310799 | 57.76421 | 93.13712 | 70.21369 | 29.89771 | 0.763712 | 1.803435 | 0.321057 |
| 7 | 60795000 | 60798999 | 52.33979 | 84.7775 | 64.7259 | 27.70046 | 0.752497 | 1.677941 | 0.321068 |
| 5 | 144576600 | 144599599 | 267.4565 | 378.4274 | 569.9592 | 235.8888 | -0.40532 | 4.403948 | 0.321076 |
| 2 | 89097600 | 89102999 | 86.21585 | 135.7256 | 124.7493 | 51.75877 | 0.443439 | 2.515668 | 0.321096 |
| 5 | 120753800 | 120755399 | 18.95134 | 32.07418 | 59.07974 | 25.86166 | -0.49663 | 1.062219 | 0.321096 |
| 6 | 59726800 | 59729199 | 24.00413 | 42.03674 | 31.32295 | 13.82755 | 0.809778 | 0.692323 | 0.321105 |
| 9 | 8277400 | 8279399 | 13.25437 | 24.53446 | 20.36879 | 8.572137 | 0.706464 | 0.198999 | 0.321105 |
| 16 | 95615200 | 95620199 | 48.36793 | 76.88454 | 133.3207 | 55.6305 | -0.45912 | 2.25317 | 0.321111 |
| 17 | 59449800 | 59450199 | 4.516385 | 8.814807 | 22.2861 | 7.908452 | -0.66448 | -0.47582 | 0.321135 |
| 7 | 5539400 | 5540599 | 31.39595 | 52.27408 | 49.22747 | 21.37997 | 0.467561 | 1.249554 | 0.321144 |
| 10 | 77778000 | 77786599 | 110.9057 | 167.8244 | 269.2666 | 111.383 | -0.40413 | 3.287062 | 0.321149 |
| 7 | 33655400 | 33669799 | 334.2134 | 475.414 | 411.0457 | 169.9678 | 0.402869 | 4.327686 | 0.32115 |
| 1 | 103968600 | 103969799 | 15.4961 | 28.4437 | 24.60217 | 10.47294 | 0.636045 | 0.274006 | 0.321168 |
| 3 | 12099400 | 12102599 | 40.11487 | 65.55392 | 60.92395 | 26.19192 | 0.479502 | 1.505121 | 0.321178 |
| 15 | 96096400 | 96098399 | 10.07129 | 18.34928 | 36.02039 | 15.48323 | -0.5482 | 0.30114 | 0.32118 |
| 6 | 132383800 | 132385599 | 70.97012 | 112.5159 | 103.4931 | 43.07079 | 0.450483 | 2.341043 | 0.321183 |
| 5 | 138365600 | 138367599 | 26.45535 | 45.88314 | 36.47885 | 16.00534 | 0.716076 | 0.846787 | 0.321184 |
| 1 | 127596000 | 127597999 | 29.19417 | 49.0139 | 45.91133 | 20.0446 | 0.474955 | 1.151325 | 0.321191 |
| 5 | 94365800 | 94366199 | 9.900647 | 19.16008 | 18.65494 | 7.110241 | 0.602496 | -0.17333 | 0.321195 |
| X | 61345200 | 61346799 | 23.9238 | 41.72458 | 34.41245 | 15.14513 | 0.664199 | 0.759715 | 0.321198 |

| | | | | | | | | | |
|---|---|---|---|---|---|---|---|---|---|
| 1 | 153812600 | 153813199 | 11.57639 | 22.22525 | 20.10131 | 7.877257 | 0.670828 | -0.06032 | 0.3212 |
| 13 | 86738200 | 86739199 | 14.7833 | 25.32676 | 57.40356 | 25.17391 | -0.79218 | 0.935413 | 0.321211 |
| 5 | 111452600 | 111453799 | 15.70523 | 28.27281 | 25.71026 | 11.21849 | 0.541072 | 0.387827 | 0.321215 |
| 6 | 69038800 | 69040599 | 14.91354 | 28.06431 | 21.67115 | 8.881954 | 0.839903 | 0.118119 | 0.321228 |
| 1 | 85846000 | 85851399 | 50.78535 | 80.62979 | 140.1308 | 58.40151 | -0.46413 | 2.290772 | 0.321243 |
| 16 | 88562800 | 88565399 | 40.33831 | 65.83506 | 61.36688 | 26.37008 | 0.473977 | 1.52477 | 0.321263 |
| 18 | 15072000 | 15073399 | 9.814792 | 17.71032 | 35.88141 | 15.61065 | -0.60462 | 0.355695 | 0.321264 |
| 12 | 6205600 | 6208199 | 26.50759 | 45.75066 | 38.10617 | 16.71687 | 0.647807 | 0.890429 | 0.321265 |
| 9 | 77562200 | 77563399 | 5.893509 | 11.2366 | 28.2943 | 11.2903 | -0.81771 | -0.03632 | 0.321276 |
| 14 | 52601200 | 52607199 | 92.83031 | 145.2582 | 135.4614 | 56.19506 | 0.415852 | 2.634025 | 0.321279 |
| 16 | 89382400 | 89383399 | 7.646104 | 14.0329 | 33.02343 | 14.06029 | -0.79723 | 0.23731 | 0.321279 |
| 7 | 132081800 | 132085399 | 15.29665 | 26.54524 | 49.51475 | 21.72532 | -0.5086 | 0.809621 | 0.321283 |
| 16 | 16324400 | 16326199 | 35.49134 | 56.76352 | 116.6884 | 49.10106 | -0.69438 | 2.043986 | 0.321291 |
| 2 | 44360600 | 44362799 | 24.83223 | 42.63971 | 37.7441 | 16.55544 | 0.558786 | 0.925988 | 0.321294 |
| 2 | 146751600 | 146754399 | 35.93439 | 59.74888 | 50.94391 | 22.10501 | 0.610412 | 1.280376 | 0.321316 |
| 3 | 27166400 | 27167999 | 11.27908 | 20.00002 | 39.32083 | 17.38035 | -0.57751 | 0.481275 | 0.321318 |
| 7 | 125089400 | 125092199 | 11.00681 | 19.44409 | 39.91073 | 17.65618 | -0.64133 | 0.522506 | 0.32133 |
| 10 | 54331000 | 54332999 | 14.467 | 25.21788 | 47.34308 | 20.82898 | -0.51714 | 0.738694 | 0.321346 |
| 12 | 83501000 | 83503399 | 15.80728 | 27.16216 | 53.22307 | 23.34538 | -0.5825 | 0.904364 | 0.321352 |
| 1 | 172095600 | 172098999 | 22.05146 | 38.33356 | 35.65632 | 15.66035 | 0.49094 | 0.795613 | 0.321352 |
| X | 26024000 | 26025999 | 28.60848 | 49.10047 | 38.77393 | 17.00649 | 0.723027 | 0.952201 | 0.321401 |
| X | 108746000 | 108747999 | 29.04479 | 49.26465 | 43.15877 | 18.90495 | 0.571684 | 1.062339 | 0.321405 |
| 17 | 20135800 | 20139399 | 103.2938 | 160.0447 | 151.8498 | 63.0742 | 0.379973 | 2.810992 | 0.321415 |
| 9 | 35018800 | 35021199 | 10.08004 | 17.85475 | 39.89693 | 17.64061 | -0.76163 | 0.485566 | 0.321418 |
| 7 | 17029800 | 17050399 | 362.4614 | 512.9861 | 429.0651 | 177.5693 | 0.44333 | 4.403198 | 0.321437 |
| 5 | 100369000 | 100371799 | 45.27912 | 72.06906 | 129.3345 | 54.05586 | -0.50654 | 2.209185 | 0.321446 |
| 8 | 73808200 | 73808799 | 9.066536 | 17.70631 | 14.42248 | 5.064429 | 0.963432 | -0.47648 | 0.321471 |
| 15 | 66925200 | 66925799 | 9.066536 | 17.70631 | 14.42248 | 5.064429 | 0.963432 | -0.47648 | 0.321471 |
| 11 | 79036400 | 79037399 | 6.352568 | 11.96143 | 29.64802 | 12.13897 | -0.82694 | 0.062625 | 0.321473 |
| X | 24562000 | 24564199 | 55.19632 | 88.94394 | 72.80442 | 30.97664 | 0.644311 | 1.820518 | 0.321485 |
| 11 | 60913000 | 60914999 | 16.7531 | 28.31756 | 59.50133 | 26.03612 | -0.68589 | 1.064966 | 0.321485 |
| 6 | 11030800 | 11031999 | 11.565 | 22.08116 | 20.48138 | 8.148252 | 0.614158 | -0.01315 | 0.321498 |
| 7 | 142088800 | 142092399 | 16.6605 | 28.40656 | 55.8488 | 24.48265 | -0.58919 | 0.982705 | 0.321499 |
| 5 | 57813000 | 57815999 | 25.52798 | 44.67406 | 30.69554 | 13.54257 | 0.926224 | 0.710778 | 0.321511 |
| 1 | 8734200 | 8734999 | 15.88568 | 28.58271 | 26.18521 | 11.43263 | 0.529903 | 0.393257 | 0.321518 |
| 18 | 69927000 | 69929399 | 10.59835 | 18.9761 | 38.88415 | 17.04737 | -0.62877 | 0.413233 | 0.321522 |
| 15 | 87488200 | 87496999 | 63.23551 | 99.05416 | 171.1678 | 70.99986 | -0.47172 | 2.605086 | 0.32153 |
| 8 | 77687800 | 77689199 | 12.47418 | 23.77808 | 20.98245 | 8.376568 | 0.681459 | 0.009028 | 0.321531 |
| 1 | 4172800 | 4173599 | 6.10918 | 11.83576 | 26.8127 | 10.16695 | -0.59719 | -0.20755 | 0.321544 |
| 11 | 76741800 | 76743999 | 12.04084 | 21.39865 | 43.52594 | 19.20277 | -0.62477 | 0.478467 | 0.321551 |
| 7 | 121850600 | 121852599 | 9.54296 | 17.11325 | 38.93014 | 17.045 | -0.77719 | 0.39593 | 0.321554 |
| 7 | 85147600 | 85150399 | 42.00548 | 68.6405 | 55.4562 | 23.84089 | 0.68092 | 1.505911 | 0.321574 |
| 11 | 55518600 | 55520799 | 19.52797 | 34.34451 | 31.17532 | 13.80393 | 0.526701 | 0.603843 | 0.321591 |
| 1 | 85130600 | 85138799 | 246.7345 | 359.9365 | 322.3129 | 132.8094 | 0.388211 | 3.950027 | 0.321612 |
| 3 | 51113800 | 51118799 | 35.81894 | 57.64224 | 103.634 | 44.00892 | -0.49088 | 1.842932 | 0.321616 |
| 16 | 10216600 | 10218399 | 10.26047 | 18.53397 | 36.39342 | 15.81264 | -0.55869 | 0.353315 | 0.32162 |
| 9 | 73770000 | 73770799 | 15.68098 | 28.51468 | 25.61541 | 11.0373 | 0.570102 | 0.330034 | 0.321681 |

| | | | | | | | | | |
|---|---|---|---|---|---|---|---|---|---|
| 8 | 93682200 | 93684599 | 11.3305 | 19.93202 | 41.97384 | 18.58448 | -0.68071 | 0.573646 | 0.321682 |
| 6 | 16487800 | 16489399 | 23.20952 | 40.94295 | 28.72006 | 12.7091 | 0.898388 | 0.590725 | 0.32169 |
| 10 | 63875200 | 63877999 | 20.53295 | 34.55505 | 67.00966 | 29.28267 | -0.57124 | 1.121208 | 0.32169 |
| 8 | 112177400 | 112179199 | 8.909602 | 16.10369 | 35.4224 | 15.37895 | -0.72029 | 0.338789 | 0.321707 |
| 15 | 60385400 | 60386599 | 15.96175 | 29.06188 | 22.15784 | 9.532433 | 0.806709 | 0.317749 | 0.32171 |
| 4 | 118837400 | 118839999 | 16.76767 | 28.33677 | 58.89959 | 25.77132 | -0.67017 | 1.068391 | 0.321717 |
| X | 88863400 | 88864599 | 23.34288 | 41.64483 | 24.75493 | 10.88587 | 1.143319 | 0.493777 | 0.321725 |
| 9 | 119226400 | 119229399 | 14.53361 | 25.06593 | 53.22734 | 23.34988 | -0.69685 | 0.864427 | 0.321735 |
| 11 | 108429000 | 108431399 | 15.3411 | 26.70699 | 59.4197 | 26.12096 | -0.76158 | 0.777992 | 0.32174 |
| 17 | 58781400 | 58782199 | 14.76248 | 28.05282 | 20.15206 | 8.098035 | 0.967515 | 0.04915 | 0.321743 |
| 5 | 141082400 | 141085599 | 22.58737 | 37.58602 | 67.87577 | 29.46788 | -0.46875 | 1.248248 | 0.321744 |
| 8 | 127494000 | 127496999 | 25.75797 | 42.2586 | 78.39946 | 33.75773 | -0.51205 | 1.432038 | 0.321746 |
| 6 | 136498200 | 136503799 | 45.19596 | 72.29339 | 138.0836 | 58.11376 | -0.58867 | 2.054734 | 0.32175 |
| 5 | 78973400 | 78975999 | 19.68538 | 34.71226 | 27.73354 | 12.27353 | 0.710846 | 0.567255 | 0.321759 |
| 11 | 5700800 | 5704599 | 43.04892 | 70.4646 | 58.43252 | 25.13591 | 0.64515 | 1.486663 | 0.321759 |
| 7 | 99418400 | 99421399 | 32.41106 | 53.91673 | 50.87103 | 22.08079 | 0.464786 | 1.263905 | 0.321775 |
| 11 | 65681200 | 65682799 | 7.72564 | 14.38565 | 31.08083 | 12.96799 | -0.65363 | 0.147625 | 0.321779 |
| 19 | 56010000 | 56015799 | 49.14744 | 78.32862 | 131.0052 | 54.85342 | -0.4031 | 2.170191 | 0.321783 |
| 1 | 89427200 | 89433999 | 50.18409 | 79.61296 | 139.3213 | 58.01875 | -0.47514 | 2.314003 | 0.321784 |
| 9 | 99125000 | 99125999 | 6.791211 | 12.81196 | 29.15509 | 11.89692 | -0.70129 | 0.050946 | 0.321815 |
| X | 32671200 | 32672999 | 13.67853 | 25.37986 | 22.92852 | 9.618228 | 0.589164 | 0.197994 | 0.321816 |
| 7 | 104267000 | 104268799 | 19.23947 | 32.6337 | 59.14688 | 25.90837 | -0.47334 | 1.043404 | 0.321817 |
| 15 | 32477800 | 32479399 | 22.52272 | 39.83322 | 28.17292 | 12.46061 | 0.887858 | 0.558407 | 0.321819 |
| 3 | 14535800 | 14539199 | 49.25424 | 79.46672 | 73.69249 | 31.40492 | 0.466666 | 1.77403 | 0.321835 |
| 11 | 90314000 | 90317599 | 36.30984 | 58.42793 | 103.0302 | 43.74787 | -0.46311 | 1.846111 | 0.321836 |
| 7 | 62328200 | 62330999 | 32.28782 | 55.88714 | 31.72159 | 13.91722 | 1.199346 | 0.884362 | 0.32186 |
| 6 | 86996000 | 87000399 | 35.67056 | 57.53079 | 99.41277 | 42.34688 | -0.42988 | 1.774982 | 0.321865 |
| 19 | 57379400 | 57381999 | 15.14351 | 26.16004 | 51.10157 | 22.41845 | -0.57638 | 0.850831 | 0.321865 |
| 4 | 93782200 | 93783799 | 16.01712 | 29.51469 | 24.05174 | 10.18456 | 0.726912 | 0.255828 | 0.32187 |
| 1 | 192365600 | 192367599 | 10.2295 | 18.11203 | 41.42187 | 18.31638 | -0.79537 | 0.492727 | 0.321878 |
| 9 | 36359600 | 36361599 | 23.39892 | 41.33145 | 26.79113 | 11.8524 | 1.011962 | 0.572265 | 0.32188 |
| 12 | 3575200 | 3578799 | 51.77878 | 83.34037 | 77.77851 | 33.03754 | 0.453388 | 1.849522 | 0.321893 |
| 1 | 117667400 | 117669199 | 16.81414 | 31.79421 | 19.32169 | 7.845844 | 1.193838 | 0.098824 | 0.3219 |
| 12 | 38017600 | 38018599 | 7.015321 | 13.45145 | 29.78595 | 11.71513 | -0.61544 | -0.06953 | 0.321927 |
| 2 | 26689200 | 26693399 | 70.65532 | 112.711 | 94.18861 | 39.48844 | 0.597689 | 2.151309 | 0.321951 |
| 1 | 167111200 | 167112999 | 11.02569 | 19.39709 | 43.00225 | 19.03957 | -0.75453 | 0.57199 | 0.321955 |
| 17 | 38890200 | 38892599 | 19.21484 | 35.86993 | 19.51413 | 8.107155 | 1.325846 | 0.199508 | 0.321957 |
| 19 | 34895600 | 34895999 | 3.125648 | 6.113281 | 18.41131 | 6.221578 | -0.84595 | -0.68191 | 0.321965 |
| X | 8641800 | 8643599 | 34.87405 | 58.34396 | 47.55289 | 20.71521 | 0.675207 | 1.21061 | 0.322018 |
| X | 114738600 | 114741199 | 26.85334 | 47.04698 | 29.65705 | 13.09699 | 1.050077 | 0.701016 | 0.322023 |
| 17 | 90902400 | 90904599 | 27.3328 | 46.63185 | 41.98971 | 18.4081 | 0.533017 | 1.000596 | 0.322025 |
| 8 | 59570800 | 59572599 | 23.50943 | 40.61267 | 37.77523 | 16.57214 | 0.489443 | 0.868517 | 0.32203 |
| 7 | 116501800 | 116503199 | 10.54275 | 18.6505 | 39.40628 | 17.42811 | -0.68196 | 0.503118 | 0.322032 |
| 10 | 120027800 | 120029399 | 14.91565 | 25.98412 | 50.03453 | 22.0047 | -0.55385 | 0.751838 | 0.322034 |
| 6 | 116277800 | 116279199 | 13.73984 | 26.02076 | 21.30517 | 8.623287 | 0.771258 | 0.065027 | 0.322045 |
| 6 | 133067200 | 133070399 | 35.45458 | 58.29709 | 54.1358 | 23.33219 | 0.484881 | 1.411031 | 0.322062 |
| 12 | 27388800 | 27389799 | 7.101969 | 13.32916 | 30.14799 | 12.41172 | -0.70357 | 0.08792 | 0.322062 |

| | | | | | | | | | |
|---|---|---|---|---|---|---|---|---|---|
| 7 | 100406200 | 100408399 | 19.37865 | 34.74245 | 26.73333 | 11.70736 | 0.77713 | 0.440691 | 0.322063 |
| 1 | 140643400 | 140643999 | 10.24447 | 19.84317 | 18.65798 | 7.081787 | 0.658075 | -0.18293 | 0.322068 |
| 14 | 8348600 | 8349599 | 7.188161 | 13.26676 | 33.07463 | 13.97486 | -0.87232 | 0.198267 | 0.322084 |
| 7 | 49864600 | 49866199 | 25.02257 | 42.96436 | 39.89104 | 17.49707 | 0.490311 | 0.931855 | 0.322087 |
| 7 | 26179800 | 26185399 | 89.03117 | 140.4694 | 110.0275 | 45.66668 | 0.676308 | 2.47158 | 0.322092 |
| 16 | 5527000 | 5528799 | 17.08317 | 30.86553 | 25.87268 | 11.23809 | 0.661844 | 0.376363 | 0.322094 |
| 12 | 101298400 | 101301599 | 37.93532 | 62.78667 | 52.18139 | 22.57025 | 0.646717 | 1.336755 | 0.322096 |
| 9 | 121399200 | 121400399 | 16.06315 | 29.12362 | 26.12314 | 11.29766 | 0.568946 | 0.351111 | 0.32211 |
| 7 | 12811400 | 12813999 | 21.23732 | 35.54894 | 65.17306 | 28.40063 | -0.49048 | 1.177183 | 0.322117 |
| 2 | 88199000 | 88200999 | 20.68308 | 36.15289 | 32.88281 | 14.49315 | 0.523639 | 0.706441 | 0.322123 |
| X | 152549000 | 152551999 | 36.83876 | 61.14792 | 51.51102 | 22.32471 | 0.627645 | 1.303551 | 0.322123 |
| 4 | 141422200 | 141422999 | 4.564406 | 8.869961 | 24.55704 | 9.144702 | -0.85816 | -0.29614 | 0.322126 |
| 9 | 26250400 | 26255199 | 61.49617 | 98.75764 | 79.66017 | 33.72564 | 0.659413 | 1.935117 | 0.322137 |
| 15 | 8546200 | 8550599 | 56.74353 | 89.40169 | 156.8105 | 65.06794 | -0.48668 | 2.48877 | 0.32216 |
| X | 32879400 | 32881999 | 21.99908 | 39.62579 | 24.0039 | 10.46643 | 1.124498 | 0.427363 | 0.322184 |
| 16 | 48998400 | 49001199 | 39.20763 | 64.13286 | 59.84385 | 25.73657 | 0.474639 | 1.487699 | 0.322212 |
| 7 | 9121200 | 9124799 | 29.47165 | 51.68306 | 26.1552 | 11.55975 | 1.365046 | 0.695841 | 0.322221 |
| 2 | 150232800 | 150234399 | 31.00825 | 52.65985 | 42.45261 | 18.60258 | 0.691124 | 1.056162 | 0.322228 |
| 4 | 86805400 | 86806599 | 11.91161 | 22.77843 | 20.4794 | 8.117228 | 0.663634 | -0.02132 | 0.322231 |
| 2 | 85097000 | 85097599 | 4.107497 | 8.011447 | 21.44691 | 7.6894 | -0.75853 | -0.44633 | 0.322255 |
| 2 | 52028000 | 52029999 | 29.43113 | 50.16244 | 41.95156 | 18.38992 | 0.639076 | 1.018666 | 0.322267 |
| 2 | 28472200 | 28473399 | 6.1487 | 11.76441 | 27.8129 | 11.00693 | -0.71579 | -0.05809 | 0.322277 |
| 1 | 53035400 | 53037399 | 18.56986 | 33.93416 | 23.07412 | 9.881407 | 0.974673 | 0.318406 | 0.322282 |
| 3 | 87466800 | 87467999 | 16.27412 | 29.15959 | 26.27804 | 11.51362 | 0.550181 | 0.422706 | 0.322297 |
| 5 | 50829600 | 50830399 | 13.5988 | 25.94433 | 20.32005 | 8.09724 | 0.854325 | 0.012124 | 0.322313 |
| X | 54654400 | 54657399 | 47.26827 | 78.20792 | 44.95446 | 19.43994 | 1.176989 | 1.364083 | 0.322323 |
| 7 | 91828800 | 91831399 | 18.17263 | 30.70434 | 59.00684 | 25.81652 | -0.55763 | 1.074265 | 0.322326 |
| 14 | 105993800 | 105997599 | 54.54215 | 87.48927 | 83.0591 | 35.10312 | 0.422832 | 1.960289 | 0.32234 |
| 11 | 105594600 | 105596999 | 36.2038 | 60.37958 | 48.70794 | 21.17657 | 0.690055 | 1.246255 | 0.322341 |
| 7 | 38831800 | 38838599 | 112.9149 | 174.9597 | 150.7172 | 62.60372 | 0.519233 | 2.814874 | 0.322342 |
| 17 | 15181000 | 15181599 | 5.706439 | 10.98902 | 26.37561 | 10.2462 | -0.7128 | -0.13158 | 0.322357 |
| 14 | 68247000 | 68248199 | 19.51429 | 35.59321 | 23.6954 | 10.17129 | 1.003302 | 0.338831 | 0.322359 |
| 4 | 139370600 | 139372599 | 8.367833 | 15.01468 | 36.94449 | 16.16764 | -0.88845 | 0.38261 | 0.322369 |
| X | 148811800 | 148815999 | 51.40112 | 83.26417 | 65.3538 | 27.97094 | 0.712766 | 1.673538 | 0.322379 |
| 8 | 26078000 | 26079399 | 14.92283 | 27.12386 | 24.80548 | 10.69432 | 0.544125 | 0.326001 | 0.322386 |
| 14 | 39620000 | 39621199 | 14.92283 | 27.12386 | 24.80548 | 10.69432 | 0.544125 | 0.326001 | 0.322386 |
| 18 | 75398800 | 75399999 | 13.1016 | 22.77024 | 48.42629 | 21.26591 | -0.69655 | 0.776232 | 0.322394 |
| 8 | 3637200 | 3640799 | 22.0865 | 36.35888 | 73.32043 | 31.61632 | -0.62978 | 1.37958 | 0.322399 |
| 7 | 39620600 | 39623599 | 23.56547 | 39.13905 | 70.18612 | 30.43089 | -0.45893 | 1.274714 | 0.322412 |
| 11 | 13765600 | 13766199 | 8.91046 | 17.33352 | 17.23236 | 6.36372 | 0.615383 | -0.29495 | 0.322417 |
| 1 | 155064200 | 155064799 | 5.666976 | 11.00653 | 25.19533 | 9.424061 | -0.59306 | -0.26414 | 0.322437 |
| 2 | 113579400 | 113580599 | 11.21214 | 19.68023 | 42.58277 | 18.83476 | -0.71973 | 0.609129 | 0.322443 |
| 6 | 63846200 | 63848399 | 18.96669 | 33.89826 | 27.25363 | 11.96655 | 0.711574 | 0.457108 | 0.322457 |
| 13 | 112040400 | 112042199 | 11.63172 | 20.45545 | 42.47457 | 18.80737 | -0.66082 | 0.579264 | 0.322461 |
| 10 | 91779400 | 91780599 | 12.75059 | 24.41115 | 19.21839 | 7.592036 | 0.857591 | -0.02784 | 0.322492 |
| 4 | 32325800 | 32327999 | 12.57279 | 22.19157 | 41.91812 | 18.54739 | -0.52285 | 0.545236 | 0.322517 |
| 3 | 35592200 | 35593799 | 15.31831 | 28.74401 | 21.54288 | 8.872076 | 0.876977 | 0.138319 | 0.322526 |

| | | | | | | | | | |
|---|---|---|---|---|---|---|---|---|---|
| 10 | 96034200 | 96035799 | 20.01545 | 35.16259 | 31.67459 | 14.02017 | 0.537397 | 0.622216 | 0.322526 |
| 10 | 76982400 | 76984399 | 10.56086 | 18.62609 | 40.95589 | 18.12442 | -0.74105 | 0.538859 | 0.322531 |
| 14 | 78892000 | 78903599 | 135.8319 | 203.1542 | 318.4725 | 131.3273 | -0.38556 | 3.521903 | 0.322531 |
| 11 | 4544200 | 4544999 | 7.942804 | 14.57361 | 33.00354 | 14.05684 | -0.74304 | 0.239225 | 0.322534 |
| 7 | 124605400 | 124606999 | 17.15363 | 30.40121 | 26.26656 | 11.61258 | 0.601607 | 0.49668 | 0.322553 |
| 13 | 22818400 | 22821799 | 92.37984 | 145.0627 | 128.6833 | 53.37346 | 0.4918 | 2.571203 | 0.322558 |
| 5 | 93654800 | 93655399 | 9.709714 | 18.85617 | 17.59029 | 6.585849 | 0.687357 | -0.23896 | 0.322589 |
| 1 | 83253400 | 83257399 | 35.02795 | 56.4017 | 99.78322 | 42.40895 | -0.46661 | 1.823515 | 0.322608 |
| 1 | 59599600 | 59603599 | 39.90937 | 64.07341 | 109.4708 | 46.32712 | -0.42173 | 1.925491 | 0.322646 |
| 8 | 36542800 | 36543999 | 13.76172 | 25.64007 | 23.34085 | 9.729064 | 0.584982 | 0.174138 | 0.322685 |
| 15 | 33711200 | 33713199 | 28.76004 | 48.42716 | 44.26862 | 19.34826 | 0.510054 | 1.120785 | 0.322697 |
| 4 | 7418000 | 7419199 | 18.90761 | 33.85162 | 27.34574 | 11.98937 | 0.706321 | 0.447229 | 0.322708 |
| 8 | 100395400 | 100397999 | 23.88967 | 41.55739 | 35.97603 | 15.80754 | 0.594176 | 0.795538 | 0.322709 |
| 10 | 76986000 | 76989999 | 20.45293 | 33.81993 | 71.24389 | 30.79552 | -0.69089 | 1.318475 | 0.322728 |
| X | 133326400 | 133329199 | 16.05703 | 27.83592 | 51.68619 | 22.67736 | -0.50257 | 0.823602 | 0.322736 |
| 12 | 66455000 | 66455799 | 11.25665 | 21.58266 | 20.01929 | 7.875132 | 0.629063 | -0.05339 | 0.322754 |
| 15 | 8647000 | 8649799 | 20.04325 | 33.82424 | 62.4838 | 27.32779 | -0.50116 | 1.092345 | 0.322759 |
| 10 | 53378400 | 53379999 | 14.74007 | 26.81275 | 23.85731 | 10.27543 | 0.584386 | 0.318021 | 0.322765 |
| 1 | 82563800 | 82565399 | 8.077245 | 14.88317 | 32.52922 | 13.77523 | -0.68594 | 0.216415 | 0.322776 |
| 12 | 103445000 | 103447999 | 21.03094 | 34.93334 | 67.54946 | 29.2899 | -0.56715 | 1.265631 | 0.322787 |
| 6 | 29612000 | 29612999 | 14.25935 | 26.79277 | 22.27576 | 9.153126 | 0.73123 | 0.123196 | 0.3228 |
| 10 | 108433200 | 108434199 | 23.91317 | 41.37878 | 37.13474 | 16.29198 | 0.541252 | 0.853879 | 0.322803 |
| 17 | 37732800 | 37734399 | 34.00984 | 56.30453 | 52.51531 | 22.71883 | 0.480932 | 1.326063 | 0.32282 |
| 2 | 132053400 | 132059199 | 39.96623 | 63.88773 | 122.6647 | 51.57835 | -0.59705 | 2.06266 | 0.322824 |
| 3 | 65941800 | 65943199 | 16.00046 | 29.07623 | 25.72092 | 11.09202 | 0.591666 | 0.338039 | 0.322834 |
| 11 | 18690400 | 18692599 | 21.74752 | 36.3 | 66.2432 | 28.81542 | -0.4838 | 1.210446 | 0.322839 |
| 7 | 49764600 | 49768399 | 28.5254 | 46.23652 | 96.16543 | 41.1768 | -0.68887 | 1.637825 | 0.322858 |
| 9 | 99099800 | 99100799 | 5.702361 | 10.96044 | 26.29121 | 10.26895 | -0.7187 | -0.11179 | 0.322866 |
| 16 | 32482200 | 32483399 | 4.484209 | 8.613706 | 27.88301 | 10.91008 | -1.14923 | -0.11663 | 0.322881 |
| 12 | 115849400 | 115852399 | 41.98083 | 68.80092 | 53.31533 | 22.94656 | 0.74329 | 1.465554 | 0.322887 |
| 19 | 5819400 | 5823999 | 27.43412 | 44.80507 | 80.37245 | 34.5456 | -0.46725 | 1.499277 | 0.322907 |
| X | 118349000 | 118351999 | 44.75067 | 73.98727 | 43.34828 | 18.73573 | 1.149396 | 1.363226 | 0.322915 |
| 2 | 167072200 | 167074599 | 13.25078 | 23.04262 | 47.89703 | 21.04102 | -0.66359 | 0.768216 | 0.322933 |
| X | 33630600 | 33632399 | 27.13936 | 46.99698 | 36.06813 | 15.8244 | 0.7665 | 0.864628 | 0.322946 |
| 14 | 32574600 | 32575399 | 5.109966 | 9.911456 | 24.41607 | 9.207889 | -0.70845 | -0.2439 | 0.322953 |
| 14 | 108135800 | 108136599 | 11.56996 | 21.85505 | 19.97808 | 8.130313 | 0.606787 | 0.058546 | 0.322956 |
| 14 | 50443400 | 50446399 | 82.38496 | 129.558 | 121.1635 | 50.26411 | 0.417299 | 2.534682 | 0.322962 |
| 17 | 62254200 | 62255599 | 18.06146 | 31.90227 | 29.23533 | 12.93381 | 0.515386 | 0.545346 | 0.32297 |
| 5 | 73663200 | 73664799 | 14.19577 | 26.30633 | 23.75519 | 9.982144 | 0.587652 | 0.20951 | 0.322983 |
| 7 | 27194200 | 27196999 | 18.39704 | 30.86035 | 62.15761 | 27.12303 | -0.62551 | 1.143888 | 0.322988 |
| 5 | 37183800 | 37191199 | 38.55045 | 61.64185 | 124.2686 | 52.27354 | -0.6668 | 2.055125 | 0.323006 |
| 17 | 22472600 | 22476399 | 31.04737 | 50.47983 | 93.55971 | 40.14821 | -0.51901 | 1.578351 | 0.32302 |
| 16 | 19374600 | 19378399 | 75.11668 | 118.7655 | 113.655 | 47.22254 | 0.390178 | 2.412831 | 0.323035 |
| 1 | 67128800 | 67129399 | 7.844779 | 14.66813 | 31.16316 | 12.91145 | -0.62174 | 0.120521 | 0.32305 |
| 8 | 35272000 | 35278799 | 35.38443 | 56.48337 | 132.0439 | 55.41073 | -0.8827 | 2.106193 | 0.323052 |
| 15 | 86069000 | 86081199 | 193.1683 | 278.2506 | 474.4892 | 195.8094 | -0.55847 | 4.137593 | 0.323057 |
| 5 | 89658800 | 89661199 | 45.22622 | 73.11456 | 69.62381 | 29.74794 | 0.431777 | 1.706748 | 0.323057 |

| | | | | | | | | | |
|---|---|---|---|---|---|---|---|---|---|
| 1 | 186565600 | 186566999 | 11.49261 | 20.19781 | 41.98987 | 18.5922 | -0.66268 | 0.586592 | 0.323059 |
| X | 114994200 | 114995599 | 16.40774 | 31.11995 | 18.79336 | 7.582527 | 1.210586 | 0.073579 | 0.323059 |
| 5 | 90332000 | 90333399 | 14.90802 | 27.3892 | 24.73083 | 10.51466 | 0.575829 | 0.265333 | 0.323062 |
| 9 | 83662600 | 83663199 | 7.349212 | 13.94301 | 29.32719 | 11.82536 | -0.57415 | 0.015149 | 0.323073 |
| 16 | 25343800 | 25344799 | 14.12832 | 26.50895 | 21.00595 | 8.65212 | 0.79661 | 0.125606 | 0.32308 |
| 16 | 7606800 | 7608999 | 17.08405 | 31.22891 | 23.1574 | 9.913149 | 0.850576 | 0.304329 | 0.323082 |
| 2 | 164545600 | 164550599 | 37.13243 | 59.74192 | 103.0415 | 43.74239 | -0.43154 | 1.852953 | 0.323101 |
| 19 | 36144800 | 36146399 | 10.28232 | 18.64247 | 36.46822 | 15.77178 | -0.54872 | 0.333804 | 0.323119 |
| 2 | 75467400 | 75470199 | 19.49572 | 32.65679 | 63.9168 | 27.86561 | -0.58442 | 1.163035 | 0.323122 |
| 13 | 12120200 | 12122799 | 20.32134 | 33.55659 | 75.81808 | 32.74193 | -0.79191 | 1.3358 | 0.323134 |
| 2 | 87212000 | 87213399 | 16.87926 | 30.57662 | 25.35616 | 10.97793 | 0.6802 | 0.358849 | 0.323153 |
| 9 | 46754000 | 46756599 | 13.04949 | 22.92413 | 42.98908 | 19.02773 | -0.51489 | 0.604654 | 0.323155 |
| 2 | 109741200 | 109744199 | 30.645 | 51.42579 | 46.9658 | 20.49897 | 0.511361 | 1.161499 | 0.323158 |
| 9 | 35977800 | 35980199 | 32.9959 | 56.10899 | 40.16888 | 17.6052 | 0.861956 | 1.049789 | 0.323182 |
| 5 | 57158200 | 57160199 | 24.27211 | 42.46411 | 32.62297 | 14.38944 | 0.766233 | 0.713581 | 0.323183 |
| 8 | 119738000 | 119742799 | 35.88709 | 57.60436 | 106.8317 | 45.19667 | -0.54009 | 1.926055 | 0.323206 |
| 15 | 5570400 | 5572599 | 10.91419 | 19.48766 | 40.4194 | 17.7618 | -0.64864 | 0.435169 | 0.323208 |
| X | 49995400 | 49997799 | 21.92965 | 39.33913 | 25.01358 | 10.94878 | 1.051341 | 0.452574 | 0.323213 |
| Y | 1988800 | 1991199 | 19.40518 | 34.31885 | 17.49716 | 7.738858 | 1.356208 | 0.513063 | 0.323224 |
| 4 | 96488000 | 96488799 | 5.974392 | 11.54843 | 27.84332 | 10.67009 | -0.70139 | -0.17248 | 0.323227 |
| 18 | 32567600 | 32568999 | 8.482121 | 15.7327 | 32.07986 | 13.46346 | -0.57757 | 0.179081 | 0.323241 |
| 7 | 34449000 | 34449799 | 10.57707 | 20.29742 | 19.6412 | 7.707041 | 0.572404 | -0.06782 | 0.323243 |
| 6 | 107660800 | 107661599 | 9.78991 | 18.94689 | 18.69567 | 7.123802 | 0.584016 | -0.17462 | 0.323257 |
| 7 | 84741600 | 84746199 | 50.43876 | 81.91489 | 61.97265 | 26.57399 | 0.769085 | 1.620143 | 0.323268 |
| 4 | 22254000 | 22256999 | 35.17493 | 58.17348 | 53.73307 | 23.22914 | 0.494749 | 1.339478 | 0.32327 |
| 18 | 37206600 | 37207199 | 22.45995 | 39.02932 | 36.20129 | 15.89602 | 0.495009 | 0.804398 | 0.323277 |
| 12 | 92616800 | 92617999 | 16.16229 | 27.80581 | 52.67675 | 23.1066 | -0.53352 | 0.896835 | 0.323286 |
| 2 | 8220600 | 8222799 | 39.05863 | 63.82117 | 60.32418 | 25.93308 | 0.455063 | 1.502868 | 0.323317 |
| 9 | 31282800 | 31283199 | 3.877944 | 7.57816 | 19.99099 | 6.932943 | -0.69207 | -0.56915 | 0.32332 |
| 7 | 11468600 | 11470199 | 33.46072 | 55.78095 | 50.32191 | 21.87238 | 0.529308 | 1.244656 | 0.323328 |
| 11 | 58618000 | 58620399 | 13.19329 | 23.07566 | 45.70584 | 20.15563 | -0.59362 | 0.678871 | 0.32333 |
| 14 | 21758200 | 21760199 | 14.4665 | 25.16836 | 48.42835 | 21.28062 | -0.55286 | 0.767207 | 0.323351 |
| 7 | 127906400 | 127909599 | 18.43578 | 30.84901 | 68.10162 | 29.67046 | -0.75732 | 1.177607 | 0.323383 |
| 8 | 111004800 | 111005399 | 10.43514 | 20.06879 | 19.02831 | 7.419918 | 0.609321 | -0.093 | 0.323385 |
| 7 | 98495400 | 98501399 | 32.47311 | 52.25846 | 100.5834 | 42.71414 | -0.58889 | 1.838266 | 0.323389 |
| 16 | 51248800 | 51249399 | 8.153001 | 15.30002 | 31.24978 | 12.8678 | -0.55796 | 0.101641 | 0.323389 |
| 3 | 111704200 | 111706799 | 11.95135 | 23.00892 | 15.85072 | 6.162222 | 1.070714 | -0.09407 | 0.323393 |
| 8 | 106418800 | 106422599 | 39.55262 | 63.15743 | 120.0617 | 50.40567 | -0.58437 | 2.098426 | 0.323405 |
| 9 | 20296200 | 20297599 | 15.37913 | 26.36691 | 53.17683 | 23.32169 | -0.62437 | 0.924574 | 0.323413 |
| 11 | 107296000 | 107298599 | 18.77545 | 31.73387 | 59.63582 | 26.09446 | -0.52557 | 1.072455 | 0.323418 |
| 17 | 39147600 | 39153399 | 86.51761 | 137.2679 | 101.9692 | 42.46844 | 0.759347 | 2.332816 | 0.323424 |
| 1 | 190484400 | 190493599 | 89.58767 | 137.2116 | 225.9158 | 93.7605 | -0.42722 | 3.033624 | 0.323437 |
| 4 | 53176600 | 53179599 | 23.04297 | 38.32382 | 70.85769 | 30.7512 | -0.50279 | 1.256423 | 0.32344 |
| 18 | 46598200 | 46598999 | 6.581368 | 12.63286 | 27.42177 | 10.75224 | -0.58188 | -0.08752 | 0.323442 |
| 1 | 93681600 | 93683799 | 14.39468 | 24.85415 | 55.37521 | 24.29294 | -0.76546 | 0.857859 | 0.323469 |
| 10 | 83358000 | 83360599 | 18.94574 | 31.96219 | 60.72304 | 26.5552 | -0.54138 | 1.090031 | 0.323507 |
| 6 | 64704200 | 64707999 | 33.23605 | 53.54845 | 99.83147 | 42.4663 | -0.54095 | 1.800933 | 0.323507 |

| | | | | | | | | | |
|---|---|---|---|---|---|---|---|---|---|
| 1 | 30665600 | 30668799 | 44.87356 | 73.43201 | 57.39956 | 24.68921 | 0.729896 | 1.493924 | 0.323527 |
| X | 108683000 | 108685199 | 26.44932 | 46.3249 | 29.52817 | 13.03669 | 1.033935 | 0.701691 | 0.323529 |
| 2 | 28944000 | 28945199 | 6.853006 | 12.75154 | 31.07687 | 12.9822 | -0.82709 | 0.145318 | 0.323535 |
| 19 | 7710600 | 7718399 | 93.74861 | 147.3094 | 126.3823 | 52.42359 | 0.540663 | 2.557747 | 0.32354 |
| 7 | 23259400 | 23263999 | 69.91616 | 111.813 | 86.40904 | 36.35605 | 0.714235 | 2.073397 | 0.323544 |
| 15 | 67197000 | 67199399 | 17.77141 | 31.76109 | 27.38422 | 12.02418 | 0.611235 | 0.449044 | 0.323549 |
| 3 | 129010400 | 129012599 | 22.10796 | 36.33751 | 77.38912 | 33.34877 | -0.70935 | 1.39956 | 0.323549 |
| 1 | 51481600 | 51483399 | 19.39779 | 34.85777 | 25.55065 | 11.1675 | 0.84844 | 0.423311 | 0.323551 |
| 8 | 71387000 | 71390199 | 17.74729 | 29.81643 | 63.86832 | 27.89558 | -0.71412 | 1.124818 | 0.323554 |
| 7 | 75593000 | 75593799 | 6.560433 | 12.63688 | 28.22266 | 10.95549 | -0.60959 | -0.1211 | 0.323559 |
| 7 | 11590200 | 11604199 | 233.2191 | 345.2593 | 263.7917 | 108.6375 | 0.637163 | 3.716006 | 0.323563 |
| 6 | 85977000 | 85979199 | 32.57042 | 54.50308 | 48.63787 | 21.19106 | 0.544774 | 1.199994 | 0.323566 |
| 1 | 49727600 | 49728399 | 11.84581 | 22.82217 | 19.53988 | 7.580755 | 0.762427 | -0.09152 | 0.32357 |
| X | 88747400 | 88749199 | 22.37862 | 39.80981 | 26.52989 | 11.69475 | 0.977207 | 0.508453 | 0.323578 |
| 7 | 12325200 | 12344799 | 281.2452 | 407.9942 | 337.3132 | 139.0711 | 0.495543 | 4.037559 | 0.323586 |
| 7 | 36317400 | 36320799 | 28.09663 | 45.80486 | 82.02136 | 35.2313 | -0.46657 | 1.527425 | 0.323593 |
| 12 | 4273800 | 4278399 | 34.19147 | 55.25699 | 96.24141 | 41.10316 | -0.43786 | 1.712371 | 0.323602 |
| 4 | 59222200 | 59223999 | 8.451266 | 15.84578 | 32.3553 | 13.34292 | -0.55958 | 0.111609 | 0.323612 |
| 12 | 76469800 | 76470799 | 7.181945 | 13.7222 | 28.98926 | 11.51586 | -0.56131 | -0.03596 | 0.323615 |
| 17 | 42448000 | 42450599 | 25.23885 | 43.56035 | 38.74756 | 16.99823 | 0.553382 | 0.886319 | 0.323641 |
| 19 | 4062200 | 4063399 | 5.872674 | 11.22506 | 27.23978 | 10.81108 | -0.7574 | -0.0553 | 0.323642 |
| 11 | 12870000 | 12873199 | 48.21652 | 77.82324 | 73.14337 | 31.18681 | 0.447988 | 1.760055 | 0.323644 |
| 2 | 144320800 | 144321999 | 6.315219 | 12.03337 | 27.73273 | 11.08035 | -0.69239 | -0.02755 | 0.32365 |
| 14 | 120746200 | 120747399 | 6.315219 | 12.03337 | 27.73273 | 11.08035 | -0.69239 | -0.02755 | 0.32365 |
| 1 | 4090200 | 4092799 | 21.40242 | 38.44673 | 23.69832 | 10.36092 | 1.096457 | 0.435874 | 0.323658 |
| 12 | 11156600 | 11157799 | 13.94024 | 26.39972 | 20.65424 | 8.360207 | 0.835983 | 0.064507 | 0.32366 |
| 2 | 168411000 | 168415799 | 64.19932 | 100.1848 | 179.2699 | 74.38946 | -0.52626 | 2.67361 | 0.323663 |
| 3 | 7856000 | 7857599 | 18.82772 | 33.91054 | 26.37298 | 11.50063 | 0.766031 | 0.409638 | 0.323664 |
| 1 | 194109400 | 194110599 | 14.69196 | 26.88451 | 24.71789 | 10.56651 | 0.544236 | 0.285174 | 0.323666 |
| 12 | 110699600 | 110702199 | 13.18166 | 23.12569 | 44.05362 | 19.47711 | -0.5376 | 0.627056 | 0.323675 |
| 4 | 134650200 | 134651999 | 6.848879 | 12.76981 | 31.63527 | 13.17002 | -0.84698 | 0.131392 | 0.323677 |
| 18 | 27432200 | 27437599 | 65.53496 | 104.457 | 98.52087 | 41.25777 | 0.422126 | 2.173288 | 0.323678 |
| 18 | 66590400 | 66592799 | 17.41443 | 29.31439 | 62.77977 | 27.43638 | -0.71366 | 1.105587 | 0.323691 |
| 4 | 113467200 | 113469799 | 36.20187 | 60.96294 | 43.0327 | 18.80854 | 0.881844 | 1.147772 | 0.323692 |
| 1 | 54675000 | 54675999 | 6.463727 | 12.3058 | 29.99105 | 12.00438 | -0.77574 | -0.01788 | 0.323693 |
| X | 54787000 | 54788199 | 17.16622 | 31.29479 | 23.74735 | 10.19996 | 0.81552 | 0.326034 | 0.323714 |
| 1 | 164600200 | 164601999 | 26.12479 | 44.97717 | 35.63575 | 15.63192 | 0.718896 | 0.907707 | 0.323715 |
| 18 | 37471800 | 37473599 | 19.3339 | 35.18997 | 24.23388 | 10.42988 | 0.95237 | 0.350222 | 0.323732 |
| 3 | 57584200 | 57585199 | 14.36927 | 26.49979 | 23.40349 | 9.899079 | 0.613203 | 0.238621 | 0.323732 |
| 4 | 21050000 | 21051999 | 25.16611 | 44.12786 | 30.16104 | 13.32898 | 0.933752 | 0.678297 | 0.32374 |
| 8 | 75987800 | 75990399 | 43.39533 | 70.31483 | 66.86827 | 28.62487 | 0.436852 | 1.652889 | 0.323765 |
| 14 | 92061600 | 92067799 | 119.0228 | 183.0239 | 169.0388 | 70.19674 | 0.410465 | 2.959766 | 0.323773 |
| 18 | 35135600 | 35136599 | 8.305796 | 15.60976 | 32.41986 | 13.31631 | -0.57914 | 0.096067 | 0.323775 |
| 16 | 93257800 | 93259599 | 8.660424 | 15.9423 | 33.02338 | 14.00291 | -0.61104 | 0.225217 | 0.323791 |
| 7 | 100371400 | 100371999 | 2.937745 | 5.740278 | 19.2815 | 6.704163 | -1.03696 | -0.57689 | 0.323796 |
| 17 | 45434800 | 45436799 | 19.06636 | 32.50516 | 61.45388 | 26.93954 | -0.53319 | 0.999284 | 0.323801 |
| 1 | 37972400 | 37974599 | 20.77799 | 36.35297 | 33.37058 | 14.71983 | 0.510459 | 0.693642 | 0.323808 |

| | | | | | | | | | |
|---|---|---|---|---|---|---|---|---|---|
| 8 | 101123000 | 101126599 | 75.92853 | 120.3015 | 112.9488 | 46.97901 | 0.420073 | 2.36423 | 0.32382 |
| 5 | 111454600 | 111455599 | 6.596961 | 12.59351 | 27.79438 | 11.06159 | -0.62586 | -0.03844 | 0.323823 |
| 15 | 79974600 | 79978999 | 23.14101 | 37.79799 | 79.40334 | 34.1305 | -0.6941 | 1.490071 | 0.323823 |
| 8 | 121014200 | 121017599 | 18.70291 | 31.00552 | 69.43257 | 30.06571 | -0.77833 | 1.283047 | 0.323833 |
| 13 | 108350600 | 108352799 | 15.01574 | 25.89496 | 51.08398 | 22.40956 | -0.59095 | 0.866718 | 0.323835 |
| 11 | 8997800 | 8999799 | 10.33392 | 18.45361 | 37.35988 | 16.41579 | -0.6133 | 0.427699 | 0.323836 |
| 17 | 18928600 | 18937799 | 139.5437 | 213.9525 | 167.1821 | 69.39787 | 0.648246 | 3.024274 | 0.323836 |
| 17 | 87896200 | 87902999 | 54.38669 | 86.34431 | 144.6157 | 60.26843 | -0.41095 | 2.296141 | 0.32385 |
| 11 | 115618600 | 115622999 | 25.6103 | 42.02389 | 76.54289 | 32.96082 | -0.48543 | 1.429346 | 0.323868 |
| 13 | 17533000 | 17533599 | 6.755724 | 12.90977 | 29.49187 | 11.71182 | -0.672 | -0.03663 | 0.32387 |
| 7 | 104372600 | 104375999 | 49.144 | 79.71417 | 59.51753 | 25.49813 | 0.786397 | 1.639233 | 0.323885 |
| 5 | 112016400 | 112028599 | 209.617 | 301.2246 | 482.5478 | 199.2065 | -0.47179 | 4.17398 | 0.323886 |
| 2 | 91934400 | 91937199 | 28.71621 | 48.86852 | 41.93433 | 18.38008 | 0.601667 | 1.027287 | 0.323898 |
| 19 | 8357800 | 8358799 | 18.69116 | 32.90186 | 30.25403 | 13.39363 | 0.508641 | 0.58767 | 0.323906 |
| 17 | 45918200 | 45919199 | 12.89422 | 24.44959 | 21.38735 | 8.633902 | 0.679817 | 0.048817 | 0.323908 |
| 7 | 82584800 | 82586599 | 17.79368 | 32.95661 | 21.67654 | 9.115088 | 1.04081 | 0.236792 | 0.323912 |
| 10 | 116259800 | 116265999 | 84.71226 | 130.1023 | 218.5965 | 90.76114 | -0.45337 | 2.976627 | 0.323925 |
| 13 | 75958600 | 75958999 | 3.596548 | 7.012005 | 23.12143 | 8.3342 | -1.06277 | -0.42991 | 0.323941 |
| 2 | 82312800 | 82316399 | 50.65754 | 81.69042 | 74.54277 | 31.74703 | 0.48905 | 1.78971 | 0.323946 |
| 3 | 107066200 | 107068399 | 16.03872 | 27.33197 | 57.45258 | 25.18463 | -0.68546 | 0.985896 | 0.323973 |
| 5 | 91144600 | 91146199 | 16.3636 | 29.91861 | 25.03372 | 10.71338 | 0.678219 | 0.303519 | 0.323996 |
| 1 | 111641800 | 111643999 | 29.42652 | 51.0509 | 32.89923 | 14.43489 | 1.01757 | 0.851866 | 0.324002 |
| 17 | 53356800 | 53357799 | 15.34986 | 28.05135 | 25.14045 | 10.76566 | 0.579147 | 0.298525 | 0.32403 |
| 1 | 65853200 | 65856799 | 67.78074 | 107.6812 | 102.5503 | 42.76654 | 0.403259 | 2.277682 | 0.324031 |
| 12 | 115842600 | 115844399 | 27.21001 | 46.79912 | 35.63594 | 15.63153 | 0.775727 | 0.922072 | 0.324055 |
| 10 | 79093400 | 79095199 | 28.98063 | 48.77433 | 45.33948 | 19.8133 | 0.485968 | 1.127461 | 0.324069 |
| 2 | 151129200 | 151130599 | 14.71458 | 28.21822 | 17.97211 | 7.065811 | 1.167331 | -0.02536 | 0.324097 |
| 11 | 87289000 | 87289999 | 6.411974 | 12.37732 | 27.11636 | 10.45817 | -0.57286 | -0.14557 | 0.324103 |
| 19 | 32506800 | 32509399 | 13.98815 | 24.42416 | 46.26262 | 20.37512 | -0.5296 | 0.710788 | 0.324117 |
| 9 | 59126200 | 59131599 | 33.37882 | 53.85191 | 96.13021 | 40.96561 | -0.4761 | 1.759592 | 0.324129 |
| X | 8594800 | 8600199 | 107.8777 | 168.0603 | 138.7735 | 57.59817 | 0.586254 | 2.720662 | 0.324142 |
| 5 | 143518600 | 143520799 | 10.27313 | 18.51981 | 36.36781 | 15.84102 | -0.56109 | 0.365248 | 0.324149 |
| 12 | 113794400 | 113797999 | 78.52675 | 124.4655 | 111.0699 | 46.21257 | 0.493758 | 2.355142 | 0.32415 |
| 9 | 111745400 | 111746199 | 11.86674 | 22.45583 | 19.57764 | 7.937223 | 0.679853 | 0.049808 | 0.324157 |
| 15 | 28128200 | 28130399 | 18.52946 | 31.65243 | 58.716 | 25.74344 | -0.50544 | 0.975147 | 0.324164 |
| X | 48656000 | 48656599 | 10.0867 | 19.47208 | 19.17342 | 7.392521 | 0.570943 | -0.13089 | 0.324166 |
| 15 | 19958600 | 19961199 | 38.83617 | 63.92001 | 51.58661 | 22.24274 | 0.687231 | 1.404975 | 0.324168 |
| 17 | 22745800 | 22747599 | 24.23259 | 42.54185 | 29.57603 | 13.08367 | 0.909139 | 0.652809 | 0.324173 |
| 6 | 98424800 | 98426799 | 22.51467 | 39.70069 | 29.155 | 12.90264 | 0.832198 | 0.590109 | 0.324182 |
| 4 | 142703400 | 142706399 | 28.84469 | 46.72897 | 87.82173 | 37.58926 | -0.54367 | 1.644604 | 0.324183 |
| 6 | 42111400 | 42112399 | 10.95363 | 19.60361 | 37.86367 | 16.60612 | -0.544 | 0.419529 | 0.324185 |
| 10 | 3949400 | 3950999 | 10.85304 | 19.19367 | 39.4063 | 17.42917 | -0.64094 | 0.509026 | 0.324213 |
| 19 | 10080400 | 10082799 | 12.06951 | 21.36038 | 40.34328 | 17.84102 | -0.52114 | 0.50904 | 0.324226 |
| 1 | 99070600 | 99072799 | 18.18748 | 32.95414 | 24.64664 | 10.66762 | 0.828474 | 0.365664 | 0.324232 |
| 18 | 21077000 | 21080399 | 18.90927 | 32.03536 | 59.13546 | 25.894 | -0.49975 | 1.051742 | 0.324265 |
| 5 | 25383000 | 25384999 | 14.31071 | 24.93278 | 47.35021 | 20.82547 | -0.53375 | 0.745085 | 0.324266 |
| 3 | 8384000 | 8384799 | 15.99101 | 28.82602 | 25.79235 | 11.23539 | 0.56605 | 0.383832 | 0.324284 |

| | | | | | | | | | |
|---|---|---|---|---|---|---|---|---|---|
| 3 | 6558800 | 6560399 | 15.90558 | 29.74538 | 22.05198 | 9.133829 | 0.886244 | 0.165958 | 0.324329 |
| 3 | 73847800 | 73849999 | 18.43348 | 32.70571 | 26.7159 | 11.80285 | 0.682546 | 0.495255 | 0.324338 |
| 1 | 19304000 | 19307999 | 30.38687 | 49.31554 | 89.57853 | 38.39184 | -0.49237 | 1.60897 | 0.324346 |
| 5 | 71083200 | 71084199 | 10.50129 | 20.19024 | 19.60175 | 7.649811 | 0.57463 | -0.08727 | 0.324349 |
| 5 | 103960400 | 103963599 | 22.64714 | 37.44216 | 71.57971 | 30.93726 | -0.55147 | 1.32575 | 0.324358 |
| 6 | 121484000 | 121491399 | 54.7322 | 86.74363 | 145.6557 | 60.59752 | -0.41704 | 2.346773 | 0.324383 |
| 5 | 32628000 | 32630399 | 14.1406 | 24.78342 | 47.24708 | 20.8715 | -0.53884 | 0.653349 | 0.324385 |
| X | 8327400 | 8332599 | 103.6593 | 163.2783 | 110.4463 | 45.82779 | 0.885826 | 2.516714 | 0.324385 |
| 8 | 126343800 | 126357999 | 146.1747 | 216.8179 | 349.1612 | 143.816 | -0.43532 | 3.673343 | 0.32439 |
| 10 | 19253800 | 19257599 | 60.44658 | 96.59292 | 91.85553 | 38.60466 | 0.414257 | 2.087718 | 0.324391 |
| 10 | 47548600 | 47550799 | 18.84136 | 33.25297 | 30.09215 | 13.31502 | 0.532929 | 0.559768 | 0.324404 |
| X | 5590400 | 5592999 | 37.28246 | 62.65149 | 38.75858 | 16.92969 | 1.071329 | 1.164642 | 0.324449 |
| 13 | 114448200 | 114448999 | 12.19829 | 23.05452 | 21.40115 | 8.699393 | 0.586023 | 0.063991 | 0.32447 |
| 17 | 19745200 | 19749399 | 53.55419 | 86.7096 | 64.36332 | 27.53711 | 0.792452 | 1.686578 | 0.32448 |
| 4 | 132439200 | 132440399 | 8.778211 | 16.34175 | 32.70306 | 13.65029 | -0.54485 | 0.161183 | 0.324483 |
| 17 | 76034200 | 76035599 | 20.36226 | 36.27027 | 26.48126 | 11.66044 | 0.846937 | 0.484678 | 0.324495 |
| 3 | 25900800 | 25901399 | 3.935344 | 7.682209 | 20.38734 | 7.211265 | -0.72738 | -0.49912 | 0.324495 |
| 13 | 8781200 | 8782999 | 11.92181 | 21.06252 | 40.37043 | 17.8592 | -0.5432 | 0.527329 | 0.324509 |
| 3 | 7160800 | 7163399 | 24.42835 | 42.82252 | 30.739 | 13.58125 | 0.863379 | 0.678836 | 0.324509 |
| 2 | 87124400 | 87126599 | 21.77844 | 37.99239 | 33.8616 | 14.90502 | 0.552561 | 0.7412 | 0.324514 |
| 13 | 35818000 | 35820599 | 28.96901 | 49.79741 | 38.62558 | 16.94264 | 0.74909 | 0.938224 | 0.32452 |
| 6 | 51200000 | 51203999 | 31.87755 | 51.67082 | 89.99501 | 38.53517 | -0.43356 | 1.638451 | 0.324558 |
| 1 | 190048200 | 190050599 | 22.53112 | 37.61424 | 69.7955 | 30.36386 | -0.50789 | 1.212975 | 0.324561 |
| 9 | 37455800 | 37457599 | 11.54735 | 20.17744 | 45.24818 | 19.93769 | -0.7712 | 0.689454 | 0.324589 |
| 6 | 90346000 | 90347799 | 12.47172 | 21.7242 | 47.87525 | 21.05359 | -0.74727 | 0.739436 | 0.32459 |
| 10 | 91708600 | 91709799 | 8.597453 | 16.07877 | 32.8465 | 13.60417 | -0.56546 | 0.129566 | 0.324592 |
| 5 | 135858200 | 135865399 | 46.02718 | 73.32384 | 127.8626 | 53.50542 | -0.46387 | 2.179593 | 0.324639 |
| 5 | 49681600 | 49683999 | 22.25974 | 36.98723 | 69.56319 | 30.17028 | -0.52738 | 1.263989 | 0.324651 |
| 12 | 42298200 | 42298799 | 5.810358 | 11.28356 | 25.79472 | 9.65831 | -0.59256 | -0.25663 | 0.324661 |
| 7 | 17312800 | 17314199 | 19.24004 | 34.28709 | 24.99598 | 11.00217 | 0.84981 | 0.473831 | 0.324662 |
| 10 | 16669000 | 16669399 | 4.645215 | 9.065938 | 22.27992 | 7.910945 | -0.62465 | -0.47107 | 0.32467 |
| 17 | 18585000 | 18586199 | 37.33091 | 61.12054 | 56.83219 | 24.45007 | 0.480612 | 1.470434 | 0.324678 |
| 4 | 101701800 | 101711199 | 152.6517 | 231.5527 | 200.817 | 83.13102 | 0.485653 | 3.242909 | 0.324682 |
| 1 | 139781000 | 139783199 | 21.52361 | 37.99012 | 28.59561 | 12.65262 | 0.797128 | 0.571474 | 0.324691 |
| 6 | 117091800 | 117093599 | 9.921498 | 18.04829 | 35.46013 | 15.27206 | -0.5512 | 0.307928 | 0.324693 |
| 1 | 178145800 | 178146999 | 8.031532 | 14.9124 | 31.56496 | 13.22869 | -0.62943 | 0.168533 | 0.324702 |
| 1 | 96374000 | 96375999 | 21.62845 | 37.77356 | 34.44241 | 15.17149 | 0.520129 | 0.727395 | 0.324709 |
| 6 | 56401800 | 56403199 | 8.143548 | 15.31648 | 31.31317 | 12.84518 | -0.55498 | 0.087083 | 0.324711 |
| 9 | 22422400 | 22423799 | 7.659855 | 14.41838 | 31.91232 | 13.07308 | -0.66748 | 0.076108 | 0.324747 |
| 5 | 127780600 | 127790799 | 103.1876 | 156.9414 | 251.7957 | 104.3179 | -0.39797 | 3.180484 | 0.324751 |
| 8 | 13552600 | 13554799 | 9.990501 | 17.90419 | 38.90996 | 17.04548 | -0.71228 | 0.403035 | 0.324777 |
| 10 | 19724600 | 19728599 | 35.26563 | 56.53642 | 112.9545 | 47.69666 | -0.64945 | 1.965164 | 0.324798 |
| 8 | 36934800 | 36936999 | 23.02959 | 40.10472 | 35.5637 | 15.63702 | 0.559863 | 0.777157 | 0.324825 |
| 9 | 29131200 | 29133399 | 23.81235 | 39.25855 | 72.57196 | 31.31335 | -0.50387 | 1.364477 | 0.324827 |
| 13 | 106444600 | 106446399 | 17.18632 | 31.32669 | 24.26712 | 10.42518 | 0.784929 | 0.323983 | 0.324829 |
| X | 14308600 | 14310399 | 45.13689 | 74.40863 | 50.60458 | 21.83401 | 0.934411 | 1.404399 | 0.324846 |
| X | 112643800 | 112645399 | 8.160696 | 15.37016 | 33.0515 | 13.52628 | -0.62459 | 0.083009 | 0.32485 |

| | | | | | | | | | |
|---|---|---|---|---|---|---|---|---|---|
| 14 | 34497400 | 34508199 | 93.06593 | 142.4906 | 228.7908 | 94.94625 | -0.39143 | 3.041551 | 0.324853 |
| 14 | 124143000 | 124144599 | 20.80287 | 36.50698 | 32.13493 | 14.21113 | 0.570559 | 0.64389 | 0.324859 |
| 18 | 32593200 | 32594399 | 10.8208 | 19.17187 | 40.36244 | 17.8455 | -0.67627 | 0.487422 | 0.324868 |
| 15 | 8547600 | 8549599 | 46.16107 | 73.26343 | 138.0802 | 57.52758 | -0.58127 | 2.299168 | 0.324885 |
| 10 | 93627400 | 93630199 | 21.99973 | 38.36128 | 35.04411 | 15.4212 | 0.517291 | 0.752711 | 0.324896 |
| 1 | 190377400 | 190378999 | 22.2207 | 37.13801 | 66.89027 | 29.12074 | -0.46501 | 1.198591 | 0.324928 |
| 12 | 33196400 | 33199999 | 22.50741 | 37.1514 | 73.79787 | 31.8606 | -0.6068 | 1.345119 | 0.324936 |
| X | 125181600 | 125191999 | 148.2182 | 227.0062 | 163.8533 | 68.00282 | 0.761398 | 3.04826 | 0.324938 |
| 16 | 10830600 | 10831799 | 7.738106 | 14.43144 | 31.63949 | 13.16613 | -0.6711 | 0.141286 | 0.324945 |
| 12 | 25808200 | 25810999 | 25.65332 | 44.55691 | 35.87871 | 15.74958 | 0.697953 | 0.82458 | 0.324953 |
| 18 | 61041400 | 61046999 | 29.85775 | 48.20117 | 93.51629 | 39.8837 | -0.59482 | 1.735965 | 0.324963 |
| 8 | 43533600 | 43536999 | 43.74668 | 69.73524 | 126.4426 | 52.9578 | -0.51905 | 2.15628 | 0.324971 |
| 13 | 57880000 | 57883999 | 33.21217 | 53.70742 | 93.20221 | 39.83681 | -0.43155 | 1.691573 | 0.324972 |
| 7 | 13363000 | 13367599 | 60.53735 | 97.74276 | 64.52549 | 27.5244 | 0.957066 | 1.769546 | 0.324984 |
| 7 | 84920200 | 84923799 | 47.49543 | 77.08372 | 65.04365 | 27.87723 | 0.611143 | 1.629499 | 0.324984 |
| 12 | 102964600 | 102965399 | 5.685318 | 11.00735 | 26.12833 | 9.937891 | -0.66822 | -0.20508 | 0.324996 |
| 11 | 66896400 | 66899799 | 25.16863 | 41.22041 | 86.26616 | 37.11786 | -0.68643 | 1.458138 | 0.325042 |
| 14 | 3096200 | 3098799 | 43.26006 | 70.65749 | 60.59546 | 26.04584 | 0.594644 | 1.519177 | 0.325057 |
| 2 | 161069000 | 161069999 | 6.470751 | 12.29823 | 28.23248 | 11.34123 | -0.69403 | -0.00788 | 0.325059 |
| 4 | 136126000 | 136131399 | 33.88827 | 54.58625 | 99.15853 | 42.16545 | -0.50401 | 1.809754 | 0.325066 |
| 9 | 59012000 | 59012399 | 4.917564 | 9.579508 | 23.07571 | 8.409328 | -0.62966 | -0.36941 | 0.325067 |
| 11 | 92987800 | 92992199 | 40.94823 | 65.43685 | 119.9756 | 50.42509 | -0.53111 | 2.07347 | 0.325067 |
| 9 | 7879600 | 7887399 | 112.9756 | 174.0368 | 164.2963 | 68.2441 | 0.380869 | 2.921956 | 0.325076 |
| 4 | 112213000 | 112221199 | 199.2191 | 296.1014 | 265.8016 | 109.5038 | 0.41004 | 3.642035 | 0.325087 |
| 18 | 82609000 | 82613399 | 24.61362 | 40.09675 | 81.79042 | 35.12255 | -0.65464 | 1.531506 | 0.325102 |
| 9 | 80936600 | 80940199 | 36.49777 | 61.14649 | 45.29493 | 19.74913 | 0.812451 | 1.196954 | 0.325111 |
| 4 | 144258000 | 144261199 | 41.38598 | 68.11569 | 55.05784 | 23.73929 | 0.685026 | 1.414307 | 0.325138 |
| 6 | 10672200 | 10673599 | 15.91299 | 29.49954 | 21.77128 | 9.143839 | 0.876699 | 0.215381 | 0.325144 |
| 7 | 13345200 | 13356999 | 161.0084 | 244.1497 | 198.8488 | 82.30829 | 0.575847 | 3.252867 | 0.325146 |
| 16 | 11097000 | 11098999 | 11.24768 | 20.08598 | 38.33624 | 16.84434 | -0.52897 | 0.434782 | 0.325175 |
| 12 | 84147600 | 84149399 | 9.022669 | 16.41084 | 34.44056 | 14.83563 | -0.64555 | 0.300652 | 0.325182 |
| 17 | 40019000 | 40026999 | 158.1919 | 239.305 | 210.3303 | 86.99534 | 0.46311 | 3.300662 | 0.325193 |
| 17 | 18198800 | 18201799 | 54.45114 | 87.45769 | 82.21186 | 34.81429 | 0.439301 | 1.920546 | 0.325195 |
| X | 107120800 | 107121399 | 9.303188 | 18.11933 | 16.60276 | 6.065548 | 0.745095 | -0.33083 | 0.325195 |
| 11 | 29974200 | 29976399 | 19.62293 | 32.97825 | 61.77623 | 26.98446 | -0.52135 | 1.12741 | 0.3252 |
| 5 | 127140000 | 127142599 | 25.3407 | 44.52178 | 28.4084 | 12.57539 | 1.032222 | 0.646008 | 0.325203 |
| 10 | 100064200 | 100067999 | 30.43887 | 49.25864 | 91.23606 | 39.01974 | -0.52427 | 1.668386 | 0.325223 |
| 2 | 66464600 | 66466199 | 12.77257 | 22.26073 | 46.82241 | 20.59785 | -0.68018 | 0.731966 | 0.325241 |
| 8 | 106052200 | 106053399 | 7.123876 | 13.30929 | 30.6045 | 12.69758 | -0.73617 | 0.119258 | 0.325245 |
| 2 | 97712000 | 97714199 | 27.24668 | 46.9465 | 38.65745 | 16.95783 | 0.663818 | 0.909921 | 0.325272 |
| X | 166916000 | 166917599 | 17.49437 | 33.2959 | 17.72139 | 7.097898 | 1.400677 | 0.059271 | 0.325273 |
| Y | 88675200 | 88675999 | 8.750022 | 17.0538 | 9.305852 | 3.368963 | 1.494331 | -0.39942 | 0.325276 |
| 1 | 117392800 | 117402399 | 117.6927 | 181.4985 | 161.1536 | 66.94035 | 0.470394 | 2.903327 | 0.325278 |
| 3 | 30656200 | 30657199 | 9.504231 | 17.23155 | 34.92392 | 15.10412 | -0.59928 | 0.325124 | 0.325279 |
| 4 | 40947600 | 40948799 | 9.504231 | 17.23155 | 34.92392 | 15.10412 | -0.59928 | 0.325124 | 0.325279 |
| 17 | 55455600 | 55456999 | 19.80738 | 34.89189 | 30.68866 | 13.58409 | 0.572965 | 0.588646 | 0.325281 |
| 19 | 29565800 | 29567799 | 28.27226 | 48.71856 | 37.64472 | 16.51363 | 0.755341 | 0.914154 | 0.325289 |

| | | | | | | | | | |
|---|---|---|---|---|---|---|---|---|---|
| 14 | 100520800 | 100523999 | 20.39502 | 33.53213 | 77.67505 | 33.47624 | -0.82994 | 1.392178 | 0.325294 |
| 7 | 16948800 | 16951399 | 57.78132 | 92.47547 | 88.4512 | 37.25467 | 0.408302 | 2.036973 | 0.325296 |
| 7 | 63587000 | 63590399 | 21.07786 | 34.47313 | 78.70042 | 33.84448 | -0.81233 | 1.465393 | 0.325301 |
| 6 | 104946200 | 104946999 | 5.389366 | 10.46883 | 25.2085 | 9.418213 | -0.66402 | -0.2728 | 0.325311 |
| 10 | 29022400 | 29024599 | 20.26303 | 33.58176 | 70.26172 | 30.41842 | -0.68083 | 1.290127 | 0.325319 |
| 8 | 17582200 | 17586199 | 29.70111 | 47.93587 | 94.71102 | 40.37976 | -0.62149 | 1.742946 | 0.325324 |
| 12 | 12912800 | 12913399 | 5.35949 | 10.42467 | 24.71698 | 9.135803 | -0.62646 | -0.30884 | 0.325348 |
| 15 | 38467800 | 38474199 | 33.13733 | 53.24792 | 106.9504 | 45.3199 | -0.65301 | 1.887405 | 0.325348 |
| 12 | 115896400 | 115897399 | 13.05967 | 25.22205 | 17.68731 | 6.80597 | 1.058242 | -0.11058 | 0.32535 |
| 7 | 109720800 | 109722599 | 10.27506 | 18.0915 | 41.01552 | 18.15692 | -0.7856 | 0.557982 | 0.325358 |
| 5 | 116733800 | 116735199 | 9.934042 | 18.03365 | 35.42912 | 15.29835 | -0.5535 | 0.320273 | 0.32538 |
| 17 | 41724800 | 41726599 | 16.74423 | 31.44544 | 19.67067 | 8.089801 | 1.136972 | 0.141111 | 0.325389 |
| 16 | 79226600 | 79227799 | 17.60083 | 32.61023 | 21.68453 | 9.114336 | 1.025566 | 0.233316 | 0.325394 |
| 13 | 67700000 | 67701999 | 15.0912 | 26.15711 | 52.76325 | 23.14979 | -0.62166 | 0.820609 | 0.325402 |
| 11 | 11316800 | 11318599 | 19.12082 | 33.80677 | 29.42823 | 13.01658 | 0.589531 | 0.540476 | 0.325404 |
| 4 | 118044800 | 118049799 | 48.01022 | 76.4419 | 129.0499 | 53.96216 | -0.41804 | 2.199833 | 0.325416 |
| 14 | 75812600 | 75814199 | 16.13091 | 29.24122 | 26.17797 | 11.32392 | 0.571622 | 0.353458 | 0.325424 |
| Y | 16614200 | 16614999 | 13.59072 | 25.46007 | 12.86963 | 5.315034 | 1.436149 | 0.107521 | 0.325426 |
| 13 | 31138400 | 31140199 | 16.89524 | 30.65489 | 26.01953 | 11.24247 | 0.648964 | 0.352862 | 0.325427 |
| 4 | 85431000 | 85433399 | 12.77824 | 21.94478 | 55.78313 | 24.46788 | -0.95633 | 0.89975 | 0.325432 |
| 2 | 12101000 | 12103999 | 39.21416 | 64.70281 | 54.43691 | 23.49641 | 0.628784 | 1.3781 | 0.325435 |
| 17 | 59881600 | 59882799 | 15.73296 | 28.66485 | 25.64948 | 11.02516 | 0.578045 | 0.320045 | 0.325442 |
| 6 | 21456200 | 21457799 | 19.99232 | 36.11982 | 24.5038 | 10.6442 | 0.965179 | 0.395503 | 0.325462 |
| 7 | 29109800 | 29113999 | 21.18248 | 34.78722 | 75.86599 | 32.68084 | -0.74381 | 1.408518 | 0.325468 |
| X | 54646200 | 54647799 | 17.26079 | 31.1121 | 25.72708 | 11.20826 | 0.679167 | 0.394313 | 0.325476 |
| 5 | 135509800 | 135521999 | 94.55222 | 144.2915 | 237.4934 | 98.48295 | -0.43087 | 3.109284 | 0.325483 |
| 3 | 54641800 | 54644199 | 16.91998 | 28.57274 | 59.48562 | 26.02182 | -0.67266 | 1.074197 | 0.325492 |
| 4 | 139527200 | 139530199 | 16.91998 | 28.57274 | 59.48562 | 26.02182 | -0.67266 | 1.074197 | 0.325492 |
| 9 | 96899600 | 96901399 | 16.91998 | 28.57274 | 59.48562 | 26.02182 | -0.67266 | 1.074197 | 0.325492 |
| 17 | 41590800 | 41592599 | 23.40819 | 40.9741 | 32.51679 | 14.34895 | 0.719427 | 0.698767 | 0.325508 |
| 1 | 15721200 | 15722799 | 10.1526 | 18.20226 | 36.85532 | 16.13974 | -0.61015 | 0.3995 | 0.325514 |
| 3 | 47832400 | 47834199 | 19.12765 | 34.43601 | 26.29828 | 11.47419 | 0.791608 | 0.413668 | 0.325538 |
| 8 | 21673600 | 21675599 | 19.12765 | 34.43601 | 26.29828 | 11.47419 | 0.791608 | 0.413668 | 0.325538 |
| 2 | 172263400 | 172265799 | 16.25586 | 27.55952 | 58.44245 | 25.59603 | -0.69886 | 1.035508 | 0.325566 |
| 17 | 78009600 | 78011999 | 19.15375 | 33.94363 | 27.68904 | 12.24163 | 0.683772 | 0.511481 | 0.325579 |
| 16 | 5188800 | 5189999 | 8.597196 | 16.00831 | 33.26995 | 13.88172 | -0.59885 | 0.157674 | 0.325587 |
| 10 | 119641400 | 119649199 | 81.70362 | 126.0537 | 214.0683 | 88.91999 | -0.46375 | 2.886667 | 0.325595 |
| X | 26685200 | 26689999 | 40.49204 | 67.54071 | 45.36764 | 19.72604 | 0.953391 | 1.256355 | 0.325598 |
| 7 | 84725600 | 84728399 | 63.05529 | 100.7467 | 93.17414 | 39.15061 | 0.454129 | 2.095009 | 0.325608 |
| 5 | 149100400 | 149101999 | 9.152247 | 16.71549 | 33.96012 | 14.55323 | -0.59422 | 0.276493 | 0.32561 |
| 8 | 117241800 | 117245599 | 63.46922 | 101.3129 | 94.28513 | 39.5648 | 0.443674 | 2.125152 | 0.325632 |
| 4 | 75070200 | 75072399 | 26.88236 | 46.33424 | 38.65825 | 16.95836 | 0.645001 | 0.905099 | 0.325641 |
| 3 | 44359600 | 44361999 | 30.68442 | 51.63352 | 46.3884 | 20.27069 | 0.535005 | 1.136282 | 0.325659 |
| 11 | 99827200 | 99828799 | 10.2571 | 18.60446 | 35.93952 | 15.53506 | -0.53032 | 0.329125 | 0.32566 |
| 17 | 64503800 | 64507799 | 32.0794 | 51.79345 | 94.97384 | 40.51113 | -0.51354 | 1.735199 | 0.325666 |
| 11 | 44409200 | 44411199 | 13.28611 | 23.03581 | 48.95843 | 21.48065 | -0.69601 | 0.808364 | 0.325669 |
| 9 | 89360200 | 89360999 | 11.1718 | 21.3418 | 20.5811 | 8.178686 | 0.560198 | -0.01936 | 0.325676 |

| | | | | | | | | | |
|---|---|---|---|---|---|---|---|---|---|
| 5 | 56427000 | 56429199 | 13.49015 | 25.37664 | 22.39801 | 9.185456 | 0.647991 | 0.110071 | 0.325686 |
| 6 | 12152400 | 12153599 | 11.80846 | 22.43829 | 20.37039 | 8.186593 | 0.631989 | 0.020501 | 0.325698 |
| 14 | 107396600 | 107397599 | 11.80846 | 22.43829 | 20.37039 | 8.186593 | 0.631989 | 0.020501 | 0.325698 |
| 2 | 163597200 | 163597799 | 4.52015 | 8.811027 | 21.98434 | 7.94412 | -0.66891 | -0.41072 | 0.325714 |
| 3 | 41098800 | 41099399 | 4.52015 | 8.811027 | 21.98434 | 7.94412 | -0.66891 | -0.41072 | 0.325714 |
| 8 | 81059800 | 81062399 | 23.3098 | 40.70378 | 33.66971 | 14.8297 | 0.659851 | 0.735678 | 0.325716 |
| 11 | 24608600 | 24610199 | 17.06723 | 31.87001 | 21.3515 | 8.866158 | 1.028312 | 0.183348 | 0.325742 |
| X | 91751400 | 91753199 | 17.06723 | 31.87001 | 21.3515 | 8.866158 | 1.028312 | 0.183348 | 0.325742 |
| 2 | 46897400 | 46900199 | 29.52387 | 47.71001 | 98.03186 | 41.8637 | -0.67579 | 1.708205 | 0.325764 |
| 15 | 21719000 | 21722799 | 57.98162 | 92.75334 | 88.03544 | 37.0554 | 0.418667 | 2.049493 | 0.325781 |
| 4 | 67328400 | 67328999 | 12.16627 | 23.10015 | 20.88392 | 8.406842 | 0.635988 | 0.030888 | 0.325805 |
| 9 | 57614600 | 57616599 | 12.89928 | 22.58094 | 44.13286 | 19.47619 | -0.57426 | 0.66224 | 0.32581 |
| 17 | 40841400 | 40844599 | 27.21218 | 44.64025 | 79.74788 | 34.337 | -0.45779 | 1.438057 | 0.325815 |
| 13 | 60645200 | 60648799 | 22.70286 | 37.87766 | 69.19756 | 30.09206 | -0.48545 | 1.219971 | 0.325817 |
| 16 | 5334200 | 5343799 | 73.54728 | 114.6346 | 185.6936 | 77.06671 | -0.38424 | 2.70012 | 0.325833 |
| 5 | 14275600 | 14276599 | 5.984547 | 11.53991 | 27.04337 | 10.46558 | -0.67394 | -0.14049 | 0.325833 |
| 17 | 26471600 | 26472399 | 5.984547 | 11.53991 | 27.04337 | 10.46558 | -0.67394 | -0.14049 | 0.325833 |
| X | 71916600 | 71917999 | 19.78885 | 35.45813 | 26.42811 | 11.57912 | 0.822715 | 0.448388 | 0.325841 |
| 15 | 31449400 | 31452199 | 13.55637 | 23.21277 | 54.69949 | 23.98706 | -0.84811 | 0.930124 | 0.325846 |
| 5 | 138060200 | 138061399 | 10.64997 | 18.9371 | 38.84386 | 17.12929 | -0.63593 | 0.458283 | 0.325851 |
| 10 | 61064000 | 61066999 | 19.49506 | 32.67872 | 62.80196 | 27.39176 | -0.55812 | 1.155107 | 0.325855 |
| 17 | 38087400 | 38088999 | 18.50273 | 32.7951 | 28.94337 | 12.79578 | 0.571131 | 0.509547 | 0.325855 |
| 7 | 23480400 | 23500999 | 423.0663 | 591.9536 | 472.3477 | 195.8407 | 0.494646 | 4.574091 | 0.325856 |
| X | 79758200 | 79758799 | 5.054364 | 9.853227 | 25.1818 | 9.089036 | -0.70217 | -0.39635 | 0.325859 |
| 5 | 83900400 | 83902799 | 23.70587 | 41.05328 | 37.71459 | 16.54678 | 0.507967 | 0.847858 | 0.32586 |
| 17 | 23636000 | 23641599 | 95.24512 | 149.8297 | 122.3888 | 50.7745 | 0.612612 | 2.536236 | 0.325864 |
| 19 | 4530200 | 4531599 | 7.81968 | 14.81359 | 30.98882 | 12.53463 | -0.57043 | 0.034747 | 0.325882 |
| 2 | 146059200 | 146061599 | 19.96494 | 33.49753 | 62.29461 | 27.18645 | -0.5109 | 1.146537 | 0.325883 |
| 8 | 97407200 | 97407799 | 4.352354 | 8.495289 | 21.00793 | 7.44539 | -0.63009 | -0.48447 | 0.325888 |
| 6 | 7280600 | 7295799 | 208.9619 | 309.0582 | 280.8472 | 115.6603 | 0.385843 | 3.727851 | 0.325892 |
| 16 | 85147400 | 85149599 | 20.13734 | 35.37534 | 31.62221 | 13.99652 | 0.548385 | 0.622953 | 0.325894 |
| 16 | 85387400 | 85389399 | 20.13734 | 35.37534 | 31.62221 | 13.99652 | 0.548385 | 0.622953 | 0.325894 |
| 5 | 76198800 | 76201399 | 30.22076 | 51.52628 | 41.35768 | 18.1301 | 0.698164 | 1.017541 | 0.325907 |
| 17 | 93320200 | 93322399 | 36.34344 | 59.91453 | 55.42556 | 23.91442 | 0.491788 | 1.380852 | 0.325922 |
| 12 | 22292800 | 22294199 | 21.59561 | 40.51446 | 15.42758 | 6.355871 | 1.844769 | 0.181076 | 0.325926 |
| 4 | 138190000 | 138192999 | 11.01349 | 19.41754 | 40.94049 | 18.11888 | -0.68109 | 0.545421 | 0.325928 |
| 5 | 43965200 | 43965799 | 5.395899 | 10.46304 | 25.01805 | 9.44766 | -0.66827 | -0.23518 | 0.325928 |
| 6 | 107210600 | 107212999 | 22.16123 | 38.71601 | 34.39722 | 15.15453 | 0.557466 | 0.72618 | 0.325949 |
| 15 | 57054600 | 57055999 | 7.045541 | 13.50939 | 35.31537 | 13.88993 | -0.85304 | -0.0544 | 0.325952 |
| 5 | 79187000 | 79188799 | 11.91058 | 22.86859 | 18.76674 | 7.35267 | 0.809096 | -0.06411 | 0.32597 |
| 6 | 64235000 | 64236399 | 21.07947 | 37.09402 | 31.01192 | 13.73004 | 0.645137 | 0.607205 | 0.325972 |
| 7 | 27144600 | 27149199 | 29.96237 | 48.49554 | 89.89781 | 38.45209 | -0.52521 | 1.664083 | 0.325983 |
| 1 | 115279800 | 115281399 | 15.92644 | 29.2762 | 24.63136 | 10.46461 | 0.677891 | 0.27032 | 0.325989 |
| 2 | 81483800 | 81485399 | 21.83286 | 38.02692 | 35.13329 | 15.44937 | 0.500948 | 0.76801 | 0.326016 |
| 1 | 150329800 | 150331799 | 13.23408 | 23.11938 | 45.22284 | 19.92361 | -0.57548 | 0.699645 | 0.326029 |
| 12 | 22447000 | 22447399 | 8.196916 | 15.95639 | 14.09943 | 5.178028 | 0.790772 | -0.33551 | 0.326031 |
| 1 | 35453000 | 35454399 | 9.619106 | 17.58531 | 34.49444 | 14.76417 | -0.54277 | 0.276209 | 0.326039 |

| | | | | | | | | | |
|---|---|---|---|---|---|---|---|---|---|
| Y | 20332800 | 20333399 | 7.938844 | 15.50134 | 8.787935 | 3.095033 | 1.474511 | -0.50908 | 0.326043 |
| 5 | 38433200 | 38435599 | 27.02036 | 43.9273 | 84.39528 | 36.21407 | -0.57069 | 1.566699 | 0.326059 |
| 12 | 59725000 | 59726199 | 13.22194 | 25.2397 | 20.33589 | 8.093801 | 0.815192 | 0.004372 | 0.326067 |
| 13 | 53969800 | 53971599 | 14.02947 | 24.38535 | 47.90544 | 21.03855 | -0.58263 | 0.779107 | 0.326069 |
| 7 | 16354800 | 16356599 | 9.480924 | 17.26466 | 34.44676 | 14.81647 | -0.57171 | 0.298075 | 0.326073 |
| 12 | 79685600 | 79686799 | 9.480924 | 17.26466 | 34.44676 | 14.81647 | -0.57171 | 0.298075 | 0.326073 |
| 3 | 16350000 | 16353399 | 41.40169 | 68.65743 | 48.83907 | 21.153 | 0.870701 | 1.323751 | 0.326079 |
| 9 | 27210200 | 27212999 | 28.82799 | 46.73054 | 86.91583 | 37.21832 | -0.5279 | 1.630653 | 0.326089 |
| 17 | 18164200 | 18167199 | 43.86123 | 71.30352 | 58.68409 | 25.1811 | 0.649134 | 1.58514 | 0.326092 |
| 3 | 41675000 | 41675999 | 10.57798 | 20.43903 | 18.49559 | 7.100971 | 0.696765 | -0.14343 | 0.326101 |
| 6 | 134416400 | 134417999 | 8.423438 | 15.3367 | 34.46629 | 14.82763 | -0.74233 | 0.289857 | 0.326102 |
| 17 | 66676400 | 66677199 | 5.545476 | 10.71448 | 25.3622 | 9.73991 | -0.67676 | -0.17362 | 0.326128 |
| X | 129217800 | 129219399 | 17.05347 | 31.92321 | 20.81229 | 8.606116 | 1.072014 | 0.166768 | 0.326136 |
| 18 | 54336800 | 54337399 | 5.524319 | 10.73738 | 25.31237 | 9.411603 | -0.62729 | -0.28767 | 0.326138 |
| 19 | 12911600 | 12912599 | 10.92186 | 21.22935 | 17.0964 | 6.354312 | 0.908172 | -0.2481 | 0.326141 |
| 15 | 86586200 | 86591599 | 58.89785 | 92.54353 | 164.1899 | 68.10585 | -0.50625 | 2.543956 | 0.326152 |
| 5 | 62050000 | 62051599 | 13.51144 | 23.75225 | 45.05108 | 19.94971 | -0.53133 | 0.599044 | 0.326179 |
| 6 | 136404400 | 136405799 | 9.241803 | 17.01437 | 35.73877 | 15.15208 | -0.63141 | 0.233142 | 0.326185 |
| 9 | 20819000 | 20823999 | 23.46849 | 38.3196 | 79.52611 | 34.17895 | -0.67702 | 1.494731 | 0.326188 |
| 6 | 73071400 | 73074399 | 28.56054 | 48.58754 | 43.06805 | 18.8765 | 0.555072 | 1.032724 | 0.326213 |
| 4 | 48627800 | 48630199 | 27.00092 | 43.96028 | 82.17667 | 35.28096 | -0.52974 | 1.544821 | 0.326213 |
| 1 | 6514600 | 6515799 | 10.09066 | 19.42112 | 17.9961 | 7.002054 | 0.645054 | -0.10695 | 0.326214 |
| 9 | 36367000 | 36367599 | 10.09066 | 19.42112 | 17.9961 | 7.002054 | 0.645054 | -0.10695 | 0.326214 |
| 18 | 42962000 | 42963199 | 18.1549 | 32.35623 | 24.60818 | 10.83065 | 0.789077 | 0.464461 | 0.326226 |
| 15 | 103508800 | 103510599 | 11.08427 | 19.87048 | 37.85629 | 16.57959 | -0.52306 | 0.408902 | 0.326228 |
| 17 | 60723200 | 60728399 | 60.58594 | 95.59961 | 155.8864 | 64.70369 | -0.37995 | 2.458883 | 0.326237 |
| 18 | 36568200 | 36571199 | 31.8546 | 53.10967 | 49.76874 | 21.63355 | 0.474574 | 1.236509 | 0.326238 |
| 6 | 103191800 | 103195199 | 64.83088 | 103.1953 | 98.70505 | 41.25776 | 0.399858 | 2.215243 | 0.326242 |
| 1 | 79590400 | 79594599 | 44.84218 | 73.09742 | 61.26424 | 26.3144 | 0.625705 | 1.54968 | 0.32626 |
| 10 | 87327400 | 87330799 | 29.69068 | 47.94274 | 93.74003 | 39.99127 | -0.60562 | 1.727821 | 0.326266 |
| 3 | 52873800 | 52875199 | 8.53361 | 15.63986 | 33.49159 | 14.28676 | -0.66491 | 0.250313 | 0.326273 |
| 3 | 69629000 | 69630199 | 8.53361 | 15.63986 | 33.49159 | 14.28676 | -0.66491 | 0.250313 | 0.326273 |
| 12 | 50963800 | 50967599 | 38.55862 | 64.10448 | 49.32945 | 21.40269 | 0.757888 | 1.287706 | 0.326275 |
| 8 | 19250600 | 19252399 | 20.57906 | 37.06053 | 25.00909 | 10.90747 | 0.969235 | 0.419067 | 0.326278 |
| X | 53236800 | 53239199 | 28.72228 | 50.03838 | 31.99047 | 14.06947 | 1.02983 | 0.795798 | 0.326283 |
| 15 | 77452400 | 77455999 | 45.35616 | 73.30321 | 69.63651 | 29.74215 | 0.434734 | 1.714864 | 0.326284 |
| 7 | 62150600 | 62158399 | 112.0485 | 173.8645 | 147.5999 | 61.29802 | 0.541887 | 2.789906 | 0.326293 |
| 4 | 143927000 | 143934399 | 129.0919 | 198.8074 | 158.8445 | 65.97702 | 0.621003 | 2.934375 | 0.326304 |
| 16 | 48086400 | 48090599 | 43.01839 | 68.8464 | 117.9314 | 49.66947 | -0.43091 | 2.029912 | 0.326321 |
| 18 | 66593800 | 66595399 | 9.909736 | 18.13935 | 35.58486 | 15.20543 | -0.54146 | 0.272569 | 0.326325 |
| 7 | 24110800 | 24117199 | 50.15598 | 79.22479 | 150.3401 | 62.43462 | -0.59713 | 2.426432 | 0.326342 |
| 9 | 92860600 | 92863799 | 41.91062 | 68.54536 | 60.00679 | 25.80543 | 0.566358 | 1.496656 | 0.326365 |
| 10 | 115529000 | 115531799 | 20.27443 | 33.93274 | 66.82482 | 29.11801 | -0.59341 | 1.175734 | 0.326369 |
| 1 | 12174800 | 12175399 | 8.893981 | 17.36994 | 14.42834 | 5.063385 | 0.936142 | -0.48132 | 0.326381 |
| 16 | 94760600 | 94761999 | 6.887124 | 13.13922 | 28.35746 | 11.30059 | -0.59587 | -0.03057 | 0.326383 |
| 19 | 53023400 | 53027799 | 22.58367 | 37.11449 | 74.36975 | 32.04586 | -0.62163 | 1.402883 | 0.326389 |
| 7 | 13680200 | 13688199 | 106.6314 | 166.8589 | 128.0347 | 53.11401 | 0.696923 | 2.642234 | 0.326392 |

| | | | | | | | | | |
|---|---|---|---|---|---|---|---|---|---|
| X | 3420400 | 3423199 | 47.61039 | 77.13739 | 60.27961 | 25.80262 | 0.719321 | 1.661222 | 0.326405 |
| 12 | 20416600 | 20419199 | 55.37261 | 88.78566 | 83.16602 | 35.12682 | 0.441514 | 1.976008 | 0.326408 |
| 3 | 68776000 | 68778399 | 19.88265 | 33.10697 | 67.66475 | 29.38885 | -0.64664 | 1.231795 | 0.326413 |
| 14 | 60230400 | 60234999 | 27.75655 | 45.34032 | 80.43145 | 34.57337 | -0.45103 | 1.497466 | 0.32642 |
| 9 | 113865000 | 113866599 | 7.927643 | 14.57724 | 33.54333 | 14.24482 | -0.76319 | 0.226193 | 0.326428 |
| 3 | 145804800 | 145813399 | 107.4878 | 163.2762 | 257.3915 | 106.5961 | -0.3741 | 3.208032 | 0.326433 |
| 3 | 13279200 | 13280799 | 18.93297 | 33.71702 | 27.7635 | 12.22729 | 0.674331 | 0.479351 | 0.326434 |
| 1 | 122694800 | 122695799 | 11.50682 | 22.42257 | 13.93191 | 5.056204 | 1.306693 | -0.33379 | 0.326444 |
| 18 | 37698000 | 37698599 | 22.19783 | 38.6228 | 32.76942 | 14.40056 | 0.623297 | 0.781776 | 0.326469 |
| 6 | 124935600 | 124938399 | 12.27482 | 21.50882 | 43.59973 | 19.25683 | -0.62657 | 0.641787 | 0.326473 |
| 12 | 19509600 | 19532799 | 504.8151 | 693.9328 | 521.3132 | 216.7452 | 0.555863 | 4.844 | 0.326478 |
| 3 | 129941600 | 129942799 | 8.172575 | 15.16831 | 32.09088 | 13.4563 | -0.62955 | 0.172073 | 0.326492 |
| 4 | 137719800 | 137721799 | 11.50386 | 20.19197 | 42.54629 | 18.81839 | -0.68197 | 0.608535 | 0.326493 |
| 3 | 34751400 | 34755799 | 39.77326 | 63.75335 | 110.8846 | 46.81199 | -0.45013 | 1.975558 | 0.326517 |
| 3 | 65435000 | 65437199 | 14.30501 | 24.52464 | 53.13649 | 23.30394 | -0.72717 | 0.921264 | 0.32653 |
| 4 | 14632400 | 14633799 | 11.7718 | 20.82063 | 39.86074 | 17.6298 | -0.54082 | 0.51386 | 0.326535 |
| 15 | 77533000 | 77533799 | 5.374127 | 10.44703 | 24.06395 | 8.936541 | -0.59183 | -0.29552 | 0.326544 |
| 4 | 44719800 | 44722799 | 18.23141 | 31.11411 | 56.46631 | 24.75521 | -0.47405 | 0.981048 | 0.326557 |
| 7 | 14130800 | 14136399 | 118.469 | 183.6958 | 144.0397 | 59.82586 | 0.655544 | 2.803822 | 0.326562 |
| 4 | 101417200 | 101419199 | 10.42083 | 18.81801 | 36.38743 | 15.81593 | -0.53698 | 0.356522 | 0.326565 |
| 13 | 90702800 | 90705599 | 14.77803 | 25.80269 | 55.53536 | 24.45866 | -0.71353 | 0.726353 | 0.326576 |
| 2 | 22352800 | 22354399 | 35.52333 | 59.50144 | 46.41795 | 20.2361 | 0.738054 | 1.197184 | 0.326586 |
| 11 | 51822600 | 51824399 | 14.62985 | 25.49896 | 47.34463 | 20.82824 | -0.50128 | 0.74131 | 0.326594 |
| 1 | 51907000 | 51910999 | 54.47436 | 86.21034 | 145.7089 | 60.56978 | -0.42815 | 2.382737 | 0.326596 |
| 7 | 17074800 | 17098999 | 563.5833 | 767.1741 | 627.0029 | 261.0873 | 0.420092 | 4.997886 | 0.326598 |
| X | 67890800 | 67893199 | 25.35645 | 44.42696 | 30.74221 | 13.57687 | 0.916141 | 0.692528 | 0.3266 |
| 8 | 100667400 | 100668399 | 12.87447 | 24.82533 | 18.91871 | 7.321661 | 0.932303 | -0.08992 | 0.326616 |
| 16 | 12749000 | 12751399 | 15.47445 | 26.88009 | 50.04165 | 21.96804 | -0.50561 | 0.798587 | 0.326626 |
| 6 | 14847600 | 14849199 | 21.38455 | 37.9509 | 27.56584 | 12.17584 | 0.851581 | 0.517199 | 0.326633 |
| 7 | 61216800 | 61220399 | 56.32365 | 90.43329 | 82.90916 | 35.09071 | 0.474728 | 1.93482 | 0.326636 |
| 9 | 45082000 | 45086599 | 36.49199 | 58.80179 | 101.0323 | 42.98507 | -0.42325 | 1.8039 | 0.326668 |
| 7 | 41663800 | 41665799 | 31.33927 | 53.98981 | 35.7409 | 15.67836 | 0.977305 | 0.926068 | 0.326672 |
| 18 | 75282200 | 75283199 | 6.519803 | 12.24948 | 29.62457 | 12.17441 | -0.79674 | 0.075824 | 0.326677 |
| 1 | 39406600 | 39410399 | 21.08941 | 35.26378 | 65.12539 | 28.36079 | -0.50098 | 1.188204 | 0.326682 |
| 4 | 119226000 | 119230199 | 31.56454 | 51.14573 | 89.19271 | 38.18187 | -0.43582 | 1.644869 | 0.326718 |
| 7 | 59115400 | 59129599 | 163.93 | 247.298 | 219.7815 | 90.82492 | 0.443707 | 3.357397 | 0.326729 |
| 15 | 21753800 | 21757599 | 50.71009 | 82.11849 | 66.44921 | 28.43305 | 0.668473 | 1.678733 | 0.326736 |
| 16 | 47485200 | 47486599 | 21.19757 | 37.62928 | 27.57093 | 12.17531 | 0.839305 | 0.5142 | 0.326757 |
| 1 | 79075400 | 79077799 | 25.75141 | 42.41651 | 75.66408 | 32.63488 | -0.45323 | 1.382775 | 0.326768 |
| 11 | 102967000 | 102975199 | 57.40791 | 90.26045 | 165.1093 | 68.49292 | -0.54957 | 2.529867 | 0.326771 |
| 19 | 25847200 | 25848999 | 23.10951 | 38.31593 | 69.43577 | 30.06989 | -0.47419 | 1.293375 | 0.326778 |
| 7 | 17931000 | 17943799 | 186.0615 | 279.298 | 229.1794 | 94.58088 | 0.552495 | 3.458019 | 0.326789 |
| 7 | 31474000 | 31474999 | 11.80771 | 22.8462 | 18.56479 | 7.082377 | 0.860174 | -0.14803 | 0.326796 |
| 4 | 103349000 | 103351199 | 19.31644 | 33.90983 | 31.29629 | 13.84419 | 0.50258 | 0.628783 | 0.326796 |
| 14 | 42507000 | 42509399 | 30.66516 | 52.11462 | 41.3171 | 18.10758 | 0.715113 | 1.04579 | 0.326806 |
| X | 26139600 | 26142799 | 48.01271 | 79.57728 | 41.49344 | 17.96473 | 1.317147 | 1.343742 | 0.326809 |
| 8 | 95895600 | 95897199 | 11.29374 | 20.02694 | 38.82937 | 17.16244 | -0.55735 | 0.48133 | 0.326811 |

| | | | | | | | | | |
|---|---|---|---|---|---|---|---|---|---|
| 13 | 44311800 | 44313199 | 8.111459 | 15.2897 | 31.96934 | 13.06563 | -0.58321 | 0.07242 | 0.326817 |
| 17 | 47514000 | 47517399 | 12.92184 | 22.53419 | 46.29194 | 20.37206 | -0.64618 | 0.723673 | 0.326831 |
| 5 | 37287600 | 37293799 | 41.06182 | 65.7288 | 117.5991 | 49.54411 | -0.49324 | 2.020684 | 0.326856 |
| 2 | 156314000 | 156319199 | 23.6107 | 38.59603 | 79.87264 | 34.34223 | -0.67184 | 1.476801 | 0.32686 |
| 1 | 148088000 | 148090399 | 29.90085 | 51.31967 | 38.56391 | 16.9144 | 0.794213 | 0.954929 | 0.326879 |
| 5 | 119345800 | 119356799 | 99.48193 | 151.5487 | 246.0109 | 101.9703 | -0.41296 | 3.146204 | 0.326902 |
| 12 | 11709800 | 11713199 | 28.42229 | 48.49722 | 41.37945 | 18.14077 | 0.610414 | 1.002284 | 0.326916 |
| 18 | 65788000 | 65793199 | 40.23321 | 64.58135 | 110.1026 | 46.58138 | -0.41897 | 1.931784 | 0.326917 |
| 4 | 48599200 | 48604999 | 59.20078 | 93.63162 | 155.3433 | 64.55424 | -0.40221 | 2.402123 | 0.326925 |
| 15 | 59605000 | 59606599 | 6.450986 | 12.31825 | 27.80026 | 11.05745 | -0.65709 | -0.04233 | 0.326932 |
| 12 | 21310000 | 21317799 | 43.61697 | 69.45407 | 129.1402 | 54.00756 | -0.55693 | 2.191708 | 0.326968 |
| 1 | 32583600 | 32587599 | 32.18888 | 53.59138 | 50.32898 | 21.85762 | 0.471506 | 1.25287 | 0.326974 |
| 8 | 19726600 | 19728599 | 48.04442 | 77.60891 | 62.47236 | 26.66503 | 0.672295 | 1.729756 | 0.326975 |
| 7 | 107574000 | 107575399 | 21.3818 | 37.61818 | 30.95409 | 13.70501 | 0.667812 | 0.610794 | 0.326977 |
| 8 | 45473000 | 45481399 | 207.3664 | 298.2952 | 477.3497 | 197.029 | -0.46879 | 4.158795 | 0.327001 |
| 5 | 104980200 | 104988199 | 189.3111 | 282.824 | 249.1229 | 102.7077 | 0.444094 | 3.54837 | 0.327005 |
| 13 | 60843800 | 60847199 | 25.93534 | 42.5305 | 77.1299 | 33.20477 | -0.47956 | 1.438165 | 0.327006 |
| X | 25749200 | 25750799 | 27.38217 | 46.78767 | 41.00319 | 17.97783 | 0.572434 | 0.98865 | 0.327016 |
| 7 | 33382800 | 33396999 | 265.1114 | 387.1468 | 313.2299 | 129.0528 | 0.535972 | 3.936325 | 0.327025 |
| 18 | 8273200 | 8275399 | 16.23828 | 28.17365 | 58.91697 | 25.85052 | -0.6733 | 0.825359 | 0.327042 |
| 3 | 131592200 | 131594599 | 16.56286 | 28.67317 | 59.45171 | 26.08338 | -0.66173 | 0.847138 | 0.327044 |
| 11 | 93763400 | 93765599 | 27.38867 | 47.15941 | 38.03852 | 16.68598 | 0.693195 | 0.913856 | 0.327045 |
| 2 | 98709600 | 98714199 | 38.35374 | 63.52048 | 52.1664 | 22.57389 | 0.663941 | 1.331608 | 0.327045 |
| 11 | 57690400 | 57693599 | 15.17689 | 25.4394 | 64.59363 | 28.17255 | -0.9582 | 1.14375 | 0.327051 |
| 5 | 144581000 | 144596999 | 208.7236 | 302.0674 | 457.2844 | 188.5927 | -0.38014 | 4.069327 | 0.327056 |
| X | 21691600 | 21693799 | 27.98857 | 49.0767 | 27.99103 | 12.36994 | 1.193649 | 0.692607 | 0.327057 |
| 6 | 22039200 | 22041199 | 12.85585 | 22.68323 | 42.94135 | 19.00146 | -0.52641 | 0.550909 | 0.327072 |
| 10 | 81941200 | 81948199 | 172.5564 | 261.6716 | 188.0166 | 77.82554 | 0.756559 | 3.255184 | 0.327107 |
| 1 | 75256600 | 75259199 | 35.07202 | 57.99633 | 54.25502 | 23.45284 | 0.476471 | 1.341776 | 0.32711 |
| 16 | 71551200 | 71553199 | 25.54925 | 44.30201 | 37.0362 | 16.24978 | 0.643813 | 0.844503 | 0.327116 |
| 14 | 42405600 | 42407199 | 17.24651 | 31.49882 | 24.31019 | 10.41781 | 0.792605 | 0.313903 | 0.327117 |
| 17 | 72552800 | 72554799 | 9.445403 | 17.31553 | 34.01045 | 14.50287 | -0.54084 | 0.257651 | 0.327124 |
| 17 | 38610000 | 38616199 | 81.53366 | 129.6757 | 97.23128 | 40.59562 | 0.749297 | 2.25763 | 0.327128 |
| 8 | 61303600 | 61304599 | 14.44273 | 24.8416 | 55.82803 | 24.48873 | -0.77889 | 0.895792 | 0.327128 |
| 11 | 49478200 | 49481599 | 22.61773 | 37.08476 | 77.13758 | 33.20607 | -0.67676 | 1.432366 | 0.327137 |
| 1 | 186811000 | 186814799 | 46.52038 | 75.57443 | 64.0637 | 27.47684 | 0.605771 | 1.606054 | 0.327139 |
| 19 | 34168200 | 34170399 | 13.38412 | 23.30158 | 47.88541 | 21.0513 | -0.64712 | 0.751155 | 0.327148 |
| 5 | 77635200 | 77636799 | 12.7398 | 22.34468 | 43.04917 | 19.02884 | -0.5536 | 0.62868 | 0.327155 |
| 5 | 16087200 | 16088999 | 10.35747 | 18.42851 | 37.86975 | 16.69094 | -0.63773 | 0.452311 | 0.327157 |
| 7 | 99629000 | 99629799 | 6.91678 | 13.10574 | 28.71815 | 11.6113 | -0.63588 | 0.019069 | 0.327159 |
| 6 | 68220400 | 68228199 | 236.2519 | 348.084 | 265.5228 | 109.3448 | 0.633037 | 3.79385 | 0.327165 |
| 9 | 97155000 | 97160999 | 58.59341 | 92.6783 | 151.9374 | 63.14143 | -0.3849 | 2.400254 | 0.327184 |
| 17 | 38081400 | 38083599 | 28.6343 | 48.57591 | 38.46296 | 16.84882 | 0.716903 | 1.052325 | 0.327195 |
| 15 | 54690400 | 54691999 | 7.797833 | 14.77447 | 31.01603 | 12.54157 | -0.57532 | 0.032089 | 0.3272 |
| 14 | 50353400 | 50356599 | 43.05292 | 69.91842 | 65.39437 | 28.04078 | 0.463801 | 1.606212 | 0.327234 |
| 6 | 142185400 | 142187399 | 21.85077 | 38.0618 | 35.01562 | 15.39856 | 0.506955 | 0.764805 | 0.327237 |
| 6 | 145260400 | 145264399 | 36.14402 | 58.26579 | 100.0497 | 42.59124 | -0.42164 | 1.789624 | 0.327258 |

| | | | | | | | | | |
|---|---|---|---|---|---|---|---|---|---|
| 6 | 112153800 | 112157199 | 42.67907 | 68.35463 | 117.7671 | 49.65514 | -0.43799 | 2.00608 | 0.327271 |
| 14 | 3604400 | 3606399 | 36.06519 | 59.17693 | 52.50918 | 22.61107 | 0.549127 | 1.436477 | 0.327276 |
| 12 | 110157400 | 110164599 | 47.69619 | 75.78542 | 136.3961 | 56.89023 | -0.51331 | 2.264237 | 0.327285 |
| 1 | 179630200 | 179631799 | 11.33066 | 19.95284 | 40.95449 | 18.12941 | -0.6432 | 0.561489 | 0.3273 |
| 16 | 3664800 | 3665999 | 8.834006 | 16.25371 | 33.02083 | 14.01135 | -0.5837 | 0.231078 | 0.327306 |
| 2 | 100063000 | 100064999 | 24.22067 | 42.58359 | 28.41015 | 12.5804 | 0.968243 | 0.629122 | 0.32731 |
| 5 | 137317000 | 137321799 | 29.67456 | 47.97719 | 90.44842 | 38.64984 | -0.55099 | 1.69068 | 0.327328 |
| 6 | 111135800 | 111137799 | 25.88558 | 44.99151 | 34.7049 | 15.24095 | 0.759657 | 0.813685 | 0.327337 |
| 15 | 33900000 | 33901399 | 9.476983 | 17.18009 | 35.93868 | 15.54532 | -0.64518 | 0.325089 | 0.327338 |
| 18 | 26959800 | 26963799 | 35.66609 | 59.48714 | 49.24401 | 21.41062 | 0.652955 | 1.245767 | 0.327342 |
| 7 | 24936600 | 24937999 | 8.996414 | 16.43589 | 33.96481 | 14.54969 | -0.61826 | 0.273161 | 0.327353 |
| 1 | 124017600 | 124019399 | 14.9053 | 27.5266 | 23.57931 | 9.95458 | 0.658268 | 0.232754 | 0.327365 |
| 14 | 20160000 | 20162399 | 29.13943 | 49.51703 | 43.03704 | 18.86121 | 0.583066 | 1.043902 | 0.327402 |
| 13 | 46667200 | 46671399 | 27.47674 | 44.72659 | 82.88906 | 35.58439 | -0.51752 | 1.548333 | 0.32741 |
| 7 | 3151200 | 3158799 | 70.07763 | 111.5614 | 93.40811 | 39.05069 | 0.591773 | 2.214658 | 0.327413 |
| 10 | 26323000 | 26324599 | 15.1093 | 27.82233 | 24.09519 | 10.21269 | 0.638584 | 0.251808 | 0.327445 |
| 15 | 70371400 | 70371999 | 9.740819 | 18.81101 | 18.65351 | 7.180193 | 0.563025 | -0.14211 | 0.327454 |
| 5 | 39384000 | 39384599 | 11.03297 | 21.1954 | 20.11983 | 7.870161 | 0.603878 | -0.07231 | 0.32746 |
| 19 | 10502200 | 10503399 | 8.687425 | 15.95319 | 32.99842 | 14.03867 | -0.61219 | 0.240737 | 0.32746 |
| 7 | 10641200 | 10645799 | 90.44883 | 141.3561 | 121.8347 | 50.54862 | 0.527998 | 2.653514 | 0.327469 |
| 1 | 9506800 | 9508399 | 19.89309 | 34.949 | 31.72648 | 14.0436 | 0.526348 | 0.62148 | 0.327489 |
| 6 | 132148200 | 132151999 | 74.25727 | 118.0422 | 104.2248 | 43.49136 | 0.513115 | 2.266416 | 0.327494 |
| 7 | 31653000 | 31658599 | 20.49972 | 36.53435 | 24.67239 | 10.85913 | 0.95944 | 0.480455 | 0.327505 |
| 15 | 99440000 | 99445599 | 34.12396 | 54.76715 | 108.8941 | 46.0601 | -0.64055 | 1.927754 | 0.327506 |
| 11 | 94258000 | 94259599 | 9.252666 | 17.15591 | 34.82176 | 14.62117 | -0.57187 | 0.193087 | 0.327516 |
| 12 | 42527800 | 42529399 | 9.252666 | 17.15591 | 34.82176 | 14.62117 | -0.57187 | 0.193087 | 0.327516 |
| 7 | 72176800 | 72177799 | 10.95497 | 19.54079 | 37.83983 | 16.64236 | -0.55056 | 0.43995 | 0.327522 |
| 13 | 53666400 | 53668999 | 33.55711 | 55.94394 | 50.33535 | 21.8788 | 0.533086 | 1.243606 | 0.327532 |
| 17 | 49263200 | 49269999 | 37.07762 | 59.33565 | 114.4156 | 48.18507 | -0.60137 | 2.024886 | 0.327534 |
| 7 | 32054600 | 32061199 | 146.7586 | 222.2428 | 207.8764 | 86.00891 | 0.374677 | 3.276376 | 0.327553 |
| 17 | 86473800 | 86476399 | 14.69406 | 24.98653 | 57.31585 | 25.12033 | -0.8112 | 1.002207 | 0.327553 |
| 7 | 120532400 | 120534199 | 7.59808 | 14.08543 | 31.55073 | 13.25105 | -0.71253 | 0.174189 | 0.327557 |
| 17 | 18614600 | 18619599 | 79.69509 | 126.6451 | 101.8946 | 42.50516 | 0.646579 | 2.281251 | 0.327562 |
| 8 | 111094200 | 111094999 | 4.785267 | 9.311724 | 22.91665 | 8.441277 | -0.67425 | -0.33338 | 0.327563 |
| 3 | 110322200 | 110323999 | 21.25836 | 37.00775 | 33.20145 | 14.59527 | 0.543074 | 0.768781 | 0.327568 |
| 12 | 92070800 | 92072399 | 20.76998 | 37.39294 | 25.00126 | 10.90822 | 0.982166 | 0.42223 | 0.327581 |
| 12 | 88111800 | 88116399 | 55.43662 | 89.31516 | 74.47803 | 31.68049 | 0.617223 | 1.823959 | 0.327584 |
| 7 | 116539600 | 116541199 | 14.21512 | 26.0128 | 22.21285 | 9.495216 | 0.649986 | 0.274358 | 0.3276 |
| 10 | 121375600 | 121380799 | 59.44876 | 93.68443 | 157.9389 | 65.53803 | -0.4294 | 2.484444 | 0.327623 |
| 1 | 178431400 | 178432199 | 8.661172 | 16.05256 | 32.604 | 13.69765 | -0.57287 | 0.187593 | 0.327624 |
| X | 10812800 | 10815199 | 33.72088 | 57.36031 | 35.95016 | 15.75663 | 1.053157 | 1.046255 | 0.32763 |
| 3 | 15820400 | 15821999 | 18.78269 | 33.57863 | 27.25831 | 11.96584 | 0.697938 | 0.454026 | 0.327631 |
| X | 108063600 | 108067799 | 40.5532 | 68.39096 | 37.94579 | 16.59258 | 1.227937 | 1.14691 | 0.327654 |
| 17 | 17413400 | 17423599 | 257.3796 | 378.6203 | 278.2187 | 114.5731 | 0.684839 | 3.822556 | 0.327658 |
| 10 | 58467400 | 58471199 | 29.60939 | 48.01145 | 87.91719 | 37.65537 | -0.50513 | 1.625225 | 0.327679 |
| 4 | 79175400 | 79177799 | 21.28174 | 37.33516 | 32.23998 | 14.25365 | 0.598245 | 0.653549 | 0.327721 |
| 2 | 177169600 | 177175799 | 118.9036 | 182.6549 | 171.3588 | 71.15024 | 0.386796 | 2.979296 | 0.327737 |

| | | | | | | | | | |
|---|---|---|---|---|---|---|---|---|---|
| 7 | 41083800 | 41088599 | 66.67088 | 106.8752 | 80.833 | 34.12863 | 0.749066 | 1.995627 | 0.327741 |
| X | 85360000 | 85363399 | 22.78141 | 37.949 | 69.55112 | 30.21582 | -0.48996 | 1.240076 | 0.327745 |
| 1 | 5238800 | 5244599 | 170.2265 | 254.955 | 235.4744 | 97.12834 | 0.379051 | 3.497372 | 0.327758 |
| 9 | 25753800 | 25756799 | 27.82312 | 48.40545 | 32.93399 | 14.47195 | 0.940044 | 0.808409 | 0.32776 |
| 5 | 115368800 | 115371799 | 19.17733 | 32.12846 | 64.46812 | 28.10872 | -0.62024 | 1.160349 | 0.327787 |
| 8 | 8760800 | 8762199 | 12.30575 | 21.50973 | 44.70886 | 19.70546 | -0.66246 | 0.685665 | 0.327789 |
| 3 | 151776800 | 151777399 | 7.227542 | 13.65491 | 29.19494 | 11.8767 | -0.60843 | 0.044495 | 0.327808 |
| 19 | 57377800 | 57378599 | 7.227542 | 13.65491 | 29.19494 | 11.8767 | -0.60843 | 0.044495 | 0.327808 |
| 11 | 64869600 | 64870799 | 6.292191 | 12.06117 | 27.95058 | 11.00302 | -0.68134 | -0.07876 | 0.327816 |
| 15 | 45855400 | 45856799 | 18.45427 | 33.41316 | 25.21124 | 10.92196 | 0.815158 | 0.373546 | 0.327818 |
| 7 | 133283000 | 133293599 | 124.8647 | 187.5313 | 298.1244 | 123.051 | -0.40031 | 3.440333 | 0.32782 |
| 13 | 4378600 | 4380399 | 16.54881 | 28.61738 | 52.76283 | 23.14798 | -0.4932 | 0.849524 | 0.327821 |
| 18 | 9108200 | 9112399 | 66.56598 | 106.5167 | 87.27743 | 36.75147 | 0.630847 | 2.052166 | 0.327844 |
| 5 | 147211000 | 147214799 | 39.09806 | 62.74266 | 109.0233 | 46.10621 | -0.44681 | 1.938034 | 0.32785 |
| 6 | 81835800 | 81836799 | 12.87242 | 23.97736 | 22.37404 | 9.329901 | 0.548962 | 0.165122 | 0.327851 |
| 2 | 37096600 | 37100199 | 65.3576 | 104.0496 | 100.1226 | 41.85911 | 0.391432 | 2.211504 | 0.327868 |
| 19 | 23854200 | 23856599 | 24.4899 | 42.22451 | 38.77554 | 17.00998 | 0.507248 | 0.891088 | 0.327869 |
| 3 | 113112000 | 113113799 | 31.23718 | 52.18991 | 49.19349 | 21.41188 | 0.466127 | 1.2126 | 0.327871 |
| 4 | 135765400 | 135765999 | 5.507362 | 10.69586 | 25.70772 | 9.62056 | -0.66247 | -0.25936 | 0.327892 |
| 12 | 93557000 | 93557799 | 8.279254 | 15.07699 | 33.97892 | 14.61456 | -0.74572 | 0.28894 | 0.327896 |
| 5 | 128962600 | 128963999 | 10.50927 | 18.65658 | 39.35323 | 17.37771 | -0.67742 | 0.468437 | 0.327902 |
| 9 | 78349800 | 78351599 | 17.15207 | 29.20791 | 55.83712 | 24.47504 | -0.54917 | 0.997019 | 0.327915 |
| 7 | 25496000 | 25503999 | 49.16685 | 78.1149 | 135.3735 | 56.45843 | -0.459 | 2.268456 | 0.327918 |
| 7 | 49032600 | 49035799 | 17.21075 | 29.60596 | 54.36137 | 23.84547 | -0.48892 | 0.901622 | 0.327919 |
| 5 | 52616600 | 52617799 | 12.36262 | 23.40947 | 21.39856 | 8.663501 | 0.613096 | 0.052679 | 0.327919 |
| 3 | 154019000 | 154022399 | 32.86384 | 53.17724 | 92.21335 | 39.43461 | -0.42961 | 1.676172 | 0.32792 |
| 2 | 164705200 | 164715599 | 65.86862 | 102.2851 | 202.9821 | 84.28862 | -0.68094 | 2.762676 | 0.327925 |
| 7 | 5342400 | 5350999 | 115.5806 | 179.3685 | 144.5899 | 60.04682 | 0.616644 | 2.789652 | 0.327931 |
| 7 | 81982600 | 81989399 | 51.59166 | 81.95434 | 138.2795 | 57.66121 | -0.42074 | 2.275372 | 0.327935 |
| X | 32790200 | 32797799 | 205.864 | 305.6995 | 263.9429 | 108.728 | 0.46495 | 3.659665 | 0.327968 |
| 9 | 7487200 | 7496999 | 143.2698 | 217.1317 | 204.0109 | 84.43069 | 0.369165 | 3.259033 | 0.327974 |
| 10 | 126218400 | 126220199 | 11.2867 | 19.99297 | 39.33484 | 17.39193 | -0.57912 | 0.492124 | 0.327977 |
| 3 | 116553400 | 116555399 | 30.33495 | 49.03588 | 92.66442 | 39.58738 | -0.55471 | 1.699822 | 0.327977 |
| 1 | 39688200 | 39690599 | 17.56203 | 29.59169 | 60.58532 | 26.48517 | -0.64884 | 1.096494 | 0.327982 |
| 7 | 85274600 | 85282599 | 98.75669 | 155.0776 | 124.8852 | 51.79817 | 0.631168 | 2.573597 | 0.327988 |
| 1 | 189553800 | 189556399 | 18.44828 | 31.27041 | 58.00084 | 25.40113 | -0.50652 | 1.045402 | 0.327992 |
| 11 | 80307200 | 80308599 | 21.56476 | 37.64039 | 34.07269 | 15.00314 | 0.530602 | 0.735963 | 0.327998 |
| 7 | 97363800 | 97364599 | 10.52898 | 20.39043 | 18.61125 | 7.070262 | 0.699179 | -0.17741 | 0.328006 |
| 7 | 131024000 | 131027199 | 27.06356 | 46.21556 | 42.00745 | 18.41726 | 0.519752 | 0.991941 | 0.32801 |
| 9 | 19382800 | 19383799 | 11.33139 | 21.93156 | 18.62113 | 7.09334 | 0.799541 | -0.1577 | 0.328018 |
| 17 | 17486800 | 17492399 | 134.7891 | 207.457 | 155.849 | 64.72748 | 0.709173 | 2.948418 | 0.328022 |
| 4 | 80829000 | 80830599 | 11.79967 | 20.72061 | 42.51464 | 18.81163 | -0.64392 | 0.602154 | 0.328026 |
| 2 | 23750800 | 23752599 | 25.04044 | 43.53185 | 35.9074 | 15.77115 | 0.66351 | 0.809604 | 0.328042 |
| 16 | 59285000 | 59287199 | 25.37755 | 43.65283 | 38.71666 | 16.98294 | 0.556498 | 0.914413 | 0.328052 |
| 9 | 41992600 | 41994999 | 16.50061 | 28.13829 | 55.84837 | 24.48276 | -0.60278 | 0.980423 | 0.328054 |
| 1 | 102463000 | 102463399 | 10.58978 | 20.55665 | 18.12517 | 6.809047 | 0.763931 | -0.21425 | 0.328077 |
| 7 | 13446000 | 13451799 | 105.5833 | 165.292 | 117.7432 | 48.84245 | 0.804531 | 2.631043 | 0.328092 |

| | | | | | | | | | |
|---|---|---|---|---|---|---|---|---|---|
| 2 | 58315000 | 58316599 | 16.52472 | 28.24298 | 54.75795 | 24.0091 | -0.56843 | 0.961324 | 0.328094 |
| 1 | 91816600 | 91823799 | 65.82765 | 103.2071 | 168.7981 | 70.01043 | -0.39145 | 2.590888 | 0.328098 |
| 7 | 31888800 | 31889799 | 19.9786 | 35.03401 | 32.29627 | 14.27337 | 0.504295 | 0.65154 | 0.32813 |
| 8 | 14981400 | 14985199 | 17.98121 | 30.03503 | 64.73673 | 28.17289 | -0.72286 | 1.190919 | 0.328156 |
| 6 | 128303800 | 128305399 | 10.56188 | 18.64584 | 39.96257 | 17.68143 | -0.70347 | 0.528543 | 0.328176 |
| 19 | 39934600 | 39939999 | 84.30117 | 132.9504 | 122.2786 | 50.74551 | 0.444487 | 2.479607 | 0.328185 |
| 16 | 97818600 | 97827799 | 212.3318 | 313.55 | 284.7507 | 117.2659 | 0.384627 | 3.754323 | 0.328214 |
| 17 | 20752800 | 20759399 | 81.09137 | 128.9164 | 99.06578 | 41.34196 | 0.713291 | 2.269426 | 0.328222 |
| 17 | 93506600 | 93511599 | 48.83452 | 77.54616 | 139.6211 | 58.19929 | -0.51479 | 2.283724 | 0.328231 |
| 12 | 88015400 | 88021599 | 160.6047 | 241.6005 | 221.7192 | 91.55448 | 0.393879 | 3.410501 | 0.328244 |
| 5 | 109843200 | 109848799 | 94.5797 | 148.0562 | 135.069 | 56.02992 | 0.447989 | 2.628925 | 0.328247 |
| 18 | 12566600 | 12569199 | 14.47254 | 25.16376 | 48.43427 | 21.27519 | -0.55337 | 0.776008 | 0.328255 |
| 14 | 13746800 | 13748599 | 16.87688 | 30.19705 | 27.58735 | 12.10165 | 0.52928 | 0.437259 | 0.328275 |
| 8 | 48371600 | 48375999 | 31.61349 | 51.04526 | 103.3293 | 44.07966 | -0.65576 | 1.736311 | 0.328276 |
| 1 | 49672400 | 49673799 | 17.481 | 32.0624 | 23.8232 | 10.15195 | 0.852687 | 0.291972 | 0.328283 |
| 5 | 27184400 | 27186199 | 12.87286 | 22.60393 | 42.99624 | 19.02384 | -0.53539 | 0.609717 | 0.328283 |
| 7 | 14836400 | 14865399 | 533.1648 | 733.3749 | 542.6239 | 225.5826 | 0.578753 | 4.835478 | 0.328311 |
| 12 | 22701800 | 22704399 | 28.73263 | 48.69926 | 38.94516 | 17.05544 | 0.702556 | 1.059923 | 0.328315 |
| 12 | 65850000 | 65851199 | 26.78061 | 46.12141 | 35.55496 | 15.59664 | 0.758326 | 0.907472 | 0.328315 |
| 10 | 15248400 | 15249999 | 10.24543 | 18.62181 | 35.97386 | 15.5097 | -0.52796 | 0.316843 | 0.328347 |
| 13 | 67830600 | 67831999 | 9.596463 | 17.54705 | 34.49086 | 14.75934 | -0.5457 | 0.273806 | 0.328354 |
| 4 | 148365600 | 148374999 | 68.8799 | 106.807 | 194.4306 | 80.75502 | -0.55819 | 2.78746 | 0.328356 |
| X | 24238400 | 24240599 | 50.89944 | 82.19892 | 69.7723 | 29.77064 | 0.595605 | 1.750065 | 0.328359 |
| 10 | 82289400 | 82290599 | 19.54307 | 34.53548 | 29.98243 | 13.26309 | 0.593196 | 0.550577 | 0.32836 |
| 17 | 70924400 | 70926199 | 16.17827 | 29.58908 | 25.04026 | 10.7118 | 0.662331 | 0.300181 | 0.32837 |
| 17 | 87904000 | 87904399 | 4.103012 | 8.015123 | 21.22126 | 7.41621 | -0.70975 | -0.5398 | 0.328371 |
| 8 | 115130000 | 115131799 | 10.0525 | 18.30069 | 38.17316 | 16.42469 | -0.63644 | 0.308225 | 0.328375 |
| 1 | 103378200 | 103379799 | 19.69542 | 34.88321 | 28.18414 | 12.46197 | 0.697281 | 0.522527 | 0.328378 |
| 9 | 73213600 | 73214999 | 17.94004 | 30.47691 | 57.47851 | 25.18943 | -0.53034 | 1.022661 | 0.328392 |
| 18 | 81794800 | 81797399 | 22.00453 | 38.40096 | 34.366 | 15.13055 | 0.546781 | 0.738568 | 0.328397 |
| 3 | 102903000 | 102903599 | 5.537215 | 10.72343 | 24.93238 | 9.471338 | -0.63623 | -0.21257 | 0.328412 |
| X | 89796600 | 89798399 | 21.49608 | 39.83342 | 17.45669 | 7.335757 | 1.622247 | 0.255673 | 0.328427 |
| 11 | 20607400 | 20610199 | 23.26224 | 38.50567 | 71.17427 | 30.78775 | -0.50291 | 1.312801 | 0.328427 |
| X | 125592600 | 125593599 | 14.20294 | 26.94535 | 20.70824 | 8.350417 | 0.866476 | 0.055792 | 0.328439 |
| 19 | 7787600 | 7790999 | 43.32897 | 71.30955 | 53.96089 | 23.26575 | 0.779898 | 1.420443 | 0.328439 |
| 18 | 85190600 | 85195799 | 78.90142 | 124.7613 | 115.5866 | 48.02741 | 0.436977 | 2.415174 | 0.32846 |
| 12 | 56177600 | 56178599 | 13.32507 | 24.85075 | 22.79315 | 9.485092 | 0.575866 | 0.161924 | 0.32847 |
| 16 | 15924800 | 15928799 | 67.08438 | 106.8026 | 100.9975 | 42.22703 | 0.416559 | 2.211216 | 0.328485 |
| Y | 4006400 | 4010399 | 32.86155 | 54.76027 | 33.70492 | 14.64614 | 1.078684 | 1.226775 | 0.328493 |
| 10 | 47787600 | 47787999 | 8.105726 | 15.833 | 14.51076 | 5.078332 | 0.799405 | -0.50236 | 0.328527 |
| X | 106104000 | 106104799 | 12.36931 | 24.01359 | 16.78392 | 6.299789 | 1.096656 | -0.20073 | 0.328539 |
| 13 | 47363200 | 47364399 | 10.28208 | 19.72202 | 17.07414 | 6.708839 | 0.729602 | -0.07532 | 0.32854 |
| 7 | 19313600 | 19315399 | 9.755645 | 17.36781 | 37.93208 | 16.71021 | -0.72422 | 0.446794 | 0.328556 |
| X | 5070400 | 5073199 | 20.93531 | 37.85307 | 23.93235 | 10.38581 | 1.067185 | 0.397378 | 0.328564 |
| 12 | 78008400 | 78009999 | 13.40718 | 23.34243 | 46.83824 | 20.59133 | -0.61266 | 0.75136 | 0.328589 |
| 13 | 62738400 | 62747199 | 143.1132 | 217.2707 | 202.0295 | 83.65447 | 0.386212 | 3.223973 | 0.32859 |
| X | 140201800 | 140202599 | 13.46922 | 25.85616 | 19.3202 | 7.573762 | 0.943166 | -0.04302 | 0.3286 |

| | | | | | | | | | |
|---|---|---|---|---|---|---|---|---|---|
| 15 | 103847400 | 103849999 | 34.6522 | 57.81231 | 49.24798 | 21.41614 | 0.611681 | 1.237574 | 0.328601 |
| X | 74601400 | 74605799 | 46.46947 | 76.19277 | 54.62125 | 23.51326 | 0.855608 | 1.472982 | 0.32861 |
| 15 | 33133000 | 33134999 | 37.1239 | 60.8087 | 57.98999 | 24.95288 | 0.444464 | 1.464418 | 0.328617 |
| 7 | 42690400 | 42697999 | 80.16205 | 127.7134 | 89.72326 | 37.55003 | 0.845658 | 2.198965 | 0.328617 |
| 13 | 56361200 | 56364999 | 23.28571 | 38.48408 | 71.63533 | 30.95362 | -0.51317 | 1.331794 | 0.328619 |
| 3 | 91400000 | 91400599 | 18.96357 | 33.43144 | 30.25784 | 13.39131 | 0.532301 | 0.574347 | 0.328634 |
| 10 | 120037000 | 120038999 | 12.14062 | 21.20096 | 48.9067 | 21.54136 | -0.81262 | 0.701446 | 0.328643 |
| 6 | 14413000 | 14413999 | 16.09371 | 28.85038 | 25.80862 | 11.30331 | 0.561138 | 0.417361 | 0.328657 |
| 10 | 46632000 | 46634999 | 31.18434 | 52.76183 | 44.13852 | 19.31948 | 0.637786 | 1.089867 | 0.328673 |
| 1 | 137115000 | 137115599 | 11.17263 | 21.62726 | 18.54775 | 7.060823 | 0.785412 | -0.1635 | 0.328679 |
| X | 146722600 | 146724999 | 51.64106 | 83.0777 | 73.53799 | 31.21399 | 0.528734 | 1.865887 | 0.32868 |
| Y | 5330200 | 5331999 | 10.30964 | 19.93239 | 11.42267 | 4.373306 | 1.352562 | -0.17974 | 0.328681 |
| 13 | 91264600 | 91265799 | 18.35345 | 32.8867 | 27.35908 | 11.98684 | 0.664745 | 0.437911 | 0.328683 |
| 4 | 46571200 | 46572199 | 16.59693 | 30.47501 | 24.55168 | 10.44646 | 0.738513 | 0.281435 | 0.328688 |
| 7 | 23445000 | 23447599 | 50.42879 | 81.38123 | 73.32241 | 31.25686 | 0.508581 | 1.769132 | 0.328695 |
| 7 | 139979200 | 139981199 | 20.42691 | 34.30457 | 62.36324 | 27.23239 | -0.47831 | 1.137762 | 0.328695 |
| 8 | 22211400 | 22214599 | 46.22687 | 74.93243 | 65.70306 | 28.13425 | 0.554176 | 1.648038 | 0.328712 |
| 11 | 12087000 | 12089999 | 33.636 | 56.62074 | 44.69365 | 19.53227 | 0.721067 | 1.141419 | 0.328727 |
| 2 | 51891000 | 51892599 | 11.62572 | 20.48669 | 40.92448 | 18.11361 | -0.60398 | 0.554849 | 0.32873 |
| 15 | 44650400 | 44651599 | 7.748317 | 14.35935 | 31.54668 | 13.25512 | -0.68518 | 0.177682 | 0.328732 |
| 2 | 15422000 | 15424999 | 42.23965 | 69.15915 | 58.94238 | 25.35817 | 0.60592 | 1.481903 | 0.328755 |
| 6 | 85854400 | 85855599 | 14.53293 | 27.29413 | 22.31555 | 9.176707 | 0.754553 | 0.129447 | 0.328763 |
| 14 | 104220800 | 104222599 | 10.5619 | 19.06148 | 37.41745 | 16.27567 | -0.55945 | 0.36199 | 0.328769 |
| 13 | 3285000 | 3288199 | 36.38622 | 59.98153 | 53.80812 | 23.21597 | 0.535914 | 1.378959 | 0.328779 |
| 5 | 132829600 | 132834599 | 59.57033 | 95.42968 | 87.61416 | 36.9545 | 0.468847 | 2.009284 | 0.328783 |
| 7 | 13935800 | 13936999 | 19.73708 | 35.09773 | 28.30445 | 12.48089 | 0.703206 | 0.495852 | 0.328783 |
| 17 | 11648600 | 11652399 | 54.10427 | 86.83981 | 83.09462 | 35.15071 | 0.41253 | 1.941638 | 0.328799 |
| 7 | 30002800 | 30006399 | 16.82965 | 28.73761 | 54.77316 | 24.01399 | -0.54416 | 0.970097 | 0.328809 |
| 6 | 43939400 | 43939999 | 5.978544 | 11.54496 | 27.13815 | 10.44834 | -0.67137 | -0.15789 | 0.32881 |
| 7 | 95972400 | 95975799 | 56.45932 | 89.23303 | 149.2238 | 61.98919 | -0.4144 | 2.41627 | 0.328814 |
| 18 | 80608000 | 80610999 | 23.74033 | 38.86869 | 78.36275 | 33.71267 | -0.63275 | 1.453731 | 0.328815 |
| 1 | 136636800 | 136639799 | 21.52484 | 35.64996 | 69.80013 | 30.20514 | -0.58552 | 1.301118 | 0.328849 |
| 3 | 137929800 | 137933999 | 31.25737 | 50.5716 | 89.98267 | 38.47719 | -0.46678 | 1.675103 | 0.328853 |
| 13 | 86083200 | 86083999 | 13.05897 | 22.85826 | 44.13469 | 19.4754 | -0.55679 | 0.664962 | 0.328877 |
| 3 | 26655400 | 26658999 | 31.92778 | 51.55064 | 93.62047 | 39.93582 | -0.49955 | 1.734013 | 0.328895 |
| 12 | 46512600 | 46514599 | 24.71209 | 43.10883 | 34.28056 | 15.08915 | 0.716559 | 0.75991 | 0.328902 |
| 3 | 73062600 | 73063599 | 15.52446 | 29.48946 | 18.21053 | 7.323792 | 1.182076 | 0.052964 | 0.32891 |
| 15 | 93416400 | 93417999 | 8.192268 | 15.12167 | 34.14671 | 14.42564 | -0.73106 | 0.206624 | 0.328947 |
| X | 112112600 | 112114199 | 18.18999 | 33.5685 | 22.85447 | 9.658017 | 0.986773 | 0.263393 | 0.328967 |
| 8 | 21213000 | 21213599 | 9.994624 | 19.50795 | 14.2621 | 5.056364 | 1.105033 | -0.43169 | 0.328974 |
| 19 | 9653800 | 9656799 | 47.47624 | 76.75768 | 70.35473 | 30.06309 | 0.486884 | 1.710471 | 0.328994 |
| 18 | 19665600 | 19669199 | 25.37372 | 44.25444 | 33.58409 | 14.78079 | 0.783534 | 0.763358 | 0.328996 |
| 2 | 110002800 | 110005599 | 19.66576 | 32.89496 | 65.01742 | 28.32011 | -0.59853 | 1.179511 | 0.329043 |
| 7 | 19009400 | 19013999 | 29.0168 | 46.94359 | 90.13276 | 38.53957 | -0.57636 | 1.672128 | 0.329044 |
| 10 | 86629200 | 86633399 | 38.06303 | 60.85769 | 116.5143 | 49.00624 | -0.59252 | 2.054669 | 0.329064 |
| 8 | 119977000 | 119978599 | 13.66454 | 23.97805 | 44.56655 | 19.70751 | -0.50227 | 0.627567 | 0.329081 |
| 17 | 26788800 | 26791799 | 18.06779 | 30.33374 | 62.74855 | 27.39553 | -0.66393 | 1.132859 | 0.329098 |

| | | | | | | | | | |
|---|---|---|---|---|---|---|---|---|---|
| 1 | 100912600 | 100916199 | 46.83548 | 76.09093 | 58.8786 | 25.25409 | 0.737093 | 1.602962 | 0.329101 |
| 10 | 97596400 | 97597399 | 13.6134 | 25.37797 | 22.73021 | 9.46553 | 0.609012 | 0.166506 | 0.329103 |
| 16 | 81460800 | 81461999 | 13.2055 | 23.15427 | 43.56105 | 19.25004 | -0.51956 | 0.636904 | 0.32911 |
| 10 | 31124400 | 31127599 | 33.70879 | 54.37243 | 96.06031 | 40.92433 | -0.46157 | 1.766771 | 0.329127 |
| 3 | 41377200 | 41379799 | 14.62299 | 25.40506 | 49.51001 | 21.7368 | -0.57141 | 0.78898 | 0.329128 |
| 14 | 13305600 | 13306599 | 6.594382 | 12.58489 | 28.93451 | 11.52249 | -0.68569 | -0.0355 | 0.329128 |
| X | 25421000 | 25428399 | 126.0193 | 194.8513 | 147.3142 | 61.19087 | 0.705019 | 2.859882 | 0.329138 |
| 2 | 169045600 | 169053399 | 89.03242 | 136.9702 | 217.5235 | 90.33668 | -0.37027 | 2.944979 | 0.329159 |
| X | 90649400 | 90652799 | 43.53686 | 71.26108 | 58.3434 | 25.0973 | 0.663538 | 1.490137 | 0.32916 |
| 13 | 52355200 | 52363199 | 69.21425 | 108.1332 | 176.3766 | 73.18034 | -0.39154 | 2.656133 | 0.329167 |
| 11 | 77667600 | 77670399 | 14.93795 | 25.62799 | 53.66245 | 23.53596 | -0.67787 | 0.920148 | 0.329189 |
| 16 | 91926600 | 91937199 | 73.02765 | 113.447 | 188.753 | 78.37534 | -0.42652 | 2.757222 | 0.329191 |
| X | 125101600 | 125103999 | 35.94749 | 60.36245 | 40.4479 | 17.66123 | 0.956542 | 1.168875 | 0.329195 |
| 7 | 15363200 | 15368799 | 120.3729 | 185.1764 | 164.0392 | 68.1244 | 0.47105 | 2.953232 | 0.329204 |
| 17 | 39753400 | 39754799 | 22.17394 | 39.31965 | 26.90056 | 11.89019 | 0.936709 | 0.525646 | 0.329216 |
| 5 | 52845600 | 52848399 | 28.51367 | 46.20315 | 87.37116 | 37.40264 | -0.55227 | 1.63802 | 0.329223 |
| 10 | 67577600 | 67579599 | 11.64094 | 20.47446 | 41.46345 | 18.35915 | -0.62467 | 0.577321 | 0.329234 |
| 12 | 114546400 | 114556199 | 143.2096 | 219.4463 | 166.8432 | 69.25172 | 0.687084 | 3.036445 | 0.32924 |
| 10 | 67756400 | 67758799 | 17.8833 | 30.58308 | 55.94881 | 24.53242 | -0.48511 | 0.959573 | 0.329243 |
| 17 | 38851800 | 38853799 | 51.20274 | 82.2947 | 76.30429 | 32.34381 | 0.46024 | 1.89423 | 0.329257 |
| 6 | 74412400 | 74413199 | 10.46527 | 20.2452 | 18.55603 | 7.084876 | 0.686134 | -0.16238 | 0.329258 |
| 7 | 26069000 | 26075999 | 109.0566 | 170.9718 | 120.5849 | 50.01397 | 0.82045 | 2.610236 | 0.329262 |
| 11 | 74026400 | 74034799 | 150.4074 | 227.7482 | 209.7254 | 86.77151 | 0.397082 | 3.278997 | 0.329264 |
| 8 | 51449000 | 51450999 | 18.02381 | 31.78697 | 28.33981 | 12.54148 | 0.554011 | 0.559886 | 0.329271 |
| 8 | 111457800 | 111459399 | 9.334079 | 16.96702 | 34.43198 | 14.84312 | -0.59809 | 0.307217 | 0.32928 |
| 15 | 98941600 | 98942999 | 9.334079 | 16.96702 | 34.43198 | 14.84312 | -0.59809 | 0.307217 | 0.32928 |
| 6 | 71504000 | 71505799 | 13.21897 | 23.09458 | 45.21643 | 19.92165 | -0.57688 | 0.697664 | 0.329285 |
| X | 147809600 | 147812999 | 26.53221 | 46.24915 | 32.96956 | 14.50524 | 0.873285 | 0.775904 | 0.32929 |
| 7 | 121443200 | 121444799 | 10.55018 | 19.07423 | 37.98756 | 16.48693 | -0.57817 | 0.352311 | 0.329294 |
| X | 66957400 | 66959199 | 19.16979 | 35.32947 | 22.15912 | 9.381201 | 1.102327 | 0.27656 | 0.3293 |
| 18 | 54803400 | 54805399 | 28.74547 | 49.16692 | 40.24437 | 17.64657 | 0.670575 | 0.980941 | 0.329315 |
| 2 | 45953600 | 45955599 | 14.30255 | 24.99851 | 46.24194 | 20.38012 | -0.49549 | 0.697601 | 0.329318 |
| 1 | 166206400 | 166207799 | 10.24527 | 18.10598 | 41.44244 | 18.33234 | -0.79758 | 0.514096 | 0.329334 |
| 9 | 112063200 | 112065399 | 16.35092 | 27.88783 | 55.29701 | 24.24139 | -0.60127 | 0.978265 | 0.329345 |
| 2 | 23450400 | 23453199 | 27.15468 | 47.60059 | 29.00599 | 12.81531 | 1.098621 | 0.693021 | 0.329366 |
| 14 | 13331800 | 13340199 | 108.1179 | 167.4306 | 154.4756 | 64.16503 | 0.419707 | 2.823224 | 0.329375 |
| 7 | 6578400 | 6582799 | 78.02527 | 123.1216 | 116.4826 | 48.34919 | 0.404424 | 2.462463 | 0.329389 |
| 14 | 11118400 | 11120999 | 30.10166 | 51.90675 | 36.31109 | 15.92894 | 0.898276 | 0.91271 | 0.329391 |
| 8 | 85157400 | 85158799 | 7.531876 | 14.19023 | 29.68312 | 12.14138 | -0.5841 | 0.066932 | 0.329419 |
| 2 | 156161200 | 156162999 | 17.20948 | 30.49011 | 27.27113 | 12.0575 | 0.551833 | 0.503113 | 0.329435 |
| X | 33729800 | 33734399 | 77.30197 | 123.3218 | 88.26619 | 37.00876 | 0.82089 | 2.154777 | 0.329441 |
| 11 | 4186200 | 4191999 | 55.35211 | 86.96511 | 162.7481 | 67.50708 | -0.58315 | 2.545104 | 0.329472 |
| 6 | 121398000 | 121404399 | 34.52746 | 55.62568 | 98.80064 | 42.02405 | -0.47134 | 1.804989 | 0.329481 |
| 7 | 4956200 | 4957399 | 8.744409 | 15.8759 | 34.45645 | 14.87619 | -0.69546 | 0.311198 | 0.329488 |
| 3 | 11900600 | 11902599 | 21.73801 | 38.01192 | 33.79183 | 14.89879 | 0.556512 | 0.707706 | 0.329529 |
| 7 | 62350600 | 62371399 | 708.365 | 943.2515 | 778.3359 | 325.1636 | 0.374361 | 5.347697 | 0.329533 |
| 7 | 86226000 | 86226999 | 13.12107 | 24.69702 | 22.4101 | 9.181177 | 0.609399 | 0.102715 | 0.329538 |

| | | | | | | | | | |
|---|---|---|---|---|---|---|---|---|---|
| 1 | 141603600 | 141605399 | 17.20717 | 31.29007 | 24.79599 | 10.68417 | 0.749666 | 0.338685 | 0.329566 |
| 7 | 140199400 | 140208799 | 120.9632 | 187.0611 | 151.6414 | 62.98826 | 0.604649 | 2.854536 | 0.32958 |
| 1 | 133375400 | 133377399 | 12.38956 | 21.81554 | 41.92266 | 18.55833 | -0.54901 | 0.569074 | 0.329585 |
| 18 | 76253400 | 76255999 | 26.78055 | 45.4839 | 42.66284 | 18.69517 | 0.473527 | 1.040928 | 0.329592 |
| 7 | 16181400 | 16181999 | 5.266366 | 10.20042 | 25.52151 | 9.688393 | -0.7404 | -0.2179 | 0.329597 |
| 14 | 113596800 | 113598399 | 22.5027 | 39.40357 | 29.26879 | 12.91968 | 0.81451 | 0.684538 | 0.329603 |
| 10 | 42588400 | 42592199 | 18.54298 | 31.70208 | 58.16143 | 25.50201 | -0.48932 | 0.967638 | 0.329603 |
| 1 | 193166800 | 193168799 | 9.279863 | 16.97874 | 34.50352 | 14.75242 | -0.5926 | 0.267114 | 0.329605 |
| 13 | 65898600 | 65901799 | 32.54198 | 55.3818 | 40.8024 | 17.88402 | 0.821001 | 1.043562 | 0.329607 |
| 10 | 55614400 | 55616799 | 21.9318 | 38.56845 | 30.94995 | 13.70441 | 0.703624 | 0.619301 | 0.329617 |
| 3 | 110995000 | 110995599 | 5.505774 | 10.6999 | 25.8167 | 9.609507 | -0.66089 | -0.27848 | 0.329632 |
| 4 | 35997400 | 35998199 | 10.80715 | 20.69531 | 20.07066 | 7.923337 | 0.560735 | -0.04615 | 0.32964 |
| 1 | 128743200 | 128744599 | 7.329055 | 13.92209 | 32.3128 | 12.99495 | -0.71243 | 0.010846 | 0.329641 |
| X | 47121800 | 47123999 | 29.22561 | 50.06252 | 37.33824 | 16.37423 | 0.804825 | 0.971608 | 0.329645 |
| 6 | 73195800 | 73198999 | 24.08122 | 39.77679 | 72.89008 | 31.48377 | -0.49074 | 1.340824 | 0.329653 |
| X | 148484600 | 148489599 | 85.32274 | 135.0308 | 112.0958 | 46.59718 | 0.596094 | 2.398954 | 0.329658 |
| 8 | 53555200 | 53558399 | 54.03165 | 86.73313 | 82.10552 | 34.73832 | 0.42818 | 1.936801 | 0.329661 |
| 4 | 75076200 | 75079799 | 61.3645 | 98.39694 | 84.20717 | 35.56863 | 0.571646 | 1.981213 | 0.329683 |
| 13 | 81866600 | 81867599 | 5.814502 | 11.28744 | 30.29746 | 11.37811 | -0.82724 | -0.23738 | 0.329685 |
| 7 | 114025800 | 114028599 | 30.78025 | 51.59811 | 48.0668 | 20.96551 | 0.482959 | 1.175744 | 0.329687 |
| 4 | 53931400 | 53935599 | 30.0682 | 48.84478 | 85.99203 | 36.88057 | -0.44603 | 1.588408 | 0.329714 |
| 4 | 13622000 | 13624599 | 26.88556 | 46.95745 | 31.22953 | 13.75672 | 0.973113 | 0.748553 | 0.329716 |
| 10 | 107329200 | 107330999 | 10.51612 | 18.67344 | 38.35595 | 16.93372 | -0.63889 | 0.46634 | 0.329734 |
| 12 | 27885000 | 27886999 | 16.7855 | 30.8113 | 24.54466 | 10.44806 | 0.754235 | 0.284815 | 0.329755 |
| 13 | 119132400 | 119134399 | 18.29716 | 31.28596 | 60.8658 | 26.68808 | -0.57354 | 0.970419 | 0.329759 |
| 17 | 19155000 | 19159199 | 111.3941 | 171.3861 | 158.6331 | 65.88047 | 0.407935 | 2.945182 | 0.329764 |
| X | 32107400 | 32113799 | 293.1571 | 426.1452 | 286.6974 | 118.1776 | 0.795604 | 4.007962 | 0.329776 |
| 14 | 45075800 | 45078999 | 20.89633 | 35.21984 | 64.14207 | 28.04242 | -0.48075 | 1.109155 | 0.32978 |
| 12 | 17295400 | 17311799 | 159.3749 | 235.5237 | 367.3724 | 151.2923 | -0.39431 | 3.737931 | 0.329782 |
| 8 | 47426400 | 47431599 | 60.48258 | 96.99742 | 84.42485 | 35.66875 | 0.547572 | 1.976754 | 0.329784 |
| 4 | 156561400 | 156562199 | 15.31018 | 27.69604 | 25.16328 | 10.91322 | 0.547839 | 0.355368 | 0.329804 |
| X | 108418400 | 108422199 | 55.16999 | 88.88737 | 73.99597 | 31.47636 | 0.61974 | 1.825486 | 0.329809 |
| 5 | 10608000 | 10610999 | 37.80374 | 61.8465 | 59.09044 | 25.41406 | 0.440835 | 1.482988 | 0.329832 |
| 11 | 73689600 | 73691199 | 19.55243 | 34.77881 | 28.30904 | 12.48006 | 0.690078 | 0.492828 | 0.329835 |
| 7 | 112397400 | 112399799 | 12.53582 | 22.10995 | 41.90254 | 18.54324 | -0.52817 | 0.55137 | 0.32984 |
| 10 | 13614000 | 13615999 | 10.85074 | 19.22646 | 38.88488 | 17.19195 | -0.61818 | 0.487593 | 0.32984 |
| 4 | 115761200 | 115763399 | 11.83959 | 20.70087 | 44.70309 | 19.70772 | -0.71731 | 0.6776 | 0.329864 |
| 6 | 87615200 | 87617599 | 13.99388 | 24.4188 | 46.27393 | 20.37204 | -0.53034 | 0.719965 | 0.329869 |
| X | 14267200 | 14268599 | 22.55061 | 39.63975 | 30.50503 | 13.50856 | 0.763626 | 0.627866 | 0.329871 |
| 1 | 186165400 | 186165999 | 7.028233 | 13.43455 | 28.42772 | 11.28076 | -0.56222 | -0.0427 | 0.329875 |
| 11 | 27652200 | 27656599 | 55.17518 | 89.17279 | 66.7093 | 28.49208 | 0.778645 | 1.731461 | 0.329875 |
| 1 | 88721600 | 88728599 | 61.36823 | 96.41289 | 162.5189 | 67.4116 | -0.43263 | 2.548719 | 0.329875 |
| 7 | 30987600 | 30993199 | 91.23524 | 142.9135 | 134.4462 | 55.76786 | 0.404429 | 2.615447 | 0.329884 |
| 10 | 40405600 | 40410399 | 63.83493 | 101.9364 | 94.44823 | 39.65514 | 0.450613 | 2.112296 | 0.329892 |
| 14 | 9871200 | 9874199 | 15.10058 | 26.24411 | 48.97559 | 21.50698 | -0.50908 | 0.786123 | 0.329898 |
| 9 | 102632400 | 102637199 | 26.94349 | 43.99882 | 80.95659 | 34.79525 | -0.50393 | 1.499251 | 0.329905 |
| X | 33198800 | 33208399 | 38.03664 | 63.18955 | 46.03802 | 19.96436 | 0.836431 | 1.292168 | 0.329929 |

| | | | | | | | | | |
|---|---|---|---|---|---|---|---|---|---|
| 7 | 37244600 | 37246999 | 23.01429 | 39.8143 | 37.26429 | 16.3487 | 0.480892 | 0.852953 | 0.329938 |
| 7 | 33007000 | 33016199 | 206.899 | 306.4819 | 273.174 | 112.5025 | 0.415808 | 3.702662 | 0.329949 |
| 2 | 91852800 | 91854799 | 10.49099 | 18.69594 | 37.84749 | 16.65866 | -0.61502 | 0.441983 | 0.329974 |
| 7 | 127830400 | 127831599 | 10.49099 | 18.69594 | 37.84749 | 16.65866 | -0.61502 | 0.441983 | 0.329974 |
| 7 | 42387000 | 42393199 | 108.7312 | 167.948 | 159.237 | 66.14369 | 0.377573 | 2.870894 | 0.329981 |
| 1 | 155443200 | 155444199 | 11.87049 | 22.44807 | 21.04932 | 8.545836 | 0.573665 | 0.057976 | 0.329984 |
| 10 | 15299400 | 15300999 | 15.71713 | 28.45468 | 26.15691 | 11.33295 | 0.53207 | 0.356729 | 0.329985 |
| 5 | 60909400 | 60910399 | 12.13549 | 23.03582 | 21.50367 | 8.661008 | 0.58982 | 0.035203 | 0.329987 |
| 13 | 81858600 | 81860399 | 15.58392 | 26.57369 | 55.2557 | 24.22285 | -0.66951 | 0.977845 | 0.329989 |
| 12 | 63214800 | 63215999 | 12.12608 | 22.92663 | 20.83396 | 8.462127 | 0.617316 | 0.058636 | 0.329998 |
| 17 | 3411400 | 3413799 | 13.2081 | 23.10363 | 44.66473 | 19.698 | -0.55868 | 0.677054 | 0.330026 |
| X | 67803200 | 67805999 | 33.19293 | 55.96365 | 44.7245 | 19.55516 | 0.703466 | 1.128944 | 0.33003 |
| X | 20598000 | 20600399 | 26.11773 | 46.07943 | 26.26719 | 11.62335 | 1.196032 | 0.60263 | 0.330045 |
| 1 | 102290000 | 102293399 | 64.57384 | 102.9255 | 98.14254 | 41.0996 | 0.406405 | 2.172437 | 0.33006 |
| 14 | 10956200 | 10957599 | 12.20474 | 21.6139 | 41.35889 | 18.28775 | -0.53968 | 0.503334 | 0.3301 |
| 9 | 98627000 | 98630599 | 35.84499 | 57.63918 | 103.0569 | 43.71439 | -0.4843 | 1.868116 | 0.33012 |
| 13 | 57306200 | 57307799 | 24.85853 | 43.22033 | 35.90655 | 15.77186 | 0.653228 | 0.807056 | 0.330128 |
| X | 156572800 | 156574999 | 28.48943 | 49.27936 | 36.39734 | 15.96829 | 0.821144 | 0.879317 | 0.33014 |
| X | 57683400 | 57685199 | 24.51402 | 42.85604 | 33.09307 | 14.5878 | 0.758848 | 0.726145 | 0.330143 |
| 10 | 82957000 | 82958599 | 12.5998 | 22.13981 | 42.49301 | 18.81221 | -0.54802 | 0.598353 | 0.33015 |
| 18 | 25720600 | 25728199 | 49.03382 | 77.81087 | 138.6676 | 57.76338 | -0.50093 | 2.304037 | 0.330155 |
| 2 | 37121800 | 37122399 | 7.167962 | 13.69142 | 30.17658 | 11.99504 | -0.62323 | -0.03301 | 0.33016 |
| 10 | 30076000 | 30076799 | 7.167962 | 13.69142 | 30.17658 | 11.99504 | -0.62323 | -0.03301 | 0.33016 |
| 10 | 104215600 | 104216799 | 7.167962 | 13.69142 | 30.17658 | 11.99504 | -0.62323 | -0.03301 | 0.33016 |
| X | 166442000 | 166442799 | 11.60734 | 22.53941 | 17.49424 | 6.554398 | 0.949711 | -0.21318 | 0.330174 |
| 8 | 22121000 | 22123399 | 23.44828 | 41.10323 | 30.74252 | 13.58246 | 0.804262 | 0.672657 | 0.330177 |
| 11 | 99457000 | 99457599 | 4.648822 | 9.061444 | 21.97982 | 7.947086 | -0.62926 | -0.40613 | 0.330179 |
| 5 | 119521600 | 119528199 | 45.98811 | 73.31536 | 128.8426 | 53.97028 | -0.47392 | 2.155296 | 0.330187 |
| 3 | 97968800 | 97971599 | 10.79259 | 19.22226 | 38.33399 | 16.88105 | -0.59414 | 0.448043 | 0.330198 |
| 1 | 99329600 | 99330599 | 11.93732 | 23.0414 | 18.95616 | 7.313575 | 0.826592 | -0.11035 | 0.330204 |
| 9 | 88106400 | 88109999 | 24.27716 | 40.18097 | 72.8437 | 31.50735 | -0.47499 | 1.320528 | 0.330205 |
| 15 | 9544600 | 9545799 | 9.428592 | 17.43089 | 35.29429 | 14.87885 | -0.57255 | 0.212535 | 0.330207 |
| 19 | 7801000 | 7804199 | 38.96041 | 63.79789 | 59.79927 | 25.72817 | 0.469137 | 1.473702 | 0.33021 |
| 14 | 106235600 | 106236999 | 14.46398 | 26.74771 | 22.51429 | 9.487496 | 0.685018 | 0.218396 | 0.330215 |
| 1 | 31475600 | 31477199 | 19.02276 | 32.42971 | 62.62749 | 27.45394 | -0.56382 | 0.999027 | 0.330222 |
| 11 | 41701800 | 41702599 | 4.644257 | 9.06713 | 23.05463 | 8.138478 | -0.66592 | -0.48993 | 0.330234 |
| 2 | 73630400 | 73631599 | 11.8579 | 22.38037 | 20.90671 | 8.523185 | 0.573336 | 0.068769 | 0.330245 |
| 14 | 74506400 | 74508399 | 13.03505 | 24.89004 | 20.34381 | 8.092083 | 0.795363 | 0.00049 | 0.330256 |
| 17 | 8478600 | 8480999 | 26.67539 | 43.5051 | 80.99618 | 34.79597 | -0.52208 | 1.520311 | 0.33026 |
| X | 33056800 | 33060999 | 38.5479 | 65.92736 | 30.4518 | 13.35257 | 1.492307 | 1.012578 | 0.330264 |
| 8 | 55905400 | 55908599 | 42.0589 | 68.4525 | 63.75415 | 27.36816 | 0.47233 | 1.567555 | 0.33027 |
| 5 | 144328000 | 144329999 | 17.16559 | 29.22801 | 55.82262 | 24.46849 | -0.54779 | 0.998728 | 0.330281 |
| 19 | 53129800 | 53131799 | 18.4395 | 30.82837 | 63.66278 | 27.7213 | -0.6613 | 1.182755 | 0.330303 |
| 1 | 8705000 | 8707199 | 18.98392 | 33.86163 | 27.78914 | 12.22217 | 0.680512 | 0.470487 | 0.330321 |
| 9 | 19630600 | 19634999 | 59.76672 | 95.95453 | 81.82493 | 34.628 | 0.578886 | 1.940904 | 0.330322 |
| 11 | 3410800 | 3412599 | 13.45075 | 25.0283 | 22.74704 | 9.502329 | 0.58449 | 0.176791 | 0.330327 |
| 2 | 150152000 | 150154999 | 28.28093 | 45.98046 | 86.56048 | 37.14428 | -0.54141 | 1.568114 | 0.330336 |

| | | | | | | | | | |
|---|---|---|---|---|---|---|---|---|---|
| 13 | 66037600 | 66040199 | 52.66642 | 84.58367 | 81.12285 | 34.34851 | 0.410249 | 1.920881 | 0.330359 |
| X | 45319400 | 45322199 | 35.53088 | 59.26409 | 49.23915 | 21.40907 | 0.647707 | 1.2453 | 0.330361 |
| 1 | 44328000 | 44336599 | 170.8539 | 256.2599 | 237.3466 | 97.93 | 0.376849 | 3.471034 | 0.330363 |
| 5 | 16915400 | 16916599 | 15.78968 | 28.96939 | 24.53496 | 10.4508 | 0.66551 | 0.27833 | 0.330372 |
| 4 | 46543000 | 46544199 | 8.185697 | 15.19375 | 31.56018 | 13.23248 | -0.60287 | 0.172043 | 0.330374 |
| 2 | 37203000 | 37204599 | 26.62375 | 45.7724 | 36.14236 | 15.85348 | 0.723408 | 0.922756 | 0.330398 |
| 17 | 19435600 | 19436599 | 16.1403 | 29.8143 | 23.24982 | 9.812375 | 0.79391 | 0.244169 | 0.330399 |
| 13 | 27890200 | 27890999 | 13.04132 | 25.30948 | 16.81771 | 6.325867 | 1.166763 | -0.18489 | 0.330401 |
| 14 | 92317000 | 92320199 | 22.94764 | 40.3735 | 29.64007 | 13.12319 | 0.831406 | 0.616752 | 0.330405 |
| 6 | 130794600 | 130811999 | 352.1369 | 503.4746 | 370.0383 | 152.8812 | 0.644539 | 4.25513 | 0.330406 |
| 15 | 28564000 | 28566799 | 28.22965 | 48.86392 | 35.18939 | 15.43866 | 0.857464 | 0.868464 | 0.330408 |
| 7 | 40768800 | 40775199 | 130.452 | 199.2856 | 185.0697 | 76.76271 | 0.395265 | 3.094323 | 0.330416 |
| 2 | 108042200 | 108042999 | 15.29091 | 27.71803 | 25.12888 | 10.87035 | 0.55326 | 0.343277 | 0.330435 |
| 19 | 33263600 | 33264199 | 6.905602 | 13.14293 | 28.27224 | 11.3239 | -0.59719 | -0.01205 | 0.330439 |
| 8 | 91086600 | 91088399 | 17.71907 | 30.43172 | 55.43976 | 24.31732 | -0.47811 | 0.920093 | 0.330448 |
| 3 | 121085000 | 121085799 | 5.553432 | 10.74022 | 26.62807 | 10.18002 | -0.73759 | -0.18817 | 0.330448 |
| 14 | 84075800 | 84083999 | 107.3256 | 166.0835 | 156.0169 | 64.80552 | 0.39296 | 2.836243 | 0.330452 |
| 13 | 62084000 | 62086999 | 79.78087 | 126.3372 | 113.6326 | 47.25058 | 0.481293 | 2.38026 | 0.330464 |
| 13 | 67385800 | 67396199 | 174.8079 | 262.2417 | 237.2475 | 97.89421 | 0.410969 | 3.467904 | 0.330472 |
| 17 | 59004800 | 59006599 | 20.02294 | 35.66231 | 27.6983 | 12.19737 | 0.758378 | 0.486554 | 0.33048 |
| 3 | 115185200 | 115187799 | 15.65403 | 27.23706 | 53.8414 | 23.6607 | -0.59167 | 0.780722 | 0.330488 |
| 13 | 61663200 | 61664399 | 26.13757 | 44.86344 | 40.41043 | 17.72439 | 0.53388 | 0.941598 | 0.330495 |
| 14 | 104105400 | 104107599 | 22.91314 | 40.48344 | 27.37723 | 12.11109 | 0.951346 | 0.567731 | 0.330518 |
| X | 142828400 | 142830999 | 35.96688 | 59.66194 | 52.60835 | 22.79007 | 0.561865 | 1.308682 | 0.33053 |
| 2 | 171881400 | 171890999 | 70.17135 | 108.7989 | 206.0462 | 85.58071 | -0.61532 | 2.791501 | 0.330551 |
| 7 | 139266600 | 139272399 | 31.60102 | 51.05877 | 91.95935 | 39.26267 | -0.48639 | 1.712658 | 0.330553 |
| 5 | 101805400 | 101807799 | 16.96683 | 28.53889 | 67.12296 | 29.32759 | -0.8484 | 1.115868 | 0.330578 |
| 4 | 113740000 | 113741399 | 21.46334 | 37.64996 | 32.2411 | 14.25293 | 0.610269 | 0.656319 | 0.330584 |
| 10 | 65002800 | 65003199 | 10.35946 | 20.20006 | 15.43493 | 5.553668 | 1.024022 | -0.37215 | 0.330587 |
| 1 | 147199800 | 147202599 | 24.52246 | 42.57221 | 35.92336 | 15.7644 | 0.630566 | 0.824689 | 0.330591 |
| 3 | 111808200 | 111811399 | 34.18631 | 57.9763 | 40.72049 | 17.83605 | 0.889172 | 1.077909 | 0.330603 |
| 15 | 88476000 | 88487799 | 140.8075 | 209.8166 | 333.6481 | 137.4904 | -0.41102 | 3.591496 | 0.330609 |
| 13 | 79804400 | 79805799 | 13.8124 | 25.68057 | 23.24456 | 9.723401 | 0.588857 | 0.186393 | 0.33064 |
| 11 | 15639600 | 15640799 | 7.806623 | 14.76246 | 30.36295 | 12.32797 | -0.5511 | 0.044539 | 0.330645 |
| 11 | 15453000 | 15454999 | 19.23283 | 35.09733 | 23.63452 | 10.13883 | 0.987277 | 0.333135 | 0.330645 |
| 16 | 55676200 | 55678599 | 24.29823 | 41.79456 | 38.83138 | 17.03324 | 0.489787 | 0.911063 | 0.330649 |
| 14 | 42640800 | 42646199 | 66.75483 | 107.0282 | 80.7461 | 34.10085 | 0.752976 | 1.991262 | 0.330683 |
| 13 | 98190200 | 98191199 | 15.3 | 27.72808 | 25.31458 | 10.95386 | 0.543286 | 0.347071 | 0.330689 |
| 7 | 3096000 | 3149799 | 1010.506 | 1309.228 | 1036.467 | 434.3386 | 0.394739 | 5.791318 | 0.330697 |
| 9 | 100067200 | 100068399 | 15.15705 | 28.23985 | 23.23916 | 9.686635 | 0.729718 | 0.186182 | 0.330698 |
| 7 | 13654600 | 13657599 | 54.25101 | 87.68495 | 67.69885 | 28.91719 | 0.733361 | 1.729223 | 0.330742 |
| X | 23453200 | 23455199 | 24.60722 | 43.37006 | 27.92659 | 12.36017 | 1.020607 | 0.606103 | 0.330746 |
| 3 | 49557000 | 49559399 | 27.43537 | 47.27697 | 38.03902 | 16.6866 | 0.697002 | 0.907618 | 0.330755 |
| 15 | 29038600 | 29039799 | 5.956767 | 11.56571 | 27.01848 | 10.13182 | -0.62568 | -0.24596 | 0.330758 |
| 9 | 117452400 | 117453799 | 20.4392 | 36.16497 | 29.37017 | 12.98895 | 0.689698 | 0.542127 | 0.330781 |
| 13 | 13763200 | 13765599 | 13.18072 | 23.04818 | 47.82842 | 21.08741 | -0.66067 | 0.684493 | 0.330783 |
| 14 | 102616800 | 102617799 | 16.71188 | 30.3266 | 22.20462 | 9.592385 | 0.861123 | 0.34314 | 0.330792 |

| | | | | | | | | | |
|---|---|---|---|---|---|---|---|---|---|
| 7 | 9321800 | 9323799 | 30.56947 | 51.93104 | 38.84537 | 17.02352 | 0.798774 | 1.049277 | 0.33081 |
| 7 | 78865000 | 78869399 | 39.2479 | 63.04558 | 107.7727 | 45.64597 | -0.42157 | 1.907017 | 0.33082 |
| 8 | 101123200 | 101126599 | 75.60187 | 119.8022 | 112.9431 | 46.98123 | 0.41434 | 2.360219 | 0.330822 |
| 19 | 40258200 | 40261599 | 23.1317 | 38.23395 | 71.01644 | 30.68864 | -0.51003 | 1.330068 | 0.330827 |
| 2 | 65025400 | 65026799 | 10.59698 | 19.16113 | 36.94038 | 16.03007 | -0.53109 | 0.352361 | 0.330835 |
| 10 | 20748200 | 20749999 | 10.59698 | 19.16113 | 36.94038 | 16.03007 | -0.53109 | 0.352361 | 0.330835 |
| 13 | 75107400 | 75108799 | 10.59698 | 19.16113 | 36.94038 | 16.03007 | -0.53109 | 0.352361 | 0.330835 |
| 12 | 115089600 | 115092599 | 75.29649 | 119.5128 | 106.3305 | 44.30275 | 0.500049 | 2.313457 | 0.330841 |
| 12 | 17866600 | 17871399 | 50.92512 | 82.62943 | 62.11159 | 26.61568 | 0.777111 | 1.636971 | 0.330842 |
| 10 | 118195400 | 118196399 | 13.81019 | 26.28037 | 20.19368 | 8.090511 | 0.874606 | 0.030031 | 0.330864 |
| 16 | 90395600 | 90398599 | 33.15462 | 55.61378 | 46.95368 | 20.48823 | 0.62433 | 1.174537 | 0.33087 |
| 2 | 15122400 | 15125599 | 21.3958 | 37.91255 | 28.11702 | 12.43114 | 0.820658 | 0.530072 | 0.330874 |
| 4 | 139669200 | 139676199 | 77.02605 | 118.99 | 200.3219 | 83.20945 | -0.44974 | 2.861411 | 0.330878 |
| 1 | 171533200 | 171534799 | 13.41286 | 24.95104 | 23.36058 | 9.763107 | 0.541673 | 0.180848 | 0.330883 |
| 16 | 97904400 | 97908999 | 241.718 | 357.9174 | 232.9006 | 95.91582 | 0.868928 | 3.711383 | 0.330895 |
| X | 122284200 | 122285199 | 21.96302 | 38.85186 | 25.32866 | 11.20213 | 1.004837 | 0.547575 | 0.330899 |
| 13 | 64710400 | 64715199 | 60.85851 | 97.54166 | 85.7673 | 36.20282 | 0.53188 | 1.994725 | 0.330914 |
| 17 | 22596400 | 22601599 | 113.8171 | 176.8371 | 142.8988 | 59.33419 | 0.614403 | 2.768276 | 0.33092 |
| X | 25384200 | 25391999 | 103.6964 | 163.0635 | 115.6756 | 47.98711 | 0.815453 | 2.550266 | 0.330927 |
| 7 | 5002400 | 5007999 | 55.94783 | 89.68816 | 85.88039 | 36.2611 | 0.409439 | 1.983238 | 0.330939 |
| 8 | 25671800 | 25679999 | 69.09487 | 107.3712 | 185.9454 | 77.20593 | -0.48386 | 2.75049 | 0.330943 |
| X | 39865600 | 39866399 | 11.07425 | 21.41139 | 19.12965 | 7.322885 | 0.719692 | -0.14395 | 0.330963 |
| 6 | 95452000 | 95454599 | 33.29531 | 55.20239 | 52.5195 | 22.74363 | 0.452503 | 1.306561 | 0.330967 |
| 18 | 82533200 | 82533799 | 4.105168 | 8.01336 | 21.01706 | 7.439818 | -0.71274 | -0.49398 | 0.330977 |
| 7 | 10861800 | 10863399 | 22.89867 | 40.48554 | 28.17064 | 12.46037 | 0.911159 | 0.564274 | 0.330978 |
| 7 | 12351600 | 12354799 | 60.19944 | 96.38312 | 88.79702 | 37.42131 | 0.462893 | 2.027382 | 0.330992 |
| 19 | 14574200 | 14576599 | 14.25138 | 24.59227 | 50.53571 | 22.16947 | -0.64921 | 0.858299 | 0.331011 |
| 4 | 124498800 | 124502599 | 26.13185 | 42.73969 | 78.61196 | 33.80664 | -0.50178 | 1.474277 | 0.331026 |
| 10 | 63257000 | 63259199 | 13.52093 | 23.58455 | 46.80652 | 20.60159 | -0.59677 | 0.723042 | 0.331044 |
| 14 | 55350400 | 55355999 | 38.46937 | 61.30051 | 134.49 | 56.29993 | -0.79407 | 2.166656 | 0.33105 |
| 13 | 54362600 | 54363999 | 6.560158 | 12.2392 | 31.11244 | 12.93854 | -0.88232 | 0.124211 | 0.331078 |
| 12 | 52213600 | 52216799 | 35.94524 | 59.93049 | 48.717 | 21.17652 | 0.679023 | 1.247531 | 0.331081 |
| 9 | 19003800 | 19009999 | 93.04583 | 145.4928 | 137.4139 | 57.00918 | 0.39677 | 2.646997 | 0.331085 |
| 10 | 89521400 | 89522399 | 8.074917 | 14.84708 | 33.01638 | 14.02228 | -0.71423 | 0.227146 | 0.331108 |
| 15 | 96725800 | 96734199 | 60.43135 | 95.25316 | 157.0437 | 65.16838 | -0.39705 | 2.480995 | 0.331135 |
| 17 | 78752600 | 78754199 | 12.40724 | 21.84721 | 41.42085 | 18.3361 | -0.52952 | 0.569148 | 0.331136 |
| 4 | 119378200 | 119379999 | 9.309603 | 16.99771 | 33.95544 | 14.55677 | -0.57036 | 0.279822 | 0.331148 |
| 11 | 36116200 | 36117999 | 13.33295 | 23.41666 | 44.01675 | 19.47877 | -0.51848 | 0.610118 | 0.331149 |
| 13 | 58492200 | 58498799 | 25.31535 | 41.45356 | 78.07639 | 33.5916 | -0.53483 | 1.455154 | 0.331182 |
| 5 | 136923600 | 136925599 | 12.21801 | 21.59897 | 40.85728 | 18.07237 | -0.52409 | 0.522571 | 0.331197 |
| 19 | 58301600 | 58310599 | 77.57568 | 120.61 | 196.5985 | 81.6231 | -0.39609 | 2.746241 | 0.331201 |
| 1 | 101169600 | 101172599 | 46.01603 | 74.60576 | 61.46597 | 26.32384 | 0.644127 | 1.643498 | 0.331202 |
| 9 | 122333000 | 122333799 | 4.622163 | 8.987963 | 23.39201 | 8.665853 | -0.76103 | -0.31174 | 0.331204 |
| 18 | 52598600 | 52599199 | 5.557913 | 10.73706 | 26.53634 | 10.19725 | -0.74007 | -0.17041 | 0.331216 |
| 6 | 104794800 | 104797399 | 22.48184 | 39.46922 | 31.40866 | 13.89465 | 0.715357 | 0.64609 | 0.331218 |
| 9 | 121672200 | 121673199 | 15.1184 | 27.41382 | 24.84095 | 10.74162 | 0.554644 | 0.341502 | 0.33123 |
| 4 | 127137600 | 127139399 | 8.212143 | 14.76555 | 36.94533 | 16.14009 | -0.91079 | 0.36775 | 0.331239 |

| | | | | | | | | | |
|---|---|---|---|---|---|---|---|---|---|
| 1 | 102474400 | 102476399 | 24.57942 | 42.96772 | 33.14222 | 14.60871 | 0.760513 | 0.728131 | 0.331257 |
| 16 | 93253600 | 93256999 | 21.21774 | 35.12882 | 69.71746 | 30.16195 | -0.605 | 1.304751 | 0.331261 |
| 1 | 125270000 | 125280399 | 124.1434 | 186.8864 | 291.6228 | 120.4431 | -0.37049 | 3.394246 | 0.331265 |
| 4 | 115497200 | 115497799 | 6.112771 | 11.83047 | 26.10795 | 9.947689 | -0.56621 | -0.1925 | 0.331277 |
| 5 | 73155800 | 73157399 | 12.88662 | 22.64204 | 42.46847 | 18.80012 | -0.51516 | 0.599838 | 0.331283 |
| 15 | 96111600 | 96112399 | 9.359701 | 16.90514 | 35.39726 | 15.38177 | -0.6506 | 0.346107 | 0.331291 |
| 1 | 89040400 | 89041599 | 8.506189 | 15.66334 | 33.02869 | 13.99913 | -0.63615 | 0.221799 | 0.331302 |
| 18 | 37637200 | 37640199 | 93.03968 | 145.0732 | 132.9997 | 55.19441 | 0.436514 | 2.696658 | 0.331308 |
| 10 | 81951200 | 81955799 | 99.34839 | 156.2923 | 119.326 | 49.50419 | 0.710036 | 2.538378 | 0.331327 |
| 2 | 55700600 | 55701399 | 10.90948 | 19.60181 | 37.3531 | 16.32777 | -0.52149 | 0.391929 | 0.331328 |
| 12 | 106601400 | 106602999 | 10.90948 | 19.60181 | 37.3531 | 16.32777 | -0.52149 | 0.391929 | 0.331328 |
| 4 | 63580200 | 63583599 | 21.7346 | 36.27809 | 66.84262 | 29.07599 | -0.49766 | 1.210307 | 0.331333 |
| 7 | 10471600 | 10472799 | 24.07053 | 42.14716 | 31.89298 | 14.07754 | 0.787729 | 0.696259 | 0.331362 |
| 3 | 24281800 | 24283799 | 23.50945 | 41.30786 | 30.20463 | 13.37146 | 0.836769 | 0.637178 | 0.33137 |
| X | 99981800 | 99983999 | 22.84045 | 40.84959 | 24.79966 | 10.88278 | 1.115089 | 0.475685 | 0.331374 |
| 10 | 60727800 | 60731599 | 17.06238 | 28.83421 | 59.52497 | 26.04479 | -0.66053 | 1.067663 | 0.331391 |
| 3 | 33116000 | 33116999 | 12.10746 | 23.09207 | 20.9965 | 8.372174 | 0.639981 | 0.001299 | 0.331398 |
| 4 | 150777000 | 150780399 | 21.42398 | 35.7933 | 65.1065 | 28.33762 | -0.47911 | 1.198733 | 0.331414 |
| X | 125753400 | 125762599 | 120.5174 | 188.0164 | 123.7903 | 51.36928 | 0.914152 | 2.701822 | 0.331423 |
| X | 33313400 | 33318399 | 61.52767 | 99.02221 | 72.80449 | 30.92047 | 0.796637 | 1.868434 | 0.331428 |
| X | 121083600 | 121085599 | 19.27632 | 35.42647 | 22.76148 | 9.672986 | 1.064163 | 0.294363 | 0.331435 |
| 2 | 165328200 | 165329599 | 7.810711 | 14.29623 | 34.53266 | 14.75714 | -0.83933 | 0.253613 | 0.331449 |
| 13 | 117344400 | 117345399 | 8.620735 | 15.98352 | 32.61148 | 13.6939 | -0.57925 | 0.182605 | 0.331469 |
| 18 | 14703000 | 14704399 | 8.620735 | 15.98352 | 32.61148 | 13.6939 | -0.57925 | 0.182605 | 0.331469 |
| 17 | 68238200 | 68242199 | 41.21392 | 66.09392 | 112.3578 | 47.46813 | -0.41645 | 1.96261 | 0.331477 |
| 14 | 89490400 | 89492199 | 19.6626 | 35.31018 | 26.11788 | 11.42181 | 0.835049 | 0.430974 | 0.331492 |
| 6 | 49171400 | 49174399 | 35.34624 | 56.99008 | 97.96976 | 41.71556 | -0.42293 | 1.782922 | 0.331495 |
| 18 | 16939800 | 16942999 | 24.20059 | 39.31864 | 86.65189 | 37.16748 | -0.76885 | 1.574031 | 0.331532 |
| 4 | 136581800 | 136588199 | 43.11755 | 68.70852 | 126.412 | 52.91959 | -0.54059 | 2.167236 | 0.331532 |
| 13 | 97983600 | 97984999 | 19.61972 | 34.59853 | 30.63797 | 13.55874 | 0.563759 | 0.575536 | 0.331541 |
| 12 | 46893800 | 46894799 | 10.72639 | 20.69172 | 19.65599 | 7.59809 | 0.618745 | -0.11436 | 0.331542 |
| 4 | 41800400 | 41801599 | 6.584971 | 12.59291 | 29.02081 | 11.50126 | -0.68283 | -0.05157 | 0.331545 |
| 1 | 3430200 | 3432399 | 12.21062 | 21.6758 | 41.37624 | 18.27181 | -0.53444 | 0.483845 | 0.331578 |
| 5 | 135422400 | 135423599 | 6.125923 | 11.46136 | 31.15312 | 12.89368 | -0.97244 | 0.101777 | 0.331584 |
| 12 | 98329600 | 98330599 | 15.90171 | 28.55938 | 26.23259 | 11.47095 | 0.524677 | 0.404916 | 0.331606 |
| 9 | 83981800 | 83982399 | 11.3321 | 21.60126 | 19.91275 | 7.949367 | 0.618484 | -0.00673 | 0.331615 |
| 3 | 144425000 | 144426399 | 10.28301 | 18.08899 | 41.51961 | 18.38343 | -0.80398 | 0.56812 | 0.331628 |
| 12 | 71656400 | 71657999 | 19.47164 | 34.44947 | 29.37058 | 12.98939 | 0.619733 | 0.535799 | 0.331629 |
| 3 | 84568200 | 84574999 | 51.30473 | 81.37215 | 141.1129 | 58.75137 | -0.4624 | 2.321363 | 0.331649 |
| 1 | 25445200 | 25448799 | 34.38045 | 58.24027 | 41.29183 | 18.08016 | 0.875656 | 1.088744 | 0.331651 |
| 9 | 25547800 | 25549999 | 19.4716 | 32.72946 | 61.19998 | 26.73412 | -0.51871 | 1.125441 | 0.331652 |
| 15 | 13945600 | 13947999 | 25.49951 | 44.4961 | 33.62574 | 14.80378 | 0.789723 | 0.758254 | 0.331675 |
| 5 | 126816800 | 126819799 | 25.80324 | 43.89829 | 39.54706 | 17.33331 | 0.531754 | 1.017003 | 0.331692 |
| 5 | 140689800 | 140691999 | 12.45666 | 21.74394 | 45.75928 | 20.14986 | -0.6806 | 0.706699 | 0.3317 |
| 13 | 65773600 | 65783599 | 73.04553 | 116.7357 | 78.37997 | 32.94514 | 0.915577 | 2.094288 | 0.331702 |
| 17 | 72500400 | 72502599 | 12.36084 | 21.8425 | 42.41171 | 18.76149 | -0.56198 | 0.529618 | 0.331702 |
| 6 | 148352800 | 148354199 | 13.80184 | 26.25766 | 20.84052 | 8.354248 | 0.827979 | 0.034381 | 0.331726 |

| | | | | | | | | | |
|---|---|---|---|---|---|---|---|---|---|
| X | 15656000 | 15657799 | 22.49357 | 40.4865 | 23.90342 | 10.43228 | 1.16043 | 0.434541 | 0.33173 |
| 7 | 38500000 | 38507199 | 96.67056 | 151.7654 | 127.5669 | 52.90883 | 0.569188 | 2.577944 | 0.331745 |
| X | 125476400 | 125476999 | 13.34556 | 24.93802 | 22.93353 | 9.511735 | 0.576337 | 0.151988 | 0.331763 |
| 8 | 3884200 | 3888199 | 47.39418 | 77.54546 | 54.15875 | 23.294 | 0.891712 | 1.498385 | 0.331773 |
| 7 | 128497800 | 128498799 | 13.48919 | 25.20427 | 22.78009 | 9.44946 | 0.600567 | 0.151502 | 0.331779 |
| 18 | 63087800 | 63090199 | 19.10773 | 32.31132 | 59.08622 | 25.85782 | -0.48627 | 1.069855 | 0.33179 |
| 4 | 92051200 | 92053199 | 26.81647 | 47.02218 | 28.35021 | 12.52883 | 1.113684 | 0.684266 | 0.331792 |
| 7 | 14477400 | 14503799 | 539.5753 | 740.6655 | 559.3697 | 232.6198 | 0.546183 | 4.86757 | 0.3318 |
| 4 | 151585600 | 151590399 | 44.27931 | 70.49437 | 128.2945 | 53.63876 | -0.52637 | 2.199387 | 0.331831 |
| 18 | 8579600 | 8580199 | 5.076174 | 9.888274 | 23.11555 | 8.426177 | -0.58782 | -0.36795 | 0.33184 |
| 16 | 3315600 | 3320799 | 54.15786 | 87.82337 | 61.53862 | 26.35878 | 0.87747 | 1.657055 | 0.331844 |
| 16 | 4688600 | 4689799 | 6.729828 | 12.86942 | 28.44243 | 11.27332 | -0.62296 | -0.05051 | 0.331849 |
| 8 | 80783600 | 80785399 | 10.04755 | 17.88654 | 37.87434 | 16.68552 | -0.68029 | 0.446225 | 0.331852 |
| 18 | 68491000 | 68492599 | 19.72176 | 34.51863 | 30.95379 | 13.65833 | 0.544012 | 0.677264 | 0.331872 |
| 14 | 25793200 | 25796199 | 24.3602 | 41.85514 | 39.40049 | 17.2823 | 0.470697 | 0.922491 | 0.331875 |
| 8 | 61040000 | 61041399 | 17.82111 | 32.15209 | 26.34746 | 11.46513 | 0.69277 | 0.391437 | 0.331882 |
| 3 | 112264600 | 112265999 | 19.36819 | 34.46045 | 28.31363 | 12.47923 | 0.676853 | 0.4898 | 0.331883 |
| 13 | 55900400 | 55901399 | 4.526897 | 8.804183 | 22.85706 | 8.456272 | -0.75647 | -0.3224 | 0.331946 |
| Y | 21878000 | 21878199 | 1.869916 | 3.657991 | 17.25302 | 5.71262 | -1.44716 | -0.77653 | 0.331949 |
| 5 | 46133800 | 46135799 | 15.19325 | 27.48792 | 25.22982 | 10.94038 | 0.53347 | 0.354478 | 0.331963 |
| 12 | 69903800 | 69905999 | 15.19325 | 27.48792 | 25.22982 | 10.94038 | 0.53347 | 0.354478 | 0.331963 |
| 1 | 51611600 | 51612599 | 11.54007 | 22.11933 | 19.96528 | 7.860755 | 0.666761 | -0.04824 | 0.331999 |
| 14 | 18923800 | 18924199 | 11.54007 | 22.11933 | 19.96528 | 7.860755 | 0.666761 | -0.04824 | 0.331999 |
| 14 | 50437000 | 50441199 | 61.31924 | 97.78658 | 89.30247 | 37.42335 | 0.469334 | 2.149947 | 0.332034 |
| 15 | 92976000 | 92978399 | 21.00043 | 36.77881 | 33.31169 | 14.70624 | 0.529716 | 0.679459 | 0.332041 |
| 5 | 20674000 | 20678399 | 53.73774 | 86.65813 | 73.64929 | 31.36677 | 0.591427 | 1.795375 | 0.332049 |
| 16 | 49070800 | 49071999 | 10.89491 | 19.70821 | 39.62667 | 17.18662 | -0.5912 | 0.355969 | 0.332056 |
| 11 | 15262000 | 15262599 | 11.31435 | 21.74789 | 17.45915 | 6.818642 | 0.845799 | -0.08172 | 0.332068 |
| 17 | 9564600 | 9565799 | 16.49864 | 30.37806 | 23.97569 | 10.16323 | 0.771637 | 0.263623 | 0.332073 |
| 1 | 46758400 | 46760999 | 26.06004 | 43.00148 | 81.8358 | 35.32346 | -0.54541 | 1.364858 | 0.332099 |
| 9 | 16653400 | 16654399 | 13.94992 | 26.76436 | 19.35143 | 7.597881 | 0.988564 | -0.03227 | 0.332109 |
| 1 | 4857000 | 4859599 | 41.76256 | 67.997 | 63.71011 | 27.35318 | 0.464125 | 1.56184 | 0.332113 |
| 3 | 64492400 | 64496599 | 65.21075 | 103.9543 | 98.51134 | 41.26153 | 0.415526 | 2.168707 | 0.33212 |
| 2 | 168645000 | 168650599 | 34.21399 | 55.09296 | 99.69495 | 42.37353 | -0.49906 | 1.820678 | 0.332149 |
| 4 | 51691000 | 51691599 | 6.715385 | 12.88386 | 29.21398 | 11.47093 | -0.64714 | -0.08036 | 0.332187 |
| 10 | 78601400 | 78604599 | 36.00257 | 60.1519 | 48.12144 | 20.9453 | 0.702031 | 1.227945 | 0.332195 |
| 14 | 44943200 | 44944999 | 10.72322 | 19.34686 | 37.41039 | 16.27866 | -0.53821 | 0.365187 | 0.332198 |
| 17 | 19337800 | 19351999 | 16.52785 | 30.00181 | 22.20757 | 9.589999 | 0.845815 | 0.339883 | 0.332207 |
| 10 | 19197400 | 19198799 | 6.881663 | 13.11908 | 30.66932 | 12.24104 | -0.71286 | -0.02201 | 0.332207 |
| X | 26317400 | 26319799 | 22.2403 | 39.97474 | 23.70665 | 10.35957 | 1.15249 | 0.439206 | 0.332212 |
| 4 | 52922800 | 52923199 | 4.444807 | 8.671351 | 22.68199 | 8.107695 | -0.72074 | -0.44395 | 0.332218 |
| 4 | 106326800 | 106329199 | 9.671044 | 17.45712 | 35.3888 | 15.38951 | -0.60485 | 0.352519 | 0.33224 |
| 7 | 35030400 | 35035199 | 29.01485 | 46.94037 | 90.79657 | 38.82337 | -0.58704 | 1.671987 | 0.332242 |
| 7 | 12425400 | 12428199 | 33.09323 | 55.58651 | 42.14016 | 18.40367 | 0.77913 | 1.15754 | 0.332246 |
| 18 | 35797600 | 35798599 | 6.76961 | 12.83037 | 28.72395 | 11.60697 | -0.66588 | 0.015291 | 0.332267 |
| 14 | 37345600 | 37346799 | 17.38654 | 31.16345 | 22.18981 | 9.719578 | 0.888061 | 0.418501 | 0.332271 |
| 17 | 32351000 | 32352799 | 8.190602 | 15.23503 | 31.59353 | 13.20597 | -0.59701 | 0.160801 | 0.332283 |

| | | | | | | | | | |
|---|---|---|---|---|---|---|---|---|---|
| 15 | 75840400 | 75841799 | 9.333168 | 16.92946 | 34.91748 | 15.09235 | -0.62399 | 0.319522 | 0.332284 |
| 12 | 18292800 | 18306399 | 307.4885 | 445.4178 | 325.1471 | 134.0802 | 0.673018 | 4.061306 | 0.332303 |
| X | 137772400 | 137772999 | 9.838474 | 19.10364 | 18.21954 | 6.828879 | 0.655054 | -0.22985 | 0.332305 |
| 6 | 42914000 | 42915199 | 12.90176 | 24.78271 | 19.34409 | 7.569186 | 0.882999 | -0.05499 | 0.332323 |
| 9 | 107872200 | 107874199 | 9.109327 | 16.32929 | 36.41705 | 15.94989 | -0.74839 | 0.393759 | 0.332335 |
| 12 | 20162600 | 20164199 | 23.25995 | 40.57266 | 29.02618 | 12.77576 | 0.868385 | 0.74565 | 0.332356 |
| 2 | 80354600 | 80355999 | 14.96672 | 27.43626 | 24.63412 | 10.50513 | 0.580613 | 0.276718 | 0.332362 |
| 16 | 94363000 | 94364399 | 19.03488 | 33.56636 | 29.47689 | 13.04497 | 0.57543 | 0.567458 | 0.332364 |
| 4 | 120738400 | 120740599 | 10.69 | 18.9282 | 38.88388 | 17.19395 | -0.64108 | 0.493239 | 0.332394 |
| 3 | 67315600 | 67317599 | 26.18484 | 44.86924 | 40.91237 | 17.94212 | 0.515903 | 0.955895 | 0.332407 |
| X | 116809200 | 116810399 | 18.88777 | 35.89319 | 16.40105 | 6.590245 | 1.614453 | 0.078896 | 0.332445 |
| 12 | 109339000 | 109341999 | 24.78262 | 40.3407 | 88.13613 | 37.83682 | -0.75428 | 1.546916 | 0.33245 |
| 5 | 3621600 | 3625199 | 35.98951 | 59.32426 | 55.87132 | 24.10558 | 0.466017 | 1.383664 | 0.332457 |
| 9 | 96625200 | 96627199 | 21.63673 | 37.65245 | 34.56061 | 15.18921 | 0.510067 | 0.776203 | 0.332474 |
| 4 | 9841800 | 9847799 | 47.8525 | 76.05018 | 135.3556 | 56.46841 | -0.49696 | 2.257843 | 0.332479 |
| 9 | 38553200 | 38554999 | 26.99227 | 46.05388 | 42.06453 | 18.441 | 0.512647 | 1.000646 | 0.332484 |
| 18 | 87713400 | 87714999 | 24.51771 | 42.55119 | 33.54203 | 14.7171 | 0.728487 | 0.827514 | 0.332496 |
| 11 | 9752200 | 9754399 | 19.94473 | 34.91545 | 32.41597 | 14.30579 | 0.494179 | 0.678273 | 0.332499 |
| 17 | 63186400 | 63191399 | 33.17637 | 53.23478 | 116.3792 | 49.20993 | -0.77751 | 1.936581 | 0.332515 |
| 5 | 42901800 | 42903999 | 19.00928 | 34.87469 | 22.57305 | 9.615753 | 1.050727 | 0.299846 | 0.332518 |
| 7 | 28037600 | 28039399 | 12.09393 | 21.28269 | 41.43439 | 18.3443 | -0.56789 | 0.573623 | 0.332548 |
| 5 | 66966400 | 66971999 | 37.09635 | 59.72838 | 105.5449 | 44.85357 | -0.46513 | 1.829223 | 0.332572 |
| 16 | 90489600 | 90490399 | 7.066246 | 13.3853 | 28.71234 | 11.61564 | -0.60607 | 0.022844 | 0.33258 |
| 11 | 69955200 | 69958199 | 13.0725 | 22.81053 | 45.76112 | 20.14601 | -0.61202 | 0.715323 | 0.332584 |
| 8 | 128339400 | 128341599 | 9.984154 | 18.17757 | 37.08757 | 15.95615 | -0.60402 | 0.309708 | 0.332589 |
| 14 | 12283600 | 12284399 | 12.60507 | 23.63958 | 22.43644 | 9.248801 | 0.537955 | 0.120716 | 0.332605 |
| 1 | 124854800 | 124857999 | 34.20114 | 57.44367 | 45.82099 | 20.01035 | 0.705957 | 1.164121 | 0.332605 |
| X | 111335200 | 111340399 | 66.21838 | 106.0938 | 76.65796 | 32.33987 | 0.813964 | 2.01059 | 0.332612 |
| 1 | 114298600 | 114301999 | 47.22034 | 76.27832 | 72.04232 | 30.75007 | 0.442312 | 1.734087 | 0.332614 |
| 6 | 108696000 | 108698999 | 23.52113 | 39.10975 | 74.4197 | 32.29175 | -0.54424 | 1.262727 | 0.332618 |
| 14 | 82909600 | 82910399 | 11.97938 | 22.94518 | 20.53444 | 8.101248 | 0.676645 | -0.0347 | 0.332621 |
| 7 | 26227600 | 26233999 | 30.49062 | 51.37045 | 41.6197 | 18.19365 | 0.683661 | 1.120896 | 0.332624 |
| 13 | 61633200 | 61635599 | 43.10908 | 70.88073 | 54.03421 | 23.28807 | 0.768668 | 1.428788 | 0.332625 |
| 4 | 143304800 | 143309999 | 151.1764 | 229.334 | 200.8142 | 83.13194 | 0.4719 | 3.242898 | 0.332626 |
| 2 | 123931800 | 123933999 | 17.78223 | 29.80799 | 68.78091 | 29.99982 | -0.82106 | 1.154484 | 0.33265 |
| 10 | 69397200 | 69398399 | 10.40993 | 18.91054 | 36.49539 | 15.74523 | -0.5272 | 0.322462 | 0.332664 |
| 16 | 80004200 | 80005399 | 12.96775 | 24.42242 | 21.79578 | 8.920846 | 0.63393 | 0.094825 | 0.332673 |
| 2 | 18044000 | 18046399 | 15.63892 | 26.52862 | 57.31047 | 25.10457 | -0.72533 | 1.028589 | 0.332683 |
| 6 | 28178000 | 28178599 | 6.601581 | 12.60039 | 27.77683 | 11.05819 | -0.62421 | -0.03572 | 0.332687 |
| 2 | 80750400 | 80752399 | 18.42438 | 33.34914 | 25.84666 | 11.20139 | 0.776583 | 0.377303 | 0.332689 |
| 14 | 110748600 | 110750199 | 18.42438 | 33.34914 | 25.84666 | 11.20139 | 0.776583 | 0.377303 | 0.332689 |
| 7 | 62890200 | 62892199 | 23.69748 | 41.53005 | 31.31481 | 13.83249 | 0.792717 | 0.679373 | 0.332689 |
| 6 | 49408200 | 49408799 | 6.745375 | 12.87326 | 27.78849 | 11.06573 | -0.59481 | -0.03456 | 0.3327 |
| 14 | 111736000 | 111738199 | 35.91572 | 59.45767 | 48.26134 | 20.87799 | 0.680686 | 1.325653 | 0.332709 |
| 8 | 74152800 | 74156999 | 62.66284 | 99.9178 | 96.36058 | 40.37475 | 0.390655 | 2.156838 | 0.332712 |
| 6 | 116690600 | 116691399 | 5.813197 | 11.26482 | 26.13545 | 9.899889 | -0.62889 | -0.21329 | 0.332748 |
| 11 | 116653600 | 116658599 | 43.26107 | 68.98188 | 124.5405 | 52.18285 | -0.51238 | 2.145859 | 0.332754 |

| | | | | | | | | | |
|---|---|---|---|---|---|---|---|---|---|
| 3 | 54018600 | 54020399 | 9.57479 | 17.65775 | 35.23072 | 14.90176 | -0.55503 | 0.226809 | 0.33276 |
| 11 | 117054400 | 117055999 | 16.62591 | 28.47368 | 53.74446 | 23.56858 | -0.52957 | 0.939692 | 0.332762 |
| X | 32082200 | 32083199 | 31.397 | 52.79885 | 39.62814 | 17.31325 | 0.793562 | 1.139234 | 0.332767 |
| 3 | 61945200 | 61946199 | 13.69567 | 24.00296 | 44.61076 | 19.70644 | -0.5021 | 0.649083 | 0.332778 |
| 17 | 86234800 | 86241799 | 43.6301 | 69.44559 | 129.9611 | 54.31907 | -0.56689 | 2.205771 | 0.332783 |
| 2 | 174103800 | 174106199 | 16.41241 | 27.86692 | 56.80653 | 24.89092 | -0.64187 | 1.023565 | 0.332787 |
| 11 | 49602800 | 49604999 | 11.65323 | 20.46582 | 42.01496 | 18.59234 | -0.64459 | 0.599424 | 0.332791 |
| 4 | 109058600 | 109059599 | 5.556294 | 10.70271 | 25.80003 | 10.015 | -0.71741 | -0.13579 | 0.332799 |
| 11 | 77540400 | 77541199 | 5.556294 | 10.70271 | 25.80003 | 10.015 | -0.71741 | -0.13579 | 0.332799 |
| 15 | 5229200 | 5229999 | 5.556294 | 10.70271 | 25.80003 | 10.015 | -0.71741 | -0.13579 | 0.332799 |
| 11 | 88803200 | 88803799 | 14.02881 | 26.07428 | 18.76344 | 7.852754 | 0.916896 | 0.179248 | 0.332821 |
| 1 | 93594200 | 93597199 | 16.24482 | 27.56636 | 57.91661 | 25.37286 | -0.68546 | 1.026699 | 0.33283 |
| 4 | 107735400 | 107737399 | 10.39374 | 18.40583 | 38.92319 | 17.21092 | -0.68237 | 0.49164 | 0.332833 |
| 18 | 3167600 | 3169399 | 26.70609 | 45.94339 | 39.31997 | 17.24764 | 0.60802 | 0.923448 | 0.332857 |
| 16 | 45088400 | 45089199 | 6.264772 | 12.10835 | 26.5942 | 10.19684 | -0.56802 | -0.16741 | 0.332865 |
| 3 | 96826200 | 96827799 | 13.54117 | 23.60558 | 46.29494 | 20.36869 | -0.57969 | 0.732221 | 0.332867 |
| 6 | 125762200 | 125764399 | 10.09423 | 18.31893 | 35.42342 | 15.30165 | -0.5311 | 0.323534 | 0.332869 |
| 11 | 33377800 | 33378799 | 9.165694 | 16.66494 | 34.9435 | 15.0591 | -0.64493 | 0.303925 | 0.332883 |
| 2 | 169190600 | 169192599 | 16.05908 | 27.90888 | 53.33844 | 23.4223 | -0.54351 | 0.796448 | 0.33291 |
| 5 | 93289800 | 93291799 | 10.28007 | 18.57045 | 35.88609 | 15.59106 | -0.5355 | 0.35334 | 0.332911 |
| 7 | 26389000 | 26390399 | 24.17905 | 41.8948 | 37.65363 | 16.5203 | 0.539568 | 0.845046 | 0.332913 |
| 5 | 3563800 | 3567599 | 28.74597 | 48.27633 | 45.39527 | 19.82316 | 0.469431 | 1.146791 | 0.332938 |
| 2 | 64268200 | 64268999 | 6.87496 | 13.19021 | 28.00705 | 10.99656 | -0.55253 | -0.07942 | 0.332991 |
| 8 | 20514400 | 20514999 | 13.58877 | 25.05214 | 22.43265 | 9.492682 | 0.592774 | 0.230676 | 0.333004 |
| 8 | 71565800 | 71567799 | 18.86377 | 33.97738 | 26.2439 | 11.44354 | 0.775773 | 0.408201 | 0.33303 |
| 2 | 102974000 | 102976599 | 15.40041 | 26.77289 | 50.05394 | 21.98448 | -0.51176 | 0.784189 | 0.333033 |
| 1 | 184186800 | 184187999 | 8.441344 | 15.72259 | 33.27818 | 13.87844 | -0.62448 | 0.154127 | 0.33304 |
| X | 31960200 | 31966199 | 48.97811 | 80.16658 | 54.04831 | 23.24962 | 0.942975 | 1.501278 | 0.333043 |
| 9 | 77237400 | 77242399 | 75.1771 | 118.9013 | 114.2762 | 47.48878 | 0.384364 | 2.405693 | 0.333052 |
| 7 | 71955800 | 71956799 | 5.838208 | 11.26265 | 25.87552 | 10.00107 | -0.64329 | -0.14772 | 0.333076 |
| 3 | 124754000 | 124755399 | 13.25271 | 24.72557 | 22.2309 | 9.244923 | 0.605019 | 0.15681 | 0.333077 |
| 6 | 29177000 | 29182999 | 39.44718 | 63.22716 | 110.7899 | 46.76807 | -0.46093 | 1.976716 | 0.333084 |
| X | 56497800 | 56498599 | 14.02068 | 26.11066 | 23.38186 | 9.753673 | 0.60784 | 0.180215 | 0.333088 |
| 19 | 57446800 | 57451599 | 31.84928 | 51.21766 | 102.496 | 43.48095 | -0.64626 | 1.860716 | 0.33309 |
| 6 | 95836000 | 95836599 | 11.1399 | 21.39878 | 20.07815 | 7.856018 | 0.619931 | -0.0711 | 0.333095 |
| 7 | 11807400 | 11818599 | 154.0892 | 234.7659 | 182.8517 | 75.79402 | 0.645798 | 3.155334 | 0.333121 |
| 2 | 27536000 | 27540999 | 35.82935 | 57.69282 | 100.0412 | 42.51933 | -0.43774 | 1.824173 | 0.333125 |
| 7 | 131032800 | 131033599 | 12.68416 | 23.74789 | 21.02908 | 8.693401 | 0.634389 | 0.129733 | 0.333128 |
| 8 | 86413000 | 86416199 | 87.02874 | 136.7323 | 129.387 | 53.66903 | 0.399305 | 2.557188 | 0.333136 |
| 7 | 31193200 | 31194199 | 13.99218 | 26.61225 | 20.83259 | 8.356069 | 0.847029 | 0.03821 | 0.333137 |
| 1 | 48333800 | 48334999 | 15.7663 | 29.70562 | 21.04673 | 8.608059 | 0.9655 | 0.116691 | 0.333143 |
| 7 | 18254400 | 18255999 | 29.21422 | 49.37696 | 45.28427 | 19.81471 | 0.505599 | 1.092932 | 0.333156 |
| 12 | 85691000 | 85694799 | 29.11581 | 47.27822 | 85.07609 | 36.47669 | -0.47804 | 1.594767 | 0.333168 |
| 13 | 43654000 | 43655599 | 15.58761 | 27.09581 | 49.51196 | 21.74517 | -0.47884 | 0.78685 | 0.333171 |
| 13 | 112184400 | 112185799 | 10.66257 | 18.92762 | 38.84136 | 17.15251 | -0.63795 | 0.469303 | 0.333193 |
| 1 | 12738400 | 12739199 | 8.810986 | 16.28989 | 32.56156 | 13.72593 | -0.55362 | 0.202076 | 0.333222 |
| 7 | 19502000 | 19508599 | 39.85978 | 63.61545 | 123.6321 | 51.86717 | -0.61703 | 2.11322 | 0.333239 |

| | | | | | | | | | |
|---|---|---|---|---|---|---|---|---|---|
| 7 | 5466800 | 5469399 | 41.16506 | 68.12474 | 50.56286 | 21.86852 | 0.80941 | 1.346555 | 0.333258 |
| 7 | 66383200 | 66384999 | 15.37927 | 26.34479 | 53.7101 | 23.55396 | -0.64016 | 0.932285 | 0.333259 |
| 3 | 89423200 | 89424599 | 28.36591 | 47.98167 | 43.75378 | 19.14968 | 0.513952 | 1.083621 | 0.333264 |
| 10 | 76422200 | 76423999 | 10.9993 | 19.45118 | 39.38755 | 17.42108 | -0.62127 | 0.509816 | 0.333274 |
| 4 | 13377000 | 13378399 | 16.08197 | 28.86897 | 26.70958 | 11.68436 | 0.513902 | 0.410298 | 0.333275 |
| 10 | 116704400 | 116708999 | 28.5896 | 46.54893 | 83.1448 | 35.6972 | -0.46429 | 1.54726 | 0.333288 |
| 12 | 23544800 | 23546599 | 31.94838 | 53.08019 | 47.12339 | 20.43634 | 0.552272 | 1.274741 | 0.333297 |
| 16 | 81838600 | 81839999 | 17.15519 | 30.9827 | 26.43195 | 11.48702 | 0.636237 | 0.380976 | 0.333304 |
| 9 | 57248600 | 57252199 | 16.11677 | 27.76429 | 51.63294 | 22.64965 | -0.50659 | 0.881758 | 0.333308 |
| 7 | 27594200 | 27595999 | 19.45049 | 34.74002 | 27.17563 | 11.93894 | 0.750307 | 0.464094 | 0.333322 |
| 16 | 82874200 | 82875999 | 19.45049 | 34.74002 | 27.17563 | 11.93894 | 0.750307 | 0.464094 | 0.333322 |
| 12 | 11937600 | 11940799 | 40.30299 | 66.0389 | 59.34918 | 25.54148 | 0.530048 | 1.469022 | 0.333322 |
| X | 146717400 | 146719999 | 21.2034 | 38.00254 | 25.42666 | 11.13793 | 0.977306 | 0.452292 | 0.333324 |
| 13 | 66641600 | 66653199 | 272.6337 | 393.7285 | 352.7071 | 145.5036 | 0.373489 | 4.103202 | 0.333333 |
| 9 | 36193000 | 36205199 | 170.5893 | 257.8634 | 206.1762 | 85.25676 | 0.598613 | 3.317478 | 0.333337 |
| 3 | 93820800 | 93825399 | 26.2606 | 45.00663 | 40.34141 | 17.69197 | 0.540469 | 0.953071 | 0.333347 |
| 11 | 50627200 | 50629199 | 23.80536 | 39.26775 | 72.04585 | 31.09329 | -0.49277 | 1.357471 | 0.333351 |
| 3 | 121219400 | 121222399 | 16.29651 | 27.53435 | 59.45984 | 26.01469 | -0.72517 | 1.067178 | 0.333352 |
| 7 | 13695800 | 13712599 | 460.2712 | 635.6181 | 544.4702 | 226.2078 | 0.373278 | 4.772889 | 0.333355 |
| 12 | 58646800 | 58648999 | 15.5593 | 27.06024 | 50.59499 | 22.22787 | -0.51184 | 0.779792 | 0.333356 |
| 8 | 114016200 | 114017399 | 12.28748 | 21.61377 | 41.97545 | 18.58548 | -0.56445 | 0.582693 | 0.333357 |
| 5 | 110627200 | 110630199 | 31.86873 | 51.20407 | 104.2269 | 44.15914 | -0.6723 | 1.890218 | 0.333365 |
| 6 | 91801600 | 91806599 | 33.71302 | 54.32126 | 99.39442 | 42.28526 | -0.51388 | 1.798913 | 0.333391 |
| 16 | 28745000 | 28748999 | 20.22326 | 33.61288 | 67.57807 | 29.31504 | -0.62312 | 1.254417 | 0.333391 |
| 5 | 50526200 | 50526599 | 2.739273 | 5.358618 | 18.55912 | 6.168732 | -1.02228 | -0.74711 | 0.333422 |
| 18 | 89633800 | 89636399 | 46.51227 | 75.29366 | 62.48651 | 26.73038 | 0.631591 | 1.675745 | 0.33344 |
| 4 | 154761600 | 154768999 | 48.28583 | 76.54139 | 149.7491 | 62.31226 | -0.63689 | 2.338583 | 0.333461 |
| 11 | 85650800 | 85652799 | 8.583653 | 15.58683 | 35.46493 | 15.30817 | -0.76344 | 0.307905 | 0.333475 |
| 6 | 96838600 | 96839999 | 14.88605 | 27.74765 | 23.19824 | 9.662076 | 0.707739 | 0.180096 | 0.333489 |
| 7 | 41049400 | 41050999 | 14.88605 | 27.74765 | 23.19824 | 9.662076 | 0.707739 | 0.180096 | 0.333489 |
| 8 | 101642200 | 101644399 | 26.54326 | 43.69496 | 78.16926 | 33.70696 | -0.45779 | 1.392658 | 0.333495 |
| 15 | 74822200 | 74824399 | 19.24675 | 32.57054 | 59.1232 | 25.87982 | -0.47568 | 1.063276 | 0.333518 |
| 17 | 54517400 | 54518799 | 16.13641 | 30.03018 | 22.45679 | 9.378664 | 0.864245 | 0.198294 | 0.333518 |
| 4 | 103272800 | 103276999 | 47.15953 | 75.21055 | 125.2513 | 52.49267 | -0.39586 | 2.14501 | 0.333527 |
| 6 | 85871800 | 85873799 | 15.48903 | 26.65058 | 52.15162 | 22.87633 | -0.58014 | 0.889998 | 0.33354 |
| 17 | 83553800 | 83565799 | 108.3292 | 163.6137 | 275.4641 | 113.8782 | -0.47591 | 3.32499 | 0.333544 |
| 2 | 35900600 | 35904999 | 32.45707 | 52.27109 | 98.57549 | 41.90611 | -0.55819 | 1.813979 | 0.333544 |
| 18 | 13182800 | 13185799 | 21.66764 | 37.9097 | 33.82876 | 14.922 | 0.551136 | 0.699661 | 0.333548 |
| 1 | 59951000 | 59953399 | 16.46415 | 28.20111 | 53.74403 | 23.5687 | -0.54334 | 0.937352 | 0.333556 |
| 5 | 131402000 | 131403199 | 13.52282 | 25.1525 | 23.30627 | 9.742438 | 0.556169 | 0.181858 | 0.333577 |
| 6 | 64704600 | 64707999 | 29.83651 | 48.20047 | 94.10982 | 40.17296 | -0.60284 | 1.713734 | 0.333609 |
| 13 | 62225600 | 62229399 | 84.56344 | 132.9424 | 126.5315 | 52.48887 | 0.391636 | 2.541495 | 0.333622 |
| 15 | 83220600 | 83222999 | 14.62103 | 25.45197 | 47.89123 | 21.05227 | -0.52065 | 0.759109 | 0.333639 |
| 2 | 112209400 | 112209999 | 6.188349 | 11.76534 | 28.79023 | 11.55729 | -0.78472 | -0.01274 | 0.333662 |
| 8 | 100393400 | 100394399 | 11.68362 | 22.51886 | 18.90177 | 7.324618 | 0.791744 | -0.10014 | 0.333674 |
| X | 8189400 | 8194199 | 114.6 | 178.4719 | 134.5504 | 55.84822 | 0.717034 | 2.729954 | 0.33368 |
| 7 | 118088800 | 118091399 | 22.72451 | 36.99137 | 88.23383 | 37.87907 | -0.88044 | 1.543087 | 0.333681 |

| | | | | | | | | | |
|---|---|---|---|---|---|---|---|---|---|
| 19 | 3582800 | 3584199 | 13.08473 | 22.92689 | 44.118 | 19.48493 | -0.55182 | 0.649854 | 0.333699 |
| 7 | 77669800 | 77671599 | 15.34999 | 28.66139 | 22.60281 | 9.386857 | 0.794626 | 0.171785 | 0.333709 |
| 13 | 65833600 | 65841399 | 127.6216 | 199.0128 | 116.7121 | 48.43452 | 1.080407 | 2.712883 | 0.333711 |
| 1 | 119671400 | 119672799 | 34.07119 | 54.96708 | 97.08626 | 41.37121 | -0.46087 | 1.764388 | 0.333715 |
| 19 | 48475000 | 48476999 | 19.07143 | 33.6906 | 29.42439 | 13.01711 | 0.584267 | 0.548826 | 0.33373 |
| 5 | 118757800 | 118760399 | 14.19557 | 24.70137 | 47.36765 | 20.81684 | -0.5477 | 0.764182 | 0.333739 |
| X | 20786600 | 20787599 | 22.74863 | 39.8354 | 31.3806 | 13.85222 | 0.730346 | 0.69082 | 0.333745 |
| 3 | 49770800 | 49772399 | 16.65883 | 30.44492 | 24.96565 | 10.69017 | 0.706549 | 0.307782 | 0.333759 |
| 6 | 99051800 | 99052999 | 20.48282 | 35.82091 | 33.46652 | 14.75671 | 0.485229 | 0.699623 | 0.333771 |
| 6 | 145253600 | 145254399 | 14.31288 | 24.98565 | 46.26576 | 20.37368 | -0.49708 | 0.716095 | 0.333779 |
| 19 | 51729600 | 51730599 | 16.56228 | 29.51311 | 26.30351 | 11.57836 | 0.561512 | 0.458279 | 0.333797 |
| 8 | 107811400 | 107817999 | 51.47676 | 81.73604 | 136.6817 | 56.96998 | -0.40843 | 2.288804 | 0.333811 |
| 12 | 84828200 | 84828999 | 6.145147 | 11.783 | 26.79026 | 10.53676 | -0.65187 | -0.08299 | 0.333814 |
| 10 | 119006600 | 119008599 | 8.731918 | 15.85506 | 34.9453 | 15.08507 | -0.71783 | 0.308829 | 0.333838 |
| 7 | 32234600 | 32238599 | 29.5068 | 51.24771 | 33.51923 | 14.71343 | 0.996657 | 0.841664 | 0.333853 |
| 13 | 43862600 | 43864399 | 17.92622 | 31.91526 | 27.88955 | 12.28569 | 0.588799 | 0.474383 | 0.33387 |
| X | 103687400 | 103689199 | 18.88058 | 34.31043 | 24.71294 | 10.65717 | 0.886065 | 0.35588 | 0.33388 |
| 17 | 38467000 | 38468399 | 20.35216 | 35.83538 | 30.68676 | 13.58447 | 0.611159 | 0.597232 | 0.333887 |
| 17 | 88371400 | 88373999 | 17.93344 | 30.07325 | 62.10605 | 27.09518 | -0.66141 | 1.145928 | 0.333902 |
| 10 | 24098200 | 24100199 | 12.42963 | 21.73536 | 46.24125 | 20.38807 | -0.69621 | 0.6787 | 0.333912 |
| 16 | 65113000 | 65114399 | 14.98355 | 28.33244 | 20.54873 | 8.344265 | 0.940528 | 0.084374 | 0.333921 |
| 1 | 170341400 | 170347599 | 59.70143 | 94.33287 | 155.3615 | 64.53048 | -0.39274 | 2.424998 | 0.333947 |
| 1 | 113179200 | 113182599 | 31.61634 | 53.89327 | 40.77814 | 17.87571 | 0.782975 | 1.024928 | 0.333949 |
| 1 | 48747200 | 48749799 | 41.38817 | 67.49181 | 53.19667 | 22.85225 | 0.714369 | 1.530895 | 0.333958 |
| X | 5163200 | 5171999 | 98.95355 | 156.2872 | 106.4276 | 44.19586 | 0.879162 | 2.45463 | 0.333963 |
| 7 | 17045000 | 17050599 | 95.89129 | 149.8557 | 138.5661 | 57.49061 | 0.426674 | 2.658943 | 0.333971 |
| 7 | 15490400 | 15492999 | 49.78853 | 80.5034 | 69.20486 | 29.5752 | 0.57929 | 1.709197 | 0.333972 |
| 7 | 14427400 | 14430799 | 70.47495 | 112.0065 | 106.0472 | 44.24269 | 0.412253 | 2.269607 | 0.333995 |
| 7 | 120012600 | 120013199 | 5.824069 | 11.26234 | 25.4208 | 9.72364 | -0.60324 | -0.18309 | 0.334002 |
| 13 | 24926800 | 24927599 | 5.824069 | 11.26234 | 25.4208 | 9.72364 | -0.60324 | -0.18309 | 0.334002 |
| X | 31848000 | 31857999 | 212.1526 | 314.4424 | 265.5075 | 109.3514 | 0.494578 | 3.693884 | 0.334006 |
| 11 | 57808800 | 57812399 | 23.92336 | 39.56308 | 71.14751 | 30.75634 | -0.4635 | 1.326242 | 0.334054 |
| 3 | 26400800 | 26403199 | 24.52361 | 40.17413 | 76.7463 | 33.02526 | -0.5549 | 1.448092 | 0.334077 |
| 3 | 111485600 | 111487799 | 34.55966 | 57.40376 | 51.43234 | 22.30104 | 0.539057 | 1.290052 | 0.334082 |
| 2 | 178422000 | 178426199 | 38.32524 | 61.40097 | 112.0709 | 47.27614 | -0.52046 | 1.989808 | 0.334125 |
| 1 | 31403200 | 31404999 | 16.53407 | 30.67119 | 22.97222 | 9.639547 | 0.857064 | 0.221424 | 0.334132 |
| 12 | 60144600 | 60145799 | 16.53407 | 30.67119 | 22.97222 | 9.639547 | 0.857064 | 0.221424 | 0.334132 |
| 14 | 39936400 | 39937799 | 20.30187 | 35.90718 | 28.10771 | 12.43165 | 0.741893 | 0.540642 | 0.334134 |
| X | 102021400 | 102022999 | 16.32736 | 30.88799 | 19.95261 | 8.094137 | 1.107802 | 0.092956 | 0.334142 |
| 18 | 48897200 | 48899399 | 18.2534 | 32.64516 | 27.95492 | 12.26759 | 0.621799 | 0.450632 | 0.33416 |
| 15 | 87636000 | 87644199 | 64.5459 | 101.0706 | 172.2394 | 71.44695 | -0.45209 | 2.612074 | 0.334165 |
| 1 | 112786600 | 112788999 | 17.54102 | 31.58868 | 26.93683 | 11.74856 | 0.633656 | 0.401421 | 0.334181 |
| 14 | 73784000 | 73786399 | 12.75216 | 22.27039 | 45.74563 | 20.14989 | -0.64584 | 0.700815 | 0.33419 |
| 3 | 145618600 | 145619999 | 7.085105 | 13.36113 | 29.15246 | 11.90525 | -0.64233 | 0.055751 | 0.334191 |
| 17 | 45337400 | 45337999 | 7.085105 | 13.36113 | 29.15246 | 11.90525 | -0.64233 | 0.055751 | 0.334191 |
| 16 | 82402200 | 82402599 | 4.965528 | 9.667554 | 24.90882 | 9.136394 | -0.736 | -0.34514 | 0.334209 |
| 6 | 93654200 | 93656399 | 28.33751 | 45.96617 | 85.79847 | 36.75796 | -0.53209 | 1.615521 | 0.334219 |

| | | | | | | | | | |
|---|---|---|---|---|---|---|---|---|---|
| 2 | 21210400 | 21211199 | 7.26829 | 13.57434 | 31.13382 | 12.92485 | -0.73351 | 0.122892 | 0.33422 |
| 3 | 103875200 | 103876199 | 7.26829 | 13.57434 | 31.13382 | 12.92485 | -0.73351 | 0.122892 | 0.33422 |
| 11 | 121311800 | 121312799 | 3.889707 | 7.550616 | 24.92297 | 9.362764 | -1.11971 | -0.27045 | 0.334224 |
| 18 | 23192800 | 23193399 | 6.581232 | 12.61211 | 27.37165 | 10.78272 | -0.58825 | -0.07374 | 0.33424 |
| 7 | 94885200 | 94886599 | 19.29284 | 34.453 | 27.87044 | 12.24582 | 0.702294 | 0.466866 | 0.334253 |
| 12 | 119491200 | 119493399 | 19.57308 | 33.18481 | 59.69976 | 26.14703 | -0.46267 | 1.049361 | 0.334288 |
| 16 | 88664000 | 88666399 | 15.93998 | 27.72011 | 52.76588 | 23.18045 | -0.53756 | 0.787189 | 0.334304 |
| 13 | 80987000 | 80988599 | 20.75303 | 36.60803 | 29.79302 | 13.18402 | 0.684851 | 0.575683 | 0.334316 |
| 13 | 97207600 | 97208599 | 9.531305 | 17.13838 | 36.37426 | 15.88987 | -0.67513 | 0.375305 | 0.334371 |
| 11 | 86762800 | 86763999 | 8.639539 | 16.01953 | 32.07434 | 13.46705 | -0.55188 | 0.182579 | 0.334376 |
| 6 | 130085800 | 130088599 | 43.14498 | 70.02049 | 66.345 | 28.43421 | 0.444261 | 1.621709 | 0.334377 |
| X | 122450000 | 122456999 | 64.83081 | 104.246 | 73.09661 | 31.00477 | 0.86333 | 1.900001 | 0.334388 |
| 14 | 8057400 | 8059399 | 19.22507 | 34.05027 | 29.43603 | 13.01551 | 0.600488 | 0.523579 | 0.33439 |
| 10 | 75437600 | 75441999 | 36.02434 | 57.87177 | 105 | 44.47644 | -0.50713 | 1.899618 | 0.33441 |
| 13 | 77757400 | 77759999 | 28.511 | 48.50821 | 43.05199 | 18.86961 | 0.553251 | 1.031027 | 0.334435 |
| 12 | 3366400 | 3368599 | 15.99355 | 29.26058 | 25.0472 | 10.71011 | 0.646344 | 0.29684 | 0.334455 |
| 19 | 11104400 | 11106599 | 15.99355 | 29.26058 | 25.0472 | 10.71011 | 0.646344 | 0.29684 | 0.334455 |
| 16 | 82628400 | 82631399 | 32.62412 | 55.79788 | 37.33108 | 16.36906 | 0.960319 | 0.994383 | 0.334461 |
| 11 | 32323200 | 32325599 | 14.37341 | 24.87412 | 49.4971 | 21.71574 | -0.60222 | 0.830469 | 0.334462 |
| 7 | 14364600 | 14368799 | 56.89958 | 91.49509 | 78.96844 | 33.49657 | 0.564248 | 1.887003 | 0.334467 |
| 4 | 89192800 | 89196199 | 34.16815 | 55.15051 | 96.59483 | 41.18781 | -0.44801 | 1.747275 | 0.334469 |
| 5 | 67239600 | 67243399 | 55.7761 | 89.41565 | 85.8784 | 36.26197 | 0.405141 | 1.981973 | 0.334512 |
| 6 | 28418000 | 28419399 | 8.759001 | 15.83618 | 35.4263 | 15.37494 | -0.74413 | 0.33557 | 0.334517 |
| 1 | 188891400 | 188892199 | 8.299872 | 15.53942 | 31.19684 | 12.89675 | -0.53802 | 0.117083 | 0.334534 |
| 12 | 102148000 | 102155799 | 105.5455 | 160.0223 | 260.7761 | 107.9365 | -0.42418 | 3.246085 | 0.334537 |
| 11 | 52396000 | 52399399 | 32.52653 | 52.68685 | 91.20999 | 39.03579 | -0.42577 | 1.65373 | 0.334538 |
| 7 | 31225600 | 31238199 | 224.0932 | 331.8003 | 271.8444 | 111.954 | 0.536679 | 3.712214 | 0.334542 |
| 15 | 60864400 | 60865599 | 7.799346 | 14.71319 | 31.42181 | 12.82309 | -0.61178 | 0.062317 | 0.334548 |
| 8 | 108723600 | 108729199 | 37.47333 | 60.20859 | 104.8422 | 44.41929 | -0.44772 | 1.896877 | 0.334554 |
| 4 | 137424400 | 137426799 | 9.171176 | 16.28874 | 38.43418 | 16.96517 | -0.83723 | 0.456789 | 0.334556 |
| 2 | 36702400 | 36702999 | 10.28537 | 19.71837 | 18.54143 | 7.29594 | 0.609242 | -0.06664 | 0.334571 |
| 14 | 29151600 | 29152999 | 7.996319 | 14.91081 | 32.19484 | 13.40057 | -0.65052 | 0.140851 | 0.334572 |
| 15 | 38360600 | 38362199 | 27.1054 | 46.52896 | 40.33916 | 17.69326 | 0.588792 | 0.944353 | 0.334581 |
| 12 | 105500600 | 105508599 | 56.2839 | 88.0962 | 181.7746 | 75.40557 | -0.72811 | 2.617864 | 0.334587 |
| 18 | 3198000 | 3198399 | 12.75444 | 23.78233 | 20.5098 | 8.537467 | 0.664202 | 0.15383 | 0.334595 |
| 10 | 119934800 | 119937199 | 18.14138 | 32.16054 | 29.12577 | 12.87595 | 0.534362 | 0.507146 | 0.334598 |
| 8 | 126319600 | 126320599 | 18.86287 | 33.11986 | 28.3511 | 12.54336 | 0.610508 | 0.617039 | 0.334599 |
| 14 | 67319000 | 67322999 | 21.91135 | 36.50178 | 67.93314 | 29.51398 | -0.51208 | 1.232589 | 0.334622 |
| 13 | 18830600 | 18835399 | 52.99879 | 85.80479 | 64.95567 | 27.78922 | 0.764072 | 1.686439 | 0.334631 |
| 3 | 85954200 | 85955599 | 23.46971 | 40.80623 | 36.60253 | 16.07771 | 0.543184 | 0.802207 | 0.334633 |
| 7 | 103052600 | 103053599 | 9.760892 | 17.91359 | 35.10515 | 14.94727 | -0.53611 | 0.256357 | 0.334676 |
| 5 | 93867000 | 93867599 | 15.07545 | 27.75149 | 24.71377 | 10.47925 | 0.598406 | 0.25575 | 0.334699 |
| 9 | 25340600 | 25342599 | 20.19928 | 35.33816 | 32.37502 | 14.2799 | 0.513225 | 0.689121 | 0.3347 |
| 11 | 49139600 | 49141999 | 13.84364 | 24.09544 | 47.8904 | 21.05011 | -0.59925 | 0.756993 | 0.334701 |
| 6 | 13969600 | 13970999 | 16.31614 | 30.94303 | 19.41324 | 7.834336 | 1.155974 | 0.075213 | 0.334702 |
| 5 | 96430600 | 96432199 | 21.38016 | 38.3878 | 24.87196 | 10.87857 | 1.02482 | 0.441774 | 0.334705 |
| 9 | 108410800 | 108412399 | 8.494055 | 15.71799 | 32.04394 | 13.4913 | -0.58059 | 0.192636 | 0.334705 |

| | | | | | | | | | |
|---|---|---|---|---|---|---|---|---|---|
| 11 | 24174000 | 24177399 | 17.53659 | 30.04944 | 55.97591 | 24.54821 | -0.51069 | 0.939425 | 0.334715 |
| 8 | 19131800 | 19137199 | 50.42623 | 81.96146 | 60.8918 | 26.12625 | 0.796291 | 1.603426 | 0.334719 |
| 2 | 82613200 | 82614399 | 15.78846 | 29.16976 | 24.16192 | 10.19476 | 0.70735 | 0.240734 | 0.334722 |
| 16 | 38736200 | 38737599 | 9.462773 | 17.23466 | 34.43852 | 14.80964 | -0.57382 | 0.295698 | 0.334736 |
| 1 | 89507600 | 89511399 | 20.43581 | 33.83052 | 70.1913 | 30.36464 | -0.66889 | 1.304008 | 0.334742 |
| 15 | 95424600 | 95426799 | 20.36157 | 33.87684 | 67.6793 | 29.37935 | -0.61395 | 1.242459 | 0.334745 |
| 11 | 50343200 | 50344999 | 14.83796 | 27.53146 | 23.6733 | 9.930495 | 0.65956 | 0.207481 | 0.334752 |
| 19 | 8210800 | 8211999 | 14.83796 | 27.53146 | 23.6733 | 9.930495 | 0.65956 | 0.207481 | 0.334752 |
| 3 | 7628000 | 7628599 | 20.67222 | 36.10569 | 33.5705 | 14.78798 | 0.492168 | 0.720844 | 0.334753 |
| 15 | 33668400 | 33669599 | 6.420465 | 12.34977 | 27.54311 | 10.73526 | -0.61301 | -0.11109 | 0.334758 |
| 18 | 53989400 | 53989999 | 10.69782 | 20.48786 | 20.11668 | 7.939367 | 0.543605 | -0.04734 | 0.334761 |
| 6 | 102548600 | 102551999 | 38.90361 | 64.03715 | 56.61969 | 24.41385 | 0.556027 | 1.407762 | 0.334786 |
| 8 | 75939400 | 75943999 | 32.61131 | 52.51897 | 99.34481 | 42.23255 | -0.56261 | 1.814143 | 0.334787 |
| X | 68288800 | 68290799 | 27.91944 | 48.0286 | 38.74442 | 16.99515 | 0.692996 | 0.928726 | 0.334797 |
| 1 | 175376200 | 175379199 | 18.70296 | 31.7601 | 60.86007 | 26.66809 | -0.55344 | 1.030772 | 0.334826 |
| 17 | 38866600 | 38869599 | 43.03291 | 70.50369 | 57.9037 | 24.91816 | 0.659553 | 1.474852 | 0.334832 |
| X | 77370400 | 77372399 | 22.01333 | 38.90938 | 28.60559 | 12.65346 | 0.831834 | 0.560791 | 0.334835 |
| 6 | 110841400 | 110841999 | 6.431493 | 12.34982 | 26.86391 | 10.52215 | -0.58339 | -0.09442 | 0.334838 |
| 8 | 114859400 | 114861399 | 12.60573 | 22.01743 | 45.22948 | 19.92412 | -0.64583 | 0.698246 | 0.334857 |
| 2 | 151090000 | 151092999 | 26.88221 | 46.60959 | 36.40598 | 15.97323 | 0.741377 | 0.850537 | 0.334883 |
| 2 | 181330800 | 181333599 | 12.82639 | 22.22622 | 50.00656 | 21.94014 | -0.7781 | 0.811031 | 0.334893 |
| 14 | 29646000 | 29647799 | 18.29446 | 33.35628 | 24.20446 | 10.39448 | 0.879107 | 0.331385 | 0.334894 |
| 13 | 65617400 | 65618599 | 22.16066 | 38.49818 | 35.27385 | 15.48599 | 0.512626 | 0.806295 | 0.334914 |
| 5 | 73562200 | 73565599 | 23.90365 | 39.12688 | 86.0059 | 36.99723 | -0.75737 | 1.461646 | 0.334932 |
| 2 | 84034200 | 84037399 | 47.71981 | 77.75943 | 58.66366 | 25.19398 | 0.776673 | 1.558443 | 0.334937 |
| 18 | 64341000 | 64342599 | 8.621346 | 15.93404 | 34.73255 | 14.64762 | -0.67884 | 0.203643 | 0.334939 |
| 4 | 81080200 | 81081799 | 17.31008 | 31.67906 | 24.35572 | 10.41003 | 0.800705 | 0.303725 | 0.334954 |
| 3 | 144142600 | 144144999 | 12.44999 | 21.73014 | 46.79988 | 20.60679 | -0.71395 | 0.708474 | 0.334961 |
| 1 | 76077000 | 76077799 | 13.28913 | 25.46594 | 19.86547 | 7.826655 | 0.874919 | -0.02776 | 0.334979 |
| 11 | 15409200 | 15409999 | 11.50039 | 21.80768 | 21.04904 | 8.496474 | 0.53934 | 0.035271 | 0.334988 |
| 7 | 17262400 | 17263599 | 17.42336 | 31.72588 | 25.46303 | 10.95313 | 0.733315 | 0.335846 | 0.334993 |
| 5 | 77553600 | 77558199 | 64.67612 | 101.4541 | 166.5665 | 69.08119 | -0.39637 | 2.581095 | 0.334999 |
| X | 65397800 | 65400599 | 31.26548 | 54.4807 | 29.56294 | 13.00359 | 1.26515 | 0.801946 | 0.335004 |
| 7 | 31465000 | 31466799 | 22.89615 | 39.95525 | 31.36392 | 13.80844 | 0.735108 | 0.740595 | 0.335025 |
| 12 | 105732200 | 105739399 | 69.99674 | 108.967 | 185.8509 | 77.14614 | -0.45989 | 2.718942 | 0.335028 |
| X | 162127600 | 162129799 | 20.17204 | 36.33897 | 25.66994 | 11.19128 | 0.904157 | 0.416435 | 0.335034 |
| 2 | 166466800 | 166469999 | 23.59698 | 38.60399 | 77.50931 | 33.33566 | -0.62745 | 1.465477 | 0.335044 |
| 12 | 104902400 | 104903399 | 17.49209 | 31.21196 | 27.984 | 12.30411 | 0.553897 | 0.458561 | 0.335045 |
| 4 | 14679000 | 14682199 | 65.51737 | 102.6951 | 168.2383 | 69.78004 | -0.39411 | 2.595305 | 0.33505 |
| 8 | 76459200 | 76460999 | 25.21075 | 44.21065 | 29.51851 | 13.04616 | 0.967206 | 0.675025 | 0.335063 |
| 2 | 143510800 | 143511199 | 3.230899 | 6.31887 | 17.53266 | 5.743442 | -0.68851 | -0.79272 | 0.335064 |
| 8 | 92578000 | 92579399 | 8.54156 | 15.61164 | 35.02608 | 14.99621 | -0.73576 | 0.267945 | 0.335095 |
| 9 | 58942600 | 58944399 | 11.77028 | 20.73786 | 41.42176 | 18.33434 | -0.60404 | 0.557658 | 0.335096 |
| 18 | 66044800 | 66046999 | 11.77028 | 20.73786 | 41.42176 | 18.33434 | -0.60404 | 0.557658 | 0.335096 |
| 2 | 32839000 | 32846199 | 46.43574 | 73.74558 | 135.7186 | 56.57914 | -0.54612 | 2.27863 | 0.335104 |
| X | 4321000 | 4322999 | 17.82937 | 32.75361 | 23.92342 | 10.17254 | 0.879739 | 0.28741 | 0.335105 |
| X | 89071400 | 89073199 | 17.82937 | 32.75361 | 23.92342 | 10.17254 | 0.879739 | 0.28741 | 0.335105 |

| | | | | | | | | | |
|---|---|---|---|---|---|---|---|---|---|
| X | 125461400 | 125468799 | 89.04262 | 141.1478 | 103.8398 | 43.20883 | 0.772082 | 2.36106 | 0.335117 |
| X | 127218600 | 127223199 | 59.20387 | 95.4919 | 68.89312 | 29.34869 | 0.827611 | 1.798137 | 0.335118 |
| 10 | 75803800 | 75810199 | 60.57623 | 95.10687 | 175.6617 | 72.85699 | -0.56508 | 2.563055 | 0.335123 |
| 7 | 76382000 | 76382599 | 6.333079 | 12.01179 | 28.17835 | 11.36892 | -0.73012 | 0.006504 | 0.335133 |
| 17 | 91137600 | 91138599 | 11.84215 | 22.39865 | 20.89701 | 8.480642 | 0.580797 | 0.053787 | 0.335133 |
| 3 | 34701800 | 34705799 | 38.71424 | 61.85728 | 118.9168 | 49.96913 | -0.59955 | 2.0762 | 0.335142 |
| 7 | 21049800 | 21052999 | 26.04156 | 45.90018 | 26.75945 | 11.84349 | 1.16325 | 0.612722 | 0.335147 |
| X | 32930600 | 32933199 | 42.18736 | 69.32411 | 49.41309 | 21.29108 | 0.865081 | 1.434012 | 0.335155 |
| 12 | 92805200 | 92806199 | 7.673698 | 14.57072 | 31.06359 | 12.50354 | -0.59114 | 0.018282 | 0.335179 |
| 8 | 94670600 | 94671199 | 4.817392 | 9.361348 | 23.38587 | 8.710363 | -0.71111 | -0.29386 | 0.335195 |
| 12 | 6959400 | 6960799 | 15.50256 | 28.73222 | 23.59641 | 9.913418 | 0.723821 | 0.21895 | 0.3352 |
| 4 | 120460200 | 120462399 | 10.71236 | 18.89266 | 40.44891 | 17.90018 | -0.70277 | 0.539698 | 0.335236 |
| 4 | 112421600 | 112431799 | 186.6785 | 279.2822 | 245.1401 | 101.0932 | 0.450991 | 3.52195 | 0.335274 |
| 4 | 113516200 | 113517599 | 24.99502 | 44.70081 | 21.80923 | 9.570888 | 1.428031 | 0.484564 | 0.335276 |
| 5 | 147043200 | 147052999 | 119.7164 | 179.7667 | 296.3986 | 122.3327 | -0.45313 | 3.443062 | 0.33529 |
| 9 | 70155400 | 70157399 | 9.89601 | 18.07815 | 35.53807 | 15.22495 | -0.5471 | 0.282873 | 0.335311 |
| 11 | 72545800 | 72547199 | 14.05337 | 26.91278 | 19.1726 | 7.567923 | 1.002001 | -0.01644 | 0.335317 |
| 1 | 54964200 | 54967199 | 22.39154 | 37.35631 | 66.85239 | 29.07126 | -0.45574 | 1.218318 | 0.335331 |
| 5 | 76590200 | 76592199 | 22.58603 | 39.2436 | 36.67709 | 16.10371 | 0.484062 | 0.806828 | 0.335337 |
| 11 | 15193400 | 15194999 | 17.58316 | 31.67384 | 26.31329 | 11.47239 | 0.671263 | 0.39772 | 0.335339 |
| 3 | 150738000 | 150739999 | 10.43165 | 18.9667 | 43.71856 | 18.84076 | -0.78127 | 0.330716 | 0.335363 |
| 10 | 4192400 | 4194599 | 18.71539 | 31.79341 | 58.62674 | 25.69246 | -0.49809 | 1.026235 | 0.335371 |
| 6 | 63673000 | 63673799 | 11.54406 | 22.3127 | 19.07499 | 7.312706 | 0.780656 | -0.13438 | 0.335385 |
| 1 | 47734600 | 47734999 | 9.035326 | 17.53918 | 17.72721 | 6.65955 | 0.56922 | -0.23285 | 0.335403 |
| 8 | 74738400 | 74739399 | 14.23506 | 27.30823 | 18.61723 | 7.312064 | 1.071516 | -0.03159 | 0.335415 |
| 2 | 118318400 | 118322399 | 25.24227 | 41.56483 | 74.51394 | 32.13454 | -0.46056 | 1.383333 | 0.335418 |
| 17 | 20015800 | 20017799 | 32.45994 | 54.80685 | 44.70354 | 19.55455 | 0.674139 | 1.113486 | 0.335422 |
| 3 | 22860800 | 22864399 | 50.57703 | 81.72687 | 64.25184 | 27.43621 | 0.706886 | 1.728331 | 0.335432 |
| 12 | 13801600 | 13803799 | 17.91081 | 30.60559 | 55.3776 | 24.28038 | -0.46942 | 0.967363 | 0.335434 |
| 19 | 5626000 | 5628399 | 27.00748 | 46.33107 | 36.72558 | 16.10747 | 0.71735 | 0.944926 | 0.335438 |
| 3 | 84205800 | 84206999 | 13.51672 | 25.30624 | 22.92043 | 9.475545 | 0.601859 | 0.141456 | 0.335439 |
| 15 | 95912200 | 95915199 | 52.85506 | 84.86724 | 80.22473 | 33.95644 | 0.430682 | 1.926026 | 0.335446 |
| 5 | 104903600 | 104912799 | 224.0632 | 330.0811 | 294.1775 | 121.1449 | 0.408514 | 3.795273 | 0.335464 |
| 2 | 132931400 | 132932999 | 15.41366 | 28.49592 | 24.17538 | 10.19136 | 0.673901 | 0.233826 | 0.335474 |
| 12 | 22839600 | 22841199 | 50.90431 | 81.80029 | 75.91652 | 32.17103 | 0.458549 | 1.897457 | 0.335481 |
| 17 | 29759200 | 29760799 | 10.62282 | 18.95751 | 38.34285 | 16.8567 | -0.61274 | 0.433764 | 0.33549 |
| 10 | 114007800 | 114012799 | 59.5915 | 95.62972 | 82.00504 | 34.68023 | 0.570041 | 1.952717 | 0.335503 |
| 5 | 10410800 | 10414999 | 76.19119 | 120.5341 | 115.3481 | 47.94005 | 0.390839 | 2.401246 | 0.335514 |
| 4 | 146130600 | 146133999 | 44.88546 | 74.93973 | 40.27044 | 17.5202 | 1.274116 | 1.25758 | 0.335518 |
| 8 | 33845200 | 33847799 | 36.81815 | 60.62736 | 56.00156 | 24.1512 | 0.493275 | 1.396073 | 0.335531 |
| 19 | 30364000 | 30368399 | 65.86807 | 105.283 | 83.88849 | 35.26892 | 0.669337 | 2.082075 | 0.335536 |
| 15 | 31827400 | 31828599 | 10.19908 | 18.18338 | 37.38317 | 16.44845 | -0.63655 | 0.438139 | 0.335562 |
| 15 | 36496400 | 36496999 | 4.81286 | 9.365768 | 22.95143 | 8.451525 | -0.66851 | -0.3364 | 0.335573 |
| X | 115397400 | 115397999 | 13.22242 | 24.77146 | 22.46825 | 9.278321 | 0.601199 | 0.134179 | 0.335577 |
| 1 | 30581200 | 30582599 | 13.04576 | 23.01009 | 43.47619 | 19.2395 | -0.52378 | 0.557997 | 0.335584 |
| 3 | 10530600 | 10531799 | 11.12143 | 19.76383 | 38.32562 | 16.90912 | -0.55579 | 0.46521 | 0.335592 |
| 3 | 138829400 | 138830599 | 11.12143 | 19.76383 | 38.32562 | 16.90912 | -0.55579 | 0.46521 | 0.335592 |

| | | | | | | | | | |
|---|---|---|---|---|---|---|---|---|---|
| 12 | 102243200 | 102246799 | 36.55959 | 60.35475 | 48.83811 | 21.0868 | 0.684864 | 1.359322 | 0.335601 |
| 4 | 112032200 | 112033999 | 29.07729 | 49.70171 | 39.09917 | 17.1435 | 0.727364 | 0.985294 | 0.335605 |
| 19 | 13242400 | 13248199 | 145.6014 | 220.5235 | 207.7182 | 85.94753 | 0.364763 | 3.272979 | 0.335606 |
| 5 | 148009400 | 148017599 | 91.13144 | 139.4532 | 228.278 | 94.7216 | -0.41986 | 3.051851 | 0.335643 |
| 6 | 70821000 | 70822199 | 6.461196 | 12.30813 | 27.72778 | 11.08482 | -0.66052 | -0.0237 | 0.335653 |
| 1 | 64159600 | 64161999 | 13.55366 | 23.56586 | 47.90056 | 21.04064 | -0.63154 | 0.771433 | 0.335658 |
| 18 | 90563200 | 90564599 | 10.91268 | 19.53662 | 37.83706 | 16.58995 | -0.54786 | 0.41501 | 0.335673 |
| 7 | 141788200 | 141792799 | 87.04428 | 136.5468 | 131.5055 | 54.53989 | 0.372248 | 2.587097 | 0.335688 |
| 13 | 61745000 | 61746599 | 28.34533 | 48.79059 | 38.65633 | 16.95682 | 0.718998 | 0.924328 | 0.335689 |
| 8 | 118532800 | 118535199 | 12.58492 | 22.06231 | 43.06794 | 19.02933 | -0.57247 | 0.635657 | 0.335691 |
| 18 | 37066600 | 37070199 | 45.1602 | 74.34016 | 52.33756 | 22.568 | 0.883822 | 1.42119 | 0.335706 |
| 12 | 87884000 | 87894599 | 101.5579 | 158.4914 | 139.5763 | 57.918 | 0.495399 | 2.686255 | 0.335722 |
| 13 | 40588000 | 40589399 | 7.836679 | 14.72179 | 30.17334 | 12.40257 | -0.5608 | 0.089046 | 0.335753 |
| 4 | 121586400 | 121588399 | 41.98705 | 68.37296 | 63.66892 | 27.33702 | 0.473164 | 1.561029 | 0.33576 |
| 13 | 66603600 | 66604599 | 14.01023 | 27.20407 | 15.49473 | 5.807575 | 1.390905 | -0.18981 | 0.335781 |
| 13 | 99218600 | 99221199 | 15.01305 | 25.86714 | 51.6163 | 22.64246 | -0.60772 | 0.874776 | 0.335822 |
| 7 | 141788000 | 141792799 | 90.25222 | 141.2638 | 135.4985 | 56.20886 | 0.375594 | 2.628463 | 0.335833 |
| 17 | 74241200 | 74242999 | 11.33383 | 19.97691 | 40.44898 | 17.90232 | -0.62301 | 0.551269 | 0.335835 |
| 17 | 93736400 | 93736999 | 5.657252 | 11.00138 | 25.9918 | 9.624395 | -0.6245 | -0.2965 | 0.335844 |
| 7 | 33376600 | 33377999 | 36.54123 | 60.39413 | 54.24576 | 23.44034 | 0.534998 | 1.353723 | 0.335851 |
| 18 | 90073200 | 90077599 | 42.86824 | 70.33815 | 56.78953 | 24.45402 | 0.684949 | 1.455938 | 0.33586 |
| 16 | 56162600 | 56163799 | 16.14089 | 29.39148 | 25.52351 | 10.97876 | 0.620195 | 0.325172 | 0.335879 |
| X | 57049200 | 57051399 | 20.72451 | 37.4723 | 24.59611 | 10.67373 | 1.013691 | 0.397914 | 0.335886 |
| 17 | 6226000 | 6227599 | 17.31131 | 31.18267 | 27.06178 | 11.79942 | 0.608899 | 0.399686 | 0.335911 |
| 3 | 8237400 | 8241199 | 83.48284 | 131.2163 | 126.8496 | 52.61939 | 0.368966 | 2.545677 | 0.335921 |
| 7 | 93959800 | 93962199 | 30.9726 | 52.01103 | 47.49916 | 20.74069 | 0.511455 | 1.157994 | 0.335947 |
| 17 | 63549400 | 63555399 | 72.78147 | 112.8543 | 191.0084 | 79.3339 | -0.45327 | 2.78982 | 0.335952 |
| 15 | 97116400 | 97124399 | 73.428 | 114.3078 | 185.836 | 77.14 | -0.39084 | 2.721028 | 0.335955 |
| 9 | 12364200 | 12370999 | 129.9308 | 198.5875 | 184.6059 | 76.57906 | 0.394383 | 3.083906 | 0.335958 |
| 16 | 14774400 | 14776999 | 16.44528 | 28.21928 | 53.22579 | 23.3448 | -0.5281 | 0.920015 | 0.335963 |
| 8 | 21416600 | 21424199 | 141.4263 | 214.7704 | 201.4156 | 83.40733 | 0.374246 | 3.217983 | 0.335974 |
| 1 | 117916000 | 117923999 | 199.2826 | 295.9908 | 269.0527 | 110.8353 | 0.391055 | 3.654242 | 0.336002 |
| X | 109437200 | 109438399 | 17.36701 | 31.34839 | 26.6023 | 11.56884 | 0.643548 | 0.387936 | 0.336007 |
| 10 | 90978400 | 90980199 | 10.15477 | 18.16932 | 37.8601 | 16.60809 | -0.65321 | 0.413186 | 0.336022 |
| 3 | 28346800 | 28348399 | 9.916321 | 17.99612 | 36.47329 | 15.7551 | -0.59871 | 0.322595 | 0.336025 |
| 12 | 50960000 | 50960999 | 9.916321 | 17.99612 | 36.47329 | 15.7551 | -0.59871 | 0.322595 | 0.336025 |
| 9 | 8885600 | 8885999 | 3.93065 | 7.687328 | 20.38546 | 6.920582 | -0.67325 | -0.64633 | 0.336073 |
| 14 | 121185600 | 121191999 | 58.78491 | 92.65218 | 157.9375 | 65.53872 | -0.44518 | 2.480771 | 0.336086 |
| 11 | 112825800 | 112831399 | 46.62601 | 74.46238 | 125.8538 | 52.84765 | -0.41501 | 2.100859 | 0.336108 |
| 8 | 3939000 | 3941199 | 39.92443 | 65.68224 | 56.6517 | 24.42211 | 0.591415 | 1.417157 | 0.336114 |
| 1 | 6231000 | 6239599 | 143.2864 | 217.193 | 204.1008 | 84.47232 | 0.369141 | 3.255374 | 0.336122 |
| 13 | 83840600 | 83843199 | 32.05639 | 53.65199 | 47.54639 | 20.71772 | 0.554682 | 1.193384 | 0.336134 |
| 6 | 135284400 | 135287399 | 24.40104 | 40.43498 | 71.86258 | 31.10899 | -0.44631 | 1.304983 | 0.336144 |
| 3 | 49586800 | 49587399 | 5.400119 | 10.45925 | 24.93758 | 9.467745 | -0.67145 | -0.21678 | 0.336171 |
| 8 | 34628200 | 34628999 | 5.400119 | 10.45925 | 24.93758 | 9.467745 | -0.67145 | -0.21678 | 0.336171 |
| 1 | 144222800 | 144224199 | 19.95204 | 35.70115 | 25.39987 | 11.14159 | 0.88759 | 0.451832 | 0.336187 |
| 7 | 136524400 | 136525599 | 14.94183 | 27.32123 | 25.26206 | 10.81004 | 0.535236 | 0.293325 | 0.336202 |

| | | | | | | | | | |
|---|---|---|---|---|---|---|---|---|---|
| 4 | 38814800 | 38815999 | 12.71481 | 24.42883 | 19.35204 | 7.567661 | 0.862587 | -0.05899 | 0.336218 |
| 17 | 19108000 | 19113399 | 96.52598 | 151.8899 | 120.6232 | 50.044 | 0.65355 | 2.53063 | 0.336219 |
| 4 | 78607000 | 78608399 | 12.65187 | 24.22031 | 19.82654 | 7.833976 | 0.801721 | -0.02616 | 0.336227 |
| 6 | 41223000 | 41224399 | 23.11889 | 40.41708 | 27.0116 | 11.90703 | 0.966306 | 0.709967 | 0.336236 |
| 4 | 38208200 | 38209799 | 18.63736 | 32.81014 | 30.19713 | 13.3682 | 0.507312 | 0.585566 | 0.336236 |
| 12 | 73811600 | 73813199 | 8.185282 | 15.15631 | 32.05417 | 13.48386 | -0.63232 | 0.185681 | 0.336241 |
| 3 | 153098600 | 153100799 | 33.90241 | 56.04949 | 53.65085 | 23.18804 | 0.443438 | 1.340545 | 0.336243 |
| 9 | 96546000 | 96550999 | 49.6353 | 78.92506 | 133.1135 | 55.5623 | -0.41871 | 2.243874 | 0.336261 |
| 7 | 140560200 | 140564199 | 20.40846 | 34.32745 | 61.84971 | 27.02349 | -0.4654 | 1.122728 | 0.336268 |
| X | 131669800 | 131672199 | 32.65058 | 54.84164 | 46.94218 | 20.50051 | 0.604608 | 1.160814 | 0.336269 |
| 13 | 119125200 | 119126399 | 11.57455 | 20.65426 | 38.84494 | 17.07862 | -0.50849 | 0.443125 | 0.336279 |
| 2 | 70964400 | 70966799 | 10.56754 | 18.61632 | 42.4888 | 18.80713 | -0.79538 | 0.553242 | 0.336295 |
| 2 | 79271200 | 79272399 | 14.46144 | 27.09665 | 22.79262 | 9.409948 | 0.709198 | 0.1438 | 0.336302 |
| 19 | 53752400 | 53757599 | 41.54569 | 66.62184 | 112.363 | 47.46584 | -0.40516 | 1.965162 | 0.336337 |
| 6 | 85045400 | 85046799 | 9.717414 | 17.36762 | 37.87869 | 16.6317 | -0.71943 | 0.417558 | 0.336345 |
| 5 | 109684000 | 109687199 | 29.45067 | 50.6709 | 37.40604 | 16.40815 | 0.820288 | 0.929311 | 0.33636 |
| 12 | 22368600 | 22374399 | 24.07768 | 41.69929 | 33.0947 | 14.51989 | 0.718065 | 0.842427 | 0.336373 |
| 7 | 14154400 | 14156599 | 44.03865 | 71.55659 | 64.76416 | 27.78124 | 0.51181 | 1.599931 | 0.33638 |
| 9 | 26618000 | 26621199 | 58.98573 | 94.37854 | 89.67961 | 37.75817 | 0.417376 | 2.047164 | 0.336384 |
| 6 | 29210400 | 29212599 | 11.30855 | 19.97298 | 39.89289 | 17.64902 | -0.6024 | 0.526216 | 0.336386 |
| 17 | 8030200 | 8031799 | 8.237819 | 15.1082 | 32.98895 | 14.05911 | -0.6916 | 0.243561 | 0.336413 |
| 10 | 3383600 | 3387399 | 55.62785 | 89.17876 | 85.80061 | 36.22961 | 0.402644 | 1.981704 | 0.336415 |
| 5 | 125341800 | 125349999 | 51.12826 | 80.71729 | 153.4457 | 63.70706 | -0.60035 | 2.439282 | 0.336415 |
| 14 | 51556800 | 51564999 | 164.9567 | 247.9049 | 231.5377 | 95.58384 | 0.367307 | 3.431691 | 0.336439 |
| 2 | 81294400 | 81298399 | 45.08788 | 72.94191 | 69.69966 | 29.80217 | 0.427866 | 1.689512 | 0.336447 |
| 6 | 125999400 | 126002799 | 34.54051 | 55.63087 | 98.56368 | 41.90596 | -0.46825 | 1.814393 | 0.336448 |
| 6 | 125999600 | 126002799 | 34.54051 | 55.63087 | 98.56368 | 41.90596 | -0.46825 | 1.814393 | 0.336448 |
| 3 | 45802600 | 45803999 | 19.18729 | 35.28056 | 22.02723 | 9.354561 | 1.105457 | 0.287996 | 0.336466 |
| 5 | 75734400 | 75737999 | 25.01188 | 40.92943 | 77.91036 | 33.51134 | -0.55057 | 1.463704 | 0.336468 |
| 13 | 53733600 | 53736799 | 39.97334 | 65.59396 | 57.79865 | 24.88948 | 0.559165 | 1.447489 | 0.336497 |
| 1 | 172491200 | 172492999 | 13.81484 | 24.15488 | 45.70761 | 20.15084 | -0.52809 | 0.687651 | 0.336501 |
| 9 | 31797800 | 31799999 | 13.81484 | 24.15488 | 45.70761 | 20.15084 | -0.52809 | 0.687651 | 0.336501 |
| 13 | 102381600 | 102382199 | 5.230117 | 10.1872 | 24.39425 | 8.903207 | -0.62463 | -0.35978 | 0.336504 |
| 17 | 75295600 | 75298999 | 47.42086 | 76.50182 | 73.40879 | 31.28036 | 0.417331 | 1.773293 | 0.336508 |
| 4 | 26346200 | 26346999 | 5.677225 | 10.99319 | 24.92719 | 9.474935 | -0.60115 | -0.20836 | 0.336512 |
| 9 | 53075400 | 53076199 | 5.677225 | 10.99319 | 24.92719 | 9.474935 | -0.60115 | -0.20836 | 0.336512 |
| 5 | 6938200 | 6941799 | 81.90442 | 128.8672 | 121.9095 | 50.57688 | 0.40131 | 2.52477 | 0.336531 |
| 6 | 52331400 | 52332999 | 18.27956 | 32.33702 | 29.6663 | 13.12045 | 0.51467 | 0.531452 | 0.336544 |
| 8 | 22200200 | 22203999 | 59.25163 | 95.01079 | 78.12621 | 33.00102 | 0.62918 | 1.97672 | 0.336545 |
| 7 | 59936600 | 59944999 | 148.8731 | 226.073 | 185.8225 | 76.94952 | 0.564301 | 3.21942 | 0.336548 |
| 7 | 60582400 | 60584999 | 44.9818 | 72.8016 | 69.05989 | 29.53798 | 0.438957 | 1.681375 | 0.336572 |
| 3 | 91545600 | 91551599 | 91.31488 | 143.7754 | 123.3656 | 51.18569 | 0.542715 | 2.516883 | 0.336586 |
| 4 | 139339000 | 139344999 | 35.80792 | 57.28489 | 122.2315 | 51.45265 | -0.74769 | 2.03704 | 0.336601 |
| 7 | 41768200 | 41771199 | 55.16972 | 88.4929 | 84.63876 | 35.76887 | 0.412095 | 1.964311 | 0.336607 |
| 9 | 19808200 | 19814199 | 131.6193 | 200.5345 | 189.8456 | 78.6934 | 0.364515 | 3.148199 | 0.336619 |
| 5 | 119266600 | 119268799 | 12.79243 | 22.26672 | 46.84287 | 20.59022 | -0.6805 | 0.75184 | 0.336624 |
| X | 127826600 | 127830399 | 36.80906 | 62.07242 | 41.86505 | 18.30598 | 0.947318 | 1.136417 | 0.336639 |

| | | | | | | | | | |
|---|---|---|---|---|---|---|---|---|---|
| 1 | 111873000 | 111875399 | 23.7811 | 41.34893 | 35.94479 | 15.78911 | 0.588064 | 0.800445 | 0.336671 |
| X | 58881200 | 58882599 | 14.04274 | 26.94505 | 18.62557 | 7.310847 | 1.052477 | -0.03557 | 0.336698 |
| 3 | 130593000 | 130594799 | 10.07463 | 18.43078 | 36.11374 | 15.44327 | -0.54058 | 0.278326 | 0.3367 |
| 4 | 126361400 | 126362999 | 7.518637 | 13.45535 | 37.99365 | 16.66384 | -1.08767 | 0.396019 | 0.336714 |
| 18 | 78461000 | 78465199 | 31.0166 | 52.43762 | 44.69499 | 19.55848 | 0.610884 | 1.096248 | 0.336716 |
| 14 | 71208800 | 71210999 | 13.0387 | 22.87926 | 43.01933 | 19.02291 | -0.51875 | 0.622386 | 0.336719 |
| 12 | 101061600 | 101067599 | 162.6831 | 244.4277 | 228.4332 | 94.29599 | 0.366087 | 3.435861 | 0.336725 |
| 12 | 31948800 | 31950399 | 20.2659 | 35.76223 | 26.27569 | 11.62665 | 0.831589 | 0.56157 | 0.336764 |
| 12 | 20245800 | 20252999 | 174.2199 | 260.4644 | 241.2881 | 99.49136 | 0.37235 | 3.531797 | 0.336774 |
| 8 | 75093000 | 75093599 | 5.368153 | 10.45522 | 25.01788 | 9.14043 | -0.62499 | -0.35193 | 0.336788 |
| 4 | 147044400 | 147047399 | 77.40146 | 123.0768 | 92.99804 | 38.81816 | 0.737821 | 2.257081 | 0.336796 |
| 13 | 54758400 | 54759399 | 5.84186 | 11.20167 | 27.30912 | 10.74024 | -0.75082 | -0.08249 | 0.336815 |
| 1 | 172508200 | 172509799 | 16.26813 | 29.82403 | 24.45841 | 10.4292 | 0.710545 | 0.286014 | 0.336816 |
| 2 | 73424400 | 73430199 | 55.93057 | 88.46474 | 148.047 | 61.52407 | -0.41457 | 2.397075 | 0.336831 |
| 1 | 27033600 | 27035799 | 21.58941 | 38.20759 | 28.67939 | 12.68295 | 0.802753 | 0.545777 | 0.336837 |
| 11 | 12847600 | 12849999 | 20.44636 | 36.68666 | 25.44803 | 11.1368 | 0.926367 | 0.439828 | 0.336851 |
| 16 | 18817000 | 18818199 | 17.53743 | 30.10574 | 54.89448 | 24.07748 | -0.47952 | 0.922193 | 0.336889 |
| 10 | 115371600 | 115372399 | 8.70427 | 15.8744 | 34.47008 | 14.79844 | -0.69134 | 0.281541 | 0.336905 |
| X | 24247400 | 24253599 | 62.12024 | 99.99868 | 65.49566 | 27.82556 | 0.963379 | 1.85748 | 0.336906 |
| X | 38134200 | 38138399 | 49.7309 | 80.53587 | 66.66707 | 28.52707 | 0.63585 | 1.678611 | 0.336909 |
| 17 | 18531800 | 18532199 | 10.85461 | 20.85717 | 20.126 | 7.867799 | 0.581171 | -0.07631 | 0.33692 |
| 1 | 59478600 | 59482399 | 31.57462 | 51.0699 | 90.65787 | 38.75753 | -0.46385 | 1.681381 | 0.336922 |
| 4 | 142771800 | 142772199 | 3.755178 | 7.342659 | 18.95523 | 6.472776 | -0.6409 | -0.63051 | 0.336928 |
| 7 | 6023400 | 6031399 | 144.9237 | 220.482 | 195.7267 | 81.09565 | 0.455521 | 3.181978 | 0.336949 |
| X | 62618000 | 62623799 | 88.94584 | 140.5294 | 114.0381 | 47.3645 | 0.626952 | 2.441849 | 0.336949 |
| 7 | 10052200 | 10069599 | 318.8557 | 456.8426 | 380.8005 | 157.2762 | 0.466029 | 4.222935 | 0.336952 |
| 9 | 113758800 | 113761799 | 15.65688 | 26.88512 | 53.21377 | 23.34081 | -0.59718 | 0.909948 | 0.336967 |
| 7 | 5398800 | 5407799 | 30.74296 | 52.83282 | 33.80252 | 14.82694 | 1.025568 | 0.942224 | 0.336969 |
| 7 | 25813400 | 25813999 | 17.21069 | 30.48862 | 27.74312 | 12.26648 | 0.527064 | 0.505234 | 0.336973 |
| 6 | 63232200 | 63235599 | 29.8264 | 48.24048 | 89.47832 | 38.25221 | -0.527 | 1.679864 | 0.337006 |
| 18 | 60118800 | 60121799 | 30.39122 | 52.16205 | 37.36813 | 16.38995 | 0.862741 | 0.953954 | 0.337015 |
| 14 | 51450400 | 51453999 | 59.67613 | 95.42666 | 91.18899 | 38.36148 | 0.408297 | 2.064711 | 0.337056 |
| 3 | 118927000 | 118928999 | 22.98719 | 40.24114 | 33.12676 | 14.61933 | 0.666894 | 0.696571 | 0.337056 |
| 1 | 94086600 | 94087199 | 4.948942 | 9.616301 | 23.38184 | 8.71383 | -0.67316 | -0.28951 | 0.337059 |
| 1 | 190868200 | 190870399 | 16.22978 | 28.10842 | 51.13697 | 22.43573 | -0.47348 | 0.832811 | 0.337064 |
| 4 | 52378000 | 52378999 | 23.62397 | 40.94762 | 37.74491 | 16.56053 | 0.503422 | 0.840675 | 0.33708 |
| 12 | 115237400 | 115241999 | 83.85642 | 132.6285 | 114.9485 | 47.76814 | 0.533327 | 2.412869 | 0.337089 |
| 1 | 85562200 | 85567599 | 135.3846 | 206.9142 | 183.4697 | 76.107 | 0.462442 | 3.08654 | 0.3371 |
| 12 | 113346400 | 113347799 | 13.77065 | 26.02287 | 21.1898 | 8.615838 | 0.773112 | 0.07848 | 0.337122 |
| 10 | 22533600 | 22535599 | 16.77036 | 28.76794 | 53.22659 | 23.34455 | -0.50052 | 0.924739 | 0.337127 |
| 15 | 12805000 | 12807999 | 19.03755 | 32.29928 | 59.73812 | 26.16934 | -0.50251 | 1.039385 | 0.337128 |
| 14 | 3136000 | 3161399 | 541.3968 | 738.8276 | 613.7193 | 255.4566 | 0.40033 | 4.958258 | 0.337132 |
| 2 | 174572600 | 174574399 | 14.53347 | 25.14696 | 49.49781 | 21.71594 | -0.58661 | 0.832949 | 0.337136 |
| 7 | 54946200 | 54950599 | 37.16238 | 61.86946 | 49.30075 | 21.41056 | 0.707762 | 1.267435 | 0.33714 |
| 11 | 3050000 | 3077799 | 1448.537 | 1819.568 | 1389.022 | 583.8771 | 0.40232 | 6.294019 | 0.337148 |
| 1 | 24761600 | 24762799 | 14.02038 | 27.12444 | 17.21616 | 6.571224 | 1.212495 | -0.12621 | 0.337162 |
| 14 | 99434000 | 99436799 | 42.42068 | 69.07391 | 63.78807 | 27.38743 | 0.485055 | 1.562132 | 0.337175 |

| | | | | | | | | | |
|---|---|---|---|---|---|---|---|---|---|
| 14 | 20438600 | 20439199 | 3.990771 | 7.780317 | 21.96212 | 7.920122 | -0.84196 | -0.42604 | 0.337185 |
| 1 | 20728200 | 20729199 | 8.32685 | 15.40851 | 33.11447 | 13.94211 | -0.65652 | 0.19169 | 0.337187 |
| 12 | 18285200 | 18286199 | 35.75847 | 58.7369 | 51.42509 | 22.15434 | 0.569022 | 1.422278 | 0.337191 |
| 4 | 141756000 | 141761599 | 40.78636 | 65.23671 | 115.2692 | 48.50864 | -0.47644 | 2.045212 | 0.337215 |
| 14 | 31302600 | 31303799 | 11.7052 | 20.47015 | 45.74822 | 20.17201 | -0.76616 | 0.679294 | 0.337221 |
| X | 24431600 | 24433199 | 19.57679 | 35.97317 | 22.01072 | 9.355889 | 1.13355 | 0.294774 | 0.337232 |
| 5 | 115340600 | 115343199 | 13.1694 | 23.0885 | 45.11671 | 19.93569 | -0.57424 | 0.638688 | 0.33724 |
| 3 | 123639400 | 123640999 | 20.23046 | 35.84919 | 28.11892 | 12.43187 | 0.740104 | 0.520739 | 0.337251 |
| 7 | 120633000 | 120634799 | 11.27402 | 19.96763 | 40.3458 | 17.83944 | -0.6176 | 0.494187 | 0.337255 |
| 16 | 19126800 | 19127799 | 15.60084 | 28.24766 | 26.21745 | 11.35741 | 0.518512 | 0.355837 | 0.337266 |
| 14 | 12300000 | 12301999 | 25.86315 | 44.21566 | 41.50172 | 18.19719 | 0.473793 | 0.975968 | 0.337267 |
| 8 | 61819600 | 61820199 | 5.98048 | 11.53469 | 25.87016 | 10.00502 | -0.60962 | -0.14364 | 0.337272 |
| 10 | 101438200 | 101439199 | 9.251426 | 17.15803 | 34.27046 | 14.38464 | -0.54842 | 0.190566 | 0.337289 |
| 17 | 15524600 | 15527599 | 27.3983 | 44.39933 | 87.15913 | 37.31405 | -0.60594 | 1.634249 | 0.337296 |
| X | 54562200 | 54566399 | 84.0113 | 132.4192 | 116.7218 | 48.43595 | 0.505125 | 2.491499 | 0.33733 |
| 4 | 152922000 | 152931799 | 101.9536 | 154.8412 | 253.6583 | 105.0442 | -0.42971 | 3.209923 | 0.337332 |
| 16 | 9568800 | 9569999 | 14.35133 | 24.86108 | 49.50314 | 21.71906 | -0.60292 | 0.820159 | 0.337344 |
| 8 | 25969000 | 25970399 | 11.40994 | 20.37029 | 38.33243 | 16.84648 | -0.50887 | 0.43786 | 0.337378 |
| 15 | 16524400 | 16527399 | 21.80154 | 38.12681 | 33.19006 | 14.63471 | 0.586546 | 0.704508 | 0.337382 |
| 9 | 12684200 | 12686599 | 33.34463 | 55.35226 | 52.01535 | 22.54446 | 0.470343 | 1.290672 | 0.337391 |
| 10 | 42833200 | 42834399 | 6.614782 | 12.56768 | 28.78936 | 11.57328 | -0.69228 | -0.00397 | 0.337393 |
| 9 | 3690200 | 3696199 | 36.90841 | 61.81673 | 41.54049 | 18.10887 | 0.952547 | 1.199195 | 0.337405 |
| 5 | 70595600 | 70596599 | 13.06846 | 24.27619 | 22.88741 | 9.585762 | 0.529383 | 0.184965 | 0.337415 |
| 18 | 48087200 | 48088599 | 18.51813 | 32.98738 | 28.39535 | 12.50265 | 0.610995 | 0.476836 | 0.337419 |
| 8 | 53555400 | 53558399 | 53.70898 | 86.22927 | 82.09524 | 34.74261 | 0.420249 | 1.931407 | 0.33743 |
| 17 | 17519400 | 17535599 | 452.8484 | 630.3597 | 494.6439 | 205.2498 | 0.51129 | 4.65256 | 0.337446 |
| 8 | 26544200 | 26545799 | 17.15892 | 30.74493 | 27.48334 | 12.04173 | 0.561567 | 0.429813 | 0.337446 |
| 7 | 12553800 | 12564399 | 158.5489 | 240.4779 | 200.3325 | 82.92836 | 0.543565 | 3.246886 | 0.337459 |
| 2 | 94743400 | 94745199 | 13.39475 | 23.32236 | 46.83588 | 20.59117 | -0.61388 | 0.74943 | 0.337463 |
| 3 | 60353000 | 60354599 | 14.93356 | 27.63023 | 24.24301 | 10.20886 | 0.62683 | 0.22592 | 0.337487 |
| 8 | 69187000 | 69189599 | 15.51574 | 26.69429 | 51.60567 | 22.63679 | -0.56256 | 0.891919 | 0.337489 |
| 2 | 172371400 | 172374199 | 17.08059 | 28.81278 | 61.14637 | 26.73953 | -0.70039 | 1.084603 | 0.337495 |
| 12 | 68085800 | 68086799 | 13.06659 | 22.96666 | 42.98009 | 19.03249 | -0.51187 | 0.596732 | 0.337495 |
| 1 | 29048400 | 29049799 | 19.59392 | 34.97719 | 27.80113 | 12.21933 | 0.727168 | 0.470762 | 0.337496 |
| 12 | 61162400 | 61164199 | 19.59392 | 34.97719 | 27.80113 | 12.21933 | 0.727168 | 0.470762 | 0.337496 |
| 1 | 113428200 | 113431199 | 46.36092 | 75.64231 | 59.68423 | 25.6443 | 0.713466 | 1.539631 | 0.3375 |
| 2 | 146062600 | 146066199 | 36.86162 | 59.11511 | 109.1364 | 46.10821 | -0.53513 | 1.955929 | 0.337505 |
| 2 | 112271800 | 112273399 | 8.753047 | 16.3819 | 35.76241 | 14.79327 | -0.65937 | 0.131995 | 0.337514 |
| 4 | 105775600 | 105777999 | 21.53509 | 37.726 | 33.26118 | 14.6875 | 0.56842 | 0.678124 | 0.337517 |
| 10 | 95638400 | 95640599 | 17.23414 | 29.71812 | 54.374 | 23.85277 | -0.48305 | 0.880966 | 0.337521 |
| 12 | 22334800 | 22336199 | 24.89472 | 42.99175 | 31.21445 | 13.69376 | 0.845172 | 0.869114 | 0.337531 |
| 3 | 134974200 | 134975799 | 8.313222 | 15.41934 | 33.16367 | 13.91843 | -0.65436 | 0.178134 | 0.337531 |
| 7 | 39725800 | 39727999 | 12.51599 | 22.13152 | 42.93413 | 18.99003 | -0.56034 | 0.524075 | 0.337534 |
| 7 | 24173600 | 24175399 | 9.671456 | 17.4223 | 35.8701 | 15.63865 | -0.62964 | 0.364404 | 0.337575 |
| 6 | 87106800 | 87108799 | 21.154 | 36.93107 | 34.52859 | 15.20586 | 0.484132 | 0.730574 | 0.337579 |
| 10 | 24354600 | 24356799 | 13.49508 | 23.64289 | 44.60962 | 19.69994 | -0.52389 | 0.651911 | 0.337602 |
| 17 | 15641400 | 15643599 | 22.85964 | 39.48667 | 36.8246 | 16.15505 | 0.485666 | 0.86537 | 0.337603 |

| | | | | | | | | | |
|---|---|---|---|---|---|---|---|---|---|
| 1 | 176096400 | 176097599 | 14.64438 | 26.79604 | 24.83157 | 10.61586 | 0.533091 | 0.286646 | 0.337622 |
| 15 | 95985800 | 95988999 | 31.59026 | 52.63789 | 50.30142 | 21.85696 | 0.446483 | 1.243249 | 0.337624 |
| 1 | 34625000 | 34629399 | 32.99744 | 52.99009 | 105.6404 | 44.72358 | -0.6433 | 1.907759 | 0.337628 |
| 17 | 23201800 | 23207999 | 79.59057 | 126.3885 | 104.91 | 43.7317 | 0.600649 | 2.3014 | 0.337628 |
| 6 | 59402800 | 59403599 | 5.038304 | 9.685969 | 29.14844 | 11.37853 | -1.0431 | -0.11718 | 0.337634 |
| 2 | 119107600 | 119108799 | 7.563338 | 14.12054 | 30.59273 | 12.7076 | -0.65244 | 0.127466 | 0.337651 |
| 2 | 69098600 | 69099599 | 11.23354 | 21.61927 | 19.5034 | 7.590038 | 0.683096 | -0.08956 | 0.337659 |
| 14 | 71810400 | 71814399 | 48.0761 | 78.49124 | 57.94182 | 24.90148 | 0.809995 | 1.535981 | 0.337666 |
| 13 | 112866000 | 112869199 | 33.27575 | 53.57137 | 99.02319 | 42.07574 | -0.52995 | 1.827092 | 0.337672 |
| 5 | 94869600 | 94869999 | 13.98347 | 26.02691 | 20.27128 | 8.464049 | 0.806405 | 0.174643 | 0.337707 |
| 7 | 57932400 | 57933199 | 16.42505 | 29.32274 | 26.31688 | 11.5662 | 0.55306 | 0.446254 | 0.337712 |
| 2 | 47494400 | 47496799 | 32.3257 | 53.99079 | 49.74022 | 21.64536 | 0.499071 | 1.221193 | 0.33772 |
| 5 | 27022000 | 27023799 | 14.05556 | 24.33211 | 49.4957 | 21.71535 | -0.63373 | 0.825501 | 0.337732 |
| 3 | 60909600 | 60911799 | 18.43119 | 30.8132 | 65.99976 | 28.73818 | -0.71403 | 1.182204 | 0.337743 |
| 3 | 20212400 | 20219399 | 84.83569 | 133.6425 | 124.7867 | 51.7788 | 0.421472 | 2.502693 | 0.337746 |
| 3 | 121819600 | 121820199 | 5.824626 | 11.2768 | 25.53353 | 9.712711 | -0.60068 | -0.20396 | 0.337747 |
| 1 | 80360000 | 80363199 | 26.47039 | 43.2683 | 78.5013 | 33.75192 | -0.48263 | 1.484289 | 0.337747 |
| 1 | 29376000 | 29377999 | 23.54795 | 41.57692 | 27.94736 | 12.36497 | 0.959822 | 0.580672 | 0.337747 |
| 9 | 87587400 | 87588199 | 5.710685 | 10.687 | 32.22029 | 13.33 | -1.1202 | 0.098733 | 0.337755 |
| 12 | 21178800 | 21184599 | 35.69201 | 57.40224 | 101.7473 | 43.16879 | -0.47151 | 1.863046 | 0.33776 |
| X | 33675400 | 33676199 | 19.63403 | 34.8506 | 28.89993 | 12.76317 | 0.661779 | 0.508081 | 0.337766 |
| 15 | 98521400 | 98523399 | 18.01843 | 31.95084 | 28.70319 | 12.68699 | 0.546211 | 0.504215 | 0.337792 |
| 12 | 3110800 | 3114399 | 44.86962 | 72.58102 | 66.54481 | 28.45097 | 0.487295 | 1.691738 | 0.337795 |
| 5 | 135076400 | 135079799 | 26.87458 | 43.66835 | 83.56137 | 35.84506 | -0.56549 | 1.576322 | 0.337805 |
| X | 126526400 | 126528799 | 21.24442 | 37.84633 | 27.05922 | 11.91368 | 0.877351 | 0.492424 | 0.337809 |
| X | 117749200 | 117753799 | 50.76157 | 82.89939 | 55.24065 | 23.74308 | 0.95757 | 1.53647 | 0.337812 |
| 1 | 124260000 | 124260599 | 5.054279 | 9.853426 | 24.53872 | 8.852637 | -0.66444 | -0.39977 | 0.337821 |
| 5 | 132647400 | 132648599 | 16.83997 | 29.89902 | 26.27725 | 11.60231 | 0.578517 | 0.481995 | 0.337827 |
| 9 | 103376000 | 103382399 | 54.62309 | 86.4272 | 146.4875 | 60.88685 | -0.43247 | 2.387555 | 0.337831 |
| 15 | 3100200 | 3113599 | 160.9169 | 243.2799 | 213.1467 | 88.14342 | 0.466942 | 3.313203 | 0.337832 |
| 3 | 101795000 | 101796399 | 7.728827 | 14.35959 | 31.57452 | 13.22112 | -0.68312 | 0.161502 | 0.337833 |
| 14 | 47101400 | 47102999 | 7.728827 | 14.35959 | 31.57452 | 13.22112 | -0.68312 | 0.161502 | 0.337833 |
| 13 | 61869800 | 61880199 | 202.9625 | 302.8558 | 248.053 | 102.2428 | 0.547405 | 3.572748 | 0.337835 |
| 5 | 64348200 | 64349799 | 29.4314 | 49.45167 | 46.96361 | 20.51282 | 0.455184 | 1.146052 | 0.337838 |
| 3 | 107730600 | 107732999 | 12.28077 | 21.57222 | 41.99913 | 18.58847 | -0.56848 | 0.600708 | 0.337847 |
| 15 | 85093200 | 85101399 | 120.0869 | 181.058 | 283.6038 | 117.1794 | -0.37399 | 3.363047 | 0.337864 |
| 10 | 70263400 | 70267799 | 30.0479 | 48.86833 | 85.6479 | 36.75681 | -0.4382 | 1.567654 | 0.337881 |
| X | 125494200 | 125497199 | 36.08802 | 60.32657 | 47.56485 | 20.70995 | 0.722945 | 1.223219 | 0.337883 |
| 5 | 118922800 | 118935199 | 158.8143 | 234.1575 | 372.2102 | 153.2809 | -0.42466 | 3.775859 | 0.337908 |
| 17 | 34075800 | 34078199 | 33.34578 | 55.58426 | 45.47695 | 19.76539 | 0.669767 | 1.240317 | 0.337914 |
| 6 | 105602800 | 105603999 | 14.49933 | 25.30448 | 46.79857 | 20.60491 | -0.49531 | 0.722576 | 0.337923 |
| 6 | 14439200 | 14443399 | 37.57647 | 62.67912 | 47.04346 | 20.45502 | 0.793733 | 1.246095 | 0.337939 |
| 15 | 72961200 | 72963199 | 9.071018 | 16.341 | 38.46899 | 16.77128 | -0.82206 | 0.363063 | 0.337949 |
| 8 | 97495200 | 97498399 | 24.46453 | 43.08627 | 27.84582 | 12.32627 | 1.014808 | 0.612005 | 0.337956 |
| 18 | 73689600 | 73690399 | 7.504877 | 14.2923 | 30.02457 | 12.01009 | -0.56208 | -0.00669 | 0.337965 |
| X | 102501000 | 102501999 | 7.504877 | 14.2923 | 30.02457 | 12.01009 | -0.56208 | -0.00669 | 0.337965 |
| 12 | 107070200 | 107071799 | 9.946979 | 18.16476 | 35.53815 | 15.2318 | -0.54037 | 0.287667 | 0.337968 |

| | | | | | | | | | |
|---|---|---|---|---|---|---|---|---|---|
| 15 | 75818400 | 75820999 | 13.99957 | 24.41349 | 46.28509 | 20.36897 | -0.53108 | 0.729078 | 0.33797 |
| 7 | 33176200 | 33182799 | 127.3711 | 196.3512 | 157.6765 | 65.49618 | 0.614801 | 2.914706 | 0.338001 |
| 15 | 90542200 | 90543799 | 15.46195 | 28.25254 | 25.08048 | 10.74172 | 0.592572 | 0.299456 | 0.338013 |
| 16 | 53208800 | 53211599 | 37.80319 | 62.10467 | 57.13399 | 24.61531 | 0.497801 | 1.427399 | 0.338018 |
| 1 | 81185800 | 81187799 | 14.28478 | 25.02061 | 47.25927 | 20.86596 | -0.52557 | 0.663669 | 0.338039 |
| X | 125880000 | 125882999 | 39.1112 | 65.22673 | 47.05185 | 20.45622 | 0.850928 | 1.256508 | 0.338041 |
| 15 | 99749200 | 99752399 | 20.74189 | 34.78141 | 63.49809 | 27.70176 | -0.48443 | 1.155062 | 0.338056 |
| 1 | 66214000 | 66214999 | 7.368328 | 13.91924 | 29.19943 | 11.88151 | -0.58178 | 0.045596 | 0.338069 |
| 5 | 81522200 | 81527599 | 38.94881 | 62.66862 | 109.0143 | 46.28258 | -0.44373 | 1.856232 | 0.338081 |
| 16 | 10640000 | 10641999 | 20.27791 | 34.01693 | 62.32551 | 27.19705 | -0.48951 | 1.149112 | 0.338109 |
| 3 | 3282000 | 3288399 | 97.05023 | 152.3802 | 117.7053 | 48.81947 | 0.691054 | 2.573555 | 0.33811 |
| 5 | 51532600 | 51533399 | 5.957903 | 11.56474 | 29.47326 | 11.07653 | -0.75295 | -0.23035 | 0.338118 |
| 1 | 100672400 | 100675599 | 31.16849 | 53.0156 | 41.87365 | 18.35276 | 0.720788 | 1.042278 | 0.338118 |
| 3 | 69459800 | 69461999 | 13.56652 | 23.55845 | 48.43796 | 21.2598 | -0.6482 | 0.790667 | 0.33813 |
| 5 | 6733400 | 6739399 | 24.85527 | 43.00132 | 34.13108 | 14.97412 | 0.71774 | 0.857672 | 0.33815 |
| 10 | 40301400 | 40302599 | 7.658629 | 14.41945 | 31.35251 | 12.83866 | -0.64151 | 0.073436 | 0.338162 |
| 4 | 70570400 | 70573199 | 49.66905 | 80.04749 | 75.98676 | 32.33653 | 0.43118 | 1.806273 | 0.338175 |
| 7 | 38201600 | 38212599 | 312.0957 | 447.9312 | 377.757 | 155.9769 | 0.451681 | 4.196623 | 0.338176 |
| 7 | 5497800 | 5500799 | 35.11111 | 58.89146 | 46.39788 | 20.24481 | 0.723846 | 1.18258 | 0.338177 |
| 16 | 90911200 | 90917399 | 57.15199 | 90.19728 | 151.9567 | 63.07833 | -0.42673 | 2.451687 | 0.338184 |
| 14 | 49774600 | 49775199 | 15.27144 | 27.74032 | 25.09836 | 10.82897 | 0.558504 | 0.33107 | 0.338187 |
| 14 | 50576000 | 50578599 | 22.74431 | 39.81753 | 30.11784 | 13.29189 | 0.78836 | 0.689786 | 0.338189 |
| 11 | 6241400 | 6242599 | 7.755052 | 14.33752 | 32.03963 | 13.50609 | -0.71323 | 0.191234 | 0.338191 |
| 7 | 59850800 | 59855399 | 62.76089 | 101.0103 | 71.64051 | 30.42757 | 0.848461 | 1.866138 | 0.338192 |
| 19 | 9735800 | 9737799 | 23.67005 | 41.34266 | 34.26963 | 15.09619 | 0.657042 | 0.736964 | 0.33821 |
| X | 74395600 | 74396199 | 9.832481 | 19.1191 | 17.69452 | 6.55658 | 0.71286 | -0.27123 | 0.338233 |
| 17 | 38306200 | 38307999 | 27.59409 | 46.63543 | 44.28001 | 19.37487 | 0.455749 | 1.087265 | 0.338236 |
| 6 | 131050400 | 131057999 | 119.6093 | 185.3954 | 145.6154 | 60.48366 | 0.652737 | 2.812223 | 0.33825 |
| 7 | 135697200 | 135698399 | 9.306085 | 16.95442 | 34.44353 | 14.80618 | -0.59719 | 0.292396 | 0.338269 |
| 17 | 63356200 | 63358399 | 20.39325 | 34.30138 | 62.43081 | 27.27727 | -0.47998 | 1.122592 | 0.338281 |
| 6 | 136356200 | 136364199 | 64.43257 | 100.7994 | 188.5943 | 78.23865 | -0.58774 | 2.630993 | 0.338306 |
| 10 | 33122000 | 33123999 | 24.44582 | 42.45224 | 37.09818 | 16.2831 | 0.580512 | 0.824057 | 0.338318 |
| 12 | 114584400 | 114587999 | 73.11383 | 116.1075 | 106.1993 | 44.27063 | 0.460892 | 2.296621 | 0.338325 |
| 18 | 53497800 | 53500599 | 20.19023 | 35.73329 | 29.31072 | 12.96202 | 0.675408 | 0.537009 | 0.338333 |
| 1 | 144614800 | 144615399 | 7.09026 | 13.33984 | 29.65405 | 12.1578 | -0.67393 | 0.070636 | 0.33837 |
| 7 | 89250000 | 89250999 | 7.38007 | 13.87609 | 30.19055 | 12.39136 | -0.64473 | 0.078051 | 0.338385 |
| 7 | 106007600 | 106008599 | 13.68012 | 25.37621 | 23.42193 | 9.828681 | 0.558055 | 0.200517 | 0.338391 |
| X | 147717800 | 147720199 | 23.55696 | 41.3956 | 29.56738 | 13.09049 | 0.870279 | 0.633846 | 0.338391 |
| 1 | 99927400 | 99929199 | 17.1221 | 31.3451 | 24.36348 | 10.4088 | 0.785472 | 0.300379 | 0.338398 |
| 7 | 23098600 | 23110999 | 278.8937 | 405.8507 | 323.6274 | 133.3852 | 0.552039 | 3.990358 | 0.338408 |
| 19 | 46950200 | 46951199 | 7.714499 | 14.39864 | 30.58788 | 12.71172 | -0.62481 | 0.131045 | 0.338409 |
| 7 | 16602600 | 16604199 | 7.424746 | 13.82607 | 32.6984 | 13.64172 | -0.78354 | 0.145695 | 0.338436 |
| 8 | 96586600 | 96588599 | 19.08259 | 33.89922 | 28.91283 | 12.76014 | 0.622028 | 0.499059 | 0.33844 |
| 7 | 31442000 | 31446599 | 83.57635 | 131.3613 | 125.6111 | 52.10556 | 0.384686 | 2.546851 | 0.338447 |
| 4 | 112673000 | 112684399 | 26.53059 | 45.6963 | 35.63558 | 15.63206 | 0.741869 | 0.906276 | 0.338447 |
| 5 | 9869400 | 9873799 | 55.68897 | 89.75315 | 72.9548 | 31.04799 | 0.654603 | 1.812266 | 0.338458 |
| 4 | 143518800 | 143523599 | 89.09019 | 140.4944 | 120.7136 | 50.09566 | 0.542497 | 2.483699 | 0.338466 |

| | | | | | | | | | |
|---|---|---|---|---|---|---|---|---|---|
| 2 | 56188400 | 56188999 | 6.940361 | 13.08982 | 29.15031 | 11.90174 | -0.67104 | 0.05466 | 0.338482 |
| 5 | 129702600 | 129703599 | 7.046696 | 13.50625 | 30.29045 | 11.92714 | -0.63461 | -0.06111 | 0.338492 |
| 15 | 59035800 | 59036799 | 21.28308 | 37.05125 | 33.55788 | 14.75207 | 0.529267 | 0.767408 | 0.338501 |
| 10 | 28905400 | 28907199 | 20.72398 | 34.37785 | 68.04764 | 29.47892 | -0.60093 | 1.279942 | 0.338501 |
| 7 | 85575400 | 85577599 | 28.08292 | 48.17621 | 36.18204 | 15.86908 | 0.794935 | 0.949574 | 0.33851 |
| 4 | 113173400 | 113180799 | 165.5734 | 249.5711 | 222.6691 | 91.99571 | 0.437051 | 3.373625 | 0.338511 |
| 3 | 22569400 | 22571799 | 24.68491 | 43.15751 | 32.42574 | 14.2944 | 0.798013 | 0.723644 | 0.338512 |
| 11 | 26411600 | 26413199 | 24.68491 | 43.15751 | 32.42574 | 14.2944 | 0.798013 | 0.723644 | 0.338512 |
| 16 | 13827400 | 13828999 | 9.942723 | 18.08801 | 34.94298 | 15.04817 | -0.52672 | 0.307941 | 0.338525 |
| 1 | 8077600 | 8077999 | 9.911899 | 19.20847 | 17.20156 | 6.513955 | 0.731977 | -0.19618 | 0.338528 |
| 2 | 87859400 | 87860399 | 15.74077 | 29.38188 | 21.94172 | 9.117125 | 0.871866 | 0.174829 | 0.338561 |
| X | 153234200 | 153238199 | 84.74709 | 134.713 | 96.92386 | 40.44354 | 0.807988 | 2.276736 | 0.338565 |
| 3 | 78665800 | 78667399 | 8.426489 | 15.80192 | 32.37375 | 13.34681 | -0.56426 | 0.109031 | 0.338568 |
| Y | 4230200 | 4235799 | 44.7901 | 72.87443 | 35.49503 | 15.23541 | 1.404108 | 1.554031 | 0.338586 |
| 11 | 107829600 | 107832399 | 40.10265 | 66.30882 | 52.7656 | 22.80751 | 0.709185 | 1.35608 | 0.338586 |
| 13 | 52135800 | 52139599 | 29.03109 | 46.9439 | 89.78117 | 38.37217 | -0.57139 | 1.684788 | 0.338593 |
| 7 | 128604400 | 128606999 | 16.23497 | 27.98676 | 52.73251 | 23.13247 | -0.52534 | 0.875801 | 0.338605 |
| 16 | 22055200 | 22057199 | 16.23497 | 27.98676 | 52.73251 | 23.13247 | -0.52534 | 0.875801 | 0.338605 |
| 1 | 19131800 | 19135199 | 36.00646 | 60.13491 | 48.13139 | 20.94447 | 0.701272 | 1.2302 | 0.338616 |
| 2 | 168048200 | 168050599 | 12.43573 | 21.73391 | 46.25374 | 20.38398 | -0.69668 | 0.688068 | 0.338634 |
| 2 | 156197000 | 156208199 | 162.2241 | 237.9835 | 390.816 | 160.9695 | -0.47851 | 3.857499 | 0.338644 |
| 7 | 5654000 | 5656599 | 28.17913 | 49.37195 | 28.57451 | 12.6194 | 1.172755 | 0.706783 | 0.338653 |
| 16 | 45830000 | 45831199 | 7.737802 | 14.00748 | 36.49415 | 15.81348 | -0.96093 | 0.322263 | 0.338658 |
| 6 | 89177400 | 89178599 | 8.10246 | 14.82005 | 35.04367 | 14.98952 | -0.81001 | 0.260318 | 0.338666 |
| 17 | 20748200 | 20751999 | 80.04801 | 126.6273 | 115.7284 | 48.09653 | 0.457076 | 2.406149 | 0.33867 |
| 6 | 73915800 | 73917999 | 14.35332 | 26.36181 | 23.09631 | 9.823927 | 0.618181 | 0.256878 | 0.338679 |
| 5 | 38655800 | 38659599 | 22.15262 | 36.69778 | 69.321 | 30.00236 | -0.53395 | 1.29996 | 0.338695 |
| 8 | 126690200 | 126693199 | 24.08761 | 39.83037 | 71.14987 | 30.75506 | -0.45387 | 1.328116 | 0.338696 |
| 6 | 126872600 | 126873599 | 7.027272 | 13.39894 | 29.50703 | 11.77291 | -0.62669 | -0.02378 | 0.338703 |
| 15 | 51593000 | 51594799 | 13.19572 | 23.16625 | 42.99973 | 19.02219 | -0.50022 | 0.61532 | 0.338705 |
| X | 152902800 | 152903599 | 13.86168 | 26.54621 | 19.84153 | 7.831534 | 0.933828 | -0.01593 | 0.338706 |
| 15 | 37840000 | 37844199 | 61.8288 | 98.79896 | 93.16597 | 39.15387 | 0.426318 | 2.089214 | 0.338724 |
| 12 | 35107600 | 35109399 | 25.11679 | 43.40387 | 38.14051 | 16.73256 | 0.571162 | 0.872723 | 0.338724 |
| 7 | 140190200 | 140190799 | 5.519021 | 10.71313 | 24.47748 | 9.197298 | -0.59573 | -0.24977 | 0.338732 |
| 17 | 39662200 | 39662999 | 15.10849 | 27.67888 | 25.24077 | 10.77297 | 0.557736 | 0.283973 | 0.338732 |
| X | 34776800 | 34782399 | 118.5617 | 184.3086 | 137.6414 | 57.14663 | 0.728669 | 2.76196 | 0.338735 |
| 13 | 62238600 | 62247799 | 206.2242 | 304.539 | 279.6127 | 115.15 | 0.36887 | 3.752453 | 0.33874 |
| 1 | 65246000 | 65251999 | 86.11189 | 135.5793 | 125.7966 | 52.19414 | 0.429986 | 2.513823 | 0.338744 |
| 7 | 138379000 | 138381799 | 15.64825 | 26.86328 | 54.83995 | 24.05361 | -0.64176 | 0.913154 | 0.338746 |
| 1 | 63875200 | 63875799 | 4.655327 | 9.053317 | 22.85344 | 8.459786 | -0.71707 | -0.31799 | 0.338761 |
| 18 | 5899200 | 5902599 | 37.50785 | 61.49207 | 58.56708 | 25.21065 | 0.446813 | 1.455686 | 0.33877 |
| 10 | 61195600 | 61197399 | 20.57316 | 34.12637 | 67.96622 | 29.44288 | -0.60972 | 1.280965 | 0.338774 |
| 7 | 17826400 | 17828199 | 32.27512 | 55.09098 | 39.63078 | 17.37465 | 0.85583 | 1.014204 | 0.338774 |
| 9 | 102594400 | 102596599 | 12.75074 | 22.24984 | 47.32793 | 20.83623 | -0.69625 | 0.71367 | 0.338787 |
| 13 | 21853000 | 21855399 | 26.40331 | 46.08953 | 32.38487 | 14.26066 | 0.894166 | 0.755409 | 0.338792 |
| 4 | 44357600 | 44359399 | 17.9235 | 30.64926 | 55.37604 | 24.28111 | -0.46714 | 0.961444 | 0.338796 |
| 2 | 75999800 | 76000799 | 15.69277 | 29.42099 | 22.17119 | 9.144339 | 0.867884 | 0.149746 | 0.338816 |

| | | | | | | | | | |
|---|---|---|---|---|---|---|---|---|---|
| 5 | 64001600 | 64003599 | 14.04212 | 26.19398 | 19.82195 | 8.245487 | 0.852117 | 0.16074 | 0.338823 |
| 9 | 32606800 | 32608199 | 9.875567 | 17.65013 | 36.8798 | 16.19322 | -0.65774 | 0.418483 | 0.338825 |
| 3 | 154033400 | 154042199 | 75.68008 | 116.6668 | 225.0715 | 93.4909 | -0.6484 | 2.900941 | 0.338832 |
| 17 | 40045200 | 40048599 | 64.06222 | 102.1827 | 97.32617 | 40.79785 | 0.409091 | 2.148924 | 0.338836 |
| 15 | 72416800 | 72424199 | 63.66515 | 99.46404 | 175.4209 | 72.7838 | -0.50411 | 2.652777 | 0.338847 |
| 1 | 85239200 | 85244799 | 196.5134 | 293.7542 | 243.7215 | 100.4921 | 0.531127 | 3.538738 | 0.338851 |
| 14 | 42912200 | 42919799 | 156.3568 | 236.8828 | 206.1357 | 85.30063 | 0.479311 | 3.269097 | 0.338871 |
| 7 | 40018600 | 40021799 | 25.36595 | 43.42333 | 40.97913 | 17.96998 | 0.466295 | 0.961466 | 0.338877 |
| 3 | 86747800 | 86750999 | 50.35957 | 81.05817 | 77.98115 | 33.12485 | 0.409727 | 1.847536 | 0.338881 |
| 2 | 122064200 | 122065799 | 8.127532 | 14.80776 | 33.97141 | 14.6026 | -0.7709 | 0.28326 | 0.3389 |
| 5 | 7514800 | 7518999 | 54.1662 | 87.05108 | 80.26527 | 34.01905 | 0.468101 | 1.8998 | 0.338904 |
| 6 | 122204000 | 122204999 | 12.73462 | 22.30932 | 43.59447 | 19.25072 | -0.57401 | 0.647996 | 0.338915 |
| 2 | 146422400 | 146426199 | 19.52287 | 33.19742 | 65.42937 | 28.6763 | -0.59374 | 1.026807 | 0.338939 |
| 4 | 86257600 | 86261399 | 32.42944 | 52.28219 | 96.79245 | 41.20915 | -0.52976 | 1.779701 | 0.338953 |
| 14 | 42103800 | 42110999 | 83.61953 | 132.1674 | 117.3059 | 48.73165 | 0.49831 | 2.42766 | 0.338954 |
| 15 | 38903200 | 38904599 | 8.59595 | 16.00963 | 32.72099 | 13.64741 | -0.5744 | 0.155113 | 0.338959 |
| 2 | 157586600 | 157589799 | 23.38466 | 38.8902 | 69.10337 | 29.98884 | -0.44559 | 1.257762 | 0.338972 |
| 7 | 57712000 | 57714399 | 17.13576 | 31.68113 | 22.83186 | 9.62438 | 0.907485 | 0.243696 | 0.339011 |
| 6 | 59760400 | 59760799 | 9.583204 | 18.67244 | 15.97921 | 5.806143 | 0.849193 | -0.3486 | 0.339011 |
| 18 | 29098200 | 29099199 | 12.34361 | 23.31682 | 22.00664 | 8.95554 | 0.561249 | 0.071976 | 0.339032 |
| 3 | 59192800 | 59195399 | 27.37254 | 44.43402 | 91.07005 | 39.03789 | -0.66569 | 1.601552 | 0.339038 |
| 9 | 18966600 | 18974199 | 109.6911 | 170.2363 | 149.1716 | 61.94913 | 0.496422 | 2.785606 | 0.339043 |
| 7 | 23713000 | 23724399 | 169.1592 | 256.3105 | 196.9843 | 81.5169 | 0.658565 | 3.270486 | 0.339046 |
| 8 | 59540800 | 59543799 | 20.25801 | 36.35885 | 25.45337 | 11.13652 | 0.913394 | 0.436703 | 0.339063 |
| 6 | 20424000 | 20425199 | 14.11982 | 26.03965 | 24.34176 | 10.29599 | 0.531528 | 0.237666 | 0.339071 |
| 1 | 116805600 | 116809799 | 65.84172 | 104.8426 | 101.1178 | 42.28761 | 0.388446 | 2.205264 | 0.339075 |
| 13 | 19974400 | 19975999 | 14.2487 | 26.59348 | 22.64709 | 9.412198 | 0.683499 | 0.164298 | 0.339077 |
| 4 | 20090800 | 20091199 | 4.230472 | 8.247205 | 21.37084 | 7.712118 | -0.72029 | -0.42025 | 0.339091 |
| 4 | 80058000 | 80059799 | 17.35726 | 31.26719 | 26.94343 | 11.747 | 0.618955 | 0.39825 | 0.339101 |
| 1 | 97073800 | 97075599 | 23.18787 | 40.86662 | 29.08268 | 12.87197 | 0.877027 | 0.599493 | 0.339116 |
| 7 | 31421200 | 31423799 | 24.87476 | 43.45563 | 33.00257 | 14.53988 | 0.782616 | 0.7374 | 0.339133 |
| 8 | 14986200 | 14988399 | 6.921678 | 12.72323 | 33.03821 | 14.0367 | -0.9361 | 0.220339 | 0.339141 |
| 11 | 36318200 | 36320399 | 12.74562 | 22.30145 | 44.14854 | 19.47537 | -0.59263 | 0.669027 | 0.339141 |
| 17 | 33354800 | 33357399 | 29.23987 | 47.60872 | 83.20955 | 35.72529 | -0.433 | 1.548821 | 0.339151 |
| 17 | 69463400 | 69467599 | 27.1079 | 44.22701 | 83.31068 | 35.79515 | -0.5387 | 1.513296 | 0.33916 |
| 17 | 92924600 | 92925199 | 9.643678 | 18.80779 | 15.46027 | 5.549461 | 0.922348 | -0.39076 | 0.339163 |
| 12 | 19639000 | 19677799 | 1202.293 | 1533.165 | 1208.104 | 507.0457 | 0.378477 | 6.050111 | 0.339164 |
| 3 | 68555000 | 68558799 | 33.84578 | 54.7182 | 93.97191 | 40.1527 | -0.417 | 1.699622 | 0.339186 |
| 7 | 22370800 | 22371399 | 18.71523 | 33.54997 | 21.3049 | 9.330797 | 1.052658 | 0.426677 | 0.339192 |
| 19 | 53504600 | 53520199 | 98.48376 | 148.4761 | 311.1769 | 128.5697 | -0.79383 | 3.360968 | 0.339194 |
| 12 | 16578200 | 16581599 | 26.21545 | 43.05525 | 78.61768 | 33.86704 | -0.48859 | 1.419548 | 0.339212 |
| 14 | 10758600 | 10759799 | 16.97865 | 30.22123 | 27.30398 | 12.03009 | 0.541159 | 0.470935 | 0.339222 |
| 12 | 119900800 | 119901999 | 7.835875 | 14.78402 | 30.27171 | 12.35035 | -0.55051 | 0.062175 | 0.339239 |
| 15 | 71472800 | 71475399 | 26.0759 | 44.80103 | 40.36302 | 17.70447 | 0.533766 | 0.931938 | 0.339242 |
| 8 | 19477000 | 19491199 | 66.83971 | 106.7081 | 86.62189 | 36.35633 | 0.640659 | 2.119911 | 0.339247 |
| 7 | 6591800 | 6593999 | 31.32244 | 52.16426 | 47.09511 | 20.45698 | 0.528279 | 1.246638 | 0.339276 |
| 1 | 13394200 | 13395599 | 8.113796 | 15.22438 | 32.3905 | 13.34058 | -0.61726 | 0.101752 | 0.339279 |

| | | | | | | | | | |
|---|---|---|---|---|---|---|---|---|---|
| 5 | 38824800 | 38836399 | 209.6244 | 302.2746 | 465.066 | 191.8938 | -0.40863 | 4.123138 | 0.339283 |
| 2 | 100769000 | 100770599 | 19.88618 | 35.92389 | 25.08649 | 10.8989 | 0.923658 | 0.396913 | 0.33929 |
| 1 | 65664800 | 65665599 | 19.1101 | 33.5921 | 30.84816 | 13.6605 | 0.509884 | 0.607541 | 0.339291 |
| 10 | 66814800 | 66817399 | 21.36155 | 35.47641 | 73.81088 | 32.00111 | -0.67257 | 1.273508 | 0.339293 |
| 3 | 3751800 | 3755399 | 50.09087 | 80.77983 | 67.4645 | 28.73394 | 0.616581 | 1.784187 | 0.339295 |
| 9 | 94858200 | 94859599 | 15.89498 | 29.3625 | 24.10499 | 10.17265 | 0.719868 | 0.24171 | 0.339325 |
| 10 | 35017800 | 35018599 | 6.559748 | 12.63772 | 27.63264 | 10.7212 | -0.57853 | -0.12407 | 0.339326 |
| 9 | 3512200 | 3515199 | 31.97909 | 54.31874 | 41.85534 | 18.3419 | 0.756057 | 1.057318 | 0.339344 |
| X | 9510600 | 9514399 | 41.37963 | 67.65126 | 60.3839 | 25.96401 | 0.538117 | 1.502839 | 0.339348 |
| 10 | 21427200 | 21427799 | 3.704265 | 7.228801 | 20.90825 | 7.436506 | -0.85769 | -0.48268 | 0.33935 |
| 18 | 12166200 | 12168199 | 10.96575 | 19.52643 | 37.83912 | 16.66618 | -0.553 | 0.451141 | 0.33936 |
| 9 | 21133400 | 21137399 | 30.49392 | 49.23941 | 92.55764 | 39.48647 | -0.54894 | 1.729911 | 0.339384 |
| 13 | 62044000 | 62046399 | 44.34163 | 71.88058 | 67.43276 | 28.87612 | 0.457086 | 1.64783 | 0.339388 |
| 13 | 33994800 | 33997599 | 34.98675 | 58.64334 | 46.9656 | 20.48381 | 0.700337 | 1.190113 | 0.339394 |
| 2 | 98026200 | 98026799 | 15.1129 | 27.42352 | 23.43644 | 10.1252 | 0.639504 | 0.334511 | 0.339395 |
| 8 | 91554600 | 91556199 | 8.59102 | 15.22234 | 38.99687 | 17.24149 | -0.95667 | 0.473209 | 0.339395 |
| X | 26043800 | 26044599 | 26.79393 | 45.9024 | 38.47151 | 16.87137 | 0.637181 | 0.961195 | 0.339398 |
| 8 | 31403800 | 31406999 | 17.90701 | 30.46798 | 58.07502 | 25.45427 | -0.54534 | 1.007795 | 0.339403 |
| 3 | 159281400 | 159284199 | 33.26446 | 55.58592 | 48.66834 | 21.18545 | 0.572146 | 1.215311 | 0.339429 |
| 16 | 84072800 | 84073999 | 12.63806 | 22.18413 | 44.60569 | 19.73031 | -0.61472 | 0.620395 | 0.339438 |
| 18 | 77546400 | 77553199 | 70.08456 | 108.8898 | 186.6601 | 77.50472 | -0.46936 | 2.75353 | 0.339448 |
| 3 | 20183800 | 20185799 | 12.23804 | 21.63493 | 40.35595 | 17.85053 | -0.50386 | 0.522636 | 0.339454 |
| 2 | 147099800 | 147101399 | 15.70734 | 28.26998 | 26.18874 | 11.43062 | 0.514196 | 0.390092 | 0.339491 |
| 1 | 148046400 | 148048599 | 30.68484 | 51.32971 | 45.94482 | 20.01328 | 0.540314 | 1.192387 | 0.339492 |
| 4 | 28365200 | 28367199 | 14.44359 | 25.24222 | 46.7719 | 20.61176 | -0.498 | 0.700183 | 0.339493 |
| 17 | 79552800 | 79553799 | 15.9322 | 28.71156 | 26.69466 | 11.63273 | 0.510548 | 0.386073 | 0.339494 |
| 7 | 117111800 | 117114199 | 20.22917 | 34.0307 | 62.42942 | 27.27795 | -0.49133 | 1.12048 | 0.339495 |
| 1 | 19582200 | 19584199 | 17.77006 | 31.80171 | 28.02427 | 12.29119 | 0.581257 | 0.443606 | 0.339499 |
| 17 | 61009800 | 61012799 | 27.20009 | 46.82348 | 39.18355 | 17.18814 | 0.640343 | 0.917831 | 0.339507 |
| X | 118902000 | 118904799 | 26.96546 | 47.70282 | 24.12832 | 10.67117 | 1.368905 | 0.577165 | 0.339518 |
| 9 | 24863200 | 24865199 | 16.26026 | 28.22458 | 52.22758 | 22.91719 | -0.49719 | 0.814566 | 0.339524 |
| 17 | 41384600 | 41393399 | 129.0181 | 197.475 | 180.6402 | 74.95881 | 0.419204 | 3.0543 | 0.339526 |
| 17 | 34877200 | 34880599 | 17.63627 | 29.5416 | 62.57894 | 27.28208 | -0.69789 | 1.15811 | 0.339541 |
| 5 | 84805800 | 84807199 | 16.35 | 30.77752 | 21.02376 | 8.611849 | 1.016217 | 0.127746 | 0.339542 |
| 9 | 89594600 | 89595799 | 12.45579 | 23.89614 | 19.29631 | 7.577669 | 0.82929 | -0.04927 | 0.339563 |
| 1 | 10439400 | 10439999 | 6.888046 | 13.16997 | 27.84832 | 11.04267 | -0.55983 | -0.04672 | 0.339566 |
| X | 8893400 | 8894999 | 22.22965 | 39.23852 | 29.35422 | 12.98813 | 0.80639 | 0.579675 | 0.339583 |
| 3 | 147011600 | 147012199 | 9.400425 | 18.16378 | 17.12679 | 6.575377 | 0.63868 | -0.15943 | 0.339596 |
| 1 | 67794800 | 67798199 | 54.76977 | 87.75175 | 80.54702 | 33.98099 | 0.469409 | 1.996878 | 0.339596 |
| 1 | 156060600 | 156062199 | 15.55443 | 28.11105 | 26.1886 | 11.37146 | 0.510908 | 0.365519 | 0.339598 |
| 6 | 135652200 | 135654199 | 11.15794 | 19.75979 | 44.56369 | 19.70488 | -0.7753 | 0.50392 | 0.339613 |
| 1 | 154118600 | 154119399 | 6.12047 | 11.81916 | 25.93644 | 9.985177 | -0.57224 | -0.15696 | 0.339619 |
| 1 | 188982000 | 188982599 | 6.12047 | 11.81916 | 25.93644 | 9.985177 | -0.57224 | -0.15696 | 0.339619 |
| 13 | 91894600 | 91896599 | 18.63515 | 31.67096 | 58.5459 | 25.65831 | -0.5014 | 1.025121 | 0.339619 |
| 7 | 86187200 | 86188999 | 20.03131 | 35.56125 | 27.57383 | 12.17593 | 0.757915 | 0.504827 | 0.33964 |
| 4 | 90114600 | 90119199 | 55.40617 | 88.99167 | 82.50049 | 34.93654 | 0.459285 | 1.921481 | 0.339654 |
| 5 | 69883000 | 69884199 | 15.01392 | 28.24952 | 21.61254 | 8.859893 | 0.852796 | 0.119186 | 0.339667 |

| | | | | | | | | | |
|---|---|---|---|---|---|---|---|---|---|
| 11 | 55547800 | 55550199 | 20.51707 | 36.19183 | 30.41453 | 13.45904 | 0.638919 | 0.576206 | 0.339669 |
| 1 | 189251200 | 189254599 | 29.55663 | 48.10007 | 84.4582 | 36.25468 | -0.4402 | 1.556815 | 0.339671 |
| 3 | 106508600 | 106508999 | 7.046927 | 13.41026 | 28.28254 | 11.33096 | -0.56951 | -0.01091 | 0.339679 |
| 14 | 113915000 | 113918199 | 61.24909 | 97.75131 | 94.46288 | 39.62923 | 0.389128 | 2.127542 | 0.339692 |
| 10 | 22560800 | 22561399 | 23.57992 | 41.05232 | 31.98186 | 14.0598 | 0.745779 | 0.777825 | 0.339709 |
| 9 | 121835800 | 121836799 | 12.40772 | 21.90859 | 40.88356 | 18.08825 | -0.50514 | 0.53818 | 0.339715 |
| 13 | 97431000 | 97433199 | 12.40772 | 21.90859 | 40.88356 | 18.08825 | -0.50514 | 0.53818 | 0.339715 |
| 1 | 193538600 | 193540799 | 26.59466 | 46.38436 | 32.96384 | 14.50791 | 0.877827 | 0.768903 | 0.339721 |
| 3 | 16828200 | 16828599 | 4.937091 | 9.630394 | 24.11768 | 8.636723 | -0.66356 | -0.43238 | 0.339724 |
| 9 | 32228600 | 32229199 | 4.752581 | 9.26239 | 23.12882 | 8.377904 | -0.6725 | -0.39164 | 0.339749 |
| 17 | 57735200 | 57738199 | 27.86035 | 47.98205 | 38.59752 | 16.9313 | 0.697234 | 0.915926 | 0.33975 |
| 17 | 22131400 | 22134199 | 27.9146 | 48.37519 | 35.19381 | 15.44115 | 0.843177 | 0.857008 | 0.339757 |
| X | 10004200 | 10004999 | 17.49536 | 32.40068 | 22.93407 | 9.645257 | 0.935926 | 0.238843 | 0.33976 |
| 3 | 30847000 | 30848199 | 9.445034 | 16.82989 | 37.41761 | 16.47114 | -0.74913 | 0.436288 | 0.33977 |
| 2 | 9003800 | 9004799 | 12.24668 | 22.98629 | 21.93985 | 9.031549 | 0.531405 | 0.110801 | 0.339772 |
| 5 | 86148200 | 86151199 | 35.75232 | 59.86662 | 46.97697 | 20.47583 | 0.729681 | 1.202204 | 0.339773 |
| 17 | 59035200 | 59036199 | 8.613541 | 16.14159 | 32.9052 | 13.58172 | -0.55828 | 0.117865 | 0.339781 |
| 2 | 53954800 | 53956799 | 14.84329 | 25.84442 | 48.4154 | 21.28556 | -0.51412 | 0.760411 | 0.339784 |
| 10 | 3506200 | 3509999 | 25.22902 | 43.5829 | 38.72045 | 16.98681 | 0.555377 | 0.878116 | 0.339792 |
| 15 | 81121000 | 81125999 | 28.16934 | 45.69758 | 86.51817 | 37.06881 | -0.55245 | 1.612576 | 0.339819 |
| 7 | 28712600 | 28715199 | 5.779898 | 10.85966 | 31.74161 | 13.04377 | -1.06705 | 0.0759 | 0.339832 |
| 12 | 4862800 | 4864199 | 6.749054 | 12.85049 | 28.29517 | 11.32282 | -0.62979 | -0.01859 | 0.339832 |
| 1 | 42239200 | 42240199 | 11.88925 | 22.34417 | 20.46389 | 8.405624 | 0.593084 | 0.095553 | 0.339838 |
| 2 | 50439600 | 50440799 | 16.05013 | 28.73174 | 25.33074 | 11.1072 | 0.58083 | 0.424331 | 0.339842 |
| 1 | 105536000 | 105541599 | 91.75623 | 143.3703 | 138.2073 | 57.34269 | 0.366473 | 2.659388 | 0.339846 |
| 5 | 143142800 | 143143399 | 6.782279 | 12.8429 | 29.76685 | 12.05115 | -0.71703 | 0.028744 | 0.339871 |
| 5 | 149449000 | 149452199 | 4.53476 | 8.796304 | 26.09894 | 9.839742 | -0.97313 | -0.24569 | 0.339873 |
| 10 | 11296600 | 11298399 | 25.94869 | 44.38963 | 41.52082 | 18.20648 | 0.478961 | 0.971805 | 0.339919 |
| 4 | 78953000 | 78954399 | 14.16908 | 26.45541 | 22.07105 | 9.166607 | 0.713541 | 0.159172 | 0.339923 |
| 5 | 32183600 | 32184199 | 3.694047 | 7.216553 | 20.46603 | 7.146606 | -0.80303 | -0.54349 | 0.339929 |
| 11 | 55186800 | 55187999 | 7.821111 | 14.69155 | 30.72744 | 12.63179 | -0.59026 | 0.089107 | 0.339948 |
| 12 | 111735600 | 111737799 | 11.47962 | 20.27026 | 39.91055 | 17.65768 | -0.58185 | 0.531282 | 0.339957 |
| 18 | 3698000 | 3705599 | 159.6395 | 240.0135 | 220.4182 | 91.00355 | 0.392148 | 3.422148 | 0.33997 |
| 5 | 120769600 | 120774399 | 45.40179 | 72.25837 | 129.9228 | 54.29559 | -0.50946 | 2.21046 | 0.339984 |
| 8 | 10407600 | 10409999 | 30.90779 | 51.4806 | 48.68294 | 21.14848 | 0.461556 | 1.24378 | 0.340002 |
| 2 | 165434800 | 165437999 | 26.82107 | 43.7058 | 81.77387 | 35.11926 | -0.53015 | 1.530662 | 0.340002 |
| 8 | 68514400 | 68516399 | 11.71533 | 20.89883 | 39.34816 | 17.3045 | -0.51046 | 0.446146 | 0.340024 |
| 7 | 11096800 | 11097799 | 35.5946 | 58.448 | 54.67523 | 23.55117 | 0.473561 | 1.427851 | 0.340033 |
| 12 | 119641600 | 119643199 | 16.93193 | 31.30483 | 23.4845 | 9.899252 | 0.850252 | 0.244228 | 0.340036 |
| 13 | 67158000 | 67165599 | 48.01152 | 77.60642 | 59.01649 | 25.21119 | 0.755259 | 1.711337 | 0.340044 |
| 1 | 194922600 | 194926399 | 22.71315 | 37.75731 | 72.59401 | 31.49438 | -0.55913 | 1.261271 | 0.340055 |
| 7 | 7412600 | 7414799 | 35.56535 | 58.40387 | 54.69563 | 23.56062 | 0.471961 | 1.426484 | 0.34006 |
| 16 | 95946200 | 95947799 | 8.340326 | 15.39784 | 33.07132 | 13.96777 | -0.65888 | 0.205104 | 0.340062 |
| 14 | 15985200 | 15988399 | 51.68894 | 83.25805 | 77.26746 | 32.85607 | 0.462794 | 1.826985 | 0.340063 |
| X | 3146200 | 3150999 | 47.48364 | 78.00609 | 50.73154 | 21.85667 | 0.996689 | 1.448386 | 0.340076 |
| 17 | 28038800 | 28039999 | 7.112442 | 13.31965 | 30.11042 | 12.44225 | -0.70705 | 0.102372 | 0.340077 |
| X | 10351000 | 10352599 | 9.613674 | 17.68566 | 35.1609 | 14.92236 | -0.55323 | 0.242404 | 0.34008 |

| | | | | | | | | | |
|---|---|---|---|---|---|---|---|---|---|
| 8 | 118323400 | 118326199 | 30.73015 | 49.8248 | 88.1329 | 37.74555 | -0.45543 | 1.629018 | 0.340095 |
| 5 | 23087000 | 23088399 | 13.37876 | 24.77559 | 23.34466 | 9.818646 | 0.525817 | 0.205544 | 0.340102 |
| 13 | 87650400 | 87652199 | 26.78044 | 46.70314 | 32.96442 | 14.50714 | 0.887639 | 0.771518 | 0.340102 |
| 10 | 59715200 | 59716999 | 7.771558 | 14.33256 | 32.0209 | 13.54367 | -0.7161 | 0.207075 | 0.340106 |
| 13 | 109808600 | 109812599 | 55.80066 | 88.27577 | 146.3639 | 60.83039 | -0.40095 | 2.39318 | 0.340118 |
| 2 | 141306400 | 141307399 | 14.16764 | 26.35607 | 20.74662 | 8.670108 | 0.79022 | 0.180728 | 0.34012 |
| 2 | 121970600 | 121972399 | 14.08547 | 25.86239 | 24.33035 | 10.35289 | 0.516202 | 0.261318 | 0.340133 |
| 7 | 38081200 | 38089199 | 104.5219 | 162.2986 | 149.8216 | 62.21423 | 0.421919 | 2.774115 | 0.34015 |
| 9 | 57038000 | 57038599 | 5.526486 | 10.73505 | 24.60191 | 9.192532 | -0.59326 | -0.27166 | 0.340154 |
| 14 | 79425800 | 79427999 | 33.23511 | 53.60509 | 95.22171 | 40.56386 | -0.46948 | 1.768001 | 0.340174 |
| 7 | 107744400 | 107747799 | 54.92202 | 88.3034 | 79.96262 | 33.91173 | 0.49491 | 1.890739 | 0.340186 |
| 6 | 122643000 | 122647599 | 24.83625 | 40.68095 | 77.43699 | 33.32062 | -0.54988 | 1.450086 | 0.340225 |
| 12 | 20621400 | 20626199 | 113.4353 | 175.574 | 143.1645 | 59.46689 | 0.597164 | 2.834066 | 0.340234 |
| 11 | 91804000 | 91804999 | 16.86102 | 30.2228 | 27.54925 | 12.06668 | 0.533926 | 0.42585 | 0.340238 |
| 2 | 108188800 | 108190399 | 13.48673 | 23.71093 | 45.04534 | 19.94681 | -0.53366 | 0.596957 | 0.34026 |
| 12 | 38882600 | 38883399 | 6.11553 | 11.82218 | 26.64001 | 10.20521 | -0.60394 | -0.17442 | 0.340261 |
| 17 | 88144600 | 88163199 | 231.2951 | 331.0581 | 504.6057 | 208.4422 | -0.4058 | 4.233192 | 0.340273 |
| 6 | 104454200 | 104455599 | 12.90921 | 24.41875 | 21.9064 | 8.888881 | 0.637291 | 0.066676 | 0.340295 |
| 12 | 17996600 | 18007799 | 235.6136 | 344.9664 | 313.1746 | 129.0064 | 0.373459 | 3.893023 | 0.340325 |
| 12 | 92726400 | 92726799 | 11.6186 | 22.57945 | 16.23757 | 6.044158 | 1.066246 | -0.24007 | 0.340331 |
| X | 52936000 | 52938999 | 54.37493 | 87.88953 | 67.55039 | 28.85559 | 0.739982 | 1.729422 | 0.340334 |
| 6 | 106746000 | 106746799 | 5.713283 | 10.98268 | 26.3115 | 10.27354 | -0.71697 | -0.11458 | 0.340358 |
| 9 | 78074000 | 78075199 | 5.86232 | 11.20395 | 27.22489 | 10.80865 | -0.75926 | -0.05257 | 0.340358 |
| 2 | 17063800 | 17066999 | 58.26465 | 93.27295 | 89.36948 | 37.65512 | 0.406192 | 2.029855 | 0.34037 |
| 7 | 13960200 | 13973399 | 163.0472 | 247.7164 | 188.9062 | 78.23776 | 0.672947 | 3.213954 | 0.340376 |
| 4 | 5258000 | 5261799 | 65.99664 | 105.1423 | 100.324 | 41.98496 | 0.404681 | 2.189058 | 0.340376 |
| 19 | 15823000 | 15825199 | 20.90185 | 36.75811 | 30.89557 | 13.68026 | 0.636952 | 0.610967 | 0.340381 |
| 14 | 82373000 | 82373399 | 9.913218 | 19.13958 | 19.17851 | 7.390138 | 0.546656 | -0.13499 | 0.340387 |
| 5 | 9655600 | 9657599 | 20.80884 | 36.73761 | 29.79939 | 13.1846 | 0.690119 | 0.567482 | 0.340401 |
| 9 | 3208000 | 3220799 | 191.4765 | 285.6995 | 254.7929 | 105.0196 | 0.42459 | 3.572002 | 0.340413 |
| 1 | 59431200 | 59436599 | 36.66712 | 58.89103 | 105.7153 | 44.76437 | -0.49218 | 1.907613 | 0.340419 |
| 17 | 87734800 | 87738399 | 24.08889 | 39.72325 | 75.7559 | 32.69022 | -0.54873 | 1.361704 | 0.340422 |
| 5 | 21991600 | 21992199 | 4.91871 | 9.577386 | 22.98912 | 8.425261 | -0.63189 | -0.34901 | 0.340439 |
| 18 | 56271200 | 56271799 | 4.77843 | 9.322152 | 22.16165 | 7.92238 | -0.58588 | -0.44436 | 0.340449 |
| 14 | 8329000 | 8331399 | 31.2691 | 53.8414 | 35.08181 | 15.389 | 0.99978 | 0.927569 | 0.340455 |
| 3 | 14470600 | 14474199 | 37.37096 | 62.48647 | 46.44099 | 20.22373 | 0.807797 | 1.221358 | 0.340463 |
| 12 | 61240000 | 61242199 | 17.32787 | 31.63382 | 24.88175 | 10.66848 | 0.765066 | 0.318679 | 0.340466 |
| 6 | 131495000 | 131496199 | 12.59018 | 23.6648 | 22.42941 | 9.210767 | 0.544238 | 0.106269 | 0.340497 |
| 10 | 11575600 | 11580399 | 70.11021 | 111.2399 | 107.9154 | 44.94651 | 0.374864 | 2.317909 | 0.340524 |
| 5 | 128541600 | 128542999 | 8.020526 | 14.8909 | 32.09639 | 13.45273 | -0.65581 | 0.168564 | 0.340527 |
| 2 | 64245200 | 64247799 | 22.36309 | 39.01468 | 34.87989 | 15.35383 | 0.548223 | 0.745691 | 0.340536 |
| 5 | 128304600 | 128306399 | 22.36309 | 39.01468 | 34.87989 | 15.35383 | 0.548223 | 0.745691 | 0.340536 |
| 13 | 61896400 | 61906999 | 214.1628 | 317.5413 | 271.8234 | 111.957 | 0.475319 | 3.686325 | 0.340549 |
| 15 | 45591200 | 45591999 | 11.06569 | 21.05433 | 19.98315 | 8.008843 | 0.572026 | 0.003975 | 0.340556 |
| 6 | 134709000 | 134710399 | 9.138366 | 16.97044 | 36.54195 | 15.31091 | -0.65417 | 0.188565 | 0.340558 |
| 18 | 53405400 | 53407599 | 18.91504 | 32.07546 | 58.03587 | 25.4198 | -0.47087 | 1.043131 | 0.340572 |
| 11 | 113890000 | 113891399 | 9.562445 | 17.13047 | 36.39406 | 15.9485 | -0.67948 | 0.400859 | 0.340573 |

| | | | | | | | | | |
|---|---|---|---|---|---|---|---|---|---|
| 17 | 42526600 | 42528199 | 16.56882 | 31.03107 | 21.43105 | 8.854682 | 0.990496 | 0.161118 | 0.340588 |
| 11 | 5880600 | 5894399 | 176.482 | 258.6087 | 398.9551 | 164.3335 | -0.38994 | 3.87652 | 0.340592 |
| 5 | 123423000 | 123430599 | 80.69878 | 123.6248 | 238.6236 | 99.03962 | -0.65612 | 3.027566 | 0.340594 |
| 4 | 113487600 | 113490399 | 36.37472 | 60.69294 | 48.70158 | 21.18004 | 0.697671 | 1.241931 | 0.340636 |
| 4 | 90346400 | 90346999 | 8.530403 | 16.5755 | 16.67612 | 6.214601 | 0.586155 | -0.27114 | 0.34064 |
| 10 | 68559200 | 68560599 | 7.444752 | 13.48576 | 35.99829 | 15.58591 | -0.9947 | 0.3158 | 0.340641 |
| 16 | 16330600 | 16332199 | 18.32931 | 33.2608 | 25.26225 | 10.91366 | 0.80831 | 0.360595 | 0.340643 |
| 7 | 15336000 | 15349399 | 350.1089 | 495.7512 | 431.0095 | 178.3593 | 0.388246 | 4.39551 | 0.340644 |
| 17 | 11537200 | 11538999 | 11.77591 | 20.76952 | 40.39966 | 17.87819 | -0.56527 | 0.54534 | 0.340662 |
| 1 | 72225400 | 72226999 | 10.24619 | 18.54998 | 35.90192 | 15.55512 | -0.53531 | 0.338953 | 0.340673 |
| 5 | 104285800 | 104286399 | 3.595141 | 7.012695 | 22.61452 | 8.099397 | -1.02258 | -0.45491 | 0.340685 |
| 17 | 94379000 | 94380599 | 15.67129 | 28.31874 | 26.12616 | 11.3461 | 0.524661 | 0.366403 | 0.340692 |
| 15 | 57847600 | 57851599 | 26.16314 | 42.70975 | 79.49397 | 34.16231 | -0.52076 | 1.502764 | 0.340694 |
| X | 43166000 | 43167799 | 26.00037 | 45.38674 | 33.155 | 14.5999 | 0.838586 | 0.756536 | 0.340705 |
| 6 | 131529600 | 131533599 | 39.62463 | 65.4585 | 53.86463 | 23.26785 | 0.660892 | 1.366105 | 0.34071 |
| 12 | 21928400 | 21930399 | 55.5818 | 89.69182 | 71.09061 | 30.30317 | 0.692956 | 1.778442 | 0.340718 |
| 3 | 128963800 | 128965199 | 7.796838 | 14.77584 | 30.44772 | 12.30603 | -0.54805 | 0.029346 | 0.340726 |
| 6 | 145793600 | 145794599 | 7.796838 | 14.77584 | 30.44772 | 12.30603 | -0.54805 | 0.029346 | 0.340726 |
| 3 | 31549400 | 31552199 | 60.3067 | 96.26732 | 92.2711 | 38.72114 | 0.401214 | 2.118279 | 0.340747 |
| 2 | 172033000 | 172034399 | 9.908405 | 18.14144 | 35.05028 | 14.97192 | -0.5192 | 0.270159 | 0.34075 |
| 14 | 20145400 | 20146999 | 9.908405 | 18.14144 | 35.05028 | 14.97192 | -0.5192 | 0.270159 | 0.34075 |
| 13 | 81171600 | 81175199 | 46.63137 | 74.40015 | 125.7815 | 52.74616 | -0.41689 | 2.129898 | 0.340756 |
| 10 | 123725800 | 123726599 | 14.74842 | 27.29659 | 24.24933 | 10.20699 | 0.609466 | 0.222451 | 0.340761 |
| 14 | 76932400 | 76936799 | 20.16193 | 33.67601 | 64.41995 | 28.03419 | -0.55146 | 1.197166 | 0.340782 |
| 6 | 23252800 | 23256999 | 30.89425 | 50.04954 | 90.46037 | 38.71879 | -0.48762 | 1.64653 | 0.340803 |
| 9 | 99616400 | 99620999 | 77.73989 | 122.7905 | 116.2768 | 48.28749 | 0.404208 | 2.439955 | 0.34082 |
| 14 | 75878400 | 75884999 | 48.30725 | 76.86076 | 132.1542 | 55.20649 | -0.44553 | 2.22237 | 0.34082 |
| 3 | 52116400 | 52119199 | 17.5597 | 30.21574 | 54.90675 | 24.08493 | -0.47386 | 0.901868 | 0.340821 |
| X | 64928600 | 64930399 | 27.18633 | 47.23567 | 34.91581 | 15.33018 | 0.821168 | 0.829124 | 0.340834 |
| 7 | 78255200 | 78255999 | 5.66296 | 10.99014 | 25.05643 | 9.424982 | -0.59419 | -0.24233 | 0.340854 |
| 13 | 69915000 | 69920999 | 61.57024 | 97.13654 | 156.8129 | 65.08273 | -0.36571 | 2.463619 | 0.340855 |
| 1 | 35154000 | 35157999 | 107.9351 | 166.6854 | 158.9472 | 66.02337 | 0.369108 | 2.874588 | 0.340862 |
| 10 | 48031600 | 48032799 | 8.738116 | 16.31719 | 36.2578 | 15.05546 | -0.68937 | 0.146172 | 0.340869 |
| 17 | 64981200 | 64981799 | 11.92588 | 22.75218 | 21.00353 | 8.369978 | 0.618968 | -0.00257 | 0.340887 |
| 15 | 76039800 | 76042399 | 16.29832 | 27.9721 | 52.6813 | 23.10632 | -0.52587 | 0.91777 | 0.340895 |
| 3 | 146769400 | 146770999 | 8.800327 | 16.19362 | 33.54488 | 14.23148 | -0.61185 | 0.228644 | 0.340918 |
| 2 | 73741000 | 73741999 | 7.570551 | 13.7438 | 34.99009 | 15.10783 | -0.92416 | 0.3021 | 0.340927 |
| 3 | 53444200 | 53445599 | 24.80391 | 42.62633 | 39.94003 | 17.51907 | 0.477523 | 0.924554 | 0.340937 |
| 11 | 111249400 | 111251599 | 41.19453 | 67.0327 | 63.48251 | 27.24962 | 0.448164 | 1.570328 | 0.34094 |
| X | 65806800 | 65809199 | 33.00359 | 55.69779 | 44.70675 | 19.55309 | 0.697218 | 1.119914 | 0.340963 |
| 8 | 36175400 | 36176399 | 6.269985 | 12.11513 | 27.73267 | 10.64687 | -0.62866 | -0.15848 | 0.340969 |
| 12 | 107880400 | 107886199 | 91.95399 | 139.7166 | 252.1976 | 104.4533 | -0.56899 | 3.198985 | 0.341006 |
| 5 | 46571600 | 46571999 | 4.224857 | 8.253094 | 20.59116 | 7.196729 | -0.62483 | -0.53858 | 0.341028 |
| 7 | 22037400 | 22037799 | 17.71606 | 31.78709 | 22.75527 | 9.958533 | 0.880839 | 0.414009 | 0.341039 |
| 5 | 137178000 | 137191399 | 368.7862 | 503.0536 | 810.4947 | 337.3856 | -0.55581 | 4.962048 | 0.341045 |
| X | 88184000 | 88185999 | 34.39583 | 58.35325 | 40.33164 | 17.66771 | 0.912486 | 1.078698 | 0.341059 |
| 5 | 8154400 | 8155399 | 15.75802 | 29.3431 | 22.47257 | 9.375984 | 0.831127 | 0.191223 | 0.341072 |

| | | | | | | | | | |
|---|---|---|---|---|---|---|---|---|---|
| 9 | 3841800 | 3843399 | 30.28328 | 50.55924 | 46.53934 | 20.24731 | 0.500302 | 1.215859 | 0.341079 |
| 12 | 92377600 | 92378999 | 14.75336 | 27.63192 | 22.7329 | 9.392022 | 0.740067 | 0.148462 | 0.341082 |
| 2 | 36400000 | 36404999 | 63.82052 | 101.9629 | 93.53238 | 39.29763 | 0.465778 | 2.097179 | 0.341084 |
| 11 | 12728000 | 12728799 | 9.999889 | 18.00017 | 35.87595 | 15.65617 | -0.5838 | 0.37306 | 0.341088 |
| 10 | 14546800 | 14550399 | 31.65379 | 53.11033 | 47.50356 | 20.73163 | 0.541349 | 1.167145 | 0.34111 |
| 7 | 33058200 | 33081799 | 450.5455 | 629.1595 | 469.7094 | 194.8067 | 0.587779 | 4.604455 | 0.341118 |
| 5 | 53241600 | 53245599 | 24.96904 | 40.94926 | 77.64979 | 33.42975 | -0.54369 | 1.434273 | 0.34112 |
| 2 | 167722200 | 167727399 | 35.24055 | 56.6742 | 111.2223 | 47.18638 | -0.61817 | 1.868194 | 0.341124 |
| 5 | 127639000 | 127642999 | 38.94131 | 62.45431 | 109.9083 | 46.43886 | -0.4661 | 1.956315 | 0.341136 |
| 5 | 3182600 | 3185199 | 18.25033 | 32.23502 | 29.05323 | 12.85332 | 0.53903 | 0.544754 | 0.34116 |
| 3 | 117077400 | 117080399 | 24.19649 | 39.73771 | 75.57046 | 32.55377 | -0.54687 | 1.417633 | 0.34118 |
| 1 | 85656400 | 85657999 | 18.05359 | 31.91471 | 29.66419 | 13.12138 | 0.495454 | 0.536921 | 0.341184 |
| 12 | 75237400 | 75239199 | 13.6525 | 23.99462 | 46.11338 | 20.41863 | -0.55039 | 0.603799 | 0.341186 |
| 2 | 73461000 | 73462599 | 8.289451 | 15.64212 | 33.74885 | 13.76463 | -0.62561 | 0.071979 | 0.341205 |
| X | 31369400 | 31372799 | 38.19435 | 64.58012 | 39.00768 | 17.07033 | 1.1059 | 1.116798 | 0.341212 |
| 1 | 51247200 | 51248399 | 7.661682 | 14.59688 | 34.25615 | 13.69091 | -0.72004 | 0.001054 | 0.341233 |
| 7 | 60405600 | 60407399 | 28.2412 | 48.30366 | 41.39795 | 18.15237 | 0.604599 | 0.981906 | 0.341234 |
| 12 | 103528600 | 103528999 | 3.931245 | 7.686685 | 19.61636 | 6.704053 | -0.62693 | -0.62499 | 0.341241 |
| 8 | 47851400 | 47854399 | 25.56654 | 42.07142 | 75.10044 | 32.37856 | -0.45481 | 1.392437 | 0.341251 |
| 3 | 13122600 | 13132399 | 189.994 | 283.2519 | 258.077 | 106.3561 | 0.392633 | 3.58704 | 0.34129 |
| 3 | 84972400 | 84975399 | 21.76991 | 38.00163 | 33.81117 | 14.88876 | 0.555108 | 0.732101 | 0.341291 |
| 2 | 118374000 | 118376599 | 12.8106 | 22.24156 | 48.43144 | 21.2544 | -0.73049 | 0.790437 | 0.34132 |
| 18 | 3081800 | 3084799 | 34.89098 | 59.94878 | 34.0186 | 14.92163 | 1.197973 | 0.963302 | 0.341325 |
| 1 | 9854000 | 9855799 | 11.11594 | 19.8226 | 37.83678 | 16.64477 | -0.53011 | 0.443016 | 0.341326 |
| 1 | 126925200 | 126926799 | 10.11691 | 18.35683 | 35.43126 | 15.30843 | -0.52852 | 0.325888 | 0.341331 |
| 12 | 106247400 | 106252799 | 65.36184 | 102.4386 | 167.9591 | 69.66514 | -0.39545 | 2.597508 | 0.341338 |
| 13 | 92274400 | 92285399 | 181.1392 | 271.6688 | 236.8143 | 97.70864 | 0.46419 | 3.473991 | 0.341339 |
| 4 | 99603200 | 99604599 | 11.63063 | 20.45584 | 41.96034 | 18.57925 | -0.64322 | 0.577246 | 0.34136 |
| 19 | 43425400 | 43426399 | 14.00545 | 25.83888 | 23.90097 | 10.10415 | 0.54731 | 0.234222 | 0.341385 |
| 17 | 19107800 | 19113399 | 98.5352 | 154.8515 | 122.9995 | 51.02127 | 0.651848 | 2.557466 | 0.341387 |
| 11 | 47878000 | 47880799 | 19.87988 | 33.09909 | 68.8293 | 29.89278 | -0.67157 | 1.233027 | 0.341397 |
| 5 | 80704800 | 80706599 | 17.87815 | 32.77974 | 23.15599 | 9.87217 | 0.924434 | 0.294805 | 0.341397 |
| 14 | 119470200 | 119470799 | 10.47749 | 20.36814 | 17.63745 | 6.548809 | 0.805176 | -0.25665 | 0.341411 |
| 14 | 61353000 | 61354199 | 14.97398 | 27.97573 | 22.61793 | 9.383803 | 0.760033 | 0.164627 | 0.341439 |
| 4 | 50101800 | 50103199 | 16.18249 | 30.32558 | 21.44703 | 8.852354 | 0.957552 | 0.153885 | 0.34144 |
| 9 | 31841600 | 31843399 | 18.69317 | 33.60703 | 26.77947 | 11.70004 | 0.729245 | 0.41894 | 0.341442 |
| 10 | 43675000 | 43677399 | 17.09825 | 30.78258 | 25.22862 | 11.00742 | 0.689789 | 0.393975 | 0.341442 |
| 13 | 68676400 | 68680399 | 55.57938 | 89.184 | 84.539 | 35.74743 | 0.425624 | 1.95199 | 0.341444 |
| 3 | 10112400 | 10115199 | 24.30507 | 42.55787 | 31.26643 | 13.80099 | 0.830034 | 0.695744 | 0.341446 |
| 5 | 132895200 | 132896999 | 11.35558 | 19.93398 | 42.04893 | 18.6002 | -0.68342 | 0.605817 | 0.341456 |
| 11 | 96092600 | 96094999 | 18.22857 | 31.06246 | 56.45337 | 24.74654 | -0.47646 | 0.994385 | 0.341468 |
| 13 | 96560800 | 96562199 | 8.646354 | 15.95671 | 32.53069 | 13.74661 | -0.58461 | 0.209514 | 0.341471 |
| 13 | 17406000 | 17407399 | 8.736893 | 16.31867 | 35.68402 | 14.81142 | -0.66589 | 0.143584 | 0.341478 |
| 10 | 78115000 | 78116799 | 9.068811 | 16.35311 | 35.90124 | 15.63256 | -0.72042 | 0.353965 | 0.341491 |
| 15 | 101220000 | 101221799 | 8.414272 | 15.35673 | 33.46632 | 14.35439 | -0.69519 | 0.27491 | 0.341497 |
| 12 | 40457400 | 40464999 | 68.06936 | 108.4292 | 92.14012 | 38.55251 | 0.571352 | 2.193129 | 0.341508 |
| 15 | 42778600 | 42779399 | 5.96714 | 11.55126 | 25.52729 | 9.716055 | -0.56669 | -0.19977 | 0.341512 |

| | | | | | | | | | |
|---|---|---|---|---|---|---|---|---|---|
| 3 | 9851200 | 9852799 | 9.086529 | 16.86636 | 33.18367 | 13.91256 | -0.52554 | 0.182012 | 0.341533 |
| 2 | 71380000 | 71383799 | 28.07471 | 45.80629 | 81.61544 | 35.06731 | -0.45858 | 1.514755 | 0.341535 |
| 17 | 58176200 | 58178599 | 25.67592 | 44.09461 | 40.36322 | 17.70432 | 0.510754 | 0.933312 | 0.341536 |
| 2 | 19913600 | 19915999 | 20.89916 | 34.60889 | 69.71349 | 30.16475 | -0.62634 | 1.300959 | 0.341537 |
| 7 | 15588000 | 15589599 | 18.06258 | 31.9005 | 29.71928 | 13.14822 | 0.49167 | 0.54742 | 0.341551 |
| 13 | 23451200 | 23454999 | 33.84801 | 54.63926 | 95.30458 | 40.64292 | -0.44188 | 1.743498 | 0.341557 |
| 8 | 99506600 | 99507399 | 9.768759 | 17.81863 | 34.4542 | 14.78865 | -0.52495 | 0.289781 | 0.341583 |
| 10 | 28928200 | 28929999 | 13.37343 | 23.50087 | 43.49294 | 19.25605 | -0.49598 | 0.602307 | 0.341597 |
| 10 | 52396000 | 52397999 | 27.27091 | 46.14172 | 43.75204 | 19.15047 | 0.457752 | 1.075996 | 0.341619 |
| 8 | 112634800 | 112637399 | 13.79992 | 23.75572 | 52.64377 | 23.09255 | -0.75854 | 0.87872 | 0.341622 |
| 8 | 35184000 | 35187599 | 70.44608 | 111.9148 | 107.1199 | 44.67189 | 0.396003 | 2.281327 | 0.341623 |
| 8 | 4283600 | 4289399 | 30.16888 | 48.70355 | 95.09677 | 40.55789 | -0.60408 | 1.735183 | 0.341626 |
| 17 | 20898600 | 20901199 | 42.01046 | 68.69254 | 60.48212 | 26.00762 | 0.55789 | 1.501791 | 0.341633 |
| 9 | 19594400 | 19595399 | 12.8304 | 24.16021 | 22.46411 | 9.196662 | 0.575402 | 0.097957 | 0.341637 |
| 8 | 100141200 | 100144199 | 27.67378 | 44.9326 | 86.70208 | 37.17137 | -0.57864 | 1.59338 | 0.341639 |
| 9 | 106469600 | 106473999 | 38.46473 | 61.73411 | 106.8901 | 45.21275 | -0.44145 | 1.933249 | 0.341644 |
| 15 | 50009200 | 50010399 | 15.70014 | 28.48283 | 26.13396 | 11.29372 | 0.537096 | 0.344614 | 0.341669 |
| 11 | 72220600 | 72222599 | 10.93714 | 19.5121 | 37.82361 | 16.63298 | -0.55203 | 0.437715 | 0.341672 |
| 13 | 84509600 | 84510599 | 8.640506 | 16.09103 | 32.15963 | 13.41551 | -0.54207 | 0.157603 | 0.34168 |
| 1 | 175813800 | 175816199 | 24.83344 | 42.68353 | 39.86589 | 17.48663 | 0.482062 | 0.921565 | 0.341684 |
| 4 | 103295600 | 103296799 | 8.519243 | 15.65213 | 32.99809 | 14.02904 | -0.63899 | 0.234911 | 0.341688 |
| 11 | 10326200 | 10328199 | 20.86857 | 36.86592 | 26.91138 | 11.90534 | 0.841817 | 0.557214 | 0.341692 |
| 12 | 23298200 | 23299199 | 29.37148 | 49.36083 | 45.39768 | 19.83005 | 0.501382 | 1.144368 | 0.341694 |
| 1 | 124985400 | 124987999 | 25.69126 | 44.15351 | 39.91534 | 17.50826 | 0.529041 | 0.928511 | 0.341709 |
| 12 | 29291600 | 29297599 | 37.30902 | 60.04712 | 101.8904 | 43.27578 | -0.40739 | 1.842458 | 0.341733 |
| 19 | 9607400 | 9607999 | 15.44557 | 27.88762 | 24.39302 | 10.60447 | 0.600425 | 0.367402 | 0.341735 |
| 17 | 19526000 | 19535799 | 129.2085 | 199.1929 | 156.2401 | 64.89951 | 0.648754 | 2.914697 | 0.34174 |
| 2 | 98060400 | 98061599 | 16.23852 | 29.76062 | 25.08443 | 10.70061 | 0.671067 | 0.289908 | 0.341759 |
| 19 | 25454600 | 25459399 | 42.36964 | 67.95728 | 115.8583 | 48.95792 | -0.42036 | 1.960988 | 0.341761 |
| 1 | 20948000 | 20950799 | 20.36303 | 34.3841 | 61.94876 | 27.09862 | -0.46515 | 1.088158 | 0.341775 |
| 7 | 86482600 | 86483799 | 14.61569 | 25.40956 | 48.97089 | 21.50926 | -0.55533 | 0.778461 | 0.341798 |
| 8 | 41692400 | 41693999 | 10.51025 | 18.35281 | 46.79794 | 20.61178 | -0.95587 | 0.693028 | 0.341799 |
| 7 | 31799800 | 31813199 | 163.6105 | 248.9395 | 181.2389 | 75.09545 | 0.741506 | 3.184514 | 0.341818 |
| 8 | 109778800 | 109780599 | 16.5076 | 30.35551 | 21.9819 | 9.334383 | 0.893265 | 0.26682 | 0.341824 |
| 19 | 47240000 | 47243199 | 23.78426 | 39.2956 | 71.02527 | 30.68331 | -0.47086 | 1.33753 | 0.341853 |
| 5 | 116249400 | 116252399 | 14.6665 | 25.37506 | 50.03516 | 21.95164 | -0.58932 | 0.833461 | 0.341854 |
| 5 | 120744600 | 120745399 | 4.367355 | 8.475582 | 23.78622 | 8.948035 | -0.8892 | -0.25962 | 0.341858 |
| 7 | 60595800 | 60603599 | 125.5694 | 193.407 | 162.2499 | 67.39262 | 0.550872 | 2.930203 | 0.341872 |
| 7 | 84489200 | 84492999 | 61.26504 | 98.15306 | 87.31699 | 36.83402 | 0.514413 | 2.009046 | 0.341873 |
| 2 | 100134400 | 100138599 | 41.28041 | 67.39471 | 62.11126 | 26.69314 | 0.490628 | 1.521824 | 0.341881 |
| 14 | 12751200 | 12751799 | 10.49068 | 20.20641 | 19.61169 | 7.613981 | 0.581926 | -0.10416 | 0.341892 |
| 17 | 20359800 | 20361599 | 26.61088 | 45.79626 | 39.18391 | 17.18815 | 0.60832 | 0.916847 | 0.341897 |
| 13 | 25969600 | 25970999 | 13.05365 | 22.86373 | 43.59811 | 19.24914 | -0.53888 | 0.653476 | 0.341908 |
| 7 | 76477800 | 76479399 | 23.09055 | 40.57598 | 30.83181 | 13.6503 | 0.781694 | 0.634888 | 0.341911 |
| 2 | 16615000 | 16615999 | 17.20988 | 31.13984 | 26.47098 | 11.47703 | 0.643608 | 0.371371 | 0.341923 |
| 12 | 62869800 | 62871199 | 17.20988 | 31.13984 | 26.47098 | 11.47703 | 0.643608 | 0.371371 | 0.341923 |
| 9 | 35432200 | 35432999 | 10.34509 | 19.97287 | 17.90599 | 6.900637 | 0.704685 | -0.13597 | 0.341927 |

| | | | | | | | | | |
|---|---|---|---|---|---|---|---|---|---|
| X | 29527600 | 29527999 | 8.035552 | 15.67021 | 15.73795 | 5.655259 | 0.636748 | -0.40494 | 0.341941 |
| 12 | 20083200 | 20098199 | 382.5323 | 538.4418 | 416.7831 | 172.6422 | 0.547131 | 4.483538 | 0.341968 |
| 9 | 114892200 | 114903199 | 106.7447 | 162.097 | 256.245 | 106.1114 | -0.37848 | 3.213717 | 0.341978 |
| 18 | 58341600 | 58344799 | 19.93287 | 33.08126 | 67.99853 | 29.46732 | -0.65509 | 1.27179 | 0.341987 |
| 16 | 5051400 | 5052199 | 4.401835 | 8.561562 | 22.86067 | 8.452761 | -0.79594 | -0.32681 | 0.341995 |
| 8 | 116432200 | 116435999 | 20.97433 | 34.5646 | 74.54656 | 32.15944 | -0.72586 | 1.362274 | 0.341997 |
| 12 | 31068000 | 31070399 | 27.84889 | 45.19343 | 85.27164 | 36.54409 | -0.54703 | 1.604905 | 0.342 |
| 4 | 112613200 | 112615799 | 41.734 | 68.70578 | 55.07633 | 23.74979 | 0.697098 | 1.412323 | 0.342003 |
| 1 | 7300000 | 7302999 | 37.06196 | 61.12151 | 56.00613 | 24.16867 | 0.505718 | 1.379443 | 0.342032 |
| 11 | 96241800 | 96244399 | 10.27691 | 18.09388 | 41.99927 | 18.5926 | -0.81977 | 0.557839 | 0.342042 |
| 13 | 50505800 | 50508199 | 23.85288 | 41.07612 | 38.85914 | 17.04602 | 0.464213 | 0.899439 | 0.342064 |
| 2 | 97689000 | 97693199 | 37.97043 | 63.00817 | 51.0356 | 22.11426 | 0.683975 | 1.307263 | 0.342068 |
| 9 | 71015600 | 71016799 | 7.660427 | 14.46226 | 29.74387 | 12.11858 | -0.55535 | 0.05313 | 0.342107 |
| 9 | 4235000 | 4236599 | 22.89678 | 39.75059 | 37.1981 | 16.32396 | 0.482123 | 0.82038 | 0.342109 |
| 7 | 7437800 | 7440599 | 34.7779 | 59.05025 | 40.14593 | 17.59077 | 0.936052 | 1.070306 | 0.34211 |
| 15 | 18267400 | 18268199 | 7.871802 | 14.61574 | 32.63704 | 13.67811 | -0.70646 | 0.167588 | 0.342118 |
| 1 | 151080200 | 151080599 | 3.039833 | 5.946139 | 16.79785 | 5.541835 | -0.72112 | -0.77069 | 0.342129 |
| 10 | 81544400 | 81545799 | 14.51234 | 26.56426 | 24.7232 | 10.56422 | 0.527161 | 0.281824 | 0.342138 |
| 6 | 37630200 | 37630999 | 12.25318 | 23.31982 | 20.3035 | 8.132735 | 0.695853 | 0.013758 | 0.342146 |
| 11 | 94212000 | 94213399 | 8.878026 | 16.15362 | 33.94325 | 14.61478 | -0.64671 | 0.297374 | 0.342158 |
| 7 | 8723400 | 8724199 | 16.94544 | 30.47864 | 23.58779 | 10.30167 | 0.771444 | 0.397203 | 0.342161 |
| 10 | 48167000 | 48167599 | 15.95557 | 28.66995 | 24.88914 | 10.87906 | 0.606183 | 0.400459 | 0.342163 |
| 14 | 44534200 | 44537999 | 45.03687 | 72.05421 | 121.7173 | 51.24125 | -0.41134 | 2.04438 | 0.34217 |
| Y | 5117600 | 5119999 | 20.05714 | 35.31881 | 18.5522 | 8.21259 | 1.31139 | 0.566287 | 0.342178 |
| 12 | 108235200 | 108236999 | 8.596125 | 15.60861 | 34.47152 | 14.8804 | -0.72008 | 0.310288 | 0.342181 |
| 13 | 44161200 | 44165199 | 31.21317 | 50.59949 | 88.80255 | 38.02765 | -0.44436 | 1.633566 | 0.342209 |
| 15 | 22920200 | 22922199 | 17.85682 | 30.61404 | 55.99053 | 24.55564 | -0.48421 | 0.936305 | 0.342224 |
| 13 | 77927400 | 77929999 | 38.52583 | 63.52462 | 55.53563 | 23.96387 | 0.573222 | 1.384714 | 0.342227 |
| 2 | 53534000 | 53534799 | 8.279278 | 15.56979 | 31.32245 | 12.85176 | -0.53236 | 0.088156 | 0.342227 |
| 12 | 113384200 | 113391199 | 71.6289 | 110.7044 | 199.3669 | 82.81253 | -0.54634 | 2.850057 | 0.342256 |
| 5 | 134737800 | 134738999 | 8.565401 | 15.65819 | 33.48487 | 14.33295 | -0.66602 | 0.267948 | 0.342271 |
| 7 | 47963600 | 47972599 | 133.8447 | 204.7523 | 181.0847 | 75.1339 | 0.467223 | 3.067499 | 0.342273 |
| 17 | 21563400 | 21566399 | 38.32814 | 61.50386 | 107.1346 | 45.3038 | -0.4504 | 1.93969 | 0.342279 |
| 13 | 60231000 | 60239799 | 69.47437 | 107.8551 | 190.6541 | 79.17263 | -0.51454 | 2.767018 | 0.34228 |
| 18 | 61095800 | 61097999 | 12.75642 | 22.29372 | 44.69908 | 19.69835 | -0.61091 | 0.689734 | 0.342309 |
| 7 | 9838200 | 9850799 | 200.9652 | 301.3734 | 222.9543 | 91.9876 | 0.700625 | 3.480647 | 0.342309 |
| 7 | 59154600 | 59155999 | 54.71139 | 87.72338 | 79.99237 | 33.78515 | 0.48017 | 1.975004 | 0.342313 |
| 7 | 19274800 | 19276799 | 10.41839 | 18.36134 | 40.48129 | 17.91702 | -0.74511 | 0.546406 | 0.342324 |
| 1 | 96395800 | 96396399 | 8.479326 | 16.48903 | 16.08328 | 5.956278 | 0.638098 | -0.29673 | 0.342336 |
| 2 | 36287200 | 36290599 | 63.66956 | 101.4414 | 98.50455 | 41.22714 | 0.379526 | 2.184172 | 0.342358 |
| 7 | 14015400 | 14020399 | 92.6622 | 146.2117 | 116.6997 | 48.4382 | 0.64934 | 2.474051 | 0.342362 |
| 9 | 26499400 | 26500599 | 10.38336 | 18.86562 | 36.49351 | 15.74092 | -0.53046 | 0.320101 | 0.342396 |
| 4 | 116349800 | 116351599 | 23.39977 | 40.74091 | 35.96731 | 15.81242 | 0.566125 | 0.780949 | 0.3424 |
| 11 | 30450600 | 30451399 | 13.90183 | 26.07118 | 22.81311 | 9.404288 | 0.654237 | 0.132918 | 0.342401 |
| 18 | 19638200 | 19641799 | 64.59575 | 103.0849 | 80.14056 | 33.6172 | 0.702299 | 2.128759 | 0.342409 |
| X | 113096200 | 113100599 | 60.34108 | 96.52344 | 90.38656 | 38.0426 | 0.438051 | 2.056728 | 0.342418 |
| 17 | 28789000 | 28793199 | 27.04027 | 44.34962 | 79.16557 | 34.08347 | -0.45676 | 1.439165 | 0.342422 |

| | | | | | | | | | |
|---|---|---|---|---|---|---|---|---|---|
| 17 | 57454800 | 57464399 | 76.86769 | 118.7424 | 199.6663 | 82.93715 | -0.44805 | 2.861598 | 0.34243 |
| 12 | 7487800 | 7489399 | 15.80949 | 28.93307 | 25.05413 | 10.70843 | 0.630225 | 0.293495 | 0.342447 |
| 13 | 51587600 | 51589399 | 10.01556 | 17.89005 | 38.37017 | 16.85579 | -0.69606 | 0.423691 | 0.342449 |
| 4 | 93815600 | 93816799 | 12.7261 | 24.40253 | 19.88939 | 7.821781 | 0.814418 | -0.03963 | 0.342475 |
| 17 | 19676800 | 19677799 | 12.7261 | 24.40253 | 19.88939 | 7.821781 | 0.814418 | -0.03963 | 0.342475 |
| 12 | 30894200 | 30896199 | 10.28284 | 18.74385 | 38.20203 | 16.41097 | -0.60133 | 0.306204 | 0.34248 |
| 13 | 38198800 | 38201599 | 22.49576 | 39.16043 | 36.11294 | 15.87478 | 0.503555 | 0.780072 | 0.342483 |
| 12 | 84627200 | 84627999 | 10.65921 | 20.7199 | 17.63097 | 6.550264 | 0.829484 | -0.25226 | 0.34249 |
| 17 | 80154800 | 80155399 | 10.65921 | 20.7199 | 17.63097 | 6.550264 | 0.829484 | -0.25226 | 0.34249 |
| 2 | 162494000 | 162495599 | 10.19569 | 18.26683 | 37.40931 | 16.39193 | -0.62667 | 0.408537 | 0.342504 |
| 16 | 82250200 | 82251799 | 15.19602 | 28.05053 | 23.51043 | 9.931742 | 0.688757 | 0.237174 | 0.342505 |
| 10 | 84923000 | 84926999 | 46.2935 | 75.0792 | 67.21078 | 28.79082 | 0.525092 | 1.641796 | 0.342505 |
| 17 | 22957400 | 22959199 | 11.60591 | 20.46986 | 41.41052 | 18.3255 | -0.62175 | 0.544432 | 0.342508 |
| 7 | 17904600 | 17922599 | 306.3398 | 440.7597 | 368.1091 | 151.9343 | 0.469279 | 4.16071 | 0.34251 |
| 18 | 79207000 | 79208999 | 16.95775 | 31.74055 | 21.41507 | 8.857012 | 1.022878 | 0.168334 | 0.342512 |
| 7 | 47207000 | 47219999 | 148.3865 | 227.1364 | 167.3288 | 69.43622 | 0.731337 | 3.059014 | 0.34252 |
| 9 | 32418600 | 32419799 | 11.27007 | 21.34287 | 20.01684 | 8.102309 | 0.576874 | 0.038598 | 0.342552 |
| 5 | 36387600 | 36397399 | 83.79518 | 128.5134 | 220.5051 | 91.53599 | -0.48515 | 3.00431 | 0.342555 |
| 11 | 24249400 | 24252399 | 11.29044 | 19.93074 | 42.96348 | 19.00943 | -0.71252 | 0.534386 | 0.342566 |
| 5 | 53191000 | 53197199 | 47.62483 | 76.05102 | 129.8987 | 54.53955 | -0.43032 | 2.10591 | 0.342598 |
| 18 | 3000000 | 3024199 | 786.5359 | 1046.111 | 766.1014 | 320.1039 | 0.544813 | 5.367786 | 0.342604 |
| 2 | 151934800 | 151937999 | 30.14804 | 48.73491 | 90.67802 | 38.73996 | -0.53239 | 1.695014 | 0.342625 |
| 3 | 56024800 | 56026999 | 26.2031 | 45.01692 | 40.33935 | 17.69406 | 0.541475 | 0.932525 | 0.342664 |
| 4 | 64160800 | 64162199 | 17.27647 | 32.1816 | 21.21363 | 8.845619 | 1.046542 | 0.199193 | 0.342665 |
| 7 | 29387400 | 29395199 | 112.8827 | 174.2473 | 161.7612 | 67.19241 | 0.407261 | 2.884518 | 0.34267 |
| 1 | 128963000 | 128963999 | 12.57863 | 23.45587 | 20.98438 | 8.734156 | 0.611659 | 0.152841 | 0.342675 |
| 4 | 16823400 | 16827999 | 47.79624 | 77.2329 | 71.17837 | 30.39364 | 0.478075 | 1.72451 | 0.342675 |
| 10 | 58616400 | 58616799 | 4.00339 | 7.822966 | 20.59783 | 7.151324 | -0.69095 | -0.56054 | 0.342683 |
| 12 | 88168400 | 88170199 | 20.60232 | 36.23015 | 31.61691 | 13.99976 | 0.583251 | 0.612833 | 0.342687 |
| 6 | 135346400 | 135348399 | 11.28955 | 19.95745 | 40.37182 | 17.85767 | -0.62028 | 0.515584 | 0.342706 |
| 14 | 50795000 | 50796399 | 24.97618 | 40.56226 | 90.03214 | 38.6058 | -0.77967 | 1.587554 | 0.342709 |
| 10 | 15054800 | 15056599 | 15.91645 | 27.52832 | 50.58662 | 22.19335 | -0.48827 | 0.84291 | 0.34271 |
| 9 | 108634400 | 108637999 | 31.90468 | 51.6097 | 90.06162 | 38.50589 | -0.43905 | 1.679998 | 0.342719 |
| 4 | 22739000 | 22741599 | 53.21754 | 85.41203 | 78.97362 | 33.40474 | 0.461836 | 1.942253 | 0.342733 |
| 17 | 57987000 | 57995599 | 90.95489 | 143.5391 | 116.51 | 48.36314 | 0.625258 | 2.467676 | 0.342735 |
| 16 | 36163800 | 36165399 | 20.48117 | 36.28004 | 28.68622 | 12.68361 | 0.728502 | 0.52801 | 0.342743 |
| X | 97153800 | 97154999 | 12.07917 | 23.03742 | 21.07787 | 8.40516 | 0.631381 | 0.002864 | 0.34275 |
| 11 | 105049200 | 105059999 | 184.9059 | 269.1989 | 422.0147 | 173.9281 | -0.42211 | 3.977769 | 0.34277 |
| 13 | 53989200 | 53989999 | 5.698684 | 10.9958 | 25.88087 | 9.997111 | -0.67712 | -0.15181 | 0.342796 |
| 7 | 86499000 | 86502199 | 32.59194 | 55.14453 | 42.98438 | 18.81443 | 0.739316 | 1.092737 | 0.3428 |
| 9 | 33637400 | 33639799 | 3.342439 | 6.530157 | 21.08103 | 7.351799 | -0.98656 | -0.55447 | 0.342816 |
| 7 | 126026400 | 126028999 | 14.39019 | 24.83292 | 50.54706 | 22.17448 | -0.63566 | 0.857081 | 0.342821 |
| 9 | 23672200 | 23676199 | 32.36956 | 55.1054 | 40.7311 | 17.85321 | 0.816293 | 1.038468 | 0.342848 |
| 12 | 56823000 | 56824399 | 14.56545 | 27.28803 | 22.74013 | 9.39031 | 0.722202 | 0.144845 | 0.342874 |
| 7 | 6216200 | 6226599 | 140.5105 | 212.9387 | 194.7151 | 80.58224 | 0.408216 | 3.258558 | 0.342875 |
| 8 | 21779800 | 21782199 | 17.93359 | 32.94057 | 23.85733 | 10.14632 | 0.891564 | 0.288358 | 0.342888 |
| 12 | 101820000 | 101820799 | 5.233958 | 10.18281 | 23.56505 | 8.69405 | -0.5898 | -0.32282 | 0.34289 |

| | | | | | | | | | |
|---|---|---|---|---|---|---|---|---|---|
| 5 | 96267400 | 96267999 | 7.105058 | 13.33586 | 29.61241 | 12.18983 | -0.67667 | 0.08786 | 0.342913 |
| 15 | 80287400 | 80290799 | 33.85888 | 54.67312 | 94.08914 | 40.14003 | -0.42199 | 1.735034 | 0.34292 |
| 7 | 66111800 | 66113999 | 18.9115 | 34.2055 | 25.76566 | 11.17663 | 0.816562 | 0.384623 | 0.342924 |
| 6 | 76499000 | 76500399 | 13.81622 | 25.5848 | 22.09592 | 9.29388 | 0.650123 | 0.204496 | 0.342938 |
| 16 | 60553800 | 60555199 | 19.67472 | 34.96895 | 25.84203 | 11.39999 | 0.828038 | 0.493832 | 0.342962 |
| 1 | 180454400 | 180456599 | 18.37267 | 32.40732 | 30.20332 | 13.36573 | 0.490402 | 0.563556 | 0.342975 |
| 10 | 127467200 | 127468599 | 7.55516 | 14.19652 | 30.16804 | 12.39509 | -0.61159 | 0.086907 | 0.342985 |
| 15 | 12737600 | 12740399 | 20.65377 | 34.42143 | 66.08785 | 28.71999 | -0.55676 | 1.223451 | 0.342989 |
| X | 73801200 | 73803999 | 60.42498 | 97.00288 | 81.54192 | 34.50364 | 0.599399 | 1.946114 | 0.343011 |
| 8 | 122384800 | 122388999 | 23.04731 | 37.90647 | 73.38227 | 31.63096 | -0.5716 | 1.393254 | 0.343015 |
| 4 | 110009200 | 110009999 | 5.149117 | 9.939589 | 26.4405 | 10.19743 | -0.84986 | -0.16235 | 0.343018 |
| 9 | 114215600 | 114219999 | 36.6816 | 58.86729 | 107.7376 | 45.56537 | -0.52142 | 1.933164 | 0.343025 |
| 6 | 80996000 | 81000199 | 44.06974 | 70.51325 | 118.7753 | 50.0091 | -0.40707 | 2.040172 | 0.343028 |
| 2 | 171186600 | 171188999 | 18.35538 | 31.45338 | 59.88777 | 26.26381 | -0.54224 | 0.947036 | 0.343039 |
| 12 | 22445800 | 22446199 | 8.284112 | 16.1314 | 16.74364 | 6.132599 | 0.564619 | -0.33297 | 0.343051 |
| 8 | 21889200 | 21891599 | 18.6427 | 34.65478 | 21.15528 | 8.852033 | 1.152877 | 0.223886 | 0.343053 |
| 3 | 53916000 | 53917999 | 17.93534 | 32.64339 | 24.74927 | 10.65212 | 0.81418 | 0.33951 | 0.343081 |
| 3 | 96249600 | 96252799 | 30.43569 | 49.30391 | 87.73132 | 37.54915 | -0.46514 | 1.647123 | 0.343085 |
| 16 | 68097000 | 68098399 | 16.54591 | 31.13688 | 21.0161 | 8.613114 | 1.032826 | 0.131423 | 0.343106 |
| 7 | 96588000 | 96589399 | 16.72808 | 30.29034 | 26.54437 | 11.49826 | 0.600848 | 0.364032 | 0.343117 |
| 5 | 128919800 | 128924799 | 43.05706 | 68.78764 | 119.0724 | 50.02572 | -0.44877 | 2.084459 | 0.343119 |
| 3 | 127534400 | 127535999 | 17.37717 | 29.95921 | 54.93558 | 24.09898 | -0.48631 | 0.883274 | 0.343144 |
| 1 | 80895200 | 80897199 | 20.28118 | 36.53103 | 25.59636 | 11.16094 | 0.915605 | 0.417299 | 0.343162 |
| 10 | 17916200 | 17917399 | 8.043303 | 14.89923 | 31.53559 | 13.25868 | -0.63265 | 0.182171 | 0.343164 |
| 18 | 64569000 | 64571199 | 12.69984 | 22.33477 | 42.43745 | 18.79077 | -0.53353 | 0.584869 | 0.343167 |
| 8 | 16654400 | 16655399 | 9.564352 | 17.67493 | 35.85275 | 15.12235 | -0.57587 | 0.216011 | 0.343187 |
| 17 | 4597000 | 4597799 | 10.19027 | 19.69265 | 19.1426 | 7.34457 | 0.595679 | -0.14708 | 0.343203 |
| 5 | 117284400 | 117286799 | 14.7457 | 25.3375 | 56.40214 | 24.73981 | -0.76568 | 0.90487 | 0.343211 |
| 8 | 21830800 | 21839999 | 116.9569 | 181.3273 | 146.2739 | 60.75502 | 0.614512 | 2.80527 | 0.343214 |
| 9 | 19167800 | 19171599 | 25.79139 | 44.71409 | 33.04623 | 14.49909 | 0.820885 | 0.843284 | 0.343216 |
| 6 | 138185800 | 138190399 | 55.83188 | 89.57195 | 84.09796 | 35.55054 | 0.439066 | 1.957158 | 0.343226 |
| 3 | 6580800 | 6582799 | 19.17168 | 35.34401 | 21.48563 | 9.092199 | 1.147225 | 0.272605 | 0.343231 |
| 19 | 33683600 | 33685399 | 16.1714 | 29.93809 | 23.51487 | 9.894317 | 0.786118 | 0.230373 | 0.343243 |
| 2 | 35405200 | 35411599 | 93.5528 | 142.5182 | 242.5269 | 100.5266 | -0.48093 | 3.144361 | 0.343252 |
| 11 | 105974600 | 105976399 | 10.64488 | 18.61692 | 47.81594 | 21.08476 | -0.96636 | 0.669463 | 0.343257 |
| 12 | 45240400 | 45243599 | 32.08205 | 51.79144 | 94.30314 | 40.21875 | -0.50359 | 1.73857 | 0.343262 |
| 1 | 82197800 | 82199599 | 9.760739 | 17.76548 | 34.94174 | 15.03859 | -0.55207 | 0.30228 | 0.343279 |
| 5 | 93186800 | 93188399 | 9.760739 | 17.76548 | 34.94174 | 15.03859 | -0.55207 | 0.30228 | 0.343279 |
| 1 | 37707600 | 37709999 | 13.25403 | 23.06433 | 46.84557 | 20.58812 | -0.63015 | 0.757669 | 0.34328 |
| 1 | 165586600 | 165589399 | 13.25403 | 23.06433 | 46.84557 | 20.58812 | -0.63015 | 0.757669 | 0.34328 |
| 2 | 129088600 | 129091599 | 17.19823 | 29.18383 | 56.84369 | 24.90272 | -0.57657 | 1.031317 | 0.343282 |
| 7 | 30331400 | 30336199 | 29.44215 | 47.76901 | 86.20364 | 36.93782 | -0.48326 | 1.612105 | 0.343283 |
| 18 | 35239200 | 35240199 | 7.068437 | 13.35995 | 29.21188 | 11.87298 | -0.63973 | 0.038122 | 0.343291 |
| 1 | 153061000 | 153062399 | 6.280942 | 11.73661 | 33.85586 | 14.04221 | -1.06077 | 0.115848 | 0.343298 |
| 1 | 7158600 | 7159799 | 10.5444 | 18.63555 | 39.9109 | 17.65467 | -0.70205 | 0.513703 | 0.3433 |
| 17 | 57894200 | 57896399 | 32.55079 | 54.14256 | 51.42928 | 22.32122 | 0.455074 | 1.265717 | 0.343313 |
| 9 | 98769600 | 98772999 | 43.35627 | 69.39392 | 116.0526 | 48.88493 | -0.39621 | 2.027268 | 0.343314 |

| | | | | | | | | | |
|---|---|---|---|---|---|---|---|---|---|
| X | 115844000 | 115845799 | 26.90402 | 46.18955 | 39.89079 | 17.49679 | 0.59445 | 0.943312 | 0.343323 |
| 12 | 100797600 | 100799599 | 15.43281 | 26.79989 | 48.98345 | 21.49953 | -0.47931 | 0.79988 | 0.343329 |
| 15 | 57673600 | 57678199 | 31.89875 | 51.54638 | 93.52917 | 39.93947 | -0.49691 | 1.708572 | 0.343333 |
| 19 | 31148400 | 31150199 | 12.86714 | 22.74618 | 46.69382 | 20.6541 | -0.64175 | 0.536173 | 0.343345 |
| 18 | 61726600 | 61729799 | 37.52719 | 61.68629 | 57.52891 | 24.79142 | 0.478572 | 1.423248 | 0.343364 |
| 11 | 111615600 | 111616399 | 9.175865 | 16.78869 | 33.99682 | 14.5355 | -0.58696 | 0.268581 | 0.343386 |
| 16 | 14857800 | 14860999 | 31.4195 | 53.68273 | 39.03832 | 17.11625 | 0.840346 | 1.000006 | 0.343397 |
| 10 | 68322200 | 68324799 | 26.04144 | 44.76448 | 39.21153 | 17.19966 | 0.574193 | 0.923833 | 0.343409 |
| 7 | 5914400 | 5933399 | 381.2003 | 538.6112 | 430.1824 | 178.0806 | 0.50747 | 4.42887 | 0.343422 |
| 10 | 38783400 | 38784799 | 14.7765 | 26.96897 | 25.29015 | 10.84714 | 0.512917 | 0.302593 | 0.34345 |
| 14 | 53688800 | 53689799 | 6.589951 | 12.67853 | 31.06756 | 12.10141 | -0.74621 | -0.09789 | 0.343463 |
| 9 | 3800000 | 3804799 | 47.61001 | 77.57109 | 60.7936 | 26.10756 | 0.72182 | 1.563607 | 0.343468 |
| 14 | 48772600 | 48774599 | 13.50987 | 23.7549 | 43.99499 | 19.48116 | -0.49694 | 0.595025 | 0.343471 |
| 2 | 37064400 | 37074199 | 170.6391 | 256.3365 | 234.6542 | 96.85946 | 0.395713 | 3.440953 | 0.343498 |
| 13 | 86273200 | 86276599 | 31.81064 | 53.422 | 47.49278 | 20.7388 | 0.550134 | 1.160128 | 0.343499 |
| 11 | 118893600 | 118894999 | 8.772965 | 16.21732 | 33.63811 | 14.18243 | -0.60731 | 0.202116 | 0.343501 |
| 11 | 63396400 | 63397399 | 5.973269 | 11.53555 | 25.91315 | 9.973554 | -0.6049 | -0.15823 | 0.343521 |
| 12 | 115556600 | 115557199 | 19.46136 | 34.22623 | 31.28249 | 13.85148 | 0.516922 | 0.605189 | 0.343523 |
| 1 | 145991000 | 145995199 | 49.20786 | 79.67868 | 66.49911 | 28.45253 | 0.623816 | 1.675517 | 0.343531 |
| 11 | 88807200 | 88814799 | 127.0505 | 190.4624 | 303.7546 | 125.3147 | -0.40729 | 3.476643 | 0.343538 |
| 5 | 22928200 | 22932399 | 49.15924 | 78.16504 | 132.1219 | 55.14627 | -0.42192 | 2.244684 | 0.343586 |
| 18 | 83084400 | 83087199 | 16.31345 | 27.86448 | 56.97501 | 24.97994 | -0.64527 | 0.964428 | 0.343613 |
| 4 | 112608400 | 112609599 | 23.60813 | 41.2374 | 34.22568 | 15.07749 | 0.655179 | 0.734991 | 0.343619 |
| 11 | 52636800 | 52641399 | 27.95355 | 45.47607 | 84.78451 | 36.39033 | -0.5268 | 1.557806 | 0.343626 |
| 1 | 31000000 | 31006199 | 80.10725 | 127.111 | 107.6171 | 44.82559 | 0.571094 | 2.324668 | 0.343644 |
| 1 | 86156600 | 86157999 | 7.735889 | 14.37374 | 31.05049 | 13.00043 | -0.65732 | 0.161468 | 0.34365 |
| 1 | 140831000 | 140832999 | 22.17117 | 38.93572 | 30.78352 | 13.62162 | 0.724512 | 0.63719 | 0.343664 |
| 2 | 158060800 | 158064599 | 22.61585 | 37.08451 | 75.93905 | 32.69118 | -0.65416 | 1.431297 | 0.343673 |
| 15 | 80695400 | 80697399 | 10.54769 | 19.15381 | 37.01816 | 15.97805 | -0.52984 | 0.325713 | 0.343685 |
| 12 | 17982000 | 17991799 | 215.5601 | 317.1772 | 289.2904 | 119.1309 | 0.373559 | 3.812903 | 0.343697 |
| 18 | 84913800 | 84916599 | 29.53963 | 49.87001 | 40.01608 | 17.50349 | 0.697539 | 1.097185 | 0.343702 |
| 12 | 109861400 | 109868799 | 82.03203 | 126.099 | 213.5151 | 88.66223 | -0.46351 | 2.957935 | 0.343706 |
| 1 | 127966200 | 127967199 | 6.729853 | 12.89427 | 27.36526 | 10.78659 | -0.55701 | -0.06979 | 0.343714 |
| 14 | 66070800 | 66071599 | 6.729853 | 12.89427 | 27.36526 | 10.78659 | -0.55701 | -0.06979 | 0.343714 |
| 16 | 81402800 | 81403599 | 6.729853 | 12.89427 | 27.36526 | 10.78659 | -0.55701 | -0.06979 | 0.343714 |
| 10 | 104636800 | 104638599 | 19.28348 | 34.3828 | 27.71788 | 12.19476 | 0.705749 | 0.474367 | 0.343719 |
| 3 | 92731000 | 92733199 | 24.07744 | 39.37047 | 78.20308 | 33.62796 | -0.61256 | 1.473125 | 0.343744 |
| 4 | 25823200 | 25823999 | 10.02354 | 19.35147 | 19.13649 | 7.375933 | 0.564931 | -0.13373 | 0.343751 |
| 5 | 139531400 | 139533599 | 17.17609 | 28.76166 | 63.09684 | 27.50236 | -0.7482 | 1.159982 | 0.343753 |
| 2 | 75937400 | 75940599 | 21.14739 | 35.23164 | 65.50095 | 28.45859 | -0.51044 | 1.229544 | 0.343757 |
| 1 | 148223400 | 148229599 | 176.6612 | 264.9798 | 219.5619 | 90.59285 | 0.537425 | 3.470576 | 0.343757 |
| 7 | 76600200 | 76601799 | 12.10111 | 21.27641 | 41.44866 | 18.35388 | -0.56929 | 0.583757 | 0.343796 |
| 18 | 78315000 | 78316199 | 13.2492 | 24.59754 | 23.3807 | 9.800172 | 0.516864 | 0.191008 | 0.343826 |
| 5 | 24234400 | 24236199 | 12.2455 | 21.52661 | 41.94622 | 18.57483 | -0.56985 | 0.586518 | 0.343834 |
| 1 | 131935000 | 131936399 | 7.866716 | 14.64656 | 31.07679 | 12.96897 | -0.62816 | 0.148648 | 0.343834 |
| 13 | 22588400 | 22591599 | 117.689 | 180.4683 | 170.6593 | 70.84258 | 0.373389 | 3.013772 | 0.343837 |
| 17 | 19974800 | 19977799 | 49.53535 | 80.34606 | 65.14426 | 27.90795 | 0.667791 | 1.645907 | 0.343843 |

| | | | | | | | | | |
|---|---|---|---|---|---|---|---|---|---|
| 17 | 12590600 | 12595799 | 28.02302 | 45.41831 | 92.45406 | 39.58514 | -0.6581 | 1.632357 | 0.343848 |
| 14 | 46297800 | 46298999 | 8.124863 | 15.34916 | 31.47058 | 12.80428 | -0.5493 | 0.057449 | 0.343856 |
| 3 | 16679000 | 16681599 | 31.79244 | 53.81757 | 43.55391 | 19.06649 | 0.685421 | 1.086683 | 0.343865 |
| 11 | 57855600 | 57860799 | 35.30131 | 56.93841 | 99.05784 | 42.19872 | -0.43957 | 1.770908 | 0.343889 |
| 17 | 62212800 | 62216399 | 37.92073 | 62.73523 | 53.81451 | 23.26964 | 0.601232 | 1.344538 | 0.343893 |
| 15 | 44904000 | 44904599 | 9.921755 | 19.12571 | 19.17574 | 7.427842 | 0.538871 | -0.11773 | 0.343899 |
| 13 | 98767400 | 98769399 | 11.7542 | 20.74975 | 40.88263 | 18.08869 | -0.58333 | 0.534845 | 0.343899 |
| 2 | 144867600 | 144869199 | 19.02962 | 33.95223 | 27.1582 | 11.94198 | 0.717342 | 0.466899 | 0.343903 |
| 16 | 10231400 | 10237599 | 40.81192 | 65.19796 | 118.1153 | 49.62052 | -0.51436 | 2.083754 | 0.343912 |
| 6 | 115668000 | 115670399 | 24.68593 | 42.72839 | 37.03481 | 16.24828 | 0.59127 | 0.854185 | 0.343916 |
| 6 | 132897400 | 132898999 | 26.29981 | 45.47578 | 38.13959 | 16.73247 | 0.638484 | 0.874608 | 0.343927 |
| 7 | 125306400 | 125307999 | 11.20165 | 19.64904 | 44.61526 | 19.72293 | -0.78946 | 0.618655 | 0.343933 |
| 11 | 33363000 | 33365199 | 22.60723 | 39.7087 | 31.45761 | 13.92204 | 0.721892 | 0.640401 | 0.343947 |
| 17 | 5541600 | 5543999 | 17.68779 | 29.92703 | 57.91781 | 25.35081 | -0.56758 | 1.059264 | 0.343961 |
| 16 | 76138800 | 76144199 | 51.32048 | 81.51466 | 135.5111 | 56.50064 | -0.39948 | 2.279113 | 0.343972 |
| 1 | 53015600 | 53017199 | 15.46468 | 28.73005 | 23.69803 | 9.924045 | 0.721085 | 0.206948 | 0.343978 |
| 7 | 7151000 | 7151799 | 15.46468 | 28.73005 | 23.69803 | 9.924045 | 0.721085 | 0.206948 | 0.343978 |
| 8 | 50462200 | 50464599 | 27.66572 | 48.05677 | 35.27201 | 15.48418 | 0.831177 | 0.833446 | 0.343978 |
| 9 | 47350200 | 47353599 | 18.02834 | 30.82571 | 56.51158 | 24.77883 | -0.48823 | 0.961764 | 0.343984 |
| 7 | 122547600 | 122550799 | 18.14044 | 30.70908 | 57.97171 | 25.37847 | -0.53189 | 1.05537 | 0.343989 |
| 13 | 66643000 | 66652599 | 257.2648 | 373.1158 | 337.4737 | 139.141 | 0.365645 | 4.0392 | 0.343995 |
| 17 | 87496600 | 87503799 | 63.18601 | 98.29119 | 196.5733 | 81.60603 | -0.69011 | 2.730854 | 0.344003 |
| 12 | 63020600 | 63021599 | 18.61356 | 32.79407 | 30.31238 | 13.41711 | 0.501742 | 0.578838 | 0.344027 |
| 6 | 66580600 | 66581799 | 12.92967 | 22.63141 | 44.15164 | 19.48249 | -0.5715 | 0.666269 | 0.344043 |
| 18 | 78500800 | 78504399 | 24.12496 | 39.7354 | 73.20558 | 31.57379 | -0.49963 | 1.376566 | 0.34406 |
| X | 118208800 | 118211799 | 52.0976 | 84.10239 | 72.46352 | 30.90402 | 0.573439 | 1.762108 | 0.344062 |
| 3 | 47555600 | 47557399 | 26.31031 | 46.05993 | 29.46622 | 13.00421 | 1.028619 | 0.707086 | 0.344075 |
| 11 | 113657200 | 113662399 | 27.84205 | 45.17464 | 90.26513 | 38.67925 | -0.62986 | 1.60869 | 0.344101 |
| 1 | 100719400 | 100719799 | 10.33506 | 19.91853 | 18.18875 | 7.049476 | 0.672404 | -0.11272 | 0.344105 |
| 11 | 34703200 | 34705799 | 20.1986 | 33.67845 | 70.49519 | 30.64426 | -0.68098 | 1.221943 | 0.344109 |
| 10 | 13319600 | 13320799 | 9.841827 | 17.65462 | 36.86099 | 16.13487 | -0.65376 | 0.393241 | 0.344115 |
| 12 | 110116000 | 110117599 | 30.88526 | 52.039 | 46.3728 | 20.27186 | 0.546809 | 1.126344 | 0.344127 |
| 5 | 37784600 | 37788399 | 42.56571 | 69.05133 | 66.35522 | 28.42977 | 0.423393 | 1.627602 | 0.344153 |
| 10 | 117782600 | 117785199 | 15.10476 | 26.30153 | 47.88844 | 21.05481 | -0.47336 | 0.757978 | 0.344158 |
| 8 | 30530600 | 30533399 | 19.78462 | 34.93776 | 28.73472 | 12.71293 | 0.670132 | 0.554914 | 0.344161 |
| 18 | 9268000 | 9269799 | 12.05406 | 21.3808 | 39.8064 | 17.59041 | -0.49917 | 0.485328 | 0.34417 |
| 6 | 7005200 | 7013999 | 166.641 | 251.1961 | 223.1818 | 92.20911 | 0.443175 | 3.37319 | 0.344171 |
| 5 | 53666000 | 53667599 | 8.607171 | 15.94499 | 34.79899 | 14.62915 | -0.67733 | 0.190182 | 0.344174 |
| 8 | 45380200 | 45382799 | 39.09966 | 64.57499 | 54.41652 | 23.50216 | 0.626648 | 1.367608 | 0.344187 |
| 14 | 35317000 | 35318199 | 10.61805 | 20.48472 | 19.69438 | 7.610789 | 0.602174 | -0.11561 | 0.344189 |
| 16 | 95658200 | 95659399 | 8.341684 | 15.4783 | 31.5554 | 13.23627 | -0.57651 | 0.17555 | 0.344221 |
| 5 | 3065000 | 3089199 | 451.6323 | 623.8812 | 538.9707 | 223.9119 | 0.361489 | 4.76773 | 0.344255 |
| 1 | 138157600 | 138161399 | 58.43504 | 93.72521 | 77.45103 | 32.72847 | 0.622492 | 1.96777 | 0.344271 |
| 1 | 30534400 | 30535799 | 11.2034 | 21.55713 | 20.13158 | 7.838823 | 0.633366 | -0.08503 | 0.344274 |
| 4 | 128875200 | 128878399 | 22.0834 | 36.36212 | 78.57478 | 33.88532 | -0.72917 | 1.37956 | 0.34428 |
| 15 | 99729400 | 99730999 | 11.23485 | 20.10253 | 37.82938 | 16.59413 | -0.50704 | 0.421221 | 0.344321 |
| 8 | 49599600 | 49600399 | 9.640723 | 17.55033 | 34.44209 | 14.82008 | -0.54833 | 0.301371 | 0.344327 |

| | | | | | | | | | |
|---|---|---|---|---|---|---|---|---|---|
| X | 148555800 | 148558599 | 45.09484 | 73.32038 | 64.50389 | 27.67993 | 0.553386 | 1.589187 | 0.344328 |
| 8 | 34931000 | 34935799 | 58.87411 | 94.26713 | 89.15761 | 37.57688 | 0.425196 | 2.023884 | 0.344335 |
| 1 | 48255800 | 48256399 | 8.698889 | 16.89046 | 16.69314 | 6.258013 | 0.604019 | -0.24779 | 0.344344 |
| 14 | 15814800 | 15817599 | 24.54913 | 42.71504 | 35.89885 | 15.77606 | 0.636743 | 0.795128 | 0.34435 |
| 17 | 34927400 | 34943799 | 121.6533 | 181.7306 | 318.3861 | 131.2612 | -0.54738 | 3.541858 | 0.34436 |
| 12 | 24401600 | 24423199 | 436.8352 | 606.5722 | 520.7595 | 216.1691 | 0.37802 | 4.688743 | 0.344369 |
| 16 | 18333800 | 18334399 | 9.189024 | 17.90532 | 16.66319 | 6.050614 | 0.730973 | -0.35273 | 0.344375 |
| 6 | 98644200 | 98646999 | 32.85122 | 54.7942 | 50.32692 | 21.88195 | 0.503427 | 1.235659 | 0.344381 |
| 11 | 112475200 | 112479799 | 31.85593 | 51.73638 | 92.31736 | 39.58634 | -0.46582 | 1.599966 | 0.344392 |
| 6 | 82189000 | 82189999 | 9.749914 | 17.93389 | 35.16044 | 14.92489 | -0.53357 | 0.243367 | 0.344406 |
| 15 | 65045400 | 65048999 | 43.5576 | 69.58875 | 118.4485 | 49.76505 | -0.42449 | 2.084721 | 0.344422 |
| 13 | 75356200 | 75359599 | 45.13505 | 72.93973 | 69.96349 | 29.87857 | 0.420731 | 1.718718 | 0.34445 |
| 12 | 118416000 | 118417799 | 16.05241 | 29.42898 | 25.09143 | 10.69892 | 0.65502 | 0.286545 | 0.344454 |
| 7 | 85043000 | 85045399 | 30.55121 | 51.98158 | 42.44854 | 18.60518 | 0.672971 | 1.038277 | 0.344459 |
| 7 | 117627800 | 117629999 | 15.92042 | 27.586 | 50.05564 | 21.96163 | -0.46951 | 0.826186 | 0.344467 |
| 7 | 60710200 | 60711199 | 12.51564 | 24.04664 | 20.0063 | 7.82305 | 0.792968 | -0.05849 | 0.344476 |
| 12 | 59821800 | 59822199 | 4.449625 | 8.658674 | 23.42921 | 8.629429 | -0.8081 | -0.33295 | 0.344489 |
| 7 | 10361800 | 10366399 | 66.73842 | 107.4066 | 68.34522 | 29.02589 | 1.004561 | 1.873713 | 0.344498 |
| 19 | 33701200 | 33702999 | 17.09941 | 31.0905 | 25.37119 | 10.9336 | 0.707578 | 0.340123 | 0.344508 |
| 5 | 82914800 | 82915799 | 13.31439 | 25.47935 | 18.51898 | 7.324984 | 0.969846 | -0.0205 | 0.344522 |
| 6 | 104651400 | 104654799 | 35.42867 | 59.64401 | 44.1227 | 19.2833 | 0.814391 | 1.147456 | 0.344531 |
| 19 | 46684400 | 46689999 | 30.12077 | 48.81771 | 87.06015 | 37.27537 | -0.46765 | 1.637206 | 0.344534 |
| 11 | 7486200 | 7489599 | 47.63718 | 76.86844 | 73.15599 | 31.18187 | 0.429526 | 1.765257 | 0.344568 |
| 14 | 22942200 | 22943599 | 8.621981 | 16.12571 | 32.27747 | 13.36699 | -0.53586 | 0.129521 | 0.344578 |
| 12 | 113178000 | 113192599 | 112.0521 | 168.3706 | 300.0949 | 123.8805 | -0.56455 | 3.427798 | 0.34458 |
| 7 | 45214000 | 45220199 | 30.46132 | 49.26296 | 89.70051 | 38.34406 | -0.50055 | 1.682382 | 0.344587 |
| 3 | 148503800 | 148506399 | 45.22984 | 72.11473 | 123.5192 | 51.74854 | -0.4367 | 2.151416 | 0.344587 |
| 7 | 68099200 | 68099599 | 3.525504 | 6.888265 | 19.33248 | 6.73393 | -0.78613 | -0.56379 | 0.344594 |
| 18 | 83872400 | 83872799 | 3.525504 | 6.888265 | 19.33248 | 6.73393 | -0.78613 | -0.56379 | 0.344594 |
| 2 | 137211400 | 137212199 | 12.90406 | 22.76414 | 42.96038 | 19.01057 | -0.52188 | 0.555171 | 0.344621 |
| 7 | 49000200 | 49001999 | 11.69816 | 20.43 | 45.2445 | 19.92936 | -0.75358 | 0.695476 | 0.344622 |
| 14 | 39476000 | 39477999 | 13.56519 | 23.53789 | 49.49852 | 21.71812 | -0.68081 | 0.800962 | 0.344635 |
| 9 | 82556600 | 82557399 | 6.412345 | 12.4033 | 27.88844 | 10.65398 | -0.59673 | -0.17493 | 0.344647 |
| 14 | 117585400 | 117587199 | 18.28901 | 32.53672 | 28.37555 | 12.50769 | 0.591107 | 0.482615 | 0.344668 |
| 6 | 96431200 | 96431999 | 4.822465 | 9.356512 | 23.82841 | 8.974152 | -0.75346 | -0.25265 | 0.344668 |
| X | 33373000 | 33380399 | 120.3088 | 187.8158 | 122.4383 | 50.80479 | 0.929173 | 2.689827 | 0.34468 |
| 4 | 134408200 | 134416399 | 76.80139 | 119.2818 | 190.0539 | 78.91823 | -0.36441 | 2.763347 | 0.344695 |
| 14 | 20228800 | 20232999 | 20.08307 | 33.34667 | 67.47494 | 29.25019 | -0.63243 | 1.26627 | 0.344715 |
| 11 | 88754000 | 88755999 | 11.88996 | 20.67757 | 48.93394 | 21.49741 | -0.84975 | 0.761366 | 0.344717 |
| 4 | 142828200 | 142832999 | 31.12869 | 50.25345 | 93.39198 | 39.83123 | -0.53293 | 1.736585 | 0.344747 |
| 13 | 5910800 | 5913399 | 16.36938 | 28.28923 | 52.76303 | 23.14762 | -0.51 | 0.85354 | 0.344755 |
| 9 | 123443600 | 123445199 | 19.72652 | 35.48489 | 26.14659 | 11.41808 | 0.842001 | 0.421707 | 0.344774 |
| X | 139177000 | 139180399 | 34.29011 | 57.13177 | 49.22707 | 21.389 | 0.595449 | 1.254364 | 0.344777 |
| 5 | 115410000 | 115410999 | 5.218613 | 9.892979 | 28.68261 | 11.58482 | -1.03396 | -0.00029 | 0.344781 |
| 14 | 101640800 | 101641799 | 12.58697 | 24.31426 | 18.33253 | 7.047299 | 0.956096 | -0.11707 | 0.344787 |
| 19 | 50373800 | 50374199 | 10.59066 | 20.70575 | 12.01339 | 4.115743 | 1.478678 | -0.54238 | 0.344802 |
| X | 32905600 | 32908199 | 20.80594 | 37.79289 | 23.44988 | 10.11797 | 1.099352 | 0.370201 | 0.344803 |

| | | | | | | | | | |
|---|---|---|---|---|---|---|---|---|---|
| 1 | 98726200 | 98728199 | 16.083 | 29.49411 | 24.46578 | 10.42779 | 0.694583 | 0.282647 | 0.344821 |
| 14 | 94554600 | 94556199 | 22.85802 | 40.0669 | 32.55059 | 14.3808 | 0.685807 | 0.67492 | 0.344825 |
| 3 | 56030400 | 56034399 | 62.13408 | 99.78625 | 79.86657 | 33.81518 | 0.670674 | 1.934464 | 0.344831 |
| 12 | 114266800 | 114271999 | 87.73199 | 138.4563 | 119.6374 | 49.66191 | 0.535187 | 2.466108 | 0.344832 |
| 7 | 122174000 | 122174599 | 4.516298 | 8.814888 | 21.6607 | 7.682744 | -0.62304 | -0.47935 | 0.344832 |
| 16 | 88504400 | 88504799 | 4.516298 | 8.814888 | 21.6607 | 7.682744 | -0.62304 | -0.47935 | 0.344832 |
| 5 | 93161400 | 93163399 | 22.5503 | 39.30872 | 35.44062 | 15.59255 | 0.53603 | 0.759171 | 0.344843 |
| 2 | 32561200 | 32567999 | 60.60936 | 95.54216 | 155.6106 | 64.57439 | -0.37937 | 2.479561 | 0.344844 |
| 5 | 146365600 | 146367199 | 9.920046 | 18.04996 | 34.93601 | 15.04183 | -0.5294 | 0.30557 | 0.344844 |
| X | 24369000 | 24370799 | 26.64965 | 46.90785 | 26.66093 | 11.80311 | 1.198914 | 0.629992 | 0.344847 |
| 1 | 127083800 | 127086199 | 13.18571 | 23.25538 | 47.76215 | 21.13645 | -0.64383 | 0.564864 | 0.344858 |
| 4 | 130268200 | 130270999 | 13.38084 | 23.48564 | 43.53406 | 19.25469 | -0.49823 | 0.622158 | 0.344903 |
| 5 | 105922400 | 105925399 | 22.40618 | 37.38111 | 66.25256 | 28.81059 | -0.44179 | 1.218458 | 0.344912 |
| 6 | 131323800 | 131325399 | 8.190759 | 15.1227 | 33.61042 | 14.1942 | -0.70787 | 0.20414 | 0.344918 |
| 13 | 57215200 | 57215999 | 16.49299 | 29.81759 | 26.51331 | 11.50747 | 0.57805 | 0.370474 | 0.34493 |
| 2 | 53778600 | 53779999 | 13.37669 | 25.12139 | 21.67886 | 8.915618 | 0.676376 | 0.115191 | 0.344941 |
| 17 | 31432600 | 31434399 | 13.04351 | 22.87424 | 43.04038 | 19.02276 | -0.51975 | 0.632177 | 0.344941 |
| 7 | 26398600 | 26399799 | 21.29191 | 37.04402 | 33.60822 | 14.76854 | 0.52678 | 0.776247 | 0.344948 |
| 2 | 102401400 | 102403399 | 13.20974 | 23.30379 | 46.13852 | 20.41688 | -0.59091 | 0.560827 | 0.344951 |
| 10 | 21012800 | 21014199 | 16.73598 | 31.04516 | 22.31448 | 9.363787 | 0.91575 | 0.220893 | 0.344968 |
| 5 | 19934800 | 19936599 | 18.81806 | 33.50892 | 28.32742 | 12.47674 | 0.636583 | 0.480697 | 0.344993 |
| 5 | 45320200 | 45320799 | 11.84687 | 22.49698 | 21.55497 | 8.674296 | 0.553702 | 0.030275 | 0.344996 |
| 8 | 104746000 | 104750199 | 37.32281 | 59.99829 | 103.0755 | 43.7041 | -0.42732 | 1.88024 | 0.345005 |
| 8 | 52460200 | 52460999 | 18.71755 | 33.26746 | 25.13045 | 11.08616 | 0.796646 | 0.486777 | 0.345012 |
| 2 | 164895600 | 164897799 | 20.84125 | 34.64779 | 67.6939 | 29.36982 | -0.58201 | 1.253049 | 0.345012 |
| 10 | 79334800 | 79336599 | 30.45953 | 51.10062 | 48.05482 | 20.97068 | 0.469365 | 1.166558 | 0.34503 |
| 12 | 5030800 | 5032599 | 19.20077 | 34.10549 | 28.85021 | 12.73366 | 0.633693 | 0.499867 | 0.345036 |
| 19 | 20461000 | 20461599 | 7.973152 | 15.04124 | 30.27953 | 12.35664 | -0.52673 | 0.063266 | 0.345038 |
| 1 | 191246400 | 191247199 | 7.319531 | 13.98018 | 29.58618 | 11.7617 | -0.5651 | -0.03195 | 0.345039 |
| 10 | 70626600 | 70627599 | 15.30178 | 27.72467 | 25.79589 | 11.16547 | 0.515805 | 0.349392 | 0.345043 |
| 2 | 81882800 | 81885199 | 18.49138 | 31.16248 | 60.112 | 26.2795 | -0.56327 | 1.097646 | 0.345048 |
| 8 | 20609600 | 20611199 | 19.26351 | 33.81901 | 31.24186 | 13.82083 | 0.501159 | 0.626719 | 0.345063 |
| 13 | 65814200 | 65819199 | 82.15455 | 130.4971 | 93.30522 | 38.90531 | 0.815971 | 2.29392 | 0.345074 |
| 18 | 8314600 | 8316399 | 21.11721 | 37.3048 | 29.86255 | 13.21093 | 0.709539 | 0.564285 | 0.345074 |
| 4 | 87537200 | 87539399 | 25.47146 | 44.51949 | 31.80185 | 14.01585 | 0.87048 | 0.731145 | 0.345086 |
| 9 | 65440200 | 65444199 | 27.45881 | 44.79559 | 80.14191 | 34.43256 | -0.46452 | 1.516067 | 0.345086 |
| 8 | 109664800 | 109669199 | 50.63866 | 80.48835 | 133.532 | 55.71473 | -0.39555 | 2.257055 | 0.345094 |
| 9 | 28181800 | 28182599 | 10.74598 | 20.6504 | 20.166 | 7.881272 | 0.564653 | -0.07753 | 0.345095 |
| 12 | 6573600 | 6576999 | 31.71585 | 53.67864 | 44.12294 | 19.31458 | 0.663087 | 1.089327 | 0.345097 |
| 1 | 72268600 | 72269599 | 12.995 | 24.24422 | 22.95608 | 9.546859 | 0.531377 | 0.158847 | 0.345115 |
| 7 | 140095800 | 140098199 | 39.44305 | 65.61334 | 48.76659 | 21.16705 | 0.807919 | 1.283809 | 0.345124 |
| 7 | 107905600 | 107908399 | 28.88371 | 48.7503 | 45.84416 | 20.05121 | 0.469509 | 1.104103 | 0.345145 |
| 15 | 81128000 | 81129999 | 26.32766 | 45.1382 | 40.95522 | 17.96169 | 0.52319 | 0.953415 | 0.345145 |
| 7 | 12291600 | 12303599 | 178.5372 | 268.9036 | 219.8079 | 90.79482 | 0.56228 | 3.392161 | 0.34515 |
| 1 | 13855200 | 13858599 | 23.57676 | 39.02066 | 71.22456 | 30.80643 | -0.48485 | 1.315127 | 0.345153 |
| 16 | 43531000 | 43533599 | 21.90174 | 36.50998 | 67.39425 | 29.29212 | -0.50031 | 1.224898 | 0.345156 |
| X | 32980400 | 32981599 | 17.44902 | 32.6351 | 21.45955 | 8.886924 | 1.058514 | 0.178098 | 0.345161 |

| | | | | | | | | | |
|---|---|---|---|---|---|---|---|---|---|
| 10 | 113858400 | 113858999 | 10.85768 | 20.9756 | 18.43793 | 7.084953 | 0.736929 | -0.13804 | 0.345177 |
| 18 | 84848200 | 84850999 | 21.84463 | 36.26182 | 68.11906 | 29.5239 | -0.52563 | 1.276645 | 0.345199 |
| 6 | 63002600 | 63003799 | 24.37142 | 42.09293 | 38.34583 | 16.82236 | 0.519424 | 0.877291 | 0.345208 |
| 1 | 93978600 | 93981599 | 18.71537 | 31.38943 | 61.61493 | 26.88358 | -0.58845 | 1.146597 | 0.345215 |
| 4 | 32425600 | 32427599 | 11.63802 | 20.50887 | 40.43205 | 17.89556 | -0.58491 | 0.55492 | 0.345218 |
| 17 | 74213200 | 74215199 | 10.55846 | 19.13837 | 36.44613 | 15.76712 | -0.51074 | 0.335605 | 0.345239 |
| 1 | 136633400 | 136635999 | 14.7822 | 25.29119 | 53.66903 | 23.53364 | -0.69785 | 0.942012 | 0.345243 |
| 8 | 22148200 | 22150599 | 18.34731 | 33.60328 | 23.72121 | 10.12816 | 0.924148 | 0.306348 | 0.345256 |
| 2 | 133954800 | 133956999 | 25.07156 | 43.73041 | 34.14913 | 15.03011 | 0.742505 | 0.761917 | 0.345275 |
| 6 | 15607000 | 15608599 | 16.26376 | 27.96329 | 52.70593 | 23.11897 | -0.52656 | 0.900062 | 0.345278 |
| 7 | 108115600 | 108118599 | 45.15705 | 73.54901 | 62.8248 | 26.97645 | 0.597579 | 1.564866 | 0.345285 |
| 1 | 192327800 | 192328999 | 15.02946 | 28.20731 | 22.14369 | 9.1166 | 0.811059 | 0.136258 | 0.345288 |
| 4 | 136646000 | 136647999 | 14.63273 | 25.44163 | 47.90548 | 21.04258 | -0.52181 | 0.776765 | 0.345294 |
| 4 | 112314000 | 112314799 | 28.09539 | 47.41582 | 45.33833 | 19.82947 | 0.445582 | 1.102094 | 0.345303 |
| X | 140304600 | 140304999 | 10.63528 | 20.44313 | 19.67922 | 7.685132 | 0.586477 | -0.08167 | 0.345311 |
| 11 | 18944200 | 18946599 | 23.52316 | 41.2466 | 31.93052 | 14.11084 | 0.755049 | 0.672335 | 0.345312 |
| 12 | 115599800 | 115602599 | 54.14939 | 87.08957 | 80.21727 | 34.03581 | 0.470958 | 1.87822 | 0.345313 |
| 9 | 34930600 | 34932399 | 9.32778 | 17.02787 | 33.96322 | 14.56335 | -0.56819 | 0.282216 | 0.345323 |
| 16 | 27035400 | 27036999 | 15.77578 | 29.30331 | 23.00142 | 9.634427 | 0.791562 | 0.207418 | 0.34533 |
| 1 | 114830000 | 114831999 | 23.61948 | 41.20821 | 34.19404 | 15.0523 | 0.655239 | 0.749947 | 0.345336 |
| 6 | 73464000 | 73465799 | 13.65534 | 23.99065 | 44.52953 | 19.7111 | -0.50036 | 0.607787 | 0.345337 |
| 14 | 21742800 | 21745199 | 24.19515 | 41.5425 | 39.42716 | 17.29356 | 0.458761 | 0.928033 | 0.345341 |
| 13 | 65733000 | 65739399 | 188.3831 | 279.9713 | 256.6752 | 105.743 | 0.379532 | 3.64159 | 0.345346 |
| 4 | 8535400 | 8536399 | 8.060709 | 14.85813 | 32.52218 | 13.7654 | -0.68799 | 0.211505 | 0.345347 |
| 11 | 96967000 | 96968599 | 8.060709 | 14.85813 | 32.52218 | 13.7654 | -0.68799 | 0.211505 | 0.345347 |
| 3 | 159271400 | 159273199 | 19.04927 | 33.71971 | 28.81208 | 12.74107 | 0.616714 | 0.525621 | 0.345365 |
| 4 | 142204000 | 142207399 | 23.56743 | 38.62764 | 76.05526 | 32.73503 | -0.59802 | 1.439109 | 0.34542 |
| 10 | 46420000 | 46430199 | 246.9826 | 360.9725 | 320.7484 | 132.141 | 0.401974 | 3.922205 | 0.345428 |
| 7 | 15566800 | 15576399 | 176.2714 | 263.2104 | 241.5127 | 99.5604 | 0.384536 | 3.554469 | 0.345467 |
| 2 | 143926600 | 143931599 | 28.99127 | 46.96219 | 88.10716 | 37.70892 | -0.54136 | 1.644798 | 0.345468 |
| 7 | 22570600 | 22578199 | 124.7914 | 194.1682 | 127.7989 | 53.053 | 0.911676 | 2.74816 | 0.345475 |
| 18 | 54481800 | 54484599 | 24.68323 | 40.86832 | 74.92057 | 32.41402 | -0.49112 | 1.315485 | 0.345477 |
| 16 | 59563200 | 59564199 | 6.714655 | 12.88463 | 28.62748 | 11.23521 | -0.61731 | -0.08327 | 0.345493 |
| 10 | 52537400 | 52539199 | 25.27324 | 43.29896 | 40.99947 | 17.98004 | 0.461674 | 0.954913 | 0.345493 |
| 16 | 17831400 | 17835999 | 14.87919 | 27.33602 | 20.68354 | 8.793992 | 0.828929 | 0.254862 | 0.345494 |
| 16 | 60729400 | 60731199 | 19.55483 | 34.3464 | 31.12728 | 13.77555 | 0.528572 | 0.620374 | 0.345498 |
| 13 | 61354800 | 61356199 | 19.10229 | 35.02967 | 23.15857 | 9.871177 | 1.019939 | 0.304687 | 0.345503 |
| 2 | 177846000 | 177848399 | 16.3506 | 27.8646 | 55.83487 | 24.47558 | -0.61661 | 0.985674 | 0.345508 |
| 7 | 13693000 | 13694599 | 27.08596 | 46.85484 | 37.52066 | 16.46114 | 0.704988 | 0.873211 | 0.345514 |
| 8 | 3106000 | 3124799 | 295.8135 | 423.9987 | 381.0592 | 157.3731 | 0.358029 | 4.214867 | 0.345528 |
| 9 | 51123400 | 51125399 | 12.61078 | 22.11787 | 43.07201 | 19.03856 | -0.56884 | 0.630002 | 0.345532 |
| 5 | 33953000 | 33959799 | 30.2114 | 52.24813 | 31.17946 | 13.67901 | 1.127351 | 0.882959 | 0.345541 |
| 13 | 43656800 | 43658799 | 13.52235 | 23.62102 | 45.2133 | 19.91837 | -0.5445 | 0.701022 | 0.345552 |
| 7 | 139346000 | 139347599 | 12.39544 | 21.77212 | 42.991 | 19.02633 | -0.58906 | 0.601289 | 0.345553 |
| 6 | 10111800 | 10112799 | 11.43503 | 22.22432 | 16.2449 | 6.043318 | 1.04363 | -0.24444 | 0.345557 |
| 9 | 124167000 | 124169999 | 73.92665 | 117.1751 | 112.7082 | 46.89454 | 0.385721 | 2.351817 | 0.345564 |
| 17 | 18926600 | 18927199 | 10.32747 | 20.09452 | 16.41206 | 6.049862 | 0.897166 | -0.28866 | 0.345567 |

| | | | | | | | | | |
|---|---|---|---|---|---|---|---|---|---|
| 3 | 111466600 | 111469199 | 31.22153 | 53.38324 | 39.63063 | 17.37696 | 0.810896 | 0.994648 | 0.345572 |
| 5 | 104613600 | 104616199 | 26.26157 | 43.06022 | 76.55374 | 32.95503 | -0.45102 | 1.440462 | 0.345586 |
| 12 | 21367600 | 21373199 | 31.77643 | 51.26026 | 95.73 | 40.78847 | -0.54127 | 1.760314 | 0.345589 |
| 13 | 66686000 | 66687999 | 20.00922 | 37.40111 | 18.92709 | 7.846102 | 1.431927 | 0.197378 | 0.345589 |
| Y | 4757800 | 4759199 | 11.3804 | 21.7598 | 12.97458 | 5.144508 | 1.249785 | -0.04945 | 0.345591 |
| 6 | 3967800 | 3968599 | 7.454842 | 13.81923 | 31.55907 | 13.25541 | -0.73977 | 0.17568 | 0.345598 |
| 6 | 27024400 | 27026799 | 26.01247 | 45.19825 | 35.83603 | 15.73468 | 0.720246 | 0.821047 | 0.345602 |
| 5 | 109492400 | 109496799 | 76.70099 | 121.8999 | 104.3402 | 43.52994 | 0.557551 | 2.276037 | 0.345623 |
| X | 33928800 | 33931999 | 59.28829 | 95.19295 | 81.3171 | 34.41668 | 0.576521 | 1.940077 | 0.345624 |
| 13 | 85694600 | 85698199 | 48.11435 | 77.89669 | 68.96207 | 29.50316 | 0.538787 | 1.68045 | 0.345645 |
| 6 | 35346800 | 35347799 | 6.169083 | 11.43695 | 32.08564 | 13.47472 | -1.03417 | 0.162878 | 0.34565 |
| 8 | 64108400 | 64108999 | 4.353289 | 8.494061 | 20.92573 | 7.462349 | -0.63265 | -0.46228 | 0.345652 |
| 14 | 113670400 | 113671799 | 8.103075 | 15.23469 | 33.04144 | 13.56269 | -0.64096 | 0.089864 | 0.345657 |
| 12 | 22875600 | 22898799 | 734.7975 | 974.8142 | 805.0603 | 336.4929 | 0.367783 | 5.407226 | 0.345674 |
| 8 | 72635000 | 72636599 | 11.36553 | 20.35722 | 39.39651 | 17.26663 | -0.54656 | 0.417284 | 0.345676 |
| 17 | 20898000 | 20900999 | 46.17372 | 75.03956 | 64.47053 | 27.65903 | 0.586983 | 1.600104 | 0.345686 |
| 7 | 62286400 | 62297399 | 341.3411 | 482.8989 | 426.8265 | 176.6547 | 0.363115 | 4.407858 | 0.345691 |
| 16 | 37702800 | 37704399 | 10.55992 | 18.62657 | 40.44889 | 17.89964 | -0.72305 | 0.536797 | 0.345696 |
| 5 | 43448400 | 43449999 | 10.41986 | 18.89398 | 35.93346 | 15.5383 | -0.50827 | 0.332377 | 0.345707 |
| 1 | 71851200 | 71851999 | 8.821964 | 16.27262 | 32.52729 | 13.75456 | -0.55686 | 0.215418 | 0.345708 |
| 4 | 113946000 | 113952799 | 133.8124 | 204.7963 | 179.3416 | 74.41851 | 0.481986 | 3.057363 | 0.345715 |
| 10 | 119643600 | 119652199 | 91.58165 | 140.3658 | 230.7802 | 95.79302 | -0.42438 | 3.019183 | 0.345729 |
| 4 | 90489000 | 90490799 | 24.63546 | 42.30708 | 39.96487 | 17.52959 | 0.465629 | 0.930097 | 0.345738 |
| 10 | 13840200 | 13843799 | 23.79233 | 38.84365 | 82.14037 | 35.30095 | -0.70427 | 1.496233 | 0.345754 |
| 10 | 119938400 | 119942199 | 46.48715 | 74.11803 | 125.6766 | 52.65102 | -0.42224 | 2.152491 | 0.345767 |
| X | 4761200 | 4764199 | 28.94354 | 50.56373 | 30.06294 | 13.24478 | 1.134119 | 0.755727 | 0.345781 |
| 9 | 83716400 | 83717199 | 4.590777 | 8.853792 | 25.78422 | 9.97355 | -0.98207 | -0.15862 | 0.345784 |
| 18 | 37408400 | 37419999 | 35.15066 | 58.45367 | 47.11084 | 20.44372 | 0.691346 | 1.272467 | 0.345792 |
| 13 | 7453400 | 7455999 | 31.45463 | 53.59037 | 40.73141 | 17.85449 | 0.776332 | 1.027213 | 0.345792 |
| 3 | 61642200 | 61642399 | 7.4078 | 14.48417 | 14.11162 | 4.824044 | 0.743103 | -0.59261 | 0.345792 |
| 4 | 112767600 | 112774199 | 95.85656 | 151.8498 | 99.54732 | 41.40137 | 0.937336 | 2.38266 | 0.345804 |
| 6 | 139900800 | 139902799 | 10.45607 | 18.77365 | 36.34142 | 15.89517 | -0.54443 | 0.391929 | 0.345808 |
| 5 | 124967600 | 124969599 | 11.36195 | 19.94394 | 42.06372 | 18.60579 | -0.68311 | 0.607881 | 0.345809 |
| 12 | 108680000 | 108680599 | 5.810171 | 11.28385 | 25.18874 | 9.42699 | -0.55781 | -0.25981 | 0.34581 |
| 8 | 89549800 | 89555599 | 56.07358 | 88.79917 | 145.174 | 60.36735 | -0.37949 | 2.368009 | 0.345815 |
| 7 | 128170600 | 128171599 | 13.42688 | 25.19856 | 22.8761 | 9.418792 | 0.602963 | 0.124588 | 0.345828 |
| 10 | 61719200 | 61720399 | 13.42688 | 25.19856 | 22.8761 | 9.418792 | 0.602963 | 0.124588 | 0.345828 |
| 5 | 135284400 | 135287999 | 41.5396 | 66.62942 | 111.9206 | 47.29771 | -0.39887 | 1.956824 | 0.345856 |
| 9 | 38045600 | 38046599 | 20.07283 | 35.28816 | 27.66317 | 12.24982 | 0.736864 | 0.604885 | 0.345862 |
| 4 | 134468400 | 134470399 | 8.601728 | 15.58969 | 34.4524 | 14.90688 | -0.72338 | 0.320216 | 0.345866 |
| 12 | 44315400 | 44316399 | 8.601728 | 15.58969 | 34.4524 | 14.90688 | -0.72338 | 0.320216 | 0.345866 |
| 8 | 104297000 | 104298399 | 12.32161 | 23.03669 | 20.88547 | 8.654614 | 0.597353 | 0.132649 | 0.345866 |
| 17 | 77858400 | 77859999 | 15.75538 | 28.77604 | 25.01322 | 10.7188 | 0.622173 | 0.303713 | 0.345876 |
| 13 | 73383000 | 73394199 | 86.13584 | 132.813 | 209.7295 | 87.118 | -0.35961 | 2.899875 | 0.345881 |
| 7 | 107972800 | 107977399 | 80.99119 | 127.7653 | 122.6121 | 50.88608 | 0.38355 | 2.471964 | 0.345897 |
| 14 | 90312400 | 90314599 | 31.26022 | 53.3177 | 40.16852 | 17.60931 | 0.7892 | 1.015934 | 0.345916 |
| 2 | 177704400 | 177706199 | 24.79523 | 42.47068 | 40.01759 | 17.54914 | 0.468737 | 0.954151 | 0.345919 |

| | | | | | | | | | |
|---|---|---|---|---|---|---|---|---|---|
| 16 | 66556000 | 66556799 | 12.86493 | 24.50983 | 20.88017 | 8.345154 | 0.730124 | 0.015174 | 0.345938 |
| 7 | 11573800 | 11582799 | 17.12728 | 31.19468 | 21.69447 | 9.329084 | 0.939216 | 0.325131 | 0.345942 |
| 6 | 108961800 | 108963799 | 14.38884 | 24.97982 | 61.57944 | 27.02322 | -0.90929 | 0.8153 | 0.345946 |
| 4 | 113768000 | 113769199 | 29.48094 | 50.01957 | 44.16945 | 19.34944 | 0.560239 | 1.061493 | 0.34595 |
| 17 | 29319200 | 29320599 | 13.9466 | 24.03155 | 51.05818 | 22.39769 | -0.69762 | 0.869725 | 0.345953 |
| 1 | 146157800 | 146159599 | 40.64202 | 66.01276 | 61.63455 | 26.43144 | 0.466521 | 1.597639 | 0.345973 |
| 3 | 19670400 | 19671599 | 8.56873 | 15.59558 | 34.97009 | 15.0512 | -0.73979 | 0.293144 | 0.346004 |
| 5 | 64876600 | 64878999 | 33.84439 | 56.66612 | 48.08438 | 20.95719 | 0.617149 | 1.197191 | 0.346011 |
| 18 | 88645000 | 88646999 | 21.19461 | 37.0968 | 33.22677 | 14.66138 | 0.545638 | 0.688328 | 0.346014 |
| 6 | 121715000 | 121717599 | 27.96463 | 47.80825 | 41.37777 | 18.14284 | 0.590323 | 0.983741 | 0.34602 |
| 3 | 7211000 | 7212599 | 20.40595 | 35.96023 | 31.0218 | 13.73049 | 0.600869 | 0.58774 | 0.346022 |
| 9 | 8192000 | 8193199 | 13.47174 | 25.86599 | 18.65056 | 7.307197 | 0.994349 | -0.04751 | 0.346029 |
| 17 | 19596400 | 19597999 | 21.45224 | 38.29261 | 25.8601 | 11.36667 | 0.960589 | 0.478648 | 0.346048 |
| 4 | 74720000 | 74721799 | 26.38816 | 45.78466 | 35.80844 | 15.71139 | 0.739572 | 0.84004 | 0.346058 |
| 3 | 138977800 | 138978199 | 9.275985 | 17.96099 | 18.1744 | 6.909477 | 0.550952 | -0.1903 | 0.346069 |
| 14 | 81397400 | 81397999 | 9.275985 | 17.96099 | 18.1744 | 6.909477 | 0.550952 | -0.1903 | 0.346069 |
| X | 4006200 | 4008799 | 30.33714 | 51.18647 | 45.81639 | 20.03705 | 0.54045 | 1.111566 | 0.346102 |
| 17 | 14363800 | 14365799 | 11.95518 | 20.99322 | 42.00365 | 18.58517 | -0.60777 | 0.602972 | 0.34611 |
| 18 | 4213600 | 4214399 | 14.04333 | 26.52079 | 21.76556 | 8.862218 | 0.760387 | 0.087551 | 0.346114 |
| 18 | 55546600 | 55548799 | 14.01407 | 24.51354 | 45.18055 | 19.92556 | -0.49013 | 0.683046 | 0.346126 |
| 8 | 35655800 | 35659599 | 29.78717 | 48.28615 | 86.53798 | 37.05696 | -0.4745 | 1.631803 | 0.346146 |
| 6 | 72011600 | 72014199 | 28.59437 | 46.67873 | 82.1301 | 35.29522 | -0.4399 | 1.510712 | 0.346157 |
| 7 | 42458800 | 42462799 | 41.66233 | 67.67869 | 65.13482 | 27.93515 | 0.423068 | 1.599281 | 0.34617 |
| X | 17853400 | 17854599 | 14.24829 | 27.275 | 19.82558 | 7.834789 | 0.972221 | -0.00806 | 0.346179 |
| 18 | 36983800 | 36990399 | 125.0122 | 191.0003 | 182.1652 | 75.56024 | 0.357 | 3.089142 | 0.346183 |
| 17 | 26934400 | 26935799 | 9.234663 | 16.64532 | 35.42111 | 15.43145 | -0.6758 | 0.359491 | 0.3462 |
| 10 | 117059800 | 117061199 | 8.350967 | 15.42245 | 32.01844 | 13.51717 | -0.60945 | 0.202598 | 0.346232 |
| 12 | 86541000 | 86546199 | 36.44328 | 58.74542 | 101.5296 | 43.21858 | -0.43109 | 1.791076 | 0.346234 |
| 6 | 30363000 | 30364999 | 9.526608 | 17.16741 | 35.38246 | 15.41925 | -0.63055 | 0.361249 | 0.346248 |
| 6 | 134617600 | 134618399 | 8.141443 | 15.25072 | 30.66061 | 12.66393 | -0.539 | 0.110846 | 0.346256 |
| X | 118044600 | 118045799 | 16.06408 | 31.0109 | 15.05656 | 5.805503 | 1.580416 | -0.07852 | 0.346259 |
| 14 | 66645400 | 66646599 | 8.734447 | 16.32163 | 34.54415 | 14.3271 | -0.61811 | 0.138402 | 0.346274 |
| 5 | 128543800 | 128549799 | 38.62052 | 61.92761 | 110.3419 | 46.60806 | -0.4843 | 1.962094 | 0.346276 |
| 17 | 48832600 | 48837199 | 31.73853 | 51.29859 | 92.32457 | 39.4361 | -0.48481 | 1.70172 | 0.346291 |
| 10 | 61873200 | 61883199 | 94.49894 | 144.4445 | 232.598 | 96.48916 | -0.39748 | 3.075554 | 0.346291 |
| 10 | 61390600 | 61392799 | 21.61795 | 36.02602 | 65.56797 | 28.49291 | -0.47988 | 1.227667 | 0.346304 |
| 10 | 42026000 | 42030999 | 36.45663 | 58.66356 | 103.8202 | 44.08269 | -0.46835 | 1.84627 | 0.346309 |
| 6 | 52104000 | 52104999 | 17.7048 | 31.69011 | 27.96737 | 12.26454 | 0.579172 | 0.441348 | 0.346311 |
| 6 | 60596200 | 60597999 | 14.30192 | 24.99926 | 45.71292 | 20.14824 | -0.4789 | 0.695708 | 0.346318 |
| 13 | 66232000 | 66233999 | 25.65048 | 44.13887 | 35.71537 | 15.66661 | 0.688549 | 0.910339 | 0.346323 |
| 7 | 5323200 | 5328599 | 75.39939 | 120.121 | 96.01612 | 40.18152 | 0.65971 | 2.197319 | 0.346373 |
| 14 | 112447600 | 112449799 | 19.35006 | 34.82781 | 26.1587 | 11.41682 | 0.815021 | 0.415394 | 0.346374 |
| 5 | 78926800 | 78931399 | 73.81519 | 117.3227 | 105.7167 | 44.10764 | 0.483642 | 2.270923 | 0.346375 |
| 1 | 165594600 | 165596799 | 23.31978 | 38.51614 | 70.88944 | 30.61802 | -0.49691 | 1.340637 | 0.346377 |
| 12 | 19683400 | 19714599 | 1259.068 | 1598.152 | 1254.605 | 526.8049 | 0.377157 | 6.125013 | 0.346384 |
| 15 | 14568200 | 14569799 | 27.38332 | 46.39876 | 44.22845 | 19.36746 | 0.450193 | 1.063802 | 0.346413 |
| 10 | 110753400 | 110756799 | 27.05034 | 44.36956 | 77.93762 | 33.55589 | -0.43355 | 1.438096 | 0.346416 |

| | | | | | | | | | |
|---|---|---|---|---|---|---|---|---|---|
| 6 | 20088600 | 20089999 | 15.93724 | 29.373 | 23.99815 | 10.15927 | 0.72333 | 0.253371 | 0.346433 |
| 8 | 22444200 | 22445799 | 15.93724 | 29.373 | 23.99815 | 10.15927 | 0.72333 | 0.253371 | 0.346433 |
| 17 | 18451400 | 18462199 | 155.415 | 235.4716 | 206.6603 | 85.5195 | 0.467119 | 3.268887 | 0.346434 |
| 7 | 31178200 | 31178999 | 23.56495 | 40.78367 | 36.35606 | 15.95044 | 0.55121 | 0.85348 | 0.346435 |
| 19 | 25170600 | 25174999 | 40.2734 | 64.45573 | 114.5069 | 48.2297 | -0.48327 | 2.024799 | 0.346452 |
| 4 | 142459600 | 142461399 | 17.25219 | 29.55508 | 53.76629 | 23.57873 | -0.47656 | 0.939502 | 0.346467 |
| 12 | 104078600 | 104079399 | 9.882914 | 18.09357 | 35.58833 | 15.20311 | -0.54511 | 0.270158 | 0.346486 |
| 4 | 49418600 | 49420999 | 29.93818 | 50.09633 | 48.09304 | 20.95316 | 0.439757 | 1.191747 | 0.346491 |
| 13 | 65724200 | 65739399 | 304.617 | 436.5972 | 385.8819 | 159.366 | 0.382055 | 4.216559 | 0.346514 |
| 19 | 46511800 | 46513799 | 12.01817 | 20.94125 | 47.34667 | 20.82557 | -0.78388 | 0.731712 | 0.346522 |
| 12 | 114968600 | 114975399 | 90.73974 | 143.3293 | 114.6117 | 47.59693 | 0.647872 | 2.448489 | 0.346533 |
| 19 | 21996400 | 21997599 | 9.417707 | 17.45176 | 35.36703 | 14.86188 | -0.57049 | 0.199094 | 0.346544 |
| X | 81203400 | 81205599 | 59.90236 | 95.78866 | 87.41227 | 36.77282 | 0.474706 | 2.066629 | 0.346545 |
| 6 | 72195200 | 72197199 | 6.733119 | 12.4722 | 31.5567 | 13.26765 | -0.88764 | 0.171836 | 0.346553 |
| 1 | 100664000 | 100667399 | 61.12872 | 98.05446 | 83.25902 | 35.18755 | 0.583531 | 1.968769 | 0.346563 |
| 15 | 86487400 | 86491199 | 33.69522 | 54.38662 | 94.55071 | 40.31569 | -0.43728 | 1.746532 | 0.346576 |
| 7 | 10074600 | 10079999 | 80.92912 | 128.8128 | 95.55547 | 39.93821 | 0.765891 | 2.233069 | 0.346577 |
| 15 | 6555200 | 6557999 | 16.09823 | 27.73525 | 51.64037 | 22.65298 | -0.50832 | 0.879944 | 0.346616 |
| 7 | 85050800 | 85053999 | 41.39647 | 67.80233 | 59.48026 | 25.59337 | 0.564533 | 1.475794 | 0.346617 |
| 7 | 12850400 | 12852599 | 25.49246 | 44.45965 | 33.53689 | 14.75963 | 0.792152 | 0.764026 | 0.346652 |
| 18 | 24359200 | 24362599 | 49.20194 | 79.58481 | 69.67633 | 29.78869 | 0.553587 | 1.699882 | 0.346666 |
| 1 | 99470000 | 99471599 | 17.95386 | 32.59964 | 25.27672 | 10.91147 | 0.779351 | 0.354088 | 0.346668 |
| 7 | 77474800 | 77476599 | 16.77387 | 30.95086 | 23.3778 | 9.881031 | 0.837447 | 0.252474 | 0.346669 |
| 17 | 15162200 | 15164199 | 14.34851 | 24.99731 | 46.82685 | 20.5946 | -0.51393 | 0.747126 | 0.346672 |
| 1 | 68942600 | 68943799 | 16.75643 | 30.90909 | 24.01667 | 10.15535 | 0.796708 | 0.256439 | 0.346674 |
| 9 | 69300600 | 69302399 | 11.08299 | 19.87242 | 37.3394 | 16.35028 | -0.50299 | 0.40667 | 0.346689 |
| 14 | 103583600 | 103584799 | 8.744779 | 15.84606 | 34.9299 | 15.11693 | -0.72036 | 0.321103 | 0.346694 |
| 5 | 69044600 | 69045799 | 15.25802 | 28.90274 | 20.6054 | 8.33589 | 0.970025 | 0.075794 | 0.346695 |
| 10 | 125837000 | 125838999 | 12.08578 | 21.28989 | 40.90817 | 18.10766 | -0.54849 | 0.561369 | 0.346706 |
| 4 | 78217000 | 78217599 | 12.64803 | 23.82387 | 22.46977 | 9.194358 | 0.555493 | 0.094268 | 0.346724 |
| 11 | 51322000 | 51330999 | 137.8063 | 204.7151 | 338.7881 | 139.5649 | -0.47263 | 3.645522 | 0.346728 |
| 13 | 67342400 | 67343799 | 7.885221 | 14.60633 | 32.59454 | 13.70399 | -0.7088 | 0.181226 | 0.346728 |
| 1 | 17553000 | 17553799 | 7.983038 | 15.02602 | 30.21946 | 12.38057 | -0.52989 | 0.078071 | 0.34674 |
| 6 | 132442600 | 132445599 | 37.26151 | 62.00245 | 49.31684 | 21.4102 | 0.710335 | 1.272586 | 0.34676 |
| 5 | 122422200 | 122423399 | 6.625069 | 12.56068 | 28.17833 | 11.37834 | -0.66754 | 0.011418 | 0.346778 |
| 2 | 99082400 | 99082999 | 10.14479 | 19.77244 | 16.11111 | 5.831711 | 0.925664 | -0.35079 | 0.346786 |
| 11 | 120563400 | 120564999 | 7.541585 | 14.17663 | 29.63929 | 12.16967 | -0.58771 | 0.081657 | 0.3468 |
| 13 | 37722600 | 37724799 | 21.93978 | 38.09528 | 35.77695 | 15.70189 | 0.477058 | 0.812271 | 0.346806 |
| 4 | 134930400 | 134933999 | 25.95613 | 42.47352 | 78.72397 | 33.86098 | -0.51232 | 1.465928 | 0.346817 |
| 18 | 74108200 | 74112999 | 41.0676 | 65.70788 | 117.9422 | 49.65637 | -0.49862 | 2.034767 | 0.34682 |
| 14 | 121270200 | 121274399 | 25.97622 | 42.08722 | 95.1909 | 40.74713 | -0.81004 | 1.638835 | 0.346821 |
| 4 | 75835800 | 75838599 | 33.96289 | 56.62211 | 45.99339 | 19.99189 | 0.680097 | 1.240293 | 0.346832 |
| 14 | 51617800 | 51622199 | 87.30167 | 134.0738 | 216.4081 | 89.85203 | -0.39557 | 2.978779 | 0.346846 |
| 5 | 109129800 | 109141999 | 209.0083 | 310.7587 | 262.5305 | 108.1599 | 0.49802 | 3.637547 | 0.346846 |
| 7 | 58321200 | 58322799 | 14.72151 | 27.45647 | 22.57315 | 9.392727 | 0.732514 | 0.172454 | 0.346855 |
| 4 | 28750800 | 28752999 | 16.58303 | 30.16514 | 26.06999 | 11.22845 | 0.626046 | 0.33647 | 0.346856 |
| 6 | 59197200 | 59197999 | 8.787423 | 16.32712 | 32.10984 | 13.44445 | -0.52333 | 0.172423 | 0.346878 |

| | | | | | | | | | |
|---|---|---|---|---|---|---|---|---|---|
| 6 | 28646200 | 28646799 | 8.319548 | 16.24932 | 15.14127 | 5.306565 | 0.774701 | -0.49063 | 0.346885 |
| 8 | 101078800 | 101079799 | 12.66945 | 24.53606 | 16.56631 | 6.292903 | 1.12925 | -0.16026 | 0.346888 |
| 8 | 110936000 | 110936599 | 8.977969 | 17.44005 | 17.69212 | 6.608896 | 0.570871 | -0.25467 | 0.346896 |
| 9 | 36056400 | 36059799 | 45.41985 | 73.43019 | 70.78302 | 30.24461 | 0.414274 | 1.706981 | 0.34691 |
| 10 | 61573600 | 61584199 | 73.7917 | 114.3491 | 192.3128 | 79.88135 | -0.44482 | 2.801166 | 0.346911 |
| 14 | 42554000 | 42557199 | 27.99663 | 48.10905 | 39.76289 | 17.44132 | 0.657757 | 0.938783 | 0.346914 |
| 12 | 119783200 | 119784799 | 17.54478 | 31.1573 | 27.4295 | 12.1088 | 0.576034 | 0.485951 | 0.346927 |
| X | 118490000 | 118493599 | 35.76196 | 60.66399 | 40.17817 | 17.59987 | 0.973688 | 1.082991 | 0.346942 |
| 1 | 32756200 | 32758199 | 20.931 | 36.70556 | 32.64796 | 14.42928 | 0.555607 | 0.656297 | 0.346948 |
| 10 | 79242800 | 79250399 | 170.2489 | 255.2721 | 238.7737 | 98.51049 | 0.362238 | 3.477659 | 0.346949 |
| 8 | 93375800 | 93376999 | 6.437053 | 12.37324 | 28.59429 | 11.1668 | -0.66549 | -0.09683 | 0.346952 |
| 1 | 44131000 | 44131599 | 4.108798 | 8.010359 | 21.3744 | 7.708947 | -0.76148 | -0.42485 | 0.346962 |
| 7 | 24801600 | 24807999 | 38.82636 | 62.14563 | 114.0917 | 48.06119 | -0.53042 | 2.020219 | 0.346978 |
| 7 | 24196400 | 24207399 | 81.31329 | 125.1864 | 210.3232 | 87.35522 | -0.45058 | 2.92684 | 0.346985 |
| 15 | 34408000 | 34414199 | 52.2356 | 82.97129 | 138.5322 | 57.76243 | -0.40571 | 2.2779 | 0.346987 |
| 3 | 38020200 | 38020999 | 9.349285 | 16.99166 | 34.44257 | 14.85097 | -0.59648 | 0.309581 | 0.347003 |
| 3 | 28442600 | 28443599 | 14.79896 | 27.32693 | 24.15164 | 10.19787 | 0.613403 | 0.234258 | 0.347011 |
| 11 | 113262000 | 113268399 | 67.59461 | 105.5564 | 175.2422 | 72.71294 | -0.41748 | 2.662869 | 0.347014 |
| X | 33646600 | 33648999 | 36.64016 | 62.07457 | 39.56565 | 17.32487 | 1.028679 | 1.095062 | 0.347038 |
| 10 | 93619000 | 93623999 | 79.84329 | 126.0672 | 119.7979 | 49.74066 | 0.398736 | 2.44835 | 0.34704 |
| 9 | 102974600 | 102974999 | 3.523022 | 6.89036 | 18.43903 | 6.240517 | -0.68068 | -0.66987 | 0.347041 |
| 1 | 36293000 | 36296799 | 33.07607 | 53.31083 | 96.8059 | 41.20107 | -0.50235 | 1.788505 | 0.347051 |
| 19 | 12824000 | 12824799 | 7.346066 | 14 | 28.89775 | 11.54164 | -0.53485 | -0.01616 | 0.347063 |
| 10 | 11487400 | 11489399 | 18.45179 | 32.92523 | 28.48096 | 12.52235 | 0.605493 | 0.467129 | 0.347064 |
| 6 | 92191000 | 92192199 | 8.854301 | 15.78744 | 38.42871 | 16.90704 | -0.87866 | 0.428411 | 0.347065 |
| 8 | 126697000 | 126697799 | 10.43938 | 20.14434 | 19.70016 | 7.608343 | 0.578542 | -0.1197 | 0.347079 |
| 14 | 48926800 | 48928799 | 16.08056 | 27.80753 | 50.58689 | 22.19335 | -0.47383 | 0.845379 | 0.347081 |
| 11 | 112679200 | 112681999 | 24.53654 | 40.142 | 79.64671 | 34.25521 | -0.61051 | 1.466081 | 0.347102 |
| 3 | 101129800 | 101132399 | 16.71363 | 28.90125 | 52.21943 | 22.90955 | -0.46414 | 0.850258 | 0.347102 |
| 5 | 140460400 | 140466599 | 47.52902 | 75.63664 | 129.0198 | 53.91105 | -0.43376 | 2.216055 | 0.347109 |
| 15 | 19421000 | 19422199 | 15.46132 | 28.37119 | 25.21766 | 10.73942 | 0.596756 | 0.277913 | 0.347114 |
| 1 | 33147400 | 33148399 | 13.82404 | 25.51242 | 23.90523 | 10.10151 | 0.52925 | 0.230782 | 0.347119 |
| 9 | 96370000 | 96371799 | 14.41541 | 26.3326 | 24.80847 | 10.62909 | 0.507203 | 0.293559 | 0.347121 |
| 7 | 123370200 | 123370999 | 17.56796 | 31.09696 | 25.95434 | 11.47739 | 0.650758 | 0.513451 | 0.347131 |
| 3 | 72755000 | 72755999 | 12.55204 | 23.4919 | 22.40016 | 9.265962 | 0.527189 | 0.132434 | 0.34714 |
| 6 | 105713800 | 105715199 | 12.55204 | 23.4919 | 22.40016 | 9.265962 | 0.527189 | 0.132434 | 0.34714 |
| 5 | 63955800 | 63956799 | 7.623814 | 14.50626 | 32.41778 | 12.99112 | -0.6537 | 0.002932 | 0.347154 |
| 17 | 22706800 | 22709799 | 23.39105 | 38.41155 | 80.17289 | 34.53509 | -0.68085 | 1.418564 | 0.347169 |
| 9 | 75401800 | 75403399 | 12.84588 | 22.62888 | 42.41941 | 18.7766 | -0.51324 | 0.567375 | 0.347175 |
| 7 | 41380800 | 41382599 | 18.14909 | 32.36638 | 23.24014 | 10.22308 | 0.872125 | 0.457925 | 0.347178 |
| 9 | 31477200 | 31478799 | 9.52654 | 17.12162 | 37.90473 | 16.56512 | -0.7363 | 0.37979 | 0.347182 |
| 13 | 86299400 | 86301199 | 23.76778 | 41.34357 | 36.52922 | 16.05017 | 0.564875 | 0.796692 | 0.347185 |
| 18 | 3949400 | 3952199 | 25.87423 | 44.54564 | 39.7901 | 17.45426 | 0.546647 | 0.911948 | 0.347186 |
| 12 | 54571000 | 54572399 | 19.31126 | 34.10628 | 25.95821 | 11.48424 | 0.781509 | 0.546317 | 0.347211 |
| 6 | 31375600 | 31376799 | 7.56879 | 14.09668 | 31.61862 | 13.18748 | -0.70719 | 0.144116 | 0.347216 |
| 6 | 3436600 | 3439399 | 37.83331 | 61.88754 | 59.62053 | 25.64094 | 0.428831 | 1.485496 | 0.347229 |
| 2 | 28207600 | 28210399 | 27.92005 | 45.60641 | 79.84941 | 34.32291 | -0.43217 | 1.497993 | 0.347236 |

| | | | | | | | | | |
|---|---|---|---|---|---|---|---|---|---|
| 8 | 46368000 | 46372199 | 35.65834 | 57.43702 | 98.98254 | 42.09018 | -0.4282 | 1.81261 | 0.347259 |
| 5 | 98319600 | 98321399 | 13.85854 | 24.21664 | 45.20928 | 19.92096 | -0.5085 | 0.699065 | 0.347279 |
| 1 | 100563600 | 100564199 | 22.02906 | 38.40936 | 25.83949 | 11.37013 | 0.956489 | 0.738234 | 0.347292 |
| 19 | 34723800 | 34726399 | 22.12847 | 36.71964 | 68.29085 | 29.59093 | -0.51137 | 1.279205 | 0.347302 |
| 14 | 15832200 | 15832799 | 27.99775 | 47.30411 | 44.7748 | 19.58967 | 0.460164 | 1.090334 | 0.347322 |
| 13 | 65707000 | 65708199 | 17.3279 | 31.48116 | 26.04522 | 11.23513 | 0.687329 | 0.349557 | 0.347329 |
| 7 | 99154200 | 99157599 | 27.19398 | 44.70479 | 79.95299 | 34.45611 | -0.4582 | 1.41197 | 0.34733 |
| 8 | 20692400 | 20699599 | 224.0944 | 322.6646 | 481.2171 | 198.5967 | -0.36583 | 4.145801 | 0.347336 |
| 8 | 26366200 | 26368999 | 18.57585 | 33.07485 | 21.81086 | 9.607213 | 0.992852 | 0.46723 | 0.347339 |
| 10 | 56010200 | 56013599 | 35.98756 | 59.84416 | 50.97016 | 22.11816 | 0.611926 | 1.27785 | 0.347367 |
| 16 | 5827000 | 5828799 | 10.45561 | 19.02526 | 41.4728 | 17.85532 | -0.70003 | 0.32368 | 0.347368 |
| X | 110732200 | 110736999 | 75.62999 | 120.3673 | 99.69318 | 41.66233 | 0.607022 | 2.232153 | 0.347391 |
| 18 | 10482800 | 10487599 | 39.64961 | 63.74945 | 116.756 | 49.51671 | -0.51924 | 1.883802 | 0.347392 |
| 14 | 68889200 | 68889999 | 11.11794 | 21.43824 | 19.5683 | 7.573105 | 0.673879 | -0.10779 | 0.347395 |
| 9 | 63998600 | 64000999 | 15.77237 | 27.28423 | 50.04696 | 21.95672 | -0.48554 | 0.840551 | 0.347397 |
| 14 | 33840800 | 33843599 | 15.77237 | 27.28423 | 50.04696 | 21.95672 | -0.48554 | 0.840551 | 0.347397 |
| 4 | 9928200 | 9930199 | 17.39066 | 30.97362 | 28.57419 | 12.58322 | 0.511523 | 0.471048 | 0.347401 |
| 17 | 22067400 | 22068599 | 11.73651 | 22.31001 | 19.79527 | 7.949094 | 0.665547 | 0.015041 | 0.347413 |
| 6 | 20924000 | 20925799 | 26.05852 | 45.07105 | 38.1104 | 16.7198 | 0.626767 | 0.870304 | 0.347416 |
| 9 | 122863600 | 122865399 | 9.215881 | 16.60861 | 36.90835 | 16.08283 | -0.73877 | 0.359411 | 0.347424 |
| 1 | 95083400 | 95085999 | 30.95493 | 52.45984 | 44.13909 | 19.32937 | 0.629636 | 1.076282 | 0.347426 |
| 18 | 80011800 | 80012999 | 16.69172 | 30.56586 | 25.12249 | 10.72894 | 0.706193 | 0.299039 | 0.347453 |
| 17 | 39152400 | 39161199 | 213.4476 | 314.7985 | 289.1563 | 119.0786 | 0.366509 | 3.77541 | 0.347456 |
| 13 | 57477000 | 57481799 | 33.67403 | 54.38613 | 94.9155 | 40.50361 | -0.44183 | 1.727877 | 0.347462 |
| 7 | 20530800 | 20540199 | 99.45771 | 154.9298 | 124.5236 | 51.68438 | 0.625034 | 2.714888 | 0.347467 |
| 12 | 34996800 | 34997999 | 6.758242 | 12.83021 | 30.48175 | 12.27389 | -0.74707 | 0.005478 | 0.34752 |
| 6 | 40917000 | 40918999 | 19.98559 | 35.02289 | 32.85051 | 14.51014 | 0.479447 | 0.663136 | 0.347537 |
| 6 | 73050200 | 73051199 | 12.43841 | 23.54411 | 21.96792 | 8.901113 | 0.582896 | 0.058064 | 0.347537 |
| 14 | 109147200 | 109151599 | 31.62116 | 50.98421 | 99.27098 | 42.26985 | -0.6022 | 1.775357 | 0.347557 |
| 1 | 16106200 | 16116199 | 151.0525 | 228.6421 | 208.9669 | 86.4478 | 0.407502 | 3.285222 | 0.347566 |
| 1 | 19394200 | 19396199 | 23.51574 | 40.76878 | 37.68513 | 16.53441 | 0.499387 | 0.836939 | 0.347589 |
| X | 31311200 | 31315999 | 40.53958 | 68.55292 | 36.74364 | 16.08008 | 1.277625 | 1.123237 | 0.347595 |
| 9 | 37852000 | 37853199 | 12.30527 | 23.2539 | 21.34769 | 8.679876 | 0.601451 | 0.065088 | 0.347597 |
| 17 | 7320200 | 7324199 | 43.80044 | 70.84276 | 67.68612 | 28.93629 | 0.427738 | 1.691458 | 0.3476 |
| 3 | 43424600 | 43427199 | 21.19119 | 37.29746 | 30.27925 | 13.40517 | 0.687161 | 0.602486 | 0.347612 |
| 3 | 45501200 | 45502199 | 10.27165 | 18.52145 | 35.85359 | 15.61249 | -0.54023 | 0.362966 | 0.34762 |
| 16 | 61977800 | 61979999 | 21.89202 | 38.37249 | 32.55318 | 14.38158 | 0.623537 | 0.668748 | 0.347624 |
| 17 | 42489000 | 42491199 | 21.89202 | 38.37249 | 32.55318 | 14.38158 | 0.623537 | 0.668748 | 0.347624 |
| 15 | 14895400 | 14897199 | 21.89749 | 38.8947 | 27.64957 | 12.20396 | 0.883368 | 0.51678 | 0.347625 |
| 1 | 70991400 | 70992399 | 10.96205 | 21.0616 | 20.08475 | 7.853835 | 0.597482 | -0.0751 | 0.347626 |
| 17 | 26451800 | 26452799 | 12.36918 | 23.26086 | 22.02074 | 9.035406 | 0.547196 | 0.101479 | 0.347627 |
| 14 | 43920600 | 43920999 | 8.604816 | 16.81737 | 13.90336 | 4.806188 | 0.961999 | -0.53817 | 0.347659 |
| 6 | 97881200 | 97884199 | 25.15142 | 43.06176 | 40.52943 | 17.77296 | 0.470254 | 0.960446 | 0.347662 |
| 6 | 29230400 | 29237599 | 37.99465 | 60.94468 | 107.58 | 45.46489 | -0.47021 | 1.951042 | 0.347671 |
| 5 | 70423600 | 70426399 | 58.31814 | 93.51042 | 78.79271 | 33.28065 | 0.593946 | 1.975643 | 0.347683 |
| 2 | 122147200 | 122153199 | 34.044 | 54.83787 | 98.64088 | 41.94619 | -0.48983 | 1.809099 | 0.3477 |
| 11 | 50736200 | 50739799 | 34.00967 | 54.84572 | 98.73 | 42.04979 | -0.48897 | 1.773359 | 0.347708 |

| | | | | | | | | | |
|---|---|---|---|---|---|---|---|---|---|
| 7 | 140917400 | 140920599 | 18.26901 | 30.58635 | 62.58552 | 27.27644 | -0.64807 | 1.166333 | 0.347724 |
| 7 | 96787600 | 96790399 | 41.05893 | 68.01642 | 50.53959 | 21.8734 | 0.807898 | 1.335784 | 0.347731 |
| 2 | 121651200 | 121653399 | 28.41922 | 47.90866 | 45.87386 | 20.05689 | 0.443507 | 1.113171 | 0.347732 |
| 5 | 132103200 | 132104399 | 5.541199 | 10.71911 | 24.86257 | 9.494487 | -0.63992 | -0.19448 | 0.347741 |
| 16 | 46239800 | 46240799 | 10.99844 | 21.26627 | 19.09641 | 7.307806 | 0.71257 | -0.14683 | 0.347742 |
| 11 | 107625000 | 107627999 | 17.00229 | 28.9578 | 55.287 | 24.23425 | -0.54721 | 0.994882 | 0.347761 |
| 5 | 119559000 | 119561399 | 8.204429 | 15.11344 | 33.56792 | 14.2203 | -0.71013 | 0.217419 | 0.347765 |
| 17 | 45449200 | 45450999 | 11.77669 | 20.70686 | 42.97962 | 19.03164 | -0.66046 | 0.582031 | 0.347775 |
| X | 120692800 | 120694799 | 23.44422 | 41.64366 | 26.42683 | 11.66389 | 1.046225 | 0.524322 | 0.347778 |
| 1 | 119428400 | 119430999 | 16.91611 | 29.06611 | 52.70711 | 23.11903 | -0.4712 | 0.909607 | 0.34778 |
| 11 | 41454000 | 41455599 | 21.4585 | 38.06784 | 28.13119 | 12.42949 | 0.826654 | 0.521523 | 0.347781 |
| 9 | 109280200 | 109284399 | 46.83622 | 75.98133 | 66.30888 | 28.41053 | 0.562075 | 1.635083 | 0.347786 |
| 7 | 11400000 | 11406799 | 105.5096 | 165.4815 | 123.9039 | 51.3883 | 0.734772 | 2.600866 | 0.347789 |
| 7 | 110204800 | 110208599 | 19.83514 | 33.59572 | 64.83776 | 28.38879 | -0.56395 | 1.060901 | 0.347803 |
| 8 | 126326200 | 126331999 | 43.07288 | 68.7562 | 120.9761 | 50.76617 | -0.47358 | 2.110389 | 0.347813 |
| X | 25469400 | 25470199 | 22.79997 | 39.48038 | 37.32668 | 16.37654 | 0.46675 | 0.845192 | 0.347821 |
| 18 | 56646800 | 56648599 | 9.132898 | 16.79609 | 33.00747 | 14.01425 | -0.53687 | 0.235452 | 0.347833 |
| 4 | 134217000 | 134222599 | 38.15392 | 61.16081 | 109.8743 | 46.38911 | -0.49662 | 1.971412 | 0.347866 |
| 9 | 20231200 | 20234399 | 58.43436 | 93.40064 | 90.37029 | 37.99382 | 0.389719 | 2.077155 | 0.347879 |
| X | 33611000 | 33611999 | 11.87463 | 23.12415 | 15.84306 | 5.799329 | 1.158133 | -0.29277 | 0.347884 |
| 12 | 115344000 | 115349199 | 103.1352 | 160.7048 | 143.2199 | 59.44185 | 0.476533 | 2.714315 | 0.347889 |
| 12 | 31549000 | 31550599 | 14.34078 | 24.8687 | 48.97694 | 21.48888 | -0.58681 | 0.809887 | 0.347912 |
| X | 145277200 | 145278599 | 16.91904 | 31.91116 | 19.78521 | 8.077473 | 1.158357 | 0.117353 | 0.347925 |
| 13 | 33644200 | 33650599 | 48.35958 | 78.64932 | 55.71782 | 23.91194 | 0.865217 | 1.585373 | 0.347939 |
| 14 | 122320400 | 122323799 | 44.38951 | 72.61537 | 57.46027 | 24.71228 | 0.711898 | 1.495716 | 0.347939 |
| 1 | 80161600 | 80165599 | 30.63762 | 49.53229 | 97.97746 | 41.86498 | -0.62033 | 1.696203 | 0.347952 |
| 2 | 172901800 | 172911599 | 95.55013 | 146.2186 | 229.7776 | 95.34439 | -0.361 | 3.05293 | 0.347957 |
| 11 | 113278400 | 113283599 | 33.12052 | 53.27728 | 109.8831 | 46.63415 | -0.68927 | 1.855628 | 0.347958 |
| 13 | 29141800 | 29143999 | 15.50688 | 26.70549 | 51.08149 | 22.40754 | -0.54699 | 0.882145 | 0.347973 |
| 7 | 69898200 | 69900399 | 13.47882 | 23.65597 | 44.55621 | 19.70562 | -0.52144 | 0.622723 | 0.347973 |
| 5 | 95125400 | 95127599 | 24.96692 | 43.4593 | 32.21909 | 14.1625 | 0.816844 | 0.786335 | 0.347987 |
| 7 | 23206400 | 23220199 | 211.7308 | 315.5532 | 248.5821 | 102.4381 | 0.601951 | 3.596889 | 0.34801 |
| 3 | 144997800 | 145001599 | 21.98838 | 36.40497 | 71.64491 | 30.99596 | -0.59304 | 1.306382 | 0.34802 |
| 11 | 48275200 | 48277399 | 12.53025 | 22.18363 | 41.35729 | 18.28816 | -0.50244 | 0.509277 | 0.34803 |
| 8 | 83276600 | 83278399 | 15.14248 | 26.31555 | 48.43923 | 21.27074 | -0.48924 | 0.788132 | 0.348033 |
| 5 | 146066600 | 146067399 | 12.54383 | 23.96252 | 21.03676 | 8.359998 | 0.694706 | -0.00404 | 0.348042 |
| 19 | 7471000 | 7472399 | 12.2956 | 21.51932 | 43.6476 | 19.25838 | -0.62732 | 0.662983 | 0.348043 |
| 4 | 92669000 | 92671599 | 40.05124 | 65.62605 | 59.25623 | 25.50142 | 0.523306 | 1.469575 | 0.348054 |
| 3 | 133199000 | 133201599 | 30.86459 | 52.73927 | 40.19307 | 17.62267 | 0.77307 | 0.999236 | 0.348063 |
| 9 | 105618400 | 105619799 | 8.340357 | 15.54194 | 31.6291 | 13.18001 | -0.56649 | 0.150464 | 0.348069 |
| 1 | 69886800 | 69888199 | 14.17001 | 24.73937 | 45.74289 | 20.14503 | -0.49488 | 0.71354 | 0.348076 |
| 7 | 74462200 | 74463799 | 21.42633 | 37.46881 | 33.96623 | 14.97633 | 0.528632 | 0.707623 | 0.348079 |
| 17 | 39831800 | 39832999 | 16.41299 | 31.35244 | 18.52632 | 7.363013 | 1.261187 | 0.027659 | 0.348106 |
| 14 | 122473600 | 122476399 | 18.47929 | 31.31642 | 57.96441 | 25.38356 | -0.50344 | 1.048739 | 0.348159 |
| 7 | 97588600 | 97589999 | 9.226236 | 17.029 | 35.25942 | 14.89716 | -0.6072 | 0.217467 | 0.348162 |
| 13 | 111391400 | 111392799 | 10.41939 | 18.81983 | 35.86969 | 15.58632 | -0.51598 | 0.354226 | 0.348164 |
| 1 | 39708600 | 39714199 | 39.58215 | 63.10913 | 131.6153 | 55.13934 | -0.72021 | 2.148751 | 0.348179 |

| | | | | | | | | | |
|---|---|---|---|---|---|---|---|---|---|
| 1 | 166064000 | 166067999 | 50.42217 | 79.87174 | 151.7145 | 63.09598 | -0.59525 | 2.359712 | 0.348194 |
| X | 119956200 | 119962799 | 61.87256 | 99.36347 | 80.2659 | 33.98281 | 0.657344 | 1.935825 | 0.3482 |
| 11 | 110962600 | 110963399 | 7.338478 | 13.95643 | 28.84271 | 11.57054 | -0.54241 | -0.00314 | 0.34821 |
| 19 | 54267800 | 54268599 | 7.338478 | 13.95643 | 28.84271 | 11.57054 | -0.54241 | -0.00314 | 0.34821 |
| 1 | 8475000 | 8478799 | 33.73678 | 57.01282 | 43.55028 | 19.05516 | 0.768438 | 1.108866 | 0.348218 |
| 17 | 17180400 | 17184199 | 81.62153 | 129.5904 | 104.5971 | 43.59344 | 0.640747 | 2.308538 | 0.348219 |
| 3 | 52414600 | 52417399 | 50.84251 | 81.70701 | 78.06589 | 33.08541 | 0.416863 | 1.897167 | 0.348221 |
| 17 | 32617800 | 32618399 | 5.213327 | 10.15909 | 23.83316 | 8.64703 | -0.58693 | -0.38071 | 0.348228 |
| 7 | 12577200 | 12588399 | 336.1476 | 478.3689 | 413.2339 | 170.8604 | 0.404766 | 4.321825 | 0.348254 |
| 15 | 86789600 | 86790799 | 6.398596 | 11.97015 | 30.09676 | 12.45889 | -0.86087 | 0.103769 | 0.348257 |
| 7 | 63476200 | 63476599 | 3.125393 | 6.113343 | 17.88633 | 5.996621 | -0.79426 | -0.71196 | 0.348287 |
| 17 | 35113000 | 35115799 | 17.44087 | 29.31907 | 60.48002 | 26.42044 | -0.65967 | 1.118217 | 0.348294 |
| 19 | 8487000 | 8496599 | 119.6865 | 184.0363 | 170.6313 | 70.85763 | 0.404855 | 2.960992 | 0.348303 |
| 18 | 44944800 | 44945599 | 7.3912 | 13.86626 | 30.14388 | 12.41872 | -0.64783 | 0.092629 | 0.348308 |
| 13 | 7393000 | 7396399 | 30.23876 | 51.73595 | 40.21786 | 17.63535 | 0.744862 | 0.985903 | 0.348322 |
| 4 | 54541600 | 54542599 | 13.45252 | 25.55037 | 20.74183 | 8.34411 | 0.790815 | 0.040598 | 0.348325 |
| 11 | 12518400 | 12521999 | 61.62705 | 98.25251 | 93.64559 | 39.22962 | 0.407392 | 2.159067 | 0.34833 |
| 14 | 3166600 | 3175999 | 122.6764 | 189.3063 | 158.4542 | 65.81886 | 0.556163 | 2.894201 | 0.348358 |
| 2 | 165614800 | 165616799 | 21.23488 | 35.55149 | 64.05254 | 27.91561 | -0.46543 | 1.174336 | 0.348377 |
| 7 | 117469400 | 117472999 | 25.87704 | 42.5926 | 75.78481 | 32.67694 | -0.45003 | 1.390288 | 0.348381 |
| 12 | 102843000 | 102845799 | 15.30089 | 26.48239 | 49.51039 | 21.72166 | -0.51268 | 0.83323 | 0.348383 |
| 12 | 36431400 | 36432199 | 6.020739 | 11.50703 | 27.25034 | 10.8181 | -0.72322 | -0.05412 | 0.348387 |
| 11 | 21197200 | 21199399 | 20.39359 | 33.87499 | 66.953 | 29.03146 | -0.59861 | 1.262666 | 0.348388 |
| 16 | 53399200 | 53403799 | 111.0501 | 170.8946 | 162.9201 | 67.66294 | 0.365604 | 2.940989 | 0.348388 |
| 18 | 36418000 | 36421199 | 18.34962 | 30.51471 | 66.78092 | 28.97972 | -0.74486 | 1.243286 | 0.34839 |
| 13 | 81967600 | 81970199 | 22.03647 | 38.82614 | 30.28549 | 13.40511 | 0.745253 | 0.597876 | 0.34839 |
| 6 | 37685600 | 37687999 | 46.6267 | 75.77041 | 63.59336 | 27.28129 | 0.620491 | 1.599207 | 0.348398 |
| 14 | 63730000 | 63732599 | 13.44966 | 23.44924 | 47.86418 | 21.06092 | -0.6368 | 0.735842 | 0.3484 |
| 16 | 90137400 | 90139999 | 30.00032 | 50.56919 | 41.12077 | 17.97822 | 0.678428 | 1.114646 | 0.348448 |
| 14 | 42615600 | 42617199 | 15.81882 | 30.08929 | 19.50676 | 7.822678 | 1.11692 | 0.051232 | 0.348468 |
| 17 | 15808800 | 15811799 | 22.04514 | 38.34262 | 36.12233 | 15.87023 | 0.472671 | 0.788681 | 0.348472 |
| 4 | 121374400 | 121384599 | 258.129 | 376.3719 | 327.2718 | 134.8598 | 0.42989 | 3.958886 | 0.348504 |
| 12 | 102434600 | 102435999 | 6.68619 | 12.52356 | 31.73157 | 13.10831 | -0.87037 | 0.099374 | 0.348511 |
| 16 | 12886800 | 12888399 | 9.602835 | 17.48338 | 34.94747 | 15.03536 | -0.57492 | 0.298986 | 0.348521 |
| 3 | 142553200 | 142554999 | 29.66572 | 50.6458 | 41.33548 | 18.12226 | 0.674396 | 1.003225 | 0.348543 |
| 3 | 114223400 | 114223999 | 7.190237 | 13.70989 | 28.34379 | 11.3084 | -0.53573 | -0.02283 | 0.348548 |
| 16 | 47732600 | 47735399 | 38.87898 | 63.53934 | 60.15845 | 25.86357 | 0.452899 | 1.502148 | 0.348564 |
| 6 | 116076800 | 116078599 | 16.54459 | 29.96724 | 26.55049 | 11.49652 | 0.585462 | 0.360801 | 0.348576 |
| 12 | 24256000 | 24259799 | 93.69744 | 145.9743 | 140.7924 | 58.43583 | 0.362431 | 2.711919 | 0.348581 |
| 3 | 43325000 | 43325599 | 10.1584 | 19.57885 | 18.69671 | 7.245518 | 0.608406 | -0.11381 | 0.348586 |
| 18 | 64864800 | 64867599 | 16.74787 | 30.67971 | 24.37903 | 10.40634 | 0.754631 | 0.293675 | 0.348593 |
| 1 | 16247800 | 16249199 | 9.397348 | 17.38526 | 34.20954 | 14.40787 | -0.53066 | 0.205062 | 0.348595 |
| 7 | 140369000 | 140369799 | 10.41631 | 20.19702 | 18.67449 | 7.053679 | 0.688655 | -0.19691 | 0.3486 |
| 18 | 12510000 | 12513799 | 53.26044 | 85.67779 | 79.6724 | 33.81489 | 0.457597 | 1.869458 | 0.348607 |
| X | 115418200 | 115422199 | 75.14642 | 119.6825 | 97.51083 | 40.79032 | 0.631769 | 2.208919 | 0.348618 |
| 3 | 71000000 | 71000599 | 4.809181 | 9.369341 | 22.52499 | 8.197561 | -0.62577 | -0.38038 | 0.348624 |
| 5 | 24976800 | 24984199 | 131.5603 | 196.1342 | 324.9217 | 133.9031 | -0.46937 | 3.581292 | 0.348631 |

| | | | | | | | | | |
|---|---|---|---|---|---|---|---|---|---|
| 1 | 65570600 | 65573599 | 20.75699 | 36.57744 | 26.75102 | 11.8403 | 0.837476 | 0.580452 | 0.34864 |
| 6 | 143213200 | 143216199 | 29.23404 | 47.52867 | 86.76794 | 37.23291 | -0.49726 | 1.570392 | 0.348645 |
| 12 | 62579600 | 62584799 | 54.77564 | 87.88089 | 84.85071 | 35.87091 | 0.398875 | 1.955981 | 0.348653 |
| 17 | 40089800 | 40097799 | 92.91278 | 146.5385 | 118.1231 | 49.0213 | 0.634561 | 2.483839 | 0.348657 |
| X | 110037200 | 110040599 | 44.84711 | 74.1059 | 49.4897 | 21.38201 | 0.961216 | 1.374395 | 0.348673 |
| 7 | 59122800 | 59126199 | 44.88841 | 72.56075 | 70.15686 | 29.97159 | 0.409677 | 1.709266 | 0.348675 |
| 13 | 61501600 | 61506599 | 67.61676 | 108.3039 | 84.13738 | 35.48011 | 0.709159 | 2.023471 | 0.348677 |
| 5 | 5651600 | 5652399 | 12.07981 | 23.47 | 16.86267 | 6.289271 | 1.065656 | -0.22515 | 0.348685 |
| 1 | 16855400 | 16858199 | 17.07063 | 29.51785 | 58.30206 | 25.57809 | -0.59226 | 0.859428 | 0.348714 |
| 10 | 52318400 | 52319999 | 12.18665 | 21.57363 | 43.46571 | 19.2208 | -0.61406 | 0.509613 | 0.348719 |
| 5 | 77989400 | 77992399 | 33.16381 | 55.30648 | 50.33675 | 21.88395 | 0.516525 | 1.238116 | 0.348733 |
| 1 | 179703800 | 179706199 | 12.38856 | 21.81628 | 41.40618 | 18.32928 | -0.53108 | 0.56704 | 0.348734 |
| 8 | 15269000 | 15275599 | 40.26012 | 64.43309 | 116.5735 | 49.09863 | -0.50958 | 2.025372 | 0.348745 |
| 8 | 52504200 | 52504599 | 8.819108 | 17.10862 | 16.16281 | 6.090748 | 0.661474 | -0.23089 | 0.348749 |
| 10 | 83572200 | 83575399 | 25.27048 | 41.62453 | 73.82937 | 31.84361 | -0.44498 | 1.380616 | 0.348757 |
| X | 26426600 | 26428799 | 24.72071 | 43.56748 | 27.85182 | 12.32723 | 1.030873 | 0.606856 | 0.34878 |
| 8 | 19182000 | 19185999 | 57.49947 | 92.23146 | 84.9442 | 35.8989 | 0.466495 | 1.965091 | 0.348781 |
| 9 | 118237400 | 118239199 | 17.05864 | 30.5074 | 28.0744 | 12.32036 | 0.518455 | 0.442561 | 0.348784 |
| X | 33824800 | 33827999 | 52.20017 | 85.00573 | 57.56213 | 24.71514 | 0.931601 | 1.579779 | 0.348789 |
| 13 | 20317000 | 20317599 | 5.096818 | 9.923631 | 23.66852 | 8.678192 | -0.62483 | -0.34729 | 0.348794 |
| 15 | 31072200 | 31072999 | 5.096818 | 9.923631 | 23.66852 | 8.678192 | -0.62483 | -0.34729 | 0.348794 |
| 11 | 13294400 | 13297799 | 37.94758 | 63.22706 | 46.50436 | 20.20628 | 0.822803 | 1.257154 | 0.348808 |
| 2 | 178113200 | 178119199 | 41.6002 | 66.43447 | 121.6775 | 51.09217 | -0.53068 | 2.094863 | 0.348812 |
| 15 | 25110600 | 25112799 | 19.43879 | 35.47272 | 22.96375 | 9.851245 | 1.043526 | 0.332599 | 0.348815 |
| 10 | 42880000 | 42881799 | 15.29276 | 26.44386 | 50.04207 | 21.9543 | -0.53046 | 0.841504 | 0.348817 |
| 9 | 99617200 | 99620399 | 68.41546 | 108.6754 | 102.676 | 42.81335 | 0.41457 | 2.283548 | 0.34882 |
| X | 117614200 | 117622999 | 94.19351 | 148.628 | 115.7529 | 48.04523 | 0.684697 | 2.475207 | 0.34882 |
| 9 | 114788200 | 114791799 | 34.01979 | 54.85561 | 97.3627 | 41.46118 | -0.46881 | 1.776776 | 0.348822 |
| 9 | 111946200 | 111948799 | 13.19103 | 23.24634 | 42.94533 | 19.00785 | -0.492 | 0.567023 | 0.348831 |
| 12 | 21725800 | 21735799 | 168.4516 | 254.3783 | 197.1911 | 81.51562 | 0.641999 | 3.335767 | 0.348842 |
| 5 | 44165000 | 44168599 | 39.50853 | 64.66528 | 59.93423 | 25.78129 | 0.484977 | 1.479244 | 0.348865 |
| 19 | 43111600 | 43113199 | 19.16308 | 33.94365 | 29.3767 | 12.98903 | 0.598854 | 0.521401 | 0.348878 |
| 2 | 156021400 | 156022399 | 8.033599 | 14.8802 | 32.05928 | 13.48015 | -0.65847 | 0.182197 | 0.348898 |
| 11 | 16909200 | 16910199 | 8.033599 | 14.8802 | 32.05928 | 13.48015 | -0.65847 | 0.182197 | 0.348898 |
| 6 | 128778800 | 128780399 | 9.144643 | 16.64837 | 37.08358 | 15.95533 | -0.73042 | 0.298559 | 0.348898 |
| 1 | 105222200 | 105223999 | 20.05923 | 35.51233 | 28.68349 | 12.68386 | 0.697547 | 0.530567 | 0.348899 |
| 4 | 63570000 | 63571199 | 12.92591 | 22.55755 | 45.24621 | 19.92088 | -0.61167 | 0.712776 | 0.348899 |
| 10 | 61531000 | 61535599 | 20.97721 | 34.55688 | 72.69654 | 31.35656 | -0.69016 | 1.366997 | 0.348902 |
| 15 | 103814400 | 103819399 | 46.88147 | 75.89425 | 63.05406 | 26.97393 | 0.630076 | 1.676447 | 0.348906 |
| X | 44768200 | 44768999 | 15.60773 | 28.64374 | 24.32353 | 10.35672 | 0.662695 | 0.277921 | 0.34892 |
| 3 | 108570200 | 108571399 | 13.37077 | 24.87742 | 22.96473 | 9.594614 | 0.562609 | 0.179809 | 0.348924 |
| X | 107083000 | 107083999 | 19.15089 | 33.72168 | 30.79713 | 13.63321 | 0.518739 | 0.58947 | 0.348935 |
| 7 | 26640000 | 26641399 | 20.10897 | 35.23777 | 32.80348 | 14.4889 | 0.490242 | 0.663849 | 0.34894 |
| X | 119578400 | 119580999 | 27.22451 | 47.84944 | 27.23184 | 12.05839 | 1.196426 | 0.650402 | 0.348952 |
| 3 | 7220200 | 7224399 | 108.4937 | 168.6844 | 134.6239 | 55.89248 | 0.633117 | 2.753093 | 0.348968 |
| 12 | 20685200 | 20689999 | 105.742 | 165.5449 | 129.0625 | 53.53778 | 0.674531 | 2.633312 | 0.348968 |
| 1 | 78867000 | 78868799 | 20.43117 | 35.90845 | 31.50904 | 13.95199 | 0.574931 | 0.616566 | 0.348994 |

| | | | | | | | | | |
|---|---|---|---|---|---|---|---|---|---|
| 14 | 74871400 | 74872599 | 13.56472 | 25.33531 | 23.34416 | 9.689976 | 0.572337 | 0.156642 | 0.349016 |
| X | 125275600 | 125278799 | 44.39217 | 72.40164 | 61.62061 | 26.4725 | 0.604185 | 1.544354 | 0.34902 |
| X | 147574000 | 147574599 | 18.72803 | 33.51515 | 24.95985 | 10.94742 | 0.822605 | 0.444421 | 0.349034 |
| 6 | 6307000 | 6308399 | 10.22021 | 18.57926 | 35.97043 | 15.50478 | -0.53104 | 0.314478 | 0.349047 |
| 15 | 45343600 | 45345399 | 32.99591 | 55.25907 | 48.62407 | 21.19569 | 0.565005 | 1.19415 | 0.349048 |
| X | 125831800 | 125834599 | 57.54667 | 93.06685 | 63.88968 | 27.31101 | 0.90343 | 1.717997 | 0.349055 |
| 1 | 36328800 | 36330999 | 11.92703 | 21.01333 | 40.90843 | 18.10725 | -0.56718 | 0.558493 | 0.34906 |
| 6 | 62721000 | 62723599 | 37.16242 | 61.63335 | 52.64022 | 22.80087 | 0.607745 | 1.314575 | 0.349063 |
| 10 | 49898600 | 49899599 | 18.61748 | 32.78624 | 28.46873 | 12.60219 | 0.591299 | 0.580905 | 0.349068 |
| 11 | 90582200 | 90584799 | 19.39292 | 32.86943 | 58.6053 | 25.66533 | -0.44982 | 1.049617 | 0.349085 |
| 16 | 55918000 | 55921999 | 50.21623 | 81.20538 | 69.73682 | 29.80748 | 0.581 | 1.707747 | 0.349088 |
| 14 | 85504800 | 85506599 | 19.51072 | 34.37677 | 31.07631 | 13.75508 | 0.533555 | 0.583587 | 0.349091 |
| X | 4802400 | 4803999 | 18.51109 | 33.96815 | 23.83373 | 10.14976 | 0.935749 | 0.298489 | 0.349107 |
| 14 | 6562400 | 6563599 | 10.88628 | 19.72364 | 40.77886 | 17.65153 | -0.62943 | 0.348524 | 0.349133 |
| 5 | 111080600 | 111082199 | 22.68852 | 39.49548 | 36.00628 | 15.82777 | 0.51992 | 0.77946 | 0.349138 |
| 2 | 117659800 | 117660599 | 15.97976 | 28.85031 | 26.72314 | 11.61837 | 0.518127 | 0.376545 | 0.349148 |
| 4 | 66660800 | 66662999 | 28.56088 | 46.15931 | 90.75385 | 38.76161 | -0.61204 | 1.699215 | 0.349153 |
| 4 | 112292000 | 112295599 | 67.42194 | 107.5719 | 96.37307 | 40.41407 | 0.49773 | 2.143757 | 0.349159 |
| 1 | 22148400 | 22149199 | 14.33888 | 25.01362 | 46.28104 | 20.37149 | -0.49597 | 0.727202 | 0.34916 |
| 5 | 53033000 | 53035399 | 21.92009 | 38.12666 | 36.15687 | 15.88581 | 0.463226 | 0.788031 | 0.349164 |
| 12 | 117254800 | 117258799 | 27.24021 | 44.52873 | 80.6067 | 34.65774 | -0.47945 | 1.484505 | 0.349191 |
| 7 | 107012600 | 107014799 | 13.51431 | 23.74751 | 43.4831 | 19.25106 | -0.48074 | 0.603049 | 0.3492 |
| 13 | 43971800 | 43973399 | 11.35532 | 19.92148 | 42.56582 | 18.81925 | -0.70197 | 0.615608 | 0.349201 |
| 16 | 3753000 | 3754599 | 8.427654 | 15.37203 | 33.99558 | 14.59223 | -0.71657 | 0.282073 | 0.349241 |
| 3 | 24616200 | 24623799 | 120.5747 | 186.5985 | 150.6013 | 62.556 | 0.611811 | 2.840455 | 0.349243 |
| 14 | 95143800 | 95144999 | 15.78038 | 28.35073 | 25.82433 | 11.28932 | 0.537026 | 0.401809 | 0.349259 |
| 12 | 26046200 | 26048399 | 14.91306 | 25.98612 | 48.41524 | 21.2959 | -0.5064 | 0.746329 | 0.349266 |
| 3 | 11188400 | 11191999 | 125.061 | 191.3807 | 168.9336 | 70.09785 | 0.470377 | 3.058011 | 0.34927 |
| 4 | 64416000 | 64416399 | 58.41204 | 95.54841 | 43.39611 | 18.66319 | 1.510432 | 1.526886 | 0.349278 |
| 9 | 89923800 | 89926199 | 29.55845 | 49.5682 | 47.52646 | 20.73464 | 0.441461 | 1.166483 | 0.349278 |
| X | 71920600 | 71924199 | 121.6672 | 187.1365 | 141.0661 | 58.5825 | 0.703475 | 2.954172 | 0.349282 |
| 12 | 105476800 | 105478999 | 13.69137 | 24.01131 | 44.06873 | 19.4779 | -0.48404 | 0.63743 | 0.349283 |
| X | 40269000 | 40270599 | 23.26756 | 40.55351 | 30.04065 | 13.21563 | 0.818303 | 0.758348 | 0.349296 |
| 7 | 109473800 | 109475599 | 9.296404 | 17.01433 | 33.4603 | 14.30028 | -0.54489 | 0.264705 | 0.349311 |
| 12 | 96392800 | 96393599 | 6.709041 | 12.92262 | 27.62461 | 10.72456 | -0.54698 | -0.12002 | 0.349314 |
| 17 | 26507400 | 26508399 | 6.709041 | 12.92262 | 27.62461 | 10.72456 | -0.54698 | -0.12002 | 0.349314 |
| 4 | 61905000 | 61907199 | 31.07291 | 51.88636 | 49.73907 | 21.6417 | 0.441894 | 1.218397 | 0.349317 |
| 15 | 99290000 | 99291799 | 8.442657 | 15.00597 | 39.46091 | 17.40748 | -0.99201 | 0.448182 | 0.349333 |
| 15 | 67543600 | 67544799 | 12.69867 | 23.86566 | 22.37907 | 9.192092 | 0.559168 | 0.107335 | 0.349348 |
| 7 | 59551200 | 59555399 | 68.1607 | 108.2851 | 105.6616 | 44.06416 | 0.368273 | 2.279039 | 0.349359 |
| 3 | 99947000 | 99948199 | 10.25533 | 18.48666 | 36.87516 | 16.06317 | -0.58376 | 0.36522 | 0.349362 |
| 7 | 43460400 | 43462999 | 17.5169 | 29.66981 | 57.39166 | 25.12872 | -0.5668 | 1.048389 | 0.349373 |
| 1 | 179767800 | 179770799 | 28.58103 | 46.53755 | 83.20899 | 35.72548 | -0.46571 | 1.545985 | 0.34938 |
| 13 | 22420400 | 22421599 | 24.10162 | 41.4157 | 39.45261 | 17.30522 | 0.453721 | 0.921313 | 0.349382 |
| 13 | 119264400 | 119265799 | 20.82918 | 36.68813 | 28.00571 | 12.39675 | 0.776121 | 0.590103 | 0.349384 |
| 4 | 147397000 | 147400199 | 45.68372 | 74.45259 | 56.68249 | 24.34444 | 0.763753 | 1.552604 | 0.349384 |
| 13 | 13276400 | 13280399 | 55.7295 | 89.4941 | 83.11801 | 35.18793 | 0.456334 | 1.927815 | 0.349387 |

| | | | | | | | | | |
|---|---|---|---|---|---|---|---|---|---|
| 7 | 127001600 | 127006199 | 52.17872 | 82.64441 | 142.2254 | 59.14604 | -0.45318 | 2.361178 | 0.349397 |
| 18 | 85858200 | 85859599 | 14.70876 | 27.09396 | 24.72625 | 10.47534 | 0.564148 | 0.24892 | 0.349397 |
| 5 | 119345000 | 119349799 | 57.13114 | 90.24255 | 149.4949 | 62.08361 | -0.40148 | 2.430703 | 0.349403 |
| 1 | 188610000 | 188610999 | 8.605804 | 16.06696 | 31.68037 | 13.15914 | -0.51793 | 0.138512 | 0.349404 |
| X | 120154000 | 120154799 | 17.9552 | 32.19611 | 26.6992 | 11.69068 | 0.670313 | 0.432347 | 0.349409 |
| 16 | 12565600 | 12566799 | 13.29567 | 25.48337 | 19.206 | 7.562653 | 0.924427 | -0.03222 | 0.34944 |
| 1 | 12854600 | 12855799 | 7.15955 | 13.70074 | 30.291 | 11.98195 | -0.62123 | -0.04912 | 0.349442 |
| 5 | 133381800 | 133384199 | 27.47032 | 46.53633 | 44.25379 | 19.37731 | 0.453671 | 1.067127 | 0.349458 |
| 3 | 105695200 | 105696199 | 17.24291 | 30.6133 | 28.65574 | 12.65289 | 0.488073 | 0.490011 | 0.349468 |
| 8 | 42670200 | 42670599 | 4.777437 | 9.323671 | 22.27373 | 7.913441 | -0.58491 | -0.46634 | 0.349489 |
| 3 | 84430200 | 84432199 | 21.64616 | 38.39034 | 28.12611 | 12.43009 | 0.838816 | 0.524514 | 0.349522 |
| 7 | 62236400 | 62240399 | 49.8146 | 82.12055 | 45.00923 | 19.41989 | 1.244595 | 1.410803 | 0.349524 |
| X | 104194600 | 104197199 | 12.74938 | 22.54803 | 48.88088 | 21.61959 | -0.71972 | 0.535347 | 0.349532 |
| 9 | 72959000 | 72960999 | 11.30074 | 19.98008 | 39.36849 | 17.41338 | -0.5823 | 0.513553 | 0.349547 |
| 16 | 95385800 | 95387599 | 11.65015 | 20.44795 | 42.53466 | 18.81264 | -0.66359 | 0.609269 | 0.349558 |
| 9 | 115243800 | 115244599 | 12.42614 | 23.31084 | 22.44163 | 9.246352 | 0.518089 | 0.117079 | 0.349582 |
| 12 | 112272800 | 112278799 | 46.64673 | 74.4109 | 124.1974 | 52.06856 | -0.3987 | 2.135845 | 0.349582 |
| 2 | 85067600 | 85068999 | 8.295678 | 15.62976 | 31.93785 | 13.06292 | -0.55094 | 0.078735 | 0.349586 |
| X | 147543000 | 147545999 | 43.52503 | 71.12359 | 60.55594 | 26.0333 | 0.605543 | 1.514444 | 0.349589 |
| 4 | 109137200 | 109138399 | 6.847735 | 12.77056 | 31.10686 | 12.94509 | -0.82226 | 0.128814 | 0.349614 |
| 10 | 81484400 | 81492999 | 235.1975 | 343.8202 | 315.9378 | 130.1574 | 0.35385 | 3.917529 | 0.349615 |
| 19 | 25172400 | 25174999 | 30.44743 | 49.27653 | 88.71993 | 37.94594 | -0.48334 | 1.666572 | 0.349639 |
| X | 13834800 | 13836599 | 24.68251 | 43.49352 | 28.50142 | 12.61513 | 0.995431 | 0.610247 | 0.349664 |
| 16 | 93938000 | 93940799 | 26.09191 | 42.86061 | 75.48514 | 32.52025 | -0.4364 | 1.417117 | 0.34967 |
| 12 | 114027000 | 114028199 | 27.25672 | 46.99608 | 36.88243 | 16.17952 | 0.732829 | 0.898788 | 0.349672 |
| 4 | 73392400 | 73393599 | 12.37883 | 23.38415 | 21.91665 | 8.91828 | 0.57098 | 0.070433 | 0.349682 |
| X | 33048000 | 33048599 | 11.86075 | 23.17916 | 11.08453 | 3.838818 | 1.738248 | -0.49215 | 0.349689 |
| 9 | 19909800 | 19912599 | 39.75925 | 65.69386 | 48.84952 | 21.10435 | 0.806619 | 1.360789 | 0.349693 |
| 2 | 146756800 | 146757199 | 4.645134 | 9.066037 | 21.65509 | 7.685307 | -0.58324 | -0.47461 | 0.3497 |
| 18 | 45186600 | 45186999 | 4.645134 | 9.066037 | 21.65509 | 7.685307 | -0.58324 | -0.47461 | 0.3497 |
| 8 | 23308800 | 23310999 | 13.52003 | 23.58494 | 46.28024 | 20.37102 | -0.58048 | 0.721182 | 0.349701 |
| 16 | 92380600 | 92381999 | 16.20286 | 29.07648 | 27.13116 | 11.87205 | 0.501353 | 0.413392 | 0.349724 |
| 14 | 98486200 | 98487599 | 22.22468 | 38.7958 | 34.99362 | 15.40954 | 0.535702 | 0.739178 | 0.349734 |
| 6 | 48727800 | 48728399 | 8.533036 | 15.71096 | 32.51282 | 13.78264 | -0.60893 | 0.224235 | 0.349737 |
| 17 | 68815600 | 68818399 | 46.1327 | 74.63475 | 60.73241 | 25.96861 | 0.659116 | 1.685494 | 0.349752 |
| 1 | 38936400 | 38937399 | 5.05824 | 9.674297 | 28.89866 | 11.44923 | -1.05236 | -0.06746 | 0.349759 |
| 4 | 90275200 | 90276599 | 14.99058 | 27.93899 | 23.14262 | 9.64095 | 0.720656 | 0.181114 | 0.349785 |
| 12 | 19183600 | 19221999 | 1265.087 | 1607.277 | 1275.373 | 535.4759 | 0.363031 | 6.110134 | 0.349788 |
| 3 | 4045400 | 4047399 | 40.2343 | 66.3144 | 50.00723 | 21.57459 | 0.785589 | 1.38942 | 0.349793 |
| 5 | 104365200 | 104369199 | 38.67806 | 63.21596 | 60.75274 | 26.11982 | 0.431476 | 1.500305 | 0.349798 |
| X | 121140400 | 121142799 | 37.7353 | 62.35556 | 54.28757 | 23.45526 | 0.579855 | 1.359491 | 0.349801 |
| 2 | 75727200 | 75728599 | 17.28472 | 30.73509 | 28.67319 | 12.64408 | 0.494229 | 0.481266 | 0.349815 |
| 6 | 48265600 | 48270399 | 106.9685 | 165.1024 | 157.2305 | 65.31047 | 0.370403 | 2.884042 | 0.349816 |
| X | 136300800 | 136301999 | 21.69642 | 38.10197 | 32.02114 | 14.16926 | 0.637054 | 0.638228 | 0.349822 |
| 15 | 7731200 | 7732399 | 12.4025 | 22.00247 | 41.90272 | 18.51419 | -0.53158 | 0.491189 | 0.349826 |
| 4 | 33291800 | 33292999 | 7.96383 | 15.05564 | 30.35473 | 12.33174 | -0.52325 | 0.048285 | 0.349831 |
| 8 | 36631000 | 36632199 | 7.96383 | 15.05564 | 30.35473 | 12.33174 | -0.52325 | 0.048285 | 0.349831 |

| | | | | | | | | | |
|---|---|---|---|---|---|---|---|---|---|
| 2 | 59196400 | 59200799 | 59.04646 | 94.32978 | 91.61469 | 38.48855 | 0.383455 | 2.09345 | 0.349842 |
| 7 | 53736400 | 53738199 | 19.73172 | 35.2995 | 26.55938 | 11.65227 | 0.807359 | 0.454567 | 0.349844 |
| 12 | 73061200 | 73063199 | 18.79367 | 32.98995 | 29.90143 | 13.22642 | 0.528555 | 0.624315 | 0.349847 |
| 2 | 89163400 | 89164799 | 12.05558 | 23.00788 | 19.77912 | 7.876062 | 0.72127 | -0.00862 | 0.349848 |
| 16 | 62966400 | 62966799 | 10.45303 | 20.11426 | 17.18553 | 6.690771 | 0.761536 | -0.1004 | 0.34985 |
| 10 | 71003600 | 71008199 | 36.61271 | 58.97797 | 101.2289 | 43.04929 | -0.42233 | 1.812299 | 0.349865 |
| 4 | 117130400 | 117132399 | 35.49387 | 58.3277 | 54.64681 | 23.54663 | 0.471811 | 1.419179 | 0.349892 |
| 10 | 126611800 | 126612199 | 4.915232 | 9.583841 | 22.66593 | 8.160372 | -0.58767 | -0.41469 | 0.349894 |
| 9 | 122575800 | 122576799 | 8.197078 | 15.17922 | 31.53119 | 13.2626 | -0.60621 | 0.185655 | 0.349899 |
| 1 | 18600800 | 18602999 | 38.5746 | 62.87883 | 59.0045 | 25.34352 | 0.463078 | 1.535319 | 0.349918 |
| 6 | 73064600 | 73068399 | 31.20076 | 50.66437 | 87.32581 | 37.4416 | -0.41611 | 1.599324 | 0.349924 |
| 1 | 66060000 | 66061199 | 18.37939 | 33.49674 | 24.77981 | 10.64728 | 0.851102 | 0.336133 | 0.349925 |
| 7 | 97577000 | 97577599 | 4.480363 | 8.748061 | 21.09723 | 7.434225 | -0.58685 | -0.50226 | 0.349928 |
| X | 3664000 | 3666799 | 23.17603 | 41.76704 | 23.25559 | 10.13368 | 1.245802 | 0.431026 | 0.34993 |
| 1 | 117894800 | 117898999 | 71.0853 | 113.0824 | 103.9695 | 43.42186 | 0.455869 | 2.241247 | 0.349939 |
| 14 | 44631200 | 44633399 | 22.08548 | 38.91838 | 29.65724 | 13.12663 | 0.778692 | 0.594485 | 0.349948 |
| 7 | 9159000 | 9160599 | 40.6642 | 66.42584 | 50.9782 | 21.91303 | 0.754522 | 1.505113 | 0.349952 |
| 8 | 11374600 | 11378999 | 21.98267 | 36.44159 | 68.78789 | 29.78593 | -0.53284 | 1.290725 | 0.34996 |
| 12 | 22118600 | 22150399 | 689.5306 | 919.6443 | 768.8262 | 321.1204 | 0.357844 | 5.32201 | 0.349963 |
| 6 | 91684800 | 91686199 | 15.70208 | 28.54639 | 25.78472 | 11.1135 | 0.562304 | 0.333859 | 0.349988 |
| X | 24611200 | 24613199 | 27.83628 | 48.77854 | 28.93662 | 12.78093 | 1.137157 | 0.701933 | 0.350012 |
| 1 | 9010800 | 9012199 | 18.72384 | 33.23699 | 28.27701 | 12.48743 | 0.624764 | 0.498193 | 0.350015 |
| 7 | 25831600 | 25835199 | 39.28537 | 64.83703 | 54.97721 | 23.73303 | 0.617669 | 1.376473 | 0.350025 |
| 19 | 25627000 | 25628399 | 10.45164 | 18.73131 | 36.83366 | 16.13541 | -0.56864 | 0.403479 | 0.350027 |
| 8 | 70124400 | 70129399 | 52.31875 | 84.87348 | 63.17279 | 27.06637 | 0.791093 | 1.646743 | 0.350035 |
| 7 | 59756400 | 59759799 | 40.18434 | 66.04601 | 57.2608 | 24.67461 | 0.58341 | 1.428895 | 0.35004 |
| 11 | 114482000 | 114484199 | 12.25119 | 21.49372 | 43.55035 | 19.25494 | -0.62608 | 0.620261 | 0.350045 |
| 1 | 165073600 | 165075599 | 13.06716 | 22.78669 | 46.28994 | 20.37066 | -0.6302 | 0.724348 | 0.350052 |
| 15 | 21852800 | 21863999 | 189.4686 | 282.1294 | 262.033 | 107.9715 | 0.363384 | 3.608198 | 0.350055 |
| 16 | 9997800 | 10006199 | 283.0607 | 407.1937 | 367.8878 | 151.8472 | 0.355657 | 4.161245 | 0.350057 |
| 4 | 137084200 | 137085199 | 7.227079 | 13.62712 | 29.15437 | 11.9086 | -0.61473 | 0.056842 | 0.35007 |
| 10 | 86396800 | 86399199 | 19.39855 | 34.36068 | 28.75155 | 12.71262 | 0.64712 | 0.520875 | 0.350076 |
| 15 | 94743200 | 94744199 | 16.63807 | 29.7763 | 27.71015 | 12.15326 | 0.503105 | 0.435575 | 0.350091 |
| 7 | 9708200 | 9710599 | 24.75124 | 43.26752 | 32.9953 | 14.54378 | 0.776737 | 0.727517 | 0.350096 |
| 13 | 56428400 | 56432799 | 28.3659 | 45.92269 | 89.67145 | 38.36212 | -0.59968 | 1.655754 | 0.350113 |
| 3 | 137587200 | 137588599 | 13.82437 | 24.27057 | 44.54991 | 19.70842 | -0.48434 | 0.620515 | 0.350124 |
| 1 | 89914600 | 89917399 | 20.55219 | 34.53658 | 62.99474 | 27.5159 | -0.48317 | 1.131364 | 0.35013 |
| 7 | 26163200 | 26170999 | 187.2533 | 279.7303 | 250.819 | 103.4056 | 0.41836 | 3.548594 | 0.350133 |
| X | 154732400 | 154734999 | 26.01788 | 45.69554 | 28.33611 | 12.53968 | 1.073238 | 0.655135 | 0.350143 |
| 2 | 18693200 | 18695599 | 21.70462 | 35.51154 | 83.59872 | 35.95588 | -0.85626 | 1.463198 | 0.350161 |
| 19 | 43986000 | 43987399 | 7.843605 | 14.6693 | 30.62524 | 12.68366 | -0.59618 | 0.117917 | 0.35018 |
| 8 | 18513200 | 18513799 | 8.132974 | 15.33495 | 30.82612 | 12.59188 | -0.52587 | 0.069667 | 0.350202 |
| 1 | 26210600 | 26212399 | 33.92215 | 56.02196 | 54.18361 | 23.40095 | 0.428418 | 1.353474 | 0.350232 |
| 14 | 42256400 | 42261999 | 51.09133 | 82.48914 | 66.68754 | 28.44568 | 0.665297 | 1.752172 | 0.350239 |
| 9 | 47967800 | 47969999 | 17.76714 | 31.56594 | 29.09344 | 12.83864 | 0.510888 | 0.490273 | 0.350239 |
| 14 | 29531800 | 29533999 | 17.76714 | 31.56594 | 29.09344 | 12.83864 | 0.510888 | 0.490273 | 0.350239 |
| 7 | 61041000 | 61045799 | 95.88482 | 149.0419 | 142.0658 | 58.98442 | 0.376876 | 2.751346 | 0.350244 |

| | | | | | | | | | |
|---|---|---|---|---|---|---|---|---|---|
| 6 | 101843600 | 101847399 | 44.32137 | 70.92406 | 118.8839 | 50.063 | -0.39987 | 2.035049 | 0.350245 |
| 11 | 20072400 | 20074399 | 12.43133 | 21.75574 | 44.14484 | 19.47692 | -0.62809 | 0.6636 | 0.350256 |
| 2 | 19801600 | 19802999 | 13.0204 | 24.34558 | 22.846 | 9.465445 | 0.548191 | 0.143284 | 0.350263 |
| 9 | 18351200 | 18352399 | 15.61466 | 29.20679 | 22.64825 | 9.377829 | 0.822311 | 0.163925 | 0.350269 |
| 2 | 19249000 | 19249799 | 13.55973 | 25.09611 | 23.83968 | 10.0348 | 0.513367 | 0.211525 | 0.350294 |
| 8 | 105724400 | 105725799 | 10.85152 | 19.15447 | 40.41718 | 17.88308 | -0.68154 | 0.530033 | 0.350304 |
| 13 | 47035800 | 47037799 | 10.14013 | 18.17899 | 37.35282 | 16.3583 | -0.63143 | 0.399464 | 0.350315 |
| 9 | 78454800 | 78455599 | 6.177504 | 11.77338 | 28.3016 | 11.30271 | -0.75269 | -0.0313 | 0.350331 |
| 16 | 60814400 | 60815999 | 14.65082 | 25.53591 | 46.81769 | 20.59662 | -0.48304 | 0.741411 | 0.350339 |
| 6 | 10379000 | 10380599 | 12.35233 | 21.85057 | 41.87895 | 18.52189 | -0.54271 | 0.516933 | 0.350339 |
| 11 | 33037600 | 33040599 | 18.42823 | 31.29488 | 56.93953 | 24.95048 | -0.47874 | 1.027792 | 0.350364 |
| 2 | 23461400 | 23462199 | 4.689068 | 9.112626 | 23.3899 | 8.7069 | -0.74916 | -0.29822 | 0.350366 |
| 12 | 116034600 | 116035199 | 4.689068 | 9.112626 | 23.3899 | 8.7069 | -0.74916 | -0.29822 | 0.350366 |
| 12 | 103496400 | 103499999 | 20.41876 | 33.8329 | 72.05446 | 31.19012 | -0.70646 | 1.293074 | 0.350368 |
| 13 | 95733400 | 95738199 | 36.01717 | 57.89261 | 102.8127 | 43.58525 | -0.47528 | 1.881351 | 0.350369 |
| X | 24115200 | 24117599 | 51.81884 | 83.33587 | 74.59313 | 31.64653 | 0.512104 | 1.87708 | 0.350377 |
| 6 | 140368800 | 140370599 | 15.83899 | 27.0893 | 56.42866 | 24.74299 | -0.67164 | 0.94994 | 0.350399 |
| 13 | 52436600 | 52439399 | 17.12833 | 29.19698 | 55.32652 | 24.25321 | -0.53632 | 0.986227 | 0.350401 |
| 10 | 122640400 | 122641199 | 16.89448 | 30.16858 | 28.11488 | 12.35372 | 0.499142 | 0.450759 | 0.350403 |
| 18 | 40327600 | 40329799 | 22.37967 | 39.80313 | 27.02818 | 11.91656 | 0.950107 | 0.510602 | 0.350423 |
| 2 | 20070800 | 20072599 | 17.5409 | 32.02232 | 24.23372 | 10.39079 | 0.820171 | 0.318147 | 0.350445 |
| 2 | 144369000 | 144370199 | 8.505853 | 15.7035 | 32.01371 | 13.52102 | -0.58379 | 0.206049 | 0.35045 |
| 7 | 55987600 | 55992399 | 51.78874 | 83.58801 | 67.23663 | 28.66789 | 0.672023 | 1.763329 | 0.350452 |
| 2 | 5181200 | 5182199 | 14.31561 | 26.64272 | 23.83581 | 9.953904 | 0.607836 | 0.187321 | 0.350455 |
| 7 | 30446600 | 30458999 | 120.3501 | 180.8733 | 291.9913 | 120.5414 | -0.42159 | 3.426074 | 0.350466 |
| 7 | 15812000 | 15813599 | 9.951368 | 17.57893 | 39.47945 | 17.46552 | -0.77004 | 0.51691 | 0.350475 |
| 5 | 113406000 | 113408199 | 11.45849 | 20.25594 | 39.36835 | 17.41384 | -0.56269 | 0.516497 | 0.350477 |
| 16 | 38310800 | 38312999 | 16.13239 | 27.86271 | 51.10273 | 22.41882 | -0.48583 | 0.859104 | 0.350483 |
| 7 | 17595200 | 17599199 | 69.33014 | 110.5722 | 96.86993 | 40.59993 | 0.529324 | 2.157675 | 0.350484 |
| Y | 2609600 | 2611599 | 21.96625 | 38.19876 | 27.7985 | 12.21185 | 0.843326 | 0.777317 | 0.350488 |
| 10 | 39419800 | 39422999 | 13.74587 | 23.77592 | 51.0829 | 22.41113 | -0.71265 | 0.832332 | 0.350494 |
| 7 | 59405200 | 59405999 | 9.975526 | 19.25856 | 19.21455 | 7.406385 | 0.552715 | -0.13215 | 0.350507 |
| 5 | 73852000 | 73853599 | 9.896511 | 18.16007 | 35.10173 | 14.94851 | -0.5167 | 0.25731 | 0.35051 |
| 6 | 93889600 | 93891599 | 10.57515 | 18.61775 | 40.98476 | 18.14372 | -0.74368 | 0.559502 | 0.350514 |
| 10 | 8203600 | 8206999 | 41.72153 | 67.88708 | 64.29189 | 27.59657 | 0.448062 | 1.570379 | 0.350533 |
| 7 | 83076800 | 83078399 | 11.24796 | 20.25284 | 39.49979 | 17.2311 | -0.55284 | 0.386154 | 0.350541 |
| 5 | 134840400 | 134841599 | 12.26334 | 21.59754 | 40.92402 | 18.11554 | -0.52849 | 0.566353 | 0.350545 |
| 4 | 74451200 | 74453999 | 25.71921 | 44.97114 | 31.79568 | 14.01804 | 0.885288 | 0.726569 | 0.35056 |
| 12 | 72116400 | 72118199 | 14.7917 | 25.30197 | 57.38932 | 25.16453 | -0.79361 | 0.949182 | 0.350573 |
| 9 | 75441600 | 75442799 | 11.57972 | 20.52735 | 39.29993 | 17.3739 | -0.53969 | 0.485079 | 0.350575 |
| 13 | 51141600 | 51143999 | 41.78822 | 67.78801 | 65.60935 | 28.10957 | 0.413066 | 1.627082 | 0.350585 |
| 2 | 75789800 | 75791599 | 8.685538 | 15.91727 | 32.96876 | 14.06498 | -0.61718 | 0.251259 | 0.350589 |
| 1 | 7247800 | 7252999 | 75.99206 | 118 | 188.6698 | 78.34596 | -0.36968 | 2.766554 | 0.350593 |
| 16 | 90079600 | 90083599 | 25.33778 | 41.42577 | 79.07618 | 34.00173 | -0.55554 | 1.47744 | 0.350607 |
| 1 | 9467000 | 9468999 | 16.41976 | 28.35562 | 51.11715 | 22.42506 | -0.46117 | 0.860333 | 0.350609 |
| 13 | 8157400 | 8159199 | 23.47479 | 40.57831 | 38.34545 | 16.82262 | 0.466976 | 0.865034 | 0.350649 |
| 15 | 59719400 | 59721199 | 23.47479 | 40.57831 | 38.34545 | 16.82262 | 0.466976 | 0.865034 | 0.350649 |

| | | | | | | | | | |
|---|---|---|---|---|---|---|---|---|---|
| 3 | 26807000 | 26808199 | 13.85497 | 25.86662 | 23.28647 | 9.672855 | 0.604721 | 0.16125 | 0.35065 |
| 6 | 35063400 | 35065399 | 15.33454 | 28.355 | 24.12642 | 10.16812 | 0.669818 | 0.231347 | 0.350655 |
| 7 | 12715600 | 12717999 | 12.92099 | 22.53459 | 45.77143 | 20.14405 | -0.62989 | 0.721817 | 0.350661 |
| 11 | 69856000 | 69858599 | 20.22207 | 33.61368 | 67.0193 | 29.07431 | -0.61112 | 1.253065 | 0.350675 |
| 14 | 41426800 | 41427799 | 10.58002 | 20.58235 | 17.03596 | 6.288598 | 0.876896 | -0.27791 | 0.350683 |
| 7 | 121855400 | 121857399 | 14.32779 | 24.91714 | 47.37503 | 20.81257 | -0.53563 | 0.771745 | 0.350684 |
| 4 | 64984200 | 64986399 | 29.2308 | 49.69582 | 43.5855 | 19.10231 | 0.570192 | 1.041539 | 0.350696 |
| X | 8533200 | 8535799 | 50.90297 | 82.57614 | 64.10015 | 27.46355 | 0.730643 | 1.645618 | 0.35072 |
| 17 | 40735800 | 40737799 | 24.52192 | 42.78609 | 34.14703 | 15.03236 | 0.71124 | 0.754045 | 0.35074 |
| 13 | 46116800 | 46119599 | 14.39982 | 24.47188 | 57.83337 | 25.34596 | -0.85416 | 1.006677 | 0.350744 |
| 4 | 56045600 | 56047199 | 14.12964 | 24.67018 | 46.26031 | 20.37357 | -0.5152 | 0.711472 | 0.350746 |
| 8 | 86161400 | 86162799 | 7.686864 | 14.478 | 29.67777 | 12.1452 | -0.55564 | 0.070627 | 0.35076 |
| 13 | 94223600 | 94226399 | 16.88978 | 28.9915 | 54.33772 | 23.83312 | -0.51907 | 0.918968 | 0.35076 |
| 17 | 31845000 | 31848199 | 36.11111 | 59.28297 | 57.46275 | 24.74956 | 0.422303 | 1.433786 | 0.350763 |
| 18 | 3187800 | 3204399 | 220.3376 | 327.5752 | 254.4565 | 104.8326 | 0.618933 | 3.641099 | 0.350767 |
| 2 | 119187400 | 119193799 | 79.79909 | 123.3223 | 199.0924 | 82.69858 | -0.38891 | 2.854764 | 0.350775 |
| 6 | 144798800 | 144800799 | 10.17205 | 18.26156 | 41.59884 | 18.19649 | -0.77812 | 0.401035 | 0.350777 |
| 7 | 27915400 | 27931799 | 209.8684 | 304.1567 | 452.7974 | 186.707 | -0.35396 | 4.04846 | 0.350788 |
| 2 | 53278400 | 53280599 | 17.15086 | 29.20925 | 55.30094 | 24.24025 | -0.5352 | 0.995436 | 0.35079 |
| 15 | 79746200 | 79748399 | 11.25763 | 20.07314 | 37.81701 | 16.63758 | -0.51154 | 0.443847 | 0.350795 |
| 13 | 90953400 | 90956799 | 33.5793 | 56.98127 | 40.69221 | 17.82692 | 0.865255 | 1.069639 | 0.350798 |
| 17 | 52963400 | 52964999 | 13.38429 | 25.36808 | 20.67934 | 8.355568 | 0.779229 | 0.053194 | 0.350805 |
| 12 | 23211400 | 23236999 | 757.8692 | 1003.995 | 831.556 | 347.6282 | 0.361578 | 5.430429 | 0.350825 |
| 6 | 111924200 | 111925399 | 8.027956 | 15.09144 | 32.39217 | 13.2992 | -0.62534 | 0.093929 | 0.350834 |
| 18 | 3074400 | 3080799 | 89.35861 | 141.856 | 98.03693 | 40.85397 | 0.864138 | 2.318223 | 0.350844 |
| 13 | 61025800 | 61028599 | 39.45274 | 64.30236 | 62.40653 | 26.80356 | 0.414519 | 1.541616 | 0.35085 |
| 11 | 103157800 | 103159399 | 7.80639 | 14.29629 | 32.99123 | 14.08774 | -0.77269 | 0.248778 | 0.350877 |
| 16 | 59062600 | 59066199 | 89.21075 | 139.6383 | 135.4988 | 56.20877 | 0.358947 | 2.626742 | 0.350886 |
| 7 | 75511400 | 75516799 | 59.3009 | 93.0021 | 165.3773 | 68.58949 | -0.51159 | 2.581605 | 0.350891 |
| 14 | 45650200 | 45650999 | 6.877137 | 13.1506 | 27.86995 | 11.04766 | -0.56296 | -0.04945 | 0.350926 |
| 7 | 35790400 | 35793599 | 16.22389 | 28.05014 | 51.13047 | 22.43171 | -0.47682 | 0.847733 | 0.350934 |
| 3 | 91326800 | 91330999 | 66.09013 | 105.4537 | 96.71202 | 40.55985 | 0.464127 | 2.140473 | 0.350938 |
| 11 | 34242000 | 34242599 | 9.907263 | 19.14928 | 19.18784 | 7.35557 | 0.553633 | -0.1525 | 0.350943 |
| X | 12102200 | 12105799 | 40.78974 | 67.22867 | 54.36861 | 23.45604 | 0.685155 | 1.398331 | 0.350947 |
| 4 | 133751000 | 133753599 | 10.60825 | 18.96845 | 37.83591 | 16.60651 | -0.59118 | 0.420229 | 0.350954 |
| 7 | 59154600 | 59162999 | 148.8806 | 226.5329 | 178.9806 | 74.16025 | 0.623634 | 3.179434 | 0.350955 |
| 6 | 112469200 | 112473399 | 23.84648 | 39.17224 | 74.64044 | 32.15639 | -0.5496 | 1.410281 | 0.350974 |
| 14 | 105561000 | 105563399 | 19.6891 | 33.41041 | 59.75872 | 26.17972 | -0.45437 | 1.040994 | 0.350979 |
| 13 | 94737800 | 94739999 | 17.22561 | 29.58779 | 53.25506 | 23.35913 | -0.46065 | 0.914245 | 0.350997 |
| 11 | 30409800 | 30410599 | 7.52152 | 14.20478 | 29.19324 | 11.88579 | -0.5531 | 0.049328 | 0.351006 |
| 3 | 17423000 | 17424599 | 21.14157 | 36.95513 | 34.43302 | 15.1767 | 0.489112 | 0.712153 | 0.35101 |
| 10 | 36382000 | 36384399 | 24.02217 | 39.42014 | 75.11722 | 32.34767 | -0.55028 | 1.424277 | 0.35102 |
| 16 | 7571400 | 7572599 | 10.54189 | 18.65117 | 38.90292 | 17.20509 | -0.66337 | 0.501013 | 0.351022 |
| 9 | 11726600 | 11728799 | 22.1109 | 39.9165 | 23.2126 | 10.09309 | 1.185178 | 0.412828 | 0.35103 |
| 14 | 111146200 | 111149799 | 34.14602 | 57.20186 | 47.5226 | 20.71936 | 0.647543 | 1.191762 | 0.35104 |
| 2 | 13378600 | 13380399 | 15.00186 | 27.62117 | 24.66454 | 10.45572 | 0.594641 | 0.2533 | 0.351045 |
| 4 | 22496800 | 22498799 | 22.2594 | 37.0155 | 67.28645 | 29.19854 | -0.47828 | 1.253899 | 0.351065 |

| | | | | | | | | | |
|---|---|---|---|---|---|---|---|---|---|
| 16 | 47999200 | 48001599 | 32.64599 | 55.22334 | 43.54393 | 19.05803 | 0.722799 | 1.095897 | 0.351065 |
| 5 | 148379600 | 148388799 | 78.26149 | 121.131 | 197.4169 | 82.00193 | -0.40088 | 2.825182 | 0.351068 |
| 17 | 51872600 | 51874799 | 17.1506 | 30.56572 | 28.0297 | 12.33675 | 0.520532 | 0.463602 | 0.351078 |
| 2 | 121778000 | 121780199 | 23.28147 | 40.37907 | 37.13708 | 16.29416 | 0.506696 | 0.830808 | 0.351078 |
| 15 | 27198200 | 27199199 | 8.585589 | 16.0956 | 32.36538 | 13.34992 | -0.53805 | 0.112664 | 0.351081 |
| 8 | 26837200 | 26838199 | 6.869813 | 13.15901 | 27.94911 | 11.02494 | -0.55953 | -0.06572 | 0.351083 |
| 12 | 77089600 | 77093199 | 37.94885 | 60.96959 | 105.0297 | 44.49556 | -0.43231 | 1.898774 | 0.351097 |
| 10 | 123099800 | 123102599 | 22.08846 | 36.35089 | 73.37485 | 31.63566 | -0.63138 | 1.382433 | 0.351101 |
| 11 | 113011000 | 113013599 | 16.53779 | 28.10973 | 56.33162 | 24.68812 | -0.61732 | 1.011831 | 0.351108 |
| 12 | 98081800 | 98083199 | 19.91719 | 35.355 | 28.14421 | 12.42888 | 0.720184 | 0.506159 | 0.35111 |
| 13 | 100260000 | 100262999 | 20.52362 | 34.56373 | 63.62029 | 27.80749 | -0.49615 | 1.112348 | 0.351119 |
| 14 | 96344800 | 96345199 | 4.67775 | 9.121478 | 23.35494 | 8.400149 | -0.70086 | -0.42363 | 0.351149 |
| 1 | 8068400 | 8072999 | 39.00874 | 63.92327 | 59.79005 | 25.73207 | 0.472608 | 1.465703 | 0.351152 |
| 14 | 39030600 | 39031799 | 13.06521 | 24.82229 | 21.39652 | 8.601957 | 0.706036 | 0.037305 | 0.35116 |
| 1 | 147257000 | 147258199 | 13.28349 | 25.51 | 18.65889 | 7.30598 | 0.974635 | -0.0515 | 0.351184 |
| 10 | 127461600 | 127463399 | 34.81846 | 57.48588 | 55.29402 | 23.87584 | 0.436341 | 1.360923 | 0.351204 |
| 3 | 57229200 | 57232199 | 18.50404 | 33.43578 | 25.09466 | 10.8989 | 0.820291 | 0.384041 | 0.351213 |
| 19 | 37665800 | 37667799 | 12.70382 | 22.38507 | 41.39341 | 18.32139 | -0.49336 | 0.562531 | 0.351214 |
| X | 118190800 | 118199199 | 105.1898 | 165.7581 | 109.1705 | 45.30087 | 0.924741 | 2.50963 | 0.351217 |
| 7 | 72032600 | 72035799 | 33.59662 | 56.36083 | 47.50077 | 20.72826 | 0.626907 | 1.176632 | 0.351233 |
| 3 | 74091200 | 74095199 | 45.42537 | 74.21608 | 59.63626 | 25.63631 | 0.688602 | 1.519369 | 0.351237 |
| 9 | 70869600 | 70871599 | 11.90403 | 21.03302 | 40.35416 | 17.85168 | -0.54468 | 0.52518 | 0.351243 |
| 8 | 5483600 | 5485799 | 28.40265 | 48.65998 | 40.82214 | 17.90227 | 0.63562 | 0.967242 | 0.351249 |
| 18 | 33172000 | 33174999 | 20.62019 | 34.75457 | 64.05132 | 28.00278 | -0.49776 | 1.109752 | 0.351258 |
| 14 | 92056400 | 92068199 | 232.5462 | 341.0768 | 310.1544 | 127.7482 | 0.373506 | 3.865523 | 0.351267 |
| X | 122736400 | 122737199 | 14.44298 | 27.64181 | 19.8176 | 7.836418 | 0.991155 | -0.00414 | 0.351273 |
| 17 | 7673200 | 7677399 | 27.79374 | 45.24054 | 82.37086 | 35.36133 | -0.49211 | 1.549213 | 0.351279 |
| 3 | 123188200 | 123188999 | 10.34868 | 19.8308 | 19.60869 | 7.725895 | 0.53574 | -0.05811 | 0.3513 |
| 5 | 55844400 | 55845199 | 10.34868 | 19.8308 | 19.60869 | 7.725895 | 0.53574 | -0.05811 | 0.3513 |
| X | 4023000 | 4024399 | 36.42524 | 59.93594 | 53.02437 | 22.86008 | 0.555075 | 1.406346 | 0.351302 |
| 10 | 111939800 | 111941999 | 40.33058 | 65.65329 | 63.47813 | 27.25174 | 0.418733 | 1.56216 | 0.351308 |
| 15 | 96979400 | 96980599 | 8.689461 | 15.88478 | 33.97418 | 14.54263 | -0.66687 | 0.266485 | 0.351315 |
| 2 | 22087000 | 22092399 | 57.37747 | 92.15335 | 82.50301 | 34.93497 | 0.509453 | 1.92748 | 0.351321 |
| 4 | 91058400 | 91058999 | 7.348239 | 13.94423 | 28.77296 | 11.59647 | -0.54605 | 0.01239 | 0.351332 |
| X | 4469000 | 4474999 | 88.47186 | 140.4718 | 87.20097 | 36.34449 | 1.018929 | 2.311618 | 0.351337 |
| 9 | 13170400 | 13172399 | 19.03984 | 33.52839 | 31.18097 | 13.80295 | 0.492598 | 0.587286 | 0.351353 |
| 3 | 49371000 | 49372999 | 22.19547 | 38.89265 | 33.16313 | 14.64731 | 0.616361 | 0.677493 | 0.351389 |
| 1 | 116944400 | 116944799 | 4.308612 | 8.42099 | 22.05489 | 7.619891 | -0.67843 | -0.57044 | 0.351393 |
| 8 | 40972800 | 40975599 | 36.92873 | 61.15683 | 53.7747 | 23.26891 | 0.565746 | 1.33026 | 0.351401 |
| 7 | 4529400 | 4531199 | 12.89861 | 22.58145 | 43.61527 | 19.24899 | -0.55726 | 0.660314 | 0.351405 |
| 14 | 96493400 | 96493999 | 9.251006 | 17.99412 | 17.10731 | 6.322823 | 0.677108 | -0.29073 | 0.351406 |
| X | 126716200 | 126719999 | 61.35391 | 98.51093 | 81.35424 | 34.43315 | 0.625207 | 1.94133 | 0.351412 |
| 9 | 98547600 | 98552199 | 42.92963 | 68.51191 | 120.5563 | 50.57367 | -0.47404 | 2.117744 | 0.351415 |
| 13 | 79736800 | 79737799 | 7.32858 | 13.96883 | 28.92146 | 11.54735 | -0.5391 | -0.01886 | 0.351416 |
| 15 | 73863800 | 73874999 | 66.64975 | 103.2125 | 194.6506 | 80.85267 | -0.6106 | 2.811014 | 0.351424 |
| 8 | 47604400 | 47607799 | 49.26541 | 80.25386 | 59.7664 | 25.66476 | 0.795081 | 1.56895 | 0.351425 |
| 5 | 62453800 | 62456799 | 56.13887 | 90.18637 | 82.55934 | 34.97168 | 0.477817 | 1.917235 | 0.351427 |

| | | | | | | | | | |
|---|---|---|---|---|---|---|---|---|---|
| 10 | 5120600 | 5121599 | 10.26063 | 19.95126 | 16.35073 | 6.059543 | 0.885007 | -0.27215 | 0.35143 |
| 5 | 72716200 | 72717399 | 9.937273 | 17.59039 | 38.9223 | 17.21191 | -0.7477 | 0.491321 | 0.351446 |
| 7 | 6174800 | 6176999 | 12.41931 | 21.76119 | 43.59083 | 19.25229 | -0.60962 | 0.642504 | 0.35145 |
| 4 | 109394400 | 109396199 | 16.94199 | 29.03341 | 53.21874 | 23.33923 | -0.48732 | 0.934937 | 0.351477 |
| 5 | 129375200 | 129378599 | 19.12433 | 32.2884 | 59.05517 | 25.83186 | -0.48661 | 1.083955 | 0.351479 |
| 5 | 7515600 | 7519199 | 50.51675 | 81.36066 | 75.61669 | 32.14881 | 0.460453 | 1.825706 | 0.351494 |
| 8 | 77482200 | 77483799 | 10.74357 | 19.49079 | 38.07724 | 16.45492 | -0.54634 | 0.335999 | 0.351507 |
| 12 | 20047000 | 20050399 | 51.40611 | 82.83416 | 61.68307 | 26.24374 | 0.780359 | 1.80776 | 0.351509 |
| 1 | 81959400 | 81963199 | 54.80346 | 87.99001 | 83.89639 | 35.50703 | 0.418193 | 1.933523 | 0.35151 |
| 7 | 36369000 | 36373599 | 43.882 | 70.01516 | 122.8845 | 51.53303 | -0.47074 | 2.125783 | 0.351517 |
| 16 | 72985800 | 72986399 | 11.98981 | 22.92146 | 21.05804 | 8.353382 | 0.631843 | -0.01574 | 0.351522 |
| 5 | 151059600 | 151060999 | 12.78746 | 24.09001 | 21.80215 | 8.918977 | 0.61441 | 0.091134 | 0.351524 |
| 3 | 72421000 | 72421799 | 8.112553 | 15.22558 | 31.83188 | 13.10535 | -0.59171 | 0.099114 | 0.351529 |
| 7 | 23685000 | 23692399 | 81.29311 | 129.1026 | 104.0468 | 43.37538 | 0.643274 | 2.300712 | 0.351532 |
| 11 | 34981600 | 34983199 | 9.546317 | 17.6085 | 35.24189 | 14.90273 | -0.55939 | 0.224348 | 0.351544 |
| 1 | 102889800 | 102891399 | 18.08395 | 30.87148 | 55.36559 | 24.27322 | -0.45688 | 0.977279 | 0.351545 |
| 12 | 88111600 | 88113999 | 32.29911 | 54.24267 | 46.93314 | 20.49451 | 0.58901 | 1.159076 | 0.351545 |
| 12 | 109894400 | 109897999 | 26.71089 | 43.88175 | 76.86413 | 33.11574 | -0.42858 | 1.417058 | 0.351548 |
| 7 | 31345400 | 31349599 | 57.28809 | 92.13492 | 78.62033 | 33.35683 | 0.580972 | 1.884924 | 0.351572 |
| 19 | 9756600 | 9758999 | 29.07878 | 49.63169 | 41.33622 | 18.12223 | 0.645202 | 1.002171 | 0.351584 |
| 8 | 46585800 | 46588399 | 35.75427 | 60.26613 | 43.58249 | 19.0543 | 0.847133 | 1.139299 | 0.351588 |
| 7 | 45193400 | 45195999 | 14.7876 | 25.78928 | 46.80657 | 20.59924 | -0.4685 | 0.733039 | 0.351614 |
| 7 | 69986000 | 69988199 | 12.86644 | 22.56559 | 43.54646 | 19.24779 | -0.55614 | 0.629321 | 0.351622 |
| 9 | 39270000 | 39271599 | 17.25352 | 29.77116 | 54.92103 | 24.09322 | -0.49466 | 0.876319 | 0.351627 |
| 3 | 79474200 | 79475999 | 10.14744 | 17.82508 | 42.06259 | 18.61676 | -0.84416 | 0.587481 | 0.351635 |
| 5 | 86766400 | 86768199 | 19.94507 | 35.87774 | 25.49418 | 11.13325 | 0.893844 | 0.421175 | 0.351649 |
| 4 | 92069200 | 92070599 | 16.22652 | 30.1858 | 23.03802 | 9.627919 | 0.834444 | 0.20336 | 0.351653 |
| 6 | 74203800 | 74204199 | 6.276461 | 12.0753 | 27.55057 | 10.73173 | -0.64481 | -0.11512 | 0.351667 |
| 2 | 97862200 | 97864999 | 33.83402 | 56.14533 | 52.58758 | 22.78706 | 0.475029 | 1.296295 | 0.351676 |
| 4 | 97275600 | 97276999 | 16.90408 | 29.14739 | 52.1929 | 22.89593 | -0.45204 | 0.87746 | 0.351679 |
| 8 | 35649400 | 35652599 | 38.46225 | 61.7858 | 104.8548 | 44.41249 | -0.411 | 1.90483 | 0.351679 |
| 15 | 91365800 | 91366999 | 14.76722 | 27.46906 | 23.35964 | 9.76356 | 0.679765 | 0.19436 | 0.351686 |
| X | 17720000 | 17723399 | 65.84132 | 104.9424 | 97.43371 | 40.80092 | 0.44476 | 2.175762 | 0.351697 |
| 17 | 14717200 | 14718199 | 6.899817 | 13.1888 | 28.38527 | 11.26461 | -0.58593 | -0.04114 | 0.351712 |
| 1 | 144712000 | 144713999 | 48.4625 | 78.07297 | 75.17307 | 31.97256 | 0.410034 | 1.815203 | 0.351717 |
| 4 | 10873600 | 10875199 | 9.862986 | 17.66221 | 36.37187 | 15.94384 | -0.63521 | 0.404827 | 0.351719 |
| 11 | 102475800 | 102478199 | 26.43841 | 44.88036 | 42.68231 | 18.70087 | 0.453643 | 1.043754 | 0.351729 |
| 5 | 57119000 | 57122799 | 18.65782 | 33.15273 | 25.99101 | 11.46829 | 0.743058 | 0.488869 | 0.351738 |
| 6 | 48061800 | 48063799 | 14.35569 | 24.93231 | 47.91523 | 21.03221 | -0.55116 | 0.792899 | 0.351744 |
| 18 | 70558000 | 70559199 | 6.257328 | 12.12124 | 26.80471 | 10.16995 | -0.56344 | -0.20332 | 0.351751 |
| 11 | 58996400 | 58999599 | 28.3368 | 45.96688 | 85.21406 | 36.50868 | -0.52217 | 1.614438 | 0.351761 |
| 7 | 48183800 | 48184799 | 26.10892 | 44.39804 | 39.62613 | 17.36733 | 0.545169 | 1.024339 | 0.351761 |
| 12 | 18490000 | 18492399 | 55.44509 | 88.99462 | 82.27681 | 34.80639 | 0.462151 | 1.94457 | 0.351775 |
| 5 | 6170200 | 6172199 | 26.91791 | 46.93221 | 33.5836 | 14.77751 | 0.868033 | 0.777248 | 0.35178 |
| X | 43738600 | 43740799 | 26.91791 | 46.93221 | 33.5836 | 14.77751 | 0.868033 | 0.777248 | 0.35178 |
| X | 25551800 | 25556399 | 59.96411 | 96.89587 | 63.51043 | 27.1198 | 0.968657 | 1.744523 | 0.351794 |
| 12 | 103765400 | 103770199 | 111.0382 | 171.6376 | 159.2805 | 66.16152 | 0.409312 | 2.858918 | 0.3518 |

| | | | | | | | | | |
|---|---|---|---|---|---|---|---|---|---|
| 13 | 115942600 | 115943799 | 7.381107 | 13.94126 | 29.18919 | 11.88116 | -0.57913 | 0.048229 | 0.351814 |
| 5 | 90238000 | 90238799 | 11.36593 | 21.56415 | 20.44353 | 8.242804 | 0.566035 | 0.02718 | 0.351818 |
| 14 | 48514200 | 48515599 | 17.92262 | 31.71528 | 29.66727 | 13.12008 | 0.48689 | 0.525547 | 0.35184 |
| 12 | 102630000 | 102631399 | 16.83163 | 30.06114 | 27.74948 | 12.19153 | 0.513097 | 0.449976 | 0.351848 |
| 19 | 29657200 | 29674799 | 408.1786 | 555.1916 | 827.563 | 344.6437 | -0.44782 | 5.00789 | 0.351852 |
| 11 | 73399600 | 73400399 | 14.16257 | 26.11981 | 24.23644 | 10.25074 | 0.541965 | 0.236177 | 0.351866 |
| 1 | 16902200 | 16904199 | 32.14003 | 53.78087 | 44.34611 | 19.32073 | 0.658381 | 1.194515 | 0.351874 |
| 17 | 38859600 | 38865199 | 53.8819 | 87.59069 | 58.164 | 24.95762 | 0.958002 | 1.608066 | 0.351885 |
| 12 | 114581800 | 114583399 | 35.76682 | 59.98978 | 45.26588 | 19.75057 | 0.785839 | 1.18216 | 0.351899 |
| 11 | 103237200 | 103244399 | 55.40649 | 87.86703 | 142.2581 | 59.22094 | -0.36362 | 2.329143 | 0.3519 |
| 9 | 36003600 | 36007799 | 79.84784 | 125.8088 | 122.602 | 50.8735 | 0.360093 | 2.495124 | 0.351904 |
| 18 | 11253000 | 11255199 | 15.58404 | 28.07114 | 25.10254 | 10.94527 | 0.565574 | 0.380954 | 0.351928 |
| 5 | 59861800 | 59864799 | 28.14229 | 48.32162 | 39.68415 | 17.40634 | 0.66661 | 0.944263 | 0.35193 |
| 15 | 13131600 | 13132199 | 4.273121 | 8.329932 | 22.59281 | 8.15845 | -0.78756 | -0.4165 | 0.351933 |
| 14 | 48406000 | 48407799 | 12.94852 | 24.22081 | 22.28335 | 9.226112 | 0.577162 | 0.138124 | 0.351953 |
| 3 | 21081400 | 21081799 | 4.350343 | 8.497931 | 20.58605 | 7.199268 | -0.58342 | -0.53371 | 0.351967 |
| 8 | 105677400 | 105677799 | 4.350343 | 8.497931 | 20.58605 | 7.199268 | -0.58342 | -0.53371 | 0.351967 |
| 14 | 109186000 | 109186399 | 4.350343 | 8.497931 | 20.58605 | 7.199268 | -0.58342 | -0.53371 | 0.351967 |
| 15 | 14432200 | 14432599 | 4.350343 | 8.497931 | 20.58605 | 7.199268 | -0.58342 | -0.53371 | 0.351967 |
| 14 | 77892200 | 77892799 | 13.33903 | 25.81519 | 17.86758 | 6.808874 | 1.090956 | -0.13698 | 0.351967 |
| 2 | 118435200 | 118436999 | 20.68196 | 36.29337 | 32.6884 | 14.45541 | 0.537661 | 0.645133 | 0.351994 |
| 5 | 23510400 | 23511399 | 20.68196 | 36.29337 | 32.6884 | 14.45541 | 0.537661 | 0.645133 | 0.351994 |
| 11 | 59211000 | 59212599 | 16.4288 | 29.31703 | 27.26125 | 11.98552 | 0.501823 | 0.450634 | 0.352003 |
| 4 | 53439800 | 53441399 | 10.45754 | 18.35197 | 44.10486 | 19.50465 | -0.87074 | 0.606863 | 0.352008 |
| 13 | 80218800 | 80220599 | 17.59633 | 32.58309 | 22.86802 | 9.619322 | 0.947783 | 0.239825 | 0.352013 |
| 1 | 41506800 | 41507599 | 7.815308 | 14.75067 | 29.73673 | 12.12261 | -0.52742 | 0.056853 | 0.352014 |
| 8 | 110999600 | 111001799 | 11.4185 | 20.21516 | 41.89508 | 18.526 | -0.65441 | 0.501335 | 0.352022 |
| 12 | 99237200 | 99240399 | 15.95368 | 27.54037 | 50.5713 | 22.18535 | -0.4877 | 0.861363 | 0.352027 |
| 9 | 124181600 | 124182599 | 12.76642 | 22.50197 | 42.9771 | 19.02125 | -0.53945 | 0.564734 | 0.35203 |
| 9 | 18955800 | 18961599 | 105.0076 | 162.5628 | 156.0011 | 64.79882 | 0.362677 | 2.827981 | 0.352035 |
| 10 | 78676400 | 78676999 | 13.46202 | 24.86671 | 23.42279 | 9.885521 | 0.522951 | 0.221407 | 0.352062 |
| 2 | 83407000 | 83407999 | 8.827397 | 16.17112 | 33.47603 | 14.28874 | -0.61714 | 0.254629 | 0.352071 |
| 5 | 34281200 | 34283399 | 17.15886 | 29.33533 | 54.78344 | 24.02084 | -0.51455 | 0.961162 | 0.352071 |
| 7 | 15651000 | 15658599 | 58.23729 | 91.50797 | 165.627 | 68.7022 | -0.53499 | 2.54267 | 0.352088 |
| 7 | 44084200 | 44088199 | 23.50249 | 38.66827 | 74.20743 | 31.99124 | -0.55896 | 1.388748 | 0.3521 |
| 8 | 58366600 | 58367599 | 8.030836 | 15.00681 | 31.14399 | 12.91692 | -0.58885 | 0.129223 | 0.352106 |
| 13 | 99038800 | 99043599 | 46.81648 | 74.79721 | 124.0321 | 52.1129 | -0.38689 | 2.088846 | 0.352123 |
| 3 | 82147800 | 82148799 | 7.805634 | 14.76382 | 29.80562 | 12.09608 | -0.52379 | 0.041818 | 0.352138 |
| 17 | 15474400 | 15475799 | 11.26475 | 19.97367 | 39.81763 | 17.60063 | -0.59759 | 0.481219 | 0.352143 |
| 8 | 53075400 | 53076599 | 13.84045 | 25.43174 | 23.78835 | 10.11185 | 0.525329 | 0.253057 | 0.352146 |
| X | 42468200 | 42470599 | 31.60757 | 53.39866 | 45.2495 | 19.79685 | 0.619283 | 1.105868 | 0.352163 |
| 5 | 34205200 | 34205799 | 4.925843 | 9.564221 | 23.85688 | 8.945031 | -0.71707 | -0.26425 | 0.352163 |
| 11 | 95038000 | 95042799 | 29.07756 | 46.90655 | 95.5982 | 40.73229 | -0.66696 | 1.756436 | 0.352164 |
| 15 | 63902200 | 63906599 | 19.91195 | 33.10336 | 66.38508 | 28.80227 | -0.61936 | 1.250794 | 0.35217 |
| 2 | 70167600 | 70168799 | 8.395113 | 15.35712 | 33.47215 | 14.31506 | -0.69287 | 0.259794 | 0.352189 |
| 7 | 5364000 | 5371999 | 178.8842 | 267.7302 | 244.2216 | 100.7224 | 0.396029 | 3.513611 | 0.352206 |
| 11 | 73329800 | 73335999 | 106.1611 | 165.596 | 138.7787 | 57.58987 | 0.566325 | 2.698018 | 0.352227 |

| | | | | | | | | | |
|---|---|---|---|---|---|---|---|---|---|
| 14 | 109378800 | 109379599 | 6.254194 | 12.12666 | 28.16973 | 10.64059 | -0.62799 | -0.21542 | 0.352234 |
| 3 | 66009400 | 66011199 | 22.97829 | 39.86249 | 37.16929 | 16.3084 | 0.487042 | 0.827671 | 0.352249 |
| X | 77040800 | 77041399 | 11.4459 | 22.09723 | 19.65081 | 7.574555 | 0.717145 | -0.11521 | 0.352252 |
| 17 | 26565200 | 26568799 | 17.54691 | 29.99214 | 56.49582 | 24.77123 | -0.5273 | 0.962495 | 0.352297 |
| 7 | 44267400 | 44276999 | 103.2882 | 156.7936 | 254.0186 | 105.1784 | -0.41425 | 3.220078 | 0.352304 |
| 6 | 122778400 | 122780199 | 14.29041 | 24.86225 | 49.49945 | 21.75142 | -0.60196 | 0.766341 | 0.352308 |
| 5 | 127826200 | 127839199 | 130.2066 | 195.2682 | 304.8121 | 125.762 | -0.37589 | 3.469777 | 0.352308 |
| 13 | 52140800 | 52141999 | 15.00984 | 27.44062 | 25.31413 | 10.83477 | 0.538452 | 0.295729 | 0.352311 |
| 7 | 95056400 | 95056599 | 8.072336 | 15.7892 | 11.63386 | 3.828649 | 1.188205 | -0.71774 | 0.352315 |
| 7 | 17721000 | 17724399 | 66.10156 | 106.0586 | 79.67795 | 33.68733 | 0.76028 | 1.967881 | 0.352323 |
| X | 147660000 | 147662599 | 40.2063 | 65.68807 | 61.39058 | 26.3903 | 0.471263 | 1.510223 | 0.352326 |
| 15 | 38476800 | 38484599 | 61.6716 | 96.93329 | 160.6038 | 66.62116 | -0.4073 | 2.539299 | 0.352332 |
| 16 | 29017400 | 29020199 | 16.88013 | 28.99961 | 54.34928 | 23.83922 | -0.51876 | 0.911035 | 0.352363 |
| 2 | 113647200 | 113652599 | 49.47199 | 78.66112 | 132.3617 | 55.24552 | -0.41545 | 2.245126 | 0.352393 |
| 10 | 40514200 | 40515799 | 10.15116 | 18.20355 | 36.3465 | 15.91407 | -0.5899 | 0.397265 | 0.352419 |
| 7 | 115814000 | 115816199 | 29.61725 | 49.86628 | 46.94125 | 20.51583 | 0.467974 | 1.129391 | 0.352439 |
| 10 | 88720800 | 88723799 | 24.14651 | 39.7151 | 73.64034 | 31.74245 | -0.50968 | 1.394708 | 0.35244 |
| X | 157227000 | 157231199 | 42.1227 | 69.35844 | 55.07343 | 23.75038 | 0.710896 | 1.411725 | 0.352441 |
| 5 | 140703800 | 140706199 | 26.09001 | 42.80447 | 76.02399 | 32.73525 | -0.44926 | 1.432057 | 0.352442 |
| X | 25801800 | 25805399 | 35.43217 | 60.2821 | 39.03651 | 17.1096 | 1.006357 | 1.055502 | 0.352446 |
| 2 | 164364800 | 164370999 | 95.22548 | 145.2912 | 236.1679 | 97.92896 | -0.41307 | 3.113222 | 0.352449 |
| 10 | 112762800 | 112765199 | 21.74512 | 36.30264 | 65.11998 | 28.3301 | -0.45909 | 1.207657 | 0.352464 |
| X | 151314400 | 151315399 | 18.30128 | 33.20004 | 25.89537 | 11.19134 | 0.770062 | 0.364377 | 0.352469 |
| 15 | 35408200 | 35409599 | 11.41327 | 20.44481 | 38.36724 | 16.81407 | -0.50206 | 0.417367 | 0.352475 |
| X | 167746400 | 167748799 | 27.30433 | 47.88594 | 29.07006 | 12.84884 | 1.104129 | 0.687532 | 0.3525 |
| 4 | 135565600 | 135568399 | 22.07538 | 36.3645 | 72.28238 | 31.17739 | -0.60868 | 1.370003 | 0.352502 |
| 4 | 21644400 | 21645999 | 25.45672 | 44.00432 | 38.72082 | 16.9873 | 0.569407 | 0.87416 | 0.352509 |
| 16 | 68000400 | 68002799 | 41.721 | 67.73495 | 65.2321 | 27.96498 | 0.421312 | 1.609183 | 0.352517 |
| 9 | 112351400 | 112352999 | 9.575626 | 17.50784 | 35.01848 | 14.98631 | -0.57094 | 0.273804 | 0.352525 |
| 7 | 10377000 | 10381199 | 64.26745 | 103.4758 | 68.92343 | 29.29029 | 0.939516 | 1.855461 | 0.352538 |
| X | 49686200 | 49689999 | 15.93441 | 27.73268 | 55.54229 | 24.41246 | -0.61056 | 0.778657 | 0.352542 |
| 2 | 153003000 | 153003999 | 5.274891 | 10.19421 | 25.37152 | 9.732195 | -0.74699 | -0.18189 | 0.352549 |
| 10 | 14580800 | 14581399 | 4.839761 | 9.416254 | 25.41328 | 9.373033 | -0.80976 | -0.32629 | 0.35255 |
| 3 | 43204200 | 43205999 | 16.75367 | 29.99276 | 26.97679 | 11.82844 | 0.551902 | 0.43053 | 0.352562 |
| X | 77272200 | 77276799 | 59.5322 | 95.70381 | 79.05198 | 33.51953 | 0.627052 | 1.900287 | 0.352564 |
| 3 | 61369000 | 61370399 | 16.61509 | 28.04484 | 59.46632 | 26.00971 | -0.69902 | 1.076908 | 0.352599 |
| 7 | 35077800 | 35081599 | 21.64435 | 36.02303 | 66.03718 | 28.67308 | -0.49021 | 1.243113 | 0.3526 |
| 9 | 81183200 | 81184999 | 24.19442 | 42.33445 | 32.41658 | 14.3002 | 0.770727 | 0.708272 | 0.352611 |
| 7 | 47905200 | 47910399 | 75.88183 | 120.5454 | 106.2711 | 44.3161 | 0.51442 | 2.286873 | 0.352619 |
| 2 | 40375400 | 40378799 | 37.94339 | 61.0242 | 103.067 | 43.72986 | -0.40199 | 1.865647 | 0.352628 |
| 14 | 46505600 | 46506399 | 8.158979 | 15.37418 | 31.92314 | 13.05407 | -0.57342 | 0.077653 | 0.352629 |
| 9 | 25062200 | 25063599 | 11.50314 | 20.19224 | 42.03601 | 18.59389 | -0.66461 | 0.606553 | 0.352639 |
| 5 | 70678400 | 70678999 | 11.64643 | 22.50924 | 19.03215 | 7.298291 | 0.795792 | -0.13311 | 0.352651 |
| 17 | 66077800 | 66080399 | 19.24483 | 32.4891 | 60.21641 | 26.33914 | -0.50582 | 1.084208 | 0.352657 |
| 6 | 130528600 | 130536599 | 150.4429 | 229.9376 | 172.6567 | 71.62221 | 0.702125 | 3.089602 | 0.352659 |
| 7 | 21143800 | 21145599 | 23.17415 | 40.0467 | 36.89094 | 16.18436 | 0.503558 | 0.86507 | 0.352683 |
| 2 | 150968800 | 150974199 | 68.65569 | 109.4543 | 96.9958 | 40.631 | 0.512206 | 2.167487 | 0.352691 |

| | | | | | | | | | |
|---|---|---|---|---|---|---|---|---|---|
| 5 | 141111600 | 141112599 | 8.054519 | 14.91726 | 31.53857 | 13.26325 | -0.63108 | 0.184658 | 0.352702 |
| 9 | 119918000 | 119921999 | 35.31199 | 56.90668 | 99.29186 | 42.25135 | -0.4452 | 1.796182 | 0.352702 |
| 2 | 70734800 | 70736599 | 12.42037 | 21.83144 | 41.44906 | 18.35495 | -0.53249 | 0.58943 | 0.352715 |
| 4 | 104658400 | 104659199 | 6.255887 | 12.1055 | 26.73777 | 10.19534 | -0.56895 | -0.18812 | 0.352722 |
| 10 | 65740200 | 65742399 | 19.36844 | 35.17058 | 24.62605 | 10.62967 | 0.925717 | 0.363299 | 0.352723 |
| 10 | 20377000 | 20380199 | 21.526 | 36.09137 | 65.27412 | 28.47415 | -0.47101 | 1.159922 | 0.352729 |
| 11 | 15684000 | 15685599 | 19.3987 | 34.38608 | 29.38007 | 12.98863 | 0.617737 | 0.515827 | 0.352741 |
| 3 | 110327200 | 110328999 | 18.13728 | 32.63495 | 26.79388 | 11.69807 | 0.686983 | 0.409479 | 0.352764 |
| 2 | 118575800 | 118577399 | 10.23094 | 17.8332 | 47.33768 | 20.82611 | -1.0137 | 0.719081 | 0.352773 |
| 3 | 33587800 | 33589799 | 33.5143 | 55.8857 | 45.99872 | 19.99679 | 0.661125 | 1.237373 | 0.352786 |
| 11 | 107451000 | 107452599 | 8.840553 | 16.19629 | 32.96431 | 14.06906 | -0.59251 | 0.254626 | 0.352794 |
| 6 | 69905800 | 69907399 | 15.53403 | 28.37214 | 25.63853 | 10.98659 | 0.566705 | 0.304242 | 0.352797 |
| 7 | 69862400 | 69863799 | 9.40821 | 17.36935 | 33.61008 | 14.19532 | -0.50949 | 0.215934 | 0.3528 |
| 13 | 112296000 | 112296799 | 5.404266 | 10.45553 | 24.86724 | 9.490726 | -0.67505 | -0.19865 | 0.352817 |
| 1 | 98657000 | 98658399 | 15.24454 | 28.11891 | 24.70118 | 10.44546 | 0.620777 | 0.246078 | 0.352819 |
| 7 | 43111200 | 43118999 | 50.65518 | 79.58684 | 176.0276 | 73.01546 | -0.82388 | 2.542837 | 0.352819 |
| 14 | 102166600 | 102168999 | 20.34805 | 33.87212 | 68.31077 | 29.66402 | -0.62751 | 1.236068 | 0.352826 |
| 1 | 44311200 | 44314599 | 60.80659 | 97.75803 | 76.68349 | 32.51791 | 0.701483 | 1.897254 | 0.352829 |
| X | 60765800 | 60767599 | 24.00547 | 41.70869 | 34.73912 | 15.25253 | 0.649774 | 0.8145 | 0.352838 |
| 12 | 115792400 | 115796599 | 81.30219 | 128.2336 | 123.373 | 51.20039 | 0.379711 | 2.475562 | 0.352842 |
| 15 | 96474200 | 96485399 | 104.9741 | 159.7271 | 250.1956 | 103.6683 | -0.36291 | 3.172026 | 0.352847 |
| 7 | 16844800 | 16846999 | 36.81399 | 60.37817 | 58.56253 | 25.21287 | 0.420831 | 1.44869 | 0.352849 |
| 2 | 49950200 | 49950599 | 3.932604 | 7.685214 | 19.42824 | 6.732346 | -0.63131 | -0.57611 | 0.352851 |
| 2 | 53959800 | 53961399 | 14.46931 | 26.86508 | 23.68628 | 9.927057 | 0.624505 | 0.200475 | 0.352864 |
| 17 | 9636000 | 9638199 | 21.68481 | 37.86608 | 34.99058 | 15.41145 | 0.500994 | 0.731238 | 0.352873 |
| 1 | 166514800 | 166516599 | 11.8566 | 21.13969 | 40.38105 | 17.76665 | -0.53178 | 0.451344 | 0.352881 |
| 7 | 40704800 | 40710199 | 102.9219 | 160.4883 | 131.1586 | 54.43007 | 0.602158 | 2.700461 | 0.352882 |
| 13 | 28725200 | 28727599 | 16.03571 | 28.76341 | 25.05784 | 10.96902 | 0.599234 | 0.409166 | 0.352894 |
| 6 | 24597400 | 24598199 | 6.890325 | 13.11205 | 29.42753 | 11.79006 | -0.65907 | -0.01185 | 0.352895 |
| 11 | 35234200 | 35237399 | 24.11694 | 39.74294 | 72.67815 | 31.35324 | -0.48865 | 1.369622 | 0.35293 |
| 4 | 93213600 | 93216999 | 35.43758 | 58.79884 | 53.15965 | 23.03292 | 0.525923 | 1.304269 | 0.352943 |
| 3 | 52832000 | 52832999 | 13.74645 | 26.43271 | 18.71012 | 7.299311 | 1.026754 | -0.05727 | 0.352952 |
| 9 | 11799400 | 11800599 | 13.74645 | 26.43271 | 18.71012 | 7.299311 | 1.026754 | -0.05727 | 0.352952 |
| 8 | 67199400 | 67200999 | 14.65562 | 26.93915 | 24.686 | 10.48836 | 0.55522 | 0.259501 | 0.352973 |
| 4 | 31753400 | 31754799 | 10.30853 | 18.48075 | 36.34391 | 15.91656 | -0.56833 | 0.400393 | 0.352985 |
| 9 | 95062000 | 95064599 | 13.05537 | 22.99034 | 42.44219 | 18.78792 | -0.49128 | 0.574466 | 0.353006 |
| 12 | 6133600 | 6134999 | 13.05537 | 22.99034 | 42.44219 | 18.78792 | -0.49128 | 0.574466 | 0.353006 |
| 10 | 11729200 | 11732799 | 45.97471 | 74.45461 | 69.29076 | 29.65024 | 0.46729 | 1.670967 | 0.353009 |
| 2 | 73422400 | 73423599 | 6.870976 | 12.75554 | 31.57161 | 13.23433 | -0.85337 | 0.15917 | 0.353014 |
| 12 | 116830200 | 116831799 | 11.89969 | 21.08815 | 39.29904 | 17.37456 | -0.50113 | 0.491063 | 0.353015 |
| 15 | 77242600 | 77243999 | 5.357412 | 10.19468 | 29.88683 | 11.9748 | -1.04046 | -0.02478 | 0.353015 |
| 3 | 121541000 | 121545399 | 71.95879 | 112.113 | 183.238 | 76.05225 | -0.39754 | 2.707698 | 0.353028 |
| 8 | 22887800 | 22888799 | 7.075255 | 13.37371 | 28.66233 | 11.64801 | -0.61051 | 0.038041 | 0.353037 |
| 14 | 41780000 | 41793799 | 207.9511 | 309.7091 | 255.639 | 105.3388 | 0.533732 | 3.608004 | 0.353043 |
| 11 | 21208600 | 21209199 | 5.958788 | 11.56167 | 25.72462 | 9.686445 | -0.5618 | -0.23649 | 0.353058 |
| 11 | 97917000 | 97925999 | 69.29748 | 108.0362 | 178.0152 | 73.87741 | -0.40851 | 2.693749 | 0.353062 |
| 7 | 55699600 | 55700799 | 8.53046 | 15.67379 | 32.48518 | 13.80972 | -0.61421 | 0.234901 | 0.353105 |

| | | | | | | | | | |
|---|---|---|---|---|---|---|---|---|---|
| 6 | 7005800 | 7008999 | 55.95614 | 89.93342 | 81.53627 | 34.56093 | 0.492522 | 1.90243 | 0.353118 |
| 5 | 26638000 | 26641199 | 33.69062 | 54.53846 | 94.83161 | 40.56712 | -0.43297 | 1.670822 | 0.353125 |
| 14 | 95457400 | 95459999 | 18.95114 | 33.93137 | 27.21819 | 11.93291 | 0.716278 | 0.445809 | 0.353137 |
| 2 | 37342000 | 37342999 | 5.84504 | 11.25548 | 25.81138 | 10.02814 | -0.64759 | -0.13049 | 0.353153 |
| 4 | 62883400 | 62884399 | 5.84504 | 11.25548 | 25.81138 | 10.02814 | -0.64759 | -0.13049 | 0.353153 |
| 4 | 138093400 | 138094199 | 11.82918 | 22.27554 | 21.03873 | 8.61268 | 0.553124 | 0.085106 | 0.353189 |
| 11 | 27580400 | 27581399 | 14.92231 | 28.50044 | 18.37714 | 7.306151 | 1.134719 | 0.0119 | 0.3532 |
| 6 | 7669200 | 7669999 | 5.233259 | 10.12459 | 24.82041 | 9.473274 | -0.71725 | -0.19708 | 0.353205 |
| 11 | 74225000 | 74225999 | 12.44991 | 23.27429 | 20.5352 | 8.510878 | 0.636531 | 0.135161 | 0.35321 |
| 14 | 42198000 | 42201399 | 30.21741 | 51.11079 | 45.25087 | 19.80459 | 0.556308 | 1.090216 | 0.353217 |
| 13 | 90596000 | 90596399 | 7.284569 | 14.22409 | 9.071364 | 2.869276 | 1.438754 | -0.87556 | 0.353227 |
| 10 | 10118200 | 10119799 | 8.109202 | 15.29266 | 30.8437 | 12.59553 | -0.53056 | 0.067047 | 0.353234 |
| 5 | 38808000 | 38810799 | 20.68304 | 36.55242 | 29.80606 | 13.18489 | 0.683117 | 0.556294 | 0.353246 |
| 15 | 32823000 | 32827999 | 41.54373 | 68.53186 | 52.3284 | 22.58427 | 0.768122 | 1.383526 | 0.353252 |
| 4 | 94661400 | 94662999 | 14.76791 | 27.59125 | 23.25527 | 9.647919 | 0.70069 | 0.165154 | 0.353255 |
| 13 | 112368400 | 112376799 | 62.75416 | 98.76566 | 160.0378 | 66.39783 | -0.37377 | 2.512839 | 0.353256 |
| 4 | 81797000 | 81798999 | 27.97039 | 48.57871 | 35.23214 | 15.46521 | 0.848278 | 0.836604 | 0.353261 |
| 19 | 49349200 | 49350599 | 12.19798 | 21.69475 | 40.87356 | 18.02079 | -0.51404 | 0.470714 | 0.353267 |
| 16 | 37342200 | 37344999 | 40.58059 | 66.39527 | 54.81997 | 23.57814 | 0.650953 | 1.485508 | 0.353283 |
| X | 19153800 | 19157799 | 47.36401 | 77.61526 | 55.20298 | 23.75796 | 0.866685 | 1.482858 | 0.353307 |
| 3 | 115884800 | 115885799 | 13.52537 | 25.41012 | 20.26379 | 8.328373 | 0.7906 | 0.112444 | 0.353309 |
| 8 | 3154000 | 3157199 | 48.52675 | 78.1217 | 74.7855 | 31.77542 | 0.417154 | 1.837966 | 0.353335 |
| 11 | 73388800 | 73390599 | 27.45962 | 46.41073 | 44.2912 | 19.3801 | 0.448465 | 1.086743 | 0.353345 |
| 13 | 25193800 | 25195199 | 13.18597 | 24.87213 | 22.45435 | 9.165651 | 0.621054 | 0.090642 | 0.353348 |
| 1 | 100713600 | 100714199 | 12.38957 | 23.14927 | 21.9237 | 9.094478 | 0.533755 | 0.140396 | 0.353348 |
| 1 | 117008200 | 117010199 | 11.86396 | 21.21011 | 42.03802 | 18.45354 | -0.58247 | 0.437653 | 0.353358 |
| 9 | 96096800 | 96097399 | 9.550682 | 18.54678 | 18.26006 | 6.83824 | 0.61092 | -0.23553 | 0.353362 |
| X | 33488400 | 33488799 | 8.314592 | 16.26289 | 10.16375 | 3.343445 | 1.419743 | -0.72211 | 0.353362 |
| 10 | 94034600 | 94035399 | 5.532792 | 10.72824 | 24.43222 | 9.227428 | -0.59858 | -0.23405 | 0.353368 |
| 7 | 13675000 | 13677199 | 37.1488 | 61.19106 | 57.04817 | 24.60582 | 0.480193 | 1.3961 | 0.353396 |
| 14 | 83788400 | 83792399 | 43.01498 | 70.40873 | 59.03871 | 25.39664 | 0.629145 | 1.486518 | 0.353403 |
| 6 | 100382400 | 100389199 | 57.33578 | 90.51482 | 165.4805 | 68.70269 | -0.54419 | 2.447239 | 0.353404 |
| 5 | 137705800 | 137706999 | 7.346299 | 13.90637 | 29.82872 | 12.08456 | -0.6085 | 0.030548 | 0.353419 |
| X | 49393200 | 49398999 | 61.72242 | 99.15041 | 79.70819 | 33.76106 | 0.664776 | 1.926969 | 0.353431 |
| 1 | 182839800 | 182841599 | 10.34087 | 18.52884 | 36.88454 | 16.16059 | -0.586 | 0.407161 | 0.353434 |
| 11 | 58130000 | 58131199 | 9.444082 | 17.31742 | 33.48481 | 14.27387 | -0.51796 | 0.255226 | 0.353434 |
| 4 | 3223000 | 3233799 | 585.2777 | 791.875 | 668.9573 | 278.8184 | 0.363541 | 5.09384 | 0.353435 |
| 8 | 24647400 | 24649999 | 17.69836 | 29.9418 | 57.89952 | 25.34201 | -0.56638 | 1.060918 | 0.353446 |
| 14 | 113750800 | 113752599 | 21.62835 | 38.01117 | 31.39764 | 13.90225 | 0.661739 | 0.624275 | 0.353448 |
| 7 | 85901800 | 85904399 | 24.93109 | 43.74225 | 30.72672 | 13.58592 | 0.894537 | 0.669178 | 0.353465 |
| 17 | 28207800 | 28209599 | 9.191818 | 16.64416 | 34.90548 | 15.11994 | -0.64988 | 0.328487 | 0.353485 |
| 14 | 79670600 | 79673399 | 25.5638 | 42.29089 | 78.92513 | 34.12685 | -0.51691 | 1.332454 | 0.353487 |
| 7 | 15582600 | 15585599 | 35.38949 | 58.51149 | 46.12968 | 19.93819 | 0.722453 | 1.336933 | 0.353492 |
| 7 | 18560600 | 18562999 | 20.60056 | 34.46232 | 64.55902 | 28.12236 | -0.52148 | 1.180684 | 0.353504 |
| 4 | 50466600 | 50467999 | 16.80967 | 31.25183 | 22.36652 | 9.356815 | 0.925191 | 0.209873 | 0.353514 |
| 4 | 99247200 | 99248999 | 16.64591 | 29.96167 | 26.98795 | 11.77824 | 0.55442 | 0.397296 | 0.353524 |
| X | 148478000 | 148480999 | 39.61165 | 65.70952 | 46.06268 | 19.95481 | 0.891795 | 1.31402 | 0.353534 |

| | | | | | | | | | |
|---|---|---|---|---|---|---|---|---|---|
| X | 116078200 | 116080199 | 21.46733 | 39.15115 | 21.74047 | 9.333417 | 1.262773 | 0.349445 | 0.353545 |
| 8 | 102972600 | 102973599 | 8.758439 | 15.84569 | 38.03554 | 16.49363 | -0.84445 | 0.337358 | 0.353558 |
| 13 | 60712800 | 60714199 | 15.56872 | 28.30727 | 26.17311 | 11.27923 | 0.528695 | 0.331473 | 0.353559 |
| 6 | 146274400 | 146277599 | 26.22812 | 43.02002 | 79.10489 | 34.05816 | -0.49942 | 1.435815 | 0.353585 |
| 11 | 89450200 | 89452799 | 21.52429 | 37.76098 | 31.98219 | 14.13978 | 0.625789 | 0.651614 | 0.353586 |
| 9 | 27252800 | 27255199 | 15.57654 | 27.11154 | 48.96485 | 21.52234 | -0.46187 | 0.767457 | 0.353596 |
| 4 | 115696200 | 115697999 | 7.959767 | 14.55642 | 36.10553 | 15.44752 | -0.8789 | 0.261783 | 0.353602 |
| 14 | 20757400 | 20758199 | 5.140055 | 9.944823 | 26.02358 | 9.929906 | -0.81172 | -0.20087 | 0.353628 |
| 16 | 92367800 | 92369999 | 21.4428 | 35.74136 | 65.6003 | 28.51059 | -0.49203 | 1.224161 | 0.353641 |
| 6 | 98513800 | 98514799 | 13.31103 | 25.39994 | 20.92554 | 8.335307 | 0.782552 | 0.009916 | 0.353648 |
| Y | 28757000 | 28758199 | 11.09246 | 21.25176 | 12.98979 | 5.121016 | 1.222032 | -0.07083 | 0.353661 |
| X | 62112000 | 62113799 | 26.24906 | 45.71739 | 34.7672 | 15.2881 | 0.780542 | 0.790217 | 0.353668 |
| 1 | 166433400 | 166436399 | 25.49494 | 44.51693 | 33.57233 | 14.78628 | 0.792734 | 0.749188 | 0.353682 |
| 7 | 10910200 | 10936599 | 675.7041 | 903.738 | 755.759 | 315.5414 | 0.361453 | 5.277313 | 0.353698 |
| 2 | 15308000 | 15310999 | 35.49731 | 58.78285 | 53.75931 | 23.26131 | 0.50916 | 1.325556 | 0.353709 |
| X | 25915400 | 25916999 | 31.9572 | 53.83486 | 45.32391 | 19.81219 | 0.628593 | 1.133605 | 0.35371 |
| X | 90921800 | 90924399 | 33.19016 | 57.38166 | 32.87591 | 14.42314 | 1.185796 | 0.901308 | 0.353718 |
| 1 | 16025800 | 16026599 | 13.2795 | 24.55005 | 22.46006 | 9.468443 | 0.565929 | 0.21297 | 0.353725 |
| 7 | 18311000 | 18314999 | 62.88268 | 100.2455 | 97.94451 | 41.02546 | 0.371546 | 2.165303 | 0.35373 |
| 1 | 32770000 | 32771799 | 16.00096 | 29.55417 | 24.04689 | 10.15004 | 0.732348 | 0.242684 | 0.353732 |
| 1 | 168572600 | 168574599 | 29.48446 | 49.80916 | 45.83766 | 20.05385 | 0.500699 | 1.099875 | 0.353734 |
| 8 | 35671400 | 35673199 | 10.43163 | 18.80435 | 35.84835 | 15.616 | -0.51861 | 0.366155 | 0.353735 |
| 13 | 102523000 | 102524799 | 36.71702 | 59.17881 | 124.9363 | 53.17311 | -0.7195 | 1.80679 | 0.353763 |
| 7 | 14225000 | 14238599 | 202.2802 | 303.2342 | 224.6837 | 92.69352 | 0.697887 | 3.487982 | 0.35378 |
| 4 | 103419800 | 103422199 | 12.73919 | 22.34531 | 42.5339 | 18.80229 | -0.53625 | 0.626714 | 0.353796 |
| 7 | 34755000 | 34757599 | 12.73919 | 22.34531 | 42.5339 | 18.80229 | -0.53625 | 0.626714 | 0.353796 |
| 7 | 46969200 | 46979199 | 167.0846 | 251.2816 | 231.8832 | 95.74472 | 0.38557 | 3.418085 | 0.353798 |
| 7 | 63321400 | 63322199 | 7.056191 | 13.39828 | 28.22307 | 11.36006 | -0.57364 | 0.004693 | 0.353805 |
| 14 | 120355600 | 120357199 | 11.03695 | 19.86044 | 37.87657 | 16.5358 | -0.52156 | 0.383345 | 0.353807 |
| 3 | 96136200 | 96137999 | 9.11175 | 16.76027 | 33.00435 | 14.00943 | -0.53972 | 0.23301 | 0.353814 |
| 15 | 27493600 | 27495199 | 9.11175 | 16.76027 | 33.00435 | 14.00943 | -0.53972 | 0.23301 | 0.353814 |
| 17 | 46222200 | 46225599 | 46.24891 | 73.67355 | 126.5495 | 52.95245 | -0.4422 | 2.18293 | 0.353829 |
| 3 | 3210200 | 3224199 | 185.5994 | 278.5423 | 232.8282 | 96.08102 | 0.525535 | 3.463077 | 0.353869 |
| 3 | 3289800 | 3291599 | 17.55646 | 32.75088 | 21.92538 | 9.119849 | 1.027737 | 0.1956 | 0.35387 |
| 7 | 52791200 | 52793599 | 21.53032 | 37.73993 | 33.20683 | 14.67082 | 0.571257 | 0.668551 | 0.353871 |
| 7 | 67089000 | 67090999 | 13.81409 | 24.15537 | 45.18068 | 19.91979 | -0.51139 | 0.68575 | 0.35389 |
| 17 | 54661600 | 54662799 | 14.10616 | 25.80396 | 23.2383 | 9.938523 | 0.573307 | 0.278228 | 0.353895 |
| 7 | 15498800 | 15500199 | 40.25103 | 65.39024 | 61.70162 | 26.46395 | 0.451549 | 1.591945 | 0.353909 |
| 16 | 55793800 | 55795399 | 11.81048 | 20.71515 | 42.5581 | 18.81211 | -0.64572 | 0.621896 | 0.35391 |
| X | 54723400 | 54727199 | 54.89967 | 88.50563 | 72.83902 | 31.00983 | 0.63727 | 1.807374 | 0.353912 |
| 15 | 93009400 | 93015399 | 41.75765 | 66.85276 | 112.9383 | 47.59084 | -0.4103 | 2.017436 | 0.353918 |
| 19 | 47864000 | 47865799 | 9.708493 | 17.39022 | 36.37395 | 15.94115 | -0.65734 | 0.401712 | 0.353927 |
| 4 | 98219200 | 98221599 | 22.53541 | 39.33601 | 34.83377 | 15.3386 | 0.561964 | 0.739243 | 0.353928 |
| 3 | 129176400 | 129179999 | 39.7045 | 63.4768 | 124.7958 | 52.48319 | -0.63091 | 2.06277 | 0.353961 |
| 2 | 169679400 | 169687399 | 51.32358 | 80.96674 | 164.4274 | 68.24129 | -0.69635 | 2.459134 | 0.353961 |
| 1 | 185483600 | 185483999 | 3.148176 | 6.158419 | 17.32287 | 5.766102 | -0.72659 | -0.73402 | 0.354015 |
| 1 | 113766400 | 113768599 | 19.38398 | 35.11586 | 25.15543 | 10.89026 | 0.89034 | 0.377687 | 0.354015 |

| | | | | | | | | | |
|---|---|---|---|---|---|---|---|---|---|
| 18 | 19579400 | 19581199 | 19.38398 | 35.11586 | 25.15543 | 10.89026 | 0.89034 | 0.377687 | 0.354015 |
| 2 | 98219600 | 98220399 | 15.37797 | 26.34572 | 53.17853 | 23.32107 | -0.62578 | 0.930643 | 0.354016 |
| 2 | 71918400 | 71919599 | 13.00911 | 24.15022 | 20.00728 | 8.386919 | 0.713805 | 0.180922 | 0.354017 |
| 6 | 114769400 | 114770599 | 6.593706 | 12.61303 | 27.28555 | 10.80516 | -0.59027 | -0.05466 | 0.35403 |
| 4 | 103228800 | 103232599 | 24.80605 | 43.10221 | 36.43557 | 15.99797 | 0.628183 | 0.815598 | 0.354035 |
| 17 | 20363000 | 20366599 | 94.38696 | 147.9623 | 121.4213 | 50.36125 | 0.602121 | 2.602772 | 0.354066 |
| 6 | 140393600 | 140394399 | 4.684355 | 9.116353 | 22.95599 | 8.448321 | -0.70667 | -0.34085 | 0.35407 |
| 12 | 116878000 | 116878799 | 4.684355 | 9.116353 | 22.95599 | 8.448321 | -0.70667 | -0.34085 | 0.35407 |
| 15 | 57376600 | 57377799 | 15.07802 | 27.63659 | 24.57343 | 10.48115 | 0.594289 | 0.277663 | 0.35407 |
| 5 | 134132600 | 134142199 | 99.54099 | 151.8128 | 239.3914 | 99.25614 | -0.36981 | 3.122796 | 0.354084 |
| 14 | 45894400 | 45896199 | 13.55677 | 23.54119 | 48.96741 | 21.492 | -0.66496 | 0.79056 | 0.354085 |
| X | 116246600 | 116248199 | 18.59312 | 33.11541 | 28.14444 | 12.39392 | 0.62948 | 0.480519 | 0.354093 |
| 9 | 10628000 | 10629399 | 8.260775 | 15.59722 | 32.62944 | 13.29145 | -0.57994 | 0.063954 | 0.354102 |
| 16 | 35461800 | 35468399 | 49.28627 | 78.07576 | 151.1889 | 62.87876 | -0.62295 | 2.357935 | 0.35412 |
| 11 | 89887000 | 89890199 | 29.28927 | 47.48862 | 87.25942 | 37.37145 | -0.51018 | 1.625752 | 0.354136 |
| 3 | 77938800 | 77940599 | 24.68055 | 40.52351 | 75.62786 | 32.57538 | -0.51994 | 1.421587 | 0.354142 |
| 2 | 24029400 | 24031599 | 32.36624 | 54.85518 | 39.29015 | 17.20626 | 0.861025 | 1.075603 | 0.354146 |
| 14 | 108345600 | 108346799 | 13.16156 | 24.94006 | 21.95438 | 8.875166 | 0.669285 | 0.058006 | 0.354148 |
| 4 | 82977800 | 82979399 | 16.10345 | 29.45771 | 24.98623 | 10.68598 | 0.659179 | 0.297774 | 0.354152 |
| 7 | 58761400 | 58762799 | 23.90303 | 41.153 | 39.39961 | 17.28298 | 0.447021 | 0.902932 | 0.354153 |
| 13 | 114202200 | 114203599 | 10.7144 | 19.29141 | 37.89105 | 16.53009 | -0.56315 | 0.377006 | 0.354173 |
| 5 | 27491000 | 27493199 | 16.99136 | 30.62653 | 26.95665 | 11.74389 | 0.589203 | 0.391897 | 0.354177 |
| 16 | 22446600 | 22448599 | 15.98137 | 27.55741 | 51.09204 | 22.41303 | -0.50181 | 0.873117 | 0.354199 |
| 13 | 68219200 | 68225799 | 179.451 | 268.0676 | 251.428 | 103.6561 | 0.353428 | 3.548472 | 0.354204 |
| 5 | 82405400 | 82405799 | 3.816364 | 7.453165 | 19.87515 | 6.977943 | -0.72462 | -0.53031 | 0.354214 |
| 3 | 41146600 | 41147799 | 7.995062 | 14.91185 | 31.65317 | 13.17004 | -0.62561 | 0.138274 | 0.354214 |
| 1 | 31581000 | 31582799 | 17.34813 | 30.84952 | 28.55404 | 12.59096 | 0.505429 | 0.47987 | 0.354235 |
| 11 | 10984400 | 10984799 | 5.101561 | 9.919167 | 23.484 | 8.707032 | -0.62893 | -0.30754 | 0.354244 |
| 1 | 184441200 | 184442999 | 16.21403 | 28.0038 | 52.21033 | 22.90473 | -0.5096 | 0.857651 | 0.35427 |
| 15 | 97421400 | 97423799 | 29.77499 | 50.04576 | 47.4925 | 20.74594 | 0.45627 | 1.144951 | 0.354272 |
| 7 | 83753000 | 83754599 | 11.41159 | 20.30637 | 38.30425 | 16.8808 | -0.51514 | 0.458002 | 0.354283 |
| 2 | 121243800 | 121244999 | 7.024771 | 13.00656 | 31.54673 | 13.27164 | -0.82801 | 0.17632 | 0.354288 |
| 4 | 53238000 | 53239199 | 7.024771 | 13.00656 | 31.54673 | 13.27164 | -0.82801 | 0.17632 | 0.354288 |
| 7 | 114159600 | 114160599 | 8.619328 | 15.985 | 32.07587 | 13.4643 | -0.55498 | 0.180083 | 0.354296 |
| 7 | 86177800 | 86178799 | 20.48844 | 36.01313 | 26.80131 | 11.86855 | 0.811529 | 0.60862 | 0.354306 |
| 3 | 23231400 | 23233799 | 20.54935 | 36.10878 | 32.11894 | 14.2196 | 0.555531 | 0.622724 | 0.35431 |
| 12 | 113503200 | 113510599 | 92.45848 | 145.0839 | 132.0548 | 54.76683 | 0.453961 | 2.58463 | 0.354335 |
| 10 | 37199200 | 37200999 | 15.11357 | 27.71718 | 23.43955 | 9.98996 | 0.666708 | 0.271348 | 0.354347 |
| X | 125143400 | 125144599 | 17.092 | 32.54129 | 18.28153 | 7.31692 | 1.324205 | 0.054383 | 0.354348 |
| 16 | 71587400 | 71587999 | 16.76272 | 30.16488 | 23.12638 | 10.0959 | 0.785326 | 0.391781 | 0.354352 |
| 12 | 32059200 | 32061999 | 34.15094 | 56.31397 | 53.58139 | 23.12114 | 0.451514 | 1.372715 | 0.354358 |
| 14 | 51450600 | 51453999 | 57.36239 | 91.83811 | 88.83625 | 37.43617 | 0.392642 | 2.024498 | 0.354362 |
| 7 | 33514400 | 33517999 | 100.6226 | 156.6935 | 141.9342 | 58.91364 | 0.452423 | 2.724706 | 0.354363 |
| 5 | 4262400 | 4265399 | 27.64469 | 47.09492 | 41.40273 | 18.14863 | 0.567036 | 1.01219 | 0.354376 |
| 1 | 60156200 | 60158399 | 17.89804 | 30.77641 | 56.55271 | 24.8064 | -0.49013 | 0.911689 | 0.354381 |
| 1 | 187067400 | 187071999 | 24.5924 | 40.11283 | 80.29357 | 34.49566 | -0.62621 | 1.511488 | 0.354388 |
| 1 | 133488400 | 133490399 | 18.91534 | 33.54658 | 29.44604 | 13.0136 | 0.579444 | 0.509048 | 0.354389 |

| | | | | | | | | | |
|---|---|---|---|---|---|---|---|---|---|
| 1 | 183835400 | 183837799 | 18.91534 | 33.54658 | 29.44604 | 13.0136 | 0.579444 | 0.509048 | 0.354389 |
| 5 | 37732800 | 37736799 | 27.63157 | 44.98016 | 82.36976 | 35.36189 | -0.50032 | 1.547572 | 0.354391 |
| 12 | 67607400 | 67608599 | 17.41852 | 29.38682 | 63.80477 | 27.90321 | -0.73309 | 1.090579 | 0.354416 |
| 5 | 27994600 | 27997199 | 13.33212 | 22.98526 | 51.04859 | 22.39385 | -0.76106 | 0.861834 | 0.354428 |
| 1 | 102250200 | 102251599 | 19.77366 | 35.5056 | 26.04538 | 11.39119 | 0.846777 | 0.431826 | 0.354437 |
| 4 | 5078000 | 5078999 | 17.13163 | 30.38011 | 27.81155 | 12.29228 | 0.519166 | 0.49581 | 0.354451 |
| 5 | 134104400 | 134108999 | 42.65834 | 68.29652 | 120.3085 | 50.69859 | -0.47061 | 2.017325 | 0.354454 |
| 4 | 86596400 | 86597199 | 8.595196 | 16.08197 | 31.74058 | 13.13586 | -0.5151 | 0.124354 | 0.354458 |
| 15 | 59575000 | 59575999 | 8.595196 | 16.08197 | 31.74058 | 13.13586 | -0.5151 | 0.124354 | 0.354458 |
| 14 | 5664200 | 5666599 | 12.16561 | 21.24057 | 48.36877 | 21.30189 | -0.79371 | 0.70746 | 0.354459 |
| 5 | 131522800 | 131529799 | 64.57013 | 101.53 | 164.0992 | 68.07467 | -0.37114 | 2.532752 | 0.354464 |
| 13 | 95470200 | 95473199 | 28.89227 | 47.24139 | 83.07902 | 35.72407 | -0.43758 | 1.487935 | 0.354483 |
| 5 | 132982200 | 132983799 | 11.35449 | 20.3695 | 39.95267 | 17.48715 | -0.56454 | 0.408006 | 0.354484 |
| 6 | 96643000 | 96645599 | 24.10213 | 42.03913 | 34.15157 | 15.0311 | 0.685652 | 0.755867 | 0.354499 |
| 8 | 21558400 | 21565399 | 50.28149 | 81.39138 | 62.33672 | 26.66537 | 0.747046 | 1.683131 | 0.354505 |
| 8 | 105310800 | 105312999 | 35.88025 | 59.55831 | 52.63686 | 22.81302 | 0.558615 | 1.299713 | 0.354505 |
| X | 67139000 | 67141399 | 23.10166 | 41.45988 | 24.27093 | 10.61975 | 1.170097 | 0.45624 | 0.354536 |
| 12 | 22200200 | 22208599 | 145.4591 | 219.944 | 205.6796 | 85.05963 | 0.373159 | 3.305634 | 0.354539 |
| 14 | 3393600 | 3398999 | 69.37271 | 110.7862 | 84.9284 | 35.66115 | 0.723616 | 2.114969 | 0.354542 |
| 1 | 182566200 | 182567399 | 7.657403 | 14.42051 | 30.79638 | 12.60593 | -0.61525 | 0.070762 | 0.354544 |
| 5 | 147229600 | 147231599 | 19.19273 | 32.1189 | 63.86899 | 27.82818 | -0.60719 | 1.172104 | 0.354552 |
| 7 | 59567600 | 59572399 | 76.85627 | 122.1155 | 105.5195 | 44.01066 | 0.543552 | 2.283656 | 0.354572 |
| 3 | 3793600 | 3794399 | 15.02999 | 27.15836 | 24.33023 | 10.5669 | 0.566842 | 0.358666 | 0.354585 |
| 11 | 68710200 | 68711199 | 6.138494 | 11.79273 | 26.29645 | 10.28601 | -0.61662 | -0.10255 | 0.354589 |
| 10 | 77639000 | 77654999 | 170.3896 | 250.5407 | 383.5821 | 157.9672 | -0.37425 | 3.821817 | 0.354592 |
| 10 | 84480400 | 84481599 | 15.46897 | 28.07051 | 26.25144 | 11.34092 | 0.510216 | 0.342813 | 0.354602 |
| 9 | 115892800 | 115897199 | 24.71401 | 40.87142 | 71.86236 | 31.06579 | -0.43113 | 1.32815 | 0.354604 |
| X | 126656000 | 126657999 | 29.30595 | 50.12312 | 40.39035 | 17.71051 | 0.693274 | 0.987824 | 0.354619 |
| 12 | 50694600 | 50696399 | 14.49039 | 25.15016 | 48.45086 | 21.25878 | -0.55486 | 0.802061 | 0.354632 |
| 11 | 11290400 | 11293999 | 36.30182 | 58.48248 | 99.22875 | 42.20426 | -0.40555 | 1.808812 | 0.354633 |
| 16 | 88587000 | 88589599 | 26.09631 | 44.6891 | 40.89117 | 17.9319 | 0.510678 | 0.960121 | 0.354643 |
| 18 | 35669800 | 35671599 | 17.29706 | 30.59499 | 26.3381 | 11.6488 | 0.605766 | 0.518926 | 0.354645 |
| 14 | 10202200 | 10203999 | 33.4007 | 55.49038 | 52.55134 | 22.78954 | 0.459287 | 1.282789 | 0.35467 |
| 14 | 65662200 | 65664599 | 15.93403 | 27.50934 | 50.57691 | 22.18788 | -0.4895 | 0.85953 | 0.354715 |
| X | 4585200 | 4594799 | 132.7716 | 205.2786 | 141.5297 | 58.78816 | 0.837817 | 2.864471 | 0.354715 |
| 3 | 120114000 | 120122799 | 124.1019 | 190.3644 | 176.5461 | 73.28955 | 0.401816 | 3.009595 | 0.354718 |
| 7 | 72104000 | 72104599 | 6.40753 | 12.38356 | 26.62543 | 10.21157 | -0.53829 | -0.16613 | 0.354732 |
| X | 105237600 | 105238599 | 6.40753 | 12.38356 | 26.62543 | 10.21157 | -0.53829 | -0.16613 | 0.354732 |
| 13 | 66011400 | 66016999 | 155.1754 | 233.7037 | 206.2273 | 85.18382 | 0.451869 | 3.387324 | 0.354738 |
| 17 | 37944800 | 37956799 | 236.8247 | 347.2463 | 311.2086 | 128.1849 | 0.394045 | 3.871001 | 0.354754 |
| 5 | 8238400 | 8244599 | 61.02045 | 97.74569 | 88.02337 | 37.12305 | 0.496542 | 2.012943 | 0.354761 |
| 10 | 35813400 | 35814599 | 11.24571 | 21.59996 | 20.03518 | 7.841361 | 0.635806 | -0.06989 | 0.354767 |
| 8 | 125917400 | 125919199 | 12.88603 | 22.64277 | 41.95186 | 18.57264 | -0.49752 | 0.597837 | 0.354778 |
| 11 | 94955000 | 94957199 | 12.88603 | 22.64277 | 41.95186 | 18.57264 | -0.49752 | 0.597837 | 0.354778 |
| 7 | 85115400 | 85118199 | 54.02973 | 87.2561 | 70.56014 | 30.10768 | 0.665381 | 1.752523 | 0.35478 |
| 8 | 54595000 | 54596799 | 18.7193 | 33.26359 | 25.59518 | 11.29317 | 0.77001 | 0.488927 | 0.354803 |
| 15 | 71694600 | 71694999 | 10.523 | 20.15725 | 20.1219 | 7.936678 | 0.520693 | -0.05128 | 0.354804 |

| | | | | | | | | | |
|---|---|---|---|---|---|---|---|---|---|
| 7 | 15111200 | 15115599 | 97.37692 | 153.8781 | 106.629 | 44.29126 | 0.854653 | 2.440801 | 0.354805 |
| 1 | 183388800 | 183391599 | 18.95081 | 32.02951 | 58.52517 | 25.6083 | -0.48516 | 1.073244 | 0.354807 |
| 18 | 50254200 | 50256599 | 18.41002 | 31.31715 | 56.42518 | 24.73072 | -0.46437 | 1.011529 | 0.35481 |
| 10 | 37426600 | 37427799 | 21.78426 | 38.13438 | 33.7325 | 14.88635 | 0.563633 | 0.690948 | 0.354818 |
| X | 89451200 | 89452799 | 21.78426 | 38.13438 | 33.7325 | 14.88635 | 0.563633 | 0.690948 | 0.354818 |
| 6 | 34129800 | 34132599 | 13.65087 | 23.92918 | 44.0679 | 19.4699 | -0.48906 | 0.643061 | 0.354819 |
| 6 | 39921400 | 39922799 | 11.76817 | 20.77734 | 39.87454 | 17.64216 | -0.54536 | 0.532838 | 0.354825 |
| 12 | 84084200 | 84090599 | 54.87737 | 86.61229 | 162.4308 | 67.42807 | -0.58115 | 2.45045 | 0.354837 |
| 3 | 138388400 | 138392999 | 75.64022 | 117.5152 | 187.9408 | 78.03715 | -0.36945 | 2.757033 | 0.354852 |
| 3 | 11752000 | 11753399 | 18.79285 | 33.10114 | 30.79778 | 13.63269 | 0.492178 | 0.583743 | 0.354865 |
| 1 | 112646800 | 112647399 | 8.435404 | 16.48667 | 13.90927 | 4.805243 | 0.933729 | -0.54314 | 0.354871 |
| 5 | 5484600 | 5485999 | 8.094728 | 15.24272 | 34.88018 | 14.27974 | -0.71492 | 0.083197 | 0.354874 |
| 8 | 31640600 | 31641399 | 5.966143 | 11.54855 | 25.49285 | 9.706015 | -0.56505 | -0.19688 | 0.354875 |
| 7 | 30394200 | 30395799 | 7.816179 | 14.27488 | 33.50864 | 14.36031 | -0.80024 | 0.268776 | 0.354877 |
| 6 | 135229200 | 135239199 | 96.29397 | 147.5612 | 230.3808 | 95.62481 | -0.35003 | 3.025302 | 0.354882 |
| 9 | 20549600 | 20550799 | 7.551691 | 14.13239 | 30.09909 | 12.45226 | -0.62325 | 0.110654 | 0.35489 |
| 15 | 101421000 | 101422399 | 7.551691 | 14.13239 | 30.09909 | 12.45226 | -0.62325 | 0.110654 | 0.35489 |
| 1 | 165461600 | 165464199 | 12.43485 | 21.73398 | 45.72956 | 20.15424 | -0.68029 | 0.686173 | 0.35489 |
| 16 | 85571000 | 85571599 | 11.1724 | 21.4653 | 19.44683 | 7.604829 | 0.670335 | -0.0756 | 0.354911 |
| 3 | 148521800 | 148523799 | 20.27002 | 33.5865 | 73.56595 | 31.84444 | -0.74673 | 1.296428 | 0.354914 |
| 1 | 82488200 | 82489199 | 13.79772 | 24.16798 | 45.13302 | 19.92752 | -0.50918 | 0.657302 | 0.354922 |
| 5 | 99866200 | 99868199 | 12.10089 | 21.45309 | 40.34929 | 17.83749 | -0.51402 | 0.491848 | 0.354928 |
| 4 | 147326400 | 147329999 | 43.85593 | 71.92569 | 56.84811 | 24.47449 | 0.715318 | 1.463732 | 0.354935 |
| 2 | 3054800 | 3056199 | 33.85268 | 55.98487 | 53.59305 | 23.16809 | 0.443417 | 1.338566 | 0.354948 |
| 8 | 34621600 | 34622399 | 5.083694 | 9.877439 | 23.37781 | 8.7173 | -0.63532 | -0.28515 | 0.354952 |
| 15 | 21656400 | 21656799 | 17.1623 | 32.05991 | 20.06536 | 8.326649 | 1.125913 | 0.177685 | 0.354963 |
| 6 | 131520400 | 131524399 | 9.703997 | 18.9585 | 11.98638 | 4.181238 | 1.332285 | -0.49774 | 0.354965 |
| 6 | 89258800 | 89260799 | 22.46035 | 37.24314 | 68.26301 | 29.56395 | -0.4905 | 1.289104 | 0.354965 |
| 18 | 63784200 | 63793399 | 96.61724 | 146.73 | 254.7485 | 105.4943 | -0.5135 | 3.209141 | 0.35498 |
| 8 | 27563400 | 27564799 | 9.675653 | 17.39231 | 36.86874 | 16.11027 | -0.67384 | 0.378432 | 0.35499 |
| X | 26151800 | 26154399 | 24.8503 | 44.02111 | 25.6376 | 11.33558 | 1.167198 | 0.551005 | 0.355015 |
| 17 | 93332200 | 93333799 | 21.06334 | 37.10871 | 30.28765 | 13.40627 | 0.680034 | 0.5916 | 0.355015 |
| X | 20896000 | 20903199 | 169.2101 | 256.7154 | 191.2721 | 79.18401 | 0.704789 | 3.245359 | 0.35506 |
| 7 | 5300600 | 5313199 | 226.2257 | 334.204 | 283.497 | 116.7501 | 0.483528 | 3.748949 | 0.355062 |
| 2 | 144322600 | 144322999 | 9.376157 | 18.20917 | 18.26504 | 6.836145 | 0.584951 | -0.23985 | 0.355064 |
| 1 | 191879200 | 191880599 | 7.680853 | 14.40039 | 30.67897 | 12.65295 | -0.62043 | 0.099962 | 0.355071 |
| 4 | 103030800 | 103032199 | 7.680853 | 14.40039 | 30.67897 | 12.65295 | -0.62043 | 0.099962 | 0.355071 |
| 19 | 3128000 | 3134599 | 100.4257 | 156.1112 | 148.3917 | 61.61025 | 0.380965 | 2.754205 | 0.35508 |
| 12 | 20556600 | 20569199 | 217.4136 | 323.2479 | 237.7412 | 97.94677 | 0.697796 | 3.638152 | 0.355084 |
| 4 | 94733400 | 94733999 | 9.519557 | 17.25629 | 34.93527 | 15.11236 | -0.59773 | 0.327471 | 0.355096 |
| 6 | 145366000 | 145367799 | 14.60834 | 25.41411 | 48.43134 | 21.28139 | -0.53905 | 0.767862 | 0.35511 |
| 17 | 43327200 | 43328799 | 9.848838 | 17.31364 | 42.54485 | 18.83989 | -0.90229 | 0.573764 | 0.355137 |
| 7 | 60974400 | 60976199 | 24.89434 | 43.37537 | 34.11572 | 15.00542 | 0.731957 | 0.773983 | 0.355145 |
| 15 | 96996600 | 96997799 | 12.09184 | 22.85105 | 21.9718 | 8.933248 | 0.535513 | 0.065604 | 0.355146 |
| X | 74297000 | 74298599 | 19.0595 | 35.226 | 21.54768 | 9.087195 | 1.141884 | 0.25854 | 0.355151 |
| 5 | 25412600 | 25414199 | 15.09151 | 28.45683 | 21.66808 | 8.850271 | 0.863899 | 0.107202 | 0.355155 |
| 18 | 64468000 | 64468599 | 8.820251 | 17.10726 | 17.17356 | 6.477959 | 0.573131 | -0.22461 | 0.355166 |

| | | | | | | | | | |
|---|---|---|---|---|---|---|---|---|---|
| 12 | 113516000 | 113522599 | 97.0323 | 152.4177 | 126.1851 | 52.3395 | 0.591659 | 2.566049 | 0.35517 |
| 10 | 62143400 | 62144599 | 15.79879 | 28.53219 | 26.72782 | 11.61602 | 0.502297 | 0.373344 | 0.35517 |
| 3 | 48228000 | 48228999 | 12.46674 | 23.48934 | 22.47543 | 9.192056 | 0.535409 | 0.090574 | 0.355178 |
| 18 | 81078600 | 81081399 | 43.78159 | 70.79958 | 68.26341 | 29.17756 | 0.414358 | 1.695847 | 0.35518 |
| 1 | 191116000 | 191121599 | 60.30183 | 94.52381 | 177.2266 | 73.50779 | -0.58851 | 2.593097 | 0.355182 |
| 1 | 121129200 | 121131599 | 26.95808 | 45.78431 | 43.11988 | 18.89531 | 0.467565 | 1.040193 | 0.355185 |
| 7 | 57377600 | 57378999 | 14.81291 | 25.70151 | 48.45209 | 21.25895 | -0.52384 | 0.8071 | 0.355191 |
| 12 | 102118800 | 102121999 | 15.91091 | 27.43712 | 52.73225 | 23.13258 | -0.55374 | 0.870944 | 0.355193 |
| 3 | 144869000 | 144870399 | 13.88885 | 25.56768 | 24.31379 | 10.30936 | 0.504309 | 0.244917 | 0.355206 |
| 4 | 113912000 | 113912599 | 23.38786 | 40.24047 | 36.93229 | 16.20035 | 0.507569 | 0.903394 | 0.355231 |
| 8 | 108321600 | 108324199 | 28.37796 | 47.84409 | 45.86614 | 20.05414 | 0.44179 | 1.111549 | 0.355237 |
| 14 | 56835800 | 56838599 | 13.96212 | 24.39587 | 46.2473 | 20.37738 | -0.53076 | 0.699564 | 0.355251 |
| X | 26061000 | 26064199 | 42.15713 | 69.23835 | 57.2691 | 24.67063 | 0.650759 | 1.444279 | 0.355256 |
| 12 | 110486800 | 110491399 | 27.66073 | 44.98978 | 82.06266 | 35.21926 | -0.49544 | 1.561349 | 0.355259 |
| 3 | 58321800 | 58324999 | 22.81728 | 36.99669 | 95.81196 | 41.03559 | -1.00374 | 1.618258 | 0.355262 |
| 10 | 128373400 | 128374599 | 15.01576 | 27.175 | 25.23379 | 10.93798 | 0.517145 | 0.351252 | 0.355289 |
| 11 | 66498400 | 66499799 | 13.67972 | 24.03388 | 44.01056 | 19.48164 | -0.48086 | 0.607868 | 0.355311 |
| 3 | 115508200 | 115509199 | 7.461615 | 13.80475 | 31.53208 | 13.27938 | -0.74333 | 0.184271 | 0.355319 |
| 9 | 25392600 | 25394799 | 17.16835 | 29.18974 | 55.81146 | 24.4607 | -0.54973 | 1.011859 | 0.355334 |
| 4 | 119173800 | 119174599 | 4.232975 | 8.244564 | 21.81915 | 7.968488 | -0.7659 | -0.37512 | 0.355337 |
| 10 | 104269400 | 104270799 | 25.40017 | 43.91128 | 38.69321 | 16.97524 | 0.567382 | 0.872318 | 0.355362 |
| 1 | 92718200 | 92719399 | 6.156828 | 11.44384 | 31.58914 | 13.21953 | -1.00728 | 0.146589 | 0.355378 |
| 7 | 66180400 | 66182799 | 15.68569 | 26.88996 | 52.65666 | 23.09338 | -0.58208 | 0.925862 | 0.355397 |
| 9 | 96265000 | 96266999 | 14.4773 | 25.20696 | 46.83297 | 20.58974 | -0.50232 | 0.754798 | 0.355402 |
| 2 | 42807200 | 42811399 | 44.81933 | 72.95141 | 63.91117 | 27.43641 | 0.560513 | 1.573805 | 0.355405 |
| 14 | 85417600 | 85420399 | 15.56029 | 26.9988 | 50.06611 | 21.96714 | -0.50038 | 0.810784 | 0.35541 |
| 4 | 126039400 | 126047199 | 39.05309 | 62.34497 | 121.6381 | 51.05031 | -0.62221 | 2.103524 | 0.355433 |
| 17 | 36525600 | 36529599 | 26.09495 | 42.75433 | 77.72213 | 33.44718 | -0.48355 | 1.449227 | 0.355439 |
| 6 | 112153600 | 112157199 | 43.65311 | 69.87285 | 118.3414 | 49.85288 | -0.41437 | 2.025766 | 0.355443 |
| 5 | 130965800 | 130972399 | 46.35053 | 74.05775 | 122.4351 | 51.44682 | -0.38241 | 2.086492 | 0.355447 |
| 17 | 62246400 | 62246999 | 6.40563 | 12.36466 | 27.74905 | 10.70387 | -0.60802 | -0.14541 | 0.355454 |
| 15 | 94799200 | 94800999 | 18.67358 | 33.1242 | 26.04928 | 11.51054 | 0.73725 | 0.499764 | 0.355461 |
| 7 | 42078800 | 42080799 | 21.86011 | 38.48111 | 30.84375 | 13.65464 | 0.705754 | 0.607167 | 0.355463 |
| 15 | 90184600 | 90186799 | 21.86011 | 38.48111 | 30.84375 | 13.65464 | 0.705754 | 0.607167 | 0.355463 |
| 1 | 143468600 | 143469199 | 4.5152 | 8.815981 | 21.77439 | 7.674327 | -0.62221 | -0.50188 | 0.355465 |
| 18 | 24841600 | 24843799 | 17.90716 | 30.09513 | 60.5306 | 26.44046 | -0.62341 | 1.121333 | 0.355478 |
| 12 | 103605800 | 103608199 | 17.50097 | 29.61425 | 59.60421 | 26.08984 | -0.62409 | 1.056685 | 0.355491 |
| 14 | 41330200 | 41339599 | 88.64922 | 140.2747 | 110.4411 | 45.90593 | 0.672229 | 2.40328 | 0.355509 |
| 18 | 86728000 | 86731799 | 30.97531 | 49.97567 | 95.64138 | 40.75885 | -0.5762 | 1.754218 | 0.355519 |
| 17 | 84234400 | 84237399 | 17.95146 | 30.54866 | 55.85386 | 24.48106 | -0.48528 | 1.006606 | 0.355521 |
| 17 | 68048200 | 68048599 | 125.3076 | 194.7062 | 124.3943 | 51.65016 | 0.953023 | 2.767822 | 0.355526 |
| X | 90669000 | 90670799 | 22.86276 | 40.0989 | 32.1392 | 14.20614 | 0.705336 | 0.666713 | 0.355532 |
| 12 | 82741000 | 82742799 | 13.19047 | 23.17269 | 42.45782 | 18.7932 | -0.48161 | 0.603398 | 0.355534 |
| 11 | 109449800 | 109451999 | 19.15372 | 33.5875 | 31.41552 | 13.88412 | 0.48348 | 0.645554 | 0.355548 |
| 5 | 125772200 | 125786799 | 123.6078 | 186.1562 | 291.9519 | 120.5923 | -0.37724 | 3.38553 | 0.355556 |
| 15 | 94390800 | 94392199 | 10.13662 | 18.51475 | 38.80658 | 16.62852 | -0.63904 | 0.295113 | 0.355557 |
| 9 | 103005000 | 103009399 | 31.06068 | 50.3689 | 86.94607 | 37.24172 | -0.42005 | 1.626481 | 0.35558 |

| | | | | | | | | | |
|---|---|---|---|---|---|---|---|---|---|
| 13 | 89687200 | 89688999 | 10.00063 | 17.89832 | 37.86341 | 16.60579 | -0.67468 | 0.410081 | 0.355593 |
| X | 24423400 | 24424599 | 12.64684 | 24.04484 | 21.05435 | 8.451446 | 0.686174 | 0.030018 | 0.355597 |
| 10 | 37957600 | 37959199 | 18.97048 | 34.55197 | 24.10909 | 10.36736 | 0.934021 | 0.342185 | 0.355602 |
| 6 | 10879400 | 10881399 | 21.99819 | 38.62647 | 31.9698 | 14.14486 | 0.65896 | 0.641754 | 0.355618 |
| 1 | 181076800 | 181079399 | 18.5664 | 31.60321 | 56.43978 | 24.73838 | -0.45154 | 1.00637 | 0.355629 |
| 12 | 52438800 | 52442199 | 18.5664 | 31.60321 | 56.43978 | 24.73838 | -0.45154 | 1.00637 | 0.355629 |
| 9 | 73814000 | 73814599 | 13.94647 | 25.97076 | 23.84854 | 9.94945 | 0.57152 | 0.180243 | 0.355634 |
| 15 | 59570600 | 59572399 | 12.40708 | 21.76673 | 43.03323 | 19.02594 | -0.59078 | 0.621073 | 0.355643 |
| 3 | 30701000 | 30703199 | 12.75127 | 22.27071 | 45.2249 | 19.92158 | -0.62935 | 0.698934 | 0.355644 |
| 6 | 32845800 | 32848599 | 22.93072 | 40.11835 | 33.64743 | 14.84163 | 0.640099 | 0.705868 | 0.355645 |
| 7 | 41528200 | 41533199 | 79.24 | 125.7178 | 108.6782 | 45.26139 | 0.540906 | 2.329188 | 0.355659 |
| X | 25975400 | 25976799 | 18.49333 | 33.11926 | 27.97449 | 12.26211 | 0.64268 | 0.444657 | 0.355661 |
| 2 | 114506800 | 114507999 | 17.20477 | 31.48255 | 24.9342 | 10.65902 | 0.758072 | 0.30519 | 0.355682 |
| 1 | 94336000 | 94337199 | 17.78652 | 31.40146 | 28.69134 | 12.69438 | 0.519232 | 0.546525 | 0.355685 |
| 5 | 84165800 | 84168399 | 21.41799 | 37.52549 | 33.16628 | 14.64712 | 0.56475 | 0.674072 | 0.355688 |
| 10 | 125673600 | 125675199 | 13.04828 | 22.86927 | 43.06113 | 19.02256 | -0.52073 | 0.641894 | 0.355704 |
| 13 | 101546600 | 101548599 | 21.30151 | 37.55401 | 30.29258 | 13.40662 | 0.697354 | 0.586385 | 0.35571 |
| 1 | 23383600 | 23383999 | 7.665913 | 14.9576 | 15.70494 | 5.600547 | 0.582114 | -0.43905 | 0.355711 |
| 10 | 13595000 | 13595399 | 7.665913 | 14.9576 | 15.70494 | 5.600547 | 0.582114 | -0.43905 | 0.355711 |
| 19 | 16694200 | 16694599 | 7.665913 | 14.9576 | 15.70494 | 5.600547 | 0.582114 | -0.43905 | 0.355711 |
| 11 | 118948600 | 118950399 | 15.23736 | 26.08477 | 53.16056 | 23.31174 | -0.6397 | 0.937859 | 0.355716 |
| 9 | 121944600 | 121947399 | 11.76931 | 20.73852 | 40.9087 | 18.10683 | -0.58601 | 0.555614 | 0.355735 |
| 4 | 128725600 | 128727399 | 15.15514 | 27.57744 | 25.79797 | 11.10641 | 0.513032 | 0.324018 | 0.355749 |
| 18 | 26682400 | 26682799 | 3.570659 | 6.984663 | 18.58094 | 6.201059 | -0.65252 | -0.71373 | 0.355752 |
| 7 | 35955000 | 35956999 | 12.34567 | 21.85699 | 42.40102 | 18.74967 | -0.55964 | 0.508358 | 0.355754 |
| 1 | 148442200 | 148444399 | 20.31558 | 36.92093 | 24.05545 | 10.37266 | 1.029989 | 0.365116 | 0.355765 |
| 14 | 88878400 | 88880199 | 20.39181 | 35.62486 | 33.47923 | 14.7501 | 0.476728 | 0.71423 | 0.355771 |
| 1 | 139306800 | 139307599 | 5.827634 | 11.25684 | 25.35288 | 9.74764 | -0.60711 | -0.16537 | 0.355789 |
| 3 | 11198600 | 11199799 | 25.96901 | 44.18715 | 40.63121 | 17.80872 | 0.502455 | 1.019558 | 0.355794 |
| 19 | 53105200 | 53107199 | 22.6489 | 40.21367 | 26.26398 | 11.5962 | 1.004286 | 0.519525 | 0.355801 |
| 7 | 27463800 | 27466199 | 15.71534 | 26.81053 | 57.46792 | 25.1936 | -0.71329 | 0.973791 | 0.355807 |
| 5 | 103385000 | 103388999 | 55.77931 | 89.58277 | 82.00242 | 34.72077 | 0.47742 | 1.927203 | 0.355812 |
| 2 | 31006200 | 31008399 | 16.45123 | 28.25739 | 52.15577 | 22.87695 | -0.49667 | 0.910549 | 0.355816 |
| 16 | 65051600 | 65053599 | 22.17418 | 38.68234 | 35.5087 | 15.62993 | 0.510391 | 0.748155 | 0.355817 |
| 1 | 127612800 | 127614999 | 12.19834 | 21.56363 | 41.36381 | 18.29652 | -0.54426 | 0.522507 | 0.355825 |
| 6 | 43928000 | 43929799 | 22.86313 | 40.01517 | 31.90224 | 14.07632 | 0.712625 | 0.694828 | 0.355826 |
| 6 | 123010600 | 123014199 | 47.60306 | 77.15621 | 67.90462 | 29.07503 | 0.548777 | 1.656387 | 0.35583 |
| 7 | 139925000 | 139925799 | 5.37831 | 10.44133 | 23.90448 | 8.972732 | -0.59716 | -0.25723 | 0.355837 |
| 8 | 78939800 | 78940399 | 5.37831 | 10.44133 | 23.90448 | 8.972732 | -0.59716 | -0.25723 | 0.355837 |
| 5 | 22485600 | 22486599 | 6.73658 | 12.88514 | 27.29782 | 10.81287 | -0.56106 | -0.05348 | 0.355843 |
| 5 | 29400400 | 29401199 | 6.73658 | 12.88514 | 27.29782 | 10.81287 | -0.56106 | -0.05348 | 0.355843 |
| 18 | 88224600 | 88225199 | 9.316943 | 18.09486 | 18.23587 | 6.823133 | 0.578258 | -0.24281 | 0.355844 |
| 8 | 65105000 | 65107399 | 19.09471 | 34.16978 | 27.84147 | 12.21178 | 0.693587 | 0.452535 | 0.355857 |
| 9 | 32654400 | 32656399 | 11.31397 | 19.92021 | 42.46464 | 18.79852 | -0.6979 | 0.56338 | 0.355869 |
| 6 | 69183200 | 69184799 | 15.54521 | 28.74405 | 23.48815 | 9.899355 | 0.727517 | 0.230818 | 0.355881 |
| 7 | 109422400 | 109425599 | 19.9555 | 33.50852 | 61.21117 | 26.72711 | -0.4852 | 1.136912 | 0.355886 |
| X | 146598200 | 146601199 | 19.2807 | 34.49307 | 27.83676 | 12.21256 | 0.707136 | 0.455628 | 0.355886 |

| | | | | | | | | | |
|---|---|---|---|---|---|---|---|---|---|
| 6 | 137685600 | 137686599 | 12.06821 | 23.06868 | 21.09182 | 8.369556 | 0.638504 | -0.013 | 0.355888 |
| 7 | 92250000 | 92255599 | 67.56429 | 107.8238 | 96.00992 | 40.27489 | 0.506892 | 2.136616 | 0.355897 |
| 2 | 50914400 | 50916599 | 17.20842 | 29.14539 | 62.28921 | 27.27255 | -0.71024 | 1.052275 | 0.355902 |
| 16 | 25392200 | 25393199 | 12.52104 | 22.19688 | 44.00494 | 19.45376 | -0.59005 | 0.498334 | 0.355929 |
| 6 | 56958400 | 56959399 | 22.52433 | 39.0907 | 36.317 | 15.93465 | 0.492586 | 0.823485 | 0.355935 |
| 1 | 92230400 | 92232599 | 18.1494 | 30.69981 | 57.95802 | 25.36635 | -0.53202 | 1.062502 | 0.355942 |
| 19 | 41025600 | 41029799 | 24.84108 | 41.14593 | 74.96129 | 32.44049 | -0.48213 | 1.311427 | 0.355942 |
| 1 | 138475600 | 138478799 | 26.36977 | 43.32963 | 77.50044 | 33.39258 | -0.45857 | 1.413353 | 0.355946 |
| 2 | 170065400 | 170067999 | 12.40638 | 21.8482 | 40.90796 | 18.10864 | -0.51148 | 0.567113 | 0.355957 |
| 2 | 167204000 | 167213799 | 60.68025 | 94.95831 | 182.1059 | 75.54454 | -0.62293 | 2.623109 | 0.35597 |
| 3 | 111535600 | 111537399 | 22.267 | 38.80351 | 34.89156 | 15.34815 | 0.539743 | 0.759802 | 0.355979 |
| 14 | 87501800 | 87502599 | 11.58536 | 21.83674 | 20.01151 | 8.17925 | 0.597951 | 0.073439 | 0.355993 |
| 19 | 59243200 | 59245799 | 22.88164 | 39.98605 | 34.22271 | 15.08059 | 0.611037 | 0.724349 | 0.355998 |
| 18 | 37381000 | 37391799 | 156.5688 | 237.9911 | 193.2728 | 80.05805 | 0.582847 | 3.20229 | 0.355999 |
| 9 | 12764000 | 12764999 | 12.14985 | 23.00708 | 22.01984 | 8.914628 | 0.547642 | 0.053195 | 0.356004 |
| 14 | 82408800 | 82409799 | 12.14985 | 23.00708 | 22.01984 | 8.914628 | 0.547642 | 0.053195 | 0.356004 |
| 6 | 132221800 | 132224599 | 24.50988 | 42.8186 | 30.51394 | 13.44355 | 0.873952 | 0.731888 | 0.356015 |
| 2 | 28351800 | 28356399 | 26.70652 | 43.41209 | 83.10174 | 35.65751 | -0.56553 | 1.567419 | 0.35602 |
| 4 | 43769200 | 43771399 | 17.08067 | 29.34413 | 52.70732 | 23.11901 | -0.45757 | 0.911988 | 0.35604 |
| 4 | 112558600 | 112559599 | 20.40124 | 35.71842 | 33.45193 | 14.76431 | 0.481764 | 0.682099 | 0.356047 |
| X | 108005600 | 108006199 | 14.78979 | 27.38089 | 23.3959 | 9.843977 | 0.666071 | 0.221437 | 0.356057 |
| 10 | 76146200 | 76149799 | 26.03823 | 42.43806 | 80.24285 | 34.46408 | -0.54501 | 1.527793 | 0.356061 |
| 4 | 37141000 | 37142799 | 19.16597 | 34.29164 | 27.8998 | 12.23907 | 0.695647 | 0.454791 | 0.356102 |
| 10 | 8253600 | 8256399 | 30.09565 | 51.58821 | 39.65699 | 17.39205 | 0.761319 | 0.968333 | 0.356129 |
| 18 | 58820400 | 58822399 | 25.36594 | 43.77146 | 39.24244 | 17.21523 | 0.542011 | 0.888632 | 0.356139 |
| 10 | 86703000 | 86705199 | 14.80179 | 25.84475 | 46.78858 | 20.60673 | -0.46466 | 0.716736 | 0.356149 |
| 7 | 17272000 | 17275999 | 71.14177 | 112.7253 | 108.1741 | 44.99469 | 0.388671 | 2.358011 | 0.356152 |
| 13 | 59970400 | 59972599 | 11.12864 | 19.71831 | 38.8255 | 17.1658 | -0.5801 | 0.487015 | 0.356153 |
| 4 | 73672600 | 73675599 | 27.65135 | 48.04055 | 34.61837 | 15.19897 | 0.857439 | 0.828527 | 0.356203 |
| 1 | 102007000 | 102007799 | 13.39678 | 25.26344 | 22.35947 | 9.131303 | 0.648605 | 0.092806 | 0.356216 |
| 3 | 64670400 | 64672199 | 33.48796 | 55.71592 | 51.97292 | 22.56123 | 0.481223 | 1.267468 | 0.356223 |
| 17 | 88532600 | 88534199 | 20.56918 | 36.66976 | 27.64774 | 12.16543 | 0.801774 | 0.48464 | 0.35623 |
| 4 | 46521000 | 46524399 | 34.58391 | 55.59846 | 101.6653 | 43.10966 | -0.51697 | 1.87467 | 0.35624 |
| 16 | 15348600 | 15349799 | 36.62444 | 60.4673 | 53.63873 | 23.16101 | 0.552861 | 1.366865 | 0.356247 |
| 5 | 4451400 | 4456599 | 44.45708 | 72.20614 | 59.86257 | 25.67069 | 0.637434 | 1.603404 | 0.356294 |
| 1 | 85607600 | 85610399 | 18.07232 | 30.82984 | 55.36713 | 24.27252 | -0.45905 | 0.983121 | 0.356301 |
| 10 | 64870400 | 64870999 | 16.5798 | 29.96931 | 26.2508 | 11.39602 | 0.599814 | 0.374277 | 0.35631 |
| 3 | 121830200 | 121830999 | 12.89582 | 24.16905 | 22.8938 | 9.44779 | 0.539388 | 0.128025 | 0.356314 |
| 10 | 43838000 | 43838799 | 12.89582 | 24.16905 | 22.8938 | 9.44779 | 0.539388 | 0.128025 | 0.356314 |
| 11 | 22940400 | 22941599 | 6.931431 | 13.10341 | 28.65969 | 11.64401 | -0.63898 | 0.036936 | 0.356319 |
| 8 | 113230800 | 113235199 | 41.5066 | 67.672 | 62.73858 | 26.94939 | 0.480676 | 1.54027 | 0.356342 |
| 11 | 68558000 | 68559799 | 23.91083 | 41.50836 | 37.6533 | 16.52304 | 0.526809 | 0.826824 | 0.35635 |
| 2 | 16790800 | 16794599 | 38.32592 | 64.03124 | 47.01269 | 20.46222 | 0.825406 | 1.234354 | 0.35635 |
| X | 150798800 | 150800199 | 19.66791 | 35.31012 | 26.45314 | 11.57107 | 0.816992 | 0.436114 | 0.356354 |
| 17 | 65766800 | 65773999 | 73.70376 | 116.9214 | 100.0135 | 41.64815 | 0.555914 | 2.324189 | 0.356354 |
| 7 | 111784200 | 111790399 | 46.55163 | 74.07744 | 128.6738 | 53.76263 | -0.45996 | 2.216605 | 0.356356 |
| X | 119922200 | 119923399 | 18.60304 | 33.07553 | 28.18713 | 12.43093 | 0.624174 | 0.491523 | 0.356369 |

| | | | | | | | | | |
|---|---|---|---|---|---|---|---|---|---|
| 7 | 32796600 | 32802199 | 138.3848 | 210.219 | 196.8047 | 81.50576 | 0.377131 | 3.211097 | 0.356414 |
| 13 | 30465600 | 30471399 | 66.04997 | 105.456 | 95.35176 | 40.02506 | 0.485539 | 2.120598 | 0.356421 |
| 3 | 144758200 | 144759399 | 17.47715 | 30.89431 | 28.215 | 12.48046 | 0.520595 | 0.530244 | 0.356431 |
| 1 | 4953000 | 4955799 | 29.38161 | 50.5884 | 37.97192 | 16.65674 | 0.796638 | 0.924217 | 0.356443 |
| 7 | 19003800 | 19006799 | 24.35626 | 39.95534 | 75.01364 | 32.29861 | -0.52905 | 1.430525 | 0.356445 |
| 5 | 105870600 | 105872199 | 14.4273 | 25.19777 | 47.31449 | 20.84193 | -0.51723 | 0.70935 | 0.356456 |
| 13 | 120057000 | 120068199 | 2275.506 | 2675.268 | 4105.134 | 1740.826 | -0.70201 | 7.343439 | 0.356461 |
| 7 | 18477200 | 18479199 | 26.67436 | 46.05965 | 38.68992 | 16.97319 | 0.635871 | 0.889448 | 0.356467 |
| 1 | 35887000 | 35889999 | 19.67532 | 32.85961 | 68.40752 | 29.76743 | -0.67325 | 1.197845 | 0.356485 |
| 15 | 67493800 | 67494799 | 11.49094 | 22.41248 | 14.06527 | 5.042757 | 1.308573 | -0.37392 | 0.356492 |
| 15 | 41022000 | 41023199 | 13.1074 | 24.44523 | 23.40965 | 9.741348 | 0.514327 | 0.162447 | 0.356498 |
| 11 | 66069600 | 66070999 | 18.85527 | 34.00263 | 23.77977 | 10.35359 | 0.919587 | 0.396854 | 0.3565 |
| 7 | 139467400 | 139469199 | 10.28418 | 18.50746 | 35.83659 | 15.64369 | -0.54291 | 0.374805 | 0.356503 |
| 13 | 4386000 | 4389399 | 53.575 | 86.04288 | 82.74666 | 35.03606 | 0.406323 | 1.921544 | 0.35651 |
| 1 | 140239600 | 140241599 | 21.91137 | 38.57717 | 30.21973 | 13.37798 | 0.738624 | 0.603793 | 0.356536 |
| 8 | 12789400 | 12799999 | 94.52208 | 144.8132 | 227.4581 | 94.40725 | -0.35898 | 3.028649 | 0.356547 |
| X | 74340200 | 74341999 | 29.49686 | 50.48549 | 40.82228 | 17.90093 | 0.688396 | 0.98119 | 0.356564 |
| 14 | 78891600 | 78903399 | 138.204 | 206.6103 | 317.4694 | 130.9024 | -0.35727 | 3.53045 | 0.356575 |
| 16 | 71370400 | 71371199 | 11.15111 | 21.37871 | 20.60337 | 8.106645 | 0.574205 | -0.05169 | 0.356576 |
| 3 | 53558800 | 53565799 | 133.5566 | 197.5862 | 379.8691 | 156.4406 | -0.69427 | 3.717622 | 0.356584 |
| 15 | 82909200 | 82913199 | 19.40733 | 32.35232 | 64.7981 | 28.164 | -0.61742 | 1.21719 | 0.356597 |
| 12 | 51502000 | 51504199 | 19.93273 | 35.32497 | 28.68965 | 12.68395 | 0.690235 | 0.519082 | 0.356599 |
| X | 53197600 | 53203999 | 98.29111 | 155.357 | 106.187 | 44.11261 | 0.874686 | 2.438068 | 0.356603 |
| 1 | 61410400 | 61411199 | 13.5333 | 25.39502 | 22.82677 | 9.40052 | 0.616795 | 0.125644 | 0.356636 |
| 16 | 26383200 | 26384799 | 15.18779 | 27.6972 | 25.66592 | 11.01945 | 0.528867 | 0.310134 | 0.356645 |
| 7 | 20519800 | 20529599 | 140.5755 | 215.2963 | 173.4101 | 71.96876 | 0.603325 | 3.046313 | 0.356658 |
| 12 | 108261800 | 108265799 | 29.75065 | 48.29056 | 85.80951 | 36.78075 | -0.46046 | 1.603353 | 0.356667 |
| 8 | 73796000 | 73801199 | 60.82793 | 97.55466 | 85.10803 | 35.95929 | 0.544277 | 1.976456 | 0.356683 |
| 5 | 120403000 | 120404799 | 8.865298 | 16.16806 | 33.44501 | 14.35839 | -0.6215 | 0.282481 | 0.356716 |
| 16 | 47860600 | 47863199 | 30.16991 | 51.6737 | 39.63052 | 17.37931 | 0.764443 | 0.974865 | 0.356726 |
| 1 | 55276000 | 55277999 | 13.53971 | 23.57381 | 46.83516 | 20.59033 | -0.59853 | 0.750093 | 0.356746 |
| 7 | 11962400 | 11974399 | 201.9971 | 301.9095 | 241.8125 | 99.70232 | 0.581768 | 3.542682 | 0.356747 |
| 5 | 143803200 | 143803999 | 5.853016 | 11.26831 | 26.96419 | 10.48279 | -0.71 | -0.12738 | 0.356767 |
| 1 | 165828800 | 165831199 | 14.93314 | 25.54801 | 53.68008 | 23.53838 | -0.6837 | 0.942587 | 0.356782 |
| 3 | 135640200 | 135642399 | 18.59246 | 31.56869 | 56.94076 | 24.94979 | -0.46625 | 1.030018 | 0.356802 |
| 3 | 136375000 | 136377999 | 26.08558 | 42.37634 | 84.016 | 36.03384 | -0.61683 | 1.57917 | 0.356803 |
| 1 | 78473600 | 78474199 | 11.02162 | 20.96514 | 20.59207 | 8.257336 | 0.522693 | 0.008316 | 0.356803 |
| 7 | 5934200 | 5945599 | 164.7794 | 249.9531 | 196.7829 | 81.46146 | 0.625169 | 3.247093 | 0.356823 |
| 8 | 61859800 | 61860599 | 7.165928 | 13.69371 | 29.01802 | 11.52353 | -0.56561 | -0.03868 | 0.356845 |
| 8 | 67603800 | 67605199 | 14.28605 | 27.16712 | 20.11649 | 8.07138 | 0.925675 | 0.038821 | 0.356872 |
| 11 | 47325000 | 47326799 | 21.24969 | 37.2214 | 32.05476 | 14.15328 | 0.601857 | 0.67368 | 0.356892 |
| 13 | 103646000 | 103646799 | 11.86587 | 22.77013 | 20.59779 | 8.082921 | 0.668559 | -0.05223 | 0.356892 |
| 13 | 3394800 | 3396399 | 20.94445 | 37.31829 | 27.63737 | 12.1665 | 0.827059 | 0.490731 | 0.356895 |
| 1 | 150997800 | 150999799 | 25.66164 | 44.11947 | 35.75524 | 15.68368 | 0.686085 | 0.918329 | 0.35691 |
| 2 | 129773800 | 129776399 | 17.03526 | 30.18245 | 27.74331 | 12.26619 | 0.512634 | 0.502253 | 0.356913 |
| 15 | 33063800 | 33067599 | 80.63223 | 127.143 | 123.6596 | 51.31753 | 0.363768 | 2.480258 | 0.356917 |
| 14 | 122712800 | 122715199 | 24.79879 | 40.69434 | 77.11977 | 33.20995 | -0.54247 | 1.425845 | 0.356921 |

| | | | | | | | | | |
|---|---|---|---|---|---|---|---|---|---|
| 9 | 56601600 | 56602999 | 12.08171 | 21.34431 | 39.85809 | 17.63267 | -0.50578 | 0.528176 | 0.356926 |
| 6 | 82135200 | 82139199 | 36.487 | 58.69895 | 100.9597 | 42.85384 | -0.42762 | 1.85382 | 0.356929 |
| 9 | 110632000 | 110635799 | 22.40704 | 37.2995 | 66.78129 | 28.9998 | -0.45635 | 1.242048 | 0.356939 |
| 10 | 15345600 | 15346399 | 13.36762 | 23.38213 | 44.14657 | 19.46912 | -0.52469 | 0.677846 | 0.356952 |
| 13 | 102373200 | 102376399 | 26.8063 | 46.58233 | 35.83311 | 15.73441 | 0.763783 | 0.82447 | 0.35697 |
| 12 | 4843600 | 4857999 | 274.6109 | 386.1631 | 590.0394 | 244.4466 | -0.43503 | 4.494932 | 0.356983 |
| X | 32229200 | 32233999 | 58.77015 | 95.18782 | 59.98655 | 25.67114 | 1.028542 | 1.689537 | 0.357011 |
| 12 | 41862000 | 41863599 | 15.95883 | 29.18813 | 25.66431 | 10.97855 | 0.60768 | 0.300712 | 0.357012 |
| 16 | 95631800 | 95633999 | 10.02083 | 17.90782 | 36.85936 | 16.18571 | -0.63637 | 0.419326 | 0.357013 |
| 14 | 95292600 | 95292999 | 9.282223 | 17.95272 | 18.17915 | 6.947968 | 0.542833 | -0.17234 | 0.357025 |
| 5 | 111707800 | 111710999 | 27.74437 | 44.89118 | 88.59299 | 37.885 | -0.61565 | 1.667179 | 0.35703 |
| 9 | 62080000 | 62081999 | 21.02738 | 35.37245 | 63.04273 | 27.54569 | -0.44983 | 1.124079 | 0.35703 |
| 3 | 144274600 | 144276799 | 13.13838 | 22.74112 | 50.01038 | 21.94106 | -0.74571 | 0.822633 | 0.357034 |
| 18 | 53550000 | 53551599 | 8.554377 | 15.6761 | 32.98872 | 14.07447 | -0.63961 | 0.252727 | 0.357037 |
| 1 | 21974200 | 21975399 | 14.7449 | 27.16885 | 24.10842 | 10.20933 | 0.604562 | 0.244971 | 0.357052 |
| 6 | 122316200 | 122317599 | 8.325372 | 15.4097 | 32.57984 | 13.71227 | -0.63267 | 0.189185 | 0.357102 |
| 8 | 72973400 | 72976999 | 21.2797 | 35.08002 | 71.5645 | 30.87752 | -0.64551 | 1.359518 | 0.357104 |
| 1 | 176640800 | 176642599 | 14.14511 | 24.25637 | 53.68463 | 23.5448 | -0.75781 | 0.917014 | 0.357107 |
| 3 | 33768800 | 33770199 | 10.91478 | 19.77646 | 40.2242 | 17.41013 | -0.60573 | 0.348547 | 0.357113 |
| 10 | 67136000 | 67137799 | 10.69643 | 18.90361 | 39.40533 | 17.43115 | -0.66324 | 0.514552 | 0.357123 |
| 4 | 107015600 | 107017599 | 10.09284 | 18.32076 | 34.90381 | 15.07272 | -0.50945 | 0.321196 | 0.357134 |
| 13 | 14732000 | 14732999 | 10.09284 | 18.32076 | 34.90381 | 15.07272 | -0.50945 | 0.321196 | 0.357134 |
| 13 | 68374000 | 68381199 | 151.6106 | 230.4108 | 197.9377 | 81.98573 | 0.501582 | 3.205241 | 0.357141 |
| 14 | 42031200 | 42036599 | 32.73621 | 55.14635 | 41.48079 | 18.13218 | 0.790362 | 1.128828 | 0.357149 |
| 1 | 18125800 | 18129799 | 30.40371 | 49.36287 | 85.2826 | 36.5616 | -0.41965 | 1.600212 | 0.357161 |
| 9 | 74760400 | 74761999 | 13.90691 | 24.05707 | 49.4933 | 21.71381 | -0.65019 | 0.83144 | 0.357165 |
| 7 | 5267600 | 5272199 | 67.6274 | 108.2608 | 86.33439 | 36.37607 | 0.670544 | 2.041116 | 0.357173 |
| 7 | 94470600 | 94472199 | 13.28134 | 24.54717 | 22.94064 | 9.674413 | 0.535029 | 0.215463 | 0.357178 |
| 5 | 54428400 | 54430599 | 21.24195 | 37.48289 | 30.35795 | 13.43312 | 0.692126 | 0.577478 | 0.357183 |
| 3 | 135128600 | 135129599 | 4.45288 | 8.62924 | 26.05855 | 9.868413 | -1.00524 | -0.23432 | 0.357205 |
| X | 67144200 | 67148199 | 57.68472 | 93.06025 | 70.56236 | 30.06756 | 0.756346 | 1.790439 | 0.357214 |
| 10 | 82258800 | 82259399 | 14.36308 | 26.41345 | 24.74821 | 10.50828 | 0.524072 | 0.255145 | 0.357225 |
| 17 | 56513200 | 56514799 | 10.20927 | 18.1705 | 37.3897 | 16.47502 | -0.63923 | 0.449318 | 0.357226 |
| 7 | 67888800 | 67891599 | 15.52802 | 26.64665 | 52.12542 | 22.86227 | -0.58029 | 0.915744 | 0.357228 |
| X | 23829400 | 23829999 | 18.28089 | 33.09177 | 26.0342 | 11.28166 | 0.755462 | 0.377868 | 0.357236 |
| 17 | 54993800 | 54995999 | 28.1541 | 48.29425 | 40.24237 | 17.64978 | 0.645534 | 0.953738 | 0.357246 |
| 11 | 32861600 | 32865599 | 22.98437 | 37.93855 | 71.51571 | 30.87525 | -0.53143 | 1.345549 | 0.357248 |
| 15 | 76139800 | 76144599 | 33.65718 | 54.4266 | 94.19242 | 40.25961 | -0.42768 | 1.691253 | 0.357272 |
| 1 | 63896400 | 63898799 | 12.5095 | 21.72265 | 51.58115 | 22.6392 | -0.85485 | 0.791195 | 0.357275 |
| 7 | 41365800 | 41371799 | 84.5366 | 133.9039 | 111.0318 | 46.17534 | 0.598751 | 2.376856 | 0.357303 |
| 5 | 119141400 | 119144999 | 18.48207 | 31.14632 | 60.68236 | 26.52873 | -0.57764 | 1.097512 | 0.357321 |
| 18 | 88495800 | 88496199 | 12.75105 | 23.91684 | 21.7649 | 8.969757 | 0.598006 | 0.117615 | 0.357328 |
| 15 | 86827200 | 86833599 | 52.55751 | 83.47962 | 138.6588 | 57.81313 | -0.39828 | 2.279162 | 0.357341 |
| 12 | 119005000 | 119005999 | 6.493449 | 12.28459 | 29.22389 | 11.85454 | -0.7578 | 0.025771 | 0.357345 |
| 5 | 56222400 | 56226199 | 44.32646 | 72.24042 | 62.39029 | 26.79586 | 0.582295 | 1.552039 | 0.357367 |
| 6 | 48770800 | 48772599 | 12.23097 | 21.64541 | 39.82734 | 17.61297 | -0.48363 | 0.509895 | 0.357369 |
| X | 113239600 | 113240399 | 21.40867 | 37.52813 | 33.25307 | 14.69172 | 0.56122 | 0.667837 | 0.357372 |

| | | | | | | | | | |
|---|---|---|---|---|---|---|---|---|---|
| 16 | 3270400 | 3302999 | 515.4455 | 709.2417 | 559.436 | 232.5594 | 0.486976 | 4.829838 | 0.357391 |
| 3 | 147816400 | 147818199 | 25.55846 | 44.5678 | 34.10586 | 15.00856 | 0.771524 | 0.767817 | 0.357405 |
| 3 | 96268000 | 96271599 | 69.41456 | 110.2389 | 107.3479 | 44.75191 | 0.370734 | 2.289777 | 0.357405 |
| 14 | 91810800 | 91811399 | 14.22936 | 26.52332 | 20.77551 | 8.653149 | 0.801241 | 0.169443 | 0.357407 |
| 12 | 88277400 | 88282199 | 53.97022 | 86.92739 | 77.12135 | 32.79109 | 0.527498 | 1.831838 | 0.357411 |
| 17 | 55298600 | 55301199 | 58.65387 | 93.98845 | 87.96228 | 37.1175 | 0.441719 | 2.000298 | 0.35742 |
| 9 | 21083200 | 21084599 | 7.25546 | 13.61423 | 29.60829 | 12.19409 | -0.64743 | 0.091507 | 0.357424 |
| Y | 10591200 | 10596399 | 22.28422 | 38.97455 | 18.60916 | 8.206097 | 1.447577 | 0.67782 | 0.357449 |
| 5 | 141903400 | 141905599 | 14.80634 | 25.83637 | 46.799 | 20.60313 | -0.46552 | 0.725896 | 0.35745 |
| 1 | 48124000 | 48126199 | 23.48941 | 41.38928 | 29.01294 | 12.84165 | 0.898558 | 0.603115 | 0.357454 |
| 8 | 65443600 | 65445599 | 27.31068 | 46.67083 | 41.9498 | 18.39304 | 0.535907 | 0.985486 | 0.357459 |
| 3 | 7882600 | 7883999 | 27.35353 | 46.56584 | 41.97336 | 18.39771 | 0.531003 | 1.018378 | 0.357464 |
| 9 | 124165800 | 124170199 | 90.52498 | 142.0773 | 132.4751 | 54.94252 | 0.419424 | 2.580262 | 0.357466 |
| 12 | 115275000 | 115276399 | 23.10918 | 40.47342 | 33.1681 | 14.6433 | 0.673444 | 0.691148 | 0.357495 |
| 7 | 62163200 | 62172999 | 148.4634 | 226.2529 | 187.7395 | 77.82608 | 0.554827 | 3.147602 | 0.357505 |
| 17 | 18581000 | 18594399 | 187.5486 | 281.6398 | 228.9709 | 94.50468 | 0.566344 | 3.451427 | 0.357525 |
| 16 | 69012600 | 69014599 | 35.27284 | 59.86718 | 40.12209 | 17.57824 | 0.956632 | 1.075252 | 0.357528 |
| 8 | 72151000 | 72158199 | 59.18943 | 93.05533 | 163.1981 | 67.69952 | -0.48893 | 2.531923 | 0.357531 |
| 10 | 119591800 | 119594599 | 18.05139 | 30.92109 | 54.87175 | 24.06332 | -0.44097 | 0.944932 | 0.357532 |
| 10 | 117214200 | 117214599 | 5.095175 | 9.925155 | 23.77743 | 8.668966 | -0.62387 | -0.36766 | 0.357546 |
| 12 | 8258000 | 8258599 | 5.095175 | 9.925155 | 23.77743 | 8.668966 | -0.62387 | -0.36766 | 0.357546 |
| 7 | 31164800 | 31181199 | 293.3834 | 423.6751 | 357.8225 | 147.6148 | 0.458376 | 4.105728 | 0.357547 |
| 9 | 30572200 | 30577999 | 82.71915 | 130.4872 | 123.0385 | 51.06249 | 0.408877 | 2.47302 | 0.357548 |
| 1 | 103232600 | 103234599 | 30.4098 | 51.05312 | 48.0422 | 20.97449 | 0.468355 | 1.158303 | 0.357567 |
| 15 | 77287600 | 77288799 | 6.775295 | 12.8604 | 29.27948 | 11.79402 | -0.685 | 0.010638 | 0.357577 |
| 11 | 110286200 | 110290399 | 28.28762 | 46.00227 | 82.90195 | 35.57744 | -0.47828 | 1.563929 | 0.357596 |
| 2 | 37004600 | 37006399 | 9.427139 | 16.965 | 60.32909 | 26.3356 | -1.41528 | 0.415145 | 0.357618 |
| 7 | 68548000 | 68549399 | 21.7183 | 38.28297 | 29.66122 | 13.1276 | 0.754998 | 0.588739 | 0.35763 |
| 17 | 31034200 | 31034999 | 10.30048 | 20.0106 | 18.19406 | 6.792206 | 0.728353 | -0.2387 | 0.357641 |
| 10 | 116568200 | 116569399 | 16.86911 | 30.02334 | 28.13656 | 12.39794 | 0.488348 | 0.470828 | 0.357644 |
| 15 | 9353400 | 9358199 | 60.08959 | 96.24836 | 88.84698 | 37.46671 | 0.4608 | 2.014709 | 0.357645 |
| X | 50827800 | 50829999 | 31.88549 | 55.28916 | 32.94274 | 14.45376 | 1.13036 | 0.870567 | 0.35765 |
| 3 | 114272400 | 114273599 | 11.80778 | 22.57818 | 21.06512 | 8.351182 | 0.610511 | -0.01964 | 0.357664 |
| 10 | 65838400 | 65838999 | 5.808023 | 11.28719 | 25.29801 | 9.417108 | -0.55649 | -0.27893 | 0.35767 |
| 6 | 146755400 | 146757199 | 18.21911 | 31.19966 | 55.4161 | 24.30149 | -0.44234 | 0.948905 | 0.357684 |
| 1 | 150068400 | 150070199 | 27.97762 | 47.4627 | 44.17099 | 19.34934 | 0.484661 | 1.054901 | 0.357687 |
| 13 | 42804800 | 42805799 | 5.991339 | 11.53416 | 26.38941 | 10.26049 | -0.64582 | -0.12629 | 0.357696 |
| 1 | 105926000 | 105926999 | 7.020443 | 13.44476 | 27.94173 | 11.02856 | -0.52916 | -0.06177 | 0.357714 |
| 8 | 28979000 | 28979799 | 7.020443 | 13.44476 | 27.94173 | 11.02856 | -0.52916 | -0.06177 | 0.357714 |
| 9 | 30897000 | 30899399 | 14.13808 | 24.71711 | 45.18422 | 19.91809 | -0.47848 | 0.691128 | 0.357738 |
| 14 | 67216800 | 67218199 | 14.13808 | 24.71711 | 45.18422 | 19.91809 | -0.47848 | 0.691128 | 0.357738 |
| 14 | 19419400 | 19421199 | 8.479981 | 14.99583 | 40.51025 | 17.9177 | -1.03418 | 0.495985 | 0.357765 |
| 8 | 85094000 | 85096799 | 18.40218 | 31.45235 | 55.93464 | 24.52507 | -0.44455 | 0.968257 | 0.357766 |
| 8 | 46349800 | 46350799 | 12.88039 | 24.47802 | 21.40445 | 8.60004 | 0.686201 | 0.033491 | 0.357784 |
| 9 | 49271200 | 49273199 | 11.72999 | 20.76769 | 40.3287 | 17.83317 | -0.5609 | 0.500935 | 0.357797 |
| 5 | 64718800 | 64721199 | 15.21332 | 27.71191 | 23.46766 | 10.09014 | 0.657179 | 0.314592 | 0.357801 |
| 12 | 90639000 | 90641799 | 27.03408 | 46.74021 | 36.89036 | 16.18435 | 0.725509 | 0.874843 | 0.357801 |

| | | | | | | | | | |
|---|---|---|---|---|---|---|---|---|---|
| X | 67453600 | 67454599 | 14.53822 | 27.82143 | 19.76288 | 7.816925 | 1.003807 | -0.00297 | 0.357803 |
| 17 | 94599800 | 94604599 | 52.13323 | 84.37798 | 66.6496 | 28.50757 | 0.702484 | 1.691104 | 0.357812 |
| 1 | 68642600 | 68643599 | 9.23317 | 18.0179 | 14.22494 | 5.061626 | 0.989827 | -0.43189 | 0.357818 |
| 5 | 104529200 | 104533999 | 29.35268 | 47.85113 | 86.02003 | 36.94848 | -0.47231 | 1.530988 | 0.357867 |
| 13 | 18413800 | 18414599 | 6.430226 | 12.36782 | 26.911 | 10.49065 | -0.57704 | -0.10844 | 0.357867 |
| 2 | 82773600 | 82774399 | 13.51418 | 23.6785 | 43.57321 | 19.24373 | -0.48787 | 0.650057 | 0.357868 |
| 6 | 78240200 | 78242399 | 21.9521 | 38.76307 | 29.10464 | 12.87674 | 0.801041 | 0.571127 | 0.357881 |
| 12 | 47805400 | 47807199 | 16.0434 | 30.29041 | 20.43662 | 8.325479 | 1.040077 | 0.104115 | 0.357889 |
| 13 | 69144600 | 69149199 | 29.11919 | 47.22434 | 88.57444 | 37.94146 | -0.53943 | 1.620374 | 0.35789 |
| 4 | 58238200 | 58239799 | 16.39399 | 30.03434 | 25.18664 | 10.74967 | 0.67809 | 0.294747 | 0.357898 |
| 4 | 153224400 | 153262799 | 522.2889 | 699.5334 | 976.5498 | 407.6415 | -0.37565 | 5.253463 | 0.357912 |
| 2 | 14082400 | 14085399 | 27.66872 | 46.94898 | 44.18205 | 19.35548 | 0.46868 | 1.052156 | 0.357913 |
| 2 | 125033600 | 125035599 | 20.39624 | 35.73047 | 33.39946 | 14.74841 | 0.484464 | 0.672581 | 0.357926 |
| 18 | 27548800 | 27550999 | 17.99032 | 32.51403 | 26.3241 | 11.43 | 0.712094 | 0.382779 | 0.357938 |
| 2 | 53080600 | 53082599 | 14.33903 | 24.49553 | 55.78317 | 24.45767 | -0.79974 | 0.955211 | 0.357938 |
| 10 | 44684000 | 44685599 | 20.96578 | 37.44099 | 26.4533 | 11.62303 | 0.89621 | 0.473875 | 0.357959 |
| 9 | 19767200 | 19769599 | 31.32484 | 53.66095 | 38.4703 | 16.87075 | 0.861374 | 0.976689 | 0.357966 |
| 11 | 20747600 | 20749399 | 18.68384 | 33.10287 | 29.61348 | 13.09308 | 0.551708 | 0.518221 | 0.35797 |
| 12 | 76866800 | 76869799 | 20.2135 | 33.61972 | 66.49148 | 28.85735 | -0.59942 | 1.245504 | 0.35798 |
| 19 | 29649800 | 29653999 | 28.43728 | 46.27586 | 82.43847 | 35.38696 | -0.46101 | 1.554496 | 0.357986 |
| 7 | 31383800 | 31414399 | 659.3447 | 885.451 | 727.1987 | 303.4352 | 0.393415 | 5.21274 | 0.358 |
| 19 | 19151800 | 19153399 | 20.17068 | 35.78195 | 28.70215 | 12.68215 | 0.708826 | 0.513487 | 0.358001 |
| 4 | 113947200 | 113947999 | 19.69714 | 34.63657 | 30.84913 | 13.65993 | 0.55408 | 0.608105 | 0.358016 |
| 3 | 42478600 | 42479999 | 13.36513 | 25.71054 | 18.72696 | 7.29686 | 0.987356 | -0.06532 | 0.358018 |
| 3 | 15313800 | 15314599 | 10.45111 | 20.26682 | 18.02952 | 6.806419 | 0.743931 | -0.20165 | 0.358024 |
| 9 | 123070800 | 123071999 | 14.00683 | 26.26473 | 22.76183 | 9.38518 | 0.667673 | 0.133969 | 0.358031 |
| 8 | 118243000 | 118244999 | 9.82783 | 17.66482 | 36.35334 | 15.88584 | -0.63135 | 0.379331 | 0.358035 |
| 16 | 88761200 | 88766199 | 111.0411 | 170.9063 | 164.4902 | 68.31639 | 0.352044 | 2.938178 | 0.358056 |
| 7 | 10961600 | 11000599 | 920.9139 | 1202.009 | 972.5634 | 407.2573 | 0.373942 | 5.670803 | 0.358059 |
| 1 | 34930400 | 34935399 | 34.56698 | 55.56812 | 102.5368 | 43.47564 | -0.53018 | 1.875328 | 0.35806 |
| X | 37463800 | 37465599 | 19.6379 | 34.92494 | 28.065 | 12.37424 | 0.7087 | 0.496501 | 0.358068 |
| 13 | 50127400 | 50130799 | 21.824 | 36.23894 | 67.07679 | 29.07846 | -0.50436 | 1.271006 | 0.358069 |
| 2 | 148889600 | 148892399 | 45.09806 | 72.92483 | 70.77377 | 30.24862 | 0.404859 | 1.700646 | 0.358079 |
| 3 | 59907800 | 59910199 | 25.3317 | 43.84364 | 38.11276 | 16.72118 | 0.587159 | 0.860436 | 0.358084 |
| 6 | 90044400 | 90046599 | 20.77763 | 36.56624 | 31.56814 | 13.97604 | 0.599128 | 0.605628 | 0.358099 |
| 13 | 18167400 | 18169199 | 10.88365 | 19.50999 | 39.412 | 17.26157 | -0.60747 | 0.407932 | 0.358102 |
| 12 | 84767600 | 84769399 | 16.39941 | 29.36198 | 27.65538 | 12.12479 | 0.486082 | 0.430213 | 0.358107 |
| 10 | 95185200 | 95190199 | 69.15604 | 108.2932 | 172.199 | 71.42996 | -0.35226 | 2.615141 | 0.358126 |
| 7 | 67999600 | 68006599 | 65.04217 | 101.8943 | 167.5399 | 69.49428 | -0.4 | 2.604588 | 0.358137 |
| 5 | 109449200 | 109451999 | 41.13518 | 67.75932 | 55.59337 | 23.97864 | 0.664049 | 1.404922 | 0.358138 |
| X | 150282200 | 150283999 | 24.40643 | 43.56476 | 24.08221 | 10.58543 | 1.247839 | 0.496634 | 0.358184 |
| 11 | 119570200 | 119572599 | 10.75667 | 19.37425 | 36.351 | 15.85134 | -0.49674 | 0.374796 | 0.358205 |
| 7 | 6572400 | 6574999 | 51.53672 | 82.82075 | 79.1232 | 33.53238 | 0.416914 | 1.897415 | 0.358217 |
| 15 | 56319800 | 56320199 | 3.261076 | 6.378044 | 18.36683 | 6.21602 | -0.78194 | -0.6683 | 0.358228 |
| 13 | 60485200 | 60489399 | 39.08096 | 62.78368 | 105.9247 | 44.86976 | -0.40247 | 1.902984 | 0.358234 |
| 6 | 6705400 | 6706599 | 6.850053 | 13.18171 | 30.66737 | 11.93835 | -0.67246 | -0.10399 | 0.358238 |
| 5 | 63389000 | 63390199 | 17.44946 | 30.87444 | 27.87242 | 12.32661 | 0.537927 | 0.52108 | 0.358247 |

| | | | | | | | | | |
|---|---|---|---|---|---|---|---|---|---|
| 2 | 177550200 | 177553199 | 45.98857 | 74.5611 | 62.07207 | 26.58338 | 0.629133 | 1.643355 | 0.358261 |
| 12 | 114254800 | 114257399 | 54.44138 | 87.47616 | 81.37596 | 34.48 | 0.455035 | 1.910099 | 0.358269 |
| Y | 41108200 | 41110799 | 23.00685 | 40.08894 | 20.73402 | 9.119958 | 1.333095 | 0.739972 | 0.358287 |
| 5 | 77325800 | 77327199 | 10.55626 | 19.23736 | 38.20202 | 16.4162 | -0.56461 | 0.308047 | 0.358287 |
| 6 | 18971800 | 18973199 | 27.71903 | 47.35526 | 37.84752 | 16.59401 | 0.704734 | 0.986551 | 0.358295 |
| X | 153013600 | 153014599 | 19.31962 | 34.36486 | 28.93253 | 12.75488 | 0.641768 | 0.493356 | 0.358313 |
| 3 | 108628200 | 108630199 | 9.629475 | 17.45955 | 34.89716 | 15.09155 | -0.57958 | 0.32368 | 0.358316 |
| 11 | 88529600 | 88531399 | 9.629475 | 17.45955 | 34.89716 | 15.09155 | -0.57958 | 0.32368 | 0.358316 |
| 19 | 4495800 | 4497799 | 9.629475 | 17.45955 | 34.89716 | 15.09155 | -0.57958 | 0.32368 | 0.358316 |
| X | 89636200 | 89637799 | 21.51566 | 39.83569 | 19.21341 | 8.083355 | 1.484627 | 0.264669 | 0.358317 |
| 8 | 15690200 | 15693399 | 40.13265 | 64.4512 | 109.8697 | 46.51713 | -0.41797 | 1.918514 | 0.358318 |
| 13 | 37803800 | 37805599 | 9.26912 | 16.5805 | 37.39333 | 16.40681 | -0.76659 | 0.408309 | 0.358324 |
| 2 | 98953800 | 98955399 | 15.15024 | 27.75681 | 25.13322 | 10.72631 | 0.567735 | 0.282496 | 0.35833 |
| 1 | 144935000 | 144936999 | 22.49993 | 39.23745 | 33.72998 | 14.84386 | 0.604327 | 0.748466 | 0.358347 |
| 9 | 8172400 | 8174599 | 42.46155 | 68.90776 | 66.87663 | 28.66095 | 0.409642 | 1.621253 | 0.358349 |
| 10 | 22540200 | 22541999 | 12.50294 | 22.08453 | 43.9927 | 19.46246 | -0.59907 | 0.536641 | 0.358384 |
| X | 109546800 | 109552599 | 74.60086 | 118.8094 | 98.77037 | 41.31515 | 0.602677 | 2.209385 | 0.358393 |
| 1 | 5183600 | 5185199 | 10.94866 | 20.85756 | 17.61421 | 7.040896 | 0.742981 | -0.01115 | 0.358405 |
| 7 | 31685200 | 31694199 | 114.9618 | 177.788 | 157.1245 | 65.26577 | 0.480223 | 2.851198 | 0.358423 |
| 12 | 56838600 | 56839799 | 10.64096 | 18.91628 | 39.85655 | 17.57963 | -0.67483 | 0.460392 | 0.358429 |
| 3 | 11119200 | 11124799 | 108.806 | 167.8673 | 161.5246 | 67.09432 | 0.355039 | 2.892976 | 0.358436 |
| 10 | 42696400 | 42697199 | 4.659268 | 9.048418 | 23.30166 | 8.722911 | -0.76047 | -0.27605 | 0.358438 |
| 9 | 9451600 | 9455199 | 51.89209 | 83.78859 | 72.56476 | 30.95576 | 0.56639 | 1.755169 | 0.358441 |
| 15 | 22108800 | 22109399 | 8.913579 | 17.3266 | 17.77185 | 6.602775 | 0.562848 | -0.27222 | 0.358442 |
| 12 | 115562800 | 115574199 | 159.6088 | 242.9734 | 183.977 | 76.24148 | 0.685515 | 3.173201 | 0.358448 |
| 14 | 94624200 | 94625799 | 21.39734 | 37.30192 | 34.66422 | 15.25173 | 0.492773 | 0.753298 | 0.358448 |
| 3 | 39841800 | 39844799 | 28.81687 | 46.70523 | 88.30612 | 37.80932 | -0.55178 | 1.632656 | 0.358455 |
| 13 | 66610800 | 66633599 | 477.8316 | 658.5858 | 561.6448 | 233.4155 | 0.376849 | 4.803256 | 0.358457 |
| 4 | 57510000 | 57511799 | 7.853757 | 14.75145 | 30.16549 | 12.40279 | -0.55761 | 0.091637 | 0.358459 |
| 17 | 57984400 | 57984999 | 7.529096 | 14.15548 | 29.64505 | 12.16866 | -0.59009 | 0.079062 | 0.358463 |
| 1 | 84641000 | 84641799 | 6.101923 | 11.84647 | 27.70043 | 10.39521 | -0.62851 | -0.24144 | 0.358463 |
| 6 | 59201400 | 59202399 | 22.92407 | 39.89505 | 36.55909 | 16.06812 | 0.512518 | 0.785772 | 0.358463 |
| 8 | 91566800 | 91569799 | 17.32559 | 29.40446 | 56.89284 | 24.92541 | -0.5671 | 1.028497 | 0.35847 |
| 15 | 43904200 | 43906199 | 27.19435 | 46.43071 | 42.51746 | 18.6406 | 0.509022 | 0.994103 | 0.358474 |
| X | 45234600 | 45237199 | 24.4251 | 43.00519 | 29.00192 | 12.83873 | 0.953758 | 0.617402 | 0.358489 |
| X | 67145200 | 67147799 | 42.24748 | 69.51262 | 55.56157 | 23.95342 | 0.701029 | 1.422695 | 0.35849 |
| X | 122006600 | 122008999 | 33.76016 | 57.08365 | 43.58937 | 19.07544 | 0.769045 | 1.106276 | 0.35849 |
| 3 | 128166800 | 128168399 | 9.212895 | 17.12175 | 35.47704 | 14.84947 | -0.59842 | 0.179481 | 0.358497 |
| X | 153107400 | 153111599 | 68.37336 | 109.5101 | 84.06329 | 35.44609 | 0.726304 | 2.02678 | 0.35851 |
| 9 | 18901800 | 18906399 | 122.6213 | 187.5672 | 180.4581 | 74.87247 | 0.345756 | 3.065045 | 0.358511 |
| 5 | 134424000 | 134425399 | 14.21262 | 24.72938 | 47.36886 | 20.81648 | -0.54605 | 0.766103 | 0.358516 |
| 10 | 81494000 | 81496799 | 66.56563 | 106.6526 | 84.19098 | 35.51913 | 0.686622 | 2.012407 | 0.358522 |
| 12 | 42197600 | 42198799 | 9.907073 | 18.14352 | 34.51828 | 14.73967 | -0.49671 | 0.267746 | 0.358527 |
| X | 15811800 | 15814199 | 40.19335 | 66.40444 | 53.86198 | 23.26826 | 0.681634 | 1.367304 | 0.358546 |
| 5 | 121172600 | 121175799 | 26.28558 | 42.99262 | 78.09175 | 33.58335 | -0.48371 | 1.473483 | 0.358555 |
| 9 | 46236200 | 46237199 | 5.385566 | 10.43143 | 24.34055 | 9.239556 | -0.63969 | -0.21698 | 0.358561 |
| 4 | 51064400 | 51068599 | 49.43517 | 79.82722 | 73.34798 | 31.29277 | 0.481341 | 1.749671 | 0.358564 |

| | | | | | | | | | |
|---|---|---|---|---|---|---|---|---|---|
| 6 | 130064200 | 130065399 | 15.79558 | 30.1972 | 18.41046 | 7.303068 | 1.218153 | 0.014391 | 0.358574 |
| 4 | 64925000 | 64926399 | 21.08731 | 37.05786 | 30.27123 | 13.40275 | 0.6775 | 0.618349 | 0.358576 |
| 5 | 130015600 | 130020199 | 23.30951 | 38.50504 | 70.36781 | 30.39592 | -0.4867 | 1.337921 | 0.35859 |
| 10 | 8510200 | 8513599 | 71.08782 | 112.6349 | 108.3869 | 45.08222 | 0.384627 | 2.358449 | 0.358608 |
| 3 | 86457600 | 86458199 | 4.557088 | 8.874514 | 23.64424 | 8.651927 | -0.78003 | -0.3622 | 0.358609 |
| 1 | 134178200 | 134179599 | 18.0275 | 31.78238 | 29.77272 | 13.17651 | 0.482789 | 0.56604 | 0.358609 |
| 19 | 56618200 | 56620999 | 20.27216 | 33.98945 | 62.33403 | 27.19177 | -0.49088 | 1.154206 | 0.358632 |
| 8 | 22482400 | 22485999 | 24.67116 | 40.4142 | 78.07074 | 33.59461 | -0.57104 | 1.448202 | 0.358642 |
| 17 | 19300400 | 19301599 | 17.99719 | 33.21497 | 23.38148 | 9.879812 | 0.939001 | 0.262427 | 0.358645 |
| 2 | 168963200 | 168976999 | 184.4447 | 268.7065 | 420.1498 | 173.1475 | -0.41743 | 3.967133 | 0.358651 |
| 18 | 54537600 | 54538799 | 6.419769 | 12.35052 | 26.96653 | 10.50542 | -0.58192 | -0.11404 | 0.358652 |
| 8 | 121325800 | 121328799 | 29.25362 | 47.64495 | 82.5554 | 35.44818 | -0.42022 | 1.545218 | 0.358653 |
| 16 | 78172800 | 78174999 | 14.12632 | 24.27116 | 52.10424 | 22.85455 | -0.71343 | 0.897933 | 0.358655 |
| 9 | 14695400 | 14696399 | 6.154309 | 11.7817 | 26.75207 | 10.5702 | -0.65658 | -0.0693 | 0.358669 |
| 11 | 18461600 | 18462199 | 8.660062 | 16.05461 | 32.07321 | 13.46996 | -0.54875 | 0.185074 | 0.358678 |
| 14 | 43039800 | 43042999 | 24.34666 | 42.4543 | 34.71695 | 15.27738 | 0.676223 | 0.762281 | 0.358684 |
| 9 | 36878400 | 36881199 | 13.06822 | 22.84865 | 44.1695 | 19.47218 | -0.55849 | 0.683854 | 0.358688 |
| 9 | 36287000 | 36294199 | 97.69702 | 152.6219 | 138.5744 | 57.4962 | 0.452548 | 2.667758 | 0.358703 |
| 18 | 37939000 | 37942199 | 24.79316 | 40.33062 | 88.00619 | 37.77174 | -0.7532 | 1.556899 | 0.358713 |
| 10 | 78316400 | 78320199 | 23.79249 | 39.2161 | 72.67544 | 31.35472 | -0.50768 | 1.365967 | 0.358719 |
| 10 | 86663000 | 86665999 | 25.71859 | 42.32028 | 75.15507 | 32.40168 | -0.4474 | 1.392851 | 0.358723 |
| 6 | 24196800 | 24199399 | 12.84067 | 22.70417 | 46.15162 | 20.41341 | -0.62758 | 0.531928 | 0.358752 |
| 17 | 19270200 | 19271199 | 12.85676 | 24.89007 | 17.83544 | 6.786139 | 1.042925 | -0.14828 | 0.35876 |
| 12 | 20360200 | 20368399 | 281.717 | 405.6764 | 366.2444 | 151.1457 | 0.3582 | 4.146541 | 0.358763 |
| 12 | 91106400 | 91109799 | 39.44606 | 63.22815 | 109.5586 | 46.25141 | -0.44473 | 1.974992 | 0.358766 |
| 5 | 18485200 | 18488799 | 31.75704 | 53.37516 | 47.47877 | 20.7397 | 0.549283 | 1.151818 | 0.358768 |
| 2 | 155314600 | 155315999 | 7.270094 | 13.58168 | 30.07514 | 12.47917 | -0.68219 | 0.120251 | 0.358769 |
| 6 | 101084000 | 101084799 | 7.270094 | 13.58168 | 30.07514 | 12.47917 | -0.68219 | 0.120251 | 0.358769 |
| 14 | 13938000 | 13938999 | 10.12983 | 18.27221 | 35.36129 | 15.39129 | -0.53925 | 0.3598 | 0.358793 |
| 15 | 46249600 | 46251599 | 18.2138 | 33.52628 | 23.25274 | 9.8607 | 0.956427 | 0.277377 | 0.358796 |
| 2 | 8147200 | 8147999 | 13.96798 | 26.69588 | 20.32321 | 8.06824 | 0.89969 | 0.004088 | 0.358799 |
| 12 | 102493600 | 102496599 | 22.69806 | 37.88353 | 66.91904 | 29.10792 | -0.43705 | 1.214412 | 0.358801 |
| 7 | 62908400 | 62910399 | 24.50875 | 42.36711 | 38.722 | 16.98784 | 0.514819 | 0.86831 | 0.358815 |
| 6 | 70330000 | 70330999 | 6.733132 | 12.92148 | 27.41531 | 10.75612 | -0.54999 | -0.08353 | 0.358828 |
| 2 | 4086600 | 4088199 | 12.03078 | 21.41203 | 39.83277 | 17.55178 | -0.49523 | 0.463353 | 0.358831 |
| 3 | 13535800 | 13538799 | 26.00053 | 42.45027 | 78.91495 | 33.91522 | -0.51886 | 1.499913 | 0.358852 |
| X | 77118400 | 77121199 | 30.80831 | 53.69539 | 30.01016 | 13.20206 | 1.222688 | 0.796653 | 0.358853 |
| 18 | 37461200 | 37464199 | 41.91638 | 68.66323 | 59.50976 | 25.60754 | 0.582058 | 1.475659 | 0.358868 |
| 16 | 60966800 | 60974399 | 87.95473 | 139.2868 | 108.276 | 45.02415 | 0.691474 | 2.383713 | 0.358869 |
| 6 | 29115800 | 29117799 | 13.18023 | 23.12673 | 43.00854 | 19.01753 | -0.50302 | 0.623115 | 0.35887 |
| 8 | 21615200 | 21628999 | 223.9191 | 331.6058 | 273.7321 | 112.7317 | 0.526115 | 3.709094 | 0.358874 |
| 19 | 39546600 | 39547799 | 21.94805 | 38.30686 | 30.38846 | 13.38032 | 0.719892 | 0.730823 | 0.358875 |
| 2 | 39828400 | 39829599 | 16.20618 | 29.84567 | 24.56108 | 10.40849 | 0.712079 | 0.261703 | 0.358889 |
| X | 66385000 | 66387399 | 18.79071 | 33.65802 | 22.72691 | 9.960386 | 0.964033 | 0.435612 | 0.358891 |
| 1 | 41511600 | 41513199 | 17.95283 | 32.31257 | 26.8004 | 11.69706 | 0.672677 | 0.406318 | 0.358897 |
| 18 | 11323600 | 11326199 | 32.23949 | 54.63454 | 43.54501 | 19.06897 | 0.707472 | 1.080164 | 0.358913 |
| 17 | 64863200 | 64865799 | 14.12312 | 24.73309 | 45.13647 | 19.92573 | -0.47609 | 0.662769 | 0.358922 |

| | | | | | | | | | |
|---|---|---|---|---|---|---|---|---|---|
| 8 | 63751800 | 63754799 | 34.24661 | 57.96023 | 40.119 | 17.56172 | 0.909932 | 1.092114 | 0.358948 |
| 14 | 32614600 | 32617599 | 16.01105 | 27.00852 | 61.6146 | 26.94427 | -0.80423 | 1.081307 | 0.358949 |
| 15 | 77979200 | 77980999 | 12.29046 | 21.52419 | 43.1165 | 19.0344 | -0.60941 | 0.651502 | 0.358953 |
| 7 | 32257600 | 32261999 | 61.95494 | 99.26309 | 79.36755 | 33.48303 | 0.668138 | 2.006611 | 0.358959 |
| 7 | 6371600 | 6372799 | 7.658992 | 14.52277 | 29.31268 | 11.83321 | -0.51652 | 0.02273 | 0.358966 |
| 7 | 57794400 | 57797999 | 51.14425 | 82.17461 | 79.91254 | 33.8577 | 0.391036 | 1.904364 | 0.358972 |
| 11 | 21942800 | 21944599 | 24.98806 | 40.94875 | 76.33644 | 32.85356 | -0.51957 | 1.444039 | 0.358976 |
| 14 | 57361600 | 57363199 | 10.08323 | 18.41569 | 35.00108 | 14.99946 | -0.49883 | 0.28621 | 0.358985 |
| 11 | 78524800 | 78525999 | 12.25745 | 23.20799 | 21.97496 | 8.898752 | 0.562533 | 0.054303 | 0.359025 |
| 12 | 93060600 | 93061599 | 7.633175 | 14.56452 | 30.73209 | 12.24335 | -0.56364 | -0.01853 | 0.35905 |
| 18 | 77441800 | 77442199 | 4.104017 | 8.014294 | 20.5012 | 7.20771 | -0.66815 | -0.52028 | 0.359072 |
| 10 | 21168600 | 21169799 | 15.29464 | 28.14733 | 24.60093 | 10.43526 | 0.624678 | 0.257672 | 0.359076 |
| 9 | 77226200 | 77226999 | 6.79266 | 12.81275 | 29.14806 | 11.89786 | -0.70091 | 0.053567 | 0.359083 |
| 6 | 56993400 | 56995999 | 40.91847 | 67.10767 | 53.78864 | 23.15725 | 0.695164 | 1.455908 | 0.359085 |
| 10 | 56494800 | 56496799 | 17.24187 | 31.77933 | 23.93974 | 10.13374 | 0.839954 | 0.264257 | 0.359098 |
| 1 | 188739200 | 188740399 | 9.874253 | 17.708 | 36.40283 | 15.93796 | -0.63124 | 0.397824 | 0.3591 |
| 7 | 137962800 | 137963799 | 5.987042 | 11.52653 | 25.80656 | 10.03228 | -0.61404 | -0.12645 | 0.35911 |
| 8 | 94650400 | 94654599 | 35.3715 | 56.84667 | 103.2464 | 43.76007 | -0.50785 | 1.884788 | 0.359122 |
| 3 | 64273400 | 64274799 | 15.48214 | 26.69229 | 50.56214 | 22.18046 | -0.53274 | 0.870482 | 0.359127 |
| 1 | 166258200 | 166259999 | 10.46714 | 18.76004 | 36.34133 | 15.91905 | -0.54693 | 0.403517 | 0.359129 |
| 5 | 24741200 | 24743399 | 15.18007 | 26.10493 | 52.14403 | 22.87263 | -0.6098 | 0.89325 | 0.359133 |
| 5 | 72176600 | 72178399 | 10.77175 | 19.43212 | 36.90307 | 16.06028 | -0.51217 | 0.367529 | 0.359152 |
| 5 | 49935000 | 49935599 | 8.569247 | 16.68394 | 17.25249 | 6.327542 | 0.567839 | -0.32372 | 0.359163 |
| 6 | 127543400 | 127544599 | 14.14192 | 24.66073 | 46.28286 | 20.36747 | -0.51661 | 0.729752 | 0.359165 |
| X | 32785600 | 32801199 | 293.6652 | 424.471 | 348.3819 | 143.7027 | 0.500942 | 4.092842 | 0.359177 |
| 7 | 8425000 | 8431399 | 87.36976 | 138.9105 | 94.05704 | 39.25536 | 0.895705 | 2.272286 | 0.359187 |
| 1 | 113279800 | 113282599 | 24.69509 | 40.94014 | 71.99453 | 31.17553 | -0.43109 | 1.29987 | 0.359187 |
| 4 | 3223000 | 3236599 | 700.0369 | 933.6778 | 776.0206 | 324.1243 | 0.366287 | 5.322073 | 0.359191 |
| 16 | 20481800 | 20484199 | 11.80889 | 20.69845 | 43.59418 | 19.25915 | -0.68154 | 0.633525 | 0.359195 |
| 12 | 72880800 | 72882599 | 11.45014 | 20.19812 | 41.41092 | 18.32513 | -0.64087 | 0.541532 | 0.359202 |
| 17 | 58848400 | 58853399 | 52.73657 | 85.35241 | 65.6023 | 28.05911 | 0.741754 | 1.691543 | 0.359205 |
| 16 | 95733800 | 95734999 | 13.00844 | 24.21721 | 22.97349 | 9.588926 | 0.524568 | 0.172734 | 0.359214 |
| 7 | 36703400 | 36706199 | 22.87329 | 39.46983 | 37.36177 | 16.3902 | 0.463964 | 0.875315 | 0.359238 |
| 4 | 74444200 | 74449999 | 141.5879 | 214.7572 | 203.871 | 84.39372 | 0.355273 | 3.241769 | 0.359242 |
| 8 | 35183600 | 35187599 | 73.92736 | 117.1739 | 113.3438 | 47.15807 | 0.377578 | 2.352495 | 0.359253 |
| 2 | 39372800 | 39374199 | 9.378321 | 16.91211 | 38.49813 | 16.76223 | -0.77235 | 0.362171 | 0.359264 |
| 7 | 45114400 | 45116199 | 8.55365 | 15.60444 | 34.47475 | 14.79484 | -0.71579 | 0.278224 | 0.359271 |
| X | 14646400 | 14647599 | 21.31446 | 37.68784 | 28.99398 | 12.82433 | 0.767254 | 0.556128 | 0.35928 |
| 4 | 111827400 | 111829199 | 20.34965 | 35.83993 | 30.34583 | 13.43288 | 0.627126 | 0.590312 | 0.359306 |
| 16 | 19096600 | 19098399 | 22.90725 | 39.7323 | 37.74536 | 16.56107 | 0.460271 | 0.830676 | 0.359315 |
| 9 | 19367000 | 19371799 | 55.0452 | 88.61145 | 78.1603 | 33.20711 | 0.534931 | 1.851296 | 0.359315 |
| X | 164845800 | 164848199 | 25.26297 | 44.25759 | 31.48447 | 13.90319 | 0.876642 | 0.697094 | 0.359322 |
| 4 | 40289400 | 40297199 | 124.2532 | 190.117 | 182.3968 | 75.68177 | 0.350136 | 3.059988 | 0.35933 |
| 2 | 150335000 | 150338599 | 46.8448 | 76.07103 | 65.24211 | 27.97402 | 0.588437 | 1.615027 | 0.359331 |
| 14 | 24536800 | 24538599 | 7.613008 | 14.07779 | 31.53332 | 13.28896 | -0.71576 | 0.19022 | 0.359332 |
| 17 | 36419400 | 36422599 | 22.83521 | 37.65548 | 73.18971 | 31.5827 | -0.57596 | 1.357647 | 0.359335 |
| 10 | 28993800 | 28995199 | 24.52776 | 42.47257 | 38.25965 | 16.78587 | 0.536263 | 0.854857 | 0.359336 |

| | | | | | | | | | |
|---|---|---|---|---|---|---|---|---|---|
| 12 | 117035000 | 117037199 | 30.52188 | 50.97291 | 46.53912 | 20.25106 | 0.51206 | 1.214534 | 0.359343 |
| 8 | 125663600 | 125666599 | 26.43494 | 43.26722 | 77.62091 | 33.3894 | -0.46513 | 1.463405 | 0.359346 |
| 13 | 94266800 | 94268999 | 9.223349 | 16.66186 | 34.92359 | 15.17253 | -0.65149 | 0.345241 | 0.359358 |
| 17 | 20477200 | 20480399 | 40.30434 | 67.4393 | 42.45855 | 18.49781 | 1.046239 | 1.219332 | 0.359378 |
| 5 | 151089600 | 151090999 | 8.450685 | 15.71398 | 32.127 | 13.43592 | -0.57792 | 0.162865 | 0.35938 |
| 13 | 72637800 | 72640599 | 27.25836 | 46.91288 | 39.73806 | 17.43125 | 0.622863 | 0.921311 | 0.3594 |
| 6 | 79806600 | 79807199 | 15.06439 | 27.60362 | 24.85206 | 10.60418 | 0.576607 | 0.283036 | 0.359403 |
| 1 | 21232600 | 21233599 | 6.14001 | 11.79698 | 26.31615 | 10.29036 | -0.61718 | -0.10534 | 0.359417 |
| 10 | 82708000 | 82709199 | 6.14001 | 11.79698 | 26.31615 | 10.29036 | -0.61718 | -0.10534 | 0.359417 |
| 14 | 33731200 | 33732999 | 16.89403 | 30.38766 | 27.53669 | 12.02581 | 0.545278 | 0.405012 | 0.359438 |
| 19 | 44701400 | 44703999 | 36.49341 | 58.73925 | 99.89311 | 42.4325 | -0.41042 | 1.837896 | 0.359483 |
| 5 | 67620400 | 67621999 | 18.87804 | 32.12322 | 57.00083 | 24.98364 | -0.44241 | 1.01075 | 0.359495 |
| 7 | 82958000 | 82959599 | 19.42407 | 34.32318 | 30.5375 | 13.50879 | 0.558038 | 0.551878 | 0.359496 |
| 3 | 144509400 | 144510799 | 12.85794 | 23.99643 | 22.85964 | 9.501444 | 0.522986 | 0.153762 | 0.35951 |
| 7 | 23071800 | 23072999 | 29.65209 | 50.34356 | 44.15185 | 19.34565 | 0.570121 | 1.056488 | 0.359514 |
| 18 | 65637800 | 65642399 | 30.45117 | 49.2606 | 89.24881 | 38.15959 | -0.49297 | 1.6756 | 0.359533 |
| 17 | 50249000 | 50252999 | 43.83687 | 70.09288 | 118.205 | 49.72095 | -0.40984 | 2.058537 | 0.35954 |
| 1 | 36068600 | 36070999 | 22.00509 | 36.43935 | 68.67893 | 29.71656 | -0.53073 | 1.304181 | 0.359547 |
| 17 | 38560800 | 38562199 | 25.70591 | 44.4296 | 38.66442 | 16.96249 | 0.585228 | 0.875386 | 0.359555 |
| 1 | 93669200 | 93671599 | 16.76133 | 29.7898 | 27.7662 | 12.24918 | 0.495117 | 0.478873 | 0.359569 |
| 1 | 187468800 | 187469799 | 5.252436 | 10.21039 | 25.32674 | 9.406111 | -0.69866 | -0.29642 | 0.359588 |
| 5 | 147583200 | 147584399 | 6.573505 | 12.62132 | 26.88105 | 10.5317 | -0.5539 | -0.09321 | 0.359607 |
| X | 149005200 | 149013599 | 146.7267 | 223.8812 | 184.5135 | 76.51096 | 0.566017 | 3.123084 | 0.359627 |
| 15 | 101753400 | 101755399 | 9.502969 | 17.23333 | 34.4134 | 14.87893 | -0.5777 | 0.322793 | 0.359648 |
| 3 | 83342200 | 83342799 | 7.933759 | 15.02356 | 32.18701 | 13.03054 | -0.60629 | 0.038679 | 0.359649 |
| 4 | 152003200 | 152008199 | 60.45903 | 95.30595 | 154.6947 | 64.1944 | -0.37441 | 2.479487 | 0.359649 |
| 1 | 67909800 | 67910999 | 11.68265 | 22.42327 | 20.60493 | 8.08081 | 0.646851 | -0.05621 | 0.359657 |
| 7 | 38664000 | 38666399 | 23.43255 | 40.90133 | 34.9136 | 15.3736 | 0.614906 | 0.746595 | 0.35966 |
| 13 | 103825600 | 103826199 | 11.87896 | 23.0974 | 16.98704 | 6.296703 | 1.040902 | -0.24698 | 0.359666 |
| 13 | 41822600 | 41825399 | 13.82001 | 24.27751 | 44.53121 | 19.71017 | -0.48335 | 0.610603 | 0.359692 |
| Y | 20742200 | 20744799 | 13.01407 | 24.44826 | 13.41968 | 5.515943 | 1.323894 | 0.086285 | 0.359737 |
| 5 | 146000200 | 146001799 | 31.5918 | 53.21575 | 45.80584 | 20.02244 | 0.596569 | 1.129021 | 0.359738 |
| 6 | 100297400 | 100299799 | 15.12765 | 26.13539 | 51.10866 | 22.42163 | -0.57796 | 0.848991 | 0.359758 |
| 13 | 106192200 | 106192799 | 11.42357 | 20.29324 | 38.30151 | 16.90375 | -0.51736 | 0.469049 | 0.359761 |
| 15 | 24366200 | 24366599 | 8.665619 | 16.90349 | 16.24419 | 5.816501 | 0.703884 | -0.40739 | 0.359763 |
| 1 | 191658800 | 191661399 | 18.97463 | 31.96565 | 59.57057 | 26.04003 | -0.51369 | 1.102561 | 0.359785 |
| 4 | 66594200 | 66596199 | 27.65668 | 47.73131 | 38.01048 | 16.67486 | 0.712278 | 0.895641 | 0.359788 |
| 6 | 134224200 | 134227399 | 20.381 | 33.47379 | 79.63364 | 34.30505 | -0.86878 | 1.409559 | 0.359799 |
| 7 | 128650600 | 128652799 | 11.04765 | 19.84788 | 37.33723 | 16.32236 | -0.50318 | 0.392792 | 0.359802 |
| 8 | 92792000 | 92795799 | 22.49996 | 37.16926 | 70.93251 | 30.64071 | -0.549 | 1.334306 | 0.359806 |
| 7 | 4542000 | 4547399 | 36.46045 | 58.78468 | 98.99794 | 42.15161 | -0.39342 | 1.785181 | 0.359807 |
| 10 | 64772800 | 64775399 | 22.00463 | 38.59438 | 31.93118 | 14.11436 | 0.659418 | 0.657871 | 0.359811 |
| X | 148933600 | 148941999 | 126.8772 | 196.9489 | 135.0516 | 56.08355 | 0.850017 | 2.789743 | 0.359816 |
| 18 | 3124200 | 3129399 | 51.02306 | 83.79495 | 48.47338 | 20.88108 | 1.165007 | 1.461097 | 0.359826 |
| 7 | 59138800 | 59141199 | 31.02906 | 51.73075 | 49.75814 | 21.62827 | 0.43705 | 1.23441 | 0.359833 |
| 17 | 81371600 | 81372599 | 6.365941 | 12.01598 | 30.26007 | 12.33409 | -0.84558 | 0.042566 | 0.359834 |
| 6 | 149181800 | 149182799 | 22.88614 | 39.81787 | 35.42204 | 15.56563 | 0.554923 | 0.785843 | 0.359845 |

| | | | | | | | | | |
|---|---|---|---|---|---|---|---|---|---|
| 12 | 24294200 | 24303999 | 202.495 | 299.8295 | 279.6398 | 115.1645 | 0.349856 | 3.708685 | 0.359857 |
| 7 | 131491400 | 131492799 | 15.23464 | 26.55712 | 48.41142 | 21.29959 | -0.47501 | 0.742552 | 0.359866 |
| 14 | 16507400 | 16509199 | 14.31752 | 24.8504 | 49.51345 | 21.72516 | -0.60335 | 0.801205 | 0.359868 |
| 5 | 138908200 | 138910999 | 19.6113 | 33.06862 | 59.60923 | 26.0642 | -0.46581 | 1.097248 | 0.359871 |
| 16 | 49951200 | 49954599 | 41.14051 | 67.56088 | 57.32609 | 24.69398 | 0.613915 | 1.440912 | 0.359877 |
| 7 | 137602200 | 137604599 | 38.81214 | 63.69484 | 59.29836 | 25.53637 | 0.480154 | 1.446566 | 0.359878 |
| 3 | 136374400 | 136378199 | 28.86696 | 46.67671 | 90.59119 | 38.71646 | -0.59265 | 1.684141 | 0.359913 |
| 6 | 45420200 | 45422199 | 13.70842 | 24.06644 | 44.56577 | 19.71505 | -0.49684 | 0.621847 | 0.359922 |
| 4 | 52820200 | 52822199 | 19.35206 | 32.9964 | 60.38626 | 26.47179 | -0.48645 | 0.997217 | 0.359925 |
| 17 | 20319200 | 20332199 | 162.8474 | 247.4251 | 190.5399 | 78.91556 | 0.6589 | 3.213842 | 0.359929 |
| 10 | 56141400 | 56141999 | 5.395513 | 10.46337 | 24.43742 | 9.223934 | -0.63388 | -0.23831 | 0.359943 |
| 11 | 10440600 | 10441199 | 10.09633 | 19.45475 | 19.69892 | 7.638932 | 0.523293 | -0.11069 | 0.359952 |
| X | 47966200 | 47967199 | 10.63758 | 19.27903 | 40.22407 | 17.40464 | -0.64174 | 0.346755 | 0.359953 |
| X | 120367000 | 120373199 | 69.56103 | 111.4688 | 81.39962 | 34.3494 | 0.799049 | 2.012218 | 0.359958 |
| 17 | 13253000 | 13258799 | 29.66689 | 47.94389 | 94.3268 | 40.28448 | -0.61325 | 1.702163 | 0.359965 |
| 2 | 54546600 | 54547599 | 11.58704 | 21.83475 | 20.49676 | 8.380929 | 0.563008 | 0.076122 | 0.359989 |
| 13 | 66272600 | 66273399 | 10.93231 | 21.32587 | 14.08544 | 5.040813 | 1.237578 | -0.38797 | 0.360002 |
| 7 | 35654400 | 35657999 | 31.09001 | 50.35285 | 87.73741 | 37.54571 | -0.43523 | 1.653303 | 0.360012 |
| 4 | 149510400 | 149511399 | 4.585706 | 8.856963 | 25.28511 | 9.726367 | -0.94598 | -0.17913 | 0.360019 |
| 8 | 8750000 | 8751599 | 17.31137 | 29.52555 | 54.77617 | 24.01304 | -0.50566 | 0.982822 | 0.360039 |
| 7 | 9755400 | 9758599 | 63.26572 | 101.2221 | 90.7723 | 38.21581 | 0.500675 | 2.053483 | 0.360052 |
| 16 | 15345800 | 15347599 | 44.46062 | 71.90315 | 70.08537 | 29.95928 | 0.398835 | 1.695935 | 0.360061 |
| 1 | 17161600 | 17162399 | 7.877908 | 14.63561 | 31.04406 | 13.00017 | -0.63158 | 0.16248 | 0.360069 |
| 16 | 17731000 | 17733599 | 18.22814 | 30.58303 | 62.75839 | 27.38858 | -0.6524 | 1.140088 | 0.360072 |
| 14 | 44170400 | 44170799 | 6.652619 | 12.99872 | 13.67337 | 4.757236 | 0.61209 | -0.54553 | 0.360073 |
| 10 | 84869400 | 84876999 | 63.93926 | 100.0553 | 169.2562 | 70.21424 | -0.44217 | 2.623968 | 0.36008 |
| 17 | 13350000 | 13357599 | 42.26442 | 67.45096 | 121.9108 | 51.14261 | -0.51261 | 2.116192 | 0.36008 |
| 2 | 102230400 | 102231799 | 10.44681 | 18.34844 | 42.05197 | 18.60991 | -0.80239 | 0.591041 | 0.360081 |
| 5 | 13669000 | 13669999 | 9.085243 | 16.86833 | 32.6409 | 13.6802 | -0.50133 | 0.179481 | 0.360088 |
| 17 | 20851800 | 20858199 | 73.01925 | 116.6378 | 89.35145 | 37.53121 | 0.725036 | 2.113984 | 0.360089 |
| 8 | 96548800 | 96552199 | 26.92767 | 44.01219 | 79.32007 | 34.10294 | -0.47319 | 1.485306 | 0.360091 |
| 19 | 21887200 | 21888399 | 6.424316 | 12.35996 | 26.37155 | 10.272 | -0.5482 | -0.11419 | 0.360098 |
| X | 109321600 | 109326799 | 78.85556 | 125.3032 | 104.2324 | 43.47765 | 0.598474 | 2.282668 | 0.360105 |
| 2 | 92457600 | 92459599 | 10.19315 | 18.14381 | 37.85397 | 16.67803 | -0.65925 | 0.447051 | 0.360109 |
| 10 | 116785600 | 116786799 | 10.19315 | 18.14381 | 37.85397 | 16.67803 | -0.65925 | 0.447051 | 0.360109 |
| 4 | 17536200 | 17537799 | 18.59963 | 33.72656 | 25.82394 | 11.16656 | 0.796026 | 0.368514 | 0.360111 |
| 4 | 26162800 | 26165399 | 36.36187 | 59.99138 | 56.00128 | 24.17075 | 0.479207 | 1.372112 | 0.360119 |
| 2 | 93448800 | 93450199 | 6.292029 | 12.06215 | 26.78525 | 10.54103 | -0.61887 | -0.07902 | 0.360129 |
| 17 | 29660600 | 29661799 | 6.292029 | 12.06215 | 26.78525 | 10.54103 | -0.61887 | -0.07902 | 0.360129 |
| 19 | 16647800 | 16650999 | 39.14421 | 62.73668 | 109.1776 | 46.08202 | -0.45114 | 1.979133 | 0.360145 |
| 4 | 140185600 | 140197799 | 110.8148 | 166.9659 | 281.962 | 116.4801 | -0.48328 | 3.374053 | 0.360168 |
| 10 | 9727200 | 9727999 | 11.2256 | 19.66771 | 46.74113 | 20.64201 | -0.85448 | 0.646406 | 0.360169 |
| 3 | 62029600 | 62030399 | 18.40246 | 33.06209 | 21.31362 | 9.317287 | 1.032877 | 0.41116 | 0.360179 |
| 8 | 60463600 | 60464799 | 14.43218 | 26.30689 | 24.84018 | 10.67119 | 0.50155 | 0.306108 | 0.360194 |
| X | 22948400 | 22950399 | 24.18725 | 42.67682 | 28.51539 | 12.61799 | 0.968214 | 0.593548 | 0.360198 |
| 7 | 144625400 | 144626599 | 8.554633 | 15.64508 | 32.96411 | 14.10251 | -0.6444 | 0.263141 | 0.360208 |
| 6 | 133803600 | 133804799 | 8.256378 | 15.52565 | 31.90023 | 13.09035 | -0.56303 | 0.088217 | 0.360219 |

| | | | | | | | | | |
|---|---|---|---|---|---|---|---|---|---|
| 6 | 133517400 | 133518199 | 11.2891 | 21.64901 | 19.38457 | 7.62128 | 0.67981 | -0.05785 | 0.360223 |
| 14 | 51081000 | 51082999 | 9.951872 | 17.5814 | 39.46069 | 17.4569 | -0.76925 | 0.514762 | 0.360236 |
| 7 | 105458800 | 105461399 | 17.47544 | 29.65352 | 57.45097 | 25.16858 | -0.56907 | 1.030588 | 0.360238 |
| 1 | 46891800 | 46893599 | 14.43969 | 24.77685 | 53.70552 | 23.55513 | -0.7278 | 0.912066 | 0.360263 |
| 11 | 22776000 | 22777999 | 18.65227 | 31.45717 | 59.01985 | 25.8079 | -0.52329 | 1.09148 | 0.360276 |
| 7 | 27932600 | 27936399 | 59.93916 | 95.76318 | 93.52629 | 39.29586 | 0.375561 | 2.093408 | 0.36028 |
| 5 | 92774000 | 92775599 | 15.5741 | 27.04419 | 48.98772 | 21.50087 | -0.46645 | 0.800522 | 0.360284 |
| 7 | 51938200 | 51940399 | 30.32664 | 51.0564 | 42.16138 | 18.42631 | 0.656227 | 1.127107 | 0.360291 |
| 4 | 100638400 | 100639999 | 8.300826 | 15.47684 | 31.10291 | 12.9486 | -0.54777 | 0.142828 | 0.360293 |
| 12 | 107880200 | 107885999 | 91.92534 | 139.7535 | 248.4894 | 102.9351 | -0.54657 | 3.186929 | 0.360317 |
| 2 | 105056200 | 105057199 | 4.946702 | 9.618954 | 22.87541 | 8.475087 | -0.6336 | -0.31219 | 0.360337 |
| 6 | 143175400 | 143176999 | 9.544856 | 17.61028 | 34.69441 | 14.66614 | -0.5364 | 0.221872 | 0.360359 |
| 8 | 119160000 | 119162399 | 14.97263 | 26.12145 | 47.33111 | 20.83485 | -0.46605 | 0.730394 | 0.360382 |
| 18 | 38342200 | 38348399 | 42.53042 | 67.68712 | 136.5896 | 57.08009 | -0.67565 | 2.211694 | 0.360382 |
| 4 | 13015000 | 13018799 | 32.7967 | 56.16394 | 37.90039 | 16.62038 | 0.948322 | 0.985972 | 0.360402 |
| 3 | 94379400 | 94381999 | 10.49305 | 18.66687 | 38.33768 | 16.89936 | -0.63732 | 0.453085 | 0.360403 |
| 19 | 8595400 | 8596999 | 12.26536 | 21.54814 | 41.47221 | 18.35741 | -0.55203 | 0.596634 | 0.360406 |
| 7 | 39163400 | 39168599 | 59.73255 | 94.43724 | 151.4381 | 62.91946 | -0.3536 | 2.411445 | 0.360412 |
| 14 | 33440000 | 33441799 | 10.72605 | 19.28061 | 37.3455 | 16.31847 | -0.54466 | 0.386489 | 0.360424 |
| 11 | 115644000 | 115647399 | 25.30237 | 43.41801 | 40.97578 | 17.97196 | 0.466795 | 0.941081 | 0.360443 |
| 5 | 95637800 | 95641399 | 61.37766 | 98.58881 | 79.9945 | 33.87842 | 0.651268 | 1.927157 | 0.360464 |
| 15 | 81947800 | 81950599 | 17.93731 | 30.06411 | 62.11267 | 27.08919 | -0.66202 | 1.15102 | 0.360471 |
| 10 | 54483800 | 54484399 | 6.87432 | 13.1912 | 27.43497 | 10.76674 | -0.52215 | -0.08233 | 0.360487 |
| 2 | 73516600 | 73517399 | 10.00981 | 17.98507 | 35.87575 | 15.68217 | -0.5866 | 0.384805 | 0.360505 |
| X | 9137600 | 9141199 | 31.14336 | 53.284 | 39.73794 | 17.42492 | 0.804609 | 0.991136 | 0.360509 |
| 10 | 111045800 | 111046999 | 13.38286 | 25.00292 | 23.35053 | 9.68784 | 0.55355 | 0.153059 | 0.360512 |
| 4 | 11075800 | 11076999 | 7.357867 | 13.93224 | 28.71182 | 11.62506 | -0.55 | 0.02773 | 0.36052 |
| 16 | 70357600 | 70361399 | 26.52547 | 43.67832 | 76.47029 | 32.9784 | -0.42655 | 1.387512 | 0.360526 |
| 17 | 9386000 | 9387599 | 9.122215 | 16.69966 | 33.98324 | 14.51889 | -0.59376 | 0.261351 | 0.360569 |
| 10 | 123392800 | 123394399 | 18.45359 | 32.87775 | 28.33662 | 12.47508 | 0.609235 | 0.474612 | 0.36058 |
| 2 | 36463400 | 36464399 | 19.80865 | 34.71629 | 32.84932 | 14.51083 | 0.466856 | 0.66039 | 0.360582 |
| 9 | 41801000 | 41802599 | 8.006398 | 14.52802 | 34.46406 | 14.88971 | -0.82362 | 0.307197 | 0.360604 |
| 5 | 56054400 | 56059199 | 75.90475 | 120.6739 | 104.2024 | 43.48667 | 0.545334 | 2.264724 | 0.360606 |
| 18 | 89567200 | 89568599 | 17.67887 | 31.52318 | 28.64695 | 12.60303 | 0.53422 | 0.467444 | 0.360632 |
| 1 | 33196400 | 33196999 | 5.514793 | 10.67747 | 25.4413 | 9.675408 | -0.67175 | -0.20369 | 0.360635 |
| 1 | 151919200 | 151922999 | 23.04138 | 38.29031 | 69.69046 | 30.22782 | -0.48024 | 1.264578 | 0.360649 |
| 7 | 5622000 | 5629999 | 136.4052 | 210.2466 | 151.5221 | 62.93979 | 0.770576 | 2.921387 | 0.360672 |
| 15 | 90108000 | 90109999 | 12.19527 | 21.62452 | 40.30554 | 17.81566 | -0.50116 | 0.488271 | 0.36068 |
| 10 | 10955000 | 10958999 | 33.20386 | 53.60082 | 93.94529 | 40.06458 | -0.44875 | 1.741154 | 0.360683 |
| 5 | 146642800 | 146654799 | 89.07432 | 135.5106 | 268.433 | 111.2191 | -0.70152 | 3.172535 | 0.360687 |
| 3 | 13535600 | 13538599 | 25.83919 | 42.1903 | 78.91388 | 33.91576 | -0.5276 | 1.498224 | 0.360694 |
| 9 | 3937400 | 3938199 | 17.17302 | 30.50028 | 27.82401 | 12.28189 | 0.525508 | 0.487115 | 0.360699 |
| 13 | 96883400 | 96885799 | 23.92698 | 39.59226 | 69.99825 | 30.27177 | -0.43892 | 1.319181 | 0.360728 |
| 4 | 33121800 | 33125999 | 32.16998 | 52.19146 | 90.40663 | 38.73694 | -0.42453 | 1.6202 | 0.360731 |
| 19 | 29406200 | 29406999 | 7.318542 | 13.98144 | 29.00964 | 11.527 | -0.53613 | -0.03479 | 0.360735 |
| 10 | 95141600 | 95148199 | 40.87411 | 65.52498 | 112.1884 | 47.37017 | -0.42735 | 1.972915 | 0.360757 |
| 13 | 12991200 | 12991999 | 12.1537 | 23.31528 | 20.55065 | 8.071544 | 0.704377 | -0.04706 | 0.360769 |

| | | | | | | | | | |
|---|---|---|---|---|---|---|---|---|---|
| 2 | 144817200 | 144819399 | 22.08315 | 39.24129 | 26.28099 | 11.59571 | 0.968841 | 0.510509 | 0.36077 |
| 3 | 128356200 | 128357399 | 14.52079 | 27.40392 | 21.69401 | 8.845733 | 0.810156 | 0.096076 | 0.360795 |
| 18 | 23245600 | 23250199 | 71.42329 | 113.1772 | 110.4777 | 45.95431 | 0.364136 | 2.356835 | 0.360799 |
| 4 | 74497800 | 74502599 | 66.40733 | 105.9868 | 95.65667 | 40.13172 | 0.487568 | 2.13099 | 0.360829 |
| 5 | 87536200 | 87538199 | 29.62084 | 50.75805 | 39.63076 | 17.3801 | 0.738813 | 0.967876 | 0.360831 |
| 17 | 87515800 | 87517799 | 16.70368 | 28.48752 | 60.15531 | 26.37106 | -0.69176 | 0.987283 | 0.360833 |
| 12 | 23995600 | 24000999 | 100.6066 | 156.5308 | 134.3085 | 55.75571 | 0.529504 | 2.73633 | 0.360861 |
| 5 | 94848000 | 94849999 | 25.97402 | 45.40588 | 32.37065 | 14.26866 | 0.873466 | 0.733116 | 0.360863 |
| 13 | 66457400 | 66475399 | 414.6233 | 577.0561 | 503.7187 | 209.0207 | 0.357403 | 4.65359 | 0.360864 |
| 17 | 26770800 | 26774199 | 14.96751 | 26.04988 | 47.36498 | 20.81831 | -0.47137 | 0.764381 | 0.360865 |
| 9 | 45094400 | 45097399 | 23.12446 | 38.26632 | 69.34167 | 29.98891 | -0.47432 | 1.315823 | 0.360866 |
| 6 | 92744600 | 92746199 | 21.47428 | 38.00158 | 29.24695 | 12.9341 | 0.766905 | 0.547088 | 0.360884 |
| 1 | 8380400 | 8382599 | 19.58285 | 34.90324 | 27.64513 | 12.16667 | 0.730712 | 0.478241 | 0.360886 |
| 8 | 21980200 | 21980799 | 24.32931 | 41.78556 | 37.00341 | 16.23067 | 0.558538 | 0.92508 | 0.360904 |
| 10 | 47516200 | 47517399 | 13.8716 | 26.13981 | 22.29589 | 9.116578 | 0.700058 | 0.101286 | 0.360909 |
| 4 | 138547600 | 138552399 | 47.61653 | 75.96232 | 125.2738 | 52.52401 | -0.38134 | 2.135097 | 0.360934 |
| 14 | 118622800 | 118624399 | 8.463909 | 15.70182 | 32.08174 | 13.46087 | -0.58042 | 0.176583 | 0.360939 |
| 7 | 60349600 | 60360799 | 134.2871 | 206.3752 | 165.0909 | 68.55181 | 0.616969 | 2.976675 | 0.360963 |
| 2 | 175002400 | 175008999 | 133.8552 | 205.354 | 171.9099 | 71.36626 | 0.549581 | 3.01021 | 0.360964 |
| 17 | 68363800 | 68365399 | 13.83807 | 24.1365 | 45.75929 | 20.14002 | -0.53101 | 0.724592 | 0.360964 |
| 6 | 86471000 | 86472399 | 13.3563 | 23.34889 | 44.67486 | 19.69252 | -0.54389 | 0.687145 | 0.360987 |
| 8 | 47826800 | 47831599 | 71.20908 | 110.6842 | 183.8284 | 76.32412 | -0.4233 | 2.746934 | 0.361022 |
| 16 | 12927400 | 12928399 | 8.702748 | 15.87543 | 33.95247 | 14.57186 | -0.66924 | 0.279159 | 0.361023 |
| 7 | 18751000 | 18759199 | 60.05383 | 94.33406 | 161.4624 | 66.97211 | -0.4548 | 2.550197 | 0.361027 |
| 1 | 110740600 | 110741599 | 18.31018 | 32.50372 | 25.58626 | 11.29886 | 0.736423 | 0.49162 | 0.361037 |
| 18 | 54128200 | 54129599 | 9.407893 | 17.47064 | 36.0081 | 15.08761 | -0.59175 | 0.188041 | 0.361059 |
| 2 | 71396200 | 71397799 | 16.77044 | 28.82104 | 52.16326 | 22.88061 | -0.46835 | 0.907345 | 0.361062 |
| 4 | 84116600 | 84119399 | 13.40935 | 23.27136 | 49.49793 | 21.71798 | -0.69711 | 0.798441 | 0.36108 |
| X | 8685200 | 8685799 | 13.5304 | 25.27019 | 23.43713 | 9.729234 | 0.563167 | 0.158162 | 0.361084 |
| 2 | 101913600 | 101914799 | 19.22533 | 34.67813 | 26.21225 | 11.40933 | 0.80844 | 0.402825 | 0.361084 |
| 14 | 11123200 | 11123799 | 4.810571 | 9.368004 | 22.44329 | 8.214131 | -0.62805 | -0.35987 | 0.361106 |
| 16 | 86212000 | 86213799 | 22.50307 | 39.93984 | 27.49519 | 12.14363 | 0.928729 | 0.524385 | 0.361112 |
| 6 | 29001200 | 29003399 | 14.90555 | 25.93873 | 49.50552 | 21.75739 | -0.54127 | 0.765886 | 0.361117 |
| 11 | 30958800 | 30964599 | 81.50711 | 129.4699 | 102.7396 | 42.83953 | 0.66586 | 2.29359 | 0.361118 |
| 19 | 24865800 | 24867599 | 14.46545 | 25.2194 | 46.28603 | 20.36607 | -0.48456 | 0.735 | 0.361143 |
| 7 | 10770600 | 10784199 | 285.1797 | 412.4632 | 354.5724 | 146.2442 | 0.435044 | 4.082443 | 0.361147 |
| 8 | 43704400 | 43711199 | 56.81936 | 91.31754 | 81.60939 | 34.59013 | 0.513166 | 1.905177 | 0.361147 |
| 11 | 39837400 | 39839599 | 19.22329 | 32.11075 | 63.76937 | 27.75246 | -0.60523 | 1.193186 | 0.361149 |
| 7 | 17290800 | 17293399 | 38.65932 | 63.9819 | 47.72408 | 20.64162 | 0.802347 | 1.339208 | 0.36115 |
| 12 | 112956400 | 112958199 | 10.63408 | 18.94892 | 37.8275 | 16.65136 | -0.59519 | 0.442812 | 0.361152 |
| 10 | 6340200 | 6341199 | 13.18759 | 25.32391 | 19.28167 | 7.551486 | 0.917173 | -0.0498 | 0.36116 |
| 1 | 128304400 | 128304999 | 9.743336 | 17.85373 | 33.99544 | 14.50528 | -0.49709 | 0.261968 | 0.361185 |
| 13 | 49142600 | 49143999 | 10.23176 | 18.5662 | 35.40987 | 15.29818 | -0.5116 | 0.32444 | 0.361187 |
| 18 | 37741400 | 37744199 | 45.8711 | 74.28129 | 69.44254 | 29.71356 | 0.460658 | 1.670801 | 0.361192 |
| 12 | 11939200 | 11940799 | 28.80483 | 48.57056 | 46.40267 | 20.28959 | 0.446794 | 1.113641 | 0.361202 |
| 17 | 45619200 | 45630799 | 131.4341 | 197.2192 | 303.1141 | 125.0802 | -0.35279 | 3.459145 | 0.361213 |
| 8 | 70148000 | 70150399 | 42.32663 | 68.73917 | 66.3287 | 28.44149 | 0.418941 | 1.606627 | 0.361218 |

| | | | | | | | | | |
|---|---|---|---|---|---|---|---|---|---|
| 12 | 115874200 | 115874999 | 15.19598 | 28.32131 | 22.49626 | 9.371968 | 0.78041 | 0.180586 | 0.361224 |
| X | 4030200 | 4032799 | 31.1937 | 53.47054 | 39.09502 | 17.1456 | 0.83337 | 0.972979 | 0.361226 |
| X | 26128200 | 26133399 | 59.45981 | 96.12534 | 63.35923 | 27.07136 | 0.961383 | 1.732254 | 0.361241 |
| 12 | 23153400 | 23187399 | 1150.105 | 1470.459 | 1176.795 | 493.7788 | 0.359912 | 6.007238 | 0.361292 |
| 19 | 49651600 | 49655199 | 76.40809 | 121.0527 | 102.7635 | 42.74111 | 0.565075 | 2.362888 | 0.36131 |
| 7 | 28059200 | 28062599 | 21.60847 | 35.58531 | 72.13719 | 31.11104 | -0.63705 | 1.373022 | 0.361319 |
| 1 | 101028200 | 101031999 | 33.26687 | 55.87471 | 46.36885 | 20.24965 | 0.649005 | 1.160257 | 0.361324 |
| 18 | 34116800 | 34121999 | 34.05009 | 54.83343 | 98.48687 | 41.86492 | -0.48816 | 1.81738 | 0.361344 |
| 4 | 69191400 | 69192199 | 10.12207 | 19.66544 | 18.20054 | 6.790654 | 0.703658 | -0.24305 | 0.361348 |
| 7 | 45518800 | 45522399 | 23.39181 | 38.38161 | 75.01026 | 32.30015 | -0.58667 | 1.424047 | 0.361351 |
| 18 | 37745200 | 37753199 | 112.9453 | 175.361 | 146.5632 | 60.86201 | 0.56506 | 2.779196 | 0.361355 |
| 17 | 5859400 | 5863199 | 32.54493 | 52.57764 | 92.94521 | 39.67751 | -0.4598 | 1.716792 | 0.361358 |
| 9 | 111100000 | 111101799 | 12.77777 | 22.25802 | 46.31135 | 20.36627 | -0.66463 | 0.739135 | 0.361373 |
| 4 | 36207600 | 36208599 | 17.69635 | 31.87382 | 26.24512 | 11.44443 | 0.683786 | 0.398588 | 0.361376 |
| 13 | 66205000 | 66205599 | 29.1775 | 48.96409 | 43.87848 | 19.15169 | 0.538737 | 1.156489 | 0.361386 |
| 18 | 27735000 | 27736999 | 26.1633 | 45.61238 | 33.49096 | 14.73591 | 0.830838 | 0.772528 | 0.361394 |
| 7 | 14747400 | 14751199 | 82.7512 | 130.6294 | 120.618 | 50.07549 | 0.439892 | 2.458208 | 0.361411 |
| 8 | 71723400 | 71726199 | 17.52576 | 29.63716 | 57.92109 | 25.34817 | -0.58171 | 1.062546 | 0.361419 |
| 7 | 18566200 | 18567599 | 20.08336 | 35.2316 | 32.83689 | 14.51714 | 0.488617 | 0.64788 | 0.361426 |
| 11 | 55350800 | 55354999 | 20.67233 | 34.39163 | 67.16003 | 29.15263 | -0.58121 | 1.243522 | 0.36143 |
| 4 | 152851400 | 152865399 | 142.8967 | 212.7166 | 333.1052 | 137.2527 | -0.39021 | 3.60934 | 0.361441 |
| 2 | 59166600 | 59166999 | 3.45387 | 6.753875 | 18.32839 | 6.249749 | -0.70874 | -0.64112 | 0.361464 |
| 13 | 56899800 | 56901799 | 18.92702 | 31.58153 | 64.23557 | 27.93618 | -0.63954 | 1.204958 | 0.361464 |
| 14 | 65927200 | 65929199 | 16.76146 | 28.73154 | 53.22818 | 23.34386 | -0.50258 | 0.930821 | 0.361467 |
| 18 | 68557400 | 68560999 | 40.772 | 66.27268 | 64.63298 | 27.73235 | 0.404771 | 1.584008 | 0.361474 |
| 9 | 54984400 | 54991599 | 50.26324 | 79.50543 | 145.5902 | 60.50489 | -0.54417 | 2.391522 | 0.361477 |
| 10 | 62066200 | 62067999 | 12.07784 | 21.29765 | 40.38236 | 17.87127 | -0.52881 | 0.549006 | 0.36149 |
| 12 | 17444600 | 17446399 | 6.587446 | 12.24979 | 34.28057 | 14.33562 | -1.02712 | 0.151053 | 0.361494 |
| 16 | 90996200 | 90997399 | 13.137 | 24.39262 | 23.43497 | 9.82106 | 0.501871 | 0.190005 | 0.36153 |
| 9 | 41087600 | 41090599 | 19.8626 | 33.74701 | 60.3239 | 26.43691 | -0.45341 | 1.031889 | 0.361535 |
| 15 | 58724200 | 58725999 | 13.34172 | 23.40777 | 43.01028 | 19.0167 | -0.48573 | 0.625898 | 0.361545 |
| 8 | 73821600 | 73823599 | 17.7215 | 31.65666 | 28.49744 | 12.51806 | 0.549103 | 0.454848 | 0.361545 |
| 10 | 66619000 | 66621999 | 22.69834 | 37.83974 | 67.47086 | 29.32631 | -0.45052 | 1.226991 | 0.361547 |
| 6 | 123093200 | 123095799 | 29.06227 | 49.37185 | 44.13488 | 19.34173 | 0.542634 | 1.048152 | 0.361555 |
| 9 | 103181800 | 103182999 | 8.123145 | 15.21542 | 31.21365 | 12.89381 | -0.56835 | 0.110898 | 0.361567 |
| 12 | 49390600 | 49391599 | 17.40776 | 29.98982 | 53.81655 | 23.60756 | -0.45541 | 0.889822 | 0.361568 |
| 17 | 35037800 | 35047799 | 120.3038 | 180.6097 | 295.2211 | 121.8397 | -0.44092 | 3.447112 | 0.361569 |
| 6 | 26988400 | 26989799 | 15.86267 | 29.45136 | 23.57968 | 9.88273 | 0.762739 | 0.212453 | 0.36157 |
| 4 | 69986200 | 69989199 | 17.04826 | 28.87875 | 62.28785 | 27.27327 | -0.72341 | 1.050079 | 0.361575 |
| 14 | 59265200 | 59267199 | 39.80368 | 65.74169 | 54.43464 | 23.51096 | 0.651946 | 1.369233 | 0.36159 |
| 4 | 62827600 | 62828599 | 12.48345 | 24.14312 | 18.40662 | 7.036384 | 0.947945 | -0.13584 | 0.361602 |
| 7 | 85418600 | 85419599 | 26.71585 | 45.45046 | 41.64862 | 18.25441 | 0.507429 | 1.026005 | 0.361607 |
| 10 | 113310200 | 113310799 | 4.391998 | 8.569496 | 21.56532 | 7.691867 | -0.66435 | -0.46193 | 0.361608 |
| 10 | 115295400 | 115295799 | 4.391998 | 8.569496 | 21.56532 | 7.691867 | -0.66435 | -0.46193 | 0.361608 |
| 18 | 56694800 | 56695999 | 8.435267 | 15.79094 | 31.19971 | 12.90066 | -0.5156 | 0.118133 | 0.361613 |
| 3 | 106325600 | 106328199 | 26.33501 | 46.11927 | 30.62264 | 13.51855 | 0.975392 | 0.704829 | 0.361616 |
| 1 | 109521800 | 109524799 | 41.72 | 68.57913 | 56.76501 | 24.46234 | 0.650157 | 1.431291 | 0.361617 |

| | | | | | | | | | |
|---|---|---|---|---|---|---|---|---|---|
| 2 | 36494600 | 36497799 | 38.48894 | 64.32112 | 45.88135 | 19.97323 | 0.866831 | 1.229441 | 0.361626 |
| 6 | 13887000 | 13888399 | 18.28229 | 32.87777 | 27.4148 | 11.97472 | 0.664504 | 0.416321 | 0.361626 |
| 12 | 70343400 | 70345399 | 18.28229 | 32.87777 | 27.4148 | 11.97472 | 0.664504 | 0.416321 | 0.361626 |
| 14 | 97087600 | 97089599 | 22.54554 | 37.75501 | 67.52439 | 29.43326 | -0.45493 | 1.179569 | 0.361629 |
| 1 | 160408400 | 160411799 | 33.48908 | 56.54105 | 44.68128 | 19.54446 | 0.719647 | 1.117785 | 0.36165 |
| 13 | 73273000 | 73273399 | 10.75389 | 20.77416 | 19.04508 | 7.319793 | 0.676755 | -0.13628 | 0.361654 |
| 13 | 15574800 | 15581399 | 57.71032 | 91.26613 | 149.4833 | 62.11627 | -0.38376 | 2.401801 | 0.36166 |
| 16 | 10807000 | 10809399 | 24.47109 | 40.20539 | 74.79892 | 32.22708 | -0.51515 | 1.40913 | 0.361674 |
| 1 | 13608600 | 13610799 | 21.0522 | 36.91668 | 32.59902 | 14.40718 | 0.565951 | 0.657016 | 0.361675 |
| 7 | 98768600 | 98770599 | 12.75259 | 22.54029 | 42.95948 | 19.00287 | -0.5349 | 0.533483 | 0.361701 |
| 10 | 61741600 | 61745599 | 43.71423 | 69.88795 | 116.4408 | 48.97011 | -0.39258 | 2.062966 | 0.361707 |
| 17 | 55084200 | 55084799 | 8.278194 | 15.57139 | 30.76816 | 12.61933 | -0.50613 | 0.085494 | 0.361714 |
| 8 | 120623800 | 120625399 | 9.968904 | 17.95067 | 35.84634 | 15.63641 | -0.58645 | 0.368459 | 0.361714 |
| 8 | 60964200 | 60967199 | 29.57656 | 51.22434 | 35.14508 | 15.41948 | 0.927297 | 0.870968 | 0.361721 |
| 10 | 116828600 | 116829999 | 9.307196 | 16.91609 | 35.45192 | 15.28513 | -0.64481 | 0.307192 | 0.361768 |
| 14 | 112204400 | 112206599 | 23.69388 | 41.12523 | 36.51889 | 16.02365 | 0.557251 | 0.824303 | 0.361776 |
| 17 | 25262600 | 25263999 | 7.267114 | 13.57523 | 30.60244 | 12.69937 | -0.70825 | 0.120299 | 0.361788 |
| 13 | 46310000 | 46311199 | 14.34501 | 26.94279 | 22.8481 | 9.394758 | 0.702273 | 0.128435 | 0.361792 |
| 3 | 57938600 | 57940199 | 9.396006 | 17.38722 | 33.66427 | 14.17328 | -0.50706 | 0.202561 | 0.361805 |
| 8 | 87552400 | 87553399 | 9.396006 | 17.38722 | 33.66427 | 14.17328 | -0.50706 | 0.202561 | 0.361805 |
| 2 | 158719600 | 158720799 | 6.298836 | 12.05527 | 27.31546 | 10.80078 | -0.65501 | -0.06525 | 0.361808 |
| 10 | 42016600 | 42017599 | 5.975206 | 11.54124 | 25.37273 | 9.757873 | -0.57335 | -0.16409 | 0.361813 |
| 10 | 93336800 | 93337599 | 5.975206 | 11.54124 | 25.37273 | 9.757873 | -0.57335 | -0.16409 | 0.361813 |
| 15 | 6709200 | 6714799 | 63.62531 | 99.97424 | 162.076 | 67.22925 | -0.37628 | 2.546265 | 0.361813 |
| 10 | 75690600 | 75699999 | 78.35025 | 120.6641 | 207.8131 | 86.31652 | -0.48593 | 2.919261 | 0.361815 |
| 2 | 113628800 | 113630199 | 9.124496 | 16.74611 | 32.97262 | 14.03874 | -0.54269 | 0.246039 | 0.361829 |
| 19 | 59957400 | 59958599 | 16.75953 | 29.98396 | 28.13766 | 12.34414 | 0.49076 | 0.438627 | 0.361829 |
| 10 | 69154400 | 69157999 | 29.54672 | 48.13915 | 82.80812 | 35.56123 | -0.40945 | 1.53964 | 0.361836 |
| 12 | 23087400 | 23113999 | 661.281 | 883.9703 | 729.4165 | 304.5655 | 0.379946 | 5.285439 | 0.361836 |
| 17 | 50036200 | 50037999 | 21.80217 | 38.09399 | 34.9053 | 15.37971 | 0.513134 | 0.722734 | 0.361843 |
| 6 | 33054800 | 33059199 | 45.82289 | 73.98192 | 71.26564 | 30.3979 | 0.413027 | 1.745347 | 0.361855 |
| 3 | 89621800 | 89622999 | 4.854911 | 9.407353 | 24.9258 | 9.444046 | -0.8192 | -0.23076 | 0.361879 |
| 1 | 188469000 | 188469399 | 3.377914 | 6.607035 | 19.05721 | 6.409602 | -0.77643 | -0.68518 | 0.361883 |
| 12 | 73954000 | 73955599 | 10.84297 | 19.15924 | 39.89328 | 17.64761 | -0.66187 | 0.517425 | 0.361919 |
| 4 | 68871200 | 68873399 | 18.36151 | 30.874 | 60.60083 | 26.47512 | -0.5884 | 1.117636 | 0.361966 |
| 4 | 8303800 | 8305799 | 20.76147 | 37.16138 | 25.91268 | 11.36347 | 0.916868 | 0.457368 | 0.361975 |
| 15 | 79749400 | 79758399 | 90.10163 | 138.4056 | 217.3532 | 90.24725 | -0.35578 | 2.974773 | 0.361986 |
| 4 | 112788400 | 112792399 | 72.10231 | 114.8386 | 100.0662 | 41.86146 | 0.535051 | 2.204895 | 0.361993 |
| 11 | 8666800 | 8669399 | 14.76717 | 25.81719 | 46.763 | 20.61423 | -0.46538 | 0.696227 | 0.361996 |
| 10 | 58746200 | 58748599 | 27.23267 | 46.6315 | 40.82354 | 17.90214 | 0.57417 | 0.96519 | 0.361996 |
| 5 | 117961400 | 117964799 | 23.29138 | 38.45856 | 71.60579 | 30.92149 | -0.51364 | 1.342029 | 0.361997 |
| 19 | 49339800 | 49342999 | 27.98958 | 47.76034 | 42.48289 | 18.62456 | 0.550629 | 1.002076 | 0.362003 |
| X | 119666000 | 119666799 | 13.05701 | 24.745 | 21.99888 | 8.890838 | 0.655624 | 0.056896 | 0.36201 |
| 12 | 24209400 | 24210599 | 54.52569 | 87.39926 | 79.86836 | 33.7173 | 0.476561 | 1.983815 | 0.362013 |
| 9 | 23433800 | 23435999 | 20.36543 | 36.07288 | 29.25366 | 12.93467 | 0.692209 | 0.529329 | 0.362016 |
| 14 | 41083400 | 41090399 | 76.49258 | 121.625 | 102.8011 | 42.9077 | 0.576325 | 2.260415 | 0.362032 |
| 13 | 34459800 | 34463599 | 20.87887 | 35.00267 | 65.24307 | 28.4587 | -0.51437 | 1.158297 | 0.362033 |

| | | | | | | | | | |
|---|---|---|---|---|---|---|---|---|---|
| 5 | 21984400 | 21985199 | 6.412225 | 12.35822 | 27.05553 | 10.48581 | -0.57882 | -0.13111 | 0.362043 |
| 18 | 30104600 | 30105999 | 13.17606 | 25.35228 | 18.73538 | 7.295635 | 0.967396 | -0.06935 | 0.36205 |
| 15 | 64813800 | 64814399 | 4.39953 | 8.563454 | 22.35343 | 8.215653 | -0.75551 | -0.35003 | 0.362056 |
| 7 | 16928800 | 16953399 | 417.2721 | 582.0099 | 501.7727 | 208.1426 | 0.378483 | 4.620578 | 0.362069 |
| 17 | 35422600 | 35424999 | 13.32166 | 22.99021 | 50.52767 | 22.16601 | -0.74571 | 0.851889 | 0.362073 |
| 3 | 98686200 | 98687399 | 19.56714 | 33.13267 | 58.59275 | 25.65231 | -0.43809 | 1.058988 | 0.362101 |
| 7 | 26977600 | 26982599 | 28.98432 | 46.97611 | 86.40385 | 36.99454 | -0.51209 | 1.633428 | 0.362108 |
| 17 | 19792200 | 19793399 | 15.11143 | 29.01358 | 18.65113 | 7.307891 | 1.159209 | -0.02925 | 0.362116 |
| 7 | 139944000 | 139945799 | 22.2413 | 38.80051 | 35.43279 | 15.59679 | 0.517761 | 0.746872 | 0.362118 |
| Y | 41076800 | 41079199 | 16.86581 | 30.4157 | 15.41083 | 6.709591 | 1.380282 | 0.359197 | 0.362129 |
| 18 | 17767200 | 17768199 | 8.34896 | 15.52517 | 31.58828 | 13.21012 | -0.57029 | 0.164332 | 0.362131 |
| 3 | 38769200 | 38771399 | 14.02771 | 24.43079 | 46.31045 | 20.36349 | -0.53094 | 0.749047 | 0.362133 |
| 7 | 10704800 | 10706599 | 46.22249 | 74.82206 | 60.12826 | 25.72094 | 0.677887 | 1.67547 | 0.36214 |
| 9 | 48301800 | 48304599 | 13.65893 | 23.873 | 45.1942 | 19.91796 | -0.52869 | 0.692417 | 0.362161 |
| 6 | 11625200 | 11627799 | 25.51013 | 43.774 | 40.91095 | 17.94351 | 0.480716 | 0.940471 | 0.362174 |
| 3 | 125955400 | 125956799 | 12.80632 | 23.85815 | 22.31934 | 9.304502 | 0.545468 | 0.162593 | 0.36218 |
| 3 | 86987200 | 86992999 | 33.95632 | 54.51791 | 105.9303 | 44.83127 | -0.60674 | 1.915366 | 0.36218 |
| 7 | 83992600 | 83995599 | 23.71535 | 39.48876 | 69.72902 | 30.28537 | -0.43651 | 1.245879 | 0.36218 |
| 1 | 148049600 | 148053199 | 40.13282 | 65.88155 | 58.84681 | 25.34506 | 0.539879 | 1.445839 | 0.362185 |
| 4 | 146428800 | 146429799 | 23.08075 | 40.52842 | 28.28092 | 12.51301 | 0.903836 | 0.641275 | 0.362187 |
| 1 | 117066400 | 117067999 | 19.73867 | 35.09311 | 28.80365 | 12.70329 | 0.677759 | 0.498002 | 0.362188 |
| 12 | 25828000 | 25829199 | 20.1297 | 35.35612 | 32.27587 | 14.28386 | 0.518408 | 0.628352 | 0.362188 |
| X | 78303600 | 78304599 | 18.17673 | 32.32663 | 29.10646 | 12.83333 | 0.54535 | 0.487569 | 0.362198 |
| 14 | 8017400 | 8018599 | 23.11827 | 39.97543 | 38.3453 | 16.82273 | 0.445552 | 0.860114 | 0.362212 |
| 17 | 53606200 | 53608199 | 22.79132 | 39.49684 | 37.80442 | 16.58654 | 0.449285 | 0.838518 | 0.362217 |
| 9 | 72842000 | 72843999 | 22.27213 | 38.57017 | 35.3626 | 15.51487 | 0.510937 | 0.839165 | 0.36222 |
| 13 | 50340800 | 50341999 | 5.689054 | 10.97784 | 25.29829 | 9.770483 | -0.64606 | -0.15198 | 0.362236 |
| 9 | 5243400 | 5244399 | 16.56823 | 29.50511 | 27.72143 | 12.20899 | 0.485236 | 0.464798 | 0.362243 |
| 7 | 39075800 | 39077399 | 30.12418 | 50.99195 | 45.25461 | 19.81081 | 0.552912 | 1.084277 | 0.36225 |
| 4 | 108590600 | 108593199 | 34.02937 | 56.7815 | 50.3398 | 21.89295 | 0.554288 | 1.235417 | 0.362258 |
| 16 | 24205600 | 24207999 | 25.74353 | 44.57788 | 38.11285 | 16.72147 | 0.611202 | 0.858889 | 0.362265 |
| 11 | 89856600 | 89858999 | 21.95319 | 38.47435 | 33.11402 | 14.62763 | 0.602883 | 0.672722 | 0.362278 |
| 5 | 14530000 | 14532799 | 36.49121 | 60.23634 | 55.47403 | 23.95047 | 0.498814 | 1.365522 | 0.362282 |
| 13 | 55053800 | 55055799 | 13.06606 | 22.89903 | 43.07344 | 19.02702 | -0.5192 | 0.643929 | 0.362283 |
| 5 | 48033000 | 48033999 | 7.492655 | 14.24571 | 28.83543 | 11.57437 | -0.51338 | 0.000687 | 0.362294 |
| 16 | 19184000 | 19185199 | 17.53822 | 30.97401 | 27.87068 | 12.33041 | 0.54155 | 0.540829 | 0.362301 |
| 5 | 58655600 | 58657799 | 19.0123 | 33.57712 | 30.53534 | 13.50931 | 0.526217 | 0.554341 | 0.362306 |
| 1 | 59133400 | 59134399 | 7.97385 | 14.97759 | 30.17554 | 12.40612 | -0.5367 | 0.090115 | 0.36231 |
| 5 | 69614600 | 69615199 | 7.97385 | 14.97759 | 30.17554 | 12.40612 | -0.5367 | 0.090115 | 0.36231 |
| 11 | 93710400 | 93712199 | 14.61278 | 26.62147 | 25.32165 | 10.8855 | 0.490444 | 0.311779 | 0.36232 |
| 11 | 105071000 | 105073399 | 17.93479 | 30.44627 | 57.48382 | 25.18631 | -0.53199 | 1.028286 | 0.362326 |
| 7 | 27979400 | 27981199 | 12.36546 | 21.83794 | 40.85353 | 18.0743 | -0.50879 | 0.533895 | 0.362341 |
| 2 | 52631600 | 52633799 | 17.39668 | 29.80092 | 53.77765 | 23.5836 | -0.46502 | 0.94008 | 0.362369 |
| 7 | 60014400 | 60017599 | 23.91272 | 42.70283 | 24.6856 | 10.84652 | 1.184135 | 0.491871 | 0.36237 |
| 19 | 29658400 | 29674999 | 403.1247 | 548.8208 | 817.0747 | 340.2275 | -0.44473 | 4.993356 | 0.362373 |
| 9 | 3109400 | 3146799 | 793.7727 | 1048.849 | 856.3146 | 358.0923 | 0.378594 | 5.472279 | 0.36239 |
| 1 | 178100200 | 178104399 | 41.02164 | 65.75508 | 115.5052 | 48.76343 | -0.46445 | 1.976505 | 0.362391 |

| | | | | | | | | | |
|---|---|---|---|---|---|---|---|---|---|
| 12 | 87634200 | 87637599 | 28.07126 | 48.4894 | 36.84958 | 16.16602 | 0.779652 | 0.887699 | 0.362396 |
| X | 93870600 | 93872599 | 20.95733 | 37.26767 | 28.17441 | 12.42355 | 0.795921 | 0.50386 | 0.362413 |
| 3 | 32299400 | 32300999 | 16.01889 | 29.51033 | 24.5686 | 10.40707 | 0.695865 | 0.258287 | 0.362415 |
| 13 | 8216600 | 8217799 | 12.2744 | 22.94753 | 21.01301 | 8.708052 | 0.583354 | 0.134165 | 0.36244 |
| 1 | 118041000 | 118042999 | 51.53447 | 82.86097 | 80.74133 | 34.24441 | 0.389436 | 1.883618 | 0.362446 |
| 13 | 67621200 | 67621599 | 10.84511 | 20.63974 | 20.09254 | 8.048477 | 0.536786 | 0.001692 | 0.362458 |
| 17 | 13178200 | 13179799 | 9.785653 | 17.69625 | 35.89221 | 15.57286 | -0.60383 | 0.341365 | 0.362483 |
| 14 | 24125400 | 24126399 | 20.86752 | 36.59055 | 33.2498 | 14.69402 | 0.52498 | 0.659574 | 0.362498 |
| 9 | 26090800 | 26091999 | 15.46886 | 28.80997 | 23.06984 | 9.622172 | 0.767733 | 0.189223 | 0.3625 |
| 10 | 129349400 | 129353199 | 36.53786 | 60.83854 | 50.40114 | 21.89044 | 0.651867 | 1.265313 | 0.362507 |
| 19 | 13136000 | 13138799 | 26.88342 | 46.60556 | 36.93228 | 16.20408 | 0.720601 | 0.852288 | 0.362508 |
| 7 | 102987200 | 102988199 | 13.97098 | 25.65561 | 24.38823 | 10.37563 | 0.501583 | 0.260364 | 0.362513 |
| 13 | 102484000 | 102484999 | 8.468756 | 15.81667 | 31.13876 | 12.92644 | -0.51478 | 0.134888 | 0.362531 |
| 1 | 184560800 | 184565999 | 35.1847 | 56.6947 | 97.06952 | 41.29846 | -0.41815 | 1.800481 | 0.362567 |
| 7 | 11046600 | 11056999 | 153.531 | 233.0824 | 201.1785 | 83.30162 | 0.493544 | 3.22681 | 0.362574 |
| 2 | 110381000 | 110383799 | 26.04447 | 44.87115 | 39.76503 | 17.44365 | 0.558218 | 0.906307 | 0.362595 |
| 4 | 28812800 | 28815799 | 26.82322 | 43.36396 | 93.09564 | 39.77719 | -0.73812 | 1.686822 | 0.362596 |
| 17 | 70481400 | 70484199 | 26.71996 | 45.30485 | 43.71272 | 19.14548 | 0.432789 | 1.055138 | 0.36261 |
| 3 | 41286600 | 41289399 | 28.75996 | 49.13875 | 41.33553 | 18.12345 | 0.631099 | 0.991772 | 0.362621 |
| 14 | 115317400 | 115319399 | 17.80452 | 32.18806 | 26.33126 | 11.42875 | 0.697577 | 0.37957 | 0.362622 |
| 13 | 56858400 | 56861799 | 27.52757 | 45.15833 | 78.63658 | 33.85873 | -0.42099 | 1.43779 | 0.362629 |
| 14 | 75666600 | 75672199 | 50.11584 | 79.75922 | 130.1867 | 54.40427 | -0.3702 | 2.215857 | 0.362631 |
| 13 | 13186000 | 13186799 | 15.39851 | 28.5359 | 24.17359 | 10.15759 | 0.679305 | 0.220469 | 0.362633 |
| 4 | 89744000 | 89746599 | 25.18293 | 44.03605 | 32.98978 | 14.54523 | 0.802313 | 0.725607 | 0.362643 |
| 5 | 139678800 | 139680199 | 23.32891 | 38.52841 | 70.84374 | 30.59674 | -0.49551 | 1.342065 | 0.362649 |
| 6 | 132539000 | 132539799 | 12.03087 | 22.94959 | 20.95835 | 8.354198 | 0.633901 | -0.00142 | 0.362664 |
| 8 | 66628000 | 66629199 | 12.03087 | 22.94959 | 20.95835 | 8.354198 | 0.633901 | -0.00142 | 0.362664 |
| 13 | 73615200 | 73615999 | 5.591308 | 10.71145 | 26.75511 | 10.54999 | -0.78999 | -0.08231 | 0.36267 |
| X | 25875200 | 25881599 | 77.2457 | 122.8911 | 90.99595 | 38.00734 | 0.767782 | 2.243807 | 0.362672 |
| 9 | 66950600 | 66951399 | 5.860396 | 11.26253 | 26.88771 | 10.50642 | -0.71349 | -0.11042 | 0.362677 |
| 1 | 15004600 | 15006599 | 16.4643 | 30.46605 | 23.44558 | 9.871453 | 0.814664 | 0.234824 | 0.362691 |
| 11 | 88306600 | 88308199 | 8.788963 | 16.2563 | 32.03207 | 13.49467 | -0.53261 | 0.197099 | 0.362695 |
| 2 | 125500400 | 125500799 | 8.793401 | 17.10639 | 17.27092 | 6.376187 | 0.594045 | -0.29636 | 0.362704 |
| 17 | 32142400 | 32145599 | 16.71884 | 28.34065 | 57.38603 | 25.13233 | -0.6325 | 1.037378 | 0.362706 |
| 9 | 8851200 | 8851799 | 6.108461 | 11.83682 | 25.61348 | 9.702823 | -0.53011 | -0.21376 | 0.362712 |
| X | 44324000 | 44333199 | 178.093 | 266.4474 | 245.7817 | 101.3585 | 0.37944 | 3.520365 | 0.362715 |
| 16 | 57206000 | 57207599 | 8.001692 | 14.87488 | 34.88853 | 14.59766 | -0.77559 | 0.165014 | 0.362718 |
| 2 | 37145400 | 37147999 | 45.73122 | 74.03617 | 69.93339 | 29.91804 | 0.445435 | 1.676121 | 0.362753 |
| 12 | 59994000 | 59994999 | 22.99085 | 37.96824 | 70.32623 | 30.37221 | -0.50607 | 1.340028 | 0.36276 |
| 3 | 138277600 | 138286399 | 91.4432 | 140.4251 | 221.4379 | 91.93956 | -0.36207 | 2.980937 | 0.362772 |
| 14 | 9278200 | 9279999 | 14.14306 | 24.71179 | 45.19941 | 19.91555 | -0.47926 | 0.700445 | 0.362779 |
| 5 | 147325000 | 147328799 | 19.93767 | 33.67993 | 60.18792 | 26.3317 | -0.45327 | 1.083721 | 0.362781 |
| 17 | 26506600 | 26527599 | 273.4939 | 387.2207 | 561.3927 | 232.3186 | -0.34941 | 4.394823 | 0.362792 |
| 13 | 65106400 | 65110599 | 65.01324 | 103.7325 | 87.85912 | 36.84563 | 0.578635 | 2.137342 | 0.362805 |
| 10 | 26619600 | 26620799 | 12.25666 | 21.51966 | 41.9922 | 18.57795 | -0.5719 | 0.606514 | 0.362823 |
| 12 | 22725400 | 22725599 | 9.337047 | 18.05938 | 17.7113 | 6.768055 | 0.589249 | -0.17252 | 0.362832 |
| 4 | 43991800 | 43994199 | 21.09948 | 35.2875 | 63.421 | 27.62196 | -0.46183 | 1.185516 | 0.36285 |

| | | | | | | | | | |
|---|---|---|---|---|---|---|---|---|---|
| 16 | 42112000 | 42112799 | 6.151123 | 11.77427 | 26.73697 | 10.56765 | -0.65668 | -0.06656 | 0.362858 |
| 3 | 144761400 | 144763199 | 18.64389 | 33.73794 | 25.71226 | 11.14666 | 0.80022 | 0.379074 | 0.362899 |
| 10 | 19559400 | 19562999 | 34.89704 | 57.51378 | 55.87338 | 24.10475 | 0.42124 | 1.381276 | 0.362899 |
| 5 | 123169000 | 123171399 | 23.7722 | 41.11525 | 38.80451 | 17.0243 | 0.468885 | 0.86154 | 0.362936 |
| 18 | 34208000 | 34209199 | 8.583004 | 16.02334 | 32.23735 | 13.39198 | -0.54718 | 0.138554 | 0.362941 |
| 12 | 102378400 | 102384999 | 59.57791 | 93.56641 | 161.5662 | 67.01327 | -0.46773 | 2.554453 | 0.362945 |
| 3 | 45544400 | 45545999 | 22.17545 | 39.31443 | 27.40271 | 12.11407 | 0.909839 | 0.527768 | 0.362952 |
| X | 78076400 | 78078999 | 28.90153 | 50.16839 | 35.23571 | 15.46145 | 0.894383 | 0.849172 | 0.362968 |
| X | 4394400 | 4403399 | 116.6734 | 182.7941 | 116.2358 | 48.21334 | 0.969001 | 2.626042 | 0.362992 |
| 7 | 32264000 | 32266599 | 28.66644 | 49.58474 | 36.28431 | 15.91869 | 0.834373 | 0.8787 | 0.362993 |
| 1 | 119015400 | 119018399 | 17.17974 | 29.10707 | 58.50696 | 25.61928 | -0.62209 | 1.04373 | 0.362994 |
| 9 | 30338200 | 30340599 | 17.17974 | 29.10707 | 58.50696 | 25.61928 | -0.62209 | 1.04373 | 0.362994 |
| 1 | 192471400 | 192473799 | 11.75322 | 20.75042 | 40.36845 | 17.86077 | -0.56503 | 0.532772 | 0.363002 |
| 1 | 88762600 | 88762999 | 11.65279 | 22.49404 | 19.57286 | 7.548614 | 0.747093 | -0.11271 | 0.363036 |
| 14 | 76514000 | 76517999 | 42.2588 | 67.53724 | 116.3511 | 48.90791 | -0.44128 | 2.07219 | 0.363052 |
| 15 | 34024600 | 34025599 | 12.03753 | 23.23347 | 18.90698 | 7.296585 | 0.841561 | -0.10911 | 0.363053 |
| 4 | 93865600 | 93869799 | 38.25487 | 63.55901 | 51.00708 | 22.12085 | 0.697461 | 1.295446 | 0.363067 |
| 2 | 151082200 | 151088999 | 88.64252 | 138.7751 | 133.9418 | 55.56179 | 0.366872 | 2.622163 | 0.363088 |
| 2 | 151622800 | 151624799 | 22.0878 | 36.75664 | 66.749 | 28.97974 | -0.47676 | 1.24436 | 0.363091 |
| 5 | 119373600 | 119378199 | 39.56987 | 63.55527 | 106.5537 | 45.1217 | -0.3938 | 1.910691 | 0.363091 |
| Y | 6621800 | 6626799 | 33.31878 | 55.66015 | 28.67957 | 12.48195 | 1.33383 | 1.195606 | 0.363101 |
| 10 | 53330000 | 53331799 | 19.86955 | 33.73526 | 59.75366 | 26.18189 | -0.44029 | 1.037628 | 0.363116 |
| 17 | 54700400 | 54704199 | 65.50168 | 104.6096 | 87.32893 | 36.67179 | 0.600932 | 2.109229 | 0.363135 |
| 6 | 38248000 | 38253799 | 42.3437 | 67.92178 | 114.806 | 48.51987 | -0.40783 | 1.957225 | 0.363148 |
| X | 125276200 | 125278799 | 40.05393 | 65.77903 | 58.21419 | 25.07695 | 0.55345 | 1.441491 | 0.363151 |
| 11 | 32592200 | 32593799 | 12.40352 | 21.80368 | 41.43319 | 18.34778 | -0.53381 | 0.587343 | 0.363157 |
| 7 | 20772800 | 20773999 | 15.18508 | 28.14825 | 24.27867 | 10.19786 | 0.654098 | 0.218489 | 0.363173 |
| X | 25652200 | 25670799 | 251.0512 | 370.6985 | 265.6806 | 109.4124 | 0.725903 | 3.761014 | 0.363192 |
| 3 | 137555400 | 137559999 | 47.6839 | 76.10546 | 125.7516 | 52.75918 | -0.38336 | 2.122378 | 0.363221 |
| 17 | 83101200 | 83101799 | 6.115116 | 11.82259 | 26.05143 | 9.974984 | -0.57119 | -0.17747 | 0.363228 |
| 6 | 128037600 | 128039599 | 20.56857 | 34.72259 | 61.34401 | 26.83206 | -0.43691 | 1.092082 | 0.363234 |
| 10 | 32880000 | 32881799 | 26.08171 | 44.92937 | 39.34652 | 17.25999 | 0.575386 | 0.90945 | 0.363245 |
| 3 | 11189000 | 11191999 | 119.4995 | 183.29 | 163.1104 | 67.71125 | 0.461236 | 3.010222 | 0.363259 |
| 11 | 68441000 | 68443399 | 10.1661 | 18.16169 | 37.34563 | 16.40362 | -0.63541 | 0.422372 | 0.363301 |
| 1 | 35207800 | 35208599 | 5.812932 | 11.28898 | 25.84635 | 9.674699 | -0.59483 | -0.25959 | 0.363306 |
| 3 | 129556600 | 129557599 | 15.48899 | 29.11405 | 22.18311 | 9.109325 | 0.856991 | 0.131778 | 0.363312 |
| 5 | 59396600 | 59397199 | 5.510779 | 10.72666 | 24.26097 | 8.915282 | -0.55182 | -0.33056 | 0.363321 |
| 13 | 85078000 | 85078399 | 5.510779 | 10.72666 | 24.26097 | 8.915282 | -0.55182 | -0.33056 | 0.363321 |
| 7 | 56098800 | 56102199 | 33.15232 | 55.10671 | 52.51482 | 22.78197 | 0.450399 | 1.277245 | 0.363332 |
| 3 | 65772400 | 65775999 | 35.20727 | 56.62943 | 100.4462 | 42.62413 | -0.47226 | 1.858181 | 0.363336 |
| 1 | 155432600 | 155434799 | 9.929337 | 17.59251 | 39.3892 | 17.41518 | -0.76431 | 0.480455 | 0.363339 |
| X | 123030000 | 123035399 | 167.7413 | 252.4852 | 225.9794 | 93.32841 | 0.430735 | 3.402822 | 0.363345 |
| 4 | 128474200 | 128484199 | 195.8509 | 284.1131 | 435.594 | 179.5916 | -0.39504 | 4.032843 | 0.36335 |
| 18 | 44314600 | 44317199 | 20.41727 | 36.61646 | 26.60242 | 11.64682 | 0.859806 | 0.445706 | 0.363363 |
| 1 | 35026000 | 35027999 | 11.9188 | 21.02033 | 40.38263 | 17.87086 | -0.54756 | 0.54611 | 0.363365 |
| 18 | 76685000 | 76691799 | 52.34357 | 82.90471 | 141.5872 | 58.88036 | -0.44219 | 2.361498 | 0.363381 |
| 7 | 26023400 | 26045999 | 569.447 | 774.7845 | 636.4609 | 265.0453 | 0.412038 | 5.005691 | 0.363387 |

| | | | | | | | | | |
|---|---|---|---|---|---|---|---|---|---|
| 9 | 62982800 | 62985599 | 23.76371 | 40.95176 | 39.39928 | 17.28333 | 0.440276 | 0.894255 | 0.363387 |
| 16 | 87393200 | 87394799 | 18.1801 | 31.18456 | 56.00519 | 24.5631 | -0.45794 | 0.933173 | 0.363388 |
| 18 | 43528800 | 43529999 | 14.49823 | 27.42021 | 21.81063 | 8.852959 | 0.809326 | 0.082771 | 0.363392 |
| 15 | 79793200 | 79795599 | 19.22779 | 32.54577 | 58.59567 | 25.65065 | -0.4639 | 1.060095 | 0.363392 |
| 6 | 103896400 | 103898199 | 20.02869 | 35.79167 | 27.81795 | 12.21568 | 0.760308 | 0.467963 | 0.363414 |
| 11 | 50783200 | 50786399 | 61.63332 | 98.64348 | 90.49483 | 38.11713 | 0.468449 | 2.041629 | 0.363421 |
| 1 | 147519600 | 147521199 | 14.30341 | 26.36298 | 24.83868 | 10.51445 | 0.519452 | 0.243564 | 0.363442 |
| 6 | 47034000 | 47034999 | 14.30341 | 26.36298 | 24.83868 | 10.51445 | 0.519452 | 0.243564 | 0.363442 |
| 18 | 37572400 | 37573999 | 19.39501 | 34.69372 | 27.77282 | 12.18565 | 0.718596 | 0.456464 | 0.363443 |
| 2 | 149435600 | 149437599 | 25.90185 | 44.3087 | 42.02409 | 18.42714 | 0.458983 | 0.971678 | 0.363448 |
| 9 | 116936000 | 116937399 | 8.329811 | 15.49558 | 31.06112 | 12.98112 | -0.54831 | 0.159266 | 0.363457 |
| 10 | 43131400 | 43132199 | 8.329811 | 15.49558 | 31.06112 | 12.98112 | -0.54831 | 0.159266 | 0.363457 |
| 10 | 114963200 | 114964799 | 18.54342 | 32.92556 | 28.28154 | 12.4866 | 0.611224 | 0.495186 | 0.363458 |
| 7 | 97114400 | 97119599 | 53.03738 | 85.24205 | 82.02955 | 34.76983 | 0.406703 | 1.899331 | 0.363459 |
| 7 | 110106000 | 110109399 | 53.67206 | 86.18686 | 83.66492 | 35.41744 | 0.392625 | 1.927456 | 0.363472 |
| 3 | 83421000 | 83421999 | 8.100574 | 15.23712 | 31.90864 | 13.08733 | -0.58974 | 0.084544 | 0.363473 |
| 7 | 40958000 | 40973999 | 208.5626 | 309.9507 | 266.3038 | 109.7093 | 0.473067 | 3.645364 | 0.363488 |
| 14 | 92252000 | 92252599 | 9.401644 | 18.16242 | 18.14093 | 6.971373 | 0.554852 | -0.15337 | 0.363513 |
| 7 | 75424000 | 75424799 | 12.64708 | 23.56883 | 22.00322 | 9.167881 | 0.549517 | 0.160516 | 0.363545 |
| 10 | 28789800 | 28791599 | 22.61785 | 39.62576 | 33.06912 | 14.60361 | 0.647187 | 0.681647 | 0.363546 |
| 7 | 7973000 | 7974399 | 34.80843 | 57.29626 | 51.40716 | 22.16626 | 0.534995 | 1.394816 | 0.363554 |
| 18 | 66415400 | 66417999 | 16.5467 | 28.45697 | 54.90738 | 24.0848 | -0.5603 | 0.899947 | 0.363576 |
| 17 | 19227000 | 19228199 | 18.96249 | 35.0424 | 22.15088 | 9.343143 | 1.094845 | 0.260048 | 0.363591 |
| 13 | 66747600 | 66752199 | 165.7725 | 248.8182 | 232.385 | 95.90134 | 0.365742 | 3.45681 | 0.363593 |
| 7 | 136975600 | 136977799 | 13.0147 | 22.90483 | 41.9075 | 18.55347 | -0.47889 | 0.578412 | 0.363645 |
| 2 | 68376600 | 68385599 | 199.5827 | 288.0972 | 457.8518 | 188.8909 | -0.45386 | 4.106453 | 0.363647 |
| 7 | 142998600 | 143002199 | 24.72101 | 40.41961 | 77.51658 | 33.33179 | -0.56206 | 1.477496 | 0.36365 |
| 2 | 10673600 | 10673999 | 8.607534 | 16.81091 | 15.08899 | 5.293082 | 0.826545 | -0.48416 | 0.363653 |
| 19 | 11943000 | 11943999 | 7.057332 | 13.37045 | 28.7243 | 11.61673 | -0.60817 | 0.020188 | 0.363656 |
| 2 | 149206200 | 149207599 | 11.12845 | 21.52107 | 17.82513 | 6.816248 | 0.828227 | -0.15325 | 0.36366 |
| 11 | 72143800 | 72146599 | 20.59086 | 34.46978 | 64.02242 | 27.90071 | -0.50918 | 1.172712 | 0.363682 |
| 16 | 3702000 | 3702999 | 9.392299 | 16.89477 | 35.39949 | 15.45915 | -0.656 | 0.372202 | 0.363685 |
| 9 | 114879200 | 114887599 | 59.55808 | 93.52959 | 164.8443 | 68.37239 | -0.49732 | 2.555062 | 0.363713 |
| 14 | 79411200 | 79414199 | 40.75631 | 65.26572 | 111.7881 | 47.12381 | -0.42973 | 2.007355 | 0.363728 |
| 5 | 134867600 | 134882399 | 160.5227 | 237.5451 | 360.8512 | 148.609 | -0.35431 | 3.715347 | 0.363732 |
| 7 | 3170800 | 3172399 | 21.12943 | 36.79511 | 35.17325 | 15.46519 | 0.451966 | 0.766535 | 0.363736 |
| 5 | 134763000 | 134765399 | 18.81461 | 31.726 | 59.02107 | 25.80721 | -0.51109 | 1.093624 | 0.363741 |
| 16 | 94852800 | 94854599 | 26.9138 | 46.77969 | 34.61589 | 15.20208 | 0.81934 | 0.81839 | 0.363762 |
| 4 | 88648200 | 88651199 | 41.91638 | 67.97874 | 66.22933 | 28.36991 | 0.403224 | 1.632032 | 0.363791 |
| Y | 67891400 | 67893999 | 13.54414 | 25.2494 | 14.45637 | 6.019837 | 1.249156 | 0.142613 | 0.363792 |
| 12 | 50915200 | 50918799 | 36.4293 | 60.37644 | 53.75811 | 23.27416 | 0.547792 | 1.320096 | 0.363805 |
| 3 | 31873400 | 31878199 | 41.81737 | 68.62325 | 58.36823 | 25.13527 | 0.610105 | 1.454064 | 0.363807 |
| 5 | 64968400 | 64970399 | 9.067625 | 16.34246 | 37.42982 | 16.30877 | -0.78204 | 0.358506 | 0.363808 |
| 3 | 83476800 | 83478999 | 14.43917 | 25.18885 | 46.80088 | 20.60046 | -0.50219 | 0.725821 | 0.36381 |
| 11 | 100227400 | 100229999 | 25.39828 | 41.78296 | 75.10451 | 32.37653 | -0.46495 | 1.394852 | 0.363815 |
| 7 | 84665000 | 84665599 | 9.65989 | 18.77085 | 18.24666 | 6.79804 | 0.635519 | -0.25313 | 0.363832 |
| 13 | 76212600 | 76212999 | 2.947019 | 5.759692 | 19.30382 | 6.67312 | -1.02392 | -0.59271 | 0.36385 |

| | | | | | | | | | |
|---|---|---|---|---|---|---|---|---|---|
| 11 | 29925400 | 29928799 | 43.28482 | 70.47413 | 57.52063 | 24.69562 | 0.662674 | 1.562915 | 0.363854 |
| 6 | 147114400 | 147115999 | 16.40448 | 29.73128 | 25.93735 | 11.2226 | 0.608101 | 0.353798 | 0.363871 |
| 14 | 50190800 | 50193599 | 36.1566 | 59.50345 | 57.51508 | 24.79782 | 0.427732 | 1.404742 | 0.363909 |
| 15 | 8764200 | 8766799 | 28.219 | 48.07429 | 38.81239 | 17.0132 | 0.689531 | 1.012783 | 0.363922 |
| 6 | 52434200 | 52436199 | 26.14265 | 42.72931 | 77.9251 | 33.50325 | -0.49015 | 1.483627 | 0.363959 |
| 13 | 72115200 | 72117799 | 17.52607 | 30.11939 | 53.81779 | 23.60634 | -0.45007 | 0.910927 | 0.363968 |
| X | 117430600 | 117432599 | 34.29962 | 57.5881 | 46.93189 | 20.49078 | 0.675188 | 1.170186 | 0.363973 |
| 12 | 33551400 | 33554999 | 29.08201 | 49.29571 | 44.13871 | 19.33038 | 0.540116 | 1.06531 | 0.363989 |
| 12 | 26446800 | 26451199 | 23.5212 | 39.07104 | 70.88889 | 30.73859 | -0.47582 | 1.271606 | 0.364002 |
| X | 34594400 | 34597799 | 27.87324 | 48.84834 | 28.2768 | 12.49058 | 1.172146 | 0.698682 | 0.364006 |
| 5 | 86595400 | 86596599 | 15.90695 | 29.7409 | 22.58317 | 9.357589 | 0.851482 | 0.168552 | 0.364038 |
| 15 | 64792000 | 64795199 | 28.41834 | 45.89376 | 92.24498 | 39.35956 | -0.64509 | 1.721873 | 0.364058 |
| 9 | 89880800 | 89885599 | 29.84585 | 48.64619 | 85.42887 | 36.6921 | -0.43885 | 1.534827 | 0.364067 |
| 14 | 80029000 | 80034199 | 47.71312 | 76.97887 | 74.75292 | 31.85608 | 0.400267 | 1.771609 | 0.364072 |
| 18 | 56759200 | 56761999 | 32.4631 | 53.77868 | 49.82842 | 21.56609 | 0.490415 | 1.309008 | 0.364097 |
| 1 | 8134000 | 8136999 | 41.64254 | 68.71676 | 53.94571 | 23.28542 | 0.728495 | 1.383955 | 0.364109 |
| 13 | 45987800 | 45988799 | 12.4081 | 21.99041 | 40.85266 | 18.06122 | -0.49661 | 0.497773 | 0.36412 |
| 18 | 81380400 | 81381999 | 12.4081 | 21.99041 | 40.85266 | 18.06122 | -0.49661 | 0.497773 | 0.36412 |
| 14 | 105771600 | 105773199 | 19.06082 | 34.53518 | 25.86782 | 11.19674 | 0.826813 | 0.377333 | 0.364141 |
| 6 | 17636800 | 17637999 | 7.911613 | 14.58791 | 32.5269 | 13.76147 | -0.71407 | 0.208072 | 0.364167 |
| X | 57890000 | 57892799 | 45.09372 | 73.04699 | 66.69394 | 28.54455 | 0.49518 | 1.664704 | 0.364169 |
| 1 | 40501000 | 40503799 | 19.19283 | 32.53447 | 59.74583 | 26.16583 | -0.49234 | 1.047151 | 0.36418 |
| 12 | 84233000 | 84234799 | 16.07501 | 27.35516 | 55.23406 | 24.20938 | -0.62773 | 0.999639 | 0.36418 |
| 15 | 87369600 | 87373399 | 28.35316 | 45.93871 | 86.78829 | 37.15053 | -0.55102 | 1.63888 | 0.36418 |
| 14 | 121235600 | 121238399 | 23.3289 | 38.46137 | 71.38698 | 30.80202 | -0.50968 | 1.362167 | 0.364182 |
| 6 | 122615400 | 122622999 | 76.05833 | 118.3848 | 186.7021 | 77.50128 | -0.34721 | 2.725213 | 0.364194 |
| 5 | 15678000 | 15678199 | 8.06537 | 15.74211 | 15.03707 | 5.332477 | 0.723286 | -0.45675 | 0.364201 |
| 9 | 74758200 | 74759199 | 8.923066 | 16.57686 | 32.10763 | 13.44605 | -0.50185 | 0.173433 | 0.364206 |
| 4 | 4792000 | 4793799 | 12.30098 | 21.47633 | 45.23926 | 19.92329 | -0.68192 | 0.70217 | 0.364242 |
| 15 | 23205600 | 23207599 | 16.99091 | 30.88965 | 25.94367 | 11.18193 | 0.666248 | 0.341561 | 0.364242 |
| 10 | 43385400 | 43388199 | 38.11541 | 60.81589 | 120.9851 | 50.73826 | -0.651 | 2.120472 | 0.364244 |
| 11 | 7595600 | 7597199 | 18.75519 | 34.75307 | 22.28664 | 9.365909 | 1.078179 | 0.24485 | 0.364252 |
| 9 | 60446200 | 60447599 | 10.1428 | 17.81722 | 41.57453 | 18.401 | -0.82788 | 0.587558 | 0.36426 |
| 14 | 114730200 | 114731199 | 14.87525 | 27.45558 | 24.71459 | 10.4419 | 0.586623 | 0.239204 | 0.364263 |
| 5 | 66869400 | 66871399 | 10.23953 | 18.72655 | 36.10556 | 15.44639 | -0.51787 | 0.281671 | 0.364268 |
| 7 | 36323400 | 36324799 | 19.28823 | 34.15485 | 29.99178 | 13.26181 | 0.578006 | 0.527865 | 0.364278 |
| 4 | 82673200 | 82676399 | 14.85299 | 25.5987 | 51.08512 | 22.40965 | -0.60774 | 0.870652 | 0.364285 |
| X | 4259800 | 4262399 | 23.03386 | 41.17497 | 25.29662 | 11.10553 | 1.097653 | 0.480956 | 0.364302 |
| X | 104298800 | 104299399 | 7.167122 | 13.74429 | 28.60205 | 11.24534 | -0.52588 | -0.07135 | 0.364317 |
| 3 | 100192800 | 100193599 | 6.7528 | 12.88545 | 27.77139 | 11.06251 | -0.59263 | -0.03184 | 0.364322 |
| 8 | 3564400 | 3567999 | 22.66831 | 37.42511 | 70.89641 | 30.61255 | -0.53846 | 1.341987 | 0.364336 |
| 14 | 59130400 | 59131999 | 15.93093 | 27.1242 | 55.73459 | 24.4298 | -0.65264 | 0.996631 | 0.364342 |
| 2 | 165221000 | 165225199 | 105.9291 | 160.1046 | 269.2515 | 111.3353 | -0.47337 | 3.310487 | 0.364355 |
| 13 | 67228800 | 67229999 | 12.23789 | 21.57669 | 40.38208 | 17.87167 | -0.5102 | 0.551898 | 0.36439 |
| 7 | 84593200 | 84594599 | 14.72511 | 27.71137 | 22.214 | 9.103526 | 0.786416 | 0.117103 | 0.364402 |
| 6 | 132004200 | 132006999 | 48.78992 | 79.57932 | 59.20147 | 25.43383 | 0.797861 | 1.550335 | 0.364403 |
| 1 | 144959800 | 144960999 | 14.41608 | 26.95713 | 22.64063 | 9.379229 | 0.707039 | 0.15386 | 0.364409 |

| | | | | | | | | | |
|---|---|---|---|---|---|---|---|---|---|
| 5 | 100583400 | 100585199 | 19.42271 | 33.99981 | 31.00641 | 13.68312 | 0.51986 | 0.673843 | 0.36441 |
| 16 | 18901200 | 18902599 | 20.42467 | 34.30731 | 61.25816 | 26.75303 | -0.45247 | 1.134847 | 0.36441 |
| 14 | 66168200 | 66169799 | 12.893 | 22.58627 | 43.07978 | 19.02308 | -0.53922 | 0.648798 | 0.364415 |
| 11 | 90057800 | 90060999 | 26.55462 | 45.00059 | 43.20947 | 18.92201 | 0.43973 | 1.058588 | 0.364418 |
| 12 | 115357400 | 115359599 | 26.23855 | 46.1375 | 28.40388 | 12.57728 | 1.083958 | 0.641655 | 0.364427 |
| 7 | 60673600 | 60677999 | 59.33916 | 95.37283 | 79.34233 | 33.63209 | 0.616371 | 1.904705 | 0.364445 |
| 2 | 31444000 | 31444999 | 16.5854 | 28.63693 | 51.11743 | 22.42507 | -0.44704 | 0.862781 | 0.364447 |
| 8 | 8639200 | 8639999 | 11.50173 | 21.80538 | 21.56218 | 8.707695 | 0.50407 | 0.038019 | 0.364468 |
| 19 | 17532600 | 17533399 | 11.50173 | 21.80538 | 21.56218 | 8.707695 | 0.50407 | 0.038019 | 0.364468 |
| X | 52936200 | 52939199 | 52.38355 | 84.76938 | 67.50418 | 28.86964 | 0.690655 | 1.698103 | 0.364515 |
| 17 | 94584200 | 94585599 | 15.6271 | 28.88252 | 24.06202 | 10.1474 | 0.699342 | 0.235784 | 0.364531 |
| 5 | 58474400 | 58474999 | 24.55744 | 41.98404 | 39.03408 | 17.11523 | 0.48751 | 0.968186 | 0.36454 |
| 1 | 191730800 | 191731999 | 10.17962 | 18.24871 | 36.38141 | 15.93367 | -0.58771 | 0.401808 | 0.364544 |
| 2 | 177249000 | 177251599 | 25.02485 | 43.27041 | 38.77649 | 17.01188 | 0.543309 | 0.870475 | 0.364577 |
| 5 | 127422200 | 127423999 | 9.321232 | 16.9068 | 35.42442 | 15.31261 | -0.64686 | 0.319506 | 0.364595 |
| 18 | 77134600 | 77135199 | 4.819523 | 9.359329 | 23.32391 | 8.733558 | -0.71461 | -0.27469 | 0.364604 |
| 3 | 125254400 | 125255199 | 11.20064 | 20.07587 | 38.35065 | 16.79838 | -0.5273 | 0.409716 | 0.364606 |
| 11 | 116471400 | 116472799 | 9.149734 | 16.30816 | 38.40936 | 16.90381 | -0.83185 | 0.432307 | 0.364645 |
| 14 | 9286800 | 9288999 | 22.75872 | 39.55995 | 37.15319 | 16.31697 | 0.477112 | 0.802495 | 0.364647 |
| 6 | 86748200 | 86748799 | 5.074785 | 9.890279 | 22.60692 | 8.189517 | -0.54759 | -0.39206 | 0.364656 |
| 7 | 3641200 | 3644799 | 18.3634 | 30.84337 | 63.37922 | 27.67843 | -0.65445 | 1.128559 | 0.364658 |
| 7 | 29357000 | 29357799 | 13.65741 | 25.16122 | 23.44973 | 9.93245 | 0.53457 | 0.238065 | 0.364672 |
| 9 | 70161800 | 70163199 | 13.65741 | 25.16122 | 23.44973 | 9.93245 | 0.53457 | 0.238065 | 0.364672 |
| 5 | 140508400 | 140517199 | 134.0256 | 199.9229 | 320.2026 | 131.9729 | -0.41995 | 3.571182 | 0.364672 |
| 3 | 92592800 | 92593599 | 6.264063 | 12.09576 | 25.95967 | 9.996786 | -0.54119 | -0.15569 | 0.364674 |
| 4 | 64545600 | 64546399 | 7.013232 | 13.45423 | 28.02999 | 11.00853 | -0.526 | -0.07822 | 0.364681 |
| 13 | 66673600 | 66676599 | 40.37634 | 67.61009 | 43.59436 | 19.00157 | 1.012019 | 1.214937 | 0.364698 |
| X | 8335400 | 8338399 | 40.37634 | 67.61009 | 43.59436 | 19.00157 | 1.012019 | 1.214937 | 0.364698 |
| 10 | 12110400 | 12111199 | 11.85951 | 22.3785 | 21.41572 | 8.73474 | 0.538158 | 0.071465 | 0.364731 |
| 5 | 25128800 | 25131199 | 15.27391 | 26.51333 | 48.45499 | 21.26222 | -0.4792 | 0.804192 | 0.364732 |
| 6 | 65358000 | 65359399 | 9.799263 | 17.72607 | 34.87477 | 15.12586 | -0.55961 | 0.339077 | 0.364734 |
| 2 | 118242000 | 118245599 | 27.92665 | 45.57148 | 79.74094 | 34.26376 | -0.43234 | 1.512208 | 0.364751 |
| 7 | 13812400 | 13822799 | 128.5472 | 196.6684 | 181.5117 | 75.31287 | 0.405881 | 3.064412 | 0.364753 |
| 6 | 66820600 | 66821399 | 5.834698 | 11.28097 | 27.2503 | 10.43176 | -0.70263 | -0.17968 | 0.36476 |
| 3 | 133007000 | 133009399 | 52.49562 | 84.23971 | 81.33981 | 34.39712 | 0.39914 | 1.946143 | 0.364765 |
| 2 | 111003000 | 111007599 | 36.27963 | 60.56208 | 49.2438 | 21.42206 | 0.67866 | 1.238037 | 0.36477 |
| 5 | 16658400 | 16661599 | 68.79393 | 109.4528 | 104.6053 | 43.69597 | 0.400314 | 2.23497 | 0.364779 |
| 14 | 5098400 | 5101999 | 36.7235 | 60.42865 | 57.62475 | 24.84376 | 0.447076 | 1.406392 | 0.364795 |
| 17 | 61908800 | 61910599 | 32.453 | 54.60149 | 46.91973 | 20.50195 | 0.599031 | 1.145237 | 0.364796 |
| 5 | 28114400 | 28114999 | 6.469465 | 12.29662 | 27.68137 | 11.11824 | -0.66535 | -0.00798 | 0.364803 |
| 6 | 5476200 | 5476999 | 12.19962 | 22.90781 | 21.06187 | 8.663819 | 0.586266 | 0.107045 | 0.364804 |
| X | 104569800 | 104575599 | 69.49584 | 111.4805 | 77.92479 | 32.93526 | 0.863773 | 1.980387 | 0.364806 |
| 14 | 41358400 | 41359199 | 11.0151 | 21.48602 | 14.75927 | 5.286252 | 1.181356 | -0.38133 | 0.36481 |
| 3 | 4709400 | 4711999 | 32.76215 | 55.46882 | 42.96275 | 18.8087 | 0.748494 | 1.087848 | 0.364818 |
| 7 | 31439200 | 31440799 | 26.27543 | 44.70839 | 43.16186 | 18.9179 | 0.432414 | 1.022822 | 0.364837 |
| 2 | 55325600 | 55328199 | 25.02738 | 40.92551 | 77.17153 | 33.18489 | -0.53769 | 1.474444 | 0.364838 |
| 7 | 41889400 | 41894999 | 79.80616 | 126.8788 | 101.8405 | 42.5018 | 0.650621 | 2.267907 | 0.364856 |

| | | | | | | | | | |
|---|---|---|---|---|---|---|---|---|---|
| 13 | 103024000 | 103027599 | 27.12942 | 44.23953 | 80.14704 | 34.42946 | -0.48304 | 1.520373 | 0.364859 |
| 4 | 115125200 | 115126399 | 14.26075 | 26.78903 | 22.80617 | 9.374851 | 0.696854 | 0.12584 | 0.364867 |
| 10 | 119324400 | 119329799 | 60.87139 | 96.13394 | 153.8108 | 63.87038 | -0.3516 | 2.437616 | 0.364868 |
| 2 | 85792400 | 85795999 | 53.82797 | 86.49201 | 82.78747 | 35.07894 | 0.41399 | 1.907657 | 0.364875 |
| 12 | 78719600 | 78721399 | 28.03489 | 47.14732 | 41.77905 | 18.25325 | 0.554798 | 1.128171 | 0.364882 |
| 1 | 131605800 | 131608799 | 28.60634 | 46.53156 | 82.38002 | 35.3565 | -0.45249 | 1.561106 | 0.364895 |
| 11 | 32061800 | 32066799 | 53.38998 | 84.79156 | 138.0255 | 57.5421 | -0.36939 | 2.283677 | 0.36491 |
| 11 | 21187800 | 21189799 | 11.45089 | 20.26391 | 38.84346 | 17.17803 | -0.54227 | 0.503738 | 0.36491 |
| 11 | 52020600 | 52021999 | 8.05494 | 14.88625 | 31.51172 | 13.29048 | -0.63606 | 0.195663 | 0.364915 |
| 9 | 3296200 | 3298599 | 26.53184 | 45.69288 | 39.73885 | 17.43195 | 0.585122 | 0.911718 | 0.364931 |
| 17 | 20512600 | 20514199 | 30.07581 | 50.91549 | 45.24451 | 19.80702 | 0.551055 | 1.082624 | 0.364939 |
| 13 | 18047200 | 18053799 | 91.80097 | 144.3457 | 127.9558 | 53.08091 | 0.494292 | 2.544032 | 0.364984 |
| 2 | 39304800 | 39306599 | 18.40193 | 32.77698 | 28.95421 | 12.74983 | 0.573826 | 0.478184 | 0.36499 |
| 1 | 81544400 | 81545199 | 10.03943 | 19.5592 | 15.91094 | 5.7898 | 0.919473 | -0.33803 | 0.364993 |
| 8 | 72706800 | 72709399 | 39.31125 | 64.35066 | 60.93946 | 26.21498 | 0.454189 | 1.479534 | 0.365019 |
| X | 109155600 | 109156399 | 12.61105 | 24.17959 | 20.56922 | 8.091376 | 0.753467 | -0.03635 | 0.365034 |
| X | 81890000 | 81891599 | 21.33489 | 37.4715 | 32.17478 | 14.23866 | 0.606362 | 0.636034 | 0.365051 |
| 12 | 95986600 | 95988799 | 23.13004 | 40.39092 | 34.18593 | 15.05732 | 0.626912 | 0.734897 | 0.365053 |
| 13 | 67749600 | 67750999 | 27.68901 | 47.422 | 41.37611 | 18.14473 | 0.579121 | 0.967272 | 0.365059 |
| 14 | 91401000 | 91407799 | 22.77119 | 39.76813 | 31.1912 | 13.73911 | 0.736239 | 0.72753 | 0.365068 |
| 16 | 44293000 | 44294799 | 14.93323 | 26.10303 | 47.30018 | 20.848 | -0.46599 | 0.700803 | 0.365088 |
| 2 | 84728800 | 84740399 | 250.9695 | 356.8173 | 530.223 | 219.2667 | -0.37889 | 4.332483 | 0.365118 |
| 6 | 95955800 | 95958199 | 16.36175 | 28.3613 | 51.1528 | 22.44326 | -0.46086 | 0.824976 | 0.365119 |
| 12 | 91716600 | 91719199 | 18.05425 | 30.85265 | 54.85012 | 24.04923 | -0.44412 | 0.966316 | 0.365136 |
| 7 | 141754200 | 141756399 | 17.94314 | 30.51357 | 55.85543 | 24.4804 | -0.48716 | 1.012306 | 0.365143 |
| 2 | 58721000 | 58723399 | 13.33575 | 22.98976 | 54.21311 | 23.782 | -0.8472 | 0.867522 | 0.365145 |
| 2 | 41670400 | 41671599 | 11.92807 | 22.4995 | 21.97486 | 8.969854 | 0.508158 | 0.076782 | 0.365152 |
| 2 | 9802200 | 9802999 | 5.977568 | 11.52776 | 25.84591 | 9.998919 | -0.6091 | -0.14082 | 0.365154 |
| 7 | 40951400 | 40957199 | 86.77619 | 136.9033 | 122.1709 | 50.70521 | 0.488375 | 2.473722 | 0.36516 |
| 4 | 23260400 | 23263999 | 99.00102 | 153.4671 | 145.6501 | 60.49719 | 0.380099 | 2.800205 | 0.365166 |
| 14 | 60029800 | 60032999 | 18.57692 | 31.46037 | 59.13849 | 25.89235 | -0.52591 | 1.052862 | 0.365175 |
| 8 | 16574000 | 16574399 | 11.96973 | 22.96722 | 20.55825 | 8.069621 | 0.683078 | -0.05103 | 0.365179 |
| 4 | 63849800 | 63850999 | 19.46798 | 34.20831 | 31.82756 | 14.09049 | 0.491105 | 0.617189 | 0.36519 |
| 5 | 115042000 | 115049199 | 70.88299 | 110.6123 | 177.5354 | 73.66988 | -0.36962 | 2.677285 | 0.365224 |
| 5 | 138804800 | 138808799 | 24.74782 | 40.37515 | 79.02314 | 33.95256 | -0.5936 | 1.50961 | 0.365235 |
| 10 | 120486000 | 120488199 | 24.97512 | 42.88207 | 40.38705 | 17.7146 | 0.469755 | 0.931731 | 0.365277 |
| 8 | 97055800 | 97056999 | 8.43016 | 15.88109 | 31.95236 | 13.0717 | -0.52919 | 0.079817 | 0.365283 |
| 2 | 12624000 | 12627399 | 41.71951 | 67.97785 | 63.82007 | 27.40788 | 0.461977 | 1.550539 | 0.365292 |
| 1 | 109135800 | 109137999 | 22.24899 | 38.82433 | 34.2815 | 15.09256 | 0.565973 | 0.739918 | 0.365292 |
| 13 | 117529000 | 117530599 | 17.67242 | 32.39471 | 24.33215 | 10.37701 | 0.836402 | 0.297752 | 0.365298 |
| 2 | 149141600 | 149145199 | 68.91207 | 109.9013 | 97.74667 | 40.96526 | 0.507533 | 2.159406 | 0.365342 |
| 13 | 84552600 | 84553199 | 4.566512 | 8.868725 | 23.89902 | 8.942926 | -0.82578 | -0.28014 | 0.365353 |
| 3 | 25202800 | 25204599 | 37.65136 | 61.79812 | 58.22265 | 25.07454 | 0.463011 | 1.43902 | 0.365362 |
| 1 | 30711400 | 30715999 | 38.07157 | 62.54052 | 58.77885 | 25.32311 | 0.467022 | 1.431461 | 0.365381 |
| 17 | 67419000 | 67419999 | 6.170889 | 11.7642 | 27.19706 | 10.85492 | -0.69547 | -0.03422 | 0.365387 |
| 11 | 6632400 | 6634599 | 29.42196 | 49.46792 | 47.49034 | 20.74697 | 0.43964 | 1.140753 | 0.365412 |
| 3 | 116244800 | 116246599 | 17.76699 | 31.57911 | 28.42454 | 12.53896 | 0.544982 | 0.484522 | 0.365412 |

| | | | | | | | | | |
|---|---|---|---|---|---|---|---|---|---|
| 5 | 73737600 | 73742999 | 45.65106 | 72.97614 | 119.6899 | 50.32843 | -0.37016 | 2.06894 | 0.365416 |
| 12 | 35323200 | 35325999 | 28.66799 | 49.29239 | 39.09734 | 17.1497 | 0.71699 | 0.933933 | 0.365416 |
| 6 | 50902000 | 50905199 | 23.72599 | 40.85491 | 39.39957 | 17.28305 | 0.436614 | 0.900527 | 0.365417 |
| 10 | 33468800 | 33469799 | 7.502093 | 14.23229 | 28.76621 | 11.60047 | -0.51714 | 0.016185 | 0.365418 |
| 10 | 114467600 | 114468799 | 7.880272 | 14.75935 | 31.18649 | 12.88429 | -0.60991 | 0.113931 | 0.365431 |
| 6 | 24663600 | 24665599 | 16.83199 | 28.26 | 65.31252 | 28.51559 | -0.82312 | 1.132954 | 0.365432 |
| 9 | 6910400 | 6912399 | 28.11807 | 47.54274 | 44.74911 | 19.58285 | 0.468228 | 1.083821 | 0.365432 |
| 11 | 10120800 | 10123599 | 27.5213 | 47.87949 | 34.03291 | 14.95467 | 0.877262 | 0.808803 | 0.365448 |
| 15 | 101903800 | 101908399 | 36.37154 | 58.46043 | 102.5463 | 43.4704 | -0.45752 | 1.884295 | 0.365462 |
| 18 | 23289600 | 23291399 | 16.32208 | 28.03427 | 52.13987 | 22.86994 | -0.50755 | 0.911745 | 0.365475 |
| X | 78465000 | 78466999 | 24.839 | 43.58206 | 30.46288 | 13.46962 | 0.901876 | 0.669883 | 0.365476 |
| 1 | 42723200 | 42726799 | 25.36439 | 41.40811 | 79.43281 | 34.13671 | -0.56412 | 1.499505 | 0.36548 |
| 13 | 78384000 | 78384799 | 8.741042 | 16.3137 | 32.22955 | 13.39525 | -0.52164 | 0.14213 | 0.365481 |
| 10 | 81111000 | 81113399 | 19.77382 | 32.82359 | 67.43931 | 29.22885 | -0.65443 | 1.268521 | 0.365483 |
| 5 | 100895200 | 100895999 | 12.20101 | 22.90505 | 21.54891 | 8.867678 | 0.55285 | 0.109687 | 0.365548 |
| 10 | 105709800 | 105712999 | 16.49318 | 28.55365 | 66.22795 | 29.05637 | -0.82285 | 0.856223 | 0.365561 |
| 7 | 49294000 | 49297199 | 40.97004 | 66.58535 | 65.16583 | 27.95812 | 0.399556 | 1.587982 | 0.365579 |
| X | 46064800 | 46066999 | 33.39768 | 56.12141 | 42.62806 | 18.61968 | 0.776338 | 1.154862 | 0.365587 |
| 13 | 12770800 | 12776799 | 77.09907 | 122.9992 | 89.60926 | 37.57233 | 0.795381 | 2.154285 | 0.365621 |
| 2 | 163865800 | 163871199 | 38.79678 | 62.17917 | 109.9854 | 46.42222 | -0.47465 | 1.978168 | 0.36564 |
| X | 122383600 | 122386199 | 25.42839 | 44.40385 | 33.52447 | 14.76585 | 0.791084 | 0.74722 | 0.365644 |
| 5 | 44238000 | 44241199 | 24.02462 | 41.54673 | 38.75026 | 17.00044 | 0.485799 | 0.862775 | 0.365645 |
| 6 | 144479000 | 144482199 | 31.524 | 51.1734 | 87.33556 | 37.43722 | -0.40231 | 1.606099 | 0.365646 |
| 7 | 32074800 | 32079799 | 45.96084 | 75.19851 | 52.33153 | 22.50791 | 0.896861 | 1.495249 | 0.365665 |
| 18 | 62405600 | 62406999 | 15.2531 | 28.345 | 23.80461 | 9.964847 | 0.695942 | 0.204947 | 0.365671 |
| 10 | 70622600 | 70623999 | 14.27865 | 26.83008 | 22.17429 | 9.110835 | 0.739503 | 0.12166 | 0.365674 |
| 11 | 110280800 | 110282999 | 23.25806 | 40.57579 | 35.35401 | 15.56248 | 0.585228 | 0.751636 | 0.36568 |
| 10 | 53456200 | 53457599 | 6.058744 | 11.47299 | 28.14818 | 11.39546 | -0.7986 | 0.014314 | 0.365704 |
| 3 | 88239600 | 88241799 | 12.05178 | 20.93375 | 48.42134 | 21.25612 | -0.81708 | 0.779646 | 0.365706 |
| 15 | 72373200 | 72378999 | 42.4129 | 67.71933 | 120.4437 | 50.56038 | -0.48872 | 2.101576 | 0.365728 |
| 13 | 64309200 | 64329199 | 319.3873 | 445.4314 | 645.8946 | 267.9311 | -0.37156 | 4.620041 | 0.365741 |
| 9 | 118513000 | 118515799 | 16.76002 | 28.90432 | 51.6471 | 22.65662 | -0.44885 | 0.875148 | 0.365753 |
| 2 | 59464400 | 59466799 | 27.23412 | 46.53221 | 42.51654 | 18.64128 | 0.512363 | 0.988266 | 0.365755 |
| 2 | 122635400 | 122639399 | 57.25957 | 90.52657 | 147.2875 | 61.19469 | -0.37448 | 2.408829 | 0.365757 |
| 10 | 34270000 | 34272199 | 15.99318 | 27.41119 | 52.66625 | 23.09719 | -0.55485 | 0.928796 | 0.365765 |
| 14 | 49762000 | 49763599 | 15.99318 | 27.41119 | 52.66625 | 23.09719 | -0.55485 | 0.928796 | 0.365765 |
| 15 | 100263000 | 100268199 | 35.49636 | 57.21955 | 96.77789 | 41.19637 | -0.39989 | 1.788629 | 0.365774 |
| 15 | 36955600 | 36957199 | 18.13908 | 30.71046 | 57.43084 | 25.1426 | -0.51833 | 1.053841 | 0.365775 |
| 8 | 84784400 | 84786399 | 12.04069 | 21.3342 | 39.79659 | 17.59516 | -0.50276 | 0.493992 | 0.365798 |
| 7 | 17924000 | 17928799 | 64.71871 | 103.9639 | 77.82168 | 32.96214 | 0.767522 | 1.930324 | 0.365814 |
| 15 | 56797000 | 56798999 | 17.42175 | 31.13311 | 28.56063 | 12.54177 | 0.52236 | 0.450935 | 0.365823 |
| 17 | 68231600 | 68232799 | 7.483084 | 14.25936 | 28.91364 | 11.551 | -0.50995 | -0.01501 | 0.36583 |
| X | 33403400 | 33406399 | 47.79433 | 78.7723 | 49.54355 | 21.37424 | 1.046713 | 1.41266 | 0.365832 |
| 7 | 61784800 | 61788199 | 63.09162 | 100.4831 | 96.49322 | 40.36452 | 0.394986 | 2.194123 | 0.36584 |
| 3 | 111630600 | 111633599 | 36.25239 | 60.65223 | 43.20399 | 18.82118 | 0.868826 | 1.20985 | 0.365858 |
| X | 31510600 | 31512599 | 19.4287 | 34.95527 | 26.72642 | 11.66957 | 0.789178 | 0.419945 | 0.365871 |
| 2 | 128971400 | 128976199 | 59.96594 | 94.45744 | 155.4598 | 64.50555 | -0.39525 | 2.493745 | 0.365879 |

| | | | | | | | | | |
|---|---|---|---|---|---|---|---|---|---|
| 1 | 81687000 | 81688999 | 17.13321 | 30.58821 | 27.99168 | 12.30185 | 0.524942 | 0.452424 | 0.365889 |
| 4 | 86116400 | 86117799 | 12.0654 | 21.43808 | 39.31512 | 17.34791 | -0.47596 | 0.474442 | 0.365895 |
| 16 | 17257000 | 17266199 | 261.5408 | 378.5758 | 346.108 | 142.7379 | 0.347386 | 4.068135 | 0.3659 |
| 4 | 71700400 | 71704399 | 55.61577 | 89.28197 | 84.65917 | 35.8224 | 0.425915 | 1.938877 | 0.3659 |
| 12 | 35819000 | 35820199 | 11.17311 | 21.33946 | 21.10864 | 8.392359 | 0.52317 | -0.01653 | 0.365926 |
| 19 | 39355000 | 39359799 | 140.9766 | 213.5945 | 202.8634 | 83.94811 | 0.353266 | 3.264303 | 0.365932 |
| 9 | 55747200 | 55747599 | 8.578745 | 16.66783 | 17.24337 | 6.431 | 0.545595 | -0.26496 | 0.365946 |
| 19 | 51103200 | 51103599 | 8.578745 | 16.66783 | 17.24337 | 6.431 | 0.545595 | -0.26496 | 0.365946 |
| 13 | 29725800 | 29726199 | 12.1473 | 22.76261 | 21.03887 | 8.6846 | 0.574592 | 0.118871 | 0.365948 |
| 7 | 47449400 | 47451999 | 40.85015 | 66.69338 | 62.57255 | 26.89155 | 0.464967 | 1.521845 | 0.365961 |
| 4 | 115451400 | 115451999 | 5.516892 | 10.71663 | 23.974 | 8.956431 | -0.55785 | -0.27189 | 0.365968 |
| 8 | 122935000 | 122935799 | 5.516892 | 10.71663 | 23.974 | 8.956431 | -0.55785 | -0.27189 | 0.365968 |
| 18 | 3710000 | 3712599 | 57.04175 | 91.47357 | 86.47923 | 36.53324 | 0.428393 | 1.97333 | 0.365983 |
| 17 | 72074000 | 72075399 | 10.36913 | 18.88378 | 36.00543 | 15.48514 | -0.50704 | 0.305361 | 0.365983 |
| 17 | 60863800 | 60864599 | 19.15683 | 33.58351 | 29.56234 | 13.062 | 0.570677 | 0.647433 | 0.365988 |
| 1 | 167455000 | 167456399 | 13.66615 | 23.91463 | 44.12303 | 19.46583 | -0.49167 | 0.671804 | 0.365989 |
| 1 | 139865600 | 139866999 | 16.85995 | 30.45524 | 27.00178 | 11.73189 | 0.581288 | 0.379185 | 0.365997 |
| 3 | 129254600 | 129259599 | 62.92341 | 99.04327 | 158.7355 | 65.85847 | -0.35782 | 2.51072 | 0.366026 |
| 2 | 168278600 | 168286799 | 105.1873 | 159.4768 | 257.4458 | 106.5576 | -0.41059 | 3.246327 | 0.36604 |
| 12 | 42016400 | 42018799 | 16.16079 | 27.61833 | 54.79683 | 24.02597 | -0.60178 | 0.9574 | 0.366043 |
| 4 | 135249400 | 135253799 | 20.72043 | 34.39261 | 66.92553 | 29.00483 | -0.57629 | 1.272661 | 0.366052 |
| 18 | 85779200 | 85782999 | 48.74387 | 78.46313 | 75.2522 | 31.96893 | 0.414306 | 1.842452 | 0.366059 |
| X | 90034000 | 90034999 | 15.37325 | 28.4877 | 24.2723 | 10.19975 | 0.671227 | 0.221975 | 0.36608 |
| X | 146710800 | 146713399 | 26.18331 | 45.54976 | 35.21908 | 15.47547 | 0.756435 | 0.804472 | 0.366111 |
| 5 | 104840200 | 104844799 | 125.5471 | 191.6428 | 178.7351 | 74.12185 | 0.388211 | 3.108156 | 0.366112 |
| 7 | 12888000 | 12891199 | 40.59961 | 66.56289 | 58.71368 | 25.27409 | 0.557274 | 1.464218 | 0.366118 |
| 17 | 57252800 | 57262799 | 75.49382 | 117.7472 | 189.29 | 78.55169 | -0.37254 | 2.690136 | 0.366121 |
| 18 | 74165400 | 74169799 | 46.8542 | 74.62825 | 125.8728 | 52.66001 | -0.41615 | 2.186066 | 0.366133 |
| 10 | 15110600 | 15111399 | 11.45179 | 20.37141 | 38.33772 | 16.90011 | -0.51186 | 0.462435 | 0.366145 |
| 3 | 32654800 | 32660999 | 41.75728 | 66.74313 | 116.4483 | 48.95648 | -0.45934 | 2.067671 | 0.36615 |
| 6 | 65880800 | 65883599 | 20.64046 | 36.66239 | 28.08825 | 12.39746 | 0.775721 | 0.507324 | 0.366165 |
| 7 | 11471600 | 11485999 | 241.9458 | 356.8981 | 281.1285 | 115.7729 | 0.588363 | 3.775871 | 0.366167 |
| 7 | 72966800 | 72978799 | 102.1882 | 155.5843 | 248.3239 | 102.9118 | -0.38924 | 3.158123 | 0.366197 |
| 13 | 82341600 | 82343399 | 23.65117 | 41.41376 | 33.02514 | 14.57754 | 0.712521 | 0.695977 | 0.366206 |
| 6 | 12091200 | 12092599 | 24.45325 | 42.28042 | 34.62976 | 15.19262 | 0.672391 | 0.860945 | 0.366211 |
| 11 | 32481600 | 32486799 | 27.48067 | 44.72305 | 81.66451 | 35.05584 | -0.49643 | 1.551078 | 0.366212 |
| 6 | 112448600 | 112450399 | 14.68511 | 25.33569 | 51.08937 | 22.41222 | -0.6224 | 0.860019 | 0.366221 |
| 13 | 63486200 | 63487999 | 10.87976 | 19.7354 | 38.07183 | 16.45515 | -0.5285 | 0.336901 | 0.366224 |
| 4 | 128934000 | 128935599 | 10.25749 | 18.53728 | 35.35967 | 15.35446 | -0.51649 | 0.348715 | 0.366228 |
| 16 | 13595800 | 13598999 | 41.90779 | 67.92297 | 65.58536 | 28.08131 | 0.415285 | 1.643887 | 0.366229 |
| 4 | 88606800 | 88607399 | 10.77204 | 20.59564 | 19.65837 | 7.7957 | 0.578077 | -0.03128 | 0.366243 |
| 16 | 30417600 | 30420799 | 21.48389 | 35.72509 | 66.01469 | 28.64623 | -0.50177 | 1.252099 | 0.366248 |
| 10 | 61430400 | 61432599 | 11.05369 | 19.39643 | 42.06755 | 18.60228 | -0.72328 | 0.610115 | 0.366258 |
| 9 | 87287800 | 87291999 | 32.19824 | 53.69081 | 50.86555 | 22.11244 | 0.458874 | 1.236926 | 0.366267 |
| 4 | 20778000 | 20779199 | 10.07088 | 17.8679 | 37.88679 | 16.73838 | -0.68503 | 0.46833 | 0.366267 |
| 16 | 49431600 | 49433399 | 26.08524 | 45.88191 | 27.74985 | 12.28693 | 1.109391 | 0.635481 | 0.366293 |
| 16 | 15937400 | 15945599 | 118.9104 | 184.2666 | 148.8963 | 61.84734 | 0.611551 | 2.81609 | 0.36631 |

| | | | | | | | | | |
|---|---|---|---|---|---|---|---|---|---|
| 11 | 47269800 | 47270999 | 8.183905 | 15.15762 | 31.52821 | 13.25803 | -0.60808 | 0.183168 | 0.366333 |
| 5 | 138316000 | 138316599 | 15.94957 | 28.68054 | 27.21314 | 11.88741 | 0.479159 | 0.400067 | 0.366351 |
| 17 | 48130200 | 48132799 | 13.2364 | 23.05103 | 46.31271 | 20.36359 | -0.61451 | 0.745004 | 0.366351 |
| 6 | 105099800 | 105103599 | 42.63695 | 70.03736 | 56.27669 | 24.24472 | 0.692438 | 1.440722 | 0.366354 |
| X | 3751400 | 3753799 | 45.47663 | 74.05718 | 63.44444 | 27.24086 | 0.593195 | 1.568002 | 0.366367 |
| 1 | 191751200 | 191752999 | 13.86737 | 24.27346 | 44.6469 | 19.70154 | -0.48713 | 0.670973 | 0.36637 |
| 12 | 23899200 | 23900199 | 20.846 | 36.64676 | 28.61622 | 12.6719 | 0.742572 | 0.612291 | 0.366371 |
| 2 | 17063800 | 17074399 | 177.2771 | 265.508 | 245.6888 | 101.341 | 0.376277 | 3.500248 | 0.36639 |
| 13 | 78683600 | 78685199 | 22.19994 | 39.22847 | 29.11028 | 12.87742 | 0.818339 | 0.56579 | 0.366394 |
| 14 | 21457600 | 21458799 | 13.49979 | 25.04337 | 23.92539 | 10.03904 | 0.508688 | 0.19954 | 0.366397 |
| 7 | 50484800 | 50486999 | 25.67313 | 44.5082 | 37.5279 | 16.46542 | 0.631451 | 0.846883 | 0.366416 |
| 10 | 17071800 | 17072399 | 4.676206 | 9.122666 | 23.48617 | 8.396583 | -0.70093 | -0.44521 | 0.366429 |
| 4 | 62539200 | 62541799 | 11.96238 | 20.96703 | 43.08083 | 19.032 | -0.64599 | 0.63433 | 0.366437 |
| 3 | 155961800 | 155964399 | 23.92086 | 41.28948 | 39.34787 | 17.26163 | 0.454401 | 0.880383 | 0.36644 |
| 2 | 102353200 | 102354799 | 14.96991 | 27.89283 | 23.76748 | 9.905858 | 0.67986 | 0.185192 | 0.366442 |
| 5 | 109247600 | 109248399 | 14.96991 | 27.89283 | 23.76748 | 9.905858 | 0.67986 | 0.185192 | 0.366442 |
| 13 | 50325800 | 50328799 | 36.67067 | 59.01235 | 100.3366 | 42.60808 | -0.41048 | 1.845354 | 0.366463 |
| 15 | 30073800 | 30074799 | 29.28105 | 49.00605 | 45.98449 | 20.03692 | 0.472599 | 1.186003 | 0.366467 |
| 16 | 56987000 | 56987799 | 5.561945 | 10.73421 | 25.88623 | 9.99316 | -0.71111 | -0.1559 | 0.366488 |
| 4 | 117156600 | 117157799 | 7.243462 | 13.59302 | 30.14918 | 12.41484 | -0.67606 | 0.088986 | 0.366499 |
| 14 | 122521000 | 122522199 | 7.243462 | 13.59302 | 30.14918 | 12.41484 | -0.67606 | 0.088986 | 0.366499 |
| 8 | 36313000 | 36315199 | 20.87852 | 36.43602 | 34.49165 | 15.1859 | 0.466138 | 0.733129 | 0.366575 |
| 16 | 31397600 | 31399399 | 16.31219 | 28.11513 | 51.62468 | 22.64637 | -0.48808 | 0.879673 | 0.366599 |
| 6 | 73584800 | 73585799 | 16.48609 | 29.92203 | 26.64532 | 11.50828 | 0.580682 | 0.35014 | 0.366603 |
| 13 | 110739000 | 110739999 | 22.33182 | 39.25064 | 28.99386 | 12.83876 | 0.821763 | 0.620717 | 0.366609 |
| X | 31729600 | 31732799 | 24.94834 | 44.96428 | 21.4533 | 9.347492 | 1.467052 | 0.439683 | 0.366629 |
| X | 54405600 | 54409599 | 51.30959 | 83.66206 | 57.4911 | 24.69612 | 0.911753 | 1.561294 | 0.366636 |
| 13 | 91608200 | 91610599 | 18.85339 | 31.62185 | 62.24323 | 27.15822 | -0.59254 | 1.145376 | 0.36665 |
| 1 | 132257800 | 132260599 | 12.59533 | 22.00077 | 45.22322 | 19.92221 | -0.64679 | 0.696268 | 0.366652 |
| 3 | 75910800 | 75911399 | 12.55174 | 23.59165 | 22.52182 | 9.249315 | 0.534169 | 0.107801 | 0.366673 |
| 17 | 95162200 | 95164199 | 22.82111 | 39.87609 | 34.82281 | 15.34379 | 0.582209 | 0.727525 | 0.366697 |
| 10 | 7339200 | 7344999 | 62.32503 | 97.06841 | 192.7244 | 79.99382 | -0.67829 | 2.709255 | 0.366703 |
| 7 | 36538800 | 36542199 | 26.58859 | 43.66048 | 76.06726 | 32.76595 | -0.42107 | 1.423669 | 0.366723 |
| 17 | 76146000 | 76147599 | 16.72169 | 29.67982 | 27.28099 | 12.04864 | 0.513965 | 0.485472 | 0.366729 |
| X | 80704200 | 80705799 | 26.7238 | 46.66338 | 32.95222 | 14.5133 | 0.88715 | 0.754762 | 0.36678 |
| 7 | 17515600 | 17534999 | 375.8432 | 531.8965 | 427.4588 | 176.9062 | 0.50087 | 4.40514 | 0.36679 |
| 17 | 19926000 | 19926999 | 17.21644 | 31.1249 | 24.04901 | 10.43815 | 0.779604 | 0.371833 | 0.366798 |
| 15 | 93592800 | 93595599 | 20.03739 | 33.83043 | 60.26308 | 26.36034 | -0.4488 | 1.086344 | 0.366806 |
| 7 | 9940200 | 9947199 | 91.61461 | 145.3337 | 99.80163 | 41.55968 | 0.872357 | 2.341438 | 0.366809 |
| 8 | 93169800 | 93173199 | 23.69215 | 39.3335 | 70.28474 | 30.46512 | -0.45392 | 1.278312 | 0.366823 |
| 11 | 116384200 | 116393799 | 51.53795 | 81.17312 | 169.0487 | 70.14309 | -0.73443 | 2.492595 | 0.366854 |
| 17 | 54690200 | 54693199 | 32.27442 | 54.28793 | 46.36178 | 20.25667 | 0.607854 | 1.144143 | 0.366862 |
| X | 120826800 | 120827599 | 15.4774 | 28.32948 | 25.73209 | 10.99599 | 0.562204 | 0.293089 | 0.366867 |
| 13 | 66865800 | 66886599 | 420.0617 | 584.9872 | 510.0444 | 211.6252 | 0.359992 | 4.644142 | 0.366872 |
| 17 | 18835400 | 18853599 | 292.9831 | 424.8987 | 335.9443 | 138.5113 | 0.559546 | 4.04301 | 0.366876 |
| Y | 3903600 | 3905399 | 12.7131 | 23.94646 | 13.43122 | 5.495885 | 1.298124 | 0.066847 | 0.366878 |
| Y | 13419200 | 13421599 | 12.7131 | 23.94646 | 13.43122 | 5.495885 | 1.298124 | 0.066847 | 0.366878 |

| | | | | | | | | | |
|---|---|---|---|---|---|---|---|---|---|
| 9 | 61571400 | 61572199 | 6.01322 | 11.51395 | 26.75703 | 10.56594 | -0.68899 | -0.07324 | 0.366888 |
| 9 | 36100800 | 36102199 | 24.78479 | 42.79343 | 37.867 | 16.61213 | 0.561167 | 0.882295 | 0.366899 |
| 4 | 65531400 | 65533399 | 41.04401 | 65.42861 | 123.3981 | 51.68445 | -0.57562 | 2.154542 | 0.366901 |
| 1 | 8257400 | 8260799 | 47.28892 | 76.31958 | 74.0328 | 31.56259 | 0.402379 | 1.762195 | 0.366912 |
| 12 | 99645400 | 99647199 | 16.87767 | 31.04774 | 24.47894 | 10.38938 | 0.771922 | 0.272883 | 0.366941 |
| 13 | 54712400 | 54713999 | 9.122272 | 16.31877 | 37.90358 | 16.62988 | -0.80885 | 0.407423 | 0.366951 |
| 14 | 18142000 | 18143799 | 15.05724 | 27.52194 | 25.70617 | 11.00524 | 0.520203 | 0.296652 | 0.366958 |
| 19 | 47394200 | 47397199 | 35.61267 | 57.04959 | 110.2804 | 46.51396 | -0.60319 | 1.990144 | 0.366976 |
| 5 | 101018000 | 101018599 | 16.43067 | 29.31418 | 27.73923 | 12.19783 | 0.476559 | 0.452821 | 0.366981 |
| 12 | 70392000 | 70394799 | 23.68218 | 38.92612 | 73.49279 | 31.67015 | -0.536 | 1.401862 | 0.367002 |
| 1 | 66894600 | 66900399 | 41.74602 | 69.06262 | 46.65419 | 20.17326 | 0.94476 | 1.350402 | 0.367044 |
| 6 | 49393000 | 49394399 | 16.7124 | 30.3251 | 26.53015 | 11.46217 | 0.605627 | 0.351938 | 0.367057 |
| 13 | 118285600 | 118288399 | 56.62016 | 90.58233 | 87.9877 | 37.03931 | 0.385482 | 2.047059 | 0.367067 |
| 17 | 34342800 | 34343799 | 6.116299 | 11.82528 | 25.43929 | 9.739028 | -0.53609 | -0.17765 | 0.367075 |
| X | 33144600 | 33148399 | 30.9963 | 54.05654 | 29.41734 | 12.94631 | 1.26098 | 0.788543 | 0.367094 |
| 3 | 38656400 | 38657599 | 15.15834 | 28.23566 | 23.76027 | 9.90775 | 0.697269 | 0.188739 | 0.367096 |
| X | 88657600 | 88660399 | 29.35394 | 50.73034 | 36.88362 | 16.18119 | 0.84354 | 0.891458 | 0.367103 |
| 5 | 4804800 | 4809799 | 18.36008 | 30.16722 | 81.10916 | 34.94686 | -1.04442 | 1.399735 | 0.367142 |
| 13 | 119599200 | 119599999 | 5.109605 | 9.911782 | 23.84672 | 8.988832 | -0.67389 | -0.24704 | 0.367203 |
| 12 | 93465400 | 93465999 | 5.238429 | 10.17775 | 23.40355 | 8.729248 | -0.59504 | -0.28378 | 0.367246 |
| 9 | 50207600 | 50208999 | 14.55615 | 27.09105 | 23.10591 | 9.648696 | 0.67584 | 0.185281 | 0.367271 |
| 11 | 30989000 | 30995599 | 74.65641 | 119.099 | 92.67558 | 38.8565 | 0.700482 | 2.153792 | 0.367275 |
| 7 | 32876400 | 32879799 | 42.58377 | 70.77822 | 46.54682 | 20.19181 | 0.983759 | 1.300247 | 0.367292 |
| 3 | 118107800 | 118110799 | 33.86252 | 56.80195 | 47.4978 | 20.72582 | 0.63818 | 1.177614 | 0.36732 |
| 18 | 83695800 | 83696199 | 3.414152 | 6.674491 | 18.88624 | 6.482829 | -0.77881 | -0.62076 | 0.367322 |
| 1 | 7557800 | 7560399 | 24.81288 | 40.79586 | 82.98406 | 35.76437 | -0.64322 | 1.407756 | 0.367322 |
| 4 | 109830800 | 109836199 | 64.59059 | 100.6451 | 182.3197 | 75.66976 | -0.54565 | 2.699718 | 0.367341 |
| X | 97649400 | 97650399 | 17.55425 | 31.66917 | 27.09088 | 11.78931 | 0.631436 | 0.393409 | 0.367376 |
| 15 | 56509800 | 56511399 | 17.98853 | 31.83295 | 29.55136 | 13.06874 | 0.497701 | 0.524168 | 0.367379 |
| 4 | 120775600 | 120780599 | 65.84143 | 104.7752 | 103.1577 | 43.10256 | 0.357769 | 2.226081 | 0.36738 |
| 3 | 75854000 | 75854799 | 13.05332 | 22.82398 | 44.16029 | 19.46911 | -0.55982 | 0.681859 | 0.3674 |
| 12 | 69939200 | 69940399 | 9.074002 | 16.88551 | 32.69768 | 13.65748 | -0.49866 | 0.16574 | 0.36742 |
| 16 | 58001000 | 58002199 | 8.184457 | 15.19534 | 31.03526 | 13.00786 | -0.57827 | 0.16951 | 0.367428 |
| 11 | 117801600 | 117805399 | 25.88082 | 42.1606 | 80.24834 | 34.46022 | -0.55507 | 1.533703 | 0.367438 |
| 1 | 146229400 | 146231399 | 27.32235 | 46.2303 | 44.23757 | 19.36318 | 0.444566 | 1.074736 | 0.36744 |
| 18 | 80248800 | 80250599 | 9.619528 | 17.51904 | 33.91607 | 14.58613 | -0.52846 | 0.296629 | 0.367455 |
| 7 | 60219800 | 60228799 | 112.3824 | 174.488 | 148.1358 | 61.51504 | 0.542499 | 2.780126 | 0.367455 |
| 1 | 155716600 | 155718199 | 17.10924 | 31.19185 | 24.20305 | 10.3953 | 0.782606 | 0.321599 | 0.367457 |
| 11 | 120205600 | 120207999 | 13.24715 | 23.02417 | 47.90378 | 21.03701 | -0.66508 | 0.775084 | 0.367466 |
| 2 | 84728800 | 84738599 | 237.3404 | 338.8254 | 508.4148 | 210.1035 | -0.38698 | 4.271912 | 0.367477 |
| 13 | 56063800 | 56065399 | 12.1709 | 21.23395 | 45.26915 | 19.92745 | -0.69899 | 0.714609 | 0.367485 |
| 9 | 41888200 | 41889399 | 17.47983 | 31.01417 | 29.19404 | 12.8975 | 0.479603 | 0.497355 | 0.36749 |
| 13 | 85157600 | 85158799 | 17.80016 | 31.50048 | 29.72463 | 13.14529 | 0.474457 | 0.524757 | 0.367492 |
| 12 | 21808000 | 21815399 | 128.526 | 198.9212 | 144.2875 | 59.93335 | 0.765805 | 2.843453 | 0.367494 |
| 12 | 109275000 | 109276799 | 16.98755 | 30.43309 | 28.15399 | 12.3372 | 0.512456 | 0.432607 | 0.367499 |
| 4 | 120262200 | 120263199 | 4.961258 | 9.601733 | 24.27343 | 9.249302 | -0.7589 | -0.20851 | 0.367512 |
| 8 | 36136600 | 36137799 | 19.66423 | 35.30536 | 26.62378 | 11.6452 | 0.807167 | 0.433209 | 0.367513 |

| | | | | | | | | | |
|---|---|---|---|---|---|---|---|---|---|
| 17 | 22712800 | 22717999 | 62.59578 | 100.0712 | 93.30804 | 39.2395 | 0.443296 | 2.077127 | 0.36754 |
| X | 161471200 | 161472799 | 16.67826 | 31.14613 | 22.55099 | 9.363003 | 0.917533 | 0.182884 | 0.367545 |
| 17 | 6148200 | 6148999 | 10.55896 | 20.33207 | 20.18831 | 7.844337 | 0.548477 | -0.09838 | 0.367585 |
| 12 | 115753400 | 115754799 | 21.49969 | 37.57094 | 34.94359 | 15.3982 | 0.491739 | 0.719383 | 0.367587 |
| 12 | 5460000 | 5462599 | 16.19911 | 27.61439 | 54.75111 | 24.00107 | -0.60127 | 0.981931 | 0.367613 |
| 12 | 71306400 | 71309999 | 26.99232 | 45.86932 | 43.64431 | 19.12738 | 0.452942 | 1.036142 | 0.367637 |
| 7 | 38243600 | 38246199 | 44.03123 | 71.57856 | 65.41275 | 28.06815 | 0.498503 | 1.591277 | 0.367652 |
| 4 | 75164000 | 75165799 | 24.03675 | 41.97004 | 34.7869 | 15.32105 | 0.657146 | 0.744129 | 0.367678 |
| 5 | 124981800 | 124984399 | 17.81605 | 30.17317 | 57.42852 | 25.14402 | -0.54363 | 1.049455 | 0.36768 |
| 2 | 116010600 | 116010999 | 7.597858 | 14.81552 | 12.57147 | 4.521979 | 0.875102 | -0.42292 | 0.367695 |
| X | 81450400 | 81452599 | 29.45683 | 50.33846 | 37.23831 | 16.32727 | 0.815931 | 0.991018 | 0.367704 |
| 11 | 11634400 | 11636399 | 18.56847 | 31.09021 | 62.70102 | 27.32817 | -0.62738 | 1.164181 | 0.367733 |
| 15 | 38477000 | 38479199 | 19.67945 | 32.94453 | 62.18628 | 27.10077 | -0.53223 | 1.170529 | 0.367738 |
| 3 | 93737600 | 93743399 | 125.9306 | 192.6106 | 183.7842 | 76.25078 | 0.357565 | 3.067777 | 0.367746 |
| 8 | 55979400 | 55982399 | 52.6646 | 84.51461 | 82.59572 | 34.93057 | 0.381842 | 1.94542 | 0.367749 |
| 1 | 191746400 | 191752999 | 65.58258 | 103.1448 | 163.5815 | 67.86186 | -0.34361 | 2.52968 | 0.367758 |
| 6 | 26952200 | 26952599 | 4.611385 | 9.009367 | 23.97341 | 8.359541 | -0.71433 | -0.52906 | 0.367769 |
| 15 | 102922600 | 102924399 | 10.13792 | 18.21619 | 35.83822 | 15.66509 | -0.56702 | 0.383384 | 0.367781 |
| 4 | 132169400 | 132171599 | 26.37354 | 45.84355 | 35.79284 | 15.71957 | 0.742397 | 0.817498 | 0.367792 |
| X | 114323600 | 114324599 | 18.25572 | 32.64005 | 24.63357 | 10.81265 | 0.802869 | 0.446511 | 0.367843 |
| 11 | 47546000 | 47547799 | 11.88036 | 21.05333 | 39.79718 | 17.59489 | -0.52172 | 0.491005 | 0.367856 |
| 1 | 120722600 | 120723999 | 8.424145 | 15.80487 | 31.26184 | 12.87826 | -0.51291 | 0.103764 | 0.367859 |
| 9 | 38262800 | 38263199 | 9.035652 | 17.53857 | 18.26349 | 6.864281 | 0.525851 | -0.22969 | 0.367859 |
| 7 | 85287400 | 85292199 | 53.4175 | 86.45587 | 66.64867 | 28.50699 | 0.737538 | 1.694992 | 0.367864 |
| 4 | 30534600 | 30537399 | 21.02948 | 37.61623 | 26.48375 | 11.62037 | 0.902658 | 0.464916 | 0.367871 |
| 14 | 97402400 | 97407399 | 43.7015 | 70.91867 | 67.14945 | 28.77748 | 0.445131 | 1.62297 | 0.367884 |
| 13 | 68648200 | 68650799 | 45.17397 | 73.04325 | 70.35023 | 30.0653 | 0.415711 | 1.700936 | 0.367891 |
| 1 | 14308600 | 14310399 | 9.42131 | 17.27882 | 33.48061 | 14.26854 | -0.52089 | 0.252803 | 0.367901 |
| 18 | 74367400 | 74402599 | 676.0741 | 885.7842 | 1219.056 | 510.2752 | -0.38681 | 5.607946 | 0.367917 |
| 6 | 19926600 | 19927599 | 12.10963 | 23.42619 | 18.423 | 7.03416 | 0.905052 | -0.14418 | 0.36792 |
| 15 | 85743600 | 85745199 | 7.6962 | 14.46308 | 29.63437 | 12.17363 | -0.55937 | 0.085324 | 0.367922 |
| 12 | 84717000 | 84719399 | 18.60358 | 31.51032 | 57.46437 | 25.16054 | -0.48223 | 1.051593 | 0.367931 |
| 5 | 147057600 | 147066399 | 88.0813 | 135.462 | 214.2705 | 88.98168 | -0.36488 | 2.9504 | 0.367932 |
| 7 | 31437600 | 31448199 | 155.529 | 234.731 | 219.3752 | 90.66997 | 0.371742 | 3.347192 | 0.36794 |
| X | 122367000 | 122369599 | 17.09601 | 31.7594 | 22.94545 | 9.609422 | 0.910762 | 0.218431 | 0.367943 |
| 7 | 67021600 | 67023999 | 15.02102 | 25.86157 | 51.08141 | 22.40731 | -0.59325 | 0.881207 | 0.367946 |
| 5 | 30295800 | 30298399 | 22.43173 | 37.43941 | 66.20829 | 28.79895 | -0.43856 | 1.215137 | 0.367953 |
| X | 60489200 | 60489599 | 9.185992 | 17.96594 | 12.71856 | 4.306709 | 1.209548 | -0.60079 | 0.367965 |
| 6 | 66512000 | 66512799 | 6.768642 | 12.83134 | 28.18249 | 11.38291 | -0.63787 | 0.012542 | 0.367979 |
| 11 | 121127400 | 121128599 | 6.768642 | 12.83134 | 28.18249 | 11.38291 | -0.63787 | 0.012542 | 0.367979 |
| 8 | 64746600 | 64747799 | 8.634426 | 16.2065 | 34.09684 | 14.03569 | -0.60031 | 0.111317 | 0.36799 |
| 5 | 131320800 | 131323199 | 16.95147 | 28.97735 | 53.73776 | 23.56212 | -0.50462 | 0.958052 | 0.367999 |
| 13 | 54817000 | 54821399 | 26.60611 | 43.54396 | 76.98667 | 33.11562 | -0.44422 | 1.465276 | 0.368011 |
| 7 | 31995400 | 31996199 | 19.47658 | 34.18462 | 31.45727 | 13.91322 | 0.507005 | 0.634966 | 0.368015 |
| 10 | 14818200 | 14820599 | 19.97318 | 35.04266 | 32.74009 | 14.47585 | 0.485009 | 0.643778 | 0.368025 |
| 13 | 117598600 | 117600199 | 19.97318 | 35.04266 | 32.74009 | 14.47585 | 0.485009 | 0.643778 | 0.368025 |
| 2 | 53101600 | 53103799 | 24.33834 | 42.19106 | 38.23182 | 16.77423 | 0.528045 | 0.843962 | 0.368051 |

| | | | | | | | | | |
|---|---|---|---|---|---|---|---|---|---|
| X | 29318800 | 29320999 | 47.63794 | 77.16649 | 66.34529 | 28.39482 | 0.581666 | 1.670445 | 0.368056 |
| 8 | 5319800 | 5322599 | 23.33809 | 40.38139 | 38.34497 | 16.82291 | 0.460301 | 0.856102 | 0.368059 |
| X | 69851400 | 69853799 | 33.15179 | 55.6212 | 48.04805 | 20.96865 | 0.591462 | 1.175019 | 0.368064 |
| 3 | 37722000 | 37724599 | 44.20273 | 70.1567 | 146.4687 | 61.02364 | -0.72864 | 2.297171 | 0.368066 |
| 5 | 93622000 | 93624799 | 41.82954 | 68.26374 | 62.26323 | 26.75452 | 0.505106 | 1.52717 | 0.368077 |
| 6 | 145516800 | 145518799 | 11.24958 | 19.98438 | 39.31195 | 17.34954 | -0.5769 | 0.468128 | 0.368079 |
| 16 | 70063200 | 70067599 | 42.85455 | 69.60007 | 66.01389 | 28.30757 | 0.443725 | 1.604473 | 0.368089 |
| 19 | 7956800 | 7958799 | 28.25982 | 48.78676 | 38.00985 | 16.67473 | 0.743869 | 0.896608 | 0.368094 |
| 10 | 22630200 | 22632599 | 24.49744 | 42.93422 | 31.82534 | 14.05863 | 0.81735 | 0.684745 | 0.36813 |
| 3 | 91312000 | 91315399 | 53.6043 | 86.38257 | 76.36323 | 32.49303 | 0.533604 | 1.816699 | 0.368133 |
| 1 | 9769600 | 9771399 | 13.89872 | 24.04239 | 50.02397 | 21.94661 | -0.66648 | 0.831328 | 0.368151 |
| 3 | 92397600 | 92398599 | 11.39734 | 21.53013 | 20.47474 | 8.331486 | 0.550225 | 0.057522 | 0.368171 |
| 15 | 92511200 | 92512599 | 11.39734 | 21.53013 | 20.47474 | 8.331486 | 0.550225 | 0.057522 | 0.368171 |
| 18 | 88515200 | 88516399 | 11.39734 | 21.53013 | 20.47474 | 8.331486 | 0.550225 | 0.057522 | 0.368171 |
| 1 | 71758000 | 71758599 | 5.214436 | 10.15716 | 23.11002 | 8.429018 | -0.54983 | -0.36342 | 0.368182 |
| 16 | 93376200 | 93378399 | 15.47501 | 26.62921 | 52.15987 | 22.88002 | -0.58154 | 0.888194 | 0.3682 |
| 7 | 132914200 | 132918999 | 36.69553 | 58.91239 | 104.5802 | 44.25599 | -0.47679 | 1.921367 | 0.368206 |
| 13 | 98935600 | 98939399 | 55.50283 | 89.06267 | 85.51881 | 36.16267 | 0.407114 | 1.952749 | 0.36821 |
| X | 113571200 | 113573399 | 45.52239 | 74.41601 | 59.59719 | 25.62472 | 0.694001 | 1.512933 | 0.36823 |
| 7 | 5546200 | 5550799 | 60.35644 | 97.12874 | 75.94337 | 32.24947 | 0.707896 | 1.868667 | 0.368248 |
| 8 | 23240000 | 23244999 | 38.31828 | 61.57636 | 104.2231 | 44.1676 | -0.40654 | 1.894026 | 0.368254 |
| 1 | 102401800 | 102403399 | 16.25417 | 30.59375 | 21.6217 | 8.858434 | 0.967436 | 0.12934 | 0.368261 |
| 13 | 19806800 | 19809599 | 18.66349 | 31.35988 | 63.98851 | 27.9492 | -0.64437 | 1.125842 | 0.368269 |
| 10 | 49037200 | 49037799 | 9.79042 | 18.94591 | 19.24035 | 7.33491 | 0.542155 | -0.17155 | 0.368281 |
| 8 | 55868000 | 55870599 | 33.04111 | 53.39904 | 91.21557 | 38.95779 | -0.40981 | 1.707149 | 0.368289 |
| 8 | 62796200 | 62799399 | 50.04767 | 80.53495 | 78.98299 | 33.53761 | 0.381564 | 1.855839 | 0.36832 |
| 15 | 73926400 | 73932199 | 46.06666 | 73.28924 | 128.9686 | 53.86904 | -0.47901 | 2.22338 | 0.368324 |
| 7 | 27680400 | 27686999 | 135.9278 | 207.5587 | 188.0052 | 77.9723 | 0.430714 | 3.105339 | 0.368325 |
| 17 | 73952200 | 73954399 | 12.21624 | 21.6009 | 39.82466 | 17.61479 | -0.48704 | 0.518385 | 0.368341 |
| 13 | 81172000 | 81174999 | 44.4887 | 71.10275 | 119.3846 | 50.18478 | -0.40427 | 2.073444 | 0.368346 |
| 6 | 87280600 | 87281599 | 11.1351 | 19.8536 | 37.85344 | 16.65434 | -0.52852 | 0.445248 | 0.368346 |
| X | 61124200 | 61128999 | 56.6201 | 90.91845 | 82.68403 | 35.00079 | 0.486541 | 1.933613 | 0.368351 |
| 7 | 23323000 | 23324999 | 78.20758 | 123.3673 | 109.0689 | 45.26482 | 0.501653 | 2.469515 | 0.368363 |
| 1 | 47596800 | 47598399 | 22.38708 | 39.1789 | 33.64465 | 14.84414 | 0.606136 | 0.697766 | 0.368367 |
| 3 | 118936800 | 118938799 | 29.07353 | 49.8445 | 39.63101 | 17.3809 | 0.712775 | 0.960864 | 0.36838 |
| 11 | 8703200 | 8705999 | 32.21131 | 53.64943 | 49.25648 | 21.39705 | 0.503956 | 1.246077 | 0.36841 |
| 4 | 118045800 | 118049799 | 39.45923 | 63.27906 | 107.7867 | 45.53597 | -0.41927 | 1.960553 | 0.368424 |
| 4 | 133407600 | 133408999 | 7.607562 | 14.09782 | 31.55569 | 13.25976 | -0.71147 | 0.179173 | 0.368444 |
| 3 | 25472800 | 25475799 | 40.73375 | 66.5037 | 54.75256 | 23.53083 | 0.652935 | 1.515865 | 0.368446 |
| 2 | 72831200 | 72844799 | 144.5716 | 215.4421 | 328.0416 | 135.1812 | -0.34834 | 3.590932 | 0.36845 |
| 2 | 7799600 | 7800399 | 4.702143 | 9.102527 | 24.27992 | 9.240996 | -0.83412 | -0.21689 | 0.36846 |
| 10 | 77848400 | 77853599 | 38.01779 | 60.97314 | 116.0959 | 49.05325 | -0.57951 | 1.958818 | 0.368465 |
| 10 | 101441400 | 101443999 | 21.21783 | 37.25103 | 31.9735 | 14.14482 | 0.606639 | 0.638337 | 0.368484 |
| 3 | 29095000 | 29095999 | 12.74167 | 24.05102 | 22.42976 | 9.141968 | 0.576135 | 0.080584 | 0.368485 |
| 1 | 11566400 | 11570199 | 40.51305 | 67.1697 | 50.52964 | 21.88204 | 0.790227 | 1.324163 | 0.368495 |
| 8 | 13807200 | 13809399 | 10.26753 | 18.47643 | 36.33369 | 15.86182 | -0.56531 | 0.374774 | 0.368497 |
| 13 | 66424600 | 66441199 | 354.4914 | 507.1923 | 368.8255 | 152.3636 | 0.660881 | 4.245235 | 0.368502 |

| | | | | | | | | | |
|---|---|---|---|---|---|---|---|---|---|
| X | 79149600 | 79150599 | 17.59254 | 31.94628 | 26.09314 | 11.26271 | 0.705298 | 0.355179 | 0.368515 |
| 9 | 61020000 | 61024599 | 33.22873 | 53.65419 | 92.58431 | 39.49672 | -0.42586 | 1.734974 | 0.368523 |
| 7 | 63537600 | 63541799 | 21.90318 | 35.79065 | 80.08945 | 34.4305 | -0.78462 | 1.478919 | 0.368531 |
| 15 | 11821000 | 11822399 | 19.56311 | 36.02115 | 22.66184 | 9.606061 | 1.096777 | 0.285834 | 0.368534 |
| 14 | 34287600 | 34290999 | 24.26119 | 39.60922 | 78.56026 | 33.76238 | -0.61199 | 1.49693 | 0.368534 |
| 9 | 37054800 | 37059599 | 26.89922 | 43.6429 | 86.9619 | 37.26256 | -0.62582 | 1.607036 | 0.368538 |
| 6 | 124270000 | 124273599 | 57.90793 | 93.12237 | 73.03589 | 30.98294 | 0.702171 | 1.884828 | 0.368543 |
| 17 | 18713200 | 18726999 | 218.1634 | 322.5469 | 284.9909 | 117.3663 | 0.425819 | 3.735719 | 0.368543 |
| 3 | 25905800 | 25907199 | 15.09681 | 28.47462 | 21.01508 | 8.579335 | 0.9088 | 0.102973 | 0.368571 |
| 5 | 85290600 | 85293199 | 19.99916 | 35.4351 | 29.25591 | 12.93486 | 0.666626 | 0.523383 | 0.368575 |
| 5 | 76950800 | 76953199 | 26.32695 | 44.93719 | 42.60322 | 18.67778 | 0.459161 | 0.994833 | 0.368589 |
| 3 | 146390600 | 146391399 | 5.444321 | 10.46823 | 26.87965 | 10.48795 | -0.81542 | -0.11967 | 0.368601 |
| 12 | 63507600 | 63511199 | 14.66314 | 27.17913 | 21.67859 | 9.106476 | 0.76586 | 0.207427 | 0.368601 |
| 10 | 17597600 | 17600399 | 23.778 | 38.57336 | 91.01759 | 38.9963 | -0.86922 | 1.605853 | 0.368601 |
| 4 | 22200400 | 22201399 | 13.50408 | 23.69028 | 43.01202 | 19.01588 | -0.46856 | 0.628679 | 0.368604 |
| 12 | 52230400 | 52232399 | 19.70823 | 34.88893 | 29.87519 | 13.21098 | 0.61387 | 0.532408 | 0.368611 |
| 1 | 60457600 | 60459399 | 20.21681 | 35.6655 | 30.97487 | 13.70675 | 0.591774 | 0.574601 | 0.368618 |
| X | 150449400 | 150451199 | 17.66569 | 32.87046 | 22.39028 | 9.353653 | 0.997889 | 0.212875 | 0.368624 |
| 18 | 24666400 | 24668799 | 47.24779 | 76.15493 | 68.46261 | 29.13933 | 0.509755 | 1.79687 | 0.368644 |
| 4 | 134613800 | 134617999 | 38.11577 | 61.22974 | 104.3369 | 44.19366 | -0.41688 | 1.903087 | 0.368666 |
| 1 | 37468600 | 37470199 | 11.63179 | 20.48127 | 40.43086 | 17.89794 | -0.58733 | 0.563081 | 0.368684 |
| 11 | 119650200 | 119651199 | 13.4638 | 24.86364 | 23.91374 | 10.09624 | 0.492657 | 0.223891 | 0.36871 |
| X | 88653200 | 88655399 | 38.02356 | 63.79122 | 44.69823 | 19.50604 | 0.892413 | 1.188814 | 0.368725 |
| X | 153232000 | 153233199 | 29.0953 | 49.12753 | 46.37566 | 20.28619 | 0.464059 | 1.102416 | 0.368729 |
| 14 | 122456600 | 122457599 | 19.75005 | 34.78597 | 31.62639 | 13.99957 | 0.52524 | 0.5906 | 0.368734 |
| 13 | 54747000 | 54748199 | 14.94307 | 27.09118 | 25.80394 | 11.16048 | 0.482858 | 0.342906 | 0.368735 |
| 9 | 48405400 | 48407799 | 14.82137 | 25.69251 | 48.45186 | 21.25826 | -0.5246 | 0.815644 | 0.368743 |
| 5 | 10140200 | 10140799 | 4.680615 | 9.119267 | 22.52998 | 8.194664 | -0.66408 | -0.38492 | 0.368743 |
| 15 | 51860200 | 51861799 | 9.120229 | 16.81518 | 32.51438 | 13.75757 | -0.51009 | 0.219831 | 0.368751 |
| X | 134108800 | 134112399 | 17.57076 | 30.28479 | 54.92083 | 24.09232 | -0.47043 | 0.887515 | 0.368757 |
| 2 | 34691800 | 34692999 | 6.071255 | 11.49343 | 29.22765 | 11.83955 | -0.85127 | 0.01714 | 0.368769 |
| 9 | 98608800 | 98609599 | 5.103878 | 9.917028 | 23.40805 | 8.725943 | -0.63171 | -0.28812 | 0.368776 |
| 7 | 40824800 | 40836599 | 170.5516 | 257.7856 | 210.2801 | 86.95154 | 0.569669 | 3.319712 | 0.368782 |
| 17 | 64177800 | 64178599 | 14.94357 | 27.31771 | 25.76108 | 11.02696 | 0.506708 | 0.29572 | 0.368789 |
| 1 | 121128200 | 121131599 | 32.31781 | 53.82992 | 50.88503 | 22.10529 | 0.462058 | 1.248451 | 0.368809 |
| 10 | 54398800 | 54400999 | 16.18012 | 29.58507 | 25.55066 | 10.93371 | 0.632884 | 0.302576 | 0.368818 |
| 7 | 118417000 | 118418199 | 7.747105 | 14.36075 | 31.02797 | 13.03266 | -0.66087 | 0.175162 | 0.368835 |
| 16 | 64707600 | 64711799 | 58.19214 | 93.44884 | 83.26393 | 35.24976 | 0.516099 | 1.932864 | 0.368856 |
| 14 | 14761400 | 14762399 | 12.60288 | 22.31418 | 44.01956 | 19.46507 | -0.58339 | 0.515645 | 0.368893 |
| 5 | 31994800 | 31997199 | 19.53694 | 34.32756 | 32.22186 | 14.26438 | 0.478307 | 0.617845 | 0.368909 |
| 1 | 153824200 | 153826199 | 20.00459 | 34.99273 | 33.50796 | 14.77789 | 0.449871 | 0.69354 | 0.368916 |
| 3 | 125730800 | 125732999 | 16.10077 | 27.69095 | 52.1697 | 22.88395 | -0.52583 | 0.89599 | 0.368926 |
| 1 | 78452400 | 78452799 | 5.076068 | 9.888427 | 22.515 | 8.203366 | -0.54944 | -0.3713 | 0.368927 |
| 4 | 89316000 | 89316599 | 5.076068 | 9.888427 | 22.515 | 8.203366 | -0.54944 | -0.3713 | 0.368927 |
| 18 | 32689200 | 32689799 | 5.076068 | 9.888427 | 22.515 | 8.203366 | -0.54944 | -0.3713 | 0.368927 |
| 10 | 91420400 | 91423599 | 35.50603 | 57.23369 | 96.69661 | 41.16043 | -0.39836 | 1.789741 | 0.368964 |
| 4 | 152815600 | 152818599 | 31.25002 | 50.59975 | 87.74395 | 37.5418 | -0.42863 | 1.658284 | 0.368972 |

| | | | | | | | | | |
|---|---|---|---|---|---|---|---|---|---|
| 4 | 76446000 | 76447599 | 19.80013 | 36.17788 | 23.59089 | 10.10426 | 1.034869 | 0.330633 | 0.368983 |
| 1 | 88904000 | 88906399 | 33.22173 | 53.62043 | 93.12153 | 39.70445 | -0.43581 | 1.746787 | 0.368985 |
| 12 | 24470800 | 24471599 | 30.21055 | 50.5942 | 48.08162 | 20.95919 | 0.454349 | 1.185391 | 0.368991 |
| 10 | 86788800 | 86789399 | 4.83954 | 9.416379 | 24.79503 | 9.140662 | -0.77379 | -0.32957 | 0.368996 |
| X | 87306400 | 87309399 | 52.60363 | 84.99424 | 70.96615 | 30.29989 | 0.620124 | 1.737254 | 0.369013 |
| 17 | 13226400 | 13227799 | 10.4404 | 18.34953 | 41.52294 | 18.38624 | -0.78412 | 0.578998 | 0.369017 |
| 10 | 47682800 | 47683399 | 10.72013 | 20.45064 | 20.1293 | 8.02259 | 0.52745 | -0.01521 | 0.369023 |
| 2 | 177280400 | 177283599 | 26.38601 | 45.53195 | 39.18751 | 17.19114 | 0.600693 | 0.893234 | 0.369036 |
| 7 | 9207200 | 9208399 | 12.62663 | 24.20995 | 20.49327 | 8.06102 | 0.760191 | -0.03793 | 0.369042 |
| 11 | 15433800 | 15439399 | 62.5678 | 100.2931 | 84.61791 | 35.72409 | 0.591549 | 1.991727 | 0.369069 |
| 3 | 142223600 | 142225799 | 26.46733 | 43.22192 | 79.09119 | 33.99368 | -0.49596 | 1.49718 | 0.369076 |
| X | 17489800 | 17492399 | 54.07975 | 86.7683 | 83.12976 | 35.14577 | 0.410137 | 1.954878 | 0.369081 |
| 5 | 104927800 | 104933199 | 73.55661 | 117.6209 | 85.58106 | 36.00157 | 0.800859 | 2.082128 | 0.369091 |
| 8 | 92736600 | 92739199 | 15.13731 | 26.20695 | 48.98144 | 21.48978 | -0.51217 | 0.829 | 0.369105 |
| 13 | 73543400 | 73545599 | 12.74377 | 22.34029 | 42.55571 | 18.80265 | -0.53728 | 0.636464 | 0.369109 |
| 4 | 86475400 | 86477599 | 14.9564 | 25.94886 | 48.45474 | 21.26008 | -0.51023 | 0.807736 | 0.369111 |
| 4 | 119349200 | 119350399 | 5.882493 | 11.24509 | 27.2553 | 10.81375 | -0.75568 | -0.05802 | 0.369119 |
| 3 | 32173000 | 32174599 | 8.348365 | 15.36497 | 35.17172 | 14.91768 | -0.75494 | 0.225775 | 0.369125 |
| 1 | 148665400 | 148667199 | 17.50321 | 31.85765 | 26.02525 | 11.20096 | 0.707565 | 0.340576 | 0.36913 |
| 6 | 28427800 | 28428199 | 3.524822 | 6.888852 | 18.81434 | 6.506517 | -0.73776 | -0.59137 | 0.369136 |
| 16 | 83114600 | 83114999 | 3.524822 | 6.888852 | 18.81434 | 6.506517 | -0.73776 | -0.59137 | 0.369136 |
| 11 | 104359600 | 104360599 | 7.76585 | 14.32008 | 33.05581 | 13.984 | -0.76373 | 0.207111 | 0.36914 |
| 1 | 145279400 | 145281799 | 22.28918 | 39.14143 | 30.72467 | 13.59513 | 0.734773 | 0.63792 | 0.369151 |
| 10 | 119249200 | 119251399 | 16.93281 | 28.99654 | 53.22033 | 23.33856 | -0.48938 | 0.94097 | 0.369156 |
| 14 | 8750600 | 8752599 | 16.20273 | 28.01297 | 51.678 | 22.6719 | -0.4941 | 0.847618 | 0.36917 |
| 15 | 46447600 | 46449399 | 14.10915 | 24.74817 | 46.15527 | 20.40303 | -0.50736 | 0.637701 | 0.369174 |
| 9 | 26213400 | 26217199 | 40.11808 | 65.92049 | 58.3223 | 25.12934 | 0.554142 | 1.433685 | 0.369175 |
| 6 | 123157800 | 123159399 | 18.99302 | 33.61101 | 29.93192 | 13.237 | 0.557354 | 0.531206 | 0.369177 |
| 12 | 106247200 | 106252799 | 66.5225 | 104.1961 | 167.9555 | 69.66785 | -0.37158 | 2.608508 | 0.369178 |
| 6 | 81993800 | 81998799 | 37.12085 | 59.78422 | 99.85455 | 42.45218 | -0.38345 | 1.819814 | 0.36919 |
| 3 | 61928800 | 61929999 | 9.282875 | 17.03142 | 32.96736 | 14.04251 | -0.51868 | 0.249421 | 0.369197 |
| 4 | 12906000 | 12907599 | 9.282875 | 17.03142 | 32.96736 | 14.04251 | -0.51868 | 0.249421 | 0.369197 |
| 9 | 33042400 | 33044199 | 26.06969 | 44.62598 | 42.00464 | 18.41961 | 0.470042 | 0.966314 | 0.36923 |
| 16 | 55078000 | 55079399 | 11.8715 | 22.92089 | 17.66916 | 6.807933 | 0.920437 | -0.1208 | 0.369234 |
| 7 | 56497600 | 56499199 | 16.98559 | 29.07455 | 53.72382 | 23.55854 | -0.49903 | 0.947877 | 0.369239 |
| 18 | 21733000 | 21735199 | 35.84219 | 60.03814 | 47.52265 | 20.71826 | 0.717216 | 1.200217 | 0.369243 |
| 14 | 79392800 | 79393799 | 12.19983 | 23.05234 | 21.92361 | 8.915929 | 0.550739 | 0.0667 | 0.369247 |
| 5 | 105627200 | 105629999 | 41.63648 | 66.47649 | 118.4517 | 49.7208 | -0.49141 | 2.103052 | 0.369257 |
| 2 | 11103000 | 11106399 | 39.45773 | 63.20015 | 110.9584 | 46.78976 | -0.46492 | 1.998521 | 0.369265 |
| 17 | 40424400 | 40428799 | 52.93794 | 84.09411 | 135.721 | 56.59513 | -0.35668 | 2.276618 | 0.36929 |
| 15 | 96462800 | 96466599 | 30.721 | 49.81391 | 87.60972 | 37.52367 | -0.44708 | 1.626724 | 0.369291 |
| 3 | 45913800 | 45915399 | 15.48584 | 28.77075 | 23.59423 | 9.879928 | 0.729199 | 0.205437 | 0.369298 |
| 12 | 62101000 | 62103999 | 23.36121 | 41.24863 | 28.45645 | 12.59013 | 0.922554 | 0.579828 | 0.369299 |
| 5 | 66143800 | 66146599 | 24.51609 | 42.03827 | 40.45845 | 17.74403 | 0.438415 | 0.942135 | 0.3693 |
| 9 | 110636800 | 110637799 | 10.4799 | 20.2229 | 19.62964 | 7.580716 | 0.588778 | -0.12128 | 0.369303 |
| 14 | 100587200 | 100588399 | 10.4799 | 20.2229 | 19.62964 | 7.580716 | 0.588778 | -0.12128 | 0.369303 |
| 4 | 104641000 | 104641999 | 10.56708 | 19.00004 | 37.34963 | 16.31652 | -0.56563 | 0.383332 | 0.369309 |

| | | | | | | | | | |
|---|---|---|---|---|---|---|---|---|---|
| X | 90688600 | 90691399 | 23.64377 | 41.56202 | 30.76874 | 13.62529 | 0.81915 | 0.633425 | 0.369331 |
| 14 | 32358800 | 32364199 | 50.75557 | 80.81715 | 130.9705 | 54.76911 | -0.35911 | 2.200595 | 0.369332 |
| 15 | 22462600 | 22463999 | 20.92481 | 36.71668 | 32.59103 | 14.4114 | 0.558504 | 0.646582 | 0.369337 |
| 6 | 56999800 | 57000399 | 8.848067 | 17.2425 | 16.67494 | 6.047754 | 0.677391 | -0.36187 | 0.369339 |
| 11 | 41750800 | 41751399 | 12.61256 | 23.89902 | 18.9122 | 7.645852 | 0.821813 | 0.045428 | 0.36934 |
| 10 | 116843200 | 116846199 | 26.92025 | 44.01847 | 78.79325 | 33.88161 | -0.46297 | 1.478872 | 0.36936 |
| 13 | 77814000 | 77822399 | 86.94702 | 136.4192 | 132.2508 | 54.84799 | 0.362953 | 2.582961 | 0.36936 |
| 14 | 62603600 | 62604799 | 8.042961 | 14.93469 | 31.04237 | 13.00841 | -0.60298 | 0.168502 | 0.369362 |
| 7 | 28157200 | 28160399 | 22.82905 | 37.72383 | 69.75175 | 30.13667 | -0.50346 | 1.332474 | 0.369362 |
| 15 | 88764000 | 88767199 | 25.43569 | 41.75021 | 75.44101 | 32.49056 | -0.47379 | 1.423779 | 0.369372 |
| 15 | 97855400 | 97857799 | 14.81362 | 25.75377 | 47.37982 | 20.81047 | -0.48838 | 0.77943 | 0.369373 |
| 16 | 62137200 | 62138599 | 15.28032 | 26.65197 | 47.86648 | 21.06739 | -0.45343 | 0.735558 | 0.369382 |
| 11 | 11061600 | 11066799 | 51.6019 | 81.96886 | 134.8413 | 56.22631 | -0.38423 | 2.27563 | 0.3694 |
| 10 | 127349200 | 127352999 | 27.3648 | 44.41697 | 85.33378 | 36.57626 | -0.57275 | 1.598908 | 0.369402 |
| 19 | 29499200 | 29500599 | 11.16634 | 19.68862 | 39.92644 | 17.66981 | -0.62509 | 0.544203 | 0.369407 |
| 14 | 15650800 | 15654399 | 35.9212 | 59.61679 | 53.18463 | 23.04788 | 0.545137 | 1.302514 | 0.369414 |
| 10 | 129548200 | 129549799 | 15.16221 | 28.71832 | 21.20902 | 8.581939 | 0.919438 | 0.077421 | 0.369414 |
| 8 | 11485600 | 11487199 | 20.14213 | 35.19162 | 33.56698 | 14.78954 | 0.455495 | 0.712857 | 0.369419 |
| 17 | 25332000 | 25334999 | 15.36667 | 26.44262 | 51.07126 | 22.4025 | -0.56109 | 0.889635 | 0.369428 |
| 5 | 67683000 | 67685799 | 31.57722 | 52.44861 | 50.85883 | 22.05198 | 0.425271 | 1.278781 | 0.369462 |
| 2 | 129286000 | 129289599 | 48.91276 | 78.8199 | 76.86368 | 32.70457 | 0.392235 | 1.809747 | 0.369462 |
| 1 | 94739200 | 94740999 | 22.4857 | 39.42868 | 31.87943 | 14.08863 | 0.692447 | 0.66411 | 0.369483 |
| 14 | 41175600 | 41176599 | 22.65721 | 39.54694 | 34.79322 | 15.32035 | 0.571273 | 0.739919 | 0.369509 |
| 1 | 162844800 | 162846599 | 20.19209 | 35.27947 | 33.12005 | 14.59278 | 0.478393 | 0.712955 | 0.36953 |
| 16 | 67675800 | 67677199 | 19.21912 | 34.06434 | 26.51679 | 11.72304 | 0.751357 | 0.51087 | 0.36954 |
| 17 | 17158200 | 17162399 | 44.74395 | 72.56281 | 68.28339 | 29.24931 | 0.453161 | 1.639309 | 0.369545 |
| 7 | 104167800 | 104168999 | 11.74838 | 21.02025 | 39.92656 | 17.51487 | -0.52046 | 0.430234 | 0.369551 |
| 14 | 76317800 | 76319799 | 14.47807 | 25.34627 | 45.67603 | 20.15826 | -0.45793 | 0.672167 | 0.369594 |
| 4 | 3412600 | 3414399 | 19.37165 | 33.91506 | 30.469 | 13.44751 | 0.541316 | 0.669905 | 0.369599 |
| 11 | 115004400 | 115006799 | 14.19071 | 24.77381 | 45.74815 | 20.14648 | -0.49299 | 0.71551 | 0.369599 |
| 18 | 54115600 | 54118799 | 35.17078 | 56.77324 | 95.3866 | 40.67669 | -0.388 | 1.747604 | 0.369604 |
| 7 | 8530600 | 8542199 | 156.6573 | 239.0994 | 175.2237 | 72.66345 | 0.735592 | 3.120365 | 0.369606 |
| 13 | 72771200 | 72773999 | 28.23955 | 46.09335 | 79.85944 | 34.31774 | -0.41786 | 1.509116 | 0.369622 |
| 8 | 21649400 | 21654199 | 37.42941 | 62.10396 | 52.67338 | 22.82229 | 0.617798 | 1.309458 | 0.369622 |
| 3 | 92097800 | 92100199 | 26.75266 | 45.53466 | 43.65605 | 19.13498 | 0.442418 | 1.022055 | 0.369632 |
| 10 | 90710800 | 90712199 | 14.66588 | 26.7196 | 25.2063 | 10.83521 | 0.502104 | 0.310319 | 0.369635 |
| 12 | 7328400 | 7329999 | 31.62907 | 52.89216 | 43.32796 | 18.86956 | 0.667841 | 1.198587 | 0.369655 |
| 5 | 113229800 | 113233599 | 37.6215 | 60.46512 | 103.2702 | 43.77263 | -0.4193 | 1.886597 | 0.369656 |
| 5 | 112113400 | 112116599 | 27.34137 | 44.43696 | 83.23464 | 35.7109 | -0.53448 | 1.571383 | 0.369666 |
| 17 | 69381600 | 69385999 | 37.63556 | 60.39958 | 107.5052 | 45.46864 | -0.48129 | 1.933205 | 0.369669 |
| 5 | 128330800 | 128355599 | 243.8217 | 348.6693 | 512.7535 | 211.8382 | -0.35548 | 4.247358 | 0.3697 |
| X | 3246600 | 3253399 | 79.7128 | 126.46 | 109.5345 | 45.61513 | 0.53799 | 2.331313 | 0.369709 |
| 8 | 55837000 | 55839399 | 20.69913 | 36.25332 | 33.85947 | 14.94883 | 0.485654 | 0.677952 | 0.369716 |
| 4 | 141705400 | 141707599 | 18.19207 | 30.68042 | 63.92613 | 27.95312 | -0.67408 | 1.094791 | 0.369718 |
| 11 | 50355400 | 50376599 | 265.6706 | 377.1277 | 545.8805 | 225.8011 | -0.3433 | 4.356197 | 0.36972 |
| X | 156552400 | 156552799 | 5.072244 | 9.893949 | 23.46428 | 8.403589 | -0.58569 | -0.43118 | 0.369733 |
| 1 | 17622800 | 17623399 | 16.80343 | 29.90482 | 28.25768 | 12.45187 | 0.476623 | 0.472226 | 0.369758 |

| | | | | | | | | | |
|---|---|---|---|---|---|---|---|---|---|
| 1 | 113071600 | 113073599 | 34.13062 | 56.34694 | 55.27346 | 23.86599 | 0.408073 | 1.359306 | 0.369787 |
| 11 | 70909800 | 70910799 | 5.998522 | 11.52753 | 26.3216 | 10.28637 | -0.64978 | -0.10934 | 0.369795 |
| 7 | 29193600 | 29196799 | 21.58565 | 36.0656 | 63.99183 | 27.85353 | -0.44336 | 1.197946 | 0.369808 |
| 8 | 95526400 | 95529599 | 18.17013 | 30.70742 | 57.92557 | 25.34527 | -0.53086 | 1.071241 | 0.36981 |
| 7 | 59993600 | 59998799 | 44.16145 | 72.53316 | 56.25144 | 24.23266 | 0.743458 | 1.448031 | 0.369816 |
| 13 | 3089200 | 3091799 | 40.60017 | 65.99851 | 64.63206 | 27.73279 | 0.398903 | 1.582392 | 0.369824 |
| 12 | 72936800 | 72938599 | 11.67949 | 20.87027 | 39.88282 | 17.51457 | -0.53052 | 0.434829 | 0.369873 |
| 14 | 21800200 | 21801399 | 6.525611 | 12.26239 | 29.62887 | 12.17282 | -0.79549 | 0.073235 | 0.369875 |
| 15 | 74206200 | 74215799 | 72.20124 | 111.4008 | 202.1894 | 83.98568 | -0.55943 | 2.881861 | 0.369881 |
| 1 | 65657800 | 65665799 | 108.1351 | 167.4001 | 157.4753 | 65.41115 | 0.391353 | 2.829752 | 0.369886 |
| 8 | 122929600 | 122934199 | 22.56169 | 37.1239 | 72.86101 | 31.41205 | -0.59111 | 1.385214 | 0.369888 |
| 4 | 146793400 | 146798599 | 155.5683 | 236.3911 | 196.8408 | 81.52783 | 0.546363 | 3.209673 | 0.369915 |
| 7 | 62927800 | 62932999 | 59.1455 | 94.70214 | 89.40502 | 37.68144 | 0.427834 | 2.023434 | 0.369916 |
| 11 | 90280000 | 90283799 | 18.61158 | 31.06273 | 64.23214 | 27.93897 | -0.66328 | 1.200944 | 0.369925 |
| 16 | 18937000 | 18943199 | 83.41258 | 131.7872 | 118.9605 | 49.40896 | 0.47353 | 2.438244 | 0.369932 |
| 9 | 41010400 | 41012799 | 11.29623 | 19.95302 | 39.87516 | 17.64085 | -0.60325 | 0.52407 | 0.369938 |
| 7 | 31641800 | 31647399 | 76.9472 | 122.7867 | 89.04592 | 37.34884 | 0.802348 | 2.146701 | 0.369973 |
| 10 | 13694000 | 13698199 | 31.37996 | 53.14232 | 43.53654 | 19.06138 | 0.667819 | 1.080667 | 0.369981 |
| 4 | 123531800 | 123532999 | 20.03995 | 35.1133 | 33.397 | 14.7498 | 0.459517 | 0.667106 | 0.369993 |
| 2 | 99411800 | 99414599 | 26.93158 | 46.76579 | 36.36683 | 15.96142 | 0.74809 | 0.835451 | 0.369996 |
| 7 | 89400200 | 89404599 | 51.45605 | 83.21369 | 68.85526 | 29.43221 | 0.634522 | 1.707344 | 0.369999 |
| 7 | 32856600 | 32859399 | 81.65135 | 128.305 | 116.9344 | 48.5048 | 0.453621 | 2.548019 | 0.370007 |
| 4 | 53630800 | 53631999 | 7.875717 | 14.2629 | 34.96523 | 15.14332 | -0.87304 | 0.318536 | 0.370009 |
| 4 | 94767200 | 94768999 | 10.03244 | 18.3206 | 36.06819 | 15.45921 | -0.54983 | 0.286197 | 0.37002 |
| 5 | 110198000 | 110201399 | 61.74295 | 99.09604 | 82.75117 | 35.00316 | 0.608533 | 1.953273 | 0.370039 |
| 9 | 17625400 | 17625999 | 9.336545 | 18.06027 | 17.21095 | 6.573693 | 0.631047 | -0.17558 | 0.37004 |
| X | 154661600 | 154662199 | 5.968275 | 11.53662 | 26.53188 | 10.16746 | -0.63223 | -0.16997 | 0.37006 |
| X | 125578400 | 125580799 | 30.26876 | 52.43412 | 35.18626 | 15.43765 | 0.959319 | 0.873772 | 0.370063 |
| 12 | 33159400 | 33161199 | 10.05636 | 18.23932 | 37.52687 | 16.22236 | -0.62119 | 0.328167 | 0.370068 |
| 1 | 149388800 | 149391999 | 43.26881 | 69.98137 | 67.68647 | 28.93597 | 0.410081 | 1.690281 | 0.37012 |
| 18 | 82891800 | 82896999 | 26.94027 | 44.00148 | 79.20906 | 34.04639 | -0.47224 | 1.495751 | 0.370128 |
| 1 | 18462200 | 18464599 | 24.91401 | 43.33673 | 36.42335 | 16.00349 | 0.636653 | 0.802057 | 0.370163 |
| 7 | 62281000 | 62297399 | 437.4226 | 606.1003 | 532.6958 | 221.1974 | 0.341126 | 4.720884 | 0.370175 |
| 5 | 138776200 | 138781999 | 44.07384 | 70.21276 | 127.4015 | 53.31322 | -0.52109 | 2.176684 | 0.370184 |
| 13 | 94898800 | 94899799 | 7.506549 | 14.17866 | 29.2036 | 11.88619 | -0.55612 | 0.046696 | 0.370185 |
| 7 | 56664200 | 56669799 | 52.92288 | 85.3633 | 74.76902 | 31.85291 | 0.548406 | 1.788184 | 0.370187 |
| 2 | 36287400 | 36290799 | 62.67891 | 99.89535 | 98.48358 | 41.23682 | 0.358208 | 2.172834 | 0.370205 |
| 7 | 106118600 | 106120199 | 25.11755 | 42.96194 | 41.56067 | 18.22376 | 0.430573 | 0.96964 | 0.37021 |
| 8 | 25012600 | 25014399 | 8.827137 | 16.21165 | 32.46924 | 13.81133 | -0.56595 | 0.239256 | 0.370228 |
| 6 | 144441400 | 144444999 | 31.37161 | 50.84586 | 88.2156 | 37.77057 | -0.42811 | 1.63755 | 0.370228 |
| 19 | 39747200 | 39747999 | 17.76251 | 31.37716 | 29.77712 | 13.17325 | 0.465294 | 0.543722 | 0.37023 |
| 3 | 152818000 | 152819399 | 7.835619 | 14.72322 | 29.63444 | 12.17626 | -0.53433 | 0.086394 | 0.370233 |
| 5 | 3581600 | 3582399 | 5.824234 | 11.2772 | 24.95213 | 9.486999 | -0.56693 | -0.20705 | 0.370234 |
| 5 | 68595200 | 68596399 | 14.15987 | 27.06195 | 19.65447 | 7.803167 | 0.966568 | 0.003572 | 0.370238 |
| 17 | 27643200 | 27649599 | 95.63983 | 145.6656 | 240.538 | 99.69565 | -0.43783 | 3.149779 | 0.370238 |
| 13 | 61785800 | 61797599 | 247.3418 | 362.9744 | 305.0513 | 125.6421 | 0.487902 | 3.858986 | 0.370241 |
| X | 25355800 | 25359599 | 89.51936 | 141.9031 | 104.8033 | 43.60968 | 0.766476 | 2.362743 | 0.370257 |

| | | | | | | | | | |
|---|---|---|---|---|---|---|---|---|---|
| 9 | 121719400 | 121722799 | 23.63805 | 38.94792 | 72.62956 | 31.33029 | -0.51683 | 1.369842 | 0.370267 |
| 5 | 77247400 | 77249399 | 15.81323 | 28.64543 | 24.92026 | 10.78924 | 0.611375 | 0.349207 | 0.370267 |
| X | 8549400 | 8553799 | 74.81522 | 119.3907 | 91.84077 | 38.52387 | 0.717552 | 2.145304 | 0.37027 |
| 5 | 77921000 | 77924599 | 31.90624 | 53.14601 | 51.40471 | 22.33148 | 0.429094 | 1.24843 | 0.370295 |
| 18 | 16164800 | 16166999 | 18.21976 | 32.46132 | 28.95855 | 12.74883 | 0.559928 | 0.475139 | 0.370296 |
| 3 | 100838600 | 100840999 | 14.1486 | 24.70578 | 45.21326 | 19.9137 | -0.48013 | 0.709694 | 0.370318 |
| 15 | 22450800 | 22454999 | 100.7975 | 157.2865 | 142.0632 | 58.95066 | 0.458827 | 2.688824 | 0.370321 |
| 17 | 30168200 | 30170399 | 12.38756 | 21.81702 | 40.89187 | 18.10119 | -0.513 | 0.565003 | 0.37035 |
| 8 | 92692000 | 92695799 | 30.61254 | 49.60655 | 87.06461 | 37.27275 | -0.44488 | 1.641873 | 0.370361 |
| 10 | 18138600 | 18140199 | 26.21999 | 44.65426 | 43.12982 | 18.90523 | 0.43189 | 1.013718 | 0.370395 |
| 10 | 8393000 | 8395799 | 18.50026 | 30.78241 | 66.88713 | 29.03701 | -0.7346 | 1.23601 | 0.370398 |
| 6 | 68994600 | 68996399 | 16.02239 | 29.60292 | 23.40959 | 9.876882 | 0.773362 | 0.238706 | 0.370409 |
| 4 | 69566800 | 69567799 | 11.34073 | 21.69619 | 21.11838 | 8.360019 | 0.5519 | -0.02864 | 0.370441 |
| 1 | 149943000 | 149944799 | 11.7357 | 21.05034 | 42.65951 | 18.67419 | -0.61148 | 0.418451 | 0.370442 |
| 7 | 39777400 | 39784799 | 78.53138 | 124.2489 | 107.7128 | 44.76901 | 0.533334 | 2.3981 | 0.370442 |
| 10 | 123631000 | 123632199 | 15.33577 | 28.69842 | 22.60732 | 9.353534 | 0.800428 | 0.15776 | 0.370464 |
| 3 | 11759000 | 11764399 | 53.20907 | 85.95747 | 69.56259 | 29.69393 | 0.664761 | 1.74004 | 0.370511 |
| 7 | 62036200 | 62037599 | 18.22239 | 32.06278 | 29.8862 | 13.23194 | 0.488941 | 0.589322 | 0.370518 |
| 14 | 22161600 | 22162799 | 10.71683 | 20.60069 | 19.50918 | 7.617874 | 0.60901 | -0.08483 | 0.370539 |
| 4 | 99547800 | 99552399 | 33.87701 | 54.58044 | 97.51046 | 41.4782 | -0.47963 | 1.801609 | 0.370542 |
| 5 | 10705800 | 10706999 | 14.13028 | 26.50637 | 21.53458 | 8.8734 | 0.76043 | 0.128236 | 0.370564 |
| 2 | 128971200 | 128976199 | 60.30055 | 94.95346 | 155.9754 | 64.71677 | -0.39284 | 2.500546 | 0.370572 |
| 5 | 14344800 | 14347399 | 27.08455 | 44.52056 | 79.12861 | 34.09943 | -0.44922 | 1.414459 | 0.370577 |
| 17 | 71452600 | 71454999 | 18.38851 | 31.56477 | 58.76853 | 25.77656 | -0.50959 | 0.931623 | 0.370584 |
| 10 | 30757000 | 30757599 | 6.434409 | 12.33535 | 26.81777 | 10.55321 | -0.58926 | -0.08058 | 0.370585 |
| 8 | 61823000 | 61827599 | 30.57289 | 49.54039 | 88.00765 | 37.6754 | -0.46243 | 1.640375 | 0.370597 |
| 4 | 112243800 | 112246599 | 47.75889 | 77.066 | 73.27001 | 31.23108 | 0.430992 | 1.764442 | 0.370602 |
| 13 | 26979400 | 26980399 | 13.32683 | 24.84828 | 23.30656 | 9.702532 | 0.543345 | 0.164494 | 0.370622 |
| X | 149083200 | 149084199 | 20.943 | 36.63216 | 34.00605 | 14.99685 | 0.494528 | 0.701549 | 0.370623 |
| 7 | 76221000 | 76222399 | 16.80591 | 30.56276 | 25.95059 | 11.18059 | 0.650937 | 0.338287 | 0.370632 |
| 1 | 52271200 | 52272799 | 14.2665 | 24.57529 | 50.53083 | 22.16615 | -0.65057 | 0.874684 | 0.370638 |
| 11 | 86653800 | 86654599 | 11.11448 | 21.30804 | 20.67537 | 8.135457 | 0.564846 | -0.05012 | 0.370659 |
| 11 | 82532600 | 82534399 | 26.606 | 45.05186 | 42.72872 | 18.70698 | 0.457458 | 1.065874 | 0.37066 |
| 3 | 5795400 | 5797599 | 24.91383 | 42.88238 | 40.38668 | 17.71583 | 0.470542 | 0.910865 | 0.370687 |
| 1 | 117743800 | 117752199 | 168.6847 | 254.5169 | 220.2875 | 91.03923 | 0.482098 | 3.354983 | 0.370689 |
| 1 | 35028800 | 35029799 | 6.268066 | 12.10267 | 25.93077 | 9.988802 | -0.53878 | -0.15285 | 0.370693 |
| 3 | 10455600 | 10457799 | 15.55205 | 27.00552 | 49.52738 | 21.73754 | -0.48432 | 0.800413 | 0.370702 |
| 10 | 57399000 | 57399999 | 17.46936 | 31.58641 | 27.02159 | 11.72639 | 0.633795 | 0.379219 | 0.370702 |
| 13 | 8054600 | 8057199 | 31.59078 | 53.99052 | 40.16672 | 17.61139 | 0.807776 | 1.000854 | 0.370721 |
| 18 | 6240800 | 6242999 | 15.15566 | 26.18817 | 49.50594 | 21.71862 | -0.52907 | 0.847492 | 0.370729 |
| 1 | 28683800 | 28684799 | 11.57879 | 20.52813 | 38.78716 | 17.1468 | -0.52069 | 0.482945 | 0.370757 |
| 1 | 48358600 | 48359399 | 9.506362 | 18.38408 | 18.08191 | 6.91457 | 0.583163 | -0.16977 | 0.370784 |
| 15 | 36130200 | 36133399 | 31.57145 | 54.04732 | 39.65657 | 17.39004 | 0.828061 | 0.986946 | 0.370801 |
| 7 | 19078400 | 19079999 | 8.273265 | 15.0714 | 33.95766 | 14.59897 | -0.74537 | 0.284186 | 0.370808 |
| 19 | 39345200 | 39346999 | 42.5509 | 69.33238 | 63.89947 | 27.44196 | 0.48858 | 1.552477 | 0.370825 |
| 9 | 78735200 | 78737799 | 25.2349 | 43.62627 | 29.27465 | 12.8432 | 0.958691 | 0.858855 | 0.370843 |
| 18 | 39119600 | 39120199 | 8.815077 | 17.1135 | 17.69308 | 6.64089 | 0.537528 | -0.2401 | 0.37086 |

| | | | | | | | | | |
|---|---|---|---|---|---|---|---|---|---|
| 1 | 8231400 | 8234999 | 36.05124 | 59.71517 | 49.32684 | 21.34711 | 0.655612 | 1.32221 | 0.370864 |
| 7 | 9934600 | 9939399 | 63.75491 | 102.4113 | 78.50463 | 33.24935 | 0.733182 | 1.92925 | 0.370883 |
| 6 | 58677600 | 58679999 | 29.77409 | 50.20099 | 46.92248 | 20.51639 | 0.478224 | 1.117782 | 0.370884 |
| 19 | 5084400 | 5086399 | 10.62697 | 19.04136 | 36.33874 | 15.92154 | -0.5257 | 0.406638 | 0.370905 |
| 16 | 47018800 | 47020999 | 31.56922 | 52.94127 | 43.28173 | 18.88301 | 0.670663 | 1.170003 | 0.37091 |
| 4 | 56691200 | 56692999 | 16.03057 | 28.99602 | 26.75203 | 11.60386 | 0.526024 | 0.366931 | 0.370919 |
| 1 | 146027200 | 146030999 | 32.28406 | 54.74958 | 43.5392 | 19.0706 | 0.710737 | 1.074689 | 0.370927 |
| 15 | 38742400 | 38744199 | 12.71132 | 22.32594 | 41.95866 | 18.56875 | -0.51804 | 0.602927 | 0.370935 |
| 7 | 13783200 | 13786799 | 60.65619 | 97.54004 | 70.5813 | 29.94084 | 0.818059 | 1.887799 | 0.370936 |
| 7 | 84254600 | 84256199 | 11.28 | 20.04445 | 37.81665 | 16.68538 | -0.51639 | 0.466079 | 0.370936 |
| 18 | 19517600 | 19521999 | 87.93318 | 137.6214 | 135.265 | 56.11251 | 0.340336 | 2.628609 | 0.370941 |
| 8 | 121595800 | 121603999 | 59.16064 | 92.71643 | 181.1559 | 75.1391 | -0.64816 | 2.595654 | 0.370949 |
| 11 | 83007200 | 83008199 | 8.644961 | 15.95817 | 32.00498 | 13.51981 | -0.56072 | 0.207031 | 0.370961 |
| 15 | 34253800 | 34257199 | 23.67682 | 39.39062 | 69.79271 | 30.29571 | -0.44149 | 1.253835 | 0.370965 |
| 12 | 23091400 | 23107999 | 476.6123 | 654.4172 | 544.1101 | 226.2641 | 0.407905 | 4.861033 | 0.370973 |
| 2 | 166115800 | 166122199 | 64.34853 | 100.8654 | 166.9355 | 69.23949 | -0.40878 | 2.593246 | 0.370976 |
| X | 88174400 | 88188399 | 186.1017 | 280.4494 | 210.6721 | 87.03383 | 0.684786 | 3.38417 | 0.371025 |
| 11 | 6852600 | 6854599 | 11.11773 | 19.7272 | 38.30767 | 16.9229 | -0.55914 | 0.473991 | 0.371026 |
| 1 | 89054800 | 89061999 | 49.80615 | 79.18132 | 131.6161 | 54.92662 | -0.39802 | 2.249575 | 0.371046 |
| 13 | 74465000 | 74465799 | 5.579171 | 10.61699 | 27.67429 | 11.08759 | -0.87072 | -0.0229 | 0.371053 |
| 7 | 127440000 | 127441999 | 10.60978 | 18.60926 | 43.09939 | 19.05145 | -0.81758 | 0.615549 | 0.371059 |
| 7 | 38231600 | 38248399 | 239.4088 | 351.5491 | 306.5431 | 126.2516 | 0.435593 | 3.848491 | 0.37106 |
| 10 | 95100200 | 95101199 | 6.708374 | 12.92355 | 27.04054 | 10.49266 | -0.51556 | -0.12298 | 0.371086 |
| 16 | 95548400 | 95549399 | 13.58092 | 25.3047 | 23.86127 | 9.945001 | 0.534576 | 0.17315 | 0.371095 |
| 6 | 91647200 | 91649199 | 11.61002 | 20.54242 | 38.84331 | 17.17849 | -0.52275 | 0.506699 | 0.371106 |
| Y | 8704000 | 8705799 | 13.24124 | 24.74775 | 14.46791 | 5.998687 | 1.224223 | 0.123918 | 0.371107 |
| 7 | 13806400 | 13811399 | 62.62648 | 100.8322 | 72.64781 | 30.87236 | 0.826527 | 1.856622 | 0.371115 |
| 4 | 88172200 | 88175999 | 42.72315 | 68.33418 | 114.6526 | 48.24879 | -0.40195 | 2.047255 | 0.371147 |
| 5 | 41794800 | 41795399 | 6.256987 | 12.12183 | 26.20252 | 9.936662 | -0.53012 | -0.20643 | 0.371152 |
| 1 | 177899000 | 177901199 | 24.21525 | 42.06247 | 33.39577 | 14.66038 | 0.718143 | 0.81265 | 0.371161 |
| 13 | 59227000 | 59230199 | 20.7708 | 34.87367 | 61.78866 | 26.97738 | -0.4413 | 1.142079 | 0.371165 |
| 7 | 62018800 | 62019399 | 12.37054 | 23.25792 | 22.52668 | 9.246694 | 0.513983 | 0.104135 | 0.371169 |
| 15 | 30578600 | 30579799 | 16.57911 | 28.57836 | 51.11287 | 22.42193 | -0.45037 | 0.877364 | 0.371189 |
| 7 | 108894400 | 108896599 | 12.59857 | 22.02159 | 44.18086 | 19.47277 | -0.61178 | 0.685158 | 0.371192 |
| 5 | 141319800 | 141320599 | 5.872068 | 11.22575 | 26.7 | 10.59185 | -0.72794 | -0.05815 | 0.371204 |
| 16 | 32334800 | 32339399 | 23.60692 | 38.59697 | 77.51584 | 33.33114 | -0.62848 | 1.473404 | 0.371207 |
| 5 | 9913600 | 9915199 | 17.8576 | 31.63361 | 29.55529 | 13.0678 | 0.48908 | 0.512686 | 0.371211 |
| 5 | 92915600 | 92919799 | 30.5374 | 49.54932 | 89.65833 | 38.42003 | -0.48715 | 1.612885 | 0.371213 |
| 1 | 84539600 | 84543799 | 41.52045 | 67.83982 | 61.70893 | 26.52782 | 0.510209 | 1.509785 | 0.371217 |
| 5 | 144313200 | 144326799 | 126.4341 | 189.9515 | 295.5864 | 122.0143 | -0.36908 | 3.433615 | 0.371217 |
| 4 | 21847600 | 21848599 | 9.007471 | 16.42528 | 33.42988 | 14.35415 | -0.59854 | 0.28341 | 0.371219 |
| 6 | 13644800 | 13649199 | 59.91643 | 96.29673 | 79.88397 | 33.85969 | 0.6204 | 1.907789 | 0.371234 |
| 5 | 92940800 | 92944599 | 34.80892 | 56.25541 | 95.35613 | 40.72558 | -0.39879 | 1.711637 | 0.371251 |
| 17 | 15594600 | 15598199 | 21.37845 | 35.37887 | 69.16375 | 29.91304 | -0.58339 | 1.309831 | 0.371252 |
| 17 | 84318400 | 84322999 | 31.30945 | 50.4966 | 94.14851 | 40.10403 | -0.53919 | 1.765976 | 0.371274 |
| 15 | 47052600 | 47053399 | 9.332518 | 18.06735 | 18.74438 | 7.131901 | 0.514428 | -0.18431 | 0.371287 |
| X | 54752600 | 54755399 | 33.75357 | 57.11922 | 43.56884 | 19.07119 | 0.770645 | 1.099256 | 0.37129 |

| | | | | | | | | | |
|---|---|---|---|---|---|---|---|---|---|
| 5 | 23896200 | 23922199 | 835.0464 | 1073.616 | 1478.059 | 619.8525 | -0.41454 | 5.915542 | 0.371303 |
| 8 | 24034800 | 24036199 | 18.27248 | 32.35174 | 30.10425 | 13.3119 | 0.494713 | 0.522917 | 0.371311 |
| 9 | 33828800 | 33830799 | 13.36183 | 23.34517 | 44.69067 | 19.69054 | -0.54463 | 0.696468 | 0.371343 |
| 18 | 4416800 | 4419199 | 24.12077 | 41.5481 | 39.91481 | 17.50924 | 0.442252 | 0.900773 | 0.371364 |
| 1 | 189481400 | 189483799 | 14.36268 | 24.88443 | 48.44834 | 21.25637 | -0.57048 | 0.818514 | 0.37137 |
| 7 | 68398400 | 68399399 | 5.148714 | 9.939806 | 25.8746 | 9.974511 | -0.81813 | -0.16535 | 0.371375 |
| 3 | 65663200 | 65663799 | 9.488343 | 17.31569 | 33.95888 | 14.56704 | -0.54432 | 0.285539 | 0.37141 |
| 10 | 130010800 | 130011799 | 10.55376 | 19.24212 | 37.11855 | 15.94094 | -0.52236 | 0.303304 | 0.371434 |
| 9 | 96835400 | 96837999 | 13.55881 | 23.56276 | 47.38342 | 20.80678 | -0.61618 | 0.778391 | 0.371435 |
| 5 | 149453200 | 149454999 | 10.6492 | 19.01044 | 36.85008 | 16.19623 | -0.55117 | 0.43166 | 0.371446 |
| 18 | 37200400 | 37208599 | 112.6536 | 175.586 | 137.2865 | 56.97687 | 0.66545 | 2.714393 | 0.371448 |
| 4 | 135932800 | 135934799 | 7.703729 | 14.38088 | 30.58958 | 12.70893 | -0.6266 | 0.128503 | 0.371453 |
| X | 144817200 | 144821599 | 76.4156 | 120.8287 | 116.6984 | 48.48896 | 0.377023 | 2.413883 | 0.371473 |
| 3 | 13734800 | 13736599 | 21.92104 | 38.18527 | 34.93615 | 15.36388 | 0.514784 | 0.763533 | 0.371473 |
| 13 | 45996000 | 45997799 | 12.92765 | 24.11699 | 23.41521 | 9.738887 | 0.495141 | 0.158882 | 0.371482 |
| 18 | 16527800 | 16528999 | 12.92765 | 24.11699 | 23.41521 | 9.738887 | 0.495141 | 0.158882 | 0.371482 |
| 11 | 52594000 | 52598599 | 32.77079 | 52.8099 | 103.878 | 44.20103 | -0.61776 | 1.796423 | 0.371484 |
| 17 | 92632200 | 92634799 | 23.91277 | 42.08477 | 28.93107 | 12.80848 | 0.92582 | 0.61756 | 0.371489 |
| 13 | 90588200 | 90592199 | 38.57872 | 62.12098 | 105.8118 | 44.9731 | -0.41206 | 1.829399 | 0.371494 |
| 16 | 75434800 | 75435999 | 17.83931 | 31.86361 | 28.4373 | 12.4928 | 0.561326 | 0.455681 | 0.371497 |
| 12 | 106224000 | 106231599 | 53.22292 | 84.59045 | 136.6681 | 57.01952 | -0.35751 | 2.261232 | 0.371506 |
| 1 | 113621800 | 113622599 | 11.28143 | 21.54362 | 21.06721 | 8.37812 | 0.539094 | -0.01537 | 0.371528 |
| 7 | 112506000 | 112506999 | 5.536849 | 10.72383 | 24.36167 | 9.250098 | -0.60229 | -0.21567 | 0.371535 |
| 16 | 50002800 | 50003599 | 5.536849 | 10.72383 | 24.36167 | 9.250098 | -0.60229 | -0.21567 | 0.371535 |
| 17 | 3607600 | 3608399 | 5.536849 | 10.72383 | 24.36167 | 9.250098 | -0.60229 | -0.21567 | 0.371535 |
| 2 | 6349400 | 6350999 | 14.68116 | 25.55356 | 47.36433 | 20.81907 | -0.49885 | 0.763064 | 0.371541 |
| X | 118289200 | 118292999 | 54.79872 | 88.26296 | 76.9278 | 32.7094 | 0.553122 | 1.834628 | 0.371554 |
| 5 | 127840000 | 127842999 | 39.04248 | 63.93426 | 60.4659 | 26.01532 | 0.456244 | 1.472614 | 0.371568 |
| 1 | 65847600 | 65849799 | 27.65659 | 46.96726 | 44.1637 | 19.35229 | 0.469865 | 1.044911 | 0.371572 |
| 12 | 51152400 | 51153799 | 13.76088 | 25.75786 | 22.71827 | 9.39537 | 0.638898 | 0.142016 | 0.371596 |
| 4 | 90306200 | 90308199 | 27.93246 | 47.10559 | 45.87848 | 20.06116 | 0.419082 | 1.108387 | 0.371605 |
| 5 | 88304800 | 88305399 | 10.19132 | 19.69099 | 14.68045 | 5.637461 | 0.974703 | -0.15677 | 0.371607 |
| 14 | 123714200 | 123716199 | 20.66176 | 36.12485 | 34.47129 | 15.19655 | 0.454891 | 0.706104 | 0.371608 |
| 9 | 62084400 | 62086199 | 9.693645 | 17.43018 | 35.37679 | 15.45421 | -0.6108 | 0.376235 | 0.371613 |
| 5 | 101923800 | 101928399 | 23.993 | 39.4317 | 74.1691 | 31.95977 | -0.53074 | 1.403918 | 0.371614 |
| 3 | 133291800 | 133294199 | 14.23126 | 24.56573 | 50.02022 | 21.94355 | -0.63604 | 0.852918 | 0.371622 |
| 16 | 29887200 | 29889799 | 13.19067 | 23.11797 | 43.0506 | 19.0173 | -0.50493 | 0.642612 | 0.371625 |
| 7 | 35133600 | 35135599 | 15.53107 | 26.6749 | 51.59664 | 22.63165 | -0.56385 | 0.907931 | 0.371657 |
| 1 | 137146400 | 137149799 | 19.87857 | 33.51075 | 61.35639 | 26.82607 | -0.48842 | 1.099501 | 0.371663 |
| 7 | 10382000 | 10394199 | 181.1358 | 273.3959 | 209.1058 | 86.42734 | 0.660773 | 3.35129 | 0.3717 |
| 8 | 101714200 | 101719599 | 116.4825 | 178.7742 | 173.7218 | 72.12247 | 0.335115 | 2.996252 | 0.371703 |
| 17 | 12298400 | 12303199 | 29.10781 | 47.30492 | 82.84854 | 35.54361 | -0.43789 | 1.576963 | 0.371735 |
| 9 | 51765200 | 51766799 | 13.04928 | 23.00318 | 41.9134 | 18.55004 | -0.4719 | 0.562167 | 0.371736 |
| 18 | 56591600 | 56592599 | 10.19897 | 18.21548 | 36.89442 | 16.20912 | -0.61351 | 0.426904 | 0.37174 |
| 11 | 41111600 | 41115599 | 45.88177 | 74.18193 | 71.52224 | 30.5631 | 0.41408 | 1.705675 | 0.371747 |
| 14 | 48090200 | 48093999 | 50.85171 | 81.69342 | 77.30169 | 32.74504 | 0.430177 | 1.907954 | 0.371758 |
| X | 147630600 | 147633799 | 52.4855 | 84.66056 | 74.58766 | 31.77698 | 0.540068 | 1.787249 | 0.371764 |

| | | | | | | | | | |
|---|---|---|---|---|---|---|---|---|---|
| 15 | 33102000 | 33105199 | 55.23276 | 88.84449 | 80.27653 | 34.06791 | 0.498886 | 1.876514 | 0.371774 |
| 18 | 38367200 | 38368399 | 8.170575 | 15.26955 | 30.61019 | 12.69326 | -0.53918 | 0.127666 | 0.3718 |
| 3 | 83787400 | 83790199 | 20.82831 | 35.19548 | 64.26779 | 28.11928 | -0.4845 | 1.084981 | 0.371801 |
| 3 | 28116000 | 28116999 | 10.58388 | 19.18561 | 35.92739 | 15.54154 | -0.48641 | 0.335626 | 0.371804 |
| 4 | 58184000 | 58185399 | 10.58388 | 19.18561 | 35.92739 | 15.54154 | -0.48641 | 0.335626 | 0.371804 |
| 2 | 40188600 | 40190999 | 33.08224 | 54.77048 | 53.62604 | 23.19982 | 0.411044 | 1.322268 | 0.371809 |
| 8 | 21703400 | 21705999 | 37.89688 | 62.79733 | 53.26578 | 23.05811 | 0.617544 | 1.325526 | 0.371809 |
| 1 | 85949400 | 85950399 | 7.114362 | 13.32413 | 29.58357 | 12.22317 | -0.68078 | 0.102317 | 0.371836 |
| 2 | 72306200 | 72307599 | 7.114362 | 13.32413 | 29.58357 | 12.22317 | -0.68078 | 0.102317 | 0.371836 |
| 9 | 48703000 | 48704199 | 10.53205 | 19.0939 | 36.44171 | 15.76171 | -0.51382 | 0.333259 | 0.371837 |
| 7 | 26097400 | 26116599 | 289.0503 | 417.6573 | 359.2196 | 148.1798 | 0.432941 | 4.096746 | 0.371864 |
| 5 | 3241200 | 3243999 | 38.22981 | 62.97304 | 57.17816 | 24.66211 | 0.518196 | 1.397566 | 0.371864 |
| X | 113797800 | 113804799 | 66.64763 | 106.6455 | 88.96642 | 37.46176 | 0.604972 | 2.054005 | 0.371869 |
| 2 | 8261800 | 8262599 | 6.48908 | 12.27979 | 28.14175 | 11.40875 | -0.70306 | 0.022956 | 0.371869 |
| 6 | 127047800 | 127050199 | 15.28439 | 26.56329 | 47.9118 | 21.03969 | -0.46008 | 0.786985 | 0.371873 |
| 6 | 136738200 | 136740199 | 20.11017 | 33.78928 | 60.68133 | 26.50433 | -0.4607 | 1.130788 | 0.371888 |
| 18 | 27771400 | 27772199 | 11.08783 | 21.37606 | 20.19292 | 7.819161 | 0.624516 | -0.10322 | 0.371895 |
| 13 | 61720400 | 61729399 | 229.0812 | 336.1107 | 310.8065 | 128.0141 | 0.349801 | 3.859922 | 0.371935 |
| 10 | 59683400 | 59686399 | 14.04409 | 24.34055 | 48.44683 | 21.25593 | -0.60209 | 0.813514 | 0.371951 |
| 8 | 122325800 | 122330599 | 26.05413 | 42.39595 | 81.99299 | 35.19519 | -0.57919 | 1.549996 | 0.371954 |
| 10 | 62863200 | 62864999 | 11.5403 | 20.56415 | 38.80757 | 17.08247 | -0.51475 | 0.449867 | 0.371965 |
| 15 | 92647000 | 92649199 | 12.17669 | 21.58184 | 42.39784 | 18.74409 | -0.57711 | 0.494606 | 0.371967 |
| 4 | 22339800 | 22340399 | 5.669494 | 10.97748 | 24.89754 | 9.466653 | -0.60139 | -0.20547 | 0.371974 |
| 18 | 76940400 | 76942399 | 15.96106 | 27.40482 | 52.6893 | 23.11061 | -0.55549 | 0.911247 | 0.371985 |
| 14 | 41724000 | 41726199 | 16.45585 | 30.15082 | 24.96542 | 10.65373 | 0.695986 | 0.291788 | 0.37199 |
| X | 26267600 | 26269199 | 15.75451 | 29.75027 | 21.05518 | 8.574236 | 0.972247 | 0.102004 | 0.372022 |
| 1 | 52348400 | 52350599 | 20.83405 | 36.44329 | 34.4237 | 15.18163 | 0.469459 | 0.699454 | 0.372032 |
| 11 | 120045800 | 120046999 | 19.42428 | 34.08963 | 32.38457 | 14.32222 | 0.46122 | 0.636852 | 0.372042 |
| 7 | 6333200 | 6336199 | 25.02292 | 40.92052 | 77.22792 | 33.20983 | -0.53888 | 1.473118 | 0.372051 |
| 17 | 76516000 | 76518799 | 24.73821 | 43.01707 | 35.83063 | 15.73988 | 0.649419 | 0.803245 | 0.372053 |
| 10 | 66681200 | 66683199 | 15.45626 | 26.92018 | 48.41748 | 21.29069 | -0.45568 | 0.75801 | 0.372062 |
| 12 | 76768800 | 76779599 | 94.17609 | 144.354 | 226.2238 | 93.90276 | -0.35516 | 3.018505 | 0.372071 |
| X | 120468200 | 120471799 | 52.56999 | 84.99161 | 69.1077 | 29.52957 | 0.659299 | 1.720742 | 0.372072 |
| 7 | 26310000 | 26312199 | 20.15495 | 35.94134 | 28.28776 | 12.44411 | 0.740501 | 0.482119 | 0.372076 |
| X | 84553400 | 84555399 | 20.984 | 36.81624 | 29.24638 | 12.93116 | 0.717959 | 0.644631 | 0.372081 |
| 8 | 115204000 | 115205599 | 19.63695 | 33.03166 | 60.10274 | 26.26066 | -0.4794 | 1.119401 | 0.372097 |
| 2 | 107762800 | 107763199 | 6.255487 | 12.10597 | 26.14041 | 9.962773 | -0.53583 | -0.19119 | 0.372097 |
| 13 | 58922000 | 58922999 | 6.255487 | 12.10597 | 26.14041 | 9.962773 | -0.53583 | -0.19119 | 0.372097 |
| 8 | 43658600 | 43660399 | 17.41447 | 31.18239 | 27.71575 | 12.15026 | 0.569593 | 0.438869 | 0.372105 |
| 5 | 56416800 | 56420799 | 5.166962 | 9.856032 | 29.35477 | 11.70172 | -1.0555 | -0.0419 | 0.372114 |
| 18 | 24345200 | 24346799 | 16.25008 | 29.85896 | 24.45213 | 10.39423 | 0.715732 | 0.273179 | 0.372116 |
| 5 | 69854400 | 69858999 | 75.31672 | 118.9379 | 114.5756 | 47.57594 | 0.379261 | 2.441769 | 0.372139 |
| 9 | 111862400 | 111863799 | 12.08488 | 21.29074 | 40.39753 | 17.88122 | -0.53031 | 0.559327 | 0.372145 |
| 19 | 46135200 | 46137799 | 20.419 | 33.84622 | 74.84963 | 32.40861 | -0.76057 | 1.292041 | 0.372165 |
| 13 | 40749000 | 40750799 | 9.772218 | 17.75367 | 34.3968 | 14.83817 | -0.53256 | 0.312318 | 0.372165 |
| 14 | 54459600 | 54463599 | 16.82123 | 28.27314 | 61.02064 | 26.65539 | -0.72473 | 1.116469 | 0.372167 |
| 1 | 72022800 | 72024599 | 10.37559 | 18.96828 | 36.64261 | 15.68363 | -0.52127 | 0.285007 | 0.372183 |

| | | | | | | | | | |
|---|---|---|---|---|---|---|---|---|---|
| 3 | 39824800 | 39825199 | 2.935437 | 5.74238 | 16.74161 | 5.562798 | -0.77529 | -0.74853 | 0.372186 |
| X | 31984400 | 31988999 | 95.48512 | 149.5653 | 134.678 | 55.86422 | 0.467471 | 2.61958 | 0.372195 |
| 12 | 119960400 | 119960999 | 10.89517 | 19.62134 | 36.33303 | 15.8459 | -0.47811 | 0.37567 | 0.372203 |
| 8 | 47236000 | 47236599 | 4.52004 | 8.811134 | 21.39562 | 7.72769 | -0.62935 | -0.41414 | 0.372212 |
| 7 | 60185000 | 60188599 | 41.43519 | 67.35451 | 61.62058 | 26.44041 | 0.496888 | 1.585918 | 0.372216 |
| 6 | 84249800 | 84254199 | 33.67783 | 54.4296 | 92.03842 | 39.31066 | -0.3952 | 1.707342 | 0.372234 |
| 10 | 84457600 | 84458799 | 9.084534 | 16.79107 | 32.55097 | 13.72653 | -0.51045 | 0.20405 | 0.372245 |
| 13 | 14457400 | 14458199 | 10.7086 | 20.46985 | 20.6363 | 8.188679 | 0.498604 | -0.02833 | 0.372292 |
| 18 | 58387400 | 58387799 | 10.7086 | 20.46985 | 20.6363 | 8.188679 | 0.498604 | -0.02833 | 0.372292 |
| 10 | 7905600 | 7907399 | 16.61044 | 29.62066 | 28.21809 | 12.41344 | 0.466536 | 0.458067 | 0.372293 |
| 12 | 113827000 | 113827399 | 10.35909 | 19.81656 | 19.61951 | 7.768505 | 0.52737 | -0.04184 | 0.372294 |
| 3 | 124946600 | 124947999 | 16.08696 | 29.78741 | 23.46164 | 9.869377 | 0.782243 | 0.227884 | 0.372318 |
| 5 | 116087200 | 116096599 | 119.9337 | 180.3669 | 287.4095 | 118.6709 | -0.40198 | 3.412993 | 0.372322 |
| 15 | 35381800 | 35383199 | 23.79131 | 41.54908 | 34.74407 | 15.30394 | 0.644396 | 0.739513 | 0.372347 |
| 13 | 65383600 | 65385599 | 28.64846 | 46.51458 | 83.64568 | 35.86069 | -0.47716 | 1.596346 | 0.372348 |
| 19 | 23727200 | 23728599 | 8.157789 | 15.18235 | 31.06679 | 12.97369 | -0.57721 | 0.153211 | 0.372349 |
| 4 | 51088000 | 51089399 | 15.58715 | 28.6725 | 24.53539 | 10.41396 | 0.654287 | 0.262037 | 0.372369 |
| 10 | 44728600 | 44729799 | 7.84206 | 14.85371 | 30.96238 | 12.528 | -0.56548 | 0.037405 | 0.372381 |
| 17 | 87962200 | 87972599 | 106.5165 | 162.2566 | 250.0356 | 103.637 | -0.33802 | 3.149716 | 0.372389 |
| 7 | 13919600 | 13922599 | 69.28161 | 110.1739 | 104.9771 | 43.82168 | 0.403923 | 2.252864 | 0.372406 |
| 3 | 77587600 | 77589799 | 16.72671 | 31.01009 | 23.49027 | 9.864909 | 0.840072 | 0.227427 | 0.372412 |
| 9 | 38182200 | 38184799 | 35.04974 | 58.30128 | 52.04743 | 22.59008 | 0.544332 | 1.27334 | 0.372414 |
| 17 | 19764800 | 19767999 | 66.95311 | 106.7176 | 99.85899 | 41.81823 | 0.433547 | 2.175919 | 0.372417 |
| 9 | 24844400 | 24844999 | 10.25441 | 18.60844 | 34.89848 | 15.0761 | -0.48723 | 0.324457 | 0.372442 |
| 5 | 111053200 | 111054999 | 13.85631 | 24.29817 | 44.07049 | 19.47704 | -0.46705 | 0.640198 | 0.372461 |
| 11 | 57813200 | 57817799 | 26.31937 | 42.95105 | 79.55814 | 34.1859 | -0.51417 | 1.507006 | 0.372461 |
| 6 | 10358600 | 10361799 | 48.50202 | 78.39346 | 71.86278 | 30.69604 | 0.486202 | 1.719293 | 0.372468 |
| 1 | 188687000 | 188689599 | 22.02093 | 36.91076 | 64.71796 | 28.2265 | -0.42641 | 1.164683 | 0.372486 |
| X | 29313000 | 29313399 | 13.77963 | 25.78661 | 22.95945 | 9.499071 | 0.625676 | 0.147633 | 0.372498 |
| 4 | 37916200 | 37919599 | 33.98836 | 57.2337 | 45.22686 | 19.76737 | 0.719522 | 1.140305 | 0.372553 |
| 19 | 5497400 | 5500799 | 27.34257 | 44.45196 | 82.00477 | 35.18854 | -0.5122 | 1.566738 | 0.372554 |
| 12 | 76696800 | 76702999 | 92.075 | 141.0884 | 225.5075 | 93.60135 | -0.38392 | 3.023878 | 0.372573 |
| 12 | 113905000 | 113906199 | 17.30331 | 32.0472 | 23.46597 | 9.868043 | 0.88742 | 0.237868 | 0.372579 |
| 12 | 5389800 | 5392599 | 17.40289 | 29.70587 | 55.92204 | 24.51698 | -0.52668 | 0.973836 | 0.372594 |
| 18 | 43443800 | 43445399 | 10.88842 | 19.71988 | 36.95954 | 16.00606 | -0.48886 | 0.344409 | 0.372596 |
| 4 | 117741600 | 117744999 | 16.13065 | 27.66832 | 52.6825 | 23.10576 | -0.54176 | 0.921522 | 0.372609 |
| 1 | 10319000 | 10322599 | 25.73766 | 44.56085 | 38.20083 | 16.75999 | 0.607407 | 0.86198 | 0.372616 |
| X | 91201200 | 91202999 | 21.98478 | 39.87128 | 22.72802 | 9.825861 | 1.218998 | 0.385898 | 0.372619 |
| 8 | 23311800 | 23315599 | 58.89844 | 94.09008 | 92.74802 | 38.96277 | 0.362032 | 2.094806 | 0.37262 |
| 4 | 16187600 | 16192999 | 49.38037 | 80.2365 | 63.16791 | 27.0966 | 0.712456 | 1.607933 | 0.372675 |
| 1 | 21096800 | 21101199 | 26.00118 | 42.9057 | 76.65835 | 33.08912 | -0.45456 | 1.359234 | 0.372678 |
| 10 | 54997800 | 54998399 | 10.2824 | 20.06052 | 14.78663 | 5.283239 | 1.083367 | -0.40009 | 0.372697 |
| X | 4283200 | 4286199 | 43.52721 | 70.81695 | 56.73819 | 24.3532 | 0.688728 | 1.576894 | 0.372711 |
| 13 | 63710800 | 63714599 | 26.00565 | 42.07775 | 88.81754 | 37.98093 | -0.71238 | 1.664192 | 0.372716 |
| 8 | 3584600 | 3588399 | 19.12705 | 32.24605 | 59.04426 | 25.8157 | -0.48833 | 1.096281 | 0.372724 |
| 4 | 84217600 | 84218599 | 13.02327 | 24.91659 | 20.36552 | 8.060607 | 0.801755 | -0.01545 | 0.372725 |
| 2 | 117080200 | 117080999 | 5.27027 | 10.19756 | 24.8719 | 9.486969 | -0.71031 | -0.20283 | 0.372733 |

| | | | | | | | | | |
|---|---|---|---|---|---|---|---|---|---|
| 2 | 53555200 | 53556399 | 14.74003 | 27.66934 | 22.74077 | 9.35943 | 0.74583 | 0.134195 | 0.372734 |
| X | 144720400 | 144722199 | 17.98203 | 32.04473 | 29.07626 | 12.7983 | 0.53588 | 0.473496 | 0.372745 |
| 6 | 81909200 | 81910799 | 21.30912 | 37.28697 | 34.3353 | 15.14699 | 0.505973 | 0.696167 | 0.372747 |
| 2 | 174037600 | 174042599 | 33.26891 | 53.51509 | 102.0739 | 43.31888 | -0.57673 | 1.85267 | 0.372768 |
| 13 | 19131000 | 19133199 | 18.67236 | 31.39018 | 60.6271 | 26.48493 | -0.56523 | 1.120251 | 0.372801 |
| 2 | 174904000 | 174904599 | 4.754446 | 9.257862 | 22.95473 | 8.410317 | -0.6771 | -0.35046 | 0.372803 |
| X | 20959200 | 20964399 | 62.45422 | 100.5268 | 73.75549 | 31.33041 | 0.799889 | 1.865222 | 0.372805 |
| 11 | 10117000 | 10118199 | 24.00721 | 39.49959 | 72.91339 | 31.43368 | -0.50295 | 1.39161 | 0.372814 |
| 19 | 61319600 | 61319799 | 18.02193 | 33.2374 | 20.03631 | 8.474363 | 1.159536 | 0.257731 | 0.372821 |
| 15 | 92580600 | 92581799 | 7.185488 | 13.67227 | 28.29259 | 11.33792 | -0.54293 | -0.00976 | 0.372835 |
| 17 | 23362000 | 23362199 | 3.754345 | 7.343949 | 18.6286 | 6.218957 | -0.58646 | -0.71189 | 0.372837 |
| 3 | 156508600 | 156510399 | 19.56503 | 34.27448 | 32.50403 | 14.35305 | 0.463775 | 0.666622 | 0.372837 |
| X | 64571800 | 64574799 | 41.50901 | 68.72472 | 51.66015 | 22.35001 | 0.791257 | 1.34505 | 0.372845 |
| 10 | 120985400 | 120987399 | 17.65594 | 29.96475 | 62.37741 | 27.32816 | -0.67237 | 1.035563 | 0.372871 |
| 4 | 98810400 | 98812799 | 14.40793 | 24.82003 | 50.54121 | 22.17073 | -0.63674 | 0.873474 | 0.372913 |
| 3 | 53167400 | 53167999 | 6.104437 | 11.84275 | 25.71719 | 9.689431 | -0.52779 | -0.23222 | 0.372914 |
| X | 61942400 | 61944599 | 23.70074 | 42.51068 | 23.57572 | 10.32054 | 1.247008 | 0.461937 | 0.372925 |
| 17 | 54431800 | 54436199 | 94.60256 | 147.6798 | 142.4684 | 59.11632 | 0.364351 | 2.677831 | 0.372932 |
| 17 | 39314200 | 39316599 | 40.75613 | 66.79469 | 60.01348 | 25.82954 | 0.530503 | 1.467395 | 0.372937 |
| 3 | 95166400 | 95168999 | 14.89423 | 26.00203 | 48.39433 | 21.31181 | -0.50466 | 0.719093 | 0.372958 |
| 7 | 51271400 | 51277799 | 48.24942 | 76.96199 | 128.1615 | 53.72514 | -0.39555 | 2.140072 | 0.372964 |
| 15 | 19572600 | 19576399 | 66.50148 | 105.9005 | 102.6557 | 42.93488 | 0.38127 | 2.203996 | 0.372974 |
| 16 | 59129800 | 59132599 | 40.96295 | 67.71719 | 52.27998 | 22.59435 | 0.752643 | 1.358803 | 0.372987 |
| 1 | 121185600 | 121186999 | 14.40942 | 26.872 | 23.78911 | 9.900189 | 0.626717 | 0.174529 | 0.372993 |
| 2 | 173527800 | 173529799 | 11.398 | 20.32125 | 37.79694 | 16.63025 | -0.49334 | 0.444677 | 0.372995 |
| 3 | 101046600 | 101048399 | 15.0075 | 27.37687 | 25.6714 | 11.01755 | 0.512207 | 0.306823 | 0.373002 |
| 5 | 22524000 | 22525599 | 15.15117 | 27.87852 | 25.2704 | 10.72173 | 0.572524 | 0.260649 | 0.373003 |
| 5 | 148657600 | 148661199 | 32.06588 | 51.81878 | 91.06822 | 38.89083 | -0.45084 | 1.707137 | 0.373003 |
| 3 | 79003000 | 79004199 | 12.44075 | 23.85862 | 20.50134 | 8.059288 | 0.739462 | -0.0419 | 0.373036 |
| 3 | 112488400 | 112489599 | 12.44075 | 23.85862 | 20.50134 | 8.059288 | 0.739462 | -0.0419 | 0.373036 |
| 10 | 40662400 | 40663399 | 15.4992 | 29.69229 | 18.49742 | 7.293879 | 1.195322 | -0.00667 | 0.373038 |
| 5 | 75204400 | 75209199 | 28.16251 | 45.68817 | 87.24331 | 37.38051 | -0.56474 | 1.611207 | 0.373047 |
| 1 | 125524800 | 125529399 | 61.89342 | 99.09929 | 89.73862 | 37.82045 | 0.487801 | 2.028841 | 0.373047 |
| 3 | 142882200 | 142884999 | 38.84242 | 64.03318 | 56.13018 | 24.21827 | 0.569252 | 1.388219 | 0.373052 |
| 4 | 105456200 | 105457599 | 15.23536 | 28.25162 | 23.55225 | 9.8879 | 0.70286 | 0.212965 | 0.373061 |
| 4 | 138052800 | 138059599 | 54.30095 | 85.5214 | 165.0544 | 68.48539 | -0.62485 | 2.495394 | 0.373081 |
| 14 | 35117000 | 35118199 | 11.91264 | 22.77655 | 21.02396 | 8.337104 | 0.625287 | -0.01848 | 0.373082 |
| 2 | 168894800 | 168898999 | 44.03248 | 70.30583 | 119.6295 | 50.21914 | -0.42496 | 2.103085 | 0.373083 |
| 9 | 6939600 | 6940799 | 11.9246 | 23.07122 | 18.4312 | 7.033051 | 0.883323 | -0.14835 | 0.373097 |
| 10 | 25348800 | 25351999 | 25.73242 | 41.89123 | 80.70889 | 34.64965 | -0.5732 | 1.543265 | 0.373105 |
| 4 | 62358800 | 62363799 | 47.77725 | 77.04641 | 74.24608 | 31.62107 | 0.410531 | 1.78352 | 0.373116 |
| 16 | 78809000 | 78810599 | 17.09851 | 30.93475 | 27.03456 | 11.72306 | 0.603861 | 0.372788 | 0.373118 |
| 13 | 109411000 | 109412399 | 21.36765 | 37.66268 | 30.85547 | 13.65631 | 0.675036 | 0.590544 | 0.373133 |
| 15 | 39073600 | 39074599 | 6.88927 | 13.11295 | 28.86454 | 11.55908 | -0.63066 | -0.01464 | 0.373145 |
| 13 | 62402800 | 62413799 | 226.4178 | 332.8421 | 305.4131 | 125.7775 | 0.363592 | 3.829722 | 0.373145 |
| 5 | 90029800 | 90032599 | 18.37112 | 30.86688 | 60.58162 | 26.46067 | -0.58831 | 1.124445 | 0.373147 |
| 4 | 45303800 | 45306599 | 30.19888 | 49.25687 | 85.41727 | 36.6967 | -0.41995 | 1.526759 | 0.373169 |

| | | | | | | | | | |
|---|---|---|---|---|---|---|---|---|---|
| 16 | 45353000 | 45356199 | 24.14844 | 39.68305 | 74.79622 | 32.22858 | -0.5338 | 1.405561 | 0.373175 |
| 6 | 4923800 | 4924999 | 25.09308 | 42.85376 | 41.58279 | 18.23118 | 0.42585 | 0.983154 | 0.373222 |
| 9 | 74726400 | 74727799 | 10.11141 | 18.24218 | 35.34674 | 15.38158 | -0.54096 | 0.357486 | 0.373229 |
| 12 | 112027000 | 112032399 | 50.8939 | 80.58312 | 152.6379 | 63.4657 | -0.59175 | 2.369812 | 0.373236 |
| 9 | 22226600 | 22231399 | 79.85266 | 125.8548 | 124.1107 | 51.50159 | 0.343337 | 2.489407 | 0.373241 |
| 13 | 31730400 | 31732199 | 8.938575 | 16.63932 | 32.15334 | 13.42216 | -0.49474 | 0.162167 | 0.373255 |
| 9 | 19825800 | 19826399 | 10.92762 | 20.88448 | 20.5473 | 8.157705 | 0.532395 | -0.02599 | 0.373272 |
| 17 | 25023200 | 25023999 | 5.135201 | 9.947621 | 25.52724 | 9.684879 | -0.77588 | -0.22211 | 0.373281 |
| 4 | 147271600 | 147280199 | 204.3201 | 301.9773 | 283.0159 | 116.5525 | 0.340367 | 3.739197 | 0.373293 |
| 6 | 70951600 | 70953399 | 11.91269 | 20.98756 | 41.40721 | 18.32811 | -0.58647 | 0.558422 | 0.373296 |
| 6 | 49536200 | 49537399 | 11.18548 | 21.44868 | 19.97923 | 7.856774 | 0.623186 | -0.05616 | 0.373315 |
| 2 | 35380400 | 35386999 | 64.46367 | 100.76 | 171.2915 | 71.06693 | -0.45058 | 2.644524 | 0.373318 |
| 12 | 36399400 | 36401399 | 26.62836 | 45.67705 | 41.43658 | 18.17355 | 0.523529 | 0.950187 | 0.373349 |
| 13 | 107072600 | 107074199 | 9.293103 | 16.56435 | 36.94151 | 16.25728 | -0.75312 | 0.433273 | 0.373359 |
| 15 | 7139000 | 7140199 | 12.68259 | 23.88986 | 22.37992 | 9.157011 | 0.564928 | 0.09273 | 0.373361 |
| 13 | 75168000 | 75169799 | 19.89962 | 34.93376 | 32.78054 | 14.50082 | 0.478835 | 0.635321 | 0.373363 |
| 10 | 127512600 | 127513399 | 5.545114 | 10.71486 | 24.80232 | 9.520402 | -0.64402 | -0.17665 | 0.373364 |
| 10 | 122606600 | 122608999 | 16.14283 | 27.73357 | 59.79298 | 26.22701 | -0.72006 | 0.915015 | 0.373374 |
| X | 14451200 | 14451999 | 20.40625 | 35.96121 | 27.19556 | 12.03693 | 0.789914 | 0.581648 | 0.373379 |
| 3 | 15547600 | 15549399 | 21.33323 | 37.95009 | 26.30372 | 11.59507 | 0.9204 | 0.498438 | 0.373399 |
| 3 | 130574400 | 130576599 | 13.32715 | 23.36811 | 43.54063 | 19.24139 | -0.50584 | 0.635583 | 0.37342 |
| 3 | 22171800 | 22173799 | 20.94478 | 36.68078 | 33.77248 | 14.90947 | 0.506097 | 0.679366 | 0.373439 |
| X | 44323800 | 44333199 | 179.1183 | 267.8662 | 248.1677 | 102.3339 | 0.372606 | 3.528591 | 0.373449 |
| 10 | 124913800 | 124914799 | 8.131979 | 15.33669 | 30.27192 | 12.36056 | -0.4992 | 0.066978 | 0.373457 |
| 6 | 72740800 | 72743999 | 21.81194 | 36.20566 | 67.12257 | 29.09095 | -0.50674 | 1.274155 | 0.373476 |
| 5 | 38475400 | 38478999 | 44.7386 | 71.22818 | 128.2415 | 53.61931 | -0.51082 | 2.197584 | 0.373486 |
| 11 | 121795400 | 121796399 | 19.1115 | 33.58873 | 31.82505 | 14.09179 | 0.464941 | 0.611547 | 0.373498 |
| 1 | 163135200 | 163137799 | 16.70587 | 28.73314 | 53.81753 | 23.60681 | -0.51755 | 0.898983 | 0.373509 |
| 12 | 17902800 | 17916799 | 329.6059 | 469.1754 | 416.7269 | 172.2975 | 0.365 | 4.317854 | 0.37351 |
| 4 | 37436600 | 37438599 | 26.87453 | 46.35614 | 35.55556 | 15.59765 | 0.766094 | 0.894918 | 0.373514 |
| 4 | 27351600 | 27352199 | 11.41854 | 22.18953 | 17.52435 | 6.525824 | 0.932646 | -0.23639 | 0.373516 |
| 7 | 81478400 | 81479799 | 8.546212 | 15.2508 | 36.96638 | 16.25226 | -0.87143 | 0.420163 | 0.373527 |
| 6 | 36050200 | 36052399 | 11.41107 | 20.55727 | 42.23122 | 18.41061 | -0.62692 | 0.388907 | 0.373539 |
| 1 | 130463400 | 130464399 | 8.115336 | 14.82105 | 32.97598 | 14.13226 | -0.7239 | 0.268256 | 0.373541 |
| 5 | 142131400 | 142144999 | 145.7528 | 217.3675 | 329.8404 | 135.9392 | -0.3424 | 3.577069 | 0.373551 |
| 8 | 69540400 | 69541999 | 10.28021 | 18.46589 | 36.32849 | 15.88372 | -0.56741 | 0.386501 | 0.373557 |
| X | 23698800 | 23704799 | 107.1091 | 166.9383 | 139.6988 | 57.97831 | 0.567533 | 2.713458 | 0.373586 |
| 10 | 81156200 | 81158599 | 15.57945 | 26.61278 | 53.66313 | 23.52803 | -0.62485 | 0.961293 | 0.373587 |
| X | 9231600 | 9235199 | 49.437 | 80.06707 | 68.61014 | 29.36045 | 0.586186 | 1.675455 | 0.37359 |
| 18 | 6893200 | 6898199 | 83.72993 | 131.5523 | 129.4478 | 53.69438 | 0.343004 | 2.554812 | 0.373616 |
| 2 | 84638200 | 84642999 | 45.37104 | 72.38304 | 121.0168 | 50.74201 | -0.40095 | 2.131385 | 0.373632 |
| 19 | 30196600 | 30198799 | 19.04504 | 33.73257 | 29.93614 | 13.23655 | 0.562855 | 0.52272 | 0.373634 |
| 13 | 66800600 | 66811999 | 148.9501 | 227.7297 | 174.9789 | 72.59094 | 0.66933 | 3.083686 | 0.373637 |
| 7 | 52602000 | 52602999 | 16.56316 | 29.70674 | 26.82633 | 11.74409 | 0.548206 | 0.421114 | 0.373638 |
| 1 | 78634800 | 78636799 | 12.43054 | 21.75599 | 43.62841 | 19.25029 | -0.61116 | 0.661678 | 0.373641 |
| 8 | 63914000 | 63915799 | 19.56941 | 34.73421 | 29.33999 | 12.95804 | 0.635172 | 0.508018 | 0.373641 |
| 4 | 14481800 | 14483399 | 31.16597 | 52.66059 | 45.80503 | 20.04082 | 0.581767 | 1.104179 | 0.373652 |

| | | | | | | | | | |
|---|---|---|---|---|---|---|---|---|---|
| X | 25495400 | 25500799 | 68.48021 | 110.1055 | 71.27001 | 30.22654 | 0.978341 | 1.906643 | 0.373653 |
| 13 | 62628200 | 62632999 | 84.61792 | 133.4714 | 123.9224 | 51.42879 | 0.431059 | 2.476788 | 0.373665 |
| 1 | 110768200 | 110769599 | 14.90723 | 28.12484 | 21.0244 | 8.578104 | 0.891152 | 0.099265 | 0.373666 |
| 15 | 12543000 | 12546199 | 23.6159 | 38.94626 | 73.3421 | 31.64963 | -0.53046 | 1.358265 | 0.37367 |
| 17 | 81216400 | 81217399 | 15.76823 | 27.35621 | 48.9881 | 21.49407 | -0.45012 | 0.813519 | 0.373687 |
| 7 | 23176200 | 23184799 | 141.2942 | 217.1103 | 160.4006 | 66.60278 | 0.731437 | 2.982474 | 0.373695 |
| X | 48567200 | 48569999 | 16.42035 | 28.53099 | 52.77974 | 23.17404 | -0.49668 | 0.801643 | 0.373706 |
| 7 | 17725400 | 17727199 | 48.2494 | 77.6868 | 74.07887 | 31.482 | 0.423064 | 1.833416 | 0.373736 |
| 9 | 55283200 | 55284399 | 14.67593 | 26.90857 | 25.20273 | 10.74469 | 0.520397 | 0.274824 | 0.373744 |
| 2 | 51820400 | 51820999 | 6.583496 | 12.59426 | 27.88363 | 11.04008 | -0.62403 | -0.05728 | 0.373764 |
| 15 | 5055600 | 5055999 | 4.403915 | 8.55985 | 22.80206 | 8.477003 | -0.79968 | -0.3072 | 0.373783 |
| 2 | 123228800 | 123229599 | 5.809984 | 11.28414 | 24.58984 | 9.198538 | -0.52264 | -0.263 | 0.37379 |
| 5 | 47097200 | 47098599 | 20.30045 | 35.52308 | 33.91502 | 14.96198 | 0.454064 | 0.689446 | 0.3738 |
| 15 | 100947800 | 100953599 | 56.02717 | 88.36761 | 151.5602 | 62.89811 | -0.45319 | 2.464847 | 0.373839 |
| 14 | 51014000 | 51016199 | 15.25858 | 26.6015 | 47.34046 | 20.8297 | -0.44037 | 0.740808 | 0.373857 |
| 7 | 13172400 | 13175199 | 48.04355 | 77.47025 | 75.83697 | 32.29561 | 0.387814 | 1.788106 | 0.373877 |
| 3 | 57972600 | 57974199 | 16.33133 | 29.46713 | 27.03658 | 11.76358 | 0.530793 | 0.381465 | 0.373892 |
| 6 | 140263200 | 140266199 | 84.74062 | 132.8735 | 129.6361 | 53.76709 | 0.353083 | 2.593276 | 0.373914 |
| 8 | 21452000 | 21460199 | 110.015 | 171.341 | 141.6283 | 58.78557 | 0.584454 | 2.72655 | 0.373915 |
| 13 | 65058000 | 65058999 | 12.7254 | 24.45388 | 18.68388 | 7.302331 | 0.914483 | -0.0635 | 0.373919 |
| 17 | 83324600 | 83329599 | 30.59873 | 49.49642 | 91.24863 | 39.01297 | -0.51812 | 1.676696 | 0.373947 |
| 1 | 33852600 | 33856199 | 29.04089 | 47.30982 | 84.05247 | 36.09416 | -0.45607 | 1.539443 | 0.373951 |
| 2 | 6462400 | 6468199 | 52.54297 | 83.52112 | 136.7818 | 57.07471 | -0.37684 | 2.256148 | 0.373964 |
| 8 | 115877200 | 115880799 | 24.73203 | 40.76089 | 74.03019 | 31.93794 | -0.4788 | 1.369538 | 0.373968 |
| 15 | 71457000 | 71457999 | 9.289114 | 17.18859 | 33.65684 | 14.17109 | -0.52295 | 0.204054 | 0.37397 |
| 5 | 130653200 | 130654999 | 12.04573 | 21.27966 | 40.85466 | 18.07372 | -0.54585 | 0.528045 | 0.37398 |
| 1 | 5426200 | 5429399 | 28.89493 | 49.04862 | 44.68801 | 19.57943 | 0.515254 | 1.05473 | 0.374009 |
| 15 | 77590400 | 77591399 | 15.36476 | 27.83381 | 25.85277 | 11.19259 | 0.518188 | 0.351735 | 0.374012 |
| 13 | 43938200 | 43940599 | 15.30653 | 26.0169 | 57.90718 | 25.37862 | -0.76815 | 1.007667 | 0.374015 |
| 6 | 140539600 | 140542199 | 15.00495 | 25.82334 | 53.76905 | 23.58598 | -0.66937 | 0.886041 | 0.374019 |
| 9 | 20093800 | 20097799 | 61.90013 | 99.18599 | 87.62667 | 36.97341 | 0.52465 | 2.004812 | 0.374021 |
| 8 | 79655200 | 79657599 | 18.94128 | 31.96599 | 59.07967 | 25.83931 | -0.50171 | 1.085535 | 0.374026 |
| 12 | 118799800 | 118803199 | 23.33843 | 38.85784 | 69.8167 | 30.32182 | -0.46153 | 1.24367 | 0.374026 |
| 17 | 5616400 | 5617999 | 9.403574 | 16.85309 | 36.37811 | 15.93575 | -0.70207 | 0.395471 | 0.37403 |
| 6 | 39846200 | 39850599 | 41.44487 | 66.18393 | 119.5944 | 50.2148 | -0.51129 | 2.095606 | 0.374031 |
| 18 | 34475000 | 34477399 | 16.49818 | 28.35 | 52.67033 | 23.10295 | -0.50584 | 0.909623 | 0.374038 |
| 1 | 12427200 | 12429199 | 20.82011 | 37.1669 | 27.67128 | 12.16203 | 0.821053 | 0.47883 | 0.374042 |
| 14 | 50607600 | 50609399 | 15.2284 | 26.49922 | 48.43269 | 21.28522 | -0.47905 | 0.767409 | 0.374063 |
| 10 | 47088400 | 47090599 | 21.89854 | 38.88826 | 28.14877 | 12.42653 | 0.857289 | 0.518917 | 0.374074 |
| 8 | 125909200 | 125912199 | 18.86829 | 32.13584 | 56.47072 | 24.75311 | -0.42821 | 1.001741 | 0.37408 |
| 2 | 21485000 | 21489599 | 46.60412 | 75.49771 | 69.48129 | 29.73868 | 0.483772 | 1.665466 | 0.374096 |
| 2 | 69908400 | 69910199 | 13.82759 | 24.26548 | 43.51656 | 19.24409 | -0.45089 | 0.626408 | 0.374111 |
| 6 | 128953800 | 128955399 | 13.82759 | 24.26548 | 43.51656 | 19.24409 | -0.45089 | 0.626408 | 0.374111 |
| 4 | 22826000 | 22828199 | 29.20473 | 49.39466 | 45.80672 | 20.04747 | 0.489645 | 1.087406 | 0.374119 |
| 18 | 36481400 | 36482399 | 6.744494 | 12.87446 | 27.23909 | 10.84182 | -0.56542 | -0.03739 | 0.374139 |
| 9 | 18094200 | 18095599 | 25.97088 | 44.34348 | 42.60278 | 18.67806 | 0.440111 | 0.990255 | 0.374142 |
| 13 | 48642800 | 48644799 | 36.39749 | 58.37361 | 106.8313 | 45.13769 | -0.52244 | 1.954505 | 0.374152 |

| | | | | | | | | | |
|---|---|---|---|---|---|---|---|---|---|
| Y | 11854200 | 11855399 | 12.41414 | 23.44388 | 13.44388 | 5.473991 | 1.272368 | 0.047144 | 0.374152 |
| 4 | 26038600 | 26041599 | 35.69669 | 60.63592 | 38.94633 | 17.06733 | 1.017672 | 1.068252 | 0.374167 |
| 7 | 59051800 | 59077799 | 427.8051 | 597.2155 | 494.9352 | 205.2789 | 0.436721 | 4.608075 | 0.374168 |
| 11 | 113055800 | 113056599 | 4.460883 | 8.625453 | 25.90449 | 9.913032 | -1.01158 | -0.198 | 0.374168 |
| 1 | 117525000 | 117529999 | 70.08102 | 111.9136 | 92.41055 | 38.80106 | 0.61655 | 2.117276 | 0.374169 |
| 19 | 58397800 | 58399999 | 25.19056 | 43.12475 | 41.49998 | 18.19843 | 0.438224 | 0.960768 | 0.374174 |
| 17 | 12216600 | 12217799 | 12.12775 | 22.92473 | 21.35459 | 8.677637 | 0.58125 | 0.06135 | 0.374175 |
| 4 | 26423600 | 26424399 | 14.56792 | 26.48855 | 25.29597 | 10.90075 | 0.482413 | 0.321797 | 0.374188 |
| 4 | 90471400 | 90473599 | 24.13983 | 42.17112 | 32.38453 | 14.26747 | 0.766465 | 0.730802 | 0.374189 |
| 4 | 112517600 | 112528599 | 265.8677 | 385.2202 | 351.9664 | 145.135 | 0.34968 | 4.054469 | 0.374212 |
| X | 57817600 | 57820199 | 41.81554 | 68.09109 | 64.81718 | 27.82965 | 0.441456 | 1.561646 | 0.374221 |
| 7 | 129157000 | 129157799 | 7.656177 | 14.42158 | 30.24407 | 12.37496 | -0.58871 | 0.068085 | 0.374229 |
| 10 | 11233000 | 11235999 | 43.73093 | 70.87482 | 68.68979 | 29.40958 | 0.40989 | 1.649836 | 0.374255 |
| 16 | 83030000 | 83031799 | 12.09095 | 21.47558 | 39.84377 | 17.58663 | -0.49286 | 0.478798 | 0.374266 |
| 18 | 86556600 | 86558199 | 18.02564 | 32.43064 | 27.36247 | 11.94738 | 0.647846 | 0.410872 | 0.374277 |
| 5 | 134097400 | 134105999 | 79.21816 | 123.1548 | 198.0021 | 82.20674 | -0.37634 | 2.748378 | 0.374299 |
| 15 | 46677200 | 46681999 | 46.33482 | 75.0256 | 69.5455 | 29.75567 | 0.47275 | 1.673012 | 0.374315 |
| 8 | 129820800 | 129825999 | 62.076 | 99.69527 | 80.61671 | 34.13399 | 0.655932 | 1.933299 | 0.374322 |
| 15 | 10882400 | 10885799 | 51.94995 | 83.88997 | 72.18977 | 30.79955 | 0.57572 | 1.751059 | 0.374331 |
| 1 | 34838200 | 34839799 | 7.581839 | 14.0756 | 34.276 | 14.36743 | -0.83101 | 0.168168 | 0.374331 |
| 6 | 3186200 | 3187399 | 16.25999 | 29.70493 | 22.67068 | 9.712002 | 0.809345 | 0.303228 | 0.374333 |
| 14 | 63352800 | 63354399 | 10.88476 | 19.56476 | 36.82179 | 16.09042 | -0.50342 | 0.387388 | 0.374335 |
| 19 | 13176600 | 13183599 | 91.81738 | 145.1289 | 113.2571 | 47.05039 | 0.683737 | 2.432776 | 0.37435 |
| 9 | 75316400 | 75318599 | 11.20829 | 19.65401 | 42.58504 | 18.82004 | -0.72201 | 0.622634 | 0.374355 |
| 16 | 12240200 | 12242599 | 26.46792 | 45.32652 | 42.00395 | 18.41992 | 0.492577 | 0.965012 | 0.37436 |
| 6 | 116240400 | 116242199 | 10.27016 | 18.52309 | 35.34185 | 15.38516 | -0.51917 | 0.360681 | 0.374366 |
| 12 | 20958400 | 20972999 | 524.9356 | 715.5184 | 618.2378 | 257.3843 | 0.341793 | 4.976127 | 0.374369 |
| 10 | 113896200 | 113897799 | 16.67718 | 30.13425 | 27.00822 | 11.73023 | 0.566073 | 0.37598 | 0.374377 |
| 12 | 36665000 | 36668199 | 63.61535 | 101.4682 | 98.28377 | 41.19829 | 0.384863 | 2.149759 | 0.374377 |
| 19 | 59931400 | 59932399 | 6.633891 | 12.55044 | 28.13732 | 11.41347 | -0.67232 | 0.026713 | 0.374392 |
| 19 | 39773800 | 39775399 | 23.46308 | 40.39193 | 38.34747 | 16.8214 | 0.45899 | 0.899195 | 0.374402 |
| 5 | 62453600 | 62456599 | 54.32954 | 87.34819 | 81.90502 | 34.73398 | 0.444622 | 1.890248 | 0.374406 |
| 11 | 63400800 | 63402199 | 7.562226 | 14.12167 | 30.06604 | 12.48401 | -0.62694 | 0.124876 | 0.37442 |
| 12 | 23416200 | 23416999 | 18.85684 | 33.43335 | 26.51809 | 11.7224 | 0.724612 | 0.504891 | 0.374424 |
| 1 | 99022000 | 99022799 | 12.2024 | 22.9023 | 22.04233 | 9.074328 | 0.519752 | 0.112327 | 0.374424 |
| 17 | 20292600 | 20294799 | 41.92026 | 68.52344 | 61.70334 | 26.5297 | 0.525234 | 1.505404 | 0.374429 |
| 6 | 48777600 | 48778799 | 9.570936 | 17.66412 | 33.60306 | 14.19855 | -0.4855 | 0.21938 | 0.374447 |
| 8 | 81281400 | 81282799 | 9.570936 | 17.66412 | 33.60306 | 14.19855 | -0.4855 | 0.21938 | 0.374447 |
| 9 | 100900400 | 100901599 | 12.72475 | 22.42251 | 40.89334 | 18.09996 | -0.47339 | 0.562603 | 0.374473 |
| 15 | 19557200 | 19559599 | 42.27852 | 68.83598 | 64.90145 | 27.8645 | 0.455101 | 1.56402 | 0.374484 |
| 18 | 37170200 | 37181399 | 144.127 | 219.7221 | 190.8502 | 79.12253 | 0.489323 | 3.140229 | 0.374491 |
| 17 | 22448400 | 22457999 | 182.0011 | 266.5446 | 398.165 | 164.0138 | -0.34428 | 3.88613 | 0.374494 |
| 16 | 6082200 | 6083799 | 10.74141 | 20.57177 | 20.02936 | 7.90443 | 0.555285 | -0.04891 | 0.374494 |
| 5 | 75360600 | 75361799 | 7.786315 | 14.7263 | 30.38338 | 12.33228 | -0.55543 | 0.041896 | 0.374504 |
| 9 | 118463800 | 118465599 | 28.53353 | 48.12411 | 45.8849 | 20.06458 | 0.449701 | 1.111127 | 0.374521 |
| 4 | 147380000 | 147386799 | 111.0635 | 173.1623 | 137.5483 | 57.08281 | 0.64302 | 2.707503 | 0.374528 |
| 8 | 21794600 | 21796399 | 28.61281 | 48.65998 | 44.14635 | 19.34859 | 0.521623 | 1.037545 | 0.374532 |

| | | | | | | | | | |
|---|---|---|---|---|---|---|---|---|---|
| 19 | 24876600 | 24878599 | 20.90287 | 34.60029 | 69.72082 | 30.15906 | -0.62692 | 1.305471 | 0.374546 |
| 10 | 69059200 | 69061599 | 14.81103 | 25.59359 | 51.64357 | 22.65644 | -0.62297 | 0.845801 | 0.374553 |
| 7 | 133701400 | 133703199 | 16.24069 | 28.03247 | 50.58329 | 22.19052 | -0.46272 | 0.862588 | 0.374558 |
| 7 | 67481200 | 67487399 | 60.71311 | 95.88916 | 152.3761 | 63.27635 | -0.34171 | 2.436143 | 0.374566 |
| 5 | 143717200 | 143718799 | 7.18547 | 13.26845 | 32.0356 | 13.5266 | -0.82558 | 0.193294 | 0.374572 |
| 11 | 70127800 | 70129199 | 7.18547 | 13.26845 | 32.0356 | 13.5266 | -0.82558 | 0.193294 | 0.374572 |
| 17 | 75295400 | 75298999 | 48.73012 | 78.48657 | 77.29108 | 32.86274 | 0.377268 | 1.824805 | 0.374585 |
| 17 | 12814800 | 12818199 | 45.42242 | 72.38521 | 124.1323 | 51.96905 | -0.43921 | 2.168799 | 0.374596 |
| 6 | 60222600 | 60223799 | 12.38819 | 21.94123 | 39.81251 | 17.60359 | -0.46318 | 0.502136 | 0.374597 |
| 1 | 103208800 | 103209199 | 13.30359 | 24.74514 | 23.91899 | 9.998279 | 0.495842 | 0.182383 | 0.374597 |
| X | 108652000 | 108654999 | 34.73519 | 58.34918 | 46.35889 | 20.24712 | 0.711676 | 1.165129 | 0.374601 |
| 16 | 3506600 | 3508199 | 17.42778 | 29.87661 | 53.2282 | 23.34406 | -0.44636 | 0.934159 | 0.374608 |
| 3 | 64153000 | 64154999 | 25.58304 | 43.51892 | 40.127 | 17.58734 | 0.498415 | 1.019176 | 0.374619 |
| 13 | 104083000 | 104086799 | 78.18767 | 123.2389 | 120.3457 | 49.9397 | 0.357531 | 2.486301 | 0.374635 |
| 1 | 109893200 | 109893799 | 10.70787 | 20.73201 | 19.1706 | 7.290706 | 0.679163 | -0.17012 | 0.374644 |
| 9 | 120846200 | 120854199 | 83.50338 | 127.9011 | 237.2172 | 98.45229 | -0.59878 | 3.031399 | 0.374666 |
| 1 | 54596200 | 54597999 | 10.9802 | 19.43972 | 38.84389 | 17.17667 | -0.60163 | 0.494836 | 0.374671 |
| 12 | 64153200 | 64154799 | 15.85984 | 29.52902 | 22.40804 | 9.351038 | 0.843917 | 0.192234 | 0.374687 |
| 8 | 74133600 | 74136399 | 36.15537 | 60.55239 | 47.52393 | 20.71658 | 0.729438 | 1.202743 | 0.374707 |
| 3 | 64663000 | 64666399 | 40.36883 | 66.77803 | 52.80287 | 22.83048 | 0.71833 | 1.351109 | 0.374733 |
| 2 | 47488200 | 47491599 | 33.7988 | 56.80548 | 46.91856 | 20.49485 | 0.655923 | 1.158826 | 0.374737 |
| Y | 20273200 | 20275199 | 13.13862 | 24.55337 | 15.00045 | 6.220599 | 1.161212 | 0.125475 | 0.37474 |
| 16 | 27471800 | 27475199 | 26.75365 | 46.42752 | 36.93502 | 16.20578 | 0.715299 | 0.843211 | 0.374749 |
| 13 | 104245200 | 104246799 | 10.77387 | 18.87036 | 44.6661 | 19.72042 | -0.84899 | 0.641574 | 0.374751 |
| X | 4495000 | 4506399 | 138.028 | 213.7718 | 131.5423 | 54.63914 | 1.003573 | 2.834497 | 0.374761 |
| 17 | 66408000 | 66409199 | 6.050449 | 11.5063 | 28.79609 | 11.55296 | -0.8162 | -0.01657 | 0.374771 |
| 7 | 75629600 | 75631799 | 14.15321 | 24.85205 | 46.13423 | 20.41219 | -0.50059 | 0.622184 | 0.374771 |
| 7 | 25813200 | 25828599 | 257.4981 | 374.3796 | 340.6712 | 140.423 | 0.360381 | 4.002364 | 0.374813 |
| 4 | 146077000 | 146087799 | 359.5239 | 506.4665 | 454.3228 | 188.1668 | 0.335639 | 4.470973 | 0.374822 |
| 2 | 74829800 | 74831199 | 6.354697 | 12.02273 | 29.77488 | 12.07668 | -0.81537 | 0.024744 | 0.374828 |
| 1 | 77196400 | 77197999 | 6.899914 | 13.10424 | 28.79322 | 11.5845 | -0.63393 | 0.00098 | 0.374836 |
| 13 | 94787600 | 94788599 | 6.899914 | 13.10424 | 28.79322 | 11.5845 | -0.63393 | 0.00098 | 0.374836 |
| 18 | 4924000 | 4925599 | 7.511401 | 14.21909 | 28.70558 | 11.62923 | -0.52121 | 0.031494 | 0.374839 |
| 13 | 54187800 | 54192999 | 45.49146 | 73.46167 | 70.93758 | 30.26585 | 0.409892 | 1.741363 | 0.374845 |
| 9 | 39767000 | 39769799 | 23.6755 | 41.10707 | 37.62147 | 16.51091 | 0.514035 | 0.822422 | 0.374866 |
| 5 | 38335000 | 38337599 | 17.17524 | 29.13036 | 56.88024 | 24.91498 | -0.58027 | 1.03355 | 0.374872 |
| 13 | 31740400 | 31744199 | 44.06995 | 70.2148 | 126.8792 | 53.10325 | -0.51494 | 2.172728 | 0.374899 |
| 1 | 124608000 | 124610599 | 24.5084 | 42.82156 | 33.52148 | 14.76979 | 0.7391 | 0.733959 | 0.374927 |
| 2 | 151078600 | 151079999 | 24.29545 | 40.10145 | 71.5169 | 30.87513 | -0.45167 | 1.351614 | 0.374941 |
| 6 | 46509800 | 46511399 | 12.82563 | 24.59104 | 19.83895 | 7.804085 | 0.828483 | -0.03844 | 0.374947 |
| 11 | 91645000 | 91646999 | 39.62132 | 64.8776 | 61.00396 | 26.24604 | 0.464561 | 1.476342 | 0.37495 |
| 6 | 104414000 | 104414799 | 11.95869 | 22.45517 | 21.99813 | 9.049103 | 0.49504 | 0.10609 | 0.37495 |
| 12 | 8677400 | 8680199 | 51.54228 | 82.76012 | 78.76487 | 33.339 | 0.420942 | 1.923496 | 0.37496 |
| 4 | 145440000 | 145443999 | 96.52804 | 151.4087 | 121.6248 | 50.44257 | 0.633536 | 2.59222 | 0.374974 |
| 19 | 4233600 | 4236199 | 12.63612 | 21.98526 | 46.84768 | 20.58681 | -0.69905 | 0.756219 | 0.374987 |
| 9 | 112203400 | 112205799 | 17.59854 | 30.32187 | 56.00271 | 24.56664 | -0.4968 | 0.892722 | 0.374994 |
| 12 | 21900000 | 21927399 | 862.8038 | 1128.669 | 928.1876 | 388.5857 | 0.353697 | 5.634317 | 0.375013 |

| | | | | | | | | | |
|---|---|---|---|---|---|---|---|---|---|
| 18 | 86516800 | 86517399 | 6.399566 | 12.39475 | 27.45834 | 10.42587 | -0.56749 | -0.19891 | 0.37502 |
| 1 | 32833400 | 32835199 | 25.38267 | 41.8968 | 72.38253 | 31.24925 | -0.40608 | 1.352944 | 0.375032 |
| 10 | 59451400 | 59452799 | 8.059308 | 14.85927 | 32.00203 | 13.54061 | -0.66436 | 0.209032 | 0.375043 |
| 4 | 146088800 | 146090999 | 33.35131 | 56.35709 | 44.67469 | 19.54669 | 0.715221 | 1.110315 | 0.375051 |
| Y | 40977600 | 40980199 | 24.28035 | 42.21785 | 19.18194 | 8.425966 | 1.518453 | 0.772886 | 0.375061 |
| 18 | 66102600 | 66104599 | 14.00122 | 24.53699 | 44.0812 | 19.47152 | -0.4534 | 0.650604 | 0.375082 |
| 19 | 4190400 | 4197599 | 46.38671 | 73.87804 | 125.4228 | 52.46623 | -0.42565 | 2.188547 | 0.375083 |
| X | 33382600 | 33384599 | 18.78189 | 33.68416 | 27.87766 | 12.20495 | 0.673013 | 0.437211 | 0.375095 |
| 3 | 109799200 | 109800399 | 10.81563 | 19.20216 | 37.82832 | 16.70306 | -0.57917 | 0.468022 | 0.3751 |
| 8 | 42370400 | 42371399 | 13.15486 | 25.40439 | 17.6207 | 6.782204 | 1.073257 | -0.10933 | 0.375106 |
| 1 | 18392200 | 18395599 | 28.52413 | 46.68374 | 80.31231 | 34.54633 | -0.40497 | 1.470649 | 0.375106 |
| 11 | 29114400 | 29116799 | 19.47468 | 34.08355 | 32.02821 | 14.1315 | 0.476891 | 0.679693 | 0.375109 |
| 13 | 94342400 | 94345599 | 24.56758 | 40.61706 | 70.63082 | 30.52707 | -0.41524 | 1.330999 | 0.37511 |
| 2 | 166481400 | 166483199 | 24.885 | 41.13263 | 71.2537 | 30.79152 | -0.40977 | 1.334595 | 0.375112 |
| 6 | 95733000 | 95734599 | 14.606 | 25.52738 | 45.71097 | 20.14511 | -0.44887 | 0.699079 | 0.375116 |
| 17 | 24842800 | 24846199 | 15.48502 | 26.6235 | 52.15309 | 22.87645 | -0.5819 | 0.896239 | 0.375127 |
| X | 134203000 | 134203999 | 6.251001 | 12.1322 | 29.61046 | 11.13437 | -0.6927 | -0.22767 | 0.37514 |
| 1 | 44140800 | 44141999 | 7.006016 | 13.03093 | 32.1261 | 13.42979 | -0.84513 | 0.14893 | 0.375145 |
| 5 | 145805200 | 145812799 | 156.6004 | 236.8974 | 215.3404 | 89.06757 | 0.414594 | 3.299956 | 0.37518 |
| 9 | 89887600 | 89891399 | 27.41388 | 44.78244 | 79.32731 | 34.09927 | -0.44892 | 1.494319 | 0.375196 |
| 5 | 115438200 | 115440599 | 11.97596 | 21.00366 | 42.0644 | 18.59257 | -0.60899 | 0.624754 | 0.375206 |
| 8 | 70106800 | 70107799 | 7.367366 | 13.92044 | 28.6589 | 11.65621 | -0.55425 | 0.042885 | 0.375212 |
| 7 | 12738400 | 12744999 | 49.55729 | 78.52339 | 141.1968 | 58.73003 | -0.51594 | 2.348972 | 0.375221 |
| 8 | 19370200 | 19372399 | 24.22709 | 42.24394 | 35.27931 | 15.52442 | 0.646046 | 0.763456 | 0.375237 |
| 14 | 35098200 | 35102199 | 54.10927 | 87.17578 | 76.69435 | 32.62278 | 0.540088 | 1.822926 | 0.375258 |
| 17 | 78632200 | 78633999 | 13.38044 | 23.41816 | 43.62077 | 19.24667 | -0.5052 | 0.668476 | 0.375264 |
| 1 | 139531200 | 139533199 | 19.69138 | 34.68582 | 31.5753 | 13.97669 | 0.523405 | 0.58849 | 0.375274 |
| 15 | 77493000 | 77493799 | 5.408343 | 10.45188 | 24.80647 | 9.516477 | -0.67906 | -0.18078 | 0.375286 |
| 5 | 91051600 | 91053599 | 11.19846 | 20.14987 | 37.34739 | 16.30558 | -0.48063 | 0.384248 | 0.375291 |
| 10 | 45159600 | 45165999 | 47.32053 | 75.35329 | 127.3784 | 53.27255 | -0.41973 | 2.193715 | 0.37532 |
| 8 | 11210200 | 11213599 | 27.12653 | 44.29729 | 78.45465 | 33.71979 | -0.44897 | 1.500206 | 0.375325 |
| 10 | 26294000 | 26296999 | 22.26319 | 38.75923 | 36.58943 | 16.0852 | 0.469883 | 0.777384 | 0.375327 |
| 10 | 125739000 | 125740599 | 10.08735 | 17.85545 | 38.42292 | 16.9912 | -0.70769 | 0.492177 | 0.375332 |
| X | 38792800 | 38794599 | 19.33158 | 35.97623 | 21.18708 | 8.848487 | 1.206734 | 0.223409 | 0.375344 |
| 7 | 40949600 | 40950599 | 13.94257 | 26.32784 | 22.35416 | 9.103819 | 0.711595 | 0.089161 | 0.375352 |
| X | 127319600 | 127321199 | 16.22035 | 30.906 | 19.62155 | 7.840229 | 1.151869 | 0.045033 | 0.375364 |
| X | 45554000 | 45556399 | 21.48293 | 38.96917 | 24.00963 | 10.37725 | 1.108155 | 0.384623 | 0.375368 |
| 10 | 58762800 | 58767199 | 25.26991 | 41.45492 | 78.40056 | 33.75718 | -0.53975 | 1.430848 | 0.3754 |
| 11 | 110214600 | 110216199 | 9.430928 | 17.33639 | 32.99295 | 14.01666 | -0.49164 | 0.239818 | 0.375409 |
| X | 60226800 | 60229199 | 53.22159 | 86.42469 | 61.53651 | 26.3848 | 0.856256 | 1.628817 | 0.375421 |
| 3 | 37693600 | 37695399 | 12.93232 | 22.49331 | 48.42085 | 21.27379 | -0.71394 | 0.762975 | 0.375426 |
| 16 | 35972200 | 35973599 | 15.38209 | 28.9922 | 21.60292 | 8.830134 | 0.893956 | 0.111977 | 0.375455 |
| 16 | 83536000 | 83537399 | 14.73847 | 26.97103 | 24.60131 | 10.51592 | 0.555456 | 0.283702 | 0.375458 |
| 8 | 95810800 | 95812799 | 10.02961 | 17.9244 | 36.37622 | 15.97289 | -0.6159 | 0.419367 | 0.375459 |
| 8 | 27850800 | 27852399 | 12.26203 | 21.51636 | 42.01546 | 18.57895 | -0.5729 | 0.616398 | 0.375466 |
| 8 | 48549000 | 48552199 | 19.09204 | 32.18182 | 61.30558 | 26.80309 | -0.54538 | 1.09737 | 0.375483 |
| 10 | 39486600 | 39490999 | 25.27439 | 41.46843 | 76.60056 | 32.98422 | -0.50574 | 1.428558 | 0.37549 |

| | | | | | | | | | |
|---|---|---|---|---|---|---|---|---|---|
| 6 | 126619000 | 126623399 | 33.81184 | 54.27337 | 112.1603 | 47.45119 | -0.69589 | 1.926203 | 0.375506 |
| 8 | 65649000 | 65651799 | 4.488847 | 8.734295 | 24.64945 | 9.084751 | -0.8703 | -0.32939 | 0.375512 |
| 10 | 85702200 | 85704599 | 36.92044 | 60.5768 | 59.20956 | 25.49571 | 0.409945 | 1.443525 | 0.375517 |
| 17 | 87547400 | 87549999 | 34.55801 | 55.62898 | 98.17162 | 41.70659 | -0.46351 | 1.831933 | 0.375549 |
| 19 | 7937200 | 7938599 | 30.4662 | 50.83029 | 49.73062 | 21.62649 | 0.412625 | 1.2262 | 0.37555 |
| 6 | 29649000 | 29649999 | 5.399736 | 10.45958 | 24.36634 | 9.246436 | -0.6375 | -0.21988 | 0.375561 |
| 9 | 25311800 | 25312599 | 5.399736 | 10.45958 | 24.36634 | 9.246436 | -0.6375 | -0.21988 | 0.375561 |
| 1 | 28661800 | 28662199 | 4.77833 | 9.322305 | 21.55016 | 7.700033 | -0.54512 | -0.44785 | 0.375576 |
| 2 | 43012800 | 43013399 | 4.77833 | 9.322305 | 21.55016 | 7.700033 | -0.54512 | -0.44785 | 0.375576 |
| 6 | 59434400 | 59435199 | 12.74598 | 23.89591 | 22.52857 | 9.291998 | 0.547253 | 0.125876 | 0.375578 |
| 1 | 147516200 | 147517799 | 22.26734 | 38.57759 | 37.32608 | 16.37653 | 0.433653 | 0.837732 | 0.375605 |
| 7 | 23503400 | 23508799 | 75.22345 | 120.0601 | 91.01742 | 38.18658 | 0.738795 | 2.14025 | 0.375609 |
| 4 | 133655400 | 133656999 | 7.067422 | 13.36093 | 28.6704 | 11.64755 | -0.61219 | 0.035404 | 0.375613 |
| 10 | 22610600 | 22611199 | 6.92238 | 13.11722 | 28.16854 | 11.38748 | -0.60637 | 0.018981 | 0.375626 |
| 1 | 58431600 | 58432999 | 8.808531 | 15.80189 | 35.92043 | 15.72247 | -0.77513 | 0.385259 | 0.375628 |
| 4 | 88932600 | 88933199 | 6.10152 | 11.84706 | 26.44783 | 9.915838 | -0.56084 | -0.24778 | 0.375631 |
| 1 | 22161600 | 22163399 | 13.50205 | 23.76794 | 42.93946 | 19.00948 | -0.46063 | 0.580871 | 0.375635 |
| 1 | 174080200 | 174082399 | 19.11393 | 32.35197 | 57.9924 | 25.38631 | -0.4575 | 1.061355 | 0.375637 |
| 5 | 134667400 | 134668599 | 8.541967 | 15.66009 | 32.46765 | 13.84406 | -0.6178 | 0.247883 | 0.375654 |
| 15 | 77289800 | 77293399 | 19.51698 | 32.61142 | 65.55768 | 28.53629 | -0.62289 | 1.190349 | 0.375678 |
| 13 | 106709600 | 106710799 | 8.59346 | 16.01228 | 31.63272 | 13.18334 | -0.5247 | 0.149983 | 0.375679 |
| 2 | 170795400 | 170797799 | 16.73599 | 28.86953 | 51.11867 | 22.42497 | -0.43577 | 0.871617 | 0.375683 |
| 8 | 91859400 | 91862799 | 23.12112 | 38.24456 | 69.34959 | 29.98355 | -0.47536 | 1.320292 | 0.375712 |
| 7 | 85131600 | 85137399 | 85.67011 | 136.2659 | 94.66289 | 39.5278 | 0.85938 | 2.257999 | 0.375714 |
| 15 | 10291000 | 10292399 | 16.27345 | 30.03418 | 24.60993 | 10.39867 | 0.721361 | 0.251072 | 0.375751 |
| X | 65785200 | 65787999 | 28.32966 | 49.29119 | 34.02935 | 14.95413 | 0.919246 | 0.81224 | 0.375754 |
| 14 | 83084200 | 83084799 | 8.490304 | 15.67767 | 32.01005 | 13.5161 | -0.58593 | 0.203582 | 0.375759 |
| 9 | 83393600 | 83394999 | 14.55162 | 27.32378 | 22.74848 | 9.357891 | 0.727875 | 0.130549 | 0.375786 |
| 2 | 108251200 | 108253999 | 25.04262 | 41.30495 | 73.54635 | 31.73966 | -0.44984 | 1.360493 | 0.375786 |
| 1 | 82492800 | 82494599 | 20.18015 | 36.22511 | 25.3808 | 11.10321 | 0.912224 | 0.434425 | 0.375789 |
| 12 | 53209400 | 53210599 | 9.998634 | 18.00208 | 35.37083 | 15.43126 | -0.563 | 0.370789 | 0.375792 |
| 5 | 145019800 | 145025599 | 38.42102 | 61.78148 | 103.1694 | 43.76214 | -0.38599 | 1.871569 | 0.375793 |
| X | 16202000 | 16206599 | 45.73649 | 75.66172 | 48.34587 | 20.90426 | 1.024833 | 1.36373 | 0.375795 |
| 15 | 96869600 | 96877399 | 79.25324 | 122.64 | 197.0517 | 81.85035 | -0.3806 | 2.829342 | 0.375806 |
| 14 | 17786400 | 17788399 | 15.99614 | 28.81485 | 27.2368 | 11.87512 | 0.486471 | 0.390651 | 0.375831 |
| 10 | 47493000 | 47494799 | 16.86292 | 30.21971 | 28.04375 | 12.28551 | 0.508074 | 0.428074 | 0.375846 |
| 7 | 66637400 | 66642799 | 55.57474 | 88.014 | 143.1134 | 59.51217 | -0.37162 | 2.365824 | 0.375847 |
| 3 | 157597600 | 157598999 | 15.58671 | 28.27381 | 26.68952 | 11.53546 | 0.496309 | 0.346043 | 0.37587 |
| 1 | 5238600 | 5244399 | 169.7712 | 254.1599 | 241.2401 | 99.49798 | 0.339091 | 3.505775 | 0.375905 |
| 9 | 37597200 | 37598399 | 8.687932 | 15.88581 | 33.45526 | 14.31619 | -0.64439 | 0.264083 | 0.375917 |
| 15 | 32253600 | 32255399 | 29.0079 | 49.49646 | 42.44681 | 18.6087 | 0.603174 | 1.006525 | 0.37592 |
| X | 45794600 | 45796999 | 31.15597 | 53.20298 | 40.8464 | 17.90819 | 0.762512 | 1.009487 | 0.375941 |
| 13 | 96491000 | 96491999 | 8.75573 | 16.3776 | 31.73316 | 13.13921 | -0.48921 | 0.127959 | 0.375964 |
| 13 | 48070600 | 48072199 | 17.39668 | 31.46065 | 26.96901 | 11.70094 | 0.630971 | 0.37689 | 0.375966 |
| 8 | 94846800 | 94850199 | 18.87679 | 32.12483 | 56.45633 | 24.74522 | -0.42851 | 1.009176 | 0.375973 |
| 17 | 22580000 | 22581799 | 10.9719 | 19.41996 | 40.35934 | 17.84773 | -0.65836 | 0.499004 | 0.375984 |
| 1 | 188407600 | 188408199 | 5.977691 | 11.54565 | 25.97197 | 9.989913 | -0.60697 | -0.16392 | 0.375997 |

| | | | | | | | | | |
|---|---|---|---|---|---|---|---|---|---|
| 6 | 22980800 | 22981599 | 14.65098 | 27.4372 | 23.31332 | 9.633587 | 0.693732 | 0.150031 | 0.376015 |
| 3 | 96898400 | 96899999 | 15.1239 | 27.40414 | 26.28575 | 11.37658 | 0.472181 | 0.348472 | 0.376015 |
| 8 | 103599200 | 103601199 | 19.03948 | 34.00815 | 28.31734 | 12.43983 | 0.660899 | 0.463741 | 0.376026 |
| 17 | 7391200 | 7393199 | 9.3443 | 16.9192 | 34.39242 | 14.89888 | -0.60517 | 0.329404 | 0.37603 |
| 11 | 113312800 | 113316999 | 32.865 | 53.17565 | 89.75667 | 38.38217 | -0.39085 | 1.67657 | 0.37604 |
| 11 | 12904400 | 12905599 | 15.22261 | 27.87719 | 25.69061 | 10.97073 | 0.541997 | 0.287323 | 0.37604 |
| 1 | 71599400 | 71600999 | 21.39575 | 37.37072 | 34.90934 | 15.37809 | 0.485334 | 0.724639 | 0.376053 |
| 17 | 18619800 | 18628799 | 172.0726 | 259.7846 | 215.7219 | 89.16929 | 0.542561 | 3.343836 | 0.376074 |
| 9 | 88883600 | 88885999 | 18.37149 | 32.84565 | 28.39318 | 12.46334 | 0.607929 | 0.45367 | 0.376077 |
| 3 | 124540600 | 124541399 | 10.24499 | 19.8421 | 19.21878 | 7.298217 | 0.614948 | -0.17984 | 0.376097 |
| 5 | 106707200 | 106709599 | 18.68333 | 31.82524 | 57.03297 | 24.99984 | -0.45643 | 0.999349 | 0.376099 |
| 6 | 97354400 | 97356999 | 28.98713 | 49.02022 | 40.56738 | 17.75375 | 0.653306 | 1.082569 | 0.37612 |
| 2 | 29014000 | 29015799 | 9.188067 | 16.6762 | 33.91284 | 14.64719 | -0.60281 | 0.313941 | 0.376142 |
| 13 | 24774200 | 24775799 | 14.7308 | 26.82655 | 25.75419 | 11.07656 | 0.476623 | 0.315062 | 0.37616 |
| 5 | 133243400 | 133244799 | 8.099324 | 15.23833 | 31.34737 | 12.85203 | -0.56369 | 0.08188 | 0.376165 |
| 11 | 30903800 | 30904799 | 5.747603 | 10.98766 | 27.80947 | 11.03238 | -0.81762 | -0.05908 | 0.376166 |
| 17 | 19414400 | 19422599 | 107.6071 | 168.125 | 134.8251 | 55.93985 | 0.631629 | 2.669944 | 0.376195 |
| X | 87402800 | 87407799 | 49.51025 | 80.67751 | 59.59934 | 25.59589 | 0.80706 | 1.567873 | 0.376197 |
| 8 | 21457200 | 21459599 | 36.86044 | 61.01666 | 54.36558 | 23.51738 | 0.546656 | 1.334578 | 0.376211 |
| 13 | 6517400 | 6519999 | 29.1507 | 49.0171 | 47.49479 | 20.74959 | 0.426382 | 1.139749 | 0.376224 |
| 4 | 12997400 | 12999999 | 30.68115 | 52.72545 | 37.90262 | 16.62538 | 0.858158 | 0.945834 | 0.37623 |
| 10 | 3532400 | 3537199 | 56.29061 | 90.08778 | 88.73179 | 37.37174 | 0.366037 | 2.035823 | 0.376231 |
| 1 | 156878400 | 156879999 | 11.27983 | 19.96367 | 39.33271 | 17.39397 | -0.58172 | 0.500714 | 0.376248 |
| 4 | 73854800 | 73856599 | 22.76133 | 39.77428 | 34.78136 | 15.32626 | 0.580199 | 0.725545 | 0.376257 |
| 4 | 137781200 | 137782199 | 5.966765 | 11.55172 | 24.94644 | 9.490419 | -0.53296 | -0.20285 | 0.37631 |
| 17 | 21583600 | 21584199 | 5.966765 | 11.55172 | 24.94644 | 9.490419 | -0.53296 | -0.20285 | 0.37631 |
| 18 | 43030800 | 43033799 | 26.16116 | 42.71161 | 78.33917 | 33.6675 | -0.49951 | 1.500438 | 0.376314 |
| 6 | 146028400 | 146030999 | 18.87913 | 32.02128 | 58.07255 | 25.43749 | -0.47426 | 1.039784 | 0.376314 |
| 6 | 68328400 | 68330399 | 24.17163 | 42.52494 | 29.50512 | 13.06357 | 0.912472 | 0.624505 | 0.376347 |
| 7 | 81145000 | 81148399 | 22.47424 | 37.2285 | 68.19738 | 29.51491 | -0.48978 | 1.301128 | 0.376358 |
| 11 | 110640200 | 110642199 | 14.67365 | 25.3415 | 50.56321 | 22.18208 | -0.60689 | 0.850039 | 0.376359 |
| 10 | 24314000 | 24315599 | 24.72838 | 42.28415 | 41.045 | 17.99723 | 0.425568 | 0.969311 | 0.376369 |
| 11 | 26160000 | 26160999 | 6.283325 | 12.06917 | 26.89385 | 10.52417 | -0.61708 | -0.10119 | 0.376392 |
| 16 | 70512200 | 70512799 | 6.283325 | 12.06917 | 26.89385 | 10.52417 | -0.61708 | -0.10119 | 0.376392 |
| X | 55104800 | 55111799 | 101.9591 | 160.1368 | 124.4749 | 51.62929 | 0.682392 | 2.573464 | 0.376409 |
| 5 | 136878400 | 136879999 | 15.90316 | 27.60972 | 48.99084 | 21.49936 | -0.43684 | 0.805578 | 0.376415 |
| 5 | 104213800 | 104217999 | 43.68671 | 69.8407 | 116.6542 | 49.0566 | -0.39628 | 2.063595 | 0.376423 |
| 3 | 85790800 | 85791799 | 6.893635 | 13.20164 | 27.8976 | 11.0118 | -0.55244 | -0.06021 | 0.376439 |
| 13 | 54498200 | 54502399 | 49.20546 | 78.13356 | 133.8003 | 55.77183 | -0.44257 | 2.285119 | 0.376451 |
| 7 | 5247400 | 5257599 | 161.8052 | 244.9084 | 210.7265 | 87.17462 | 0.494452 | 3.290174 | 0.376471 |
| 9 | 114051800 | 114053199 | 9.74747 | 17.93846 | 34.08336 | 14.45775 | -0.48781 | 0.238462 | 0.376478 |
| 3 | 94523400 | 94524399 | 7.007836 | 13.41767 | 29.12548 | 11.50305 | -0.59277 | -0.05878 | 0.376494 |
| 19 | 49238800 | 49242199 | 31.12541 | 52.52717 | 45.79162 | 20.02746 | 0.578386 | 1.112085 | 0.37652 |
| 18 | 12161200 | 12163199 | 23.70166 | 40.74599 | 39.42634 | 17.29393 | 0.431319 | 0.914759 | 0.376545 |
| 6 | 83030200 | 83032399 | 14.56679 | 25.06744 | 51.06956 | 22.40179 | -0.63781 | 0.883884 | 0.376572 |
| 17 | 24938400 | 24941799 | 29.46038 | 47.76551 | 85.36205 | 36.55855 | -0.47015 | 1.626526 | 0.376575 |
| 10 | 126761200 | 126768999 | 39.50411 | 63.10719 | 121.3279 | 50.96839 | -0.6 | 2.082348 | 0.37659 |

| | | | | | | | | | |
|---|---|---|---|---|---|---|---|---|---|
| 9 | 105954000 | 105955599 | 11.74541 | 20.80044 | 38.80447 | 17.15826 | -0.50302 | 0.496762 | 0.376593 |
| 1 | 85156400 | 85161199 | 178.0866 | 267.7965 | 230.4911 | 95.16695 | 0.485877 | 3.422853 | 0.376594 |
| 3 | 136192400 | 136192999 | 6.430871 | 12.3507 | 26.30524 | 10.29835 | -0.55248 | -0.09734 | 0.376598 |
| 9 | 118116000 | 118116999 | 6.430871 | 12.3507 | 26.30524 | 10.29835 | -0.55248 | -0.09734 | 0.376598 |
| 14 | 48683600 | 48684199 | 6.430871 | 12.3507 | 26.30524 | 10.29835 | -0.55248 | -0.09734 | 0.376598 |
| 7 | 55248000 | 55249399 | 24.336 | 39.95029 | 74.02527 | 31.88256 | -0.50977 | 1.419891 | 0.376602 |
| 6 | 20300200 | 20301799 | 20.76427 | 36.58521 | 30.85407 | 13.65673 | 0.633134 | 0.59001 | 0.376602 |
| 4 | 76367600 | 76369799 | 15.65224 | 26.95757 | 50.55952 | 22.17861 | -0.51875 | 0.881054 | 0.376635 |
| 3 | 54822000 | 54823599 | 16.18896 | 27.62235 | 54.22914 | 23.774 | -0.58688 | 0.972777 | 0.376646 |
| 18 | 76511200 | 76513399 | 16.02331 | 27.34029 | 55.30975 | 24.24778 | -0.63006 | 0.971963 | 0.37666 |
| 17 | 30060400 | 30062799 | 19.20537 | 32.12434 | 62.71525 | 27.31706 | -0.58067 | 1.17737 | 0.376668 |
| 13 | 17827600 | 17828599 | 13.15107 | 24.63035 | 23.44806 | 9.687894 | 0.53105 | 0.136839 | 0.376713 |
| 4 | 124430800 | 124432599 | 10.57422 | 18.61822 | 40.48107 | 17.92033 | -0.72581 | 0.557466 | 0.376725 |
| 11 | 114422200 | 114430399 | 60.55339 | 95.08795 | 161.3289 | 66.9139 | -0.4425 | 2.557017 | 0.37673 |
| 15 | 83633400 | 83634399 | 7.422233 | 13.82764 | 31.62427 | 13.1835 | -0.73452 | 0.140566 | 0.37674 |
| X | 145023800 | 145026399 | 22.99529 | 40.78676 | 27.41001 | 12.1127 | 0.962725 | 0.531166 | 0.376751 |
| 1 | 94669800 | 94670999 | 7.345192 | 14.00147 | 28.33695 | 11.31231 | -0.50597 | -0.01897 | 0.376753 |
| 16 | 47279200 | 47279599 | 6.56133 | 12.66534 | 26.52526 | 10.23131 | -0.50844 | -0.14447 | 0.376764 |
| 2 | 172369200 | 172372799 | 24.77658 | 40.69279 | 77.85914 | 33.54052 | -0.55579 | 1.41472 | 0.376766 |
| 2 | 35767000 | 35770999 | 19.5264 | 32.63379 | 63.27578 | 27.54655 | -0.57086 | 1.18781 | 0.376776 |
| 12 | 5130000 | 5137999 | 58.05007 | 91.73502 | 150.2786 | 62.42319 | -0.38488 | 2.419742 | 0.376792 |
| 1 | 47544600 | 47546799 | 22.07034 | 39.08235 | 28.55076 | 12.62454 | 0.841958 | 0.541878 | 0.3768 |
| 1 | 166396400 | 166399599 | 35.81044 | 57.66779 | 98.99831 | 42.08183 | -0.42311 | 1.82009 | 0.376812 |
| 14 | 83088400 | 83089999 | 18.82534 | 34.55919 | 21.90643 | 9.323719 | 1.081169 | 0.292567 | 0.37682 |
| 14 | 21100400 | 21103199 | 13.37498 | 23.30565 | 46.30033 | 20.36544 | -0.59835 | 0.73669 | 0.376821 |
| 13 | 67594400 | 67595399 | 13.38488 | 25.17589 | 22.96758 | 9.420766 | 0.600551 | 0.111661 | 0.376828 |
| Y | 20958800 | 20961399 | 12.40783 | 23.36399 | 15.03559 | 6.1557 | 1.10154 | 0.071627 | 0.376853 |
| 6 | 130412000 | 130414799 | 39.79703 | 65.76371 | 54.44277 | 23.52263 | 0.652259 | 1.363455 | 0.376854 |
| 18 | 4538400 | 4545599 | 129.3848 | 197.3716 | 190.5489 | 79.00913 | 0.337671 | 3.123136 | 0.376861 |
| 7 | 47472000 | 47474599 | 50.46927 | 81.27824 | 78.34136 | 33.30369 | 0.40793 | 1.830408 | 0.376869 |
| 9 | 73838400 | 73840999 | 31.68651 | 51.41217 | 87.28044 | 37.40017 | -0.39536 | 1.616047 | 0.376878 |
| 7 | 6016800 | 6022599 | 90.30529 | 142.6402 | 116.8925 | 48.54367 | 0.612512 | 2.448157 | 0.37688 |
| 13 | 16794400 | 16796999 | 21.1826 | 35.65955 | 62.50664 | 27.31749 | -0.42588 | 1.117881 | 0.376883 |
| 6 | 52512800 | 52514999 | 37.68154 | 61.57376 | 60.09498 | 25.83496 | 0.409049 | 1.500682 | 0.376886 |
| X | 77788600 | 77790199 | 42.6084 | 69.81814 | 58.83976 | 25.32251 | 0.62258 | 1.475693 | 0.376895 |
| 11 | 86698000 | 86698799 | 4.412155 | 8.553249 | 23.76152 | 8.984125 | -0.88244 | -0.24389 | 0.376896 |
| 11 | 15953000 | 15954599 | 19.30672 | 34.61168 | 27.17493 | 11.90127 | 0.748083 | 0.440644 | 0.376946 |
| 12 | 20427600 | 20488999 | 1371.026 | 1729.322 | 1383.286 | 581.2127 | 0.340918 | 6.227271 | 0.376948 |
| 6 | 147201400 | 147209799 | 100.8679 | 153.8348 | 238.8447 | 99.02918 | -0.34745 | 3.123235 | 0.376949 |
| 13 | 29886400 | 29887599 | 6.197128 | 11.75891 | 28.19303 | 11.36466 | -0.76063 | 4.56E-05 | 0.376954 |
| X | 87935400 | 87937799 | 20.17591 | 36.05066 | 27.44325 | 12.05229 | 0.790194 | 0.470264 | 0.376973 |
| X | 97183800 | 97185999 | 23.89998 | 42.00452 | 30.82791 | 13.65026 | 0.831649 | 0.638324 | 0.376982 |
| X | 4948000 | 4948999 | 18.09263 | 34.23344 | 18.6367 | 7.557547 | 1.352892 | 0.104803 | 0.376982 |
| 13 | 67385600 | 67395599 | 165.0708 | 248.3769 | 232.0266 | 95.8167 | 0.368552 | 3.407945 | 0.376984 |
| 10 | 4080000 | 4082799 | 16.92323 | 29.05708 | 52.16449 | 22.88043 | -0.45696 | 0.915888 | 0.376986 |
| 3 | 17847600 | 17851399 | 34.01203 | 54.88068 | 94.56755 | 40.3065 | -0.42506 | 1.755833 | 0.376992 |
| 6 | 131762800 | 131766199 | 49.67621 | 80.42205 | 69.06176 | 29.54493 | 0.582593 | 1.685019 | 0.376997 |

| | | | | | | | | | |
|---|---|---|---|---|---|---|---|---|---|
| X | 21091200 | 21093199 | 31.49716 | 53.16494 | 44.78051 | 19.5864 | 0.628034 | 1.115195 | 0.377028 |
| 1 | 139642400 | 139645399 | 40.14558 | 64.21323 | 123.3517 | 51.91165 | -0.5968 | 2.045982 | 0.377042 |
| X | 47800000 | 47802599 | 33.12137 | 55.25043 | 49.74395 | 21.62987 | 0.532233 | 1.238609 | 0.377046 |
| X | 147618800 | 147628399 | 140.6548 | 215.0693 | 181.4158 | 75.26312 | 0.534921 | 3.079093 | 0.377065 |
| 7 | 125043800 | 125045999 | 16.04419 | 27.8512 | 49.53203 | 21.73521 | -0.44019 | 0.807992 | 0.377075 |
| 13 | 102571000 | 102573199 | 15.69863 | 26.9059 | 56.94995 | 24.97581 | -0.694 | 0.932968 | 0.377089 |
| 1 | 36752000 | 36753599 | 18.44479 | 32.46818 | 30.85036 | 13.65763 | 0.461676 | 0.588189 | 0.377089 |
| 4 | 143938200 | 143942599 | 63.17317 | 101.2404 | 86.24834 | 36.39981 | 0.577239 | 2.001038 | 0.3771 |
| 12 | 16497000 | 16498399 | 15.87134 | 29.30973 | 24.7249 | 10.43867 | 0.680702 | 0.245687 | 0.377117 |
| 14 | 7878400 | 7879199 | 5.096612 | 9.923819 | 23.06428 | 8.452587 | -0.58709 | -0.35061 | 0.377138 |
| 2 | 165513600 | 165517599 | 29.29113 | 47.56044 | 83.7759 | 35.91785 | -0.44735 | 1.596649 | 0.377155 |
| 7 | 41550000 | 41553599 | 70.07399 | 111.4342 | 106.0148 | 44.25498 | 0.406137 | 2.253345 | 0.377161 |
| 2 | 64510000 | 64510199 | 4.223919 | 8.25406 | 20.07208 | 6.965265 | -0.57883 | -0.56582 | 0.377168 |
| 8 | 125206200 | 125210799 | 32.84596 | 53.14678 | 90.49875 | 38.70054 | -0.40346 | 1.675242 | 0.377171 |
| 4 | 144487600 | 144491999 | 35.74742 | 59.67854 | 48.69675 | 21.18517 | 0.673501 | 1.232893 | 0.377179 |
| 12 | 86772600 | 86773399 | 7.046 | 13.41147 | 27.73285 | 11.10552 | -0.54062 | -0.0137 | 0.377179 |
| 7 | 21016400 | 21032199 | 229.9548 | 337.8558 | 306.955 | 126.4169 | 0.377135 | 3.838915 | 0.377182 |
| 7 | 31343000 | 31344599 | 19.96916 | 36.66062 | 22.51261 | 9.58114 | 1.127198 | 0.303957 | 0.377195 |
| 15 | 59873600 | 59874999 | 17.12492 | 31.64293 | 23.99954 | 10.12449 | 0.833773 | 0.250164 | 0.3772 |
| 4 | 115507000 | 115510799 | 29.91073 | 48.57465 | 84.10178 | 36.06203 | -0.4224 | 1.593228 | 0.377208 |
| X | 22227400 | 22228799 | 23.5622 | 41.05069 | 34.13408 | 15.01268 | 0.652505 | 0.773782 | 0.377228 |
| 6 | 68437800 | 68439199 | 11.79264 | 20.75397 | 46.13536 | 20.42528 | -0.75839 | 0.578079 | 0.377232 |
| 7 | 30502000 | 30512799 | 107.2933 | 162.9024 | 255.2322 | 105.6866 | -0.36583 | 3.216962 | 0.377261 |
| 11 | 104750200 | 104755199 | 81.88353 | 128.7985 | 127.1215 | 52.7373 | 0.339911 | 2.530675 | 0.37728 |
| 14 | 10184800 | 10185999 | 6.823691 | 12.78622 | 30.11939 | 12.43389 | -0.76495 | 0.095125 | 0.377303 |
| 7 | 14241000 | 14247399 | 200.0946 | 296.1801 | 277.574 | 114.3133 | 0.342455 | 3.713811 | 0.377312 |
| 12 | 25849600 | 25850799 | 12.17689 | 22.95301 | 22.5268 | 9.206419 | 0.499986 | 0.085795 | 0.377316 |
| 6 | 35936800 | 35938399 | 15.58848 | 29.29724 | 22.12748 | 9.088484 | 0.869175 | 0.132837 | 0.377335 |
| 6 | 24222600 | 24223199 | 8.928869 | 16.65673 | 32.21113 | 13.39782 | -0.49167 | 0.148242 | 0.377346 |
| 19 | 26987000 | 26987599 | 4.182271 | 8.163676 | 20.38017 | 7.217258 | -0.64148 | -0.4896 | 0.377354 |
| 2 | 68420200 | 68420799 | 4.806193 | 9.372213 | 22.10718 | 7.948327 | -0.58284 | -0.42588 | 0.377358 |
| 9 | 50686400 | 50686999 | 4.806193 | 9.372213 | 22.10718 | 7.948327 | -0.58284 | -0.42588 | 0.377358 |
| 5 | 109503200 | 109503799 | 18.26149 | 32.35537 | 26.94311 | 11.91237 | 0.654767 | 0.511678 | 0.377361 |
| X | 38222800 | 38229199 | 70.97049 | 113.6671 | 82.52223 | 34.79501 | 0.806568 | 2.031057 | 0.377367 |
| 1 | 108023400 | 108024199 | 6.267932 | 12.08266 | 27.06304 | 10.48239 | -0.61072 | -0.13517 | 0.377376 |
| 16 | 16322400 | 16326199 | 49.90803 | 79.13885 | 138.1065 | 57.48656 | -0.47174 | 2.328355 | 0.377379 |
| 3 | 87877000 | 87883199 | 29.41206 | 47.41427 | 97.92524 | 41.68763 | -0.68725 | 1.776958 | 0.377388 |
| 1 | 35926800 | 35937399 | 140.9792 | 210.4841 | 324.6787 | 133.8411 | -0.36458 | 3.554998 | 0.377392 |
| 13 | 65804600 | 65806599 | 39.45277 | 65.57575 | 44.91066 | 19.48237 | 0.925376 | 1.288762 | 0.377416 |
| 7 | 55603400 | 55603799 | 14.0109 | 25.57782 | 23.35119 | 10.01248 | 0.551499 | 0.289953 | 0.377422 |
| 16 | 60141200 | 60141599 | 6.131738 | 11.80264 | 25.80157 | 10.03635 | -0.58067 | -0.12241 | 0.377427 |
| 2 | 10701400 | 10702799 | 10.78171 | 19.51629 | 38.02792 | 16.47964 | -0.54506 | 0.350484 | 0.377442 |
| 14 | 100974800 | 100975599 | 6.165863 | 11.75283 | 27.18635 | 10.85414 | -0.69626 | -0.03152 | 0.377458 |
| 18 | 42770600 | 42771399 | 6.958139 | 13.05608 | 31.79022 | 13.09314 | -0.81018 | 0.089617 | 0.377459 |
| 14 | 119854800 | 119856199 | 11.37987 | 20.28368 | 38.81186 | 17.08051 | -0.5344 | 0.446811 | 0.377499 |
| 14 | 110163600 | 110166799 | 33.06077 | 55.47791 | 48.04339 | 20.96886 | 0.587845 | 1.171862 | 0.377502 |
| 11 | 54225800 | 54229999 | 23.1895 | 38.16407 | 72.39304 | 31.21277 | -0.542 | 1.384174 | 0.377502 |

| | | | | | | | | | |
|---|---|---|---|---|---|---|---|---|---|
| 11 | 34111400 | 34112799 | 16.14329 | 28.97432 | 27.07193 | 11.8445 | 0.499472 | 0.411114 | 0.377535 |
| 7 | 32145600 | 32149199 | 37.79617 | 63.37745 | 45.25985 | 19.74478 | 0.865106 | 1.193203 | 0.377546 |
| 3 | 105707800 | 105708799 | 7.865554 | 14.64771 | 30.5499 | 12.74421 | -0.60305 | 0.146086 | 0.377547 |
| 12 | 74391200 | 74394599 | 26.86847 | 44.12322 | 76.28941 | 32.86264 | -0.41012 | 1.421745 | 0.377566 |
| 6 | 124228600 | 124229999 | 7.71342 | 14.39991 | 30.06159 | 12.48819 | -0.59932 | 0.128458 | 0.377572 |
| 7 | 101222800 | 101223599 | 7.71342 | 14.39991 | 30.06159 | 12.48819 | -0.59932 | 0.128458 | 0.377572 |
| 15 | 100937000 | 100938799 | 22.20167 | 36.67958 | 69.61942 | 30.07524 | -0.54108 | 1.333727 | 0.377597 |
| 16 | 50257400 | 50258799 | 12.68225 | 22.34908 | 41.37913 | 18.31474 | -0.4952 | 0.560414 | 0.377602 |
| 5 | 87806000 | 87807199 | 16.9105 | 30.35496 | 28.06639 | 12.27835 | 0.51474 | 0.418857 | 0.377613 |
| 14 | 50160600 | 50161799 | 15.92137 | 28.53091 | 27.23792 | 11.933 | 0.467471 | 0.420923 | 0.377638 |
| 19 | 24007000 | 24008999 | 14.06313 | 24.32635 | 48.97028 | 21.48432 | -0.61893 | 0.832175 | 0.37764 |
| 9 | 37889200 | 37890399 | 18.45079 | 32.45691 | 29.94441 | 13.25835 | 0.503645 | 0.594265 | 0.377662 |
| 8 | 127429000 | 127429999 | 6.887414 | 13.17094 | 27.29194 | 10.81691 | -0.53012 | -0.04957 | 0.377677 |
| 14 | 49865800 | 49866399 | 6.887414 | 13.17094 | 27.29194 | 10.81691 | -0.53012 | -0.04957 | 0.377677 |
| 11 | 30979200 | 30981999 | 54.26538 | 87.38745 | 77.79877 | 33.0707 | 0.52219 | 1.837497 | 0.377688 |
| 10 | 77580400 | 77588399 | 62.22276 | 97.16535 | 187.6383 | 77.85665 | -0.63535 | 2.662017 | 0.377694 |
| 5 | 138169200 | 138171999 | 16.67648 | 28.36932 | 55.82447 | 24.46751 | -0.59096 | 1.003418 | 0.377722 |
| 1 | 64364600 | 64367999 | 35.77236 | 59.79227 | 48.11716 | 20.94896 | 0.69342 | 1.220255 | 0.377728 |
| 4 | 57022200 | 57024399 | 16.62342 | 28.47625 | 52.67673 | 23.1008 | -0.50052 | 0.936414 | 0.37774 |
| 18 | 70838200 | 70839399 | 12.44373 | 21.80464 | 42.58697 | 18.8094 | -0.57308 | 0.642677 | 0.37775 |
| 14 | 3064200 | 3064799 | 13.15347 | 24.31985 | 22.82568 | 9.62107 | 0.529093 | 0.21051 | 0.377751 |
| 17 | 20215400 | 20217399 | 15.68306 | 29.08763 | 21.54944 | 9.044602 | 0.872514 | 0.212386 | 0.377753 |
| 2 | 15320000 | 15320599 | 18.88697 | 33.17929 | 31.83054 | 14.0889 | 0.447043 | 0.616609 | 0.377753 |
| 15 | 31304800 | 31310799 | 53.78181 | 85.15976 | 142.9957 | 59.45799 | -0.41812 | 2.368359 | 0.377772 |
| X | 80778400 | 80779599 | 14.24093 | 27.30947 | 19.84727 | 7.803396 | 0.979208 | -0.02422 | 0.377773 |
| 1 | 6322600 | 6324799 | 28.56418 | 48.92517 | 39.06935 | 17.13327 | 0.706231 | 0.964763 | 0.377784 |
| 18 | 24662000 | 24665599 | 29.24335 | 47.53903 | 83.68737 | 35.90969 | -0.44498 | 1.571007 | 0.377785 |
| 7 | 64932400 | 64934599 | 15.22212 | 26.50442 | 48.42622 | 21.29052 | -0.4785 | 0.758536 | 0.377791 |
| 2 | 101887600 | 101889599 | 9.107527 | 16.3244 | 37.39657 | 16.3803 | -0.78739 | 0.393689 | 0.377806 |
| 9 | 88798800 | 88801199 | 13.79052 | 24.23814 | 45.57251 | 20.17981 | -0.51899 | 0.602539 | 0.37781 |
| 1 | 141728200 | 141729399 | 19.78975 | 34.68287 | 32.49889 | 14.35594 | 0.481049 | 0.661799 | 0.377819 |
| 11 | 11271800 | 11274199 | 29.48254 | 50.89592 | 37.41039 | 16.41174 | 0.827525 | 0.901949 | 0.377849 |
| 3 | 158936600 | 158939199 | 23.94625 | 41.68266 | 35.85862 | 15.76233 | 0.603174 | 0.785495 | 0.377868 |
| 8 | 92568600 | 92570999 | 16.60063 | 28.61245 | 51.10651 | 22.41907 | -0.44847 | 0.879183 | 0.377869 |
| 4 | 131106000 | 131109399 | 22.84987 | 37.68316 | 70.77029 | 30.53988 | -0.5264 | 1.356803 | 0.377878 |
| 17 | 39242800 | 39245999 | 34.91829 | 59.58479 | 38.43599 | 16.85043 | 1.012479 | 1.026766 | 0.3779 |
| 8 | 10315600 | 10316999 | 7.197578 | 13.75305 | 28.396 | 11.27849 | -0.52792 | -0.03607 | 0.377904 |
| 2 | 117146000 | 117148199 | 31.97981 | 54.89336 | 37.90029 | 16.62347 | 0.915918 | 0.962357 | 0.377907 |
| 12 | 48530600 | 48535199 | 60.43917 | 96.78685 | 90.0073 | 37.94322 | 0.449777 | 2.022471 | 0.377928 |
| 4 | 145236200 | 145244999 | 131.1775 | 201.3371 | 172.9463 | 71.80098 | 0.512935 | 3.00039 | 0.377957 |
| 3 | 159003600 | 159005599 | 27.53854 | 46.725 | 44.72506 | 19.59297 | 0.444262 | 1.053162 | 0.377957 |
| 14 | 106400800 | 106402199 | 32.06492 | 53.76272 | 43.74511 | 19.08298 | 0.677394 | 1.171828 | 0.377968 |
| 17 | 74536600 | 74538399 | 14.01575 | 25.97446 | 24.27406 | 10.19925 | 0.538536 | 0.208535 | 0.377968 |
| 1 | 103387200 | 103388199 | 27.96453 | 47.05534 | 45.38983 | 19.83417 | 0.43287 | 1.12753 | 0.377972 |
| 14 | 101966600 | 101968599 | 14.45862 | 24.76817 | 53.6874 | 23.54404 | -0.72816 | 0.927799 | 0.377978 |
| 14 | 52356600 | 52357999 | 7.254769 | 13.5845 | 30.10882 | 12.4442 | -0.67928 | 0.103425 | 0.377994 |
| X | 54430400 | 54433399 | 31.88585 | 55.2824 | 33.4664 | 14.68346 | 1.10751 | 0.872311 | 0.377997 |

| | | | | | | | | | |
|---|---|---|---|---|---|---|---|---|---|
| 4 | 112685800 | 112691799 | 101.6958 | 158.5333 | 143.8814 | 59.71237 | 0.450794 | 2.706736 | 0.378011 |
| 8 | 92529400 | 92534799 | 43.77337 | 69.72954 | 124.9565 | 52.28532 | -0.50319 | 2.178904 | 0.378015 |
| 4 | 65593800 | 65596199 | 20.5196 | 36.16714 | 30.29296 | 13.40741 | 0.643081 | 0.582985 | 0.378017 |
| 10 | 58474200 | 58477599 | 20.85536 | 34.63822 | 67.07236 | 29.08102 | -0.56923 | 1.26403 | 0.378034 |
| 12 | 85587800 | 85588599 | 4.752506 | 9.262582 | 22.52862 | 8.156586 | -0.63414 | -0.39504 | 0.37805 |
| 2 | 98774200 | 98776599 | 28.27859 | 48.7183 | 39.12729 | 17.16387 | 0.699668 | 0.916193 | 0.378051 |
| 14 | 34763200 | 34768599 | 31.88598 | 51.6105 | 88.48759 | 37.84971 | -0.41284 | 1.666508 | 0.378052 |
| 7 | 85237000 | 85239399 | 25.37473 | 44.53592 | 30.72113 | 13.58739 | 0.920672 | 0.667083 | 0.378057 |
| 8 | 37581000 | 37585399 | 97.51715 | 151.7014 | 148.013 | 61.44625 | 0.344169 | 2.739235 | 0.378059 |
| 3 | 159277200 | 159278999 | 22.45495 | 39.55278 | 30.16168 | 13.35102 | 0.777513 | 0.602246 | 0.378063 |
| 8 | 92144600 | 92146999 | 16.31205 | 31.07673 | 19.55699 | 7.816552 | 1.163923 | 0.046168 | 0.378096 |
| 6 | 131899600 | 131901999 | 32.93657 | 55.23226 | 48.60402 | 21.20454 | 0.564826 | 1.179437 | 0.378103 |
| 5 | 38921800 | 38922999 | 6.600777 | 12.60168 | 27.23049 | 10.83556 | -0.59495 | -0.03855 | 0.378125 |
| 17 | 66065200 | 66074599 | 67.34136 | 104.9694 | 191.4925 | 79.47254 | -0.55538 | 2.696763 | 0.37813 |
| 19 | 38466600 | 38472199 | 28.38229 | 46.3058 | 81.74265 | 35.12132 | -0.44529 | 1.515512 | 0.378135 |
| 3 | 7567800 | 7569199 | 9.953977 | 17.96348 | 35.35164 | 15.37801 | -0.5629 | 0.354288 | 0.378153 |
| 4 | 69894600 | 69895999 | 19.50892 | 34.89359 | 27.70795 | 12.15833 | 0.73003 | 0.4573 | 0.378175 |
| 10 | 57100200 | 57101799 | 12.82356 | 24.53496 | 21.01225 | 8.316132 | 0.735373 | -0.01494 | 0.378195 |
| 7 | 62256400 | 62264799 | 120.5576 | 185.8455 | 164.7169 | 68.42064 | 0.472572 | 2.912403 | 0.378198 |
| 11 | 21702400 | 21703599 | 13.63895 | 25.59 | 22.77629 | 9.381764 | 0.630538 | 0.126699 | 0.37822 |
| 3 | 3239800 | 3247399 | 56.52624 | 90.81084 | 76.16477 | 32.26409 | 0.603845 | 1.913838 | 0.37824 |
| 10 | 48075600 | 48077199 | 22.17599 | 38.95858 | 31.90988 | 14.12444 | 0.673918 | 0.634706 | 0.37824 |
| 7 | 9170200 | 9171599 | 21.66356 | 37.87761 | 34.01124 | 14.99415 | 0.542385 | 0.7123 | 0.378261 |
| 3 | 87226000 | 87226599 | 13.61158 | 25.01788 | 23.90916 | 10.1595 | 0.495149 | 0.251141 | 0.37827 |
| 2 | 51824600 | 51826199 | 10.397 | 18.7827 | 35.85888 | 15.57819 | -0.51831 | 0.351903 | 0.378292 |
| 12 | 38796800 | 38799999 | 33.73345 | 54.32006 | 97.72194 | 41.53593 | -0.49057 | 1.818857 | 0.378295 |
| 11 | 120237200 | 120239199 | 19.51412 | 32.2732 | 69.37945 | 29.99323 | -0.71992 | 1.313532 | 0.378297 |
| 15 | 68200600 | 68202799 | 20.74563 | 36.40593 | 32.59008 | 14.41222 | 0.546323 | 0.643803 | 0.378317 |
| 13 | 84613600 | 84616399 | 29.88448 | 51.0908 | 41.31259 | 18.11427 | 0.688074 | 0.992131 | 0.378317 |
| 10 | 71214600 | 71218199 | 27.74677 | 45.3028 | 79.2739 | 34.06989 | -0.43185 | 1.502898 | 0.378333 |
| 14 | 42004200 | 42016599 | 138.8533 | 213.0372 | 169.1969 | 70.24073 | 0.625537 | 3.0112 | 0.378335 |
| 13 | 103845200 | 103860799 | 238.9577 | 341.214 | 507.2037 | 209.5947 | -0.37307 | 4.268457 | 0.378348 |
| 16 | 46140200 | 46144399 | 68.19324 | 108.5085 | 104.9189 | 43.83337 | 0.383691 | 2.231726 | 0.378362 |
| 9 | 115216200 | 115218599 | 19.62612 | 32.94626 | 61.21525 | 26.7242 | -0.50964 | 1.137881 | 0.378369 |
| 16 | 93288200 | 93291599 | 21.91027 | 36.5345 | 65.10648 | 28.30279 | -0.44962 | 1.220891 | 0.378388 |
| 13 | 61467000 | 61470999 | 58.02099 | 93.79022 | 64.6827 | 27.63339 | 0.896022 | 1.730244 | 0.378403 |
| 13 | 67772000 | 67773799 | 21.03279 | 37.16824 | 29.11315 | 12.87853 | 0.740638 | 0.556545 | 0.378407 |
| 17 | 17123000 | 17126399 | 37.52856 | 62.09972 | 54.39281 | 23.52317 | 0.571211 | 1.340647 | 0.378411 |
| 10 | 129992000 | 129993399 | 9.722416 | 17.98488 | 38.17803 | 16.08139 | -0.6398 | 0.216088 | 0.378419 |
| 8 | 101715000 | 101719199 | 109.869 | 169.1799 | 164.9999 | 68.53241 | 0.333456 | 2.929103 | 0.378423 |
| 7 | 139772000 | 139773399 | 11.54795 | 20.61878 | 37.79899 | 16.61061 | -0.47136 | 0.4365 | 0.378427 |
| 2 | 177418600 | 177422599 | 36.71787 | 61.32318 | 48.68751 | 21.1865 | 0.712912 | 1.233233 | 0.378441 |
| 2 | 135484200 | 135485199 | 14.83982 | 27.5277 | 24.19445 | 10.15272 | 0.627789 | 0.21 | 0.378442 |
| 5 | 119362800 | 119365399 | 22.59835 | 37.61094 | 66.14864 | 28.72155 | -0.43066 | 1.245661 | 0.378449 |
| 6 | 60515000 | 60515799 | 10.7624 | 20.76155 | 19.5825 | 7.569286 | 0.628462 | -0.11595 | 0.37845 |
| 15 | 58822600 | 58824799 | 9.096109 | 16.34148 | 35.4104 | 15.48028 | -0.70522 | 0.377628 | 0.37845 |
| 6 | 132329200 | 132335199 | 72.53081 | 115.2548 | 99.14822 | 41.35757 | 0.550088 | 2.274032 | 0.378453 |

| | | | | | | | | | |
|---|---|---|---|---|---|---|---|---|---|
| 17 | 92929400 | 92929999 | 11.04348 | 21.49667 | 16.39986 | 6.024971 | 0.999195 | -0.29058 | 0.378454 |
| X | 26119200 | 26120399 | 16.09544 | 30.81469 | 18.47094 | 7.296906 | 1.248172 | 0.005136 | 0.378455 |
| 1 | 3143200 | 3144399 | 12.09102 | 23.47593 | 16.73567 | 6.276036 | 1.069045 | -0.208 | 0.378476 |
| 7 | 17783600 | 17806799 | 397.543 | 563.0211 | 405.5224 | 167.8072 | 0.659932 | 4.395484 | 0.378518 |
| 3 | 64118600 | 64119999 | 13.42532 | 25.54997 | 21.40411 | 8.574042 | 0.751452 | 0.029499 | 0.378519 |
| 12 | 19020600 | 19035399 | 371.001 | 523.1864 | 414.3138 | 171.5664 | 0.516957 | 4.455529 | 0.378524 |
| 15 | 64062400 | 64063799 | 5.521485 | 10.74044 | 24.20031 | 8.940309 | -0.55395 | -0.31371 | 0.378531 |
| 10 | 6357200 | 6358799 | 18.33108 | 33.25591 | 25.77353 | 11.13793 | 0.779097 | 0.36292 | 0.378544 |
| 8 | 79161000 | 79165199 | 42.00002 | 66.92579 | 123.8646 | 51.8511 | -0.54914 | 2.167083 | 0.378581 |
| 16 | 91450800 | 91453399 | 25.47308 | 41.7524 | 75.18695 | 32.36174 | -0.46963 | 1.442818 | 0.378588 |
| 19 | 35668600 | 35669199 | 13.64521 | 25.18406 | 24.40505 | 10.31179 | 0.481058 | 0.229817 | 0.37862 |
| 6 | 137578800 | 137582399 | 22.85508 | 38.16673 | 66.37979 | 28.88345 | -0.41471 | 1.208621 | 0.37862 |
| 3 | 51391200 | 51392399 | 8.20937 | 14.76683 | 35.9382 | 15.69451 | -0.87059 | 0.363222 | 0.378624 |
| 3 | 111661800 | 111663199 | 26.96694 | 45.55225 | 39.67972 | 17.35931 | 0.579585 | 1.083375 | 0.378624 |
| 7 | 62895800 | 62901199 | 54.01937 | 86.81639 | 82.8474 | 35.1142 | 0.418687 | 1.902229 | 0.378627 |
| 14 | 53601000 | 53603599 | 31.77832 | 52.79151 | 51.94073 | 22.5237 | 0.404399 | 1.278768 | 0.378628 |
| 16 | 61242000 | 61244199 | 17.93504 | 32.93473 | 24.37643 | 10.37073 | 0.860107 | 0.290785 | 0.378636 |
| 1 | 85270800 | 85273599 | 51.90739 | 84.01668 | 61.76085 | 26.41751 | 0.806024 | 1.689023 | 0.378655 |
| 14 | 19753200 | 19754199 | 12.80498 | 24.21916 | 22.48472 | 9.125484 | 0.58809 | 0.068295 | 0.378666 |
| 5 | 149563600 | 149564999 | 7.804645 | 14.76518 | 29.25238 | 11.86607 | -0.49618 | 0.039093 | 0.378679 |
| 14 | 15154200 | 15155799 | 8.261958 | 15.08693 | 32.96094 | 14.12804 | -0.69812 | 0.269192 | 0.378685 |
| 19 | 48301200 | 48302999 | 18.0935 | 32.03977 | 30.1049 | 13.31178 | 0.48084 | 0.51995 | 0.378694 |
| 16 | 44027000 | 44028599 | 5.617855 | 10.73397 | 30.06376 | 11.94223 | -0.9647 | -0.0516 | 0.3787 |
| 7 | 18880800 | 18884199 | 13.61484 | 23.51461 | 50.01267 | 21.94061 | -0.69842 | 0.844948 | 0.378702 |
| 3 | 9134400 | 9135599 | 5.851668 | 11.24826 | 25.75578 | 10.05768 | -0.65225 | -0.1135 | 0.378704 |
| 5 | 130372000 | 130376799 | 67.94768 | 106.4725 | 169.6339 | 70.36107 | -0.35432 | 2.601276 | 0.378706 |
| 7 | 3706800 | 3709799 | 13.8775 | 24.18701 | 45.78164 | 20.14029 | -0.52867 | 0.737557 | 0.378721 |
| 12 | 69954600 | 69955999 | 13.8775 | 24.18701 | 45.78164 | 20.14029 | -0.52867 | 0.737557 | 0.378721 |
| 6 | 20295400 | 20296599 | 14.50433 | 25.38878 | 46.21158 | 20.39226 | -0.47222 | 0.676103 | 0.378724 |
| 10 | 53949200 | 53952399 | 26.26475 | 42.62311 | 85.65082 | 36.70687 | -0.63764 | 1.600983 | 0.378728 |
| 7 | 84722200 | 84728399 | 96.74953 | 151.7011 | 132.9493 | 55.14136 | 0.507657 | 2.601909 | 0.378729 |
| 6 | 64849600 | 64849999 | 7.509015 | 14.64573 | 15.70784 | 5.632735 | 0.544717 | -0.42205 | 0.378738 |
| 7 | 29486600 | 29487399 | 14.51326 | 26.37672 | 22.45446 | 9.681971 | 0.646702 | 0.317673 | 0.37874 |
| 17 | 24751200 | 24752999 | 8.149797 | 14.7847 | 33.97294 | 14.6819 | -0.77793 | 0.31047 | 0.378756 |
| 10 | 113250400 | 113250799 | 2.83581 | 5.545843 | 16.27232 | 5.316626 | -0.76271 | -0.81365 | 0.378763 |
| 7 | 45740000 | 45741399 | 13.1553 | 23.1483 | 42.41416 | 18.77843 | -0.48115 | 0.58119 | 0.378767 |
| 7 | 19807600 | 19831999 | 394.1085 | 552.7094 | 477.8696 | 198.0591 | 0.382296 | 4.538987 | 0.378769 |
| 5 | 105666600 | 105668199 | 12.75037 | 22.27102 | 44.70635 | 19.6942 | -0.61273 | 0.697051 | 0.378777 |
| X | 18863200 | 18865999 | 26.36894 | 46.07954 | 33.04533 | 14.5616 | 0.865204 | 0.743605 | 0.378792 |
| 17 | 41061400 | 41066399 | 64.96042 | 103.596 | 99.05304 | 41.51097 | 0.403245 | 2.155161 | 0.378845 |
| 19 | 36681600 | 36686999 | 40.6916 | 65.30104 | 110.3182 | 46.65508 | -0.40627 | 1.939197 | 0.378846 |
| 4 | 27151600 | 27153199 | 22.41953 | 39.65945 | 27.89603 | 12.3376 | 0.895553 | 0.552708 | 0.378857 |
| 8 | 21979800 | 21986399 | 92.02481 | 144.8744 | 125.3101 | 51.99163 | 0.531014 | 2.52052 | 0.378863 |
| 1 | 183800400 | 183801399 | 8.088441 | 15.24884 | 31.99345 | 13.07182 | -0.58803 | 0.069803 | 0.37888 |
| 17 | 23865600 | 23866799 | 7.679951 | 14.03105 | 34.51195 | 14.78373 | -0.86766 | 0.263046 | 0.378925 |
| 12 | 115403200 | 115411199 | 166.1005 | 250.2708 | 227.0892 | 93.81237 | 0.412268 | 3.381266 | 0.378925 |
| 7 | 60368600 | 60370199 | 21.94647 | 38.26454 | 36.02588 | 15.85222 | 0.473909 | 0.754452 | 0.378925 |

| | | | | | | | | | |
|---|---|---|---|---|---|---|---|---|---|
| 7 | 15530600 | 15554799 | 717.3757 | 953.3663 | 806.1814 | 336.8874 | 0.336199 | 5.37922 | 0.378931 |
| 7 | 26104000 | 26107199 | 63.02955 | 100.801 | 92.57107 | 38.94943 | 0.465733 | 2.065861 | 0.37894 |
| 7 | 16684200 | 16690399 | 45.30088 | 72.33141 | 122.9011 | 51.5929 | -0.42295 | 2.104258 | 0.378983 |
| 11 | 74089200 | 74091199 | 26.86733 | 45.50953 | 44.24259 | 19.37172 | 0.421889 | 1.064375 | 0.378991 |
| 5 | 119551400 | 119552799 | 22.08365 | 38.64691 | 34.33258 | 15.14734 | 0.557616 | 0.699589 | 0.378993 |
| 1 | 25054200 | 25057399 | 38.11501 | 61.18963 | 105.9317 | 44.8261 | -0.44077 | 1.923001 | 0.379003 |
| X | 146770600 | 146772999 | 18.02061 | 32.22903 | 28.57123 | 12.53798 | 0.572241 | 0.451161 | 0.379006 |
| 16 | 80151800 | 80152399 | 11.06368 | 21.13313 | 21.14892 | 8.406134 | 0.507051 | -0.01769 | 0.379012 |
| 2 | 160472400 | 160475799 | 44.2696 | 71.58574 | 70.66948 | 30.20434 | 0.380317 | 1.698807 | 0.379025 |
| 1 | 172705200 | 172707399 | 16.53156 | 29.75248 | 27.5456 | 12.02332 | 0.514973 | 0.398691 | 0.37905 |
| 7 | 42252200 | 42262399 | 249.0601 | 364.9413 | 311.1372 | 128.1609 | 0.465126 | 3.883212 | 0.379053 |
| 13 | 61525400 | 61529399 | 62.70553 | 100.6162 | 82.91918 | 35.06113 | 0.627117 | 1.96173 | 0.379059 |
| 6 | 76409000 | 76411799 | 24.70562 | 43.09806 | 34.66961 | 15.26052 | 0.699856 | 0.758521 | 0.379066 |
| 1 | 166439400 | 166443799 | 45.27336 | 74.0209 | 58.62569 | 25.20849 | 0.710071 | 1.506594 | 0.379084 |
| 1 | 81474000 | 81475399 | 15.43535 | 28.43935 | 21.74207 | 9.208003 | 0.818522 | 0.247842 | 0.379084 |
| 17 | 88919200 | 88930599 | 121.6995 | 183.3929 | 283.2817 | 117.0299 | -0.35456 | 3.37323 | 0.379092 |
| 18 | 37658600 | 37664799 | 148.4802 | 227.6113 | 148.1591 | 61.49978 | 0.911324 | 3.028932 | 0.379102 |
| 5 | 145187800 | 145189399 | 14.25422 | 25.99047 | 24.8439 | 10.66869 | 0.48431 | 0.3028 | 0.379115 |
| X | 5823600 | 5825599 | 29.5125 | 51.53541 | 30.58213 | 13.46986 | 1.136883 | 0.76552 | 0.379123 |
| 10 | 69365800 | 69366399 | 4.938158 | 9.629118 | 22.10164 | 7.950977 | -0.54452 | -0.42125 | 0.379127 |
| 12 | 88266200 | 88276199 | 134.1581 | 205.2719 | 182.0038 | 75.51865 | 0.463797 | 3.062739 | 0.37914 |
| X | 120418800 | 120420999 | 29.95292 | 51.99872 | 34.62007 | 15.19228 | 0.971186 | 0.852102 | 0.379141 |
| 16 | 15461800 | 15465399 | 23.14944 | 38.32008 | 79.73291 | 34.49064 | -0.67323 | 1.319967 | 0.379149 |
| 4 | 87019000 | 87021799 | 16.36524 | 28.29281 | 51.13804 | 22.43498 | -0.46472 | 0.848352 | 0.37915 |
| 7 | 27835600 | 27837799 | 16.36524 | 28.29281 | 51.13804 | 22.43498 | -0.46472 | 0.848352 | 0.37915 |
| 17 | 20910000 | 20911199 | 26.59968 | 46.03238 | 34.36914 | 15.07869 | 0.805579 | 0.862503 | 0.379155 |
| 17 | 40804400 | 40807799 | 28.75825 | 48.96277 | 43.00273 | 18.84908 | 0.568395 | 1.02475 | 0.379166 |
| 19 | 37865800 | 37867599 | 11.96126 | 21.01842 | 40.9772 | 18.14159 | -0.57041 | 0.591065 | 0.379207 |
| 16 | 48845000 | 48851799 | 88.47483 | 138.8994 | 132.9953 | 55.16031 | 0.381493 | 2.572852 | 0.379224 |
| 18 | 88122000 | 88122399 | 3.522528 | 6.890629 | 18.00187 | 5.999095 | -0.62626 | -0.72635 | 0.379258 |
| 16 | 68069000 | 68072199 | 32.24601 | 54.88497 | 41.83168 | 18.33535 | 0.772207 | 1.041055 | 0.379278 |
| 2 | 5261000 | 5263799 | 36.18595 | 60.08828 | 52.65659 | 22.82747 | 0.570844 | 1.296025 | 0.379279 |
| 12 | 6382600 | 6384999 | 28.33896 | 48.85075 | 37.37468 | 16.39535 | 0.769385 | 0.906889 | 0.379279 |
| 15 | 21917400 | 21919199 | 40.28232 | 65.48862 | 64.59014 | 27.71586 | 0.388785 | 1.580414 | 0.379286 |
| 10 | 82444000 | 82448999 | 44.44575 | 72.60393 | 61.27435 | 26.33895 | 0.617859 | 1.520511 | 0.37929 |
| 1 | 89174400 | 89176399 | 27.0955 | 46.3365 | 42.5155 | 18.64216 | 0.506539 | 0.980089 | 0.379299 |
| 5 | 145842000 | 145847199 | 93.40556 | 146.6506 | 131.0982 | 54.37396 | 0.480518 | 2.573937 | 0.379299 |
| 15 | 39522600 | 39523799 | 8.825877 | 16.17234 | 32.95634 | 14.06218 | -0.59421 | 0.252209 | 0.3793 |
| 7 | 78990800 | 78993199 | 19.23179 | 32.50035 | 59.13413 | 25.87377 | -0.47918 | 1.07416 | 0.379308 |
| X | 126055800 | 126059399 | 33.19791 | 57.10431 | 35.61008 | 15.62024 | 1.062709 | 0.946955 | 0.379314 |
| 14 | 58623800 | 58626199 | 15.08485 | 26.20512 | 48.44931 | 21.26992 | -0.49579 | 0.79113 | 0.37933 |
| 14 | 106328400 | 106333199 | 47.45511 | 75.78997 | 123.4338 | 51.83484 | -0.36152 | 2.10015 | 0.379339 |
| 4 | 121334400 | 121341399 | 58.49505 | 94.14674 | 69.93649 | 29.7048 | 0.781821 | 1.85748 | 0.37935 |
| 3 | 96362000 | 96363199 | 7.473292 | 13.79467 | 31.51462 | 13.31352 | -0.7468 | 0.197707 | 0.379365 |
| 3 | 82455200 | 82456399 | 8.874676 | 16.11228 | 34.93132 | 15.08137 | -0.69449 | 0.30974 | 0.379379 |
| 4 | 8728200 | 8731199 | 14.77671 | 24.9722 | 61.0082 | 26.69396 | -0.90242 | 1.060384 | 0.379383 |
| 3 | 37862400 | 37865399 | 35.64014 | 59.02213 | 54.82157 | 23.7217 | 0.486921 | 1.328045 | 0.379386 |

| | | | | | | | | | |
|---|---|---|---|---|---|---|---|---|---|
| 5 | 31931200 | 31932799 | 20.57563 | 36.03838 | 33.90241 | 14.96827 | 0.475337 | 0.677245 | 0.379388 |
| 1 | 68607200 | 68609399 | 17.89506 | 30.78275 | 54.90665 | 24.08465 | -0.44723 | 0.90666 | 0.379397 |
| 17 | 51652600 | 51653599 | 7.517018 | 14.16788 | 29.15065 | 11.91554 | -0.55986 | 0.061636 | 0.379405 |
| 10 | 10326600 | 10328399 | 16.75017 | 30.77116 | 23.20517 | 9.867088 | 0.833253 | 0.274554 | 0.379407 |
| 9 | 113231800 | 113232399 | 6.54497 | 12.65474 | 30.48647 | 11.66838 | -0.69908 | -0.16431 | 0.37942 |
| 9 | 120662800 | 120668399 | 62.92021 | 98.57261 | 165.9233 | 68.82353 | -0.43374 | 2.602772 | 0.379469 |
| 19 | 12980000 | 12985199 | 69.60728 | 110.9007 | 100.8649 | 42.21452 | 0.473819 | 2.190684 | 0.379472 |
| 18 | 84098400 | 84098999 | 2.741853 | 5.357756 | 19.286 | 6.697734 | -1.13433 | -0.5867 | 0.379478 |
| 9 | 20360800 | 20362999 | 25.46792 | 44.03042 | 38.63714 | 16.95067 | 0.573303 | 0.871091 | 0.37949 |
| 2 | 98119600 | 98120799 | 12.93951 | 24.35671 | 22.94165 | 9.39801 | 0.555997 | 0.101699 | 0.379493 |
| 1 | 132331800 | 132333399 | 14.84629 | 27.03344 | 25.69617 | 11.05336 | 0.490634 | 0.31598 | 0.379533 |
| 1 | 86353000 | 86355999 | 16.94154 | 28.98755 | 53.21219 | 23.33344 | -0.48978 | 0.948724 | 0.379541 |
| 19 | 51557800 | 51559399 | 19.40532 | 34.16601 | 31.01803 | 13.7313 | 0.52691 | 0.589548 | 0.379545 |
| 4 | 45331200 | 45332999 | 16.55321 | 28.68898 | 50.60485 | 22.20275 | -0.42886 | 0.827587 | 0.379563 |
| 11 | 54361600 | 54363799 | 21.55885 | 37.95432 | 31.40846 | 13.90438 | 0.660009 | 0.605588 | 0.379568 |
| 12 | 36114400 | 36116199 | 21.55885 | 37.95432 | 31.40846 | 13.90438 | 0.660009 | 0.605588 | 0.379568 |
| X | 71838200 | 71840399 | 21.45588 | 38.34088 | 27.18117 | 11.93637 | 0.892259 | 0.476938 | 0.37958 |
| 12 | 21816400 | 21819199 | 99.877 | 154.826 | 145.2677 | 60.3383 | 0.396605 | 2.801387 | 0.379581 |
| 8 | 4686400 | 4688199 | 21.51438 | 37.80612 | 32.11697 | 14.21924 | 0.621683 | 0.628955 | 0.379582 |
| 9 | 38587200 | 38588799 | 17.14235 | 31.14608 | 26.55738 | 11.45513 | 0.644103 | 0.34863 | 0.379583 |
| 3 | 121288200 | 121290399 | 27.57262 | 46.81655 | 44.72002 | 19.59504 | 0.44729 | 1.047603 | 0.379598 |
| 2 | 36664000 | 36674399 | 153.7149 | 233.6262 | 197.1151 | 81.64679 | 0.527655 | 3.205207 | 0.379608 |
| 10 | 109545200 | 109546999 | 14.36351 | 26.6654 | 24.25942 | 10.16894 | 0.57949 | 0.202002 | 0.379612 |
| 16 | 33404000 | 33405799 | 11.69078 | 20.43282 | 44.20382 | 19.48127 | -0.71986 | 0.682388 | 0.379624 |
| 3 | 88559200 | 88561199 | 11.68928 | 20.44184 | 43.68616 | 19.2624 | -0.70221 | 0.673091 | 0.379628 |
| 9 | 62138000 | 62145399 | 58.0413 | 91.36889 | 168.0234 | 69.71297 | -0.55582 | 2.506985 | 0.379639 |
| 3 | 75097200 | 75098399 | 14.43904 | 27.63487 | 19.71544 | 7.795538 | 0.997796 | -0.00572 | 0.379647 |
| X | 73596800 | 73600399 | 78.86102 | 124.7815 | 117.635 | 48.89522 | 0.412669 | 2.399364 | 0.379647 |
| 7 | 48516800 | 48518599 | 8.173906 | 15.13418 | 32.58569 | 13.70877 | -0.65836 | 0.185706 | 0.37965 |
| 1 | 27288200 | 27288599 | 9.388508 | 18.18016 | 17.23478 | 6.550078 | 0.645554 | -0.19067 | 0.379651 |
| 17 | 63805600 | 63808599 | 24.44654 | 40.24479 | 72.64105 | 31.32364 | -0.47046 | 1.383199 | 0.379654 |
| 18 | 79539000 | 79539599 | 5.509032 | 10.72952 | 24.37992 | 8.908253 | -0.55102 | -0.35076 | 0.379663 |
| 7 | 38310800 | 38312799 | 20.07343 | 35.25671 | 32.72605 | 14.48289 | 0.494412 | 0.628267 | 0.379675 |
| 3 | 11811800 | 11813799 | 22.50023 | 39.40141 | 33.02732 | 14.57943 | 0.640771 | 0.687109 | 0.379683 |
| 14 | 118720800 | 118724799 | 30.03212 | 48.81326 | 86.9116 | 37.29022 | -0.46157 | 1.575974 | 0.379705 |
| 3 | 51470400 | 51472999 | 16.2775 | 27.91526 | 53.22698 | 23.34423 | -0.54387 | 0.923759 | 0.379716 |
| 14 | 23040200 | 23041399 | 20.8691 | 36.37338 | 34.21234 | 15.05091 | 0.47538 | 0.752607 | 0.379724 |
| 6 | 37436400 | 37436999 | 5.807835 | 11.28748 | 24.68882 | 9.185996 | -0.52087 | -0.28215 | 0.379731 |
| 7 | 73767000 | 73768599 | 8.769958 | 16.22001 | 32.56339 | 13.71982 | -0.55973 | 0.197121 | 0.379746 |
| 17 | 49522400 | 49522999 | 5.081567 | 9.880499 | 22.87135 | 8.478451 | -0.59571 | -0.30779 | 0.379752 |
| 15 | 84240200 | 84243599 | 32.55447 | 52.58661 | 91.60646 | 39.09983 | -0.43885 | 1.720351 | 0.379753 |
| 17 | 37520000 | 37523799 | 20.01491 | 33.46376 | 68.3078 | 29.74479 | -0.64485 | 1.190203 | 0.379763 |
| 7 | 59375200 | 59377199 | 26.74134 | 45.2749 | 39.6946 | 17.3779 | 0.570445 | 1.064381 | 0.37978 |
| 13 | 76102000 | 76102799 | 10.65568 | 19.05211 | 36.86488 | 16.18114 | -0.54718 | 0.422565 | 0.379794 |
| X | 25964800 | 25968199 | 33.9833 | 58.84937 | 31.07054 | 13.63178 | 1.303466 | 0.888258 | 0.379798 |
| 7 | 31129600 | 31163999 | 826.3147 | 1088.837 | 885.3271 | 370.3456 | 0.380534 | 5.51693 | 0.3798 |
| X | 80863200 | 80868999 | 66.4846 | 106.6895 | 79.99309 | 33.82844 | 0.763428 | 1.965872 | 0.379803 |

| | | | | | | | | | |
|---|---|---|---|---|---|---|---|---|---|
| 7 | 38422400 | 38426199 | 74.49209 | 118.5527 | 102.4191 | 42.7997 | 0.546226 | 2.231338 | 0.379806 |
| 4 | 90968200 | 90969599 | 8.910905 | 16.46171 | 34.72735 | 14.65451 | -0.63271 | 0.208084 | 0.379819 |
| 16 | 8939400 | 8940799 | 13.86908 | 25.72564 | 23.14638 | 9.713608 | 0.593865 | 0.19851 | 0.379821 |
| 6 | 139072000 | 139073599 | 9.401746 | 17.48251 | 38.39166 | 16.05583 | -0.68082 | 0.184515 | 0.379825 |
| 10 | 3167800 | 3177199 | 165.4941 | 251.408 | 190.3247 | 78.8228 | 0.683374 | 3.21799 | 0.379837 |
| 6 | 95093600 | 95095599 | 10.45536 | 18.34078 | 42.11068 | 18.61643 | -0.80483 | 0.612931 | 0.379842 |
| 11 | 71848000 | 71851799 | 50.05285 | 80.60612 | 78.67553 | 33.44484 | 0.389831 | 1.831545 | 0.379851 |
| 6 | 15446000 | 15447399 | 14.43338 | 26.53105 | 25.30132 | 10.74957 | 0.498258 | 0.26003 | 0.379868 |
| 13 | 37296400 | 37297999 | 14.43338 | 26.53105 | 25.30132 | 10.74957 | 0.498258 | 0.26003 | 0.379868 |
| X | 82901200 | 82906599 | 52.0787 | 84.66165 | 60.92246 | 26.1415 | 0.842438 | 1.605776 | 0.37987 |
| 9 | 38441400 | 38445799 | 67.89657 | 108.4116 | 95.17923 | 39.95643 | 0.528066 | 2.120716 | 0.37991 |
| 14 | 115792600 | 115794199 | 14.02246 | 24.5852 | 44.59667 | 19.70666 | -0.46724 | 0.644919 | 0.379912 |
| 6 | 66670800 | 66672399 | 19.72127 | 34.62417 | 32.77938 | 14.50154 | 0.466087 | 0.632528 | 0.379927 |
| 6 | 66447200 | 66447999 | 7.195563 | 13.66126 | 28.22951 | 11.36556 | -0.54675 | 0.005825 | 0.379941 |
| 3 | 95565200 | 95566999 | 10.71841 | 18.8884 | 39.96194 | 17.68747 | -0.68611 | 0.548009 | 0.379979 |
| 3 | 7039800 | 7040999 | 11.44759 | 21.89858 | 21.07928 | 8.346785 | 0.567296 | -0.02747 | 0.379979 |
| 5 | 37762000 | 37762799 | 11.44759 | 21.89858 | 21.07928 | 8.346785 | 0.567296 | -0.02747 | 0.379979 |
| 14 | 45722400 | 45723399 | 11.44759 | 21.89858 | 21.07928 | 8.346785 | 0.567296 | -0.02747 | 0.379979 |
| 10 | 16205800 | 16207799 | 22.78953 | 39.97327 | 32.4406 | 14.34021 | 0.687201 | 0.663249 | 0.37998 |
| 8 | 89834200 | 89836599 | 13.17951 | 23.12724 | 42.48932 | 18.78916 | -0.48553 | 0.621142 | 0.379985 |
| 14 | 5744400 | 5748199 | 51.59562 | 82.13261 | 132.1502 | 55.24184 | -0.34919 | 2.210706 | 0.379986 |
| 19 | 42677000 | 42678999 | 9.616206 | 17.43033 | 35.926 | 15.54223 | -0.62422 | 0.325992 | 0.380007 |
| 15 | 23699600 | 23705599 | 56.92261 | 91.78483 | 72.55799 | 30.89797 | 0.695468 | 1.801332 | 0.380047 |
| 8 | 38279000 | 38281199 | 32.06897 | 55.36816 | 35.084 | 15.39122 | 1.040763 | 0.909836 | 0.380062 |
| 10 | 39376400 | 39380199 | 29.44435 | 47.79901 | 83.847 | 35.94267 | -0.44168 | 1.6006 | 0.380068 |
| 4 | 141171800 | 141172999 | 7.656901 | 14.0278 | 32.49642 | 13.86935 | -0.77738 | 0.245408 | 0.380082 |
| 17 | 51299600 | 51301199 | 19.67201 | 35.52875 | 26.24278 | 11.40456 | 0.843103 | 0.399732 | 0.380084 |
| 19 | 23064800 | 23067199 | 21.53924 | 37.40654 | 32.83297 | 14.41146 | 0.574212 | 0.803264 | 0.380091 |
| 1 | 75592200 | 75593199 | 10.76828 | 19.29245 | 36.31797 | 15.9139 | -0.5063 | 0.40749 | 0.380142 |
| X | 120900800 | 120904599 | 36.90722 | 62.84643 | 37.25352 | 16.32926 | 1.133495 | 1.05125 | 0.380143 |
| X | 23789800 | 23792399 | 36.72475 | 61.3161 | 49.23604 | 21.4211 | 0.696719 | 1.238741 | 0.380171 |
| 7 | 142130400 | 142136999 | 86.3746 | 132.5547 | 215.8691 | 89.61938 | -0.40919 | 2.992212 | 0.380172 |
| 8 | 89831200 | 89833199 | 12.85503 | 22.7681 | 42.403 | 18.749 | -0.50077 | 0.508718 | 0.380185 |
| 14 | 115353200 | 115353599 | 7.675544 | 13.97042 | 36.57169 | 15.73984 | -0.96052 | 0.29471 | 0.380196 |
| 6 | 88910400 | 88911399 | 5.823739 | 11.26285 | 24.85324 | 9.502019 | -0.57013 | -0.18615 | 0.380219 |
| 2 | 166066200 | 166069199 | 30.50816 | 49.64409 | 85.91218 | 36.8776 | -0.41944 | 1.560088 | 0.380224 |
| 4 | 74151000 | 74154199 | 24.47639 | 40.21724 | 73.60111 | 31.71205 | -0.49142 | 1.408041 | 0.380238 |
| 7 | 17328000 | 17339399 | 172.5332 | 260.2263 | 203.2807 | 84.00084 | 0.629445 | 3.362442 | 0.380238 |
| 5 | 100130800 | 100135199 | 36.59222 | 58.97109 | 100.9364 | 42.95248 | -0.41754 | 1.797716 | 0.380247 |
| 2 | 165520800 | 165522599 | 23.01137 | 37.9457 | 70.77152 | 30.53912 | -0.51649 | 1.35864 | 0.380251 |
| 7 | 9887800 | 9889999 | 21.21789 | 38.32647 | 25.00961 | 10.8666 | 1.020947 | 0.407477 | 0.38026 |
| 17 | 32220200 | 32223599 | 33.25731 | 55.04974 | 54.1806 | 23.43663 | 0.403544 | 1.325459 | 0.380264 |
| 5 | 39069200 | 39069999 | 9.209193 | 17.04581 | 34.22787 | 14.40564 | -0.55917 | 0.199107 | 0.380272 |
| 8 | 84116200 | 84117199 | 7.461542 | 13.80365 | 32.0484 | 13.49811 | -0.76714 | 0.184297 | 0.380281 |
| 3 | 72760400 | 72762199 | 12.41804 | 21.96854 | 40.35818 | 17.84902 | -0.4815 | 0.517124 | 0.380284 |
| 13 | 67675600 | 67677599 | 15.75038 | 27.25252 | 49.5195 | 21.72546 | -0.47195 | 0.836958 | 0.380291 |
| 4 | 105702000 | 105706399 | 28.82307 | 46.73596 | 93.03579 | 39.84738 | -0.62534 | 1.627216 | 0.380297 |

| | | | | | | | | | |
|---|---|---|---|---|---|---|---|---|---|
| 5 | 116767600 | 116772999 | 32.07142 | 51.78604 | 93.25641 | 39.78398 | -0.4873 | 1.731036 | 0.380317 |
| 2 | 174232600 | 174235399 | 29.19223 | 47.74957 | 82.67671 | 35.55589 | -0.41485 | 1.482269 | 0.380337 |
| 5 | 31728400 | 31729599 | 6.623785 | 12.56015 | 28.18782 | 11.37851 | -0.66803 | 0.008757 | 0.38034 |
| 7 | 9215800 | 9217399 | 18.6684 | 33.91692 | 25.86853 | 11.15877 | 0.803926 | 0.35875 | 0.38035 |
| Y | 67874600 | 67876599 | 15.91717 | 28.91632 | 15.43168 | 6.657293 | 1.314296 | 0.310451 | 0.380354 |
| X | 148657800 | 148661599 | 76.34575 | 121.1718 | 110.3473 | 45.97393 | 0.466413 | 2.315504 | 0.38036 |
| 2 | 133103400 | 133103999 | 7.302215 | 13.54905 | 31.03904 | 13.02031 | -0.74344 | 0.164663 | 0.380385 |
| 8 | 129760000 | 129762999 | 25.6478 | 44.57063 | 36.36138 | 15.9667 | 0.67925 | 0.817759 | 0.380389 |
| 13 | 12409800 | 12410799 | 6.800233 | 12.80159 | 29.66388 | 12.1499 | -0.73232 | 0.06327 | 0.380406 |
| 14 | 11158600 | 11160599 | 12.83655 | 22.6383 | 41.36761 | 18.30685 | -0.47593 | 0.552978 | 0.380413 |
| 7 | 56515000 | 56516999 | 21.35975 | 37.42576 | 33.11836 | 14.62672 | 0.562943 | 0.672044 | 0.380424 |
| 6 | 87933800 | 87944599 | 138.8234 | 207.9596 | 315.648 | 130.2029 | -0.33693 | 3.492645 | 0.380433 |
| 12 | 32381800 | 32384399 | 33.47459 | 55.26547 | 54.13135 | 23.37331 | 0.410242 | 1.356554 | 0.380435 |
| 3 | 61781400 | 61782399 | 7.174644 | 13.79892 | 30.58854 | 11.92603 | -0.60496 | -0.09313 | 0.380445 |
| 7 | 30188400 | 30196199 | 57.6634 | 90.92163 | 151.9583 | 63.06091 | -0.41626 | 2.471895 | 0.380488 |
| 2 | 97568600 | 97572799 | 46.75345 | 75.833 | 67.93224 | 29.1021 | 0.524231 | 1.640407 | 0.380503 |
| 2 | 76773000 | 76776399 | 53.40681 | 85.82633 | 82.5636 | 34.99061 | 0.407008 | 1.903948 | 0.380513 |
| 1 | 136602000 | 136604999 | 17.04518 | 29.39261 | 51.66303 | 22.66353 | -0.42534 | 0.876363 | 0.380522 |
| 15 | 61856800 | 61860199 | 26.26657 | 42.62425 | 86.8878 | 37.23583 | -0.65832 | 1.601931 | 0.380535 |
| 11 | 80806400 | 80808199 | 31.90798 | 51.6518 | 88.34065 | 37.78993 | -0.40912 | 1.665477 | 0.380536 |
| 6 | 66466400 | 66466999 | 20.37518 | 35.57265 | 33.17298 | 14.6092 | 0.487925 | 0.724818 | 0.38054 |
| 13 | 96678800 | 96680199 | 11.23804 | 19.646 | 44.20675 | 19.4863 | -0.77624 | 0.67632 | 0.380567 |
| 17 | 68570400 | 68571399 | 11.44941 | 21.66222 | 21.52088 | 8.728368 | 0.491842 | 0.050549 | 0.380577 |
| 4 | 147415400 | 147418199 | 31.11929 | 54.488 | 26.66624 | 11.7702 | 1.413359 | 0.729716 | 0.380593 |
| 9 | 91872800 | 91873399 | 5.812708 | 11.2892 | 25.23504 | 9.441453 | -0.55986 | -0.26277 | 0.380594 |
| 12 | 34111600 | 34112399 | 5.812708 | 11.2892 | 25.23504 | 9.441453 | -0.55986 | -0.26277 | 0.380594 |
| 3 | 10302600 | 10303399 | 7.58402 | 14.0996 | 30.53441 | 12.77422 | -0.66002 | 0.1554 | 0.380617 |
| 16 | 87323600 | 87324999 | 9.326616 | 17.02984 | 33.4478 | 14.33801 | -0.54577 | 0.279827 | 0.380619 |
| 19 | 9470200 | 9472199 | 30.25488 | 50.86677 | 48.02494 | 20.98032 | 0.463637 | 1.14435 | 0.380624 |
| 2 | 101911600 | 101912599 | 15.16315 | 28.3142 | 23.14118 | 9.609476 | 0.743519 | 0.17079 | 0.380645 |
| 8 | 62202000 | 62203599 | 17.60389 | 31.22444 | 29.622 | 13.09004 | 0.468085 | 0.500294 | 0.380656 |
| 10 | 129836600 | 129842199 | 86.1573 | 136.2682 | 115.8584 | 48.14635 | 0.560979 | 2.414388 | 0.380676 |
| 17 | 94917200 | 94917999 | 15.45498 | 27.87168 | 26.76439 | 11.65258 | 0.46509 | 0.378831 | 0.380679 |
| 11 | 52226000 | 52227199 | 8.56043 | 16.04735 | 32.94728 | 13.5915 | -0.56827 | 0.112716 | 0.380687 |
| 1 | 120741600 | 120742599 | 9.655572 | 17.43632 | 34.86621 | 15.1545 | -0.58487 | 0.347905 | 0.380698 |
| 17 | 6858600 | 6863799 | 22.50463 | 37.18022 | 69.75501 | 30.13387 | -0.52445 | 1.333159 | 0.380718 |
| 7 | 78876200 | 78877399 | 8.959458 | 16.60203 | 32.06332 | 13.47422 | -0.50108 | 0.189565 | 0.380727 |
| 1 | 152189200 | 152191399 | 27.93942 | 47.5193 | 43.58577 | 19.10304 | 0.505817 | 1.03152 | 0.380738 |
| 6 | 133141200 | 133145199 | 43.68666 | 71.96741 | 53.49132 | 23.07268 | 0.806163 | 1.407872 | 0.380741 |
| 12 | 115221600 | 115223799 | 44.52068 | 72.14128 | 69.12816 | 29.59319 | 0.426016 | 1.657256 | 0.380746 |
| 7 | 33500400 | 33502799 | 46.22554 | 75.1283 | 65.21848 | 27.98115 | 0.572132 | 1.597821 | 0.380748 |
| 4 | 142181800 | 142193999 | 114.8788 | 173.0964 | 283.1288 | 116.9634 | -0.43721 | 3.373611 | 0.380755 |
| 11 | 57779400 | 57780399 | 16.79275 | 30.14696 | 28.12397 | 12.30238 | 0.5021 | 0.418002 | 0.380759 |
| 12 | 114011800 | 114012799 | 16.79275 | 30.14696 | 28.12397 | 12.30238 | 0.5021 | 0.418002 | 0.380759 |
| 5 | 75348400 | 75350999 | 16.13213 | 27.74596 | 51.09155 | 22.41106 | -0.49284 | 0.896228 | 0.380763 |
| 7 | 103059000 | 103059799 | 4.93954 | 9.627478 | 22.00908 | 7.964506 | -0.54634 | -0.40005 | 0.380764 |
| 13 | 50791400 | 50796199 | 50.7101 | 80.35992 | 139.4081 | 57.99168 | -0.4641 | 2.347047 | 0.380767 |

| | | | | | | | | | |
|---|---|---|---|---|---|---|---|---|---|
| 3 | 91185200 | 91186399 | 25.02614 | 42.68372 | 41.07041 | 18.00465 | 0.437643 | 0.993206 | 0.380818 |
| 1 | 98305000 | 98306399 | 22.58064 | 39.46239 | 34.78035 | 15.32709 | 0.568918 | 0.722879 | 0.380819 |
| 7 | 117370000 | 117373399 | 38.08949 | 61.38346 | 106.7249 | 45.4151 | -0.44016 | 1.803474 | 0.380824 |
| 1 | 24356600 | 24359199 | 13.95647 | 24.52351 | 46.65114 | 20.65294 | -0.53581 | 0.609364 | 0.380863 |
| 16 | 3441200 | 3449599 | 141.3965 | 215.2825 | 197.2487 | 81.74524 | 0.410863 | 3.165596 | 0.380878 |
| 12 | 18459200 | 18488999 | 1208.381 | 1538.01 | 1232.639 | 517.4406 | 0.351273 | 6.080629 | 0.380895 |
| 7 | 23391600 | 23394399 | 63.32387 | 100.9739 | 97.83508 | 40.99371 | 0.38398 | 2.159069 | 0.380899 |
| 13 | 104455400 | 104455999 | 6.411706 | 12.40461 | 26.68642 | 10.18499 | -0.53206 | -0.18106 | 0.380921 |
| 14 | 93097000 | 93097599 | 6.411706 | 12.40461 | 26.68642 | 10.18499 | -0.53206 | -0.18106 | 0.380921 |
| 7 | 35459200 | 35464999 | 57.27004 | 90.56358 | 145.9927 | 60.66364 | -0.36091 | 2.403626 | 0.380932 |
| X | 127387800 | 127393199 | 56.48302 | 91.16781 | 71.18675 | 30.35346 | 0.714911 | 1.773905 | 0.380943 |
| 3 | 49781600 | 49783199 | 17.99243 | 32.39334 | 26.16803 | 11.41688 | 0.710649 | 0.402624 | 0.380947 |
| 19 | 29164200 | 29166199 | 18.19014 | 32.2951 | 29.46314 | 13.00937 | 0.524858 | 0.497045 | 0.380958 |
| 5 | 104658000 | 104659399 | 15.60625 | 28.49187 | 26.19432 | 11.23074 | 0.541612 | 0.309022 | 0.380961 |
| 7 | 19209200 | 19217199 | 50.29727 | 79.82677 | 138.3756 | 57.64967 | -0.46087 | 2.300127 | 0.380972 |
| 10 | 91115000 | 91121999 | 70.10247 | 109.7292 | 175.1562 | 72.65976 | -0.35831 | 2.622175 | 0.380974 |
| 1 | 134433800 | 134434399 | 7.408784 | 13.86984 | 29.57554 | 12.22835 | -0.62396 | 0.106988 | 0.380994 |
| 17 | 37177200 | 37179399 | 13.8074 | 24.11491 | 45.7294 | 20.14361 | -0.53126 | 0.704278 | 0.38101 |
| 5 | 127588800 | 127591199 | 15.58749 | 26.97516 | 49.51913 | 21.72541 | -0.48659 | 0.834479 | 0.381029 |
| 5 | 124450200 | 124450999 | 20.68082 | 36.09015 | 31.25641 | 13.76129 | 0.594106 | 0.729821 | 0.38103 |
| Y | 4340000 | 4341599 | 11.21043 | 21.4392 | 12.97622 | 5.142131 | 1.229078 | -0.05336 | 0.381032 |
| 15 | 97873600 | 97878999 | 66.4945 | 103.7335 | 174.8516 | 72.55841 | -0.4405 | 2.680475 | 0.381048 |
| X | 102014600 | 102016399 | 19.53124 | 36.337 | 21.17821 | 8.849227 | 1.221093 | 0.226934 | 0.381051 |
| 5 | 138017800 | 138019999 | 8.617921 | 15.98649 | 31.54363 | 13.23627 | -0.53045 | 0.177558 | 0.381055 |
| 14 | 86520000 | 86521999 | 17.28752 | 31.62 | 25.50195 | 10.90828 | 0.73155 | 0.309995 | 0.381057 |
| 7 | 31030800 | 31035399 | 72.58922 | 115.7406 | 96.48108 | 40.42276 | 0.600523 | 2.168362 | 0.381057 |
| 19 | 20120800 | 20125399 | 38.59687 | 62.13047 | 103.0213 | 43.76694 | -0.3739 | 1.838972 | 0.381071 |
| 1 | 130332400 | 130336199 | 47.5541 | 77.15877 | 66.93525 | 28.68242 | 0.570914 | 1.634178 | 0.381076 |
| 5 | 73489600 | 73491199 | 8.493974 | 15.63978 | 33.55712 | 14.22442 | -0.6614 | 0.221812 | 0.381083 |
| 10 | 71680600 | 71683799 | 38.00894 | 62.65323 | 56.65183 | 24.44237 | 0.524548 | 1.387443 | 0.381105 |
| 7 | 79735200 | 79736799 | 11.12771 | 19.71898 | 38.31835 | 16.94117 | -0.56109 | 0.484888 | 0.381125 |
| 13 | 21028800 | 21030399 | 9.44729 | 17.25107 | 33.42378 | 14.3268 | -0.5265 | 0.278366 | 0.381151 |
| 2 | 83729000 | 83730199 | 9.917464 | 18.12728 | 33.95402 | 14.53635 | -0.47693 | 0.278088 | 0.381153 |
| 14 | 88284600 | 88285399 | 9.917464 | 18.12728 | 33.95402 | 14.53635 | -0.47693 | 0.278088 | 0.381153 |
| 4 | 8722400 | 8723199 | 4.850268 | 9.410117 | 24.42549 | 9.200566 | -0.78173 | -0.25243 | 0.381153 |
| 13 | 25231600 | 25239399 | 89.81547 | 137.8726 | 216.9631 | 90.07692 | -0.3595 | 2.987827 | 0.381153 |
| 5 | 42993400 | 42993799 | 10.33743 | 19.91278 | 19.72782 | 7.657351 | 0.55358 | -0.10381 | 0.381194 |
| 1 | 133108800 | 133111199 | 13.38395 | 22.94403 | 52.5758 | 23.05792 | -0.80741 | 0.918661 | 0.381199 |
| 17 | 31605000 | 31607599 | 13.24235 | 23.04743 | 46.32288 | 20.36032 | -0.61512 | 0.753926 | 0.3812 |
| 5 | 150998800 | 150999399 | 12.85942 | 24.43187 | 22.02605 | 8.854619 | 0.642191 | 0.037822 | 0.381209 |
| 1 | 27794000 | 27794599 | 11.52395 | 21.76741 | 20.0994 | 8.180267 | 0.592684 | 0.060084 | 0.38121 |
| 5 | 146585600 | 146587799 | 14.92964 | 25.97178 | 47.37099 | 20.81481 | -0.47604 | 0.769383 | 0.381213 |
| 1 | 63287400 | 63289199 | 23.87946 | 41.03985 | 39.94004 | 17.5191 | 0.422966 | 0.918812 | 0.38122 |
| 10 | 56196600 | 56198599 | 20.71705 | 37.13953 | 26.52585 | 11.61713 | 0.883912 | 0.449664 | 0.381224 |
| 3 | 15969600 | 15971399 | 25.66804 | 43.67967 | 42.10547 | 18.45504 | 0.434529 | 1.01692 | 0.381233 |
| 8 | 45482200 | 45499999 | 281.7511 | 396.5884 | 579.9069 | 240.2119 | -0.37022 | 4.481014 | 0.381234 |
| 7 | 88284400 | 88285199 | 17.19654 | 30.90328 | 24.59947 | 10.75142 | 0.730471 | 0.407153 | 0.381242 |

| | | | | | | | | | |
|---|---|---|---|---|---|---|---|---|---|
| X | 26457000 | 26463799 | 105.477 | 165.6647 | 121.6506 | 50.45792 | 0.764382 | 2.575046 | 0.381247 |
| 16 | 97257200 | 97265999 | 81.13874 | 125.7915 | 195.4468 | 81.17943 | -0.33019 | 2.796886 | 0.381263 |
| 11 | 68170800 | 68172399 | 7.236517 | 13.72364 | 29.8295 | 12.03821 | -0.62143 | 0.022115 | 0.381274 |
| 7 | 108971000 | 108971799 | 7.995666 | 14.95282 | 30.08589 | 12.4615 | -0.54333 | 0.118913 | 0.381278 |
| 4 | 11190200 | 11192399 | 16.04833 | 27.36204 | 54.20745 | 23.76307 | -0.60002 | 0.979777 | 0.381279 |
| 13 | 59483000 | 59493399 | 83.63014 | 129.0897 | 203.4175 | 84.49564 | -0.35534 | 2.877938 | 0.381318 |
| 1 | 170715600 | 170716999 | 13.68973 | 25.50427 | 23.81173 | 9.926441 | 0.548449 | 0.174168 | 0.381327 |
| 16 | 84227200 | 84227999 | 13.68973 | 25.50427 | 23.81173 | 9.926441 | 0.548449 | 0.174168 | 0.381327 |
| 17 | 9093600 | 9096599 | 16.74412 | 28.85806 | 51.11321 | 22.42197 | -0.43644 | 0.879788 | 0.381332 |
| 1 | 74885400 | 74886199 | 6.729199 | 12.89517 | 26.80599 | 10.56108 | -0.52663 | -0.07267 | 0.381345 |
| 12 | 31684000 | 31684999 | 6.729199 | 12.89517 | 26.80599 | 10.56108 | -0.52663 | -0.07267 | 0.381345 |
| 14 | 27244000 | 27248399 | 44.74042 | 72.77643 | 65.12155 | 27.9495 | 0.529394 | 1.582146 | 0.381354 |
| 7 | 42741800 | 42744199 | 51.01844 | 82.80022 | 63.83043 | 27.35694 | 0.741141 | 1.634381 | 0.381364 |
| 4 | 53876400 | 53877199 | 10.29291 | 18.45572 | 36.32648 | 15.90677 | -0.56959 | 0.398122 | 0.381367 |
| 14 | 21851800 | 21853599 | 8.552083 | 15.60531 | 33.95678 | 14.56819 | -0.69368 | 0.27584 | 0.38137 |
| 9 | 87174200 | 87175799 | 11.93288 | 22.61675 | 20.8395 | 8.420579 | 0.603827 | 0.039739 | 0.381391 |
| 6 | 122066400 | 122068599 | 35.03609 | 58.70982 | 48.068 | 20.96096 | 0.668639 | 1.19453 | 0.381395 |
| 5 | 107354200 | 107356599 | 16.43189 | 28.27973 | 51.10263 | 22.41636 | -0.46563 | 0.891175 | 0.381401 |
| 3 | 60985000 | 60987399 | 24.47446 | 42.64477 | 35.24663 | 15.50307 | 0.660798 | 0.773722 | 0.381406 |
| 17 | 92767000 | 92769399 | 34.65387 | 57.08265 | 56.33968 | 24.30046 | 0.398199 | 1.388761 | 0.381438 |
| X | 23695600 | 23705399 | 151.4252 | 230.6965 | 186.6516 | 77.36803 | 0.590893 | 3.156616 | 0.381441 |
| 4 | 83492800 | 83494799 | 18.50908 | 33.27807 | 27.29001 | 11.92249 | 0.688075 | 0.41804 | 0.381443 |
| 15 | 85445200 | 85447199 | 19.67734 | 32.92413 | 62.19325 | 27.09505 | -0.53329 | 1.175531 | 0.381457 |
| 7 | 19719200 | 19727599 | 115.3065 | 177.029 | 172.5491 | 71.63891 | 0.331105 | 2.988941 | 0.381462 |
| 11 | 86449200 | 86450199 | 6.150446 | 11.79386 | 27.39083 | 10.77113 | -0.68299 | -0.08561 | 0.381473 |
| 16 | 23081800 | 23085799 | 25.64254 | 41.97279 | 76.29588 | 32.82014 | -0.48395 | 1.460378 | 0.381488 |
| 14 | 83867600 | 83870199 | 18.57273 | 31.81347 | 67.83367 | 29.74756 | -0.70516 | 0.960277 | 0.381493 |
| 13 | 80828600 | 80831799 | 34.32732 | 55.50348 | 92.92008 | 39.70953 | -0.38009 | 1.696389 | 0.381498 |
| 2 | 89106200 | 89107399 | 14.75011 | 27.29264 | 24.76549 | 10.42793 | 0.578698 | 0.224949 | 0.381505 |
| 9 | 109557600 | 109560599 | 32.8319 | 55.12844 | 48.03409 | 20.97311 | 0.579062 | 1.165794 | 0.381511 |
| 7 | 49273400 | 49275199 | 12.02967 | 21.29127 | 40.83258 | 18.05717 | -0.54327 | 0.506809 | 0.381514 |
| 4 | 153036600 | 153040399 | 29.15534 | 47.21848 | 86.86853 | 37.17329 | -0.5134 | 1.648883 | 0.381533 |
| 14 | 67309000 | 67310599 | 9.08024 | 16.34535 | 35.38689 | 15.44205 | -0.70236 | 0.363565 | 0.381537 |
| 5 | 112187600 | 112190599 | 15.52686 | 26.56063 | 56.97157 | 24.98144 | -0.7137 | 0.946921 | 0.381538 |
| 3 | 3607200 | 3609599 | 20.70312 | 37.19134 | 25.98063 | 11.35855 | 0.917333 | 0.435996 | 0.381539 |
| 3 | 6907200 | 6908599 | 20.70312 | 37.19134 | 25.98063 | 11.35855 | 0.917333 | 0.435996 | 0.381539 |
| 1 | 46299000 | 46301799 | 45.0803 | 73.48385 | 62.96835 | 27.04612 | 0.593814 | 1.552271 | 0.381558 |
| 13 | 66724400 | 66728999 | 103.4744 | 161.4484 | 141.5329 | 58.73119 | 0.50175 | 2.692014 | 0.381562 |
| 5 | 123399200 | 123400799 | 8.649813 | 16.00517 | 31.51399 | 13.26936 | -0.53077 | 0.193606 | 0.381577 |
| 1 | 25867600 | 25870399 | 21.43036 | 35.69954 | 66.76766 | 29.00713 | -0.51913 | 1.230198 | 0.381577 |
| 13 | 73994000 | 73996599 | 21.43036 | 35.69954 | 66.76766 | 29.00713 | -0.51913 | 1.230198 | 0.381577 |
| 19 | 25804400 | 25805399 | 10.38272 | 19.99696 | 20.19454 | 7.841969 | 0.52501 | -0.10243 | 0.381591 |
| 4 | 60115000 | 60116199 | 24.83509 | 42.56001 | 41.04152 | 17.99889 | 0.435564 | 0.951422 | 0.381591 |
| 6 | 133911200 | 133912399 | 9.248483 | 16.61074 | 35.40871 | 15.48306 | -0.68194 | 0.380778 | 0.381606 |
| 19 | 12191400 | 12193999 | 32.00412 | 54.25252 | 39.81238 | 17.43604 | 0.826163 | 1.072745 | 0.381625 |
| 7 | 15211200 | 15216999 | 103.1317 | 160.5762 | 135.3086 | 56.16478 | 0.556402 | 2.726188 | 0.38163 |
| X | 166946800 | 166947599 | 5.955875 | 11.57055 | 27.72187 | 10.34806 | -0.65539 | -0.25876 | 0.381641 |

| | | | | | | | | | |
|---|---|---|---|---|---|---|---|---|---|
| 7 | 27901400 | 27906199 | 31.85874 | 51.64076 | 87.69114 | 37.5489 | -0.39706 | 1.63645 | 0.381674 |
| X | 58831800 | 58834799 | 42.84166 | 69.93883 | 52.58973 | 22.60057 | 0.783339 | 1.522416 | 0.381693 |
| 3 | 13094000 | 13094599 | 11.83216 | 22.27064 | 22.03409 | 9.027364 | 0.485603 | 0.090447 | 0.381694 |
| 6 | 3646200 | 3648599 | 17.39285 | 29.46393 | 62.78431 | 27.49107 | -0.70589 | 1.05382 | 0.381734 |
| 8 | 106015400 | 106018199 | 17.42072 | 28.97369 | 72.87671 | 31.62779 | -0.94502 | 1.243261 | 0.38174 |
| 6 | 117689400 | 117694199 | 42.22146 | 67.5317 | 114.6525 | 48.2487 | -0.41896 | 2.046176 | 0.381741 |
| 8 | 25248600 | 25252999 | 75.66864 | 119.4538 | 115.6176 | 48.00051 | 0.372074 | 2.450218 | 0.381754 |
| 17 | 18866800 | 18871399 | 56.77827 | 91.59625 | 72.35663 | 30.8314 | 0.697333 | 1.789351 | 0.381756 |
| 10 | 42585800 | 42586399 | 9.932682 | 19.35092 | 16.55889 | 6.026806 | 0.847365 | -0.33613 | 0.381768 |
| X | 72659600 | 72663599 | 20.25397 | 34.36402 | 64.19391 | 28.12152 | -0.51686 | 1.050541 | 0.381772 |
| 4 | 96268600 | 96271999 | 33.41048 | 56.07295 | 46.93241 | 20.48586 | 0.636748 | 1.168878 | 0.381774 |
| 15 | 82377800 | 82381399 | 38.87675 | 62.5039 | 107.8458 | 45.73473 | -0.43343 | 1.87783 | 0.381795 |
| 4 | 130345000 | 130349599 | 24.92462 | 41.00101 | 73.84713 | 31.83359 | -0.46785 | 1.39395 | 0.381804 |
| 18 | 44811400 | 44811799 | 3.636706 | 7.113303 | 17.90729 | 6.016668 | -0.58383 | -0.69458 | 0.381804 |
| 13 | 12167800 | 12169599 | 10.0567 | 18.29632 | 34.92331 | 15.03872 | -0.50971 | 0.306487 | 0.381804 |
| 17 | 65946800 | 65954599 | 58.36098 | 92.23097 | 151.9698 | 63.12728 | -0.39319 | 2.418734 | 0.381807 |
| 17 | 6780400 | 6780599 | 10.82362 | 20.76028 | 20.19423 | 7.935312 | 0.563298 | -0.05817 | 0.381813 |
| 4 | 14678200 | 14681999 | 66.66757 | 104.4838 | 166.5608 | 69.08522 | -0.35494 | 2.598186 | 0.381823 |
| 9 | 33229600 | 33231199 | 17.51688 | 31.8069 | 26.54424 | 11.45831 | 0.674261 | 0.355153 | 0.381829 |
| 8 | 14354200 | 14357599 | 184.4088 | 267.6507 | 430.1557 | 177.34 | -0.46228 | 4.02442 | 0.381837 |
| X | 40117600 | 40118399 | 13.61501 | 25.3747 | 22.99044 | 9.577906 | 0.592798 | 0.17207 | 0.381841 |
| X | 89340600 | 89349199 | 123.4277 | 190.7405 | 153.1997 | 63.63578 | 0.617238 | 2.869832 | 0.381845 |
| 11 | 107210200 | 107211199 | 8.151033 | 15.30344 | 30.16186 | 12.41005 | -0.50585 | 0.096358 | 0.381851 |
| 10 | 84393600 | 84397399 | 46.87166 | 75.52548 | 71.90507 | 30.59162 | 0.426642 | 1.806012 | 0.381857 |
| 10 | 85259600 | 85265199 | 43.01232 | 68.91062 | 112.8584 | 47.60884 | -0.36444 | 1.995501 | 0.381862 |
| 8 | 95743000 | 95745799 | 11.40169 | 19.90413 | 45.25379 | 19.92834 | -0.79137 | 0.699403 | 0.381867 |
| 11 | 117261800 | 117265599 | 31.48503 | 50.76549 | 94.43538 | 40.21162 | -0.53637 | 1.776019 | 0.381868 |
| X | 62169800 | 62170799 | 12.48088 | 23.6253 | 21.57609 | 8.741713 | 0.614286 | 0.059541 | 0.381871 |
| 8 | 56300200 | 56302999 | 20.19701 | 34.31027 | 67.1396 | 29.4231 | -0.58366 | 1.040411 | 0.381872 |
| 6 | 7962800 | 7964799 | 12.70878 | 22.53806 | 46.18157 | 20.41067 | -0.63718 | 0.50121 | 0.381876 |
| 13 | 48211600 | 48213599 | 24.75746 | 42.54038 | 40.95338 | 17.96348 | 0.438533 | 0.926097 | 0.381891 |
| 16 | 78343200 | 78347199 | 41.35704 | 66.38471 | 109.8747 | 46.4843 | -0.37636 | 1.933126 | 0.381898 |
| 8 | 114045200 | 114047999 | 21.14658 | 37.17093 | 31.95883 | 14.15241 | 0.604243 | 0.619891 | 0.381908 |
| 5 | 88386600 | 88389999 | 35.58085 | 59.78183 | 45.79293 | 20.00125 | 0.76422 | 1.16582 | 0.381928 |
| 9 | 30662200 | 30663199 | 12.31959 | 23.71832 | 20.04368 | 7.791891 | 0.778316 | -0.07923 | 0.381997 |
| 10 | 127745600 | 127746199 | 5.099009 | 9.921563 | 22.97727 | 8.468264 | -0.5894 | -0.33054 | 0.382021 |
| X | 77939400 | 77940399 | 16.58386 | 30.97506 | 22.499 | 9.33875 | 0.9131 | 0.180332 | 0.382027 |
| 13 | 56867400 | 56868199 | 17.42915 | 31.38002 | 23.677 | 10.32973 | 0.809068 | 0.39642 | 0.382031 |
| 13 | 119071200 | 119073599 | 27.24076 | 46.34293 | 44.17179 | 19.36028 | 0.450717 | 1.028663 | 0.382037 |
| 9 | 119049800 | 119056199 | 60.45897 | 95.56327 | 156.8981 | 65.18024 | -0.38783 | 2.419557 | 0.382047 |
| 13 | 21842600 | 21843999 | 9.850476 | 17.70737 | 35.37467 | 15.45682 | -0.58831 | 0.379395 | 0.382059 |
| 8 | 129756400 | 129758799 | 22.19947 | 37.29257 | 68.16634 | 29.7719 | -0.48634 | 1.145181 | 0.38206 |
| 1 | 96386600 | 96388399 | 17.84098 | 31.63143 | 29.67338 | 13.11748 | 0.483927 | 0.505481 | 0.382061 |
| 6 | 128332200 | 128333399 | 9.008571 | 16.4631 | 32.94264 | 14.10646 | -0.57145 | 0.270766 | 0.382078 |
| X | 80848600 | 80850399 | 18.54418 | 34.5422 | 21.88554 | 9.126152 | 1.10396 | 0.213348 | 0.382091 |
| 4 | 52287000 | 52289199 | 23.95863 | 41.91859 | 33.55165 | 14.79994 | 0.707236 | 0.71069 | 0.382094 |
| 3 | 145757400 | 145759199 | 10.49142 | 18.66783 | 37.83052 | 16.6729 | -0.61793 | 0.450918 | 0.382099 |

| | | | | | | | | | |
|---|---|---|---|---|---|---|---|---|---|
| 8 | 66042600 | 66044199 | 17.10427 | 29.26139 | 53.21262 | 23.33333 | -0.47631 | 0.951049 | 0.382102 |
| 8 | 56649600 | 56651799 | 18.60805 | 34.24021 | 22.57366 | 9.577201 | 1.02838 | 0.280245 | 0.382131 |
| 7 | 133309400 | 133313199 | 26.09035 | 42.89901 | 74.37516 | 32.05501 | -0.41324 | 1.404948 | 0.382134 |
| 19 | 37719200 | 37721599 | 17.89111 | 30.63331 | 53.77876 | 23.58318 | -0.42555 | 0.947092 | 0.382135 |
| 5 | 40757800 | 40758799 | 6.868072 | 13.1304 | 29.03476 | 11.5166 | -0.62509 | -0.04649 | 0.382152 |
| 7 | 4044400 | 4046599 | 24.18288 | 42.36725 | 31.77615 | 14.03251 | 0.800382 | 0.687485 | 0.382161 |
| 9 | 64287000 | 64290799 | 40.21948 | 64.56812 | 107.1094 | 45.32441 | -0.37934 | 1.926311 | 0.382178 |
| 4 | 86679600 | 86682999 | 43.25785 | 70.06587 | 69.08982 | 29.56772 | 0.384216 | 1.661443 | 0.382225 |
| 4 | 95855000 | 95856199 | 9.110316 | 16.76184 | 32.47909 | 13.78177 | -0.51619 | 0.230556 | 0.382232 |
| 7 | 99580600 | 99582199 | 9.110316 | 16.76184 | 32.47909 | 13.78177 | -0.51619 | 0.230556 | 0.382232 |
| 14 | 71200400 | 71201599 | 9.110316 | 16.76184 | 32.47909 | 13.78177 | -0.51619 | 0.230556 | 0.382232 |
| 15 | 61958600 | 61959999 | 9.110316 | 16.76184 | 32.47909 | 13.78177 | -0.51619 | 0.230556 | 0.382232 |
| 1 | 110251200 | 110253599 | 39.07553 | 64.56701 | 49.4222 | 21.35242 | 0.764897 | 1.356596 | 0.382249 |
| 9 | 18832400 | 18833199 | 18.48245 | 32.4823 | 30.79445 | 13.63515 | 0.463972 | 0.604897 | 0.382253 |
| 2 | 61865200 | 61865799 | 11.48013 | 22.12329 | 20.1141 | 7.79782 | 0.677326 | -0.09667 | 0.382293 |
| 11 | 19425000 | 19425799 | 4.555454 | 8.875483 | 23.13766 | 8.414843 | -0.74092 | -0.38609 | 0.382298 |
| X | 130883800 | 130885399 | 16.97152 | 31.68036 | 22.48208 | 9.341085 | 0.945367 | 0.187477 | 0.3823 |
| X | 31984400 | 31994799 | 151.8359 | 230.9838 | 194.0387 | 80.39478 | 0.535018 | 3.186122 | 0.382304 |
| 16 | 89534200 | 89534999 | 5.381622 | 10.43681 | 23.83833 | 8.996299 | -0.6011 | -0.23852 | 0.382305 |
| X | 147457200 | 147459599 | 28.92972 | 49.46207 | 41.31345 | 18.11476 | 0.641429 | 0.986477 | 0.382308 |
| X | 68010400 | 68016599 | 92.0669 | 144.5976 | 117.9504 | 48.92298 | 0.612851 | 2.566284 | 0.382322 |
| 2 | 152018600 | 152022799 | 26.08928 | 42.37408 | 83.37575 | 35.75409 | -0.60615 | 1.582947 | 0.382325 |
| 16 | 38611000 | 38612399 | 9.112647 | 16.91606 | 32.63509 | 13.68122 | -0.4971 | 0.181986 | 0.382337 |
| 13 | 60684800 | 60686599 | 9.969482 | 17.99251 | 34.85778 | 15.16208 | -0.54016 | 0.354313 | 0.382348 |
| 2 | 86406600 | 86407399 | 12.26512 | 23.38014 | 21.60335 | 8.624004 | 0.615932 | 0.00954 | 0.382375 |
| 1 | 149172000 | 149172999 | 8.339428 | 15.54375 | 31.09694 | 12.95326 | -0.5415 | 0.147894 | 0.382385 |
| 12 | 78751200 | 78751999 | 8.339428 | 15.54375 | 31.09694 | 12.95326 | -0.5415 | 0.147894 | 0.382385 |
| 3 | 101284000 | 101285999 | 9.882958 | 17.69231 | 36.40904 | 15.96445 | -0.63421 | 0.40935 | 0.382389 |
| 10 | 119481000 | 119486399 | 36.26536 | 58.46677 | 99.77978 | 42.48134 | -0.41267 | 1.785621 | 0.382402 |
| 10 | 31515000 | 31516399 | 15.20824 | 26.10936 | 51.59718 | 22.63138 | -0.59481 | 0.909343 | 0.382407 |
| 10 | 16397200 | 16398799 | 8.09796 | 15.30738 | 30.36597 | 12.33943 | -0.50057 | 0.049387 | 0.382423 |
| 5 | 110088400 | 110098799 | 208.1922 | 311.2063 | 238.4864 | 98.33321 | 0.645624 | 3.541698 | 0.382441 |
| 5 | 42249400 | 42250199 | 8.323766 | 15.45186 | 31.02779 | 13.00716 | -0.55434 | 0.170518 | 0.382445 |
| 17 | 95135400 | 95138799 | 50.67688 | 81.9588 | 70.21765 | 30.01694 | 0.584603 | 1.707468 | 0.382462 |
| X | 77421200 | 77424599 | 65.16765 | 104.4187 | 86.1814 | 36.36197 | 0.622602 | 2.010479 | 0.382475 |
| 2 | 97959600 | 97961399 | 27.41366 | 47.32494 | 37.98023 | 16.66173 | 0.701164 | 0.891382 | 0.382481 |
| 1 | 67012400 | 67013399 | 11.9379 | 23.04013 | 19.52415 | 7.536351 | 0.78363 | -0.10736 | 0.382482 |
| 6 | 97640400 | 97641799 | 11.9379 | 23.04013 | 19.52415 | 7.536351 | 0.78363 | -0.10736 | 0.382482 |
| 17 | 28750800 | 28753199 | 16.13986 | 27.66102 | 52.67538 | 23.10109 | -0.54212 | 0.929377 | 0.382492 |
| 7 | 53013800 | 53015799 | 36.4587 | 59.93439 | 58.65181 | 25.27611 | 0.40932 | 1.419162 | 0.382508 |
| 1 | 34723000 | 34724399 | 8.196475 | 15.14578 | 31.50206 | 13.28884 | -0.61125 | 0.196651 | 0.382532 |
| 14 | 110006600 | 110009999 | 38.10225 | 62.38745 | 60.90233 | 26.20183 | 0.410611 | 1.472766 | 0.38254 |
| 13 | 51206400 | 51207599 | 6.876432 | 13.15143 | 27.30988 | 10.8205 | -0.53309 | -0.0523 | 0.382554 |
| 13 | 112653000 | 112657399 | 58.80867 | 92.69641 | 152.8703 | 63.4366 | -0.39741 | 2.479352 | 0.38257 |
| 17 | 60610600 | 60612799 | 31.48894 | 52.34654 | 51.37609 | 22.28936 | 0.408039 | 1.27038 | 0.382597 |
| 1 | 86746800 | 86747799 | 8.340486 | 15.48005 | 31.03085 | 13.01171 | -0.55191 | 0.17302 | 0.3826 |
| X | 77083000 | 77088399 | 47.89029 | 78.03863 | 55.64864 | 23.89929 | 0.857804 | 1.559165 | 0.382602 |

| | | | | | | | | | |
|---|---|---|---|---|---|---|---|---|---|
| 18 | 3698000 | 3712399 | 252.922 | 367.5885 | 340.7094 | 140.4443 | 0.333297 | 4.00746 | 0.38261 |
| 10 | 23354000 | 23355399 | 16.58017 | 29.89466 | 27.58236 | 12.01182 | 0.52208 | 0.389245 | 0.38262 |
| 3 | 69223200 | 69226399 | 25.88375 | 42.15332 | 80.25159 | 34.45802 | -0.55568 | 1.537471 | 0.38263 |
| 6 | 98792600 | 98793999 | 13.28965 | 24.77224 | 23.91575 | 9.960725 | 0.501723 | 0.168576 | 0.382636 |
| 3 | 112930400 | 112933999 | 29.0195 | 49.46977 | 43.00117 | 18.8503 | 0.583564 | 1.015588 | 0.382647 |
| 14 | 42090200 | 42095399 | 67.30145 | 107.292 | 99.99293 | 41.8847 | 0.439606 | 2.169076 | 0.382665 |
| 18 | 50385400 | 50386599 | 10.23025 | 18.56788 | 34.8887 | 15.06854 | -0.48989 | 0.322103 | 0.382715 |
| 2 | 137959200 | 137961199 | 28.17963 | 48.46556 | 39.65867 | 17.39607 | 0.672284 | 0.93025 | 0.382716 |
| 18 | 90197600 | 90199199 | 16.12562 | 29.70288 | 24.51066 | 10.38446 | 0.708274 | 0.259249 | 0.382729 |
| 8 | 71872600 | 71877399 | 72.32705 | 114.6196 | 113.7331 | 47.31269 | 0.340676 | 2.356307 | 0.38273 |
| 4 | 26185200 | 26188399 | 48.1256 | 77.52329 | 75.7048 | 32.19454 | 0.389555 | 1.817508 | 0.38273 |
| 2 | 117336400 | 117338399 | 43.57336 | 69.74919 | 115.1119 | 48.49637 | -0.37694 | 2.025373 | 0.382731 |
| 7 | 19003400 | 19006999 | 27.50266 | 44.7238 | 81.42331 | 34.94233 | -0.49291 | 1.563559 | 0.382731 |
| X | 24573200 | 24576199 | 59.18612 | 95.10096 | 80.27476 | 34.01391 | 0.595018 | 1.916995 | 0.382734 |
| 7 | 13680200 | 13680799 | 21.85345 | 37.98264 | 33.35285 | 14.64703 | 0.573819 | 0.795954 | 0.382747 |
| 12 | 114249800 | 114252399 | 57.72509 | 92.66522 | 84.58212 | 35.78807 | 0.480723 | 1.942172 | 0.382748 |
| 5 | 26755000 | 26757799 | 18.18925 | 31.10709 | 54.87511 | 24.06167 | -0.43285 | 0.959217 | 0.382752 |
| 9 | 75594600 | 75599599 | 66.99525 | 106.9275 | 96.81048 | 40.61707 | 0.483074 | 2.132121 | 0.382753 |
| X | 112794000 | 112795999 | 17.66666 | 32.29302 | 25.48734 | 10.91091 | 0.761848 | 0.316664 | 0.382756 |
| 1 | 111185800 | 111189399 | 48.01552 | 77.30446 | 75.36502 | 32.02512 | 0.391028 | 1.834763 | 0.382785 |
| X | 33956400 | 33963599 | 96.83074 | 152.8204 | 114.2628 | 47.43165 | 0.743683 | 2.473116 | 0.382802 |
| 13 | 94458200 | 94461199 | 36.87487 | 60.64417 | 58.19719 | 25.08468 | 0.43766 | 1.413612 | 0.382808 |
| 3 | 112314400 | 112315999 | 20.18191 | 35.50059 | 32.06924 | 14.19927 | 0.533704 | 0.607079 | 0.38281 |
| 1 | 94041600 | 94045999 | 46.85477 | 75.9455 | 68.53908 | 29.3473 | 0.512643 | 1.653504 | 0.382822 |
| 14 | 3823800 | 3824799 | 8.730754 | 16.25501 | 33.27493 | 13.88667 | -0.57756 | 0.158693 | 0.382826 |
| 19 | 9350400 | 9350799 | 8.147948 | 15.92981 | 13.98538 | 4.792881 | 0.887455 | -0.57402 | 0.382838 |
| 9 | 63915800 | 63922799 | 43.15928 | 68.67958 | 139.2307 | 58.17397 | -0.68251 | 2.215338 | 0.382846 |
| 13 | 39784200 | 39786399 | 22.6227 | 39.25896 | 37.16973 | 16.30877 | 0.465182 | 0.82265 | 0.382857 |
| 13 | 28579800 | 28583199 | 29.33686 | 49.61536 | 45.7994 | 20.04392 | 0.496263 | 1.087929 | 0.382861 |
| 4 | 139648800 | 139649999 | 7.189333 | 13.71124 | 27.78728 | 11.08112 | -0.50653 | -0.02565 | 0.382882 |
| 11 | 116764200 | 116765399 | 7.189333 | 13.71124 | 27.78728 | 11.08112 | -0.50653 | -0.02565 | 0.382882 |
| 1 | 103563400 | 103564799 | 13.56873 | 25.63614 | 22.37118 | 9.100068 | 0.673757 | 0.081706 | 0.382887 |
| 5 | 31124800 | 31128199 | 14.12328 | 24.2591 | 54.24796 | 23.79465 | -0.77236 | 0.901038 | 0.382892 |
| 10 | 125660000 | 125666399 | 53.7676 | 85.46442 | 137.8207 | 57.50595 | -0.35466 | 2.260164 | 0.382926 |
| 6 | 5607200 | 5608399 | 26.09818 | 44.28076 | 42.70149 | 18.70645 | 0.433637 | 1.04657 | 0.382927 |
| 6 | 68432400 | 68436199 | 49.11549 | 79.45412 | 69.82703 | 29.85565 | 0.548242 | 1.696347 | 0.382951 |
| X | 116856200 | 116857999 | 38.22562 | 62.97959 | 57.07072 | 24.618 | 0.521231 | 1.397101 | 0.382963 |
| 13 | 59557600 | 59560999 | 33.87337 | 56.19706 | 53.68746 | 23.25978 | 0.446573 | 1.300421 | 0.382966 |
| 3 | 12018800 | 12019999 | 17.69263 | 31.48608 | 29.17175 | 12.85509 | 0.504889 | 0.480693 | 0.382989 |
| 18 | 61859000 | 61859799 | 5.41903 | 10.44225 | 25.25428 | 9.792535 | -0.72054 | -0.14282 | 0.382992 |
| 17 | 13041200 | 13042599 | 30.29744 | 51.54321 | 43.5548 | 19.08986 | 0.623836 | 1.040316 | 0.382997 |
| 11 | 54080000 | 54082799 | 29.24136 | 49.32635 | 46.9192 | 20.51793 | 0.453057 | 1.111374 | 0.383021 |
| 10 | 57241200 | 57242399 | 17.88593 | 31.76672 | 29.20542 | 12.89157 | 0.51437 | 0.494681 | 0.383033 |
| X | 4288400 | 4291599 | 25.9175 | 44.66048 | 35.71449 | 15.66689 | 0.705921 | 0.900257 | 0.383039 |
| 17 | 43699800 | 43700999 | 7.814321 | 14.75201 | 29.19032 | 11.89439 | -0.50008 | 0.054152 | 0.383079 |
| 13 | 30069200 | 30071399 | 58.54975 | 93.49875 | 86.32262 | 36.24468 | 0.455777 | 2.103857 | 0.383086 |
| 5 | 74005600 | 74008199 | 17.69917 | 30.38962 | 53.26584 | 23.36352 | -0.42266 | 0.919581 | 0.383097 |

| | | | | | | | | | |
|---|---|---|---|---|---|---|---|---|---|
| 11 | 16443800 | 16444599 | 12.69851 | 24.02071 | 22.52703 | 9.1403 | 0.574087 | 0.067195 | 0.383149 |
| 11 | 110130600 | 110131599 | 12.69851 | 24.02071 | 22.52703 | 9.1403 | 0.574087 | 0.067195 | 0.383149 |
| 9 | 71484400 | 71485399 | 12.02064 | 22.52746 | 21.41274 | 8.837474 | 0.534236 | 0.116347 | 0.383157 |
| X | 142447000 | 142447799 | 10.99582 | 21.00676 | 21.11479 | 8.389804 | 0.500951 | -0.02042 | 0.383159 |
| 2 | 11843000 | 11843599 | 11.13162 | 21.26442 | 20.66446 | 8.212326 | 0.549568 | -0.0178 | 0.383177 |
| 13 | 68614000 | 68619999 | 123.5866 | 188.7199 | 177.2485 | 73.51163 | 0.378508 | 3.09867 | 0.383189 |
| 4 | 37657800 | 37658399 | 10.0708 | 18.43756 | 34.51138 | 14.74304 | -0.47381 | 0.271104 | 0.383198 |
| 17 | 24998400 | 25000199 | 10.0708 | 18.43756 | 34.51138 | 14.74304 | -0.47381 | 0.271104 | 0.383198 |
| 13 | 104724600 | 104729999 | 28.21013 | 46.09967 | 79.50394 | 34.17966 | -0.41006 | 1.491063 | 0.383198 |
| 6 | 61064400 | 61065799 | 10.35704 | 18.82115 | 36.49202 | 15.73678 | -0.53371 | 0.317739 | 0.383209 |
| 18 | 3793400 | 3795999 | 30.74849 | 52.55181 | 40.14206 | 17.6006 | 0.769753 | 0.995767 | 0.383222 |
| 3 | 101663600 | 101665399 | 19.48204 | 34.23323 | 32.17105 | 14.24263 | 0.476555 | 0.615766 | 0.383239 |
| 11 | 92715600 | 92716599 | 21.70906 | 37.80647 | 36.17859 | 15.90775 | 0.450494 | 0.771001 | 0.383259 |
| 9 | 51568200 | 51569599 | 18.53134 | 32.70862 | 30.31593 | 13.4138 | 0.498892 | 0.559593 | 0.383297 |
| X | 125343000 | 125345999 | 46.20725 | 75.24441 | 63.44435 | 27.24048 | 0.616068 | 1.570839 | 0.383309 |
| 8 | 48863400 | 48864399 | 22.77377 | 39.35711 | 38.37375 | 16.83491 | 0.42192 | 0.863491 | 0.383325 |
| 6 | 144696000 | 144698199 | 14.29936 | 25.08162 | 45.09502 | 19.93423 | -0.45466 | 0.638597 | 0.383336 |
| 19 | 3358800 | 3361599 | 22.84913 | 39.36734 | 36.89545 | 16.18487 | 0.477828 | 0.888103 | 0.383337 |
| 9 | 118056400 | 118058799 | 11.40729 | 20.37412 | 37.30605 | 16.38959 | -0.46925 | 0.433466 | 0.383354 |
| 3 | 85812400 | 85814999 | 24.8235 | 42.69836 | 40.36358 | 17.70534 | 0.464932 | 0.915269 | 0.383372 |
| 12 | 42826200 | 42829399 | 41.87848 | 68.28559 | 63.69302 | 27.36043 | 0.472113 | 1.543086 | 0.383375 |
| 13 | 66509000 | 66521399 | 269.7644 | 391.0794 | 312.8352 | 128.9897 | 0.542172 | 4.045991 | 0.383384 |
| 14 | 63699800 | 63701999 | 12.40552 | 21.84919 | 40.3973 | 17.88218 | -0.49329 | 0.565076 | 0.383406 |
| 1 | 83786600 | 83787799 | 7.631313 | 14.56754 | 29.56833 | 11.76849 | -0.50674 | -0.02418 | 0.383432 |
| 7 | 128125000 | 128126199 | 8.90671 | 16.10598 | 34.4114 | 14.93129 | -0.67793 | 0.334174 | 0.383455 |
| 2 | 43246200 | 43246999 | 14.03125 | 25.78106 | 23.0051 | 9.779563 | 0.593297 | 0.255487 | 0.383531 |
| 1 | 127081400 | 127082799 | 8.133719 | 15.16171 | 32.74431 | 13.63735 | -0.6518 | 0.144453 | 0.383552 |
| 5 | 59253200 | 59254399 | 15.60609 | 28.03322 | 26.77739 | 11.7075 | 0.4687 | 0.403158 | 0.383554 |
| 19 | 29624400 | 29626799 | 13.5154 | 23.62651 | 44.15653 | 19.46368 | -0.51013 | 0.687933 | 0.38358 |
| 8 | 92998800 | 92999399 | 4.517286 | 8.813901 | 20.95792 | 7.475975 | -0.58327 | -0.46073 | 0.38358 |
| 18 | 32692200 | 32693199 | 5.852284 | 11.26881 | 26.39525 | 10.25662 | -0.67868 | -0.13034 | 0.383587 |
| 5 | 88323400 | 88324999 | 15.41109 | 28.89949 | 22.66437 | 9.344135 | 0.810974 | 0.146157 | 0.383595 |
| 14 | 26359000 | 26360199 | 6.873922 | 12.73219 | 31.04225 | 13.05252 | -0.83455 | 0.172806 | 0.383599 |
| 5 | 9132600 | 9134199 | 9.893835 | 18.16428 | 34.03089 | 14.48259 | -0.47117 | 0.252446 | 0.383604 |
| 7 | 67480400 | 67481799 | 9.893835 | 18.16428 | 34.03089 | 14.48259 | -0.47117 | 0.252446 | 0.383604 |
| 13 | 102393400 | 102394999 | 9.893835 | 18.16428 | 34.03089 | 14.48259 | -0.47117 | 0.252446 | 0.383604 |
| 13 | 93229400 | 93235599 | 60.21155 | 95.07885 | 151.4691 | 62.89362 | -0.34555 | 2.439884 | 0.383614 |
| 17 | 36534400 | 36536599 | 16.05721 | 27.77361 | 50.0577 | 21.96135 | -0.46043 | 0.841794 | 0.383618 |
| 12 | 16799800 | 16803999 | 21.23937 | 35.58738 | 62.37392 | 27.19772 | -0.42576 | 1.165024 | 0.383621 |
| 12 | 77563400 | 77564999 | 11.53035 | 20.72674 | 38.38892 | 16.77416 | -0.48046 | 0.395085 | 0.383628 |
| 19 | 39641000 | 39643599 | 21.84809 | 39.18209 | 25.94668 | 11.35983 | 0.992773 | 0.454745 | 0.383635 |
| 19 | 25243600 | 25247599 | 29.41877 | 47.80862 | 82.97885 | 35.5986 | -0.42499 | 1.577686 | 0.383662 |
| 7 | 144532400 | 144535199 | 38.13188 | 61.28575 | 102.5724 | 43.47726 | -0.3901 | 1.887862 | 0.383696 |
| 13 | 76091200 | 76092199 | 8.258766 | 15.08015 | 33.45946 | 14.34285 | -0.72061 | 0.269191 | 0.383708 |
| 2 | 152226200 | 152227799 | 21.36085 | 37.39686 | 34.32813 | 15.15046 | 0.510518 | 0.688747 | 0.383717 |
| 10 | 40365400 | 40367999 | 23.8912 | 39.64303 | 68.4596 | 29.6632 | -0.4048 | 1.285268 | 0.383722 |
| 1 | 69999000 | 70001799 | 28.81942 | 49.69012 | 37.94126 | 16.644 | 0.772406 | 0.908961 | 0.383739 |

| | | | | | | | | | |
|---|---|---|---|---|---|---|---|---|---|
| 9 | 86166800 | 86168999 | 30.98248 | 52.33064 | 45.78499 | 20.02974 | 0.573252 | 1.104614 | 0.383764 |
| 8 | 117541000 | 117542199 | 6.713194 | 12.88615 | 27.46929 | 10.77026 | -0.55658 | -0.08908 | 0.383766 |
| 9 | 49396200 | 49398199 | 10.40773 | 18.36294 | 39.9442 | 17.67538 | -0.72531 | 0.531828 | 0.383775 |
| 2 | 178273400 | 178276199 | 21.34489 | 35.40301 | 67.6144 | 29.28848 | -0.54957 | 1.281877 | 0.383776 |
| 11 | 108656400 | 108658199 | 23.21471 | 40.68743 | 32.39802 | 14.31199 | 0.714489 | 0.677054 | 0.383787 |
| X | 153604400 | 153605399 | 12.80511 | 24.06814 | 21.56392 | 8.855763 | 0.625393 | 0.10876 | 0.383802 |
| 16 | 64799600 | 64801799 | 10.50665 | 18.80224 | 43.68912 | 19.16224 | -0.80937 | 0.42769 | 0.383806 |
| 11 | 57985600 | 57989599 | 19.43926 | 32.76552 | 59.06756 | 25.82424 | -0.46596 | 1.098934 | 0.383817 |
| 2 | 36848200 | 36857399 | 116.2142 | 179.633 | 158.803 | 65.96317 | 0.479233 | 2.862285 | 0.383839 |
| 2 | 38105000 | 38106999 | 19.45765 | 35.41251 | 24.01814 | 10.33801 | 0.973798 | 0.349663 | 0.383847 |
| 9 | 122381600 | 122382399 | 7.503191 | 14.29548 | 28.8904 | 11.54548 | -0.5053 | -0.0123 | 0.383848 |
| 18 | 69657400 | 69658199 | 7.503191 | 14.29548 | 28.8904 | 11.54548 | -0.5053 | -0.0123 | 0.383848 |
| 5 | 79426800 | 79429399 | 23.93726 | 41.68533 | 36.44757 | 16.02564 | 0.580035 | 0.781678 | 0.383871 |
| 14 | 11307600 | 11308999 | 6.459546 | 12.3096 | 27.19851 | 10.86463 | -0.63207 | -0.0292 | 0.383877 |
| 1 | 40504600 | 40506199 | 9.344156 | 16.95442 | 33.90926 | 14.65113 | -0.57927 | 0.317202 | 0.383877 |
| 11 | 107576200 | 107577599 | 9.344156 | 16.95442 | 33.90926 | 14.65113 | -0.57927 | 0.317202 | 0.383877 |
| 12 | 79218800 | 79220599 | 9.344156 | 16.95442 | 33.90926 | 14.65113 | -0.57927 | 0.317202 | 0.383877 |
| 18 | 82592800 | 82593999 | 12.44088 | 21.85299 | 41.49069 | 18.36309 | -0.53241 | 0.601527 | 0.383879 |
| 14 | 72763600 | 72766599 | 29.59127 | 48.06134 | 83.44774 | 35.78461 | -0.42628 | 1.590169 | 0.383883 |
| 4 | 142492200 | 142495399 | 15.84821 | 27.47732 | 52.7984 | 23.16543 | -0.55188 | 0.820877 | 0.383896 |
| 17 | 87700200 | 87705399 | 33.07541 | 53.37563 | 92.89481 | 39.59952 | -0.43914 | 1.751943 | 0.383906 |
| 2 | 57082000 | 57083199 | 15.40335 | 28.34354 | 24.54284 | 10.41255 | 0.637728 | 0.258632 | 0.383915 |
| 19 | 39936600 | 39939999 | 63.00137 | 100.4645 | 99.01439 | 41.49061 | 0.359489 | 2.15568 | 0.383916 |
| 18 | 70146200 | 70147399 | 17.63782 | 31.16241 | 29.35403 | 12.98562 | 0.476144 | 0.540866 | 0.38392 |
| 11 | 20322200 | 20329999 | 78.49794 | 121.2914 | 196.9025 | 81.78901 | -0.39708 | 2.857085 | 0.383929 |
| 2 | 22357600 | 22358399 | 14.7871 | 25.67856 | 47.92084 | 21.03312 | -0.50917 | 0.794828 | 0.383937 |
| 6 | 56690600 | 56692599 | 14.7871 | 25.67856 | 47.92084 | 21.03312 | -0.50917 | 0.794828 | 0.383937 |
| 18 | 74573400 | 74577199 | 36.54443 | 58.57566 | 110.1011 | 46.47865 | -0.56191 | 1.971692 | 0.383954 |
| 7 | 10270200 | 10274199 | 50.62484 | 81.71246 | 74.55402 | 31.78933 | 0.490753 | 1.765072 | 0.383967 |
| 2 | 51378800 | 51379199 | 8.317135 | 16.25538 | 14.57655 | 5.036819 | 0.847145 | -0.54109 | 0.383985 |
| 9 | 71335600 | 71335999 | 8.317135 | 16.25538 | 14.57655 | 5.036819 | 0.847145 | -0.54109 | 0.383985 |
| 6 | 104671400 | 104672399 | 12.48227 | 23.45945 | 22.99408 | 9.444375 | 0.496044 | 0.107855 | 0.383986 |
| 14 | 55404000 | 55407999 | 34.29261 | 55.04905 | 111.7394 | 47.2783 | -0.66993 | 1.924379 | 0.38399 |
| 6 | 17174800 | 17175399 | 7.166445 | 13.7453 | 28.02206 | 11.01199 | -0.49573 | -0.07425 | 0.384004 |
| X | 126858000 | 126861199 | 87.39307 | 137.6951 | 110.2776 | 45.75945 | 0.64279 | 2.499644 | 0.384005 |
| 13 | 72824600 | 72826199 | 12.26226 | 21.48972 | 43.07248 | 19.02614 | -0.61044 | 0.637806 | 0.384036 |
| 9 | 90379800 | 90380399 | 5.505505 | 10.70063 | 24.61065 | 9.151934 | -0.59091 | -0.28493 | 0.38404 |
| 19 | 55528000 | 55528599 | 5.505505 | 10.70063 | 24.61065 | 9.151934 | -0.59091 | -0.28493 | 0.38404 |
| 3 | 94119800 | 94127599 | 139.4044 | 212.8736 | 187.6905 | 77.8423 | 0.469608 | 3.105158 | 0.384047 |
| 8 | 5490400 | 5492599 | 33.14412 | 55.33875 | 50.32859 | 21.89666 | 0.517549 | 1.225423 | 0.384053 |
| 5 | 96732600 | 96733199 | 9.913768 | 19.13868 | 19.71706 | 7.601292 | 0.506269 | -0.13199 | 0.38406 |
| 1 | 96471600 | 96473799 | 20.69494 | 36.29749 | 32.59715 | 14.4088 | 0.541708 | 0.651488 | 0.38407 |
| 13 | 40925400 | 40929199 | 55.56386 | 89.13345 | 86.2996 | 36.47594 | 0.394638 | 1.961122 | 0.384071 |
| 1 | 112278000 | 112278999 | 8.257742 | 15.60174 | 30.91994 | 12.57761 | -0.50053 | 0.055838 | 0.384091 |
| 6 | 105548600 | 105549999 | 8.882019 | 16.63279 | 34.06256 | 14.07649 | -0.56672 | 0.125205 | 0.384098 |
| 5 | 129697400 | 129701199 | 21.21933 | 35.13739 | 67.99294 | 29.41929 | -0.56855 | 1.302266 | 0.384132 |
| 18 | 36034000 | 36038399 | 52.95182 | 85.72597 | 67.18565 | 28.74256 | 0.714163 | 1.689276 | 0.384132 |

| | | | | | | | | | |
|---|---|---|---|---|---|---|---|---|---|
| 6 | 78496200 | 78500799 | 36.92907 | 61.74312 | 48.11011 | 20.94963 | 0.739855 | 1.222591 | 0.384134 |
| 10 | 125000600 | 125001799 | 7.623448 | 14.45024 | 31.05383 | 12.54551 | -0.60799 | 0.025652 | 0.384141 |
| 3 | 14643400 | 14644999 | 10.12436 | 18.22945 | 35.33319 | 15.41378 | -0.54369 | 0.369362 | 0.38415 |
| 19 | 20816400 | 20818199 | 25.67431 | 44.09749 | 36.29394 | 15.9195 | 0.663615 | 0.927931 | 0.38416 |
| 15 | 87322400 | 87326799 | 45.5426 | 73.78852 | 69.47776 | 29.74028 | 0.451054 | 1.659651 | 0.384164 |
| X | 148906200 | 148910199 | 51.24191 | 82.79959 | 72.04144 | 30.76165 | 0.560923 | 1.73302 | 0.384172 |
| 18 | 90384600 | 90385799 | 23.24181 | 40.38592 | 37.64319 | 16.52863 | 0.487918 | 0.810012 | 0.384182 |
| 10 | 25655400 | 25659999 | 34.13029 | 56.4545 | 55.30183 | 23.90986 | 0.41027 | 1.337453 | 0.384182 |
| 6 | 13336200 | 13339599 | 37.09545 | 62.36984 | 44.09339 | 19.26298 | 0.879536 | 1.163661 | 0.384208 |
| 12 | 103875200 | 103878799 | 48.90787 | 78.91903 | 75.51495 | 32.18744 | 0.42177 | 1.769816 | 0.384217 |
| X | 25222800 | 25225999 | 22.94347 | 40.84425 | 26.97217 | 11.88244 | 0.990926 | 0.508757 | 0.384232 |
| 3 | 100829000 | 100830599 | 9.217416 | 16.62389 | 34.88601 | 15.18729 | -0.65544 | 0.352586 | 0.384242 |
| 1 | 94882000 | 94883999 | 23.0986 | 40.5307 | 32.48583 | 14.36479 | 0.705184 | 0.660424 | 0.38425 |
| X | 4590600 | 4592599 | 37.86791 | 62.7037 | 52.03422 | 22.51368 | 0.649093 | 1.336666 | 0.384253 |
| 18 | 9917000 | 9917799 | 14.85582 | 26.84537 | 24.80252 | 10.77116 | 0.522674 | 0.357753 | 0.384256 |
| 9 | 69565400 | 69565999 | 3.702072 | 7.230208 | 19.87872 | 6.975049 | -0.76756 | -0.53514 | 0.384267 |
| 2 | 89206000 | 89207999 | 23.41806 | 41.2674 | 28.94678 | 12.81212 | 0.897571 | 0.601002 | 0.384272 |
| 15 | 88030600 | 88033199 | 18.61775 | 31.42988 | 59.07716 | 25.84067 | -0.52595 | 1.081225 | 0.38429 |
| 16 | 52473400 | 52473799 | 3.23057 | 6.319279 | 16.82312 | 5.551948 | -0.63877 | -0.7688 | 0.384294 |
| 10 | 123744000 | 123745399 | 17.69055 | 30.40088 | 53.27555 | 23.36846 | -0.42215 | 0.911639 | 0.384297 |
| 7 | 105411000 | 105412599 | 13.36696 | 23.38272 | 43.62883 | 19.242 | -0.50769 | 0.675936 | 0.384302 |
| 5 | 43004000 | 43006399 | 25.94614 | 44.22835 | 43.14375 | 18.91273 | 0.417878 | 1.005163 | 0.384316 |
| X | 26820800 | 26826199 | 56.41695 | 91.5174 | 60.53262 | 25.93428 | 0.960939 | 1.655178 | 0.384321 |
| 2 | 129365800 | 129371999 | 52.45963 | 83.22189 | 136.879 | 57.00095 | -0.38583 | 2.315804 | 0.384326 |
| 7 | 80397800 | 80399999 | 14.35628 | 24.93181 | 52.17737 | 22.90217 | -0.67378 | 0.798518 | 0.384332 |
| 11 | 44904200 | 44904999 | 14.24903 | 27.27173 | 20.38138 | 8.058408 | 0.931827 | -0.0052 | 0.38434 |
| 16 | 58861200 | 58862399 | 10.52153 | 19.19145 | 36.58322 | 15.70275 | -0.50491 | 0.298542 | 0.384346 |
| 1 | 152055400 | 152059199 | 45.05329 | 72.843 | 71.41512 | 30.51763 | 0.390031 | 1.703839 | 0.384348 |
| 5 | 109252200 | 109261599 | 247.2115 | 364.3109 | 284.0568 | 116.9776 | 0.601752 | 3.792678 | 0.384362 |
| 2 | 4022200 | 4024199 | 14.12168 | 24.67613 | 45.19289 | 19.91353 | -0.48119 | 0.698462 | 0.384363 |
| 6 | 145010400 | 145015999 | 37.43979 | 60.14131 | 107.583 | 45.56582 | -0.48693 | 1.90308 | 0.384363 |
| 4 | 150199000 | 150201999 | 19.44418 | 32.84776 | 58.52877 | 25.60618 | -0.44901 | 1.079744 | 0.384391 |
| 16 | 31555200 | 31557599 | 12.53078 | 22.27009 | 41.38811 | 18.25909 | -0.49506 | 0.478924 | 0.384403 |
| 4 | 115551000 | 115551599 | 7.353925 | 13.98677 | 28.26991 | 11.33911 | -0.51003 | -0.00324 | 0.384408 |
| 11 | 12121000 | 12121599 | 9.499177 | 18.39099 | 18.08221 | 6.878216 | 0.590738 | -0.18769 | 0.384409 |
| 4 | 60444200 | 60445999 | 20.8285 | 36.45455 | 34.37446 | 15.16728 | 0.471921 | 0.690055 | 0.384429 |
| 5 | 24689400 | 24692599 | 13.19035 | 22.72152 | 52.10262 | 22.85564 | -0.80736 | 0.869174 | 0.384443 |
| 17 | 13064800 | 13065799 | 8.112881 | 14.81471 | 33.97716 | 14.56361 | -0.76807 | 0.268254 | 0.384456 |
| 5 | 146695400 | 146703399 | 63.26309 | 99.51106 | 158.9841 | 65.95588 | -0.35403 | 2.524244 | 0.384459 |
| 18 | 79860000 | 79863399 | 48.13713 | 78.25133 | 64.16815 | 27.53386 | 0.654381 | 1.598668 | 0.384462 |
| 2 | 156866800 | 156873999 | 70.13931 | 108.8365 | 186.5233 | 77.46265 | -0.47043 | 2.77562 | 0.38448 |
| 18 | 48584400 | 48585999 | 19.31158 | 33.91648 | 32.22766 | 14.26158 | 0.460709 | 0.622879 | 0.384493 |
| 1 | 139237400 | 139238199 | 4.651299 | 9.058357 | 21.83729 | 7.986397 | -0.63514 | -0.36465 | 0.384504 |
| 9 | 86760200 | 86763199 | 21.53267 | 35.63282 | 69.20827 | 29.93156 | -0.57406 | 1.31027 | 0.384522 |
| 5 | 141857800 | 141861799 | 38.09008 | 61.25467 | 102.5539 | 43.50655 | -0.38954 | 1.867942 | 0.384534 |
| 17 | 50201200 | 50247999 | 470.9607 | 629.3973 | 1080.038 | 450.9823 | -0.67651 | 5.287978 | 0.38455 |
| 18 | 13639800 | 13641199 | 11.57144 | 20.58962 | 37.79084 | 16.65536 | -0.47591 | 0.458875 | 0.384563 |

| | | | | | | | | | |
|---|---|---|---|---|---|---|---|---|---|
| 10 | 85771400 | 85775599 | 48.86822 | 78.73945 | 77.52426 | 32.98038 | 0.378239 | 1.812675 | 0.384568 |
| 12 | 107394200 | 107398599 | 35.29768 | 57.0302 | 95.97715 | 40.97681 | -0.38889 | 1.720411 | 0.384589 |
| 5 | 125681000 | 125697999 | 208.2917 | 301.7941 | 444.015 | 183.0919 | -0.33732 | 4.051834 | 0.384592 |
| 3 | 85949800 | 85950799 | 12.80233 | 23.93826 | 23.45957 | 9.719348 | 0.486328 | 0.143806 | 0.384593 |
| 3 | 3160400 | 3180599 | 270.9086 | 393.7052 | 315.2889 | 129.9665 | 0.544002 | 4.009294 | 0.384595 |
| 19 | 54981800 | 54985599 | 22.15788 | 38.50374 | 37.25034 | 16.35648 | 0.434508 | 0.804789 | 0.384599 |
| 4 | 7869000 | 7869999 | 10.08356 | 18.26966 | 34.87049 | 15.09365 | -0.51439 | 0.331075 | 0.384606 |
| 5 | 149810000 | 149814399 | 42.76378 | 69.53718 | 66.03093 | 28.33624 | 0.443568 | 1.583474 | 0.384608 |
| 1 | 68621600 | 68623799 | 14.42145 | 27.21369 | 22.28798 | 9.089866 | 0.761386 | 0.097683 | 0.384616 |
| 2 | 55045800 | 55046399 | 12.14786 | 23.21824 | 21.12533 | 8.385632 | 0.645268 | -0.01026 | 0.384623 |
| 2 | 177299800 | 177300199 | 5.518849 | 10.71342 | 23.89999 | 8.976138 | -0.56082 | -0.25294 | 0.384636 |
| 14 | 105942600 | 105943399 | 10.607 | 20.5123 | 19.7385 | 7.548528 | 0.614927 | -0.1505 | 0.384637 |
| 17 | 19399000 | 19400399 | 19.55416 | 35.10403 | 27.19945 | 11.89806 | 0.768278 | 0.434592 | 0.384641 |
| X | 32833000 | 32834599 | 20.79448 | 37.8537 | 23.42399 | 10.07954 | 1.105654 | 0.358021 | 0.384691 |
| X | 130082400 | 130084999 | 23.04264 | 40.45944 | 32.02043 | 14.16697 | 0.723432 | 0.65004 | 0.384693 |
| 6 | 63968200 | 63971399 | 39.51545 | 65.33608 | 54.40958 | 23.51765 | 0.643826 | 1.356792 | 0.384705 |
| 7 | 17948400 | 17980999 | 566.1055 | 770.9202 | 635.3955 | 264.5654 | 0.40853 | 4.991586 | 0.384706 |
| 8 | 4689600 | 4698199 | 122.3237 | 188.0851 | 172.0314 | 71.43872 | 0.424405 | 2.962472 | 0.384709 |
| 3 | 33221200 | 33223199 | 21.2934 | 37.57491 | 30.24203 | 13.38174 | 0.701028 | 0.576112 | 0.384721 |
| 8 | 124362000 | 124367599 | 67.71111 | 105.9145 | 169.8559 | 70.46596 | -0.36574 | 2.633309 | 0.384736 |
| 9 | 38827200 | 38833799 | 74.54673 | 118.8737 | 95.3988 | 39.97489 | 0.65541 | 2.168084 | 0.384738 |
| 5 | 55565800 | 55566799 | 12.00462 | 23.2514 | 18.5001 | 7.02345 | 0.896253 | -0.16332 | 0.384789 |
| 13 | 9881400 | 9884799 | 38.65389 | 63.26475 | 61.44135 | 26.42933 | 0.417759 | 1.480462 | 0.384827 |
| 18 | 5725800 | 5726399 | 13.6909 | 25.67612 | 23.43434 | 9.660052 | 0.594161 | 0.133443 | 0.38488 |
| X | 145177000 | 145178799 | 21.42116 | 38.59242 | 25.52805 | 11.12826 | 0.998518 | 0.424645 | 0.384889 |
| 14 | 112184400 | 112184799 | 4.55696 | 8.874582 | 23.04386 | 8.428273 | -0.74253 | -0.36553 | 0.384893 |
| 19 | 16396000 | 16397399 | 14.50901 | 26.79699 | 24.728 | 10.43834 | 0.551884 | 0.232316 | 0.384899 |
| 17 | 9934000 | 9935799 | 19.47916 | 35.78011 | 23.12914 | 9.836392 | 1.054466 | 0.298683 | 0.384914 |
| 3 | 85952600 | 85956599 | 48.52206 | 78.35748 | 73.99401 | 31.57086 | 0.442046 | 1.745431 | 0.384924 |
| 12 | 64032800 | 64035199 | 28.98723 | 49.44781 | 43.03255 | 18.86508 | 0.582087 | 1.011001 | 0.384938 |
| 6 | 37593000 | 37597799 | 46.76046 | 74.69927 | 124.3768 | 52.24937 | -0.39297 | 2.090997 | 0.384956 |
| 4 | 68347200 | 68350799 | 20.74452 | 37.03807 | 27.60906 | 12.13299 | 0.819344 | 0.476591 | 0.384973 |
| 2 | 108778400 | 108779599 | 13.81589 | 26.21308 | 21.90826 | 8.832868 | 0.746867 | 0.055271 | 0.384985 |
| 3 | 109091600 | 109092599 | 13.81589 | 26.21308 | 21.90826 | 8.832868 | 0.746867 | 0.055271 | 0.384985 |
| 11 | 24866600 | 24867799 | 13.81589 | 26.21308 | 21.90826 | 8.832868 | 0.746867 | 0.055271 | 0.384985 |
| 10 | 127894200 | 127895599 | 10.09817 | 17.85258 | 38.98214 | 17.24274 | -0.72909 | 0.50931 | 0.384988 |
| 7 | 5562000 | 5576199 | 233.6361 | 344.6276 | 288.9772 | 119.0051 | 0.498161 | 3.775717 | 0.385012 |
| 5 | 100740600 | 100742999 | 40.94892 | 66.80484 | 63.76595 | 27.39699 | 0.439412 | 1.532181 | 0.385056 |
| 6 | 78952400 | 78957599 | 104.3705 | 161.4594 | 158.4523 | 65.81722 | 0.329593 | 2.841 | 0.385067 |
| 1 | 92190000 | 92194599 | 34.34239 | 55.45215 | 92.97354 | 39.66252 | -0.38451 | 1.73628 | 0.385076 |
| 10 | 101287400 | 101287999 | 8.684132 | 15.91864 | 32.45388 | 13.84066 | -0.5941 | 0.248835 | 0.385083 |
| 3 | 131449200 | 131450399 | 15.40873 | 29.5167 | 19.12724 | 7.543894 | 1.138934 | -0.00498 | 0.385084 |
| 19 | 19486000 | 19486599 | 7.347267 | 13.94546 | 28.22325 | 11.36963 | -0.51765 | 0.009627 | 0.385084 |
| X | 125810000 | 125810999 | 18.55461 | 32.91736 | 29.62429 | 13.08884 | 0.544053 | 0.506641 | 0.385125 |
| 14 | 21001000 | 21004799 | 42.57367 | 69.76087 | 59.4417 | 25.5815 | 0.606742 | 1.475549 | 0.385133 |
| 9 | 56655800 | 56659199 | 28.80912 | 48.42073 | 47.50043 | 20.7492 | 0.408585 | 1.142317 | 0.385135 |
| 3 | 144940200 | 144940599 | 4.804712 | 9.373641 | 22.21267 | 7.938228 | -0.58168 | -0.4475 | 0.385143 |

| | | | | | | | | | |
|---|---|---|---|---|---|---|---|---|---|
| 12 | 37652800 | 37653399 | 4.804712 | 9.373641 | 22.21267 | 7.938228 | -0.58168 | -0.4475 | 0.385143 |
| 13 | 100358000 | 100362999 | 58.22249 | 92.06064 | 147.2886 | 61.19908 | -0.35015 | 2.406228 | 0.385145 |
| 4 | 53269400 | 53271599 | 15.61679 | 26.94995 | 50.04133 | 21.95263 | -0.50382 | 0.861172 | 0.385172 |
| 19 | 32348400 | 32350199 | 15.61989 | 27.21673 | 48.9488 | 21.53007 | -0.45559 | 0.753501 | 0.385184 |
| 8 | 4181800 | 4188599 | 43.6993 | 69.7185 | 124.9958 | 52.413 | -0.50153 | 2.12756 | 0.385213 |
| 1 | 136497200 | 136499199 | 14.26031 | 24.87498 | 48.40244 | 21.30443 | -0.56866 | 0.726804 | 0.385224 |
| 14 | 45766600 | 45768199 | 11.93342 | 21.0081 | 40.41379 | 17.89111 | -0.55048 | 0.566705 | 0.38523 |
| X | 81430600 | 81434399 | 46.07356 | 76.02834 | 50.66018 | 21.8672 | 0.964325 | 1.394126 | 0.385236 |
| 2 | 123094800 | 123096399 | 14.45216 | 26.89463 | 23.68677 | 9.893045 | 0.629754 | 0.186819 | 0.385246 |
| 6 | 107878000 | 107878599 | 6.441566 | 12.32796 | 26.75703 | 10.58116 | -0.59346 | -0.06418 | 0.385246 |
| 4 | 60483200 | 60484799 | 12.62183 | 23.87937 | 22.49228 | 9.123208 | 0.568063 | 0.064544 | 0.385248 |
| Y | 10608000 | 10610199 | 22.9204 | 39.8814 | 22.30806 | 9.80352 | 1.220708 | 0.762641 | 0.38525 |
| 11 | 6655800 | 6665799 | 132.5835 | 197.7557 | 320.0291 | 131.8993 | -0.43499 | 3.572027 | 0.385251 |
| 7 | 79906400 | 79911399 | 40.25189 | 64.49028 | 110.2047 | 46.49177 | -0.42554 | 1.990182 | 0.385269 |
| X | 78135400 | 78140199 | 84.59502 | 133.8855 | 101.0163 | 41.99256 | 0.733798 | 2.396416 | 0.385291 |
| 7 | 5017400 | 5018999 | 9.707071 | 17.39134 | 35.87164 | 15.71822 | -0.63708 | 0.399484 | 0.385303 |
| 16 | 52321600 | 52322599 | 16.23389 | 29.2291 | 27.61073 | 12.04539 | 0.486666 | 0.394657 | 0.385316 |
| 7 | 127002200 | 127005999 | 49.03185 | 77.86325 | 133.2847 | 55.56072 | -0.4419 | 2.282542 | 0.385324 |
| 13 | 3499000 | 3501599 | 44.97459 | 73.23509 | 58.33994 | 25.0486 | 0.697704 | 1.563187 | 0.385331 |
| 12 | 51180400 | 51181999 | 17.76042 | 30.26228 | 54.80427 | 24.0235 | -0.47129 | 0.992016 | 0.385337 |
| 7 | 59978200 | 59982599 | 51.9543 | 85.01127 | 53.05462 | 22.81956 | 1.053747 | 1.511275 | 0.38534 |
| X | 90887800 | 90890199 | 34.25526 | 57.29768 | 44.86356 | 19.5413 | 0.732851 | 1.207328 | 0.385346 |
| 13 | 120598600 | 120599799 | 19.47558 | 34.08255 | 32.51685 | 14.34618 | 0.455101 | 0.681609 | 0.38536 |
| 2 | 17946800 | 17948799 | 10.91935 | 19.48347 | 37.80757 | 16.62368 | -0.55351 | 0.435479 | 0.385374 |
| 19 | 25482400 | 25484399 | 15.78956 | 26.78911 | 56.26242 | 24.64686 | -0.68477 | 1.026031 | 0.385393 |
| 5 | 87419400 | 87421399 | 19.75837 | 34.7648 | 32.17198 | 14.244 | 0.499271 | 0.602744 | 0.385398 |
| 11 | 57485400 | 57490599 | 39.37421 | 63.32289 | 106.7758 | 45.30088 | -0.39977 | 1.869171 | 0.385399 |
| 18 | 37903200 | 37911799 | 170.8936 | 257.3465 | 227.6594 | 94.03362 | 0.448129 | 3.394546 | 0.385408 |
| 10 | 86428800 | 86429999 | 10.76293 | 19.24796 | 36.80865 | 16.15357 | -0.5305 | 0.418904 | 0.385412 |
| 3 | 87715800 | 87719599 | 32.26899 | 52.01321 | 96.12243 | 40.91344 | -0.52751 | 1.784652 | 0.385417 |
| 19 | 39340400 | 39348199 | 195.9904 | 293.3446 | 241.3772 | 99.55005 | 0.544707 | 3.514815 | 0.385417 |
| 11 | 40800200 | 40802399 | 31.06171 | 50.45373 | 85.40556 | 36.62639 | -0.38966 | 1.593304 | 0.385427 |
| 1 | 126611400 | 126614599 | 19.81441 | 33.61649 | 60.91718 | 26.68565 | -0.47323 | 1.042946 | 0.385442 |
| 13 | 69884200 | 69895599 | 93.83177 | 143.5501 | 230.96 | 95.829 | -0.39525 | 3.056977 | 0.385444 |
| 11 | 25030200 | 25031199 | 18.07252 | 31.8848 | 26.5558 | 11.7511 | 0.652076 | 0.553786 | 0.38545 |
| 11 | 6754600 | 6759799 | 58.30914 | 93.64971 | 83.47552 | 35.34679 | 0.515763 | 1.928279 | 0.385472 |
| 4 | 57897200 | 57899599 | 14.47757 | 25.34731 | 45.14703 | 19.92607 | -0.44112 | 0.670242 | 0.385475 |
| 2 | 70673600 | 70675399 | 10.09226 | 18.50535 | 35.62717 | 15.18781 | -0.51195 | 0.265538 | 0.385482 |
| 17 | 88212400 | 88215199 | 25.38209 | 41.85982 | 72.34685 | 31.22042 | -0.40703 | 1.363039 | 0.385485 |
| 12 | 4283600 | 4285199 | 11.16167 | 19.71929 | 38.90254 | 17.20988 | -0.58434 | 0.52127 | 0.385489 |
| 2 | 98353000 | 98354599 | 26.87893 | 46.60379 | 33.20756 | 14.56991 | 0.873137 | 0.845347 | 0.385499 |
| 3 | 106532200 | 106533999 | 19.74256 | 34.80492 | 31.5802 | 13.97654 | 0.528598 | 0.580391 | 0.385505 |
| X | 117818400 | 117821399 | 38.78589 | 65.22919 | 42.97556 | 18.77614 | 0.980963 | 1.167739 | 0.385513 |
| 3 | 52136400 | 52137999 | 29.11013 | 48.71699 | 47.03344 | 20.49317 | 0.43165 | 1.186799 | 0.385514 |
| 17 | 71136000 | 71140799 | 32.0482 | 51.81664 | 90.7495 | 38.78024 | -0.44502 | 1.691455 | 0.385516 |
| 9 | 25010000 | 25013999 | 46.44239 | 75.09708 | 72.1023 | 30.8156 | 0.42031 | 1.704552 | 0.385533 |
| 13 | 63857400 | 63859399 | 14.27676 | 24.91037 | 46.26938 | 20.36899 | -0.50171 | 0.721323 | 0.385539 |

| | | | | | | | | | |
|---|---|---|---|---|---|---|---|---|---|
| 7 | 45952000 | 45953399 | 11.14673 | 19.77645 | 37.84398 | 16.71963 | -0.53793 | 0.476242 | 0.385546 |
| 13 | 66244800 | 66247199 | 44.00897 | 71.84636 | 56.66533 | 24.35231 | 0.714626 | 1.524564 | 0.385549 |
| X | 154404400 | 154408199 | 47.50327 | 77.08255 | 64.79589 | 27.76734 | 0.616423 | 1.635278 | 0.38556 |
| 13 | 68437400 | 68456999 | 480.6446 | 660.8554 | 579.5858 | 240.9645 | 0.332919 | 4.840401 | 0.385563 |
| 5 | 119168400 | 119172999 | 34.17761 | 55.16353 | 93.43556 | 39.83862 | -0.39984 | 1.747768 | 0.385567 |
| 18 | 67272800 | 67274399 | 9.458568 | 17.19421 | 34.40675 | 14.83128 | -0.57822 | 0.305767 | 0.385567 |
| 6 | 5073200 | 5074599 | 18.57118 | 33.82733 | 25.28747 | 10.87302 | 0.835505 | 0.341786 | 0.385578 |
| 10 | 3580600 | 3582199 | 15.78572 | 28.342 | 27.25639 | 11.92185 | 0.458585 | 0.408544 | 0.38559 |
| 13 | 65549800 | 65550999 | 28.18621 | 47.52979 | 41.28112 | 18.05046 | 0.583875 | 1.107296 | 0.385612 |
| 1 | 32479400 | 32483199 | 34.45739 | 57.72597 | 48.62312 | 21.19976 | 0.627804 | 1.194795 | 0.385616 |
| 18 | 37349800 | 37351599 | 14.61463 | 26.61838 | 25.81123 | 11.09933 | 0.46252 | 0.314145 | 0.385668 |
| 3 | 66911400 | 66911799 | 3.881636 | 7.571185 | 20.80415 | 7.455477 | -0.79277 | -0.44821 | 0.385706 |
| X | 29332200 | 29333199 | 22.11541 | 38.5302 | 35.67006 | 15.68826 | 0.498177 | 0.767397 | 0.385717 |
| 2 | 76781600 | 76783799 | 21.3136 | 35.40816 | 67.74744 | 29.37879 | -0.55205 | 1.262081 | 0.38572 |
| 1 | 67922800 | 67928999 | 59.11397 | 95.09491 | 78.46957 | 33.30478 | 0.629699 | 1.879779 | 0.385741 |
| 3 | 18651000 | 18654199 | 93.62029 | 146.024 | 143.5035 | 59.55098 | 0.336734 | 2.691849 | 0.385744 |
| X | 25518000 | 25520599 | 23.45496 | 42.4366 | 22.75446 | 9.866575 | 1.304361 | 0.410797 | 0.385772 |
| X | 8898200 | 8901399 | 27.16405 | 46.83688 | 38.77349 | 17.00909 | 0.656508 | 0.906088 | 0.385782 |
| 8 | 123499200 | 123500199 | 4.986719 | 9.651655 | 24.29505 | 9.254639 | -0.753 | -0.21139 | 0.385786 |
| X | 58611000 | 58613999 | 38.33501 | 63.13218 | 55.30355 | 23.8513 | 0.569762 | 1.403161 | 0.385829 |
| 10 | 28944000 | 28945599 | 18.14641 | 32.1696 | 29.44113 | 13.01535 | 0.519086 | 0.505696 | 0.385843 |
| 1 | 104024600 | 104026399 | 34.85614 | 57.47652 | 55.91407 | 24.12767 | 0.419536 | 1.374177 | 0.385867 |
| 13 | 111708800 | 111718599 | 96.07386 | 147.038 | 228.8625 | 94.96733 | -0.34705 | 3.049971 | 0.385872 |
| 17 | 13734600 | 13736799 | 31.95422 | 53.31019 | 51.38763 | 22.34635 | 0.433981 | 1.232565 | 0.385877 |
| 8 | 41179200 | 41181599 | 26.96324 | 46.42254 | 39.68673 | 17.40894 | 0.609647 | 0.915311 | 0.385878 |
| 7 | 13788800 | 13798199 | 165.6333 | 249.8913 | 222.4717 | 91.93941 | 0.441327 | 3.356743 | 0.385891 |
| 6 | 130954800 | 130963399 | 160.4664 | 242.4004 | 220.198 | 91.03588 | 0.413799 | 3.329949 | 0.385904 |
| 16 | 58228200 | 58229799 | 12.87697 | 22.55954 | 43.0676 | 19.01867 | -0.54059 | 0.646769 | 0.385911 |
| 10 | 17897800 | 17902999 | 54.41244 | 86.44773 | 138.372 | 57.70815 | -0.34461 | 2.274598 | 0.385921 |
| 5 | 122555800 | 122557599 | 14.2369 | 24.64037 | 52.67355 | 23.10941 | -0.7051 | 0.833751 | 0.385924 |
| 7 | 84833400 | 84840199 | 127.0688 | 195.0046 | 176.3545 | 73.21458 | 0.438601 | 3.002498 | 0.385932 |
| 12 | 23759200 | 23794199 | 1154.925 | 1475.21 | 1188.537 | 498.7521 | 0.348856 | 6.022795 | 0.385944 |
| 2 | 68709000 | 68709799 | 14.5669 | 27.2843 | 23.27185 | 9.613719 | 0.688434 | 0.147458 | 0.385949 |
| 18 | 18948400 | 18951999 | 37.78272 | 62.74067 | 47.1351 | 20.43402 | 0.792237 | 1.296033 | 0.38596 |
| 5 | 100418800 | 100422799 | 48.64123 | 77.48043 | 125.9233 | 52.68663 | -0.36261 | 2.18535 | 0.385978 |
| 14 | 42290400 | 42291399 | 18.32384 | 32.95422 | 27.29509 | 11.92191 | 0.673984 | 0.414889 | 0.385981 |
| 18 | 84112600 | 84113399 | 4.442812 | 8.633778 | 25.15575 | 9.366694 | -0.93123 | -0.29728 | 0.385989 |
| 12 | 19333200 | 19340799 | 193.6345 | 286.9049 | 260.9576 | 107.4741 | 0.386924 | 3.695284 | 0.386011 |
| 14 | 36604600 | 36605199 | 4.779565 | 9.320418 | 21.46317 | 7.715375 | -0.54737 | -0.42621 | 0.386014 |
| 6 | 92258000 | 92259599 | 11.08626 | 19.9514 | 36.85461 | 16.08757 | -0.47535 | 0.383411 | 0.386022 |
| 7 | 10478600 | 10484599 | 43.09705 | 70.34708 | 57.09887 | 24.53728 | 0.673198 | 1.525984 | 0.386028 |
| 17 | 12897000 | 12899999 | 39.80819 | 64.68434 | 62.74639 | 26.9162 | 0.412063 | 1.588201 | 0.386031 |
| 7 | 119899200 | 119901799 | 24.64865 | 40.48787 | 81.76305 | 35.22417 | -0.63334 | 1.417363 | 0.386051 |
| X | 11953400 | 11958399 | 28.29018 | 46.30685 | 86.47972 | 37.201 | -0.52294 | 1.475015 | 0.386053 |
| 8 | 79193200 | 79195199 | 21.49554 | 37.61551 | 33.69303 | 14.86444 | 0.545546 | 0.693649 | 0.386068 |
| 7 | 26983600 | 26986599 | 17.36462 | 29.37281 | 57.38193 | 25.11433 | -0.58112 | 1.058846 | 0.386077 |
| 12 | 105968000 | 105970999 | 29.24354 | 47.5582 | 82.51113 | 35.4101 | -0.42361 | 1.565092 | 0.386078 |

| | | | | | | | | | |
|---|---|---|---|---|---|---|---|---|---|
| 11 | 107158000 | 107165599 | 54.05989 | 85.87343 | 136.9768 | 57.11683 | -0.33985 | 2.278358 | 0.386083 |
| 4 | 15015200 | 15016599 | 16.16887 | 30.36918 | 21.44844 | 8.818833 | 0.963848 | 0.139617 | 0.386085 |
| 1 | 185213600 | 185215199 | 6.942998 | 13.06959 | 29.12462 | 11.93019 | -0.67638 | 0.064283 | 0.386085 |
| 2 | 122607000 | 122607799 | 6.942998 | 13.06959 | 29.12462 | 11.93019 | -0.67638 | 0.064283 | 0.386085 |
| 15 | 102043800 | 102044999 | 6.942998 | 13.06959 | 29.12462 | 11.93019 | -0.67638 | 0.064283 | 0.386085 |
| 9 | 22105200 | 22115599 | 163.2419 | 245.7737 | 231.1986 | 95.49025 | 0.359289 | 3.396494 | 0.386089 |
| 8 | 129828400 | 129832199 | 52.064 | 83.94967 | 76.39388 | 32.531 | 0.492859 | 1.79669 | 0.386102 |
| 16 | 14250000 | 14251399 | 6.58751 | 12.60468 | 26.75163 | 10.58526 | -0.56215 | -0.06026 | 0.38611 |
| 8 | 67805600 | 67810799 | 70.40985 | 111.6111 | 109.9393 | 45.7467 | 0.351594 | 2.345288 | 0.386111 |
| X | 73314600 | 73317599 | 39.46239 | 63.43863 | 105.9696 | 44.93207 | -0.38692 | 1.884507 | 0.386125 |
| 2 | 68041600 | 68042599 | 8.487844 | 15.78509 | 31.0559 | 12.98517 | -0.52209 | 0.162798 | 0.386127 |
| 1 | 48127400 | 48128799 | 12.92917 | 24.37779 | 19.39649 | 7.923377 | 0.801058 | 0.082971 | 0.38614 |
| 2 | 7598000 | 7599199 | 22.37164 | 39.00029 | 31.96944 | 14.06604 | 0.67279 | 0.752653 | 0.386149 |
| 4 | 153395800 | 153409999 | 152.9548 | 226.5391 | 348.6744 | 143.5955 | -0.37205 | 3.700067 | 0.386159 |
| 5 | 65264600 | 65265599 | 6.665993 | 12.52546 | 29.09823 | 11.95598 | -0.73922 | 0.074243 | 0.386173 |
| 17 | 34188400 | 34189999 | 27.7336 | 46.8517 | 45.85276 | 20.0605 | 0.412176 | 1.092304 | 0.386179 |
| 7 | 78580000 | 78580999 | 12.02384 | 22.57377 | 22.04639 | 9.0715 | 0.499318 | 0.108679 | 0.386179 |
| 9 | 84164800 | 84165599 | 12.02384 | 22.57377 | 22.04639 | 9.0715 | 0.499318 | 0.108679 | 0.386179 |
| 14 | 61944000 | 61944799 | 12.02384 | 22.57377 | 22.04639 | 9.0715 | 0.499318 | 0.108679 | 0.386179 |
| 13 | 66912200 | 66920999 | 154.7641 | 234.085 | 194.7231 | 80.53627 | 0.542272 | 3.301792 | 0.386198 |
| 9 | 41348000 | 41352599 | 58.32182 | 93.86843 | 77.5572 | 32.94186 | 0.628721 | 1.86124 | 0.386222 |
| 1 | 101007800 | 101010599 | 27.40308 | 47.33653 | 38.57413 | 16.9226 | 0.679501 | 0.887961 | 0.386228 |
| 2 | 51342600 | 51343199 | 4.817202 | 9.361519 | 22.81744 | 8.494672 | -0.67517 | -0.29708 | 0.386228 |
| 17 | 31588800 | 31589999 | 11.51123 | 22.0561 | 21.13505 | 8.329943 | 0.580202 | -0.0409 | 0.386233 |
| 9 | 120219200 | 120221399 | 13.67612 | 23.90516 | 44.15834 | 19.46289 | -0.49335 | 0.690617 | 0.386235 |
| 6 | 52776400 | 52777199 | 7.019751 | 13.44568 | 27.37435 | 10.79944 | -0.49898 | -0.06465 | 0.386242 |
| 12 | 19358800 | 19367199 | 218.0888 | 320.188 | 295.3338 | 121.6363 | 0.35433 | 3.847989 | 0.386247 |
| 4 | 43201400 | 43203799 | 29.28634 | 49.87856 | 42.98569 | 18.84211 | 0.595686 | 1.024114 | 0.386273 |
| 14 | 6959600 | 6960399 | 14.82993 | 25.7316 | 47.92508 | 21.02773 | -0.50636 | 0.807209 | 0.386275 |
| 7 | 143012600 | 143020399 | 178.2375 | 259.4881 | 416.4086 | 171.6178 | -0.45581 | 3.977131 | 0.386303 |
| 2 | 28283200 | 28285399 | 12.74832 | 22.33533 | 42.57724 | 18.80296 | -0.53831 | 0.646141 | 0.386311 |
| 6 | 135124800 | 135126799 | 11.04165 | 19.40123 | 41.00918 | 18.15477 | -0.68634 | 0.586227 | 0.386314 |
| 10 | 44044200 | 44046599 | 22.48059 | 37.20148 | 68.73408 | 29.72649 | -0.50223 | 1.312902 | 0.386327 |
| X | 45447800 | 45451399 | 35.38455 | 58.36108 | 56.37373 | 24.32835 | 0.429941 | 1.376142 | 0.386335 |
| 10 | 70232200 | 70235999 | 61.78106 | 98.59934 | 96.32797 | 40.41111 | 0.373374 | 2.12734 | 0.386338 |
| 13 | 116025000 | 116026999 | 15.50043 | 28.00527 | 26.79108 | 11.63732 | 0.47272 | 0.369256 | 0.386338 |
| Y | 5992800 | 5995599 | 18.76588 | 33.20126 | 18.02008 | 7.969514 | 1.266762 | 0.505513 | 0.386346 |
| 5 | 41050800 | 41056199 | 69.23707 | 110.4 | 99.01198 | 41.48536 | 0.495258 | 2.165562 | 0.386351 |
| 12 | 14543800 | 14544999 | 21.43638 | 37.43543 | 35.06784 | 15.44629 | 0.481487 | 0.729878 | 0.386362 |
| 4 | 146123200 | 146125399 | 48.43421 | 78.29226 | 64.64047 | 27.61487 | 0.636998 | 1.714648 | 0.386364 |
| 19 | 13537400 | 13539399 | 33.6332 | 56.13904 | 50.34056 | 21.89786 | 0.537862 | 1.229897 | 0.386406 |
| 1 | 78777400 | 78780399 | 34.50365 | 55.69392 | 93.51053 | 39.87441 | -0.3871 | 1.746335 | 0.386412 |
| X | 15828200 | 15836799 | 260.2255 | 378.618 | 337.5052 | 139.1076 | 0.391086 | 3.992111 | 0.386415 |
| 7 | 60064200 | 60065599 | 27.06767 | 46.05504 | 43.17721 | 18.92458 | 0.474631 | 1.027833 | 0.386421 |
| 7 | 67496000 | 67501799 | 46.66955 | 74.36384 | 124.414 | 52.07846 | -0.40385 | 2.172228 | 0.386429 |
| 15 | 100501200 | 100504199 | 13.5695 | 23.58168 | 46.86411 | 20.57893 | -0.5991 | 0.778497 | 0.386431 |
| 7 | 27628200 | 27629199 | 6.126994 | 11.81144 | 27.65689 | 10.71119 | -0.67425 | -0.13626 | 0.38645 |

| | | | | | | | | | |
|---|---|---|---|---|---|---|---|---|---|
| 18 | 9583800 | 9590199 | 91.55578 | 143.9397 | 129.288 | 53.63255 | 0.475159 | 2.547595 | 0.386473 |
| 1 | 149389200 | 149391999 | 41.13331 | 66.72235 | 65.52493 | 28.07238 | 0.392052 | 1.626768 | 0.386474 |
| 7 | 143466400 | 143471199 | 35.49116 | 57.25551 | 95.2593 | 40.59049 | -0.37492 | 1.765525 | 0.386493 |
| 9 | 62822800 | 62823599 | 16.37427 | 30.66 | 22.62374 | 9.350929 | 0.89558 | 0.164255 | 0.386499 |
| X | 128338400 | 128339999 | 16.37427 | 30.66 | 22.62374 | 9.350929 | 0.89558 | 0.164255 | 0.386499 |
| 18 | 80232200 | 80234199 | 12.20115 | 21.61779 | 39.27958 | 17.36669 | -0.46503 | 0.494852 | 0.386505 |
| 7 | 11057800 | 11067799 | 148.3153 | 225.3752 | 186.3891 | 77.19947 | 0.556235 | 3.206154 | 0.386505 |
| 13 | 93430000 | 93431399 | 9.105493 | 16.32846 | 35.89814 | 15.71771 | -0.7277 | 0.389252 | 0.386513 |
| 7 | 96600400 | 96601199 | 8.172492 | 15.21068 | 30.54027 | 12.75244 | -0.54963 | 0.153174 | 0.386522 |
| 12 | 114204400 | 114209799 | 119.8559 | 183.7267 | 175.7454 | 72.95035 | 0.356474 | 3.022514 | 0.386533 |
| 5 | 49680400 | 49683999 | 29.85082 | 48.61244 | 84.12386 | 36.12027 | -0.41857 | 1.546412 | 0.386537 |
| X | 119291200 | 119296399 | 68.24174 | 108.8431 | 96.46282 | 40.43277 | 0.51281 | 2.156887 | 0.386544 |
| 2 | 31643200 | 31644999 | 21.17271 | 37.47148 | 29.81355 | 13.18442 | 0.719301 | 0.545449 | 0.386548 |
| 6 | 53944400 | 53948199 | 29.60255 | 47.98 | 89.1841 | 38.18605 | -0.52727 | 1.633174 | 0.386553 |
| 9 | 18072400 | 18072599 | 6.515252 | 12.7421 | 12.45791 | 4.078476 | 0.792854 | -0.76071 | 0.386568 |
| 14 | 44839800 | 44841599 | 15.58351 | 27.18917 | 48.93896 | 21.54359 | -0.45664 | 0.733492 | 0.386587 |
| 4 | 140658600 | 140668199 | 112.3468 | 169.5404 | 273.8027 | 113.1616 | -0.41692 | 3.342609 | 0.386617 |
| 7 | 14660000 | 14661999 | 38.54527 | 64.14582 | 49.87941 | 21.65611 | 0.74296 | 1.276762 | 0.386639 |
| 1 | 166815200 | 166815599 | 5.993433 | 11.51838 | 25.75123 | 10.06189 | -0.61881 | -0.10949 | 0.38666 |
| 5 | 21857200 | 21860599 | 17.77195 | 30.17836 | 56.9398 | 24.95008 | -0.531 | 1.024506 | 0.386683 |
| 10 | 22810400 | 22813199 | 31.13974 | 52.86694 | 43.51649 | 19.06699 | 0.661161 | 1.059788 | 0.386702 |
| 5 | 105046200 | 105052599 | 97.45624 | 152.7166 | 134.2261 | 55.67438 | 0.502815 | 2.611761 | 0.386729 |
| 13 | 108545600 | 108545999 | 8.866719 | 17.21211 | 18.26802 | 6.862001 | 0.499322 | -0.23399 | 0.386746 |
| 14 | 24885400 | 24886199 | 11.31101 | 21.49734 | 21.57329 | 8.666351 | 0.489473 | 0.018837 | 0.386763 |
| 8 | 20664600 | 20664999 | 8.287412 | 16.12557 | 17.28311 | 6.369475 | 0.510707 | -0.30968 | 0.386796 |
| 7 | 96310200 | 96311199 | 6.602647 | 12.57797 | 27.75032 | 11.09313 | -0.63128 | -0.02524 | 0.386802 |
| 1 | 159287200 | 159288999 | 30.33807 | 50.70406 | 49.71113 | 21.6416 | 0.409578 | 1.208061 | 0.386811 |
| 4 | 52774600 | 52775399 | 12.42773 | 23.30837 | 22.95409 | 9.461245 | 0.485112 | 0.11971 | 0.386818 |
| 17 | 13749800 | 13751599 | 11.20882 | 20.13624 | 36.814 | 16.09442 | -0.46225 | 0.39369 | 0.386819 |
| 4 | 37957400 | 37963399 | 79.85382 | 126.7233 | 109.3094 | 45.53561 | 0.544383 | 2.32216 | 0.386825 |
| 7 | 14505000 | 14532799 | 624.258 | 845.2527 | 660.4764 | 275.2517 | 0.477131 | 5.082885 | 0.386837 |
| 11 | 3716600 | 3722399 | 41.08668 | 65.72913 | 113.8157 | 47.90943 | -0.44706 | 2.038772 | 0.386839 |
| 8 | 98734800 | 98737399 | 24.9475 | 43.54002 | 34.66417 | 15.26253 | 0.714836 | 0.75407 | 0.386846 |
| 17 | 70119600 | 70122999 | 40.50078 | 66.55355 | 56.2484 | 24.23641 | 0.619915 | 1.429049 | 0.386847 |
| 4 | 73305000 | 73308799 | 63.84356 | 102.0917 | 92.75405 | 39.02042 | 0.481042 | 2.069969 | 0.386852 |
| 7 | 14014800 | 14020199 | 96.34323 | 151.6137 | 123.926 | 51.41476 | 0.612103 | 2.530692 | 0.386852 |
| 8 | 111211400 | 111213599 | 20.38284 | 33.89818 | 65.3122 | 28.34369 | -0.56176 | 1.247743 | 0.386863 |
| 7 | 33529200 | 33540599 | 383.2722 | 536.4982 | 479.4827 | 198.786 | 0.331761 | 4.566238 | 0.386864 |
| 19 | 28524400 | 28526199 | 16.72956 | 30.03845 | 28.07033 | 12.27722 | 0.499707 | 0.415721 | 0.386868 |
| X | 60482200 | 60484599 | 29.0976 | 49.94523 | 40.28986 | 17.67154 | 0.692432 | 0.954617 | 0.386868 |
| 2 | 168212000 | 168227199 | 197.4589 | 287.388 | 425.67 | 175.4401 | -0.34069 | 3.983018 | 0.386874 |
| 5 | 148335800 | 148337199 | 11.01513 | 19.46154 | 38.92297 | 17.21871 | -0.60383 | 0.520486 | 0.386885 |
| 6 | 82138600 | 82139199 | 11.69467 | 20.46027 | 43.68909 | 19.27086 | -0.70083 | 0.667647 | 0.386891 |
| Y | 6588200 | 6590599 | 15.60269 | 28.41611 | 15.43876 | 6.639514 | 1.291501 | 0.293831 | 0.38691 |
| 13 | 62362200 | 62363599 | 29.46317 | 49.39703 | 48.60086 | 21.20029 | 0.404384 | 1.170439 | 0.386917 |
| 10 | 113165800 | 113166599 | 9.854495 | 18.12717 | 35.16727 | 14.92676 | -0.5187 | 0.241887 | 0.386925 |
| 4 | 121548600 | 121553999 | 108.9862 | 168.6442 | 159.3037 | 66.17077 | 0.384908 | 2.838338 | 0.386926 |

| | | | | | | | | | |
|---|---|---|---|---|---|---|---|---|---|
| 8 | 122426400 | 122430199 | 47.76017 | 75.81819 | 134.1123 | 55.88682 | -0.48961 | 2.29228 | 0.386932 |
| 6 | 50322200 | 50325199 | 16.46758 | 28.28601 | 51.61752 | 22.64087 | -0.48024 | 0.910669 | 0.38694 |
| 5 | 73959000 | 73962199 | 30.37387 | 49.4329 | 83.34636 | 35.77816 | -0.38173 | 1.557757 | 0.386942 |
| 12 | 73860200 | 73862999 | 14.13708 | 24.62157 | 46.82741 | 20.59097 | -0.53593 | 0.747676 | 0.386944 |
| 2 | 78335000 | 78335599 | 6.905449 | 13.11853 | 27.6858 | 11.13216 | -0.57497 | -0.00189 | 0.386947 |
| 5 | 93779600 | 93780999 | 44.06539 | 71.36354 | 68.32407 | 29.23689 | 0.426526 | 1.667903 | 0.386961 |
| 6 | 6862600 | 6865399 | 29.43396 | 47.76145 | 85.09464 | 36.46585 | -0.46469 | 1.608702 | 0.386963 |
| 4 | 24468800 | 24473199 | 50.90472 | 82.23532 | 71.62642 | 30.57513 | 0.558969 | 1.73685 | 0.386985 |
| 13 | 3313400 | 3344199 | 62.71414 | 100.5255 | 78.44029 | 33.1149 | 0.703985 | 1.990742 | 0.386986 |
| 3 | 50804000 | 50809399 | 40.77447 | 65.24844 | 125.3357 | 52.78015 | -0.59607 | 2.032954 | 0.387008 |
| 3 | 10285400 | 10289199 | 55.31941 | 88.64854 | 87.92677 | 37.10463 | 0.358129 | 1.994624 | 0.387014 |
| 2 | 116805600 | 116807199 | 7.171786 | 13.27697 | 31.53945 | 13.27101 | -0.79866 | 0.177318 | 0.387017 |
| 13 | 50618600 | 50619399 | 7.171786 | 13.27697 | 31.53945 | 13.27101 | -0.79866 | 0.177318 | 0.387017 |
| 10 | 128877800 | 128879399 | 36.78008 | 60.17311 | 53.52522 | 23.02071 | 0.543718 | 1.476694 | 0.387024 |
| 11 | 83653800 | 83654999 | 16.87058 | 30.53003 | 27.14916 | 11.76907 | 0.579351 | 0.371015 | 0.387039 |
| 16 | 67414000 | 67418199 | 49.49652 | 78.61675 | 133.2429 | 55.55435 | -0.42731 | 2.27654 | 0.387059 |
| 8 | 120732800 | 120735199 | 11.60265 | 20.55128 | 38.3179 | 16.94251 | -0.50198 | 0.493842 | 0.387062 |
| 8 | 43464800 | 43467399 | 20.96522 | 36.85765 | 31.96045 | 14.1527 | 0.592084 | 0.617065 | 0.387078 |
| 18 | 19557000 | 19559599 | 27.3615 | 46.43219 | 44.72398 | 19.59346 | 0.43526 | 1.050951 | 0.387079 |
| 3 | 43441400 | 43442199 | 15.7073 | 29.45756 | 21.40516 | 8.823652 | 0.919938 | 0.144054 | 0.387079 |
| X | 159377000 | 159377999 | 13.33546 | 25.83867 | 17.88824 | 6.779365 | 1.097914 | -0.15485 | 0.387091 |
| 15 | 53377000 | 53379399 | 21.33377 | 37.46978 | 32.52816 | 14.39513 | 0.590443 | 0.63461 | 0.387095 |
| 18 | 75309400 | 75311999 | 19.17712 | 32.59915 | 57.56697 | 25.22973 | -0.43577 | 1.020792 | 0.387101 |
| X | 5288200 | 5295399 | 81.23446 | 129.6401 | 86.23242 | 36.17357 | 0.926981 | 2.14999 | 0.387133 |
| 3 | 48032600 | 48033199 | 6.573839 | 12.64499 | 26.36559 | 10.27584 | -0.51602 | -0.11016 | 0.38715 |
| 11 | 79100400 | 79103799 | 20.93316 | 35.10182 | 61.78179 | 26.95285 | -0.43175 | 1.155966 | 0.387154 |
| 11 | 79762400 | 79766399 | 30.04933 | 48.96036 | 82.80745 | 35.56156 | -0.38504 | 1.540833 | 0.3872 |
| 10 | 129288200 | 129289599 | 21.56033 | 37.70056 | 30.85501 | 13.60375 | 0.675282 | 0.702965 | 0.38721 |
| 13 | 56536400 | 56541199 | 55.12272 | 87.16539 | 145.4775 | 60.44866 | -0.41094 | 2.40195 | 0.387244 |
| 5 | 85970400 | 85971199 | 12.47099 | 23.24202 | 22.49385 | 9.371383 | 0.497951 | 0.159527 | 0.387251 |
| 2 | 51683400 | 51686199 | 36.47055 | 58.77071 | 97.48254 | 41.47834 | -0.3724 | 1.800084 | 0.387265 |
| 7 | 12727400 | 12728599 | 9.051054 | 16.7585 | 34.23611 | 14.4028 | -0.58344 | 0.19563 | 0.387266 |
| 5 | 140653000 | 140657199 | 61.04034 | 95.43583 | 169.9968 | 70.52778 | -0.51752 | 2.638604 | 0.38727 |
| 12 | 13558200 | 13559399 | 11.07578 | 19.97219 | 36.88215 | 16.05962 | -0.47264 | 0.3716 | 0.387283 |
| 14 | 99759000 | 99759999 | 10.2582 | 18.68978 | 34.96095 | 15.02786 | -0.4788 | 0.302084 | 0.387292 |
| 12 | 5153000 | 5157799 | 30.05569 | 48.7955 | 86.00946 | 36.87191 | -0.4486 | 1.599238 | 0.387301 |
| 18 | 50773400 | 50774399 | 12.00418 | 22.99316 | 20.45371 | 8.06891 | 0.684898 | -0.03628 | 0.387306 |
| 9 | 9607800 | 9609599 | 31.98494 | 53.02335 | 51.38495 | 22.25043 | 0.425992 | 1.303595 | 0.387308 |
| 15 | 25819200 | 25821799 | 40.08096 | 65.73812 | 60.00318 | 25.83341 | 0.508231 | 1.45639 | 0.387308 |
| 11 | 50643600 | 50647399 | 21.98958 | 36.419 | 68.76218 | 29.7556 | -0.53331 | 1.30127 | 0.387315 |
| 4 | 134275400 | 134278999 | 40.28444 | 64.49981 | 110.7857 | 46.68995 | -0.43401 | 2.011948 | 0.38732 |
| 19 | 15900800 | 15903399 | 19.33059 | 32.41615 | 61.15204 | 26.67895 | -0.53142 | 1.148714 | 0.38733 |
| 6 | 131935800 | 131939199 | 68.20482 | 108.4107 | 107.1973 | 44.72751 | 0.349875 | 2.265 | 0.387344 |
| 7 | 25849000 | 25858399 | 111.474 | 172.6923 | 155.5461 | 64.60956 | 0.454766 | 2.818905 | 0.387358 |
| 2 | 57332800 | 57334399 | 19.7634 | 34.78203 | 31.51725 | 13.95345 | 0.529451 | 0.597279 | 0.387384 |
| 11 | 71123400 | 71125199 | 11.63871 | 20.5505 | 39.40538 | 17.43429 | -0.54385 | 0.532127 | 0.387392 |
| 3 | 148342600 | 148345799 | 33.01752 | 53.45759 | 89.36264 | 38.23222 | -0.37603 | 1.663255 | 0.387397 |

| | | | | | | | | | |
|---|---|---|---|---|---|---|---|---|---|
| 17 | 33095800 | 33100399 | 43.21664 | 69.12655 | 114.1336 | 48.03341 | -0.37874 | 2.046555 | 0.387405 |
| 5 | 92408000 | 92412599 | 22.62695 | 37.43906 | 69.95357 | 30.25132 | -0.51845 | 1.314504 | 0.387412 |
| 8 | 111572600 | 111573999 | 5.776939 | 10.96889 | 28.19626 | 11.35003 | -0.85827 | -0.00869 | 0.387428 |
| X | 8020800 | 8022799 | 26.04757 | 45.00851 | 37.29697 | 16.36245 | 0.655734 | 0.881257 | 0.387434 |
| 1 | 26987400 | 26989399 | 17.78619 | 32.10429 | 25.65196 | 11.15587 | 0.729231 | 0.385399 | 0.387448 |
| 4 | 152571400 | 152606199 | 562.8875 | 736.6155 | 1345.41 | 563.2336 | -0.79677 | 5.626492 | 0.387452 |
| X | 32201200 | 32207599 | 64.40092 | 103.5166 | 76.43157 | 32.40203 | 0.788365 | 1.912451 | 0.387488 |
| 6 | 20161000 | 20163399 | 21.15035 | 37.44116 | 29.12427 | 12.87897 | 0.751622 | 0.539799 | 0.387493 |
| 18 | 21919600 | 21922599 | 32.85004 | 55.2605 | 46.90772 | 20.49569 | 0.616594 | 1.146744 | 0.387499 |
| 14 | 117054400 | 117056399 | 25.87108 | 45.02161 | 34.60629 | 15.20939 | 0.764812 | 0.796275 | 0.387501 |
| 13 | 118351800 | 118352599 | 12.86562 | 22.61004 | 41.42065 | 18.33962 | -0.48136 | 0.593752 | 0.387516 |
| 14 | 121212200 | 121217399 | 43.53362 | 69.50791 | 120.2549 | 50.48026 | -0.44895 | 2.10282 | 0.387529 |
| 10 | 32882600 | 32884399 | 34.91884 | 58.06321 | 47.64998 | 20.67646 | 0.665385 | 1.274398 | 0.387536 |
| 16 | 79912800 | 79913799 | 16.08174 | 28.87476 | 26.52163 | 11.60003 | 0.524207 | 0.406594 | 0.387574 |
| 2 | 40859800 | 40861999 | 12.98047 | 22.94454 | 42.92235 | 18.98619 | -0.50843 | 0.530741 | 0.387584 |
| X | 31764400 | 31766199 | 30.20084 | 51.8616 | 39.03692 | 17.12253 | 0.791877 | 0.95169 | 0.387613 |
| 4 | 17828800 | 17829199 | 4.644009 | 9.067431 | 21.14039 | 7.451624 | -0.53975 | -0.50066 | 0.38762 |
| 14 | 81723400 | 81726399 | 35.2367 | 59.13077 | 46.92889 | 20.48277 | 0.71328 | 1.177886 | 0.38762 |
| 12 | 53217600 | 53219799 | 28.1908 | 48.43602 | 40.21725 | 17.64055 | 0.651039 | 0.93983 | 0.387667 |
| 4 | 115538400 | 115544399 | 47.58025 | 77.21652 | 66.92934 | 28.68409 | 0.572387 | 1.630646 | 0.387682 |
| 8 | 16692200 | 16694799 | 25.70388 | 44.69229 | 36.35517 | 15.96941 | 0.683507 | 0.81101 | 0.387694 |
| 9 | 26616400 | 26617199 | 13.80635 | 25.31074 | 23.93808 | 10.20742 | 0.506575 | 0.267445 | 0.387699 |
| 11 | 118620200 | 118622599 | 22.35722 | 36.91993 | 69.67186 | 30.08853 | -0.53286 | 1.33856 | 0.387713 |
| 12 | 114644000 | 114645199 | 18.95059 | 33.46309 | 31.14059 | 13.7775 | 0.493202 | 0.557779 | 0.38774 |
| 19 | 8734000 | 8735199 | 7.0861 | 13.3311 | 30.20203 | 12.38391 | -0.7016 | 0.070703 | 0.387747 |
| 1 | 22912600 | 22914599 | 19.105 | 32.36672 | 57.46302 | 25.16162 | -0.44361 | 1.052669 | 0.387791 |
| 5 | 147991600 | 147993599 | 17.35823 | 29.84315 | 52.73463 | 23.13151 | -0.43395 | 0.905196 | 0.387792 |
| 14 | 62282000 | 62283599 | 19.43481 | 34.68766 | 28.48369 | 12.52101 | 0.68053 | 0.473531 | 0.387802 |
| 13 | 17590800 | 17591999 | 16.3791 | 30.22498 | 24.55417 | 10.37706 | 0.733391 | 0.252041 | 0.387812 |
| 10 | 122143800 | 122145999 | 26.0259 | 44.13673 | 42.17272 | 18.47209 | 0.446797 | 1.048906 | 0.387814 |
| X | 33591600 | 33594999 | 34.98991 | 60.35707 | 32.17166 | 14.11321 | 1.288801 | 0.927138 | 0.387816 |
| 3 | 65886600 | 65887799 | 12.91058 | 24.41593 | 22.44129 | 9.110205 | 0.601973 | 0.069394 | 0.38782 |
| 17 | 58835000 | 58837999 | 27.02875 | 46.21229 | 37.74894 | 16.55183 | 0.673391 | 0.974424 | 0.387831 |
| 18 | 65971800 | 65985599 | 131.0731 | 196.8885 | 300.209 | 123.917 | -0.33996 | 3.437827 | 0.387836 |
| 8 | 16184200 | 16184999 | 11.76998 | 22.54453 | 21.15715 | 8.346458 | 0.609098 | -0.03419 | 0.387847 |
| 16 | 56059000 | 56061399 | 15.86656 | 28.82932 | 26.61291 | 11.47846 | 0.530157 | 0.337997 | 0.387855 |
| 3 | 147067800 | 147068199 | 5.557079 | 10.73762 | 25.38813 | 9.746813 | -0.67523 | -0.17647 | 0.387864 |
| 10 | 70117200 | 70120799 | 32.06279 | 51.88032 | 88.35233 | 37.78306 | -0.40355 | 1.673817 | 0.387866 |
| 8 | 7405600 | 7406199 | 11.79901 | 22.883 | 18.61807 | 7.025272 | 0.873029 | -0.18409 | 0.387867 |
| 8 | 98418600 | 98419399 | 7.460137 | 14.29226 | 30.99544 | 12.2205 | -0.58923 | -0.05465 | 0.387874 |
| 13 | 49306400 | 49311799 | 38.44808 | 61.72937 | 104.707 | 44.31332 | -0.41123 | 1.921026 | 0.387884 |
| X | 33550800 | 33561199 | 34.10941 | 57.8863 | 37.59023 | 16.46844 | 1.001943 | 1.069838 | 0.387887 |
| 12 | 4004000 | 4005399 | 11.91786 | 21.02109 | 39.87257 | 17.64473 | -0.52917 | 0.544051 | 0.387894 |
| X | 29428000 | 29430399 | 28.21936 | 47.95857 | 38.48526 | 16.86651 | 0.697872 | 1.036183 | 0.387903 |
| 16 | 4501800 | 4502599 | 6.419073 | 12.35128 | 26.39558 | 10.278 | -0.55047 | -0.11699 | 0.387914 |
| 4 | 82985200 | 82985599 | 5.816082 | 11.27468 | 24.42181 | 9.234426 | -0.52845 | -0.22556 | 0.387951 |
| 8 | 125148000 | 125148799 | 5.816082 | 11.27468 | 24.42181 | 9.234426 | -0.52845 | -0.22556 | 0.387951 |

| | | | | | | | | | |
|---|---|---|---|---|---|---|---|---|---|
| 13 | 63001800 | 63011399 | 86.01293 | 132.1416 | 213.7056 | 88.73427 | -0.39799 | 2.970324 | 0.387955 |
| 9 | 9758200 | 9760199 | 16.45198 | 28.56697 | 52.76699 | 23.15876 | -0.49456 | 0.812165 | 0.387955 |
| 10 | 56604600 | 56606599 | 18.38764 | 32.80912 | 28.92432 | 12.71782 | 0.578131 | 0.467076 | 0.387964 |
| 5 | 140518600 | 140521799 | 35.83107 | 57.77155 | 96.24724 | 40.98189 | -0.37779 | 1.783156 | 0.387971 |
| 15 | 101719400 | 101720199 | 10.06396 | 19.29428 | 19.15556 | 7.537175 | 0.531976 | -0.06614 | 0.387975 |
| 2 | 147018600 | 147021199 | 15.60635 | 27.00837 | 48.98571 | 21.4914 | -0.46919 | 0.834592 | 0.387993 |
| 1 | 191626400 | 191627999 | 10.78799 | 19.32464 | 36.33615 | 15.92403 | -0.50463 | 0.409755 | 0.388002 |
| 7 | 29514400 | 29530399 | 132.8388 | 198.7172 | 310.7719 | 128.1568 | -0.38184 | 3.516999 | 0.388003 |
| X | 52810000 | 52810999 | 17.18329 | 30.71373 | 28.67638 | 12.59027 | 0.497414 | 0.449265 | 0.388003 |
| 7 | 27547600 | 27549399 | 17.89754 | 31.73321 | 29.24479 | 12.9275 | 0.509545 | 0.505543 | 0.388004 |
| 11 | 19037600 | 19039799 | 19.6976 | 35.10392 | 27.57631 | 12.1375 | 0.742353 | 0.479062 | 0.388014 |
| 1 | 59579800 | 59581799 | 17.90642 | 30.07507 | 61.10356 | 26.6842 | -0.63802 | 1.126391 | 0.388016 |
| 18 | 37787000 | 37790999 | 75.30785 | 118.9502 | 118.0591 | 49.02792 | 0.336514 | 2.436997 | 0.388016 |
| 13 | 103164400 | 103168999 | 39.57168 | 63.55326 | 105.2411 | 44.56061 | -0.37612 | 1.912756 | 0.388021 |
| 9 | 82010000 | 82011599 | 16.43773 | 29.91166 | 25.96442 | 11.17792 | 0.619947 | 0.331728 | 0.38803 |
| 7 | 62122000 | 62126199 | 47.69645 | 77.80899 | 60.81281 | 26.1277 | 0.727044 | 1.547288 | 0.388044 |
| 6 | 23797600 | 23800399 | 36.01868 | 59.78087 | 53.21468 | 23.06152 | 0.548254 | 1.30149 | 0.388047 |
| 9 | 35228200 | 35230599 | 18.43918 | 31.24062 | 56.92203 | 24.92519 | -0.48093 | 1.047862 | 0.388048 |
| 4 | 148735600 | 148740399 | 42.94893 | 68.52659 | 119.5167 | 50.12087 | -0.46162 | 2.125429 | 0.388049 |
| 12 | 28239200 | 28241999 | 46.93869 | 75.70213 | 74.99987 | 31.94645 | 0.370864 | 1.781596 | 0.38805 |
| 14 | 84818800 | 84819599 | 8.625521 | 16.03918 | 31.04997 | 12.98536 | -0.49937 | 0.163813 | 0.388072 |
| 9 | 60644200 | 60652199 | 63.59951 | 99.53272 | 168.2359 | 69.79032 | -0.44091 | 2.621472 | 0.388074 |
| X | 25413000 | 25414199 | 25.77811 | 44.75634 | 37.52607 | 16.4687 | 0.63993 | 0.833668 | 0.388076 |
| 15 | 29141200 | 29141999 | 6.853899 | 13.22296 | 27.72331 | 10.71313 | -0.51296 | -0.13313 | 0.38808 |
| 11 | 59369600 | 59371999 | 12.2118 | 21.76236 | 39.85559 | 17.54268 | -0.4716 | 0.457507 | 0.388096 |
| 5 | 62288400 | 62289399 | 10.30015 | 18.60783 | 35.38517 | 15.37231 | -0.51221 | 0.353365 | 0.388098 |
| X | 34732600 | 34737199 | 58.55332 | 94.42472 | 64.77339 | 27.58693 | 0.899832 | 1.799486 | 0.388103 |
| 3 | 97037000 | 97038999 | 17.73819 | 32.20055 | 26.42648 | 11.41111 | 0.697819 | 0.356951 | 0.388105 |
| X | 81463400 | 81468799 | 100.3449 | 157.0535 | 135.0301 | 56.01481 | 0.53323 | 2.636344 | 0.388107 |
| 4 | 112944200 | 112945399 | 26.61929 | 45.70075 | 41.41528 | 18.16546 | 0.525182 | 0.942355 | 0.38812 |
| 1 | 60939000 | 60941599 | 19.66171 | 32.93995 | 68.98551 | 30.07836 | -0.68182 | 1.166323 | 0.388126 |
| 1 | 139546800 | 139548599 | 16.90974 | 31.43372 | 22.29857 | 9.330289 | 0.937504 | 0.210874 | 0.388128 |
| 2 | 51603600 | 51606599 | 28.73374 | 48.84206 | 43.5569 | 19.0895 | 0.546097 | 1.039312 | 0.388128 |
| 18 | 28732200 | 28734399 | 23.25192 | 40.59107 | 35.30829 | 15.54861 | 0.587689 | 0.742563 | 0.388128 |
| 19 | 8474000 | 8476799 | 44.82828 | 72.40546 | 69.15828 | 29.51444 | 0.425949 | 1.733606 | 0.388132 |
| 1 | 183373800 | 183374399 | 10.0692 | 19.50414 | 19.22549 | 7.296386 | 0.59062 | -0.18405 | 0.388135 |
| 12 | 39884600 | 39885399 | 10.0692 | 19.50414 | 19.22549 | 7.296386 | 0.59062 | -0.18405 | 0.388135 |
| 1 | 85285000 | 85285599 | 15.69291 | 28.11949 | 24.41393 | 10.69645 | 0.603596 | 0.413664 | 0.388143 |
| 9 | 51482000 | 51482599 | 6.417012 | 12.37025 | 25.87771 | 10.02279 | -0.51229 | -0.13424 | 0.388147 |
| 13 | 71096800 | 71097399 | 6.417012 | 12.37025 | 25.87771 | 10.02279 | -0.51229 | -0.13424 | 0.388147 |
| 3 | 17352200 | 17354799 | 24.6713 | 42.92225 | 36.38661 | 15.98915 | 0.624261 | 0.797583 | 0.388153 |
| 11 | 30964600 | 30968199 | 45.72904 | 73.16499 | 125.6038 | 52.88086 | -0.43452 | 2.045559 | 0.388158 |
| 3 | 13959600 | 13960999 | 14.92711 | 27.48076 | 24.86998 | 10.54426 | 0.575778 | 0.256277 | 0.388168 |
| 6 | 88166600 | 88168999 | 19.65133 | 32.9256 | 61.7111 | 26.90821 | -0.5221 | 1.159365 | 0.388177 |
| 7 | 95852400 | 95854199 | 10.6375 | 18.91793 | 38.82854 | 17.12024 | -0.63681 | 0.456071 | 0.388179 |
| X | 72107000 | 72110799 | 19.99835 | 34.00254 | 60.36861 | 26.45806 | -0.4435 | 1.025104 | 0.388184 |
| 3 | 55173200 | 55173799 | 9.66314 | 18.76343 | 18.78796 | 7.041187 | 0.585454 | -0.23099 | 0.388187 |

| | | | | | | | | | |
|---|---|---|---|---|---|---|---|---|---|
| 1 | 117276800 | 117278199 | 16.78154 | 30.13989 | 27.45668 | 12.0061 | 0.536306 | 0.412054 | 0.388238 |
| 16 | 78775200 | 78775799 | 11.59433 | 22.59086 | 15.19994 | 5.523552 | 1.192295 | -0.32393 | 0.388243 |
| 8 | 61435600 | 61436799 | 9.20038 | 16.69561 | 33.9273 | 14.65666 | -0.60178 | 0.316294 | 0.388251 |
| 9 | 5899000 | 5900999 | 18.40328 | 32.54619 | 30.59601 | 13.53244 | 0.479403 | 0.536425 | 0.388258 |
| 11 | 32068200 | 32072399 | 22.84003 | 38.18663 | 66.42756 | 28.92576 | -0.41503 | 1.19572 | 0.388274 |
| 6 | 37714400 | 37715599 | 15.20286 | 29.18626 | 18.58631 | 7.284539 | 1.172049 | -0.02806 | 0.38828 |
| 2 | 18605400 | 18607999 | 21.04836 | 36.74859 | 34.95346 | 15.39348 | 0.45935 | 0.728505 | 0.388281 |
| 13 | 62089200 | 62110799 | 514.9426 | 704.339 | 612.341 | 254.7917 | 0.337958 | 4.921803 | 0.388283 |
| 16 | 57305000 | 57306799 | 8.422973 | 15.80634 | 30.71122 | 12.64646 | -0.48681 | 0.101126 | 0.38829 |
| 2 | 148995600 | 148997199 | 23.76011 | 42.06191 | 27.96935 | 12.3682 | 0.976889 | 0.555803 | 0.388297 |
| 6 | 73530600 | 73533799 | 25.79664 | 42.30769 | 85.91765 | 36.98965 | -0.64236 | 1.442362 | 0.388297 |
| 17 | 33311200 | 33311999 | 7.012536 | 13.45516 | 27.45451 | 10.77731 | -0.49548 | -0.08112 | 0.388302 |
| 3 | 20971000 | 20972999 | 16.32892 | 28.32635 | 52.26173 | 22.93035 | -0.4933 | 0.818114 | 0.388311 |
| 9 | 95958400 | 95962999 | 27.03961 | 44.4195 | 75.70439 | 32.61537 | -0.3892 | 1.418158 | 0.388327 |
| 3 | 39920600 | 39921999 | 16.80518 | 29.90166 | 28.73893 | 12.66618 | 0.452054 | 0.474389 | 0.388345 |
| 17 | 40426000 | 40428799 | 40.59921 | 65.01752 | 110.4552 | 46.56563 | -0.41785 | 2.005401 | 0.388358 |
| 7 | 85720600 | 85721199 | 19.83907 | 34.67253 | 32.62907 | 14.37953 | 0.474913 | 0.705728 | 0.388366 |
| 5 | 113288200 | 113290199 | 22.48114 | 37.22124 | 68.16396 | 29.49023 | -0.48941 | 1.307099 | 0.388379 |
| 12 | 31791400 | 31794999 | 52.13288 | 84.1229 | 74.68752 | 31.83471 | 0.529573 | 1.77583 | 0.388382 |
| 11 | 31261400 | 31263199 | 17.1895 | 31.52607 | 24.93169 | 10.6251 | 0.763327 | 0.292497 | 0.388401 |
| 7 | 14642600 | 14663999 | 458.2751 | 635.9364 | 521.8204 | 216.631 | 0.442342 | 4.699735 | 0.388404 |
| 4 | 79408400 | 79409399 | 22.13243 | 38.55708 | 36.58239 | 16.08879 | 0.462751 | 0.767848 | 0.388419 |
| 5 | 149401000 | 149402799 | 22.13243 | 38.55708 | 36.58239 | 16.08879 | 0.462751 | 0.767848 | 0.388419 |
| 3 | 5752200 | 5752999 | 10.57888 | 20.294 | 20.69266 | 8.127547 | 0.496123 | -0.06202 | 0.388436 |
| 15 | 37590600 | 37593399 | 35.32123 | 56.96834 | 95.19087 | 40.54936 | -0.38151 | 1.771527 | 0.388444 |
| 6 | 92200400 | 92202399 | 17.16022 | 30.39476 | 28.63727 | 12.66219 | 0.476935 | 0.507291 | 0.388446 |
| 6 | 131879400 | 131889799 | 218.9452 | 323.9097 | 284.3297 | 117.0948 | 0.436111 | 3.725295 | 0.38845 |
| 5 | 34130000 | 34131799 | 19.54338 | 34.31531 | 32.77823 | 14.50226 | 0.453255 | 0.629732 | 0.388465 |
| 14 | 122172600 | 122174599 | 31.62259 | 52.83295 | 50.82454 | 22.12158 | 0.436851 | 1.215942 | 0.38847 |
| 7 | 114145600 | 114147599 | 22.9085 | 38.31985 | 68.60888 | 29.88484 | -0.45646 | 1.194578 | 0.388482 |
| 3 | 4259800 | 4261199 | 14.0956 | 26.73207 | 21.9363 | 8.852375 | 0.772197 | 0.061735 | 0.388486 |
| 4 | 127040400 | 127042599 | 20.3359 | 35.90272 | 31.03314 | 13.73046 | 0.599114 | 0.568531 | 0.388487 |
| 7 | 15088000 | 15092199 | 110.5174 | 173.5477 | 116.4822 | 48.31299 | 0.893688 | 2.580278 | 0.388488 |
| 9 | 11955400 | 11956799 | 22.12038 | 39.134 | 28.47731 | 12.59444 | 0.846932 | 0.551074 | 0.388491 |
| 3 | 44218400 | 44219799 | 17.95664 | 32.06966 | 28.37632 | 12.46716 | 0.573496 | 0.456513 | 0.388492 |
| 12 | 23458800 | 23460199 | 27.96788 | 47.47706 | 37.9881 | 16.64418 | 0.701902 | 1.043603 | 0.388498 |
| 14 | 17657400 | 17660199 | 17.94351 | 31.72595 | 29.54813 | 13.06956 | 0.492509 | 0.53261 | 0.388499 |
| 12 | 119614800 | 119615199 | 8.993939 | 17.41811 | 18.22238 | 6.921693 | 0.504681 | -0.19581 | 0.388545 |
| 13 | 86412000 | 86412799 | 11.94079 | 21.04052 | 39.90794 | 17.66391 | -0.52956 | 0.55652 | 0.38857 |
| 11 | 22236400 | 22237599 | 10.97167 | 19.4454 | 38.3185 | 16.94073 | -0.58106 | 0.481896 | 0.388576 |
| 10 | 32188200 | 32191399 | 65.21566 | 104.1158 | 88.15638 | 37.00061 | 0.579989 | 2.120395 | 0.388576 |
| 1 | 64655400 | 64656999 | 10.61506 | 19.05799 | 35.83 | 15.67226 | -0.50255 | 0.392826 | 0.388577 |
| 12 | 105035000 | 105037799 | 22.03512 | 36.80527 | 65.21451 | 28.38022 | -0.44146 | 1.202183 | 0.388588 |
| 17 | 21245400 | 21248599 | 53.4856 | 87.50901 | 51.36694 | 22.09294 | 1.141801 | 1.515233 | 0.388598 |
| 5 | 125041000 | 125067199 | 253.1369 | 354.9774 | 658.0219 | 272.729 | -0.71777 | 4.535893 | 0.388602 |
| 16 | 9576400 | 9576999 | 4.841842 | 9.414974 | 24.08618 | 8.924255 | -0.73906 | -0.31308 | 0.388613 |
| 7 | 48273800 | 48276999 | 52.81041 | 84.82736 | 83.72026 | 35.45621 | 0.369254 | 1.917041 | 0.388635 |

| | | | | | | | | | |
|---|---|---|---|---|---|---|---|---|---|
| 5 | 142686000 | 142692199 | 40.97564 | 65.40932 | 119.7021 | 50.23254 | -0.53015 | 2.107292 | 0.388644 |
| 12 | 115792200 | 115796599 | 82.83208 | 130.4702 | 127.4005 | 52.86252 | 0.356854 | 2.503937 | 0.388645 |
| 17 | 83570600 | 83573599 | 18.79002 | 31.66238 | 60.16013 | 26.29969 | -0.54162 | 1.098673 | 0.388659 |
| 2 | 145515800 | 145520199 | 36.42672 | 58.70797 | 100.3232 | 42.69495 | -0.4151 | 1.795334 | 0.388664 |
| 7 | 43751400 | 43757399 | 53.94442 | 85.38473 | 144.1629 | 59.93178 | -0.42648 | 2.377448 | 0.388676 |
| 12 | 69205400 | 69206199 | 6.652569 | 12.52896 | 28.59973 | 11.70557 | -0.70919 | 0.059439 | 0.388676 |
| 7 | 9851600 | 9875399 | 395.51 | 557.9496 | 440.4854 | 182.3932 | 0.521925 | 4.45241 | 0.388701 |
| 15 | 97438600 | 97446599 | 81.59641 | 126.5913 | 199.5807 | 82.88761 | -0.35043 | 2.784865 | 0.388703 |
| 5 | 33411800 | 33414599 | 27.83302 | 47.28018 | 44.72785 | 19.60118 | 0.461277 | 1.045925 | 0.388706 |
| 11 | 20150800 | 20152199 | 11.51349 | 20.18676 | 42.08104 | 18.59541 | -0.66649 | 0.626211 | 0.388708 |
| 17 | 66142600 | 66143999 | 5.751177 | 10.94705 | 27.16058 | 10.87601 | -0.80033 | -0.02724 | 0.38871 |
| 16 | 91265000 | 91266199 | 16.53573 | 30.28383 | 25.53328 | 10.9026 | 0.669566 | 0.296614 | 0.388725 |
| 7 | 21093600 | 21096599 | 22.49303 | 39.99178 | 27.45764 | 12.10921 | 0.93395 | 0.513556 | 0.388728 |
| 2 | 174729400 | 174730599 | 7.226 | 13.62828 | 28.61929 | 11.68541 | -0.58755 | 0.054151 | 0.388736 |
| 14 | 30032200 | 30035999 | 17.86939 | 30.59924 | 53.78976 | 23.5882 | -0.42746 | 0.945333 | 0.388762 |
| 10 | 119030400 | 119032799 | 29.20262 | 47.6923 | 80.68169 | 34.67823 | -0.38299 | 1.503577 | 0.388766 |
| 12 | 105186800 | 105192799 | 61.81444 | 97.16968 | 159.066 | 65.98428 | -0.38979 | 2.536264 | 0.38877 |
| 1 | 140226600 | 140227999 | 20.56687 | 36.09587 | 32.58914 | 14.41303 | 0.534065 | 0.641021 | 0.388777 |
| 3 | 107006800 | 107013799 | 51.62296 | 81.8472 | 140.3692 | 58.42055 | -0.44692 | 2.331774 | 0.38881 |
| 10 | 36933800 | 36935399 | 18.42566 | 32.99637 | 28.41968 | 12.4578 | 0.61456 | 0.444608 | 0.38881 |
| 6 | 23517600 | 23519799 | 17.5092 | 31.66017 | 26.90897 | 11.67672 | 0.643012 | 0.377773 | 0.388818 |
| 2 | 98231400 | 98232199 | 8.45825 | 15.93137 | 31.9298 | 13.06614 | -0.52375 | 0.08243 | 0.38883 |
| 14 | 93593000 | 93593999 | 8.45825 | 15.93137 | 31.9298 | 13.06614 | -0.52375 | 0.08243 | 0.38883 |
| X | 89667800 | 89673599 | 82.7068 | 131.0777 | 101.0574 | 42.04251 | 0.704287 | 2.361412 | 0.388832 |
| 6 | 40213200 | 40214799 | 11.73124 | 20.72788 | 41.37541 | 18.30343 | -0.60167 | 0.524268 | 0.388836 |
| 16 | 59269800 | 59272199 | 14.33 | 25.11904 | 45.1194 | 19.93405 | -0.45321 | 0.650348 | 0.388842 |
| 17 | 19682000 | 19683599 | 34.49617 | 56.87643 | 56.35346 | 24.31609 | 0.393174 | 1.376994 | 0.388846 |
| 8 | 21893400 | 21898799 | 45.3658 | 74.0787 | 55.60436 | 23.89838 | 0.786084 | 1.524108 | 0.388849 |
| 4 | 112333600 | 112339399 | 97.12601 | 152.1337 | 135.4358 | 56.17861 | 0.483908 | 2.620633 | 0.388857 |
| 13 | 95440800 | 95442199 | 13.88881 | 25.80471 | 24.32193 | 10.18409 | 0.529938 | 0.193981 | 0.388866 |
| 8 | 58361600 | 58363599 | 21.53293 | 38.4137 | 26.9432 | 11.8487 | 0.905913 | 0.48524 | 0.388867 |
| 7 | 33100400 | 33110399 | 233.5464 | 341.4005 | 319.6932 | 131.7047 | 0.326602 | 3.917556 | 0.388873 |
| 8 | 22974400 | 22976599 | 23.72497 | 40.71599 | 39.47985 | 17.31601 | 0.427888 | 0.930603 | 0.388903 |
| 1 | 182514800 | 182518199 | 33.1109 | 54.73102 | 54.14933 | 23.40056 | 0.395891 | 1.34122 | 0.388916 |
| 5 | 109300600 | 109304799 | 54.67139 | 88.13364 | 75.64319 | 32.20177 | 0.576709 | 1.806579 | 0.38892 |
| 11 | 33699000 | 33702799 | 29.82022 | 48.23796 | 87.78664 | 37.5333 | -0.49939 | 1.675557 | 0.388921 |
| 10 | 67103600 | 67105399 | 9.972888 | 18.20882 | 35.53879 | 15.23548 | -0.53698 | 0.290063 | 0.388926 |
| 7 | 59273800 | 59275399 | 19.52882 | 35.51609 | 25.24884 | 10.87767 | 0.905897 | 0.35819 | 0.388935 |
| 17 | 20231200 | 20244399 | 242.6507 | 356.3919 | 305.5573 | 125.8438 | 0.460285 | 3.845489 | 0.388937 |
| 7 | 84683600 | 84688799 | 52.28556 | 84.63889 | 61.18221 | 26.17231 | 0.830359 | 1.687639 | 0.388979 |
| 15 | 26055200 | 26056599 | 14.86087 | 28.22245 | 21.29469 | 8.568313 | 0.895821 | 0.057365 | 0.388982 |
| 1 | 75094800 | 75097599 | 14.66121 | 25.4613 | 47.39385 | 20.80251 | -0.50534 | 0.794293 | 0.388985 |
| 11 | 29534600 | 29536599 | 19.27057 | 35.50098 | 22.60334 | 9.574481 | 1.080161 | 0.279971 | 0.388996 |
| 1 | 170889600 | 170894599 | 123.4953 | 184.2355 | 319.8861 | 131.8453 | -0.53618 | 3.567085 | 0.389004 |
| 6 | 95728800 | 95731799 | 27.71755 | 47.17442 | 44.14575 | 19.34954 | 0.477202 | 1.026335 | 0.389006 |
| 13 | 5661400 | 5661799 | 4.309748 | 8.418484 | 20.53961 | 7.185202 | -0.59298 | -0.53048 | 0.389021 |
| 10 | 82745200 | 82746999 | 11.72523 | 20.82144 | 38.27661 | 16.89803 | -0.47996 | 0.472881 | 0.38903 |

| | | | | | | | | | |
|---|---|---|---|---|---|---|---|---|---|
| 4 | 113254600 | 113266599 | 79.93333 | 126.8412 | 100.4393 | 41.83786 | 0.667524 | 2.321815 | 0.389035 |
| 16 | 44008000 | 44009399 | 18.45199 | 32.51453 | 30.63565 | 13.55977 | 0.474088 | 0.574533 | 0.38904 |
| 17 | 8910400 | 8918799 | 106.3354 | 164.4936 | 159.5298 | 66.2648 | 0.34662 | 2.842238 | 0.389049 |
| 19 | 55799600 | 55801199 | 11.05856 | 19.90968 | 36.32709 | 15.84919 | -0.45729 | 0.378844 | 0.389056 |
| 8 | 61832400 | 61835399 | 28.55094 | 46.16713 | 96.44612 | 41.22033 | -0.69856 | 1.685942 | 0.389066 |
| 7 | 62225800 | 62235599 | 137.2602 | 210.1988 | 178.3929 | 74.03509 | 0.527845 | 3.047093 | 0.389082 |
| 3 | 153656200 | 153659399 | 26.36356 | 43.30655 | 76.92267 | 33.13947 | -0.44875 | 1.416273 | 0.389096 |
| 2 | 149323200 | 149323799 | 7.618633 | 14.51262 | 29.50061 | 11.79416 | -0.51476 | -0.01105 | 0.389132 |
| 6 | 14873800 | 14875599 | 16.64093 | 30.20087 | 26.47798 | 11.43705 | 0.602659 | 0.349584 | 0.389136 |
| 18 | 65774200 | 65775799 | 9.218828 | 17.11399 | 32.64009 | 13.68243 | -0.48093 | 0.180487 | 0.389151 |
| X | 134574400 | 134576599 | 21.92117 | 38.26858 | 35.53874 | 15.65025 | 0.493879 | 0.737721 | 0.389153 |
| 9 | 108053800 | 108054399 | 10.16769 | 19.55781 | 19.21729 | 7.489901 | 0.559941 | -0.09387 | 0.389163 |
| 6 | 15553000 | 15559399 | 114.1549 | 175.2504 | 169.5219 | 70.38152 | 0.341993 | 2.989197 | 0.389169 |
| 13 | 53944000 | 53947199 | 38.8761 | 63.54593 | 61.88156 | 26.6061 | 0.412585 | 1.500447 | 0.389178 |
| 13 | 18090600 | 18092799 | 67.7575 | 105.8598 | 171.8473 | 71.3009 | -0.38463 | 2.655124 | 0.389183 |
| 1 | 145899200 | 145900799 | 19.32016 | 34.08594 | 26.26304 | 11.62156 | 0.763636 | 0.560227 | 0.389189 |
| 17 | 33418200 | 33418799 | 5.373748 | 10.39509 | 24.81668 | 9.477383 | -0.68011 | -0.1929 | 0.38919 |
| 2 | 117946200 | 117951799 | 36.80762 | 59.15183 | 103.0948 | 43.69271 | -0.44862 | 1.887775 | 0.389195 |
| 1 | 53527200 | 53529399 | 18.52514 | 33.23841 | 27.82597 | 12.17777 | 0.656812 | 0.431814 | 0.389197 |
| 14 | 21541600 | 21543799 | 18.52514 | 33.23841 | 27.82597 | 12.17777 | 0.656812 | 0.431814 | 0.389197 |
| 2 | 36609400 | 36621199 | 189.8495 | 283.3724 | 258.0119 | 106.3538 | 0.39516 | 3.565097 | 0.389207 |
| 4 | 86323600 | 86326399 | 18.07371 | 31.88293 | 30.7482 | 13.6066 | 0.441403 | 0.561949 | 0.389211 |
| 7 | 65364800 | 65369799 | 33.22882 | 53.63192 | 92.44917 | 39.41781 | -0.42506 | 1.746927 | 0.389214 |
| X | 97143600 | 97146199 | 22.89432 | 40.30597 | 30.72613 | 13.60228 | 0.777815 | 0.612085 | 0.389221 |
| 14 | 112526000 | 112528399 | 34.74173 | 57.1302 | 50.34978 | 21.70129 | 0.560137 | 1.404947 | 0.389233 |
| 17 | 20748400 | 20752199 | 78.55505 | 124.314 | 117.5164 | 48.84924 | 0.408871 | 2.396224 | 0.38924 |
| X | 4796400 | 4798999 | 25.75635 | 45.07736 | 32.513 | 14.34539 | 0.856896 | 0.716467 | 0.38924 |
| 17 | 79043200 | 79044599 | 11.01912 | 19.4324 | 39.44258 | 17.45507 | -0.62616 | 0.541376 | 0.389278 |
| 4 | 113133600 | 113138399 | 76.80695 | 122.3357 | 96.51196 | 40.37638 | 0.678254 | 2.205176 | 0.389285 |
| 7 | 9687000 | 9694199 | 84.17671 | 134.1104 | 88.98638 | 37.24197 | 0.927959 | 2.203931 | 0.389298 |
| X | 52975000 | 52980399 | 65.5119 | 104.7177 | 89.38101 | 37.58016 | 0.570386 | 2.086644 | 0.389314 |
| 11 | 108776200 | 108776599 | 4.227898 | 8.249913 | 20.34422 | 7.247187 | -0.63258 | -0.47049 | 0.389314 |
| 15 | 16184000 | 16186599 | 32.87651 | 54.72211 | 43.91528 | 19.06908 | 0.697515 | 1.25128 | 0.389333 |
| 14 | 52839800 | 52841199 | 25.72432 | 43.73128 | 42.65993 | 18.69654 | 0.417248 | 1.027548 | 0.389336 |
| 10 | 117211800 | 117212799 | 5.865164 | 11.23335 | 26.20463 | 10.34016 | -0.69293 | -0.0773 | 0.389337 |
| 1 | 140674400 | 140675199 | 6.698275 | 12.90116 | 28.25554 | 10.97127 | -0.58233 | -0.11986 | 0.389337 |
| 6 | 29635600 | 29639399 | 30.05903 | 50.34775 | 49.15745 | 21.43012 | 0.415413 | 1.181987 | 0.38934 |
| Y | 88849800 | 88850599 | 11.82121 | 22.43795 | 13.4727 | 5.427045 | 1.220042 | 0.006918 | 0.389343 |
| 16 | 47731800 | 47735599 | 42.70774 | 69.30875 | 67.36025 | 28.86866 | 0.407677 | 1.622826 | 0.389349 |
| 1 | 60241000 | 60242999 | 16.6295 | 30.37471 | 26.09984 | 11.18346 | 0.639122 | 0.314027 | 0.389349 |
| 6 | 57314200 | 57318999 | 63.5938 | 102.2401 | 76.33248 | 32.36956 | 0.772675 | 1.902027 | 0.389349 |
| X | 54886000 | 54898599 | 24.75569 | 43.34855 | 29.00592 | 12.80362 | 0.964457 | 0.696927 | 0.389354 |
| 12 | 99961800 | 99963799 | 9.586183 | 17.49826 | 33.9301 | 14.55052 | -0.52828 | 0.281688 | 0.389368 |
| Y | 6392600 | 6395799 | 17.64024 | 31.60792 | 15.39838 | 6.746643 | 1.430562 | 0.403241 | 0.389372 |
| 15 | 71608600 | 71611599 | 25.43064 | 41.35466 | 87.35637 | 37.48857 | -0.70676 | 1.563558 | 0.389376 |
| 2 | 64579600 | 64581199 | 19.04147 | 32.29536 | 57.52724 | 25.19832 | -0.44844 | 1.040451 | 0.389384 |
| 5 | 42533200 | 42533599 | 9.336043 | 18.06116 | 16.72097 | 6.383473 | 0.673173 | -0.17864 | 0.389394 |

| | | | | | | | | | |
|---|---|---|---|---|---|---|---|---|---|
| Y | 10446400 | 10447399 | 8.65393 | 16.83699 | 11.50552 | 4.243773 | 1.151095 | -0.3263 | 0.389428 |
| 17 | 94258000 | 94259199 | 9.434625 | 17.41933 | 33.07077 | 13.96622 | -0.48187 | 0.215919 | 0.389459 |
| 8 | 109367200 | 109369999 | 16.47637 | 28.12894 | 54.79929 | 24.02507 | -0.57578 | 0.967913 | 0.389468 |
| 15 | 94328000 | 94328999 | 8.253948 | 15.52834 | 30.78223 | 12.62157 | -0.51065 | 0.082893 | 0.389476 |
| 13 | 37105800 | 37109399 | 72.1843 | 114.2462 | 111.3074 | 46.27116 | 0.365536 | 2.386676 | 0.389476 |
| 10 | 87833000 | 87834399 | 16.47032 | 28.19511 | 52.6693 | 23.09629 | -0.51471 | 0.941857 | 0.389478 |
| 11 | 109754600 | 109757399 | 15.73065 | 27.27161 | 48.9925 | 21.49553 | -0.455 | 0.818301 | 0.38948 |
| 14 | 62855200 | 62856599 | 11.23207 | 20.10607 | 36.80118 | 16.13735 | -0.46684 | 0.416787 | 0.389481 |
| 2 | 103998000 | 103998799 | 6.304818 | 12.04601 | 26.69845 | 10.6064 | -0.62914 | -0.04917 | 0.389491 |
| 11 | 17911800 | 17912599 | 5.824973 | 11.28761 | 27.49492 | 10.41502 | -0.70022 | -0.2158 | 0.389491 |
| 7 | 131141400 | 131151799 | 32.87614 | 52.71018 | 116.2405 | 49.09062 | -0.79142 | 1.962167 | 0.389493 |
| 9 | 88708600 | 88711799 | 23.94767 | 41.08229 | 40.01488 | 17.55013 | 0.421375 | 0.93639 | 0.389493 |
| 6 | 48097000 | 48100599 | 29.17397 | 47.62808 | 82.57722 | 35.48823 | -0.41876 | 1.508322 | 0.389517 |
| 2 | 25031200 | 25032399 | 11.43998 | 20.20341 | 40.36914 | 17.86002 | -0.60324 | 0.526931 | 0.389529 |
| 2 | 119424600 | 119425399 | 3.643935 | 7.104954 | 20.75723 | 7.474283 | -0.8865 | -0.43592 | 0.389537 |
| 13 | 18837600 | 18839999 | 22.88961 | 40.2305 | 30.65251 | 13.57309 | 0.777577 | 0.628649 | 0.389547 |
| 1 | 139338200 | 139338999 | 22.69913 | 39.17084 | 37.86684 | 16.61179 | 0.433742 | 0.874588 | 0.389562 |
| 18 | 61217000 | 61219999 | 24.41359 | 40.46272 | 70.09973 | 30.34946 | -0.40954 | 1.302487 | 0.389562 |
| 10 | 124111600 | 124112799 | 13.44922 | 24.88859 | 24.39555 | 10.27046 | 0.468357 | 0.213039 | 0.389575 |
| 2 | 27708000 | 27712399 | 36.12291 | 58.23395 | 97.10567 | 41.33995 | -0.37936 | 1.786756 | 0.389577 |
| 1 | 54903800 | 54906199 | 11.94919 | 20.97156 | 42.0118 | 18.58043 | -0.6096 | 0.610817 | 0.389632 |
| 19 | 50007600 | 50009999 | 22.78261 | 39.51345 | 38.28895 | 16.79954 | 0.431802 | 0.831805 | 0.389645 |
| 12 | 111094600 | 111106199 | 95.97462 | 146.6844 | 229.587 | 95.23774 | -0.35664 | 3.07807 | 0.389651 |
| 2 | 97600800 | 97601399 | 10.28386 | 19.91966 | 18.58348 | 7.053205 | 0.668731 | -0.18453 | 0.389668 |
| 18 | 37731000 | 37732199 | 22.55903 | 39.29271 | 36.51647 | 16.05786 | 0.492405 | 0.771764 | 0.389674 |
| 14 | 15803400 | 15806199 | 18.84366 | 33.24922 | 31.08512 | 13.75503 | 0.486033 | 0.563888 | 0.389686 |
| 1 | 31037200 | 31037999 | 7.685891 | 14.47956 | 29.13958 | 11.92022 | -0.52875 | 0.067948 | 0.389693 |
| 13 | 66075600 | 66083599 | 228.6537 | 335.9655 | 260.2839 | 107.1953 | 0.60693 | 3.835851 | 0.389698 |
| 4 | 98639800 | 98642999 | 42.06029 | 68.30631 | 66.19778 | 28.38525 | 0.41271 | 1.607156 | 0.389713 |
| 1 | 75123000 | 75124399 | 7.705417 | 14.44839 | 29.59766 | 12.20421 | -0.56335 | 0.09985 | 0.38972 |
| 14 | 124487800 | 124490599 | 37.8073 | 61.93472 | 60.86207 | 26.18991 | 0.401372 | 1.466646 | 0.389738 |
| 4 | 151565000 | 151565999 | 12.83025 | 23.88688 | 23.47545 | 9.797471 | 0.473963 | 0.171748 | 0.389745 |
| 4 | 16577800 | 16579199 | 15.86875 | 29.09424 | 25.61209 | 10.91993 | 0.609234 | 0.285596 | 0.389752 |
| 15 | 4488000 | 4489599 | 17.07777 | 30.70293 | 28.03231 | 12.24737 | 0.534105 | 0.410419 | 0.389754 |
| 12 | 20041600 | 20045999 | 105.607 | 163.5709 | 137.8107 | 57.24283 | 0.55079 | 2.817004 | 0.389762 |
| 2 | 77414400 | 77418199 | 51.73843 | 83.45536 | 75.8389 | 32.30997 | 0.495457 | 1.785495 | 0.389793 |
| 7 | 40919400 | 40939399 | 301.1234 | 433.1906 | 377.271 | 155.7194 | 0.408634 | 4.162671 | 0.389796 |
| X | 153045000 | 153048399 | 62.03661 | 99.52157 | 84.78181 | 35.83704 | 0.579042 | 1.969627 | 0.389798 |
| 3 | 59588800 | 59592199 | 41.46745 | 68.18016 | 56.8143 | 24.48604 | 0.640617 | 1.426761 | 0.389798 |
| X | 70093200 | 70096599 | 41.5478 | 68.08957 | 56.97799 | 24.52269 | 0.632886 | 1.472007 | 0.389803 |
| 2 | 171031200 | 171033199 | 14.25166 | 24.98566 | 49.41242 | 21.83364 | -0.59173 | 0.650164 | 0.389806 |
| 14 | 5038600 | 5040599 | 19.4306 | 32.93037 | 58.02136 | 25.40892 | -0.43269 | 1.051416 | 0.389827 |
| 13 | 65714600 | 65722399 | 114.1023 | 178.5388 | 124.8488 | 51.7927 | 0.830659 | 2.646851 | 0.389858 |
| 7 | 13649200 | 13668399 | 309.6388 | 446.4872 | 358.97 | 148.1172 | 0.527286 | 4.129134 | 0.389871 |
| X | 109555800 | 109556999 | 15.38857 | 28.86184 | 22.33758 | 9.206926 | 0.830845 | 0.146606 | 0.389876 |
| 7 | 52650800 | 52651599 | 11.64047 | 22.21169 | 21.59217 | 8.601273 | 0.54556 | -0.00478 | 0.389877 |
| 6 | 105387600 | 105388999 | 15.57122 | 28.91627 | 24.16914 | 10.12691 | 0.701452 | 0.210476 | 0.389884 |

| | | | | | | | | | |
|---|---|---|---|---|---|---|---|---|---|
| 19 | 9521600 | 9522199 | 7.520584 | 14.2061 | 28.65316 | 11.66054 | -0.52558 | 0.04662 | 0.389889 |
| 1 | 34586800 | 34589199 | 23.5472 | 41.1303 | 34.70111 | 15.28678 | 0.63158 | 0.734889 | 0.389916 |
| 16 | 60509600 | 60512799 | 30.30995 | 51.07392 | 47.46546 | 20.75004 | 0.486458 | 1.125652 | 0.389917 |
| 17 | 39581800 | 39584399 | 27.4498 | 46.71602 | 38.92999 | 17.06334 | 0.643834 | 1.019743 | 0.389919 |
| 16 | 45279000 | 45281199 | 25.53496 | 44.34116 | 33.90335 | 14.88031 | 0.772342 | 0.825415 | 0.389929 |
| X | 125961000 | 125962399 | 21.10387 | 37.35316 | 29.75494 | 13.15826 | 0.717562 | 0.543282 | 0.389953 |
| 1 | 118184000 | 118185599 | 23.79636 | 40.89516 | 39.98857 | 17.54036 | 0.416198 | 0.920011 | 0.389959 |
| 11 | 49055000 | 49058599 | 43.27073 | 70.92795 | 59.04875 | 25.41616 | 0.64031 | 1.470977 | 0.389968 |
| 10 | 4215600 | 4217199 | 11.96157 | 20.96719 | 42.56793 | 18.80662 | -0.62876 | 0.632377 | 0.389969 |
| 7 | 5265400 | 5266799 | 20.35709 | 36.09528 | 29.19956 | 12.90716 | 0.696312 | 0.518606 | 0.389982 |
| X | 33408600 | 33415199 | 182.7813 | 276.344 | 199.1988 | 82.37629 | 0.748267 | 3.318619 | 0.389994 |
| 7 | 123174800 | 123176999 | 18.48958 | 32.47513 | 30.8567 | 13.65571 | 0.460803 | 0.61486 | 0.389997 |
| 2 | 166376400 | 166378999 | 27.0264 | 43.94261 | 81.3001 | 34.88751 | -0.51622 | 1.564893 | 0.390005 |
| 2 | 65261000 | 65261599 | 6.101318 | 11.84736 | 25.83099 | 9.680029 | -0.52634 | -0.25096 | 0.390048 |
| 5 | 24859400 | 24865399 | 48.81302 | 77.65773 | 128.7601 | 53.78122 | -0.39333 | 2.225891 | 0.390065 |
| 18 | 11692800 | 11693399 | 5.22842 | 10.18911 | 23.26476 | 8.434222 | -0.54774 | -0.38718 | 0.390069 |
| 14 | 65426200 | 65428399 | 22.85653 | 39.77304 | 37.1412 | 16.32282 | 0.485489 | 0.788848 | 0.39008 |
| 6 | 73378000 | 73382199 | 46.10499 | 74.62421 | 71.05871 | 30.39433 | 0.433553 | 1.68246 | 0.390083 |
| 9 | 121013600 | 121028999 | 245.8369 | 350.72 | 511.2365 | 211.2933 | -0.34617 | 4.282093 | 0.390089 |
| 10 | 79243000 | 79247599 | 88.99352 | 139.2971 | 136.5519 | 56.64567 | 0.344268 | 2.62815 | 0.390095 |
| 17 | 5859600 | 5863199 | 32.53653 | 52.5842 | 91.29056 | 38.99042 | -0.43312 | 1.704813 | 0.390102 |
| 12 | 13103000 | 13106399 | 20.12379 | 33.68382 | 62.85443 | 27.38961 | -0.51581 | 1.171852 | 0.390103 |
| 15 | 90436600 | 90436799 | 3.636414 | 7.113562 | 18.10249 | 5.989723 | -0.57949 | -0.74821 | 0.39011 |
| 4 | 111952800 | 111955199 | 44.87923 | 72.72707 | 69.42288 | 29.7208 | 0.431579 | 1.654667 | 0.390117 |
| 4 | 152932800 | 152936599 | 36.5413 | 58.61591 | 105.9831 | 44.79287 | -0.50467 | 1.947258 | 0.390125 |
| 5 | 62012200 | 62017599 | 97.72775 | 151.8452 | 149.6781 | 62.14875 | 0.328061 | 2.760062 | 0.390139 |
| 17 | 37722800 | 37726799 | 47.56093 | 76.60478 | 75.34488 | 32.03607 | 0.379077 | 1.821527 | 0.390141 |
| 3 | 124080600 | 124081999 | 15.87941 | 29.19671 | 24.46786 | 10.39178 | 0.683495 | 0.266393 | 0.390142 |
| 13 | 92182200 | 92183799 | 16.41902 | 30.52788 | 23.55941 | 9.854569 | 0.817553 | 0.209421 | 0.390142 |
| 19 | 6423800 | 6426599 | 15.38551 | 26.34006 | 52.64196 | 23.08445 | -0.61164 | 0.936801 | 0.390154 |
| 7 | 39739400 | 39742999 | 18.55744 | 31.8496 | 58.76875 | 25.77632 | -0.49671 | 0.933985 | 0.390162 |
| 9 | 34471800 | 34474199 | 15.42446 | 26.88775 | 47.34198 | 20.82894 | -0.42506 | 0.743425 | 0.390163 |
| 4 | 118624000 | 118624599 | 14.06754 | 25.98187 | 21.5735 | 9.10663 | 0.703904 | 0.224293 | 0.390171 |
| 17 | 38523800 | 38545199 | 26.68608 | 45.65701 | 35.38739 | 15.5172 | 0.74913 | 0.966366 | 0.390177 |
| 6 | 89043200 | 89045599 | 27.80391 | 45.26003 | 80.50283 | 34.56022 | -0.45837 | 1.548369 | 0.390181 |
| 7 | 39503000 | 39506799 | 32.96754 | 55.05077 | 50.32635 | 21.89741 | 0.510101 | 1.223418 | 0.390245 |
| 7 | 42565400 | 42566399 | 22.73651 | 39.26141 | 38.37434 | 16.83476 | 0.418144 | 0.869913 | 0.390251 |
| 7 | 142594200 | 142596799 | 13.99998 | 24.37097 | 46.31356 | 20.35881 | -0.53475 | 0.75411 | 0.390283 |
| 10 | 81974800 | 81978999 | 89.36537 | 140.6659 | 127.5779 | 52.93148 | 0.462524 | 2.522094 | 0.390288 |
| 10 | 45422400 | 45427199 | 55.58332 | 89.54409 | 76.35044 | 32.47305 | 0.585017 | 1.827131 | 0.390295 |
| X | 64462200 | 64465399 | 32.59399 | 56.12591 | 36.30711 | 15.92651 | 1.01049 | 0.937746 | 0.390298 |
| 7 | 38435000 | 38436599 | 22.41587 | 40.25 | 25.38138 | 11.10078 | 1.063953 | 0.450478 | 0.390323 |
| 17 | 7040800 | 7042799 | 9.754201 | 17.83868 | 33.4368 | 14.30496 | -0.47725 | 0.272345 | 0.390334 |
| 17 | 95020800 | 95023399 | 23.7425 | 41.40235 | 35.95004 | 15.82038 | 0.590247 | 0.76263 | 0.39034 |
| 8 | 54197800 | 54198399 | 9.562248 | 18.51616 | 16.73887 | 6.362305 | 0.713514 | -0.18956 | 0.39034 |
| 9 | 54081800 | 54082399 | 4.91856 | 9.577665 | 22.40516 | 8.207309 | -0.59431 | -0.35233 | 0.390343 |
| 10 | 94957600 | 94959999 | 18.36349 | 31.52139 | 55.45517 | 24.32326 | -0.42811 | 0.927787 | 0.390368 |

| | | | | | | | | | |
|---|---|---|---|---|---|---|---|---|---|
| 14 | 94127400 | 94129399 | 16.30934 | 27.92394 | 52.66883 | 23.09638 | -0.52855 | 0.939523 | 0.390381 |
| 17 | 12149600 | 12152999 | 29.86454 | 48.65471 | 82.28596 | 35.3364 | -0.38507 | 1.540527 | 0.390386 |
| 3 | 32679800 | 32680999 | 17.52144 | 29.23274 | 68.43398 | 29.75987 | -0.84155 | 1.20724 | 0.39039 |
| 3 | 8365400 | 8370399 | 54.29696 | 87.42056 | 78.8849 | 33.52252 | 0.502424 | 1.844563 | 0.390397 |
| 19 | 18713800 | 18714399 | 5.106164 | 9.914954 | 23.34218 | 8.747677 | -0.63496 | -0.26901 | 0.390408 |
| 1 | 148342400 | 148343399 | 13.92592 | 25.51376 | 24.84986 | 10.60536 | 0.463394 | 0.273227 | 0.390411 |
| 16 | 50488400 | 50490799 | 24.80212 | 40.76246 | 73.02852 | 31.46862 | -0.46065 | 1.406211 | 0.390418 |
| 3 | 42774400 | 42776199 | 24.797 | 42.641 | 40.95316 | 17.9638 | 0.442179 | 0.919941 | 0.39044 |
| 4 | 145247800 | 145249999 | 31.12473 | 52.70308 | 44.64967 | 19.54795 | 0.619635 | 1.082657 | 0.390453 |
| 6 | 37592600 | 37597599 | 48.07777 | 76.73378 | 126.6587 | 53.13944 | -0.38191 | 2.120947 | 0.390454 |
| 13 | 58943800 | 58947199 | 43.60397 | 70.53609 | 70.07853 | 29.96254 | 0.371691 | 1.688374 | 0.390461 |
| 13 | 50454800 | 50455799 | 9.48056 | 16.80784 | 37.92466 | 16.76558 | -0.77468 | 0.469447 | 0.390486 |
| 18 | 37831600 | 37844399 | 321.3774 | 457.9678 | 412.4947 | 170.5161 | 0.346454 | 4.301732 | 0.390499 |
| 1 | 190855800 | 190859599 | 21.01337 | 34.87547 | 68.8079 | 29.81864 | -0.59631 | 1.273142 | 0.390535 |
| 13 | 80189400 | 80189999 | 8.299637 | 15.47814 | 30.56922 | 12.72151 | -0.52235 | 0.140252 | 0.390558 |
| 12 | 113782200 | 113786399 | 65.0199 | 104.1014 | 88.63592 | 37.35362 | 0.576456 | 2.033172 | 0.3906 |
| 13 | 62062200 | 62071799 | 197.8158 | 295.4079 | 252.4705 | 104.0795 | 0.487079 | 3.557391 | 0.390603 |
| 5 | 13295600 | 13297199 | 17.55102 | 31.87063 | 26.43377 | 11.40985 | 0.682983 | 0.353695 | 0.390617 |
| 11 | 33675600 | 33679199 | 27.46647 | 44.76661 | 79.45357 | 34.1253 | -0.45398 | 1.528941 | 0.390617 |
| 8 | 14536000 | 14538799 | 17.20877 | 29.76579 | 52.77347 | 23.15282 | -0.43688 | 0.851054 | 0.390639 |
| 2 | 56420200 | 56420599 | 4.349431 | 8.499126 | 20.06698 | 6.967708 | -0.5374 | -0.5609 | 0.390662 |
| 17 | 85395200 | 85398599 | 16.62576 | 28.37573 | 55.89169 | 24.50343 | -0.59173 | 0.971689 | 0.390667 |
| 9 | 121091600 | 121093599 | 13.54279 | 23.69999 | 43.62261 | 19.2459 | -0.48808 | 0.671191 | 0.390671 |
| 7 | 92431600 | 92434599 | 17.52208 | 30.19569 | 52.75278 | 23.14113 | -0.41686 | 0.885219 | 0.390676 |
| 11 | 73365800 | 73371599 | 78.63006 | 124.412 | 118.5059 | 49.25638 | 0.397729 | 2.399467 | 0.39068 |
| 18 | 58201000 | 58203199 | 18.51753 | 31.55141 | 56.48606 | 24.76065 | -0.45494 | 0.995496 | 0.390688 |
| 14 | 48397200 | 48399799 | 30.65685 | 49.48265 | 91.90725 | 39.18873 | -0.53237 | 1.741208 | 0.390691 |
| 3 | 72659400 | 72667999 | 196.7407 | 291.4355 | 276.2214 | 113.7575 | 0.32723 | 3.700239 | 0.390694 |
| 19 | 19130800 | 19133199 | 34.18912 | 56.70561 | 47.68129 | 20.6538 | 0.630148 | 1.300253 | 0.390695 |
| 2 | 87654600 | 87658199 | 33.38258 | 55.96819 | 48.60941 | 21.20402 | 0.583683 | 1.182476 | 0.390708 |
| 8 | 117573400 | 117579999 | 86.39686 | 132.9654 | 208.6734 | 86.66574 | -0.35275 | 2.935777 | 0.390716 |
| 7 | 43802400 | 43805599 | 50.76238 | 81.85435 | 76.7379 | 32.67927 | 0.450076 | 1.793843 | 0.390718 |
| X | 3746400 | 3754399 | 97.07575 | 152.8642 | 121.678 | 50.48628 | 0.651023 | 2.51894 | 0.390726 |
| 16 | 19818800 | 19819399 | 11.26639 | 21.41666 | 21.13842 | 8.488202 | 0.514198 | 0.017615 | 0.390742 |
| 16 | 32629400 | 32631799 | 14.47505 | 25.27026 | 45.21643 | 19.91234 | -0.44781 | 0.715003 | 0.390745 |
| 2 | 129183800 | 129186999 | 74.70459 | 118.2144 | 117.0161 | 48.63995 | 0.342472 | 2.39405 | 0.390755 |
| 7 | 23047800 | 23062599 | 335.3374 | 476.2327 | 426.5258 | 176.4174 | 0.349663 | 4.352091 | 0.390759 |
| 5 | 85740800 | 85741199 | 7.959843 | 15.49228 | 16.22917 | 5.96832 | 0.546185 | -0.32516 | 0.390765 |
| 10 | 66572600 | 66574599 | 35.41626 | 58.10167 | 56.30978 | 24.2458 | 0.422102 | 1.439298 | 0.390773 |
| X | 11345400 | 11349199 | 19.10421 | 32.60107 | 59.76051 | 26.19925 | -0.48849 | 0.990428 | 0.390786 |
| 7 | 27706800 | 27710199 | 20.37344 | 34.27958 | 61.34711 | 26.80651 | -0.45571 | 1.118042 | 0.390804 |
| 11 | 107027000 | 107032799 | 40.9014 | 65.52436 | 110.3105 | 46.52935 | -0.40419 | 1.994203 | 0.390807 |
| 4 | 29268600 | 29270799 | 33.12485 | 54.91225 | 54.17124 | 23.4567 | 0.400361 | 1.308614 | 0.390817 |
| 14 | 53029400 | 53031399 | 14.13261 | 24.79543 | 44.03471 | 19.46969 | -0.43694 | 0.63189 | 0.390819 |
| 8 | 104137200 | 104138199 | 8.045117 | 14.87082 | 31.50642 | 13.28493 | -0.63727 | 0.19319 | 0.390821 |
| 2 | 108155400 | 108158799 | 25.74478 | 44.57424 | 38.6393 | 16.95237 | 0.59134 | 0.860628 | 0.390822 |
| 9 | 19392600 | 19403999 | 175.8643 | 266.0128 | 204.3219 | 84.50758 | 0.65731 | 3.306951 | 0.390845 |

| | | | | | | | | | |
|---|---|---|---|---|---|---|---|---|---|
| 13 | 120358000 | 120360399 | 20.63893 | 34.54851 | 62.24498 | 27.12524 | -0.46527 | 1.174686 | 0.390854 |
| 3 | 138197800 | 138198599 | 10.88593 | 20.79945 | 21.15465 | 8.403402 | 0.484575 | -0.02159 | 0.390878 |
| 1 | 7696800 | 7702599 | 55.11381 | 88.93445 | 72.45796 | 30.88504 | 0.65326 | 1.77731 | 0.390892 |
| 5 | 142219400 | 142236399 | 186.6301 | 272.4401 | 412.014 | 169.759 | -0.36652 | 3.936537 | 0.390908 |
| 16 | 38428600 | 38431799 | 29.53149 | 48.27606 | 83.21206 | 35.77844 | -0.40896 | 1.492121 | 0.390909 |
| 9 | 62344800 | 62350399 | 72.30638 | 112.0882 | 188.6146 | 78.34271 | -0.4452 | 2.793707 | 0.390913 |
| 7 | 47123800 | 47126599 | 56.22888 | 90.06706 | 87.63564 | 36.9575 | 0.38505 | 2.010105 | 0.390916 |
| 8 | 94655400 | 94656999 | 11.66762 | 20.50401 | 41.00916 | 18.15693 | -0.6069 | 0.589575 | 0.39092 |
| X | 31755800 | 31760599 | 73.8917 | 117.4124 | 95.57565 | 39.86567 | 0.629627 | 2.277692 | 0.390926 |
| 10 | 118818600 | 118821399 | 35.04182 | 56.44884 | 96.75373 | 41.14759 | -0.42019 | 1.809616 | 0.390927 |
| 4 | 135342600 | 135358399 | 126.1471 | 188.0082 | 348.2298 | 143.5022 | -0.63065 | 3.586624 | 0.390946 |
| 7 | 60617400 | 60621999 | 70.45624 | 112.7422 | 86.77058 | 36.53821 | 0.721051 | 2.057028 | 0.390966 |
| 7 | 62895200 | 62896999 | 24.41283 | 41.92559 | 40.97494 | 17.97264 | 0.416696 | 0.929264 | 0.390976 |
| 5 | 117296000 | 117297399 | 8.593792 | 15.58076 | 34.42514 | 14.88855 | -0.72305 | 0.315525 | 0.390979 |
| 13 | 17860000 | 17861799 | 14.1247 | 24.80739 | 45.05823 | 19.93977 | -0.46932 | 0.616142 | 0.390983 |
| 7 | 20581800 | 20586599 | 33.146 | 56.90538 | 36.71868 | 16.10544 | 1.013143 | 0.964186 | 0.390992 |
| 4 | 82518200 | 82520399 | 27.77957 | 47.92081 | 39.12963 | 17.16558 | 0.676229 | 0.90269 | 0.390993 |
| 13 | 95702600 | 95705799 | 30.51172 | 49.70415 | 84.18542 | 36.15079 | -0.38734 | 1.544222 | 0.390996 |
| 15 | 82199600 | 82200999 | 11.01882 | 19.41383 | 39.94388 | 17.6802 | -0.6463 | 0.551657 | 0.391004 |
| 7 | 10502600 | 10531999 | 623.9815 | 841.939 | 698.0675 | 291.0732 | 0.386565 | 5.13767 | 0.391026 |
| 9 | 120448200 | 120463199 | 169.5022 | 249.9208 | 371.6898 | 153.0666 | -0.32868 | 3.776821 | 0.391026 |
| 14 | 48209000 | 48211399 | 32.66142 | 52.9616 | 89.15579 | 38.18656 | -0.38407 | 1.631034 | 0.391045 |
| 5 | 39664000 | 39664799 | 12.21535 | 21.60186 | 39.31184 | 17.38756 | -0.46829 | 0.51629 | 0.391049 |
| 8 | 34864200 | 34866599 | 12.21535 | 21.60186 | 39.31184 | 17.38756 | -0.46829 | 0.51629 | 0.391049 |
| 16 | 82360000 | 82362799 | 26.61327 | 45.31657 | 43.08343 | 18.88464 | 0.454497 | 1.01537 | 0.391053 |
| 1 | 166291800 | 166292799 | 5.523454 | 10.65628 | 25.81166 | 9.974145 | -0.71707 | -0.14741 | 0.391055 |
| 6 | 143166200 | 143170599 | 27.92533 | 45.14119 | 90.89419 | 38.83158 | -0.64605 | 1.690879 | 0.391058 |
| 7 | 14322200 | 14332599 | 211.8263 | 312.4288 | 291.749 | 120.1464 | 0.342832 | 3.773111 | 0.391059 |
| 12 | 7897600 | 7901399 | 32.61627 | 54.73852 | 48.58974 | 21.20891 | 0.552314 | 1.170245 | 0.391086 |
| 10 | 39359200 | 39359799 | 10.54922 | 20.35076 | 20.21315 | 7.81256 | 0.555037 | -0.11546 | 0.391092 |
| 3 | 48190800 | 48191599 | 10.61072 | 20.39519 | 20.09796 | 7.849475 | 0.552027 | -0.08312 | 0.391099 |
| 8 | 22852600 | 22854399 | 22.73495 | 39.87348 | 33.05458 | 14.61036 | 0.656802 | 0.66653 | 0.391108 |
| 5 | 101153200 | 101154999 | 13.35294 | 23.4576 | 41.94155 | 18.56433 | -0.44647 | 0.604219 | 0.391126 |
| 17 | 20040200 | 20047399 | 95.49173 | 150.8976 | 111.577 | 46.34635 | 0.761133 | 2.441244 | 0.391126 |
| 4 | 91897000 | 91898599 | 14.92649 | 27.7578 | 23.61682 | 9.875587 | 0.67785 | 0.194884 | 0.391135 |
| 13 | 32757600 | 32758599 | 9.04584 | 16.83586 | 32.70753 | 13.65782 | -0.50318 | 0.163219 | 0.391136 |
| 5 | 149616000 | 149629999 | 145.9474 | 217.7971 | 331.4394 | 136.6153 | -0.34569 | 3.564997 | 0.391151 |
| 2 | 36450200 | 36452399 | 30.41477 | 52.00804 | 41.35463 | 18.13301 | 0.71233 | 0.994357 | 0.391152 |
| 3 | 147576400 | 147577199 | 10.32296 | 18.50757 | 35.85721 | 15.70274 | -0.5467 | 0.40043 | 0.391166 |
| 4 | 151816800 | 151818399 | 10.32296 | 18.50757 | 35.85721 | 15.70274 | -0.5467 | 0.40043 | 0.391166 |
| 5 | 91038600 | 91039599 | 10.68121 | 18.89604 | 39.36904 | 17.41153 | -0.66206 | 0.501746 | 0.391167 |
| 10 | 119653600 | 119655799 | 10.68121 | 18.89604 | 39.36904 | 17.41153 | -0.66206 | 0.501746 | 0.391167 |
| 18 | 50136200 | 50137399 | 9.393056 | 17.30917 | 33.02922 | 13.98533 | -0.4922 | 0.224249 | 0.391178 |
| 15 | 103882600 | 103885599 | 47.42332 | 76.67567 | 64.02104 | 27.35829 | 0.621194 | 1.705803 | 0.391184 |
| X | 116695400 | 116698599 | 35.04125 | 59.8322 | 38.46773 | 16.8652 | 1.017434 | 1.022742 | 0.391193 |
| 2 | 172176400 | 172188199 | 141.1779 | 210.227 | 326.9216 | 134.7094 | -0.37973 | 3.603216 | 0.391199 |
| 18 | 47576800 | 47578199 | 18.00206 | 32.82147 | 25.31068 | 10.87025 | 0.791896 | 0.331899 | 0.391215 |

| | | | | | | | | | |
|---|---|---|---|---|---|---|---|---|---|
| X | 140814600 | 140816599 | 24.13109 | 42.81459 | 26.83205 | 11.8542 | 1.063246 | 0.536238 | 0.391223 |
| 1 | 97135200 | 97138399 | 29.4141 | 49.92467 | 44.10263 | 19.32234 | 0.559573 | 1.055331 | 0.391225 |
| 6 | 133529800 | 133532799 | 75.50879 | 119.3332 | 119.0207 | 49.44098 | 0.330093 | 2.424818 | 0.391232 |
| 4 | 132848000 | 132850599 | 12.23702 | 21.57764 | 39.87215 | 17.64559 | -0.4918 | 0.549844 | 0.391278 |
| 4 | 10470600 | 10476199 | 55.29934 | 89.17525 | 72.55474 | 30.9001 | 0.654125 | 1.794742 | 0.391281 |
| 2 | 70945600 | 70948999 | 23.62183 | 39.05724 | 69.31274 | 29.95939 | -0.44438 | 1.327362 | 0.391283 |
| 3 | 116045000 | 116046999 | 21.3029 | 35.43339 | 66.08462 | 28.68174 | -0.51511 | 1.247145 | 0.391288 |
| 14 | 27272600 | 27274599 | 18.63866 | 31.44117 | 58.49778 | 25.58133 | -0.51126 | 1.088355 | 0.391288 |
| 4 | 41745000 | 41749999 | 61.18187 | 97.7209 | 95.83568 | 40.24999 | 0.36912 | 2.103036 | 0.391318 |
| 18 | 77624800 | 77628599 | 50.38411 | 81.01569 | 80.53209 | 34.15832 | 0.360844 | 1.88027 | 0.391329 |
| 16 | 51783600 | 51786399 | 29.7979 | 51.37494 | 37.94 | 16.64343 | 0.820431 | 0.914773 | 0.391337 |
| 15 | 46649600 | 46651199 | 21.15946 | 37.4165 | 29.68902 | 13.13157 | 0.722567 | 0.55245 | 0.391347 |
| 2 | 7001000 | 7002199 | 15.27646 | 28.1792 | 24.5987 | 10.40141 | 0.629631 | 0.244585 | 0.39137 |
| 8 | 33955600 | 33956799 | 9.207658 | 17.04734 | 33.67984 | 14.17005 | -0.5355 | 0.196601 | 0.39137 |
| 6 | 116592600 | 116594799 | 19.24275 | 32.57552 | 57.48667 | 25.16622 | -0.43506 | 1.058701 | 0.391379 |
| 1 | 158936400 | 158938399 | 12.3418 | 21.86049 | 40.29848 | 17.81824 | -0.48605 | 0.499896 | 0.391384 |
| 4 | 133298200 | 133302599 | 31.89442 | 51.60113 | 88.35585 | 37.78093 | -0.41155 | 1.675686 | 0.391405 |
| 7 | 59778200 | 59783399 | 65.05724 | 104.1153 | 90.19468 | 37.98652 | 0.550854 | 2.047279 | 0.391411 |
| X | 80426000 | 80430399 | 59.04193 | 95.4041 | 66.31096 | 28.31505 | 0.884226 | 1.745337 | 0.391413 |
| X | 4508800 | 4510399 | 23.77118 | 41.69651 | 32.43731 | 14.33916 | 0.748061 | 0.669447 | 0.391416 |
| 7 | 36114800 | 36118599 | 21.06347 | 35.254 | 63.50133 | 27.67018 | -0.46509 | 1.175869 | 0.391419 |
| 17 | 49418200 | 49420999 | 22.21557 | 37.05896 | 65.1691 | 28.33743 | -0.43055 | 1.216998 | 0.391422 |
| 12 | 24791000 | 24792999 | 13.50345 | 23.69098 | 42.4928 | 18.78755 | -0.45106 | 0.62671 | 0.391432 |
| 11 | 7158200 | 7164399 | 40.66968 | 64.89287 | 120.4287 | 50.50652 | -0.55095 | 2.120813 | 0.391457 |
| 14 | 53050000 | 53050999 | 12.33249 | 23.34127 | 22.54123 | 9.135354 | 0.533473 | 0.059686 | 0.391463 |
| 1 | 27140200 | 27142599 | 22.14627 | 39.05374 | 30.17372 | 13.3533 | 0.759339 | 0.588433 | 0.391469 |
| 12 | 22730400 | 22731999 | 50.00333 | 80.39165 | 71.03759 | 30.12477 | 0.530064 | 1.880556 | 0.391497 |
| X | 25429200 | 25431599 | 23.63898 | 41.59173 | 30.71795 | 13.6003 | 0.823016 | 0.623458 | 0.391551 |
| 4 | 113119600 | 113125399 | 82.85735 | 130.8255 | 122.4816 | 50.85481 | 0.420209 | 2.452904 | 0.391557 |
| 3 | 104544200 | 104547399 | 18.27518 | 30.90991 | 59.10577 | 25.86801 | -0.55054 | 1.062766 | 0.391577 |
| 8 | 72712000 | 72712799 | 17.36637 | 30.7783 | 28.82163 | 12.74218 | 0.486349 | 0.50525 | 0.3916 |
| 13 | 68514200 | 68524199 | 216.4948 | 319.5472 | 294.536 | 121.2951 | 0.362595 | 3.762576 | 0.391601 |
| 13 | 89741400 | 89743399 | 28.06052 | 48.51545 | 36.81658 | 16.15199 | 0.781954 | 0.879473 | 0.391631 |
| 18 | 19994800 | 19995799 | 15.68959 | 26.91619 | 56.42181 | 24.7461 | -0.67984 | 0.923466 | 0.391634 |
| 13 | 75854200 | 75857999 | 23.59735 | 38.95796 | 72.27814 | 31.20863 | -0.50859 | 1.34417 | 0.391653 |
| 14 | 51508400 | 51510199 | 10.95129 | 19.80785 | 39.08279 | 16.95324 | -0.56401 | 0.358327 | 0.39169 |
| 6 | 113052800 | 113055199 | 14.78823 | 25.63721 | 48.98496 | 21.49221 | -0.54352 | 0.813662 | 0.391699 |
| 16 | 33853600 | 33855999 | 14.78823 | 25.63721 | 48.98496 | 21.49221 | -0.54352 | 0.813662 | 0.391699 |
| 16 | 8723800 | 8724399 | 4.646071 | 9.064871 | 20.95341 | 7.478759 | -0.54354 | -0.45603 | 0.391701 |
| 8 | 11047600 | 11048799 | 7.482129 | 14.26073 | 28.34985 | 11.32042 | -0.48095 | -0.01781 | 0.391705 |
| 4 | 57709600 | 57711199 | 10.96098 | 19.5906 | 36.84183 | 16.17616 | -0.50673 | 0.4265 | 0.391707 |
| 2 | 72703600 | 72705399 | 10.97083 | 19.42036 | 39.84513 | 17.61993 | -0.63983 | 0.496891 | 0.391726 |
| 3 | 15109800 | 15112599 | 29.58327 | 50.31352 | 44.10695 | 19.33141 | 0.570937 | 1.040253 | 0.39174 |
| 3 | 127630600 | 127631599 | 7.077402 | 13.35161 | 28.62448 | 11.68089 | -0.6165 | 0.050438 | 0.391747 |
| 14 | 53054600 | 53055399 | 6.565548 | 12.69803 | 27.87989 | 10.65701 | -0.56347 | -0.17077 | 0.39175 |
| 13 | 27483800 | 27484599 | 10.71412 | 20.71823 | 19.70645 | 7.538346 | 0.630927 | -0.14922 | 0.391756 |
| 15 | 38975600 | 38976799 | 10.30558 | 18.44583 | 36.32764 | 15.93098 | -0.57185 | 0.409638 | 0.391765 |

| | | | | | | | | | |
|---|---|---|---|---|---|---|---|---|---|
| 7 | 6071800 | 6079199 | 129.3418 | 198.0008 | 183.8015 | 76.27333 | 0.398207 | 3.052163 | 0.391771 |
| 14 | 43242800 | 43243799 | 18.38407 | 32.39001 | 31.23593 | 13.82587 | 0.440764 | 0.577928 | 0.39179 |
| 6 | 88051000 | 88052399 | 9.387386 | 17.24969 | 34.59064 | 14.70314 | -0.56762 | 0.242278 | 0.391794 |
| 6 | 117571000 | 117571399 | 3.813785 | 7.455314 | 18.91032 | 6.500963 | -0.62574 | -0.60905 | 0.391808 |
| X | 5583600 | 5589599 | 81.95713 | 130.9315 | 81.1463 | 34.09092 | 1.030422 | 2.118468 | 0.391817 |
| 2 | 161696400 | 161696999 | 4.480294 | 8.748177 | 20.48752 | 7.215793 | -0.54415 | -0.50585 | 0.391832 |
| 8 | 87292200 | 87294199 | 13.23554 | 23.05145 | 45.79333 | 20.13627 | -0.59825 | 0.743171 | 0.391846 |
| 6 | 17955600 | 17958399 | 24.54898 | 42.70342 | 37.04683 | 16.27783 | 0.591229 | 0.801913 | 0.391851 |
| 17 | 22360200 | 22362199 | 40.46372 | 66.22583 | 61.61765 | 26.50501 | 0.479432 | 1.484814 | 0.391856 |
| 5 | 93951200 | 93952399 | 22.15595 | 38.73123 | 30.43167 | 13.41393 | 0.733972 | 0.711109 | 0.391874 |
| 5 | 12598400 | 12599199 | 10.12846 | 18.274 | 34.85356 | 15.16587 | -0.51806 | 0.357512 | 0.391876 |
| 13 | 65913000 | 65921599 | 109.556 | 171.2588 | 119.6052 | 49.62237 | 0.831384 | 2.658623 | 0.391887 |
| 13 | 75758800 | 75762999 | 39.07964 | 62.67839 | 110.1987 | 46.56406 | -0.4647 | 1.954573 | 0.391889 |
| 3 | 97206600 | 97207799 | 11.39157 | 20.274 | 38.28779 | 16.87128 | -0.51679 | 0.455784 | 0.391893 |
| 5 | 97292600 | 97293399 | 11.70491 | 22.23431 | 21.47817 | 8.637722 | 0.542377 | 0.024913 | 0.391912 |
| X | 3730800 | 3733399 | 27.56861 | 48.79666 | 23.83731 | 10.54131 | 1.418891 | 0.57413 | 0.391913 |
| 7 | 108074000 | 108083599 | 176.3169 | 264.5048 | 242.6688 | 100.1311 | 0.390783 | 3.469047 | 0.391921 |
| 7 | 60926200 | 60927399 | 19.7263 | 34.84567 | 30.98384 | 13.70714 | 0.558678 | 0.557651 | 0.391933 |
| 1 | 49012400 | 49019799 | 113.6343 | 175.9633 | 156.083 | 64.83272 | 0.476373 | 2.827581 | 0.391945 |
| 6 | 118544600 | 118546999 | 22.52247 | 37.7979 | 69.33374 | 30.26313 | -0.49142 | 1.157377 | 0.391952 |
| 13 | 98238600 | 98243399 | 45.80746 | 73.22052 | 118.4436 | 49.79893 | -0.35039 | 2.071019 | 0.391957 |
| 10 | 51114800 | 51115799 | 11.72255 | 20.88871 | 37.78986 | 16.63462 | -0.45399 | 0.450793 | 0.391972 |
| 5 | 119581200 | 119583599 | 19.9078 | 33.48666 | 60.74713 | 26.54207 | -0.4752 | 1.118672 | 0.391975 |
| 1 | 137181000 | 137181799 | 13.42056 | 25.61235 | 20.22844 | 8.055364 | 0.842685 | 0.006679 | 0.391977 |
| 13 | 65524200 | 65530199 | 181.8566 | 270.6561 | 244.7033 | 100.8255 | 0.401494 | 3.615247 | 0.391989 |
| 7 | 12598600 | 12607599 | 102.3956 | 159.9478 | 139.8343 | 58.01952 | 0.506963 | 2.670836 | 0.392031 |
| 17 | 84023400 | 84028799 | 56.58383 | 89.51928 | 144.4353 | 60.03035 | -0.36164 | 2.391536 | 0.392039 |
| 6 | 30905000 | 30906399 | 19.50309 | 34.39144 | 27.6682 | 12.24457 | 0.701369 | 0.569283 | 0.39204 |
| 9 | 33666800 | 33667399 | 7.817032 | 14.81837 | 29.30542 | 11.83715 | -0.48802 | 0.026515 | 0.392058 |
| 14 | 78572000 | 78572799 | 7.817032 | 14.81837 | 29.30542 | 11.83715 | -0.48802 | 0.026515 | 0.392058 |
| 10 | 41068000 | 41075799 | 68.7306 | 106.9052 | 193.3676 | 80.27645 | -0.5455 | 2.735265 | 0.392066 |
| 8 | 73656200 | 73659399 | 60.48117 | 96.60149 | 95.20437 | 39.9847 | 0.362061 | 2.103659 | 0.3921 |
| 19 | 57487800 | 57488999 | 16.52081 | 29.56975 | 28.08524 | 12.31661 | 0.473722 | 0.433271 | 0.392112 |
| 17 | 22045800 | 22048999 | 39.9157 | 66.25814 | 51.05733 | 22.12874 | 0.755773 | 1.310254 | 0.392115 |
| 12 | 64200000 | 64204999 | 78.86154 | 124.314 | 123.8722 | 51.40377 | 0.328548 | 2.485364 | 0.392128 |
| 17 | 73574000 | 73575399 | 17.99631 | 32.00764 | 29.60213 | 13.05122 | 0.506911 | 0.486697 | 0.39214 |
| X | 109435600 | 109436399 | 16.04716 | 29.44232 | 19.39464 | 8.261044 | 1.026116 | 0.270304 | 0.392145 |
| 17 | 89144800 | 89146199 | 22.62167 | 39.37139 | 37.10777 | 16.30996 | 0.472168 | 0.784356 | 0.392157 |
| X | 34609800 | 34612199 | 16.2719 | 31.15732 | 17.19014 | 6.787929 | 1.366647 | 0.001758 | 0.392161 |
| 2 | 59927400 | 59929199 | 17.89152 | 30.47992 | 55.91119 | 24.50835 | -0.48988 | 0.994031 | 0.392165 |
| 15 | 52907000 | 52909799 | 17.64483 | 30.38582 | 56.06885 | 24.59527 | -0.49581 | 0.894169 | 0.392167 |
| 14 | 42547200 | 42549799 | 21.55234 | 39.40797 | 20.57469 | 8.804409 | 1.353713 | 0.333201 | 0.392175 |
| 3 | 96011400 | 96012399 | 5.983149 | 11.51772 | 25.78688 | 10.02789 | -0.61401 | -0.12364 | 0.392177 |
| 3 | 100139800 | 100141799 | 35.49382 | 58.32774 | 57.98138 | 24.98348 | 0.386585 | 1.421036 | 0.392177 |
| 10 | 108250800 | 108251599 | 7.309841 | 13.99246 | 28.52767 | 11.27492 | -0.50383 | -0.05378 | 0.39218 |
| 5 | 94395400 | 94396599 | 11.62698 | 21.98728 | 22.03095 | 8.944607 | 0.478324 | 0.057021 | 0.392184 |
| 16 | 8588400 | 8590199 | 13.56257 | 23.73139 | 44.14966 | 19.4762 | -0.50337 | 0.675116 | 0.39219 |

| | | | | | | | | | |
|---|---|---|---|---|---|---|---|---|---|
| 15 | 5325000 | 5325799 | 13.46559 | 24.86057 | 24.40971 | 10.30923 | 0.462674 | 0.226373 | 0.392193 |
| 16 | 78233600 | 78238799 | 39.2023 | 63.00902 | 107.7748 | 45.68544 | -0.42141 | 1.887354 | 0.39221 |
| 6 | 59625200 | 59627399 | 22.36887 | 39.23938 | 33.0528 | 14.61208 | 0.633825 | 0.661002 | 0.39221 |
| 8 | 21973600 | 21977999 | 52.72433 | 85.64447 | 61.08539 | 26.19752 | 0.854167 | 1.619605 | 0.392238 |
| 6 | 132767200 | 132774799 | 160.0818 | 241.6521 | 223.477 | 92.3713 | 0.387182 | 3.343455 | 0.392238 |
| 6 | 49162200 | 49164599 | 28.6335 | 48.1331 | 47.49938 | 20.74972 | 0.400052 | 1.140218 | 0.392255 |
| 13 | 23912400 | 23914199 | 13.06552 | 22.89984 | 42.55939 | 18.80115 | -0.50189 | 0.64198 | 0.392267 |
| 15 | 22318400 | 22319399 | 14.26739 | 25.97204 | 23.93083 | 10.29761 | 0.535192 | 0.310529 | 0.39227 |
| 15 | 80094800 | 80096799 | 15.93005 | 27.12586 | 55.20554 | 24.19813 | -0.6388 | 0.995049 | 0.39227 |
| 7 | 27372800 | 27375999 | 21.95747 | 36.48683 | 66.06445 | 28.65385 | -0.47264 | 1.261145 | 0.392273 |
| 16 | 87420600 | 87430399 | 106.3831 | 162.1096 | 252.8121 | 104.7986 | -0.35484 | 3.142192 | 0.392278 |
| 1 | 165021200 | 165022199 | 8.839177 | 16.16133 | 32.93539 | 14.09485 | -0.5973 | 0.265024 | 0.392283 |
| 2 | 171466200 | 171467599 | 8.839177 | 16.16133 | 32.93539 | 14.09485 | -0.5973 | 0.265024 | 0.392283 |
| 6 | 100758600 | 100762199 | 48.98813 | 78.81241 | 77.68338 | 32.97512 | 0.373914 | 1.859747 | 0.392297 |
| 13 | 10938800 | 10941599 | 27.25774 | 46.03401 | 45.34247 | 19.83545 | 0.402957 | 1.092972 | 0.392297 |
| 4 | 90421000 | 90422999 | 25.5591 | 44.04665 | 40.31733 | 17.68607 | 0.511831 | 0.902576 | 0.392299 |
| 6 | 146478600 | 146480199 | 12.73308 | 22.25394 | 45.2048 | 19.92198 | -0.62982 | 0.687654 | 0.392303 |
| 8 | 54792400 | 54794199 | 17.15319 | 31.53025 | 25.0421 | 10.63957 | 0.760513 | 0.281162 | 0.392331 |
| 7 | 141686400 | 141689599 | 18.26355 | 30.9475 | 57.47937 | 25.17034 | -0.50853 | 1.04553 | 0.392335 |
| X | 24379400 | 24385999 | 77.9157 | 124.2356 | 93.07235 | 38.99554 | 0.754406 | 2.175305 | 0.392337 |
| 3 | 15922600 | 15926999 | 46.73646 | 76.55117 | 57.44096 | 24.71625 | 0.789415 | 1.488696 | 0.39238 |
| 1 | 12114600 | 12115599 | 11.46946 | 22.22841 | 17.26173 | 6.534649 | 0.933831 | -0.18382 | 0.392382 |
| 3 | 69280200 | 69280999 | 11.14247 | 21.59155 | 19.21733 | 7.279924 | 0.739467 | -0.17682 | 0.392401 |
| 18 | 33009800 | 33010799 | 15.63595 | 28.18294 | 27.24614 | 11.87028 | 0.454821 | 0.384293 | 0.392406 |
| 6 | 122931600 | 122934799 | 29.61233 | 50.55377 | 41.85188 | 18.34864 | 0.653906 | 1.003096 | 0.392431 |
| Y | 39174200 | 39176799 | 15.07639 | 27.52105 | 16.50192 | 7.076273 | 1.153255 | 0.27993 | 0.392434 |
| 7 | 33409000 | 33409399 | 10.52399 | 20.15575 | 20.64774 | 8.148023 | 0.48297 | -0.04841 | 0.392454 |
| 6 | 119599800 | 119602399 | 38.09407 | 62.40068 | 60.93085 | 26.21883 | 0.41048 | 1.467434 | 0.392487 |
| 6 | 112536200 | 112543199 | 54.2414 | 85.97615 | 140.2481 | 58.35484 | -0.37516 | 2.341959 | 0.392495 |
| 7 | 4843800 | 4848999 | 43.35518 | 69.27486 | 118.2424 | 49.68812 | -0.42829 | 2.078992 | 0.392506 |
| 3 | 88120800 | 88124599 | 27.76594 | 44.88524 | 90.23123 | 38.55024 | -0.64362 | 1.689516 | 0.392516 |
| 8 | 15776000 | 15777799 | 16.35107 | 29.43646 | 27.55178 | 12.02151 | 0.499656 | 0.395525 | 0.392519 |
| X | 74525000 | 74527799 | 29.88953 | 52.25643 | 29.39869 | 12.95985 | 1.213462 | 0.748854 | 0.392542 |
| 8 | 39987200 | 39990399 | 27.56437 | 45.11132 | 77.15904 | 33.18951 | -0.39661 | 1.467645 | 0.392542 |
| 16 | 49960400 | 49968799 | 116.5965 | 176.04 | 273.2355 | 112.9447 | -0.3591 | 3.333441 | 0.392545 |
| 4 | 141979800 | 141982599 | 28.53349 | 46.67088 | 79.02839 | 33.98657 | -0.38277 | 1.478866 | 0.392546 |
| 15 | 75038000 | 75040999 | 25.9676 | 42.07843 | 92.0766 | 39.41761 | -0.76224 | 1.635917 | 0.39255 |
| 16 | 34949000 | 34950999 | 12.89232 | 22.58679 | 42.56556 | 18.79721 | -0.52192 | 0.646856 | 0.392555 |
| 5 | 16454200 | 16455199 | 11.2346 | 21.61758 | 20.0594 | 7.810285 | 0.64209 | -0.08661 | 0.392571 |
| 7 | 108228800 | 108230199 | 11.2346 | 21.61758 | 20.0594 | 7.810285 | 0.64209 | -0.08661 | 0.392571 |
| 12 | 55445200 | 55445999 | 7.247785 | 13.60519 | 29.08342 | 11.96993 | -0.62222 | 0.086282 | 0.392582 |
| 6 | 125540200 | 125541999 | 9.182814 | 16.62687 | 35.4068 | 15.33823 | -0.6721 | 0.328468 | 0.392594 |
| 11 | 24312200 | 24314199 | 9.182814 | 16.62687 | 35.4068 | 15.33823 | -0.6721 | 0.328468 | 0.392594 |
| 14 | 35749800 | 35750999 | 6.870221 | 13.25064 | 28.27563 | 10.93558 | -0.53902 | -0.12731 | 0.392607 |
| 17 | 53580000 | 53581799 | 19.02525 | 32.4995 | 56.51339 | 24.77773 | -0.41247 | 0.975612 | 0.392611 |
| 14 | 39786600 | 39789999 | 24.48981 | 42.61118 | 36.3858 | 15.9899 | 0.613846 | 0.795019 | 0.392616 |
| 17 | 54228000 | 54229399 | 20.02987 | 35.78604 | 28.32023 | 12.43851 | 0.734233 | 0.470155 | 0.392632 |

| | | | | | | | | | |
|---|---|---|---|---|---|---|---|---|---|
| 4 | 121230000 | 121236999 | 133.3019 | 203.5713 | 187.6993 | 77.8474 | 0.405215 | 3.100539 | 0.392633 |
| 2 | 155953000 | 155953999 | 7.265938 | 13.57612 | 30.07493 | 12.47566 | -0.68274 | 0.117703 | 0.392638 |
| 14 | 31717200 | 31721399 | 47.19911 | 76.02431 | 73.59883 | 31.29504 | 0.40193 | 1.818909 | 0.392695 |
| 4 | 146698400 | 146700599 | 15.8322 | 29.53375 | 23.06057 | 9.592353 | 0.806976 | 0.182519 | 0.392713 |
| 19 | 16529400 | 16531399 | 20.1389 | 35.37253 | 32.62882 | 14.44015 | 0.503283 | 0.626929 | 0.39272 |
| 4 | 100432400 | 100435599 | 41.68979 | 69.01054 | 52.23156 | 22.59417 | 0.781394 | 1.348226 | 0.392772 |
| 16 | 75954200 | 75957199 | 31.5083 | 52.53559 | 50.85901 | 22.10833 | 0.427766 | 1.235293 | 0.392777 |
| 15 | 20247600 | 20248799 | 14.48769 | 26.68718 | 25.33938 | 10.73485 | 0.507566 | 0.249448 | 0.392781 |
| 19 | 44475600 | 44477799 | 18.7189 | 31.83965 | 55.89979 | 24.50026 | -0.42718 | 1.012755 | 0.392791 |
| 19 | 39202400 | 39206199 | 66.46267 | 106.0501 | 97.89478 | 41.057 | 0.454759 | 2.139249 | 0.392799 |
| 7 | 60654600 | 60656199 | 18.9108 | 34.58217 | 24.85721 | 10.63539 | 0.896466 | 0.322647 | 0.392806 |
| 2 | 83295600 | 83296599 | 11.58301 | 20.5753 | 37.79122 | 16.67936 | -0.47832 | 0.469917 | 0.392807 |
| 8 | 116445400 | 116447199 | 10.59521 | 19.02544 | 35.81186 | 15.66219 | -0.50427 | 0.390544 | 0.392809 |
| 17 | 4045800 | 4047799 | 34.36004 | 56.63222 | 55.75728 | 24.05529 | 0.402105 | 1.381002 | 0.392827 |
| 18 | 46037800 | 46040199 | 18.52703 | 33.39298 | 26.12831 | 11.38369 | 0.757738 | 0.400417 | 0.392829 |
| 18 | 82356400 | 82357799 | 12.58361 | 22.06337 | 42.04213 | 18.57846 | -0.53774 | 0.631744 | 0.392835 |
| X | 113587600 | 113589799 | 34.41745 | 57.94745 | 39.49242 | 17.26022 | 0.932365 | 1.141095 | 0.392848 |
| 14 | 106917000 | 106917999 | 8.145157 | 15.19438 | 30.57484 | 12.71771 | -0.54859 | 0.136685 | 0.392852 |
| 6 | 57778000 | 57778799 | 11.34908 | 21.43826 | 20.58739 | 8.377919 | 0.536556 | 0.059059 | 0.392856 |
| X | 141497800 | 141498199 | 3.339891 | 6.533212 | 16.81967 | 5.554237 | -0.59169 | -0.76347 | 0.392866 |
| 7 | 42267600 | 42269199 | 38.54268 | 63.89771 | 53.23825 | 23.05373 | 0.643373 | 1.323677 | 0.392867 |
| 14 | 71639800 | 71642599 | 25.35744 | 41.9802 | 71.98143 | 31.14015 | -0.3949 | 1.317952 | 0.392877 |
| 7 | 44026000 | 44032599 | 48.09448 | 76.74711 | 124.4358 | 52.19398 | -0.35641 | 2.126153 | 0.392893 |
| 6 | 129822800 | 129823399 | 18.89885 | 33.16021 | 31.94643 | 14.12583 | 0.441085 | 0.636324 | 0.392903 |
| 3 | 13084000 | 13086799 | 23.95649 | 42.01727 | 32.43811 | 14.33828 | 0.759048 | 0.672208 | 0.39292 |
| 6 | 86373600 | 86374399 | 5.859715 | 11.26311 | 26.32705 | 10.28238 | -0.68253 | -0.11335 | 0.39292 |
| 10 | 117854800 | 117856999 | 21.66518 | 37.62453 | 36.20789 | 15.89263 | 0.441903 | 0.807745 | 0.392928 |
| 2 | 144881600 | 144882999 | 15.2899 | 27.74735 | 26.7505 | 11.55559 | 0.466637 | 0.341889 | 0.392934 |
| 12 | 83200600 | 83205799 | 35.03064 | 56.41873 | 96.76562 | 41.13986 | -0.42153 | 1.815716 | 0.392941 |
| 10 | 49506800 | 49509599 | 38.50065 | 63.83285 | 53.2525 | 23.06111 | 0.641519 | 1.322218 | 0.392966 |
| 6 | 76930600 | 76931999 | 11.38429 | 20.33631 | 37.28911 | 16.37993 | -0.47124 | 0.431222 | 0.39297 |
| 6 | 90283000 | 90287199 | 57.51573 | 90.67541 | 151.5637 | 62.89596 | -0.41657 | 2.475033 | 0.392972 |
| 4 | 118625400 | 118626799 | 14.39348 | 26.72933 | 23.63751 | 9.904082 | 0.620303 | 0.197968 | 0.392978 |
| 1 | 64998400 | 65001199 | 43.97195 | 71.61501 | 58.13331 | 24.96148 | 0.670852 | 1.560612 | 0.39298 |
| 5 | 10828400 | 10829999 | 19.90544 | 34.92028 | 33.336 | 14.73807 | 0.454192 | 0.64707 | 0.392992 |
| 7 | 22345200 | 22345999 | 12.45691 | 23.89419 | 19.85584 | 7.801235 | 0.787629 | -0.04637 | 0.392992 |
| 7 | 42884000 | 42891799 | 210.8155 | 310.6622 | 292.8591 | 120.6024 | 0.327984 | 3.788523 | 0.392994 |
| 9 | 93327000 | 93329599 | 20.29621 | 36.04733 | 29.29074 | 12.92929 | 0.691413 | 0.509187 | 0.393004 |
| 14 | 76662000 | 76664199 | 29.13509 | 47.28749 | 83.65448 | 35.85484 | -0.45409 | 1.604709 | 0.393026 |
| 10 | 8259800 | 8261199 | 19.53079 | 35.61633 | 24.06793 | 10.33373 | 0.981419 | 0.339725 | 0.393035 |
| 9 | 8220200 | 8220599 | 10.79752 | 20.7095 | 20.61662 | 8.102086 | 0.529263 | -0.05962 | 0.393046 |
| 11 | 75893600 | 75895399 | 9.814933 | 17.75546 | 34.38072 | 14.91048 | -0.53651 | 0.339098 | 0.393048 |
| 16 | 61611600 | 61615199 | 35.60798 | 59.26869 | 52.04463 | 22.59907 | 0.568164 | 1.268338 | 0.393053 |
| 5 | 106212200 | 106215799 | 47.07396 | 75.88644 | 75.54555 | 32.16038 | 0.363186 | 1.794472 | 0.393057 |
| 6 | 68543200 | 68544399 | 16.8544 | 30.04283 | 28.58487 | 12.57933 | 0.467812 | 0.461899 | 0.393065 |
| 3 | 102842600 | 102846399 | 45.87816 | 73.18451 | 120.9144 | 50.69184 | -0.38402 | 2.135732 | 0.393069 |
| 3 | 80984600 | 80984999 | 7.54536 | 14.70953 | 15.75802 | 5.686473 | 0.539004 | -0.3976 | 0.39307 |

| | | | | | | | | | |
|---|---|---|---|---|---|---|---|---|---|
| 4 | 20031400 | 20033999 | 22.18218 | 38.99022 | 32.00864 | 14.17454 | 0.670777 | 0.628217 | 0.393075 |
| 10 | 9218600 | 9219999 | 10.55126 | 19.24688 | 36.04405 | 15.47011 | -0.47924 | 0.298552 | 0.393081 |
| 7 | 6524000 | 6526199 | 36.19581 | 59.42384 | 58.10523 | 25.0266 | 0.4097 | 1.432682 | 0.393081 |
| 15 | 96132600 | 96132999 | 4.350267 | 8.498029 | 19.97371 | 6.981626 | -0.53949 | -0.53735 | 0.393081 |
| 19 | 40718400 | 40718799 | 4.350267 | 8.498029 | 19.97371 | 6.981626 | -0.53949 | -0.53735 | 0.393081 |
| 1 | 12719000 | 12720399 | 17.02014 | 31.53919 | 23.421 | 9.842143 | 0.867806 | 0.231898 | 0.393093 |
| 1 | 187553600 | 187558599 | 38.23394 | 61.54016 | 102.5073 | 43.5423 | -0.38063 | 1.840177 | 0.393109 |
| 9 | 117778200 | 117779599 | 13.90268 | 26.44015 | 21.33996 | 8.560853 | 0.802882 | 0.038292 | 0.393111 |
| 15 | 45635600 | 45637999 | 24.1884 | 42.0884 | 33.37912 | 14.66897 | 0.720034 | 0.789726 | 0.393118 |
| 11 | 117363800 | 117370599 | 63.81896 | 99.34965 | 180.6086 | 74.97009 | -0.55176 | 2.715821 | 0.393123 |
| 3 | 144461000 | 144462999 | 15.56104 | 27.05812 | 47.90803 | 21.0449 | -0.43349 | 0.779542 | 0.393139 |
| X | 151639800 | 151644799 | 55.1364 | 88.8653 | 76.00545 | 32.34755 | 0.581428 | 1.814794 | 0.393145 |
| X | 151861400 | 151863399 | 24.33754 | 42.92879 | 29.15215 | 12.90056 | 0.94497 | 0.599847 | 0.393146 |
| 4 | 64443600 | 64445199 | 14.78883 | 27.63145 | 23.15725 | 9.606596 | 0.70861 | 0.16363 | 0.393156 |
| 1 | 17295600 | 17298999 | 43.95289 | 71.80828 | 61.81685 | 26.57342 | 0.589454 | 1.517918 | 0.393163 |
| 1 | 3121400 | 3125999 | 79.79856 | 125.8898 | 124.414 | 51.63512 | 0.341274 | 2.469951 | 0.393176 |
| 17 | 86515600 | 86517599 | 12.57632 | 22.18507 | 40.38356 | 17.87044 | -0.46996 | 0.549451 | 0.393192 |
| 17 | 65263000 | 65266999 | 27.73678 | 46.99472 | 44.18004 | 19.34589 | 0.470021 | 1.063731 | 0.393195 |
| 5 | 72231200 | 72231999 | 16.50024 | 29.80535 | 24.11135 | 10.47609 | 0.712751 | 0.370937 | 0.393224 |
| 15 | 14857800 | 14858999 | 14.6373 | 27.55077 | 22.1682 | 9.082161 | 0.78118 | 0.114493 | 0.393225 |
| 11 | 76448600 | 76450399 | 14.10921 | 24.6362 | 46.25648 | 20.37276 | -0.51712 | 0.709501 | 0.393243 |
| 11 | 73337800 | 73341199 | 35.34294 | 58.21194 | 57.47884 | 24.79077 | 0.397535 | 1.392879 | 0.393273 |
| 4 | 8037000 | 8037999 | 16.97174 | 30.98649 | 25.68581 | 11.01198 | 0.690414 | 0.31976 | 0.393279 |
| 3 | 146430000 | 146433999 | 43.96901 | 71.91815 | 60.71463 | 26.11066 | 0.618708 | 1.501279 | 0.393293 |
| 17 | 63367200 | 63367599 | 4.225813 | 8.252089 | 19.8972 | 6.996061 | -0.58358 | -0.51906 | 0.393333 |
| 5 | 77163200 | 77173799 | 113.9212 | 172.4268 | 264.996 | 109.6243 | -0.34181 | 3.282522 | 0.393369 |
| 2 | 150976800 | 150979799 | 37.69855 | 62.49097 | 53.77052 | 23.28235 | 0.59703 | 1.322918 | 0.393375 |
| 1 | 98539800 | 98540799 | 13.04158 | 24.42838 | 23.49338 | 9.704535 | 0.516863 | 0.135793 | 0.393382 |
| 7 | 8232800 | 8233599 | 12.61279 | 23.99609 | 14.98778 | 6.007779 | 1.170917 | 0.006351 | 0.393394 |
| 18 | 68913200 | 68916799 | 55.09971 | 88.45998 | 85.2601 | 36.08067 | 0.402531 | 1.93586 | 0.393398 |
| 18 | 79673400 | 79675399 | 16.9934 | 30.77211 | 25.79388 | 11.17255 | 0.66429 | 0.362053 | 0.393402 |
| 5 | 8970800 | 8973799 | 21.53327 | 36.32234 | 65.35263 | 28.57858 | -0.46333 | 1.10521 | 0.393442 |
| 1 | 37175600 | 37177199 | 15.24314 | 27.61015 | 26.71905 | 11.56938 | 0.458953 | 0.351685 | 0.393449 |
| 12 | 119776800 | 119778599 | 15.24314 | 27.61015 | 26.71905 | 11.56938 | 0.458953 | 0.351685 | 0.393449 |
| 14 | 30473400 | 30475799 | 20.78159 | 37.92259 | 22.88245 | 9.816537 | 1.144482 | 0.343353 | 0.393452 |
| X | 26619600 | 26627799 | 88.69121 | 140.9087 | 97.98475 | 40.86479 | 0.85637 | 2.296114 | 0.393458 |
| 17 | 32477000 | 32478599 | 11.02396 | 19.96288 | 36.94874 | 16.00386 | -0.47113 | 0.345304 | 0.393465 |
| 6 | 69438200 | 69439199 | 6.583927 | 12.69655 | 29.39202 | 11.37226 | -0.65558 | -0.12384 | 0.393465 |
| 4 | 147467000 | 147469399 | 73.64999 | 116.871 | 101.7418 | 42.38054 | 0.531023 | 2.315262 | 0.393474 |
| 14 | 37120400 | 37121999 | 16.34926 | 29.95194 | 25.54112 | 10.90118 | 0.653746 | 0.293259 | 0.393475 |
| 9 | 37356600 | 37358399 | 14.10707 | 24.75023 | 44.55428 | 19.69915 | -0.45649 | 0.631805 | 0.393476 |
| 10 | 44315800 | 44318399 | 15.77163 | 27.18819 | 50.57365 | 22.18543 | -0.50675 | 0.871697 | 0.393478 |
| 8 | 13352400 | 13357199 | 36.94361 | 59.50497 | 98.72182 | 41.97694 | -0.37358 | 1.815414 | 0.39348 |
| 14 | 40112400 | 40114199 | 23.96288 | 41.6535 | 36.9907 | 16.24572 | 0.557385 | 0.80658 | 0.39348 |
| 2 | 164184400 | 164185999 | 17.47355 | 31.46533 | 27.38281 | 11.94394 | 0.604321 | 0.401383 | 0.393519 |
| 12 | 96304000 | 96305999 | 17.47355 | 31.46533 | 27.38281 | 11.94394 | 0.604321 | 0.401383 | 0.393519 |
| 6 | 125649000 | 125651399 | 19.34588 | 32.93338 | 57.03425 | 24.99915 | -0.4074 | 1.008392 | 0.393522 |

| | | | | | | | | | |
|---|---|---|---|---|---|---|---|---|---|
| 15 | 98223600 | 98224399 | 6.099422 | 11.85016 | 27.21632 | 10.15419 | -0.59507 | -0.26357 | 0.393528 |
| 1 | 162455400 | 162457599 | 22.74818 | 37.82191 | 66.16771 | 28.70972 | -0.42317 | 1.257049 | 0.393528 |
| 2 | 66994200 | 66994799 | 9.472228 | 17.18364 | 34.38779 | 14.86131 | -0.58068 | 0.318093 | 0.393534 |
| 12 | 64123400 | 64123999 | 12.92259 | 23.91176 | 21.99097 | 9.259337 | 0.559646 | 0.203538 | 0.393536 |
| 19 | 56513200 | 56517399 | 41.76609 | 67.99065 | 65.4391 | 28.09368 | 0.425278 | 1.56522 | 0.39355 |
| X | 36137800 | 36138999 | 20.84091 | 36.68675 | 29.83028 | 13.20599 | 0.685545 | 0.603103 | 0.393579 |
| 14 | 4267600 | 4269399 | 24.70362 | 41.03624 | 71.46186 | 30.9876 | -0.41678 | 1.277229 | 0.393582 |
| 3 | 102663000 | 102664799 | 9.838797 | 17.65666 | 35.84507 | 15.68458 | -0.61308 | 0.388758 | 0.393619 |
| 7 | 122269800 | 122271399 | 12.31346 | 21.47142 | 45.26528 | 19.91919 | -0.68323 | 0.720497 | 0.393619 |
| 3 | 21743000 | 21743599 | 10.77669 | 20.88816 | 19.23195 | 7.276639 | 0.692519 | -0.1853 | 0.393619 |
| 16 | 67624800 | 67626399 | 9.077459 | 16.98707 | 34.59521 | 14.31329 | -0.55994 | 0.133994 | 0.393666 |
| 7 | 15458800 | 15466399 | 135.4581 | 206.6814 | 188.9957 | 78.36907 | 0.416161 | 3.121565 | 0.3937 |
| 17 | 20027600 | 20032799 | 65.10341 | 104.5522 | 79.53637 | 33.67403 | 0.743797 | 1.940492 | 0.393706 |
| X | 78946000 | 78948599 | 42.39155 | 69.49949 | 59.46586 | 25.59767 | 0.60105 | 1.468461 | 0.393709 |
| 14 | 13656000 | 13656999 | 13.99155 | 26.29555 | 22.77163 | 9.35328 | 0.673069 | 0.119586 | 0.39373 |
| 6 | 60241400 | 60242799 | 10.05353 | 19.53302 | 18.17203 | 6.777921 | 0.696261 | -0.24603 | 0.393739 |
| 7 | 11549600 | 11565999 | 409.9307 | 571.5743 | 507.5472 | 210.5492 | 0.335387 | 4.625842 | 0.393741 |
| 17 | 18186200 | 18187799 | 19.77284 | 34.99582 | 30.43478 | 13.45901 | 0.591586 | 0.53668 | 0.39376 |
| 4 | 122503600 | 122515799 | 55.95325 | 89.84489 | 76.86332 | 32.53533 | 0.574459 | 1.929915 | 0.393773 |
| 5 | 70818200 | 70824399 | 117.7894 | 180.8268 | 176.5293 | 73.29064 | 0.328746 | 2.992091 | 0.393782 |
| 6 | 96599600 | 96600799 | 13.22389 | 23.18863 | 42.53936 | 18.8 | -0.48323 | 0.635031 | 0.3938 |
| 5 | 97315400 | 97315799 | 3.755136 | 7.342724 | 18.35313 | 6.263955 | -0.59397 | -0.63432 | 0.39381 |
| 8 | 45696000 | 45697999 | 15.52167 | 28.16089 | 26.6385 | 11.51083 | 0.49341 | 0.343691 | 0.393823 |
| 2 | 64813200 | 64814799 | 18.98381 | 33.62725 | 29.87923 | 13.21114 | 0.561071 | 0.520542 | 0.393823 |
| 10 | 22410200 | 22414799 | 78.99039 | 124.8363 | 119.7098 | 49.7274 | 0.386721 | 2.425826 | 0.393845 |
| 8 | 27334000 | 27337399 | 19.25492 | 32.08672 | 64.21937 | 27.90613 | -0.61635 | 1.220159 | 0.393855 |
| 14 | 45787800 | 45792599 | 21.90678 | 36.13199 | 75.73188 | 32.68257 | -0.68425 | 1.357126 | 0.393857 |
| 6 | 16117400 | 16118799 | 9.303107 | 16.9571 | 33.41007 | 14.35334 | -0.55265 | 0.287649 | 0.393889 |
| 1 | 133902200 | 133904199 | 11.04996 | 19.78442 | 37.30277 | 16.35427 | -0.50937 | 0.413638 | 0.393901 |
| 13 | 64228200 | 64229999 | 11.04996 | 19.78442 | 37.30277 | 16.35427 | -0.50937 | 0.413638 | 0.393901 |
| 18 | 81882800 | 81884799 | 11.04996 | 19.78442 | 37.30277 | 16.35427 | -0.50937 | 0.413638 | 0.393901 |
| 6 | 124373600 | 124377399 | 43.92921 | 70.04363 | 121.2026 | 50.78064 | -0.45132 | 2.148339 | 0.393912 |
| 14 | 45736200 | 45737999 | 9.581269 | 17.64713 | 33.02362 | 13.99253 | -0.46476 | 0.230127 | 0.393918 |
| 3 | 22921200 | 22924599 | 38.61013 | 63.05886 | 62.49483 | 26.86166 | 0.38645 | 1.510264 | 0.393925 |
| 15 | 16951400 | 16951799 | 3.818943 | 7.462628 | 20.55113 | 7.13296 | -0.75329 | -0.5622 | 0.393954 |
| 16 | 28631000 | 28634199 | 39.95421 | 64.09884 | 106.8234 | 45.15687 | -0.38711 | 1.948671 | 0.393963 |
| 3 | 159379800 | 159382199 | 18.71106 | 31.43703 | 64.94328 | 28.36547 | -0.66196 | 1.13157 | 0.393968 |
| 6 | 84302600 | 84303599 | 7.099769 | 13.33118 | 29.0875 | 11.96577 | -0.65102 | 0.08263 | 0.393987 |
| 11 | 60122800 | 60124399 | 7.099769 | 13.33118 | 29.0875 | 11.96577 | -0.65102 | 0.08263 | 0.393987 |
| 7 | 85620000 | 85623999 | 44.56495 | 72.69446 | 62.80677 | 26.98357 | 0.582711 | 1.54323 | 0.393989 |
| 1 | 85581000 | 85582399 | 22.24109 | 38.46561 | 37.35887 | 16.39006 | 0.427708 | 0.852648 | 0.394009 |
| 7 | 59551400 | 59559799 | 148.3592 | 224.2714 | 216.8019 | 89.65203 | 0.325014 | 3.312703 | 0.394014 |
| 17 | 87274400 | 87276599 | 11.14056 | 19.74229 | 37.83665 | 16.72811 | -0.54108 | 0.484944 | 0.394019 |
| 8 | 74716000 | 74717799 | 18.7351 | 33.22888 | 29.94761 | 13.23461 | 0.541577 | 0.508183 | 0.394021 |
| 7 | 20861000 | 20862799 | 36.06742 | 60.37245 | 48.08744 | 20.95508 | 0.708195 | 1.207552 | 0.394032 |
| 16 | 53473800 | 53476399 | 17.37161 | 29.96844 | 52.2107 | 22.90408 | -0.41251 | 0.87475 | 0.394034 |
| 18 | 15445800 | 15448399 | 23.06606 | 39.74553 | 38.91351 | 17.07019 | 0.415095 | 0.891002 | 0.394035 |

| | | | | | | | | | |
|---|---|---|---|---|---|---|---|---|---|
| 4 | 101903800 | 101908399 | 44.92022 | 71.5016 | 126.5405 | 52.89198 | -0.4864 | 2.205204 | 0.394047 |
| 5 | 105011000 | 105016799 | 101.2154 | 158.0891 | 139.5947 | 57.92066 | 0.492492 | 2.673255 | 0.394059 |
| 7 | 8077200 | 8079999 | 14.13162 | 26.16411 | 21.24517 | 8.938251 | 0.73864 | 0.207936 | 0.394059 |
| 8 | 25129000 | 25129999 | 9.35678 | 16.90776 | 34.38467 | 14.93304 | -0.60813 | 0.341505 | 0.39406 |
| 14 | 76116400 | 76119999 | 30.87455 | 49.68872 | 105.3734 | 44.75194 | -0.72831 | 1.837535 | 0.394064 |
| 3 | 103032600 | 103033799 | 13.25599 | 24.94869 | 22.26528 | 9.123258 | 0.632545 | 0.10225 | 0.394077 |
| 2 | 11978400 | 11979599 | 15.2707 | 26.67117 | 47.31382 | 20.84183 | -0.43552 | 0.715405 | 0.394083 |
| 7 | 99416400 | 99417199 | 7.380256 | 13.94268 | 28.6517 | 11.65702 | -0.55172 | 0.045519 | 0.394087 |
| X | 3873800 | 3875999 | 23.17305 | 40.57962 | 33.72538 | 14.88758 | 0.653341 | 0.695088 | 0.394091 |
| X | 93231600 | 93232199 | 11.05501 | 21.3161 | 19.71548 | 7.630875 | 0.655872 | -0.10461 | 0.394102 |
| 13 | 95017600 | 95022599 | 43.88394 | 70.03152 | 119.7487 | 50.23146 | -0.43286 | 2.119452 | 0.394105 |
| 17 | 28754000 | 28766599 | 200.7885 | 289.2726 | 495.3232 | 204.4028 | -0.56421 | 4.136198 | 0.394107 |
| 13 | 62574600 | 62580199 | 91.0223 | 142.6781 | 136.5054 | 56.618 | 0.380917 | 2.603268 | 0.394122 |
| 6 | 144128000 | 144129999 | 19.43444 | 34.13093 | 32.178 | 14.23913 | 0.471925 | 0.623613 | 0.394124 |
| 5 | 103430600 | 103435799 | 41.01118 | 65.81124 | 110.4196 | 46.69506 | -0.39646 | 1.940768 | 0.394128 |
| 17 | 33965800 | 33968999 | 19.30493 | 32.46565 | 59.56055 | 26.02194 | -0.49124 | 1.11893 | 0.394144 |
| 4 | 103193200 | 103194799 | 14.96072 | 27.28345 | 26.27769 | 11.28194 | 0.473693 | 0.310663 | 0.394183 |
| 10 | 58897800 | 58898599 | 6.260406 | 12.11591 | 25.51482 | 9.722754 | -0.49925 | -0.19138 | 0.394193 |
| X | 113241200 | 113242199 | 27.36603 | 46.25026 | 44.31441 | 19.38996 | 0.442778 | 1.087864 | 0.394208 |
| 2 | 120648400 | 120652399 | 60.89069 | 97.56866 | 88.69459 | 37.4241 | 0.483532 | 2.003636 | 0.394213 |
| 3 | 18576800 | 18577999 | 11.20431 | 21.55515 | 20.68414 | 8.057972 | 0.593796 | -0.08208 | 0.394233 |
| 6 | 44745400 | 44747599 | 25.18149 | 43.29449 | 40.88855 | 17.93537 | 0.466174 | 0.921715 | 0.394237 |
| 6 | 143888600 | 143889999 | 13.61347 | 25.01505 | 24.39666 | 10.37021 | 0.465673 | 0.253583 | 0.394254 |
| 12 | 18431000 | 18452799 | 728.9796 | 966.8129 | 816.449 | 341.2653 | 0.335211 | 5.41217 | 0.394259 |
| 6 | 29639200 | 29643399 | 35.84345 | 57.65786 | 99.49749 | 42.22705 | -0.43254 | 1.855616 | 0.394263 |
| 3 | 14804600 | 14806599 | 16.41509 | 30.31805 | 22.74186 | 9.599644 | 0.848329 | 0.241623 | 0.394306 |
| 17 | 89073800 | 89075999 | 23.71944 | 40.82843 | 38.28827 | 16.79538 | 0.476559 | 0.900391 | 0.394321 |
| 16 | 13726000 | 13732199 | 78.24851 | 123.9423 | 115.3908 | 47.98787 | 0.431903 | 2.374565 | 0.394325 |
| 14 | 8839600 | 8840199 | 8.47568 | 16.49237 | 16.61391 | 6.121434 | 0.598474 | -0.31315 | 0.394337 |
| 4 | 122434800 | 122437799 | 53.1391 | 85.67465 | 76.11288 | 32.40716 | 0.527264 | 1.800104 | 0.394342 |
| 13 | 4390200 | 4390999 | 15 | 27.62465 | 19.86823 | 8.419618 | 0.905316 | 0.241812 | 0.394343 |
| 4 | 70942000 | 70943399 | 16.89289 | 31.47822 | 21.76296 | 9.069246 | 0.978896 | 0.194659 | 0.394355 |
| 6 | 125226000 | 125231599 | 74.25841 | 115.5933 | 180.9549 | 75.1146 | -0.33643 | 2.722513 | 0.394365 |
| 7 | 140365800 | 140368199 | 20.42006 | 34.23918 | 61.81009 | 26.96409 | -0.46824 | 1.153259 | 0.394371 |
| 11 | 91465800 | 91468799 | 42.65246 | 67.95939 | 123.248 | 51.58665 | -0.52002 | 2.170799 | 0.394377 |
| 7 | 102910600 | 102915799 | 84.89287 | 133.4865 | 129.4721 | 53.71005 | 0.364639 | 2.538239 | 0.394401 |
| 17 | 38301400 | 38304399 | 41.20802 | 67.68254 | 58.44471 | 25.17754 | 0.588815 | 1.440616 | 0.394401 |
| X | 65992200 | 65997399 | 68.75951 | 110.0685 | 86.87366 | 36.60109 | 0.685378 | 2.045093 | 0.394407 |
| 8 | 33819600 | 33821799 | 17.98579 | 30.93964 | 55.49916 | 24.34457 | -0.45547 | 0.905336 | 0.394414 |
| 18 | 47909000 | 47912799 | 51.8161 | 83.53287 | 70.43486 | 29.98554 | 0.60225 | 1.799073 | 0.394417 |
| 8 | 34220800 | 34222799 | 12.2365 | 21.49842 | 41.95339 | 18.5711 | -0.57211 | 0.594477 | 0.394421 |
| 14 | 6166200 | 6171399 | 56.13415 | 90.10139 | 85.88195 | 36.33197 | 0.418186 | 1.944546 | 0.394424 |
| 1 | 194963000 | 194965399 | 36.11259 | 59.7215 | 49.32352 | 21.3222 | 0.655741 | 1.341351 | 0.394425 |
| 12 | 23266600 | 23267799 | 30.97681 | 51.59035 | 45.02202 | 19.55689 | 0.577118 | 1.24348 | 0.39443 |
| 13 | 6562600 | 6564399 | 20.96745 | 37.43589 | 26.96055 | 11.84806 | 0.868607 | 0.47606 | 0.394435 |
| 7 | 113364000 | 113367599 | 23.88346 | 39.12634 | 77.37709 | 33.29702 | -0.60472 | 1.445017 | 0.394448 |
| 9 | 78181000 | 78181999 | 12.38658 | 21.81779 | 40.37981 | 17.87411 | -0.49477 | 0.562964 | 0.394451 |

| | | | | | | | | | |
|---|---|---|---|---|---|---|---|---|---|
| X | 150734000 | 150736999 | 23.39417 | 41.29167 | 29.66757 | 13.12693 | 0.864277 | 0.587447 | 0.394455 |
| 4 | 128655200 | 128658199 | 46.6596 | 75.36774 | 73.52265 | 31.37936 | 0.395667 | 1.736977 | 0.394459 |
| 13 | 36931800 | 36933399 | 13.317 | 23.51026 | 42.92438 | 18.9918 | -0.4742 | 0.547085 | 0.394461 |
| X | 65153400 | 65155199 | 20.81279 | 37.60673 | 25.67295 | 11.15037 | 0.956753 | 0.404817 | 0.394494 |
| 11 | 60649000 | 60650999 | 11.23579 | 20.04159 | 37.28951 | 16.40039 | -0.49342 | 0.439427 | 0.3945 |
| 13 | 49964000 | 49964799 | 11.23579 | 20.04159 | 37.28951 | 16.40039 | -0.49342 | 0.439427 | 0.3945 |
| 8 | 103042600 | 103043999 | 13.05404 | 22.9933 | 41.40724 | 18.32893 | -0.45548 | 0.570394 | 0.39451 |
| 1 | 152140600 | 152142199 | 8.787604 | 16.25792 | 31.50522 | 13.26812 | -0.50829 | 0.194598 | 0.394512 |
| 14 | 48152800 | 48153799 | 8.787604 | 16.25792 | 31.50522 | 13.26812 | -0.50829 | 0.194598 | 0.394512 |
| 3 | 133031000 | 133035799 | 33.65633 | 54.37969 | 93.32838 | 39.84727 | -0.41705 | 1.714821 | 0.394518 |
| 7 | 84057800 | 84060999 | 20.55283 | 34.49911 | 62.42807 | 27.25227 | -0.47174 | 1.141816 | 0.394525 |
| 8 | 114613200 | 114614199 | 11.58263 | 22.0543 | 21.53812 | 8.618475 | 0.533152 | 0.008239 | 0.394528 |
| 12 | 18267800 | 18282799 | 490.8473 | 673.6017 | 590.9228 | 245.7516 | 0.329763 | 4.870306 | 0.394532 |
| 7 | 108003400 | 108011599 | 121.248 | 186.1487 | 177.052 | 73.50834 | 0.366366 | 2.991834 | 0.394541 |
| 10 | 116185800 | 116186999 | 10.24139 | 18.62782 | 34.40383 | 14.8195 | -0.46256 | 0.309786 | 0.394554 |
| X | 137255000 | 137257999 | 40.30644 | 66.03025 | 61.06609 | 26.27796 | 0.488706 | 1.47261 | 0.394559 |
| X | 102931000 | 102932199 | 9.955502 | 18.25702 | 35.62944 | 15.18603 | -0.53108 | 0.264589 | 0.394572 |
| 2 | 170924400 | 170925799 | 12.9954 | 23.00711 | 43.99489 | 19.4545 | -0.53906 | 0.515852 | 0.394576 |
| 4 | 91642800 | 91646399 | 106.3274 | 163.9904 | 160.5819 | 66.70286 | 0.329442 | 2.897245 | 0.394578 |
| 7 | 30939000 | 30945599 | 156.1697 | 235.5231 | 226.1088 | 93.43115 | 0.332103 | 3.36196 | 0.394589 |
| 11 | 30978400 | 30981999 | 60.54678 | 97.11342 | 86.02347 | 36.35157 | 0.522495 | 1.972037 | 0.394589 |
| 10 | 95104000 | 95106199 | 11.90212 | 21.03455 | 39.32944 | 17.39756 | -0.50748 | 0.521008 | 0.394594 |
| 4 | 131113400 | 131116799 | 46.30839 | 73.94773 | 119.7545 | 50.27931 | -0.35358 | 2.102402 | 0.3946 |
| 7 | 38382600 | 38384199 | 33.48309 | 55.46085 | 54.72087 | 23.68135 | 0.400067 | 1.319695 | 0.394609 |
| 16 | 20707600 | 20710199 | 26.70538 | 43.82461 | 76.24402 | 32.83334 | -0.41944 | 1.429571 | 0.39461 |
| 13 | 52109600 | 52112399 | 14.47025 | 25.2143 | 45.77275 | 20.13347 | -0.46888 | 0.742177 | 0.39461 |
| X | 126485600 | 126492399 | 69.0989 | 110.9768 | 75.37487 | 31.91568 | 0.907146 | 1.942235 | 0.39462 |
| 10 | 87825200 | 87826799 | 20.9477 | 36.92812 | 31.47359 | 13.92946 | 0.618459 | 0.588191 | 0.394635 |
| 7 | 12756000 | 12759399 | 28.39249 | 46.34458 | 79.33783 | 34.09401 | -0.40058 | 1.508337 | 0.394648 |
| X | 102499000 | 102499999 | 12.897 | 22.78082 | 41.38792 | 18.31006 | -0.46653 | 0.538536 | 0.394649 |
| 6 | 129917200 | 129922799 | 125.6258 | 191.6333 | 184.924 | 76.67642 | 0.338303 | 3.121263 | 0.39466 |
| 3 | 126390000 | 126391599 | 11.02107 | 19.39777 | 40.95351 | 18.13083 | -0.68405 | 0.563883 | 0.394665 |
| 17 | 42077600 | 42078999 | 16.5758 | 28.52575 | 51.10902 | 22.41912 | -0.45343 | 0.891774 | 0.394666 |
| 17 | 18194000 | 18201799 | 107.2382 | 166.3544 | 154.3846 | 64.11833 | 0.413173 | 2.792784 | 0.394667 |
| 10 | 3050000 | 3078599 | 945.1002 | 1228.475 | 1016.168 | 425.6956 | 0.336124 | 5.738248 | 0.394684 |
| 15 | 10261200 | 10261799 | 7.495213 | 14.19061 | 28.71679 | 11.63002 | -0.52451 | 0.028844 | 0.394712 |
| 17 | 21921200 | 21921999 | 7.495213 | 14.19061 | 28.71679 | 11.63002 | -0.52451 | 0.028844 | 0.394712 |
| 12 | 112050200 | 112054999 | 38.69201 | 61.91213 | 120.5037 | 50.74084 | -0.61506 | 2.032037 | 0.394718 |
| 1 | 49918400 | 49919399 | 13.43536 | 25.04753 | 23.25333 | 9.68241 | 0.557695 | 0.165517 | 0.394736 |
| 19 | 35976800 | 35977799 | 12.43978 | 23.54146 | 22.49985 | 9.120933 | 0.54786 | 0.06079 | 0.39474 |
| 11 | 68770400 | 68773199 | 15.0798 | 26.26423 | 46.8245 | 20.59007 | -0.44312 | 0.752388 | 0.394741 |
| 1 | 92244600 | 92246399 | 14.45319 | 25.0996 | 49.5159 | 21.7337 | -0.58907 | 0.793219 | 0.394745 |
| 15 | 55989600 | 55990999 | 10.8814 | 19.64021 | 35.84097 | 15.58784 | -0.45457 | 0.361519 | 0.394749 |
| 12 | 21026600 | 21036799 | 113.7918 | 174.7888 | 170.7218 | 70.88503 | 0.328627 | 2.978539 | 0.394753 |
| 12 | 48454000 | 48454999 | 12.22064 | 21.74518 | 39.32121 | 17.32956 | -0.45458 | 0.466444 | 0.394764 |
| 17 | 51595800 | 51597799 | 17.26915 | 29.58823 | 52.68552 | 23.10502 | -0.44567 | 0.937984 | 0.394765 |
| 1 | 35901800 | 35906399 | 31.75492 | 51.25327 | 94.91032 | 40.46744 | -0.52821 | 1.743087 | 0.394801 |

| | | | | | | | | | |
|---|---|---|---|---|---|---|---|---|---|
| 4 | 98214200 | 98216399 | 21.93991 | 38.2773 | 35.98345 | 15.83985 | 0.476143 | 0.745423 | 0.394802 |
| 3 | 135115200 | 135115799 | 13.10895 | 24.44247 | 23.92174 | 9.958334 | 0.482687 | 0.165018 | 0.394817 |
| 1 | 124276000 | 124276799 | 8.297727 | 15.54257 | 30.11594 | 12.44018 | -0.4862 | 0.111847 | 0.394835 |
| 4 | 64166200 | 64167199 | 8.297727 | 15.54257 | 30.11594 | 12.44018 | -0.4862 | 0.111847 | 0.394835 |
| 4 | 47154800 | 47158999 | 35.16724 | 58.5645 | 51.49847 | 22.369 | 0.566085 | 1.260046 | 0.394837 |
| X | 24106600 | 24108199 | 20.2555 | 36.46518 | 26.7388 | 11.66692 | 0.849978 | 0.423255 | 0.394854 |
| 16 | 92065800 | 92068199 | 20.572 | 34.55657 | 60.73364 | 26.52482 | -0.42972 | 1.133889 | 0.394864 |
| 17 | 22323000 | 22327999 | 77.95268 | 123.4642 | 116.4858 | 48.44138 | 0.412629 | 2.375105 | 0.394866 |
| 2 | 61632800 | 61636399 | 32.52926 | 52.58991 | 90.82911 | 38.80995 | -0.42512 | 1.694797 | 0.394873 |
| 7 | 113046400 | 113047399 | 5.965251 | 11.54486 | 26.03747 | 9.921273 | -0.59644 | -0.19091 | 0.394909 |
| 1 | 130524400 | 130530199 | 57.92171 | 91.57606 | 146.1285 | 60.71402 | -0.34646 | 2.408666 | 0.39492 |
| 6 | 22516000 | 22518399 | 21.08774 | 36.86051 | 34.90093 | 15.38274 | 0.46599 | 0.712055 | 0.394923 |
| 9 | 38546600 | 38552399 | 186.2288 | 276.9251 | 264.2826 | 108.8832 | 0.322391 | 3.63157 | 0.394942 |
| 4 | 122974800 | 122976999 | 15.30953 | 26.42892 | 49.5084 | 21.71907 | -0.51632 | 0.856398 | 0.394978 |
| 10 | 42376400 | 42379199 | 17.60647 | 29.98081 | 54.79282 | 24.01724 | -0.48456 | 0.997226 | 0.394987 |
| 11 | 70611200 | 70613199 | 10.94584 | 19.46405 | 37.80327 | 16.6698 | -0.55759 | 0.457813 | 0.39499 |
| 18 | 71039000 | 71040799 | 10.94584 | 19.46405 | 37.80327 | 16.6698 | -0.55759 | 0.457813 | 0.39499 |
| 2 | 30013800 | 30014999 | 5.695204 | 10.96985 | 25.24662 | 9.801284 | -0.65113 | -0.1347 | 0.395001 |
| 10 | 76171600 | 76173999 | 14.76288 | 25.65126 | 48.44619 | 21.2714 | -0.52636 | 0.786044 | 0.39501 |
| 12 | 114266200 | 114268999 | 46.14019 | 75.00058 | 65.71282 | 28.19604 | 0.559026 | 1.596187 | 0.395019 |
| 17 | 94303800 | 94305599 | 17.45614 | 32.57829 | 21.19667 | 8.81008 | 1.068879 | 0.18896 | 0.395038 |
| 9 | 57988400 | 57991199 | 22.343 | 38.58367 | 36.35847 | 15.95043 | 0.470644 | 0.86456 | 0.395041 |
| 7 | 15053600 | 15068399 | 521.2566 | 716.6217 | 571.4226 | 237.5744 | 0.470079 | 4.843333 | 0.395049 |
| 14 | 73802400 | 73805599 | 28.3282 | 48.40255 | 42.44597 | 18.61037 | 0.571351 | 0.991561 | 0.395051 |
| 7 | 120094800 | 120095799 | 15.87642 | 27.64629 | 48.96477 | 21.52857 | -0.4338 | 0.761824 | 0.395057 |
| 4 | 122479600 | 122482599 | 52.51126 | 84.35878 | 83.63829 | 35.42879 | 0.362944 | 1.912607 | 0.395058 |
| 16 | 23332600 | 23333599 | 8.899633 | 16.60872 | 31.67449 | 13.16585 | -0.47111 | 0.143121 | 0.395058 |
| 1 | 179373200 | 179374599 | 8.126244 | 14.80803 | 33.96126 | 14.59502 | -0.77046 | 0.280881 | 0.395079 |
| 7 | 39171000 | 39173399 | 26.08253 | 45.27812 | 37.49578 | 16.45387 | 0.657703 | 0.836816 | 0.39508 |
| 1 | 5577400 | 5580599 | 34.05266 | 57.0673 | 48.61803 | 21.20208 | 0.61141 | 1.189131 | 0.395081 |
| 9 | 57055000 | 57055999 | 5.84386 | 11.23198 | 25.73989 | 10.05475 | -0.65339 | -0.11071 | 0.395112 |
| 18 | 63624800 | 63629399 | 33.6478 | 54.46592 | 91.38328 | 39.09029 | -0.38165 | 1.668959 | 0.39513 |
| 17 | 19612600 | 19614799 | 23.12814 | 40.87638 | 29.04425 | 12.84772 | 0.880821 | 0.569978 | 0.395133 |
| 10 | 129378400 | 129380999 | 39.02657 | 63.86586 | 53.12895 | 22.85255 | 0.640333 | 1.477059 | 0.395151 |
| 16 | 8170000 | 8172199 | 26.55826 | 44.9958 | 44.25314 | 19.37762 | 0.405251 | 1.061651 | 0.395159 |
| 3 | 127732600 | 127734599 | 22.4721 | 37.20899 | 68.2049 | 29.50944 | -0.49076 | 1.305646 | 0.395187 |
| 7 | 41207000 | 41215999 | 148.0243 | 224.0603 | 214.3174 | 88.66147 | 0.341886 | 3.285163 | 0.39519 |
| 4 | 122970400 | 122973799 | 19.0301 | 32.3585 | 56.45819 | 24.74436 | -0.41835 | 1.017115 | 0.395205 |
| 1 | 98055000 | 98056799 | 14.24001 | 26.50029 | 24.31299 | 10.1543 | 0.571458 | 0.187346 | 0.395223 |
| 19 | 53801600 | 53806199 | 27.55855 | 44.65341 | 86.06521 | 36.84232 | -0.57973 | 1.635945 | 0.395223 |
| 6 | 69473000 | 69474399 | 16.73938 | 30.57143 | 26.04431 | 11.16148 | 0.651194 | 0.314952 | 0.395231 |
| 7 | 22175600 | 22176599 | 34.75204 | 57.61572 | 47.68489 | 20.64953 | 0.652933 | 1.307446 | 0.395248 |
| X | 15361800 | 15362999 | 17.76402 | 33.04009 | 22.97614 | 9.604954 | 0.96766 | 0.217936 | 0.395266 |
| X | 148445000 | 148448199 | 27.12092 | 46.48785 | 35.83751 | 15.71694 | 0.757313 | 0.952978 | 0.39527 |
| 9 | 84320200 | 84321399 | 13.49619 | 25.80807 | 20.29791 | 8.045756 | 0.854761 | -0.00646 | 0.395272 |
| 18 | 27595600 | 27597599 | 16.5938 | 28.7037 | 50.58675 | 22.1935 | -0.42812 | 0.846284 | 0.3953 |
| 13 | 25015400 | 25017599 | 12.28057 | 21.48346 | 43.64492 | 19.24997 | -0.62978 | 0.668454 | 0.395318 |

| | | | | | | | | | |
|---|---|---|---|---|---|---|---|---|---|
| 7 | 11255600 | 11256799 | 12.77036 | 24.65993 | 19.55819 | 7.530673 | 0.882193 | -0.10532 | 0.395322 |
| 16 | 57076600 | 57078399 | 24.54515 | 40.64745 | 69.59357 | 30.11328 | -0.39269 | 1.310654 | 0.395335 |
| 10 | 21084400 | 21088799 | 43.65695 | 70.67731 | 70.10984 | 29.9933 | 0.374989 | 1.673634 | 0.395335 |
| 6 | 43837800 | 43839199 | 14.29617 | 27.1057 | 21.84452 | 8.819968 | 0.797105 | 0.063957 | 0.395344 |
| 7 | 111634000 | 111635399 | 14.29617 | 27.1057 | 21.84452 | 8.819968 | 0.797105 | 0.063957 | 0.395344 |
| 3 | 97065800 | 97066999 | 7.242275 | 13.59392 | 29.61443 | 12.18988 | -0.64981 | 0.086343 | 0.395348 |
| 14 | 112329000 | 112330199 | 14.41631 | 27.27657 | 21.12114 | 8.56558 | 0.848304 | 0.07585 | 0.395354 |
| 6 | 10595000 | 10596799 | 24.72593 | 43.04093 | 36.38037 | 15.99193 | 0.628572 | 0.790729 | 0.395363 |
| X | 61972800 | 61976599 | 57.67633 | 92.49993 | 83.97289 | 35.47953 | 0.487021 | 1.974158 | 0.395371 |
| X | 119053600 | 119055199 | 26.41221 | 45.69661 | 38.69278 | 16.97546 | 0.624985 | 0.871742 | 0.395373 |
| 7 | 33126600 | 33164399 | 772.9734 | 1029.18 | 776.712 | 324.4896 | 0.502989 | 5.350535 | 0.395381 |
| 8 | 91388200 | 91391799 | 42.10974 | 68.29343 | 67.95153 | 29.10798 | 0.372952 | 1.633314 | 0.395384 |
| 6 | 40074200 | 40074999 | 10.2989 | 19.70368 | 19.57213 | 7.747209 | 0.522818 | -0.04459 | 0.395386 |
| 9 | 3928200 | 3928799 | 10.2989 | 19.70368 | 19.57213 | 7.747209 | 0.522818 | -0.04459 | 0.395386 |
| 4 | 91583200 | 91584599 | 16.0435 | 28.74044 | 27.66159 | 12.12193 | 0.45547 | 0.423995 | 0.395386 |
| 1 | 104583200 | 104584599 | 21.2243 | 36.9711 | 33.96393 | 14.93621 | 0.508876 | 0.758405 | 0.395388 |
| 6 | 28057200 | 28060199 | 20.39252 | 34.6257 | 64.24463 | 28.15029 | -0.50708 | 1.043833 | 0.395412 |
| 8 | 29995000 | 29997599 | 20.57391 | 34.7126 | 60.22146 | 26.33669 | -0.41077 | 1.096064 | 0.395417 |
| 8 | 73227800 | 73229799 | 10.21507 | 18.11729 | 38.84996 | 17.16904 | -0.70178 | 0.47123 | 0.395422 |
| 8 | 81751200 | 81751799 | 9.87508 | 19.202 | 17.6247 | 6.530503 | 0.723992 | -0.27284 | 0.395426 |
| 7 | 10594800 | 10596599 | 36.89743 | 60.52468 | 59.06069 | 25.42906 | 0.412254 | 1.448546 | 0.395444 |
| 2 | 16274800 | 16279399 | 28.41536 | 48.91457 | 35.38863 | 15.52346 | 0.84937 | 0.918038 | 0.395465 |
| 3 | 46838600 | 46839799 | 14.18364 | 26.51571 | 23.8926 | 9.902164 | 0.606317 | 0.158506 | 0.395475 |
| 18 | 46331200 | 46332999 | 17.19853 | 29.09601 | 57.93726 | 25.35848 | -0.60862 | 1.056532 | 0.395495 |
| 15 | 62642000 | 62644799 | 30.67733 | 49.9043 | 84.66441 | 36.338 | -0.39115 | 1.564444 | 0.395498 |
| 4 | 150919200 | 150922799 | 20.34447 | 33.92094 | 63.79563 | 27.73158 | -0.52697 | 1.216984 | 0.395527 |
| 2 | 18000600 | 18001799 | 12.25575 | 21.60871 | 39.89059 | 17.65407 | -0.49036 | 0.551969 | 0.395531 |
| 2 | 119505000 | 119506399 | 9.429889 | 16.83357 | 37.88245 | 16.65015 | -0.76538 | 0.422742 | 0.395533 |
| 11 | 113502600 | 113503799 | 6.73709 | 12.86229 | 27.25327 | 10.84395 | -0.56746 | -0.04011 | 0.395538 |
| 1 | 145104400 | 145106599 | 21.71699 | 38.035 | 34.28005 | 15.13417 | 0.536825 | 0.684711 | 0.395539 |
| 10 | 16307000 | 16308799 | 23.51498 | 41.28229 | 31.80689 | 14.07062 | 0.761764 | 0.652945 | 0.395542 |
| 13 | 74704800 | 74707199 | 27.38797 | 46.80509 | 41.91144 | 18.37629 | 0.541304 | 0.984313 | 0.395548 |
| 10 | 9610400 | 9612599 | 15.9517 | 28.91128 | 27.17814 | 11.75915 | 0.501325 | 0.354858 | 0.395552 |
| 11 | 17903400 | 17903999 | 6.426453 | 12.34361 | 26.32477 | 10.30264 | -0.55432 | -0.10012 | 0.395555 |
| 3 | 155546000 | 155546599 | 7.968634 | 14.93376 | 30.67461 | 12.66001 | -0.56912 | 0.10465 | 0.395557 |
| 5 | 78150800 | 78151799 | 13.04786 | 22.9266 | 41.43934 | 18.34539 | -0.46196 | 0.598659 | 0.395572 |
| 8 | 86833600 | 86836199 | 18.64583 | 31.58725 | 66.30477 | 29.03276 | -0.68454 | 1.059582 | 0.395594 |
| 11 | 9391200 | 9392799 | 11.52906 | 20.72888 | 37.86265 | 16.5412 | -0.4603 | 0.39283 | 0.395639 |
| 10 | 58478400 | 58480399 | 14.61854 | 25.45396 | 46.30559 | 20.35843 | -0.47208 | 0.753633 | 0.395647 |
| X | 34919800 | 34920999 | 25.43791 | 44.00556 | 38.77639 | 17.01208 | 0.5677 | 0.868952 | 0.395648 |
| 5 | 149905000 | 149905399 | 4.389627 | 8.571343 | 21.15161 | 7.446688 | -0.6195 | -0.51027 | 0.395651 |
| 18 | 47789000 | 47792799 | 59.87026 | 95.73462 | 93.64592 | 39.39362 | 0.374612 | 2.066656 | 0.395651 |
| 8 | 117831000 | 117837399 | 50.32036 | 79.92364 | 132.7309 | 55.33975 | -0.39805 | 2.278088 | 0.395656 |
| 1 | 189723000 | 189724999 | 18.4014 | 31.38512 | 54.82819 | 24.03587 | -0.41936 | 0.986092 | 0.395671 |
| 7 | 86490600 | 86493799 | 69.41763 | 110.2954 | 108.373 | 45.20043 | 0.358433 | 2.276454 | 0.395674 |
| 9 | 26120600 | 26122199 | 16.62559 | 30.81885 | 24.07668 | 10.11286 | 0.795823 | 0.228948 | 0.395686 |
| 12 | 22960200 | 22968199 | 150.4812 | 228.2109 | 209.0404 | 86.52959 | 0.406591 | 3.247747 | 0.395688 |

| | | | | | | | | | |
|---|---|---|---|---|---|---|---|---|---|
| 13 | 33125800 | 33132399 | 52.54674 | 83.61254 | 134.3453 | 56.12778 | -0.34785 | 2.224471 | 0.395693 |
| 2 | 139519000 | 139520999 | 13.58098 | 23.57204 | 47.39895 | 20.79738 | -0.61617 | 0.797629 | 0.395696 |
| 4 | 70898200 | 70900199 | 24.57327 | 43.0035 | 33.54683 | 14.80058 | 0.744205 | 0.711431 | 0.395698 |
| 16 | 38370200 | 38374399 | 30.85461 | 50.1549 | 84.5418 | 36.27541 | -0.38262 | 1.575949 | 0.395717 |
| 7 | 10489000 | 10494999 | 113.3653 | 176.2112 | 146.0571 | 60.6417 | 0.578277 | 2.761679 | 0.395754 |
| 5 | 21878200 | 21881799 | 22.14154 | 36.32222 | 75.41912 | 32.4732 | -0.67392 | 1.423445 | 0.395782 |
| 1 | 73768400 | 73770999 | 25.46295 | 44.27327 | 36.95141 | 16.23129 | 0.646677 | 0.811514 | 0.395783 |
| 13 | 57540600 | 57542199 | 8.93397 | 16.5621 | 31.53266 | 13.24326 | -0.48013 | 0.184554 | 0.395824 |
| 12 | 35903800 | 35906199 | 21.79438 | 36.27757 | 64.9597 | 28.20776 | -0.45642 | 1.240844 | 0.395831 |
| 4 | 146739400 | 146740199 | 11.99319 | 22.91348 | 17.49481 | 6.95059 | 0.895037 | -0.01963 | 0.39584 |
| 12 | 17102400 | 17103999 | 10.15864 | 18.28477 | 35.36696 | 15.43526 | -0.54084 | 0.373963 | 0.395843 |
| 9 | 16596200 | 16597399 | 10.10103 | 18.4849 | 35.04242 | 14.97872 | -0.49253 | 0.275924 | 0.395843 |
| 10 | 114007400 | 114010999 | 48.26593 | 78.04749 | 70.87091 | 30.2902 | 0.50057 | 1.70444 | 0.395864 |
| 9 | 63088600 | 63089999 | 7.05527 | 13.39948 | 27.68046 | 11.13646 | -0.54506 | 0.001923 | 0.395881 |
| 11 | 19869600 | 19870799 | 7.05527 | 13.39948 | 27.68046 | 11.13646 | -0.54506 | 0.001923 | 0.395881 |
| 11 | 89405400 | 89407599 | 14.58742 | 25.48109 | 45.71912 | 20.14101 | -0.45183 | 0.706373 | 0.395884 |
| 13 | 119347400 | 119349199 | 22.91125 | 39.9223 | 36.47735 | 16.04473 | 0.516837 | 0.768053 | 0.395888 |
| 7 | 23567000 | 23577399 | 132.3505 | 201.9847 | 173.7702 | 72.05936 | 0.504329 | 3.113231 | 0.395888 |
| 5 | 119668000 | 119669199 | 8.107091 | 15.29542 | 29.73459 | 12.13172 | -0.47661 | 0.061663 | 0.395896 |
| 7 | 14147200 | 14150999 | 61.02725 | 98.02501 | 82.07842 | 34.76046 | 0.605992 | 1.92915 | 0.395908 |
| 12 | 21986600 | 22023599 | 1289.953 | 1632.685 | 1306.856 | 548.9038 | 0.34498 | 6.171252 | 0.395917 |
| 14 | 21828200 | 21829799 | 15.57557 | 28.07909 | 27.19256 | 11.8443 | 0.45245 | 0.381984 | 0.395917 |
| 17 | 32996800 | 32999599 | 17.16996 | 29.65508 | 52.2292 | 22.91301 | -0.42783 | 0.86225 | 0.395919 |
| 17 | 53458800 | 53462399 | 94.01055 | 147.8738 | 126.2941 | 52.39403 | 0.548316 | 2.537161 | 0.395919 |
| 7 | 43895400 | 43896599 | 9.668425 | 17.42492 | 34.85759 | 15.1884 | -0.58774 | 0.359847 | 0.395946 |
| 13 | 47463200 | 47464999 | 19.74636 | 34.49694 | 32.09965 | 14.14222 | 0.491001 | 0.709144 | 0.395952 |
| 15 | 14813800 | 14814599 | 12.41812 | 23.44877 | 22.56341 | 9.189404 | 0.532711 | 0.077315 | 0.396002 |
| 7 | 14298200 | 14305799 | 164.6165 | 247.952 | 211.1251 | 87.20966 | 0.503504 | 3.386919 | 0.396007 |
| 11 | 88883800 | 88885999 | 10.10345 | 18.30678 | 34.3694 | 14.87565 | -0.49047 | 0.331094 | 0.396009 |
| 10 | 37361800 | 37363199 | 13.34949 | 25.28344 | 22.48458 | 9.099181 | 0.653389 | 0.064512 | 0.396013 |
| 17 | 89747800 | 89748399 | 5.96583 | 11.54913 | 24.9168 | 9.482138 | -0.53153 | -0.19996 | 0.396013 |
| 5 | 89373600 | 89375799 | 18.54363 | 32.72671 | 31.13861 | 13.77812 | 0.460979 | 0.560218 | 0.396017 |
| 9 | 81745400 | 81746399 | 7.724457 | 14.41817 | 30.06293 | 12.49218 | -0.59762 | 0.131 | 0.396018 |
| 16 | 65609000 | 65610199 | 16.81274 | 30.85501 | 24.94829 | 10.62282 | 0.732369 | 0.285755 | 0.396022 |
| X | 17699000 | 17701399 | 70.48332 | 111.9856 | 107.0108 | 44.63106 | 0.398549 | 2.280375 | 0.396024 |
| 12 | 36658000 | 36659799 | 19.59044 | 34.67835 | 30.43577 | 13.45903 | 0.578525 | 0.533723 | 0.39603 |
| 12 | 85762400 | 85763799 | 21.41946 | 37.33461 | 36.06486 | 15.86642 | 0.436961 | 0.755599 | 0.396038 |
| 1 | 65010000 | 65013799 | 22.85095 | 38.22803 | 65.84976 | 28.68512 | -0.4009 | 1.18936 | 0.396059 |
| 19 | 39480800 | 39482999 | 28.73212 | 49.11956 | 38.26228 | 16.77678 | 0.741769 | 0.985029 | 0.396081 |
| 1 | 112764400 | 112768199 | 40.39915 | 66.02349 | 56.44936 | 24.26891 | 0.599593 | 1.501739 | 0.396089 |
| 17 | 83651800 | 83654799 | 38.11782 | 61.30563 | 101.1537 | 42.91897 | -0.36834 | 1.865287 | 0.396089 |
| 8 | 19066400 | 19067599 | 22.9367 | 39.78338 | 36.85258 | 16.16936 | 0.496681 | 0.832145 | 0.396093 |
| Y | 20638800 | 20640199 | 13.36981 | 24.93152 | 14.45768 | 6.017478 | 1.231378 | 0.139052 | 0.396096 |
| 5 | 115352800 | 115355799 | 14.45695 | 25.17533 | 46.30404 | 20.35912 | -0.48783 | 0.751039 | 0.396108 |
| 6 | 138160800 | 138161599 | 6.572829 | 12.62215 | 26.31887 | 10.30647 | -0.52282 | -0.09612 | 0.396115 |
| 10 | 95524800 | 95525999 | 6.572829 | 12.62215 | 26.31887 | 10.30647 | -0.52282 | -0.09612 | 0.396115 |
| 4 | 13996800 | 13997799 | 13.07228 | 24.27069 | 23.88868 | 10.01219 | 0.466925 | 0.190029 | 0.396115 |

| | | | | | | | | | |
|---|---|---|---|---|---|---|---|---|---|
| 4 | 27100800 | 27101799 | 13.07228 | 24.27069 | 23.88868 | 10.01219 | 0.466925 | 0.190029 | 0.396115 |
| 5 | 130861600 | 130869199 | 70.84499 | 110.6553 | 174.4644 | 72.38868 | -0.34288 | 2.660454 | 0.396123 |
| 1 | 117379200 | 117382399 | 36.51108 | 61.1079 | 47.53214 | 20.71156 | 0.742285 | 1.209134 | 0.396134 |
| 8 | 109515000 | 109527599 | 147.6829 | 216.5212 | 418.7813 | 172.4944 | -0.71526 | 3.866126 | 0.396146 |
| 4 | 7691400 | 7695999 | 62.87653 | 100.7214 | 88.55277 | 37.34777 | 0.531167 | 2.01667 | 0.396154 |
| 4 | 152872200 | 152889999 | 252.3328 | 358.0564 | 535.1097 | 221.3601 | -0.38995 | 4.360859 | 0.39616 |
| 16 | 35410400 | 35415599 | 35.6301 | 57.37921 | 100.3792 | 42.67079 | -0.45024 | 1.818421 | 0.39616 |
| 4 | 156292200 | 156295199 | 34.1713 | 56.35272 | 54.77733 | 23.63812 | 0.420723 | 1.370031 | 0.396181 |
| 1 | 121689000 | 121690599 | 16.43438 | 30.47449 | 24.08482 | 10.11173 | 0.779662 | 0.225461 | 0.396191 |
| 7 | 54666200 | 54668799 | 13.57084 | 23.71086 | 50.5113 | 22.25539 | -0.69817 | 0.707273 | 0.396195 |
| 10 | 14730800 | 14732199 | 15.2936 | 28.33805 | 24.74788 | 10.40062 | 0.635616 | 0.221988 | 0.396207 |
| 18 | 60395800 | 60399399 | 33.0524 | 55.49856 | 48.03065 | 20.97157 | 0.588735 | 1.165198 | 0.39621 |
| 3 | 33558600 | 33561199 | 27.98231 | 45.93675 | 82.46754 | 35.51884 | -0.46451 | 1.431296 | 0.396217 |
| 7 | 102263400 | 102264199 | 5.514723 | 10.7202 | 23.46892 | 8.716465 | -0.5191 | -0.29437 | 0.396217 |
| 4 | 142127400 | 142133599 | 66.14266 | 105.249 | 104.6115 | 43.70685 | 0.344026 | 2.228001 | 0.396223 |
| 14 | 63629600 | 63630799 | 9.379542 | 17.32403 | 33.07519 | 13.96107 | -0.48974 | 0.210981 | 0.396235 |
| 7 | 122142000 | 122144599 | 17.90123 | 30.47234 | 55.89787 | 24.50094 | -0.49006 | 1.001492 | 0.396248 |
| 12 | 18705000 | 18758399 | 1832.152 | 2255.145 | 1777.296 | 748.8415 | 0.326623 | 6.61935 | 0.396252 |
| 4 | 92704000 | 92705599 | 21.52592 | 37.9852 | 29.60985 | 13.10201 | 0.746814 | 0.575588 | 0.396259 |
| 10 | 12946600 | 12947399 | 10.89242 | 20.93022 | 20.04928 | 7.837365 | 0.591222 | -0.07789 | 0.396262 |
| X | 146552600 | 146555999 | 42.88531 | 70.11056 | 61.66514 | 26.51461 | 0.559254 | 1.507425 | 0.396274 |
| 19 | 8219800 | 8227999 | 125.9713 | 193.0785 | 181.1267 | 75.18291 | 0.384434 | 3.026596 | 0.396288 |
| 15 | 101092400 | 101094399 | 17.46375 | 29.69223 | 55.30505 | 24.23877 | -0.51217 | 1.011387 | 0.3963 |
| 12 | 119516000 | 119522599 | 362.3307 | 512.4021 | 404.0685 | 167.2459 | 0.52712 | 4.414418 | 0.396304 |
| 17 | 19780200 | 19784799 | 64.52179 | 103.4166 | 86.15597 | 36.36879 | 0.609678 | 2.000161 | 0.396306 |
| 16 | 25942000 | 25942399 | 10.06231 | 19.66829 | 12.45868 | 4.293422 | 1.344324 | -0.53125 | 0.396308 |
| 8 | 79168000 | 79170799 | 14.92369 | 26.11793 | 46.73897 | 20.62172 | -0.44804 | 0.680132 | 0.396312 |
| 14 | 10925000 | 10928399 | 20.65561 | 34.99262 | 67.74691 | 29.6644 | -0.56852 | 1.065206 | 0.396316 |
| 4 | 58700800 | 58702399 | 14.89793 | 25.99868 | 46.7925 | 20.60152 | -0.45649 | 0.72265 | 0.396318 |
| 1 | 34637600 | 34645599 | 178.2315 | 261.307 | 389.3998 | 160.3924 | -0.33931 | 3.868597 | 0.396349 |
| 18 | 51787400 | 51790999 | 39.09851 | 65.74076 | 42.96057 | 18.76836 | 0.992675 | 1.170327 | 0.396381 |
| 13 | 66208200 | 66220799 | 351.1942 | 494.931 | 442.5944 | 183.2971 | 0.340711 | 4.464768 | 0.396383 |
| X | 113575400 | 113579199 | 48.38976 | 78.58183 | 65.7382 | 28.18724 | 0.624362 | 1.620585 | 0.396387 |
| 3 | 16687400 | 16690999 | 69.464 | 110.4212 | 107.926 | 45.03526 | 0.366691 | 2.263241 | 0.396392 |
| 15 | 22493000 | 22494399 | 19.74027 | 35.0885 | 29.30605 | 12.92717 | 0.652586 | 0.500151 | 0.396394 |
| 16 | 76082400 | 76084199 | 11.51187 | 20.66389 | 37.83485 | 16.55204 | -0.46521 | 0.402185 | 0.3964 |
| 1 | 25641800 | 25645399 | 31.53527 | 53.01004 | 48.0205 | 20.97726 | 0.523079 | 1.149585 | 0.396404 |
| 1 | 109177400 | 109179799 | 32.81505 | 54.64342 | 52.50775 | 22.80636 | 0.438564 | 1.256275 | 0.396416 |
| 15 | 20073200 | 20074199 | 11.27647 | 21.66317 | 19.39598 | 7.587027 | 0.686613 | -0.07439 | 0.396427 |
| 10 | 108178000 | 108180399 | 17.82174 | 30.59028 | 54.92214 | 24.08939 | -0.45739 | 0.923366 | 0.396433 |
| 12 | 101128600 | 101132199 | 58.46723 | 93.65316 | 90.26817 | 38.06812 | 0.398651 | 2.012353 | 0.396435 |
| 1 | 167444000 | 167446199 | 10.47667 | 18.67727 | 37.32286 | 16.42266 | -0.59622 | 0.437571 | 0.396438 |
| 15 | 98237800 | 98238999 | 10.47667 | 18.67727 | 37.32286 | 16.42266 | -0.59622 | 0.437571 | 0.396438 |
| 14 | 91098000 | 91098399 | 7.380726 | 14.40714 | 15.75836 | 5.579404 | 0.53312 | -0.46745 | 0.39644 |
| 17 | 65398800 | 65399999 | 11.24353 | 20.09126 | 36.79945 | 16.1605 | -0.46927 | 0.428181 | 0.396441 |
| 15 | 5082400 | 5085199 | 32.70764 | 55.06697 | 46.90088 | 20.4979 | 0.611806 | 1.139449 | 0.396485 |
| 3 | 85186600 | 85189799 | 15.40893 | 26.76459 | 47.92065 | 21.03609 | -0.44968 | 0.794421 | 0.39649 |

| | | | | | | | | | |
|---|---|---|---|---|---|---|---|---|---|
| 17 | 74636400 | 74636799 | 3.524124 | 6.889453 | 18.29394 | 6.280139 | -0.68792 | -0.61951 | 0.39649 |
| 13 | 19381400 | 19383599 | 19.85092 | 34.82679 | 33.28978 | 14.71841 | 0.452274 | 0.645019 | 0.396494 |
| X | 112183600 | 112185399 | 19.10433 | 34.92502 | 24.84974 | 10.63641 | 0.910706 | 0.325978 | 0.396507 |
| 1 | 72322600 | 72323999 | 7.185353 | 13.63795 | 29.35397 | 11.82548 | -0.60633 | 0.008683 | 0.396508 |
| 4 | 61827800 | 61831799 | 65.11847 | 103.5725 | 102.7924 | 42.91999 | 0.345468 | 2.240744 | 0.396508 |
| 1 | 18780600 | 18783399 | 25.24449 | 44.08388 | 34.04681 | 14.99617 | 0.758492 | 0.74644 | 0.396511 |
| 13 | 23533400 | 23536599 | 38.79613 | 64.2079 | 54.40707 | 23.53204 | 0.618911 | 1.34347 | 0.396513 |
| 5 | 131606200 | 131609399 | 19.76779 | 33.23154 | 60.15482 | 26.27857 | -0.47209 | 1.123518 | 0.396535 |
| X | 109036800 | 109045399 | 116.5239 | 180.6109 | 150.4607 | 62.49633 | 0.567876 | 2.81231 | 0.396536 |
| 10 | 119998800 | 120002399 | 24.51281 | 40.56928 | 71.98936 | 31.13691 | -0.4443 | 1.317479 | 0.396541 |
| 7 | 15369800 | 15388399 | 349.578 | 497.799 | 410.0901 | 169.5424 | 0.474126 | 4.313586 | 0.396555 |
| 6 | 10307000 | 10308999 | 27.73387 | 48.08929 | 36.86182 | 16.17326 | 0.768422 | 0.854463 | 0.396559 |
| 19 | 10858600 | 10864199 | 46.98581 | 75.14525 | 128.7431 | 54.1714 | -0.43229 | 2.058861 | 0.39657 |
| X | 109298000 | 109302999 | 75.87404 | 120.763 | 92.38292 | 38.61057 | 0.721513 | 2.228232 | 0.39657 |
| 12 | 15494800 | 15495799 | 14.56785 | 26.9562 | 20.83458 | 8.772398 | 0.808814 | 0.214867 | 0.396582 |
| 6 | 10784800 | 10787599 | 31.88599 | 51.85126 | 97.95285 | 42.03572 | -0.54636 | 1.580735 | 0.396582 |
| 4 | 80994600 | 80995399 | 6.913206 | 13.13126 | 27.67659 | 11.13205 | -0.57317 | 0.000789 | 0.396589 |
| 1 | 186930200 | 186932199 | 15.08655 | 26.32536 | 46.26552 | 20.3683 | -0.42222 | 0.72539 | 0.396607 |
| 2 | 100105800 | 100106599 | 10.69173 | 20.76777 | 18.11312 | 6.769669 | 0.785808 | -0.23164 | 0.396612 |
| 17 | 32569000 | 32569799 | 4.994958 | 9.64545 | 24.74777 | 9.528683 | -0.79501 | -0.17265 | 0.396621 |
| 3 | 91281400 | 91282199 | 11.4628 | 22.04518 | 20.70343 | 8.073116 | 0.623683 | -0.07525 | 0.396634 |
| 15 | 27630200 | 27631199 | 7.876588 | 14.63691 | 30.5221 | 12.77714 | -0.60674 | 0.159939 | 0.396654 |
| 10 | 50029400 | 50030599 | 19.4197 | 34.09949 | 32.82513 | 14.5235 | 0.442189 | 0.629 | 0.396658 |
| 5 | 100988200 | 100992199 | 45.35445 | 73.45388 | 70.02483 | 29.96548 | 0.432677 | 1.667247 | 0.396661 |
| 11 | 27755800 | 27756599 | 9.452491 | 18.27596 | 19.23599 | 7.363307 | 0.485368 | -0.16208 | 0.396662 |
| 2 | 160278000 | 160282399 | 24.37319 | 40.2825 | 71.87712 | 31.05871 | -0.4524 | 1.333271 | 0.396663 |
| X | 117304200 | 117307599 | 41.88518 | 68.75041 | 58.9076 | 25.37003 | 0.599686 | 1.452567 | 0.396671 |
| 10 | 110940200 | 110943799 | 22.75019 | 37.78561 | 66.70946 | 28.92383 | -0.43642 | 1.269224 | 0.396686 |
| 9 | 90741200 | 90741599 | 8.076997 | 15.72915 | 16.71077 | 6.115176 | 0.531904 | -0.34052 | 0.396693 |
| 12 | 108807000 | 108808999 | 14.03332 | 24.38158 | 46.86683 | 20.57687 | -0.55138 | 0.784241 | 0.396697 |
| 10 | 91770600 | 91771599 | 13.08985 | 24.24552 | 23.90315 | 10.05075 | 0.461286 | 0.20358 | 0.396732 |
| 6 | 124623000 | 124624799 | 14.16278 | 24.83218 | 44.06403 | 19.47193 | -0.43568 | 0.643684 | 0.396755 |
| 1 | 30377200 | 30378999 | 19.54511 | 34.53087 | 30.9848 | 13.70714 | 0.545673 | 0.554731 | 0.396761 |
| 2 | 97913800 | 97916199 | 19.54511 | 34.53087 | 30.9848 | 13.70714 | 0.545673 | 0.554731 | 0.396761 |
| 8 | 21554000 | 21557599 | 33.30848 | 56.53842 | 42.94146 | 18.81399 | 0.776861 | 1.070351 | 0.396773 |
| 5 | 100053200 | 100055599 | 22.04455 | 36.01032 | 84.00524 | 36.10976 | -0.84482 | 1.487548 | 0.396781 |
| 4 | 115300600 | 115301399 | 21.35906 | 37.18212 | 36.21715 | 15.92072 | 0.424988 | 0.774693 | 0.3968 |
| 15 | 37469800 | 37471999 | 25.32453 | 43.81318 | 39.24557 | 17.21796 | 0.54407 | 0.866986 | 0.396811 |
| 17 | 20847000 | 20848999 | 46.94144 | 76.0049 | 52.56664 | 22.49048 | 0.894159 | 1.667054 | 0.396834 |
| 11 | 43888400 | 43889399 | 11.89443 | 22.3382 | 21.94939 | 9.026527 | 0.490861 | 0.1035 | 0.396836 |
| 6 | 13069400 | 13070799 | 7.401042 | 13.8576 | 29.57532 | 12.22489 | -0.62518 | 0.104424 | 0.396837 |
| 1 | 61607200 | 61608799 | 11.39651 | 20.32284 | 37.28621 | 16.40264 | -0.47351 | 0.442493 | 0.396848 |
| 13 | 79620800 | 79621799 | 13.22108 | 24.42782 | 23.86735 | 10.06969 | 0.470622 | 0.217968 | 0.396869 |
| X | 34890200 | 34894599 | 69.54247 | 111.1532 | 91.36105 | 38.40915 | 0.624612 | 2.092171 | 0.396881 |
| 7 | 29817400 | 29818799 | 19.61681 | 34.36308 | 32.5527 | 14.3738 | 0.465397 | 0.668647 | 0.396883 |
| 14 | 91934400 | 91935199 | 6.998472 | 13.47368 | 28.23736 | 10.97747 | -0.5208 | -0.11177 | 0.396897 |
| Y | 8694800 | 8699199 | 32.92387 | 54.92773 | 30.74776 | 13.3709 | 1.214916 | 1.21107 | 0.396912 |

| | | | | | | | | | |
|---|---|---|---|---|---|---|---|---|---|
| 14 | 83434800 | 83435199 | 7.226452 | 14.09713 | 14.19754 | 5.079248 | 0.638307 | -0.43867 | 0.396921 |
| 14 | 15064600 | 15066999 | 10.84957 | 19.15543 | 39.40439 | 17.43416 | -0.64486 | 0.525885 | 0.396932 |
| 17 | 13332000 | 13344599 | 74.52326 | 115.3353 | 197.4139 | 82.00073 | -0.4716 | 2.824561 | 0.396947 |
| 4 | 80331600 | 80333399 | 20.33658 | 35.64638 | 33.8057 | 14.93502 | 0.463619 | 0.662902 | 0.396957 |
| 17 | 43096400 | 43097999 | 13.94794 | 25.9675 | 24.36725 | 10.16983 | 0.540042 | 0.182797 | 0.396997 |
| 7 | 105033000 | 105034599 | 9.64106 | 17.44919 | 34.36936 | 14.89672 | -0.5606 | 0.333535 | 0.397002 |
| 5 | 15811600 | 15813599 | 25.80876 | 44.01286 | 42.5709 | 18.66225 | 0.430069 | 0.998421 | 0.39702 |
| 3 | 127382000 | 127383599 | 18.3556 | 32.99691 | 27.97333 | 12.22265 | 0.640538 | 0.420861 | 0.397069 |
| 12 | 23328200 | 23331199 | 34.77824 | 57.91111 | 47.08807 | 20.45205 | 0.678678 | 1.256157 | 0.397092 |
| 1 | 107176800 | 107181999 | 60.08427 | 96.2602 | 89.6088 | 37.80012 | 0.449039 | 2.008325 | 0.397096 |
| 2 | 171988000 | 171990599 | 21.69395 | 36.35999 | 63.01116 | 27.4808 | -0.40956 | 1.162878 | 0.397098 |
| 16 | 97340000 | 97343599 | 34.53892 | 57.27515 | 54.27199 | 23.50564 | 0.458285 | 1.307924 | 0.397098 |
| 8 | 18822400 | 18824199 | 12.54729 | 22.09761 | 40.37973 | 17.87461 | -0.47656 | 0.565836 | 0.397128 |
| 5 | 21989400 | 21990399 | 7.680253 | 14.43883 | 29.10458 | 11.94798 | -0.53537 | 0.080072 | 0.397134 |
| 4 | 146351600 | 146353999 | 90.60245 | 143.1235 | 117.237 | 48.6891 | 0.613254 | 2.445886 | 0.397156 |
| 7 | 61522800 | 61523599 | 12.72317 | 23.9557 | 23.02908 | 9.429687 | 0.52755 | 0.099552 | 0.397156 |
| 9 | 73995200 | 73996399 | 12.72317 | 23.9557 | 23.02908 | 9.429687 | 0.52755 | 0.099552 | 0.397156 |
| 17 | 6367000 | 6368399 | 8.158392 | 15.14478 | 32.09348 | 13.45402 | -0.63185 | 0.169572 | 0.397157 |
| 8 | 15305200 | 15311999 | 68.1136 | 106.7483 | 167.6731 | 69.54668 | -0.33366 | 2.598716 | 0.397162 |
| 2 | 155616400 | 155617999 | 14.35787 | 24.85613 | 48.45048 | 21.25679 | -0.57247 | 0.825118 | 0.397166 |
| 5 | 53590200 | 53592199 | 12.99457 | 23.00838 | 43.45882 | 19.21699 | -0.52129 | 0.513741 | 0.397197 |
| 13 | 65025200 | 65030399 | 97.33941 | 151.0204 | 143.4812 | 59.58873 | 0.379515 | 2.785041 | 0.397203 |
| 12 | 44264400 | 44265399 | 6.255088 | 12.10644 | 25.5493 | 9.73281 | -0.50231 | -0.19427 | 0.397229 |
| 16 | 42491800 | 42493199 | 9.547255 | 17.70307 | 33.71701 | 14.15626 | -0.48064 | 0.192541 | 0.39723 |
| 6 | 60009000 | 60013199 | 89.01993 | 139.7424 | 134.167 | 55.64561 | 0.377475 | 2.575603 | 0.397242 |
| X | 64028600 | 64030799 | 21.35724 | 38.31924 | 26.54029 | 11.61526 | 0.928807 | 0.44989 | 0.397251 |
| 17 | 44173000 | 44174199 | 11.17304 | 20.10307 | 37.86319 | 16.53151 | -0.50383 | 0.384215 | 0.397252 |
| 3 | 92125000 | 92125399 | 9.18872 | 17.95074 | 15.13315 | 5.284412 | 0.922587 | -0.48926 | 0.397255 |
| 12 | 18807400 | 18835799 | 962.6472 | 1248.212 | 1034.433 | 433.4521 | 0.329873 | 5.77792 | 0.397259 |
| 13 | 3098400 | 3099799 | 18.20583 | 32.26805 | 29.99677 | 13.26156 | 0.4966 | 0.51001 | 0.397259 |
| 19 | 33926400 | 33926799 | 5.403887 | 10.45586 | 24.30502 | 9.271744 | -0.64154 | -0.20172 | 0.397268 |
| 17 | 5327000 | 5330199 | 16.98682 | 28.94111 | 54.23321 | 23.77297 | -0.52032 | 0.989997 | 0.397272 |
| 17 | 18456600 | 18459199 | 46.79082 | 75.46444 | 74.93992 | 31.92136 | 0.367511 | 1.781284 | 0.397278 |
| 6 | 117509200 | 117510399 | 9.852975 | 18.12895 | 34.62347 | 14.6909 | -0.49582 | 0.239435 | 0.397279 |
| 1 | 90406400 | 90408199 | 19.11721 | 33.57566 | 32.37524 | 14.32708 | 0.439811 | 0.62348 | 0.397292 |
| 13 | 114805000 | 114807999 | 29.33991 | 49.95872 | 43.53987 | 19.08469 | 0.579566 | 1.027392 | 0.397351 |
| 8 | 14094400 | 14097599 | 47.29001 | 76.31727 | 75.20868 | 32.06169 | 0.379562 | 1.764181 | 0.397385 |
| 15 | 91342600 | 91345399 | 13.49497 | 23.77978 | 42.40866 | 18.77066 | -0.44148 | 0.568621 | 0.397388 |
| 1 | 109677400 | 109680399 | 26.91505 | 46.52527 | 37.9851 | 16.66453 | 0.67686 | 0.877675 | 0.397392 |
| 16 | 13087000 | 13090399 | 37.19926 | 59.68081 | 113.5989 | 48.0232 | -0.57844 | 1.946152 | 0.397393 |
| 16 | 5217400 | 5220399 | 22.72906 | 37.9395 | 65.15755 | 28.34372 | -0.3965 | 1.213299 | 0.397404 |
| 11 | 12078400 | 12080199 | 17.98477 | 30.59921 | 55.82493 | 24.46798 | -0.48213 | 1.01001 | 0.397407 |
| 7 | 31575200 | 31575799 | 16.6402 | 29.57149 | 28.30467 | 12.48781 | 0.456974 | 0.480211 | 0.397413 |
| 1 | 103238200 | 103241999 | 45.08133 | 74.41694 | 51.77018 | 22.35153 | 0.902397 | 1.388488 | 0.397417 |
| 3 | 20653600 | 20655799 | 17.5825 | 31.93728 | 25.80405 | 11.1339 | 0.72069 | 0.349903 | 0.397419 |
| 7 | 59064400 | 59066199 | 26.53389 | 45.02806 | 44.21868 | 19.37213 | 0.407457 | 1.047253 | 0.397425 |
| 5 | 63530600 | 63532199 | 13.85767 | 26.1716 | 22.31496 | 9.084862 | 0.705804 | 0.086524 | 0.397443 |

| | | | | | | | | | |
|---|---|---|---|---|---|---|---|---|---|
| 3 | 51825400 | 51829399 | 34.8233 | 56.20722 | 93.59207 | 39.90674 | -0.37523 | 1.748111 | 0.397446 |
| X | 77749200 | 77752799 | 34.45569 | 58.53445 | 41.83131 | 18.33223 | 0.864464 | 1.067799 | 0.397453 |
| 12 | 44268400 | 44269199 | 6.112057 | 11.83152 | 24.94076 | 9.493843 | -0.49916 | -0.19865 | 0.397471 |
| 18 | 6254800 | 6257999 | 47.72957 | 76.88026 | 76.5738 | 32.56098 | 0.36104 | 1.820701 | 0.397479 |
| X | 122611400 | 122613599 | 17.56892 | 32.29341 | 23.74733 | 10.08989 | 0.870404 | 0.280095 | 0.39748 |
| 8 | 21596400 | 21599799 | 51.66975 | 83.36585 | 75.10821 | 32.00921 | 0.508289 | 1.779032 | 0.397486 |
| 1 | 84902000 | 84904399 | 23.05332 | 38.27995 | 67.92042 | 29.44404 | -0.44368 | 1.272814 | 0.397495 |
| 13 | 76626200 | 76627399 | 12.4682 | 23.48652 | 22.99809 | 9.409601 | 0.501823 | 0.09325 | 0.397495 |
| 5 | 17248000 | 17250799 | 21.29645 | 37.64718 | 30.3139 | 13.40877 | 0.701484 | 0.558842 | 0.397502 |
| 1 | 66092600 | 66093799 | 11.61674 | 22.50337 | 19.16888 | 7.272438 | 0.800089 | -0.16705 | 0.397506 |
| 16 | 9983000 | 9984599 | 27.97236 | 48.19558 | 39.68619 | 17.40909 | 0.663835 | 0.914877 | 0.397509 |
| 5 | 127314800 | 127317799 | 29.65704 | 49.65067 | 49.15373 | 21.42203 | 0.395543 | 1.187086 | 0.39751 |
| 9 | 15268600 | 15270799 | 13.81832 | 24.28022 | 42.94987 | 19.014 | -0.43122 | 0.604613 | 0.397512 |
| 18 | 59862200 | 59864799 | 51.83001 | 81.75206 | 161.0202 | 66.82451 | -0.65244 | 2.463859 | 0.397512 |
| 1 | 119029200 | 119031599 | 15.31451 | 26.36053 | 51.62615 | 22.6461 | -0.58123 | 0.884101 | 0.397515 |
| 12 | 45004000 | 45005199 | 12.33109 | 23.07883 | 23.00863 | 9.517354 | 0.463663 | 0.132954 | 0.397522 |
| 17 | 12833000 | 12835399 | 13.80735 | 24.21898 | 43.00282 | 19.00961 | -0.43662 | 0.632182 | 0.397523 |
| 2 | 60325400 | 60328999 | 19.35247 | 32.81174 | 58.09494 | 25.44438 | -0.43981 | 1.044739 | 0.397549 |
| Y | 86578000 | 86579399 | 11.52587 | 21.93434 | 13.48752 | 5.403212 | 1.192892 | -0.01362 | 0.39756 |
| 8 | 92111200 | 92112799 | 7.980664 | 14.92378 | 30.62413 | 12.68586 | -0.57194 | 0.118962 | 0.397561 |
| 19 | 59683200 | 59684599 | 7.980664 | 14.92378 | 30.62413 | 12.68586 | -0.57194 | 0.118962 | 0.397561 |
| 6 | 26691600 | 26692399 | 12.30259 | 23.12856 | 22.99974 | 9.441987 | 0.475873 | 0.10419 | 0.397564 |
| 11 | 92147600 | 92148199 | 12.30259 | 23.12856 | 22.99974 | 9.441987 | 0.475873 | 0.10419 | 0.397564 |
| 12 | 75432600 | 75436399 | 31.86231 | 51.63678 | 87.03423 | 37.26242 | -0.38659 | 1.640082 | 0.397573 |
| 12 | 13622400 | 13624799 | 20.07903 | 35.24241 | 33.28381 | 14.72122 | 0.469604 | 0.640084 | 0.397574 |
| 9 | 11944200 | 11948999 | 71.47843 | 113.3921 | 112.5483 | 46.86833 | 0.34169 | 2.325135 | 0.397605 |
| 5 | 24599200 | 24599799 | 4.157847 | 8.097903 | 23.61222 | 8.628005 | -0.90655 | -0.37265 | 0.397613 |
| 5 | 145933200 | 145943199 | 140.7506 | 214.8404 | 189.5539 | 78.60762 | 0.468159 | 3.114115 | 0.397617 |
| 12 | 14979400 | 14979999 | 5.38138 | 10.47594 | 24.32069 | 8.929285 | -0.58861 | -0.33807 | 0.397639 |
| 7 | 133825000 | 133827799 | 19.59242 | 32.58479 | 65.27519 | 28.32832 | -0.61775 | 1.244165 | 0.397646 |
| 8 | 43533400 | 43536399 | 43.56185 | 69.48967 | 121.3183 | 50.86129 | -0.46344 | 2.131838 | 0.397646 |
| 10 | 19905400 | 19911399 | 65.33879 | 104.407 | 94.58627 | 39.75037 | 0.484038 | 2.094051 | 0.397668 |
| 17 | 18635400 | 18638399 | 42.64465 | 70.51007 | 46.66403 | 20.16978 | 0.974262 | 1.359728 | 0.397674 |
| 12 | 17178400 | 17179599 | 10.76892 | 20.53971 | 20.5493 | 8.193369 | 0.503097 | -0.01395 | 0.397689 |
| 7 | 61631800 | 61634199 | 22.95695 | 40.34309 | 31.22601 | 13.82537 | 0.755063 | 0.632724 | 0.397695 |
| 16 | 61426800 | 61428999 | 22.30032 | 39.14836 | 32.43095 | 14.34622 | 0.657648 | 0.647247 | 0.397724 |
| X | 125767800 | 125768599 | 20.12587 | 35.36826 | 32.71892 | 14.48647 | 0.499283 | 0.620426 | 0.397729 |
| 19 | 40066800 | 40067599 | 5.957214 | 11.56682 | 26.34484 | 9.877579 | -0.58862 | -0.24618 | 0.397733 |
| 2 | 70323000 | 70323799 | 5.532423 | 10.72863 | 23.8594 | 9.006735 | -0.56383 | -0.23718 | 0.397739 |
| 10 | 40037200 | 40037999 | 5.665965 | 10.98433 | 24.39721 | 9.222577 | -0.56344 | -0.22688 | 0.397758 |
| 2 | 148085600 | 148085999 | 4.391907 | 8.56956 | 20.96244 | 7.473194 | -0.62306 | -0.46543 | 0.397764 |
| 19 | 39514600 | 39515799 | 18.1679 | 33.10353 | 25.93598 | 11.14733 | 0.768715 | 0.339028 | 0.397771 |
| 6 | 73618400 | 73619399 | 10.87599 | 20.81773 | 21.16876 | 8.36768 | 0.491372 | -0.03767 | 0.397772 |
| 3 | 159535800 | 159536799 | 13.61603 | 25.52481 | 19.88123 | 8.201321 | 0.820454 | 0.126796 | 0.397775 |
| 16 | 59349200 | 59350999 | 18.38656 | 30.90497 | 64.38656 | 28.12594 | -0.67409 | 1.125903 | 0.397778 |
| 12 | 15292800 | 15293199 | 6.849573 | 13.22974 | 28.4463 | 10.9413 | -0.5428 | -0.14752 | 0.397781 |
| 14 | 44214800 | 44216599 | 9.746249 | 17.94074 | 33.54896 | 14.22617 | -0.46457 | 0.236005 | 0.397789 |

| | | | | | | | | | |
|---|---|---|---|---|---|---|---|---|---|
| 7 | 123069600 | 123071599 | 18.09826 | 32.54878 | 27.92399 | 12.19652 | 0.623704 | 0.41542 | 0.397789 |
| 2 | 53448800 | 53451599 | 30.53104 | 51.30118 | 43.2455 | 18.88903 | 0.626544 | 1.145728 | 0.397802 |
| 2 | 82590200 | 82591599 | 15.90969 | 28.91623 | 25.99656 | 11.20848 | 0.568218 | 0.334197 | 0.397814 |
| 15 | 36006400 | 36008999 | 26.18377 | 44.33543 | 41.68758 | 18.25098 | 0.469819 | 1.062263 | 0.397817 |
| 2 | 23001200 | 23001999 | 9.923869 | 19.3702 | 14.19299 | 5.030302 | 1.101493 | -0.43489 | 0.397828 |
| 5 | 8328600 | 8330399 | 26.58472 | 46.4271 | 32.84952 | 14.46936 | 0.884244 | 0.750827 | 0.397831 |
| 13 | 112055200 | 112057199 | 15.90644 | 27.43968 | 51.11825 | 22.42481 | -0.50878 | 0.865819 | 0.397853 |
| 2 | 157718000 | 157719999 | 8.930457 | 16.08044 | 35.39041 | 15.43792 | -0.7256 | 0.360389 | 0.397877 |
| 5 | 89022800 | 89023799 | 9.083112 | 16.79271 | 32.01914 | 13.49759 | -0.4863 | 0.201555 | 0.397893 |
| 9 | 62009000 | 62009799 | 7.159361 | 13.75589 | 28.11986 | 10.99474 | -0.49283 | -0.09087 | 0.397894 |
| 1 | 188270800 | 188273799 | 43.17012 | 69.88495 | 69.43417 | 29.70292 | 0.372625 | 1.673787 | 0.397907 |
| 16 | 26375400 | 26376199 | 6.753391 | 12.86248 | 27.18869 | 10.87335 | -0.57007 | -0.0215 | 0.397918 |
| 6 | 132130600 | 132134599 | 58.1593 | 93.55255 | 79.75471 | 33.84676 | 0.582622 | 1.879641 | 0.397931 |
| 10 | 83379400 | 83380199 | 6.165538 | 11.78234 | 27.26067 | 10.82503 | -0.69063 | -0.05295 | 0.397932 |
| 1 | 104763200 | 104763999 | 17.04766 | 30.16829 | 28.75824 | 12.71796 | 0.459763 | 0.517204 | 0.397938 |
| 2 | 87681600 | 87683799 | 24.73185 | 42.42261 | 41.49737 | 18.20016 | 0.415154 | 0.941728 | 0.397945 |
| 9 | 7489000 | 7496999 | 129.1749 | 196.8018 | 192.0456 | 79.60432 | 0.32077 | 3.147942 | 0.397947 |
| 9 | 20113400 | 20115199 | 32.75774 | 54.37652 | 53.62761 | 23.2428 | 0.400886 | 1.291401 | 0.397949 |
| 13 | 103362000 | 103364199 | 17.85302 | 31.47583 | 30.25787 | 13.39113 | 0.445669 | 0.565202 | 0.397952 |
| 7 | 84822800 | 84827199 | 84.52059 | 132.9925 | 129.7423 | 53.82698 | 0.357062 | 2.525854 | 0.397953 |
| X | 4404200 | 4406599 | 25.5497 | 45.03366 | 28.35894 | 12.55139 | 1.052686 | 0.61233 | 0.397956 |
| 14 | 67118000 | 67120799 | 13.80236 | 24.22457 | 42.98114 | 19.00953 | -0.43559 | 0.622375 | 0.397977 |
| 4 | 143074400 | 143082399 | 62.5 | 98.08355 | 164.8904 | 68.38554 | -0.42999 | 2.570146 | 0.397979 |
| 16 | 47814200 | 47817599 | 39.82529 | 65.93463 | 52.79837 | 22.84186 | 0.700234 | 1.339618 | 0.39799 |
| 14 | 121302600 | 121306999 | 63.18002 | 99.25686 | 160.3358 | 66.50587 | -0.37129 | 2.55041 | 0.397998 |
| 2 | 100676400 | 100677799 | 20.18597 | 36.44086 | 25.5133 | 11.09369 | 0.919133 | 0.403264 | 0.397998 |
| 5 | 109097000 | 109104399 | 87.60129 | 138.8696 | 107.7992 | 44.85855 | 0.694798 | 2.358357 | 0.398003 |
| 18 | 56071000 | 56076199 | 62.84206 | 101.0315 | 77.89917 | 33.03391 | 0.726319 | 1.902281 | 0.398036 |
| X | 4567400 | 4569599 | 33.83768 | 57.20356 | 44.64837 | 19.53772 | 0.737508 | 1.108162 | 0.398042 |
| 17 | 27122800 | 27134599 | 183.1698 | 267.6914 | 402.0103 | 165.6196 | -0.35485 | 3.919082 | 0.39808 |
| 5 | 148517400 | 148523399 | 66.24967 | 104.0017 | 164.5918 | 68.26459 | -0.34261 | 2.56623 | 0.398097 |
| 6 | 66353400 | 66353799 | 3.523003 | 6.890364 | 17.83483 | 6.032929 | -0.63228 | -0.67374 | 0.398109 |
| 2 | 23139400 | 23140999 | 18.99423 | 33.91821 | 28.93647 | 12.7147 | 0.625998 | 0.467342 | 0.39811 |
| 4 | 115199600 | 115201999 | 18.99423 | 33.91821 | 28.93647 | 12.7147 | 0.625998 | 0.467342 | 0.39811 |
| 7 | 41587000 | 41588399 | 21.23941 | 37.21345 | 33.67676 | 14.87285 | 0.530812 | 0.67327 | 0.398135 |
| 8 | 104233800 | 104235799 | 21.23941 | 37.21345 | 33.67676 | 14.87285 | 0.530812 | 0.67327 | 0.398135 |
| 8 | 25047800 | 25049799 | 17.22763 | 29.65029 | 51.64994 | 22.65646 | -0.41298 | 0.895011 | 0.398142 |
| 1 | 121254400 | 121255999 | 10.42124 | 18.71316 | 36.82408 | 16.10479 | -0.56819 | 0.389613 | 0.398156 |
| 2 | 26951200 | 26957599 | 37.5852 | 60.09704 | 121.9682 | 51.30419 | -0.67644 | 2.054314 | 0.398157 |
| 12 | 21401000 | 21404999 | 45.87376 | 74.36725 | 63.14023 | 27.03877 | 0.600711 | 1.646363 | 0.39817 |
| 10 | 27741200 | 27744599 | 67.56881 | 107.9931 | 84.74044 | 35.62321 | 0.691276 | 2.086964 | 0.398174 |
| 11 | 14468000 | 14470199 | 21.53641 | 37.96362 | 30.17223 | 13.35367 | 0.718466 | 0.587903 | 0.398176 |
| 14 | 23359000 | 23360999 | 15.06181 | 26.28179 | 46.79397 | 20.60074 | -0.44099 | 0.725291 | 0.398196 |
| 7 | 41919800 | 41924999 | 65.99335 | 105.8432 | 83.10418 | 35.11481 | 0.696082 | 1.981293 | 0.3982 |
| 1 | 93888200 | 93890599 | 35.5868 | 58.64042 | 57.58028 | 24.83932 | 0.405768 | 1.386664 | 0.398206 |
| X | 26386200 | 26392199 | 24.83715 | 43.30203 | 26.65969 | 11.73045 | 1.084022 | 0.758049 | 0.398216 |
| 3 | 88062000 | 88063999 | 8.810657 | 15.8049 | 36.40376 | 15.9347 | -0.79429 | 0.385226 | 0.39822 |

| | | | | | | | | | |
|---|---|---|---|---|---|---|---|---|---|
| 6 | 3786600 | 3790999 | 115.0459 | 176.432 | 168.5311 | 69.96005 | 0.358975 | 3.0106 | 0.39823 |
| 7 | 44294800 | 44295799 | 6.030191 | 11.47187 | 27.15333 | 10.88568 | -0.73434 | -0.01957 | 0.398236 |
| X | 155104800 | 155107199 | 41.27964 | 67.42437 | 61.96103 | 26.63302 | 0.495262 | 1.517656 | 0.39824 |
| 5 | 147160400 | 147165799 | 38.43651 | 61.6633 | 108.238 | 45.75412 | -0.46188 | 1.94507 | 0.398257 |
| 1 | 55492400 | 55493999 | 12.68897 | 22.34328 | 40.87531 | 18.0946 | -0.47839 | 0.568631 | 0.398267 |
| 6 | 48797000 | 48798599 | 9.964505 | 18.12876 | 34.43276 | 14.8273 | -0.50214 | 0.307953 | 0.398267 |
| 4 | 39200200 | 39201599 | 18.37499 | 32.40383 | 31.18745 | 13.80188 | 0.444082 | 0.567662 | 0.398276 |
| 7 | 7923400 | 7925599 | 49.67137 | 79.83653 | 76.17646 | 32.29144 | 0.419052 | 1.890061 | 0.398283 |
| 7 | 48358400 | 48360999 | 29.97079 | 50.64515 | 45.32127 | 19.82965 | 0.541011 | 1.100606 | 0.398283 |
| 15 | 79747800 | 79758399 | 97.58644 | 149.2254 | 229.2849 | 95.12384 | -0.3294 | 3.067986 | 0.398286 |
| 5 | 55898800 | 55900599 | 23.24909 | 40.37095 | 38.20013 | 16.76597 | 0.466212 | 0.820401 | 0.398296 |
| 10 | 69353600 | 69355999 | 16.2663 | 29.173 | 28.16752 | 12.33171 | 0.451918 | 0.420135 | 0.398302 |
| 7 | 103920800 | 103922999 | 14.68064 | 25.55476 | 46.83957 | 20.58948 | -0.48275 | 0.761242 | 0.398315 |
| X | 116227000 | 116231599 | 58.60571 | 94.18923 | 81.67906 | 34.62006 | 0.556556 | 1.908854 | 0.398316 |
| 8 | 111985000 | 111986199 | 8.966013 | 16.42291 | 33.46493 | 14.28655 | -0.59487 | 0.255577 | 0.39833 |
| 3 | 70528200 | 70529399 | 12.79425 | 24.61915 | 19.98535 | 7.780996 | 0.833706 | -0.06993 | 0.398336 |
| 8 | 10235200 | 10236199 | 13.65756 | 25.77283 | 19.83348 | 8.088426 | 0.850949 | 0.085704 | 0.398344 |
| 2 | 60688200 | 60694799 | 59.99421 | 94.97231 | 153.2494 | 63.71326 | -0.36113 | 2.380934 | 0.398345 |
| 2 | 33585000 | 33594399 | 83.1045 | 128.3708 | 201.5179 | 83.70626 | -0.34907 | 2.864095 | 0.39835 |
| 10 | 129277400 | 129279999 | 27.20799 | 46.40483 | 43.59693 | 19.11223 | 0.471971 | 1.004815 | 0.398379 |
| 8 | 91456200 | 91457999 | 12.72331 | 22.36255 | 40.92386 | 18.12144 | -0.47974 | 0.591 | 0.398381 |
| 5 | 101569000 | 101570999 | 14.71557 | 25.73111 | 50.55564 | 22.28914 | -0.58243 | 0.696182 | 0.398412 |
| 12 | 87731600 | 87736399 | 58.01784 | 91.78886 | 147.4041 | 61.26456 | -0.35489 | 2.392136 | 0.398417 |
| 7 | 23604600 | 23624999 | 303.8609 | 438.6654 | 358.3474 | 147.8375 | 0.505952 | 4.111435 | 0.398438 |
| 10 | 27734800 | 27737599 | 24.54382 | 42.72866 | 36.37955 | 15.99269 | 0.618146 | 0.788153 | 0.398441 |
| 12 | 21297200 | 21305599 | 57.73043 | 91.18473 | 150.4159 | 62.46458 | -0.39544 | 2.431117 | 0.398443 |
| 19 | 8370400 | 8373799 | 38.20522 | 63.02406 | 56.66661 | 24.4565 | 0.533094 | 1.379295 | 0.398474 |
| X | 122516600 | 122517999 | 16.81428 | 30.50469 | 26.24777 | 11.34262 | 0.629659 | 0.355771 | 0.398477 |
| 18 | 40351800 | 40352599 | 5.576425 | 10.72362 | 25.76486 | 10.04927 | -0.71958 | -0.12153 | 0.398478 |
| 10 | 7767400 | 7769199 | 25.26751 | 43.71923 | 39.22002 | 17.20681 | 0.541913 | 0.865137 | 0.398482 |
| X | 55027400 | 55032999 | 19.63744 | 34.84093 | 26.05128 | 11.50948 | 0.809966 | 0.506111 | 0.39849 |
| 3 | 42474000 | 42475199 | 13.65461 | 23.92547 | 43.04351 | 19.01041 | -0.45545 | 0.64883 | 0.39849 |
| 4 | 113221000 | 113223799 | 41.69514 | 68.17183 | 62.16715 | 26.73146 | 0.507331 | 1.503223 | 0.398507 |
| 8 | 22290000 | 22290999 | 11.35304 | 21.67206 | 21.63666 | 8.611562 | 0.508608 | -0.00982 | 0.398523 |
| Y | 5828600 | 5829799 | 10.63684 | 20.42458 | 13.00697 | 5.094884 | 1.171911 | -0.09653 | 0.398539 |
| 9 | 4195400 | 4196799 | 15.10647 | 28.00051 | 24.75511 | 10.39901 | 0.618449 | 0.218504 | 0.398565 |
| 3 | 72180200 | 72182999 | 26.52786 | 45.94968 | 38.02207 | 16.68179 | 0.658218 | 0.859203 | 0.398582 |
| 1 | 188192800 | 188194199 | 13.51644 | 25.00831 | 24.44098 | 10.2922 | 0.472453 | 0.215522 | 0.398588 |
| 15 | 96519400 | 96523599 | 48.11236 | 76.7921 | 123.1547 | 51.67229 | -0.34032 | 2.119757 | 0.398602 |
| 6 | 141002600 | 141005799 | 51.40009 | 82.77548 | 79.46983 | 33.78234 | 0.413603 | 1.834305 | 0.398619 |
| 9 | 83378800 | 83380799 | 11.45167 | 20.31001 | 37.81822 | 16.71343 | -0.49916 | 0.480057 | 0.39862 |
| 9 | 5688600 | 5689999 | 21.14484 | 36.8483 | 35.01158 | 15.40103 | 0.460586 | 0.754399 | 0.398622 |
| 2 | 138936800 | 138941399 | 63.77048 | 101.7792 | 98.18196 | 41.1917 | 0.391732 | 2.130351 | 0.398631 |
| 1 | 41076200 | 41077599 | 12.16776 | 21.74035 | 42.56246 | 18.6931 | -0.56543 | 0.443776 | 0.398639 |
| 7 | 122671200 | 122672399 | 7.3569 | 13.93346 | 28.16906 | 11.40003 | -0.5219 | 0.024992 | 0.398656 |
| 13 | 63626400 | 63628599 | 22.18087 | 38.40061 | 37.2955 | 16.36276 | 0.427949 | 0.842428 | 0.398658 |
| 1 | 163294800 | 163296599 | 21.07422 | 36.88184 | 34.29136 | 15.12966 | 0.49205 | 0.691455 | 0.398661 |

| | | | | | | | | | |
|---|---|---|---|---|---|---|---|---|---|
| X | 89574000 | 89574799 | 15.18646 | 28.14345 | 24.79677 | 10.41918 | 0.623213 | 0.220999 | 0.39868 |
| 1 | 57543600 | 57546199 | 29.80341 | 50.73503 | 38.87175 | 17.03818 | 0.764761 | 1.028536 | 0.398683 |
| X | 25345600 | 25350399 | 156.3471 | 236.4391 | 216.5204 | 89.54667 | 0.403538 | 3.307327 | 0.398687 |
| 6 | 100475400 | 100478199 | 16.92489 | 28.96201 | 53.22192 | 23.3379 | -0.49133 | 0.946971 | 0.398699 |
| 8 | 75775800 | 75778999 | 16.92489 | 28.96201 | 53.22192 | 23.3379 | -0.49133 | 0.946971 | 0.398699 |
| 15 | 3195200 | 3195999 | 17.90914 | 31.6988 | 28.40689 | 12.56211 | 0.548828 | 0.522784 | 0.3987 |
| 17 | 29056000 | 29063199 | 84.64779 | 130.54 | 202.7454 | 84.21685 | -0.33548 | 2.895966 | 0.398703 |
| 6 | 51730000 | 51732799 | 22.17948 | 38.4593 | 37.35585 | 16.39003 | 0.428412 | 0.830509 | 0.398715 |
| 9 | 103614800 | 103616399 | 17.04398 | 29.39423 | 51.1258 | 22.42794 | -0.41019 | 0.874658 | 0.398724 |
| 17 | 48465400 | 48469599 | 56.22169 | 89.99193 | 89.65516 | 37.76923 | 0.349834 | 2.030208 | 0.398733 |
| 5 | 17737000 | 17738799 | 10.85952 | 19.77184 | 39.28618 | 16.90148 | -0.56668 | 0.316993 | 0.398755 |
| 17 | 78872400 | 78873999 | 12.35605 | 21.84682 | 39.80652 | 17.60698 | -0.47026 | 0.519172 | 0.398764 |
| 6 | 79148000 | 79149999 | 17.62517 | 32.15441 | 25.32618 | 10.8684 | 0.762237 | 0.325288 | 0.398764 |
| 3 | 57790200 | 57793399 | 33.69178 | 56.27741 | 50.85029 | 22.12966 | 0.526988 | 1.225723 | 0.398773 |
| 5 | 30682800 | 30687199 | 28.8277 | 46.69278 | 88.18077 | 37.73777 | -0.55099 | 1.64551 | 0.398773 |
| X | 94660600 | 94662399 | 24.09073 | 41.60803 | 39.84335 | 17.47932 | 0.447631 | 0.876326 | 0.398775 |
| 14 | 3726800 | 3728399 | 24.5886 | 42.91538 | 35.30785 | 15.54634 | 0.667777 | 0.753908 | 0.398789 |
| 7 | 64176800 | 64178199 | 9.759798 | 18.03157 | 35.27766 | 14.88489 | -0.5242 | 0.219451 | 0.398822 |
| 1 | 39558200 | 39559199 | 15.80064 | 28.31768 | 27.2992 | 11.95874 | 0.453653 | 0.420073 | 0.398829 |
| 13 | 49752600 | 49753999 | 13.93774 | 24.47016 | 44.53401 | 19.70177 | -0.4722 | 0.619183 | 0.398849 |
| 4 | 151973400 | 151981799 | 77.79712 | 120.3545 | 195.3715 | 81.15253 | -0.39566 | 2.833479 | 0.398854 |
| 17 | 41656800 | 41657999 | 22.25098 | 38.45275 | 36.89009 | 16.18401 | 0.44517 | 0.859233 | 0.398865 |
| 2 | 170453800 | 170456199 | 29.5509 | 48.19934 | 81.09785 | 34.84084 | -0.37647 | 1.523541 | 0.398872 |
| 2 | 162530800 | 162532399 | 16.05387 | 28.91364 | 27.61654 | 12.0434 | 0.471122 | 0.391486 | 0.398911 |
| 14 | 46301400 | 46302999 | 16.30397 | 29.93549 | 25.64481 | 10.91236 | 0.650444 | 0.281963 | 0.398924 |
| 9 | 3609000 | 3615799 | 71.93754 | 114.6619 | 99.33157 | 41.59714 | 0.544591 | 2.180892 | 0.398942 |
| 14 | 37850200 | 37851799 | 16.10469 | 27.72907 | 50.57098 | 22.18345 | -0.47871 | 0.884671 | 0.398945 |
| 14 | 3779000 | 3782599 | 55.04647 | 88.20234 | 87.01433 | 36.714 | 0.365727 | 1.997873 | 0.398961 |
| 18 | 74565800 | 74567399 | 15.93598 | 28.70482 | 27.67418 | 12.06669 | 0.457952 | 0.390616 | 0.398988 |
| 2 | 21141800 | 21147399 | 68.23991 | 107.1006 | 166.9737 | 69.25033 | -0.32128 | 2.572885 | 0.398998 |
| 7 | 7837600 | 7840399 | 23.68637 | 41.28687 | 31.48297 | 13.85087 | 0.776754 | 0.762061 | 0.399 |
| 3 | 46173400 | 46176399 | 32.03329 | 53.6454 | 39.73609 | 17.32103 | 0.812597 | 1.182815 | 0.399034 |
| 7 | 48051600 | 48053399 | 50.36219 | 80.91102 | 77.81886 | 32.96648 | 0.40681 | 1.905927 | 0.399038 |
| 3 | 8943200 | 8949799 | 87.72968 | 135.3947 | 212.1087 | 88.10588 | -0.34711 | 2.884859 | 0.399059 |
| 3 | 33759000 | 33760399 | 22.10881 | 38.47793 | 36.0093 | 15.82693 | 0.482211 | 0.77863 | 0.399085 |
| X | 38454400 | 38456599 | 19.71022 | 36.93344 | 19.52893 | 8.061762 | 1.373967 | 0.181373 | 0.399089 |
| 6 | 10937600 | 10939999 | 22.88408 | 39.77294 | 37.64117 | 16.53 | 0.466029 | 0.80493 | 0.399118 |
| X | 120455200 | 120457199 | 21.96336 | 38.84794 | 29.68944 | 13.13108 | 0.776691 | 0.555851 | 0.399119 |
| 7 | 33352200 | 33365599 | 276.1229 | 399.238 | 361.0032 | 148.9028 | 0.361679 | 4.086681 | 0.399121 |
| 3 | 4568200 | 4573199 | 83.41203 | 131.1743 | 128.98 | 53.5068 | 0.345085 | 2.534848 | 0.399123 |
| 10 | 113974000 | 113977999 | 48.61083 | 78.3723 | 76.09109 | 32.39706 | 0.399375 | 1.792461 | 0.399124 |
| 18 | 66938000 | 66939599 | 20.35432 | 35.83113 | 32.02329 | 14.17677 | 0.549475 | 0.599848 | 0.399129 |
| 3 | 41517600 | 41520199 | 20.46289 | 34.20251 | 62.81654 | 27.34844 | -0.49293 | 1.188983 | 0.399136 |
| 9 | 124160200 | 124164599 | 82.11136 | 130.0685 | 114.6355 | 47.67516 | 0.510928 | 2.375012 | 0.39914 |
| 3 | 116505400 | 116506199 | 7.591839 | 14.08002 | 31.0273 | 13.02345 | -0.68869 | 0.169171 | 0.399145 |
| 5 | 27970000 | 27971399 | 7.591839 | 14.08002 | 31.0273 | 13.02345 | -0.68869 | 0.169171 | 0.399145 |
| 17 | 88463400 | 88465399 | 29.72956 | 49.74827 | 49.1666 | 21.42117 | 0.397953 | 1.190986 | 0.399149 |

| | | | | | | | | | |
|---|---|---|---|---|---|---|---|---|---|
| 18 | 9342800 | 9344799 | 22.67112 | 39.76401 | 33.00551 | 14.58944 | 0.654927 | 0.664486 | 0.399167 |
| 4 | 26064000 | 26067399 | 43.3985 | 71.01877 | 60.60697 | 26.06889 | 0.603658 | 1.49448 | 0.399168 |
| 16 | 24018000 | 24020599 | 14.43739 | 25.18999 | 45.74298 | 20.13693 | -0.46923 | 0.7221 | 0.399171 |
| 19 | 45306800 | 45308999 | 9.41763 | 17.35564 | 32.50046 | 13.76016 | -0.46491 | 0.224236 | 0.399172 |
| 2 | 40375200 | 40378599 | 37.44387 | 60.26433 | 99.63957 | 42.31994 | -0.37002 | 1.8409 | 0.399183 |
| 10 | 41259400 | 41260399 | 10.05327 | 18.37317 | 33.94222 | 14.53362 | -0.4574 | 0.279024 | 0.399185 |
| 11 | 6708400 | 6710199 | 13.5082 | 23.6854 | 42.51523 | 18.78813 | -0.45213 | 0.636476 | 0.39919 |
| 14 | 122430400 | 122438399 | 78.15503 | 121.2394 | 190.551 | 79.13877 | -0.34606 | 2.783915 | 0.399195 |
| 2 | 140237000 | 140238399 | 11.64132 | 20.43371 | 42.52113 | 18.80762 | -0.66423 | 0.607207 | 0.399228 |
| 19 | 40519400 | 40521999 | 24.22716 | 42.00194 | 38.69854 | 16.97905 | 0.504099 | 0.84198 | 0.399237 |
| 7 | 3508200 | 3509799 | 19.52646 | 34.16692 | 33.05551 | 14.58215 | 0.435114 | 0.68553 | 0.399248 |
| 16 | 64581200 | 64585799 | 53.08407 | 85.54871 | 77.55595 | 33.00304 | 0.497415 | 1.814403 | 0.399267 |
| 5 | 3070400 | 3078799 | 254.7123 | 368.9679 | 332.7019 | 137.1955 | 0.368343 | 4.056276 | 0.399277 |
| 8 | 14024200 | 14025999 | 15.1957 | 26.18186 | 50.02325 | 21.94364 | -0.54513 | 0.875731 | 0.399278 |
| 8 | 71116000 | 71118599 | 15.1957 | 26.18186 | 50.02325 | 21.94364 | -0.54513 | 0.875731 | 0.399278 |
| 2 | 149968000 | 149970999 | 90.84016 | 141.869 | 140.9311 | 58.47577 | 0.322636 | 2.668529 | 0.399287 |
| 16 | 42026600 | 42028399 | 17.38198 | 30.8024 | 29.61456 | 13.09304 | 0.448215 | 0.505927 | 0.399292 |
| 13 | 31416200 | 31419799 | 35.90727 | 59.58334 | 52.68733 | 22.82964 | 0.557751 | 1.301787 | 0.399322 |
| 3 | 111598800 | 111600799 | 18.42095 | 30.87259 | 65.9257 | 28.7509 | -0.70952 | 1.158711 | 0.399326 |
| 9 | 109960600 | 109961799 | 7.377745 | 13.87817 | 29.11436 | 11.94026 | -0.5915 | 0.072726 | 0.39935 |
| 5 | 138339600 | 138341399 | 10.86548 | 19.14649 | 39.9438 | 17.67964 | -0.66612 | 0.548775 | 0.399353 |
| 12 | 4967000 | 4968199 | 10.86548 | 19.14649 | 39.9438 | 17.67964 | -0.66612 | 0.548775 | 0.399353 |
| 1 | 3419000 | 3421999 | 27.49808 | 47.36191 | 39.66035 | 17.39757 | 0.639518 | 0.914489 | 0.399354 |
| 1 | 60219400 | 60220199 | 7.397933 | 13.88303 | 29.07933 | 11.97409 | -0.59361 | 0.089931 | 0.39936 |
| 6 | 95017600 | 95018799 | 7.397933 | 13.88303 | 29.07933 | 11.97409 | -0.59361 | 0.089931 | 0.39936 |
| 6 | 72652200 | 72656799 | 26.94025 | 43.97014 | 80.38083 | 34.54109 | -0.49515 | 1.505829 | 0.399364 |
| 7 | 41523600 | 41526799 | 29.18786 | 50.83219 | 32.81591 | 14.42941 | 1.015658 | 0.802051 | 0.39937 |
| X | 91280200 | 91287599 | 94.62957 | 148.2425 | 136.6266 | 56.67182 | 0.434099 | 2.617065 | 0.399375 |
| 10 | 98937800 | 98940399 | 31.01359 | 50.39763 | 85.14114 | 36.52477 | -0.38625 | 1.582329 | 0.399387 |
| 2 | 150084200 | 150090399 | 117.4865 | 181.4128 | 162.2495 | 67.39513 | 0.461359 | 2.880343 | 0.39939 |
| 5 | 28377400 | 28381799 | 31.27257 | 50.48774 | 94.29983 | 40.22049 | -0.54014 | 1.734478 | 0.399393 |
| 18 | 80468400 | 80472399 | 24.78567 | 40.75897 | 72.5538 | 31.2717 | -0.45108 | 1.398028 | 0.399398 |
| 3 | 153661800 | 153664999 | 36.81671 | 60.58787 | 58.69669 | 25.30693 | 0.424435 | 1.407451 | 0.399398 |
| 1 | 11749200 | 11751599 | 31.47312 | 53.44493 | 44.08403 | 19.31698 | 0.658259 | 1.061355 | 0.399407 |
| 13 | 51137200 | 51140599 | 26.51853 | 45.1841 | 43.62667 | 19.12374 | 0.432453 | 1.010675 | 0.399409 |
| 7 | 105997400 | 106000999 | 44.49057 | 72.94217 | 57.48542 | 24.74537 | 0.719277 | 1.469213 | 0.399443 |
| X | 24558600 | 24566199 | 110.6449 | 172.4479 | 138.7666 | 57.5915 | 0.623972 | 2.715661 | 0.399452 |
| 12 | 105250200 | 105267599 | 174.695 | 255.6897 | 395.6 | 162.9634 | -0.39574 | 3.894444 | 0.399463 |
| 12 | 5639400 | 5640599 | 17.57116 | 30.12434 | 52.70221 | 23.11349 | -0.42023 | 0.933094 | 0.399486 |
| 14 | 38132600 | 38134799 | 20.93729 | 36.91876 | 30.80404 | 13.63247 | 0.648903 | 0.582666 | 0.399489 |
| 16 | 16607000 | 16608199 | 7.420976 | 13.82842 | 31.09246 | 12.95685 | -0.70965 | 0.137997 | 0.399497 |
| 14 | 121315800 | 121320399 | 35.12009 | 56.69035 | 95.71167 | 40.81432 | -0.39507 | 1.746258 | 0.39951 |
| 1 | 72047400 | 72049399 | 12.85045 | 22.62428 | 40.87503 | 18.09501 | -0.46052 | 0.571495 | 0.399529 |
| 5 | 108979400 | 108982799 | 36.30931 | 60.10314 | 55.41243 | 23.9699 | 0.497564 | 1.335438 | 0.399531 |
| 14 | 22184000 | 22185799 | 20.44862 | 36.09572 | 27.87741 | 12.33463 | 0.7604 | 0.563789 | 0.399534 |
| 1 | 171172000 | 171173999 | 33.87422 | 56.60065 | 45.95574 | 20.00371 | 0.680684 | 1.217438 | 0.39954 |
| X | 23689800 | 23692399 | 44.97345 | 73.00758 | 65.76455 | 28.19102 | 0.517438 | 1.622222 | 0.399544 |

| | | | | | | | | | |
|---|---|---|---|---|---|---|---|---|---|
| 3 | 149868600 | 149870399 | 14.85876 | 27.22823 | 25.70377 | 10.96682 | 0.508326 | 0.280607 | 0.399547 |
| 5 | 14486400 | 14487199 | 8.466094 | 15.91382 | 31.29899 | 12.85208 | -0.50063 | 0.094424 | 0.399558 |
| 9 | 27396400 | 27397799 | 20.00286 | 36.34875 | 25.29322 | 10.90783 | 0.935852 | 0.367088 | 0.399579 |
| 4 | 7445800 | 7447399 | 15.10774 | 27.23283 | 24.91285 | 10.85276 | 0.534182 | 0.375182 | 0.399592 |
| 6 | 97219200 | 97220199 | 6.696863 | 12.93905 | 27.24569 | 10.46338 | -0.51067 | -0.15769 | 0.399601 |
| 7 | 135424200 | 135425999 | 15.71809 | 28.5019 | 26.79086 | 11.58446 | 0.5022 | 0.350743 | 0.399608 |
| 1 | 11604400 | 11607799 | 39.26284 | 63.02896 | 104.8088 | 44.34729 | -0.38287 | 1.926532 | 0.399636 |
| X | 43603800 | 43605399 | 26.70936 | 45.93164 | 40.88775 | 17.93514 | 0.551356 | 0.928462 | 0.399659 |
| 1 | 84075200 | 84077199 | 20.69979 | 36.25169 | 34.36616 | 15.17155 | 0.464258 | 0.67988 | 0.399661 |
| 10 | 121585000 | 121589799 | 45.49371 | 72.23392 | 142.4855 | 59.38735 | -0.64634 | 2.286653 | 0.399662 |
| 17 | 4427200 | 4427799 | 4.655161 | 9.053523 | 22.28887 | 8.24691 | -0.68052 | -0.32125 | 0.399676 |
| 9 | 45077600 | 45078999 | 11.51242 | 20.59034 | 38.31428 | 16.81292 | -0.4915 | 0.425121 | 0.399676 |
| 5 | 57386400 | 57387399 | 7.174262 | 13.68459 | 27.80436 | 11.08434 | -0.5101 | -0.02836 | 0.399728 |
| X | 139409000 | 139412399 | 14.72652 | 25.7545 | 51.0535 | 22.51177 | -0.59483 | 0.700009 | 0.399733 |
| X | 117902600 | 117903999 | 16.53546 | 30.57441 | 24.65196 | 10.39016 | 0.74719 | 0.243791 | 0.399748 |
| 10 | 59554200 | 59554999 | 8.336106 | 15.4385 | 31.00142 | 13.03766 | -0.55764 | 0.18414 | 0.399757 |
| 3 | 131353200 | 131354399 | 12.81752 | 24.15329 | 21.16756 | 8.65534 | 0.660635 | 0.086921 | 0.399773 |
| 16 | 76200400 | 76202399 | 11.6881 | 20.93699 | 37.81523 | 16.57225 | -0.44732 | 0.416842 | 0.399798 |
| 10 | 12769600 | 12770399 | 5.134789 | 9.947842 | 24.94796 | 9.460569 | -0.74225 | -0.22521 | 0.399806 |
| 10 | 61985400 | 61990599 | 35.80133 | 57.82004 | 95.84158 | 40.89714 | -0.36774 | 1.73368 | 0.399814 |
| 5 | 93632200 | 93634599 | 22.50246 | 39.39744 | 30.55155 | 13.48426 | 0.752518 | 0.686975 | 0.399819 |
| 19 | 60590400 | 60592199 | 15.23225 | 26.55965 | 46.80582 | 20.59644 | -0.42634 | 0.73701 | 0.39983 |
| 10 | 91875800 | 91876799 | 9.766072 | 17.82231 | 33.41445 | 14.33372 | -0.48008 | 0.285016 | 0.399839 |
| X | 54760400 | 54762199 | 19.55266 | 35.3913 | 26.29729 | 11.39672 | 0.837154 | 0.386976 | 0.399844 |
| 7 | 32973400 | 32980999 | 161.3742 | 244.5708 | 207.1446 | 85.72771 | 0.518758 | 3.264341 | 0.399859 |
| 4 | 105097600 | 105098599 | 8.995858 | 16.47974 | 32.44576 | 13.84898 | -0.54494 | 0.255571 | 0.399859 |
| 19 | 53744800 | 53746999 | 20.04292 | 33.87506 | 58.60754 | 25.64428 | -0.40674 | 1.076429 | 0.399867 |
| 17 | 85523600 | 85525599 | 17.055 | 30.15978 | 27.87161 | 12.32761 | 0.50397 | 0.523641 | 0.399871 |
| X | 148686000 | 148688799 | 46.85296 | 76.42369 | 61.90748 | 26.59683 | 0.675401 | 1.547625 | 0.399876 |
| 17 | 19517600 | 19522399 | 41.65221 | 69.77706 | 43.00271 | 18.74886 | 1.077026 | 1.214793 | 0.399877 |
| 8 | 100749200 | 100751799 | 34.4102 | 57.02966 | 54.80575 | 23.72761 | 0.437989 | 1.314788 | 0.399891 |
| 12 | 92178200 | 92180399 | 19.74674 | 33.25069 | 59.6426 | 26.06778 | -0.45888 | 1.108352 | 0.399891 |
| 1 | 4724000 | 4724799 | 5.81952 | 11.28211 | 24.45213 | 9.242963 | -0.52935 | -0.22847 | 0.399895 |
| 1 | 22596200 | 22596799 | 5.81952 | 11.28211 | 24.45213 | 9.242963 | -0.52935 | -0.22847 | 0.399895 |
| 6 | 24383000 | 24385599 | 23.71957 | 39.42752 | 67.37424 | 29.22838 | -0.38943 | 1.263784 | 0.399913 |
| 11 | 113489800 | 113500599 | 114.9686 | 173.9571 | 267.3588 | 110.5907 | -0.34226 | 3.289059 | 0.399917 |
| 12 | 14643200 | 14646399 | 38.96129 | 63.37513 | 61.09003 | 26.2191 | 0.422304 | 1.568192 | 0.39993 |
| 10 | 51625000 | 51626599 | 21.55433 | 38.19233 | 27.90937 | 12.3394 | 0.841609 | 0.529477 | 0.399935 |
| 10 | 15446600 | 15449599 | 43.5369 | 69.66944 | 113.2748 | 47.70164 | -0.35591 | 2.033225 | 0.399942 |
| X | 113900800 | 113902799 | 48.05252 | 77.80037 | 69.98113 | 29.94021 | 0.515983 | 1.681191 | 0.39995 |
| 12 | 76930400 | 76931999 | 14.27129 | 25.96663 | 24.87911 | 10.71211 | 0.478569 | 0.315248 | 0.399952 |
| 18 | 13617200 | 13618999 | 11.17276 | 20.18395 | 37.42336 | 16.25804 | -0.47655 | 0.360537 | 0.399978 |
| 11 | 80032400 | 80033199 | 5.832652 | 11.26843 | 24.8145 | 9.533147 | -0.57425 | -0.17124 | 0.399985 |
| X | 77434000 | 77434999 | 20.02096 | 35.08783 | 31.57979 | 13.94985 | 0.539005 | 0.662694 | 0.40001 |
| 7 | 18063800 | 18066199 | 71.1237 | 112.838 | 109.8963 | 45.76752 | 0.369108 | 2.323598 | 0.400012 |
| 15 | 29345600 | 29348399 | 34.61869 | 57.8879 | 47.55417 | 20.70926 | 0.663709 | 1.210088 | 0.400019 |
| 1 | 3966600 | 3968999 | 20.62571 | 36.70001 | 28.05525 | 12.36474 | 0.780274 | 0.496411 | 0.400025 |

| | | | | | | | | | |
|---|---|---|---|---|---|---|---|---|---|
| 8 | 34639400 | 34643199 | 57.71932 | 92.36152 | 91.66977 | 38.60158 | 0.354794 | 2.041779 | 0.400031 |
| 1 | 161558600 | 161559599 | 7.164268 | 13.69559 | 27.87773 | 11.06017 | -0.50662 | -0.04437 | 0.400036 |
| X | 77757400 | 77766399 | 106.2984 | 166.2871 | 133.7838 | 55.50069 | 0.628352 | 2.647929 | 0.400054 |
| 8 | 110430600 | 110433199 | 13.20216 | 23.07053 | 44.17011 | 19.46368 | -0.54476 | 0.691905 | 0.400058 |
| 16 | 14842000 | 14844999 | 30.76715 | 51.78373 | 48.01364 | 20.98224 | 0.489718 | 1.137094 | 0.400067 |
| 3 | 93934400 | 93937799 | 54.74135 | 88.4924 | 69.02742 | 29.4911 | 0.71886 | 1.725666 | 0.400069 |
| 5 | 75575800 | 75579399 | 44.00348 | 71.31221 | 69.7048 | 29.84282 | 0.397584 | 1.653672 | 0.400081 |
| 1 | 25008000 | 25009199 | 15.02706 | 27.71821 | 25.3177 | 10.70593 | 0.565082 | 0.246608 | 0.400087 |
| 8 | 52715200 | 52715999 | 12.54046 | 23.61869 | 23.03479 | 9.427287 | 0.507435 | 0.095864 | 0.400108 |
| 4 | 150801800 | 150805199 | 22.70631 | 37.42818 | 70.63405 | 30.47482 | -0.53362 | 1.363522 | 0.400125 |
| 10 | 15810800 | 15814599 | 88.74104 | 138.7263 | 137.705 | 57.13146 | 0.324809 | 2.648816 | 0.400126 |
| 5 | 83420200 | 83421999 | 24.12811 | 42.29218 | 32.34167 | 14.28821 | 0.772583 | 0.6812 | 0.400145 |
| 8 | 101324800 | 101326799 | 47.59561 | 76.84965 | 63.42023 | 27.05404 | 0.63576 | 1.742834 | 0.400158 |
| 15 | 19309600 | 19311599 | 21.54691 | 37.39845 | 36.28208 | 15.91966 | 0.430289 | 0.815695 | 0.400166 |
| 17 | 39818600 | 39820199 | 34.35845 | 56.87462 | 55.28726 | 23.91597 | 0.421417 | 1.330441 | 0.400175 |
| 1 | 181695200 | 181699799 | 50.09968 | 79.77966 | 127.5554 | 53.34734 | -0.33952 | 2.196472 | 0.400181 |
| 12 | 55916600 | 55919799 | 42.95293 | 69.88299 | 66.06754 | 28.3576 | 0.450467 | 1.576494 | 0.400202 |
| 11 | 93580000 | 93583999 | 50.08114 | 80.72814 | 77.75625 | 33.09829 | 0.410583 | 1.802747 | 0.400241 |
| 15 | 27496800 | 27498799 | 10.43486 | 18.70424 | 36.81787 | 16.1263 | -0.57009 | 0.401211 | 0.400247 |
| 15 | 74216600 | 74233199 | 174.045 | 254.737 | 393.9976 | 162.3034 | -0.39528 | 3.894591 | 0.400262 |
| 1 | 76742200 | 76743199 | 9.768344 | 17.70915 | 34.88108 | 15.08692 | -0.55884 | 0.324583 | 0.400263 |
| 10 | 41394600 | 41396399 | 9.768344 | 17.70915 | 34.88108 | 15.08692 | -0.55884 | 0.324583 | 0.400263 |
| 13 | 51925200 | 51927399 | 17.66241 | 29.92665 | 62.91953 | 27.55197 | -0.68677 | 1.04989 | 0.400265 |
| 18 | 30093400 | 30095199 | 11.83871 | 21.24994 | 41.03906 | 17.95427 | -0.54191 | 0.410321 | 0.400268 |
| 11 | 23491000 | 23496399 | 46.36396 | 75.15658 | 69.07538 | 29.57885 | 0.486513 | 1.65112 | 0.400273 |
| 1 | 86706800 | 86708199 | 16.44134 | 31.00935 | 21.6215 | 8.827644 | 0.990725 | 0.118462 | 0.400273 |
| 9 | 56454200 | 56455599 | 9.147813 | 16.6791 | 33.41436 | 14.34974 | -0.57619 | 0.284339 | 0.400283 |
| 18 | 34491200 | 34493399 | 14.46467 | 25.28507 | 44.66294 | 19.68632 | -0.42916 | 0.694601 | 0.40032 |
| 2 | 178861000 | 178862799 | 11.53459 | 20.63541 | 37.29125 | 16.36038 | -0.44915 | 0.422952 | 0.40032 |
| 3 | 97244600 | 97246399 | 8.961828 | 16.06418 | 37.91025 | 16.60449 | -0.83 | 0.392797 | 0.400333 |
| 11 | 61573600 | 61576999 | 22.5674 | 37.11968 | 72.24945 | 31.14355 | -0.57934 | 1.389581 | 0.400357 |
| 6 | 28434200 | 28436399 | 14.96699 | 25.98856 | 47.39605 | 20.80158 | -0.47604 | 0.797468 | 0.400378 |
| 13 | 62502400 | 62506799 | 93.78161 | 147.4479 | 128.6965 | 53.38758 | 0.516525 | 2.546891 | 0.400388 |
| 12 | 33029600 | 33030999 | 14.53164 | 27.03181 | 24.25691 | 10.13798 | 0.602343 | 0.191886 | 0.400389 |
| X | 33691000 | 33700799 | 190.6783 | 283.8082 | 261.2085 | 107.627 | 0.375912 | 3.61709 | 0.400448 |
| 4 | 96043200 | 96045999 | 37.30675 | 60.90974 | 57.27649 | 24.61569 | 0.461708 | 1.509603 | 0.400453 |
| 2 | 161449000 | 161452199 | 26.22323 | 44.80078 | 43.07914 | 18.88781 | 0.43894 | 0.985408 | 0.400469 |
| 7 | 57692800 | 57694799 | 25.84628 | 45.26722 | 31.71827 | 14.00323 | 0.898191 | 0.702531 | 0.400472 |
| 12 | 24028600 | 24031599 | 37.64379 | 62.23415 | 50.48233 | 21.8184 | 0.681522 | 1.348149 | 0.400477 |
| 10 | 44943000 | 44943799 | 5.395128 | 10.46371 | 23.86407 | 9.003172 | -0.59911 | -0.24144 | 0.400478 |
| 13 | 96142200 | 96143999 | 11.33576 | 20.39025 | 37.85833 | 16.53322 | -0.48351 | 0.387378 | 0.400503 |
| 13 | 99057800 | 99059399 | 12.82818 | 22.64703 | 40.83662 | 18.06811 | -0.45624 | 0.540509 | 0.400512 |
| 6 | 100710400 | 100725399 | 209.6517 | 304.044 | 446.4232 | 184.0625 | -0.33311 | 4.0382 | 0.400513 |
| 1 | 101351800 | 101353399 | 14.49264 | 27.4776 | 21.19128 | 8.556621 | 0.859686 | 0.063436 | 0.400532 |
| X | 134872800 | 134875799 | 26.82672 | 46.49294 | 38.05279 | 16.69555 | 0.674179 | 0.857025 | 0.400534 |
| 17 | 93305400 | 93305799 | 8.97828 | 17.43963 | 18.24676 | 6.819311 | 0.526031 | -0.25148 | 0.400544 |
| X | 102234800 | 102244399 | 132.9256 | 203.8169 | 176.1317 | 73.11344 | 0.503192 | 3.022104 | 0.400548 |

| | | | | | | | | | |
|---|---|---|---|---|---|---|---|---|---|
| 7 | 30197200 | 30208199 | 86.19284 | 131.8544 | 239.9439 | 99.55498 | -0.57282 | 3.057945 | 0.40055 |
| 1 | 117652800 | 117654399 | 35.33541 | 58.1208 | 57.99868 | 25.00065 | 0.381562 | 1.409542 | 0.400565 |
| 6 | 79790000 | 79792199 | 24.63909 | 43.03635 | 34.65656 | 15.26689 | 0.698544 | 0.741654 | 0.400569 |
| 8 | 97844600 | 97847199 | 27.57705 | 46.97569 | 44.14468 | 19.35042 | 0.47136 | 1.018388 | 0.400579 |
| X | 118357800 | 118358799 | 15.82693 | 28.83819 | 24.89553 | 10.70009 | 0.630038 | 0.321542 | 0.400583 |
| 9 | 90156600 | 90158999 | 14.61604 | 25.51533 | 45.21142 | 19.90962 | -0.43386 | 0.715686 | 0.400591 |
| 15 | 47238600 | 47250799 | 148.8043 | 225.0181 | 215.9427 | 89.30598 | 0.335929 | 3.305032 | 0.400593 |
| 13 | 69478000 | 69480599 | 18.16985 | 31.1292 | 54.35407 | 23.83662 | -0.41773 | 0.94211 | 0.400602 |
| 14 | 90417400 | 90418799 | 9.394337 | 17.49678 | 34.41333 | 14.353 | -0.51941 | 0.166658 | 0.400615 |
| 6 | 19861000 | 19863199 | 21.12908 | 37.21238 | 31.35874 | 13.88128 | 0.634056 | 0.59777 | 0.400621 |
| 12 | 87595000 | 87599399 | 48.81597 | 77.67862 | 127.5996 | 53.31186 | -0.37957 | 2.219091 | 0.400625 |
| 3 | 127419200 | 127422799 | 16.66582 | 28.50886 | 58.04376 | 25.45158 | -0.63867 | 0.956252 | 0.400631 |
| 12 | 19380800 | 19386599 | 75.13021 | 118.5973 | 114.9909 | 47.73899 | 0.369471 | 2.45054 | 0.40064 |
| 2 | 149068200 | 149075999 | 271.1984 | 391.4234 | 364.6582 | 150.4456 | 0.316127 | 4.111734 | 0.400645 |
| 17 | 83933600 | 83934399 | 10.20547 | 18.68165 | 34.50297 | 14.74209 | -0.45492 | 0.272046 | 0.400664 |
| 8 | 16464800 | 16469399 | 42.50445 | 69.41665 | 62.36563 | 26.80577 | 0.527592 | 1.518467 | 0.400668 |
| 11 | 103446000 | 103448999 | 24.52303 | 40.78708 | 70.94583 | 30.79034 | -0.41498 | 1.261631 | 0.400673 |
| 19 | 13224800 | 13228999 | 72.31819 | 115.1087 | 103.9009 | 43.42712 | 0.483277 | 2.225113 | 0.400688 |
| 6 | 136007200 | 136008199 | 16.0075 | 29.40463 | 25.70449 | 10.93093 | 0.622167 | 0.27763 | 0.400716 |
| Y | 29097200 | 29099999 | 14.97607 | 27.41505 | 15.45352 | 6.602643 | 1.244873 | 0.26001 | 0.400722 |
| 10 | 42606000 | 42608199 | 17.87501 | 30.82575 | 55.50058 | 24.34704 | -0.45998 | 0.884102 | 0.400735 |
| 11 | 3132600 | 3150199 | 23776.02 | 23606.79 | 14822.03 | 6459.859 | 0.339482 | 10.04072 | 0.400749 |
| 10 | 90811600 | 90812599 | 6.593111 | 12.61398 | 26.73719 | 10.58297 | -0.56038 | -0.05752 | 0.40075 |
| 2 | 124476400 | 124479799 | 18.88345 | 32.06306 | 56.43233 | 24.72708 | -0.43108 | 1.029547 | 0.400759 |
| 7 | 44614600 | 44618199 | 20.20006 | 33.64281 | 64.29455 | 27.9289 | -0.54999 | 1.229027 | 0.400771 |
| 10 | 81597800 | 81599799 | 16.65563 | 30.80191 | 23.96499 | 10.09826 | 0.798198 | 0.240778 | 0.400778 |
| 3 | 57643000 | 57644199 | 8.956571 | 16.06572 | 35.90004 | 15.71498 | -0.75083 | 0.38612 | 0.400808 |
| 10 | 84172200 | 84175599 | 25.15911 | 41.61709 | 73.20644 | 31.65186 | -0.43182 | 1.326166 | 0.400842 |
| 15 | 18724600 | 18727199 | 22.49172 | 39.66524 | 28.95877 | 12.81548 | 0.840457 | 0.586624 | 0.400853 |
| 3 | 27961400 | 27962799 | 16.44894 | 29.72212 | 27.62536 | 11.99812 | 0.514078 | 0.37651 | 0.400855 |
| 6 | 47332600 | 47333799 | 7.815165 | 14.27645 | 33.00493 | 14.1402 | -0.77804 | 0.266381 | 0.400868 |
| X | 126194400 | 126197999 | 46.94618 | 77.30758 | 52.37899 | 22.58926 | 0.939218 | 1.422453 | 0.400871 |
| 1 | 118906200 | 118910399 | 26.03816 | 42.40609 | 80.37338 | 34.51069 | -0.54926 | 1.536563 | 0.400877 |
| 18 | 90457800 | 90459399 | 19.63697 | 34.84111 | 29.89611 | 13.2078 | 0.612408 | 0.512348 | 0.400906 |
| 5 | 147651600 | 147654599 | 22.81941 | 37.65182 | 72.7116 | 31.38429 | -0.56635 | 1.349201 | 0.400909 |
| 10 | 116001000 | 116002599 | 17.39909 | 31.90998 | 24.28318 | 10.34893 | 0.81822 | 0.291964 | 0.400921 |
| 13 | 104073800 | 104075799 | 16.6272 | 29.59293 | 28.74173 | 12.66455 | 0.437225 | 0.471353 | 0.400939 |
| 3 | 30066200 | 30067399 | 6.582759 | 12.59493 | 27.32253 | 10.81279 | -0.59414 | -0.06014 | 0.400958 |
| 16 | 78841400 | 78841799 | 6.582759 | 12.59493 | 27.32253 | 10.81279 | -0.59414 | -0.06014 | 0.400958 |
| 9 | 21852000 | 21853599 | 9.770723 | 17.75518 | 33.88187 | 14.61169 | -0.5105 | 0.30997 | 0.400992 |
| 6 | 63986400 | 63989199 | 25.78467 | 44.46308 | 40.31781 | 17.68661 | 0.525548 | 0.89871 | 0.401009 |
| 17 | 41706400 | 41709799 | 38.01172 | 62.53448 | 58.30707 | 25.13555 | 0.479284 | 1.412272 | 0.401033 |
| 8 | 28218600 | 28220199 | 11.35036 | 20.45755 | 37.37495 | 16.28393 | -0.45803 | 0.375633 | 0.401059 |
| 10 | 14484800 | 14491399 | 135.9086 | 207.1924 | 177.1233 | 73.4316 | 0.51226 | 3.135476 | 0.401061 |
| X | 74224400 | 74225399 | 14.15804 | 26.5932 | 23.38717 | 9.615447 | 0.650222 | 0.127424 | 0.401063 |
| 3 | 40269800 | 40270199 | 9.464801 | 18.25752 | 18.71249 | 7.233976 | 0.510243 | -0.13011 | 0.401065 |
| 7 | 96206200 | 96209199 | 20.87814 | 35.03964 | 61.87706 | 27.00904 | -0.43662 | 1.144535 | 0.401081 |

| | | | | | | | | | |
|---|---|---|---|---|---|---|---|---|---|
| 14 | 60571200 | 60572799 | 16.48765 | 29.91761 | 27.14865 | 11.72923 | 0.553345 | 0.352469 | 0.401082 |
| 14 | 3056800 | 3092199 | 598.1037 | 811.5545 | 657.3763 | 273.871 | 0.428306 | 5.04909 | 0.401094 |
| 5 | 87415000 | 87416799 | 21.3559 | 37.41005 | 34.27791 | 15.13577 | 0.513089 | 0.679254 | 0.401094 |
| 7 | 25691600 | 25700399 | 132 | 202.0415 | 183.2871 | 76.0574 | 0.431211 | 3.056548 | 0.401095 |
| 18 | 38675000 | 38676599 | 10.47455 | 18.71056 | 36.32943 | 15.95883 | -0.55313 | 0.424138 | 0.401101 |
| 1 | 162582200 | 162582999 | 6.263669 | 12.09622 | 25.38687 | 9.77159 | -0.50848 | -0.15871 | 0.401102 |
| 15 | 101973200 | 101974999 | 11.78952 | 20.79645 | 39.40442 | 17.43727 | -0.52737 | 0.543349 | 0.401118 |
| 19 | 54474400 | 54478199 | 48.17799 | 76.87559 | 124.3985 | 52.17372 | -0.35365 | 2.129999 | 0.40113 |
| 7 | 61282200 | 61287999 | 58.25647 | 93.80906 | 76.91808 | 32.69354 | 0.640601 | 1.847 | 0.401139 |
| 3 | 38943200 | 38947199 | 43.54611 | 69.5778 | 116.0623 | 48.76988 | -0.39523 | 2.080565 | 0.401144 |
| 5 | 74061600 | 74065599 | 36.50593 | 58.72203 | 99.49931 | 42.22626 | -0.40628 | 1.858113 | 0.401153 |
| 17 | 12815000 | 12818399 | 46.06434 | 73.44395 | 122.2707 | 51.22421 | -0.39578 | 2.153184 | 0.401169 |
| 8 | 21641000 | 21648399 | 77.44916 | 123.4125 | 95.93227 | 40.15812 | 0.700236 | 2.193125 | 0.401177 |
| 3 | 94319800 | 94320999 | 8.472588 | 15.30133 | 34.4353 | 14.96564 | -0.75372 | 0.338913 | 0.401195 |
| X | 90577000 | 90579199 | 27.97693 | 48.67519 | 35.21327 | 15.47394 | 0.852185 | 0.813899 | 0.401197 |
| 4 | 16683600 | 16685599 | 21.54679 | 37.9423 | 30.7329 | 13.60465 | 0.690705 | 0.600107 | 0.401217 |
| 6 | 61109600 | 61110399 | 15.57895 | 28.55479 | 22.71234 | 9.687336 | 0.754433 | 0.279567 | 0.401263 |
| 14 | 49828800 | 49829599 | 5.254559 | 10.20888 | 24.02029 | 8.960769 | -0.6288 | -0.28354 | 0.401263 |
| X | 148914800 | 148922399 | 116.1508 | 179.9049 | 154.2619 | 64.0762 | 0.52549 | 2.824514 | 0.401263 |
| 17 | 36073600 | 36078199 | 25.24741 | 41.50833 | 73.85497 | 31.82929 | -0.45065 | 1.401741 | 0.401265 |
| X | 30894200 | 30895599 | 36.91064 | 61.80867 | 42.61086 | 18.57325 | 0.915802 | 1.202032 | 0.40128 |
| 15 | 25754400 | 25756999 | 20.8018 | 34.69638 | 63.88858 | 27.77851 | -0.49663 | 1.213559 | 0.401284 |
| 12 | 91612200 | 91620599 | 144.7338 | 214.9596 | 335.1831 | 138.0768 | -0.3871 | 3.649905 | 0.401298 |
| 2 | 86451000 | 86456999 | 138.2408 | 210.1483 | 202.9541 | 84.07023 | 0.333119 | 3.196025 | 0.401306 |
| 6 | 3875400 | 3875799 | 5.073367 | 9.892327 | 22.09611 | 7.953629 | -0.50632 | -0.41661 | 0.401318 |
| 10 | 36140200 | 36141799 | 31.43096 | 52.13479 | 50.31331 | 21.795 | 0.43203 | 1.293747 | 0.401335 |
| 19 | 14059200 | 14061399 | 32.60283 | 54.43394 | 51.40149 | 22.36318 | 0.463561 | 1.226316 | 0.401337 |
| 6 | 11510000 | 11510399 | 7.556752 | 14.75289 | 12.75981 | 4.506424 | 0.869242 | -0.48846 | 0.401348 |
| 13 | 44081200 | 44084999 | 34.99411 | 56.43811 | 94.51307 | 40.25822 | -0.38475 | 1.771944 | 0.401368 |
| 10 | 11577000 | 11579599 | 53.44163 | 85.65792 | 84.75335 | 35.7772 | 0.362002 | 1.983823 | 0.401371 |
| 3 | 37385800 | 37386799 | 6.27504 | 12.07665 | 26.40199 | 10.27434 | -0.58223 | -0.12102 | 0.401388 |
| 1 | 122246400 | 122247599 | 12.5671 | 22.29782 | 40.85391 | 18.04851 | -0.47588 | 0.489916 | 0.401399 |
| 13 | 33343400 | 33343999 | 5.803598 | 11.29407 | 25.5536 | 9.407436 | -0.5554 | -0.31808 | 0.401409 |
| 5 | 130721600 | 130724399 | 26.43976 | 45.19626 | 43.07829 | 18.8883 | 0.451747 | 0.981842 | 0.401411 |
| 4 | 3471400 | 3474999 | 61.2615 | 98.05993 | 83.11331 | 35.01382 | 0.58261 | 2.035314 | 0.40143 |
| 4 | 119176400 | 119178599 | 11.01173 | 19.43935 | 38.92254 | 17.22139 | -0.60601 | 0.528869 | 0.401448 |
| 2 | 165768000 | 165771999 | 55.18246 | 86.69722 | 159.2942 | 66.07426 | -0.55671 | 2.5438 | 0.401455 |
| 10 | 123703600 | 123704999 | 15.70683 | 28.62391 | 25.48129 | 10.94967 | 0.58536 | 0.3162 | 0.401463 |
| 6 | 89279400 | 89281999 | 17.91404 | 30.54725 | 54.27881 | 23.79462 | -0.4438 | 0.985187 | 0.40147 |
| 10 | 73600000 | 73601599 | 23.12559 | 37.81742 | 81.93166 | 35.23242 | -0.73731 | 1.473787 | 0.40147 |
| 7 | 72688600 | 72690199 | 8.643149 | 15.91741 | 32.49494 | 13.7707 | -0.5897 | 0.220307 | 0.401475 |
| 7 | 55442000 | 55444399 | 11.56187 | 20.49758 | 39.28168 | 17.36551 | -0.54116 | 0.482889 | 0.401485 |
| 7 | 26328000 | 26332399 | 57.26615 | 92.06753 | 81.58554 | 34.59741 | 0.525997 | 1.89636 | 0.401527 |
| 17 | 30700800 | 30703199 | 18.32394 | 31.13503 | 56.88623 | 24.93049 | -0.48456 | 1.024993 | 0.401529 |
| 1 | 16814600 | 16816399 | 13.48627 | 25.07197 | 24.42335 | 10.21805 | 0.483749 | 0.188452 | 0.401533 |
| 11 | 54161800 | 54163999 | 13.9915 | 24.42082 | 44.70427 | 19.68413 | -0.48057 | 0.714365 | 0.401537 |
| 12 | 33543800 | 33545199 | 14.67452 | 27.15788 | 24.72202 | 10.40696 | 0.574243 | 0.222464 | 0.401558 |

| | | | | | | | | | |
|---|---|---|---|---|---|---|---|---|---|
| 13 | 67011400 | 67013999 | 22.20805 | 39.19599 | 30.18163 | 13.35453 | 0.764685 | 0.580286 | 0.40158 |
| X | 125350600 | 125352399 | 39.79621 | 66.2776 | 49.29578 | 21.41314 | 0.806987 | 1.274589 | 0.401586 |
| 1 | 36163200 | 36166199 | 34.19518 | 55.12498 | 94.73149 | 40.32802 | -0.42269 | 1.783894 | 0.401592 |
| 19 | 7016800 | 7020999 | 20.42821 | 34.23218 | 61.79213 | 26.94598 | -0.4681 | 1.159906 | 0.401595 |
| 4 | 57341400 | 57345599 | 35.65438 | 57.55911 | 94.94122 | 40.49151 | -0.36133 | 1.745488 | 0.401609 |
| 10 | 52217400 | 52220399 | 20.55198 | 34.57914 | 60.22034 | 26.31396 | -0.41649 | 1.118819 | 0.40161 |
| 2 | 5587200 | 5587599 | 4.108691 | 8.010443 | 20.80082 | 7.498496 | -0.72183 | -0.42828 | 0.401626 |
| 12 | 52052800 | 52053399 | 4.108691 | 8.010443 | 20.80082 | 7.498496 | -0.72183 | -0.42828 | 0.401626 |
| 9 | 75517800 | 75518999 | 7.269037 | 13.5828 | 29.55429 | 12.25833 | -0.65653 | 0.117653 | 0.401645 |
| X | 16092600 | 16096799 | 42.76223 | 71.20981 | 46.50355 | 20.19891 | 0.994011 | 1.281677 | 0.401648 |
| 1 | 45762400 | 45765799 | 38.9246 | 62.63274 | 102.2618 | 43.41885 | -0.35237 | 1.851875 | 0.401649 |
| X | 146421200 | 146431199 | 234.3697 | 344.991 | 277.6301 | 114.3296 | 0.554625 | 3.808896 | 0.40165 |
| 5 | 150832400 | 150834399 | 18.94291 | 33.4812 | 31.09258 | 13.75366 | 0.496683 | 0.547324 | 0.401693 |
| Y | 2605400 | 2607999 | 18.03115 | 31.85963 | 21.56926 | 9.541663 | 0.949631 | 0.523912 | 0.401704 |
| 11 | 67974800 | 67976199 | 9.143322 | 16.85465 | 32.51726 | 13.76224 | -0.50693 | 0.222289 | 0.401724 |
| 14 | 79449800 | 79452999 | 24.34897 | 39.93729 | 73.9304 | 31.83014 | -0.50887 | 1.430827 | 0.40175 |
| 1 | 121454600 | 121457199 | 27.21578 | 44.16579 | 83.26523 | 35.6841 | -0.5458 | 1.603795 | 0.401752 |
| 2 | 83731000 | 83731999 | 7.174199 | 13.64682 | 29.43582 | 11.80331 | -0.60353 | -0.00687 | 0.401752 |
| 4 | 134230800 | 134232199 | 10.71948 | 19.43363 | 35.39173 | 15.30765 | -0.44639 | 0.334199 | 0.401755 |
| 3 | 17437600 | 17438199 | 12.21422 | 22.87892 | 21.57759 | 8.912978 | 0.545052 | 0.124065 | 0.401763 |
| 5 | 95448200 | 95450799 | 31.60018 | 54.30758 | 37.8679 | 16.61018 | 0.90201 | 0.949804 | 0.401767 |
| 3 | 60408200 | 60409199 | 10.10987 | 18.24367 | 34.8372 | 15.15543 | -0.51971 | 0.355197 | 0.401789 |
| 7 | 49697000 | 49698599 | 10.10987 | 18.24367 | 34.8372 | 15.15543 | -0.51971 | 0.355197 | 0.401789 |
| 14 | 33529200 | 33531599 | 17.82401 | 30.1662 | 56.87744 | 24.89762 | -0.53012 | 1.055086 | 0.40179 |
| 6 | 82062600 | 82064599 | 26.05736 | 44.84219 | 40.84452 | 17.91656 | 0.518487 | 0.917704 | 0.401799 |
| 1 | 191529200 | 191531599 | 26.2554 | 43.09661 | 74.29188 | 31.99595 | -0.40601 | 1.426065 | 0.401804 |
| 17 | 19401400 | 19405599 | 55.89132 | 90.24128 | 71.33452 | 30.42888 | 0.69772 | 1.762955 | 0.401812 |
| 10 | 122458000 | 122458799 | 10.67868 | 20.51994 | 20.67629 | 8.081994 | 0.519299 | -0.07739 | 0.401813 |
| 16 | 49904200 | 49905799 | 16.16468 | 29.16151 | 27.70137 | 12.05538 | 0.481049 | 0.384309 | 0.401865 |
| 2 | 78211600 | 78213399 | 21.58618 | 37.85511 | 33.70875 | 14.89831 | 0.554112 | 0.662903 | 0.401868 |
| 1 | 12105800 | 12113399 | 97.45636 | 151.2412 | 149.6524 | 62.14925 | 0.32122 | 2.780589 | 0.401874 |
| 13 | 74869600 | 74872399 | 28.84333 | 49.03354 | 44.69227 | 19.58726 | 0.514904 | 1.043028 | 0.401885 |
| 4 | 119052200 | 119053399 | 6.654516 | 12.53468 | 28.60109 | 11.70289 | -0.70864 | 0.056832 | 0.401891 |
| 12 | 119488800 | 119489799 | 10.70482 | 20.61487 | 19.52448 | 7.584338 | 0.615728 | -0.10169 | 0.401893 |
| 1 | 176479200 | 176480399 | 14.07708 | 25.67536 | 24.84762 | 10.6662 | 0.466929 | 0.299489 | 0.401893 |
| 15 | 98994000 | 98996199 | 16.08089 | 27.2987 | 57.39398 | 25.14684 | -0.68646 | 1.021241 | 0.401895 |
| 9 | 118431200 | 118432799 | 7.726112 | 14.3619 | 30.52614 | 12.77324 | -0.63367 | 0.156415 | 0.401895 |
| 7 | 47141200 | 47142999 | 21.87979 | 38.39701 | 32.43667 | 14.34461 | 0.629513 | 0.649392 | 0.401898 |
| 6 | 146683800 | 146686799 | 26.05078 | 42.92276 | 72.8834 | 31.43961 | -0.38212 | 1.377325 | 0.401899 |
| 7 | 40912200 | 40914999 | 37.28689 | 61.89706 | 53.22488 | 23.06886 | 0.598072 | 1.30431 | 0.401918 |
| 11 | 11735000 | 11736599 | 7.666763 | 14.41248 | 29.64559 | 12.17148 | -0.56482 | 0.080135 | 0.40192 |
| 13 | 71719800 | 71720999 | 13.40065 | 24.74527 | 24.36193 | 10.28645 | 0.458934 | 0.223904 | 0.401922 |
| 12 | 51164200 | 51166999 | 29.0652 | 47.2716 | 83.81231 | 35.96946 | -0.45478 | 1.563179 | 0.40193 |
| 2 | 114779600 | 114782599 | 27.36579 | 47.1237 | 40.29155 | 17.67434 | 0.609646 | 0.918592 | 0.401949 |
| 1 | 184621400 | 184624399 | 24.15181 | 39.69455 | 73.02788 | 31.46864 | -0.49883 | 1.403235 | 0.401972 |
| 19 | 9645200 | 9647599 | 22.33012 | 38.88549 | 31.54444 | 13.86954 | 0.68764 | 0.766763 | 0.401986 |
| 9 | 19338400 | 19350999 | 198.6208 | 297.0066 | 246.3902 | 101.5993 | 0.531753 | 3.532872 | 0.401991 |

| | | | | | | | | | |
|---|---|---|---|---|---|---|---|---|---|
| 1 | 167743600 | 167748799 | 27.22385 | 46.76443 | 37.19802 | 16.31611 | 0.712683 | 0.935001 | 0.401997 |
| 1 | 101933400 | 101935399 | 17.83199 | 32.74154 | 24.95961 | 10.62059 | 0.817723 | 0.292265 | 0.402011 |
| 18 | 56578600 | 56581199 | 12.73703 | 22.27688 | 43.64004 | 19.24409 | -0.57762 | 0.674562 | 0.402018 |
| 12 | 60121000 | 60121999 | 15.59109 | 28.52657 | 26.1871 | 11.19631 | 0.54637 | 0.296427 | 0.402028 |
| 12 | 86593200 | 86595599 | 17.57676 | 30.05855 | 53.22358 | 23.33744 | -0.4381 | 0.956264 | 0.402057 |
| 18 | 76612000 | 76615199 | 32.04044 | 51.86171 | 87.96764 | 37.62795 | -0.39735 | 1.665795 | 0.402069 |
| 8 | 62598200 | 62599799 | 16.21725 | 29.25796 | 27.59449 | 12.00804 | 0.491217 | 0.382871 | 0.402076 |
| 12 | 25120200 | 25125599 | 42.20688 | 67.59301 | 111.0504 | 46.81749 | -0.36965 | 2.006682 | 0.402092 |
| 5 | 85461200 | 85461999 | 13.03189 | 22.8862 | 41.44771 | 18.34074 | -0.46484 | 0.606557 | 0.402104 |
| 8 | 115194600 | 115199199 | 37.83214 | 60.61481 | 110.2259 | 46.50232 | -0.51499 | 1.986147 | 0.402105 |
| 11 | 79714200 | 79715199 | 8.938261 | 16.49212 | 31.99046 | 13.522 | -0.51369 | 0.211456 | 0.402119 |
| 4 | 143117200 | 143122199 | 57.87459 | 91.55448 | 145.7259 | 60.56446 | -0.34217 | 2.394138 | 0.402124 |
| 1 | 28148200 | 28149199 | 16.23925 | 29.02532 | 28.18449 | 12.37467 | 0.440934 | 0.440683 | 0.402145 |
| 12 | 78326800 | 78330199 | 53.10772 | 85.63865 | 69.5684 | 29.62719 | 0.656348 | 1.792502 | 0.402164 |
| 17 | 38691600 | 38696599 | 73.18297 | 116.4565 | 103.2114 | 43.12426 | 0.509279 | 2.235475 | 0.402166 |
| 16 | 96630800 | 96631399 | 4.967373 | 9.666121 | 22.98231 | 8.465222 | -0.62631 | -0.33498 | 0.402172 |
| 17 | 74775000 | 74775799 | 4.967373 | 9.666121 | 22.98231 | 8.465222 | -0.62631 | -0.33498 | 0.402172 |
| 5 | 104656000 | 104656799 | 6.139356 | 11.79771 | 25.76237 | 10.06896 | -0.58592 | -0.10827 | 0.402188 |
| 1 | 76651600 | 76652599 | 12.95512 | 24.32492 | 23.4603 | 9.651444 | 0.517258 | 0.118875 | 0.402236 |
| 14 | 83019200 | 83021399 | 23.65563 | 40.96009 | 38.72475 | 16.9899 | 0.466638 | 0.849486 | 0.402238 |
| 13 | 62089200 | 62111999 | 526.0421 | 718.2637 | 626.3723 | 260.7014 | 0.330972 | 4.949281 | 0.40224 |
| 6 | 7027000 | 7030799 | 66.8951 | 106.674 | 90.23927 | 37.81371 | 0.579541 | 2.158641 | 0.402241 |
| 17 | 57647800 | 57649799 | 27.38525 | 44.80799 | 77.79986 | 33.46239 | -0.41848 | 1.469756 | 0.402258 |
| 17 | 38732400 | 38735999 | 43.50902 | 71.67669 | 54.06662 | 23.32109 | 0.784967 | 1.407312 | 0.402279 |
| 7 | 5562000 | 5562999 | 20.06477 | 35.06408 | 32.1955 | 14.18747 | 0.510334 | 0.708498 | 0.402289 |
| 13 | 67888000 | 67894199 | 94.23465 | 147.0179 | 143.6331 | 59.60325 | 0.345476 | 2.687492 | 0.402294 |
| 9 | 114904000 | 114907199 | 34.03282 | 54.97972 | 91.75498 | 39.16788 | -0.37697 | 1.721544 | 0.402298 |
| 5 | 94813600 | 94813999 | 17.81409 | 31.69761 | 25.60671 | 11.28547 | 0.70117 | 0.473795 | 0.40231 |
| 7 | 60732800 | 60733599 | 16.93893 | 30.30308 | 27.34483 | 11.99693 | 0.547207 | 0.440977 | 0.402319 |
| 4 | 72879200 | 72880399 | 9.714875 | 17.99876 | 34.81103 | 14.63231 | -0.50392 | 0.201062 | 0.402325 |
| 10 | 90707800 | 90709799 | 24.93707 | 42.83666 | 41.47599 | 18.19255 | 0.430207 | 0.930133 | 0.402332 |
| 2 | 52434400 | 52436199 | 18.93699 | 33.70987 | 28.76569 | 12.6732 | 0.622806 | 0.483678 | 0.402337 |
| 11 | 15962400 | 15962999 | 5.838552 | 11.27831 | 25.97813 | 9.986481 | -0.64009 | -0.16804 | 0.402341 |
| 18 | 62368000 | 62370599 | 21.87469 | 38.43166 | 32.47256 | 14.37426 | 0.629304 | 0.633152 | 0.402356 |
| 13 | 106274000 | 106277399 | 40.56815 | 65.89991 | 65.17715 | 27.95289 | 0.383737 | 1.595655 | 0.402361 |
| 16 | 25927000 | 25928199 | 14.28089 | 25.02522 | 44.06492 | 19.46309 | -0.4247 | 0.652036 | 0.402362 |
| 6 | 129941400 | 129942999 | 27.99702 | 47.42679 | 45.82653 | 20.06573 | 0.430716 | 1.071143 | 0.40237 |
| 8 | 61277200 | 61277799 | 12.24975 | 21.52369 | 40.94486 | 18.1261 | -0.53535 | 0.592554 | 0.402393 |
| 8 | 116417800 | 116420399 | 18.48118 | 31.7171 | 56.5881 | 24.81977 | -0.44842 | 0.929906 | 0.402393 |
| 6 | 43872800 | 43873999 | 12.18753 | 21.57248 | 39.79268 | 17.597 | -0.48731 | 0.505555 | 0.402421 |
| 17 | 33143600 | 33144799 | 11.47833 | 20.23507 | 39.42325 | 17.44775 | -0.56767 | 0.547933 | 0.402424 |
| 10 | 38728000 | 38729599 | 15.96591 | 28.66381 | 27.07532 | 11.84324 | 0.484037 | 0.407978 | 0.402437 |
| 9 | 106198200 | 106200599 | 14.31404 | 24.92953 | 45.7831 | 20.13164 | -0.48558 | 0.748544 | 0.40244 |
| 10 | 48273200 | 48277199 | 52.88307 | 85.23002 | 77.61694 | 33.03164 | 0.491005 | 1.811891 | 0.402456 |
| 15 | 97198800 | 97199599 | 15.42891 | 27.72256 | 26.77976 | 11.70583 | 0.452791 | 0.400012 | 0.402457 |
| 2 | 23377200 | 23381399 | 68.81574 | 109.741 | 99.29086 | 41.60893 | 0.482752 | 2.160611 | 0.402458 |
| 6 | 48596200 | 48600999 | 39.16593 | 62.66654 | 111.8666 | 47.09981 | -0.49059 | 2.031794 | 0.402468 |

| | | | | | | | | | |
|---|---|---|---|---|---|---|---|---|---|
| 10 | 106375800 | 106377999 | 31.18775 | 53.4999 | 39.60115 | 17.3686 | 0.815819 | 0.966869 | 0.402478 |
| 14 | 7713400 | 7714199 | 10.04055 | 18.39014 | 33.97925 | 14.50718 | -0.45472 | 0.266277 | 0.40248 |
| 16 | 72936200 | 72940199 | 66.76261 | 106.4369 | 92.3535 | 38.68668 | 0.542595 | 2.165308 | 0.402481 |
| 8 | 82687000 | 82688199 | 8.637018 | 16.02306 | 31.01829 | 13.01457 | -0.50276 | 0.17753 | 0.402493 |
| 9 | 8205400 | 8206399 | 8.637018 | 16.02306 | 31.01829 | 13.01457 | -0.50276 | 0.17753 | 0.402493 |
| 15 | 60043800 | 60044599 | 8.637018 | 16.02306 | 31.01829 | 13.01457 | -0.50276 | 0.17753 | 0.402493 |
| 17 | 35576200 | 35579199 | 26.24787 | 43.04982 | 74.8767 | 32.23692 | -0.41934 | 1.435474 | 0.402495 |
| 10 | 52108000 | 52110399 | 15.71369 | 27.28773 | 48.45702 | 21.26851 | -0.43795 | 0.799382 | 0.4025 |
| 18 | 11091200 | 11092799 | 9.118963 | 16.81709 | 31.98976 | 13.53094 | -0.4862 | 0.217357 | 0.40251 |
| 13 | 45938800 | 45941799 | 20.03488 | 34.01877 | 58.65081 | 25.69931 | -0.40145 | 1.036747 | 0.402559 |
| 4 | 34919000 | 34920399 | 7.952452 | 15.07326 | 29.31536 | 11.84419 | -0.46464 | 0.027634 | 0.402568 |
| 7 | 83459400 | 83461599 | 19.12346 | 33.4805 | 31.05842 | 13.70763 | 0.495385 | 0.670416 | 0.402573 |
| 3 | 33979000 | 33979599 | 11.65551 | 21.9446 | 22.03854 | 9.024687 | 0.464717 | 0.086762 | 0.402574 |
| 3 | 140937600 | 140939999 | 26.54476 | 45.66311 | 40.822 | 17.90647 | 0.54523 | 0.923079 | 0.402586 |
| 12 | 43446200 | 43446799 | 8.122094 | 15.35401 | 29.79022 | 12.10378 | -0.46832 | 0.049311 | 0.402586 |
| 7 | 42322400 | 42335399 | 313.621 | 449.8284 | 389.249 | 160.7383 | 0.413626 | 4.206426 | 0.402602 |
| 7 | 20883600 | 20884599 | 40.65364 | 65.99547 | 59.33053 | 25.43311 | 0.520259 | 1.607755 | 0.402604 |
| 1 | 129017800 | 129020999 | 53.73623 | 86.31414 | 84.24758 | 35.67932 | 0.385244 | 1.919907 | 0.402619 |
| 17 | 60896000 | 60897199 | 14.93 | 28.55411 | 20.30551 | 8.045339 | 1.000082 | 0.007713 | 0.402627 |
| 11 | 69354000 | 69357199 | 25.57529 | 41.62921 | 80.77187 | 34.6748 | -0.58353 | 1.54409 | 0.402637 |
| X | 31638000 | 31640399 | 55.86365 | 90.5091 | 63.96127 | 27.37854 | 0.864106 | 1.679022 | 0.402646 |
| 17 | 8569200 | 8570999 | 8.451574 | 15.00295 | 38.46808 | 16.98751 | -0.9567 | 0.454922 | 0.402651 |
| 4 | 137934600 | 137936999 | 10.62182 | 18.92615 | 38.32185 | 16.86965 | -0.61573 | 0.442765 | 0.402654 |
| 9 | 111489000 | 111492399 | 23.792 | 41.54669 | 35.26455 | 15.53235 | 0.62295 | 0.74138 | 0.40266 |
| 14 | 104710400 | 104712199 | 17.74344 | 30.23639 | 54.81694 | 24.02925 | -0.47287 | 0.9903 | 0.402665 |
| X | 25824200 | 25828799 | 28.94766 | 50.96638 | 25.47075 | 11.2756 | 1.38373 | 0.640065 | 0.402679 |
| 17 | 29412400 | 29415199 | 18.28992 | 30.99319 | 56.3661 | 24.68307 | -0.47819 | 1.045792 | 0.402687 |
| 5 | 142073200 | 142078399 | 41.36463 | 66.44588 | 108.9615 | 46.14958 | -0.36175 | 1.910348 | 0.40269 |
| 6 | 123786800 | 123787599 | 13.17536 | 25.29064 | 20.57343 | 8.066342 | 0.82199 | -0.0408 | 0.40269 |
| 2 | 53544200 | 53545399 | 16.01993 | 28.60748 | 27.20425 | 11.95338 | 0.470112 | 0.442248 | 0.402716 |
| 8 | 34301400 | 34303199 | 16.11569 | 27.89476 | 49.51516 | 21.72379 | -0.43816 | 0.837228 | 0.402721 |
| 4 | 16709600 | 16711999 | 28.15845 | 48.52169 | 39.07118 | 17.13937 | 0.695905 | 0.912575 | 0.402783 |
| 11 | 115438600 | 115440999 | 28.15845 | 48.52169 | 39.07118 | 17.13937 | 0.695905 | 0.912575 | 0.402783 |
| 14 | 52894400 | 52896599 | 21.04519 | 37.56037 | 27.53484 | 12.10433 | 0.842931 | 0.480479 | 0.402784 |
| 7 | 144249600 | 144251199 | 18.09478 | 30.91932 | 54.28621 | 23.80183 | -0.42609 | 0.968182 | 0.402801 |
| 5 | 147012600 | 147016199 | 19.04981 | 31.50103 | 75.57642 | 32.66916 | -0.87789 | 1.317743 | 0.402814 |
| 7 | 85469600 | 85470199 | 6.400081 | 12.37 | 26.66452 | 10.22333 | -0.54193 | -0.16899 | 0.402821 |
| 7 | 9587000 | 9587199 | 7.447965 | 14.53872 | 14.75277 | 5.221351 | 0.642194 | -0.46817 | 0.402828 |
| 16 | 96633200 | 96635399 | 22.19148 | 38.90574 | 33.62088 | 14.85597 | 0.597123 | 0.669846 | 0.402844 |
| 7 | 137217000 | 137221399 | 36.86057 | 59.10472 | 107.221 | 45.28887 | -0.51013 | 1.959772 | 0.402862 |
| 17 | 23782800 | 23783799 | 8.310712 | 15.46606 | 30.53296 | 12.75201 | -0.52588 | 0.154194 | 0.40289 |
| 16 | 30361000 | 30377199 | 151.1755 | 224.6521 | 338.132 | 139.2995 | -0.3354 | 3.640953 | 0.40291 |
| 5 | 38661800 | 38663999 | 11.29908 | 20.02017 | 37.83664 | 16.72862 | -0.52111 | 0.487933 | 0.402918 |
| 6 | 68826200 | 68828399 | 30.68243 | 52.72097 | 38.43918 | 16.86067 | 0.837811 | 0.947492 | 0.40292 |
| 1 | 179874200 | 179877599 | 42.90993 | 70.55361 | 56.83858 | 24.49676 | 0.689285 | 1.431358 | 0.402921 |
| 1 | 78286400 | 78289199 | 10.04928 | 17.56521 | 49.39081 | 21.76867 | -1.09607 | 0.679787 | 0.402935 |
| 4 | 138905400 | 138912599 | 55.27557 | 87.55393 | 140.804 | 58.55646 | -0.35554 | 2.362183 | 0.402935 |

| | | | | | | | | | |
|---|---|---|---|---|---|---|---|---|---|
| 2 | 24306600 | 24307399 | 5.412352 | 10.44829 | 24.75477 | 9.544877 | -0.68346 | -0.16317 | 0.402942 |
| 9 | 56324400 | 56325399 | 5.412352 | 10.44829 | 24.75477 | 9.544877 | -0.68346 | -0.16317 | 0.402942 |
| 3 | 24147600 | 24149199 | 13.59819 | 25.39885 | 23.24833 | 9.649538 | 0.581598 | 0.155123 | 0.402944 |
| 2 | 76726400 | 76732399 | 67.04956 | 105.4391 | 164.9306 | 68.42043 | -0.32389 | 2.532969 | 0.402944 |
| 17 | 94746200 | 94752799 | 78.92189 | 125.2671 | 108.2431 | 45.09963 | 0.542083 | 2.313631 | 0.402957 |
| 5 | 109164200 | 109164999 | 18.365 | 32.25757 | 30.91074 | 13.68007 | 0.448694 | 0.614123 | 0.402964 |
| 15 | 57939000 | 57940399 | 13.22866 | 23.05545 | 45.26336 | 19.91191 | -0.58118 | 0.732333 | 0.402968 |
| 17 | 43614800 | 43615799 | 5.833807 | 11.28154 | 26.06488 | 9.968472 | -0.63746 | -0.18578 | 0.402973 |
| 12 | 29538400 | 29540599 | 30.042 | 50.62794 | 47.47063 | 20.75298 | 0.47373 | 1.124633 | 0.402978 |
| 1 | 41404400 | 41404799 | 5.074678 | 9.890433 | 22.00408 | 7.967317 | -0.5082 | -0.39546 | 0.402984 |
| 1 | 136889400 | 136889999 | 5.074678 | 9.890433 | 22.00408 | 7.967317 | -0.5082 | -0.39546 | 0.402984 |
| 7 | 62144400 | 62145399 | 29.50579 | 49.3241 | 48.10368 | 20.94475 | 0.417172 | 1.201737 | 0.402998 |
| 12 | 23728400 | 23731399 | 41.31066 | 67.23092 | 65.8912 | 28.28496 | 0.398937 | 1.568952 | 0.403001 |
| 4 | 102752800 | 102754399 | 30.78747 | 52.12247 | 45.77473 | 20.0394 | 0.568011 | 1.085775 | 0.403006 |
| Y | 89255600 | 89256999 | 10.43587 | 20.12129 | 11.94684 | 4.62468 | 1.287236 | -0.14041 | 0.403016 |
| 4 | 8517000 | 8520399 | 57.2322 | 91.6435 | 90.65469 | 38.21123 | 0.360667 | 2.020568 | 0.403035 |
| 15 | 100242400 | 100247399 | 32.04088 | 51.90948 | 86.9029 | 37.19737 | -0.37726 | 1.647524 | 0.403036 |
| 12 | 112791200 | 112793599 | 10.73852 | 18.86914 | 42.51523 | 18.82196 | -0.77823 | 0.584508 | 0.40304 |
| 7 | 5267400 | 5269999 | 41.47537 | 67.74388 | 63.29296 | 27.20551 | 0.471411 | 1.516822 | 0.403042 |
| 12 | 5222000 | 5227399 | 31.9198 | 51.52028 | 93.04177 | 39.67162 | -0.49205 | 1.742687 | 0.403065 |
| 11 | 109212400 | 109214999 | 7.313455 | 13.5364 | 35.82133 | 15.07702 | -0.95383 | 0.189821 | 0.403067 |
| 15 | 97395200 | 97396999 | 20.29083 | 35.37959 | 34.16644 | 15.03449 | 0.43748 | 0.742813 | 0.403075 |
| 8 | 66834000 | 66836799 | 34.87484 | 57.74818 | 47.71334 | 20.6443 | 0.655151 | 1.318948 | 0.403081 |
| 9 | 24200000 | 24201399 | 7.109457 | 13.31622 | 30.11368 | 12.44017 | -0.70755 | 0.099806 | 0.403089 |
| 4 | 100630400 | 100632599 | 25.84688 | 43.96684 | 43.14702 | 18.91097 | 0.408744 | 1.022306 | 0.403139 |
| 12 | 109573400 | 109585599 | 102.6404 | 155.4474 | 278.5373 | 115.2483 | -0.56233 | 3.26863 | 0.403147 |
| 3 | 134345200 | 134346599 | 24.12896 | 42.43845 | 30.65321 | 13.57263 | 0.854762 | 0.629866 | 0.403162 |
| 18 | 19027400 | 19029799 | 21.54389 | 37.54996 | 36.02986 | 15.85061 | 0.446574 | 0.756265 | 0.403163 |
| 14 | 61364800 | 61365399 | 5.213127 | 10.15944 | 22.60139 | 8.192265 | -0.50956 | -0.38749 | 0.403163 |
| X | 25895600 | 25899999 | 71.70011 | 114.3975 | 95.8652 | 40.20099 | 0.59396 | 2.149275 | 0.403191 |
| 13 | 19386800 | 19388799 | 25.46523 | 43.56845 | 42.58312 | 18.67249 | 0.415845 | 0.969843 | 0.403195 |
| 17 | 5735400 | 5736599 | 10.63937 | 19.16193 | 36.38088 | 15.86546 | -0.51354 | 0.376226 | 0.403196 |
| 12 | 113494800 | 113496799 | 29.38679 | 49.27723 | 48.60029 | 21.20228 | 0.400861 | 1.167291 | 0.403205 |
| 9 | 26845800 | 26846599 | 5.705104 | 10.99017 | 25.26297 | 9.804437 | -0.64948 | -0.13751 | 0.403206 |
| 12 | 56119400 | 56120199 | 16.10008 | 30.08599 | 23.11115 | 9.584993 | 0.833921 | 0.174728 | 0.403206 |
| 10 | 117881400 | 117886799 | 34.45805 | 55.68998 | 94.22799 | 40.24528 | -0.39616 | 1.708747 | 0.40324 |
| 15 | 8739600 | 8740199 | 5.812484 | 11.28942 | 24.63075 | 9.211047 | -0.52445 | -0.26596 | 0.403242 |
| 11 | 114548400 | 114550399 | 16.40036 | 28.38404 | 49.52367 | 21.72745 | -0.41355 | 0.838476 | 0.403242 |
| 10 | 81587600 | 81590199 | 44.26516 | 72.70909 | 48.98958 | 21.10499 | 0.945625 | 1.438856 | 0.403257 |
| X | 23219000 | 23220999 | 20.63796 | 37.13154 | 26.72475 | 11.66868 | 0.876125 | 0.429568 | 0.403262 |
| X | 78232800 | 78234799 | 32.22737 | 54.5403 | 45.22628 | 19.79711 | 0.650558 | 1.094597 | 0.403263 |
| 9 | 8003000 | 8004999 | 16.95205 | 28.90728 | 54.80215 | 24.02407 | -0.53691 | 0.980633 | 0.403274 |
| 9 | 71732400 | 71734199 | 11.87842 | 21.05489 | 38.76644 | 17.13837 | -0.48377 | 0.486747 | 0.40328 |
| 2 | 131630600 | 131633199 | 41.94772 | 68.58514 | 62.27752 | 26.77897 | 0.513456 | 1.502675 | 0.403301 |
| 14 | 46015200 | 46015999 | 8.603788 | 16.00137 | 31.04966 | 12.98166 | -0.50262 | 0.161298 | 0.40334 |
| 18 | 58864600 | 58865799 | 16.07275 | 29.18818 | 26.76187 | 11.55052 | 0.539598 | 0.345079 | 0.403341 |
| 14 | 42388200 | 42389399 | 25.96423 | 44.05598 | 42.71753 | 18.71379 | 0.425808 | 1.046033 | 0.403347 |

| | | | | | | | | | |
|---|---|---|---|---|---|---|---|---|---|
| 3 | 47869400 | 47871199 | 15.88671 | 29.05431 | 26.12673 | 11.17737 | 0.575267 | 0.300702 | 0.403362 |
| 1 | 183548400 | 183550399 | 11.9334 | 21.04903 | 39.38467 | 17.42873 | -0.50946 | 0.54412 | 0.403376 |
| 14 | 63318800 | 63322799 | 23.65131 | 38.93787 | 71.97085 | 31.03538 | -0.50455 | 1.379926 | 0.403411 |
| 10 | 124924000 | 124926399 | 22.56678 | 39.27856 | 37.07524 | 16.29636 | 0.47 | 0.782435 | 0.40342 |
| 10 | 10016800 | 10018799 | 23.00335 | 40.538 | 30.10335 | 13.32394 | 0.815889 | 0.600692 | 0.403457 |
| 18 | 31742400 | 31743399 | 6.887356 | 13.21471 | 27.40885 | 10.76001 | -0.51832 | -0.07955 | 0.403461 |
| 11 | 29603000 | 29606599 | 42.09156 | 68.58478 | 64.97532 | 27.90411 | 0.448976 | 1.552357 | 0.403465 |
| 8 | 125490200 | 125494399 | 21.90475 | 36.50727 | 65.1151 | 28.29773 | -0.45089 | 1.225706 | 0.403473 |
| 17 | 66887800 | 66889799 | 24.01084 | 39.88428 | 68.60422 | 29.74787 | -0.39903 | 1.272046 | 0.40349 |
| 19 | 3187200 | 3193399 | 118.8391 | 182.2414 | 179.1209 | 74.35561 | 0.317789 | 3.013441 | 0.403501 |
| 4 | 6551000 | 6552799 | 28.31077 | 48.08585 | 44.67255 | 19.57621 | 0.487232 | 1.046398 | 0.403504 |
| 4 | 10038600 | 10039199 | 8.689288 | 16.90413 | 17.76369 | 6.557228 | 0.536095 | -0.29913 | 0.403504 |
| 7 | 11264600 | 11267599 | 47.17156 | 76.31939 | 71.60334 | 30.61808 | 0.454371 | 1.693805 | 0.403506 |
| X | 80010200 | 80011999 | 17.92844 | 32.31688 | 27.57591 | 12.01309 | 0.634085 | 0.40205 | 0.403512 |
| 7 | 40525600 | 40530199 | 36.17359 | 59.62714 | 57.11755 | 24.64317 | 0.441539 | 1.382811 | 0.403521 |
| X | 33920800 | 33923199 | 19.58579 | 35.86764 | 24.24234 | 10.35221 | 0.986621 | 0.31863 | 0.403535 |
| 18 | 36014800 | 36016399 | 18.4834 | 32.48143 | 31.28368 | 13.85085 | 0.441302 | 0.606899 | 0.403553 |
| 1 | 121719600 | 121721799 | 15.70144 | 27.29893 | 48.44496 | 21.27913 | -0.43686 | 0.781995 | 0.403555 |
| 7 | 82025000 | 82027799 | 18.30637 | 31.44519 | 57.15894 | 25.07151 | -0.47502 | 0.921296 | 0.403572 |
| X | 146562200 | 146565599 | 49.01933 | 79.56352 | 60.10894 | 25.76381 | 0.770315 | 1.626929 | 0.40358 |
| 12 | 71355200 | 71357999 | 16.75883 | 28.73395 | 52.15956 | 22.87565 | -0.47323 | 0.927525 | 0.403607 |
| 1 | 97087000 | 97089199 | 20.60862 | 36.45932 | 29.69285 | 13.13204 | 0.68538 | 0.543626 | 0.403688 |
| 10 | 79527800 | 79532599 | 36.85443 | 59.1326 | 105.0362 | 44.40909 | -0.4787 | 1.939522 | 0.403698 |
| 6 | 10752000 | 10753799 | 20.10298 | 36.7351 | 22.31085 | 9.554753 | 1.136421 | 0.328638 | 0.403713 |
| 3 | 52601400 | 52603399 | 15.02011 | 27.16997 | 26.20259 | 11.36467 | 0.462273 | 0.355869 | 0.403725 |
| 7 | 29611200 | 29616399 | 25.70628 | 44.21918 | 36.75939 | 16.1244 | 0.649701 | 0.915493 | 0.40373 |
| 8 | 10738200 | 10740799 | 14.14097 | 24.78284 | 43.54927 | 19.23718 | -0.42169 | 0.649375 | 0.403737 |
| 3 | 58072800 | 58074399 | 8.60356 | 16.07014 | 30.59986 | 12.70047 | -0.46692 | 0.133333 | 0.403743 |
| 5 | 113795600 | 113798799 | 25.70292 | 43.75914 | 43.14571 | 18.91169 | 0.402173 | 1.014367 | 0.403745 |
| 2 | 41234200 | 41236799 | 42.17297 | 69.18291 | 59.52059 | 25.62778 | 0.593479 | 1.459986 | 0.403748 |
| 6 | 4505200 | 4506599 | 11.52256 | 20.65045 | 37.30011 | 16.33983 | -0.44699 | 0.411499 | 0.403779 |
| 7 | 51068000 | 51070599 | 19.82779 | 33.26131 | 66.7078 | 29.10907 | -0.61952 | 1.152501 | 0.403801 |
| 7 | 18021200 | 18030599 | 210.6794 | 311.1214 | 293.2892 | 120.7823 | 0.330866 | 3.753717 | 0.403807 |
| 16 | 44326800 | 44328199 | 8.749968 | 16.30296 | 31.08722 | 12.95727 | -0.47403 | 0.15097 | 0.403842 |
| 7 | 55937000 | 55938199 | 14.46905 | 26.4255 | 25.18734 | 10.79401 | 0.490055 | 0.294481 | 0.403853 |
| 7 | 9648000 | 9651799 | 34.77087 | 58.74969 | 44.06838 | 19.2808 | 0.794586 | 1.114677 | 0.403861 |
| 4 | 147380200 | 147383399 | 62.71955 | 100.5784 | 85.73687 | 36.21951 | 0.577453 | 1.981 | 0.403866 |
| 8 | 41348400 | 41349999 | 11.0891 | 19.75123 | 37.29236 | 16.42174 | -0.51567 | 0.44757 | 0.403882 |
| 14 | 122023800 | 122024999 | 8.156572 | 15.18351 | 30.53816 | 12.74805 | -0.55201 | 0.150653 | 0.403897 |
| 8 | 61586200 | 61589599 | 48.7934 | 78.70573 | 76.47214 | 32.58017 | 0.399156 | 1.781162 | 0.403924 |
| 10 | 79608000 | 79611599 | 38.41877 | 61.7333 | 103.4659 | 43.84203 | -0.39254 | 1.894046 | 0.403939 |
| 8 | 63404400 | 63405199 | 6.953926 | 13.06109 | 29.09159 | 11.96162 | -0.68 | 0.078975 | 0.403943 |
| 19 | 29876000 | 29879799 | 52.80392 | 84.83785 | 83.85184 | 35.52467 | 0.367605 | 1.909294 | 0.403955 |
| 2 | 10748200 | 10749399 | 6.897366 | 13.249 | 29.19257 | 11.42283 | -0.60035 | -0.08467 | 0.403961 |
| 8 | 109791800 | 109793799 | 20.44868 | 36.13397 | 30.93136 | 13.68299 | 0.613451 | 0.558896 | 0.403965 |
| 12 | 78664200 | 78664799 | 16.73223 | 29.66699 | 25.95381 | 11.47277 | 0.584111 | 0.489949 | 0.403975 |
| 9 | 14756600 | 14758799 | 18.39245 | 31.24451 | 56.42863 | 24.72909 | -0.46814 | 1.022862 | 0.403982 |

| | | | | | | | | | |
|---|---|---|---|---|---|---|---|---|---|
| 7 | 31849400 | 31858799 | 33.13908 | 56.0317 | 40.32985 | 17.64885 | 0.853868 | 1.098721 | 0.404 |
| 2 | 17849400 | 17853799 | 29.48703 | 48.14524 | 83.88896 | 36.05362 | -0.4258 | 1.507543 | 0.404017 |
| 13 | 92317600 | 92318399 | 14.10597 | 26.68636 | 22.45212 | 9.107542 | 0.730008 | 0.079546 | 0.404018 |
| 11 | 18573200 | 18573799 | 5.884887 | 11.16988 | 27.6447 | 11.13639 | -0.8039 | -0.00231 | 0.404018 |
| 19 | 7916200 | 7917799 | 35.63353 | 58.47831 | 58.52621 | 25.20384 | 0.376064 | 1.437748 | 0.404035 |
| 4 | 117165800 | 117167199 | 13.13573 | 24.33796 | 23.29665 | 9.791652 | 0.503742 | 0.199572 | 0.404039 |
| 7 | 122641400 | 122645999 | 37.27184 | 60.06947 | 98.41599 | 41.88279 | -0.35446 | 1.797519 | 0.404061 |
| 14 | 50148400 | 50149999 | 18.23962 | 33.00887 | 26.87109 | 11.64793 | 0.705909 | 0.378689 | 0.404072 |
| X | 90163000 | 90166599 | 68.58967 | 109.2874 | 102.0531 | 42.71612 | 0.435971 | 2.18903 | 0.404091 |
| 6 | 11166200 | 11167999 | 19.64763 | 35.64247 | 25.70821 | 11.11044 | 0.882213 | 0.373553 | 0.404093 |
| 8 | 4893400 | 4895199 | 20.37521 | 36.04643 | 30.31971 | 13.40928 | 0.63917 | 0.544152 | 0.404111 |
| 5 | 100413800 | 100422999 | 80.75175 | 124.4225 | 218.8437 | 90.90418 | -0.51576 | 2.912137 | 0.404113 |
| 12 | 18009200 | 18017599 | 211.5687 | 311.3534 | 292.6512 | 120.515 | 0.330397 | 3.810369 | 0.404115 |
| 16 | 97762800 | 97764599 | 16.39652 | 27.83278 | 55.77404 | 24.43661 | -0.61738 | 1.022712 | 0.404151 |
| 11 | 79423600 | 79425399 | 9.09602 | 16.77786 | 31.98504 | 13.52556 | -0.48926 | 0.214897 | 0.404154 |
| X | 65000000 | 65001799 | 19.15078 | 35.45299 | 22.79637 | 9.592301 | 1.073142 | 0.254291 | 0.404158 |
| 9 | 68770000 | 68772199 | 45.98694 | 74.32649 | 73.02119 | 31.18984 | 0.386492 | 1.717899 | 0.404171 |
| 1 | 37765000 | 37769999 | 41.04022 | 65.86881 | 107.1291 | 45.31507 | -0.35133 | 1.93561 | 0.404185 |
| 10 | 67728800 | 67730199 | 8.516317 | 15.65456 | 31.96147 | 13.57911 | -0.59222 | 0.230018 | 0.404203 |
| 7 | 111840000 | 111842199 | 14.1402 | 24.66186 | 45.23406 | 19.90802 | -0.48356 | 0.726043 | 0.40423 |
| 14 | 50437400 | 50440999 | 57.01907 | 91.12282 | 84.73657 | 35.61661 | 0.44658 | 2.078158 | 0.404254 |
| 9 | 123952400 | 123960599 | 163.6748 | 248.009 | 206.6353 | 85.51164 | 0.542194 | 3.269184 | 0.404273 |
| 1 | 105831600 | 105832799 | 11.47979 | 21.96435 | 20.45441 | 8.094941 | 0.614961 | -0.03189 | 0.404281 |
| 7 | 110106800 | 110109399 | 50.3923 | 81.07849 | 80.772 | 34.2905 | 0.358761 | 1.861372 | 0.404306 |
| X | 77334200 | 77337199 | 44.78676 | 73.38458 | 52.81047 | 22.72732 | 0.849701 | 1.476093 | 0.404325 |
| 2 | 56636800 | 56637599 | 11.13129 | 21.63234 | 18.16417 | 6.761482 | 0.845795 | -0.23891 | 0.404327 |
| 12 | 114237800 | 114259799 | 104.5606 | 163.2378 | 127.8029 | 53.03055 | 0.665298 | 2.678843 | 0.40434 |
| 13 | 110222000 | 110224199 | 31.67077 | 53.21489 | 48.57085 | 21.2149 | 0.512285 | 1.155824 | 0.404347 |
| 1 | 13624800 | 13626199 | 20.59417 | 35.89704 | 35.12538 | 15.45335 | 0.418495 | 0.749789 | 0.40435 |
| 4 | 145547400 | 145547799 | 11.37023 | 21.77143 | 18.39311 | 7.263145 | 0.757605 | -0.04452 | 0.404351 |
| X | 80761600 | 80763999 | 34.21689 | 57.4357 | 48.03664 | 20.97001 | 0.637971 | 1.174162 | 0.404354 |
| 6 | 7181800 | 7184199 | 19.17371 | 33.72372 | 32.17076 | 14.24451 | 0.455388 | 0.60219 | 0.404356 |
| 18 | 82844800 | 82847799 | 16.13851 | 27.66205 | 52.14416 | 22.86834 | -0.52746 | 0.927731 | 0.404365 |
| X | 63687800 | 63704999 | 29.0216 | 49.92663 | 35.04743 | 15.37349 | 0.892734 | 0.928341 | 0.404368 |
| 15 | 103692200 | 103692999 | 12.70895 | 22.379 | 40.37969 | 17.87512 | -0.45847 | 0.568704 | 0.404372 |
| 4 | 82494400 | 82496199 | 18.6632 | 33.68693 | 26.74577 | 11.63172 | 0.738658 | 0.39554 | 0.404372 |
| X | 47826800 | 47827799 | 13.17678 | 24.74626 | 22.05076 | 9.068376 | 0.631645 | 0.118726 | 0.404374 |
| 1 | 10107600 | 10108599 | 5.162069 | 9.931842 | 25.76015 | 10.0328 | -0.8267 | -0.13081 | 0.404398 |
| 13 | 61650600 | 61652799 | 33.65605 | 56.77628 | 45.20291 | 19.7677 | 0.70882 | 1.123963 | 0.4044 |
| 8 | 115032600 | 115034399 | 15.58895 | 28.06681 | 26.06594 | 11.37199 | 0.51078 | 0.385493 | 0.404417 |
| 4 | 90157600 | 90163599 | 130.1031 | 199.3999 | 180.4895 | 74.91781 | 0.435887 | 3.028789 | 0.404422 |
| 7 | 31648200 | 31650399 | 30.27351 | 52.28311 | 36.7698 | 16.13092 | 0.891039 | 0.900602 | 0.404439 |
| 7 | 13764800 | 13782399 | 275.4529 | 401.3438 | 327.6432 | 135.0228 | 0.519895 | 3.971833 | 0.404445 |
| 6 | 143268200 | 143270799 | 21.25115 | 35.50078 | 63.43968 | 27.61028 | -0.4536 | 1.197417 | 0.404447 |
| 1 | 121820400 | 121820799 | 9.215654 | 17.84956 | 16.74363 | 6.356552 | 0.66215 | -0.19798 | 0.404468 |
| 4 | 54248200 | 54250199 | 19.02539 | 34.04398 | 28.28901 | 12.40862 | 0.665287 | 0.452485 | 0.404474 |
| 7 | 12365600 | 12368199 | 44.82187 | 72.50745 | 70.43703 | 30.1162 | 0.404013 | 1.688859 | 0.404484 |

| | | | | | | | | | |
|---|---|---|---|---|---|---|---|---|---|
| 15 | 92544400 | 92545199 | 12.61181 | 24.17689 | 21.12068 | 8.3124 | 0.714761 | -0.03346 | 0.404493 |
| 16 | 9466800 | 9469199 | 19.66041 | 34.54044 | 32.67499 | 14.46354 | 0.467084 | 0.620576 | 0.404516 |
| 18 | 3814400 | 3819199 | 71.35864 | 113.0377 | 112.9751 | 46.9851 | 0.329787 | 2.365976 | 0.404535 |
| 5 | 43227400 | 43230599 | 30.56201 | 49.63496 | 83.58084 | 35.84006 | -0.38231 | 1.593672 | 0.40454 |
| X | 26838400 | 26839199 | 20.43538 | 36.28265 | 29.40339 | 12.9826 | 0.69513 | 0.513588 | 0.404562 |
| 18 | 11168200 | 11174199 | 55.76163 | 89.6083 | 83.3632 | 35.32817 | 0.455151 | 1.906685 | 0.40457 |
| X | 8637400 | 8640599 | 55.47447 | 89.85124 | 64.94034 | 27.7914 | 0.831373 | 1.685919 | 0.404583 |
| 8 | 121859200 | 121867199 | 71.27071 | 110.2139 | 208.7834 | 86.7237 | -0.61863 | 2.84021 | 0.404583 |
| 18 | 24286400 | 24289599 | 35.04908 | 56.43713 | 97.12567 | 41.28096 | -0.42675 | 1.822978 | 0.404585 |
| 2 | 152525600 | 152530599 | 59.76057 | 94.20341 | 153.0211 | 63.49975 | -0.37553 | 2.478002 | 0.404612 |
| 15 | 52737000 | 52738199 | 12.37329 | 21.74763 | 42.95679 | 19.02184 | -0.58964 | 0.589198 | 0.404616 |
| 4 | 125864800 | 125868799 | 28.41413 | 46.34482 | 79.167 | 34.01122 | -0.39822 | 1.519914 | 0.404617 |
| 14 | 72758000 | 72758999 | 9.895038 | 19.16925 | 19.23221 | 7.294556 | 0.566092 | -0.18827 | 0.404618 |
| 3 | 112960000 | 112962999 | 50.39413 | 81.16492 | 77.93472 | 33.13536 | 0.413564 | 1.826185 | 0.404621 |
| 13 | 74787400 | 74788599 | 8.506785 | 15.63209 | 32.99367 | 14.02368 | -0.64058 | 0.232476 | 0.404644 |
| 16 | 44648000 | 44649599 | 10.6257 | 19.04311 | 35.83322 | 15.69725 | -0.50526 | 0.404408 | 0.404646 |
| 5 | 95647600 | 95650399 | 59.43202 | 95.1742 | 91.06774 | 38.39038 | 0.408715 | 2.021303 | 0.404666 |
| 5 | 144026200 | 144031399 | 49.25987 | 78.47128 | 126.4909 | 52.92919 | -0.35069 | 2.183093 | 0.404695 |
| 8 | 69992800 | 69994999 | 23.14132 | 39.93281 | 39.42398 | 17.29487 | 0.403584 | 0.880049 | 0.404701 |
| 15 | 73314400 | 73316799 | 12.83637 | 22.54089 | 43.49503 | 19.24527 | -0.55614 | 0.607567 | 0.404705 |
| 9 | 11259000 | 11260199 | 15.1098 | 27.4289 | 26.7556 | 11.55338 | 0.450135 | 0.338632 | 0.404711 |
| 3 | 11589600 | 11590199 | 13.47109 | 24.8512 | 21.56528 | 9.117647 | 0.638845 | 0.224756 | 0.404713 |
| X | 123123000 | 123125799 | 25.09771 | 43.6467 | 33.38372 | 14.66593 | 0.772053 | 0.802536 | 0.404714 |
| 8 | 103977800 | 103980199 | 23.01087 | 40.50877 | 30.66649 | 13.57603 | 0.787724 | 0.612833 | 0.40472 |
| 14 | 34696800 | 34699599 | 18.78841 | 31.66338 | 59.61305 | 26.06145 | -0.52842 | 1.097185 | 0.404751 |
| 1 | 133058600 | 133060999 | 15.47206 | 26.63069 | 51.09517 | 22.41316 | -0.55172 | 0.884819 | 0.404751 |
| X | 117082800 | 117085199 | 19.00635 | 35.73455 | 19.62227 | 8.054273 | 1.326337 | 0.155226 | 0.404754 |
| 17 | 39343000 | 39347999 | 71.79167 | 114.9037 | 85.28971 | 35.92554 | 0.773521 | 2.051792 | 0.404759 |
| 6 | 112897400 | 112902799 | 42.3814 | 67.72205 | 118.0258 | 49.60035 | -0.45822 | 2.076238 | 0.404776 |
| 9 | 46572000 | 46575999 | 29.95044 | 48.5317 | 85.30858 | 36.51992 | -0.44713 | 1.639418 | 0.404776 |
| 11 | 58605600 | 58606399 | 8.84346 | 17.25003 | 16.13253 | 5.778107 | 0.740958 | -0.40749 | 0.404793 |
| 4 | 22477400 | 22480199 | 22.89803 | 38.00672 | 67.88769 | 29.42163 | -0.45333 | 1.276987 | 0.404799 |
| 2 | 168509200 | 168526799 | 128.3991 | 190.405 | 366.4341 | 150.9268 | -0.69255 | 3.676328 | 0.404828 |
| 5 | 91497400 | 91499399 | 11.50975 | 20.76067 | 39.53748 | 17.20968 | -0.51686 | 0.376092 | 0.404832 |
| 2 | 50273200 | 50274399 | 12.80448 | 24.58973 | 20.5186 | 8.03451 | 0.786743 | -0.05037 | 0.404833 |
| 17 | 40001600 | 40009999 | 103.5466 | 161.7828 | 139.172 | 57.74334 | 0.530499 | 2.667355 | 0.404836 |
| X | 19069400 | 19073599 | 36.92412 | 62.2229 | 44.08743 | 19.27364 | 0.876496 | 1.145554 | 0.404837 |
| 8 | 3139200 | 3139999 | 14.60058 | 26.44441 | 24.89269 | 10.77951 | 0.498667 | 0.343918 | 0.404849 |
| 11 | 16020800 | 16021599 | 14.60058 | 26.44441 | 24.89269 | 10.77951 | 0.498667 | 0.343918 | 0.404849 |
| 8 | 36411200 | 36412999 | 12.85791 | 22.61717 | 40.88971 | 18.10475 | -0.46197 | 0.58166 | 0.404894 |
| 8 | 64360000 | 64363399 | 42.35685 | 67.90508 | 109.9101 | 46.41145 | -0.34634 | 1.973471 | 0.404902 |
| 15 | 65045200 | 65048599 | 44.36092 | 70.87852 | 116.2738 | 48.85758 | -0.37121 | 2.08213 | 0.40491 |
| 16 | 37179600 | 37181399 | 33.2768 | 55.67502 | 50.28921 | 21.90802 | 0.527377 | 1.205869 | 0.404918 |
| X | 125109800 | 125122799 | 65.54317 | 104.8096 | 84.84422 | 35.6929 | 0.647244 | 2.070402 | 0.404945 |
| 2 | 36253200 | 36258399 | 169.683 | 254.767 | 240.0128 | 99.05748 | 0.353643 | 3.451917 | 0.404954 |
| 1 | 164360800 | 164363999 | 33.69303 | 54.38887 | 92.15781 | 39.30076 | -0.40013 | 1.742533 | 0.404954 |
| 6 | 14302200 | 14306599 | 109.7134 | 168.8628 | 164.3541 | 68.25988 | 0.335897 | 2.9372 | 0.404957 |

| | | | | | | | | | |
|---|---|---|---|---|---|---|---|---|---|
| 13 | 4265600 | 4267199 | 27.93132 | 47.87258 | 37.64656 | 16.5102 | 0.728547 | 0.956987 | 0.40496 |
| 19 | 9480400 | 9483799 | 27.93132 | 47.87258 | 37.64656 | 16.5102 | 0.728547 | 0.956987 | 0.40496 |
| 15 | 87340000 | 87341599 | 8.744279 | 16.39595 | 31.24695 | 12.8848 | -0.46071 | 0.111039 | 0.404968 |
| X | 9408000 | 9409599 | 27.31053 | 46.94309 | 40.38596 | 17.71477 | 0.600328 | 0.935724 | 0.404975 |
| 17 | 38675600 | 38690599 | 215.9846 | 319.6443 | 285.1714 | 117.4419 | 0.413219 | 3.719268 | 0.404975 |
| 2 | 17089200 | 17090399 | 28.82916 | 48.38484 | 48.05191 | 20.97472 | 0.390936 | 1.157216 | 0.404978 |
| 4 | 24461800 | 24465799 | 51.32472 | 82.95195 | 71.65401 | 30.60505 | 0.571679 | 1.725575 | 0.404985 |
| 13 | 10972400 | 10974399 | 25.11618 | 43.91867 | 33.46134 | 14.75295 | 0.77805 | 0.725506 | 0.404992 |
| 7 | 96784400 | 96784999 | 9.143264 | 17.77081 | 18.26462 | 6.792834 | 0.557898 | -0.26623 | 0.404998 |
| 6 | 76710200 | 76711599 | 17.03555 | 31.09857 | 25.98037 | 11.14025 | 0.678663 | 0.319196 | 0.404998 |
| 7 | 88979800 | 88980999 | 16.63976 | 29.77938 | 28.02528 | 12.29146 | 0.486761 | 0.434114 | 0.405012 |
| 8 | 88744600 | 88750399 | 35.49427 | 56.76336 | 122.0388 | 51.34623 | -0.75914 | 2.046759 | 0.405026 |
| 9 | 36209400 | 36211599 | 20.86443 | 36.97012 | 29.82068 | 13.18442 | 0.700281 | 0.531062 | 0.405033 |
| 12 | 100609000 | 100611999 | 31.42186 | 53.20922 | 45.24146 | 19.80738 | 0.614596 | 1.087019 | 0.405036 |
| 17 | 27123400 | 27134599 | 180.281 | 263.8233 | 395.224 | 162.8102 | -0.34944 | 3.897709 | 0.40504 |
| X | 53022800 | 53024799 | 47.72421 | 77.04492 | 73.3777 | 31.29525 | 0.429304 | 1.755658 | 0.405046 |
| 12 | 29576200 | 29578999 | 17.18297 | 29.52066 | 53.27577 | 23.36837 | -0.4645 | 0.91067 | 0.405073 |
| 13 | 23585400 | 23586599 | 18.93998 | 33.26774 | 32.37396 | 14.32774 | 0.426622 | 0.620674 | 0.405084 |
| 18 | 51940000 | 51941999 | 27.39253 | 46.87986 | 38.32291 | 16.80482 | 0.672648 | 0.970016 | 0.405095 |
| 5 | 41936000 | 41939999 | 33.81541 | 54.67099 | 91.72707 | 39.19527 | -0.38339 | 1.696491 | 0.4051 |
| 9 | 115185000 | 115187999 | 30.7166 | 49.79923 | 87.02286 | 37.26815 | -0.43804 | 1.629163 | 0.4051 |
| 16 | 51794800 | 51796799 | 26.74114 | 45.58244 | 43.15898 | 18.91937 | 0.460791 | 1.010201 | 0.4051 |
| 4 | 97798400 | 97799999 | 15.62764 | 28.03328 | 26.15605 | 11.44913 | 0.501007 | 0.408798 | 0.405102 |
| 17 | 12357400 | 12363799 | 57.52113 | 90.65251 | 164.1383 | 68.11106 | -0.53219 | 2.483546 | 0.405104 |
| X | 22249000 | 22251199 | 27.39841 | 46.67873 | 44.14456 | 19.35061 | 0.462274 | 1.016128 | 0.405113 |
| 2 | 38815000 | 38817399 | 16.74256 | 28.6364 | 57.04196 | 25.01204 | -0.60749 | 0.952536 | 0.405118 |
| 13 | 53423600 | 53428999 | 92.61356 | 145.5346 | 128.8259 | 53.4376 | 0.495661 | 2.554938 | 0.405136 |
| 5 | 145553200 | 145558799 | 148.541 | 225.112 | 212.7837 | 88.0603 | 0.360462 | 3.26163 | 0.405144 |
| 1 | 38712400 | 38713999 | 15.23075 | 28.57773 | 22.02673 | 9.072411 | 0.836537 | 0.138362 | 0.405156 |
| 1 | 89292400 | 89294199 | 21.57975 | 36.16752 | 62.36948 | 27.20044 | -0.40239 | 1.164097 | 0.405185 |
| 4 | 149175800 | 149177799 | 11.59619 | 20.69826 | 37.83025 | 16.62885 | -0.46708 | 0.440981 | 0.405186 |
| 9 | 78458600 | 78461999 | 20.14378 | 34.17257 | 59.80586 | 26.19819 | -0.42315 | 1.044282 | 0.405191 |
| 9 | 4498000 | 4499999 | 23.69379 | 41.0748 | 33.67854 | 14.77648 | 0.671641 | 0.833857 | 0.405194 |
| 10 | 113773200 | 113774599 | 14.56494 | 26.70189 | 25.76302 | 10.98534 | 0.477739 | 0.276298 | 0.40521 |
| 7 | 20885800 | 20890799 | 47.75263 | 77.97323 | 59.80467 | 25.70328 | 0.755028 | 1.53257 | 0.405212 |
| 9 | 25010800 | 25013399 | 40.30657 | 65.69743 | 64.76499 | 27.81705 | 0.392082 | 1.544345 | 0.405221 |
| 12 | 73098800 | 73102599 | 28.23801 | 46.04766 | 79.81075 | 34.2851 | -0.41938 | 1.521919 | 0.405224 |
| 18 | 50752600 | 50754799 | 22.54303 | 39.68988 | 31.40649 | 13.9033 | 0.724451 | 0.611861 | 0.40523 |
| 14 | 33676800 | 33678599 | 18.79323 | 31.30584 | 64.12354 | 27.85802 | -0.64955 | 1.22356 | 0.405256 |
| 19 | 8379200 | 8380799 | 18.184 | 33.63949 | 22.7137 | 9.566692 | 1.001708 | 0.248716 | 0.405265 |
| 7 | 42420400 | 42425799 | 89.86016 | 140.3165 | 139.008 | 57.67879 | 0.326404 | 2.670615 | 0.405269 |
| 4 | 20354800 | 20357199 | 21.63653 | 38.43159 | 28.59588 | 12.62153 | 0.817813 | 0.515848 | 0.405278 |
| 6 | 115127000 | 115128999 | 18.45805 | 33.40383 | 26.23212 | 11.3711 | 0.7573 | 0.378157 | 0.4053 |
| 2 | 14313800 | 14314399 | 4.680491 | 9.119355 | 21.93699 | 7.975198 | -0.62521 | -0.38829 | 0.405305 |
| 5 | 133940400 | 133940999 | 4.680491 | 9.119355 | 21.93699 | 7.975198 | -0.62521 | -0.38829 | 0.405305 |
| 6 | 130936200 | 130950599 | 223.7677 | 333.1622 | 249.3954 | 102.7628 | 0.674228 | 3.616304 | 0.405309 |
| 7 | 12119000 | 12119799 | 30.69489 | 51.13576 | 47.09282 | 20.46018 | 0.499725 | 1.241761 | 0.405312 |

| | | | | | | | | | |
|---|---|---|---|---|---|---|---|---|---|
| 13 | 33237400 | 33242599 | 62.01601 | 98.79076 | 97.94885 | 40.98531 | 0.349289 | 2.187549 | 0.405336 |
| 6 | 70896600 | 70898999 | 13.51089 | 23.5897 | 44.69939 | 19.68605 | -0.5301 | 0.706409 | 0.405348 |
| 1 | 15780600 | 15781799 | 10.2364 | 18.50242 | 35.34736 | 15.34574 | -0.51862 | 0.346389 | 0.405363 |
| 11 | 63090800 | 63093199 | 47.54283 | 76.70287 | 75.93338 | 32.35845 | 0.372512 | 1.772141 | 0.405384 |
| 4 | 124848400 | 124851399 | 17.07108 | 29.20706 | 53.76903 | 23.57747 | -0.49401 | 0.949174 | 0.405395 |
| 16 | 90338600 | 90340399 | 9.718697 | 17.38168 | 35.88052 | 15.74535 | -0.63968 | 0.410967 | 0.405413 |
| 5 | 45092200 | 45093599 | 16.88372 | 30.56999 | 26.36711 | 11.42249 | 0.623278 | 0.363474 | 0.405423 |
| 8 | 74274000 | 74275799 | 18.57358 | 33.92409 | 24.1086 | 10.33084 | 0.910858 | 0.323171 | 0.405435 |
| 10 | 38775600 | 38777999 | 28.87227 | 48.92251 | 45.78518 | 20.04925 | 0.476593 | 1.070505 | 0.405438 |
| 19 | 16648400 | 16650999 | 38.14153 | 61.24718 | 103.2982 | 43.72795 | -0.40271 | 1.915862 | 0.405438 |
| 13 | 89386600 | 89387399 | 6.137909 | 11.79359 | 25.74669 | 10.06611 | -0.58554 | -0.10548 | 0.405441 |
| 5 | 114685200 | 114699399 | 104.68 | 157.5927 | 302.0605 | 124.7554 | -0.66688 | 3.390254 | 0.405446 |
| 1 | 7846400 | 7848199 | 12.0624 | 21.54046 | 39.36567 | 17.29696 | -0.46674 | 0.443283 | 0.405446 |
| 15 | 25165600 | 25166599 | 10.97346 | 20.82574 | 21.06453 | 8.487834 | 0.474367 | 0.023894 | 0.405459 |
| 1 | 3740400 | 3744399 | 30.09588 | 48.77956 | 85.50692 | 36.61156 | -0.44283 | 1.633852 | 0.405462 |
| 6 | 131667800 | 131669999 | 43.58973 | 70.9041 | 66.15743 | 28.394 | 0.469167 | 1.58047 | 0.405471 |
| 8 | 19229600 | 19235999 | 81.56178 | 128.798 | 122.6464 | 50.92685 | 0.395932 | 2.449012 | 0.405488 |
| X | 24006400 | 24008599 | 28.48219 | 49.95253 | 28.85294 | 12.75306 | 1.175471 | 0.693134 | 0.40549 |
| 3 | 16798000 | 16801799 | 38.091 | 63.43316 | 50.41991 | 21.90044 | 0.711411 | 1.269054 | 0.40551 |
| 1 | 156326400 | 156329999 | 31.3505 | 50.89086 | 85.56013 | 36.6757 | -0.38073 | 1.604814 | 0.40553 |
| 5 | 6099200 | 6099999 | 14.50767 | 26.3903 | 24.4611 | 10.53533 | 0.52597 | 0.318548 | 0.405534 |
| X | 5696800 | 5700599 | 64.63296 | 103.8435 | 78.96126 | 33.4529 | 0.745232 | 1.926198 | 0.405536 |
| 19 | 44182200 | 44186999 | 30.56569 | 49.54601 | 86.2983 | 36.95315 | -0.43355 | 1.632464 | 0.405543 |
| 9 | 117868600 | 117870199 | 10.39072 | 18.37094 | 38.90215 | 17.20714 | -0.68556 | 0.506573 | 0.405547 |
| 18 | 37519400 | 37520399 | 36.16722 | 59.2014 | 56.75706 | 24.41535 | 0.436477 | 1.470319 | 0.40555 |
| 17 | 57115200 | 57129599 | 80.38749 | 123.3604 | 217.7109 | 90.38342 | -0.52509 | 2.990532 | 0.405551 |
| X | 36007200 | 36009199 | 20.13982 | 35.7099 | 29.50045 | 13.0402 | 0.666526 | 0.52059 | 0.40556 |
| 7 | 38094600 | 38097199 | 40.93145 | 66.4903 | 66.32275 | 28.44428 | 0.371512 | 1.597446 | 0.405582 |
| 3 | 67125600 | 67126799 | 9.929707 | 18.03864 | 33.87725 | 14.61521 | -0.48793 | 0.313244 | 0.405597 |
| 14 | 112845200 | 112846199 | 14.30313 | 27.12592 | 21.20082 | 8.55538 | 0.841292 | 0.059653 | 0.405628 |
| 13 | 108592600 | 108595999 | 59.41108 | 94.90292 | 95.28675 | 40.02564 | 0.335429 | 2.098144 | 0.405638 |
| 8 | 87949200 | 87950599 | 7.072973 | 13.33951 | 29.70952 | 12.13081 | -0.67226 | 0.053207 | 0.405643 |
| X | 5056800 | 5058199 | 46.94691 | 76.64369 | 60.87392 | 26.1611 | 0.704612 | 1.534219 | 0.40566 |
| 7 | 69579000 | 69579999 | 9.230228 | 17.09909 | 32.05286 | 13.47484 | -0.45906 | 0.191558 | 0.405684 |
| 4 | 151517000 | 151531399 | 114.9814 | 173.1788 | 279.4364 | 115.4251 | -0.41812 | 3.382006 | 0.405687 |
| 11 | 89950200 | 89951199 | 11.74696 | 22.41206 | 21.55325 | 8.588054 | 0.56049 | -0.00363 | 0.40569 |
| 6 | 109682400 | 109683399 | 12.33978 | 23.66317 | 21.10072 | 8.296718 | 0.686342 | -0.0402 | 0.4057 |
| 17 | 25234200 | 25236199 | 9.712685 | 17.37007 | 36.36037 | 15.95659 | -0.65992 | 0.410927 | 0.405708 |
| 15 | 86476600 | 86478199 | 11.80288 | 20.74377 | 40.46985 | 17.91966 | -0.57138 | 0.58623 | 0.405738 |
| 6 | 76286400 | 76288599 | 25.67794 | 44.77042 | 34.5869 | 15.21902 | 0.757878 | 0.770322 | 0.40574 |
| X | 44732800 | 44736799 | 60.8663 | 97.43558 | 83.86259 | 35.3337 | 0.560609 | 2.031764 | 0.405776 |
| 19 | 13709600 | 13710199 | 10.08476 | 19.57856 | 18.79991 | 7.05602 | 0.643916 | -0.21942 | 0.405785 |
| 5 | 117311400 | 117312999 | 9.694043 | 17.46498 | 34.89344 | 15.21067 | -0.58602 | 0.36446 | 0.405792 |
| 14 | 101777400 | 101778799 | 8.979235 | 16.45228 | 32.43527 | 13.84107 | -0.54679 | 0.253155 | 0.405795 |
| 15 | 19376000 | 19378799 | 15.51142 | 26.97839 | 51.15433 | 22.47435 | -0.5321 | 0.777773 | 0.405806 |
| 9 | 49050000 | 49052199 | 21.82493 | 38.75565 | 28.59023 | 12.62194 | 0.829942 | 0.518852 | 0.405806 |
| 13 | 58292400 | 58294999 | 38.12562 | 62.4437 | 61.46762 | 26.44825 | 0.398784 | 1.469968 | 0.405832 |

| | | | | | | | | | |
|---|---|---|---|---|---|---|---|---|---|
| 1 | 114771800 | 114772399 | 5.659654 | 11.01606 | 24.30627 | 8.934413 | -0.5173 | -0.32913 | 0.405835 |
| 7 | 10150800 | 10152799 | 28.39309 | 48.32543 | 44.17074 | 19.36054 | 0.511142 | 1.030806 | 0.40584 |
| 18 | 19916800 | 19918199 | 15.49594 | 29.12823 | 22.07906 | 9.065999 | 0.864161 | 0.130236 | 0.405843 |
| 17 | 19096400 | 19105399 | 134.3711 | 206.206 | 173.6116 | 72.07561 | 0.541679 | 3.005781 | 0.405865 |
| 10 | 66245200 | 66247999 | 34.85603 | 57.66938 | 55.40264 | 23.95759 | 0.438258 | 1.335189 | 0.40588 |
| 13 | 60473600 | 60477199 | 44.78859 | 72.32451 | 72.09335 | 30.75779 | 0.364108 | 1.740739 | 0.405887 |
| 16 | 87859000 | 87860399 | 8.264378 | 15.51684 | 30.1709 | 12.41297 | -0.48681 | 0.094836 | 0.40589 |
| 12 | 93269200 | 93271399 | 15.54988 | 27.14957 | 48.93499 | 21.55136 | -0.45858 | 0.722392 | 0.405896 |
| 18 | 3814800 | 3819199 | 68.7154 | 109.0234 | 108.3243 | 45.11541 | 0.340391 | 2.318679 | 0.405896 |
| 16 | 54181200 | 54183399 | 20.51368 | 36.24547 | 30.98437 | 13.7067 | 0.615434 | 0.561044 | 0.4059 |
| 7 | 15393800 | 15415599 | 430.0695 | 601.1048 | 489.8993 | 203.1519 | 0.462594 | 4.590522 | 0.405905 |
| 2 | 23593000 | 23594399 | 33.50023 | 55.42847 | 55.25459 | 23.89425 | 0.385161 | 1.332834 | 0.40592 |
| 19 | 44813800 | 44814599 | 4.982307 | 9.654975 | 23.79247 | 9.011298 | -0.7147 | -0.23275 | 0.405922 |
| 8 | 15950600 | 15951799 | 9.380962 | 17.40933 | 32.63303 | 13.6855 | -0.45653 | 0.183993 | 0.405925 |
| 5 | 41019600 | 41020599 | 7.008986 | 13.51697 | 28.31169 | 10.95257 | -0.51306 | -0.12606 | 0.405941 |
| 15 | 12321200 | 12322399 | 7.923163 | 14.5799 | 31.98546 | 13.56943 | -0.69356 | 0.218828 | 0.405943 |
| 10 | 129038200 | 129043999 | 69.92944 | 111.1081 | 109.1109 | 45.50799 | 0.35923 | 2.277332 | 0.405943 |
| 14 | 76651400 | 76656399 | 31.03916 | 49.6577 | 126.7246 | 53.34672 | -1.00527 | 2.030439 | 0.405956 |
| 2 | 181137800 | 181139599 | 8.322415 | 15.41206 | 31.52048 | 13.25723 | -0.58426 | 0.184166 | 0.405972 |
| 1 | 190278000 | 190280799 | 20.42217 | 34.31037 | 60.15982 | 26.27582 | -0.42631 | 1.131928 | 0.405997 |
| 7 | 93983600 | 93984799 | 14.14647 | 26.39371 | 23.75419 | 9.877994 | 0.603778 | 0.168425 | 0.405997 |
| 9 | 96563400 | 96564799 | 10.43062 | 19.0987 | 42.32515 | 18.07795 | -0.71617 | 0.288458 | 0.406003 |
| 5 | 72752000 | 72753399 | 16.21528 | 28.89208 | 27.72654 | 12.20571 | 0.455247 | 0.458702 | 0.40601 |
| 15 | 7910000 | 7911599 | 16.21528 | 28.89208 | 27.72654 | 12.20571 | 0.455247 | 0.458702 | 0.40601 |
| 8 | 92852400 | 92854399 | 10.70431 | 19.36995 | 35.35434 | 15.3272 | -0.45186 | 0.343995 | 0.406024 |
| 3 | 139527000 | 139527999 | 11.90306 | 21.23352 | 38.32099 | 16.8529 | -0.44955 | 0.447073 | 0.406025 |
| 7 | 47375600 | 47375999 | 5.6751 | 11.01615 | 24.54339 | 9.221728 | -0.56068 | -0.25136 | 0.406037 |
| X | 66184800 | 66188199 | 45.38281 | 73.53183 | 69.76007 | 29.86177 | 0.440289 | 1.661392 | 0.406047 |
| 13 | 65942000 | 65943799 | 17.012 | 32.29597 | 20.00235 | 8.054492 | 1.17752 | 0.076862 | 0.406053 |
| 2 | 98747400 | 98749599 | 24.26134 | 42.39724 | 33.473 | 14.75026 | 0.726815 | 0.729318 | 0.406084 |
| 11 | 5178800 | 5179999 | 17.84928 | 31.61114 | 30.21541 | 13.36032 | 0.456435 | 0.518362 | 0.406088 |
| 12 | 114407400 | 114412399 | 61.45365 | 98.99967 | 73.27421 | 31.16401 | 0.788755 | 1.837778 | 0.406112 |
| 1 | 72434600 | 72438799 | 31.73943 | 51.29781 | 89.87704 | 38.38849 | -0.44621 | 1.702101 | 0.406112 |
| 5 | 85266200 | 85267199 | 13.03496 | 24.81467 | 22.04598 | 8.825438 | 0.668425 | 0.026177 | 0.406113 |
| 5 | 104093800 | 104105199 | 140.7737 | 210.1863 | 318.6779 | 131.3685 | -0.33967 | 3.554057 | 0.406129 |
| 15 | 88273400 | 88279799 | 66.41216 | 104.306 | 162.9054 | 67.56928 | -0.32304 | 2.556994 | 0.406135 |
| 10 | 62447000 | 62452399 | 65.60593 | 104.5361 | 101.8544 | 42.63458 | 0.374776 | 2.183394 | 0.406137 |
| 11 | 22684000 | 22698599 | 135.1805 | 202.3938 | 306.89 | 126.5771 | -0.33617 | 3.501414 | 0.406167 |
| 2 | 95966800 | 95967599 | 8.09722 | 15.39764 | 31.79358 | 12.75742 | -0.54194 | 0.011638 | 0.406189 |
| 9 | 57384200 | 57392999 | 72.32444 | 112.0596 | 203.5516 | 84.54968 | -0.55598 | 2.804444 | 0.406189 |
| 7 | 68030800 | 68032999 | 12.5365 | 22.05828 | 40.89001 | 18.10382 | -0.49775 | 0.575961 | 0.406196 |
| 9 | 83764200 | 83765999 | 12.5365 | 22.05828 | 40.89001 | 18.10382 | -0.49775 | 0.575961 | 0.406196 |
| 7 | 71199000 | 71201599 | 24.55296 | 42.47715 | 35.17103 | 15.43035 | 0.657015 | 0.858051 | 0.406199 |
| 15 | 85212800 | 85213999 | 15.22223 | 27.68784 | 26.34096 | 11.34616 | 0.488502 | 0.328743 | 0.406202 |
| 10 | 55991000 | 55993599 | 16.68999 | 28.79621 | 51.13221 | 22.43105 | -0.43983 | 0.867964 | 0.406209 |
| X | 116872400 | 116875399 | 46.93388 | 76.26342 | 65.59237 | 28.13718 | 0.585266 | 1.606356 | 0.406212 |
| 6 | 89065000 | 89073599 | 84.93942 | 130.7079 | 208.0784 | 86.41729 | -0.37343 | 2.937083 | 0.406221 |

| | | | | | | | | | |
|---|---|---|---|---|---|---|---|---|---|
| 2 | 114380200 | 114381799 | 17.09741 | 31.75359 | 23.47735 | 9.835732 | 0.877307 | 0.220956 | 0.406229 |
| 7 | 103805400 | 103805799 | 11.81922 | 22.29195 | 22.53747 | 9.201441 | 0.458535 | 0.078384 | 0.40623 |
| 5 | 3887800 | 3889399 | 20.032 | 36.31714 | 25.69213 | 11.11213 | 0.909341 | 0.380033 | 0.406236 |
| X | 54380000 | 54388799 | 115.314 | 180.2689 | 127.7511 | 53.00169 | 0.810427 | 2.663942 | 0.406258 |
| 4 | 114353400 | 114354399 | 7.335452 | 14.0179 | 27.84944 | 11.05906 | -0.47274 | -0.03774 | 0.406261 |
| 18 | 17557800 | 17558599 | 7.335452 | 14.0179 | 27.84944 | 11.05906 | -0.47274 | -0.03774 | 0.406261 |
| 3 | 67033600 | 67037399 | 58.69608 | 94.48701 | 77.61047 | 32.9727 | 0.6375 | 1.85697 | 0.406262 |
| 7 | 20559400 | 20571399 | 194.0939 | 289.8393 | 255.0849 | 105.1565 | 0.444741 | 3.558331 | 0.406266 |
| 1 | 48858800 | 48860599 | 24.22787 | 42.24169 | 35.80286 | 15.75396 | 0.624804 | 0.765297 | 0.406285 |
| 19 | 45174400 | 45178799 | 63.74812 | 100.2068 | 160.9418 | 66.76222 | -0.36238 | 2.537597 | 0.406315 |
| 3 | 4320000 | 4322799 | 31.67979 | 54.31959 | 37.25884 | 16.3407 | 0.924946 | 0.967421 | 0.406315 |
| 19 | 11243000 | 11244199 | 11.41481 | 21.51146 | 20.99293 | 8.582963 | 0.507412 | 0.075104 | 0.406318 |
| 1 | 23275400 | 23276199 | 13.78426 | 25.90651 | 23.40189 | 9.611821 | 0.612913 | 0.12011 | 0.406334 |
| 17 | 29097200 | 29125599 | 272.0937 | 380.1027 | 671.5206 | 278.4896 | -0.654 | 4.59381 | 0.406337 |
| 11 | 9033400 | 9040199 | 75.26726 | 119.0624 | 118.7552 | 49.35396 | 0.331098 | 2.402055 | 0.406353 |
| 11 | 43427600 | 43429599 | 8.444679 | 15.33915 | 33.98364 | 14.66319 | -0.72393 | 0.30703 | 0.406356 |
| 17 | 24457600 | 24464399 | 51.73521 | 82.23863 | 132.4607 | 55.27267 | -0.35284 | 2.254248 | 0.406368 |
| 14 | 42590600 | 42591999 | 23.62291 | 41.2021 | 32.20212 | 14.17284 | 0.741194 | 0.75158 | 0.406371 |
| 1 | 155965000 | 155965999 | 11.79507 | 22.11543 | 21.04539 | 8.678742 | 0.533873 | 0.111612 | 0.406372 |
| 10 | 123628400 | 123630199 | 19.78097 | 34.97231 | 30.98821 | 13.70672 | 0.564194 | 0.549331 | 0.406375 |
| 4 | 29779200 | 29779799 | 9.611688 | 18.62383 | 19.26993 | 7.302447 | 0.523388 | -0.19381 | 0.406381 |
| 15 | 42074800 | 42075399 | 8.457336 | 15.93341 | 31.369 | 12.83153 | -0.49767 | 0.079753 | 0.406387 |
| 8 | 24768000 | 24768999 | 5.996996 | 11.5405 | 28.1491 | 10.95707 | -0.73962 | -0.12023 | 0.406388 |
| 2 | 129061600 | 129063599 | 12.84347 | 22.78875 | 44.56178 | 19.68616 | -0.56963 | 0.495612 | 0.406408 |
| 16 | 84784000 | 84785399 | 10.08074 | 18.42008 | 33.94859 | 14.5396 | -0.45407 | 0.281424 | 0.406411 |
| 14 | 118650800 | 118653799 | 21.86675 | 36.57879 | 63.54235 | 27.67867 | -0.41298 | 1.184518 | 0.406438 |
| X | 67923000 | 67923999 | 13.46419 | 25.32081 | 23.00622 | 9.4393 | 0.606213 | 0.114266 | 0.406444 |
| 5 | 95486200 | 95488399 | 28.18111 | 48.30105 | 37.05903 | 16.25254 | 0.763892 | 0.956502 | 0.406448 |
| 10 | 129341600 | 129344799 | 68.16417 | 108.2797 | 108.798 | 45.36754 | 0.325922 | 2.282592 | 0.406457 |
| 8 | 50696000 | 50697199 | 7.15656 | 13.70417 | 27.96037 | 11.03886 | -0.50341 | -0.06059 | 0.406459 |
| 5 | 23580000 | 23581999 | 12.99131 | 22.93159 | 40.83598 | 18.06837 | -0.43838 | 0.543422 | 0.406461 |
| 15 | 34978600 | 34981199 | 32.45252 | 55.9453 | 36.18922 | 15.87535 | 1.010744 | 0.926138 | 0.406477 |
| 2 | 65068200 | 65069799 | 12.10433 | 21.24337 | 41.50301 | 18.35507 | -0.57359 | 0.611631 | 0.406489 |
| 1 | 101942000 | 101947999 | 80.06708 | 127.3862 | 100.912 | 42.15208 | 0.670642 | 2.24659 | 0.406498 |
| 9 | 108182400 | 108184199 | 10.6148 | 18.6087 | 42.61112 | 18.82761 | -0.80122 | 0.623369 | 0.406503 |
| 1 | 50834200 | 50835599 | 9.35368 | 17.35298 | 33.73575 | 14.15431 | -0.50953 | 0.186545 | 0.40651 |
| 2 | 10321000 | 10323599 | 15.84863 | 27.53278 | 49.53069 | 21.74508 | -0.45663 | 0.794941 | 0.406514 |
| 5 | 17207000 | 17207399 | 4.056893 | 7.91944 | 19.80397 | 7.005622 | -0.64238 | -0.49791 | 0.406521 |
| 5 | 112863000 | 112863599 | 4.056893 | 7.91944 | 19.80397 | 7.005622 | -0.64238 | -0.49791 | 0.406521 |
| 16 | 84058000 | 84059199 | 13.81122 | 26.69586 | 16.36864 | 6.273958 | 1.254706 | -0.1191 | 0.406523 |
| 9 | 31473400 | 31476199 | 18.83992 | 32.17268 | 55.42188 | 24.29855 | -0.39904 | 0.975968 | 0.406531 |
| 18 | 13595000 | 13596199 | 13.14676 | 25.10667 | 19.62973 | 7.805858 | 0.858558 | -0.00275 | 0.406535 |
| 9 | 115266200 | 115269599 | 62.78286 | 99.9876 | 96.27552 | 40.27134 | 0.39109 | 2.19715 | 0.406539 |
| 1 | 44405000 | 44408399 | 55.49909 | 88.98431 | 88.21011 | 37.25464 | 0.359852 | 1.978502 | 0.406552 |
| 10 | 61184200 | 61187399 | 30.16606 | 48.72001 | 90.26865 | 38.52017 | -0.52781 | 1.721775 | 0.40656 |
| 2 | 164931400 | 164932799 | 8.830902 | 16.32785 | 32.03571 | 13.50298 | -0.5266 | 0.202056 | 0.406577 |
| 8 | 73651800 | 73654599 | 88.5311 | 138.3758 | 135.1697 | 56.08054 | 0.347771 | 2.652663 | 0.406586 |

| | | | | | | | | | |
|---|---|---|---|---|---|---|---|---|---|
| 11 | 14761000 | 14761999 | 14.58734 | 26.46229 | 25.81199 | 11.15549 | 0.449375 | 0.336407 | 0.406604 |
| 13 | 15691600 | 15693999 | 11.36121 | 20.44151 | 36.82104 | 16.07658 | -0.43967 | 0.38515 | 0.406643 |
| 12 | 108347200 | 108349199 | 12.36435 | 21.75062 | 42.94788 | 19.01576 | -0.58871 | 0.579102 | 0.406645 |
| 15 | 55525600 | 55529399 | 32.78736 | 55.05302 | 48.58081 | 21.21167 | 0.5608 | 1.165655 | 0.406656 |
| 15 | 87835800 | 87836799 | 10.67276 | 20.31282 | 20.60158 | 8.251442 | 0.478149 | 0.000645 | 0.406666 |
| 4 | 153192200 | 153197599 | 52.57251 | 83.54736 | 132.9934 | 55.48017 | -0.33626 | 2.263283 | 0.40667 |
| 3 | 19241600 | 19242799 | 11.16803 | 19.71235 | 38.92269 | 17.22197 | -0.58608 | 0.531785 | 0.40669 |
| 2 | 71750400 | 71752399 | 19.81195 | 33.6795 | 58.70535 | 25.72872 | -0.4171 | 1.023829 | 0.406693 |
| 18 | 72590000 | 72592999 | 41.1059 | 66.62662 | 64.32646 | 27.54331 | 0.415524 | 1.641087 | 0.406694 |
| 9 | 7772200 | 7774799 | 13.47583 | 23.65801 | 42.45594 | 18.78163 | -0.45191 | 0.614806 | 0.406697 |
| 7 | 79090600 | 79092799 | 19.65906 | 34.37551 | 33.5567 | 14.79473 | 0.422233 | 0.697519 | 0.406718 |
| 13 | 34851200 | 34853599 | 28.02494 | 47.20917 | 46.43131 | 20.29607 | 0.405022 | 1.12001 | 0.406736 |
| X | 153599600 | 153602999 | 37.65306 | 62.91579 | 48.67861 | 21.18916 | 0.750065 | 1.23204 | 0.406737 |
| 13 | 62112800 | 62118399 | 109.5941 | 169.1502 | 165.4969 | 68.74418 | 0.331432 | 2.885901 | 0.406742 |
| 6 | 34349800 | 34352799 | 20.07967 | 33.34923 | 65.80946 | 28.53306 | -0.59632 | 1.262238 | 0.406744 |
| 1 | 101264000 | 101266999 | 32.86858 | 54.66354 | 53.04115 | 23.01859 | 0.424514 | 1.271533 | 0.406746 |
| 13 | 3208800 | 3210599 | 32.86858 | 54.66354 | 53.04115 | 23.01859 | 0.424514 | 1.271533 | 0.406746 |
| 2 | 25910800 | 25912399 | 7.595579 | 14.08795 | 30.51633 | 12.80775 | -0.66367 | 0.16914 | 0.406751 |
| 17 | 22613000 | 22617199 | 72.19749 | 114.8014 | 107.3798 | 44.82103 | 0.430472 | 2.257628 | 0.406756 |
| X | 113093400 | 113099799 | 81.10411 | 128.1745 | 120.2927 | 49.96908 | 0.417747 | 2.430644 | 0.40676 |
| 15 | 89505400 | 89507999 | 26.53631 | 43.1853 | 80.13858 | 34.40056 | -0.51956 | 1.550342 | 0.406772 |
| 13 | 3914800 | 3916799 | 12.43004 | 21.78361 | 42.063 | 18.58012 | -0.55676 | 0.638696 | 0.406795 |
| 15 | 33204000 | 33205799 | 25.37673 | 44.50518 | 31.81385 | 14.06154 | 0.869578 | 0.68065 | 0.40682 |
| 12 | 59651200 | 59654199 | 47.18101 | 76.26136 | 72.87678 | 31.13077 | 0.426406 | 1.716197 | 0.40683 |
| 14 | 13713800 | 13715199 | 14.58362 | 26.9228 | 25.28843 | 10.68138 | 0.52598 | 0.237265 | 0.406833 |
| 13 | 97892800 | 97893999 | 6.250684 | 12.1117 | 25.64658 | 9.718039 | -0.49989 | -0.21244 | 0.406836 |
| 11 | 13905400 | 13906999 | 17.21504 | 31.7951 | 24.57417 | 10.37328 | 0.806252 | 0.255116 | 0.40685 |
| 10 | 59312200 | 59315799 | 42.56324 | 69.80455 | 59.49248 | 25.61279 | 0.606898 | 1.464832 | 0.406853 |
| 4 | 37337600 | 37339399 | 19.61297 | 34.61921 | 30.92613 | 13.68369 | 0.551468 | 0.563772 | 0.406889 |
| 10 | 45857000 | 45858399 | 8.707387 | 16.35444 | 33.02874 | 13.58055 | -0.54086 | 0.101917 | 0.4069 |
| 7 | 20525200 | 20529599 | 76.1195 | 120.9416 | 108.1708 | 45.11525 | 0.493848 | 2.283621 | 0.406902 |
| 17 | 22609800 | 22610799 | 24.77279 | 42.24772 | 39.51642 | 17.32327 | 0.478251 | 0.990224 | 0.406902 |
| 5 | 62701800 | 62702599 | 14.44114 | 26.79746 | 24.82327 | 10.41167 | 0.553148 | 0.207031 | 0.40693 |
| 6 | 132369600 | 132371799 | 24.44609 | 43.20576 | 28.4618 | 12.59007 | 0.98952 | 0.577946 | 0.406934 |
| 4 | 138225000 | 138229999 | 49.10588 | 78.196 | 126.6932 | 52.98569 | -0.35864 | 2.195416 | 0.406945 |
| 13 | 8317600 | 8320199 | 50.20444 | 80.835 | 79.8772 | 33.94639 | 0.371753 | 1.837842 | 0.406954 |
| 2 | 23376400 | 23381999 | 79.90239 | 126.7267 | 111.91 | 46.59189 | 0.509735 | 2.338376 | 0.406972 |
| 5 | 91203600 | 91205199 | 15.71724 | 29.38795 | 23.12821 | 9.582466 | 0.800298 | 0.167552 | 0.406974 |
| 1 | 39492600 | 39495599 | 20.74713 | 34.71249 | 63.47621 | 27.65318 | -0.48677 | 1.178333 | 0.406984 |
| 1 | 182587800 | 182595199 | 42.83848 | 68.19599 | 134.6398 | 56.28652 | -0.64374 | 2.202058 | 0.407007 |
| 1 | 11962600 | 11964999 | 21.45052 | 37.96433 | 29.05875 | 12.85055 | 0.774611 | 0.54354 | 0.407021 |
| 17 | 71355800 | 71357399 | 19.21532 | 33.65921 | 30.63051 | 13.52485 | 0.523048 | 0.662982 | 0.407024 |
| 13 | 67760600 | 67765399 | 76.4315 | 121.5745 | 104.0427 | 43.4462 | 0.559011 | 2.249828 | 0.407029 |
| 8 | 14355400 | 14357599 | 177.0342 | 257.8891 | 411.6054 | 169.6277 | -0.44716 | 3.968212 | 0.40704 |
| 17 | 59320000 | 59320399 | 3.986808 | 7.782807 | 20.42271 | 7.221607 | -0.71201 | -0.5023 | 0.407044 |
| 12 | 91297400 | 91299599 | 14.64046 | 25.02973 | 56.83574 | 24.9207 | -0.79531 | 0.952945 | 0.407051 |
| 19 | 39743600 | 39748399 | 51.7483 | 83.55055 | 74.29708 | 31.69188 | 0.528225 | 1.757849 | 0.407058 |

| | | | | | | | | | |
|---|---|---|---|---|---|---|---|---|---|
| 14 | 11390200 | 11392599 | 28.41413 | 48.51147 | 42.42892 | 18.60172 | 0.574953 | 0.997951 | 0.407077 |
| 4 | 153054600 | 153066999 | 108.32 | 164.4432 | 255.2121 | 105.6748 | -0.35228 | 3.219225 | 0.407091 |
| 4 | 24283600 | 24284999 | 45.80941 | 73.89682 | 70.71324 | 30.12895 | 0.421214 | 1.772306 | 0.407096 |
| 4 | 4096400 | 4096999 | 5.257098 | 10.20711 | 23.94009 | 8.978295 | -0.63124 | -0.26448 | 0.407104 |
| 4 | 77514800 | 77515599 | 12.39302 | 23.14502 | 22.92349 | 9.516649 | 0.468649 | 0.145577 | 0.407139 |
| 2 | 3872800 | 3874599 | 14.78623 | 25.67912 | 47.39348 | 20.80271 | -0.49321 | 0.793044 | 0.407147 |
| X | 26104600 | 26104999 | 12.84627 | 24.12363 | 23.5031 | 9.666956 | 0.503124 | 0.117816 | 0.407166 |
| 2 | 87309600 | 87310399 | 8.598196 | 16.7073 | 16.5739 | 6.176299 | 0.605507 | -0.27274 | 0.407168 |
| 1 | 7465800 | 7469199 | 27.89921 | 45.15156 | 95.38132 | 40.79445 | -0.71338 | 1.665259 | 0.407171 |
| 19 | 4763800 | 4767799 | 27.02168 | 43.93987 | 80.78433 | 34.6676 | -0.50705 | 1.562514 | 0.407186 |
| 3 | 32816800 | 32819599 | 16.83401 | 29.16832 | 51.16077 | 22.44648 | -0.42093 | 0.8306 | 0.407188 |
| 10 | 64590600 | 64592599 | 13.5403 | 23.89917 | 46.68104 | 20.65484 | -0.57122 | 0.558322 | 0.407196 |
| 12 | 104007600 | 104013599 | 45.88358 | 73.10258 | 123.1219 | 51.52894 | -0.41362 | 2.175826 | 0.407198 |
| 4 | 40528600 | 40529799 | 7.961941 | 15.14904 | 30.60318 | 12.26494 | -0.50874 | 0.004929 | 0.40723 |
| 5 | 131431400 | 131435399 | 39.23199 | 63.09416 | 102.8201 | 43.62261 | -0.3506 | 1.868379 | 0.407267 |
| 13 | 77125800 | 77126999 | 13.38269 | 24.7701 | 24.34981 | 10.2486 | 0.46432 | 0.210554 | 0.407278 |
| 10 | 120379000 | 120379799 | 14.00077 | 25.60213 | 23.97205 | 10.2572 | 0.517481 | 0.283566 | 0.407284 |
| 15 | 40067000 | 40069199 | 27.30915 | 46.76804 | 42.49648 | 18.63594 | 0.520821 | 0.968812 | 0.407291 |
| 7 | 12340000 | 12340599 | 11.47221 | 21.63134 | 19.60152 | 8.006736 | 0.614998 | 0.069668 | 0.407292 |
| 8 | 52708800 | 52709999 | 13.12063 | 24.20172 | 22.97922 | 9.717067 | 0.509529 | 0.225306 | 0.407303 |
| 7 | 7776400 | 7777599 | 12.8478 | 24.46528 | 22.05408 | 8.823403 | 0.648298 | 0.022334 | 0.407308 |
| Y | 1763400 | 1765599 | 17.2767 | 30.89759 | 16.97717 | 7.450335 | 1.257025 | 0.416559 | 0.407313 |
| 18 | 18968600 | 18970599 | 20.94556 | 36.67934 | 34.28308 | 15.1339 | 0.484493 | 0.681291 | 0.407317 |
| 16 | 46141400 | 46143799 | 53.48533 | 85.78968 | 86.21499 | 36.43407 | 0.340818 | 1.962612 | 0.407323 |
| 6 | 28829800 | 28834199 | 31.19344 | 50.2028 | 99.2957 | 42.17179 | -0.62797 | 1.83243 | 0.407328 |
| 9 | 40204800 | 40205999 | 8.870195 | 16.24467 | 32.98131 | 14.0833 | -0.5891 | 0.259456 | 0.407332 |
| 17 | 52401600 | 52403599 | 11.91266 | 21.21397 | 38.31451 | 16.87462 | -0.45205 | 0.458244 | 0.407334 |
| 11 | 63515800 | 63517799 | 16.12531 | 29.41897 | 26.02151 | 11.1664 | 0.595973 | 0.31511 | 0.407341 |
| 3 | 114644200 | 114645799 | 17.23085 | 31.53194 | 24.81588 | 10.60768 | 0.766982 | 0.303713 | 0.407348 |
| 15 | 19926400 | 19928799 | 25.54226 | 44.32011 | 38.05821 | 16.69854 | 0.605476 | 0.839551 | 0.407353 |
| 10 | 33561400 | 33568599 | 77.52005 | 122.6136 | 107.9227 | 44.84976 | 0.511111 | 2.403869 | 0.40736 |
| 11 | 85785600 | 85793199 | 60.45632 | 95.20532 | 154.1691 | 63.96787 | -0.37216 | 2.498887 | 0.407378 |
| 10 | 15453600 | 15455599 | 15.67566 | 29.10612 | 24.11649 | 10.10679 | 0.713645 | 0.211473 | 0.407384 |
| 7 | 59560800 | 59564399 | 48.3901 | 78.81367 | 62.07353 | 26.65372 | 0.714338 | 1.570253 | 0.407408 |
| 1 | 156590600 | 156595799 | 86.44177 | 133.3382 | 203.9291 | 84.70845 | -0.31305 | 2.89239 | 0.407432 |
| 15 | 93954400 | 93956399 | 32.21166 | 52.22719 | 87.0208 | 37.26942 | -0.36936 | 1.632712 | 0.407456 |
| 6 | 144227600 | 144229399 | 18.02375 | 30.96343 | 53.27576 | 23.36832 | -0.3959 | 0.916398 | 0.407467 |
| 8 | 106562000 | 106562999 | 6.282618 | 12.06985 | 26.33066 | 10.29881 | -0.58598 | -0.10412 | 0.407474 |
| 9 | 102339000 | 102340799 | 11.09111 | 19.8601 | 36.3123 | 15.91867 | -0.46491 | 0.413719 | 0.407486 |
| 7 | 103007600 | 103008599 | 13.99041 | 24.47913 | 43.60632 | 19.23593 | -0.44122 | 0.675309 | 0.407487 |
| 14 | 95554400 | 95556599 | 17.76989 | 32.10824 | 27.46266 | 11.92781 | 0.633247 | 0.385603 | 0.407488 |
| X | 31953600 | 31955199 | 22.12451 | 39.6513 | 26.51772 | 11.61639 | 0.97822 | 0.46234 | 0.407491 |
| 3 | 61352000 | 61352799 | 16.06295 | 27.90294 | 48.45274 | 21.27079 | -0.40571 | 0.797696 | 0.407501 |
| X | 148830800 | 148833399 | 34.58004 | 57.9788 | 43.71947 | 19.07159 | 0.786876 | 1.182438 | 0.407507 |
| 9 | 61025400 | 61026799 | 8.825776 | 16.21319 | 31.95333 | 13.58727 | -0.54245 | 0.236815 | 0.407513 |
| 7 | 108815600 | 108821599 | 35.33248 | 56.89033 | 97.54718 | 41.45678 | -0.42162 | 1.823339 | 0.40752 |
| 6 | 23089400 | 23091199 | 19.00222 | 33.58292 | 31.144 | 13.77666 | 0.498685 | 0.549475 | 0.40753 |

| | | | | | | | | | |
|---|---|---|---|---|---|---|---|---|---|
| 11 | 89281200 | 89283199 | 14.80822 | 25.62518 | 48.98199 | 21.48971 | -0.54463 | 0.8306 | 0.407589 |
| 7 | 61054600 | 61070799 | 306.0139 | 438.6136 | 395.0843 | 163.1657 | 0.35476 | 4.216269 | 0.407597 |
| 4 | 51383400 | 51386399 | 20.00745 | 34.06863 | 60.38875 | 26.47006 | -0.44085 | 1.012033 | 0.407608 |
| X | 98499400 | 98502799 | 15.79904 | 27.47543 | 48.96683 | 21.51171 | -0.44278 | 0.782675 | 0.407613 |
| 12 | 114705200 | 114716399 | 176.9991 | 266.8146 | 222.4038 | 91.88806 | 0.535199 | 3.375404 | 0.407613 |
| 2 | 85725200 | 85726599 | 12.15603 | 21.66996 | 39.836 | 17.5293 | -0.47692 | 0.45305 | 0.407624 |
| 1 | 36502600 | 36504399 | 13.89432 | 24.21467 | 45.78674 | 20.14167 | -0.52713 | 0.739502 | 0.407639 |
| 14 | 105908400 | 105908799 | 12.66441 | 23.53985 | 23.00406 | 9.626693 | 0.479028 | 0.179432 | 0.407642 |
| 1 | 141957400 | 141959399 | 22.40618 | 39.38892 | 31.22646 | 13.82782 | 0.720689 | 0.624266 | 0.407654 |
| 10 | 54375200 | 54376399 | 13.31632 | 24.8631 | 20.31298 | 8.436496 | 0.74396 | 0.14903 | 0.407657 |
| 16 | 17707600 | 17710799 | 18.00496 | 30.3917 | 57.93315 | 25.33935 | -0.54603 | 1.07988 | 0.407675 |
| 8 | 105421600 | 105422799 | 13.20368 | 23.15638 | 42.01122 | 18.56874 | -0.46735 | 0.631024 | 0.407695 |
| 12 | 56150000 | 56150599 | 4.396135 | 8.566189 | 21.33327 | 7.745449 | -0.67222 | -0.39769 | 0.407704 |
| 17 | 19706400 | 19710199 | 68.54501 | 109.1631 | 94.70956 | 39.61566 | 0.541119 | 2.200859 | 0.407707 |
| 13 | 18078200 | 18078799 | 5.950759 | 11.57686 | 28.13467 | 10.38369 | -0.66126 | -0.30052 | 0.407723 |
| 7 | 101226400 | 101227399 | 17.01659 | 28.93996 | 54.73225 | 23.98827 | -0.53389 | 1.008173 | 0.407729 |
| X | 91620800 | 91622799 | 24.6917 | 43.40845 | 30.64661 | 13.57094 | 0.88732 | 0.638339 | 0.407737 |
| 4 | 99290000 | 99293999 | 28.08123 | 45.76116 | 80.92243 | 34.75412 | -0.44897 | 1.531436 | 0.407754 |
| 8 | 27547800 | 27552799 | 28.8642 | 46.68416 | 87.57707 | 37.43937 | -0.54326 | 1.676759 | 0.407758 |
| 15 | 97930200 | 97932999 | 16.06683 | 27.71527 | 50.58405 | 22.19044 | -0.47932 | 0.866545 | 0.407763 |
| 6 | 84789200 | 84791199 | 20.11892 | 33.74343 | 60.6561 | 26.46521 | -0.46204 | 1.149386 | 0.40777 |
| 8 | 41598000 | 41599199 | 7.793854 | 14.77995 | 28.76709 | 11.61052 | -0.46439 | 0.0211 | 0.407788 |
| 9 | 77803800 | 77804599 | 7.793854 | 14.77995 | 28.76709 | 11.61052 | -0.46439 | 0.0211 | 0.407788 |
| 3 | 4349200 | 4351399 | 17.66115 | 32.06651 | 26.37373 | 11.38576 | 0.6948 | 0.354595 | 0.407792 |
| 6 | 87458000 | 87461399 | 25.65413 | 41.97675 | 75.61592 | 32.52341 | -0.47127 | 1.465845 | 0.407812 |
| 15 | 29389200 | 29391199 | 18.43671 | 33.15344 | 27.23157 | 11.8953 | 0.685803 | 0.415749 | 0.40785 |
| 15 | 38090200 | 38090799 | 5.962246 | 11.55735 | 24.44643 | 9.246284 | -0.4953 | -0.22423 | 0.407855 |
| 15 | 62534800 | 62535399 | 5.962246 | 11.55735 | 24.44643 | 9.246284 | -0.4953 | -0.22423 | 0.407855 |
| 14 | 70109200 | 70111599 | 15.72956 | 27.21936 | 49.52348 | 21.72728 | -0.47384 | 0.835103 | 0.407861 |
| 3 | 12838400 | 12840199 | 17.53009 | 32.39047 | 22.68285 | 9.569482 | 0.947529 | 0.249098 | 0.407909 |
| 11 | 41271600 | 41272799 | 15.87049 | 27.19665 | 56.39165 | 24.73068 | -0.66436 | 0.935437 | 0.407918 |
| 13 | 108778600 | 108779199 | 8.961367 | 17.46291 | 16.63498 | 6.034972 | 0.6982 | -0.36039 | 0.407923 |
| 12 | 85420800 | 85421599 | 3.419402 | 6.671022 | 20.25167 | 7.237155 | -0.93149 | -0.47037 | 0.407953 |
| X | 152876000 | 152879199 | 85.14589 | 134.7711 | 115.5317 | 48.02909 | 0.550001 | 2.395783 | 0.407953 |
| 13 | 63364400 | 63368199 | 46.26714 | 74.81209 | 63.57207 | 27.16913 | 0.595983 | 1.698902 | 0.407955 |
| 1 | 3692200 | 3692799 | 15.93896 | 28.50551 | 27.76243 | 12.18326 | 0.437207 | 0.434549 | 0.407965 |
| 15 | 16220600 | 16221199 | 10.54687 | 20.12121 | 20.64231 | 8.226111 | 0.468212 | -0.01626 | 0.407996 |
| 9 | 116090000 | 116090999 | 11.80993 | 22.42613 | 21.63578 | 8.707403 | 0.544123 | 0.03183 | 0.408 |
| 3 | 12948600 | 12952799 | 48.19753 | 77.62624 | 77.88282 | 33.11278 | 0.350353 | 1.824609 | 0.408001 |
| 19 | 38217200 | 38218599 | 14.72652 | 27.70739 | 22.75534 | 9.329158 | 0.751264 | 0.119761 | 0.408015 |
| X | 23986600 | 23987999 | 24.84405 | 43.2699 | 32.36881 | 14.23338 | 0.804157 | 0.78004 | 0.408018 |
| 15 | 98136800 | 98141199 | 42.02831 | 67.47684 | 110.0505 | 46.57405 | -0.3548 | 1.927912 | 0.408028 |
| X | 148500200 | 148503199 | 22.85981 | 40.06179 | 30.10556 | 13.29827 | 0.797762 | 0.674803 | 0.408036 |
| 5 | 16870200 | 16872799 | 51.23453 | 82.48531 | 71.2841 | 30.29009 | 0.564503 | 1.838012 | 0.40804 |
| 17 | 37569400 | 37572399 | 23.42337 | 40.9233 | 34.09231 | 15.02073 | 0.649669 | 0.729064 | 0.408051 |
| 1 | 127132600 | 127134599 | 14.5793 | 25.04561 | 51.59416 | 22.63071 | -0.65431 | 0.899832 | 0.408053 |
| 4 | 153590000 | 153604599 | 189.0663 | 274.7181 | 425.8846 | 175.5587 | -0.40874 | 4.007072 | 0.408066 |

| | | | | | | | | | |
|---|---|---|---|---|---|---|---|---|---|
| 9 | 93498800 | 93501999 | 28.95205 | 50.25333 | 35.68471 | 15.65793 | 0.878545 | 0.849685 | 0.408082 |
| 19 | 13240400 | 13249599 | 186.2928 | 277.8948 | 262.1561 | 108.0492 | 0.34325 | 3.576346 | 0.408085 |
| 6 | 119517600 | 119521799 | 25.94342 | 42.50142 | 75.30297 | 32.40462 | -0.44661 | 1.449403 | 0.40809 |
| 2 | 148999400 | 149000799 | 17.56691 | 31.7012 | 26.80265 | 11.65964 | 0.648099 | 0.388212 | 0.408127 |
| Y | 2240800 | 2244399 | 27.66286 | 47.1025 | 26.02415 | 11.40707 | 1.233927 | 0.99586 | 0.408129 |
| 9 | 86313800 | 86316799 | 28.07644 | 47.76462 | 44.68043 | 19.58322 | 0.477692 | 1.032391 | 0.408132 |
| 13 | 58816400 | 58817399 | 9.534961 | 17.28117 | 34.94692 | 15.12074 | -0.59618 | 0.329817 | 0.408151 |
| X | 128232200 | 128233599 | 16.29508 | 30.36588 | 23.72294 | 9.885129 | 0.804736 | 0.196254 | 0.408187 |
| 16 | 60702600 | 60706199 | 47.17959 | 76.18521 | 74.41375 | 31.74761 | 0.393333 | 1.744601 | 0.408194 |
| 8 | 56599200 | 56599999 | 6.767674 | 12.83231 | 27.64579 | 11.16097 | -0.60957 | 0.00979 | 0.408206 |
| 11 | 19958400 | 19980399 | 299.8104 | 415.7602 | 721.9326 | 299.7378 | -0.63976 | 4.705675 | 0.408213 |
| 7 | 33086200 | 33089799 | 74.11073 | 117.6469 | 110.7386 | 46.14575 | 0.419055 | 2.309035 | 0.408217 |
| 19 | 58832000 | 58838399 | 51.73683 | 81.82669 | 153.8195 | 63.92104 | -0.58209 | 2.397204 | 0.408221 |
| 6 | 124966400 | 124969199 | 51.47848 | 81.85688 | 131.328 | 54.81771 | -0.3467 | 2.246013 | 0.408239 |
| 16 | 73849600 | 73850599 | 11.14918 | 21.14868 | 21.57344 | 8.702066 | 0.460905 | 0.030445 | 0.408242 |
| X | 33243000 | 33259399 | 248.791 | 363.3873 | 320.1003 | 131.8807 | 0.413639 | 3.933204 | 0.408278 |
| 15 | 97761000 | 97763199 | 12.94169 | 22.54334 | 45.28513 | 19.91484 | -0.61404 | 0.740003 | 0.408292 |
| 15 | 70785000 | 70786399 | 22.24272 | 38.46353 | 37.86471 | 16.61191 | 0.408265 | 0.854381 | 0.408332 |
| 7 | 64146200 | 64153799 | 69.67245 | 108.4814 | 191.8444 | 79.63189 | -0.51186 | 2.717489 | 0.408343 |
| 1 | 127604200 | 127608599 | 39.93563 | 63.9963 | 108.254 | 45.68266 | -0.41054 | 1.98342 | 0.40835 |
| 10 | 121007800 | 121013799 | 48.49876 | 77.081 | 131.0364 | 54.66918 | -0.43077 | 2.256208 | 0.408361 |
| 11 | 10623800 | 10624999 | 18.1923 | 31.24407 | 53.81814 | 23.60602 | -0.39758 | 0.920438 | 0.408369 |
| 12 | 100179400 | 100182199 | 31.58533 | 52.64573 | 51.94177 | 22.57406 | 0.400528 | 1.241083 | 0.408373 |
| 8 | 92613800 | 92617399 | 31.51975 | 51.10095 | 87.16385 | 37.32832 | -0.40325 | 1.63104 | 0.408405 |
| 8 | 92614000 | 92617399 | 31.51975 | 51.10095 | 87.16385 | 37.32832 | -0.40325 | 1.63104 | 0.408405 |
| 2 | 94630600 | 94631599 | 14.40221 | 26.35293 | 25.77769 | 11.02034 | 0.45538 | 0.285696 | 0.408426 |
| 16 | 44389600 | 44392399 | 47.74342 | 76.92941 | 77.26889 | 32.87317 | 0.349571 | 1.810302 | 0.408447 |
| 9 | 61698600 | 61703199 | 46.29266 | 73.99931 | 118.654 | 49.89048 | -0.33765 | 2.070206 | 0.408452 |
| 1 | 24631800 | 24634999 | 36.94194 | 59.45605 | 99.63995 | 42.31964 | -0.38949 | 1.839759 | 0.408461 |
| 2 | 84888200 | 84890599 | 19.13096 | 32.18135 | 59.58135 | 26.03305 | -0.50441 | 1.11521 | 0.408498 |
| 3 | 148343800 | 148346799 | 31.51638 | 51.21623 | 85.14017 | 36.52518 | -0.363 | 1.583518 | 0.408504 |
| 1 | 55227800 | 55231599 | 25.34872 | 41.82644 | 73.06646 | 31.53786 | -0.42216 | 1.355733 | 0.408505 |
| 8 | 41586200 | 41587399 | 7.195787 | 13.22298 | 32.0249 | 13.61193 | -0.83625 | 0.22468 | 0.408519 |
| 14 | 47253600 | 47254399 | 3.937244 | 7.680108 | 20.26625 | 7.257047 | -0.73501 | -0.4548 | 0.408531 |
| 4 | 3391600 | 3394199 | 30.41257 | 51.048 | 49.12918 | 21.44646 | 0.436044 | 1.161197 | 0.408541 |
| 13 | 80013600 | 80015999 | 45.82749 | 74.33848 | 68.90798 | 29.52225 | 0.475207 | 1.638034 | 0.408549 |
| 2 | 41269200 | 41270399 | 8.477336 | 15.80245 | 30.5634 | 12.72875 | -0.49301 | 0.146349 | 0.408552 |
| 18 | 7371600 | 7372199 | 8.477336 | 15.80245 | 30.5634 | 12.72875 | -0.49301 | 0.146349 | 0.408552 |
| 4 | 54622000 | 54622999 | 13.51404 | 25.70891 | 21.9901 | 8.815952 | 0.720853 | 0.034982 | 0.408552 |
| 5 | 148278600 | 148280799 | 14.19182 | 24.62132 | 47.40172 | 20.79803 | -0.55396 | 0.803947 | 0.408553 |
| 17 | 34614800 | 34619199 | 28.05327 | 45.78189 | 80.56771 | 34.62006 | -0.44053 | 1.509842 | 0.408558 |
| 3 | 18651200 | 18655599 | 110.0849 | 169.8721 | 165.7714 | 68.85827 | 0.334949 | 2.88962 | 0.408558 |
| 13 | 7682800 | 7683399 | 9.135684 | 16.7337 | 32.43083 | 13.84507 | -0.52271 | 0.256519 | 0.408564 |
| 9 | 92862200 | 92863799 | 29.00351 | 49.16955 | 45.79721 | 20.05757 | 0.483548 | 1.068345 | 0.40857 |
| X | 67719200 | 67722999 | 40.75771 | 67.75804 | 49.31723 | 21.39685 | 0.838091 | 1.298122 | 0.408571 |
| 4 | 35864600 | 35867999 | 37.92364 | 61.04762 | 99.88744 | 42.43643 | -0.35469 | 1.835808 | 0.408579 |
| 16 | 18586400 | 18587399 | 5.974837 | 11.54169 | 24.80988 | 9.536906 | -0.54046 | -0.16712 | 0.408589 |

| | | | | | | | | | |
|---|---|---|---|---|---|---|---|---|---|
| 17 | 15747000 | 15747799 | 5.974837 | 11.54169 | 24.80988 | 9.536906 | -0.54046 | -0.16712 | 0.408589 |
| 12 | 115220000 | 115223999 | 60.59567 | 97.1336 | 87.64725 | 37.004 | 0.494886 | 1.993302 | 0.408597 |
| 2 | 168541400 | 168545199 | 37.35397 | 59.89034 | 106.5244 | 44.9883 | -0.48188 | 1.963729 | 0.408601 |
| X | 126308600 | 126318599 | 90.33839 | 143.1275 | 97.2614 | 40.45342 | 0.885707 | 2.373567 | 0.408603 |
| 5 | 134480600 | 134482399 | 18.73629 | 31.81886 | 55.87545 | 24.4844 | -0.42776 | 1.027451 | 0.408606 |
| 18 | 46730000 | 46731799 | 12.12429 | 21.29003 | 41.00484 | 18.14313 | -0.55292 | 0.603869 | 0.408611 |
| 2 | 73071600 | 73073399 | 11.26674 | 20.32851 | 39.01857 | 16.97736 | -0.52739 | 0.374367 | 0.408617 |
| 4 | 68585800 | 68587999 | 20.48631 | 34.26232 | 68.37921 | 29.78158 | -0.61248 | 1.188247 | 0.408627 |
| 11 | 30965400 | 30967999 | 40.94957 | 65.78662 | 110.3236 | 46.73419 | -0.39377 | 1.908602 | 0.408652 |
| 17 | 18860800 | 18865599 | 61.56942 | 98.95278 | 80.49214 | 34.11717 | 0.648673 | 1.91209 | 0.40866 |
| 2 | 93152200 | 93156399 | 28.00674 | 45.47316 | 89.50769 | 38.37417 | -0.60725 | 1.597333 | 0.408676 |
| 10 | 94262000 | 94264799 | 20.59876 | 34.49298 | 61.77566 | 26.92649 | -0.45678 | 1.168573 | 0.408698 |
| 18 | 50458400 | 50458999 | 18.543 | 32.55746 | 29.07563 | 12.86366 | 0.549944 | 0.618887 | 0.408699 |
| 1 | 55966600 | 55967199 | 9.682628 | 18.698 | 19.14757 | 7.371545 | 0.516414 | -0.14195 | 0.408704 |
| 4 | 126405000 | 126406399 | 12.27161 | 23.08592 | 21.82468 | 8.949344 | 0.549167 | 0.094742 | 0.408739 |
| 13 | 39806800 | 39807999 | 12.27161 | 23.08592 | 21.82468 | 8.949344 | 0.549167 | 0.094742 | 0.408739 |
| 4 | 151583400 | 151584599 | 14.49776 | 26.40888 | 25.86732 | 11.12168 | 0.448317 | 0.313226 | 0.408746 |
| 12 | 71745800 | 71746799 | 18.74574 | 33.20561 | 30.49475 | 13.48329 | 0.513812 | 0.521067 | 0.40875 |
| 12 | 52855200 | 52858199 | 25.72061 | 43.98552 | 41.44072 | 18.1709 | 0.468371 | 0.970739 | 0.40877 |
| 2 | 178922600 | 178925599 | 25.42672 | 41.78098 | 73.18516 | 31.53355 | -0.42839 | 1.409224 | 0.408778 |
| 13 | 87504000 | 87505599 | 22.64055 | 39.86895 | 30.67093 | 13.57718 | 0.764791 | 0.607129 | 0.408779 |
| 5 | 6851400 | 6856199 | 69.2793 | 110.2455 | 96.5839 | 40.35564 | 0.52591 | 2.22781 | 0.4088 |
| 11 | 9094600 | 9098799 | 60.17308 | 96.5223 | 86.09529 | 36.38667 | 0.51264 | 1.968477 | 0.408805 |
| 10 | 74587000 | 74588399 | 21.70417 | 37.81828 | 36.64713 | 16.11921 | 0.432506 | 0.763923 | 0.408808 |
| 13 | 42336800 | 42337999 | 13.51689 | 23.74323 | 41.94329 | 18.56353 | -0.42915 | 0.607034 | 0.40883 |
| X | 38230200 | 38235199 | 63.51637 | 101.7503 | 88.79539 | 37.45227 | 0.541924 | 2.016071 | 0.408834 |
| 2 | 129452800 | 129454199 | 13.41605 | 24.94554 | 24.38237 | 10.19806 | 0.479079 | 0.185936 | 0.408838 |
| 3 | 11232600 | 11233199 | 5.091974 | 9.928104 | 23.39386 | 8.426369 | -0.58449 | -0.41286 | 0.408845 |
| 3 | 18539600 | 18540199 | 9.953967 | 19.30467 | 19.28964 | 7.278767 | 0.579132 | -0.20363 | 0.408863 |
| 1 | 34348600 | 34350399 | 15.39733 | 26.84455 | 46.80732 | 20.59568 | -0.41107 | 0.739632 | 0.408865 |
| 13 | 108034400 | 108036399 | 19.3428 | 34.0469 | 32.12499 | 14.22209 | 0.471529 | 0.594948 | 0.408879 |
| 8 | 21718200 | 21726999 | 102.9773 | 160.5669 | 144.9287 | 60.14502 | 0.458958 | 2.703305 | 0.408894 |
| 1 | 118786600 | 118790599 | 28.62098 | 46.46606 | 85.14829 | 36.50263 | -0.50448 | 1.595944 | 0.408895 |
| X | 15686400 | 15686999 | 11.688 | 22.25736 | 20.65614 | 8.263634 | 0.607364 | 0.009613 | 0.408897 |
| 5 | 126654200 | 126657599 | 22.31593 | 37.34962 | 66.47672 | 28.96637 | -0.44802 | 1.181548 | 0.408913 |
| 7 | 96864600 | 96865599 | 11.89274 | 22.40208 | 20.13117 | 8.236883 | 0.625293 | 0.082368 | 0.408938 |
| 2 | 51373000 | 51374199 | 7.708079 | 14.35436 | 31.61713 | 13.18956 | -0.68163 | 0.14514 | 0.408952 |
| 9 | 4374800 | 4376999 | 15.40834 | 26.83062 | 46.82754 | 20.5886 | -0.41259 | 0.757578 | 0.408953 |
| 1 | 91945600 | 91946599 | 7.302249 | 14.0021 | 28.62776 | 11.25848 | -0.5012 | -0.07015 | 0.408954 |
| 17 | 40676600 | 40678399 | 20.37033 | 36.01201 | 29.68862 | 13.13205 | 0.667415 | 0.54906 | 0.408978 |
| 7 | 97858600 | 97860399 | 27.79565 | 47.37411 | 44.14384 | 19.35091 | 0.48367 | 1.014927 | 0.408986 |
| 18 | 67220000 | 67222999 | 17.08448 | 28.8106 | 58.95509 | 25.77891 | -0.648 | 1.085596 | 0.408988 |
| 4 | 11007000 | 11008599 | 11.50955 | 20.20535 | 41.04025 | 18.15484 | -0.62911 | 0.604648 | 0.408994 |
| 6 | 3603600 | 3609999 | 83.0724 | 131.9659 | 104.4438 | 43.5533 | 0.669796 | 2.29505 | 0.408997 |
| 19 | 39159600 | 39160999 | 11.16122 | 20.19934 | 38.00401 | 16.47327 | -0.4955 | 0.350825 | 0.409006 |
| 18 | 75471400 | 75476399 | 45.157 | 72.20878 | 115.9159 | 48.76294 | -0.33873 | 2.057921 | 0.409009 |
| 1 | 107054200 | 107055599 | 12.52696 | 22.18831 | 39.27852 | 17.36728 | -0.42776 | 0.500815 | 0.409018 |

| | | | | | | | | | |
|---|---|---|---|---|---|---|---|---|---|
| 17 | 69694400 | 69696399 | 18.3746 | 30.9208 | 58.92732 | 25.75026 | -0.54578 | 1.111196 | 0.409018 |
| 2 | 80074600 | 80075599 | 7.464631 | 14.28581 | 28.51928 | 11.27829 | -0.47447 | -0.04985 | 0.409019 |
| 12 | 88546400 | 88547199 | 7.464631 | 14.28581 | 28.51928 | 11.27829 | -0.47447 | -0.04985 | 0.409019 |
| 7 | 10241400 | 10244999 | 51.44111 | 83.00211 | 75.82145 | 32.31577 | 0.488477 | 1.776325 | 0.409034 |
| 7 | 91623200 | 91624399 | 11.90485 | 21.08209 | 38.29583 | 16.93346 | -0.46477 | 0.49762 | 0.409039 |
| 12 | 35201400 | 35203199 | 9.217435 | 17.11581 | 32.09846 | 13.45066 | -0.45638 | 0.177954 | 0.409046 |
| 14 | 12738200 | 12738799 | 13.49161 | 25.20055 | 18.89952 | 7.844088 | 0.867506 | 0.144849 | 0.409049 |
| 5 | 146737400 | 146739999 | 29.58056 | 50.24251 | 40.37575 | 17.69328 | 0.695628 | 1.046719 | 0.409083 |
| 8 | 62591600 | 62591999 | 13.16368 | 24.3455 | 22.05087 | 9.290973 | 0.581002 | 0.209489 | 0.409084 |
| 4 | 74334800 | 74337199 | 23.05778 | 40.30979 | 34.69194 | 15.29247 | 0.603045 | 0.719663 | 0.409092 |
| 3 | 19340000 | 19341599 | 10.49249 | 19.13727 | 37.12906 | 15.93831 | -0.53044 | 0.298532 | 0.409096 |
| 8 | 55551600 | 55552199 | 9.660725 | 18.88918 | 13.25279 | 4.530761 | 1.210122 | -0.56013 | 0.409111 |
| 3 | 22300800 | 22308199 | 218.6916 | 321.2388 | 304.2189 | 125.2917 | 0.317025 | 3.840793 | 0.409116 |
| 18 | 37262400 | 37264399 | 17.84526 | 32.68249 | 25.47832 | 10.87896 | 0.782128 | 0.307405 | 0.409122 |
| 14 | 72658000 | 72664399 | 85.76827 | 135.5589 | 119.4078 | 49.60395 | 0.509191 | 2.43027 | 0.409157 |
| X | 96323200 | 96325199 | 22.93574 | 40.9034 | 26.9343 | 11.84793 | 0.996442 | 0.497773 | 0.409158 |
| 13 | 4813800 | 4816999 | 28.01577 | 47.71022 | 44.11935 | 19.3389 | 0.494397 | 1.021745 | 0.409171 |
| 14 | 33223800 | 33229599 | 39.88353 | 64.02016 | 106.4113 | 45.01978 | -0.38242 | 1.929427 | 0.409172 |
| 2 | 173814000 | 173818799 | 55.75031 | 89.3139 | 89.31915 | 37.67606 | 0.345783 | 2.003423 | 0.409223 |
| 17 | 17283200 | 17284999 | 32.08133 | 53.51397 | 50.33305 | 21.88562 | 0.469196 | 1.23098 | 0.409223 |
| 8 | 85715800 | 85717999 | 10.841 | 19.16021 | 38.88193 | 17.19942 | -0.62476 | 0.513247 | 0.409224 |
| 14 | 16288000 | 16289799 | 10.841 | 19.16021 | 38.88193 | 17.19942 | -0.62476 | 0.513247 | 0.409224 |
| 7 | 39132400 | 39133199 | 10.63692 | 20.43928 | 20.73852 | 8.106777 | 0.509755 | -0.07581 | 0.409244 |
| 8 | 15845400 | 15846599 | 15.63664 | 28.3659 | 26.5811 | 11.48783 | 0.506676 | 0.344591 | 0.409247 |
| 1 | 92356400 | 92358199 | 18.53518 | 32.56787 | 31.82805 | 14.0902 | 0.420415 | 0.610978 | 0.409258 |
| 13 | 76227200 | 76229999 | 18.53518 | 32.56787 | 31.82805 | 14.0902 | 0.420415 | 0.610978 | 0.409258 |
| 9 | 19636200 | 19640799 | 76.39803 | 120.8042 | 119.6364 | 49.71044 | 0.340923 | 2.41184 | 0.409267 |
| 2 | 84623600 | 84627399 | 67.49931 | 107.2357 | 108.0591 | 45.06456 | 0.321941 | 2.278523 | 0.409287 |
| 1 | 23975800 | 23977199 | 27.34355 | 46.32496 | 45.27728 | 19.82591 | 0.414174 | 1.065199 | 0.40929 |
| 6 | 139354200 | 139356199 | 23.10824 | 40.44528 | 34.11296 | 15.05212 | 0.632005 | 0.701097 | 0.40933 |
| X | 30898400 | 30900799 | 24.23295 | 43.07533 | 26.20398 | 11.55858 | 1.107317 | 0.524125 | 0.409342 |
| 17 | 11192400 | 11193599 | 8.11299 | 15.36994 | 29.86686 | 12.07965 | -0.46478 | 0.034155 | 0.409355 |
| 1 | 109109600 | 109110199 | 5.211891 | 10.1616 | 22.70478 | 8.181457 | -0.50822 | -0.40855 | 0.409359 |
| 7 | 50662600 | 50662999 | 5.211891 | 10.1616 | 22.70478 | 8.181457 | -0.50822 | -0.40855 | 0.409359 |
| 5 | 59236400 | 59237399 | 11.70944 | 22.47515 | 20.53361 | 8.05237 | 0.654735 | -0.05779 | 0.409361 |
| 2 | 114485000 | 114485599 | 4.555325 | 8.875552 | 22.53497 | 8.19177 | -0.70245 | -0.38946 | 0.409364 |
| 11 | 94072000 | 94073799 | 8.387008 | 15.38462 | 31.96103 | 13.61765 | -0.61986 | 0.242079 | 0.409366 |
| 5 | 109952000 | 109956199 | 38.01009 | 62.43408 | 53.73344 | 23.14859 | 0.593617 | 1.428687 | 0.409374 |
| 5 | 76475600 | 76479399 | 26.56172 | 43.75849 | 74.61339 | 32.18397 | -0.38821 | 1.382289 | 0.409378 |
| X | 153280400 | 153284799 | 71.87038 | 114.6932 | 95.40676 | 40.0206 | 0.604909 | 2.143058 | 0.409392 |
| 18 | 63435000 | 63435799 | 11.1153 | 21.306 | 21.21128 | 8.350403 | 0.527385 | -0.04723 | 0.409403 |
| 2 | 160772800 | 160774799 | 18.22987 | 30.97319 | 56.42785 | 24.72948 | -0.48063 | 1.02063 | 0.40941 |
| 5 | 57124000 | 57128799 | 76.5676 | 120.9987 | 120.2935 | 49.96806 | 0.334667 | 2.427992 | 0.409414 |
| X | 3246600 | 3250999 | 58.40376 | 93.72636 | 85.97198 | 36.35944 | 0.47317 | 1.954816 | 0.409421 |
| 10 | 65505600 | 65507799 | 28.98769 | 49.77462 | 40.16637 | 17.61789 | 0.691838 | 0.948093 | 0.409428 |
| 2 | 94727600 | 94728599 | 10.82285 | 20.80251 | 19.46074 | 7.601048 | 0.625919 | -0.08361 | 0.409451 |
| 8 | 78336800 | 78337599 | 5.383716 | 10.47382 | 23.60057 | 8.709552 | -0.55258 | -0.3214 | 0.409455 |

| | | | | | | | | | |
|---|---|---|---|---|---|---|---|---|---|
| 3 | 9252800 | 9257399 | 35.12609 | 56.68484 | 94.96319 | 40.48115 | -0.38434 | 1.753939 | 0.409459 |
| 7 | 17340200 | 17375399 | 666.8638 | 898.9849 | 688.6526 | 287.1873 | 0.499398 | 5.152743 | 0.409463 |
| 14 | 49526000 | 49526599 | 3.932553 | 7.685266 | 18.83328 | 6.522891 | -0.58607 | -0.57981 | 0.409474 |
| 2 | 7511400 | 7515199 | 35.36743 | 56.91255 | 107.5467 | 45.66691 | -0.56267 | 1.846709 | 0.409475 |
| 16 | 56259400 | 56260599 | 13.54081 | 25.70241 | 21.88095 | 8.813696 | 0.721385 | 0.048799 | 0.409485 |
| 7 | 32303000 | 32318999 | 221.5983 | 327.972 | 282.1128 | 116.1824 | 0.465935 | 3.719252 | 0.409487 |
| 10 | 16239600 | 16242199 | 27.74774 | 47.39908 | 42.44671 | 18.61032 | 0.541118 | 0.990517 | 0.409496 |
| 7 | 20556200 | 20557199 | 18.48382 | 32.86017 | 28.73008 | 12.67148 | 0.587391 | 0.492863 | 0.409498 |
| 8 | 92002800 | 92003399 | 12.70422 | 23.70605 | 23.51327 | 9.775553 | 0.465268 | 0.156828 | 0.409508 |
| 7 | 23295000 | 23295599 | 29.99023 | 50.01517 | 47.08032 | 20.46852 | 0.468183 | 1.227513 | 0.40951 |
| 8 | 8739600 | 8740599 | 7.253587 | 13.5854 | 29.57977 | 12.2208 | -0.65328 | 0.100805 | 0.409519 |
| 4 | 22548000 | 22549199 | 12.53357 | 22.17885 | 39.29451 | 17.37766 | -0.42947 | 0.511542 | 0.409529 |
| 14 | 66461000 | 66463199 | 12.53357 | 22.17885 | 39.29451 | 17.37766 | -0.42947 | 0.511542 | 0.409529 |
| 8 | 8811400 | 8814599 | 22.47308 | 38.92837 | 37.74544 | 16.56045 | 0.430449 | 0.838871 | 0.409536 |
| 6 | 138397400 | 138400799 | 19.99222 | 33.38261 | 64.45776 | 28.04538 | -0.56493 | 1.198515 | 0.409545 |
| 3 | 90843000 | 90845399 | 32.07994 | 54.19036 | 45.20312 | 19.77586 | 0.641814 | 1.106301 | 0.409547 |
| 8 | 74367400 | 74368999 | 34.13075 | 57.42159 | 42.00076 | 18.35231 | 0.830557 | 1.145594 | 0.409566 |
| X | 157108400 | 157110799 | 32.29619 | 53.99478 | 49.71533 | 21.64801 | 0.499928 | 1.212492 | 0.409587 |
| 4 | 144351000 | 144362599 | 170.0893 | 255.9707 | 233.6099 | 96.47483 | 0.402381 | 3.406259 | 0.409622 |
| 9 | 67072400 | 67074999 | 26.33149 | 43.33329 | 76.53627 | 32.9986 | -0.43958 | 1.39278 | 0.409627 |
| 5 | 67609600 | 67612399 | 16.81972 | 29.18638 | 52.82189 | 23.17633 | -0.46557 | 0.818878 | 0.409631 |
| 18 | 3404800 | 3414599 | 116.8374 | 180.9587 | 154.2617 | 64.07615 | 0.533852 | 2.826037 | 0.409656 |
| 14 | 118488400 | 118491399 | 25.93188 | 44.42459 | 42.53085 | 18.65153 | 0.445823 | 0.959649 | 0.409657 |
| 17 | 22723000 | 22732199 | 107.8201 | 166.6519 | 160.5891 | 66.70511 | 0.354951 | 2.857345 | 0.409658 |
| 3 | 16685400 | 16691199 | 90.3322 | 141.9187 | 132.693 | 55.04015 | 0.416567 | 2.560082 | 0.40966 |
| 3 | 146635800 | 146637599 | 12.34453 | 21.80298 | 40.83854 | 18.06734 | -0.5106 | 0.531751 | 0.409711 |
| X | 122679000 | 122684999 | 91.51245 | 144.145 | 114.2503 | 47.40616 | 0.65745 | 2.508136 | 0.409728 |
| 5 | 78849400 | 78851199 | 22.74353 | 39.83483 | 34.19153 | 15.09572 | 0.60694 | 0.689549 | 0.409728 |
| 4 | 112593000 | 112594599 | 34.39331 | 56.94301 | 55.36689 | 23.95352 | 0.421037 | 1.326536 | 0.409752 |
| 1 | 118111600 | 118112999 | 16.97596 | 30.46086 | 28.61726 | 12.52337 | 0.491623 | 0.42337 | 0.409761 |
| 7 | 18254200 | 18279999 | 410.8228 | 573.1051 | 508.362 | 210.8708 | 0.337659 | 4.617898 | 0.409766 |
| 18 | 58460200 | 58461399 | 10.59032 | 18.9818 | 36.30305 | 15.90159 | -0.52877 | 0.402102 | 0.409786 |
| 17 | 11819800 | 11823199 | 39.42706 | 65.40076 | 52.69224 | 22.82619 | 0.691712 | 1.319923 | 0.409796 |
| 18 | 7033600 | 7035599 | 19.81776 | 35.11489 | 29.76331 | 13.1591 | 0.628926 | 0.522523 | 0.409812 |
| 7 | 7291800 | 7294999 | 20.21834 | 34.08538 | 59.13165 | 25.8538 | -0.4108 | 1.099441 | 0.409816 |
| 6 | 39653600 | 39654199 | 9.993001 | 19.22256 | 19.73503 | 7.690807 | 0.497186 | -0.09496 | 0.40982 |
| 14 | 112230600 | 112231999 | 12.80742 | 22.66891 | 41.85812 | 18.5105 | -0.48897 | 0.512974 | 0.409831 |
| 1 | 92858600 | 92862599 | 56.84607 | 90.09687 | 142.5531 | 59.30873 | -0.33134 | 2.34687 | 0.409832 |
| 12 | 93178600 | 93178999 | 4.775375 | 9.326832 | 21.88871 | 7.674017 | -0.54231 | -0.51513 | 0.409834 |
| 18 | 17091600 | 17096399 | 38.95524 | 62.75084 | 108.956 | 46.3347 | -0.43902 | 1.820703 | 0.409837 |
| 5 | 67154200 | 67155199 | 6.712464 | 12.88692 | 26.89804 | 10.5412 | -0.52569 | -0.09199 | 0.40984 |
| 18 | 37621600 | 37625599 | 109.3588 | 168.4898 | 167.0849 | 69.40379 | 0.31009 | 2.918555 | 0.409841 |
| 8 | 88703400 | 88707399 | 27.9423 | 45.55211 | 78.99352 | 33.93041 | -0.42038 | 1.526411 | 0.409857 |
| 11 | 66943200 | 66943999 | 4.814862 | 9.363801 | 22.30977 | 8.256888 | -0.63478 | -0.31984 | 0.409862 |
| 11 | 121771800 | 121775599 | 72.32645 | 114.4462 | 114.5603 | 47.61811 | 0.326304 | 2.392159 | 0.409872 |
| 10 | 26155800 | 26156399 | 4.444687 | 8.671701 | 21.4749 | 7.668739 | -0.641 | -0.45094 | 0.409894 |
| 6 | 3649200 | 3652199 | 30.66384 | 51.48654 | 49.12175 | 21.44762 | 0.448593 | 1.159762 | 0.409923 |

| | | | | | | | | | |
|---|---|---|---|---|---|---|---|---|---|
| 2 | 78596800 | 78597799 | 8.040728 | 15.05553 | 31.18113 | 12.88838 | -0.58176 | 0.11755 | 0.409948 |
| 19 | 44083400 | 44085799 | 26.00439 | 45.2595 | 36.34336 | 15.97414 | 0.702283 | 0.799954 | 0.409948 |
| 12 | 18027800 | 18039999 | 164.9033 | 247.9918 | 236.6672 | 97.71904 | 0.337082 | 3.418768 | 0.409965 |
| 14 | 79537400 | 79539599 | 16.44722 | 27.79776 | 57.31402 | 25.07826 | -0.65858 | 1.063292 | 0.409967 |
| 7 | 34306600 | 34309999 | 28.494 | 46.25559 | 82.9394 | 35.55325 | -0.4732 | 1.599729 | 0.40998 |
| 13 | 52198800 | 52205199 | 58.33485 | 91.84097 | 155.3071 | 64.43788 | -0.43487 | 2.502348 | 0.409989 |
| 4 | 43649600 | 43650599 | 28.86269 | 48.45325 | 42.86933 | 18.71548 | 0.557123 | 1.151016 | 0.41001 |
| 18 | 54347200 | 54348399 | 16.31884 | 29.48658 | 24.11527 | 10.47384 | 0.697418 | 0.36773 | 0.410014 |
| 2 | 82508000 | 82510199 | 29.14786 | 49.19902 | 41.62839 | 18.20765 | 0.621342 | 1.100128 | 0.41002 |
| 15 | 65337400 | 65338999 | 22.56085 | 39.51884 | 34.19055 | 15.09656 | 0.595527 | 0.686822 | 0.410033 |
| 11 | 7012800 | 7014399 | 13.03171 | 22.80207 | 43.61454 | 19.23982 | -0.54341 | 0.668517 | 0.410033 |
| 11 | 27312400 | 27313999 | 27.3087 | 46.21562 | 45.84837 | 20.06935 | 0.392786 | 1.076633 | 0.410047 |
| 7 | 43803000 | 43805399 | 46.5239 | 75.3195 | 70.99961 | 30.37419 | 0.448427 | 1.679909 | 0.410053 |
| X | 148876800 | 148880599 | 50.81574 | 81.71138 | 72.63525 | 30.80972 | 0.521816 | 1.879192 | 0.410079 |
| 17 | 33345400 | 33348799 | 36.84276 | 60.65394 | 58.69174 | 25.3089 | 0.426349 | 1.403244 | 0.410106 |
| 3 | 134660200 | 134662399 | 19.8892 | 35.91198 | 26.10656 | 11.34893 | 0.865499 | 0.40148 | 0.410116 |
| 3 | 158585400 | 158588799 | 56.73075 | 90.95084 | 78.25616 | 33.04613 | 0.563571 | 1.979047 | 0.410127 |
| 1 | 31827800 | 31829199 | 16.9198 | 31.34942 | 24.01212 | 10.09223 | 0.823397 | 0.233416 | 0.410134 |
| 4 | 47095000 | 47105799 | 88.55864 | 135.3919 | 243.4074 | 100.978 | -0.55601 | 3.070699 | 0.410145 |
| 5 | 149640800 | 149641599 | 12.16175 | 22.73975 | 21.07664 | 8.732575 | 0.566388 | 0.133124 | 0.410154 |
| 2 | 42834000 | 42834999 | 17.9644 | 31.64114 | 29.94393 | 13.2548 | 0.467811 | 0.577649 | 0.410156 |
| X | 33458200 | 33460399 | 29.02579 | 50.57846 | 33.43844 | 14.70836 | 0.98168 | 0.795672 | 0.410164 |
| 5 | 53458800 | 53461599 | 26.50151 | 43.71115 | 74.28159 | 32.05694 | -0.38273 | 1.365409 | 0.410168 |
| 7 | 29281000 | 29288399 | 22.97699 | 40.03668 | 32.86429 | 14.45547 | 0.670689 | 0.762877 | 0.410168 |
| 3 | 99906400 | 99906999 | 5.672642 | 10.99915 | 23.85473 | 9.010301 | -0.5287 | -0.23293 | 0.410174 |
| 15 | 79927800 | 79932999 | 38.95753 | 62.69448 | 103.3824 | 43.9035 | -0.36639 | 1.848884 | 0.410196 |
| 2 | 74568600 | 74569799 | 7.215309 | 13.63978 | 28.12814 | 11.42855 | -0.55531 | 0.036429 | 0.410198 |
| 14 | 70305200 | 70306599 | 7.215309 | 13.63978 | 28.12814 | 11.42855 | -0.55531 | 0.036429 | 0.410198 |
| 8 | 86646400 | 86647199 | 6.965462 | 13.05199 | 29.06711 | 11.9993 | -0.68405 | 0.096068 | 0.410212 |
| 7 | 8167000 | 8167399 | 10.38528 | 20.26963 | 13.48438 | 4.783994 | 1.237427 | -0.42735 | 0.410217 |
| 5 | 111709000 | 111715599 | 71.97699 | 112.0002 | 179.8135 | 74.64496 | -0.37322 | 2.727316 | 0.410236 |
| 9 | 119836600 | 119837599 | 18.22172 | 32.19921 | 30.31847 | 13.41184 | 0.476882 | 0.545462 | 0.410242 |
| X | 88032400 | 88034199 | 18.06954 | 33.50309 | 23.43425 | 9.839988 | 0.95465 | 0.238431 | 0.410247 |
| 9 | 97241000 | 97245599 | 44.51772 | 71.2505 | 114.2927 | 48.14134 | -0.33623 | 2.029786 | 0.410263 |
| 6 | 47659200 | 47659399 | 2.524378 | 4.935819 | 15.70124 | 5.085352 | -0.86081 | -0.84519 | 0.410265 |
| X | 44783800 | 44785799 | 19.79088 | 36.14096 | 24.76127 | 10.6129 | 0.963634 | 0.336867 | 0.410291 |
| X | 101993800 | 101997399 | 18.37649 | 31.59086 | 56.0185 | 24.57188 | -0.43905 | 0.915381 | 0.410305 |
| 3 | 84931400 | 84934799 | 20.17447 | 33.61835 | 66.08977 | 28.71851 | -0.59077 | 1.222298 | 0.410323 |
| 7 | 10593200 | 10597799 | 61.44652 | 98.43344 | 88.67419 | 37.40055 | 0.496159 | 2.014841 | 0.410328 |
| 7 | 5546000 | 5549999 | 53.58969 | 86.58338 | 71.72761 | 30.62314 | 0.63135 | 1.740412 | 0.410332 |
| 14 | 46340200 | 46342599 | 26.40651 | 45.99946 | 33.97694 | 14.94221 | 0.822141 | 0.784576 | 0.410348 |
| 13 | 110489800 | 110490999 | 10.47678 | 18.74818 | 35.84145 | 15.72065 | -0.52921 | 0.412784 | 0.410351 |
| 2 | 132261800 | 132264399 | 26.34158 | 44.88965 | 43.62652 | 19.12392 | 0.423086 | 1.00841 | 0.410389 |
| 3 | 95712200 | 95713599 | 11.10563 | 19.78162 | 36.80742 | 16.20754 | -0.49449 | 0.447618 | 0.410402 |
| 7 | 5642000 | 5649799 | 135.3547 | 207.5665 | 161.3501 | 66.9786 | 0.655926 | 3.017799 | 0.410407 |
| 5 | 70607600 | 70609399 | 23.16149 | 40.29427 | 37.62985 | 16.53514 | 0.485361 | 0.793874 | 0.41041 |
| 1 | 71270400 | 71271999 | 16.56977 | 30.19819 | 26.57073 | 11.41703 | 0.602476 | 0.326477 | 0.410425 |

| | | | | | | | | | |
|---|---|---|---|---|---|---|---|---|---|
| 1 | 111142600 | 111144399 | 21.77941 | 38.14522 | 34.19408 | 15.09651 | 0.544489 | 0.683396 | 0.410448 |
| 11 | 121772400 | 121775399 | 66.4923 | 105.6003 | 104.3396 | 43.49873 | 0.34979 | 2.286532 | 0.410452 |
| X | 142594000 | 142595599 | 20.82091 | 38.86217 | 19.42092 | 8.070523 | 1.447373 | 0.214237 | 0.410473 |
| 7 | 41564000 | 41567599 | 67.94893 | 108.2611 | 102.387 | 42.85293 | 0.417601 | 2.190283 | 0.41048 |
| 17 | 89037400 | 89039799 | 23.81743 | 39.74863 | 70.50401 | 30.66575 | -0.44308 | 1.22095 | 0.410481 |
| 5 | 96633000 | 96634199 | 17.39017 | 30.95952 | 29.23224 | 12.87764 | 0.478073 | 0.476838 | 0.410485 |
| 15 | 10429800 | 10431399 | 16.23358 | 29.47095 | 27.10708 | 11.70413 | 0.5344 | 0.346859 | 0.410496 |
| 8 | 115486400 | 115488199 | 20.48517 | 35.81691 | 34.97639 | 15.41952 | 0.421688 | 0.705315 | 0.410502 |
| 3 | 138027800 | 138031999 | 45.46344 | 72.64578 | 118.257 | 49.69641 | -0.36001 | 2.080513 | 0.410521 |
| 11 | 102873000 | 102876199 | 19.53888 | 32.65531 | 62.12155 | 27.04441 | -0.54335 | 1.188085 | 0.410533 |
| 18 | 22429400 | 22431199 | 13.0963 | 22.85362 | 48.38125 | 21.30041 | -0.68923 | 0.719422 | 0.410542 |
| 7 | 33778800 | 33780799 | 25.46946 | 44.02741 | 39.16581 | 17.18254 | 0.553643 | 0.872814 | 0.410555 |
| 1 | 115037800 | 115039399 | 17.43055 | 32.20261 | 23.28261 | 9.82414 | 0.901781 | 0.250604 | 0.410577 |
| 8 | 96392400 | 96394199 | 25.78163 | 42.36278 | 73.65349 | 31.7349 | -0.41766 | 1.412613 | 0.410587 |
| 3 | 10149600 | 10153399 | 30.65694 | 49.49744 | 90.71883 | 38.69677 | -0.51269 | 1.732748 | 0.410599 |
| 11 | 17884200 | 17889599 | 65.20117 | 104.357 | 90.35576 | 38.0602 | 0.551789 | 2.044003 | 0.410599 |
| 10 | 113659400 | 113660199 | 15.45955 | 28.2562 | 20.35194 | 8.713427 | 0.89276 | 0.288156 | 0.410605 |
| 13 | 61555000 | 61560599 | 95.3286 | 148.8557 | 143.7707 | 59.65497 | 0.362993 | 2.674188 | 0.410609 |
| 18 | 7988200 | 7989999 | 10.2907 | 18.86407 | 40.06503 | 17.086 | -0.65331 | 0.277866 | 0.410611 |
| 9 | 98565200 | 98568399 | 29.47896 | 47.75795 | 92.9113 | 39.76858 | -0.59346 | 1.646347 | 0.410628 |
| 10 | 23733200 | 23734599 | 13.73471 | 25.6958 | 23.86854 | 9.879576 | 0.563982 | 0.14875 | 0.410666 |
| 18 | 73202400 | 73203799 | 13.73471 | 25.6958 | 23.86854 | 9.879576 | 0.563982 | 0.14875 | 0.410666 |
| 12 | 19093600 | 19094999 | 78.07594 | 123.1446 | 108.135 | 44.8765 | 0.511309 | 2.472039 | 0.410678 |
| 6 | 54793400 | 54794599 | 10.42244 | 18.75837 | 35.30697 | 15.41034 | -0.5022 | 0.373409 | 0.410706 |
| 8 | 105470000 | 105471799 | 10.42244 | 18.75837 | 35.30697 | 15.41034 | -0.5022 | 0.373409 | 0.410706 |
| 18 | 84856600 | 84857599 | 10.42244 | 18.75837 | 35.30697 | 15.41034 | -0.5022 | 0.373409 | 0.410706 |
| 6 | 12305200 | 12306199 | 21.2914 | 37.29619 | 34.36244 | 15.1728 | 0.505272 | 0.68057 | 0.41073 |
| 7 | 85566200 | 85574199 | 101.0368 | 158.0599 | 137.8764 | 57.19838 | 0.511829 | 2.644107 | 0.410739 |
| 4 | 29219600 | 29221999 | 30.31491 | 51.76245 | 37.71662 | 16.5359 | 0.837521 | 1.002354 | 0.410742 |
| 8 | 36897800 | 36899799 | 14.12106 | 24.73512 | 43.55804 | 19.23262 | -0.4248 | 0.656968 | 0.410752 |
| 18 | 67426000 | 67428399 | 17.62336 | 30.10395 | 53.7289 | 23.55685 | -0.44968 | 0.969027 | 0.410761 |
| 7 | 45764400 | 45768599 | 31.785 | 51.26332 | 93.12166 | 39.66637 | -0.50173 | 1.766072 | 0.410776 |
| 1 | 184485200 | 184488399 | 42.62603 | 69.11352 | 69.11284 | 29.6 | 0.365416 | 1.640156 | 0.41078 |
| 7 | 114526200 | 114527199 | 15.86142 | 27.21063 | 51.59289 | 22.62815 | -0.53543 | 0.920594 | 0.410793 |
| 6 | 125954000 | 125955599 | 10.18065 | 18.54948 | 36.01225 | 15.47872 | -0.53252 | 0.299682 | 0.410795 |
| 5 | 109919400 | 109926799 | 198.2235 | 295.0723 | 268.3304 | 110.5628 | 0.393397 | 3.615341 | 0.410804 |
| 3 | 15912000 | 15918599 | 60.00184 | 96.4327 | 81.57172 | 34.57439 | 0.592322 | 1.909744 | 0.410806 |
| X | 25448800 | 25450399 | 47.50325 | 76.86993 | 67.8633 | 29.02269 | 0.542221 | 1.691624 | 0.410813 |
| 9 | 13167400 | 13169599 | 13.29585 | 23.45605 | 42.39142 | 18.75868 | -0.46002 | 0.553291 | 0.410819 |
| Y | 90154000 | 90156399 | 19.10922 | 33.70002 | 19.07106 | 8.439751 | 1.205357 | 0.544975 | 0.410821 |
| 12 | 40087200 | 40088399 | 8.129696 | 14.49129 | 37.47576 | 16.49177 | -0.96529 | 0.424438 | 0.410821 |
| 15 | 94584800 | 94586199 | 18.18363 | 31.96162 | 30.84767 | 13.66033 | 0.438329 | 0.601327 | 0.410827 |
| 12 | 15719200 | 15721199 | 30.30671 | 51.28003 | 46.33487 | 20.2812 | 0.527086 | 1.089486 | 0.410829 |
| 13 | 58304600 | 58306799 | 15.45406 | 26.67684 | 49.51217 | 21.72069 | -0.50308 | 0.857012 | 0.410834 |
| 7 | 70635000 | 70637999 | 31.48719 | 53.29291 | 45.1974 | 19.78506 | 0.618105 | 1.088533 | 0.410844 |
| 3 | 18643200 | 18644799 | 22.26529 | 39.37347 | 29.62663 | 13.10387 | 0.798861 | 0.559566 | 0.41085 |
| 17 | 20249800 | 20250799 | 14.36604 | 27.38487 | 19.51172 | 7.789089 | 0.986787 | 0.021631 | 0.410857 |

| | | | | | | | | | |
|---|---|---|---|---|---|---|---|---|---|
| 13 | 112490800 | 112492599 | 11.37656 | 20.28626 | 37.78035 | 16.6207 | -0.49516 | 0.442446 | 0.410877 |
| 12 | 86901000 | 86903399 | 13.01997 | 22.89887 | 40.88956 | 18.10522 | -0.44427 | 0.584505 | 0.410878 |
| 9 | 18761800 | 18762399 | 9.425253 | 18.31693 | 18.22646 | 6.784463 | 0.602807 | -0.26048 | 0.410901 |
| 10 | 57979200 | 57983399 | 1623.136 | 1971.715 | 2660.663 | 1122.951 | -0.46853 | 6.82795 | 0.410909 |
| 4 | 12828600 | 12829799 | 19.84507 | 34.83745 | 33.23612 | 14.70214 | 0.454989 | 0.635235 | 0.410915 |
| 18 | 38429600 | 38430199 | 5.526067 | 10.73548 | 23.42442 | 8.744372 | -0.52162 | -0.27806 | 0.41092 |
| 3 | 51480000 | 51483199 | 24.8972 | 41.1564 | 70.06469 | 30.27954 | -0.38469 | 1.333444 | 0.410931 |
| 5 | 85548800 | 85551199 | 21.79035 | 38.46471 | 30.81085 | 13.63252 | 0.708215 | 0.577912 | 0.41095 |
| 5 | 109735000 | 109737199 | 20.65678 | 36.13401 | 34.93057 | 15.40547 | 0.436209 | 0.698662 | 0.410951 |
| 18 | 90329600 | 90333799 | 61.38269 | 98.57533 | 82.38768 | 34.88035 | 0.608265 | 1.934575 | 0.410952 |
| 7 | 41873200 | 41877999 | 91.00438 | 142.795 | 133.5431 | 55.38368 | 0.414857 | 2.586865 | 0.410959 |
| X | 5099800 | 5103599 | 43.54571 | 70.5441 | 69.51568 | 29.75369 | 0.385425 | 1.660761 | 0.410965 |
| 10 | 126431800 | 126438999 | 93.57476 | 146.0538 | 143.6371 | 59.60217 | 0.336457 | 2.67949 | 0.410966 |
| 18 | 54030600 | 54031799 | 13.90749 | 25.77073 | 24.84011 | 10.4388 | 0.494142 | 0.210054 | 0.410976 |
| 8 | 26652400 | 26652799 | 9.802243 | 17.77273 | 33.883 | 14.65303 | -0.51155 | 0.324618 | 0.410986 |
| 1 | 144712200 | 144716999 | 84.44196 | 132.7052 | 131.3188 | 54.47229 | 0.335174 | 2.54927 | 0.410998 |
| 7 | 35778200 | 35781599 | 31.21415 | 50.64673 | 84.71252 | 36.30032 | -0.3739 | 1.614204 | 0.411017 |
| 7 | 25609800 | 25612399 | 16.83542 | 29.09428 | 51.14564 | 22.43805 | -0.42495 | 0.853893 | 0.411021 |
| 6 | 122583400 | 122586799 | 26.78352 | 43.77018 | 76.8334 | 33.03206 | -0.43538 | 1.486778 | 0.411028 |
| 18 | 77728800 | 77729999 | 8.725368 | 16.4264 | 32.52682 | 13.32156 | -0.50786 | 0.08726 | 0.411031 |
| 6 | 123113000 | 123116599 | 43.19696 | 69.30302 | 114.2664 | 48.30512 | -0.37173 | 1.953534 | 0.411041 |
| 8 | 4778000 | 4779599 | 22.06524 | 38.16786 | 37.35859 | 16.39002 | 0.416592 | 0.85018 | 0.411048 |
| 12 | 43974800 | 43976199 | 15.25719 | 26.60355 | 46.27855 | 20.36462 | -0.40762 | 0.737111 | 0.411049 |
| Y | 4055200 | 4059399 | 20.16556 | 35.42294 | 19.08823 | 8.448377 | 1.273862 | 0.597277 | 0.411052 |
| 6 | 70299400 | 70300599 | 9.681529 | 17.94032 | 34.82585 | 14.63483 | -0.50909 | 0.198571 | 0.411054 |
| 6 | 129894400 | 129901199 | 203.9617 | 301.9656 | 285.1271 | 117.4241 | 0.331913 | 3.710428 | 0.411058 |
| 18 | 7625400 | 7627199 | 12.70443 | 22.33074 | 40.90554 | 18.11427 | -0.48122 | 0.588914 | 0.41107 |
| 2 | 68351400 | 68352799 | 15.32136 | 28.73606 | 22.6184 | 9.322518 | 0.805904 | 0.143574 | 0.411079 |
| 5 | 130632600 | 130635599 | 29.0433 | 48.80844 | 48.03104 | 20.98021 | 0.404014 | 1.143182 | 0.411099 |
| X | 3056800 | 3060199 | 49.66918 | 80.29856 | 72.38556 | 30.9286 | 0.510754 | 1.71723 | 0.41113 |
| 14 | 114423000 | 114429199 | 81.55515 | 128.8009 | 113.0095 | 46.92776 | 0.513964 | 2.443569 | 0.411138 |
| 17 | 72383600 | 72387199 | 19.33697 | 33.02101 | 57.07994 | 25.02546 | -0.40404 | 0.980082 | 0.411145 |
| 3 | 71676400 | 71677799 | 12.20829 | 23.61563 | 18.37229 | 7.01675 | 0.919851 | -0.1429 | 0.411147 |
| 9 | 87614600 | 87616199 | 11.26071 | 20.0175 | 37.29205 | 16.44884 | -0.49805 | 0.461714 | 0.411151 |
| 12 | 101253600 | 101254999 | 12.33997 | 22.02204 | 40.3887 | 17.75604 | -0.47259 | 0.449333 | 0.41117 |
| 18 | 47916200 | 47917799 | 12.33997 | 22.02204 | 40.3887 | 17.75604 | -0.47259 | 0.449333 | 0.41117 |
| 17 | 39246800 | 39248399 | 15.55199 | 29.43334 | 21.73907 | 8.811118 | 0.917322 | 0.087665 | 0.411176 |
| 5 | 76890400 | 76891799 | 14.72932 | 27.05581 | 25.75276 | 10.95195 | 0.499906 | 0.266815 | 0.411183 |
| 8 | 78430000 | 78431399 | 14.72932 | 27.05581 | 25.75276 | 10.95195 | 0.499906 | 0.266815 | 0.411183 |
| 7 | 120990600 | 120995199 | 30.80371 | 49.73771 | 91.01732 | 38.82742 | -0.51038 | 1.729631 | 0.411185 |
| 2 | 122015800 | 122017599 | 24.09729 | 41.67289 | 31.74224 | 13.9259 | 0.776742 | 0.853952 | 0.411207 |
| 1 | 101134200 | 101135399 | 18.04679 | 31.92549 | 30.59745 | 13.53225 | 0.451829 | 0.530532 | 0.411219 |
| 9 | 38804600 | 38805799 | 13.66721 | 24.05786 | 42.41567 | 18.77772 | -0.42559 | 0.581701 | 0.411237 |
| 3 | 91041600 | 91045799 | 81.36695 | 128.7697 | 117.9074 | 49.0129 | 0.454872 | 2.397268 | 0.411245 |
| 4 | 140228200 | 140230599 | 16.96175 | 28.90081 | 54.79107 | 24.01767 | -0.53711 | 0.988155 | 0.411245 |
| 6 | 129870200 | 129876199 | 115.3929 | 178.1261 | 165.0718 | 68.56757 | 0.409823 | 2.886 | 0.411253 |
| 8 | 39342200 | 39343599 | 10.52972 | 19.17962 | 34.938 | 15.02275 | -0.4413 | 0.303926 | 0.411277 |

| | | | | | | | | | |
|---|---|---|---|---|---|---|---|---|---|
| 13 | 61057400 | 61057999 | 9.333025 | 18.06645 | 19.28435 | 7.340913 | 0.473014 | -0.18123 | 0.411281 |
| 7 | 25999800 | 26008999 | 251.4865 | 364.9843 | 342.1022 | 141.0392 | 0.315181 | 4.028203 | 0.411283 |
| 7 | 40767000 | 40767999 | 14.8452 | 27.07625 | 26.3306 | 11.30284 | 0.460115 | 0.30974 | 0.41136 |
| 8 | 23697800 | 23698999 | 10.05146 | 17.86979 | 37.35883 | 16.47682 | -0.66317 | 0.452956 | 0.411376 |
| 1 | 109910000 | 109911599 | 16.2457 | 27.5551 | 56.84573 | 24.90023 | -0.65925 | 1.029199 | 0.411382 |
| 7 | 26296400 | 26306999 | 159.1817 | 239.9162 | 229.8138 | 94.94537 | 0.334533 | 3.375021 | 0.411399 |
| X | 17437800 | 17439599 | 20.63462 | 36.53155 | 29.98989 | 13.2615 | 0.674709 | 0.540135 | 0.411401 |
| X | 80411400 | 80414199 | 33.56322 | 58.11626 | 32.77159 | 14.37805 | 1.209036 | 0.889752 | 0.411401 |
| 15 | 34985200 | 34988999 | 42.42854 | 69.2781 | 63.41323 | 27.25396 | 0.500564 | 1.523014 | 0.411402 |
| 17 | 20944400 | 20947199 | 34.26501 | 58.26686 | 41.80982 | 18.32439 | 0.858792 | 1.058333 | 0.411405 |
| 2 | 21197400 | 21199599 | 19.52353 | 34.30109 | 32.37098 | 14.32936 | 0.470747 | 0.621262 | 0.411407 |
| 11 | 117474800 | 117482999 | 76.40901 | 118.8515 | 183.4854 | 76.17356 | -0.31724 | 2.736371 | 0.411408 |
| 8 | 27427000 | 27430999 | 36.93201 | 59.46442 | 99.2662 | 42.18594 | -0.38314 | 1.826555 | 0.41141 |
| 7 | 129266800 | 129267999 | 7.236425 | 13.61721 | 28.58648 | 11.71665 | -0.59144 | 0.06898 | 0.411415 |
| 11 | 4586800 | 4588199 | 8.106816 | 15.18032 | 32.33174 | 13.36241 | -0.62299 | 0.113568 | 0.411421 |
| 19 | 51684400 | 51685799 | 15.28887 | 28.82138 | 21.55265 | 8.807054 | 0.88895 | 0.109354 | 0.411446 |
| 2 | 15198000 | 15201199 | 78.982 | 124.2741 | 116.4125 | 48.2966 | 0.415528 | 2.520064 | 0.41145 |
| 19 | 16254800 | 16256399 | 14.14164 | 24.71321 | 44.15442 | 19.4574 | -0.44554 | 0.696668 | 0.411452 |
| 11 | 14050000 | 14052399 | 29.01576 | 49.96249 | 39.04094 | 17.12572 | 0.73887 | 0.922782 | 0.411455 |
| 1 | 97104600 | 97106399 | 22.87563 | 39.78805 | 33.07222 | 14.5304 | 0.652574 | 0.788759 | 0.411461 |
| 14 | 46663200 | 46667199 | 18.13931 | 30.83801 | 62.97522 | 27.60021 | -0.64458 | 1.023566 | 0.411469 |
| 1 | 121608200 | 121610399 | 25.42733 | 43.32944 | 43.15949 | 18.91911 | 0.387757 | 1.005825 | 0.411476 |
| 8 | 67055600 | 67057199 | 15.54858 | 27.07148 | 47.35878 | 20.82056 | -0.41604 | 0.760066 | 0.411479 |
| 9 | 19599000 | 19602999 | 54.18038 | 87.02315 | 85.07751 | 36.02802 | 0.382812 | 1.92165 | 0.411486 |
| 12 | 41428600 | 41430599 | 14.80846 | 25.93256 | 46.19634 | 20.39312 | -0.44139 | 0.670022 | 0.411489 |
| 7 | 42460400 | 42462799 | 28.54413 | 47.94952 | 46.98419 | 20.52029 | 0.410258 | 1.146608 | 0.411496 |
| 2 | 73933200 | 73933799 | 10.61068 | 20.20231 | 20.04794 | 8.023278 | 0.510138 | -0.00485 | 0.411505 |
| 14 | 40723000 | 40727199 | 18.49104 | 30.78882 | 66.39842 | 28.83828 | -0.72375 | 1.226861 | 0.41151 |
| 10 | 22344200 | 22350599 | 132.9233 | 202.2207 | 197.8908 | 81.99179 | 0.31501 | 3.178371 | 0.411512 |
| 14 | 76509000 | 76517599 | 89.84251 | 138.2436 | 210.5334 | 87.43608 | -0.30964 | 2.940936 | 0.411522 |
| 11 | 71981200 | 71983999 | 11.15748 | 19.6955 | 38.90184 | 17.21243 | -0.5866 | 0.529647 | 0.411523 |
| 12 | 71403200 | 71404199 | 15.34194 | 27.84329 | 26.64428 | 11.50893 | 0.477158 | 0.340437 | 0.411526 |
| 9 | 46509000 | 46514599 | 46.49412 | 74.16027 | 120.7346 | 50.61001 | -0.36296 | 2.138295 | 0.411528 |
| 4 | 52438600 | 52439599 | 6.530032 | 12.21336 | 29.57802 | 12.24869 | -0.80712 | 0.107321 | 0.411534 |
| 12 | 86584200 | 86587199 | 23.68831 | 39.42332 | 66.90139 | 29.04756 | -0.37935 | 1.248531 | 0.411546 |
| 7 | 85188600 | 85196599 | 28.96366 | 49.59898 | 37.6133 | 16.49446 | 0.780569 | 0.969227 | 0.411578 |
| 5 | 119065400 | 119068599 | 27.60514 | 45.05971 | 77.94179 | 33.49417 | -0.41543 | 1.50667 | 0.41158 |
| 14 | 99263200 | 99266199 | 40.64923 | 66.20463 | 63.7861 | 27.38887 | 0.424233 | 1.553171 | 0.411589 |
| 19 | 39823000 | 39823999 | 17.50692 | 30.95857 | 29.83585 | 13.19643 | 0.443859 | 0.531631 | 0.411598 |
| 8 | 65773400 | 65778199 | 40.93422 | 65.58551 | 127.4655 | 53.77605 | -0.61085 | 1.995678 | 0.411604 |
| 3 | 3341800 | 3352399 | 116.3847 | 180.3306 | 153.6301 | 63.81367 | 0.535191 | 2.817105 | 0.411604 |
| 11 | 118868800 | 118870799 | 12.05354 | 21.32143 | 38.80172 | 17.16102 | -0.46813 | 0.511239 | 0.411607 |
| 2 | 65954200 | 65955599 | 11.45678 | 20.25766 | 38.35901 | 16.96658 | -0.52511 | 0.512308 | 0.411609 |
| 5 | 59418200 | 59418799 | 16.80693 | 29.89849 | 29.22366 | 12.8821 | 0.427717 | 0.47655 | 0.411633 |
| 6 | 87210800 | 87212199 | 12.92842 | 23.90417 | 23.43459 | 9.877494 | 0.466901 | 0.211034 | 0.411635 |
| 2 | 99782200 | 99783799 | 18.00671 | 32.16768 | 27.75223 | 12.19023 | 0.609834 | 0.452909 | 0.411642 |
| 10 | 68900400 | 68901599 | 16.39561 | 30.64098 | 21.85033 | 9.060708 | 0.940186 | 0.172639 | 0.411653 |

| | | | | | | | | | |
|---|---|---|---|---|---|---|---|---|---|
| 9 | 84256400 | 84258999 | 22.66607 | 39.86277 | 31.90636 | 14.12769 | 0.707407 | 0.624647 | 0.411657 |
| 3 | 5983400 | 5984999 | 21.02138 | 37.4331 | 28.70787 | 12.64408 | 0.776555 | 0.497279 | 0.411657 |
| 2 | 90115800 | 90116799 | 16.32334 | 29.07526 | 27.84704 | 12.26208 | 0.458012 | 0.463152 | 0.411707 |
| 5 | 16164200 | 16165599 | 16.47749 | 29.84387 | 27.00322 | 11.69344 | 0.555116 | 0.360774 | 0.411726 |
| 10 | 32817000 | 32818599 | 24.33337 | 42.61197 | 33.58511 | 14.8258 | 0.729471 | 0.700634 | 0.411728 |
| 19 | 45241000 | 45243799 | 26.88547 | 44.17409 | 74.49205 | 32.09546 | -0.37382 | 1.415293 | 0.411777 |
| 1 | 121210200 | 121211999 | 15.6836 | 27.31536 | 48.96576 | 21.5314 | -0.45115 | 0.757299 | 0.411785 |
| 6 | 130573200 | 130575399 | 30.74302 | 52.51531 | 41.8303 | 18.34006 | 0.709546 | 1.003482 | 0.411796 |
| 10 | 13334400 | 13336999 | 39.65071 | 65.33651 | 56.75768 | 24.48439 | 0.581991 | 1.395768 | 0.411811 |
| 9 | 19003200 | 19005399 | 33.6208 | 55.49156 | 55.8094 | 24.09329 | 0.372154 | 1.360812 | 0.411824 |
| 5 | 148674600 | 148682799 | 73.49858 | 114.2171 | 185.0318 | 76.82632 | -0.3877 | 2.750684 | 0.41183 |
| X | 41215800 | 41219799 | 71.0805 | 113.2569 | 101.4868 | 42.4795 | 0.495388 | 2.191128 | 0.411846 |
| 18 | 50463200 | 50464599 | 16.72824 | 31.00475 | 24.0207 | 10.09129 | 0.807468 | 0.229935 | 0.411867 |
| 4 | 30632000 | 30635399 | 37.92781 | 63.08729 | 51.55366 | 22.37544 | 0.671529 | 1.282165 | 0.411876 |
| 15 | 77333800 | 77335199 | 8.951503 | 16.47899 | 31.96186 | 13.55183 | -0.51674 | 0.224684 | 0.411912 |
| 5 | 109589800 | 109594399 | 59.88736 | 96.14067 | 84.42119 | 35.72246 | 0.536563 | 1.943709 | 0.411912 |
| 1 | 118122400 | 118124199 | 25.17667 | 43.48597 | 39.74531 | 17.43634 | 0.51456 | 0.879714 | 0.411916 |
| 1 | 34608600 | 34610399 | 26.26816 | 44.72118 | 44.19586 | 19.37198 | 0.398899 | 1.018575 | 0.411921 |
| 4 | 130129400 | 130134599 | 44.01681 | 70.31893 | 116.9047 | 49.11301 | -0.39065 | 2.085533 | 0.411925 |
| 16 | 75581600 | 75583799 | 15.55802 | 28.60279 | 25.67485 | 10.90595 | 0.585166 | 0.268409 | 0.41193 |
| 17 | 17843600 | 17846199 | 29.0354 | 46.93083 | 97.66525 | 41.71881 | -0.69376 | 1.699199 | 0.411933 |
| 12 | 115703600 | 115707399 | 53.91251 | 86.84799 | 77.9076 | 33.13287 | 0.511844 | 1.826483 | 0.41195 |
| 12 | 21692000 | 21712799 | 879.0379 | 1147.449 | 959.0562 | 401.5976 | 0.327232 | 5.668941 | 0.411958 |
| X | 147346200 | 147349399 | 20.90051 | 37.09245 | 29.1544 | 12.87735 | 0.738416 | 0.516798 | 0.411966 |
| 10 | 96276400 | 96278199 | 16.80165 | 30.40345 | 27.60251 | 11.9665 | 0.549461 | 0.370985 | 0.411966 |
| 13 | 18097600 | 18103799 | 64.47344 | 103.3001 | 87.95691 | 37.10768 | 0.577621 | 2.011666 | 0.411985 |
| 2 | 127779200 | 127780199 | 11.4897 | 21.826 | 22.09515 | 8.885427 | 0.475657 | 0.025424 | 0.411999 |
| 4 | 72871000 | 72871799 | 11.4897 | 21.826 | 22.09515 | 8.885427 | 0.475657 | 0.025424 | 0.411999 |
| 16 | 90163400 | 90168999 | 44.55866 | 71.02392 | 122.0664 | 51.11927 | -0.44206 | 2.158898 | 0.412019 |
| 16 | 10993200 | 10994799 | 11.91061 | 20.9887 | 40.38062 | 17.87289 | -0.55018 | 0.554332 | 0.412023 |
| 8 | 34619000 | 34619999 | 7.336523 | 13.9589 | 27.73366 | 11.1152 | -0.4847 | -0.00872 | 0.412033 |
| 8 | 115582600 | 115585599 | 19.47769 | 32.7556 | 65.6528 | 28.68348 | -0.61859 | 1.125165 | 0.412034 |
| 7 | 43291200 | 43296599 | 45.9477 | 73.42116 | 119.0959 | 50.05051 | -0.35488 | 2.080516 | 0.412035 |
| 2 | 154433800 | 154436999 | 21.53198 | 35.28138 | 82.67224 | 35.57874 | -0.84854 | 1.441663 | 0.412036 |
| 5 | 53184000 | 53188199 | 29.53878 | 48.04922 | 83.94215 | 36.0272 | -0.43339 | 1.561841 | 0.41204 |
| X | 117521000 | 117525799 | 60.55578 | 96.88669 | 92.29066 | 38.85526 | 0.413764 | 2.051805 | 0.412044 |
| 13 | 61830400 | 61836399 | 114.2369 | 176.2162 | 167.0119 | 69.3737 | 0.37666 | 2.898069 | 0.412054 |
| 3 | 50195200 | 50196199 | 11.43714 | 21.91668 | 21.10606 | 8.315757 | 0.573288 | -0.04366 | 0.412072 |
| 7 | 113541800 | 113543599 | 33.34605 | 55.01114 | 52.99968 | 22.87441 | 0.433811 | 1.365097 | 0.412074 |
| 3 | 138287200 | 138292199 | 41.8444 | 67.99638 | 66.00015 | 28.313 | 0.411189 | 1.591246 | 0.412077 |
| 18 | 51998200 | 51999999 | 13.19389 | 23.16839 | 41.44947 | 18.33997 | -0.4473 | 0.609364 | 0.412089 |
| X | 11639800 | 11641199 | 14.21907 | 26.83235 | 22.92198 | 9.348032 | 0.701335 | 0.098207 | 0.412112 |
| 1 | 8839600 | 8844999 | 66.63426 | 106.8624 | 83.26199 | 35.17714 | 0.707011 | 1.984375 | 0.41212 |
| 11 | 9684600 | 9685999 | 14.9662 | 26.05191 | 46.31249 | 20.35783 | -0.43892 | 0.76074 | 0.412124 |
| 17 | 25001400 | 25003199 | 9.702223 | 17.82588 | 33.50444 | 14.24398 | -0.47484 | 0.244184 | 0.412139 |
| 18 | 47142600 | 47145199 | 30.9997 | 52.00047 | 44.31367 | 19.33658 | 0.61095 | 1.165097 | 0.41214 |
| 8 | 115854600 | 115870399 | 164.055 | 242.6073 | 358.2806 | 147.5492 | -0.31451 | 3.726946 | 0.412167 |

| | | | | | | | | | |
|---|---|---|---|---|---|---|---|---|---|
| 18 | 5698200 | 5699799 | 11.45054 | 20.22727 | 38.86116 | 17.19198 | -0.54658 | 0.522835 | 0.412175 |
| X | 66947800 | 66950999 | 37.04491 | 62.48066 | 43.50322 | 19.02325 | 0.901619 | 1.137744 | 0.412199 |
| 16 | 5397800 | 5400399 | 21.84714 | 36.70294 | 62.49761 | 27.29657 | -0.38432 | 1.138323 | 0.412217 |
| 19 | 8870000 | 8870799 | 5.373757 | 10.44753 | 22.89207 | 8.493321 | -0.5189 | -0.302 | 0.412226 |
| 14 | 115362200 | 115364999 | 33.33359 | 55.60507 | 51.44595 | 22.3701 | 0.492915 | 1.236545 | 0.412226 |
| 4 | 47207400 | 47210399 | 16.66601 | 28.35433 | 55.83898 | 24.47406 | -0.59212 | 1.001716 | 0.41223 |
| 8 | 23261800 | 23262799 | 5.707703 | 10.9534 | 25.70026 | 10.08376 | -0.69306 | -0.09794 | 0.412232 |
| 15 | 57755200 | 57756399 | 7.066408 | 13.36191 | 28.13342 | 11.42415 | -0.58436 | 0.032684 | 0.412251 |
| 10 | 47659800 | 47660599 | 9.723826 | 17.98226 | 33.64939 | 14.17944 | -0.45908 | 0.209499 | 0.41226 |
| 12 | 95946000 | 95949599 | 31.27295 | 53.20618 | 43.50936 | 19.07004 | 0.670977 | 1.04303 | 0.412282 |
| 17 | 74162000 | 74163199 | 19.38746 | 34.41331 | 29.84707 | 13.18133 | 0.597294 | 0.507152 | 0.412299 |
| 3 | 11039200 | 11040399 | 17.68679 | 31.16248 | 29.82924 | 13.20316 | 0.451452 | 0.570552 | 0.412316 |
| 6 | 44241600 | 44243399 | 23.43567 | 41.36629 | 29.5423 | 13.07142 | 0.872827 | 0.586078 | 0.412331 |
| 11 | 8768800 | 8772399 | 19.47679 | 33.26524 | 59.89371 | 26.25945 | -0.46267 | 0.981085 | 0.412332 |
| 5 | 20897200 | 20900399 | 60.03428 | 95.88059 | 96.09476 | 40.35442 | 0.337732 | 2.104239 | 0.41234 |
| 17 | 33212000 | 33214399 | 16.07691 | 27.81179 | 48.98969 | 21.49288 | -0.42735 | 0.84017 | 0.412352 |
| 8 | 117669800 | 117672399 | 12.15269 | 21.20828 | 44.73566 | 19.69616 | -0.68391 | 0.706853 | 0.412357 |
| 9 | 32259000 | 32261399 | 17.19397 | 29.70092 | 51.13846 | 22.4346 | -0.39517 | 0.860644 | 0.412359 |
| 10 | 105469200 | 105471599 | 12.83949 | 22.79594 | 42.95823 | 18.96775 | -0.51599 | 0.489193 | 0.412361 |
| 15 | 60287400 | 60288799 | 9.092371 | 16.95687 | 32.20374 | 13.40127 | -0.4663 | 0.151809 | 0.41237 |
| 14 | 48645200 | 48647199 | 14.59978 | 25.53467 | 44.63963 | 19.6848 | -0.41438 | 0.685933 | 0.412376 |
| 6 | 145119600 | 145122999 | 24.04995 | 39.38452 | 75.55015 | 32.50576 | -0.56105 | 1.449874 | 0.412376 |
| 12 | 31999200 | 32000999 | 13.49071 | 23.23666 | 51.58432 | 22.62821 | -0.76086 | 0.870501 | 0.412394 |
| 13 | 3089200 | 3101999 | 60.33869 | 96.47984 | 83.84164 | 35.26732 | 0.544976 | 2.065703 | 0.412404 |
| 11 | 87627600 | 87628999 | 7.253344 | 13.57472 | 31.73091 | 13.12627 | -0.75697 | 0.108711 | 0.412423 |
| 13 | 74676200 | 74679199 | 143.1633 | 211.2138 | 361.5519 | 148.8927 | -0.53059 | 3.764361 | 0.412431 |
| 1 | 188926400 | 188928599 | 22.03987 | 36.88617 | 62.97743 | 27.44094 | -0.38808 | 1.180161 | 0.412436 |
| 9 | 69164800 | 69166999 | 14.62521 | 25.57761 | 44.64789 | 19.68743 | -0.41217 | 0.687931 | 0.412449 |
| 9 | 121014800 | 121028999 | 239.7508 | 343.011 | 494.7619 | 204.39 | -0.32687 | 4.240149 | 0.412454 |
| 13 | 112287000 | 112287799 | 5.083524 | 9.877682 | 22.81035 | 8.501727 | -0.59941 | -0.28837 | 0.412464 |
| 7 | 38093400 | 38097199 | 54.45412 | 87.45235 | 85.23686 | 36.08892 | 0.386988 | 1.925029 | 0.412469 |
| 10 | 116232400 | 116235399 | 21.87509 | 36.49952 | 65.761 | 28.59903 | -0.46546 | 1.212859 | 0.412479 |
| 5 | 126657600 | 126660399 | 30.82917 | 52.15872 | 45.76398 | 20.03069 | 0.569258 | 1.090065 | 0.412482 |
| 12 | 43080000 | 43081199 | 13.46698 | 24.81786 | 23.77341 | 10.06542 | 0.495175 | 0.233164 | 0.412489 |
| 5 | 141708400 | 141711799 | 31.25376 | 50.54317 | 88.15213 | 37.68088 | -0.43863 | 1.683295 | 0.412522 |
| 19 | 29852000 | 29852599 | 11.25219 | 21.77519 | 19.72228 | 7.517748 | 0.70588 | -0.15445 | 0.412534 |
| X | 91573000 | 91574599 | 18.67022 | 34.50038 | 23.28675 | 9.822982 | 1.000919 | 0.260586 | 0.412541 |
| 11 | 97027000 | 97031999 | 27.42894 | 44.38091 | 87.42625 | 37.38613 | -0.61301 | 1.665436 | 0.412543 |
| 2 | 168970000 | 168976999 | 86.13281 | 132.4427 | 210.4264 | 87.38327 | -0.37145 | 2.952618 | 0.41255 |
| 14 | 89728000 | 89730399 | 31.02688 | 53.07622 | 41.2894 | 18.10499 | 0.743864 | 0.992491 | 0.412561 |
| 10 | 33854800 | 33857399 | 49.86118 | 80.21313 | 80.71117 | 34.25798 | 0.344151 | 1.864248 | 0.412565 |
| 1 | 104135600 | 104137999 | 20.57364 | 36.28855 | 30.80785 | 13.63333 | 0.624139 | 0.576894 | 0.412578 |
| 15 | 72676400 | 72685599 | 170.3655 | 249.6849 | 385.7616 | 158.8974 | -0.39186 | 3.873477 | 0.41261 |
| 19 | 13404000 | 13406599 | 26.70444 | 45.62313 | 37.91136 | 16.62203 | 0.648765 | 0.982088 | 0.412643 |
| 14 | 64792600 | 64793799 | 12.85814 | 22.7626 | 40.31208 | 17.8262 | -0.4287 | 0.51099 | 0.412645 |
| 7 | 32241800 | 32243199 | 29.96048 | 50.65554 | 40.59158 | 17.7632 | 0.699699 | 1.089656 | 0.412665 |
| 4 | 100174600 | 100175399 | 8.437122 | 15.86941 | 30.21106 | 12.39206 | -0.45283 | 0.086507 | 0.41267 |

| | | | | | | | | | |
|---|---|---|---|---|---|---|---|---|---|
| 6 | 48129000 | 48129799 | 8.437122 | 15.86941 | 30.21106 | 12.39206 | -0.45283 | 0.086507 | 0.41267 |
| 11 | 63711400 | 63713199 | 16.37353 | 29.7143 | 27.20242 | 11.75059 | 0.540954 | 0.35157 | 0.41268 |
| 13 | 62960600 | 62965999 | 95.80051 | 148.7437 | 145.9186 | 60.59425 | 0.33418 | 2.771375 | 0.412685 |
| 13 | 70319600 | 70321199 | 9.743097 | 17.78355 | 33.4042 | 14.32614 | -0.48271 | 0.282624 | 0.412687 |
| 4 | 25300200 | 25300799 | 5.096411 | 9.924003 | 22.46866 | 8.230362 | -0.54891 | -0.35393 | 0.412717 |
| 2 | 93899200 | 93900799 | 18.93989 | 33.75963 | 29.41305 | 12.94397 | 0.594279 | 0.478421 | 0.412719 |
| 4 | 66492600 | 66496599 | 36.34151 | 59.76176 | 59.1815 | 25.508 | 0.392442 | 1.415824 | 0.412736 |
| 19 | 29908200 | 29909199 | 8.821393 | 17.1059 | 18.22775 | 6.882329 | 0.486359 | -0.21834 | 0.41274 |
| 7 | 47969000 | 47972999 | 62.88423 | 100.3133 | 99.70463 | 41.80039 | 0.347867 | 2.144941 | 0.412758 |
| 6 | 55086400 | 55087799 | 16.16053 | 28.93447 | 28.26025 | 12.38991 | 0.434138 | 0.43074 | 0.412763 |
| 1 | 191746000 | 191749999 | 46.9632 | 75.01895 | 119.534 | 50.211 | -0.32968 | 2.092756 | 0.412767 |
| 3 | 16567400 | 16570199 | 34.3505 | 57.13714 | 48.18632 | 20.91399 | 0.626164 | 1.268408 | 0.412776 |
| 8 | 15370600 | 15372799 | 20.63183 | 36.18034 | 33.71092 | 14.89887 | 0.488937 | 0.656724 | 0.412778 |
| 4 | 78234400 | 78235999 | 16.84462 | 31.04974 | 25.10948 | 10.62768 | 0.738586 | 0.263828 | 0.412786 |
| 10 | 36046800 | 36048399 | 23.36302 | 40.83126 | 35.29557 | 15.55464 | 0.596859 | 0.728189 | 0.412786 |
| 6 | 24390200 | 24393199 | 27.1064 | 44.24318 | 78.57858 | 33.76714 | -0.45354 | 1.505336 | 0.412793 |
| 16 | 74160000 | 74160599 | 9.036622 | 15.94582 | 42.5654 | 18.83479 | -1.01805 | 0.53092 | 0.412829 |
| 2 | 5164400 | 5166399 | 16.43112 | 28.23267 | 51.09985 | 22.41403 | -0.46845 | 0.905422 | 0.412836 |
| 2 | 36898800 | 36902599 | 54.98644 | 88.33536 | 83.11073 | 35.20546 | 0.438412 | 1.917812 | 0.412843 |
| 3 | 157652400 | 157653999 | 9.548116 | 17.11028 | 36.35818 | 15.92848 | -0.67979 | 0.396327 | 0.412893 |
| 3 | 56116000 | 56117999 | 16.17679 | 27.69208 | 56.95276 | 24.97462 | -0.65299 | 0.946015 | 0.412918 |
| 8 | 60277200 | 60277799 | 5.231268 | 10.18588 | 22.46363 | 8.233311 | -0.51207 | -0.34944 | 0.412924 |
| 11 | 11969600 | 11979599 | 177.1975 | 260.0011 | 384.6718 | 158.4369 | -0.3278 | 3.85568 | 0.412948 |
| 15 | 31288400 | 31289199 | 7.866287 | 14.89074 | 32.06627 | 12.99099 | -0.61354 | 0.04553 | 0.412967 |
| X | 32217000 | 32223799 | 80.44696 | 127.7958 | 94.42337 | 39.37494 | 0.768815 | 2.291196 | 0.412979 |
| 9 | 78654000 | 78655999 | 20.17116 | 35.31586 | 34.41188 | 15.18809 | 0.424778 | 0.681208 | 0.412983 |
| 16 | 57686800 | 57688599 | 15.28294 | 28.46539 | 23.60999 | 9.846474 | 0.717413 | 0.188235 | 0.412991 |
| 11 | 63203000 | 63205399 | 20.06309 | 33.80279 | 59.11452 | 25.84148 | -0.42238 | 1.10423 | 0.412994 |
| 4 | 21254000 | 21254599 | 9.094883 | 17.63525 | 17.2648 | 6.520433 | 0.608197 | -0.21449 | 0.413004 |
| 18 | 40232400 | 40234999 | 25.66254 | 44.3076 | 39.71904 | 17.4246 | 0.542284 | 0.885206 | 0.413016 |
| 18 | 85457600 | 85459399 | 21.68923 | 39.33324 | 23.84862 | 10.31099 | 1.130625 | 0.386401 | 0.413019 |
| 4 | 64669800 | 64671199 | 12.86492 | 22.61058 | 40.90729 | 18.11351 | -0.4634 | 0.591746 | 0.413034 |
| 7 | 58774800 | 58775599 | 9.768816 | 18.77024 | 18.69213 | 7.307893 | 0.536351 | -0.09084 | 0.413058 |
| 19 | 3296800 | 3298199 | 27.81055 | 46.73737 | 44.89476 | 19.61035 | 0.438847 | 1.137551 | 0.413061 |
| 7 | 4874600 | 4875799 | 6.418315 | 12.39124 | 25.92511 | 9.99243 | -0.50552 | -0.14874 | 0.413069 |
| 11 | 63117400 | 63118799 | 6.418315 | 12.39124 | 25.92511 | 9.99243 | -0.50552 | -0.14874 | 0.413069 |
| 4 | 140995200 | 141000599 | 43.85173 | 70.05728 | 116.9026 | 49.11418 | -0.39595 | 2.084351 | 0.413074 |
| 2 | 55865400 | 55868199 | 22.86378 | 40.04393 | 34.14824 | 15.07611 | 0.616238 | 0.690253 | 0.413086 |
| 5 | 22757400 | 22760399 | 40.07107 | 64.22168 | 109.2466 | 46.11062 | -0.41842 | 1.97889 | 0.413114 |
| 7 | 12108400 | 12113399 | 47.04954 | 77.08745 | 57.4921 | 24.74144 | 0.798343 | 1.485506 | 0.41312 |
| 5 | 60684600 | 60687999 | 82.88574 | 130.8893 | 113.3462 | 47.06521 | 0.532737 | 2.446965 | 0.413149 |
| 10 | 50688600 | 50691199 | 43.89437 | 70.31036 | 114.4319 | 48.2586 | -0.35574 | 2.004596 | 0.413165 |
| 17 | 68156600 | 68157799 | 8.285701 | 15.3997 | 34.33345 | 14.3694 | -0.70331 | 0.167006 | 0.413177 |
| 7 | 19429800 | 19431199 | 17.94065 | 30.51642 | 54.78279 | 24.01073 | -0.45907 | 1.009167 | 0.413181 |
| 7 | 107263200 | 107265799 | 15.07667 | 26.21078 | 47.37971 | 20.81027 | -0.46328 | 0.778824 | 0.4132 |
| 12 | 12142400 | 12143199 | 13.19575 | 23.07388 | 43.635 | 19.23814 | -0.52705 | 0.680654 | 0.413211 |
| 12 | 113609400 | 113613399 | 40.35474 | 66.34799 | 58.88559 | 25.3783 | 0.549963 | 1.428425 | 0.413218 |

| | | | | | | | | | |
|---|---|---|---|---|---|---|---|---|---|
| 10 | 90123600 | 90126199 | 31.30602 | 52.39623 | 49.72881 | 21.66872 | 0.456128 | 1.193524 | 0.413223 |
| 12 | 26346800 | 26348799 | 15.76329 | 27.45311 | 47.8869 | 21.0565 | -0.41174 | 0.759439 | 0.413227 |
| 2 | 105778000 | 105779599 | 13.32559 | 23.36894 | 42.4996 | 18.78376 | -0.471 | 0.631664 | 0.413228 |
| 1 | 85034600 | 85042399 | 153.778 | 231.6212 | 224.7874 | 92.85259 | 0.314853 | 3.385458 | 0.413236 |
| 7 | 17937200 | 17938399 | 20.2553 | 35.4233 | 34.42708 | 15.18036 | 0.428481 | 0.698737 | 0.413238 |
| 17 | 59155800 | 59156799 | 20.2553 | 35.4233 | 34.42708 | 15.18036 | 0.428481 | 0.698737 | 0.413238 |
| 6 | 25023800 | 25024399 | 5.516719 | 10.71692 | 23.39456 | 8.735869 | -0.5221 | -0.27509 | 0.413254 |
| 14 | 121826600 | 121830199 | 17.54689 | 29.57568 | 60.08673 | 26.26978 | -0.63778 | 1.091758 | 0.413261 |
| 15 | 36135000 | 36142999 | 75.72191 | 117.4178 | 204.6156 | 84.98511 | -0.49531 | 2.792062 | 0.413274 |
| 4 | 146043800 | 146047199 | 28.32568 | 48.55529 | 41.90758 | 18.37911 | 0.594975 | 0.963958 | 0.413284 |
| X | 114762400 | 114770799 | 228.2505 | 336.5972 | 294.7513 | 121.3826 | 0.43529 | 3.779587 | 0.413304 |
| 7 | 103462800 | 103463399 | 4.515024 | 8.816146 | 20.52847 | 7.228032 | -0.53648 | -0.50905 | 0.413312 |
| 16 | 35933200 | 35933599 | 4.515024 | 8.816146 | 20.52847 | 7.228032 | -0.53648 | -0.50905 | 0.413312 |
| 12 | 22794200 | 22806999 | 343.8252 | 485.7512 | 444.1967 | 183.8852 | 0.312104 | 4.42785 | 0.41333 |
| 5 | 20898400 | 20900399 | 50.98838 | 81.94784 | 82.4151 | 34.93261 | 0.34317 | 1.895993 | 0.413346 |
| 16 | 77713000 | 77715999 | 24.02383 | 41.11542 | 40.01878 | 17.54849 | 0.421834 | 0.956788 | 0.413351 |
| 8 | 24141000 | 24142999 | 18.51538 | 32.77388 | 30.49146 | 13.48378 | 0.494726 | 0.526605 | 0.413359 |
| 3 | 66964400 | 66966399 | 31.55764 | 52.9645 | 49.13604 | 21.44804 | 0.488834 | 1.165899 | 0.413369 |
| 3 | 158280400 | 158282799 | 16.39921 | 29.97787 | 25.43314 | 10.88697 | 0.657915 | 0.304438 | 0.413375 |
| 16 | 71045400 | 71047599 | 13.96586 | 24.39311 | 44.68159 | 19.68279 | -0.48145 | 0.70318 | 0.413381 |
| 11 | 90577200 | 90580399 | 21.91975 | 36.46767 | 66.193 | 28.73328 | -0.47612 | 1.245765 | 0.413386 |
| 9 | 19053600 | 19055999 | 48.54672 | 78.23334 | 78.1083 | 33.23617 | 0.358421 | 1.807755 | 0.413399 |
| 11 | 120978400 | 120980799 | 12.40467 | 21.85018 | 39.88897 | 17.65676 | -0.47496 | 0.563037 | 0.413437 |
| 3 | 6334400 | 6337399 | 54.58992 | 87.55864 | 87.21024 | 36.85286 | 0.353616 | 1.964677 | 0.413478 |
| 13 | 13105800 | 13106199 | 18.27397 | 32.36969 | 24.73223 | 10.93539 | 0.777692 | 0.506827 | 0.413481 |
| 1 | 154772400 | 154777599 | 42.89598 | 69.6199 | 68.21537 | 29.2369 | 0.396029 | 1.619698 | 0.413487 |
| 4 | 11722000 | 11725799 | 43.00574 | 69.87071 | 67.60704 | 28.99841 | 0.415514 | 1.600683 | 0.413493 |
| 2 | 124740400 | 124741999 | 23.22765 | 38.60904 | 66.21065 | 28.72319 | -0.39453 | 1.261407 | 0.413502 |
| 14 | 16310200 | 16312199 | 22.98147 | 39.98547 | 37.62883 | 16.53583 | 0.474351 | 0.791308 | 0.413513 |
| 14 | 43859800 | 43860599 | 23.96264 | 41.05868 | 38.06029 | 16.69125 | 0.492318 | 0.945443 | 0.413516 |
| 1 | 151896400 | 151901599 | 44.97947 | 71.87088 | 117.4208 | 49.3435 | -0.36526 | 2.080511 | 0.413519 |
| X | 125347400 | 125349399 | 33.30084 | 55.77743 | 49.16164 | 21.43189 | 0.562676 | 1.195759 | 0.413539 |
| 4 | 138390600 | 138393599 | 25.75794 | 42.40491 | 72.69169 | 31.34586 | -0.39622 | 1.387802 | 0.413563 |
| 4 | 60317600 | 60317999 | 8.877283 | 17.19499 | 17.74861 | 6.736613 | 0.525323 | -0.20022 | 0.413583 |
| 4 | 45859200 | 45861799 | 24.75852 | 40.84866 | 70.48366 | 30.42181 | -0.40435 | 1.356111 | 0.413604 |
| 17 | 42598200 | 42599599 | 20.33084 | 35.65745 | 33.75383 | 14.91958 | 0.466232 | 0.653241 | 0.413629 |
| 6 | 137363000 | 137363599 | 6.267732 | 12.10325 | 25.36246 | 9.765264 | -0.50627 | -0.15586 | 0.413642 |
| 9 | 80694200 | 80696399 | 27.04408 | 46.45732 | 41.35346 | 18.13882 | 0.551029 | 0.938018 | 0.413648 |
| 16 | 19860800 | 19862599 | 29.90803 | 50.42313 | 40.14462 | 17.55245 | 0.708871 | 1.112784 | 0.413649 |
| 10 | 118575600 | 118577999 | 19.31371 | 32.90814 | 57.06704 | 25.01533 | -0.40917 | 0.999233 | 0.413652 |
| 19 | 61157600 | 61161399 | 64.02858 | 101.8947 | 102.657 | 42.8959 | 0.324689 | 2.222919 | 0.413653 |
| 13 | 98771800 | 98774199 | 17.33222 | 29.399 | 55.80858 | 24.44617 | -0.53971 | 1.032672 | 0.413661 |
| X | 120934000 | 120936599 | 26.98134 | 47.09615 | 34.04628 | 14.99178 | 0.853545 | 0.763078 | 0.413666 |
| 7 | 34232200 | 34244199 | 107.6496 | 163.795 | 250.781 | 103.9103 | -0.33006 | 3.17272 | 0.413673 |
| 7 | 7259800 | 7262199 | 21.28761 | 38.12206 | 27.01892 | 11.84362 | 0.894171 | 0.461194 | 0.413686 |
| 16 | 41093200 | 41095199 | 18.41241 | 33.03267 | 28.39329 | 12.42716 | 0.618935 | 0.433195 | 0.413712 |
| 12 | 117509800 | 117513599 | 21.69914 | 36.46049 | 61.91277 | 27.04252 | -0.38029 | 1.136362 | 0.413714 |

| | | | | | | | | | |
|---|---|---|---|---|---|---|---|---|---|
| Y | 6417600 | 6420199 | 15.87262 | 28.70127 | 17.00022 | 7.37686 | 1.160007 | 0.340935 | 0.413715 |
| 1 | 82932400 | 82934199 | 13.35593 | 23.39356 | 42.55741 | 18.78991 | -0.47132 | 0.653053 | 0.413718 |
| 14 | 50799800 | 50803799 | 55.83781 | 89.73367 | 83.8203 | 35.52358 | 0.449362 | 1.906823 | 0.413722 |
| 13 | 78535000 | 78538999 | 46.43579 | 75.73281 | 62.53048 | 26.86314 | 0.647765 | 1.547939 | 0.413727 |
| 9 | 46491400 | 46492599 | 29.71484 | 50.65064 | 43.56754 | 19.09844 | 0.598748 | 1.021662 | 0.413736 |
| 11 | 73910600 | 73914199 | 43.38659 | 71.5538 | 54.56374 | 23.54649 | 0.770016 | 1.396404 | 0.413738 |
| X | 167916000 | 167920599 | 32.24069 | 52.38822 | 97.73605 | 41.92583 | -0.52921 | 1.59501 | 0.413739 |
| 17 | 80680000 | 80683399 | 28.10952 | 45.83434 | 78.99107 | 33.93196 | -0.41123 | 1.524271 | 0.413744 |
| 2 | 21829600 | 21832799 | 32.10329 | 53.39476 | 53.05411 | 23.02539 | 0.390346 | 1.266912 | 0.413758 |
| 3 | 136147800 | 136148799 | 7.949502 | 15.00542 | 29.19324 | 11.89857 | -0.47638 | 0.055247 | 0.41376 |
| 3 | 157129000 | 157129799 | 7.949502 | 15.00542 | 29.19324 | 11.89857 | -0.47638 | 0.055247 | 0.41376 |
| 15 | 21949400 | 21950399 | 13.85739 | 25.444 | 24.93392 | 10.6095 | 0.457902 | 0.261845 | 0.413775 |
| X | 15176000 | 15177599 | 16.48888 | 30.71798 | 23.71526 | 9.886856 | 0.821185 | 0.1998 | 0.413783 |
| 2 | 98765000 | 98766399 | 11.05716 | 19.91182 | 35.81288 | 15.62028 | -0.43638 | 0.376574 | 0.413791 |
| 12 | 115245200 | 115253199 | 146.566 | 222.2537 | 191.4431 | 79.24331 | 0.495113 | 3.247206 | 0.413802 |
| X | 61965600 | 61969799 | 60.47938 | 97.3081 | 78.11865 | 33.16409 | 0.669585 | 1.876661 | 0.413805 |
| 13 | 68668400 | 68675399 | 105.59 | 163.645 | 157.2263 | 65.30736 | 0.362177 | 2.807718 | 0.413809 |
| 12 | 4669000 | 4670599 | 19.94865 | 35.33591 | 30.38358 | 13.43416 | 0.608318 | 0.528997 | 0.413821 |
| 18 | 78205200 | 78206599 | 9.255375 | 17.06637 | 31.97962 | 13.52913 | -0.46501 | 0.218333 | 0.413829 |
| 10 | 93844000 | 93845399 | 13.64584 | 25.13304 | 24.26755 | 10.28267 | 0.48299 | 0.23905 | 0.413833 |
| 18 | 59535200 | 59535799 | 5.219615 | 10.14819 | 22.86728 | 8.481818 | -0.55796 | -0.3034 | 0.41384 |
| 17 | 20308000 | 20309999 | 30.39093 | 51.24001 | 47.46406 | 20.75307 | 0.491233 | 1.122011 | 0.413857 |
| 17 | 36002000 | 36007199 | 35.31222 | 56.90638 | 96.19264 | 40.93184 | -0.39955 | 1.79554 | 0.413863 |
| 2 | 74244400 | 74246599 | 21.52577 | 37.51155 | 36.64603 | 16.11984 | 0.420854 | 0.76132 | 0.413867 |
| 17 | 24569800 | 24571999 | 16.61214 | 28.53615 | 51.0948 | 22.41164 | -0.45298 | 0.909593 | 0.413868 |
| 3 | 50549200 | 50553599 | 49.28213 | 80.01937 | 65.96203 | 28.28019 | 0.645343 | 1.623937 | 0.413872 |
| 3 | 60598800 | 60599399 | 5.95215 | 11.57431 | 25.41162 | 9.41313 | -0.52048 | -0.29395 | 0.413878 |
| 7 | 13907600 | 13919199 | 182.4269 | 273.8865 | 238.4328 | 98.404 | 0.467431 | 3.455246 | 0.413879 |
| 7 | 14108600 | 14120799 | 212.7638 | 316.1169 | 270.248 | 111.3314 | 0.479951 | 3.651017 | 0.413885 |
| 13 | 115738200 | 115741199 | 33.66968 | 55.84554 | 54.73289 | 23.70872 | 0.409838 | 1.305116 | 0.413886 |
| 7 | 16723400 | 16724199 | 8.447107 | 15.85271 | 30.15713 | 12.41686 | -0.45612 | 0.10107 | 0.413887 |
| 7 | 136433600 | 136436999 | 22.40378 | 37.24065 | 66.77133 | 28.96698 | -0.45858 | 1.257696 | 0.413897 |
| 7 | 14181200 | 14191799 | 141.4242 | 217.7661 | 154.4557 | 64.15274 | 0.792508 | 2.942259 | 0.413909 |
| 7 | 140008600 | 140009799 | 23.18064 | 40.51652 | 35.29455 | 15.55549 | 0.585771 | 0.72552 | 0.413911 |
| 1 | 45771600 | 45774999 | 47.31207 | 76.27069 | 76.0095 | 32.35811 | 0.361671 | 1.795983 | 0.413916 |
| 3 | 90138600 | 90139599 | 7.229628 | 13.60349 | 29.11924 | 11.9364 | -0.61983 | 0.069047 | 0.413921 |
| 5 | 33492600 | 33493799 | 7.229628 | 13.60349 | 29.11924 | 11.9364 | -0.61983 | 0.069047 | 0.413921 |
| 3 | 128300400 | 128302399 | 20.5025 | 35.98287 | 28.82983 | 12.7504 | 0.705699 | 0.636375 | 0.413929 |
| 2 | 18140400 | 18142399 | 16.20835 | 28.00812 | 50.05885 | 21.96121 | -0.44871 | 0.850754 | 0.413935 |
| 19 | 31811400 | 31812399 | 14.85938 | 27.04445 | 26.34984 | 11.34128 | 0.45496 | 0.322174 | 0.413978 |
| X | 147318600 | 147320399 | 13.06712 | 24.51735 | 19.89184 | 8.193496 | 0.76358 | 0.115888 | 0.413978 |
| 8 | 50292200 | 50293799 | 22.50695 | 38.93617 | 38.4006 | 16.84722 | 0.40581 | 0.853944 | 0.41398 |
| 18 | 37178600 | 37179399 | 28.79842 | 48.20337 | 47.03373 | 20.49546 | 0.416408 | 1.184273 | 0.413992 |
| 5 | 88397800 | 88399799 | 17.39842 | 31.45625 | 27.47709 | 11.92496 | 0.603723 | 0.379187 | 0.413993 |
| 11 | 27554000 | 27555199 | 17.39842 | 31.45625 | 27.47709 | 11.92496 | 0.603723 | 0.379187 | 0.413993 |
| 10 | 69759600 | 69761999 | 10.80905 | 19.15506 | 39.83126 | 17.60991 | -0.65848 | 0.48311 | 0.413996 |
| 3 | 93919400 | 93921399 | 19.90494 | 33.56808 | 58.56232 | 25.60743 | -0.41882 | 1.095294 | 0.414006 |

| | | | | | | | | | |
|---|---|---|---|---|---|---|---|---|---|
| 13 | 62434400 | 62436999 | 57.88779 | 92.67364 | 88.72239 | 37.38476 | 0.407406 | 2.025363 | 0.41402 |
| 10 | 119261800 | 119262199 | 4.132247 | 8.076707 | 20.70098 | 7.142638 | -0.64446 | -0.57958 | 0.414026 |
| X | 23712200 | 23715999 | 42.71122 | 71.1308 | 46.49904 | 20.19801 | 0.99253 | 1.280176 | 0.41403 |
| 12 | 27592200 | 27593399 | 12.23447 | 23.55719 | 20.01124 | 7.776668 | 0.771047 | -0.08205 | 0.414036 |
| 11 | 70256400 | 70259399 | 39.9163 | 65.70313 | 52.70835 | 22.72718 | 0.695945 | 1.408472 | 0.414064 |
| 4 | 109344400 | 109345599 | 10.24169 | 18.55497 | 34.35074 | 14.86986 | -0.47065 | 0.331994 | 0.414082 |
| 8 | 89860200 | 89862199 | 10.24169 | 18.55497 | 34.35074 | 14.86986 | -0.47065 | 0.331994 | 0.414082 |
| 9 | 118088600 | 118089799 | 10.24169 | 18.55497 | 34.35074 | 14.86986 | -0.47065 | 0.331994 | 0.414082 |
| 12 | 48318800 | 48320599 | 18.75765 | 32.97097 | 32.31634 | 14.31079 | 0.416258 | 0.607898 | 0.414082 |
| 5 | 138620800 | 138621999 | 7.778039 | 14.31831 | 31.48631 | 13.35073 | -0.69615 | 0.215397 | 0.4141 |
| 15 | 18717200 | 18718599 | 23.4533 | 40.84979 | 37.00168 | 16.27109 | 0.52931 | 0.777666 | 0.414103 |
| 5 | 127693200 | 127699599 | 48.12094 | 76.61614 | 126.99 | 53.09885 | -0.39164 | 2.199008 | 0.414111 |
| 12 | 17058400 | 17065799 | 73.39864 | 113.9073 | 201.9823 | 83.88124 | -0.51947 | 2.776494 | 0.41413 |
| 8 | 102887600 | 102891799 | 35.81268 | 59.41417 | 53.7873 | 23.30296 | 0.523953 | 1.305463 | 0.414148 |
| 9 | 35535000 | 35535799 | 18.23457 | 31.00943 | 55.34621 | 24.25723 | -0.45088 | 1.011834 | 0.414157 |
| 3 | 71218400 | 71219599 | 11.0243 | 20.96084 | 21.62132 | 8.678276 | 0.451291 | 0.013882 | 0.414162 |
| 5 | 148426200 | 148432999 | 57.898 | 91.48327 | 147.3068 | 61.18489 | -0.3602 | 2.422336 | 0.414177 |
| 18 | 46805000 | 46805399 | 6.269105 | 12.08973 | 25.31971 | 9.795952 | -0.5124 | -0.1413 | 0.414182 |
| 12 | 84754200 | 84754999 | 5.388706 | 10.46937 | 23.42939 | 8.741232 | -0.55712 | -0.2824 | 0.414193 |
| 14 | 99520800 | 99522199 | 5.388706 | 10.46937 | 23.42939 | 8.741232 | -0.55712 | -0.2824 | 0.414193 |
| 8 | 10797800 | 10799599 | 18.51801 | 32.59063 | 31.72627 | 14.04596 | 0.426643 | 0.590887 | 0.414193 |
| 3 | 91559600 | 91562999 | 44.75664 | 73.49088 | 56.9094 | 24.5133 | 0.74543 | 1.452216 | 0.414193 |
| 8 | 24038600 | 24038999 | 9.835723 | 19.04364 | 16.48929 | 6.270626 | 0.77314 | -0.18854 | 0.414204 |
| 17 | 85404400 | 85407799 | 47.01597 | 76.35663 | 66.4567 | 28.4969 | 0.567421 | 1.613988 | 0.41421 |
| 6 | 61287800 | 61289199 | 21.64626 | 37.55864 | 36.85122 | 16.16894 | 0.414066 | 0.821224 | 0.414221 |
| 3 | 12997800 | 13003199 | 104.5467 | 162.3192 | 141.3834 | 58.71116 | 0.505518 | 2.775468 | 0.414244 |
| X | 5303600 | 5308399 | 57.17166 | 92.37427 | 69.10729 | 29.50591 | 0.778188 | 1.745271 | 0.414256 |
| 4 | 53208800 | 53210599 | 11.00566 | 19.89743 | 36.37649 | 15.78799 | -0.45541 | 0.352724 | 0.414258 |
| 17 | 20955400 | 20957999 | 28.39257 | 49.45944 | 33.97026 | 14.93933 | 0.926787 | 0.796631 | 0.414265 |
| 7 | 104430600 | 104432799 | 15.88036 | 27.64084 | 47.89607 | 21.0528 | -0.4024 | 0.767059 | 0.414268 |
| 8 | 69241800 | 69244799 | 37.1251 | 61.56067 | 54.34422 | 23.5361 | 0.560195 | 1.317071 | 0.414287 |
| 3 | 6014800 | 6019199 | 27.89649 | 45.81856 | 79.20143 | 34.11928 | -0.40968 | 1.426128 | 0.414309 |
| 8 | 104432800 | 104434799 | 26.48853 | 46.12276 | 35.73517 | 15.71124 | 0.753687 | 0.795161 | 0.414312 |
| 3 | 41649000 | 41650599 | 17.33079 | 31.47107 | 27.06019 | 11.68008 | 0.631477 | 0.354228 | 0.414328 |
| 9 | 11922400 | 11924999 | 42.25877 | 68.87571 | 65.01369 | 27.92332 | 0.454488 | 1.548971 | 0.414336 |
| 1 | 57014600 | 57017599 | 23.01163 | 37.91714 | 70.78002 | 30.53291 | -0.51822 | 1.36726 | 0.414337 |
| 3 | 22986200 | 22987999 | 16.18976 | 27.75477 | 56.94948 | 24.97612 | -0.64926 | 0.933956 | 0.414342 |
| 8 | 61308400 | 61310399 | 15.33728 | 28.69448 | 23.14528 | 9.57994 | 0.766093 | 0.160361 | 0.414346 |
| 6 | 123672000 | 123674599 | 22.87661 | 39.99701 | 34.69102 | 15.29334 | 0.591878 | 0.716985 | 0.414347 |
| 1 | 189733200 | 189737799 | 28.12705 | 45.788 | 88.90932 | 38.16975 | -0.58467 | 1.552448 | 0.414368 |
| 3 | 51097400 | 51098999 | 9.584623 | 17.49962 | 33.40922 | 14.32283 | -0.50566 | 0.279299 | 0.414372 |
| 5 | 85533000 | 85534599 | 16.53726 | 30.66094 | 24.02928 | 10.09034 | 0.791402 | 0.22645 | 0.414379 |
| 10 | 101281400 | 101283399 | 16.53726 | 30.66094 | 24.02928 | 10.09034 | 0.791402 | 0.22645 | 0.414379 |
| X | 121243600 | 121247199 | 46.3954 | 76.50118 | 51.81703 | 22.35935 | 0.940355 | 1.404556 | 0.414386 |
| 15 | 89094400 | 89096599 | 9.626401 | 17.46222 | 33.87198 | 14.63955 | -0.53597 | 0.319015 | 0.414402 |
| 7 | 40709400 | 40710199 | 14.20632 | 26.12072 | 25.40356 | 10.78905 | 0.470711 | 0.258115 | 0.414404 |
| 13 | 76652800 | 76656599 | 47.50641 | 76.58127 | 75.36618 | 32.08277 | 0.379708 | 1.797541 | 0.414405 |

| | | | | | | | | | |
|---|---|---|---|---|---|---|---|---|---|
| 13 | 93716800 | 93720199 | 56.52075 | 90.47277 | 90.53566 | 38.14144 | 0.343493 | 2.031791 | 0.414415 |
| 9 | 22342800 | 22344399 | 9.535204 | 17.72299 | 33.78384 | 14.13832 | -0.47852 | 0.178903 | 0.41443 |
| 6 | 128662600 | 128664599 | 24.14204 | 39.78503 | 70.82682 | 30.55482 | -0.44992 | 1.37005 | 0.414454 |
| 16 | 26480000 | 26480599 | 11.09952 | 21.01639 | 21.02148 | 8.511959 | 0.484102 | 0.040301 | 0.414477 |
| 7 | 5504600 | 5514599 | 139.1811 | 213.6123 | 170.2778 | 70.69271 | 0.620617 | 3.005096 | 0.414488 |
| 6 | 26257400 | 26259199 | 16.31408 | 30.41095 | 22.98504 | 9.5725 | 0.852179 | 0.19065 | 0.414491 |
| 8 | 41165800 | 41169599 | 61.35793 | 97.85505 | 98.62846 | 41.33548 | 0.327388 | 2.150279 | 0.414492 |
| 16 | 56818000 | 56819599 | 22.44991 | 39.04947 | 37.66247 | 16.54765 | 0.438986 | 0.792374 | 0.414501 |
| 5 | 93837400 | 93852799 | 254.6265 | 373.3518 | 309.423 | 127.4493 | 0.506883 | 3.872587 | 0.414512 |
| 4 | 156193600 | 156194799 | 10.14829 | 18.20611 | 35.33662 | 15.46638 | -0.54885 | 0.392789 | 0.41453 |
| 8 | 105776200 | 105778599 | 17.00111 | 29.37575 | 51.14566 | 22.43795 | -0.41116 | 0.856351 | 0.414531 |
| 2 | 124208400 | 124210399 | 21.2997 | 38.06614 | 27.56092 | 12.10196 | 0.861954 | 0.474637 | 0.414533 |
| 1 | 135157400 | 135158399 | 7.502348 | 14.29707 | 28.33012 | 11.31622 | -0.47643 | -0.01511 | 0.414549 |
| 13 | 61566600 | 61570999 | 101.21 | 157.1453 | 148.626 | 61.71837 | 0.386902 | 2.775818 | 0.414569 |
| 7 | 66633400 | 66634599 | 14.61647 | 26.61529 | 26.30505 | 11.31511 | 0.434881 | 0.316507 | 0.414587 |
| 4 | 4158800 | 4162599 | 39.41889 | 65.07687 | 56.15323 | 24.24571 | 0.592553 | 1.373704 | 0.414587 |
| X | 61973800 | 61976599 | 48.23346 | 77.77795 | 73.66615 | 31.37192 | 0.435413 | 1.789869 | 0.414589 |
| 12 | 104791800 | 104794599 | 15.86096 | 27.14447 | 52.11674 | 22.85348 | -0.55407 | 0.942167 | 0.41459 |
| 10 | 122683600 | 122685799 | 9.598248 | 17.44181 | 34.91142 | 15.05564 | -0.5792 | 0.309032 | 0.414597 |
| 8 | 3916000 | 3929799 | 155.2415 | 234.9259 | 217.3422 | 89.90569 | 0.389647 | 3.292657 | 0.414602 |
| X | 114658400 | 114660799 | 21.65759 | 38.84281 | 26.46261 | 11.58512 | 0.9521 | 0.453851 | 0.414614 |
| 17 | 91099000 | 91101599 | 19.05035 | 33.55106 | 31.02048 | 13.73156 | 0.50085 | 0.583825 | 0.414621 |
| 12 | 38576400 | 38578399 | 21.16927 | 35.62177 | 61.39852 | 26.8297 | -0.40171 | 1.12017 | 0.414629 |
| 18 | 87975800 | 87987199 | 114.9157 | 177.157 | 168.1884 | 69.86264 | 0.373555 | 2.909138 | 0.414654 |
| 10 | 116808600 | 116810799 | 24.52572 | 40.15478 | 75.60809 | 32.52765 | -0.53444 | 1.453741 | 0.414657 |
| 6 | 131542000 | 131543199 | 19.11596 | 33.81421 | 31.044 | 13.72996 | 0.513561 | 0.539728 | 0.414663 |
| 11 | 59910400 | 59912199 | 11.1644 | 19.66919 | 39.92551 | 17.67227 | -0.62702 | 0.552421 | 0.414684 |
| X | 44545800 | 44547599 | 29.09457 | 50.44876 | 31.73909 | 13.92564 | 1.052229 | 0.856272 | 0.414694 |
| 16 | 58297800 | 58299399 | 18.45486 | 32.91664 | 29.47969 | 12.96613 | 0.555308 | 0.47149 | 0.4147 |
| 17 | 38301000 | 38311999 | 119.386 | 184.1321 | 166.0369 | 68.96876 | 0.448273 | 2.903051 | 0.414703 |
| 8 | 33852400 | 33855199 | 43.21419 | 70.40328 | 59.31853 | 25.47335 | 0.617541 | 1.553743 | 0.414716 |
| 5 | 109140400 | 109144399 | 63.15267 | 100.8371 | 97.46606 | 40.91658 | 0.389544 | 2.114956 | 0.414737 |
| 1 | 57014800 | 57017799 | 22.51258 | 37.15983 | 69.72483 | 30.10209 | -0.52472 | 1.343372 | 0.414763 |
| 4 | 123689600 | 123692599 | 19.41572 | 33.03841 | 57.52856 | 25.21497 | -0.41523 | 1.015077 | 0.414771 |
| X | 153105400 | 153106599 | 21.77228 | 38.15194 | 34.27269 | 15.13766 | 0.54156 | 0.677206 | 0.414778 |
| 17 | 23654400 | 23655199 | 12.26402 | 23.86103 | 16.29485 | 6.009693 | 1.152509 | -0.26242 | 0.41478 |
| 4 | 80719200 | 80722599 | 49.34807 | 79.3326 | 78.57059 | 33.31588 | 0.36573 | 1.884215 | 0.41478 |
| 16 | 45338800 | 45340199 | 10.92642 | 19.74961 | 36.40643 | 15.80556 | -0.4672 | 0.356492 | 0.414782 |
| 4 | 17267400 | 17268999 | 16.13163 | 30.57984 | 21.18584 | 8.557531 | 1.013301 | 0.081217 | 0.414785 |
| 16 | 19395800 | 19396999 | 19.35105 | 33.86331 | 33.05413 | 14.58276 | 0.422341 | 0.682822 | 0.414812 |
| 5 | 77916800 | 77918799 | 30.72132 | 51.56739 | 41.22898 | 17.99802 | 0.702685 | 1.155444 | 0.414815 |
| 14 | 71357800 | 71360999 | 31.2558 | 50.5565 | 87.55785 | 37.43346 | -0.42821 | 1.678884 | 0.414825 |
| 10 | 35148200 | 35148799 | 5.518819 | 10.74335 | 23.69658 | 8.701042 | -0.51538 | -0.3369 | 0.414846 |
| X | 51912400 | 51914399 | 30.51016 | 52.35994 | 40.165 | 17.61651 | 0.764641 | 0.960892 | 0.414856 |
| 18 | 56575200 | 56577599 | 14.26309 | 25.04752 | 45.5882 | 20.17232 | -0.47235 | 0.618865 | 0.41487 |
| 2 | 101508600 | 101509399 | 4.399346 | 8.563599 | 21.78692 | 8.003641 | -0.71803 | -0.35334 | 0.414874 |
| 1 | 126014600 | 126018599 | 25.64211 | 42.44121 | 73.87127 | 31.95239 | -0.41657 | 1.321299 | 0.414875 |

| | | | | | | | | | |
|---|---|---|---|---|---|---|---|---|---|
| 7 | 143820400 | 143822599 | 12.59378 | 22.09447 | 41.525 | 18.35925 | -0.51786 | 0.62205 | 0.414881 |
| 15 | 91669800 | 91672799 | 25.46914 | 41.31889 | 89.84363 | 38.49477 | -0.7515 | 1.610385 | 0.414892 |
| 5 | 93122200 | 93124399 | 14.46168 | 25.37889 | 45.07931 | 19.93561 | -0.43725 | 0.631597 | 0.414918 |
| 8 | 101286000 | 101286799 | 10.1283 | 19.38452 | 19.06974 | 7.540254 | 0.538089 | -0.05137 | 0.414934 |
| 5 | 35465800 | 35469999 | 33.52379 | 54.09371 | 93.29158 | 39.76344 | -0.42626 | 1.753819 | 0.41494 |
| 12 | 79962200 | 79964399 | 16.68368 | 28.94385 | 50.07815 | 21.97233 | -0.40087 | 0.81795 | 0.414945 |
| 5 | 23896200 | 23918599 | 813.8203 | 1049.169 | 1416.021 | 593.6802 | -0.38198 | 5.871247 | 0.414964 |
| 6 | 123701200 | 123702799 | 22.88132 | 40.36329 | 30.11406 | 13.32602 | 0.809743 | 0.589727 | 0.414969 |
| 12 | 32590400 | 32595599 | 62.45343 | 99.91839 | 92.27047 | 38.8443 | 0.458347 | 2.052523 | 0.414975 |
| 12 | 117824400 | 117825999 | 11.94779 | 20.99766 | 40.45235 | 17.91133 | -0.55339 | 0.586977 | 0.414979 |
| 2 | 74778000 | 74779599 | 8.276026 | 15.5746 | 29.67112 | 12.15974 | -0.45279 | 0.080163 | 0.414992 |
| 13 | 26466800 | 26468199 | 8.276026 | 15.5746 | 29.67112 | 12.15974 | -0.45279 | 0.080163 | 0.414992 |
| 3 | 41301000 | 41302199 | 8.265542 | 15.59014 | 29.72697 | 12.13556 | -0.4496 | 0.065377 | 0.414999 |
| 3 | 90161200 | 90162199 | 7.864393 | 14.64886 | 30.0268 | 12.52121 | -0.57769 | 0.143521 | 0.415001 |
| 10 | 43909200 | 43910399 | 17.73981 | 31.62009 | 29.52428 | 12.99307 | 0.494872 | 0.470391 | 0.415002 |
| 13 | 65025000 | 65030399 | 97.86646 | 151.7325 | 146.2943 | 60.76334 | 0.357537 | 2.797519 | 0.415018 |
| 3 | 121361200 | 121367399 | 54.35417 | 86.3519 | 136.8202 | 57.05882 | -0.33002 | 2.274264 | 0.415026 |
| 6 | 7265000 | 7265799 | 12.47671 | 23.77207 | 22.07028 | 8.819336 | 0.607491 | 0.014636 | 0.415027 |
| 3 | 114207200 | 114208199 | 12.40684 | 23.87932 | 20.64096 | 8.030801 | 0.745126 | -0.07346 | 0.41503 |
| 4 | 50861600 | 50862999 | 15.93875 | 27.13092 | 60.02506 | 26.3099 | -0.75901 | 1.003529 | 0.415032 |
| 11 | 31915400 | 31916599 | 11.43465 | 20.23939 | 38.31646 | 16.94427 | -0.52436 | 0.499458 | 0.415038 |
| 10 | 28882800 | 28883999 | 15.11528 | 28.42972 | 22.09621 | 9.063846 | 0.829351 | 0.122907 | 0.415039 |
| 1 | 103424000 | 103425599 | 23.02894 | 40.43568 | 32.37465 | 14.32377 | 0.706628 | 0.648614 | 0.415068 |
| 7 | 13790600 | 13791399 | 33.25292 | 55.00939 | 52.99458 | 22.91408 | 0.434352 | 1.334094 | 0.41507 |
| 2 | 117901600 | 117903799 | 16.73467 | 28.9493 | 50.05781 | 21.96148 | -0.40078 | 0.845168 | 0.415076 |
| 14 | 91633800 | 91635199 | 19.40828 | 34.12418 | 32.71842 | 14.48803 | 0.447982 | 0.609125 | 0.415087 |
| 17 | 83583400 | 83587399 | 22.7909 | 37.76995 | 67.11084 | 29.05366 | -0.44585 | 1.296301 | 0.415089 |
| 7 | 13863600 | 13869199 | 108.9541 | 168.9958 | 155.038 | 64.39086 | 0.429661 | 2.79643 | 0.415092 |
| 13 | 66812800 | 66824199 | 224.0497 | 334.0709 | 241.593 | 99.56559 | 0.725916 | 3.590736 | 0.415105 |
| X | 164608400 | 164610799 | 45.91062 | 74.33745 | 70.25783 | 30.06007 | 0.444862 | 1.676317 | 0.415109 |
| 6 | 4745800 | 4749599 | 89.59252 | 136.7892 | 229.3435 | 95.11405 | -0.45693 | 3.097605 | 0.41512 |
| 13 | 77773800 | 77795799 | 463.2109 | 627.5699 | 853.3441 | 355.5961 | -0.32105 | 5.070869 | 0.415129 |
| 9 | 19174400 | 19186199 | 152.5324 | 232.0791 | 196.4337 | 81.39085 | 0.524332 | 3.182013 | 0.415138 |
| 10 | 22663000 | 22663999 | 15.3663 | 28.12277 | 26.29017 | 11.2363 | 0.520851 | 0.294556 | 0.415154 |
| 1 | 4621200 | 4621999 | 6.402628 | 12.39044 | 25.54255 | 9.735974 | -0.46952 | -0.19008 | 0.415157 |
| 6 | 109542200 | 109543199 | 6.402628 | 12.39044 | 25.54255 | 9.735974 | -0.46952 | -0.19008 | 0.415157 |
| 8 | 119635200 | 119638599 | 23.16777 | 38.6861 | 65.85322 | 28.65302 | -0.38383 | 1.209711 | 0.415174 |
| X | 120747800 | 120750399 | 31.46421 | 52.76674 | 48.59735 | 21.20252 | 0.499451 | 1.176431 | 0.415175 |
| 15 | 100195000 | 100199999 | 37.28518 | 59.90898 | 103.1963 | 43.72483 | -0.43204 | 1.893598 | 0.415185 |
| 17 | 40451200 | 40452399 | 19.14968 | 33.59256 | 32.83774 | 14.5171 | 0.420068 | 0.641705 | 0.415191 |
| 6 | 12554600 | 12555999 | 12.0797 | 21.50118 | 38.82365 | 17.10573 | -0.45212 | 0.463457 | 0.415195 |
| 4 | 8515600 | 8520999 | 68.75073 | 109.316 | 107.4902 | 44.86518 | 0.358388 | 2.254399 | 0.415227 |
| X | 19747600 | 19750399 | 26.09596 | 45.78692 | 30.53861 | 13.50296 | 0.968888 | 0.674782 | 0.415233 |
| X | 44052200 | 44054199 | 18.79712 | 34.45749 | 24.91785 | 10.62633 | 0.89123 | 0.309108 | 0.415235 |
| X | 65298000 | 65299399 | 16.54801 | 31.12579 | 22.09775 | 9.063832 | 0.959541 | 0.13673 | 0.415254 |
| 1 | 89892000 | 89893199 | 8.207101 | 15.16646 | 30.99248 | 13.07265 | -0.58555 | 0.196629 | 0.415259 |
| 1 | 90852400 | 90854599 | 18.43873 | 32.47958 | 31.77997 | 14.06726 | 0.419889 | 0.582076 | 0.415293 |

| | | | | | | | | | |
|---|---|---|---|---|---|---|---|---|---|
| 14 | 79587400 | 79588999 | 18.43873 | 32.47958 | 31.77997 | 14.06726 | 0.419889 | 0.582076 | 0.415293 |
| 10 | 36480600 | 36481999 | 18.51358 | 33.95521 | 24.87345 | 10.59996 | 0.873297 | 0.303322 | 0.415301 |
| 4 | 150264200 | 150271399 | 70.58098 | 110.0473 | 176.8194 | 73.38014 | -0.37213 | 2.691574 | 0.415364 |
| Y | 84079400 | 84080599 | 9.312901 | 18.07841 | 11.46503 | 4.306273 | 1.233681 | -0.25546 | 0.415371 |
| 8 | 56555600 | 56555999 | 5.969269 | 11.54285 | 24.84858 | 9.505791 | -0.5355 | -0.18199 | 0.415375 |
| 9 | 14661600 | 14663799 | 19.28384 | 32.48737 | 58.50728 | 25.57543 | -0.46453 | 1.102235 | 0.415375 |
| X | 114582800 | 114583399 | 15.09197 | 27.51958 | 25.88117 | 11.11332 | 0.507969 | 0.313063 | 0.415392 |
| 10 | 48707600 | 48708799 | 16.95003 | 29.22414 | 51.63984 | 22.65325 | -0.43287 | 0.879399 | 0.415399 |
| 7 | 29482400 | 29484199 | 13.07812 | 22.88085 | 43.13369 | 19.02601 | -0.52228 | 0.67262 | 0.415407 |
| 1 | 85492000 | 85493399 | 26.42999 | 45.21723 | 43.06178 | 18.88232 | 0.45313 | 0.974201 | 0.415409 |
| 9 | 94268000 | 94270199 | 25.65586 | 44.55019 | 37.46192 | 16.44197 | 0.635687 | 0.829901 | 0.415414 |
| 9 | 83297800 | 83300399 | 35.67008 | 58.95814 | 56.00966 | 24.20362 | 0.454271 | 1.348808 | 0.415416 |
| 10 | 86332200 | 86334999 | 16.88271 | 29.04749 | 51.11574 | 22.42192 | -0.42768 | 0.894774 | 0.415425 |
| 5 | 75576800 | 75579399 | 37.49932 | 61.31409 | 61.84272 | 26.59234 | 0.362329 | 1.492047 | 0.415426 |
| X | 124958800 | 124962799 | 49.91366 | 80.54234 | 67.91908 | 28.94994 | 0.603699 | 1.768793 | 0.415431 |
| 8 | 5273200 | 5277399 | 59.92104 | 95.87747 | 93.24647 | 39.26134 | 0.383929 | 2.047717 | 0.41544 |
| 13 | 61507600 | 61513399 | 75.25061 | 119.9487 | 97.7674 | 40.94498 | 0.632457 | 2.182094 | 0.415448 |
| 8 | 55799600 | 55803999 | 86.30308 | 132.6361 | 209.8958 | 87.15675 | -0.36627 | 2.963274 | 0.415459 |
| 1 | 11875600 | 11876399 | 9.680754 | 17.4861 | 37.99796 | 16.51061 | -0.70379 | 0.354338 | 0.415469 |
| 19 | 34335800 | 34337199 | 16.13325 | 30.51555 | 21.05566 | 8.544696 | 1.012954 | 0.094636 | 0.415493 |
| 7 | 103075800 | 103076999 | 19.39002 | 33.96606 | 33.4894 | 14.7875 | 0.407883 | 0.667755 | 0.415497 |
| 6 | 42055600 | 42064399 | 168.0616 | 252.7995 | 216.5616 | 89.42299 | 0.493044 | 3.413915 | 0.415503 |
| 1 | 152930200 | 152931799 | 7.515883 | 14.16905 | 28.61339 | 11.6913 | -0.53258 | 0.05895 | 0.415533 |
| 2 | 135693800 | 135694799 | 11.65515 | 22.18391 | 22.10882 | 8.853978 | 0.503283 | 0.013733 | 0.415541 |
| 10 | 75457400 | 75458399 | 11.65515 | 22.18391 | 22.10882 | 8.853978 | 0.503283 | 0.013733 | 0.415541 |
| 10 | 99433600 | 99435999 | 29.04879 | 47.58509 | 82.71597 | 35.59344 | -0.4193 | 1.463298 | 0.415541 |
| 16 | 26174400 | 26175199 | 10.3595 | 20.0376 | 19.72088 | 7.534872 | 0.583581 | -0.15755 | 0.415542 |
| 6 | 145268000 | 145270399 | 25.06926 | 41.37507 | 70.57293 | 30.46577 | -0.38765 | 1.352613 | 0.415561 |
| 7 | 59684400 | 59687599 | 45.87425 | 74.83195 | 56.72536 | 24.37104 | 0.770904 | 1.540366 | 0.415579 |
| 12 | 8690800 | 8691399 | 5.103673 | 9.917203 | 22.83516 | 8.508404 | -0.59552 | -0.29134 | 0.415579 |
| 5 | 70356000 | 70357199 | 19.53121 | 34.33832 | 32.67006 | 14.46678 | 0.459008 | 0.609868 | 0.41558 |
| 10 | 76692400 | 76694199 | 11.40317 | 20.26442 | 37.77475 | 16.66614 | -0.4993 | 0.464689 | 0.415591 |
| 9 | 41902000 | 41903799 | 8.647643 | 16.07771 | 31.05068 | 12.98923 | -0.49607 | 0.166327 | 0.41562 |
| 17 | 32173200 | 32175799 | 36.29645 | 58.48857 | 96.27955 | 40.96484 | -0.3615 | 1.799823 | 0.415645 |
| 16 | 67912400 | 67914399 | 21.96592 | 38.44267 | 34.76155 | 15.3372 | 0.532065 | 0.697485 | 0.415664 |
| 7 | 11009000 | 11011399 | 43.05155 | 70.32725 | 57.17192 | 24.57561 | 0.671711 | 1.514376 | 0.415664 |
| 7 | 63190400 | 63195599 | 47.45602 | 75.69274 | 121.8038 | 51.05655 | -0.34626 | 2.139932 | 0.415668 |
| 16 | 84210800 | 84212399 | 19.78267 | 34.75388 | 32.62683 | 14.44174 | 0.478001 | 0.621313 | 0.415675 |
| X | 25928200 | 25931199 | 52.63483 | 85.03539 | 72.70173 | 31.03655 | 0.585872 | 1.741443 | 0.415677 |
| 11 | 98017400 | 98019399 | 16.75139 | 29.80449 | 29.16928 | 12.85638 | 0.42592 | 0.474334 | 0.415693 |
| 14 | 88337200 | 88338999 | 16.56175 | 29.87553 | 26.91415 | 11.71392 | 0.5566 | 0.383218 | 0.415706 |
| 9 | 102953200 | 102957199 | 31.20403 | 50.60862 | 85.31225 | 36.54576 | -0.38573 | 1.622374 | 0.415721 |
| X | 89728800 | 89730999 | 32.88604 | 55.16953 | 49.14124 | 21.44431 | 0.547423 | 1.176913 | 0.415738 |
| 17 | 58265400 | 58268999 | 42.0791 | 68.71893 | 63.41142 | 27.25483 | 0.489104 | 1.519648 | 0.415767 |
| 5 | 21721600 | 21723999 | 16.01817 | 27.81016 | 48.99785 | 21.50282 | -0.42675 | 0.80433 | 0.41579 |
| 14 | 121663400 | 121667399 | 43.51025 | 69.67294 | 111.9771 | 47.20056 | -0.33817 | 2.011887 | 0.415815 |
| 11 | 85810800 | 85812999 | 14.4451 | 25.13905 | 46.83764 | 20.58569 | -0.50652 | 0.759837 | 0.415824 |

| | | | | | | | | | |
|---|---|---|---|---|---|---|---|---|---|
| 12 | 40454600 | 40456199 | 21.19687 | 37.0935 | 30.78947 | 13.58366 | 0.65494 | 0.688193 | 0.415826 |
| 6 | 6510400 | 6511599 | 9.2186 | 17.03655 | 32.54517 | 13.72662 | -0.48974 | 0.205035 | 0.415858 |
| 3 | 116552000 | 116555399 | 42.81188 | 68.37781 | 117.9182 | 49.52219 | -0.44371 | 2.094812 | 0.415869 |
| X | 48346800 | 48349199 | 37.09081 | 61.59928 | 53.1692 | 23.05189 | 0.592736 | 1.300312 | 0.415879 |
| 3 | 33104400 | 33106199 | 11.07084 | 20.10447 | 39.19646 | 16.91758 | -0.54234 | 0.334961 | 0.415883 |
| Y | 4225200 | 4226999 | 13.29259 | 24.72706 | 15.51048 | 6.482424 | 1.114595 | 0.157077 | 0.415896 |
| 3 | 104977400 | 104980399 | 17.567 | 30.0702 | 52.69662 | 23.10813 | -0.42303 | 0.946914 | 0.415913 |
| 3 | 16777400 | 16779799 | 35.50928 | 59.54009 | 47.49551 | 20.71952 | 0.70595 | 1.186911 | 0.415921 |
| 3 | 8574200 | 8577599 | 25.23486 | 41.60566 | 71.10883 | 30.68262 | -0.39108 | 1.365807 | 0.415925 |
| 7 | 65846400 | 65850999 | 31.77672 | 51.25124 | 93.87139 | 39.987 | -0.5136 | 1.764809 | 0.415927 |
| 4 | 111129000 | 111132199 | 52.3602 | 83.98079 | 81.60238 | 34.48217 | 0.389237 | 1.961774 | 0.415944 |
| 10 | 93681400 | 93687999 | 83.91638 | 129.6331 | 198.7335 | 82.54963 | -0.31484 | 2.863553 | 0.415984 |
| 14 | 50960800 | 50965199 | 85.53844 | 134.1375 | 134.8676 | 55.9363 | 0.309827 | 2.592405 | 0.415988 |
| 8 | 75514400 | 75519399 | 91.56905 | 143.1752 | 141.4418 | 58.68063 | 0.331934 | 2.647739 | 0.416001 |
| 12 | 99084800 | 99091399 | 51.54687 | 82.05819 | 130.591 | 54.593 | -0.33335 | 2.208135 | 0.416001 |
| 2 | 69466000 | 69468599 | 33.63764 | 55.46777 | 55.77503 | 24.06776 | 0.371997 | 1.372286 | 0.416021 |
| 13 | 61032200 | 61034199 | 25.36274 | 44.28849 | 34.0014 | 14.97577 | 0.767013 | 0.747116 | 0.416023 |
| 7 | 5914800 | 5920599 | 127.891 | 196.3511 | 176.1131 | 73.11869 | 0.450968 | 2.994928 | 0.416028 |
| X | 44125400 | 44128599 | 44.31889 | 73.12814 | 53.43012 | 23.0622 | 0.831827 | 1.393059 | 0.416033 |
| 13 | 62199800 | 62211999 | 311.4639 | 448.1608 | 374.6296 | 154.624 | 0.468066 | 4.160434 | 0.416069 |
| 7 | 137457600 | 137458399 | 4.556831 | 8.87465 | 22.45224 | 8.208031 | -0.70461 | -0.36886 | 0.416092 |
| 5 | 106610600 | 106618799 | 62.7471 | 98.25559 | 169.0719 | 70.13293 | -0.46601 | 2.609885 | 0.416092 |
| 1 | 9416600 | 9419399 | 13.99794 | 24.64612 | 48.28693 | 21.37592 | -0.57715 | 0.589401 | 0.416105 |
| X | 54477600 | 54482599 | 55.27278 | 89.7737 | 60.5318 | 25.95788 | 0.934928 | 1.628823 | 0.416113 |
| 5 | 150387200 | 150388599 | 8.873269 | 16.55843 | 32.23289 | 13.39944 | -0.50086 | 0.143162 | 0.416119 |
| 15 | 10716000 | 10719199 | 40.60239 | 65.86205 | 64.95752 | 27.82957 | 0.385897 | 1.624502 | 0.416122 |
| 14 | 87128800 | 87129599 | 12.91871 | 23.91682 | 21.05736 | 8.86006 | 0.622907 | 0.198529 | 0.416124 |
| 14 | 49223000 | 49224399 | 6.150546 | 11.77511 | 26.19648 | 10.3491 | -0.62661 | -0.06943 | 0.416125 |
| 8 | 100279400 | 100280799 | 15.54117 | 28.1312 | 27.1542 | 11.76761 | 0.461759 | 0.358122 | 0.416131 |
| 16 | 58551400 | 58553199 | 9.215337 | 16.97734 | 34.65656 | 14.67949 | -0.58956 | 0.225768 | 0.416131 |
| 1 | 189632800 | 189635799 | 27.1697 | 44.22443 | 79.74378 | 34.23345 | -0.47808 | 1.548227 | 0.416132 |
| 1 | 98029400 | 98031199 | 16.08888 | 29.78317 | 23.99049 | 10.09543 | 0.749741 | 0.230384 | 0.416148 |
| 12 | 30012400 | 30014199 | 15.46048 | 26.63719 | 50.56892 | 22.18299 | -0.53619 | 0.875009 | 0.416151 |
| 3 | 56761200 | 56762199 | 10.36472 | 18.88939 | 34.41786 | 14.789 | -0.441 | 0.298254 | 0.416151 |
| 6 | 131037400 | 131038399 | 15.41693 | 28.48717 | 25.21568 | 10.63743 | 0.612336 | 0.238798 | 0.416152 |
| 12 | 49845200 | 49845799 | 5.958449 | 11.56264 | 24.49582 | 9.217681 | -0.49009 | -0.23985 | 0.416155 |
| 15 | 65627600 | 65628999 | 14.51652 | 26.55857 | 25.72365 | 10.99922 | 0.469252 | 0.286633 | 0.416171 |
| 7 | 5973800 | 5996599 | 409.8799 | 575.2593 | 473.4199 | 196.1919 | 0.454551 | 4.528487 | 0.416176 |
| 11 | 82166400 | 82169999 | 16.50621 | 28.0733 | 58.56356 | 25.66757 | -0.6751 | 1.00563 | 0.416205 |
| 5 | 108535800 | 108537399 | 11.73483 | 20.7639 | 38.80232 | 17.16025 | -0.50602 | 0.505325 | 0.416205 |
| 16 | 76088400 | 76092599 | 55.57295 | 87.61479 | 150.5132 | 62.46019 | -0.45597 | 2.473548 | 0.416215 |
| 16 | 78096600 | 78099999 | 24.51824 | 40.56456 | 70.8019 | 30.61617 | -0.42056 | 1.320788 | 0.41622 |
| 7 | 86288800 | 86292599 | 42.86086 | 70.63214 | 55.67901 | 24.01992 | 0.721827 | 1.404762 | 0.41623 |
| 12 | 11061200 | 11064199 | 22.18614 | 38.70366 | 35.91183 | 15.80749 | 0.49483 | 0.746769 | 0.416255 |
| 18 | 44357600 | 44360999 | 42.52691 | 68.19982 | 109.463 | 46.24545 | -0.33368 | 1.963738 | 0.416256 |
| 17 | 38969800 | 38983399 | 209.5703 | 312.522 | 254.1957 | 104.7621 | 0.55663 | 3.585638 | 0.416271 |
| 2 | 180351200 | 180352799 | 11.40022 | 19.90354 | 44.23983 | 19.48304 | -0.7587 | 0.697621 | 0.416285 |

| | | | | | | | | | |
|---|---|---|---|---|---|---|---|---|---|
| 12 | 57771200 | 57772199 | 10.57827 | 19.11008 | 34.83674 | 15.13423 | -0.45184 | 0.35284 | 0.416293 |
| 14 | 119336200 | 119337599 | 7.5411 | 13.77495 | 35.55746 | 15.23495 | -0.93717 | 0.264504 | 0.416297 |
| 9 | 20410000 | 20410999 | 11.08141 | 21.50501 | 17.98801 | 6.767313 | 0.836499 | -0.20594 | 0.416332 |
| 1 | 111295800 | 111296799 | 10.03616 | 18.49895 | 34.66938 | 14.67474 | -0.4661 | 0.232175 | 0.416338 |
| 9 | 121161000 | 121167999 | 39.11536 | 62.70748 | 107.1012 | 45.22428 | -0.4237 | 1.969 | 0.416343 |
| 7 | 12796200 | 12799999 | 55.25237 | 88.68716 | 77.27803 | 32.69275 | 0.547416 | 1.939825 | 0.416355 |
| 1 | 139283600 | 139283999 | 4.938044 | 9.629251 | 21.49597 | 7.729354 | -0.50405 | -0.42469 | 0.416366 |
| 4 | 116677200 | 116680199 | 18.52611 | 31.15363 | 58.95337 | 25.7561 | -0.53574 | 1.116942 | 0.416369 |
| 11 | 69349200 | 69351399 | 23.9109 | 39.57817 | 68.3393 | 29.56079 | -0.40489 | 1.313816 | 0.416369 |
| 9 | 38529000 | 38532799 | 89.75501 | 140.6427 | 138.0799 | 57.27309 | 0.343304 | 2.609256 | 0.41637 |
| 5 | 122108800 | 122111599 | 22.21505 | 36.65137 | 69.54908 | 30.01813 | -0.54099 | 1.349722 | 0.416377 |
| 2 | 148058600 | 148059199 | 5.664111 | 11.01023 | 23.49851 | 8.729829 | -0.48361 | -0.29298 | 0.416379 |
| 10 | 63133400 | 63134199 | 5.664111 | 11.01023 | 23.49851 | 8.729829 | -0.48361 | -0.29298 | 0.416379 |
| 12 | 60741600 | 60742999 | 15.31662 | 28.10898 | 25.63442 | 10.91514 | 0.559849 | 0.275481 | 0.416395 |
| 1 | 183155600 | 183156599 | 20.82922 | 36.45303 | 34.88427 | 15.39124 | 0.450717 | 0.691968 | 0.416408 |
| 13 | 79733400 | 79734199 | 10.70203 | 20.74489 | 19.2081 | 7.265261 | 0.684557 | -0.18822 | 0.416411 |
| 13 | 50260200 | 50262199 | 16.68323 | 28.86642 | 50.06808 | 21.96613 | -0.40523 | 0.841463 | 0.416416 |
| 4 | 27757400 | 27760399 | 32.52917 | 55.08246 | 43.49182 | 19.04117 | 0.720829 | 1.085534 | 0.416426 |
| 19 | 57659400 | 57660799 | 8.077864 | 14.86091 | 31.48008 | 13.35951 | -0.64342 | 0.222227 | 0.416448 |
| 6 | 128851200 | 128852399 | 15.09837 | 28.47084 | 21.56124 | 8.806071 | 0.871389 | 0.105654 | 0.416456 |
| 19 | 29292800 | 29297199 | 55.89304 | 89.80506 | 84.2174 | 35.68186 | 0.443369 | 1.913702 | 0.416456 |
| 3 | 58624200 | 58626399 | 28.13549 | 47.81644 | 45.23761 | 19.8258 | 0.461256 | 1.042962 | 0.41646 |
| 5 | 109549200 | 109552799 | 36.82091 | 61.83224 | 46.36302 | 20.24467 | 0.795026 | 1.177837 | 0.416481 |
| 19 | 43204800 | 43205799 | 7.806073 | 14.70652 | 29.14584 | 11.92204 | -0.50721 | 0.066421 | 0.416498 |
| 14 | 26967400 | 26969399 | 10.06417 | 18.23344 | 35.37834 | 15.31462 | -0.53822 | 0.331056 | 0.416506 |
| 16 | 33667000 | 33668399 | 11.75601 | 20.84219 | 37.79179 | 16.7068 | -0.46144 | 0.483864 | 0.416509 |
| 7 | 38143000 | 38147999 | 128.9832 | 196.5551 | 194.1677 | 80.48952 | 0.303393 | 3.143542 | 0.416511 |
| 3 | 57745800 | 57747999 | 16.95747 | 30.81219 | 27.07411 | 11.67706 | 0.601056 | 0.347703 | 0.416512 |
| 16 | 20884400 | 20886199 | 31.60272 | 53.35769 | 42.01574 | 18.37865 | 0.724653 | 1.10643 | 0.416522 |
| 7 | 85975000 | 85977399 | 20.20126 | 35.70558 | 30.87909 | 13.6592 | 0.598805 | 0.55382 | 0.416523 |
| 13 | 6671600 | 6673599 | 27.64542 | 46.55319 | 46.43347 | 20.29481 | 0.384793 | 1.121013 | 0.416526 |
| 7 | 55252800 | 55255399 | 24.60998 | 43.1557 | 31.70706 | 14.01303 | 0.829988 | 0.675876 | 0.416537 |
| 3 | 63261200 | 63262799 | 15.44317 | 28.54449 | 24.59428 | 10.37092 | 0.651172 | 0.234815 | 0.416554 |
| 7 | 17600000 | 17609999 | 169.2504 | 255.9078 | 213.4215 | 88.26023 | 0.538184 | 3.3138 | 0.416576 |
| 19 | 8031400 | 8038399 | 90.72185 | 142.9953 | 124.4407 | 51.63899 | 0.523552 | 2.494902 | 0.41658 |
| X | 24519200 | 24533199 | 180.6445 | 272.3512 | 218.493 | 90.27598 | 0.590583 | 3.374198 | 0.416585 |
| 18 | 46651800 | 46654199 | 17.72059 | 30.61622 | 55.518 | 24.35606 | -0.46966 | 0.867075 | 0.416589 |
| 1 | 157275800 | 157279799 | 23.348 | 38.84809 | 66.90638 | 29.04514 | -0.40064 | 1.249331 | 0.416596 |
| 10 | 62043200 | 62049999 | 40.78277 | 65.14413 | 119.169 | 50.05529 | -0.52854 | 2.085781 | 0.416609 |
| 1 | 144576600 | 144578399 | 20.43909 | 35.71831 | 34.98339 | 15.41603 | 0.417421 | 0.712614 | 0.416618 |
| 9 | 5549000 | 5550999 | 14.97093 | 27.43012 | 25.64971 | 10.94575 | 0.521653 | 0.281549 | 0.416638 |
| 15 | 95256400 | 95258999 | 35.17444 | 58.16301 | 56.44673 | 24.39947 | 0.423666 | 1.345404 | 0.416638 |
| 17 | 7049400 | 7050999 | 13.22857 | 23.08614 | 44.21094 | 19.46529 | -0.54519 | 0.712367 | 0.416669 |
| X | 117480600 | 117482399 | 34.96005 | 57.81822 | 54.6903 | 23.64299 | 0.460658 | 1.343534 | 0.416675 |
| 3 | 103098800 | 103101599 | 44.75561 | 72.30389 | 72.87512 | 31.10944 | 0.348903 | 1.727395 | 0.416679 |
| 9 | 22642600 | 22643399 | 14.64419 | 25.54667 | 45.23178 | 19.90987 | -0.43277 | 0.726795 | 0.416686 |
| 19 | 9168200 | 9170199 | 19.4604 | 33.14455 | 58.74238 | 25.74887 | -0.44062 | 1.004536 | 0.416688 |

| | | | | | | | | | |
|---|---|---|---|---|---|---|---|---|---|
| 7 | 25366800 | 25370399 | 21.98645 | 36.43833 | 67.0694 | 29.03656 | -0.49657 | 1.292777 | 0.416705 |
| 5 | 30238600 | 30239999 | 17.39441 | 30.96723 | 29.56119 | 13.0225 | 0.46214 | 0.475381 | 0.416723 |
| 13 | 61599400 | 61604599 | 63.56487 | 102.2506 | 76.29107 | 32.37815 | 0.774607 | 1.886208 | 0.416725 |
| 19 | 37151200 | 37152999 | 12.4134 | 21.79604 | 40.97273 | 18.12775 | -0.51825 | 0.605344 | 0.416744 |
| 9 | 9390400 | 9391399 | 17.82146 | 31.36946 | 29.82802 | 13.20516 | 0.46046 | 0.581515 | 0.416759 |
| 13 | 52257800 | 52259199 | 10.39015 | 18.8569 | 34.36483 | 14.84346 | -0.44607 | 0.323027 | 0.416782 |
| 13 | 70254000 | 70255799 | 9.405012 | 17.29626 | 32.46276 | 13.78327 | -0.47125 | 0.234929 | 0.416787 |
| 6 | 52245200 | 52248399 | 17.41037 | 29.63618 | 58.64209 | 25.70374 | -0.59905 | 1.000903 | 0.416788 |
| 17 | 37929600 | 37932599 | 40.73772 | 66.49966 | 63.64131 | 27.34934 | 0.436274 | 1.525081 | 0.416793 |
| 6 | 83360600 | 83362999 | 14.29399 | 25.091 | 44.01672 | 19.46989 | -0.41935 | 0.624858 | 0.416794 |
| 12 | 12378400 | 12380399 | 14.29399 | 25.091 | 44.01672 | 19.46989 | -0.41935 | 0.624858 | 0.416794 |
| 11 | 117801600 | 117817599 | 112.0699 | 167.8091 | 324.9646 | 134.0346 | -0.68868 | 3.496918 | 0.416821 |
| 1 | 84628000 | 84628999 | 6.622793 | 12.56101 | 27.65065 | 11.15652 | -0.63972 | 0.006002 | 0.416835 |
| 2 | 70339400 | 70342399 | 17.33547 | 29.93975 | 51.68657 | 22.67493 | -0.39909 | 0.862979 | 0.416844 |
| 4 | 40093800 | 40094799 | 7.194564 | 13.66237 | 27.68403 | 11.14075 | -0.51803 | 0.003057 | 0.416845 |
| 15 | 26994800 | 26995399 | 7.194564 | 13.66237 | 27.68403 | 11.14075 | -0.51803 | 0.003057 | 0.416845 |
| 10 | 66490000 | 66493799 | 47.19204 | 76.16706 | 75.44348 | 32.1661 | 0.372383 | 1.759647 | 0.416866 |
| 1 | 182078400 | 182084599 | 58.92336 | 92.9536 | 150.6955 | 62.54139 | -0.37186 | 2.461965 | 0.416867 |
| 16 | 93624000 | 93627399 | 24.55084 | 40.63972 | 68.99044 | 29.84362 | -0.38048 | 1.315291 | 0.416876 |
| 6 | 58480200 | 58481399 | 10.39145 | 18.73412 | 36.33982 | 15.82775 | -0.54367 | 0.363817 | 0.416877 |
| 14 | 84463400 | 84464799 | 20.89224 | 36.57927 | 28.45136 | 12.55761 | 0.74833 | 0.675182 | 0.416879 |
| 5 | 14340400 | 14344199 | 49.49822 | 80.01055 | 72.92916 | 31.15503 | 0.494573 | 1.721228 | 0.416894 |
| 15 | 81350200 | 81351199 | 7.411907 | 13.84806 | 29.54738 | 12.25801 | -0.629 | 0.118696 | 0.416904 |
| 5 | 61573800 | 61576399 | 25.49562 | 43.50739 | 43.15721 | 18.92028 | 0.394063 | 0.994267 | 0.416908 |
| 3 | 154061800 | 154062599 | 6.410765 | 12.35962 | 25.90219 | 10.02898 | -0.51484 | -0.13706 | 0.416912 |
| 1 | 71238800 | 71239999 | 12.95081 | 24.65834 | 22.01581 | 8.810436 | 0.661542 | 0.023473 | 0.416916 |
| 9 | 17063800 | 17065199 | 8.773738 | 16.27446 | 31.00978 | 13.01342 | -0.48043 | 0.178534 | 0.416922 |
| 5 | 61191800 | 61192199 | 4.776303 | 9.325411 | 21.13477 | 7.454096 | -0.50001 | -0.49587 | 0.416933 |
| 13 | 120217000 | 120217799 | 8.540676 | 15.6616 | 31.95773 | 13.62204 | -0.59457 | 0.245456 | 0.416972 |
| 6 | 44535600 | 44536999 | 10.47775 | 18.74727 | 40.47349 | 17.75444 | -0.70408 | 0.421297 | 0.416995 |
| 13 | 59267600 | 59278399 | 161.7811 | 238.5494 | 362.6687 | 149.3517 | -0.36032 | 3.774742 | 0.416998 |
| 9 | 29041600 | 29044999 | 33.31192 | 55.58502 | 51.96424 | 22.59959 | 0.478033 | 1.235698 | 0.417004 |
| 9 | 117674800 | 117679599 | 27.1263 | 44.24229 | 78.41049 | 33.68566 | -0.45123 | 1.51692 | 0.417013 |
| 1 | 157438400 | 157441799 | 43.40174 | 69.36421 | 127.6385 | 53.65296 | -0.53624 | 2.076153 | 0.417015 |
| 5 | 52702400 | 52705799 | 21.12021 | 35.26325 | 62.80241 | 27.32374 | -0.44866 | 1.203518 | 0.417026 |
| 14 | 111098200 | 111100799 | 38.68011 | 63.3987 | 54.82318 | 23.59666 | 0.585595 | 1.457168 | 0.4171 |
| 7 | 42032400 | 42033999 | 35.24996 | 58.10995 | 56.88588 | 24.54416 | 0.410458 | 1.382683 | 0.417111 |
| 13 | 114675800 | 114676999 | 8.733784 | 16.41283 | 31.31562 | 12.86445 | -0.45805 | 0.096561 | 0.417116 |
| 16 | 80411800 | 80412199 | 8.733784 | 16.41283 | 31.31562 | 12.86445 | -0.45805 | 0.096561 | 0.417116 |
| 7 | 59152600 | 59153799 | 26.86324 | 46.15426 | 41.35368 | 18.13901 | 0.541654 | 0.935648 | 0.417125 |
| 2 | 50572600 | 50575199 | 21.46142 | 37.66156 | 30.09117 | 13.30698 | 0.709605 | 0.645086 | 0.417135 |
| 17 | 18922800 | 18923599 | 11.20374 | 21.841 | 15.95545 | 5.759326 | 1.084097 | -0.3487 | 0.417145 |
| 18 | 65243800 | 65247799 | 40.48166 | 66.07083 | 64.29643 | 27.62922 | 0.412031 | 1.525792 | 0.41717 |
| 12 | 115154400 | 115160399 | 76.76888 | 122.3308 | 97.8906 | 40.97896 | 0.658496 | 2.19357 | 0.417189 |
| 4 | 95335400 | 95337999 | 23.04834 | 38.31815 | 66.24489 | 28.74175 | -0.40615 | 1.257965 | 0.417191 |
| 7 | 11033000 | 11045799 | 154.1815 | 235.2615 | 182.7628 | 75.78539 | 0.651263 | 3.12509 | 0.417216 |
| 5 | 67560800 | 67561199 | 8.457165 | 16.44233 | 17.8147 | 6.609174 | 0.486584 | -0.28239 | 0.41722 |

| | | | | | | | | | |
|---|---|---|---|---|---|---|---|---|---|
| 7 | 23578200 | 23581999 | 84.08045 | 132.58 | 126.3891 | 52.45022 | 0.39264 | 2.482805 | 0.417236 |
| 4 | 90340600 | 90341999 | 15.56598 | 27.05276 | 47.38433 | 20.8081 | -0.41803 | 0.786485 | 0.41726 |
| 13 | 92666800 | 92667999 | 9.224767 | 16.60353 | 35.36614 | 15.4353 | -0.67934 | 0.36444 | 0.417293 |
| 11 | 121753800 | 121759799 | 100.4178 | 156.8514 | 142.3293 | 59.055 | 0.453258 | 2.670927 | 0.417302 |
| 16 | 59014400 | 59016199 | 31.80944 | 53.87187 | 40.44166 | 17.70659 | 0.793555 | 1.082001 | 0.417302 |
| X | 153105000 | 153114199 | 142.8855 | 219.1805 | 168.9575 | 70.13942 | 0.668348 | 3.017373 | 0.417309 |
| 3 | 95212200 | 95214199 | 22.69385 | 39.21721 | 38.42961 | 16.8594 | 0.414773 | 0.864716 | 0.417315 |
| 7 | 11819400 | 11832199 | 157.7369 | 240.3226 | 186.7685 | 77.41747 | 0.648969 | 3.156621 | 0.417322 |
| X | 29324000 | 29333799 | 102.5472 | 161.4489 | 118.8444 | 49.30933 | 0.763498 | 2.526862 | 0.417334 |
| 7 | 81692400 | 81693799 | 14.69362 | 27.12215 | 25.23921 | 10.66258 | 0.539052 | 0.238239 | 0.417394 |
| 9 | 60924000 | 60928999 | 41.42568 | 66.35528 | 109.411 | 46.14042 | -0.37447 | 2.000048 | 0.417398 |
| 5 | 77412400 | 77414399 | 18.41253 | 31.22767 | 56.40842 | 24.70802 | -0.46847 | 1.037433 | 0.417398 |
| 5 | 28191400 | 28192199 | 6.404323 | 12.36589 | 25.98285 | 10.00838 | -0.5116 | -0.15443 | 0.417401 |
| 1 | 49809000 | 49809399 | 4.93663 | 9.630929 | 21.58846 | 7.715928 | -0.5022 | -0.4463 | 0.417403 |
| 1 | 66698600 | 66698999 | 4.93663 | 9.630929 | 21.58846 | 7.715928 | -0.5022 | -0.4463 | 0.417403 |
| 14 | 44946400 | 44946799 | 4.93663 | 9.630929 | 21.58846 | 7.715928 | -0.5022 | -0.4463 | 0.417403 |
| 9 | 102374200 | 102375399 | 9.49913 | 17.1629 | 34.36399 | 14.92632 | -0.58608 | 0.342395 | 0.417431 |
| X | 72921200 | 72925999 | 21.19234 | 35.51732 | 68.33721 | 29.8026 | -0.55981 | 1.171445 | 0.417436 |
| 3 | 74666200 | 74668199 | 18.53775 | 33.53387 | 26.80209 | 11.62432 | 0.731676 | 0.382791 | 0.417437 |
| 4 | 35186000 | 35189199 | 32.17843 | 53.502 | 52.53656 | 22.79556 | 0.407256 | 1.269199 | 0.417445 |
| 1 | 78243800 | 78244799 | 12.32474 | 23.22956 | 22.3928 | 9.153086 | 0.525014 | 0.085339 | 0.41745 |
| 7 | 80799400 | 80803399 | 58.15935 | 91.619 | 152.9891 | 63.48105 | -0.41602 | 2.490247 | 0.417461 |
| 1 | 14780600 | 14782199 | 14.94686 | 27.08508 | 26.78036 | 11.58972 | 0.428683 | 0.347552 | 0.417462 |
| 8 | 42110400 | 42111799 | 23.98941 | 41.0366 | 39.00536 | 17.10343 | 0.455823 | 0.959493 | 0.417474 |
| 7 | 31535200 | 31540399 | 59.5229 | 95.90742 | 74.41413 | 31.6575 | 0.721163 | 1.828456 | 0.417481 |
| 13 | 66782000 | 66784599 | 24.38751 | 43.50007 | 25.76173 | 11.33046 | 1.148729 | 0.504564 | 0.417497 |
| 10 | 43527000 | 43529999 | 20.12059 | 34.19998 | 62.09739 | 27.21507 | -0.47593 | 1.028643 | 0.417528 |
| 1 | 18262400 | 18264199 | 24.88926 | 43.17663 | 34.4173 | 15.10093 | 0.712203 | 0.832938 | 0.41755 |
| 4 | 101297200 | 101299999 | 21.88804 | 37.8725 | 36.85515 | 16.16929 | 0.425004 | 0.845971 | 0.417575 |
| 15 | 18660400 | 18663799 | 41.87027 | 68.77799 | 58.50174 | 25.20331 | 0.610575 | 1.440857 | 0.417589 |
| 3 | 7975000 | 7975999 | 14.63026 | 25.44361 | 46.32666 | 20.35223 | -0.47347 | 0.77135 | 0.417595 |
| 16 | 55832600 | 55834599 | 18.85547 | 33.06397 | 32.44325 | 14.33813 | 0.414724 | 0.645981 | 0.417596 |
| 6 | 111100600 | 111102399 | 14.34954 | 27.01427 | 22.77176 | 9.326437 | 0.715014 | 0.112406 | 0.417623 |
| 2 | 163215600 | 163219999 | 55.64971 | 87.89687 | 146.4063 | 60.79896 | -0.4094 | 2.430264 | 0.417628 |
| 14 | 124448200 | 124451599 | 78.33031 | 123.8566 | 121.0336 | 50.29035 | 0.360127 | 2.415198 | 0.417656 |
| X | 42948800 | 42951599 | 25.94056 | 45.48007 | 31.81597 | 14.05883 | 0.900653 | 0.688901 | 0.417683 |
| 12 | 57975600 | 57976399 | 14.0251 | 26.22806 | 23.81513 | 9.864483 | 0.595667 | 0.153387 | 0.417685 |
| 12 | 6443600 | 6446599 | 23.26368 | 40.84786 | 31.75095 | 14.04783 | 0.749016 | 0.648094 | 0.417689 |
| 12 | 49578400 | 49580999 | 20.5277 | 36.16416 | 31.97787 | 14.15448 | 0.565204 | 0.592566 | 0.417715 |
| 3 | 38264000 | 38264399 | 4.516124 | 8.815051 | 20.44135 | 7.243133 | -0.53871 | -0.48644 | 0.417719 |
| 2 | 22027200 | 22028799 | 20.19037 | 35.48302 | 28.74956 | 12.72944 | 0.689545 | 0.61306 | 0.41772 |
| 7 | 43228800 | 43230599 | 12.05946 | 21.26962 | 39.85398 | 17.63671 | -0.51175 | 0.544822 | 0.417728 |
| 3 | 137843600 | 137845799 | 15.62159 | 28.38637 | 23.15845 | 9.987569 | 0.707366 | 0.328237 | 0.417739 |
| 6 | 42621800 | 42622599 | 7.011841 | 13.45609 | 26.88432 | 10.54839 | -0.46462 | -0.08403 | 0.417747 |
| 7 | 55087800 | 55090199 | 18.66778 | 32.05618 | 56.58694 | 24.82038 | -0.43295 | 0.926162 | 0.417785 |
| X | 33693800 | 33700199 | 161.2919 | 242.3949 | 230.0172 | 94.95588 | 0.34439 | 3.432038 | 0.417785 |
| 9 | 114512000 | 114514799 | 23.83032 | 39.67674 | 67.53671 | 29.33157 | -0.38375 | 1.244113 | 0.4178 |

| | | | | | | | | | |
|---|---|---|---|---|---|---|---|---|---|
| X | 126672400 | 126674199 | 26.90404 | 46.52277 | 39.19019 | 17.19341 | 0.632217 | 0.879004 | 0.417824 |
| 12 | 115180400 | 115186999 | 89.61512 | 141.892 | 111.4002 | 46.31929 | 0.677016 | 2.390701 | 0.417843 |
| 18 | 26897600 | 26897999 | 3.876216 | 7.581625 | 18.51768 | 6.232693 | -0.54299 | -0.68031 | 0.417843 |
| 4 | 15541400 | 15543799 | 23.7567 | 41.37196 | 36.97012 | 16.25567 | 0.54874 | 0.780913 | 0.417844 |
| 7 | 54031000 | 54035199 | 60.89249 | 97.23266 | 97.34203 | 40.86714 | 0.339032 | 2.111674 | 0.417862 |
| 4 | 58333000 | 58333999 | 11.00544 | 21.25155 | 20.19958 | 7.778099 | 0.622866 | -0.12328 | 0.417864 |
| 6 | 79109600 | 79109999 | 11.00544 | 21.25155 | 20.19958 | 7.778099 | 0.622866 | -0.12328 | 0.417864 |
| 1 | 112621400 | 112624399 | 39.01104 | 64.64088 | 53.33137 | 23.08585 | 0.657371 | 1.328475 | 0.417877 |
| 1 | 61299200 | 61300799 | 9.368886 | 17.42715 | 32.69117 | 13.66419 | -0.45406 | 0.170288 | 0.417881 |
| 7 | 25921400 | 25927999 | 118.3551 | 182.019 | 174.4388 | 72.44041 | 0.357339 | 2.957558 | 0.417888 |
| 10 | 117516000 | 117520599 | 29.10386 | 47.23288 | 85.15818 | 36.49711 | -0.48159 | 1.604301 | 0.417893 |
| 16 | 53400000 | 53404399 | 109.5803 | 168.6061 | 166.655 | 69.21256 | 0.313334 | 2.943979 | 0.417894 |
| 1 | 32585400 | 32586599 | 15.49199 | 27.83732 | 26.6571 | 11.65171 | 0.465158 | 0.398587 | 0.417911 |
| 4 | 118596200 | 118599399 | 26.99919 | 44.35881 | 76.45066 | 32.93867 | -0.40526 | 1.415271 | 0.417915 |
| X | 101499000 | 101499799 | 8.639176 | 16.19449 | 31.77817 | 13.10987 | -0.50276 | 0.11519 | 0.417931 |
| 1 | 47262000 | 47263599 | 15.06867 | 27.11349 | 24.89652 | 10.8697 | 0.526702 | 0.384672 | 0.417946 |
| 2 | 180929400 | 180930199 | 9.132836 | 16.69104 | 32.91619 | 14.09424 | -0.55088 | 0.269304 | 0.417954 |
| 14 | 100483400 | 100488399 | 65.28249 | 102.6215 | 160.0178 | 66.37915 | -0.31974 | 2.53933 | 0.417954 |
| 7 | 74719600 | 74720599 | 11.84069 | 20.72979 | 42.64745 | 18.82218 | -0.64747 | 0.654993 | 0.417957 |
| 15 | 34457000 | 34458199 | 14.26062 | 26.28278 | 24.93615 | 10.55643 | 0.509639 | 0.245053 | 0.41797 |
| 10 | 8509600 | 8516799 | 105.4367 | 162.8205 | 162.4465 | 67.47694 | 0.303863 | 2.874 | 0.417995 |
| X | 34885200 | 34897199 | 160.4136 | 244.2376 | 187.3978 | 77.66283 | 0.666635 | 3.171518 | 0.418009 |
| 9 | 83113800 | 83114799 | 12.00645 | 22.74257 | 21.94388 | 8.878066 | 0.536434 | 0.047879 | 0.418016 |
| 18 | 85646000 | 85647199 | 12.00645 | 22.74257 | 21.94388 | 8.878066 | 0.536434 | 0.047879 | 0.418016 |
| 13 | 115671600 | 115673199 | 14.93682 | 27.62191 | 25.27899 | 10.65265 | 0.565675 | 0.230907 | 0.418031 |
| 17 | 48135400 | 48138599 | 31.56584 | 51.04532 | 88.88905 | 37.99343 | -0.43649 | 1.685 | 0.418037 |
| 10 | 73830400 | 73832399 | 16.91032 | 29.21816 | 50.58822 | 22.19338 | -0.403 | 0.857691 | 0.418042 |
| 16 | 4350800 | 4353399 | 14.10898 | 24.68533 | 44.10927 | 19.45695 | -0.4458 | 0.675919 | 0.418084 |
| 18 | 81513800 | 81515999 | 14.10898 | 24.68533 | 44.10927 | 19.45695 | -0.4458 | 0.675919 | 0.418084 |
| X | 112129400 | 112132199 | 44.46916 | 73.0831 | 50.5952 | 21.80162 | 0.907074 | 1.43799 | 0.418085 |
| 7 | 61478200 | 61483999 | 89.35487 | 140.1234 | 136.9045 | 56.78105 | 0.351146 | 2.594971 | 0.418094 |
| 2 | 18060800 | 18062799 | 13.58642 | 23.51287 | 49.49562 | 21.71511 | -0.68307 | 0.824625 | 0.4181 |
| 16 | 82473600 | 82475199 | 39.21204 | 63.91695 | 63.99231 | 27.48573 | 0.369898 | 1.541802 | 0.41812 |
| 7 | 14605200 | 14625799 | 423.1559 | 590.6486 | 503.0634 | 208.6543 | 0.397093 | 4.610523 | 0.418137 |
| 9 | 59180400 | 59181999 | 14.33091 | 26.07045 | 24.93757 | 10.73968 | 0.480844 | 0.317619 | 0.418178 |
| 10 | 18089000 | 18089999 | 7.184251 | 13.63886 | 28.79592 | 11.59526 | -0.57809 | 0.005918 | 0.418197 |
| 9 | 74246600 | 74247599 | 16.20031 | 28.07471 | 48.9965 | 21.49703 | -0.41358 | 0.823934 | 0.418221 |
| 14 | 9385000 | 9386599 | 15.99741 | 29.25289 | 26.07249 | 11.15611 | 0.58775 | 0.301634 | 0.418226 |
| 3 | 126798800 | 126800199 | 9.441441 | 17.32121 | 32.44242 | 13.82015 | -0.47151 | 0.25037 | 0.418227 |
| 11 | 62645400 | 62649199 | 22.34787 | 36.92927 | 75.49561 | 32.61869 | -0.64799 | 1.333939 | 0.41823 |
| 3 | 104106000 | 104109199 | 47.73326 | 75.94072 | 127.7126 | 53.34447 | -0.41372 | 2.226038 | 0.418233 |
| 9 | 24833400 | 24837399 | 35.26174 | 58.59116 | 52.65683 | 22.83769 | 0.53457 | 1.284893 | 0.418253 |
| 2 | 111662400 | 111663399 | 20.29405 | 35.72832 | 31.97319 | 14.15433 | 0.547582 | 0.597745 | 0.418258 |
| 4 | 91539800 | 91541399 | 25.08957 | 43.32692 | 39.86638 | 17.48935 | 0.505035 | 0.883993 | 0.41826 |
| 12 | 118483200 | 118485799 | 17.7583 | 31.29311 | 29.22991 | 12.93749 | 0.486542 | 0.567157 | 0.418265 |
| 6 | 8453600 | 8454599 | 13.80424 | 26.25025 | 21.94024 | 8.803568 | 0.752745 | 0.03994 | 0.418273 |
| 15 | 56555600 | 56557199 | 10.57122 | 18.6146 | 40.46408 | 17.91249 | -0.72555 | 0.555358 | 0.418275 |

| | | | | | | | | | |
|---|---|---|---|---|---|---|---|---|---|
| 14 | 52820600 | 52822199 | 23.84434 | 40.96825 | 40.52406 | 17.77512 | 0.399614 | 0.923466 | 0.418276 |
| X | 147802400 | 147805799 | 54.65803 | 87.72251 | 83.78201 | 35.43773 | 0.415154 | 1.947588 | 0.418279 |
| 8 | 101276600 | 101279199 | 37.67759 | 62.3023 | 55.5444 | 24.00949 | 0.545685 | 1.349524 | 0.418282 |
| 7 | 38447200 | 38451799 | 46.81865 | 76.80063 | 56.394 | 24.28112 | 0.821361 | 1.469338 | 0.418294 |
| 14 | 110933000 | 110933599 | 6.553139 | 12.67881 | 26.12578 | 9.96894 | -0.47041 | -0.18283 | 0.418305 |
| 13 | 74675800 | 74679199 | 146.748 | 216.2648 | 365.1129 | 150.3574 | -0.51214 | 3.783024 | 0.41831 |
| 19 | 4991600 | 4994199 | 14.44857 | 24.7808 | 51.57879 | 22.62152 | -0.66947 | 0.915118 | 0.418328 |
| 17 | 17078000 | 17079399 | 22.97448 | 39.61608 | 39.44957 | 17.30572 | 0.391005 | 0.885786 | 0.418366 |
| 7 | 9741000 | 9773799 | 560.888 | 766.9745 | 602.0197 | 250.5078 | 0.484962 | 4.927551 | 0.418371 |
| 7 | 23373800 | 23376199 | 39.51032 | 65.3554 | 54.9868 | 23.77413 | 0.629112 | 1.352251 | 0.418383 |
| 12 | 66333400 | 66334199 | 6.557649 | 12.64015 | 25.89583 | 10.03257 | -0.48313 | -0.13299 | 0.418407 |
| 7 | 43435600 | 43437799 | 13.65695 | 23.98833 | 41.92934 | 18.55675 | -0.414 | 0.607773 | 0.418414 |
| 13 | 93510000 | 93511399 | 12.7547 | 24.13397 | 21.82462 | 8.849888 | 0.626254 | 0.061377 | 0.418431 |
| 5 | 44307200 | 44309399 | 37.2709 | 60.95638 | 61.34475 | 26.38071 | 0.365779 | 1.486516 | 0.418434 |
| 7 | 122335200 | 122336799 | 13.95363 | 25.47165 | 23.76285 | 10.1898 | 0.520246 | 0.289945 | 0.418438 |
| 6 | 72490800 | 72493199 | 16.10774 | 27.91007 | 48.98703 | 21.49277 | -0.42165 | 0.827039 | 0.418449 |
| 8 | 38674400 | 38675999 | 16.65936 | 30.28372 | 27.13455 | 11.69674 | 0.573605 | 0.34353 | 0.418451 |
| 9 | 10921400 | 10922599 | 23.17894 | 40.53642 | 34.85078 | 15.36723 | 0.604752 | 0.717657 | 0.418457 |
| 7 | 8415600 | 8422199 | 74.14059 | 118.9414 | 74.55379 | 31.51456 | 1.020844 | 1.982387 | 0.418458 |
| 15 | 33935200 | 33939199 | 46.33302 | 75.22353 | 67.9045 | 29.11089 | 0.514504 | 1.621539 | 0.418458 |
| 14 | 25782600 | 25784399 | 8.052458 | 14.84563 | 32.51918 | 13.7607 | -0.68903 | 0.209049 | 0.418459 |
| X | 115682200 | 115683599 | 18.90061 | 34.64247 | 24.85591 | 10.60188 | 0.902186 | 0.310044 | 0.418464 |
| 6 | 58856200 | 58857999 | 13.33871 | 23.32127 | 44.14549 | 19.46138 | -0.52853 | 0.683253 | 0.418464 |
| 3 | 84305800 | 84307599 | 20.9621 | 36.47918 | 36.22047 | 15.91891 | 0.397345 | 0.776448 | 0.418466 |
| 8 | 24232400 | 24234799 | 13.19404 | 23.24121 | 40.87662 | 18.09385 | -0.42159 | 0.569116 | 0.418469 |
| 12 | 59338200 | 59340999 | 12.42121 | 21.83087 | 46.12974 | 20.42609 | -0.68636 | 0.597449 | 0.418482 |
| 4 | 32050200 | 32053799 | 22.64838 | 37.4595 | 68.13906 | 29.45849 | -0.47983 | 1.319398 | 0.418501 |
| 18 | 17266000 | 17266999 | 7.638822 | 14.55535 | 28.34255 | 11.32415 | -0.45202 | -0.01396 | 0.418506 |
| 9 | 19309600 | 19315599 | 72.13231 | 115.3168 | 89.7976 | 37.76304 | 0.702748 | 2.087659 | 0.418508 |
| 2 | 88718800 | 88719199 | 5.966391 | 11.55218 | 24.37246 | 9.267608 | -0.49885 | -0.20594 | 0.418519 |
| 12 | 63094400 | 63095799 | 12.47088 | 21.91448 | 42.00806 | 18.59838 | -0.54603 | 0.597688 | 0.418522 |
| 3 | 11546600 | 11547999 | 18.12303 | 30.93559 | 54.80037 | 24.02526 | -0.43908 | 0.979136 | 0.418532 |
| 17 | 70259000 | 70259599 | 5.515858 | 10.74659 | 26.54377 | 9.654386 | -0.66523 | -0.36424 | 0.418549 |
| 2 | 177480200 | 177482399 | 10.76038 | 19.20896 | 37.29939 | 16.39366 | -0.55404 | 0.430212 | 0.418554 |
| 17 | 80061400 | 80061999 | 4.686439 | 9.114695 | 22.31335 | 8.253476 | -0.67286 | -0.32425 | 0.418572 |
| 1 | 48257200 | 48258599 | 13.96339 | 26.05971 | 23.7618 | 9.876286 | 0.585597 | 0.164856 | 0.418578 |
| 18 | 50944800 | 50945599 | 8.436417 | 15.98045 | 34.59044 | 13.99421 | -0.62009 | 0.048394 | 0.418601 |
| 1 | 143113200 | 143114199 | 11.54659 | 21.98003 | 22.14519 | 8.866206 | 0.488206 | 0.012593 | 0.418602 |
| 4 | 114011000 | 114011599 | 11.54659 | 21.98003 | 22.14519 | 8.866206 | 0.488206 | 0.012593 | 0.418602 |
| 8 | 64231000 | 64232199 | 14.45958 | 25.29231 | 44.12232 | 19.45851 | -0.41126 | 0.683329 | 0.418605 |
| 5 | 70885600 | 70887799 | 19.07301 | 33.67326 | 31.58965 | 13.97668 | 0.481484 | 0.560618 | 0.418616 |
| X | 118169600 | 118175399 | 91.47152 | 143.923 | 128.1069 | 53.14815 | 0.489109 | 2.532391 | 0.418668 |
| 5 | 42895600 | 42896399 | 21.33931 | 37.27959 | 30.43798 | 13.41008 | 0.678675 | 0.714975 | 0.418686 |
| 5 | 45774200 | 45784199 | 147.7488 | 224.2957 | 207.9657 | 86.11104 | 0.390274 | 3.226486 | 0.41875 |
| 4 | 86672400 | 86673199 | 11.62802 | 22.23534 | 21.61959 | 8.570011 | 0.551494 | -0.02071 | 0.418751 |
| 11 | 55266000 | 55267399 | 8.168426 | 15.17226 | 30.50881 | 12.77955 | -0.55549 | 0.16447 | 0.418755 |
| 14 | 17346400 | 17347399 | 8.168426 | 15.17226 | 30.50881 | 12.77955 | -0.55549 | 0.16447 | 0.418755 |

| | | | | | | | | | |
|---|---|---|---|---|---|---|---|---|---|
| 7 | 41417600 | 41420799 | 38.57162 | 64.32096 | 49.82962 | 21.66262 | 0.748337 | 1.257434 | 0.418761 |
| 4 | 112781400 | 112786999 | 124.411 | 190.9389 | 179.0901 | 74.351 | 0.386116 | 3.000077 | 0.418764 |
| 4 | 14482200 | 14485399 | 46.01672 | 75.01937 | 63.54846 | 27.29649 | 0.610531 | 1.554158 | 0.418803 |
| 17 | 39550400 | 39554199 | 41.64645 | 68.93826 | 52.8161 | 22.84691 | 0.763867 | 1.348082 | 0.418815 |
| 7 | 61959000 | 61962799 | 31.21897 | 53.34464 | 41.84949 | 18.34892 | 0.73154 | 1.003982 | 0.418849 |
| 14 | 20182400 | 20184199 | 16.0507 | 29.1479 | 27.11363 | 11.70246 | 0.518574 | 0.3436 | 0.418859 |
| 4 | 19740600 | 19742599 | 18.24226 | 32.24353 | 29.98433 | 13.26337 | 0.49448 | 0.538416 | 0.418865 |
| 3 | 69063600 | 69064399 | 6.551085 | 12.64769 | 25.9762 | 10.01163 | -0.47973 | -0.15032 | 0.41887 |
| 19 | 43372800 | 43377399 | 30.93427 | 49.99684 | 90.39371 | 38.60952 | -0.49143 | 1.700131 | 0.418871 |
| 1 | 34216200 | 34219599 | 40.96081 | 65.57192 | 110.5982 | 46.59952 | -0.4081 | 2.018829 | 0.41889 |
| X | 121898800 | 121903199 | 91.14789 | 142.9954 | 134.9621 | 55.97301 | 0.401454 | 2.589909 | 0.418891 |
| 1 | 39635200 | 39638399 | 25.43989 | 41.81835 | 72.55896 | 31.26871 | -0.41449 | 1.405197 | 0.418894 |
| 3 | 139725000 | 139728999 | 55.32953 | 88.56013 | 88.48221 | 37.27285 | 0.345656 | 2.031375 | 0.418912 |
| X | 32244800 | 32255399 | 166.2412 | 253.0333 | 183.3643 | 75.98426 | 0.748641 | 3.17849 | 0.418916 |
| 1 | 167513800 | 167522599 | 53.33452 | 83.94851 | 170.1244 | 70.58352 | -0.69588 | 2.506461 | 0.418921 |
| 7 | 6057600 | 6080799 | 403.1304 | 564.0486 | 496.7522 | 205.9597 | 0.352322 | 4.573741 | 0.418948 |
| 7 | 40335200 | 40339199 | 40.00378 | 65.07188 | 65.68349 | 28.19072 | 0.356017 | 1.573317 | 0.418971 |
| 4 | 28461400 | 28461999 | 7.888749 | 14.62505 | 30.5022 | 12.80976 | -0.61026 | 0.173642 | 0.418987 |
| 11 | 104169200 | 104171199 | 11.28966 | 20.03207 | 37.31993 | 16.4867 | -0.49949 | 0.474911 | 0.419004 |
| 1 | 117778600 | 117781199 | 41.19349 | 67.15662 | 64.43175 | 27.67596 | 0.431286 | 1.541664 | 0.419013 |
| 6 | 25624400 | 25624999 | 19.66542 | 34.36725 | 33.60765 | 14.81013 | 0.419748 | 0.70683 | 0.419015 |
| 13 | 99261400 | 99264199 | 16.0119 | 27.81572 | 48.99296 | 21.50867 | -0.42624 | 0.795693 | 0.419045 |
| 7 | 16049600 | 16053199 | 16.48263 | 28.26412 | 51.60766 | 22.63385 | -0.48146 | 0.926481 | 0.419054 |
| 17 | 18216800 | 18218199 | 32.66188 | 53.97603 | 53.5522 | 23.13873 | 0.391865 | 1.343041 | 0.419056 |
| 4 | 3455800 | 3468799 | 311.8301 | 444.5047 | 412.3623 | 170.4522 | 0.304347 | 4.296038 | 0.419063 |
| 2 | 165269200 | 165273199 | 26.78842 | 43.73143 | 77.94004 | 33.49478 | -0.45835 | 1.502108 | 0.419077 |
| 8 | 4401800 | 4402999 | 13.88906 | 26.10012 | 23.3577 | 9.595769 | 0.625905 | 0.121172 | 0.419096 |
| 17 | 75873200 | 75874399 | 13.52622 | 25.14729 | 24.33443 | 10.18013 | 0.493088 | 0.186944 | 0.419098 |
| 2 | 148152200 | 148153999 | 11.06838 | 19.89467 | 35.80386 | 15.64569 | -0.43903 | 0.388321 | 0.419098 |
| 5 | 87422200 | 87425199 | 45.87337 | 74.36786 | 69.61113 | 29.8102 | 0.460327 | 1.648732 | 0.419101 |
| 9 | 96638000 | 96640399 | 18.2951 | 31.0275 | 55.82497 | 24.45222 | -0.46265 | 1.038701 | 0.419102 |
| 16 | 47422400 | 47423799 | 15.7463 | 29.29959 | 24.17162 | 10.0975 | 0.723446 | 0.200383 | 0.419126 |
| 4 | 86259200 | 86261599 | 25.64029 | 41.97481 | 75.14804 | 32.32817 | -0.46196 | 1.457991 | 0.419128 |
| 14 | 121043000 | 121046199 | 35.71405 | 57.39492 | 100.5709 | 42.62363 | -0.45626 | 1.886956 | 0.419128 |
| 18 | 18013200 | 18014399 | 15.4734 | 27.85564 | 26.61748 | 11.61426 | 0.469859 | 0.386889 | 0.419138 |
| 14 | 90800600 | 90801799 | 17.09247 | 30.26659 | 29.72624 | 13.14439 | 0.417293 | 0.512902 | 0.41915 |
| X | 55019600 | 55021999 | 40.50349 | 66.25449 | 62.69995 | 26.96517 | 0.454462 | 1.493775 | 0.419165 |
| 8 | 90947200 | 90949799 | 55.03057 | 88.68518 | 73.74113 | 31.37945 | 0.621777 | 1.811837 | 0.419167 |
| 17 | 37941800 | 37942599 | 16.79103 | 29.74625 | 28.82069 | 12.74283 | 0.437117 | 0.504897 | 0.419177 |
| 18 | 58937400 | 58937799 | 4.11132 | 8.008252 | 21.25994 | 7.756196 | -0.76891 | -0.38298 | 0.419181 |
| 9 | 86171200 | 86171999 | 11.06491 | 21.13066 | 21.67922 | 8.621061 | 0.470786 | -0.01486 | 0.419181 |
| 16 | 95591600 | 95593599 | 21.66599 | 36.51261 | 62.55211 | 27.34629 | -0.39288 | 1.1105 | 0.419182 |
| 18 | 67698600 | 67700199 | 8.896421 | 16.53469 | 31.57263 | 13.21403 | -0.48094 | 0.168383 | 0.419186 |
| 3 | 31645800 | 31647199 | 14.36159 | 26.5202 | 25.38399 | 10.71773 | 0.499566 | 0.235315 | 0.419205 |
| X | 159145400 | 159147399 | 21.04196 | 37.61189 | 27.82083 | 12.21434 | 0.8317 | 0.474396 | 0.419217 |
| 3 | 25278400 | 25279799 | 11.88563 | 23.12828 | 16.31101 | 6.008431 | 1.107921 | -0.27131 | 0.419224 |
| 7 | 55171200 | 55174199 | 41.96976 | 68.4099 | 63.97577 | 27.4783 | 0.467919 | 1.544956 | 0.419228 |

| | | | | | | | | | |
|---|---|---|---|---|---|---|---|---|---|
| 4 | 64393600 | 64396199 | 33.35816 | 56.22753 | 46.32533 | 20.2535 | 0.659583 | 1.131167 | 0.419237 |
| 5 | 57625000 | 57625399 | 8.491818 | 16.4779 | 16.16828 | 6.085541 | 0.608735 | -0.23943 | 0.419243 |
| 13 | 16426600 | 16429399 | 27.15358 | 46.75551 | 39.07175 | 17.13923 | 0.642293 | 0.913016 | 0.419244 |
| 12 | 114195800 | 114199999 | 61.56848 | 98.70534 | 87.82731 | 37.08708 | 0.515231 | 1.988656 | 0.419248 |
| 4 | 63475000 | 63479799 | 22.17349 | 36.32047 | 78.23907 | 33.66616 | -0.72773 | 1.44306 | 0.419269 |
| 15 | 36384200 | 36385199 | 7.599697 | 14.53567 | 30.30103 | 12.00335 | -0.53915 | -0.0401 | 0.419277 |
| 3 | 108983000 | 108983999 | 14.27157 | 24.96737 | 44.111 | 19.45611 | -0.42953 | 0.678625 | 0.419298 |
| 4 | 81117200 | 81119199 | 30.95896 | 52.44726 | 45.76137 | 20.03755 | 0.577389 | 1.08086 | 0.419304 |
| X | 4999400 | 5001599 | 35.36978 | 59.86512 | 38.06265 | 16.66191 | 1.032163 | 1.096467 | 0.419305 |
| 6 | 124955000 | 124956399 | 13.98477 | 26.20996 | 23.91267 | 9.868243 | 0.593311 | 0.140744 | 0.419306 |
| X | 54345000 | 54351399 | 93.75281 | 147.4456 | 128.8499 | 53.45328 | 0.515117 | 2.542375 | 0.419306 |
| 18 | 32821000 | 32824199 | 32.9341 | 55.40469 | 46.89849 | 20.49197 | 0.620587 | 1.145644 | 0.419307 |
| 2 | 169879400 | 169882199 | 16.757 | 28.78334 | 51.10068 | 22.41418 | -0.44081 | 0.910184 | 0.419314 |
| X | 110020000 | 110022399 | 29.68389 | 51.64532 | 33.35245 | 14.65629 | 1.015069 | 0.81838 | 0.419323 |
| Y | 42326000 | 42328999 | 20.18629 | 35.27532 | 26.25728 | 11.57038 | 0.811381 | 0.696563 | 0.419328 |
| 3 | 15084200 | 15084799 | 5.811984 | 11.28104 | 23.92015 | 8.991304 | -0.4898 | -0.24729 | 0.419347 |
| 14 | 70800600 | 70801399 | 5.811984 | 11.28104 | 23.92015 | 8.991304 | -0.4898 | -0.24729 | 0.419347 |
| 1 | 107136200 | 107137599 | 19.2244 | 33.86089 | 26.26246 | 11.62515 | 0.753173 | 0.576556 | 0.419355 |
| 3 | 42593000 | 42594999 | 19.08562 | 33.67217 | 30.97774 | 13.70784 | 0.508972 | 0.565424 | 0.419364 |
| 14 | 5859800 | 5860799 | 7.648058 | 14.5404 | 28.27228 | 11.34961 | -0.45587 | 0.00173 | 0.41939 |
| 2 | 64142800 | 64143999 | 7.064413 | 13.38768 | 27.63608 | 11.16989 | -0.5498 | 0.017356 | 0.419402 |
| 11 | 73367200 | 73371399 | 59.93844 | 95.76447 | 96.06804 | 40.36526 | 0.337001 | 2.093215 | 0.419409 |
| 6 | 120525400 | 120528799 | 21.69285 | 36.31742 | 63.0017 | 27.45725 | -0.41102 | 1.17451 | 0.419443 |
| 1 | 38489200 | 38490199 | 6.903906 | 13.14553 | 27.18377 | 10.87772 | -0.53936 | -0.01765 | 0.419446 |
| 2 | 157464400 | 157464999 | 6.435369 | 12.35793 | 26.28642 | 10.29434 | -0.55064 | -0.09456 | 0.419466 |
| 7 | 41874000 | 41886999 | 191.3076 | 286.8491 | 238.3026 | 98.31627 | 0.533248 | 3.482515 | 0.419467 |
| 13 | 69564600 | 69568999 | 45.71406 | 73.70283 | 69.58986 | 29.62948 | 0.439562 | 1.78603 | 0.419467 |
| 5 | 50711800 | 50712399 | 11.59191 | 21.88284 | 21.1242 | 8.608106 | 0.527583 | 0.068124 | 0.419468 |
| 2 | 177707000 | 177708799 | 22.09247 | 38.9897 | 30.75189 | 13.60702 | 0.730273 | 0.581577 | 0.419469 |
| 5 | 52443400 | 52446599 | 22.73037 | 38.06731 | 65.79234 | 28.6796 | -0.40567 | 1.178907 | 0.419473 |
| 9 | 80352400 | 80354799 | 51.07536 | 82.38348 | 69.14153 | 29.45559 | 0.609893 | 1.782867 | 0.41948 |
| X | 31591600 | 31596199 | 39.00146 | 65.27751 | 46.97323 | 20.46832 | 0.854325 | 1.219352 | 0.419484 |
| 1 | 183167800 | 183171399 | 31.32357 | 50.98833 | 85.4493 | 36.68346 | -0.37286 | 1.557172 | 0.419504 |
| X | 25716800 | 25717399 | 19.41413 | 34.36723 | 30.59942 | 13.53132 | 0.558092 | 0.534317 | 0.419504 |
| X | 113243400 | 113244199 | 15.92263 | 29.11314 | 26.27185 | 11.24287 | 0.570256 | 0.304581 | 0.419512 |
| 4 | 152819400 | 152826199 | 50.90259 | 80.95465 | 130.7608 | 54.59128 | -0.35621 | 2.238452 | 0.419523 |
| 1 | 118059000 | 118067799 | 135.692 | 207.4719 | 187.2152 | 77.66833 | 0.437674 | 3.078885 | 0.419529 |
| 3 | 144641600 | 144642799 | 20.88257 | 36.29488 | 33.79213 | 14.83986 | 0.489583 | 0.789904 | 0.419545 |
| 4 | 38002400 | 38004599 | 26.93309 | 46.28222 | 40.77647 | 17.88592 | 0.56582 | 0.932787 | 0.419555 |
| 3 | 90675400 | 90678599 | 50.36417 | 81.55085 | 69.40784 | 29.69703 | 0.595673 | 1.680923 | 0.419558 |
| 18 | 61211600 | 61220599 | 86.98219 | 133.664 | 211.1545 | 87.67799 | -0.3639 | 2.965537 | 0.419563 |
| X | 150191000 | 150193799 | 32.42961 | 56.14365 | 33.88878 | 14.86812 | 1.111225 | 0.886698 | 0.419592 |
| 12 | 59650800 | 59654199 | 49.66659 | 80.13477 | 76.36015 | 32.54322 | 0.427563 | 1.773506 | 0.419632 |
| 1 | 19440600 | 19442999 | 19.88169 | 33.30303 | 73.36872 | 31.98616 | -0.75456 | 1.176025 | 0.419636 |
| 17 | 18422200 | 18433999 | 178.1769 | 268.9479 | 215.916 | 89.244 | 0.591005 | 3.349378 | 0.419641 |
| 8 | 25640600 | 25647199 | 60.77682 | 95.75654 | 152.6268 | 63.33195 | -0.34881 | 2.489382 | 0.419646 |
| 5 | 141814600 | 141816999 | 32.14005 | 53.23393 | 53.56782 | 23.18165 | 0.371747 | 1.315279 | 0.419649 |

| | | | | | | | | | |
|---|---|---|---|---|---|---|---|---|---|
| 10 | 19105600 | 19107599 | 16.56657 | 29.69368 | 28.6024 | 12.52844 | 0.454765 | 0.426334 | 0.419654 |
| 1 | 82816600 | 82817999 | 11.95894 | 20.9906 | 41.01069 | 18.13909 | -0.57353 | 0.608917 | 0.419679 |
| 13 | 53158800 | 53161199 | 15.78238 | 27.3339 | 48.45704 | 21.25995 | -0.4361 | 0.828761 | 0.419687 |
| X | 4407400 | 4413399 | 55.10404 | 90.20718 | 49.66089 | 21.3636 | 1.234585 | 1.511377 | 0.419702 |
| X | 32823800 | 32827199 | 65.66056 | 105.5907 | 66.83239 | 28.3518 | 1.01087 | 1.894537 | 0.419712 |
| 7 | 12212000 | 12220599 | 114.0567 | 175.4512 | 173.2677 | 71.95647 | 0.314347 | 2.951387 | 0.419714 |
| 8 | 53258000 | 53261999 | 32.78378 | 55.63759 | 42.90969 | 18.79902 | 0.754738 | 1.068109 | 0.419719 |
| 1 | 100781400 | 100790399 | 100.5513 | 156.8435 | 145.0653 | 60.19873 | 0.424204 | 2.69458 | 0.41972 |
| X | 144624600 | 144626799 | 24.19657 | 42.59275 | 30.6613 | 13.57395 | 0.860084 | 0.62196 | 0.419726 |
| 2 | 23180400 | 23192599 | 216.8556 | 318.8337 | 303.9101 | 125.1573 | 0.30891 | 3.826413 | 0.419745 |
| 3 | 84139800 | 84141799 | 19.0044 | 33.59097 | 30.98767 | 13.70716 | 0.506124 | 0.54595 | 0.419754 |
| 6 | 146686600 | 146689799 | 39.85554 | 65.13153 | 56.44987 | 24.26865 | 0.579936 | 1.500541 | 0.419766 |
| 8 | 22143600 | 22146799 | 23.2693 | 40.82958 | 32.33093 | 14.29611 | 0.722444 | 0.659815 | 0.419771 |
| 1 | 109153800 | 109154599 | 10.14764 | 19.76011 | 16.43719 | 6.020669 | 0.879029 | -0.31299 | 0.41978 |
| 3 | 53749200 | 53750799 | 12.36957 | 21.89455 | 38.7828 | 17.15044 | -0.429 | 0.506472 | 0.419783 |
| 13 | 45646800 | 45648399 | 12.36957 | 21.89455 | 38.7828 | 17.15044 | -0.429 | 0.506472 | 0.419783 |
| 16 | 19040400 | 19043399 | 35.32697 | 59.47101 | 45.19963 | 19.75378 | 0.775507 | 1.147179 | 0.419799 |
| 1 | 97697000 | 97698599 | 14.45609 | 25.1759 | 45.77965 | 20.12961 | -0.47141 | 0.749209 | 0.419801 |
| 2 | 152608400 | 152610399 | 11.21711 | 20.00729 | 37.78414 | 16.61863 | -0.51496 | 0.439381 | 0.41981 |
| 1 | 181952000 | 181952999 | 7.045057 | 13.38212 | 27.6898 | 11.13663 | -0.54731 | -0.00075 | 0.419818 |
| 1 | 111118200 | 111120199 | 25.56614 | 44.54146 | 35.73164 | 15.71519 | 0.703723 | 0.782233 | 0.419821 |
| 8 | 120313200 | 120314799 | 15.70806 | 27.23533 | 48.99584 | 21.49706 | -0.45703 | 0.816428 | 0.419863 |
| 13 | 66764200 | 66766799 | 40.37333 | 66.30103 | 59.53837 | 25.64695 | 0.53238 | 1.441047 | 0.419907 |
| 3 | 113133600 | 113137599 | 58.92853 | 94.50773 | 88.01107 | 37.18543 | 0.450262 | 1.974217 | 0.419919 |
| 5 | 47174000 | 47174599 | 12.24593 | 23.44011 | 21.65643 | 8.560257 | 0.628732 | -0.02228 | 0.419927 |
| X | 25473600 | 25475599 | 27.33276 | 47.26171 | 34.5 | 15.13572 | 0.837928 | 0.875436 | 0.419927 |
| 15 | 96881400 | 96883199 | 16.44745 | 29.48344 | 28.65807 | 12.55165 | 0.441942 | 0.425485 | 0.419937 |
| 9 | 6289200 | 6290999 | 12.93279 | 24.23928 | 22.79905 | 9.408065 | 0.549242 | 0.1265 | 0.419947 |
| 12 | 115694000 | 115702799 | 158.2531 | 239.3426 | 219.0293 | 90.58679 | 0.404632 | 3.305271 | 0.419956 |
| 14 | 115335400 | 115336599 | 7.486858 | 14.32671 | 28.49045 | 11.27057 | -0.46908 | -0.04711 | 0.419964 |
| 8 | 13996400 | 13999599 | 26.88706 | 43.66041 | 88.70063 | 38.03152 | -0.65255 | 1.592383 | 0.419989 |
| 12 | 22322200 | 22325999 | 118.8374 | 182.8033 | 140.443 | 58.32458 | 0.676203 | 2.950578 | 0.419994 |
| 15 | 82447400 | 82450399 | 17.98963 | 30.93472 | 53.29629 | 23.37802 | -0.39755 | 0.906635 | 0.420018 |
| 8 | 32407600 | 32408999 | 11.01788 | 19.41439 | 39.44217 | 17.45773 | -0.62803 | 0.54961 | 0.42002 |
| X | 23380600 | 23381999 | 15.45615 | 28.6284 | 24.83183 | 10.44128 | 0.645119 | 0.226974 | 0.42002 |
| 14 | 101264600 | 101265999 | 15.54891 | 29.43517 | 21.08576 | 8.541627 | 0.961389 | 0.083384 | 0.420022 |
| 10 | 54830800 | 54832799 | 18.89608 | 32.00606 | 56.95851 | 24.93912 | -0.44713 | 1.051125 | 0.420024 |
| 2 | 15318000 | 15322199 | 44.79565 | 72.76208 | 67.84478 | 29.09547 | 0.468505 | 1.608457 | 0.420036 |
| 7 | 21055000 | 21059599 | 36.99902 | 62.00755 | 47.50381 | 20.71672 | 0.764168 | 1.19834 | 0.420046 |
| 8 | 52513800 | 52515199 | 16.37413 | 29.41016 | 28.23989 | 12.34788 | 0.461235 | 0.413152 | 0.420048 |
| 17 | 11169800 | 11170799 | 10.55975 | 19.2309 | 34.94353 | 15.02848 | -0.43778 | 0.306304 | 0.420079 |
| 9 | 21772400 | 21773799 | 15.60794 | 28.48797 | 26.70308 | 11.45239 | 0.51354 | 0.311403 | 0.420089 |
| 12 | 28600000 | 28605199 | 42.84812 | 68.61462 | 111.256 | 46.89865 | -0.35085 | 2.009688 | 0.420109 |
| 6 | 67579800 | 67581799 | 21.06779 | 36.89199 | 34.241 | 15.11485 | 0.494517 | 0.681997 | 0.420111 |
| 3 | 31605600 | 31606999 | 15.13421 | 28.31508 | 23.78427 | 9.844104 | 0.708299 | 0.160889 | 0.420129 |
| 9 | 20552800 | 20556799 | 27.30129 | 44.10565 | 98.1538 | 41.9002 | -0.79107 | 1.711837 | 0.42013 |
| 4 | 93510200 | 93514599 | 46.91304 | 75.66975 | 76.2895 | 32.50076 | 0.345891 | 1.778176 | 0.420149 |

| | | | | | | | | | |
|---|---|---|---|---|---|---|---|---|---|
| 7 | 64673400 | 64676599 | 44.4822 | 72.12907 | 69.88951 | 29.93447 | 0.410919 | 1.643052 | 0.420164 |
| 11 | 100733000 | 100734399 | 12.38668 | 21.94377 | 38.78527 | 17.14857 | -0.42532 | 0.497895 | 0.420168 |
| 15 | 33492800 | 33494199 | 12.38668 | 21.94377 | 38.78527 | 17.14857 | -0.42532 | 0.497895 | 0.420168 |
| 1 | 9832000 | 9833999 | 13.97842 | 25.61126 | 24.73916 | 10.55737 | 0.475095 | 0.271754 | 0.420168 |
| 10 | 119288400 | 119291399 | 19.76905 | 32.82347 | 66.36598 | 28.76838 | -0.63134 | 1.264358 | 0.420181 |
| 18 | 56825600 | 56826399 | 16.38318 | 29.60459 | 23.71665 | 10.29995 | 0.727255 | 0.367759 | 0.4202 |
| 12 | 103781200 | 103797199 | 201.3586 | 292.0976 | 434.8801 | 179.2976 | -0.35274 | 4.03306 | 0.420206 |
| 5 | 100563600 | 100564399 | 5.77601 | 10.96952 | 27.66283 | 11.13016 | -0.83018 | -0.01146 | 0.420211 |
| X | 32359800 | 32361399 | 31.34879 | 52.94805 | 46.88995 | 20.51298 | 0.555926 | 1.109331 | 0.420217 |
| X | 31850200 | 31856199 | 161.7509 | 243.4645 | 227.3288 | 93.88528 | 0.36968 | 3.402186 | 0.420264 |
| 17 | 73564200 | 73566799 | 15.70104 | 27.19069 | 50.60542 | 22.20247 | -0.50627 | 0.828313 | 0.420272 |
| 7 | 119700400 | 119702799 | 20.82979 | 34.63994 | 66.05033 | 28.66314 | -0.54695 | 1.247484 | 0.420274 |
| 11 | 57447000 | 57455599 | 68.56674 | 106.6567 | 194.7575 | 80.85275 | -0.55911 | 2.733406 | 0.42028 |
| 5 | 74934400 | 74936399 | 13.17562 | 23.27293 | 40.83879 | 18.06713 | -0.41681 | 0.538015 | 0.420282 |
| 3 | 120943600 | 120946399 | 37.70649 | 62.46142 | 54.89329 | 23.75728 | 0.566561 | 1.331482 | 0.420284 |
| 8 | 3911000 | 3911799 | 15.32239 | 27.68537 | 27.28423 | 11.85057 | 0.430116 | 0.368344 | 0.420287 |
| 18 | 84219000 | 84220799 | 21.31605 | 37.27202 | 35.40779 | 15.61055 | 0.461262 | 0.709323 | 0.42029 |
| 6 | 135967400 | 135972399 | 45.07555 | 72.96257 | 71.59067 | 30.6233 | 0.390477 | 1.678937 | 0.420312 |
| 14 | 51568600 | 51569199 | 14.7943 | 26.73274 | 24.93515 | 10.82949 | 0.509148 | 0.359192 | 0.420342 |
| X | 80942200 | 80943199 | 18.06641 | 32.42286 | 28.56486 | 12.4987 | 0.584039 | 0.430644 | 0.420353 |
| 3 | 11460600 | 11461599 | 13.91519 | 25.67662 | 24.19205 | 10.22489 | 0.520593 | 0.230278 | 0.42036 |
| 4 | 92842000 | 92844599 | 35.34077 | 58.46992 | 50.41848 | 21.80193 | 0.593684 | 1.333572 | 0.420378 |
| 14 | 42091800 | 42093999 | 42.38983 | 68.71329 | 68.58894 | 29.37018 | 0.367652 | 1.643297 | 0.420382 |
| 10 | 92353600 | 92355799 | 32.66758 | 54.65247 | 50.29667 | 21.91045 | 0.500459 | 1.20316 | 0.420384 |
| 16 | 61460000 | 61460399 | 9.637476 | 18.58026 | 19.7474 | 7.646272 | 0.456085 | -0.12007 | 0.420394 |
| 18 | 72658200 | 72661199 | 35.3501 | 59.40271 | 46.33736 | 20.23993 | 0.738061 | 1.163594 | 0.420396 |
| X | 135743800 | 135745799 | 18.57315 | 33.66029 | 26.96085 | 11.66608 | 0.730991 | 0.374602 | 0.420399 |
| 9 | 25668400 | 25670799 | 30.22788 | 49.11358 | 82.46481 | 35.3728 | -0.37757 | 1.583384 | 0.420427 |
| 6 | 116775200 | 116777999 | 37.01746 | 61.24314 | 55.98726 | 24.20976 | 0.509711 | 1.344243 | 0.420436 |
| 1 | 86286000 | 86288399 | 14.3798 | 24.79834 | 50.55437 | 22.17723 | -0.63818 | 0.861656 | 0.420452 |
| 13 | 100807400 | 100809799 | 19.6222 | 34.5907 | 32.13339 | 14.22153 | 0.494822 | 0.581674 | 0.420465 |
| 9 | 18719800 | 18721399 | 18.55749 | 33.87606 | 25.28039 | 10.84043 | 0.840428 | 0.3294 | 0.420475 |
| 19 | 13407400 | 13412399 | 49.06919 | 79.70401 | 65.4489 | 28.06808 | 0.651391 | 1.615209 | 0.420477 |
| 16 | 80064400 | 80067799 | 36.9723 | 59.52659 | 97.72032 | 41.52623 | -0.35909 | 1.829005 | 0.42048 |
| 15 | 94300800 | 94304399 | 21.2078 | 35.12627 | 67.51477 | 29.21698 | -0.55887 | 1.298039 | 0.420483 |
| 11 | 83127400 | 83129999 | 27.45309 | 47.18066 | 41.3533 | 18.13899 | 0.573405 | 0.93665 | 0.420502 |
| 8 | 114088400 | 114089799 | 24.75886 | 42.61171 | 40.47805 | 17.75586 | 0.458295 | 0.913278 | 0.420504 |
| 13 | 66778000 | 66781199 | 76.94916 | 121.8652 | 106.9531 | 44.47537 | 0.516752 | 2.374536 | 0.420524 |
| 16 | 33135400 | 33138599 | 30.06993 | 48.96041 | 82.03873 | 35.22265 | -0.37229 | 1.551037 | 0.420527 |
| 17 | 25875600 | 25879999 | 55.0108 | 86.46785 | 157.2325 | 65.22304 | -0.54122 | 2.534785 | 0.420532 |
| 12 | 68770800 | 68771999 | 14.9353 | 26.09981 | 45.1862 | 19.91184 | -0.40041 | 0.702541 | 0.420536 |
| 16 | 95391600 | 95393399 | 24.51894 | 42.05789 | 41.55621 | 18.22599 | 0.400797 | 0.942337 | 0.42058 |
| 3 | 11795200 | 11799799 | 33.25689 | 55.73719 | 48.6187 | 21.20321 | 0.577416 | 1.184357 | 0.420593 |
| 4 | 62850000 | 62852399 | 30.41233 | 50.9171 | 50.25042 | 21.90052 | 0.399982 | 1.188926 | 0.420596 |
| 15 | 78511800 | 78515199 | 26.45187 | 43.31619 | 75.19937 | 32.35429 | -0.41751 | 1.457302 | 0.420625 |
| 19 | 59154400 | 59158599 | 37.51401 | 60.13936 | 106.5317 | 44.98278 | -0.47623 | 1.96771 | 0.420628 |
| 8 | 77552400 | 77555199 | 26.03118 | 45.13722 | 38.64148 | 16.9541 | 0.609778 | 0.850052 | 0.420628 |

| | | | | | | | | | |
|---|---|---|---|---|---|---|---|---|---|
| 3 | 15331000 | 15336199 | 62.47351 | 99.88458 | 94.36351 | 39.68833 | 0.424529 | 2.073222 | 0.420638 |
| 2 | 129985000 | 129990399 | 44.89275 | 71.52292 | 123.0107 | 51.48137 | -0.4438 | 2.174643 | 0.420647 |
| 1 | 26574600 | 26576399 | 18.7633 | 34.15816 | 25.79824 | 11.10041 | 0.820184 | 0.347427 | 0.420656 |
| 15 | 82950000 | 82952799 | 28.45979 | 47.84297 | 48.03228 | 20.98276 | 0.375311 | 1.13958 | 0.420664 |
| 2 | 7571200 | 7572199 | 26.38368 | 44.80648 | 43.74388 | 19.16845 | 0.416042 | 1.042412 | 0.420667 |
| 1 | 39578800 | 39580199 | 19.32639 | 34.04491 | 32.22734 | 14.26515 | 0.467466 | 0.588256 | 0.420686 |
| 18 | 38677600 | 38691799 | 141.0732 | 211.0259 | 315.6618 | 130.1722 | -0.31777 | 3.519366 | 0.420724 |
| 18 | 24323800 | 24326199 | 27.12635 | 46.56041 | 41.33282 | 18.12895 | 0.554681 | 0.944686 | 0.420737 |
| 12 | 42467600 | 42469199 | 12.23614 | 21.57858 | 39.36461 | 17.42058 | -0.47326 | 0.547788 | 0.420763 |
| 15 | 68130000 | 68131599 | 14.29495 | 24.94669 | 44.69968 | 19.68024 | -0.4499 | 0.717692 | 0.420769 |
| 11 | 25979200 | 25980199 | 12.00446 | 22.88808 | 22.12182 | 8.827029 | 0.551791 | 0.005772 | 0.420781 |
| 18 | 5938000 | 5939399 | 13.55148 | 25.36043 | 23.87582 | 9.877684 | 0.545252 | 0.145146 | 0.420783 |
| 4 | 112405200 | 112414599 | 136.2463 | 209.5566 | 164.2302 | 68.20201 | 0.647445 | 2.961702 | 0.420786 |
| 7 | 49798600 | 49799199 | 5.215639 | 10.15507 | 22.42973 | 8.22339 | -0.51398 | -0.3464 | 0.420797 |
| 5 | 145507200 | 145509399 | 33.9332 | 57.2613 | 45.75807 | 20.01233 | 0.703566 | 1.123932 | 0.4208 |
| 5 | 22927600 | 22932399 | 53.76321 | 85.26954 | 136.5021 | 56.83043 | -0.34718 | 2.323725 | 0.420807 |
| 2 | 124651600 | 124654399 | 18.46725 | 31.73591 | 58.29283 | 25.56922 | -0.48992 | 0.919068 | 0.420808 |
| 5 | 150898800 | 150901999 | 27.6747 | 45.41136 | 76.35786 | 32.88115 | -0.37048 | 1.433332 | 0.420814 |
| 15 | 32214600 | 32215399 | 12.68575 | 23.48542 | 21.93099 | 9.227705 | 0.538278 | 0.197579 | 0.420815 |
| 17 | 46626000 | 46635799 | 96.72932 | 147.5901 | 231.789 | 96.11388 | -0.36343 | 3.112482 | 0.420845 |
| 18 | 40845800 | 40849799 | 56.02294 | 89.97731 | 85.38343 | 36.15432 | 0.425616 | 1.926336 | 0.420853 |
| X | 5685400 | 5687199 | 15.4862 | 28.66186 | 21.72707 | 9.145422 | 0.837009 | 0.223839 | 0.420867 |
| 2 | 162987600 | 162991999 | 36.98419 | 59.42051 | 101.6298 | 43.056 | -0.42192 | 1.896161 | 0.420871 |
| 8 | 18033600 | 18035199 | 19.8135 | 34.89482 | 32.02772 | 14.17745 | 0.51146 | 0.591276 | 0.420892 |
| 10 | 65536800 | 65541399 | 49.75942 | 80.32616 | 75.73498 | 32.29837 | 0.443704 | 1.759798 | 0.4209 |
| 7 | 106371200 | 106374399 | 42.13904 | 68.88536 | 61.84768 | 26.59237 | 0.529479 | 1.503541 | 0.420901 |
| X | 34821800 | 34838599 | 223.39 | 332.7823 | 248.0129 | 102.1989 | 0.681325 | 3.607191 | 0.420905 |
| 7 | 83387800 | 83390199 | 13.04055 | 22.83254 | 42.57532 | 18.79182 | -0.50677 | 0.657166 | 0.420911 |
| 3 | 40827400 | 40829199 | 15.04701 | 27.3198 | 26.70584 | 11.52931 | 0.447184 | 0.336275 | 0.420925 |
| X | 49511000 | 49511999 | 7.860064 | 14.91153 | 32.16808 | 12.97209 | -0.61025 | 0.030278 | 0.420925 |
| 13 | 81111000 | 81113199 | 17.41164 | 29.78442 | 52.68914 | 23.10362 | -0.43669 | 0.952319 | 0.420929 |
| 7 | 35402200 | 35403599 | 6.601645 | 12.57885 | 27.20417 | 10.8697 | -0.60205 | -0.02805 | 0.420932 |
| 6 | 52511000 | 52515199 | 54.60396 | 87.47391 | 88.39158 | 37.28253 | 0.330774 | 2.003064 | 0.420935 |
| 1 | 5153000 | 5153799 | 8.286407 | 15.55926 | 29.62278 | 12.18605 | -0.45622 | 0.094778 | 0.420937 |
| 1 | 16734600 | 16735999 | 8.286407 | 15.55926 | 29.62278 | 12.18605 | -0.45622 | 0.094778 | 0.420937 |
| 16 | 67182200 | 67182999 | 11.7241 | 22.68124 | 19.67436 | 7.510614 | 0.765554 | -0.1448 | 0.420947 |
| X | 91813400 | 91815399 | 22.12948 | 38.48032 | 36.29734 | 15.94478 | 0.471074 | 0.795671 | 0.420948 |
| 3 | 20908200 | 20910199 | 19.20705 | 32.52188 | 57.52184 | 25.18321 | -0.43836 | 1.055381 | 0.420965 |
| 18 | 20731200 | 20733199 | 19.48758 | 33.2493 | 57.08168 | 25.02447 | -0.3944 | 0.988211 | 0.420969 |
| 7 | 14729200 | 14753199 | 407.1303 | 573.2724 | 450.0187 | 186.3981 | 0.527414 | 4.480484 | 0.420982 |
| 6 | 135150000 | 135150799 | 15.29488 | 27.54894 | 26.13672 | 11.39689 | 0.480721 | 0.381448 | 0.420983 |
| 19 | 56536200 | 56537399 | 6.34575 | 12.01441 | 27.61792 | 11.18486 | -0.70609 | 0.01646 | 0.420985 |
| 5 | 25854200 | 25855199 | 9.818331 | 19.06534 | 18.11843 | 6.788648 | 0.65953 | -0.23443 | 0.420991 |
| 13 | 10010800 | 10011999 | 13.28054 | 25.09607 | 22.96489 | 9.338092 | 0.606309 | 0.079586 | 0.421005 |
| 18 | 77770200 | 77771999 | 9.17694 | 16.73028 | 32.92919 | 14.14342 | -0.55057 | 0.286729 | 0.421007 |
| 15 | 54207000 | 54209399 | 17.95717 | 30.83588 | 56.6045 | 24.82808 | -0.48925 | 0.921026 | 0.421018 |
| 15 | 103853400 | 103855199 | 11.15729 | 20.1191 | 37.37588 | 16.27467 | -0.48157 | 0.370137 | 0.421024 |

| | | | | | | | | | |
|---|---|---|---|---|---|---|---|---|---|
| 9 | 74803800 | 74806599 | 15.73028 | 27.41673 | 47.34358 | 20.82767 | -0.39719 | 0.746709 | 0.421027 |
| 5 | 41923000 | 41926799 | 28.38083 | 46.33092 | 78.81487 | 33.87071 | -0.39138 | 1.505872 | 0.42103 |
| 8 | 53240800 | 53241399 | 10.01771 | 19.35964 | 19.70335 | 7.558884 | 0.530026 | -0.14819 | 0.421049 |
| 10 | 108300200 | 108300999 | 10.01771 | 19.35964 | 19.70335 | 7.558884 | 0.530026 | -0.14819 | 0.421049 |
| 1 | 22842200 | 22845999 | 27.90126 | 45.47885 | 84.04416 | 36.098 | -0.51215 | 1.527842 | 0.421049 |
| 15 | 58760200 | 58761799 | 7.595777 | 13.75766 | 35.98641 | 15.58314 | -0.96599 | 0.316702 | 0.421053 |
| 2 | 95787400 | 95787999 | 13.23804 | 24.7287 | 24.00182 | 9.960537 | 0.498438 | 0.156135 | 0.421056 |
| 9 | 79950600 | 79956799 | 72.11108 | 114.4949 | 111.6876 | 46.5503 | 0.367897 | 2.300125 | 0.421057 |
| X | 63675800 | 63676199 | 5.056112 | 9.849326 | 22.47637 | 8.191745 | -0.55232 | -0.36824 | 0.421068 |
| 13 | 33726800 | 33733199 | 84.46452 | 133.7582 | 115.0985 | 47.8609 | 0.545102 | 2.383686 | 0.421069 |
| 4 | 8690000 | 8692399 | 15.40304 | 26.32705 | 52.62863 | 23.07557 | -0.61233 | 0.952266 | 0.421088 |
| 15 | 98172000 | 98173799 | 24.51183 | 42.36157 | 39.77154 | 17.44815 | 0.47613 | 0.871756 | 0.421109 |
| 7 | 35870200 | 35872199 | 25.34319 | 43.75393 | 40.34262 | 17.69814 | 0.501935 | 0.886921 | 0.421112 |
| Y | 89196000 | 89197999 | 16.64401 | 29.89435 | 16.98224 | 7.425592 | 1.212775 | 0.385533 | 0.421113 |
| 3 | 94048200 | 94056199 | 120.4441 | 186.5833 | 152.3215 | 63.27016 | 0.596434 | 2.823233 | 0.421115 |
| 14 | 88275000 | 88278799 | 53.5288 | 85.7881 | 85.05358 | 35.89788 | 0.358927 | 1.988651 | 0.421136 |
| 5 | 142059400 | 142061399 | 12.10991 | 21.24068 | 41.5272 | 18.35666 | -0.57459 | 0.621467 | 0.421172 |
| 18 | 10856600 | 10861799 | 67.47996 | 107.3297 | 107.1547 | 44.74212 | 0.33695 | 2.243993 | 0.42118 |
| X | 126182200 | 126182599 | 10.89207 | 20.89625 | 19.21648 | 7.546234 | 0.644925 | -0.06119 | 0.4212 |
| 10 | 40633600 | 40636399 | 20.85397 | 35.16165 | 59.72387 | 26.1138 | -0.38047 | 1.100926 | 0.42121 |
| 13 | 117566200 | 117566599 | 4.819333 | 9.359498 | 22.76493 | 8.5203 | -0.67917 | -0.27787 | 0.421211 |
| 12 | 56333200 | 56333799 | 5.399354 | 10.45991 | 23.80233 | 9.028079 | -0.60319 | -0.22298 | 0.421213 |
| 17 | 90217800 | 90218599 | 7.773838 | 14.80747 | 28.92103 | 11.56376 | -0.45759 | -0.01 | 0.421216 |
| X | 118322600 | 118324199 | 20.50943 | 36.68456 | 28.21912 | 12.38221 | 0.775825 | 0.465928 | 0.421216 |
| X | 98150400 | 98152199 | 19.99506 | 35.51842 | 29.85971 | 13.17858 | 0.642772 | 0.507513 | 0.421254 |
| 10 | 111545400 | 111546399 | 14.40916 | 27.2499 | 22.31674 | 9.059688 | 0.767177 | 0.082835 | 0.421258 |
| 1 | 37713600 | 37719599 | 32.80686 | 52.75058 | 99.68282 | 42.27944 | -0.56398 | 1.865017 | 0.421271 |
| 6 | 40882600 | 40884399 | 20.37163 | 34.49861 | 58.63272 | 25.67065 | -0.38102 | 1.061131 | 0.421276 |
| 16 | 73458400 | 73459999 | 14.27682 | 26.74598 | 23.86166 | 9.854252 | 0.62447 | 0.145411 | 0.421282 |
| 13 | 47106200 | 47107599 | 9.074965 | 16.73787 | 32.5099 | 13.7488 | -0.51633 | 0.214912 | 0.421295 |
| 2 | 93669600 | 93670799 | 6.941861 | 13.07049 | 28.59467 | 11.70851 | -0.64946 | 0.061609 | 0.421315 |
| X | 5633400 | 5637199 | 62.49223 | 100.1638 | 88.29914 | 37.27362 | 0.528274 | 1.997801 | 0.421315 |
| 1 | 159608600 | 159612199 | 28.67816 | 48.25268 | 47.48269 | 20.74824 | 0.404096 | 1.130726 | 0.421316 |
| 2 | 166081800 | 166085399 | 38.45181 | 61.80025 | 101.449 | 43.0014 | -0.3622 | 1.887659 | 0.421317 |
| 8 | 13991000 | 13992999 | 21.56836 | 36.13298 | 61.82467 | 26.95557 | -0.39115 | 1.16771 | 0.421321 |
| 6 | 132197600 | 132200399 | 35.14511 | 59.17434 | 45.19849 | 19.75424 | 0.768353 | 1.145069 | 0.421326 |
| 3 | 106579000 | 106580999 | 34.66318 | 56.02818 | 91.58812 | 39.12378 | -0.34629 | 1.706329 | 0.421332 |
| 4 | 47947000 | 47949599 | 12.70985 | 22.32681 | 40.93208 | 18.1168 | -0.48238 | 0.598937 | 0.421356 |
| 11 | 94668200 | 94671599 | 19.24894 | 32.42336 | 60.76052 | 26.5591 | -0.52184 | 1.103317 | 0.421356 |
| X | 111385000 | 111386199 | 20.71399 | 36.48227 | 32.1264 | 14.22101 | 0.571167 | 0.598895 | 0.421374 |
| 3 | 37117000 | 37118799 | 24.10431 | 39.8087 | 69.33412 | 29.94459 | -0.41757 | 1.341786 | 0.421393 |
| 2 | 119050400 | 119052799 | 17.53776 | 30.04812 | 52.71257 | 23.11686 | -0.42437 | 0.937379 | 0.421399 |
| 13 | 76337400 | 76341999 | 36.96055 | 59.65986 | 99.51572 | 42.43525 | -0.37773 | 1.753171 | 0.421407 |
| 19 | 11182800 | 11183999 | 12.84403 | 22.69938 | 39.80499 | 17.60789 | -0.41549 | 0.527992 | 0.421414 |
| 12 | 93283400 | 93284599 | 10.42353 | 18.98521 | 34.95519 | 15.03111 | -0.45642 | 0.305384 | 0.421448 |
| 3 | 51303000 | 51305399 | 11.96226 | 21.05749 | 39.94533 | 17.68396 | -0.53019 | 0.568883 | 0.421458 |
| 11 | 115470800 | 115472199 | 7.701223 | 14.59193 | 30.38504 | 12.28552 | -0.56274 | 0.033549 | 0.421463 |

| | | | | | | | | | |
|---|---|---|---|---|---|---|---|---|---|
| 6 | 128713000 | 128715999 | 22.92172 | 37.9879 | 67.79396 | 29.35001 | -0.45221 | 1.295142 | 0.421477 |
| 16 | 43606600 | 43607999 | 17.97069 | 32.07408 | 29.54453 | 12.98747 | 0.51551 | 0.46454 | 0.421478 |
| 6 | 115317800 | 115318999 | 16.69428 | 29.66283 | 28.36385 | 12.51562 | 0.458356 | 0.482418 | 0.421503 |
| 12 | 105386600 | 105388799 | 11.22657 | 19.64585 | 45.71131 | 20.16679 | -0.82462 | 0.659152 | 0.42151 |
| X | 25370200 | 25376599 | 169.5802 | 257.0701 | 178.7029 | 73.96268 | 0.803806 | 3.261036 | 0.421522 |
| 4 | 146629000 | 146631599 | 86.48504 | 136.3219 | 115.8874 | 48.08984 | 0.557324 | 2.490879 | 0.42153 |
| 6 | 57792200 | 57792999 | 5.536482 | 10.72422 | 23.79817 | 9.031805 | -0.56801 | -0.21876 | 0.421536 |
| 9 | 118620800 | 118622799 | 13.83703 | 24.18589 | 43.64226 | 19.23507 | -0.45968 | 0.691404 | 0.421556 |
| 1 | 48866000 | 48868799 | 28.47331 | 49.29866 | 36.78431 | 16.13853 | 0.806655 | 0.869667 | 0.421599 |
| 4 | 105365600 | 105368999 | 35.18021 | 56.79154 | 93.50546 | 39.87728 | -0.35875 | 1.745745 | 0.421602 |
| 2 | 60866400 | 60868599 | 29.61747 | 50.44786 | 44.09215 | 19.32728 | 0.575578 | 1.027237 | 0.421602 |
| 12 | 119992200 | 119992799 | 259.8137 | 384.7401 | 203.3801 | 83.75843 | 1.168604 | 3.710671 | 0.421608 |
| 5 | 87299000 | 87300999 | 20.51572 | 35.95401 | 34.31531 | 15.15789 | 0.45449 | 0.667591 | 0.421612 |
| 6 | 25291200 | 25291799 | 6.866521 | 13.13179 | 27.89233 | 11.05294 | -0.56605 | -0.05218 | 0.421625 |
| 9 | 89818200 | 89819199 | 10.29899 | 18.6101 | 34.87576 | 15.14656 | -0.49093 | 0.351065 | 0.421626 |
| X | 7688000 | 7692799 | 72.55327 | 115.7928 | 92.54705 | 38.82569 | 0.662631 | 2.14223 | 0.421629 |
| 3 | 27147200 | 27150599 | 46.4322 | 75.63966 | 62.6222 | 26.89124 | 0.642682 | 1.562048 | 0.421634 |
| 3 | 50172000 | 50172599 | 5.669863 | 11.02054 | 24.74256 | 9.197893 | -0.55762 | -0.28951 | 0.421643 |
| 13 | 36858400 | 36859799 | 13.61505 | 25.24057 | 24.89305 | 10.45443 | 0.462026 | 0.205573 | 0.42165 |
| 2 | 99399600 | 99402799 | 59.1595 | 94.56173 | 94.72028 | 39.82388 | 0.339847 | 2.07915 | 0.421658 |
| 1 | 94277200 | 94280199 | 31.71033 | 53.8263 | 44.64064 | 19.55852 | 0.650462 | 1.066785 | 0.421664 |
| 9 | 7497800 | 7499199 | 16.7733 | 30.36658 | 26.93776 | 11.67138 | 0.582948 | 0.364894 | 0.421664 |
| 7 | 25691600 | 25696799 | 91.66623 | 143.4057 | 140.412 | 58.2501 | 0.345404 | 2.639349 | 0.421684 |
| 10 | 3890000 | 3893399 | 19.33715 | 32.41062 | 60.5886 | 26.42464 | -0.51832 | 1.153937 | 0.421689 |
| 14 | 101043200 | 101043799 | 5.129848 | 9.950691 | 24.44782 | 9.216957 | -0.70485 | -0.24682 | 0.421691 |
| 7 | 93653800 | 93655799 | 22.47813 | 39.48064 | 32.37244 | 14.32651 | 0.67235 | 0.640227 | 0.421697 |
| 12 | 115290200 | 115294199 | 79.95439 | 126.6045 | 117.5927 | 48.89582 | 0.43492 | 2.382468 | 0.421698 |
| 10 | 43568600 | 43568999 | 9.327396 | 18.07637 | 19.30552 | 7.309514 | 0.479463 | -0.19936 | 0.421707 |
| 18 | 37248800 | 37249599 | 14.67905 | 26.72445 | 26.35429 | 11.33883 | 0.437978 | 0.318884 | 0.421737 |
| 12 | 57931800 | 57933799 | 24.08457 | 42.35394 | 31.28083 | 13.85094 | 0.82285 | 0.633217 | 0.421744 |
| 8 | 51459000 | 51460799 | 22.00868 | 38.33982 | 37.12981 | 16.3291 | 0.433325 | 0.769015 | 0.421748 |
| 4 | 79206200 | 79209599 | 29.64631 | 50.29272 | 45.19964 | 19.79985 | 0.534826 | 1.062294 | 0.421749 |
| 12 | 11541600 | 11548999 | 63.48378 | 101.4781 | 95.17157 | 40.01605 | 0.434734 | 2.083018 | 0.421752 |
| 7 | 32948400 | 32950799 | 69.45817 | 110.1849 | 107.3947 | 44.72132 | 0.367869 | 2.324503 | 0.421752 |
| 13 | 55867000 | 55868599 | 14.93775 | 26.02222 | 45.7723 | 20.13001 | -0.42365 | 0.748057 | 0.421753 |
| X | 125118400 | 125120599 | 42.4252 | 68.91859 | 66.80758 | 28.65276 | 0.412743 | 1.603009 | 0.421759 |
| 5 | 116460400 | 116472999 | 124.5705 | 187.1862 | 292.8642 | 120.8965 | -0.37664 | 3.429122 | 0.421762 |
| Y | 5550400 | 5552999 | 24.01096 | 41.46807 | 25.46276 | 11.17052 | 1.085528 | 0.853066 | 0.421768 |
| 13 | 44099200 | 44104999 | 49.40156 | 78.82983 | 126.7931 | 53.17982 | -0.34496 | 2.129269 | 0.42177 |
| 1 | 53617400 | 53619199 | 13.68462 | 25.61937 | 22.6709 | 9.373139 | 0.634304 | 0.139449 | 0.42178 |
| 2 | 66715000 | 66716599 | 16.82669 | 30.52124 | 26.98189 | 11.66271 | 0.590151 | 0.355162 | 0.421791 |
| 12 | 67511600 | 67512199 | 7.004509 | 13.46574 | 26.96595 | 10.52699 | -0.46114 | -0.10074 | 0.421794 |
| 13 | 11153800 | 11155999 | 22.98366 | 40.25235 | 34.10456 | 15.05635 | 0.625491 | 0.690957 | 0.421803 |
| 16 | 20998400 | 20999799 | 21.10938 | 36.67061 | 35.80727 | 15.71983 | 0.421141 | 0.801505 | 0.421838 |
| 1 | 88798600 | 88803399 | 39.51373 | 63.49912 | 108.5009 | 45.98223 | -0.42023 | 1.892766 | 0.421895 |
| 11 | 68595000 | 68596999 | 13.34859 | 23.5503 | 41.36912 | 18.30612 | -0.41899 | 0.553433 | 0.4219 |
| 7 | 25120200 | 25122599 | 12.45111 | 21.76785 | 43.16483 | 19.02792 | -0.59489 | 0.680665 | 0.421938 |

| | | | | | | | | | |
|---|---|---|---|---|---|---|---|---|---|
| 13 | 78962200 | 78963599 | 17.65755 | 31.1938 | 30.15237 | 13.33909 | 0.438781 | 0.541507 | 0.421943 |
| 4 | 109893400 | 109894999 | 12.81329 | 22.59933 | 41.35357 | 18.3003 | -0.47795 | 0.550851 | 0.421968 |
| 1 | 101811000 | 101814399 | 42.64081 | 70.28161 | 55.70021 | 24.03082 | 0.714202 | 1.40155 | 0.421975 |
| X | 4524200 | 4527799 | 42.02534 | 70.32662 | 44.14652 | 19.23452 | 1.050647 | 1.228144 | 0.421979 |
| 1 | 143216600 | 143218199 | 11.5393 | 20.8201 | 38.4702 | 16.73896 | -0.47299 | 0.373839 | 0.42199 |
| X | 5394200 | 5402799 | 100.2869 | 158.4489 | 109.228 | 45.36683 | 0.86196 | 2.449006 | 0.421992 |
| 1 | 117638200 | 117640599 | 23.72019 | 41.60281 | 33.05171 | 14.6095 | 0.717998 | 0.672727 | 0.421994 |
| 4 | 119001400 | 119002799 | 13.46738 | 24.8575 | 24.91053 | 10.52442 | 0.433004 | 0.228853 | 0.422007 |
| 17 | 92886400 | 92888199 | 24.6228 | 42.28566 | 40.91051 | 17.94437 | 0.431155 | 0.928655 | 0.422028 |
| 14 | 81590600 | 81591199 | 11.33977 | 21.45265 | 22.08156 | 8.957657 | 0.44095 | 0.052168 | 0.422031 |
| 10 | 125970800 | 125974799 | 31.93254 | 51.55993 | 99.09179 | 42.27144 | -0.58103 | 1.735443 | 0.422036 |
| 2 | 100726400 | 100728999 | 19.772 | 34.72943 | 33.27433 | 14.72631 | 0.448948 | 0.626714 | 0.422046 |
| 19 | 3135400 | 3137599 | 28.05476 | 47.6362 | 39.52639 | 17.31945 | 0.649716 | 1.04435 | 0.422057 |
| 5 | 19052600 | 19058199 | 103.9686 | 160.5408 | 159.4199 | 66.21976 | 0.310563 | 2.874864 | 0.422062 |
| 9 | 53001000 | 53003399 | 19.90272 | 35.38289 | 28.61605 | 12.62112 | 0.698679 | 0.497414 | 0.422063 |
| 7 | 92820400 | 92822599 | 21.95018 | 38.90561 | 29.06797 | 12.85086 | 0.810228 | 0.532536 | 0.422079 |
| 7 | 17271400 | 17275399 | 68.6752 | 108.9107 | 108.7583 | 45.27574 | 0.33251 | 2.331357 | 0.422104 |
| 1 | 39297800 | 39298799 | 15.3922 | 27.77005 | 25.92704 | 11.28213 | 0.506152 | 0.375702 | 0.422132 |
| 5 | 41869800 | 41870999 | 14.94479 | 27.96896 | 23.79245 | 9.84256 | 0.690719 | 0.157287 | 0.42215 |
| 5 | 110626600 | 110630599 | 38.36263 | 61.39621 | 110.4319 | 46.51179 | -0.50108 | 2.023355 | 0.422153 |
| 7 | 3376400 | 3382199 | 42.66199 | 68.29374 | 113.7593 | 47.92991 | -0.39023 | 2.019667 | 0.422153 |
| 7 | 27004400 | 27016799 | 98.89161 | 150.8991 | 233.5008 | 96.82517 | -0.34201 | 3.111524 | 0.422177 |
| 12 | 86704400 | 86705599 | 14.07909 | 25.67281 | 25.33385 | 10.87822 | 0.4387 | 0.301865 | 0.422193 |
| 6 | 59669800 | 59671399 | 21.41212 | 37.39348 | 28.94917 | 12.7513 | 0.754945 | 0.717267 | 0.422213 |
| X | 148728200 | 148735999 | 90.72162 | 143.0555 | 111.2825 | 46.18131 | 0.68564 | 2.48621 | 0.422217 |
| 1 | 147564200 | 147565799 | 11.49746 | 20.60441 | 37.80741 | 16.56278 | -0.46975 | 0.411459 | 0.422225 |
| 7 | 34528800 | 34529799 | 11.38489 | 21.54346 | 21.47734 | 8.707879 | 0.487059 | 0.0479 | 0.422245 |
| 8 | 60207200 | 60208399 | 13.86329 | 26.54018 | 20.96267 | 8.282215 | 0.853534 | -0.0102 | 0.422275 |
| 12 | 6562800 | 6563399 | 10.5408 | 19.08429 | 34.84207 | 15.09499 | -0.45159 | 0.33845 | 0.422277 |
| 15 | 91156200 | 91157399 | 10.5408 | 19.08429 | 34.84207 | 15.09499 | -0.45159 | 0.33845 | 0.422277 |
| 11 | 57730800 | 57733399 | 12.17862 | 21.57986 | 39.261 | 17.35816 | -0.46691 | 0.492669 | 0.422286 |
| 14 | 112292800 | 112293999 | 14.90875 | 28.12144 | 21.57042 | 8.805025 | 0.853699 | 0.10195 | 0.422315 |
| 5 | 150983000 | 150987199 | 43.7366 | 69.88477 | 115.2318 | 48.42346 | -0.37848 | 2.080327 | 0.422325 |
| 1 | 103637600 | 103637999 | 8.675706 | 16.92358 | 16.13896 | 5.776959 | 0.713758 | -0.41217 | 0.422337 |
| 9 | 59657600 | 59659199 | 10.35655 | 18.40628 | 36.8565 | 16.25967 | -0.60153 | 0.456855 | 0.422345 |
| X | 153454400 | 153456799 | 34.94855 | 58.28564 | 51.42065 | 22.35585 | 0.56146 | 1.247426 | 0.422349 |
| 15 | 67087200 | 67088999 | 12.98725 | 22.82077 | 42.42779 | 18.78344 | -0.50273 | 0.596421 | 0.422359 |
| 17 | 26509200 | 26527799 | 259.7247 | 369.8052 | 522.7519 | 216.124 | -0.30474 | 4.311586 | 0.422365 |
| 14 | 125034600 | 125039399 | 77.64455 | 122.9259 | 118.6024 | 49.31137 | 0.379927 | 2.384922 | 0.422379 |
| 8 | 19732200 | 19732799 | 19.81696 | 34.86065 | 32.7206 | 14.48751 | 0.478893 | 0.606854 | 0.422389 |
| 7 | 26462800 | 26463999 | 16.32509 | 29.07227 | 28.32318 | 12.47396 | 0.43335 | 0.465323 | 0.422403 |
| 18 | 83717800 | 83719199 | 20.04222 | 35.01891 | 33.97748 | 14.9709 | 0.430861 | 0.710779 | 0.422419 |
| 11 | 41911200 | 41915399 | 40.81201 | 67.53318 | 52.84217 | 22.85262 | 0.733445 | 1.347992 | 0.422421 |
| 8 | 25938800 | 25939599 | 6.696798 | 12.90272 | 27.07355 | 10.50202 | -0.51953 | -0.1258 | 0.422425 |
| 15 | 67535800 | 67536999 | 10.67451 | 19.50624 | 36.63182 | 15.68758 | -0.48136 | 0.289277 | 0.422453 |
| 17 | 32724200 | 32726199 | 7.630955 | 14.05197 | 31.4896 | 13.34642 | -0.72278 | 0.211977 | 0.422454 |
| 15 | 81628400 | 81630999 | 12.88177 | 22.55673 | 42.5735 | 18.79256 | -0.52416 | 0.654437 | 0.422463 |

| | | | | | | | | | |
|---|---|---|---|---|---|---|---|---|---|
| 6 | 82698800 | 82701599 | 22.87227 | 39.34326 | 36.47261 | 15.99859 | 0.493087 | 0.90255 | 0.422469 |
| 18 | 5705600 | 5707599 | 21.5485 | 38.35683 | 27.99504 | 12.33417 | 0.847105 | 0.500594 | 0.422477 |
| 4 | 148053400 | 148054399 | 9.260113 | 16.997 | 32.43909 | 13.80938 | -0.49794 | 0.244561 | 0.422504 |
| X | 159165600 | 159168999 | 40.49923 | 66.74531 | 57.27543 | 24.7094 | 0.59977 | 1.398591 | 0.422507 |
| 6 | 52263400 | 52266599 | 17.66 | 30.52579 | 54.44435 | 23.88538 | -0.44576 | 0.859946 | 0.422521 |
| 12 | 39485800 | 39486199 | 11.30269 | 21.3029 | 21.0497 | 8.604301 | 0.490008 | 0.07402 | 0.422525 |
| 4 | 5055600 | 5056999 | 10.25092 | 18.8279 | 36.22163 | 15.40595 | -0.50857 | 0.258448 | 0.42253 |
| 14 | 120950600 | 120950999 | 10.68543 | 20.2962 | 20.11626 | 8.091302 | 0.505908 | 0.013431 | 0.422582 |
| 2 | 17328800 | 17330799 | 17.69476 | 30.56276 | 53.31889 | 23.39109 | -0.41416 | 0.866615 | 0.422593 |
| 16 | 32416600 | 32417799 | 7.204784 | 13.73859 | 27.77085 | 11.07817 | -0.50292 | -0.02294 | 0.422599 |
| 7 | 137998200 | 138001199 | 21.58801 | 35.2667 | 86.39803 | 37.13912 | -0.91523 | 1.485822 | 0.422606 |
| 1 | 182705600 | 182720399 | 198.2579 | 288.4986 | 419.2192 | 172.7846 | -0.31336 | 3.985868 | 0.42261 |
| 4 | 17308400 | 17311799 | 32.28337 | 53.37468 | 52.47661 | 22.68108 | 0.404961 | 1.335292 | 0.422618 |
| 10 | 52469200 | 52471799 | 22.16884 | 37.20998 | 63.07209 | 27.53262 | -0.37777 | 1.149096 | 0.422641 |
| 15 | 31813200 | 31816599 | 15.04389 | 26.29904 | 46.76055 | 20.61187 | -0.43886 | 0.697613 | 0.422669 |
| 9 | 76596400 | 76598199 | 24.18255 | 40.10598 | 68.55532 | 29.6942 | -0.39021 | 1.290747 | 0.422673 |
| X | 3057600 | 3065399 | 89.78268 | 142.0282 | 114.7146 | 47.67548 | 0.635139 | 2.413683 | 0.422682 |
| 8 | 25540600 | 25542599 | 14.1703 | 24.68222 | 45.26691 | 19.9062 | -0.48351 | 0.746046 | 0.422706 |
| 14 | 50874600 | 50877399 | 29.49261 | 50.12359 | 44.63507 | 19.56176 | 0.548197 | 1.045733 | 0.422717 |
| 1 | 119943200 | 119945199 | 12.67573 | 22.50112 | 39.78645 | 17.56997 | -0.42459 | 0.484245 | 0.422717 |
| 8 | 84031600 | 84032999 | 12.67573 | 22.50112 | 39.78645 | 17.56997 | -0.42459 | 0.484245 | 0.422717 |
| 17 | 14108200 | 14109399 | 11.29108 | 19.9888 | 37.83562 | 16.73104 | -0.52389 | 0.496549 | 0.422738 |
| 5 | 116753600 | 116755399 | 13.79158 | 24.23662 | 42.4126 | 18.77836 | -0.41577 | 0.600531 | 0.422738 |
| 7 | 30343800 | 30345599 | 8.733161 | 15.52616 | 37.94053 | 16.7333 | -0.88662 | 0.445549 | 0.422744 |
| 18 | 32436200 | 32440399 | 52.04906 | 83.68302 | 83.20725 | 35.28699 | 0.360223 | 1.887369 | 0.422758 |
| 11 | 110088200 | 110090999 | 24.46985 | 42.25511 | 39.77051 | 17.44727 | 0.472297 | 0.878158 | 0.422777 |
| 10 | 18873800 | 18877199 | 36.60709 | 60.06174 | 60.30895 | 25.96894 | 0.371185 | 1.443792 | 0.422782 |
| X | 79510000 | 79510799 | 19.58061 | 34.38908 | 30.54494 | 13.51696 | 0.557928 | 0.625286 | 0.422785 |
| 15 | 97147400 | 97151999 | 48.96757 | 78.01589 | 124.2469 | 51.99965 | -0.33307 | 2.178934 | 0.422788 |
| 2 | 176794400 | 176794799 | 11.53738 | 20.71518 | 36.80621 | 16.09842 | -0.42172 | 0.399979 | 0.42279 |
| 8 | 34202600 | 34206599 | 43.54904 | 69.62413 | 113.8582 | 47.88425 | -0.3657 | 2.061971 | 0.422807 |
| 5 | 131087600 | 131090599 | 44.11884 | 72.41594 | 56.94292 | 24.52378 | 0.723127 | 1.453723 | 0.422808 |
| 4 | 66415800 | 66417399 | 12.86559 | 24.57417 | 20.25503 | 8.051413 | 0.783657 | -0.00497 | 0.422827 |
| 9 | 71121600 | 71124599 | 16.24974 | 27.56371 | 63.40798 | 27.77523 | -0.81597 | 1.034912 | 0.422839 |
| 4 | 50475600 | 50476999 | 15.46607 | 28.24619 | 26.098 | 11.18438 | 0.534656 | 0.304231 | 0.422847 |
| 3 | 121371600 | 121372999 | 13.44761 | 24.6811 | 22.9949 | 9.790097 | 0.529676 | 0.255839 | 0.422854 |
| 10 | 65286000 | 65287199 | 15.17748 | 28.67824 | 21.63307 | 8.798125 | 0.882431 | 0.093524 | 0.422872 |
| 11 | 13478000 | 13480199 | 21.24992 | 38.29785 | 25.39306 | 11.06438 | 0.99532 | 0.420035 | 0.422877 |
| X | 151468800 | 151469799 | 15.10295 | 27.85297 | 25.36028 | 10.72668 | 0.569491 | 0.249071 | 0.422879 |
| 3 | 54692600 | 54693799 | 10.61378 | 19.05977 | 35.3241 | 15.4482 | -0.4818 | 0.390579 | 0.42288 |
| 6 | 128506200 | 128506999 | 6.418378 | 12.35203 | 25.83049 | 10.05307 | -0.51867 | -0.11994 | 0.422881 |
| 6 | 70372200 | 70373999 | 19.40061 | 34.38116 | 30.38684 | 13.43436 | 0.569048 | 0.52007 | 0.422891 |
| 1 | 190861000 | 190867199 | 65.51035 | 103.1132 | 160.8905 | 66.75243 | -0.31927 | 2.512436 | 0.422896 |
| 7 | 85178000 | 85187799 | 195.5312 | 291.3554 | 266.4326 | 109.792 | 0.386679 | 3.597774 | 0.422904 |
| 1 | 49250000 | 49251999 | 26.78245 | 45.21782 | 44.83946 | 19.61363 | 0.393284 | 1.093637 | 0.422905 |
| 9 | 55456000 | 55457399 | 10.52734 | 19.09897 | 34.86717 | 15.06726 | -0.44916 | 0.326268 | 0.42294 |
| 15 | 30037200 | 30039799 | 28.25261 | 47.88633 | 40.50845 | 17.74013 | 0.621798 | 1.059595 | 0.422947 |

| | | | | | | | | | |
|---|---|---|---|---|---|---|---|---|---|
| 9 | 23440000 | 23441399 | 12.85172 | 22.57343 | 41.42913 | 18.335 | -0.48404 | 0.601673 | 0.422961 |
| X | 102168400 | 102170199 | 19.13468 | 35.53082 | 21.58982 | 9.047184 | 1.15797 | 0.234417 | 0.422972 |
| 1 | 115286400 | 115287999 | 20.14401 | 35.30934 | 33.93547 | 14.99253 | 0.444621 | 0.663508 | 0.422973 |
| 6 | 10560200 | 10562799 | 30.07027 | 51.67941 | 39.57551 | 17.3592 | 0.767381 | 0.944902 | 0.423037 |
| 3 | 112192600 | 112193799 | 13.6715 | 26.17857 | 20.97181 | 8.280773 | 0.834035 | -0.01413 | 0.423038 |
| 12 | 65829400 | 65834199 | 66.01699 | 105.1067 | 104.8743 | 43.84997 | 0.339311 | 2.210834 | 0.423048 |
| X | 3162200 | 3175799 | 336.9374 | 478.8334 | 417.9701 | 172.8587 | 0.387735 | 4.342818 | 0.423053 |
| 11 | 59236800 | 59238999 | 15.78644 | 27.21765 | 49.51208 | 21.71984 | -0.47459 | 0.870108 | 0.423059 |
| 12 | 103615600 | 103617199 | 18.21352 | 34.35677 | 20.34001 | 8.30189 | 1.22528 | 0.130258 | 0.423064 |
| 5 | 19782200 | 19782599 | 4.643927 | 9.067531 | 20.52343 | 7.230567 | -0.49666 | -0.50424 | 0.42307 |
| 11 | 75865000 | 75869199 | 34.87193 | 56.28527 | 93.87426 | 40.02667 | -0.37755 | 1.750455 | 0.423072 |
| 15 | 34037000 | 34037999 | 12.99758 | 22.92416 | 40.33153 | 17.84804 | -0.42141 | 0.551761 | 0.423086 |
| 4 | 146301000 | 146304399 | 82.63006 | 131.22 | 106.3894 | 44.35003 | 0.634578 | 2.303586 | 0.423106 |
| 9 | 12506400 | 12508399 | 23.64253 | 40.97973 | 34.6284 | 15.19317 | 0.628258 | 0.835499 | 0.423108 |
| 4 | 134917000 | 134929599 | 94.86495 | 144.1246 | 265.1217 | 109.8026 | -0.5964 | 3.201842 | 0.42311 |
| X | 118760200 | 118766799 | 81.67628 | 129.6512 | 108.2288 | 45.09796 | 0.591999 | 2.316847 | 0.423117 |
| 13 | 72428000 | 72429399 | 8.948951 | 16.72757 | 32.26257 | 13.3739 | -0.4838 | 0.136716 | 0.423123 |
| 9 | 24583200 | 24587999 | 48.39747 | 78.5467 | 67.42149 | 28.89607 | 0.586532 | 1.633415 | 0.42313 |
| 12 | 21837400 | 21843199 | 98.07108 | 153.0518 | 145.0698 | 60.19748 | 0.389456 | 2.684674 | 0.423143 |
| 3 | 62589000 | 62590599 | 20.15255 | 36.14989 | 27.05427 | 11.84238 | 0.817323 | 0.442539 | 0.423144 |
| 12 | 70338600 | 70341399 | 24.48257 | 40.73278 | 69.878 | 30.33358 | -0.39507 | 1.255921 | 0.423146 |
| 1 | 65316800 | 65317199 | 4.653053 | 9.056161 | 21.78065 | 8.01058 | -0.63911 | -0.34442 | 0.423161 |
| 1 | 80317400 | 80318999 | 9.561358 | 17.104 | 36.36244 | 15.9539 | -0.68193 | 0.407827 | 0.423162 |
| 7 | 130467400 | 130468399 | 9.561358 | 17.104 | 36.36244 | 15.9539 | -0.68193 | 0.407827 | 0.423162 |
| 1 | 34558200 | 34559999 | 15.64416 | 26.92641 | 50.03479 | 21.948 | -0.50564 | 0.885655 | 0.423168 |
| 1 | 101046200 | 101048399 | 25.59424 | 44.80311 | 32.24727 | 14.23069 | 0.859625 | 0.709135 | 0.42317 |
| 10 | 29495600 | 29496199 | 7.18843 | 13.71258 | 27.23577 | 10.85606 | -0.47702 | -0.02847 | 0.423187 |
| 9 | 38321000 | 38330999 | 203.7224 | 301.5689 | 287.8451 | 118.5433 | 0.316149 | 3.713236 | 0.423187 |
| 13 | 110068200 | 110069599 | 13.49776 | 24.80574 | 24.45274 | 10.39216 | 0.450633 | 0.252625 | 0.423197 |
| 6 | 132094400 | 132095399 | 13.0243 | 24.91431 | 20.92748 | 8.28708 | 0.762021 | -0.0126 | 0.423209 |
| 3 | 146887400 | 146888599 | 9.75209 | 17.34751 | 36.88563 | 16.26025 | -0.68679 | 0.446934 | 0.42321 |
| 8 | 72786000 | 72790199 | 60.65444 | 96.80047 | 97.19543 | 40.77453 | 0.333934 | 2.126889 | 0.423235 |
| 10 | 74872600 | 74873599 | 5.556662 | 10.7379 | 24.82373 | 9.525639 | -0.64229 | -0.1795 | 0.423238 |
| 4 | 134425600 | 134429599 | 21.73033 | 36.22062 | 65.70836 | 28.55756 | -0.47531 | 1.223579 | 0.423239 |
| 16 | 95567000 | 95569599 | 14.72918 | 25.78552 | 45.66214 | 20.15189 | -0.433 | 0.671564 | 0.42325 |
| 17 | 36541800 | 36542999 | 24.24523 | 41.64425 | 40.03716 | 17.56139 | 0.440473 | 0.928336 | 0.423251 |
| 4 | 148542800 | 148545999 | 16.82246 | 28.64571 | 54.23435 | 23.77244 | -0.53525 | 0.993499 | 0.423257 |
| 2 | 9885800 | 9888599 | 26.70223 | 43.41512 | 80.78547 | 34.66655 | -0.52449 | 1.562953 | 0.423262 |
| 16 | 70574000 | 70575999 | 15.2043 | 26.17075 | 54.27021 | 23.80593 | -0.66326 | 0.88928 | 0.423263 |
| 9 | 114790600 | 114799799 | 91.35528 | 139.7541 | 224.3086 | 93.06812 | -0.39181 | 3.057172 | 0.423284 |
| 8 | 46470200 | 46471399 | 13.28081 | 25.49111 | 20.45949 | 8.023427 | 0.840295 | -0.04119 | 0.423305 |
| 9 | 113923000 | 113926999 | 34.04512 | 54.82349 | 96.69011 | 41.09905 | -0.46197 | 1.817555 | 0.423311 |
| 14 | 61108600 | 61113199 | 32.38394 | 52.30013 | 91.61668 | 39.09359 | -0.44724 | 1.725404 | 0.423321 |
| 2 | 99863600 | 99865599 | 21.88409 | 38.18441 | 31.96114 | 14.07027 | 0.642903 | 0.737691 | 0.42333 |
| 8 | 56023600 | 56025599 | 18.96817 | 34.43796 | 26.31357 | 11.36046 | 0.800491 | 0.365237 | 0.42334 |
| 17 | 14806800 | 14808999 | 18.80204 | 31.74072 | 57.41377 | 25.11303 | -0.47064 | 1.082144 | 0.423344 |
| 9 | 33037600 | 33038599 | 12.02647 | 21.34847 | 38.25092 | 16.89195 | -0.44351 | 0.476707 | 0.423351 |

| | | | | | | | | | |
|---|---|---|---|---|---|---|---|---|---|
| 10 | 94218600 | 94223599 | 51.20365 | 81.50989 | 130.5932 | 54.59212 | -0.34307 | 2.208207 | 0.423356 |
| 19 | 24426800 | 24428799 | 15.75031 | 27.3847 | 47.37873 | 20.81117 | -0.40024 | 0.782208 | 0.423369 |
| X | 48440200 | 48440999 | 9.444047 | 17.40112 | 32.50372 | 13.76499 | -0.46139 | 0.226691 | 0.423372 |
| 5 | 142550600 | 142555999 | 60.19605 | 95.09715 | 148.6116 | 61.7212 | -0.31731 | 2.428685 | 0.423378 |
| 2 | 146854200 | 146855799 | 10.06421 | 17.8544 | 38.8527 | 17.16647 | -0.72265 | 0.468225 | 0.423388 |
| 1 | 148598800 | 148599399 | 14.72948 | 27.33956 | 18.93146 | 7.937669 | 0.970683 | 0.194804 | 0.423403 |
| 7 | 133872600 | 133875799 | 27.79937 | 47.17713 | 45.79806 | 20.06426 | 0.424082 | 1.054586 | 0.423436 |
| 18 | 33824400 | 33825599 | 11.49491 | 21.94577 | 20.98076 | 8.348185 | 0.570415 | -0.01305 | 0.423441 |
| 4 | 4147400 | 4149599 | 15.76725 | 27.23613 | 48.98696 | 21.49072 | -0.45774 | 0.851915 | 0.423447 |
| 7 | 124114400 | 124114999 | 8.443321 | 16.47038 | 16.8071 | 6.015595 | 0.618018 | -0.41175 | 0.423469 |
| 15 | 60249000 | 60249599 | 6.109782 | 11.82792 | 24.97088 | 9.502306 | -0.50135 | -0.20154 | 0.423482 |
| 17 | 55752000 | 55754799 | 25.17503 | 41.56762 | 72.52489 | 31.31776 | -0.42027 | 1.346825 | 0.423486 |
| 14 | 73857400 | 73859399 | 26.34196 | 45.03837 | 43.04712 | 18.87494 | 0.447739 | 0.97836 | 0.423498 |
| 13 | 113323000 | 113329199 | 66.02372 | 105.095 | 105.314 | 44.02079 | 0.332796 | 2.217823 | 0.423521 |
| 19 | 6651000 | 6655399 | 29.36733 | 47.43166 | 90.19241 | 38.48935 | -0.5651 | 1.718808 | 0.423521 |
| X | 122671600 | 122675399 | 34.47247 | 57.40213 | 51.95753 | 22.56707 | 0.524619 | 1.263161 | 0.423525 |
| 10 | 75623600 | 75625999 | 12.83723 | 22.70874 | 39.79032 | 17.59808 | -0.41386 | 0.517392 | 0.423536 |
| 6 | 100447200 | 100451999 | 55.83103 | 88.59422 | 140.2611 | 58.42519 | -0.33054 | 2.310238 | 0.423541 |
| 16 | 8929200 | 8929999 | 14.87342 | 27.0131 | 26.37329 | 11.38128 | 0.449629 | 0.334486 | 0.423542 |
| 1 | 14010800 | 14013599 | 31.89726 | 51.64257 | 86.64057 | 37.06697 | -0.3812 | 1.661125 | 0.42356 |
| 3 | 91273000 | 91275799 | 25.33921 | 44.06421 | 36.88302 | 16.20266 | 0.642467 | 0.807703 | 0.423588 |
| 14 | 45805200 | 45808999 | 32.67381 | 52.8913 | 88.1808 | 37.72143 | -0.37245 | 1.664926 | 0.423598 |
| 15 | 19829800 | 19830599 | 6.249668 | 12.13456 | 25.8233 | 9.682698 | -0.49237 | -0.24665 | 0.423622 |
| 8 | 71270400 | 71273399 | 43.40479 | 69.48763 | 112.6267 | 47.45736 | -0.35072 | 2.02079 | 0.42363 |
| 9 | 114594600 | 114596599 | 20.20118 | 33.62835 | 64.30103 | 27.92316 | -0.55077 | 1.2338 | 0.423647 |
| 5 | 147846600 | 147848999 | 20.20759 | 34.09782 | 58.59964 | 25.62817 | -0.39723 | 1.090989 | 0.42365 |
| 11 | 58741000 | 58743199 | 23.71697 | 40.93296 | 39.9381 | 17.5205 | 0.420606 | 0.882704 | 0.423669 |
| 6 | 98966600 | 98968399 | 11.74258 | 20.75813 | 38.82091 | 17.17169 | -0.50762 | 0.515978 | 0.423671 |
| 4 | 57955600 | 57957999 | 17.90375 | 30.07638 | 60.01623 | 26.21234 | -0.61212 | 1.123468 | 0.423673 |
| 15 | 18444000 | 18447399 | 52.01085 | 83.59205 | 83.75728 | 35.50207 | 0.348537 | 1.89798 | 0.423675 |
| X | 15133800 | 15135199 | 16.9513 | 31.78411 | 22.61056 | 9.323756 | 0.950725 | 0.160964 | 0.42368 |
| 14 | 71278400 | 71279199 | 4.563793 | 8.870309 | 22.82459 | 8.487659 | -0.75128 | -0.30581 | 0.423689 |
| 10 | 36696000 | 36697599 | 10.43341 | 18.96343 | 34.92686 | 15.05835 | -0.45932 | 0.317785 | 0.423691 |
| 2 | 14381000 | 14383599 | 20.52328 | 35.75327 | 34.74971 | 15.28265 | 0.428223 | 0.758191 | 0.423695 |
| 10 | 129205400 | 129210799 | 75.87591 | 119.9114 | 119.8288 | 49.77626 | 0.327274 | 2.424281 | 0.423717 |
| 10 | 82213400 | 82218799 | 116.7959 | 181.8889 | 139.0857 | 57.73084 | 0.696574 | 2.730548 | 0.42372 |
| 7 | 56541000 | 56545199 | 34.75792 | 58.51797 | 46.32759 | 20.24725 | 0.716899 | 1.146495 | 0.423721 |
| 7 | 4205600 | 4210799 | 48.08345 | 77.7494 | 72.96772 | 31.18481 | 0.453067 | 1.708088 | 0.423734 |
| 2 | 57308000 | 57315799 | 90.21693 | 138.7328 | 211.715 | 87.91927 | -0.3133 | 2.952904 | 0.423736 |
| 5 | 114943600 | 114945799 | 13.02493 | 22.80484 | 43.07759 | 19.01327 | -0.52545 | 0.657078 | 0.423749 |
| 17 | 83938200 | 83940399 | 20.89901 | 34.60226 | 68.03646 | 29.43537 | -0.59161 | 1.301536 | 0.423757 |
| 2 | 93119200 | 93128999 | 96.33192 | 147.2003 | 228.2632 | 94.68408 | -0.34348 | 3.082033 | 0.423763 |
| 3 | 23791800 | 23792199 | 3.453366 | 6.754762 | 17.28048 | 5.80122 | -0.60401 | -0.7003 | 0.423768 |
| X | 61652400 | 61653599 | 15.49999 | 29.68866 | 19.06373 | 7.520928 | 1.151395 | -0.0038 | 0.423771 |
| 15 | 100728200 | 100729999 | 15.46973 | 26.70444 | 48.98347 | 21.48876 | -0.4861 | 0.857127 | 0.423791 |
| 13 | 63711400 | 63714599 | 23.49263 | 38.31445 | 78.27899 | 33.62871 | -0.65566 | 1.510458 | 0.423802 |
| 18 | 50351800 | 50353599 | 17.70522 | 30.38168 | 52.18199 | 22.88701 | -0.39359 | 0.924166 | 0.423808 |

| | | | | | | | | | |
|---|---|---|---|---|---|---|---|---|---|
| 6 | 116603200 | 116604399 | 11.68854 | 20.44288 | 46.76978 | 20.62404 | -0.80023 | 0.675113 | 0.423816 |
| 1 | 33945200 | 33949199 | 25.27531 | 41.09841 | 86.84161 | 37.26645 | -0.70731 | 1.563242 | 0.423824 |
| X | 15828800 | 15838599 | 271.6943 | 393.8453 | 354.699 | 146.2528 | 0.37112 | 4.048704 | 0.42384 |
| 4 | 91685400 | 91689799 | 53.11995 | 85.43167 | 82.5585 | 35.02785 | 0.401811 | 1.877783 | 0.423849 |
| 9 | 17663800 | 17664799 | 14.28989 | 25.01402 | 43.5795 | 19.23116 | -0.40938 | 0.669246 | 0.423871 |
| 10 | 66416800 | 66418799 | 19.91401 | 33.55735 | 58.5473 | 25.59485 | -0.41895 | 1.102231 | 0.423881 |
| 4 | 23111800 | 23114199 | 25.22728 | 43.06973 | 41.64185 | 18.25663 | 0.431069 | 0.988348 | 0.423894 |
| 16 | 55928200 | 55929599 | 27.65832 | 48.29503 | 33.42197 | 14.72018 | 0.916238 | 0.760749 | 0.423909 |
| 5 | 145312400 | 145315599 | 50.1349 | 80.69521 | 80.08786 | 34.01892 | 0.36488 | 1.849877 | 0.423958 |
| 4 | 124217400 | 124220399 | 32.68691 | 52.95858 | 87.04234 | 37.25834 | -0.35072 | 1.647843 | 0.42396 |
| 6 | 94675600 | 94677999 | 14.61841 | 25.40747 | 46.85767 | 20.5776 | -0.49204 | 0.780012 | 0.423977 |
| 14 | 51383000 | 51383599 | 5.510423 | 10.72724 | 23.04756 | 8.46125 | -0.4769 | -0.33717 | 0.423978 |
| 14 | 115607000 | 115607599 | 5.510423 | 10.72724 | 23.04756 | 8.46125 | -0.4769 | -0.33717 | 0.423978 |
| 19 | 52097400 | 52098799 | 27.91309 | 47.13605 | 46.92079 | 20.52527 | 0.387755 | 1.097465 | 0.423983 |
| 5 | 106076800 | 106081199 | 61.73215 | 97.35048 | 151.8745 | 63.02724 | -0.3169 | 2.471684 | 0.424006 |
| X | 124967400 | 124970399 | 31.26207 | 54.58134 | 29.33628 | 12.91976 | 1.278952 | 0.775192 | 0.424013 |
| 4 | 135257200 | 135261199 | 44.40394 | 70.75683 | 122.3755 | 51.22821 | -0.45157 | 2.168209 | 0.424032 |
| 5 | 94412800 | 94415199 | 21.47303 | 37.63412 | 30.71442 | 13.56803 | 0.679229 | 0.66641 | 0.424056 |
| 5 | 105311800 | 105313399 | 19.0401 | 32.40533 | 55.37247 | 24.27031 | -0.3881 | 1.00828 | 0.424074 |
| 5 | 147035200 | 147039999 | 30.62775 | 49.48891 | 89.51809 | 38.22304 | -0.4925 | 1.7074 | 0.424077 |
| X | 144816200 | 144821599 | 83.12377 | 130.8589 | 128.3608 | 53.25722 | 0.349734 | 2.516079 | 0.424081 |
| 15 | 20651600 | 20653799 | 31.42 | 52.74848 | 44.30288 | 19.34182 | 0.631853 | 1.159072 | 0.424093 |
| 10 | 36040800 | 36042999 | 18.38258 | 33.48601 | 25.81453 | 11.09873 | 0.791444 | 0.340853 | 0.424104 |
| 14 | 112655600 | 112657599 | 21.81734 | 38.30678 | 32.99388 | 14.59726 | 0.601751 | 0.642915 | 0.424109 |
| 1 | 137388600 | 137389399 | 6.399642 | 12.37041 | 26.07245 | 9.991564 | -0.50902 | -0.17203 | 0.424117 |
| 1 | 134297000 | 134300399 | 36.85181 | 60.67834 | 52.62273 | 22.69334 | 0.583984 | 1.395593 | 0.424136 |
| X | 83446000 | 83447599 | 23.46758 | 41.41238 | 30.23519 | 13.37866 | 0.84123 | 0.591631 | 0.424136 |
| 1 | 46704800 | 46705599 | 13.06735 | 24.36392 | 24.00896 | 9.995336 | 0.473062 | 0.166533 | 0.424152 |
| 5 | 101357200 | 101358799 | 16.12271 | 27.71222 | 50.56508 | 22.1797 | -0.47991 | 0.900771 | 0.424156 |
| 13 | 37106400 | 37109399 | 67.05324 | 106.4477 | 103.2354 | 43.02081 | 0.37607 | 2.299438 | 0.42416 |
| 7 | 38796400 | 38810799 | 355.7617 | 503.4275 | 447.3411 | 185.1375 | 0.355826 | 4.405145 | 0.424165 |
| X | 121866400 | 121874999 | 105.0484 | 165.9034 | 105.4586 | 43.79211 | 0.978317 | 2.461894 | 0.424171 |
| 11 | 31748200 | 31749199 | 12.73424 | 23.60823 | 23.4144 | 9.833112 | 0.453942 | 0.194091 | 0.424175 |
| 15 | 87517000 | 87526999 | 72.0038 | 112.4553 | 173.2963 | 71.90467 | -0.31004 | 2.662814 | 0.424175 |
| 18 | 81470800 | 81471799 | 6.566424 | 12.60916 | 26.90491 | 10.53761 | -0.55648 | -0.09598 | 0.424187 |
| 17 | 57881200 | 57881799 | 9.810009 | 17.67796 | 34.83584 | 15.18121 | -0.56645 | 0.360726 | 0.42419 |
| 18 | 65543400 | 65544599 | 13.38877 | 24.83443 | 24.47754 | 10.27241 | 0.463902 | 0.201063 | 0.424205 |
| X | 152997000 | 152997799 | 21.35266 | 37.37933 | 34.45412 | 15.20497 | 0.504735 | 0.692072 | 0.424217 |
| 12 | 23734800 | 23736599 | 3.453566 | 6.754373 | 20.26881 | 6.872467 | -0.84554 | -0.65496 | 0.424218 |
| 11 | 67019800 | 67021199 | 13.82687 | 24.19716 | 43.08652 | 19.0094 | -0.44063 | 0.670681 | 0.424234 |
| 12 | 24219800 | 24245999 | 673.5715 | 898.6509 | 777.8712 | 324.8839 | 0.308132 | 5.316631 | 0.424239 |
| 2 | 17744200 | 17746799 | 12.9904 | 22.93264 | 40.31762 | 17.8386 | -0.41988 | 0.541353 | 0.424255 |
| 7 | 113007600 | 113009999 | 12.9904 | 22.93264 | 40.31762 | 17.8386 | -0.41988 | 0.541353 | 0.424255 |
| 5 | 88504000 | 88504799 | 11.10415 | 19.91563 | 36.33 | 15.90473 | -0.46016 | 0.404474 | 0.424265 |
| X | 125240000 | 125241199 | 22.25401 | 39.03606 | 33.24321 | 14.69567 | 0.618321 | 0.663659 | 0.424271 |
| 1 | 35730000 | 35733199 | 32.4289 | 53.78248 | 54.16306 | 23.46037 | 0.370705 | 1.300982 | 0.424282 |
| 12 | 49260000 | 49263399 | 31.84626 | 54.12057 | 43.47958 | 19.05526 | 0.696178 | 1.054176 | 0.424287 |

| | | | | | | | | | |
|---|---|---|---|---|---|---|---|---|---|
| 11 | 68974400 | 68975799 | 13.34858 | 23.35373 | 43.10131 | 19.01151 | -0.49206 | 0.67202 | 0.424288 |
| X | 102084200 | 102087199 | 39.1149 | 65.20241 | 49.83685 | 21.66063 | 0.767731 | 1.263365 | 0.424295 |
| X | 126352800 | 126364399 | 111.3284 | 173.7118 | 124.0378 | 51.47107 | 0.797434 | 2.693766 | 0.424298 |
| 1 | 185372200 | 185373199 | 9.539063 | 17.61751 | 32.53202 | 13.73303 | -0.44196 | 0.211946 | 0.424303 |
| 7 | 133786000 | 133788599 | 29.82093 | 50.10009 | 49.11189 | 21.45036 | 0.409615 | 1.150917 | 0.424314 |
| 8 | 19151600 | 19154399 | 19.25525 | 34.19694 | 29.23154 | 12.90366 | 0.618423 | 0.500614 | 0.42434 |
| 4 | 92214000 | 92215599 | 14.16235 | 26.66984 | 22.77998 | 9.325077 | 0.696652 | 0.108722 | 0.424356 |
| 6 | 134547000 | 134549999 | 19.96779 | 33.49437 | 67.34155 | 29.38462 | -0.62315 | 1.152859 | 0.424357 |
| 11 | 77563800 | 77564799 | 15.20708 | 28.59068 | 22.68828 | 9.31316 | 0.798978 | 0.128162 | 0.424367 |
| 5 | 148057600 | 148058999 | 10.86315 | 19.76537 | 35.9837 | 15.49351 | -0.44177 | 0.315236 | 0.424369 |
| 9 | 20073400 | 20080199 | 82.73821 | 130.9479 | 117.6568 | 48.91022 | 0.482167 | 2.395763 | 0.424375 |
| 17 | 84480400 | 84485599 | 48.49143 | 77.10569 | 128.8408 | 53.77852 | -0.4053 | 2.242025 | 0.424375 |
| Y | 4355000 | 4357599 | 15.4487 | 28.00003 | 17.53259 | 7.585683 | 1.083195 | 0.32609 | 0.424382 |
| X | 112943600 | 112945399 | 19.69719 | 35.40682 | 27.79744 | 12.14631 | 0.750633 | 0.429884 | 0.42439 |
| 1 | 184098800 | 184101599 | 41.69027 | 68.02585 | 63.96271 | 27.48323 | 0.461185 | 1.530249 | 0.424401 |
| 19 | 42466800 | 42469199 | 13.94641 | 24.46198 | 42.99182 | 19.00414 | -0.42198 | 0.632907 | 0.424416 |
| 7 | 104245600 | 104247199 | 15.98204 | 28.62271 | 28.26311 | 12.38831 | 0.418656 | 0.427632 | 0.424419 |
| 5 | 135772600 | 135785599 | 136.0327 | 203.6158 | 306.0421 | 126.2209 | -0.32385 | 3.506446 | 0.424433 |
| 8 | 90490000 | 90491999 | 22.44896 | 38.87067 | 38.85775 | 17.04825 | 0.386584 | 0.845459 | 0.424449 |
| 14 | 12376200 | 12378399 | 19.09769 | 34.16032 | 28.85201 | 12.65964 | 0.641687 | 0.456943 | 0.424468 |
| 7 | 13847800 | 13851799 | 62.36151 | 99.74967 | 93.79706 | 39.47484 | 0.431953 | 2.060148 | 0.424487 |
| 6 | 71075200 | 71076199 | 8.106052 | 15.29679 | 29.18613 | 11.90249 | -0.44918 | 0.058967 | 0.424507 |
| 15 | 96535600 | 96539999 | 25.32934 | 41.45469 | 75.04213 | 32.27927 | -0.47808 | 1.461203 | 0.424508 |
| 4 | 149224600 | 149225799 | 13.87052 | 25.41731 | 24.47431 | 10.44225 | 0.480338 | 0.272285 | 0.424511 |
| 3 | 59402000 | 59402599 | 6.848801 | 13.23095 | 27.23729 | 10.46629 | -0.47906 | -0.15356 | 0.424527 |
| X | 78469400 | 78470799 | 17.75494 | 33.09567 | 22.98426 | 9.572749 | 0.973756 | 0.204278 | 0.424531 |
| 7 | 119009800 | 119012199 | 20.00882 | 33.98645 | 57.56978 | 25.22832 | -0.37606 | 1.031956 | 0.424551 |
| 19 | 52048800 | 52050399 | 16.7245 | 30.61193 | 26.04421 | 11.12947 | 0.655835 | 0.302367 | 0.424552 |
| 4 | 78339200 | 78340199 | 8.887331 | 16.62553 | 31.18753 | 12.91098 | -0.44275 | 0.126397 | 0.424557 |
| 1 | 34916600 | 34917999 | 9.267818 | 17.05051 | 31.95211 | 13.55936 | -0.46829 | 0.231512 | 0.424563 |
| 10 | 109337800 | 109338799 | 11.52982 | 22.13507 | 20.54168 | 8.050642 | 0.633104 | -0.06178 | 0.424566 |
| 10 | 54841400 | 54843399 | 20.12492 | 36.56782 | 25.09548 | 10.82357 | 0.955331 | 0.366522 | 0.424566 |
| 9 | 121547600 | 121549399 | 8.746344 | 15.85729 | 33.43106 | 14.45869 | -0.65552 | 0.31646 | 0.424577 |
| 7 | 42413400 | 42419399 | 58.05117 | 93.25644 | 83.75639 | 35.47763 | 0.505241 | 1.92001 | 0.424586 |
| 5 | 68613000 | 68615399 | 25.80486 | 45.01029 | 35.15097 | 15.47123 | 0.742458 | 0.767092 | 0.424595 |
| 1 | 70645200 | 70647599 | 20.69771 | 34.9315 | 59.16674 | 25.87768 | -0.37635 | 1.091955 | 0.424615 |
| 9 | 65136600 | 65137799 | 8.629417 | 15.97443 | 30.98657 | 13.0395 | -0.50887 | 0.188617 | 0.42462 |
| 2 | 12000000 | 12003999 | 74.07053 | 117.2427 | 117.5291 | 48.85987 | 0.32459 | 2.387628 | 0.42463 |
| 8 | 92685800 | 92686799 | 8.704916 | 16.35745 | 31.89418 | 13.10366 | -0.48951 | 0.096619 | 0.424638 |
| 3 | 31147400 | 31150199 | 18.1435 | 30.67606 | 56.88403 | 24.89291 | -0.50627 | 1.064916 | 0.424652 |
| 6 | 103045200 | 103046199 | 8.753552 | 16.38109 | 30.642 | 12.6775 | -0.43777 | 0.122743 | 0.424653 |
| 17 | 50479400 | 50489999 | 110.0282 | 166.772 | 255.8651 | 105.8946 | -0.33757 | 3.252436 | 0.424659 |
| 1 | 31791800 | 31792599 | 12.48245 | 23.22455 | 21.56513 | 9.009908 | 0.554176 | 0.168265 | 0.424667 |
| 5 | 4780200 | 4781399 | 14.91453 | 26.04895 | 45.19388 | 19.90792 | -0.40357 | 0.709819 | 0.424673 |
| 19 | 12827400 | 12827999 | 10.16931 | 19.55423 | 20.25231 | 7.901058 | 0.483165 | -0.08799 | 0.424676 |
| 7 | 3966400 | 3966999 | 5.210628 | 10.1638 | 22.82102 | 8.174035 | -0.50748 | -0.42998 | 0.424679 |
| 15 | 43531400 | 43531999 | 5.210628 | 10.1638 | 22.82102 | 8.174035 | -0.50748 | -0.42998 | 0.424679 |

| | | | | | | | | | |
|---|---|---|---|---|---|---|---|---|---|
| 18 | 10439200 | 10442999 | 25.23489 | 41.86587 | 73.70467 | 31.9323 | -0.43259 | 1.295905 | 0.424681 |
| 5 | 124609400 | 124612199 | 14.10044 | 24.6405 | 45.18626 | 19.91162 | -0.48312 | 0.696478 | 0.424688 |
| 15 | 64994200 | 64996399 | 25.33551 | 43.32385 | 42.55037 | 18.65735 | 0.408664 | 0.97227 | 0.424695 |
| 13 | 25066400 | 25070199 | 47.28477 | 75.52976 | 119.6453 | 50.25507 | -0.32131 | 2.09418 | 0.424706 |
| X | 47429600 | 47432599 | 39.29213 | 65.07616 | 51.50565 | 22.28833 | 0.717244 | 1.338626 | 0.424713 |
| 5 | 142210600 | 142221599 | 110.8118 | 166.6775 | 307.3688 | 126.918 | -0.61234 | 3.408317 | 0.424718 |
| 8 | 80914600 | 80916399 | 19.26905 | 32.54257 | 57.44816 | 25.13123 | -0.43565 | 1.079961 | 0.424725 |
| 5 | 68079600 | 68080799 | 17.63457 | 31.3866 | 25.60901 | 11.28393 | 0.687092 | 0.470758 | 0.424725 |
| 6 | 65139000 | 65139999 | 15.25242 | 27.41296 | 26.78214 | 11.70416 | 0.436757 | 0.396862 | 0.424733 |
| 15 | 99368200 | 99369999 | 8.429348 | 15.353 | 32.9683 | 14.17679 | -0.67613 | 0.287501 | 0.424754 |
| 17 | 23790200 | 23800599 | 119.2119 | 179.237 | 284.0379 | 117.271 | -0.39434 | 3.417279 | 0.424757 |
| 15 | 15037600 | 15039399 | 17.34803 | 31.04626 | 25.58563 | 11.22167 | 0.677292 | 0.435455 | 0.424758 |
| 10 | 50262800 | 50265999 | 30.05947 | 51.4186 | 41.82993 | 18.34193 | 0.679498 | 0.988439 | 0.424776 |
| 13 | 20761200 | 20762799 | 15.66759 | 28.537 | 26.61004 | 11.44243 | 0.518431 | 0.322392 | 0.424777 |
| 17 | 22489200 | 22493799 | 85.60836 | 134.6498 | 119.6153 | 49.62294 | 0.491546 | 2.532426 | 0.424787 |
| 12 | 19464200 | 19496599 | 1166.786 | 1488.653 | 1227.827 | 515.2969 | 0.313366 | 6.041441 | 0.424796 |
| 16 | 91173200 | 91176399 | 23.45966 | 38.761 | 68.7183 | 29.68759 | -0.44302 | 1.334486 | 0.424811 |
| Y | 4538000 | 4542799 | 31.04885 | 52.08254 | 29.17213 | 12.72941 | 1.213718 | 1.144853 | 0.42483 |
| 14 | 102156400 | 102158399 | 16.51452 | 28.52715 | 50.6005 | 22.1986 | -0.43795 | 0.855552 | 0.424837 |
| 1 | 161615400 | 161616399 | 6.96472 | 13.05272 | 29.06487 | 11.99508 | -0.68386 | 0.093498 | 0.424868 |
| 18 | 79963000 | 79964799 | 17.14422 | 31.29288 | 25.92244 | 11.11721 | 0.690555 | 0.320118 | 0.424869 |
| 3 | 151406200 | 151411199 | 42.18539 | 68.54967 | 57.98597 | 24.87483 | 0.609143 | 1.588746 | 0.424872 |
| 7 | 66637200 | 66642599 | 56.88036 | 90.05797 | 142.0921 | 59.07959 | -0.32854 | 2.372818 | 0.424893 |
| X | 112179200 | 112180199 | 15.16471 | 28.31048 | 23.6771 | 9.835924 | 0.710079 | 0.173369 | 0.424896 |
| 6 | 9757200 | 9757999 | 12.69584 | 24.42793 | 20.59767 | 8.02116 | 0.77929 | -0.06821 | 0.424905 |
| X | 66456600 | 66460199 | 40.69774 | 68.40004 | 42.37644 | 18.51073 | 1.069365 | 1.179373 | 0.424905 |
| 12 | 115999000 | 116003799 | 52.41288 | 84.15638 | 84.27269 | 35.66924 | 0.347683 | 1.927604 | 0.424949 |
| 15 | 77190200 | 77191599 | 11.50277 | 20.23767 | 39.98314 | 17.70261 | -0.58873 | 0.572599 | 0.424956 |
| 13 | 69542800 | 69544999 | 13.63305 | 23.88914 | 43.03108 | 19.00589 | -0.45729 | 0.646798 | 0.424962 |
| 5 | 6652600 | 6653199 | 10.28252 | 19.76664 | 20.75015 | 8.101492 | 0.462576 | -0.08384 | 0.424963 |
| X | 3165800 | 3175799 | 272.7317 | 394.6791 | 351.4932 | 144.9635 | 0.384854 | 4.074074 | 0.424963 |
| 5 | 42860000 | 42862799 | 28.35818 | 45.97403 | 91.40439 | 39.14356 | -0.62369 | 1.631188 | 0.424974 |
| 10 | 104670000 | 104673199 | 70.13014 | 111.1052 | 111.0891 | 46.20356 | 0.329331 | 2.361035 | 0.424989 |
| 4 | 118787600 | 118788399 | 7.933129 | 15.10331 | 29.48293 | 11.80094 | -0.45819 | -0.00335 | 0.425013 |
| 10 | 12816600 | 12818599 | 14.04988 | 24.5156 | 45.24486 | 19.91854 | -0.49212 | 0.722124 | 0.425017 |
| 17 | 22906800 | 22912999 | 126.0105 | 193.9843 | 166.9171 | 69.32621 | 0.513728 | 2.942623 | 0.425021 |
| 10 | 112685600 | 112686799 | 16.73135 | 30.03516 | 28.57002 | 12.49805 | 0.474051 | 0.41796 | 0.425024 |
| 14 | 54955600 | 54957399 | 16.73135 | 30.03516 | 28.57002 | 12.49805 | 0.474051 | 0.41796 | 0.425024 |
| 10 | 51758600 | 51761599 | 41.63289 | 67.94349 | 57.5253 | 24.71872 | 0.612283 | 1.526204 | 0.425037 |
| 13 | 11775800 | 11776999 | 7.462416 | 14.22535 | 28.37029 | 11.32489 | -0.48542 | -0.02052 | 0.42504 |
| 18 | 80212800 | 80214599 | 13.50875 | 23.63166 | 43.10312 | 19.01075 | -0.47513 | 0.674726 | 0.425041 |
| 11 | 108954400 | 108957799 | 28.25405 | 47.65008 | 45.84273 | 20.0458 | 0.43662 | 1.106678 | 0.425042 |
| 2 | 132747600 | 132748999 | 10.05186 | 18.37504 | 33.41808 | 14.30485 | -0.43462 | 0.276626 | 0.425049 |
| 3 | 138765400 | 138766199 | 10.05186 | 18.37504 | 33.41808 | 14.30485 | -0.43462 | 0.276626 | 0.425049 |
| 13 | 89951600 | 89953599 | 24.10388 | 41.78538 | 39.24967 | 17.22082 | 0.476364 | 0.843106 | 0.425087 |
| 4 | 73362800 | 73363799 | 6.728545 | 12.89606 | 26.25226 | 10.33796 | -0.49592 | -0.07556 | 0.425126 |
| 14 | 79553400 | 79555199 | 8.143933 | 15.19555 | 30.04449 | 12.4923 | -0.52291 | 0.134104 | 0.425127 |

| | | | | | | | | | |
|---|---|---|---|---|---|---|---|---|---|
| 8 | 52049200 | 52052399 | 42.88997 | 70.28171 | 60.21196 | 25.91339 | 0.598885 | 1.473903 | 0.425138 |
| X | 42916400 | 42919999 | 61.78051 | 99.05109 | 87.34075 | 36.88492 | 0.528396 | 1.989256 | 0.425148 |
| 3 | 85139800 | 85140599 | 6.564981 | 12.63153 | 25.82459 | 10.0568 | -0.48709 | -0.1159 | 0.425159 |
| 5 | 59364400 | 59364799 | 6.564981 | 12.63153 | 25.82459 | 10.0568 | -0.48709 | -0.1159 | 0.425159 |
| 9 | 38546600 | 38555399 | 229.8009 | 336.1551 | 321.0483 | 132.2573 | 0.299115 | 3.906718 | 0.425163 |
| 15 | 79545600 | 79546399 | 12.58857 | 23.50404 | 22.57119 | 9.374574 | 0.512939 | 0.150634 | 0.425164 |
| 5 | 100453400 | 100454799 | 8.298448 | 15.47944 | 30.03928 | 12.49617 | -0.49667 | 0.137674 | 0.425166 |
| 4 | 24576400 | 24578399 | 30.18867 | 50.70476 | 43.26298 | 18.89333 | 0.60914 | 1.14828 | 0.425176 |
| 2 | 105772600 | 105773999 | 12.29084 | 23.27198 | 21.88957 | 8.863558 | 0.571767 | 0.052755 | 0.425178 |
| 1 | 89980000 | 89983199 | 20.51694 | 34.12757 | 65.45354 | 28.40869 | -0.55526 | 1.243705 | 0.425191 |
| 15 | 57631600 | 57633199 | 10.29995 | 18.54118 | 34.84215 | 15.2041 | -0.49939 | 0.372566 | 0.425199 |
| 3 | 14951000 | 14952999 | 11.64777 | 20.44847 | 41.00692 | 18.14045 | -0.61097 | 0.603319 | 0.425203 |
| 16 | 87087600 | 87088799 | 25.08895 | 43.1177 | 36.25615 | 15.90321 | 0.632922 | 0.919017 | 0.425209 |
| 1 | 24746000 | 24747799 | 16.5246 | 28.51912 | 50.59526 | 22.19569 | -0.43846 | 0.863811 | 0.42521 |
| 18 | 10048800 | 10054399 | 61.4097 | 96.31625 | 175.3804 | 72.73771 | -0.54572 | 2.582539 | 0.425212 |
| 2 | 5501400 | 5503799 | 20.24848 | 35.43163 | 34.37808 | 15.16608 | 0.430827 | 0.689352 | 0.425222 |
| 2 | 111909000 | 111914999 | 70.95957 | 112.8811 | 107.0276 | 44.69835 | 0.411533 | 2.242462 | 0.425241 |
| X | 153341400 | 153345599 | 91.24064 | 142.5723 | 139.5186 | 57.88645 | 0.344893 | 2.660047 | 0.425257 |
| 7 | 3229400 | 3232799 | 55.17424 | 88.28599 | 85.73552 | 36.101 | 0.386122 | 2.037228 | 0.425259 |
| 3 | 87920200 | 87922199 | 19.06628 | 32.36881 | 55.87783 | 24.48304 | -0.4032 | 1.031901 | 0.425259 |
| 11 | 106106800 | 106108399 | 9.37221 | 16.55003 | 39.97822 | 17.68741 | -0.87497 | 0.515691 | 0.425272 |
| 14 | 116808200 | 116809999 | 30.90912 | 51.98854 | 43.74913 | 19.11482 | 0.629096 | 1.139455 | 0.425283 |
| 14 | 99336400 | 99337599 | 8.307281 | 15.42411 | 31.02507 | 13.00276 | -0.55675 | 0.168015 | 0.425291 |
| 7 | 136364400 | 136367999 | 20.8074 | 35.10531 | 61.45254 | 26.87479 | -0.42385 | 1.095199 | 0.425303 |
| 15 | 96941400 | 96942999 | 11.64885 | 20.90297 | 38.34563 | 16.78009 | -0.46833 | 0.405282 | 0.425342 |
| X | 7803200 | 7806399 | 41.54399 | 68.42354 | 56.04447 | 24.17183 | 0.666463 | 1.410725 | 0.425355 |
| 12 | 21690200 | 21712799 | 895.7369 | 1167.419 | 984.409 | 412.2806 | 0.312233 | 5.69441 | 0.425356 |
| 7 | 9980400 | 9986999 | 188.6656 | 280.2057 | 266.6788 | 109.8568 | 0.324737 | 3.65411 | 0.425363 |
| 14 | 54286800 | 54288199 | 14.21985 | 26.89941 | 22.32633 | 9.058226 | 0.748717 | 0.079089 | 0.425379 |
| 13 | 15122600 | 15123999 | 11.16972 | 20.18798 | 36.3694 | 15.79084 | -0.43467 | 0.35594 | 0.425385 |
| 16 | 88916000 | 88918399 | 24.17464 | 42.11389 | 36.96551 | 16.25706 | 0.574547 | 0.779159 | 0.425393 |
| 17 | 22494600 | 22499799 | 65.21962 | 104.4231 | 89.87482 | 37.87682 | 0.560949 | 2.033479 | 0.425401 |
| 4 | 38508000 | 38512799 | 66.45785 | 105.7653 | 105.5335 | 44.10061 | 0.338648 | 2.223854 | 0.425401 |
| 5 | 50260000 | 50260599 | 7.817553 | 14.6951 | 29.10222 | 11.94956 | -0.51053 | 0.081145 | 0.425429 |
| 10 | 41573000 | 41573799 | 7.083247 | 13.36349 | 28.09276 | 11.46235 | -0.58781 | 0.050363 | 0.425441 |
| 13 | 60343200 | 60344599 | 27.83694 | 45.19555 | 82.4851 | 35.36231 | -0.49849 | 1.593378 | 0.425461 |
| X | 54998400 | 55007199 | 121.0255 | 187.2064 | 156.6371 | 65.06324 | 0.55935 | 2.851181 | 0.425467 |
| 16 | 54998600 | 55000799 | 20.91853 | 36.95973 | 30.19568 | 13.35788 | 0.680147 | 0.55992 | 0.425472 |
| 6 | 87741400 | 87743199 | 10.55412 | 19.06981 | 34.82188 | 15.12445 | -0.45418 | 0.350516 | 0.42548 |
| 5 | 21248200 | 21251399 | 27.81266 | 47.29514 | 44.6699 | 19.57791 | 0.463686 | 1.034101 | 0.42548 |
| 9 | 119245800 | 119247999 | 20.32407 | 33.9926 | 61.6529 | 26.85285 | -0.47481 | 1.182271 | 0.425491 |
| 4 | 35047400 | 35047799 | 9.463259 | 18.47978 | 14.283 | 5.021572 | 1.03563 | -0.46844 | 0.425523 |
| X | 148871800 | 148874999 | 25.42574 | 45.00811 | 26.64691 | 11.78354 | 1.143353 | 0.564071 | 0.425538 |
| 17 | 88777600 | 88780199 | 26.90823 | 46.26721 | 41.35397 | 18.13956 | 0.5454 | 0.929518 | 0.425539 |
| 1 | 7815000 | 7816799 | 23.59612 | 40.8071 | 39.84456 | 17.48055 | 0.420031 | 0.862536 | 0.425557 |
| 7 | 31826400 | 31836599 | 190.8701 | 283.7617 | 268.1899 | 110.4904 | 0.336094 | 3.63751 | 0.42557 |
| 12 | 76384000 | 76386199 | 10.14052 | 18.15054 | 37.33257 | 16.37156 | -0.63435 | 0.408711 | 0.425578 |

| | | | | | | | | | |
|---|---|---|---|---|---|---|---|---|---|
| X | 15368400 | 15369999 | 17.94227 | 33.51733 | 22.44626 | 9.314475 | 1.029981 | 0.191465 | 0.425607 |
| 2 | 97624200 | 97624599 | 5.052014 | 9.858575 | 22.8943 | 8.147604 | -0.54667 | -0.45335 | 0.425616 |
| 6 | 141692200 | 141694999 | 21.15342 | 37.15387 | 33.03914 | 14.62155 | 0.555893 | 0.633792 | 0.425632 |
| 15 | 41812200 | 41813599 | 9.106156 | 16.83649 | 31.49505 | 13.2754 | -0.45858 | 0.201534 | 0.425633 |
| 17 | 45506400 | 45507999 | 20.05916 | 35.00389 | 34.07468 | 15.00172 | 0.426023 | 0.72915 | 0.425633 |
| 3 | 40822600 | 40824799 | 17.01812 | 30.37314 | 29.04235 | 12.767 | 0.461748 | 0.455971 | 0.425637 |
| 4 | 90185200 | 90186999 | 20.75371 | 36.39266 | 33.66654 | 14.87877 | 0.499187 | 0.657442 | 0.42565 |
| 2 | 39848800 | 39849999 | 12.88203 | 24.19172 | 23.42339 | 9.633536 | 0.511796 | 0.116284 | 0.425681 |
| 2 | 105706200 | 105706799 | 12.84747 | 23.81486 | 23.35856 | 9.811505 | 0.469534 | 0.195089 | 0.425716 |
| X | 34813000 | 34816999 | 63.5798 | 102.1665 | 80.17404 | 33.97397 | 0.700097 | 1.920079 | 0.425754 |
| 9 | 11101000 | 11104799 | 46.758 | 75.52571 | 74.87195 | 31.95471 | 0.372463 | 1.737018 | 0.425755 |
| 10 | 127603000 | 127605399 | 27.4444 | 46.43705 | 46.3734 | 20.29808 | 0.383257 | 1.078677 | 0.425772 |
| 17 | 64061600 | 64062399 | 7.637246 | 14.49013 | 28.22225 | 11.37872 | -0.46401 | 0.014553 | 0.425784 |
| 4 | 60102800 | 60105399 | 20.88279 | 35.17931 | 59.68522 | 26.08994 | -0.37883 | 1.109463 | 0.425791 |
| 11 | 35685800 | 35690199 | 35.04111 | 56.30107 | 103.8675 | 44.00675 | -0.53085 | 1.891655 | 0.425796 |
| 7 | 11709600 | 11719199 | 140.5799 | 215.2744 | 178.8793 | 74.23631 | 0.558273 | 3.04923 | 0.425797 |
| 17 | 86886600 | 86888799 | 21.53985 | 37.65984 | 34.55273 | 15.23237 | 0.511456 | 0.713502 | 0.425811 |
| 3 | 71024800 | 71025799 | 14.9604 | 28.00591 | 23.16235 | 9.577418 | 0.731301 | 0.153153 | 0.425815 |
| 3 | 78683600 | 78684599 | 9.018411 | 16.41481 | 32.91107 | 14.16414 | -0.57911 | 0.293569 | 0.425818 |
| 17 | 42325600 | 42325999 | 6.762165 | 12.85071 | 27.14626 | 10.90734 | -0.57502 | -0.0058 | 0.425819 |
| 14 | 3404400 | 3407999 | 60.1238 | 95.93818 | 97.08394 | 40.71848 | 0.322468 | 2.132757 | 0.425825 |
| 15 | 82050600 | 82052799 | 9.328744 | 16.93344 | 33.38764 | 14.41832 | -0.55846 | 0.312509 | 0.425827 |
| X | 15415800 | 15425999 | 171.6065 | 259.9753 | 203.2049 | 84.08671 | 0.634016 | 3.27542 | 0.425838 |
| 14 | 35354600 | 35356999 | 22.45456 | 39.26293 | 35.2905 | 15.55889 | 0.540733 | 0.714818 | 0.425866 |
| 15 | 8769400 | 8770799 | 15.40801 | 27.95145 | 27.19554 | 11.75322 | 0.452965 | 0.34512 | 0.425877 |
| 3 | 49510400 | 49511799 | 16.49706 | 29.8107 | 27.51962 | 11.95138 | 0.523731 | 0.375067 | 0.425886 |
| 6 | 113974400 | 113976199 | 15.84793 | 27.60087 | 48.42679 | 21.29407 | -0.4203 | 0.758065 | 0.425904 |
| 4 | 49529000 | 49531399 | 51.73499 | 83.40854 | 70.59321 | 30.05594 | 0.596979 | 1.795739 | 0.425918 |
| 5 | 114283200 | 114285599 | 14.92794 | 25.97309 | 46.31566 | 20.3532 | -0.44354 | 0.765757 | 0.425922 |
| 3 | 6552400 | 6552999 | 10.06315 | 19.51525 | 19.26121 | 7.270088 | 0.596109 | -0.2023 | 0.425944 |
| 10 | 7556600 | 7557799 | 8.373714 | 15.3983 | 31.46282 | 13.36031 | -0.59255 | 0.226603 | 0.425946 |
| 10 | 88292600 | 88297399 | 41.62816 | 66.48462 | 115.556 | 48.52744 | -0.45507 | 2.09231 | 0.425953 |
| 10 | 41723400 | 41725799 | 27.03827 | 45.76863 | 45.33683 | 19.84684 | 0.394999 | 1.074059 | 0.425981 |
| 10 | 119519800 | 119520599 | 4.107281 | 8.011615 | 20.28767 | 7.266773 | -0.67756 | -0.45327 | 0.425991 |
| 13 | 34992200 | 34999199 | 79.86447 | 126.0765 | 124.6307 | 51.74224 | 0.34166 | 2.455349 | 0.426008 |
| X | 103920800 | 103922599 | 19.25405 | 34.85751 | 26.87924 | 11.64612 | 0.784249 | 0.385175 | 0.426009 |
| 19 | 35957600 | 35959799 | 18.8729 | 33.64573 | 29.36006 | 12.91887 | 0.59207 | 0.476193 | 0.426026 |
| 8 | 96839600 | 96842999 | 28.64074 | 48.42442 | 46.34806 | 20.28231 | 0.444181 | 1.087812 | 0.426031 |
| 9 | 96992600 | 96993599 | 6.786903 | 12.81046 | 28.63492 | 11.67188 | -0.6749 | 0.043002 | 0.426031 |
| 4 | 72117200 | 72120199 | 24.37612 | 39.5379 | 91.46291 | 39.1829 | -0.84087 | 1.61215 | 0.426056 |
| 19 | 3183400 | 3193399 | 145.8708 | 221.0869 | 214.0637 | 88.59272 | 0.325926 | 3.259325 | 0.426064 |
| 8 | 39270800 | 39271399 | 9.032803 | 17.54384 | 18.833 | 7.0439 | 0.488636 | -0.24532 | 0.426081 |
| 7 | 32267800 | 32272999 | 41.36453 | 68.2801 | 55.62962 | 24.01748 | 0.675176 | 1.381199 | 0.426088 |
| 19 | 58880200 | 58882199 | 20.36016 | 35.52075 | 34.5356 | 15.20238 | 0.427714 | 0.734323 | 0.426107 |
| 1 | 78508200 | 78509399 | 8.486828 | 15.78677 | 30.53065 | 12.76067 | -0.497 | 0.160248 | 0.426133 |
| 7 | 27638800 | 27643599 | 48.22744 | 77.89099 | 75.11425 | 32.05375 | 0.412024 | 1.742419 | 0.426146 |
| X | 112865600 | 112869399 | 71.00731 | 113.1321 | 101.7265 | 42.57555 | 0.490301 | 2.194293 | 0.42615 |

| | | | | | | | | | |
|---|---|---|---|---|---|---|---|---|---|
| X | 26107000 | 26109799 | 50.75647 | 81.98815 | 74.26975 | 31.69966 | 0.50236 | 1.742609 | 0.426156 |
| 2 | 116298400 | 116300999 | 22.84203 | 40.24522 | 31.30231 | 13.85519 | 0.7495 | 0.60536 | 0.426157 |
| 14 | 18069000 | 18069799 | 10.29886 | 19.89928 | 19.66162 | 7.549393 | 0.571044 | -0.14278 | 0.426171 |
| 1 | 15688200 | 15690999 | 29.87662 | 51.11409 | 41.83 | 18.3422 | 0.670983 | 0.986129 | 0.426172 |
| 9 | 110741400 | 110742999 | 7.330958 | 13.53529 | 32.02769 | 13.52572 | -0.79713 | 0.194279 | 0.426173 |
| 4 | 47126000 | 47127599 | 22.62028 | 39.02315 | 38.93906 | 17.08201 | 0.388171 | 0.879224 | 0.426174 |
| 9 | 19328800 | 19337599 | 114.1031 | 177.1292 | 150.8925 | 62.66138 | 0.537393 | 2.784726 | 0.426178 |
| 2 | 7576400 | 7578399 | 18.71497 | 33.71343 | 26.63012 | 11.61019 | 0.743568 | 0.405858 | 0.42623 |
| 7 | 41683600 | 41684599 | 22.36171 | 38.97057 | 37.0934 | 16.31723 | 0.458232 | 0.765294 | 0.426242 |
| 7 | 10240200 | 10244999 | 59.18441 | 95.00774 | 85.80218 | 36.30424 | 0.496071 | 1.946002 | 0.426246 |
| 11 | 115732000 | 115736799 | 35.7197 | 57.36056 | 102.06 | 43.20645 | -0.47963 | 1.910948 | 0.426249 |
| 3 | 52604600 | 52606799 | 27.36685 | 47.11953 | 40.82167 | 17.90681 | 0.5907 | 0.920273 | 0.426252 |
| 7 | 126980600 | 126986799 | 83.07935 | 126.9832 | 241.8305 | 100.318 | -0.63932 | 3.074883 | 0.426262 |
| 1 | 188826000 | 188828599 | 15.5453 | 26.95765 | 48.99561 | 21.49708 | -0.4717 | 0.813921 | 0.426267 |
| 16 | 5257600 | 5258799 | 12.94099 | 24.3537 | 23.47287 | 9.619494 | 0.522551 | 0.104364 | 0.426277 |
| 13 | 86643000 | 86644999 | 20.59522 | 35.89583 | 35.62949 | 15.6743 | 0.397963 | 0.75163 | 0.426285 |
| 7 | 31795400 | 31798999 | 43.90627 | 71.94579 | 59.69611 | 25.69119 | 0.644931 | 1.476434 | 0.426311 |
| 11 | 94982200 | 94983199 | 17.01613 | 30.14841 | 27.47185 | 12.14618 | 0.525382 | 0.503697 | 0.426314 |
| 6 | 112941200 | 112942799 | 15.20173 | 27.47728 | 26.85277 | 11.65791 | 0.442673 | 0.365161 | 0.426315 |
| 18 | 76265400 | 76268599 | 46.15168 | 73.70275 | 117.7667 | 49.44972 | -0.33413 | 2.099767 | 0.42632 |
| 13 | 64309000 | 64322599 | 273.5156 | 387.14 | 549.015 | 227.2072 | -0.31798 | 4.398759 | 0.426325 |
| 1 | 179647400 | 179648399 | 25.08439 | 41.43455 | 70.55601 | 30.47598 | -0.38512 | 1.343868 | 0.426334 |
| 7 | 26470600 | 26472999 | 34.40867 | 56.64077 | 56.92763 | 24.54709 | 0.371668 | 1.395283 | 0.426364 |
| 8 | 87195800 | 87198399 | 15.68813 | 27.31107 | 47.89459 | 21.05448 | -0.41963 | 0.76255 | 0.426369 |
| 6 | 17315800 | 17316799 | 6.146238 | 11.79035 | 25.71181 | 10.09965 | -0.5907 | -0.09155 | 0.42638 |
| 12 | 40914200 | 40915599 | 16.9463 | 30.08536 | 29.6897 | 13.10992 | 0.412234 | 0.490478 | 0.426382 |
| 15 | 82322200 | 82323199 | 9.301618 | 16.95845 | 32.89775 | 14.12901 | -0.53006 | 0.285273 | 0.426397 |
| X | 77698000 | 77701399 | 37.61668 | 62.52305 | 52.6205 | 22.8267 | 0.629151 | 1.292599 | 0.426441 |
| 10 | 43700200 | 43712799 | 134.7314 | 201.6239 | 310.6464 | 128.1144 | -0.35984 | 3.510011 | 0.426471 |
| 9 | 43965600 | 43967999 | 15.05789 | 26.36297 | 47.80578 | 21.09929 | -0.46715 | 0.675196 | 0.426488 |
| 9 | 89722400 | 89725999 | 21.49255 | 36.12637 | 61.96393 | 27.06782 | -0.39475 | 1.131066 | 0.4265 |
| 5 | 150307000 | 150309399 | 16.86263 | 29.06726 | 50.59098 | 22.193 | -0.41132 | 0.876874 | 0.426512 |
| Y | 9464600 | 9465399 | 9.040787 | 17.56349 | 11.47858 | 4.285045 | 1.19879 | -0.28012 | 0.426513 |
| 17 | 41061800 | 41066399 | 61.84372 | 98.77507 | 97.25917 | 40.84611 | 0.363292 | 2.105014 | 0.426526 |
| 19 | 40107800 | 40108799 | 8.389423 | 15.84863 | 30.93296 | 12.58609 | -0.47916 | 0.056936 | 0.426528 |
| 17 | 23858600 | 23864799 | 58.04834 | 91.3609 | 153.6008 | 63.7273 | -0.42684 | 2.509316 | 0.426553 |
| 18 | 59807200 | 59808199 | 8.095875 | 15.31013 | 29.2512 | 11.87544 | -0.44546 | 0.043951 | 0.426553 |
| 3 | 54073200 | 54075599 | 19.42597 | 34.31836 | 31.54204 | 13.95382 | 0.511246 | 0.55603 | 0.42656 |
| 17 | 16516000 | 16517399 | 15.9149 | 28.7835 | 27.29915 | 11.84287 | 0.486368 | 0.368347 | 0.426572 |
| 14 | 121235800 | 121238999 | 26.77075 | 43.83285 | 75.25578 | 32.37683 | -0.40162 | 1.45943 | 0.426575 |
| 1 | 13627200 | 13627999 | 11.4579 | 20.22164 | 38.88096 | 17.2039 | -0.54821 | 0.533337 | 0.426583 |
| 14 | 81418600 | 81419599 | 12.16937 | 23.09569 | 21.95668 | 8.847057 | 0.562775 | 0.036407 | 0.426599 |
| 9 | 23110200 | 23111799 | 15.90353 | 28.76278 | 27.64371 | 11.99243 | 0.467043 | 0.366873 | 0.426602 |
| 5 | 110143200 | 110144799 | 15.92532 | 27.42753 | 50.57548 | 22.18561 | -0.49453 | 0.880448 | 0.426603 |
| 7 | 108523800 | 108528399 | 31.60611 | 51.05328 | 97.94684 | 41.80457 | -0.57784 | 1.722823 | 0.426634 |
| 16 | 79934600 | 79936399 | 15.08934 | 26.20174 | 47.39568 | 20.80178 | -0.46441 | 0.79622 | 0.426646 |
| 1 | 15304600 | 15305799 | 22.20739 | 38.37174 | 37.86546 | 16.61183 | 0.404547 | 0.860842 | 0.426648 |

| | | | | | | | | | |
|---|---|---|---|---|---|---|---|---|---|
| 8 | 36186200 | 36189199 | 41.72748 | 67.79923 | 67.07175 | 28.77046 | 0.383706 | 1.594345 | 0.426656 |
| 13 | 3587400 | 3590399 | 20.64907 | 34.40604 | 64.42399 | 27.99304 | -0.52072 | 1.22412 | 0.426659 |
| 5 | 67523000 | 67530799 | 87.86369 | 137.7296 | 137.4349 | 57.0034 | 0.320295 | 2.601 | 0.426667 |
| X | 19925000 | 19928799 | 49.48755 | 80.24714 | 67.30596 | 28.82836 | 0.618715 | 1.653494 | 0.426671 |
| 7 | 119548400 | 119552199 | 37.07026 | 60.64046 | 60.81129 | 26.15322 | 0.371059 | 1.482296 | 0.426672 |
| 18 | 21546800 | 21551399 | 46.86596 | 74.9766 | 127.9613 | 53.86585 | -0.42625 | 2.050145 | 0.426683 |
| 15 | 47429200 | 47431999 | 30.03471 | 51.86364 | 37.90994 | 16.63103 | 0.835633 | 0.902804 | 0.426685 |
| 4 | 13010200 | 13011799 | 17.2393 | 31.67439 | 25.6102 | 10.88755 | 0.734396 | 0.286035 | 0.426687 |
| 18 | 9605400 | 9607199 | 21.61843 | 38.08641 | 31.92395 | 14.13056 | 0.642059 | 0.599597 | 0.426691 |
| 4 | 150570200 | 150570999 | 5.544753 | 10.71525 | 24.24927 | 9.303722 | -0.61096 | -0.17968 | 0.426715 |
| 7 | 140256200 | 140260599 | 69.41822 | 110.6603 | 100.679 | 42.16876 | 0.47419 | 2.172798 | 0.426722 |
| 19 | 9112000 | 9113199 | 7.528797 | 14.12589 | 29.60764 | 12.19561 | -0.59552 | 0.091054 | 0.426735 |
| 16 | 17254000 | 17266599 | 287.9153 | 412.9715 | 385.641 | 159.2476 | 0.303814 | 4.203892 | 0.426764 |
| 2 | 153065400 | 153068599 | 17.65246 | 29.8837 | 56.89276 | 24.9063 | -0.54405 | 1.051238 | 0.426766 |
| 5 | 119366400 | 119367999 | 17.65246 | 29.8837 | 56.89276 | 24.9063 | -0.54405 | 1.051238 | 0.426766 |
| 1 | 152947000 | 152948799 | 13.32566 | 23.3265 | 43.58841 | 19.23533 | -0.50999 | 0.662438 | 0.426777 |
| 10 | 119726200 | 119729599 | 63.54408 | 101.1669 | 102.8269 | 42.99233 | 0.312642 | 2.208909 | 0.426808 |
| 10 | 59922600 | 59925199 | 14.03256 | 24.38224 | 46.34748 | 20.34989 | -0.5353 | 0.782449 | 0.426824 |
| 10 | 102864000 | 102868199 | 29.8875 | 48.55624 | 83.11673 | 35.65011 | -0.40547 | 1.583662 | 0.42683 |
| 3 | 127945200 | 127945799 | 8.926782 | 16.66047 | 31.12532 | 12.93615 | -0.44121 | 0.143081 | 0.426834 |
| X | 87369000 | 87370599 | 16.49835 | 30.66054 | 24.23338 | 10.14366 | 0.783004 | 0.215986 | 0.426849 |
| 14 | 112718800 | 112719399 | 14.57195 | 26.48312 | 26.27514 | 11.32952 | 0.42707 | 0.326493 | 0.42686 |
| 3 | 23584400 | 23586599 | 14.0959 | 24.76218 | 43.98476 | 19.4683 | -0.43734 | 0.610086 | 0.426872 |
| 4 | 86659600 | 86661599 | 33.99713 | 56.33982 | 47.69063 | 20.64587 | 0.620466 | 1.308889 | 0.426874 |
| 11 | 61305600 | 61306599 | 15.38977 | 27.60678 | 26.76518 | 11.71569 | 0.446148 | 0.409323 | 0.426881 |
| 7 | 50766400 | 50767399 | 13.36469 | 24.6269 | 23.99468 | 10.16049 | 0.47121 | 0.236136 | 0.426882 |
| 9 | 80475400 | 80476999 | 17.10342 | 31.08266 | 26.39673 | 11.38249 | 0.649845 | 0.344809 | 0.426885 |
| X | 147648000 | 147649799 | 32.83404 | 55.12139 | 49.13194 | 21.44985 | 0.546391 | 1.168685 | 0.426903 |
| X | 93360400 | 93362999 | 15.08606 | 26.41471 | 46.72731 | 20.62488 | -0.43147 | 0.673364 | 0.426905 |
| 4 | 146344600 | 146346399 | 37.37435 | 61.97835 | 48.25898 | 20.90165 | 0.740614 | 1.310016 | 0.426905 |
| 12 | 114482800 | 114491399 | 184.8903 | 276.468 | 255.7336 | 105.4515 | 0.374739 | 3.531352 | 0.426914 |
| 10 | 15459600 | 15460199 | 9.544433 | 18.51844 | 18.70808 | 7.045049 | 0.565719 | -0.21824 | 0.426943 |
| 14 | 77794800 | 77796199 | 10.05357 | 18.29938 | 33.87982 | 14.58064 | -0.46534 | 0.301767 | 0.426946 |
| X | 24222800 | 24229599 | 130.8154 | 201.2543 | 168.0732 | 69.80037 | 0.556121 | 2.956109 | 0.426979 |
| 13 | 4664200 | 4664799 | 6.131691 | 11.80589 | 25.26572 | 9.819928 | -0.5495 | -0.12817 | 0.426997 |
| 19 | 39505400 | 39508399 | 49.64085 | 80.73396 | 63.77713 | 27.3708 | 0.708602 | 1.594891 | 0.427003 |
| 2 | 129731600 | 129732799 | 18.65236 | 33.19452 | 30.00277 | 13.22113 | 0.540331 | 0.487768 | 0.427005 |
| 14 | 100626200 | 100628199 | 39.79828 | 65.15215 | 55.32322 | 23.79964 | 0.611154 | 1.476791 | 0.42702 |
| 15 | 60830000 | 60832399 | 20.13978 | 35.32291 | 33.88463 | 14.97759 | 0.447288 | 0.653847 | 0.427022 |
| 18 | 26349600 | 26350199 | 3.523406 | 6.890066 | 17.77108 | 6.054777 | -0.6365 | -0.64825 | 0.427025 |
| 19 | 13546200 | 13550599 | 56.45098 | 90.71366 | 84.04676 | 35.61647 | 0.461004 | 1.909005 | 0.427045 |
| 17 | 23842400 | 23843199 | 6.439592 | 12.32353 | 27.32886 | 10.80893 | -0.6249 | -0.06406 | 0.427057 |
| 12 | 116249200 | 116252799 | 42.33425 | 68.72373 | 68.59939 | 29.40638 | 0.369634 | 1.615816 | 0.427076 |
| 2 | 21009800 | 21011599 | 12.03675 | 21.40393 | 37.76199 | 16.65159 | -0.41977 | 0.465765 | 0.42708 |
| 17 | 39807000 | 39816599 | 140.4315 | 215.8497 | 163.6054 | 67.93633 | 0.694861 | 2.97621 | 0.427104 |
| 7 | 122679600 | 122683999 | 27.39552 | 44.88236 | 75.46778 | 32.47616 | -0.37133 | 1.452976 | 0.42711 |
| 14 | 104368800 | 104371199 | 35.4595 | 58.74793 | 49.30309 | 21.34008 | 0.632874 | 1.318997 | 0.427118 |

| | | | | | | | | | |
|---|---|---|---|---|---|---|---|---|---|
| Y | 3849600 | 3853399 | 19.75981 | 34.71098 | 20.13141 | 8.910249 | 1.168601 | 0.596972 | 0.427124 |
| 6 | 112965400 | 112965799 | 3.984461 | 7.784258 | 19.98736 | 6.973834 | -0.66369 | -0.55196 | 0.427128 |
| 10 | 49762600 | 49765199 | 26.36534 | 45.93913 | 35.72809 | 15.71477 | 0.748341 | 0.78567 | 0.427163 |
| 15 | 90970800 | 90973799 | 21.35409 | 35.42597 | 72.12371 | 31.24646 | -0.64151 | 1.283341 | 0.427181 |
| 1 | 114190600 | 114194799 | 50.44864 | 82.06173 | 62.7145 | 26.91643 | 0.756417 | 1.593334 | 0.42719 |
| 19 | 39616200 | 39618399 | 31.72643 | 53.27421 | 49.12851 | 21.45147 | 0.497422 | 1.1613 | 0.4272 |
| 15 | 3611400 | 3613199 | 11.2067 | 19.65419 | 41.5717 | 18.37455 | -0.68737 | 0.618706 | 0.427208 |
| 15 | 52781200 | 52782999 | 53.0403 | 85.15052 | 85.78693 | 36.30156 | 0.338712 | 1.935499 | 0.427212 |
| 10 | 61422000 | 61426399 | 34.36295 | 55.40285 | 93.53239 | 39.82442 | -0.39688 | 1.780246 | 0.427217 |
| 4 | 99008400 | 99009599 | 6.048474 | 11.50761 | 27.70603 | 11.10529 | -0.75946 | -0.02215 | 0.427224 |
| 17 | 22523400 | 22527999 | 67.25431 | 107.2834 | 100.0391 | 41.94003 | 0.439791 | 2.150559 | 0.427236 |
| 2 | 36332400 | 36335599 | 35.19508 | 58.83449 | 45.38622 | 19.76149 | 0.754245 | 1.216235 | 0.427252 |
| 4 | 115282200 | 115282799 | 10.81086 | 20.49148 | 19.59819 | 7.917933 | 0.551349 | 0.027192 | 0.427257 |
| 16 | 94033200 | 94033999 | 10.81086 | 20.49148 | 19.59819 | 7.917933 | 0.551349 | 0.027192 | 0.427257 |
| 3 | 11691800 | 11693199 | 13.8915 | 26.17829 | 22.20079 | 9.077113 | 0.708015 | 0.099744 | 0.427268 |
| 19 | 42861600 | 42864199 | 13.21994 | 23.39674 | 44.00449 | 19.46006 | -0.5153 | 0.523156 | 0.427281 |
| 15 | 33886200 | 33886799 | 10.80224 | 19.25239 | 36.33249 | 15.99058 | -0.51415 | 0.441555 | 0.427295 |
| 7 | 33734400 | 33740399 | 113.2618 | 174.0985 | 173.7485 | 72.14875 | 0.298377 | 2.965156 | 0.427315 |
| 4 | 130819800 | 130826399 | 58.94339 | 93.01318 | 148.6219 | 61.69023 | -0.35062 | 2.455197 | 0.427321 |
| 18 | 17799200 | 17808999 | 151.8241 | 230.5512 | 206.2885 | 85.42418 | 0.441956 | 3.221966 | 0.427331 |
| 14 | 73214400 | 73218199 | 38.26015 | 61.0951 | 124.8027 | 52.39791 | -0.68792 | 2.097282 | 0.427334 |
| 2 | 78817800 | 78819199 | 14.60124 | 26.88579 | 25.80256 | 10.93691 | 0.491564 | 0.25287 | 0.427335 |
| 10 | 127111800 | 127115799 | 27.06995 | 44.37664 | 74.88357 | 32.23327 | -0.37607 | 1.44424 | 0.427346 |
| 17 | 33036800 | 33039999 | 30.46344 | 51.14469 | 48.584 | 21.21143 | 0.454745 | 1.157076 | 0.427356 |
| 14 | 27262600 | 27264399 | 19.40236 | 35.05041 | 26.70552 | 11.60208 | 0.798755 | 0.396498 | 0.427376 |
| 16 | 45302600 | 45305399 | 30.28526 | 51.66864 | 42.37344 | 18.57632 | 0.66733 | 1.012085 | 0.427387 |
| 5 | 151453600 | 151459799 | 131.4362 | 200.2932 | 197.5571 | 81.89452 | 0.305607 | 3.14491 | 0.427396 |
| 1 | 84240800 | 84242599 | 20.29501 | 33.56088 | 73.40606 | 31.73149 | -0.7449 | 1.320223 | 0.427397 |
| 3 | 7267400 | 7268199 | 23.85387 | 40.94934 | 40.54736 | 17.78465 | 0.397909 | 0.931372 | 0.427397 |
| 6 | 58269000 | 58271399 | 26.90111 | 45.88575 | 44.14325 | 19.35188 | 0.437934 | 1.003573 | 0.427405 |
| 1 | 3111200 | 3115799 | 62.39552 | 99.96604 | 89.56957 | 37.7853 | 0.504121 | 2.010126 | 0.427417 |
| 5 | 78528000 | 78531399 | 34.14499 | 56.67806 | 54.77313 | 23.73892 | 0.430132 | 1.297325 | 0.427418 |
| 18 | 87097800 | 87104799 | 123.6666 | 189.4954 | 165.7036 | 68.77842 | 0.485406 | 3.028915 | 0.427421 |
| 3 | 104995600 | 104999599 | 21.65342 | 35.89078 | 68.83317 | 29.80267 | -0.55574 | 1.29 | 0.427423 |
| 6 | 19458600 | 19460799 | 22.70725 | 40.17783 | 29.62976 | 13.10376 | 0.828124 | 0.5571 | 0.427426 |
| X | 57442800 | 57444999 | 34.31223 | 57.28273 | 50.84433 | 22.1195 | 0.552688 | 1.234797 | 0.427437 |
| 7 | 13279800 | 13283199 | 54.20537 | 87.05926 | 85.60959 | 36.25079 | 0.374358 | 1.92358 | 0.427448 |
| 6 | 90403400 | 90405999 | 15.48826 | 26.6861 | 49.50729 | 21.71731 | -0.50292 | 0.875242 | 0.427463 |
| 12 | 35420400 | 35421599 | 10.51375 | 19.11384 | 34.89737 | 15.0413 | -0.4469 | 0.313968 | 0.427491 |
| X | 82588200 | 82592799 | 44.28245 | 72.82763 | 56.33298 | 24.2813 | 0.747921 | 1.431977 | 0.427517 |
| 11 | 105196200 | 105198399 | 18.00171 | 30.8535 | 53.2695 | 23.36174 | -0.40144 | 0.936482 | 0.427528 |
| 16 | 20987800 | 20988799 | 7.332348 | 13.91927 | 28.23566 | 11.37095 | -0.52086 | 0.006956 | 0.427537 |
| 9 | 105904000 | 105906399 | 29.38462 | 49.39861 | 48.56779 | 21.21683 | 0.405335 | 1.142077 | 0.427544 |
| 4 | 125940600 | 125941399 | 7.241087 | 13.59483 | 29.08387 | 11.96681 | -0.62331 | 0.083698 | 0.427545 |
| 16 | 20406800 | 20410599 | 33.20806 | 53.58036 | 92.54261 | 39.44052 | -0.42848 | 1.75538 | 0.42755 |
| 1 | 162778400 | 162779799 | 19.27477 | 34.19336 | 23.83549 | 10.53161 | 0.910533 | 0.500285 | 0.42756 |
| 17 | 69580200 | 69582199 | 11.4325 | 20.28339 | 37.29506 | 16.47717 | -0.48083 | 0.475721 | 0.427562 |

| | | | | | | | | | |
|---|---|---|---|---|---|---|---|---|---|
| 12 | 23032200 | 23032399 | 9.385361 | 18.18746 | 19.32857 | 7.320586 | 0.486419 | -0.19645 | 0.427566 |
| 5 | 118642000 | 118644999 | 12.52432 | 23.77741 | 19.82667 | 7.983079 | 0.751593 | 0.032969 | 0.427567 |
| 19 | 45685600 | 45689999 | 35.68782 | 57.37344 | 99.78588 | 42.31302 | -0.44489 | 1.874077 | 0.427567 |
| 1 | 41884000 | 41884999 | 13.61314 | 25.82898 | 22.53691 | 9.085895 | 0.685572 | 0.055791 | 0.427568 |
| 4 | 61573000 | 61573999 | 13.61314 | 25.82898 | 22.53691 | 9.085895 | 0.685572 | 0.055791 | 0.427568 |
| 10 | 5772000 | 5774599 | 23.02369 | 37.8803 | 73.63581 | 31.74474 | -0.57746 | 1.386316 | 0.427568 |
| 18 | 60620600 | 60624199 | 21.02223 | 35.33008 | 60.80482 | 26.56018 | -0.39961 | 1.130218 | 0.42758 |
| 17 | 27916400 | 27920999 | 78.07445 | 120.4087 | 199.0779 | 82.69342 | -0.42553 | 2.892828 | 0.4276 |
| 10 | 65033400 | 65037799 | 48.34665 | 78.17746 | 72.54048 | 31.00349 | 0.469459 | 1.706546 | 0.4276 |
| 7 | 58447000 | 58447999 | 5.391148 | 10.46724 | 23.35957 | 8.761483 | -0.56011 | -0.26334 | 0.427601 |
| 15 | 63362600 | 63365599 | 41.91522 | 68.13065 | 67.10669 | 28.79367 | 0.390497 | 1.587434 | 0.42761 |
| 2 | 60559400 | 60562399 | 18.3458 | 31.55316 | 54.38669 | 23.85645 | -0.39814 | 0.908565 | 0.427612 |
| 4 | 144893600 | 144896199 | 21.71112 | 37.57747 | 36.35443 | 15.94976 | 0.433502 | 0.841754 | 0.427615 |
| 6 | 8914800 | 8915799 | 7.971548 | 14.98022 | 29.09694 | 11.95327 | -0.48329 | 0.084811 | 0.427616 |
| 5 | 10198400 | 10202599 | 60.0836 | 96.17023 | 92.86591 | 39.11992 | 0.39474 | 2.037502 | 0.427616 |
| 7 | 107765400 | 107771799 | 101.7645 | 158.6792 | 146.2393 | 60.68875 | 0.428961 | 2.701921 | 0.427625 |
| 8 | 119425000 | 119429599 | 45.78521 | 73.18309 | 117.3013 | 49.31691 | -0.33721 | 2.07096 | 0.427627 |
| 9 | 83820800 | 83824999 | 69.88552 | 110.9843 | 110.1905 | 45.93588 | 0.342758 | 2.291114 | 0.427632 |
| 15 | 22532000 | 22533599 | 20.52087 | 36.63104 | 28.75445 | 12.63859 | 0.745147 | 0.479305 | 0.427639 |
| 19 | 45228600 | 45233599 | 41.26004 | 65.93657 | 115.7362 | 48.64479 | -0.4683 | 2.071742 | 0.427642 |
| 13 | 30552600 | 30554599 | 28.5475 | 48.14286 | 47.46317 | 20.75411 | 0.401536 | 1.111233 | 0.427643 |
| 6 | 107305800 | 107306799 | 8.456422 | 15.93544 | 30.81188 | 12.59861 | -0.4713 | 0.077073 | 0.427643 |
| 10 | 49658400 | 49659799 | 8.555918 | 15.62076 | 32.43541 | 13.90608 | -0.62559 | 0.271065 | 0.427648 |
| 4 | 146229400 | 146230399 | 21.15615 | 38.03997 | 26.51371 | 11.58266 | 0.921444 | 0.435285 | 0.427655 |
| 2 | 165822800 | 165827799 | 24.86603 | 41.04975 | 71.28184 | 30.77702 | -0.41342 | 1.347779 | 0.427666 |
| 1 | 17738000 | 17740599 | 34.61932 | 57.51554 | 54.24924 | 23.52603 | 0.465131 | 1.28814 | 0.427678 |
| 1 | 4548800 | 4551199 | 13.77397 | 24.1315 | 44.06892 | 19.46118 | -0.47712 | 0.651399 | 0.427689 |
| 10 | 95884000 | 95885399 | 8.743184 | 16.39771 | 30.69724 | 12.65314 | -0.43462 | 0.108408 | 0.427701 |
| 5 | 139629200 | 139632999 | 31.15314 | 50.24769 | 92.21707 | 39.28451 | -0.51632 | 1.763832 | 0.427708 |
| 5 | 109323600 | 109329999 | 102.7443 | 160.0824 | 146.8748 | 60.95954 | 0.434543 | 2.717025 | 0.427716 |
| 6 | 28133200 | 28136999 | 30.16336 | 48.71093 | 90.34695 | 38.54876 | -0.5295 | 1.723884 | 0.427743 |
| X | 111315000 | 111316399 | 16.96017 | 30.06321 | 29.72896 | 13.1429 | 0.408077 | 0.501339 | 0.427755 |
| 7 | 25895000 | 25896199 | 22.56524 | 39.10575 | 38.80569 | 17.02586 | 0.397375 | 0.837663 | 0.427769 |
| 4 | 54414400 | 54414799 | 5.508816 | 10.72988 | 23.14374 | 8.44852 | -0.47515 | -0.35744 | 0.427781 |
| 13 | 112653600 | 112657399 | 56.11634 | 88.72528 | 143.5584 | 59.6474 | -0.3664 | 2.405782 | 0.427792 |
| 2 | 49072600 | 49073799 | 12.18668 | 23.33386 | 21.00471 | 8.295329 | 0.666502 | -0.02947 | 0.427815 |
| 15 | 46424800 | 46426599 | 27.40374 | 46.85421 | 38.86589 | 17.0416 | 0.651459 | 0.979294 | 0.427816 |
| 1 | 95731000 | 95734599 | 59.83304 | 95.58727 | 95.72984 | 40.2179 | 0.339253 | 2.09456 | 0.427824 |
| 12 | 73659200 | 73660999 | 22.05166 | 38.24005 | 37.38607 | 16.40333 | 0.419088 | 0.829878 | 0.427836 |
| 8 | 29032800 | 29041399 | 107.8193 | 167.0697 | 158.4012 | 65.79541 | 0.381106 | 2.81322 | 0.427838 |
| 12 | 107694400 | 107698999 | 44.37175 | 70.95269 | 114.1547 | 48.02167 | -0.34194 | 2.057548 | 0.427843 |
| 4 | 22736800 | 22741599 | 65.61913 | 104.439 | 104.7136 | 43.76286 | 0.331848 | 2.221985 | 0.427857 |
| 6 | 41168800 | 41170399 | 11.8971 | 21.09107 | 37.76856 | 16.69673 | -0.44365 | 0.484671 | 0.427861 |
| 12 | 73630200 | 73633199 | 22.25101 | 36.99421 | 65.6089 | 28.46668 | -0.44281 | 1.254531 | 0.42789 |
| 13 | 66149600 | 66156199 | 137.4692 | 210.3828 | 166.382 | 69.04011 | 0.628838 | 3.059924 | 0.427906 |
| 2 | 162437400 | 162438999 | 20.53413 | 33.74614 | 78.02532 | 33.621 | -0.82742 | 1.400731 | 0.427918 |
| 6 | 128588000 | 128591199 | 60.32349 | 96.67952 | 89.6616 | 37.84386 | 0.455092 | 1.997976 | 0.427919 |

| | | | | | | | | | |
|---|---|---|---|---|---|---|---|---|---|
| 10 | 59239000 | 59240599 | 10.82904 | 19.26022 | 36.86291 | 16.25238 | -0.53598 | 0.457093 | 0.427922 |
| 18 | 44180200 | 44180799 | 13.19329 | 24.53738 | 24.47531 | 10.23267 | 0.450697 | 0.183918 | 0.427941 |
| 2 | 145497200 | 145498399 | 7.494168 | 14.19173 | 28.17128 | 11.40382 | -0.49629 | 0.026108 | 0.427954 |
| 9 | 24370200 | 24371399 | 8.417404 | 15.90254 | 30.34853 | 12.34658 | -0.44652 | 0.056863 | 0.427975 |
| 4 | 109333000 | 109334599 | 12.01841 | 21.2992 | 39.2616 | 17.3579 | -0.48564 | 0.489681 | 0.427975 |
| 18 | 20438600 | 20441599 | 24.30861 | 42.62986 | 32.28172 | 14.26775 | 0.786638 | 0.674354 | 0.427994 |
| 17 | 33579600 | 33581799 | 23.19683 | 38.60448 | 65.73776 | 28.54186 | -0.38428 | 1.246144 | 0.428 |
| 1 | 181139200 | 181142199 | 32.85111 | 53.11876 | 89.11312 | 38.08461 | -0.38298 | 1.690191 | 0.428025 |
| 7 | 23074400 | 23075399 | 23.91207 | 41.18918 | 39.98674 | 17.5408 | 0.427296 | 0.901384 | 0.42803 |
| 7 | 23257400 | 23273799 | 255.6084 | 374.9987 | 307.3606 | 126.5955 | 0.523634 | 3.864796 | 0.428043 |
| 14 | 124011400 | 124013999 | 20.25819 | 35.83728 | 30.88416 | 13.65907 | 0.604369 | 0.545461 | 0.428058 |
| 8 | 118649200 | 118651399 | 17.15055 | 29.54533 | 52.76315 | 23.14544 | -0.44861 | 0.884877 | 0.428065 |
| 6 | 69624000 | 69626599 | 22.76009 | 39.76147 | 35.93202 | 15.83053 | 0.533101 | 0.732574 | 0.428077 |
| 15 | 77501400 | 77505599 | 23.91814 | 39.51124 | 69.41951 | 29.98671 | -0.43014 | 1.33707 | 0.428099 |
| 3 | 79593000 | 79597999 | 61.60777 | 98.6035 | 91.70733 | 38.62835 | 0.448686 | 2.039516 | 0.428099 |
| 13 | 103468400 | 103472599 | 32.73187 | 52.79203 | 93.5144 | 39.83521 | -0.46549 | 1.765922 | 0.4281 |
| 14 | 95616000 | 95618399 | 24.2374 | 41.62595 | 41.49597 | 18.2012 | 0.388229 | 0.928555 | 0.428106 |
| 10 | 46560800 | 46561799 | 18.13436 | 32.18678 | 26.52916 | 11.71673 | 0.670284 | 0.492894 | 0.42811 |
| 12 | 93530200 | 93532399 | 26.27596 | 45.55591 | 38.58533 | 16.9294 | 0.625039 | 0.851304 | 0.428119 |
| 7 | 59509800 | 59513399 | 48.8807 | 78.97625 | 67.57898 | 28.85243 | 0.584736 | 1.729622 | 0.428122 |
| 7 | 27612600 | 27615399 | 33.77674 | 57.46672 | 41.76854 | 18.30707 | 0.840368 | 1.05049 | 0.428146 |
| 8 | 43294200 | 43296799 | 26.00125 | 45.23038 | 36.27693 | 15.9397 | 0.703813 | 0.805514 | 0.428153 |
| 13 | 56330200 | 56331599 | 18.70765 | 31.85308 | 54.83203 | 24.03425 | -0.39861 | 1.002187 | 0.428164 |
| 7 | 58692400 | 58697999 | 88.45344 | 139.6239 | 111.813 | 46.41886 | 0.644968 | 2.461132 | 0.428175 |
| 6 | 10808800 | 10812599 | 44.70803 | 72.29947 | 72.04382 | 30.79437 | 0.366963 | 1.69735 | 0.428178 |
| 13 | 33136000 | 33136599 | 98.82483 | 153.8489 | 135.6194 | 56.29497 | 0.491263 | 2.726301 | 0.428186 |
| 5 | 72861800 | 72864399 | 35.12009 | 57.72469 | 58.56642 | 25.23762 | 0.357276 | 1.417328 | 0.428188 |
| 1 | 87635600 | 87640399 | 40.05168 | 64.30663 | 104.6495 | 44.29177 | -0.3515 | 1.921434 | 0.428192 |
| X | 118343800 | 118345799 | 21.3583 | 38.3127 | 27.05167 | 11.84127 | 0.901021 | 0.452113 | 0.428257 |
| 8 | 57623800 | 57624399 | 8.01121 | 14.98288 | 29.5575 | 12.24128 | -0.51464 | 0.118864 | 0.428263 |
| 7 | 26291400 | 26293999 | 30.56962 | 51.11196 | 50.81219 | 22.12665 | 0.389523 | 1.203826 | 0.42828 |
| 1 | 37299800 | 37303399 | 29.38796 | 47.89583 | 79.81838 | 34.28132 | -0.36346 | 1.533566 | 0.428291 |
| 17 | 53712000 | 53714799 | 18.43897 | 32.90248 | 28.80959 | 12.6668 | 0.587804 | 0.465667 | 0.428292 |
| 15 | 39722000 | 39723599 | 16.18706 | 28.02549 | 49.53162 | 21.73119 | -0.43204 | 0.832278 | 0.428303 |
| 9 | 100951600 | 100953199 | 19.95482 | 35.07204 | 29.16303 | 12.91344 | 0.652223 | 0.610155 | 0.428334 |
| 5 | 118055200 | 118059799 | 32.56329 | 52.56314 | 91.3349 | 38.94789 | -0.43641 | 1.741169 | 0.428357 |
| 6 | 14805400 | 14809999 | 47.4592 | 77.13522 | 64.8854 | 27.83876 | 0.617449 | 1.598456 | 0.428376 |
| 12 | 114816800 | 114823199 | 143.4115 | 217.1944 | 213.5118 | 88.34318 | 0.30311 | 3.274331 | 0.428391 |
| 17 | 38490200 | 38499399 | 124.0134 | 192.2589 | 146.2675 | 60.75276 | 0.698904 | 2.810179 | 0.428422 |
| 4 | 82662200 | 82665199 | 27.86206 | 45.5628 | 78.44804 | 33.73433 | -0.40698 | 1.478725 | 0.428426 |
| 6 | 50255200 | 50257399 | 17.36145 | 29.98525 | 51.13854 | 22.43452 | -0.38153 | 0.863095 | 0.428426 |
| 3 | 90915800 | 90920799 | 63.94739 | 102.2398 | 94.89495 | 39.91113 | 0.450006 | 2.07701 | 0.428432 |
| 14 | 39193000 | 39197799 | 39.01837 | 62.38555 | 124.0873 | 52.19197 | -0.64756 | 2.056651 | 0.428439 |
| 14 | 71481600 | 71482399 | 17.06596 | 30.14727 | 28.40444 | 12.56594 | 0.475644 | 0.536282 | 0.428447 |
| 18 | 63519000 | 63520799 | 16.58352 | 28.51788 | 50.57403 | 22.18395 | -0.4389 | 0.898136 | 0.428448 |
| Y | 5430600 | 5433199 | 14.76807 | 27.10938 | 14.93178 | 6.355693 | 1.281655 | 0.241321 | 0.428455 |
| 19 | 14953800 | 14954999 | 13.51212 | 23.75126 | 41.40227 | 18.33426 | -0.41001 | 0.595032 | 0.428465 |

| | | | | | | | | | |
|---|---|---|---|---|---|---|---|---|---|
| 10 | 130410400 | 130417999 | 128.1392 | 197.2292 | 168.3095 | 69.90295 | 0.525487 | 2.946008 | 0.428469 |
| 4 | 21054600 | 21056599 | 19.00715 | 32.52993 | 55.44626 | 24.31356 | -0.3833 | 0.957044 | 0.428472 |
| 4 | 13044000 | 13045599 | 13.50253 | 23.86839 | 44.52692 | 19.69579 | -0.50441 | 0.537417 | 0.428475 |
| 9 | 19148600 | 19149599 | 17.51458 | 30.94286 | 29.89021 | 13.22297 | 0.440032 | 0.542166 | 0.428475 |
| 19 | 53835200 | 53840199 | 32.8428 | 53.15006 | 88.11702 | 37.68642 | -0.36475 | 1.671057 | 0.428492 |
| 5 | 31514200 | 31515399 | 8.968975 | 16.58511 | 31.50452 | 13.27694 | -0.4802 | 0.200546 | 0.428512 |
| 8 | 67792800 | 67795399 | 32.37062 | 52.59529 | 89.30482 | 38.3069 | -0.39374 | 1.592035 | 0.428512 |
| 18 | 33002800 | 33003599 | 7.315135 | 14.05225 | 28.11136 | 10.99803 | -0.46266 | -0.08687 | 0.428512 |
| 11 | 75825200 | 75827199 | 9.345642 | 16.89059 | 34.36786 | 14.92247 | -0.60885 | 0.33918 | 0.428513 |
| 12 | 115175200 | 115178799 | 16.32197 | 29.87851 | 22.69583 | 9.697126 | 0.818774 | 0.292998 | 0.428528 |
| 7 | 84922600 | 84924399 | 32.32642 | 53.84024 | 53.04255 | 23.04147 | 0.40269 | 1.253241 | 0.428535 |
| 16 | 68722200 | 68725799 | 37.28267 | 61.45248 | 57.22293 | 24.68816 | 0.482222 | 1.385683 | 0.42855 |
| X | 20927600 | 20930399 | 27.51831 | 47.24057 | 41.92565 | 18.3892 | 0.555221 | 0.94796 | 0.428555 |
| 19 | 52807200 | 52810199 | 37.34349 | 60.12455 | 99.962 | 42.47918 | -0.37733 | 1.831714 | 0.428558 |
| 13 | 72411400 | 72412599 | 9.880273 | 18.0273 | 33.91114 | 14.55023 | -0.48539 | 0.28594 | 0.428566 |
| 3 | 91062200 | 91066799 | 94.71837 | 148.4538 | 137.525 | 57.04159 | 0.427247 | 2.608129 | 0.428567 |
| 12 | 19987200 | 20040799 | 1774.466 | 2188.25 | 1752.093 | 738.0498 | 0.306541 | 6.589468 | 0.428577 |
| 9 | 12213800 | 12214199 | 8.125982 | 15.80166 | 17.2861 | 6.402556 | 0.474912 | -0.29443 | 0.428606 |
| 7 | 89409600 | 89411399 | 12.18912 | 21.63147 | 38.24718 | 16.89407 | -0.42468 | 0.479717 | 0.428613 |
| 4 | 22463200 | 22464199 | 13.52367 | 23.58258 | 44.7286 | 19.68347 | -0.53162 | 0.724717 | 0.428624 |
| 7 | 33463200 | 33467999 | 97.95322 | 153.7295 | 132.5317 | 54.96567 | 0.532358 | 2.586635 | 0.428628 |
| 18 | 5730200 | 5732399 | 26.93712 | 46.79138 | 37.48646 | 16.45635 | 0.705469 | 0.833588 | 0.42864 |
| 2 | 31666800 | 31669399 | 17.95719 | 30.73331 | 54.28263 | 23.80332 | -0.43426 | 0.95399 | 0.428646 |
| 14 | 99573000 | 99573799 | 6.266421 | 12.08387 | 25.90856 | 10.0254 | -0.54671 | -0.14114 | 0.428649 |
| 1 | 103385400 | 103389199 | 45.19822 | 73.28796 | 69.89005 | 29.93405 | 0.433825 | 1.645802 | 0.42866 |
| 2 | 89689200 | 89691799 | 46.60747 | 75.60656 | 62.24641 | 26.66898 | 0.645857 | 1.633005 | 0.428665 |
| 14 | 99447800 | 99449999 | 13.49338 | 23.78248 | 41.36509 | 18.30791 | -0.40537 | 0.564532 | 0.42867 |
| 17 | 45243200 | 45244599 | 10.32175 | 18.50932 | 35.35556 | 15.48029 | -0.52614 | 0.398195 | 0.428679 |
| 3 | 78089200 | 78091599 | 20.73261 | 36.42734 | 32.47486 | 14.37602 | 0.552165 | 0.624114 | 0.428681 |
| 2 | 53177000 | 53177399 | 5.070901 | 9.895889 | 22.31809 | 7.936219 | -0.50443 | -0.46004 | 0.428689 |
| 3 | 96475400 | 96480199 | 33.22237 | 53.56967 | 93.44739 | 39.79375 | -0.44382 | 1.774493 | 0.428692 |
| 2 | 63013800 | 63015599 | 13.71467 | 23.9098 | 44.74565 | 19.68795 | -0.51227 | 0.731244 | 0.428701 |
| 16 | 51060400 | 51062599 | 17.82873 | 31.67595 | 25.20568 | 11.12283 | 0.721654 | 0.482639 | 0.428722 |
| 9 | 116579000 | 116583399 | 36.97999 | 60.57465 | 61.36154 | 26.40396 | 0.357618 | 1.465855 | 0.428724 |
| 5 | 145355600 | 145359599 | 93.70841 | 146.0927 | 146.4796 | 60.78998 | 0.307313 | 2.701168 | 0.42873 |
| 11 | 47681600 | 47682199 | 4.972213 | 9.662439 | 22.83915 | 8.50504 | -0.63229 | -0.29569 | 0.428741 |
| 6 | 103192200 | 103195399 | 60.40171 | 96.32257 | 97.55048 | 40.87931 | 0.32037 | 2.151913 | 0.428753 |
| 4 | 152730000 | 152739999 | 94.20457 | 142.7911 | 274.6936 | 113.688 | -0.66367 | 3.249587 | 0.42876 |
| 3 | 51467000 | 51468999 | 14.00521 | 24.36737 | 45.80313 | 20.12783 | -0.51909 | 0.761168 | 0.428769 |
| 2 | 120325200 | 120328599 | 18.34929 | 31.02704 | 57.36689 | 25.10496 | -0.50194 | 1.068114 | 0.42877 |
| 9 | 6784200 | 6786199 | 17.78926 | 31.3337 | 30.42219 | 13.46641 | 0.43104 | 0.576816 | 0.428779 |
| 7 | 26244000 | 26245799 | 23.86492 | 41.895 | 31.11993 | 13.76627 | 0.814106 | 0.65315 | 0.428794 |
| 13 | 44765000 | 44766599 | 12.0132 | 21.43565 | 37.77145 | 16.60754 | -0.41518 | 0.443481 | 0.428811 |
| 5 | 114073000 | 114074399 | 16.95582 | 29.01597 | 52.14066 | 22.86383 | -0.45903 | 0.947218 | 0.428829 |
| 2 | 126614000 | 126615799 | 11.8922 | 21.25575 | 37.81402 | 16.6025 | -0.4273 | 0.433601 | 0.428854 |
| 15 | 58936800 | 58940599 | 54.0612 | 85.34043 | 146.7933 | 60.94256 | -0.45635 | 2.443184 | 0.42886 |
| 13 | 85129200 | 85129999 | 14.36601 | 26.2282 | 25.89765 | 11.10324 | 0.43946 | 0.2998 | 0.428863 |

| | | | | | | | | | |
|---|---|---|---|---|---|---|---|---|---|
| 4 | 95973400 | 95973999 | 4.963222 | 9.669243 | 22.56035 | 8.212699 | -0.58398 | -0.3789 | 0.428872 |
| 9 | 118020400 | 118022399 | 13.0736 | 22.88765 | 42.60225 | 18.80179 | -0.50405 | 0.661201 | 0.428873 |
| 4 | 8093200 | 8094799 | 19.12787 | 33.55144 | 32.97839 | 14.57823 | 0.412346 | 0.64508 | 0.428873 |
| 5 | 109473600 | 109476399 | 45.96693 | 74.59605 | 62.19959 | 26.65645 | 0.628089 | 1.624398 | 0.428877 |
| 5 | 8293000 | 8293599 | 6.254688 | 12.10691 | 24.9647 | 9.505546 | -0.46839 | -0.19735 | 0.428891 |
| 13 | 62384200 | 62396399 | 153.7457 | 233.6757 | 201.677 | 83.53665 | 0.494991 | 3.2045 | 0.428891 |
| 3 | 110787000 | 110788999 | 16.40178 | 29.83011 | 27.095 | 11.67253 | 0.554484 | 0.337889 | 0.428899 |
| 1 | 20161200 | 20161999 | 28.59685 | 47.85615 | 45.99036 | 20.0382 | 0.438263 | 1.184374 | 0.428916 |
| 7 | 121133200 | 121134599 | 8.116723 | 15.28285 | 29.13441 | 11.92915 | -0.45267 | 0.073807 | 0.428917 |
| 9 | 119421400 | 119422599 | 8.116723 | 15.28285 | 29.13441 | 11.92915 | -0.45267 | 0.073807 | 0.428917 |
| 18 | 65537200 | 65540799 | 32.46632 | 53.64039 | 53.01256 | 22.90189 | 0.397425 | 1.344289 | 0.428951 |
| 4 | 122996800 | 123002999 | 52.48958 | 83.14719 | 138.6476 | 57.66183 | -0.40759 | 2.357547 | 0.428959 |
| 10 | 9861600 | 9863799 | 21.25078 | 37.45008 | 31.92756 | 14.13131 | 0.617841 | 0.593861 | 0.428979 |
| 3 | 68261800 | 68264199 | 30.7476 | 51.66393 | 49.11653 | 21.45327 | 0.453646 | 1.151957 | 0.428981 |
| 16 | 3430600 | 3454999 | 324.9966 | 464.0793 | 410.4387 | 169.6075 | 0.375765 | 4.272183 | 0.428987 |
| X | 14549400 | 14553199 | 46.19294 | 74.97529 | 65.81125 | 28.20773 | 0.554496 | 1.628511 | 0.428989 |
| X | 148838600 | 148840799 | 27.18058 | 47.14807 | 37.39767 | 16.40856 | 0.719372 | 0.848728 | 0.428992 |
| 7 | 106256800 | 106259799 | 36.84831 | 60.93416 | 56.53974 | 24.4406 | 0.488261 | 1.349517 | 0.429003 |
| X | 164391400 | 164392999 | 17.24189 | 32.39602 | 22.12051 | 9.08973 | 1.013588 | 0.150313 | 0.429029 |
| 5 | 16901000 | 16901799 | 12.83037 | 22.71821 | 39.77627 | 17.58853 | -0.41224 | 0.506704 | 0.42903 |
| 7 | 34471200 | 34471799 | 14.63565 | 26.58333 | 23.05585 | 9.948682 | 0.619559 | 0.324742 | 0.429036 |
| 7 | 101407800 | 101409799 | 11.77328 | 20.77207 | 38.88096 | 17.20492 | -0.50982 | 0.539155 | 0.429042 |
| 2 | 35416400 | 35418799 | 31.40671 | 50.81302 | 87.10564 | 37.24685 | -0.41316 | 1.673608 | 0.429048 |
| 3 | 47918400 | 47923199 | 44.66474 | 72.82227 | 63.62812 | 27.33167 | 0.565995 | 1.547531 | 0.429055 |
| 3 | 5891800 | 5894799 | 20.47657 | 34.73012 | 59.25371 | 25.95473 | -0.38655 | 1.04716 | 0.429055 |
| X | 125008800 | 125013399 | 44.27224 | 72.65865 | 52.26376 | 22.5074 | 0.851255 | 1.454876 | 0.429056 |
| 2 | 162851800 | 162853599 | 15.07743 | 27.48607 | 26.72856 | 11.48085 | 0.459376 | 0.313956 | 0.429058 |
| 7 | 42120400 | 42125399 | 56.18075 | 90.78409 | 70.19298 | 29.97408 | 0.730955 | 1.740741 | 0.429085 |
| 11 | 107731400 | 107732399 | 12.56678 | 23.55581 | 23.54827 | 9.715469 | 0.462849 | 0.127496 | 0.429106 |
| 3 | 148893000 | 148893999 | 6.90755 | 13.08365 | 30.41227 | 12.30541 | -0.72216 | 0.021986 | 0.429107 |
| X | 80335600 | 80337799 | 37.93088 | 63.75482 | 45.20961 | 19.74878 | 0.87524 | 1.171402 | 0.429109 |
| 2 | 8142800 | 8144999 | 22.72036 | 37.95418 | 64.06051 | 27.88004 | -0.37152 | 1.204087 | 0.429111 |
| 4 | 71128600 | 71130599 | 22.11267 | 38.84671 | 32.37097 | 14.32834 | 0.649135 | 0.634619 | 0.429117 |
| 6 | 133537600 | 133541599 | 92.50819 | 144.7483 | 140.6581 | 58.35111 | 0.356495 | 2.635216 | 0.429126 |
| 8 | 125002400 | 125003799 | 19.01355 | 31.97946 | 58.95262 | 25.75727 | -0.49807 | 1.118052 | 0.429132 |
| 9 | 96705400 | 96712199 | 59.50702 | 93.22159 | 175.5768 | 72.82805 | -0.59569 | 2.603158 | 0.429138 |
| 10 | 8371600 | 8374999 | 31.12699 | 51.80271 | 52.46771 | 22.78053 | 0.362869 | 1.256811 | 0.429148 |
| 2 | 179370800 | 179373199 | 23.35438 | 38.88365 | 65.76614 | 28.5623 | -0.37453 | 1.241864 | 0.429163 |
| 5 | 130555000 | 130556599 | 17.23299 | 30.65466 | 28.43664 | 12.53578 | 0.502337 | 0.475444 | 0.429176 |
| 10 | 73384200 | 73386599 | 24.99192 | 42.57971 | 41.66488 | 18.26368 | 0.413356 | 1.005162 | 0.429188 |
| 6 | 33756800 | 33758999 | 25.85048 | 45.24077 | 32.8252 | 14.48278 | 0.848176 | 0.715752 | 0.429197 |
| 12 | 6583200 | 6583799 | 10.73567 | 20.42483 | 21.15286 | 8.479025 | 0.447399 | 0.006131 | 0.4292 |
| 11 | 98624400 | 98625999 | 23.9457 | 41.24299 | 40.95329 | 17.96473 | 0.394711 | 0.901742 | 0.429231 |
| 7 | 85330200 | 85339799 | 127.2267 | 196.1167 | 164.1451 | 68.18315 | 0.555071 | 2.916574 | 0.429242 |
| 7 | 33709400 | 33716799 | 123.9384 | 189.6108 | 185.6154 | 77.01497 | 0.320929 | 3.061992 | 0.429242 |
| 16 | 31679600 | 31681399 | 22.6577 | 39.45528 | 36.68648 | 16.13023 | 0.491856 | 0.778826 | 0.429256 |
| 13 | 57680400 | 57682799 | 16.24707 | 28.02568 | 49.51791 | 21.72273 | -0.43263 | 0.867387 | 0.42928 |

| | | | | | | | | | |
|---|---|---|---|---|---|---|---|---|---|
| 13 | 85742200 | 85744399 | 25.75189 | 44.32089 | 41.3958 | 18.15844 | 0.482462 | 0.916394 | 0.42928 |
| 2 | 37056000 | 37060599 | 89.21038 | 139.8147 | 138.3369 | 57.37861 | 0.332315 | 2.605344 | 0.429281 |
| 4 | 128678200 | 128682399 | 20.23758 | 33.61055 | 65.25895 | 28.29383 | -0.573 | 1.262833 | 0.429287 |
| 14 | 62162600 | 62163599 | 11.40448 | 21.57174 | 22.11975 | 8.976072 | 0.446201 | 0.054815 | 0.429291 |
| X | 121656000 | 121657599 | 22.7925 | 40.02049 | 32.99057 | 14.59627 | 0.664845 | 0.649134 | 0.4293 |
| 2 | 41300400 | 41301199 | 6.260135 | 12.08898 | 25.9895 | 10.00513 | -0.54362 | -0.15854 | 0.429306 |
| X | 153523800 | 153525399 | 24.19048 | 42.05614 | 38.12842 | 16.74714 | 0.527868 | 0.8091 | 0.429319 |
| 1 | 59158200 | 59159799 | 10.20092 | 18.601 | 33.89967 | 14.5564 | -0.44066 | 0.292571 | 0.429338 |
| 17 | 73175200 | 73177599 | 22.77605 | 39.73859 | 36.42877 | 16.03652 | 0.512293 | 0.749413 | 0.429342 |
| 6 | 68202400 | 68204999 | 29.72204 | 50.99572 | 40.70464 | 17.85286 | 0.70742 | 0.959161 | 0.429348 |
| 9 | 70226000 | 70228199 | 9.796301 | 17.72903 | 33.85825 | 14.67635 | -0.51632 | 0.334451 | 0.429354 |
| 12 | 16776800 | 16779399 | 13.66978 | 23.91104 | 43.10492 | 19.00999 | -0.4583 | 0.677429 | 0.429355 |
| 6 | 17370000 | 17371399 | 12.42933 | 23.30591 | 23.47224 | 9.678654 | 0.452502 | 0.122339 | 0.429374 |
| 10 | 11647200 | 11648599 | 17.71917 | 32.08124 | 27.56189 | 11.94074 | 0.629336 | 0.375113 | 0.429386 |
| 1 | 112490400 | 112492599 | 31.8342 | 53.02985 | 51.97415 | 22.57989 | 0.410064 | 1.246779 | 0.429392 |
| 7 | 27796400 | 27805199 | 97.72758 | 149.1467 | 231.6997 | 96.08186 | -0.3475 | 3.107907 | 0.429393 |
| 3 | 135531000 | 135532599 | 9.355319 | 16.90906 | 33.88252 | 14.71066 | -0.58661 | 0.339201 | 0.429398 |
| 7 | 7301200 | 7303999 | 27.27105 | 44.12945 | 87.49443 | 37.41781 | -0.6222 | 1.66272 | 0.429429 |
| 15 | 102312400 | 102315199 | 12.20981 | 21.55384 | 39.32678 | 17.40033 | -0.47307 | 0.535236 | 0.429434 |
| 6 | 3207800 | 3208399 | 18.55039 | 32.65191 | 30.47794 | 13.49294 | 0.487325 | 0.59037 | 0.429443 |
| X | 85712800 | 85714399 | 21.11357 | 38.12431 | 26.10503 | 11.34767 | 0.951594 | 0.411166 | 0.429451 |
| 6 | 68716000 | 68716999 | 14.1081 | 25.82638 | 25.83613 | 11.03857 | 0.423893 | 0.281407 | 0.429473 |
| 14 | 3936800 | 3938599 | 12.24192 | 21.62916 | 38.84116 | 17.18251 | -0.44956 | 0.526941 | 0.429474 |
| 14 | 73131000 | 73131599 | 5.712023 | 10.98395 | 25.21532 | 9.836038 | -0.65442 | -0.12045 | 0.429506 |
| 17 | 66022200 | 66023199 | 13.44412 | 24.99829 | 24.0064 | 10.04042 | 0.504866 | 0.187419 | 0.429507 |
| X | 121796400 | 121797799 | 15.81407 | 30.21345 | 20.21559 | 8.031539 | 1.083724 | 0.024468 | 0.429514 |
| 9 | 120142400 | 120143799 | 7.670478 | 14.57755 | 28.80975 | 11.57457 | -0.47997 | 0.007194 | 0.429515 |
| 6 | 146350000 | 146352199 | 33.62225 | 55.48951 | 56.36585 | 24.33217 | 0.357819 | 1.362087 | 0.429583 |
| 4 | 81317800 | 81319799 | 24.59098 | 40.76999 | 70.09267 | 30.35322 | -0.39846 | 1.299851 | 0.429589 |
| 8 | 72125800 | 72126599 | 6.004929 | 11.52165 | 25.7163 | 10.09544 | -0.62319 | -0.09552 | 0.429606 |
| 6 | 136570600 | 136571999 | 13.82904 | 25.50204 | 25.40505 | 10.74651 | 0.440298 | 0.238202 | 0.429625 |
| 2 | 64600600 | 64603599 | 31.55921 | 51.12803 | 85.58771 | 36.63385 | -0.37714 | 1.646628 | 0.429648 |
| 6 | 9650800 | 9652999 | 37.09997 | 61.04958 | 52.03488 | 22.43402 | 0.608617 | 1.404537 | 0.429652 |
| 3 | 153749600 | 153751199 | 10.31918 | 18.94817 | 37.35035 | 15.89184 | -0.54429 | 0.259406 | 0.429669 |
| 18 | 76767400 | 76769199 | 21.26768 | 35.5925 | 61.7568 | 26.90844 | -0.41118 | 1.178327 | 0.429688 |
| X | 125797600 | 125799599 | 28.75464 | 49.22145 | 42.47878 | 18.62751 | 0.594867 | 0.979711 | 0.42969 |
| 3 | 154957600 | 154958199 | 6.107683 | 11.84948 | 25.07766 | 9.446872 | -0.49017 | -0.23243 | 0.4297 |
| 3 | 140576400 | 140577399 | 10.77483 | 18.85376 | 46.24746 | 20.40187 | -0.89992 | 0.662972 | 0.429703 |
| 7 | 72619800 | 72620399 | 9.943037 | 19.09128 | 20.21856 | 7.920986 | 0.445388 | -0.07808 | 0.429717 |
| 17 | 80684400 | 80685599 | 13.90513 | 25.89913 | 23.70941 | 9.887941 | 0.575907 | 0.176202 | 0.429722 |
| 10 | 46398000 | 46400599 | 27.98277 | 48.15815 | 40.77631 | 17.88667 | 0.623468 | 0.926248 | 0.429728 |
| 9 | 8627800 | 8628999 | 8.581052 | 16.10206 | 30.15782 | 12.41979 | -0.43424 | 0.102132 | 0.429729 |
| 5 | 93687000 | 93688199 | 8.974727 | 17.4445 | 18.2706 | 6.791102 | 0.531616 | -0.2706 | 0.429734 |
| X | 121899800 | 121903199 | 84.43949 | 132.9863 | 128.5675 | 53.34592 | 0.371115 | 2.508617 | 0.429737 |
| 12 | 112407400 | 112417799 | 124.2737 | 186.3739 | 294.0661 | 121.3289 | -0.39132 | 3.468028 | 0.429739 |
| 17 | 37732400 | 37743599 | 135.2287 | 207.3056 | 178.6561 | 74.15924 | 0.506917 | 3.026063 | 0.429751 |
| 16 | 78727600 | 78729199 | 12.74858 | 22.27165 | 43.67595 | 19.24233 | -0.57914 | 0.693281 | 0.429773 |

| | | | | | | | | | |
|---|---|---|---|---|---|---|---|---|---|
| 1 | 72677200 | 72681399 | 38.58487 | 62.11194 | 99.97443 | 42.47328 | -0.33107 | 1.83725 | 0.429782 |
| 17 | 25279800 | 25281399 | 8.149713 | 15.24003 | 29.55133 | 12.24129 | -0.49042 | 0.119909 | 0.429812 |
| X | 65279600 | 65280399 | 16.76846 | 30.2794 | 27.71809 | 12.04837 | 0.535371 | 0.384359 | 0.429815 |
| 14 | 12199600 | 12201199 | 24.10226 | 39.84477 | 68.79661 | 29.73278 | -0.40494 | 1.330244 | 0.429823 |
| 6 | 82357000 | 82358799 | 14.24481 | 24.99094 | 44.54722 | 19.6954 | -0.44242 | 0.632531 | 0.429828 |
| 13 | 57012000 | 57012799 | 13.26126 | 24.66538 | 24.01122 | 10.03754 | 0.485893 | 0.183892 | 0.429829 |
| 13 | 90842200 | 90847599 | 80.13479 | 127.0365 | 114.5938 | 47.68715 | 0.478474 | 2.352953 | 0.429861 |
| 12 | 20165000 | 20167999 | 50.0393 | 80.82427 | 63.00347 | 26.88868 | 0.718451 | 1.738378 | 0.429864 |
| 12 | 24855200 | 24858399 | 28.61009 | 48.31475 | 47.45295 | 20.75828 | 0.407067 | 1.100593 | 0.429877 |
| 14 | 110085600 | 110086999 | 11.62723 | 22.21139 | 19.20085 | 7.630505 | 0.715399 | -0.02194 | 0.429892 |
| 9 | 114813200 | 114817799 | 40.7644 | 65.18945 | 113.5325 | 47.7652 | -0.45592 | 2.049435 | 0.429896 |
| 4 | 96799600 | 96801599 | 26.29121 | 45.02015 | 43.0607 | 18.88321 | 0.44709 | 0.965942 | 0.429911 |
| 1 | 186436000 | 186438599 | 14.76896 | 25.30135 | 51.57547 | 22.61799 | -0.63975 | 0.927682 | 0.429912 |
| 9 | 112186600 | 112187999 | 13.63235 | 23.94659 | 41.91364 | 18.55076 | -0.41604 | 0.605699 | 0.429914 |
| 5 | 145979400 | 145998799 | 483.6052 | 663.4948 | 594.5524 | 247.271 | 0.298756 | 4.873972 | 0.429922 |
| 6 | 106121800 | 106123799 | 22.76637 | 40.15812 | 30.68742 | 13.5799 | 0.775389 | 0.591013 | 0.429931 |
| 19 | 5354800 | 5361199 | 58.63223 | 92.01838 | 171.6381 | 71.18451 | -0.57967 | 2.569317 | 0.429937 |
| 4 | 115500200 | 115505199 | 35.11163 | 56.69806 | 94.04211 | 40.12188 | -0.36891 | 1.734509 | 0.429942 |
| 6 | 81717600 | 81719799 | 17.22771 | 30.84322 | 29.0344 | 12.73 | 0.486796 | 0.438729 | 0.429948 |
| 12 | 35177600 | 35179399 | 18.99512 | 34.12153 | 27.72671 | 12.12238 | 0.700457 | 0.427427 | 0.429969 |
| 1 | 15288800 | 15291199 | 24.62172 | 42.43314 | 40.31872 | 17.6867 | 0.458096 | 0.896781 | 0.429981 |
| X | 8295600 | 8296199 | 18.81507 | 33.04962 | 32.42186 | 14.34941 | 0.415104 | 0.619938 | 0.43 |
| 6 | 128440200 | 128443599 | 29.69013 | 48.43369 | 80.05069 | 34.39285 | -0.35065 | 1.52032 | 0.430003 |
| 8 | 114705800 | 114707199 | 15.41328 | 26.90654 | 46.25682 | 20.36972 | -0.39053 | 0.721531 | 0.430009 |
| 8 | 84681800 | 84683199 | 5.624425 | 10.68525 | 27.13317 | 10.90924 | -0.83865 | -0.01537 | 0.430016 |
| 18 | 28307000 | 28309399 | 34.73117 | 58.14711 | 44.82363 | 19.53522 | 0.755238 | 1.198887 | 0.43002 |
| 2 | 21249200 | 21252199 | 21.52402 | 35.66451 | 73.24233 | 31.70446 | -0.65416 | 1.298501 | 0.43003 |
| 2 | 63227600 | 63228799 | 14.55962 | 25.19055 | 52.67447 | 23.10958 | -0.67351 | 0.838709 | 0.430044 |
| 7 | 53470400 | 53470999 | 11.2261 | 21.20779 | 20.97925 | 8.535807 | 0.493819 | 0.056497 | 0.430082 |
| 7 | 77280000 | 77280799 | 5.54253 | 10.74784 | 24.46352 | 9.236332 | -0.59791 | -0.23697 | 0.430083 |
| 1 | 34216600 | 34219599 | 38.9884 | 62.52984 | 105.6692 | 44.64775 | -0.40766 | 1.957118 | 0.430087 |
| 6 | 129681400 | 129682399 | 13.04289 | 24.42511 | 24.01186 | 9.922644 | 0.484914 | 0.138407 | 0.430126 |
| 3 | 49683400 | 49684999 | 23.32669 | 41.12756 | 30.68067 | 13.57816 | 0.80975 | 0.599656 | 0.430126 |
| 11 | 73015200 | 73019199 | 29.37595 | 47.82928 | 80.99678 | 34.77499 | -0.38759 | 1.54713 | 0.430141 |
| 11 | 32556400 | 32558599 | 14.75521 | 25.83343 | 44.60661 | 19.68756 | -0.39665 | 0.669731 | 0.430148 |
| X | 49030400 | 49031599 | 15.64835 | 29.61203 | 21.68686 | 8.792263 | 0.928945 | 0.088761 | 0.430164 |
| 8 | 3694000 | 3697399 | 22.39734 | 37.27555 | 65.10431 | 28.26767 | -0.4207 | 1.242695 | 0.430172 |
| 17 | 83343800 | 83345799 | 11.08751 | 19.7525 | 36.78644 | 16.19612 | -0.49577 | 0.445393 | 0.430185 |
| 19 | 9928800 | 9930599 | 20.33732 | 35.64491 | 34.31419 | 15.15868 | 0.442121 | 0.664845 | 0.430189 |
| 3 | 59277000 | 59278799 | 26.94921 | 45.51725 | 45.87567 | 20.06928 | 0.369935 | 1.091799 | 0.430191 |
| 11 | 19571200 | 19573599 | 49.16488 | 79.07675 | 79.99257 | 33.94288 | 0.336114 | 1.868827 | 0.430199 |
| 7 | 62335800 | 62337399 | 21.37687 | 37.14539 | 36.33784 | 15.95549 | 0.418608 | 0.800965 | 0.430204 |
| 9 | 4092200 | 4092999 | 16.01308 | 28.77755 | 28.24619 | 12.34489 | 0.430133 | 0.406862 | 0.430204 |
| 16 | 95940800 | 95943399 | 27.99903 | 47.12265 | 47.49397 | 20.75486 | 0.369783 | 1.128198 | 0.430221 |
| 15 | 92737400 | 92738799 | 10.03486 | 18.50127 | 34.12588 | 14.43969 | -0.44287 | 0.229706 | 0.430227 |
| 9 | 100427200 | 100429399 | 15.59955 | 26.55099 | 60.66157 | 26.58868 | -0.80549 | 0.999501 | 0.430243 |
| 12 | 115898600 | 115901999 | 50.97438 | 82.63967 | 66.73749 | 28.58017 | 0.672827 | 1.656202 | 0.430253 |

| | | | | | | | | | |
|---|---|---|---|---|---|---|---|---|---|
| 1 | 3949800 | 3950399 | 10.14713 | 19.60503 | 20.30871 | 7.799857 | 0.503795 | -0.13943 | 0.430257 |
| 18 | 54197200 | 54197799 | 10.14713 | 19.60503 | 20.30871 | 7.799857 | 0.503795 | -0.13943 | 0.430257 |
| 2 | 15059200 | 15060799 | 12.39727 | 21.85949 | 39.36447 | 17.42103 | -0.45477 | 0.550683 | 0.430286 |
| 10 | 89820800 | 89828199 | 88.61806 | 136.5809 | 206.2078 | 85.65535 | -0.29537 | 2.908553 | 0.430288 |
| X | 25977600 | 25981199 | 39.7955 | 67.30567 | 38.91067 | 17.02928 | 1.168917 | 1.120619 | 0.430297 |
| 5 | 86939000 | 86941799 | 18.11931 | 31.18694 | 56.63986 | 24.84536 | -0.47317 | 0.901265 | 0.430299 |
| 3 | 104501000 | 104501599 | 8.864603 | 16.20597 | 32.44918 | 13.88876 | -0.57173 | 0.267427 | 0.430312 |
| 15 | 14394600 | 14396999 | 37.17555 | 60.81345 | 61.83769 | 26.59476 | 0.351128 | 1.484692 | 0.430314 |
| 17 | 85529800 | 85532199 | 26.67218 | 43.85605 | 74.696 | 32.1934 | -0.38777 | 1.403829 | 0.430317 |
| 7 | 110229400 | 110231799 | 12.85752 | 22.57016 | 41.45425 | 18.33689 | -0.48509 | 0.611601 | 0.430319 |
| 2 | 162565200 | 162566599 | 18.05043 | 31.75397 | 31.33577 | 13.87427 | 0.406905 | 0.592531 | 0.430321 |
| 7 | 141175000 | 141176799 | 11.46347 | 20.25062 | 38.38029 | 16.97915 | -0.52691 | 0.522899 | 0.430329 |
| X | 136446200 | 136447599 | 16.82446 | 31.01101 | 25.20696 | 10.66968 | 0.731396 | 0.265324 | 0.43033 |
| 8 | 7027200 | 7027799 | 9.341995 | 18.05069 | 17.24421 | 6.62077 | 0.620564 | -0.15784 | 0.430335 |
| 4 | 50057000 | 50058999 | 16.29189 | 29.7782 | 26.01003 | 11.13554 | 0.616139 | 0.305899 | 0.430336 |
| 1 | 3381400 | 3382199 | 10.98062 | 20.81671 | 18.64671 | 7.530866 | 0.645766 | 0.025479 | 0.430339 |
| 7 | 84973000 | 84984799 | 163.3804 | 248.882 | 183.4091 | 76.02142 | 0.725403 | 3.161847 | 0.430339 |
| X | 26027200 | 26029999 | 29.82653 | 52.34954 | 27.15362 | 12.00957 | 1.329796 | 0.687389 | 0.430353 |
| 12 | 31115600 | 31116399 | 12.37516 | 23.99117 | 18.52069 | 7.00108 | 0.945305 | -0.17294 | 0.43036 |
| 12 | 23436400 | 23438399 | 46.67493 | 75.50086 | 56.79746 | 24.28342 | 0.77195 | 1.688496 | 0.430368 |
| 7 | 38029800 | 38043599 | 185.572 | 279.0333 | 229.7967 | 94.87847 | 0.549401 | 3.425426 | 0.430375 |
| 10 | 7705600 | 7706799 | 8.434944 | 15.72855 | 30.5642 | 12.72207 | -0.49955 | 0.141282 | 0.430397 |
| 10 | 89279000 | 89280199 | 8.434944 | 15.72855 | 30.5642 | 12.72207 | -0.49955 | 0.141282 | 0.430397 |
| 1 | 26706000 | 26707599 | 21.7695 | 38.43735 | 30.12794 | 13.32971 | 0.739292 | 0.572346 | 0.4304 |
| 13 | 68260600 | 68277599 | 399.1459 | 557.4089 | 505.7316 | 209.7449 | 0.306638 | 4.601935 | 0.430416 |
| 14 | 27839000 | 27842199 | 35.53592 | 59.24499 | 52.01209 | 22.60811 | 0.568471 | 1.25218 | 0.430422 |
| 1 | 52556200 | 52557999 | 16.97277 | 30.76764 | 27.58956 | 11.93438 | 0.569253 | 0.362181 | 0.430433 |
| 14 | 32729600 | 32730999 | 8.786245 | 16.25954 | 30.98186 | 13.0432 | -0.48373 | 0.192093 | 0.430437 |
| 1 | 12726000 | 12728199 | 5.087938 | 9.871348 | 29.29384 | 11.03374 | -0.97325 | -0.23446 | 0.430444 |
| 14 | 54586000 | 54590199 | 62.89017 | 99.13788 | 153.3065 | 63.61829 | -0.30465 | 2.480506 | 0.43045 |
| 2 | 147214600 | 147216799 | 21.48249 | 36.13703 | 61.9841 | 27.0829 | -0.39476 | 1.124262 | 0.430451 |
| 13 | 104751400 | 104754799 | 25.37199 | 41.8718 | 70.66926 | 30.50589 | -0.37249 | 1.353467 | 0.430452 |
| 5 | 131138400 | 131145599 | 75.01387 | 118.7487 | 118.8478 | 49.41083 | 0.327023 | 2.385525 | 0.430459 |
| 19 | 39661000 | 39661599 | 14.37857 | 26.20145 | 25.43772 | 10.93192 | 0.461568 | 0.309939 | 0.430468 |
| 8 | 124930600 | 124933999 | 32.0226 | 51.93385 | 85.38196 | 36.57439 | -0.34977 | 1.625477 | 0.430487 |
| 16 | 80812400 | 80817999 | 128.9493 | 196.2212 | 189.3675 | 78.46588 | 0.335334 | 3.172069 | 0.430488 |
| 1 | 169884800 | 169888799 | 40.83679 | 66.32694 | 67.34072 | 28.87778 | 0.345881 | 1.601774 | 0.430503 |
| 5 | 96636600 | 96639199 | 34.53959 | 55.7559 | 92.67532 | 39.5219 | -0.37243 | 1.745542 | 0.430536 |
| 19 | 38862400 | 38863599 | 10.59819 | 20.25728 | 21.19872 | 8.413651 | 0.445026 | -0.02665 | 0.430543 |
| 3 | 49090800 | 49092799 | 44.02477 | 71.34287 | 69.95762 | 29.94987 | 0.392876 | 1.652653 | 0.430554 |
| 3 | 58432400 | 58433599 | 10.75396 | 19.30937 | 35.30154 | 15.43977 | -0.46243 | 0.391441 | 0.430555 |
| 10 | 89284000 | 89285199 | 11.36796 | 20.53164 | 36.89433 | 16.03305 | -0.43175 | 0.363771 | 0.430574 |
| 12 | 50952200 | 50953999 | 17.36909 | 30.06937 | 52.23896 | 22.91893 | -0.40741 | 0.84342 | 0.430578 |
| 6 | 87096600 | 87099199 | 18.53775 | 31.48957 | 56.44503 | 24.73586 | -0.45736 | 1.023406 | 0.430586 |
| 14 | 113919200 | 113924999 | 56.72551 | 91.46314 | 75.33651 | 32.07942 | 0.63628 | 1.804698 | 0.430594 |
| X | 67171000 | 67173799 | 56.01601 | 90.22206 | 78.25644 | 33.2735 | 0.560056 | 1.838043 | 0.430595 |
| 3 | 84091200 | 84092599 | 18.37614 | 32.40208 | 31.68432 | 14.02209 | 0.421246 | 0.569712 | 0.430596 |

| | | | | | | | | | |
|---|---|---|---|---|---|---|---|---|---|
| 16 | 20951400 | 20952199 | 18.37614 | 32.40208 | 31.68432 | 14.02209 | 0.421246 | 0.569712 | 0.430596 |
| 7 | 12073000 | 12084599 | 154.026 | 233.6389 | 209.8032 | 86.85116 | 0.435495 | 3.245169 | 0.430604 |
| 19 | 19573800 | 19574799 | 17.83076 | 31.67296 | 25.66796 | 11.32882 | 0.695262 | 0.484787 | 0.430607 |
| 8 | 79718800 | 79720399 | 16.92797 | 30.89857 | 26.5544 | 11.38805 | 0.638029 | 0.320651 | 0.430609 |
| 4 | 111020800 | 111027199 | 69.58449 | 110.6372 | 108.3352 | 45.21604 | 0.364359 | 2.256569 | 0.430615 |
| 3 | 65401400 | 65402599 | 7.924242 | 15.11719 | 30.1634 | 12.02134 | -0.48419 | -0.01632 | 0.43062 |
| 10 | 48875600 | 48880399 | 64.89627 | 103.432 | 102.6905 | 42.99987 | 0.348118 | 2.175913 | 0.430636 |
| X | 11869400 | 11874599 | 26.29809 | 43.42011 | 84.65365 | 36.55414 | -0.58005 | 1.362897 | 0.430654 |
| 2 | 97459400 | 97461199 | 22.34424 | 37.36674 | 63.57922 | 27.68968 | -0.38318 | 1.18907 | 0.430656 |
| 2 | 59335000 | 59339199 | 41.89946 | 68.15506 | 66.60685 | 28.58695 | 0.40251 | 1.576483 | 0.430662 |
| 14 | 45546400 | 45547999 | 14.32119 | 24.97519 | 44.72231 | 19.68164 | -0.44901 | 0.728772 | 0.430672 |
| 18 | 37799800 | 37911599 | 4405.678 | 4955.309 | 5663.145 | 2418.942 | -0.34209 | 8.040143 | 0.430677 |
| 12 | 113377800 | 113379799 | 26.24673 | 44.61294 | 44.73766 | 19.60692 | 0.377549 | 1.033167 | 0.430691 |
| 1 | 129018000 | 129020999 | 51.59929 | 82.98512 | 82.51773 | 35.01001 | 0.360695 | 1.877655 | 0.43071 |
| 17 | 37834200 | 37835599 | 16.64292 | 30.19717 | 26.98915 | 11.66138 | 0.574783 | 0.351913 | 0.430722 |
| 18 | 34152800 | 34156799 | 34.99449 | 56.52514 | 91.99281 | 39.2621 | -0.34106 | 1.726954 | 0.430764 |
| 9 | 34113600 | 34114799 | 7.990845 | 15.01403 | 29.09481 | 11.95571 | -0.48002 | 0.087403 | 0.430767 |
| 11 | 41937200 | 41937599 | 5.249157 | 10.21277 | 23.60663 | 8.706746 | -0.58834 | -0.32583 | 0.430787 |
| 16 | 87085800 | 87086599 | 21.88934 | 38.17597 | 31.04764 | 13.66416 | 0.684207 | 0.743051 | 0.43079 |
| 9 | 55531800 | 55532599 | 12.67967 | 24.07026 | 19.4363 | 7.827332 | 0.797877 | 0.035565 | 0.430791 |
| 2 | 72415000 | 72415999 | 13.90259 | 25.93808 | 24.45773 | 10.17095 | 0.537459 | 0.17047 | 0.430791 |
| 1 | 9805400 | 9806999 | 14.0366 | 24.66744 | 44.53195 | 19.71687 | -0.46037 | 0.607608 | 0.430796 |
| 6 | 84647400 | 84653599 | 61.73631 | 97.66238 | 159.9363 | 66.46999 | -0.38323 | 2.401074 | 0.430837 |
| 6 | 66070200 | 66076999 | 71.67661 | 114.1897 | 102.429 | 42.86505 | 0.493657 | 2.195984 | 0.430839 |
| 2 | 109328600 | 109329399 | 9.866043 | 18.03974 | 33.94234 | 14.52486 | -0.48326 | 0.273282 | 0.430849 |
| X | 14831000 | 14835599 | 58.82256 | 95.07107 | 67.39493 | 28.78628 | 0.856252 | 1.740197 | 0.430859 |
| 13 | 22825200 | 22832599 | 154.0271 | 233.1882 | 198.1893 | 81.99402 | 0.512453 | 3.283102 | 0.430887 |
| 10 | 67973600 | 67975599 | 13.63764 | 23.94153 | 41.93805 | 18.55218 | -0.41715 | 0.615621 | 0.430906 |
| 4 | 134837200 | 134844199 | 70.77751 | 110.2969 | 175.1756 | 72.70355 | -0.35598 | 2.701034 | 0.430913 |
| 6 | 27667600 | 27668799 | 8.550713 | 16.14539 | 31.49148 | 12.82667 | -0.47956 | 0.06337 | 0.430926 |
| 5 | 109740000 | 109742599 | 30.62156 | 50.93333 | 51.36716 | 22.29417 | 0.368962 | 1.260628 | 0.430931 |
| 6 | 79882600 | 79883399 | 15.81718 | 28.29081 | 27.82225 | 12.20839 | 0.423436 | 0.433708 | 0.430944 |
| 11 | 19388000 | 19388999 | 15.81718 | 28.29081 | 27.82225 | 12.20839 | 0.423436 | 0.433708 | 0.430944 |
| 6 | 142848600 | 142849599 | 12.09464 | 22.96258 | 21.37642 | 8.606348 | 0.593639 | 0.030972 | 0.430973 |
| 4 | 130978200 | 130981599 | 18.47865 | 31.12712 | 59.5936 | 26.04934 | -0.55253 | 1.09984 | 0.430983 |
| 15 | 27036400 | 27037599 | 15.57286 | 28.9117 | 24.69995 | 10.35303 | 0.669724 | 0.213001 | 0.430997 |
| 5 | 34086000 | 34088999 | 29.79916 | 49.99967 | 49.67079 | 21.67777 | 0.390478 | 1.16417 | 0.431043 |
| 14 | 110503000 | 110503599 | 6.107742 | 11.83788 | 24.44074 | 9.249608 | -0.46143 | -0.21999 | 0.43106 |
| 14 | 41993000 | 41993799 | 44.66565 | 72.11687 | 67.88769 | 28.95877 | 0.446309 | 1.743282 | 0.431084 |
| 5 | 37482600 | 37484399 | 21.97801 | 38.03467 | 35.43906 | 15.5481 | 0.487597 | 0.84446 | 0.431089 |
| 5 | 83268000 | 83269199 | 16.599 | 29.63962 | 29.15904 | 12.81301 | 0.421434 | 0.451237 | 0.431095 |
| 13 | 50992000 | 50992599 | 8.591813 | 16.08677 | 30.10805 | 12.44563 | -0.43751 | 0.116517 | 0.431104 |
| 17 | 85147600 | 85148799 | 8.591813 | 16.08677 | 30.10805 | 12.44563 | -0.43751 | 0.116517 | 0.431104 |
| 14 | 84176200 | 84177199 | 8.707412 | 16.02401 | 32.51406 | 13.79254 | -0.5812 | 0.2301 | 0.431121 |
| 11 | 3596600 | 3600999 | 41.86194 | 67.20979 | 107.4838 | 45.48785 | -0.32651 | 1.926866 | 0.431144 |
| 18 | 39125400 | 39128599 | 21.11411 | 35.16752 | 65.61278 | 28.50274 | -0.5156 | 1.229787 | 0.431157 |
| 5 | 59596800 | 59598599 | 29.26166 | 47.60415 | 80.63425 | 34.60899 | -0.3887 | 1.56056 | 0.431204 |

| | | | | | | | | | |
|---|---|---|---|---|---|---|---|---|---|
| 4 | 32545200 | 32546199 | 7.155813 | 13.70503 | 27.38966 | 10.80836 | -0.47308 | -0.06346 | 0.431214 |
| 5 | 67887000 | 67887999 | 7.155813 | 13.70503 | 27.38966 | 10.80836 | -0.47308 | -0.06346 | 0.431214 |
| 16 | 68939000 | 68939999 | 7.155813 | 13.70503 | 27.38966 | 10.80836 | -0.47308 | -0.06346 | 0.431214 |
| Y | 6591400 | 6593999 | 16.99036 | 30.39054 | 17.50217 | 7.679652 | 1.189899 | 0.414828 | 0.431228 |
| 4 | 132773200 | 132774999 | 6.856135 | 12.76514 | 30.04151 | 12.53674 | -0.77602 | 0.137679 | 0.431245 |
| 7 | 136661600 | 136664399 | 21.2104 | 35.50173 | 62.97542 | 27.4419 | -0.44308 | 1.174933 | 0.431247 |
| 17 | 4447000 | 4450999 | 46.30185 | 73.92589 | 118.4076 | 49.70296 | -0.33796 | 2.106625 | 0.431259 |
| 6 | 3649400 | 3652399 | 29.15371 | 49.16426 | 48.00395 | 20.99045 | 0.415487 | 1.115463 | 0.431263 |
| 5 | 32735600 | 32736399 | 12.9925 | 24.01204 | 23.97619 | 10.11185 | 0.440113 | 0.216003 | 0.431269 |
| 11 | 118937400 | 118939799 | 19.45181 | 32.68952 | 59.5885 | 26.02856 | -0.48213 | 1.124644 | 0.431282 |
| 19 | 38446200 | 38448399 | 24.55437 | 42.47414 | 35.67377 | 15.65086 | 0.636484 | 0.859784 | 0.431293 |
| 8 | 15803600 | 15807199 | 76.66341 | 120.843 | 120.7466 | 50.10641 | 0.324511 | 2.4838 | 0.431302 |
| 3 | 54468000 | 54470599 | 28.40927 | 48.23087 | 45.77053 | 20.05456 | 0.456687 | 1.052855 | 0.431306 |
| X | 92004200 | 92005399 | 26.1897 | 44.73142 | 43.65101 | 19.13812 | 0.417782 | 0.989918 | 0.431313 |
| 15 | 99841000 | 99844999 | 62.23531 | 97.14862 | 171.469 | 71.15026 | -0.50612 | 2.668037 | 0.431315 |
| 2 | 167869200 | 167872399 | 39.90745 | 64.07565 | 103.8541 | 43.95574 | -0.34567 | 1.921289 | 0.431316 |
| 9 | 60605200 | 60608799 | 22.6512 | 37.88927 | 64.76021 | 28.20834 | -0.38972 | 1.187903 | 0.431329 |
| X | 116033200 | 116035599 | 26.97334 | 47.17741 | 32.8291 | 14.47712 | 0.908309 | 0.731891 | 0.431333 |
| 15 | 20505200 | 20507599 | 22.10556 | 38.93051 | 31.86738 | 14.10655 | 0.675862 | 0.60607 | 0.431341 |
| 1 | 87780000 | 87782999 | 15.49283 | 26.59441 | 52.14836 | 22.87289 | -0.58383 | 0.910402 | 0.431353 |
| 17 | 17365000 | 17366799 | 31.50802 | 55.22867 | 25.97105 | 11.47389 | 1.470479 | 0.714431 | 0.431358 |
| 9 | 77431200 | 77434399 | 17.23808 | 29.52004 | 52.16242 | 22.87469 | -0.43482 | 0.940611 | 0.431363 |
| 8 | 59672200 | 59674199 | 19.7024 | 33.29122 | 58.06351 | 25.40431 | -0.41836 | 1.075995 | 0.431372 |
| 16 | 18941600 | 18943199 | 29.83643 | 50.62931 | 45.76522 | 20.04929 | 0.526647 | 1.063326 | 0.431396 |
| 10 | 22030000 | 22030599 | 5.369226 | 10.45375 | 22.4586 | 8.236265 | -0.47535 | -0.34496 | 0.431413 |
| 17 | 90223400 | 90223799 | 5.369226 | 10.45375 | 22.4586 | 8.236265 | -0.47535 | -0.34496 | 0.431413 |
| 5 | 74228600 | 74229999 | 11.54847 | 20.55631 | 37.26335 | 16.41779 | -0.45784 | 0.454489 | 0.431415 |
| 9 | 61211600 | 61213399 | 20.49194 | 35.80553 | 35.53046 | 15.6554 | 0.398682 | 0.716479 | 0.431435 |
| 7 | 57766000 | 57771199 | 48.63794 | 78.49058 | 76.57247 | 32.64202 | 0.394076 | 1.767993 | 0.431437 |
| 15 | 40310600 | 40311599 | 9.578026 | 17.56947 | 32.4286 | 13.81674 | -0.45084 | 0.251324 | 0.431438 |
| 9 | 7388600 | 7392399 | 48.57892 | 78.5293 | 73.57533 | 31.43352 | 0.455054 | 1.716925 | 0.431472 |
| 15 | 76883200 | 76886399 | 31.85469 | 52.77373 | 52.44694 | 22.70028 | 0.389731 | 1.311772 | 0.431476 |
| 6 | 90705000 | 90709199 | 19.96021 | 33.42034 | 61.74896 | 26.91314 | -0.50157 | 1.166952 | 0.431477 |
| 9 | 36057000 | 36059599 | 40.46864 | 65.74178 | 66.80715 | 28.65308 | 0.344823 | 1.59623 | 0.431483 |
| 2 | 59667000 | 59669999 | 23.94914 | 39.46404 | 70.97089 | 30.61595 | -0.46461 | 1.368302 | 0.431487 |
| 17 | 92823800 | 92827599 | 32.87457 | 55.61988 | 45.17941 | 19.7749 | 0.680126 | 1.097209 | 0.43152 |
| 7 | 41703200 | 41706799 | 37.25256 | 63.02464 | 42.38332 | 18.55085 | 0.951757 | 1.113568 | 0.431535 |
| 9 | 12941400 | 12941999 | 10.16036 | 19.73991 | 18.13563 | 6.766204 | 0.713518 | -0.24466 | 0.431545 |
| 12 | 78455800 | 78458399 | 24.58916 | 42.61397 | 35.13041 | 15.41335 | 0.663799 | 0.843142 | 0.43155 |
| 15 | 19573000 | 19576199 | 60.72495 | 96.95534 | 97.87483 | 41.08495 | 0.326898 | 2.114658 | 0.43155 |
| 5 | 114458800 | 114462199 | 18.83376 | 32.06179 | 55.94318 | 24.52093 | -0.41819 | 1.00419 | 0.431551 |
| 13 | 94247200 | 94250599 | 37.14546 | 59.88917 | 97.70294 | 41.60028 | -0.34764 | 1.786926 | 0.431554 |
| 8 | 116114400 | 116116199 | 12.09999 | 21.32033 | 39.40347 | 17.44074 | -0.49233 | 0.557355 | 0.431572 |
| 4 | 75548400 | 75550199 | 18.98495 | 33.62522 | 30.38413 | 13.4347 | 0.536872 | 0.522655 | 0.43158 |
| 12 | 19464200 | 19492399 | 1109.654 | 1421.379 | 1181.088 | 495.4909 | 0.308382 | 5.976917 | 0.431586 |
| 1 | 54447000 | 54449799 | 23.84713 | 43.2111 | 22.70199 | 9.825324 | 1.33531 | 0.405361 | 0.431586 |
| 3 | 78955200 | 78956599 | 8.073459 | 15.26339 | 30.38616 | 12.34365 | -0.50548 | 0.046748 | 0.431587 |

| | | | | | | | | | |
|---|---|---|---|---|---|---|---|---|---|
| 13 | 107470000 | 107473399 | 36.85575 | 60.77807 | 57.73257 | 24.91311 | 0.453888 | 1.379999 | 0.431617 |
| 13 | 55681800 | 55684799 | 18.77324 | 31.77483 | 56.37401 | 24.67774 | -0.44268 | 1.057885 | 0.431619 |
| X | 107732000 | 107734999 | 33.42332 | 56.70687 | 44.06278 | 19.30254 | 0.744126 | 1.076767 | 0.43162 |
| 15 | 46833800 | 46835199 | 18.81057 | 33.59225 | 29.44542 | 12.93768 | 0.587153 | 0.46646 | 0.431627 |
| X | 164449600 | 164450599 | 15.39351 | 28.04348 | 26.8148 | 11.52755 | 0.483027 | 0.322022 | 0.431642 |
| 2 | 20385000 | 20386399 | 23.25015 | 40.18559 | 39.86794 | 17.4905 | 0.396964 | 0.865911 | 0.431646 |
| 12 | 33289600 | 33291799 | 24.45592 | 42.71752 | 35.18023 | 15.49783 | 0.66632 | 0.740872 | 0.431653 |
| 5 | 148991600 | 148993399 | 12.04836 | 21.38835 | 37.76173 | 16.67526 | -0.4222 | 0.476762 | 0.431666 |
| 11 | 120798800 | 120799999 | 10.90415 | 19.60904 | 35.29691 | 15.41782 | -0.43889 | 0.382906 | 0.431686 |
| 4 | 45972000 | 45973199 | 7.225006 | 13.62934 | 28.08926 | 11.46377 | -0.56003 | 0.051457 | 0.431736 |
| 7 | 39924600 | 39925799 | 7.225006 | 13.62934 | 28.08926 | 11.46377 | -0.56003 | 0.051457 | 0.431736 |
| 3 | 68807200 | 68810199 | 25.20359 | 43.1257 | 42.61612 | 18.68714 | 0.400124 | 0.968703 | 0.431746 |
| 4 | 146076800 | 146094599 | 435.3657 | 603.6067 | 543.6678 | 225.7322 | 0.306636 | 4.711965 | 0.431754 |
| 8 | 38516200 | 38523399 | 72.33463 | 115.5162 | 93.39298 | 39.21536 | 0.647018 | 2.119796 | 0.431765 |
| 1 | 85139600 | 85149799 | 250.6595 | 366.2228 | 330.7856 | 136.2789 | 0.377903 | 3.928037 | 0.431788 |
| 19 | 16771200 | 16773999 | 43.59854 | 70.61752 | 70.88299 | 30.33507 | 0.358621 | 1.663058 | 0.43181 |
| 7 | 55347200 | 55350599 | 36.98957 | 61.0054 | 58.21621 | 25.12297 | 0.447368 | 1.381197 | 0.431831 |
| 14 | 28576600 | 28577199 | 10.68683 | 20.29435 | 20.61315 | 8.29506 | 0.470199 | 0.0162 | 0.431852 |
| 8 | 43519600 | 43522799 | 33.39003 | 55.38577 | 54.6941 | 23.69312 | 0.398998 | 1.304226 | 0.431861 |
| 7 | 32805200 | 32806799 | 19.9976 | 35.93264 | 27.73024 | 12.12099 | 0.774881 | 0.433879 | 0.431884 |
| 15 | 46076400 | 46080799 | 55.17006 | 88.4921 | 88.34209 | 37.33305 | 0.350388 | 1.965812 | 0.431891 |
| 10 | 14866600 | 14867399 | 6.777277 | 12.82296 | 27.60812 | 11.19649 | -0.61436 | 0.025096 | 0.431894 |
| 10 | 85890400 | 85893399 | 31.54101 | 51.12592 | 85.1916 | 36.4787 | -0.36982 | 1.634969 | 0.431903 |
| 1 | 124829200 | 124830799 | 13.82865 | 26.23815 | 21.82556 | 8.798919 | 0.753367 | 0.053711 | 0.431908 |
| 4 | 34233800 | 34234799 | 8.55952 | 15.98212 | 32.24933 | 13.39325 | -0.55129 | 0.136009 | 0.431911 |
| 13 | 64431800 | 64432799 | 11.78596 | 22.70994 | 19.41869 | 7.530805 | 0.763959 | -0.09594 | 0.431918 |
| 11 | 65902000 | 65904999 | 23.28585 | 40.77595 | 34.06134 | 15.03551 | 0.64585 | 0.694381 | 0.431941 |
| 14 | 58608600 | 58611199 | 35.4586 | 58.29103 | 59.15546 | 25.49329 | 0.357022 | 1.416596 | 0.431961 |
| 8 | 12092800 | 12094999 | 10.68753 | 18.9305 | 37.38416 | 16.52963 | -0.58418 | 0.486878 | 0.431967 |
| 13 | 98951000 | 98952399 | 10.68753 | 18.9305 | 37.38416 | 16.52963 | -0.58418 | 0.486878 | 0.431967 |
| 13 | 79655800 | 79658799 | 37.73711 | 61.62167 | 61.89259 | 26.60101 | 0.367006 | 1.509702 | 0.431985 |
| 6 | 102102000 | 102105799 | 33.62484 | 56.15314 | 50.89369 | 22.14544 | 0.522488 | 1.224915 | 0.431988 |
| 2 | 94484800 | 94486599 | 27.69923 | 47.66528 | 39.60662 | 17.37352 | 0.650307 | 0.921884 | 0.432006 |
| 19 | 43313000 | 43314399 | 20.78624 | 36.53577 | 33.12497 | 14.66863 | 0.528171 | 0.621672 | 0.432014 |
| X | 65576400 | 65580399 | 42.96684 | 70.93726 | 54.52277 | 23.54242 | 0.759049 | 1.383749 | 0.432026 |
| 7 | 47401000 | 47421399 | 371.9651 | 525.7215 | 449.7691 | 186.1795 | 0.408763 | 4.423823 | 0.432042 |
| 7 | 11638200 | 11642599 | 58.34847 | 93.78093 | 83.10819 | 35.2286 | 0.525409 | 1.906206 | 0.432052 |
| 6 | 57597000 | 57599199 | 18.3422 | 32.91214 | 28.33977 | 12.40201 | 0.616442 | 0.430919 | 0.432062 |
| 1 | 80762400 | 80763799 | 8.229935 | 15.55498 | 30.37695 | 12.34702 | -0.47863 | 0.050489 | 0.432067 |
| 15 | 25901200 | 25903799 | 25.06825 | 41.28192 | 71.64547 | 30.89865 | -0.41384 | 1.379503 | 0.43207 |
| 1 | 27547600 | 27549199 | 13.13512 | 24.53755 | 19.83934 | 8.232528 | 0.759869 | 0.14286 | 0.432073 |
| 12 | 105273200 | 105280599 | 88.11169 | 135.8516 | 206.1427 | 85.62815 | -0.30221 | 2.901039 | 0.432083 |
| 9 | 88017600 | 88018199 | 6.586843 | 12.60549 | 26.20509 | 10.36407 | -0.53179 | -0.06312 | 0.432091 |
| 7 | 25989600 | 25992799 | 57.46675 | 92.11975 | 89.84839 | 37.9344 | 0.38286 | 1.987738 | 0.432092 |
| 13 | 62238600 | 62250999 | 239.2865 | 348.8238 | 332.7335 | 137.1134 | 0.296115 | 3.960299 | 0.432105 |
| 13 | 108054600 | 108056999 | 23.00864 | 39.68098 | 39.37374 | 17.27253 | 0.396055 | 0.882749 | 0.432107 |
| 19 | 53690000 | 53691799 | 13.13939 | 23.10439 | 41.88998 | 18.5489 | -0.46645 | 0.587194 | 0.432112 |

| | | | | | | | | | |
|---|---|---|---|---|---|---|---|---|---|
| 7 | 42185400 | 42194199 | 112.3513 | 174.4744 | 151.2997 | 62.82701 | 0.512153 | 2.776764 | 0.432128 |
| 8 | 56644000 | 56645599 | 17.0026 | 31.18538 | 25.46338 | 10.85069 | 0.71757 | 0.291547 | 0.432138 |
| 7 | 7782000 | 7783799 | 30.38697 | 51.09501 | 43.76383 | 19.1195 | 0.603665 | 1.139925 | 0.432143 |
| 13 | 60168800 | 60171599 | 17.89825 | 31.57446 | 31.24063 | 13.82442 | 0.404636 | 0.561109 | 0.432158 |
| 7 | 17561000 | 17563799 | 47.257 | 76.96527 | 64.10344 | 27.52496 | 0.633901 | 1.571827 | 0.43217 |
| 8 | 57251200 | 57251799 | 11.93287 | 22.59669 | 22.62701 | 9.160023 | 0.483248 | 0.053711 | 0.432171 |
| X | 60966400 | 60970599 | 38.03934 | 63.70735 | 47.51494 | 20.71261 | 0.802759 | 1.209043 | 0.432177 |
| X | 148485400 | 148489599 | 74.41219 | 118.3327 | 107.266 | 44.7772 | 0.475761 | 2.257777 | 0.432197 |
| 8 | 15546200 | 15551999 | 47.15911 | 76.14064 | 75.01325 | 31.99715 | 0.380686 | 1.748826 | 0.432206 |
| 4 | 7299600 | 7303199 | 57.72035 | 92.29483 | 91.91241 | 38.6632 | 0.348818 | 2.066401 | 0.43221 |
| 5 | 148311400 | 148317399 | 73.40938 | 113.9005 | 184.9753 | 76.82077 | -0.393 | 2.777976 | 0.432219 |
| 3 | 111688200 | 111688999 | 12.40875 | 23.175 | 22.57488 | 9.37151 | 0.493033 | 0.147053 | 0.432221 |
| 7 | 9876400 | 9884599 | 121.9205 | 189.6634 | 138.6627 | 57.56461 | 0.760081 | 2.75223 | 0.432233 |
| 2 | 15800400 | 15801199 | 11.86739 | 22.8602 | 20.02849 | 7.773784 | 0.728359 | -0.09015 | 0.43224 |
| 2 | 88993200 | 88998399 | 76.65035 | 121.6315 | 113.0216 | 47.07892 | 0.437099 | 2.321609 | 0.432243 |
| 13 | 66899200 | 66904799 | 81.64975 | 129.1159 | 120.9174 | 50.24417 | 0.421521 | 2.415769 | 0.432243 |
| X | 14379800 | 14383199 | 82.78833 | 130.7934 | 121.8195 | 50.59494 | 0.428345 | 2.439928 | 0.432244 |
| 6 | 105297000 | 105299199 | 33.06731 | 54.63493 | 52.4414 | 22.65561 | 0.439494 | 1.347049 | 0.432249 |
| 6 | 54990400 | 54992399 | 18.80569 | 32.97871 | 32.39411 | 14.31715 | 0.413127 | 0.643937 | 0.432256 |
| X | 146572000 | 146573799 | 20.00746 | 35.13727 | 33.32179 | 14.74527 | 0.463743 | 0.631588 | 0.432262 |
| 4 | 143823200 | 143827199 | 73.93182 | 117.7328 | 102.3349 | 42.80065 | 0.538344 | 2.210875 | 0.432271 |
| 3 | 96989800 | 97000799 | 140.0676 | 209.8249 | 311.9081 | 128.6604 | -0.30686 | 3.492314 | 0.432274 |
| 13 | 51382000 | 51383999 | 16.63914 | 28.45964 | 52.13397 | 22.85996 | -0.48677 | 0.950295 | 0.432289 |
| 3 | 96850400 | 96851199 | 12.57509 | 24.31354 | 19.60523 | 7.504394 | 0.866371 | -0.12686 | 0.432289 |
| 9 | 123608400 | 123614399 | 73.1781 | 116.2899 | 109.8488 | 45.82364 | 0.415435 | 2.276386 | 0.432298 |
| 7 | 86587800 | 86592599 | 52.74234 | 84.85752 | 81.53802 | 34.61448 | 0.410692 | 1.864159 | 0.432309 |
| 9 | 36187400 | 36190599 | 40.50927 | 66.31019 | 61.75965 | 26.56754 | 0.478057 | 1.481212 | 0.43232 |
| 7 | 94461800 | 94463399 | 12.05121 | 21.46288 | 37.78278 | 16.639 | -0.41524 | 0.456896 | 0.432326 |
| 13 | 113241600 | 113244799 | 22.35228 | 37.35808 | 63.55842 | 27.67029 | -0.38302 | 1.195558 | 0.432336 |
| 7 | 31170600 | 31171999 | 21.82305 | 38.29054 | 33.56514 | 14.84141 | 0.576541 | 0.654651 | 0.432345 |
| 5 | 124067800 | 124069599 | 8.296655 | 15.54415 | 29.58133 | 12.21457 | -0.45989 | 0.109225 | 0.432347 |
| 3 | 9673800 | 9676399 | 15.30264 | 26.33959 | 52.16788 | 22.88372 | -0.59745 | 0.883983 | 0.432364 |
| 7 | 27968800 | 27971599 | 13.40325 | 23.38574 | 48.38568 | 21.30035 | -0.65641 | 0.722742 | 0.43237 |
| 7 | 136835000 | 136836999 | 10.55284 | 19.14639 | 34.35952 | 14.84676 | -0.42434 | 0.326285 | 0.432374 |
| 15 | 59003600 | 59006799 | 17.20919 | 29.67798 | 50.59059 | 22.19313 | -0.38126 | 0.875382 | 0.432378 |
| 1 | 172770000 | 172770399 | 7.998273 | 15.57524 | 17.32874 | 6.344498 | 0.46608 | -0.33558 | 0.432387 |
| X | 33298000 | 33299599 | 14.42613 | 26.83268 | 24.83167 | 10.38007 | 0.558084 | 0.193422 | 0.432434 |
| 5 | 140520000 | 140520999 | 22.474 | 37.25324 | 66.50932 | 28.79776 | -0.45265 | 1.292626 | 0.432435 |
| 3 | 34595800 | 34598599 | 39.20607 | 64.449 | 59.49315 | 25.63839 | 0.493273 | 1.425877 | 0.432438 |
| 13 | 72045600 | 72047999 | 33.99649 | 57.61171 | 44.04498 | 19.28788 | 0.767373 | 1.08768 | 0.432446 |
| 10 | 130415400 | 130417999 | 55.45978 | 88.90017 | 89.44169 | 37.76115 | 0.338115 | 1.985611 | 0.432447 |
| 15 | 11173000 | 11175199 | 24.92589 | 41.04903 | 70.97861 | 30.61154 | -0.40846 | 1.379228 | 0.432457 |
| 5 | 6309800 | 6312399 | 43.6219 | 70.51062 | 71.63475 | 30.60136 | 0.338364 | 1.709525 | 0.432486 |
| 13 | 61295000 | 61297799 | 49.31624 | 79.75949 | 72.47481 | 30.9822 | 0.499921 | 1.705239 | 0.432512 |
| 13 | 50027400 | 50034199 | 85.69192 | 135.0523 | 127.9692 | 53.10245 | 0.400809 | 2.495132 | 0.432533 |
| 7 | 139799800 | 139805999 | 70.60938 | 109.6261 | 183.7589 | 76.30835 | -0.43789 | 2.765478 | 0.432534 |
| 14 | 120995000 | 120996199 | 7.01347 | 13.41229 | 27.32172 | 10.82805 | -0.50624 | -0.05113 | 0.432539 |

| | | | | | | | | | |
|---|---|---|---|---|---|---|---|---|---|
| 7 | 47344000 | 47366399 | 304.415 | 439.889 | 357.1153 | 147.3105 | 0.516305 | 4.097427 | 0.43254 |
| 10 | 109054200 | 109054599 | 5.81475 | 11.28709 | 23.95049 | 8.999885 | -0.49093 | -0.25023 | 0.432544 |
| 5 | 135487000 | 135489399 | 20.30957 | 33.95387 | 61.68662 | 26.86031 | -0.47727 | 1.185681 | 0.432545 |
| 9 | 46698200 | 46700999 | 21.88153 | 36.56367 | 62.94348 | 27.39897 | -0.39991 | 1.196094 | 0.432559 |
| 10 | 73425800 | 73429199 | 27.19629 | 46.68839 | 40.75324 | 17.87499 | 0.578894 | 0.94184 | 0.432575 |
| 12 | 22193000 | 22196999 | 123.7928 | 189.8568 | 155.5683 | 64.57948 | 0.580094 | 3.012198 | 0.432579 |
| 9 | 118463400 | 118469999 | 61.54233 | 97.24424 | 150.2938 | 62.42641 | -0.30112 | 2.425721 | 0.432583 |
| 11 | 53735600 | 53740599 | 25.22131 | 41.5296 | 72.26808 | 31.16578 | -0.41777 | 1.381175 | 0.432599 |
| 3 | 50617600 | 50620199 | 19.45023 | 34.11243 | 33.29455 | 14.71637 | 0.422252 | 0.647158 | 0.432609 |
| 11 | 22741800 | 22744599 | 17.8299 | 30.8104 | 54.99662 | 24.12745 | -0.44703 | 0.864128 | 0.432618 |
| 9 | 75381200 | 75383199 | 12.89362 | 22.69879 | 40.39757 | 17.88388 | -0.43878 | 0.573675 | 0.432618 |
| 18 | 65563400 | 65564799 | 11.81351 | 20.68408 | 43.62098 | 19.25331 | -0.68347 | 0.650781 | 0.432633 |
| 13 | 22606200 | 22607999 | 18.15031 | 30.77131 | 61.25675 | 26.82861 | -0.60816 | 1.046353 | 0.43264 |
| 10 | 119393800 | 119395999 | 23.34907 | 40.17273 | 40.01157 | 17.55107 | 0.390128 | 0.908458 | 0.432649 |
| 7 | 27480400 | 27486599 | 127.5731 | 195.1195 | 188.7837 | 78.3247 | 0.337521 | 3.070081 | 0.432653 |
| 17 | 33215600 | 33220799 | 71.36368 | 113.5571 | 106.4728 | 44.48457 | 0.428075 | 2.234153 | 0.432657 |
| 17 | 19229000 | 19248999 | 347.7509 | 497.3687 | 386.5467 | 159.6933 | 0.56448 | 4.250713 | 0.43266 |
| 15 | 41145800 | 41147599 | 11.3936 | 20.39403 | 36.28763 | 15.91293 | -0.42629 | 0.417676 | 0.432677 |
| 18 | 89878000 | 89881199 | 72.28344 | 114.71 | 100.2929 | 41.77974 | 0.524885 | 2.311481 | 0.432682 |
| 1 | 102895200 | 102897199 | 20.08485 | 35.45206 | 29.67906 | 13.1319 | 0.644432 | 0.562726 | 0.432702 |
| 16 | 70763400 | 70765599 | 27.99546 | 47.61897 | 45.23648 | 19.82666 | 0.455534 | 1.035106 | 0.432705 |
| 3 | 151808400 | 151809399 | 11.62075 | 21.79492 | 21.04865 | 8.675817 | 0.51326 | 0.107976 | 0.432706 |
| X | 61581200 | 61582999 | 24.76021 | 42.89196 | 39.29816 | 17.24171 | 0.512045 | 0.854302 | 0.432713 |
| 4 | 66066400 | 66067999 | 17.44144 | 30.12185 | 53.82799 | 23.61435 | -0.44862 | 0.868944 | 0.432737 |
| 11 | 13711000 | 13713399 | 21.2393 | 35.93664 | 65.91683 | 28.85216 | -0.49083 | 1.079274 | 0.432759 |
| 6 | 11504000 | 11504599 | 6.259018 | 12.10175 | 24.88763 | 9.525865 | -0.47169 | -0.17941 | 0.432764 |
| 6 | 99411800 | 99412599 | 6.259018 | 12.10175 | 24.88763 | 9.525865 | -0.47169 | -0.17941 | 0.432764 |
| 4 | 112263200 | 112272199 | 133.9204 | 206.1382 | 164.4658 | 68.30707 | 0.622519 | 2.946452 | 0.432779 |
| 3 | 38296000 | 38297799 | 19.61978 | 35.27357 | 27.74157 | 12.1202 | 0.748153 | 0.427594 | 0.43278 |
| 11 | 73637800 | 73642599 | 55.09336 | 89.04969 | 70.23132 | 30.00034 | 0.702801 | 1.730685 | 0.432799 |
| 11 | 5878800 | 5888599 | 147.5523 | 219.9296 | 323.3162 | 133.2367 | -0.29742 | 3.587533 | 0.432816 |
| 7 | 140836400 | 140837399 | 6.165088 | 11.75395 | 26.65448 | 10.6368 | -0.66716 | -0.03433 | 0.432832 |
| 18 | 58991000 | 58993599 | 24.44094 | 42.23172 | 40.34459 | 17.69945 | 0.451178 | 0.874703 | 0.432837 |
| 7 | 108043200 | 108045399 | 37.19317 | 61.15058 | 59.36725 | 25.58599 | 0.420822 | 1.418279 | 0.432845 |
| X | 16556200 | 16559199 | 62.42223 | 99.91534 | 91.63193 | 38.60112 | 0.469077 | 2.04141 | 0.432845 |
| X | 91280600 | 91283999 | 41.54486 | 67.81703 | 64.33438 | 27.64715 | 0.448903 | 1.526893 | 0.432846 |
| 1 | 23777600 | 23781399 | 57.17124 | 91.85506 | 84.59318 | 35.83898 | 0.469379 | 1.916246 | 0.432852 |
| 8 | 3121400 | 3122599 | 19.89119 | 34.69864 | 33.1351 | 14.58485 | 0.45358 | 0.731902 | 0.432853 |
| 14 | 121279800 | 121282999 | 25.98565 | 42.8827 | 73.1667 | 31.58303 | -0.38822 | 1.355927 | 0.432859 |
| X | 17245200 | 17248999 | 36.79216 | 61.61096 | 48.63169 | 21.19776 | 0.721289 | 1.208299 | 0.432865 |
| 1 | 113582600 | 113584599 | 18.55054 | 33.77125 | 26.43786 | 11.37545 | 0.768924 | 0.347962 | 0.432866 |
| 2 | 68484000 | 68484799 | 9.512379 | 17.15272 | 34.35883 | 14.96122 | -0.58901 | 0.354375 | 0.43289 |
| 15 | 24604200 | 24606599 | 19.08416 | 32.46998 | 55.88437 | 24.49405 | -0.39848 | 1.013258 | 0.432902 |
| 7 | 43219200 | 43224399 | 39.71272 | 63.91649 | 104.0067 | 44.17532 | -0.34711 | 1.845236 | 0.432907 |
| 14 | 6556000 | 6559599 | 36.59524 | 58.94418 | 98.20137 | 41.756 | -0.3796 | 1.813613 | 0.432932 |
| X | 77074400 | 77077399 | 36.45453 | 60.59311 | 53.13688 | 23.05115 | 0.570015 | 1.289462 | 0.432933 |
| 15 | 85880200 | 85882399 | 10.53318 | 18.6418 | 37.88085 | 16.75199 | -0.62577 | 0.494627 | 0.432935 |

| | | | | | | | | | |
|---|---|---|---|---|---|---|---|---|---|
| 15 | 57570800 | 57571999 | 7.647377 | 14.47792 | 28.16508 | 11.40788 | -0.46808 | 0.029874 | 0.432943 |
| 2 | 73838600 | 73841599 | 34.13333 | 55.18381 | 91.20615 | 38.9716 | -0.36179 | 1.697294 | 0.432949 |
| 6 | 131767000 | 131772399 | 94.33928 | 149.0042 | 116.7552 | 48.48533 | 0.677039 | 2.452522 | 0.432952 |
| 4 | 4917200 | 4918999 | 17.28009 | 30.94528 | 28.42129 | 12.45837 | 0.522221 | 0.435096 | 0.432954 |
| 13 | 61952000 | 61952399 | 13.01389 | 23.97084 | 21.95022 | 9.299856 | 0.559264 | 0.228653 | 0.432981 |
| 14 | 41322400 | 41323399 | 28.20248 | 47.35824 | 46.99225 | 20.51906 | 0.392179 | 1.149157 | 0.433012 |
| 3 | 66305800 | 66306799 | 9.216043 | 17.11764 | 31.56013 | 13.22044 | -0.43158 | 0.17542 | 0.433012 |
| 11 | 104534200 | 104538599 | 24.15661 | 39.70553 | 71.84172 | 30.95853 | -0.4748 | 1.402142 | 0.433013 |
| X | 78109000 | 78111199 | 25.86246 | 45.16998 | 34.56814 | 15.22726 | 0.771723 | 0.749101 | 0.433016 |
| 13 | 65660600 | 65663199 | 52.20959 | 84.69706 | 66.07642 | 28.31059 | 0.723461 | 1.648268 | 0.433042 |
| 16 | 62882600 | 62883999 | 14.83007 | 27.05495 | 26.18084 | 11.23528 | 0.467106 | 0.305886 | 0.433043 |
| 3 | 6222600 | 6233399 | 177.1056 | 264.5745 | 257.8831 | 106.3181 | 0.298021 | 3.547661 | 0.433044 |
| 19 | 54956400 | 54958199 | 14.74779 | 27.02167 | 26.26554 | 11.20833 | 0.466327 | 0.282078 | 0.433054 |
| 6 | 134011800 | 134013199 | 10.1601 | 18.19508 | 35.34501 | 15.49314 | -0.55151 | 0.404343 | 0.433078 |
| 17 | 12619800 | 12630399 | 85.6256 | 131.7229 | 209.0397 | 86.81286 | -0.36927 | 2.94311 | 0.433092 |
| 10 | 41827000 | 41831599 | 47.82125 | 76.18146 | 123.1758 | 51.54377 | -0.35502 | 2.180836 | 0.433113 |
| 8 | 17385600 | 17390999 | 86.75036 | 136.0351 | 136.93 | 56.79183 | 0.308179 | 2.594363 | 0.433132 |
| 17 | 45654000 | 45656199 | 42.48586 | 68.78959 | 70.00607 | 29.95153 | 0.33831 | 1.668712 | 0.433152 |
| 3 | 7218200 | 7219399 | 16.63365 | 30.03958 | 28.12234 | 12.22226 | 0.503131 | 0.382004 | 0.43316 |
| 19 | 17206000 | 17207199 | 16.63365 | 30.03958 | 28.12234 | 12.22226 | 0.503131 | 0.382004 | 0.43316 |
| 7 | 106759400 | 106760399 | 7.173251 | 13.68572 | 27.24924 | 10.8579 | -0.48044 | -0.03118 | 0.43317 |
| 12 | 52039800 | 52042399 | 16.53165 | 28.63932 | 48.99694 | 21.497 | -0.38508 | 0.828926 | 0.433174 |
| 2 | 36401800 | 36405199 | 49.99079 | 80.55262 | 79.03593 | 33.62614 | 0.383301 | 1.812807 | 0.433203 |
| 5 | 73879200 | 73879999 | 7.526838 | 14.1578 | 28.57822 | 11.72152 | -0.53632 | 0.073737 | 0.433213 |
| 1 | 108524800 | 108527999 | 37.28391 | 60.17003 | 99.52935 | 42.43035 | -0.366 | 1.759272 | 0.433217 |
| 14 | 52792400 | 52795599 | 42.38045 | 68.02128 | 109.1671 | 46.17861 | -0.33214 | 1.938275 | 0.433227 |
| 6 | 75431600 | 75434399 | 23.21477 | 40.4416 | 36.96747 | 16.25789 | 0.516177 | 0.773142 | 0.433232 |
| 17 | 12433600 | 12435999 | 14.64079 | 25.39288 | 47.40298 | 20.79839 | -0.50988 | 0.807734 | 0.433261 |
| X | 53095800 | 53098199 | 28.23222 | 48.21736 | 43.59464 | 19.11333 | 0.527503 | 0.998843 | 0.433262 |
| 2 | 7456800 | 7457399 | 8.936806 | 16.4936 | 31.4665 | 13.29596 | -0.48949 | 0.208975 | 0.433275 |
| X | 31416800 | 31430799 | 808.7977 | 1074.112 | 788.1968 | 329.4018 | 0.539748 | 5.392294 | 0.433303 |
| 12 | 16455400 | 16459199 | 48.90065 | 79.14427 | 71.48466 | 30.57513 | 0.509511 | 1.688692 | 0.433328 |
| 2 | 71353200 | 71354199 | 6.27433 | 12.07733 | 25.83639 | 10.04935 | -0.55042 | -0.12397 | 0.433328 |
| 5 | 19481400 | 19481999 | 6.27433 | 12.07733 | 25.83639 | 10.04935 | -0.55042 | -0.12397 | 0.433328 |
| 12 | 87518400 | 87519199 | 6.27433 | 12.07733 | 25.83639 | 10.04935 | -0.55042 | -0.12397 | 0.433328 |
| 12 | 114495000 | 114502999 | 101.0201 | 159.0952 | 120.0881 | 49.8274 | 0.727675 | 2.518455 | 0.433334 |
| X | 17816000 | 17817599 | 22.67083 | 39.84002 | 28.07042 | 12.42816 | 0.890217 | 0.628556 | 0.433344 |
| 14 | 15109800 | 15112399 | 15.12757 | 27.39768 | 27.26944 | 11.80939 | 0.418569 | 0.353107 | 0.433366 |
| 15 | 74239600 | 74241599 | 22.11518 | 36.73158 | 65.51563 | 28.40656 | -0.45109 | 1.266401 | 0.433372 |
| 19 | 39523000 | 39523399 | 11.49052 | 21.60663 | 20.59312 | 8.450876 | 0.537344 | 0.089745 | 0.433376 |
| 12 | 19253000 | 19284399 | 1020.172 | 1315.388 | 1095.68 | 459.3677 | 0.314397 | 5.869531 | 0.43338 |
| 5 | 116004000 | 116008399 | 28.63621 | 46.51512 | 81.44435 | 34.92825 | -0.43816 | 1.585846 | 0.433399 |
| 5 | 118309200 | 118314999 | 50.85418 | 80.18628 | 163.0994 | 67.68445 | -0.69917 | 2.46874 | 0.433407 |
| 5 | 21259600 | 21261599 | 15.1368 | 26.1649 | 48.45735 | 21.25882 | -0.4995 | 0.842187 | 0.433414 |
| 6 | 10438400 | 10440199 | 16.88257 | 30.18784 | 29.06988 | 12.75889 | 0.453193 | 0.443879 | 0.433422 |
| 5 | 110221600 | 110226999 | 147.7482 | 225.304 | 191.8057 | 79.52369 | 0.518602 | 3.136324 | 0.433424 |
| 7 | 128592200 | 128596199 | 44.90368 | 72.54033 | 73.51708 | 31.38193 | 0.340908 | 1.728733 | 0.433425 |

| | | | | | | | | | |
|---|---|---|---|---|---|---|---|---|---|
| 7 | 9950400 | 9953999 | 51.98791 | 84.41714 | 64.4458 | 27.63119 | 0.755886 | 1.627957 | 0.433435 |
| 19 | 11430400 | 11431799 | 13.78123 | 25.72724 | 23.76942 | 9.874579 | 0.567265 | 0.161284 | 0.433452 |
| 7 | 28190000 | 28193399 | 23.00691 | 38.40681 | 64.69891 | 28.14583 | -0.36879 | 1.21132 | 0.433468 |
| X | 67207600 | 67210399 | 14.95073 | 27.95081 | 21.18804 | 8.780704 | 0.853932 | 0.15838 | 0.433478 |
| 15 | 65496400 | 65497599 | 13.37429 | 25.20801 | 23.50923 | 9.609255 | 0.572928 | 0.099711 | 0.433479 |
| 8 | 102605200 | 102608399 | 40.32772 | 66.64123 | 55.60275 | 24.02395 | 0.641015 | 1.365792 | 0.433479 |
| 12 | 85452600 | 85459399 | 96.33059 | 149.4376 | 147.3247 | 61.18603 | 0.326101 | 2.786503 | 0.433489 |
| 1 | 37683200 | 37688199 | 41.1024 | 65.77211 | 110.6219 | 46.58293 | -0.40467 | 2.030434 | 0.4335 |
| 16 | 87580600 | 87583399 | 18.9902 | 32.29336 | 57.03656 | 24.99801 | -0.43595 | 1.01529 | 0.433513 |
| 8 | 18186400 | 18188599 | 24.66025 | 42.4403 | 41.47571 | 18.19346 | 0.417424 | 0.913035 | 0.433527 |
| 7 | 17540600 | 17567799 | 488.4193 | 676.9806 | 523.1661 | 217.2291 | 0.527149 | 4.71822 | 0.433533 |
| 17 | 16903200 | 16906399 | 34.32529 | 57.32836 | 50.89902 | 22.1491 | 0.552139 | 1.226553 | 0.433542 |
| 16 | 20419000 | 20421599 | 16.81185 | 28.65365 | 53.71208 | 23.54524 | -0.52074 | 0.984418 | 0.433576 |
| X | 81592000 | 81593999 | 22.69657 | 39.65285 | 35.49276 | 15.63759 | 0.546934 | 0.731993 | 0.43358 |
| 18 | 14687200 | 14688799 | 12.44121 | 21.77593 | 46.75057 | 20.6289 | -0.70915 | 0.665712 | 0.433582 |
| Y | 2502000 | 2503999 | 14.27714 | 26.2072 | 16.51915 | 7.031711 | 1.088941 | 0.24237 | 0.433585 |
| 5 | 17519200 | 17523199 | 60.51213 | 96.74927 | 95.06635 | 39.98429 | 0.367923 | 2.07119 | 0.43359 |
| 2 | 94626800 | 94629199 | 14.6557 | 25.72809 | 49.928 | 22.08619 | -0.56446 | 0.63458 | 0.433609 |
| 10 | 79161800 | 79162599 | 12.89422 | 24.75488 | 21.11635 | 8.275461 | 0.754366 | -0.04467 | 0.433636 |
| 3 | 13947200 | 13951199 | 58.68485 | 94.02241 | 82.67977 | 34.87958 | 0.531117 | 2.002519 | 0.433654 |
| 2 | 110074200 | 110076399 | 25.86283 | 45.08385 | 35.1164 | 15.45009 | 0.746074 | 0.774765 | 0.433671 |
| 5 | 40951400 | 40953199 | 19.05664 | 33.70824 | 27.6066 | 12.20977 | 0.677441 | 0.533487 | 0.433683 |
| 19 | 31801800 | 31803599 | 19.05664 | 33.70824 | 27.6066 | 12.20977 | 0.677441 | 0.533487 | 0.433683 |
| 4 | 99184000 | 99185399 | 9.48417 | 17.17438 | 33.86602 | 14.66878 | -0.56156 | 0.327982 | 0.433693 |
| 13 | 62494600 | 62507199 | 239.8921 | 353.6945 | 293.3283 | 120.7963 | 0.513446 | 3.783893 | 0.433693 |
| 16 | 33439800 | 33440399 | 8.949328 | 17.31324 | 18.1874 | 6.942923 | 0.491761 | -0.18068 | 0.433718 |
| 17 | 18623000 | 18628399 | 115.1457 | 178.0134 | 162.252 | 67.39577 | 0.435406 | 2.857809 | 0.433726 |
| 7 | 41451400 | 41454199 | 36.16298 | 60.81198 | 46.32995 | 20.24107 | 0.772092 | 1.16171 | 0.433751 |
| 6 | 117014600 | 117015799 | 15.99633 | 28.5974 | 28.3013 | 12.42367 | 0.414014 | 0.439006 | 0.433762 |
| 7 | 102558200 | 102558999 | 8.679104 | 16.01936 | 31.49565 | 13.30672 | -0.53195 | 0.209478 | 0.433764 |
| 6 | 122530600 | 122531199 | 3.701989 | 7.230254 | 19.29679 | 6.767516 | -0.72432 | -0.53875 | 0.433769 |
| 6 | 37013600 | 37015799 | 15.84677 | 27.53406 | 48.45409 | 21.27462 | -0.42494 | 0.791363 | 0.43377 |
| 17 | 33453000 | 33453799 | 18.69378 | 33.38737 | 29.50119 | 12.96098 | 0.575811 | 0.46563 | 0.433794 |
| 1 | 10037000 | 10039399 | 14.58416 | 25.07319 | 54.78283 | 24.02992 | -0.73868 | 0.897461 | 0.433796 |
| 13 | 58130000 | 58134999 | 37.16352 | 59.63732 | 104.0926 | 44.02002 | -0.45323 | 1.934521 | 0.4338 |
| 14 | 42043200 | 42046999 | 43.8579 | 71.40778 | 59.92525 | 25.72809 | 0.622657 | 1.563903 | 0.433801 |
| 13 | 24574000 | 24575399 | 10.41506 | 18.82526 | 34.33154 | 14.90482 | -0.45178 | 0.347325 | 0.433808 |
| Y | 4347600 | 4351199 | 30.40315 | 51.08032 | 29.16361 | 12.7378 | 1.18616 | 1.126199 | 0.43381 |
| 10 | 7349000 | 7352399 | 32.98145 | 55.78192 | 45.74815 | 20.02179 | 0.666362 | 1.102009 | 0.433814 |
| 14 | 33699400 | 33702999 | 26.85891 | 44.10086 | 74.55532 | 32.1136 | -0.37785 | 1.422199 | 0.433818 |
| 10 | 34621800 | 34623999 | 14.94203 | 27.60925 | 22.26265 | 9.391857 | 0.74613 | 0.229271 | 0.433834 |
| 6 | 25453400 | 25455399 | 19.6559 | 34.75766 | 30.88252 | 13.65946 | 0.56022 | 0.545018 | 0.433846 |
| X | 76377200 | 76379399 | 38.78141 | 63.56439 | 60.82131 | 26.17828 | 0.440059 | 1.464639 | 0.43386 |
| 6 | 137854600 | 137856599 | 20.83799 | 36.66515 | 32.47775 | 14.37933 | 0.562004 | 0.60831 | 0.433867 |
| 14 | 108574200 | 108579599 | 64.95901 | 102.2239 | 159.9026 | 66.3407 | -0.32309 | 2.518071 | 0.43387 |
| 14 | 32593600 | 32594999 | 9.257363 | 16.94905 | 33.44743 | 14.2877 | -0.54967 | 0.259884 | 0.433873 |
| 2 | 152016400 | 152017399 | 16.56711 | 28.59367 | 49.5205 | 21.72423 | -0.40372 | 0.864043 | 0.433874 |

| | | | | | | | | | |
|---|---|---|---|---|---|---|---|---|---|
| 8 | 42885800 | 42889199 | 25.04448 | 43.09387 | 41.41625 | 18.16728 | 0.441318 | 0.914818 | 0.433885 |
| 8 | 126978600 | 126980199 | 13.15996 | 23.14375 | 40.87232 | 18.09779 | -0.4285 | 0.585255 | 0.433894 |
| 13 | 40809600 | 40810799 | 8.263171 | 15.51817 | 29.62751 | 12.18461 | -0.46014 | 0.09219 | 0.433912 |
| 2 | 31754800 | 31762399 | 54.15702 | 85.60643 | 144.0787 | 59.85616 | -0.42344 | 2.409854 | 0.433914 |
| 6 | 130994800 | 131010399 | 195.3506 | 292.8301 | 239.5665 | 98.83237 | 0.555012 | 3.487452 | 0.433921 |
| 1 | 164625400 | 164626799 | 9.540781 | 17.53921 | 32.97985 | 14.01014 | -0.47464 | 0.238342 | 0.433927 |
| 6 | 68164000 | 68172999 | 130.8041 | 200.345 | 184.283 | 76.48249 | 0.412002 | 3.04241 | 0.433939 |
| 5 | 43976000 | 43976799 | 6.290178 | 12.06399 | 25.70731 | 10.10385 | -0.55838 | -0.08758 | 0.433945 |
| 4 | 53260600 | 53263599 | 25.1756 | 42.88651 | 42.62313 | 18.68351 | 0.390839 | 1.006011 | 0.433965 |
| 13 | 17647400 | 17651199 | 36.46244 | 61.19553 | 47.48013 | 20.72504 | 0.74593 | 1.183214 | 0.43397 |
| 7 | 32183000 | 32192999 | 15.89357 | 29.0393 | 23.6421 | 10.1265 | 0.716769 | 0.301347 | 0.433976 |
| 6 | 47919000 | 47920599 | 9.431494 | 17.26787 | 32.41066 | 13.84482 | -0.47754 | 0.260826 | 0.433979 |
| 15 | 58027200 | 58029199 | 14.92224 | 27.28506 | 25.6059 | 10.95735 | 0.513578 | 0.291846 | 0.433995 |
| 8 | 112310200 | 112312199 | 22.07114 | 38.30554 | 38.28931 | 16.80005 | 0.387339 | 0.821768 | 0.43402 |
| 10 | 43499200 | 43500199 | 8.095576 | 15.24197 | 29.68582 | 12.15636 | -0.4838 | 0.073873 | 0.434033 |
| 12 | 21743000 | 21780199 | 993.8476 | 1285.334 | 1082.87 | 453.8644 | 0.3024 | 5.820172 | 0.434035 |
| 6 | 91449400 | 91451999 | 15.59764 | 26.5426 | 54.717 | 23.98273 | -0.65762 | 0.995321 | 0.434036 |
| X | 65858600 | 65860599 | 23.66881 | 42.01537 | 27.27872 | 12.04645 | 1.012814 | 0.530186 | 0.434037 |
| 12 | 35981600 | 35982999 | 12.58284 | 22.25934 | 39.84395 | 17.62171 | -0.44422 | 0.517977 | 0.434071 |
| 1 | 46688600 | 46689999 | 21.48037 | 37.40859 | 37.1647 | 16.34145 | 0.3966 | 0.77013 | 0.434072 |
| 17 | 88282000 | 88283199 | 7.459052 | 13.80698 | 30.50647 | 12.83863 | -0.69489 | 0.179256 | 0.434073 |
| 7 | 39120400 | 39127399 | 118.7655 | 183.1501 | 167.7042 | 69.66137 | 0.426019 | 2.906083 | 0.434084 |
| 3 | 122412200 | 122414199 | 13.55017 | 23.56751 | 45.81434 | 20.12838 | -0.56735 | 0.764202 | 0.43409 |
| 16 | 15692000 | 15693399 | 13.89065 | 25.80137 | 24.84304 | 10.40599 | 0.498993 | 0.196513 | 0.434094 |
| 14 | 90107800 | 90109599 | 21.09679 | 36.68673 | 36.73393 | 16.13715 | 0.385154 | 0.787693 | 0.434098 |
| 13 | 29420400 | 29423199 | 22.8632 | 39.75362 | 38.21324 | 16.78573 | 0.443818 | 0.801197 | 0.434117 |
| 15 | 19130800 | 19131399 | 8.318979 | 16.17742 | 16.62143 | 6.154938 | 0.563764 | -0.29788 | 0.43412 |
| 16 | 22606800 | 22609199 | 20.93999 | 35.01408 | 61.74103 | 26.88703 | -0.43437 | 1.185754 | 0.434123 |
| 3 | 3295200 | 3302599 | 106.7444 | 165.8065 | 153.6302 | 63.79249 | 0.416951 | 2.770768 | 0.434143 |
| 6 | 66024400 | 66025799 | 9.932252 | 18.19537 | 33.46779 | 14.28497 | -0.44763 | 0.265331 | 0.434158 |
| 3 | 103767000 | 103768599 | 11.96185 | 22.4506 | 23.01179 | 9.473593 | 0.42924 | 0.111372 | 0.434183 |
| 2 | 90118800 | 90120999 | 19.82355 | 34.87661 | 32.57741 | 14.42374 | 0.485742 | 0.603411 | 0.434188 |
| 8 | 87528400 | 87530599 | 16.37895 | 28.50968 | 49.5104 | 21.76286 | -0.4056 | 0.771405 | 0.43419 |
| 5 | 68807800 | 68809999 | 11.7442 | 21.03067 | 37.84083 | 16.588 | -0.44195 | 0.421369 | 0.434206 |
| 5 | 132600800 | 132602999 | 27.03583 | 46.27235 | 43.01422 | 18.86208 | 0.487916 | 0.972304 | 0.434218 |
| 15 | 68846400 | 68847999 | 11.03342 | 19.79836 | 36.279 | 15.87592 | -0.46656 | 0.397648 | 0.434232 |
| 12 | 69984800 | 69986399 | 10.15054 | 18.65858 | 35.70471 | 15.16943 | -0.50016 | 0.248697 | 0.434234 |
| X | 77268200 | 77275599 | 122.4635 | 188.7306 | 167.3211 | 69.50248 | 0.471867 | 2.919842 | 0.43424 |
| 10 | 82187800 | 82196599 | 141.3241 | 214.2291 | 209.7896 | 86.82812 | 0.309762 | 3.254001 | 0.434265 |
| 3 | 3480800 | 3484599 | 40.02739 | 67.32837 | 41.77976 | 18.25461 | 1.066992 | 1.168473 | 0.434267 |
| 1 | 121309800 | 121311399 | 15.09641 | 27.70527 | 26.25305 | 11.17833 | 0.505124 | 0.276011 | 0.434274 |
| 11 | 19678400 | 19679799 | 15.09641 | 27.70527 | 26.25305 | 11.17833 | 0.505124 | 0.276011 | 0.434274 |
| 15 | 20341800 | 20342399 | 7.151478 | 13.76771 | 27.63632 | 10.74504 | -0.45915 | -0.11068 | 0.434279 |
| 10 | 85285800 | 85289999 | 27.72725 | 45.37966 | 76.00271 | 32.69259 | -0.3663 | 1.46699 | 0.43428 |
| 5 | 125341400 | 125345799 | 32.77874 | 52.7659 | 96.7237 | 41.09265 | -0.51812 | 1.8271 | 0.434294 |
| 11 | 105017000 | 105018599 | 11.41208 | 20.2181 | 38.78506 | 17.14833 | -0.54295 | 0.488621 | 0.434322 |
| 7 | 17832400 | 17844799 | 185.9286 | 278.5119 | 246.1133 | 101.5195 | 0.44295 | 3.499716 | 0.434323 |

| | | | | | | | | | |
|---|---|---|---|---|---|---|---|---|---|
| X | 125890400 | 125896399 | 82.00335 | 130.5245 | 98.91932 | 41.345 | 0.735112 | 2.234838 | 0.434364 |
| 17 | 23581200 | 23584999 | 48.53751 | 78.74452 | 61.7419 | 26.45457 | 0.716243 | 1.636194 | 0.434385 |
| 16 | 63683600 | 63685199 | 8.159398 | 15.2889 | 29.58518 | 12.21358 | -0.48333 | 0.10817 | 0.434396 |
| 10 | 40039400 | 40042999 | 28.33689 | 46.3443 | 79.15027 | 34.03765 | -0.39524 | 1.47845 | 0.434411 |
| 11 | 29604400 | 29606599 | 32.44573 | 53.9762 | 53.61679 | 23.27313 | 0.390775 | 1.265992 | 0.434412 |
| 18 | 68065800 | 68067799 | 11.99231 | 20.94551 | 44.20576 | 19.47287 | -0.68458 | 0.693087 | 0.434417 |
| 5 | 97966200 | 97967999 | 23.52454 | 41.23933 | 33.47927 | 14.79233 | 0.686813 | 0.678167 | 0.434431 |
| 4 | 37969200 | 37975199 | 58.19041 | 93.24478 | 90.12539 | 38.02963 | 0.395256 | 1.999065 | 0.434442 |
| 10 | 36998400 | 36998999 | 8.523899 | 16.58359 | 17.76915 | 6.555404 | 0.508971 | -0.30357 | 0.434457 |
| 13 | 67191600 | 67192399 | 18.87894 | 33.7088 | 29.49653 | 12.96193 | 0.58958 | 0.468701 | 0.43446 |
| 8 | 70110800 | 70113199 | 36.62946 | 60.9592 | 52.63255 | 22.85187 | 0.592454 | 1.273748 | 0.43446 |
| X | 25534200 | 25536599 | 31.57411 | 54.03183 | 41.26681 | 18.09562 | 0.770388 | 0.9918 | 0.434462 |
| X | 148407800 | 148408799 | 18.77439 | 33.93527 | 27.47017 | 11.92578 | 0.712832 | 0.392075 | 0.434484 |
| 7 | 60332000 | 60332799 | 19.75561 | 34.55313 | 31.23089 | 13.77227 | 0.532943 | 0.692005 | 0.434484 |
| 9 | 81167400 | 81172399 | 61.82831 | 98.72875 | 97.80395 | 41.06211 | 0.354225 | 2.111624 | 0.434486 |
| 16 | 80376000 | 80376999 | 12.5569 | 23.293 | 22.92557 | 9.620203 | 0.465718 | 0.188075 | 0.434501 |
| 13 | 105430400 | 105432599 | 12.66402 | 22.36577 | 39.80256 | 17.60925 | -0.43712 | 0.533443 | 0.434518 |
| 16 | 46537800 | 46538799 | 17.53972 | 31.10244 | 30.22104 | 13.35773 | 0.433647 | 0.503816 | 0.434522 |
| 11 | 73519000 | 73522399 | 69.08212 | 109.7197 | 110.4802 | 46.06139 | 0.32263 | 2.289461 | 0.434531 |
| 10 | 38700200 | 38703599 | 61.17604 | 97.66254 | 98.07065 | 41.15937 | 0.334271 | 2.118736 | 0.434533 |
| 5 | 7989800 | 7995799 | 97.99578 | 151.964 | 153.0002 | 63.54669 | 0.295369 | 2.794218 | 0.434539 |
| X | 164229600 | 164231599 | 27.26516 | 46.52389 | 43.65219 | 19.13717 | 0.474059 | 1.003642 | 0.43455 |
| 3 | 153663000 | 153664799 | 28.63276 | 48.27079 | 48.00798 | 20.99024 | 0.388972 | 1.115965 | 0.434578 |
| X | 162351200 | 162351599 | 9.434118 | 18.4548 | 11.48826 | 3.799028 | 1.422051 | -0.67824 | 0.43459 |
| 2 | 36990400 | 36996999 | 85.73932 | 134.8069 | 132.6066 | 55.00981 | 0.34428 | 2.542573 | 0.434595 |
| 16 | 84969400 | 84971999 | 22.9392 | 37.998 | 67.15938 | 29.06531 | -0.43832 | 1.301281 | 0.434599 |
| 5 | 138979800 | 138983399 | 24.11283 | 39.34941 | 77.99301 | 33.51314 | -0.61151 | 1.501823 | 0.4346 |
| 7 | 84696400 | 84701799 | 74.03597 | 117.6628 | 109.4221 | 45.64943 | 0.438067 | 2.274948 | 0.434604 |
| 7 | 129550400 | 129553199 | 25.98714 | 44.60267 | 41.92699 | 18.38948 | 0.47244 | 0.941225 | 0.434621 |
| 1 | 136430200 | 136431199 | 15.45874 | 27.86532 | 27.74096 | 12.08488 | 0.412783 | 0.383387 | 0.434623 |
| 14 | 18289800 | 18290399 | 4.102754 | 8.015314 | 19.38132 | 6.763058 | -0.57782 | -0.55074 | 0.434636 |
| 7 | 60152600 | 60156999 | 51.70679 | 83.67173 | 70.07433 | 29.96816 | 0.618037 | 1.69635 | 0.434639 |
| 5 | 150324800 | 150331599 | 47.89774 | 76.48321 | 123.1521 | 51.70344 | -0.34541 | 2.10499 | 0.434643 |
| 3 | 23234800 | 23238599 | 49.14265 | 79.10359 | 79.75918 | 33.88269 | 0.342205 | 1.84371 | 0.434647 |
| 3 | 123301800 | 123302599 | 5.274107 | 10.19475 | 24.2566 | 9.295751 | -0.68113 | -0.18796 | 0.434662 |
| 4 | 138590200 | 138591199 | 5.328648 | 10.21075 | 27.31311 | 10.7625 | -0.88728 | -0.08732 | 0.434678 |
| 6 | 70289400 | 70292199 | 28.65758 | 49.29187 | 40.75266 | 17.87599 | 0.657584 | 0.934 | 0.434692 |
| 2 | 83192600 | 83193199 | 10.58872 | 18.98294 | 35.7941 | 15.67586 | -0.50822 | 0.39987 | 0.434704 |
| 5 | 148597200 | 148604199 | 64.71761 | 101.8351 | 157.3304 | 65.27283 | -0.30532 | 2.517722 | 0.434713 |
| 12 | 109931400 | 109935399 | 28.9516 | 46.97944 | 83.85715 | 35.93594 | -0.46733 | 1.606573 | 0.434738 |
| 5 | 40307000 | 40309599 | 27.70868 | 47.19167 | 44.66799 | 19.57953 | 0.460956 | 1.020515 | 0.434743 |
| 3 | 99061200 | 99063199 | 19.33564 | 34.31218 | 30.45998 | 13.45502 | 0.563813 | 0.51068 | 0.434743 |
| 9 | 12665000 | 12666199 | 8.55537 | 16.05277 | 30.71445 | 12.65048 | -0.46523 | 0.102187 | 0.434744 |
| 19 | 50665200 | 50667399 | 15.72099 | 27.17943 | 50.0581 | 21.96121 | -0.4917 | 0.843365 | 0.434751 |
| 5 | 80400200 | 80401399 | 16.93336 | 29.9394 | 28.8781 | 12.77324 | 0.442477 | 0.527041 | 0.434779 |
| 16 | 95894600 | 95896599 | 16.93336 | 29.9394 | 28.8781 | 12.77324 | 0.442477 | 0.527041 | 0.434779 |
| 15 | 38432600 | 38435999 | 54.35232 | 87.05834 | 88.0002 | 37.10938 | 0.330081 | 2.00812 | 0.434782 |

| | | | | | | | | | |
|---|---|---|---|---|---|---|---|---|---|
| 5 | 21183400 | 21185199 | 14.62075 | 25.50966 | 44.70286 | 19.67893 | -0.41798 | 0.722975 | 0.434804 |
| 1 | 190352000 | 190353199 | 19.81047 | 34.88105 | 32.67819 | 14.46609 | 0.482071 | 0.596763 | 0.434805 |
| 17 | 13989200 | 13989799 | 10.27334 | 19.7848 | 20.77397 | 8.068726 | 0.469152 | -0.1007 | 0.434809 |
| 1 | 138453400 | 138454999 | 13.18469 | 24.97911 | 22.41761 | 9.066098 | 0.640751 | 0.059188 | 0.434821 |
| 3 | 105986800 | 105988999 | 28.97636 | 48.87297 | 48.00283 | 20.99097 | 0.406978 | 1.113327 | 0.434828 |
| 17 | 56173400 | 56175399 | 37.65975 | 62.42095 | 54.919 | 23.77813 | 0.564996 | 1.324088 | 0.434831 |
| 13 | 103271800 | 103274799 | 30.85992 | 51.57324 | 51.34131 | 22.35048 | 0.387621 | 1.211883 | 0.434844 |
| 16 | 24346600 | 24348199 | 24.59315 | 42.21798 | 42.01968 | 18.43041 | 0.390406 | 0.934946 | 0.434846 |
| 19 | 10621000 | 10622599 | 8.60255 | 16.00268 | 30.5197 | 12.75521 | -0.47734 | 0.158748 | 0.434849 |
| X | 19334400 | 19340399 | 72.75894 | 115.8129 | 104.9595 | 43.87101 | 0.477509 | 2.226006 | 0.434849 |
| X | 23960400 | 23960999 | 17.27332 | 31.50756 | 26.63979 | 11.43437 | 0.660881 | 0.328744 | 0.434853 |
| 13 | 9428400 | 9429799 | 16.49347 | 30.26589 | 26.15645 | 11.13885 | 0.637074 | 0.287765 | 0.434875 |
| 4 | 120494800 | 120497199 | 19.4621 | 33.20921 | 57.09906 | 25.0323 | -0.39658 | 0.986485 | 0.434875 |
| 12 | 37157400 | 37159799 | 18.26153 | 30.57163 | 61.01479 | 26.59067 | -0.61228 | 1.163951 | 0.434881 |
| 1 | 161919800 | 161920399 | 9.102441 | 17.64573 | 18.73062 | 7.081862 | 0.489752 | -0.20964 | 0.434882 |
| 8 | 59796000 | 59796599 | 10.19941 | 19.47799 | 19.64046 | 7.810171 | 0.495601 | -0.02968 | 0.434899 |
| 9 | 89963200 | 89968799 | 45.16429 | 72.35518 | 117.1165 | 49.40184 | -0.34763 | 2.001328 | 0.434899 |
| X | 126206200 | 126208599 | 25.05785 | 44.17242 | 29.45721 | 13.03714 | 0.970568 | 0.609905 | 0.434908 |
| 6 | 10848000 | 10849799 | 17.03812 | 31.48028 | 24.479 | 10.32777 | 0.797803 | 0.250169 | 0.434925 |
| 10 | 67660000 | 67662399 | 14.65702 | 25.72494 | 47.21544 | 20.88251 | -0.48431 | 0.634523 | 0.434931 |
| 7 | 48782600 | 48788399 | 39.92275 | 63.98734 | 107.2117 | 45.25534 | -0.39649 | 1.977032 | 0.434939 |
| 6 | 29306600 | 29307199 | 9.253889 | 18.00946 | 18.37333 | 6.761157 | 0.582996 | -0.29923 | 0.434952 |
| 1 | 104150000 | 104151999 | 18.8062 | 33.99466 | 27.36126 | 11.87768 | 0.720929 | 0.390627 | 0.434957 |
| 5 | 117359800 | 117362199 | 23.74388 | 39.34911 | 67.2327 | 29.10633 | -0.3896 | 1.298808 | 0.434959 |
| 7 | 85168800 | 85187599 | 429.9081 | 597.0163 | 536.203 | 222.5762 | 0.313122 | 4.687255 | 0.43496 |
| 17 | 20269200 | 20278599 | 334.1485 | 472.4658 | 428.5886 | 177.4 | 0.324881 | 4.415335 | 0.43496 |
| 14 | 23855200 | 23858199 | 18.22565 | 30.94031 | 56.97593 | 24.96384 | -0.49615 | 1.027816 | 0.434972 |
| 5 | 110257800 | 110258599 | 7.451536 | 14.23694 | 28.45478 | 11.30413 | -0.48248 | -0.03646 | 0.434999 |
| 9 | 90419800 | 90421399 | 13.81878 | 24.10855 | 44.70966 | 19.68096 | -0.49934 | 0.718915 | 0.435026 |
| 2 | 12001000 | 12004199 | 64.9181 | 103.2473 | 104.6777 | 43.70336 | 0.314678 | 2.24441 | 0.435026 |
| 14 | 77830000 | 77835199 | 37.74202 | 60.80115 | 98.4454 | 41.86959 | -0.33823 | 1.810967 | 0.435029 |
| 3 | 99516800 | 99520199 | 40.56148 | 66.36607 | 62.65947 | 26.95023 | 0.458084 | 1.491164 | 0.43506 |
| 16 | 22796600 | 22799399 | 44.75391 | 72.64025 | 62.54206 | 26.80668 | 0.58222 | 1.620214 | 0.435077 |
| 6 | 139890800 | 139891799 | 7.162062 | 13.82418 | 30.89974 | 11.92167 | -0.60273 | -0.12727 | 0.435079 |
| 18 | 84997600 | 84999399 | 16.186 | 28.1663 | 48.43657 | 21.28727 | -0.39156 | 0.772112 | 0.435079 |
| 10 | 126577800 | 126581199 | 31.69202 | 51.40552 | 84.78898 | 36.32445 | -0.35425 | 1.621254 | 0.435084 |
| 4 | 113536000 | 113543599 | 91.79649 | 145.0824 | 116.6168 | 48.44383 | 0.641059 | 2.435243 | 0.435101 |
| 13 | 71498200 | 71498799 | 6.100916 | 11.84795 | 24.61768 | 9.2167 | -0.45604 | -0.25731 | 0.435103 |
| 19 | 37432200 | 37434599 | 22.93394 | 38.14395 | 65.56578 | 28.45534 | -0.39775 | 1.254446 | 0.435104 |
| 5 | 74768000 | 74769399 | 8.130138 | 15.3399 | 29.17705 | 11.90261 | -0.44486 | 0.061592 | 0.435117 |
| 1 | 180071000 | 180073399 | 11.49201 | 20.29501 | 38.92341 | 17.22045 | -0.54391 | 0.529271 | 0.435157 |
| 10 | 117150200 | 117153199 | 19.50547 | 32.64929 | 61.12408 | 26.63582 | -0.52043 | 1.169082 | 0.435158 |
| 15 | 33575200 | 33577799 | 29.11599 | 49.43935 | 45.76047 | 20.05121 | 0.492607 | 1.05448 | 0.435167 |
| X | 53793600 | 53794999 | 21.60041 | 38.16717 | 27.32285 | 12.08684 | 0.869983 | 0.560534 | 0.435172 |
| 11 | 106862800 | 106863799 | 9.566035 | 17.16905 | 35.42536 | 15.4995 | -0.63562 | 0.389348 | 0.4352 |
| X | 148538200 | 148541199 | 21.41086 | 38.26053 | 27.49233 | 12.07298 | 0.872674 | 0.475465 | 0.435201 |
| 2 | 68112400 | 68113599 | 19.63394 | 34.40823 | 33.85247 | 14.95328 | 0.410841 | 0.661528 | 0.435203 |

| | | | | | | | | | |
|---|---|---|---|---|---|---|---|---|---|
| 11 | 22425600 | 22427199 | 24.38694 | 42.05555 | 35.17295 | 15.42969 | 0.64159 | 0.883684 | 0.43522 |
| 9 | 112796000 | 112799599 | 34.8305 | 58.26614 | 49.76702 | 21.67858 | 0.607761 | 1.210527 | 0.435223 |
| 14 | 95084000 | 95085799 | 25.25804 | 44.03781 | 32.15104 | 14.14709 | 0.839121 | 0.766141 | 0.435228 |
| 15 | 80943800 | 80945399 | 12.16407 | 21.66051 | 38.25879 | 16.85084 | -0.4204 | 0.457669 | 0.435247 |
| 4 | 150089200 | 150090799 | 14.77214 | 25.68934 | 46.32461 | 20.35082 | -0.45966 | 0.772003 | 0.435251 |
| 1 | 43207200 | 43208199 | 6.884249 | 13.14254 | 26.70015 | 10.62507 | -0.50728 | -0.03905 | 0.435265 |
| 7 | 127615800 | 127621399 | 45.3198 | 72.47825 | 114.7659 | 48.28779 | -0.31881 | 2.053998 | 0.435276 |
| 6 | 140253200 | 140258799 | 87.56903 | 137.0194 | 138.6095 | 57.50129 | 0.298559 | 2.632393 | 0.435276 |
| 10 | 56119000 | 56120599 | 14.99154 | 27.40008 | 26.16507 | 11.20261 | 0.488264 | 0.296643 | 0.435286 |
| 8 | 85938000 | 85941799 | 56.08558 | 90.62351 | 70.30452 | 30.01881 | 0.72598 | 1.741689 | 0.435302 |
| X | 121263000 | 121270999 | 90.60379 | 143.0308 | 121.2733 | 50.34853 | 0.56281 | 2.463211 | 0.435313 |
| 9 | 51818000 | 51818599 | 4.352205 | 8.495482 | 19.82443 | 7.019047 | -0.54566 | -0.49157 | 0.435316 |
| 17 | 62614200 | 62614799 | 4.055953 | 7.920752 | 19.28755 | 6.77635 | -0.59546 | -0.52427 | 0.435327 |
| 8 | 120384200 | 120385599 | 18.50361 | 31.56409 | 54.86718 | 24.05325 | -0.41228 | 0.983204 | 0.435335 |
| 8 | 19542000 | 19545799 | 60.32828 | 96.83459 | 77.74449 | 32.89006 | 0.665288 | 1.946186 | 0.435347 |
| 15 | 88469000 | 88492199 | 272.6693 | 386.0412 | 549.4528 | 227.3787 | -0.32285 | 4.394798 | 0.435347 |
| 4 | 141252600 | 141256399 | 32.67105 | 52.94916 | 86.58246 | 37.06988 | -0.34294 | 1.640885 | 0.435351 |
| Y | 6633400 | 6634599 | 11.04191 | 21.12125 | 12.97785 | 5.139756 | 1.208229 | -0.05727 | 0.43536 |
| 4 | 101339200 | 101340999 | 16.01297 | 29.84695 | 24.22179 | 10.08911 | 0.750471 | 0.192727 | 0.435381 |
| 7 | 107807000 | 107808999 | 19.51899 | 34.5888 | 30.33078 | 13.40974 | 0.58025 | 0.520866 | 0.435387 |
| 10 | 74579800 | 74583999 | 36.78824 | 61.11254 | 53.77236 | 23.31754 | 0.565076 | 1.295285 | 0.435391 |
| 10 | 77674600 | 77684399 | 116.8981 | 176.7119 | 265.4144 | 109.7542 | -0.31023 | 3.30885 | 0.4354 |
| 11 | 116938600 | 116947399 | 86.91628 | 133.7197 | 207.3586 | 86.11572 | -0.33586 | 2.942162 | 0.435439 |
| 9 | 34946000 | 34947399 | 9.426741 | 16.83928 | 35.89484 | 15.7676 | -0.68672 | 0.416181 | 0.435447 |
| 10 | 122047600 | 122048599 | 13.76219 | 25.38349 | 25.3629 | 10.72596 | 0.436125 | 0.235741 | 0.435451 |
| 10 | 124677400 | 124679599 | 16.03423 | 27.7951 | 48.46366 | 21.26344 | -0.41215 | 0.819676 | 0.435467 |
| 12 | 64799400 | 64801599 | 17.89435 | 30.52035 | 53.75412 | 23.56502 | -0.43109 | 0.981869 | 0.435469 |
| 7 | 10642400 | 10645799 | 80.04028 | 125.8925 | 112.2383 | 46.56255 | 0.486379 | 2.528237 | 0.435503 |
| 14 | 88926000 | 88929599 | 33.03302 | 55.54625 | 48.0109 | 20.97519 | 0.59058 | 1.151767 | 0.435508 |
| 14 | 32892200 | 32898599 | 43.4856 | 69.67014 | 111.198 | 46.90969 | -0.32728 | 1.994725 | 0.435522 |
| 5 | 134204400 | 134205799 | 8.86267 | 16.17101 | 32.42957 | 13.91411 | -0.57638 | 0.277714 | 0.435536 |
| 10 | 120898800 | 120900399 | 16.48507 | 28.49406 | 52.23947 | 22.91805 | -0.48533 | 0.850586 | 0.435552 |
| 2 | 74439600 | 74442399 | 30.40692 | 51.51489 | 45.19048 | 19.7879 | 0.569543 | 1.07544 | 0.435558 |
| 14 | 42441800 | 42445199 | 60.11433 | 96.28546 | 82.99649 | 34.99816 | 0.559374 | 2.012443 | 0.435585 |
| 11 | 66089600 | 66089999 | 9.393332 | 18.16888 | 17.2644 | 6.596204 | 0.635117 | -0.1727 | 0.435586 |
| 1 | 186238600 | 186241999 | 25.39217 | 41.78819 | 72.22523 | 31.14013 | -0.40829 | 1.388622 | 0.435601 |
| 1 | 6769800 | 6771199 | 17.91641 | 31.54854 | 31.33798 | 13.87259 | 0.39804 | 0.581639 | 0.435617 |
| 6 | 113639600 | 113642199 | 24.54091 | 40.54511 | 70.12231 | 30.29292 | -0.40751 | 1.337109 | 0.435631 |
| 4 | 79392000 | 79392999 | 16.09488 | 28.67462 | 28.26791 | 12.4451 | 0.416636 | 0.460051 | 0.43567 |
| 15 | 17945400 | 17946199 | 14.10162 | 25.88592 | 24.49385 | 10.42581 | 0.507952 | 0.265229 | 0.435723 |
| 7 | 25701800 | 25715599 | 235.2405 | 344.3872 | 325.3343 | 134.0162 | 0.315978 | 3.895146 | 0.435727 |
| 18 | 3205400 | 3222799 | 81.82241 | 129.892 | 98.45891 | 41.02972 | 0.731002 | 2.310391 | 0.435729 |
| 6 | 132007800 | 132009599 | 27.59359 | 48.1145 | 34.53632 | 15.19757 | 0.863599 | 0.780478 | 0.435738 |
| 3 | 112160800 | 112161599 | 9.499841 | 18.39036 | 18.631 | 7.090406 | 0.547238 | -0.1846 | 0.435744 |
| X | 116014800 | 116015799 | 13.18848 | 24.86522 | 23.51625 | 9.60726 | 0.553426 | 0.09601 | 0.435747 |
| 7 | 79826400 | 79828799 | 19.07483 | 32.35692 | 55.86857 | 24.47293 | -0.40353 | 1.039168 | 0.435763 |
| 18 | 50783000 | 50783999 | 33.24537 | 55.33888 | 43.34889 | 18.82374 | 0.732216 | 1.249797 | 0.435767 |

| | | | | | | | | | |
|---|---|---|---|---|---|---|---|---|---|
| 10 | 46413200 | 46416399 | 29.29926 | 49.9573 | 44.09125 | 19.32849 | 0.56178 | 1.017 | 0.435776 |
| 7 | 118069200 | 118078599 | 73.65902 | 114.4997 | 199.0097 | 82.62645 | -0.48878 | 2.74819 | 0.435796 |
| 1 | 50198400 | 50201399 | 51.11407 | 82.28633 | 72.34005 | 30.73604 | 0.53976 | 1.840685 | 0.4358 |
| X | 24000000 | 24005599 | 73.9359 | 118.2342 | 88.15002 | 37.08532 | 0.765861 | 2.081811 | 0.4358 |
| 16 | 64006000 | 64007799 | 19.47537 | 34.24328 | 32.62245 | 14.44544 | 0.457109 | 0.60778 | 0.435815 |
| 8 | 48066000 | 48067999 | 19.18435 | 32.64905 | 55.3866 | 24.2764 | -0.37774 | 1.00883 | 0.435816 |
| 12 | 23534600 | 23538399 | 48.73388 | 78.64726 | 67.24411 | 28.66631 | 0.583978 | 1.763113 | 0.435834 |
| 3 | 37224200 | 37225599 | 19.40137 | 33.94674 | 33.59194 | 14.81831 | 0.402752 | 0.686741 | 0.435872 |
| 6 | 130190800 | 130194399 | 43.55466 | 71.22566 | 61.91498 | 26.62487 | 0.576386 | 1.502811 | 0.435879 |
| 1 | 45204200 | 45206999 | 47.26348 | 75.15629 | 138.6518 | 57.87609 | -0.54783 | 2.243796 | 0.435879 |
| 2 | 66841800 | 66842999 | 11.40535 | 21.85158 | 21.71797 | 8.561537 | 0.52779 | -0.03921 | 0.435893 |
| 18 | 10173400 | 10175599 | 18.43237 | 31.21107 | 56.38721 | 24.68668 | -0.46877 | 1.051845 | 0.435895 |
| Y | 10821000 | 10821399 | 2.452983 | 4.797417 | 17.71958 | 6.013329 | -1.1382 | -0.67891 | 0.435906 |
| 5 | 145825200 | 145827199 | 57.09085 | 91.46896 | 90.86384 | 38.3324 | 0.355567 | 2.003235 | 0.435909 |
| 11 | 17247400 | 17248399 | 11.74361 | 20.85937 | 37.28373 | 16.45618 | -0.43926 | 0.470785 | 0.435911 |
| 12 | 80666200 | 80667599 | 12.94611 | 23.88138 | 23.4636 | 9.921003 | 0.46056 | 0.224343 | 0.435914 |
| 5 | 106275600 | 106276599 | 8.530649 | 15.85148 | 32.1098 | 13.44277 | -0.56496 | 0.172914 | 0.435925 |
| 10 | 50065600 | 50066399 | 6.095481 | 11.84226 | 27.93014 | 10.42226 | -0.6338 | -0.26334 | 0.435945 |
| 12 | 103765000 | 103803399 | 504.2014 | 680.0804 | 907.6161 | 378.469 | -0.30053 | 5.141726 | 0.435955 |
| 6 | 147204600 | 147212199 | 88.71754 | 136.4531 | 208.0214 | 86.3877 | -0.3116 | 2.948798 | 0.435967 |
| 9 | 51760000 | 51761599 | 10.78647 | 19.22734 | 36.31021 | 15.97863 | -0.51512 | 0.439326 | 0.435978 |
| 14 | 67190600 | 67192199 | 10.78647 | 19.22734 | 36.31021 | 15.97863 | -0.51512 | 0.439326 | 0.435978 |
| 2 | 104134800 | 104137599 | 14.88205 | 25.90387 | 48.43108 | 21.2883 | -0.51167 | 0.760325 | 0.435996 |
| 6 | 134846200 | 134849199 | 22.25875 | 36.98782 | 65.58135 | 28.4445 | -0.44251 | 1.260707 | 0.436001 |
| 4 | 33655600 | 33656599 | 11.03663 | 20.90513 | 20.46615 | 8.280548 | 0.515634 | 0.03491 | 0.436022 |
| X | 149148600 | 149152799 | 134.9615 | 205.8388 | 186.4521 | 77.30695 | 0.429338 | 3.130298 | 0.436048 |
| 3 | 61270400 | 61273399 | 25.52125 | 41.31959 | 94.90133 | 40.60109 | -0.83312 | 1.652846 | 0.436049 |
| 2 | 36968400 | 36975799 | 68.76933 | 109.4959 | 96.30294 | 40.26974 | 0.521143 | 2.209089 | 0.436051 |
| 13 | 72357600 | 72360199 | 13.64451 | 23.8818 | 43.07252 | 19.00576 | -0.45907 | 0.665954 | 0.436055 |
| 14 | 99962200 | 99964799 | 57.22667 | 91.89302 | 78.44798 | 33.20795 | 0.577584 | 1.93039 | 0.436067 |
| 5 | 135523000 | 135524999 | 17.7313 | 30.24695 | 53.75371 | 23.56514 | -0.44398 | 0.979584 | 0.436068 |
| 19 | 11854800 | 11857799 | 25.41414 | 41.82178 | 71.55635 | 30.85093 | -0.39374 | 1.390156 | 0.436092 |
| 7 | 85246400 | 85257199 | 144.4985 | 220.7921 | 185.7902 | 77.06547 | 0.537644 | 3.09435 | 0.436116 |
| 1 | 19519800 | 19520599 | 13.37031 | 23.43263 | 42.03625 | 18.5691 | -0.45116 | 0.643504 | 0.436159 |
| 7 | 64359200 | 64361199 | 15.72296 | 28.0878 | 27.27057 | 11.9787 | 0.440683 | 0.438333 | 0.436164 |
| 8 | 30525800 | 30527199 | 15.01599 | 25.86591 | 56.48458 | 24.77784 | -0.73752 | 0.881916 | 0.43618 |
| 4 | 60267600 | 60275799 | 190.9995 | 283.2872 | 274.8404 | 113.2033 | 0.295239 | 3.677664 | 0.436182 |
| 6 | 62331600 | 62333999 | 24.88081 | 43.50369 | 34.59988 | 15.25392 | 0.716464 | 0.727909 | 0.436184 |
| X | 5776400 | 5778999 | 26.58156 | 46.07495 | 38.55598 | 16.9164 | 0.642324 | 0.85442 | 0.436197 |
| 14 | 84665200 | 84666599 | 15.4745 | 27.83872 | 27.76482 | 12.12383 | 0.408021 | 0.395142 | 0.436198 |
| 14 | 104824600 | 104824999 | 8.038431 | 15.6628 | 14.76519 | 5.363789 | 0.713839 | -0.36973 | 0.4362 |
| 18 | 73959400 | 73960399 | 8.447981 | 15.95437 | 30.88914 | 12.5764 | -0.46797 | 0.062205 | 0.436205 |
| 7 | 106548200 | 106549399 | 15.78928 | 28.33618 | 28.23057 | 12.35244 | 0.407531 | 0.413024 | 0.436209 |
| X | 94494800 | 94496599 | 24.02858 | 42.00445 | 35.21139 | 15.52066 | 0.64095 | 0.727656 | 0.436213 |
| 6 | 20632600 | 20635799 | 38.9785 | 64.55316 | 53.91486 | 23.32999 | 0.639707 | 1.334256 | 0.436213 |
| 2 | 11557000 | 11561399 | 31.5317 | 51.13645 | 84.72621 | 36.29283 | -0.36097 | 1.624436 | 0.436239 |
| 5 | 101552800 | 101553399 | 8.876065 | 16.64096 | 31.25174 | 12.88952 | -0.4401 | 0.112095 | 0.436241 |

| | | | | | | | | | |
|---|---|---|---|---|---|---|---|---|---|
| 13 | 86617000 | 86617999 | 8.876065 | 16.64096 | 31.25174 | 12.88952 | -0.4401 | 0.112095 | 0.436241 |
| 18 | 9355400 | 9358799 | 105.8601 | 163.2205 | 162.3674 | 67.44462 | 0.306394 | 2.903123 | 0.436265 |
| 7 | 12226400 | 12228199 | 43.93972 | 72.67731 | 52.29245 | 22.60814 | 0.854134 | 1.363806 | 0.43631 |
| 16 | 44350800 | 44355599 | 31.13841 | 50.26331 | 98.0911 | 41.82912 | -0.60352 | 1.742518 | 0.436337 |
| 3 | 90923600 | 90924799 | 22.79549 | 39.48875 | 39.34901 | 17.26403 | 0.391332 | 0.843767 | 0.436351 |
| 17 | 20013600 | 20017599 | 49.93813 | 81.07381 | 66.41972 | 28.47364 | 0.654098 | 1.629916 | 0.436357 |
| 15 | 54819400 | 54820799 | 12.64738 | 22.45438 | 39.77311 | 17.56171 | -0.42704 | 0.482045 | 0.436359 |
| 14 | 49709600 | 49710799 | 16.34894 | 30.76926 | 21.44757 | 8.788477 | 0.98651 | 0.128858 | 0.436362 |
| 6 | 71382400 | 71385399 | 39.88128 | 65.79167 | 56.76628 | 24.49996 | 0.592423 | 1.383378 | 0.436379 |
| 3 | 3756200 | 3763999 | 176.9695 | 264.2081 | 254.38 | 104.8613 | 0.314937 | 3.559536 | 0.436379 |
| 4 | 66826800 | 66829999 | 36.76323 | 61.06532 | 52.70562 | 22.8535 | 0.592699 | 1.293225 | 0.436397 |
| X | 57083000 | 57085399 | 28.83169 | 49.32681 | 37.78056 | 16.56656 | 0.766241 | 0.979267 | 0.436397 |
| 5 | 150978800 | 150980799 | 18.2924 | 31.07946 | 61.86579 | 27.11265 | -0.60796 | 1.028322 | 0.436423 |
| 14 | 39846400 | 39847399 | 14.18212 | 26.33043 | 24.78987 | 10.39035 | 0.530423 | 0.201023 | 0.436438 |
| 6 | 75597800 | 75600799 | 31.6612 | 51.4425 | 84.62201 | 36.29817 | -0.34812 | 1.586411 | 0.436459 |
| X | 66568600 | 66570799 | 29.1943 | 50.28474 | 39.71094 | 17.41978 | 0.723942 | 0.924885 | 0.436473 |
| 7 | 62278800 | 62279399 | 21.64027 | 37.56841 | 37.8299 | 16.60101 | 0.376871 | 0.816209 | 0.436491 |
| 10 | 113565200 | 113567999 | 45.40026 | 73.66291 | 61.63043 | 26.40895 | 0.622935 | 1.625667 | 0.436495 |
| 7 | 54073400 | 54075199 | 27.01965 | 46.19341 | 43.60794 | 19.12072 | 0.465618 | 0.984492 | 0.436505 |
| 14 | 121671000 | 121673599 | 15.73466 | 27.40971 | 46.82056 | 20.59066 | -0.38165 | 0.753843 | 0.436511 |
| 14 | 90940200 | 90940799 | 10.74775 | 20.64673 | 21.25094 | 8.313458 | 0.488004 | -0.07168 | 0.436527 |
| 10 | 54087400 | 54088399 | 10.99792 | 20.00324 | 36.50873 | 15.72679 | -0.44596 | 0.318505 | 0.436531 |
| X | 92337200 | 92338599 | 8.339159 | 15.76137 | 34.37504 | 13.97217 | -0.63636 | 0.065022 | 0.436532 |
| 6 | 81481000 | 81482399 | 11.11215 | 19.8283 | 36.32015 | 15.96914 | -0.47039 | 0.43645 | 0.436537 |
| 1 | 146144000 | 146144599 | 11.85769 | 22.79143 | 21.19109 | 8.277039 | 0.63542 | -0.06587 | 0.436539 |
| 7 | 10997200 | 10999599 | 36.13946 | 59.32987 | 60.26888 | 25.95824 | 0.35485 | 1.43578 | 0.436546 |
| 13 | 95376400 | 95379399 | 34.31665 | 56.71427 | 56.96997 | 24.61656 | 0.374042 | 1.349041 | 0.436557 |
| 1 | 182081000 | 182084599 | 39.43476 | 63.21841 | 106.5874 | 45.00616 | -0.40513 | 1.969673 | 0.436559 |
| 2 | 22904200 | 22905799 | 8.933683 | 16.44507 | 32.47979 | 13.7727 | -0.54306 | 0.224697 | 0.436563 |
| 14 | 118762200 | 118764599 | 25.44226 | 41.78855 | 72.51818 | 31.24049 | -0.41518 | 1.414894 | 0.43657 |
| 8 | 78263400 | 78265799 | 20.68293 | 36.08597 | 36.12478 | 15.90252 | 0.385914 | 0.739316 | 0.43658 |
| 9 | 6831000 | 6833199 | 27.82371 | 46.71795 | 45.94762 | 20.06178 | 0.404915 | 1.147209 | 0.436607 |
| 14 | 52166800 | 52168599 | 7.875269 | 14.6382 | 30.00398 | 12.55586 | -0.58166 | 0.157395 | 0.436623 |
| 14 | 41116600 | 41118999 | 22.44836 | 40.22685 | 25.78056 | 11.29461 | 1.038965 | 0.462645 | 0.436627 |
| 4 | 69725000 | 69726999 | 15.69662 | 27.37701 | 47.3328 | 20.83199 | -0.39891 | 0.735726 | 0.43663 |
| 9 | 16363000 | 16364399 | 10.37515 | 18.40992 | 40.40534 | 17.84929 | -0.73457 | 0.474114 | 0.436655 |
| 9 | 79950600 | 79951799 | 19.86395 | 34.80305 | 34.40236 | 15.19312 | 0.40416 | 0.66824 | 0.436694 |
| 16 | 21986600 | 21988799 | 19.86395 | 34.80305 | 34.40236 | 15.19312 | 0.40416 | 0.66824 | 0.436694 |
| 10 | 4303400 | 4307799 | 54.67576 | 86.3923 | 144.2185 | 59.90227 | -0.41212 | 2.420297 | 0.436705 |
| 3 | 15879600 | 15881199 | 20.49383 | 36.01655 | 31.90997 | 14.12885 | 0.560982 | 0.617223 | 0.436706 |
| 3 | 93837000 | 93838199 | 14.80009 | 26.72426 | 26.35454 | 11.45609 | 0.428411 | 0.366078 | 0.436709 |
| 12 | 87722800 | 87725399 | 68.82887 | 109.1471 | 110.7439 | 46.09933 | 0.309476 | 2.333624 | 0.436722 |
| 17 | 38896400 | 38902199 | 74.89914 | 119.7368 | 87.42668 | 36.76455 | 0.795398 | 2.090389 | 0.436723 |
| 10 | 120249800 | 120251199 | 9.980114 | 17.98022 | 34.34826 | 14.97318 | -0.522 | 0.363937 | 0.436738 |
| 7 | 18748000 | 18748999 | 7.589318 | 14.11034 | 30.02511 | 12.55981 | -0.63438 | 0.155365 | 0.43676 |
| 9 | 119667200 | 119668199 | 8.398296 | 15.76262 | 30.72185 | 12.64731 | -0.49117 | 0.098542 | 0.436762 |
| 9 | 93221200 | 93222999 | 17.32541 | 29.95315 | 51.15282 | 22.44148 | -0.38322 | 0.852963 | 0.436765 |

| | | | | | | | | | |
|---|---|---|---|---|---|---|---|---|---|
| 16 | 92902200 | 92904799 | 17.32541 | 29.95315 | 51.15282 | 22.44148 | -0.38322 | 0.852963 | 0.436765 |
| 15 | 11791600 | 11791999 | 6.39488 | 12.37501 | 26.17296 | 9.977896 | -0.5069 | -0.1899 | 0.436775 |
| 17 | 18362800 | 18370599 | 104.3224 | 162.5464 | 147.5387 | 61.23482 | 0.450086 | 2.716509 | 0.436775 |
| 1 | 170929400 | 170930999 | 7.589821 | 14.0717 | 32.07463 | 13.46902 | -0.73793 | 0.171726 | 0.436775 |
| 13 | 64765800 | 64767599 | 17.72505 | 31.64947 | 29.49825 | 12.96236 | 0.498894 | 0.459231 | 0.43678 |
| 11 | 116889400 | 116890799 | 14.42115 | 26.32381 | 26.29318 | 11.27552 | 0.422489 | 0.300717 | 0.436782 |
| 3 | 41277800 | 41278999 | 16.66315 | 29.96606 | 28.64816 | 12.51343 | 0.468428 | 0.407796 | 0.436788 |
| 19 | 54254400 | 54255999 | 17.64177 | 32.39447 | 25.49282 | 10.84647 | 0.772214 | 0.291312 | 0.436839 |
| X | 77720800 | 77722599 | 31.13864 | 52.29478 | 48.57632 | 21.2123 | 0.487109 | 1.160546 | 0.43684 |
| 12 | 3644600 | 3645399 | 5.815748 | 11.2752 | 23.85006 | 9.013871 | -0.49373 | -0.22868 | 0.436879 |
| 3 | 138383800 | 138385799 | 14.16667 | 26.114 | 25.25809 | 10.69029 | 0.481722 | 0.241113 | 0.436928 |
| 5 | 126223600 | 126226999 | 31.75254 | 52.94387 | 52.49692 | 22.82062 | 0.393347 | 1.238141 | 0.43696 |
| 17 | 83387200 | 83389399 | 20.32902 | 35.47939 | 35.6157 | 15.68144 | 0.381973 | 0.732182 | 0.436975 |
| 6 | 80997000 | 81000399 | 41.25592 | 66.19895 | 106.673 | 45.10547 | -0.33838 | 1.944767 | 0.436982 |
| 1 | 5216800 | 5218199 | 12.69253 | 23.73043 | 24.01879 | 9.952839 | 0.439973 | 0.145366 | 0.436982 |
| 14 | 6504200 | 6509199 | 61.05615 | 97.59281 | 96.14394 | 40.4221 | 0.363778 | 2.080787 | 0.436998 |
| 3 | 91293600 | 91297399 | 43.52394 | 71.31077 | 60.72581 | 26.13302 | 0.607538 | 1.478983 | 0.437024 |
| 7 | 38266600 | 38268599 | 69.65663 | 110.3584 | 105.0711 | 43.70125 | 0.399887 | 2.35919 | 0.437033 |
| 15 | 23462800 | 23463999 | 16.58887 | 30.60362 | 24.4392 | 10.33235 | 0.757362 | 0.253979 | 0.437037 |
| 10 | 17605200 | 17607199 | 14.75043 | 25.83998 | 44.58877 | 19.6893 | -0.39573 | 0.660177 | 0.437057 |
| 4 | 112570800 | 112571399 | 8.678224 | 16.91998 | 16.70876 | 6.020463 | 0.655648 | -0.38739 | 0.437066 |
| 13 | 106759600 | 106761199 | 19.94098 | 35.35971 | 30.33636 | 13.40895 | 0.612111 | 0.51827 | 0.437091 |
| 3 | 144766400 | 144768399 | 12.97928 | 23.03208 | 44.01816 | 19.44433 | -0.53673 | 0.494281 | 0.43712 |
| 5 | 91221600 | 91223199 | 12.88599 | 22.59218 | 41.52007 | 18.34683 | -0.48585 | 0.633274 | 0.437154 |
| Y | 36423800 | 36425599 | 16.08941 | 29.0998 | 15.9473 | 6.918025 | 1.271303 | 0.338104 | 0.437179 |
| X | 76975000 | 76978199 | 45.77535 | 75.06089 | 58.55371 | 25.20698 | 0.734164 | 1.474049 | 0.437181 |
| X | 88088400 | 88089599 | 12.00511 | 23.24987 | 19.08836 | 7.250371 | 0.850748 | -0.16023 | 0.437188 |
| 17 | 40845400 | 40848599 | 30.68807 | 51.7844 | 47.43877 | 20.74728 | 0.50713 | 1.113391 | 0.437233 |
| 19 | 27679800 | 27682999 | 25.8839 | 42.1578 | 85.53941 | 36.72982 | -0.6473 | 1.538894 | 0.437248 |
| 7 | 61861400 | 61861999 | 11.51216 | 22.05413 | 21.68722 | 8.55174 | 0.54248 | -0.03802 | 0.437273 |
| 6 | 4746000 | 4749599 | 89.07083 | 136.0717 | 225.9883 | 93.73518 | -0.4426 | 3.085398 | 0.437284 |
| 15 | 10713600 | 10715199 | 12.13225 | 21.20055 | 44.69951 | 19.69669 | -0.6832 | 0.686363 | 0.437313 |
| 7 | 6153800 | 6159399 | 37.28016 | 59.89493 | 103.2027 | 43.72122 | -0.43265 | 1.89646 | 0.437316 |
| 5 | 73954000 | 73955599 | 16.55105 | 29.7197 | 28.57422 | 12.49698 | 0.458898 | 0.414824 | 0.437325 |
| 14 | 20967600 | 20968799 | 9.391164 | 17.5029 | 32.74751 | 13.64233 | -0.44633 | 0.158966 | 0.437328 |
| 8 | 45056600 | 45057599 | 10.02985 | 17.90044 | 35.86641 | 15.76713 | -0.59879 | 0.426264 | 0.437337 |
| 4 | 30355000 | 30357799 | 20.46726 | 34.74279 | 59.26751 | 25.96723 | -0.38632 | 1.039916 | 0.437366 |
| 14 | 114439000 | 114441399 | 24.95812 | 42.98125 | 41.43619 | 18.1765 | 0.437156 | 0.908003 | 0.437404 |
| 17 | 62677400 | 62679999 | 17.83262 | 32.43672 | 26.36213 | 11.35491 | 0.713974 | 0.345649 | 0.437409 |
| 5 | 39967600 | 39969799 | 31.37662 | 50.98 | 83.76668 | 35.93121 | -0.34645 | 1.587085 | 0.437422 |
| 7 | 42399200 | 42402399 | 79.87911 | 125.6159 | 127.0394 | 52.7016 | 0.30453 | 2.533736 | 0.437436 |
| 5 | 22804000 | 22806399 | 25.77365 | 44.77191 | 38.0186 | 16.69032 | 0.62175 | 0.826887 | 0.437448 |
| 13 | 65988000 | 65990399 | 30.92181 | 52.13258 | 43.1813 | 18.8802 | 0.651959 | 1.119072 | 0.437454 |
| 6 | 81129400 | 81130599 | 12.85209 | 24.53638 | 21.46324 | 8.540508 | 0.697692 | 0.002255 | 0.437465 |
| 9 | 18145200 | 18147399 | 19.61526 | 34.61073 | 32.09044 | 14.19987 | 0.498058 | 0.571389 | 0.437481 |
| 12 | 48561600 | 48563199 | 14.71836 | 27.36489 | 24.77799 | 10.36489 | 0.588454 | 0.197951 | 0.437488 |
| 4 | 78477200 | 78479199 | 17.01969 | 30.98563 | 27.11738 | 11.66721 | 0.609138 | 0.337802 | 0.437502 |

| | | | | | | | | | |
|---|---|---|---|---|---|---|---|---|---|
| 18 | 79817600 | 79818599 | 13.64521 | 24.96765 | 23.97814 | 10.25107 | 0.481954 | 0.276875 | 0.437507 |
| 7 | 139593400 | 139597199 | 92.76994 | 144.6456 | 143.6354 | 59.60865 | 0.321352 | 2.699237 | 0.437515 |
| 5 | 61337400 | 61338199 | 17.3308 | 30.64816 | 30.31999 | 13.41048 | 0.406283 | 0.530779 | 0.43753 |
| 3 | 75863000 | 75864599 | 12.31873 | 21.46922 | 44.7656 | 19.69273 | -0.66748 | 0.727719 | 0.43754 |
| 1 | 133330200 | 133332999 | 14.59188 | 25.47705 | 44.67995 | 19.67747 | -0.41905 | 0.711843 | 0.437564 |
| 7 | 9055000 | 9055599 | 24.51234 | 41.90575 | 38.61926 | 16.93331 | 0.500286 | 0.969487 | 0.437576 |
| 17 | 39235600 | 39239599 | 42.0795 | 69.30816 | 56.83955 | 24.51514 | 0.664604 | 1.408472 | 0.437612 |
| 5 | 94003000 | 94008199 | 39.82441 | 64.9888 | 64.79212 | 27.84077 | 0.377075 | 1.526667 | 0.437619 |
| 7 | 142217600 | 142224799 | 127.5834 | 191.5785 | 289.6589 | 119.5509 | -0.32824 | 3.443931 | 0.437622 |
| 7 | 105357800 | 105359599 | 15.7569 | 27.24608 | 48.46028 | 21.26033 | -0.44137 | 0.841858 | 0.43763 |
| 6 | 146975600 | 146983799 | 103.8064 | 157.61 | 247.3059 | 102.4048 | -0.36794 | 3.215751 | 0.437631 |
| 3 | 40580000 | 40584399 | 28.86264 | 47.10858 | 78.87726 | 33.8952 | -0.36877 | 1.50962 | 0.437637 |
| 2 | 39666600 | 39670599 | 35.33213 | 58.35383 | 57.57087 | 24.86482 | 0.399884 | 1.360032 | 0.437669 |
| 1 | 95057400 | 95059599 | 21.0793 | 36.81373 | 36.07952 | 15.89242 | 0.416558 | 0.728314 | 0.43767 |
| 2 | 168931800 | 168938599 | 78.66274 | 121.9823 | 188.286 | 78.20237 | -0.32043 | 2.790272 | 0.437673 |
| 10 | 49780800 | 49782199 | 29.65163 | 49.90606 | 46.96436 | 20.52367 | 0.468507 | 1.13532 | 0.437674 |
| 6 | 133243200 | 133245399 | 11.47262 | 20.21238 | 51.01672 | 22.58154 | -0.94037 | 0.570235 | 0.437691 |
| 6 | 57345000 | 57347399 | 24.64488 | 42.75365 | 38.61784 | 16.94397 | 0.532704 | 0.837271 | 0.437714 |
| 11 | 74141800 | 74145399 | 37.22488 | 61.30905 | 58.83312 | 25.37442 | 0.438537 | 1.397479 | 0.437724 |
| 15 | 78745400 | 78751199 | 35.03037 | 56.39902 | 96.02082 | 40.803 | -0.41152 | 1.826034 | 0.437755 |
| 6 | 74818400 | 74819199 | 5.672271 | 11.01851 | 24.03976 | 8.980691 | -0.52305 | -0.27348 | 0.437788 |
| 7 | 7648400 | 7651399 | 24.45201 | 43.11777 | 30.10409 | 13.32261 | 0.904959 | 0.604686 | 0.437805 |
| 4 | 47385800 | 47387399 | 15.98371 | 27.85056 | 51.10767 | 22.47959 | -0.48454 | 0.759037 | 0.437806 |
| 13 | 66269800 | 66270399 | 17.01624 | 32.54007 | 19.05574 | 7.552857 | 1.277698 | 0.026405 | 0.437807 |
| 13 | 60755200 | 60758399 | 16.72824 | 28.81561 | 50.05017 | 21.95529 | -0.40851 | 0.882711 | 0.43781 |
| 10 | 41037600 | 41039199 | 16.60289 | 30.11561 | 27.60342 | 11.9312 | 0.53847 | 0.355693 | 0.437812 |
| 8 | 52451200 | 52453399 | 24.2669 | 42.49493 | 34.0579 | 15.03431 | 0.705397 | 0.700537 | 0.437819 |
| 3 | 20369400 | 20370399 | 14.30676 | 26.35564 | 25.85628 | 10.95284 | 0.460744 | 0.248488 | 0.437834 |
| 2 | 9893800 | 9895999 | 19.0317 | 32.25993 | 56.98757 | 24.95751 | -0.43651 | 1.044464 | 0.437843 |
| 13 | 101677000 | 101678599 | 10.51037 | 18.70721 | 36.36986 | 16.02506 | -0.55781 | 0.448846 | 0.437855 |
| 7 | 132929200 | 132930799 | 16.8757 | 29.96466 | 29.31657 | 12.94348 | 0.424899 | 0.489727 | 0.437868 |
| 19 | 58088600 | 58095399 | 45.03118 | 71.73739 | 124.6045 | 52.14229 | -0.45819 | 2.176937 | 0.437871 |
| 3 | 3991400 | 3995599 | 39.10283 | 64.51785 | 56.21534 | 24.26382 | 0.578374 | 1.377668 | 0.437884 |
| 7 | 11375600 | 11378999 | 49.39625 | 79.61486 | 76.6666 | 32.62956 | 0.410753 | 1.806652 | 0.437891 |
| 9 | 12768000 | 12770399 | 20.05829 | 35.07659 | 35.05155 | 15.4549 | 0.388674 | 0.701243 | 0.437893 |
| 4 | 84935600 | 84936999 | 14.04604 | 26.5141 | 22.85207 | 9.313372 | 0.689082 | 0.092942 | 0.437899 |
| 8 | 81090600 | 81093199 | 19.09614 | 32.17901 | 59.09101 | 25.83251 | -0.49257 | 1.098347 | 0.437933 |
| 2 | 137569000 | 137570199 | 7.437197 | 14.25233 | 30.44087 | 11.99211 | -0.56651 | -0.06029 | 0.437952 |
| 4 | 19787800 | 19790399 | 33.28308 | 56.33271 | 45.17563 | 19.77555 | 0.698603 | 1.096155 | 0.437967 |
| 11 | 27904000 | 27907999 | 50.05602 | 80.6886 | 68.28733 | 29.06812 | 0.596832 | 1.798735 | 0.437973 |
| 17 | 58280600 | 58281199 | 11.55855 | 20.49971 | 38.25974 | 16.90845 | -0.50281 | 0.478616 | 0.437974 |
| 18 | 64135000 | 64137599 | 19.9482 | 33.99787 | 61.01871 | 26.74809 | -0.45864 | 1.002744 | 0.437976 |
| 1 | 174287800 | 174290399 | 29.02618 | 49.10273 | 42.11501 | 18.43292 | 0.601934 | 1.081696 | 0.437976 |
| 1 | 100665600 | 100668799 | 60.7963 | 97.3133 | 92.1476 | 38.81863 | 0.42296 | 2.036163 | 0.437978 |
| 9 | 14374400 | 14375599 | 10.6563 | 18.93205 | 36.82974 | 16.2529 | -0.56049 | 0.460699 | 0.437983 |
| 7 | 105351800 | 105354799 | 18.33323 | 31.41169 | 53.27016 | 23.36143 | -0.37575 | 0.941172 | 0.437984 |
| 2 | 21634400 | 21638599 | 47.6911 | 77.29416 | 69.82059 | 29.89404 | 0.511201 | 1.657508 | 0.437984 |

| | | | | | | | | | |
|---|---|---|---|---|---|---|---|---|---|
| 2 | 43057400 | 43057999 | 12.29725 | 22.89119 | 21.3789 | 8.924694 | 0.546281 | 0.160702 | 0.437991 |
| 4 | 26066000 | 26067599 | 24.33369 | 41.9676 | 40.93262 | 17.9564 | 0.420881 | 0.892187 | 0.438 |
| 10 | 77576200 | 77579399 | 20.48657 | 34.16234 | 63.30798 | 27.52187 | -0.50579 | 1.214545 | 0.438004 |
| 2 | 84624600 | 84627199 | 58.23486 | 92.98402 | 94.16758 | 39.53143 | 0.322352 | 2.109408 | 0.438013 |
| 15 | 6768400 | 6769199 | 8.764501 | 16.46783 | 31.29992 | 12.8616 | -0.45265 | 0.099156 | 0.438016 |
| 9 | 120356000 | 120358599 | 15.54214 | 27.01314 | 47.38325 | 20.80851 | -0.42015 | 0.784582 | 0.438024 |
| 9 | 6286000 | 6286399 | 4.477853 | 8.752179 | 20.16837 | 6.959572 | -0.49461 | -0.57993 | 0.438024 |
| 11 | 120102000 | 120105199 | 35.28115 | 57.05341 | 93.90284 | 40.13649 | -0.35539 | 1.693898 | 0.438073 |
| 7 | 17763000 | 17782199 | 289.7842 | 420.762 | 340.6346 | 140.4273 | 0.527116 | 4.025615 | 0.43809 |
| 13 | 11294200 | 11294799 | 5.843058 | 11.27521 | 25.33565 | 9.784413 | -0.61081 | -0.15355 | 0.438111 |
| 8 | 117502600 | 117504399 | 21.96575 | 38.05406 | 38.37175 | 16.8353 | 0.374331 | 0.838115 | 0.438133 |
| 2 | 173576200 | 173578399 | 18.27447 | 32.3475 | 31.09952 | 13.75266 | 0.447701 | 0.527129 | 0.438144 |
| 10 | 11313600 | 11316199 | 19.28575 | 32.94026 | 57.11486 | 25.04118 | -0.40846 | 0.976615 | 0.438149 |
| 5 | 129583800 | 129585599 | 10.07701 | 18.22402 | 34.83764 | 15.11438 | -0.5189 | 0.340864 | 0.438158 |
| 5 | 126070600 | 126071599 | 18.25414 | 32.38989 | 30.51294 | 13.47951 | 0.478413 | 0.503518 | 0.438177 |
| X | 92342000 | 92344599 | 18.98687 | 32.37308 | 59.69611 | 26.1692 | -0.49714 | 0.999113 | 0.438177 |
| 9 | 12850000 | 12852599 | 25.64763 | 44.18308 | 40.43216 | 17.73624 | 0.512254 | 0.909222 | 0.438178 |
| 18 | 37793000 | 37797799 | 124.9962 | 190.9295 | 187.8156 | 77.89961 | 0.312166 | 3.095679 | 0.438203 |
| 15 | 103837800 | 103844599 | 89.25047 | 141.284 | 113.2392 | 47.07877 | 0.646995 | 2.395472 | 0.438209 |
| 4 | 53019600 | 53020199 | 8.643079 | 16.80328 | 17.71468 | 6.573667 | 0.524273 | -0.2826 | 0.438224 |
| X | 19337600 | 19340399 | 34.37907 | 57.31764 | 52.53387 | 22.83525 | 0.506553 | 1.249003 | 0.43823 |
| 15 | 74968600 | 74971599 | 37.33505 | 61.02478 | 62.37593 | 26.81831 | 0.342828 | 1.496751 | 0.43826 |
| 7 | 41937000 | 41947999 | 22.17514 | 38.92817 | 29.34147 | 12.97943 | 0.793132 | 0.641081 | 0.438279 |
| 5 | 151593200 | 151599399 | 109.1178 | 168.3723 | 167.5659 | 69.60373 | 0.306602 | 2.890693 | 0.438291 |
| 1 | 100109200 | 100110999 | 20.2663 | 35.81292 | 31.44059 | 13.90817 | 0.577204 | 0.558048 | 0.438307 |
| 18 | 35763800 | 35771199 | 70.95758 | 112.7314 | 100.1418 | 41.76334 | 0.503558 | 2.279629 | 0.438311 |
| 10 | 107837200 | 107837599 | 8.783808 | 17.00041 | 16.66887 | 6.350327 | 0.59329 | -0.19407 | 0.438312 |
| 16 | 81488600 | 81490799 | 37.72795 | 61.67197 | 62.3386 | 26.80305 | 0.358975 | 1.497564 | 0.438331 |
| 8 | 99143800 | 99145199 | 16.84744 | 31.13823 | 24.48796 | 10.32692 | 0.782045 | 0.246719 | 0.438336 |
| 10 | 83621600 | 83624799 | 23.65052 | 38.99248 | 69.6942 | 30.07189 | -0.4554 | 1.36203 | 0.438341 |
| 2 | 74392800 | 74395199 | 20.72641 | 34.949 | 59.13021 | 25.85476 | -0.37476 | 1.100546 | 0.438353 |
| 14 | 43768200 | 43770599 | 57.39739 | 90.7483 | 144.3277 | 59.96623 | -0.34166 | 2.406941 | 0.438369 |
| 18 | 82269600 | 82272399 | 22.99619 | 39.69498 | 39.34838 | 17.26188 | 0.397731 | 0.874564 | 0.438395 |
| 2 | 133393000 | 133396999 | 26.12506 | 42.36921 | 86.76408 | 37.16639 | -0.66572 | 1.613483 | 0.438458 |
| 4 | 15866600 | 15870599 | 44.11204 | 72.14123 | 61.91367 | 26.62485 | 0.59487 | 1.504015 | 0.438463 |
| 4 | 25106000 | 25108599 | 24.05226 | 40.03182 | 67.4259 | 29.2767 | -0.36851 | 1.250598 | 0.438466 |
| 3 | 23949400 | 23951999 | 27.47288 | 47.12104 | 42.46014 | 18.62232 | 0.533168 | 0.95565 | 0.438517 |
| 12 | 14629600 | 14632599 | 39.66321 | 65.19567 | 57.3692 | 24.72231 | 0.562031 | 1.423982 | 0.43852 |
| 3 | 41094800 | 41097799 | 41.49114 | 67.89085 | 61.83639 | 26.59674 | 0.509776 | 1.488866 | 0.438533 |
| 2 | 93565000 | 93566399 | 9.241495 | 17.0844 | 31.48521 | 13.2737 | -0.43754 | 0.202522 | 0.438552 |
| 8 | 110773400 | 110774399 | 9.241495 | 17.0844 | 31.48521 | 13.2737 | -0.43754 | 0.202522 | 0.438552 |
| 7 | 42386600 | 42412399 | 470.6153 | 647.4608 | 583.9341 | 242.7489 | 0.293493 | 4.830696 | 0.438572 |
| 5 | 70963200 | 70964199 | 18.03224 | 32.55862 | 28.01373 | 12.17821 | 0.624013 | 0.393412 | 0.438594 |
| 14 | 105699600 | 105699999 | 9.770512 | 18.98404 | 18.79577 | 7.007901 | 0.607961 | -0.24855 | 0.438601 |
| 3 | 123071000 | 123072999 | 15.8183 | 27.55661 | 51.15272 | 22.49647 | -0.50137 | 0.756355 | 0.438604 |
| 1 | 117337400 | 117338799 | 20.37616 | 36.04378 | 30.82954 | 13.63508 | 0.61507 | 0.546244 | 0.438608 |
| 11 | 92798800 | 92799999 | 11.87252 | 21.29617 | 37.83684 | 16.5628 | -0.42251 | 0.410728 | 0.438608 |

| | | | | | | | | | |
|---|---|---|---|---|---|---|---|---|---|
| 18 | 74989200 | 74993599 | 40.53429 | 65.08476 | 104.8314 | 44.37214 | -0.33661 | 1.920495 | 0.438613 |
| 10 | 100355600 | 100356999 | 12.05262 | 21.32229 | 38.29371 | 16.93594 | -0.44906 | 0.509137 | 0.438615 |
| 1 | 70445400 | 70446599 | 10.41802 | 18.71534 | 35.79712 | 15.65004 | -0.52703 | 0.385114 | 0.438616 |
| 5 | 93943200 | 93944399 | 16.26742 | 30.53445 | 22.59103 | 9.298844 | 0.896271 | 0.147477 | 0.438622 |
| 9 | 60207000 | 60209799 | 44.44239 | 72.33647 | 66.28083 | 28.45348 | 0.495907 | 1.573764 | 0.438633 |
| 3 | 91319200 | 91321999 | 29.94885 | 50.68479 | 46.87525 | 20.51899 | 0.493626 | 1.086653 | 0.438681 |
| 17 | 75856400 | 75859799 | 22.07042 | 36.80709 | 63.43547 | 27.579 | -0.40152 | 1.218731 | 0.438704 |
| 5 | 142222600 | 142234199 | 145.4762 | 216.8652 | 324.3503 | 133.6658 | -0.32207 | 3.584155 | 0.438718 |
| 12 | 28935400 | 28936999 | 17.47286 | 31.05495 | 29.48026 | 13.00515 | 0.468638 | 0.48499 | 0.43872 |
| 5 | 7299400 | 7300599 | 16.45067 | 29.71815 | 28.12883 | 12.22044 | 0.487696 | 0.378802 | 0.438729 |
| 15 | 63654200 | 63655799 | 18.78361 | 34.20541 | 25.15487 | 10.81952 | 0.858492 | 0.343607 | 0.438737 |
| 10 | 115479200 | 115483199 | 28.34009 | 48.58734 | 41.84956 | 18.35389 | 0.597858 | 0.961007 | 0.438752 |
| 15 | 60314400 | 60317799 | 48.22416 | 77.8975 | 75.16542 | 32.08187 | 0.411413 | 1.738027 | 0.438762 |
| 16 | 97562800 | 97569199 | 43.63094 | 69.79058 | 116.8297 | 49.16938 | -0.39861 | 2.045018 | 0.438765 |
| 18 | 76995400 | 76997399 | 17.04312 | 29.47618 | 50.06325 | 21.96372 | -0.37513 | 0.84826 | 0.438769 |
| 1 | 37632400 | 37633399 | 13.59878 | 25.27108 | 24.89455 | 10.42092 | 0.466994 | 0.191995 | 0.438769 |
| 9 | 8930400 | 8931799 | 13.59878 | 25.27108 | 24.89455 | 10.42092 | 0.466994 | 0.191995 | 0.438769 |
| 4 | 17748000 | 17750399 | 19.05606 | 34.28896 | 27.75859 | 12.11902 | 0.707281 | 0.41814 | 0.438771 |
| 5 | 146007400 | 146011399 | 81.16064 | 127.5732 | 129.4441 | 53.69602 | 0.299281 | 2.543336 | 0.438782 |
| 14 | 34727800 | 34731599 | 24.33892 | 40.38486 | 68.01908 | 29.47178 | -0.3689 | 1.285282 | 0.438789 |
| 4 | 48042600 | 48046199 | 28.80517 | 46.72094 | 83.66983 | 35.84386 | -0.47263 | 1.615787 | 0.438796 |
| 7 | 46623600 | 46625799 | 20.37042 | 34.50074 | 58.08432 | 25.43148 | -0.3674 | 1.059603 | 0.438805 |
| 17 | 55523600 | 55527399 | 36.81908 | 60.85473 | 57.11758 | 24.68168 | 0.471695 | 1.355199 | 0.438819 |
| 7 | 108113200 | 108114399 | 15.04771 | 28.15798 | 23.74545 | 9.825242 | 0.702832 | 0.158324 | 0.438838 |
| 5 | 121110200 | 121116199 | 43.86579 | 69.99204 | 119.8674 | 50.27054 | -0.43535 | 2.123359 | 0.438841 |
| 19 | 40179000 | 40181199 | 16.3194 | 28.2715 | 50.61903 | 22.20868 | -0.45059 | 0.827961 | 0.438843 |
| 3 | 143419200 | 143421599 | 23.81647 | 41.46965 | 37.52775 | 16.49925 | 0.530649 | 0.78466 | 0.438847 |
| 7 | 30528800 | 30532399 | 41.40751 | 66.30015 | 109.3389 | 46.08249 | -0.3754 | 2.011558 | 0.438851 |
| 5 | 74780600 | 74781199 | 5.081397 | 9.880744 | 22.30262 | 8.263723 | -0.55892 | -0.31104 | 0.438874 |
| 13 | 42224800 | 42225399 | 5.081397 | 9.880744 | 22.30262 | 8.263723 | -0.55892 | -0.31104 | 0.438874 |
| 14 | 81349000 | 81350799 | 14.95033 | 26.1689 | 45.14509 | 19.92128 | -0.3953 | 0.676388 | 0.438891 |
| X | 164516600 | 164519399 | 44.4372 | 72.63016 | 56.11245 | 24.12496 | 0.745541 | 1.510025 | 0.438894 |
| 11 | 13967400 | 13968599 | 7.907792 | 15.05403 | 30.08768 | 12.04477 | -0.49245 | -0.00326 | 0.438896 |
| 4 | 113846800 | 113848999 | 35.25998 | 59.09767 | 49.16525 | 21.44234 | 0.645669 | 1.194439 | 0.43891 |
| 1 | 167926600 | 167930399 | 49.64461 | 80.08744 | 76.96568 | 32.79501 | 0.415093 | 1.777647 | 0.438912 |
| 15 | 100221600 | 100225199 | 21.87227 | 36.50147 | 64.63642 | 28.1132 | -0.44057 | 1.210076 | 0.438913 |
| 5 | 104135600 | 104138199 | 21.0752 | 35.41235 | 60.18302 | 26.28705 | -0.3814 | 1.13355 | 0.438931 |
| 15 | 81376200 | 81383399 | 72.95998 | 113.5567 | 177.859 | 73.83099 | -0.33731 | 2.724294 | 0.43896 |
| X | 67170800 | 67179199 | 146.2957 | 222.8678 | 195.6882 | 81.11225 | 0.4729 | 3.156597 | 0.438962 |
| 12 | 12418600 | 12420999 | 12.35274 | 21.91605 | 38.24344 | 16.89618 | -0.406 | 0.482723 | 0.43898 |
| 5 | 91678200 | 91678799 | 9.848799 | 19.07749 | 19.84684 | 7.531689 | 0.514054 | -0.18357 | 0.438988 |
| 15 | 51800200 | 51802599 | 23.78268 | 41.02201 | 39.81869 | 17.4678 | 0.427663 | 0.88479 | 0.439 |
| 12 | 7453200 | 7457999 | 35.61026 | 59.26229 | 53.16833 | 23.08429 | 0.537273 | 1.271052 | 0.439016 |
| X | 122894200 | 122897199 | 26.17131 | 45.28065 | 39.72204 | 17.42703 | 0.574083 | 0.870889 | 0.439023 |
| 2 | 160264600 | 160267199 | 13.1667 | 23.13775 | 40.89233 | 18.10476 | -0.42985 | 0.595319 | 0.439028 |
| 2 | 158541800 | 158543999 | 16.60469 | 28.4539 | 51.62263 | 22.63916 | -0.47254 | 0.931365 | 0.439049 |
| 3 | 113823800 | 113825999 | 14.94024 | 25.845 | 50.58596 | 22.19319 | -0.57891 | 0.834596 | 0.439053 |

| | | | | | | | | | |
|---|---|---|---|---|---|---|---|---|---|
| 16 | 77541400 | 77542799 | 12.68036 | 23.51327 | 23.03269 | 9.670051 | 0.472334 | 0.19307 | 0.439059 |
| 7 | 126806600 | 126808599 | 10.62934 | 18.95228 | 36.31251 | 15.97607 | -0.53567 | 0.436264 | 0.43909 |
| 2 | 29195200 | 29196399 | 8.127487 | 15.26884 | 29.09167 | 11.95698 | -0.45623 | 0.088473 | 0.439093 |
| 11 | 83801000 | 83802199 | 8.127487 | 15.26884 | 29.09167 | 11.95698 | -0.45623 | 0.088473 | 0.439093 |
| 19 | 13684200 | 13685199 | 16.24921 | 28.85361 | 27.82183 | 12.28305 | 0.445658 | 0.480784 | 0.439094 |
| 16 | 48099200 | 48106199 | 67.83324 | 106.5336 | 169.6842 | 70.38059 | -0.35141 | 2.558428 | 0.439095 |
| 16 | 80561400 | 80561799 | 11.02045 | 21.0547 | 20.70717 | 8.227105 | 0.532948 | -0.01896 | 0.439105 |
| 17 | 22704400 | 22706599 | 42.09635 | 68.68236 | 58.70741 | 25.22434 | 0.598309 | 1.530934 | 0.439113 |
| 7 | 51450800 | 51452199 | 18.55437 | 32.54295 | 32.00091 | 14.1452 | 0.411668 | 0.640533 | 0.439117 |
| X | 151369600 | 151375199 | 52.67393 | 83.69442 | 134.5826 | 56.13365 | -0.35105 | 2.270681 | 0.43912 |
| 16 | 38160400 | 38160999 | 11.76322 | 22.23316 | 22.62287 | 9.195203 | 0.45524 | 0.065023 | 0.439135 |
| 7 | 31070600 | 31089399 | 435.3781 | 605.3524 | 530.7563 | 220.2734 | 0.349625 | 4.669016 | 0.439149 |
| 17 | 18678600 | 18680199 | 26.64121 | 45.64333 | 43.04444 | 18.87688 | 0.467506 | 0.962878 | 0.439158 |
| 8 | 113755000 | 113756199 | 8.291857 | 15.6373 | 29.71552 | 12.13457 | -0.44486 | 0.067998 | 0.439168 |
| 1 | 13691600 | 13694399 | 34.27167 | 56.48552 | 57.49762 | 24.80593 | 0.354099 | 1.380571 | 0.439195 |
| X | 54636600 | 54641999 | 69.165 | 110.7407 | 87.96747 | 37.0733 | 0.676458 | 2.038635 | 0.439216 |
| 12 | 26372200 | 26374799 | 14.58416 | 25.55317 | 44.0584 | 19.45797 | -0.39456 | 0.655497 | 0.439227 |
| 1 | 156918400 | 156919999 | 17.1962 | 30.35793 | 27.01066 | 11.95087 | 0.55751 | 0.541312 | 0.439233 |
| Y | 5434000 | 5436599 | 13.42506 | 24.91307 | 15.50074 | 6.501121 | 1.122441 | 0.171838 | 0.439263 |
| 13 | 49805800 | 49806999 | 7.679158 | 14.44013 | 28.57375 | 11.72545 | -0.50836 | 0.077411 | 0.439263 |
| 17 | 38342400 | 38346999 | 52.20738 | 84.13881 | 71.1992 | 30.2994 | 0.596609 | 1.80714 | 0.439264 |
| 13 | 33602400 | 33604399 | 20.83515 | 36.68919 | 32.58156 | 14.423 | 0.558944 | 0.601674 | 0.439267 |
| 15 | 91083400 | 91083999 | 10.85335 | 19.59764 | 35.31152 | 15.34969 | -0.43595 | 0.356908 | 0.439274 |
| X | 25631000 | 25648999 | 217.839 | 324.9011 | 248.4958 | 102.4159 | 0.645536 | 3.58502 | 0.439281 |
| 12 | 22493600 | 22498399 | 43.54301 | 71.25522 | 55.58312 | 23.90797 | 0.732868 | 1.489847 | 0.439306 |
| 1 | 143033800 | 143035599 | 20.32898 | 35.89075 | 31.38217 | 13.88487 | 0.582397 | 0.567088 | 0.439309 |
| 11 | 65687200 | 65689999 | 21.33546 | 37.23096 | 36.56501 | 16.09808 | 0.41338 | 0.740666 | 0.439325 |
| X | 21482600 | 21485599 | 27.88049 | 48.21711 | 39.1081 | 17.15746 | 0.686636 | 0.881797 | 0.439349 |
| 11 | 88806400 | 88811999 | 113.246 | 171.0427 | 266.1821 | 110.0416 | -0.36246 | 3.32531 | 0.439352 |
| 14 | 53748200 | 53748799 | 7.147984 | 13.71376 | 27.47393 | 10.78784 | -0.46989 | -0.07992 | 0.439356 |
| 1 | 118044000 | 118047399 | 65.01195 | 103.8198 | 97.92117 | 41.11546 | 0.425035 | 2.113425 | 0.43936 |
| 4 | 148679600 | 148688799 | 74.57207 | 115.6572 | 187.5042 | 77.87581 | -0.39098 | 2.78532 | 0.439364 |
| 14 | 4329400 | 4329799 | 7.762428 | 15.11061 | 15.6655 | 5.753138 | 0.56236 | -0.33596 | 0.439402 |
| 3 | 46161800 | 46164199 | 23.7179 | 41.47376 | 34.6025 | 15.25613 | 0.647609 | 0.719114 | 0.439402 |
| 5 | 106692800 | 106694199 | 20.33588 | 35.46893 | 35.65845 | 15.69403 | 0.379764 | 0.74127 | 0.43942 |
| 7 | 143457800 | 143459999 | 12.92829 | 22.6341 | 42.61379 | 18.80748 | -0.52036 | 0.660493 | 0.439445 |
| 19 | 37740200 | 37744599 | 48.56394 | 77.44811 | 124.2643 | 52.07694 | -0.34231 | 2.145729 | 0.43945 |
| 3 | 52899800 | 52902999 | 21.22274 | 35.52606 | 61.83728 | 26.94771 | -0.41582 | 1.173648 | 0.439454 |
| 18 | 36609200 | 36610199 | 9.971248 | 19.26766 | 20.31492 | 7.797626 | 0.479266 | -0.14356 | 0.439464 |
| 1 | 34598600 | 34600799 | 17.24733 | 29.56099 | 51.62926 | 22.64243 | -0.41783 | 0.932932 | 0.439472 |
| 7 | 69524400 | 69526199 | 13.90397 | 26.43659 | 21.89777 | 8.788861 | 0.765152 | 0.041068 | 0.439472 |
| 5 | 114700200 | 114704399 | 50.96492 | 81.02218 | 129.2044 | 53.91924 | -0.33829 | 2.251106 | 0.439475 |
| 5 | 95579400 | 95579799 | 7.861262 | 15.36502 | 15.24626 | 5.262785 | 0.704356 | -0.549 | 0.439479 |
| 7 | 3160000 | 3165399 | 61.83398 | 98.57301 | 99.1334 | 41.52306 | 0.329941 | 2.165957 | 0.439491 |
| 16 | 80556600 | 80561799 | 36.9011 | 60.66099 | 49.91291 | 21.50997 | 0.658666 | 1.412174 | 0.439498 |
| 6 | 39832600 | 39833399 | 10.32174 | 18.64353 | 35.39835 | 15.38142 | -0.51007 | 0.355688 | 0.439509 |
| 8 | 13481800 | 13483599 | 13.21271 | 23.14531 | 42.05736 | 18.5709 | -0.46957 | 0.650401 | 0.439538 |

| | | | | | | | | | |
|---|---|---|---|---|---|---|---|---|---|
| 7 | 47088400 | 47089199 | 20.11392 | 35.15444 | 34.63722 | 15.26509 | 0.409032 | 0.710647 | 0.439543 |
| 13 | 68937200 | 68939999 | 28.17565 | 47.28339 | 47.52612 | 20.74393 | 0.373661 | 1.155722 | 0.439552 |
| 1 | 192299800 | 192303999 | 37.23857 | 60.01343 | 97.9816 | 41.69574 | -0.34951 | 1.797427 | 0.439564 |
| 5 | 104841000 | 104844799 | 116.4382 | 178.3401 | 172.3602 | 71.53403 | 0.34069 | 3.036831 | 0.439569 |
| 17 | 34029200 | 34030599 | 7.89874 | 14.25108 | 34.46586 | 14.99718 | -0.85787 | 0.340422 | 0.439572 |
| 13 | 68886200 | 68888199 | 15.7186 | 27.35016 | 46.83789 | 20.58335 | -0.3855 | 0.769693 | 0.439596 |
| 8 | 101081200 | 101083799 | 27.84014 | 48.43985 | 35.66137 | 15.67139 | 0.826979 | 0.811875 | 0.439602 |
| 1 | 28954200 | 28956799 | 24.15733 | 42.44292 | 31.9034 | 14.12257 | 0.797427 | 0.647188 | 0.439612 |
| 13 | 15637200 | 15639999 | 21.20241 | 35.11992 | 67.55459 | 29.23569 | -0.56 | 1.296581 | 0.439622 |
| 1 | 33453400 | 33454999 | 20.65665 | 36.35153 | 32.4795 | 14.37967 | 0.54966 | 0.605466 | 0.439638 |
| 12 | 54201400 | 54203399 | 19.78184 | 34.96967 | 31.4944 | 13.93095 | 0.540764 | 0.551414 | 0.439639 |
| 8 | 60829800 | 60832199 | 25.30835 | 44.35427 | 32.85899 | 14.51491 | 0.818298 | 0.69206 | 0.439653 |
| X | 75403200 | 75406799 | 57.72955 | 92.75866 | 82.33028 | 34.88386 | 0.522678 | 1.916739 | 0.439682 |
| 1 | 53424600 | 53429599 | 66.0845 | 103.9259 | 159.5308 | 66.18075 | -0.29669 | 2.53123 | 0.439691 |
| 2 | 149994000 | 149996399 | 28.17062 | 47.88324 | 45.20809 | 19.8131 | 0.46416 | 1.040128 | 0.439723 |
| 2 | 118324000 | 118327399 | 18.89973 | 32.09515 | 55.33781 | 24.24852 | -0.40142 | 1.028122 | 0.439786 |
| 2 | 158352600 | 158355199 | 17.41762 | 29.34156 | 57.82921 | 25.27855 | -0.59397 | 1.09709 | 0.439814 |
| 1 | 19721000 | 19722999 | 15.31541 | 26.58276 | 56.04116 | 24.58955 | -0.68457 | 0.815618 | 0.439817 |
| 13 | 66899400 | 66904799 | 81.14996 | 128.3501 | 120.9111 | 50.24639 | 0.413245 | 2.410929 | 0.439821 |
| 18 | 16091600 | 16092799 | 11.03151 | 20.06076 | 36.50905 | 15.73045 | -0.44194 | 0.320873 | 0.439837 |
| X | 75402000 | 75407199 | 71.12302 | 113.3711 | 99.18251 | 41.5385 | 0.530567 | 2.179706 | 0.43985 |
| 7 | 25793400 | 25800399 | 119.8746 | 184.2604 | 176.9432 | 73.47521 | 0.353868 | 2.968572 | 0.439857 |
| 13 | 51073200 | 51077199 | 35.56835 | 58.90234 | 55.9886 | 24.22643 | 0.453667 | 1.327758 | 0.439858 |
| 4 | 78042000 | 78043999 | 20.85854 | 37.22948 | 28.05199 | 12.33104 | 0.80358 | 0.479598 | 0.439858 |
| 18 | 5587600 | 5588999 | 14.5365 | 27.28133 | 23.90904 | 9.843882 | 0.65366 | 0.137355 | 0.439886 |
| 11 | 50755200 | 50756399 | 22.06925 | 36.84822 | 62.89486 | 27.36416 | -0.38761 | 1.206093 | 0.439899 |
| 18 | 42502400 | 42504399 | 23.0919 | 39.91864 | 38.93706 | 17.08219 | 0.421532 | 0.864914 | 0.439909 |
| 17 | 17513600 | 17514999 | 20.5263 | 35.70481 | 33.74078 | 14.82491 | 0.468281 | 0.776032 | 0.439914 |
| 18 | 18558200 | 18561799 | 32.16741 | 54.60228 | 44.03304 | 19.29232 | 0.690633 | 1.065706 | 0.439919 |
| 5 | 19581400 | 19582999 | 21.45988 | 37.44651 | 36.53182 | 16.08304 | 0.422934 | 0.741341 | 0.439926 |
| 19 | 46634000 | 46635399 | 6.797902 | 12.80314 | 28.59876 | 11.70444 | -0.67873 | 0.057918 | 0.439927 |
| 9 | 36559800 | 36562799 | 46.18263 | 74.44727 | 74.96152 | 31.91049 | 0.346728 | 1.792609 | 0.43993 |
| 17 | 87267600 | 87273399 | 29.60004 | 47.65208 | 106.2302 | 45.13576 | -0.79969 | 1.824184 | 0.439952 |
| 15 | 103827800 | 103828399 | 13.438 | 24.69424 | 24.89354 | 10.58047 | 0.418305 | 0.252629 | 0.439964 |
| 3 | 147484600 | 147486399 | 21.83633 | 38.70149 | 29.64264 | 13.10508 | 0.774622 | 0.533859 | 0.439966 |
| 6 | 69635200 | 69638399 | 33.405 | 55.49801 | 54.14719 | 23.48174 | 0.416529 | 1.286184 | 0.439974 |
| X | 141615200 | 141619999 | 25.39258 | 42.09309 | 71.94879 | 31.15399 | -0.3902 | 1.30442 | 0.439977 |
| 17 | 29686600 | 29688199 | 7.865972 | 14.61987 | 30.51757 | 12.77204 | -0.60818 | 0.15743 | 0.43999 |
| 16 | 59041000 | 59041999 | 17.17327 | 30.38156 | 29.66661 | 13.12045 | 0.425 | 0.522198 | 0.440021 |
| 5 | 95654200 | 95666199 | 202.7595 | 302.2535 | 262.9799 | 108.3745 | 0.458993 | 3.591384 | 0.440029 |
| 1 | 181677800 | 181681399 | 23.95653 | 39.45855 | 70.92829 | 30.59178 | -0.4642 | 1.373988 | 0.440034 |
| 11 | 31024000 | 31027999 | 44.7484 | 71.46983 | 132.4248 | 55.61352 | -0.54724 | 2.09979 | 0.440045 |
| 6 | 59106200 | 59108399 | 19.30775 | 32.57639 | 64.56871 | 28.2379 | -0.6023 | 1.095748 | 0.440084 |
| 15 | 76451200 | 76453799 | 26.76958 | 45.86063 | 43.02893 | 18.86999 | 0.474816 | 0.963452 | 0.440125 |
| 14 | 37223200 | 37227599 | 34.6238 | 55.95333 | 91.75229 | 39.1838 | -0.35115 | 1.710615 | 0.440136 |
| 2 | 44797000 | 44798399 | 23.55283 | 40.89155 | 34.62459 | 15.19508 | 0.625787 | 0.819716 | 0.440137 |
| 5 | 147687800 | 147692599 | 43.3668 | 69.30696 | 114.3977 | 48.08577 | -0.37969 | 2.072582 | 0.440139 |

| | | | | | | | | | |
|---|---|---|---|---|---|---|---|---|---|
| 17 | 89792400 | 89795599 | 31.32478 | 52.60158 | 49.66893 | 21.68791 | 0.463464 | 1.161261 | 0.440167 |
| 5 | 130651800 | 130652399 | 5.381295 | 10.43725 | 23.27762 | 8.780448 | -0.56622 | -0.24165 | 0.440179 |
| 13 | 35100400 | 35101199 | 5.381295 | 10.43725 | 23.27762 | 8.780448 | -0.56622 | -0.24165 | 0.440179 |
| 17 | 68178400 | 68180399 | 11.7447 | 20.72062 | 39.85454 | 17.63591 | -0.54915 | 0.539019 | 0.440194 |
| 6 | 140822000 | 140824399 | 23.7507 | 41.38743 | 36.9301 | 16.24446 | 0.550896 | 0.772031 | 0.440202 |
| 6 | 10623400 | 10626999 | 31.18771 | 53.20965 | 42.35396 | 18.56782 | 0.71027 | 1.01598 | 0.440218 |
| 5 | 19388000 | 19390199 | 22.01525 | 38.32426 | 37.68343 | 16.56523 | 0.411403 | 0.779706 | 0.440225 |
| 6 | 59175200 | 59177599 | 23.96233 | 39.70217 | 69.36249 | 30.02368 | -0.42159 | 1.305831 | 0.440233 |
| 10 | 32836200 | 32837599 | 13.02723 | 24.1796 | 24.48706 | 10.26728 | 0.425897 | 0.194061 | 0.44024 |
| 1 | 167488000 | 167489799 | 11.77079 | 20.70822 | 40.41313 | 17.89316 | -0.57154 | 0.571936 | 0.440256 |
| 4 | 31552000 | 31554199 | 15.35702 | 27.643 | 27.16735 | 11.85586 | 0.429478 | 0.388285 | 0.440285 |
| X | 63621000 | 63623399 | 32.77484 | 54.89809 | 45.39211 | 19.78896 | 0.654455 | 1.188556 | 0.440295 |
| 4 | 147253400 | 147255199 | 21.08298 | 37.08368 | 29.10003 | 12.88473 | 0.735872 | 0.609203 | 0.440329 |
| 14 | 22377000 | 22378799 | 20.84763 | 36.41575 | 36.04277 | 15.87805 | 0.402371 | 0.723684 | 0.440334 |
| 1 | 118284400 | 118286399 | 16.9005 | 30.64215 | 27.54299 | 11.91154 | 0.56592 | 0.359834 | 0.440359 |
| 17 | 48882800 | 48885999 | 33.81017 | 54.76959 | 89.11583 | 38.14133 | -0.33647 | 1.653582 | 0.440369 |
| X | 81203000 | 81209999 | 96.9573 | 151.8068 | 135.3403 | 56.14139 | 0.481375 | 2.629162 | 0.4404 |
| X | 89448200 | 89449399 | 20.63607 | 36.11551 | 35.00196 | 15.44334 | 0.432656 | 0.692577 | 0.440441 |
| 2 | 140012000 | 140012999 | 7.375824 | 13.91004 | 28.08467 | 11.46825 | -0.53126 | 0.055169 | 0.440447 |
| 14 | 45409600 | 45411999 | 23.43991 | 40.46499 | 40.41117 | 17.72817 | 0.387147 | 0.878319 | 0.440451 |
| 10 | 17790800 | 17791999 | 9.913245 | 17.5992 | 37.36963 | 16.50122 | -0.68664 | 0.460972 | 0.440455 |
| 1 | 173314600 | 173315799 | 19.59711 | 34.4561 | 28.68255 | 12.70091 | 0.650755 | 0.602491 | 0.440457 |
| 17 | 48782400 | 48783999 | 12.36751 | 21.97583 | 38.26521 | 16.88402 | -0.40156 | 0.473973 | 0.440476 |
| 6 | 131813600 | 131816199 | 33.35376 | 55.64506 | 52.50971 | 22.83429 | 0.464591 | 1.238605 | 0.440478 |
| 14 | 46585600 | 46588599 | 39.52689 | 65.22044 | 56.76359 | 24.50093 | 0.580053 | 1.379712 | 0.440478 |
| 1 | 117939000 | 117943599 | 90.48956 | 141.1018 | 142.0876 | 58.96557 | 0.3013 | 2.694703 | 0.440491 |
| 3 | 138910400 | 138912599 | 25.03572 | 41.47052 | 69.06374 | 29.88933 | -0.35283 | 1.309066 | 0.440494 |
| 16 | 30327600 | 30329799 | 11.16339 | 19.66967 | 39.42243 | 17.44919 | -0.6087 | 0.550376 | 0.440494 |
| 8 | 125874400 | 125876799 | 18.21324 | 31.0293 | 54.29282 | 23.79902 | -0.42193 | 0.99381 | 0.440508 |
| 17 | 56507400 | 56511999 | 28.91273 | 47.01987 | 81.34436 | 34.90928 | -0.4195 | 1.563292 | 0.440513 |
| X | 21627600 | 21631399 | 27.9979 | 48.10531 | 36.68776 | 16.09241 | 0.773281 | 0.936713 | 0.440539 |
| 7 | 18691200 | 18701999 | 78.28691 | 120.749 | 200.03 | 83.08887 | -0.42823 | 2.889543 | 0.440564 |
| 4 | 70486600 | 70487399 | 8.283247 | 15.65438 | 29.78253 | 12.10764 | -0.44091 | 0.053053 | 0.440566 |
| 12 | 106312600 | 106314399 | 31.28537 | 52.00482 | 53.02191 | 23.00239 | 0.353232 | 1.27099 | 0.440574 |
| 4 | 57978400 | 57982599 | 53.88331 | 85.65831 | 133.0592 | 55.5257 | -0.30058 | 2.255115 | 0.440591 |
| 17 | 33769400 | 33770399 | 7.701055 | 14.41438 | 29.04274 | 12.01175 | -0.54328 | 0.109038 | 0.440592 |
| 12 | 115230800 | 115236599 | 97.49822 | 152.5037 | 141.042 | 58.51246 | 0.427407 | 2.64587 | 0.4406 |
| 11 | 121505000 | 121506399 | 23.20027 | 39.98953 | 39.98591 | 17.54079 | 0.38501 | 0.891726 | 0.440626 |
| 9 | 5279000 | 5281399 | 42.85718 | 69.36208 | 70.49134 | 30.14995 | 0.339757 | 1.67861 | 0.440634 |
| 2 | 88126400 | 88126799 | 7.197512 | 13.69913 | 27.1828 | 10.88649 | -0.48159 | -0.01266 | 0.44064 |
| 17 | 72125000 | 72129399 | 49.08371 | 79.09558 | 79.29893 | 33.74088 | 0.35236 | 1.810836 | 0.440643 |
| 9 | 35616600 | 35617799 | 9.642631 | 17.4913 | 38.07831 | 16.45796 | -0.70165 | 0.327788 | 0.440653 |
| 2 | 89931800 | 89933999 | 29.70211 | 49.83479 | 49.66487 | 21.67463 | 0.385938 | 1.165257 | 0.440675 |
| 7 | 38390600 | 38398599 | 128.9423 | 197.6022 | 168.7028 | 70.02445 | 0.520064 | 3.028837 | 0.440684 |
| 3 | 89561800 | 89563999 | 13.94638 | 24.40504 | 43.58454 | 19.22829 | -0.445 | 0.671294 | 0.440684 |
| 2 | 32017800 | 32018399 | 3.815204 | 7.454148 | 18.77519 | 6.542408 | -0.63316 | -0.56092 | 0.440689 |
| 7 | 37612400 | 37612999 | 7.606001 | 14.52801 | 28.44646 | 11.30749 | -0.45379 | -0.03256 | 0.44069 |

| | | | | | | | | | |
|---|---|---|---|---|---|---|---|---|---|
| 4 | 79801000 | 79801799 | 12.25546 | 23.41052 | 22.17324 | 8.812518 | 0.585968 | -0.00343 | 0.440693 |
| 19 | 30587200 | 30588399 | 11.79893 | 22.59663 | 21.64732 | 8.543483 | 0.578602 | -0.0329 | 0.4407 |
| 11 | 3998000 | 4001199 | 72.12998 | 114.0638 | 105.8321 | 43.97621 | 0.434929 | 2.404939 | 0.440714 |
| X | 91081000 | 91085799 | 61.87981 | 99.33189 | 85.03893 | 35.9798 | 0.572988 | 1.950547 | 0.440715 |
| 6 | 80938600 | 80939399 | 12.30409 | 23.12593 | 23.52141 | 9.660003 | 0.443097 | 0.106843 | 0.440717 |
| 2 | 168864200 | 168867399 | 33.01615 | 53.40394 | 87.98748 | 37.61717 | -0.35644 | 1.681725 | 0.440741 |
| 14 | 42882800 | 42885399 | 23.17687 | 41.16039 | 27.8756 | 12.30543 | 0.952671 | 0.525522 | 0.440756 |
| 6 | 131544000 | 131546199 | 22.13226 | 37.19759 | 63.70004 | 27.82529 | -0.39252 | 1.13548 | 0.440771 |
| 8 | 34898200 | 34900399 | 29.23237 | 49.07623 | 49.12281 | 21.44622 | 0.379572 | 1.154906 | 0.440771 |
| 7 | 18360600 | 18365199 | 66.56734 | 106.1286 | 93.05668 | 38.98276 | 0.527532 | 2.164053 | 0.440783 |
| 2 | 144021000 | 144025799 | 34.63444 | 55.97058 | 91.07787 | 38.89583 | -0.34008 | 1.710756 | 0.440796 |
| 13 | 94721800 | 94725599 | 32.30932 | 52.45632 | 86.42282 | 37.05019 | -0.35137 | 1.603337 | 0.440804 |
| 2 | 16893400 | 16897799 | 47.39614 | 76.63563 | 73.50561 | 31.41342 | 0.421668 | 1.707609 | 0.440818 |
| 1 | 67581600 | 67582599 | 11.60702 | 20.50301 | 38.358 | 16.96949 | -0.50843 | 0.523681 | 0.440829 |
| 14 | 44482400 | 44489599 | 86.52934 | 132.7776 | 229.4162 | 95.24212 | -0.49463 | 2.995622 | 0.440852 |
| 1 | 85964600 | 85966599 | 11.05993 | 19.77581 | 36.27501 | 15.92161 | -0.47094 | 0.420634 | 0.440879 |
| 7 | 4151400 | 4153399 | 17.81089 | 30.15072 | 56.35628 | 24.67115 | -0.51764 | 1.0519 | 0.440893 |
| 7 | 31837400 | 31848399 | 142.2321 | 217.9131 | 176.9175 | 73.4306 | 0.592297 | 3.040345 | 0.440906 |
| 7 | 47395800 | 47400199 | 52.53239 | 84.80507 | 75.08271 | 32.02143 | 0.534309 | 1.762713 | 0.440914 |
| 18 | 75183600 | 75184799 | 8.28072 | 15.44537 | 30.57021 | 12.71843 | -0.52534 | 0.137716 | 0.44093 |
| 5 | 67573800 | 67575999 | 12.64844 | 22.3802 | 39.77334 | 17.58972 | -0.43412 | 0.512272 | 0.440943 |
| 16 | 53409000 | 53410199 | 16.62314 | 30.72266 | 25.21974 | 10.636 | 0.720984 | 0.248755 | 0.440943 |
| 2 | 87015600 | 87016799 | 12.39717 | 23.62423 | 22.04152 | 8.804928 | 0.600624 | 0.011926 | 0.440945 |
| 16 | 78189400 | 78190999 | 13.25373 | 25.16201 | 22.48562 | 9.053825 | 0.652508 | 0.046638 | 0.440958 |
| 6 | 61157000 | 61158199 | 10.16135 | 18.1646 | 35.83209 | 15.73049 | -0.5752 | 0.415754 | 0.440958 |
| X | 113898400 | 113903399 | 70.89101 | 113.0574 | 99.09242 | 41.52933 | 0.528639 | 2.1631 | 0.440964 |
| 5 | 88887000 | 88890399 | 42.84956 | 69.46102 | 70.17781 | 30.05135 | 0.350356 | 1.646283 | 0.440975 |
| 17 | 57728600 | 57731599 | 30.49528 | 49.81446 | 83.94721 | 36.08473 | -0.37727 | 1.504753 | 0.440979 |
| 12 | 23194600 | 23197199 | 56.46518 | 91.25836 | 69.74192 | 29.78764 | 0.747986 | 1.735952 | 0.440995 |
| 12 | 118202800 | 118205999 | 44.51534 | 73.11608 | 58.00911 | 24.99009 | 0.710735 | 1.449764 | 0.441009 |
| 6 | 58694600 | 58695599 | 13.48789 | 25.06853 | 24.93856 | 10.43735 | 0.453238 | 0.19099 | 0.441012 |
| 2 | 63435800 | 63438599 | 103.6441 | 160.7434 | 137.3849 | 57.05997 | 0.531578 | 2.791355 | 0.441012 |
| 6 | 17064200 | 17065799 | 12.36184 | 21.90436 | 38.25353 | 16.91275 | -0.40803 | 0.493588 | 0.441013 |
| 10 | 114774400 | 114775599 | 12.30713 | 23.39698 | 22.59386 | 9.071052 | 0.545243 | 0.029143 | 0.441024 |
| 18 | 80072000 | 80073799 | 14.27083 | 24.96794 | 43.588 | 19.22665 | -0.41235 | 0.676713 | 0.441061 |
| 17 | 94102400 | 94103399 | 15.01922 | 27.59005 | 20.38893 | 8.670115 | 0.862959 | 0.257294 | 0.441074 |
| 7 | 72830200 | 72837399 | 67.24698 | 105.1339 | 182.513 | 75.7209 | -0.48057 | 2.644028 | 0.441085 |
| 17 | 47538800 | 47542799 | 53.67493 | 84.89858 | 141.5075 | 58.80696 | -0.4087 | 2.394579 | 0.441093 |
| 10 | 125624800 | 125626199 | 17.82443 | 31.50983 | 30.59469 | 13.53294 | 0.432698 | 0.536004 | 0.441104 |
| 7 | 36026600 | 36030199 | 28.24602 | 47.49596 | 48.02934 | 20.98313 | 0.364916 | 1.135878 | 0.441106 |
| 16 | 44978400 | 44979799 | 11.44959 | 20.22794 | 38.35801 | 16.96899 | -0.52774 | 0.520751 | 0.44111 |
| 5 | 105786800 | 105788399 | 13.8228 | 24.14833 | 43.64978 | 19.23081 | -0.46224 | 0.698726 | 0.441118 |
| 3 | 128817000 | 128820199 | 59.26317 | 93.63013 | 146.4072 | 60.80788 | -0.31807 | 2.426995 | 0.441125 |
| 6 | 131833400 | 131838399 | 96.735 | 150.5913 | 150.3595 | 62.41377 | 0.311714 | 2.727968 | 0.441128 |
| 5 | 134054800 | 134058399 | 22.501 | 37.14315 | 70.3803 | 30.38645 | -0.53887 | 1.341807 | 0.441178 |
| 9 | 11168800 | 11169999 | 17.18359 | 30.37106 | 29.72127 | 13.14711 | 0.421377 | 0.532786 | 0.441186 |
| 12 | 97566000 | 97567599 | 12.46968 | 22.18251 | 39.7951 | 17.54229 | -0.4437 | 0.46803 | 0.441186 |

| | | | | | | | | | |
|---|---|---|---|---|---|---|---|---|---|
| 16 | 36054200 | 36061399 | 112.5364 | 174.182 | 159.9823 | 66.45255 | 0.425637 | 2.836583 | 0.441226 |
| 4 | 151063800 | 151072399 | 74.78573 | 115.4715 | 213.7016 | 88.76732 | -0.58675 | 2.87016 | 0.441229 |
| X | 31698400 | 31703399 | 76.67529 | 122.1851 | 98.96037 | 41.42841 | 0.641177 | 2.19343 | 0.441246 |
| 8 | 8544000 | 8547799 | 28.28998 | 45.97422 | 81.68525 | 35.04364 | -0.45866 | 1.573982 | 0.441258 |
| 15 | 39060000 | 39062799 | 24.16229 | 39.31899 | 88.07039 | 37.80441 | -0.79043 | 1.549092 | 0.441278 |
| 12 | 23248000 | 23258999 | 260.4145 | 376.5712 | 356.556 | 147.0661 | 0.295434 | 4.083141 | 0.441283 |
| 8 | 67244000 | 67244799 | 14.92302 | 28.59139 | 20.33918 | 8.017778 | 1.006303 | -0.0083 | 0.441288 |
| X | 157277800 | 157281199 | 33.6187 | 57.41108 | 40.06066 | 17.56377 | 0.899695 | 1.0156 | 0.441301 |
| 16 | 95204400 | 95206999 | 18.52284 | 31.54645 | 54.84439 | 24.04017 | -0.41282 | 0.998216 | 0.441311 |
| X | 5446000 | 5446799 | 18.63884 | 32.92712 | 29.89066 | 13.22333 | 0.529331 | 0.551384 | 0.441314 |
| 5 | 51377000 | 51379599 | 35.0765 | 58.7737 | 49.17837 | 21.44427 | 0.637316 | 1.194735 | 0.441316 |
| 3 | 103111200 | 103114199 | 23.80371 | 39.65834 | 67.60148 | 29.3718 | -0.38583 | 1.236329 | 0.44133 |
| 14 | 36783400 | 36785799 | 20.80394 | 35.16377 | 59.79425 | 26.1641 | -0.38199 | 1.078303 | 0.441333 |
| 12 | 41832200 | 41833399 | 14.10716 | 26.74883 | 22.40725 | 9.046126 | 0.741808 | 0.062918 | 0.441334 |
| 19 | 8227000 | 8227799 | 18.54887 | 32.55012 | 32.43399 | 14.3429 | 0.392635 | 0.632729 | 0.441354 |
| 2 | 15457000 | 15458599 | 21.81671 | 37.8177 | 37.26376 | 16.34946 | 0.40752 | 0.829009 | 0.441383 |
| X | 36327200 | 36329599 | 22.34811 | 39.42642 | 31.31599 | 13.85763 | 0.720064 | 0.588651 | 0.44143 |
| 17 | 14575000 | 14576799 | 14.15507 | 25.75221 | 25.41926 | 10.94625 | 0.435823 | 0.316689 | 0.441431 |
| 7 | 14306600 | 14342199 | 602.2035 | 813.8485 | 703.628 | 293.3202 | 0.328486 | 5.11194 | 0.441432 |
| 18 | 70620200 | 70622799 | 43.1652 | 70.49741 | 63.0229 | 27.08851 | 0.534762 | 1.519904 | 0.441446 |
| 1 | 30825200 | 30827199 | 13.61653 | 24.04519 | 45.05381 | 19.93294 | -0.51142 | 0.548616 | 0.441449 |
| X | 117907800 | 117910999 | 39.68582 | 65.25906 | 58.14103 | 25.05918 | 0.544611 | 1.426532 | 0.441462 |
| 5 | 56792600 | 56794399 | 22.70672 | 39.4657 | 38.16212 | 16.75904 | 0.435132 | 0.804176 | 0.441515 |
| 15 | 103869400 | 103881199 | 197.6554 | 294.3258 | 269.8636 | 111.1983 | 0.381978 | 3.609956 | 0.44153 |
| 17 | 92122600 | 92123799 | 15.22816 | 27.63362 | 27.20135 | 11.75124 | 0.436584 | 0.341867 | 0.441531 |
| 1 | 80734000 | 80736999 | 20.04172 | 33.70967 | 61.35011 | 26.80498 | -0.47992 | 1.119043 | 0.441545 |
| 12 | 51648600 | 51650799 | 13.22299 | 23.38813 | 42.41612 | 18.75988 | -0.46331 | 0.527485 | 0.441554 |
| 12 | 72454000 | 72456199 | 19.23116 | 32.09513 | 62.65461 | 27.2519 | -0.58054 | 1.201725 | 0.441562 |
| X | 90180000 | 90187599 | 80.68734 | 128.2403 | 105.6998 | 44.0988 | 0.612016 | 2.28002 | 0.441564 |
| 10 | 125654200 | 125657199 | 26.21931 | 44.9836 | 42.47795 | 18.63055 | 0.465858 | 0.947005 | 0.441591 |
| X | 107443200 | 107444399 | 16.16472 | 29.6823 | 25.79331 | 10.9743 | 0.630412 | 0.2825 | 0.441599 |
| X | 76315200 | 76318599 | 28.77159 | 50.06821 | 35.08399 | 15.41916 | 0.897971 | 0.814164 | 0.441607 |
| 4 | 122548600 | 122549599 | 16.51643 | 29.77799 | 28.03418 | 12.20948 | 0.492991 | 0.389211 | 0.441609 |
| 12 | 114991800 | 114999199 | 110.6049 | 172.6274 | 139.2657 | 57.78851 | 0.621852 | 2.690063 | 0.441623 |
| Y | 1893600 | 1896599 | 22.16007 | 38.5676 | 23.34383 | 10.26023 | 1.107593 | 0.75722 | 0.441627 |
| 2 | 87952800 | 87954799 | 23.45659 | 38.79236 | 75.68923 | 32.72039 | -0.58083 | 1.327996 | 0.441631 |
| 16 | 64925200 | 64926599 | 9.428274 | 17.34022 | 31.94723 | 13.56314 | -0.44436 | 0.23492 | 0.441632 |
| 4 | 16148000 | 16149799 | 17.8084 | 32.17959 | 27.35304 | 11.87937 | 0.642055 | 0.384157 | 0.441643 |
| 15 | 20354400 | 20356799 | 22.2233 | 39.13555 | 31.81383 | 14.08301 | 0.685737 | 0.606821 | 0.441643 |
| 1 | 189780800 | 189783199 | 26.21183 | 44.88214 | 43.09139 | 18.8966 | 0.441697 | 0.967124 | 0.441672 |
| 13 | 66762800 | 66766799 | 52.7501 | 85.22059 | 73.41428 | 31.34025 | 0.574935 | 1.741625 | 0.441684 |
| X | 103876000 | 103878799 | 37.44764 | 62.24235 | 47.66248 | 20.67598 | 0.76493 | 1.286799 | 0.441696 |
| 14 | 11081400 | 11084399 | 29.01226 | 47.55198 | 79.29671 | 34.13022 | -0.35923 | 1.453408 | 0.441702 |
| X | 153775600 | 153779799 | 57.57593 | 92.78781 | 76.5138 | 32.55965 | 0.633843 | 1.820957 | 0.441702 |
| 13 | 88476200 | 88477799 | 22.40723 | 39.38617 | 31.74069 | 14.05585 | 0.697086 | 0.626265 | 0.44173 |
| 12 | 18041000 | 18080599 | 1026.455 | 1327.322 | 1102.189 | 461.9678 | 0.323064 | 5.822855 | 0.441731 |
| 19 | 18467000 | 18467999 | 14.39955 | 25.21722 | 44.5948 | 19.68663 | -0.43101 | 0.662258 | 0.441754 |

| | | | | | | | | | |
|---|---|---|---|---|---|---|---|---|---|
| 11 | 24113400 | 24114999 | 17.33185 | 29.42627 | 54.73873 | 23.98444 | -0.51044 | 1.023967 | 0.441782 |
| 4 | 22275800 | 22277999 | 22.60951 | 39.43455 | 35.83985 | 15.77388 | 0.524497 | 0.750737 | 0.441783 |
| 3 | 72005000 | 72005799 | 15.8471 | 28.43531 | 28.28319 | 12.37717 | 0.409829 | 0.415301 | 0.441802 |
| 5 | 67304400 | 67306799 | 18.76386 | 33.16585 | 31.59536 | 13.97596 | 0.460072 | 0.546474 | 0.441813 |
| 3 | 145614400 | 145615599 | 10.83537 | 20.65393 | 21.63807 | 8.638546 | 0.435684 | -0.00561 | 0.441813 |
| 7 | 12565200 | 12565999 | 10.83537 | 20.65393 | 21.63807 | 8.638546 | 0.435684 | -0.00561 | 0.441813 |
| 7 | 42328000 | 42340999 | 267.4959 | 389.9711 | 334.4915 | 137.8369 | 0.449403 | 3.962529 | 0.441816 |
| 15 | 35795600 | 35797399 | 18.85653 | 33.43367 | 30.39134 | 13.43429 | 0.528966 | 0.511113 | 0.441818 |
| 8 | 25249000 | 25252999 | 72.67229 | 114.8699 | 114.4404 | 47.54241 | 0.331905 | 2.417716 | 0.441847 |
| 1 | 14795000 | 14797599 | 31.8682 | 53.33476 | 50.82982 | 22.14785 | 0.450199 | 1.197509 | 0.441848 |
| X | 19380800 | 19386999 | 70.54578 | 112.8653 | 89.64973 | 37.74103 | 0.675354 | 2.064017 | 0.44186 |
| 6 | 134850200 | 134852199 | 10.67776 | 19.32532 | 35.34366 | 15.31918 | -0.45466 | 0.34166 | 0.441862 |
| 6 | 105007400 | 105010799 | 56.36586 | 90.17635 | 91.34424 | 38.45279 | 0.325095 | 2.046148 | 0.441864 |
| 11 | 121837400 | 121840399 | 25.83898 | 44.85956 | 37.99569 | 16.67423 | 0.625296 | 0.834188 | 0.441872 |
| 2 | 120624200 | 120627799 | 62.73186 | 100.2432 | 96.96816 | 40.75248 | 0.389641 | 2.091585 | 0.441891 |
| 5 | 101286200 | 101289199 | 24.12035 | 41.99241 | 37.49799 | 16.4846 | 0.549744 | 0.787895 | 0.441904 |
| 5 | 32423800 | 32426199 | 21.72736 | 38.03129 | 34.71893 | 15.32048 | 0.518304 | 0.692769 | 0.441908 |
| 9 | 65341200 | 65343599 | 14.14543 | 24.65776 | 44.72754 | 19.67847 | -0.46767 | 0.733261 | 0.44191 |
| 15 | 22576200 | 22579199 | 39.19104 | 64.06515 | 63.66447 | 27.37336 | 0.383481 | 1.500829 | 0.441921 |
| 5 | 109837200 | 109839799 | 22.3879 | 37.28407 | 64.57034 | 28.04748 | -0.40852 | 1.235063 | 0.441975 |
| 10 | 45156400 | 45158799 | 19.33739 | 34.27231 | 30.33197 | 13.40985 | 0.567066 | 0.517886 | 0.44198 |
| 7 | 17595200 | 17605999 | 205.256 | 305.4183 | 269.8765 | 111.1953 | 0.434274 | 3.624124 | 0.441984 |
| X | 164256800 | 164258799 | 22.91921 | 41.03163 | 26.3522 | 11.5543 | 1.035326 | 0.473309 | 0.441986 |
| 17 | 7436800 | 7438399 | 11.15369 | 19.76607 | 37.36063 | 16.51757 | -0.52103 | 0.485001 | 0.441988 |
| 5 | 105018000 | 105036799 | 518.759 | 709.0743 | 625.7429 | 260.3962 | 0.315401 | 4.932848 | 0.441988 |
| 1 | 35230200 | 35236799 | 82.64866 | 129.963 | 131.5871 | 54.58775 | 0.302797 | 2.536959 | 0.441989 |
| 12 | 115728000 | 115730999 | 37.33431 | 61.75321 | 56.5815 | 24.46285 | 0.506425 | 1.34784 | 0.441991 |
| 3 | 20277200 | 20279399 | 14.12764 | 24.62637 | 45.24111 | 19.90423 | -0.48598 | 0.733161 | 0.442006 |
| 10 | 76515200 | 76519599 | 36.62421 | 58.90366 | 99.85467 | 42.36807 | -0.40727 | 1.861092 | 0.442011 |
| X | 94814600 | 94816999 | 38.20124 | 63.25926 | 55.48712 | 24.00859 | 0.569306 | 1.336998 | 0.442024 |
| 14 | 108574800 | 108579199 | 58.79289 | 92.9481 | 146.5153 | 60.87295 | -0.32891 | 2.411779 | 0.442027 |
| 5 | 109516800 | 109519999 | 36.85546 | 61.34063 | 52.10447 | 22.62394 | 0.615894 | 1.271808 | 0.442032 |
| 11 | 32138200 | 32141599 | 14.27205 | 24.52909 | 50.53861 | 22.16802 | -0.65432 | 0.893505 | 0.442032 |
| 15 | 58952000 | 58954599 | 16.40451 | 27.82719 | 55.23338 | 24.19468 | -0.60368 | 1.028449 | 0.442033 |
| 4 | 28656800 | 28657599 | 11.98561 | 22.64303 | 23.06664 | 9.384748 | 0.452017 | 0.069924 | 0.442045 |
| X | 148999000 | 149001799 | 47.8453 | 77.64187 | 68.60047 | 29.39895 | 0.544745 | 1.634386 | 0.442058 |
| 16 | 73634200 | 73637799 | 31.00478 | 52.90603 | 42.35406 | 18.5681 | 0.702066 | 1.013705 | 0.44206 |
| 5 | 149207600 | 149209799 | 21.01068 | 35.34165 | 60.2708 | 26.33392 | -0.3864 | 1.121946 | 0.442067 |
| 5 | 28237000 | 28239399 | 19.22206 | 32.55172 | 56.41928 | 24.70153 | -0.40916 | 1.053983 | 0.442068 |
| 15 | 66336200 | 66336799 | 5.071918 | 9.894419 | 21.58292 | 7.718487 | -0.46396 | -0.44162 | 0.442088 |
| 1 | 7707800 | 7710399 | 34.6787 | 57.88129 | 51.98988 | 22.6146 | 0.535546 | 1.237451 | 0.442096 |
| 3 | 23235000 | 23238599 | 47.99296 | 77.33178 | 78.11529 | 33.23343 | 0.341381 | 1.809673 | 0.44212 |
| 7 | 84985600 | 84989399 | 88.8091 | 139.8871 | 117.5114 | 48.75932 | 0.5737 | 2.506357 | 0.44214 |
| 16 | 32019000 | 32021999 | 26.68315 | 43.9167 | 72.95003 | 31.45382 | -0.35116 | 1.391687 | 0.44215 |
| 9 | 29974200 | 29976199 | 13.78833 | 24.17475 | 42.49083 | 18.77615 | -0.42208 | 0.63792 | 0.442166 |
| 1 | 7865200 | 7867199 | 21.84675 | 38.5246 | 31.25921 | 13.83375 | 0.689065 | 0.588879 | 0.442173 |
| 19 | 8266600 | 8270799 | 86.29941 | 134.9229 | 133.894 | 55.54975 | 0.325308 | 2.645689 | 0.44221 |

| | | | | | | | | | |
|---|---|---|---|---|---|---|---|---|---|
| 2 | 47430800 | 47437799 | 72.486 | 114.8458 | 116.912 | 48.62717 | 0.303517 | 2.362701 | 0.442221 |
| 11 | 70223800 | 70225599 | 18.76284 | 32.88875 | 31.9083 | 14.09708 | 0.430882 | 0.649634 | 0.442237 |
| 17 | 21352000 | 21355799 | 30.07731 | 48.81386 | 82.81822 | 35.49487 | -0.39405 | 1.605883 | 0.442243 |
| 5 | 7992000 | 7995799 | 80.90806 | 127.0965 | 127.7194 | 52.97626 | 0.312528 | 2.556082 | 0.442247 |
| 12 | 52986200 | 52987399 | 7.982536 | 14.96772 | 29.0605 | 11.98311 | -0.48698 | 0.099337 | 0.44225 |
| 15 | 76646000 | 76647599 | 7.982536 | 14.96772 | 29.0605 | 11.98311 | -0.48698 | 0.099337 | 0.44225 |
| 4 | 26315600 | 26318999 | 41.18894 | 68.05581 | 54.56569 | 23.57382 | 0.698251 | 1.367218 | 0.442254 |
| 16 | 30465400 | 30469199 | 29.65691 | 48.57183 | 83.57208 | 35.95888 | -0.40475 | 1.466071 | 0.442259 |
| 19 | 14285000 | 14286599 | 13.65442 | 24.08203 | 41.87622 | 18.53172 | -0.4047 | 0.559156 | 0.442265 |
| 1 | 30725600 | 30730399 | 81.75584 | 128.672 | 129.9229 | 53.90353 | 0.30773 | 2.520048 | 0.442267 |
| 7 | 94167800 | 94174199 | 43.31129 | 71.44556 | 48.37787 | 20.87901 | 0.940906 | 1.387273 | 0.442267 |
| 8 | 9440200 | 9440799 | 5.525598 | 10.70241 | 23.77769 | 9.026876 | -0.56955 | -0.21587 | 0.442273 |
| 10 | 79009000 | 79009599 | 5.525598 | 10.70241 | 23.77769 | 9.026876 | -0.56955 | -0.21587 | 0.442273 |
| 17 | 43621200 | 43622199 | 7.970293 | 15.04565 | 28.64202 | 11.67229 | -0.44475 | 0.055174 | 0.442307 |
| X | 94511600 | 94513199 | 33.76956 | 56.21648 | 53.05063 | 23.03792 | 0.464649 | 1.264246 | 0.442325 |
| 8 | 112621000 | 112622199 | 12.91365 | 22.85123 | 41.38142 | 18.30061 | -0.46082 | 0.519544 | 0.442329 |
| 8 | 76841200 | 76842799 | 14.90082 | 27.40385 | 22.75601 | 9.661654 | 0.697373 | 0.255567 | 0.442354 |
| 9 | 3469000 | 3469999 | 9.410547 | 17.4655 | 32.08878 | 13.45594 | -0.42749 | 0.183967 | 0.442358 |
| 6 | 11707400 | 11710199 | 24.80107 | 43.11255 | 37.45142 | 16.44855 | 0.58907 | 0.810528 | 0.442381 |
| 3 | 77672000 | 77673799 | 21.18617 | 37.33967 | 31.87723 | 14.10877 | 0.615852 | 0.591746 | 0.442387 |
| 3 | 39311000 | 39313399 | 14.93471 | 26.10073 | 44.66004 | 19.68122 | -0.38352 | 0.700652 | 0.4424 |
| 13 | 103165400 | 103168799 | 35.80718 | 57.71712 | 94.46236 | 40.20758 | -0.35266 | 1.789577 | 0.442402 |
| 13 | 36118800 | 36120599 | 17.63994 | 33.04126 | 22.52755 | 9.305754 | 1.009551 | 0.172859 | 0.44241 |
| 5 | 63961800 | 63966799 | 68.84298 | 107.697 | 167.297 | 69.40357 | -0.3197 | 2.631349 | 0.442416 |
| 13 | 57964200 | 57969999 | 58.25302 | 92.13172 | 143.8031 | 59.7583 | -0.31423 | 2.40059 | 0.442426 |
| 13 | 62415000 | 62426399 | 218.7517 | 324.304 | 279.5424 | 115.1347 | 0.465167 | 3.690506 | 0.442445 |
| 4 | 55396600 | 55404999 | 72.65222 | 113.4747 | 173.065 | 71.80826 | -0.29505 | 2.661958 | 0.442446 |
| 13 | 62364400 | 62365999 | 39.1676 | 63.82182 | 65.17627 | 27.99063 | 0.340995 | 1.546566 | 0.442449 |
| 10 | 41827400 | 41831799 | 46.5092 | 74.16928 | 119.7958 | 50.20242 | -0.35185 | 2.144978 | 0.442461 |
| 18 | 46461600 | 46463799 | 22.30121 | 38.92606 | 32.29907 | 14.22222 | 0.655364 | 0.734438 | 0.442463 |
| 2 | 53136600 | 53138399 | 18.70603 | 33.29184 | 29.89548 | 13.17325 | 0.549579 | 0.486366 | 0.442464 |
| 5 | 94924600 | 94925999 | 21.0447 | 37.2069 | 30.77036 | 13.61033 | 0.663132 | 0.555838 | 0.442496 |
| 17 | 12489600 | 12491599 | 19.88408 | 33.12781 | 63.16036 | 27.4421 | -0.54653 | 1.223827 | 0.442507 |
| 2 | 23600400 | 23603599 | 45.10879 | 74.61444 | 51.1847 | 22.12992 | 0.922758 | 1.365586 | 0.442513 |
| X | 76944000 | 76950599 | 70.7588 | 113.3827 | 84.10302 | 35.48755 | 0.776448 | 2.016119 | 0.442521 |
| 10 | 6583800 | 6584199 | 5.518607 | 10.74358 | 23.08785 | 8.473457 | -0.47741 | -0.34021 | 0.442535 |
| 13 | 94540400 | 94541599 | 19.71042 | 34.45804 | 34.59385 | 15.24928 | 0.38201 | 0.703303 | 0.442538 |
| 11 | 113431600 | 113435799 | 49.33342 | 78.52573 | 125.805 | 52.58405 | -0.34307 | 2.210281 | 0.442559 |
| 5 | 109665000 | 109667199 | 29.49827 | 49.78771 | 47.44422 | 20.75106 | 0.450427 | 1.106957 | 0.442562 |
| 10 | 83000600 | 83002399 | 19.10647 | 32.50089 | 56.40958 | 24.72382 | -0.41058 | 1.016528 | 0.442574 |
| 5 | 128554200 | 128578399 | 245.6839 | 351.2859 | 505.7507 | 208.9494 | -0.32506 | 4.249022 | 0.442581 |
| 18 | 64893800 | 64897199 | 36.25644 | 59.60309 | 59.81314 | 25.77681 | 0.373131 | 1.418466 | 0.442583 |
| 10 | 121863000 | 121865599 | 18.73158 | 31.73632 | 56.41572 | 24.70356 | -0.44551 | 1.047394 | 0.442615 |
| 9 | 106411000 | 106412399 | 20.90058 | 36.56731 | 29.46576 | 12.99736 | 0.697609 | 0.688475 | 0.442618 |
| 17 | 88607800 | 88609999 | 16.84811 | 29.15055 | 49.53229 | 21.73104 | -0.37569 | 0.842199 | 0.442636 |
| 14 | 91917400 | 91919199 | 17.68034 | 32.79466 | 23.98659 | 10.06641 | 0.891143 | 0.233966 | 0.442642 |
| 2 | 17851400 | 17852999 | 15.85834 | 27.58991 | 47.36933 | 20.81529 | -0.38925 | 0.772162 | 0.442643 |

| | | | | | | | | | |
|---|---|---|---|---|---|---|---|---|---|
| 11 | 113190200 | 113191799 | 17.3542 | 29.84728 | 51.12261 | 22.42467 | -0.38899 | 0.900165 | 0.442658 |
| 19 | 43098200 | 43099199 | 20.69575 | 36.51873 | 28.43134 | 12.58064 | 0.748791 | 0.570884 | 0.442662 |
| 17 | 18361000 | 18365399 | 66.85483 | 106.5848 | 102.3029 | 42.85495 | 0.397259 | 2.168204 | 0.442667 |
| 3 | 112959800 | 112962999 | 50.89805 | 81.93038 | 80.23268 | 34.08455 | 0.384247 | 1.845327 | 0.442676 |
| 3 | 132344200 | 132344799 | 5.957647 | 11.56308 | 23.94482 | 9.003086 | -0.4568 | -0.24594 | 0.442686 |
| X | 112864600 | 112869599 | 79.99886 | 126.94 | 111.326 | 46.37081 | 0.520377 | 2.326912 | 0.442697 |
| 7 | 34537800 | 34541199 | 29.42036 | 49.63326 | 46.90067 | 20.51045 | 0.462465 | 1.109149 | 0.442699 |
| 10 | 25539800 | 25541399 | 19.31054 | 34.39416 | 29.9085 | 13.17044 | 0.596444 | 0.486679 | 0.44272 |
| 10 | 116291400 | 116293999 | 26.92171 | 43.98139 | 78.16537 | 33.60063 | -0.45359 | 1.490788 | 0.44274 |
| 15 | 6503600 | 6509599 | 58.48598 | 94.04116 | 82.14796 | 34.84262 | 0.546863 | 1.89289 | 0.442744 |
| 14 | 120338000 | 120338599 | 11.22461 | 21.50982 | 21.7252 | 8.559223 | 0.505505 | -0.04316 | 0.442779 |
| 5 | 93711600 | 93720199 | 124.5902 | 193.3097 | 145.0932 | 60.25772 | 0.719286 | 2.797689 | 0.442781 |
| 1 | 162875600 | 162876999 | 14.7446 | 27.13962 | 25.88466 | 10.97729 | 0.500211 | 0.257771 | 0.442793 |
| 15 | 51944000 | 51948999 | 94.48992 | 146.8459 | 145.028 | 60.21601 | 0.325544 | 2.755475 | 0.442797 |
| 4 | 69997800 | 70001199 | 31.27385 | 52.73558 | 47.99335 | 20.98533 | 0.516599 | 1.122912 | 0.442801 |
| 11 | 94865600 | 94867399 | 8.255758 | 15.08188 | 32.44125 | 13.89805 | -0.67549 | 0.264402 | 0.442818 |
| 5 | 143741400 | 143744199 | 20.41354 | 34.28009 | 59.61997 | 26.03351 | -0.41462 | 1.135636 | 0.442828 |
| 3 | 49331000 | 49333199 | 17.47512 | 30.17912 | 51.69494 | 22.67856 | -0.38794 | 0.863593 | 0.442836 |
| 17 | 14218800 | 14219599 | 6.114338 | 11.83513 | 24.9116 | 9.485745 | -0.49701 | -0.19576 | 0.442857 |
| 17 | 52941200 | 52943199 | 17.48597 | 32.44495 | 23.9957 | 10.06566 | 0.875666 | 0.230474 | 0.442859 |
| 10 | 52186800 | 52189399 | 21.94821 | 38.26107 | 37.05886 | 16.30502 | 0.433145 | 0.758136 | 0.442864 |
| 7 | 73982200 | 73989199 | 65.64402 | 103.4144 | 161.1891 | 66.88395 | -0.31677 | 2.49497 | 0.442864 |
| 3 | 94505200 | 94508599 | 28.015 | 45.86579 | 76.80344 | 33.04136 | -0.36584 | 1.462305 | 0.442887 |
| 2 | 146697600 | 146698599 | 7.054349 | 13.40068 | 27.14281 | 10.91507 | -0.51618 | -0.00085 | 0.442917 |
| 16 | 84631200 | 84634599 | 20.96493 | 34.56527 | 76.82857 | 33.15047 | -0.76893 | 1.358714 | 0.442917 |
| X | 5010600 | 5014599 | 96.62393 | 152.561 | 116.7676 | 48.48216 | 0.710497 | 2.462856 | 0.442918 |
| Y | 38491600 | 38493199 | 12.38674 | 23.21912 | 15.54664 | 6.413822 | 1.03603 | 0.100863 | 0.442919 |
| 4 | 100866000 | 100869999 | 69.30753 | 109.8865 | 112.1169 | 46.6625 | 0.3012 | 2.339277 | 0.442923 |
| 9 | 70959400 | 70960399 | 9.823974 | 18.1637 | 34.20436 | 14.41736 | -0.46876 | 0.210503 | 0.442939 |
| 15 | 31184800 | 31190799 | 42.17454 | 67.54713 | 110.7256 | 46.68674 | -0.36627 | 2.002898 | 0.442956 |
| 2 | 124135200 | 124137199 | 28.59807 | 49.19477 | 40.73001 | 17.86611 | 0.655546 | 0.932206 | 0.443003 |
| 2 | 173324600 | 173325599 | 10.08896 | 18.51323 | 34.02466 | 14.48981 | -0.44417 | 0.258258 | 0.443008 |
| 8 | 108291200 | 108295199 | 26.53121 | 43.63226 | 72.85854 | 31.4039 | -0.35915 | 1.401083 | 0.443011 |
| X | 89345800 | 89347199 | 33.66879 | 56.05668 | 51.92647 | 22.55175 | 0.491388 | 1.262567 | 0.443037 |
| 13 | 34927400 | 34932799 | 69.14912 | 107.3192 | 196.9893 | 81.80694 | -0.56915 | 2.773328 | 0.443047 |
| 1 | 192153600 | 192156599 | 22.87814 | 38.02244 | 66.25149 | 28.73795 | -0.41749 | 1.260698 | 0.44305 |
| 16 | 20868800 | 20872599 | 73.69708 | 117.3102 | 104.836 | 43.82145 | 0.497755 | 2.225239 | 0.443052 |
| 7 | 14214600 | 14215999 | 23.33501 | 40.62583 | 32.57232 | 14.31936 | 0.704588 | 0.778329 | 0.443054 |
| 10 | 50768000 | 50771799 | 24.07007 | 39.37252 | 75.39873 | 32.4263 | -0.55956 | 1.465876 | 0.443065 |
| 8 | 60985200 | 60987399 | 29.75557 | 50.48959 | 45.78211 | 20.05639 | 0.522185 | 1.064465 | 0.443073 |
| 7 | 58816200 | 58817799 | 19.09833 | 33.50454 | 32.7898 | 14.49666 | 0.418334 | 0.639655 | 0.443083 |
| 10 | 106594200 | 106595599 | 13.27705 | 24.79681 | 24.44184 | 10.14645 | 0.475673 | 0.157202 | 0.443086 |
| 4 | 126347200 | 126348199 | 7.172074 | 13.7369 | 26.80589 | 10.57638 | -0.43831 | -0.06358 | 0.443088 |
| 16 | 13931400 | 13932399 | 7.172074 | 13.7369 | 26.80589 | 10.57638 | -0.43831 | -0.06358 | 0.443088 |
| 6 | 96792400 | 96793599 | 21.40227 | 37.53316 | 33.84166 | 14.95724 | 0.536337 | 0.663511 | 0.443105 |
| X | 18016600 | 18018999 | 31.67891 | 53.96181 | 43.49278 | 19.06463 | 0.692072 | 1.034517 | 0.443111 |
| 3 | 11291200 | 11292199 | 12.66787 | 23.53406 | 23.99379 | 10.04797 | 0.417519 | 0.184509 | 0.443115 |

| | | | | | | | | | |
|---|---|---|---|---|---|---|---|---|---|
| 7 | 30579000 | 30582399 | 25.56657 | 42.07129 | 71.60594 | 30.8719 | -0.38622 | 1.39057 | 0.443125 |
| 2 | 82326000 | 82326599 | 5.053941 | 9.854223 | 22.05238 | 7.94028 | -0.50867 | -0.4135 | 0.443154 |
| 2 | 156670000 | 156680599 | 202.6904 | 294.2263 | 425.2813 | 175.3279 | -0.30914 | 4.023406 | 0.443159 |
| 6 | 37764600 | 37766799 | 32.57511 | 53.91264 | 54.71674 | 23.66615 | 0.359329 | 1.325142 | 0.44319 |
| 3 | 130541000 | 130545199 | 48.64943 | 78.62436 | 73.70393 | 31.47869 | 0.453864 | 1.722682 | 0.443191 |
| 15 | 72614600 | 72615599 | 7.183478 | 13.67451 | 27.1922 | 10.88673 | -0.48457 | -0.01535 | 0.443216 |
| 3 | 138482400 | 138484799 | 14.08531 | 24.70278 | 43.50638 | 19.22973 | -0.42508 | 0.635594 | 0.443221 |
| 12 | 58403400 | 58405199 | 13.60494 | 23.97435 | 42.38481 | 18.75998 | -0.42909 | 0.567266 | 0.443231 |
| 6 | 129824400 | 129828399 | 72.30812 | 114.6519 | 115.6391 | 48.13174 | 0.317894 | 2.340638 | 0.443233 |
| 4 | 106541200 | 106542399 | 7.234365 | 13.64493 | 28.08848 | 11.46712 | -0.55845 | 0.054077 | 0.443247 |
| 15 | 79030000 | 79031399 | 7.234365 | 13.64493 | 28.08848 | 11.46712 | -0.55845 | 0.054077 | 0.443247 |
| 6 | 23278400 | 23281999 | 31.50554 | 52.56277 | 52.46281 | 22.81421 | 0.383914 | 1.232401 | 0.443258 |
| 13 | 7672200 | 7674599 | 35.02076 | 57.68071 | 57.58632 | 24.83702 | 0.381584 | 1.389725 | 0.443277 |
| 7 | 38768000 | 38778799 | 134.7128 | 207.4881 | 161.7167 | 67.17109 | 0.656907 | 2.934232 | 0.44328 |
| 5 | 10870600 | 10870999 | 8.754758 | 16.97653 | 18.30025 | 6.909981 | 0.470441 | -0.21675 | 0.443313 |
| 5 | 65072200 | 65076999 | 64.58743 | 102.8235 | 104.7963 | 43.81309 | 0.308648 | 2.21232 | 0.443319 |
| 13 | 32646400 | 32647199 | 6.281912 | 12.07052 | 25.77331 | 10.07595 | -0.55455 | -0.10705 | 0.443342 |
| 10 | 128350800 | 128353799 | 18.15426 | 30.69472 | 56.33172 | 24.65133 | -0.49132 | 1.065048 | 0.443364 |
| 8 | 74268600 | 74271999 | 50.89444 | 81.84137 | 83.05239 | 35.23051 | 0.331185 | 1.880064 | 0.443368 |
| 9 | 96546000 | 96548199 | 21.52527 | 36.09196 | 61.34063 | 26.75911 | -0.38154 | 1.156499 | 0.44339 |
| 18 | 58761600 | 58762399 | 18.63171 | 33.61207 | 27.88442 | 12.13505 | 0.675237 | 0.40154 | 0.443399 |
| X | 18812000 | 18812799 | 12.98403 | 24.9746 | 20.55221 | 8.01065 | 0.812777 | -0.06296 | 0.443405 |
| 4 | 112753200 | 112754199 | 12.72809 | 24.39797 | 21.01749 | 8.273565 | 0.733895 | -0.03386 | 0.443417 |
| 3 | 73758000 | 73759999 | 22.81789 | 39.5387 | 38.75302 | 17.00269 | 0.415046 | 0.838917 | 0.443434 |
| 7 | 59318800 | 59329599 | 189.5246 | 282.467 | 242.6865 | 100.0089 | 0.476884 | 3.59193 | 0.443434 |
| 8 | 92814400 | 92817199 | 13.4587 | 23.74532 | 41.34041 | 18.2933 | -0.40629 | 0.552072 | 0.443453 |
| 2 | 88132400 | 88132999 | 13.35294 | 24.64964 | 24.95958 | 10.54318 | 0.418447 | 0.227876 | 0.443453 |
| 13 | 67824600 | 67826799 | 26.11939 | 45.48623 | 36.26404 | 15.94514 | 0.712621 | 0.791816 | 0.44346 |
| 5 | 139969200 | 139981199 | 141.7835 | 211.9389 | 312.7413 | 128.9502 | -0.29907 | 3.532562 | 0.443461 |
| 1 | 50095800 | 50097799 | 20.94891 | 36.49812 | 36.64868 | 16.11874 | 0.381315 | 0.760514 | 0.443463 |
| 14 | 63843400 | 63846199 | 21.88069 | 36.13275 | 68.50716 | 29.5854 | -0.53958 | 1.336237 | 0.443464 |
| 13 | 70167400 | 70169999 | 10.91535 | 19.59382 | 35.29894 | 15.44216 | -0.44157 | 0.394582 | 0.44348 |
| 6 | 144784400 | 144785399 | 8.36037 | 15.41234 | 30.9648 | 13.10406 | -0.56481 | 0.210952 | 0.443491 |
| 4 | 17442400 | 17445799 | 29.56039 | 50.11627 | 45.195 | 19.79414 | 0.52985 | 1.066137 | 0.443504 |
| 16 | 95686000 | 95689199 | 39.43452 | 63.50519 | 104.5742 | 44.45335 | -0.3632 | 1.828403 | 0.443527 |
| 11 | 6683200 | 6690599 | 79.93901 | 124.077 | 187.4814 | 77.85731 | -0.28868 | 2.775199 | 0.443559 |
| 5 | 23604600 | 23606999 | 14.28444 | 25.02074 | 43.03796 | 19.00262 | -0.39104 | 0.657769 | 0.443569 |
| 8 | 46387400 | 46388199 | 5.41357 | 10.48601 | 24.96918 | 9.476759 | -0.66972 | -0.21968 | 0.44357 |
| 18 | 65215800 | 65218799 | 26.65146 | 43.06882 | 95.52386 | 40.79338 | -0.78563 | 1.701981 | 0.443571 |
| 9 | 25709000 | 25713599 | 46.99011 | 76.10143 | 70.87285 | 30.32794 | 0.466289 | 1.670831 | 0.443578 |
| 7 | 96836600 | 96837799 | 10.36456 | 18.81378 | 34.35305 | 14.83508 | -0.44879 | 0.320672 | 0.443578 |
| 13 | 76652000 | 76656799 | 55.61136 | 89.17559 | 87.96304 | 37.15743 | 0.367217 | 1.976095 | 0.443595 |
| X | 121023600 | 121025399 | 21.89833 | 38.34992 | 30.35229 | 13.39932 | 0.723466 | 0.68153 | 0.443608 |
| 14 | 58590000 | 58592199 | 24.74579 | 42.40024 | 42.56453 | 18.66657 | 0.37768 | 0.95281 | 0.443614 |
| 1 | 75727800 | 75729199 | 15.71198 | 28.86763 | 26.27732 | 11.17145 | 0.564481 | 0.275725 | 0.443625 |
| 19 | 57855000 | 57857999 | 25.84147 | 42.71663 | 70.82222 | 30.60657 | -0.3467 | 1.335784 | 0.443648 |
| X | 100565400 | 100567799 | 13.67976 | 24.14123 | 42.93191 | 18.99662 | -0.43657 | 0.555039 | 0.443651 |

| | | | | | | | | | |
|---|---|---|---|---|---|---|---|---|---|
| 12 | 51737600 | 51739399 | 14.33288 | 24.81494 | 48.98031 | 21.4893 | -0.5906 | 0.823147 | 0.443685 |
| X | 92900000 | 92902999 | 13.00458 | 23.09252 | 41.89118 | 18.49426 | -0.46116 | 0.487917 | 0.443687 |
| 12 | 105877400 | 105880599 | 32.81879 | 53.17544 | 87.25256 | 37.34997 | -0.34825 | 1.645774 | 0.443716 |
| 7 | 62430800 | 62434999 | 70.84562 | 112.4949 | 112.4061 | 46.85362 | 0.333394 | 2.297355 | 0.44372 |
| 10 | 122004600 | 122008999 | 42.67464 | 68.45554 | 108.4984 | 45.85709 | -0.31508 | 1.954787 | 0.443732 |
| 13 | 56818200 | 56819599 | 8.944748 | 16.54757 | 30.97715 | 13.04689 | -0.45878 | 0.195567 | 0.443738 |
| 16 | 93163000 | 93164399 | 8.944748 | 16.54757 | 30.97715 | 13.04689 | -0.45878 | 0.195567 | 0.443738 |
| 16 | 23467200 | 23469799 | 17.16285 | 30.39218 | 29.61217 | 13.09387 | 0.428628 | 0.511521 | 0.443758 |
| 4 | 43867800 | 43870799 | 24.91219 | 42.7146 | 42.54689 | 18.65986 | 0.389033 | 0.947369 | 0.44376 |
| 15 | 37270000 | 37272999 | 19.98963 | 34.10124 | 59.31555 | 26.00351 | -0.41328 | 0.993911 | 0.44377 |
| 9 | 39099800 | 39100799 | 37.71728 | 61.5118 | 61.62217 | 26.47264 | 0.369536 | 1.529919 | 0.443777 |
| 5 | 104557400 | 104558999 | 13.82831 | 24.35992 | 42.40689 | 18.77099 | -0.40697 | 0.57437 | 0.443799 |
| 7 | 15328200 | 15331599 | 41.94214 | 68.22291 | 66.29918 | 28.45466 | 0.410649 | 1.580006 | 0.443803 |
| 13 | 49031400 | 49032399 | 5.976837 | 11.54633 | 24.8307 | 9.541917 | -0.54112 | -0.16997 | 0.443807 |
| 10 | 61108000 | 61111799 | 20.41361 | 34.21372 | 61.26667 | 26.71981 | -0.45661 | 1.156897 | 0.443815 |
| 18 | 88172200 | 88174199 | 22.04342 | 38.75021 | 28.76583 | 12.73825 | 0.81498 | 0.618581 | 0.443822 |
| X | 80304600 | 80308399 | 61.45748 | 98.38286 | 91.36888 | 38.49683 | 0.451149 | 2.036962 | 0.443849 |
| 4 | 130130400 | 130134399 | 37.63154 | 60.4557 | 101.1195 | 42.8348 | -0.38996 | 1.89952 | 0.443849 |
| 5 | 115769200 | 115771199 | 10.58746 | 18.61028 | 40.5006 | 17.935 | -0.72814 | 0.575724 | 0.443856 |
| 5 | 68376000 | 68383999 | 82.08551 | 126.8767 | 194.2048 | 80.66828 | -0.31195 | 2.852242 | 0.443865 |
| 6 | 37813200 | 37814599 | 14.62793 | 27.12805 | 25.34083 | 10.63729 | 0.540343 | 0.213046 | 0.443872 |
| 12 | 67711600 | 67718399 | 57.71238 | 91.60014 | 146.5746 | 61.06878 | -0.3456 | 2.307764 | 0.443876 |
| 2 | 150196800 | 150200999 | 86.21173 | 135.6305 | 132.0547 | 54.78512 | 0.359724 | 2.530905 | 0.443879 |
| 11 | 10883800 | 10884199 | 7.638632 | 14.87764 | 16.22173 | 5.924903 | 0.497551 | -0.35432 | 0.443902 |
| 9 | 19339200 | 19342999 | 72.02391 | 114.7345 | 103.0683 | 43.12813 | 0.491439 | 2.198952 | 0.443903 |
| 5 | 136678200 | 136679999 | 18.43974 | 32.47759 | 32.27546 | 14.28693 | 0.397521 | 0.58411 | 0.443904 |
| 7 | 84821200 | 84826999 | 99.71535 | 155.3468 | 149.3482 | 61.98709 | 0.36708 | 2.717003 | 0.443918 |
| 1 | 102539800 | 102541599 | 18.59906 | 32.70487 | 31.67108 | 14.02396 | 0.433597 | 0.599628 | 0.443969 |
| 12 | 57564400 | 57566999 | 14.42907 | 25.25671 | 43.57016 | 19.22644 | -0.39527 | 0.669951 | 0.443977 |
| 13 | 76309400 | 76310799 | 19.02612 | 33.37449 | 33.38271 | 14.75754 | 0.387126 | 0.642944 | 0.443982 |
| 3 | 137026600 | 137028999 | 12.83638 | 22.70991 | 39.27444 | 17.36952 | -0.39497 | 0.515291 | 0.443983 |
| 7 | 72904800 | 72905599 | 5.523476 | 10.73825 | 22.91694 | 8.504862 | -0.48211 | -0.3006 | 0.444008 |
| 13 | 4596200 | 4596599 | 4.178613 | 8.170132 | 18.98296 | 6.492214 | -0.49362 | -0.61868 | 0.444045 |
| 10 | 111250600 | 111254799 | 44.61018 | 71.4058 | 114.9037 | 48.40571 | -0.34064 | 2.02521 | 0.444067 |
| 4 | 153512400 | 153516599 | 53.11706 | 84.20338 | 136.4265 | 56.77105 | -0.36519 | 2.337554 | 0.444077 |
| 3 | 79581400 | 79583199 | 18.34807 | 31.3332 | 53.78443 | 23.58067 | -0.39389 | 0.971855 | 0.444093 |
| 2 | 149277200 | 149283199 | 87.44309 | 137.7289 | 129.2387 | 53.62504 | 0.414191 | 2.507602 | 0.444097 |
| 4 | 59224800 | 59227399 | 20.8559 | 35.06004 | 60.2516 | 26.32069 | -0.39747 | 1.12665 | 0.444098 |
| 14 | 84060400 | 84061799 | 15.90097 | 27.61266 | 47.39554 | 20.80247 | -0.38904 | 0.800241 | 0.444103 |
| 3 | 82810200 | 82813999 | 34.19714 | 55.12313 | 93.46369 | 39.78364 | -0.40349 | 1.786173 | 0.44411 |
| X | 25911000 | 25917799 | 73.38688 | 117.2935 | 89.92657 | 37.80405 | 0.724805 | 2.098726 | 0.444112 |
| 5 | 95122200 | 95125599 | 48.51059 | 78.68678 | 68.26576 | 29.246 | 0.5704 | 1.642087 | 0.44412 |
| 8 | 59581600 | 59587399 | 62.09696 | 98.97896 | 101.28 | 42.4143 | 0.304788 | 2.170165 | 0.444129 |
| 12 | 48855000 | 48857399 | 30.39547 | 52.84918 | 34.51296 | 15.1601 | 0.999139 | 0.833417 | 0.444136 |
| 7 | 138239800 | 138241599 | 16.38999 | 28.39928 | 48.46238 | 21.26259 | -0.38129 | 0.826543 | 0.444171 |
| 6 | 5977600 | 5979399 | 15.48908 | 28.02869 | 27.75612 | 12.03288 | 0.424912 | 0.361877 | 0.444174 |
| 7 | 41787400 | 41790399 | 42.60312 | 69.84124 | 60.75234 | 26.15064 | 0.577236 | 1.468962 | 0.444178 |

| | | | | | | | | | |
|---|---|---|---|---|---|---|---|---|---|
| 17 | 56645400 | 56649199 | 17.28809 | 29.40441 | 54.78257 | 24.0105 | -0.51247 | 1.005873 | 0.44418 |
| 5 | 142822400 | 142824599 | 13.52051 | 23.73719 | 41.45299 | 18.33844 | -0.41259 | 0.614969 | 0.4442 |
| 10 | 93250400 | 93254999 | 36.8 | 60.75586 | 51.51053 | 22.24173 | 0.61788 | 1.364114 | 0.444211 |
| X | 25439400 | 25442399 | 25.35013 | 44.46239 | 31.64526 | 13.98805 | 0.875647 | 0.67728 | 0.444212 |
| 17 | 41914600 | 41921399 | 76.46635 | 120.9336 | 121.4341 | 50.46192 | 0.321187 | 2.408725 | 0.444265 |
| 1 | 182469600 | 182479599 | 89.73628 | 137.9535 | 209.4966 | 86.99457 | -0.30656 | 2.958424 | 0.444267 |
| 9 | 116752600 | 116755999 | 24.68072 | 40.96095 | 68.01429 | 29.47415 | -0.34838 | 1.284532 | 0.444268 |
| 1 | 7247200 | 7253199 | 81.50558 | 126.2409 | 190.9843 | 79.32972 | -0.29281 | 2.813711 | 0.444268 |
| 7 | 28047600 | 28053599 | 28.43337 | 46.23308 | 81.86561 | 35.13097 | -0.45313 | 1.564484 | 0.444269 |
| 1 | 18014200 | 18016199 | 23.42741 | 40.42775 | 39.29688 | 17.23913 | 0.425763 | 0.878219 | 0.444274 |
| 3 | 49690400 | 49690999 | 6.263275 | 12.09668 | 24.82052 | 9.549085 | -0.47541 | -0.16172 | 0.444287 |
| 8 | 23284000 | 23287199 | 21.4434 | 35.71413 | 63.91109 | 27.7626 | -0.45557 | 1.231138 | 0.444307 |
| 8 | 22045600 | 22049999 | 29.14737 | 50.71082 | 35.0852 | 15.41752 | 0.916239 | 0.81928 | 0.444319 |
| 9 | 86542800 | 86545999 | 24.81037 | 41.23198 | 69.29068 | 30.0554 | -0.36554 | 1.269333 | 0.444326 |
| 14 | 14763600 | 14771199 | 86.89997 | 136.195 | 138.0922 | 57.27709 | 0.29708 | 2.603821 | 0.444328 |
| 6 | 35903600 | 35904199 | 6.556728 | 12.6729 | 25.45059 | 9.757394 | -0.43997 | -0.16807 | 0.444337 |
| 2 | 129635400 | 129639599 | 35.46071 | 58.74508 | 49.84477 | 21.5733 | 0.61708 | 1.320309 | 0.444347 |
| 14 | 124930400 | 124931599 | 19.29128 | 33.89326 | 33.31697 | 14.7482 | 0.412128 | 0.620357 | 0.444356 |
| 1 | 119864200 | 119866399 | 12.84318 | 22.70054 | 39.29091 | 17.38008 | -0.39667 | 0.525905 | 0.444363 |
| 3 | 104307400 | 104308599 | 15.68093 | 28.10022 | 27.83905 | 12.19629 | 0.414474 | 0.421449 | 0.444376 |
| 8 | 7331000 | 7331799 | 10.78966 | 19.26716 | 35.82378 | 15.74102 | -0.49119 | 0.428103 | 0.444386 |
| 14 | 76466600 | 76467799 | 12.43626 | 22.02233 | 39.84673 | 17.61937 | -0.45904 | 0.506491 | 0.444386 |
| 13 | 4450400 | 4454199 | 55.00475 | 88.60473 | 79.11748 | 33.6459 | 0.518456 | 1.830696 | 0.44441 |
| 15 | 58994800 | 58996799 | 19.15419 | 33.71183 | 32.77073 | 14.50823 | 0.428867 | 0.597569 | 0.444417 |
| 19 | 33567000 | 33567399 | 4.944244 | 9.621885 | 21.79704 | 8.023447 | -0.55537 | -0.33854 | 0.444421 |
| 19 | 36713800 | 36714399 | 4.944244 | 9.621885 | 21.79704 | 8.023447 | -0.55537 | -0.33854 | 0.444421 |
| 19 | 43554000 | 43554799 | 4.944244 | 9.621885 | 21.79704 | 8.023447 | -0.55537 | -0.33854 | 0.444421 |
| 1 | 3914800 | 3923399 | 100.7911 | 157.1511 | 147.6767 | 61.28585 | 0.400849 | 2.702873 | 0.444437 |
| 1 | 109084000 | 109085199 | 11.2293 | 21.39349 | 21.55219 | 8.613896 | 0.49004 | 0.000542 | 0.444465 |
| 12 | 111243600 | 111247599 | 38.587 | 62.10882 | 99.2882 | 42.17531 | -0.32141 | 1.840382 | 0.444476 |
| 12 | 118264600 | 118266799 | 17.36905 | 29.00916 | 72.02908 | 31.34418 | -0.92647 | 1.194624 | 0.444529 |
| 18 | 46276800 | 46278399 | 9.819763 | 17.32046 | 39.47945 | 17.47075 | -0.79239 | 0.530715 | 0.44456 |
| 7 | 121594800 | 121598399 | 25.70955 | 42.36219 | 71.16532 | 30.699 | -0.36668 | 1.37418 | 0.444565 |
| 8 | 90082200 | 90083599 | 11.3813 | 20.33952 | 36.26888 | 15.92613 | -0.43081 | 0.426817 | 0.444565 |
| 9 | 20563200 | 20563999 | 5.679192 | 11.0126 | 23.88044 | 9.017163 | -0.52859 | -0.23585 | 0.444565 |
| X | 66185000 | 66188199 | 42.89633 | 69.65208 | 68.59337 | 29.40875 | 0.389395 | 1.613434 | 0.444581 |
| 19 | 40556800 | 40561199 | 50.26014 | 80.08635 | 124.9168 | 52.29002 | -0.30289 | 2.175617 | 0.444585 |
| 15 | 85056400 | 85057799 | 9.629058 | 17.65222 | 32.96396 | 14.0566 | -0.46816 | 0.258596 | 0.444588 |
| 7 | 23373000 | 23389199 | 258.5885 | 378.5851 | 317.7621 | 130.8966 | 0.486523 | 3.897554 | 0.444596 |
| 1 | 108845200 | 108847399 | 20.25622 | 35.42223 | 34.93533 | 15.40348 | 0.40739 | 0.700636 | 0.444598 |
| 14 | 71531800 | 71532599 | 7.327353 | 14.11521 | 30.7264 | 11.9248 | -0.57276 | -0.10602 | 0.444601 |
| 1 | 7184600 | 7185599 | 7.609409 | 14.60319 | 30.51889 | 11.97367 | -0.52954 | -0.07002 | 0.444601 |
| 16 | 30817600 | 30820199 | 25.01677 | 41.40352 | 69.08234 | 29.87965 | -0.35567 | 1.318055 | 0.444602 |
| 11 | 117099000 | 117100799 | 11.76553 | 20.7799 | 38.358 | 16.97 | -0.48926 | 0.526608 | 0.444605 |
| 8 | 104678200 | 104679599 | 8.894957 | 16.53619 | 31.03698 | 12.98526 | -0.45587 | 0.16584 | 0.444615 |
| 12 | 20910200 | 20927399 | 472.5823 | 649.0412 | 582.5622 | 242.2449 | 0.297876 | 4.857346 | 0.444629 |
| 17 | 48081400 | 48084799 | 53.97326 | 85.43401 | 139.8179 | 58.12669 | -0.38148 | 2.376584 | 0.444631 |

| | | | | | | | | | |
|---|---|---|---|---|---|---|---|---|---|
| 1 | 99745000 | 99746999 | 24.46406 | 42.44722 | 38.61859 | 16.94439 | 0.52238 | 0.834765 | 0.444646 |
| 5 | 37814000 | 37817199 | 25.24353 | 41.48957 | 73.2854 | 31.57984 | -0.44033 | 1.404685 | 0.444649 |
| 9 | 8165600 | 8170599 | 53.09915 | 85.24154 | 86.09522 | 36.42977 | 0.334995 | 1.935494 | 0.444657 |
| 5 | 95461400 | 95461799 | 8.955057 | 17.47373 | 16.8295 | 5.996935 | 0.706414 | -0.4193 | 0.444664 |
| 13 | 6760600 | 6762199 | 13.71508 | 26.08481 | 21.9073 | 8.787428 | 0.746048 | 0.037244 | 0.444664 |
| 8 | 120841200 | 120843399 | 16.6898 | 28.93509 | 48.99955 | 21.49864 | -0.37018 | 0.822944 | 0.444676 |
| 1 | 123971800 | 123974799 | 25.90094 | 42.49469 | 74.98063 | 32.28612 | -0.43985 | 1.429238 | 0.44468 |
| 4 | 16029200 | 16029799 | 6.245796 | 12.11754 | 25.1511 | 9.474164 | -0.46317 | -0.23402 | 0.444697 |
| 14 | 113303600 | 113305799 | 25.27558 | 43.13796 | 43.66597 | 19.14353 | 0.364949 | 0.991854 | 0.444723 |
| 15 | 59790400 | 59791999 | 16.03438 | 29.17992 | 27.10895 | 11.66952 | 0.522801 | 0.331328 | 0.444732 |
| 1 | 128441000 | 128443599 | 19.6445 | 34.56677 | 32.57884 | 14.42395 | 0.472933 | 0.600567 | 0.444736 |
| 14 | 13803200 | 13804599 | 10.85344 | 19.67871 | 34.85512 | 15.07341 | -0.40646 | 0.332773 | 0.44477 |
| 19 | 37723600 | 37724999 | 10.85344 | 19.67871 | 34.85512 | 15.07341 | -0.40646 | 0.332773 | 0.44477 |
| 15 | 57175400 | 57175799 | 12.18056 | 22.71009 | 22.55329 | 9.389613 | 0.461895 | 0.154945 | 0.444778 |
| 13 | 106562000 | 106563999 | 23.34154 | 41.02592 | 32.36036 | 14.32966 | 0.728124 | 0.635856 | 0.44478 |
| 9 | 36140600 | 36147199 | 79.90442 | 126.2815 | 122.739 | 50.98585 | 0.36733 | 2.42923 | 0.444785 |
| 12 | 5477000 | 5477799 | 11.22758 | 21.24297 | 22.12497 | 8.972974 | 0.424555 | 0.051062 | 0.44481 |
| X | 31296600 | 31299599 | 53.95114 | 87.74568 | 52.36818 | 22.47465 | 1.11203 | 1.599499 | 0.444825 |
| 2 | 34661400 | 34662799 | 6.908329 | 13.09744 | 27.64684 | 11.16423 | -0.58096 | 0.010915 | 0.444825 |
| 8 | 101520800 | 101521599 | 6.908329 | 13.09744 | 27.64684 | 11.16423 | -0.58096 | 0.010915 | 0.444825 |
| 4 | 3481400 | 3489199 | 184.3592 | 274.6396 | 268.1582 | 110.5002 | 0.291847 | 3.600376 | 0.444835 |
| 1 | 107160600 | 107163199 | 40.7736 | 66.63648 | 57.56635 | 24.74927 | 0.584729 | 1.50358 | 0.44485 |
| 3 | 53025400 | 53028999 | 25.16756 | 41.64824 | 69.69939 | 30.14411 | -0.36002 | 1.319798 | 0.444851 |
| 18 | 75680600 | 75683999 | 25.16756 | 41.64824 | 69.69939 | 30.14411 | -0.36002 | 1.319798 | 0.444851 |
| 18 | 77107000 | 77108599 | 15.01452 | 27.42896 | 26.81498 | 11.48774 | 0.454357 | 0.302904 | 0.444854 |
| 1 | 113548400 | 113552799 | 29.34958 | 49.59327 | 42.6434 | 18.65774 | 0.598246 | 1.092939 | 0.444856 |
| 5 | 82782400 | 82784599 | 21.65591 | 38.3452 | 30.20906 | 13.35804 | 0.733688 | 0.543653 | 0.444863 |
| 13 | 31329800 | 31330399 | 5.514549 | 10.72049 | 22.88755 | 8.496531 | -0.48246 | -0.29761 | 0.444867 |
| 3 | 153985600 | 153987999 | 13.01104 | 22.85274 | 40.91734 | 18.10816 | -0.44851 | 0.602492 | 0.444871 |
| 14 | 67042600 | 67045399 | 20.73453 | 34.78869 | 60.17928 | 26.26324 | -0.40689 | 1.146397 | 0.444872 |
| 8 | 73892000 | 73895199 | 37.50766 | 62.48436 | 50.98469 | 22.15082 | 0.673675 | 1.263014 | 0.444884 |
| 3 | 134209400 | 134209999 | 4.64599 | 9.064969 | 20.3622 | 7.264204 | -0.50186 | -0.45953 | 0.444888 |
| 4 | 4090000 | 4092599 | 20.70099 | 36.81221 | 29.13395 | 12.84654 | 0.730142 | 0.502927 | 0.444907 |
| 6 | 143434600 | 143435999 | 14.92066 | 25.929 | 46.85854 | 20.5761 | -0.46295 | 0.783211 | 0.444911 |
| 2 | 151910600 | 151912799 | 12.90863 | 22.67096 | 40.97026 | 18.1303 | -0.46159 | 0.605899 | 0.444923 |
| 18 | 11239600 | 11242399 | 21.54421 | 36.03346 | 62.404 | 27.18005 | -0.40856 | 1.184136 | 0.444935 |
| 5 | 96935400 | 96937399 | 20.3485 | 35.44969 | 35.24419 | 15.50037 | 0.395664 | 0.757423 | 0.444953 |
| 6 | 37757800 | 37761599 | 92.19967 | 144.2604 | 141.7023 | 58.78452 | 0.340932 | 2.636009 | 0.444967 |
| 1 | 88644800 | 88646799 | 16.82251 | 28.70796 | 58.03002 | 25.44064 | -0.62892 | 0.977998 | 0.444971 |
| 9 | 21378600 | 21380999 | 18.54147 | 31.57871 | 54.29362 | 23.79878 | -0.39679 | 0.998344 | 0.444983 |
| 12 | 86089200 | 86090799 | 34.87613 | 57.27726 | 57.8774 | 24.93215 | 0.362694 | 1.427679 | 0.444999 |
| 15 | 67605200 | 67608799 | 34.8808 | 56.12325 | 106.1478 | 45.06564 | -0.56411 | 1.848815 | 0.445003 |
| 17 | 20315000 | 20318199 | 54.25178 | 86.88679 | 88.24741 | 37.20678 | 0.322976 | 2.010448 | 0.445006 |
| 3 | 135142800 | 135146199 | 20.74512 | 34.35278 | 73.67779 | 31.87943 | -0.71662 | 1.303742 | 0.445009 |
| 19 | 45171600 | 45172199 | 5.37142 | 10.45073 | 22.38346 | 8.25504 | -0.47836 | -0.32494 | 0.445022 |
| 8 | 93816600 | 93817399 | 7.456563 | 14.29743 | 28.03601 | 11.02702 | -0.44147 | -0.06908 | 0.445022 |
| 13 | 104995600 | 104996999 | 13.65743 | 25.85713 | 22.3551 | 9.053846 | 0.692359 | 0.067826 | 0.445029 |

| | | | | | | | | | |
|---|---|---|---|---|---|---|---|---|---|
| 7 | 13669200 | 13673199 | 38.7823 | 64.21208 | 47.74635 | 20.65717 | 0.806704 | 1.330399 | 0.44503 |
| 4 | 113664200 | 113668999 | 53.61308 | 86.65912 | 71.81131 | 30.6762 | 0.631652 | 1.728286 | 0.445041 |
| 19 | 48946000 | 48946599 | 10.17764 | 19.53539 | 20.24829 | 7.940024 | 0.475274 | -0.07131 | 0.445046 |
| 5 | 18874400 | 18875599 | 18.68626 | 33.0952 | 31.10175 | 13.75209 | 0.480787 | 0.524565 | 0.445047 |
| 7 | 54960000 | 54962399 | 18.68626 | 33.0952 | 31.10175 | 13.75209 | 0.480787 | 0.524565 | 0.445047 |
| 10 | 68801200 | 68801599 | 9.39507 | 18.16476 | 19.30198 | 7.388911 | 0.472256 | -0.16049 | 0.445063 |
| 1 | 65809600 | 65811399 | 24.16969 | 42.13184 | 36.92531 | 16.24582 | 0.576785 | 0.770252 | 0.445067 |
| 9 | 24739200 | 24739799 | 6.249479 | 12.1349 | 25.21373 | 9.449699 | -0.45746 | -0.24982 | 0.445098 |
| 7 | 118998400 | 119001399 | 29.90416 | 50.28107 | 49.11001 | 21.45486 | 0.414849 | 1.143072 | 0.445102 |
| 2 | 64301600 | 64302999 | 10.36892 | 18.9796 | 33.9655 | 14.51374 | -0.40977 | 0.272991 | 0.445105 |
| 8 | 83755000 | 83756599 | 17.09535 | 30.66502 | 29.06423 | 12.72249 | 0.478627 | 0.426453 | 0.445106 |
| 5 | 136739800 | 136744199 | 27.50672 | 44.70891 | 80.21444 | 34.41694 | -0.47236 | 1.569947 | 0.445131 |
| 1 | 112556600 | 112557599 | 11.39018 | 21.74362 | 21.56836 | 8.583831 | 0.517524 | -0.01138 | 0.445155 |
| 11 | 14498000 | 14499199 | 11.39018 | 21.74362 | 21.56836 | 8.583831 | 0.517524 | -0.01138 | 0.445155 |
| 19 | 28219400 | 28219799 | 4.48432 | 8.741623 | 20.79155 | 7.508167 | -0.59847 | -0.41446 | 0.445158 |
| 17 | 26440000 | 26442599 | 18.97946 | 31.96048 | 57.93013 | 25.31975 | -0.47375 | 1.105013 | 0.44516 |
| 3 | 79918400 | 79920799 | 27.78368 | 47.70369 | 41.28993 | 18.1106 | 0.591093 | 0.9445 | 0.445169 |
| X | 55642600 | 55643999 | 16.81652 | 30.49397 | 27.23881 | 11.77802 | 0.575415 | 0.36042 | 0.445174 |
| 5 | 126140600 | 126145599 | 60.78863 | 97.4318 | 89.90397 | 37.95019 | 0.462481 | 1.996109 | 0.445196 |
| 5 | 93414000 | 93416599 | 17.62274 | 29.9649 | 54.23835 | 23.77142 | -0.47099 | 1.010495 | 0.445198 |
| 17 | 22914200 | 22922799 | 161.6304 | 243.9554 | 226.8907 | 93.77816 | 0.378834 | 3.347603 | 0.445206 |
| 12 | 51020600 | 51022399 | 17.67918 | 31.17037 | 30.74664 | 13.60736 | 0.408662 | 0.564364 | 0.445213 |
| 2 | 135526200 | 135528199 | 18.13513 | 31.10272 | 53.83217 | 23.60976 | -0.40489 | 0.930877 | 0.445228 |
| 19 | 39560800 | 39561799 | 26.52823 | 45.17859 | 44.73459 | 19.60863 | 0.39623 | 1.018408 | 0.445247 |
| 2 | 7768000 | 7770199 | 20.11983 | 33.68655 | 61.20146 | 26.67399 | -0.47735 | 1.167577 | 0.445251 |
| 1 | 35947000 | 35947799 | 10.24019 | 18.55664 | 33.83899 | 14.64401 | -0.44869 | 0.32967 | 0.445271 |
| 5 | 135372400 | 135373799 | 10.24019 | 18.55664 | 33.83899 | 14.64401 | -0.44869 | 0.32967 | 0.445271 |
| 1 | 103243200 | 103244799 | 15.43859 | 28.88032 | 23.09174 | 9.559844 | 0.778281 | 0.161396 | 0.44528 |
| 1 | 190710200 | 190711799 | 11.28284 | 19.9335 | 39.34775 | 17.40681 | -0.58551 | 0.51984 | 0.445284 |
| 4 | 73611000 | 73612799 | 20.20912 | 36.15362 | 28.79618 | 12.63359 | 0.726055 | 0.464207 | 0.445301 |
| 7 | 25966200 | 25967799 | 20.20912 | 36.15362 | 28.79618 | 12.63359 | 0.726055 | 0.464207 | 0.445301 |
| 16 | 35961200 | 35966199 | 65.23721 | 103.9016 | 104.7463 | 43.81767 | 0.325004 | 2.199387 | 0.445315 |
| 17 | 20333200 | 20334199 | 13.96012 | 26.72299 | 20.92034 | 8.267756 | 0.865698 | -0.00903 | 0.445315 |
| 13 | 83410800 | 83411399 | 9.466636 | 18.25477 | 19.75229 | 7.643508 | 0.431196 | -0.12414 | 0.445317 |
| 13 | 28231600 | 28236199 | 43.32815 | 70.92689 | 60.84363 | 26.17387 | 0.596426 | 1.486898 | 0.44532 |
| 12 | 112315200 | 112317199 | 17.74067 | 30.23875 | 53.74467 | 23.55965 | -0.44429 | 0.98712 | 0.445321 |
| 8 | 121792600 | 121796599 | 34.33037 | 55.49894 | 89.7917 | 38.36416 | -0.33116 | 1.700249 | 0.445325 |
| 2 | 49673800 | 49677999 | 35.6102 | 57.41125 | 95.9783 | 40.86353 | -0.38293 | 1.782537 | 0.445347 |
| 2 | 121626600 | 121628799 | 15.39586 | 26.93596 | 47.27983 | 20.85491 | -0.42033 | 0.688159 | 0.445361 |
| 10 | 33063400 | 33065399 | 30.07167 | 50.54172 | 49.10729 | 21.45098 | 0.42243 | 1.149461 | 0.445365 |
| 2 | 53635000 | 53636999 | 21.32921 | 37.48185 | 32.98382 | 14.60339 | 0.570894 | 0.626706 | 0.445368 |
| 3 | 45213400 | 45217199 | 32.20106 | 53.47446 | 53.63602 | 23.25305 | 0.376611 | 1.284143 | 0.445369 |
| 9 | 35992800 | 35996999 | 52.56293 | 84.98293 | 64.61085 | 27.60923 | 0.755996 | 1.715981 | 0.44538 |
| 13 | 119090800 | 119093399 | 32.52435 | 55.46794 | 42.35268 | 18.56666 | 0.770054 | 1.026132 | 0.445399 |
| 19 | 9074800 | 9078399 | 22.57258 | 37.60583 | 63.97387 | 27.79502 | -0.38278 | 1.232403 | 0.445403 |
| 13 | 28425400 | 28427999 | 33.30257 | 56.26007 | 46.30617 | 20.25891 | 0.661152 | 1.113177 | 0.445407 |
| 10 | 22192600 | 22193599 | 12.77611 | 24.6713 | 19.48431 | 7.501855 | 0.887793 | -0.10693 | 0.445422 |

| | | | | | | | | | |
|---|---|---|---|---|---|---|---|---|---|
| 5 | 50710200 | 50713599 | 27.89586 | 47.52947 | 44.70071 | 19.5945 | 0.470343 | 1.01993 | 0.445441 |
| 2 | 23653200 | 23653799 | 10.31974 | 19.95843 | 20.33673 | 7.774307 | 0.533714 | -0.1529 | 0.445454 |
| 11 | 70963400 | 70967199 | 104.181 | 160.7276 | 158.353 | 65.7769 | 0.320951 | 2.891104 | 0.445461 |
| 3 | 92048200 | 92050199 | 22.83742 | 39.59397 | 39.34911 | 17.2645 | 0.395415 | 0.837193 | 0.445464 |
| 8 | 97905800 | 97908599 | 24.75573 | 43.34054 | 34.0147 | 15.0122 | 0.735513 | 0.706663 | 0.445468 |
| 1 | 118436000 | 118438399 | 12.24135 | 21.52908 | 39.90753 | 17.67045 | -0.49781 | 0.578459 | 0.445469 |
| 3 | 90744000 | 90744599 | 11.73317 | 22.0373 | 22.10162 | 9.087761 | 0.462206 | 0.103963 | 0.445472 |
| 9 | 121242000 | 121245399 | 41.24536 | 66.035 | 109.2461 | 46.03753 | -0.38009 | 2.013914 | 0.445473 |
| 11 | 27572400 | 27573399 | 13.91789 | 26.39736 | 22.41695 | 9.044659 | 0.722947 | 0.059134 | 0.445473 |
| 17 | 94454400 | 94457599 | 26.2231 | 45.43941 | 37.96326 | 16.65618 | 0.644413 | 0.852998 | 0.445482 |
| 7 | 14675200 | 14697799 | 435.9667 | 607.4349 | 516.6634 | 214.3552 | 0.39642 | 4.637971 | 0.445548 |
| 2 | 171193600 | 171196199 | 13.95106 | 24.45765 | 42.49253 | 18.77534 | -0.40542 | 0.640682 | 0.445548 |
| 8 | 16572400 | 16572999 | 9.928214 | 19.36066 | 16.00796 | 5.754022 | 0.911923 | -0.38114 | 0.445552 |
| 5 | 140348800 | 140354799 | 51.11752 | 81.27421 | 128.8014 | 53.75728 | -0.32916 | 2.247566 | 0.445573 |
| 10 | 86136000 | 86139599 | 15.40942 | 26.33842 | 52.09687 | 22.84244 | -0.59704 | 0.95239 | 0.445577 |
| 15 | 45754200 | 45756199 | 26.2972 | 45.77054 | 35.66269 | 15.67527 | 0.745415 | 0.798181 | 0.445578 |
| 5 | 28315600 | 28316999 | 10.37844 | 18.80042 | 34.33167 | 14.8639 | -0.45128 | 0.332893 | 0.445587 |
| 6 | 52282600 | 52285799 | 19.7617 | 33.17833 | 60.71638 | 26.50866 | -0.48784 | 1.135337 | 0.445616 |
| 2 | 89392800 | 89394399 | 25.67427 | 43.98294 | 43.0909 | 18.89712 | 0.412705 | 0.960133 | 0.445625 |
| 3 | 93558000 | 93559399 | 10.97864 | 19.84742 | 36.8973 | 16.01522 | -0.47963 | 0.352699 | 0.445631 |
| 15 | 21866000 | 21871399 | 70.73635 | 113.0924 | 91.58577 | 38.51857 | 0.646397 | 2.084001 | 0.445639 |
| X | 13710800 | 13713999 | 56.11907 | 90.50746 | 74.89419 | 31.90114 | 0.630077 | 1.799169 | 0.445644 |
| 19 | 27035000 | 27037399 | 38.6574 | 63.36896 | 62.09842 | 26.72937 | 0.405751 | 1.461659 | 0.445646 |
| 7 | 135125000 | 135127999 | 21.84402 | 37.92171 | 37.74345 | 16.56295 | 0.393491 | 0.815598 | 0.445653 |
| 18 | 6400000 | 6403199 | 41.75561 | 67.91876 | 67.14303 | 28.81671 | 0.385945 | 1.577236 | 0.445656 |
| 10 | 73279000 | 73280399 | 13.0008 | 24.6376 | 22.42671 | 9.064428 | 0.621147 | 0.055415 | 0.445657 |
| 15 | 19095600 | 19096599 | 16.73252 | 28.8744 | 49.52093 | 21.72431 | -0.38974 | 0.866484 | 0.445662 |
| 16 | 60875200 | 60879599 | 58.36148 | 93.44466 | 92.2255 | 38.86883 | 0.363823 | 2.024385 | 0.445682 |
| 3 | 110381000 | 110381399 | 4.350191 | 8.498127 | 19.37329 | 6.768369 | -0.49509 | -0.54099 | 0.4457 |
| 1 | 133793200 | 133795399 | 17.00151 | 30.39718 | 29.01375 | 12.73543 | 0.465724 | 0.444717 | 0.4457 |
| 19 | 39947000 | 39948799 | 28.7172 | 48.20449 | 48.58442 | 21.20926 | 0.369644 | 1.153909 | 0.445742 |
| 1 | 81515800 | 81516999 | 13.30542 | 24.50975 | 24.92863 | 10.56061 | 0.408989 | 0.238608 | 0.445745 |
| 3 | 7103600 | 7104599 | 19.56324 | 34.49031 | 28.21049 | 12.48509 | 0.677593 | 0.573454 | 0.445754 |
| 15 | 98640800 | 98642999 | 18.24238 | 31.0021 | 54.79768 | 24.01553 | -0.43703 | 1.017636 | 0.445781 |
| 7 | 17106800 | 17113999 | 86.23347 | 137.29 | 92.77743 | 38.79057 | 0.900567 | 2.229256 | 0.445781 |
| 15 | 22644000 | 22645599 | 18.85383 | 34.40169 | 25.20759 | 10.81523 | 0.866061 | 0.33363 | 0.445789 |
| 10 | 46444400 | 46448999 | 40.91959 | 67.10446 | 61.22431 | 26.35764 | 0.50877 | 1.460799 | 0.445792 |
| 9 | 95470200 | 95470999 | 10.07592 | 19.27742 | 20.1799 | 7.987355 | 0.448186 | -0.04409 | 0.445815 |
| 19 | 20450400 | 20453999 | 31.11788 | 52.06238 | 51.35594 | 22.37258 | 0.400714 | 1.19991 | 0.445824 |
| 8 | 61616800 | 61617999 | 12.59546 | 23.93542 | 22.54977 | 9.061112 | 0.579438 | 0.03409 | 0.445827 |
| 16 | 30038400 | 30042799 | 33.00902 | 53.36531 | 89.12141 | 38.08007 | -0.37673 | 1.695041 | 0.445842 |
| 12 | 65736000 | 65737999 | 16.35001 | 28.30649 | 48.99754 | 21.49684 | -0.40212 | 0.833017 | 0.445847 |
| 7 | 107362400 | 107365199 | 19.69346 | 34.6814 | 32.5845 | 14.42409 | 0.477926 | 0.592551 | 0.445862 |
| 15 | 28539800 | 28540999 | 13.39437 | 23.45805 | 42.58467 | 18.80002 | -0.46813 | 0.6571 | 0.445874 |
| 4 | 113080200 | 113089199 | 108.3741 | 169.7615 | 131.1276 | 54.39127 | 0.688545 | 2.626424 | 0.445885 |
| 9 | 25388000 | 25389599 | 8.950018 | 16.48039 | 31.44262 | 13.32712 | -0.49274 | 0.222222 | 0.445888 |
| 2 | 9543400 | 9544799 | 13.96684 | 24.59967 | 42.9201 | 18.99882 | -0.41036 | 0.577164 | 0.445893 |

| | | | | | | | | | |
|---|---|---|---|---|---|---|---|---|---|
| 9 | 12341600 | 12342599 | 10.5694 | 19.21278 | 34.39312 | 14.82612 | -0.41846 | 0.316348 | 0.445896 |
| 15 | 45784200 | 45785199 | 9.374396 | 17.53544 | 34.60162 | 14.32605 | -0.51563 | 0.138641 | 0.445912 |
| 5 | 151501400 | 151501999 | 14.2217 | 26.08308 | 25.9153 | 11.04425 | 0.436525 | 0.27346 | 0.445936 |
| 2 | 27962400 | 27964799 | 28.41143 | 46.40883 | 77.53074 | 33.32618 | -0.36465 | 1.498405 | 0.44594 |
| 16 | 74497800 | 74499599 | 15.12565 | 26.34489 | 45.77901 | 20.13073 | -0.40612 | 0.752594 | 0.44594 |
| 5 | 60877800 | 60879599 | 22.7944 | 40.30036 | 28.91853 | 12.79123 | 0.866682 | 0.562748 | 0.445945 |
| 17 | 55613800 | 55615399 | 21.39216 | 37.21434 | 37.17668 | 16.33551 | 0.388457 | 0.783948 | 0.445948 |
| 13 | 66319200 | 66321999 | 32.0757 | 54.816 | 41.80838 | 18.3311 | 0.772089 | 1.006164 | 0.445965 |
| 17 | 85760400 | 85763199 | 21.91716 | 36.46944 | 65.07193 | 28.24989 | -0.45144 | 1.243039 | 0.445975 |
| 10 | 35005000 | 35008599 | 41.98496 | 68.27473 | 67.63524 | 29.02532 | 0.382724 | 1.582491 | 0.44598 |
| 19 | 39350000 | 39360599 | 222.8417 | 327.1341 | 315.3967 | 129.9002 | 0.290374 | 3.85074 | 0.445997 |
| 9 | 76655200 | 76656199 | 15.71484 | 27.44155 | 47.31515 | 20.83987 | -0.39479 | 0.719429 | 0.446009 |
| 7 | 52728800 | 52731199 | 39.99154 | 65.09263 | 66.33531 | 28.47696 | 0.342888 | 1.564516 | 0.446033 |
| 3 | 11780600 | 11782399 | 19.94161 | 34.91956 | 34.3687 | 15.17108 | 0.410301 | 0.676469 | 0.446039 |
| 3 | 21559000 | 21564999 | 148.2942 | 224.4402 | 219.5738 | 90.83237 | 0.309245 | 3.288674 | 0.446045 |
| 1 | 15336800 | 15338199 | 18.07594 | 31.87942 | 31.73515 | 14.044 | 0.395752 | 0.566058 | 0.446045 |
| 4 | 137527400 | 137529399 | 12.68541 | 22.29922 | 40.88818 | 18.10643 | -0.48268 | 0.586827 | 0.446051 |
| 14 | 88814400 | 88816999 | 23.04999 | 40.14005 | 32.8212 | 14.43113 | 0.676128 | 0.770512 | 0.446055 |
| 15 | 50396600 | 50397999 | 13.13422 | 23.24579 | 40.80243 | 18.0439 | -0.41621 | 0.514684 | 0.446073 |
| 2 | 34862200 | 34864599 | 19.12301 | 32.25063 | 57.42058 | 25.10848 | -0.44798 | 1.091802 | 0.446075 |
| 13 | 87840800 | 87841399 | 6.250357 | 12.09686 | 26.1808 | 9.97501 | -0.5391 | -0.19408 | 0.446077 |
| 3 | 106231800 | 106232799 | 21.15577 | 36.83835 | 36.07558 | 15.86122 | 0.41711 | 0.766985 | 0.446078 |
| 8 | 60438800 | 60441199 | 22.7854 | 39.50802 | 39.325 | 17.25398 | 0.393164 | 0.835322 | 0.446086 |
| 4 | 43336200 | 43336999 | 15.27072 | 28.83595 | 22.86443 | 9.311521 | 0.809868 | 0.103114 | 0.446098 |
| 6 | 92232400 | 92235399 | 22.04866 | 37.0563 | 62.46585 | 27.286 | -0.36975 | 1.136849 | 0.446107 |
| 12 | 114789800 | 114790999 | 43.17947 | 69.79283 | 69.70309 | 29.77473 | 0.362944 | 1.709942 | 0.446118 |
| 17 | 7776800 | 7778199 | 15.65552 | 28.37987 | 27.73906 | 11.99899 | 0.445681 | 0.353041 | 0.446154 |
| 16 | 56011000 | 56012199 | 18.02894 | 32.1642 | 26.51347 | 11.65928 | 0.674394 | 0.461201 | 0.446161 |
| 1 | 179539200 | 179545399 | 68.12342 | 106.2768 | 171.6533 | 71.23108 | -0.37897 | 2.680461 | 0.446171 |
| 7 | 41178000 | 41180799 | 48.32036 | 77.87313 | 78.43915 | 33.37972 | 0.345819 | 1.807239 | 0.446185 |
| 7 | 62213800 | 62215599 | 68.70881 | 109.0346 | 111.7669 | 46.55843 | 0.295743 | 2.313442 | 0.446198 |
| 17 | 24239200 | 24241599 | 15.83252 | 27.17316 | 50.55478 | 22.17364 | -0.50818 | 0.913721 | 0.446199 |
| 8 | 74989000 | 74992599 | 33.19668 | 56.09998 | 45.73856 | 20.01262 | 0.67475 | 1.108405 | 0.446205 |
| 8 | 118915400 | 118916799 | 12.09011 | 21.28567 | 39.40285 | 17.44329 | -0.49517 | 0.565503 | 0.446225 |
| 4 | 130598400 | 130601399 | 25.0669 | 42.63578 | 42.67556 | 18.70418 | 0.380367 | 1.020282 | 0.446225 |
| 4 | 78572600 | 78573599 | 13.86389 | 26.55614 | 19.06413 | 7.520062 | 0.991321 | -0.02196 | 0.446236 |
| 7 | 13177200 | 13178999 | 46.16135 | 74.42381 | 74.65895 | 31.78767 | 0.352378 | 1.790532 | 0.446268 |
| 7 | 22303200 | 22303799 | 16.66944 | 29.92625 | 24.66558 | 10.78937 | 0.679698 | 0.409323 | 0.446274 |
| 19 | 45315800 | 45317999 | 14.6418 | 25.43342 | 46.34696 | 20.34594 | -0.47482 | 0.788828 | 0.44628 |
| 10 | 25786000 | 25787399 | 14.51913 | 26.93005 | 25.38447 | 10.65358 | 0.527677 | 0.212052 | 0.446293 |
| 2 | 83343600 | 83344799 | 13.01846 | 23.06503 | 40.29943 | 17.81773 | -0.40887 | 0.50322 | 0.446297 |
| 1 | 49456400 | 49460599 | 77.29396 | 121.9724 | 124.5687 | 51.70656 | 0.294253 | 2.461455 | 0.446319 |
| 13 | 65627200 | 65629199 | 22.06514 | 40.06361 | 24.47879 | 10.56737 | 1.121278 | 0.384641 | 0.446325 |
| 7 | 85803600 | 85804999 | 19.33472 | 34.31673 | 26.12948 | 11.54037 | 0.784275 | 0.502305 | 0.446349 |
| 13 | 113816400 | 113817799 | 21.78088 | 37.83981 | 35.87795 | 15.75157 | 0.463562 | 0.806154 | 0.446352 |
| 4 | 98826800 | 98831399 | 26.29531 | 43.40179 | 74.42673 | 32.13106 | -0.39548 | 1.354982 | 0.446358 |
| X | 116538400 | 116541399 | 32.38457 | 55.82698 | 36.68655 | 16.0935 | 0.988056 | 0.926013 | 0.446361 |

| | | | | | | | | | |
|---|---|---|---|---|---|---|---|---|---|
| 15 | 91018800 | 91019799 | 9.75654 | 18.03941 | 33.64285 | 14.18027 | -0.45435 | 0.211978 | 0.446374 |
| 5 | 146929800 | 146936599 | 55.1999 | 87.56267 | 136.842 | 56.9814 | -0.31232 | 2.319497 | 0.446386 |
| 18 | 67475800 | 67477399 | 13.3447 | 23.4046 | 41.4844 | 18.33648 | -0.43398 | 0.629786 | 0.446417 |
| 13 | 90237200 | 90238199 | 9.134264 | 16.73524 | 31.91815 | 13.62161 | -0.49934 | 0.254101 | 0.446435 |
| 17 | 93608000 | 93609199 | 18.59913 | 33.35113 | 28.89745 | 12.6531 | 0.607065 | 0.438542 | 0.446436 |
| 2 | 110870600 | 110871599 | 10.20271 | 18.79992 | 34.66105 | 14.67761 | -0.44306 | 0.235602 | 0.446439 |
| 4 | 113243000 | 113245599 | 35.79106 | 59.74162 | 51.46919 | 22.38899 | 0.595494 | 1.239116 | 0.446445 |
| 2 | 100948800 | 100951199 | 29.44575 | 47.74017 | 84.32935 | 36.11536 | -0.45346 | 1.625455 | 0.446445 |
| 7 | 62208800 | 62210399 | 30.89309 | 51.86635 | 49.67594 | 21.68824 | 0.44298 | 1.160615 | 0.446477 |
| 7 | 5357400 | 5397799 | 684.5875 | 918.1522 | 749.6292 | 312.8552 | 0.400021 | 5.234124 | 0.446485 |
| 9 | 82330400 | 82331199 | 9.739179 | 17.85946 | 32.42336 | 13.82043 | -0.42757 | 0.2547 | 0.446487 |
| 17 | 72168000 | 72169799 | 23.91838 | 40.98664 | 41.55776 | 18.2252 | 0.363384 | 0.947371 | 0.446491 |
| Y | 41202000 | 41208399 | 40.43608 | 66.07964 | 37.03647 | 15.92248 | 1.206625 | 1.483236 | 0.446505 |
| 1 | 93601200 | 93605599 | 34.47895 | 55.75566 | 89.99872 | 38.46733 | -0.32736 | 1.691552 | 0.446515 |
| 9 | 113902200 | 113905999 | 35.1641 | 56.63242 | 95.81727 | 40.73536 | -0.40197 | 1.814964 | 0.446548 |
| 3 | 145999400 | 145999799 | 5.657509 | 11.01887 | 23.80309 | 8.69575 | -0.47891 | -0.35264 | 0.446557 |
| 2 | 126621800 | 126622999 | 7.073433 | 13.37607 | 27.59933 | 11.20572 | -0.55483 | 0.032601 | 0.44659 |
| 1 | 23035800 | 23037999 | 20.56919 | 36.09232 | 30.15282 | 13.33315 | 0.645515 | 0.640954 | 0.446594 |
| 12 | 23836400 | 23857799 | 596.1314 | 802.6366 | 698.5485 | 291.3687 | 0.313502 | 5.170725 | 0.446595 |
| 12 | 8641600 | 8650599 | 136.0763 | 203.1158 | 313.1374 | 129.0775 | -0.36407 | 3.55825 | 0.446607 |
| 14 | 3230800 | 3249799 | 393.4544 | 551.9204 | 446.7943 | 185.1728 | 0.477544 | 4.534795 | 0.446657 |
| 7 | 59557000 | 59558599 | 50.10564 | 80.50473 | 78.20406 | 33.13317 | 0.3926 | 1.904455 | 0.446668 |
| 1 | 30035400 | 30036199 | 12.36872 | 21.89562 | 38.2731 | 16.92464 | -0.40986 | 0.504361 | 0.446671 |
| 17 | 78791800 | 78793399 | 11.71979 | 20.89253 | 36.77176 | 16.18076 | -0.41413 | 0.446433 | 0.446683 |
| 1 | 145608200 | 145612399 | 47.29021 | 76.75724 | 68.64304 | 29.42175 | 0.527624 | 1.628428 | 0.446689 |
| X | 104102400 | 104105999 | 19.63425 | 33.56449 | 58.79699 | 25.78048 | -0.42302 | 0.972548 | 0.446711 |
| 7 | 38963200 | 38963999 | 16.01858 | 28.97011 | 23.25808 | 10.09017 | 0.725114 | 0.359029 | 0.446724 |
| 13 | 62821800 | 62829399 | 170.0133 | 256.4865 | 226.3441 | 93.53824 | 0.453945 | 3.358304 | 0.446741 |
| 13 | 66036600 | 66043799 | 95.29762 | 149.0863 | 141.96 | 58.89231 | 0.385581 | 2.641458 | 0.446757 |
| 13 | 14706800 | 14708399 | 12.5159 | 22.13131 | 38.77998 | 17.15264 | -0.41409 | 0.517929 | 0.446766 |
| 15 | 84413600 | 84415399 | 12.5159 | 22.13131 | 38.77998 | 17.15264 | -0.41409 | 0.517929 | 0.446766 |
| 7 | 13331000 | 13343999 | 192.5061 | 287.9334 | 251.4238 | 103.6798 | 0.458185 | 3.528577 | 0.446791 |
| 16 | 35455000 | 35457599 | 12.76172 | 22.32072 | 42.64019 | 18.80364 | -0.54131 | 0.674745 | 0.44682 |
| 16 | 18418800 | 18421599 | 16.45084 | 28.14434 | 52.67952 | 23.09963 | -0.5178 | 0.946097 | 0.446847 |
| 19 | 50822800 | 50823599 | 7.927305 | 15.03121 | 28.77443 | 11.61646 | -0.44117 | 0.022223 | 0.446852 |
| 5 | 101814600 | 101817399 | 18.08708 | 30.68927 | 55.86638 | 24.47388 | -0.47956 | 1.031457 | 0.446876 |
| 7 | 128572000 | 128574999 | 24.85873 | 41.16746 | 68.50119 | 29.64096 | -0.35165 | 1.310365 | 0.446883 |
| 7 | 72137800 | 72139199 | 10.32515 | 18.94035 | 34.5455 | 14.71827 | -0.43416 | 0.260164 | 0.446886 |
| X | 26860000 | 26865999 | 69.23567 | 111.0908 | 80.97269 | 34.23641 | 0.803925 | 1.974027 | 0.446888 |
| 7 | 12887800 | 12888999 | 21.91545 | 38.04002 | 37.3839 | 16.40363 | 0.411916 | 0.820722 | 0.446949 |
| 8 | 46905800 | 46909199 | 55.70406 | 89.57603 | 83.38884 | 35.37351 | 0.455399 | 1.887029 | 0.446968 |
| 13 | 117592200 | 117595599 | 40.38011 | 64.9423 | 103.7012 | 43.99811 | -0.32128 | 1.869742 | 0.446976 |
| X | 121150200 | 121157799 | 128.2528 | 196.8563 | 177.2474 | 73.58704 | 0.445142 | 3.000768 | 0.446989 |
| 7 | 114316400 | 114317599 | 9.403512 | 17.29785 | 31.93919 | 13.55634 | -0.44741 | 0.232478 | 0.446991 |
| 3 | 13234600 | 13236999 | 32.60128 | 54.59588 | 50.82403 | 22.15409 | 0.483981 | 1.194182 | 0.446997 |
| 1 | 107139400 | 107141199 | 18.04792 | 31.92369 | 31.09652 | 13.7533 | 0.428456 | 0.532628 | 0.447003 |
| 5 | 109115400 | 109119599 | 59.74696 | 95.69489 | 91.92421 | 38.76149 | 0.403378 | 2.016455 | 0.447015 |

| | | | | | | | | | |
|---|---|---|---|---|---|---|---|---|---|
| 13 | 56859400 | 56861799 | 23.14995 | 38.70915 | 64.7678 | 28.20502 | -0.35909 | 1.194 | 0.447016 |
| 6 | 41375000 | 41379599 | 46.21953 | 75.27047 | 64.81521 | 27.82989 | 0.585845 | 1.568376 | 0.447022 |
| 4 | 134214600 | 134215799 | 7.486226 | 14.3279 | 27.91961 | 11.03946 | -0.43927 | -0.04997 | 0.447026 |
| 2 | 4960800 | 4963599 | 24.36132 | 40.36027 | 67.9262 | 29.40049 | -0.36796 | 1.303359 | 0.447031 |
| 12 | 21633800 | 21643599 | 126.2748 | 193.6382 | 182.7886 | 75.8778 | 0.375977 | 3.017231 | 0.447039 |
| 4 | 150325000 | 150326799 | 6.912363 | 13.13255 | 27.14182 | 10.91187 | -0.54443 | -0.00199 | 0.447062 |
| X | 114018400 | 114020999 | 32.29526 | 54.5406 | 46.89008 | 20.51235 | 0.598572 | 1.114752 | 0.447068 |
| 2 | 39624000 | 39631599 | 86.63984 | 136.8854 | 121.8928 | 50.62672 | 0.493123 | 2.44028 | 0.447076 |
| 4 | 56726000 | 56731199 | 47.90365 | 76.06977 | 141.3028 | 58.90169 | -0.55967 | 2.283958 | 0.447077 |
| 5 | 70977200 | 70979199 | 18.47802 | 33.35486 | 27.26776 | 11.85776 | 0.696784 | 0.39463 | 0.447103 |
| 17 | 19730800 | 19736199 | 88.50043 | 139.7762 | 110.1056 | 45.72296 | 0.669348 | 2.448287 | 0.447103 |
| 12 | 19390600 | 19404199 | 223.6122 | 329.1787 | 279.6423 | 115.1583 | 0.47665 | 3.805892 | 0.447144 |
| 13 | 33655400 | 33657399 | 34.15108 | 57.1905 | 44.84663 | 19.54839 | 0.730685 | 1.195396 | 0.447148 |
| 6 | 7603800 | 7606399 | 21.01022 | 35.4514 | 59.1904 | 25.88636 | -0.35572 | 1.094621 | 0.447148 |
| 1 | 9142400 | 9144999 | 37.82501 | 62.00394 | 62.02668 | 26.69838 | 0.376076 | 1.459666 | 0.447185 |
| 17 | 88747000 | 88749599 | 17.48061 | 29.67717 | 54.75173 | 23.98972 | -0.49862 | 1.024509 | 0.447219 |
| 14 | 80115200 | 80117399 | 16.35973 | 28.24277 | 50.05999 | 21.96107 | -0.43707 | 0.859651 | 0.447232 |
| 15 | 86707000 | 86709999 | 38.51378 | 63.54912 | 57.28642 | 24.72665 | 0.529514 | 1.377103 | 0.447241 |
| 7 | 39162600 | 39168199 | 62.05042 | 98.02417 | 150.8188 | 62.63707 | -0.29491 | 2.431898 | 0.447246 |
| 15 | 94404400 | 94406399 | 22.58314 | 37.74567 | 64.01837 | 27.87134 | -0.37847 | 1.198698 | 0.447246 |
| 2 | 11441000 | 11441399 | 9.473595 | 18.24431 | 18.22974 | 7.08087 | 0.540419 | -0.11593 | 0.447248 |
| 2 | 132824400 | 132824999 | 5.805263 | 11.29149 | 23.5824 | 8.717989 | -0.44612 | -0.30813 | 0.447276 |
| 13 | 67243800 | 67244599 | 7.77597 | 14.89364 | 30.37347 | 11.98381 | -0.50203 | -0.04964 | 0.447288 |
| 12 | 107298800 | 107303199 | 33.67134 | 54.36759 | 91.18587 | 38.89895 | -0.38502 | 1.733867 | 0.447292 |
| 12 | 84104800 | 84106399 | 10.11394 | 18.29298 | 33.85048 | 14.68393 | -0.47176 | 0.340907 | 0.447305 |
| 4 | 118754800 | 118758999 | 33.18868 | 53.71403 | 87.45165 | 37.40402 | -0.33852 | 1.670849 | 0.447325 |
| 18 | 3733600 | 3739999 | 99.43465 | 154.7584 | 137.5266 | 57.08884 | 0.479341 | 2.730426 | 0.447329 |
| 18 | 72884400 | 72888599 | 41.22388 | 68.14286 | 54.55406 | 23.5758 | 0.700449 | 1.362855 | 0.447339 |
| 4 | 78296000 | 78297599 | 11.71362 | 20.98823 | 36.79517 | 16.12163 | -0.4042 | 0.414658 | 0.44735 |
| 14 | 103763800 | 103766199 | 21.42091 | 37.52015 | 31.21408 | 13.78173 | 0.651686 | 0.675887 | 0.447352 |
| 3 | 59403600 | 59404999 | 15.39581 | 26.84642 | 45.74687 | 20.1312 | -0.378 | 0.735933 | 0.447356 |
| 10 | 43304200 | 43309399 | 46.94623 | 74.97356 | 118.1933 | 49.63068 | -0.31469 | 2.099469 | 0.447365 |
| 7 | 43674600 | 43676199 | 15.59952 | 28.65153 | 23.02954 | 9.795106 | 0.74195 | 0.267664 | 0.447389 |
| 15 | 31781400 | 31785999 | 58.53773 | 92.86587 | 149.0847 | 62.08497 | -0.35097 | 2.322751 | 0.447392 |
| 15 | 7839000 | 7843599 | 36.26259 | 58.05164 | 122.9594 | 51.80882 | -0.73608 | 2.01438 | 0.447393 |
| Y | 2313000 | 2315599 | 18.28009 | 32.2733 | 20.63215 | 9.128591 | 1.031323 | 0.531833 | 0.447393 |
| 10 | 81505200 | 81506599 | 22.02316 | 38.37404 | 32.51489 | 14.30163 | 0.625179 | 0.758245 | 0.447398 |
| 14 | 35721000 | 35724399 | 28.24477 | 49.1212 | 33.89353 | 14.8902 | 0.919795 | 0.815849 | 0.447401 |
| 17 | 4289000 | 4291999 | 50.65812 | 82.12482 | 61.32824 | 26.26317 | 0.785429 | 1.654066 | 0.447402 |
| 1 | 27544400 | 27546599 | 20.49908 | 36.73293 | 28.19413 | 12.35225 | 0.780451 | 0.454631 | 0.447402 |
| 3 | 47864600 | 47867799 | 41.55683 | 67.87878 | 63.98982 | 27.50532 | 0.458518 | 1.515752 | 0.447404 |
| 7 | 42071400 | 42076599 | 68.51879 | 109.9553 | 80.70901 | 34.1319 | 0.794022 | 1.966986 | 0.447412 |
| 2 | 7588600 | 7592399 | 54.61472 | 87.63628 | 86.8689 | 36.73275 | 0.361305 | 1.953737 | 0.447419 |
| 16 | 48405400 | 48408599 | 27.14211 | 46.06663 | 45.24573 | 19.82292 | 0.407195 | 1.048606 | 0.447433 |
| 7 | 76868000 | 76868399 | 5.403507 | 10.45619 | 23.74997 | 9.055699 | -0.60769 | -0.20479 | 0.447477 |
| X | 127853200 | 127853999 | 15.47151 | 28.12755 | 26.38685 | 11.37286 | 0.508174 | 0.334389 | 0.447482 |
| 13 | 24409400 | 24414999 | 107.2153 | 165.1892 | 166.1273 | 69.00614 | 0.289873 | 2.917451 | 0.44749 |

| | | | | | | | | | |
|---|---|---|---|---|---|---|---|---|---|
| 5 | 112595800 | 112597399 | 13.05643 | 22.86067 | 42.07815 | 18.57262 | -0.48803 | 0.657255 | 0.447495 |
| 4 | 9135600 | 9136799 | 11.90021 | 21.03608 | 38.31442 | 16.94777 | -0.46966 | 0.516828 | 0.447507 |
| 6 | 10840600 | 10842199 | 19.01709 | 33.53809 | 32.23285 | 14.26435 | 0.44632 | 0.574381 | 0.44752 |
| 18 | 45306600 | 45311199 | 33.39174 | 56.49754 | 45.74446 | 20.0225 | 0.684852 | 1.100963 | 0.447526 |
| 13 | 98598000 | 98598399 | 7.799845 | 15.18107 | 16.75773 | 6.162057 | 0.471326 | -0.32633 | 0.447533 |
| 16 | 31254200 | 31263399 | 83.27851 | 128.6056 | 196.8315 | 81.75974 | -0.31281 | 2.868955 | 0.447535 |
| 2 | 152942600 | 152944999 | 14.81355 | 25.82322 | 45.24698 | 19.9074 | -0.41788 | 0.738492 | 0.447547 |
| 12 | 44306600 | 44310199 | 49.95831 | 80.43263 | 80.80249 | 34.33586 | 0.34787 | 1.841303 | 0.447566 |
| 10 | 22842800 | 22844199 | 15.74521 | 28.93018 | 26.18826 | 11.1328 | 0.572312 | 0.274239 | 0.447574 |
| X | 8903400 | 8905599 | 20.90972 | 36.72558 | 33.27401 | 14.7254 | 0.529327 | 0.635735 | 0.447575 |
| 4 | 54068200 | 54070999 | 39.80686 | 65.13283 | 63.27268 | 27.2142 | 0.417072 | 1.488103 | 0.447598 |
| 15 | 24219600 | 24221199 | 17.58558 | 31.77728 | 27.98101 | 12.15195 | 0.591615 | 0.384689 | 0.447617 |
| 1 | 103392400 | 103392999 | 10.47795 | 20.36661 | 18.81295 | 6.992037 | 0.71155 | -0.25019 | 0.447627 |
| 3 | 41885200 | 41886399 | 16.05646 | 27.91288 | 47.3806 | 20.81007 | -0.37292 | 0.785414 | 0.447629 |
| 7 | 31668200 | 31673199 | 79.06438 | 125.5868 | 99.19347 | 41.35992 | 0.672555 | 2.291933 | 0.447632 |
| 5 | 24910600 | 24912999 | 12.14778 | 21.34143 | 41.01497 | 18.15491 | -0.54965 | 0.598048 | 0.447638 |
| 13 | 43107600 | 43112399 | 33.31022 | 54.0922 | 92.02012 | 39.45564 | -0.39726 | 1.604674 | 0.447648 |
| 4 | 141925800 | 141928199 | 26.75666 | 43.85309 | 74.20542 | 31.93829 | -0.38016 | 1.446288 | 0.447669 |
| 4 | 81155400 | 81156399 | 12.36019 | 23.28053 | 23.56531 | 9.643448 | 0.454207 | 0.094846 | 0.447675 |
| 12 | 114940800 | 114941999 | 14.91763 | 28.05182 | 23.29343 | 9.558487 | 0.734488 | 0.126085 | 0.447694 |
| 11 | 54022800 | 54024799 | 14.29412 | 24.94735 | 44.17949 | 19.45225 | -0.43302 | 0.715825 | 0.447713 |
| 8 | 87385800 | 87386599 | 14.3416 | 25.00907 | 44.73013 | 19.68424 | -0.44726 | 0.730725 | 0.447728 |
| 9 | 8477400 | 8477999 | 10.20081 | 19.47627 | 20.14233 | 8.013562 | 0.458701 | -0.02685 | 0.447733 |
| 9 | 27595800 | 27596799 | 10.20081 | 19.47627 | 20.14233 | 8.013562 | 0.458701 | -0.02685 | 0.447733 |
| 17 | 87580800 | 87581599 | 14.40417 | 26.34991 | 26.28167 | 11.23963 | 0.427087 | 0.288098 | 0.447742 |
| 17 | 62061800 | 62063399 | 17.19781 | 31.1677 | 27.48095 | 11.89116 | 0.592993 | 0.363967 | 0.447788 |
| 1 | 4757800 | 4759799 | 25.12699 | 41.29427 | 71.89353 | 30.97887 | -0.41949 | 1.407316 | 0.447797 |
| 7 | 15315600 | 15331599 | 158.2253 | 241.9173 | 152.0451 | 63.07713 | 0.958949 | 3.084037 | 0.447812 |
| 7 | 17298600 | 17301999 | 44.67595 | 72.41714 | 63.63515 | 27.24841 | 0.551066 | 1.645927 | 0.447818 |
| 2 | 115996200 | 115997599 | 12.45185 | 21.72893 | 44.2178 | 19.46735 | -0.63243 | 0.708313 | 0.447824 |
| 8 | 12419600 | 12420399 | 8.896885 | 16.71444 | 31.30768 | 12.86757 | -0.43215 | 0.100222 | 0.447826 |
| 10 | 35203400 | 35204199 | 8.896885 | 16.71444 | 31.30768 | 12.86757 | -0.43215 | 0.100222 | 0.447826 |
| 4 | 48101400 | 48102599 | 8.798611 | 16.24484 | 30.95944 | 13.07481 | -0.48722 | 0.20551 | 0.447844 |
| 3 | 29108600 | 29109399 | 7.630163 | 14.44437 | 28.72145 | 11.63486 | -0.49992 | 0.029957 | 0.44786 |
| 10 | 50688000 | 50691199 | 47.84431 | 76.4357 | 122.0356 | 51.2706 | -0.33237 | 2.091976 | 0.447875 |
| 1 | 118003400 | 118005399 | 48.6019 | 78.29207 | 79.64876 | 33.87272 | 0.330714 | 1.821151 | 0.447876 |
| 7 | 11940000 | 11945199 | 93.38858 | 145.6118 | 146.9989 | 61.00441 | 0.297597 | 2.699179 | 0.447886 |
| 11 | 82955800 | 82961799 | 42.0518 | 67.4781 | 107.0502 | 45.26713 | -0.31587 | 1.9444 | 0.447893 |
| 2 | 8534400 | 8535799 | 14.32382 | 26.08267 | 23.07133 | 9.923999 | 0.59427 | 0.308179 | 0.447902 |
| 14 | 4232800 | 4233599 | 10.42321 | 19.92697 | 21.20403 | 8.410756 | 0.421879 | -0.03056 | 0.447912 |
| 17 | 56492800 | 56498799 | 31.81141 | 51.25754 | 93.71408 | 39.86905 | -0.51259 | 1.795652 | 0.447916 |
| 10 | 51776000 | 51776799 | 11.0858 | 21.0317 | 21.53394 | 8.683298 | 0.455773 | 0.027775 | 0.447918 |
| 16 | 25460000 | 25461599 | 19.46499 | 34.48784 | 30.94727 | 13.68272 | 0.547223 | 0.524371 | 0.447931 |
| 4 | 52535800 | 52536599 | 7.802668 | 14.7679 | 28.15889 | 11.41194 | -0.44006 | 0.033636 | 0.447931 |
| 16 | 55124600 | 55126799 | 33.8675 | 55.95418 | 56.90207 | 24.58182 | 0.356337 | 1.352138 | 0.447934 |
| X | 28808600 | 28809399 | 17.27556 | 31.28313 | 24.55711 | 10.63682 | 0.75885 | 0.364477 | 0.447936 |
| 2 | 87066600 | 87070399 | 59.63446 | 95.22374 | 97.40621 | 40.89402 | 0.308022 | 2.111106 | 0.447937 |

| | | | | | | | | | |
|---|---|---|---|---|---|---|---|---|---|
| 2 | 52426600 | 52428199 | 19.88399 | 33.64714 | 56.96576 | 24.93505 | -0.37548 | 1.064262 | 0.447943 |
| 8 | 13835200 | 13836599 | 8.824414 | 16.21474 | 31.44072 | 13.36476 | -0.51873 | 0.234371 | 0.447947 |
| 17 | 50836000 | 50837599 | 12.86007 | 22.53301 | 42.53845 | 18.78715 | -0.52459 | 0.642796 | 0.447947 |
| 8 | 3325200 | 3330599 | 36.82952 | 59.15219 | 101.382 | 42.92981 | -0.42551 | 1.906499 | 0.44796 |
| 10 | 93008800 | 93009799 | 13.38466 | 24.76736 | 24.85983 | 10.46693 | 0.434072 | 0.213055 | 0.447986 |
| 12 | 88071600 | 88079799 | 107.4038 | 165.8992 | 165.635 | 68.8013 | 0.303118 | 2.870559 | 0.448004 |
| 13 | 62047200 | 62053799 | 102.3056 | 159.5477 | 146.8976 | 60.96049 | 0.430534 | 2.700151 | 0.448018 |
| 7 | 48110600 | 48113199 | 34.88837 | 57.87677 | 55.32484 | 23.96796 | 0.44579 | 1.307789 | 0.448021 |
| 1 | 163292200 | 163293199 | 10.43812 | 18.86362 | 34.34821 | 14.91534 | -0.44963 | 0.349652 | 0.448031 |
| 3 | 4649400 | 4650599 | 33.22107 | 54.84617 | 50.83545 | 21.94998 | 0.489713 | 1.355229 | 0.448035 |
| 10 | 117382200 | 117387999 | 54.70758 | 86.71826 | 138.7008 | 57.71314 | -0.34675 | 2.340533 | 0.448037 |
| X | 66804600 | 66806199 | 25.70187 | 44.79177 | 36.40078 | 16.01259 | 0.685368 | 0.782264 | 0.448043 |
| 17 | 66191600 | 66193799 | 18.5428 | 32.50414 | 29.03173 | 12.82657 | 0.549628 | 0.642816 | 0.448043 |
| 9 | 31121400 | 31124399 | 18.03371 | 30.36985 | 57.88976 | 25.3026 | -0.54609 | 1.100759 | 0.448071 |
| 3 | 152831200 | 152833599 | 16.86575 | 29.12851 | 49.52661 | 21.72736 | -0.37712 | 0.858849 | 0.448077 |
| 7 | 47991600 | 47991999 | 12.15248 | 23.00182 | 23.08286 | 9.353784 | 0.478561 | 0.058654 | 0.448079 |
| 18 | 38086400 | 38092799 | 42.8446 | 68.16549 | 136.854 | 57.16546 | -0.66881 | 2.221116 | 0.44808 |
| 15 | 32934800 | 32936199 | 8.608543 | 16.15098 | 30.15099 | 12.42044 | -0.42967 | 0.104715 | 0.44809 |
| 18 | 37629400 | 37632999 | 117.4435 | 179.8227 | 177.5733 | 73.69221 | 0.309315 | 3.044112 | 0.448092 |
| 17 | 58710800 | 58712199 | 19.07449 | 34.23285 | 25.32644 | 11.08148 | 0.83438 | 0.430037 | 0.448098 |
| 3 | 114483000 | 114483599 | 11.55649 | 20.78379 | 36.8298 | 16.08633 | -0.41643 | 0.390607 | 0.448112 |
| 3 | 9254800 | 9257199 | 23.18868 | 38.57347 | 65.74689 | 28.53692 | -0.3857 | 1.250869 | 0.448118 |
| 4 | 74729800 | 74731799 | 17.82686 | 31.44749 | 31.34219 | 13.86945 | 0.394301 | 0.56247 | 0.448138 |
| 19 | 9601600 | 9606399 | 20.02635 | 35.17712 | 29.10565 | 12.88165 | 0.659266 | 0.61877 | 0.448139 |
| 13 | 33845000 | 33850199 | 62.27899 | 99.73105 | 83.31298 | 35.12007 | 0.604163 | 2.022018 | 0.448148 |
| 13 | 35313000 | 35315999 | 11.71728 | 22.46191 | 18.88698 | 7.433435 | 0.768806 | -0.05005 | 0.44816 |
| 5 | 109303600 | 109304599 | 18.44606 | 32.67787 | 31.05168 | 13.72927 | 0.46493 | 0.519413 | 0.448165 |
| 18 | 5423400 | 5424599 | 10.724 | 20.4442 | 21.67588 | 8.651359 | 0.419083 | -0.00676 | 0.448166 |
| 18 | 16093600 | 16094199 | 10.1449 | 19.54533 | 20.17161 | 7.825688 | 0.494857 | -0.1093 | 0.448184 |
| 15 | 84168400 | 84169399 | 9.555278 | 17.68985 | 32.0404 | 13.48141 | -0.41069 | 0.198522 | 0.448229 |
| 4 | 136785200 | 136786399 | 11.16458 | 21.12702 | 22.0872 | 8.954744 | 0.419363 | 0.048413 | 0.448241 |
| 10 | 39606800 | 39609399 | 16.89646 | 28.22491 | 67.93814 | 29.56731 | -0.88158 | 1.189825 | 0.448247 |
| 13 | 75357200 | 75359399 | 40.18983 | 65.24442 | 66.60119 | 28.55041 | 0.337453 | 1.610028 | 0.448254 |
| 3 | 23738200 | 23743199 | 85.27184 | 133.4071 | 132.2303 | 54.85568 | 0.327803 | 2.632766 | 0.448255 |
| 19 | 11370200 | 11370599 | 4.918471 | 9.577835 | 21.83021 | 7.992934 | -0.55638 | -0.35566 | 0.44826 |
| 3 | 44490800 | 44494199 | 39.03049 | 64.14186 | 60.48333 | 26.06192 | 0.462542 | 1.431638 | 0.448263 |
| 8 | 50303400 | 50304999 | 17.24083 | 31.38015 | 27.01127 | 11.6253 | 0.632447 | 0.339623 | 0.448271 |
| 12 | 76429800 | 76433399 | 34.01057 | 55.99157 | 55.28502 | 23.8399 | 0.397238 | 1.390759 | 0.448281 |
| 1 | 107155400 | 107156999 | 16.83439 | 29.25655 | 50.61985 | 22.22864 | -0.40061 | 0.796418 | 0.448306 |
| 9 | 31561000 | 31562599 | 8.737832 | 16.24869 | 31.04337 | 12.9817 | -0.48077 | 0.162313 | 0.448315 |
| 9 | 56688000 | 56690399 | 17.42916 | 29.81493 | 52.14947 | 22.86701 | -0.42036 | 0.952443 | 0.448323 |
| 12 | 61035600 | 61037399 | 17.68946 | 31.15998 | 30.79773 | 13.63246 | 0.405328 | 0.574629 | 0.448328 |
| 2 | 7956800 | 7957999 | 8.431772 | 15.79532 | 29.57712 | 12.21542 | -0.43717 | 0.110279 | 0.448339 |
| 3 | 127419000 | 127420199 | 8.431772 | 15.79532 | 29.57712 | 12.21542 | -0.43717 | 0.110279 | 0.448339 |
| 11 | 121738400 | 121739199 | 8.431772 | 15.79532 | 29.57712 | 12.21542 | -0.43717 | 0.110279 | 0.448339 |
| 8 | 25690800 | 25691999 | 9.990675 | 17.93198 | 34.83788 | 15.24189 | -0.54979 | 0.387409 | 0.448343 |
| 1 | 117944400 | 117947399 | 66.0872 | 105.365 | 102.5323 | 42.95329 | 0.377519 | 2.166902 | 0.448352 |

| | | | | | | | | | |
|---|---|---|---|---|---|---|---|---|---|
| 10 | 54402600 | 54406399 | 89.1858 | 139.7618 | 139.1137 | 57.70141 | 0.323584 | 2.606921 | 0.448365 |
| 16 | 22201400 | 22204199 | 19.56739 | 32.96107 | 59.11585 | 25.8405 | -0.4587 | 1.103125 | 0.448368 |
| 17 | 64758400 | 64759199 | 10.32943 | 19.85721 | 20.68967 | 8.077423 | 0.472892 | -0.08541 | 0.448379 |
| 2 | 14735600 | 14738399 | 22.96994 | 40.01499 | 37.55434 | 16.51596 | 0.478386 | 0.773648 | 0.448386 |
| 4 | 127682800 | 127683599 | 7.576525 | 14.0911 | 30.01184 | 12.5474 | -0.6356 | 0.150339 | 0.448386 |
| 4 | 22083200 | 22084999 | 23.40705 | 40.52422 | 39.82226 | 17.47134 | 0.41113 | 0.851724 | 0.448388 |
| 2 | 39746200 | 39749599 | 74.22523 | 117.4099 | 119.7449 | 49.76439 | 0.298976 | 2.403414 | 0.448414 |
| 8 | 46324400 | 46326799 | 18.77493 | 32.93895 | 32.97568 | 14.5795 | 0.385978 | 0.639532 | 0.448416 |
| 10 | 7656000 | 7658199 | 16.56856 | 28.08638 | 61.18347 | 26.79556 | -0.73761 | 1.0404 | 0.448426 |
| 8 | 85557400 | 85559399 | 10.96764 | 19.42154 | 38.31691 | 16.94294 | -0.58334 | 0.490543 | 0.448447 |
| 18 | 74364600 | 74366599 | 24.01588 | 42.06222 | 34.01152 | 15.0159 | 0.692583 | 0.6958 | 0.448478 |
| 14 | 53936600 | 53938799 | 31.87045 | 54.22877 | 44.05155 | 19.30786 | 0.680609 | 1.045679 | 0.448492 |
| 6 | 78184800 | 78186399 | 20.22703 | 35.63777 | 32.5388 | 14.4025 | 0.519274 | 0.590976 | 0.448493 |
| 16 | 28401800 | 28402799 | 8.490167 | 15.72279 | 30.47902 | 12.81911 | -0.50716 | 0.185111 | 0.448497 |
| 6 | 69124200 | 69126199 | 23.40951 | 40.71901 | 38.09524 | 16.7379 | 0.48268 | 0.796977 | 0.448519 |
| 4 | 129825600 | 129830399 | 34.86107 | 56.06981 | 99.41852 | 42.18503 | -0.47186 | 1.857358 | 0.448521 |
| 14 | 8298600 | 8300999 | 39.97526 | 65.86169 | 58.40005 | 25.19133 | 0.552446 | 1.401293 | 0.448536 |
| 7 | 136778200 | 136783399 | 54.14438 | 86.77857 | 89.04647 | 37.58569 | 0.309456 | 1.988528 | 0.448557 |
| 7 | 8386600 | 8388399 | 51.49861 | 82.71802 | 78.73606 | 33.34398 | 0.421375 | 1.916967 | 0.448578 |
| 18 | 73508200 | 73513399 | 45.30405 | 72.53779 | 114.3583 | 48.19667 | -0.31065 | 2.017906 | 0.448589 |
| 9 | 121686000 | 121687399 | 7.591455 | 13.7327 | 33.98751 | 14.74052 | -0.88765 | 0.321941 | 0.44859 |
| 8 | 17790800 | 17792799 | 15.58511 | 27.18637 | 46.27004 | 20.36606 | -0.37617 | 0.733297 | 0.44859 |
| 12 | 93263600 | 93265599 | 18.13713 | 32.77707 | 26.13297 | 11.348 | 0.733739 | 0.382214 | 0.448612 |
| 12 | 22225600 | 22226999 | 24.46999 | 42.25484 | 36.1798 | 15.87202 | 0.608262 | 0.874376 | 0.448621 |
| 4 | 125961600 | 125964599 | 19.01643 | 31.9751 | 63.89204 | 27.91294 | -0.61408 | 1.124455 | 0.448637 |
| 14 | 71635200 | 71637599 | 19.92517 | 35.11328 | 32.58878 | 14.42402 | 0.495932 | 0.587349 | 0.448647 |
| 14 | 28681800 | 28684799 | 22.30698 | 37.46335 | 63.11923 | 27.56318 | -0.36912 | 1.142873 | 0.448656 |
| 13 | 93213600 | 93215599 | 23.41781 | 40.97166 | 35.24024 | 15.54456 | 0.604023 | 0.71162 | 0.448688 |
| 7 | 31863400 | 31868199 | 64.64134 | 103.2194 | 98.6129 | 41.40104 | 0.406428 | 2.117043 | 0.44869 |
| 18 | 4939800 | 4940999 | 7.944657 | 15.1849 | 30.83134 | 12.23782 | -0.50385 | -0.02671 | 0.448691 |
| 15 | 68239200 | 68240599 | 12.24655 | 21.79233 | 38.81986 | 17.10791 | -0.4329 | 0.466497 | 0.448697 |
| 7 | 109037400 | 109038599 | 9.131291 | 16.69224 | 32.40235 | 13.87004 | -0.52789 | 0.266904 | 0.448705 |
| 16 | 84499600 | 84500799 | 9.131291 | 16.69224 | 32.40235 | 13.87004 | -0.52789 | 0.266904 | 0.448705 |
| 10 | 119339200 | 119359199 | 295.9632 | 417.4533 | 580.6966 | 240.457 | -0.29524 | 4.451157 | 0.448707 |
| 11 | 102719000 | 102720399 | 17.4981 | 31.20281 | 29.47688 | 12.96747 | 0.478369 | 0.465107 | 0.448713 |
| 12 | 119309800 | 119310399 | 4.651197 | 9.058482 | 21.27078 | 7.775557 | -0.59681 | -0.36799 | 0.448745 |
| 3 | 57770600 | 57772399 | 17.62324 | 31.85474 | 27.36098 | 11.87829 | 0.627411 | 0.380951 | 0.448754 |
| 6 | 53977600 | 53978399 | 12.05206 | 23.11458 | 21.70409 | 8.530629 | 0.613053 | -0.04246 | 0.448794 |
| 8 | 48687600 | 48695199 | 116.1204 | 175.1955 | 269.5873 | 111.4115 | -0.34753 | 3.34789 | 0.448796 |
| X | 15064400 | 15066599 | 42.23469 | 69.57558 | 49.97187 | 21.55463 | 0.855508 | 1.404674 | 0.448825 |
| 1 | 44796400 | 44798199 | 15.96503 | 29.17174 | 23.22478 | 9.946956 | 0.749049 | 0.301358 | 0.448837 |
| X | 121470200 | 121470799 | 15.55251 | 28.44729 | 26.79202 | 11.4597 | 0.509399 | 0.3003 | 0.448844 |
| 2 | 89141000 | 89145599 | 45.23875 | 73.0964 | 74.20801 | 31.68513 | 0.338768 | 1.723854 | 0.448845 |
| 15 | 9813600 | 9815199 | 15.77276 | 28.5433 | 27.04042 | 11.72196 | 0.488188 | 0.359903 | 0.448861 |
| 9 | 5552800 | 5556599 | 81.61686 | 128.347 | 130.1753 | 54.00244 | 0.300364 | 2.534674 | 0.448868 |
| 15 | 8584600 | 8587199 | 36.05653 | 59.11133 | 59.72164 | 25.70738 | 0.361844 | 1.450859 | 0.448874 |
| 4 | 22689800 | 22691199 | 26.54178 | 44.79798 | 44.31042 | 19.38044 | 0.396874 | 1.093704 | 0.448875 |

| | | | | | | | | | |
|---|---|---|---|---|---|---|---|---|---|
| X | 159352400 | 159357199 | 69.59217 | 111.5933 | 82.576 | 34.88127 | 0.781151 | 1.994015 | 0.448889 |
| 3 | 77208600 | 77209599 | 12.99386 | 24.94376 | 21.08176 | 8.264198 | 0.76703 | -0.04347 | 0.448893 |
| X | 124926600 | 124928999 | 17.22842 | 31.72818 | 25.61902 | 10.85684 | 0.739557 | 0.273117 | 0.448895 |
| 12 | 23114800 | 23187599 | 2285.586 | 2758.467 | 2176.946 | 919.6098 | 0.295782 | 6.931138 | 0.448909 |
| 7 | 128089400 | 128090999 | 33.47145 | 55.61007 | 54.7545 | 23.7456 | 0.403346 | 1.285011 | 0.448912 |
| X | 47935200 | 47935799 | 9.147388 | 17.75489 | 17.80235 | 6.691958 | 0.580571 | -0.22696 | 0.448919 |
| 9 | 27702000 | 27706799 | 33.81559 | 54.62436 | 91.46905 | 39.04175 | -0.38199 | 1.720973 | 0.44894 |
| 14 | 16147800 | 16150599 | 37.92713 | 61.87563 | 62.9632 | 27.05218 | 0.347338 | 1.523983 | 0.448942 |
| 12 | 18031000 | 18039599 | 123.6207 | 189.0739 | 188.5704 | 78.23617 | 0.293773 | 3.068029 | 0.44895 |
| 1 | 37084800 | 37088599 | 30.74357 | 51.67229 | 44.29141 | 19.34746 | 0.602582 | 1.145585 | 0.448954 |
| 9 | 77094600 | 77096199 | 15.73789 | 28.77724 | 26.78558 | 11.46178 | 0.525883 | 0.303639 | 0.448961 |
| 15 | 3775400 | 3776399 | 9.053979 | 16.82641 | 31.0311 | 12.98855 | -0.4311 | 0.169363 | 0.448973 |
| 14 | 74954400 | 74956399 | 21.65763 | 36.45872 | 60.86597 | 26.60035 | -0.35576 | 1.1182 | 0.448986 |
| 18 | 24295000 | 24300399 | 69.77414 | 110.6953 | 112.9581 | 47.04192 | 0.301861 | 2.321406 | 0.448991 |
| 12 | 21737800 | 21785399 | 1211.598 | 1542.211 | 1287.054 | 540.2793 | 0.292993 | 6.079506 | 0.448995 |
| 4 | 97331200 | 97333199 | 19.49723 | 35.12615 | 27.79093 | 12.11475 | 0.741728 | 0.415116 | 0.449007 |
| 5 | 6239200 | 6239599 | 8.534785 | 16.57011 | 18.27699 | 6.857496 | 0.44562 | -0.24259 | 0.449016 |
| 17 | 70036000 | 70037199 | 8.880593 | 16.74407 | 33.77394 | 13.79675 | -0.53144 | 0.081573 | 0.449046 |
| 2 | 148972400 | 148974399 | 27.07602 | 46.88237 | 37.92765 | 16.64018 | 0.690479 | 0.863543 | 0.449072 |
| 12 | 113663400 | 113666399 | 37.56702 | 61.57731 | 61.52754 | 26.48286 | 0.3777 | 1.458058 | 0.449073 |
| 3 | 27651800 | 27653599 | 15.44163 | 28.06791 | 23.16207 | 9.985321 | 0.691286 | 0.324964 | 0.449125 |
| 10 | 83549200 | 83552399 | 24.04525 | 39.84569 | 67.31565 | 29.14079 | -0.37338 | 1.299695 | 0.449141 |
| 7 | 97051200 | 97053399 | 28.25027 | 47.60871 | 48.01077 | 20.98915 | 0.368987 | 1.116977 | 0.449146 |
| 5 | 141709200 | 141711599 | 27.10618 | 44.27966 | 76.26388 | 32.7824 | -0.40844 | 1.492915 | 0.449151 |
| 8 | 88466400 | 88467799 | 12.31107 | 22.8966 | 23.01191 | 9.620841 | 0.43983 | 0.172381 | 0.449152 |
| 7 | 11559000 | 11565599 | 151.5244 | 228.8368 | 224.3965 | 92.76449 | 0.303257 | 3.333336 | 0.449155 |
| 10 | 63505800 | 63507199 | 10.19021 | 18.70467 | 33.48369 | 14.25357 | -0.40626 | 0.254351 | 0.449178 |
| 19 | 24575000 | 24577599 | 22.27873 | 39.3077 | 31.263 | 13.83392 | 0.718149 | 0.586523 | 0.4492 |
| 7 | 72842600 | 72845599 | 37.74178 | 61.68829 | 62.52346 | 26.88153 | 0.354932 | 1.49659 | 0.44923 |
| 2 | 113840800 | 113842399 | 13.30682 | 23.52536 | 40.81195 | 18.05208 | -0.39998 | 0.528263 | 0.449235 |
| 17 | 20876400 | 20879399 | 75.0528 | 119.0538 | 100.2413 | 41.74038 | 0.578605 | 2.32902 | 0.449242 |
| 15 | 100992400 | 100993599 | 7.056754 | 13.43287 | 27.17886 | 10.88209 | -0.50886 | -0.0138 | 0.449243 |
| 3 | 103448400 | 103450199 | 21.68193 | 36.32772 | 61.90658 | 26.99338 | -0.3854 | 1.165062 | 0.449265 |
| 17 | 14312600 | 14315599 | 19.44056 | 32.76438 | 66.88952 | 29.24309 | -0.64499 | 1.105917 | 0.449274 |
| 5 | 130851600 | 130852999 | 9.537621 | 17.61933 | 31.99898 | 13.50334 | -0.41773 | 0.209458 | 0.449313 |
| 5 | 4508200 | 4510799 | 24.12995 | 41.36811 | 42.07573 | 18.45303 | 0.359019 | 0.945329 | 0.449316 |
| 14 | 123863600 | 123864599 | 9.496922 | 17.19849 | 33.38005 | 14.45721 | -0.53855 | 0.328 | 0.449321 |
| 3 | 93131400 | 93133799 | 21.84795 | 36.6461 | 61.36132 | 26.77417 | -0.36013 | 1.153979 | 0.449335 |
| 4 | 16990400 | 16997399 | 135.3674 | 205.5507 | 205.1302 | 84.94328 | 0.284543 | 3.217982 | 0.449355 |
| 19 | 12520000 | 12526799 | 90.24238 | 140.7263 | 141.1969 | 58.59534 | 0.306609 | 2.694451 | 0.449362 |
| 13 | 81086000 | 81087599 | 8.476314 | 15.80414 | 30.03654 | 12.50451 | -0.46744 | 0.143776 | 0.449375 |
| 3 | 4809200 | 4810399 | 24.53382 | 42.51712 | 35.59741 | 15.61824 | 0.641504 | 0.843028 | 0.449408 |
| 13 | 43257600 | 43258799 | 28.05815 | 47.28586 | 47.48507 | 20.75945 | 0.37512 | 1.117711 | 0.449409 |
| 2 | 170533800 | 170538599 | 45.44387 | 72.3113 | 125.251 | 52.32916 | -0.45591 | 2.217423 | 0.449411 |
| 12 | 26165000 | 26166799 | 18.16588 | 31.13367 | 52.73023 | 23.12517 | -0.37382 | 0.937185 | 0.449413 |
| 6 | 130073200 | 130076399 | 41.58653 | 68.50725 | 57.38855 | 24.75362 | 0.634148 | 1.405691 | 0.449423 |
| 6 | 131522200 | 131526199 | 37.29502 | 61.18408 | 61.48077 | 26.47237 | 0.370083 | 1.447814 | 0.449441 |

| | | | | | | | | | |
|---|---|---|---|---|---|---|---|---|---|
| 5 | 86459600 | 86461599 | 23.91249 | 40.86951 | 39.08885 | 17.13884 | 0.446786 | 0.96834 | 0.449445 |
| 11 | 115040000 | 115041599 | 20.20871 | 34.05032 | 58.58994 | 25.61266 | -0.39909 | 1.103256 | 0.449448 |
| 14 | 4807600 | 4810599 | 20.8166 | 34.66444 | 63.84651 | 27.73164 | -0.49699 | 1.231029 | 0.449463 |
| 4 | 37729200 | 37733399 | 38.65085 | 64.49414 | 49.8327 | 21.67267 | 0.752058 | 1.250091 | 0.449478 |
| 12 | 113838400 | 113843199 | 69.05053 | 109.8708 | 107.6443 | 44.97092 | 0.364699 | 2.2339 | 0.449481 |
| 8 | 79506400 | 79509799 | 41.53523 | 67.59865 | 67.00073 | 28.76159 | 0.382807 | 1.570396 | 0.449489 |
| 7 | 33005200 | 33026999 | 398.0737 | 559.1346 | 476.9595 | 197.6343 | 0.403958 | 4.516655 | 0.449499 |
| 5 | 148210800 | 148213399 | 19.1567 | 32.68408 | 54.88004 | 24.05924 | -0.36246 | 0.984795 | 0.449507 |
| 10 | 33853400 | 33858599 | 65.32736 | 103.906 | 106.5326 | 44.48239 | 0.29864 | 2.241772 | 0.449542 |
| 3 | 121531600 | 121533799 | 13.48642 | 23.71093 | 40.87176 | 18.0986 | -0.39388 | 0.590934 | 0.449542 |
| 1 | 166200800 | 166202199 | 13.0361 | 24.81118 | 22.59794 | 9.05076 | 0.632199 | 0.028966 | 0.449562 |
| X | 32206200 | 32207599 | 22.65817 | 39.45275 | 37.19725 | 16.35394 | 0.471886 | 0.780647 | 0.449635 |
| 18 | 6431400 | 6432799 | 12.15276 | 21.60241 | 38.75005 | 17.0925 | -0.44413 | 0.468687 | 0.449644 |
| 14 | 37115400 | 37116399 | 16.47626 | 28.32921 | 50.56021 | 22.17778 | -0.44795 | 0.901135 | 0.449649 |
| 9 | 70809000 | 70811399 | 23.12474 | 38.24084 | 67.62441 | 29.23253 | -0.43926 | 1.32229 | 0.449651 |
| 10 | 40045200 | 40048599 | 22.50054 | 37.64932 | 63.04204 | 27.46549 | -0.36015 | 1.183172 | 0.449665 |
| X | 8662400 | 8665399 | 73.59006 | 117.6375 | 90.42929 | 38.02421 | 0.721236 | 2.092994 | 0.44967 |
| 10 | 56898600 | 56901399 | 27.57913 | 46.57884 | 45.29896 | 19.81668 | 0.421142 | 1.091255 | 0.449685 |
| X | 146512800 | 146513599 | 16.72755 | 30.07692 | 28.69744 | 12.5367 | 0.471214 | 0.410088 | 0.449702 |
| 13 | 66477000 | 66478599 | 16.4031 | 31.44416 | 19.10145 | 7.518137 | 1.234321 | -0.00135 | 0.44972 |
| 3 | 38948800 | 38950799 | 21.01143 | 36.79556 | 34.1976 | 15.09647 | 0.492522 | 0.679978 | 0.449728 |
| 6 | 67605000 | 67606799 | 18.52362 | 32.97615 | 29.9006 | 13.17249 | 0.535865 | 0.483332 | 0.44973 |
| 7 | 53217600 | 53219799 | 36.19522 | 59.23997 | 60.63809 | 26.0837 | 0.342119 | 1.474666 | 0.449737 |
| 5 | 140588400 | 140590599 | 18.10794 | 30.8953 | 53.73186 | 23.55592 | -0.41276 | 0.981775 | 0.449753 |
| X | 38213600 | 38217799 | 68.35214 | 109.2505 | 83.57909 | 35.13748 | 0.727936 | 2.088813 | 0.449787 |
| 1 | 127317000 | 127320399 | 29.36269 | 49.38125 | 49.1028 | 21.45311 | 0.38917 | 1.141064 | 0.449827 |
| 6 | 3266600 | 3269199 | 35.72882 | 58.58909 | 59.53094 | 25.62808 | 0.353898 | 1.449666 | 0.44983 |
| 11 | 30691000 | 30694399 | 40.46636 | 65.67542 | 67.26777 | 28.83036 | 0.332211 | 1.615393 | 0.449843 |
| 4 | 128755800 | 128757599 | 8.465525 | 15.75326 | 30.00372 | 12.53219 | -0.4745 | 0.155177 | 0.449849 |
| 13 | 7064600 | 7066399 | 19.17754 | 33.91593 | 31.59768 | 13.97544 | 0.492449 | 0.54398 | 0.449851 |
| 2 | 15584200 | 15589799 | 65.85353 | 105.2391 | 96.00851 | 40.35326 | 0.474121 | 2.092383 | 0.449853 |
| 6 | 38034000 | 38044399 | 109.5442 | 166.8213 | 251.1418 | 104.0866 | -0.30473 | 3.155858 | 0.449854 |
| 16 | 47809400 | 47811199 | 16.69215 | 30.70062 | 25.537 | 10.84126 | 0.69486 | 0.274368 | 0.449856 |
| X | 31287000 | 31293999 | 99.86053 | 157.0156 | 127.1591 | 52.75028 | 0.624514 | 2.550015 | 0.449861 |
| 15 | 59045400 | 59047999 | 18.9681 | 32.3668 | 55.43783 | 24.30405 | -0.39103 | 0.982381 | 0.449894 |
| 9 | 19429800 | 19434399 | 56.8127 | 91.33491 | 83.62748 | 35.4613 | 0.478812 | 1.896154 | 0.449895 |
| 8 | 113872600 | 113874599 | 21.19448 | 37.31537 | 32.43246 | 14.35751 | 0.589588 | 0.603926 | 0.4499 |
| 13 | 61321600 | 61327799 | 112.811 | 175.154 | 152.2008 | 63.20302 | 0.508988 | 2.781752 | 0.449911 |
| 7 | 38528000 | 38534199 | 77.5246 | 122.8637 | 117.2148 | 48.764 | 0.397386 | 2.359925 | 0.449913 |
| 17 | 5717000 | 5719199 | 15.69582 | 27.37794 | 46.79712 | 20.59732 | -0.38249 | 0.733871 | 0.449922 |
| 8 | 54423800 | 54424399 | 10.40422 | 19.95973 | 21.23635 | 8.340135 | 0.435285 | -0.06309 | 0.449923 |
| 1 | 178164200 | 178165999 | 12.75859 | 22.36831 | 41.55102 | 18.36015 | -0.50105 | 0.634594 | 0.449936 |
| 6 | 86981400 | 86983199 | 17.1615 | 30.32826 | 29.89051 | 13.22231 | 0.411354 | 0.536308 | 0.44994 |
| 12 | 70714600 | 70717999 | 47.68839 | 76.74583 | 77.53873 | 32.92892 | 0.338912 | 1.847989 | 0.449949 |
| 1 | 139650600 | 139652999 | 27.17392 | 46.57295 | 43.02817 | 18.8703 | 0.497119 | 0.962141 | 0.449953 |
| 17 | 54431400 | 54436199 | 95.31093 | 148.598 | 148.3101 | 61.54925 | 0.313966 | 2.700798 | 0.449957 |
| 2 | 15088000 | 15089999 | 25.97127 | 45.03232 | 39.14518 | 17.17508 | 0.587784 | 0.849936 | 0.449959 |

| | | | | | | | | | |
|---|---|---|---|---|---|---|---|---|---|
| 5 | 144913400 | 144923599 | 99.88648 | 152.7667 | 228.1678 | 94.66386 | -0.28838 | 3.065241 | 0.449979 |
| 8 | 14050600 | 14055399 | 53.81727 | 85.14603 | 154.7562 | 64.32133 | -0.53319 | 2.390067 | 0.449979 |
| 9 | 7062400 | 7065199 | 3.231555 | 6.318695 | 22.10137 | 7.55027 | -1.07458 | -0.62842 | 0.449986 |
| 11 | 70172800 | 70175599 | 15.85233 | 27.52966 | 47.92577 | 21.0359 | -0.40946 | 0.798241 | 0.449992 |
| 18 | 29691400 | 29693199 | 15.77569 | 28.9951 | 25.56996 | 10.86541 | 0.60999 | 0.270317 | 0.449996 |
| 8 | 45613800 | 45619999 | 73.57304 | 115.1337 | 176.8122 | 73.34855 | -0.30287 | 2.62903 | 0.450003 |
| 18 | 90200000 | 90203199 | 34.02195 | 57.36115 | 46.30973 | 20.24825 | 0.688775 | 1.131191 | 0.450005 |
| 18 | 5924200 | 5927799 | 34.48476 | 57.6391 | 51.42377 | 22.38833 | 0.545203 | 1.221293 | 0.450009 |
| 19 | 61244600 | 61247399 | 22.18727 | 37.24487 | 61.92809 | 27.03504 | -0.35006 | 1.147783 | 0.45003 |
| 18 | 41519000 | 41522399 | 30.98626 | 52.9039 | 42.93236 | 18.82247 | 0.682708 | 1.010692 | 0.450039 |
| 6 | 59559800 | 59561799 | 32.01977 | 52.96942 | 51.90172 | 22.44154 | 0.409943 | 1.328691 | 0.450043 |
| 5 | 89456400 | 89458199 | 23.34686 | 38.94209 | 64.68043 | 28.12311 | -0.34845 | 1.221703 | 0.450061 |
| X | 54726000 | 54727199 | 27.63787 | 46.83281 | 44.80805 | 19.62161 | 0.445015 | 1.068159 | 0.450084 |
| X | 34592200 | 34593199 | 12.65228 | 24.57391 | 18.0299 | 6.743107 | 1.032428 | -0.20574 | 0.450105 |
| 3 | 18297000 | 18297999 | 5.692847 | 11.00081 | 24.26202 | 9.311098 | -0.57494 | -0.1784 | 0.450109 |
| 4 | 32492800 | 32500199 | 70.95649 | 112.6635 | 112.6292 | 46.94348 | 0.332596 | 2.298864 | 0.450111 |
| 14 | 80207600 | 80210599 | 6.743167 | 12.90322 | 31.37075 | 12.41104 | -0.75669 | -0.04694 | 0.450112 |
| X | 153083800 | 153084799 | 22.67817 | 39.46037 | 38.17726 | 16.77753 | 0.434603 | 0.789896 | 0.450119 |
| 8 | 124361800 | 124367399 | 68.34573 | 106.9703 | 165.9993 | 68.86184 | -0.31769 | 2.622747 | 0.450135 |
| 12 | 52848400 | 52850199 | 15.38458 | 26.86057 | 45.72119 | 20.1358 | -0.37628 | 0.717692 | 0.450159 |
| 7 | 7732600 | 7735199 | 34.80947 | 59.28425 | 40.60592 | 17.79896 | 0.925887 | 1.044455 | 0.450195 |
| 12 | 112408600 | 112417999 | 118.3013 | 178.1206 | 277.0793 | 114.4415 | -0.36577 | 3.388954 | 0.450228 |
| 18 | 16219400 | 16225599 | 79.62384 | 126.0329 | 119.8315 | 49.81708 | 0.400794 | 2.391461 | 0.450232 |
| X | 121221000 | 121222799 | 23.04067 | 40.47449 | 32.98382 | 14.59884 | 0.681372 | 0.644179 | 0.450233 |
| 6 | 66293600 | 66294599 | 8.416429 | 15.90418 | 29.79022 | 12.11383 | -0.41914 | 0.054153 | 0.450233 |
| 5 | 143613600 | 143617599 | 27.73826 | 45.36484 | 75.89976 | 32.6397 | -0.36554 | 1.477585 | 0.450252 |
| 11 | 73627000 | 73628799 | 19.84041 | 34.78199 | 33.97631 | 15.01233 | 0.421313 | 0.66004 | 0.450264 |
| 4 | 71881000 | 71883999 | 33.69551 | 56.03431 | 48.72506 | 21.14434 | 0.582126 | 1.269623 | 0.450265 |
| 13 | 82258200 | 82262799 | 56.12146 | 89.88264 | 91.61012 | 38.62553 | 0.317972 | 2.013899 | 0.450269 |
| 4 | 140323200 | 140325799 | 51.14968 | 81.25878 | 130.4279 | 54.39158 | -0.34865 | 2.272376 | 0.450288 |
| 6 | 75041800 | 75043199 | 12.02359 | 21.42161 | 37.25382 | 16.40095 | -0.39754 | 0.452496 | 0.450321 |
| 2 | 173807600 | 173811399 | 29.31539 | 47.95544 | 79.2678 | 34.091 | -0.34798 | 1.478186 | 0.450326 |
| 19 | 56071200 | 56074999 | 43.44212 | 70.68112 | 59.93311 | 25.72487 | 0.607016 | 1.571748 | 0.450338 |
| 8 | 100378000 | 100381199 | 25.18845 | 44.00583 | 35.13545 | 15.48005 | 0.710884 | 0.742338 | 0.450354 |
| 10 | 46928200 | 46929199 | 14.56568 | 26.31976 | 25.82155 | 11.21439 | 0.436543 | 0.358253 | 0.450375 |
| X | 104309400 | 104310999 | 21.96768 | 39.05614 | 29.18766 | 12.87261 | 0.812482 | 0.514554 | 0.450383 |
| 7 | 128584000 | 128588199 | 44.65223 | 72.14093 | 73.92648 | 31.56054 | 0.325144 | 1.726985 | 0.450384 |
| 14 | 99525600 | 99526399 | 10.68823 | 20.2925 | 21.11724 | 8.501909 | 0.434835 | 0.018966 | 0.450402 |
| 9 | 10696800 | 10699199 | 19.89569 | 34.94258 | 33.73588 | 14.92909 | 0.437907 | 0.629433 | 0.450411 |
| Y | 86739600 | 86741799 | 13.83688 | 25.60858 | 14.95976 | 6.297142 | 1.208797 | 0.188394 | 0.450414 |
| 19 | 45188200 | 45199799 | 123.3491 | 184.5552 | 323.5796 | 133.4433 | -0.54642 | 3.513137 | 0.450433 |
| 5 | 5848200 | 5851599 | 40.41802 | 66.20588 | 62.29123 | 26.80393 | 0.463868 | 1.474053 | 0.450457 |
| 3 | 82495800 | 82498799 | 33.24943 | 55.28555 | 54.21495 | 23.52465 | 0.409228 | 1.274108 | 0.450466 |
| 1 | 146680400 | 146681999 | 12.99737 | 24.3791 | 20.80444 | 8.57379 | 0.690377 | 0.118572 | 0.450471 |
| 19 | 33804600 | 33805999 | 13.49415 | 25.36606 | 23.98729 | 9.849117 | 0.547758 | 0.118034 | 0.450472 |
| 17 | 57868400 | 57874199 | 64.52823 | 103.2048 | 94.2486 | 39.6591 | 0.473921 | 2.065987 | 0.450497 |
| 9 | 30543600 | 30545999 | 13.9328 | 24.36533 | 44.64152 | 19.68371 | -0.48183 | 0.682548 | 0.4505 |

| | | | | | | | | | |
|---|---|---|---|---|---|---|---|---|---|
| 9 | 26770800 | 26773199 | 16.37335 | 28.34751 | 48.46351 | 21.26256 | -0.38422 | 0.833129 | 0.450523 |
| 14 | 42205800 | 42207399 | 27.28965 | 46.60157 | 37.44354 | 16.41631 | 0.697076 | 0.989105 | 0.450528 |
| 5 | 145449600 | 145457399 | 115.3913 | 179.0139 | 153.3928 | 63.70578 | 0.528245 | 2.795766 | 0.450551 |
| 3 | 12726400 | 12729799 | 41.98541 | 68.09787 | 69.63421 | 29.83073 | 0.333747 | 1.63268 | 0.450553 |
| 2 | 23799800 | 23802399 | 30.53701 | 51.79774 | 45.73463 | 20.0335 | 0.560325 | 1.067663 | 0.450579 |
| 1 | 105398800 | 105401799 | 32.18492 | 54.33639 | 46.86882 | 20.50101 | 0.593691 | 1.114056 | 0.450591 |
| 6 | 118757200 | 118758599 | 9.550708 | 17.60297 | 31.96341 | 13.5306 | -0.42069 | 0.22274 | 0.450597 |
| 5 | 117453000 | 117457999 | 33.70145 | 54.31442 | 94.66394 | 40.28527 | -0.44345 | 1.790068 | 0.450603 |
| 4 | 113654200 | 113657199 | 55.54778 | 89.25191 | 85.42943 | 36.19632 | 0.413927 | 1.912862 | 0.450608 |
| 4 | 112520000 | 112531799 | 260.1103 | 377.632 | 354.4951 | 146.1431 | 0.313481 | 4.024594 | 0.450609 |
| 15 | 45427200 | 45427799 | 11.77442 | 22.2067 | 22.61961 | 9.235642 | 0.448089 | 0.079922 | 0.450614 |
| 2 | 82761800 | 82767999 | 68.38759 | 109.2053 | 97.32634 | 40.86294 | 0.506584 | 2.118961 | 0.450616 |
| 18 | 35138400 | 35140599 | 20.29198 | 35.45953 | 35.13771 | 15.48341 | 0.400846 | 0.715205 | 0.450621 |
| 18 | 83196000 | 83200199 | 48.19349 | 76.54435 | 130.8704 | 54.56317 | -0.43994 | 2.275795 | 0.450624 |
| 4 | 46553400 | 46554799 | 9.107449 | 16.76498 | 31.43815 | 13.33096 | -0.46843 | 0.225641 | 0.450629 |
| 8 | 36328400 | 36331199 | 28.71715 | 46.82581 | 78.71044 | 33.81159 | -0.37536 | 1.523334 | 0.450644 |
| 16 | 32294000 | 32296399 | 19.88499 | 34.70378 | 34.07355 | 15.00226 | 0.413705 | 0.72651 | 0.450649 |
| 6 | 115970400 | 115972199 | 15.97497 | 27.51128 | 49.50952 | 21.71797 | -0.45937 | 0.882494 | 0.450651 |
| 11 | 39936200 | 39937399 | 15.25043 | 26.48079 | 46.87069 | 20.57104 | -0.43317 | 0.79697 | 0.450653 |
| 5 | 63961600 | 63966599 | 70.01823 | 109.4351 | 168.9664 | 70.10273 | -0.31197 | 2.648359 | 0.450658 |
| 18 | 67201000 | 67217799 | 287.8731 | 406.3519 | 565.2947 | 234.0499 | -0.29424 | 4.439452 | 0.450678 |
| 7 | 108225000 | 108227199 | 24.2184 | 42.02076 | 39.20087 | 17.19987 | 0.486431 | 0.835265 | 0.450679 |
| 17 | 50815200 | 50817199 | 13.71391 | 23.83557 | 45.82703 | 20.12509 | -0.55159 | 0.775551 | 0.450705 |
| 10 | 117771400 | 117773799 | 17.05473 | 29.31701 | 50.57998 | 22.1863 | -0.39951 | 0.903512 | 0.450709 |
| 19 | 33571400 | 33573199 | 47.19574 | 75.24986 | 120.8375 | 50.62565 | -0.34378 | 2.152862 | 0.450713 |
| 13 | 35943800 | 35948799 | 52.89012 | 84.20787 | 131.572 | 55.01335 | -0.30667 | 2.205984 | 0.450724 |
| 12 | 21559400 | 21580999 | 254.8819 | 363.0021 | 521.6981 | 215.6791 | -0.32824 | 4.307108 | 0.450726 |
| 14 | 15050600 | 15051599 | 7.618928 | 14.4544 | 28.78977 | 11.60919 | -0.49657 | 0.014643 | 0.450726 |
| 6 | 41348000 | 41349399 | 19.85128 | 35.03568 | 30.8155 | 13.63506 | 0.573605 | 0.56531 | 0.450734 |
| 17 | 64972600 | 64973999 | 11.63409 | 20.91881 | 37.83936 | 16.53031 | -0.44649 | 0.391403 | 0.450742 |
| 19 | 46998800 | 47005199 | 46.06643 | 73.26018 | 140.5378 | 58.66962 | -0.60398 | 2.237911 | 0.450744 |
| X | 14697000 | 14699199 | 29.96469 | 51.89222 | 37.35091 | 16.38722 | 0.858604 | 0.877624 | 0.450752 |
| 1 | 5531400 | 5532199 | 11.2471 | 21.79068 | 19.77007 | 7.495136 | 0.710727 | -0.17241 | 0.450803 |
| 5 | 68102600 | 68103399 | 11.2471 | 21.79068 | 19.77007 | 7.495136 | 0.710727 | -0.17241 | 0.450803 |
| 7 | 41774400 | 41778199 | 88.72463 | 138.7641 | 138.7541 | 57.564 | 0.31479 | 2.642876 | 0.450806 |
| 3 | 115790800 | 115793399 | 38.67079 | 63.23053 | 63.66173 | 27.37468 | 0.364917 | 1.495719 | 0.45083 |
| 14 | 25131200 | 25136599 | 39.73244 | 63.88777 | 101.5236 | 43.06162 | -0.31464 | 1.874566 | 0.450834 |
| 6 | 124413600 | 124416199 | 15.4211 | 26.7018 | 47.93221 | 21.02953 | -0.45422 | 0.826757 | 0.450858 |
| 7 | 22539400 | 22559599 | 531.9533 | 730.0392 | 592.1901 | 246.2772 | 0.442778 | 4.871096 | 0.450864 |
| 14 | 52661600 | 52662799 | 11.60085 | 20.47273 | 38.85984 | 17.19476 | -0.52985 | 0.534113 | 0.450881 |
| 9 | 81134800 | 81138999 | 83.30578 | 130.9161 | 132.7585 | 55.06931 | 0.299877 | 2.549457 | 0.450893 |
| 5 | 81670600 | 81673399 | 17.63462 | 30.47306 | 52.24834 | 22.92176 | -0.38915 | 0.859495 | 0.450896 |
| 9 | 77450600 | 77452799 | 15.5313 | 26.59451 | 51.59703 | 22.62656 | -0.56899 | 0.934105 | 0.450899 |
| 5 | 133000000 | 133001799 | 19.68046 | 33.57371 | 56.51607 | 24.77623 | -0.36611 | 0.990622 | 0.450908 |
| 16 | 55725000 | 55727399 | 18.20777 | 30.58147 | 60.60971 | 26.46868 | -0.6024 | 1.125935 | 0.450909 |
| 1 | 18229200 | 18229599 | 5.074572 | 9.890586 | 21.4108 | 7.748786 | -0.46837 | -0.39886 | 0.450917 |
| 16 | 93076000 | 93076599 | 5.074572 | 9.890586 | 21.4108 | 7.748786 | -0.46837 | -0.39886 | 0.450917 |

| | | | | | | | | | |
|---|---|---|---|---|---|---|---|---|---|
| X | 153129200 | 153132999 | 64.71987 | 103.4392 | 95.80196 | 40.27294 | 0.452579 | 2.089651 | 0.450919 |
| 19 | 47837800 | 47840199 | 14.12174 | 24.81184 | 42.41589 | 18.77663 | -0.3822 | 0.606175 | 0.450924 |
| 14 | 8236600 | 8237599 | 8.399085 | 15.93342 | 31.1085 | 12.54598 | -0.46826 | 0.029256 | 0.450932 |
| 19 | 61146200 | 61146599 | 5.955892 | 11.56737 | 23.99283 | 8.97614 | -0.45203 | -0.26186 | 0.450933 |
| 4 | 147291200 | 147296199 | 72.85243 | 115.1223 | 111.6199 | 46.36409 | 0.3707 | 2.424157 | 0.450957 |
| 1 | 174541000 | 174541599 | 5.113206 | 9.90841 | 23.23617 | 8.801547 | -0.64378 | -0.23164 | 0.450972 |
| 19 | 21543400 | 21543999 | 5.113206 | 9.90841 | 23.23617 | 8.801547 | -0.64378 | -0.23164 | 0.450972 |
| 4 | 66001000 | 66001599 | 11.04084 | 21.50405 | 16.4247 | 5.999071 | 1.004846 | -0.31056 | 0.45098 |
| X | 169275800 | 169276199 | 4.974599 | 9.660717 | 22.78242 | 8.529111 | -0.63597 | -0.2765 | 0.450986 |
| 12 | 113592000 | 113600399 | 94.81443 | 148.1764 | 144.2995 | 59.86923 | 0.352044 | 2.659448 | 0.45099 |
| 1 | 180458400 | 180459999 | 10.74101 | 19.32507 | 34.79222 | 15.19113 | -0.43873 | 0.37748 | 0.450992 |
| 4 | 17018000 | 17019199 | 19.94814 | 34.77096 | 33.72025 | 14.83517 | 0.431402 | 0.744753 | 0.450996 |
| 2 | 162322800 | 162323599 | 11.56002 | 21.95125 | 22.65759 | 9.118915 | 0.447026 | 0.030873 | 0.450996 |
| 6 | 35568600 | 35570199 | 15.51299 | 29.0799 | 23.15522 | 9.552434 | 0.788362 | 0.149823 | 0.450996 |
| 7 | 55245000 | 55249399 | 43.99111 | 70.45731 | 111.0241 | 46.81337 | -0.30939 | 2.005726 | 0.451007 |
| 12 | 22820800 | 22836799 | 561.9734 | 760.462 | 673.4096 | 280.6674 | 0.295813 | 5.090741 | 0.451009 |
| X | 125533800 | 125542999 | 125.9473 | 194.4273 | 160.5687 | 66.69703 | 0.57596 | 2.890759 | 0.45102 |
| 2 | 75810200 | 75812199 | 14.49624 | 25.41181 | 44.0675 | 19.47021 | -0.40266 | 0.649204 | 0.451038 |
| 8 | 50810200 | 50811599 | 15.63879 | 28.36298 | 27.08651 | 11.70978 | 0.479232 | 0.346918 | 0.451065 |
| 2 | 23412800 | 23413799 | 11.49192 | 22.0978 | 21.20758 | 8.273346 | 0.591597 | -0.0739 | 0.451093 |
| 7 | 103654800 | 103655999 | 9.514737 | 17.75697 | 34.51653 | 14.34673 | -0.49876 | 0.153757 | 0.4511 |
| 16 | 32305200 | 32308399 | 15.35604 | 25.99047 | 61.5909 | 26.96127 | -0.85855 | 1.050913 | 0.451107 |
| 7 | 19604200 | 19608399 | 34.17984 | 55.13919 | 92.03394 | 39.21537 | -0.37955 | 1.761933 | 0.451114 |
| 16 | 21471200 | 21479999 | 86.15682 | 132.8089 | 202.1663 | 83.97651 | -0.30702 | 2.90392 | 0.451119 |
| 9 | 81392200 | 81393199 | 8.000583 | 14.99911 | 29.06155 | 11.98686 | -0.4841 | 0.101912 | 0.451125 |
| 9 | 79666600 | 79667399 | 5.715171 | 11.00895 | 25.83751 | 10.02902 | -0.68001 | -0.13735 | 0.451138 |
| 17 | 92982600 | 92984199 | 16.25448 | 29.19851 | 28.29485 | 12.37074 | 0.448249 | 0.412294 | 0.451138 |
| 7 | 18258600 | 18260599 | 31.87019 | 52.80887 | 53.06212 | 22.96939 | 0.373848 | 1.3077 | 0.451141 |
| 3 | 61549600 | 61551399 | 18.64717 | 31.49674 | 62.339 | 27.27194 | -0.60011 | 1.081126 | 0.45116 |
| 9 | 19135600 | 19143399 | 181.5016 | 271.1079 | 231.656 | 95.50311 | 0.487655 | 3.549025 | 0.451201 |
| 18 | 45952000 | 45954199 | 30.82201 | 51.38767 | 52.45064 | 22.7992 | 0.351743 | 1.237093 | 0.451202 |
| 19 | 7802200 | 7803999 | 30.82201 | 51.38767 | 52.45064 | 22.7992 | 0.351743 | 1.237093 | 0.451202 |
| X | 89616000 | 89621199 | 89.3999 | 140.7335 | 130.1723 | 54.00854 | 0.434316 | 2.520936 | 0.45121 |
| 3 | 157237800 | 157240599 | 22.43996 | 37.33145 | 77.10696 | 33.4698 | -0.66208 | 1.257115 | 0.451224 |
| 19 | 10320800 | 10325599 | 43.17799 | 69.04256 | 113.6394 | 47.80365 | -0.37476 | 2.054957 | 0.451259 |
| 8 | 26370000 | 26372599 | 26.04871 | 44.86141 | 41.3554 | 18.14108 | 0.501447 | 0.911368 | 0.451268 |
| 14 | 6821000 | 6824199 | 58.92364 | 94.25804 | 94.76135 | 39.88508 | 0.335796 | 2.054411 | 0.45127 |
| 13 | 99968600 | 99972599 | 60.45817 | 96.56928 | 97.84524 | 41.09657 | 0.322243 | 2.102542 | 0.451273 |
| 14 | 36017800 | 36018999 | 13.80831 | 25.95272 | 23.32064 | 9.577823 | 0.620215 | 0.118574 | 0.451297 |
| 3 | 99342400 | 99345199 | 33.79389 | 54.72254 | 89.12823 | 38.13602 | -0.33842 | 1.660469 | 0.451303 |
| 9 | 106438000 | 106439999 | 17.91151 | 30.55014 | 53.21001 | 23.32652 | -0.41501 | 0.981995 | 0.451311 |
| 10 | 81386000 | 81388399 | 16.95887 | 28.90265 | 53.723 | 23.54992 | -0.50865 | 0.984976 | 0.451317 |
| X | 149092800 | 149094599 | 25.13023 | 43.95819 | 34.593 | 15.25628 | 0.731708 | 0.723311 | 0.451351 |
| 8 | 34185000 | 34187599 | 19.6801 | 32.94387 | 66.08337 | 28.79816 | -0.61974 | 1.173927 | 0.451358 |
| 5 | 19069600 | 19071599 | 26.06975 | 44.08817 | 43.78054 | 19.16014 | 0.391321 | 1.074932 | 0.451361 |
| 2 | 181586400 | 181590399 | 52.68222 | 85.38471 | 67.83918 | 29.04602 | 0.695982 | 1.667776 | 0.451374 |
| 8 | 75344200 | 75347199 | 38.90581 | 64.17348 | 57.83792 | 24.96099 | 0.529622 | 1.383446 | 0.451386 |

| | | | | | | | | | |
|---|---|---|---|---|---|---|---|---|---|
| 6 | 5212400 | 5214999 | 27.17711 | 44.8386 | 75.49316 | 32.58384 | -0.3692 | 1.36524 | 0.451396 |
| 13 | 35987800 | 35989399 | 19.86277 | 33.93056 | 60.42917 | 26.49459 | -0.44692 | 0.983162 | 0.45142 |
| 3 | 110157000 | 110158999 | 18.85648 | 31.61973 | 60.03965 | 26.19293 | -0.54076 | 1.146284 | 0.45143 |
| 15 | 8581600 | 8589599 | 91.78799 | 143.1565 | 144.9295 | 60.14316 | 0.293828 | 2.691943 | 0.451439 |
| X | 59528800 | 59530199 | 13.55885 | 25.19567 | 24.97616 | 10.45577 | 0.45821 | 0.193503 | 0.451442 |
| X | 104836000 | 104838199 | 17.51362 | 31.45013 | 29.081 | 12.71794 | 0.515084 | 0.423483 | 0.451444 |
| 16 | 8739400 | 8742799 | 30.64872 | 49.97255 | 82.00623 | 35.22663 | -0.34092 | 1.529903 | 0.451482 |
| 14 | 34821000 | 34823999 | 21.33392 | 35.84645 | 60.83191 | 26.57033 | -0.37942 | 1.132821 | 0.451484 |
| 5 | 42345600 | 42346799 | 7.637897 | 14.55684 | 27.78402 | 11.09573 | -0.42271 | -0.01676 | 0.451488 |
| X | 41214800 | 41219799 | 79.89512 | 126.7003 | 114.4568 | 47.64661 | 0.476869 | 2.343559 | 0.45152 |
| 4 | 23485000 | 23488399 | 39.75157 | 65.026 | 63.31448 | 27.22968 | 0.413489 | 1.490718 | 0.451532 |
| 2 | 145451800 | 145453799 | 14.11935 | 24.67757 | 43.62447 | 19.22605 | -0.43032 | 0.69279 | 0.451541 |
| 12 | 109875000 | 109885799 | 134.1399 | 200.6991 | 301.7495 | 124.4404 | -0.32482 | 3.514065 | 0.451546 |
| 15 | 43346800 | 43351399 | 51.038 | 82.6434 | 70.43963 | 30.13889 | 0.593656 | 1.684381 | 0.451549 |
| 7 | 45506000 | 45508799 | 15.08465 | 25.81544 | 51.57843 | 22.6174 | -0.6112 | 0.938391 | 0.451551 |
| 7 | 7296800 | 7303999 | 90.0898 | 137.7043 | 224.1632 | 92.99001 | -0.4131 | 3.073671 | 0.451559 |
| 4 | 112952000 | 112962399 | 58.25235 | 93.57288 | 77.49563 | 32.82226 | 0.621852 | 1.921093 | 0.451594 |
| 6 | 5676000 | 5677399 | 24.65523 | 42.12288 | 42.06284 | 18.44225 | 0.384627 | 0.976695 | 0.451601 |
| 14 | 48221800 | 48223599 | 24.65523 | 42.12288 | 42.06284 | 18.44225 | 0.384627 | 0.976695 | 0.451601 |
| 17 | 20267800 | 20286599 | 447.771 | 618.6946 | 562.9861 | 233.88 | 0.286922 | 4.764356 | 0.451603 |
| 3 | 150879400 | 150880199 | 19.87603 | 34.78115 | 35.00394 | 15.44265 | 0.378429 | 0.689164 | 0.451644 |
| 4 | 5305000 | 5306199 | 19.87603 | 34.78115 | 35.00394 | 15.44265 | 0.378429 | 0.689164 | 0.451644 |
| 8 | 75730400 | 75732999 | 14.25605 | 25.05621 | 43.96592 | 19.4683 | -0.41979 | 0.602925 | 0.451649 |
| 15 | 79488800 | 79489799 | 12.42361 | 21.78829 | 41.024 | 18.13238 | -0.52051 | 0.625052 | 0.451654 |
| 8 | 30558800 | 30561199 | 23.45897 | 41.12695 | 33.4314 | 14.77196 | 0.68489 | 0.676132 | 0.451655 |
| 13 | 61837400 | 61840599 | 52.6983 | 84.92329 | 77.94872 | 33.16807 | 0.479457 | 1.812518 | 0.451655 |
| 3 | 14598800 | 14599599 | 7.381217 | 13.85299 | 29.6129 | 12.19183 | -0.62319 | 0.087409 | 0.451676 |
| 10 | 104419800 | 104422599 | 30.7939 | 51.89844 | 48.5468 | 21.22388 | 0.477092 | 1.126505 | 0.451686 |
| 12 | 23487200 | 23488599 | 36.25049 | 59.71332 | 50.94073 | 21.97189 | 0.607921 | 1.388776 | 0.451688 |
| 13 | 62966800 | 62970999 | 56.14118 | 90.00056 | 90.20845 | 38.09009 | 0.343693 | 1.984163 | 0.451707 |
| 4 | 143553000 | 143555599 | 48.05354 | 77.5279 | 76.21662 | 32.47973 | 0.38262 | 1.775323 | 0.451708 |
| 1 | 4047200 | 4049999 | 35.99187 | 59.35636 | 57.70509 | 24.90177 | 0.420533 | 1.376554 | 0.451716 |
| 7 | 82877600 | 82878999 | 15.21499 | 28.19033 | 25.23498 | 10.60638 | 0.599945 | 0.222002 | 0.451721 |
| 8 | 61874800 | 61876599 | 11.31248 | 20.51389 | 37.51298 | 16.21998 | -0.45234 | 0.339624 | 0.451729 |
| 8 | 93340200 | 93346199 | 51.91201 | 82.4173 | 133.3569 | 55.57756 | -0.36118 | 2.290769 | 0.45173 |
| 16 | 19054000 | 19056999 | 30.64548 | 51.30803 | 43.32683 | 18.88316 | 0.624121 | 1.182955 | 0.451732 |
| 5 | 18846800 | 18850599 | 29.62763 | 50.51999 | 43.49398 | 19.06675 | 0.597404 | 1.01536 | 0.451742 |
| 14 | 52957800 | 52958399 | 13.15175 | 23.09722 | 41.41568 | 18.32691 | -0.4507 | 0.605224 | 0.451753 |
| X | 110118000 | 110118399 | 10.10391 | 19.50576 | 18.77289 | 7.237294 | 0.604698 | -0.12946 | 0.451766 |
| 19 | 15818200 | 15819199 | 19.33195 | 32.46477 | 58.96455 | 25.7478 | -0.47682 | 1.13265 | 0.451779 |
| 12 | 22784600 | 22786999 | 74.79805 | 118.1508 | 112.5494 | 46.74097 | 0.395729 | 2.43627 | 0.45181 |
| 6 | 75205800 | 75206999 | 14.01003 | 26.25832 | 23.83334 | 9.834847 | 0.600501 | 0.139214 | 0.451813 |
| 5 | 109963200 | 109964599 | 16.89743 | 30.98874 | 26.04431 | 11.10007 | 0.676091 | 0.293022 | 0.451829 |
| 7 | 41225000 | 41229399 | 37.77668 | 62.10139 | 60.47958 | 26.06386 | 0.416268 | 1.423318 | 0.451831 |
| 6 | 12301400 | 12306799 | 62.99568 | 100.9188 | 90.64591 | 38.23431 | 0.500447 | 2.014883 | 0.451852 |
| 8 | 50306600 | 50308599 | 14.59223 | 25.63187 | 46.65408 | 20.64828 | -0.47246 | 0.618546 | 0.451859 |
| 18 | 79964400 | 79966599 | 15.01745 | 26.25246 | 47.29696 | 20.84806 | -0.45786 | 0.697517 | 0.451868 |

| | | | | | | | | | |
|---|---|---|---|---|---|---|---|---|---|
| 9 | 92334800 | 92335799 | 11.48419 | 20.22917 | 42.99499 | 19.03156 | -0.69321 | 0.562259 | 0.451888 |
| 19 | 12282200 | 12284199 | 26.78034 | 45.83474 | 43.57878 | 19.10963 | 0.45556 | 0.972787 | 0.451898 |
| 6 | 98252400 | 98254199 | 13.87534 | 24.14273 | 45.29527 | 19.90534 | -0.51621 | 0.760613 | 0.451905 |
| 17 | 92934800 | 92936399 | 16.79596 | 30.13968 | 29.12187 | 12.74355 | 0.451358 | 0.422475 | 0.451927 |
| 3 | 82901400 | 82903199 | 11.45796 | 20.30101 | 37.33634 | 16.50753 | -0.482 | 0.488793 | 0.451938 |
| 2 | 47084400 | 47089199 | 80.80336 | 127.5292 | 126.4064 | 52.4733 | 0.337515 | 2.461127 | 0.451939 |
| 8 | 47913000 | 47914799 | 31.28849 | 51.90564 | 53.01205 | 22.96624 | 0.350553 | 1.293898 | 0.451944 |
| 1 | 170517400 | 170520799 | 27.18068 | 44.64309 | 74.02741 | 31.89041 | -0.34986 | 1.419023 | 0.451954 |
| 15 | 33792600 | 33802399 | 137.9999 | 209.9337 | 205.0198 | 84.94431 | 0.3179 | 3.181816 | 0.451971 |
| 9 | 103330000 | 103331599 | 9.378266 | 17.41331 | 31.55388 | 13.22365 | -0.40719 | 0.178932 | 0.451977 |
| 18 | 56431000 | 56434799 | 39.38291 | 64.72843 | 60.07115 | 25.88557 | 0.485494 | 1.428886 | 0.451977 |
| 12 | 47732600 | 47734599 | 27.75058 | 46.82638 | 47.46501 | 20.7586 | 0.361663 | 1.103783 | 0.451997 |
| 19 | 9308800 | 9310599 | 23.30199 | 40.72401 | 35.77883 | 15.76741 | 0.573503 | 0.728088 | 0.452007 |
| 7 | 43811800 | 43820399 | 114.9712 | 177.1212 | 172.3418 | 71.58655 | 0.337352 | 2.922716 | 0.452012 |
| 10 | 59750400 | 59752999 | 12.8795 | 22.52691 | 43.11573 | 19.01349 | -0.54429 | 0.673354 | 0.452019 |
| 4 | 112385400 | 112387399 | 34.32927 | 57.09858 | 54.22449 | 23.53377 | 0.455451 | 1.274973 | 0.452026 |
| 8 | 3123400 | 3124399 | 13.14592 | 24.4651 | 23.07251 | 9.635622 | 0.532683 | 0.177783 | 0.452047 |
| 5 | 80107600 | 80109199 | 15.36996 | 26.87901 | 46.75101 | 20.61411 | -0.40724 | 0.693642 | 0.452049 |
| 3 | 94269200 | 94272599 | 39.20825 | 62.98078 | 104.2825 | 44.16661 | -0.37568 | 1.905209 | 0.452075 |
| 12 | 32327600 | 32329199 | 10.58824 | 18.94368 | 36.79435 | 16.14145 | -0.55258 | 0.413551 | 0.452082 |
| 10 | 128983600 | 128987199 | 34.30691 | 55.57331 | 90.3852 | 38.68413 | -0.33589 | 1.655682 | 0.452086 |
| 10 | 117820800 | 117822399 | 11.5339 | 20.56993 | 36.75467 | 16.16742 | -0.43555 | 0.441138 | 0.452098 |
| 13 | 109932000 | 109932799 | 6.732605 | 12.96726 | 26.97725 | 10.47499 | -0.5075 | -0.11735 | 0.452105 |
| 7 | 11348200 | 11355199 | 40.64197 | 66.36872 | 57.63 | 24.76948 | 0.576501 | 1.512593 | 0.452108 |
| 18 | 79525600 | 79526599 | 11.64117 | 22.52237 | 19.64978 | 7.498801 | 0.757417 | -0.14769 | 0.452126 |
| 13 | 8610600 | 8611799 | 14.39483 | 26.1692 | 26.27988 | 11.32724 | 0.410036 | 0.323219 | 0.452154 |
| 2 | 64093400 | 64094599 | 14.26355 | 26.78154 | 23.88342 | 9.825584 | 0.629406 | 0.13112 | 0.452155 |
| 7 | 96193600 | 96195199 | 17.2204 | 31.26697 | 27.62531 | 11.92573 | 0.592442 | 0.35566 | 0.452167 |
| 7 | 12719200 | 12723799 | 29.36317 | 47.89706 | 78.82527 | 33.86552 | -0.34454 | 1.519791 | 0.452169 |
| 5 | 138403200 | 138404599 | 15.71648 | 30.08578 | 20.30059 | 8.022306 | 1.078861 | 0.007441 | 0.452171 |
| 16 | 21519800 | 21521399 | 13.7701 | 24.26178 | 42.38365 | 18.76002 | -0.41202 | 0.570143 | 0.452178 |
| 8 | 29021600 | 29022199 | 11.76261 | 22.56639 | 21.74073 | 8.537485 | 0.577593 | -0.04763 | 0.452195 |
| 8 | 3442600 | 3445399 | 17.36765 | 29.89693 | 50.58771 | 22.19064 | -0.37118 | 0.892236 | 0.45221 |
| 12 | 113566000 | 113567399 | 17.04136 | 30.56658 | 29.16646 | 12.7678 | 0.469088 | 0.427877 | 0.452217 |
| 18 | 56666600 | 56667199 | 3.759375 | 7.336383 | 19.73365 | 7.01812 | -0.75219 | -0.48185 | 0.452228 |
| 17 | 69530400 | 69533599 | 32.32451 | 52.47087 | 85.10632 | 36.4806 | -0.32911 | 1.607206 | 0.452238 |
| 17 | 53383000 | 53384799 | 18.68262 | 33.42732 | 29.47823 | 12.93133 | 0.580094 | 0.454383 | 0.452251 |
| 17 | 82209600 | 82211399 | 16.76604 | 30.20633 | 28.5804 | 12.45855 | 0.485084 | 0.396969 | 0.452255 |
| 18 | 18434800 | 18443999 | 167.6654 | 251.676 | 245.6171 | 101.3638 | 0.302452 | 3.456034 | 0.452258 |
| 16 | 81437000 | 81439599 | 26.12181 | 45.1523 | 40.25218 | 17.65911 | 0.55066 | 0.879044 | 0.452278 |
| 16 | 38628200 | 38630799 | 24.41977 | 40.32717 | 68.81587 | 29.71948 | -0.38818 | 1.342788 | 0.452288 |
| 7 | 55506400 | 55507399 | 8.589247 | 16.18748 | 30.26528 | 12.37286 | -0.42312 | 0.075475 | 0.452293 |
| 2 | 14608800 | 14610599 | 13.7486 | 23.77427 | 48.44917 | 21.25548 | -0.63668 | 0.832027 | 0.452304 |
| X | 20912000 | 20914399 | 32.90194 | 55.9347 | 43.53877 | 19.08175 | 0.741869 | 1.058659 | 0.452319 |
| 18 | 57721000 | 57721799 | 6.579559 | 12.63576 | 25.74747 | 10.08113 | -0.48918 | -0.09626 | 0.452328 |
| 2 | 53522000 | 53523399 | 21.7893 | 37.75473 | 38.3716 | 16.83532 | 0.36303 | 0.835623 | 0.452329 |
| 17 | 94293000 | 94295799 | 26.70472 | 43.85539 | 73.34868 | 31.59621 | -0.36227 | 1.419344 | 0.452343 |

| | | | | | | | | | |
|---|---|---|---|---|---|---|---|---|---|
| 3 | 145224200 | 145227199 | 13.94324 | 24.54092 | 42.38953 | 18.76658 | -0.39636 | 0.583209 | 0.452345 |
| X | 91891200 | 91893399 | 34.40415 | 58.01083 | 46.87691 | 20.49681 | 0.687565 | 1.134408 | 0.452352 |
| 8 | 41306600 | 41309199 | 30.05922 | 48.81132 | 82.41525 | 35.33648 | -0.38638 | 1.593897 | 0.452354 |
| 2 | 105489600 | 105492399 | 28.49364 | 46.18504 | 85.95363 | 36.80424 | -0.52892 | 1.628749 | 0.452366 |
| 12 | 21098000 | 21101599 | 80.94362 | 127.378 | 129.4349 | 53.7002 | 0.298455 | 2.522011 | 0.452393 |
| 15 | 65665800 | 65667799 | 15.34403 | 27.84038 | 27.14763 | 11.72988 | 0.449882 | 0.342768 | 0.452397 |
| 19 | 58029400 | 58029799 | 10.21826 | 19.7181 | 20.30756 | 7.842433 | 0.50498 | -0.1192 | 0.452408 |
| 13 | 52627600 | 52629599 | 19.25443 | 33.76691 | 33.92911 | 14.99606 | 0.380672 | 0.649719 | 0.45243 |
| 17 | 21269800 | 21284999 | 162.6555 | 240.5374 | 351.879 | 144.9129 | -0.30087 | 3.726489 | 0.452432 |
| X | 118320600 | 118321799 | 16.31964 | 30.33195 | 24.6819 | 10.32977 | 0.741158 | 0.213487 | 0.452438 |
| 15 | 101899200 | 101901599 | 18.94965 | 31.99196 | 56.8939 | 24.88613 | -0.44625 | 1.081161 | 0.452439 |
| 3 | 127904200 | 127907599 | 36.92022 | 59.47432 | 97.06948 | 41.28144 | -0.34981 | 1.810348 | 0.452444 |
| 8 | 92958400 | 92959799 | 6.423066 | 12.36174 | 25.25589 | 9.827875 | -0.48465 | -0.12009 | 0.452461 |
| 9 | 105019600 | 105023399 | 33.37066 | 53.85939 | 90.82492 | 38.72393 | -0.39351 | 1.746322 | 0.452484 |
| 7 | 109820600 | 109821599 | 6.746119 | 12.85345 | 26.65835 | 10.65282 | -0.54215 | -0.027 | 0.452485 |
| X | 39598400 | 39599999 | 20.97138 | 38.13111 | 25.24223 | 10.87785 | 1.00816 | 0.371257 | 0.452486 |
| 14 | 10418800 | 10421399 | 25.71069 | 44.21015 | 41.9271 | 18.39099 | 0.460257 | 0.92425 | 0.452487 |
| 6 | 75735000 | 75735599 | 9.312299 | 18.08767 | 17.56323 | 6.566099 | 0.631404 | -0.25057 | 0.452498 |
| 7 | 136737000 | 136738199 | 12.23527 | 21.57952 | 38.85953 | 17.19666 | -0.45457 | 0.54573 | 0.452514 |
| 2 | 82871400 | 82872199 | 18.20645 | 31.14825 | 52.70689 | 23.11165 | -0.37284 | 0.954469 | 0.452515 |
| X | 61796200 | 61797999 | 19.74478 | 35.42081 | 28.21768 | 12.35094 | 0.727895 | 0.442177 | 0.452542 |
| 12 | 74229200 | 74233799 | 50.74339 | 81.60957 | 83.04419 | 35.23397 | 0.327487 | 1.875799 | 0.452549 |
| 5 | 127036000 | 127041799 | 113.7859 | 174.7995 | 175.5824 | 72.90432 | 0.288378 | 2.976768 | 0.452574 |
| 10 | 59158200 | 59159799 | 11.601 | 20.55321 | 37.31171 | 16.49684 | -0.46347 | 0.489596 | 0.452578 |
| 12 | 22306400 | 22307999 | 31.71548 | 53.0062 | 42.34404 | 18.43437 | 0.70405 | 1.205333 | 0.45259 |
| 7 | 108241800 | 108245799 | 50.33972 | 81.48915 | 70.60734 | 30.2042 | 0.569558 | 1.686415 | 0.452608 |
| 7 | 53547600 | 53551399 | 40.91611 | 67.25733 | 59.58683 | 25.67832 | 0.552371 | 1.431281 | 0.452619 |
| 7 | 22505200 | 22508199 | 54.4806 | 88.31459 | 65.72949 | 28.1463 | 0.790222 | 1.666414 | 0.452645 |
| 10 | 95072200 | 95072999 | 6.406393 | 12.38515 | 24.882 | 9.529283 | -0.439 | -0.17526 | 0.452658 |
| 3 | 136988600 | 136989199 | 4.106041 | 8.012633 | 19.77276 | 7.036065 | -0.63209 | -0.47872 | 0.452681 |
| 5 | 86994200 | 86996599 | 20.72782 | 36.7305 | 30.2159 | 13.35898 | 0.671943 | 0.528812 | 0.452698 |
| 14 | 8246600 | 8247599 | 11.34598 | 20.46403 | 35.81049 | 15.58859 | -0.39533 | 0.368792 | 0.452702 |
| 17 | 20304000 | 20317999 | 188.4924 | 281.927 | 255.0232 | 105.1639 | 0.40728 | 3.527791 | 0.45271 |
| 1 | 178132600 | 178136999 | 38.41444 | 62.59308 | 63.50657 | 27.27332 | 0.35031 | 1.542787 | 0.452712 |
| 12 | 88076200 | 88078199 | 36.34084 | 59.39755 | 60.02792 | 25.80845 | 0.359121 | 1.493616 | 0.45272 |
| 16 | 17788800 | 17790199 | 8.181978 | 15.19852 | 29.99663 | 12.56378 | -0.52836 | 0.164437 | 0.452732 |
| 6 | 63096600 | 63097799 | 19.88237 | 34.97284 | 33.13141 | 14.66964 | 0.465436 | 0.607518 | 0.452733 |
| 1 | 78361000 | 78362199 | 10.88324 | 19.72924 | 34.86598 | 15.08146 | -0.40331 | 0.33512 | 0.452741 |
| 14 | 18591200 | 18593399 | 18.33796 | 32.4607 | 27.13204 | 11.99813 | 0.648526 | 0.519566 | 0.452744 |
| 16 | 48317000 | 48321399 | 50.09154 | 79.78995 | 124.6031 | 52.13365 | -0.30514 | 2.186382 | 0.452746 |
| 13 | 59339400 | 59340999 | 11.1551 | 20.20745 | 35.87996 | 15.53411 | -0.4112 | 0.341564 | 0.452764 |
| 7 | 85014200 | 85016599 | 38.37423 | 62.87394 | 55.3369 | 23.81402 | 0.56002 | 1.461601 | 0.45277 |
| X | 89210000 | 89213999 | 54.11831 | 87.19996 | 77.24661 | 32.86294 | 0.530419 | 1.823868 | 0.452789 |
| 1 | 53267800 | 53269399 | 21.05943 | 36.86855 | 35.39065 | 15.61918 | 0.446332 | 0.689167 | 0.452826 |
| 3 | 113693200 | 113695399 | 25.10874 | 43.35554 | 40.84947 | 17.92057 | 0.470846 | 0.884363 | 0.452833 |
| 10 | 51425000 | 51426199 | 9.0672 | 16.81109 | 30.99545 | 13.01553 | -0.43407 | 0.183033 | 0.452854 |
| Y | 88024400 | 88026599 | 12.08634 | 22.71556 | 15.55909 | 6.390486 | 1.008591 | 0.081631 | 0.452854 |

| | | | | | | | | | |
|---|---|---|---|---|---|---|---|---|---|
| 17 | 64329000 | 64329799 | 6.410036 | 12.36032 | 25.3345 | 9.804376 | -0.4823 | -0.14005 | 0.452861 |
| 2 | 151511800 | 151513199 | 17.02728 | 28.95861 | 54.19155 | 23.75132 | -0.51863 | 1.008301 | 0.45287 |
| 3 | 16921000 | 16922399 | 12.57824 | 23.68511 | 23.48679 | 9.615528 | 0.482891 | 0.096998 | 0.452885 |
| 3 | 63931600 | 63933199 | 22.93439 | 39.68517 | 39.86854 | 17.49137 | 0.379273 | 0.854515 | 0.452886 |
| 4 | 150628800 | 150630599 | 7.065394 | 13.3629 | 27.60112 | 11.20286 | -0.55625 | 0.029961 | 0.452905 |
| 4 | 49460000 | 49462599 | 21.61871 | 37.79326 | 35.88886 | 15.81982 | 0.461804 | 0.714586 | 0.45291 |
| 17 | 53044400 | 53045599 | 24.09364 | 41.15296 | 41.10988 | 18.02406 | 0.384119 | 0.977091 | 0.452919 |
| X | 81334200 | 81337999 | 38.43967 | 64.09161 | 50.94714 | 22.14642 | 0.711327 | 1.259715 | 0.452921 |
| 7 | 18591600 | 18592599 | 7.178953 | 13.72666 | 26.74339 | 10.60349 | -0.4426 | -0.04733 | 0.452972 |
| 7 | 64847600 | 64848999 | 21.4117 | 37.26497 | 37.26429 | 16.3786 | 0.387246 | 0.780249 | 0.452974 |
| 4 | 96973000 | 96974399 | 16.02329 | 29.35166 | 26.73313 | 11.41158 | 0.559435 | 0.295261 | 0.452975 |
| 16 | 22173000 | 22174199 | 8.566127 | 16.04131 | 30.11199 | 12.44381 | -0.44164 | 0.113951 | 0.452987 |
| 7 | 38428600 | 38434199 | 54.83802 | 88.85935 | 66.29285 | 28.37939 | 0.786284 | 1.675211 | 0.452995 |
| 10 | 100764200 | 100765999 | 17.85491 | 30.76685 | 51.6714 | 22.66671 | -0.36016 | 0.886662 | 0.453007 |
| 15 | 22450000 | 22454999 | 107.4269 | 166.7917 | 155.4575 | 64.55559 | 0.407948 | 2.778952 | 0.453014 |
| X | 26681200 | 26683799 | 23.90171 | 41.83733 | 30.61167 | 13.50809 | 0.836195 | 0.700552 | 0.453043 |
| X | 26043600 | 26053199 | 102.6417 | 161.3521 | 124.5874 | 51.68192 | 0.692976 | 2.557978 | 0.45305 |
| X | 22407000 | 22408199 | 18.04509 | 33.69564 | 23.03764 | 9.56649 | 0.999727 | 0.196594 | 0.45306 |
| 2 | 171991600 | 171995999 | 51.10807 | 81.28676 | 127.868 | 53.3884 | -0.31796 | 2.236955 | 0.453063 |
| 15 | 23297200 | 23300199 | 26.39723 | 43.53814 | 72.91978 | 31.46893 | -0.36175 | 1.366684 | 0.453087 |
| 4 | 131258200 | 131260199 | 11.88124 | 21.27828 | 37.3065 | 16.35235 | -0.40473 | 0.419996 | 0.453094 |
| 19 | 33934000 | 33936799 | 22.08056 | 38.65807 | 35.31168 | 15.58496 | 0.517615 | 0.693995 | 0.453097 |
| 9 | 122329800 | 122332199 | 17.87046 | 30.73981 | 51.65808 | 22.65965 | -0.36155 | 0.902805 | 0.453117 |
| 6 | 76048800 | 76050199 | 24.30331 | 41.52435 | 42.13484 | 18.47391 | 0.361724 | 0.976736 | 0.453135 |
| 14 | 55770600 | 55771999 | 13.18848 | 23.07759 | 42.58499 | 18.78649 | -0.49182 | 0.667395 | 0.453137 |
| 7 | 107972800 | 107981199 | 107.0313 | 165.6611 | 162.2835 | 67.40843 | 0.33276 | 2.833785 | 0.45316 |
| 1 | 21955600 | 21956399 | 6.398027 | 12.39691 | 25.04555 | 9.491328 | -0.43268 | -0.21132 | 0.453162 |
| 1 | 150394800 | 150396599 | 21.18243 | 37.08248 | 35.35302 | 15.60208 | 0.456131 | 0.689869 | 0.453174 |
| 8 | 19546600 | 19555599 | 30.6889 | 51.62552 | 44.2631 | 19.34022 | 0.602204 | 1.137212 | 0.453187 |
| 12 | 23651200 | 23653399 | 18.87406 | 34.18239 | 27.40985 | 11.87046 | 0.728546 | 0.381032 | 0.453195 |
| 14 | 113524000 | 113526799 | 20.37193 | 35.74567 | 33.87869 | 14.98048 | 0.46466 | 0.648946 | 0.453214 |
| 3 | 90890600 | 90892999 | 54.59151 | 87.59903 | 87.55057 | 37.021 | 0.349432 | 1.953603 | 0.453226 |
| 7 | 16347400 | 16349799 | 14.92719 | 25.92502 | 46.86756 | 20.57241 | -0.46352 | 0.791903 | 0.453242 |
| 3 | 106760600 | 106761399 | 26.70566 | 45.14749 | 45.36181 | 19.85003 | 0.374508 | 1.083864 | 0.453277 |
| 14 | 8406600 | 8408199 | 10.17559 | 18.55365 | 34.41747 | 14.7799 | -0.4663 | 0.292563 | 0.453283 |
| 12 | 61280600 | 61281199 | 5.809611 | 11.28472 | 23.41449 | 8.750662 | -0.45108 | -0.26938 | 0.453289 |
| 16 | 59722600 | 59725399 | 23.5379 | 41.18021 | 35.20083 | 15.52668 | 0.612885 | 0.712314 | 0.453301 |
| 18 | 63189200 | 63190399 | 12.18387 | 22.86149 | 22.91776 | 9.438941 | 0.460311 | 0.113489 | 0.453318 |
| 1 | 148509000 | 148510599 | 16.27724 | 29.5518 | 27.00369 | 11.65871 | 0.543681 | 0.345404 | 0.453321 |
| 2 | 11637600 | 11638599 | 12.0176 | 22.97121 | 21.53808 | 8.547411 | 0.601684 | -0.01447 | 0.453323 |
| 13 | 17835000 | 17835999 | 12.46096 | 23.25734 | 23.98529 | 9.968021 | 0.40961 | 0.153292 | 0.453366 |
| 9 | 7389400 | 7392399 | 42.45161 | 69.22496 | 65.63063 | 28.19348 | 0.448691 | 1.542939 | 0.453384 |
| 6 | 57263000 | 57265399 | 34.16594 | 56.74137 | 49.28939 | 21.36967 | 0.583458 | 1.286073 | 0.453406 |
| 12 | 94888000 | 94889799 | 19.24001 | 34.77812 | 26.0733 | 11.31885 | 0.822392 | 0.389559 | 0.453415 |
| 4 | 39049600 | 39050599 | 32.90135 | 54.33977 | 52.43138 | 22.64524 | 0.43194 | 1.351001 | 0.453423 |
| 11 | 7741000 | 7744199 | 46.9911 | 75.73487 | 76.85592 | 32.7082 | 0.335137 | 1.800385 | 0.453438 |
| 16 | 12146400 | 12149399 | 15.69325 | 28.74594 | 23.61518 | 10.08118 | 0.707086 | 0.285337 | 0.453449 |

| | | | | | | | | | |
|---|---|---|---|---|---|---|---|---|---|
| 12 | 116481000 | 116483199 | 15.49828 | 27.04834 | 48.40243 | 21.31131 | -0.44837 | 0.725746 | 0.453459 |
| 7 | 67361200 | 67363199 | 19.77258 | 34.72773 | 33.77937 | 14.94883 | 0.427245 | 0.6287 | 0.453479 |
| 13 | 86659200 | 86659999 | 15.66862 | 28.12277 | 28.28606 | 12.37556 | 0.394033 | 0.412168 | 0.453482 |
| 7 | 74389200 | 74389599 | 15.73326 | 29.33915 | 24.19091 | 10.06837 | 0.728417 | 0.186635 | 0.453485 |
| 11 | 23615400 | 23616599 | 10.17179 | 18.18417 | 35.35618 | 15.52092 | -0.55426 | 0.415794 | 0.453493 |
| 16 | 78918000 | 78918599 | 6.118558 | 11.81705 | 25.99615 | 10.00188 | -0.57581 | -0.16265 | 0.453509 |
| 7 | 17267800 | 17269799 | 40.34283 | 65.47348 | 67.17774 | 28.79131 | 0.329693 | 1.616271 | 0.453519 |
| 4 | 3453800 | 3470599 | 356.7404 | 502.5634 | 467.3507 | 193.5614 | 0.28376 | 4.470433 | 0.453534 |
| 13 | 67044600 | 67048199 | 39.19309 | 64.31233 | 61.58197 | 26.51845 | 0.439614 | 1.45086 | 0.453547 |
| 1 | 48909000 | 48910199 | 18.95248 | 33.45851 | 32.14278 | 14.22157 | 0.447398 | 0.561912 | 0.453556 |
| 15 | 37095800 | 37100799 | 56.49226 | 90.56058 | 80.17866 | 33.85347 | 0.522271 | 1.98107 | 0.453574 |
| 8 | 12491200 | 12493999 | 20.67976 | 36.15666 | 34.85822 | 15.36725 | 0.439971 | 0.704707 | 0.453618 |
| 8 | 126837800 | 126839799 | 18.06619 | 31.73467 | 31.44628 | 13.91781 | 0.400597 | 0.612566 | 0.453623 |
| 13 | 62957600 | 62965999 | 115.8961 | 177.6575 | 179.1085 | 74.34638 | 0.280734 | 3.020325 | 0.453648 |
| X | 54579000 | 54584199 | 93.94152 | 147.0151 | 127.7643 | 53.00134 | 0.517614 | 2.632882 | 0.453653 |
| X | 38712400 | 38713599 | 17.2135 | 32.50891 | 20.94188 | 8.526138 | 1.107317 | 0.114385 | 0.453665 |
| X | 78718600 | 78720799 | 28.90631 | 49.33618 | 44.11606 | 19.34091 | 0.543248 | 1.007856 | 0.45368 |
| 10 | 71478000 | 71479599 | 16.61919 | 30.07523 | 28.11698 | 12.18855 | 0.50745 | 0.370068 | 0.453684 |
| 18 | 53007600 | 53010199 | 32.65809 | 52.50036 | 106.6469 | 45.2194 | -0.66845 | 1.874983 | 0.45369 |
| 16 | 92666400 | 92670399 | 28.07042 | 45.83475 | 76.95808 | 33.07553 | -0.37229 | 1.501621 | 0.453694 |
| 12 | 72319200 | 72321199 | 16.43266 | 28.19047 | 51.09729 | 22.41078 | -0.47096 | 0.919505 | 0.453699 |
| 11 | 74025400 | 74042599 | 290.3853 | 417.7088 | 388.2452 | 160.2511 | 0.314683 | 4.162888 | 0.453699 |
| 13 | 102329600 | 102330399 | 12.84791 | 23.85456 | 24.49181 | 10.26471 | 0.406646 | 0.190554 | 0.453703 |
| 8 | 46423800 | 46425199 | 29.00571 | 48.54671 | 49.6863 | 21.65033 | 0.347725 | 1.188431 | 0.453708 |
| 5 | 64004400 | 64005199 | 7.959262 | 14.99425 | 28.60111 | 11.70037 | -0.45246 | 0.067437 | 0.453748 |
| 18 | 8935200 | 8936199 | 9.25857 | 16.99841 | 31.91993 | 13.58376 | -0.4743 | 0.242125 | 0.453797 |
| 19 | 39484200 | 39492999 | 112.2752 | 174.6121 | 148.178 | 61.51724 | 0.544968 | 2.75134 | 0.453805 |
| 1 | 185508400 | 185509599 | 12.117 | 21.34838 | 39.42354 | 17.44993 | -0.49113 | 0.559471 | 0.453835 |
| X | 126802200 | 126805599 | 44.7177 | 73.28997 | 60.2405 | 25.92766 | 0.658659 | 1.477914 | 0.453838 |
| 8 | 105734600 | 105738999 | 52.95729 | 84.87197 | 86.40209 | 36.46689 | 0.320794 | 1.987284 | 0.453848 |
| 9 | 32659000 | 32661599 | 12.12448 | 21.39866 | 39.40539 | 17.43585 | -0.48606 | 0.540873 | 0.453859 |
| 10 | 52326800 | 52327999 | 17.43389 | 30.73112 | 28.52849 | 12.62624 | 0.495852 | 0.562895 | 0.453859 |
| 3 | 18505200 | 18506399 | 12.5585 | 23.91871 | 22.64759 | 9.057396 | 0.578495 | 0.020143 | 0.45387 |
| 2 | 89259200 | 89262399 | 40.24049 | 66.97261 | 49.90172 | 21.66707 | 0.804355 | 1.280959 | 0.45387 |
| 8 | 93435600 | 93439799 | 41.50959 | 68.47347 | 56.79115 | 24.51002 | 0.649277 | 1.38986 | 0.45388 |
| X | 126022400 | 126029799 | 76.33273 | 122.0871 | 86.12921 | 36.24353 | 0.845697 | 2.079587 | 0.453885 |
| 12 | 77573200 | 77574399 | 13.31834 | 23.37356 | 41.44269 | 18.32811 | -0.43454 | 0.617922 | 0.453915 |
| 8 | 4158600 | 4162799 | 42.31183 | 68.87001 | 67.17938 | 28.83942 | 0.405882 | 1.56979 | 0.453921 |
| 4 | 16985800 | 16995399 | 188.7667 | 280.1185 | 272.6272 | 112.2975 | 0.291355 | 3.668083 | 0.453929 |
| 4 | 86433800 | 86435799 | 20.04596 | 35.32407 | 32.5404 | 14.40278 | 0.506577 | 0.588106 | 0.453938 |
| 8 | 24729800 | 24731799 | 20.35571 | 35.6095 | 35.47722 | 15.64735 | 0.39293 | 0.697174 | 0.45394 |
| 17 | 62683800 | 62684399 | 5.661603 | 11.0135 | 22.99134 | 8.490513 | -0.44403 | -0.31582 | 0.453941 |
| 8 | 107403400 | 107404399 | 6.527276 | 12.24224 | 28.56438 | 11.77329 | -0.74861 | 0.085108 | 0.453957 |
| 7 | 55139600 | 55141999 | 77.02903 | 121.3381 | 122.1843 | 50.6986 | 0.312612 | 2.497733 | 0.45399 |
| 1 | 136952600 | 136956399 | 40.04727 | 65.22279 | 65.39174 | 28.07808 | 0.366951 | 1.553815 | 0.453996 |
| 8 | 106291600 | 106294199 | 14.27649 | 24.53719 | 50.01472 | 21.93824 | -0.63879 | 0.893623 | 0.454019 |
| 1 | 53556800 | 53557799 | 11.24518 | 21.46398 | 22.2097 | 8.841534 | 0.457243 | -0.00815 | 0.454027 |

| | | | | | | | | | |
|---|---|---|---|---|---|---|---|---|---|
| 10 | 99178600 | 99180999 | 32.43037 | 54.0861 | 46.53109 | 20.23009 | 0.597457 | 1.231621 | 0.45403 |
| 17 | 36937200 | 36941599 | 49.72923 | 80.10372 | 80.51627 | 34.23896 | 0.347948 | 1.824468 | 0.454036 |
| 1 | 98307600 | 98308999 | 16.14131 | 30.51594 | 22.26247 | 9.043744 | 0.932287 | 0.101613 | 0.45405 |
| X | 64128400 | 64131399 | 44.88929 | 73.60706 | 52.22558 | 22.48265 | 0.870522 | 1.468221 | 0.454058 |
| 1 | 14511000 | 14513399 | 19.66621 | 32.91381 | 68.40353 | 29.80537 | -0.67077 | 1.176496 | 0.45407 |
| 3 | 89740200 | 89741799 | 15.25944 | 26.42627 | 47.93167 | 21.0294 | -0.46906 | 0.824267 | 0.454108 |
| 3 | 53545400 | 53545999 | 6.097209 | 11.84051 | 25.26739 | 9.461508 | -0.49497 | -0.25709 | 0.45415 |
| 7 | 6665000 | 6666599 | 11.49421 | 20.60735 | 36.76729 | 16.10134 | -0.42911 | 0.407 | 0.45417 |
| 5 | 135196800 | 135197799 | 7.214321 | 13.64088 | 27.59631 | 11.20731 | -0.5272 | 0.033709 | 0.45419 |
| 3 | 93368000 | 93369999 | 17.16481 | 30.73564 | 28.98326 | 12.70583 | 0.484358 | 0.436468 | 0.454205 |
| X | 92180000 | 92182199 | 21.07836 | 37.6626 | 28.71308 | 12.61012 | 0.787888 | 0.477359 | 0.454219 |
| 9 | 22816400 | 22817799 | 11.11301 | 19.73098 | 36.7999 | 16.24838 | -0.50065 | 0.467554 | 0.454223 |
| 11 | 112587200 | 112589599 | 21.16093 | 35.8299 | 60.3556 | 26.42376 | -0.36832 | 1.066199 | 0.454233 |
| 12 | 3420000 | 3422199 | 18.12413 | 31.88468 | 31.23072 | 13.82767 | 0.417516 | 0.591185 | 0.454234 |
| 14 | 25297200 | 25299599 | 16.87448 | 29.11763 | 49.52443 | 21.72581 | -0.37785 | 0.867095 | 0.454237 |
| 13 | 22818200 | 22824399 | 127.0595 | 195.0966 | 178.6166 | 74.15925 | 0.421509 | 2.992665 | 0.454266 |
| 2 | 172983000 | 172984399 | 14.28152 | 24.95859 | 43.62623 | 19.22526 | -0.41411 | 0.695469 | 0.45428 |
| 5 | 78238400 | 78240999 | 28.21555 | 48.37067 | 42.40753 | 18.5985 | 0.572412 | 0.962068 | 0.454281 |
| X | 150178200 | 150179199 | 15.00215 | 27.60667 | 25.41648 | 10.78247 | 0.550656 | 0.261178 | 0.45429 |
| 5 | 131774600 | 131776799 | 14.42192 | 25.20118 | 44.13922 | 19.4495 | -0.41709 | 0.698049 | 0.45431 |
| 14 | 35380200 | 35382599 | 26.05422 | 45.10645 | 38.52562 | 16.90246 | 0.612475 | 0.860634 | 0.454314 |
| 13 | 111501000 | 111504799 | 29.79454 | 48.70321 | 81.69661 | 35.12609 | -0.36993 | 1.490649 | 0.454332 |
| 6 | 60023800 | 60024599 | 12.4757 | 23.23482 | 23.98878 | 10.00585 | 0.403877 | 0.167218 | 0.454334 |
| 9 | 7582000 | 7582599 | 12.4757 | 23.23482 | 23.98878 | 10.00585 | 0.403877 | 0.167218 | 0.454334 |
| 11 | 66457800 | 66460199 | 15.69562 | 27.24438 | 47.93115 | 21.03188 | -0.42465 | 0.804333 | 0.454339 |
| 11 | 22288200 | 22292199 | 29.41254 | 49.52323 | 42.16067 | 18.42665 | 0.612371 | 1.121748 | 0.454377 |
| 19 | 4705000 | 4705799 | 6.546156 | 12.65332 | 25.47986 | 9.765531 | -0.44379 | -0.17093 | 0.45438 |
| 12 | 67994600 | 67998399 | 44.16185 | 71.35673 | 73.28244 | 31.29015 | 0.322196 | 1.722679 | 0.454406 |
| 7 | 18207800 | 18221199 | 292.8381 | 422.2193 | 346.2994 | 142.8909 | 0.498367 | 4.127924 | 0.454419 |
| 15 | 33994400 | 33997999 | 40.19672 | 64.66063 | 103.2542 | 43.82149 | -0.32096 | 1.862395 | 0.454432 |
| 2 | 122777000 | 122778799 | 17.96036 | 31.64825 | 31.83065 | 14.08869 | 0.380597 | 0.575574 | 0.454432 |
| 1 | 45123600 | 45127799 | 49.37899 | 79.65435 | 77.97327 | 33.22189 | 0.388483 | 1.780112 | 0.454457 |
| 7 | 62052400 | 62053399 | 12.91158 | 23.97451 | 24.03424 | 10.07194 | 0.441148 | 0.19053 | 0.454475 |
| 4 | 17962200 | 17965199 | 36.5176 | 60.71657 | 53.23339 | 23.09791 | 0.570261 | 1.282932 | 0.454476 |
| 19 | 9701000 | 9702399 | 22.0565 | 38.32645 | 38.22128 | 16.78251 | 0.390811 | 0.804569 | 0.454482 |
| 7 | 105148400 | 105149999 | 14.69525 | 26.69123 | 26.86774 | 11.59372 | 0.40583 | 0.33356 | 0.454503 |
| 5 | 75609000 | 75612799 | 26.39692 | 42.8956 | 87.64594 | 37.59941 | -0.65978 | 1.575596 | 0.454512 |
| 6 | 20618000 | 20618999 | 12.93084 | 24.1118 | 24.44541 | 10.17532 | 0.432195 | 0.164028 | 0.454512 |
| 7 | 141765600 | 141768999 | 25.31882 | 41.42702 | 75.7264 | 32.57062 | -0.49241 | 1.463714 | 0.45452 |
| 4 | 32972800 | 32983599 | 134.423 | 204.4008 | 204.2456 | 84.61218 | 0.284323 | 3.189562 | 0.454523 |
| 10 | 24719600 | 24721399 | 19.47119 | 34.80166 | 29.35112 | 12.88698 | 0.643181 | 0.46531 | 0.454531 |
| Y | 75174200 | 75174799 | 2.452765 | 4.797419 | 17.20323 | 5.793165 | -1.0858 | -0.70868 | 0.454565 |
| 13 | 51395000 | 51396999 | 19.50904 | 33.21767 | 55.39965 | 24.28306 | -0.35316 | 1.005917 | 0.454578 |
| 18 | 90510200 | 90516999 | 123.0692 | 187.9144 | 188.8488 | 78.32126 | 0.280877 | 3.10181 | 0.454579 |
| 9 | 17285200 | 17288399 | 34.49232 | 56.99373 | 57.53489 | 24.85739 | 0.366901 | 1.352187 | 0.454586 |
| 5 | 59184000 | 59184799 | 16.96124 | 30.06151 | 30.2178 | 13.35933 | 0.384514 | 0.503465 | 0.454592 |
| 6 | 121938800 | 121940199 | 15.12415 | 27.95193 | 25.79598 | 10.87919 | 0.552883 | 0.236853 | 0.454594 |

| | | | | | | | | | |
|---|---|---|---|---|---|---|---|---|---|
| 3 | 112086000 | 112087599 | 16.90014 | 30.15945 | 29.59653 | 13.01176 | 0.42447 | 0.457327 | 0.4546 |
| 2 | 65625200 | 65629599 | 95.13737 | 144.6843 | 258.8823 | 107.2509 | -0.55529 | 3.181921 | 0.454614 |
| 4 | 92771400 | 92773399 | 14.73118 | 27.05235 | 26.2672 | 11.17459 | 0.470999 | 0.269248 | 0.454618 |
| 1 | 29469200 | 29472999 | 33.77268 | 56.29869 | 53.10679 | 23.08223 | 0.465116 | 1.247688 | 0.454645 |
| 19 | 56238800 | 56243599 | 28.17359 | 45.69442 | 82.89399 | 35.51027 | -0.49122 | 1.615621 | 0.454653 |
| 9 | 111139000 | 111142199 | 36.58723 | 58.97525 | 96.07177 | 40.89333 | -0.34596 | 1.789959 | 0.454656 |
| 17 | 15409200 | 15410799 | 8.399846 | 15.3395 | 32.9333 | 14.1153 | -0.67346 | 0.267734 | 0.45466 |
| 2 | 87832000 | 87833199 | 13.0515 | 24.76231 | 19.42706 | 7.832838 | 0.837762 | 0.043161 | 0.454666 |
| 1 | 35390800 | 35392999 | 15.88199 | 27.56535 | 47.40247 | 20.7987 | -0.39183 | 0.806963 | 0.454667 |
| 3 | 49274400 | 49275999 | 22.68674 | 39.70842 | 35.23632 | 15.54808 | 0.55914 | 0.700816 | 0.454682 |
| 12 | 118834200 | 118835999 | 16.34119 | 29.0449 | 28.84434 | 12.72414 | 0.40426 | 0.478528 | 0.454682 |
| 5 | 98382600 | 98386599 | 36.1478 | 59.39673 | 58.74627 | 25.31203 | 0.393728 | 1.421555 | 0.454684 |
| 11 | 18034000 | 18035199 | 15.3702 | 27.79292 | 26.01519 | 11.28862 | 0.504387 | 0.358535 | 0.454687 |
| X | 4414200 | 4422399 | 95.74814 | 151.4121 | 112.6229 | 46.79788 | 0.753441 | 2.426753 | 0.45471 |
| 19 | 10925600 | 10926799 | 6.011899 | 11.51523 | 25.67349 | 10.12845 | -0.62821 | -0.07899 | 0.454716 |
| 16 | 47654200 | 47657599 | 32.8147 | 54.97535 | 50.81602 | 22.15621 | 0.494175 | 1.191246 | 0.454718 |
| 7 | 31685800 | 31689599 | 50.75814 | 81.66457 | 82.56492 | 35.05008 | 0.337472 | 1.863943 | 0.45473 |
| 7 | 121462800 | 121464799 | 11.41557 | 20.36208 | 36.29366 | 15.96253 | -0.43165 | 0.440353 | 0.454738 |
| 2 | 22507400 | 22515799 | 70.19589 | 108.8162 | 206.4908 | 85.7678 | -0.61842 | 2.794075 | 0.454774 |
| 3 | 7185600 | 7186199 | 4.776226 | 9.32553 | 20.51838 | 7.233105 | -0.45693 | -0.49944 | 0.45478 |
| 3 | 18638400 | 18639399 | 10.05045 | 18.37691 | 32.89687 | 14.07748 | -0.41161 | 0.274226 | 0.454802 |
| 9 | 8899800 | 8901999 | 23.82965 | 39.1611 | 72.46033 | 31.22273 | -0.50713 | 1.402601 | 0.454808 |
| 6 | 53614800 | 53615999 | 7.320385 | 13.93033 | 27.74673 | 11.11683 | -0.48819 | -0.01141 | 0.454812 |
| 5 | 11242200 | 11243599 | 20.12896 | 35.06355 | 33.28422 | 14.63741 | 0.462092 | 0.754497 | 0.454819 |
| 19 | 27300000 | 27301599 | 19.88196 | 33.14347 | 62.60803 | 27.21207 | -0.5332 | 1.217634 | 0.454834 |
| 2 | 20596800 | 20598399 | 16.64561 | 29.56258 | 29.74918 | 13.13205 | 0.38456 | 0.486657 | 0.454847 |
| 18 | 29821000 | 29821799 | 12.76866 | 23.56758 | 22.98541 | 9.711215 | 0.471916 | 0.218424 | 0.454857 |
| 16 | 43695400 | 43697199 | 12.54543 | 22.0994 | 39.36283 | 17.42366 | -0.43965 | 0.561758 | 0.454863 |
| 1 | 169891200 | 169893799 | 16.00711 | 27.89461 | 47.88156 | 21.06215 | -0.38874 | 0.749884 | 0.45488 |
| 15 | 93035400 | 93036799 | 11.12253 | 19.81268 | 36.32835 | 15.99593 | -0.47326 | 0.447666 | 0.454887 |
| 14 | 106265200 | 106267599 | 32.88598 | 55.17576 | 49.70013 | 21.68976 | 0.531246 | 1.174111 | 0.454893 |
| 18 | 48796400 | 48797199 | 11.0389 | 20.93741 | 22.13468 | 8.930341 | 0.409615 | 0.032078 | 0.454901 |
| 3 | 77835000 | 77835999 | 10.01117 | 19.36881 | 19.73532 | 7.531404 | 0.535452 | -0.1659 | 0.454903 |
| 1 | 106227400 | 106232799 | 39.22751 | 64.1039 | 55.27115 | 23.76172 | 0.587412 | 1.499247 | 0.454906 |
| 11 | 113003600 | 113005799 | 13.33603 | 23.36188 | 42.02913 | 18.5588 | -0.45539 | 0.649088 | 0.454931 |
| 17 | 57690200 | 57691599 | 33.33209 | 54.9421 | 56.28424 | 24.28342 | 0.345011 | 1.377002 | 0.454944 |
| 16 | 95356800 | 95359599 | 23.02119 | 38.38954 | 64.0917 | 27.86286 | -0.35583 | 1.222681 | 0.454947 |
| 14 | 102218400 | 102220399 | 16.25932 | 28.34856 | 50.57569 | 22.25437 | -0.44405 | 0.749552 | 0.454953 |
| 10 | 60462200 | 60476199 | 163.9756 | 240.9986 | 371.2469 | 152.8917 | -0.38373 | 3.827892 | 0.454961 |
| X | 8829000 | 8832199 | 48.39332 | 78.56702 | 59.3047 | 25.42377 | 0.771953 | 1.622991 | 0.454978 |
| 7 | 61719200 | 61725599 | 105.8176 | 163.7861 | 144.9991 | 60.2289 | 0.478669 | 2.830374 | 0.454992 |
| 9 | 19124200 | 19128799 | 60.77339 | 97.17331 | 96.11869 | 40.43062 | 0.358482 | 2.070875 | 0.455 |
| X | 25340600 | 25350399 | 219.45 | 325.3226 | 277.6469 | 114.3534 | 0.479432 | 3.692041 | 0.455068 |
| 18 | 12834600 | 12838599 | 49.10818 | 79.22884 | 77.83739 | 33.17011 | 0.383531 | 1.776335 | 0.455087 |
| 4 | 38507200 | 38512999 | 71.27274 | 113.114 | 113.708 | 47.37148 | 0.323974 | 2.312395 | 0.455089 |
| 1 | 73511200 | 73513599 | 22.88413 | 39.98039 | 35.78415 | 15.76594 | 0.546745 | 0.729985 | 0.455096 |
| 16 | 95580400 | 95584799 | 30.82084 | 50.23166 | 81.94859 | 35.19637 | -0.3329 | 1.536641 | 0.455105 |

| | | | | | | | | | |
|---|---|---|---|---|---|---|---|---|---|
| 10 | 3055000 | 3078799 | 818.8179 | 1075.455 | 928.6558 | 388.6117 | 0.289101 | 5.569155 | 0.455139 |
| 7 | 77950600 | 77952999 | 17.14925 | 29.60713 | 50.6069 | 22.20103 | -0.38493 | 0.863514 | 0.455153 |
| X | 29413000 | 29415799 | 39.08616 | 65.02373 | 52.11289 | 22.62049 | 0.699567 | 1.285888 | 0.455177 |
| 12 | 36663800 | 36668599 | 71.38882 | 113.2459 | 114.7529 | 47.78473 | 0.311845 | 2.327231 | 0.455186 |
| 4 | 152374400 | 152376599 | 19.19181 | 32.15207 | 60.00263 | 26.1614 | -0.51581 | 1.158646 | 0.455194 |
| 2 | 151290400 | 151302199 | 134.2514 | 205.0913 | 193.0875 | 80.08845 | 0.375538 | 3.095898 | 0.455194 |
| 4 | 88856000 | 88856999 | 9.940704 | 18.02582 | 33.35583 | 14.42302 | -0.46876 | 0.32319 | 0.455207 |
| 7 | 34801600 | 34803599 | 21.86722 | 37.95229 | 38.37037 | 16.83562 | 0.371105 | 0.82238 | 0.455208 |
| 4 | 65074000 | 65074799 | 6.511601 | 12.27188 | 28.08118 | 11.48001 | -0.71165 | 0.05045 | 0.455209 |
| 9 | 19205400 | 19206799 | 16.33549 | 29.46065 | 28.04096 | 12.20783 | 0.4776 | 0.386026 | 0.45526 |
| 13 | 53006400 | 53007799 | 18.506 | 31.31569 | 56.84486 | 24.88221 | -0.47544 | 1.061665 | 0.455265 |
| 19 | 20820800 | 20821799 | 11.09055 | 21.07882 | 22.17731 | 8.908567 | 0.422145 | 0.019348 | 0.455273 |
| 17 | 23522000 | 23523799 | 16.74546 | 30.94596 | 25.07623 | 10.57676 | 0.739088 | 0.248227 | 0.455274 |
| 11 | 32977000 | 32981599 | 37.1125 | 59.91087 | 97.16243 | 41.4369 | -0.33707 | 1.749496 | 0.455291 |
| 3 | 4065800 | 4071999 | 79.40653 | 125.5629 | 111.929 | 46.50816 | 0.492539 | 2.412978 | 0.455293 |
| X | 122580200 | 122585599 | 45.60265 | 74.74139 | 59.69082 | 25.69122 | 0.700085 | 1.480054 | 0.455298 |
| 15 | 21857600 | 21863199 | 135.5563 | 205.7638 | 205.8623 | 85.23727 | 0.280479 | 3.225428 | 0.4553 |
| X | 131282600 | 131283999 | 24.737 | 42.45167 | 42.07415 | 18.45384 | 0.396512 | 0.940278 | 0.4553 |
| 12 | 16814000 | 16816799 | 26.05067 | 42.96818 | 71.20029 | 30.72737 | -0.34635 | 1.363523 | 0.455309 |
| 10 | 76359400 | 76361199 | 16.42194 | 29.9237 | 27.2325 | 11.70386 | 0.554176 | 0.329391 | 0.455347 |
| 6 | 144241600 | 144245599 | 30.31477 | 49.52079 | 81.57955 | 35.06729 | -0.34457 | 1.502277 | 0.455351 |
| 6 | 7016800 | 7020999 | 51.70558 | 83.31915 | 79.83228 | 33.96696 | 0.417568 | 1.815333 | 0.455368 |
| 6 | 19125200 | 19126199 | 7.14416 | 13.77869 | 27.74701 | 10.7318 | -0.45672 | -0.1278 | 0.455399 |
| 5 | 90965800 | 90967199 | 14.03479 | 24.55382 | 43.6281 | 19.24358 | -0.43743 | 0.679319 | 0.455422 |
| 15 | 12657600 | 12659799 | 16.46964 | 28.63313 | 50.07686 | 21.99525 | -0.41604 | 0.787619 | 0.45543 |
| 13 | 68472200 | 68496599 | 519.389 | 710.4429 | 627.0078 | 260.8932 | 0.316311 | 4.921551 | 0.455438 |
| X | 71893000 | 71896399 | 36.08104 | 60.35958 | 50.31231 | 21.91644 | 0.642952 | 1.217361 | 0.45544 |
| 6 | 112620400 | 112621599 | 13.06865 | 24.36071 | 24.51953 | 10.21189 | 0.442241 | 0.169101 | 0.455463 |
| X | 116053000 | 116053799 | 16.96518 | 30.55597 | 28.22402 | 12.3078 | 0.519535 | 0.401615 | 0.455469 |
| 7 | 67812600 | 67815599 | 24.17508 | 40.32726 | 67.57346 | 29.38271 | -0.36116 | 1.2259 | 0.45547 |
| 1 | 119185800 | 119191199 | 69.46404 | 110.3533 | 111.328 | 46.42621 | 0.32024 | 2.281679 | 0.455475 |
| 6 | 73081200 | 73084399 | 21.73296 | 36.40759 | 61.27471 | 26.71526 | -0.3674 | 1.168336 | 0.455486 |
| X | 146456200 | 146469199 | 323.7355 | 463.6356 | 396.714 | 163.8597 | 0.427661 | 4.229985 | 0.455496 |
| 8 | 18612200 | 18613399 | 7.652501 | 14.42477 | 28.61167 | 11.69313 | -0.50737 | 0.060038 | 0.455499 |
| 18 | 44357800 | 44361199 | 43.17377 | 69.21681 | 108.8692 | 45.97402 | -0.30504 | 1.973385 | 0.455501 |
| 13 | 65619400 | 65625799 | 168.2926 | 251.6226 | 242.2309 | 99.88166 | 0.317039 | 3.529829 | 0.455504 |
| X | 118244800 | 118250199 | 57.88128 | 93.19417 | 79.01395 | 33.57878 | 0.59225 | 1.852047 | 0.455527 |
| 8 | 60688400 | 60690999 | 32.50529 | 54.65768 | 49.11334 | 21.45668 | 0.53481 | 1.152685 | 0.45553 |
| 17 | 40726400 | 40727599 | 14.61053 | 26.78685 | 25.66368 | 10.94221 | 0.48768 | 0.274821 | 0.455553 |
| 7 | 61142400 | 61149799 | 95.63238 | 150.8027 | 122.0921 | 50.66735 | 0.628103 | 2.489907 | 0.45556 |
| 4 | 153234800 | 153260799 | 341.6009 | 475.2513 | 649.4576 | 269.5112 | -0.28958 | 4.65756 | 0.455568 |
| 14 | 11029600 | 11030999 | 22.08399 | 38.28707 | 38.83183 | 17.03753 | 0.366137 | 0.832041 | 0.455623 |
| 7 | 119683600 | 119685599 | 18.74519 | 32.94966 | 29.52306 | 13.07378 | 0.544704 | 0.598951 | 0.45564 |
| 9 | 105209600 | 105210999 | 9.892737 | 18.01639 | 33.36722 | 14.3503 | -0.46518 | 0.296106 | 0.455642 |
| 15 | 95432400 | 95433999 | 9.892737 | 18.01639 | 33.36722 | 14.3503 | -0.46518 | 0.296106 | 0.455642 |
| 5 | 133921400 | 133924399 | 24.84741 | 41.233 | 68.01674 | 29.47295 | -0.33892 | 1.286456 | 0.455643 |
| 2 | 123588400 | 123590799 | 18.02888 | 32.57769 | 26.71454 | 11.60222 | 0.693387 | 0.383626 | 0.455643 |

| | | | | | | | | | |
|---|---|---|---|---|---|---|---|---|---|
| 10 | 45431200 | 45431999 | 13.74125 | 25.23413 | 24.98713 | 10.6302 | 0.443264 | 0.260891 | 0.45565 |
| 5 | 145984000 | 145998199 | 378.8335 | 530.3254 | 490.7183 | 203.4417 | 0.281776 | 4.565047 | 0.455652 |
| 1 | 39695400 | 39698999 | 32.19134 | 53.4863 | 53.64176 | 23.26395 | 0.376824 | 1.278034 | 0.455663 |
| 10 | 11481400 | 11484999 | 34.38993 | 57.34761 | 52.5788 | 22.85777 | 0.506005 | 1.244917 | 0.455671 |
| 2 | 111956000 | 111957799 | 22.06348 | 38.42938 | 37.67724 | 16.56825 | 0.415674 | 0.772776 | 0.455679 |
| 15 | 33994200 | 33997999 | 41.18401 | 66.19962 | 105.5852 | 44.76256 | -0.32058 | 1.88795 | 0.455686 |
| 14 | 121308600 | 121311399 | 27.71256 | 45.31147 | 76.59225 | 32.93449 | -0.3806 | 1.480063 | 0.455686 |
| 1 | 102200400 | 102201599 | 10.37295 | 19.09776 | 35.74573 | 15.15402 | -0.46593 | 0.243943 | 0.45569 |
| 6 | 101429600 | 101431599 | 25.94807 | 42.49714 | 74.10768 | 31.88648 | -0.42387 | 1.452373 | 0.455696 |
| 18 | 30628000 | 30628599 | 6.893968 | 13.13153 | 26.65348 | 10.65707 | -0.51194 | -0.02315 | 0.455702 |
| 16 | 41351800 | 41353999 | 15.56308 | 26.99599 | 47.40335 | 20.79864 | -0.42201 | 0.810499 | 0.455714 |
| 19 | 40246600 | 40248799 | 20.18428 | 35.29293 | 35.52095 | 15.66038 | 0.378355 | 0.703834 | 0.455718 |
| 18 | 87492200 | 87492999 | 8.244757 | 15.62112 | 29.31684 | 11.8547 | -0.41483 | 0.03253 | 0.45574 |
| 19 | 29875600 | 29877799 | 38.96932 | 63.50744 | 65.21637 | 28.00925 | 0.33313 | 1.54363 | 0.455746 |
| 15 | 66254000 | 66254599 | 5.823451 | 11.278 | 23.81022 | 9.044157 | -0.49832 | -0.21324 | 0.45576 |
| 12 | 11498600 | 11498999 | 4.514936 | 8.816228 | 19.92224 | 7.011103 | -0.49287 | -0.51264 | 0.45578 |
| 8 | 27209600 | 27210799 | 14.50548 | 26.64505 | 26.36146 | 11.21058 | 0.44468 | 0.267344 | 0.455791 |
| 6 | 19127800 | 19128399 | 11.73402 | 20.95083 | 36.79025 | 16.16744 | -0.40946 | 0.437428 | 0.455797 |
| 10 | 43701400 | 43707999 | 74.37136 | 115.8134 | 183.3356 | 76.09849 | -0.35211 | 2.714697 | 0.455797 |
| 3 | 17434000 | 17438199 | 46.67111 | 75.81864 | 68.02785 | 29.17688 | 0.524044 | 1.610271 | 0.455801 |
| 3 | 79635400 | 79636599 | 7.863231 | 14.65 | 29.50764 | 12.30001 | -0.55209 | 0.140954 | 0.45581 |
| 18 | 50571200 | 50573399 | 35.42517 | 59.65646 | 46.31199 | 20.24199 | 0.745095 | 1.146533 | 0.455812 |
| 2 | 49762800 | 49763999 | 7.916537 | 15.04381 | 28.85058 | 11.5932 | -0.43791 | 0.006782 | 0.455832 |
| 5 | 75820600 | 75821599 | 9.522391 | 17.17176 | 33.88007 | 14.75168 | -0.56703 | 0.3544 | 0.455833 |
| 14 | 78544400 | 78546599 | 20.64603 | 34.42718 | 62.75873 | 27.28257 | -0.48214 | 1.215155 | 0.455838 |
| 4 | 81192600 | 81193399 | 10.10349 | 19.59571 | 18.62171 | 7.024957 | 0.650379 | -0.20704 | 0.45588 |
| 1 | 149373600 | 149374599 | 19.15236 | 33.99506 | 26.6017 | 11.74835 | 0.745082 | 0.501439 | 0.455889 |
| 10 | 128583600 | 128584599 | 7.840085 | 14.67279 | 29.03495 | 12.01105 | -0.51792 | 0.110088 | 0.455914 |
| 3 | 12069000 | 12070599 | 18.75018 | 34.20852 | 25.79655 | 11.06956 | 0.824849 | 0.335082 | 0.455918 |
| 7 | 110879400 | 110881799 | 16.23474 | 28.28194 | 48.4205 | 21.2948 | -0.38496 | 0.758201 | 0.45592 |
| 1 | 8174400 | 8180399 | 93.32156 | 146.4567 | 136.3439 | 56.54654 | 0.42162 | 2.583526 | 0.455923 |
| 18 | 78843600 | 78846199 | 14.09032 | 24.69897 | 43.00239 | 18.99879 | -0.40853 | 0.643354 | 0.455941 |
| 13 | 55509800 | 55510799 | 6.322093 | 12.02508 | 26.62457 | 10.67765 | -0.63953 | -0.01738 | 0.455947 |
| 1 | 130172400 | 130175399 | 35.73013 | 59.08711 | 51.5086 | 22.26629 | 0.578001 | 1.34135 | 0.45595 |
| 15 | 72826000 | 72827599 | 16.2227 | 28.11772 | 47.93262 | 21.03035 | -0.37971 | 0.822289 | 0.455952 |
| 16 | 42650000 | 42651599 | 19.32112 | 34.74268 | 28.32126 | 12.37164 | 0.696927 | 0.425837 | 0.455968 |
| 3 | 37813200 | 37815599 | 22.79148 | 38.25113 | 65.41855 | 28.55505 | -0.39069 | 1.153346 | 0.455968 |
| 3 | 37763800 | 37765199 | 12.67297 | 22.50557 | 38.753 | 17.10761 | -0.38614 | 0.479955 | 0.455984 |
| 3 | 116770400 | 116771999 | 20.34408 | 36.77138 | 26.65729 | 11.57334 | 0.870805 | 0.400664 | 0.455984 |
| 1 | 139312800 | 139314199 | 9.233054 | 17.31556 | 34.11453 | 14.06295 | -0.50818 | 0.120713 | 0.455995 |
| 12 | 93580600 | 93581199 | 4.936515 | 9.631062 | 20.98048 | 7.49498 | -0.46061 | -0.44979 | 0.456001 |
| 5 | 6510400 | 6517199 | 67.1065 | 107.3412 | 91.99794 | 38.71857 | 0.565502 | 2.064051 | 0.456007 |
| 4 | 61529200 | 61531199 | 14.98332 | 28.30563 | 22.84013 | 9.293035 | 0.785738 | 0.09675 | 0.456021 |
| X | 148170000 | 148172599 | 14.98332 | 28.30563 | 22.84013 | 9.293035 | 0.785738 | 0.09675 | 0.456021 |
| 2 | 113270400 | 113272999 | 15.54977 | 26.60162 | 51.5863 | 22.62018 | -0.56845 | 0.943625 | 0.456028 |
| 19 | 21177200 | 21178799 | 20.92674 | 36.68358 | 34.82143 | 15.38474 | 0.462364 | 0.667456 | 0.456036 |
| 12 | 58257600 | 58261799 | 30.11671 | 48.40561 | 110.3434 | 46.77378 | -0.83473 | 1.877841 | 0.456038 |

| | | | | | | | | | |
|---|---|---|---|---|---|---|---|---|---|
| 10 | 106645600 | 106646399 | 10.19143 | 19.69082 | 20.25951 | 7.780593 | 0.513052 | -0.14102 | 0.456052 |
| X | 43609400 | 43613199 | 43.44576 | 71.5199 | 57.37016 | 24.73835 | 0.696192 | 1.420906 | 0.456068 |
| 14 | 102088200 | 102089599 | 16.03875 | 28.93779 | 28.10261 | 12.22816 | 0.449305 | 0.381962 | 0.456073 |
| X | 31645800 | 31646799 | 38.05324 | 63.63412 | 49.22341 | 21.43685 | 0.750385 | 1.226193 | 0.456095 |
| 17 | 75205200 | 75206799 | 12.03942 | 21.48351 | 37.27486 | 16.38851 | -0.39282 | 0.443529 | 0.456113 |
| 9 | 47926000 | 47927999 | 13.64275 | 24.10346 | 42.38809 | 18.7517 | -0.41994 | 0.540384 | 0.456129 |
| 7 | 17210000 | 17213199 | 40.75044 | 66.34102 | 66.28933 | 28.45928 | 0.371457 | 1.564905 | 0.456132 |
| 3 | 46880200 | 46883599 | 37.75082 | 62.78935 | 52.68539 | 22.86575 | 0.633484 | 1.282917 | 0.456149 |
| 10 | 14123800 | 14124599 | 10.56748 | 20.61529 | 14.70898 | 5.260225 | 1.128205 | -0.39387 | 0.456157 |
| 4 | 39136200 | 39137599 | 23.99302 | 41.79859 | 37.49055 | 16.48823 | 0.543484 | 0.778392 | 0.456159 |
| 14 | 43051800 | 43054199 | 20.29524 | 35.71761 | 33.13416 | 14.66934 | 0.495924 | 0.605239 | 0.456185 |
| 16 | 37131600 | 37132399 | 8.34354 | 15.61966 | 30.64179 | 12.66938 | -0.5046 | 0.1196 | 0.456196 |
| 7 | 86352600 | 86356599 | 61.25458 | 97.74165 | 100.0187 | 41.94882 | 0.306377 | 2.134592 | 0.456203 |
| 3 | 87929000 | 87942599 | 105.657 | 160.2862 | 272.6507 | 112.8715 | -0.48531 | 3.234647 | 0.456208 |
| 17 | 90069800 | 90071599 | 12.78026 | 22.69809 | 42.9524 | 18.96004 | -0.52186 | 0.48479 | 0.456218 |
| 5 | 4143600 | 4145999 | 21.81884 | 38.32657 | 33.5954 | 14.86901 | 0.576678 | 0.638465 | 0.456237 |
| 5 | 31357400 | 31357999 | 34.75685 | 57.19733 | 51.8873 | 22.37096 | 0.519057 | 1.399817 | 0.456281 |
| 11 | 10616400 | 10617999 | 16.38631 | 29.86578 | 27.09436 | 11.64108 | 0.558664 | 0.325542 | 0.456286 |
| 1 | 12245600 | 12248799 | 31.96093 | 53.2696 | 51.9867 | 22.59301 | 0.416169 | 1.240965 | 0.4563 |
| 6 | 134843000 | 134843799 | 8.430188 | 15.68196 | 31.59268 | 13.20378 | -0.55621 | 0.157801 | 0.456312 |
| 2 | 143604600 | 143607199 | 20.68598 | 36.54454 | 31.3342 | 13.86186 | 0.610864 | 0.562518 | 0.456329 |
| 5 | 95622000 | 95623399 | 17.92343 | 31.74744 | 29.93742 | 13.23699 | 0.475608 | 0.513418 | 0.456342 |
| 7 | 60801400 | 60807799 | 77.12502 | 122.6693 | 106.0673 | 44.28748 | 0.544066 | 2.253242 | 0.456345 |
| X | 54545600 | 54548799 | 31.72906 | 54.08599 | 43.49206 | 19.06541 | 0.695568 | 1.028827 | 0.456347 |
| 7 | 59033800 | 59044399 | 228.5981 | 335.033 | 322.575 | 132.8704 | 0.29007 | 3.877043 | 0.456348 |
| 17 | 19445000 | 19460799 | 241.6252 | 355.9128 | 299.5377 | 123.3522 | 0.490978 | 3.799279 | 0.456363 |
| 15 | 19525200 | 19526999 | 25.07568 | 43.2196 | 41.39761 | 18.16003 | 0.446642 | 0.900533 | 0.456366 |
| 6 | 29237000 | 29238399 | 13.48532 | 24.82687 | 25.42378 | 10.7788 | 0.398424 | 0.244485 | 0.45638 |
| 6 | 123228200 | 123229199 | 11.82514 | 22.65115 | 21.02181 | 8.292066 | 0.624332 | -0.03735 | 0.456387 |
| 8 | 3479400 | 3487399 | 50.22658 | 79.94102 | 127.4805 | 53.27849 | -0.33656 | 2.21204 | 0.456398 |
| 11 | 74499800 | 74503199 | 25.71499 | 42.29523 | 72.23288 | 31.13588 | -0.39143 | 1.396444 | 0.456399 |
| 7 | 46021800 | 46030799 | 69.11793 | 107.7352 | 177.6986 | 73.74817 | -0.4102 | 2.695233 | 0.456402 |
| 12 | 25237200 | 25239199 | 19.35896 | 33.85277 | 34.09321 | 15.03202 | 0.37747 | 0.696035 | 0.456424 |
| 14 | 37805000 | 37805999 | 8.705767 | 16.45812 | 33.3304 | 13.54088 | -0.53028 | 0.06025 | 0.456429 |
| 18 | 9469400 | 9474999 | 84.37663 | 132.7332 | 132.7266 | 55.06332 | 0.32121 | 2.531195 | 0.456447 |
| 13 | 91888600 | 91890999 | 15.40013 | 26.7178 | 47.4029 | 20.79855 | -0.43683 | 0.807986 | 0.456454 |
| 5 | 9836600 | 9837799 | 9.030181 | 16.85413 | 31.12914 | 12.93686 | -0.42512 | 0.14157 | 0.456456 |
| 7 | 19834800 | 19835999 | 15.22842 | 28.00108 | 26.3379 | 11.18529 | 0.518614 | 0.267984 | 0.456475 |
| 5 | 120728200 | 120728999 | 10.74844 | 20.40368 | 21.15921 | 8.521581 | 0.439636 | 0.021642 | 0.456482 |
| 9 | 114254200 | 114260799 | 62.3637 | 97.89781 | 158.2368 | 65.62855 | -0.37303 | 2.564709 | 0.456521 |
| 14 | 60099800 | 60102599 | 24.36245 | 40.64027 | 69.85547 | 30.3752 | -0.39783 | 1.22861 | 0.456528 |
| X | 121504400 | 121507999 | 37.59619 | 62.96661 | 48.63128 | 21.19955 | 0.752579 | 1.208873 | 0.45653 |
| 3 | 92344200 | 92344799 | 6.691655 | 12.94605 | 26.15727 | 9.983675 | -0.44301 | -0.18154 | 0.456533 |
| 13 | 19993800 | 19995999 | 18.4853 | 32.47969 | 32.27218 | 14.28662 | 0.396555 | 0.610896 | 0.456548 |
| 13 | 27347600 | 27347999 | 5.660385 | 10.99523 | 23.39007 | 8.739184 | -0.48589 | -0.27075 | 0.456549 |
| 3 | 103369400 | 103370399 | 7.979954 | 15.03076 | 28.59871 | 11.70264 | -0.44892 | 0.070048 | 0.45656 |
| 10 | 43664600 | 43665399 | 7.161021 | 13.82641 | 29.03818 | 11.18741 | -0.5114 | -0.13628 | 0.456562 |

| | | | | | | | | | |
|---|---|---|---|---|---|---|---|---|---|
| 2 | 150073800 | 150077799 | 111.2613 | 171.9468 | 164.493 | 68.32675 | 0.365274 | 2.864116 | 0.456568 |
| 4 | 16804800 | 16806199 | 14.8245 | 27.55878 | 24.731 | 10.34739 | 0.600945 | 0.198956 | 0.456574 |
| 4 | 6102400 | 6103199 | 13.09594 | 24.46569 | 24.44872 | 10.14429 | 0.456535 | 0.15362 | 0.456575 |
| X | 89466800 | 89468399 | 20.20545 | 35.73852 | 31.49702 | 13.93055 | 0.572243 | 0.548931 | 0.456586 |
| 13 | 68915200 | 68916599 | 10.84388 | 19.48506 | 38.37158 | 16.77216 | -0.56901 | 0.389574 | 0.456606 |
| 11 | 9099400 | 9100799 | 19.29079 | 32.06187 | 69.60729 | 30.19454 | -0.73349 | 1.25559 | 0.45661 |
| 16 | 47996200 | 47998399 | 25.30325 | 43.69634 | 40.25191 | 17.65849 | 0.503214 | 0.882038 | 0.456614 |
| 16 | 30802800 | 30804999 | 13.17618 | 23.08412 | 42.02735 | 18.55954 | -0.4725 | 0.646345 | 0.45662 |
| 2 | 171232000 | 171232999 | 12.28323 | 21.56865 | 40.50256 | 17.92055 | -0.51653 | 0.604687 | 0.456622 |
| 13 | 9544400 | 9545399 | 7.625947 | 14.50381 | 27.7333 | 11.12441 | -0.43113 | -0.00374 | 0.456641 |
| X | 52887600 | 52889399 | 24.03407 | 42.03469 | 35.28343 | 15.55887 | 0.639152 | 0.721686 | 0.456641 |
| 5 | 57050200 | 57051999 | 22.20239 | 37.07202 | 62.95615 | 27.39153 | -0.38035 | 1.205032 | 0.456644 |
| 15 | 96114800 | 96116999 | 22.20239 | 37.07202 | 62.95615 | 27.39153 | -0.38035 | 1.205032 | 0.456644 |
| 7 | 133288600 | 133293399 | 74.9359 | 116.5841 | 178.5616 | 74.12718 | -0.30555 | 2.732094 | 0.456644 |
| 6 | 133590400 | 133592999 | 29.30155 | 50.54064 | 38.41438 | 16.85175 | 0.779079 | 0.908002 | 0.456646 |
| 1 | 42221400 | 42223199 | 16.42383 | 29.21182 | 27.58104 | 12.15982 | 0.476923 | 0.469213 | 0.456658 |
| 7 | 51931600 | 51934999 | 33.67807 | 56.94499 | 45.16378 | 19.76445 | 0.714433 | 1.104744 | 0.456668 |
| 5 | 131157600 | 131161399 | 35.91934 | 58.0105 | 94.27229 | 40.22737 | -0.33926 | 1.731926 | 0.45667 |
| 7 | 23224200 | 23239199 | 343.8112 | 487.8589 | 437.9529 | 181.1692 | 0.345107 | 4.365076 | 0.456704 |
| 13 | 98775400 | 98779399 | 36.08299 | 58.24657 | 94.74554 | 40.40406 | -0.34143 | 1.74638 | 0.456711 |
| 9 | 114468400 | 114470399 | 15.07353 | 26.34236 | 44.65356 | 19.67776 | -0.37013 | 0.701342 | 0.456727 |
| 9 | 73722400 | 73724599 | 17.04576 | 31.19533 | 25.34813 | 10.8342 | 0.721247 | 0.302769 | 0.456733 |
| 3 | 148525000 | 148527799 | 21.35481 | 35.98912 | 59.72786 | 26.1118 | -0.34714 | 1.107335 | 0.456734 |
| 8 | 117224200 | 117225599 | 13.987 | 24.56315 | 41.93274 | 18.55511 | -0.38011 | 0.613392 | 0.456745 |
| 7 | 87779000 | 87780399 | 13.49675 | 23.64112 | 42.03091 | 18.55805 | -0.43838 | 0.651829 | 0.456746 |
| 15 | 93492200 | 93494599 | 14.11994 | 24.63026 | 44.70514 | 19.67712 | -0.46853 | 0.722229 | 0.456752 |
| 7 | 10126800 | 10128399 | 30.1248 | 50.79789 | 48.54316 | 21.22572 | 0.446404 | 1.119606 | 0.456757 |
| 3 | 23619200 | 23628999 | 197.6152 | 292.9132 | 285.4273 | 117.5563 | 0.288081 | 3.690422 | 0.456779 |
| 17 | 14233000 | 14234799 | 18.95313 | 31.56364 | 68.58106 | 29.78947 | -0.73449 | 1.230236 | 0.456803 |
| X | 43061400 | 43064199 | 23.97003 | 42.11484 | 32.41505 | 14.34934 | 0.763431 | 0.646368 | 0.456805 |
| 1 | 54544200 | 54544799 | 5.666953 | 11.02303 | 24.24099 | 8.957799 | -0.52006 | -0.31229 | 0.45683 |
| 2 | 118257000 | 118257599 | 10.65857 | 20.55376 | 15.56941 | 6.024399 | 0.941497 | -0.12388 | 0.456831 |
| 5 | 22933400 | 22934799 | 17.51112 | 31.39514 | 28.95008 | 12.67786 | 0.517508 | 0.431273 | 0.456835 |
| 7 | 19014800 | 19018199 | 19.75071 | 33.21392 | 59.08463 | 25.81368 | -0.44703 | 1.118978 | 0.456845 |
| 7 | 42557600 | 42576199 | 338.5147 | 481.1631 | 433.2125 | 179.1569 | 0.343346 | 4.339789 | 0.456857 |
| 6 | 56900400 | 56901399 | 11.86172 | 21.21366 | 36.76997 | 16.13616 | -0.38953 | 0.426937 | 0.456862 |
| 4 | 40301000 | 40302399 | 10.64987 | 19.43718 | 34.96647 | 14.99755 | -0.42116 | 0.292327 | 0.456889 |
| 19 | 13172200 | 13174599 | 36.23553 | 59.85408 | 57.70019 | 24.92304 | 0.433151 | 1.35966 | 0.456896 |
| 14 | 14638800 | 14639599 | 10.90012 | 21.04724 | 20.79976 | 8.010892 | 0.567067 | -0.12152 | 0.456915 |
| 15 | 45526200 | 45527199 | 7.597396 | 14.53855 | 28.54008 | 11.28952 | -0.45102 | -0.04867 | 0.45692 |
| 15 | 46885000 | 46888399 | 37.3659 | 62.09632 | 53.25981 | 23.10149 | 0.601825 | 1.291139 | 0.456937 |
| X | 109143400 | 109144399 | 14.94924 | 27.84019 | 24.90442 | 10.38863 | 0.609445 | 0.191823 | 0.456937 |
| X | 62023200 | 62024399 | 15.95379 | 30.09683 | 22.78931 | 9.299842 | 0.873314 | 0.115337 | 0.456938 |
| 9 | 101172600 | 101173399 | 12.56752 | 23.28163 | 21.06196 | 8.854231 | 0.584847 | 0.191565 | 0.45694 |
| X | 8494600 | 8499599 | 81.7649 | 130.0192 | 104.1521 | 43.47531 | 0.653811 | 2.265308 | 0.45697 |
| 15 | 75964400 | 75965799 | 15.75455 | 27.37795 | 46.85944 | 20.5764 | -0.38481 | 0.78914 | 0.456981 |
| 5 | 120893600 | 120900399 | 57.64364 | 90.86362 | 150.1623 | 62.3132 | -0.40038 | 2.476687 | 0.456993 |

| | | | | | | | | | |
|---|---|---|---|---|---|---|---|---|---|
| 12 | 107616600 | 107620199 | 27.23723 | 44.51074 | 77.06239 | 33.13026 | -0.4156 | 1.486618 | 0.457027 |
| 4 | 105451000 | 105452799 | 14.06181 | 24.42832 | 50.54683 | 22.19099 | -0.65724 | 0.791875 | 0.457031 |
| 5 | 5644000 | 5645399 | 10.67508 | 19.40591 | 34.36732 | 14.81403 | -0.40323 | 0.314895 | 0.45704 |
| 10 | 74178200 | 74179599 | 10.67508 | 19.40591 | 34.36732 | 14.81403 | -0.40323 | 0.314895 | 0.45704 |
| X | 26395600 | 26397799 | 33.39877 | 56.1588 | 42.15096 | 18.41483 | 0.793593 | 1.150952 | 0.457057 |
| 1 | 85003200 | 85008399 | 224.4836 | 328.5485 | 315.5366 | 129.9824 | 0.291798 | 3.89769 | 0.457063 |
| 10 | 126303000 | 126303599 | 14.01187 | 25.82425 | 25.90829 | 10.96802 | 0.429377 | 0.244098 | 0.457072 |
| 14 | 92529400 | 92532399 | 26.35522 | 45.37902 | 41.33465 | 18.13185 | 0.518573 | 0.914366 | 0.457092 |
| X | 120959600 | 120961799 | 31.60334 | 53.21744 | 48.55352 | 21.22135 | 0.513006 | 1.138213 | 0.457093 |
| 16 | 16470400 | 16471199 | 13.2229 | 23.19044 | 41.51588 | 18.35004 | -0.44811 | 0.631108 | 0.4571 |
| 2 | 84657200 | 84658199 | 6.556946 | 12.64095 | 25.32869 | 9.808041 | -0.45061 | -0.13597 | 0.457111 |
| 5 | 141488200 | 141489199 | 6.556946 | 12.64095 | 25.32869 | 9.808041 | -0.45061 | -0.13597 | 0.457111 |
| 5 | 6097000 | 6098399 | 19.24901 | 33.77755 | 33.87849 | 14.98123 | 0.383231 | 0.639979 | 0.457129 |
| 15 | 4989800 | 4992199 | 28.32868 | 47.57748 | 47.50254 | 20.74389 | 0.383152 | 1.145246 | 0.457131 |
| 11 | 68176800 | 68179599 | 17.4463 | 29.6454 | 55.32009 | 24.24423 | -0.51503 | 1.015365 | 0.457136 |
| 13 | 61098800 | 61101799 | 36.65727 | 60.53316 | 52.11281 | 22.50512 | 0.595833 | 1.359707 | 0.45714 |
| X | 128649800 | 128651599 | 23.9029 | 41.98103 | 27.52701 | 12.18164 | 0.993485 | 0.644149 | 0.457153 |
| 15 | 95981800 | 95986199 | 34.84242 | 56.14146 | 94.07922 | 40.02446 | -0.38727 | 1.798995 | 0.457184 |
| 1 | 121101000 | 121103799 | 29.4466 | 49.56323 | 49.10057 | 21.45751 | 0.394518 | 1.133165 | 0.45719 |
| 4 | 112203600 | 112221999 | 381.9129 | 536.7712 | 481.6756 | 199.5691 | 0.331805 | 4.506385 | 0.457197 |
| X | 4465000 | 4466399 | 30.64032 | 51.9554 | 46.29769 | 20.27806 | 0.547139 | 1.072533 | 0.457216 |
| 4 | 66659200 | 66662199 | 40.09315 | 64.31445 | 104.4864 | 44.16137 | -0.35062 | 1.950932 | 0.45723 |
| 7 | 144316200 | 144319799 | 26.08072 | 42.78901 | 73.15356 | 31.4992 | -0.39439 | 1.43007 | 0.457237 |
| 8 | 32314200 | 32314799 | 10.47709 | 20.06212 | 21.24515 | 8.388334 | 0.435151 | -0.04396 | 0.457242 |
| 17 | 85679600 | 85682599 | 17.66108 | 29.87814 | 56.344 | 24.66103 | -0.5304 | 1.056857 | 0.457244 |
| X | 157676600 | 157677599 | 13.98956 | 26.03958 | 24.49126 | 10.22517 | 0.536746 | 0.186837 | 0.457289 |
| 6 | 65232800 | 65233199 | 7.406622 | 14.48674 | 14.15089 | 4.766485 | 0.75782 | -0.64308 | 0.457315 |
| 17 | 19079000 | 19085199 | 98.96265 | 154.8232 | 141.298 | 58.61755 | 0.446753 | 2.643216 | 0.457316 |
| 3 | 17170400 | 17172999 | 26.10678 | 44.26828 | 45.30808 | 19.84464 | 0.348009 | 1.054276 | 0.45732 |
| 10 | 104676600 | 104680399 | 35.66511 | 58.82931 | 58.69598 | 25.33153 | 0.383387 | 1.37634 | 0.457379 |
| 5 | 108206800 | 108209999 | 36.74718 | 60.39146 | 60.41826 | 26.03424 | 0.377379 | 1.423669 | 0.457379 |
| 7 | 27679800 | 27692399 | 254.1436 | 372.3373 | 318.7965 | 131.317 | 0.458799 | 3.886172 | 0.457386 |
| 11 | 82399200 | 82400799 | 11.8188 | 21.28156 | 37.87131 | 16.51987 | -0.42152 | 0.385134 | 0.457418 |
| X | 155467600 | 155469799 | 28.25069 | 48.88499 | 38.6337 | 16.94962 | 0.724208 | 0.879798 | 0.457431 |
| 10 | 48905600 | 48908799 | 43.29944 | 71.69621 | 51.7334 | 22.38324 | 0.850094 | 1.348178 | 0.457443 |
| 3 | 114059400 | 114061199 | 21.24849 | 37.55213 | 31.43977 | 13.90754 | 0.645491 | 0.564373 | 0.457449 |
| X | 62731600 | 62736199 | 60.65176 | 97.38205 | 85.25163 | 36.08188 | 0.54123 | 1.943374 | 0.457452 |
| 12 | 106878600 | 106879999 | 10.95566 | 19.45673 | 36.80218 | 16.24578 | -0.52061 | 0.464536 | 0.45747 |
| 15 | 99084000 | 99085199 | 6.59424 | 12.59675 | 26.16073 | 10.39668 | -0.53675 | -0.04694 | 0.457484 |
| 14 | 112092000 | 112095199 | 30.00047 | 52.07401 | 35.60093 | 15.62134 | 0.933055 | 0.854374 | 0.457509 |
| 7 | 130191600 | 130193199 | 7.417456 | 13.85933 | 29.03433 | 12.04383 | -0.60226 | 0.118647 | 0.457521 |
| 8 | 27249400 | 27254399 | 54.81469 | 87.92551 | 86.99107 | 36.76429 | 0.363363 | 1.961329 | 0.45754 |
| 5 | 71856800 | 71858799 | 14.44318 | 25.13986 | 45.7865 | 20.12579 | -0.47381 | 0.756196 | 0.457546 |
| 13 | 16981200 | 16982399 | 14.76122 | 26.60536 | 26.33191 | 11.47248 | 0.421059 | 0.375647 | 0.457558 |
| 16 | 89854800 | 89855399 | 17.89354 | 31.77956 | 25.98617 | 11.47076 | 0.682093 | 0.485541 | 0.457578 |
| 3 | 51286000 | 51286799 | 5.683687 | 11.00871 | 23.81485 | 9.0406 | -0.53241 | -0.21745 | 0.457584 |
| 7 | 110182800 | 110183599 | 5.683687 | 11.00871 | 23.81485 | 9.0406 | -0.53241 | -0.21745 | 0.457584 |

| | | | | | | | | | |
|---|---|---|---|---|---|---|---|---|---|
| 5 | 60788000 | 60788399 | 3.754258 | 7.344104 | 17.45914 | 5.777413 | -0.48223 | -0.74692 | 0.457614 |
| 5 | 122817000 | 122819399 | 17.66449 | 29.85634 | 57.42359 | 25.12652 | -0.55884 | 1.065352 | 0.457625 |
| 6 | 56913800 | 56914599 | 8.396996 | 15.76385 | 30.17209 | 12.41611 | -0.46468 | 0.095901 | 0.457636 |
| 4 | 113574000 | 113578399 | 60.41665 | 97.29542 | 77.03948 | 32.74898 | 0.691039 | 1.849433 | 0.457637 |
| 8 | 127447200 | 127458599 | 85.93066 | 131.8346 | 235.2091 | 97.64457 | -0.54106 | 2.998713 | 0.457643 |
| 12 | 20881600 | 20909399 | 738.6208 | 980.5378 | 845.6487 | 353.4287 | 0.306235 | 5.396796 | 0.457645 |
| 9 | 56823400 | 56826399 | 19.45448 | 32.74845 | 57.95256 | 25.32675 | -0.43935 | 1.109773 | 0.457661 |
| 13 | 34623200 | 34625199 | 11.54688 | 20.55775 | 36.75652 | 16.19165 | -0.43789 | 0.452318 | 0.457729 |
| 14 | 123693000 | 123696199 | 38.37394 | 63.36145 | 57.73506 | 24.93135 | 0.514217 | 1.371718 | 0.457733 |
| 1 | 24848400 | 24851399 | 30.47697 | 52.14808 | 42.37086 | 18.57956 | 0.681382 | 0.988792 | 0.457765 |
| 16 | 5011000 | 5013999 | 16.74959 | 28.70057 | 51.62955 | 22.64209 | -0.46038 | 0.931945 | 0.457778 |
| 2 | 40151400 | 40154399 | 32.76419 | 54.94979 | 47.49628 | 20.72283 | 0.590437 | 1.172657 | 0.457785 |
| 14 | 52758800 | 52759599 | 12.04436 | 21.33039 | 37.76639 | 16.69917 | -0.42802 | 0.4963 | 0.457793 |
| 7 | 42585400 | 42586799 | 31.80757 | 53.25394 | 51.35895 | 22.38388 | 0.43316 | 1.195038 | 0.457794 |
| 4 | 149903800 | 149907599 | 20.05814 | 33.85551 | 57.48986 | 25.14503 | -0.37993 | 1.087472 | 0.457804 |
| 5 | 12039200 | 12043799 | 55.26041 | 88.41071 | 90.31576 | 38.02161 | 0.312986 | 2.044172 | 0.457807 |
| 18 | 4541400 | 4544799 | 83.65254 | 131.1721 | 132.7327 | 55.05122 | 0.300546 | 2.593164 | 0.457811 |
| 5 | 42293000 | 42294399 | 10.06075 | 18.23585 | 34.34222 | 14.85732 | -0.49478 | 0.326406 | 0.457833 |
| 10 | 129192400 | 129202599 | 227.2387 | 333.1598 | 321.5375 | 132.44 | 0.287121 | 3.871483 | 0.457844 |
| 15 | 29284200 | 29285799 | 11.01184 | 20.10045 | 37.70407 | 16.1691 | -0.48102 | 0.300706 | 0.457846 |
| 2 | 99902600 | 99904799 | 25.38198 | 43.73761 | 41.37703 | 18.15088 | 0.464396 | 0.903547 | 0.457848 |
| 18 | 21644800 | 21645399 | 4.518472 | 8.812704 | 20.3022 | 7.282507 | -0.54504 | -0.44242 | 0.457867 |
| 18 | 17351000 | 17351599 | 5.65925 | 11.01659 | 23.08178 | 8.476174 | -0.44186 | -0.33574 | 0.45787 |
| 16 | 48259200 | 48260599 | 9.558429 | 17.80544 | 33.2911 | 13.88093 | -0.4464 | 0.165083 | 0.457882 |
| 18 | 11285200 | 11289399 | 53.85108 | 85.53398 | 133.9063 | 55.83145 | -0.31302 | 2.279362 | 0.457891 |
| 5 | 145645800 | 145651799 | 128.7417 | 196.8582 | 190.7774 | 79.14737 | 0.334887 | 3.075737 | 0.457893 |
| 7 | 109783400 | 109783799 | 5.803022 | 11.29497 | 23.68331 | 8.706331 | -0.44442 | -0.32797 | 0.457929 |
| 10 | 8645600 | 8648799 | 31.1972 | 50.61547 | 83.60788 | 35.825 | -0.356 | 1.614359 | 0.457935 |
| 10 | 36006400 | 36008799 | 26.74642 | 46.68192 | 35.18236 | 15.49117 | 0.793793 | 0.765444 | 0.457936 |
| 13 | 88904800 | 88908999 | 56.1114 | 88.7777 | 156.2085 | 64.9257 | -0.48646 | 2.39183 | 0.457964 |
| 7 | 79473600 | 79478399 | 39.25859 | 63.09121 | 101.0265 | 42.81669 | -0.3266 | 1.891328 | 0.457991 |
| 9 | 22386800 | 22387399 | 5.981693 | 11.54228 | 24.77179 | 9.567735 | -0.54512 | -0.15245 | 0.457993 |
| 13 | 105386200 | 105386999 | 5.981693 | 11.54228 | 24.77179 | 9.567735 | -0.54512 | -0.15245 | 0.457993 |
| X | 19270400 | 19272999 | 24.08652 | 42.96122 | 26.8625 | 11.81506 | 1.070891 | 0.505037 | 0.457998 |
| X | 5699200 | 5700999 | 30.84942 | 52.16014 | 47.42566 | 20.75404 | 0.518084 | 1.099012 | 0.458011 |
| 8 | 44093200 | 44094199 | 18.43556 | 32.70211 | 26.6673 | 11.78359 | 0.685347 | 0.500289 | 0.45802 |
| X | 78446800 | 78448399 | 16.72475 | 31.60046 | 21.58005 | 8.777385 | 1.025086 | 0.108549 | 0.458045 |
| 7 | 41657000 | 41662199 | 68.06066 | 109.1728 | 83.4706 | 35.2765 | 0.734577 | 1.980724 | 0.458047 |
| 2 | 174997000 | 174999799 | 60.96169 | 97.4102 | 97.56836 | 41.00212 | 0.339401 | 2.091683 | 0.458077 |
| 1 | 60382200 | 60383199 | 8.193735 | 15.14837 | 30.47055 | 12.84488 | -0.56247 | 0.191659 | 0.45809 |
| 4 | 19966400 | 19968199 | 20.04206 | 35.10891 | 34.91135 | 15.41561 | 0.395647 | 0.672912 | 0.458092 |
| 9 | 5605600 | 5606399 | 11.59662 | 22.17107 | 22.22618 | 8.815279 | 0.507378 | -0.01629 | 0.458122 |
| 7 | 105992600 | 105993799 | 16.40508 | 29.3533 | 29.13299 | 12.77952 | 0.410424 | 0.436852 | 0.458124 |
| 14 | 94885800 | 94887799 | 52.53567 | 84.24447 | 86.36871 | 36.48455 | 0.311649 | 1.970014 | 0.458127 |
| 8 | 86313600 | 86317399 | 24.95868 | 41.34132 | 72.16173 | 31.22925 | -0.42057 | 1.308356 | 0.458146 |
| 10 | 49178000 | 49188199 | 51.36476 | 83.22888 | 61.88368 | 26.4913 | 0.791029 | 1.665993 | 0.45816 |
| 6 | 67151400 | 67153199 | 20.45407 | 35.86943 | 34.22214 | 15.1251 | 0.454915 | 0.655916 | 0.458162 |

| | | | | | | | | | |
|---|---|---|---|---|---|---|---|---|---|
| 8 | 33341800 | 33344199 | 30.28934 | 49.03898 | 92.93149 | 39.75819 | -0.55657 | 1.660952 | 0.458165 |
| X | 24120000 | 24123799 | 48.8969 | 78.98689 | 74.92602 | 31.9811 | 0.436094 | 1.740553 | 0.458166 |
| 3 | 50870600 | 50872399 | 15.45752 | 26.63867 | 49.51263 | 21.71979 | -0.50568 | 0.871608 | 0.458178 |
| 7 | 9976600 | 9979199 | 35.90688 | 60.75191 | 44.02874 | 19.27142 | 0.844161 | 1.108305 | 0.458201 |
| X | 33619000 | 33619799 | 13.0489 | 25.2534 | 19.00238 | 7.241726 | 0.971008 | -0.13794 | 0.458206 |
| 11 | 14340000 | 14340599 | 9.820399 | 19.15012 | 16.67005 | 5.993237 | 0.838669 | -0.3792 | 0.458212 |
| 7 | 20944000 | 20951999 | 66.87725 | 106.7267 | 90.69888 | 38.04665 | 0.574082 | 2.132985 | 0.458212 |
| 6 | 35314200 | 35315599 | 13.17403 | 24.32722 | 24.96427 | 10.54054 | 0.399727 | 0.22443 | 0.458234 |
| 7 | 19637200 | 19651399 | 175.1637 | 256.5338 | 385.0107 | 158.5951 | -0.35102 | 3.88472 | 0.458241 |
| 4 | 78906400 | 78907399 | 18.15556 | 31.85912 | 30.49686 | 13.48597 | 0.450212 | 0.627026 | 0.458264 |
| 13 | 66541400 | 66555999 | 202.9848 | 303.5327 | 225.5615 | 93.01056 | 0.690425 | 3.532645 | 0.458266 |
| 10 | 110984600 | 110985999 | 16.28476 | 29.14121 | 29.18711 | 12.80207 | 0.397516 | 0.436011 | 0.458281 |
| 2 | 93121800 | 93131799 | 101.7775 | 155.1101 | 235.724 | 97.71858 | -0.31742 | 3.137511 | 0.458294 |
| 19 | 56451800 | 56452999 | 10.92647 | 19.57886 | 35.30429 | 15.46772 | -0.44434 | 0.406153 | 0.458296 |
| 17 | 20180800 | 20190999 | 144.8505 | 221.2565 | 188.6328 | 78.23854 | 0.518416 | 3.102065 | 0.458315 |
| 4 | 135460000 | 135462599 | 32.01708 | 51.9412 | 84.26393 | 36.10382 | -0.33017 | 1.618557 | 0.458319 |
| 10 | 95840800 | 95843399 | 33.67536 | 56.52785 | 44.22526 | 19.30643 | 0.733819 | 1.166901 | 0.458319 |
| 10 | 79204000 | 79205599 | 15.3388 | 28.69054 | 23.68833 | 9.808635 | 0.732222 | 0.162959 | 0.458352 |
| 7 | 122401800 | 122402799 | 13.08064 | 24.33087 | 24.52036 | 10.24936 | 0.436307 | 0.182908 | 0.458352 |
| 10 | 73517200 | 73518399 | 16.46719 | 29.68052 | 28.64038 | 12.47806 | 0.457425 | 0.392921 | 0.458364 |
| 2 | 128941600 | 128943599 | 14.11646 | 24.81986 | 42.39675 | 18.7737 | -0.38095 | 0.596159 | 0.458382 |
| 2 | 52399800 | 52400599 | 6.579218 | 12.61456 | 25.7092 | 10.11328 | -0.4961 | -0.08241 | 0.458386 |
| 5 | 46442400 | 46446199 | 91.89149 | 142.9584 | 140.6735 | 58.39879 | 0.33198 | 2.734387 | 0.458389 |
| 9 | 19876600 | 19881599 | 127.1461 | 193.7692 | 194.9667 | 80.82242 | 0.276982 | 3.142029 | 0.458421 |
| 9 | 36346400 | 36350599 | 37.5675 | 62.37538 | 52.7438 | 22.86351 | 0.62214 | 1.299901 | 0.458425 |
| 12 | 40080400 | 40082999 | 22.76076 | 40.18639 | 30.63991 | 13.5563 | 0.779109 | 0.580726 | 0.458481 |
| 9 | 122567400 | 122569599 | 28.54553 | 46.73135 | 76.69033 | 32.99179 | -0.33682 | 1.466173 | 0.458489 |
| 6 | 146436400 | 146438399 | 18.52955 | 31.15047 | 63.89346 | 27.91194 | -0.65171 | 1.12334 | 0.458493 |
| 1 | 54644000 | 54644999 | 8.50043 | 15.66845 | 30.95026 | 13.10029 | -0.5409 | 0.211929 | 0.458522 |
| X | 152742200 | 152746599 | 59.73421 | 95.73415 | 90.55928 | 38.22218 | 0.42659 | 1.998621 | 0.458525 |
| 13 | 75285800 | 75288199 | 22.51993 | 39.34569 | 36.41431 | 16.04399 | 0.498803 | 0.729813 | 0.458528 |
| 7 | 39044600 | 39049999 | 111.3901 | 171.8731 | 153.0037 | 63.55486 | 0.467249 | 2.890083 | 0.458535 |
| 8 | 48358400 | 48363799 | 77.25098 | 119.1325 | 212.2013 | 88.14467 | -0.53296 | 2.894705 | 0.45854 |
| 1 | 136598200 | 136600999 | 18.17258 | 31.06739 | 53.25442 | 23.35035 | -0.39167 | 0.960291 | 0.458551 |
| 6 | 149483400 | 149488199 | 111.652 | 174.7554 | 126.3729 | 52.42338 | 0.782618 | 2.638476 | 0.458552 |
| 1 | 174614800 | 174615399 | 16.15573 | 28.75295 | 26.50092 | 11.67746 | 0.512495 | 0.464697 | 0.458563 |
| 2 | 147311400 | 147314799 | 29.704 | 48.28995 | 82.70953 | 35.49066 | -0.40549 | 1.569866 | 0.458564 |
| 19 | 13435200 | 13438999 | 79.58787 | 125.6952 | 110.4795 | 45.87737 | 0.511413 | 2.440121 | 0.458596 |
| 5 | 10122000 | 10122599 | 24.32206 | 41.52453 | 36.10978 | 15.83126 | 0.583314 | 0.975615 | 0.458604 |
| X | 5250200 | 5273999 | 334.2904 | 480.276 | 370.8477 | 153.0995 | 0.580324 | 4.184795 | 0.458613 |
| 11 | 107246600 | 107247999 | 7.675005 | 14.49702 | 28.11235 | 11.44175 | -0.46931 | 0.047643 | 0.458615 |
| 6 | 131198000 | 131203399 | 73.55973 | 117.1468 | 103.0668 | 43.11005 | 0.520971 | 2.208552 | 0.458626 |
| 8 | 69808800 | 69818599 | 203.1718 | 304.4805 | 237.1068 | 97.81303 | 0.62583 | 3.493848 | 0.458629 |
| X | 139796200 | 139799399 | 42.58887 | 69.10577 | 68.99683 | 29.56687 | 0.368724 | 1.626954 | 0.458644 |
| 16 | 22807000 | 22809799 | 38.33994 | 62.47702 | 64.52368 | 27.71099 | 0.324894 | 1.543489 | 0.458679 |
| 10 | 128204000 | 128205199 | 7.688742 | 14.39368 | 29.03938 | 12.00705 | -0.54512 | 0.106475 | 0.458682 |
| 9 | 94950200 | 94951999 | 26.11548 | 44.38706 | 45.27636 | 19.84291 | 0.353051 | 1.034183 | 0.458686 |

| | | | | | | | | | |
|---|---|---|---|---|---|---|---|---|---|
| 16 | 33328000 | 33329199 | 11.19278 | 20.37597 | 38.17385 | 16.42559 | -0.48238 | 0.318856 | 0.458692 |
| 8 | 53108400 | 53113999 | 80.94339 | 127.5088 | 129.2102 | 53.61416 | 0.303561 | 2.500138 | 0.458711 |
| 3 | 157116000 | 157117199 | 14.30984 | 25.16774 | 43.97761 | 19.4725 | -0.41335 | 0.59706 | 0.458711 |
| 4 | 147143200 | 147143999 | 16.91431 | 32.6107 | 16.89566 | 6.541324 | 1.483641 | -0.05162 | 0.458712 |
| 6 | 122851200 | 122852199 | 7.631168 | 14.05141 | 31.99342 | 13.5612 | -0.74598 | 0.211993 | 0.458716 |
| X | 8560000 | 8566399 | 108.0646 | 168.5594 | 143.7517 | 59.65374 | 0.541122 | 2.700835 | 0.458738 |
| 11 | 63088800 | 63092999 | 64.04704 | 102.1339 | 102.3191 | 42.87655 | 0.335974 | 2.15747 | 0.458741 |
| 3 | 111711400 | 111712399 | 10.93188 | 21.11096 | 20.17518 | 7.766211 | 0.615242 | -0.12613 | 0.458774 |
| 3 | 4275400 | 4278199 | 30.80152 | 51.70553 | 50.21873 | 21.92333 | 0.422927 | 1.163151 | 0.458815 |
| 11 | 17874800 | 17882199 | 80.17624 | 124.8748 | 205.9922 | 85.5005 | -0.41131 | 2.719053 | 0.458823 |
| 5 | 84035400 | 84037399 | 15.08439 | 26.41779 | 45.12292 | 19.92057 | -0.38108 | 0.66758 | 0.458854 |
| 17 | 78090200 | 78091599 | 12.67523 | 22.60814 | 40.37744 | 17.75918 | -0.43493 | 0.455464 | 0.458864 |
| 18 | 74400600 | 74402599 | 18.73933 | 31.66525 | 58.58824 | 25.63422 | -0.50329 | 1.07156 | 0.458909 |
| 14 | 66963800 | 66966999 | 43.00003 | 68.89354 | 109.1447 | 46.04487 | -0.31653 | 1.99304 | 0.458934 |
| 3 | 50045000 | 50046999 | 21.29798 | 37.64083 | 31.33594 | 13.86152 | 0.653527 | 0.562998 | 0.458939 |
| 10 | 99268200 | 99269999 | 18.65719 | 32.95219 | 31.54149 | 13.95426 | 0.452714 | 0.55265 | 0.458949 |
| X | 64503600 | 64505599 | 18.86495 | 34.82852 | 24.45629 | 10.32911 | 0.943315 | 0.270478 | 0.458957 |
| 2 | 65052200 | 65053599 | 8.442774 | 15.78157 | 29.53956 | 12.24493 | -0.44081 | 0.124551 | 0.458966 |
| 5 | 135543000 | 135545999 | 44.52154 | 71.07402 | 116.4247 | 48.86147 | -0.37048 | 2.108418 | 0.458969 |
| X | 71139600 | 71140999 | 22.43235 | 39.41712 | 33.07816 | 14.64413 | 0.63931 | 0.637101 | 0.458975 |
| 6 | 42096800 | 42099999 | 41.53373 | 67.71824 | 65.99796 | 28.34976 | 0.408867 | 1.544337 | 0.458978 |
| 10 | 3224000 | 3227799 | 61.46737 | 98.28176 | 87.25627 | 36.70218 | 0.514088 | 2.068403 | 0.458985 |
| 1 | 141112400 | 141115199 | 34.17535 | 55.25045 | 89.12206 | 38.07994 | -0.32677 | 1.698889 | 0.459009 |
| 6 | 130523200 | 130527799 | 56.37113 | 91.2336 | 67.47958 | 28.86176 | 0.797121 | 1.702444 | 0.459012 |
| 1 | 187215400 | 187217199 | 13.4257 | 23.70294 | 42.90475 | 18.98299 | -0.46195 | 0.545721 | 0.459022 |
| 16 | 8779600 | 8785599 | 39.02883 | 62.75157 | 102.9286 | 43.65268 | -0.36043 | 1.874724 | 0.459028 |
| 3 | 96634600 | 96636399 | 11.58339 | 20.52388 | 37.28696 | 16.48559 | -0.46462 | 0.487412 | 0.459042 |
| 8 | 107903800 | 107905599 | 24.0096 | 41.74273 | 38.65349 | 16.97804 | 0.497464 | 0.808344 | 0.459075 |
| 7 | 4814000 | 4815999 | 15.91403 | 27.07629 | 53.65258 | 23.51586 | -0.60073 | 0.998383 | 0.45908 |
| 10 | 49105400 | 49107799 | 20.89908 | 37.27753 | 29.25343 | 12.86624 | 0.744979 | 0.487599 | 0.459091 |
| 10 | 83476600 | 83478199 | 8.455932 | 15.83799 | 29.57607 | 12.21837 | -0.43335 | 0.112848 | 0.459092 |
| 6 | 140347400 | 140351799 | 35.73738 | 57.65277 | 98.9467 | 42.16175 | -0.41972 | 1.764329 | 0.459097 |
| 2 | 96028400 | 96030799 | 29.52828 | 50.04981 | 42.0474 | 18.41424 | 0.631782 | 1.066905 | 0.459109 |
| 6 | 122227400 | 122228599 | 6.600643 | 12.57973 | 26.66322 | 10.64857 | -0.57253 | -0.03086 | 0.45911 |
| 8 | 46712600 | 46713999 | 12.02456 | 21.62743 | 39.46037 | 17.23406 | -0.4585 | 0.397386 | 0.459124 |
| 3 | 34634400 | 34635399 | 11.7092 | 22.08213 | 23.09793 | 9.432085 | 0.40969 | 0.079987 | 0.459126 |
| 15 | 37832600 | 37837399 | 90.58492 | 141.7568 | 142.2287 | 59.00192 | 0.310545 | 2.632795 | 0.459133 |
| X | 164246800 | 164251999 | 52.52774 | 84.49297 | 81.90956 | 34.76072 | 0.397599 | 1.874731 | 0.459134 |
| 13 | 62933000 | 62934799 | 17.31522 | 29.88873 | 50.60697 | 22.20095 | -0.37138 | 0.865958 | 0.459146 |
| 6 | 87923400 | 87938199 | 221.6805 | 319.9007 | 453.562 | 187.1284 | -0.28971 | 4.11228 | 0.459152 |
| X | 127279800 | 127280799 | 15.93349 | 29.55241 | 25.30146 | 10.61923 | 0.665598 | 0.224025 | 0.459156 |
| X | 18870800 | 18872599 | 27.75693 | 47.63148 | 42.47665 | 18.63029 | 0.548257 | 0.95374 | 0.459163 |
| 2 | 15439400 | 15441199 | 19.94208 | 34.83663 | 34.68235 | 15.27946 | 0.394095 | 0.717226 | 0.459163 |
| X | 12350600 | 12354599 | 21.72908 | 36.73198 | 72.3061 | 31.64006 | -0.59257 | 1.093723 | 0.459211 |
| 6 | 57310000 | 57313199 | 67.14303 | 106.845 | 106.8064 | 44.62626 | 0.335873 | 2.230008 | 0.459213 |
| 3 | 120721000 | 120722599 | 14.11045 | 24.37343 | 53.1673 | 23.32463 | -0.73461 | 0.853416 | 0.459246 |
| 17 | 9646400 | 9647799 | 9.524686 | 17.5538 | 32.48865 | 13.75423 | -0.44836 | 0.222754 | 0.459255 |

| | | | | | | | | | |
|---|---|---|---|---|---|---|---|---|---|
| 4 | 146315800 | 146316399 | 19.13325 | 34.58114 | 26.65877 | 11.57469 | 0.782366 | 0.390972 | 0.459266 |
| 2 | 74336200 | 74337599 | 10.78556 | 19.32848 | 35.32824 | 15.4768 | -0.46357 | 0.4053 | 0.459274 |
| X | 155065600 | 155068399 | 27.80543 | 47.75709 | 41.94405 | 18.39778 | 0.570403 | 0.946029 | 0.459274 |
| 5 | 69578000 | 69587399 | 202.1975 | 298.7117 | 292.718 | 120.5476 | 0.275507 | 3.746027 | 0.45928 |
| 7 | 44777800 | 44781799 | 21.09072 | 34.84458 | 67.85749 | 29.31434 | -0.57797 | 1.328501 | 0.459285 |
| 6 | 53821400 | 53822399 | 8.285329 | 15.56085 | 29.0864 | 11.96069 | -0.42938 | 0.092131 | 0.459329 |
| 19 | 8266400 | 8270799 | 87.33007 | 136.3999 | 137.3502 | 56.98943 | 0.303174 | 2.664044 | 0.459335 |
| 7 | 16133000 | 16136399 | 18.52498 | 31.13498 | 58.41905 | 25.51849 | -0.52355 | 1.120689 | 0.459335 |
| 14 | 36551000 | 36553399 | 21.1548 | 35.63739 | 59.75442 | 26.12015 | -0.36197 | 1.108868 | 0.45934 |
| 1 | 142238000 | 142241199 | 62.48472 | 99.94592 | 94.59683 | 39.81095 | 0.422538 | 2.060272 | 0.459365 |
| 7 | 82005000 | 82006599 | 14.68001 | 26.90246 | 26.22214 | 11.18698 | 0.462496 | 0.279659 | 0.459367 |
| 13 | 31739600 | 31744799 | 53.71919 | 85.12962 | 140.2616 | 58.34757 | -0.38987 | 2.347106 | 0.459371 |
| 3 | 60590400 | 60593399 | 40.82858 | 66.26342 | 68.41576 | 29.32242 | 0.320696 | 1.617174 | 0.459376 |
| 1 | 117497000 | 117499599 | 29.72631 | 50.49546 | 45.73156 | 20.04093 | 0.523888 | 1.052745 | 0.459387 |
| 7 | 61972000 | 61979399 | 92.32208 | 145.7589 | 121.1578 | 50.30326 | 0.591522 | 2.461017 | 0.459394 |
| 6 | 70168400 | 70170799 | 21.30499 | 37.29564 | 35.31508 | 15.58484 | 0.465866 | 0.690571 | 0.459404 |
| 19 | 61158400 | 61161399 | 59.26158 | 94.53819 | 96.82407 | 40.59497 | 0.304756 | 2.13931 | 0.459404 |
| 13 | 57784800 | 57786799 | 10.48655 | 18.67073 | 36.32341 | 16.00031 | -0.55877 | 0.444405 | 0.459406 |
| 9 | 95138800 | 95140799 | 17.44889 | 30.13319 | 52.24832 | 22.92131 | -0.40549 | 0.863477 | 0.459424 |
| 8 | 26771000 | 26771399 | 6.109365 | 11.82832 | 24.39252 | 9.277765 | -0.46703 | -0.20463 | 0.459425 |
| 3 | 135269000 | 135270799 | 20.67946 | 34.79942 | 59.71694 | 26.09531 | -0.39533 | 1.116244 | 0.459426 |
| 9 | 101962800 | 101964999 | 18.91433 | 31.98974 | 56.39525 | 24.68146 | -0.43363 | 1.063882 | 0.459445 |
| 9 | 19290400 | 19302799 | 189.4888 | 284.0258 | 246.3234 | 101.6187 | 0.470818 | 3.488089 | 0.459457 |
| 15 | 72481400 | 72483599 | 17.22886 | 29.48292 | 52.164 | 22.87404 | -0.43686 | 0.946613 | 0.459457 |
| 13 | 19749400 | 19752599 | 28.84118 | 47.16329 | 77.34276 | 33.25887 | -0.33689 | 1.481371 | 0.459461 |
| 4 | 103071200 | 103072599 | 9.240336 | 16.60581 | 34.39499 | 15.03239 | -0.6403 | 0.373984 | 0.459465 |
| 12 | 76450200 | 76453199 | 22.24931 | 36.2616 | 86.09237 | 36.97601 | -0.87251 | 1.522187 | 0.459466 |
| 12 | 115339000 | 115342199 | 56.77146 | 91.44212 | 79.18961 | 33.67203 | 0.562381 | 1.837478 | 0.459467 |
| 10 | 93014400 | 93016399 | 23.41645 | 38.9363 | 65.60733 | 28.46824 | -0.36914 | 1.258814 | 0.459468 |
| 16 | 12498800 | 12500399 | 21.96723 | 38.05172 | 38.88341 | 17.05972 | 0.355169 | 0.839864 | 0.459469 |
| 5 | 124010800 | 124011999 | 8.265647 | 15.68917 | 29.94385 | 12.06182 | -0.43391 | 0.022616 | 0.459476 |
| 17 | 73183000 | 73184799 | 26.44115 | 44.74738 | 45.8558 | 20.0726 | 0.346117 | 1.072831 | 0.459489 |
| 15 | 37832800 | 37837399 | 89.57123 | 140.2593 | 141.0634 | 58.51469 | 0.307803 | 2.62098 | 0.459489 |
| 15 | 31322200 | 31323799 | 16.48202 | 29.23634 | 29.3067 | 12.95005 | 0.389155 | 0.492412 | 0.459494 |
| 10 | 59836400 | 59837599 | 7.393359 | 13.84547 | 29.57549 | 12.2216 | -0.6264 | 0.10186 | 0.459502 |
| 7 | 60852400 | 60852799 | 5.663911 | 11.01049 | 22.91197 | 8.507908 | -0.44671 | -0.29622 | 0.459507 |
| 10 | 126389000 | 126389599 | 5.663911 | 11.01049 | 22.91197 | 8.507908 | -0.44671 | -0.29622 | 0.459507 |
| 10 | 70781000 | 70783399 | 11.88318 | 21.00544 | 38.79989 | 17.1626 | -0.48996 | 0.516764 | 0.459508 |
| 17 | 49952000 | 49953799 | 11.87277 | 21.19628 | 36.76784 | 16.15906 | -0.39206 | 0.438264 | 0.459512 |
| X | 59405400 | 59407199 | 22.90839 | 40.83461 | 28.0861 | 12.35969 | 0.933738 | 0.503519 | 0.459512 |
| 4 | 150665400 | 150665999 | 6.278542 | 12.08474 | 25.24549 | 9.821149 | -0.51573 | -0.12132 | 0.459519 |
| X | 122394400 | 122394999 | 7.661656 | 14.59661 | 28.32329 | 11.32013 | -0.44712 | -0.01125 | 0.459539 |
| 10 | 16066200 | 16068799 | 15.90586 | 29.01186 | 27.15955 | 11.65815 | 0.514511 | 0.318014 | 0.459549 |
| 8 | 7213200 | 7214199 | 8.149065 | 15.30686 | 29.08993 | 11.95959 | -0.45266 | 0.091063 | 0.45955 |
| 5 | 145754800 | 145767599 | 259.0483 | 378.7363 | 327.1849 | 134.7898 | 0.44307 | 3.918685 | 0.45957 |
| 17 | 82747200 | 82749599 | 24.86455 | 42.63734 | 42.53008 | 18.65264 | 0.386988 | 0.9456 | 0.459574 |
| 11 | 73748400 | 73752999 | 41.48301 | 68.82974 | 45.48224 | 19.7065 | 0.97669 | 1.314812 | 0.459583 |

| | | | | | | | | | |
|---|---|---|---|---|---|---|---|---|---|
| 10 | 51676000 | 51677799 | 28.58018 | 48.37318 | 41.11906 | 17.99983 | 0.614882 | 1.075431 | 0.459592 |
| 5 | 18872400 | 18875399 | 33.11761 | 55.79341 | 47.99937 | 20.98212 | 0.597498 | 1.138919 | 0.459622 |
| 14 | 40687000 | 40689799 | 18.77319 | 31.70232 | 56.90958 | 24.89495 | -0.45974 | 1.077364 | 0.459625 |
| 15 | 62406600 | 62407999 | 19.57152 | 35.04796 | 28.7597 | 12.60697 | 0.68398 | 0.452666 | 0.459625 |
| 3 | 98867600 | 98868999 | 15.79569 | 28.95832 | 26.08567 | 11.12287 | 0.576033 | 0.28559 | 0.459633 |
| 17 | 18756000 | 18759999 | 69.84874 | 111.506 | 97.6342 | 40.97524 | 0.531605 | 2.12908 | 0.459634 |
| 3 | 120118800 | 120122199 | 65.33135 | 103.9005 | 107.0633 | 44.69699 | 0.291231 | 2.245556 | 0.459644 |
| 7 | 11503800 | 11514999 | 164.4726 | 250.0736 | 194.2553 | 80.47161 | 0.647257 | 3.199911 | 0.459652 |
| 2 | 8207600 | 8210599 | 38.75895 | 63.14274 | 63.84161 | 27.41536 | 0.355205 | 1.549497 | 0.459678 |
| 3 | 133017400 | 133019599 | 16.8313 | 29.09819 | 49.53298 | 21.73081 | -0.37856 | 0.848706 | 0.459687 |
| 1 | 5552800 | 5562199 | 135.8595 | 207.1414 | 199.2004 | 82.5865 | 0.342725 | 3.135845 | 0.459689 |
| 14 | 25780000 | 25780799 | 5.958131 | 11.56323 | 23.91494 | 8.994709 | -0.45493 | -0.24301 | 0.459693 |
| 6 | 39487800 | 39489199 | 8.905923 | 16.69807 | 30.69026 | 12.65649 | -0.40884 | 0.112044 | 0.459698 |
| X | 31308000 | 31310399 | 22.09139 | 39.43935 | 27.40357 | 12.04359 | 0.920069 | 0.485494 | 0.459705 |
| X | 143470600 | 143472999 | 19.70069 | 33.62809 | 57.0632 | 25.01728 | -0.37749 | 0.986399 | 0.459706 |
| 5 | 109306400 | 109308799 | 54.95079 | 87.94723 | 90.37782 | 38.06728 | 0.304999 | 2.03305 | 0.459721 |
| 1 | 33637400 | 33641999 | 55.3581 | 87.86662 | 135.5013 | 56.45724 | -0.2923 | 2.301801 | 0.459765 |
| 13 | 67377000 | 67384599 | 96.2819 | 150.0215 | 149.2183 | 61.93171 | 0.318233 | 2.711094 | 0.459771 |
| 4 | 116324600 | 116326199 | 15.74769 | 29.29538 | 24.70831 | 10.32577 | 0.691321 | 0.202926 | 0.459779 |
| 3 | 25179200 | 25179599 | 4.310458 | 8.416943 | 19.86085 | 6.989552 | -0.55277 | -0.51104 | 0.459782 |
| 1 | 10613200 | 10615399 | 25.25448 | 43.17286 | 43.63627 | 19.13305 | 0.367417 | 0.976676 | 0.459783 |
| 10 | 69603400 | 69604399 | 16.182 | 29.12031 | 28.70874 | 12.53266 | 0.424915 | 0.400615 | 0.459785 |
| 9 | 83817000 | 83819199 | 24.15282 | 41.94327 | 38.6133 | 16.94808 | 0.505581 | 0.823083 | 0.459798 |
| 3 | 38738600 | 38738999 | 4.935412 | 9.632362 | 21.07276 | 7.481567 | -0.45875 | -0.47183 | 0.459803 |
| 8 | 63223000 | 63224399 | 21.35055 | 37.30603 | 29.77472 | 13.11937 | 0.711164 | 0.70982 | 0.459835 |
| 1 | 137277600 | 137278199 | 13.33441 | 24.83764 | 20.83538 | 8.688443 | 0.701545 | 0.165554 | 0.459878 |
| 14 | 42358200 | 42359599 | 21.02508 | 36.53896 | 36.83682 | 16.17594 | 0.375358 | 0.797667 | 0.459891 |
| 12 | 104578800 | 104582599 | 33.66153 | 54.42189 | 88.60028 | 37.8589 | -0.34002 | 1.695513 | 0.459903 |
| X | 164192400 | 164194799 | 39.27514 | 64.52868 | 60.42062 | 26.0323 | 0.472579 | 1.436153 | 0.459914 |
| 6 | 7586400 | 7589399 | 30.99067 | 51.81572 | 51.89818 | 22.59949 | 0.378817 | 1.207316 | 0.459915 |
| X | 14434800 | 14436599 | 18.81768 | 34.315 | 26.47851 | 11.3691 | 0.791656 | 0.34131 | 0.459926 |
| 9 | 34019800 | 34021799 | 8.696515 | 15.98681 | 35.13813 | 14.92744 | -0.69775 | 0.239918 | 0.459934 |
| 12 | 60785800 | 60786599 | 10.7385 | 20.66624 | 21.28669 | 8.284379 | 0.493838 | -0.08843 | 0.459937 |
| 2 | 159498000 | 159498799 | 6.940726 | 13.07138 | 28.06927 | 11.48886 | -0.62228 | 0.058932 | 0.459942 |
| 17 | 26332000 | 26332999 | 6.940726 | 13.07138 | 28.06927 | 11.48886 | -0.62228 | 0.058932 | 0.459942 |
| 1 | 3176200 | 3177799 | 15.46924 | 28.7158 | 25.28025 | 10.59803 | 0.626631 | 0.214527 | 0.459949 |
| 10 | 69196000 | 69198799 | 42.24335 | 68.35222 | 69.44525 | 29.69727 | 0.33981 | 1.679747 | 0.459953 |
| 3 | 10033800 | 10034199 | 4.444637 | 8.671849 | 20.88668 | 7.455046 | -0.60052 | -0.45443 | 0.459957 |
| 15 | 46364800 | 46365399 | 7.641177 | 14.87509 | 15.722 | 5.77228 | 0.535546 | -0.3374 | 0.459978 |
| 6 | 118583000 | 118585399 | 11.90765 | 21.02969 | 38.33545 | 16.96022 | -0.47139 | 0.527388 | 0.459995 |
| 14 | 67421800 | 67426399 | 36.59729 | 58.9426 | 96.9145 | 41.20386 | -0.36078 | 1.81584 | 0.459997 |
| 10 | 101779400 | 101780799 | 14.74532 | 25.65985 | 46.31046 | 20.35316 | -0.46085 | 0.761258 | 0.460001 |
| 15 | 103888400 | 103891199 | 29.55024 | 50.65299 | 41.80911 | 18.33588 | 0.659069 | 0.967862 | 0.460016 |
| 4 | 61321400 | 61328599 | 119.4947 | 182.7242 | 176.9805 | 73.42401 | 0.335728 | 3.071197 | 0.460027 |
| 4 | 37757400 | 37759799 | 30.54065 | 50.81564 | 51.88059 | 22.52217 | 0.351408 | 1.258323 | 0.460046 |
| 3 | 72654200 | 72669999 | 267.7542 | 386.9473 | 369.7459 | 152.5188 | 0.281378 | 4.092879 | 0.460055 |
| 14 | 62344800 | 62346799 | 17.29072 | 31.40569 | 26.90702 | 11.61055 | 0.636612 | 0.35041 | 0.460061 |

| | | | | | | | | | |
|---|---|---|---|---|---|---|---|---|---|
| 17 | 27080000 | 27082399 | 18.33735 | 31.34415 | 53.25478 | 23.35022 | -0.37896 | 0.962604 | 0.460063 |
| 3 | 158577800 | 158581399 | 41.46183 | 67.35529 | 66.00782 | 28.31028 | 0.396986 | 1.595285 | 0.460076 |
| 7 | 14626600 | 14641599 | 259.6717 | 383.7036 | 234.379 | 96.52254 | 0.957319 | 3.748869 | 0.46008 |
| 19 | 38278400 | 38279999 | 14.74496 | 25.84744 | 44.04075 | 19.45815 | -0.37756 | 0.648602 | 0.4601 |
| 12 | 65958200 | 65959599 | 14.93846 | 27.61775 | 25.80365 | 10.87758 | 0.535642 | 0.233397 | 0.460106 |
| 3 | 23746600 | 23747399 | 15.3611 | 28.03196 | 23.69249 | 10.16369 | 0.661648 | 0.30394 | 0.460106 |
| 12 | 32830400 | 32832399 | 18.14683 | 32.91579 | 26.81565 | 11.59086 | 0.7073 | 0.36432 | 0.460107 |
| X | 8604600 | 8612999 | 156.7775 | 238.6449 | 195.3041 | 80.93368 | 0.573397 | 3.175522 | 0.460125 |
| 14 | 24679800 | 24680599 | 6.254833 | 12.09324 | 25.4932 | 9.759223 | -0.50782 | -0.17923 | 0.460127 |
| 16 | 7935000 | 7936199 | 9.685064 | 17.8437 | 32.48212 | 13.75742 | -0.42502 | 0.22618 | 0.460131 |
| 2 | 164746800 | 164750199 | 39.59642 | 63.58031 | 102.4515 | 43.3663 | -0.33714 | 1.918939 | 0.460153 |
| 12 | 114322800 | 114328799 | 100.108 | 157.2031 | 130.7362 | 54.22515 | 0.584844 | 2.574874 | 0.460161 |
| 10 | 122911600 | 122912799 | 6.726943 | 12.87242 | 26.21101 | 10.36915 | -0.5028 | -0.06194 | 0.460163 |
| 19 | 4320400 | 4324199 | 22.58668 | 37.10552 | 71.5517 | 30.82667 | -0.56662 | 1.405057 | 0.460182 |
| 16 | 73125600 | 73126799 | 12.80953 | 24.0537 | 23.58047 | 9.699413 | 0.494484 | 0.119348 | 0.460207 |
| 6 | 129856000 | 129861999 | 99.74667 | 156.3469 | 135.5034 | 56.20263 | 0.523318 | 2.608455 | 0.46022 |
| 4 | 153005400 | 153042199 | 400.1375 | 545.1712 | 862.364 | 359.0432 | -0.53106 | 4.981478 | 0.46022 |
| 7 | 85761200 | 85762999 | 34.85019 | 57.53928 | 58.08714 | 25.08221 | 0.366791 | 1.362961 | 0.460224 |
| 8 | 55370600 | 55372199 | 20.35183 | 34.39803 | 57.53587 | 25.17555 | -0.35819 | 1.076097 | 0.460249 |
| 12 | 63336200 | 63339599 | 77.74311 | 122.3913 | 122.354 | 50.76516 | 0.322421 | 2.510314 | 0.46025 |
| 1 | 37579200 | 37580599 | 8.65479 | 15.94793 | 30.94629 | 13.10425 | -0.51581 | 0.215359 | 0.460253 |
| 13 | 28557800 | 28558799 | 8.65479 | 15.94793 | 30.94629 | 13.10425 | -0.51581 | 0.215359 | 0.460253 |
| 16 | 27809400 | 27812199 | 24.07863 | 42.19455 | 34.00375 | 15.01922 | 0.697438 | 0.688328 | 0.460258 |
| 14 | 18060400 | 18062399 | 16.18492 | 28.92677 | 28.61971 | 12.56629 | 0.413924 | 0.440636 | 0.460301 |
| 13 | 51545200 | 51547799 | 21.88824 | 36.55681 | 62.36784 | 27.13993 | -0.38694 | 1.201144 | 0.460308 |
| 17 | 39442400 | 39446399 | 63.72768 | 101.8801 | 96.23662 | 40.47068 | 0.424575 | 2.079006 | 0.460308 |
| 3 | 44456600 | 44459399 | 31.78736 | 53.48442 | 49.10851 | 21.4587 | 0.503756 | 1.144278 | 0.460324 |
| 12 | 88583000 | 88585599 | 18.79911 | 33.32917 | 26.6627 | 11.78687 | 0.712449 | 0.50628 | 0.460332 |
| 7 | 52551200 | 52552799 | 32.07932 | 53.45965 | 53.00565 | 23.03386 | 0.39351 | 1.247564 | 0.460349 |
| 12 | 64427200 | 64428599 | 15.65007 | 28.83475 | 25.63222 | 10.85563 | 0.60204 | 0.256365 | 0.46038 |
| 11 | 114730000 | 114733399 | 34.17621 | 55.32493 | 89.4408 | 38.26124 | -0.32808 | 1.669372 | 0.460399 |
| 8 | 30170400 | 30171399 | 14.38277 | 27.32804 | 21.83383 | 8.776028 | 0.814927 | 0.049826 | 0.460413 |
| 13 | 77206600 | 77208599 | 17.66203 | 31.54209 | 29.44737 | 12.93822 | 0.496544 | 0.45699 | 0.460419 |
| X | 127449000 | 127450799 | 16.73405 | 30.40657 | 27.68979 | 11.94243 | 0.549967 | 0.348236 | 0.460437 |
| 18 | 76015000 | 76018599 | 29.17979 | 47.77095 | 78.02654 | 33.567 | -0.33004 | 1.467473 | 0.460451 |
| 4 | 136930000 | 136932199 | 28.43037 | 46.26497 | 79.53329 | 34.14 | -0.4097 | 1.552586 | 0.46046 |
| 11 | 112975000 | 112976999 | 11.47242 | 20.21053 | 38.92212 | 17.22836 | -0.55149 | 0.55409 | 0.460461 |
| 18 | 80564400 | 80570799 | 48.12794 | 76.22702 | 149.909 | 62.34073 | -0.64543 | 2.359717 | 0.460473 |
| 16 | 12350800 | 12353599 | 23.9835 | 39.91496 | 66.41277 | 28.83559 | -0.35101 | 1.24809 | 0.460474 |
| 15 | 38920000 | 38921799 | 20.47885 | 34.60615 | 58.67126 | 25.67088 | -0.3777 | 1.077999 | 0.460484 |
| 11 | 73834600 | 73839399 | 13.3487 | 25.36779 | 19.72983 | 7.927331 | 0.853868 | 0.033937 | 0.460492 |
| X | 147693000 | 147694999 | 14.98667 | 27.84237 | 25.33915 | 10.61112 | 0.580184 | 0.206516 | 0.460493 |
| 8 | 122106600 | 122107799 | 9.324287 | 17.0338 | 32.42611 | 13.89167 | -0.50031 | 0.275041 | 0.460554 |
| 8 | 59821600 | 59822799 | 16.25912 | 28.99756 | 29.19823 | 12.84334 | 0.387072 | 0.45631 | 0.460581 |
| 10 | 88535400 | 88542999 | 79.22305 | 123.148 | 184.709 | 76.68905 | -0.27638 | 2.749131 | 0.460593 |
| 5 | 80099000 | 80099799 | 11.49106 | 21.82368 | 22.62956 | 9.104648 | 0.440658 | 0.028192 | 0.460603 |
| 7 | 7124000 | 7125199 | 7.7895 | 14.67427 | 29.69895 | 12.14972 | -0.53773 | 0.06649 | 0.460633 |

| | | | | | | | | | |
|---|---|---|---|---|---|---|---|---|---|
| 15 | 34556200 | 34557999 | 18.84538 | 33.83975 | 28.92221 | 12.64909 | 0.627929 | 0.432479 | 0.460649 |
| 3 | 36529400 | 36531199 | 13.85881 | 24.08648 | 45.82393 | 20.12323 | -0.53653 | 0.776201 | 0.460657 |
| 8 | 4842600 | 4845999 | 35.75528 | 59.64589 | 52.0436 | 22.63024 | 0.577234 | 1.245299 | 0.460674 |
| 19 | 7985200 | 7990799 | 107.1272 | 165.2116 | 165.7073 | 68.83189 | 0.294764 | 2.898508 | 0.460678 |
| 6 | 125280600 | 125287199 | 58.22894 | 91.15744 | 176.1144 | 73.05617 | -0.63311 | 2.614914 | 0.46068 |
| 17 | 69745200 | 69748399 | 22.28989 | 36.94158 | 66.59073 | 28.82953 | -0.46658 | 1.292345 | 0.460696 |
| 6 | 104196400 | 104204199 | 180.8446 | 263.5763 | 402.7689 | 165.9772 | -0.38369 | 3.960233 | 0.460697 |
| 14 | 63580200 | 63582599 | 18.47286 | 31.17739 | 56.87807 | 24.87687 | -0.48294 | 1.081705 | 0.460701 |
| 14 | 50738000 | 50740799 | 37.3276 | 61.864 | 54.98175 | 23.80359 | 0.550391 | 1.322321 | 0.460749 |
| 1 | 131299800 | 131301599 | 18.75583 | 33.18339 | 31.55019 | 13.95331 | 0.463381 | 0.535927 | 0.460751 |
| X | 95983600 | 95991599 | 111.6608 | 173.0187 | 158.9643 | 66.02932 | 0.426399 | 2.815785 | 0.460773 |
| 4 | 44466800 | 44472399 | 44.31874 | 70.77842 | 117.6566 | 49.40628 | -0.39103 | 2.095163 | 0.46078 |
| 2 | 125792400 | 125796999 | 53.43085 | 86.24782 | 75.68851 | 32.27608 | 0.546895 | 1.767782 | 0.460781 |
| 9 | 5662400 | 5664199 | 22.61975 | 39.61989 | 34.62394 | 15.28723 | 0.581015 | 0.687433 | 0.460802 |
| 3 | 157400600 | 157401599 | 8.579919 | 16.10367 | 29.61531 | 12.19157 | -0.40758 | 0.099492 | 0.460805 |
| 13 | 95510800 | 95511999 | 8.579919 | 16.10367 | 29.61531 | 12.19157 | -0.40758 | 0.099492 | 0.460805 |
| 14 | 111788800 | 111791599 | 23.61922 | 40.76256 | 40.93346 | 17.95714 | 0.379143 | 0.882467 | 0.460809 |
| 9 | 15404400 | 15405799 | 11.43103 | 20.28496 | 36.79533 | 16.25357 | -0.46116 | 0.473579 | 0.460825 |
| 5 | 109477600 | 109487399 | 115.8664 | 180.2646 | 145.6789 | 60.47636 | 0.61582 | 2.747643 | 0.460842 |
| X | 33244800 | 33251999 | 164.8723 | 247.5703 | 234.0914 | 96.61638 | 0.348481 | 3.449486 | 0.460842 |
| 10 | 65560400 | 65561599 | 13.68647 | 25.36163 | 25.44704 | 10.6936 | 0.436831 | 0.210581 | 0.460851 |
| X | 125916000 | 125920599 | 24.51225 | 42.86606 | 31.06146 | 13.69578 | 0.85013 | 0.718585 | 0.460854 |
| 19 | 39974200 | 39976199 | 20.58759 | 36.60673 | 29.70688 | 13.10033 | 0.694147 | 0.504266 | 0.460856 |
| 1 | 113647800 | 113651999 | 62.40979 | 100.1787 | 83.33995 | 35.25885 | 0.614157 | 1.948835 | 0.460858 |
| 6 | 115678400 | 115680199 | 19.94089 | 33.5256 | 58.50113 | 25.55687 | -0.41927 | 1.122826 | 0.460863 |
| 11 | 90997400 | 90997799 | 3.700874 | 7.230966 | 18.77825 | 6.539563 | -0.67611 | -0.5658 | 0.460864 |
| 17 | 21849000 | 21850799 | 11.98922 | 21.57207 | 38.92271 | 16.99379 | -0.44225 | 0.392832 | 0.460871 |
| 10 | 87143400 | 87144199 | 9.511058 | 17.65288 | 32.08822 | 13.45483 | -0.41243 | 0.182468 | 0.460877 |
| 12 | 64047400 | 64048199 | 9.511058 | 17.65288 | 32.08822 | 13.45483 | -0.41243 | 0.182468 | 0.460877 |
| 4 | 103136800 | 103140599 | 35.32335 | 58.85735 | 53.14226 | 23.09114 | 0.528115 | 1.257696 | 0.460878 |
| 11 | 33835000 | 33836799 | 15.70001 | 28.28002 | 28.27776 | 12.32973 | 0.405763 | 0.391144 | 0.460886 |
| X | 4490000 | 4491199 | 27.70895 | 47.84591 | 35.04792 | 15.37545 | 0.832594 | 0.890273 | 0.460889 |
| 17 | 20920800 | 20930599 | 75.88045 | 120.1831 | 108.3138 | 45.04315 | 0.478611 | 2.371162 | 0.46091 |
| X | 89901800 | 89903999 | 23.64849 | 41.85157 | 29.56804 | 13.07595 | 0.889865 | 0.561302 | 0.460913 |
| 16 | 14542800 | 14544799 | 19.45544 | 34.47942 | 30.27539 | 13.385 | 0.578302 | 0.518682 | 0.460918 |
| X | 33118800 | 33120799 | 20.52964 | 36.92449 | 27.80644 | 12.14434 | 0.811003 | 0.433209 | 0.460928 |
| 10 | 119884200 | 119888199 | 40.2904 | 65.40898 | 67.30638 | 28.85313 | 0.325919 | 1.609004 | 0.460946 |
| 3 | 22920600 | 22924199 | 36.44244 | 59.69004 | 61.3625 | 26.4036 | 0.336372 | 1.464657 | 0.460947 |
| 13 | 62440400 | 62460999 | 567.4093 | 767.4919 | 686.8262 | 286.2779 | 0.280015 | 5.097643 | 0.460947 |
| 18 | 32174200 | 32176399 | 53.46378 | 85.69018 | 88.35974 | 37.29743 | 0.302475 | 1.9863 | 0.460967 |
| 4 | 88012400 | 88013399 | 9.132782 | 16.65464 | 33.40309 | 14.34173 | -0.57778 | 0.281954 | 0.460975 |
| 13 | 58248600 | 58253199 | 39.56823 | 63.61928 | 100.8315 | 42.76372 | -0.31097 | 1.876266 | 0.461005 |
| 3 | 8235000 | 8241399 | 107.4431 | 165.8254 | 167.4283 | 69.54646 | 0.286064 | 2.885549 | 0.461015 |
| 4 | 151679000 | 151681599 | 18.45043 | 31.26772 | 55.29551 | 24.21485 | -0.43792 | 1.04492 | 0.461034 |
| 3 | 21066800 | 21072399 | 62.56462 | 99.66669 | 102.9844 | 43.09355 | 0.289831 | 2.189137 | 0.461058 |
| 17 | 19787000 | 19790199 | 53.86143 | 86.95781 | 74.71653 | 31.86867 | 0.577571 | 1.762415 | 0.461064 |
| 10 | 89456800 | 89458199 | 9.914239 | 18.05667 | 32.86697 | 14.13407 | -0.44005 | 0.296115 | 0.461065 |

| | | | | | | | | | |
|---|---|---|---|---|---|---|---|---|---|
| 15 | 98060800 | 98062399 | 9.914239 | 18.05667 | 32.86697 | 14.13407 | -0.44005 | 0.296115 | 0.461065 |
| 13 | 45604400 | 45604999 | 5.193621 | 10.13738 | 25.01728 | 8.865736 | -0.6284 | -0.46016 | 0.461077 |
| 17 | 53487400 | 53490799 | 51.93433 | 83.46025 | 84.67067 | 35.88373 | 0.330478 | 1.90265 | 0.461094 |
| 18 | 40328600 | 40331399 | 27.89223 | 47.31878 | 46.34851 | 20.30334 | 0.41125 | 1.059476 | 0.461096 |
| 4 | 143793800 | 143795799 | 17.82934 | 33.39536 | 21.85779 | 9.029389 | 1.067694 | 0.172326 | 0.461105 |
| 6 | 133866800 | 133870999 | 43.8788 | 71.86408 | 60.86979 | 26.19065 | 0.614991 | 1.482803 | 0.461108 |
| 5 | 125869800 | 125871999 | 19.47288 | 34.46406 | 31.50119 | 13.93067 | 0.520199 | 0.537138 | 0.461116 |
| 16 | 16748200 | 16749199 | 8.932032 | 16.44615 | 31.95498 | 13.54554 | -0.51922 | 0.222236 | 0.461123 |
| 6 | 117377000 | 117378799 | 13.96405 | 24.39408 | 43.64091 | 19.2264 | -0.44748 | 0.699415 | 0.461124 |
| 18 | 25221600 | 25224199 | 13.96405 | 24.39408 | 43.64091 | 19.2264 | -0.44748 | 0.699415 | 0.461124 |
| 13 | 68574800 | 68579199 | 120.637 | 185.0501 | 159.1737 | 66.07776 | 0.510304 | 3.007216 | 0.461135 |
| 11 | 120626200 | 120626999 | 9.960384 | 19.06709 | 19.07399 | 7.537465 | 0.514869 | -0.0553 | 0.461135 |
| 1 | 82293800 | 82294799 | 9.544846 | 17.11162 | 35.35236 | 15.48234 | -0.63902 | 0.39186 | 0.461153 |
| 4 | 138699600 | 138701199 | 7.887436 | 14.62633 | 29.98899 | 12.58987 | -0.5854 | 0.171119 | 0.461162 |
| 14 | 57901800 | 57902999 | 7.887436 | 14.62633 | 29.98899 | 12.58987 | -0.5854 | 0.171119 | 0.461162 |
| 6 | 94773200 | 94774599 | 18.36835 | 33.533 | 25.81376 | 11.06823 | 0.795959 | 0.328461 | 0.461168 |
| 16 | 7630200 | 7633199 | 37.93632 | 61.13499 | 97.21119 | 41.36509 | -0.31155 | 1.801238 | 0.461175 |
| 12 | 60718600 | 60721599 | 33.05498 | 55.16714 | 52.53839 | 22.85214 | 0.451331 | 1.232035 | 0.461199 |
| 10 | 61693200 | 61702399 | 111.8846 | 169.2341 | 260.7056 | 107.8273 | -0.34601 | 3.294772 | 0.461203 |
| 18 | 88887800 | 88889399 | 15.19354 | 26.18464 | 48.98001 | 21.48617 | -0.51459 | 0.87233 | 0.461205 |
| 5 | 18883600 | 18883999 | 4.44142 | 8.680695 | 20.74886 | 7.165723 | -0.54705 | -0.57292 | 0.461217 |
| 11 | 99248400 | 99249799 | 13.94411 | 25.70415 | 25.86889 | 10.94857 | 0.425007 | 0.24164 | 0.46122 |
| 10 | 82263400 | 82268799 | 102.9649 | 159.7203 | 158.3936 | 65.78366 | 0.317542 | 2.792093 | 0.461224 |
| 5 | 148883600 | 148884399 | 12.92097 | 24.25413 | 24.0612 | 9.90326 | 0.476592 | 0.123046 | 0.461224 |
| 14 | 61754800 | 61757799 | 26.7552 | 43.88842 | 73.67531 | 31.72167 | -0.36818 | 1.435595 | 0.461225 |
| 4 | 59309600 | 59310799 | 9.080269 | 16.796 | 30.96559 | 13.04444 | -0.43725 | 0.196557 | 0.461236 |
| 19 | 47825200 | 47827199 | 14.7072 | 25.73583 | 45.1386 | 19.91351 | -0.41923 | 0.677115 | 0.461238 |
| 13 | 65397800 | 65401999 | 94.25215 | 149.0804 | 113.6061 | 47.21187 | 0.71881 | 2.419347 | 0.46125 |
| 2 | 55217200 | 55218799 | 18.58606 | 33.59707 | 27.98899 | 12.15003 | 0.671679 | 0.39116 | 0.461258 |
| 1 | 13191600 | 13194199 | 23.1567 | 39.75401 | 39.55586 | 17.34982 | 0.390915 | 0.926482 | 0.461263 |
| 4 | 69293200 | 69294599 | 13.18028 | 23.26495 | 39.8194 | 17.6186 | -0.38134 | 0.544334 | 0.461274 |
| 12 | 37723800 | 37725199 | 13.18028 | 23.26495 | 39.8194 | 17.6186 | -0.38134 | 0.544334 | 0.461274 |
| 11 | 21426200 | 21427599 | 26.36201 | 44.73268 | 45.27578 | 19.83475 | 0.364018 | 1.048129 | 0.461277 |
| 1 | 121017000 | 121018999 | 20.75454 | 36.39129 | 34.18079 | 15.10505 | 0.477368 | 0.659394 | 0.461281 |
| 12 | 101155600 | 101156199 | 6.245339 | 12.10091 | 26.29347 | 9.964821 | -0.53753 | -0.21224 | 0.461282 |
| 14 | 33924400 | 33926799 | 15.56167 | 27.13266 | 45.74836 | 20.13047 | -0.36282 | 0.738556 | 0.461287 |
| 5 | 32737400 | 32743399 | 98.06747 | 152.6768 | 151.5421 | 62.90398 | 0.320329 | 2.727651 | 0.461289 |
| 12 | 8165000 | 8166199 | 9.21465 | 17.11947 | 31.02524 | 12.99182 | -0.40652 | 0.172883 | 0.461291 |
| 3 | 142415600 | 142420999 | 71.78508 | 111.7587 | 177.2629 | 73.58049 | -0.35517 | 2.717645 | 0.461294 |
| 14 | 18315600 | 18318199 | 21.42439 | 37.17431 | 37.83269 | 16.59958 | 0.361465 | 0.820366 | 0.461316 |
| 8 | 13886800 | 13889599 | 39.1474 | 63.85592 | 65.28922 | 28.04988 | 0.340322 | 1.531354 | 0.461321 |
| 1 | 7737200 | 7739799 | 29.56285 | 49.82134 | 43.2278 | 18.89823 | 0.585192 | 1.117428 | 0.461322 |
| 17 | 39699200 | 39709799 | 146.9295 | 224.8387 | 179.9121 | 74.65272 | 0.611842 | 3.066145 | 0.461369 |
| 12 | 68351200 | 68352799 | 14.17319 | 25.93423 | 26.38185 | 11.27841 | 0.399392 | 0.286224 | 0.46137 |
| 17 | 19644000 | 19668199 | 435.1649 | 606.807 | 514.114 | 213.2707 | 0.403236 | 4.62585 | 0.461398 |
| 11 | 13278000 | 13279799 | 22.65542 | 39.76707 | 32.27612 | 14.27631 | 0.686897 | 0.649425 | 0.461401 |
| 4 | 109468200 | 109469799 | 12.22365 | 22.94775 | 21.78254 | 8.963695 | 0.539154 | 0.106688 | 0.46144 |

| | | | | | | | | | |
|---|---|---|---|---|---|---|---|---|---|
| X | 152992400 | 152997999 | 77.50498 | 123.0872 | 110.633 | 46.12131 | 0.486151 | 2.300264 | 0.461445 |
| 17 | 29013800 | 29015799 | 22.64904 | 38.00474 | 63.10754 | 27.54295 | -0.3482 | 1.153754 | 0.461476 |
| 12 | 36494600 | 36496599 | 33.23787 | 55.04548 | 55.76984 | 24.13253 | 0.361959 | 1.322888 | 0.461477 |
| 4 | 141698200 | 141703799 | 70.53295 | 110.1565 | 169.8465 | 70.47337 | -0.31084 | 2.662004 | 0.461481 |
| 13 | 65486600 | 65496999 | 230.9422 | 341.0635 | 293.8258 | 121.0033 | 0.460802 | 3.755732 | 0.461486 |
| X | 77198200 | 77198999 | 21.89533 | 38.33727 | 34.91603 | 15.41152 | 0.521958 | 0.692687 | 0.461489 |
| 11 | 27413800 | 27414999 | 10.79884 | 19.65538 | 35.98029 | 15.48512 | -0.44947 | 0.310491 | 0.461528 |
| 10 | 43810800 | 43812599 | 12.53035 | 22.11566 | 38.81797 | 17.17583 | -0.41751 | 0.538992 | 0.461529 |
| 1 | 75329200 | 75330999 | 13.64412 | 24.00607 | 40.85665 | 18.08921 | -0.37518 | 0.583676 | 0.461562 |
| 8 | 19209000 | 19211199 | 15.85749 | 27.59073 | 46.83689 | 20.5824 | -0.37294 | 0.770349 | 0.461564 |
| 9 | 63306000 | 63306999 | 8.078957 | 14.83152 | 31.95689 | 13.60094 | -0.67127 | 0.232885 | 0.461568 |
| 4 | 48750800 | 48752799 | 12.65164 | 22.54002 | 39.29646 | 17.30057 | -0.40131 | 0.460035 | 0.46157 |
| 6 | 146264000 | 146265599 | 12.65164 | 22.54002 | 39.29646 | 17.30057 | -0.40131 | 0.460035 | 0.46157 |
| 1 | 168528800 | 168530199 | 15.76209 | 28.88786 | 26.70026 | 11.38967 | 0.53893 | 0.289484 | 0.461574 |
| 10 | 18407200 | 18409199 | 15.29663 | 26.44013 | 47.93501 | 21.02964 | -0.46889 | 0.842921 | 0.461601 |
| 18 | 7815800 | 7818999 | 31.50292 | 52.91779 | 44.79323 | 19.56292 | 0.620564 | 1.152725 | 0.461601 |
| 8 | 72009800 | 72012999 | 21.96934 | 36.47489 | 64.89399 | 28.12933 | -0.44747 | 1.27073 | 0.461613 |
| 1 | 78165800 | 78167999 | 19.09474 | 32.20958 | 57.45634 | 25.12568 | -0.45076 | 1.088544 | 0.461623 |
| 3 | 70755800 | 70757399 | 16.79286 | 31.26512 | 24.09467 | 10.05546 | 0.822211 | 0.205378 | 0.461624 |
| 9 | 18312400 | 18316599 | 44.8489 | 72.69165 | 70.6927 | 30.26865 | 0.405032 | 1.650914 | 0.461628 |
| 10 | 68357000 | 68360199 | 38.89678 | 63.6106 | 63.62078 | 27.35872 | 0.374655 | 1.494826 | 0.461641 |
| 3 | 22010400 | 22011999 | 11.64694 | 20.83308 | 37.77571 | 16.57515 | -0.45435 | 0.423754 | 0.461642 |
| 7 | 41715200 | 41728399 | 174.4774 | 263.6024 | 218.0595 | 90.15575 | 0.548931 | 3.330554 | 0.461651 |
| 7 | 31294800 | 31306599 | 191.3687 | 286.4321 | 251.5421 | 103.7427 | 0.450881 | 3.515252 | 0.461689 |
| 3 | 159007400 | 159014799 | 113.4863 | 174.2623 | 176.5695 | 73.30973 | 0.27538 | 2.985634 | 0.461691 |
| 9 | 120467600 | 120471999 | 45.84887 | 73.14201 | 118.0132 | 49.47947 | -0.34979 | 2.132412 | 0.461693 |
| 1 | 185278800 | 185281199 | 10.7638 | 19.36294 | 34.81273 | 15.20216 | -0.43677 | 0.379776 | 0.461694 |
| X | 21267400 | 21269399 | 19.36812 | 34.55203 | 29.93724 | 13.1653 | 0.60301 | 0.477836 | 0.461768 |
| 4 | 23768400 | 23770399 | 21.08923 | 37.06012 | 33.59162 | 14.87249 | 0.528522 | 0.627226 | 0.461773 |
| Y | 1361800 | 1364199 | 12.59374 | 23.61213 | 14.4823 | 5.972586 | 1.16155 | 0.097721 | 0.461774 |
| 3 | 24286400 | 24290399 | 64.0856 | 102.1356 | 103.6457 | 43.39735 | 0.316551 | 2.176818 | 0.461777 |
| 18 | 5552400 | 5555199 | 29.28411 | 50.73642 | 37.92308 | 16.63848 | 0.804532 | 0.871318 | 0.461794 |
| 15 | 97111000 | 97115399 | 40.04763 | 64.40811 | 101.7763 | 43.18193 | -0.30617 | 1.868247 | 0.461805 |
| 9 | 46830400 | 46831999 | 14.91171 | 27.51715 | 25.13775 | 10.62369 | 0.56496 | 0.240185 | 0.46181 |
| 1 | 182446000 | 182448999 | 32.05581 | 51.81085 | 88.17768 | 37.66425 | -0.40435 | 1.700848 | 0.461834 |
| 5 | 20901400 | 20903399 | 16.14688 | 29.3809 | 27.05602 | 11.6486 | 0.535217 | 0.332239 | 0.461835 |
| 11 | 29196600 | 29199599 | 20.49941 | 34.58649 | 58.64236 | 25.64601 | -0.37788 | 1.092023 | 0.461846 |
| 8 | 129892000 | 129893399 | 17.9386 | 31.89672 | 30.5473 | 13.47089 | 0.456521 | 0.488793 | 0.461853 |
| 8 | 47644000 | 47647599 | 55.45884 | 87.79189 | 139.5546 | 58.01938 | -0.33947 | 2.376444 | 0.46186 |
| 14 | 36322800 | 36323999 | 10.37362 | 20.01281 | 20.77963 | 8.030585 | 0.491588 | -0.11655 | 0.461883 |
| Y | 4042200 | 4044999 | 21.35643 | 37.25556 | 22.80351 | 10.03742 | 1.091704 | 0.719209 | 0.461885 |
| 14 | 13788800 | 13789799 | 18.07094 | 31.72884 | 30.54561 | 13.51408 | 0.442116 | 0.618504 | 0.461886 |
| X | 128510200 | 128517799 | 92.50651 | 145.2977 | 123.1226 | 51.06871 | 0.558049 | 2.565375 | 0.461895 |
| 16 | 50820000 | 50822199 | 25.12345 | 42.9513 | 43.64983 | 19.13908 | 0.35961 | 0.97611 | 0.4619 |
| 1 | 65788000 | 65788599 | 4.311997 | 8.413582 | 20.31224 | 7.24222 | -0.60233 | -0.46263 | 0.461901 |
| X | 43327600 | 43332799 | 51.98491 | 84.74426 | 60.47445 | 25.97555 | 0.85725 | 1.567268 | 0.461907 |
| 1 | 102947000 | 102949399 | 18.75838 | 33.75948 | 28.33939 | 12.37011 | 0.655494 | 0.416376 | 0.461922 |

| | | | | | | | | | |
|---|---|---|---|---|---|---|---|---|---|
| 8 | 42362000 | 42363199 | 12.49393 | 22.23601 | 38.25022 | 16.85466 | -0.38287 | 0.463755 | 0.461957 |
| 5 | 82791400 | 82793799 | 22.93618 | 40.10213 | 34.60678 | 15.25651 | 0.599046 | 0.715685 | 0.461961 |
| 5 | 146327800 | 146331799 | 40.22331 | 64.45297 | 108.6464 | 45.84356 | -0.40565 | 1.984649 | 0.461971 |
| X | 44126000 | 44128199 | 37.47539 | 62.67787 | 49.76839 | 21.67639 | 0.712809 | 1.224149 | 0.462005 |
| 18 | 70641000 | 70643599 | 15.77086 | 27.43737 | 46.84297 | 20.58383 | -0.381 | 0.77353 | 0.462008 |
| 10 | 126573400 | 126575799 | 24.0381 | 41.36236 | 41.45636 | 18.18488 | 0.380989 | 0.910258 | 0.462021 |
| 11 | 3652000 | 3658799 | 46.35563 | 73.77452 | 125.2428 | 52.33791 | -0.42675 | 2.210738 | 0.462028 |
| 10 | 123169000 | 123170999 | 13.49457 | 23.88558 | 42.92116 | 18.98054 | -0.44987 | 0.520528 | 0.462047 |
| 1 | 122605200 | 122605999 | 13.00124 | 22.99809 | 39.27392 | 17.36981 | -0.37693 | 0.518245 | 0.46205 |
| 6 | 132087400 | 132090399 | 42.49528 | 69.36647 | 65.07797 | 27.96641 | 0.464831 | 1.529083 | 0.462062 |
| 10 | 92938200 | 92939999 | 13.79166 | 24.32266 | 41.86273 | 18.52597 | -0.39006 | 0.559923 | 0.462115 |
| 2 | 23703600 | 23704199 | 7.013757 | 13.50848 | 26.4482 | 10.27019 | -0.42132 | -0.11784 | 0.462118 |
| 13 | 106407000 | 106407999 | 11.16667 | 20.19201 | 35.32462 | 15.32816 | -0.39197 | 0.351334 | 0.462138 |
| 1 | 3286600 | 3287799 | 20.25278 | 35.34534 | 35.68823 | 15.7131 | 0.373702 | 0.733468 | 0.462155 |
| 10 | 45123800 | 45125799 | 15.0924 | 26.40351 | 44.62756 | 19.68404 | -0.36595 | 0.684625 | 0.46216 |
| 8 | 21812600 | 21822199 | 23.60212 | 40.99884 | 34.61932 | 15.19782 | 0.629865 | 0.813012 | 0.46219 |
| 17 | 72032800 | 72043799 | 79.1811 | 122.2884 | 213.465 | 88.66893 | -0.50223 | 2.867285 | 0.462195 |
| 8 | 46802200 | 46806999 | 67.53903 | 106.1564 | 160.735 | 66.67565 | -0.27753 | 2.542292 | 0.462233 |
| 4 | 63147000 | 63149599 | 14.88991 | 25.68691 | 48.97408 | 21.48422 | -0.54159 | 0.862919 | 0.462242 |
| 5 | 94483000 | 94484199 | 14.99557 | 28.25915 | 23.35645 | 9.549471 | 0.745463 | 0.114155 | 0.462243 |
| 16 | 15739600 | 15740999 | 15.69949 | 28.91145 | 26.28564 | 11.13989 | 0.569409 | 0.262767 | 0.462254 |
| 18 | 51532800 | 51538799 | 171.6305 | 256.5037 | 253.3157 | 104.4437 | 0.279668 | 3.538219 | 0.462265 |
| X | 127096200 | 127096799 | 13.01779 | 24.48831 | 24.03946 | 9.858453 | 0.495647 | 0.109631 | 0.46228 |
| 5 | 80451200 | 80452999 | 21.13811 | 36.96173 | 35.92236 | 15.83771 | 0.428518 | 0.708534 | 0.462291 |
| 12 | 114099600 | 114105199 | 80.21977 | 127.3223 | 112.0802 | 46.69657 | 0.51532 | 2.319091 | 0.462299 |
| X | 152424800 | 152426599 | 15.76271 | 28.69136 | 27.30631 | 11.7515 | 0.488771 | 0.330966 | 0.462307 |
| 6 | 61112600 | 61112999 | 4.675766 | 9.122969 | 20.99047 | 7.489733 | -0.53737 | -0.45918 | 0.462307 |
| 6 | 144431800 | 144435599 | 56.61313 | 90.73996 | 90.91705 | 38.37824 | 0.343883 | 1.990497 | 0.462336 |
| 17 | 55512600 | 55516999 | 43.31139 | 70.13309 | 71.89895 | 30.76361 | 0.327874 | 1.669115 | 0.462337 |
| 2 | 169020400 | 169025399 | 34.93675 | 56.39404 | 94.58722 | 40.33449 | -0.38562 | 1.744874 | 0.462338 |
| 15 | 93030400 | 93031799 | 13.43854 | 23.69014 | 41.33646 | 18.29458 | -0.41001 | 0.560292 | 0.462338 |
| 1 | 31449400 | 31451199 | 24.82979 | 43.32518 | 36.28192 | 15.97264 | 0.642243 | 0.758959 | 0.462369 |
| 13 | 51208600 | 51212399 | 33.29122 | 53.90728 | 89.12908 | 38.13571 | -0.36008 | 1.65929 | 0.462369 |
| 2 | 41755600 | 41757199 | 14.89053 | 27.73662 | 24.78936 | 10.33699 | 0.610748 | 0.187776 | 0.462383 |
| 19 | 12265600 | 12267599 | 17.30503 | 31.55927 | 27.05962 | 11.61739 | 0.640394 | 0.329644 | 0.462446 |
| 2 | 119656400 | 119659199 | 17.17722 | 29.82568 | 51.1709 | 22.45799 | -0.38861 | 0.811915 | 0.46245 |
| 12 | 113680000 | 113683399 | 52.50007 | 84.37876 | 85.11022 | 36.07581 | 0.338959 | 1.898837 | 0.462498 |
| 5 | 115929400 | 115931799 | 24.9776 | 42.6038 | 43.69406 | 19.15492 | 0.346016 | 0.996541 | 0.462501 |
| 8 | 48985000 | 48986999 | 13.8491 | 24.43352 | 43.45984 | 19.23124 | -0.437 | 0.560001 | 0.462503 |
| 1 | 48205800 | 48210999 | 54.81035 | 87.89754 | 89.49199 | 37.8075 | 0.321714 | 1.971444 | 0.462506 |
| 1 | 61479400 | 61480399 | 9.180981 | 17.16379 | 31.73268 | 13.15099 | -0.42338 | 0.133642 | 0.462515 |
| 10 | 18592200 | 18592999 | 6.242391 | 12.12159 | 25.25976 | 9.464071 | -0.46168 | -0.25277 | 0.462519 |
| 19 | 53061800 | 53065399 | 34.51174 | 56.66901 | 57.87911 | 24.93106 | 0.347179 | 1.428255 | 0.462529 |
| 7 | 60227400 | 60228799 | 26.49221 | 44.79312 | 45.87607 | 20.07594 | 0.346943 | 1.081561 | 0.462544 |
| 2 | 100801000 | 100801399 | 7.387886 | 14.39672 | 15.77735 | 5.723652 | 0.499808 | -0.38122 | 0.462551 |
| 3 | 36729200 | 36730399 | 11.78607 | 22.12715 | 23.01703 | 9.471054 | 0.408652 | 0.107724 | 0.46256 |
| 16 | 43864200 | 43868199 | 95.80672 | 149.3636 | 131.6707 | 54.64422 | 0.492798 | 2.700123 | 0.462575 |

| | | | | | | | | | |
|---|---|---|---|---|---|---|---|---|---|
| 6 | 143514000 | 143515999 | 20.9569 | 35.527 | 59.83486 | 26.20531 | -0.36802 | 1.053646 | 0.462582 |
| 9 | 20144200 | 20146999 | 49.92116 | 80.2578 | 82.70846 | 35.07291 | 0.308625 | 1.885977 | 0.462583 |
| 7 | 57625800 | 57629999 | 50.88412 | 81.711 | 84.27237 | 35.67526 | 0.305448 | 1.923638 | 0.462598 |
| 9 | 94473400 | 94474199 | 6.566206 | 12.65736 | 25.30908 | 9.803662 | -0.44769 | -0.13315 | 0.4626 |
| 16 | 49931400 | 49933599 | 25.854 | 42.69863 | 70.17702 | 30.30877 | -0.33423 | 1.346158 | 0.462605 |
| 19 | 4761400 | 4762799 | 12.56724 | 22.03935 | 41.00809 | 18.12477 | -0.50359 | 0.625777 | 0.46262 |
| 13 | 61755800 | 61761799 | 108.5404 | 168.8103 | 151.4504 | 62.87545 | 0.464838 | 2.750043 | 0.462624 |
| X | 125347000 | 125355399 | 121.0679 | 187.7125 | 150.7448 | 62.61141 | 0.621096 | 2.809766 | 0.462625 |
| X | 17511200 | 17515599 | 48.36006 | 78.52354 | 68.13679 | 29.21295 | 0.571544 | 1.623713 | 0.462625 |
| 6 | 141539800 | 141543399 | 49.32281 | 79.73803 | 74.1643 | 31.68833 | 0.465712 | 1.717314 | 0.462626 |
| 12 | 69257000 | 69257799 | 6.268443 | 12.09052 | 24.76223 | 9.575574 | -0.4797 | -0.14429 | 0.462635 |
| 7 | 42379200 | 42385199 | 76.10705 | 120.9498 | 109.7462 | 45.78294 | 0.473453 | 2.278605 | 0.462644 |
| 1 | 31076400 | 31079199 | 36.7616 | 60.27853 | 61.45281 | 26.45468 | 0.348984 | 1.452025 | 0.462657 |
| 18 | 3873000 | 3875999 | 54.61854 | 87.39636 | 88.33648 | 37.19594 | 0.328511 | 2.038738 | 0.46266 |
| 4 | 4137400 | 4140999 | 26.72877 | 43.77873 | 74.89542 | 32.22634 | -0.39636 | 1.45287 | 0.462671 |
| 13 | 95707400 | 95709599 | 22.04705 | 38.47679 | 37.04619 | 16.31101 | 0.441919 | 0.743982 | 0.462684 |
| 5 | 130125600 | 130127399 | 11.5682 | 20.66466 | 36.28635 | 15.93957 | -0.40898 | 0.43215 | 0.462688 |
| 3 | 88678600 | 88679999 | 7.537995 | 14.29603 | 28.73751 | 11.5964 | -0.50955 | 0.02152 | 0.462689 |
| 9 | 4082600 | 4085399 | 27.44536 | 46.72247 | 44.65751 | 19.57423 | 0.446783 | 1.022253 | 0.462711 |
| 17 | 33449400 | 33451999 | 57.18524 | 91.69547 | 87.44057 | 36.93433 | 0.415863 | 1.980254 | 0.462729 |
| 7 | 106069600 | 106070799 | 12.24114 | 21.63031 | 38.33656 | 16.95889 | -0.43062 | 0.524856 | 0.462736 |
| 10 | 84168400 | 84169599 | 15.68258 | 28.09716 | 28.32223 | 12.41021 | 0.389439 | 0.423673 | 0.462743 |
| 9 | 21076000 | 21080199 | 25.32952 | 41.41214 | 76.82745 | 33.03477 | -0.51451 | 1.475341 | 0.462743 |
| 1 | 171549400 | 171557599 | 196.8617 | 291.8797 | 284.7399 | 117.2766 | 0.286841 | 3.685421 | 0.462764 |
| 17 | 67850600 | 67858199 | 92.76048 | 141.6111 | 246.7807 | 102.3432 | -0.51277 | 3.101262 | 0.462769 |
| 13 | 72200600 | 72202199 | 18.32734 | 32.94606 | 28.31757 | 12.37308 | 0.620524 | 0.419387 | 0.46279 |
| 5 | 133685600 | 133688199 | 14.79448 | 25.85851 | 44.14383 | 19.45555 | -0.38014 | 0.698069 | 0.462795 |
| 17 | 11311600 | 11312199 | 5.246277 | 10.21489 | 23.09999 | 8.46803 | -0.5489 | -0.34917 | 0.462801 |
| 11 | 60472800 | 60473999 | 11.26296 | 19.63909 | 48.90157 | 21.51898 | -0.92217 | 0.72094 | 0.462806 |
| 3 | 43588800 | 43591799 | 37.17268 | 61.67877 | 54.40224 | 23.57158 | 0.561435 | 1.308584 | 0.462817 |
| 7 | 27080600 | 27087799 | 53.84027 | 85.54455 | 133.6112 | 55.72648 | -0.30921 | 2.269699 | 0.462829 |
| 5 | 135477000 | 135493399 | 132.0019 | 196.6045 | 341.6018 | 140.7614 | -0.53937 | 3.599153 | 0.462839 |
| 1 | 170903400 | 170906999 | 96.37903 | 146.7652 | 235.0382 | 97.41095 | -0.39386 | 3.152749 | 0.462843 |
| 12 | 56994600 | 56995399 | 10.74768 | 20.78333 | 19.64703 | 7.515269 | 0.639241 | -0.15082 | 0.462855 |
| 7 | 84534400 | 84537999 | 37.90207 | 61.10898 | 97.60045 | 41.55658 | -0.31714 | 1.786386 | 0.462882 |
| 3 | 36337600 | 36340799 | 51.87053 | 83.34082 | 85.35896 | 36.16481 | 0.316386 | 1.908222 | 0.462888 |
| 2 | 101016600 | 101019799 | 28.00725 | 48.39602 | 39.02296 | 17.11967 | 0.694734 | 0.887575 | 0.462897 |
| 8 | 101328600 | 101330999 | 20.76945 | 34.33959 | 72.89245 | 31.50576 | -0.70186 | 1.323112 | 0.462899 |
| 7 | 7138600 | 7140799 | 19.39151 | 33.96348 | 34.48876 | 15.22681 | 0.365539 | 0.671619 | 0.462906 |
| 6 | 60529200 | 60530999 | 43.20376 | 70.19672 | 68.76692 | 29.49724 | 0.397821 | 1.600875 | 0.462911 |
| 3 | 4035000 | 4037999 | 30.82026 | 53.29893 | 36.69897 | 16.10051 | 0.921835 | 0.891547 | 0.462913 |
| 7 | 14094200 | 14096999 | 44.00815 | 71.79067 | 65.16926 | 27.9992 | 0.511619 | 1.541529 | 0.462934 |
| 5 | 96197400 | 96199799 | 50.46985 | 81.27735 | 72.32332 | 30.74435 | 0.522946 | 1.829124 | 0.462941 |
| 13 | 61763200 | 61769999 | 139.804 | 213.2387 | 197.0639 | 81.7081 | 0.400531 | 3.12892 | 0.462944 |
| 2 | 16964200 | 16967599 | 37.94125 | 62.53464 | 53.19767 | 22.95098 | 0.612297 | 1.387947 | 0.462947 |
| 7 | 65336200 | 65337199 | 7.239901 | 13.59573 | 28.55763 | 11.7457 | -0.59654 | 0.081051 | 0.462952 |
| 18 | 19407000 | 19408199 | 12.61009 | 23.90436 | 23.06297 | 9.315665 | 0.53886 | 0.052155 | 0.46296 |

| | | | | | | | | | |
|---|---|---|---|---|---|---|---|---|---|
| 7 | 15217800 | 15219199 | 9.86902 | 18.32673 | 34.32151 | 14.38116 | -0.45444 | 0.188471 | 0.462967 |
| 2 | 46274400 | 46279399 | 116.1378 | 178.3307 | 178.679 | 74.18539 | 0.291512 | 2.988159 | 0.462973 |
| 4 | 74967800 | 74970599 | 27.47851 | 47.18435 | 41.84919 | 18.35603 | 0.556066 | 0.943209 | 0.46299 |
| 5 | 43532600 | 43533799 | 9.337658 | 17.29008 | 33.13314 | 13.93778 | -0.49171 | 0.195086 | 0.463017 |
| 4 | 50687400 | 50688599 | 13.7853 | 25.14801 | 25.39514 | 10.89862 | 0.406192 | 0.297627 | 0.463019 |
| 5 | 141157200 | 141157999 | 6.267398 | 12.10383 | 24.80064 | 9.544434 | -0.47341 | -0.15888 | 0.463022 |
| 18 | 66693800 | 66695799 | 17.06677 | 29.21013 | 52.16358 | 22.87415 | -0.45018 | 0.944284 | 0.463029 |
| 2 | 88547200 | 88548199 | 16.07549 | 29.17901 | 27.76351 | 11.99009 | 0.485863 | 0.349746 | 0.463036 |
| 3 | 50303000 | 50304399 | 13.13811 | 25.07212 | 21.41192 | 8.527505 | 0.730827 | 0.007287 | 0.463036 |
| 2 | 181011600 | 181012599 | 7.293063 | 14.01383 | 27.5598 | 10.77345 | -0.43736 | -0.09258 | 0.46304 |
| 9 | 15493600 | 15496599 | 21.37165 | 35.96195 | 59.69446 | 26.08499 | -0.34749 | 1.121083 | 0.46305 |
| 4 | 131573600 | 131576399 | 12.39643 | 21.86053 | 38.85946 | 17.19714 | -0.43608 | 0.548627 | 0.463055 |
| 3 | 104764400 | 104765999 | 11.03994 | 20.04385 | 35.41033 | 15.28774 | -0.40111 | 0.328486 | 0.463094 |
| 7 | 23063400 | 23075199 | 89.06723 | 140.7059 | 108.9675 | 45.256 | 0.694125 | 2.440654 | 0.463097 |
| 11 | 119517800 | 119520399 | 20.60418 | 34.48788 | 60.65912 | 26.43296 | -0.43073 | 1.1723 | 0.463106 |
| X | 164617600 | 164621399 | 68.02016 | 108.7878 | 93.35547 | 39.28228 | 0.563503 | 2.071961 | 0.463114 |
| Y | 89140200 | 89141599 | 19.51119 | 34.60637 | 27.34319 | 12.08336 | 0.730383 | 0.508119 | 0.463121 |
| 13 | 41403000 | 41403999 | 11.74272 | 20.80305 | 37.28687 | 16.48605 | -0.44531 | 0.490398 | 0.46317 |
| 14 | 120811200 | 120812999 | 13.87592 | 24.14239 | 49.47932 | 21.74329 | -0.64332 | 0.766291 | 0.463179 |
| 10 | 26457400 | 26459999 | 24.60209 | 42.58116 | 40.30227 | 17.68179 | 0.46499 | 0.860577 | 0.463184 |
| X | 136470400 | 136474999 | 23.19121 | 38.92447 | 69.36948 | 30.28083 | -0.44993 | 1.159009 | 0.463188 |
| 6 | 56207000 | 56207599 | 7.66745 | 14.45396 | 28.07617 | 11.47292 | -0.47694 | 0.059967 | 0.463188 |
| 6 | 97635800 | 97636799 | 7.66745 | 14.45396 | 28.07617 | 11.47292 | -0.47694 | 0.059967 | 0.463188 |
| 19 | 22442200 | 22442799 | 7.66745 | 14.45396 | 28.07617 | 11.47292 | -0.47694 | 0.059967 | 0.463188 |
| 8 | 5039800 | 5045399 | 58.99965 | 94.53518 | 91.72253 | 38.69963 | 0.389661 | 2.004278 | 0.463193 |
| 11 | 14378200 | 14378999 | 9.943954 | 19.0898 | 20.74504 | 8.131202 | 0.407778 | -0.07516 | 0.4632 |
| X | 123104800 | 123113399 | 119.8395 | 185.5454 | 157.1814 | 65.28902 | 0.542537 | 2.833792 | 0.4632 |
| 8 | 70448400 | 70449599 | 8.347795 | 15.39197 | 30.95422 | 13.09633 | -0.56618 | 0.208495 | 0.463219 |
| 15 | 77345400 | 77347999 | 26.18522 | 45.05379 | 37.71095 | 16.54192 | 0.639886 | 0.91623 | 0.463219 |
| 17 | 44758000 | 44759599 | 12.96457 | 23.05496 | 42.98492 | 18.9539 | -0.49941 | 0.479049 | 0.46322 |
| 17 | 22283200 | 22284599 | 24.54145 | 42.19599 | 42.03695 | 18.43913 | 0.389573 | 0.922101 | 0.463246 |
| X | 44231400 | 44233399 | 19.4176 | 36.21174 | 21.10156 | 8.78171 | 1.225661 | 0.210762 | 0.463264 |
| 9 | 5146400 | 5147399 | 14.81497 | 26.89881 | 27.31129 | 11.79052 | 0.392932 | 0.336804 | 0.463272 |
| 11 | 51553000 | 51554999 | 13.48527 | 23.65024 | 41.46945 | 18.32929 | -0.41854 | 0.630511 | 0.463287 |
| 10 | 25671600 | 25675199 | 36.0328 | 60.06841 | 52.06946 | 22.63187 | 0.586674 | 1.252523 | 0.463294 |
| 17 | 40985600 | 40988599 | 46.99268 | 76.18219 | 63.10071 | 27.02212 | 0.63637 | 1.64995 | 0.463301 |
| 1 | 61707800 | 61711599 | 30.82355 | 51.27385 | 52.99542 | 23.00226 | 0.333682 | 1.261956 | 0.463306 |
| 5 | 118997000 | 118998199 | 6.587687 | 12.6478 | 26.2814 | 10.29851 | -0.51798 | -0.09057 | 0.463311 |
| 17 | 11315400 | 11317199 | 19.40771 | 33.93595 | 34.13552 | 15.04991 | 0.379273 | 0.698029 | 0.463325 |
| 12 | 66720400 | 66721799 | 11.64916 | 20.99219 | 36.82779 | 16.04926 | -0.40014 | 0.377414 | 0.463344 |
| 3 | 157255800 | 157259199 | 40.67379 | 66.84497 | 59.62215 | 25.69126 | 0.542528 | 1.432263 | 0.463347 |
| 7 | 6551200 | 6555799 | 66.75881 | 106.2311 | 107.5393 | 44.93109 | 0.317682 | 2.22995 | 0.463353 |
| 3 | 159111800 | 159114199 | 22.30154 | 39.14434 | 33.46609 | 14.80216 | 0.612404 | 0.651188 | 0.463354 |
| 1 | 181604200 | 181607799 | 34.43818 | 55.77437 | 91.14204 | 39.00208 | -0.343 | 1.659326 | 0.463357 |
| 19 | 53584600 | 53585999 | 11.48195 | 20.7019 | 36.31223 | 15.81404 | -0.39926 | 0.371952 | 0.463358 |
| 1 | 31043400 | 31046599 | 56.06659 | 89.86307 | 90.42045 | 38.16805 | 0.337789 | 1.991411 | 0.463377 |
| 6 | 73081800 | 73084199 | 18.90966 | 32.13825 | 54.79785 | 24.01574 | -0.38525 | 1.020927 | 0.463383 |

| | | | | | | | | | |
|---|---|---|---|---|---|---|---|---|---|
| 8 | 100643800 | 100644199 | 7.784145 | 15.19923 | 15.69748 | 5.528809 | 0.620712 | -0.48197 | 0.463411 |
| 18 | 23641200 | 23643599 | 17.51624 | 31.00112 | 30.59961 | 13.53196 | 0.409753 | 0.521669 | 0.46344 |
| 16 | 53479800 | 53481999 | 23.00188 | 39.99702 | 38.09977 | 16.73648 | 0.456715 | 0.798678 | 0.463449 |
| 10 | 70761000 | 70774399 | 143.7918 | 214.6251 | 315.4646 | 130.0359 | -0.29534 | 3.560242 | 0.463465 |
| 1 | 8814000 | 8816799 | 31.84439 | 53.24319 | 46.49498 | 20.24664 | 0.575939 | 1.204135 | 0.463467 |
| 8 | 14128200 | 14129599 | 7.162783 | 13.69729 | 26.75733 | 10.60557 | -0.4463 | -0.05007 | 0.463469 |
| 8 | 128115600 | 128116199 | 10.01352 | 19.4529 | 19.39279 | 7.240429 | 0.596744 | -0.23801 | 0.46348 |
| 14 | 111318800 | 111321999 | 40.12925 | 65.42883 | 65.9553 | 28.33152 | 0.360286 | 1.539906 | 0.463506 |
| 2 | 11413400 | 11415599 | 13.16366 | 23.09077 | 41.46596 | 18.33081 | -0.4528 | 0.624962 | 0.463513 |
| 3 | 122206400 | 122208999 | 13.16366 | 23.09077 | 41.46596 | 18.33081 | -0.4528 | 0.624962 | 0.463513 |
| 13 | 102119200 | 102119999 | 10.88262 | 21.08997 | 19.78701 | 7.492469 | 0.664261 | -0.18087 | 0.46352 |
| 8 | 70183800 | 70187799 | 39.30884 | 64.35571 | 63.26543 | 27.21718 | 0.400479 | 1.478984 | 0.463542 |
| 13 | 53101400 | 53102799 | 16.04076 | 28.71706 | 28.80333 | 12.62911 | 0.395979 | 0.432111 | 0.463556 |
| 17 | 20087600 | 20102599 | 185.8466 | 280.0713 | 222.2663 | 91.8242 | 0.60564 | 3.383601 | 0.463567 |
| 13 | 77048600 | 77052599 | 41.57663 | 68.0174 | 63.31223 | 27.22958 | 0.478399 | 1.496035 | 0.463568 |
| 19 | 21857000 | 21857799 | 16.67928 | 29.95313 | 27.37578 | 11.97178 | 0.532159 | 0.416374 | 0.463579 |
| 4 | 81231600 | 81233799 | 17.1111 | 30.63126 | 29.59153 | 12.97541 | 0.449605 | 0.440099 | 0.463623 |
| 7 | 59832000 | 59834199 | 25.1047 | 43.62227 | 38.57673 | 16.93837 | 0.563476 | 0.820435 | 0.463624 |
| 15 | 55983400 | 55986399 | 25.1047 | 43.62227 | 38.57673 | 16.93837 | 0.563476 | 0.820435 | 0.463624 |
| 10 | 121062600 | 121064599 | 16.8222 | 28.59953 | 55.30482 | 24.23854 | -0.56611 | 1.008097 | 0.463624 |
| 10 | 118818000 | 118820599 | 35.35067 | 56.98309 | 93.54328 | 39.81802 | -0.35692 | 1.788789 | 0.463626 |
| 19 | 9631000 | 9640399 | 101.0514 | 157.1815 | 153.5036 | 63.72671 | 0.342698 | 2.744241 | 0.463635 |
| 13 | 73633000 | 73634999 | 29.36235 | 49.20099 | 49.69284 | 21.66912 | 0.36667 | 1.175201 | 0.463645 |
| 9 | 115689200 | 115692799 | 42.8251 | 68.70609 | 108.0637 | 45.68265 | -0.30381 | 1.950427 | 0.463659 |
| 7 | 131494200 | 131498599 | 57.37962 | 91.92527 | 91.75663 | 38.70497 | 0.348554 | 2.007471 | 0.463668 |
| 5 | 33200800 | 33202799 | 22.24389 | 38.33211 | 37.95849 | 16.65129 | 0.398816 | 0.885454 | 0.463685 |
| 18 | 34960400 | 34962199 | 10.56576 | 19.05716 | 34.29873 | 14.93015 | -0.43542 | 0.360184 | 0.463689 |
| 5 | 105876600 | 105881599 | 33.95275 | 54.53558 | 109.129 | 46.21517 | -0.64838 | 1.903653 | 0.4637 |
| 2 | 45572600 | 45572999 | 11.13433 | 21.62094 | 19.31079 | 7.234932 | 0.749156 | -0.21364 | 0.463711 |
| 10 | 56361200 | 56362999 | 11.32951 | 20.39732 | 35.77222 | 15.60727 | -0.40068 | 0.378319 | 0.463711 |
| 1 | 89457600 | 89462199 | 39.726 | 63.77511 | 103.2547 | 43.69246 | -0.34435 | 1.924697 | 0.463726 |
| X | 80884000 | 80887399 | 22.21894 | 38.75064 | 36.1634 | 15.91558 | 0.486828 | 0.755195 | 0.463732 |
| 15 | 95920200 | 95926199 | 61.564 | 98.34535 | 98.39487 | 41.33322 | 0.34057 | 2.100277 | 0.463737 |
| 18 | 72749400 | 72756799 | 102.4277 | 160.6013 | 134.3207 | 55.71025 | 0.575011 | 2.605918 | 0.463738 |
| 8 | 55990800 | 55994199 | 38.62448 | 62.08493 | 100.2324 | 42.4928 | -0.338 | 1.884283 | 0.463747 |
| 1 | 180602400 | 180604199 | 15.23562 | 26.55573 | 45.23199 | 19.89927 | -0.37741 | 0.740503 | 0.46375 |
| 13 | 104150600 | 104153799 | 27.16549 | 44.66046 | 73.54235 | 31.69417 | -0.3393 | 1.406727 | 0.463751 |
| 1 | 117673000 | 117687199 | 244.9805 | 359.1592 | 321.3214 | 132.3513 | 0.396403 | 3.874208 | 0.463755 |
| 19 | 36323600 | 36326999 | 27.91476 | 45.52018 | 78.0123 | 33.51242 | -0.40315 | 1.520242 | 0.463772 |
| 5 | 71137000 | 71137799 | 13.11101 | 24.21535 | 24.92068 | 10.51947 | 0.395743 | 0.221959 | 0.463785 |
| 3 | 21412200 | 21413199 | 14.19291 | 25.72613 | 24.83655 | 10.7447 | 0.463137 | 0.335092 | 0.463794 |
| 2 | 13588200 | 13588999 | 12.53644 | 23.98478 | 22.17571 | 8.777486 | 0.62573 | -0.01432 | 0.463805 |
| 10 | 130425400 | 130430999 | 285.0281 | 414.5173 | 304.9236 | 125.6845 | 0.667471 | 3.999082 | 0.463806 |
| 1 | 8025800 | 8027399 | 13.78297 | 25.77471 | 24.50946 | 10.15245 | 0.529969 | 0.155438 | 0.463822 |
| 5 | 132723600 | 132724599 | 20.52908 | 35.82804 | 35.50558 | 15.63282 | 0.400332 | 0.731574 | 0.463829 |
| 5 | 144572400 | 144575799 | 32.32985 | 52.4325 | 84.92877 | 36.38045 | -0.32836 | 1.625097 | 0.463833 |
| 8 | 92995800 | 92997399 | 27.66805 | 47.14799 | 45.22449 | 19.82432 | 0.442007 | 1.017538 | 0.463837 |

| | | | | | | | | | |
|---|---|---|---|---|---|---|---|---|---|
| 7 | 24128800 | 24133999 | 74.10422 | 114.8478 | 185.071 | 76.87326 | -0.38308 | 2.799174 | 0.463853 |
| 3 | 158659200 | 158661399 | 19.99219 | 35.19162 | 33.18185 | 14.68983 | 0.472704 | 0.601641 | 0.463872 |
| 5 | 59529000 | 59531199 | 19.99879 | 35.92685 | 28.24492 | 12.34825 | 0.748096 | 0.436118 | 0.463875 |
| 6 | 48570200 | 48579799 | 139.7421 | 205.9867 | 387.0917 | 159.4088 | -0.66632 | 3.778852 | 0.463876 |
| 1 | 43324800 | 43329599 | 39.38793 | 65.16224 | 55.61887 | 24.04977 | 0.608387 | 1.349234 | 0.463878 |
| 4 | 71752400 | 71752999 | 14.48601 | 26.431 | 26.84072 | 11.51643 | 0.398375 | 0.30549 | 0.463898 |
| 7 | 42577000 | 42579599 | 30.93738 | 53.70792 | 35.01752 | 15.36672 | 1.001216 | 0.85668 | 0.463899 |
| 12 | 110394000 | 110395799 | 24.73663 | 40.92263 | 68.35002 | 29.55431 | -0.35712 | 1.323237 | 0.463902 |
| 1 | 119835800 | 119838399 | 22.53704 | 37.70768 | 62.42757 | 27.19655 | -0.34378 | 1.184866 | 0.463906 |
| 11 | 119403000 | 119404399 | 14.33148 | 24.96234 | 44.75113 | 19.67907 | -0.45082 | 0.746813 | 0.46392 |
| 2 | 177106800 | 177107999 | 7.027084 | 13.55265 | 27.67398 | 10.70411 | -0.47605 | -0.12622 | 0.463928 |
| 1 | 18143400 | 18145399 | 25.16009 | 41.70268 | 68.61124 | 29.70637 | -0.33531 | 1.300746 | 0.46393 |
| X | 127505600 | 127509399 | 71.74756 | 114.5865 | 85.40392 | 35.86425 | 0.764262 | 2.116017 | 0.463931 |
| 1 | 96499400 | 96500599 | 11.69552 | 20.92651 | 36.26212 | 15.90683 | -0.38871 | 0.421626 | 0.463938 |
| 16 | 88547000 | 88548599 | 11.69552 | 20.92651 | 36.26212 | 15.90683 | -0.38871 | 0.421626 | 0.463938 |
| 11 | 22544800 | 22547399 | 27.63338 | 45.50724 | 75.52422 | 32.57179 | -0.3496 | 1.387865 | 0.463941 |
| 15 | 102533000 | 102534599 | 12.09731 | 21.27937 | 39.42436 | 17.45592 | -0.49686 | 0.575687 | 0.46397 |
| 5 | 24137600 | 24138799 | 22.84109 | 39.8172 | 37.5469 | 16.51964 | 0.471656 | 0.764061 | 0.463971 |
| 8 | 6542600 | 6545999 | 45.52998 | 73.72416 | 65.29367 | 27.93698 | 0.538401 | 1.669592 | 0.463976 |
| 16 | 3563600 | 3573599 | 132.209 | 202.7578 | 165.5125 | 68.7072 | 0.585513 | 3.015613 | 0.463976 |
| 7 | 133913800 | 133915399 | 17.68733 | 31.26174 | 31.29722 | 13.8442 | 0.388919 | 0.540834 | 0.463997 |
| 5 | 53298600 | 53299999 | 14.72059 | 26.63964 | 25.95836 | 11.25267 | 0.448328 | 0.353223 | 0.464005 |
| 7 | 137856000 | 137856999 | 4.810336 | 9.368219 | 21.28961 | 7.784565 | -0.55111 | -0.36653 | 0.46401 |
| 11 | 101028200 | 101029999 | 11.49049 | 20.79258 | 36.92184 | 16.00675 | -0.4126 | 0.352658 | 0.464012 |
| 17 | 19786000 | 19790999 | 80.17014 | 127.5028 | 104.7811 | 43.74575 | 0.617198 | 2.258629 | 0.464013 |
| 18 | 37112800 | 37115599 | 71.97056 | 113.9037 | 111.9078 | 46.51984 | 0.353431 | 2.388738 | 0.464044 |
| 4 | 151008600 | 151011399 | 22.37364 | 37.26332 | 64.60361 | 28.06337 | -0.41009 | 1.233551 | 0.464069 |
| 14 | 14254000 | 14256199 | 13.33858 | 22.98186 | 58.05135 | 25.46531 | -0.94648 | 0.873983 | 0.464082 |
| 7 | 51558000 | 51560799 | 21.7412 | 38.00597 | 35.85281 | 15.80343 | 0.471272 | 0.715275 | 0.464085 |
| 9 | 52770800 | 52773599 | 25.09119 | 43.00932 | 43.07517 | 18.89112 | 0.381126 | 0.951355 | 0.464094 |
| 16 | 87546600 | 87547999 | 16.07455 | 28.55105 | 27.82388 | 12.28121 | 0.430625 | 0.477769 | 0.464097 |
| 5 | 149158000 | 149161399 | 50.49929 | 80.19259 | 130.4865 | 54.39363 | -0.36889 | 2.283772 | 0.464105 |
| 10 | 126116400 | 126118999 | 21.63908 | 37.75179 | 37.05056 | 16.3094 | 0.414327 | 0.745829 | 0.464107 |
| 14 | 107505600 | 107506799 | 13.33902 | 23.41021 | 40.94727 | 18.10925 | -0.41492 | 0.618004 | 0.464109 |
| 4 | 135997800 | 136005599 | 85.1453 | 130.8619 | 224.4436 | 93.19807 | -0.48224 | 2.963351 | 0.464127 |
| 11 | 84509000 | 84509999 | 13.12536 | 24.68748 | 24.00214 | 9.845305 | 0.509097 | 0.110699 | 0.464137 |
| 7 | 132180200 | 132182599 | 15.541 | 27.16219 | 46.22956 | 20.37408 | -0.37604 | 0.703759 | 0.464171 |
| 6 | 83601800 | 83604399 | 26.6169 | 43.55399 | 74.58251 | 32.07943 | -0.39837 | 1.467166 | 0.464171 |
| 17 | 19197000 | 19208399 | 127.2086 | 196.7116 | 155.8898 | 64.75293 | 0.637299 | 2.859485 | 0.464183 |
| 9 | 39072200 | 39075199 | 72.04396 | 114.3359 | 103.1035 | 42.95283 | 0.48045 | 2.311165 | 0.464193 |
| 4 | 103156400 | 103158799 | 12.01633 | 21.30057 | 38.23266 | 16.90219 | -0.44726 | 0.48542 | 0.464213 |
| 19 | 56125800 | 56133599 | 88.52824 | 138.9642 | 137.2885 | 56.93993 | 0.336239 | 2.576623 | 0.464224 |
| X | 34790600 | 34794399 | 44.25096 | 73.21887 | 52.31191 | 22.62226 | 0.864303 | 1.360277 | 0.464252 |
| 15 | 6718200 | 6720199 | 17.67988 | 31.27834 | 31.25003 | 13.8207 | 0.392341 | 0.53027 | 0.464258 |
| 18 | 8099200 | 8100799 | 21.48124 | 37.40695 | 37.68017 | 16.56722 | 0.376735 | 0.771954 | 0.464327 |
| 3 | 106026000 | 106028199 | 22.74354 | 39.51987 | 38.79263 | 17.03328 | 0.413686 | 0.811016 | 0.464329 |
| 9 | 20250600 | 20255999 | 69.72586 | 111.1998 | 101.2704 | 42.44239 | 0.473454 | 2.160627 | 0.46433 |

| | | | | | | | | | |
|---|---|---|---|---|---|---|---|---|---|
| 18 | 24139200 | 24143399 | 37.38484 | 61.50578 | 59.96104 | 25.84903 | 0.415213 | 1.411579 | 0.464341 |
| 13 | 85477200 | 85479199 | 19.82491 | 33.20476 | 66.14301 | 28.83428 | -0.60974 | 1.167832 | 0.464347 |
| 2 | 53973200 | 53976599 | 27.42749 | 44.75217 | 77.42796 | 33.26861 | -0.41619 | 1.510235 | 0.464348 |
| 1 | 31973800 | 31977599 | 58.46471 | 93.5716 | 94.19702 | 39.67619 | 0.334675 | 2.038722 | 0.464353 |
| 14 | 29319600 | 29320599 | 13.36654 | 24.79261 | 24.85864 | 10.43268 | 0.438881 | 0.199582 | 0.464368 |
| 2 | 32002200 | 32007799 | 59.42031 | 93.79849 | 146.5593 | 60.84481 | -0.31797 | 2.446832 | 0.464376 |
| 15 | 48777400 | 48779199 | 17.66673 | 30.61834 | 54.47678 | 23.90153 | -0.44147 | 0.836679 | 0.464386 |
| 9 | 64914800 | 64916599 | 19.07792 | 33.46071 | 33.92784 | 14.99676 | 0.367629 | 0.646952 | 0.464413 |
| 15 | 89841200 | 89842799 | 17.00667 | 29.03529 | 58.00999 | 25.43277 | -0.61196 | 0.976167 | 0.464462 |
| 9 | 71392600 | 71393799 | 7.012711 | 13.41306 | 26.7636 | 10.60182 | -0.4759 | -0.05398 | 0.464469 |
| 6 | 42023400 | 42026999 | 40.92825 | 65.79038 | 105.7329 | 44.82706 | -0.33144 | 1.888556 | 0.464481 |
| 17 | 10480600 | 10482399 | 26.74844 | 46.65116 | 35.75507 | 15.73607 | 0.769586 | 0.776279 | 0.464502 |
| 5 | 112710200 | 112714199 | 23.32359 | 38.44927 | 69.13345 | 29.82856 | -0.46402 | 1.361412 | 0.464504 |
| 2 | 161475400 | 161477199 | 13.61681 | 25.23728 | 25.41002 | 10.67532 | 0.431996 | 0.208087 | 0.464507 |
| 17 | 94427600 | 94428599 | 12.95208 | 24.65515 | 22.5728 | 9.037688 | 0.624973 | 0.026264 | 0.464522 |
| 8 | 32861800 | 32863199 | 14.80421 | 27.90694 | 23.36525 | 9.548192 | 0.7275 | 0.110459 | 0.464528 |
| 4 | 47979600 | 47980599 | 8.106339 | 15.2316 | 29.09384 | 11.95455 | -0.45976 | 0.085881 | 0.464553 |
| 14 | 74292400 | 74293599 | 10.93808 | 19.63146 | 35.31936 | 15.4532 | -0.43966 | 0.396862 | 0.464553 |
| 7 | 144691400 | 144694199 | 37.2165 | 61.1566 | 60.88907 | 26.23599 | 0.384319 | 1.428434 | 0.464572 |
| 16 | 35729400 | 35731199 | 18.55175 | 33.7651 | 26.95821 | 11.60293 | 0.740434 | 0.350312 | 0.464591 |
| 3 | 46030000 | 46030399 | 5.67229 | 10.99961 | 23.28997 | 8.792731 | -0.4936 | -0.23606 | 0.464605 |
| 13 | 33861200 | 33863199 | 21.97231 | 38.70299 | 31.76429 | 14.0604 | 0.672069 | 0.601855 | 0.464607 |
| 18 | 46316400 | 46318199 | 15.70559 | 27.45648 | 47.83459 | 21.08574 | -0.40964 | 0.70281 | 0.464616 |
| 16 | 57215200 | 57216999 | 16.71522 | 29.68782 | 29.63955 | 13.08305 | 0.395823 | 0.485265 | 0.46462 |
| 15 | 68520000 | 68520599 | 5.962376 | 11.55838 | 24.47726 | 9.255014 | -0.49702 | -0.22714 | 0.464626 |
| 5 | 67239000 | 67243399 | 58.27393 | 93.18141 | 95.5759 | 40.20512 | 0.3063 | 2.066131 | 0.464661 |
| 6 | 3050000 | 3055799 | 102.7602 | 160.6744 | 127.6111 | 52.94257 | 0.646387 | 2.653199 | 0.464667 |
| 16 | 3516400 | 3518999 | 18.01111 | 30.74198 | 54.32484 | 23.81662 | -0.4358 | 0.974736 | 0.46467 |
| 1 | 150061400 | 150065999 | 56.69989 | 91.25238 | 81.47195 | 34.60157 | 0.517032 | 1.862431 | 0.464672 |
| 13 | 61636600 | 61637799 | 18.06008 | 33.5574 | 24.08266 | 10.08281 | 0.921078 | 0.229029 | 0.464681 |
| 1 | 102166800 | 102180799 | 136.0901 | 209.2915 | 168.0837 | 69.80116 | 0.612067 | 2.964296 | 0.464694 |
| 10 | 82155800 | 82160199 | 51.58845 | 83.23227 | 77.68401 | 33.10571 | 0.457374 | 1.779597 | 0.464731 |
| 7 | 13885800 | 13886399 | 23.01068 | 39.55426 | 38.05414 | 16.69198 | 0.439686 | 0.912964 | 0.46477 |
| 7 | 7930400 | 7932199 | 29.21676 | 49.22694 | 42.20921 | 18.45165 | 0.602209 | 1.117092 | 0.464772 |
| 12 | 103909200 | 103926999 | 192.2824 | 280.416 | 407.0287 | 167.7221 | -0.30872 | 3.951719 | 0.464785 |
| 6 | 24012000 | 24012799 | 11.24782 | 21.59662 | 21.14968 | 8.285697 | 0.556767 | -0.06406 | 0.464789 |
| 3 | 10374800 | 10377399 | 27.4677 | 46.5414 | 46.89187 | 20.53371 | 0.370572 | 1.067626 | 0.464809 |
| 12 | 48404200 | 48406199 | 17.64427 | 31.61661 | 29.55596 | 12.94913 | 0.4976 | 0.438032 | 0.464816 |
| 1 | 105881400 | 105884399 | 30.35363 | 50.96464 | 50.20635 | 21.9209 | 0.402546 | 1.16008 | 0.464856 |
| 2 | 154491600 | 154493599 | 21.05101 | 36.66786 | 37.22463 | 16.36977 | 0.365643 | 0.766215 | 0.464867 |
| 14 | 50777000 | 50783999 | 96.03822 | 150.7464 | 134.0716 | 55.6054 | 0.487623 | 2.580587 | 0.46488 |
| 3 | 47772000 | 47775399 | 37.16803 | 61.02687 | 61.46945 | 26.47687 | 0.367133 | 1.437889 | 0.464882 |
| 7 | 38549400 | 38553799 | 54.87954 | 88.584 | 74.78897 | 31.89143 | 0.60253 | 1.769997 | 0.464892 |
| 18 | 20495800 | 20497199 | 12.86335 | 23.82641 | 24.50366 | 10.303 | 0.400968 | 0.204104 | 0.464905 |
| 14 | 96944800 | 96946399 | 20.1663 | 35.53404 | 32.4915 | 14.3813 | 0.517164 | 0.588863 | 0.464912 |
| X | 79577200 | 79578799 | 27.11835 | 46.49442 | 38.32051 | 16.80621 | 0.661275 | 0.953398 | 0.464917 |
| 17 | 27118200 | 27119599 | 10.44901 | 18.84557 | 34.34075 | 14.94946 | -0.453 | 0.361616 | 0.464932 |

| | | | | | | | | | |
|---|---|---|---|---|---|---|---|---|---|
| X | 146583400 | 146584999 | 21.50608 | 37.84683 | 33.03785 | 14.62683 | 0.583235 | 0.612994 | 0.46497 |
| 5 | 43674600 | 43675199 | 5.826979 | 11.25785 | 24.24194 | 9.311707 | -0.54138 | -0.17142 | 0.464972 |
| X | 148662400 | 148667399 | 81.41746 | 128.8129 | 120.7831 | 50.20103 | 0.420317 | 2.404849 | 0.464987 |
| 13 | 64656600 | 64663599 | 85.21688 | 134.0304 | 133.5007 | 55.38316 | 0.326642 | 2.535546 | 0.464994 |
| 1 | 21078200 | 21079599 | 8.556765 | 15.98429 | 31.15107 | 12.92711 | -0.50046 | 0.130829 | 0.465011 |
| 12 | 28559600 | 28561799 | 13.93952 | 24.46813 | 41.92578 | 18.54459 | -0.38555 | 0.619151 | 0.465021 |
| 4 | 29357800 | 29359999 | 33.42969 | 55.59372 | 54.14484 | 23.49668 | 0.419177 | 1.275412 | 0.465022 |
| 6 | 128982400 | 128983799 | 21.54898 | 37.26914 | 36.38986 | 15.96494 | 0.42006 | 0.847651 | 0.465024 |
| 2 | 15189800 | 15201799 | 146.9725 | 222.0994 | 222.5212 | 92.00714 | 0.273027 | 3.318976 | 0.465048 |
| 17 | 39559600 | 39562599 | 29.65852 | 51.36734 | 37.92084 | 16.63733 | 0.822296 | 0.876269 | 0.465052 |
| 6 | 56276200 | 56277199 | 11.48326 | 20.26507 | 38.42457 | 17.00218 | -0.52798 | 0.53554 | 0.46507 |
| 6 | 116765600 | 116766399 | 7.370169 | 13.95952 | 27.62636 | 11.17883 | -0.49081 | 0.024906 | 0.46507 |
| 4 | 36222200 | 36222799 | 8.795016 | 16.3151 | 30.49131 | 12.7946 | -0.45234 | 0.178506 | 0.465081 |
| 4 | 143461000 | 143465599 | 105.5245 | 163.2273 | 163.7018 | 67.99777 | 0.298197 | 2.843786 | 0.465087 |
| X | 126579000 | 126580399 | 24.99257 | 43.11305 | 41.41828 | 18.16957 | 0.442673 | 0.893632 | 0.465089 |
| X | 34983600 | 34990999 | 170.0645 | 255.2413 | 242.4438 | 100.0505 | 0.341305 | 3.460181 | 0.465103 |
| 6 | 65173000 | 65174799 | 22.67262 | 39.47509 | 38.14207 | 16.76858 | 0.436528 | 0.781089 | 0.465143 |
| 16 | 5939800 | 5941799 | 21.07718 | 35.47024 | 59.64761 | 26.06536 | -0.3661 | 1.120069 | 0.465146 |
| 4 | 118504000 | 118505399 | 5.985808 | 11.52805 | 24.70854 | 9.596088 | -0.55014 | -0.13237 | 0.465165 |
| 8 | 60815200 | 60816799 | 19.05572 | 33.91036 | 29.21079 | 12.87277 | 0.60896 | 0.486654 | 0.465172 |
| 9 | 59950400 | 59953199 | 43.92576 | 70.91435 | 71.99373 | 30.70588 | 0.337354 | 1.745383 | 0.465204 |
| 14 | 53366000 | 53367399 | 16.92109 | 31.0417 | 25.41492 | 10.82728 | 0.713815 | 0.289121 | 0.465206 |
| 16 | 38910000 | 38911599 | 15.47623 | 27.83579 | 28.25565 | 12.34041 | 0.382539 | 0.397401 | 0.46521 |
| 2 | 78426800 | 78429799 | 25.43684 | 41.74908 | 72.52695 | 31.23459 | -0.41716 | 1.423153 | 0.465222 |
| 4 | 23942400 | 23942999 | 8.384466 | 15.77557 | 30.23304 | 12.39415 | -0.46217 | 0.081303 | 0.465243 |
| Y | 86914600 | 86915199 | 8.390459 | 16.31305 | 12.59376 | 4.671289 | 0.970035 | -0.30591 | 0.465244 |
| 10 | 85765800 | 85768399 | 16.2311 | 28.10652 | 47.93352 | 21.03016 | -0.38057 | 0.830755 | 0.46526 |
| 10 | 118866600 | 118869199 | 16.2311 | 28.10652 | 47.93352 | 21.03016 | -0.38057 | 0.830755 | 0.46526 |
| 3 | 87388000 | 87389399 | 17.52432 | 30.84925 | 30.36611 | 13.44314 | 0.410871 | 0.57993 | 0.465269 |
| 3 | 45906800 | 45908399 | 22.94713 | 39.84591 | 39.27157 | 17.23603 | 0.407677 | 0.820789 | 0.465299 |
| 3 | 96989000 | 97000599 | 143.5063 | 214.7251 | 314.1583 | 129.5592 | -0.28547 | 3.514597 | 0.465305 |
| 7 | 126574400 | 126576199 | 10.9942 | 19.48868 | 36.86423 | 16.29723 | -0.52193 | 0.479875 | 0.465312 |
| 10 | 75027400 | 75027799 | 9.151304 | 17.75824 | 19.34212 | 7.27891 | 0.459473 | -0.22205 | 0.465325 |
| 1 | 83205800 | 83208999 | 47.18028 | 75.93116 | 75.72096 | 32.15883 | 0.357772 | 1.845108 | 0.46533 |
| 4 | 71215400 | 71218599 | 63.38902 | 101.0712 | 102.6593 | 43.01124 | 0.315925 | 2.161106 | 0.465331 |
| 1 | 6219200 | 6221999 | 35.76144 | 59.29054 | 50.40144 | 21.82625 | 0.614458 | 1.312027 | 0.465335 |
| 13 | 13621600 | 13626199 | 28.93876 | 47.43028 | 82.5635 | 35.5359 | -0.42111 | 1.452689 | 0.46535 |
| 13 | 61954400 | 61970799 | 363.2341 | 514.6936 | 443.1461 | 183.3606 | 0.403287 | 4.386371 | 0.46535 |
| 2 | 104372000 | 104375599 | 38.59746 | 63.16793 | 63.23215 | 27.19896 | 0.374122 | 1.48136 | 0.465354 |
| 2 | 13178800 | 13181199 | 18.65659 | 31.79781 | 54.869 | 24.05239 | -0.40194 | 0.991288 | 0.465373 |
| 12 | 111874000 | 111875799 | 14.8393 | 25.63672 | 47.93544 | 21.02951 | -0.51332 | 0.845708 | 0.46538 |
| 10 | 63417800 | 63421199 | 22.49133 | 37.65995 | 62.50216 | 27.24216 | -0.34739 | 1.175197 | 0.465384 |
| 5 | 104367800 | 104369199 | 20.11973 | 35.14002 | 35.18088 | 15.49717 | 0.386029 | 0.721764 | 0.465393 |
| 5 | 128726200 | 128728199 | 15.74649 | 27.20655 | 48.46189 | 21.26053 | -0.44378 | 0.848349 | 0.465394 |
| 7 | 11020000 | 11023599 | 92.97294 | 144.5287 | 140.3029 | 58.2517 | 0.350667 | 2.749131 | 0.465433 |
| 10 | 47269000 | 47270799 | 23.17682 | 40.52999 | 35.77002 | 15.77167 | 0.566998 | 0.718148 | 0.465434 |
| 1 | 177464000 | 177468599 | 31.55383 | 51.05416 | 86.65701 | 37.05667 | -0.39884 | 1.671712 | 0.465437 |

| | | | | | | | | | |
|---|---|---|---|---|---|---|---|---|---|
| 6 | 126476600 | 126478999 | 19.76103 | 33.20615 | 59.06812 | 25.80055 | -0.447 | 1.125786 | 0.465464 |
| 9 | 3489000 | 3495199 | 76.57881 | 121.2384 | 109.3697 | 45.47263 | 0.476754 | 2.382004 | 0.465482 |
| 17 | 53438400 | 53439999 | 18.24462 | 32.66704 | 29.43908 | 12.90636 | 0.549455 | 0.445931 | 0.465483 |
| 14 | 59347800 | 59348999 | 7.43325 | 13.8294 | 29.50942 | 12.33015 | -0.63729 | 0.146766 | 0.465517 |
| 4 | 119396000 | 119397599 | 9.482593 | 17.17556 | 33.36131 | 14.44592 | -0.53962 | 0.325661 | 0.465532 |
| 6 | 120435600 | 120441999 | 35.46598 | 56.78093 | 118.6982 | 50.01929 | -0.71697 | 2.010639 | 0.465535 |
| 16 | 31329000 | 31330599 | 9.278957 | 17.11103 | 31.46384 | 13.30968 | -0.43764 | 0.218319 | 0.465545 |
| 7 | 34353000 | 34355999 | 22.28172 | 36.96887 | 65.49619 | 28.37729 | -0.44153 | 1.279065 | 0.465557 |
| 7 | 144442800 | 144446199 | 24.03682 | 39.89637 | 66.23879 | 28.70675 | -0.34816 | 1.280382 | 0.465571 |
| 15 | 86075400 | 86082599 | 173.7632 | 253.7964 | 393.9572 | 162.3099 | -0.40333 | 3.927035 | 0.465578 |
| X | 32843800 | 32847799 | 31.31496 | 54.35141 | 35.01859 | 15.36506 | 1.018279 | 0.861685 | 0.465589 |
| 3 | 15369400 | 15372599 | 58.06523 | 92.77124 | 95.79772 | 40.25143 | 0.295312 | 2.09045 | 0.46559 |
| 18 | 70456600 | 70460199 | 57.18863 | 90.58273 | 140.0176 | 58.22896 | -0.29851 | 2.362954 | 0.465592 |
| 6 | 85406000 | 85407599 | 7.976259 | 15.1194 | 28.74595 | 11.61248 | -0.43167 | 0.027549 | 0.465598 |
| 7 | 110666000 | 110669399 | 32.03742 | 53.96372 | 43.67348 | 19.09007 | 0.685225 | 1.130583 | 0.465604 |
| 9 | 46504000 | 46505799 | 15.45849 | 26.7155 | 47.93573 | 21.02985 | -0.45408 | 0.845384 | 0.465618 |
| 10 | 24206200 | 24209999 | 33.73005 | 55.97644 | 55.35746 | 23.98837 | 0.396892 | 1.297919 | 0.465619 |
| 3 | 121537000 | 121540199 | 34.3536 | 55.5695 | 89.83861 | 38.40972 | -0.32914 | 1.685675 | 0.465619 |
| 12 | 98886000 | 98887199 | 5.967135 | 11.55441 | 24.39815 | 9.274452 | -0.50013 | -0.20883 | 0.46562 |
| 4 | 124097200 | 124099999 | 35.49059 | 58.16157 | 60.12854 | 25.87841 | 0.328597 | 1.455859 | 0.465632 |
| 2 | 165052200 | 165055199 | 21.10541 | 35.24786 | 61.72992 | 26.86161 | -0.42453 | 1.199182 | 0.465639 |
| 8 | 79234600 | 79236599 | 16.92077 | 28.96534 | 51.62516 | 22.63837 | -0.44727 | 0.942075 | 0.465663 |
| 7 | 131772200 | 131773399 | 14.88771 | 26.07949 | 44.58185 | 19.68562 | -0.38231 | 0.660889 | 0.465663 |
| 17 | 63951400 | 63952799 | 19.29853 | 34.43354 | 29.88752 | 13.14162 | 0.600522 | 0.475601 | 0.465664 |
| 2 | 15660000 | 15660599 | 5.950193 | 11.57231 | 24.22672 | 8.962918 | -0.45109 | -0.30346 | 0.46567 |
| 3 | 87578600 | 87580599 | 16.82519 | 29.27111 | 51.71422 | 22.72411 | -0.43052 | 0.782543 | 0.465675 |
| 1 | 94214600 | 94218799 | 68.08298 | 108.5394 | 103.0331 | 43.15893 | 0.41315 | 2.171287 | 0.465718 |
| 4 | 20301000 | 20302999 | 17.2649 | 29.86677 | 53.36922 | 23.41428 | -0.44829 | 0.84786 | 0.465756 |
| 15 | 42058000 | 42061799 | 59.46491 | 95.19226 | 93.91573 | 39.5694 | 0.364039 | 2.033616 | 0.465769 |
| 10 | 117348000 | 117348599 | 6.249879 | 12.11265 | 24.46486 | 9.261291 | -0.4308 | -0.21866 | 0.465791 |
| 18 | 18106400 | 18107199 | 6.249879 | 12.11265 | 24.46486 | 9.261291 | -0.4308 | -0.21866 | 0.465791 |
| 13 | 29150800 | 29151799 | 14.43038 | 26.12542 | 26.33064 | 11.40806 | 0.400163 | 0.347632 | 0.465803 |
| 3 | 33628800 | 33631999 | 41.478 | 67.86782 | 63.42679 | 27.28058 | 0.472774 | 1.492061 | 0.465816 |
| 10 | 45884800 | 45886799 | 19.05088 | 33.34003 | 33.58913 | 14.81953 | 0.376943 | 0.681326 | 0.465831 |
| X | 119963800 | 119966199 | 50.17276 | 80.81015 | 69.3162 | 29.47198 | 0.576038 | 1.823759 | 0.465832 |
| 2 | 74415000 | 74416799 | 22.12595 | 38.83368 | 34.1206 | 15.09084 | 0.573298 | 0.653765 | 0.465844 |
| 10 | 103631200 | 103635999 | 85.16682 | 131.7495 | 196.1746 | 81.48614 | -0.27127 | 2.838154 | 0.465848 |
| 14 | 101495800 | 101496599 | 7.203991 | 13.74015 | 27.22291 | 10.85446 | -0.47356 | -0.02576 | 0.465856 |
| 13 | 53119800 | 53123199 | 42.22005 | 67.63409 | 107.9554 | 45.53311 | -0.32754 | 1.997156 | 0.465884 |
| 15 | 84089800 | 84091199 | 9.475037 | 17.34126 | 32.43356 | 13.86147 | -0.47265 | 0.265649 | 0.465896 |
| 7 | 11838400 | 11858399 | 312.3479 | 450.065 | 373.1648 | 153.9891 | 0.481836 | 4.139985 | 0.465899 |
| 7 | 17465800 | 17468199 | 43.88743 | 71.64713 | 64.57653 | 27.75208 | 0.522647 | 1.531088 | 0.465909 |
| 16 | 86220000 | 86221999 | 20.13608 | 33.88826 | 69.66462 | 30.44316 | -0.65502 | 1.125042 | 0.465909 |
| 17 | 19964800 | 19969599 | 49.63536 | 80.57355 | 67.74281 | 29.03852 | 0.616629 | 1.630365 | 0.465927 |
| 3 | 121308000 | 121311999 | 30.94483 | 50.51451 | 87.14672 | 37.45077 | -0.41177 | 1.515571 | 0.465929 |
| 19 | 13119400 | 13122599 | 66.37405 | 105.4885 | 109.1878 | 45.55272 | 0.283847 | 2.266942 | 0.465988 |
| 1 | 103116400 | 103118999 | 22.20955 | 39.18929 | 31.20972 | 13.81009 | 0.716249 | 0.584394 | 0.466007 |

| | | | | | | | | | |
|---|---|---|---|---|---|---|---|---|---|
| 7 | 58185400 | 58185999 | 13.69346 | 25.66878 | 24.49463 | 10.1052 | 0.5294 | 0.138691 | 0.466024 |
| 17 | 49078400 | 49083999 | 37.13761 | 59.5984 | 105.6759 | 44.69259 | -0.47586 | 1.932213 | 0.466033 |
| 17 | 84274400 | 84281799 | 71.93375 | 112.4615 | 172.2352 | 71.45681 | -0.29997 | 2.643608 | 0.466034 |
| 8 | 100251400 | 100254599 | 37.88772 | 62.65956 | 56.68151 | 24.50365 | 0.524792 | 1.349568 | 0.466046 |
| X | 43040000 | 43042599 | 22.10958 | 39.77284 | 26.45285 | 11.55153 | 0.989101 | 0.43948 | 0.466053 |
| 2 | 169906600 | 169909799 | 51.2139 | 82.20505 | 84.74601 | 35.85221 | 0.305326 | 1.937578 | 0.466054 |
| 18 | 80434400 | 80442599 | 88.51504 | 135.4586 | 218.8164 | 90.79701 | -0.40064 | 3.048956 | 0.46608 |
| 1 | 150148600 | 150149399 | 7.664326 | 14.4146 | 28.57368 | 11.72217 | -0.51083 | 0.074814 | 0.466088 |
| 8 | 53422400 | 53425199 | 22.19256 | 38.90158 | 34.65585 | 15.31128 | 0.55344 | 0.673734 | 0.466089 |
| 19 | 5006000 | 5008199 | 17.8782 | 30.65152 | 51.64236 | 22.64872 | -0.36611 | 0.932733 | 0.466104 |
| 7 | 37647400 | 37649599 | 21.50938 | 38.71621 | 27.12125 | 11.8362 | 0.915386 | 0.433714 | 0.466118 |
| 5 | 77163000 | 77164599 | 11.9341 | 20.9765 | 39.9271 | 17.68083 | -0.53615 | 0.582882 | 0.466125 |
| X | 54669800 | 54676599 | 203.7579 | 300.6307 | 287.204 | 118.2751 | 0.310428 | 3.767807 | 0.466125 |
| 2 | 101387800 | 101388999 | 15.0979 | 28.1224 | 24.15592 | 10.07315 | 0.66737 | 0.187247 | 0.466136 |
| 6 | 5330200 | 5332199 | 30.99794 | 51.45399 | 52.98778 | 22.96508 | 0.338724 | 1.286942 | 0.466159 |
| 10 | 71196600 | 71198399 | 10.51551 | 19.04186 | 34.82981 | 15.08635 | -0.45421 | 0.336111 | 0.466175 |
| 1 | 114989600 | 114992999 | 43.98257 | 71.35877 | 70.50919 | 30.21232 | 0.383501 | 1.631182 | 0.466192 |
| 14 | 27044200 | 27046199 | 13.64197 | 24.10492 | 41.86009 | 18.51768 | -0.40177 | 0.538304 | 0.466217 |
| 12 | 60757200 | 60763399 | 110.6271 | 170.0709 | 172.0746 | 71.45494 | 0.278742 | 2.96218 | 0.466226 |
| 6 | 8955800 | 8959199 | 30.43052 | 49.26455 | 85.33087 | 36.50484 | -0.4272 | 1.657875 | 0.466229 |
| 16 | 90509000 | 90515999 | 131.7882 | 197.33 | 297.3251 | 122.6339 | -0.32694 | 3.499315 | 0.466237 |
| X | 79292200 | 79293399 | 20.78819 | 36.33634 | 35.96402 | 15.85109 | 0.402347 | 0.712468 | 0.466239 |
| 19 | 48801800 | 48802999 | 8.590685 | 16.08837 | 29.57147 | 12.21906 | -0.4111 | 0.1139 | 0.466244 |
| 6 | 129951200 | 129952599 | 18.99356 | 33.35457 | 33.4128 | 14.78477 | 0.385069 | 0.626718 | 0.466262 |
| 8 | 54650800 | 54653199 | 35.02407 | 57.50587 | 59.57834 | 25.66216 | 0.326595 | 1.43175 | 0.466267 |
| 2 | 122049400 | 122050999 | 11.55974 | 20.5457 | 36.76125 | 16.21695 | -0.44031 | 0.463402 | 0.466275 |
| 9 | 88481200 | 88482199 | 9.576587 | 17.57121 | 31.91025 | 13.59128 | -0.4272 | 0.248893 | 0.466297 |
| 12 | 77874000 | 77875199 | 15.40991 | 27.94819 | 27.69986 | 11.97477 | 0.426157 | 0.347448 | 0.466308 |
| 16 | 93889400 | 93896599 | 58.54701 | 92.45393 | 147.1965 | 61.12074 | -0.3446 | 2.436463 | 0.466324 |
| 13 | 111861400 | 111862799 | 12.42178 | 21.75957 | 41.56245 | 18.3522 | -0.5412 | 0.644377 | 0.46633 |
| 5 | 96891000 | 96891599 | 6.541969 | 12.62967 | 26.59288 | 10.25093 | -0.51603 | -0.15015 | 0.466348 |
| 2 | 87389600 | 87392199 | 20.84858 | 36.47028 | 30.85024 | 13.60615 | 0.62779 | 0.692196 | 0.46636 |
| 8 | 9201800 | 9203399 | 12.90027 | 22.77337 | 39.85667 | 17.63468 | -0.41312 | 0.542785 | 0.466368 |
| 7 | 11047400 | 11048999 | 29.52312 | 49.41464 | 50.76726 | 22.12189 | 0.342258 | 1.190859 | 0.466378 |
| 5 | 144468200 | 144471799 | 31.67269 | 51.42867 | 83.26287 | 35.69854 | -0.32603 | 1.598793 | 0.466385 |
| 19 | 20310600 | 20313399 | 23.3372 | 41.05491 | 32.31623 | 14.30835 | 0.731528 | 0.625905 | 0.466392 |
| 2 | 81289600 | 81293199 | 75.74429 | 119.4383 | 118.2799 | 49.08515 | 0.337819 | 2.476385 | 0.466404 |
| 17 | 28298800 | 28302399 | 22.94379 | 37.97037 | 67.13459 | 29.03599 | -0.43895 | 1.311746 | 0.46641 |
| 17 | 29865600 | 29866999 | 13.42284 | 25.30747 | 22.81284 | 9.319644 | 0.621615 | 0.093946 | 0.466415 |
| 3 | 51816000 | 51817199 | 8.261963 | 15.51951 | 29.08818 | 11.9581 | -0.43319 | 0.089541 | 0.466427 |
| 7 | 47163800 | 47173399 | 157.7024 | 238.6479 | 220.4652 | 91.19732 | 0.391732 | 3.293727 | 0.466427 |
| 11 | 81149400 | 81151199 | 20.97354 | 36.83879 | 33.55311 | 14.84929 | 0.521486 | 0.632973 | 0.466437 |
| 2 | 173167800 | 173169999 | 21.91084 | 36.57027 | 62.31817 | 27.10696 | -0.38522 | 1.2091 | 0.466444 |
| 2 | 6136000 | 6140599 | 45.77012 | 73.23152 | 114.8343 | 48.34754 | -0.30408 | 2.039897 | 0.466473 |
| X | 28693600 | 28693999 | 14.18542 | 26.51019 | 19.91818 | 8.259644 | 0.866274 | 0.151824 | 0.466479 |
| 6 | 22434200 | 22437599 | 52.68858 | 84.54606 | 87.39391 | 36.9551 | 0.300928 | 1.94971 | 0.466485 |
| 12 | 97887400 | 97889199 | 18.08374 | 31.79168 | 30.73778 | 13.6113 | 0.435748 | 0.597154 | 0.466489 |

| | | | | | | | | | |
|---|---|---|---|---|---|---|---|---|---|
| 2 | 168110400 | 168112799 | 24.84613 | 40.664 | 74.47185 | 32.03401 | -0.49481 | 1.458821 | 0.466518 |
| 8 | 41391200 | 41393799 | 21.21916 | 35.88773 | 60.84872 | 26.63048 | -0.37771 | 1.07813 | 0.466531 |
| 5 | 34069600 | 34074199 | 50.19262 | 80.85658 | 80.71545 | 34.32762 | 0.357992 | 1.822336 | 0.466555 |
| 16 | 22038800 | 22044199 | 69.17383 | 110.3581 | 100.2707 | 42.04386 | 0.477329 | 2.148226 | 0.466561 |
| 7 | 40742600 | 40748999 | 119.8727 | 183.691 | 184.7728 | 76.69405 | 0.283622 | 3.028498 | 0.466573 |
| 14 | 96336000 | 96337599 | 10.571 | 19.33023 | 35.0211 | 14.98503 | -0.42872 | 0.283555 | 0.466578 |
| 1 | 181467000 | 181469199 | 14.90758 | 26.05682 | 44.12648 | 19.4483 | -0.36872 | 0.696774 | 0.466604 |
| 8 | 112872400 | 112874199 | 14.90758 | 26.05682 | 44.12648 | 19.4483 | -0.36872 | 0.696774 | 0.466604 |
| 6 | 34317400 | 34319199 | 12.9336 | 24.36875 | 20.87628 | 8.538172 | 0.693762 | 0.091014 | 0.466609 |
| 15 | 19567000 | 19570599 | 36.84352 | 60.79006 | 58.77887 | 25.37029 | 0.428686 | 1.376331 | 0.466616 |
| 11 | 58149000 | 58151799 | 20.20473 | 34.10109 | 57.50903 | 25.153 | -0.37005 | 1.08799 | 0.466653 |
| 14 | 66380200 | 66381799 | 11.23973 | 20.27129 | 35.84157 | 15.60318 | -0.40999 | 0.370226 | 0.466655 |
| 3 | 47506800 | 47510999 | 29.16019 | 49.86642 | 42.93301 | 18.8252 | 0.597949 | 0.987747 | 0.466663 |
| 7 | 56854800 | 56856999 | 18.63676 | 32.9828 | 30.9419 | 13.68346 | 0.482644 | 0.529497 | 0.466682 |
| X | 30973800 | 30975999 | 19.0409 | 35.34284 | 22.73719 | 9.533067 | 1.075862 | 0.238461 | 0.466683 |
| 8 | 3802600 | 3807199 | 85.2717 | 133.965 | 117.4388 | 48.71266 | 0.509379 | 2.554503 | 0.466701 |
| 11 | 50692600 | 50712199 | 294.6321 | 416.1595 | 574.895 | 237.9995 | -0.2832 | 4.430998 | 0.466707 |
| 15 | 74912200 | 74914599 | 30.13593 | 50.13878 | 51.35283 | 22.292 | 0.346784 | 1.256255 | 0.466708 |
| 4 | 145216200 | 145217799 | 23.34108 | 40.91867 | 34.00057 | 15.02296 | 0.653408 | 0.677362 | 0.466717 |
| X | 117521800 | 117526799 | 58.75081 | 94.02912 | 93.23529 | 39.27088 | 0.356491 | 2.040387 | 0.46675 |
| X | 4541800 | 4551799 | 169.0755 | 255.7621 | 194.0854 | 80.27554 | 0.674831 | 3.30339 | 0.466769 |
| 3 | 37194600 | 37195999 | 14.46368 | 25.37514 | 42.98789 | 19.00612 | -0.36916 | 0.633513 | 0.466788 |
| 9 | 54217200 | 54219399 | 30.47558 | 50.71078 | 52.44517 | 22.76832 | 0.332837 | 1.256682 | 0.466792 |
| 1 | 179750400 | 179751799 | 11.36744 | 20.35478 | 35.75935 | 15.67635 | -0.40778 | 0.413195 | 0.466796 |
| 19 | 30428200 | 30430199 | 23.75449 | 40.6782 | 41.57825 | 18.23319 | 0.35168 | 0.952799 | 0.466799 |
| 11 | 11970200 | 11979999 | 180.5588 | 264.8235 | 381.2667 | 157.0385 | -0.28905 | 3.86243 | 0.466822 |
| 7 | 41860200 | 41860799 | 22.70856 | 39.03905 | 36.02053 | 15.80001 | 0.499705 | 0.906489 | 0.466831 |
| 5 | 143754600 | 143766199 | 190.1946 | 276.4708 | 414.0511 | 170.6734 | -0.35836 | 4.000983 | 0.466854 |
| 18 | 9554800 | 9557199 | 21.19135 | 35.77573 | 59.18606 | 25.88857 | -0.34247 | 1.091422 | 0.46688 |
| 4 | 127455800 | 127457799 | 15.09569 | 26.24854 | 45.8044 | 20.12062 | -0.4125 | 0.775324 | 0.46689 |
| 14 | 48114600 | 48116799 | 18.45684 | 32.44517 | 32.85944 | 14.55103 | 0.369327 | 0.60633 | 0.466901 |
| 9 | 14321000 | 14323199 | 22.91549 | 40.15921 | 34.00679 | 15.02148 | 0.62619 | 0.679415 | 0.466901 |
| 13 | 41465200 | 41465599 | 4.677244 | 9.121837 | 20.90925 | 7.506 | -0.53975 | -0.43743 | 0.466905 |
| 15 | 27278800 | 27280399 | 25.75679 | 43.96779 | 43.69276 | 19.15558 | 0.391591 | 0.994064 | 0.466911 |
| 5 | 115308000 | 115309399 | 14.42836 | 25.25737 | 43.04818 | 18.99731 | -0.3779 | 0.668029 | 0.466914 |
| 13 | 13977800 | 13983799 | 47.94557 | 76.3213 | 126.2594 | 52.77841 | -0.3892 | 2.204841 | 0.466921 |
| 3 | 9015800 | 9017199 | 18.89963 | 32.22362 | 54.29135 | 23.79986 | -0.36738 | 0.991337 | 0.466921 |
| 4 | 119611200 | 119614599 | 39.19107 | 64.649 | 58.3538 | 25.18445 | 0.527552 | 1.385068 | 0.466925 |
| 1 | 102522800 | 102525599 | 25.47538 | 44.22627 | 39.14556 | 17.17835 | 0.562124 | 0.835823 | 0.466929 |
| 17 | 89498200 | 89498599 | 4.266236 | 8.334073 | 20.63974 | 7.211526 | -0.61483 | -0.54181 | 0.466936 |
| 16 | 48813800 | 48814999 | 7.711265 | 14.40243 | 29.02111 | 12.04666 | -0.54759 | 0.123276 | 0.466941 |
| 5 | 10197800 | 10202599 | 63.21205 | 100.9656 | 98.77887 | 41.48729 | 0.372638 | 2.105849 | 0.466945 |
| 7 | 130293200 | 130295799 | 16.77963 | 28.67786 | 52.14346 | 22.86271 | -0.4763 | 0.956814 | 0.466945 |
| X | 87726200 | 87728799 | 26.18267 | 45.74354 | 35.13228 | 15.47881 | 0.766737 | 0.74844 | 0.466953 |
| 17 | 70424800 | 70425399 | 12.05381 | 22.51993 | 23.07536 | 9.573639 | 0.421096 | 0.142473 | 0.466953 |
| 2 | 111689600 | 111692999 | 51.24256 | 82.29791 | 84.93578 | 35.96362 | 0.3047 | 1.920497 | 0.466965 |
| 7 | 104285000 | 104287399 | 27.05476 | 46.32636 | 43.59196 | 19.11614 | 0.470618 | 0.970924 | 0.466969 |

| | | | | | | | | | |
|---|---|---|---|---|---|---|---|---|---|
| 1 | 193212200 | 193213199 | 14.05677 | 25.90831 | 25.82269 | 10.93104 | 0.438619 | 0.242609 | 0.466975 |
| 19 | 60568800 | 60569999 | 9.353859 | 16.91037 | 33.38325 | 14.48966 | -0.5649 | 0.336895 | 0.466976 |
| 3 | 57288200 | 57288999 | 17.93538 | 31.69262 | 31.6935 | 14.02016 | 0.390259 | 0.544475 | 0.466988 |
| 15 | 84232400 | 84233799 | 21.67205 | 37.6857 | 38.26821 | 16.81078 | 0.365007 | 0.793939 | 0.466998 |
| X | 5063200 | 5065399 | 32.52925 | 55.49143 | 41.74775 | 18.30187 | 0.791307 | 1.021669 | 0.467016 |
| 9 | 91208000 | 91209399 | 13.8307 | 25.49858 | 25.9142 | 10.96571 | 0.411279 | 0.240671 | 0.467021 |
| 1 | 184612600 | 184615999 | 26.94213 | 44.01792 | 75.6889 | 32.53693 | -0.40593 | 1.490046 | 0.467031 |
| 2 | 147871400 | 147872599 | 10.00283 | 17.92201 | 34.84873 | 15.26943 | -0.55249 | 0.398999 | 0.467041 |
| 2 | 61008400 | 61009199 | 14.17558 | 25.74167 | 25.27867 | 10.91102 | 0.440743 | 0.325199 | 0.467042 |
| 4 | 40488000 | 40490199 | 15.03609 | 26.3062 | 45.14194 | 19.91169 | -0.38784 | 0.682528 | 0.467046 |
| 2 | 3562600 | 3566199 | 33.1522 | 56.23019 | 44.01163 | 19.27839 | 0.733524 | 1.075585 | 0.467059 |
| 15 | 28429200 | 28431199 | 25.67948 | 44.76541 | 35.64736 | 15.68395 | 0.714337 | 0.773995 | 0.467064 |
| 12 | 12378800 | 12384399 | 60.02636 | 94.92676 | 144.7062 | 60.13051 | -0.28033 | 2.403885 | 0.467074 |
| 6 | 140520400 | 140523399 | 32.04637 | 51.87756 | 85.53889 | 36.5923 | -0.35642 | 1.662782 | 0.467078 |
| 17 | 20050200 | 20053799 | 47.94393 | 77.96305 | 66.37934 | 28.48609 | 0.600686 | 1.596359 | 0.467088 |
| 8 | 17693400 | 17693999 | 12.2622 | 22.77446 | 21.54922 | 9.027459 | 0.524106 | 0.175975 | 0.467134 |
| 18 | 23245800 | 23250599 | 70.977 | 112.3959 | 116.1368 | 48.29191 | 0.281434 | 2.373366 | 0.467157 |
| 7 | 34810800 | 34813599 | 21.29792 | 35.43691 | 63.85894 | 27.72214 | -0.46562 | 1.241699 | 0.467174 |
| X | 64652600 | 64654999 | 27.62546 | 47.83405 | 39.05807 | 17.13614 | 0.677299 | 0.86919 | 0.467182 |
| X | 95723600 | 95731599 | 93.96976 | 147.6541 | 133.3203 | 55.30136 | 0.467024 | 2.561052 | 0.46719 |
| 12 | 14725600 | 14726599 | 9.018169 | 16.86818 | 31.18897 | 12.9143 | -0.42247 | 0.127441 | 0.467198 |
| 11 | 44441400 | 44443399 | 30.72628 | 51.19625 | 52.42455 | 22.77893 | 0.347148 | 1.244437 | 0.467214 |
| 19 | 9379200 | 9379799 | 13.26154 | 24.37774 | 24.89815 | 10.57784 | 0.399962 | 0.249232 | 0.467226 |
| 13 | 81876000 | 81876799 | 16.35193 | 29.02672 | 27.94557 | 12.34101 | 0.447789 | 0.485186 | 0.467233 |
| 3 | 14554000 | 14554799 | 7.285966 | 14.02287 | 28.25965 | 10.99581 | -0.46617 | -0.1065 | 0.467264 |
| 13 | 53212600 | 53215399 | 21.27505 | 37.15187 | 36.49031 | 16.07235 | 0.413287 | 0.729576 | 0.467283 |
| 16 | 90363600 | 90365399 | 18.29037 | 32.75778 | 28.82065 | 12.63236 | 0.583964 | 0.442298 | 0.467287 |
| X | 52976200 | 52979399 | 49.35673 | 79.67576 | 72.16176 | 30.77434 | 0.501631 | 1.761505 | 0.467297 |
| 16 | 50154400 | 50157799 | 60.70085 | 96.69177 | 95.04822 | 39.76934 | 0.361595 | 2.186945 | 0.467305 |
| 8 | 59582000 | 59587199 | 59.11307 | 94.35841 | 97.76753 | 41.02553 | 0.288994 | 2.120884 | 0.467322 |
| 1 | 125037800 | 125042599 | 55.14182 | 88.746 | 81.93888 | 34.80054 | 0.468716 | 1.859119 | 0.467343 |
| X | 79319800 | 79323799 | 41.144 | 68.53952 | 49.87263 | 21.66715 | 0.838512 | 1.276236 | 0.467346 |
| 4 | 129173400 | 129174999 | 9.774794 | 17.99452 | 33.01926 | 14.00025 | -0.43717 | 0.235996 | 0.467359 |
| 17 | 47264600 | 47265399 | 5.373572 | 10.44778 | 22.31865 | 8.276667 | -0.48186 | -0.30524 | 0.467369 |
| 3 | 10206600 | 10210999 | 27.65256 | 45.36477 | 75.83628 | 32.65342 | -0.36225 | 1.434847 | 0.467388 |
| 2 | 90416200 | 90418799 | 24.29779 | 41.67939 | 42.57954 | 18.67484 | 0.352797 | 0.941508 | 0.467448 |
| 4 | 86791600 | 86792999 | 9.120216 | 16.75078 | 31.42074 | 13.36439 | -0.47199 | 0.238727 | 0.46746 |
| 9 | 34327200 | 34328399 | 9.120216 | 16.75078 | 31.42074 | 13.36439 | -0.47199 | 0.238727 | 0.46746 |
| 10 | 43606600 | 43608399 | 20.88544 | 36.81307 | 32.44361 | 14.35918 | 0.570265 | 0.590152 | 0.467474 |
| 6 | 132418600 | 132419599 | 13.33927 | 25.31472 | 23.06796 | 9.295823 | 0.623733 | 0.052073 | 0.467511 |
| 19 | 3095400 | 3100399 | 80.15745 | 126.6491 | 112.4444 | 46.70344 | 0.497424 | 2.433092 | 0.467516 |
| 7 | 117973600 | 117975999 | 27.74542 | 47.1088 | 45.21363 | 19.81118 | 0.440358 | 1.046867 | 0.467525 |
| 3 | 91785200 | 91789599 | 48.25832 | 77.92072 | 76.79005 | 32.75795 | 0.380392 | 1.750949 | 0.467527 |
| 10 | 50468600 | 50469999 | 6.896767 | 13.10691 | 27.15403 | 10.90975 | -0.54773 | -0.00734 | 0.467528 |
| 8 | 73174600 | 73176999 | 14.92015 | 25.97959 | 45.25573 | 19.89707 | -0.4098 | 0.753233 | 0.467532 |
| 3 | 24657200 | 24657799 | 6.254288 | 12.10738 | 24.38688 | 9.28108 | -0.43409 | -0.20043 | 0.467533 |
| 8 | 15670200 | 15671199 | 24.28883 | 41.4175 | 40.64884 | 17.81954 | 0.409227 | 0.992899 | 0.467533 |

| | | | | | | | | | |
|---|---|---|---|---|---|---|---|---|---|
| 10 | 106848400 | 106849799 | 15.19335 | 28.63124 | 22.71125 | 9.286018 | 0.803995 | 0.113629 | 0.467535 |
| 10 | 58633400 | 58637399 | 31.699 | 51.31109 | 86.85662 | 37.15379 | -0.39437 | 1.662855 | 0.467548 |
| X | 88274200 | 88275399 | 20.27059 | 35.8504 | 31.54696 | 13.9529 | 0.574472 | 0.551091 | 0.467571 |
| 6 | 132655400 | 132671399 | 191.9412 | 286.0363 | 270.705 | 111.5537 | 0.337315 | 3.595364 | 0.467572 |
| 3 | 93945800 | 93947199 | 12.7195 | 23.8344 | 23.94765 | 9.885629 | 0.454519 | 0.129642 | 0.467579 |
| 5 | 99171800 | 99172799 | 8.67125 | 15.97534 | 30.95754 | 13.11227 | -0.51393 | 0.217812 | 0.46758 |
| X | 135584600 | 135588799 | 61.69731 | 98.67621 | 95.5701 | 40.21465 | 0.389248 | 2.065014 | 0.46758 |
| 13 | 66298600 | 66316999 | 275.2027 | 401.9311 | 322.8465 | 133.0133 | 0.546581 | 3.936683 | 0.467594 |
| 1 | 139653800 | 139654199 | 8.545023 | 16.55733 | 17.24883 | 6.535983 | 0.514344 | -0.21177 | 0.467617 |
| 4 | 125028000 | 125029199 | 7.387722 | 13.93019 | 28.08462 | 11.47142 | -0.52921 | 0.057788 | 0.46763 |
| 6 | 87173600 | 87173999 | 7.387722 | 13.93019 | 28.08462 | 11.47142 | -0.52921 | 0.057788 | 0.46763 |
| 1 | 94327000 | 94327799 | 14.21396 | 25.85474 | 25.47434 | 10.97242 | 0.438333 | 0.319059 | 0.467631 |
| 4 | 150321000 | 150322799 | 8.946651 | 16.43607 | 31.9296 | 13.57625 | -0.52211 | 0.235342 | 0.467649 |
| 13 | 73667800 | 73669399 | 8.946651 | 16.43607 | 31.9296 | 13.57625 | -0.52211 | 0.235342 | 0.467649 |
| 6 | 10988800 | 10991799 | 25.77892 | 42.577 | 76.26901 | 32.94111 | -0.45823 | 1.346056 | 0.467662 |
| 1 | 48407400 | 48410199 | 31.20897 | 50.85022 | 84.11592 | 36.12359 | -0.35311 | 1.544587 | 0.467695 |
| X | 25446200 | 25471399 | 381.5666 | 540.6416 | 426.7134 | 176.5628 | 0.528642 | 4.388875 | 0.467705 |
| 1 | 10908800 | 10911399 | 24.28103 | 42.40527 | 36.34486 | 16.00927 | 0.609113 | 0.745224 | 0.467721 |
| 7 | 19130400 | 19138199 | 63.3434 | 98.96837 | 180.2144 | 74.7721 | -0.55007 | 2.652687 | 0.467725 |
| 1 | 125939600 | 125941999 | 19.19703 | 33.74004 | 32.06743 | 14.19947 | 0.460197 | 0.608782 | 0.467726 |
| 19 | 13157800 | 13163199 | 62.27473 | 99.8972 | 87.56141 | 37.00838 | 0.537897 | 1.971648 | 0.467735 |
| 7 | 16508400 | 16516399 | 59.35425 | 92.95174 | 175.2926 | 72.7126 | -0.5979 | 2.6087 | 0.467736 |
| 2 | 172005000 | 172010399 | 60.79112 | 96.24085 | 149.0817 | 61.98281 | -0.30221 | 2.378842 | 0.467759 |
| 4 | 83515400 | 83517999 | 22.3717 | 38.73015 | 39.42189 | 17.29567 | 0.360688 | 0.848436 | 0.467762 |
| 1 | 36067600 | 36070599 | 25.76282 | 42.2759 | 72.52949 | 31.23312 | -0.39928 | 1.426686 | 0.467804 |
| 14 | 106368000 | 106368799 | 16.19347 | 28.87129 | 26.05031 | 11.46175 | 0.544537 | 0.453619 | 0.467809 |
| 11 | 13351800 | 13353199 | 9.170451 | 17.17773 | 32.36634 | 13.36917 | -0.44678 | 0.122021 | 0.467814 |
| 7 | 85343800 | 85345599 | 25.25537 | 44.11575 | 35.70356 | 15.72878 | 0.691466 | 0.746183 | 0.467821 |
| 13 | 115237800 | 115240599 | 16.53171 | 28.51328 | 49.52487 | 21.7257 | -0.40818 | 0.868605 | 0.467844 |
| 10 | 42418200 | 42419999 | 12.82865 | 22.72055 | 38.74324 | 17.13092 | -0.37415 | 0.502472 | 0.467888 |
| 4 | 57782000 | 57783399 | 8.93535 | 16.49508 | 30.94599 | 13.07152 | -0.46505 | 0.206492 | 0.467904 |
| 8 | 13015000 | 13021599 | 107.0666 | 162.8727 | 241.6959 | 100.1394 | -0.28525 | 3.177449 | 0.467923 |
| 4 | 119452400 | 119454999 | 23.18361 | 40.88294 | 30.56635 | 13.52706 | 0.806505 | 0.595414 | 0.467935 |
| 2 | 97879600 | 97882799 | 44.70485 | 72.63612 | 62.13342 | 26.65195 | 0.592931 | 1.598578 | 0.467961 |
| 4 | 47982000 | 47982599 | 6.255964 | 12.12362 | 24.43504 | 9.252935 | -0.42775 | -0.21575 | 0.467965 |
| 10 | 8717400 | 8720599 | 26.05422 | 42.84498 | 71.61462 | 30.86688 | -0.36057 | 1.400179 | 0.467971 |
| 14 | 85379400 | 85384599 | 81.78736 | 128.9637 | 128.7685 | 53.43763 | 0.325919 | 2.481857 | 0.467975 |
| 9 | 19007400 | 19009999 | 44.94707 | 72.69511 | 73.74369 | 31.52524 | 0.34146 | 1.696442 | 0.467978 |
| 15 | 96950800 | 96954799 | 87.93744 | 134.5804 | 235.5539 | 97.74313 | -0.51622 | 3.049994 | 0.467981 |
| 5 | 71428800 | 71431399 | 29.35321 | 50.1404 | 43.49267 | 19.06888 | 0.586997 | 0.999236 | 0.467983 |
| 3 | 113640200 | 113641799 | 20.13165 | 34.15187 | 60.77322 | 26.62192 | -0.44691 | 1.049649 | 0.46801 |
| 7 | 62467400 | 62468399 | 20.24472 | 35.87864 | 27.33544 | 12.08519 | 0.782128 | 0.520073 | 0.468015 |
| X | 4928400 | 4934399 | 179.2996 | 271.2269 | 208.2679 | 86.14144 | 0.657806 | 3.307138 | 0.468029 |
| 19 | 19342600 | 19344999 | 25.55859 | 44.23534 | 40.30064 | 17.68106 | 0.519876 | 0.866452 | 0.468034 |
| 10 | 18069200 | 18071399 | 17.33211 | 29.75706 | 52.19703 | 22.89458 | -0.42383 | 0.915871 | 0.468086 |
| 6 | 69913200 | 69914399 | 18.72558 | 32.85186 | 33.42549 | 14.77818 | 0.362653 | 0.639446 | 0.468121 |
| 1 | 185677000 | 185678799 | 15.38945 | 26.85439 | 45.20578 | 19.90197 | -0.36037 | 0.724983 | 0.468122 |

| | | | | | | | | | |
|---|---|---|---|---|---|---|---|---|---|
| 13 | 93818000 | 93818799 | 6.113875 | 11.8238 | 24.32417 | 9.300328 | -0.47067 | -0.18662 | 0.468122 |
| 7 | 32331600 | 32335399 | 48.25041 | 78.04056 | 73.66047 | 31.4905 | 0.445268 | 1.701604 | 0.468143 |
| 5 | 61109800 | 61110799 | 11.19844 | 21.67517 | 19.04956 | 7.255087 | 0.749189 | -0.16203 | 0.468144 |
| 17 | 19135400 | 19142799 | 83.9136 | 133.496 | 100.6467 | 42.03686 | 0.741808 | 2.252485 | 0.468145 |
| X | 3237000 | 3245399 | 120.1989 | 186.3508 | 152.7202 | 63.43264 | 0.591752 | 2.810442 | 0.468146 |
| 7 | 11935200 | 11950599 | 218.9686 | 323.6691 | 294.9348 | 121.4612 | 0.381093 | 3.739882 | 0.468169 |
| 3 | 74173800 | 74180399 | 111.6837 | 172.8462 | 162.6113 | 67.54462 | 0.390946 | 2.837732 | 0.468189 |
| 6 | 67991600 | 67994799 | 31.07141 | 52.15405 | 50.22919 | 21.92669 | 0.435021 | 1.16414 | 0.468197 |
| 4 | 108814600 | 108817599 | 33.90743 | 56.14379 | 56.46523 | 24.43026 | 0.372459 | 1.324937 | 0.4682 |
| 7 | 43945000 | 43948799 | 42.02925 | 68.41135 | 66.73234 | 28.6477 | 0.405857 | 1.566934 | 0.468201 |
| 14 | 38207600 | 38208399 | 13.09371 | 24.29844 | 24.52892 | 10.28745 | 0.430362 | 0.196564 | 0.468207 |
| 18 | 56213000 | 56214199 | 13.10479 | 22.8832 | 47.8452 | 21.07296 | -0.67119 | 0.710337 | 0.468213 |
| 16 | 84040200 | 84044999 | 61.82896 | 97.67631 | 148.6205 | 61.72473 | -0.27885 | 2.429931 | 0.468245 |
| 1 | 20814000 | 20814599 | 6.581286 | 12.59628 | 26.21636 | 10.36517 | -0.53339 | -0.06586 | 0.468276 |
| 4 | 17309200 | 17311799 | 29.12905 | 48.5954 | 48.65328 | 21.15689 | 0.379465 | 1.221694 | 0.468279 |
| 11 | 40987600 | 40990799 | 38.14196 | 63.01673 | 56.73219 | 24.50857 | 0.531556 | 1.360627 | 0.468333 |
| 9 | 79253600 | 79254799 | 9.84955 | 17.81181 | 34.41268 | 14.93091 | -0.5334 | 0.343762 | 0.468372 |
| 18 | 83043200 | 83045399 | 12.84597 | 22.50627 | 43.56386 | 19.23856 | -0.56062 | 0.644654 | 0.468374 |
| 3 | 25192200 | 25193399 | 19.4118 | 33.92904 | 33.20487 | 14.63447 | 0.41873 | 0.703561 | 0.468376 |
| 10 | 14281400 | 14284599 | 36.63638 | 61.15584 | 51.48604 | 22.39706 | 0.628674 | 1.241229 | 0.468377 |
| 7 | 85755800 | 85763399 | 102.4792 | 159.9747 | 145.6003 | 60.41595 | 0.44826 | 2.683438 | 0.46838 |
| 2 | 150850000 | 150851999 | 14.55542 | 25.44584 | 44.64028 | 19.67773 | -0.41952 | 0.691314 | 0.468403 |
| 16 | 83859000 | 83861999 | 30.52819 | 51.94553 | 45.17515 | 19.80036 | 0.582576 | 1.04161 | 0.468404 |
| 13 | 14295000 | 14296199 | 13.55305 | 25.35744 | 24.40927 | 10.10239 | 0.512957 | 0.14775 | 0.468411 |
| 9 | 19210800 | 19213999 | 37.04115 | 61.16994 | 58.21931 | 25.14305 | 0.451577 | 1.366829 | 0.468424 |
| 3 | 96396400 | 96398799 | 30.92961 | 51.45224 | 53.02269 | 23.01471 | 0.33791 | 1.260916 | 0.468435 |
| 6 | 104193400 | 104204399 | 224.5523 | 322.2288 | 469.2245 | 193.7494 | -0.33626 | 4.195303 | 0.468437 |
| 13 | 67479600 | 67481999 | 16.20834 | 28.2252 | 47.35583 | 20.82159 | -0.35597 | 0.761529 | 0.468438 |
| 2 | 165572800 | 165579399 | 44.02775 | 70.38134 | 112.733 | 47.4033 | -0.33601 | 2.065507 | 0.46845 |
| 8 | 19729400 | 19732799 | 53.25088 | 85.84719 | 78.93299 | 33.60403 | 0.477661 | 1.805191 | 0.468458 |
| 18 | 6912200 | 6913399 | 14.22534 | 26.03102 | 26.28783 | 11.23742 | 0.409709 | 0.284752 | 0.46846 |
| 17 | 72428800 | 72430199 | 14.84302 | 27.25407 | 26.21871 | 11.156 | 0.484017 | 0.270199 | 0.46848 |
| 18 | 65345400 | 65346199 | 5.968958 | 11.54343 | 24.2884 | 9.287071 | -0.50206 | -0.18504 | 0.468507 |
| 13 | 61472600 | 61480199 | 89.99619 | 142.5808 | 111.828 | 46.51127 | 0.679162 | 2.377567 | 0.468509 |
| X | 126624600 | 126626999 | 22.11268 | 38.89804 | 32.93689 | 14.58335 | 0.626676 | 0.620162 | 0.46852 |
| 17 | 56979000 | 56980999 | 11.91647 | 20.98493 | 39.38223 | 17.43333 | -0.51481 | 0.560528 | 0.468525 |
| 16 | 65830600 | 65831599 | 18.34499 | 32.44984 | 26.26525 | 11.61649 | 0.694307 | 0.526005 | 0.468546 |
| 13 | 67030800 | 67036399 | 78.53015 | 125.082 | 100.2294 | 41.93235 | 0.655847 | 2.209076 | 0.468549 |
| 10 | 23891400 | 23892599 | 24.04864 | 41.69306 | 35.63543 | 15.63511 | 0.61198 | 0.83741 | 0.46856 |
| 1 | 10308800 | 10309799 | 7.345323 | 13.94791 | 27.13816 | 10.92232 | -0.45994 | 0.004095 | 0.468562 |
| 5 | 104873600 | 104886399 | 33.08798 | 55.94441 | 40.8375 | 17.87092 | 0.833621 | 1.098586 | 0.468568 |
| 8 | 125902400 | 125907599 | 38.56136 | 61.96749 | 101.815 | 43.1476 | -0.36378 | 1.890557 | 0.468576 |
| 17 | 24942600 | 24943599 | 4.990323 | 9.648921 | 23.70266 | 9.07151 | -0.72436 | -0.19645 | 0.468617 |
| 17 | 38993400 | 38998199 | 114.6237 | 176.6821 | 155.5844 | 64.62719 | 0.4818 | 2.911318 | 0.468665 |
| X | 37993800 | 37995599 | 21.72783 | 37.9316 | 35.64923 | 15.69905 | 0.476844 | 0.738333 | 0.46867 |
| 9 | 77299800 | 77301199 | 9.986494 | 18.46297 | 33.64745 | 14.1839 | -0.42184 | 0.211471 | 0.468682 |
| 12 | 8875000 | 8879999 | 32.86624 | 53.0868 | 89.3141 | 38.11823 | -0.38885 | 1.721369 | 0.468686 |

| | | | | | | | | | |
|---|---|---|---|---|---|---|---|---|---|
| 1 | 184807600 | 184810599 | 29.76445 | 48.3562 | 91.2591 | 39.14243 | -0.54594 | 1.589124 | 0.468692 |
| 13 | 60536600 | 60537999 | 16.2547 | 28.01766 | 48.99025 | 21.49076 | -0.41785 | 0.873866 | 0.468716 |
| 9 | 58674600 | 58674999 | 12.56632 | 22.07895 | 39.93442 | 17.67811 | -0.46286 | 0.594197 | 0.468726 |
| 1 | 148425600 | 148429399 | 57.42389 | 91.75552 | 94.99972 | 39.92166 | 0.291661 | 2.086784 | 0.468767 |
| 7 | 112661000 | 112662999 | 11.6871 | 20.80086 | 37.24007 | 16.40901 | -0.44015 | 0.455312 | 0.468772 |
| 15 | 52741400 | 52744799 | 35.89469 | 60.09226 | 50.30265 | 21.92246 | 0.636811 | 1.208858 | 0.468773 |
| 7 | 21759600 | 21760999 | 39.98587 | 65.29665 | 50.42074 | 21.67085 | 0.745348 | 1.507553 | 0.468833 |
| 4 | 87671400 | 87675199 | 41.72182 | 66.98029 | 105.3632 | 44.58582 | -0.30284 | 1.928764 | 0.468837 |
| 3 | 114348200 | 114349599 | 18.57606 | 32.87039 | 31.55114 | 13.95331 | 0.4498 | 0.532976 | 0.468855 |
| 3 | 69532800 | 69534999 | 15.3838 | 26.86151 | 45.19165 | 19.90365 | -0.35947 | 0.715819 | 0.468878 |
| 13 | 26516600 | 26518999 | 18.44883 | 32.46031 | 32.81631 | 14.52931 | 0.372353 | 0.596276 | 0.468898 |
| 11 | 48131600 | 48131999 | 4.643845 | 9.067631 | 19.91774 | 7.0137 | -0.45307 | -0.50783 | 0.468913 |
| 2 | 55207200 | 55210199 | 26.86957 | 46.64964 | 37.9325 | 16.64669 | 0.683915 | 0.838872 | 0.468935 |
| 13 | 109809600 | 109813799 | 59.73667 | 94.36956 | 145.3826 | 60.37949 | -0.29673 | 2.428962 | 0.468944 |
| 12 | 117752200 | 117753199 | 7.656252 | 14.52717 | 27.66673 | 11.15315 | -0.43136 | 0.014463 | 0.468946 |
| 17 | 46307600 | 46308799 | 7.656252 | 14.52717 | 27.66673 | 11.15315 | -0.43136 | 0.014463 | 0.468946 |
| 3 | 115681400 | 115682199 | 5.103467 | 9.917378 | 22.27065 | 8.294188 | -0.55897 | -0.29455 | 0.468946 |
| 12 | 109777600 | 109781399 | 38.39195 | 62.63894 | 64.55038 | 27.73492 | 0.329285 | 1.527029 | 0.468993 |
| 12 | 73362600 | 73365199 | 20.65807 | 35.06587 | 58.14303 | 25.47437 | -0.34543 | 1.040717 | 0.469006 |
| 8 | 54598600 | 54601799 | 38.37505 | 62.42401 | 61.50792 | 26.39883 | 0.390806 | 1.567853 | 0.46901 |
| 12 | 4736200 | 4739999 | 28.10958 | 45.81072 | 77.78596 | 33.40797 | -0.39037 | 1.530866 | 0.46903 |
| 3 | 8408800 | 8410399 | 18.01402 | 31.97759 | 30.46711 | 13.45438 | 0.462445 | 0.498318 | 0.469074 |
| 5 | 113987000 | 113988799 | 16.38998 | 27.87328 | 53.65428 | 23.51574 | -0.55919 | 1.005138 | 0.469117 |
| 4 | 148495000 | 148497999 | 25.19524 | 41.75475 | 68.52879 | 29.66763 | -0.33177 | 1.303669 | 0.469119 |
| 17 | 39891000 | 39893799 | 31.02718 | 53.62514 | 37.87747 | 16.61724 | 0.88517 | 0.900492 | 0.46915 |
| 13 | 61096000 | 61097799 | 28.99363 | 48.47116 | 48.10563 | 20.94648 | 0.391914 | 1.196427 | 0.469161 |
| 11 | 82617800 | 82620199 | 17.34317 | 30.01558 | 50.0732 | 21.96854 | -0.34921 | 0.842987 | 0.469174 |
| X | 154797200 | 154800999 | 35.96632 | 60.92112 | 42.87009 | 18.77069 | 0.886472 | 1.097112 | 0.469178 |
| 12 | 24474600 | 24478599 | 133.0648 | 201.9687 | 197.1466 | 81.62696 | 0.315928 | 3.225943 | 0.469187 |
| 13 | 65604600 | 65626199 | 368.7796 | 518.1946 | 481.8623 | 199.6535 | 0.280118 | 4.509105 | 0.469189 |
| 8 | 96505200 | 96508599 | 51.32392 | 82.84612 | 69.52023 | 29.6454 | 0.611289 | 1.760896 | 0.469198 |
| 2 | 4472000 | 4476799 | 60.42677 | 96.7205 | 94.18795 | 39.6774 | 0.382617 | 2.038068 | 0.469198 |
| 7 | 28741800 | 28744399 | 18.07832 | 30.73154 | 54.26243 | 23.78147 | -0.43545 | 1.013832 | 0.469204 |
| X | 147631200 | 147633799 | 45.05352 | 73.04859 | 71.00985 | 30.4122 | 0.406156 | 1.647598 | 0.469205 |
| 6 | 24527000 | 24527999 | 9.453789 | 17.19778 | 32.86847 | 14.15556 | -0.51139 | 0.298699 | 0.469225 |
| 5 | 113558800 | 113562999 | 22.38235 | 37.25668 | 64.57859 | 28.04245 | -0.40977 | 1.239823 | 0.469256 |
| 14 | 12961400 | 12962999 | 19.25549 | 33.95026 | 32.27924 | 14.28557 | 0.461738 | 0.579391 | 0.469313 |
| 5 | 26536200 | 26537799 | 21.91163 | 38.04976 | 38.36472 | 16.83855 | 0.375104 | 0.815697 | 0.469315 |
| 9 | 54125800 | 54127999 | 11.74225 | 20.86125 | 36.78164 | 16.23174 | -0.41947 | 0.468633 | 0.469325 |
| 7 | 42067600 | 42070199 | 47.58446 | 76.97736 | 73.59817 | 31.46805 | 0.426979 | 1.69674 | 0.469329 |
| 16 | 19528000 | 19530999 | 25.78801 | 42.32095 | 80.60752 | 34.71302 | -0.54969 | 1.431434 | 0.469363 |
| 6 | 46800400 | 46803399 | 18.84475 | 33.24749 | 27.73682 | 12.27369 | 0.649545 | 0.559835 | 0.469368 |
| X | 34694800 | 34696599 | 19.22478 | 35.93344 | 21.79175 | 9.034231 | 1.173175 | 0.197497 | 0.469376 |
| 13 | 15341800 | 15342999 | 15.16986 | 27.8278 | 26.80929 | 11.42202 | 0.48088 | 0.280863 | 0.469389 |
| 17 | 15924600 | 15926599 | 16.79418 | 28.29838 | 62.11532 | 27.15417 | -0.7488 | 1.096767 | 0.46941 |
| 17 | 18735600 | 18736999 | 29.90294 | 50.06388 | 50.7631 | 22.12384 | 0.361162 | 1.189968 | 0.469432 |
| 13 | 37094800 | 37096599 | 14.7209 | 26.49709 | 25.36664 | 11.07049 | 0.467192 | 0.380601 | 0.469437 |

| | | | | | | | | | |
|---|---|---|---|---|---|---|---|---|---|
| 7 | 59375200 | 59382199 | 80.10141 | 126.3727 | 110.2444 | 45.75527 | 0.521038 | 2.466921 | 0.469447 |
| 6 | 37585600 | 37590399 | 42.9832 | 68.85217 | 109.7774 | 46.29708 | -0.32609 | 1.999198 | 0.469454 |
| 11 | 5956200 | 5960799 | 92.78064 | 140.148 | 288.1515 | 119.128 | -0.76379 | 3.321416 | 0.469455 |
| 11 | 101744800 | 101747399 | 18.98581 | 32.48155 | 53.81418 | 23.59719 | -0.34257 | 0.958124 | 0.469475 |
| 2 | 124881000 | 124882799 | 22.25851 | 38.98862 | 35.26542 | 15.57095 | 0.531705 | 0.687243 | 0.469478 |
| 11 | 31902600 | 31903199 | 4.964642 | 9.668185 | 21.88637 | 8.00872 | -0.54732 | -0.36175 | 0.469503 |
| 2 | 39529000 | 39530599 | 14.613 | 27.16609 | 25.35404 | 10.60779 | 0.54496 | 0.199484 | 0.469503 |
| X | 14379400 | 14386999 | 132.2238 | 203.04 | 176.0654 | 73.1007 | 0.499848 | 2.993781 | 0.46953 |
| 2 | 23807600 | 23811399 | 52.57315 | 84.269 | 85.34917 | 36.03074 | 0.328427 | 1.981056 | 0.469537 |
| 12 | 114769600 | 114772199 | 44.70678 | 72.49454 | 71.0068 | 30.41341 | 0.395405 | 1.644478 | 0.46955 |
| X | 54418200 | 54424799 | 84.77602 | 134.9241 | 98.68384 | 41.24078 | 0.786153 | 2.24131 | 0.469556 |
| 12 | 101057800 | 101069199 | 214.9129 | 316.1251 | 309.4053 | 127.4167 | 0.27141 | 3.81892 | 0.469578 |
| 6 | 3794400 | 3794799 | 11.55231 | 21.71923 | 20.65126 | 8.477105 | 0.540641 | 0.092344 | 0.469581 |
| 13 | 30314200 | 30315599 | 12.32345 | 22.05001 | 38.30636 | 16.80358 | -0.39255 | 0.429226 | 0.469583 |
| 14 | 113926800 | 113928399 | 17.05013 | 30.9187 | 26.86597 | 11.61628 | 0.614427 | 0.356955 | 0.469587 |
| 3 | 53335000 | 53336999 | 18.5504 | 34.45127 | 23.36206 | 9.787756 | 1.001254 | 0.232991 | 0.469593 |
| 14 | 121301400 | 121306399 | 70.54391 | 110.1672 | 169.5681 | 70.35828 | -0.3084 | 2.663315 | 0.469601 |
| 5 | 101836000 | 101839999 | 29.81726 | 48.71451 | 79.16481 | 34.03094 | -0.3248 | 1.497721 | 0.469612 |
| 2 | 111352600 | 111354799 | 27.43993 | 46.77143 | 45.21468 | 19.82044 | 0.430812 | 1.013562 | 0.469615 |
| 2 | 54026400 | 54027599 | 18.10322 | 31.83663 | 32.36641 | 14.33152 | 0.364054 | 0.598654 | 0.469615 |
| 8 | 74162800 | 74164199 | 8.234665 | 15.63624 | 29.39921 | 11.83301 | -0.41163 | 0.017176 | 0.469619 |
| 12 | 52253400 | 52255799 | 35.61704 | 57.05074 | 115.7907 | 48.83043 | -0.67357 | 1.995156 | 0.469621 |
| Y | 74452000 | 74453799 | 9.513646 | 18.37583 | 13.59604 | 5.228759 | 0.982376 | -0.16616 | 0.469633 |
| 19 | 60889400 | 60890199 | 10.60226 | 20.52445 | 20.35194 | 7.745413 | 0.578492 | -0.16525 | 0.469635 |
| 1 | 21553600 | 21555199 | 15.45907 | 28.09106 | 27.73819 | 11.96267 | 0.43377 | 0.3376 | 0.46966 |
| 19 | 42116600 | 42117799 | 7.252644 | 13.64327 | 28.58633 | 11.72308 | -0.58873 | 0.074182 | 0.469666 |
| 5 | 97920800 | 97923399 | 25.90791 | 44.26834 | 37.37526 | 16.38716 | 0.625843 | 0.975408 | 0.469676 |
| 13 | 57189400 | 57192399 | 31.39139 | 50.92178 | 83.41442 | 35.73758 | -0.34419 | 1.619615 | 0.469679 |
| X | 30996800 | 30999999 | 37.10752 | 61.30605 | 57.61554 | 24.88973 | 0.469839 | 1.362621 | 0.46968 |
| 9 | 12798600 | 12800999 | 19.89337 | 35.19855 | 31.39731 | 13.8855 | 0.554972 | 0.541661 | 0.469691 |
| 5 | 70273600 | 70275199 | 22.39463 | 38.96148 | 33.11628 | 14.55212 | 0.620557 | 0.782976 | 0.469693 |
| 5 | 23586600 | 23587199 | 6.725299 | 12.93581 | 25.79974 | 10.05644 | -0.45247 | -0.10907 | 0.469696 |
| 8 | 20053600 | 20054399 | 6.725299 | 12.93581 | 25.79974 | 10.05644 | -0.45247 | -0.10907 | 0.469696 |
| 7 | 108410400 | 108411399 | 19.94558 | 33.6076 | 57.96609 | 25.34009 | -0.40238 | 1.103986 | 0.4697 |
| 11 | 54742600 | 54743599 | 11.60067 | 20.6119 | 36.80724 | 16.24159 | -0.43754 | 0.467817 | 0.469705 |
| 7 | 117119800 | 117121799 | 23.40252 | 40.43085 | 40.45465 | 17.74765 | 0.384697 | 0.87315 | 0.469709 |
| 7 | 26685800 | 26686599 | 12.87694 | 23.80171 | 24.02908 | 10.13227 | 0.424654 | 0.21502 | 0.46973 |
| 17 | 33251600 | 33252399 | 5.957531 | 11.56243 | 24.5672 | 9.238671 | -0.49438 | -0.24573 | 0.469738 |
| 7 | 62486800 | 62492399 | 86.11037 | 136.1181 | 121.5187 | 50.48433 | 0.490044 | 2.429159 | 0.469759 |
| 3 | 142429800 | 142430599 | 7.908759 | 15.14145 | 31.61593 | 12.48945 | -0.53782 | -0.04271 | 0.469801 |
| 5 | 36087800 | 36090599 | 13.50205 | 23.63687 | 42.05382 | 18.55919 | -0.4394 | 0.661399 | 0.469805 |
| 5 | 96022400 | 96023599 | 15.03873 | 27.12941 | 27.36867 | 11.91106 | 0.394829 | 0.376168 | 0.469815 |
| 11 | 7525400 | 7527199 | 24.52574 | 41.89898 | 42.6021 | 18.67859 | 0.358666 | 0.97774 | 0.469855 |
| 11 | 99008800 | 99012599 | 26.14737 | 43.15526 | 71.43302 | 30.83747 | -0.34457 | 1.353943 | 0.469865 |
| 4 | 132340800 | 132343799 | 56.08655 | 89.94047 | 89.90291 | 37.97874 | 0.348125 | 1.973907 | 0.469875 |
| 14 | 50222400 | 50223199 | 13.33096 | 24.47026 | 23.05986 | 9.815981 | 0.513638 | 0.254885 | 0.469892 |
| 2 | 112940800 | 112941799 | 8.153827 | 15.14794 | 30.50391 | 12.77429 | -0.55752 | 0.161963 | 0.469918 |

| | | | | | | | | | |
|---|---|---|---|---|---|---|---|---|---|
| 4 | 44694400 | 44695599 | 10.42583 | 18.88405 | 33.85076 | 14.65783 | -0.42451 | 0.335254 | 0.469941 |
| 1 | 164228800 | 164232399 | 31.12964 | 52.23091 | 49.69417 | 21.68767 | 0.452499 | 1.166692 | 0.469953 |
| 18 | 38931600 | 38932799 | 6.434821 | 12.35896 | 25.73761 | 10.07455 | -0.51959 | -0.09748 | 0.469954 |
| 3 | 129828800 | 129829799 | 14.5978 | 25.62273 | 43.48106 | 19.23196 | -0.3717 | 0.626403 | 0.469956 |
| 9 | 22497200 | 22498599 | 22.92553 | 39.77416 | 33.63248 | 14.75663 | 0.627506 | 0.821965 | 0.469987 |
| 16 | 33829800 | 33842199 | 206.3584 | 298.7075 | 432.4699 | 178.3538 | -0.31551 | 4.065526 | 0.469993 |
| X | 67829800 | 67835199 | 66.83176 | 107.283 | 82.7119 | 34.99623 | 0.72387 | 1.956905 | 0.470006 |
| 8 | 101007600 | 101008599 | 16.91303 | 30.35026 | 24.73652 | 10.82439 | 0.695538 | 0.414737 | 0.470018 |
| 10 | 129841400 | 129841999 | 12.55566 | 23.58232 | 24.07351 | 9.898429 | 0.436688 | 0.115745 | 0.470047 |
| 6 | 37585400 | 37590199 | 43.14908 | 69.11579 | 109.7798 | 46.29593 | -0.32067 | 2.000445 | 0.470052 |
| 18 | 85081000 | 85085599 | 46.15253 | 74.85292 | 69.79967 | 29.90051 | 0.466355 | 1.640006 | 0.47007 |
| 18 | 55974200 | 55977199 | 34.22801 | 55.12662 | 102.7037 | 43.66509 | -0.54063 | 1.816339 | 0.470084 |
| 15 | 28024200 | 28026799 | 20.10845 | 33.67025 | 61.22723 | 26.6866 | -0.47868 | 1.166012 | 0.470102 |
| X | 54702400 | 54709599 | 100.0702 | 157.6481 | 122.3935 | 50.78601 | 0.68747 | 2.513729 | 0.470109 |
| 7 | 124651800 | 124653799 | 18.34394 | 32.90809 | 28.85183 | 12.62841 | 0.590399 | 0.433149 | 0.470115 |
| 17 | 22854400 | 22868199 | 224.5746 | 331.6479 | 297.5384 | 122.5322 | 0.402282 | 3.755747 | 0.470123 |
| 13 | 74623000 | 74625799 | 24.45156 | 40.71749 | 68.22285 | 29.63251 | -0.3611 | 1.244092 | 0.470138 |
| 18 | 74855200 | 74856799 | 12.70377 | 22.29101 | 40.9663 | 18.11469 | -0.48593 | 0.616698 | 0.470143 |
| 18 | 63634600 | 63637399 | 14.74915 | 25.84168 | 43.53349 | 19.22579 | -0.3612 | 0.6563 | 0.470157 |
| 1 | 117303000 | 117305399 | 22.01602 | 38.32237 | 38.20158 | 16.79212 | 0.391699 | 0.781522 | 0.47016 |
| 8 | 21341400 | 21341799 | 4.351172 | 8.496838 | 19.30601 | 6.788871 | -0.49865 | -0.51784 | 0.470161 |
| 14 | 29496800 | 29498599 | 19.0029 | 32.45748 | 53.79521 | 23.58575 | -0.34347 | 0.973442 | 0.470165 |
| 3 | 28128800 | 28131999 | 31.69941 | 52.80836 | 53.56127 | 23.27026 | 0.36087 | 1.249739 | 0.470168 |
| 2 | 176947200 | 176951199 | 33.2311 | 55.39149 | 47.07337 | 20.45885 | 0.615094 | 1.239895 | 0.470172 |
| 2 | 105692400 | 105697799 | 53.44702 | 86.34433 | 74.07347 | 31.62064 | 0.580893 | 1.744132 | 0.470178 |
| 18 | 54396600 | 54398199 | 14.88855 | 26.97779 | 27.22597 | 11.78346 | 0.399129 | 0.347528 | 0.47019 |
| 17 | 22212200 | 22220199 | 153.2935 | 233.3642 | 198.1208 | 82.10328 | 0.52061 | 3.172674 | 0.470206 |
| 12 | 116614800 | 116616599 | 14.74009 | 25.85412 | 44.02112 | 19.45915 | -0.37657 | 0.638892 | 0.47021 |
| X | 164570200 | 164575199 | 52.23854 | 85.05734 | 61.66943 | 26.47744 | 0.83321 | 1.583944 | 0.47021 |
| X | 53003000 | 53006199 | 46.84926 | 77.19904 | 54.09873 | 23.33746 | 0.891129 | 1.415986 | 0.47023 |
| 19 | 23596600 | 23597599 | 8.707124 | 15.89581 | 31.92923 | 13.69015 | -0.57803 | 0.272002 | 0.47023 |
| 6 | 38566800 | 38567999 | 17.71539 | 31.93481 | 28.43869 | 12.38799 | 0.572615 | 0.399666 | 0.470232 |
| 8 | 91372000 | 91374399 | 28.32274 | 47.75262 | 48.5462 | 21.22617 | 0.357438 | 1.114806 | 0.470237 |
| 12 | 49447800 | 49448799 | 8.338091 | 15.48353 | 29.99295 | 12.56774 | -0.502 | 0.167953 | 0.470238 |
| 15 | 17903200 | 17903599 | 8.338091 | 15.48353 | 29.99295 | 12.56774 | -0.502 | 0.167953 | 0.470238 |
| 19 | 4765200 | 4767799 | 20.8153 | 34.76337 | 61.64406 | 26.82425 | -0.44235 | 1.199767 | 0.470246 |
| 4 | 122555800 | 122558399 | 56.28842 | 90.09123 | 93.14705 | 39.23563 | 0.296276 | 2.03456 | 0.470246 |
| 8 | 51758400 | 51760999 | 25.82639 | 44.91537 | 37.97494 | 16.68326 | 0.628154 | 0.811522 | 0.470249 |
| 10 | 53246200 | 53250999 | 72.94841 | 116.1546 | 105.2365 | 44.00781 | 0.478489 | 2.215492 | 0.470257 |
| 11 | 74054600 | 74060399 | 66.55027 | 105.8839 | 108.1795 | 45.18946 | 0.304187 | 2.234058 | 0.470259 |
| 12 | 40495400 | 40498399 | 21.35901 | 35.38118 | 66.44036 | 28.75454 | -0.5254 | 1.295848 | 0.470264 |
| 18 | 66878600 | 66879199 | 11.82219 | 22.38983 | 22.66482 | 9.172963 | 0.468164 | 0.052602 | 0.470286 |
| Y | 20336800 | 20338999 | 13.86792 | 25.60038 | 15.48096 | 6.539289 | 1.15588 | 0.204307 | 0.470291 |
| 12 | 103221200 | 103223999 | 16.40043 | 28.214 | 50.04836 | 21.95362 | -0.4392 | 0.892259 | 0.470293 |
| 16 | 79017400 | 79021199 | 74.56124 | 117.7201 | 120.95 | 50.21746 | 0.286197 | 2.44783 | 0.470297 |
| 18 | 25493800 | 25494199 | 4.833038 | 9.420186 | 22.66477 | 8.198498 | -0.6198 | -0.40432 | 0.470302 |
| 1 | 98400600 | 98401199 | 13.0104 | 23.98333 | 24.49007 | 10.3648 | 0.404411 | 0.231747 | 0.470318 |

| | | | | | | | | | |
|---|---|---|---|---|---|---|---|---|---|
| 13 | 108131000 | 108134999 | 53.73698 | 85.47908 | 131.4589 | 54.89755 | -0.28525 | 2.236121 | 0.470321 |
| X | 153276000 | 153277799 | 22.91994 | 40.29 | 32.97661 | 14.60297 | 0.675168 | 0.63356 | 0.470324 |
| 3 | 125008400 | 125010199 | 24.32401 | 39.96226 | 71.2331 | 30.69027 | -0.45354 | 1.408242 | 0.470335 |
| 7 | 22434000 | 22438399 | 81.0517 | 127.921 | 115.6404 | 48.00413 | 0.470173 | 2.459009 | 0.470341 |
| 11 | 34369800 | 34371999 | 12.71126 | 22.53362 | 38.76695 | 17.13821 | -0.38621 | 0.493051 | 0.470353 |
| 9 | 63895400 | 63897799 | 15.62982 | 27.10314 | 47.40502 | 20.79917 | -0.4163 | 0.816127 | 0.470355 |
| 1 | 53410000 | 53412399 | 57.37131 | 91.89365 | 82.25267 | 34.68548 | 0.505154 | 2.00901 | 0.470357 |
| 5 | 137675400 | 137677399 | 16.16472 | 28.73893 | 28.84665 | 12.72229 | 0.389151 | 0.475503 | 0.470362 |
| X | 77381800 | 77383399 | 41.173 | 67.19431 | 55.23842 | 23.73622 | 0.654738 | 1.524283 | 0.47037 |
| 4 | 6793800 | 6795199 | 10.33471 | 18.84258 | 33.885 | 14.55241 | -0.4218 | 0.293497 | 0.470426 |
| 5 | 133308000 | 133309199 | 14.3672 | 26.65831 | 25.31104 | 10.61731 | 0.517544 | 0.207049 | 0.470439 |
| 18 | 29686400 | 29688799 | 24.3715 | 40.6193 | 68.0823 | 29.58761 | -0.36154 | 1.237115 | 0.47044 |
| 13 | 29615600 | 29616599 | 11.62761 | 21.02312 | 38.51431 | 16.71461 | -0.45851 | 0.360464 | 0.470446 |
| 1 | 13040800 | 13042399 | 9.985884 | 17.88232 | 36.82398 | 16.14201 | -0.63561 | 0.403378 | 0.470447 |
| 11 | 7568000 | 7568399 | 7.026784 | 13.4855 | 26.30118 | 10.31798 | -0.42948 | -0.08416 | 0.470452 |
| 12 | 16657600 | 16658599 | 7.026784 | 13.4855 | 26.30118 | 10.31798 | -0.42948 | -0.08416 | 0.470452 |
| 17 | 27530800 | 27538199 | 52.09253 | 82.58997 | 137.4267 | 57.19469 | -0.40341 | 2.331832 | 0.470471 |
| 7 | 11484400 | 11486199 | 31.7444 | 53.08778 | 51.93282 | 22.61776 | 0.412693 | 1.206339 | 0.470478 |
| 7 | 134050200 | 134051399 | 11.00604 | 19.99063 | 34.87808 | 15.0501 | -0.3828 | 0.323787 | 0.470479 |
| 10 | 21397200 | 21398199 | 10.64125 | 19.28831 | 36.95909 | 15.98918 | -0.51999 | 0.33414 | 0.470481 |
| 7 | 23421400 | 23430399 | 139.9176 | 214.3851 | 180.4287 | 74.88941 | 0.54054 | 3.038515 | 0.470502 |
| 14 | 15974600 | 15976599 | 12.97399 | 22.952 | 39.25465 | 17.36097 | -0.37913 | 0.516081 | 0.470525 |
| 19 | 3132400 | 3134599 | 41.00154 | 66.4626 | 68.82852 | 29.47268 | 0.314729 | 1.642322 | 0.470527 |
| 17 | 82125200 | 82126199 | 11.49679 | 21.59818 | 22.55976 | 9.272627 | 0.404143 | 0.100361 | 0.470538 |
| 4 | 113281400 | 113282399 | 33.17457 | 54.73015 | 56.85229 | 24.53754 | 0.325496 | 1.366417 | 0.47054 |
| 8 | 91882400 | 91883999 | 21.20344 | 35.68866 | 59.14453 | 25.84682 | -0.34513 | 1.117492 | 0.470544 |
| 17 | 40052400 | 40067999 | 250.6622 | 366.9934 | 325.2405 | 133.9769 | 0.408185 | 3.896151 | 0.470544 |
| 4 | 49038200 | 49040199 | 14.90071 | 25.996 | 44.69019 | 19.67234 | -0.39057 | 0.724331 | 0.470546 |
| 10 | 33561400 | 33562799 | 39.89041 | 64.77595 | 66.55133 | 28.53476 | 0.328534 | 1.604497 | 0.47057 |
| 14 | 68519400 | 68520199 | 16.34293 | 29.04193 | 29.3242 | 12.9381 | 0.38025 | 0.48068 | 0.470578 |
| 2 | 74265600 | 74269199 | 64.20902 | 102.092 | 105.1098 | 43.87052 | 0.292141 | 2.251347 | 0.470584 |
| X | 57595000 | 57599999 | 80.35322 | 127.1727 | 118.9063 | 49.42955 | 0.424816 | 2.396663 | 0.470619 |
| 7 | 22467800 | 22479999 | 470.8525 | 646.6921 | 587.3813 | 244.2472 | 0.280825 | 4.857138 | 0.470623 |
| 12 | 19951200 | 19985999 | 1003.145 | 1302.936 | 966.8094 | 405.0485 | 0.49174 | 5.75071 | 0.47063 |
| 6 | 121888400 | 121890999 | 22.40879 | 39.45157 | 32.32743 | 14.31114 | 0.674169 | 0.611695 | 0.470701 |
| 7 | 31211600 | 31224799 | 172.5692 | 261.6427 | 202.6214 | 83.86625 | 0.648352 | 3.259577 | 0.470702 |
| 10 | 52345200 | 52346599 | 19.41158 | 34.35443 | 28.02208 | 12.3922 | 0.683542 | 0.530882 | 0.470719 |
| 18 | 48070200 | 48070799 | 5.672037 | 11.01869 | 23.44986 | 8.756221 | -0.48678 | -0.27668 | 0.470725 |
| 9 | 20701000 | 20703399 | 19.76032 | 34.48472 | 34.61298 | 15.23935 | 0.382082 | 0.727695 | 0.470726 |
| 17 | 18534800 | 18539599 | 63.00119 | 101.2014 | 83.20107 | 35.23887 | 0.632449 | 1.930137 | 0.470729 |
| 19 | 49052600 | 49054399 | 26.36956 | 44.84952 | 45.24686 | 19.83108 | 0.368981 | 1.027939 | 0.470752 |
| 6 | 120560600 | 120561799 | 14.77472 | 26.58929 | 25.43159 | 11.10128 | 0.468418 | 0.3829 | 0.470763 |
| 14 | 109684000 | 109688999 | 51.94451 | 83.70855 | 79.4951 | 33.82606 | 0.430449 | 1.812778 | 0.470763 |
| X | 115023400 | 115025199 | 23.16293 | 40.29001 | 38.67581 | 16.99295 | 0.445797 | 0.79748 | 0.470774 |
| 4 | 16986000 | 16989599 | 55.79065 | 89.19633 | 91.14581 | 38.33287 | 0.311458 | 2.066896 | 0.470797 |
| X | 163261600 | 163262199 | 12.00485 | 22.72947 | 22.65907 | 9.176006 | 0.489394 | 0.056368 | 0.470824 |
| 11 | 87938000 | 87939399 | 10.23815 | 18.1081 | 37.90329 | 16.76405 | -0.66865 | 0.499423 | 0.470827 |

| | | | | | | | | | |
|---|---|---|---|---|---|---|---|---|---|
| 14 | 89273600 | 89277799 | 81.29318 | 128.027 | 130.3798 | 54.09767 | 0.29614 | 2.505727 | 0.470846 |
| 11 | 76438400 | 76440599 | 14.60829 | 25.52449 | 43.62961 | 19.22378 | -0.38203 | 0.700817 | 0.47085 |
| 4 | 40545800 | 40548799 | 40.12519 | 65.64237 | 63.89128 | 27.47854 | 0.414108 | 1.491387 | 0.470862 |
| 15 | 13713400 | 13715399 | 52.43934 | 84.12728 | 85.74518 | 36.24558 | 0.320842 | 1.956954 | 0.470864 |
| 1 | 61629400 | 61630799 | 15.14893 | 28.26661 | 24.84016 | 10.32799 | 0.638429 | 0.180042 | 0.470866 |
| 8 | 110790600 | 110792799 | 19.20036 | 32.62882 | 54.82363 | 24.02683 | -0.36424 | 1.022017 | 0.47087 |
| 18 | 39198400 | 39199999 | 10.30738 | 18.86917 | 34.49316 | 14.73686 | -0.44038 | 0.270577 | 0.470874 |
| 14 | 75284600 | 75285399 | 3.874517 | 7.557981 | 20.86911 | 7.469948 | -0.80065 | -0.45912 | 0.47088 |
| 7 | 23447800 | 23452399 | 67.31929 | 108.1542 | 79.44921 | 33.65726 | 0.794848 | 1.931556 | 0.4709 |
| X | 140121000 | 140122999 | 22.80924 | 40.44195 | 28.94168 | 12.7935 | 0.872247 | 0.534546 | 0.470917 |
| 10 | 72073400 | 72074999 | 14.31972 | 26.6319 | 25.40367 | 10.6211 | 0.514571 | 0.194957 | 0.470937 |
| 15 | 37817000 | 37821999 | 67.13701 | 106.9285 | 105.6604 | 44.20067 | 0.353857 | 2.200197 | 0.470956 |
| 7 | 93540800 | 93541599 | 11.23452 | 21.48777 | 22.24066 | 8.810452 | 0.463138 | -0.02411 | 0.470961 |
| 1 | 188400800 | 188401199 | 5.075857 | 9.888732 | 21.34159 | 7.768451 | -0.47142 | -0.37803 | 0.470966 |
| 14 | 52639200 | 52641999 | 25.24383 | 43.19053 | 43.62145 | 19.12783 | 0.368658 | 0.969021 | 0.470975 |
| 7 | 57157200 | 57160799 | 32.34751 | 52.36325 | 86.33606 | 36.93239 | -0.35644 | 1.663319 | 0.471006 |
| 16 | 43893000 | 43895399 | 47.52645 | 76.62573 | 77.76513 | 33.11076 | 0.335694 | 1.79411 | 0.47101 |
| 2 | 122637200 | 122639399 | 44.8616 | 71.58816 | 116.7525 | 48.97119 | -0.36478 | 2.121541 | 0.471012 |
| 9 | 59393000 | 59394399 | 10.49702 | 18.69028 | 35.85086 | 15.79151 | -0.53828 | 0.444453 | 0.471016 |
| X | 54856000 | 54861399 | 76.13189 | 121.4899 | 84.33542 | 35.37486 | 0.865369 | 2.143914 | 0.471021 |
| 6 | 133207600 | 133209199 | 14.36707 | 26.97243 | 23.84113 | 9.810332 | 0.641743 | 0.132175 | 0.471031 |
| 8 | 67493600 | 67497399 | 51.846 | 83.4722 | 82.02533 | 34.85574 | 0.379638 | 1.843084 | 0.471038 |
| 10 | 41225200 | 41227599 | 16.63521 | 28.92026 | 49.53387 | 21.75641 | -0.38603 | 0.788375 | 0.471042 |
| 15 | 52640400 | 52641999 | 12.87831 | 22.95679 | 45.23466 | 19.90544 | -0.57663 | 0.469216 | 0.471044 |
| X | 77269600 | 77271399 | 45.87623 | 74.20239 | 73.68523 | 31.50166 | 0.37225 | 1.698468 | 0.471045 |
| 6 | 69468200 | 69472199 | 56.99989 | 91.12932 | 94.35359 | 39.68177 | 0.292521 | 2.06991 | 0.471071 |
| 13 | 65671000 | 65695599 | 659.8853 | 882.8752 | 778.8531 | 325.1687 | 0.284829 | 5.27207 | 0.471077 |
| X | 32288800 | 32289999 | 14.61823 | 27.15267 | 22.26995 | 9.328284 | 0.729046 | 0.197904 | 0.47111 |
| 2 | 151161800 | 151165799 | 33.39085 | 56.35974 | 46.85951 | 20.49555 | 0.64655 | 1.120495 | 0.471147 |
| 9 | 19822400 | 19824599 | 39.57076 | 64.75944 | 63.3479 | 27.24853 | 0.407267 | 1.48373 | 0.471161 |
| 3 | 42039600 | 42040799 | 13.18962 | 24.29972 | 24.97906 | 10.58003 | 0.394054 | 0.23764 | 0.471191 |
| 17 | 33841200 | 33842999 | 25.4608 | 43.15749 | 42.24577 | 18.50137 | 0.412112 | 1.052304 | 0.471199 |
| X | 155164400 | 155165999 | 15.02988 | 27.39128 | 27.32852 | 11.7429 | 0.422433 | 0.317789 | 0.471208 |
| 4 | 37297200 | 37303399 | 74.02777 | 117.1288 | 120.7207 | 50.1767 | 0.284132 | 2.396856 | 0.471209 |
| X | 25685000 | 25692599 | 102.0982 | 159.3769 | 144.5493 | 59.97995 | 0.453291 | 2.68461 | 0.47121 |
| 12 | 21574600 | 21586999 | 108.572 | 165.405 | 251.2566 | 104.1463 | -0.31722 | 3.147239 | 0.471261 |
| X | 26054600 | 26064399 | 107.7324 | 168.1612 | 142.8391 | 59.27069 | 0.547689 | 2.688294 | 0.471262 |
| 11 | 73622800 | 73629199 | 64.22362 | 102.8066 | 92.84583 | 39.11676 | 0.491324 | 2.041202 | 0.471293 |
| 11 | 28136600 | 28140399 | 35.26449 | 58.36835 | 57.08528 | 24.69198 | 0.412619 | 1.333955 | 0.471294 |
| 17 | 72928000 | 72929999 | 28.37541 | 47.52552 | 47.55454 | 20.73072 | 0.380232 | 1.177679 | 0.471319 |
| 4 | 149297200 | 149300599 | 20.0193 | 33.90547 | 56.45137 | 24.71639 | -0.35142 | 1.056124 | 0.471343 |
| 6 | 43864400 | 43865599 | 12.69258 | 22.39865 | 38.81775 | 17.17624 | -0.39933 | 0.541902 | 0.471346 |
| X | 92332000 | 92334599 | 35.42225 | 58.54809 | 57.52818 | 24.8597 | 0.405845 | 1.352713 | 0.471358 |
| 7 | 20778600 | 20780999 | 39.49877 | 64.7211 | 55.31345 | 23.80466 | 0.602383 | 1.465341 | 0.471359 |
| 2 | 112563600 | 112565999 | 21.26123 | 37.17564 | 35.88668 | 15.82148 | 0.438195 | 0.709226 | 0.471386 |
| 12 | 58163600 | 58165599 | 35.99001 | 58.93324 | 61.26198 | 26.35737 | 0.32024 | 1.467476 | 0.471404 |
| 7 | 31582200 | 31583399 | 18.37646 | 32.58955 | 31.11109 | 13.74993 | 0.459074 | 0.510038 | 0.471415 |

| | | | | | | | | | |
|---|---|---|---|---|---|---|---|---|---|
| 9 | 11162600 | 11163799 | 7.290086 | 13.95856 | 27.97885 | 11.0491 | -0.47864 | -0.0594 | 0.471422 |
| 12 | 55466400 | 55468599 | 17.13323 | 28.79082 | 63.74689 | 27.84431 | -0.76154 | 1.122804 | 0.471427 |
| 14 | 36690200 | 36690999 | 9.715304 | 17.46722 | 34.91726 | 15.25487 | -0.58874 | 0.378558 | 0.47143 |
| 6 | 78953000 | 78957399 | 97.80536 | 151.7826 | 155.6177 | 64.62658 | 0.270041 | 2.781665 | 0.471457 |
| 17 | 76908200 | 76910199 | 22.70135 | 39.66171 | 28.61014 | 12.60531 | 0.856784 | 0.720835 | 0.47146 |
| 16 | 80812000 | 80817999 | 130.8593 | 198.8716 | 197.5842 | 81.83899 | 0.291936 | 3.199291 | 0.471465 |
| 16 | 65856800 | 65859199 | 46.81681 | 75.49448 | 77.82031 | 33.14156 | 0.313531 | 1.787296 | 0.471482 |
| 4 | 27049400 | 27051399 | 29.59547 | 49.32138 | 50.27065 | 21.84564 | 0.353762 | 1.235097 | 0.471487 |
| 6 | 141902000 | 141903599 | 22.91982 | 39.90442 | 34.01161 | 14.95243 | 0.616666 | 0.776695 | 0.471488 |
| 14 | 85117400 | 85119199 | 19.63389 | 35.21815 | 28.79246 | 12.60365 | 0.690721 | 0.443556 | 0.471493 |
| 2 | 168999000 | 169000599 | 10.96193 | 19.66689 | 35.84363 | 15.68753 | -0.45843 | 0.40138 | 0.471502 |
| X | 154847000 | 154847999 | 23.9388 | 41.11697 | 37.59336 | 16.48946 | 0.512732 | 0.924979 | 0.471507 |
| 6 | 133779200 | 133781799 | 35.80851 | 59.18313 | 50.93615 | 22.01021 | 0.596395 | 1.348767 | 0.471531 |
| 17 | 49389800 | 49394199 | 30.35737 | 49.45736 | 79.94575 | 34.33137 | -0.31999 | 1.540934 | 0.471579 |
| 17 | 48157800 | 48160199 | 20.01091 | 33.80934 | 57.53862 | 25.17394 | -0.38312 | 1.077173 | 0.471583 |
| 6 | 124371000 | 124377399 | 67.58436 | 106.0925 | 160.6813 | 66.64233 | -0.27933 | 2.567937 | 0.471583 |
| 5 | 77168400 | 77175199 | 87.12243 | 134.3326 | 200.6324 | 83.33924 | -0.27931 | 2.899618 | 0.471597 |
| 10 | 78962000 | 78963799 | 18.8122 | 32.14007 | 53.80578 | 23.59088 | -0.35786 | 0.969403 | 0.471608 |
| 6 | 69778400 | 69781999 | 46.77404 | 75.58213 | 75.56027 | 32.26528 | 0.361031 | 1.725932 | 0.471608 |
| 5 | 59517600 | 59519999 | 23.81247 | 40.86745 | 42.0741 | 18.45391 | 0.341807 | 0.934559 | 0.471612 |
| 17 | 68566200 | 68567999 | 15.32133 | 28.38339 | 25.18765 | 10.5885 | 0.612119 | 0.22299 | 0.471613 |
| 7 | 11880200 | 11885399 | 55.79543 | 89.84162 | 80.4286 | 34.18319 | 0.514024 | 1.84448 | 0.471614 |
| 7 | 114347000 | 114347599 | 4.686243 | 9.114844 | 21.75201 | 8.042055 | -0.63566 | -0.32751 | 0.471617 |
| 7 | 66157600 | 66158799 | 17.23089 | 31.81765 | 25.00484 | 10.55811 | 0.78184 | 0.25609 | 0.471618 |
| Y | 42190200 | 42191399 | 10.58073 | 20.06825 | 19.06007 | 7.689987 | 0.562794 | 0.017993 | 0.471623 |
| 4 | 134038000 | 134040599 | 24.57657 | 40.60487 | 68.28778 | 29.50083 | -0.36717 | 1.337536 | 0.471643 |
| 7 | 40790400 | 40800199 | 117.3907 | 182.527 | 146.9883 | 61.02538 | 0.620236 | 2.7587 | 0.471649 |
| 11 | 108894400 | 108898599 | 40.52653 | 65.09279 | 103.1584 | 43.6849 | -0.31272 | 1.910093 | 0.471656 |
| 2 | 142037800 | 142039799 | 24.30444 | 41.83651 | 42.00094 | 18.42397 | 0.378787 | 0.90992 | 0.47167 |
| 1 | 106788600 | 106790399 | 14.24711 | 24.92794 | 43.57706 | 19.22281 | -0.41436 | 0.674707 | 0.471686 |
| 5 | 7482200 | 7485599 | 48.88139 | 78.71811 | 80.92481 | 34.40008 | 0.315087 | 1.830818 | 0.471694 |
| 13 | 56367200 | 56367999 | 5.254099 | 10.20916 | 22.85644 | 8.518621 | -0.55629 | -0.28995 | 0.471695 |
| X | 61141000 | 61142799 | 27.29457 | 46.68528 | 43.62107 | 19.1272 | 0.480565 | 0.982659 | 0.4717 |
| 4 | 145331200 | 145337199 | 122.6984 | 188.5278 | 179.2691 | 74.43708 | 0.367623 | 2.977089 | 0.471713 |
| 9 | 99239400 | 99241399 | 20.88395 | 34.60275 | 66.99221 | 28.99828 | -0.56927 | 1.291455 | 0.471751 |
| 8 | 47916600 | 47918799 | 22.50683 | 37.59619 | 63.01111 | 27.42272 | -0.36145 | 1.20109 | 0.471757 |
| 5 | 92629600 | 92635399 | 68.33769 | 108.3708 | 111.889 | 46.57707 | 0.284407 | 2.332764 | 0.471791 |
| 7 | 48080400 | 48082799 | 37.42324 | 61.48604 | 60.8577 | 26.22062 | 0.392711 | 1.43158 | 0.471796 |
| 7 | 133004600 | 133007799 | 19.29896 | 32.92498 | 54.89341 | 24.06499 | -0.3523 | 0.985355 | 0.471797 |
| 1 | 108655800 | 108656399 | 7.022092 | 13.40404 | 26.70505 | 10.6298 | -0.47997 | -0.03789 | 0.471804 |
| 4 | 12361600 | 12362799 | 7.022092 | 13.40404 | 26.70505 | 10.6298 | -0.47997 | -0.03789 | 0.471804 |
| 1 | 108778800 | 108783599 | 80.72439 | 127.0417 | 131.0858 | 54.38561 | 0.276455 | 2.520175 | 0.471824 |
| 7 | 11893800 | 11904799 | 133.9355 | 206.15 | 167.6494 | 69.62876 | 0.594912 | 2.947986 | 0.471824 |
| 17 | 83477400 | 83482599 | 46.54067 | 74.18145 | 131.4456 | 55.04542 | -0.48611 | 2.164632 | 0.47183 |
| 17 | 22064400 | 22065599 | 15.75512 | 28.0536 | 26.90054 | 11.8487 | 0.455936 | 0.45856 | 0.471845 |
| 14 | 66481600 | 66482999 | 11.44864 | 20.22861 | 37.85743 | 16.74715 | -0.50874 | 0.518666 | 0.471847 |
| 1 | 193762600 | 193762999 | 5.413133 | 10.48624 | 24.39305 | 9.253659 | -0.63555 | -0.22278 | 0.471868 |

| | | | | | | | | | |
|---|---|---|---|---|---|---|---|---|---|
| 7 | 38792200 | 38810999 | 409.751 | 573.176 | 505.4086 | 209.5565 | 0.350214 | 4.577536 | 0.471872 |
| 8 | 119967400 | 119976199 | 75.63729 | 117.4975 | 182.1834 | 75.64779 | -0.32495 | 2.757755 | 0.471878 |
| 8 | 34091600 | 34093199 | 9.903913 | 17.95695 | 34.3475 | 14.85404 | -0.51677 | 0.323157 | 0.47188 |
| X | 33670400 | 33671799 | 17.96147 | 33.39111 | 23.39096 | 9.786417 | 0.955993 | 0.222467 | 0.471889 |
| 9 | 72068600 | 72069199 | 4.647229 | 9.063425 | 20.29869 | 7.285491 | -0.50539 | -0.43776 | 0.47189 |
| 13 | 33245400 | 33247999 | 11.96223 | 21.34259 | 38.78185 | 17.05322 | -0.45963 | 0.443386 | 0.471894 |
| 13 | 61641600 | 61643999 | 34.45719 | 58.08704 | 47.43355 | 20.73877 | 0.672495 | 1.137504 | 0.471895 |
| 17 | 69832200 | 69833199 | 7.172242 | 13.68686 | 26.69927 | 10.63373 | -0.45047 | -0.03401 | 0.471907 |
| 4 | 57697200 | 57698999 | 12.7018 | 22.55315 | 38.76966 | 17.11727 | -0.38377 | 0.482157 | 0.471907 |
| 10 | 84320600 | 84323199 | 20.11037 | 33.75274 | 67.4016 | 29.42138 | -0.61343 | 1.146662 | 0.471909 |
| 5 | 11525400 | 11526399 | 25.95693 | 44.34225 | 37.91133 | 16.6219 | 0.607765 | 0.978755 | 0.471925 |
| 1 | 14719000 | 14719999 | 8.249352 | 15.72151 | 31.31757 | 12.50803 | -0.48459 | -0.00296 | 0.471933 |
| 7 | 61110200 | 61112999 | 37.48829 | 62.84169 | 48.05371 | 20.95951 | 0.766823 | 1.197798 | 0.471961 |
| 16 | 62018000 | 62021399 | 24.89759 | 43.3856 | 36.82666 | 16.19887 | 0.622631 | 0.777443 | 0.471965 |
| 1 | 65450400 | 65452999 | 20.83571 | 37.09674 | 29.73347 | 13.09751 | 0.7131 | 0.498564 | 0.471977 |
| 17 | 49518800 | 49520199 | 13.01028 | 22.85317 | 40.40622 | 17.88313 | -0.43041 | 0.600498 | 0.471981 |
| 11 | 95704800 | 95707399 | 21.46771 | 36.15265 | 60.33345 | 26.37064 | -0.35525 | 1.112987 | 0.472007 |
| 5 | 122321600 | 122326799 | 54.1615 | 85.58857 | 143.1934 | 59.47962 | -0.41519 | 2.416539 | 0.472008 |
| 10 | 54348600 | 54349999 | 10.97517 | 19.8505 | 35.84336 | 15.54858 | -0.43723 | 0.348089 | 0.472016 |
| X | 148996600 | 148997999 | 24.29299 | 42.33723 | 37.51655 | 16.50361 | 0.561014 | 0.775967 | 0.472018 |
| 3 | 7203200 | 7204799 | 23.49375 | 40.36904 | 41.55414 | 18.22699 | 0.342363 | 0.922043 | 0.472024 |
| 2 | 108940200 | 108940799 | 12.18359 | 22.70532 | 23.53906 | 9.80763 | 0.399331 | 0.160083 | 0.472035 |
| X | 66011800 | 66015199 | 39.1942 | 64.4621 | 60.53798 | 26.09424 | 0.468854 | 1.422381 | 0.472036 |
| 2 | 50972600 | 50975799 | 22.65755 | 37.93354 | 62.50447 | 27.24103 | -0.33703 | 1.177246 | 0.472043 |
| X | 132804400 | 132806599 | 21.33191 | 38.24646 | 28.20343 | 12.35057 | 0.838508 | 0.457982 | 0.472048 |
| 13 | 48445200 | 48451199 | 50.89443 | 80.98915 | 126.7463 | 52.95821 | -0.30976 | 2.218767 | 0.47205 |
| 9 | 80721200 | 80722999 | 23.59007 | 41.02755 | 34.52466 | 15.16781 | 0.634914 | 0.795662 | 0.472051 |
| 7 | 11494000 | 11502799 | 127.4848 | 196.0399 | 158.397 | 65.7777 | 0.60323 | 2.962383 | 0.472054 |
| 15 | 39548000 | 39548999 | 8.853098 | 16.67258 | 31.40332 | 12.85395 | -0.43559 | 0.082994 | 0.472056 |
| 5 | 3814000 | 3817199 | 37.68021 | 61.79432 | 59.56124 | 25.64205 | 0.429893 | 1.455411 | 0.472059 |
| 7 | 54100600 | 54101199 | 6.24539 | 12.11802 | 24.55363 | 9.24459 | -0.42797 | -0.23717 | 0.472064 |
| 1 | 79225600 | 79226999 | 13.66763 | 25.74365 | 24.00477 | 9.822932 | 0.571765 | 0.107154 | 0.472069 |
| 2 | 20050800 | 20052199 | 13.66763 | 25.74365 | 24.00477 | 9.822932 | 0.571765 | 0.107154 | 0.472069 |
| X | 145831200 | 145831599 | 8.949794 | 17.49005 | 15.13699 | 5.244601 | 0.894125 | -0.52082 | 0.472179 |
| 14 | 118674200 | 118677599 | 28.4434 | 46.28765 | 88.39185 | 37.94301 | -0.561 | 1.55705 | 0.47218 |
| 7 | 131782200 | 131783199 | 12.31474 | 21.96542 | 37.74692 | 16.60202 | -0.3796 | 0.44738 | 0.472189 |
| 3 | 136963600 | 136965599 | 20.1174 | 34.20357 | 57.07137 | 25.01288 | -0.35428 | 1.021922 | 0.472229 |
| 18 | 34319800 | 34326399 | 82.67001 | 130.3743 | 128.8898 | 53.48899 | 0.3404 | 2.480189 | 0.47223 |
| 2 | 73090800 | 73096999 | 50.81535 | 80.68687 | 143.2058 | 59.69005 | -0.49421 | 2.288743 | 0.472236 |
| 17 | 20678600 | 20683799 | 39.09977 | 64.9693 | 47.12212 | 20.43743 | 0.842951 | 1.294255 | 0.472237 |
| 3 | 15240200 | 15240599 | 3.814444 | 7.454767 | 18.25438 | 6.315399 | -0.58343 | -0.58845 | 0.472245 |
| 6 | 139838600 | 139840599 | 16.28582 | 28.09417 | 55.98692 | 24.56063 | -0.60564 | 0.875147 | 0.472261 |
| 11 | 16335600 | 16336599 | 18.44259 | 31.38089 | 54.24882 | 23.77734 | -0.40472 | 1.008641 | 0.472272 |
| 1 | 19173200 | 19175399 | 21.51426 | 36.28038 | 59.74561 | 26.12448 | -0.33595 | 1.102547 | 0.472277 |
| 7 | 30091000 | 30095599 | 27.43456 | 44.76388 | 76.79514 | 32.99675 | -0.40398 | 1.510378 | 0.472279 |
| 16 | 89117400 | 89120399 | 27.07222 | 46.90022 | 39.06122 | 17.13779 | 0.64897 | 0.861759 | 0.472293 |
| 7 | 129988600 | 129991199 | 18.03646 | 30.94393 | 51.64777 | 22.65234 | -0.35247 | 0.927234 | 0.472299 |

| | | | | | | | | | |
|---|---|---|---|---|---|---|---|---|---|
| 12 | 10548000 | 10553399 | 66.71665 | 106.3247 | 104.4496 | 43.73158 | 0.363258 | 2.180674 | 0.472308 |
| 18 | 34183400 | 34184599 | 6.703224 | 12.89629 | 25.83835 | 10.06491 | -0.45886 | -0.11467 | 0.47232 |
| 14 | 31372200 | 31373599 | 11.5649 | 20.76568 | 36.30629 | 15.87907 | -0.39843 | 0.400016 | 0.472326 |
| 12 | 92108200 | 92109799 | 18.56371 | 33.70622 | 27.47547 | 11.86095 | 0.707957 | 0.364847 | 0.47234 |
| 18 | 58200000 | 58203199 | 22.46469 | 37.63759 | 63.10857 | 27.51674 | -0.36218 | 1.168471 | 0.472349 |
| 11 | 27990400 | 27992799 | 29.13829 | 49.70528 | 39.79509 | 17.44579 | 0.701859 | 1.00662 | 0.472366 |
| 15 | 100716200 | 100717599 | 13.96527 | 24.60259 | 41.87027 | 18.53325 | -0.37447 | 0.573091 | 0.472371 |
| 6 | 17785800 | 17788199 | 20.62847 | 36.38646 | 30.69555 | 13.58354 | 0.63314 | 0.575531 | 0.472378 |
| X | 159166400 | 159168399 | 27.84642 | 47.69371 | 43.54908 | 19.09772 | 0.513863 | 0.971344 | 0.472389 |
| 15 | 98823400 | 98824399 | 8.738182 | 16.52414 | 32.13215 | 13.04615 | -0.47102 | 0.057462 | 0.472389 |
| 11 | 40143400 | 40144399 | 8.225457 | 15.65007 | 30.06791 | 12.05101 | -0.43731 | 0.004421 | 0.472406 |
| 11 | 100792200 | 100793199 | 8.225457 | 15.65007 | 30.06791 | 12.05101 | -0.43731 | 0.004421 | 0.472406 |
| 15 | 52132800 | 52135599 | 35.61148 | 57.4864 | 92.10349 | 39.2782 | -0.31957 | 1.744637 | 0.47241 |
| 9 | 97496800 | 97497199 | 3.754503 | 7.343702 | 17.29926 | 5.812947 | -0.48891 | -0.6933 | 0.472412 |
| 12 | 49322800 | 49325999 | 31.81709 | 53.96692 | 45.14901 | 19.77674 | 0.637798 | 1.071556 | 0.472412 |
| 7 | 63346600 | 63348799 | 18.15184 | 31.08688 | 52.73172 | 23.12446 | -0.37621 | 0.943215 | 0.47242 |
| 2 | 144630400 | 144631199 | 10.93058 | 20.88063 | 21.62059 | 8.59208 | 0.457981 | -0.02032 | 0.47242 |
| 10 | 37936200 | 37940199 | 90.87408 | 141.5616 | 144.5301 | 59.98803 | 0.280327 | 2.712209 | 0.472424 |
| 8 | 42528400 | 42531399 | 22.25958 | 39.39694 | 30.08823 | 13.30577 | 0.777968 | 0.551134 | 0.472424 |
| 1 | 53594600 | 53595799 | 9.603413 | 17.8431 | 33.75804 | 14.13471 | -0.46794 | 0.183926 | 0.472453 |
| 16 | 5450400 | 5454999 | 54.84656 | 87.83424 | 91.11208 | 38.41118 | 0.292506 | 2.015094 | 0.47246 |
| 13 | 90949000 | 90949999 | 14.47065 | 26.46005 | 26.83172 | 11.48153 | 0.402888 | 0.292903 | 0.472472 |
| 9 | 116480000 | 116481799 | 11.81333 | 22.43247 | 21.97255 | 8.843035 | 0.521391 | 0.028806 | 0.472478 |
| 5 | 114835000 | 114838599 | 27.93947 | 45.55558 | 77.26357 | 33.18948 | -0.38821 | 1.522969 | 0.472481 |
| 5 | 106332600 | 106333999 | 12.68594 | 23.88481 | 23.44608 | 9.600967 | 0.497015 | 0.098072 | 0.472482 |
| 16 | 3482400 | 3486199 | 46.10037 | 74.54145 | 74.45965 | 31.82168 | 0.363249 | 1.705669 | 0.472491 |
| 12 | 18340600 | 18345599 | 94.45994 | 148.2732 | 136.086 | 56.44107 | 0.442344 | 2.581522 | 0.472498 |
| 8 | 26507600 | 26508199 | 10.1781 | 19.71184 | 19.76925 | 7.508519 | 0.56456 | -0.17965 | 0.47252 |
| 3 | 90830600 | 90835799 | 55.75647 | 90.15503 | 69.81781 | 29.83801 | 0.729666 | 1.721084 | 0.47255 |
| 5 | 56410000 | 56412199 | 24.13886 | 41.93676 | 39.18973 | 17.20566 | 0.484222 | 0.819414 | 0.472563 |
| 7 | 106093600 | 106098599 | 59.3853 | 95.64696 | 77.90101 | 33.12145 | 0.650624 | 1.842113 | 0.472575 |
| 3 | 22991600 | 22993199 | 16.29385 | 29.77442 | 26.53153 | 11.36232 | 0.587211 | 0.308288 | 0.472648 |
| 16 | 36297000 | 36300399 | 34.35845 | 55.40788 | 91.81761 | 39.10549 | -0.36973 | 1.773056 | 0.472657 |
| 1 | 127290600 | 127294599 | 31.21606 | 50.64421 | 82.89252 | 35.51752 | -0.34284 | 1.615729 | 0.472675 |
| 15 | 14910600 | 14913599 | 29.83725 | 49.92735 | 51.32094 | 22.35947 | 0.341497 | 1.194796 | 0.472694 |
| 1 | 65014800 | 65015999 | 9.725389 | 17.87862 | 31.92866 | 13.56355 | -0.40046 | 0.239288 | 0.472699 |
| 9 | 7866600 | 7873799 | 55.0393 | 87.3223 | 134.9149 | 56.18823 | -0.29561 | 2.314315 | 0.4727 |
| 4 | 150231600 | 150236399 | 54.91897 | 87.09094 | 135.3586 | 56.34681 | -0.3048 | 2.328648 | 0.4727 |
| 2 | 72651000 | 72652399 | 22.47467 | 39.49335 | 29.32966 | 12.98514 | 0.814462 | 0.6283 | 0.472702 |
| 14 | 83305400 | 83307199 | 18.03785 | 32.32675 | 28.26032 | 12.37991 | 0.593547 | 0.434707 | 0.472709 |
| 2 | 13621800 | 13623999 | 30.31763 | 50.50681 | 52.43922 | 22.78287 | 0.327211 | 1.243734 | 0.472723 |
| X | 78787800 | 78791399 | 41.53211 | 68.1882 | 59.92332 | 25.81025 | 0.563463 | 1.451078 | 0.472729 |
| 3 | 101999600 | 102001399 | 15.80927 | 28.77061 | 27.72295 | 11.93365 | 0.470593 | 0.331859 | 0.472743 |
| 5 | 98042000 | 98044199 | 28.15537 | 48.21226 | 43.53486 | 19.09114 | 0.529815 | 0.974242 | 0.472745 |
| 3 | 83593800 | 83594599 | 9.920471 | 17.9923 | 33.33853 | 14.41236 | -0.47063 | 0.320842 | 0.472748 |
| 3 | 151973400 | 151974799 | 9.920471 | 17.9923 | 33.33853 | 14.41236 | -0.47063 | 0.320842 | 0.472748 |
| 14 | 71357600 | 71360399 | 29.73843 | 48.32098 | 80.65369 | 34.59733 | -0.36893 | 1.580125 | 0.472748 |

| | | | | | | | | | |
|---|---|---|---|---|---|---|---|---|---|
| Y | 2545200 | 2547199 | 13.68579 | 25.19443 | 17.05544 | 7.231504 | 0.990861 | 0.22302 | 0.472762 |
| 15 | 10328200 | 10329399 | 9.698248 | 18.02949 | 32.67481 | 13.67057 | -0.40573 | 0.177343 | 0.472765 |
| 15 | 16173000 | 16174599 | 22.11429 | 38.59521 | 36.98148 | 16.28281 | 0.448765 | 0.742696 | 0.472765 |
| 2 | 93407200 | 93408999 | 16.53521 | 28.71836 | 47.92456 | 21.03665 | -0.34863 | 0.801626 | 0.472775 |
| 14 | 112057400 | 112057999 | 12.90937 | 24.2875 | 24.07588 | 9.871228 | 0.482062 | 0.108563 | 0.472783 |
| 1 | 166852800 | 166857599 | 56.24174 | 90.16501 | 90.4536 | 38.1957 | 0.342464 | 1.98244 | 0.472822 |
| 5 | 88367200 | 88370399 | 33.09317 | 55.39795 | 51.39356 | 22.39682 | 0.488938 | 1.200165 | 0.472847 |
| 5 | 50049600 | 50051199 | 12.48874 | 23.57851 | 22.93673 | 9.34447 | 0.516005 | 0.076928 | 0.472851 |
| 13 | 58934400 | 58946999 | 106.3723 | 162.1617 | 241.8591 | 100.2693 | -0.29003 | 3.132754 | 0.472867 |
| 11 | 117980400 | 117983599 | 48.82835 | 77.58306 | 128.922 | 53.77191 | -0.39837 | 2.26436 | 0.472878 |
| 10 | 36879800 | 36882999 | 42.15952 | 69.07667 | 61.37832 | 26.41405 | 0.546312 | 1.473357 | 0.472885 |
| 3 | 16432800 | 16437799 | 94.97881 | 147.5203 | 151.076 | 62.73259 | 0.272619 | 2.76571 | 0.472901 |
| 8 | 43747200 | 43750199 | 38.55117 | 62.84914 | 65.08775 | 27.95777 | 0.321377 | 1.538737 | 0.472925 |
| 3 | 100336600 | 100339199 | 17.76523 | 30.25708 | 53.19397 | 23.31679 | -0.42872 | 0.994741 | 0.472934 |
| 2 | 68454200 | 68456999 | 39.43552 | 64.62115 | 62.78953 | 27.02271 | 0.41776 | 1.467525 | 0.472937 |
| 7 | 141747400 | 141749599 | 31.90043 | 53.27782 | 52.50164 | 22.84632 | 0.402201 | 1.221078 | 0.472967 |
| 5 | 112618400 | 112619399 | 11.76437 | 22.23044 | 23.14249 | 9.410802 | 0.421913 | 0.067733 | 0.472971 |
| 7 | 127216200 | 127218399 | 16.53015 | 28.72668 | 47.91727 | 21.04133 | -0.34792 | 0.792958 | 0.472974 |
| 16 | 36038400 | 36043799 | 69.20074 | 110.228 | 95.68694 | 40.03488 | 0.540603 | 2.195673 | 0.472978 |
| 1 | 55444600 | 55446399 | 13.37124 | 23.30687 | 44.22606 | 19.45713 | -0.53231 | 0.729312 | 0.472978 |
| 1 | 120415800 | 120417399 | 12.89591 | 22.58351 | 41.56806 | 18.35015 | -0.48801 | 0.652592 | 0.472999 |
| 1 | 74262200 | 74267999 | 40.82291 | 65.62789 | 105.1448 | 44.58445 | -0.32682 | 1.882248 | 0.47301 |
| 11 | 50352200 | 50366799 | 224.0581 | 323.4865 | 452.3452 | 186.6146 | -0.26908 | 4.105591 | 0.473029 |
| 2 | 57317000 | 57317999 | 14.01802 | 26.58397 | 22.37468 | 9.029978 | 0.735253 | 0.060247 | 0.473031 |
| 15 | 81539200 | 81543199 | 43.73124 | 70.73231 | 72.51389 | 31.00067 | 0.32632 | 1.691681 | 0.473036 |
| Y | 29274600 | 29276599 | 16.73371 | 29.8781 | 18.5472 | 8.150447 | 1.08112 | 0.426675 | 0.473062 |
| 11 | 108118000 | 108119199 | 6.742732 | 12.87685 | 26.15627 | 10.401 | -0.50573 | -0.04305 | 0.473064 |
| 3 | 109020400 | 109023199 | 35.78101 | 59.3975 | 55.42996 | 24.02465 | 0.480391 | 1.301342 | 0.473065 |
| 3 | 5110600 | 5111399 | 14.06966 | 25.51802 | 23.96598 | 10.36031 | 0.503493 | 0.329534 | 0.473068 |
| 3 | 105499800 | 105501999 | 15.02099 | 26.24979 | 45.69971 | 20.13828 | -0.4086 | 0.70115 | 0.473069 |
| 14 | 96240200 | 96241399 | 10.64832 | 19.43907 | 34.43417 | 14.76476 | -0.39871 | 0.289942 | 0.473073 |
| 16 | 91044400 | 91045799 | 12.53747 | 22.10796 | 38.83779 | 17.18775 | -0.41924 | 0.549396 | 0.473075 |
| 12 | 17631000 | 17636799 | 51.75636 | 82.24352 | 129.3464 | 53.95432 | -0.31896 | 2.264274 | 0.473077 |
| 1 | 127015400 | 127018199 | 21.31125 | 37.28217 | 35.87906 | 15.82497 | 0.442629 | 0.701886 | 0.47311 |
| 12 | 54772600 | 54775599 | 33.86121 | 55.96712 | 57.47731 | 24.83748 | 0.342237 | 1.347607 | 0.473131 |
| 8 | 55901000 | 55908399 | 103.7442 | 161.0432 | 157.6561 | 65.47043 | 0.336946 | 2.780034 | 0.473132 |
| 12 | 20599800 | 20600799 | 17.08367 | 30.69057 | 25.62226 | 11.20148 | 0.661938 | 0.410788 | 0.473163 |
| 14 | 78527400 | 78531999 | 42.14945 | 67.59321 | 107.8085 | 45.54515 | -0.32466 | 1.962008 | 0.473163 |
| X | 23852200 | 23855599 | 40.4631 | 68.70876 | 36.59406 | 16.03483 | 1.286886 | 1.089085 | 0.473164 |
| 2 | 68732400 | 68733399 | 11.21619 | 21.26307 | 22.65177 | 9.14861 | 0.39733 | 0.038613 | 0.473168 |
| X | 63319200 | 63321799 | 28.44559 | 48.14904 | 46.90864 | 20.53468 | 0.418911 | 1.081873 | 0.473171 |
| 5 | 145999600 | 146013799 | 228.4441 | 335.9954 | 313.427 | 129.0737 | 0.340554 | 3.821699 | 0.473173 |
| 2 | 39934200 | 39935799 | 12.60554 | 22.12806 | 45.11224 | 19.95514 | -0.63476 | 0.618249 | 0.473176 |
| 6 | 132238400 | 132239599 | 19.84088 | 34.78013 | 34.47721 | 15.23266 | 0.400215 | 0.661987 | 0.473189 |
| X | 54455200 | 54456799 | 29.33729 | 50.36795 | 41.88599 | 18.37195 | 0.648791 | 0.955875 | 0.473213 |
| 19 | 25486000 | 25490599 | 44.1492 | 70.63687 | 111.7356 | 47.04176 | -0.31663 | 2.037877 | 0.473215 |
| 7 | 70406400 | 70409199 | 33.52508 | 55.62735 | 55.83835 | 24.19408 | 0.375489 | 1.301296 | 0.47323 |

| | | | | | | | | | |
|---|---|---|---|---|---|---|---|---|---|
| X | 125015200 | 125018599 | 76.6008 | 121.7391 | 109.9593 | 45.87356 | 0.480083 | 2.277935 | 0.473257 |
| X | 31607000 | 31610399 | 22.15041 | 39.43976 | 29.15728 | 12.84307 | 0.829219 | 0.506667 | 0.473269 |
| 3 | 132321600 | 132322999 | 15.01834 | 28.23753 | 23.24733 | 9.541673 | 0.746392 | 0.127147 | 0.473304 |
| 4 | 95215600 | 95216999 | 15.01834 | 28.23753 | 23.24733 | 9.541673 | 0.746392 | 0.127147 | 0.473304 |
| 11 | 44882200 | 44883599 | 7.781916 | 14.79636 | 27.72658 | 11.1282 | -0.40289 | 8.61E-05 | 0.473304 |
| 14 | 4396000 | 4403199 | 26.06957 | 44.62635 | 38.41321 | 16.84474 | 0.598536 | 0.962632 | 0.473306 |
| 16 | 83195600 | 83198399 | 47.41974 | 76.42638 | 79.09315 | 33.66069 | 0.30695 | 1.803122 | 0.473338 |
| 17 | 27981600 | 27989399 | 75.52938 | 117.1974 | 182.4257 | 75.7612 | -0.33181 | 2.777806 | 0.47335 |
| 2 | 22048600 | 22049999 | 19.00696 | 33.40128 | 33.22341 | 14.70959 | 0.395027 | 0.613383 | 0.473353 |
| 7 | 51042000 | 51049399 | 101.7347 | 158.0982 | 155.7064 | 64.65014 | 0.329512 | 2.760357 | 0.473364 |
| 7 | 53585600 | 53587999 | 27.06773 | 46.03945 | 45.77249 | 20.06154 | 0.390048 | 1.031117 | 0.473435 |
| 14 | 120258600 | 120260599 | 15.52104 | 27.02987 | 46.29243 | 20.35475 | -0.38553 | 0.754481 | 0.473453 |
| 7 | 14121600 | 14138999 | 306.7177 | 440.9941 | 388.7272 | 160.4619 | 0.390447 | 4.171416 | 0.47347 |
| 16 | 91848400 | 91851599 | 50.27997 | 79.94251 | 127.4217 | 53.18145 | -0.33744 | 2.246016 | 0.473471 |
| 16 | 18938200 | 18939799 | 30.44306 | 50.74241 | 50.78191 | 22.07014 | 0.380155 | 1.239384 | 0.473473 |
| 4 | 5241800 | 5245599 | 39.50647 | 65.62244 | 52.20157 | 22.63505 | 0.710073 | 1.301553 | 0.473487 |
| 13 | 98241200 | 98244799 | 40.99716 | 65.92089 | 105.2216 | 44.62963 | -0.3211 | 1.87673 | 0.473489 |
| 11 | 78879400 | 78882199 | 50.53896 | 81.24241 | 84.12615 | 35.6686 | 0.301511 | 1.890014 | 0.473495 |
| 3 | 58599000 | 58601999 | 20.40221 | 34.22171 | 60.18329 | 26.26044 | -0.43063 | 1.147351 | 0.473521 |
| 14 | 85759400 | 85761599 | 29.84828 | 49.68512 | 48.66936 | 21.13433 | 0.410988 | 1.248525 | 0.473528 |
| 11 | 120434000 | 120435999 | 9.874678 | 17.63186 | 35.37278 | 15.54391 | -0.60018 | 0.420988 | 0.47354 |
| 5 | 105056200 | 105058399 | 37.43969 | 61.76126 | 58.8695 | 25.40725 | 0.449134 | 1.379432 | 0.473576 |
| 1 | 151944400 | 151946399 | 13.95982 | 24.51883 | 41.39844 | 18.32084 | -0.36434 | 0.60933 | 0.473593 |
| 8 | 76533200 | 76534599 | 13.61745 | 23.96129 | 40.83978 | 18.08142 | -0.37729 | 0.581578 | 0.473598 |
| X | 49547600 | 49550599 | 37.05174 | 62.07269 | 49.18465 | 21.44469 | 0.715724 | 1.204167 | 0.473607 |
| 3 | 51416400 | 51422599 | 42.78442 | 68.22089 | 129.4524 | 54.23992 | -0.5841 | 2.147164 | 0.473632 |
| 16 | 88760800 | 88766399 | 112.9771 | 173.524 | 176.7074 | 73.36951 | 0.268409 | 2.980557 | 0.473642 |
| 5 | 127084000 | 127085799 | 17.61409 | 30.4173 | 51.70332 | 22.68219 | -0.37693 | 0.864206 | 0.473646 |
| 1 | 90468000 | 90472799 | 59.10664 | 94.50933 | 96.37457 | 40.5389 | 0.314635 | 2.068529 | 0.473675 |
| 12 | 19988200 | 20005599 | 707.6362 | 939.1287 | 819.6577 | 342.5785 | 0.2886 | 5.401222 | 0.473705 |
| 7 | 31307800 | 31339199 | 481.8261 | 667.5921 | 541.3002 | 224.7721 | 0.457315 | 4.72333 | 0.473741 |
| 5 | 23304800 | 23307799 | 20.66734 | 34.80981 | 59.18323 | 25.86902 | -0.38194 | 1.107894 | 0.473753 |
| X | 123114400 | 123119799 | 97.31406 | 152.5856 | 137.5174 | 57.03601 | 0.467359 | 2.603437 | 0.473794 |
| 10 | 42466200 | 42472399 | 41.34881 | 65.92163 | 128.5449 | 53.84821 | -0.62355 | 2.151215 | 0.473795 |
| 5 | 33141200 | 33142399 | 10.68425 | 19.25845 | 35.79728 | 15.59505 | -0.48278 | 0.365602 | 0.473796 |
| X | 163724600 | 163726399 | 24.1796 | 40.31875 | 66.41054 | 28.86979 | -0.33647 | 1.229474 | 0.473799 |
| 14 | 45966200 | 45968399 | 17.15441 | 29.67376 | 49.53653 | 21.73275 | -0.35036 | 0.845295 | 0.473827 |
| 15 | 92721000 | 92722399 | 20.9884 | 35.55642 | 58.69102 | 25.6991 | -0.33907 | 1.059423 | 0.473831 |
| 7 | 142311000 | 142311599 | 4.483125 | 8.743565 | 20.27844 | 7.276162 | -0.55409 | -0.43929 | 0.473855 |
| 15 | 72731000 | 72731599 | 4.483125 | 8.743565 | 20.27844 | 7.276162 | -0.55409 | -0.43929 | 0.473855 |
| 11 | 41630400 | 41631999 | 17.6578 | 30.443 | 50.60203 | 22.19795 | -0.34518 | 0.879024 | 0.473879 |
| 6 | 20169800 | 20171799 | 25.75621 | 44.11996 | 43.59219 | 19.11684 | 0.400479 | 0.960582 | 0.473884 |
| 3 | 78649600 | 78650999 | 25.78225 | 42.30217 | 72.42983 | 31.18845 | -0.39645 | 1.429408 | 0.473887 |
| 7 | 55686800 | 55688399 | 15.40239 | 26.92495 | 45.1686 | 19.90963 | -0.35521 | 0.699284 | 0.473916 |
| X | 101293200 | 101296599 | 51.39128 | 82.95235 | 72.38629 | 30.86637 | 0.555025 | 1.768014 | 0.473924 |
| Y | 6993600 | 6994399 | 11.48929 | 21.70638 | 15.58605 | 6.340624 | 0.95244 | 0.042384 | 0.473932 |
| 19 | 5363200 | 5371599 | 179.0595 | 261.1721 | 398.3805 | 164.1558 | -0.38005 | 3.94844 | 0.47395 |

| | | | | | | | | | |
|---|---|---|---|---|---|---|---|---|---|
| 8 | 21831200 | 21832999 | 27.04909 | 46.24507 | 44.10269 | 19.3377 | 0.450934 | 0.98319 | 0.473966 |
| X | 65246800 | 65250799 | 54.33234 | 87.51531 | 80.46089 | 34.21366 | 0.476305 | 1.83423 | 0.473979 |
| 7 | 120223000 | 120223599 | 10.45592 | 19.8688 | 20.06926 | 8.065068 | 0.479623 | 0.006937 | 0.473986 |
| 3 | 154119800 | 154121999 | 25.18486 | 41.29211 | 74.34554 | 32.00356 | -0.46911 | 1.440806 | 0.473993 |
| X | 25545800 | 25547199 | 24.66147 | 42.55592 | 40.45465 | 17.74702 | 0.458027 | 0.890232 | 0.474018 |
| 17 | 19970600 | 19978599 | 109.2253 | 169.956 | 150.5434 | 62.49449 | 0.483774 | 2.742633 | 0.47406 |
| 15 | 97382000 | 97391399 | 124.4633 | 188.0279 | 275.6701 | 113.9722 | -0.2762 | 3.323274 | 0.474064 |
| 12 | 78108400 | 78111399 | 24.41745 | 41.91799 | 42.54541 | 18.66107 | 0.362285 | 0.934238 | 0.474095 |
| 10 | 22255400 | 22259799 | 36.64506 | 58.94604 | 97.26653 | 41.27889 | -0.36812 | 1.856565 | 0.474103 |
| 17 | 40761000 | 40762599 | 17.98662 | 32.03249 | 30.57293 | 13.46431 | 0.462764 | 0.480003 | 0.474114 |
| 8 | 5227600 | 5231999 | 39.82248 | 65.38192 | 60.69765 | 26.14387 | 0.484409 | 1.442282 | 0.474124 |
| X | 75054800 | 75056599 | 24.70548 | 42.79064 | 40.32499 | 17.69214 | 0.471499 | 0.855929 | 0.474127 |
| 2 | 76443600 | 76444599 | 7.771826 | 14.81026 | 27.79316 | 11.10229 | -0.39905 | -0.01561 | 0.474199 |
| 16 | 76088000 | 76092599 | 59.88208 | 94.16208 | 153.8523 | 63.82416 | -0.38669 | 2.529092 | 0.47421 |
| 9 | 99511000 | 99512599 | 10.10688 | 18.16627 | 35.80352 | 15.64568 | -0.56957 | 0.378809 | 0.474227 |
| 2 | 79567200 | 79568999 | 23.18168 | 40.51218 | 36.33982 | 16.01411 | 0.543707 | 0.729284 | 0.474229 |
| 9 | 5317800 | 5318399 | 5.84234 | 11.2757 | 24.77656 | 9.56383 | -0.57807 | -0.15654 | 0.474235 |
| X | 76604600 | 76605399 | 16.06338 | 29.21873 | 27.75828 | 11.95618 | 0.490495 | 0.337512 | 0.474245 |
| 13 | 91223000 | 91225399 | 19.75823 | 35.03772 | 30.84932 | 13.635 | 0.57495 | 0.517407 | 0.474252 |
| 15 | 42218200 | 42219999 | 19.75823 | 35.03772 | 30.84932 | 13.635 | 0.57495 | 0.517407 | 0.474252 |
| 16 | 92893800 | 92895999 | 14.95944 | 25.99715 | 45.81923 | 20.12009 | -0.4268 | 0.783424 | 0.474263 |
| 14 | 72853200 | 72854799 | 27.26108 | 46.47393 | 45.21458 | 19.82065 | 0.421669 | 1.011294 | 0.474266 |
| 7 | 32160600 | 32163799 | 26.38355 | 45.8372 | 38.54787 | 16.92402 | 0.63575 | 0.82956 | 0.474269 |
| 4 | 11117200 | 11118599 | 9.745464 | 18.06604 | 33.1553 | 13.92521 | -0.42761 | 0.196012 | 0.474277 |
| 9 | 20698600 | 20700199 | 19.52121 | 33.1361 | 55.3688 | 24.25889 | -0.35643 | 1.033652 | 0.474287 |
| 5 | 93400600 | 93403199 | 20.70579 | 36.12936 | 36.01887 | 15.85686 | 0.39173 | 0.736073 | 0.474303 |
| 1 | 45994400 | 45996999 | 25.89269 | 44.72188 | 40.78512 | 17.8925 | 0.517743 | 0.884703 | 0.474304 |
| 8 | 111651000 | 111653199 | 13.47359 | 23.65955 | 40.90413 | 18.09868 | -0.39829 | 0.608849 | 0.474307 |
| 16 | 11361600 | 11363199 | 15.42377 | 26.65773 | 48.46081 | 21.26027 | -0.47294 | 0.843427 | 0.474321 |
| 8 | 28955600 | 28957799 | 21.39647 | 37.56866 | 34.11684 | 15.09438 | 0.525832 | 0.642618 | 0.474339 |
| 3 | 9161800 | 9167599 | 55.9035 | 88.49184 | 139.5447 | 58.01393 | -0.32799 | 2.376807 | 0.474341 |
| 7 | 128614400 | 128618599 | 56.21576 | 90.09937 | 91.11474 | 38.4593 | 0.330499 | 1.992536 | 0.474342 |
| 17 | 20591400 | 20593399 | 22.98761 | 40.82672 | 28.3944 | 12.53382 | 0.91458 | 0.524654 | 0.474345 |
| 7 | 106689400 | 106691799 | 24.38522 | 42.90813 | 31.68931 | 14.03015 | 0.823161 | 0.628823 | 0.474353 |
| 8 | 112394800 | 112397399 | 28.18277 | 46.07131 | 77.28908 | 33.23177 | -0.36994 | 1.483726 | 0.474353 |
| 3 | 47017800 | 47019199 | 13.66601 | 25.14505 | 25.9226 | 10.99889 | 0.388026 | 0.25035 | 0.474374 |
| 1 | 129258400 | 129260199 | 24.98145 | 43.25313 | 38.55706 | 16.9163 | 0.550996 | 0.854136 | 0.474377 |
| 3 | 59251600 | 59252199 | 4.942109 | 9.624431 | 21.28611 | 7.787731 | -0.51299 | -0.36203 | 0.47438 |
| 10 | 45733400 | 45733999 | 4.942109 | 9.624431 | 21.28611 | 7.787731 | -0.51299 | -0.36203 | 0.47438 |
| 1 | 109760600 | 109762199 | 18.78494 | 32.99898 | 32.72538 | 14.48458 | 0.399274 | 0.616444 | 0.474382 |
| 12 | 106322000 | 106328399 | 68.1489 | 106.6357 | 166.5506 | 69.09229 | -0.32727 | 2.625849 | 0.474393 |
| 4 | 20004200 | 20005799 | 23.47353 | 40.40474 | 41.51484 | 18.21072 | 0.345473 | 0.906073 | 0.474397 |
| 19 | 12160400 | 12163799 | 28.88904 | 49.06124 | 46.29303 | 20.28548 | 0.464911 | 1.05213 | 0.4744 |
| 8 | 113456600 | 113460399 | 46.90761 | 75.54178 | 78.15513 | 33.22414 | 0.305986 | 1.825935 | 0.474406 |
| 19 | 12405200 | 12408199 | 23.31467 | 40.33578 | 39.79837 | 17.46049 | 0.405027 | 0.856374 | 0.474409 |
| 3 | 7206800 | 7209599 | 40.84604 | 66.69176 | 64.53588 | 27.73609 | 0.420494 | 1.519187 | 0.474414 |
| 5 | 125647000 | 125656799 | 89.06697 | 136.5619 | 235.6868 | 97.83459 | -0.49389 | 3.012064 | 0.474422 |

| | | | | | | | | | |
|---|---|---|---|---|---|---|---|---|---|
| 7 | 100595400 | 100597599 | 11.77289 | 20.73562 | 38.90072 | 17.2196 | -0.5141 | 0.557728 | 0.474423 |
| 11 | 97880600 | 97883399 | 11.77289 | 20.73562 | 38.90072 | 17.2196 | -0.5141 | 0.557728 | 0.474423 |
| 4 | 51792800 | 51794199 | 13.70544 | 25.75658 | 23.9082 | 9.820941 | 0.57362 | 0.120159 | 0.474431 |
| 8 | 19527000 | 19527999 | 13.70544 | 25.75658 | 23.9082 | 9.820941 | 0.57362 | 0.120159 | 0.474431 |
| 1 | 84659400 | 84667799 | 83.08166 | 128.3923 | 194.421 | 80.75819 | -0.29666 | 2.855403 | 0.474439 |
| X | 159284800 | 159285999 | 8.921839 | 16.7925 | 31.9269 | 13.08084 | -0.44977 | 0.090883 | 0.474442 |
| 6 | 79879200 | 79884399 | 46.74725 | 75.38812 | 78.00736 | 33.22451 | 0.308153 | 1.783944 | 0.474446 |
| 5 | 98601600 | 98602399 | 11.91503 | 21.02336 | 38.35699 | 16.97287 | -0.47312 | 0.537863 | 0.474474 |
| 1 | 103470200 | 103471399 | 38.75237 | 63.06664 | 64.33747 | 27.61786 | 0.341289 | 1.564361 | 0.474474 |
| 3 | 42434000 | 42436799 | 31.81494 | 54.48414 | 41.20011 | 18.06744 | 0.784789 | 0.985289 | 0.474498 |
| 8 | 64119000 | 64119599 | 5.664237 | 11.0254 | 24.35972 | 8.951324 | -0.51936 | -0.33217 | 0.474502 |
| 18 | 37707400 | 37709599 | 78.74555 | 123.8215 | 123.7891 | 51.35261 | 0.32104 | 2.534686 | 0.474517 |
| 10 | 102781600 | 102781999 | 5.114885 | 9.959181 | 23.71648 | 8.693129 | -0.62284 | -0.35033 | 0.474522 |
| 2 | 108187200 | 108187999 | 10.4384 | 20.06476 | 20.65491 | 8.066069 | 0.4896 | -0.0842 | 0.474528 |
| 13 | 93360000 | 93362599 | 31.02574 | 51.74212 | 52.99336 | 23.03927 | 0.346831 | 1.235663 | 0.474532 |
| 4 | 140606200 | 140619999 | 166.2029 | 244.9604 | 361.4044 | 148.8303 | -0.31765 | 3.782259 | 0.474536 |
| 8 | 20014800 | 20017199 | 37.28061 | 60.02609 | 96.08035 | 40.83241 | -0.3226 | 1.826759 | 0.474551 |
| 2 | 10710600 | 10711399 | 6.851958 | 13.17969 | 25.832 | 10.06847 | -0.42816 | -0.11064 | 0.474558 |
| 13 | 80732800 | 80733199 | 8.696736 | 16.89354 | 18.84403 | 7.039876 | 0.435157 | -0.25398 | 0.47456 |
| X | 78213000 | 78218799 | 113.5214 | 174.9622 | 170.1493 | 70.67723 | 0.338605 | 2.915464 | 0.474577 |
| 16 | 15350800 | 15352999 | 47.47404 | 76.76316 | 74.57456 | 31.87109 | 0.403331 | 1.708685 | 0.474579 |
| 1 | 129847800 | 129849199 | 11.67005 | 21.06833 | 36.87291 | 16.03221 | -0.39449 | 0.367857 | 0.474581 |
| 14 | 42279000 | 42281599 | 16.79195 | 30.42978 | 24.6362 | 10.66125 | 0.715465 | 0.357115 | 0.474594 |
| 3 | 156025800 | 156028199 | 26.19929 | 45.52206 | 38.547 | 16.92481 | 0.625874 | 0.827031 | 0.4746 |
| 2 | 163486400 | 163487199 | 6.020517 | 11.48403 | 26.13133 | 10.41708 | -0.67065 | -0.04102 | 0.474613 |
| 3 | 52377200 | 52384999 | 83.64379 | 131.5942 | 132.9756 | 55.16736 | 0.306204 | 2.527973 | 0.474613 |
| 2 | 52909000 | 52911199 | 26.20273 | 44.68103 | 44.68935 | 19.59081 | 0.381932 | 1.003555 | 0.474685 |
| 12 | 119553600 | 119557999 | 47.43593 | 76.39959 | 79.39424 | 33.75537 | 0.299763 | 1.824868 | 0.474698 |
| 14 | 89432400 | 89434799 | 30.71891 | 51.439 | 44.89939 | 19.57046 | 0.576556 | 1.18379 | 0.474706 |
| 6 | 136508400 | 136509999 | 10.76786 | 19.55476 | 35.43644 | 15.29423 | -0.43707 | 0.326673 | 0.474707 |
| 7 | 31036400 | 31040999 | 69.68314 | 111.4548 | 91.72535 | 38.59993 | 0.623835 | 2.069894 | 0.474724 |
| 2 | 62420800 | 62422199 | 9.520938 | 17.74665 | 32.20002 | 13.41185 | -0.40225 | 0.157421 | 0.474745 |
| 10 | 122200000 | 122200799 | 9.520938 | 17.74665 | 32.20002 | 13.41185 | -0.40225 | 0.157421 | 0.474745 |
| 11 | 30282600 | 30283799 | 9.520938 | 17.74665 | 32.20002 | 13.41185 | -0.40225 | 0.157421 | 0.474745 |
| 8 | 103672400 | 103676399 | 40.64704 | 66.90113 | 58.53214 | 25.23756 | 0.57111 | 1.413179 | 0.474775 |
| 8 | 25614000 | 25626599 | 213.4795 | 307.8072 | 447.7944 | 184.7694 | -0.32802 | 4.12412 | 0.474784 |
| 14 | 112453400 | 112453999 | 13.48734 | 24.64467 | 23.36023 | 10.00485 | 0.498624 | 0.279966 | 0.474811 |
| 15 | 29876200 | 29876999 | 10.80203 | 20.50151 | 21.58471 | 8.696445 | 0.41702 | 0.022856 | 0.474812 |
| 9 | 113364200 | 113364999 | 15.89019 | 28.78678 | 24.21923 | 10.48485 | 0.659916 | 0.350702 | 0.47482 |
| 17 | 13639000 | 13647599 | 302.1369 | 418.3182 | 720.7784 | 299.3344 | -0.63093 | 4.730248 | 0.474838 |
| 14 | 25010400 | 25012599 | 22.36727 | 39.30211 | 33.54387 | 14.8501 | 0.614985 | 0.637014 | 0.474854 |
| 1 | 103772200 | 103772599 | 6.110744 | 11.84289 | 24.41141 | 9.241438 | -0.45906 | -0.21707 | 0.474855 |
| 13 | 77724000 | 77726199 | 32.71446 | 53.05827 | 86.49139 | 37.05085 | -0.33745 | 1.628085 | 0.474873 |
| 10 | 20024400 | 20025799 | 10.25465 | 18.69682 | 33.40045 | 14.34411 | -0.41183 | 0.294965 | 0.474878 |
| 16 | 13031600 | 13032599 | 8.39681 | 15.8393 | 29.18066 | 11.91006 | -0.40021 | 0.063774 | 0.474889 |
| 13 | 6937400 | 6938399 | 6.856838 | 13.14003 | 26.83773 | 10.57296 | -0.50198 | -0.07423 | 0.474894 |
| 1 | 117862600 | 117863799 | 12.23802 | 23.10485 | 23.61979 | 9.623022 | 0.445465 | 0.078989 | 0.474911 |

| | | | | | | | | | |
|---|---|---|---|---|---|---|---|---|---|
| 6 | 69417400 | 69418999 | 17.57847 | 31.50389 | 29.50813 | 12.92639 | 0.494849 | 0.435765 | 0.474936 |
| 18 | 66523800 | 66526599 | 15.1741 | 26.47844 | 46.7902 | 20.59943 | -0.43027 | 0.724007 | 0.474938 |
| 15 | 97230400 | 97247199 | 184.8613 | 267.8118 | 467.4358 | 192.7528 | -0.58389 | 4.052087 | 0.47494 |
| 1 | 112517400 | 112518399 | 21.81204 | 37.87885 | 38.82546 | 17.04135 | 0.3514 | 0.813698 | 0.474969 |
| 7 | 42709400 | 42730199 | 362.3517 | 512.6807 | 451.8793 | 187.0201 | 0.367333 | 4.408275 | 0.474969 |
| 2 | 156471800 | 156474199 | 27.94853 | 46.9641 | 48.57019 | 21.21509 | 0.332607 | 1.141509 | 0.474973 |
| 3 | 107545200 | 107547599 | 27.20053 | 46.41893 | 44.65472 | 19.57686 | 0.438059 | 1.000494 | 0.474977 |
| 15 | 88837200 | 88839799 | 17.85901 | 30.46261 | 54.31407 | 23.81065 | -0.44877 | 0.980018 | 0.474982 |
| 16 | 65589800 | 65591399 | 16.50122 | 28.67834 | 47.91736 | 21.04226 | -0.35038 | 0.791057 | 0.475006 |
| 12 | 115611400 | 115612199 | 21.66589 | 37.69799 | 38.23396 | 16.80217 | 0.366826 | 0.78518 | 0.475013 |
| 4 | 40303400 | 40306599 | 28.71319 | 48.10871 | 49.1475 | 21.43006 | 0.350266 | 1.173708 | 0.475019 |
| 18 | 69882000 | 69885999 | 39.88433 | 64.21859 | 102.315 | 43.48221 | -0.31598 | 1.83263 | 0.475033 |
| X | 108838400 | 108840799 | 26.00648 | 45.8528 | 29.38904 | 13.00651 | 1.027636 | 0.614109 | 0.475037 |
| 7 | 104124600 | 104128599 | 48.47799 | 77.3099 | 120.3423 | 50.4323 | -0.29867 | 2.147026 | 0.475042 |
| 13 | 43966000 | 43970199 | 40.5247 | 65.16236 | 102.4912 | 43.47268 | -0.29986 | 1.875085 | 0.475044 |
| 7 | 129103600 | 129104599 | 8.845748 | 16.58686 | 31.26452 | 12.8912 | -0.44523 | 0.109518 | 0.475049 |
| 18 | 21243200 | 21245999 | 22.25165 | 38.78084 | 37.55052 | 16.51887 | 0.433541 | 0.763248 | 0.475057 |
| X | 33335800 | 33337199 | 18.1005 | 32.28831 | 30.14373 | 13.25692 | 0.495959 | 0.471194 | 0.475065 |
| 7 | 75868200 | 75870399 | 10.21727 | 18.51838 | 33.82324 | 14.63396 | -0.45091 | 0.327325 | 0.475073 |
| 3 | 66932400 | 66935999 | 51.16884 | 82.08239 | 83.36855 | 35.23661 | 0.325728 | 1.955426 | 0.475096 |
| 18 | 37675400 | 37679199 | 90.03908 | 140.9202 | 141.1974 | 58.57334 | 0.312651 | 2.631503 | 0.475097 |
| 3 | 90837400 | 90841799 | 48.36692 | 78.18282 | 66.1453 | 28.25743 | 0.601779 | 1.712171 | 0.475109 |
| 18 | 16421800 | 16423199 | 9.548923 | 17.82606 | 33.36916 | 13.86286 | -0.44391 | 0.151138 | 0.475121 |
| 4 | 144986400 | 144991799 | 45.25473 | 72.3642 | 116.442 | 48.98347 | -0.3422 | 2.05691 | 0.475133 |
| 11 | 14347600 | 14349599 | 16.16624 | 29.34689 | 27.57217 | 11.90586 | 0.50377 | 0.346836 | 0.475142 |
| 1 | 34244800 | 34246399 | 15.20164 | 26.59324 | 45.14361 | 19.91079 | -0.3723 | 0.685231 | 0.475145 |
| 7 | 55668400 | 55669599 | 9.08678 | 16.33341 | 34.87137 | 15.23779 | -0.68379 | 0.370797 | 0.475147 |
| 4 | 112621400 | 112622399 | 18.75893 | 33.15299 | 31.74261 | 14.04145 | 0.452962 | 0.549999 | 0.475154 |
| 7 | 122659600 | 122662199 | 21.62437 | 37.44074 | 38.39942 | 16.84713 | 0.349779 | 0.841562 | 0.475155 |
| 9 | 88914800 | 88916999 | 21.62437 | 37.44074 | 38.39942 | 16.84713 | 0.349779 | 0.841562 | 0.475155 |
| 7 | 144774600 | 144775399 | 11.16499 | 21.17944 | 21.1928 | 8.547878 | 0.489233 | 0.031911 | 0.475167 |
| 5 | 14457600 | 14460599 | 49.08366 | 79.1073 | 69.54724 | 29.59772 | 0.541695 | 1.800653 | 0.475223 |
| X | 87551400 | 87556399 | 55.41342 | 89.42018 | 75.64395 | 32.24411 | 0.599262 | 1.781649 | 0.475227 |
| 13 | 21013200 | 21014199 | 8.273474 | 15.50683 | 29.04939 | 11.98692 | -0.43674 | 0.104027 | 0.475234 |
| 14 | 88088000 | 88089199 | 14.09836 | 24.75928 | 41.90283 | 18.54224 | -0.3678 | 0.612058 | 0.475249 |
| X | 24090200 | 24095799 | 62.48426 | 100.8712 | 68.86503 | 29.36986 | 0.908535 | 1.779517 | 0.475276 |
| 6 | 135023800 | 135028999 | 29.41134 | 47.83855 | 79.53956 | 34.13677 | -0.36216 | 1.562383 | 0.475303 |
| 14 | 3972800 | 3975199 | 18.02105 | 30.89467 | 51.64931 | 22.65167 | -0.355 | 0.933314 | 0.475304 |
| 11 | 21579600 | 21581199 | 16.99742 | 30.61566 | 28.47115 | 12.41474 | 0.509461 | 0.398706 | 0.475312 |
| 17 | 38359800 | 38362399 | 56.31722 | 90.26524 | 91.36259 | 38.56603 | 0.329286 | 1.990821 | 0.475315 |
| 3 | 31589200 | 31590999 | 26.01528 | 45.20731 | 38.54613 | 16.92559 | 0.615941 | 0.824498 | 0.475323 |
| 2 | 118674000 | 118675399 | 18.42914 | 32.32207 | 32.53632 | 14.3814 | 0.378106 | 0.641771 | 0.475326 |
| 6 | 8695200 | 8696599 | 15.77602 | 28.35784 | 28.69415 | 12.53831 | 0.38654 | 0.403673 | 0.475326 |
| 1 | 57021600 | 57024599 | 28.00082 | 45.84883 | 76.27763 | 32.81704 | -0.35642 | 1.459772 | 0.475332 |
| 8 | 4992000 | 4996999 | 69.46201 | 110.2323 | 113.6842 | 47.35816 | 0.287003 | 2.312978 | 0.475332 |
| 19 | 39406400 | 39415999 | 126.1457 | 194.0142 | 175.4259 | 72.85105 | 0.44128 | 2.959673 | 0.475335 |
| 12 | 69438200 | 69439999 | 11.89458 | 21.16216 | 36.7725 | 16.20814 | -0.39739 | 0.46062 | 0.475342 |

| | | | | | | | | | |
|---|---|---|---|---|---|---|---|---|---|
| 17 | 15011000 | 15014399 | 44.33774 | 72.12618 | 67.90755 | 29.14592 | 0.456284 | 1.582966 | 0.475365 |
| 17 | 18046400 | 18051399 | 37.66989 | 60.35631 | 107.0759 | 45.17506 | -0.47933 | 1.984269 | 0.475369 |
| 11 | 89196800 | 89201399 | 39.55564 | 63.60195 | 100.3093 | 42.54507 | -0.30381 | 1.874287 | 0.475392 |
| 18 | 77782200 | 77784799 | 21.42193 | 37.81695 | 31.84065 | 14.08797 | 0.636655 | 0.576226 | 0.475397 |
| 4 | 89741200 | 89742799 | 14.75377 | 26.7925 | 27.26512 | 11.76795 | 0.389776 | 0.334446 | 0.475406 |
| 18 | 3872800 | 3875999 | 54.62278 | 87.39121 | 88.86323 | 37.41049 | 0.319651 | 2.043095 | 0.475433 |
| 1 | 159972600 | 159975399 | 25.17222 | 41.16098 | 75.62294 | 32.51858 | -0.50035 | 1.472589 | 0.475439 |
| 18 | 34211600 | 34216999 | 43.83929 | 70.14375 | 110.9138 | 46.69834 | -0.31602 | 2.036532 | 0.475445 |
| 12 | 25896200 | 25898399 | 18.25589 | 32.38622 | 31.01478 | 13.70363 | 0.454662 | 0.505652 | 0.47547 |
| 13 | 109460400 | 109462599 | 18.25589 | 32.38622 | 31.01478 | 13.70363 | 0.454662 | 0.505652 | 0.47547 |
| 4 | 112371200 | 112377999 | 116.0561 | 178.5963 | 174.1995 | 72.3473 | 0.332657 | 2.945843 | 0.475493 |
| 5 | 136766200 | 136784399 | 223.7157 | 321.0629 | 469.4627 | 193.8445 | -0.34207 | 4.19336 | 0.475498 |
| 17 | 37375600 | 37376999 | 7.759895 | 14.75348 | 28.29655 | 11.35861 | -0.4367 | 0.000182 | 0.475511 |
| 10 | 125791800 | 125794599 | 17.82569 | 31.3661 | 31.28412 | 13.85075 | 0.391674 | 0.587587 | 0.475527 |
| 2 | 86995800 | 87000199 | 77.33062 | 121.7859 | 124.9596 | 51.84865 | 0.285308 | 2.502379 | 0.47553 |
| 16 | 21762600 | 21762999 | 12.89144 | 24.45527 | 20.33211 | 8.20013 | 0.753815 | 0.043319 | 0.475535 |
| 1 | 116294600 | 116299399 | 50.20273 | 80.76005 | 83.1726 | 35.30113 | 0.310625 | 1.866677 | 0.475543 |
| 10 | 50918600 | 50920399 | 14.08952 | 24.69946 | 42.4802 | 18.76929 | -0.39094 | 0.641402 | 0.475545 |
| 3 | 4704000 | 4706599 | 47.1352 | 76.09616 | 77.08191 | 32.87627 | 0.34059 | 1.753859 | 0.475555 |
| 13 | 13947800 | 13948599 | 7.047687 | 13.44877 | 26.68511 | 10.62876 | -0.47428 | -0.03245 | 0.475556 |
| 2 | 49371000 | 49373199 | 35.4683 | 57.25593 | 91.35101 | 38.95789 | -0.31351 | 1.744457 | 0.475562 |
| 8 | 115078600 | 115080199 | 9.862728 | 18.04256 | 32.89265 | 14.06643 | -0.43726 | 0.268476 | 0.475568 |
| 5 | 5548200 | 5549599 | 9.212839 | 17.2245 | 31.17278 | 12.9175 | -0.39245 | 0.133598 | 0.47557 |
| 5 | 85655000 | 85656599 | 17.49835 | 30.327 | 51.16846 | 22.45003 | -0.36509 | 0.832118 | 0.47558 |
| 12 | 103270800 | 103272799 | 18.69975 | 32.73938 | 32.12162 | 14.17714 | 0.414978 | 0.67013 | 0.475587 |
| 10 | 78279200 | 78281199 | 13.70859 | 23.91803 | 43.7034 | 19.23889 | -0.47779 | 0.718432 | 0.475599 |
| 9 | 83917800 | 83918799 | 8.304734 | 15.72299 | 30.39876 | 12.31165 | -0.45906 | 0.043317 | 0.47562 |
| 2 | 81707000 | 81709599 | 36.74845 | 61.20699 | 53.16349 | 23.09443 | 0.583888 | 1.267212 | 0.475621 |
| 6 | 97532200 | 97534599 | 22.09375 | 38.13374 | 38.93831 | 17.08194 | 0.355187 | 0.871915 | 0.475623 |
| 6 | 17622800 | 17625599 | 30.23488 | 51.68346 | 43.50576 | 19.07572 | 0.630382 | 0.99849 | 0.475641 |
| 7 | 23390600 | 23404399 | 196.7345 | 294.494 | 249.1007 | 102.7378 | 0.505101 | 3.514528 | 0.47565 |
| X | 49504600 | 49507599 | 28.14698 | 47.58276 | 46.4033 | 20.3056 | 0.417447 | 1.091781 | 0.475653 |
| 13 | 66097000 | 66114599 | 369.8005 | 524.9394 | 428.9955 | 177.4523 | 0.481122 | 4.360847 | 0.475653 |
| 5 | 101861400 | 101864199 | 20.43714 | 34.64655 | 58.72614 | 25.71988 | -0.37722 | 1.049492 | 0.475667 |
| 16 | 3243600 | 3245199 | 13.95914 | 26.28143 | 23.95945 | 9.811524 | 0.60317 | 0.111913 | 0.475679 |
| 7 | 47629800 | 47632399 | 32.37647 | 53.73209 | 55.21836 | 23.9287 | 0.341719 | 1.295526 | 0.475708 |
| 2 | 177253200 | 177257399 | 26.26628 | 45.30492 | 41.33732 | 18.13399 | 0.516669 | 0.899437 | 0.475723 |
| 4 | 27449400 | 27451399 | 23.58726 | 41.36833 | 33.9935 | 15.02538 | 0.669436 | 0.672541 | 0.475737 |
| 2 | 110176000 | 110178599 | 22.71879 | 37.3929 | 76.33317 | 32.91288 | -0.64757 | 1.384651 | 0.475741 |
| 13 | 60706000 | 60709999 | 41.87284 | 67.96347 | 68.74783 | 29.46642 | 0.35025 | 1.616079 | 0.47575 |
| 18 | 7000400 | 7006999 | 46.5817 | 73.63062 | 157.2458 | 65.32566 | -0.7665 | 2.407489 | 0.475755 |
| 14 | 8890800 | 8891999 | 13.08508 | 25.11158 | 21.6805 | 8.505993 | 0.735673 | -0.03778 | 0.475758 |
| X | 53497000 | 53497999 | 13.08508 | 25.11158 | 21.6805 | 8.505993 | 0.735673 | -0.03778 | 0.475758 |
| X | 16175600 | 16178399 | 24.81044 | 43.57545 | 32.20648 | 14.25261 | 0.821513 | 0.654885 | 0.475789 |
| 4 | 145346800 | 145349199 | 57.63295 | 93.12323 | 70.46459 | 30.08693 | 0.761878 | 1.745012 | 0.475797 |
| 3 | 159260600 | 159263799 | 32.93087 | 55.45174 | 48.55123 | 21.22022 | 0.572129 | 1.14077 | 0.475805 |
| 8 | 120617600 | 120622999 | 54.00468 | 85.43801 | 139.7183 | 58.06957 | -0.381 | 2.389841 | 0.475828 |

| | | | | | | | | | |
|---|---|---|---|---|---|---|---|---|---|
| 3 | 131269600 | 131271199 | 7.903749 | 14.57973 | 32.00143 | 13.53117 | -0.6911 | 0.203136 | 0.475847 |
| 3 | 12475600 | 12478399 | 56.04614 | 89.85641 | 80.62612 | 34.04873 | 0.503234 | 1.977434 | 0.475852 |
| 7 | 70669400 | 70670599 | 21.04562 | 37.20404 | 31.28531 | 13.83844 | 0.639141 | 0.55792 | 0.475859 |
| 8 | 29776000 | 29777999 | 18.36869 | 32.35138 | 32.37053 | 14.3291 | 0.387807 | 0.585463 | 0.47586 |
| X | 88267200 | 88270799 | 29.61847 | 50.02908 | 42.67847 | 18.67098 | 0.609688 | 1.098374 | 0.475867 |
| 4 | 143378400 | 143399399 | 346.0798 | 494.9439 | 396.5947 | 163.8463 | 0.520324 | 4.251468 | 0.475877 |
| 1 | 118547000 | 118550999 | 25.35114 | 41.9903 | 68.54053 | 29.66158 | -0.32399 | 1.31009 | 0.475881 |
| 11 | 60084000 | 60086799 | 35.83079 | 57.69113 | 95.0701 | 40.40248 | -0.36449 | 1.824794 | 0.475883 |
| 14 | 40985400 | 40987599 | 10.31579 | 18.47281 | 34.84553 | 15.27494 | -0.50937 | 0.405232 | 0.475888 |
| 5 | 115129600 | 115130199 | 7.054274 | 13.37342 | 27.10742 | 10.94764 | -0.52293 | 0.011948 | 0.475897 |
| 8 | 35617200 | 35617799 | 5.951986 | 11.58157 | 25.48243 | 9.389611 | -0.51609 | -0.31061 | 0.475903 |
| 2 | 22645200 | 22646599 | 9.740077 | 17.7866 | 32.37208 | 13.87513 | -0.43682 | 0.277847 | 0.475923 |
| 9 | 60497400 | 60498799 | 14.62559 | 26.41086 | 26.83993 | 11.66615 | 0.385346 | 0.365167 | 0.475928 |
| 14 | 104117400 | 104118199 | 5.974955 | 11.55265 | 26.68796 | 10.21438 | -0.63877 | -0.18144 | 0.475933 |
| 3 | 137723600 | 137725999 | 25.21433 | 43.41182 | 41.92783 | 18.39202 | 0.434406 | 0.910881 | 0.475955 |
| 17 | 52182600 | 52186799 | 25.77696 | 42.65502 | 72.18799 | 31.21955 | -0.37621 | 1.322293 | 0.475964 |
| 5 | 65071800 | 65076999 | 66.74128 | 106.1164 | 109.4825 | 45.69477 | 0.289098 | 2.255651 | 0.47597 |
| 4 | 147367800 | 147371399 | 55.65103 | 89.53821 | 74.25832 | 31.52456 | 0.622636 | 1.865081 | 0.475984 |
| 8 | 86166000 | 86167999 | 16.3622 | 28.4402 | 47.38245 | 20.80896 | -0.34616 | 0.788616 | 0.475995 |
| 12 | 71614000 | 71614999 | 7.327391 | 14.03149 | 26.79955 | 10.58015 | -0.40837 | -0.05964 | 0.47604 |
| 5 | 109564200 | 109564999 | 14.74965 | 27.01824 | 26.77828 | 11.43105 | 0.438059 | 0.284491 | 0.476051 |
| 13 | 4468400 | 4469599 | 14.74965 | 27.01824 | 26.77828 | 11.43105 | 0.438059 | 0.284491 | 0.476051 |
| 1 | 82483600 | 82485199 | 23.33638 | 40.62137 | 33.55929 | 14.75169 | 0.661528 | 0.781964 | 0.476052 |
| 11 | 111244600 | 111245399 | 7.617799 | 14.45544 | 28.23435 | 11.37992 | -0.46792 | 0.011884 | 0.476056 |
| 10 | 69032400 | 69032999 | 11.89356 | 20.66271 | 49.96998 | 21.93848 | -0.88069 | 0.781357 | 0.476067 |
| 11 | 51868800 | 51870399 | 11.35692 | 20.44784 | 35.27394 | 15.38877 | -0.3768 | 0.378351 | 0.476069 |
| 8 | 123898800 | 123914199 | 237.8549 | 340.3574 | 483.3019 | 199.6503 | -0.30403 | 4.237301 | 0.476072 |
| 4 | 94400400 | 94401999 | 26.07554 | 44.28201 | 44.79485 | 19.62883 | 0.364993 | 1.042752 | 0.476073 |
| 3 | 55023600 | 55025199 | 14.64098 | 25.43408 | 45.82726 | 20.11882 | -0.45855 | 0.787039 | 0.476083 |
| X | 63532800 | 63533399 | 17.15329 | 30.50449 | 28.43052 | 12.53599 | 0.496018 | 0.481905 | 0.476085 |
| 1 | 139564600 | 139564799 | 4.673314 | 9.124808 | 21.19028 | 7.46695 | -0.53454 | -0.50386 | 0.476096 |
| 8 | 4907800 | 4911799 | 57.12094 | 91.28312 | 94.90283 | 39.88817 | 0.285902 | 2.082864 | 0.476124 |
| 1 | 14552200 | 14552999 | 13.08194 | 24.49074 | 19.93726 | 8.243127 | 0.754538 | 0.130213 | 0.476128 |
| 12 | 22711200 | 22711599 | 13.08194 | 24.49074 | 19.93726 | 8.243127 | 0.754538 | 0.130213 | 0.476128 |
| 7 | 124880400 | 124882799 | 18.5104 | 31.55779 | 53.77704 | 23.57427 | -0.38381 | 0.987549 | 0.476128 |
| 13 | 78327400 | 78328199 | 9.667951 | 17.39699 | 34.33347 | 14.98898 | -0.5702 | 0.367118 | 0.476136 |
| 8 | 120924000 | 120931799 | 89.84298 | 137.475 | 218.7657 | 90.77356 | -0.37915 | 3.05156 | 0.476146 |
| 6 | 68781000 | 68782399 | 15.29662 | 28.32881 | 25.80372 | 10.85208 | 0.574523 | 0.226998 | 0.476153 |
| 6 | 20139600 | 20144199 | 79.5626 | 125.2763 | 127.4397 | 52.87637 | 0.297478 | 2.50725 | 0.47617 |
| 9 | 111783800 | 111785399 | 27.51813 | 47.56919 | 35.39508 | 15.52827 | 0.81026 | 0.878267 | 0.476186 |
| 8 | 115762800 | 115763799 | 11.75302 | 22.25732 | 23.16067 | 9.374984 | 0.428257 | 0.052677 | 0.476199 |
| 1 | 27792600 | 27794999 | 3.992545 | 7.779204 | 24.27478 | 8.821745 | -0.99568 | -0.39139 | 0.476224 |
| 12 | 45249000 | 45250999 | 20.94918 | 36.89246 | 32.38585 | 14.33618 | 0.575342 | 0.598965 | 0.476225 |
| 4 | 73479600 | 73480199 | 6.312665 | 12.03651 | 26.12833 | 10.42465 | -0.60467 | -0.03599 | 0.476239 |
| 5 | 94030200 | 94030599 | 6.312665 | 12.03651 | 26.12833 | 10.42465 | -0.60467 | -0.03599 | 0.476239 |
| 9 | 19699600 | 19705399 | 105.9167 | 163.7022 | 165.8783 | 68.90213 | 0.282466 | 2.859358 | 0.476252 |
| 1 | 149048600 | 149049599 | 7.93686 | 15.02008 | 28.15988 | 11.41523 | -0.41641 | 0.034747 | 0.476261 |

| | | | | | | | | | |
|---|---|---|---|---|---|---|---|---|---|
| 6 | 29467600 | 29468599 | 7.93686 | 15.02008 | 28.15988 | 11.41523 | -0.41641 | 0.034747 | 0.476261 |
| 13 | 58190000 | 58195999 | 65.03966 | 102.3707 | 154.9988 | 64.30787 | -0.27593 | 2.512056 | 0.476265 |
| 3 | 159371600 | 159373199 | 8.190265 | 15.54642 | 35.20877 | 14.18195 | -0.67876 | 0.036315 | 0.476267 |
| 4 | 30683600 | 30684999 | 13.12713 | 23.17535 | 39.78266 | 17.6017 | -0.38562 | 0.540057 | 0.476282 |
| 3 | 53473600 | 53474599 | 8.376435 | 15.86519 | 29.89047 | 12.09538 | -0.42155 | 0.036384 | 0.476303 |
| 5 | 151222400 | 151223999 | 21.96289 | 38.23252 | 38.17307 | 16.78031 | 0.389364 | 0.779599 | 0.476311 |
| 15 | 82575600 | 82578399 | 18.37703 | 31.25727 | 54.28468 | 23.79232 | -0.41157 | 1.009242 | 0.476312 |
| 16 | 53531600 | 53533999 | 30.52918 | 51.30426 | 49.66538 | 21.69394 | 0.427601 | 1.149727 | 0.476327 |
| 5 | 41802600 | 41803599 | 7.851144 | 14.66189 | 29.01077 | 12.04488 | -0.52197 | 0.124315 | 0.476331 |
| 8 | 103809800 | 103810199 | 5.23804 | 10.17817 | 22.26714 | 8.297613 | -0.52233 | -0.2902 | 0.476341 |
| X | 63235600 | 63239799 | 45.0212 | 73.65911 | 61.79817 | 26.57927 | 0.628009 | 1.503062 | 0.476363 |
| 14 | 35770800 | 35773399 | 20.34116 | 34.41146 | 57.00285 | 24.94924 | -0.3442 | 1.067499 | 0.476374 |
| 3 | 48818200 | 48819999 | 21.0205 | 36.73378 | 36.51641 | 16.09137 | 0.396145 | 0.719009 | 0.476374 |
| 6 | 134315000 | 134316799 | 18.1529 | 31.22199 | 51.67264 | 22.66596 | -0.3397 | 0.903996 | 0.476374 |
| 15 | 37151800 | 37155799 | 28.01958 | 45.86113 | 75.59195 | 32.51668 | -0.34328 | 1.46525 | 0.476376 |
| 4 | 93934800 | 93936199 | 7.858115 | 14.91778 | 30.45849 | 12.26537 | -0.52947 | 0.021999 | 0.476381 |
| X | 155400600 | 155402199 | 23.2876 | 40.82074 | 34.6109 | 15.29131 | 0.624521 | 0.680626 | 0.476385 |
| 5 | 149305800 | 149307999 | 22.20384 | 38.67603 | 37.55694 | 16.51593 | 0.429306 | 0.770231 | 0.476418 |
| 17 | 80493800 | 80495399 | 13.32911 | 23.58775 | 40.29405 | 17.81952 | -0.37722 | 0.517708 | 0.476418 |
| 2 | 139587600 | 139587999 | 5.083355 | 9.877925 | 22.25131 | 8.289486 | -0.56315 | -0.29158 | 0.476424 |
| 10 | 69774600 | 69779399 | 42.53995 | 68.21721 | 106.8125 | 45.12213 | -0.29809 | 1.964985 | 0.476451 |
| 6 | 85955400 | 85956999 | 21.90188 | 38.15581 | 37.58582 | 16.5302 | 0.408791 | 0.766975 | 0.476494 |
| 6 | 117567400 | 117567999 | 25.79291 | 44.44259 | 32.86346 | 14.4162 | 0.81848 | 0.897589 | 0.47652 |
| X | 146445800 | 146448999 | 55.66996 | 89.6501 | 80.61236 | 34.26712 | 0.507909 | 1.843162 | 0.476528 |
| 4 | 103946800 | 103947399 | 6.107382 | 11.83841 | 23.86498 | 9.027342 | -0.42651 | -0.22311 | 0.476587 |
| 5 | 89105400 | 89108799 | 20.30167 | 34.53528 | 57.07022 | 25.01345 | -0.34024 | 1.018495 | 0.476589 |
| 18 | 9893600 | 9901399 | 103.7628 | 161.256 | 155.4556 | 64.54561 | 0.360384 | 2.761216 | 0.47659 |
| 1 | 55828000 | 55830199 | 17.41352 | 30.75911 | 31.24496 | 13.82276 | 0.367585 | 0.544106 | 0.476592 |
| 3 | 116081200 | 116082799 | 10.98739 | 19.91911 | 34.83818 | 15.06843 | -0.3886 | 0.333674 | 0.476626 |
| 4 | 151566800 | 151568799 | 15.57009 | 27.04825 | 46.33914 | 20.34362 | -0.38625 | 0.791599 | 0.476628 |
| 17 | 39863000 | 39864399 | 18.01574 | 31.06633 | 51.67876 | 22.67043 | -0.34624 | 0.880944 | 0.476642 |
| X | 138847000 | 138849999 | 28.18255 | 48.81183 | 39.11113 | 17.15956 | 0.704622 | 0.871371 | 0.47665 |
| X | 147767400 | 147770999 | 27.83484 | 48.79396 | 30.4155 | 13.43831 | 1.06607 | 0.698225 | 0.476652 |
| 13 | 89139000 | 89141399 | 24.76239 | 42.94146 | 35.59739 | 15.61854 | 0.655998 | 0.838942 | 0.476655 |
| 6 | 94529000 | 94532599 | 34.45352 | 55.79111 | 89.20113 | 38.16743 | -0.31176 | 1.66199 | 0.476659 |
| 1 | 30289800 | 30290399 | 7.157317 | 13.75896 | 26.38605 | 10.30178 | -0.39924 | -0.09966 | 0.47667 |
| 3 | 102928000 | 102928799 | 6.004265 | 11.52229 | 25.17594 | 9.878588 | -0.59198 | -0.09843 | 0.47669 |
| 9 | 113931800 | 113932599 | 11.71657 | 22.34323 | 22.70986 | 9.05821 | 0.480302 | 0.003568 | 0.4767 |
| 15 | 51944600 | 51948799 | 89.0854 | 138.9313 | 138.135 | 57.32411 | 0.319778 | 2.692389 | 0.476703 |
| 13 | 23513200 | 23515799 | 22.8912 | 39.76054 | 34.16347 | 14.99691 | 0.604816 | 0.809721 | 0.476709 |
| 3 | 128816600 | 128820199 | 61.24643 | 96.6556 | 148.1155 | 61.48264 | -0.29027 | 2.455036 | 0.47671 |
| 10 | 119983600 | 119992399 | 101.4882 | 155.3155 | 229.3464 | 95.16687 | -0.27123 | 3.052983 | 0.476734 |
| X | 4929600 | 4938399 | 305.854 | 443.2068 | 344.292 | 141.9683 | 0.583846 | 4.057229 | 0.476739 |
| 14 | 122837000 | 122837799 | 5.656424 | 11.00302 | 22.95713 | 8.480464 | -0.4431 | -0.31281 | 0.476752 |
| 11 | 94546800 | 94547799 | 5.380719 | 10.47652 | 22.49316 | 8.246429 | -0.47464 | -0.348 | 0.476755 |
| 14 | 40279000 | 40280399 | 12.69182 | 22.57382 | 38.77757 | 17.0962 | -0.38135 | 0.471169 | 0.47677 |
| 18 | 18308800 | 18312599 | 31.23102 | 53.07864 | 45.14514 | 19.78532 | 0.614245 | 1.053306 | 0.476783 |

| | | | | | | | | | |
|---|---|---|---|---|---|---|---|---|---|
| 1 | 82260000 | 82262199 | 11.66855 | 20.88163 | 36.24296 | 15.89626 | -0.391 | 0.419369 | 0.476786 |
| 2 | 117839800 | 117841199 | 13.96462 | 24.51257 | 41.42416 | 18.32295 | -0.36557 | 0.619233 | 0.476796 |
| 7 | 31447600 | 31448199 | 11.65927 | 22.00179 | 21.6631 | 8.834668 | 0.49843 | 0.073451 | 0.476801 |
| 6 | 148113000 | 148113999 | 9.243254 | 17.0127 | 31.42371 | 13.32726 | -0.44709 | 0.22661 | 0.476804 |
| 18 | 7962800 | 7972199 | 151.8995 | 229.3986 | 227.8769 | 94.20265 | 0.284578 | 3.333669 | 0.476842 |
| 13 | 24712000 | 24715199 | 22.83828 | 38.3975 | 65.38766 | 28.56547 | -0.38444 | 1.137917 | 0.476844 |
| 15 | 75395400 | 75396599 | 7.596629 | 14.53951 | 27.9622 | 11.05563 | -0.42093 | -0.05154 | 0.476855 |
| 6 | 16599000 | 16602199 | 23.10019 | 38.71915 | 69.20816 | 30.18458 | -0.45408 | 1.174959 | 0.476878 |
| 13 | 63263000 | 63265199 | 15.83916 | 28.69711 | 28.23774 | 12.22305 | 0.435451 | 0.3586 | 0.476882 |
| 12 | 21879400 | 21930599 | 1397.759 | 1756.447 | 1466.698 | 616.5032 | 0.273446 | 6.287669 | 0.476887 |
| 5 | 25838200 | 25841399 | 28.71181 | 46.91946 | 76.54256 | 32.90652 | -0.33004 | 1.49162 | 0.476888 |
| X | 30970000 | 30971599 | 25.44785 | 43.9596 | 40.89118 | 17.9391 | 0.489488 | 0.884098 | 0.476892 |
| 1 | 60599600 | 60601799 | 30.03629 | 51.23109 | 44.03786 | 19.3056 | 0.59973 | 1.015741 | 0.476911 |
| 19 | 58052800 | 58055599 | 33.19478 | 55.14988 | 55.27451 | 23.97096 | 0.377819 | 1.285448 | 0.476923 |
| 12 | 114530000 | 114532199 | 37.91119 | 62.93699 | 54.40633 | 23.58185 | 0.590453 | 1.305659 | 0.476924 |
| 8 | 112072200 | 112073399 | 8.580285 | 15.60008 | 32.43471 | 13.97846 | -0.63225 | 0.296165 | 0.476932 |
| 17 | 21012800 | 21019399 | 81.17595 | 128.8859 | 110.8973 | 46.22028 | 0.548683 | 2.309046 | 0.476935 |
| 17 | 39644600 | 39648599 | 19.46615 | 34.48533 | 27.54244 | 12.17766 | 0.714366 | 0.520275 | 0.476948 |
| 7 | 83432400 | 83432999 | 5.989213 | 11.53592 | 24.72044 | 9.597582 | -0.54988 | -0.13518 | 0.476957 |
| 9 | 60122600 | 60124799 | 12.49267 | 22.09224 | 38.75911 | 17.14309 | -0.41589 | 0.515769 | 0.476965 |
| 11 | 79073400 | 79077599 | 30.52439 | 49.62658 | 80.95392 | 34.73808 | -0.33524 | 1.570951 | 0.476972 |
| 13 | 23490800 | 23493799 | 22.98933 | 39.57687 | 39.91525 | 17.50913 | 0.372124 | 0.898847 | 0.47699 |
| 7 | 27592200 | 27593399 | 19.72332 | 34.61966 | 34.29492 | 15.16901 | 0.400997 | 0.638456 | 0.476993 |
| 8 | 73648800 | 73649599 | 12.76608 | 24.67702 | 20.1875 | 7.735744 | 0.844327 | -0.11967 | 0.477025 |
| 1 | 15325600 | 15327399 | 17.36268 | 29.73478 | 52.1781 | 22.88164 | -0.42491 | 0.939429 | 0.477089 |
| 1 | 169622200 | 169623999 | 18.78723 | 33.35899 | 30.97904 | 13.6774 | 0.499567 | 0.503809 | 0.477094 |
| 7 | 26993000 | 26995399 | 26.5496 | 45.3208 | 44.70942 | 19.60129 | 0.402051 | 0.997701 | 0.477094 |
| 12 | 19349800 | 19357999 | 164.6515 | 250.09 | 200.1734 | 82.89785 | 0.602896 | 3.221877 | 0.477115 |
| 12 | 51611800 | 51616999 | 62.77184 | 100.08 | 102.8947 | 43.10561 | 0.29833 | 2.163279 | 0.477119 |
| 14 | 117522400 | 117523599 | 34.46203 | 56.83302 | 58.64129 | 25.3057 | 0.334929 | 1.37494 | 0.477122 |
| 12 | 106624400 | 106629399 | 33.16469 | 53.65946 | 86.94013 | 37.1772 | -0.33195 | 1.675409 | 0.47713 |
| 15 | 28577200 | 28578799 | 22.37949 | 39.45564 | 30.56732 | 13.52813 | 0.755207 | 0.591993 | 0.477137 |
| 3 | 154023200 | 154023799 | 12.99606 | 24.00632 | 24.96896 | 10.53789 | 0.380844 | 0.22098 | 0.47715 |
| 15 | 8452600 | 8452999 | 5.364777 | 10.45984 | 22.12917 | 7.972447 | -0.43041 | -0.41048 | 0.477168 |
| 6 | 100759200 | 100762199 | 44.91666 | 72.46021 | 74.86923 | 31.9017 | 0.310791 | 1.769758 | 0.477171 |
| 2 | 50199200 | 50201199 | 26.73647 | 45.56419 | 45.22488 | 19.82462 | 0.39284 | 1.011949 | 0.477176 |
| 2 | 92692000 | 92694599 | 29.99207 | 51.01027 | 45.72895 | 20.04236 | 0.538835 | 1.043781 | 0.477178 |
| 18 | 39816200 | 39817799 | 20.52086 | 36.18144 | 32.44748 | 14.36002 | 0.545381 | 0.584396 | 0.477186 |
| 1 | 88818200 | 88820799 | 14.68352 | 25.69013 | 46.74126 | 20.61766 | -0.472 | 0.680849 | 0.477209 |
| 17 | 73339400 | 73342399 | 20.44782 | 34.18952 | 61.19364 | 26.64788 | -0.45584 | 1.183201 | 0.477221 |
| 18 | 44999600 | 45000799 | 14.03281 | 26.54704 | 22.891 | 9.286244 | 0.694169 | 0.078057 | 0.47723 |
| 10 | 50416800 | 50418799 | 24.14635 | 42.62343 | 30.63178 | 13.5533 | 0.86405 | 0.592856 | 0.477233 |
| 7 | 60905600 | 60911199 | 79.72113 | 126.6813 | 109.0186 | 45.47506 | 0.549598 | 2.283953 | 0.477247 |
| 7 | 23311000 | 23330399 | 65.36685 | 104.3453 | 91.30368 | 38.3154 | 0.532462 | 2.124581 | 0.477273 |
| 9 | 12165800 | 12166999 | 18.01961 | 31.72082 | 31.92656 | 14.13398 | 0.379109 | 0.587878 | 0.477273 |
| X | 83198200 | 83198599 | 11.96486 | 23.31957 | 16.54333 | 5.988938 | 1.12271 | -0.32686 | 0.477292 |
| 2 | 17524400 | 17526599 | 26.62682 | 45.30154 | 45.78848 | 20.06898 | 0.366409 | 1.027784 | 0.477299 |

| | | | | | | | | | |
|---|---|---|---|---|---|---|---|---|---|
| 12 | 40494400 | 40496799 | 18.48283 | 31.14554 | 57.40681 | 25.09624 | -0.49783 | 1.095513 | 0.47731 |
| 7 | 54036200 | 54040199 | 62.21883 | 99.59708 | 85.30864 | 35.9411 | 0.567536 | 2.035163 | 0.477311 |
| 1 | 108443600 | 108445799 | 18.40059 | 33.06501 | 28.88345 | 12.62443 | 0.597089 | 0.42392 | 0.477312 |
| X | 118744800 | 118755199 | 140.8628 | 215.2129 | 172.9548 | 71.73951 | 0.603808 | 3.095073 | 0.477315 |
| 7 | 41594600 | 41608999 | 195.216 | 292.3394 | 249.0565 | 102.7277 | 0.495296 | 3.50615 | 0.477356 |
| 19 | 3413600 | 3418999 | 63.23489 | 99.56749 | 152.5175 | 63.28146 | -0.29225 | 2.503162 | 0.477369 |
| 11 | 41399000 | 41401199 | 18.8275 | 33.75303 | 28.78309 | 12.60542 | 0.629624 | 0.440235 | 0.477373 |
| 1 | 117670000 | 117672199 | 47.05545 | 76.07608 | 67.5038 | 28.84424 | 0.533536 | 1.707805 | 0.47738 |
| 18 | 61980600 | 61982599 | 12.48457 | 22.0988 | 38.74125 | 17.13196 | -0.41429 | 0.505095 | 0.477433 |
| 9 | 114792600 | 114800399 | 68.09779 | 106.2841 | 169.754 | 70.43971 | -0.36234 | 2.671843 | 0.477442 |
| 2 | 101066600 | 101067199 | 9.327906 | 18.07547 | 19.8657 | 7.525292 | 0.437765 | -0.19625 | 0.477459 |
| 4 | 150810600 | 150813199 | 20.1093 | 33.64821 | 62.33764 | 27.15863 | -0.50549 | 1.173872 | 0.47747 |
| 12 | 40431800 | 40432399 | 9.403472 | 18.16039 | 19.73389 | 7.594391 | 0.432176 | -0.14431 | 0.477475 |
| 18 | 68911000 | 68916799 | 86.3984 | 135.5773 | 138.0908 | 57.27027 | 0.291932 | 2.58194 | 0.477531 |
| 6 | 142702200 | 142702999 | 6.711006 | 12.88845 | 25.77235 | 10.0903 | -0.46287 | -0.09783 | 0.47754 |
| 12 | 3675000 | 3675999 | 6.711006 | 12.88845 | 25.77235 | 10.0903 | -0.46287 | -0.09783 | 0.47754 |
| 15 | 25758200 | 25760599 | 17.177 | 29.58032 | 50.05902 | 21.95899 | -0.37129 | 0.886329 | 0.47754 |
| 18 | 12225600 | 12229599 | 36.479 | 58.75904 | 94.7761 | 40.30161 | -0.3329 | 1.812905 | 0.477547 |
| 4 | 136016400 | 136020999 | 51.34512 | 81.45336 | 132.8545 | 55.32471 | -0.37376 | 2.315041 | 0.477548 |
| 7 | 140232000 | 140232799 | 7.164414 | 13.74834 | 26.31328 | 10.32551 | -0.40317 | -0.08295 | 0.47755 |
| 6 | 72600600 | 72601199 | 5.518396 | 10.74381 | 22.48764 | 8.249214 | -0.43898 | -0.34352 | 0.47756 |
| 18 | 78172200 | 78174199 | 21.21496 | 35.1415 | 65.77133 | 28.4644 | -0.52053 | 1.297004 | 0.477561 |
| 10 | 89968800 | 89972199 | 79.96617 | 123.1811 | 199.2271 | 82.7529 | -0.39511 | 2.914769 | 0.477576 |
| 10 | 21970200 | 21972599 | 17.53712 | 30.97559 | 31.19378 | 13.80028 | 0.379917 | 0.544883 | 0.477588 |
| 1 | 29418200 | 29419599 | 12.01421 | 21.65108 | 39.50942 | 17.21684 | -0.45662 | 0.385647 | 0.477596 |
| 1 | 112584800 | 112588599 | 38.2972 | 63.6313 | 53.83205 | 23.34649 | 0.621595 | 1.29753 | 0.477616 |
| 4 | 75073800 | 75079999 | 86.44931 | 136.3281 | 128.4199 | 53.29365 | 0.409951 | 2.483312 | 0.477627 |
| 16 | 22122400 | 22124199 | 9.42438 | 17.22058 | 32.38066 | 13.86824 | -0.48289 | 0.271181 | 0.477634 |
| 13 | 97984000 | 97988199 | 33.86861 | 55.74689 | 57.90816 | 24.96897 | 0.324219 | 1.397511 | 0.477647 |
| 6 | 122028200 | 122029999 | 22.60795 | 39.27572 | 38.13954 | 16.74422 | 0.428918 | 0.809033 | 0.477652 |
| 16 | 61793000 | 61795399 | 18.76485 | 33.16365 | 32.10047 | 14.19973 | 0.437142 | 0.548556 | 0.477656 |
| 1 | 192017800 | 192022199 | 41.58526 | 66.54885 | 120.4232 | 50.71464 | -0.50998 | 2.031441 | 0.477658 |
| 1 | 163639600 | 163640999 | 18.93628 | 33.49737 | 27.67439 | 12.2396 | 0.665014 | 0.532702 | 0.47766 |
| X | 138771400 | 138773999 | 28.68696 | 48.71766 | 45.27921 | 19.84118 | 0.486802 | 1.054144 | 0.477671 |
| 6 | 109057400 | 109059399 | 30.51534 | 51.1293 | 51.32223 | 22.37861 | 0.375635 | 1.183346 | 0.477673 |
| 19 | 8165400 | 8168999 | 36.97779 | 61.03456 | 57.7996 | 24.95311 | 0.458665 | 1.368977 | 0.477744 |
| 5 | 148515200 | 148522999 | 84.0209 | 129.9566 | 195.515 | 81.21222 | -0.28634 | 2.839625 | 0.477747 |
| 1 | 136807600 | 136810399 | 27.39991 | 46.24939 | 47.46283 | 20.75965 | 0.343906 | 1.099489 | 0.477749 |
| 4 | 147315800 | 147324999 | 185.5413 | 279.1963 | 231.4481 | 95.57946 | 0.540878 | 3.411846 | 0.477765 |
| 2 | 171703200 | 171708999 | 47.73879 | 75.88375 | 137.4009 | 57.33248 | -0.52143 | 2.254314 | 0.477775 |
| 6 | 57838400 | 57839399 | 17.11282 | 30.23789 | 30.80553 | 13.62746 | 0.363614 | 0.538323 | 0.477795 |
| 13 | 22198000 | 22200199 | 14.11391 | 24.7426 | 41.97572 | 18.54663 | -0.3712 | 0.641488 | 0.477797 |
| 1 | 38760800 | 38767599 | 72.5956 | 112.6124 | 196.4022 | 81.56917 | -0.49604 | 2.783714 | 0.477806 |
| 19 | 39593800 | 39594399 | 6.145588 | 11.79107 | 25.1719 | 9.882842 | -0.55951 | -0.09445 | 0.477814 |
| 14 | 50348000 | 50351399 | 33.20422 | 55.86211 | 48.5589 | 21.21782 | 0.582439 | 1.148568 | 0.477863 |
| 5 | 88436600 | 88445399 | 140.9684 | 214.4564 | 206.5409 | 85.57533 | 0.338009 | 3.182096 | 0.477867 |
| 4 | 26022600 | 26025799 | 27.68195 | 47.97179 | 39.06047 | 17.13763 | 0.681604 | 0.862694 | 0.477889 |

| | | | | | | | | | |
|---|---|---|---|---|---|---|---|---|---|
| 18 | 37112200 | 37115599 | 74.98957 | 118.4354 | 118.187 | 49.07849 | 0.328607 | 2.441078 | 0.477898 |
| 9 | 109093400 | 109094999 | 18.56115 | 34.36677 | 24.53327 | 10.32098 | 0.923746 | 0.25288 | 0.477907 |
| X | 5414600 | 5420799 | 19.55387 | 34.87935 | 25.55448 | 11.23902 | 0.843633 | 0.472409 | 0.477919 |
| 7 | 108972800 | 108973399 | 11.42911 | 21.52389 | 22.13556 | 9.06047 | 0.431509 | 0.084606 | 0.477925 |
| X | 7793600 | 7798399 | 65.50307 | 104.9776 | 85.53622 | 36.10101 | 0.641501 | 2.007763 | 0.477944 |
| X | 32028600 | 32031599 | 60.55381 | 97.01657 | 80.06989 | 33.77846 | 0.622477 | 2.00661 | 0.477956 |
| 17 | 22862600 | 22867999 | 110.2632 | 170.173 | 169.7099 | 70.49422 | 0.303827 | 2.887722 | 0.477964 |
| 9 | 43789200 | 43792399 | 26.36016 | 42.90414 | 79.8649 | 34.28469 | -0.52398 | 1.544947 | 0.47797 |
| 8 | 41184200 | 41187599 | 30.95512 | 53.4158 | 37.80674 | 16.5854 | 0.881666 | 0.911897 | 0.477988 |
| 7 | 53413200 | 53413999 | 7.509742 | 14.36887 | 28.46244 | 11.26318 | -0.4636 | -0.04437 | 0.478009 |
| 14 | 103856600 | 103860999 | 33.84165 | 56.13464 | 55.38845 | 23.99384 | 0.400075 | 1.30293 | 0.47801 |
| 3 | 23617600 | 23628999 | 208.7282 | 308.1168 | 301.1288 | 124.0103 | 0.278298 | 3.76026 | 0.478033 |
| 16 | 22830600 | 22836599 | 58.00618 | 91.95341 | 139.7008 | 58.13335 | -0.27257 | 2.341976 | 0.478042 |
| 12 | 101534600 | 101536599 | 17.66478 | 31.31213 | 31.15663 | 13.77406 | 0.399192 | 0.508873 | 0.478044 |
| 8 | 50641000 | 50642399 | 8.913811 | 16.81186 | 32.58566 | 13.3043 | -0.47343 | 0.078823 | 0.478044 |
| 18 | 52676200 | 52676799 | 8.177746 | 15.25715 | 29.52716 | 12.27922 | -0.49189 | 0.1366 | 0.478046 |
| 10 | 15234400 | 15236799 | 24.53826 | 42.31444 | 41.3991 | 18.16096 | 0.416292 | 0.893275 | 0.478054 |
| 5 | 122328000 | 122333199 | 44.21162 | 70.53529 | 117.0664 | 49.0875 | -0.39033 | 2.127663 | 0.478064 |
| 1 | 37160200 | 37161599 | 7.514312 | 14.13656 | 28.57854 | 11.71841 | -0.53844 | 0.071138 | 0.478091 |
| 3 | 33480400 | 33487599 | 108.3431 | 167.8023 | 162.065 | 67.31743 | 0.353939 | 2.822517 | 0.478092 |
| 10 | 121901400 | 121902999 | 12.65524 | 22.44444 | 38.21646 | 16.88952 | -0.37118 | 0.486534 | 0.478093 |
| 1 | 119866600 | 119869599 | 24.50707 | 42.57396 | 34.96129 | 15.34866 | 0.669812 | 0.815695 | 0.478108 |
| 10 | 21601400 | 21603199 | 21.76914 | 37.78445 | 38.83149 | 17.03836 | 0.347495 | 0.820387 | 0.478109 |
| 1 | 30452200 | 30453399 | 12.96595 | 24.62008 | 23.08642 | 9.292213 | 0.584141 | 0.044475 | 0.478113 |
| 10 | 45082800 | 45083999 | 13.46061 | 25.12646 | 24.96299 | 10.37199 | 0.463366 | 0.163362 | 0.47812 |
| 2 | 13681600 | 13682399 | 8.570734 | 16.22268 | 30.42625 | 12.32909 | -0.41661 | 0.045536 | 0.478157 |
| 18 | 12359000 | 12363599 | 34.24601 | 55.1048 | 95.98498 | 40.7541 | -0.44539 | 1.843707 | 0.478161 |
| 10 | 82110800 | 82111599 | 13.23768 | 24.44021 | 25.00769 | 10.56154 | 0.403758 | 0.226899 | 0.478169 |
| 14 | 115329400 | 115332399 | 23.16158 | 38.20409 | 75.48055 | 32.57501 | -0.59933 | 1.357747 | 0.478181 |
| 8 | 34823800 | 34825799 | 15.0539 | 26.36801 | 44.58351 | 19.68471 | -0.36656 | 0.663625 | 0.478185 |
| 2 | 139541800 | 139542999 | 22.53117 | 39.3437 | 36.3506 | 16.01039 | 0.501059 | 0.735714 | 0.478217 |
| 2 | 100248800 | 100251199 | 10.86067 | 20.84591 | 18.06137 | 7.082963 | 0.731344 | -0.07136 | 0.478224 |
| 13 | 5259400 | 5259999 | 8.853556 | 17.23354 | 18.37017 | 6.749906 | 0.521682 | -0.3112 | 0.478224 |
| 7 | 120154800 | 120157599 | 35.29681 | 57.99668 | 60.25813 | 25.96312 | 0.322889 | 1.422799 | 0.478226 |
| X | 54555400 | 54561399 | 73.34213 | 117.4205 | 86.59071 | 36.48918 | 0.783426 | 2.046818 | 0.478228 |
| 18 | 50215400 | 50217799 | 19.32571 | 32.4201 | 58.97725 | 25.73622 | -0.47914 | 1.142933 | 0.478234 |
| 9 | 23194800 | 23199599 | 47.13533 | 75.92873 | 78.72973 | 33.48133 | 0.303283 | 1.819175 | 0.478235 |
| 4 | 60255600 | 60257599 | 21.42949 | 37.3194 | 37.65 | 16.55467 | 0.374483 | 0.770024 | 0.478245 |
| X | 138875400 | 138877199 | 19.05377 | 34.57009 | 27.40397 | 11.84119 | 0.7471 | 0.372242 | 0.478254 |
| 3 | 6902400 | 6903799 | 23.25402 | 40.36137 | 33.72078 | 14.79581 | 0.645055 | 0.821577 | 0.478261 |
| 5 | 46596000 | 46599199 | 37.24521 | 61.66578 | 50.99608 | 22.06282 | 0.653946 | 1.331971 | 0.478262 |
| 11 | 47751400 | 47751799 | 6.585787 | 12.62573 | 25.69832 | 10.11228 | -0.49428 | -0.07965 | 0.478263 |
| 7 | 75672200 | 75673599 | 14.35385 | 25.06223 | 44.19097 | 19.46249 | -0.42655 | 0.712545 | 0.478278 |
| 7 | 6550800 | 6555799 | 70.75734 | 112.3553 | 113.3448 | 47.24517 | 0.31964 | 2.298077 | 0.47829 |
| 4 | 113693200 | 113701399 | 128.2008 | 196.9995 | 177.0962 | 73.53569 | 0.448661 | 2.978504 | 0.478299 |
| 7 | 13859400 | 13869199 | 158.827 | 240.7847 | 213.2949 | 88.28161 | 0.454468 | 3.258492 | 0.478313 |
| 5 | 56768200 | 56769399 | 9.67447 | 17.95036 | 32.0812 | 13.45782 | -0.3886 | 0.185972 | 0.478329 |

| | | | | | | | | | |
|---|---|---|---|---|---|---|---|---|---|
| 12 | 10882600 | 10883599 | 22.45161 | 39.04591 | 38.70518 | 17.00399 | 0.399596 | 0.795977 | 0.47833 |
| 3 | 159425800 | 159428599 | 27.78918 | 47.39034 | 45.2042 | 19.81664 | 0.450153 | 1.010638 | 0.478331 |
| Y | 5010200 | 5012399 | 19.49718 | 34.20586 | 21.70489 | 9.596306 | 1.039865 | 0.619195 | 0.478345 |
| 8 | 48602000 | 48609599 | 87.80886 | 135.3284 | 202.9053 | 84.28356 | -0.28542 | 2.908095 | 0.478347 |
| 13 | 97821200 | 97823399 | 21.37308 | 37.3638 | 36.47671 | 16.07892 | 0.422128 | 0.715098 | 0.478354 |
| 17 | 26706200 | 26714799 | 109.4974 | 166.0954 | 249.6026 | 103.3269 | -0.30678 | 3.236115 | 0.478357 |
| 7 | 59748400 | 59752199 | 42.01547 | 69.47487 | 55.11939 | 23.82568 | 0.713544 | 1.362825 | 0.47839 |
| 13 | 29199600 | 29200799 | 9.53618 | 17.62114 | 31.46921 | 13.27518 | -0.39325 | 0.206968 | 0.478409 |
| 18 | 22059600 | 22060999 | 9.522967 | 17.63781 | 31.50569 | 13.24847 | -0.39029 | 0.193525 | 0.478423 |
| 10 | 119324400 | 119338199 | 157.889 | 234.6067 | 336.7202 | 138.7166 | -0.26698 | 3.643604 | 0.478428 |
| 8 | 9017800 | 9020399 | 19.02482 | 32.36459 | 54.29633 | 23.79776 | -0.3618 | 1.010839 | 0.478435 |
| 18 | 23977600 | 23981399 | 40.66784 | 66.33528 | 65.04662 | 27.94554 | 0.40044 | 1.532739 | 0.47845 |
| 5 | 83314800 | 83315199 | 4.348638 | 8.500159 | 18.92845 | 6.521077 | -0.4434 | -0.59237 | 0.47846 |
| 14 | 79095200 | 79095799 | 4.348638 | 8.500159 | 18.92845 | 6.521077 | -0.4434 | -0.59237 | 0.47846 |
| 1 | 30952000 | 30953999 | 30.60189 | 51.3131 | 43.84055 | 19.12541 | 0.607331 | 1.168824 | 0.47846 |
| 2 | 3630200 | 3632799 | 28.02102 | 47.21696 | 48.55068 | 21.22445 | 0.341078 | 1.119057 | 0.478468 |
| 16 | 47562000 | 47563399 | 16.02145 | 28.96551 | 28.0958 | 12.19409 | 0.453347 | 0.370059 | 0.478531 |
| 12 | 45271200 | 45272599 | 19.96331 | 34.89232 | 35.52681 | 15.65757 | 0.361687 | 0.708466 | 0.478551 |
| 1 | 190448400 | 190449799 | 8.597953 | 16.17098 | 29.65882 | 12.16674 | -0.39947 | 0.087519 | 0.478562 |
| 16 | 31821400 | 31822199 | 5.407589 | 10.45253 | 23.70648 | 9.085911 | -0.61258 | -0.18686 | 0.478568 |
| 9 | 122516200 | 122520599 | 22.86373 | 38.0692 | 64.05065 | 27.81839 | -0.36694 | 1.238106 | 0.478577 |
| 18 | 9418400 | 9427799 | 142.0441 | 216.2044 | 203.6943 | 84.40897 | 0.370327 | 3.172928 | 0.478577 |
| 1 | 71473000 | 71475799 | 36.97234 | 60.91149 | 59.90961 | 25.842 | 0.403289 | 1.394913 | 0.478601 |
| 16 | 3821200 | 3822599 | 8.027942 | 14.88486 | 29.9757 | 12.58662 | -0.56007 | 0.172122 | 0.478605 |
| 17 | 49351400 | 49352999 | 15.45044 | 27.69331 | 28.27168 | 12.38352 | 0.371475 | 0.418289 | 0.478611 |
| 1 | 120582000 | 120583199 | 8.758784 | 16.29237 | 29.98829 | 12.53434 | -0.4265 | 0.15973 | 0.478611 |
| 7 | 64171000 | 64172399 | 8.758784 | 16.29237 | 29.98829 | 12.53434 | -0.4265 | 0.15973 | 0.478611 |
| X | 80339000 | 80344199 | 71.24915 | 114.2229 | 72.5668 | 30.64205 | 1.000265 | 1.999019 | 0.478613 |
| 14 | 5822000 | 5824399 | 20.45654 | 34.6277 | 58.14761 | 25.45498 | -0.36383 | 1.062298 | 0.478627 |
| 6 | 68568200 | 68571399 | 81.76244 | 128.3614 | 132.7098 | 55.04188 | 0.270907 | 2.569347 | 0.478638 |
| X | 36311800 | 36313999 | 28.98229 | 49.03362 | 46.91191 | 20.53309 | 0.444979 | 1.0884 | 0.478659 |
| 9 | 49822800 | 49825399 | 20.87837 | 35.12967 | 58.57973 | 25.59738 | -0.35403 | 1.118631 | 0.478663 |
| 3 | 19948600 | 19949799 | 9.569289 | 17.12348 | 34.91541 | 15.31503 | -0.62137 | 0.405686 | 0.478664 |
| 17 | 80085400 | 80088599 | 38.94283 | 62.60328 | 99.79697 | 42.31471 | -0.31957 | 1.881051 | 0.478667 |
| 18 | 38012800 | 38013399 | 5.806922 | 11.29514 | 23.62325 | 8.730429 | -0.44826 | -0.31114 | 0.47868 |
| 5 | 80254200 | 80256999 | 35.31661 | 57.16566 | 93.33928 | 39.92618 | -0.34255 | 1.674792 | 0.478697 |
| 7 | 93365200 | 93366999 | 23.89386 | 41.54725 | 38.59829 | 16.95526 | 0.492708 | 0.804535 | 0.478699 |
| 10 | 18131800 | 18132399 | 12.28414 | 22.93395 | 23.9909 | 9.96547 | 0.389736 | 0.149716 | 0.4787 |
| 19 | 49323400 | 49325199 | 18.87058 | 33.95724 | 28.28064 | 12.34631 | 0.666642 | 0.417237 | 0.478711 |
| Y | 5610200 | 5610799 | 5.127064 | 9.636731 | 29.1043 | 11.95231 | -1.11278 | 0.058969 | 0.478718 |
| 3 | 150041800 | 150044999 | 44.8156 | 72.28268 | 71.70322 | 30.54476 | 0.369158 | 1.775325 | 0.478721 |
| 15 | 9542400 | 9542999 | 5.528974 | 10.69692 | 23.73037 | 9.056348 | -0.57457 | -0.19774 | 0.478743 |
| 14 | 84137800 | 84138399 | 8.158419 | 15.2906 | 29.05709 | 11.99088 | -0.45686 | 0.105544 | 0.478753 |
| 7 | 55268800 | 55270599 | 19.9388 | 34.01006 | 57.11727 | 25.03954 | -0.36289 | 0.991612 | 0.478778 |
| 6 | 68970000 | 68977199 | 133.2906 | 203.0005 | 203.0412 | 84.15321 | 0.284806 | 3.157557 | 0.478785 |
| 8 | 61259600 | 61261199 | 13.19965 | 23.04075 | 43.6776 | 19.23313 | -0.53069 | 0.706564 | 0.478788 |
| 12 | 29722000 | 29724399 | 13.57959 | 23.86571 | 43.43842 | 19.23365 | -0.47242 | 0.597547 | 0.478796 |

| | | | | | | | | | |
|---|---|---|---|---|---|---|---|---|---|
| 8 | 35485600 | 35488999 | 50.43875 | 81.15221 | 83.09136 | 35.2745 | 0.319294 | 1.863218 | 0.478803 |
| 18 | 19517400 | 19521599 | 83.25145 | 130.5762 | 135.2787 | 56.10572 | 0.266884 | 2.588051 | 0.478807 |
| 11 | 89074400 | 89079999 | 34.97568 | 56.51941 | 89.81279 | 38.35326 | -0.30596 | 1.712856 | 0.478817 |
| 17 | 39659200 | 39661199 | 24.24578 | 42.63138 | 31.62599 | 14.00417 | 0.81624 | 0.634607 | 0.47882 |
| 5 | 93448800 | 93451999 | 26.34487 | 45.4017 | 41.31509 | 18.12377 | 0.520215 | 0.906319 | 0.478822 |
| 3 | 73479000 | 73480799 | 17.6106 | 32.19014 | 25.84825 | 11.06559 | 0.73677 | 0.315173 | 0.478837 |
| 3 | 78547200 | 78548999 | 16.55637 | 30.23171 | 27.09097 | 11.61267 | 0.578436 | 0.316394 | 0.478841 |
| 17 | 92503600 | 92505199 | 16.55637 | 30.23171 | 27.09097 | 11.61267 | 0.578436 | 0.316394 | 0.478841 |
| 9 | 90260000 | 90261199 | 13.59522 | 23.98479 | 41.32555 | 18.28693 | -0.39147 | 0.552842 | 0.478843 |
| 5 | 132624400 | 132624799 | 10.44645 | 19.88703 | 20.6966 | 8.285173 | 0.442269 | -0.00164 | 0.478846 |
| 10 | 118298800 | 118301199 | 29.49004 | 49.62529 | 49.6437 | 21.69339 | 0.380509 | 1.136241 | 0.478858 |
| 10 | 79115400 | 79118199 | 27.09223 | 46.56418 | 42.42518 | 18.6092 | 0.517734 | 0.935228 | 0.478861 |
| 18 | 22380800 | 22381599 | 9.913547 | 18.1334 | 32.3962 | 13.85633 | -0.40807 | 0.270883 | 0.47887 |
| 1 | 181227200 | 181228799 | 7.822191 | 14.74136 | 28.07169 | 11.47676 | -0.44906 | 0.06367 | 0.478905 |
| 16 | 55523800 | 55527399 | 24.81486 | 40.89444 | 84.31219 | 36.37266 | -0.66124 | 1.380722 | 0.478908 |
| 10 | 119011400 | 119015799 | 59.87099 | 95.80851 | 94.73713 | 39.89554 | 0.360227 | 2.043928 | 0.47891 |
| 16 | 53804200 | 53807199 | 30.54032 | 49.63561 | 80.83496 | 34.6811 | -0.33323 | 1.57713 | 0.478917 |
| 6 | 8544000 | 8545199 | 18.54904 | 31.57031 | 53.74835 | 23.55849 | -0.38279 | 1.00421 | 0.478934 |
| 8 | 72683600 | 72689399 | 83.04428 | 131.4234 | 120.4218 | 50.05793 | 0.453858 | 2.399204 | 0.478937 |
| 1 | 37815800 | 37818399 | 24.79606 | 42.58698 | 36.33086 | 15.93567 | 0.612026 | 0.923985 | 0.478945 |
| 19 | 16770200 | 16773999 | 47.89041 | 77.1683 | 79.76508 | 33.93644 | 0.308299 | 1.810289 | 0.478964 |
| 14 | 22639600 | 22642599 | 15.50793 | 27.04039 | 46.26963 | 20.36052 | -0.38411 | 0.736505 | 0.478966 |
| 4 | 80719000 | 80721599 | 41.4714 | 67.14141 | 69.33321 | 29.66182 | 0.317187 | 1.667688 | 0.478974 |
| 14 | 123358800 | 123363599 | 74.34429 | 118.4326 | 103.1379 | 43.15814 | 0.536185 | 2.200691 | 0.478978 |
| X | 24162800 | 24171799 | 167.4008 | 252.9552 | 220.8927 | 91.33212 | 0.471097 | 3.326622 | 0.478978 |
| 1 | 101533800 | 101536799 | 35.63751 | 59.0071 | 50.98453 | 22.05864 | 0.590839 | 1.325385 | 0.478986 |
| 7 | 48145600 | 48147799 | 28.35166 | 48.76437 | 41.84938 | 18.35704 | 0.603972 | 0.934365 | 0.478996 |
| 5 | 127506200 | 127516399 | 115.0815 | 174.3705 | 260.3169 | 107.7252 | -0.29864 | 3.260252 | 0.479015 |
| 17 | 23791200 | 23800199 | 112.9065 | 170.504 | 263.3503 | 108.8657 | -0.35177 | 3.328299 | 0.479017 |
| 2 | 172154400 | 172161799 | 82.76636 | 128.3227 | 191.3383 | 79.4722 | -0.2708 | 2.795526 | 0.479021 |
| X | 40875400 | 40875999 | 12.09376 | 22.68532 | 23.6007 | 9.725351 | 0.407163 | 0.119175 | 0.479023 |
| 1 | 177325400 | 177327199 | 13.31062 | 23.33227 | 41.99078 | 18.55187 | -0.45599 | 0.637394 | 0.479029 |
| 8 | 4121800 | 4125599 | 14.27629 | 26.42209 | 21.38304 | 9.000826 | 0.74326 | 0.212909 | 0.47903 |
| 5 | 52405800 | 52408599 | 19.97222 | 33.96649 | 55.96043 | 24.52655 | -0.33591 | 1.018244 | 0.479032 |
| 7 | 16446200 | 16449599 | 36.12628 | 58.29082 | 92.49221 | 39.42077 | -0.30638 | 1.759796 | 0.479033 |
| 10 | 129882600 | 129883199 | 16.85352 | 29.75149 | 28.40364 | 12.56732 | 0.456274 | 0.541695 | 0.479037 |
| 1 | 114309400 | 114314999 | 54.28242 | 87.45285 | 79.78229 | 33.9352 | 0.487794 | 1.82497 | 0.479044 |
| 10 | 103545800 | 103546599 | 12.88021 | 22.55738 | 41.55184 | 18.34392 | -0.4892 | 0.650569 | 0.479071 |
| 2 | 15010400 | 15011199 | 10.54298 | 20.36279 | 20.81503 | 8.007271 | 0.520201 | -0.12975 | 0.479078 |
| 6 | 68486400 | 68487799 | 14.29268 | 26.70365 | 24.91473 | 10.3362 | 0.555117 | 0.164718 | 0.47909 |
| Y | 87983200 | 87985199 | 12.84272 | 23.90124 | 16.04277 | 6.699639 | 1.018606 | 0.151541 | 0.479098 |
| 7 | 75761000 | 75762399 | 13.53941 | 25.62449 | 23.56839 | 9.552478 | 0.603243 | 0.073728 | 0.479104 |
| 15 | 32984600 | 32990399 | 64.76991 | 103.3131 | 102.4949 | 42.96548 | 0.350438 | 2.150763 | 0.479113 |
| 10 | 65422000 | 65422599 | 11.76825 | 21.96763 | 21.53877 | 8.949083 | 0.482358 | 0.14011 | 0.479128 |
| 8 | 40323400 | 40325799 | 30.58381 | 51.6823 | 47.97424 | 20.99057 | 0.488247 | 1.102081 | 0.479133 |
| 17 | 68463600 | 68466199 | 23.21865 | 38.66657 | 64.01422 | 27.80547 | -0.34369 | 1.238802 | 0.479175 |
| 18 | 25391600 | 25393999 | 51.33334 | 81.58797 | 129.378 | 53.97851 | -0.33053 | 2.259504 | 0.47918 |

| | | | | | | | | | |
|---|---|---|---|---|---|---|---|---|---|
| X | 127951800 | 127952599 | 11.48844 | 21.9663 | 22.25508 | 8.824369 | 0.492765 | -0.01746 | 0.479181 |
| 3 | 87912800 | 87913999 | 15.18732 | 26.47071 | 46.2782 | 20.35914 | -0.41501 | 0.740301 | 0.479209 |
| 5 | 79103200 | 79104599 | 21.75596 | 38.19838 | 33.46375 | 14.80494 | 0.577323 | 0.642827 | 0.479212 |
| 13 | 120392200 | 120393999 | 21.98401 | 38.26441 | 32.08592 | 14.10336 | 0.640007 | 0.772281 | 0.479215 |
| 7 | 33214200 | 33259199 | 1071.799 | 1382.304 | 1139.533 | 477.7418 | 0.329732 | 5.866146 | 0.479219 |
| 2 | 19434200 | 19437399 | 29.07729 | 46.89284 | 98.76742 | 42.06726 | -0.71478 | 1.76798 | 0.47922 |
| 8 | 22129200 | 22130999 | 16.63997 | 28.54968 | 55.91084 | 24.5222 | -0.58212 | 0.926324 | 0.479288 |
| 18 | 85387600 | 85390399 | 19.67463 | 33.68662 | 58.23703 | 25.5382 | -0.40364 | 0.959054 | 0.479295 |
| 10 | 120598200 | 120602999 | 38.23795 | 61.46066 | 100.587 | 42.64018 | -0.35775 | 1.883177 | 0.479309 |
| 8 | 21842000 | 21848399 | 65.44753 | 105.1591 | 81.07783 | 34.35288 | 0.725394 | 1.925692 | 0.479319 |
| 1 | 148036200 | 148037799 | 14.30843 | 26.35198 | 26.37157 | 11.17501 | 0.431881 | 0.250946 | 0.479321 |
| 8 | 110830800 | 110840799 | 116.1248 | 175.5034 | 261.9104 | 108.2975 | -0.30124 | 3.312916 | 0.479329 |
| 2 | 81419400 | 81419999 | 17.39912 | 30.63062 | 30.42139 | 13.46745 | 0.39814 | 0.579175 | 0.479341 |
| 17 | 22392800 | 22394799 | 36.4139 | 59.83287 | 60.79017 | 26.19248 | 0.35514 | 1.427793 | 0.479347 |
| X | 147754400 | 147766399 | 180.1082 | 270.8931 | 237.4101 | 98.02917 | 0.460083 | 3.420118 | 0.479383 |
| 3 | 9000600 | 9001999 | 9.086031 | 16.72861 | 31.42857 | 13.32358 | -0.47102 | 0.223193 | 0.479392 |
| 7 | 52721800 | 52726599 | 71.04119 | 113.5139 | 94.60265 | 39.75472 | 0.604177 | 2.102215 | 0.479393 |
| 3 | 138387200 | 138391199 | 74.99469 | 116.9478 | 175.3886 | 72.78464 | -0.27262 | 2.69115 | 0.479396 |
| 1 | 167272400 | 167277599 | 42.58252 | 68.04563 | 112.5551 | 47.30348 | -0.3829 | 2.075648 | 0.479401 |
| 3 | 36480000 | 36485199 | 46.03955 | 74.40101 | 75.02359 | 32.0393 | 0.348727 | 1.721095 | 0.479419 |
| 18 | 51533000 | 51538799 | 169.4098 | 253.3448 | 252.1142 | 103.9611 | 0.269483 | 3.526094 | 0.479422 |
| 19 | 31090000 | 31091999 | 13.31756 | 23.37397 | 40.92901 | 18.10211 | -0.41658 | 0.615945 | 0.479427 |
| 19 | 13666600 | 13669599 | 33.25174 | 56.1733 | 46.85201 | 20.49762 | 0.642064 | 1.11304 | 0.479438 |
| 7 | 82205800 | 82206999 | 9.140407 | 16.7848 | 31.43335 | 13.37308 | -0.46973 | 0.241159 | 0.47945 |
| 18 | 80198600 | 80202199 | 46.14453 | 74.85567 | 70.76944 | 30.32063 | 0.446869 | 1.638447 | 0.479452 |
| 17 | 28290200 | 28293399 | 31.54428 | 51.1472 | 83.42561 | 35.73074 | -0.33863 | 1.628242 | 0.47946 |
| 3 | 7242800 | 7243399 | 13.29116 | 24.34122 | 23.51152 | 10.03956 | 0.474903 | 0.267756 | 0.479468 |
| 13 | 86440600 | 86442599 | 24.23639 | 42.21133 | 38.04978 | 16.73138 | 0.536352 | 0.784737 | 0.479469 |
| 9 | 32285000 | 32287799 | 15.20398 | 26.51887 | 45.21164 | 19.89895 | -0.37873 | 0.729499 | 0.479472 |
| 9 | 87889800 | 87890599 | 13.73246 | 23.8507 | 50.01841 | 21.95584 | -0.67658 | 0.791788 | 0.479491 |
| 12 | 114293800 | 114302599 | 18.04588 | 32.13146 | 27.04303 | 11.9118 | 0.642982 | 0.474493 | 0.479501 |
| 4 | 10859200 | 10860799 | 14.19519 | 25.72414 | 25.31357 | 10.95432 | 0.435451 | 0.337417 | 0.479522 |
| 1 | 60410600 | 60411199 | 7.485595 | 14.32908 | 27.3538 | 10.81055 | -0.40912 | -0.05284 | 0.479532 |
| 6 | 85938000 | 85938999 | 8.689551 | 15.86175 | 32.9135 | 14.11438 | -0.62525 | 0.271998 | 0.479533 |
| 15 | 25835200 | 25839599 | 48.87638 | 77.96407 | 122.6876 | 51.43235 | -0.314 | 2.138223 | 0.479537 |
| 13 | 24361800 | 24364399 | 15.03358 | 26.24097 | 45.19621 | 19.90137 | -0.39329 | 0.71773 | 0.479544 |
| 9 | 40692200 | 40694199 | 16.33594 | 28.25945 | 48.99836 | 21.49668 | -0.40481 | 0.839563 | 0.47955 |
| 19 | 20116200 | 20122799 | 79.35169 | 125.15 | 128.4333 | 53.30002 | 0.286666 | 2.476374 | 0.47956 |
| 4 | 126631600 | 126634599 | 21.39643 | 35.75184 | 61.80897 | 26.90453 | -0.40602 | 1.193683 | 0.479571 |
| 4 | 101574400 | 101575399 | 8.193377 | 15.18395 | 29.98236 | 12.59804 | -0.53234 | 0.178109 | 0.479578 |
| 7 | 62430800 | 62438799 | 127.0288 | 194.4604 | 189.5598 | 78.66295 | 0.327739 | 3.05182 | 0.479582 |
| 12 | 63231400 | 63234999 | 62.01342 | 98.85179 | 102.2863 | 42.83933 | 0.288777 | 2.167368 | 0.479585 |
| 18 | 55969800 | 55971799 | 23.71996 | 41.02858 | 35.25007 | 15.46494 | 0.603901 | 0.855893 | 0.479586 |
| Y | 8995000 | 8997199 | 18.87432 | 33.18785 | 21.16301 | 9.371041 | 1.032824 | 0.580215 | 0.47959 |
| 18 | 6795000 | 6798799 | 57.1949 | 91.44124 | 94.27012 | 39.64676 | 0.298697 | 2.068335 | 0.479601 |
| 12 | 44529600 | 44532399 | 24.36649 | 40.49451 | 66.29073 | 28.75424 | -0.32773 | 1.267379 | 0.479605 |
| 13 | 22298600 | 22303399 | 96.22475 | 149.3513 | 147.3274 | 61.18238 | 0.325834 | 2.77897 | 0.479615 |

| | | | | | | | | | |
|---|---|---|---|---|---|---|---|---|---|
| 7 | 23820600 | 23828599 | 86.68212 | 133.5004 | 201.3563 | 83.63506 | -0.29472 | 2.921443 | 0.479618 |
| 6 | 107990400 | 107990999 | 5.233372 | 10.18347 | 21.81257 | 8.036018 | -0.477 | -0.33266 | 0.479624 |
| 13 | 104056000 | 104058399 | 16.68491 | 28.74695 | 50.05809 | 21.95891 | -0.4122 | 0.879083 | 0.479625 |
| 19 | 57814000 | 57818599 | 44.6569 | 72.21319 | 74.16877 | 31.70062 | 0.32335 | 1.701334 | 0.479626 |
| 4 | 34495000 | 34502599 | 84.92948 | 133.3526 | 134.9573 | 55.97509 | 0.301821 | 2.567316 | 0.479641 |
| X | 91318000 | 91322999 | 54.23962 | 87.60218 | 74.51204 | 31.79727 | 0.592818 | 1.754459 | 0.479642 |
| 8 | 19747200 | 19771599 | 878.9878 | 1133.798 | 1430.463 | 599.7037 | -0.28393 | 5.862715 | 0.479648 |
| 4 | 29094200 | 29096199 | 23.22731 | 38.53765 | 65.60932 | 28.42627 | -0.38429 | 1.281582 | 0.47966 |
| 1 | 62537600 | 62539599 | 15.2444 | 27.83488 | 27.22909 | 11.6725 | 0.452681 | 0.307144 | 0.479673 |
| 12 | 17478000 | 17479399 | 8.049302 | 15.28744 | 30.00366 | 12.071 | -0.47287 | 0.013459 | 0.479677 |
| 9 | 120533800 | 120542999 | 83.31425 | 129.0097 | 192.1455 | 79.81213 | -0.27057 | 2.818539 | 0.479691 |
| X | 116479400 | 116480999 | 16.74138 | 30.84768 | 26.21349 | 11.10072 | 0.667104 | 0.267781 | 0.479719 |
| 16 | 18504600 | 18506599 | 9.499181 | 17.23868 | 32.89904 | 14.21157 | -0.51177 | 0.315786 | 0.479763 |
| 11 | 99728000 | 99728799 | 10.60044 | 19.0786 | 34.31285 | 14.97656 | -0.43675 | 0.374342 | 0.479772 |
| 9 | 36370600 | 36378599 | 175.6933 | 263.6428 | 227.7807 | 93.99496 | 0.477709 | 3.462687 | 0.479802 |
| 19 | 45174200 | 45185199 | 178.4816 | 261.4914 | 382.6107 | 157.6028 | -0.31388 | 3.879045 | 0.47981 |
| 10 | 25064000 | 25064799 | 10.87064 | 19.7519 | 34.37168 | 14.82482 | -0.37849 | 0.320534 | 0.479812 |
| 13 | 61082000 | 61084599 | 25.84733 | 44.24086 | 43.5793 | 19.10996 | 0.404626 | 0.967116 | 0.479814 |
| 10 | 49186000 | 49187999 | 36.98673 | 60.76195 | 61.49696 | 26.49477 | 0.360528 | 1.430642 | 0.479816 |
| 16 | 14227600 | 14229599 | 13.48533 | 23.5654 | 43.63604 | 19.22836 | -0.49693 | 0.691398 | 0.479836 |
| 17 | 33868600 | 33869399 | 6.42507 | 12.3451 | 25.21708 | 9.859613 | -0.49115 | -0.10597 | 0.479857 |
| 9 | 29558000 | 29559599 | 22.89533 | 38.15959 | 63.44552 | 27.57279 | -0.34983 | 1.228685 | 0.479863 |
| 15 | 16899400 | 16902599 | 30.71773 | 51.5736 | 50.2246 | 21.92823 | 0.41904 | 1.159981 | 0.479865 |
| 9 | 40044400 | 40046199 | 22.4449 | 39.28509 | 35.83276 | 15.81124 | 0.519717 | 0.701331 | 0.479866 |
| X | 77556400 | 77561599 | 76.31705 | 121.083 | 114.6966 | 47.76891 | 0.409183 | 2.327809 | 0.479875 |
| 13 | 60473800 | 60477199 | 42.98944 | 69.48252 | 72.05584 | 30.77794 | 0.308657 | 1.711379 | 0.479877 |
| 3 | 10474400 | 10478599 | 65.85231 | 105.0556 | 101.9149 | 42.7313 | 0.382925 | 2.146129 | 0.479888 |
| 7 | 11289400 | 11290799 | 46.92122 | 75.52365 | 74.95351 | 31.83864 | 0.364933 | 1.841667 | 0.47989 |
| 9 | 24233600 | 24235199 | 15.73704 | 29.0666 | 25.06293 | 10.57896 | 0.649019 | 0.241724 | 0.479921 |
| 4 | 46317800 | 46320399 | 24.82164 | 42.5884 | 43.08903 | 18.89867 | 0.36677 | 0.942387 | 0.47993 |
| 11 | 47746600 | 47749199 | 18.61376 | 33.01744 | 30.34457 | 13.40875 | 0.513456 | 0.505928 | 0.479953 |
| 10 | 36489200 | 36491999 | 45.43715 | 73.5694 | 65.16627 | 27.88123 | 0.538131 | 1.671643 | 0.479954 |
| 3 | 99519200 | 99519799 | 18.65079 | 32.71562 | 30.66805 | 13.55728 | 0.480512 | 0.638744 | 0.479957 |
| 4 | 122461000 | 122462799 | 26.86995 | 45.94616 | 44.10264 | 19.33793 | 0.44164 | 0.980883 | 0.479968 |
| 4 | 39721200 | 39723199 | 25.79899 | 44.03582 | 44.7098 | 19.60132 | 0.360631 | 0.994367 | 0.479974 |
| 14 | 8333600 | 8335599 | 29.65425 | 48.63375 | 82.48593 | 35.51202 | -0.3833 | 1.448889 | 0.479974 |
| 15 | 70337000 | 70338399 | 12.2993 | 22.91289 | 23.99397 | 10.00314 | 0.384096 | 0.16367 | 0.479979 |
| 11 | 82226600 | 82233599 | 59.29902 | 92.99821 | 172.9408 | 71.72651 | -0.57612 | 2.58252 | 0.47998 |
| 17 | 14381200 | 14384199 | 19.24235 | 32.45684 | 57.47551 | 25.13365 | -0.44028 | 1.089061 | 0.479991 |
| 19 | 55944000 | 55948999 | 50.94243 | 82.3618 | 73.91009 | 31.58238 | 0.517358 | 1.717625 | 0.479996 |
| X | 97051800 | 97054599 | 38.84618 | 63.48416 | 61.70679 | 26.529 | 0.415104 | 1.505322 | 0.480005 |
| 14 | 39326200 | 39327199 | 12.15376 | 21.76156 | 37.26867 | 16.33879 | -0.37142 | 0.421691 | 0.480021 |
| 8 | 107309000 | 107309599 | 10.99073 | 20.80374 | 22.09285 | 8.951834 | 0.397596 | 0.044654 | 0.480032 |
| 13 | 97565200 | 97566599 | 9.052544 | 16.82808 | 30.49939 | 12.76152 | -0.40578 | 0.16682 | 0.480038 |
| 10 | 95179600 | 95180199 | 9.581334 | 18.47097 | 20.25288 | 7.843279 | 0.410907 | -0.11991 | 0.480041 |
| 12 | 119402800 | 119404199 | 10.48869 | 19.23907 | 34.00061 | 14.48742 | -0.38897 | 0.261113 | 0.48005 |
| 7 | 73058200 | 73059599 | 12.39055 | 21.77362 | 39.91159 | 17.6704 | -0.48212 | 0.589291 | 0.480063 |

| | | | | | | | | | |
|---|---|---|---|---|---|---|---|---|---|
| 2 | 72817800 | 72820399 | 55.02787 | 88.16086 | 91.33485 | 38.52919 | 0.295034 | 2.002855 | 0.480076 |
| X | 151316800 | 151318399 | 20.64062 | 38.10623 | 23.19521 | 9.796673 | 1.148026 | 0.281883 | 0.480077 |
| 6 | 119456000 | 119457999 | 11.81734 | 20.69573 | 41.59429 | 18.36249 | -0.61417 | 0.645066 | 0.480082 |
| 7 | 17855200 | 17871399 | 239.8829 | 354.0258 | 293.5267 | 120.8793 | 0.515138 | 3.768353 | 0.480093 |
| 2 | 18155800 | 18156999 | 15.84899 | 29.96868 | 22.87412 | 9.288688 | 0.868141 | 0.099492 | 0.480099 |
| 9 | 121111200 | 121113799 | 29.77158 | 49.91731 | 50.77107 | 22.14786 | 0.356605 | 1.172112 | 0.480108 |
| 2 | 64532000 | 64534199 | 26.98426 | 46.18948 | 39.32404 | 17.24407 | 0.61425 | 0.96687 | 0.480132 |
| 7 | 47680400 | 47683799 | 34.43893 | 56.77282 | 58.64623 | 25.30367 | 0.333054 | 1.379233 | 0.480162 |
| 10 | 61764400 | 61769799 | 43.64525 | 69.88017 | 110.2321 | 46.45596 | -0.3115 | 2.014859 | 0.480176 |
| 18 | 37222400 | 37225199 | 40.90454 | 65.52143 | 107.0589 | 45.14948 | -0.36136 | 1.997218 | 0.480265 |
| 15 | 4055400 | 4058199 | 21.10893 | 35.19197 | 62.82407 | 27.3075 | -0.45209 | 1.21809 | 0.480268 |
| 16 | 15849600 | 15850999 | 19.61381 | 34.91352 | 30.35666 | 13.3745 | 0.596081 | 0.492693 | 0.480305 |
| X | 117003200 | 117004799 | 17.80398 | 33.67799 | 20.3753 | 8.267401 | 1.201424 | 0.1081 | 0.480308 |
| 2 | 17955000 | 17955599 | 7.676003 | 14.40458 | 28.54228 | 11.75334 | -0.51455 | 0.089419 | 0.480321 |
| 10 | 53598200 | 53599199 | 14.57621 | 27.26025 | 21.70613 | 8.990042 | 0.784077 | 0.153786 | 0.480321 |
| 9 | 18971400 | 18973999 | 42.13898 | 68.493 | 68.80466 | 29.52179 | 0.362159 | 1.589981 | 0.480322 |
| 4 | 27007200 | 27007799 | 11.04045 | 19.85621 | 34.76583 | 15.18453 | -0.39917 | 0.381514 | 0.480323 |
| 7 | 130524600 | 130526199 | 11.04045 | 19.85621 | 34.76583 | 15.18453 | -0.39917 | 0.381514 | 0.480323 |
| 6 | 15515800 | 15519199 | 37.1285 | 59.77403 | 95.92166 | 40.75789 | -0.32649 | 1.83056 | 0.480323 |
| X | 7438800 | 7443799 | 58.32425 | 93.84083 | 79.79405 | 33.87546 | 0.586829 | 1.876015 | 0.480323 |
| 6 | 22439600 | 22440199 | 5.954874 | 11.5665 | 23.43898 | 8.762166 | -0.41812 | -0.26801 | 0.480327 |
| 10 | 73991400 | 73996599 | 41.52067 | 66.77371 | 107.1517 | 45.45921 | -0.32848 | 1.873742 | 0.480332 |
| 4 | 124974400 | 124976399 | 13.66813 | 24.1689 | 42.39299 | 18.75067 | -0.41568 | 0.532001 | 0.480336 |
| 17 | 84061400 | 84064199 | 17.71124 | 30.3738 | 51.10996 | 22.41574 | -0.36422 | 0.92873 | 0.480383 |
| 3 | 16016600 | 16020199 | 44.59113 | 72.73626 | 63.74252 | 27.38547 | 0.562158 | 1.539884 | 0.480409 |
| X | 26774400 | 26791199 | 216.7827 | 323.4735 | 251.7268 | 103.7568 | 0.621159 | 3.576647 | 0.480413 |
| 6 | 145156600 | 145157199 | 5.520856 | 10.74111 | 22.40764 | 8.266232 | -0.44162 | -0.32352 | 0.480417 |
| 7 | 72059600 | 72060199 | 5.520856 | 10.74111 | 22.40764 | 8.266232 | -0.44162 | -0.32352 | 0.480417 |
| 14 | 50077800 | 50078999 | 18.70743 | 33.33832 | 30.53322 | 13.43929 | 0.522567 | 0.481122 | 0.480424 |
| 10 | 14874000 | 14876199 | 28.69925 | 48.85805 | 45.75392 | 20.05485 | 0.476302 | 1.03094 | 0.480424 |
| 1 | 179188000 | 179189199 | 10.3221 | 18.94426 | 33.47392 | 14.25199 | -0.3879 | 0.255305 | 0.480444 |
| Y | 9047600 | 9050599 | 26.68379 | 45.36329 | 29.12408 | 12.76422 | 1.018113 | 1.011181 | 0.480446 |
| 6 | 112162600 | 112163599 | 10.55824 | 20.25714 | 20.59098 | 8.084469 | 0.500355 | -0.06637 | 0.480455 |
| 17 | 48402400 | 48404599 | 24.75895 | 42.24181 | 43.68013 | 19.14922 | 0.334226 | 0.992538 | 0.480467 |
| 14 | 41400000 | 41402599 | 29.06269 | 49.19308 | 42.63285 | 18.66286 | 0.587052 | 1.077782 | 0.480491 |
| 1 | 155393400 | 155395399 | 17.78618 | 30.78184 | 52.81814 | 23.17282 | -0.38983 | 0.84703 | 0.480522 |
| 13 | 109987200 | 109990199 | 12.68291 | 24.46986 | 17.82209 | 6.886019 | 0.999195 | -0.09997 | 0.480525 |
| 1 | 71753400 | 71755599 | 14.25845 | 24.97863 | 42.50445 | 18.76923 | -0.3756 | 0.653806 | 0.480532 |
| 5 | 149203600 | 149204999 | 13.03907 | 22.83338 | 41.55368 | 18.34322 | -0.4718 | 0.653303 | 0.480535 |
| 17 | 38956400 | 38968799 | 146.3536 | 224.7681 | 166.4223 | 69.09828 | 0.727731 | 2.995681 | 0.480542 |
| 10 | 90901600 | 90902999 | 11.32205 | 21.73289 | 21.7371 | 8.522848 | 0.525691 | -0.05837 | 0.480581 |
| 14 | 5186400 | 5188599 | 28.70724 | 47.11957 | 78.10149 | 33.63619 | -0.34966 | 1.435277 | 0.480583 |
| 3 | 39655400 | 39656999 | 18.95983 | 32.01805 | 62.36799 | 27.28286 | -0.57723 | 1.083807 | 0.480589 |
| 4 | 114466400 | 114468999 | 15.39872 | 26.71879 | 46.87715 | 20.56795 | -0.4207 | 0.806215 | 0.480617 |
| 18 | 50789200 | 50790799 | 17.73205 | 31.89399 | 28.95429 | 12.64525 | 0.542625 | 0.413617 | 0.480625 |
| 10 | 121186400 | 121191399 | 50.03269 | 79.75941 | 124.4105 | 52.11019 | -0.30226 | 2.159748 | 0.480625 |
| 6 | 59903800 | 59905999 | 25.16914 | 43.20764 | 43.07314 | 18.89251 | 0.388164 | 0.939255 | 0.480631 |

| | | | | | | | | | |
|---|---|---|---|---|---|---|---|---|---|
| 1 | 170081800 | 170085599 | 58.1736 | 92.85685 | 94.17507 | 39.51687 | 0.319789 | 2.120328 | 0.480641 |
| 1 | 98383600 | 98384199 | 10.62905 | 20.18326 | 21.07486 | 8.482085 | 0.430157 | 0.01629 | 0.480657 |
| 16 | 70622400 | 70623599 | 10.62905 | 20.18326 | 21.07486 | 8.482085 | 0.430157 | 0.01629 | 0.480657 |
| 5 | 139506400 | 139509799 | 27.89842 | 45.57923 | 75.85642 | 32.60867 | -0.35901 | 1.492392 | 0.480685 |
| 13 | 88192600 | 88194199 | 17.38501 | 31.48966 | 27.98994 | 12.11972 | 0.580806 | 0.369515 | 0.480686 |
| 16 | 36356400 | 36358799 | 17.29721 | 30.59537 | 30.10106 | 13.31307 | 0.414036 | 0.525083 | 0.480696 |
| 15 | 38810000 | 38811799 | 14.10098 | 24.69097 | 42.52502 | 18.77074 | -0.39291 | 0.660652 | 0.480715 |
| 8 | 93588600 | 93591399 | 34.75622 | 57.89949 | 53.70941 | 23.3343 | 0.489228 | 1.257886 | 0.480722 |
| 13 | 66970800 | 66977999 | 244.2155 | 354.501 | 334.9346 | 138.0814 | 0.30396 | 4.024613 | 0.480725 |
| 16 | 10034200 | 10037999 | 71.4743 | 114.1068 | 96.93169 | 40.68336 | 0.575292 | 2.128759 | 0.480735 |
| 1 | 85406400 | 85416199 | 364.2368 | 513.2832 | 474.2131 | 196.3936 | 0.293627 | 4.466519 | 0.48074 |
| 13 | 18104600 | 18106999 | 23.10259 | 40.43886 | 33.97814 | 14.99371 | 0.637308 | 0.697779 | 0.480742 |
| 4 | 47262200 | 47263799 | 8.640782 | 15.96255 | 30.44853 | 12.84906 | -0.48766 | 0.199568 | 0.48075 |
| 17 | 90318000 | 90320199 | 29.51391 | 49.3102 | 51.31996 | 22.33729 | 0.323691 | 1.210155 | 0.480756 |
| 1 | 127510000 | 127511199 | 12.52788 | 23.45149 | 22.24442 | 9.198351 | 0.534592 | 0.128349 | 0.480758 |
| 17 | 52826000 | 52827799 | 24.50139 | 41.92658 | 42.56643 | 18.66543 | 0.361142 | 0.962432 | 0.480805 |
| 5 | 104722600 | 104725999 | 81.9784 | 130.1 | 112.1974 | 46.74055 | 0.54477 | 2.324347 | 0.480816 |
| X | 100000400 | 100000999 | 6.09407 | 11.85808 | 25.60057 | 9.43287 | -0.48987 | -0.31222 | 0.480847 |
| 17 | 39728200 | 39734999 | 95.41881 | 150.2358 | 128.3665 | 53.26056 | 0.548797 | 2.521308 | 0.480859 |
| X | 25770200 | 25776599 | 74.62803 | 119.4485 | 86.2308 | 36.32417 | 0.813675 | 2.055435 | 0.480865 |
| 10 | 67864400 | 67866999 | 16.63424 | 28.72802 | 51.69547 | 22.67882 | -0.4588 | 0.857774 | 0.48088 |
| 15 | 82580600 | 82582799 | 25.02904 | 43.57375 | 37.9943 | 16.70263 | 0.584 | 0.793791 | 0.480891 |
| 7 | 5631200 | 5641199 | 149.903 | 229.4837 | 176.952 | 73.43147 | 0.665699 | 3.058791 | 0.480894 |
| 1 | 133040200 | 133044199 | 30.37617 | 49.32441 | 81.8969 | 35.1134 | -0.3623 | 1.594752 | 0.480907 |
| 7 | 130264800 | 130268999 | 52.25878 | 82.99956 | 130.2841 | 54.31771 | -0.3169 | 2.279966 | 0.480926 |
| 12 | 115634800 | 115639599 | 95.04349 | 148.3053 | 148.1875 | 61.49212 | 0.31326 | 2.687215 | 0.480933 |
| 7 | 9821800 | 9834799 | 163.7032 | 248.5433 | 204.0821 | 84.50575 | 0.565573 | 3.229773 | 0.480947 |
| 6 | 47408400 | 47411599 | 17.65942 | 30.25795 | 52.73071 | 23.1249 | -0.41493 | 0.936194 | 0.480963 |
| 3 | 15416600 | 15418199 | 23.0841 | 39.72177 | 40.99538 | 17.98268 | 0.338941 | 0.90582 | 0.480998 |
| 2 | 173040600 | 173043199 | 26.05014 | 44.85819 | 41.89019 | 18.3756 | 0.482844 | 0.913052 | 0.481005 |
| X | 117530600 | 117533399 | 31.03096 | 52.45216 | 47.9735 | 20.99197 | 0.509591 | 1.102679 | 0.481028 |
| 1 | 134289000 | 134289999 | 7.50343 | 14.182 | 27.58766 | 11.21303 | -0.47232 | 0.038559 | 0.481036 |
| 6 | 49743000 | 49743999 | 7.50343 | 14.182 | 27.58766 | 11.21303 | -0.47232 | 0.038559 | 0.481036 |
| 7 | 59368400 | 59368799 | 7.074184 | 13.79334 | 15.24559 | 5.48925 | 0.497059 | -0.41489 | 0.481038 |
| 5 | 116684400 | 116690999 | 82.14621 | 127.2693 | 189.2247 | 78.59869 | -0.26746 | 2.808213 | 0.481046 |
| 8 | 10567400 | 10569599 | 13.60705 | 23.90547 | 41.37794 | 18.31581 | -0.40009 | 0.601626 | 0.481051 |
| 7 | 12711400 | 12713999 | 21.46632 | 35.67559 | 64.3956 | 27.93268 | -0.46815 | 1.255931 | 0.481057 |
| 10 | 119939600 | 119942599 | 41.52522 | 66.55968 | 106.5582 | 44.98355 | -0.33086 | 1.976597 | 0.481058 |
| 9 | 67741200 | 67742599 | 14.1292 | 25.60927 | 24.96316 | 10.80019 | 0.449477 | 0.336538 | 0.481085 |
| 9 | 4496200 | 4496999 | 16.4567 | 29.65796 | 25.04 | 10.91112 | 0.648688 | 0.385051 | 0.481094 |
| 6 | 57284600 | 57288599 | 49.28348 | 79.86226 | 71.06337 | 30.42257 | 0.532668 | 1.665446 | 0.481099 |
| 9 | 48280200 | 48281199 | 12.517 | 23.79004 | 22.5218 | 9.046882 | 0.572699 | 0.031402 | 0.48111 |
| X | 75851800 | 75853599 | 21.89262 | 38.92077 | 29.64311 | 13.07403 | 0.785169 | 0.51449 | 0.481131 |
| 13 | 23525400 | 23527799 | 27.09171 | 46.83441 | 40.18073 | 17.62783 | 0.605779 | 0.881832 | 0.481134 |
| 3 | 108133000 | 108134799 | 11.31083 | 20.00544 | 37.38482 | 16.53472 | -0.50585 | 0.50732 | 0.48115 |
| 3 | 152957600 | 152959599 | 13.11937 | 23.18324 | 39.76779 | 17.59185 | -0.38409 | 0.529557 | 0.481163 |
| 11 | 27178200 | 27179799 | 19.25511 | 33.76551 | 34.43193 | 15.2173 | 0.359483 | 0.651674 | 0.481168 |

| | | | | | | | | | |
|---|---|---|---|---|---|---|---|---|---|
| X | 135583000 | 135588799 | 70.60128 | 112.3178 | 109.0297 | 45.53819 | 0.377735 | 2.241996 | 0.481197 |
| 4 | 23548200 | 23548999 | 18.17817 | 32.31574 | 30.43938 | 13.42574 | 0.480035 | 0.490418 | 0.481215 |
| 7 | 111378200 | 111380999 | 16.45924 | 28.11248 | 53.20602 | 23.3274 | -0.53412 | 0.961347 | 0.481218 |
| 10 | 108871800 | 108872199 | 12.42382 | 23.14255 | 23.57917 | 9.831858 | 0.423543 | 0.166167 | 0.48123 |
| 11 | 35914000 | 35915799 | 32.05072 | 51.89635 | 84.94718 | 36.34525 | -0.34559 | 1.6583 | 0.48124 |
| 16 | 42831000 | 42832399 | 16.05697 | 29.77869 | 25.17151 | 10.56601 | 0.683443 | 0.223506 | 0.48126 |
| 5 | 57772800 | 57775799 | 39.66787 | 64.59102 | 66.90594 | 28.72601 | 0.319823 | 1.558649 | 0.481279 |
| 19 | 34899400 | 34900199 | 5.711394 | 10.98458 | 24.67671 | 9.6213 | -0.62269 | -0.1234 | 0.481289 |
| 3 | 124178400 | 124179599 | 12.88925 | 22.79927 | 39.31278 | 17.38711 | -0.39066 | 0.519619 | 0.481292 |
| 5 | 93730200 | 93730599 | 10.5667 | 20.45542 | 20.38688 | 7.759056 | 0.571683 | -0.16364 | 0.481299 |
| 16 | 72804400 | 72805399 | 5.369038 | 10.454 | 21.87299 | 8.017657 | -0.4368 | -0.34828 | 0.481313 |
| 6 | 109265400 | 109266999 | 11.97719 | 21.59331 | 38.9747 | 16.97505 | -0.44049 | 0.381063 | 0.48133 |
| 2 | 58537000 | 58538199 | 12.38897 | 21.86994 | 38.3352 | 16.96162 | -0.41538 | 0.536147 | 0.481347 |
| 2 | 98699600 | 98700599 | 17.25585 | 30.4321 | 30.85323 | 13.6538 | 0.36953 | 0.559943 | 0.481368 |
| 6 | 6965000 | 6970799 | 62.41343 | 99.90304 | 92.66687 | 39.0381 | 0.452708 | 2.037442 | 0.481369 |
| 6 | 60030200 | 60040999 | 196.4298 | 292.4817 | 250.9543 | 103.406 | 0.477566 | 3.609658 | 0.481373 |
| 4 | 123590600 | 123593799 | 18.11238 | 31.12418 | 52.76325 | 23.14371 | -0.37458 | 0.911744 | 0.481381 |
| 2 | 109397400 | 109397999 | 8.856964 | 16.57526 | 30.64786 | 12.67908 | -0.42146 | 0.121227 | 0.481383 |
| 3 | 54224400 | 54225599 | 8.300097 | 15.62098 | 29.12405 | 11.93414 | -0.42145 | 0.080099 | 0.481397 |
| 1 | 117649600 | 117653999 | 59.34028 | 95.09749 | 91.87778 | 38.77538 | 0.396061 | 1.999497 | 0.481415 |
| 8 | 109183800 | 109185599 | 23.75799 | 41.33403 | 38.67234 | 16.99375 | 0.482766 | 0.798338 | 0.48142 |
| 11 | 14746000 | 14747199 | 19.87676 | 34.77979 | 35.51124 | 15.66545 | 0.357705 | 0.691075 | 0.481435 |
| 6 | 145555400 | 145558199 | 15.07055 | 26.27307 | 44.70671 | 19.67057 | -0.37596 | 0.73604 | 0.481437 |
| 15 | 76817200 | 76819999 | 27.69834 | 44.93719 | 88.65416 | 37.98628 | -0.61156 | 1.612308 | 0.481448 |
| 4 | 68214800 | 68216199 | 12.25804 | 23.50513 | 21.64346 | 8.511471 | 0.639942 | -0.04008 | 0.481484 |
| 8 | 106895000 | 106896599 | 8.206178 | 15.13667 | 30.45921 | 12.88016 | -0.56625 | 0.205039 | 0.481498 |
| 7 | 4139400 | 4143199 | 19.92891 | 33.44075 | 59.60743 | 26.01551 | -0.44997 | 1.141247 | 0.481512 |
| X | 157053800 | 157058199 | 39.52729 | 65.84942 | 51.52706 | 22.38658 | 0.734024 | 1.272908 | 0.48153 |
| 3 | 61402800 | 61404799 | 19.90701 | 35.84453 | 27.65797 | 12.06515 | 0.776888 | 0.420008 | 0.481541 |
| 2 | 36638600 | 36640199 | 40.71206 | 65.97583 | 66.37391 | 28.41604 | 0.356086 | 1.646743 | 0.481543 |
| 3 | 138139000 | 138143199 | 71.36739 | 112.9663 | 117.5944 | 48.88751 | 0.270278 | 2.384447 | 0.48155 |
| 4 | 83045000 | 83047399 | 25.24186 | 44.20718 | 33.91653 | 14.97332 | 0.76805 | 0.70337 | 0.48157 |
| 12 | 22040600 | 22117599 | 1886.891 | 2314.945 | 1892.086 | 797.5311 | 0.269301 | 6.671406 | 0.481571 |
| 17 | 78931400 | 78932799 | 20.19391 | 34.11365 | 56.97814 | 24.92764 | -0.35613 | 1.07947 | 0.481585 |
| 18 | 44687200 | 44691599 | 42.01625 | 68.12377 | 70.70196 | 30.28025 | 0.311929 | 1.640772 | 0.481614 |
| X | 110178800 | 110179399 | 15.91774 | 28.77487 | 28.28992 | 12.28002 | 0.43422 | 0.372948 | 0.481619 |
| 16 | 84463400 | 84467399 | 73.3749 | 116.3351 | 117.5546 | 48.92554 | 0.31513 | 2.344636 | 0.48162 |
| 3 | 18012000 | 18013399 | 23.34515 | 40.02738 | 40.59381 | 17.80328 | 0.363284 | 0.940003 | 0.481644 |
| 15 | 91454800 | 91457799 | 18.84405 | 31.62828 | 69.02248 | 30.12945 | -0.74106 | 1.14264 | 0.481657 |
| 15 | 20622400 | 20623999 | 13.77225 | 25.93732 | 23.96779 | 9.80997 | 0.584329 | 0.108227 | 0.481662 |
| 12 | 14803000 | 14803999 | 7.002824 | 13.52818 | 26.63327 | 10.2353 | -0.41521 | -0.15246 | 0.481688 |
| 11 | 12095200 | 12097199 | 35.05535 | 56.40645 | 96.15118 | 40.82398 | -0.4143 | 1.845438 | 0.481692 |
| 1 | 129523200 | 129523799 | 6.572625 | 12.64694 | 25.25092 | 9.831833 | -0.4525 | -0.11605 | 0.4817 |
| 13 | 38291400 | 38294199 | 21.67491 | 35.93757 | 71.56741 | 30.99316 | -0.60983 | 1.290427 | 0.481703 |
| X | 156404400 | 156407799 | 28.90491 | 47.31424 | 78.37588 | 33.71527 | -0.35033 | 1.47055 | 0.481712 |
| 2 | 30637400 | 30639799 | 13.29853 | 23.45282 | 39.79763 | 17.6121 | -0.36965 | 0.553369 | 0.481731 |
| 5 | 131500600 | 131505999 | 49.93795 | 79.45383 | 125.4355 | 52.39843 | -0.32259 | 2.223395 | 0.481743 |

| | | | | | | | | | |
|---|---|---|---|---|---|---|---|---|---|
| 5 | 109774600 | 109775999 | 18.74773 | 33.20113 | 31.50538 | 13.93081 | 0.466689 | 0.525292 | 0.481751 |
| 12 | 89209000 | 89209999 | 13.15529 | 25.39144 | 20.70599 | 7.989984 | 0.839611 | -0.09111 | 0.481759 |
| 9 | 114475400 | 114477199 | 15.74851 | 27.31366 | 46.87996 | 20.56891 | -0.38922 | 0.813121 | 0.481772 |
| 2 | 94923600 | 94926599 | 44.30764 | 71.59852 | 74.55915 | 31.83885 | 0.302337 | 1.720599 | 0.481798 |
| 12 | 107373200 | 107376199 | 24.55413 | 40.53375 | 68.91208 | 29.75328 | -0.38293 | 1.34617 | 0.481802 |
| 4 | 100600800 | 100602199 | 14.47347 | 26.42069 | 26.19256 | 11.23165 | 0.4331 | 0.299252 | 0.481807 |
| 12 | 51356200 | 51366199 | 275.1272 | 388.5928 | 545.9583 | 226.0217 | -0.30761 | 4.430072 | 0.481815 |
| 19 | 11200800 | 11201399 | 11.74629 | 22.01288 | 22.12477 | 9.131645 | 0.454847 | 0.118387 | 0.481815 |
| 8 | 72826200 | 72833799 | 102.605 | 159.3095 | 157.7739 | 65.51722 | 0.320515 | 2.776179 | 0.481851 |
| 14 | 44388000 | 44390399 | 19.84816 | 33.96593 | 59.91233 | 26.27175 | -0.43268 | 0.966261 | 0.481852 |
| 17 | 72546400 | 72547599 | 19.68779 | 34.49518 | 34.90896 | 15.41715 | 0.370389 | 0.667438 | 0.481888 |
| 13 | 107398800 | 107399599 | 6.858079 | 13.27774 | 27.4309 | 10.43403 | -0.46992 | -0.18629 | 0.481891 |
| 12 | 64108000 | 64108599 | 9.711026 | 18.93654 | 17.32312 | 6.22931 | 0.767933 | -0.37726 | 0.481907 |
| 10 | 59372600 | 59376599 | 29.83551 | 48.62717 | 79.47908 | 34.13609 | -0.33553 | 1.532258 | 0.48192 |
| 8 | 105635800 | 105636799 | 9.006006 | 16.88245 | 31.25624 | 12.89413 | -0.42008 | 0.113151 | 0.481922 |
| 13 | 98399200 | 98400199 | 6.883291 | 13.14365 | 26.15828 | 10.40452 | -0.47714 | -0.04188 | 0.481927 |
| 18 | 6591200 | 6597399 | 70.20663 | 111.7132 | 108.6952 | 45.41157 | 0.374699 | 2.234319 | 0.481935 |
| 7 | 58070800 | 58072199 | 13.59202 | 23.66308 | 49.46133 | 21.74384 | -0.67131 | 0.758028 | 0.481942 |
| 10 | 42035000 | 42038199 | 60.44674 | 96.51507 | 99.54129 | 41.78689 | 0.296097 | 2.114112 | 0.481945 |
| 9 | 98673200 | 98675799 | 19.81092 | 33.75147 | 55.42672 | 24.29607 | -0.33086 | 1.001281 | 0.48195 |
| 3 | 57391600 | 57393999 | 17.64693 | 32.10107 | 26.89291 | 11.58259 | 0.670528 | 0.34472 | 0.48196 |
| 7 | 143435000 | 143435999 | 7.460292 | 14.22766 | 27.24988 | 10.86737 | -0.42617 | -0.02615 | 0.481974 |
| 7 | 20554200 | 20558399 | 42.874 | 70.09473 | 62.66759 | 26.94601 | 0.535809 | 1.509435 | 0.48198 |
| 3 | 3930200 | 3932999 | 26.33765 | 44.47348 | 45.37694 | 19.84963 | 0.352329 | 1.091947 | 0.482012 |
| 5 | 87488000 | 87489199 | 11.95365 | 21.51671 | 38.38692 | 16.75223 | -0.42569 | 0.388271 | 0.482013 |
| 12 | 81741000 | 81742599 | 10.0362 | 18.39592 | 32.40311 | 13.82019 | -0.385 | 0.259022 | 0.482025 |
| 15 | 56877400 | 56878199 | 10.0362 | 18.39592 | 32.40311 | 13.82019 | -0.385 | 0.259022 | 0.482025 |
| X | 22428200 | 22430799 | 31.28024 | 53.42415 | 43.49073 | 19.0678 | 0.678331 | 1.010511 | 0.482037 |
| 3 | 65875400 | 65878799 | 35.01765 | 57.51631 | 60.16616 | 25.91936 | 0.312931 | 1.427489 | 0.48204 |
| 15 | 100260000 | 100261799 | 11.58302 | 20.48391 | 37.81087 | 16.72364 | -0.48861 | 0.50884 | 0.48205 |
| 7 | 57521800 | 57523999 | 23.78363 | 41.22811 | 39.86852 | 17.49231 | 0.434706 | 0.844845 | 0.482061 |
| 15 | 74044400 | 74050599 | 43.16198 | 69.0542 | 111.5365 | 46.95491 | -0.34677 | 2.037633 | 0.482075 |
| 7 | 60040800 | 60041399 | 9.510987 | 18.38401 | 20.35261 | 7.800728 | 0.41154 | -0.15312 | 0.48208 |
| 6 | 98877400 | 98878199 | 6.55695 | 12.61751 | 25.84471 | 10.06135 | -0.48973 | -0.1187 | 0.482083 |
| 11 | 96903400 | 96904799 | 6.55695 | 12.61751 | 25.84471 | 10.06135 | -0.48973 | -0.1187 | 0.482083 |
| 19 | 57758800 | 57761999 | 27.74688 | 46.9652 | 47.43283 | 20.764 | 0.367033 | 1.07733 | 0.48214 |
| 5 | 123352200 | 123358999 | 88.52173 | 139.0936 | 136.0906 | 56.45048 | 0.351318 | 2.556788 | 0.482147 |
| 3 | 20559600 | 20561199 | 15.42008 | 26.62375 | 49.51882 | 21.72374 | -0.50618 | 0.853344 | 0.482176 |
| 16 | 13976600 | 13977799 | 7.210396 | 13.59695 | 29.71421 | 12.13534 | -0.64561 | 0.054298 | 0.482183 |
| 18 | 73071400 | 73073599 | 24.27293 | 41.78117 | 41.53392 | 18.2191 | 0.393046 | 0.911249 | 0.482193 |
| 7 | 44319800 | 44320799 | 11.73297 | 20.81311 | 36.76625 | 16.24576 | -0.42354 | 0.477396 | 0.482202 |
| X | 8720400 | 8724599 | 38.4334 | 64.23382 | 49.77786 | 21.67065 | 0.747869 | 1.235761 | 0.482206 |
| 10 | 75431400 | 75435599 | 47.13826 | 76.29384 | 73.52581 | 31.44875 | 0.416237 | 1.68663 | 0.482223 |
| 12 | 3385000 | 3394999 | 103.7572 | 160.4523 | 164.0236 | 68.13152 | 0.270375 | 2.848496 | 0.482238 |
| 5 | 10054200 | 10055999 | 21.96726 | 38.43913 | 35.80094 | 15.79375 | 0.489627 | 0.701304 | 0.482244 |
| 4 | 4781600 | 4784599 | 44.1848 | 72.23841 | 63.59219 | 27.34366 | 0.558038 | 1.510167 | 0.482255 |
| 1 | 102742400 | 102744199 | 16.11909 | 28.11237 | 48.94132 | 21.53922 | -0.40875 | 0.743338 | 0.482267 |

| | | | | | | | | | |
|---|---|---|---|---|---|---|---|---|---|
| X | 148571200 | 148574399 | 46.60987 | 75.72348 | 61.63515 | 26.43619 | 0.664308 | 1.605386 | 0.482309 |
| 3 | 37741000 | 37742999 | 17.34219 | 29.85993 | 50.05933 | 21.95901 | -0.35783 | 0.88874 | 0.482314 |
| 14 | 60224000 | 60224599 | 12.19464 | 22.68798 | 22.58178 | 9.435423 | 0.454604 | 0.168818 | 0.482326 |
| 11 | 104749400 | 104754999 | 82.92739 | 130.1985 | 135.1344 | 56.04798 | 0.265364 | 2.569235 | 0.482331 |
| 12 | 24746400 | 24747599 | 11.01465 | 20.77369 | 20.62283 | 8.42042 | 0.484773 | 0.066541 | 0.482338 |
| 7 | 110906600 | 110907399 | 11.8168 | 22.08715 | 23.03466 | 9.549413 | 0.39649 | 0.13633 | 0.482339 |
| 2 | 168934200 | 168941199 | 73.86276 | 114.6912 | 181.5531 | 75.39096 | -0.35527 | 2.764352 | 0.482341 |
| X | 18518800 | 18522399 | 43.65796 | 71.92302 | 57.38552 | 24.75281 | 0.704323 | 1.412923 | 0.482348 |
| 2 | 112383400 | 112384399 | 13.81606 | 24.28383 | 40.87203 | 18.0989 | -0.35973 | 0.596601 | 0.482348 |
| X | 8653800 | 8658199 | 50.43505 | 82.12778 | 63.84011 | 27.41026 | 0.733094 | 1.580509 | 0.482364 |
| 8 | 20543800 | 20544399 | 24.19584 | 41.40224 | 42.64875 | 18.70148 | 0.340412 | 0.964219 | 0.482367 |
| 9 | 114974200 | 114992599 | 361.1761 | 497.9269 | 693.5566 | 288.2256 | -0.33039 | 4.796792 | 0.482374 |
| 9 | 67580800 | 67581599 | 10.22552 | 18.65153 | 32.87267 | 14.11006 | -0.39208 | 0.2902 | 0.482375 |
| 13 | 22141000 | 22144599 | 65.25735 | 103.9379 | 105.9961 | 44.34294 | 0.308485 | 2.198603 | 0.482387 |
| 13 | 67034600 | 67041999 | 127.7981 | 195.8189 | 186.4108 | 77.37269 | 0.362965 | 3.032848 | 0.482389 |
| X | 40004400 | 40007799 | 32.59642 | 56.09399 | 38.97732 | 17.09749 | 0.907568 | 0.946105 | 0.482394 |
| 16 | 17130400 | 17133399 | 31.44199 | 52.5544 | 52.47509 | 22.84635 | 0.38326 | 1.211697 | 0.482396 |
| 6 | 52241000 | 52243599 | 22.30643 | 37.35809 | 62.5277 | 27.25686 | -0.35958 | 1.171582 | 0.482405 |
| 9 | 101323000 | 101327999 | 31.17249 | 50.59111 | 83.79839 | 35.91487 | -0.35958 | 1.607107 | 0.482431 |
| X | 74653200 | 74657199 | 45.08469 | 73.57768 | 55.97945 | 24.05458 | 0.766156 | 1.534454 | 0.48244 |
| 12 | 115497000 | 115499999 | 48.61964 | 78.9231 | 68.83418 | 29.5059 | 0.563809 | 1.629059 | 0.482458 |
| 6 | 58067200 | 58069599 | 26.63719 | 43.92959 | 72.02173 | 31.08008 | -0.33118 | 1.365074 | 0.482458 |
| 15 | 97119000 | 97122799 | 43.00536 | 68.88766 | 108.3488 | 45.69457 | -0.30647 | 1.999463 | 0.482471 |
| 5 | 96637000 | 96640399 | 39.45189 | 63.41909 | 101.0096 | 42.82639 | -0.3184 | 1.884015 | 0.4825 |
| 12 | 82295400 | 82297799 | 16.78612 | 29.07225 | 50.08429 | 21.97388 | -0.3953 | 0.83182 | 0.482519 |
| 9 | 31924800 | 31925999 | 14.80281 | 27.17192 | 26.82349 | 11.41809 | 0.446768 | 0.274111 | 0.482522 |
| 2 | 28124600 | 28129799 | 55.77717 | 88.33752 | 138.1276 | 57.43985 | -0.31519 | 2.36527 | 0.482525 |
| 9 | 3955600 | 3960599 | 63.07513 | 100.7926 | 98.3054 | 41.31499 | 0.377802 | 2.092147 | 0.482529 |
| 11 | 69810400 | 69812599 | 11.80134 | 20.71919 | 40.00262 | 17.69698 | -0.55644 | 0.602149 | 0.482535 |
| 5 | 136705000 | 136707999 | 29.75218 | 48.30696 | 81.06408 | 34.75409 | -0.37766 | 1.595833 | 0.482542 |
| 14 | 49113000 | 49117799 | 32.85038 | 53.22692 | 85.27116 | 36.50194 | -0.31375 | 1.6462 | 0.482542 |
| 12 | 18592800 | 18638399 | 1551.817 | 1933.133 | 1597.628 | 672.183 | 0.275653 | 6.427552 | 0.482545 |
| 6 | 14851200 | 14852399 | 15.93295 | 29.25818 | 26.70472 | 11.3613 | 0.559691 | 0.280057 | 0.482549 |
| 6 | 86251000 | 86254999 | 19.74457 | 33.39046 | 56.39637 | 24.68116 | -0.37203 | 1.069321 | 0.482571 |
| 4 | 10678400 | 10679799 | 9.565781 | 17.67258 | 31.47434 | 13.28068 | -0.3894 | 0.209441 | 0.482576 |
| 11 | 49859600 | 49867599 | 63.76742 | 99.78466 | 163.4306 | 67.79766 | -0.39562 | 2.623005 | 0.482591 |
| 19 | 39587000 | 39588199 | 33.94124 | 55.99542 | 58.07381 | 25.06484 | 0.327764 | 1.368169 | 0.482597 |
| 12 | 111921600 | 111923399 | 7.917946 | 14.57295 | 31.97794 | 13.56287 | -0.69388 | 0.216383 | 0.482612 |
| 13 | 88908600 | 88908999 | 7.921296 | 15.1218 | 28.42982 | 11.31424 | -0.39701 | -0.02479 | 0.482625 |
| X | 93031400 | 93032599 | 9.247529 | 17.28252 | 31.71092 | 13.1493 | -0.41279 | 0.138751 | 0.482645 |
| 16 | 4030600 | 4033199 | 14.13161 | 24.58798 | 45.79502 | 20.12462 | -0.50601 | 0.759906 | 0.482647 |
| 5 | 109960200 | 109962199 | 33.94865 | 56.49789 | 54.16286 | 23.51633 | 0.441981 | 1.268922 | 0.482659 |
| 16 | 36524600 | 36526399 | 13.78949 | 24.23882 | 40.85366 | 18.09132 | -0.3618 | 0.594515 | 0.482663 |
| 13 | 98826400 | 98827799 | 9.737794 | 17.86138 | 31.90542 | 13.59504 | -0.40394 | 0.252272 | 0.482671 |
| 17 | 95119800 | 95134199 | 301.1456 | 435.1182 | 366.8869 | 151.3434 | 0.4615 | 4.09898 | 0.482678 |
| 6 | 132171200 | 132173799 | 44.55841 | 72.92251 | 62.48744 | 26.87849 | 0.597835 | 1.496844 | 0.482689 |
| 12 | 18513000 | 18549599 | 1340.134 | 1690.359 | 1422.443 | 597.6653 | 0.267781 | 6.227092 | 0.482691 |

| | | | | | | | | | |
|---|---|---|---|---|---|---|---|---|---|
| 6 | 76297200 | 76303199 | 63.16631 | 101.2564 | 88.94447 | 37.5529 | 0.533595 | 1.993491 | 0.482697 |
| 4 | 79959600 | 79961399 | 19.27273 | 34.51883 | 29.33783 | 12.85853 | 0.633788 | 0.450936 | 0.482697 |
| 19 | 39503000 | 39508199 | 76.53295 | 121.8565 | 103.9262 | 43.45708 | 0.56549 | 2.219414 | 0.482718 |
| 1 | 15302800 | 15307199 | 45.96564 | 74.47626 | 72.36506 | 30.97691 | 0.405868 | 1.662349 | 0.482729 |
| 9 | 98746400 | 98747999 | 39.15298 | 62.67695 | 107.8628 | 45.446 | -0.43706 | 2.015264 | 0.482729 |
| 7 | 16841200 | 16847799 | 78.71369 | 124.8142 | 116.6937 | 48.57289 | 0.427236 | 2.34574 | 0.482733 |
| 7 | 14216800 | 14219999 | 43.21595 | 71.51759 | 53.9473 | 23.33202 | 0.786278 | 1.35826 | 0.482743 |
| 18 | 90084800 | 90085399 | 12.68553 | 23.50466 | 24.50805 | 10.3003 | 0.381647 | 0.20062 | 0.482747 |
| 19 | 23960000 | 23960999 | 12.68553 | 23.50466 | 24.50805 | 10.3003 | 0.381647 | 0.20062 | 0.482747 |
| 7 | 133648200 | 133649999 | 11.84921 | 21.23338 | 36.26094 | 15.88618 | -0.36653 | 0.413282 | 0.482756 |
| 6 | 136378200 | 136378999 | 13.33328 | 23.41587 | 40.40971 | 17.88166 | -0.3956 | 0.606122 | 0.482758 |
| 11 | 88137000 | 88138199 | 12.4029 | 21.74031 | 41.52241 | 18.34437 | -0.54119 | 0.632672 | 0.482759 |
| 1 | 40829400 | 40830399 | 7.324648 | 13.97382 | 26.69349 | 10.63766 | -0.42115 | -0.03013 | 0.482764 |
| 7 | 65729200 | 65730199 | 7.324648 | 13.97382 | 26.69349 | 10.63766 | -0.42115 | -0.03013 | 0.482764 |
| 15 | 12709600 | 12712799 | 26.94923 | 45.40409 | 46.95745 | 20.52702 | 0.332722 | 1.113994 | 0.482774 |
| 9 | 115202000 | 115204799 | 38.19137 | 62.6377 | 62.62783 | 26.96319 | 0.37714 | 1.454937 | 0.482793 |
| 2 | 25607400 | 25619999 | 303.9817 | 425.0366 | 600.1282 | 248.8468 | -0.32942 | 4.580947 | 0.482801 |
| 3 | 32942000 | 32945799 | 27.8872 | 47.04872 | 48.00125 | 20.99201 | 0.352292 | 1.105791 | 0.48283 |
| X | 64101200 | 64104999 | 62.84398 | 100.5987 | 92.15518 | 38.82609 | 0.470834 | 2.040492 | 0.482845 |
| 8 | 58945200 | 58945799 | 14.36922 | 26.22127 | 26.88963 | 11.53557 | 0.384578 | 0.304564 | 0.482863 |
| 4 | 5719200 | 5721399 | 19.97263 | 35.25958 | 32.55467 | 14.4038 | 0.504393 | 0.568906 | 0.482874 |
| X | 22410000 | 22412399 | 29.69588 | 51.06315 | 41.30986 | 18.12024 | 0.688826 | 0.943225 | 0.482886 |
| 11 | 99433600 | 99433999 | 4.644888 | 9.066334 | 19.84421 | 7.032251 | -0.45612 | -0.48523 | 0.482891 |
| 3 | 8424200 | 8426399 | 16.18572 | 29.46032 | 26.43512 | 11.37706 | 0.572521 | 0.328425 | 0.482903 |
| 8 | 97146800 | 97149399 | 23.30258 | 40.72194 | 36.30477 | 15.99816 | 0.552476 | 0.729968 | 0.482934 |
| 14 | 106371200 | 106374799 | 53.15698 | 85.56415 | 82.42998 | 35.01657 | 0.407832 | 1.85392 | 0.482948 |
| 9 | 95250400 | 95252399 | 11.90847 | 21.49014 | 45.6697 | 19.86699 | -0.67375 | 0.388514 | 0.482953 |
| 3 | 153779600 | 153780399 | 6.415743 | 12.37203 | 24.75744 | 9.5795 | -0.44726 | -0.14021 | 0.48296 |
| 19 | 57260200 | 57261999 | 12.34031 | 21.80567 | 38.77794 | 17.15412 | -0.43576 | 0.523424 | 0.482985 |
| X | 93679600 | 93680799 | 10.2208 | 18.90791 | 35.89496 | 15.11851 | -0.47938 | 0.216501 | 0.482988 |
| 18 | 37565800 | 37571599 | 133.2609 | 203.0848 | 172.7167 | 71.59844 | 0.519311 | 3.142548 | 0.482992 |
| 12 | 32833200 | 32833999 | 6.410144 | 12.45086 | 27.76447 | 10.37878 | -0.555 | -0.24872 | 0.482995 |
| 13 | 93716000 | 93720199 | 60.13823 | 95.98222 | 99.71809 | 41.83625 | 0.284914 | 2.126856 | 0.483 |
| 8 | 4993200 | 4996799 | 61.8222 | 98.49468 | 101.9319 | 42.65956 | 0.287744 | 2.183722 | 0.483011 |
| 11 | 17424400 | 17427199 | 32.36518 | 54.43916 | 49.11489 | 21.4594 | 0.528957 | 1.147888 | 0.483016 |
| 9 | 36678400 | 36679799 | 8.607544 | 16.15286 | 29.61138 | 12.19335 | -0.40312 | 0.102077 | 0.483048 |
| 1 | 110356800 | 110357799 | 22.0023 | 36.48392 | 65.31896 | 28.28908 | -0.45659 | 1.287171 | 0.483056 |
| 11 | 59891800 | 59893599 | 12.20707 | 21.51164 | 39.3429 | 17.4135 | -0.4775 | 0.553886 | 0.48306 |
| 12 | 102309000 | 102312599 | 27.22816 | 44.51717 | 75.95906 | 32.66161 | -0.39415 | 1.479015 | 0.483071 |
| 9 | 8354400 | 8356199 | 19.05552 | 34.06832 | 29.97684 | 13.15858 | 0.582688 | 0.46276 | 0.483081 |
| 3 | 53446400 | 53449799 | 38.70471 | 63.45732 | 62.24195 | 26.79307 | 0.404489 | 1.458014 | 0.483086 |
| 13 | 74955600 | 74957599 | 12.69945 | 22.39026 | 38.83764 | 17.18819 | -0.4011 | 0.552289 | 0.483089 |
| 3 | 106544000 | 106545799 | 21.36651 | 37.75389 | 31.89992 | 14.1116 | 0.632181 | 0.567221 | 0.483092 |
| X | 78741200 | 78743599 | 20.5118 | 36.67284 | 29.24508 | 12.8371 | 0.723786 | 0.470321 | 0.483103 |
| 7 | 48159400 | 48160999 | 22.33387 | 38.62839 | 39.93728 | 17.52129 | 0.337942 | 0.856676 | 0.483127 |
| 18 | 53285000 | 53285599 | 7.150106 | 13.76977 | 26.46853 | 10.28091 | -0.39568 | -0.1166 | 0.483138 |
| 18 | 21229000 | 21231399 | 26.69003 | 45.04339 | 46.4251 | 20.3047 | 0.33778 | 1.099249 | 0.483151 |

| | | | | | | | | | |
|---|---|---|---|---|---|---|---|---|---|
| 10 | 41833200 | 41835399 | 26.24391 | 43.23718 | 71.64013 | 30.90215 | -0.34687 | 1.379432 | 0.483153 |
| 17 | 38546600 | 38554799 | 127.1319 | 196.2866 | 161.7176 | 67.17423 | 0.579565 | 2.886576 | 0.483186 |
| 4 | 113173600 | 113191599 | 358.9704 | 507.9088 | 456.0237 | 188.7346 | 0.340715 | 4.407348 | 0.483191 |
| 6 | 14567800 | 14572399 | 32.16557 | 52.26791 | 83.43875 | 35.7936 | -0.30477 | 1.585388 | 0.483197 |
| 10 | 104669600 | 104673199 | 71.99483 | 113.8501 | 117.9128 | 48.99609 | 0.276504 | 2.407056 | 0.483199 |
| 1 | 17590400 | 17591799 | 13.33483 | 24.46468 | 24.31887 | 10.35921 | 0.435893 | 0.258274 | 0.483215 |
| 15 | 43625200 | 43627599 | 19.62477 | 34.81971 | 30.23692 | 13.35829 | 0.595056 | 0.510892 | 0.483215 |
| 3 | 109020000 | 109021799 | 23.64207 | 41.13803 | 38.61741 | 16.97108 | 0.477914 | 0.794508 | 0.483259 |
| 17 | 22340400 | 22342199 | 19.24195 | 34.39213 | 29.9716 | 13.15935 | 0.596304 | 0.465841 | 0.483287 |
| X | 65312200 | 65313999 | 18.70924 | 34.2852 | 25.91972 | 11.05849 | 0.826971 | 0.311502 | 0.483296 |
| 10 | 61201400 | 61204199 | 13.50755 | 23.55803 | 44.2032 | 19.45411 | -0.51618 | 0.720907 | 0.483303 |
| 7 | 36060000 | 36061799 | 22.08103 | 38.65593 | 35.83063 | 15.81297 | 0.496584 | 0.695912 | 0.483309 |
| 14 | 6123200 | 6124399 | 10.48193 | 20.21978 | 20.75198 | 8.022155 | 0.507623 | -0.1153 | 0.483316 |
| 3 | 44549600 | 44551399 | 18.54455 | 32.93426 | 27.49709 | 12.13818 | 0.65238 | 0.494497 | 0.483324 |
| 1 | 89780400 | 89784999 | 23.64939 | 39.37522 | 65.27644 | 28.3498 | -0.34575 | 1.241422 | 0.483329 |
| 4 | 134143600 | 134148199 | 29.6862 | 47.95782 | 94.9843 | 40.54644 | -0.62337 | 1.715632 | 0.48333 |
| 2 | 118384400 | 118385999 | 21.17585 | 36.72291 | 36.91521 | 16.19675 | 0.379127 | 0.824095 | 0.483356 |
| 4 | 81181200 | 81181999 | 14.1573 | 24.6962 | 43.68767 | 19.22965 | -0.43145 | 0.722426 | 0.483356 |
| 5 | 56521000 | 56523799 | 32.97405 | 55.32581 | 49.70605 | 21.69292 | 0.534944 | 1.173024 | 0.483361 |
| 3 | 21576800 | 21577599 | 16.73036 | 30.03719 | 24.27453 | 10.61785 | 0.708098 | 0.409362 | 0.48337 |
| 6 | 22603800 | 22604799 | 13.32376 | 24.4814 | 25.44132 | 10.81504 | 0.374607 | 0.254112 | 0.483388 |
| 2 | 88614400 | 88620399 | 65.97992 | 105.2497 | 102.0209 | 42.77039 | 0.383944 | 2.148744 | 0.483388 |
| 1 | 60716600 | 60717399 | 12.94396 | 25.11415 | 18.54002 | 6.98198 | 1.014435 | -0.17841 | 0.483396 |
| 3 | 156901200 | 156901799 | 10.046 | 19.46275 | 18.55625 | 7.036391 | 0.638448 | -0.19166 | 0.483396 |
| 15 | 37016200 | 37016799 | 5.21169 | 10.16196 | 21.48598 | 7.734793 | -0.42783 | -0.41542 | 0.483401 |
| 2 | 71605600 | 71608199 | 21.79778 | 36.27309 | 63.27355 | 27.47035 | -0.41877 | 1.242997 | 0.483422 |
| X | 23194600 | 23195999 | 19.12882 | 34.48166 | 27.57561 | 12.0125 | 0.727341 | 0.411693 | 0.483466 |
| 9 | 13348200 | 13349399 | 12.83719 | 24.49594 | 22.65254 | 9.023084 | 0.617206 | 0.009467 | 0.483477 |
| 18 | 77456600 | 77462799 | 62.72222 | 98.82669 | 150.9706 | 62.64511 | -0.28757 | 2.489113 | 0.483478 |
| 3 | 60225400 | 60226599 | 10.842 | 19.54156 | 35.27521 | 15.36883 | -0.4408 | 0.366508 | 0.483486 |
| 16 | 44329000 | 44330399 | 8.667946 | 15.89973 | 31.41971 | 13.38727 | -0.54852 | 0.241558 | 0.48349 |
| 9 | 36503400 | 36505399 | 22.4898 | 39.48835 | 34.07005 | 15.07366 | 0.599429 | 0.649565 | 0.483499 |
| 10 | 24410800 | 24411399 | 6.118308 | 11.81936 | 24.26544 | 9.325708 | -0.47474 | -0.16887 | 0.483505 |
| 3 | 3975400 | 3978999 | 102.5057 | 158.2387 | 159.6442 | 66.31304 | 0.28742 | 2.879499 | 0.483509 |
| 8 | 25873200 | 25874199 | 8.568305 | 15.97527 | 30.55881 | 12.72246 | -0.47742 | 0.142311 | 0.483509 |
| 6 | 44402200 | 44404799 | 28.43493 | 48.17701 | 47.41783 | 20.7636 | 0.404169 | 1.071839 | 0.483519 |
| 4 | 123666600 | 123667599 | 11.16781 | 21.63856 | 19.75181 | 7.485781 | 0.702154 | -0.17533 | 0.483526 |
| 14 | 48257800 | 48260199 | 16.01313 | 27.97963 | 50.57558 | 22.28453 | -0.46269 | 0.715612 | 0.483531 |
| 2 | 14815800 | 14818399 | 16.27853 | 27.56023 | 60.0091 | 26.27127 | -0.73687 | 1.050652 | 0.483537 |
| 7 | 12351400 | 12364599 | 192.7141 | 288.3029 | 255.8141 | 105.4944 | 0.435339 | 3.52423 | 0.483549 |
| 2 | 149178800 | 149179799 | 19.27108 | 33.73428 | 34.57618 | 15.25842 | 0.352258 | 0.680421 | 0.483551 |
| 9 | 6593400 | 6601799 | 240.0409 | 349.473 | 340.6321 | 140.3871 | 0.263168 | 3.980274 | 0.483551 |
| X | 77822600 | 77824599 | 32.45784 | 54.94021 | 46.85638 | 20.51178 | 0.610191 | 1.09591 | 0.483556 |
| 6 | 136612600 | 136614599 | 16.18368 | 28.16929 | 46.82992 | 20.5844 | -0.34287 | 0.766661 | 0.483576 |
| X | 8085400 | 8090399 | 51.58635 | 83.37048 | 70.95592 | 30.30485 | 0.593026 | 1.732608 | 0.483587 |
| 10 | 21087000 | 21088799 | 21.50152 | 37.21054 | 36.95157 | 16.21168 | 0.396075 | 0.844104 | 0.483595 |
| 1 | 44426400 | 44429199 | 21.27012 | 35.9094 | 60.38335 | 26.41243 | -0.36603 | 1.090038 | 0.483599 |

| | | | | | | | | | |
|---|---|---|---|---|---|---|---|---|---|
| 9 | 91431800 | 91433199 | 13.94884 | 25.48066 | 26.3547 | 11.29137 | 0.373357 | 0.293138 | 0.483625 |
| 1 | 89895200 | 89900399 | 38.99163 | 62.36104 | 111.8904 | 47.08135 | -0.49862 | 2.043406 | 0.48363 |
| 10 | 124623600 | 124626999 | 27.57506 | 47.68759 | 40.20387 | 17.63818 | 0.631078 | 0.882174 | 0.483644 |
| 9 | 35981000 | 35984599 | 41.56534 | 68.57274 | 56.86693 | 24.54383 | 0.649467 | 1.387464 | 0.483651 |
| 1 | 178233200 | 178234199 | 6.018768 | 11.50893 | 25.6385 | 10.16383 | -0.63354 | -0.06267 | 0.483655 |
| 14 | 121393400 | 121394599 | 18.96658 | 33.21334 | 33.13571 | 14.62724 | 0.39107 | 0.671221 | 0.483658 |
| 17 | 30128000 | 30129399 | 19.73478 | 34.59573 | 34.9016 | 15.42076 | 0.374894 | 0.65986 | 0.483658 |
| 7 | 14777200 | 14834999 | 1153.263 | 1481.919 | 1171.886 | 491.4794 | 0.384419 | 5.926235 | 0.48368 |
| 13 | 105982800 | 105984799 | 18.0767 | 32.3102 | 29.97365 | 13.15998 | 0.50645 | 0.456347 | 0.48369 |
| 1 | 34106400 | 34107999 | 11.64254 | 20.90955 | 36.25403 | 15.85336 | -0.38659 | 0.396312 | 0.483732 |
| 17 | 87731400 | 87733799 | 17.52449 | 30.19214 | 50.05861 | 21.95911 | -0.34169 | 0.88486 | 0.48376 |
| 3 | 12786200 | 12792999 | 62.86724 | 100.6161 | 94.3716 | 39.74876 | 0.436529 | 2.04551 | 0.483762 |
| 14 | 111580600 | 111583599 | 23.58579 | 40.98532 | 39.26195 | 17.23985 | 0.448716 | 0.814959 | 0.483785 |
| Y | 41502200 | 41504199 | 17.49461 | 30.96177 | 20.62738 | 9.122043 | 0.973413 | 0.500235 | 0.483801 |
| 1 | 46352200 | 46353199 | 23.30351 | 39.95524 | 41.08225 | 18.01747 | 0.343456 | 0.939879 | 0.483817 |
| 18 | 46963000 | 46963799 | 17.86632 | 31.56958 | 31.79059 | 14.0632 | 0.380391 | 0.545846 | 0.483826 |
| 13 | 7872000 | 7873799 | 16.75888 | 30.51836 | 27.59904 | 11.87077 | 0.562249 | 0.334493 | 0.483828 |
| 1 | 148231800 | 148232999 | 18.09659 | 32.03373 | 31.60287 | 13.97516 | 0.410662 | 0.526273 | 0.48383 |
| 5 | 117443000 | 117447799 | 38.18747 | 61.52357 | 100.0866 | 42.57244 | -0.34493 | 1.808543 | 0.483838 |
| 2 | 174797000 | 174798599 | 20.30958 | 35.3566 | 36.25045 | 15.93576 | 0.351311 | 0.76805 | 0.483852 |
| 12 | 69740200 | 69741799 | 12.67775 | 22.41676 | 38.27031 | 16.92732 | -0.37669 | 0.51878 | 0.483854 |
| 7 | 40837600 | 40850799 | 205.6462 | 305.2509 | 284.3037 | 117.1104 | 0.355145 | 3.666496 | 0.483858 |
| 4 | 56654400 | 56655599 | 9.537547 | 17.54207 | 31.92586 | 13.55313 | -0.42705 | 0.233443 | 0.483888 |
| 13 | 98566400 | 98568999 | 18.24571 | 30.96095 | 54.79335 | 24.0044 | -0.43898 | 1.030577 | 0.483906 |
| 3 | 144726400 | 144728799 | 31.0913 | 51.85786 | 53.03051 | 23.05721 | 0.348983 | 1.233087 | 0.483912 |
| 1 | 88682200 | 88686399 | 37.09026 | 59.70889 | 96.77595 | 41.11733 | -0.34096 | 1.831261 | 0.483916 |
| 2 | 33760400 | 33765999 | 39.73525 | 63.76678 | 103.0043 | 43.56241 | -0.34166 | 1.936128 | 0.483928 |
| 4 | 112493800 | 112496199 | 36.21285 | 60.35547 | 52.63888 | 22.87624 | 0.577902 | 1.255871 | 0.483941 |
| 3 | 11034800 | 11035199 | 8.839385 | 17.08423 | 17.74377 | 6.80385 | 0.502443 | -0.16722 | 0.483967 |
| 13 | 50939600 | 50943599 | 25.61672 | 42.42305 | 70.43708 | 30.47879 | -0.34862 | 1.311999 | 0.483975 |
| 8 | 108325400 | 108328399 | 20.50805 | 34.57822 | 58.08033 | 25.39513 | -0.3644 | 1.097489 | 0.483993 |
| 5 | 94510200 | 94520399 | 275.6139 | 400.1282 | 356.1247 | 146.8154 | 0.390271 | 4.026195 | 0.484002 |
| 2 | 181567000 | 181570399 | 53.22159 | 85.32611 | 88.78526 | 37.49315 | 0.289921 | 1.978561 | 0.484015 |
| 17 | 11185800 | 11187799 | 19.13013 | 33.61641 | 33.17784 | 14.68892 | 0.40617 | 0.614123 | 0.484029 |
| 5 | 92651400 | 92654399 | 19.59998 | 32.8905 | 61.27336 | 26.74347 | -0.51352 | 1.139844 | 0.484031 |
| X | 82282200 | 82283999 | 16.54484 | 31.12103 | 21.99185 | 9.019801 | 0.965931 | 0.135183 | 0.484033 |
| 17 | 94458600 | 94464599 | 69.79898 | 111.5276 | 95.14396 | 39.98147 | 0.570505 | 2.100632 | 0.484039 |
| 7 | 11760400 | 11763199 | 76.60046 | 121.2203 | 121.2974 | 50.42082 | 0.326938 | 2.395774 | 0.48405 |
| 2 | 74484600 | 74492399 | 61.90984 | 96.29821 | 195.7699 | 81.27349 | -0.71381 | 2.730816 | 0.484054 |
| 6 | 117772200 | 117773399 | 21.01411 | 36.55806 | 37.77825 | 16.60023 | 0.339945 | 0.783798 | 0.484059 |
| 8 | 14753000 | 14754799 | 11.79376 | 21.13185 | 37.26321 | 16.32669 | -0.41287 | 0.413202 | 0.484067 |
| 19 | 56641800 | 56646199 | 41.2061 | 66.00337 | 107.9615 | 45.52897 | -0.36295 | 1.997615 | 0.484083 |
| 14 | 93034600 | 93036599 | 22.85088 | 40.09316 | 33.99033 | 15.02915 | 0.624538 | 0.661483 | 0.484099 |
| 12 | 52481400 | 52484199 | 19.49994 | 33.10348 | 55.38201 | 24.26545 | -0.35822 | 1.031971 | 0.48411 |
| 15 | 23568200 | 23571599 | 61.36482 | 97.78755 | 101.8192 | 42.62538 | 0.279304 | 2.176633 | 0.484131 |
| 4 | 71045400 | 71045999 | 5.501761 | 10.70998 | 23.0085 | 8.449491 | -0.47658 | -0.33414 | 0.484166 |
| 2 | 87233200 | 87235999 | 40.67079 | 66.21458 | 67.57514 | 29.01182 | 0.341084 | 1.563347 | 0.484174 |

| | | | | | | | | | |
|---|---|---|---|---|---|---|---|---|---|
| 10 | 22138800 | 22144999 | 106.5243 | 165.6511 | 138.6206 | 57.5503 | 0.564992 | 2.747986 | 0.484209 |
| 17 | 30003400 | 30006799 | 36.37553 | 59.42237 | 61.08159 | 26.25633 | 0.334157 | 1.502483 | 0.484212 |
| 10 | 103459800 | 103461199 | 9.718683 | 17.74641 | 32.87455 | 14.09161 | -0.46242 | 0.277843 | 0.48422 |
| 4 | 15749000 | 15751999 | 48.31165 | 77.9809 | 69.74323 | 29.73909 | 0.519492 | 1.756458 | 0.484222 |
| 12 | 117560400 | 117561799 | 15.19893 | 28.22872 | 25.25168 | 10.57856 | 0.604287 | 0.208515 | 0.484237 |
| 7 | 64310200 | 64313399 | 19.97179 | 34.04349 | 55.9862 | 24.54236 | -0.33281 | 0.997658 | 0.484246 |
| 16 | 92683000 | 92684799 | 13.15966 | 23.21195 | 39.30488 | 17.39343 | -0.36639 | 0.550542 | 0.484276 |
| 5 | 125854600 | 125867399 | 159.3687 | 236.538 | 337.6719 | 139.0927 | -0.26073 | 3.663874 | 0.484276 |
| 8 | 34227200 | 34229799 | 16.01567 | 27.97569 | 48.39042 | 21.31769 | -0.39942 | 0.717318 | 0.484334 |
| 10 | 9960600 | 9960999 | 4.479235 | 8.74991 | 19.36927 | 6.771029 | -0.4538 | -0.53612 | 0.484356 |
| 12 | 101334000 | 101341199 | 81.27672 | 128.3945 | 123.6562 | 51.3556 | 0.379991 | 2.437821 | 0.484365 |
| 6 | 148344400 | 148346399 | 13.23562 | 22.99364 | 47.91092 | 21.03411 | -0.6674 | 0.780013 | 0.484369 |
| 11 | 22475000 | 22476199 | 15.59457 | 28.51852 | 27.21813 | 11.64512 | 0.489871 | 0.301222 | 0.484372 |
| 6 | 52281400 | 52281799 | 4.226748 | 8.251108 | 19.25289 | 6.809314 | -0.54407 | -0.49988 | 0.484378 |
| 11 | 57276200 | 57277399 | 14.66177 | 26.9299 | 26.22619 | 11.15432 | 0.466871 | 0.266814 | 0.484386 |
| 12 | 8345000 | 8357199 | 99.26704 | 150.0938 | 286.5637 | 118.5084 | -0.65581 | 3.300751 | 0.48439 |
| 13 | 113497800 | 113498999 | 12.25654 | 23.40721 | 22.72723 | 9.037081 | 0.549803 | -0.0006 | 0.484441 |
| 3 | 113361600 | 113362199 | 14.7729 | 25.80924 | 43.63117 | 19.22313 | -0.36616 | 0.703487 | 0.484464 |
| 16 | 20018400 | 20021199 | 25.66205 | 44.23742 | 41.92855 | 18.39303 | 0.461874 | 0.903196 | 0.484465 |
| 2 | 77061600 | 77062199 | 7.033145 | 13.47431 | 26.24146 | 10.34596 | -0.43409 | -0.06765 | 0.48447 |
| 10 | 46516600 | 46517599 | 8.65359 | 16.1583 | 30.59435 | 12.70521 | -0.45904 | 0.138428 | 0.484474 |
| 7 | 85371000 | 85372999 | 20.9188 | 36.46179 | 37.21746 | 16.37345 | 0.35793 | 0.756611 | 0.484484 |
| 4 | 131880200 | 131886799 | 62.88152 | 99.15027 | 150.2002 | 62.33137 | -0.27468 | 2.474805 | 0.484489 |
| 7 | 9998600 | 10004799 | 77.53195 | 123.6719 | 96.65933 | 40.52025 | 0.693988 | 2.163015 | 0.484493 |
| 14 | 53655200 | 53656599 | 10.34346 | 19.02286 | 33.52169 | 14.23213 | -0.38097 | 0.244754 | 0.484512 |
| 19 | 59074200 | 59077399 | 33.61171 | 54.50997 | 87.6497 | 37.54526 | -0.31791 | 1.628814 | 0.484517 |
| 15 | 87137000 | 87146199 | 82.6918 | 127.9515 | 192.589 | 79.99666 | -0.28657 | 2.831563 | 0.484519 |
| 6 | 59830400 | 59838599 | 225.7393 | 331.4734 | 319.7838 | 131.705 | 0.289812 | 3.846977 | 0.484569 |
| 17 | 76249200 | 76250799 | 17.81998 | 32.31952 | 28.023 | 12.11432 | 0.61804 | 0.366281 | 0.484574 |
| 7 | 13949600 | 13958599 | 154.2392 | 235.0982 | 173.4899 | 71.9225 | 0.724097 | 3.145182 | 0.484593 |
| 7 | 41652400 | 41654199 | 29.81441 | 50.27925 | 49.08732 | 21.46427 | 0.415634 | 1.118447 | 0.484597 |
| 3 | 90903600 | 90914799 | 160.7477 | 243.4314 | 217.7556 | 90.09691 | 0.439056 | 3.280464 | 0.484628 |
| 15 | 91876200 | 91877999 | 19.18633 | 33.8993 | 32.00202 | 14.15695 | 0.472811 | 0.553036 | 0.484636 |
| 12 | 34340400 | 34341199 | 12.06458 | 21.36677 | 37.28669 | 16.48697 | -0.40709 | 0.496359 | 0.484647 |
| 11 | 81317600 | 81322199 | 33.66656 | 54.47271 | 86.93961 | 37.17759 | -0.31026 | 1.676586 | 0.484661 |
| 9 | 19356800 | 19365599 | 105.0152 | 164.4774 | 136.6793 | 56.69491 | 0.583074 | 2.627309 | 0.484661 |
| 17 | 49780000 | 49781999 | 14.25306 | 24.87699 | 44.6841 | 19.67508 | -0.45353 | 0.713762 | 0.484674 |
| 13 | 61703400 | 61713199 | 140.8422 | 216.2253 | 173.2623 | 71.93465 | 0.613425 | 3.000477 | 0.484695 |
| 8 | 45295800 | 45299599 | 29.85705 | 50.36503 | 48.53983 | 21.22665 | 0.434165 | 1.114224 | 0.484707 |
| 16 | 47037400 | 47040399 | 35.63538 | 60.14571 | 45.72155 | 19.99757 | 0.775387 | 1.128245 | 0.484721 |
| 9 | 122672400 | 122693799 | 364.9774 | 515.0711 | 468.079 | 193.8069 | 0.319518 | 4.445283 | 0.484742 |
| 4 | 96209400 | 96210399 | 12.60648 | 23.41831 | 24.45513 | 10.24218 | 0.382873 | 0.184535 | 0.484744 |
| 9 | 83956000 | 83957199 | 8.075442 | 15.43602 | 30.26928 | 12.01215 | -0.4539 | -0.02838 | 0.484745 |
| 8 | 89423400 | 89424199 | 7.551099 | 14.1096 | 29.01881 | 12.03659 | -0.57631 | 0.11714 | 0.484758 |
| 17 | 20238200 | 20243199 | 81.39625 | 128.9933 | 117.6077 | 48.92503 | 0.462641 | 2.364928 | 0.48479 |
| X | 36799200 | 36799999 | 6.843823 | 13.23877 | 26.14942 | 9.98657 | -0.41135 | -0.17736 | 0.484803 |
| 2 | 110374800 | 110376399 | 22.01705 | 38.12904 | 39.42166 | 17.29574 | 0.338301 | 0.843472 | 0.484816 |

| | | | | | | | | | |
|---|---|---|---|---|---|---|---|---|---|
| 12 | 19872200 | 19907999 | 1562.355 | 1943.904 | 1589.019 | 668.6529 | 0.289691 | 6.446962 | 0.48484 |
| 17 | 33169200 | 33172199 | 21.99327 | 36.89176 | 60.83405 | 26.54497 | -0.33813 | 1.15303 | 0.484847 |
| 6 | 60435600 | 60437399 | 18.71331 | 33.90359 | 27.31167 | 11.82255 | 0.722163 | 0.376349 | 0.484857 |
| 7 | 47301200 | 47306199 | 81.35548 | 128.8031 | 120.2909 | 50.0138 | 0.426871 | 2.388229 | 0.484858 |
| X | 116590200 | 116591999 | 23.88228 | 42.16781 | 30.57753 | 13.52862 | 0.851215 | 0.587823 | 0.484886 |
| 13 | 120308400 | 120309999 | 13.27371 | 24.5591 | 25.41946 | 10.70431 | 0.389983 | 0.214568 | 0.484887 |
| 6 | 144416200 | 144422399 | 38.91802 | 62.54441 | 99.81523 | 42.30385 | -0.32175 | 1.889711 | 0.48492 |
| 11 | 48836200 | 48838599 | 18.84351 | 32.16797 | 53.24764 | 23.34467 | -0.34174 | 0.977138 | 0.484923 |
| X | 86013600 | 86015599 | 18.68393 | 33.42242 | 29.98732 | 13.15706 | 0.555136 | 0.456587 | 0.484926 |
| 11 | 69585600 | 69587199 | 10.42589 | 18.81155 | 33.8184 | 14.71454 | -0.43323 | 0.357015 | 0.484934 |
| 7 | 94999600 | 95001199 | 10.19787 | 18.60441 | 32.85956 | 14.10129 | -0.39507 | 0.287812 | 0.484937 |
| 17 | 8100000 | 8101199 | 10.19787 | 18.60441 | 32.85956 | 14.10129 | -0.39507 | 0.287812 | 0.484937 |
| 14 | 73351200 | 73352199 | 11.12526 | 21.28078 | 22.26866 | 8.819172 | 0.448015 | -0.02528 | 0.484942 |
| 1 | 19194000 | 19196599 | 21.14866 | 37.16542 | 33.49972 | 14.83209 | 0.536425 | 0.625884 | 0.484946 |
| 13 | 35617000 | 35618399 | 16.09117 | 28.85441 | 29.16327 | 12.76933 | 0.386153 | 0.421472 | 0.484947 |
| 15 | 21158400 | 21160199 | 16.80522 | 30.10828 | 28.99265 | 12.70406 | 0.454712 | 0.430252 | 0.484961 |
| 3 | 37248800 | 37250199 | 13.78725 | 25.14547 | 25.88368 | 11.11165 | 0.378424 | 0.300005 | 0.484972 |
| 3 | 84537200 | 84539799 | 19.18893 | 32.58656 | 54.82712 | 24.0248 | -0.36631 | 1.027651 | 0.484999 |
| 2 | 174080000 | 174086399 | 64.21362 | 101.0161 | 155.1152 | 64.35165 | -0.2968 | 2.521805 | 0.485012 |
| 17 | 91797400 | 91799199 | 17.58727 | 31.77269 | 28.49546 | 12.37909 | 0.564998 | 0.38698 | 0.485025 |
| X | 80153200 | 80156199 | 30.51318 | 52.34548 | 41.78748 | 18.32758 | 0.707272 | 0.965821 | 0.485039 |
| 4 | 95515600 | 95516199 | 7.334025 | 14.02029 | 26.73756 | 10.60743 | -0.41276 | -0.04343 | 0.485039 |
| 7 | 59761200 | 59762799 | 20.74813 | 36.4018 | 34.12951 | 15.09001 | 0.4799 | 0.649698 | 0.485047 |
| 2 | 176391600 | 176393799 | 27.34644 | 47.2255 | 40.15326 | 17.61522 | 0.618373 | 0.891264 | 0.485066 |
| 9 | 69532200 | 69535199 | 22.62234 | 37.82476 | 64.80288 | 28.21929 | -0.39315 | 1.191305 | 0.485077 |
| 12 | 19041200 | 19042199 | 26.61318 | 44.81961 | 43.34279 | 18.94455 | 0.429276 | 1.113138 | 0.485101 |
| 8 | 55840600 | 55848999 | 111.6095 | 168.9713 | 255.112 | 105.544 | -0.31587 | 3.276286 | 0.485101 |
| 5 | 147179200 | 147180399 | 10.66024 | 19.42426 | 33.87282 | 14.55827 | -0.37836 | 0.300111 | 0.48512 |
| 6 | 70936800 | 70937999 | 7.678063 | 14.44142 | 28.04739 | 11.50493 | -0.48108 | 0.074748 | 0.485126 |
| 6 | 12619200 | 12621199 | 11.50552 | 20.89225 | 38.0982 | 16.44511 | -0.44668 | 0.335358 | 0.485131 |
| 18 | 42269000 | 42270599 | 16.96231 | 30.0598 | 30.71006 | 13.57728 | 0.361167 | 0.50559 | 0.485131 |
| 7 | 44375400 | 44380599 | 48.25086 | 77.05902 | 118.6219 | 49.81243 | -0.28036 | 2.100233 | 0.485139 |
| 10 | 120358400 | 120362199 | 26.45601 | 43.67628 | 72.1014 | 31.13173 | -0.3407 | 1.351716 | 0.485158 |
| 12 | 37574400 | 37576799 | 27.94473 | 48.31039 | 40.20209 | 17.63721 | 0.649709 | 0.887081 | 0.485163 |
| 6 | 129965200 | 129968799 | 33.45386 | 56.40128 | 47.98383 | 20.98 | 0.613504 | 1.13198 | 0.485169 |
| 4 | 79654600 | 79654999 | 3.93109 | 7.686842 | 17.78463 | 6.068689 | -0.48447 | -0.63638 | 0.485179 |
| 14 | 50838000 | 50839999 | 28.50129 | 48.61367 | 45.19378 | 19.81237 | 0.487179 | 1.012253 | 0.485246 |
| 9 | 19630000 | 19641399 | 150.5524 | 228.0265 | 220.1967 | 91.11129 | 0.328922 | 3.274383 | 0.485246 |
| 10 | 64605800 | 64609199 | 37.87259 | 63.04028 | 53.2187 | 23.10938 | 0.624847 | 1.27643 | 0.485264 |
| 15 | 44030000 | 44034399 | 44.29402 | 71.82903 | 70.21835 | 30.07689 | 0.398298 | 1.643582 | 0.485267 |
| 4 | 50165800 | 50166199 | 8.839943 | 17.08356 | 18.24613 | 6.999816 | 0.461718 | -0.16419 | 0.485271 |
| 2 | 87894000 | 87895399 | 23.30864 | 40.7048 | 32.79433 | 14.44483 | 0.697879 | 0.735201 | 0.485274 |
| 9 | 99230200 | 99232999 | 19.52357 | 33.03227 | 56.44223 | 24.70476 | -0.38878 | 1.062185 | 0.485276 |
| 5 | 31746000 | 31748399 | 22.68844 | 37.89529 | 62.42723 | 27.16668 | -0.33663 | 1.203245 | 0.485295 |
| 7 | 108678600 | 108679799 | 8.710678 | 16.45015 | 30.3493 | 12.35714 | -0.39928 | 0.061691 | 0.485301 |
| 12 | 93477800 | 93479199 | 28.05323 | 47.02676 | 47.02171 | 20.50968 | 0.381248 | 1.1671 | 0.48531 |
| 13 | 62835600 | 62837199 | 29.91855 | 50.2555 | 50.20765 | 21.92735 | 0.382296 | 1.152732 | 0.485349 |

| | | | | | | | | | |
|---|---|---|---|---|---|---|---|---|---|
| 9 | 56444800 | 56447999 | 20.95777 | 35.34207 | 59.79183 | 26.14472 | -0.37481 | 1.09819 | 0.485351 |
| 7 | 10791800 | 10823599 | 713.5282 | 953.9615 | 717.1283 | 299.4282 | 0.514592 | 5.283953 | 0.485359 |
| 17 | 69476000 | 69476999 | 9.683491 | 17.84531 | 31.95211 | 13.52829 | -0.40091 | 0.223714 | 0.485372 |
| X | 64596600 | 64603199 | 68.94908 | 110.577 | 84.33994 | 35.63368 | 0.737775 | 1.989846 | 0.485373 |
| 9 | 73117000 | 73117799 | 6.403452 | 12.36671 | 24.83564 | 9.557448 | -0.44543 | -0.16045 | 0.48539 |
| 14 | 33176200 | 33178599 | 16.99578 | 29.3814 | 48.99923 | 21.49663 | -0.34907 | 0.849439 | 0.485421 |
| 11 | 61149400 | 61152199 | 16.62451 | 28.85446 | 49.00951 | 21.504 | -0.37425 | 0.810652 | 0.48544 |
| 9 | 31845400 | 31848399 | 31.20842 | 50.56363 | 84.54888 | 36.19125 | -0.37546 | 1.642456 | 0.485454 |
| 4 | 129787800 | 129789999 | 15.62881 | 27.99939 | 28.75795 | 12.60168 | 0.362384 | 0.42363 | 0.485458 |
| 3 | 69828400 | 69831999 | 29.15606 | 47.76229 | 76.96622 | 33.11971 | -0.31025 | 1.4584 | 0.485458 |
| 7 | 6082800 | 6090799 | 131.3733 | 200.9391 | 175.6095 | 72.85945 | 0.484189 | 3.068927 | 0.485459 |
| 14 | 16630000 | 16631599 | 9.807422 | 17.71768 | 33.35311 | 14.48952 | -0.49769 | 0.344218 | 0.485473 |
| 15 | 34487200 | 34487799 | 6.556368 | 12.6735 | 24.87637 | 9.532704 | -0.40651 | -0.17111 | 0.485482 |
| 16 | 84079000 | 84080399 | 14.39776 | 26.72328 | 24.69379 | 10.35402 | 0.556599 | 0.203021 | 0.485488 |
| 8 | 19109200 | 19111399 | 29.04195 | 48.93431 | 42.72487 | 18.6773 | 0.576174 | 1.116446 | 0.485494 |
| 6 | 57872200 | 57874199 | 16.12301 | 29.25975 | 28.18017 | 12.17291 | 0.468057 | 0.350622 | 0.485496 |
| 17 | 38376800 | 38378999 | 25.20785 | 42.94561 | 44.24349 | 19.39394 | 0.339331 | 1.009596 | 0.485532 |
| 10 | 17218200 | 17220199 | 18.3984 | 31.54389 | 52.71137 | 23.11759 | -0.3543 | 0.937018 | 0.485534 |
| 6 | 69396600 | 69402599 | 50.24339 | 81.88772 | 61.69597 | 26.49815 | 0.778863 | 1.564641 | 0.485536 |
| 17 | 60145400 | 60146599 | 13.60796 | 25.17362 | 24.76132 | 10.42936 | 0.462475 | 0.215028 | 0.485542 |
| 19 | 54405800 | 54406399 | 7.171402 | 13.73791 | 26.24974 | 10.35199 | -0.40747 | -0.06646 | 0.485542 |
| 15 | 4033600 | 4036399 | 54.47818 | 87.33519 | 90.4886 | 38.20879 | 0.295947 | 1.981741 | 0.485559 |
| 19 | 21877000 | 21879399 | 19.37413 | 33.99316 | 34.34232 | 15.18484 | 0.372794 | 0.642672 | 0.485567 |
| 5 | 71204800 | 71208399 | 44.83422 | 73.61915 | 59.20704 | 25.50312 | 0.691036 | 1.45436 | 0.485568 |
| 17 | 33520600 | 33525199 | 106.1871 | 163.7616 | 167.9769 | 69.77462 | 0.26246 | 2.899846 | 0.485605 |
| 2 | 89428800 | 89430999 | 27.32985 | 46.637 | 44.64357 | 19.57189 | 0.445123 | 1.001052 | 0.48561 |
| 1 | 179811400 | 179812999 | 17.71791 | 31.13157 | 30.96006 | 13.70585 | 0.395394 | 0.604938 | 0.485623 |
| 1 | 60767400 | 60773399 | 54.03963 | 86.53522 | 89.57826 | 37.76006 | 0.295373 | 2.016362 | 0.48563 |
| 3 | 116792000 | 116793199 | 8.555389 | 15.98538 | 30.60717 | 12.69647 | -0.47462 | 0.128235 | 0.485641 |
| X | 153093000 | 153098399 | 70.78903 | 113.4344 | 85.96722 | 36.27576 | 0.745594 | 2.016028 | 0.485645 |
| 5 | 94462200 | 94462799 | 13.53244 | 25.13777 | 21.35252 | 8.943064 | 0.678831 | 0.185395 | 0.485646 |
| 7 | 12918400 | 12918999 | 11.87737 | 22.19671 | 23.07943 | 9.57064 | 0.400655 | 0.138884 | 0.485662 |
| 10 | 79252800 | 79253799 | 30.58055 | 50.72629 | 50.30364 | 21.79142 | 0.392904 | 1.290208 | 0.485678 |
| 8 | 93058400 | 93060199 | 10.71543 | 19.44014 | 33.83945 | 14.62321 | -0.38066 | 0.327208 | 0.485699 |
| 13 | 14111600 | 14116399 | 72.67044 | 115.2221 | 117.5526 | 48.92636 | 0.301348 | 2.342654 | 0.485704 |
| 17 | 92679000 | 92680599 | 21.68504 | 37.65979 | 38.85034 | 17.0527 | 0.342186 | 0.813059 | 0.485706 |
| 4 | 106009000 | 106014999 | 58.94417 | 93.48334 | 145.5974 | 60.61497 | -0.30773 | 2.329602 | 0.485721 |
| 17 | 88131800 | 88138999 | 49.11402 | 78.09806 | 127.4152 | 53.1851 | -0.37082 | 2.240731 | 0.485729 |
| 7 | 59813000 | 59814999 | 11.8394 | 20.73157 | 45.18835 | 19.94687 | -0.73058 | 0.65508 | 0.485736 |
| X | 90581200 | 90583799 | 25.82433 | 45.26732 | 33.42494 | 14.76733 | 0.82316 | 0.694005 | 0.485744 |
| 19 | 13745800 | 13748799 | 8.951951 | 16.07063 | 37.44114 | 16.37824 | -0.80977 | 0.385766 | 0.485749 |
| 18 | 90301800 | 90303799 | 24.19171 | 42.05122 | 39.18307 | 17.20856 | 0.488478 | 0.812681 | 0.485751 |
| 14 | 46983000 | 46983999 | 8.806145 | 16.29751 | 30.47101 | 12.82691 | -0.45621 | 0.19207 | 0.485761 |
| 7 | 116129800 | 116131799 | 15.60145 | 28.01282 | 27.43135 | 11.99777 | 0.43295 | 0.409111 | 0.485772 |
| 18 | 19906400 | 19909199 | 56.22919 | 90.26984 | 75.37949 | 31.8975 | 0.609005 | 1.937362 | 0.485773 |
| X | 95724600 | 95726199 | 20.36892 | 35.7592 | 34.33332 | 15.18826 | 0.446022 | 0.641139 | 0.48579 |
| 7 | 40504400 | 40506999 | 21.0075 | 35.45523 | 58.09267 | 25.40811 | -0.32862 | 1.091625 | 0.485791 |

| | | | | | | | | | |
|---|---|---|---|---|---|---|---|---|---|
| 12 | 25707000 | 25708599 | 16.19628 | 29.08612 | 29.23338 | 12.78398 | 0.395562 | 0.414497 | 0.4858 |
| X | 38218800 | 38221999 | 27.0845 | 46.49735 | 43.02627 | 18.872 | 0.495232 | 0.94774 | 0.485801 |
| 13 | 31301000 | 31302199 | 10.26197 | 18.59722 | 33.34804 | 14.43046 | -0.42435 | 0.329688 | 0.485803 |
| 14 | 75903400 | 75907999 | 48.22848 | 77.08827 | 120.6738 | 50.73466 | -0.3032 | 2.07454 | 0.485825 |
| 6 | 93952000 | 93954799 | 28.28881 | 48.17945 | 45.7454 | 20.05179 | 0.456512 | 1.024733 | 0.485857 |
| 17 | 88572000 | 88580199 | 98.15398 | 150.3626 | 221.8356 | 92.07149 | -0.26881 | 3.031118 | 0.485871 |
| 5 | 61057400 | 61058199 | 9.535816 | 18.35507 | 19.67222 | 7.652006 | 0.43746 | -0.10863 | 0.485899 |
| 12 | 59679000 | 59680599 | 9.370117 | 17.54374 | 32.34002 | 13.36861 | -0.41609 | 0.128208 | 0.485902 |
| 12 | 5169400 | 5172399 | 20.11952 | 33.7128 | 59.55612 | 25.9699 | -0.43701 | 1.158243 | 0.485905 |
| 15 | 95366400 | 95368399 | 12.19756 | 21.62169 | 37.23824 | 16.46259 | -0.38781 | 0.486339 | 0.485912 |
| 18 | 20858000 | 20859799 | 11.0387 | 20.04638 | 34.88685 | 15.05724 | -0.37926 | 0.326146 | 0.485923 |
| 7 | 29504200 | 29505199 | 12.15648 | 22.74807 | 24.03577 | 9.945155 | 0.37997 | 0.134563 | 0.485929 |
| 3 | 22855400 | 22857399 | 18.85544 | 34.39607 | 25.73497 | 11.04494 | 0.835873 | 0.335999 | 0.485931 |
| 14 | 83743200 | 83745999 | 24.62888 | 42.78476 | 38.55355 | 16.92553 | 0.536365 | 0.82019 | 0.485979 |
| 14 | 52967400 | 52968199 | 16.99425 | 30.00909 | 29.94497 | 13.24848 | 0.39314 | 0.543876 | 0.486001 |
| 18 | 56214800 | 56221399 | 68.34262 | 109.1072 | 99.23574 | 41.65039 | 0.476926 | 2.126183 | 0.486003 |
| 10 | 92644800 | 92645799 | 16.66147 | 29.53653 | 30.27215 | 13.38278 | 0.356812 | 0.499674 | 0.486016 |
| 7 | 93616600 | 93618399 | 17.19973 | 30.6807 | 25.70127 | 11.30341 | 0.651217 | 0.454934 | 0.48602 |
| 9 | 42263000 | 42263999 | 6.334497 | 12.02418 | 26.59877 | 10.71331 | -0.64405 | -0.00445 | 0.486039 |
| 7 | 35290400 | 35293999 | 21.12254 | 35.29458 | 61.14818 | 26.61708 | -0.40896 | 1.194407 | 0.48604 |
| 13 | 44066400 | 44071599 | 59.82133 | 94.51634 | 147.0603 | 61.08055 | -0.31088 | 2.424488 | 0.486042 |
| 10 | 85291200 | 85295199 | 24.59352 | 40.50033 | 69.84611 | 30.1227 | -0.4047 | 1.377347 | 0.486049 |
| 2 | 72144800 | 72147399 | 14.0968 | 24.84984 | 42.90428 | 18.98857 | -0.39491 | 0.567677 | 0.486057 |
| 3 | 45143200 | 45146999 | 49.79769 | 80.22763 | 81.03531 | 34.46817 | 0.341237 | 1.820433 | 0.48606 |
| 16 | 20740200 | 20741599 | 12.64893 | 23.36115 | 22.38658 | 9.450191 | 0.497714 | 0.211005 | 0.486065 |
| X | 20889000 | 20894399 | 110.1997 | 171.1154 | 154.1464 | 64.01008 | 0.457169 | 2.779729 | 0.486088 |
| 2 | 53206600 | 53209599 | 33.47753 | 56.58539 | 46.84686 | 20.49874 | 0.652778 | 1.109859 | 0.486089 |
| 10 | 93599000 | 93602599 | 23.30495 | 38.80728 | 66.43912 | 28.85735 | -0.39208 | 1.238706 | 0.486118 |
| 1 | 12150200 | 12151399 | 17.77494 | 31.9798 | 28.3345 | 12.37174 | 0.577581 | 0.40993 | 0.486139 |
| 1 | 27551200 | 27554999 | 36.01416 | 60.37005 | 49.72967 | 21.69039 | 0.65985 | 1.194865 | 0.486144 |
| 8 | 71788800 | 71790399 | 9.595074 | 17.49016 | 32.3649 | 13.90345 | -0.46269 | 0.287134 | 0.48616 |
| 14 | 29455000 | 29456599 | 16.27633 | 29.66188 | 27.6632 | 11.88721 | 0.518987 | 0.326991 | 0.486167 |
| 15 | 15681000 | 15683599 | 31.3038 | 52.52405 | 45.35242 | 19.79305 | 0.592012 | 1.163889 | 0.486171 |
| 5 | 16960600 | 16961999 | 14.96079 | 26.14667 | 44.14559 | 19.45474 | -0.36428 | 0.700747 | 0.486179 |
| 3 | 69854000 | 69858399 | 52.76685 | 84.90865 | 83.58629 | 35.4911 | 0.376118 | 1.863021 | 0.486183 |
| 10 | 16185800 | 16187199 | 17.03471 | 30.93184 | 28.14677 | 12.14986 | 0.550212 | 0.354735 | 0.486193 |
| 6 | 9983200 | 9986799 | 29.70447 | 49.82925 | 50.75062 | 22.1458 | 0.354681 | 1.168138 | 0.486214 |
| 9 | 93242800 | 93244399 | 17.10355 | 30.92157 | 28.56303 | 12.39704 | 0.523524 | 0.379687 | 0.486217 |
| 13 | 28812600 | 28815199 | 19.36548 | 33.807 | 30.72358 | 13.52979 | 0.524807 | 0.714657 | 0.486217 |
| 7 | 31515000 | 31517799 | 37.20582 | 61.04129 | 52.55989 | 22.63177 | 0.592287 | 1.440636 | 0.486222 |
| 5 | 31242400 | 31244399 | 13.51634 | 23.67579 | 41.53513 | 18.33938 | -0.41914 | 0.651917 | 0.486223 |
| 15 | 90933200 | 90935199 | 13.95914 | 24.35447 | 44.18324 | 19.44976 | -0.4678 | 0.717731 | 0.486232 |
| 2 | 97478200 | 97479999 | 23.72036 | 41.224 | 35.03784 | 15.38619 | 0.620471 | 0.8068 | 0.486235 |
| 5 | 91374200 | 91375599 | 14.89899 | 27.16253 | 27.24342 | 11.70105 | 0.415074 | 0.313021 | 0.486244 |
| 3 | 54474400 | 54480599 | 70.66562 | 110.8593 | 168.8938 | 70.04575 | -0.28856 | 2.581937 | 0.486265 |
| 14 | 94807000 | 94814399 | 99.54316 | 156.4838 | 130.193 | 54.00736 | 0.585424 | 2.554403 | 0.48627 |
| 10 | 117955000 | 117958999 | 25.61831 | 42.46911 | 69.88229 | 30.2601 | -0.33555 | 1.300177 | 0.486272 |

| | | | | | | | | | |
|---|---|---|---|---|---|---|---|---|---|
| X | 54881600 | 54883599 | 25.86974 | 44.55496 | 41.89044 | 18.3758 | 0.473126 | 0.91065 | 0.486291 |
| X | 90317800 | 90318999 | 16.47265 | 30.81697 | 24.29025 | 10.05477 | 0.799539 | 0.174626 | 0.4863 |
| 17 | 27266600 | 27269599 | 22.72055 | 37.75137 | 65.09863 | 28.23315 | -0.40254 | 1.262215 | 0.486314 |
| 11 | 3766800 | 3768799 | 11.77999 | 21.05409 | 37.84698 | 16.61772 | -0.44207 | 0.435011 | 0.486316 |
| 2 | 125081600 | 125083799 | 18.18117 | 32.5423 | 26.41532 | 11.58405 | 0.699116 | 0.441756 | 0.486331 |
| 4 | 71235400 | 71237199 | 19.67601 | 34.96517 | 30.26888 | 13.35482 | 0.600859 | 0.502235 | 0.486333 |
| X | 88308000 | 88310999 | 27.96347 | 48.19778 | 41.33365 | 18.13198 | 0.605646 | 0.914915 | 0.486353 |
| 13 | 114385000 | 114387599 | 37.96876 | 62.34644 | 54.76046 | 23.58787 | 0.564234 | 1.434827 | 0.486363 |
| 13 | 45250800 | 45251599 | 7.297598 | 13.9515 | 27.3261 | 10.83902 | -0.45142 | -0.04604 | 0.486371 |
| 17 | 31762800 | 31767399 | 28.6337 | 46.90647 | 77.0239 | 33.1445 | -0.33737 | 1.455863 | 0.486385 |
| 4 | 38791800 | 38792599 | 11.53545 | 22.00404 | 22.71757 | 9.05588 | 0.458612 | -0.0003 | 0.486386 |
| 11 | 113206000 | 113215399 | 119.3619 | 180.3523 | 265.4031 | 109.7334 | -0.28135 | 3.318151 | 0.486407 |
| 17 | 52089400 | 52090199 | 5.256642 | 10.2074 | 22.7959 | 8.541263 | -0.55971 | -0.27083 | 0.486424 |
| 18 | 36916000 | 36916599 | 5.256642 | 10.2074 | 22.7959 | 8.541263 | -0.55971 | -0.27083 | 0.486424 |
| 3 | 120508800 | 120510399 | 13.19528 | 23.46707 | 45.71639 | 20.15699 | -0.56231 | 0.486287 | 0.486446 |
| 18 | 37367800 | 37375599 | 226.648 | 332.0515 | 325.0462 | 133.8914 | 0.265665 | 3.883067 | 0.486464 |
| 14 | 82759200 | 82766999 | 130.7006 | 200.5702 | 181.1575 | 75.20818 | 0.440391 | 3.004494 | 0.486464 |
| X | 63499200 | 63510799 | 161.8629 | 245.7432 | 205.3945 | 85.04857 | 0.539969 | 3.230249 | 0.486479 |
| 1 | 3343600 | 3350799 | 138.6606 | 211.013 | 187.8197 | 77.8261 | 0.45201 | 3.173404 | 0.486484 |
| 15 | 44730400 | 44731599 | 8.430607 | 15.79679 | 29.04417 | 11.99062 | -0.41048 | 0.107655 | 0.486496 |
| 3 | 76483000 | 76484599 | 18.52868 | 33.87938 | 26.44485 | 11.31636 | 0.778094 | 0.323149 | 0.486504 |
| 1 | 138799400 | 138801599 | 31.71053 | 51.47194 | 83.01047 | 35.58107 | -0.32089 | 1.607327 | 0.486506 |
| 5 | 75837600 | 75839799 | 19.45894 | 34.10231 | 34.35806 | 15.1769 | 0.37671 | 0.660724 | 0.486508 |
| 7 | 35072800 | 35074399 | 13.3134 | 23.43645 | 39.83105 | 17.63329 | -0.37284 | 0.573914 | 0.486512 |
| 17 | 22885200 | 22890999 | 61.75298 | 99.06742 | 88.17418 | 37.27147 | 0.515963 | 1.968682 | 0.48652 |
| 9 | 115298600 | 115299999 | 15.06847 | 27.28899 | 27.72075 | 12.00574 | 0.389136 | 0.353082 | 0.486526 |
| 1 | 109532200 | 109532799 | 10.27422 | 19.78308 | 21.32617 | 8.287309 | 0.430779 | -0.09774 | 0.486529 |
| 13 | 7779200 | 7781999 | 23.29349 | 40.48735 | 39.21027 | 17.21983 | 0.432986 | 0.808621 | 0.486544 |
| 8 | 99717800 | 99719799 | 15.89025 | 29.04638 | 27.16318 | 11.62809 | 0.518494 | 0.305488 | 0.486553 |
| 7 | 31832600 | 31834199 | 64.18124 | 102.0826 | 99.06279 | 41.36319 | 0.377877 | 2.241385 | 0.486555 |
| 10 | 38930800 | 38931799 | 6.751809 | 12.83494 | 27.15932 | 10.90557 | -0.57735 | -0.01116 | 0.486561 |
| 8 | 13348400 | 13351599 | 25.68765 | 41.9413 | 82.32995 | 35.38345 | -0.59714 | 1.50095 | 0.486564 |
| 7 | 10426200 | 10427799 | 28.20231 | 48.32871 | 43.53403 | 19.09193 | 0.533487 | 0.968288 | 0.486621 |
| 19 | 61253800 | 61255599 | 12.1354 | 21.69382 | 37.24309 | 16.34969 | -0.37632 | 0.430827 | 0.486622 |
| 7 | 106805600 | 106810999 | 113.5256 | 171.922 | 257.1139 | 106.382 | -0.30182 | 3.272169 | 0.486625 |
| 2 | 36563200 | 36569799 | 73.41015 | 117.2131 | 95.93315 | 40.2719 | 0.629111 | 2.127509 | 0.486637 |
| 8 | 76282000 | 76283199 | 14.91522 | 27.37457 | 26.77815 | 11.40088 | 0.459568 | 0.275059 | 0.486653 |
| 2 | 14321600 | 14323199 | 19.81251 | 35.18878 | 30.8799 | 13.6306 | 0.581036 | 0.508785 | 0.486655 |
| 17 | 37724200 | 37726599 | 40.1952 | 65.16672 | 65.84452 | 28.19848 | 0.350426 | 1.636303 | 0.486658 |
| 2 | 131672200 | 131678199 | 32.44694 | 53.96581 | 46.57759 | 20.21446 | 0.592827 | 1.261407 | 0.48666 |
| 5 | 127334200 | 127336799 | 21.86978 | 36.61968 | 60.75904 | 26.48262 | -0.34692 | 1.172422 | 0.486666 |
| 9 | 101169200 | 101171599 | 17.12173 | 30.22533 | 30.85539 | 13.65207 | 0.360212 | 0.548792 | 0.486667 |
| X | 149428800 | 149432199 | 21.13543 | 35.47436 | 61.19245 | 26.71159 | -0.40273 | 1.147976 | 0.486669 |
| X | 55419800 | 55424999 | 58.89242 | 94.41459 | 88.217 | 37.2513 | 0.444892 | 1.989093 | 0.486691 |
| X | 159083400 | 159087799 | 53.42005 | 85.70122 | 88.45777 | 37.39263 | 0.30269 | 1.95864 | 0.486706 |
| 14 | 82433000 | 82435599 | 29.66065 | 50.10856 | 48.52839 | 21.23127 | 0.427297 | 1.10255 | 0.486719 |
| 13 | 76601800 | 76603799 | 26.29391 | 45.01378 | 44.12648 | 19.3502 | 0.411691 | 0.969196 | 0.48673 |

| | | | | | | | | | |
|---|---|---|---|---|---|---|---|---|---|
| 5 | 151245400 | 151248399 | 28.71771 | 48.58554 | 42.65244 | 18.66869 | 0.568502 | 1.080513 | 0.48673 |
| 1 | 81719800 | 81720599 | 8.1386 | 15.3251 | 28.59382 | 11.70643 | -0.42145 | 0.07374 | 0.486749 |
| 5 | 36940400 | 36945599 | 63.33802 | 100.0043 | 150.4546 | 62.46876 | -0.26324 | 2.444936 | 0.486763 |
| 17 | 56230600 | 56231999 | 26.54398 | 44.97454 | 46.38261 | 20.31048 | 0.337088 | 1.065076 | 0.486773 |
| 12 | 118511000 | 118513599 | 13.6295 | 24.02656 | 40.31218 | 17.84128 | -0.35363 | 0.561201 | 0.486792 |
| 19 | 12112800 | 12114399 | 12.99381 | 23.00957 | 38.7427 | 17.13121 | -0.35606 | 0.505447 | 0.486796 |
| 8 | 108951200 | 108953199 | 17.69512 | 31.7094 | 29.45486 | 12.90426 | 0.506614 | 0.436611 | 0.486801 |
| X | 148555000 | 148564799 | 133.4311 | 204.4966 | 184.5351 | 76.59687 | 0.440301 | 3.031411 | 0.486836 |
| 3 | 152569000 | 152572399 | 43.68196 | 69.84512 | 111.6872 | 46.97931 | -0.33323 | 2.057058 | 0.486854 |
| 11 | 26319000 | 26320199 | 18.38542 | 32.30577 | 31.54588 | 13.96567 | 0.421845 | 0.611659 | 0.486885 |
| 19 | 7976200 | 7990799 | 230.4657 | 338.1312 | 323.2369 | 133.1341 | 0.301855 | 3.860583 | 0.486887 |
| X | 112385600 | 112387599 | 23.79638 | 42.11721 | 28.85865 | 12.762 | 0.933631 | 0.558607 | 0.48689 |
| 1 | 84051000 | 84052399 | 10.14397 | 18.5805 | 33.96638 | 14.50067 | -0.43961 | 0.264807 | 0.486897 |
| X | 115622800 | 115625199 | 25.62011 | 45.28385 | 28.24749 | 12.4951 | 1.067752 | 0.583699 | 0.4869 |
| 19 | 3198400 | 3203599 | 102.2844 | 158.0246 | 162.2848 | 67.40959 | 0.262712 | 2.864327 | 0.486903 |
| X | 3646400 | 3647999 | 24.3974 | 42.7803 | 28.57371 | 12.62758 | 0.967105 | 0.673802 | 0.486928 |
| 6 | 38642200 | 38644999 | 12.54453 | 22.10033 | 38.85813 | 17.19987 | -0.42098 | 0.559716 | 0.486928 |
| 3 | 38072800 | 38073799 | 12.8496 | 23.85148 | 24.99924 | 10.48108 | 0.376681 | 0.193083 | 0.486937 |
| 3 | 122100000 | 122102999 | 16.77049 | 28.88595 | 49.51578 | 21.72082 | -0.3895 | 0.884641 | 0.486973 |
| 6 | 84651000 | 84652999 | 33.0282 | 53.50788 | 85.79465 | 36.72242 | -0.31516 | 1.650019 | 0.486975 |
| 16 | 16519400 | 16521399 | 24.67606 | 43.28824 | 32.74647 | 14.47711 | 0.788126 | 0.672185 | 0.486985 |
| 1 | 112730200 | 112731199 | 18.42142 | 32.44505 | 31.92769 | 14.133 | 0.411823 | 0.585541 | 0.487 |
| X | 49428600 | 49431199 | 38.77663 | 63.43531 | 63.5782 | 27.34375 | 0.372 | 1.490509 | 0.487013 |
| 14 | 85379200 | 85383199 | 69.32556 | 110.1634 | 112.0538 | 46.74165 | 0.308783 | 2.274052 | 0.487049 |
| 14 | 53870400 | 53873399 | 17.84213 | 30.5704 | 51.65045 | 22.65112 | -0.37035 | 0.937019 | 0.487052 |
| 15 | 15503200 | 15505999 | 49.14837 | 79.23348 | 71.27589 | 30.34421 | 0.509117 | 1.794274 | 0.487067 |
| 7 | 52794800 | 52798199 | 39.82234 | 65.35169 | 62.26345 | 26.81309 | 0.446893 | 1.450962 | 0.487085 |
| 14 | 102260600 | 102263399 | 26.00128 | 44.42252 | 44.12922 | 19.34824 | 0.392101 | 0.984589 | 0.487095 |
| X | 21664200 | 21670799 | 88.38353 | 138.7899 | 134.9733 | 55.98248 | 0.359372 | 2.570642 | 0.487098 |
| 9 | 112409000 | 112411999 | 29.08801 | 49.51165 | 45.72881 | 20.04357 | 0.496103 | 1.032579 | 0.487103 |
| 18 | 67342800 | 67345599 | 26.00446 | 44.12336 | 45.83895 | 20.08111 | 0.326615 | 1.049739 | 0.487103 |
| 1 | 37291600 | 37292399 | 11.29035 | 21.79543 | 20.73956 | 7.996796 | 0.619283 | -0.11491 | 0.487108 |
| 3 | 115242800 | 115244799 | 19.27969 | 33.02415 | 54.93942 | 24.09294 | -0.34831 | 0.94882 | 0.487109 |
| 15 | 82460400 | 82461399 | 7.295633 | 14.0855 | 29.08273 | 11.20759 | -0.48768 | -0.13502 | 0.487119 |
| X | 59448400 | 59450799 | 38.42617 | 63.78484 | 53.72288 | 23.28383 | 0.627993 | 1.311988 | 0.487123 |
| 2 | 10251400 | 10252599 | 11.77878 | 22.47346 | 21.48046 | 8.558501 | 0.568917 | -0.00519 | 0.487132 |
| 4 | 98798800 | 98800999 | 26.06796 | 44.29975 | 45.30471 | 19.8551 | 0.349366 | 1.037019 | 0.487133 |
| 11 | 7831800 | 7832199 | 4.271168 | 8.33111 | 20.30921 | 7.276549 | -0.62448 | -0.45173 | 0.487139 |
| 10 | 29988600 | 29989999 | 13.56943 | 25.32642 | 24.92314 | 10.35763 | 0.476656 | 0.164389 | 0.487141 |
| X | 111367000 | 111371599 | 67.21841 | 107.595 | 92.30029 | 38.88406 | 0.565295 | 2.04609 | 0.487141 |
| 4 | 119510000 | 119521799 | 93.08345 | 142.9421 | 211.7938 | 87.93522 | -0.27229 | 2.981085 | 0.487154 |
| X | 107731400 | 107733399 | 28.76493 | 49.54815 | 41.29108 | 18.11296 | 0.646636 | 0.923324 | 0.487168 |
| 9 | 4914600 | 4917599 | 39.2124 | 64.55694 | 52.77392 | 22.75731 | 0.668864 | 1.401716 | 0.487182 |
| 7 | 8213400 | 8214799 | 20.14832 | 35.60235 | 28.0836 | 12.4233 | 0.730865 | 0.553192 | 0.487186 |
| 14 | 41369400 | 41371599 | 26.4647 | 45.14273 | 45.2332 | 19.82975 | 0.379446 | 1.003366 | 0.487191 |
| 2 | 147712400 | 147713199 | 11.79415 | 20.79023 | 42.46841 | 18.79575 | -0.63583 | 0.555682 | 0.487195 |
| 16 | 47864000 | 47865399 | 19.51071 | 35.066 | 28.83163 | 12.59991 | 0.68421 | 0.431227 | 0.487206 |

| | | | | | | | | | |
|---|---|---|---|---|---|---|---|---|---|
| 5 | 100011000 | 100012399 | 9.065772 | 16.81275 | 30.46892 | 12.78998 | -0.40897 | 0.180511 | 0.487207 |
| 1 | 138152600 | 138156799 | 95.28338 | 148.0841 | 153.1688 | 63.59576 | 0.258975 | 2.755581 | 0.487209 |
| 9 | 6495200 | 6496399 | 17.78586 | 32.25589 | 28.11731 | 12.15592 | 0.610501 | 0.367737 | 0.487228 |
| 7 | 12756600 | 12759199 | 25.54945 | 42.08991 | 69.98204 | 30.18598 | -0.35193 | 1.37552 | 0.487232 |
| 3 | 81687800 | 81689199 | 20.76826 | 34.93502 | 59.06124 | 25.80568 | -0.37373 | 1.122811 | 0.487248 |
| 2 | 111748200 | 111751599 | 47.0826 | 76.11427 | 75.18007 | 32.12119 | 0.379111 | 1.713353 | 0.487256 |
| 5 | 131303200 | 131305799 | 33.71755 | 55.52497 | 57.9029 | 24.97179 | 0.31889 | 1.391473 | 0.487263 |
| 4 | 131966600 | 131967599 | 11.166 | 21.12504 | 22.60638 | 9.169507 | 0.385332 | 0.051138 | 0.487274 |
| 5 | 151549000 | 151557599 | 94.71179 | 148.8358 | 134.2056 | 55.66931 | 0.469086 | 2.558784 | 0.487278 |
| 7 | 3930400 | 3933399 | 35.68391 | 59.22903 | 55.9567 | 24.2509 | 0.462714 | 1.303697 | 0.487288 |
| 3 | 102349000 | 102350799 | 16.48973 | 28.69296 | 47.90205 | 21.05149 | -0.34904 | 0.773529 | 0.487289 |
| X | 54664400 | 54678599 | 308.444 | 440.7731 | 408.6127 | 168.8321 | 0.308972 | 4.260786 | 0.487313 |
| 1 | 89940200 | 89946199 | 60.03287 | 94.91755 | 143.8116 | 59.75256 | -0.27177 | 2.408668 | 0.487324 |
| 12 | 48089800 | 48094399 | 38.70162 | 63.9056 | 56.24574 | 24.28904 | 0.564 | 1.366831 | 0.487328 |
| 14 | 87545000 | 87545799 | 5.957266 | 11.56355 | 23.3662 | 8.781247 | -0.42099 | -0.2491 | 0.487356 |
| 19 | 28181800 | 28190799 | 94.17788 | 144.7339 | 212.3259 | 88.16545 | -0.25711 | 2.966918 | 0.487382 |
| 12 | 87581200 | 87582399 | 19.988 | 34.93304 | 35.17349 | 15.50093 | 0.377927 | 0.711846 | 0.487388 |
| 8 | 34006200 | 34008399 | 15.29654 | 26.36881 | 48.98667 | 21.48925 | -0.50486 | 0.867478 | 0.487391 |
| 6 | 14127200 | 14129799 | 30.60909 | 52.33423 | 38.13465 | 16.72096 | 0.837696 | 0.991682 | 0.487418 |
| 5 | 150396000 | 150397799 | 19.54554 | 34.31108 | 34.2938 | 15.1698 | 0.388171 | 0.635667 | 0.487422 |
| 17 | 65457000 | 65457999 | 17.34204 | 30.01746 | 49.5361 | 21.73298 | -0.33357 | 0.841245 | 0.487422 |
| 2 | 33860000 | 33861999 | 19.92326 | 34.87913 | 35.50398 | 15.66907 | 0.362116 | 0.683642 | 0.487429 |
| 7 | 35547800 | 35555599 | 69.21865 | 108.5313 | 164.215 | 68.10905 | -0.27924 | 2.590106 | 0.487431 |
| 6 | 75949000 | 75949999 | 14.54965 | 26.72782 | 21.85528 | 9.293489 | 0.717915 | 0.25687 | 0.487447 |
| 5 | 83883400 | 83884399 | 9.192863 | 17.2592 | 31.86766 | 13.11223 | -0.41307 | 0.107586 | 0.487456 |
| 4 | 112581000 | 112588599 | 104.9353 | 163.9756 | 142.8508 | 59.26883 | 0.512468 | 2.666645 | 0.487472 |
| 8 | 91074000 | 91075599 | 14.16743 | 24.60478 | 45.83345 | 20.11812 | -0.5064 | 0.788101 | 0.487483 |
| 7 | 48265800 | 48278199 | 167.5471 | 252.5049 | 234.877 | 97.03518 | 0.376749 | 3.378919 | 0.487487 |
| X | 55510800 | 55514999 | 36.66172 | 61.37754 | 50.3014 | 21.92218 | 0.667316 | 1.21206 | 0.487502 |
| 10 | 119717200 | 119719599 | 20.25252 | 34.45054 | 58.21039 | 25.51311 | -0.37229 | 1.018236 | 0.487525 |
| 4 | 142219600 | 142225199 | 40.04671 | 64.19864 | 107.0652 | 45.20704 | -0.38946 | 1.969756 | 0.487542 |
| 6 | 102203000 | 102203599 | 5.678782 | 11.01294 | 23.31054 | 8.797719 | -0.49323 | -0.23898 | 0.487547 |
| 6 | 75947600 | 75947999 | 4.349328 | 8.499257 | 18.85071 | 6.53879 | -0.44648 | -0.56827 | 0.48755 |
| 11 | 70590400 | 70592599 | 13.96314 | 24.39456 | 43.12394 | 18.99966 | -0.43031 | 0.69753 | 0.487561 |
| 11 | 21755000 | 21756599 | 16.90753 | 31.39473 | 24.56459 | 10.29399 | 0.795866 | 0.222498 | 0.487581 |
| 4 | 61485200 | 61487999 | 25.82571 | 44.5984 | 41.36031 | 18.14474 | 0.493658 | 0.887639 | 0.487604 |
| 2 | 129183600 | 129190199 | 115.3737 | 177.5726 | 175.935 | 73.06955 | 0.310243 | 2.942379 | 0.487617 |
| 5 | 103370400 | 103373799 | 34.45278 | 55.79214 | 88.60298 | 37.9126 | -0.30199 | 1.660934 | 0.48765 |
| 17 | 37549200 | 37551399 | 35.60827 | 58.82411 | 52.03337 | 22.47693 | 0.556922 | 1.353864 | 0.487651 |
| 4 | 48744400 | 48746199 | 24.25151 | 41.85131 | 36.17997 | 15.8718 | 0.594252 | 0.878315 | 0.487658 |
| 9 | 3173200 | 3182599 | 190.8341 | 283.9955 | 244.7866 | 100.859 | 0.470267 | 3.618198 | 0.487659 |
| 13 | 61440000 | 61441599 | 15.88238 | 29.82244 | 23.16172 | 9.527946 | 0.827368 | 0.142823 | 0.487659 |
| 14 | 23050800 | 23052199 | 20.48328 | 36.54615 | 29.88507 | 13.14093 | 0.686227 | 0.485076 | 0.487661 |
| 10 | 7291800 | 7296199 | 51.24814 | 82.36616 | 85.26491 | 36.14108 | 0.301589 | 1.898233 | 0.487668 |
| 18 | 11760400 | 11760799 | 5.818731 | 11.28292 | 23.30592 | 8.801177 | -0.45906 | -0.23472 | 0.487669 |
| 7 | 107403400 | 107407799 | 46.82293 | 76.30865 | 65.22204 | 28.01201 | 0.597356 | 1.562981 | 0.487677 |
| 16 | 22150000 | 22155399 | 57.45206 | 92.15133 | 90.2132 | 38.1231 | 0.378529 | 1.969229 | 0.487679 |

| | | | | | | | | | |
|---|---|---|---|---|---|---|---|---|---|
| 4 | 101905000 | 101908199 | 38.80257 | 62.19078 | 104.9133 | 44.28394 | -0.40629 | 1.975956 | 0.487681 |
| 12 | 4015200 | 4017399 | 12.04343 | 21.33126 | 37.26164 | 16.4756 | -0.40856 | 0.494181 | 0.487689 |
| 14 | 8310400 | 8310999 | 12.04343 | 21.33126 | 37.26164 | 16.4756 | -0.40856 | 0.494181 | 0.487689 |
| 6 | 64952600 | 64952999 | 5.651626 | 11.01238 | 23.24495 | 8.441744 | -0.43755 | -0.37348 | 0.487689 |
| 17 | 90186600 | 90188799 | 20.72136 | 37.12287 | 28.06811 | 12.29938 | 0.801705 | 0.456312 | 0.487695 |
| 8 | 31718000 | 31722599 | 45.82151 | 73.31048 | 113.1806 | 47.64829 | -0.28169 | 2.042078 | 0.4877 |
| 7 | 29394600 | 29398399 | 34.57859 | 55.93765 | 92.91402 | 39.72679 | -0.36808 | 1.682232 | 0.487701 |
| 14 | 5815800 | 5817799 | 28.84056 | 49.60462 | 36.57709 | 16.04437 | 0.821993 | 0.930969 | 0.487722 |
| 1 | 45692200 | 45693999 | 16.8974 | 29.15779 | 48.99254 | 21.49249 | -0.36029 | 0.867211 | 0.487728 |
| 7 | 33170800 | 33212799 | 844.3891 | 1112.417 | 912.1904 | 381.5919 | 0.368017 | 5.520731 | 0.48773 |
| 6 | 89831200 | 89834199 | 37.48307 | 61.44123 | 61.24956 | 26.36349 | 0.381141 | 1.461314 | 0.48773 |
| X | 125682400 | 125683399 | 13.73081 | 26.73918 | 15.72235 | 5.754054 | 1.376892 | -0.2707 | 0.487757 |
| 1 | 40812600 | 40813999 | 8.119182 | 14.82638 | 32.50122 | 13.93078 | -0.70221 | 0.270652 | 0.487758 |
| 18 | 18216200 | 18218199 | 28.63166 | 47.81602 | 47.58783 | 20.70736 | 0.388032 | 1.208034 | 0.487777 |
| 4 | 80726400 | 80730199 | 79.11014 | 124.446 | 128.5874 | 53.34598 | 0.273952 | 2.527886 | 0.487794 |
| 7 | 6384200 | 6387199 | 24.3822 | 40.3374 | 67.26477 | 29.0856 | -0.35483 | 1.31989 | 0.487809 |
| 7 | 20164200 | 20165199 | 16.45172 | 29.71607 | 24.25573 | 10.53954 | 0.700042 | 0.372378 | 0.487813 |
| 9 | 97967600 | 97968799 | 11.43713 | 20.33091 | 36.31106 | 16.01628 | -0.43734 | 0.462635 | 0.487814 |
| 11 | 7440800 | 7442799 | 21.04987 | 36.89513 | 35.29756 | 15.59357 | 0.45108 | 0.670113 | 0.487816 |
| 3 | 159545000 | 159547399 | 14.47805 | 25.12123 | 46.36009 | 20.34108 | -0.49321 | 0.801679 | 0.487817 |
| 3 | 6691800 | 6692599 | 11.70818 | 22.36281 | 19.03142 | 7.565471 | 0.739008 | -0.01619 | 0.487831 |
| 14 | 50069000 | 50070599 | 15.97915 | 29.13309 | 27.71799 | 11.90421 | 0.490913 | 0.322772 | 0.487848 |
| 13 | 97773800 | 97775399 | 19.65084 | 33.4612 | 54.87104 | 24.05148 | -0.32889 | 1.00489 | 0.487853 |
| 18 | 80298200 | 80300399 | 15.13412 | 26.16684 | 47.41323 | 20.80092 | -0.46799 | 0.83872 | 0.487856 |
| 5 | 12099200 | 12100199 | 12.6807 | 23.75519 | 24.54869 | 10.13844 | 0.413974 | 0.133786 | 0.487867 |
| 4 | 58609200 | 58609599 | 8.462951 | 16.43057 | 18.34031 | 6.881502 | 0.429228 | -0.24099 | 0.487874 |
| 2 | 19130000 | 19132599 | 24.49923 | 42.92393 | 33.91374 | 14.9772 | 0.725794 | 0.692474 | 0.487879 |
| 4 | 31015400 | 31016999 | 16.51309 | 30.67313 | 25.21345 | 10.56083 | 0.725855 | 0.219526 | 0.487902 |
| 3 | 113431400 | 113441999 | 98.91476 | 154.2442 | 149.5135 | 62.04701 | 0.35626 | 2.699467 | 0.487914 |
| 7 | 48178000 | 48179999 | 33.76886 | 55.91177 | 57.00487 | 24.66296 | 0.352826 | 1.325805 | 0.487915 |
| 4 | 79494200 | 79495999 | 15.98425 | 29.02543 | 27.57947 | 11.90444 | 0.487921 | 0.343577 | 0.487929 |
| 7 | 106351800 | 106352599 | 11.4245 | 21.81473 | 20.94358 | 8.330647 | 0.564543 | -0.01577 | 0.487937 |
| 17 | 72460200 | 72461799 | 20.00834 | 35.1333 | 34.33096 | 15.18982 | 0.420733 | 0.635552 | 0.487976 |
| 7 | 86734200 | 86736399 | 20.14138 | 35.62426 | 31.95831 | 14.13464 | 0.546695 | 0.54886 | 0.487991 |
| 14 | 88063600 | 88065399 | 14.33373 | 26.72285 | 24.81503 | 10.33237 | 0.557495 | 0.177125 | 0.488001 |
| 10 | 21722000 | 21724599 | 20.3116 | 34.44879 | 56.48968 | 24.74368 | -0.32951 | 1.044467 | 0.48801 |
| 8 | 38039400 | 38040199 | 10.82321 | 19.19539 | 36.34549 | 16.06259 | -0.52307 | 0.472545 | 0.488016 |
| 6 | 90651200 | 90656399 | 32.99841 | 52.98968 | 109.982 | 46.55898 | -0.7014 | 1.910187 | 0.488033 |
| 1 | 95111200 | 95113199 | 23.73972 | 41.41324 | 37.47546 | 16.4956 | 0.530965 | 0.759169 | 0.488034 |
| 5 | 14198000 | 14199799 | 18.93252 | 34.13185 | 28.33351 | 12.34083 | 0.673516 | 0.407863 | 0.488043 |
| 15 | 17559600 | 17562399 | 34.50388 | 57.01178 | 49.32061 | 21.30822 | 0.5889 | 1.345059 | 0.48805 |
| 14 | 24676000 | 24677199 | 20.09612 | 35.19875 | 35.46008 | 15.65608 | 0.376974 | 0.676866 | 0.488063 |
| X | 8846600 | 8850399 | 24.73843 | 42.93402 | 38.80468 | 17.02616 | 0.532226 | 0.838774 | 0.488076 |
| 19 | 39323400 | 39326399 | 43.04951 | 70.21363 | 65.80046 | 28.26845 | 0.465557 | 1.540647 | 0.488097 |
| 16 | 95752000 | 95759999 | 70.71775 | 110.7801 | 167.4775 | 69.46876 | -0.27908 | 2.607416 | 0.488102 |
| 12 | 97581800 | 97582999 | 18.8645 | 33.0528 | 33.98362 | 15.00852 | 0.347655 | 0.661476 | 0.488105 |
| 3 | 3143200 | 3159199 | 189.2735 | 284.7875 | 230.222 | 95.07091 | 0.577014 | 3.414742 | 0.488135 |

| | | | | | | | | | |
|---|---|---|---|---|---|---|---|---|---|
| 6 | 121678000 | 121678399 | 12.54106 | 23.34736 | 24.53606 | 10.24094 | 0.377839 | 0.172302 | 0.48815 |
| 7 | 103789800 | 103793799 | 44.11315 | 70.70613 | 110.5193 | 46.65405 | -0.29648 | 1.981413 | 0.48815 |
| 4 | 99549200 | 99552399 | 28.74701 | 46.77812 | 79.01605 | 33.91741 | -0.38449 | 1.553463 | 0.488159 |
| 2 | 38228400 | 38229999 | 11.34168 | 19.90299 | 41.51787 | 18.36016 | -0.66768 | 0.607546 | 0.488171 |
| 8 | 44036800 | 44038399 | 14.25021 | 26.01377 | 26.44203 | 11.33827 | 0.397798 | 0.301256 | 0.488173 |
| 5 | 77098200 | 77099999 | 11.76843 | 20.70876 | 39.40229 | 17.44482 | -0.53497 | 0.567885 | 0.488186 |
| 9 | 121362200 | 121366599 | 43.64505 | 71.1257 | 66.85018 | 28.7107 | 0.460545 | 1.555522 | 0.488196 |
| 9 | 56513600 | 56514799 | 11.95893 | 22.26452 | 22.04997 | 9.202662 | 0.462898 | 0.160203 | 0.488206 |
| 1 | 78798000 | 78799999 | 12.04097 | 21.24137 | 39.325 | 17.4019 | -0.49443 | 0.540633 | 0.488216 |
| 2 | 179898000 | 179899599 | 12.04097 | 21.24137 | 39.325 | 17.4019 | -0.49443 | 0.540633 | 0.488216 |
| 2 | 146925600 | 146927599 | 19.42293 | 32.33957 | 68.66528 | 29.82229 | -0.7014 | 1.233142 | 0.488237 |
| 8 | 77188600 | 77192399 | 38.04705 | 63.16018 | 54.4254 | 23.5895 | 0.595027 | 1.306106 | 0.488239 |
| 12 | 103648200 | 103650199 | 14.90289 | 26.15058 | 44.53498 | 19.69323 | -0.37689 | 0.633865 | 0.488242 |
| 3 | 121619600 | 121621399 | 19.52012 | 34.56142 | 30.21227 | 13.35961 | 0.584091 | 0.527997 | 0.488243 |
| 6 | 132553600 | 132562199 | 118.259 | 184.564 | 134.0415 | 55.62105 | 0.773241 | 2.694406 | 0.48825 |
| 10 | 40277800 | 40282199 | 42.92587 | 69.65419 | 70.50563 | 30.21395 | 0.348945 | 1.625267 | 0.488258 |
| 1 | 26215400 | 26219999 | 49.4797 | 79.97569 | 74.46806 | 31.80991 | 0.463745 | 1.721078 | 0.488286 |
| 18 | 40096800 | 40098399 | 21.77248 | 36.81039 | 62.09666 | 27.17369 | -0.37056 | 1.081606 | 0.48829 |
| 7 | 117003200 | 117004999 | 10.0736 | 18.22642 | 33.81491 | 14.66205 | -0.47536 | 0.336242 | 0.488296 |
| 13 | 77951200 | 77952599 | 20.29601 | 35.7148 | 33.64354 | 14.8952 | 0.473841 | 0.607256 | 0.4883 |
| 11 | 30446400 | 30447999 | 20.62728 | 36.47019 | 31.90602 | 14.11256 | 0.582517 | 0.555515 | 0.488303 |
| 10 | 116515400 | 116517999 | 39.17728 | 63.98838 | 65.35737 | 28.09256 | 0.343131 | 1.513411 | 0.488313 |
| 5 | 96074600 | 96075599 | 15.96479 | 27.47613 | 48.98899 | 21.48921 | -0.44624 | 0.887091 | 0.488322 |
| 10 | 125351200 | 125352399 | 10.04902 | 18.3788 | 32.37889 | 13.85139 | -0.38836 | 0.271824 | 0.488333 |
| 13 | 66525400 | 66537999 | 287.3212 | 413.829 | 345.2947 | 142.4962 | 0.472116 | 4.143603 | 0.488373 |
| 18 | 53084800 | 53088399 | 24.07502 | 39.7559 | 68.29676 | 29.4943 | -0.39784 | 1.340751 | 0.488381 |
| 5 | 147045600 | 147052999 | 104.2452 | 158.2952 | 243.1444 | 100.6852 | -0.33707 | 3.213481 | 0.488391 |
| 13 | 61643000 | 61649799 | 83.59881 | 132.7716 | 108.9539 | 45.42357 | 0.617354 | 2.303111 | 0.488395 |
| 12 | 109997800 | 109998399 | 10.10852 | 19.43294 | 21.299 | 8.314824 | 0.400897 | -0.08492 | 0.488395 |
| 1 | 164157600 | 164158799 | 6.173619 | 11.77642 | 26.13618 | 10.41836 | -0.63568 | -0.04257 | 0.488411 |
| 4 | 42731200 | 42732199 | 6.173619 | 11.77642 | 26.13618 | 10.41836 | -0.63568 | -0.04257 | 0.488411 |
| 15 | 33083000 | 33083799 | 6.71867 | 12.88075 | 25.71532 | 10.11838 | -0.46725 | -0.08121 | 0.488414 |
| 4 | 35054600 | 35057199 | 21.01134 | 35.29687 | 59.70775 | 26.07763 | -0.3748 | 1.132475 | 0.488414 |
| 10 | 21773400 | 21776199 | 18.43868 | 31.774 | 53.88977 | 23.63986 | -0.37443 | 0.889555 | 0.488424 |
| 3 | 43813400 | 43815599 | 42.44454 | 68.59104 | 69.13901 | 29.52606 | 0.349282 | 1.712863 | 0.488428 |
| 11 | 73971200 | 73971999 | 10.22015 | 19.64565 | 21.27126 | 8.306277 | 0.41777 | -0.0837 | 0.488434 |
| 6 | 109178800 | 109180999 | 22.02725 | 36.81241 | 61.86413 | 26.93156 | -0.3653 | 1.193762 | 0.488438 |
| 13 | 5244200 | 5248399 | 130.5277 | 198.3136 | 197.2093 | 81.67704 | 0.290305 | 3.205035 | 0.488449 |
| 1 | 176172400 | 176174799 | 25.69409 | 43.90715 | 44.14147 | 19.35395 | 0.374989 | 0.981724 | 0.488457 |
| 14 | 24711200 | 24716799 | 43.46774 | 69.6573 | 108.9039 | 45.95776 | -0.29713 | 1.986662 | 0.488469 |
| 7 | 14755600 | 14776399 | 116.5344 | 180.4506 | 145.2257 | 60.3229 | 0.616537 | 2.827172 | 0.488486 |
| 3 | 63505400 | 63506799 | 10.83628 | 19.6156 | 34.30095 | 14.86403 | -0.39004 | 0.340262 | 0.488487 |
| 13 | 98468200 | 98470799 | 22.85788 | 38.28084 | 62.46948 | 27.23112 | -0.32308 | 1.17585 | 0.488493 |
| 1 | 182125400 | 182126999 | 15.96013 | 28.47509 | 29.25675 | 12.86498 | 0.358471 | 0.452417 | 0.488495 |
| 2 | 116349000 | 116350999 | 18.46025 | 33.22922 | 28.91849 | 12.61991 | 0.604029 | 0.41461 | 0.488499 |
| 1 | 33868000 | 33870199 | 35.14307 | 58.10123 | 58.16645 | 25.13987 | 0.378931 | 1.347843 | 0.488511 |
| 9 | 117139000 | 117142199 | 32.76193 | 54.17546 | 56.37623 | 24.36952 | 0.323262 | 1.336802 | 0.488513 |

| | | | | | | | | | |
|---|---|---|---|---|---|---|---|---|---|
| 6 | 25221000 | 25221999 | 9.184365 | 17.27403 | 33.09687 | 13.57692 | -0.4627 | 0.098355 | 0.488515 |
| 13 | 113567200 | 113568199 | 11.67078 | 20.96136 | 35.75495 | 15.63443 | -0.36297 | 0.396348 | 0.488531 |
| 6 | 89794600 | 89801599 | 77.2056 | 122.5413 | 114.5094 | 47.70987 | 0.429326 | 2.316084 | 0.488538 |
| X | 3326400 | 3328999 | 13.0416 | 24.79345 | 19.91264 | 7.994467 | 0.809445 | 0.030549 | 0.488544 |
| 10 | 121367800 | 121371999 | 33.31884 | 53.90801 | 87.0115 | 37.20727 | -0.32651 | 1.675745 | 0.488562 |
| 8 | 89909400 | 89912599 | 20.65515 | 34.82027 | 58.64914 | 25.6425 | -0.36842 | 1.099494 | 0.488595 |
| 4 | 150045800 | 150049399 | 25.37795 | 41.77372 | 71.12975 | 30.67058 | -0.38668 | 1.384731 | 0.488611 |
| 7 | 122665200 | 122666599 | 12.32553 | 21.95133 | 37.22719 | 16.3944 | -0.36185 | 0.456372 | 0.488625 |
| 10 | 66304200 | 66305399 | 15.02308 | 25.96482 | 52.15183 | 22.87955 | -0.6161 | 0.84904 | 0.488634 |
| 2 | 57963600 | 57963999 | 5.228053 | 10.18951 | 21.43822 | 7.760882 | -0.42863 | -0.39737 | 0.488645 |
| 12 | 81481600 | 81486199 | 45.28106 | 73.1793 | 75.37533 | 32.1918 | 0.318269 | 1.719321 | 0.488664 |
| 5 | 27958800 | 27960999 | 13.32325 | 23.37018 | 40.95531 | 18.10472 | -0.41772 | 0.625772 | 0.48871 |
| 18 | 90439400 | 90440399 | 17.80279 | 31.31179 | 29.12163 | 12.89448 | 0.492055 | 0.591044 | 0.48871 |
| 9 | 118165200 | 118168399 | 20.62309 | 34.84798 | 58.69268 | 25.67983 | -0.36824 | 1.078522 | 0.488731 |
| 8 | 129360200 | 129361599 | 12.21969 | 21.59688 | 37.81052 | 16.72549 | -0.41297 | 0.520617 | 0.488739 |
| 14 | 92195400 | 92201799 | 49.8032 | 80.21332 | 81.5136 | 34.65709 | 0.331997 | 1.829652 | 0.488745 |
| 10 | 63557600 | 63559599 | 27.31726 | 46.06578 | 48.01076 | 20.99322 | 0.321685 | 1.109042 | 0.488751 |
| 18 | 59295800 | 59298199 | 24.26866 | 42.13169 | 39.72868 | 17.43433 | 0.471151 | 0.830394 | 0.488757 |
| 12 | 113474800 | 113482399 | 86.73913 | 136.5084 | 132.653 | 55.03588 | 0.362887 | 2.523154 | 0.488758 |
| 10 | 28889600 | 28890599 | 29.06345 | 48.67418 | 50.23475 | 21.89806 | 0.335624 | 1.180668 | 0.48876 |
| 11 | 7043600 | 7046799 | 27.52098 | 46.91179 | 45.1953 | 19.81202 | 0.435698 | 1.012401 | 0.488767 |
| 3 | 130888800 | 130891199 | 16.81606 | 29.04881 | 49.53366 | 21.73059 | -0.38129 | 0.855174 | 0.488768 |
| 2 | 115985200 | 115987199 | 19.83904 | 35.09706 | 32.00864 | 14.15629 | 0.523185 | 0.545135 | 0.488786 |
| 18 | 76943200 | 76944999 | 15.24045 | 26.39617 | 47.93217 | 21.02969 | -0.47073 | 0.822404 | 0.488805 |
| 8 | 110947600 | 110948599 | 15.0836 | 26.32912 | 44.16181 | 19.44628 | -0.35506 | 0.717926 | 0.488806 |
| 1 | 17754400 | 17756599 | 19.79465 | 33.70029 | 55.42817 | 24.29535 | -0.33325 | 1.00702 | 0.488815 |
| 17 | 71644600 | 71650999 | 51.35842 | 81.67481 | 129.8508 | 54.20699 | -0.33349 | 2.239869 | 0.488818 |
| 2 | 158545400 | 158546199 | 11.24135 | 21.2187 | 22.63527 | 9.226037 | 0.383969 | 0.068776 | 0.48883 |
| 5 | 149152200 | 149161399 | 120.7061 | 182.2513 | 267.1336 | 110.4236 | -0.27651 | 3.333318 | 0.488831 |
| 13 | 65926800 | 65935399 | 232.4458 | 338.7221 | 309.6646 | 127.6122 | 0.356813 | 3.965079 | 0.488834 |
| 1 | 50048000 | 50048799 | 11.28864 | 21.53233 | 18.5155 | 7.381471 | 0.720355 | -0.01361 | 0.488858 |
| 1 | 34441000 | 34443399 | 19.81085 | 33.61986 | 55.94095 | 24.50782 | -0.35046 | 1.036281 | 0.48886 |
| 7 | 12399200 | 12401399 | 32.10838 | 54.42679 | 46.27178 | 20.26469 | 0.61472 | 1.080145 | 0.48887 |
| 3 | 91437000 | 91439799 | 37.29126 | 61.88348 | 55.49825 | 24.04859 | 0.537569 | 1.30962 | 0.488878 |
| 6 | 128230000 | 128232599 | 21.71248 | 36.33908 | 60.73997 | 26.4664 | -0.35751 | 1.176941 | 0.488901 |
| 13 | 66810600 | 66819999 | 177.5482 | 268.9991 | 203.6447 | 84.27092 | 0.680197 | 3.275641 | 0.488911 |
| 2 | 103913000 | 103915599 | 20.72109 | 34.89777 | 58.02912 | 25.364 | -0.3499 | 1.10978 | 0.488932 |
| 12 | 100690600 | 100691599 | 12.02347 | 21.29534 | 37.74289 | 16.68847 | -0.42953 | 0.494122 | 0.488938 |
| 4 | 120366400 | 120367599 | 14.3375 | 27.48248 | 20.94754 | 8.246472 | 0.909177 | -0.01727 | 0.488954 |
| 4 | 46421600 | 46425199 | 21.02757 | 35.32463 | 59.13723 | 25.82856 | -0.35983 | 1.13261 | 0.488957 |
| 7 | 41473200 | 41474999 | 32.51081 | 55.02477 | 41.40783 | 18.12612 | 0.789953 | 1.08932 | 0.488962 |
| 8 | 8669200 | 8671399 | 21.95715 | 36.44136 | 64.93582 | 28.14038 | -0.4498 | 1.273877 | 0.488971 |
| 9 | 60966400 | 60969799 | 33.51217 | 54.29528 | 86.52178 | 37.03533 | -0.30621 | 1.649853 | 0.489006 |
| 2 | 125531400 | 125532399 | 11.95724 | 22.26644 | 21.57381 | 9.000467 | 0.494785 | 0.157644 | 0.489011 |
| X | 34494400 | 34497999 | 60.99156 | 97.33853 | 92.28427 | 38.71426 | 0.416649 | 2.130141 | 0.489014 |
| 3 | 33655400 | 33661199 | 49.96764 | 81.21259 | 66.15291 | 28.38268 | 0.6638 | 1.606172 | 0.489016 |
| 11 | 32286400 | 32289599 | 25.88016 | 42.15651 | 85.01947 | 36.50808 | -0.63846 | 1.536477 | 0.489019 |

| | | | | | | | | | |
|---|---|---|---|---|---|---|---|---|---|
| 9 | 60139400 | 60142199 | 16.97082 | 29.33762 | 49.53695 | 21.73251 | -0.36696 | 0.849325 | 0.489042 |
| 8 | 47826000 | 47830399 | 74.78708 | 116.3131 | 177.3417 | 73.62451 | -0.2994 | 2.737871 | 0.489047 |
| 6 | 130645000 | 130651599 | 89.24879 | 140.9952 | 122.9024 | 51.04381 | 0.523772 | 2.445246 | 0.48906 |
| 3 | 42683400 | 42684399 | 12.48063 | 22.25535 | 37.74261 | 16.60388 | -0.36084 | 0.450443 | 0.489065 |
| 9 | 26590800 | 26592399 | 12.48063 | 22.25535 | 37.74261 | 16.60388 | -0.36084 | 0.450443 | 0.489065 |
| 10 | 6841800 | 6843799 | 12.48063 | 22.25535 | 37.74261 | 16.60388 | -0.36084 | 0.450443 | 0.489065 |
| 8 | 109310600 | 109313999 | 23.02898 | 38.42867 | 62.96667 | 27.38552 | -0.32888 | 1.215068 | 0.489071 |
| 19 | 42663600 | 42667599 | 43.02126 | 69.57128 | 72.72068 | 31.08344 | 0.298165 | 1.696762 | 0.489139 |
| 14 | 51248600 | 51249999 | 21.28961 | 36.96455 | 38.33938 | 16.82598 | 0.334363 | 0.812938 | 0.489167 |
| 6 | 131343800 | 131345599 | 11.19604 | 20.24632 | 34.81953 | 15.10787 | -0.36724 | 0.351358 | 0.489188 |
| 4 | 105692400 | 105694999 | 18.45609 | 32.51043 | 32.13078 | 14.22253 | 0.405304 | 0.580646 | 0.489234 |
| 14 | 114045800 | 114048199 | 20.27939 | 34.42169 | 57.06333 | 25.00101 | -0.34517 | 1.037087 | 0.489238 |
| 19 | 12772000 | 12773799 | 16.9038 | 28.93923 | 51.63158 | 22.64137 | -0.44876 | 0.940318 | 0.489243 |
| X | 73007800 | 73011599 | 39.61175 | 65.16463 | 57.51919 | 24.79557 | 0.558265 | 1.422109 | 0.48927 |
| 3 | 27130200 | 27133999 | 38.03845 | 61.30496 | 97.62542 | 41.54632 | -0.31356 | 1.79712 | 0.489275 |
| 3 | 61419200 | 61421999 | 43.86644 | 70.96086 | 73.45809 | 31.40756 | 0.312566 | 1.691754 | 0.48928 |
| 5 | 12733600 | 12735199 | 11.97409 | 21.48867 | 36.76464 | 16.08694 | -0.3681 | 0.404834 | 0.4893 |
| 13 | 76493800 | 76494599 | 11.00024 | 19.48081 | 36.89016 | 16.3118 | -0.52404 | 0.490716 | 0.489321 |
| 17 | 32407200 | 32409599 | 13.80267 | 24.11685 | 43.12233 | 19.00027 | -0.44669 | 0.694858 | 0.489336 |
| 1 | 99896000 | 99897199 | 13.70169 | 26.11931 | 21.9516 | 8.762498 | 0.751078 | 0.021699 | 0.489343 |
| 13 | 66968200 | 66977999 | 267.8539 | 385.9255 | 370.6626 | 152.958 | 0.269711 | 4.137148 | 0.489371 |
| 18 | 42408000 | 42409799 | 12.50639 | 22.03206 | 39.83223 | 17.63121 | -0.46121 | 0.559573 | 0.489383 |
| 3 | 146518000 | 146519199 | 8.550781 | 15.64959 | 31.44785 | 13.44031 | -0.57529 | 0.255915 | 0.489384 |
| 5 | 87024600 | 87025799 | 13.89654 | 26.40957 | 23.08713 | 9.275788 | 0.68702 | 0.048178 | 0.489399 |
| 2 | 177145800 | 177147999 | 58.68542 | 93.66783 | 96.70191 | 40.5735 | 0.294078 | 2.123618 | 0.489434 |
| 2 | 170748800 | 170749799 | 13.17576 | 24.32417 | 25.46745 | 10.75674 | 0.370558 | 0.226911 | 0.48944 |
| 5 | 149671800 | 149674399 | 22.55506 | 37.58647 | 62.89422 | 27.33046 | -0.35905 | 1.228135 | 0.489445 |
| X | 72457600 | 72459399 | 21.93604 | 38.62384 | 33.04102 | 14.62669 | 0.61263 | 0.610722 | 0.489469 |
| 9 | 81366800 | 81367599 | 13.98535 | 25.67593 | 24.54932 | 10.44752 | 0.493325 | 0.264277 | 0.48948 |
| 14 | 108669800 | 108670599 | 7.215299 | 13.67289 | 27.10083 | 10.95479 | -0.49165 | 0.018388 | 0.489499 |
| 16 | 17046800 | 17049599 | 28.84027 | 48.98367 | 46.28526 | 20.28273 | 0.462857 | 1.050449 | 0.489506 |
| 3 | 106285400 | 106287799 | 31.09075 | 53.62429 | 38.95285 | 17.08794 | 0.844317 | 0.918924 | 0.489517 |
| 5 | 68710800 | 68712399 | 25.28778 | 43.93192 | 39.11163 | 17.17115 | 0.553843 | 0.824748 | 0.489517 |
| 3 | 71774000 | 71774799 | 9.76266 | 18.77917 | 20.76761 | 8.093581 | 0.390293 | -0.0959 | 0.489537 |
| 6 | 31673800 | 31674799 | 7.354967 | 13.93589 | 27.09779 | 10.95634 | -0.46481 | 0.019508 | 0.489571 |
| 6 | 119431200 | 119433399 | 12.85497 | 22.61972 | 39.36246 | 17.42734 | -0.4069 | 0.575594 | 0.489583 |
| 14 | 91944600 | 91946599 | 17.60877 | 31.43668 | 26.18037 | 11.50599 | 0.660424 | 0.454288 | 0.489596 |
| 7 | 23693200 | 23699399 | 75.87698 | 120.752 | 106.5465 | 44.52207 | 0.515746 | 2.235147 | 0.489628 |
| X | 33445800 | 33449799 | 50.43627 | 82.32475 | 52.97231 | 22.76375 | 1.007294 | 1.540337 | 0.489628 |
| 4 | 27842400 | 27842799 | 4.80325 | 9.375013 | 20.46748 | 7.258794 | -0.45436 | -0.48011 | 0.489633 |
| 3 | 152979000 | 152980199 | 15.41493 | 28.81331 | 24.89187 | 10.31888 | 0.666442 | 0.172233 | 0.489643 |
| 11 | 7900600 | 7902599 | 22.39109 | 38.88924 | 39.34151 | 17.2692 | 0.370319 | 0.81635 | 0.489644 |
| 14 | 53417000 | 53419599 | 14.62352 | 25.58041 | 43.08306 | 19.00103 | -0.36073 | 0.682213 | 0.48966 |
| 8 | 4690600 | 4694199 | 72.4068 | 114.6747 | 118.3112 | 49.19924 | 0.283766 | 2.371299 | 0.489667 |
| 5 | 67043000 | 67044599 | 16.22837 | 29.787 | 26.64876 | 11.34457 | 0.587681 | 0.284384 | 0.489675 |
| 12 | 105704200 | 105706999 | 29.21351 | 47.64318 | 77.55054 | 33.31537 | -0.3289 | 1.522228 | 0.489677 |
| 4 | 147292000 | 147296199 | 70.85527 | 112.0661 | 111.084 | 46.1623 | 0.339891 | 2.401497 | 0.489694 |

| | | | | | | | | | |
|---|---|---|---|---|---|---|---|---|---|
| 4 | 155343800 | 155347599 | 46.69412 | 74.34051 | 122.2072 | 51.0953 | -0.37981 | 2.199293 | 0.489732 |
| 12 | 113317200 | 113330199 | 152.3616 | 226.2494 | 331.5941 | 136.5959 | -0.298 | 3.653529 | 0.489763 |
| 9 | 105696000 | 105696799 | 8.79726 | 16.24645 | 30.44456 | 12.85295 | -0.46264 | 0.203023 | 0.489768 |
| 16 | 71352800 | 71353399 | 8.79726 | 16.24645 | 30.44456 | 12.85295 | -0.46264 | 0.203023 | 0.489768 |
| 5 | 132658000 | 132659599 | 14.76138 | 27.49195 | 25.44127 | 10.61151 | 0.560909 | 0.190835 | 0.48978 |
| 19 | 17215400 | 17217999 | 23.34592 | 40.63624 | 38.02975 | 16.71632 | 0.482257 | 0.786289 | 0.489802 |
| 4 | 136015000 | 136019599 | 55.42042 | 87.87885 | 135.3634 | 56.34413 | -0.292 | 2.331687 | 0.489805 |
| 19 | 13536200 | 13540399 | 52.23753 | 84.35186 | 76.26807 | 32.5383 | 0.504497 | 1.755386 | 0.489814 |
| X | 115367600 | 115370799 | 24.69467 | 43.37038 | 32.78186 | 14.50674 | 0.789374 | 0.656135 | 0.489816 |
| 9 | 44144600 | 44145799 | 10.08727 | 18.32807 | 32.85283 | 14.17017 | -0.42078 | 0.311822 | 0.489818 |
| 15 | 23383400 | 23384799 | 12.3143 | 22.89213 | 24.00257 | 10.04273 | 0.378205 | 0.17747 | 0.489825 |
| 8 | 86960600 | 86961999 | 9.091926 | 16.85814 | 30.47559 | 12.79599 | -0.40549 | 0.183007 | 0.489835 |
| 3 | 12523000 | 12524599 | 23.49447 | 40.48773 | 40.91213 | 17.94697 | 0.369662 | 0.893295 | 0.489839 |
| 4 | 97226800 | 97230799 | 49.89985 | 79.43734 | 124.9371 | 52.23037 | -0.31631 | 2.204623 | 0.48984 |
| 1 | 92437600 | 92439799 | 15.23483 | 26.55657 | 44.70823 | 19.66988 | -0.36061 | 0.738658 | 0.489843 |
| 3 | 15460600 | 15462799 | 40.65802 | 66.43474 | 58.21649 | 25.02702 | 0.563969 | 1.5047 | 0.489854 |
| 17 | 45839800 | 45841999 | 13.92777 | 24.47882 | 41.35506 | 18.31194 | -0.36527 | 0.597267 | 0.489868 |
| 4 | 9626200 | 9627199 | 9.864221 | 18.211 | 31.98662 | 13.50989 | -0.3707 | 0.21636 | 0.489884 |
| 8 | 100756800 | 100758399 | 9.972047 | 18.04278 | 33.37039 | 14.46921 | -0.47043 | 0.337701 | 0.489897 |
| 8 | 56139800 | 56140199 | 6.995653 | 13.47746 | 25.90005 | 10.0491 | -0.39381 | -0.12362 | 0.489907 |
| 3 | 14334000 | 14336399 | 23.84148 | 41.70239 | 35.68881 | 15.73908 | 0.611196 | 0.717596 | 0.489908 |
| 15 | 8759400 | 8760399 | 15.38303 | 26.86244 | 44.6642 | 19.6724 | -0.34253 | 0.713944 | 0.489929 |
| 6 | 140668800 | 140675399 | 60.77011 | 96.02735 | 151.2195 | 62.812 | -0.32809 | 2.422351 | 0.489937 |
| 4 | 128851000 | 128852399 | 9.106015 | 16.76654 | 30.92271 | 13.10792 | -0.44419 | 0.22318 | 0.489938 |
| X | 113687600 | 113691399 | 35.77282 | 61.31079 | 37.758 | 16.55903 | 1.079993 | 0.992467 | 0.489986 |
| 1 | 8690600 | 8692599 | 16.69386 | 30.69585 | 26.06812 | 11.07062 | 0.664773 | 0.276808 | 0.489989 |
| 14 | 24603000 | 24604199 | 11.38751 | 21.89537 | 21.80379 | 8.507791 | 0.538629 | -0.07218 | 0.489997 |
| 3 | 8337600 | 8338799 | 8.701635 | 16.46483 | 31.01093 | 12.57508 | -0.42381 | 0.049371 | 0.490004 |
| 12 | 115335400 | 115342599 | 107.5833 | 167.3856 | 152.3712 | 63.2541 | 0.444318 | 2.743738 | 0.490019 |
| 8 | 29582200 | 29584999 | 16.71231 | 29.10147 | 49.5086 | 21.76749 | -0.37598 | 0.767709 | 0.490032 |
| 6 | 132547800 | 132551799 | 73.65596 | 116.9344 | 104.0168 | 43.34822 | 0.500609 | 2.303954 | 0.490058 |
| 7 | 15285000 | 15306999 | 336.4086 | 481.7202 | 396.8789 | 163.9312 | 0.482052 | 4.232817 | 0.49006 |
| 10 | 118213800 | 118214399 | 8.083493 | 15.32654 | 28.21289 | 11.39106 | -0.38513 | 0.023257 | 0.490075 |
| 10 | 47633400 | 47635599 | 40.72998 | 66.09664 | 68.85036 | 29.50647 | 0.307842 | 1.620367 | 0.490075 |
| 10 | 24980200 | 24985399 | 62.03752 | 97.94313 | 147.7263 | 61.33351 | -0.26696 | 2.445676 | 0.490082 |
| X | 7539200 | 7542199 | 33.54659 | 56.46671 | 45.34645 | 19.81719 | 0.696578 | 1.14853 | 0.490086 |
| 16 | 56291200 | 56293199 | 15.70527 | 29.42457 | 23.69609 | 9.784436 | 0.771183 | 0.156059 | 0.490125 |
| 14 | 110373200 | 110375199 | 23.54274 | 41.15857 | 36.29771 | 16.00062 | 0.568105 | 0.725395 | 0.490137 |
| 5 | 95586000 | 95607399 | 339.2462 | 484.1133 | 417.1834 | 172.4134 | 0.412272 | 4.282352 | 0.490141 |
| 1 | 37458200 | 37463799 | 67.4371 | 107.252 | 109.7837 | 45.83419 | 0.300868 | 2.248605 | 0.490144 |
| 12 | 36514200 | 36514999 | 7.651277 | 14.42584 | 28.07625 | 11.46977 | -0.47968 | 0.057351 | 0.490153 |
| 3 | 8448200 | 8456399 | 124.8692 | 190.2792 | 191.4528 | 79.36327 | 0.276873 | 3.142876 | 0.490154 |
| 17 | 58224000 | 58226599 | 31.66586 | 53.22882 | 50.21216 | 21.93457 | 0.464887 | 1.153461 | 0.490155 |
| 5 | 148338000 | 148341199 | 16.00832 | 27.81861 | 46.8553 | 20.57443 | -0.36189 | 0.788533 | 0.490161 |
| 17 | 64723000 | 64726999 | 47.48355 | 76.97991 | 70.50785 | 30.1947 | 0.491611 | 1.651133 | 0.490162 |
| 10 | 65698800 | 65700799 | 20.54432 | 36.40658 | 30.73233 | 13.58721 | 0.634914 | 0.527951 | 0.490166 |
| X | 54516000 | 54517599 | 59.43975 | 94.91018 | 94.94076 | 39.85737 | 0.340174 | 2.112846 | 0.490194 |

| | | | | | | | | | |
|---|---|---|---|---|---|---|---|---|---|
| 19 | 9671200 | 9673999 | 42.53455 | 69.46986 | 64.69601 | 27.808 | 0.475976 | 1.519256 | 0.490205 |
| 14 | 122869200 | 122871599 | 26.67306 | 43.92943 | 71.26905 | 30.73771 | -0.31681 | 1.382295 | 0.490211 |
| 2 | 104420600 | 104421599 | 7.196614 | 13.70046 | 26.64374 | 10.66559 | -0.45211 | -0.01545 | 0.490214 |
| 15 | 96046600 | 96047599 | 6.114044 | 11.83578 | 24.34299 | 9.26478 | -0.46315 | -0.19885 | 0.490238 |
| 10 | 8179800 | 8180799 | 7.614493 | 14.51772 | 27.24315 | 10.87106 | -0.39763 | -0.02228 | 0.490244 |
| 12 | 8893200 | 8894199 | 7.614493 | 14.51772 | 27.24315 | 10.87106 | -0.39763 | -0.02228 | 0.490244 |
| 19 | 39517200 | 39518999 | 27.61768 | 47.86562 | 39.03418 | 17.12615 | 0.679381 | 0.860831 | 0.49025 |
| 1 | 167337800 | 167339599 | 11.67156 | 20.8132 | 36.73115 | 16.15853 | -0.41795 | 0.44197 | 0.490251 |
| X | 23684400 | 23685399 | 15.71484 | 28.61666 | 26.40889 | 11.35911 | 0.533684 | 0.327714 | 0.490302 |
| 4 | 44256800 | 44257599 | 19.58586 | 34.69112 | 27.95826 | 12.36196 | 0.701234 | 0.523184 | 0.490308 |
| 8 | 3293800 | 3297599 | 36.78142 | 60.25062 | 62.53449 | 26.90891 | 0.322651 | 1.465173 | 0.49031 |
| 11 | 89008600 | 89010599 | 14.28747 | 25.10245 | 41.89549 | 18.5459 | -0.34773 | 0.606996 | 0.490316 |
| 5 | 50575200 | 50575799 | 5.073151 | 9.892637 | 20.89586 | 7.514329 | -0.42497 | -0.4235 | 0.490316 |
| 9 | 77727600 | 77728799 | 16.27688 | 28.97282 | 29.72164 | 13.09503 | 0.358765 | 0.469606 | 0.490332 |
| X | 104655600 | 104657599 | 37.27107 | 60.90766 | 62.79556 | 26.9967 | 0.330249 | 1.501484 | 0.490339 |
| 3 | 142406200 | 142407799 | 11.60471 | 20.44192 | 39.38237 | 17.43237 | -0.5523 | 0.554783 | 0.490349 |
| X | 78712400 | 78716599 | 56.77091 | 91.4412 | 79.45789 | 33.78609 | 0.557526 | 1.839468 | 0.490377 |
| 17 | 40024600 | 40027399 | 44.65106 | 72.65319 | 67.9978 | 29.18719 | 0.465085 | 1.580018 | 0.490397 |
| 4 | 63395400 | 63397599 | 16.42036 | 28.19993 | 50.04501 | 21.95101 | -0.4403 | 0.908262 | 0.490399 |
| 11 | 91126400 | 91127199 | 10.61616 | 19.00354 | 34.81782 | 15.26765 | -0.46803 | 0.409194 | 0.490404 |
| 17 | 50599600 | 50602799 | 73.37831 | 112.8989 | 219.3143 | 91.08167 | -0.66047 | 2.939027 | 0.490405 |
| 17 | 53874000 | 53875799 | 11.64979 | 20.46764 | 39.49407 | 17.48248 | -0.55551 | 0.589436 | 0.49041 |
| 3 | 91502200 | 91506999 | 65.35738 | 104.9928 | 81.78488 | 34.64217 | 0.71024 | 1.9312 | 0.490433 |
| 4 | 119544600 | 119546999 | 33.05581 | 53.4287 | 87.51649 | 37.38325 | -0.34919 | 1.70333 | 0.490451 |
| 5 | 91208800 | 91209799 | 16.27393 | 28.82592 | 27.47103 | 12.14644 | 0.460785 | 0.500371 | 0.490456 |
| 14 | 89244800 | 89246399 | 22.57481 | 39.26413 | 34.10757 | 14.98512 | 0.589276 | 0.789153 | 0.490467 |
| 1 | 51472000 | 51473199 | 17.50412 | 30.08115 | 50.59013 | 22.19056 | -0.36298 | 0.907084 | 0.49047 |
| 19 | 40667000 | 40668999 | 11.01012 | 19.75974 | 36.26236 | 15.86648 | -0.46868 | 0.395371 | 0.490484 |
| 1 | 3377000 | 3379199 | 20.34638 | 36.12312 | 30.19287 | 13.33165 | 0.650466 | 0.512065 | 0.49049 |
| 11 | 30023400 | 30024399 | 14.93225 | 25.96926 | 45.28168 | 19.89336 | -0.41134 | 0.770954 | 0.490507 |
| 10 | 50166600 | 50167799 | 11.98696 | 22.64058 | 23.59933 | 9.605955 | 0.418549 | 0.072629 | 0.49052 |
| X | 43780600 | 43782399 | 17.41561 | 33.02805 | 21.09915 | 8.514737 | 1.130629 | 0.089927 | 0.490528 |
| 8 | 57259200 | 57259599 | 8.123038 | 15.45818 | 29.44855 | 11.79796 | -0.42414 | 0.003185 | 0.49054 |
| 15 | 23568400 | 23571599 | 59.87157 | 95.48136 | 99.48723 | 41.69346 | 0.279482 | 2.149922 | 0.490547 |
| 1 | 72154600 | 72156599 | 20.64776 | 35.97584 | 37.19085 | 16.35788 | 0.339547 | 0.759082 | 0.490579 |
| 5 | 20676800 | 20678599 | 34.57479 | 57.76903 | 52.52563 | 22.86292 | 0.517998 | 1.227164 | 0.490583 |
| X | 89698200 | 89702999 | 93.73697 | 146.6086 | 143.1352 | 59.38133 | 0.349281 | 2.645988 | 0.490596 |
| 4 | 23240400 | 23241399 | 8.093796 | 15.3129 | 28.15324 | 11.41912 | -0.38912 | 0.038504 | 0.490655 |
| 6 | 117856200 | 117857199 | 8.093796 | 15.3129 | 28.15324 | 11.41912 | -0.38912 | 0.038504 | 0.490655 |
| 15 | 37531600 | 37532999 | 8.093796 | 15.3129 | 28.15324 | 11.41912 | -0.38912 | 0.038504 | 0.490655 |
| 6 | 101822400 | 101822999 | 5.80727 | 11.28837 | 22.907 | 8.51096 | -0.41147 | -0.29184 | 0.490665 |
| 8 | 121940000 | 121946399 | 68.93456 | 107.5322 | 170.2617 | 70.65427 | -0.35043 | 2.681864 | 0.490671 |
| X | 121929000 | 121936399 | 129.151 | 197.8714 | 185.3335 | 76.9237 | 0.386241 | 3.037289 | 0.490673 |
| 9 | 96379800 | 96390199 | 83.31152 | 127.7727 | 229.4096 | 95.23252 | -0.55054 | 3.004505 | 0.490728 |
| 13 | 61544600 | 61545799 | 25.75173 | 42.46169 | 70.49152 | 30.41743 | -0.34915 | 1.367136 | 0.490733 |
| Y | 89819000 | 89819599 | 9.882836 | 18.97901 | 15.16399 | 5.93778 | 0.848832 | -0.0977 | 0.490741 |
| 9 | 7096200 | 7097599 | 13.84904 | 25.42183 | 25.7878 | 10.97684 | 0.407595 | 0.262799 | 0.490752 |

| | | | | | | | | | |
|---|---|---|---|---|---|---|---|---|---|
| 14 | 32468800 | 32469599 | 4.519819 | 8.811347 | 20.24853 | 7.306397 | -0.54906 | -0.42098 | 0.490763 |
| 9 | 114465400 | 114467399 | 10.14944 | 18.14621 | 35.81047 | 15.71889 | -0.57579 | 0.413503 | 0.490766 |
| 9 | 5458800 | 5460799 | 19.6123 | 34.28809 | 35.01669 | 15.43627 | 0.357343 | 0.701216 | 0.490768 |
| 2 | 139854400 | 139855599 | 18.27488 | 32.59059 | 30.49622 | 13.41498 | 0.492225 | 0.472789 | 0.490778 |
| 10 | 15810000 | 15814199 | 88.44556 | 138.1835 | 143.0003 | 59.33197 | 0.264132 | 2.659677 | 0.490787 |
| 17 | 24215200 | 24225399 | 73.43471 | 113.8904 | 185.114 | 76.88336 | -0.3947 | 2.784737 | 0.490791 |
| 3 | 34867000 | 34869799 | 24.55154 | 41.86984 | 43.15823 | 18.91976 | 0.338675 | 0.994463 | 0.490793 |
| Y | 11857800 | 11858599 | 39.72509 | 64.84138 | 40.11325 | 17.23784 | 1.063574 | 1.501321 | 0.490819 |
| 12 | 45475400 | 45476599 | 9.941571 | 18.47012 | 37.74882 | 15.80715 | -0.579 | 0.19505 | 0.49085 |
| 13 | 120693200 | 120696199 | 44.3777 | 70.81455 | 115.9226 | 48.62169 | -0.37017 | 2.121529 | 0.490858 |
| 16 | 7993600 | 7995399 | 30.91499 | 51.21712 | 50.84554 | 22.00721 | 0.391216 | 1.30591 | 0.490862 |
| 7 | 59428400 | 59431599 | 35.06726 | 57.89564 | 48.783 | 21.06414 | 0.626752 | 1.356567 | 0.490863 |
| 1 | 100925600 | 100927599 | 25.53701 | 43.78207 | 43.6185 | 19.1297 | 0.388811 | 0.953431 | 0.490867 |
| 19 | 23867600 | 23872199 | 43.64334 | 69.53812 | 129.7275 | 54.29862 | -0.56077 | 2.174055 | 0.490878 |
| 4 | 89138400 | 89138999 | 10.22652 | 19.69425 | 20.82559 | 8.089212 | 0.459446 | -0.09912 | 0.490881 |
| 2 | 36791200 | 36793999 | 45.54707 | 73.78082 | 72.42716 | 30.99815 | 0.390795 | 1.664947 | 0.490902 |
| 5 | 148013200 | 148024199 | 122.0835 | 184.5121 | 267.0458 | 110.4216 | -0.25706 | 3.313917 | 0.490907 |
| 5 | 10839600 | 10840599 | 10.7253 | 20.44205 | 22.20551 | 8.866976 | 0.383729 | -0.00395 | 0.49094 |
| 11 | 114324600 | 114326199 | 8.823053 | 16.21628 | 30.93153 | 13.14386 | -0.49479 | 0.231925 | 0.490941 |
| 19 | 12872000 | 12876799 | 103.0775 | 159.8576 | 144.7406 | 60.11541 | 0.448416 | 2.793914 | 0.491017 |
| 8 | 37560400 | 37563599 | 30.64241 | 50.0687 | 81.04887 | 34.8424 | -0.31908 | 1.499297 | 0.491017 |
| 14 | 21993400 | 21996999 | 37.11484 | 60.9166 | 62.08159 | 26.73629 | 0.350041 | 1.441837 | 0.491033 |
| 8 | 112406800 | 112407999 | 9.775135 | 17.80982 | 32.37776 | 13.91733 | -0.43768 | 0.292822 | 0.491042 |
| 8 | 85871200 | 85882199 | 184.1534 | 274.6121 | 270.0074 | 111.278 | 0.283137 | 3.582484 | 0.491049 |
| 10 | 130242600 | 130243799 | 17.69806 | 31.48261 | 30.50335 | 13.44661 | 0.440094 | 0.483555 | 0.491062 |
| 10 | 19371400 | 19372199 | 10.73824 | 20.42057 | 22.18881 | 8.902709 | 0.377358 | 0.011701 | 0.491063 |
| 11 | 79165400 | 79165999 | 5.422491 | 10.48098 | 24.27025 | 9.303467 | -0.6432 | -0.18668 | 0.491069 |
| 10 | 120595400 | 120597799 | 18.69885 | 32.93804 | 31.52535 | 13.95454 | 0.451268 | 0.580126 | 0.491071 |
| 9 | 123148200 | 123150199 | 34.66212 | 56.16163 | 90.97616 | 38.94366 | -0.32979 | 1.652925 | 0.491072 |
| 17 | 31232800 | 31238399 | 42.74838 | 68.41165 | 109.6891 | 46.19553 | -0.33569 | 2.030352 | 0.491076 |
| 7 | 31258200 | 31264399 | 70.86356 | 112.6129 | 112.3773 | 46.87892 | 0.336384 | 2.274547 | 0.491097 |
| 9 | 120550800 | 120560399 | 106.6884 | 162.5368 | 237.1493 | 98.29904 | -0.25911 | 3.146579 | 0.491105 |
| 2 | 81289400 | 81293199 | 75.75801 | 119.4162 | 120.5693 | 50.03285 | 0.309528 | 2.48404 | 0.49112 |
| 2 | 70388600 | 70390199 | 9.199988 | 17.05495 | 30.98584 | 13.01392 | -0.41334 | 0.184033 | 0.491124 |
| 10 | 14987800 | 14989999 | 37.92444 | 62.11879 | 63.03074 | 27.12168 | 0.355231 | 1.471446 | 0.491125 |
| 2 | 149453200 | 149454199 | 12.98039 | 22.94438 | 38.75775 | 17.14404 | -0.36175 | 0.5246 | 0.491129 |
| 6 | 35653400 | 35655999 | 20.88068 | 36.55645 | 35.90487 | 15.84643 | 0.413391 | 0.68837 | 0.491144 |
| 6 | 100338800 | 100340599 | 12.49931 | 22.14924 | 37.72068 | 16.67697 | -0.37197 | 0.492264 | 0.491149 |
| 2 | 118318000 | 118319599 | 16.34467 | 29.03897 | 29.80747 | 13.15363 | 0.356461 | 0.48283 | 0.491181 |
| 7 | 19207400 | 19209799 | 16.69411 | 28.79225 | 48.99527 | 21.49335 | -0.37873 | 0.86933 | 0.491197 |
| 2 | 72833600 | 72847199 | 153.838 | 228.9834 | 326.7836 | 134.6468 | -0.25647 | 3.612554 | 0.491197 |
| 9 | 3481600 | 3487199 | 58.48951 | 94.19025 | 79.94623 | 33.98374 | 0.590976 | 1.847113 | 0.491208 |
| 6 | 18744200 | 18745399 | 16.27105 | 29.34968 | 27.99202 | 12.1838 | 0.474799 | 0.383714 | 0.491212 |
| 3 | 120123800 | 120126399 | 52.11618 | 83.82136 | 75.72392 | 32.13033 | 0.498911 | 1.872511 | 0.491214 |
| 12 | 84024600 | 84027199 | 27.64251 | 44.98336 | 78.14295 | 33.54332 | -0.42468 | 1.550948 | 0.491216 |
| 1 | 5132600 | 5132999 | 10.66363 | 20.37522 | 21.75338 | 8.640787 | 0.415508 | -0.02108 | 0.491218 |
| 3 | 47806200 | 47808399 | 19.65616 | 34.77888 | 32.00975 | 14.15635 | 0.51013 | 0.542187 | 0.491244 |

| | | | | | | | | | |
|---|---|---|---|---|---|---|---|---|---|
| 1 | 96132400 | 96133199 | 7.904531 | 15.05792 | 28.36674 | 11.33973 | -0.40577 | -0.01165 | 0.491255 |
| 10 | 87044400 | 87044799 | 4.517113 | 8.814063 | 19.78547 | 7.050877 | -0.49945 | -0.46774 | 0.491267 |
| 15 | 24033600 | 24035799 | 12.0047 | 21.67251 | 40.1096 | 17.43949 | -0.4748 | 0.376086 | 0.49127 |
| 13 | 70549200 | 70551199 | 11.45489 | 20.73676 | 36.38678 | 15.7666 | -0.39512 | 0.34801 | 0.491285 |
| 5 | 92270200 | 92273199 | 28.39216 | 47.60785 | 49.1358 | 21.43547 | 0.335541 | 1.164539 | 0.491342 |
| 4 | 102005400 | 102008199 | 24.3701 | 41.84106 | 42.52871 | 18.6539 | 0.360197 | 0.932458 | 0.491344 |
| 4 | 121109800 | 121113599 | 37.80027 | 62.11644 | 60.85528 | 26.22153 | 0.407558 | 1.431005 | 0.491345 |
| 1 | 7311400 | 7313399 | 44.16665 | 71.42666 | 74.10764 | 31.67805 | 0.308918 | 1.697989 | 0.491346 |
| 7 | 85489200 | 85491999 | 41.23512 | 67.01866 | 68.22504 | 29.27142 | 0.342889 | 1.588098 | 0.491349 |
| 8 | 56343800 | 56344999 | 19.39225 | 33.96219 | 34.99299 | 15.44843 | 0.344636 | 0.673549 | 0.491369 |
| 6 | 108848200 | 108849799 | 15.52742 | 27.09403 | 45.20058 | 19.89917 | -0.34751 | 0.725661 | 0.491415 |
| 3 | 11610800 | 11619399 | 158.9336 | 239.4439 | 237.0874 | 97.93962 | 0.286245 | 3.382959 | 0.49142 |
| 4 | 79215600 | 79218599 | 25.49661 | 44.18664 | 39.65452 | 17.39829 | 0.541741 | 0.851392 | 0.491457 |
| 17 | 85204800 | 85208599 | 40.33831 | 64.88409 | 102.1245 | 43.33787 | -0.30027 | 1.86377 | 0.491474 |
| 7 | 40385800 | 40389199 | 29.99543 | 50.0071 | 52.42768 | 22.78852 | 0.31321 | 1.234993 | 0.491495 |
| 5 | 101653600 | 101655999 | 22.21705 | 38.81917 | 36.96861 | 16.28891 | 0.457785 | 0.728346 | 0.491496 |
| 10 | 31946200 | 31947799 | 12.92135 | 24.49084 | 22.39348 | 9.048018 | 0.614922 | 0.05275 | 0.491507 |
| 4 | 65694600 | 65695399 | 12.23853 | 23.46322 | 22.26251 | 8.759752 | 0.596681 | -0.03561 | 0.491528 |
| 5 | 135607200 | 135611199 | 41.25617 | 66.04618 | 107.8335 | 45.43603 | -0.36124 | 2.01679 | 0.49153 |
| 11 | 57535200 | 57536999 | 15.25265 | 26.53672 | 45.27191 | 19.8939 | -0.37993 | 0.767278 | 0.491534 |
| 2 | 113028600 | 113030399 | 20.40778 | 34.32818 | 57.99781 | 25.34093 | -0.37283 | 1.11926 | 0.491544 |
| 15 | 74673400 | 74675599 | 29.35635 | 49.44184 | 43.75391 | 19.12454 | 0.556754 | 1.12204 | 0.491547 |
| 9 | 46269200 | 46273199 | 30 | 48.95993 | 78.41587 | 33.69567 | -0.30497 | 1.513777 | 0.49156 |
| 4 | 113597800 | 113600399 | 28.03824 | 49.42031 | 27.68755 | 12.25422 | 1.220621 | 0.627316 | 0.491579 |
| 5 | 24516400 | 24520799 | 81.6141 | 126.5873 | 187.1839 | 77.74198 | -0.25832 | 2.788419 | 0.4916 |
| 17 | 21335200 | 21336199 | 11.02776 | 20.06841 | 34.91856 | 15.03157 | -0.37657 | 0.313808 | 0.491601 |
| 4 | 90152800 | 90155399 | 29.78501 | 51.07929 | 42.35185 | 18.57461 | 0.652709 | 0.965852 | 0.491612 |
| 11 | 27771200 | 27772599 | 18.3495 | 33.48015 | 26.97331 | 11.57337 | 0.730281 | 0.33468 | 0.491613 |
| 6 | 92570200 | 92572399 | 19.3281 | 33.89429 | 34.34965 | 15.18124 | 0.368282 | 0.650351 | 0.491619 |
| 14 | 105894400 | 105895999 | 18.80059 | 33.32418 | 31.5102 | 13.93047 | 0.472307 | 0.516753 | 0.491626 |
| 1 | 110451800 | 110453199 | 17.35723 | 31.05157 | 30.1381 | 13.22236 | 0.442456 | 0.447663 | 0.491629 |
| 17 | 23194600 | 23195999 | 27.46504 | 44.78711 | 84.23023 | 36.18359 | -0.53686 | 1.524126 | 0.491659 |
| 3 | 44811800 | 44812599 | 10.39977 | 18.84463 | 33.32584 | 14.42308 | -0.40471 | 0.330589 | 0.491664 |
| 3 | 12400400 | 12402399 | 24.50802 | 42.88204 | 35.07496 | 15.47257 | 0.676144 | 0.71514 | 0.491669 |
| 12 | 102497600 | 102499199 | 13.38663 | 24.76462 | 25.37456 | 10.6874 | 0.404161 | 0.215554 | 0.491677 |
| 6 | 114415400 | 114417399 | 28.42338 | 47.57367 | 49.15422 | 21.41893 | 0.334044 | 1.184308 | 0.491698 |
| 16 | 21094400 | 21097599 | 32.88206 | 53.15268 | 86.99633 | 37.1654 | -0.34792 | 1.699636 | 0.491708 |
| 17 | 28311000 | 28321399 | 83.25035 | 128.6412 | 193.7641 | 80.48537 | -0.28909 | 2.857035 | 0.491721 |
| 18 | 25361600 | 25363199 | 12.97307 | 22.95306 | 38.73965 | 17.1328 | -0.36002 | 0.513979 | 0.491732 |
| 2 | 28357400 | 28363199 | 61.86952 | 97.57202 | 148.9068 | 61.79608 | -0.28514 | 2.469858 | 0.491735 |
| 14 | 10122200 | 10130199 | 72.98328 | 113.1102 | 186.0669 | 77.2869 | -0.41279 | 2.797004 | 0.491749 |
| 11 | 4832800 | 4835199 | 12.43365 | 21.7807 | 41.07416 | 18.13682 | -0.52272 | 0.644464 | 0.491783 |
| 1 | 5985800 | 5988199 | 24.12385 | 39.73627 | 69.18785 | 29.8418 | -0.41829 | 1.373434 | 0.491795 |
| 18 | 74387800 | 74399599 | 429.9272 | 584.0923 | 801.5652 | 333.8769 | -0.33029 | 5.026451 | 0.491799 |
| 2 | 15668400 | 15669199 | 9.230865 | 16.6298 | 36.92832 | 16.09856 | -0.7378 | 0.364022 | 0.491826 |
| 16 | 49685400 | 49687399 | 18.83874 | 33.50287 | 31.00717 | 13.67175 | 0.505784 | 0.495107 | 0.491841 |
| 17 | 42359400 | 42360999 | 13.32759 | 23.29198 | 43.63425 | 19.22914 | -0.51364 | 0.68872 | 0.491844 |

| | | | | | | | | | |
|---|---|---|---|---|---|---|---|---|---|
| 9 | 85663400 | 85665799 | 22.50093 | 37.70266 | 61.37935 | 26.76471 | -0.31967 | 1.167314 | 0.491857 |
| 12 | 113976200 | 113981799 | 67.51167 | 107.9684 | 94.34497 | 39.69544 | 0.537321 | 2.071632 | 0.491862 |
| 3 | 36218200 | 36221599 | 53.55927 | 86.26706 | 73.24546 | 31.14392 | 0.5908 | 1.830429 | 0.491871 |
| 6 | 114127000 | 114130599 | 23.3848 | 38.93854 | 64.01627 | 27.80424 | -0.33365 | 1.240776 | 0.491873 |
| 1 | 164521000 | 164522799 | 20.63343 | 35.10833 | 58.19016 | 25.50482 | -0.34449 | 1.018747 | 0.491897 |
| X | 78340200 | 78341799 | 15.31187 | 29.37666 | 20.36199 | 7.995223 | 1.048771 | -0.01658 | 0.49191 |
| 1 | 136631800 | 136632599 | 5.523268 | 10.73847 | 22.33827 | 8.286196 | -0.44478 | -0.30384 | 0.491926 |
| 12 | 16729800 | 16730399 | 5.523268 | 10.73847 | 22.33827 | 8.286196 | -0.44478 | -0.30384 | 0.491926 |
| 7 | 18068400 | 18097199 | 314.2719 | 450.4948 | 369.6568 | 152.662 | 0.489381 | 4.214705 | 0.491928 |
| 13 | 43811600 | 43817599 | 59.76365 | 94.28269 | 146.7019 | 60.88515 | -0.31266 | 2.461715 | 0.491945 |
| 1 | 185842400 | 185844799 | 8.975311 | 16.05167 | 38.44108 | 16.86942 | -0.85264 | 0.411044 | 0.491954 |
| 1 | 96103600 | 96104999 | 13.92695 | 25.87004 | 24.75697 | 10.36926 | 0.507583 | 0.195007 | 0.491955 |
| 13 | 98935200 | 98939399 | 57.65851 | 92.26593 | 95.08677 | 40.04151 | 0.300642 | 2.043282 | 0.491961 |
| 17 | 8849200 | 8851999 | 19.41833 | 33.88262 | 33.17736 | 14.60585 | 0.417623 | 0.725954 | 0.491977 |
| 13 | 38407600 | 38408599 | 19.327 | 34.0741 | 29.10845 | 12.88247 | 0.615091 | 0.572542 | 0.491988 |
| 17 | 64156400 | 64158199 | 9.392465 | 17.19415 | 33.43516 | 14.285 | -0.52889 | 0.260829 | 0.492002 |
| 5 | 100576800 | 100577999 | 7.677937 | 14.3725 | 29.56564 | 12.22602 | -0.57341 | 0.106529 | 0.49201 |
| 6 | 49933200 | 49935599 | 14.48862 | 25.25091 | 44.22217 | 19.45366 | -0.41715 | 0.738595 | 0.492025 |
| 6 | 70501000 | 70502199 | 12.87638 | 22.65536 | 39.38375 | 17.43708 | -0.40539 | 0.577693 | 0.492036 |
| 9 | 119027400 | 119035999 | 94.30801 | 144.2648 | 220.2393 | 91.37881 | -0.31968 | 3.058667 | 0.492044 |
| 18 | 60484600 | 60484999 | 10.04071 | 19.37625 | 20.85694 | 8.053663 | 0.441803 | -0.1203 | 0.492066 |
| 2 | 145707000 | 145712399 | 50.20545 | 80.05022 | 123.2426 | 51.63477 | -0.28313 | 2.153745 | 0.492074 |
| 19 | 12338400 | 12344399 | 60.79257 | 97.1757 | 76.03211 | 31.96838 | 0.695634 | 2.07317 | 0.49208 |
| 2 | 146441400 | 146443999 | 32.31658 | 52.39082 | 84.93983 | 36.37486 | -0.33022 | 1.632128 | 0.492083 |
| 4 | 24468600 | 24472799 | 46.55428 | 75.50163 | 70.93336 | 30.3853 | 0.455555 | 1.642997 | 0.492098 |
| 5 | 119870800 | 119874799 | 36.29591 | 58.39957 | 96.90435 | 41.14068 | -0.37571 | 1.846732 | 0.492098 |
| 13 | 17743000 | 17744199 | 14.70447 | 27.71588 | 23.95831 | 9.792367 | 0.681672 | 0.11206 | 0.492115 |
| 5 | 40923800 | 40924399 | 5.949978 | 11.57257 | 23.61669 | 8.733068 | -0.41387 | -0.30672 | 0.492118 |
| 13 | 65767600 | 65768599 | 45.02547 | 72.62833 | 74.00132 | 31.52774 | 0.33078 | 1.774398 | 0.492127 |
| 11 | 53319600 | 53322199 | 14.75694 | 25.7003 | 44.73283 | 19.66953 | -0.40859 | 0.748808 | 0.49215 |
| 13 | 5504200 | 5511999 | 169.1324 | 254.3643 | 242.8522 | 100.2741 | 0.336534 | 3.419205 | 0.492156 |
| 7 | 20928000 | 20930799 | 37.75049 | 61.8399 | 63.02947 | 27.12225 | 0.348837 | 1.469713 | 0.492164 |
| 5 | 42440400 | 42442599 | 22.31894 | 38.90123 | 33.41034 | 14.6983 | 0.605738 | 0.756231 | 0.492175 |
| 15 | 103479200 | 103482199 | 31.01447 | 52.07944 | 50.76105 | 22.1645 | 0.417852 | 1.16158 | 0.492175 |
| 18 | 17280200 | 17286399 | 71.72523 | 114.4455 | 99.72173 | 41.82232 | 0.537873 | 2.148275 | 0.492176 |
| 18 | 75877600 | 75880199 | 36.86802 | 60.37611 | 62.99602 | 27.1044 | 0.314943 | 1.470385 | 0.49218 |
| 1 | 102076800 | 102080599 | 50.54299 | 81.31059 | 83.84286 | 35.58762 | 0.308921 | 1.866319 | 0.492181 |
| 5 | 101772600 | 101775199 | 23.30245 | 40.35176 | 39.77616 | 17.45123 | 0.40665 | 0.84801 | 0.492213 |
| 13 | 25840400 | 25841599 | 8.093079 | 15.2444 | 28.59826 | 11.70173 | -0.42908 | 0.068521 | 0.492225 |
| 17 | 16486200 | 16486799 | 8.920912 | 16.79472 | 31.36464 | 12.84534 | -0.42362 | 0.088214 | 0.492244 |
| 7 | 35426800 | 35430199 | 31.3306 | 50.85888 | 83.27071 | 35.69449 | -0.34259 | 1.602791 | 0.492254 |
| 12 | 4837800 | 4840199 | 41.06261 | 66.58201 | 69.34688 | 29.70136 | 0.306942 | 1.637616 | 0.492257 |
| X | 26551200 | 26577599 | 389.5963 | 551.7041 | 431.3399 | 178.494 | 0.541475 | 4.397582 | 0.492296 |
| X | 150724200 | 150726399 | 21.55693 | 38.04363 | 32.45257 | 14.3596 | 0.617874 | 0.582551 | 0.492303 |
| Y | 44020000 | 44021999 | 15.42232 | 27.76227 | 19.60328 | 8.554232 | 0.903475 | 0.370403 | 0.492318 |
| 13 | 32539200 | 32543599 | 56.35552 | 90.37909 | 81.49156 | 34.42965 | 0.496706 | 1.968716 | 0.492329 |
| X | 43101200 | 43105999 | 110.6895 | 171.7941 | 156.1081 | 64.82929 | 0.444117 | 2.78798 | 0.492348 |

| | | | | | | | | | |
|---|---|---|---|---|---|---|---|---|---|
| 9 | 117956000 | 117965599 | 127.7717 | 194.8138 | 198.7876 | 82.41569 | 0.257289 | 3.133062 | 0.492359 |
| 7 | 103028800 | 103030799 | 22.13212 | 39.0982 | 30.59159 | 13.53342 | 0.742144 | 0.569858 | 0.492361 |
| 10 | 119691000 | 119692399 | 16.62087 | 30.07104 | 28.62834 | 12.41395 | 0.481114 | 0.372372 | 0.492367 |
| 1 | 66738800 | 66740799 | 14.30033 | 25.08008 | 41.96901 | 18.55035 | -0.35146 | 0.636574 | 0.49238 |
| 2 | 147513400 | 147514799 | 14.30033 | 25.08008 | 41.96901 | 18.55035 | -0.35146 | 0.636574 | 0.49238 |
| 5 | 14589400 | 14590799 | 13.349 | 23.40026 | 40.99931 | 18.1143 | -0.41731 | 0.637565 | 0.492397 |
| 18 | 52084000 | 52085799 | 14.52345 | 27.14176 | 23.66138 | 9.811973 | 0.65237 | 0.16008 | 0.492397 |
| 13 | 73071800 | 73073399 | 12.53111 | 22.18234 | 37.76567 | 16.70036 | -0.37202 | 0.505217 | 0.492413 |
| 16 | 19707000 | 19709599 | 32.36079 | 53.44251 | 55.70926 | 24.05958 | 0.320706 | 1.351709 | 0.492414 |
| 11 | 20078000 | 20083799 | 51.09113 | 82.00103 | 84.45132 | 35.72293 | 0.306593 | 1.938166 | 0.492433 |
| 8 | 81812200 | 81814799 | 26.87532 | 46.0209 | 44.1258 | 19.35031 | 0.443613 | 0.970225 | 0.492472 |
| 1 | 120920000 | 120921799 | 11.8553 | 21.02261 | 37.72463 | 16.67425 | -0.44664 | 0.480288 | 0.492483 |
| 5 | 55934200 | 55935799 | 21.79907 | 37.90191 | 38.76056 | 17.02573 | 0.354806 | 0.796385 | 0.492496 |
| 2 | 48496600 | 48497799 | 18.03759 | 32.17447 | 30.5989 | 13.45766 | 0.469259 | 0.47114 | 0.492553 |
| 18 | 24689200 | 24690599 | 14.17458 | 25.72161 | 26.41566 | 11.4119 | 0.376009 | 0.333679 | 0.492556 |
| 17 | 38413800 | 38418199 | 46.12094 | 75.38642 | 62.65404 | 26.938 | 0.640556 | 1.516422 | 0.492563 |
| 9 | 19457000 | 19460799 | 57.9867 | 92.84338 | 94.25531 | 39.72352 | 0.323171 | 2.026298 | 0.492587 |
| 3 | 53997400 | 53999799 | 22.4026 | 38.78456 | 39.86909 | 17.49188 | 0.346387 | 0.847087 | 0.492587 |
| 5 | 110147000 | 110147999 | 18.26734 | 32.36236 | 31.5575 | 13.95282 | 0.427811 | 0.518554 | 0.492592 |
| 13 | 67888000 | 67901199 | 179.7541 | 269.2452 | 256.004 | 105.5979 | 0.337268 | 3.499145 | 0.4926 |
| 14 | 120012000 | 120012999 | 12.36155 | 23.27756 | 24.09571 | 9.86447 | 0.421651 | 0.097518 | 0.492601 |
| 6 | 138318800 | 138322599 | 66.50445 | 106.3961 | 94.49093 | 39.77723 | 0.514549 | 2.061174 | 0.492605 |
| 10 | 53950000 | 53952999 | 26.34357 | 42.91124 | 78.82343 | 33.84741 | -0.50404 | 1.532614 | 0.492606 |
| 3 | 111826200 | 111828199 | 19.85796 | 35.39493 | 30.38238 | 13.37091 | 0.615679 | 0.486936 | 0.492638 |
| 3 | 40897200 | 40897999 | 6.854742 | 13.2852 | 28.19341 | 10.67165 | -0.50175 | -0.20136 | 0.492648 |
| 16 | 6500800 | 6501799 | 17.40042 | 30.62937 | 30.90313 | 13.68103 | 0.375453 | 0.581203 | 0.492658 |
| 9 | 6591400 | 6601599 | 258.3795 | 373.9284 | 364.1837 | 150.1963 | 0.255894 | 4.071358 | 0.49266 |
| 1 | 15983800 | 15988399 | 53.37006 | 85.00165 | 131.4931 | 55.0069 | -0.29176 | 2.195067 | 0.492674 |
| X | 88944400 | 88947599 | 28.21977 | 48.58372 | 41.85018 | 18.35793 | 0.598896 | 0.925812 | 0.492676 |
| X | 80759600 | 80765399 | 69.5577 | 111.0739 | 98.39315 | 41.31091 | 0.515341 | 2.122076 | 0.492694 |
| 13 | 91628000 | 91628599 | 6.866977 | 13.16236 | 25.70997 | 10.12227 | -0.43675 | -0.07725 | 0.492705 |
| 16 | 14472800 | 14476799 | 53.20979 | 84.68111 | 128.7071 | 53.7841 | -0.26754 | 2.219079 | 0.492722 |
| 16 | 21456000 | 21457799 | 14.25846 | 25.05304 | 41.87952 | 18.5398 | -0.35008 | 0.604918 | 0.492733 |
| 7 | 103264200 | 103272599 | 143.3038 | 216.9334 | 219.896 | 90.97175 | 0.258349 | 3.28347 | 0.492757 |
| 12 | 116559800 | 116563999 | 33.79645 | 54.59628 | 91.61951 | 39.10859 | -0.38503 | 1.718661 | 0.492761 |
| 17 | 22137200 | 22148799 | 335.7001 | 476.6683 | 431.1877 | 178.3506 | 0.335063 | 4.355331 | 0.492761 |
| 13 | 67361400 | 67362599 | 14.18125 | 25.89856 | 26.25142 | 11.25074 | 0.401828 | 0.295 | 0.492762 |
| 1 | 81844000 | 81845399 | 9.496808 | 17.67098 | 31.59893 | 13.20021 | -0.38486 | 0.166202 | 0.49282 |
| 4 | 141517400 | 141518999 | 9.853044 | 17.62656 | 35.33642 | 15.50317 | -0.59772 | 0.407345 | 0.492823 |
| 10 | 126976800 | 126979399 | 20.68211 | 34.8939 | 57.53078 | 25.1597 | -0.33759 | 1.092808 | 0.492826 |
| 19 | 45309800 | 45311399 | 10.64098 | 18.97791 | 35.34052 | 15.54506 | -0.4944 | 0.434122 | 0.492841 |
| 1 | 54818000 | 54820399 | 28.80232 | 48.57552 | 49.08522 | 21.46377 | 0.366127 | 1.114166 | 0.492856 |
| 2 | 27276000 | 27277799 | 19.18662 | 32.71321 | 53.77705 | 23.57442 | -0.33208 | 0.99086 | 0.492862 |
| 11 | 98430000 | 98432599 | 19.7915 | 33.86821 | 57.14655 | 25.0589 | -0.36889 | 0.962332 | 0.492867 |
| 7 | 5003800 | 5016199 | 120.879 | 182.3571 | 268.3789 | 110.9086 | -0.28342 | 3.350905 | 0.492885 |
| 14 | 124041600 | 124042799 | 9.026201 | 16.9592 | 30.75742 | 12.63912 | -0.38583 | 0.098639 | 0.492898 |
| 13 | 28623000 | 28626599 | 42.70282 | 69.85206 | 64.05056 | 27.54601 | 0.499893 | 1.499964 | 0.49291 |

| | | | | | | | | | |
|---|---|---|---|---|---|---|---|---|---|
| 10 | 106179000 | 106180999 | 41.77502 | 68.4336 | 62.98264 | 27.10227 | 0.495656 | 1.4771 | 0.492941 |
| X | 78659400 | 78660999 | 29.73438 | 50.57452 | 38.95985 | 17.07563 | 0.75685 | 1.037043 | 0.492945 |
| 11 | 45613400 | 45617199 | 29.60013 | 48.11759 | 79.94241 | 34.30238 | -0.36163 | 1.57259 | 0.492948 |
| 15 | 99918200 | 99919799 | 12.47629 | 22.17492 | 37.7124 | 16.63643 | -0.36767 | 0.470427 | 0.492976 |
| 9 | 112897400 | 112901799 | 50.256 | 80.81768 | 84.10346 | 35.67826 | 0.295023 | 1.878629 | 0.49299 |
| 6 | 3687000 | 3687799 | 6.541144 | 12.65906 | 24.98193 | 9.520304 | -0.40707 | -0.19186 | 0.492993 |
| 4 | 75073800 | 75075399 | 21.41559 | 37.50363 | 35.86358 | 15.83202 | 0.451792 | 0.687056 | 0.492995 |
| 2 | 148945600 | 148950399 | 86.9717 | 136.5194 | 137.8115 | 57.15675 | 0.305166 | 2.575259 | 0.493013 |
| 6 | 10078000 | 10082399 | 52.54315 | 84.8913 | 73.15667 | 31.23268 | 0.574475 | 1.736951 | 0.493023 |
| 2 | 51432200 | 51433599 | 12.04552 | 21.39213 | 36.75293 | 16.22413 | -0.38269 | 0.472467 | 0.493045 |
| 9 | 75385000 | 75386999 | 12.04552 | 21.39213 | 36.75293 | 16.22413 | -0.38269 | 0.472467 | 0.493045 |
| 7 | 23240200 | 23251599 | 169.4263 | 256.8528 | 207.9949 | 86.0879 | 0.583871 | 3.260964 | 0.493064 |
| 2 | 6290800 | 6292399 | 16.62819 | 28.92792 | 48.98556 | 21.52487 | -0.36956 | 0.777807 | 0.493066 |
| 4 | 67194000 | 67195799 | 19.69237 | 34.92661 | 30.79938 | 13.61106 | 0.572778 | 0.515215 | 0.493068 |
| 19 | 22700400 | 22703399 | 32.29245 | 53.36383 | 55.76521 | 24.0944 | 0.317172 | 1.343039 | 0.4931 |
| 10 | 84687400 | 84690199 | 15.52666 | 26.68036 | 49.50596 | 21.71495 | -0.5039 | 0.901252 | 0.493104 |
| 4 | 27410000 | 27418799 | 158.6077 | 238.4594 | 240.7093 | 99.3808 | 0.25581 | 3.423198 | 0.493104 |
| 13 | 75987000 | 75989999 | 15.3496 | 26.75288 | 45.74899 | 20.12601 | -0.38324 | 0.741072 | 0.493109 |
| 17 | 7564000 | 7566199 | 22.42027 | 37.36034 | 62.83182 | 27.30264 | -0.36625 | 1.229185 | 0.493134 |
| 7 | 85151200 | 85157399 | 78.59139 | 125.1375 | 103.0534 | 43.09376 | 0.615958 | 2.220817 | 0.493143 |
| 3 | 94116800 | 94128599 | 172.0766 | 259.0002 | 238.7876 | 98.61706 | 0.387932 | 3.403873 | 0.493143 |
| 2 | 97729000 | 97732399 | 36.54219 | 59.98425 | 61.93695 | 26.67541 | 0.33131 | 1.440229 | 0.493153 |
| 15 | 72762200 | 72764199 | 12.96306 | 22.89178 | 39.26796 | 17.37305 | -0.38443 | 0.535066 | 0.493154 |
| 1 | 71177000 | 71180399 | 36.92124 | 60.77591 | 60.49392 | 26.08471 | 0.385624 | 1.404615 | 0.493167 |
| 2 | 171991800 | 171996799 | 54.38671 | 86.37719 | 131.6372 | 54.88053 | -0.27438 | 2.282861 | 0.493168 |
| X | 79552200 | 79555199 | 47.02844 | 76.35188 | 61.50709 | 26.36812 | 0.678329 | 1.620693 | 0.493179 |
| 9 | 25716400 | 25717199 | 12.91644 | 23.91975 | 24.91852 | 10.48042 | 0.381927 | 0.205116 | 0.4932 |
| 12 | 92457000 | 92458199 | 12.91644 | 23.91975 | 24.91852 | 10.48042 | 0.381927 | 0.205116 | 0.4932 |
| 13 | 65051600 | 65052799 | 9.415342 | 17.28511 | 31.39957 | 13.36351 | -0.42679 | 0.243075 | 0.493235 |
| 13 | 96855800 | 96856999 | 9.415342 | 17.28511 | 31.39957 | 13.36351 | -0.42679 | 0.243075 | 0.493235 |
| 13 | 30591200 | 30596599 | 51.13215 | 82.90106 | 69.56744 | 29.79358 | 0.617736 | 1.658223 | 0.493255 |
| 5 | 114282400 | 114283999 | 12.01453 | 22.36895 | 21.93155 | 9.152608 | 0.47707 | 0.158699 | 0.493261 |
| 6 | 130391200 | 130392799 | 37.33874 | 61.43781 | 61.00558 | 26.30075 | 0.388889 | 1.412229 | 0.493262 |
| 5 | 92836600 | 92837799 | 17.24016 | 29.63154 | 50.04766 | 21.9527 | -0.36895 | 0.906019 | 0.493262 |
| 7 | 17169200 | 17176399 | 84.91511 | 134.2846 | 122.2703 | 50.80728 | 0.462096 | 2.418461 | 0.493276 |
| 6 | 131139200 | 131140799 | 26.02864 | 44.90736 | 41.31851 | 18.12589 | 0.504692 | 0.895191 | 0.493284 |
| 7 | 70656000 | 70656999 | 15.67039 | 27.93774 | 27.91414 | 12.28216 | 0.398064 | 0.453166 | 0.493299 |
| 9 | 70984600 | 70990399 | 49.75967 | 79.4065 | 121.5021 | 50.96541 | -0.27294 | 2.127127 | 0.493324 |
| 17 | 11139000 | 11140199 | 12.33092 | 23.02752 | 23.38833 | 9.710716 | 0.432266 | 0.145616 | 0.493328 |
| 10 | 31253600 | 31254599 | 14.44449 | 26.78965 | 25.87688 | 10.86132 | 0.492398 | 0.212069 | 0.49334 |
| 6 | 5324000 | 5329399 | 58.43009 | 93.2792 | 97.75985 | 41.02915 | 0.272721 | 2.116243 | 0.493349 |
| 12 | 11020400 | 11022999 | 28.56301 | 46.63046 | 76.38453 | 32.82682 | -0.33694 | 1.505522 | 0.493366 |
| 4 | 97211200 | 97211999 | 11.36229 | 21.95733 | 20.81115 | 7.985173 | 0.631813 | -0.12942 | 0.493388 |
| 8 | 90380000 | 90382599 | 17.15311 | 29.67529 | 49.00131 | 21.49802 | -0.33462 | 0.843557 | 0.493397 |
| 10 | 83295400 | 83303999 | 272.1721 | 393.5851 | 333.8634 | 137.7057 | 0.454027 | 4.082925 | 0.49341 |
| 13 | 97706600 | 97709199 | 20.45496 | 34.62921 | 57.59753 | 25.2151 | -0.35008 | 1.060772 | 0.493411 |
| X | 12141200 | 12143599 | 21.86017 | 38.57056 | 32.40111 | 14.33744 | 0.639753 | 0.586208 | 0.49342 |

| | | | | | | | | | |
|---|---|---|---|---|---|---|---|---|---|
| 14 | 4982400 | 4983399 | 13.81549 | 25.29575 | 26.39265 | 11.2733 | 0.3639 | 0.279521 | 0.493449 |
| 6 | 102744800 | 102745999 | 12.14933 | 22.9003 | 22.93086 | 9.372085 | 0.470609 | 0.08432 | 0.493477 |
| 9 | 3955400 | 3957399 | 37.48358 | 61.25475 | 64.12239 | 27.56709 | 0.308282 | 1.50065 | 0.493479 |
| X | 108862400 | 108866799 | 72.86123 | 116.2718 | 98.92895 | 41.49675 | 0.572354 | 2.145434 | 0.493502 |
| 4 | 57872000 | 57874199 | 16.91569 | 30.34456 | 29.57485 | 12.94536 | 0.43859 | 0.425559 | 0.493504 |
| 17 | 82982200 | 82984199 | 16.91569 | 30.34456 | 29.57485 | 12.94536 | 0.43859 | 0.425559 | 0.493504 |
| 13 | 113619400 | 113620399 | 7.289284 | 13.95936 | 27.40479 | 10.81722 | -0.44817 | -0.06227 | 0.493517 |
| 7 | 41112600 | 41114199 | 27.13638 | 46.2956 | 44.72093 | 19.60542 | 0.432197 | 1.006271 | 0.493527 |
| 3 | 112561400 | 112563999 | 20.99064 | 37.5803 | 28.63673 | 12.55558 | 0.790083 | 0.463881 | 0.493529 |
| 4 | 46050600 | 46058199 | 75.13695 | 117.2049 | 175.3022 | 72.74614 | -0.26841 | 2.685671 | 0.493564 |
| 5 | 132665600 | 132668999 | 43.69227 | 70.7823 | 72.52405 | 31.04157 | 0.329301 | 1.661752 | 0.49358 |
| 2 | 12619200 | 12620199 | 9.401536 | 17.37912 | 30.95548 | 13.05154 | -0.38873 | 0.203489 | 0.493597 |
| 18 | 82792200 | 82796199 | 31.3251 | 51.02652 | 81.30473 | 34.91311 | -0.29948 | 1.545513 | 0.493615 |
| 12 | 115939600 | 115947399 | 99.42237 | 155.2146 | 147.6993 | 61.28642 | 0.384184 | 2.679998 | 0.493631 |
| 5 | 100282800 | 100284799 | 16.96778 | 30.4462 | 28.96508 | 12.67558 | 0.473345 | 0.421903 | 0.493634 |
| 16 | 89915800 | 89921199 | 59.27886 | 93.46659 | 147.4717 | 61.1988 | -0.33333 | 2.47245 | 0.49365 |
| 8 | 119759800 | 119774399 | 154.6989 | 229.5447 | 333.6325 | 137.4249 | -0.28701 | 3.667418 | 0.493657 |
| 10 | 124552000 | 124553399 | 25.87307 | 42.76948 | 69.58745 | 30.07325 | -0.31957 | 1.336069 | 0.493673 |
| 9 | 119398600 | 119401399 | 22.18499 | 37.04939 | 62.42935 | 27.16529 | -0.36916 | 1.202097 | 0.493678 |
| 10 | 61359600 | 61360799 | 9.041606 | 16.76674 | 30.99165 | 13.01066 | -0.43761 | 0.180538 | 0.493685 |
| 1 | 96596800 | 96597599 | 18.64957 | 32.71775 | 33.51379 | 14.81693 | 0.353101 | 0.640772 | 0.493687 |
| 13 | 113404200 | 113407199 | 18.7824 | 32.17251 | 53.29123 | 23.37016 | -0.34205 | 0.944668 | 0.493687 |
| 4 | 144577400 | 144579999 | 33.55572 | 55.81138 | 54.80329 | 23.7848 | 0.407337 | 1.271703 | 0.493735 |
| 17 | 22016600 | 22017599 | 15.36933 | 27.59622 | 28.34343 | 12.39663 | 0.364384 | 0.408175 | 0.493737 |
| 14 | 50735800 | 50740799 | 57.19078 | 92.20062 | 77.29073 | 32.90571 | 0.610748 | 1.809641 | 0.493741 |
| 9 | 39686200 | 39686799 | 7.12372 | 13.74085 | 27.7893 | 10.74461 | -0.46271 | -0.13063 | 0.493747 |
| 7 | 98618200 | 98618999 | 11.48765 | 20.79731 | 35.86572 | 15.53966 | -0.37004 | 0.348034 | 0.493755 |
| 8 | 26888600 | 26891199 | 33.95301 | 56.1774 | 57.55813 | 24.89031 | 0.345684 | 1.334892 | 0.493765 |
| 4 | 16585200 | 16586599 | 27.63714 | 46.45058 | 48.5685 | 21.2157 | 0.316871 | 1.13891 | 0.49377 |
| 1 | 6403200 | 6405999 | 29.6259 | 51.00819 | 40.66035 | 17.83604 | 0.710279 | 0.928815 | 0.493771 |
| 13 | 69702400 | 69704599 | 11.20325 | 20.14349 | 34.76285 | 15.18671 | -0.37866 | 0.384674 | 0.493776 |
| 4 | 69346800 | 69349399 | 26.81236 | 44.23449 | 71.44053 | 30.83399 | -0.30935 | 1.361316 | 0.493776 |
| 6 | 103199800 | 103200199 | 6.262882 | 12.09714 | 24.26087 | 9.329363 | -0.44198 | -0.16474 | 0.493778 |
| 15 | 18783800 | 18785199 | 17.33791 | 30.86372 | 26.15488 | 11.52284 | 0.632914 | 0.469064 | 0.493796 |
| 3 | 33613200 | 33615799 | 19.2404 | 34.22246 | 29.71607 | 13.10078 | 0.597128 | 0.491838 | 0.493806 |
| 17 | 92554600 | 92558999 | 80.48382 | 126.4846 | 130.0222 | 53.93443 | 0.280312 | 2.548475 | 0.493809 |
| 18 | 85207200 | 85207999 | 9.044012 | 17.05524 | 32.60439 | 13.31495 | -0.4541 | 0.079908 | 0.49382 |
| 8 | 29398400 | 29401199 | 24.4436 | 42.22548 | 41.40164 | 18.16305 | 0.413725 | 0.878139 | 0.493831 |
| 4 | 74651800 | 74653199 | 19.10067 | 34.15087 | 29.87535 | 13.11344 | 0.590931 | 0.461343 | 0.493835 |
| 13 | 99969000 | 99972599 | 56.97843 | 91.1678 | 94.91767 | 39.96418 | 0.285791 | 2.045468 | 0.493845 |
| 9 | 18266400 | 18268399 | 16.52945 | 28.72783 | 47.38396 | 20.80821 | -0.33176 | 0.791165 | 0.493869 |
| 13 | 61184400 | 61192199 | 93.88988 | 147.8683 | 128.3294 | 53.24704 | 0.526609 | 2.513468 | 0.493873 |
| 6 | 41266400 | 41267199 | 6.698129 | 12.98741 | 26.34054 | 9.949733 | -0.43391 | -0.21496 | 0.493898 |
| 16 | 71876200 | 71876799 | 10.23735 | 18.63384 | 32.85842 | 14.14111 | -0.39524 | 0.302708 | 0.493899 |
| 7 | 18723600 | 18728599 | 33.31441 | 53.50928 | 108.2379 | 45.83648 | -0.66393 | 1.90342 | 0.493901 |
| 5 | 14987800 | 14988999 | 16.26105 | 28.99488 | 29.69059 | 13.06224 | 0.36276 | 0.458491 | 0.493915 |
| 10 | 45972600 | 45974399 | 14.74368 | 25.84915 | 42.98736 | 18.99519 | -0.34266 | 0.644698 | 0.493925 |

| | | | | | | | | | |
|---|---|---|---|---|---|---|---|---|---|
| 9 | 103142600 | 103143599 | 13.41925 | 24.76109 | 25.50307 | 10.77924 | 0.393527 | 0.232821 | 0.493928 |
| 4 | 152920600 | 152921199 | 7.624919 | 14.50507 | 27.18352 | 10.89877 | -0.40168 | -0.00653 | 0.493982 |
| 2 | 145973200 | 145985799 | 141.4614 | 210.8195 | 346.0203 | 142.5908 | -0.45647 | 3.589956 | 0.493999 |
| 19 | 39634200 | 39638599 | 80.37856 | 126.9523 | 126.5017 | 52.53243 | 0.330747 | 2.444339 | 0.494005 |
| 14 | 108877400 | 108882199 | 48.35257 | 78.16621 | 74.88021 | 31.99248 | 0.423122 | 1.715147 | 0.494009 |
| 17 | 75350800 | 75354599 | 56.07875 | 90.04115 | 88.21505 | 37.33707 | 0.379183 | 1.936689 | 0.494035 |
| 4 | 121066200 | 121067599 | 8.082984 | 15.42267 | 28.99441 | 11.55126 | -0.39834 | -0.01809 | 0.494066 |
| 13 | 58088400 | 58090599 | 20.71139 | 34.85822 | 58.03395 | 25.36099 | -0.3517 | 1.115039 | 0.49407 |
| X | 22263800 | 22270999 | 102.158 | 160.5744 | 114.3875 | 47.45008 | 0.808926 | 2.556062 | 0.494074 |
| 14 | 68525200 | 68526199 | 12.38472 | 22.97514 | 23.42302 | 9.828065 | 0.415291 | 0.187099 | 0.494074 |
| 14 | 106537800 | 106538599 | 12.38472 | 22.97514 | 23.42302 | 9.828065 | 0.415291 | 0.187099 | 0.494074 |
| 6 | 68664600 | 68665199 | 10.00882 | 19.18107 | 20.66485 | 8.140163 | 0.413067 | -0.06025 | 0.494083 |
| 6 | 143632000 | 143634599 | 22.48946 | 37.57023 | 63.04076 | 27.43704 | -0.36314 | 1.199549 | 0.494085 |
| 1 | 16726800 | 16727399 | 4.556574 | 8.874787 | 21.29659 | 7.778242 | -0.62758 | -0.37554 | 0.494089 |
| 4 | 108459600 | 108462399 | 20.68464 | 36.08281 | 37.14699 | 16.35075 | 0.345675 | 0.743026 | 0.4941 |
| X | 78964800 | 78965599 | 22.45939 | 39.14154 | 35.73969 | 15.72193 | 0.518163 | 0.766887 | 0.494109 |
| 9 | 62345400 | 62350199 | 71.88028 | 111.6768 | 178.1078 | 73.95394 | -0.36538 | 2.753768 | 0.494122 |
| 8 | 95946000 | 95947599 | 8.432385 | 15.73093 | 29.50366 | 12.27117 | -0.44774 | 0.136126 | 0.494147 |
| 12 | 84116200 | 84117399 | 8.432385 | 15.73093 | 29.50366 | 12.27117 | -0.44774 | 0.136126 | 0.494147 |
| 8 | 53132800 | 53133799 | 12.69474 | 24.47591 | 19.99634 | 7.7413 | 0.831763 | -0.08964 | 0.49415 |
| 16 | 93027400 | 93031399 | 43.19869 | 69.91178 | 72.86326 | 31.16345 | 0.303423 | 1.681261 | 0.494158 |
| 5 | 146119000 | 146122799 | 39.55872 | 63.63243 | 99.24396 | 42.11769 | -0.28702 | 1.86153 | 0.494166 |
| 7 | 85406200 | 85410399 | 73.71149 | 116.8124 | 119.1572 | 49.5686 | 0.300849 | 2.359214 | 0.494176 |
| 2 | 99360000 | 99361199 | 14.58712 | 26.91538 | 26.33138 | 11.12971 | 0.466908 | 0.242215 | 0.49424 |
| 13 | 84267000 | 84268399 | 14.58712 | 26.91538 | 26.33138 | 11.12971 | 0.466908 | 0.242215 | 0.49424 |
| 1 | 8324000 | 8325999 | 35.70515 | 58.90561 | 58.81036 | 25.38276 | 0.3825 | 1.372528 | 0.494269 |
| 6 | 124180800 | 124181399 | 8.624055 | 16.82322 | 16.1039 | 5.762645 | 0.70819 | -0.41532 | 0.49428 |
| 18 | 67933000 | 67933799 | 4.997843 | 9.704547 | 24.5348 | 9.195173 | -0.73789 | -0.26991 | 0.494292 |
| 10 | 82659400 | 82660799 | 13.48455 | 23.79708 | 39.80002 | 17.61076 | -0.34837 | 0.548016 | 0.494321 |
| 7 | 58306800 | 58308799 | 14.26744 | 24.87114 | 44.71371 | 19.67273 | -0.45497 | 0.731985 | 0.494326 |
| 1 | 111811000 | 111813399 | 35.30645 | 57.9828 | 60.79701 | 26.18962 | 0.309448 | 1.429523 | 0.494327 |
| 5 | 53041400 | 53044599 | 31.50616 | 51.16537 | 82.0864 | 35.19773 | -0.31274 | 1.595 | 0.494329 |
| 4 | 143222000 | 143224199 | 30.978 | 51.69114 | 51.95465 | 22.59558 | 0.373764 | 1.225744 | 0.494366 |
| 18 | 90335000 | 90341799 | 98.89186 | 153.8348 | 154.7978 | 64.26321 | 0.299672 | 2.741858 | 0.494376 |
| 13 | 44017400 | 44020999 | 25.35404 | 41.85522 | 69.00173 | 29.79247 | -0.33867 | 1.348209 | 0.494381 |
| 5 | 101053000 | 101055199 | 24.24643 | 40.24825 | 66.72056 | 28.91751 | -0.34589 | 1.28199 | 0.494383 |
| 7 | 40784000 | 40784799 | 12.77439 | 23.82206 | 24.56691 | 10.22445 | 0.408325 | 0.164519 | 0.494402 |
| 6 | 89698000 | 89701399 | 33.92037 | 56.49919 | 54.2438 | 23.5644 | 0.439778 | 1.257153 | 0.494404 |
| 2 | 124666800 | 124669999 | 29.50218 | 48.19649 | 77.29782 | 33.22756 | -0.30588 | 1.497341 | 0.494406 |
| 3 | 66932600 | 66936199 | 50.67588 | 81.30721 | 83.3583 | 35.24255 | 0.312567 | 1.94882 | 0.49441 |
| Y | 13445200 | 13445999 | 8.22961 | 16.01333 | 12.05975 | 4.443948 | 1.01372 | -0.33331 | 0.494433 |
| Y | 81115000 | 81115599 | 8.22961 | 16.01333 | 12.05975 | 4.443948 | 1.01372 | -0.33331 | 0.494433 |
| 2 | 15979000 | 15980999 | 23.62095 | 40.95404 | 40.32792 | 17.69398 | 0.408599 | 0.840994 | 0.494438 |
| 7 | 60196800 | 60206999 | 46.98949 | 76.1917 | 63.70749 | 27.28584 | 0.623031 | 1.646321 | 0.494481 |
| 2 | 80281600 | 80282799 | 11.48324 | 21.61642 | 23.06504 | 9.447871 | 0.377466 | 0.088416 | 0.494484 |
| 4 | 96357000 | 96357999 | 11.2982 | 21.64467 | 22.29755 | 8.791921 | 0.476325 | -0.03752 | 0.494514 |
| 15 | 43918800 | 43919999 | 9.845376 | 18.13786 | 31.94601 | 13.53157 | -0.37776 | 0.22714 | 0.494527 |

| | | | | | | | | | |
|---|---|---|---|---|---|---|---|---|---|
| 7 | 33621400 | 33648999 | 438.0977 | 608.9372 | 541.6346 | 224.7993 | 0.32841 | 4.674101 | 0.49455 |
| 6 | 131682200 | 131683999 | 19.77125 | 35.76046 | 27.20671 | 11.80177 | 0.802289 | 0.393336 | 0.494555 |
| 18 | 16175600 | 16176599 | 8.917971 | 16.58377 | 29.98313 | 12.53821 | -0.40139 | 0.163261 | 0.494559 |
| 15 | 21813800 | 21815599 | 14.81312 | 27.30345 | 23.15477 | 9.801141 | 0.670113 | 0.242501 | 0.494563 |
| 1 | 119452800 | 119463799 | 122.8761 | 185.09 | 273.6679 | 113.0458 | -0.29202 | 3.381112 | 0.494573 |
| 1 | 4981000 | 4982799 | 13.94399 | 24.36071 | 43.63043 | 19.22261 | -0.44916 | 0.697435 | 0.494578 |
| 17 | 47623200 | 47626399 | 26.62468 | 43.52223 | 74.48628 | 32.02611 | -0.3986 | 1.481687 | 0.49458 |
| 18 | 62884400 | 62885799 | 10.2102 | 18.78503 | 32.99648 | 14.00177 | -0.37565 | 0.24134 | 0.494581 |
| 14 | 66752600 | 66753199 | 6.393344 | 12.40351 | 24.54684 | 9.247554 | -0.39503 | -0.23289 | 0.494584 |
| 15 | 30235800 | 30236799 | 6.393344 | 12.40351 | 24.54684 | 9.247554 | -0.39503 | -0.23289 | 0.494584 |
| 1 | 115024400 | 115024999 | 10.78897 | 20.56022 | 22.23238 | 8.880623 | 0.390061 | -0.00125 | 0.494586 |
| 10 | 47084200 | 47084799 | 10.78897 | 20.56022 | 22.23238 | 8.880623 | 0.390061 | -0.00125 | 0.494586 |
| 1 | 109651800 | 109653399 | 19.47372 | 34.4614 | 32.01087 | 14.1564 | 0.496984 | 0.539237 | 0.494595 |
| 4 | 136391000 | 136394799 | 32.38953 | 52.36381 | 86.40357 | 36.9188 | -0.35936 | 1.694102 | 0.494597 |
| 9 | 9381800 | 9385799 | 77.59016 | 122.3543 | 127.3428 | 52.84646 | 0.266249 | 2.4784 | 0.494606 |
| 4 | 111390800 | 111396999 | 69.24391 | 110.0292 | 100.0352 | 41.72657 | 0.470393 | 2.272108 | 0.494616 |
| Y | 38785600 | 38786199 | 7.683742 | 14.97736 | 11.55109 | 4.175441 | 1.0045 | -0.41012 | 0.494627 |
| 16 | 49712400 | 49715399 | 29.04393 | 48.59755 | 50.75837 | 22.1137 | 0.318554 | 1.19282 | 0.49464 |
| Y | 42288800 | 42291999 | 18.88963 | 33.18844 | 21.68863 | 9.600966 | 0.99754 | 0.592284 | 0.494647 |
| 4 | 90772800 | 90773599 | 19.44797 | 34.03799 | 35.07826 | 15.47771 | 0.344434 | 0.685053 | 0.494684 |
| 14 | 40110400 | 40110799 | 5.363331 | 10.46183 | 22.23879 | 7.963892 | -0.42947 | -0.43193 | 0.494691 |
| 6 | 44219000 | 44220399 | 13.37774 | 24.86079 | 25.49875 | 10.67357 | 0.409394 | 0.192522 | 0.494698 |
| X | 22855200 | 22859399 | 70.51522 | 112.0895 | 110.5522 | 46.12998 | 0.353647 | 2.267642 | 0.494698 |
| 7 | 104300600 | 104303599 | 25.77542 | 43.92337 | 45.25365 | 19.83713 | 0.339215 | 1.009234 | 0.494704 |
| 5 | 15955400 | 15957799 | 35.41932 | 58.84141 | 55.43212 | 24.03809 | 0.466868 | 1.291421 | 0.494726 |
| X | 168062400 | 168064199 | 20.18932 | 36.25096 | 28.75661 | 12.57727 | 0.734843 | 0.441486 | 0.494734 |
| 1 | 98070200 | 98070599 | 7.330195 | 13.92131 | 27.14374 | 10.92104 | -0.46286 | 0.00142 | 0.49478 |
| 6 | 114169400 | 114170199 | 4.808826 | 9.369667 | 20.77636 | 7.550431 | -0.508 | -0.39049 | 0.494785 |
| 12 | 60796800 | 60802999 | 67.45701 | 107.9672 | 92.57022 | 38.99281 | 0.565912 | 2.048216 | 0.494791 |
| 14 | 31563000 | 31564999 | 10.71164 | 19.23958 | 35.26177 | 15.41818 | -0.46599 | 0.386875 | 0.494794 |
| 1 | 13623600 | 13628199 | 66.36226 | 105.3332 | 109.447 | 45.60174 | 0.276504 | 2.304419 | 0.494795 |
| 4 | 35131400 | 35134399 | 27.88983 | 45.6171 | 74.80441 | 32.16978 | -0.33655 | 1.475609 | 0.494811 |
| 7 | 38225000 | 38230199 | 79.65928 | 126.8723 | 101.4117 | 42.42318 | 0.659341 | 2.213502 | 0.494812 |
| 15 | 45453400 | 45454399 | 11.99861 | 22.39781 | 20.65513 | 8.581831 | 0.570499 | 0.141061 | 0.494879 |
| 3 | 46190000 | 46190599 | 8.517919 | 16.09297 | 30.37793 | 12.35766 | -0.43118 | 0.055326 | 0.494914 |
| 8 | 81922400 | 81922999 | 11.46933 | 21.86101 | 19.50358 | 7.787416 | 0.665644 | -0.00412 | 0.494915 |
| 4 | 33907800 | 33908999 | 7.469759 | 14.27846 | 26.74478 | 10.61309 | -0.38765 | -0.04225 | 0.494926 |
| 15 | 44993400 | 44994399 | 7.469759 | 14.27846 | 26.74478 | 10.61309 | -0.38765 | -0.04225 | 0.494926 |
| X | 65035400 | 65037599 | 30.4241 | 51.62565 | 46.84809 | 20.52358 | 0.52108 | 1.067501 | 0.494945 |
| 2 | 148299200 | 148302999 | 51.94103 | 82.7293 | 126.7343 | 53.0194 | -0.27759 | 2.192192 | 0.494969 |
| 6 | 122017400 | 122020399 | 38.06454 | 63.11243 | 55.5616 | 24.06176 | 0.56411 | 1.320761 | 0.494974 |
| 13 | 57230800 | 57232999 | 24.48419 | 42.13367 | 42.52835 | 18.65516 | 0.370947 | 0.913998 | 0.494978 |
| 15 | 74520000 | 74522799 | 25.05595 | 42.91187 | 43.60652 | 19.12282 | 0.360039 | 0.958989 | 0.494988 |
| 6 | 16515000 | 16517399 | 36.66282 | 60.26444 | 61.48917 | 26.49785 | 0.349208 | 1.42295 | 0.494996 |
| 4 | 34142200 | 34143199 | 6.995963 | 13.4294 | 26.33878 | 10.32598 | -0.43729 | -0.08971 | 0.494996 |
| X | 168513800 | 168515799 | 22.19549 | 38.62729 | 38.72344 | 17.01944 | 0.383573 | 0.785911 | 0.495001 |
| 10 | 20275800 | 20278799 | 23.52319 | 39.10716 | 64.60188 | 28.02868 | -0.34071 | 1.258265 | 0.495001 |

| | | | | | | | | | |
|---|---|---|---|---|---|---|---|---|---|
| 1 | 48085800 | 48089799 | 32.54202 | 55.3803 | 44.57166 | 19.53607 | 0.693974 | 1.047178 | 0.495005 |
| 14 | 44754800 | 44757999 | 23.1899 | 40.11723 | 40.34737 | 17.70131 | 0.377544 | 0.857489 | 0.495007 |
| 11 | 87830400 | 87835399 | 51.22216 | 82.33658 | 85.4051 | 36.20825 | 0.298963 | 1.893262 | 0.495026 |
| 10 | 30157800 | 30159599 | 19.89096 | 35.25868 | 31.45926 | 13.90781 | 0.555728 | 0.523908 | 0.495032 |
| X | 25755400 | 25759999 | 51.27585 | 83.27091 | 67.22841 | 28.82678 | 0.675572 | 1.626849 | 0.495033 |
| 8 | 19642600 | 19669199 | 426.191 | 593.5317 | 532.3501 | 220.8796 | 0.319195 | 4.644508 | 0.495042 |
| 7 | 63626000 | 63628599 | 23.82404 | 40.8476 | 42.61298 | 18.6888 | 0.322619 | 0.944046 | 0.495043 |
| 4 | 62839800 | 62843399 | 49.49049 | 79.57783 | 82.9853 | 35.19826 | 0.291817 | 1.879066 | 0.495081 |
| 4 | 112194200 | 112196599 | 41.70066 | 68.49487 | 61.26872 | 26.39462 | 0.538169 | 1.444782 | 0.495085 |
| X | 116654600 | 116655999 | 17.75691 | 33.08371 | 24.07087 | 10.03334 | 0.906174 | 0.209388 | 0.495088 |
| 18 | 49580000 | 49581799 | 12.54267 | 22.24819 | 37.75197 | 16.6821 | -0.36591 | 0.485784 | 0.49509 |
| 3 | 46196800 | 46198799 | 23.04723 | 40.3348 | 35.11765 | 15.49423 | 0.586351 | 0.70291 | 0.495099 |
| 1 | 104696200 | 104699799 | 23.84271 | 39.54356 | 66.85628 | 28.95888 | -0.37442 | 1.287507 | 0.495108 |
| 17 | 68164000 | 68167399 | 22.51381 | 37.58931 | 62.43355 | 27.16292 | -0.34844 | 1.206122 | 0.495123 |
| 6 | 129972600 | 129979199 | 55.57619 | 88.43725 | 159.9225 | 66.81802 | -0.5181 | 2.231852 | 0.495151 |
| 4 | 60251200 | 60254599 | 27.24685 | 46.33141 | 46.31036 | 20.29684 | 0.382289 | 1.034918 | 0.495177 |
| 9 | 74323000 | 74323799 | 10.08323 | 19.2672 | 18.72154 | 7.435697 | 0.550493 | -0.03652 | 0.495183 |
| 7 | 31348200 | 31354199 | 118.7117 | 181.9302 | 185.5963 | 77.03692 | 0.263443 | 3.026978 | 0.495186 |
| 10 | 130029600 | 130032199 | 24.9091 | 43.09698 | 40.81174 | 17.90511 | 0.46412 | 0.865399 | 0.495189 |
| 10 | 126383600 | 126384599 | 7.01045 | 13.45794 | 25.76066 | 10.09775 | -0.40186 | -0.08985 | 0.495215 |
| 19 | 25276000 | 25278199 | 15.34276 | 26.75757 | 45.73633 | 20.12829 | -0.38252 | 0.732028 | 0.495223 |
| 11 | 9364800 | 9366199 | 12.16399 | 21.74602 | 36.7446 | 16.13017 | -0.35337 | 0.430872 | 0.49524 |
| 8 | 74279800 | 74280399 | 12.52318 | 23.47867 | 23.12443 | 9.537446 | 0.484893 | 0.126398 | 0.495246 |
| 6 | 9518000 | 9520399 | 28.22363 | 48.60403 | 41.24961 | 18.09464 | 0.620256 | 0.921121 | 0.495248 |
| Y | 41056800 | 41057599 | 8.779661 | 17.04483 | 12.56586 | 4.716268 | 1.020164 | -0.26042 | 0.495285 |
| 10 | 49878400 | 49879399 | 22.10483 | 38.31326 | 38.9338 | 17.08214 | 0.363435 | 0.837018 | 0.495287 |
| 9 | 95505400 | 95506999 | 11.28656 | 20.55275 | 37.63335 | 16.18706 | -0.44938 | 0.315036 | 0.495291 |
| 8 | 43755600 | 43760599 | 67.84708 | 106.6339 | 159.7044 | 66.24758 | -0.26187 | 2.543214 | 0.495295 |
| 12 | 58360400 | 58362599 | 24.97406 | 41.1366 | 80.72853 | 34.81943 | -0.59051 | 1.384929 | 0.495307 |
| 7 | 41825000 | 41831199 | 100.0306 | 156.6166 | 140.9014 | 58.44828 | 0.468272 | 2.629449 | 0.495319 |
| 16 | 49496200 | 49497799 | 20.23627 | 35.61295 | 33.60019 | 14.87573 | 0.471583 | 0.605158 | 0.495319 |
| 2 | 153882600 | 153883799 | 14.05938 | 26.15964 | 25.3784 | 10.60235 | 0.491065 | 0.188908 | 0.495349 |
| 4 | 29514600 | 29515799 | 14.05938 | 26.15964 | 25.3784 | 10.60235 | 0.491065 | 0.188908 | 0.495349 |
| 17 | 80084800 | 80088599 | 42.05883 | 67.42802 | 105.8384 | 44.69285 | -0.30208 | 1.972439 | 0.49537 |
| 19 | 41407200 | 41411199 | 31.33549 | 50.85488 | 83.21139 | 35.66289 | -0.34199 | 1.607602 | 0.495372 |
| 3 | 3572600 | 3573799 | 11.89078 | 21.04408 | 37.28544 | 16.48875 | -0.42936 | 0.501964 | 0.495392 |
| 8 | 25406800 | 25409199 | 38.04984 | 62.17616 | 64.65436 | 27.79515 | 0.317839 | 1.504331 | 0.495403 |
| 2 | 88264600 | 88267999 | 45.20028 | 73.52269 | 67.60061 | 29.01323 | 0.490277 | 1.584026 | 0.49543 |
| 9 | 30119200 | 30121599 | 13.66424 | 24.06371 | 40.34475 | 17.85927 | -0.35304 | 0.573578 | 0.495431 |
| 6 | 70405600 | 70408199 | 44.06903 | 70.29264 | 116.6533 | 48.89969 | -0.39052 | 2.134927 | 0.495445 |
| 3 | 147876000 | 147877599 | 17.27363 | 31.44153 | 26.90995 | 11.58092 | 0.640508 | 0.338159 | 0.495475 |
| 12 | 79413000 | 79414799 | 22.80787 | 39.91585 | 35.72191 | 15.76082 | 0.546934 | 0.703425 | 0.495486 |
| 13 | 30458600 | 30462399 | 37.40548 | 61.53921 | 60.94014 | 26.27102 | 0.392752 | 1.414996 | 0.49549 |
| 10 | 99199400 | 99201799 | 32.34363 | 53.67716 | 55.79309 | 24.17763 | 0.325335 | 1.295384 | 0.495499 |
| 4 | 125699800 | 125702999 | 25.31495 | 41.93998 | 68.05289 | 29.45522 | -0.31545 | 1.305862 | 0.495502 |
| 13 | 7099400 | 7102199 | 22.91011 | 40.17377 | 33.95343 | 15.00548 | 0.628924 | 0.66982 | 0.495523 |
| 10 | 36652400 | 36657799 | 66.35712 | 104.4297 | 156.0541 | 64.74449 | -0.25718 | 2.515916 | 0.495534 |

| | | | | | | | | | |
|---|---|---|---|---|---|---|---|---|---|
| 10 | 59638600 | 59642599 | 29.64718 | 49.44318 | 51.8792 | 22.5549 | 0.31201 | 1.229635 | 0.495534 |
| 19 | 22309400 | 22310799 | 15.507 | 27.04104 | 45.73777 | 20.12754 | -0.36744 | 0.734655 | 0.495543 |
| 4 | 135093800 | 135095999 | 28.41587 | 46.34254 | 76.80133 | 32.99357 | -0.35472 | 1.520426 | 0.495543 |
| 17 | 12394800 | 12400199 | 43.18816 | 69.27437 | 107.1756 | 45.29311 | -0.2804 | 1.957894 | 0.495545 |
| 16 | 11149400 | 11151399 | 16.53452 | 28.71951 | 47.39238 | 20.80412 | -0.3325 | 0.799842 | 0.495546 |
| 2 | 97666400 | 97669599 | 31.89908 | 53.35579 | 52.45069 | 22.84582 | 0.405727 | 1.207469 | 0.495571 |
| 4 | 58486400 | 58487199 | 7.660846 | 14.59833 | 27.76829 | 11.09307 | -0.41795 | -0.01406 | 0.495579 |
| 3 | 17238200 | 17248599 | 162.7044 | 245.5245 | 232.0777 | 95.91564 | 0.35523 | 3.351828 | 0.495611 |
| 8 | 120970800 | 120971999 | 17.6012 | 29.98584 | 52.66396 | 23.08497 | -0.4272 | 0.990888 | 0.495616 |
| 17 | 92787600 | 92789199 | 13.95124 | 26.75353 | 20.96806 | 8.244411 | 0.870832 | -0.02519 | 0.49562 |
| 8 | 48253600 | 48256599 | 43.4412 | 70.12945 | 71.31161 | 30.41348 | 0.33498 | 1.7455 | 0.49563 |
| 3 | 137569600 | 137572199 | 16.27404 | 27.54846 | 61.69186 | 27.00672 | -0.77738 | 1.051919 | 0.495632 |
| 3 | 127904000 | 127907599 | 38.2128 | 61.50334 | 98.25151 | 41.73181 | -0.32051 | 1.839565 | 0.495649 |
| 4 | 3621400 | 3623199 | 14.28253 | 25.11102 | 41.87435 | 18.5425 | -0.34639 | 0.596984 | 0.495673 |
| 17 | 53338400 | 53342599 | 28.58939 | 48.24293 | 38.60633 | 16.88442 | 0.701441 | 1.097008 | 0.495674 |
| 6 | 43556000 | 43557599 | 19.42065 | 34.14699 | 29.16291 | 12.91353 | 0.613974 | 0.601666 | 0.49569 |
| 3 | 30890000 | 30893999 | 42.1674 | 67.74533 | 105.5327 | 44.70684 | -0.28756 | 1.905584 | 0.495706 |
| 4 | 127701600 | 127703199 | 13.9496 | 24.45883 | 41.45783 | 18.32051 | -0.36992 | 0.636771 | 0.495715 |
| 12 | 23001000 | 23027599 | 1055.318 | 1355.253 | 1144.71 | 480.1098 | 0.288511 | 5.934985 | 0.495726 |
| 10 | 21154200 | 21157199 | 28.35853 | 48.00013 | 47.97466 | 21.00114 | 0.382097 | 1.081604 | 0.495756 |
| Y | 41196400 | 41197999 | 12.94763 | 24.09901 | 15.5132 | 6.477339 | 1.078485 | 0.150004 | 0.495761 |
| 17 | 48715400 | 48717599 | 16.26531 | 27.56273 | 60.04801 | 26.29551 | -0.73768 | 1.041799 | 0.495767 |
| 7 | 52939600 | 52944199 | 50.27764 | 81.03341 | 79.88866 | 33.99977 | 0.376804 | 1.80603 | 0.495768 |
| 17 | 38219000 | 38223399 | 48.71372 | 78.89805 | 72.5396 | 31.04268 | 0.484923 | 1.676519 | 0.495772 |
| 4 | 111868600 | 111869399 | 14.92585 | 26.92898 | 27.75603 | 12.07785 | 0.363947 | 0.373823 | 0.495774 |
| 8 | 101877200 | 101878599 | 14.92585 | 26.92898 | 27.75603 | 12.07785 | 0.363947 | 0.373823 | 0.495774 |
| 4 | 113445000 | 113449999 | 113.5127 | 174.8375 | 175.0315 | 72.70168 | 0.296099 | 2.926953 | 0.49579 |
| 13 | 35474600 | 35477399 | 23.34791 | 39.14326 | 64.77846 | 28.25665 | -0.34331 | 1.16794 | 0.495793 |
| 4 | 90018200 | 90020599 | 19.92979 | 35.39326 | 30.82543 | 13.60779 | 0.591742 | 0.50959 | 0.495801 |
| 9 | 118023400 | 118026199 | 25.10728 | 43.25739 | 42.00057 | 18.42432 | 0.427134 | 0.907062 | 0.495807 |
| 5 | 11562000 | 11563399 | 14.1112 | 24.63456 | 43.64945 | 19.22203 | -0.43382 | 0.709338 | 0.495811 |
| 18 | 41466400 | 41472199 | 78.29175 | 123.2676 | 127.5473 | 52.92057 | 0.272925 | 2.50997 | 0.495837 |
| 9 | 20054000 | 20055999 | 36.23844 | 59.44154 | 61.8404 | 26.62548 | 0.320072 | 1.448936 | 0.495845 |
| X | 109476400 | 109477599 | 24.36454 | 42.41046 | 37.96833 | 16.68961 | 0.545985 | 0.791017 | 0.495853 |
| 6 | 49030600 | 49032399 | 17.6836 | 30.48809 | 50.06196 | 21.96105 | -0.32757 | 0.879141 | 0.495864 |
| 4 | 121124800 | 121134399 | 128.6139 | 196.5469 | 195.1443 | 80.94801 | 0.299313 | 3.087356 | 0.495873 |
| X | 4443400 | 4447799 | 95.9619 | 151.6285 | 117.8583 | 48.95347 | 0.689121 | 2.445351 | 0.495881 |
| 10 | 55417600 | 55420199 | 21.94613 | 38.48517 | 33.43322 | 14.77616 | 0.589312 | 0.661687 | 0.495893 |
| 9 | 70109800 | 70110999 | 16.9953 | 30.00743 | 30.42438 | 13.4609 | 0.370185 | 0.545948 | 0.495909 |
| 7 | 15298600 | 15299599 | 34.41961 | 56.52012 | 56.70018 | 24.42368 | 0.373069 | 1.428137 | 0.495914 |
| X | 120146800 | 120151199 | 60.83173 | 97.85455 | 81.78929 | 34.71188 | 0.611179 | 1.884927 | 0.495945 |
| Y | 4211200 | 4211599 | 7.13984 | 13.93664 | 11.03954 | 3.910068 | 0.992395 | -0.49131 | 0.495965 |
| 3 | 66182200 | 66185199 | 20.3357 | 35.64852 | 29.44295 | 13.00874 | 0.662169 | 0.655021 | 0.495975 |
| 8 | 105648600 | 105649599 | 7.146482 | 13.71549 | 26.33198 | 10.32947 | -0.4075 | -0.08573 | 0.495983 |
| 12 | 99776000 | 99780999 | 42.16495 | 68.2822 | 71.73769 | 30.69644 | 0.292721 | 1.665379 | 0.496008 |
| 13 | 46665600 | 46669999 | 34.62727 | 56.0109 | 88.34854 | 37.77202 | -0.29374 | 1.684232 | 0.496009 |
| 6 | 105571400 | 105572999 | 13.45995 | 23.83785 | 40.79086 | 18.03643 | -0.37934 | 0.509912 | 0.496011 |

| | | | | | | | | | |
|---|---|---|---|---|---|---|---|---|---|
| 8 | 18101600 | 18102399 | 16.73903 | 28.94164 | 48.46293 | 21.26144 | -0.35474 | 0.848315 | 0.496028 |
| 1 | 144270800 | 144275199 | 36.20126 | 59.79704 | 58.298 | 25.18113 | 0.416938 | 1.359516 | 0.496057 |
| 5 | 136076000 | 136079399 | 32.7278 | 52.80773 | 90.01836 | 38.36681 | -0.40948 | 1.752515 | 0.496057 |
| 10 | 90743000 | 90744999 | 11.51102 | 20.53241 | 36.73469 | 16.15645 | -0.43736 | 0.438905 | 0.496072 |
| 3 | 129830600 | 129848599 | 322.7033 | 451.4526 | 607.3013 | 251.8004 | -0.25883 | 4.573695 | 0.496087 |
| 9 | 89787800 | 89797599 | 47.41697 | 75.23776 | 145.9672 | 60.79817 | -0.62342 | 2.30667 | 0.496089 |
| 7 | 17902200 | 17903599 | 19.59495 | 35.37034 | 27.73289 | 12.06033 | 0.756783 | 0.404256 | 0.496105 |
| 13 | 22298800 | 22303399 | 95.21921 | 147.8516 | 147.3303 | 61.17981 | 0.311724 | 2.77143 | 0.496123 |
| 1 | 113688800 | 113691399 | 26.03427 | 43.2004 | 73.26854 | 31.74757 | -0.37893 | 1.295179 | 0.496132 |
| X | 161767600 | 161769999 | 27.35935 | 46.62632 | 45.76962 | 20.06344 | 0.408821 | 1.016304 | 0.496141 |
| 2 | 118845400 | 118846999 | 9.441737 | 17.40566 | 31.45931 | 13.31353 | -0.4134 | 0.221753 | 0.496145 |
| 7 | 48050000 | 48053399 | 61.44728 | 97.90733 | 102.6748 | 42.97649 | 0.268844 | 2.181688 | 0.496161 |
| 1 | 43513400 | 43514999 | 11.85989 | 21.07139 | 36.73243 | 16.20929 | -0.40332 | 0.467243 | 0.496164 |
| 14 | 54448400 | 54450399 | 14.08451 | 24.6477 | 43.0532 | 18.99448 | -0.41322 | 0.67008 | 0.496179 |
| 13 | 3386000 | 3393999 | 179.0704 | 268.3115 | 232.1135 | 95.74972 | 0.473938 | 3.491902 | 0.496206 |
| 15 | 37430800 | 37431599 | 7.033805 | 13.42734 | 26.15341 | 10.40868 | -0.44701 | -0.038 | 0.496211 |
| 16 | 90672600 | 90673399 | 7.033805 | 13.42734 | 26.15341 | 10.40868 | -0.44701 | -0.038 | 0.496211 |
| 4 | 36372000 | 36373199 | 11.93004 | 22.74458 | 22.64987 | 9.039092 | 0.508589 | 0.005867 | 0.496212 |
| X | 121352400 | 121361599 | 161.5689 | 244.4114 | 221.199 | 91.49053 | 0.420911 | 3.303102 | 0.496231 |
| 3 | 10627200 | 10627799 | 4.81229 | 9.366312 | 21.23891 | 7.809774 | -0.55531 | -0.34628 | 0.496243 |
| 13 | 25414800 | 25415799 | 11.73972 | 22.18801 | 23.04234 | 9.366355 | 0.425354 | 0.06355 | 0.496252 |
| 17 | 8276200 | 8276799 | 4.834542 | 9.419307 | 21.96689 | 7.988274 | -0.58192 | -0.38681 | 0.496262 |
| 4 | 111844200 | 111851799 | 91.93201 | 144.3708 | 136.7214 | 56.70794 | 0.397687 | 2.569327 | 0.496274 |
| 6 | 79239600 | 79242599 | 57.55116 | 92.15361 | 83.31749 | 35.11821 | 0.490187 | 2.019141 | 0.49628 |
| 14 | 3156600 | 3160599 | 105.0256 | 162.0157 | 166.2633 | 69.06273 | 0.2621 | 2.894315 | 0.496289 |
| 4 | 113385600 | 113393599 | 111.2461 | 173.1176 | 150.555 | 62.49835 | 0.510341 | 2.742243 | 0.496298 |
| 2 | 100247800 | 100249199 | 12.95806 | 24.31895 | 24.52427 | 10.09722 | 0.452411 | 0.12415 | 0.496309 |
| 13 | 66104200 | 66111599 | 153.6048 | 233.9594 | 198.1138 | 82.11311 | 0.524944 | 3.162073 | 0.496314 |
| 3 | 33566400 | 33568399 | 23.94008 | 41.98551 | 33.32878 | 14.7311 | 0.718925 | 0.672738 | 0.49632 |
| 10 | 53655400 | 53658799 | 49.71719 | 80.12728 | 80.75451 | 34.36669 | 0.345057 | 1.807841 | 0.496321 |
| 19 | 61091800 | 61094599 | 37.95115 | 62.6963 | 58.23747 | 25.15742 | 0.486649 | 1.365953 | 0.496332 |
| 17 | 13784400 | 13785599 | 9.056028 | 16.75424 | 30.95905 | 13.03841 | -0.44044 | 0.194078 | 0.496347 |
| 18 | 4010000 | 4011399 | 8.734967 | 16.32105 | 29.52664 | 12.24808 | -0.393 | 0.129184 | 0.496365 |
| 15 | 52222000 | 52223999 | 29.30289 | 49.35663 | 49.63892 | 21.69748 | 0.372874 | 1.127298 | 0.496373 |
| 18 | 90213800 | 90215199 | 24.99559 | 42.44476 | 42.74573 | 18.73048 | 0.371359 | 1.036082 | 0.496377 |
| 16 | 45095800 | 45096799 | 17.3225 | 30.87825 | 30.63761 | 13.4823 | 0.407834 | 0.470041 | 0.496378 |
| X | 107075400 | 107079199 | 44.96361 | 74.06844 | 56.91245 | 24.54798 | 0.758466 | 1.416343 | 0.496405 |
| 10 | 58055200 | 58079599 | 500.5498 | 672.5891 | 897.655 | 374.5288 | -0.30611 | 5.201697 | 0.496426 |
| 17 | 37857000 | 37857599 | 5.353392 | 10.4374 | 22.08502 | 7.958942 | -0.43036 | -0.40736 | 0.49643 |
| 8 | 105496600 | 105498599 | 23.14176 | 39.77721 | 41.55394 | 18.22708 | 0.321224 | 0.917284 | 0.496438 |
| 14 | 25785600 | 25786399 | 5.668411 | 11.00466 | 22.78404 | 8.55135 | -0.45351 | -0.25791 | 0.496454 |
| 18 | 46584000 | 46585999 | 16.64483 | 28.90956 | 47.92638 | 21.03782 | -0.33924 | 0.800375 | 0.49649 |
| 2 | 126080800 | 126081199 | 4.26737 | 8.333386 | 19.93124 | 7.005918 | -0.57261 | -0.52226 | 0.496496 |
| 8 | 115321600 | 115325999 | 44.14487 | 70.64112 | 110.5892 | 46.56956 | -0.30144 | 2.032678 | 0.496506 |
| 6 | 33056600 | 33059399 | 33.05691 | 54.71023 | 56.4289 | 24.40676 | 0.33612 | 1.327344 | 0.496507 |
| 6 | 54707000 | 54707799 | 7.15432 | 13.70674 | 26.26405 | 10.35426 | -0.41139 | -0.06922 | 0.496528 |
| X | 146450200 | 146456599 | 60.99933 | 97.97843 | 76.56256 | 32.42361 | 0.705427 | 1.927744 | 0.496534 |

| | | | | | | | | | |
|---|---|---|---|---|---|---|---|---|---|
| 2 | 80987200 | 80989399 | 20.85252 | 34.68926 | 68.7634 | 29.84733 | -0.60271 | 1.247844 | 0.496545 |
| X | 79805800 | 79807999 | 29.28151 | 49.60383 | 47.42579 | 20.76613 | 0.44597 | 1.078449 | 0.496546 |
| 10 | 64364600 | 64365599 | 9.612839 | 17.59553 | 31.90601 | 13.63384 | -0.42822 | 0.264182 | 0.49656 |
| 13 | 114005000 | 114011999 | 78.89525 | 122.1644 | 187.4975 | 77.88151 | -0.31386 | 2.816379 | 0.496569 |
| 7 | 116322600 | 116325599 | 25.5619 | 42.14581 | 69.40518 | 29.9478 | -0.3376 | 1.36567 | 0.4966 |
| 18 | 46444600 | 46446599 | 11.74183 | 20.80392 | 36.78623 | 16.26432 | -0.42576 | 0.488273 | 0.49661 |
| 3 | 129828800 | 129844599 | 323.0965 | 452.1845 | 606.9244 | 251.6284 | -0.25501 | 4.567814 | 0.496631 |
| 11 | 67423000 | 67425199 | 13.02135 | 22.84548 | 40.46133 | 17.88957 | -0.43281 | 0.620295 | 0.496633 |
| 1 | 133076000 | 133077199 | 17.51364 | 30.0718 | 50.58645 | 22.18833 | -0.36356 | 0.915039 | 0.496634 |
| 11 | 91668400 | 91669599 | 11.99506 | 21.56179 | 36.79093 | 16.0761 | -0.3628 | 0.395494 | 0.496636 |
| 2 | 38236400 | 38239199 | 21.53185 | 36.04421 | 60.76309 | 26.47992 | -0.36981 | 1.173334 | 0.49665 |
| 1 | 102048000 | 102051199 | 33.99766 | 54.9712 | 87.41159 | 37.35612 | -0.30592 | 1.693116 | 0.496696 |
| 17 | 55577000 | 55579999 | 31.84592 | 53.24077 | 47.03166 | 20.47913 | 0.559374 | 1.205541 | 0.496696 |
| 1 | 46311000 | 46313199 | 17.79191 | 30.61807 | 51.14071 | 22.43301 | -0.35277 | 0.895719 | 0.496735 |
| 8 | 40139800 | 40141199 | 13.95436 | 24.52595 | 40.85493 | 18.09073 | -0.34495 | 0.597346 | 0.496736 |
| X | 26786000 | 26789799 | 71.62383 | 114.1226 | 104.7457 | 43.84355 | 0.460837 | 2.192757 | 0.496738 |
| 10 | 90089600 | 90091399 | 26.10958 | 45.52688 | 36.17612 | 15.91916 | 0.717513 | 0.774055 | 0.496738 |
| 3 | 120189200 | 120190999 | 26.1818 | 44.57266 | 45.23608 | 19.82788 | 0.360607 | 1.018316 | 0.496769 |
| 4 | 60019400 | 60020999 | 24.59515 | 41.93498 | 43.6947 | 19.1546 | 0.323133 | 0.997774 | 0.49677 |
| 1 | 47188000 | 47189799 | 22.51762 | 37.88864 | 62.53897 | 27.3276 | -0.33946 | 1.127972 | 0.496788 |
| 17 | 59626400 | 59627199 | 11.79825 | 22.15069 | 22.64164 | 9.316112 | 0.434342 | 0.109198 | 0.496788 |
| 16 | 48064000 | 48065399 | 21.28229 | 36.83651 | 37.4187 | 16.4167 | 0.363492 | 0.841512 | 0.496798 |
| 5 | 133832800 | 133834199 | 19.42239 | 34.00076 | 34.41415 | 15.18752 | 0.370099 | 0.677803 | 0.496819 |
| 11 | 94762600 | 94766199 | 16.2372 | 27.94115 | 49.51985 | 21.72198 | -0.43781 | 0.888172 | 0.496826 |
| 1 | 112977600 | 112979799 | 30.62014 | 51.42492 | 44.84454 | 19.58292 | 0.577848 | 1.153276 | 0.496843 |
| X | 5604000 | 5605599 | 20.86503 | 38.11181 | 25.17244 | 10.78584 | 1.016455 | 0.344314 | 0.496849 |
| 10 | 103630800 | 103636199 | 89.20953 | 137.6859 | 202.0407 | 83.92381 | -0.25277 | 2.882674 | 0.496851 |
| 5 | 127381600 | 127385199 | 22.59967 | 39.28502 | 38.10427 | 16.73508 | 0.430649 | 0.800373 | 0.496857 |
| 5 | 63709000 | 63710199 | 11.34275 | 20.31385 | 35.73894 | 15.66527 | -0.40983 | 0.410933 | 0.496859 |
| 12 | 49647200 | 49647999 | 9.876528 | 18.3112 | 32.61186 | 13.69473 | -0.38453 | 0.194491 | 0.496868 |
| 13 | 90420400 | 90421799 | 19.82527 | 35.14556 | 31.40948 | 13.88554 | 0.55337 | 0.521721 | 0.496884 |
| 13 | 66954200 | 66958599 | 93.09453 | 145.117 | 149.0668 | 61.86487 | 0.272285 | 2.703868 | 0.496888 |
| 17 | 18798800 | 18800599 | 17.15916 | 32.06846 | 24.21706 | 10.04186 | 0.858913 | 0.186423 | 0.49689 |
| 17 | 20905600 | 20907199 | 16.33496 | 30.27046 | 25.73852 | 10.81557 | 0.674066 | 0.231998 | 0.496892 |
| 2 | 47690200 | 47691399 | 11.35284 | 20.56069 | 35.35757 | 15.31257 | -0.36543 | 0.344823 | 0.496895 |
| 9 | 47403400 | 47407399 | 40.94186 | 67.1693 | 62.2304 | 26.79555 | 0.487003 | 1.458782 | 0.496899 |
| 16 | 69054200 | 69055399 | 9.969104 | 18.21957 | 33.44886 | 14.32214 | -0.44794 | 0.280493 | 0.496919 |
| 11 | 99749000 | 99751199 | 10.58962 | 19.17496 | 33.84651 | 14.66149 | -0.40274 | 0.338492 | 0.496921 |
| 11 | 18147000 | 18149199 | 27.39865 | 44.87804 | 73.69436 | 31.71019 | -0.33735 | 1.454745 | 0.496924 |
| 16 | 57183000 | 57183999 | 8.349087 | 15.46756 | 29.97904 | 12.60213 | -0.5061 | 0.181599 | 0.496929 |
| 4 | 113321200 | 113335599 | 371.986 | 522.9704 | 486.997 | 201.7647 | 0.278812 | 4.50148 | 0.49693 |
| 1 | 111575800 | 111577199 | 17.34789 | 31.85311 | 26.61289 | 11.32386 | 0.686402 | 0.291838 | 0.496939 |
| 13 | 43791200 | 43791599 | 5.515069 | 10.74742 | 22.68032 | 8.225478 | -0.43599 | -0.38452 | 0.496952 |
| 3 | 16032200 | 16040399 | 48.58763 | 78.87595 | 61.76262 | 26.47586 | 0.719014 | 1.624353 | 0.496964 |
| 6 | 67627800 | 67631399 | 46.98653 | 76.19988 | 71.42129 | 30.5935 | 0.458915 | 1.64613 | 0.496981 |
| 12 | 107870000 | 107878599 | 70.48592 | 109.741 | 190.5983 | 79.11542 | -0.48592 | 2.718221 | 0.496984 |
| 12 | 114119800 | 114125799 | 137.7071 | 209.3448 | 209.681 | 86.85774 | 0.280639 | 3.196062 | 0.496984 |

| | | | | | | | | | |
|---|---|---|---|---|---|---|---|---|---|
| 10 | 35515000 | 35516799 | 18.92038 | 33.53445 | 31.45982 | 13.90838 | 0.483544 | 0.517551 | 0.496988 |
| 9 | 78180800 | 78189999 | 75.74282 | 117.821 | 178.1022 | 73.9382 | -0.28681 | 2.734244 | 0.496997 |
| 11 | 114328200 | 114331399 | 34.15031 | 55.12292 | 90.60751 | 38.63592 | -0.35655 | 1.743707 | 0.49701 |
| 17 | 93346600 | 93346999 | 5.375687 | 10.4449 | 22.26364 | 8.301043 | -0.48583 | -0.28584 | 0.49701 |
| 14 | 25619200 | 25622999 | 20.11031 | 33.72083 | 59.02841 | 25.75109 | -0.42388 | 1.150153 | 0.497014 |
| 8 | 93134600 | 93138199 | 21.51407 | 35.67351 | 64.76753 | 28.04982 | -0.47671 | 1.284674 | 0.497018 |
| 2 | 15585200 | 15589799 | 55.91812 | 89.82425 | 86.97544 | 36.83786 | 0.396838 | 1.920145 | 0.497019 |
| 12 | 65312200 | 65314199 | 28.3828 | 46.32891 | 76.4429 | 32.85001 | -0.34756 | 1.506395 | 0.497021 |
| 11 | 15809600 | 15812999 | 75.80502 | 120.1579 | 107.868 | 44.88276 | 0.485237 | 2.351743 | 0.497021 |
| X | 8568000 | 8572799 | 76.2328 | 121.2695 | 108.1867 | 45.18233 | 0.499289 | 2.248717 | 0.497028 |
| 5 | 98570200 | 98573199 | 26.8649 | 44.30875 | 71.87153 | 31.01634 | -0.31571 | 1.366828 | 0.497038 |
| 17 | 33944800 | 33947599 | 23.34158 | 40.91633 | 34.52545 | 15.25386 | 0.631336 | 0.679303 | 0.497051 |
| 8 | 54701400 | 54702199 | 17.49016 | 30.80611 | 31.44015 | 13.91708 | 0.35942 | 0.57726 | 0.497051 |
| 1 | 45591600 | 45594199 | 30.70815 | 51.59082 | 50.21774 | 21.93338 | 0.419699 | 1.153274 | 0.497076 |
| 4 | 112465800 | 112467399 | 39.92443 | 64.79063 | 64.70219 | 27.72912 | 0.368604 | 1.616884 | 0.497119 |
| 5 | 149235800 | 149239399 | 28.15995 | 46.08947 | 76.28394 | 32.81399 | -0.34931 | 1.465499 | 0.497122 |
| 3 | 99320800 | 99322799 | 27.82464 | 46.96741 | 48.53776 | 21.22996 | 0.333904 | 1.103077 | 0.49713 |
| 10 | 52142800 | 52143999 | 9.346182 | 17.36143 | 31.0177 | 12.99111 | -0.38641 | 0.173891 | 0.497183 |
| 1 | 52453800 | 52454599 | 10.54453 | 20.5474 | 17.83269 | 6.471986 | 0.831497 | -0.33308 | 0.497187 |
| 8 | 8709000 | 8712199 | 24.27625 | 39.968 | 69.80232 | 30.10067 | -0.42297 | 1.379374 | 0.497211 |
| 13 | 61347000 | 61353799 | 94.8441 | 149.3142 | 129.8018 | 53.85529 | 0.523782 | 2.522698 | 0.497212 |
| 6 | 123838400 | 123839599 | 22.48893 | 39.17822 | 31.6464 | 13.91798 | 0.693978 | 0.763657 | 0.497225 |
| 11 | 106520000 | 106525999 | 90.90543 | 139.7008 | 207.5214 | 86.1701 | -0.27516 | 2.965654 | 0.497254 |
| 17 | 90109600 | 90109799 | 4.612724 | 9.006707 | 19.8851 | 7.004324 | -0.45986 | -0.50463 | 0.497256 |
| 4 | 46845800 | 46846799 | 15.58855 | 27.17996 | 45.22792 | 19.90166 | -0.34381 | 0.738657 | 0.497259 |
| 3 | 131081800 | 131085999 | 45.20301 | 72.02311 | 120.1116 | 50.27373 | -0.39929 | 2.171313 | 0.497279 |
| 6 | 65680000 | 65681199 | 14.24484 | 26.00366 | 26.80203 | 11.49281 | 0.377489 | 0.29978 | 0.49729 |
| 6 | 13838600 | 13839199 | 4.68376 | 9.116803 | 21.24201 | 7.806406 | -0.59341 | -0.35074 | 0.497297 |
| 15 | 20954000 | 20955599 | 10.95988 | 19.95201 | 34.9064 | 15.01963 | -0.38425 | 0.309061 | 0.497301 |
| X | 116074000 | 116074999 | 17.70863 | 31.85024 | 28.70079 | 12.53507 | 0.553635 | 0.416524 | 0.497301 |
| 4 | 154728400 | 154733999 | 74.26612 | 116.0481 | 172.3716 | 71.51721 | -0.25645 | 2.654507 | 0.497328 |
| 7 | 20221000 | 20221799 | 17.57459 | 31.27817 | 25.75711 | 11.34971 | 0.67399 | 0.472157 | 0.49733 |
| 9 | 14369600 | 14370999 | 14.65486 | 27.42502 | 24.92675 | 10.31266 | 0.595826 | 0.15785 | 0.497333 |
| 17 | 8071400 | 8075799 | 38.38351 | 61.85679 | 96.93933 | 41.25051 | -0.29061 | 1.800086 | 0.497335 |
| 4 | 47054800 | 47057999 | 23.27342 | 38.47225 | 67.11265 | 29.00379 | -0.41973 | 1.325871 | 0.497339 |
| 12 | 20627000 | 20636999 | 333.2525 | 473.2334 | 383.7255 | 158.7167 | 0.492929 | 4.352184 | 0.497346 |
| 12 | 56957400 | 56960799 | 59.15036 | 94.36167 | 98.47615 | 41.28818 | 0.277718 | 2.139926 | 0.497349 |
| 5 | 119515000 | 119520799 | 54.00866 | 85.80614 | 131.5805 | 54.8757 | -0.28284 | 2.271602 | 0.497351 |
| 12 | 81390800 | 81392999 | 25.74196 | 43.83553 | 45.25471 | 19.83647 | 0.336135 | 1.014961 | 0.497361 |
| 2 | 36886200 | 36887799 | 19.02521 | 33.77428 | 30.87168 | 13.63247 | 0.522018 | 0.505419 | 0.497362 |
| X | 152880000 | 152886799 | 147.0342 | 223.678 | 205.754 | 85.24858 | 0.404174 | 3.185519 | 0.497362 |
| 12 | 27653200 | 27654799 | 14.00171 | 25.99722 | 25.32578 | 10.614 | 0.481567 | 0.200044 | 0.497367 |
| 11 | 9033000 | 9037199 | 56.21363 | 89.88675 | 93.47263 | 39.32023 | 0.286491 | 2.062216 | 0.497371 |
| 4 | 84748200 | 84749999 | 23.48039 | 41.05359 | 32.17789 | 14.18571 | 0.7374 | 0.717558 | 0.497394 |
| X | 11696200 | 11700199 | 28.73025 | 47.19922 | 80.4919 | 34.6787 | -0.39011 | 1.426459 | 0.4974 |
| 4 | 14377000 | 14380999 | 46.20373 | 74.68649 | 75.12672 | 32.09451 | 0.352706 | 1.713663 | 0.497407 |
| 10 | 122831400 | 122834599 | 20.66398 | 35.05574 | 57.0331 | 24.98372 | -0.31814 | 1.044874 | 0.497436 |

| | | | | | | | | | |
|---|---|---|---|---|---|---|---|---|---|
| 5 | 49851800 | 49854599 | 25.9642 | 45.05684 | 39.65691 | 17.40004 | 0.570128 | 0.843237 | 0.497443 |
| 16 | 4543800 | 4544999 | 10.55004 | 19.1504 | 33.33209 | 14.39425 | -0.37987 | 0.321612 | 0.49745 |
| 1 | 38560200 | 38561199 | 8.699977 | 16.36348 | 29.667 | 12.16932 | -0.38324 | 0.085992 | 0.497457 |
| 3 | 74788800 | 74789799 | 8.699977 | 16.36348 | 29.667 | 12.16932 | -0.38324 | 0.085992 | 0.497457 |
| 17 | 17064400 | 17069199 | 84.5554 | 133.1745 | 132.2936 | 54.89227 | 0.332033 | 2.507881 | 0.497491 |
| 8 | 97114000 | 97114799 | 12.78531 | 23.7366 | 24.96081 | 10.4623 | 0.37208 | 0.190578 | 0.4975 |
| 4 | 39134200 | 39139199 | 57.12608 | 91.97385 | 81.51318 | 34.63849 | 0.528342 | 1.852208 | 0.497503 |
| 1 | 191653200 | 191656199 | 33.68156 | 54.48278 | 86.80543 | 37.11309 | -0.30811 | 1.682336 | 0.497509 |
| X | 146531200 | 146535199 | 71.98151 | 114.9279 | 97.90227 | 41.09645 | 0.571442 | 2.128149 | 0.497514 |
| 1 | 155428400 | 155429999 | 18.24445 | 32.77924 | 29.55032 | 12.91763 | 0.551973 | 0.426738 | 0.497536 |
| 4 | 144054600 | 144060399 | 76.74001 | 121.6674 | 117.8766 | 49.05833 | 0.375709 | 2.347063 | 0.497559 |
| Y | 88986600 | 88989199 | 15.6756 | 28.16631 | 19.07751 | 8.338935 | 0.961395 | 0.381634 | 0.497561 |
| 14 | 68316000 | 68316799 | 8.499004 | 15.66976 | 30.4394 | 12.87966 | -0.51653 | 0.209456 | 0.497618 |
| X | 49509400 | 49510199 | 13.14932 | 24.45459 | 24.55961 | 10.26863 | 0.441111 | 0.185427 | 0.497628 |
| 2 | 171366800 | 171368799 | 11.96936 | 21.40729 | 36.73067 | 16.1182 | -0.37511 | 0.425526 | 0.497633 |
| 13 | 53059400 | 53063799 | 38.27308 | 61.47666 | 100.0099 | 42.35507 | -0.35024 | 1.904814 | 0.497656 |
| 5 | 115976200 | 115981199 | 48.39352 | 77.80048 | 80.722 | 34.22968 | 0.29882 | 1.88299 | 0.497681 |
| 1 | 156227000 | 156233999 | 83.36771 | 130.8741 | 136.2842 | 56.52397 | 0.260476 | 2.571378 | 0.497685 |
| 19 | 3121200 | 3127199 | 62.20683 | 99.4412 | 88.93397 | 37.39566 | 0.503205 | 2.077515 | 0.497687 |
| 11 | 66562000 | 66564199 | 22.8757 | 39.71748 | 39.84316 | 17.48568 | 0.382331 | 0.823811 | 0.497702 |
| 9 | 57743400 | 57745599 | 11.52526 | 20.52253 | 36.73511 | 16.18005 | -0.43948 | 0.450097 | 0.497715 |
| 11 | 38122200 | 38123599 | 16.22085 | 28.31332 | 47.858 | 21.07343 | -0.36632 | 0.729246 | 0.497718 |
| 3 | 75677600 | 75678799 | 19.38652 | 33.97198 | 34.94629 | 15.43543 | 0.346933 | 0.663984 | 0.497749 |
| 17 | 36027600 | 36030199 | 28.30178 | 47.42098 | 49.67458 | 21.66033 | 0.314268 | 1.173672 | 0.497752 |
| X | 111378200 | 111380599 | 25.87715 | 45.08888 | 37.35701 | 16.43132 | 0.657492 | 0.785302 | 0.497759 |
| X | 91030800 | 91032399 | 19.3302 | 34.41431 | 30.5665 | 13.4651 | 0.565907 | 0.492467 | 0.497764 |
| 19 | 28548600 | 28550399 | 16.04953 | 28.73279 | 29.14074 | 12.77704 | 0.379795 | 0.43065 | 0.497781 |
| 15 | 80952200 | 80954399 | 21.09025 | 36.85656 | 36.4485 | 16.06177 | 0.403523 | 0.717713 | 0.497784 |
| 16 | 73496400 | 73499799 | 35.12706 | 58.83655 | 50.85823 | 22.17158 | 0.590592 | 1.200561 | 0.497797 |
| 18 | 74467200 | 74473999 | 119.545 | 183.1812 | 186.8357 | 77.54983 | 0.263569 | 3.029791 | 0.497809 |
| 11 | 19685000 | 19686199 | 11.43647 | 21.4884 | 22.51643 | 9.252228 | 0.399725 | 0.097768 | 0.497815 |
| 3 | 15940400 | 15941399 | 20.75276 | 36.13145 | 37.29063 | 16.39397 | 0.341911 | 0.77185 | 0.49783 |
| 10 | 110245000 | 110246399 | 13.45705 | 25.54165 | 22.40532 | 9.026236 | 0.678105 | 0.048876 | 0.497834 |
| 7 | 17807600 | 17831599 | 445.2809 | 619.5307 | 531.0366 | 220.3671 | 0.383217 | 4.659926 | 0.497837 |
| 3 | 36337800 | 36340799 | 50.88244 | 81.78552 | 85.33424 | 36.17502 | 0.290379 | 1.894516 | 0.497886 |
| 10 | 106024000 | 106024799 | 16.9088 | 29.07501 | 49.51992 | 21.72152 | -0.38082 | 0.899565 | 0.497891 |
| 6 | 58400600 | 58401599 | 15.46545 | 27.01337 | 45.72443 | 20.14266 | -0.36801 | 0.714407 | 0.497892 |
| 9 | 121037200 | 121048999 | 123.8058 | 186.5649 | 272.8007 | 112.6996 | -0.27549 | 3.373571 | 0.497896 |
| X | 122631400 | 122632599 | 13.69407 | 25.66684 | 19.95 | 8.232007 | 0.823341 | 0.129375 | 0.497896 |
| 8 | 4680800 | 4685399 | 52.97595 | 85.62825 | 74.09868 | 31.65283 | 0.56928 | 1.728737 | 0.497899 |
| 5 | 32379200 | 32381399 | 25.64323 | 43.70052 | 45.26292 | 19.84128 | 0.331642 | 1.008681 | 0.497914 |
| 15 | 20513600 | 20514199 | 10.86824 | 20.60617 | 18.71579 | 7.556832 | 0.626443 | 0.024358 | 0.49792 |
| 5 | 119645200 | 119648999 | 34.50313 | 55.62745 | 91.90678 | 39.131 | -0.36587 | 1.779636 | 0.497924 |
| 7 | 97654200 | 97658199 | 35.17803 | 57.57255 | 58.90533 | 25.33793 | 0.342706 | 1.471373 | 0.497928 |
| 17 | 8428800 | 8430199 | 10.56421 | 19.05881 | 33.79371 | 14.70603 | -0.41371 | 0.357897 | 0.497929 |
| 3 | 158879400 | 158880199 | 11.75701 | 22.39522 | 19.95137 | 7.976835 | 0.665859 | 0.003693 | 0.49793 |
| 12 | 55358000 | 55358999 | 6.303233 | 12.04797 | 25.63131 | 10.17285 | -0.56913 | -0.05486 | 0.497953 |

| | | | | | | | | | |
|---|---|---|---|---|---|---|---|---|---|
| 3 | 148409800 | 148411599 | 15.42709 | 28.57533 | 25.1393 | 10.57019 | 0.624232 | 0.223978 | 0.497964 |
| 6 | 71322600 | 71326399 | 46.31683 | 74.03757 | 113.9722 | 47.9216 | -0.27891 | 2.069854 | 0.497973 |
| 3 | 24319200 | 24319599 | 6.686768 | 12.95263 | 25.66403 | 9.73888 | -0.40708 | -0.20268 | 0.497989 |
| 7 | 55020400 | 55020999 | 6.686768 | 12.95263 | 25.66403 | 9.73888 | -0.40708 | -0.20268 | 0.497989 |
| 10 | 77280800 | 77282599 | 20.56777 | 36.05875 | 35.3333 | 15.61252 | 0.416705 | 0.663898 | 0.497992 |
| 12 | 3434400 | 3445799 | 82.615 | 127.6228 | 195.5897 | 81.2438 | -0.31438 | 2.861757 | 0.498013 |
| Y | 20868400 | 20871399 | 16.53003 | 29.58 | 18.02631 | 7.906741 | 1.109264 | 0.409994 | 0.498022 |
| 7 | 46512000 | 46513399 | 9.773755 | 17.99715 | 32.49307 | 13.77241 | -0.41353 | 0.233537 | 0.498035 |
| 17 | 20128600 | 20133199 | 27.66145 | 48.00691 | 33.67455 | 14.77524 | 0.896025 | 0.842054 | 0.49804 |
| 18 | 75196400 | 75197399 | 15.19604 | 27.17999 | 26.32981 | 11.55345 | 0.44489 | 0.424656 | 0.498047 |
| 17 | 91837800 | 91838399 | 8.840502 | 17.0829 | 18.75816 | 7.199679 | 0.421344 | -0.16115 | 0.498049 |
| 1 | 171097800 | 171098599 | 5.518507 | 10.71398 | 22.76838 | 8.543199 | -0.48992 | -0.25927 | 0.498052 |
| 2 | 152660600 | 152661799 | 14.60394 | 26.63625 | 27.29785 | 11.71803 | 0.384718 | 0.308791 | 0.498067 |
| 14 | 33443600 | 33445599 | 25.27865 | 43.13288 | 44.72118 | 19.60577 | 0.330401 | 0.995048 | 0.498079 |
| 1 | 83207200 | 83208799 | 39.92468 | 64.71774 | 62.01145 | 26.55379 | 0.426551 | 1.635306 | 0.498091 |
| X | 77373400 | 77376999 | 52.40576 | 84.64841 | 75.76975 | 32.33797 | 0.519469 | 1.747108 | 0.498093 |
| 17 | 7734000 | 7735599 | 14.2981 | 25.00377 | 42.58405 | 18.77627 | -0.37671 | 0.684315 | 0.498097 |
| 15 | 42214600 | 42215599 | 13.82993 | 26.71457 | 20.10918 | 7.731223 | 0.958995 | -0.09773 | 0.498122 |
| 9 | 103697400 | 103702599 | 27.15389 | 44.76515 | 71.97732 | 31.05605 | -0.30338 | 1.37198 | 0.498137 |
| X | 113316800 | 113321999 | 64.53744 | 103.0682 | 99.75352 | 41.88839 | 0.387985 | 2.113511 | 0.498138 |
| 10 | 35745400 | 35747399 | 21.17113 | 37.38342 | 31.79312 | 14.06625 | 0.622298 | 0.571183 | 0.49814 |
| 3 | 91670400 | 91672799 | 31.54492 | 52.77669 | 52.44621 | 22.8475 | 0.390139 | 1.20341 | 0.498141 |
| 9 | 11716200 | 11716799 | 11.05331 | 20.91437 | 22.64567 | 9.1831 | 0.368971 | 0.050032 | 0.498176 |
| 1 | 38319800 | 38321599 | 9.83169 | 18.1543 | 31.98355 | 13.5052 | -0.37493 | 0.213893 | 0.498188 |
| 1 | 155402800 | 155406599 | 40.91053 | 65.51601 | 106.8811 | 45.05878 | -0.35947 | 2.004461 | 0.498203 |
| X | 119088600 | 119090599 | 36.4518 | 60.60549 | 54.82783 | 23.78928 | 0.525276 | 1.287375 | 0.498214 |
| 7 | 32992000 | 33004399 | 181.3217 | 272.6252 | 238.836 | 98.60894 | 0.460206 | 3.427735 | 0.498233 |
| 3 | 4369200 | 4371799 | 25.32714 | 43.33508 | 43.59389 | 19.11581 | 0.374324 | 0.967842 | 0.498234 |
| 5 | 7541200 | 7543999 | 22.86105 | 39.63584 | 39.80138 | 17.46295 | 0.380406 | 0.835838 | 0.498236 |
| 1 | 16649000 | 16650999 | 23.15693 | 39.90397 | 41.51478 | 18.21131 | 0.32788 | 0.894998 | 0.498239 |
| X | 111847400 | 111848999 | 16.98752 | 31.61082 | 24.62982 | 10.288 | 0.805514 | 0.211453 | 0.498254 |
| 8 | 52464800 | 52466599 | 28.0718 | 48.48977 | 40.12793 | 17.60423 | 0.657338 | 0.892836 | 0.498275 |
| 12 | 4849000 | 4857999 | 242.0431 | 344.9903 | 496.4381 | 205.2089 | -0.3289 | 4.295058 | 0.498282 |
| 7 | 66811600 | 66817999 | 70.42299 | 110.3606 | 164.3106 | 68.15245 | -0.25659 | 2.601094 | 0.498283 |
| 9 | 75615400 | 75619199 | 33.83863 | 56.3775 | 53.70781 | 23.33533 | 0.450918 | 1.252768 | 0.498285 |
| 16 | 76316000 | 76316999 | 9.088042 | 16.33999 | 33.90358 | 14.8116 | -0.64237 | 0.368566 | 0.498293 |
| 1 | 148261000 | 148262599 | 23.13402 | 40.38394 | 36.28187 | 15.97701 | 0.541176 | 0.742358 | 0.498301 |
| 5 | 136858200 | 136861199 | 42.01892 | 67.25665 | 109.6564 | 46.19378 | -0.35952 | 2.021405 | 0.498319 |
| 7 | 105953600 | 105954799 | 12.99823 | 23.10535 | 39.27141 | 17.32243 | -0.36669 | 0.477171 | 0.498321 |
| 3 | 153191400 | 153194999 | 26.61627 | 43.49451 | 85.7782 | 36.87687 | -0.6031 | 1.490897 | 0.498325 |
| 13 | 53183000 | 53186799 | 31.30264 | 50.97808 | 81.37132 | 34.93874 | -0.30228 | 1.547987 | 0.498334 |
| 6 | 20118800 | 20122599 | 69.58 | 109.123 | 162.6358 | 67.45247 | -0.25722 | 2.586932 | 0.498337 |
| 16 | 92569000 | 92571799 | 34.79642 | 56.35082 | 89.19845 | 38.1685 | -0.29723 | 1.661633 | 0.498344 |
| X | 29295000 | 29297399 | 29.97144 | 50.44936 | 43.26359 | 18.90699 | 0.602009 | 1.130143 | 0.498351 |
| 3 | 94730400 | 94731799 | 21.1914 | 36.8594 | 38.29266 | 16.82424 | 0.332288 | 0.78817 | 0.498366 |
| 11 | 90078000 | 90079799 | 10.83777 | 19.70039 | 33.84068 | 14.58819 | -0.35951 | 0.315822 | 0.498374 |
| 5 | 59596600 | 59602799 | 62.36957 | 98.47196 | 147.8555 | 61.38856 | -0.26041 | 2.444916 | 0.498399 |

| | | | | | | | | | |
|---|---|---|---|---|---|---|---|---|---|
| 12 | 3400200 | 3404399 | 27.32285 | 45.09132 | 73.2167 | 31.60553 | -0.31694 | 1.360572 | 0.498399 |
| 1 | 110972800 | 110973999 | 14.22535 | 26.39062 | 25.95655 | 10.8906 | 0.467077 | 0.21008 | 0.498406 |
| X | 120488400 | 120495599 | 84.11382 | 133.2565 | 115.9905 | 48.24165 | 0.52905 | 2.376822 | 0.498439 |
| 17 | 65967600 | 65969199 | 13.07255 | 22.88924 | 41.58616 | 18.35569 | -0.46925 | 0.657344 | 0.498441 |
| 14 | 52923000 | 52927599 | 36.58281 | 61.10812 | 51.48932 | 22.40814 | 0.627416 | 1.233344 | 0.498447 |
| X | 5519000 | 5537599 | 160.8589 | 245.2267 | 166.1747 | 68.8924 | 0.847172 | 3.144269 | 0.498482 |
| 4 | 32030600 | 32032999 | 24.92437 | 42.6743 | 42.51516 | 18.64379 | 0.388346 | 0.958189 | 0.498484 |
| 6 | 133432400 | 133433999 | 23.83622 | 41.44988 | 38.57049 | 16.94379 | 0.490339 | 0.802628 | 0.498485 |
| 19 | 57707000 | 57708199 | 15.92512 | 27.75605 | 46.28461 | 20.36168 | -0.34694 | 0.747582 | 0.498486 |
| 1 | 107483000 | 107484799 | 19.73625 | 34.38644 | 33.72424 | 14.83281 | 0.415157 | 0.749189 | 0.498488 |
| 16 | 23855800 | 23857399 | 19.17164 | 33.93414 | 32.06035 | 14.17768 | 0.472787 | 0.535495 | 0.498488 |
| 7 | 34973400 | 34982399 | 159.3742 | 235.5441 | 345.3055 | 142.2048 | -0.30473 | 3.736341 | 0.498489 |
| 11 | 107844600 | 107852599 | 64.54467 | 101.5826 | 153.435 | 63.65832 | -0.27255 | 2.513173 | 0.498501 |
| 17 | 58121000 | 58121399 | 9.839217 | 18.86313 | 20.14278 | 7.925945 | 0.427201 | -0.06709 | 0.498504 |
| 18 | 78154000 | 78155599 | 9.912787 | 18.05835 | 32.35706 | 13.91062 | -0.41718 | 0.293746 | 0.498516 |
| 3 | 5283400 | 5286799 | 22.73586 | 37.69263 | 67.30657 | 29.14581 | -0.45305 | 1.288981 | 0.498516 |
| X | 25617000 | 25619999 | 50.24553 | 81.6581 | 57.82057 | 24.80635 | 0.865305 | 1.605802 | 0.498519 |
| 18 | 60558400 | 60560799 | 20.09673 | 33.67702 | 60.14775 | 26.22907 | -0.45282 | 1.156534 | 0.498523 |
| 12 | 57697200 | 57698399 | 8.569785 | 16.22446 | 29.86135 | 12.09454 | -0.38897 | 0.042805 | 0.498529 |
| 1 | 91396000 | 91400599 | 31.11724 | 50.69749 | 82.0818 | 35.24921 | -0.32233 | 1.541159 | 0.498535 |
| 12 | 21355600 | 21356799 | 8.806295 | 16.18843 | 30.9174 | 13.13459 | -0.49654 | 0.229487 | 0.49854 |
| 3 | 92102800 | 92104399 | 20.25572 | 34.00492 | 58.5155 | 25.54642 | -0.39929 | 1.137378 | 0.498607 |
| 4 | 153109200 | 153123399 | 136.9581 | 205.018 | 300.9497 | 124.1227 | -0.28965 | 3.504549 | 0.498614 |
| 11 | 14345600 | 14346399 | 10.53062 | 20.38717 | 20.33812 | 7.737657 | 0.570001 | -0.16815 | 0.498638 |
| 5 | 74695800 | 74696799 | 6.745141 | 12.85443 | 26.12326 | 10.4341 | -0.51234 | -0.02981 | 0.498638 |
| 6 | 138513400 | 138514799 | 20.60289 | 36.55488 | 30.75178 | 13.58584 | 0.640763 | 0.519392 | 0.498639 |
| 6 | 115864800 | 115866199 | 22.11873 | 38.2765 | 37.9799 | 16.66281 | 0.397284 | 0.850346 | 0.498643 |
| 12 | 89073000 | 89073599 | 5.961939 | 11.55873 | 23.8958 | 9.030792 | -0.46183 | -0.23027 | 0.498649 |
| 9 | 82711600 | 82712799 | 7.999566 | 15.00066 | 28.53734 | 11.76604 | -0.45736 | 0.099281 | 0.498669 |
| 13 | 19579400 | 19580199 | 7.999566 | 15.00066 | 28.53734 | 11.76604 | -0.45736 | 0.099281 | 0.498669 |
| 18 | 80194000 | 80196399 | 15.34067 | 26.70196 | 46.82999 | 20.58443 | -0.41978 | 0.76069 | 0.498678 |
| 9 | 12369400 | 12370799 | 32.80919 | 54.29568 | 56.91865 | 24.61716 | 0.312776 | 1.329219 | 0.498684 |
| 18 | 65269600 | 65272999 | 21.21781 | 35.61458 | 59.10484 | 25.80121 | -0.34724 | 1.143038 | 0.4987 |
| 12 | 62208400 | 62208999 | 9.453589 | 18.27432 | 20.33357 | 7.790944 | 0.404447 | -0.15599 | 0.498702 |
| 4 | 112793400 | 112794199 | 24.83244 | 42.56546 | 43.63256 | 19.13556 | 0.347781 | 0.95184 | 0.498711 |
| 5 | 9646600 | 9649599 | 42.91064 | 70.01637 | 66.16124 | 28.42777 | 0.454115 | 1.537324 | 0.498713 |
| 13 | 3261000 | 3261999 | 28.51055 | 47.76129 | 42.9289 | 18.71657 | 0.534516 | 1.171903 | 0.498726 |
| 15 | 49161600 | 49164199 | 25.71074 | 44.21025 | 38.28788 | 16.79467 | 0.590759 | 0.920514 | 0.498736 |
| X | 148639400 | 148645399 | 76.943 | 122.7593 | 97.08332 | 40.71033 | 0.67736 | 2.156305 | 0.498755 |
| 8 | 58177600 | 58178199 | 6.851784 | 13.22628 | 25.38773 | 9.79159 | -0.38372 | -0.14512 | 0.498766 |
| 10 | 52510000 | 52514199 | 74.1829 | 117.8108 | 103.0524 | 42.96087 | 0.525257 | 2.294714 | 0.498785 |
| 8 | 66599400 | 66600399 | 13.6475 | 25.66518 | 22.69544 | 9.312011 | 0.643915 | 0.110722 | 0.49879 |
| 15 | 100836200 | 100838799 | 17.61375 | 29.90829 | 54.24352 | 23.76827 | -0.47412 | 1.021812 | 0.498792 |
| 2 | 158549600 | 158551999 | 22.17399 | 37.10035 | 61.34947 | 26.72692 | -0.34211 | 1.181373 | 0.498792 |
| 17 | 47718000 | 47721799 | 28.8163 | 47.09394 | 77.99177 | 33.53052 | -0.35168 | 1.489085 | 0.498799 |
| 19 | 41360600 | 41365799 | 32.30755 | 52.42706 | 83.93517 | 35.97037 | -0.31079 | 1.612166 | 0.498799 |
| 12 | 75409200 | 75413999 | 43.51814 | 69.86751 | 110.3444 | 46.69713 | -0.30854 | 1.930988 | 0.4988 |

| | | | | | | | | | |
|---|---|---|---|---|---|---|---|---|---|
| X | 148800000 | 148804199 | 81.06765 | 127.5136 | 125.6856 | 52.1415 | 0.341837 | 2.532096 | 0.498814 |
| X | 25685000 | 25687999 | 64.27835 | 102.5709 | 100.1104 | 41.98977 | 0.374609 | 2.13885 | 0.498844 |
| 2 | 83703800 | 83705199 | 12.97661 | 23.03623 | 38.72753 | 17.10172 | -0.35213 | 0.483778 | 0.498852 |
| 4 | 152813800 | 152814799 | 8.137006 | 15.25648 | 28.52899 | 11.76509 | -0.43319 | 0.10034 | 0.498857 |
| 14 | 95247400 | 95249199 | 16.12158 | 29.00995 | 29.1632 | 12.7318 | 0.396779 | 0.400577 | 0.498866 |
| 1 | 139596200 | 139597799 | 45.37993 | 73.49623 | 65.8634 | 28.185 | 0.521716 | 1.665685 | 0.49889 |
| 1 | 84203400 | 84207199 | 30.56713 | 49.62795 | 81.12831 | 34.78048 | -0.34003 | 1.598969 | 0.498904 |
| 3 | 91556400 | 91558599 | 42.22462 | 68.50051 | 70.73815 | 30.30844 | 0.319899 | 1.630583 | 0.498912 |
| 2 | 81120200 | 81122999 | 38.39119 | 62.94252 | 56.46648 | 24.3068 | 0.532885 | 1.454601 | 0.498913 |
| 14 | 53675800 | 53679199 | 54.75404 | 87.59664 | 90.83979 | 38.23979 | 0.291261 | 2.044875 | 0.498924 |
| 7 | 13995600 | 13996799 | 22.14463 | 38.73937 | 36.43097 | 16.06814 | 0.475902 | 0.709214 | 0.498929 |
| 7 | 32886000 | 32887999 | 35.88831 | 59.075 | 51.47463 | 22.19554 | 0.577054 | 1.396276 | 0.498931 |
| 10 | 73904400 | 73905999 | 15.01363 | 27.36802 | 27.19407 | 11.68173 | 0.428249 | 0.313942 | 0.49894 |
| 7 | 96920000 | 96923199 | 38.59203 | 63.18731 | 63.6454 | 27.38159 | 0.365583 | 1.478217 | 0.49897 |
| 2 | 97702800 | 97704399 | 22.14272 | 38.76484 | 35.75624 | 15.78046 | 0.50352 | 0.6946 | 0.498988 |
| 14 | 41495400 | 41498199 | 22.14272 | 38.76484 | 35.75624 | 15.78046 | 0.50352 | 0.6946 | 0.498988 |
| 5 | 48119200 | 48120399 | 14.45823 | 26.26712 | 27.32154 | 11.78595 | 0.358967 | 0.330273 | 0.499001 |
| 7 | 60992000 | 60995399 | 32.39758 | 54.76232 | 47.4054 | 20.7434 | 0.588625 | 1.105407 | 0.499005 |
| 3 | 145966000 | 145966999 | 6.139925 | 11.80212 | 25.22692 | 9.851732 | -0.55458 | -0.11398 | 0.499013 |
| 7 | 14715000 | 14729399 | 276.0318 | 402.0179 | 341.7946 | 140.8609 | 0.460754 | 3.978582 | 0.499014 |
| 15 | 58113000 | 58114799 | 10.49525 | 19.13406 | 33.35604 | 14.32638 | -0.37725 | 0.294431 | 0.499015 |
| X | 24614000 | 24617199 | 40.83385 | 67.91375 | 51.61864 | 22.40198 | 0.775759 | 1.292341 | 0.499038 |
| 12 | 17214000 | 17216999 | 42.15645 | 68.16141 | 70.88813 | 30.29362 | 0.305152 | 1.698509 | 0.49905 |
| 4 | 3477200 | 3489999 | 246.2603 | 358.7054 | 348.2476 | 143.5183 | 0.269577 | 3.973531 | 0.499062 |
| 17 | 14711800 | 14715799 | 27.77812 | 45.23875 | 77.7938 | 33.40279 | -0.40934 | 1.538878 | 0.499063 |
| 8 | 95420600 | 95424999 | 37.25437 | 59.97255 | 96.17464 | 40.86132 | -0.32565 | 1.831699 | 0.499076 |
| 8 | 127510600 | 127513199 | 24.99394 | 41.4815 | 66.92188 | 29.0011 | -0.30693 | 1.284239 | 0.499079 |
| 6 | 42142000 | 42144799 | 33.58411 | 55.76073 | 55.90875 | 24.23522 | 0.37714 | 1.292745 | 0.499103 |
| 3 | 108635400 | 108637999 | 25.13084 | 43.10414 | 43.60453 | 19.12423 | 0.366886 | 0.946968 | 0.499109 |
| 13 | 61248000 | 61255199 | 135.0863 | 205.852 | 184.6266 | 76.53399 | 0.442622 | 3.145058 | 0.499145 |
| 17 | 39180600 | 39183399 | 55.15025 | 88.51353 | 88.22053 | 37.31579 | 0.353762 | 1.949066 | 0.499157 |
| X | 33493200 | 33497599 | 59.09784 | 95.62421 | 66.5234 | 28.44787 | 0.885013 | 1.711968 | 0.499171 |
| 3 | 152961000 | 152961799 | 6.264911 | 12.08508 | 24.77839 | 9.57869 | -0.48126 | -0.14711 | 0.499172 |
| 12 | 115034200 | 115037199 | 60.83733 | 97.58586 | 80.603 | 34.06441 | 0.623297 | 1.968467 | 0.499173 |
| 15 | 83468000 | 83471799 | 35.79234 | 57.68268 | 93.27567 | 39.69314 | -0.33563 | 1.793836 | 0.499174 |
| 6 | 59049800 | 59056999 | 177.7721 | 264.7405 | 263.428 | 108.5475 | 0.264146 | 3.604947 | 0.499177 |
| 2 | 22071200 | 22074599 | 44.03384 | 70.30455 | 114.4546 | 48.04345 | -0.36136 | 2.103463 | 0.499178 |
| 5 | 93643200 | 93646199 | 39.64984 | 65.21121 | 54.41608 | 23.45542 | 0.638773 | 1.417383 | 0.499183 |
| 10 | 24980000 | 24985399 | 62.53744 | 98.70111 | 148.2523 | 61.54201 | -0.26134 | 2.453652 | 0.499191 |
| 8 | 81705400 | 81705799 | 7.013128 | 13.50958 | 25.87667 | 10.04332 | -0.3892 | -0.12081 | 0.499224 |
| 7 | 19632600 | 19650999 | 250.4703 | 357.2168 | 498.5865 | 206.0761 | -0.28403 | 4.286592 | 0.49925 |
| 5 | 82006000 | 82010799 | 56.71823 | 91.05172 | 88.17409 | 37.30963 | 0.395593 | 1.940717 | 0.499251 |
| 11 | 14719800 | 14720399 | 5.398105 | 10.49443 | 24.14766 | 8.962191 | -0.59067 | -0.3015 | 0.499255 |
| 4 | 113109600 | 113118399 | 125.2818 | 192.1521 | 185.5227 | 77.01478 | 0.343684 | 3.013875 | 0.499256 |
| 16 | 58132800 | 58134599 | 21.76091 | 38.19777 | 34.63723 | 15.32096 | 0.52798 | 0.650168 | 0.499274 |
| 18 | 15282800 | 15285199 | 15.47591 | 26.69817 | 47.94123 | 21.03113 | -0.45568 | 0.861939 | 0.499274 |
| 2 | 33472800 | 33476799 | 31.38114 | 50.81768 | 84.42672 | 36.12545 | -0.36684 | 1.653327 | 0.499276 |

| | | | | | | | | | |
|---|---|---|---|---|---|---|---|---|---|
| 19 | 13048600 | 13056399 | 87.5574 | 138.1162 | 127.9768 | 53.11181 | 0.434028 | 2.477816 | 0.499277 |
| 9 | 6452800 | 6454199 | 17.64331 | 31.37455 | 23.84583 | 10.51483 | 0.78825 | 0.473154 | 0.499291 |
| 11 | 100119400 | 100125399 | 56.99623 | 90.34824 | 137.527 | 57.22611 | -0.2754 | 2.341633 | 0.499295 |
| 7 | 104559800 | 104560599 | 17.30654 | 30.51517 | 31.39123 | 13.89247 | 0.348576 | 0.56412 | 0.4993 |
| 3 | 57269600 | 57270999 | 18.32936 | 31.41645 | 51.65664 | 22.65449 | -0.33121 | 0.936245 | 0.499301 |
| 11 | 4136400 | 4137999 | 8.040895 | 14.87428 | 29.96369 | 12.62141 | -0.56376 | 0.18569 | 0.499304 |
| 8 | 14459000 | 14461199 | 27.50474 | 46.33282 | 48.55296 | 21.22348 | 0.313809 | 1.119554 | 0.499409 |
| X | 81150800 | 81153999 | 48.58184 | 78.78432 | 70.88168 | 30.36158 | 0.517817 | 1.650908 | 0.499424 |
| 12 | 79485000 | 79485799 | 7.330518 | 14.10779 | 27.59249 | 10.73803 | -0.42286 | -0.10383 | 0.499426 |
| 6 | 57753200 | 57755399 | 22.70782 | 39.16114 | 40.43338 | 17.73736 | 0.338951 | 0.883122 | 0.49943 |
| 8 | 34700800 | 34703199 | 26.14375 | 43.37532 | 71.63401 | 31.03599 | -0.3407 | 1.294744 | 0.499442 |
| 6 | 71196200 | 71198199 | 9.91711 | 17.94862 | 33.81946 | 14.65856 | -0.49724 | 0.333013 | 0.499445 |
| 4 | 153847800 | 153853199 | 43.49912 | 69.72235 | 107.5578 | 45.40429 | -0.2775 | 1.980575 | 0.499455 |
| 3 | 97590600 | 97593999 | 23.44938 | 39.18698 | 64.25059 | 27.9695 | -0.32993 | 1.200026 | 0.499465 |
| 7 | 30105200 | 30106399 | 12.25324 | 22.97708 | 24.00234 | 9.896259 | 0.400198 | 0.121334 | 0.499484 |
| 5 | 105061200 | 105065599 | 70.89923 | 112.6141 | 113.7668 | 47.43518 | 0.318013 | 2.289429 | 0.499491 |
| 13 | 61137600 | 61140399 | 28.38935 | 48.69644 | 42.93134 | 18.82909 | 0.564536 | 0.958279 | 0.4995 |
| 15 | 27512800 | 27518399 | 51.68611 | 81.83209 | 150.5845 | 62.60986 | -0.55017 | 2.370932 | 0.499503 |
| X | 125909200 | 125911399 | 20.98315 | 37.16462 | 31.29889 | 13.83915 | 0.638103 | 0.538186 | 0.499516 |
| 6 | 15572400 | 15575199 | 30.29473 | 50.43918 | 52.99417 | 23.01549 | 0.310144 | 1.249852 | 0.499517 |
| 7 | 118259800 | 118260999 | 8.736373 | 16.52801 | 30.98509 | 12.56869 | -0.41733 | 0.05202 | 0.499528 |
| 16 | 16226000 | 16229999 | 34.2553 | 57.29679 | 51.98638 | 22.64356 | 0.520939 | 1.212694 | 0.499551 |
| 14 | 118090400 | 118095199 | 58.02055 | 92.7697 | 96.45113 | 40.56923 | 0.286695 | 2.069152 | 0.499565 |
| 10 | 50096200 | 50100799 | 39.96965 | 65.71303 | 61.2212 | 26.38458 | 0.480133 | 1.428407 | 0.49957 |
| 18 | 51133800 | 51137999 | 52.40485 | 82.84975 | 151.5042 | 62.94599 | -0.5428 | 2.407274 | 0.499575 |
| 6 | 66524600 | 66526599 | 18.45337 | 31.67669 | 52.21371 | 22.90158 | -0.33451 | 0.922761 | 0.499575 |
| 6 | 62629800 | 62630799 | 13.67017 | 24.05247 | 40.36095 | 17.86966 | -0.35478 | 0.583753 | 0.499628 |
| 9 | 83065800 | 83066199 | 9.872211 | 19.02001 | 20.83649 | 8.083571 | 0.410376 | -0.10723 | 0.499631 |
| 7 | 85714200 | 85721199 | 91.73258 | 144.0433 | 137.7861 | 57.14881 | 0.383135 | 2.572497 | 0.499659 |
| 10 | 124068400 | 124069599 | 14.14984 | 26.38771 | 24.82359 | 10.33083 | 0.539444 | 0.173567 | 0.499667 |
| 5 | 97853000 | 97854999 | 17.56426 | 31.53585 | 29.49007 | 12.89863 | 0.498656 | 0.424293 | 0.499674 |
| 11 | 116594400 | 116600199 | 55.32 | 87.8151 | 134.5834 | 56.08047 | -0.28321 | 2.297939 | 0.499693 |
| 4 | 86439200 | 86439999 | 5.394357 | 10.46437 | 22.74032 | 8.570933 | -0.5285 | -0.24772 | 0.49972 |
| X | 125093400 | 125095399 | 24.67412 | 42.87448 | 39.23643 | 17.22813 | 0.514432 | 0.821554 | 0.499725 |
| 2 | 170648200 | 170649799 | 10.89762 | 19.49904 | 35.26639 | 15.46992 | -0.45008 | 0.413109 | 0.499735 |
| X | 153288200 | 153303399 | 229.3983 | 340.6114 | 270.7836 | 111.5447 | 0.583853 | 3.665727 | 0.499736 |
| 10 | 118542200 | 118543599 | 9.72514 | 17.802 | 31.87383 | 13.61913 | -0.41029 | 0.262715 | 0.499736 |
| 17 | 74198000 | 74198999 | 9.72514 | 17.802 | 31.87383 | 13.61913 | -0.41029 | 0.262715 | 0.499736 |
| 19 | 17903800 | 17905399 | 12.84932 | 22.57443 | 39.89581 | 17.65987 | -0.42977 | 0.595683 | 0.49974 |
| X | 4386600 | 4390399 | 56.9278 | 91.88185 | 75.33478 | 32.12041 | 0.644577 | 1.778521 | 0.499753 |
| 12 | 41914800 | 41916199 | 37.70138 | 62.25382 | 58.39038 | 25.21513 | 0.472554 | 1.367537 | 0.499761 |
| 5 | 13107400 | 13108599 | 13.0866 | 24.48239 | 21.38679 | 8.852891 | 0.652062 | 0.138039 | 0.49977 |
| 5 | 75064000 | 75065399 | 20.42217 | 35.59212 | 36.64539 | 16.12063 | 0.345346 | 0.752699 | 0.499781 |
| 8 | 36771800 | 36773799 | 15.19656 | 26.52479 | 44.67009 | 19.66938 | -0.361 | 0.718509 | 0.499797 |
| 9 | 111955400 | 111956599 | 7.825762 | 14.64837 | 29.03157 | 12.00634 | -0.52008 | 0.107527 | 0.499824 |
| 16 | 83035000 | 83036599 | 15.7714 | 29.6865 | 23.24042 | 9.520847 | 0.820779 | 0.12735 | 0.499834 |
| 4 | 52380000 | 52384799 | 62.83488 | 100.2533 | 101.9463 | 42.75128 | 0.31526 | 2.138886 | 0.499846 |

| | | | | | | | | | |
|---|---|---|---|---|---|---|---|---|---|
| 6 | 85142000 | 85144199 | 14.31938 | 24.8827 | 45.30656 | 19.8939 | -0.47357 | 0.780156 | 0.499861 |
| 1 | 182052400 | 182054599 | 20.65235 | 34.9304 | 57.02218 | 24.95608 | -0.3232 | 1.070204 | 0.499869 |
| 4 | 97227400 | 97230999 | 48.41703 | 77.14342 | 122.0659 | 51.09112 | -0.32367 | 2.174531 | 0.499873 |
| 4 | 8980800 | 8982399 | 15.8503 | 28.4288 | 29.26713 | 12.8124 | 0.360056 | 0.419769 | 0.499874 |
| 11 | 28241800 | 28244399 | 19.64137 | 34.82689 | 31.41282 | 13.88527 | 0.540278 | 0.518734 | 0.499877 |
| 3 | 158424800 | 158427399 | 19.14702 | 33.76549 | 32.54845 | 14.40416 | 0.441771 | 0.573708 | 0.499892 |
| X | 127625800 | 127628999 | 37.46076 | 61.37349 | 62.3856 | 26.84671 | 0.352812 | 1.467729 | 0.499899 |
| 11 | 55136400 | 55139199 | 20.71605 | 36.02598 | 37.33309 | 16.40075 | 0.335912 | 0.787546 | 0.499909 |
| X | 16152600 | 16154799 | 22.54875 | 39.92801 | 30.77841 | 13.60966 | 0.765167 | 0.551337 | 0.499925 |
| 4 | 75082600 | 75083199 | 9.086897 | 17.51196 | 17.6913 | 6.85781 | 0.527025 | -0.12983 | 0.49993 |
| 10 | 54803600 | 54807799 | 42.62442 | 68.40897 | 107.1833 | 45.33553 | -0.29766 | 1.937781 | 0.499934 |
| 16 | 19579000 | 19580799 | 13.32889 | 23.36642 | 40.98131 | 18.10727 | -0.41884 | 0.635525 | 0.499976 |
| 15 | 13051200 | 13051999 | 13.29714 | 24.33397 | 24.94432 | 10.66223 | 0.388533 | 0.274988 | 0.499978 |
| 5 | 85848400 | 85849399 | 12.98721 | 24.25996 | 25.01581 | 10.38148 | 0.411469 | 0.155176 | 0.499982 |
| 3 | 41479000 | 41480199 | 17.96668 | 31.83658 | 31.60787 | 13.97449 | 0.402166 | 0.514779 | 0.499986 |
| 14 | 12732800 | 12735999 | 19.19755 | 33.68671 | 34.34115 | 15.18562 | 0.359823 | 0.639892 | 0.49999 |
| 12 | 118568400 | 118569599 | 7.939949 | 15.09268 | 27.71987 | 11.132 | -0.37486 | 0.00391 | 0.499992 |
| 6 | 90071600 | 90073199 | 20.58131 | 35.81786 | 36.82185 | 16.1836 | 0.347521 | 0.776295 | 0.499994 |
| 15 | 73300200 | 73305199 | 34.77981 | 56.14877 | 91.50076 | 39.02557 | -0.34387 | 1.739548 | 0.500012 |
| 7 | 5278000 | 5279599 | 40.89099 | 66.23449 | 65.73346 | 28.13205 | 0.375053 | 1.65656 | 0.500022 |
| 17 | 35577000 | 35588599 | 169.4708 | 250.7719 | 354.8638 | 146.1431 | -0.25213 | 3.71752 | 0.500066 |
| 5 | 77564000 | 77565799 | 13.84721 | 24.43876 | 41.8794 | 18.53059 | -0.38326 | 0.553818 | 0.500068 |
| 10 | 11373600 | 11375799 | 16.20652 | 27.2327 | 63.65779 | 27.80513 | -0.8394 | 1.120181 | 0.500068 |
| X | 24319600 | 24324999 | 63.85201 | 102.8724 | 74.23644 | 31.57727 | 0.825388 | 1.839584 | 0.500078 |
| 3 | 13157400 | 13158999 | 17.72628 | 30.29862 | 51.62901 | 22.63731 | -0.38305 | 0.959655 | 0.500108 |
| 12 | 4525200 | 4528199 | 19.01956 | 31.89833 | 65.00472 | 28.36187 | -0.64247 | 1.14957 | 0.500116 |
| 2 | 150468000 | 150469399 | 8.948177 | 16.39728 | 32.93589 | 14.05248 | -0.57379 | 0.250742 | 0.500125 |
| 11 | 26542400 | 26543999 | 11.87536 | 20.9716 | 39.3078 | 17.39058 | -0.51158 | 0.527258 | 0.500143 |
| 8 | 52050000 | 52052199 | 34.59495 | 57.74033 | 53.1338 | 23.11189 | 0.500706 | 1.237169 | 0.500147 |
| 2 | 14346400 | 14349199 | 19.2643 | 32.89582 | 54.91832 | 24.07768 | -0.35406 | 0.976053 | 0.500148 |
| 14 | 93081800 | 93083799 | 16.24526 | 29.74273 | 27.15933 | 11.60176 | 0.554825 | 0.299541 | 0.50015 |
| X | 33430200 | 33435999 | 124.345 | 192.8028 | 151.9578 | 63.1148 | 0.648161 | 2.81 | 0.500153 |
| X | 24303200 | 24308599 | 65.94419 | 105.9468 | 81.77506 | 34.64341 | 0.723641 | 1.929509 | 0.500162 |
| 1 | 94657200 | 94659999 | 20.14125 | 35.36832 | 34.16167 | 15.11552 | 0.437201 | 0.632881 | 0.500176 |
| 5 | 90152600 | 90153599 | 9.50028 | 17.78108 | 32.91098 | 13.61289 | -0.42282 | 0.131865 | 0.500177 |
| 13 | 22339400 | 22342399 | 42.36198 | 69.18621 | 58.84804 | 25.29415 | 0.60638 | 1.516282 | 0.500186 |
| X | 26592600 | 26593999 | 28.4615 | 48.62472 | 44.66358 | 19.5825 | 0.504869 | 0.998516 | 0.500192 |
| 17 | 39216800 | 39225199 | 116.3156 | 181.5722 | 137.2464 | 56.94958 | 0.715902 | 2.690279 | 0.500192 |
| X | 11399200 | 11401599 | 34.76416 | 57.79008 | 55.31945 | 23.99987 | 0.443994 | 1.284539 | 0.500196 |
| 18 | 77174200 | 77179799 | 52.61674 | 85.27827 | 69.13197 | 29.59946 | 0.667002 | 1.667489 | 0.500201 |
| 11 | 55047400 | 55048999 | 12.37565 | 21.96226 | 37.25012 | 16.45381 | -0.36486 | 0.480638 | 0.50021 |
| 4 | 118439800 | 118443199 | 36.43999 | 58.81308 | 92.11356 | 39.27323 | -0.28719 | 1.751923 | 0.500212 |
| 2 | 40186000 | 40187199 | 11.99306 | 22.91415 | 22.71705 | 9.024603 | 0.520875 | -0.00723 | 0.500234 |
| 6 | 4174000 | 4174799 | 20.22829 | 35.21706 | 35.81954 | 15.74688 | 0.362913 | 0.767496 | 0.500244 |
| 2 | 15314000 | 15316799 | 28.15323 | 46.41328 | 77.96147 | 33.63823 | -0.36632 | 1.378485 | 0.500244 |
| 4 | 105159200 | 105160799 | 13.65553 | 25.7789 | 24.04134 | 9.796497 | 0.576558 | 0.09244 | 0.500248 |
| 18 | 85158400 | 85160599 | 22.14564 | 37.24239 | 60.32135 | 26.35421 | -0.31232 | 1.128294 | 0.500298 |

| | | | | | | | | | |
|---|---|---|---|---|---|---|---|---|---|
| 11 | 113843800 | 113869599 | 398.9673 | 547.8428 | 726.7689 | 302.2182 | -0.26587 | 4.859285 | 0.50036 |
| 8 | 98602400 | 98603599 | 12.81906 | 23.68699 | 24.97366 | 10.53525 | 0.3618 | 0.217526 | 0.500373 |
| 6 | 115512600 | 115516399 | 27.42524 | 44.75383 | 76.27615 | 32.77513 | -0.39444 | 1.50792 | 0.50038 |
| 8 | 75245600 | 75247199 | 10.77558 | 19.28372 | 34.81598 | 15.2703 | -0.44718 | 0.412301 | 0.50039 |
| 6 | 88036600 | 88037999 | 8.441616 | 15.78302 | 29.01237 | 12.02173 | -0.41438 | 0.121949 | 0.500394 |
| X | 13711800 | 13714799 | 51.35203 | 83.00309 | 73.46604 | 31.38242 | 0.536835 | 1.728434 | 0.500418 |
| 12 | 76061800 | 76063399 | 8.681702 | 15.89001 | 31.41176 | 13.42396 | -0.55211 | 0.254451 | 0.50042 |
| 8 | 4906000 | 4911799 | 70.16716 | 111.1607 | 116.7575 | 48.56044 | 0.258324 | 2.363062 | 0.500439 |
| 10 | 47773000 | 47774399 | 20.12703 | 35.67273 | 31.35668 | 13.86281 | 0.577001 | 0.5255 | 0.500461 |
| 6 | 58510600 | 58513999 | 37.81371 | 61.92302 | 63.72144 | 27.41618 | 0.334834 | 1.472618 | 0.50047 |
| 13 | 61550000 | 61571599 | 378.1658 | 532.2993 | 478.0549 | 198.0223 | 0.332788 | 4.484582 | 0.50047 |
| 14 | 52609400 | 52610599 | 10.30164 | 18.41726 | 35.30737 | 15.49881 | -0.53423 | 0.414388 | 0.500501 |
| 3 | 6964600 | 6964999 | 10.27332 | 19.56286 | 20.70059 | 8.281899 | 0.419152 | -0.00549 | 0.500504 |
| 5 | 48039400 | 48039799 | 8.631398 | 16.75544 | 18.87179 | 7.088312 | 0.414984 | -0.23352 | 0.500506 |
| 1 | 169948200 | 169949599 | 11.74903 | 22.00779 | 23.112 | 9.546628 | 0.390983 | 0.12363 | 0.500543 |
| 18 | 85193600 | 85195999 | 55.71684 | 89.08945 | 91.48984 | 38.48437 | 0.304518 | 2.064308 | 0.500547 |
| 7 | 47983000 | 47985799 | 41.63973 | 67.81906 | 60.22597 | 25.86037 | 0.54163 | 1.555968 | 0.500561 |
| X | 75316000 | 75323199 | 141.0148 | 213.549 | 210.2329 | 86.98447 | 0.300932 | 3.275888 | 0.500573 |
| 16 | 58948000 | 58951599 | 46.77494 | 75.41593 | 79.07385 | 33.66904 | 0.288773 | 1.791251 | 0.500592 |
| X | 54707000 | 54709599 | 49.47935 | 80.30024 | 69.23544 | 29.67289 | 0.580084 | 1.637823 | 0.500593 |
| 6 | 44117000 | 44119799 | 38.24815 | 61.62298 | 96.01909 | 40.84386 | -0.28277 | 1.808275 | 0.500594 |
| X | 87390200 | 87390999 | 21.73813 | 37.87436 | 36.27674 | 15.95548 | 0.449401 | 0.767241 | 0.500595 |
| 8 | 4136800 | 4145399 | 134.6088 | 205.0234 | 204.6115 | 84.80588 | 0.288081 | 3.15646 | 0.500603 |
| 2 | 105683200 | 105685399 | 21.57775 | 37.67143 | 36.97873 | 16.28503 | 0.41408 | 0.734772 | 0.500625 |
| 12 | 43615400 | 43617799 | 36.83865 | 60.09953 | 62.77586 | 26.97164 | 0.30974 | 1.522576 | 0.500627 |
| 18 | 49650600 | 49652399 | 18.80532 | 33.75863 | 29.53514 | 12.92012 | 0.594321 | 0.436143 | 0.50064 |
| 7 | 72533400 | 72536599 | 24.10837 | 40.23693 | 65.97198 | 28.69578 | -0.32994 | 1.217153 | 0.500641 |
| 6 | 5652600 | 5653399 | 16.26581 | 29.40312 | 23.33023 | 10.12755 | 0.741552 | 0.364575 | 0.500647 |
| 16 | 15139400 | 15144799 | 89.52146 | 137.7567 | 203.9307 | 84.69453 | -0.26873 | 2.938915 | 0.500647 |
| 4 | 99280800 | 99283199 | 16.30095 | 28.43066 | 49.50233 | 21.78666 | -0.4092 | 0.740674 | 0.500648 |
| 10 | 27235800 | 27236599 | 7.924419 | 14.59047 | 30.47324 | 12.91793 | -0.62191 | 0.21389 | 0.500649 |
| X | 24231600 | 24234399 | 43.38415 | 71.08747 | 61.70633 | 26.55589 | 0.58008 | 1.480613 | 0.500652 |
| 17 | 22278400 | 22280399 | 34.61319 | 57.08887 | 58.52559 | 25.257 | 0.344361 | 1.376279 | 0.500653 |
| 7 | 44716400 | 44717599 | 6.87901 | 13.18393 | 25.69937 | 10.12138 | -0.4339 | -0.07449 | 0.500665 |
| 5 | 150268800 | 150274799 | 67.81297 | 107.6952 | 112.4852 | 46.89347 | 0.269768 | 2.289854 | 0.500678 |
| 10 | 53255200 | 53256199 | 9.820342 | 17.67018 | 33.83297 | 14.7712 | -0.52679 | 0.368019 | 0.500688 |
| X | 40565200 | 40565799 | 12.26306 | 23.04248 | 24.11639 | 9.908957 | 0.401878 | 0.111025 | 0.500705 |
| 1 | 128857600 | 128859199 | 12.85514 | 22.76808 | 38.25236 | 16.91378 | -0.35272 | 0.502533 | 0.50073 |
| 8 | 44215800 | 44217199 | 16.35021 | 30.30821 | 25.11405 | 10.54867 | 0.711252 | 0.227983 | 0.500762 |
| 2 | 22495400 | 22496599 | 15.80979 | 29.10308 | 26.76734 | 11.35013 | 0.552196 | 0.266178 | 0.50078 |
| 6 | 143551200 | 143555999 | 36.76029 | 59.32564 | 92.79526 | 39.56011 | -0.28544 | 1.754689 | 0.500801 |
| 12 | 113729000 | 113732399 | 46.39176 | 75.31843 | 70.20545 | 30.09718 | 0.468365 | 1.623566 | 0.500803 |
| 1 | 114157600 | 114159599 | 34.07677 | 56.55216 | 50.36717 | 21.82599 | 0.54751 | 1.295839 | 0.500804 |
| 3 | 3537000 | 3540399 | 31.44602 | 53.86328 | 41.7469 | 18.30752 | 0.749401 | 0.982268 | 0.500818 |
| 14 | 120967400 | 120971599 | 30.54267 | 49.54542 | 82.96083 | 35.54346 | -0.37579 | 1.61529 | 0.500821 |
| 6 | 143831600 | 143833799 | 30.06352 | 50.11243 | 52.40649 | 22.77695 | 0.31686 | 1.23803 | 0.500834 |
| 1 | 106601400 | 106605399 | 40.46884 | 66.21365 | 64.50945 | 27.74579 | 0.412863 | 1.490731 | 0.500846 |

| | | | | | | | | | |
|---|---|---|---|---|---|---|---|---|---|
| X | 120493400 | 120495599 | 33.87762 | 56.1989 | 54.08917 | 23.43232 | 0.436036 | 1.306576 | 0.500909 |
| 3 | 11536200 | 11537199 | 17.09807 | 30.4383 | 30.61749 | 13.48833 | 0.387029 | 0.475853 | 0.500922 |
| 18 | 66810800 | 66812199 | 12.15941 | 21.6658 | 36.72053 | 16.16478 | -0.36056 | 0.451146 | 0.500933 |
| 9 | 115393400 | 115395599 | 19.46013 | 33.966 | 34.65102 | 15.25756 | 0.358787 | 0.724374 | 0.500938 |
| 4 | 61835200 | 61843199 | 181.9673 | 271.7886 | 239.3937 | 98.69204 | 0.443832 | 3.550483 | 0.500944 |
| 14 | 96143400 | 96143999 | 5.949489 | 11.57922 | 23.67667 | 8.70898 | -0.4092 | -0.32351 | 0.50099 |
| 11 | 61714400 | 61716399 | 19.07299 | 33.46944 | 34.38182 | 15.2041 | 0.348876 | 0.639166 | 0.501034 |
| 1 | 176211000 | 176212999 | 23.04807 | 40.35329 | 35.71477 | 15.7633 | 0.562908 | 0.698733 | 0.501058 |
| 6 | 144952200 | 144954799 | 17.25803 | 29.75057 | 53.89159 | 23.64093 | -0.46898 | 0.878926 | 0.501065 |
| 9 | 5253600 | 5254199 | 9.057887 | 17.4978 | 19.30805 | 7.42097 | 0.41278 | -0.15114 | 0.501075 |
| 4 | 127781800 | 127783599 | 21.06595 | 36.58561 | 37.79701 | 16.59063 | 0.340012 | 0.806744 | 0.501088 |
| 7 | 3485600 | 3489799 | 24.33452 | 40.28098 | 68.55406 | 29.65451 | -0.38417 | 1.312234 | 0.501093 |
| 2 | 99385600 | 99388199 | 39.31443 | 63.78417 | 63.66676 | 27.28015 | 0.368901 | 1.618175 | 0.501095 |
| 2 | 151289000 | 151289999 | 16.52331 | 29.34238 | 30.291 | 13.37238 | 0.347896 | 0.487992 | 0.501116 |
| 8 | 5989800 | 5996399 | 85.13883 | 134.2627 | 129.8771 | 53.89844 | 0.371708 | 2.482596 | 0.501121 |
| 13 | 117096800 | 117098399 | 10.55735 | 19.0077 | 34.27087 | 14.95169 | -0.44019 | 0.369743 | 0.501132 |
| 13 | 62401000 | 62401999 | 40.8991 | 66.29414 | 61.869 | 26.49187 | 0.464499 | 1.640966 | 0.50115 |
| 3 | 122700800 | 122704799 | 33.56183 | 56.25756 | 51.36574 | 22.40356 | 0.511871 | 1.189315 | 0.501151 |
| 2 | 94292200 | 94293399 | 20.87647 | 34.64682 | 64.17086 | 27.80834 | -0.50524 | 1.272525 | 0.501163 |
| 1 | 29695000 | 29699999 | 103.2052 | 159.1689 | 161.048 | 66.89649 | 0.282212 | 2.89695 | 0.501171 |
| 12 | 63625400 | 63627399 | 16.63406 | 30.36052 | 27.65558 | 11.86107 | 0.554566 | 0.321167 | 0.501185 |
| 10 | 111541400 | 111543999 | 38.66797 | 63.34528 | 63.79412 | 27.45173 | 0.366086 | 1.470611 | 0.50119 |
| 18 | 75848400 | 75849599 | 6.44726 | 12.32214 | 25.63126 | 10.17549 | -0.53764 | -0.05368 | 0.501195 |
| 4 | 144407800 | 144409799 | 24.60571 | 42.83036 | 37.37285 | 16.42789 | 0.582778 | 0.790511 | 0.50121 |
| 4 | 100788400 | 100791599 | 40.97 | 66.86776 | 59.17566 | 25.42873 | 0.54871 | 1.524261 | 0.501239 |
| 12 | 73469800 | 73471199 | 16.07584 | 28.09444 | 48.38025 | 21.32031 | -0.39289 | 0.71204 | 0.501249 |
| 2 | 100753200 | 100754199 | 10.58733 | 19.09718 | 33.81454 | 14.71832 | -0.41179 | 0.360211 | 0.501249 |
| 6 | 50572600 | 50574199 | 10.58733 | 19.09718 | 33.81454 | 14.71832 | -0.41179 | 0.360211 | 0.501249 |
| 10 | 26023400 | 26026599 | 23.36107 | 40.6084 | 35.03584 | 15.38741 | 0.598944 | 0.80171 | 0.50126 |
| 10 | 24664800 | 24665999 | 9.227895 | 17.02742 | 30.92712 | 13.0716 | -0.41943 | 0.210919 | 0.501277 |
| 7 | 11410200 | 11419199 | 111.1145 | 172.6283 | 140.1642 | 58.19923 | 0.607462 | 2.767118 | 0.501297 |
| 6 | 31555000 | 31556999 | 22.25247 | 36.9671 | 64.47297 | 27.9584 | -0.41883 | 1.262678 | 0.501316 |
| 17 | 73509800 | 73511399 | 13.68806 | 25.35798 | 25.96526 | 10.91523 | 0.407344 | 0.213091 | 0.501329 |
| 10 | 95200400 | 95201599 | 14.97036 | 27.67811 | 25.72114 | 10.84207 | 0.5432 | 0.231898 | 0.501334 |
| 5 | 137665800 | 137666399 | 7.409619 | 13.85009 | 28.51373 | 11.82008 | -0.57676 | 0.113497 | 0.501395 |
| 11 | 30835200 | 30835799 | 7.409619 | 13.85009 | 28.51373 | 11.82008 | -0.57676 | 0.113497 | 0.501395 |
| 2 | 72786000 | 72787999 | 16.35154 | 29.4687 | 29.19568 | 12.72181 | 0.419532 | 0.394322 | 0.501396 |
| 10 | 111540600 | 111543199 | 41.4734 | 67.74852 | 65.65135 | 28.22042 | 0.41898 | 1.516303 | 0.501396 |
| 9 | 43821200 | 43823199 | 16.57131 | 29.68296 | 29.74969 | 13.03809 | 0.397512 | 0.434464 | 0.501406 |
| 13 | 5503200 | 5512399 | 175.7794 | 263.7415 | 251.1473 | 103.6341 | 0.337315 | 3.466213 | 0.501407 |
| 2 | 92304600 | 92308599 | 41.62701 | 67.31359 | 68.02602 | 29.0751 | 0.346654 | 1.692167 | 0.501412 |
| X | 114773400 | 114776399 | 30.2382 | 52.0774 | 40.6598 | 17.83595 | 0.740244 | 0.929817 | 0.501419 |
| 10 | 36050400 | 36053799 | 31.55316 | 52.58197 | 53.61433 | 23.29813 | 0.353187 | 1.24369 | 0.501427 |
| 2 | 78344800 | 78345399 | 12.04321 | 22.53894 | 24.07703 | 9.960104 | 0.364655 | 0.133519 | 0.501436 |
| 8 | 56393200 | 56393999 | 6.847645 | 13.23277 | 25.46651 | 9.771852 | -0.38048 | -0.16265 | 0.501439 |
| 2 | 164345800 | 164349399 | 39.21665 | 63.11333 | 98.25759 | 41.72893 | -0.28355 | 1.844832 | 0.501448 |
| 7 | 33781600 | 33823599 | 561.9582 | 750.3108 | 972.334 | 406.0607 | -0.27286 | 5.303651 | 0.501464 |

| | | | | | | | | | |
|---|---|---|---|---|---|---|---|---|---|
| 6 | 47879400 | 47885799 | 64.53158 | 100.2584 | 192.1247 | 79.77417 | -0.62997 | 2.75308 | 0.501478 |
| 12 | 119561600 | 119564599 | 34.0455 | 57.53336 | 46.27594 | 20.24763 | 0.694238 | 1.112064 | 0.501503 |
| 12 | 36371400 | 36373399 | 20.2904 | 35.53853 | 35.34753 | 15.6062 | 0.395201 | 0.676259 | 0.501504 |
| 8 | 111163600 | 111164399 | 6.864578 | 13.26324 | 26.01781 | 9.991317 | -0.40863 | -0.15676 | 0.501508 |
| 10 | 109456000 | 109456799 | 12.50051 | 23.42451 | 24.55523 | 10.13611 | 0.394039 | 0.130164 | 0.501532 |
| X | 153726000 | 153728799 | 35.98583 | 60.26596 | 50.84703 | 22.16461 | 0.625506 | 1.206883 | 0.501544 |
| 13 | 57932400 | 57937399 | 39.93376 | 63.97809 | 105.6381 | 44.56263 | -0.37612 | 1.989981 | 0.501545 |
| 13 | 56675800 | 56676399 | 3.757697 | 7.338886 | 18.7067 | 6.563192 | -0.65761 | -0.53433 | 0.501551 |
| X | 124964000 | 124965799 | 19.23218 | 34.43687 | 29.95456 | 13.13184 | 0.60045 | 0.454569 | 0.501556 |
| 9 | 18986400 | 18991799 | 70.62961 | 112.5698 | 103.9597 | 43.53133 | 0.452362 | 2.181247 | 0.501567 |
| 19 | 3124800 | 3126399 | 31.36297 | 52.68415 | 50.75937 | 22.16894 | 0.434494 | 1.159049 | 0.501571 |
| 11 | 6725800 | 6727799 | 18.75482 | 32.2027 | 53.86468 | 23.62622 | -0.35559 | 0.92279 | 0.501576 |
| 2 | 149052600 | 149054799 | 26.01581 | 45.20444 | 39.08182 | 17.15991 | 0.596013 | 0.826279 | 0.501581 |
| 6 | 37475800 | 37477399 | 26.01581 | 45.20444 | 39.08182 | 17.15991 | 0.596013 | 0.826279 | 0.501581 |
| 6 | 105969000 | 105969999 | 12.47814 | 23.20016 | 23.35059 | 9.767533 | 0.43671 | 0.174434 | 0.501588 |
| 11 | 92018200 | 92024199 | 64.64761 | 101.8943 | 153.0786 | 63.52258 | -0.26316 | 2.483509 | 0.501605 |
| 13 | 33361000 | 33361599 | 12.23568 | 21.71616 | 37.2754 | 16.46352 | -0.38182 | 0.479828 | 0.50161 |
| 5 | 144366800 | 144368999 | 15.39983 | 26.84126 | 44.70974 | 19.6692 | -0.34535 | 0.741274 | 0.501611 |
| 8 | 60475600 | 60476799 | 15.90029 | 27.69985 | 45.76394 | 20.12672 | -0.33369 | 0.752777 | 0.501619 |
| 1 | 110060400 | 110062399 | 23.89391 | 41.33992 | 40.85431 | 17.92378 | 0.402799 | 0.860765 | 0.501625 |
| 5 | 147110200 | 147113799 | 29.85756 | 48.66346 | 78.19587 | 33.58504 | -0.31099 | 1.532545 | 0.501626 |
| 3 | 9856800 | 9857199 | 9.138561 | 17.6035 | 18.73412 | 7.271558 | 0.450965 | -0.12108 | 0.501658 |
| 12 | 63787400 | 63789199 | 27.81313 | 46.73359 | 49.10503 | 21.44829 | 0.309797 | 1.142467 | 0.50169 |
| 9 | 9330600 | 9334199 | 46.02289 | 73.64768 | 114.408 | 48.17891 | -0.29028 | 2.037243 | 0.501697 |
| 5 | 45503000 | 45504799 | 15.59728 | 27.27469 | 45.69176 | 20.14588 | -0.35319 | 0.696523 | 0.501705 |
| 1 | 182942600 | 182943999 | 7.213332 | 13.64197 | 27.06931 | 10.98824 | -0.49883 | 0.030987 | 0.50171 |
| 1 | 190606600 | 190606999 | 5.961491 | 11.55828 | 23.30129 | 8.804639 | -0.42505 | -0.23047 | 0.501734 |
| 13 | 69817400 | 69822599 | 55.20023 | 87.6587 | 132.522 | 55.24279 | -0.26299 | 2.287126 | 0.50174 |
| 12 | 44841800 | 44843399 | 19.70379 | 32.88304 | 67.70347 | 29.44743 | -0.65723 | 1.207477 | 0.501744 |
| 7 | 33473400 | 33482999 | 176.2248 | 265.3262 | 236.8451 | 97.82295 | 0.434896 | 3.399405 | 0.501748 |
| 13 | 62923200 | 62932199 | 191.8098 | 286.3913 | 265.85 | 109.5915 | 0.367719 | 3.560657 | 0.501748 |
| 3 | 136325800 | 136326599 | 6.108949 | 11.82873 | 23.82122 | 9.056123 | -0.43233 | -0.20772 | 0.501752 |
| 1 | 185889000 | 185890799 | 20.79785 | 36.84216 | 31.30031 | 13.83935 | 0.625596 | 0.535219 | 0.501757 |
| 6 | 18634600 | 18636599 | 20.79785 | 36.84216 | 31.30031 | 13.83935 | 0.625596 | 0.535219 | 0.501757 |
| 13 | 99953400 | 99954799 | 16.84653 | 29.8276 | 30.81074 | 13.62426 | 0.344932 | 0.515537 | 0.50176 |
| 18 | 38490400 | 38491799 | 21.21589 | 36.81318 | 37.91761 | 16.63685 | 0.344338 | 0.820976 | 0.501771 |
| 4 | 4323000 | 4324999 | 24.16688 | 41.73562 | 36.72278 | 16.11028 | 0.569145 | 0.873036 | 0.501772 |
| 15 | 15076000 | 15079199 | 30.76942 | 50.30171 | 83.66984 | 35.97578 | -0.35773 | 1.492609 | 0.501777 |
| 10 | 108286800 | 108291599 | 36.73197 | 59.29018 | 93.62375 | 39.92219 | -0.29883 | 1.749075 | 0.501798 |
| 4 | 94586600 | 94587599 | 9.673059 | 17.95237 | 31.54154 | 13.22679 | -0.36372 | 0.183445 | 0.501813 |
| 10 | 61662600 | 61665999 | 19.16859 | 32.55545 | 54.83923 | 24.03087 | -0.36802 | 1.025966 | 0.501835 |
| 15 | 96538800 | 96541799 | 15.56188 | 26.56008 | 52.10807 | 22.84466 | -0.58576 | 0.96478 | 0.501843 |
| X | 7994000 | 7996799 | 39.96241 | 66.23689 | 54.33361 | 23.5245 | 0.66586 | 1.333504 | 0.501847 |
| 16 | 78353200 | 78362599 | 110.2404 | 168.0268 | 246.3687 | 102.1344 | -0.26566 | 3.139035 | 0.501894 |
| 19 | 39649800 | 39653199 | 36.11565 | 60.22081 | 53.18094 | 23.11844 | 0.559967 | 1.252022 | 0.501907 |
| 17 | 34532800 | 34535399 | 14.59797 | 25.37779 | 45.80636 | 20.11883 | -0.46111 | 0.774507 | 0.501911 |
| 6 | 137762400 | 137765399 | 18.63541 | 31.76491 | 54.88199 | 24.05828 | -0.40378 | 0.989571 | 0.501928 |

| | | | | | | | | | |
|---|---|---|---|---|---|---|---|---|---|
| 8 | 28050600 | 28051999 | 19.49154 | 33.17267 | 54.31923 | 23.80861 | -0.3269 | 1.008485 | 0.501936 |
| 5 | 49073800 | 49075599 | 14.75962 | 27.09124 | 26.90534 | 11.45373 | 0.438271 | 0.275592 | 0.501938 |
| 13 | 72287400 | 72289999 | 15.3719 | 26.64313 | 48.46352 | 21.26328 | -0.47306 | 0.816303 | 0.50194 |
| 15 | 37187400 | 37190199 | 31.2984 | 51.98612 | 54.66924 | 23.70434 | 0.308627 | 1.281152 | 0.501946 |
| 11 | 31424200 | 31429799 | 40.53119 | 65.11921 | 101.4001 | 42.95862 | -0.28686 | 1.900656 | 0.501957 |
| 10 | 47128600 | 47130599 | 19.22345 | 35.02603 | 26.88381 | 11.55504 | 0.798205 | 0.348791 | 0.501965 |
| 12 | 97810600 | 97812999 | 24.50223 | 40.57811 | 67.4103 | 29.16944 | -0.34932 | 1.307004 | 0.501966 |
| 17 | 22234000 | 22235199 | 21.80351 | 38.02746 | 37.66249 | 16.57571 | 0.401345 | 0.753543 | 0.501991 |
| 11 | 39196000 | 39196399 | 8.126518 | 15.80083 | 18.37074 | 6.810788 | 0.386417 | -0.28793 | 0.501994 |
| 3 | 112024000 | 112025599 | 15.52189 | 27.96098 | 28.77761 | 12.54667 | 0.36439 | 0.390239 | 0.501995 |
| 1 | 167413800 | 167415599 | 12.19916 | 21.56254 | 37.78647 | 16.71455 | -0.41438 | 0.518463 | 0.502005 |
| 3 | 16395400 | 16396399 | 21.09592 | 36.55357 | 37.38651 | 16.40312 | 0.353936 | 0.830598 | 0.502009 |
| 7 | 31997000 | 31998199 | 54.82257 | 87.67964 | 84.3206 | 35.48021 | 0.399417 | 2.053523 | 0.502016 |
| 16 | 71858800 | 71860599 | 16.16549 | 28.92478 | 29.73576 | 13.04386 | 0.36007 | 0.43738 | 0.502023 |
| 4 | 127568600 | 127571599 | 20.52262 | 34.52256 | 58.60257 | 25.60544 | -0.37973 | 1.118025 | 0.502029 |
| 11 | 56349800 | 56350999 | 22.77445 | 37.90983 | 63.88679 | 27.74227 | -0.36917 | 1.243674 | 0.502051 |
| 5 | 31788400 | 31789799 | 16.49683 | 28.29241 | 50.55281 | 22.17146 | -0.45023 | 0.924951 | 0.502082 |
| X | 28042200 | 28043799 | 19.78396 | 36.19074 | 25.81248 | 11.0392 | 0.908054 | 0.329153 | 0.502095 |
| 12 | 38779200 | 38781199 | 20.93118 | 36.37472 | 37.79003 | 16.59421 | 0.332075 | 0.797449 | 0.502102 |
| 2 | 72463800 | 72465599 | 37.6414 | 60.54326 | 97.27204 | 41.27586 | -0.32987 | 1.86178 | 0.502108 |
| X | 91656200 | 91659199 | 36.37416 | 60.34105 | 55.32431 | 23.96759 | 0.505834 | 1.313987 | 0.502128 |
| 6 | 69493200 | 69493799 | 5.800964 | 11.29818 | 23.17683 | 8.467826 | -0.40498 | -0.35147 | 0.502129 |
| 14 | 4309000 | 4310799 | 16.91402 | 29.31263 | 48.99352 | 21.49575 | -0.35139 | 0.829183 | 0.502129 |
| 1 | 53533000 | 53533799 | 7.988779 | 14.91709 | 29.00002 | 12.04291 | -0.49718 | 0.125354 | 0.502129 |
| X | 30923600 | 30924799 | 14.82382 | 28.45039 | 21.00959 | 8.24111 | 0.959598 | -0.02282 | 0.502156 |
| 5 | 96440400 | 96449399 | 179.5569 | 263.9832 | 372.4778 | 153.3955 | -0.25672 | 3.824686 | 0.502178 |
| 2 | 23762200 | 23763399 | 10.32058 | 18.94622 | 32.94229 | 14.02085 | -0.36441 | 0.252872 | 0.502204 |
| 3 | 100437600 | 100437999 | 4.675668 | 9.123035 | 20.39196 | 7.272633 | -0.49527 | -0.46268 | 0.502221 |
| 13 | 7693800 | 7698999 | 51.0246 | 82.29289 | 79.44545 | 33.84063 | 0.408149 | 1.78878 | 0.50224 |
| X | 117047000 | 117050599 | 50.85675 | 82.21525 | 74.93687 | 32.0171 | 0.494791 | 1.721679 | 0.50226 |
| 19 | 52909000 | 52909799 | 10.55107 | 20.34725 | 21.33944 | 8.256127 | 0.475806 | -0.10948 | 0.502271 |
| X | 111493000 | 111493399 | 9.629157 | 18.61372 | 19.32819 | 7.405681 | 0.504377 | -0.15346 | 0.502278 |
| 18 | 75033000 | 75035799 | 19.03738 | 32.25526 | 55.35335 | 24.23855 | -0.39487 | 1.047057 | 0.50229 |
| 12 | 103902000 | 103940599 | 450.1507 | 612.4239 | 812.1382 | 338.2077 | -0.27883 | 5.004806 | 0.502309 |
| 4 | 15513400 | 15514799 | 10.45171 | 19.03773 | 34.44047 | 14.80863 | -0.43094 | 0.305396 | 0.502311 |
| 7 | 60698000 | 60701199 | 31.78242 | 53.81984 | 47.40496 | 20.7546 | 0.563852 | 1.090374 | 0.502312 |
| 6 | 71262800 | 71265199 | 26.15681 | 45.33435 | 40.21306 | 17.64339 | 0.558608 | 0.855927 | 0.502317 |
| 10 | 95185800 | 95189799 | 66.07558 | 103.9409 | 155.7642 | 64.62054 | -0.26174 | 2.525115 | 0.502327 |
| 14 | 110199200 | 110202399 | 34.4884 | 56.62216 | 59.57913 | 25.6617 | 0.304235 | 1.430554 | 0.502329 |
| 2 | 99739800 | 99740199 | 8.376987 | 16.24926 | 18.27497 | 6.889992 | 0.411336 | -0.2282 | 0.502345 |
| 11 | 20701600 | 20703799 | 27.88576 | 47.83787 | 43.53316 | 19.09321 | 0.519074 | 0.957609 | 0.502349 |
| 12 | 101522400 | 101522799 | 10.13607 | 19.38381 | 21.24579 | 8.420072 | 0.3808 | -0.03564 | 0.502352 |
| 5 | 137855600 | 137862599 | 93.50958 | 143.7491 | 209.8231 | 87.12971 | -0.24953 | 2.960585 | 0.502356 |
| 5 | 13129800 | 13131799 | 19.15233 | 32.7652 | 53.27327 | 23.35993 | -0.31555 | 0.958821 | 0.50236 |
| 8 | 19009200 | 19011599 | 24.13957 | 41.93449 | 39.72094 | 17.43796 | 0.46479 | 0.821194 | 0.502368 |
| 5 | 105220400 | 105226399 | 82.7955 | 131.248 | 116.7558 | 48.58446 | 0.498482 | 2.358253 | 0.502401 |
| 2 | 44816600 | 44817799 | 10.967 | 19.44837 | 36.31757 | 16.05165 | -0.50337 | 0.473355 | 0.502404 |

| | | | | | | | | | |
|---|---|---|---|---|---|---|---|---|---|
| 4 | 129858400 | 129860399 | 10.967 | 19.44837 | 36.31757 | 16.05165 | -0.50337 | 0.473355 | 0.502404 |
| 5 | 93610800 | 93615199 | 68.66561 | 109.2517 | 109.6534 | 45.8064 | 0.329824 | 2.235307 | 0.502405 |
| 7 | 104852600 | 104853599 | 9.114287 | 16.70582 | 31.39381 | 13.39001 | -0.47765 | 0.249261 | 0.502408 |
| 3 | 152502200 | 152509399 | 66.37445 | 104.3787 | 156.0365 | 64.7309 | -0.25855 | 2.530816 | 0.50241 |
| 6 | 24673400 | 24673999 | 8.405802 | 15.38625 | 31.46684 | 13.44606 | -0.6001 | 0.254968 | 0.502427 |
| 6 | 66481200 | 66482999 | 17.86257 | 31.82193 | 30.47787 | 13.41876 | 0.457882 | 0.475557 | 0.502437 |
| 12 | 75905800 | 75906999 | 11.51675 | 20.65762 | 35.24614 | 15.42898 | -0.36441 | 0.40257 | 0.502438 |
| 13 | 50498000 | 50500399 | 26.55119 | 43.7805 | 71.20444 | 30.72523 | -0.31966 | 1.369027 | 0.502445 |
| 18 | 21558000 | 21560199 | 39.26157 | 64.14031 | 57.46023 | 24.70013 | 0.532052 | 1.505234 | 0.502451 |
| 8 | 129466000 | 129472999 | 107.9372 | 167.6088 | 157.7035 | 65.4868 | 0.394487 | 2.777102 | 0.502452 |
| 14 | 48316800 | 48317799 | 6.718544 | 12.94829 | 25.30377 | 9.807523 | -0.41566 | -0.12908 | 0.502456 |
| 6 | 145974200 | 145976399 | 22.06361 | 38.72365 | 34.59667 | 15.30119 | 0.549274 | 0.653675 | 0.502463 |
| 9 | 80996800 | 80997799 | 7.769883 | 14.74325 | 27.67689 | 11.1565 | -0.41118 | 0.012921 | 0.502464 |
| 2 | 76641200 | 76641999 | 10.00705 | 18.34584 | 32.39369 | 13.81273 | -0.38838 | 0.256601 | 0.502467 |
| 7 | 48399200 | 48402199 | 15.79187 | 27.1117 | 50.56752 | 22.17978 | -0.51188 | 0.908374 | 0.502483 |
| X | 32485600 | 32488399 | 37.99802 | 63.737 | 48.59907 | 21.20595 | 0.770983 | 1.194903 | 0.502489 |
| X | 26107000 | 26113399 | 96.08767 | 151.2858 | 129.27 | 53.63528 | 0.548705 | 2.522285 | 0.502489 |
| 10 | 116873000 | 116874599 | 17.13974 | 30.36593 | 31.20504 | 13.79687 | 0.35262 | 0.510694 | 0.502498 |
| 12 | 16833200 | 16834799 | 17.13974 | 30.36593 | 31.20504 | 13.79687 | 0.35262 | 0.510694 | 0.502498 |
| 10 | 83576200 | 83580799 | 82.45569 | 129.3073 | 134.653 | 55.84714 | 0.259317 | 2.590596 | 0.502499 |
| 4 | 124396000 | 124400999 | 31.9567 | 51.51497 | 100.2312 | 42.66778 | -0.60147 | 1.783508 | 0.502518 |
| 2 | 166340000 | 166343599 | 56.82175 | 90.09331 | 137.1198 | 57.07041 | -0.27486 | 2.334238 | 0.50252 |
| 15 | 47212800 | 47214199 | 14.23164 | 26.92262 | 22.24153 | 9.023181 | 0.755135 | 0.077535 | 0.502538 |
| 17 | 5535000 | 5536199 | 14.04052 | 26.05628 | 25.96323 | 10.88854 | 0.448863 | 0.20658 | 0.502573 |
| 3 | 91082800 | 91086199 | 49.91471 | 80.77049 | 73.38829 | 31.38505 | 0.500719 | 1.695189 | 0.502579 |
| 14 | 42649200 | 42652399 | 22.73004 | 39.32716 | 40.45444 | 17.74836 | 0.345352 | 0.856862 | 0.502591 |
| 17 | 86667600 | 86668799 | 11.76079 | 20.83561 | 36.81265 | 16.2763 | -0.42455 | 0.490456 | 0.502599 |
| 6 | 60873400 | 60875599 | 35.81614 | 58.92876 | 60.90628 | 26.25724 | 0.331197 | 1.408598 | 0.502607 |
| 13 | 65924200 | 65936399 | 283.2343 | 405.7722 | 378.3776 | 156.2775 | 0.304152 | 4.221315 | 0.502611 |
| 13 | 13146600 | 13149999 | 32.79081 | 54.85737 | 52.50103 | 22.87039 | 0.444208 | 1.206969 | 0.502612 |
| 11 | 86091600 | 86100399 | 107.6412 | 162.584 | 283.552 | 117.2239 | -0.52666 | 3.324156 | 0.502618 |
| 2 | 47646200 | 47648199 | 15.47103 | 26.70362 | 53.2497 | 23.36025 | -0.6063 | 0.862612 | 0.502631 |
| X | 4357400 | 4361999 | 80.94557 | 128.421 | 101.2041 | 42.14774 | 0.67416 | 2.33028 | 0.502638 |
| 13 | 74514800 | 74517799 | 29.96817 | 50.0495 | 52.43408 | 22.81657 | 0.314187 | 1.215914 | 0.502649 |
| 1 | 150868600 | 150869599 | 6.138049 | 11.79916 | 24.67336 | 9.634038 | -0.52245 | -0.11413 | 0.50267 |
| 11 | 23017400 | 23018199 | 6.138049 | 11.79916 | 24.67336 | 9.634038 | -0.52245 | -0.11413 | 0.50267 |
| 5 | 21886200 | 21892199 | 79.49426 | 123.7472 | 183.0216 | 75.97216 | -0.25455 | 2.724394 | 0.502676 |
| 19 | 59985000 | 59987999 | 32.46862 | 54.06005 | 54.72779 | 23.76881 | 0.363558 | 1.257797 | 0.50268 |
| 9 | 26317800 | 26322599 | 49.1631 | 79.09468 | 82.07503 | 34.83979 | 0.299938 | 1.863694 | 0.502693 |
| 12 | 8171800 | 8174399 | 22.17663 | 37.26228 | 60.28007 | 26.32927 | -0.31058 | 1.136679 | 0.502694 |
| 2 | 64893600 | 64895799 | 21.80863 | 36.58968 | 60.85625 | 26.55816 | -0.35051 | 1.149371 | 0.502712 |
| 18 | 22495800 | 22496799 | 8.963793 | 16.39 | 32.90987 | 14.08304 | -0.57633 | 0.263562 | 0.502712 |
| 5 | 74192800 | 74194799 | 20.34928 | 34.34726 | 56.98437 | 24.92402 | -0.34654 | 1.087008 | 0.502724 |
| 18 | 25351600 | 25353599 | 19.11476 | 34.10769 | 30.40529 | 13.36861 | 0.562263 | 0.474721 | 0.502737 |
| 4 | 81928600 | 81929199 | 6.75164 | 12.86486 | 26.12017 | 10.43623 | -0.51106 | -0.02711 | 0.502744 |
| 10 | 116585000 | 116585599 | 10.7623 | 20.38079 | 21.67106 | 8.772344 | 0.3974 | 0.0397 | 0.502748 |
| 4 | 10000600 | 10004799 | 74.26559 | 117.2733 | 122.0962 | 50.69771 | 0.267328 | 2.444182 | 0.502751 |

| | | | | | | | | | |
|---|---|---|---|---|---|---|---|---|---|
| 16 | 97896600 | 97898199 | 34.16281 | 56.1235 | 59.01331 | 25.42487 | 0.305604 | 1.421661 | 0.50277 |
| 3 | 94622400 | 94625199 | 21.51229 | 35.69326 | 70.41945 | 30.51097 | -0.59626 | 1.281267 | 0.502778 |
| 1 | 5145800 | 5147399 | 10.22422 | 18.57459 | 32.83118 | 14.16303 | -0.40095 | 0.312734 | 0.502805 |
| 1 | 145979000 | 145983799 | 102.2463 | 158.1204 | 163.8872 | 68.07487 | 0.250504 | 2.846761 | 0.502809 |
| 19 | 7909800 | 7913999 | 43.72837 | 71.37788 | 65.33145 | 28.07514 | 0.500326 | 1.530873 | 0.502815 |
| 2 | 97804200 | 97805799 | 16.99371 | 30.41794 | 30.14694 | 13.22012 | 0.412868 | 0.44147 | 0.502816 |
| X | 153141600 | 153160399 | 259.3067 | 379.326 | 327.1688 | 134.7784 | 0.446154 | 3.910901 | 0.502831 |
| 1 | 171324400 | 171326599 | 17.4732 | 30.11006 | 50.06668 | 21.96278 | -0.34586 | 0.88123 | 0.502834 |
| X | 14305800 | 14310599 | 76.47999 | 121.9654 | 99.08512 | 41.52373 | 0.637931 | 2.170814 | 0.502845 |
| 14 | 19189000 | 19192599 | 29.22595 | 48.70223 | 49.20171 | 21.38019 | 0.366489 | 1.233167 | 0.502851 |
| 18 | 19798800 | 19799599 | 18.9284 | 33.11299 | 32.70948 | 14.42676 | 0.405341 | 0.686233 | 0.502855 |
| 13 | 13473200 | 13480999 | 70.00331 | 111.6134 | 103.2903 | 43.27342 | 0.449953 | 2.168998 | 0.502907 |
| 2 | 143218600 | 143225399 | 94.26705 | 147.9904 | 137.6486 | 57.09016 | 0.423289 | 2.578719 | 0.502912 |
| 1 | 140136800 | 140139999 | 41.09591 | 67.69974 | 59.09761 | 25.4909 | 0.574884 | 1.405407 | 0.502918 |
| 4 | 147344400 | 147349799 | 61.50706 | 98.87288 | 83.21797 | 35.28303 | 0.599918 | 1.905823 | 0.502935 |
| 13 | 26729800 | 26731599 | 21.74265 | 38.22855 | 32.83569 | 14.53996 | 0.605727 | 0.621054 | 0.50294 |
| 12 | 98703400 | 98704799 | 7.857036 | 14.60221 | 31.04037 | 12.99211 | -0.63464 | 0.157465 | 0.502943 |
| 11 | 54709800 | 54712599 | 21.70349 | 36.34915 | 60.20692 | 26.24558 | -0.34445 | 1.168942 | 0.502956 |
| X | 81519400 | 81521199 | 19.01564 | 34.85041 | 24.67276 | 10.52495 | 0.920915 | 0.309495 | 0.502998 |
| 13 | 35190800 | 35199399 | 267.2017 | 377.6594 | 537.6895 | 222.5731 | -0.32576 | 4.419484 | 0.503004 |
| 12 | 91857600 | 91858399 | 12.68492 | 22.34638 | 38.83612 | 17.19032 | -0.40438 | 0.560479 | 0.503009 |
| 2 | 93225000 | 93226799 | 19.37951 | 34.15889 | 33.23555 | 14.70967 | 0.428396 | 0.584238 | 0.503011 |
| 1 | 39452600 | 39454799 | 16.33309 | 28.39387 | 46.84868 | 20.57648 | -0.33224 | 0.784914 | 0.503021 |
| 4 | 112273000 | 112276399 | 40.9474 | 67.70834 | 56.25245 | 24.31535 | 0.647206 | 1.360047 | 0.503026 |
| 17 | 38127400 | 38132599 | 46.38954 | 75.31351 | 69.86826 | 29.95224 | 0.475135 | 1.62146 | 0.503032 |
| 10 | 87774200 | 87774999 | 20.54516 | 35.717 | 36.3656 | 15.97241 | 0.361238 | 0.790127 | 0.503041 |
| 10 | 55037200 | 55038999 | 19.07551 | 33.91661 | 30.90243 | 13.62806 | 0.527964 | 0.496729 | 0.503059 |
| 16 | 68632200 | 68633399 | 10.39831 | 20.11286 | 19.66356 | 7.512627 | 0.592604 | -0.15915 | 0.50307 |
| 8 | 109877200 | 109878599 | 19.46691 | 35.05271 | 28.92568 | 12.61788 | 0.680864 | 0.421076 | 0.503096 |
| 10 | 57262400 | 57267999 | 62.9958 | 99.36906 | 149.6946 | 62.12472 | -0.26622 | 2.46633 | 0.503096 |
| 11 | 119000400 | 119004999 | 42.42545 | 67.84297 | 120.5709 | 50.72342 | -0.48518 | 2.055918 | 0.503109 |
| 18 | 5748600 | 5751199 | 22.62249 | 39.28421 | 39.36723 | 17.27857 | 0.383925 | 0.820765 | 0.50311 |
| 18 | 85147400 | 85148399 | 12.59478 | 23.77585 | 22.89363 | 9.329288 | 0.53017 | 0.078017 | 0.503114 |
| 18 | 24522800 | 24526599 | 58.02733 | 93.03243 | 91.75972 | 38.75028 | 0.36701 | 1.985059 | 0.503123 |
| 5 | 150865600 | 150867999 | 18.85846 | 32.03957 | 54.28743 | 23.79134 | -0.37638 | 1.021634 | 0.503128 |
| 8 | 25215400 | 25215999 | 4.969392 | 9.664565 | 21.76458 | 8.053508 | -0.55447 | -0.32169 | 0.503129 |
| 7 | 111510200 | 111510999 | 10.19988 | 18.69005 | 32.39748 | 13.82371 | -0.36244 | 0.262389 | 0.503134 |
| 1 | 166595600 | 166599399 | 65.45467 | 103.9472 | 102.8218 | 42.86385 | 0.348121 | 2.288111 | 0.503143 |
| 11 | 22367200 | 22368399 | 13.98912 | 25.78843 | 25.7827 | 10.91134 | 0.43432 | 0.240151 | 0.503218 |
| 16 | 38434400 | 38436799 | 26.54922 | 43.61084 | 71.57924 | 30.83723 | -0.33501 | 1.415257 | 0.503239 |
| 7 | 132495400 | 132504799 | 80.82747 | 125.2191 | 192.4368 | 79.93297 | -0.3152 | 2.807655 | 0.503283 |
| 9 | 88297000 | 88298599 | 11.86902 | 21.30357 | 36.2828 | 15.87393 | -0.36122 | 0.404014 | 0.503302 |
| X | 103655800 | 103657799 | 22.49648 | 39.44595 | 35.21065 | 15.56026 | 0.550682 | 0.672896 | 0.503303 |
| 10 | 20964600 | 20966599 | 17.51493 | 30.20605 | 49.52881 | 21.72736 | -0.32544 | 0.875008 | 0.503318 |
| 3 | 144131000 | 144133599 | 20.79778 | 35.1141 | 58.12962 | 25.42709 | -0.34347 | 1.086203 | 0.503361 |
| 14 | 121657800 | 121659199 | 19.57474 | 34.25598 | 35.5461 | 15.68263 | 0.334528 | 0.687669 | 0.503372 |
| X | 90172200 | 90172599 | 9.452937 | 18.27539 | 19.332 | 7.40276 | 0.478602 | -0.15761 | 0.503378 |

| | | | | | | | | | |
|---|---|---|---|---|---|---|---|---|---|
| 18 | 42730600 | 42732199 | 20.31058 | 35.3554 | 36.7553 | 16.15684 | 0.331377 | 0.76987 | 0.503395 |
| 6 | 3316600 | 3321999 | 38.02282 | 61.23116 | 99.25038 | 42.18962 | -0.34049 | 1.821663 | 0.503401 |
| 6 | 37706800 | 37711199 | 25.13461 | 41.79388 | 67.60664 | 29.3361 | -0.3106 | 1.263014 | 0.503411 |
| 4 | 92048400 | 92049999 | 24.65018 | 42.69391 | 40.83454 | 17.91569 | 0.45016 | 0.855813 | 0.503419 |
| 2 | 92851600 | 92855799 | 56.19096 | 88.9622 | 139.528 | 58.01206 | -0.32 | 2.372199 | 0.50342 |
| 13 | 50681400 | 50682999 | 17.8852 | 30.64162 | 51.10661 | 22.41247 | -0.35172 | 0.938905 | 0.503422 |
| 17 | 70546200 | 70547999 | 14.75708 | 25.92896 | 43.97074 | 19.46545 | -0.37077 | 0.611393 | 0.503429 |
| 15 | 56888800 | 56889399 | 10.04261 | 18.24904 | 33.33077 | 14.37423 | -0.44788 | 0.309447 | 0.503436 |
| 1 | 144347200 | 144348399 | 16.86055 | 30.03477 | 30.07251 | 13.24106 | 0.394087 | 0.468421 | 0.503439 |
| 3 | 21362600 | 21363799 | 15.79284 | 28.33036 | 29.2192 | 12.78966 | 0.357475 | 0.417495 | 0.503442 |
| 18 | 56727600 | 56729599 | 30.67628 | 50.86117 | 51.88396 | 22.46872 | 0.352233 | 1.298588 | 0.50345 |
| 13 | 40602600 | 40603999 | 7.663108 | 14.41565 | 28.04433 | 11.50048 | -0.48343 | 0.072149 | 0.503463 |
| 2 | 170772800 | 170775999 | 27.46618 | 46.27268 | 48.5522 | 21.22382 | 0.311945 | 1.117938 | 0.50349 |
| 13 | 7048000 | 7050599 | 15.69708 | 27.30258 | 45.78876 | 20.11761 | -0.35552 | 0.772971 | 0.503498 |
| X | 68034800 | 68038599 | 61.12292 | 97.78121 | 95.2842 | 40.10808 | 0.380831 | 2.057967 | 0.503501 |
| 17 | 49506000 | 49513599 | 75.93759 | 118.4947 | 176.8144 | 73.37106 | -0.26463 | 2.680023 | 0.503515 |
| 8 | 19556400 | 19557799 | 19.53691 | 34.25201 | 34.51578 | 15.25148 | 0.376665 | 0.658517 | 0.503517 |
| 18 | 27220600 | 27222399 | 15.4172 | 27.00895 | 45.64319 | 20.15548 | -0.36578 | 0.665419 | 0.503519 |
| 12 | 20655600 | 20656999 | 26.98183 | 45.88374 | 40.51722 | 17.75792 | 0.560415 | 1.026779 | 0.503545 |
| 14 | 8188000 | 8189999 | 20.67821 | 36.0945 | 37.1089 | 16.34051 | 0.34768 | 0.733934 | 0.503554 |
| 12 | 52279200 | 52281399 | 19.91882 | 35.13063 | 33.06003 | 14.63022 | 0.47645 | 0.57915 | 0.503557 |
| 17 | 50562200 | 50563799 | 16.60132 | 30.91037 | 24.64967 | 10.28604 | 0.773318 | 0.20439 | 0.503563 |
| 1 | 44157400 | 44159399 | 12.71801 | 22.23386 | 43.6375 | 19.23584 | -0.58046 | 0.679973 | 0.50357 |
| 4 | 92954000 | 92954999 | 13.60162 | 25.55869 | 24.49235 | 10.06275 | 0.527896 | 0.121745 | 0.503587 |
| 6 | 122077000 | 122078599 | 13.60162 | 25.55869 | 24.49235 | 10.06275 | 0.527896 | 0.121745 | 0.503587 |
| 3 | 120969600 | 120970799 | 11.74045 | 21.99619 | 22.98023 | 9.488918 | 0.398311 | 0.119537 | 0.503589 |
| 18 | 31027800 | 31028999 | 14.18484 | 25.70557 | 25.02416 | 10.82899 | 0.451277 | 0.338883 | 0.5036 |
| 2 | 160909800 | 160910999 | 13.01987 | 22.96939 | 38.8173 | 17.17706 | -0.36335 | 0.547712 | 0.503602 |
| 9 | 50258800 | 50260799 | 11.30088 | 20.34935 | 35.75789 | 15.59759 | -0.40339 | 0.376018 | 0.503612 |
| 17 | 90122000 | 90124199 | 17.55292 | 31.11964 | 31.05614 | 13.72912 | 0.394931 | 0.504495 | 0.503629 |
| 5 | 117470200 | 117478799 | 70.40629 | 108.648 | 211.8725 | 88.0078 | -0.66274 | 2.879049 | 0.503639 |
| 3 | 83199200 | 83201999 | 22.37342 | 37.74375 | 64.81805 | 28.34577 | -0.39631 | 1.107487 | 0.503643 |
| 13 | 106275000 | 106277399 | 32.94831 | 54.27867 | 56.29012 | 24.28012 | 0.327037 | 1.381908 | 0.503649 |
| 17 | 12494400 | 12509199 | 201.371 | 292.8465 | 412.4947 | 170.0241 | -0.26934 | 3.995242 | 0.50367 |
| 12 | 110270400 | 110278999 | 87.94144 | 135.6104 | 200.1981 | 83.1588 | -0.26242 | 2.897442 | 0.503687 |
| 5 | 150937200 | 150939399 | 42.40239 | 68.92146 | 68.94889 | 29.58377 | 0.368106 | 1.589449 | 0.503706 |
| 6 | 122826800 | 122828999 | 22.98504 | 38.33899 | 63.59486 | 27.65129 | -0.34658 | 1.218246 | 0.503706 |
| 13 | 83867800 | 83870799 | 23.12425 | 38.63398 | 63.66146 | 27.70867 | -0.33711 | 1.201036 | 0.503718 |
| Y | 8211200 | 8213999 | 21.97698 | 38.14185 | 26.47296 | 11.61543 | 0.911011 | 0.801646 | 0.503734 |
| 2 | 147180200 | 147181399 | 10.10527 | 18.24812 | 33.32477 | 14.48478 | -0.45477 | 0.348316 | 0.503741 |
| 1 | 184907200 | 184908599 | 18.08697 | 31.86209 | 32.77563 | 14.5078 | 0.348039 | 0.580409 | 0.503743 |
| 5 | 86359600 | 86369199 | 265.4011 | 382.8473 | 372.8433 | 153.8336 | 0.251395 | 4.119509 | 0.503759 |
| 13 | 3362000 | 3370999 | 102.9981 | 159.945 | 157.1616 | 65.26142 | 0.332025 | 2.772288 | 0.50376 |
| 16 | 48194000 | 48195599 | 10.65503 | 19.538 | 33.99286 | 14.49036 | -0.36696 | 0.264487 | 0.503762 |
| 12 | 95966400 | 95968199 | 19.57826 | 35.24883 | 28.87059 | 12.59581 | 0.691394 | 0.421938 | 0.503766 |
| 15 | 46428200 | 46430599 | 35.43659 | 58.32428 | 60.36576 | 26.02799 | 0.329363 | 1.40237 | 0.503774 |
| 5 | 141696400 | 141697799 | 7.814386 | 14.82316 | 27.66096 | 11.15729 | -0.40289 | 0.018258 | 0.50379 |

| | | | | | | | | | |
|---|---|---|---|---|---|---|---|---|---|
| 4 | 51698200 | 51698999 | 10.37651 | 19.7928 | 21.68736 | 8.645708 | 0.373347 | -0.0145 | 0.503797 |
| 14 | 114352400 | 114355199 | 26.50311 | 46.0196 | 38.4701 | 16.88404 | 0.644162 | 0.83561 | 0.503804 |
| 17 | 94057600 | 94060199 | 25.19993 | 43.88618 | 36.19352 | 15.91439 | 0.663805 | 0.784531 | 0.503817 |
| 8 | 66030000 | 66033599 | 35.20548 | 59.36687 | 47.41626 | 20.73316 | 0.704329 | 1.136425 | 0.503819 |
| 7 | 42192000 | 42194399 | 32.19842 | 54.36874 | 48.52772 | 21.22776 | 0.544628 | 1.115974 | 0.503821 |
| 5 | 41392600 | 41393999 | 9.387992 | 17.50902 | 31.10994 | 12.94505 | -0.37082 | 0.151251 | 0.503828 |
| X | 5188000 | 5191999 | 37.98409 | 62.72596 | 58.9045 | 25.43864 | 0.470861 | 1.36916 | 0.503835 |
| 19 | 61210400 | 61218799 | 102.316 | 154.8941 | 273.0214 | 112.9518 | -0.53912 | 3.276527 | 0.503839 |
| 2 | 27163600 | 27165999 | 29.87589 | 48.64033 | 78.61096 | 33.74992 | -0.32039 | 1.54881 | 0.503849 |
| 8 | 103973400 | 103976599 | 39.9656 | 65.1367 | 67.07332 | 28.80769 | 0.329264 | 1.545592 | 0.503867 |
| 13 | 59008800 | 59012799 | 22.49681 | 37.65358 | 61.3713 | 26.7423 | -0.32134 | 1.178913 | 0.503868 |
| 17 | 67304800 | 67309399 | 75.84053 | 119.6674 | 125.7743 | 52.20446 | 0.252782 | 2.46316 | 0.5039 |
| 18 | 90035400 | 90037999 | 34.4786 | 55.8319 | 87.86441 | 37.59556 | -0.28894 | 1.662274 | 0.503931 |
| 10 | 7538200 | 7539599 | 10.80956 | 19.20718 | 35.83705 | 15.81286 | -0.50039 | 0.459412 | 0.503933 |
| 6 | 12192400 | 12193399 | 10.33045 | 19.85578 | 21.24258 | 8.297361 | 0.434356 | -0.08248 | 0.503934 |
| X | 52666400 | 52668799 | 28.06961 | 47.11917 | 49.11175 | 21.43996 | 0.32148 | 1.154461 | 0.503939 |
| 12 | 48066800 | 48067199 | 7.734589 | 15.12934 | 14.17633 | 4.744541 | 0.825358 | -0.65864 | 0.503946 |
| 7 | 34973000 | 34982399 | 161.6212 | 238.6419 | 348.5188 | 143.5267 | -0.30051 | 3.752112 | 0.503954 |
| X | 43877000 | 43879399 | 38.44989 | 63.72845 | 47.69859 | 20.65145 | 0.797594 | 1.323473 | 0.503964 |
| 4 | 60326000 | 60330599 | 87.06655 | 136.7723 | 121.7539 | 50.50193 | 0.487182 | 2.559407 | 0.503972 |
| 5 | 119305400 | 119306399 | 11.87307 | 21.05943 | 36.73911 | 16.2352 | -0.40578 | 0.478205 | 0.503991 |
| X | 74962000 | 74964999 | 40.74351 | 67.90839 | 50.97796 | 22.15478 | 0.793705 | 1.271367 | 0.50401 |
| 14 | 11946800 | 11949199 | 19.94324 | 35.3525 | 31.35802 | 13.86297 | 0.564064 | 0.522518 | 0.504015 |
| 16 | 89800200 | 89802799 | 19.35464 | 32.72517 | 56.989 | 24.93917 | -0.41614 | 1.06698 | 0.504035 |
| 7 | 24602200 | 24604599 | 13.95637 | 24.39868 | 42.58832 | 18.77328 | -0.41209 | 0.686315 | 0.504044 |
| 3 | 83313200 | 83316199 | 46.29907 | 74.85377 | 75.59121 | 32.30027 | 0.347367 | 1.710398 | 0.504047 |
| 7 | 41408200 | 41410399 | 25.76543 | 44.83205 | 38.53068 | 16.9328 | 0.60474 | 0.805876 | 0.504056 |
| 9 | 90465600 | 90467999 | 18.54026 | 32.02169 | 57.77959 | 25.34797 | -0.46273 | 0.876111 | 0.50406 |
| 6 | 58882400 | 58885799 | 55.26424 | 87.74593 | 133.8381 | 55.78225 | -0.27601 | 2.293369 | 0.504124 |
| 7 | 42574600 | 42576199 | 33.81786 | 56.1931 | 55.89071 | 24.24058 | 0.388821 | 1.285455 | 0.504131 |
| 1 | 120934000 | 120941199 | 72.24371 | 114.8361 | 111.5392 | 46.54146 | 0.375682 | 2.267023 | 0.504166 |
| 8 | 75206000 | 75210399 | 55.08625 | 88.90479 | 76.55028 | 32.63639 | 0.573999 | 1.77555 | 0.50418 |
| 19 | 14088000 | 14088399 | 4.936427 | 9.631163 | 20.38304 | 7.278003 | -0.41856 | -0.45328 | 0.50418 |
| 10 | 61794400 | 61795999 | 10.87734 | 19.57239 | 34.26232 | 14.95846 | -0.39847 | 0.376085 | 0.504182 |
| 3 | 80172600 | 80176199 | 44.84018 | 73.03123 | 67.48965 | 28.97925 | 0.484324 | 1.566893 | 0.504186 |
| X | 117813600 | 117817599 | 35.92955 | 60.38121 | 48.5658 | 21.21381 | 0.694199 | 1.17081 | 0.504209 |
| 5 | 4532800 | 4534799 | 18.82648 | 33.02226 | 34.01123 | 15.0346 | 0.345254 | 0.645508 | 0.504236 |
| 15 | 93607600 | 93612799 | 27.99121 | 45.42927 | 82.62229 | 35.4115 | -0.494 | 1.599683 | 0.504244 |
| 10 | 118260800 | 118263199 | 28.88168 | 48.22024 | 49.18507 | 21.39865 | 0.352606 | 1.211637 | 0.504246 |
| 12 | 67683800 | 67690799 | 93.93349 | 146.5828 | 133.3228 | 55.32344 | 0.44881 | 2.682527 | 0.504246 |
| 17 | 4188400 | 4188799 | 6.726222 | 12.87318 | 25.66682 | 10.14904 | -0.47197 | -0.0648 | 0.504259 |
| 7 | 52583400 | 52585199 | 23.04847 | 40.40153 | 35.21362 | 15.55758 | 0.584985 | 0.681144 | 0.504273 |
| 9 | 113063400 | 113064399 | 10.87336 | 19.87445 | 35.00652 | 14.98698 | -0.38893 | 0.28783 | 0.504279 |
| 13 | 48642600 | 48649599 | 78.01329 | 121.4589 | 179.019 | 74.30909 | -0.24941 | 2.720892 | 0.504294 |
| 2 | 135830400 | 135831399 | 13.8091 | 24.04651 | 45.78029 | 20.1285 | -0.53744 | 0.745831 | 0.504308 |
| 6 | 76369200 | 76372199 | 32.00294 | 54.12664 | 42.04177 | 18.39965 | 0.744524 | 1.095792 | 0.50433 |
| 15 | 37170600 | 37172599 | 27.88294 | 45.16267 | 83.04356 | 35.53934 | -0.51243 | 1.643084 | 0.504331 |

| | | | | | | | | | |
|---|---|---|---|---|---|---|---|---|---|
| 13 | 120635000 | 120636599 | 12.78551 | 22.78085 | 38.75645 | 17.06135 | -0.36617 | 0.458685 | 0.504332 |
| 18 | 24165600 | 24166799 | 22.48716 | 39.05615 | 32.6911 | 14.34997 | 0.642371 | 0.806045 | 0.504343 |
| 7 | 43769000 | 43773999 | 52.83681 | 84.09046 | 128.3028 | 53.61773 | -0.27304 | 2.216683 | 0.50435 |
| 18 | 77674600 | 77677399 | 25.78404 | 42.74264 | 69.30959 | 30.01152 | -0.31448 | 1.300763 | 0.504366 |
| 13 | 87230200 | 87231999 | 15.32061 | 28.10202 | 26.67648 | 11.36671 | 0.501648 | 0.280328 | 0.504373 |
| 11 | 8688800 | 8694199 | 72.23611 | 114.85 | 101.2907 | 42.27466 | 0.515002 | 2.258523 | 0.504393 |
| 9 | 98985800 | 98989599 | 45.0691 | 71.95373 | 127.0536 | 53.32808 | -0.47849 | 2.110765 | 0.50441 |
| 17 | 18320000 | 18325799 | 69.53259 | 111.204 | 94.33989 | 39.69514 | 0.58 | 2.074747 | 0.50441 |
| 4 | 49696600 | 49697399 | 7.180947 | 13.64159 | 27.14946 | 10.91702 | -0.49153 | -0.00239 | 0.504436 |
| 13 | 3474200 | 3487399 | 379.6823 | 530.9046 | 500.4974 | 207.533 | 0.25323 | 4.581853 | 0.504444 |
| 11 | 49088400 | 49089399 | 14.71944 | 26.84348 | 27.24969 | 11.69923 | 0.398129 | 0.309714 | 0.504466 |
| 5 | 94551400 | 94554799 | 36.12434 | 59.56783 | 59.89739 | 25.84654 | 0.371989 | 1.381543 | 0.504474 |
| 3 | 3435200 | 3435799 | 13.44296 | 24.70378 | 24.55872 | 10.43792 | 0.438705 | 0.254105 | 0.504495 |
| 5 | 86756200 | 86757599 | 14.43877 | 26.5207 | 26.83771 | 11.41417 | 0.412071 | 0.267343 | 0.504499 |
| 11 | 32154600 | 32155799 | 7.192566 | 13.6646 | 26.60796 | 10.69775 | -0.45971 | -0.00249 | 0.504503 |
| 5 | 141727600 | 141730199 | 14.96928 | 26.12886 | 44.17893 | 19.45348 | -0.36647 | 0.719212 | 0.504505 |
| 7 | 8562600 | 8565199 | 44.2044 | 71.60665 | 64.62749 | 27.66019 | 0.511752 | 1.658624 | 0.504518 |
| 2 | 173461600 | 173464599 | 18.3547 | 31.27592 | 53.23181 | 23.33427 | -0.38216 | 0.991192 | 0.504527 |
| 12 | 114269800 | 114278999 | 173.209 | 260.6673 | 240.1539 | 99.17752 | 0.388775 | 3.407134 | 0.504565 |
| 15 | 25977400 | 25979999 | 25.77943 | 44.47712 | 41.89368 | 18.3782 | 0.471042 | 0.895684 | 0.504571 |
| 1 | 103170800 | 103172799 | 34.80025 | 57.21922 | 60.24971 | 25.96687 | 0.304053 | 1.413294 | 0.504572 |
| 4 | 38315800 | 38317399 | 14.25334 | 26.14865 | 25.67766 | 10.93868 | 0.453157 | 0.268079 | 0.504585 |
| 16 | 75074600 | 75076199 | 19.05392 | 33.50336 | 33.72835 | 14.93504 | 0.377973 | 0.607474 | 0.504586 |
| 7 | 114318400 | 114320399 | 10.12974 | 18.13034 | 37.83334 | 16.59187 | -0.65527 | 0.408672 | 0.504592 |
| 2 | 35314600 | 35316599 | 14.12907 | 24.7264 | 42.04592 | 18.55055 | -0.37447 | 0.670262 | 0.504605 |
| 15 | 63632000 | 63632399 | 4.911896 | 9.590092 | 20.51333 | 7.235647 | -0.41733 | -0.49465 | 0.50461 |
| 8 | 4439600 | 4442999 | 34.98106 | 56.42737 | 91.00368 | 38.77251 | -0.33024 | 1.764912 | 0.504622 |
| 6 | 76575600 | 76576599 | 12.60609 | 22.50636 | 38.77892 | 17.04234 | -0.38275 | 0.444426 | 0.504634 |
| 13 | 66422600 | 66430399 | 134.2618 | 206.7749 | 167.6749 | 69.64506 | 0.599705 | 2.93692 | 0.504646 |
| 5 | 151001400 | 151002799 | 10.17478 | 18.72805 | 32.46254 | 13.76701 | -0.35611 | 0.236439 | 0.504658 |
| 8 | 44938600 | 44940199 | 17.55613 | 31.85661 | 27.98783 | 12.0917 | 0.59959 | 0.360683 | 0.504666 |
| 1 | 67008200 | 67008999 | 11.02605 | 19.87318 | 34.26542 | 14.92899 | -0.37481 | 0.367443 | 0.504672 |
| 13 | 119314200 | 119315399 | 11.9322 | 22.29492 | 23.49072 | 9.744341 | 0.380955 | 0.139962 | 0.504673 |
| X | 153291600 | 153296999 | 95.27064 | 150.1007 | 128.0768 | 53.1449 | 0.551479 | 2.508583 | 0.504684 |
| 5 | 148357000 | 148358399 | 14.87181 | 26.09765 | 44.52606 | 19.69031 | -0.37957 | 0.63181 | 0.504685 |
| 2 | 99201600 | 99202399 | 11.46379 | 20.63659 | 35.75128 | 15.60049 | -0.38335 | 0.379191 | 0.504687 |
| 2 | 111256000 | 111259199 | 34.85205 | 58.39885 | 50.30073 | 21.93397 | 0.595646 | 1.19397 | 0.504693 |
| 14 | 106380600 | 106382999 | 23.34481 | 40.99637 | 33.97588 | 15.0341 | 0.657209 | 0.651666 | 0.504697 |
| 3 | 34566400 | 34568399 | 31.33496 | 51.84661 | 53.51648 | 23.14284 | 0.335051 | 1.325077 | 0.504708 |
| 7 | 60402000 | 60410399 | 97.30611 | 152.1092 | 146.1374 | 60.63018 | 0.371841 | 2.656693 | 0.504708 |
| 11 | 7351600 | 7352199 | 15.0852 | 27.25708 | 27.88195 | 12.11004 | 0.376824 | 0.368872 | 0.504717 |
| 3 | 51496400 | 51498999 | 18.36577 | 31.37143 | 52.16849 | 22.87266 | -0.34817 | 0.968761 | 0.504727 |
| 12 | 62384200 | 62386999 | 23.13343 | 40.37199 | 37.44107 | 16.48468 | 0.495677 | 0.749327 | 0.504735 |
| 17 | 17519400 | 17551399 | 700.1933 | 938.2953 | 766.1555 | 319.7921 | 0.398658 | 5.24774 | 0.504737 |
| 18 | 59627200 | 59629799 | 26.77348 | 44.23432 | 72.18073 | 31.18249 | -0.32394 | 1.343817 | 0.504746 |
| 13 | 112980400 | 112983599 | 50.37542 | 80.95817 | 85.19451 | 36.10778 | 0.277826 | 1.898353 | 0.504763 |
| 1 | 52326600 | 52328599 | 15.25823 | 26.5305 | 45.28482 | 19.89206 | -0.38075 | 0.776082 | 0.50477 |

| | | | | | | | | | |
|---|---|---|---|---|---|---|---|---|---|
| 13 | 61714200 | 61729399 | 302.5523 | 433.8243 | 406.2826 | 167.78 | 0.299192 | 4.210545 | 0.504773 |
| 6 | 59201400 | 59209399 | 111.709 | 173.3461 | 157.8689 | 65.56223 | 0.440711 | 2.789797 | 0.504774 |
| 17 | 35613000 | 35615199 | 15.47619 | 26.62793 | 48.98882 | 21.48896 | -0.49141 | 0.886149 | 0.504776 |
| 5 | 92132800 | 92133999 | 12.88026 | 23.79566 | 25.01631 | 10.55591 | 0.365799 | 0.219999 | 0.504793 |
| 1 | 45762600 | 45774799 | 65.44403 | 104.3899 | 93.63991 | 39.25428 | 0.495514 | 2.147445 | 0.504798 |
| 4 | 61848000 | 61849199 | 15.8367 | 28.45608 | 29.24073 | 12.78124 | 0.364198 | 0.408211 | 0.504809 |
| 4 | 121405600 | 121414799 | 199.7907 | 296.5615 | 287.0059 | 118.2236 | 0.299874 | 3.66577 | 0.504814 |
| 2 | 111563000 | 111567199 | 44.56466 | 72.13121 | 61.66484 | 26.37628 | 0.588724 | 1.674905 | 0.504845 |
| 5 | 93723600 | 93726999 | 52.75117 | 84.84435 | 76.50252 | 32.46164 | 0.501677 | 1.872739 | 0.504852 |
| 11 | 58689200 | 58693199 | 24.53171 | 40.66567 | 66.22455 | 28.67521 | -0.32052 | 1.296766 | 0.504863 |
| 7 | 64825200 | 64826399 | 8.9554 | 16.53324 | 30.44059 | 12.85683 | -0.4378 | 0.206474 | 0.504863 |
| 9 | 24653600 | 24654399 | 7.019698 | 13.49797 | 25.80688 | 10.06801 | -0.39347 | -0.10382 | 0.504894 |
| 15 | 28496200 | 28497199 | 7.019698 | 13.49797 | 25.80688 | 10.06801 | -0.39347 | -0.10382 | 0.504894 |
| 8 | 92039600 | 92040399 | 6.011239 | 11.51587 | 25.14062 | 9.913516 | -0.59738 | -0.08187 | 0.5049 |
| 15 | 96458200 | 96461399 | 24.16222 | 40.02022 | 68.58403 | 29.67999 | -0.39408 | 1.304383 | 0.504907 |
| 15 | 36407000 | 36408399 | 17.42383 | 30.745 | 31.78614 | 14.06513 | 0.34172 | 0.556599 | 0.504911 |
| 9 | 89305600 | 89306199 | 12.50874 | 23.18461 | 24.51244 | 10.2976 | 0.362163 | 0.197132 | 0.504929 |
| 9 | 35986000 | 35990999 | 58.6443 | 94.01764 | 90.89815 | 38.38399 | 0.395711 | 1.98829 | 0.504933 |
| 19 | 21779000 | 21781999 | 30.95084 | 51.42467 | 54.11248 | 23.46793 | 0.307758 | 1.275938 | 0.504935 |
| 4 | 101096400 | 101097999 | 10.0651 | 17.87225 | 35.91439 | 15.85604 | -0.60715 | 0.459739 | 0.504948 |
| 6 | 87830200 | 87831999 | 10.79389 | 19.66057 | 34.38401 | 14.78476 | -0.38308 | 0.303401 | 0.504962 |
| 8 | 37777200 | 37780199 | 51.37192 | 82.36818 | 81.85143 | 34.56985 | 0.356368 | 1.970884 | 0.504974 |
| 10 | 72327800 | 72329199 | 13.48458 | 23.71308 | 39.84925 | 17.64501 | -0.3572 | 0.586908 | 0.50498 |
| 9 | 19121000 | 19123599 | 35.65159 | 58.85865 | 59.30627 | 25.6075 | 0.369503 | 1.366139 | 0.50498 |
| 14 | 81162200 | 81164599 | 21.23093 | 37.52136 | 31.80176 | 14.06765 | 0.627706 | 0.562925 | 0.504996 |
| 13 | 24944600 | 24948999 | 28.31933 | 46.42731 | 74.63197 | 32.12571 | -0.30627 | 1.444869 | 0.505019 |
| 1 | 85457600 | 85459199 | 25.41207 | 43.50252 | 44.13929 | 19.35576 | 0.362153 | 0.965275 | 0.505047 |
| 2 | 33655600 | 33670999 | 235.386 | 336.4205 | 483.601 | 199.8115 | -0.32342 | 4.254549 | 0.505055 |
| 8 | 69952200 | 69961599 | 193.0412 | 285.864 | 285.7213 | 117.6692 | 0.250213 | 3.705784 | 0.505057 |
| 13 | 68138200 | 68143799 | 114.254 | 176.1102 | 158.3352 | 65.76983 | 0.451851 | 2.91065 | 0.505078 |
| 3 | 158864800 | 158866799 | 17.5031 | 31.48242 | 29.5706 | 12.91433 | 0.493912 | 0.414148 | 0.505082 |
| X | 65015200 | 65016599 | 17.5031 | 31.48242 | 29.5706 | 12.91433 | 0.493912 | 0.414148 | 0.505082 |
| 6 | 78795600 | 78796799 | 12.21634 | 23.28091 | 22.60934 | 9.03071 | 0.543333 | 0.010873 | 0.505098 |
| 2 | 53972400 | 53976599 | 30.88165 | 50.04167 | 83.95934 | 35.94286 | -0.38012 | 1.638284 | 0.505122 |
| 7 | 5907400 | 5913399 | 139.5485 | 213.1679 | 169.3877 | 70.25711 | 0.619882 | 3.09727 | 0.505134 |
| 3 | 56328800 | 56329199 | 7.280846 | 13.96735 | 27.49321 | 10.79832 | -0.4453 | -0.07871 | 0.505136 |
| 7 | 30628800 | 30629199 | 7.677156 | 14.9419 | 17.3299 | 6.373744 | 0.400659 | -0.32446 | 0.505145 |
| 5 | 137265600 | 137269999 | 32.52838 | 52.59046 | 86.61295 | 37.01024 | -0.35659 | 1.692087 | 0.505154 |
| 14 | 45004200 | 45008199 | 63.85182 | 101.6339 | 106.6842 | 44.59116 | 0.265878 | 2.216941 | 0.505161 |
| 3 | 36967400 | 36968399 | 17.0061 | 30.39042 | 26.11979 | 11.46991 | 0.615823 | 0.44281 | 0.505187 |
| 2 | 22209200 | 22210599 | 18.64374 | 32.09847 | 52.77231 | 23.14904 | -0.32991 | 0.898483 | 0.50521 |
| 9 | 96904400 | 96905599 | 11.87883 | 22.19432 | 23.57863 | 9.781471 | 0.369354 | 0.141478 | 0.505228 |
| 14 | 17889800 | 17890999 | 11.87883 | 22.19432 | 23.57863 | 9.781471 | 0.369354 | 0.141478 | 0.505228 |
| 16 | 35610200 | 35610999 | 11.87883 | 22.19432 | 23.57863 | 9.781471 | 0.369354 | 0.141478 | 0.505228 |
| 6 | 59206400 | 59209599 | 53.24262 | 85.70733 | 82.4618 | 35.03326 | 0.409748 | 1.849007 | 0.505242 |
| 7 | 31228400 | 31229999 | 52.31408 | 83.91507 | 85.52559 | 36.14539 | 0.32068 | 1.961436 | 0.505249 |
| 17 | 19165200 | 19169599 | 38.91382 | 65.02498 | 50.37042 | 21.92612 | 0.748397 | 1.237508 | 0.505253 |

| | | | | | | | | | |
|---|---|---|---|---|---|---|---|---|---|
| 4 | 146286000 | 146293799 | 188.4154 | 281.8642 | 259.7762 | 107.1278 | 0.380578 | 3.524145 | 0.505255 |
| 10 | 115368200 | 115372399 | 37.20902 | 59.98206 | 96.39831 | 41.03633 | -0.32637 | 1.788088 | 0.505258 |
| 18 | 34941600 | 34942399 | 5.982097 | 11.5196 | 24.69729 | 9.594848 | -0.5505 | -0.12957 | 0.505284 |
| 16 | 32150800 | 32152199 | 6.893015 | 13.13263 | 26.11885 | 10.43841 | -0.48214 | -0.02595 | 0.5053 |
| 17 | 53659600 | 53663199 | 45.70352 | 73.82411 | 76.28043 | 32.55678 | 0.312893 | 1.736637 | 0.505342 |
| X | 108347200 | 108352399 | 56.37196 | 90.73802 | 82.32162 | 34.96991 | 0.494245 | 1.859637 | 0.505357 |
| 5 | 126864600 | 126865799 | 12.14764 | 21.60675 | 37.20894 | 16.40411 | -0.38501 | 0.46221 | 0.505368 |
| 13 | 4544000 | 4545199 | 18.48648 | 32.63285 | 26.80009 | 11.85791 | 0.672339 | 0.549899 | 0.505375 |
| 10 | 109367800 | 109368399 | 4.482 | 8.745395 | 19.76356 | 7.045183 | -0.50851 | -0.46459 | 0.505384 |
| 15 | 37759800 | 37760199 | 4.482 | 8.745395 | 19.76356 | 7.045183 | -0.50851 | -0.46459 | 0.505384 |
| 8 | 9832200 | 9836199 | 37.41554 | 60.21457 | 96.71031 | 41.07168 | -0.32838 | 1.841091 | 0.50539 |
| 4 | 35275800 | 35277599 | 13.80394 | 24.41047 | 41.85202 | 18.51118 | -0.38298 | 0.530694 | 0.505398 |
| 19 | 47465800 | 47471199 | 55.05719 | 87.39545 | 132.5792 | 55.2438 | -0.26852 | 2.299397 | 0.505398 |
| 3 | 124669600 | 124671399 | 19.28601 | 33.01541 | 53.83557 | 23.60773 | -0.31962 | 0.953303 | 0.505408 |
| X | 3696000 | 3698399 | 20.77808 | 37.12438 | 29.80594 | 13.09059 | 0.713653 | 0.477836 | 0.505427 |
| X | 115597600 | 115600399 | 32.22073 | 55.85724 | 36.16604 | 15.86788 | 1.011059 | 0.878877 | 0.505441 |
| 7 | 10848400 | 10855399 | 146.7284 | 222.5853 | 216.4571 | 89.60756 | 0.320702 | 3.242439 | 0.505468 |
| 2 | 78440800 | 78441599 | 5.032595 | 9.816522 | 22.71516 | 8.136248 | -0.54911 | -0.42837 | 0.50547 |
| 1 | 169460800 | 169464199 | 28.42549 | 48.03749 | 48.52709 | 21.23279 | 0.366628 | 1.094445 | 0.505471 |
| 3 | 8737000 | 8740999 | 28.82674 | 47.11403 | 76.76895 | 33.00558 | -0.32824 | 1.488054 | 0.505485 |
| 1 | 151739200 | 151741599 | 24.2166 | 40.07103 | 66.69946 | 28.84493 | -0.3522 | 1.316703 | 0.505492 |
| 8 | 3851400 | 3859599 | 255.1654 | 369.0952 | 356.6236 | 147.0879 | 0.266684 | 4.078315 | 0.505495 |
| 6 | 27183800 | 27184799 | 12.38098 | 24.00523 | 17.83543 | 6.738388 | 1.000035 | -0.17768 | 0.505505 |
| X | 91126400 | 91127399 | 17.20157 | 30.66603 | 30.68782 | 13.50315 | 0.395738 | 0.469217 | 0.505529 |
| 18 | 75862200 | 75863599 | 16.76663 | 29.97101 | 30.1262 | 13.22605 | 0.391427 | 0.447446 | 0.505545 |
| 4 | 60335400 | 60336799 | 27.07421 | 45.45563 | 43.37429 | 18.93719 | 0.448367 | 1.141866 | 0.505547 |
| 11 | 47689400 | 47691399 | 20.18604 | 35.74339 | 31.29841 | 13.83967 | 0.581898 | 0.534796 | 0.505574 |
| 5 | 121248800 | 121253199 | 32.34192 | 52.34655 | 85.751 | 36.67816 | -0.34731 | 1.665684 | 0.505588 |
| 2 | 27107400 | 27108999 | 7.558891 | 14.12508 | 28.51046 | 11.82443 | -0.54896 | 0.117091 | 0.505597 |
| 3 | 13953600 | 13955999 | 47.43705 | 76.39773 | 80.58068 | 34.25742 | 0.278286 | 1.826773 | 0.505601 |
| 4 | 148192000 | 148195599 | 20.78857 | 34.32139 | 69.0241 | 29.80392 | -0.62441 | 1.334738 | 0.505602 |
| 12 | 115098600 | 115107399 | 110.8759 | 173.1183 | 142.4389 | 59.10284 | 0.593927 | 2.684712 | 0.505605 |
| X | 54332600 | 54335999 | 24.52019 | 43.4071 | 29.44362 | 13.02015 | 0.94844 | 0.563348 | 0.505621 |
| 2 | 74579800 | 74581199 | 8.06174 | 14.84034 | 30.94835 | 13.1229 | -0.62053 | 0.214878 | 0.505629 |
| 3 | 65577400 | 65578599 | 8.933894 | 16.49656 | 30.42905 | 12.84875 | -0.44038 | 0.204007 | 0.505643 |
| 6 | 70777200 | 70778399 | 11.72363 | 22.01356 | 22.97653 | 9.451741 | 0.403935 | 0.105136 | 0.505669 |
| 7 | 15089400 | 15096199 | 174.6299 | 264.4156 | 211.0826 | 87.33197 | 0.602957 | 3.286877 | 0.505684 |
| Y | 5258800 | 5260599 | 14.8168 | 26.87584 | 18.05913 | 7.805823 | 0.982972 | 0.317765 | 0.505684 |
| 4 | 124228000 | 124232799 | 49.71282 | 79.97602 | 83.69732 | 35.52646 | 0.287616 | 1.864593 | 0.505688 |
| 4 | 70781000 | 70782399 | 13.07165 | 23.0512 | 39.87287 | 17.64562 | -0.39688 | 0.556085 | 0.505701 |
| 9 | 12934800 | 12936399 | 22.9601 | 39.73572 | 39.79923 | 17.46104 | 0.383537 | 0.85143 | 0.505716 |
| 16 | 83567600 | 83569199 | 18.85382 | 33.2328 | 27.80336 | 12.30541 | 0.645009 | 0.570152 | 0.505724 |
| 8 | 79953800 | 79956799 | 26.08968 | 43.26944 | 72.31925 | 31.32479 | -0.358 | 1.2977 | 0.505742 |
| 18 | 63135800 | 63136399 | 12.67134 | 23.52828 | 25.00438 | 10.47858 | 0.357257 | 0.189575 | 0.50575 |
| 4 | 21258200 | 21259999 | 21.59693 | 38.08889 | 31.72643 | 14.04016 | 0.651432 | 0.585918 | 0.505751 |
| 9 | 19896400 | 19897599 | 15.99455 | 29.09171 | 28.23034 | 12.16047 | 0.459878 | 0.337491 | 0.505753 |
| X | 39957800 | 39959999 | 21.25155 | 38.10035 | 28.66608 | 12.55373 | 0.809552 | 0.45793 | 0.505775 |

| | | | | | | | | | |
|---|---|---|---|---|---|---|---|---|---|
| 4 | 83241400 | 83242799 | 9.918871 | 17.99365 | 32.83182 | 14.18911 | -0.44824 | 0.318508 | 0.505797 |
| 6 | 70660200 | 70664599 | 81.068 | 127.515 | 133.4173 | 55.3491 | 0.255828 | 2.531178 | 0.505809 |
| 4 | 119290800 | 119291999 | 8.296071 | 15.48205 | 28.99125 | 12.0509 | -0.44455 | 0.13251 | 0.505812 |
| 10 | 28877200 | 28878399 | 15.4558 | 27.04367 | 45.68063 | 20.1513 | -0.36502 | 0.686437 | 0.505819 |
| 5 | 92892600 | 92894199 | 13.29758 | 23.45387 | 39.28427 | 17.38451 | -0.35086 | 0.551312 | 0.505821 |
| 7 | 61506800 | 61509399 | 26.62828 | 45.96845 | 40.71706 | 17.86235 | 0.559441 | 0.891404 | 0.505843 |
| 1 | 89895400 | 89900399 | 38.65127 | 61.86088 | 109.1893 | 45.99132 | -0.47389 | 2.020093 | 0.505857 |
| 5 | 146933000 | 146936599 | 37.25476 | 59.9979 | 95.11311 | 40.43447 | -0.30832 | 1.818559 | 0.505859 |
| 8 | 69833400 | 69845399 | 344.9358 | 487.6061 | 462.2274 | 191.331 | 0.261069 | 4.420005 | 0.505891 |
| 18 | 56432800 | 56433999 | 17.7434 | 31.24131 | 32.32231 | 14.30856 | 0.339515 | 0.582837 | 0.505913 |
| 12 | 59132800 | 59133599 | 14.1119 | 25.82036 | 26.84393 | 11.47699 | 0.367955 | 0.286217 | 0.505916 |
| 15 | 32991600 | 32996599 | 56.17233 | 90.57388 | 77.82444 | 33.14008 | 0.575484 | 1.803214 | 0.505937 |
| 4 | 104230800 | 104232599 | 16.28716 | 29.35794 | 29.1527 | 12.70024 | 0.416359 | 0.392014 | 0.505963 |
| 17 | 93245400 | 93251999 | 108.4662 | 168.5324 | 141.8749 | 58.90859 | 0.555497 | 2.76421 | 0.505977 |
| 10 | 30448400 | 30451799 | 49.48331 | 79.58638 | 83.11364 | 35.2656 | 0.290192 | 1.871112 | 0.506002 |
| 9 | 80294600 | 80296799 | 24.44514 | 41.88364 | 42.64658 | 18.7025 | 0.357427 | 0.954604 | 0.506008 |
| 17 | 19293800 | 19299199 | 54.15908 | 87.60151 | 71.99194 | 30.77827 | 0.644695 | 1.711554 | 0.506021 |
| 6 | 109218000 | 109218399 | 4.222262 | 8.255738 | 18.40559 | 6.289604 | -0.43453 | -0.62579 | 0.506026 |
| 7 | 20657800 | 20660599 | 24.7328 | 43.33321 | 33.29042 | 14.70194 | 0.765964 | 0.691956 | 0.506029 |
| 3 | 84316600 | 84319399 | 18.40651 | 31.23258 | 54.26 | 23.77046 | -0.41235 | 1.031246 | 0.506048 |
| 3 | 93651400 | 93652399 | 12.98466 | 23.99704 | 24.32464 | 10.25911 | 0.418064 | 0.214527 | 0.50606 |
| 7 | 109469600 | 109470799 | 8.898684 | 16.84845 | 34.0029 | 13.76969 | -0.52109 | 0.054303 | 0.50607 |
| 13 | 62967000 | 62970999 | 54.17838 | 86.9416 | 89.60603 | 37.89379 | 0.305259 | 1.950619 | 0.506087 |
| 13 | 51191800 | 51203999 | 171.2682 | 252.8641 | 357.2689 | 147.13 | -0.25292 | 3.754383 | 0.506093 |
| 3 | 24467800 | 24468999 | 17.26268 | 30.77662 | 30.59136 | 13.4601 | 0.405326 | 0.467812 | 0.506094 |
| 4 | 29054600 | 29056199 | 17.34528 | 30.11922 | 50.08258 | 21.98113 | -0.34361 | 0.810887 | 0.506096 |
| 19 | 50306800 | 50308599 | 17.17941 | 29.82159 | 49.00359 | 21.50322 | -0.3266 | 0.813402 | 0.506096 |
| 10 | 120805000 | 120806199 | 11.67479 | 21.9159 | 23.57469 | 9.702282 | 0.360515 | 0.109304 | 0.506114 |
| 17 | 77096800 | 77099399 | 22.97453 | 40.35442 | 33.97429 | 15.03601 | 0.634578 | 0.646073 | 0.50613 |
| 5 | 143556800 | 143559199 | 16.83541 | 29.16635 | 47.93505 | 21.03112 | -0.3274 | 0.828542 | 0.506134 |
| 8 | 121447600 | 121451199 | 34.99565 | 56.49088 | 89.93575 | 38.35253 | -0.31043 | 1.744714 | 0.506142 |
| 4 | 140810000 | 140812999 | 23.91922 | 39.46469 | 68.83097 | 29.70873 | -0.41982 | 1.350344 | 0.506147 |
| 5 | 67379400 | 67387399 | 83.20123 | 128.6948 | 190.5503 | 79.14999 | -0.26327 | 2.838548 | 0.5062 |
| 18 | 56317400 | 56322399 | 67.41405 | 107.4894 | 104.2802 | 43.6906 | 0.382052 | 2.165968 | 0.506235 |
| 19 | 13149400 | 13156399 | 92.33617 | 145.0289 | 137.246 | 56.92674 | 0.398905 | 2.56716 | 0.506241 |
| 1 | 36295000 | 36296799 | 24.03598 | 39.81962 | 66.19652 | 28.65101 | -0.35023 | 1.301592 | 0.506297 |
| 14 | 40580800 | 40583399 | 19.92523 | 33.83093 | 57.62814 | 25.25082 | -0.38425 | 1.032514 | 0.506308 |
| 19 | 16405200 | 16407199 | 31.31424 | 52.17902 | 54.12443 | 23.51843 | 0.328549 | 1.245647 | 0.506313 |
| 13 | 5244400 | 5252399 | 176.7367 | 263.4474 | 265.44 | 109.3872 | 0.247369 | 3.588191 | 0.506323 |
| 1 | 190255000 | 190255999 | 6.694484 | 12.90487 | 25.3428 | 9.816207 | -0.42254 | -0.13472 | 0.506325 |
| 3 | 49871200 | 49872199 | 10.85121 | 20.77312 | 21.06915 | 8.325056 | 0.494882 | -0.04199 | 0.506337 |
| 2 | 53117200 | 53121599 | 73.85247 | 117.2335 | 115.6384 | 48.18612 | 0.351567 | 2.30962 | 0.506364 |
| 8 | 93736400 | 93736999 | 5.507288 | 10.7324 | 22.02676 | 7.987024 | -0.39526 | -0.38482 | 0.506369 |
| 14 | 112741200 | 112742399 | 17.49724 | 30.79423 | 31.00879 | 13.72831 | 0.378304 | 0.585407 | 0.506371 |
| 17 | 20252000 | 20254399 | 27.16516 | 47.27106 | 37.30331 | 16.39417 | 0.727269 | 0.808949 | 0.506374 |
| 10 | 67188600 | 67190799 | 15.84485 | 27.68954 | 46.22861 | 20.37136 | -0.34844 | 0.707123 | 0.506389 |
| 4 | 140805200 | 140812799 | 66.48813 | 104.1377 | 161.8722 | 67.1448 | -0.31927 | 2.607459 | 0.506408 |

| | | | | | | | | | |
|---|---|---|---|---|---|---|---|---|---|
| 8 | 68673400 | 68674199 | 11.90227 | 21.08499 | 36.78621 | 16.2648 | -0.40657 | 0.491257 | 0.506421 |
| 2 | 52298400 | 52301599 | 48.6994 | 78.95485 | 71.53565 | 30.63594 | 0.507362 | 1.655803 | 0.506451 |
| 10 | 61117000 | 61121999 | 38.69624 | 62.25152 | 99.41068 | 42.19478 | -0.32086 | 1.855535 | 0.506463 |
| 16 | 39255600 | 39256199 | 8.626359 | 16.11748 | 29.53683 | 12.25313 | -0.41115 | 0.130695 | 0.506486 |
| 3 | 127291400 | 127293999 | 25.18037 | 41.53329 | 70.15368 | 30.27588 | -0.37395 | 1.357619 | 0.506489 |
| 6 | 130885000 | 130886999 | 34.17388 | 57.80991 | 46.84385 | 20.50253 | 0.683764 | 1.10727 | 0.506502 |
| 1 | 105282600 | 105285799 | 21.49673 | 35.30135 | 78.83917 | 33.9608 | -0.77812 | 1.411325 | 0.506503 |
| 8 | 12173000 | 12176799 | 40.0746 | 65.10743 | 68.34394 | 29.31412 | 0.298289 | 1.596091 | 0.50657 |
| 1 | 125362400 | 125364199 | 13.15608 | 23.09463 | 40.41602 | 17.87788 | -0.41572 | 0.611169 | 0.506595 |
| 17 | 63848400 | 63853599 | 57.64317 | 92.39827 | 90.96536 | 38.40328 | 0.369312 | 1.987842 | 0.506611 |
| 1 | 7275800 | 7277399 | 11.56834 | 20.45742 | 38.29254 | 16.93677 | -0.50877 | 0.508779 | 0.506623 |
| 6 | 145563800 | 145565799 | 17.25086 | 29.08606 | 61.64486 | 26.95361 | -0.69846 | 1.091502 | 0.506633 |
| 2 | 111541800 | 111548399 | 59.65943 | 95.52365 | 93.89414 | 39.57259 | 0.369701 | 2.025853 | 0.506635 |
| 4 | 54654400 | 54656199 | 20.36047 | 33.92343 | 67.65467 | 29.39586 | -0.61136 | 1.228167 | 0.506638 |
| 10 | 122637800 | 122642399 | 34.12206 | 55.19466 | 87.75391 | 37.51826 | -0.30513 | 1.682016 | 0.506651 |
| 8 | 63917200 | 63919599 | 30.81604 | 51.39823 | 53.55489 | 23.28506 | 0.322053 | 1.233532 | 0.506651 |
| 11 | 111344200 | 111344599 | 5.355684 | 10.43259 | 21.91462 | 7.990025 | -0.43491 | -0.3656 | 0.506686 |
| 17 | 38151200 | 38153799 | 29.64871 | 49.80179 | 50.74498 | 22.1619 | 0.353977 | 1.153189 | 0.506708 |
| 14 | 30630000 | 30631799 | 9.931914 | 17.93936 | 33.81046 | 14.69151 | -0.49992 | 0.345085 | 0.50672 |
| 18 | 85658400 | 85660599 | 17.19461 | 31.51123 | 26.51211 | 11.31043 | 0.673487 | 0.299737 | 0.506738 |
| 2 | 66836600 | 66837799 | 8.912091 | 16.81598 | 31.44704 | 12.82753 | -0.42085 | 0.073441 | 0.50675 |
| X | 86665200 | 86665799 | 10.67446 | 20.34913 | 21.74351 | 8.678909 | 0.408279 | -0.0052 | 0.506769 |
| 2 | 116731800 | 116733999 | 16.0798 | 27.75209 | 47.94003 | 21.03088 | -0.39999 | 0.859809 | 0.506771 |
| 2 | 162859000 | 162859599 | 5.235628 | 10.1809 | 21.75751 | 8.060138 | -0.48099 | -0.31288 | 0.506777 |
| 5 | 141967800 | 141968399 | 5.235628 | 10.1809 | 21.75751 | 8.060138 | -0.48099 | -0.31288 | 0.506777 |
| 8 | 113208800 | 113210399 | 17.83932 | 30.90355 | 51.71282 | 22.68855 | -0.35338 | 0.838856 | 0.506784 |
| 18 | 37153000 | 37159999 | 131.6289 | 200.005 | 204.1116 | 84.53806 | 0.253067 | 3.204399 | 0.506787 |
| 3 | 46791200 | 46791599 | 5.207489 | 10.16932 | 23.25672 | 8.174116 | -0.50938 | -0.49662 | 0.506788 |
| 8 | 33558400 | 33560199 | 20.58807 | 36.00478 | 36.53941 | 16.11159 | 0.366478 | 0.705668 | 0.506797 |
| 13 | 120128600 | 120132399 | 31.94701 | 52.80874 | 55.20861 | 23.85882 | 0.316586 | 1.33802 | 0.506802 |
| 17 | 39008000 | 39012599 | 62.00041 | 99.46364 | 88.84153 | 37.55315 | 0.510875 | 1.972026 | 0.506806 |
| 13 | 23599600 | 23600799 | 7.340942 | 13.91143 | 27.09959 | 10.95355 | -0.46732 | 0.016851 | 0.506818 |
| 13 | 44886600 | 44887799 | 7.340942 | 13.91143 | 27.09959 | 10.95355 | -0.46732 | 0.016851 | 0.506818 |
| 17 | 89888000 | 89888399 | 8.123386 | 15.80567 | 18.39515 | 6.781861 | 0.3922 | -0.30756 | 0.506819 |
| 18 | 3994600 | 3996399 | 14.15276 | 24.85367 | 41.41735 | 18.3267 | -0.34544 | 0.61421 | 0.506827 |
| 16 | 6056400 | 6058599 | 17.1317 | 30.78807 | 28.93993 | 12.64801 | 0.491882 | 0.413487 | 0.506828 |
| 4 | 40141600 | 40144999 | 25.64438 | 41.98732 | 80.39959 | 34.58938 | -0.5587 | 1.459397 | 0.506843 |
| 6 | 142004400 | 142006599 | 20.73207 | 36.10139 | 37.16637 | 16.34104 | 0.345352 | 0.766724 | 0.50686 |
| 8 | 28191400 | 28192199 | 20.15349 | 34.39586 | 56.54798 | 24.79111 | -0.33208 | 0.988209 | 0.506865 |
| 13 | 22048800 | 22050799 | 17.35036 | 30.81448 | 31.18319 | 13.76496 | 0.376542 | 0.494221 | 0.506885 |
| 18 | 7843800 | 7846799 | 62.76161 | 99.95821 | 105.2892 | 44.04441 | 0.261817 | 2.197299 | 0.506891 |
| 14 | 116666000 | 116667799 | 17.02246 | 30.97548 | 28.15318 | 12.1203 | 0.554336 | 0.342507 | 0.506918 |
| 12 | 48859400 | 48860199 | 10.82269 | 20.67244 | 22.19924 | 8.823929 | 0.405704 | -0.01865 | 0.506921 |
| 1 | 41166600 | 41167799 | 8.954741 | 16.71375 | 30.58555 | 12.7104 | -0.41119 | 0.143041 | 0.506926 |
| 9 | 60432200 | 60433599 | 9.06834 | 16.89419 | 30.00965 | 12.51059 | -0.37252 | 0.152943 | 0.506928 |
| 13 | 98666200 | 98668399 | 26.73801 | 45.56137 | 45.76208 | 20.05992 | 0.37575 | 1.013531 | 0.506935 |
| 11 | 54713800 | 54717999 | 41.05529 | 65.99791 | 102.2405 | 43.34955 | -0.27845 | 1.885189 | 0.506937 |

| | | | | | | | | | |
|---|---|---|---|---|---|---|---|---|---|
| X | 23682400 | 23693999 | 136.9824 | 209.7123 | 184.9591 | 76.755 | 0.472086 | 3.055504 | 0.506959 |
| 9 | 76011600 | 76012999 | 14.93309 | 27.33864 | 27.28741 | 11.65745 | 0.427172 | 0.290253 | 0.506965 |
| 18 | 6489800 | 6491199 | 10.82938 | 19.16647 | 36.86364 | 16.30207 | -0.54691 | 0.494167 | 0.506978 |
| 1 | 69029800 | 69030999 | 15.56707 | 28.8072 | 26.36143 | 11.0984 | 0.566936 | 0.235457 | 0.506985 |
| 16 | 76622000 | 76623399 | 16.05941 | 29.41189 | 27.1681 | 11.60044 | 0.538745 | 0.29619 | 0.507006 |
| 2 | 39316000 | 39322599 | 69.97478 | 111.3992 | 108.0132 | 45.15745 | 0.380462 | 2.217623 | 0.507027 |
| 7 | 66598000 | 66598799 | 10.37892 | 19.08454 | 33.52197 | 14.23582 | -0.37646 | 0.247196 | 0.50703 |
| 10 | 45434800 | 45435199 | 4.94067 | 9.626133 | 20.77287 | 7.553504 | -0.46985 | -0.38594 | 0.507036 |
| 12 | 83798200 | 83801399 | 20.33384 | 34.48708 | 55.93033 | 24.49879 | -0.31357 | 1.044599 | 0.507041 |
| 10 | 93471000 | 93471599 | 6.422441 | 12.36263 | 24.70748 | 9.609799 | -0.45238 | -0.12304 | 0.507048 |
| 3 | 98569200 | 98569799 | 9.277736 | 17.1131 | 30.94645 | 13.08631 | -0.41329 | 0.215843 | 0.507068 |
| 6 | 58011600 | 58012399 | 8.892369 | 16.53885 | 29.97723 | 12.53229 | -0.40487 | 0.160746 | 0.507074 |
| 12 | 23759000 | 23833599 | 2424.321 | 2908.435 | 2337.579 | 988.2988 | 0.260649 | 7.025667 | 0.507076 |
| 7 | 113858400 | 113864199 | 71.86123 | 114.5671 | 103.0955 | 43.18734 | 0.4902 | 2.175035 | 0.507084 |
| 1 | 103216000 | 103217199 | 16.4813 | 29.42014 | 30.20434 | 13.27587 | 0.360017 | 0.454786 | 0.507085 |
| 2 | 64098400 | 64101399 | 33.38172 | 55.40618 | 55.91648 | 24.23297 | 0.367716 | 1.29541 | 0.50709 |
| 7 | 31708000 | 31712199 | 37.44458 | 62.06043 | 56.64979 | 24.52645 | 0.512009 | 1.324674 | 0.507095 |
| 17 | 65695400 | 65696399 | 10.98697 | 21.02109 | 22.22717 | 8.796926 | 0.43354 | -0.03077 | 0.507103 |
| 7 | 27666400 | 27671199 | 76.51578 | 121.1097 | 122.5197 | 50.93385 | 0.311393 | 2.390573 | 0.507143 |
| 4 | 24140800 | 24143399 | 24.55908 | 42.37208 | 41.98433 | 18.41798 | 0.398341 | 0.891697 | 0.507154 |
| 13 | 51242600 | 51245599 | 30.24463 | 50.50703 | 52.45826 | 22.8259 | 0.326584 | 1.216862 | 0.50717 |
| 1 | 93498200 | 93499999 | 37.05887 | 60.81526 | 54.19556 | 23.33852 | 0.543004 | 1.438437 | 0.507171 |
| 3 | 68405000 | 68406199 | 12.33555 | 23.07108 | 24.54888 | 10.16647 | 0.368898 | 0.140771 | 0.507179 |
| 7 | 12299000 | 12300799 | 30.72092 | 51.20763 | 53.52274 | 23.26192 | 0.317628 | 1.240897 | 0.507188 |
| 1 | 105178600 | 105179999 | 14.50682 | 26.96545 | 25.92755 | 10.84955 | 0.502197 | 0.201017 | 0.507265 |
| 14 | 120295800 | 120297399 | 18.68494 | 31.88028 | 52.7041 | 23.10538 | -0.34001 | 0.98082 | 0.507289 |
| 12 | 96901600 | 96907399 | 93.35394 | 145.4326 | 150.8872 | 62.6261 | 0.257234 | 2.713339 | 0.507301 |
| 13 | 44017200 | 44020999 | 26.0091 | 42.85937 | 70.11932 | 30.24885 | -0.32856 | 1.372498 | 0.507308 |
| 7 | 33082800 | 33114599 | 575.5827 | 781.4388 | 683.0009 | 284.5215 | 0.319432 | 5.040003 | 0.507337 |
| 6 | 130765600 | 130770799 | 56.45687 | 91.09215 | 76.62944 | 32.65871 | 0.607066 | 1.785953 | 0.50734 |
| 7 | 4754800 | 4762999 | 53.63261 | 84.4783 | 163.5191 | 67.84934 | -0.62895 | 2.491365 | 0.507345 |
| 5 | 55925400 | 55925999 | 7.674103 | 14.49847 | 27.58188 | 11.22069 | -0.44123 | 0.044933 | 0.507362 |
| 8 | 66382800 | 66383399 | 12.33119 | 22.86883 | 24.01908 | 10.08236 | 0.372461 | 0.19112 | 0.507363 |
| X | 126842200 | 126844399 | 27.39592 | 47.01043 | 36.83263 | 16.15478 | 0.734296 | 0.946545 | 0.507368 |
| 3 | 5579800 | 5583199 | 36.79605 | 60.0193 | 62.2519 | 26.74478 | 0.319682 | 1.52395 | 0.507378 |
| 1 | 38755600 | 38756199 | 5.508601 | 10.73024 | 21.94219 | 8.002291 | -0.39744 | -0.36414 | 0.507428 |
| 6 | 67105600 | 67109399 | 6.938705 | 12.71385 | 35.08306 | 14.97041 | -1.02793 | 0.243098 | 0.507447 |
| 11 | 74262400 | 74262799 | 4.480154 | 8.748408 | 19.30248 | 6.791678 | -0.45743 | -0.51303 | 0.50745 |
| 8 | 116080000 | 116081799 | 24.77493 | 42.9073 | 40.81397 | 17.90663 | 0.458003 | 0.856435 | 0.507481 |
| 12 | 28460800 | 28463199 | 34.9194 | 57.91768 | 50.93202 | 22.06211 | 0.565735 | 1.305273 | 0.507511 |
| 6 | 58101000 | 58108199 | 93.11676 | 145.5454 | 146.7806 | 60.89766 | 0.301827 | 2.655717 | 0.507515 |
| 12 | 69573400 | 69575199 | 12.79631 | 22.76566 | 38.22482 | 16.84878 | -0.34852 | 0.467612 | 0.507519 |
| 9 | 23458400 | 23459999 | 16.20652 | 28.84603 | 29.82668 | 13.14191 | 0.347514 | 0.471005 | 0.507539 |
| 13 | 15941400 | 15942999 | 16.39857 | 28.3181 | 47.93894 | 21.03076 | -0.37079 | 0.856443 | 0.507542 |
| 4 | 112006400 | 112008199 | 29.51038 | 49.93631 | 41.63377 | 18.22389 | 0.642759 | 1.082407 | 0.50755 |
| 1 | 168081800 | 168083799 | 18.82416 | 33.26145 | 32.65363 | 14.44504 | 0.41679 | 0.552784 | 0.507557 |
| 12 | 110474400 | 110476399 | 9.579961 | 17.11284 | 34.93387 | 15.34558 | -0.62444 | 0.417105 | 0.507566 |

| | | | | | | | | | |
|---|---|---|---|---|---|---|---|---|---|
| 16 | 90706200 | 90709999 | 21.50562 | 35.62538 | 66.5187 | 28.78919 | -0.51724 | 1.294832 | 0.507567 |
| 19 | 56810200 | 56810999 | 7.527838 | 14.1959 | 27.55558 | 11.2533 | -0.47491 | 0.056188 | 0.507603 |
| 6 | 94264600 | 94268199 | 43.49215 | 70.47224 | 71.13338 | 30.45084 | 0.35106 | 1.653596 | 0.507611 |
| 4 | 91257400 | 91261599 | 38.76184 | 62.32783 | 98.14483 | 41.6292 | -0.30144 | 1.871925 | 0.507612 |
| 15 | 5995000 | 5996399 | 12.0158 | 22.58829 | 24.09197 | 9.891195 | 0.375495 | 0.104764 | 0.507613 |
| 13 | 12768800 | 12771199 | 15.89875 | 27.81931 | 48.35806 | 21.33279 | -0.40641 | 0.690676 | 0.507617 |
| 15 | 14336800 | 14337999 | 13.55744 | 24.89782 | 25.84375 | 10.99389 | 0.375331 | 0.258461 | 0.507633 |
| 2 | 178254800 | 178256799 | 11.57322 | 20.74779 | 35.79472 | 15.67621 | -0.38057 | 0.409349 | 0.507661 |
| X | 102136600 | 102138199 | 23.06326 | 40.27391 | 37.03833 | 16.31356 | 0.507985 | 0.742913 | 0.507675 |
| 7 | 99196800 | 99203799 | 41.2123 | 65.64112 | 130.7074 | 54.68165 | -0.6552 | 2.182835 | 0.507688 |
| 4 | 31358200 | 31360799 | 13.29739 | 23.53937 | 39.23763 | 17.3512 | -0.34188 | 0.511324 | 0.507741 |
| 8 | 67494200 | 67497399 | 45.09995 | 73.00413 | 73.9449 | 31.63098 | 0.344767 | 1.679729 | 0.507745 |
| 2 | 164339200 | 164340399 | 6.913009 | 13.11005 | 26.58343 | 10.72833 | -0.52218 | 0.008069 | 0.507764 |
| 8 | 14343200 | 14344199 | 9.579944 | 17.50372 | 31.86462 | 13.64831 | -0.4365 | 0.272121 | 0.507766 |
| 4 | 140937200 | 140939999 | 22.06676 | 36.81102 | 61.77588 | 26.86211 | -0.36322 | 1.21457 | 0.507794 |
| 6 | 29618600 | 29622399 | 40.96706 | 66.51307 | 69.70703 | 29.8841 | 0.299683 | 1.610102 | 0.507794 |
| 7 | 6100800 | 6102599 | 32.75025 | 54.01875 | 55.65909 | 24.02042 | 0.337115 | 1.36834 | 0.507805 |
| 13 | 7930600 | 7931399 | 11.22996 | 20.18825 | 34.78391 | 15.19799 | -0.37636 | 0.386965 | 0.507808 |
| 1 | 117962200 | 117974399 | 94.3762 | 147.6989 | 131.7094 | 54.63776 | 0.480504 | 2.632546 | 0.507816 |
| 4 | 44399400 | 44401399 | 10.09929 | 18.31222 | 32.84756 | 14.20437 | -0.42416 | 0.324111 | 0.507819 |
| 12 | 19311200 | 19316399 | 148.093 | 223.5771 | 226.883 | 93.78215 | 0.253423 | 3.339578 | 0.507839 |
| 7 | 59856200 | 59857399 | 23.1604 | 40.55627 | 30.93236 | 13.6543 | 0.776323 | 0.688286 | 0.507856 |
| 6 | 46080200 | 46081999 | 42.31049 | 68.77196 | 61.27072 | 26.28928 | 0.534919 | 1.586972 | 0.507857 |
| 2 | 119598000 | 119598799 | 5.823059 | 11.2784 | 23.25025 | 8.82722 | -0.46347 | -0.21633 | 0.507859 |
| 12 | 28305600 | 28307799 | 14.48245 | 26.9459 | 24.12153 | 10.07785 | 0.605776 | 0.187854 | 0.507866 |
| 13 | 93932800 | 93935799 | 21.9672 | 36.98078 | 59.78352 | 26.12748 | -0.30952 | 1.117876 | 0.507897 |
| 4 | 107110200 | 107111399 | 8.553095 | 15.62308 | 31.43157 | 13.46763 | -0.57965 | 0.266278 | 0.50794 |
| 1 | 129948600 | 129949199 | 10.46066 | 20.09795 | 21.83587 | 8.538951 | 0.411209 | -0.07397 | 0.507975 |
| 7 | 42303000 | 42321599 | 396.6098 | 558.9387 | 466.1763 | 193.0632 | 0.441269 | 4.467754 | 0.508 |
| 13 | 50776400 | 50778599 | 13.9637 | 24.60552 | 40.82864 | 18.07135 | -0.33796 | 0.56901 | 0.508003 |
| X | 149015200 | 149018399 | 38.56313 | 64.36542 | 51.51459 | 22.40802 | 0.701477 | 1.248976 | 0.50802 |
| 3 | 137407600 | 137409599 | 19.03968 | 33.74382 | 31.40893 | 13.88604 | 0.494714 | 0.518348 | 0.508026 |
| X | 9390600 | 9391999 | 18.51108 | 33.83693 | 27.06927 | 11.58795 | 0.742682 | 0.327015 | 0.508028 |
| 9 | 60638000 | 60642999 | 58.49951 | 92.56566 | 141.0815 | 58.64147 | -0.2794 | 2.387683 | 0.50803 |
| 12 | 67328600 | 67328999 | 6.100513 | 11.84855 | 23.43353 | 8.765144 | -0.38404 | -0.26368 | 0.50805 |
| 6 | 137972400 | 137974999 | 17.69412 | 30.68237 | 51.71334 | 22.68946 | -0.3634 | 0.829767 | 0.508058 |
| 8 | 52445200 | 52445599 | 10.40243 | 19.75776 | 21.16797 | 8.515156 | 0.394353 | 0.01403 | 0.508062 |
| 1 | 112809200 | 112811999 | 25.99136 | 45.46755 | 35.0312 | 15.45041 | 0.762117 | 0.727352 | 0.50807 |
| 8 | 5622800 | 5626399 | 43.98409 | 71.35535 | 71.68502 | 30.71429 | 0.359517 | 1.633341 | 0.508103 |
| 1 | 103169200 | 103173399 | 54.83289 | 88.01702 | 89.29726 | 37.77961 | 0.328414 | 1.94254 | 0.508111 |
| 7 | 37345600 | 37348999 | 23.85566 | 39.4783 | 69.08415 | 29.87858 | -0.42424 | 1.313813 | 0.508136 |
| 18 | 3863600 | 3865599 | 13.45912 | 23.83923 | 40.26558 | 17.80374 | -0.36057 | 0.507796 | 0.508145 |
| 3 | 129255600 | 129259799 | 59.69878 | 94.33584 | 143.3171 | 59.53003 | -0.27631 | 2.421422 | 0.508153 |
| 8 | 47986200 | 47987199 | 7.492078 | 14.19398 | 27.09435 | 10.95774 | -0.43894 | 0.020627 | 0.50816 |
| 19 | 39840600 | 39843999 | 36.87813 | 61.67722 | 50.35281 | 21.93054 | 0.672782 | 1.220224 | 0.508168 |
| 14 | 53826800 | 53829199 | 23.18866 | 38.8449 | 63.05094 | 27.48909 | -0.31539 | 1.174801 | 0.50821 |
| 10 | 21250000 | 21253199 | 21.43946 | 35.7172 | 62.25368 | 27.04724 | -0.41764 | 1.227014 | 0.508219 |

| | | | | | | | | | |
|---|---|---|---|---|---|---|---|---|---|
| 3 | 97882200 | 97885199 | 23.20242 | 38.69093 | 62.94698 | 27.36484 | -0.31863 | 1.22343 | 0.508234 |
| 8 | 109801400 | 109803999 | 6.156249 | 11.77956 | 28.41742 | 11.24422 | -0.74545 | -0.0607 | 0.508243 |
| X | 78787400 | 78789399 | 22.58284 | 39.36979 | 37.18395 | 16.36068 | 0.469581 | 0.764114 | 0.50826 |
| 17 | 18435400 | 18447399 | 188.9091 | 282.735 | 257.8596 | 106.347 | 0.396313 | 3.515397 | 0.508264 |
| 9 | 56902000 | 56903399 | 9.761753 | 17.71393 | 32.8357 | 14.18547 | -0.47055 | 0.315249 | 0.508268 |
| 8 | 109130800 | 109132599 | 12.83362 | 22.58417 | 39.34124 | 17.41763 | -0.40842 | 0.573494 | 0.508329 |
| 2 | 30950400 | 30952399 | 16.372 | 28.42577 | 46.87009 | 20.56964 | -0.33139 | 0.80416 | 0.508332 |
| 16 | 45897000 | 45897799 | 6.390048 | 12.40815 | 24.64468 | 9.234577 | -0.393 | -0.25164 | 0.508335 |
| 1 | 100537200 | 100538999 | 18.14769 | 32.74724 | 28.98697 | 12.61084 | 0.58254 | 0.398848 | 0.508367 |
| 4 | 17940800 | 17942999 | 15.09712 | 26.51069 | 46.65907 | 20.6452 | -0.42419 | 0.62696 | 0.508372 |
| 6 | 68471600 | 68473599 | 32.50321 | 54.27737 | 53.60212 | 23.32325 | 0.399146 | 1.226562 | 0.508386 |
| 15 | 82354600 | 82356799 | 22.03808 | 36.94178 | 60.21866 | 26.26517 | -0.32148 | 1.161401 | 0.508388 |
| 16 | 94708200 | 94709799 | 18.18464 | 32.05775 | 32.82401 | 14.52707 | 0.355222 | 0.574402 | 0.508392 |
| 9 | 3256400 | 3266199 | 208.1556 | 306.405 | 304.583 | 125.4289 | 0.250003 | 3.806143 | 0.508403 |
| 4 | 66494800 | 66495799 | 12.03448 | 21.3401 | 36.73636 | 16.23758 | -0.38688 | 0.481207 | 0.508408 |
| 5 | 100605800 | 100608199 | 18.82272 | 32.96245 | 33.99047 | 15.00496 | 0.343411 | 0.669019 | 0.508414 |
| 8 | 25238400 | 25240199 | 11.13911 | 20.34502 | 36.06819 | 15.4559 | -0.39938 | 0.294479 | 0.508432 |
| X | 61755200 | 61758799 | 37.57461 | 63.09466 | 48.01421 | 20.96499 | 0.773748 | 1.181143 | 0.508441 |
| 8 | 34494800 | 34498599 | 33.79729 | 56.09945 | 56.45586 | 24.46778 | 0.371834 | 1.297475 | 0.508468 |
| 9 | 19922000 | 19933999 | 169.9942 | 255.3254 | 248.251 | 102.4673 | 0.3086 | 3.445333 | 0.508469 |
| X | 118614800 | 118616599 | 24.29747 | 42.31117 | 38.60685 | 16.97469 | 0.518888 | 0.788484 | 0.508478 |
| 15 | 26088600 | 26089199 | 8.406992 | 15.82646 | 28.5869 | 11.70865 | -0.37592 | 0.075901 | 0.508487 |
| 14 | 7636600 | 7637599 | 5.567202 | 10.73038 | 24.17923 | 9.369132 | -0.61871 | -0.14752 | 0.508487 |
| 5 | 9661400 | 9662399 | 13.15532 | 24.62098 | 25.02906 | 10.35343 | 0.43559 | 0.14467 | 0.508497 |
| 12 | 48176800 | 48177999 | 13.15532 | 24.62098 | 25.02906 | 10.35343 | 0.43559 | 0.14467 | 0.508497 |
| 2 | 140312800 | 140314399 | 25.65349 | 43.54202 | 45.83685 | 20.08217 | 0.307618 | 1.045311 | 0.508529 |
| 17 | 36964200 | 36965199 | 14.83594 | 26.67193 | 27.38361 | 11.96417 | 0.365942 | 0.394235 | 0.508532 |
| 5 | 126869800 | 126874999 | 51.66864 | 83.40465 | 77.4337 | 33.021 | 0.465829 | 1.763757 | 0.508545 |
| X | 149010800 | 149014399 | 55.2004 | 88.94613 | 80.44533 | 34.22522 | 0.500558 | 1.822691 | 0.508577 |
| 7 | 96281000 | 96282799 | 16.72254 | 29.53912 | 29.83347 | 13.19827 | 0.375846 | 0.536667 | 0.508582 |
| X | 31633400 | 31635599 | 46.28721 | 76.0863 | 58.11009 | 25.04194 | 0.765992 | 1.445596 | 0.508587 |
| 4 | 63316000 | 63317999 | 14.60239 | 25.46778 | 43.67159 | 19.22027 | -0.38688 | 0.726412 | 0.50859 |
| 10 | 103883400 | 103884799 | 21.34711 | 37.4332 | 35.21277 | 15.56212 | 0.475267 | 0.663955 | 0.508594 |
| 6 | 21062800 | 21063199 | 4.912815 | 9.588367 | 20.42782 | 7.251214 | -0.41974 | -0.47219 | 0.508614 |
| 3 | 54714200 | 54716999 | 20.58027 | 34.47767 | 59.60693 | 25.98864 | -0.40603 | 1.161286 | 0.508619 |
| 12 | 25126400 | 25127799 | 15.93273 | 28.69888 | 28.01386 | 12.21484 | 0.439868 | 0.389148 | 0.508619 |
| 10 | 101824400 | 101825199 | 9.85042 | 18.85204 | 20.14356 | 7.965637 | 0.419734 | -0.05075 | 0.508637 |
| 3 | 20148200 | 20149799 | 16.33656 | 29.98152 | 26.60028 | 11.32602 | 0.599346 | 0.285329 | 0.508638 |
| 4 | 113805400 | 113807399 | 29.86012 | 50.345 | 49.63301 | 21.70141 | 0.401627 | 1.121554 | 0.508649 |
| 14 | 93189200 | 93191599 | 26.78304 | 45.82817 | 44.65182 | 19.57979 | 0.420338 | 0.97603 | 0.508652 |
| 8 | 41223200 | 41229599 | 93.06284 | 145.1835 | 149.2158 | 61.92018 | 0.272397 | 2.690027 | 0.508667 |
| 1 | 16004200 | 16006199 | 16.14289 | 27.69298 | 55.36877 | 24.28418 | -0.61196 | 0.923689 | 0.50873 |
| 4 | 62452000 | 62454399 | 16.14289 | 27.69298 | 55.36877 | 24.28418 | -0.61196 | 0.923689 | 0.50873 |
| 13 | 48387000 | 48389199 | 16.14289 | 27.69298 | 55.36877 | 24.28418 | -0.61196 | 0.923689 | 0.50873 |
| 1 | 108891200 | 108891799 | 15.19201 | 27.28534 | 28.34606 | 12.39491 | 0.348204 | 0.405033 | 0.508753 |
| 9 | 14073400 | 14075199 | 34.33959 | 57.07082 | 55.9067 | 24.25068 | 0.410731 | 1.285152 | 0.508759 |
| 5 | 104071000 | 104072999 | 12.6613 | 22.6374 | 38.82036 | 17.03991 | -0.37459 | 0.437625 | 0.508763 |

| | | | | | | | | | |
|---|---|---|---|---|---|---|---|---|---|
| 11 | 19181600 | 19184799 | 30.64496 | 51.70462 | 48.53648 | 21.22649 | 0.471996 | 1.114745 | 0.508765 |
| 12 | 61963200 | 61963799 | 11.93745 | 22.31047 | 22.63386 | 9.384813 | 0.436259 | 0.138842 | 0.508783 |
| 5 | 147916800 | 147929799 | 134.1198 | 201.3033 | 290.0297 | 119.6895 | -0.25928 | 3.45363 | 0.508792 |
| 2 | 81295600 | 81298199 | 33.83444 | 55.7112 | 59.07615 | 25.47656 | 0.294755 | 1.392996 | 0.508801 |
| 7 | 36467200 | 36471399 | 45.82209 | 72.74977 | 139.8227 | 58.27304 | -0.60901 | 2.288864 | 0.508806 |
| 3 | 98602400 | 98603199 | 8.263837 | 15.69271 | 28.81626 | 11.5967 | -0.37731 | 0.017086 | 0.508829 |
| 8 | 36651000 | 36651599 | 5.807008 | 11.27924 | 23.38557 | 8.742503 | -0.44987 | -0.26642 | 0.50883 |
| 2 | 53114000 | 53121799 | 111.9572 | 173.5721 | 160.5842 | 66.69903 | 0.416991 | 2.806341 | 0.508839 |
| 7 | 128284000 | 128292999 | 119.7329 | 180.3942 | 295.4934 | 122.0684 | -0.43942 | 3.37787 | 0.508843 |
| 17 | 23945000 | 23946199 | 10.6421 | 19.07752 | 34.85342 | 15.26465 | -0.46248 | 0.402215 | 0.508844 |
| 13 | 19248600 | 19250799 | 34.80052 | 56.23444 | 88.62623 | 37.84489 | -0.29414 | 1.712621 | 0.508846 |
| 10 | 4661400 | 4663999 | 19.01493 | 32.2733 | 54.83087 | 24.02243 | -0.38035 | 1.031038 | 0.508859 |
| 19 | 51430400 | 51431599 | 9.540743 | 17.71382 | 31.01354 | 12.99837 | -0.35794 | 0.179911 | 0.508876 |
| 4 | 112012800 | 112020199 | 86.88997 | 137.7089 | 115.5584 | 48.07579 | 0.58226 | 2.366927 | 0.508878 |
| 14 | 66698000 | 66700599 | 37.93908 | 61.77558 | 62.0185 | 26.62752 | 0.364899 | 1.554204 | 0.508911 |
| 16 | 86895200 | 86896599 | 15.17141 | 27.82356 | 27.3277 | 11.64691 | 0.452832 | 0.283284 | 0.508915 |
| 5 | 133491200 | 133493799 | 18.43225 | 33.32306 | 28.41474 | 12.33298 | 0.63823 | 0.388798 | 0.508917 |
| 11 | 55358600 | 55361399 | 18.04567 | 30.75938 | 52.68864 | 23.09678 | -0.39124 | 0.986771 | 0.508961 |
| 5 | 107337000 | 107337999 | 7.920315 | 15.12333 | 27.86102 | 11.08259 | -0.36727 | -0.02762 | 0.508962 |
| 5 | 16138600 | 16141199 | 19.20619 | 32.12397 | 66.62008 | 29.0169 | -0.66762 | 1.180756 | 0.508986 |
| 14 | 100515800 | 100517999 | 26.67215 | 45.07385 | 47.46377 | 20.76745 | 0.306979 | 1.088258 | 0.508987 |
| 7 | 59353400 | 59366199 | 251.6158 | 366.1078 | 305.9267 | 126.0915 | 0.484344 | 3.988703 | 0.508988 |
| 19 | 21753800 | 21756999 | 21.4869 | 35.63315 | 66.02201 | 28.59571 | -0.50603 | 1.281454 | 0.509017 |
| 5 | 145883000 | 145886599 | 50.9596 | 82.73648 | 68.3296 | 29.29352 | 0.642595 | 1.631877 | 0.509022 |
| 10 | 40864400 | 40864799 | 4.350115 | 8.498226 | 18.78521 | 6.559633 | -0.45024 | -0.54464 | 0.509038 |
| 18 | 89403000 | 89403399 | 4.385563 | 8.574423 | 20.35792 | 6.969651 | -0.52808 | -0.6127 | 0.509052 |
| 4 | 113127200 | 113132799 | 70.12382 | 111.6351 | 98.36771 | 41.12425 | 0.518222 | 2.216084 | 0.509061 |
| 9 | 19374000 | 19376999 | 58.12962 | 92.93486 | 97.25485 | 40.90154 | 0.277141 | 2.07179 | 0.509062 |
| 9 | 119239600 | 119244599 | 83.82232 | 129.7649 | 190.2494 | 79.02466 | -0.24812 | 2.823566 | 0.509071 |
| 16 | 90938200 | 90942599 | 53.86702 | 85.60957 | 129.6519 | 54.0894 | -0.2644 | 2.261911 | 0.509084 |
| 8 | 15148000 | 15149799 | 10.69042 | 19.48047 | 33.36208 | 14.33764 | -0.35219 | 0.300121 | 0.509084 |
| 10 | 102141000 | 102142799 | 16.55204 | 29.95191 | 28.58594 | 12.39277 | 0.477658 | 0.370038 | 0.509105 |
| 12 | 98032200 | 98033999 | 16.50412 | 29.42193 | 29.58152 | 13.01663 | 0.388711 | 0.460144 | 0.509108 |
| 6 | 60369800 | 60370799 | 13.07298 | 24.08809 | 25.02855 | 10.59904 | 0.378976 | 0.236671 | 0.50912 |
| 18 | 6364400 | 6366399 | 19.26171 | 33.04899 | 54.96384 | 24.10692 | -0.34752 | 0.933195 | 0.509121 |
| 17 | 17173000 | 17231599 | 1467.197 | 1848.356 | 1441.315 | 605.6712 | 0.375881 | 6.247676 | 0.50917 |
| X | 77051200 | 77058999 | 99.04272 | 156.2905 | 120.8295 | 50.15413 | 0.695389 | 2.478957 | 0.509171 |
| 19 | 41673800 | 41674399 | 4.944063 | 9.622098 | 21.23581 | 7.813146 | -0.51729 | -0.34183 | 0.509185 |
| 18 | 63607800 | 63608399 | 10.56213 | 19.13309 | 33.32161 | 14.42666 | -0.38308 | 0.333831 | 0.509191 |
| 2 | 86996000 | 87000399 | 76.8385 | 121.0147 | 126.1135 | 52.32766 | 0.262946 | 2.501607 | 0.509192 |
| X | 38071200 | 38073599 | 28.80766 | 48.98732 | 46.31629 | 20.2995 | 0.462359 | 1.043237 | 0.509197 |
| 11 | 11291600 | 11293599 | 28.08193 | 45.77867 | 76.806 | 32.99051 | -0.37287 | 1.524714 | 0.509198 |
| 1 | 142414000 | 142415199 | 13.21725 | 24.91442 | 23.99282 | 9.805313 | 0.526807 | 0.097148 | 0.509214 |
| 13 | 88308400 | 88310999 | 83.77776 | 131.2237 | 137.0874 | 56.86377 | 0.253358 | 2.612209 | 0.509215 |
| 1 | 11304200 | 11306199 | 18.51527 | 32.44981 | 33.48279 | 14.79077 | 0.34251 | 0.654042 | 0.509228 |
| 7 | 15556200 | 15558999 | 42.33735 | 68.71739 | 70.92938 | 30.40146 | 0.321205 | 1.62017 | 0.509244 |
| 10 | 105842600 | 105843799 | 9.473966 | 17.70012 | 32.28366 | 13.39395 | -0.40538 | 0.140866 | 0.509246 |

| | | | | | | | | | |
|---|---|---|---|---|---|---|---|---|---|
| 4 | 139903800 | 139904999 | 18.6251 | 31.48663 | 55.28208 | 24.19102 | -0.42769 | 1.06689 | 0.50925 |
| 12 | 23362000 | 23367399 | 93.60424 | 146.1982 | 120.5251 | 50.00841 | 0.591421 | 2.66578 | 0.509266 |
| 11 | 68371600 | 68372799 | 7.925156 | 15.03376 | 27.67024 | 11.16028 | -0.38359 | 0.016715 | 0.509287 |
| 10 | 6521000 | 6523799 | 55.61495 | 88.87831 | 91.51871 | 38.46694 | 0.299779 | 2.079365 | 0.509287 |
| 13 | 74519400 | 74522399 | 35.76919 | 60.17797 | 48.55515 | 21.21625 | 0.689649 | 1.159307 | 0.50929 |
| 8 | 36086800 | 36090999 | 68.07644 | 107.998 | 113.7547 | 47.37198 | 0.256163 | 2.322447 | 0.509301 |
| 3 | 105038400 | 105043399 | 54.74344 | 87.60796 | 91.67943 | 38.61089 | 0.278699 | 2.035111 | 0.509317 |
| 2 | 12605000 | 12605799 | 11.11544 | 21.0287 | 22.6787 | 9.199484 | 0.374486 | 0.052681 | 0.509318 |
| 3 | 154048000 | 154048799 | 6.864196 | 13.13386 | 26.21666 | 10.37412 | -0.47499 | -0.06076 | 0.509334 |
| 8 | 53069000 | 53070799 | 16.79079 | 30.43237 | 28.625 | 12.38502 | 0.500457 | 0.363578 | 0.509335 |
| 11 | 60004400 | 60006799 | 13.63599 | 23.84318 | 42.58478 | 18.77478 | -0.44506 | 0.680936 | 0.509351 |
| 11 | 6655400 | 6661399 | 105.2441 | 159.9582 | 239.9791 | 99.40167 | -0.30205 | 3.194288 | 0.509355 |
| X | 147761400 | 147765599 | 58.50109 | 93.87308 | 89.89183 | 38.0114 | 0.41104 | 1.958338 | 0.509359 |
| 13 | 118285800 | 118289599 | 61.93772 | 98.68862 | 104.1137 | 43.57869 | 0.260242 | 2.182471 | 0.509376 |
| 12 | 63199600 | 63201199 | 16.40529 | 29.82195 | 28.12845 | 12.12513 | 0.49984 | 0.34258 | 0.50938 |
| 12 | 28297400 | 28299199 | 21.47563 | 37.41727 | 38.16353 | 16.78541 | 0.358871 | 0.764882 | 0.509391 |
| 3 | 10473200 | 10478599 | 71.96549 | 114.4027 | 111.9543 | 46.71828 | 0.364996 | 2.265132 | 0.509392 |
| 18 | 52825800 | 52828399 | 18.54843 | 31.52299 | 53.74103 | 23.55249 | -0.38509 | 1.017293 | 0.509393 |
| 15 | 83949200 | 83950999 | 34.87486 | 57.27941 | 60.69628 | 26.14727 | 0.294356 | 1.428308 | 0.509397 |
| 12 | 77764000 | 77766199 | 23.63178 | 40.92929 | 40.87607 | 17.9339 | 0.388036 | 0.851165 | 0.5094 |
| 14 | 107795200 | 107797199 | 28.14307 | 46.10309 | 75.80257 | 32.61905 | -0.33922 | 1.453613 | 0.509408 |
| 16 | 94788800 | 94790999 | 20.55251 | 35.8766 | 37.1397 | 16.3545 | 0.337834 | 0.733256 | 0.50943 |
| 6 | 132393800 | 132406599 | 36.82503 | 60.8288 | 52.67719 | 22.75366 | 0.587378 | 1.357053 | 0.509456 |
| 13 | 105764000 | 105765399 | 12.10934 | 21.72434 | 37.2681 | 16.3117 | -0.37234 | 0.407928 | 0.509467 |
| 13 | 96175600 | 96177599 | 17.57245 | 31.09033 | 31.64999 | 13.9974 | 0.365317 | 0.528039 | 0.509471 |
| 4 | 84038600 | 84040599 | 12.99482 | 22.86449 | 39.34129 | 17.41817 | -0.39079 | 0.576348 | 0.509476 |
| 16 | 18995000 | 18996199 | 11.82627 | 22.64946 | 21.58895 | 8.519929 | 0.585469 | -0.03448 | 0.509503 |
| 11 | 80670200 | 80671199 | 6.739807 | 12.84338 | 26.66549 | 10.65223 | -0.54359 | -0.0297 | 0.509508 |
| 14 | 29481200 | 29482799 | 18.52869 | 32.71535 | 32.74074 | 14.48569 | 0.388858 | 0.559526 | 0.50952 |
| 7 | 102383200 | 102385999 | 36.59115 | 59.87826 | 62.85532 | 27.03585 | 0.30572 | 1.480222 | 0.509521 |
| 11 | 45928400 | 45930399 | 14.3851 | 25.16846 | 43.56781 | 19.21812 | -0.40033 | 0.675409 | 0.509547 |
| X | 117861600 | 117863999 | 34.36402 | 57.1729 | 55.33911 | 24.02232 | 0.428096 | 1.273005 | 0.509585 |
| 2 | 22090200 | 22090799 | 21.14973 | 36.61865 | 36.45492 | 15.99401 | 0.392602 | 0.8374 | 0.509597 |
| 5 | 112915200 | 112919999 | 30.10976 | 48.79696 | 82.39783 | 35.27762 | -0.3891 | 1.635631 | 0.509603 |
| 17 | 37751400 | 37752199 | 9.027114 | 17.42713 | 17.60642 | 6.787325 | 0.533904 | -0.14997 | 0.509613 |
| 4 | 23062200 | 23062799 | 10.02146 | 19.43011 | 20.45834 | 7.729897 | 0.502739 | -0.19444 | 0.509621 |
| 7 | 42195400 | 42214399 | 313.4739 | 449.6341 | 403.5839 | 166.6568 | 0.360905 | 4.206385 | 0.509647 |
| 17 | 48945600 | 48948799 | 23.03484 | 38.33181 | 63.47098 | 27.55615 | -0.34398 | 1.244963 | 0.509668 |
| 7 | 21042200 | 21044199 | 30.29017 | 51.1239 | 43.71425 | 19.11961 | 0.6063 | 1.108337 | 0.509674 |
| 4 | 17352600 | 17354999 | 26.39394 | 44.95294 | 45.77099 | 20.06303 | 0.356035 | 1.016441 | 0.509693 |
| 14 | 17819200 | 17820599 | 8.559375 | 15.60079 | 31.91448 | 13.71156 | -0.60629 | 0.278921 | 0.509702 |
| 15 | 31082800 | 31083999 | 14.22476 | 26.07551 | 26.92236 | 11.48131 | 0.380697 | 0.2783 | 0.509706 |
| 4 | 37801800 | 37803199 | 13.69051 | 25.78773 | 23.93866 | 9.795251 | 0.577941 | 0.105612 | 0.509725 |
| 6 | 130437400 | 130443599 | 123.2043 | 188.6936 | 189.6261 | 78.69892 | 0.284368 | 3.040841 | 0.509725 |
| 3 | 82265200 | 82266799 | 9.039518 | 16.84324 | 30.00641 | 12.50578 | -0.37657 | 0.150414 | 0.509735 |
| 8 | 68913400 | 68914199 | 7.41478 | 13.88517 | 28.5425 | 11.79533 | -0.57045 | 0.104316 | 0.509746 |
| 2 | 140068400 | 140072399 | 52.50162 | 83.56304 | 127.6608 | 53.35476 | -0.27476 | 2.213763 | 0.509746 |

| | | | | | | | | | |
|---|---|---|---|---|---|---|---|---|---|
| 10 | 27674800 | 27675399 | 6.702491 | 12.89707 | 25.27605 | 9.841086 | -0.42655 | -0.11762 | 0.509757 |
| 17 | 58794600 | 58795199 | 6.702491 | 12.89707 | 25.27605 | 9.841086 | -0.42655 | -0.11762 | 0.509757 |
| 7 | 84699200 | 84701799 | 36.18801 | 59.4407 | 61.96409 | 26.69429 | 0.317933 | 1.431225 | 0.50978 |
| 5 | 52129400 | 52131199 | 13.62862 | 24.0278 | 39.79662 | 17.6127 | -0.33499 | 0.559152 | 0.509786 |
| 13 | 18134400 | 18136199 | 13.62862 | 24.0278 | 39.79662 | 17.6127 | -0.33499 | 0.559152 | 0.509786 |
| 7 | 83560000 | 83561399 | 12.98156 | 24.02965 | 25.46897 | 10.71842 | 0.35668 | 0.210108 | 0.509796 |
| 11 | 79707800 | 79708999 | 12.98156 | 24.02965 | 25.46897 | 10.71842 | 0.35668 | 0.210108 | 0.509796 |
| X | 120424600 | 120425799 | 18.23296 | 32.83222 | 24.62018 | 10.73918 | 0.818249 | 0.404212 | 0.509813 |
| 1 | 114718000 | 114722399 | 46.29897 | 75.66634 | 63.24486 | 27.19058 | 0.632242 | 1.519282 | 0.509815 |
| 8 | 113881800 | 113882599 | 12.32741 | 23.35242 | 20.46797 | 8.279523 | 0.674808 | 0.048759 | 0.509835 |
| X | 3068800 | 3071399 | 25.20305 | 44.64742 | 28.27506 | 12.50165 | 1.047288 | 0.558244 | 0.509835 |
| 11 | 118219800 | 118222199 | 25.12523 | 41.23111 | 71.29484 | 30.70191 | -0.41052 | 1.424017 | 0.50984 |
| 5 | 3907200 | 3908199 | 16.0462 | 28.58049 | 29.18244 | 12.85053 | 0.365905 | 0.462173 | 0.509911 |
| 19 | 17286400 | 17286999 | 7.66635 | 14.45526 | 27.54849 | 11.25284 | -0.44912 | 0.057278 | 0.509914 |
| 13 | 33873600 | 33875799 | 22.95229 | 40.06035 | 37.44021 | 16.48546 | 0.484581 | 0.746692 | 0.509927 |
| 14 | 3778000 | 3782999 | 65.4661 | 104.2064 | 107.9353 | 45.11592 | 0.285148 | 2.219236 | 0.509941 |
| 8 | 6369400 | 6369999 | 5.806698 | 11.29537 | 23.02067 | 8.503685 | -0.41056 | -0.31441 | 0.509948 |
| 3 | 31227400 | 31230599 | 32.15617 | 52.12376 | 84.28683 | 36.09164 | -0.32665 | 1.634211 | 0.509972 |
| 1 | 100778000 | 100778799 | 10.60044 | 20.25302 | 22.25967 | 8.843235 | 0.37349 | -0.02097 | 0.509981 |
| 9 | 77701800 | 77703399 | 17.67282 | 31.33455 | 26.66299 | 11.7863 | 0.623764 | 0.496954 | 0.510003 |
| 1 | 85541600 | 85555399 | 420.7141 | 587.055 | 523.4646 | 217.1267 | 0.330498 | 4.614693 | 0.510018 |
| 5 | 52270000 | 52273999 | 37.46521 | 60.10285 | 104.2957 | 44.08516 | -0.4454 | 1.94337 | 0.510025 |
| 8 | 28074600 | 28075799 | 16.00398 | 28.41236 | 28.89213 | 12.75777 | 0.369318 | 0.483452 | 0.510034 |
| 8 | 63448600 | 63449599 | 6.578547 | 12.61538 | 25.16673 | 9.895144 | -0.46475 | -0.0853 | 0.510057 |
| 14 | 57791200 | 57792399 | 6.578547 | 12.61538 | 25.16673 | 9.895144 | -0.46475 | -0.0853 | 0.510057 |
| 5 | 78155400 | 78156799 | 13.71977 | 25.04537 | 24.08076 | 10.32581 | 0.477623 | 0.291942 | 0.510069 |
| 10 | 53626000 | 53628199 | 20.06164 | 33.76382 | 57.47269 | 25.11553 | -0.38358 | 1.111944 | 0.510093 |
| 16 | 23131600 | 23135399 | 35.54101 | 58.24887 | 61.7803 | 26.5902 | 0.291823 | 1.456781 | 0.510113 |
| 6 | 68568200 | 68575799 | 110.5521 | 170.2037 | 175.4617 | 72.87603 | 0.25337 | 2.933965 | 0.510118 |
| X | 160421800 | 160422199 | 7.503645 | 14.38442 | 27.97725 | 11.01113 | -0.43006 | -0.06355 | 0.51012 |
| 1 | 126539200 | 126539799 | 5.957813 | 11.56382 | 23.34163 | 8.774799 | -0.41938 | -0.24616 | 0.510129 |
| 7 | 84672400 | 84678399 | 104.4622 | 162.1249 | 142.9413 | 59.36158 | 0.487571 | 2.783121 | 0.51013 |
| 17 | 7137600 | 7144399 | 73.21653 | 116.3432 | 112.3113 | 46.84778 | 0.384057 | 2.277558 | 0.510143 |
| 10 | 78329200 | 78335999 | 55.8041 | 88.53252 | 136.2955 | 56.76119 | -0.29052 | 2.315246 | 0.510147 |
| 15 | 51917600 | 51920199 | 24.74945 | 42.25498 | 44.19186 | 19.37464 | 0.318066 | 0.986587 | 0.510153 |
| 18 | 6851400 | 6861399 | 137.4535 | 209.2537 | 206.7251 | 85.66939 | 0.302132 | 3.167158 | 0.510163 |
| 2 | 47278200 | 47278399 | 7.416166 | 14.46677 | 13.77696 | 4.930774 | 0.718593 | -0.43433 | 0.510183 |
| 4 | 149925200 | 149926799 | 14.26456 | 24.91737 | 43.11522 | 18.99401 | -0.39964 | 0.700887 | 0.510189 |
| 11 | 9876200 | 9877199 | 10.90065 | 21.04609 | 21.37671 | 8.237074 | 0.527162 | -0.11852 | 0.510192 |
| 17 | 51644200 | 51645199 | 10.90065 | 21.04609 | 21.37671 | 8.237074 | 0.527162 | -0.11852 | 0.510192 |
| 4 | 78978600 | 78979399 | 8.385035 | 15.85421 | 28.69541 | 11.65408 | -0.36855 | 0.046083 | 0.51021 |
| 17 | 37422000 | 37423199 | 7.815312 | 14.91903 | 28.39088 | 11.29946 | -0.41394 | -0.02323 | 0.51022 |
| 3 | 64389800 | 64392599 | 29.51113 | 50.15872 | 46.83266 | 20.52499 | 0.48022 | 1.049188 | 0.510222 |
| 10 | 116083200 | 116084399 | 12.12403 | 22.59285 | 23.99917 | 10.00043 | 0.364151 | 0.160118 | 0.510234 |
| 15 | 44105200 | 44109599 | 58.58572 | 94.15828 | 85.24897 | 36.13384 | 0.494485 | 1.907413 | 0.510235 |
| 7 | 38554600 | 38570799 | 296.4024 | 425.6731 | 400.6765 | 165.4242 | 0.294134 | 4.187412 | 0.510238 |
| 8 | 15835000 | 15839399 | 39.1842 | 63.02246 | 100.0338 | 42.44554 | -0.31254 | 1.863489 | 0.510243 |

| | | | | | | | | | |
|---|---|---|---|---|---|---|---|---|---|
| 13 | 112535800 | 112537199 | 14.01111 | 25.58357 | 26.89932 | 11.53082 | 0.349418 | 0.297923 | 0.510243 |
| X | 8474400 | 8482599 | 170.3815 | 256.5453 | 239.0104 | 98.71898 | 0.373339 | 3.397317 | 0.510252 |
| 17 | 69253800 | 69257999 | 46.27484 | 74.06106 | 113.1468 | 47.6571 | -0.26606 | 2.031407 | 0.510255 |
| 12 | 21422800 | 21425799 | 17.88183 | 30.13417 | 61.67513 | 26.96257 | -0.64834 | 1.098434 | 0.510262 |
| 10 | 15506000 | 15506599 | 9.532274 | 18.34266 | 20.30159 | 7.903727 | 0.391016 | -0.1013 | 0.510277 |
| 11 | 101709000 | 101711599 | 29.24658 | 49.05573 | 50.76468 | 22.1504 | 0.331723 | 1.165911 | 0.51028 |
| 1 | 9069400 | 9070199 | 7.914375 | 15.04638 | 27.73173 | 11.13315 | -0.37972 | 0.001225 | 0.510292 |
| 4 | 78082600 | 78083199 | 7.914375 | 15.04638 | 27.73173 | 11.13315 | -0.37972 | 0.001225 | 0.510292 |
| 14 | 66314400 | 66315399 | 7.914375 | 15.04638 | 27.73173 | 11.13315 | -0.37972 | 0.001225 | 0.510292 |
| 17 | 53408200 | 53410599 | 26.33607 | 43.48734 | 69.6566 | 30.08046 | -0.29722 | 1.349012 | 0.510296 |
| 6 | 43107800 | 43108599 | 13.72472 | 25.0362 | 25.5036 | 10.94587 | 0.393858 | 0.299089 | 0.510299 |
| X | 126048800 | 126050799 | 22.6513 | 39.48351 | 38.15696 | 16.78729 | 0.436513 | 0.766512 | 0.510311 |
| 9 | 62165800 | 62168399 | 25.65403 | 42.59156 | 68.70087 | 29.77905 | -0.30667 | 1.283787 | 0.510325 |
| 12 | 108513400 | 108515599 | 13.89373 | 24.09121 | 50.55412 | 22.18081 | -0.67793 | 0.811404 | 0.510326 |
| X | 106780200 | 106782199 | 28.229 | 47.43842 | 49.6379 | 21.68188 | 0.315852 | 1.144555 | 0.510328 |
| 10 | 85301200 | 85304399 | 23.00978 | 38.40328 | 62.43983 | 27.15935 | -0.31777 | 1.212146 | 0.510341 |
| 15 | 31788600 | 31792199 | 37.20331 | 61.23627 | 60.56076 | 26.11283 | 0.394838 | 1.405441 | 0.510349 |
| 15 | 34837000 | 34839999 | 37.61006 | 60.47563 | 97.37235 | 41.30137 | -0.33349 | 1.869505 | 0.510357 |
| 11 | 113520600 | 113532799 | 106.5278 | 161.1817 | 276.8817 | 114.5289 | -0.50269 | 3.288438 | 0.510388 |
| 12 | 9264200 | 9265199 | 11.27454 | 20.05134 | 35.81696 | 15.79207 | -0.43713 | 0.457443 | 0.51039 |
| 5 | 4706200 | 4708999 | 22.65738 | 39.91178 | 32.8523 | 14.54185 | 0.668067 | 0.608582 | 0.510391 |
| 2 | 132174400 | 132174799 | 10.84282 | 20.45515 | 20.62622 | 8.416979 | 0.463081 | 0.062829 | 0.510392 |
| 4 | 69482200 | 69483799 | 21.69919 | 38.09181 | 34.5949 | 15.30304 | 0.52569 | 0.648109 | 0.510397 |
| 3 | 117913400 | 117917599 | 42.46586 | 69.44849 | 64.67852 | 27.81375 | 0.477265 | 1.502029 | 0.510404 |
| 10 | 51352000 | 51354399 | 17.48824 | 30.34553 | 50.0898 | 21.97757 | -0.33323 | 0.820094 | 0.510407 |
| 4 | 96817800 | 96818799 | 11.38809 | 21.54001 | 22.52467 | 9.141035 | 0.417419 | 0.053347 | 0.510412 |
| X | 153245000 | 153250799 | 94.94068 | 149.303 | 131.6079 | 54.59689 | 0.502575 | 2.54647 | 0.510413 |
| 7 | 92136200 | 92137199 | 6.874318 | 13.15392 | 25.66195 | 10.15309 | -0.44157 | -0.06088 | 0.51042 |
| 7 | 106899200 | 106901399 | 19.3865 | 33.01524 | 54.29409 | 23.79883 | -0.33284 | 1.003913 | 0.510439 |
| 6 | 106277000 | 106277599 | 5.811812 | 11.29008 | 22.86299 | 8.537962 | -0.41571 | -0.27554 | 0.510441 |
| 11 | 31111200 | 31113999 | 34.53669 | 56.71446 | 60.10467 | 25.89058 | 0.294093 | 1.429028 | 0.510483 |
| 6 | 132104600 | 132106599 | 23.37004 | 40.80375 | 37.43511 | 16.48684 | 0.511287 | 0.744842 | 0.510499 |
| 14 | 68277000 | 68278599 | 9.507025 | 17.57051 | 31.46441 | 13.26983 | -0.39707 | 0.204495 | 0.510511 |
| 17 | 43241400 | 43243399 | 12.45024 | 22.12757 | 38.208 | 16.85574 | -0.38962 | 0.470374 | 0.510523 |
| X | 5264400 | 5266999 | 40.21581 | 65.94142 | 63.35752 | 27.27445 | 0.434297 | 1.46363 | 0.51053 |
| 7 | 120930800 | 120934199 | 18.14629 | 31.32142 | 51.15274 | 22.44033 | -0.31953 | 0.871655 | 0.510534 |
| 17 | 32647800 | 32649999 | 18.34395 | 31.33733 | 52.17592 | 22.87611 | -0.34995 | 0.967023 | 0.510542 |
| 1 | 45409000 | 45411199 | 24.07186 | 42.23359 | 33.89564 | 14.98673 | 0.703256 | 0.66913 | 0.510584 |
| 5 | 123532200 | 123533799 | 9.975861 | 17.91206 | 34.31597 | 15.00884 | -0.52953 | 0.382882 | 0.510585 |
| 14 | 113299400 | 113299799 | 5.663711 | 11.01074 | 22.3338 | 8.289302 | -0.40939 | -0.29945 | 0.510587 |
| 13 | 38623600 | 38625199 | 16.85454 | 29.81326 | 30.85978 | 13.64855 | 0.341463 | 0.526194 | 0.510594 |
| 1 | 48787200 | 48788199 | 13.33368 | 23.57902 | 39.27287 | 17.3704 | -0.34124 | 0.524144 | 0.510596 |
| 2 | 47367200 | 47368799 | 20.04667 | 35.59698 | 30.77025 | 13.58467 | 0.602406 | 0.510394 | 0.510596 |
| 10 | 124542000 | 124543199 | 12.87482 | 23.92369 | 23.23397 | 9.726924 | 0.486784 | 0.179973 | 0.510607 |
| 5 | 104344200 | 104346199 | 12.50761 | 22.08009 | 38.28931 | 16.94121 | -0.40009 | 0.534785 | 0.510617 |
| 7 | 107491600 | 107495199 | 41.14906 | 67.13611 | 65.86092 | 28.29782 | 0.400144 | 1.53563 | 0.510663 |
| 9 | 61801400 | 61804199 | 16.93517 | 28.99398 | 50.56585 | 22.17409 | -0.41595 | 0.940564 | 0.510665 |

| | | | | | | | | | |
|---|---|---|---|---|---|---|---|---|---|
| 9 | 19076000 | 19079399 | 33.00391 | 55.69559 | 48.53109 | 21.2254 | 0.579158 | 1.123425 | 0.510675 |
| 17 | 23897800 | 23902799 | 36.82459 | 59.11225 | 101.2342 | 42.83197 | -0.42532 | 1.922649 | 0.510706 |
| 5 | 96991400 | 96998999 | 64.8067 | 101.2403 | 182.0125 | 75.51925 | -0.53185 | 2.655374 | 0.510714 |
| X | 162834000 | 162834999 | 9.161616 | 17.04019 | 32.67121 | 13.65776 | -0.48483 | 0.173298 | 0.510726 |
| X | 9052400 | 9055199 | 30.48351 | 52.63481 | 39.60622 | 17.37515 | 0.794093 | 0.909519 | 0.510726 |
| 13 | 117677000 | 117678399 | 18.39765 | 32.36945 | 33.27143 | 14.73089 | 0.348471 | 0.596197 | 0.51073 |
| 7 | 117875800 | 117878599 | 19.14988 | 32.52313 | 55.39279 | 24.27333 | -0.38394 | 1.025836 | 0.510736 |
| 5 | 139611400 | 139621799 | 112.9699 | 171.4985 | 247.9702 | 102.6755 | -0.25024 | 3.221343 | 0.510748 |
| 16 | 16417400 | 16418199 | 6.593333 | 12.59784 | 25.62726 | 10.17986 | -0.50645 | -0.04978 | 0.510748 |
| X | 5507600 | 5510199 | 33.52439 | 56.1595 | 51.93531 | 22.643 | 0.49351 | 1.195733 | 0.51075 |
| 4 | 74827000 | 74828799 | 17.29349 | 30.71843 | 31.13677 | 13.7426 | 0.374257 | 0.492018 | 0.510752 |
| 19 | 7644600 | 7646399 | 22.70507 | 37.92343 | 61.84461 | 26.91331 | -0.32204 | 1.203384 | 0.510761 |
| 6 | 62732800 | 62734199 | 15.91815 | 29.29854 | 26.72325 | 11.33354 | 0.563875 | 0.267137 | 0.510783 |
| 15 | 66241000 | 66242599 | 12.97325 | 24.19186 | 24.36342 | 10.14049 | 0.441558 | 0.162514 | 0.510784 |
| 7 | 59668000 | 59672799 | 57.87921 | 93.04624 | 85.53845 | 36.26979 | 0.472961 | 1.900111 | 0.510789 |
| 5 | 57511400 | 57515399 | 44.10203 | 71.46177 | 72.78992 | 31.16039 | 0.338051 | 1.65692 | 0.510792 |
| 6 | 39311000 | 39312999 | 12.67332 | 22.30537 | 39.34119 | 17.4171 | -0.42618 | 0.570637 | 0.510808 |
| 4 | 40107200 | 40108799 | 14.31265 | 25.1617 | 41.88886 | 18.54926 | -0.34398 | 0.599072 | 0.510816 |
| 13 | 50023600 | 50026199 | 57.14062 | 91.45626 | 95.16055 | 40.08442 | 0.287154 | 2.03608 | 0.510823 |
| 6 | 138312600 | 138316599 | 66.63834 | 106.188 | 105.9409 | 44.34909 | 0.340824 | 2.185371 | 0.510843 |
| 2 | 86328200 | 86332999 | 78.47289 | 123.8047 | 128.6715 | 53.40345 | 0.268788 | 2.468399 | 0.510847 |
| 11 | 34492600 | 34496399 | 35.68153 | 59.94205 | 49.12811 | 21.45701 | 0.667041 | 1.171003 | 0.510852 |
| 7 | 136303800 | 136304599 | 12.05665 | 22.51484 | 24.07845 | 9.997622 | 0.358831 | 0.147652 | 0.510864 |
| 8 | 19747400 | 19765999 | 763.7705 | 996.0536 | 1255.323 | 525.7082 | -0.26653 | 5.683625 | 0.510882 |
| 4 | 29839600 | 29841399 | 23.82646 | 41.27554 | 36.14352 | 15.85766 | 0.576919 | 0.843244 | 0.510892 |
| 7 | 13426800 | 13432199 | 86.88774 | 137.6059 | 104.955 | 43.64464 | 0.718933 | 2.381633 | 0.510897 |
| 8 | 109669800 | 109684199 | 365.4463 | 495.8347 | 890.5924 | 371.0353 | -0.72039 | 5.046531 | 0.5109 |
| 12 | 63336000 | 63339399 | 77.25927 | 121.6053 | 125.9185 | 52.24272 | 0.271543 | 2.514511 | 0.510905 |
| 1 | 111613800 | 111614799 | 11.0261 | 21.20949 | 21.82047 | 8.503886 | 0.493509 | -0.08023 | 0.51091 |
| 10 | 126282600 | 126283399 | 11.0261 | 21.20949 | 21.82047 | 8.503886 | 0.493509 | -0.08023 | 0.51091 |
| 12 | 71783400 | 71784799 | 27.563 | 46.35439 | 48.56402 | 21.21769 | 0.314109 | 1.134827 | 0.510922 |
| 2 | 168408800 | 168409599 | 12.68726 | 23.5018 | 25.01008 | 10.51499 | 0.352019 | 0.203131 | 0.510937 |
| 5 | 113136600 | 113139999 | 42.51025 | 68.15298 | 105.604 | 44.59509 | -0.28348 | 1.971385 | 0.510946 |
| X | 119951400 | 119954799 | 38.95901 | 64.75751 | 54.37272 | 23.58765 | 0.632551 | 1.296934 | 0.51095 |
| 17 | 71129400 | 71131599 | 9.688632 | 17.47357 | 36.93381 | 16.07898 | -0.66577 | 0.361702 | 0.510955 |
| 7 | 12404200 | 12408199 | 35.28201 | 58.16352 | 50.42726 | 21.75762 | 0.585034 | 1.374203 | 0.510968 |
| 11 | 8607200 | 8611599 | 35.47738 | 57.14742 | 93.5796 | 39.79459 | -0.35459 | 1.808693 | 0.510969 |
| 6 | 68668200 | 68678399 | 162.9887 | 246.6246 | 222.1694 | 91.8996 | 0.42792 | 3.297017 | 0.510988 |
| 12 | 67707000 | 67707799 | 15.30476 | 27.34826 | 27.87242 | 12.24022 | 0.371425 | 0.435775 | 0.511016 |
| 17 | 34437400 | 34454399 | 297.884 | 418.3929 | 573.7864 | 237.7571 | -0.28086 | 4.513295 | 0.511022 |
| 1 | 173362000 | 173363599 | 15.26799 | 27.94764 | 26.62387 | 11.37626 | 0.493595 | 0.290661 | 0.511026 |
| 2 | 63427600 | 63428399 | 6.843514 | 13.18938 | 25.33648 | 9.819685 | -0.39174 | -0.13066 | 0.511031 |
| 4 | 139402000 | 139412199 | 116.7194 | 176.7082 | 256.6963 | 106.1996 | -0.26025 | 3.277358 | 0.511042 |
| X | 75417000 | 75419199 | 24.28971 | 42.3851 | 37.01215 | 16.29467 | 0.582333 | 0.759415 | 0.51105 |
| 2 | 132223200 | 132223999 | 4.275599 | 8.328369 | 20.72184 | 7.558399 | -0.67676 | -0.38486 | 0.511053 |
| 2 | 121857600 | 121859999 | 13.65183 | 23.92785 | 40.98485 | 18.10583 | -0.38486 | 0.641042 | 0.511056 |
| 16 | 89458200 | 89460599 | 25.1022 | 42.9667 | 43.04648 | 18.87636 | 0.380258 | 0.962145 | 0.511068 |

| | | | | | | | | | |
|---|---|---|---|---|---|---|---|---|---|
| 3 | 15577400 | 15582399 | 39.82358 | 65.34897 | 55.94647 | 24.09201 | 0.600714 | 1.448478 | 0.511085 |
| 17 | 77164600 | 77167799 | 37.93738 | 62.86079 | 56.67981 | 24.53502 | 0.529652 | 1.328902 | 0.511101 |
| 9 | 119929800 | 119938399 | 137.7466 | 209.2749 | 211.9693 | 87.78932 | 0.26378 | 3.20885 | 0.511102 |
| 5 | 100388400 | 100391199 | 31.71232 | 53.14866 | 46.45191 | 20.25827 | 0.574649 | 1.179094 | 0.511102 |
| 9 | 3840400 | 3843799 | 44.21871 | 71.58475 | 73.60452 | 31.48769 | 0.32331 | 1.677707 | 0.511102 |
| 7 | 143189000 | 143190999 | 29.09017 | 47.32656 | 78.32313 | 33.6174 | -0.35522 | 1.558143 | 0.511103 |
| X | 121513000 | 121519199 | 97.11677 | 151.6349 | 146.7643 | 60.89765 | 0.359866 | 2.678377 | 0.511107 |
| 4 | 10622400 | 10622799 | 4.223016 | 8.254968 | 18.3292 | 6.307575 | -0.43774 | -0.6011 | 0.511137 |
| 5 | 128459400 | 128463599 | 41.00766 | 65.81402 | 103.5499 | 43.79849 | -0.30366 | 1.934832 | 0.511137 |
| 3 | 7189200 | 7190199 | 11.81071 | 22.57217 | 22.7258 | 9.022652 | 0.499498 | -0.01112 | 0.51119 |
| 4 | 114920000 | 114922599 | 17.3265 | 29.37929 | 54.21647 | 23.74631 | -0.49908 | 1.033615 | 0.51124 |
| 3 | 53408400 | 53409999 | 14.42654 | 25.19756 | 43.11679 | 18.99339 | -0.38365 | 0.703552 | 0.511245 |
| 11 | 86647200 | 86648999 | 14.42654 | 25.19756 | 43.11679 | 18.99339 | -0.38365 | 0.703552 | 0.511245 |
| 6 | 27108200 | 27110799 | 24.0212 | 42.03702 | 35.64254 | 15.72508 | 0.624518 | 0.710955 | 0.51125 |
| 17 | 47649200 | 47661399 | 175.4912 | 258.8309 | 362.9231 | 149.4569 | -0.24334 | 3.772352 | 0.51125 |
| 4 | 38554000 | 38560399 | 115.035 | 176.9155 | 159.326 | 66.16451 | 0.44697 | 2.954204 | 0.511262 |
| 6 | 58307200 | 58313199 | 149.9381 | 226.1258 | 230.9446 | 95.42949 | 0.242897 | 3.360541 | 0.511262 |
| 17 | 77014000 | 77016199 | 24.01239 | 41.2357 | 40.364 | 17.70464 | 0.414249 | 0.924224 | 0.511267 |
| 6 | 133299000 | 133300599 | 14.84101 | 28.30856 | 21.94091 | 8.746869 | 0.868886 | 0.029231 | 0.511276 |
| 13 | 37527600 | 37530799 | 22.19562 | 36.94961 | 65.76847 | 28.56098 | -0.448 | 1.237573 | 0.511276 |
| X | 84617000 | 84618199 | 19.15379 | 34.04281 | 31.09547 | 13.7176 | 0.524316 | 0.502546 | 0.511284 |
| 8 | 15808200 | 15811599 | 58.45631 | 93.58879 | 95.05347 | 40.05568 | 0.322421 | 2.028888 | 0.511292 |
| 3 | 19166000 | 19177399 | 218.3076 | 320.5784 | 318.2781 | 131.0844 | 0.248501 | 3.845259 | 0.511302 |
| 13 | 64048600 | 64050799 | 12.86896 | 22.66521 | 38.85825 | 17.20093 | -0.38491 | 0.565464 | 0.511319 |
| 18 | 9298400 | 9298799 | 4.101012 | 8.016668 | 18.33319 | 6.30511 | -0.47918 | -0.6061 | 0.511331 |
| 2 | 14033600 | 14037199 | 28.14146 | 48.56786 | 40.10036 | 17.59167 | 0.66034 | 0.899776 | 0.511341 |
| 1 | 161947600 | 161948399 | 9.936301 | 18.09797 | 32.37417 | 13.92107 | -0.41485 | 0.296124 | 0.511356 |
| 14 | 37565000 | 37567399 | 34.50092 | 56.68439 | 60.16261 | 25.92108 | 0.292212 | 1.422162 | 0.511362 |
| 11 | 38144000 | 38145199 | 15.1596 | 25.76173 | 59.38192 | 26.02451 | -0.81813 | 1.016301 | 0.511368 |
| 16 | 56798800 | 56803199 | 62.25753 | 99.42188 | 100.8159 | 42.33133 | 0.320751 | 2.11153 | 0.511369 |
| 6 | 133339600 | 133341599 | 40.59505 | 66.36611 | 58.71687 | 25.24688 | 0.550724 | 1.499006 | 0.511385 |
| 11 | 108041000 | 108041799 | 5.525648 | 10.73591 | 22.27895 | 8.308987 | -0.44841 | -0.28446 | 0.511408 |
| 1 | 33865400 | 33866199 | 9.46832 | 17.15272 | 33.34279 | 14.43474 | -0.54068 | 0.32332 | 0.511441 |
| 4 | 16688600 | 16690799 | 17.41565 | 31.74904 | 27.53375 | 11.82651 | 0.623571 | 0.332927 | 0.511444 |
| 16 | 18500200 | 18501399 | 13.38584 | 23.47569 | 41.00959 | 18.12592 | -0.41292 | 0.631918 | 0.511447 |
| 8 | 19253400 | 19254599 | 14.6195 | 25.69223 | 42.92233 | 19.00446 | -0.34919 | 0.606654 | 0.511447 |
| 10 | 89790000 | 89791799 | 20.17673 | 35.51143 | 33.55668 | 14.8562 | 0.469334 | 0.603059 | 0.511453 |
| X | 62084800 | 62088599 | 43.50067 | 71.84941 | 56.28087 | 24.30874 | 0.731835 | 1.381621 | 0.511472 |
| 12 | 57091200 | 57092399 | 10.67273 | 19.32991 | 33.79197 | 14.63365 | -0.389 | 0.33471 | 0.511508 |
| 13 | 61397600 | 61406999 | 121.2775 | 186.1152 | 185.0797 | 76.8368 | 0.301731 | 3.000987 | 0.511508 |
| 3 | 140296800 | 140298399 | 15.5962 | 28.2256 | 27.55372 | 11.94351 | 0.445208 | 0.358986 | 0.511512 |
| 1 | 116411800 | 116412399 | 6.843435 | 13.23938 | 25.55578 | 9.755166 | -0.37769 | -0.18042 | 0.511513 |
| 6 | 53972400 | 53975199 | 26.79601 | 45.61335 | 46.31577 | 20.30095 | 0.359917 | 1.024228 | 0.511557 |
| 1 | 115726800 | 115727399 | 10.71579 | 20.25653 | 21.14344 | 8.591127 | 0.419086 | 0.049459 | 0.511558 |
| X | 25921600 | 25931999 | 134.6276 | 207.587 | 163.5687 | 67.94361 | 0.642409 | 2.91548 | 0.511561 |
| 12 | 114558000 | 114563399 | 70.34239 | 111.717 | 113.0053 | 47.11232 | 0.316017 | 2.293186 | 0.511568 |
| 15 | 94150800 | 94153199 | 16.66205 | 28.97453 | 47.37556 | 20.81186 | -0.31919 | 0.783081 | 0.51158 |

| | | | | | | | | | |
|---|---|---|---|---|---|---|---|---|---|
| 5 | 85693000 | 85693799 | 10.69232 | 20.76643 | 19.31236 | 7.224799 | 0.692833 | -0.22526 | 0.511611 |
| 7 | 55280000 | 55282399 | 21.5095 | 37.54802 | 37.56347 | 16.54136 | 0.386898 | 0.737925 | 0.511615 |
| 19 | 39555400 | 39562799 | 100.6588 | 157.6025 | 141.6942 | 58.77707 | 0.469253 | 2.630818 | 0.511623 |
| 5 | 142328600 | 142332399 | 35.47629 | 57.3133 | 89.75185 | 38.31424 | -0.28534 | 1.721758 | 0.511637 |
| 5 | 46715400 | 46716999 | 14.33977 | 26.05519 | 27.37166 | 11.80571 | 0.344952 | 0.329366 | 0.511676 |
| 15 | 91848200 | 91848999 | 6.246889 | 12.08724 | 28.06051 | 10.70321 | -0.64161 | -0.18775 | 0.511678 |
| 8 | 51500200 | 51502199 | 26.45082 | 45.36904 | 38.86157 | 17.04408 | 0.606002 | 0.94742 | 0.511679 |
| 14 | 73326600 | 73328599 | 20.75907 | 36.21366 | 37.08414 | 16.32338 | 0.353225 | 0.741743 | 0.511691 |
| 3 | 13034000 | 13037399 | 36.29102 | 60.15051 | 57.14908 | 24.74318 | 0.454427 | 1.321578 | 0.511762 |
| 1 | 182524600 | 182526599 | 14.82225 | 26.69185 | 27.83265 | 12.13969 | 0.345244 | 0.384771 | 0.511794 |
| 1 | 109965000 | 109969599 | 104.7949 | 161.8443 | 167.9104 | 69.74648 | 0.247622 | 2.873051 | 0.511811 |
| 12 | 114350600 | 114361399 | 141.4772 | 216.3772 | 190.0006 | 78.82893 | 0.477287 | 3.075719 | 0.511842 |
| 13 | 88036400 | 88038599 | 22.31232 | 37.59539 | 60.90789 | 26.62609 | -0.31256 | 1.112951 | 0.511846 |
| 8 | 5748400 | 5752399 | 52.06241 | 83.57486 | 87.97096 | 37.22185 | 0.275501 | 1.937099 | 0.511856 |
| 6 | 39819600 | 39821399 | 25.5935 | 42.22454 | 69.91771 | 30.17701 | -0.34511 | 1.359706 | 0.511881 |
| 5 | 109802400 | 109803999 | 24.59478 | 42.88013 | 37.97801 | 16.71076 | 0.561772 | 0.772143 | 0.511908 |
| 13 | 75030000 | 75032799 | 31.95395 | 53.81398 | 44.23198 | 19.3332 | 0.663 | 1.133163 | 0.511915 |
| 11 | 10340000 | 10341799 | 23.17823 | 40.13397 | 40.3269 | 17.69282 | 0.379125 | 0.849129 | 0.511918 |
| X | 153129000 | 153136399 | 101.8993 | 159.4903 | 141.4287 | 58.66898 | 0.488739 | 2.638023 | 0.511918 |
| 1 | 140709400 | 140711199 | 23.36091 | 40.54618 | 40.3504 | 17.70452 | 0.393766 | 0.831246 | 0.511922 |
| 11 | 30817600 | 30820799 | 32.68527 | 54.76712 | 51.39029 | 22.40882 | 0.472496 | 1.187977 | 0.511922 |
| 17 | 37928600 | 37930799 | 33.59707 | 55.5862 | 57.43201 | 24.83507 | 0.333679 | 1.336471 | 0.51193 |
| 15 | 81093000 | 81094799 | 17.27456 | 30.40958 | 30.95597 | 13.70427 | 0.362751 | 0.580448 | 0.511931 |
| 7 | 103264400 | 103269199 | 117.4728 | 179.7991 | 185.7232 | 77.06762 | 0.244012 | 3.051862 | 0.511949 |
| 3 | 108671200 | 108673199 | 12.23222 | 21.49919 | 39.40506 | 17.44875 | -0.48198 | 0.584454 | 0.511965 |
| 18 | 27301200 | 27301799 | 11.76553 | 22.22771 | 23.66798 | 9.628982 | 0.388955 | 0.070441 | 0.51198 |
| 6 | 95086200 | 95087799 | 7.31315 | 14.05559 | 26.37918 | 10.30528 | -0.36911 | -0.09565 | 0.511981 |
| 9 | 6845800 | 6847199 | 21.22042 | 37.06073 | 32.44404 | 14.29118 | 0.57856 | 0.721801 | 0.511999 |
| 6 | 134874600 | 134875599 | 6.429404 | 12.33126 | 25.72601 | 10.11062 | -0.52874 | -0.08914 | 0.512001 |
| 4 | 73443000 | 73446599 | 51.30915 | 82.52492 | 84.68047 | 35.93165 | 0.315548 | 1.873768 | 0.512009 |
| 15 | 19323200 | 19324399 | 19.83464 | 34.85671 | 30.19442 | 13.36881 | 0.593542 | 0.613415 | 0.512014 |
| 8 | 7362200 | 7363399 | 14.61462 | 27.16198 | 25.88903 | 10.83552 | 0.514437 | 0.20202 | 0.512025 |
| 12 | 3120800 | 3137799 | 187.0489 | 280.402 | 252.1301 | 104.0166 | 0.418866 | 3.485463 | 0.512031 |
| 1 | 80671600 | 80673199 | 19.44187 | 33.06504 | 55.87062 | 24.48722 | -0.37208 | 1.01945 | 0.512037 |
| 16 | 78353400 | 78362599 | 110.2345 | 168.0374 | 245.1813 | 101.6449 | -0.25848 | 3.136607 | 0.512057 |
| 1 | 55521600 | 55526999 | 17.79367 | 30.26718 | 58.56349 | 25.66782 | -0.56684 | 1.012218 | 0.512074 |
| 19 | 40394200 | 40396799 | 17.07518 | 29.50932 | 48.99221 | 21.49334 | -0.34258 | 0.854971 | 0.512083 |
| 19 | 42668400 | 42673599 | 58.24214 | 92.14865 | 140.1995 | 58.27175 | -0.27697 | 2.391168 | 0.5121 |
| 3 | 100216400 | 100221799 | 104.836 | 161.7414 | 168.2521 | 69.8888 | 0.24264 | 2.891722 | 0.51211 |
| 15 | 9263000 | 9269199 | 63.22228 | 100.7919 | 104.4461 | 43.75182 | 0.286882 | 2.165519 | 0.51213 |
| 12 | 83997000 | 83999999 | 21.33666 | 35.42513 | 63.72099 | 27.62123 | -0.46318 | 1.267912 | 0.512151 |
| 7 | 74000200 | 74001399 | 9.738568 | 17.78813 | 31.86069 | 13.65183 | -0.41355 | 0.275454 | 0.512153 |
| 3 | 73508000 | 73509799 | 14.84539 | 27.51644 | 25.7857 | 10.83201 | 0.534805 | 0.217543 | 0.512155 |
| 14 | 103193600 | 103194599 | 13.89675 | 25.5562 | 26.34149 | 11.1846 | 0.387345 | 0.25472 | 0.51216 |
| 17 | 55491200 | 55494399 | 33.48271 | 55.58056 | 55.39209 | 24.00773 | 0.385775 | 1.293052 | 0.512174 |
| 7 | 33115400 | 33125599 | 167.6321 | 254.0157 | 213.3807 | 88.3042 | 0.530442 | 3.26989 | 0.512193 |
| 17 | 83652600 | 83654999 | 36.79002 | 59.31869 | 92.89222 | 39.55419 | -0.28864 | 1.782508 | 0.512197 |

| | | | | | | | | | |
|---|---|---|---|---|---|---|---|---|---|
| X | 26748200 | 26750799 | 33.15655 | 56.92164 | 40.03271 | 17.55898 | 0.889539 | 0.969235 | 0.5122 |
| 15 | 77630800 | 77634399 | 32.91311 | 53.07767 | 90.25228 | 38.43937 | -0.40676 | 1.770084 | 0.512204 |
| 8 | 4703800 | 4706799 | 32.30318 | 53.3509 | 56.30519 | 24.31806 | 0.302905 | 1.350141 | 0.512207 |
| Y | 8987400 | 8990199 | 25.2177 | 43.22318 | 26.00829 | 11.40618 | 1.11261 | 0.925812 | 0.512209 |
| 7 | 56573600 | 56574599 | 11.94479 | 21.03582 | 38.42481 | 17.00935 | -0.47574 | 0.560672 | 0.512219 |
| 5 | 6893600 | 6894399 | 15.79343 | 28.29261 | 27.94963 | 12.2481 | 0.418497 | 0.425982 | 0.512282 |
| 1 | 121968800 | 121971199 | 19.65828 | 33.45127 | 54.31959 | 23.80847 | -0.31492 | 1.010737 | 0.512292 |
| 1 | 46378600 | 46380399 | 14.50357 | 27.09081 | 24.83376 | 10.30702 | 0.579691 | 0.166776 | 0.512322 |
| 14 | 70356800 | 70358199 | 10.57728 | 19.04467 | 33.79058 | 14.74111 | -0.41693 | 0.369772 | 0.512365 |
| 14 | 15712200 | 15713399 | 6.769953 | 12.8403 | 26.58811 | 10.72749 | -0.55155 | 0.006943 | 0.512388 |
| 14 | 118477600 | 118478799 | 15.29071 | 27.36563 | 28.31166 | 12.41684 | 0.351134 | 0.426623 | 0.512396 |
| 15 | 86604800 | 86607799 | 16.91308 | 29.13345 | 48.99267 | 21.49139 | -0.36199 | 0.883549 | 0.5124 |
| 10 | 53664000 | 53666399 | 48.09463 | 77.32318 | 79.99564 | 33.92372 | 0.303058 | 1.881494 | 0.512411 |
| 5 | 109459200 | 109470399 | 179.3178 | 268.7315 | 255.8082 | 105.5278 | 0.336298 | 3.489694 | 0.512418 |
| 18 | 63324000 | 63325999 | 31.66099 | 51.51536 | 81.19455 | 34.85142 | -0.28496 | 1.56253 | 0.512427 |
| 1 | 41203400 | 41205599 | 18.74567 | 33.46747 | 30.41677 | 13.36747 | 0.534942 | 0.468593 | 0.51243 |
| 13 | 27542200 | 27543799 | 23.03857 | 38.53246 | 62.43407 | 27.19299 | -0.31281 | 1.190974 | 0.51243 |
| 5 | 125510200 | 125512999 | 19.9451 | 33.40706 | 60.12942 | 26.21322 | -0.46394 | 1.161098 | 0.512447 |
| 9 | 25633400 | 25635799 | 29.23552 | 48.84913 | 51.30162 | 22.33048 | 0.310724 | 1.209202 | 0.512454 |
| 6 | 13443200 | 13444199 | 12.38882 | 23.21817 | 24.59181 | 10.14902 | 0.379602 | 0.129115 | 0.512457 |
| 6 | 49013000 | 49014999 | 13.81597 | 24.10971 | 43.15948 | 19.00159 | -0.4485 | 0.713321 | 0.51248 |
| 16 | 15120000 | 15122399 | 28.9012 | 49.3564 | 45.18228 | 19.80929 | 0.509624 | 1.003528 | 0.512497 |
| 3 | 5476600 | 5478199 | 24.94836 | 42.5138 | 44.74459 | 19.61397 | 0.308597 | 1.00549 | 0.512517 |
| 8 | 18966800 | 18969399 | 19.00231 | 32.33329 | 53.77056 | 23.56773 | -0.34918 | 1.007565 | 0.51253 |
| 16 | 89431200 | 89434399 | 52.60667 | 83.56409 | 129.012 | 53.79507 | -0.2928 | 2.276352 | 0.512533 |
| 17 | 22090000 | 22093399 | 40.37045 | 65.99413 | 65.51486 | 28.16911 | 0.384959 | 1.504498 | 0.512535 |
| 13 | 22872200 | 22875599 | 33.82117 | 54.83199 | 86.18968 | 36.91108 | -0.28565 | 1.636609 | 0.51255 |
| 3 | 131528000 | 131529199 | 12.5491 | 23.73488 | 23.54675 | 9.556767 | 0.492763 | 0.067334 | 0.512551 |
| 3 | 86746200 | 86750999 | 57.62236 | 92.11146 | 97.08229 | 40.82117 | 0.266669 | 2.075112 | 0.512557 |
| 9 | 123454800 | 123455399 | 4.918381 | 9.578005 | 21.26464 | 7.782194 | -0.51809 | -0.35899 | 0.512558 |
| 6 | 24753600 | 24756599 | 24.48577 | 42.61973 | 37.93883 | 16.67622 | 0.554136 | 0.79167 | 0.512588 |
| 13 | 104078400 | 104079399 | 9.374681 | 17.20388 | 32.94414 | 14.02817 | -0.50338 | 0.245523 | 0.512597 |
| 12 | 24547600 | 24550399 | 18.13211 | 30.75488 | 59.63007 | 26.11989 | -0.57021 | 1.040055 | 0.5126 |
| 6 | 104338600 | 104339999 | 16.65964 | 29.94089 | 22.3804 | 9.780883 | 0.820796 | 0.398091 | 0.512602 |
| 4 | 149698000 | 149700799 | 39.64731 | 64.66236 | 54.17229 | 23.27259 | 0.626782 | 1.526409 | 0.512636 |
| 10 | 48150200 | 48152599 | 27.5038 | 47.34771 | 41.81272 | 18.34146 | 0.563013 | 0.921215 | 0.512647 |
| 3 | 106535000 | 106538799 | 31.13332 | 53.10949 | 43.43173 | 19.04009 | 0.671398 | 1.017437 | 0.512648 |
| 13 | 8790000 | 8791199 | 12.55697 | 23.57921 | 24.60172 | 10.11966 | 0.404918 | 0.118388 | 0.512661 |
| 15 | 89766200 | 89769399 | 32.85344 | 55.05897 | 51.91796 | 22.64152 | 0.465513 | 1.189099 | 0.512662 |
| 10 | 108134000 | 108135999 | 16.132 | 27.79816 | 53.78001 | 23.59169 | -0.56332 | 0.882231 | 0.512686 |
| 10 | 44934200 | 44936799 | 21.08001 | 35.73726 | 60.85106 | 26.65071 | -0.38364 | 1.05787 | 0.512692 |
| 4 | 16433000 | 16438199 | 87.34867 | 137.2217 | 136.5169 | 56.62575 | 0.327066 | 2.561708 | 0.512702 |
| 6 | 8072800 | 8074599 | 12.55777 | 22.14431 | 38.35695 | 16.97486 | -0.39887 | 0.549466 | 0.512705 |
| 3 | 152997400 | 152998999 | 12.69532 | 22.46963 | 37.76543 | 16.70076 | -0.35362 | 0.508183 | 0.512719 |
| 17 | 34966600 | 34969799 | 54.41222 | 87.5676 | 84.01353 | 35.67921 | 0.413401 | 1.861385 | 0.51272 |
| 4 | 80091000 | 80092799 | 21.68586 | 37.65812 | 39.36745 | 17.27878 | 0.323108 | 0.814836 | 0.512735 |
| 17 | 87464600 | 87468599 | 29.62987 | 48.369 | 79.17627 | 34.02579 | -0.3361 | 1.507664 | 0.512736 |

| | | | | | | | | | |
|---|---|---|---|---|---|---|---|---|---|
| 13 | 62644600 | 62648199 | 43.43686 | 70.68901 | 68.5367 | 29.42026 | 0.414428 | 1.574692 | 0.512743 |
| 7 | 108203400 | 108205199 | 35.14064 | 58.24369 | 50.92333 | 22.04765 | 0.574012 | 1.314637 | 0.512752 |
| 4 | 55450000 | 55451399 | 11.81792 | 21.10957 | 36.21807 | 15.91031 | -0.37606 | 0.431582 | 0.512767 |
| 1 | 142847600 | 142849599 | 13.07738 | 23.2268 | 41.86648 | 18.48001 | -0.45204 | 0.484323 | 0.512781 |
| 10 | 40403600 | 40406399 | 15.37663 | 26.87049 | 44.12201 | 19.44255 | -0.32446 | 0.702818 | 0.512783 |
| 7 | 140014400 | 140017199 | 26.00158 | 45.27135 | 37.90056 | 16.66245 | 0.642436 | 0.796608 | 0.512784 |
| 15 | 33731000 | 33737599 | 63.29666 | 101.3643 | 93.78244 | 39.53553 | 0.457189 | 2.026825 | 0.512786 |
| 9 | 59949000 | 59953199 | 54.92933 | 87.86053 | 91.97805 | 38.70863 | 0.277366 | 2.051247 | 0.512806 |
| 7 | 14354200 | 14359799 | 79.08619 | 124.3468 | 130.3845 | 54.08806 | 0.252255 | 2.537508 | 0.512811 |
| 9 | 6827200 | 6829599 | 21.79945 | 38.0997 | 36.93333 | 16.27678 | 0.432218 | 0.721033 | 0.512815 |
| 13 | 91044400 | 91046799 | 21.79945 | 38.0997 | 36.93333 | 16.27678 | 0.432218 | 0.721033 | 0.512815 |
| 12 | 17538800 | 17548199 | 60.03758 | 94.71419 | 147.4824 | 61.20885 | -0.31375 | 2.460991 | 0.51282 |
| 10 | 96081800 | 96082799 | 12.19105 | 23.54273 | 21.37726 | 8.228726 | 0.689305 | -0.10703 | 0.512828 |
| 8 | 46795400 | 46806199 | 123.1515 | 186.3621 | 269.1178 | 111.3228 | -0.25226 | 3.288636 | 0.512834 |
| 9 | 109222400 | 109224199 | 16.23235 | 30.16747 | 25.18591 | 10.54178 | 0.704157 | 0.213504 | 0.512839 |
| 16 | 97864400 | 97866599 | 52.45069 | 84.83729 | 73.34429 | 31.35619 | 0.571763 | 1.710821 | 0.512847 |
| 17 | 80240400 | 80242199 | 11.88109 | 21.11006 | 36.25469 | 15.99783 | -0.38174 | 0.467295 | 0.512849 |
| 14 | 34083400 | 34088399 | 51.29235 | 81.40202 | 130.5166 | 54.37277 | -0.34854 | 2.301668 | 0.512869 |
| 12 | 65046200 | 65048599 | 23.40999 | 40.51885 | 40.8756 | 17.93334 | 0.373349 | 0.855203 | 0.512898 |
| 12 | 56932200 | 56935199 | 22.1297 | 37.09605 | 61.4263 | 26.79221 | -0.34415 | 1.160045 | 0.512898 |
| 6 | 122022800 | 122026199 | 28.98279 | 47.67022 | 79.06527 | 34.08104 | -0.34945 | 1.409761 | 0.512904 |
| 12 | 115071400 | 115081999 | 194.7285 | 290.6 | 268.4271 | 110.6429 | 0.374173 | 3.570005 | 0.51291 |
| 7 | 84807000 | 84807799 | 17.8672 | 31.56739 | 32.2876 | 14.2834 | 0.35795 | 0.547924 | 0.512912 |
| 1 | 170447600 | 170449399 | 22.33007 | 39.03447 | 36.99408 | 16.30616 | 0.464817 | 0.723087 | 0.51293 |
| X | 34946200 | 34947999 | 22.33007 | 39.03447 | 36.99408 | 16.30616 | 0.464817 | 0.723087 | 0.51293 |
| 1 | 114176800 | 114178799 | 18.04078 | 32.71164 | 27.91786 | 12.07271 | 0.640287 | 0.368067 | 0.512944 |
| 1 | 23442000 | 23442599 | 5.652759 | 11.01016 | 22.53298 | 8.226402 | -0.40003 | -0.35627 | 0.512949 |
| 1 | 41562200 | 41562799 | 5.652759 | 11.01016 | 22.53298 | 8.226402 | -0.40003 | -0.35627 | 0.512949 |
| 18 | 17618200 | 17618599 | 5.652759 | 11.01016 | 22.53298 | 8.226402 | -0.40003 | -0.35627 | 0.512949 |
| 17 | 41608800 | 41612599 | 35.58811 | 59.32287 | 53.74005 | 23.35694 | 0.523319 | 1.253733 | 0.512958 |
| 17 | 48466000 | 48470599 | 58.76611 | 93.82658 | 99.1694 | 41.62761 | 0.260712 | 2.113451 | 0.512972 |
| 3 | 62825200 | 62827399 | 17.28425 | 30.93943 | 29.43581 | 12.90776 | 0.471274 | 0.439524 | 0.513006 |
| 13 | 58485200 | 58486999 | 10.22978 | 18.50783 | 33.3035 | 14.44145 | -0.43155 | 0.337158 | 0.513013 |
| X | 20905200 | 20907199 | 29.6786 | 50.69454 | 44.64057 | 19.57211 | 0.565507 | 1.005276 | 0.513019 |
| 5 | 81136600 | 81138399 | 19.21044 | 33.66678 | 34.94505 | 15.43618 | 0.334009 | 0.661239 | 0.513025 |
| 4 | 121438200 | 121442199 | 66.77863 | 106.5542 | 102.6246 | 43.0406 | 0.393645 | 2.14219 | 0.513042 |
| X | 53225600 | 53227399 | 26.23029 | 45.31723 | 40.89039 | 17.93883 | 0.533319 | 0.887499 | 0.513043 |
| 13 | 78875600 | 78877399 | 23.82206 | 40.71239 | 42.66701 | 18.70762 | 0.315361 | 0.973174 | 0.513047 |
| 8 | 27980200 | 27980799 | 6.253889 | 12.10785 | 23.81611 | 9.059511 | -0.39941 | -0.20351 | 0.513049 |
| 13 | 42126400 | 42127399 | 6.253889 | 12.10785 | 23.81611 | 9.059511 | -0.39941 | -0.20351 | 0.513049 |
| 4 | 145586400 | 145589399 | 50.14132 | 81.16343 | 73.37679 | 31.38761 | 0.508374 | 1.689959 | 0.513059 |
| X | 160940600 | 160943199 | 17.85902 | 30.86092 | 50.60629 | 22.20138 | -0.32497 | 0.860458 | 0.513062 |
| 10 | 120912800 | 120915999 | 41.48538 | 66.5664 | 104.882 | 44.34745 | -0.30608 | 1.942238 | 0.513071 |
| 13 | 40741000 | 40741599 | 7.558609 | 14.24258 | 28.60532 | 11.69777 | -0.52487 | 0.066795 | 0.513101 |
| 9 | 15752400 | 15757999 | 46.11241 | 73.92327 | 122.6246 | 51.77762 | -0.38184 | 1.982623 | 0.513112 |
| 5 | 118160200 | 118161599 | 8.579706 | 15.96638 | 29.98831 | 12.52489 | -0.45482 | 0.153677 | 0.513126 |
| 13 | 31508800 | 31513399 | 32.84516 | 53.23492 | 84.15534 | 36.03251 | -0.29415 | 1.639377 | 0.513139 |

| | | | | | | | | | |
|---|---|---|---|---|---|---|---|---|---|
| X | 49448600 | 49450199 | 23.25857 | 41.27211 | 29.47897 | 13.02209 | 0.875783 | 0.534106 | 0.513191 |
| 14 | 74969400 | 74970199 | 8.836948 | 16.13364 | 31.89218 | 13.67339 | -0.55538 | 0.270544 | 0.51323 |
| 1 | 133954800 | 133955799 | 14.23162 | 25.82051 | 26.46711 | 11.43668 | 0.378631 | 0.336032 | 0.51323 |
| X | 74734000 | 74735999 | 26.77767 | 46.28797 | 40.34213 | 17.69859 | 0.583345 | 0.884904 | 0.513252 |
| 2 | 113626400 | 113626999 | 7.745438 | 14.84685 | 28.04887 | 11.04164 | -0.38952 | -0.06392 | 0.513271 |
| 2 | 149109400 | 149110999 | 15.38844 | 26.94868 | 44.58682 | 19.68289 | -0.33537 | 0.669088 | 0.513278 |
| 5 | 100027400 | 100028399 | 14.47239 | 26.45673 | 27.34128 | 11.70316 | 0.375434 | 0.295298 | 0.513291 |
| 9 | 60478600 | 60479999 | 21.80647 | 37.88873 | 39.31349 | 17.26116 | 0.333877 | 0.806857 | 0.513297 |
| 9 | 88996400 | 88997799 | 17.90761 | 31.74206 | 26.7334 | 11.81863 | 0.638464 | 0.502136 | 0.51331 |
| X | 112914400 | 112915599 | 15.9018 | 29.192 | 27.31114 | 11.62417 | 0.524013 | 0.283969 | 0.513316 |
| 16 | 91525200 | 91527199 | 13.01553 | 22.80833 | 41.00343 | 18.11118 | -0.45431 | 0.639707 | 0.513318 |
| X | 8239200 | 8241399 | 32.21 | 54.42454 | 47.97765 | 20.99138 | 0.562598 | 1.111907 | 0.513321 |
| 7 | 28215200 | 28220799 | 36.57386 | 58.98681 | 93.27346 | 39.73099 | -0.30217 | 1.772344 | 0.513325 |
| 5 | 117845000 | 117849799 | 46.98922 | 74.87481 | 119.4105 | 49.98552 | -0.33482 | 2.170706 | 0.513333 |
| 15 | 46684000 | 46685799 | 18.53523 | 33.35051 | 29.47908 | 12.87058 | 0.581362 | 0.419163 | 0.513348 |
| 12 | 115303000 | 115309999 | 94.50752 | 148.0461 | 142.364 | 59.05187 | 0.372878 | 2.61869 | 0.513349 |
| 13 | 101476800 | 101478199 | 19.48042 | 34.17277 | 34.93158 | 15.44277 | 0.356045 | 0.648737 | 0.513363 |
| 8 | 14080600 | 14083599 | 26.51987 | 43.6457 | 70.61065 | 30.44448 | -0.31317 | 1.390508 | 0.513383 |
| 3 | 69329000 | 69330999 | 26.45113 | 45.00436 | 46.32326 | 20.30343 | 0.340255 | 1.027108 | 0.51339 |
| 5 | 117844800 | 117847599 | 34.36099 | 55.36662 | 92.13677 | 39.19899 | -0.37719 | 1.797157 | 0.513396 |
| 13 | 104464800 | 104465999 | 9.583678 | 17.0876 | 36.35749 | 15.99605 | -0.68596 | 0.428293 | 0.513401 |
| 10 | 116005000 | 116006999 | 23.80012 | 41.05748 | 41.98504 | 18.41822 | 0.352933 | 0.888311 | 0.513421 |
| X | 104653800 | 104658999 | 67.07176 | 106.7653 | 106.6795 | 44.59276 | 0.337007 | 2.220821 | 0.513423 |
| 6 | 107094400 | 107096399 | 18.62814 | 32.9874 | 32.06344 | 14.17773 | 0.432236 | 0.526609 | 0.513423 |
| 8 | 78341000 | 78343999 | 18.49788 | 31.56922 | 52.71223 | 23.10942 | -0.3543 | 0.976803 | 0.513446 |
| 14 | 4558200 | 4559399 | 11.72299 | 20.77244 | 36.75974 | 16.25231 | -0.42695 | 0.486089 | 0.513447 |
| 13 | 94616200 | 94620199 | 29.96812 | 49.04859 | 78.20761 | 33.64139 | -0.29564 | 1.472103 | 0.51347 |
| X | 164851000 | 164852399 | 15.59714 | 29.21067 | 24.91624 | 10.295 | 0.688444 | 0.161774 | 0.513477 |
| 1 | 51151000 | 51151599 | 5.36466 | 10.46 | 21.51814 | 7.748566 | -0.38963 | -0.41391 | 0.513491 |
| 2 | 117629400 | 117631599 | 15.52575 | 27.18398 | 45.64887 | 20.14851 | -0.35677 | 0.673649 | 0.513496 |
| 19 | 39073200 | 39074199 | 11.00552 | 20.11342 | 34.99818 | 14.98582 | -0.37172 | 0.288764 | 0.513514 |
| 11 | 73586600 | 73589599 | 36.21013 | 60.59844 | 51.43805 | 22.41217 | 0.616785 | 1.214193 | 0.513516 |
| 1 | 59520000 | 59523399 | 25.34821 | 41.41856 | 74.32913 | 31.95374 | -0.46724 | 1.483778 | 0.513524 |
| 3 | 31233200 | 31237999 | 94.52384 | 144.7633 | 215.3322 | 89.36683 | -0.2809 | 3.035512 | 0.513541 |
| 1 | 47108200 | 47110999 | 63.13136 | 100.7519 | 102.3553 | 42.93819 | 0.317054 | 2.133051 | 0.513557 |
| 11 | 8015800 | 8017599 | 11.82519 | 21.27144 | 35.75665 | 15.61487 | -0.3407 | 0.387822 | 0.513557 |
| 1 | 11448200 | 11449799 | 21.39066 | 35.79136 | 60.17006 | 26.21386 | -0.36578 | 1.177984 | 0.513564 |
| 2 | 42629400 | 42631399 | 23.29835 | 40.10479 | 42.03696 | 18.43978 | 0.316803 | 0.905355 | 0.513568 |
| 5 | 77922400 | 77924399 | 23.29835 | 40.10479 | 42.03696 | 18.43978 | 0.316803 | 0.905355 | 0.513568 |
| X | 74675400 | 74675999 | 18.50739 | 32.74762 | 31.30045 | 13.84255 | 0.456183 | 0.535802 | 0.513571 |
| 1 | 107036800 | 107037399 | 9.352748 | 16.98583 | 35.98958 | 15.53187 | -0.66093 | 0.316622 | 0.513625 |
| 3 | 18609000 | 18611799 | 26.25326 | 43.00044 | 73.07088 | 31.44145 | -0.38663 | 1.450766 | 0.513634 |
| 17 | 9947400 | 9949199 | 18.17764 | 32.07518 | 32.78308 | 14.50627 | 0.358279 | 0.564092 | 0.513639 |
| 2 | 76784800 | 76785799 | 9.762042 | 17.82783 | 31.87732 | 13.66199 | -0.41115 | 0.27785 | 0.513644 |
| 19 | 40000200 | 40009399 | 109.6446 | 169.5579 | 152.5073 | 63.34793 | 0.454845 | 2.848818 | 0.513655 |
| X | 119064200 | 119067199 | 48.54899 | 78.43635 | 76.7563 | 32.7685 | 0.391518 | 1.736231 | 0.513655 |
| 15 | 32801000 | 32801999 | 33.24016 | 54.71453 | 56.1792 | 24.22386 | 0.340986 | 1.394943 | 0.513691 |

| | | | | | | | | | |
|---|---|---|---|---|---|---|---|---|---|
| 15 | 103821200 | 103824199 | 30.49165 | 51.18431 | 51.31076 | 22.39977 | 0.377431 | 1.163469 | 0.51372 |
| 5 | 26777800 | 26779199 | 17.29042 | 30.5399 | 31.78939 | 14.06378 | 0.332538 | 0.545404 | 0.513735 |
| 3 | 76415000 | 76417799 | 39.76818 | 64.90144 | 58.55588 | 25.16172 | 0.520824 | 1.521631 | 0.513735 |
| 12 | 115038600 | 115042599 | 56.93081 | 91.78968 | 78.37164 | 33.36972 | 0.584483 | 1.807875 | 0.513765 |
| 12 | 106597400 | 106598199 | 8.074032 | 15.43848 | 29.09433 | 11.53481 | -0.39557 | -0.03407 | 0.513767 |
| 8 | 120617200 | 120622999 | 56.64765 | 89.46582 | 142.4458 | 59.15185 | -0.34449 | 2.432859 | 0.513769 |
| 13 | 41566600 | 41568799 | 26.17589 | 44.45531 | 45.7932 | 20.06669 | 0.338764 | 1.040203 | 0.513772 |
| 1 | 101238000 | 101239399 | 15.87233 | 28.81887 | 28.14565 | 12.15059 | 0.448475 | 0.34501 | 0.513781 |
| 2 | 150839600 | 150841599 | 13.05433 | 22.79184 | 42.66489 | 18.78966 | -0.51245 | 0.702131 | 0.513787 |
| 17 | 22715400 | 22716999 | 26.65086 | 45.11023 | 47.44639 | 20.76996 | 0.308727 | 1.074126 | 0.513788 |
| 5 | 66780600 | 66784399 | 28.67909 | 46.92786 | 75.01805 | 32.26663 | -0.2995 | 1.47096 | 0.513789 |
| 18 | 42739800 | 42741599 | 20.13573 | 35.10129 | 35.31198 | 15.53638 | 0.378914 | 0.749061 | 0.51379 |
| 4 | 137717800 | 137718599 | 9.11662 | 17.62921 | 19.79171 | 7.570727 | 0.393846 | -0.16734 | 0.513795 |
| 10 | 33672000 | 33674199 | 24.49714 | 42.57219 | 39.69202 | 17.42952 | 0.487578 | 0.81765 | 0.513811 |
| 2 | 18986600 | 18989599 | 26.91812 | 44.06603 | 84.24218 | 36.24017 | -0.55653 | 1.465721 | 0.513812 |
| 2 | 53132200 | 53132599 | 8.806041 | 17.07302 | 18.85766 | 7.126216 | 0.43481 | -0.21069 | 0.513819 |
| 1 | 173071000 | 173072799 | 20.70419 | 36.8022 | 30.16225 | 13.30475 | 0.679644 | 0.507221 | 0.51382 |
| 2 | 93328200 | 93330599 | 21.34244 | 36.00914 | 58.09523 | 25.4068 | -0.30641 | 1.095923 | 0.513823 |
| 5 | 12876000 | 12876599 | 6.836331 | 13.19734 | 25.41192 | 9.797831 | -0.38815 | -0.14796 | 0.513825 |
| 4 | 12401600 | 12405599 | 52.5817 | 83.59956 | 127.3758 | 53.16063 | -0.27254 | 2.249416 | 0.513842 |
| X | 80431200 | 80440599 | 154.8417 | 236.3663 | 187.8949 | 77.92096 | 0.618576 | 3.119146 | 0.513848 |
| 6 | 117559800 | 117565399 | 74.10381 | 117.4346 | 107.9771 | 44.91795 | 0.450432 | 2.356554 | 0.513851 |
| 5 | 96520600 | 96523799 | 21.94286 | 36.085 | 71.12431 | 30.65589 | -0.59756 | 1.388325 | 0.513873 |
| X | 116363000 | 116364599 | 21.24702 | 37.79185 | 30.39246 | 13.39889 | 0.707723 | 0.51039 | 0.513881 |
| X | 72700800 | 72704199 | 20.57967 | 34.97231 | 59.19566 | 25.94287 | -0.37488 | 1.033761 | 0.513881 |
| 4 | 12094400 | 12096199 | 34.83514 | 58.04102 | 48.1594 | 20.92536 | 0.649634 | 1.253478 | 0.513893 |
| 10 | 90877400 | 90879999 | 23.27759 | 40.88091 | 33.9301 | 15.01464 | 0.655038 | 0.649599 | 0.513911 |
| 11 | 19763400 | 19767599 | 59.59262 | 95.03226 | 100.4383 | 42.08944 | 0.258926 | 2.151891 | 0.513913 |
| 10 | 42894400 | 42899999 | 40.90266 | 65.4825 | 107.4145 | 45.26164 | -0.36793 | 2.01415 | 0.513916 |
| X | 80113800 | 80116199 | 28.40115 | 48.39954 | 44.7223 | 19.60426 | 0.495851 | 1.022157 | 0.513945 |
| 10 | 20751000 | 20751999 | 12.78652 | 22.29389 | 47.34003 | 20.83076 | -0.69365 | 0.728657 | 0.513974 |
| 12 | 24014800 | 24016599 | 14.504 | 26.57564 | 22.77764 | 9.719601 | 0.646746 | 0.272178 | 0.513981 |
| 6 | 79936200 | 79938199 | 21.5688 | 37.89688 | 33.40784 | 14.78914 | 0.568285 | 0.63017 | 0.513986 |
| 12 | 82140800 | 82145399 | 41.48263 | 66.62849 | 103.5245 | 43.83997 | -0.28427 | 1.911027 | 0.514011 |
| 18 | 73009400 | 73011599 | 51.34454 | 82.49308 | 84.83941 | 35.94302 | 0.310428 | 1.910207 | 0.514014 |
| 12 | 45241800 | 45243599 | 27.09447 | 44.27101 | 74.59892 | 32.06948 | -0.37671 | 1.488087 | 0.514016 |
| 13 | 81600800 | 81601399 | 9.730186 | 17.97056 | 31.46891 | 13.28418 | -0.36562 | 0.212895 | 0.514023 |
| 16 | 13572400 | 13574399 | 12.90769 | 22.67281 | 39.95914 | 17.68516 | -0.42555 | 0.601912 | 0.514025 |
| 18 | 45523200 | 45525399 | 19.66716 | 34.52933 | 29.36318 | 12.99027 | 0.620159 | 0.626456 | 0.514027 |
| 14 | 100491000 | 100494399 | 19.50636 | 34.38476 | 33.05678 | 14.63035 | 0.445418 | 0.58152 | 0.514043 |
| 13 | 46557400 | 46558999 | 12.30947 | 22.07376 | 37.27656 | 16.32139 | -0.35003 | 0.413329 | 0.514063 |
| 17 | 90631000 | 90631999 | 11.83149 | 21.09718 | 36.21906 | 15.93404 | -0.37837 | 0.442848 | 0.514066 |
| 1 | 97432000 | 97436399 | 50.08434 | 80.7206 | 81.35382 | 34.62263 | 0.345074 | 1.808551 | 0.514077 |
| X | 19924800 | 19927399 | 42.01605 | 68.7911 | 63.3016 | 27.23327 | 0.49565 | 1.487634 | 0.514087 |
| 14 | 75304400 | 75304799 | 4.96451 | 9.668275 | 21.30846 | 7.793587 | -0.50832 | -0.36508 | 0.514087 |
| 13 | 93704600 | 93707799 | 31.13953 | 51.89428 | 54.12224 | 23.5193 | 0.320729 | 1.243668 | 0.51412 |
| 1 | 171719600 | 171723199 | 52.5121 | 83.62012 | 126.6971 | 52.98741 | -0.26211 | 2.19834 | 0.514132 |

| | | | | | | | | | |
|---|---|---|---|---|---|---|---|---|---|
| 2 | 122754200 | 122755199 | 19.29272 | 33.66199 | 33.70975 | 14.83982 | 0.385409 | 0.727138 | 0.514164 |
| 2 | 112548000 | 112548599 | 9.887422 | 18.98412 | 20.29069 | 7.949699 | 0.43265 | -0.0765 | 0.51418 |
| 10 | 8645400 | 8646999 | 24.63799 | 41.87937 | 43.23867 | 18.95 | 0.335705 | 1.025869 | 0.514197 |
| 4 | 131450800 | 131454399 | 28.84414 | 47.15976 | 75.54998 | 32.48584 | -0.30341 | 1.483094 | 0.514213 |
| 7 | 85115000 | 85117999 | 54.40766 | 87.71745 | 79.80042 | 33.97601 | 0.493057 | 1.805458 | 0.51422 |
| 4 | 34102000 | 34107199 | 51.98247 | 84.06357 | 74.0678 | 31.65962 | 0.544135 | 1.713158 | 0.514249 |
| X | 143690000 | 143690399 | 8.93438 | 17.2812 | 18.82379 | 7.191924 | 0.439668 | -0.17315 | 0.51425 |
| 15 | 24670200 | 24671799 | 16.2958 | 29.77064 | 27.05675 | 11.59083 | 0.55864 | 0.310674 | 0.514262 |
| 10 | 5870800 | 5872999 | 17.03717 | 28.8687 | 54.18489 | 23.7275 | -0.52339 | 1.039773 | 0.514269 |
| 2 | 167254800 | 167256799 | 23.28598 | 39.09777 | 64.30993 | 28.07872 | -0.33462 | 1.149994 | 0.514274 |
| 8 | 96245000 | 96253799 | 85.57045 | 132.1477 | 194.2756 | 80.69805 | -0.25479 | 2.869539 | 0.514292 |
| 1 | 53376400 | 53378399 | 28.77564 | 49.52346 | 41.79143 | 18.33196 | 0.628286 | 0.929981 | 0.514303 |
| 9 | 5921800 | 5926999 | 31.40691 | 52.48647 | 47.04334 | 20.47921 | 0.538545 | 1.209015 | 0.51432 |
| 1 | 12180400 | 12181599 | 11.1689 | 20.43003 | 37.21607 | 15.91744 | -0.43648 | 0.28903 | 0.514372 |
| 15 | 97481600 | 97485599 | 40.35827 | 64.93425 | 100.2716 | 42.56871 | -0.27228 | 1.855081 | 0.514391 |
| 12 | 89562000 | 89562399 | 7.291588 | 14.01575 | 26.40201 | 10.31082 | -0.37429 | -0.09844 | 0.514396 |
| 5 | 90454600 | 90457399 | 22.28725 | 38.54693 | 40.43356 | 17.739 | 0.317112 | 0.85656 | 0.514406 |
| 5 | 59506200 | 59508799 | 38.46205 | 63.56791 | 58.39338 | 25.23234 | 0.502656 | 1.360448 | 0.514421 |
| 2 | 162075000 | 162077199 | 21.82634 | 36.80321 | 59.2182 | 25.89313 | -0.30265 | 1.10371 | 0.514422 |
| 11 | 41175800 | 41177199 | 16.64422 | 29.40327 | 29.46913 | 13.03686 | 0.387068 | 0.535956 | 0.514425 |
| 6 | 131527200 | 131528799 | 26.76867 | 45.50762 | 39.4956 | 17.30974 | 0.585203 | 1.027064 | 0.514426 |
| 3 | 54696400 | 54696999 | 6.123679 | 11.81445 | 24.20412 | 9.352592 | -0.47859 | -0.14853 | 0.514456 |
| 14 | 122739200 | 122742199 | 36.912 | 61.19835 | 56.64152 | 24.52857 | 0.492118 | 1.318977 | 0.514464 |
| 1 | 23553400 | 23554999 | 11.64175 | 20.83702 | 36.22395 | 15.88577 | -0.39329 | 0.417111 | 0.514468 |
| 15 | 52889800 | 52891199 | 11.64175 | 20.83702 | 36.22395 | 15.88577 | -0.39329 | 0.417111 | 0.514468 |
| 14 | 122271400 | 122275199 | 24.10189 | 39.81193 | 75.14148 | 32.45679 | -0.53323 | 1.343004 | 0.514469 |
| 14 | 105235200 | 105236199 | 9.667858 | 17.86204 | 31.45826 | 13.27301 | -0.37363 | 0.207953 | 0.514505 |
| 1 | 13066200 | 13069599 | 34.32009 | 55.48132 | 87.42137 | 37.35056 | -0.29313 | 1.699462 | 0.514515 |
| 10 | 61655400 | 61657399 | 18.82384 | 32.27168 | 52.20003 | 22.89324 | -0.30761 | 0.937115 | 0.514523 |
| 1 | 125029800 | 125031199 | 13.29416 | 24.76348 | 25.48084 | 10.62503 | 0.408973 | 0.176265 | 0.514545 |
| 3 | 78652000 | 78653599 | 13.29416 | 24.76348 | 25.48084 | 10.62503 | 0.408973 | 0.176265 | 0.514545 |
| X | 26351800 | 26354599 | 24.01941 | 41.71184 | 34.6631 | 15.21432 | 0.652853 | 0.81996 | 0.514568 |
| 3 | 123954200 | 123954999 | 13.75907 | 25.9712 | 24.00666 | 9.784882 | 0.588819 | 0.093525 | 0.514574 |
| 12 | 22515000 | 22517999 | 112.8838 | 173.1686 | 168.6906 | 70.02917 | 0.33104 | 3.005267 | 0.514579 |
| 2 | 69186800 | 69187799 | 12.09267 | 22.63241 | 23.99964 | 9.927424 | 0.374973 | 0.131999 | 0.514579 |
| 1 | 8788200 | 8793199 | 61.70559 | 99.07595 | 86.17885 | 36.47537 | 0.55048 | 1.941503 | 0.514597 |
| 19 | 13982400 | 13984799 | 23.69283 | 41.63503 | 32.72735 | 14.48972 | 0.732973 | 0.640052 | 0.514599 |
| 7 | 47270200 | 47271399 | 29.54214 | 49.71692 | 39.69904 | 17.34818 | 0.704558 | 1.126049 | 0.514624 |
| 7 | 23653400 | 23657199 | 34.87469 | 57.64167 | 52.03902 | 22.48729 | 0.527575 | 1.344998 | 0.51464 |
| 6 | 34932000 | 34933999 | 17.41651 | 29.77901 | 51.09445 | 22.40353 | -0.39283 | 0.95515 | 0.514655 |
| 3 | 64306400 | 64307599 | 16.37214 | 29.40385 | 24.74215 | 10.81934 | 0.650374 | 0.405325 | 0.514662 |
| 19 | 9535000 | 9541799 | 124.176 | 189.6945 | 194.9967 | 80.87986 | 0.248831 | 3.093568 | 0.514664 |
| 7 | 5886200 | 5892599 | 83.93864 | 133.1845 | 112.7935 | 46.97956 | 0.570605 | 2.330082 | 0.514672 |
| 9 | 73195600 | 73197999 | 22.23795 | 39.07669 | 33.97111 | 15.03983 | 0.588441 | 0.63485 | 0.514681 |
| 1 | 3088400 | 3091399 | 35.72769 | 58.94269 | 59.79712 | 25.80863 | 0.359536 | 1.375982 | 0.514683 |
| 3 | 78282400 | 78282799 | 3.412634 | 6.6752 | 16.71965 | 5.604887 | -0.57348 | -0.7128 | 0.514692 |
| 8 | 69748800 | 69759399 | 250.1772 | 366.9519 | 322.4672 | 132.8193 | 0.422875 | 3.867864 | 0.514697 |

| | | | | | | | | | |
|---|---|---|---|---|---|---|---|---|---|
| 10 | 3216000 | 3218999 | 28.72806 | 46.86582 | 76.33671 | 32.79758 | -0.32954 | 1.516202 | 0.514699 |
| X | 3270600 | 3272199 | 20.41144 | 35.94429 | 33.69223 | 14.91456 | 0.48146 | 0.601365 | 0.5147 |
| 9 | 57199200 | 57201399 | 28.09343 | 48.21282 | 42.35352 | 18.57641 | 0.569695 | 0.950741 | 0.514707 |
| 13 | 93413600 | 93417399 | 34.33029 | 57.48262 | 51.96322 | 22.64848 | 0.52623 | 1.202827 | 0.514715 |
| 2 | 11940800 | 11944199 | 51.70244 | 83.55298 | 75.25741 | 32.13945 | 0.511274 | 1.731374 | 0.514717 |
| 10 | 11573800 | 11574399 | 17.85681 | 33.25635 | 24.65847 | 10.28533 | 0.878448 | 0.21447 | 0.514729 |
| 5 | 107097600 | 107098999 | 9.995432 | 18.45074 | 31.97701 | 13.50832 | -0.35185 | 0.21734 | 0.51473 |
| 14 | 13815000 | 13818999 | 37.85573 | 62.22376 | 55.41516 | 23.8801 | 0.54482 | 1.420947 | 0.514736 |
| 18 | 7713200 | 7721599 | 160.4454 | 241.2212 | 244.0416 | 100.7565 | 0.252585 | 3.423668 | 0.51474 |
| 11 | 28233000 | 28240799 | 100.716 | 156.6551 | 156.1253 | 64.81538 | 0.313425 | 2.744783 | 0.514748 |
| 18 | 24097600 | 24099199 | 17.27726 | 32.12757 | 25.20319 | 10.53994 | 0.794607 | 0.220035 | 0.514748 |
| 7 | 41183600 | 41187799 | 71.92848 | 114.6944 | 102.9459 | 43.13518 | 0.494152 | 2.168595 | 0.514774 |
| 11 | 22464000 | 22466399 | 17.8516 | 30.77269 | 50.06221 | 21.96105 | -0.31427 | 0.881566 | 0.514775 |
| 10 | 21735600 | 21742399 | 47.54484 | 76.006 | 117.5148 | 49.41614 | -0.28513 | 2.06821 | 0.514796 |
| 1 | 185957800 | 185958799 | 11.54906 | 21.9748 | 23.22805 | 9.308633 | 0.418245 | 0.018173 | 0.514803 |
| 4 | 12839400 | 12842199 | 21.08426 | 37.07653 | 34.06311 | 15.08301 | 0.509355 | 0.619267 | 0.514819 |
| 6 | 95842400 | 95843799 | 14.26954 | 25.13342 | 41.85505 | 18.52728 | -0.34347 | 0.57672 | 0.514829 |
| Y | 1819000 | 1820999 | 17.0174 | 30.15671 | 20.62465 | 9.118447 | 0.936484 | 0.482691 | 0.51483 |
| 1 | 112701200 | 112704999 | 48.64312 | 78.47684 | 79.49868 | 33.87712 | 0.339369 | 1.777225 | 0.514832 |
| X | 135567000 | 135568199 | 9.460524 | 17.62234 | 33.27144 | 13.87369 | -0.46011 | 0.166593 | 0.514837 |
| Y | 84672600 | 84673599 | 10.66324 | 18.60414 | 43.76577 | 19.26599 | -0.8401 | 0.702884 | 0.514837 |
| 1 | 114665400 | 114668999 | 29.172 | 49.0424 | 49.65732 | 21.69295 | 0.363001 | 1.141302 | 0.51487 |
| 4 | 128262000 | 128265399 | 33.00302 | 53.39311 | 85.69357 | 36.64173 | -0.31844 | 1.676374 | 0.514873 |
| 19 | 52834200 | 52835999 | 18.87215 | 33.95258 | 28.80024 | 12.57549 | 0.640153 | 0.419491 | 0.514907 |
| 19 | 13997200 | 14000199 | 48.60713 | 78.13421 | 81.77055 | 34.66809 | 0.286191 | 1.888269 | 0.514919 |
| 7 | 118713400 | 118717799 | 31.98981 | 51.94405 | 82.15437 | 35.22351 | -0.29238 | 1.59855 | 0.514925 |
| 1 | 44406200 | 44408399 | 43.98624 | 71.09157 | 75.13675 | 32.09272 | 0.281289 | 1.715348 | 0.51493 |
| 3 | 49453600 | 49454999 | 12.52475 | 23.52908 | 23.9715 | 9.85306 | 0.439458 | 0.11162 | 0.51494 |
| 3 | 13023400 | 13031999 | 150.8897 | 228.6543 | 201.003 | 83.18288 | 0.464981 | 3.262762 | 0.514946 |
| 3 | 62796200 | 62796799 | 8.779466 | 17.00431 | 18.72643 | 7.110115 | 0.431347 | -0.19983 | 0.514967 |
| 5 | 5092000 | 5094599 | 24.21123 | 41.83557 | 41.38351 | 18.15525 | 0.401058 | 0.873854 | 0.51498 |
| 9 | 123023000 | 123028199 | 90.46931 | 140.8567 | 143.7594 | 59.67288 | 0.280256 | 2.72199 | 0.514996 |
| 7 | 18910400 | 18913999 | 33.71031 | 54.41314 | 88.97431 | 37.9399 | -0.34895 | 1.744169 | 0.514997 |
| 15 | 47225200 | 47227999 | 35.77513 | 59.00401 | 59.89472 | 25.84761 | 0.358479 | 1.377885 | 0.515092 |
| 10 | 37811000 | 37812199 | 13.46285 | 23.61416 | 40.93886 | 18.09687 | -0.4023 | 0.626499 | 0.515105 |
| 5 | 112602800 | 112604199 | 15.3988 | 26.77384 | 45.28008 | 19.88948 | -0.36754 | 0.776734 | 0.515105 |
| 2 | 72821400 | 72824199 | 30.29763 | 49.42562 | 79.12512 | 33.99541 | -0.30475 | 1.519524 | 0.515105 |
| 3 | 51895200 | 51896999 | 15.43607 | 26.31377 | 56.84337 | 24.91825 | -0.72441 | 0.982071 | 0.51511 |
| 2 | 114297000 | 114299599 | 20.27556 | 35.78251 | 33.01919 | 14.6103 | 0.504976 | 0.574636 | 0.515117 |
| 8 | 52220000 | 52222799 | 19.84731 | 35.43901 | 30.36731 | 13.34423 | 0.619619 | 0.475834 | 0.515124 |
| 13 | 83852400 | 83854799 | 17.12686 | 31.03722 | 27.95321 | 12.09643 | 0.562368 | 0.363939 | 0.515126 |
| 15 | 96906600 | 96910799 | 31.97514 | 52.08331 | 82.61085 | 35.47345 | -0.29298 | 1.548069 | 0.515142 |
| 9 | 72475200 | 72477599 | 27.30381 | 47.85843 | 32.69422 | 14.44392 | 0.934756 | 0.7011 | 0.515144 |
| 14 | 103993200 | 103994399 | 9.051254 | 16.82959 | 29.97169 | 12.53599 | -0.38017 | 0.164275 | 0.515153 |
| X | 67261600 | 67264199 | 27.22866 | 47.40798 | 37.29638 | 16.39687 | 0.731788 | 0.802116 | 0.515156 |
| 1 | 115816800 | 115819199 | 55.53697 | 89.03265 | 75.71358 | 31.97009 | 0.580349 | 1.977804 | 0.515165 |
| 18 | 83289400 | 83292599 | 23.00473 | 38.46291 | 61.91484 | 26.96111 | -0.30342 | 1.192959 | 0.515166 |

| | | | | | | | | | |
|---|---|---|---|---|---|---|---|---|---|
| Y | 20501600 | 20502999 | 10.21766 | 19.589 | 14.07588 | 5.538421 | 0.993512 | -0.08326 | 0.515194 |
| 4 | 55292000 | 55294799 | 18.31683 | 31.60655 | 51.69447 | 22.67778 | -0.32175 | 0.875804 | 0.515221 |
| 14 | 6098000 | 6099799 | 18.31683 | 31.60655 | 51.69447 | 22.67778 | -0.32175 | 0.875804 | 0.515221 |
| 3 | 113115800 | 113118199 | 33.16829 | 55.07185 | 56.40528 | 24.45091 | 0.346576 | 1.292732 | 0.515227 |
| 7 | 71900000 | 71900999 | 12.71479 | 22.43843 | 38.33503 | 16.96256 | -0.3787 | 0.54197 | 0.515232 |
| 2 | 131986000 | 131987799 | 12.69263 | 22.33989 | 38.85779 | 17.203 | -0.40609 | 0.570711 | 0.515234 |
| 7 | 3591000 | 3592399 | 20.22862 | 35.45522 | 27.51029 | 12.15322 | 0.751757 | 0.65281 | 0.515248 |
| 8 | 57324400 | 57326399 | 14.43 | 25.44115 | 45.03887 | 19.93267 | -0.43084 | 0.571349 | 0.515268 |
| 5 | 111459600 | 111464599 | 74.89419 | 118.2117 | 124.7692 | 51.7956 | 0.247077 | 2.455405 | 0.515272 |
| 9 | 47506000 | 47507199 | 10.16692 | 18.87115 | 34.31851 | 14.38901 | -0.41313 | 0.192981 | 0.515287 |
| 16 | 31535000 | 31537399 | 33.13471 | 55.70788 | 50.21334 | 21.93628 | 0.530373 | 1.158341 | 0.515288 |
| 1 | 89901800 | 89905599 | 27.3915 | 44.80223 | 74.2308 | 31.92315 | -0.35141 | 1.473233 | 0.515305 |
| 7 | 56116400 | 56117399 | 9.597461 | 18.64804 | 19.39577 | 7.230846 | 0.537545 | -0.24969 | 0.515331 |
| 9 | 87464200 | 87465999 | 13.70561 | 23.98361 | 47.28145 | 20.85918 | -0.58681 | 0.677852 | 0.515344 |
| 7 | 27151600 | 27152399 | 6.113461 | 11.8242 | 23.76209 | 9.081096 | -0.43643 | -0.18968 | 0.51535 |
| 7 | 65651000 | 65651599 | 6.113461 | 11.8242 | 23.76209 | 9.081096 | -0.43643 | -0.18968 | 0.51535 |
| 17 | 8580400 | 8581799 | 9.37557 | 17.41728 | 30.48846 | 12.76819 | -0.35682 | 0.173862 | 0.515356 |
| X | 23765200 | 23768999 | 39.07926 | 65.10445 | 52.63247 | 22.86715 | 0.687155 | 1.272748 | 0.515375 |
| 5 | 150984200 | 150987199 | 37.48087 | 60.27187 | 97.11506 | 41.19616 | -0.3344 | 1.868413 | 0.515403 |
| 14 | 119458000 | 119459999 | 32.59742 | 54.35639 | 54.12622 | 23.52991 | 0.387328 | 1.243679 | 0.515406 |
| 12 | 50838000 | 50839599 | 20.22121 | 35.41378 | 35.93674 | 15.865 | 0.366502 | 0.679501 | 0.515408 |
| 6 | 10843200 | 10844799 | 20.49693 | 36.24208 | 28.30842 | 12.52112 | 0.745111 | 0.547116 | 0.515416 |
| 6 | 102900600 | 102903599 | 31.38504 | 53.46792 | 44.56022 | 19.53273 | 0.643974 | 1.032 | 0.515425 |
| 5 | 93074400 | 93075999 | 8.272273 | 15.50816 | 28.52023 | 11.76397 | -0.40977 | 0.101399 | 0.515426 |
| 10 | 86782000 | 86782999 | 8.272273 | 15.50816 | 28.52023 | 11.76397 | -0.40977 | 0.101399 | 0.515426 |
| 14 | 40677400 | 40678799 | 8.272273 | 15.50816 | 28.52023 | 11.76397 | -0.40977 | 0.101399 | 0.515426 |
| 14 | 62638400 | 62642399 | 27.79217 | 45.73981 | 73.15698 | 31.54241 | -0.29679 | 1.394536 | 0.515431 |
| 16 | 84879600 | 84881199 | 13.30554 | 23.63291 | 40.29179 | 17.78438 | -0.37188 | 0.485304 | 0.515439 |
| 17 | 10374800 | 10378599 | 53.66071 | 86.05472 | 90.46386 | 38.21864 | 0.275665 | 1.969986 | 0.51544 |
| 12 | 20928600 | 20951399 | 840.4036 | 1100.044 | 974.2751 | 407.8542 | 0.247626 | 5.624301 | 0.51544 |
| 15 | 96513200 | 96517399 | 64.34154 | 101.5155 | 150.7329 | 62.56146 | -0.24513 | 2.462842 | 0.515447 |
| 10 | 3556800 | 3559199 | 44.36187 | 71.57698 | 73.99072 | 31.53406 | 0.310434 | 1.765473 | 0.515448 |
| 4 | 58779000 | 58779599 | 10.03132 | 19.40223 | 20.93128 | 7.998501 | 0.452704 | -0.1553 | 0.515477 |
| 3 | 90903600 | 90905799 | 36.07156 | 59.38434 | 60.86479 | 26.24596 | 0.343627 | 1.40338 | 0.515489 |
| 7 | 73633800 | 73635599 | 17.45872 | 31.057 | 31.1111 | 13.71446 | 0.392366 | 0.484081 | 0.515496 |
| 5 | 133522000 | 133525999 | 39.03146 | 62.80419 | 98.26934 | 41.72323 | -0.29112 | 1.849408 | 0.515501 |
| Y | 40972200 | 40974799 | 17.13447 | 30.36593 | 20.10432 | 8.888296 | 0.982984 | 0.481367 | 0.515506 |
| X | 73897600 | 73904199 | 119.2565 | 183.9539 | 154.4579 | 64.15901 | 0.551192 | 2.900663 | 0.515507 |
| X | 108861800 | 108870199 | 119.6458 | 185.5558 | 156.046 | 64.81076 | 0.554918 | 2.804295 | 0.515517 |
| 1 | 53535400 | 53536199 | 9.166711 | 17.18271 | 30.69392 | 12.6634 | -0.36885 | 0.114154 | 0.51552 |
| 7 | 50048000 | 50050199 | 19.78853 | 34.68515 | 35.45014 | 15.66129 | 0.356259 | 0.663842 | 0.515548 |
| Y | 9459400 | 9461799 | 19.96066 | 34.92714 | 22.77135 | 10.0455 | 1.001194 | 0.665865 | 0.515559 |
| 4 | 38612000 | 38612599 | 7.171232 | 13.68799 | 26.15465 | 10.4119 | -0.42017 | -0.03683 | 0.515561 |
| 10 | 3421200 | 3421799 | 14.519 | 26.17515 | 26.45416 | 11.52224 | 0.391488 | 0.373895 | 0.515599 |
| 2 | 166693600 | 166694599 | 12.96204 | 22.89265 | 38.75506 | 17.14573 | -0.36542 | 0.53299 | 0.51561 |
| 3 | 57535400 | 57535999 | 5.378843 | 10.47818 | 21.98079 | 8.009349 | -0.43347 | -0.37173 | 0.515614 |
| 13 | 99231600 | 99232199 | 6.69778 | 12.98809 | 25.73151 | 9.714931 | -0.39962 | -0.21809 | 0.515623 |

| | | | | | | | | | |
|---|---|---|---|---|---|---|---|---|---|
| 4 | 29867600 | 29870399 | 32.77498 | 55.57226 | 46.25441 | 20.2588 | 0.645212 | 1.080119 | 0.515641 |
| 6 | 49189800 | 49192799 | 18.49004 | 30.78657 | 62.49656 | 27.1433 | -0.63665 | 1.225057 | 0.51569 |
| 5 | 143754800 | 143766599 | 193.9099 | 281.7585 | 410.1479 | 169.0716 | -0.31795 | 4.007265 | 0.515721 |
| 1 | 81532000 | 81537399 | 50.28556 | 80.12195 | 122.065 | 51.09196 | -0.26906 | 2.177551 | 0.515725 |
| 7 | 63757000 | 63757999 | 14.14481 | 24.70962 | 42.6242 | 18.77837 | -0.39506 | 0.700294 | 0.515726 |
| 4 | 20634600 | 20636799 | 23.74025 | 41.41059 | 38.00364 | 16.72719 | 0.510758 | 0.761017 | 0.515728 |
| 17 | 69790600 | 69791199 | 10.63004 | 20.45596 | 21.87119 | 8.513712 | 0.440305 | -0.08671 | 0.515737 |
| 6 | 23542400 | 23544399 | 17.50752 | 31.01185 | 31.05195 | 13.72969 | 0.389616 | 0.513084 | 0.515738 |
| 1 | 53176400 | 53177999 | 12.43646 | 22.32008 | 39.39869 | 17.23844 | -0.41306 | 0.411586 | 0.515796 |
| 4 | 12987600 | 12988999 | 15.00338 | 27.79149 | 26.38604 | 11.09385 | 0.515434 | 0.225038 | 0.515804 |
| 3 | 22805000 | 22809999 | 73.03257 | 115.7942 | 118.8309 | 49.45756 | 0.29288 | 2.343986 | 0.515813 |
| 3 | 23068400 | 23069999 | 21.77006 | 37.87088 | 38.80588 | 17.05113 | 0.352122 | 0.790818 | 0.515814 |
| X | 47541600 | 47544999 | 39.51298 | 65.43616 | 54.27943 | 23.48721 | 0.649738 | 1.342573 | 0.515818 |
| 13 | 115773800 | 115775599 | 14.98758 | 27.49526 | 27.33524 | 11.64504 | 0.435841 | 0.279912 | 0.515836 |
| 9 | 12309400 | 12311799 | 23.12888 | 38.80211 | 62.58134 | 27.30983 | -0.3063 | 1.156958 | 0.515839 |
| 3 | 51653600 | 51656799 | 28.1577 | 46.28463 | 73.59703 | 31.71569 | -0.28907 | 1.411905 | 0.515843 |
| 19 | 8355000 | 8359399 | 49.53461 | 79.87268 | 80.32818 | 34.20717 | 0.348911 | 1.792836 | 0.515844 |
| 10 | 22049800 | 22051999 | 16.56932 | 29.35017 | 30.21614 | 13.36011 | 0.349828 | 0.506096 | 0.515862 |
| 17 | 46209200 | 46215599 | 63.82505 | 100.5295 | 153.0661 | 63.51177 | -0.28322 | 2.49602 | 0.515894 |
| 1 | 39073600 | 39074999 | 23.65508 | 40.40993 | 41.6594 | 18.26521 | 0.33892 | 0.97527 | 0.515894 |
| 5 | 117637600 | 117640999 | 31.31903 | 50.8683 | 82.21149 | 35.25487 | -0.32316 | 1.590914 | 0.515896 |
| 9 | 105790000 | 105791999 | 14.843 | 25.97444 | 44.60072 | 19.67685 | -0.3888 | 0.675902 | 0.515907 |
| 8 | 5688600 | 5691399 | 47.15655 | 75.18566 | 117.9415 | 49.41084 | -0.31011 | 2.151931 | 0.515945 |
| 17 | 48136000 | 48138599 | 28.53977 | 46.55122 | 77.02863 | 33.09293 | -0.35242 | 1.516919 | 0.515952 |
| X | 8447600 | 8452599 | 71.11339 | 113.4302 | 101.6897 | 42.62728 | 0.496367 | 2.159347 | 0.515968 |
| 3 | 64856400 | 64856999 | 7.314423 | 13.98672 | 26.19932 | 10.38558 | -0.38624 | -0.04901 | 0.515981 |
| 10 | 32057600 | 32059199 | 19.17732 | 32.33332 | 63.92868 | 27.95196 | -0.5987 | 1.102318 | 0.515981 |
| Y | 87047400 | 87049399 | 14.39366 | 26.17023 | 18.5901 | 8.011567 | 0.906158 | 0.302689 | 0.515994 |
| 9 | 82358000 | 82358799 | 14.085 | 24.38824 | 51.06589 | 22.40438 | -0.67509 | 0.827885 | 0.516003 |
| 15 | 90901400 | 90902399 | 6.115237 | 11.82684 | 23.74651 | 9.077998 | -0.43514 | -0.1868 | 0.516014 |
| 5 | 52787800 | 52792799 | 47.88035 | 76.41044 | 119.6122 | 50.17543 | -0.3057 | 2.121713 | 0.51602 |
| Y | 40159400 | 40160199 | 10.41633 | 19.88581 | 15.1171 | 6.015538 | 0.898381 | -0.0411 | 0.516021 |
| Y | 42321400 | 42322799 | 10.41633 | 19.88581 | 15.1171 | 6.015538 | 0.898381 | -0.0411 | 0.516021 |
| 5 | 130411200 | 130414399 | 22.61987 | 37.48665 | 64.88559 | 28.09037 | -0.40816 | 1.293658 | 0.516027 |
| 5 | 50837400 | 50840199 | 28.4417 | 46.70368 | 75.46156 | 32.50512 | -0.31274 | 1.424694 | 0.516028 |
| 2 | 128305800 | 128306599 | 6.410093 | 12.37995 | 24.25629 | 9.333021 | -0.4094 | -0.16063 | 0.516041 |
| 14 | 24294400 | 24295799 | 9.421624 | 17.17812 | 32.86604 | 14.11448 | -0.51048 | 0.283817 | 0.516052 |
| 18 | 75439600 | 75441399 | 17.07971 | 29.28894 | 50.04533 | 21.9487 | -0.38618 | 0.925635 | 0.516052 |
| 6 | 118586800 | 118588399 | 9.889506 | 18.01913 | 32.34018 | 13.90028 | -0.4195 | 0.291366 | 0.516053 |
| 12 | 64832800 | 64834199 | 7.808431 | 14.93589 | 28.47464 | 11.2777 | -0.41022 | -0.03927 | 0.516074 |
| 7 | 15248000 | 15252599 | 55.02031 | 89.11058 | 69.25584 | 29.63699 | 0.726776 | 1.686629 | 0.516094 |
| 15 | 98275200 | 98278799 | 33.34712 | 53.90409 | 86.62153 | 37.00499 | -0.32153 | 1.699598 | 0.516115 |
| 18 | 48417800 | 48418599 | 11.89109 | 22.17441 | 23.09833 | 9.613573 | 0.393892 | 0.152929 | 0.516124 |
| 9 | 22611000 | 22612399 | 18.21164 | 32.22755 | 32.20113 | 14.24058 | 0.392369 | 0.532714 | 0.516133 |
| 1 | 3761600 | 3762599 | 8.239061 | 15.47903 | 29.09039 | 11.95568 | -0.43695 | 0.086951 | 0.516141 |
| 15 | 3216600 | 3219999 | 30.99371 | 52.33978 | 49.07867 | 21.46942 | 0.473702 | 1.111021 | 0.516148 |
| 12 | 17363000 | 17366599 | 22.35525 | 37.35481 | 61.31499 | 26.6899 | -0.33143 | 1.196403 | 0.516162 |

| | | | | | | | | | |
|---|---|---|---|---|---|---|---|---|---|
| 3 | 99407200 | 99408799 | 15.4223 | 26.81291 | 45.28638 | 19.89141 | -0.3656 | 0.778642 | 0.516184 |
| 13 | 7649600 | 7656999 | 119.5151 | 183.7688 | 164.5913 | 68.34924 | 0.454675 | 2.960662 | 0.516194 |
| 18 | 66208600 | 66210599 | 13.36396 | 23.62761 | 39.80642 | 17.60713 | -0.35771 | 0.528395 | 0.516229 |
| 6 | 131107000 | 131109599 | 37.94948 | 62.17003 | 63.79412 | 27.45206 | 0.339135 | 1.467677 | 0.51623 |
| 12 | 67491000 | 67491599 | 8.418288 | 15.81224 | 28.54784 | 11.73712 | -0.37959 | 0.09055 | 0.516238 |
| 2 | 71364600 | 71365799 | 10.52674 | 19.03261 | 33.78284 | 14.66131 | -0.41285 | 0.343583 | 0.516245 |
| 2 | 97501200 | 97502799 | 20.67386 | 36.66703 | 31.31466 | 13.83889 | 0.618895 | 0.523712 | 0.516271 |
| 6 | 131726200 | 131737999 | 169.7739 | 256.2489 | 230.3534 | 95.20959 | 0.427889 | 3.349076 | 0.516274 |
| 2 | 178519000 | 178520999 | 21.05019 | 36.47821 | 37.35599 | 16.38978 | 0.352129 | 0.828731 | 0.516284 |
| 5 | 147862600 | 147867999 | 47.08909 | 75.20085 | 115.803 | 48.62616 | -0.28092 | 2.098979 | 0.516291 |
| 2 | 173774200 | 173780799 | 71.77323 | 112.2298 | 167.3778 | 69.44036 | -0.26151 | 2.640189 | 0.516336 |
| 9 | 68171800 | 68173399 | 14.55443 | 25.50847 | 42.47108 | 18.76153 | -0.3444 | 0.647632 | 0.516368 |
| 10 | 55049600 | 55053199 | 29.84705 | 50.7325 | 46.26039 | 20.27428 | 0.514179 | 1.047567 | 0.51637 |
| X | 34745400 | 34758799 | 195.3508 | 292.6809 | 250.4067 | 103.2942 | 0.48981 | 3.498104 | 0.516403 |
| 14 | 33900000 | 33907199 | 60.66646 | 96.10743 | 144.5115 | 60.10292 | -0.25856 | 2.363152 | 0.516411 |
| 15 | 89853400 | 89855399 | 15.96638 | 29.05436 | 27.58204 | 11.87411 | 0.491622 | 0.331309 | 0.516415 |
| 9 | 3079600 | 3081399 | 26.23288 | 44.31558 | 43.34088 | 18.96144 | 0.413306 | 1.087049 | 0.516423 |
| 15 | 14121200 | 14125199 | 64.0453 | 102.2166 | 102.3104 | 42.923 | 0.338558 | 2.13131 | 0.516446 |
| 11 | 53348200 | 53350399 | 16.4974 | 28.48589 | 48.99787 | 21.49525 | -0.3939 | 0.856827 | 0.516449 |
| X | 89694400 | 89702999 | 141.4226 | 215.7448 | 198.164 | 82.16752 | 0.409553 | 3.127652 | 0.516461 |
| 1 | 117980600 | 117983799 | 59.6884 | 95.26457 | 100.0516 | 41.97635 | 0.269307 | 2.126917 | 0.516462 |
| 16 | 57140000 | 57141799 | 17.92792 | 31.7058 | 32.1518 | 14.22053 | 0.370672 | 0.53603 | 0.516472 |
| 1 | 101727400 | 101734799 | 90.45985 | 142.863 | 124.2898 | 51.61172 | 0.526189 | 2.45272 | 0.516489 |
| 5 | 104921000 | 104924999 | 83.79204 | 131.4905 | 136.0205 | 56.41183 | 0.269612 | 2.5782 | 0.516493 |
| 19 | 45667000 | 45668399 | 11.65749 | 20.98025 | 35.24569 | 15.38448 | -0.33936 | 0.382393 | 0.516501 |
| X | 117058200 | 117060999 | 32.18349 | 53.95919 | 51.90094 | 22.64019 | 0.436954 | 1.182443 | 0.516525 |
| 6 | 140155000 | 140158999 | 34.29333 | 55.55415 | 86.80645 | 37.15386 | -0.27815 | 1.652771 | 0.516528 |
| 6 | 89246000 | 89249399 | 21.75199 | 36.76354 | 60.4209 | 26.43772 | -0.33307 | 1.082478 | 0.516539 |
| 1 | 109657200 | 109658399 | 17.65783 | 30.52933 | 49.53682 | 21.73255 | -0.30966 | 0.852699 | 0.51654 |
| 1 | 174242200 | 174243999 | 17.65783 | 30.52933 | 49.53682 | 21.73255 | -0.30966 | 0.852699 | 0.51654 |
| 17 | 60962200 | 60964399 | 21.03462 | 36.93792 | 34.63349 | 15.32458 | 0.479907 | 0.639006 | 0.516552 |
| 8 | 69693800 | 69696599 | 26.58483 | 45.79107 | 42.40981 | 18.60369 | 0.4948 | 0.913912 | 0.516564 |
| 14 | 46344600 | 46347799 | 38.54652 | 62.12764 | 96.40861 | 41.03176 | -0.2762 | 1.796336 | 0.516576 |
| 5 | 101228600 | 101233999 | 52.01195 | 82.79182 | 126.2671 | 52.77944 | -0.27215 | 2.209869 | 0.516598 |
| 6 | 98689200 | 98689799 | 5.817407 | 11.29267 | 23.98174 | 9.008906 | -0.49219 | -0.25317 | 0.516606 |
| 11 | 88337800 | 88338999 | 7.639015 | 14.43656 | 27.58343 | 11.21412 | -0.44718 | 0.039663 | 0.516606 |
| 2 | 56700800 | 56701999 | 20.88748 | 36.80846 | 29.47091 | 13.04408 | 0.70793 | 0.588131 | 0.516613 |
| 8 | 127839600 | 127841599 | 16.93849 | 29.36574 | 49.00982 | 21.50313 | -0.34939 | 0.822335 | 0.516627 |
| 14 | 94524000 | 94526399 | 18.70957 | 33.28402 | 30.91371 | 13.6268 | 0.50083 | 0.490681 | 0.516629 |
| 19 | 3815000 | 3819799 | 25.47358 | 41.70288 | 72.8697 | 31.34836 | -0.42699 | 1.455485 | 0.516631 |
| 7 | 41776600 | 41789599 | 179.036 | 269.9542 | 229.9876 | 95.02796 | 0.504056 | 3.371936 | 0.516638 |
| 7 | 37418000 | 37419199 | 8.905628 | 16.5253 | 29.95079 | 12.5626 | -0.40826 | 0.174483 | 0.516643 |
| 8 | 105938400 | 105940999 | 14.48506 | 25.4435 | 42.43373 | 18.78035 | -0.34663 | 0.613882 | 0.516649 |
| X | 121877200 | 121879399 | 23.61818 | 41.1994 | 38.03253 | 16.74034 | 0.50236 | 0.760348 | 0.516656 |
| 16 | 58724000 | 58727599 | 42.89373 | 69.55108 | 71.91534 | 30.80142 | 0.317381 | 1.642515 | 0.516708 |
| 9 | 17900000 | 17903399 | 30.72929 | 51.70079 | 50.2092 | 21.93924 | 0.423082 | 1.141359 | 0.51671 |
| 8 | 65027000 | 65028999 | 16.85729 | 28.69899 | 57.41959 | 25.1682 | -0.61461 | 1.000655 | 0.516717 |

| | | | | | | | | | |
|---|---|---|---|---|---|---|---|---|---|
| 16 | 92778600 | 92779199 | 12.14591 | 23.01496 | 20.47307 | 8.276739 | 0.654294 | 0.044985 | 0.516718 |
| 16 | 25567000 | 25568799 | 13.52791 | 23.72459 | 48.84931 | 21.59082 | -0.64938 | 0.642565 | 0.516725 |
| X | 8539800 | 8545399 | 74.36132 | 118.5684 | 101.9781 | 42.72701 | 0.555548 | 2.173154 | 0.516737 |
| 10 | 4860600 | 4862799 | 26.72706 | 43.89573 | 71.53195 | 30.81471 | -0.32477 | 1.418405 | 0.516737 |
| 19 | 12477800 | 12479599 | 12.51396 | 22.02812 | 39.34113 | 17.41656 | -0.44405 | 0.567777 | 0.516738 |
| 2 | 93723800 | 93725199 | 10.23719 | 18.55997 | 32.82406 | 14.1964 | -0.40413 | 0.325015 | 0.516739 |
| 2 | 16137000 | 16139799 | 24.62157 | 40.98706 | 65.91713 | 28.62505 | -0.30211 | 1.246847 | 0.516739 |
| 9 | 40157600 | 40159399 | 13.80319 | 24.41204 | 41.32441 | 18.27738 | -0.36459 | 0.528601 | 0.516752 |
| 13 | 65671800 | 65683599 | 447.2649 | 616.0406 | 568.4651 | 236.2754 | 0.262135 | 4.81253 | 0.51679 |
| 18 | 69492400 | 69493599 | 17.52357 | 29.63925 | 60.6538 | 26.54567 | -0.64794 | 1.063038 | 0.516802 |
| X | 77209600 | 77213599 | 50.84929 | 81.94717 | 81.22049 | 34.56243 | 0.369041 | 1.813323 | 0.51682 |
| 2 | 116941800 | 116942799 | 9.353015 | 17.45065 | 30.57285 | 12.7131 | -0.35002 | 0.146134 | 0.516826 |
| 10 | 30959800 | 30965199 | 46.02848 | 74.67916 | 70.97299 | 30.41137 | 0.439517 | 1.63356 | 0.516833 |
| 2 | 85724600 | 85733399 | 100.4608 | 156.4421 | 153.5531 | 63.73633 | 0.336716 | 2.723826 | 0.516868 |
| 4 | 57950800 | 57952799 | 19.47224 | 33.19654 | 53.7992 | 23.58392 | -0.31167 | 0.991976 | 0.516868 |
| 1 | 182787400 | 182789599 | 10.28614 | 18.42561 | 34.7976 | 15.24982 | -0.51107 | 0.400706 | 0.516873 |
| 6 | 144853000 | 144855599 | 21.76014 | 36.32265 | 69.06412 | 30.04333 | -0.54291 | 1.212546 | 0.516877 |
| 5 | 121532400 | 121534799 | 17.13338 | 29.69928 | 48.47026 | 21.26631 | -0.31723 | 0.824949 | 0.516877 |
| 8 | 97900200 | 97902999 | 18.24486 | 32.24051 | 27.12844 | 12.00065 | 0.637921 | 0.536439 | 0.516884 |
| 18 | 70619000 | 70622999 | 55.14895 | 88.8415 | 81.07307 | 34.48007 | 0.487232 | 1.829734 | 0.516915 |
| X | 90090400 | 90093399 | 36.51843 | 60.53644 | 56.46145 | 24.44797 | 0.481136 | 1.323048 | 0.516916 |
| 4 | 16861600 | 16864399 | 63.94724 | 102.24 | 98.10213 | 41.26028 | 0.402148 | 2.077624 | 0.516925 |
| 13 | 112864600 | 112867399 | 33.74851 | 54.42381 | 91.35553 | 38.90825 | -0.38823 | 1.774047 | 0.516934 |
| 16 | 89410800 | 89413199 | 22.37963 | 38.81677 | 39.82501 | 17.47353 | 0.349677 | 0.830204 | 0.516961 |
| 13 | 38333600 | 38336999 | 25.58464 | 42.05157 | 70.33998 | 30.31103 | -0.36187 | 1.404842 | 0.516961 |
| 14 | 37847600 | 37848199 | 9.500295 | 18.40523 | 20.42806 | 7.746752 | 0.42224 | -0.18892 | 0.516991 |
| 18 | 37083600 | 37090799 | 134.7521 | 204.5978 | 206.025 | 85.31149 | 0.271469 | 3.221256 | 0.517011 |
| 11 | 65916800 | 65918199 | 9.555384 | 17.81201 | 31.6451 | 13.17926 | -0.37172 | 0.157386 | 0.517022 |
| 13 | 98757800 | 98759399 | 9.332742 | 17.29425 | 31.51025 | 13.24125 | -0.41818 | 0.18755 | 0.517028 |
| 1 | 17808000 | 17809999 | 11.50174 | 20.90047 | 36.48476 | 15.73446 | -0.38311 | 0.328335 | 0.517049 |
| 19 | 13368200 | 13372999 | 66.44362 | 106.0035 | 93.36824 | 39.15018 | 0.522068 | 2.142771 | 0.517051 |
| 16 | 5645800 | 5650599 | 46.13055 | 74.57052 | 76.13228 | 32.52541 | 0.331404 | 1.714409 | 0.517071 |
| 7 | 62442600 | 62451399 | 111.8764 | 174.0166 | 152.7966 | 63.43333 | 0.49598 | 2.750247 | 0.517089 |
| 8 | 46867800 | 46871199 | 28.95956 | 47.58915 | 76.67325 | 33.03737 | -0.30821 | 1.4187 | 0.517106 |
| 3 | 47963600 | 47964799 | 17.05073 | 30.76112 | 29.03095 | 12.6334 | 0.49025 | 0.391519 | 0.517128 |
| 13 | 51343000 | 51345799 | 23.01753 | 38.39392 | 62.41896 | 27.14034 | -0.31762 | 1.218525 | 0.517129 |
| 2 | 47485000 | 47485599 | 10.49759 | 20.01755 | 21.74843 | 8.675736 | 0.385132 | -0.00907 | 0.51713 |
| 6 | 137830800 | 137832399 | 30.05203 | 50.26684 | 52.40819 | 22.82746 | 0.321125 | 1.201206 | 0.517144 |
| 1 | 141486200 | 141487199 | 15.15254 | 27.16977 | 28.33005 | 12.40507 | 0.341505 | 0.414316 | 0.517148 |
| 8 | 4111000 | 4115199 | 73.85851 | 116.8249 | 120.6734 | 50.14922 | 0.280583 | 2.402143 | 0.51716 |
| 7 | 40689200 | 40694199 | 124.6686 | 190.015 | 195.3982 | 81.00382 | 0.245723 | 3.139017 | 0.517169 |
| 10 | 75905800 | 75910999 | 61.48153 | 97.34803 | 146.476 | 60.90395 | -0.26015 | 2.376126 | 0.517173 |
| 14 | 118054800 | 118057599 | 20.34521 | 35.6296 | 35.90145 | 15.84891 | 0.3766 | 0.680207 | 0.517194 |
| 1 | 118029600 | 118035399 | 100.4285 | 155.9177 | 159.1964 | 66.10873 | 0.276471 | 2.776197 | 0.517218 |
| 12 | 73212200 | 73212999 | 9.908925 | 19.14657 | 20.86049 | 8.008332 | 0.431779 | -0.1434 | 0.517238 |
| 14 | 102002600 | 102002999 | 4.671944 | 9.125828 | 20.67162 | 7.233503 | -0.48991 | -0.53045 | 0.517297 |
| 13 | 62803400 | 62808399 | 58.28125 | 93.24231 | 96.51369 | 40.62997 | 0.294006 | 2.050444 | 0.517302 |

| | | | | | | | | | |
|---|---|---|---|---|---|---|---|---|---|
| X | 44309000 | 44310999 | 26.95786 | 46.27689 | 43.57503 | 19.1126 | 0.470194 | 0.94881 | 0.517313 |
| 13 | 43117800 | 43120599 | 20.97402 | 35.67257 | 58.74465 | 25.74697 | -0.3353 | 1.024805 | 0.517326 |
| 7 | 103707600 | 103708799 | 13.20874 | 24.44345 | 25.38073 | 10.68528 | 0.385523 | 0.212085 | 0.517327 |
| 7 | 47317400 | 47319399 | 32.67644 | 54.15651 | 56.91244 | 24.63977 | 0.309408 | 1.312415 | 0.517341 |
| 2 | 120238000 | 120239999 | 25.44344 | 44.72814 | 32.14069 | 14.23266 | 0.862033 | 0.645692 | 0.517344 |
| 14 | 31785000 | 31788999 | 46.09985 | 74.63484 | 74.1101 | 31.70546 | 0.373573 | 1.678523 | 0.517346 |
| 4 | 60044600 | 60048199 | 44.6544 | 72.51874 | 70.73607 | 30.33112 | 0.403371 | 1.61358 | 0.517348 |
| 9 | 63890400 | 63891999 | 9.450524 | 17.30813 | 31.40143 | 13.40799 | -0.42815 | 0.25841 | 0.517367 |
| 3 | 72112000 | 72114399 | 20.43045 | 35.8871 | 34.71288 | 15.36337 | 0.435224 | 0.63194 | 0.517384 |
| 16 | 27717200 | 27718799 | 23.14116 | 40.79214 | 32.23622 | 14.26717 | 0.726899 | 0.602893 | 0.517393 |
| X | 89249200 | 89257399 | 102.4418 | 161.4892 | 121.6279 | 50.47263 | 0.731916 | 2.502804 | 0.517399 |
| 1 | 175576400 | 175577599 | 7.307173 | 13.94262 | 26.70343 | 10.63817 | -0.42478 | -0.03284 | 0.517405 |
| X | 62185600 | 62190599 | 48.68281 | 79.46129 | 63.76909 | 27.40327 | 0.689571 | 1.54489 | 0.517413 |
| 10 | 99431000 | 99432599 | 15.24509 | 26.43565 | 46.35761 | 20.33961 | -0.42016 | 0.810493 | 0.517445 |
| 9 | 115823800 | 115826399 | 26.17626 | 44.16787 | 45.90942 | 20.078 | 0.325606 | 1.098276 | 0.517452 |
| 13 | 62162800 | 62180399 | 358.7761 | 506.4095 | 472.5284 | 195.6434 | 0.281672 | 4.442682 | 0.517491 |
| 12 | 87249800 | 87251199 | 13.73001 | 25.43694 | 25.89076 | 10.88319 | 0.415748 | 0.21159 | 0.517495 |
| 9 | 47459800 | 47461199 | 14.16842 | 24.74177 | 48.38225 | 21.3102 | -0.57539 | 0.713709 | 0.517521 |
| 7 | 94824800 | 94826999 | 25.94232 | 44.05286 | 45.32655 | 19.86141 | 0.340586 | 1.043764 | 0.517526 |
| 19 | 27062400 | 27064199 | 16.45375 | 30.40523 | 26.20813 | 11.05499 | 0.650536 | 0.248765 | 0.517537 |
| X | 58785400 | 58786799 | 19.39283 | 34.45289 | 31.4999 | 13.90104 | 0.522491 | 0.506305 | 0.517549 |
| 4 | 37271600 | 37281199 | 40.69885 | 66.4657 | 59.2935 | 25.48503 | 0.537741 | 1.513523 | 0.517551 |
| 1 | 4876200 | 4878399 | 13.53393 | 23.71461 | 46.72284 | 20.63584 | -0.58596 | 0.654113 | 0.517565 |
| 7 | 144440200 | 144441999 | 15.2222 | 26.5038 | 44.72971 | 19.66499 | -0.36435 | 0.754661 | 0.517576 |
| 16 | 53823600 | 53824399 | 13.31246 | 24.3152 | 24.49814 | 10.49756 | 0.410977 | 0.285281 | 0.517581 |
| X | 13134800 | 13136799 | 29.43382 | 50.33198 | 44.61695 | 19.56349 | 0.556139 | 0.993677 | 0.517585 |
| 5 | 113919200 | 113923599 | 38.12241 | 61.16687 | 101.8645 | 43.0679 | -0.38587 | 1.939489 | 0.517594 |
| 11 | 48666800 | 48669399 | 21.24957 | 35.5357 | 60.12671 | 26.18588 | -0.375 | 1.18403 | 0.517602 |
| 5 | 131024600 | 131025599 | 12.46721 | 22.27494 | 37.2344 | 16.35332 | -0.33849 | 0.437001 | 0.517605 |
| 7 | 60921200 | 60923199 | 23.00606 | 40.1773 | 37.43339 | 16.48842 | 0.489134 | 0.739549 | 0.517608 |
| 17 | 16574600 | 16575199 | 18.08538 | 31.71163 | 31.62173 | 13.97434 | 0.391695 | 0.642044 | 0.517609 |
| 5 | 109217600 | 109231399 | 190.8423 | 285.0456 | 266.518 | 109.8743 | 0.357765 | 3.553929 | 0.517613 |
| 9 | 29801800 | 29802399 | 9.315459 | 17.90833 | 18.75067 | 7.319983 | 0.466709 | -0.10008 | 0.517625 |
| 3 | 16668200 | 16670199 | 24.40823 | 41.93315 | 43.05811 | 18.88658 | 0.345771 | 0.928002 | 0.517629 |
| 14 | 87359800 | 87361799 | 14.67309 | 25.5 | 50.02862 | 21.96878 | -0.58083 | 0.78777 | 0.51763 |
| 6 | 53967200 | 53969199 | 16.96845 | 29.4981 | 48.45328 | 21.28111 | -0.32589 | 0.789887 | 0.517632 |
| 9 | 102680400 | 102681599 | 9.254134 | 17.15338 | 30.47087 | 12.79962 | -0.38083 | 0.186501 | 0.517653 |
| 11 | 18968400 | 18971799 | 21.53053 | 36.04533 | 60.21279 | 26.24164 | -0.35667 | 1.17191 | 0.51766 |
| X | 34841400 | 34844999 | 47.51218 | 77.0255 | 71.11632 | 30.45504 | 0.48017 | 1.656524 | 0.517663 |
| 1 | 153690600 | 153691799 | 9.66807 | 18.08565 | 33.46694 | 13.85513 | -0.42353 | 0.138077 | 0.517672 |
| 8 | 93652800 | 93655999 | 29.5961 | 50.7433 | 43.48905 | 19.07299 | 0.605172 | 0.96974 | 0.517672 |
| 3 | 11610000 | 11619399 | 162.3546 | 244.2499 | 243.6043 | 100.5934 | 0.273988 | 3.411515 | 0.517675 |
| X | 19333800 | 19336599 | 35.20479 | 58.57415 | 55.3757 | 24.03889 | 0.46198 | 1.275099 | 0.517683 |
| 13 | 71575000 | 71577199 | 21.86405 | 36.20526 | 71.40627 | 30.89634 | -0.59599 | 1.307247 | 0.517708 |
| 6 | 59353000 | 59354799 | 12.96998 | 22.88552 | 38.77406 | 17.15731 | -0.36707 | 0.543448 | 0.517711 |
| 11 | 85766400 | 85768799 | 16.73566 | 28.71101 | 50.04743 | 21.95037 | -0.41484 | 0.919142 | 0.517717 |
| 8 | 125961600 | 125963399 | 13.62254 | 23.84835 | 42.0256 | 18.5471 | -0.42579 | 0.660092 | 0.517718 |

| | | | | | | | | | |
|---|---|---|---|---|---|---|---|---|---|
| 10 | 35045800 | 35052399 | 79.46804 | 125.3951 | 129.1551 | 53.60856 | 0.281961 | 2.465413 | 0.517721 |
| 14 | 55398000 | 55399199 | 8.484574 | 15.68292 | 29.94063 | 12.62492 | -0.48804 | 0.193615 | 0.517774 |
| 11 | 13822600 | 13826399 | 53.40783 | 85.58234 | 89.66471 | 37.83606 | 0.279178 | 1.99237 | 0.517776 |
| 11 | 88812800 | 88814199 | 8.735468 | 16.52995 | 30.41696 | 12.33248 | -0.39001 | 0.049295 | 0.517776 |
| X | 48288600 | 48295399 | 76.95864 | 122.9196 | 93.91903 | 39.44492 | 0.728678 | 2.122668 | 0.517788 |
| 8 | 21724400 | 21726599 | 43.58023 | 70.6649 | 71.96786 | 30.82413 | 0.339217 | 1.644049 | 0.517789 |
| X | 108642600 | 108643399 | 16.1459 | 29.29444 | 28.31109 | 12.23304 | 0.463111 | 0.35443 | 0.5178 |
| 1 | 60136600 | 60137999 | 9.951562 | 17.89575 | 35.29647 | 15.41781 | -0.57021 | 0.373391 | 0.517805 |
| 19 | 25473600 | 25475999 | 15.1109 | 26.29333 | 44.75789 | 19.66954 | -0.37667 | 0.764807 | 0.517822 |
| 5 | 96004000 | 96004999 | 14.0527 | 25.50973 | 25.99012 | 11.22295 | 0.387974 | 0.330439 | 0.517843 |
| 4 | 146703000 | 146703799 | 9.844233 | 19.08896 | 20.46581 | 7.728042 | 0.477636 | -0.19869 | 0.517901 |
| 13 | 107248400 | 107249599 | 20.40641 | 34.32987 | 57.45683 | 25.10545 | -0.35927 | 1.117788 | 0.517905 |
| 12 | 14505600 | 14506199 | 6.267065 | 12.10441 | 24.24555 | 9.326403 | -0.44021 | -0.1619 | 0.517911 |
| 14 | 13097600 | 13098399 | 6.267065 | 12.10441 | 24.24555 | 9.326403 | -0.44021 | -0.1619 | 0.517911 |
| X | 87069200 | 87069999 | 13.31332 | 24.85364 | 25.0712 | 10.41545 | 0.442066 | 0.163854 | 0.517917 |
| 3 | 136170000 | 136171199 | 12.93621 | 23.09931 | 41.45059 | 18.21409 | -0.44102 | 0.450289 | 0.517927 |
| 6 | 131204200 | 131205799 | 12.93621 | 23.09931 | 41.45059 | 18.21409 | -0.44102 | 0.450289 | 0.517927 |
| 19 | 57813600 | 57816999 | 39.17884 | 63.80286 | 67.43561 | 28.95477 | 0.290936 | 1.556126 | 0.517986 |
| 7 | 4221600 | 4222599 | 8.874554 | 16.75506 | 30.34009 | 12.36052 | -0.37324 | 0.065417 | 0.517999 |
| 13 | 54619600 | 54620199 | 8.874554 | 16.75506 | 30.34009 | 12.36052 | -0.37324 | 0.065417 | 0.517999 |
| 1 | 47220000 | 47221199 | 18.97943 | 31.99297 | 56.31791 | 24.62269 | -0.4316 | 1.095255 | 0.518012 |
| 14 | 53799000 | 53802599 | 50.80265 | 81.51601 | 85.36406 | 36.0942 | 0.282124 | 1.94695 | 0.518036 |
| 11 | 7272200 | 7273399 | 14.89492 | 27.59445 | 26.42461 | 11.10796 | 0.503432 | 0.224051 | 0.518053 |
| 14 | 75521200 | 75522399 | 7.38487 | 13.89898 | 27.53867 | 11.28385 | -0.50812 | 0.067322 | 0.518057 |
| 6 | 103304400 | 103305399 | 9.518631 | 17.75048 | 31.10722 | 12.94662 | -0.35147 | 0.152278 | 0.518068 |
| 17 | 31375800 | 31379399 | 32.89847 | 53.02634 | 92.12133 | 39.20898 | -0.43858 | 1.784414 | 0.518076 |
| 7 | 143897800 | 143914199 | 218.3532 | 315.9964 | 436.3965 | 179.9786 | -0.24774 | 4.069842 | 0.518076 |
| 6 | 51233600 | 51236599 | 20.45889 | 34.75414 | 55.96342 | 24.52506 | -0.30329 | 1.030601 | 0.518083 |
| 11 | 75930600 | 75932999 | 18.47083 | 31.65713 | 51.66361 | 22.65744 | -0.32049 | 0.936825 | 0.518087 |
| 15 | 55122800 | 55123399 | 5.510069 | 10.72783 | 21.86848 | 8.020677 | -0.40026 | -0.3438 | 0.518092 |
| 1 | 139571800 | 139573399 | 16.22139 | 29.49676 | 28.13621 | 12.12379 | 0.48405 | 0.339307 | 0.518095 |
| 7 | 48041200 | 48046599 | 104.3879 | 161.6805 | 163.5664 | 67.94108 | 0.287115 | 2.82072 | 0.51811 |
| 16 | 16772600 | 16775599 | 38.37222 | 62.78374 | 64.80218 | 27.87177 | 0.329837 | 1.486791 | 0.518112 |
| 1 | 114589800 | 114590799 | 13.64076 | 23.93833 | 40.42123 | 17.87569 | -0.36437 | 0.619549 | 0.518116 |
| 15 | 17906200 | 17909199 | 32.49294 | 54.94972 | 47.95722 | 20.9879 | 0.57696 | 1.103192 | 0.518126 |
| 8 | 28573400 | 28575199 | 25.68302 | 41.92678 | 74.79892 | 32.14484 | -0.4597 | 1.498755 | 0.518134 |
| 10 | 44990600 | 44993199 | 24.31552 | 41.38219 | 42.21302 | 18.50233 | 0.353325 | 1.01257 | 0.518134 |
| 8 | 88519800 | 88521399 | 8.500695 | 15.70984 | 29.95383 | 12.63364 | -0.48626 | 0.196088 | 0.518155 |
| 16 | 49114000 | 49114799 | 6.563623 | 12.63319 | 24.71387 | 9.614908 | -0.42251 | -0.12181 | 0.518162 |
| 19 | 13032000 | 13036599 | 78.28311 | 123.3387 | 129.4253 | 53.70456 | 0.253454 | 2.496676 | 0.51818 |
| 17 | 37766000 | 37769599 | 31.00718 | 51.92818 | 52.43861 | 22.8584 | 0.36699 | 1.190779 | 0.518181 |
| 5 | 43641000 | 43641999 | 19.10413 | 33.51213 | 32.14755 | 14.2118 | 0.447492 | 0.644532 | 0.518182 |
| 5 | 146333800 | 146339399 | 61.82215 | 97.75194 | 145.3579 | 60.3967 | -0.24473 | 2.408711 | 0.518186 |
| 10 | 40635600 | 40638399 | 27.64152 | 46.34288 | 48.59672 | 21.19848 | 0.312736 | 1.16413 | 0.518194 |
| 11 | 51047400 | 51048799 | 14.12357 | 24.6742 | 42.61091 | 18.77333 | -0.39673 | 0.698314 | 0.518202 |
| 1 | 171560600 | 171561399 | 13.57535 | 25.15267 | 26.0048 | 10.92978 | 0.39381 | 0.2121 | 0.51822 |
| 12 | 90321600 | 90322999 | 17.73368 | 31.88992 | 29.46953 | 12.87265 | 0.516957 | 0.415868 | 0.518221 |

| | | | | | | | | | |
|---|---|---|---|---|---|---|---|---|---|
| 7 | 103651800 | 103653999 | 23.95645 | 40.93485 | 43.16994 | 18.92787 | 0.306282 | 0.975352 | 0.518233 |
| 2 | 19585800 | 19586799 | 11.17016 | 19.74166 | 40.93025 | 18.10555 | -0.65544 | 0.521198 | 0.518258 |
| 6 | 127489200 | 127490999 | 38.38731 | 62.85507 | 56.98805 | 24.51794 | 0.516931 | 1.47053 | 0.518271 |
| 8 | 79882400 | 79884199 | 15.96858 | 28.31219 | 28.40735 | 12.55765 | 0.387403 | 0.490015 | 0.518273 |
| 7 | 113965200 | 113965999 | 7.968462 | 15.2574 | 30.31935 | 11.97356 | -0.46589 | -0.04295 | 0.518278 |
| X | 88232800 | 88234599 | 20.58376 | 37.52447 | 25.65504 | 11.01959 | 0.964624 | 0.353344 | 0.518294 |
| 11 | 110227800 | 110228999 | 7.645342 | 14.4804 | 27.08911 | 10.96192 | -0.41075 | 0.0244 | 0.518304 |
| 4 | 87558000 | 87559999 | 19.29358 | 33.00498 | 53.28353 | 23.36432 | -0.30539 | 0.959393 | 0.518314 |
| 2 | 17126600 | 17128799 | 17.97137 | 31.1512 | 52.83103 | 23.17958 | -0.37255 | 0.834541 | 0.518331 |
| 3 | 105823000 | 105824399 | 10.80653 | 19.64709 | 33.82852 | 14.5797 | -0.3628 | 0.313457 | 0.518342 |
| 6 | 136763200 | 136764799 | 10.80653 | 19.64709 | 33.82852 | 14.5797 | -0.3628 | 0.313457 | 0.518342 |
| 7 | 34930600 | 34936399 | 43.79664 | 70.19135 | 107.7598 | 45.48181 | -0.27075 | 1.983795 | 0.518345 |
| 6 | 49210000 | 49213999 | 61.94424 | 98.67752 | 104.5511 | 43.74858 | 0.253708 | 2.189597 | 0.51835 |
| 13 | 12104600 | 12106399 | 15.41063 | 26.91107 | 44.14833 | 19.44538 | -0.32317 | 0.714042 | 0.518353 |
| 13 | 30393200 | 30395199 | 20.9571 | 36.3442 | 37.88996 | 16.62444 | 0.326725 | 0.823239 | 0.518359 |
| 5 | 132891400 | 132893399 | 16.46703 | 29.44673 | 30.18085 | 13.24577 | 0.363864 | 0.443521 | 0.518366 |
| 6 | 110279200 | 110280599 | 16.46703 | 29.44673 | 30.18085 | 13.24577 | 0.363864 | 0.443521 | 0.518366 |
| 4 | 38919200 | 38920199 | 15.37093 | 27.59315 | 28.83208 | 12.61264 | 0.339507 | 0.410416 | 0.518373 |
| 3 | 75277400 | 75279999 | 34.44602 | 56.78857 | 59.06143 | 25.48358 | 0.323417 | 1.381414 | 0.518379 |
| 18 | 40742200 | 40745599 | 36.42898 | 60.37696 | 57.17439 | 24.75351 | 0.459182 | 1.322018 | 0.518386 |
| 17 | 95110600 | 95118799 | 152.1554 | 230.9484 | 214.7934 | 88.93388 | 0.385661 | 3.232242 | 0.518388 |
| 4 | 31675400 | 31676999 | 18.74872 | 33.1989 | 32.01535 | 14.15664 | 0.443478 | 0.527401 | 0.518402 |
| 14 | 44535600 | 44537999 | 38.59191 | 62.1387 | 96.32878 | 40.93919 | -0.2765 | 1.828563 | 0.518412 |
| 2 | 86464800 | 86466999 | 29.7258 | 49.64475 | 52.40035 | 22.8019 | 0.303483 | 1.216501 | 0.518417 |
| 18 | 45691200 | 45695399 | 31.75724 | 51.26172 | 88.82481 | 37.87721 | -0.43237 | 1.738583 | 0.518427 |
| 8 | 71545800 | 71549399 | 32.74472 | 55.30773 | 46.84428 | 20.49322 | 0.619891 | 1.110798 | 0.518442 |
| Y | 40613800 | 40614999 | 11.02524 | 20.88805 | 15.60351 | 6.310265 | 0.903214 | 0.018584 | 0.518443 |
| 4 | 53810800 | 53811599 | 7.488859 | 14.25115 | 26.63901 | 10.67268 | -0.39675 | -0.01046 | 0.518481 |
| X | 108328800 | 108329799 | 15.06552 | 27.99644 | 19.93666 | 8.345672 | 0.932507 | 0.194776 | 0.518488 |
| 10 | 91420800 | 91423399 | 33.34919 | 53.95523 | 85.62338 | 36.61275 | -0.30212 | 1.677214 | 0.518492 |
| 1 | 140950400 | 140952999 | 34.59229 | 58.1678 | 49.12039 | 21.45986 | 0.624068 | 1.15847 | 0.518498 |
| X | 149070400 | 149072799 | 34.32148 | 57.5322 | 51.36458 | 22.40337 | 0.544156 | 1.192527 | 0.518498 |
| 3 | 42456800 | 42460399 | 44.13894 | 71.19127 | 71.64871 | 30.52153 | 0.348346 | 1.773853 | 0.518528 |
| 15 | 75412800 | 75416199 | 49.01771 | 78.90223 | 82.67748 | 35.12227 | 0.286798 | 1.845133 | 0.518552 |
| 7 | 20834800 | 20869399 | 564.1804 | 768.6891 | 653.107 | 271.9187 | 0.365435 | 4.984453 | 0.518564 |
| 11 | 45870000 | 45872199 | 20.58749 | 34.11642 | 70.69098 | 30.60236 | -0.6668 | 1.2951 | 0.518603 |
| 19 | 38404400 | 38405999 | 12.49837 | 22.15021 | 37.21264 | 16.45197 | -0.35236 | 0.490136 | 0.518631 |
| 16 | 79016800 | 79021399 | 78.09644 | 122.9463 | 129.5842 | 53.76491 | 0.246208 | 2.511983 | 0.518641 |
| 1 | 147313000 | 147314399 | 17.78179 | 31.7457 | 29.90085 | 13.14105 | 0.483519 | 0.46001 | 0.518657 |
| 16 | 66354800 | 66358799 | 26.26994 | 43.58635 | 71.72732 | 31.07724 | -0.33562 | 1.29372 | 0.518659 |
| 4 | 98160600 | 98164199 | 52.27282 | 83.84511 | 88.07444 | 37.22004 | 0.277103 | 1.962915 | 0.51866 |
| 17 | 73917000 | 73918999 | 14.55698 | 25.58553 | 42.89573 | 18.99467 | -0.35441 | 0.602487 | 0.518661 |
| 1 | 144714800 | 144719199 | 54.73853 | 88.19047 | 81.46398 | 34.6521 | 0.469936 | 1.826744 | 0.518664 |
| 7 | 5450000 | 5455599 | 65.61689 | 105.2293 | 88.15072 | 37.24172 | 0.602658 | 1.984682 | 0.518678 |
| 13 | 66557400 | 66570999 | 395.6485 | 553.5324 | 457.7721 | 189.8006 | 0.442909 | 4.574063 | 0.518682 |
| 3 | 49569600 | 49571199 | 22.08298 | 38.66265 | 35.7182 | 15.76449 | 0.501205 | 0.692587 | 0.518695 |
| 6 | 90196000 | 90198199 | 36.01226 | 59.89249 | 55.42394 | 24.05286 | 0.492691 | 1.281857 | 0.518695 |

| | | | | | | | | | |
|---|---|---|---|---|---|---|---|---|---|
| 13 | 51422000 | 51429999 | 96.39049 | 147.9215 | 214.091 | 88.88087 | -0.23922 | 2.995859 | 0.518726 |
| 6 | 79754200 | 79755599 | 22.92832 | 39.88569 | 39.17615 | 17.21381 | 0.412799 | 0.79481 | 0.518743 |
| 3 | 111445000 | 111446199 | 16.57622 | 29.43159 | 30.34118 | 13.39634 | 0.349824 | 0.490194 | 0.518756 |
| 4 | 70478800 | 70480599 | 21.7086 | 38.06566 | 34.5692 | 15.2772 | 0.525693 | 0.664155 | 0.518768 |
| 1 | 50747800 | 50752799 | 61.64376 | 98.47627 | 99.39081 | 41.75369 | 0.328026 | 2.098536 | 0.51878 |
| 16 | 35259400 | 35263199 | 47.31698 | 75.37819 | 119.9395 | 50.18956 | -0.33193 | 2.179211 | 0.51878 |
| 5 | 149622800 | 149633399 | 127.7282 | 192.8436 | 276.2087 | 114.1731 | -0.2433 | 3.336604 | 0.518822 |
| 7 | 19716600 | 19729199 | 175.5276 | 261.9965 | 265.5653 | 109.4674 | 0.240481 | 3.5633 | 0.51883 |
| 7 | 106404600 | 106405599 | 12.32142 | 23.09567 | 24.56177 | 10.13378 | 0.373933 | 0.126538 | 0.51885 |
| 12 | 8644400 | 8650799 | 121.6654 | 183.1889 | 273.1851 | 112.8336 | -0.3049 | 3.388809 | 0.518862 |
| 5 | 151593800 | 151596199 | 61.37699 | 97.77217 | 103.4093 | 43.26975 | 0.256213 | 2.188476 | 0.518878 |
| 6 | 92165800 | 92167599 | 10.14363 | 18.17614 | 34.32653 | 15.03933 | -0.51065 | 0.397623 | 0.518885 |
| 7 | 124492200 | 124496399 | 36.86937 | 60.55883 | 61.60463 | 26.53925 | 0.353053 | 1.430658 | 0.518927 |
| 7 | 113342000 | 113343799 | 16.98902 | 30.01738 | 31.34616 | 13.86678 | 0.328152 | 0.539624 | 0.518932 |
| 17 | 19075000 | 19086999 | 167.0221 | 252.2366 | 229.1963 | 94.74807 | 0.413084 | 3.337764 | 0.518934 |
| 2 | 53146200 | 53146599 | 4.937815 | 9.629515 | 20.31508 | 7.297692 | -0.42177 | -0.43159 | 0.518938 |
| 7 | 111881800 | 111882399 | 4.937815 | 9.629515 | 20.31508 | 7.297692 | -0.42177 | -0.43159 | 0.518938 |
| 5 | 84924000 | 84924799 | 18.88892 | 33.02228 | 33.68841 | 14.8503 | 0.359012 | 0.697412 | 0.518954 |
| 8 | 104538600 | 104540199 | 11.29857 | 20.43282 | 34.7862 | 15.0924 | -0.35287 | 0.349918 | 0.518973 |
| 9 | 73465200 | 73466599 | 11.29857 | 20.43282 | 34.7862 | 15.0924 | -0.35287 | 0.349918 | 0.518973 |
| 12 | 59484400 | 59489199 | 67.00113 | 106.3018 | 111.973 | 46.63427 | 0.25622 | 2.317596 | 0.518979 |
| 6 | 139559800 | 139561799 | 24.07671 | 41.30606 | 43.08943 | 18.89869 | 0.322743 | 0.939029 | 0.518983 |
| 7 | 27697600 | 27699999 | 45.64625 | 74.21053 | 69.23537 | 29.70937 | 0.468813 | 1.594363 | 0.519026 |
| 7 | 17025800 | 17028999 | 50.09254 | 80.8032 | 71.3592 | 30.40278 | 0.53667 | 1.780433 | 0.519056 |
| 9 | 80686000 | 80688199 | 17.88528 | 32.65968 | 26.94716 | 11.5504 | 0.696863 | 0.325661 | 0.51906 |
| 2 | 115939000 | 115939799 | 11.19469 | 20.24866 | 34.30529 | 14.88036 | -0.34542 | 0.349052 | 0.519061 |
| 18 | 45424400 | 45426799 | 34.48982 | 56.62038 | 60.14017 | 25.90255 | 0.290662 | 1.431774 | 0.519131 |
| 5 | 122328400 | 122333199 | 42.88136 | 68.5355 | 111.1072 | 46.70685 | -0.35365 | 2.069338 | 0.519145 |
| 18 | 21194200 | 21196999 | 21.28281 | 36.1061 | 58.73982 | 25.73148 | -0.31817 | 1.047767 | 0.519159 |
| 12 | 105843000 | 105848799 | 85.84444 | 131.8432 | 205.1333 | 85.17142 | -0.34256 | 2.975962 | 0.519162 |
| 4 | 142325800 | 142327999 | 27.23837 | 45.83995 | 48.02477 | 20.98647 | 0.314055 | 1.124826 | 0.519163 |
| 1 | 111700400 | 111711199 | 125.1021 | 192.4664 | 176.941 | 73.48324 | 0.417663 | 2.952784 | 0.519169 |
| X | 163140400 | 163140799 | 7.287393 | 14.21323 | 16.36374 | 5.869215 | 0.444538 | -0.4185 | 0.519178 |
| 12 | 62781400 | 62782199 | 9.412761 | 18.14984 | 20.25867 | 7.840746 | 0.386152 | -0.12399 | 0.519185 |
| 17 | 40925000 | 40927799 | 16.2815 | 28.43194 | 60.68401 | 26.72868 | -0.70164 | 0.739567 | 0.519191 |
| 6 | 131671200 | 131672599 | 27.59232 | 47.62467 | 41.25506 | 18.09828 | 0.59147 | 0.89894 | 0.519201 |
| X | 90162200 | 90166799 | 75.08069 | 119.1666 | 116.2876 | 48.44992 | 0.366875 | 2.31542 | 0.519218 |
| 4 | 61867600 | 61868599 | 16.75717 | 29.79601 | 30.64988 | 13.51613 | 0.353914 | 0.480809 | 0.519221 |
| 6 | 35516000 | 35517599 | 13.18556 | 23.01772 | 43.66734 | 19.22944 | -0.53185 | 0.704595 | 0.519222 |
| 7 | 52444000 | 52444999 | 24.82644 | 42.27169 | 38.2244 | 16.75472 | 0.526729 | 1.006727 | 0.519235 |
| 8 | 110711600 | 110715999 | 34.10497 | 55.24371 | 86.88014 | 37.18276 | -0.28758 | 1.653477 | 0.51924 |
| X | 130726600 | 130729399 | 23.99777 | 40.23456 | 69.43779 | 30.28993 | -0.40375 | 1.172857 | 0.519243 |
| 1 | 125800000 | 125801399 | 12.83087 | 22.64401 | 38.28884 | 16.94201 | -0.36403 | 0.540609 | 0.519248 |
| 3 | 152663800 | 152667799 | 43.00227 | 68.862 | 107.5611 | 45.34161 | -0.29701 | 2.008486 | 0.519255 |
| 8 | 79519800 | 79521999 | 28.64145 | 46.5607 | 79.12478 | 33.95195 | -0.39409 | 1.570456 | 0.519256 |
| 3 | 21542400 | 21544199 | 22.93353 | 39.52972 | 41.49603 | 18.20324 | 0.315022 | 0.890765 | 0.519258 |
| 7 | 57501200 | 57502399 | 12.77874 | 22.69934 | 38.19899 | 16.85932 | -0.3531 | 0.476415 | 0.519269 |

| | | | | | | | | | |
|---|---|---|---|---|---|---|---|---|---|
| 11 | 27307000 | 27308599 | 18.21703 | 32.73711 | 29.40608 | 12.85221 | 0.557093 | 0.423021 | 0.519347 |
| 12 | 66807400 | 66809599 | 15.39493 | 28.2259 | 27.23971 | 11.61366 | 0.477493 | 0.285171 | 0.519354 |
| 9 | 6354800 | 6355999 | 14.03841 | 24.46969 | 48.39988 | 21.29333 | -0.59196 | 0.740373 | 0.519361 |
| X | 60723200 | 60724999 | 20.12241 | 36.43863 | 27.82744 | 12.05321 | 0.798446 | 0.39161 | 0.519369 |
| 13 | 74959000 | 74961799 | 20.02657 | 33.4731 | 66.0041 | 28.73614 | -0.59503 | 1.192602 | 0.519372 |
| 3 | 21070200 | 21072199 | 33.86076 | 55.59398 | 57.8251 | 24.90664 | 0.320912 | 1.428021 | 0.519374 |
| 4 | 118846600 | 118854799 | 104.856 | 159.248 | 262.4916 | 108.702 | -0.43862 | 3.211652 | 0.519382 |
| 2 | 151001000 | 151002999 | 15.44393 | 26.94804 | 44.69565 | 19.67673 | -0.33895 | 0.727044 | 0.519388 |
| X | 128348000 | 128350399 | 24.79216 | 43.12486 | 32.58126 | 14.31539 | 0.789929 | 0.798902 | 0.519396 |
| 11 | 84644200 | 84648799 | 31.84747 | 51.65319 | 82.4465 | 35.31865 | -0.30711 | 1.622039 | 0.519417 |
| 2 | 70249600 | 70250999 | 14.25711 | 25.81337 | 26.82161 | 11.61985 | 0.356464 | 0.346715 | 0.519427 |
| 9 | 19650600 | 19651999 | 19.63869 | 35.37859 | 27.61428 | 12.03877 | 0.760825 | 0.414541 | 0.519429 |
| 16 | 57099400 | 57102599 | 26.50548 | 43.62496 | 70.65775 | 30.46566 | -0.3148 | 1.389117 | 0.519436 |
| 6 | 60361000 | 60364399 | 37.06363 | 61.10062 | 60.43985 | 26.07772 | 0.39546 | 1.389057 | 0.519455 |
| 2 | 166557400 | 166559399 | 16.79789 | 29.13126 | 47.93576 | 21.03209 | -0.3289 | 0.81818 | 0.519469 |
| 1 | 16588800 | 16590399 | 13.00281 | 22.85825 | 39.36211 | 17.43048 | -0.39242 | 0.586465 | 0.519472 |
| 5 | 26583200 | 26585999 | 59.0788 | 93.33824 | 142.5567 | 59.20841 | -0.28417 | 2.426013 | 0.519484 |
| 2 | 142036200 | 142039799 | 31.31601 | 52.30509 | 53.5711 | 23.31263 | 0.34673 | 1.220438 | 0.519486 |
| 10 | 83736600 | 83738599 | 11.28128 | 21.81364 | 17.92514 | 6.858898 | 0.839406 | -0.14749 | 0.519492 |
| X | 103243200 | 103246199 | 52.16949 | 84.61227 | 57.72159 | 24.7267 | 0.916258 | 1.650686 | 0.519495 |
| 15 | 17814000 | 17814799 | 10.29085 | 19.71228 | 21.1582 | 8.345761 | 0.416409 | -0.05223 | 0.519497 |
| 1 | 35299200 | 35305999 | 52.72146 | 83.77145 | 128.4387 | 53.57202 | -0.2824 | 2.266661 | 0.519498 |
| 12 | 9324400 | 9326799 | 23.60038 | 40.79876 | 41.4225 | 18.17261 | 0.363812 | 0.867719 | 0.519512 |
| X | 42767200 | 42774799 | 70.93774 | 112.8216 | 108.8387 | 45.44094 | 0.386298 | 2.252811 | 0.519522 |
| 16 | 20988000 | 20992999 | 66.32286 | 105.3857 | 111.1921 | 46.3795 | 0.255948 | 2.272578 | 0.519531 |
| 5 | 131197200 | 131198999 | 16.75428 | 29.64136 | 30.75685 | 13.60253 | 0.337906 | 0.521867 | 0.519533 |
| 6 | 42828200 | 42830799 | 33.67037 | 55.42169 | 59.05059 | 25.46178 | 0.287686 | 1.396809 | 0.519535 |
| 6 | 136267600 | 136272399 | 44.98045 | 72.6359 | 76.82223 | 32.77623 | 0.278871 | 1.742635 | 0.519539 |
| 11 | 113304800 | 113306199 | 14.63795 | 25.65015 | 42.51345 | 18.77735 | -0.33768 | 0.653732 | 0.519555 |
| 12 | 87819600 | 87834599 | 297.0784 | 418.2277 | 567.2974 | 234.9827 | -0.26167 | 4.478816 | 0.519572 |
| 18 | 82476200 | 82477799 | 17.84961 | 30.69195 | 50.05785 | 21.95745 | -0.31868 | 0.903987 | 0.519573 |
| 13 | 66414400 | 66417999 | 143.7082 | 216.916 | 212.0814 | 87.65866 | 0.306841 | 3.341444 | 0.519577 |
| 2 | 90113400 | 90113799 | 6.102323 | 11.82775 | 25.09559 | 9.496016 | -0.50126 | -0.2227 | 0.519587 |
| 10 | 37935200 | 37940399 | 101.6278 | 157.0031 | 163.3141 | 67.83717 | 0.244203 | 2.864965 | 0.519614 |
| 11 | 43527400 | 43528199 | 8.374241 | 15.86803 | 28.76522 | 11.62919 | -0.36504 | 0.030904 | 0.519629 |
| 5 | 89658600 | 89666599 | 138.7455 | 210.2838 | 216.2029 | 89.47758 | 0.239349 | 3.258373 | 0.519631 |
| 7 | 35620800 | 35632399 | 135.2613 | 203.0357 | 290.2012 | 119.7633 | -0.24766 | 3.452026 | 0.519638 |
| 3 | 98163200 | 98164199 | 16.44515 | 29.13199 | 30.27024 | 13.38387 | 0.336647 | 0.505298 | 0.519644 |
| X | 19359800 | 19362599 | 29.78487 | 51.51407 | 39.53451 | 17.34456 | 0.766129 | 0.890346 | 0.519655 |
| 3 | 78633000 | 78633599 | 5.516497 | 10.74587 | 21.97537 | 8.011989 | -0.39773 | -0.3672 | 0.519657 |
| 3 | 6227600 | 6233199 | 104.5376 | 161.1649 | 167.4279 | 69.54671 | 0.243777 | 2.904551 | 0.519684 |
| 2 | 134455200 | 134455799 | 8.600196 | 16.0749 | 29.00757 | 12.02558 | -0.38843 | 0.125546 | 0.519688 |
| 16 | 62606800 | 62607799 | 8.600196 | 16.0749 | 29.00757 | 12.02558 | -0.38843 | 0.125546 | 0.519688 |
| 8 | 125540800 | 125546399 | 61.51311 | 96.75471 | 152.0754 | 63.08942 | -0.3305 | 2.524207 | 0.519699 |
| 7 | 102926600 | 102933599 | 84.86823 | 133.165 | 137.2124 | 56.90536 | 0.275177 | 2.583887 | 0.519701 |
| 7 | 12941800 | 12943599 | 28.95291 | 48.31793 | 50.78872 | 22.09021 | 0.309422 | 1.218877 | 0.519719 |
| X | 54396800 | 54400799 | 36.10868 | 60.47098 | 44.79088 | 19.52454 | 0.812636 | 1.200615 | 0.519721 |

| | | | | | | | | | |
|---|---|---|---|---|---|---|---|---|---|
| 5 | 119156800 | 119158799 | 14.62296 | 25.58134 | 42.56597 | 18.77417 | -0.34331 | 0.680303 | 0.519733 |
| 1 | 142602600 | 142603799 | 11.7247 | 22.20786 | 23.07018 | 9.334652 | 0.430633 | 0.04847 | 0.519741 |
| 5 | 4116200 | 4117799 | 12.0425 | 21.33213 | 36.75954 | 16.2532 | -0.38894 | 0.492059 | 0.519744 |
| 19 | 39525200 | 39534199 | 99.33259 | 155.9626 | 135.9098 | 56.36993 | 0.51736 | 2.579726 | 0.519773 |
| 15 | 75807600 | 75811399 | 39.54481 | 64.91586 | 62.36951 | 26.8622 | 0.434951 | 1.445128 | 0.519781 |
| 5 | 144359000 | 144361399 | 15.69275 | 27.24603 | 46.34382 | 20.33849 | -0.3761 | 0.799002 | 0.519785 |
| 1 | 19119800 | 19122999 | 30.38661 | 51.78551 | 45.70929 | 20.03699 | 0.561576 | 1.027893 | 0.519787 |
| 11 | 115310400 | 115311799 | 10.88235 | 19.51021 | 34.75602 | 15.22067 | -0.4267 | 0.399384 | 0.51979 |
| 1 | 65390800 | 65391999 | 11.19322 | 19.97621 | 36.74177 | 16.15229 | -0.47662 | 0.432765 | 0.5198 |
| 1 | 174874200 | 174875999 | 25.8221 | 43.6049 | 43.83902 | 19.17715 | 0.373553 | 1.087974 | 0.519825 |
| 18 | 54291600 | 54295399 | 56.86916 | 90.27812 | 135.2205 | 56.34841 | -0.2501 | 2.299494 | 0.519833 |
| 7 | 61475000 | 61483799 | 123.1181 | 188.854 | 186.7029 | 77.50587 | 0.309703 | 3.010533 | 0.519881 |
| X | 7827800 | 7829999 | 27.42985 | 47.07353 | 42.54479 | 18.66028 | 0.529133 | 0.953773 | 0.519882 |
| X | 81442400 | 81443199 | 16.67598 | 30.42988 | 27.77979 | 11.91784 | 0.551427 | 0.325031 | 0.519895 |
| X | 40244600 | 40247799 | 54.00072 | 86.89248 | 84.26437 | 35.77805 | 0.397717 | 1.866532 | 0.519934 |
| 2 | 77976400 | 77976999 | 4.846133 | 9.412434 | 22.23 | 8.31374 | -0.63668 | -0.28401 | 0.519938 |
| 6 | 127855200 | 127856799 | 16.9806 | 30.4494 | 30.12837 | 13.19187 | 0.416699 | 0.430061 | 0.519968 |
| 4 | 41348800 | 41349599 | 5.674402 | 11.0167 | 22.80403 | 8.556272 | -0.45334 | -0.26086 | 0.519976 |
| 7 | 108723600 | 108724399 | 5.674402 | 11.0167 | 22.80403 | 8.556272 | -0.45334 | -0.26086 | 0.519976 |
| 2 | 85944200 | 85946599 | 22.43305 | 39.12445 | 36.89062 | 16.23574 | 0.471636 | 0.751811 | 0.520008 |
| 6 | 15515600 | 15519199 | 38.59846 | 62.09203 | 97.08521 | 41.20561 | -0.29049 | 1.85779 | 0.520012 |
| X | 149829200 | 149832999 | 41.8569 | 68.6619 | 62.23158 | 26.79464 | 0.518552 | 1.463486 | 0.520021 |
| 15 | 96948200 | 96949999 | 18.70893 | 32.72931 | 33.63683 | 14.83588 | 0.348363 | 0.685347 | 0.520025 |
| 8 | 43514000 | 43514399 | 8.211244 | 15.97635 | 17.88425 | 6.594384 | 0.448563 | -0.30479 | 0.520033 |
| 15 | 40412600 | 40413599 | 8.232623 | 15.63931 | 28.27348 | 11.36923 | -0.35414 | 0.011638 | 0.520036 |
| 14 | 6633200 | 6635599 | 20.1858 | 35.29025 | 36.54442 | 16.10955 | 0.337443 | 0.707622 | 0.520038 |
| 10 | 20524800 | 20527199 | 24.00739 | 40.05115 | 64.15091 | 27.89609 | -0.29624 | 1.221944 | 0.520042 |
| 4 | 40791600 | 40794999 | 45.7912 | 73.85478 | 78.40094 | 33.39709 | 0.271523 | 1.779694 | 0.520083 |
| 6 | 133840400 | 133842199 | 16.24656 | 28.08593 | 47.41333 | 20.80086 | -0.36641 | 0.845793 | 0.520103 |
| 12 | 103892000 | 103901199 | 189.4999 | 276.8362 | 390.8939 | 161.0445 | -0.2665 | 3.925715 | 0.520132 |
| 12 | 4167200 | 4170999 | 26.78531 | 44.13544 | 72.03473 | 31.07425 | -0.32523 | 1.375501 | 0.520134 |
| X | 153556200 | 153563199 | 80.41286 | 127.9495 | 105.6875 | 44.15315 | 0.610563 | 2.244653 | 0.520137 |
| 4 | 113210800 | 113217799 | 100.9109 | 158.4685 | 134.4296 | 55.75724 | 0.556299 | 2.576366 | 0.520142 |
| 4 | 80990800 | 80991399 | 8.593226 | 15.95587 | 29.96018 | 12.55554 | -0.45811 | 0.167466 | 0.520157 |
| 18 | 78538600 | 78541599 | 24.49504 | 41.79121 | 44.22902 | 19.38984 | 0.300902 | 0.993016 | 0.520161 |
| X | 142251600 | 142257399 | 79.93054 | 126.9526 | 99.33213 | 41.41446 | 0.686035 | 2.296683 | 0.520164 |
| 7 | 17872800 | 17898399 | 376.928 | 534.768 | 435.1269 | 180.0049 | 0.486627 | 4.369315 | 0.520173 |
| 5 | 21078800 | 21081199 | 28.21883 | 47.79832 | 42.64528 | 18.67228 | 0.545311 | 1.068513 | 0.520177 |
| 3 | 129209200 | 129211799 | 25.98676 | 44.49223 | 44.67326 | 19.59013 | 0.377272 | 0.967927 | 0.520209 |
| 1 | 96025600 | 96027199 | 20.82293 | 36.70196 | 32.91315 | 14.56707 | 0.545224 | 0.590074 | 0.520231 |
| 9 | 16648400 | 16650399 | 31.44301 | 52.84814 | 50.74891 | 22.16827 | 0.439319 | 1.155514 | 0.520239 |
| 6 | 42127800 | 42131399 | 41.39518 | 66.522 | 102.694 | 43.52068 | -0.27403 | 1.896631 | 0.520239 |
| 10 | 124634600 | 124635799 | 9.147253 | 17.1075 | 31.1901 | 12.91749 | -0.40274 | 0.128484 | 0.520249 |
| 11 | 116370800 | 116372199 | 14.77245 | 25.9 | 42.47821 | 18.77045 | -0.32265 | 0.644785 | 0.520275 |
| 3 | 14421000 | 14424599 | 37.17329 | 61.53149 | 57.25744 | 24.76749 | 0.484184 | 1.337506 | 0.520292 |
| 6 | 59606200 | 59608199 | 33.97278 | 56.53153 | 49.25104 | 21.38202 | 0.579503 | 1.265056 | 0.520356 |
| 7 | 30612200 | 30614599 | 15.66593 | 26.87821 | 49.51291 | 21.71609 | -0.49402 | 0.914169 | 0.520357 |

| | | | | | | | | | |
|---|---|---|---|---|---|---|---|---|---|
| 12 | 114589000 | 114599999 | 176.5894 | 264.3026 | 260.339 | 107.3701 | 0.285346 | 3.51322 | 0.520357 |
| X | 39589200 | 39590399 | 15.00708 | 28.1296 | 25.00627 | 10.31728 | 0.631045 | 0.152455 | 0.520386 |
| 11 | 27629400 | 27632999 | 29.61633 | 49.86799 | 50.19115 | 21.93665 | 0.37176 | 1.131399 | 0.520389 |
| Y | 21225600 | 21230199 | 26.23836 | 44.38883 | 34.18533 | 14.96248 | 0.756544 | 1.059039 | 0.520394 |
| 6 | 66633000 | 66636799 | 57.82833 | 92.39061 | 97.89239 | 41.13021 | 0.258216 | 2.092558 | 0.520399 |
| 6 | 93332000 | 93333599 | 10.83468 | 19.81706 | 33.94005 | 14.5179 | -0.34795 | 0.280628 | 0.520405 |
| 4 | 82697800 | 82699599 | 18.56957 | 31.93406 | 53.275 | 23.36873 | -0.35115 | 0.911197 | 0.520407 |
| 5 | 145468000 | 145470799 | 65.42749 | 104.2123 | 107.3523 | 44.91236 | 0.294019 | 2.198048 | 0.520408 |
| 15 | 32831200 | 32833999 | 40.22796 | 64.61886 | 100.4498 | 42.54346 | -0.28475 | 1.906958 | 0.520426 |
| 7 | 15128200 | 15141799 | 453.4325 | 629.6245 | 546.4452 | 226.831 | 0.362395 | 4.68997 | 0.520462 |
| 2 | 133085400 | 133087799 | 20.92225 | 37.01954 | 31.81102 | 14.06842 | 0.60861 | 0.548714 | 0.520468 |
| 7 | 11859200 | 11863799 | 24.75007 | 42.64618 | 37.19855 | 16.31787 | 0.580961 | 0.895318 | 0.520478 |
| 7 | 59562200 | 59564999 | 43.42564 | 71.11271 | 63.14129 | 27.16653 | 0.547096 | 1.485054 | 0.520479 |
| 9 | 36438200 | 36442799 | 42.07532 | 69.14434 | 60.8513 | 26.22026 | 0.561823 | 1.437931 | 0.520539 |
| Y | 38411600 | 38412999 | 11.64079 | 21.88854 | 16.0924 | 6.604228 | 0.908228 | 0.07588 | 0.520545 |
| 11 | 6408600 | 6411399 | 14.72893 | 25.86944 | 42.92031 | 18.9942 | -0.33936 | 0.615274 | 0.520546 |
| 19 | 36619400 | 36620399 | 16.19776 | 29.08249 | 29.7337 | 13.00515 | 0.370846 | 0.416737 | 0.520561 |
| 13 | 80829400 | 80831799 | 31.81375 | 51.76264 | 81.258 | 34.87834 | -0.27923 | 1.562896 | 0.520561 |
| 11 | 56380600 | 56381399 | 11.17117 | 20.28927 | 34.34915 | 14.82305 | -0.33965 | 0.324713 | 0.520565 |
| 8 | 22491600 | 22496599 | 39.07855 | 62.71604 | 101.697 | 43.01281 | -0.3471 | 1.932966 | 0.520585 |
| 13 | 100932400 | 100934199 | 16.84116 | 30.03983 | 30.69601 | 13.50057 | 0.366161 | 0.463102 | 0.520588 |
| 7 | 34860200 | 34863599 | 23.71167 | 39.20534 | 69.01406 | 29.83006 | -0.43285 | 1.323747 | 0.520599 |
| 9 | 38533600 | 38541599 | 57.21966 | 92.0029 | 76.56461 | 32.47339 | 0.616587 | 1.888916 | 0.520615 |
| 2 | 146805200 | 146810399 | 53.45538 | 85.70134 | 90.40168 | 38.17603 | 0.270246 | 1.977835 | 0.520615 |
| 1 | 179807400 | 179810399 | 32.6514 | 54.20357 | 56.38754 | 24.4401 | 0.324143 | 1.293026 | 0.520625 |
| X | 65018000 | 65019399 | 8.347639 | 15.72986 | 30.78649 | 12.58902 | -0.48834 | 0.078606 | 0.52063 |
| 4 | 7738200 | 7739999 | 16.40046 | 30.00801 | 27.7585 | 11.86786 | 0.5356 | 0.306765 | 0.520641 |
| 11 | 21758200 | 21759999 | 24.66665 | 42.52789 | 41.91251 | 18.38608 | 0.40573 | 0.895577 | 0.520658 |
| Y | 1857000 | 1857799 | 9.718226 | 18.66677 | 15.16584 | 5.93497 | 0.825643 | -0.10169 | 0.520661 |
| 3 | 89279400 | 89281399 | 12.54362 | 22.10127 | 38.35595 | 16.9772 | -0.40216 | 0.557673 | 0.520663 |
| 6 | 117848000 | 117849399 | 13.53221 | 23.71765 | 41.01911 | 18.12061 | -0.39858 | 0.64236 | 0.520672 |
| 14 | 90377000 | 90380799 | 57.31949 | 91.51115 | 96.07668 | 40.32581 | 0.270268 | 2.113548 | 0.520675 |
| 5 | 78772000 | 78773999 | 19.72125 | 34.85747 | 32.46979 | 14.36281 | 0.492223 | 0.553424 | 0.520678 |
| 1 | 47626400 | 47627199 | 11.0032 | 19.99503 | 33.83499 | 14.59127 | -0.33832 | 0.319098 | 0.520685 |
| 5 | 52599600 | 52602999 | 55.27501 | 88.57797 | 92.43334 | 39.00676 | 0.285004 | 1.994982 | 0.520689 |
| 10 | 51762800 | 51766399 | 33.16791 | 54.57428 | 57.33144 | 24.7166 | 0.30782 | 1.398873 | 0.52069 |
| 9 | 79229800 | 79232399 | 25.71772 | 44.65147 | 38.47388 | 16.88466 | 0.600576 | 0.832191 | 0.520699 |
| 18 | 14825800 | 14827399 | 22.52125 | 39.34118 | 37.4625 | 16.50401 | 0.457854 | 0.733564 | 0.520719 |
| 17 | 22397400 | 22399799 | 28.24783 | 48.83058 | 40.10596 | 17.595 | 0.668402 | 0.887075 | 0.520724 |
| 16 | 30651200 | 30653199 | 27.54436 | 46.53644 | 48.53156 | 21.23309 | 0.320906 | 1.095057 | 0.520728 |
| 3 | 50923800 | 50924999 | 14.01081 | 25.39182 | 25.49638 | 11.03246 | 0.406878 | 0.337963 | 0.520735 |
| 8 | 91560600 | 91562999 | 24.08083 | 41.05399 | 42.67132 | 18.70582 | 0.326657 | 0.995269 | 0.52074 |
| 8 | 78655600 | 78656799 | 11.81292 | 21.29093 | 35.77824 | 15.58855 | -0.33813 | 0.376044 | 0.520743 |
| 4 | 96826800 | 96832199 | 91.47579 | 142.6347 | 148.5602 | 61.65201 | 0.252636 | 2.696448 | 0.520755 |
| X | 110157600 | 110161999 | 46.07771 | 75.11727 | 55.76798 | 23.9543 | 0.800144 | 1.545399 | 0.520775 |
| 14 | 58297000 | 58299999 | 30.61311 | 51.16134 | 53.00715 | 23.07583 | 0.33012 | 1.210986 | 0.520793 |
| 16 | 7412200 | 7413799 | 21.26904 | 37.16215 | 36.97075 | 16.28969 | 0.39495 | 0.722261 | 0.520815 |

| | | | | | | | | | |
|---|---|---|---|---|---|---|---|---|---|
| 13 | 90515000 | 90517599 | 30.1667 | 51.04067 | 47.96004 | 20.99224 | 0.470726 | 1.089081 | 0.520822 |
| 3 | 87191000 | 87193599 | 25.51396 | 42.49134 | 72.24512 | 31.38177 | -0.38217 | 1.251252 | 0.520826 |
| 7 | 25804800 | 25811799 | 108.3209 | 167.5573 | 152.112 | 63.18362 | 0.441784 | 2.843341 | 0.52087 |
| 4 | 62069400 | 62070599 | 24.65962 | 41.96839 | 44.25721 | 19.39834 | 0.305552 | 1.015964 | 0.520879 |
| 16 | 63300600 | 63301599 | 9.875321 | 18.31369 | 32.07185 | 13.46324 | -0.36 | 0.191971 | 0.520896 |
| 11 | 14798200 | 14799199 | 13.40657 | 24.7372 | 25.88783 | 10.94216 | 0.370252 | 0.231341 | 0.520903 |
| 7 | 72943200 | 72944399 | 10.31708 | 18.85962 | 32.86505 | 14.06835 | -0.3736 | 0.276102 | 0.520912 |
| 5 | 96440200 | 96449199 | 179.374 | 263.7912 | 369.5835 | 152.2008 | -0.24612 | 3.820029 | 0.520955 |
| 5 | 92821600 | 92823999 | 17.3912 | 29.71402 | 51.62969 | 22.63689 | -0.41119 | 0.960956 | 0.520957 |
| 7 | 17713800 | 17720199 | 112.8028 | 174.7866 | 162.4217 | 67.46544 | 0.410031 | 2.816522 | 0.520982 |
| X | 53752200 | 53754799 | 20.79636 | 37.00874 | 30.86659 | 13.60219 | 0.655762 | 0.50428 | 0.52099 |
| 2 | 110067400 | 110067999 | 9.105022 | 17.05963 | 31.22081 | 12.8903 | -0.40388 | 0.120874 | 0.521 |
| 16 | 20584000 | 20584599 | 5.367464 | 10.45616 | 21.35971 | 7.781676 | -0.39455 | -0.37202 | 0.521013 |
| 17 | 90059200 | 90059799 | 15.8273 | 28.112 | 28.42047 | 12.54449 | 0.378076 | 0.478292 | 0.521028 |
| 15 | 48099800 | 48100999 | 13.70699 | 25.75336 | 24.45602 | 10.05002 | 0.540521 | 0.122805 | 0.521032 |
| 11 | 19509600 | 19511599 | 12.81616 | 22.59677 | 38.79391 | 17.16872 | -0.38645 | 0.550933 | 0.521045 |
| 2 | 23753800 | 23756599 | 46.61346 | 75.47615 | 66.44491 | 28.42889 | 0.547033 | 1.674062 | 0.521046 |
| 14 | 38176000 | 38180799 | 63.86012 | 101.7333 | 105.372 | 44.09434 | 0.286437 | 2.19161 | 0.521052 |
| 16 | 19482600 | 19483199 | 7.163737 | 13.74935 | 25.75482 | 10.10149 | -0.37164 | -0.08586 | 0.521062 |
| X | 80455600 | 80457599 | 24.97528 | 43.44226 | 32.09412 | 14.10113 | 0.822112 | 0.799661 | 0.521068 |
| 2 | 42813800 | 42815399 | 17.88219 | 31.59781 | 32.14781 | 14.22121 | 0.365439 | 0.544405 | 0.521091 |
| 8 | 52138800 | 52140599 | 17.88219 | 31.59781 | 32.14781 | 14.22121 | 0.365439 | 0.544405 | 0.521091 |
| 15 | 48231000 | 48231999 | 17.68915 | 31.25756 | 32.28819 | 14.28323 | 0.343834 | 0.544998 | 0.521109 |
| 2 | 10316000 | 10317199 | 15.75595 | 27.46884 | 45.22944 | 19.90096 | -0.32869 | 0.741278 | 0.521113 |
| 6 | 13480400 | 13485199 | 53.03301 | 85.2594 | 85.08459 | 36.07973 | 0.354819 | 1.889115 | 0.521119 |
| 1 | 41616400 | 41616999 | 5.804079 | 11.29813 | 23.11628 | 8.491204 | -0.40876 | -0.33431 | 0.521125 |
| 3 | 49007800 | 49010199 | 21.33628 | 37.46237 | 34.59313 | 15.30489 | 0.501802 | 0.64253 | 0.521127 |
| 1 | 7761600 | 7762199 | 11.4432 | 21.49667 | 22.64913 | 9.309963 | 0.391998 | 0.101888 | 0.521129 |
| 11 | 52077600 | 52080199 | 18.6148 | 31.4979 | 54.75912 | 23.9688 | -0.41345 | 1.05826 | 0.521132 |
| 10 | 7778400 | 7780199 | 17.30587 | 29.48936 | 52.12112 | 22.84743 | -0.43616 | 0.988187 | 0.521136 |
| 2 | 98318200 | 98320399 | 25.10609 | 43.6174 | 39.64713 | 17.40654 | 0.523977 | 0.823997 | 0.521139 |
| 5 | 145646000 | 145658399 | 213.5703 | 316.0748 | 297.8708 | 122.6719 | 0.33422 | 3.718989 | 0.521165 |
| 8 | 116724600 | 116726799 | 22.59189 | 38.92411 | 40.99506 | 17.98324 | 0.310166 | 0.892347 | 0.521213 |
| 2 | 44103400 | 44105599 | 19.94728 | 35.08411 | 33.55155 | 14.85591 | 0.451752 | 0.608182 | 0.521218 |
| 17 | 20113600 | 20118399 | 75.89014 | 120.1238 | 122.8069 | 51.05418 | 0.296287 | 2.389659 | 0.521219 |
| 15 | 32018200 | 32019399 | 9.862869 | 18.21314 | 31.45763 | 13.28187 | -0.34623 | 0.213876 | 0.521234 |
| 17 | 55457800 | 55461999 | 37.01741 | 61.34371 | 56.6777 | 24.53612 | 0.494543 | 1.323922 | 0.521243 |
| 5 | 150114000 | 150115599 | 15.41078 | 26.82741 | 44.739 | 19.6669 | -0.34718 | 0.759179 | 0.52125 |
| X | 17819200 | 17826199 | 76.29573 | 121.164 | 114.3591 | 47.67357 | 0.415709 | 2.300167 | 0.521262 |
| 2 | 17841600 | 17844999 | 22.81893 | 38.07002 | 63.03322 | 27.41077 | -0.34394 | 1.21486 | 0.521264 |
| 18 | 65698800 | 65702599 | 31.26921 | 50.97932 | 81.60927 | 35.05487 | -0.30536 | 1.531545 | 0.521271 |
| 7 | 14277800 | 14293799 | 60.67162 | 97.62291 | 71.37066 | 30.30181 | 0.804321 | 1.869866 | 0.521275 |
| 17 | 18352400 | 18354799 | 26.40245 | 45.74126 | 40.77011 | 17.88757 | 0.55161 | 0.862018 | 0.521276 |
| 1 | 36764800 | 36766799 | 16.5243 | 28.14359 | 57.43622 | 25.17632 | -0.64307 | 0.99442 | 0.521311 |
| 1 | 145291600 | 145295199 | 28.01589 | 47.89594 | 44.60733 | 19.55891 | 0.484969 | 0.987538 | 0.52132 |
| 5 | 60105000 | 60107799 | 18.44261 | 31.76866 | 51.69447 | 22.67659 | -0.31496 | 0.890885 | 0.521323 |
| 15 | 3276000 | 3276999 | 10.89183 | 19.71385 | 33.82067 | 14.65805 | -0.3625 | 0.342621 | 0.521353 |

| | | | | | | | | | |
|---|---|---|---|---|---|---|---|---|---|
| 14 | 103727600 | 103732599 | 51.24028 | 82.71565 | 76.95539 | 32.81829 | 0.462835 | 1.760775 | 0.521375 |
| 6 | 3104200 | 3106199 | 17.34926 | 31.04405 | 29.99452 | 13.15696 | 0.448912 | 0.443991 | 0.52138 |
| 14 | 98552400 | 98553999 | 19.80692 | 33.22408 | 64.51077 | 28.14916 | -0.57299 | 1.150243 | 0.521401 |
| 18 | 13472400 | 13474199 | 24.40271 | 42.68912 | 36.28537 | 16.00377 | 0.621128 | 0.721554 | 0.521411 |
| 12 | 69059400 | 69060599 | 30.2947 | 50.69323 | 51.84687 | 22.58872 | 0.348786 | 1.198436 | 0.521423 |
| 2 | 20564000 | 20565599 | 10.18515 | 18.54542 | 32.82775 | 14.12164 | -0.4006 | 0.297959 | 0.521425 |
| 6 | 149347400 | 149348399 | 6.70568 | 12.92718 | 24.76258 | 9.589913 | -0.38625 | -0.13488 | 0.521453 |
| 18 | 34639000 | 34639799 | 6.70568 | 12.92718 | 24.76258 | 9.589913 | -0.38625 | -0.13488 | 0.521453 |
| 3 | 15149600 | 15151999 | 20.58444 | 36.06972 | 35.26406 | 15.57526 | 0.419843 | 0.670151 | 0.521471 |
| 14 | 116614800 | 116616999 | 16.29447 | 28.94838 | 30.24391 | 13.34677 | 0.330983 | 0.482774 | 0.521476 |
| 14 | 13739200 | 13739999 | 13.64512 | 24.09922 | 39.78412 | 17.60098 | -0.32926 | 0.540485 | 0.52148 |
| 6 | 66983400 | 66986799 | 53.60951 | 84.58815 | 144.8878 | 60.13734 | -0.45079 | 2.454751 | 0.521502 |
| 9 | 6816800 | 6818399 | 13.75363 | 25.65676 | 25.45092 | 10.58636 | 0.464204 | 0.170532 | 0.521511 |
| 18 | 17787800 | 17794399 | 70.14098 | 111.6003 | 110.1861 | 46.02961 | 0.353497 | 2.236297 | 0.521528 |
| 14 | 44843800 | 44846199 | 25.63844 | 44.6173 | 38.47338 | 16.9091 | 0.599915 | 0.802046 | 0.521532 |
| 3 | 42565800 | 42566999 | 11.35194 | 20.37187 | 34.7419 | 15.20204 | -0.36329 | 0.397197 | 0.521542 |
| 8 | 27944200 | 27945799 | 11.35194 | 20.37187 | 34.7419 | 15.20204 | -0.36329 | 0.397197 | 0.521542 |
| 7 | 140635400 | 140637799 | 19.38231 | 32.82971 | 54.80566 | 23.99652 | -0.35517 | 1.051584 | 0.521556 |
| 8 | 40347200 | 40348399 | 10.13533 | 18.26508 | 37.4171 | 16.30511 | -0.62209 | 0.368929 | 0.521607 |
| 13 | 30117800 | 30127199 | 80.03097 | 124.4229 | 181.9983 | 75.56187 | -0.24006 | 2.746395 | 0.521607 |
| 4 | 37605800 | 37613399 | 30.76122 | 51.49587 | 45.96097 | 20.02975 | 0.544573 | 1.188192 | 0.521609 |
| 4 | 138874000 | 138881199 | 86.00758 | 134.7554 | 140.8204 | 58.41046 | 0.253245 | 2.609967 | 0.521611 |
| 4 | 10001400 | 10004799 | 67.47033 | 107.0354 | 111.3306 | 46.36202 | 0.274291 | 2.322425 | 0.521626 |
| 15 | 84244400 | 84251199 | 58.88998 | 93.0288 | 142.6877 | 59.25925 | -0.29043 | 2.428138 | 0.521627 |
| 12 | 23244200 | 23247199 | 31.45342 | 52.5324 | 53.58574 | 23.31839 | 0.352562 | 1.220912 | 0.521658 |
| 10 | 118684000 | 118685799 | 17.45104 | 30.70865 | 31.92556 | 14.13457 | 0.332311 | 0.587353 | 0.521665 |
| 19 | 13484400 | 13486999 | 15.76874 | 28.87155 | 24.16867 | 10.32361 | 0.679638 | 0.290161 | 0.521667 |
| 13 | 113504200 | 113506599 | 20.91384 | 35.00688 | 59.59456 | 25.96939 | -0.3838 | 1.171987 | 0.521668 |
| 3 | 103053200 | 103055199 | 11.7316 | 20.95532 | 35.77934 | 15.71762 | -0.36877 | 0.433033 | 0.521686 |
| 14 | 35086400 | 35087799 | 11.7316 | 20.95532 | 35.77934 | 15.71762 | -0.36877 | 0.433033 | 0.521686 |
| 3 | 48052200 | 48053999 | 14.83777 | 25.85667 | 54.93262 | 24.16366 | -0.69498 | 0.755593 | 0.521688 |
| 7 | 41411200 | 41416799 | 66.2488 | 105.9116 | 97.57465 | 41.03393 | 0.460622 | 2.083867 | 0.521689 |
| 14 | 106623000 | 106624599 | 23.15377 | 39.75856 | 42.09099 | 18.4621 | 0.301816 | 0.926879 | 0.521689 |
| 16 | 73532800 | 73534999 | 17.84817 | 31.48164 | 32.18728 | 14.24373 | 0.357397 | 0.563092 | 0.521695 |
| X | 36883000 | 36885999 | 32.49713 | 54.43077 | 51.88435 | 22.61872 | 0.450022 | 1.19576 | 0.521708 |
| 12 | 53934000 | 53935799 | 12.48251 | 22.10035 | 37.71816 | 16.67874 | -0.3756 | 0.50087 | 0.521712 |
| 13 | 78342600 | 78343999 | 12.88961 | 22.58886 | 40.53263 | 17.90402 | -0.45143 | 0.639075 | 0.521716 |
| X | 135318200 | 135320599 | 17.71582 | 30.77037 | 54.48059 | 23.91518 | -0.43386 | 0.814853 | 0.521725 |
| 6 | 135240200 | 135241799 | 17.12836 | 30.59437 | 30.62211 | 13.45238 | 0.396943 | 0.455803 | 0.521737 |
| 1 | 13839000 | 13841999 | 30.32278 | 49.43235 | 78.371 | 33.6629 | -0.29146 | 1.529876 | 0.521737 |
| 10 | 22494800 | 22496399 | 33.06818 | 55.14138 | 54.22058 | 23.57094 | 0.405363 | 1.242508 | 0.521753 |
| X | 148996400 | 149003599 | 103.2449 | 161.0296 | 149.9164 | 62.21247 | 0.414669 | 2.700286 | 0.521754 |
| 1 | 112579800 | 112580199 | 13.01753 | 23.97193 | 22.1513 | 9.388037 | 0.546388 | 0.232587 | 0.52177 |
| 3 | 144568200 | 144572999 | 53.25537 | 85.70844 | 84.39914 | 35.84454 | 0.375963 | 1.858923 | 0.521773 |
| 14 | 32083200 | 32086399 | 21.73553 | 36.31323 | 60.68602 | 26.41334 | -0.3572 | 1.19646 | 0.521795 |
| 10 | 9384400 | 9386999 | 15.88991 | 27.71854 | 45.21024 | 19.89981 | -0.31506 | 0.732891 | 0.521819 |
| 10 | 85913000 | 85918199 | 39.8481 | 64.1474 | 99.27631 | 42.17857 | -0.27453 | 1.836917 | 0.521835 |

| | | | | | | | | | |
|---|---|---|---|---|---|---|---|---|---|
| 15 | 101564000 | 101564599 | 6.927913 | 13.08156 | 27.05106 | 11.01748 | -0.56216 | 0.038614 | 0.521838 |
| X | 31653000 | 31661199 | 159.2613 | 243.3943 | 156.5349 | 64.93318 | 0.925292 | 3.094125 | 0.521857 |
| X | 161624400 | 161629399 | 33.87335 | 54.97791 | 91.0643 | 39.03076 | -0.35952 | 1.618098 | 0.521882 |
| 8 | 20490400 | 20492199 | 17.7197 | 32.59015 | 26.61662 | 11.29977 | 0.721362 | 0.285768 | 0.521889 |
| 7 | 59817200 | 59831199 | 179.391 | 269.8175 | 241.5179 | 99.72567 | 0.42963 | 3.420222 | 0.521894 |
| 17 | 27551200 | 27557399 | 49.25525 | 78.33455 | 126.4941 | 52.80978 | -0.3558 | 2.23559 | 0.5219 |
| 12 | 3895200 | 3896599 | 9.040032 | 16.76811 | 30.46256 | 12.78409 | -0.41241 | 0.178014 | 0.521927 |
| 3 | 148897800 | 148898999 | 13.65212 | 25.17101 | 26.43401 | 11.18465 | 0.364317 | 0.239708 | 0.521948 |
| X | 91063000 | 91077999 | 230.9137 | 339.7837 | 314.9662 | 129.7043 | 0.35029 | 3.816107 | 0.521949 |
| 2 | 70688600 | 70690599 | 16.84366 | 30.05716 | 30.04883 | 13.21116 | 0.397701 | 0.457276 | 0.521952 |
| 13 | 46302600 | 46305399 | 23.41712 | 40.34242 | 42.00198 | 18.42489 | 0.326688 | 0.897893 | 0.521954 |
| 17 | 54126000 | 54126399 | 10.21564 | 19.95175 | 16.31543 | 5.722632 | 0.960484 | -0.456 | 0.521959 |
| 9 | 33314000 | 33315599 | 18.46372 | 33.38986 | 27.78993 | 12.05732 | 0.673159 | 0.385055 | 0.521983 |
| 2 | 120317800 | 120320399 | 14.78855 | 25.78787 | 43.68377 | 19.22349 | -0.3693 | 0.730994 | 0.521999 |
| 7 | 11137000 | 11138399 | 14.4338 | 26.4646 | 26.18602 | 11.16391 | 0.441495 | 0.273873 | 0.522028 |
| 9 | 79959200 | 79961799 | 43.45777 | 70.29695 | 74.15708 | 31.70547 | 0.285325 | 1.690767 | 0.522033 |
| 4 | 119126600 | 119127599 | 9.736409 | 17.86329 | 31.39069 | 13.37118 | -0.38007 | 0.249843 | 0.522061 |
| 15 | 88331800 | 88333399 | 15.71628 | 28.24287 | 29.29071 | 12.80188 | 0.351121 | 0.407351 | 0.522064 |
| X | 79427400 | 79433199 | 65.91886 | 105.8587 | 84.1423 | 35.62253 | 0.680586 | 1.943563 | 0.522065 |
| 15 | 35159400 | 35163599 | 47.01823 | 75.987 | 76.67805 | 32.74831 | 0.347823 | 1.724048 | 0.522104 |
| 13 | 67397000 | 67410399 | 250.6427 | 368.3159 | 314.1526 | 129.3797 | 0.468436 | 3.839246 | 0.522107 |
| 1 | 103540400 | 103543199 | 23.34687 | 40.63446 | 38.56109 | 16.94897 | 0.462252 | 0.788101 | 0.522134 |
| 2 | 169666200 | 169668599 | 18.70329 | 32.73567 | 34.08105 | 15.0381 | 0.329757 | 0.677818 | 0.522146 |
| 2 | 80651400 | 80653199 | 16.74933 | 30.1833 | 28.43166 | 12.39008 | 0.491483 | 0.393227 | 0.522147 |
| Y | 10601600 | 10604199 | 17.33953 | 30.65952 | 21.14832 | 9.35411 | 0.923114 | 0.510063 | 0.522153 |
| 14 | 3854800 | 3856799 | 14.28104 | 25.95312 | 27.32666 | 11.78367 | 0.341778 | 0.327003 | 0.522169 |
| 8 | 79229000 | 79233599 | 42.14091 | 67.7006 | 104.398 | 44.22409 | -0.273 | 1.905582 | 0.522173 |
| 11 | 66873600 | 66875999 | 12.34829 | 21.92173 | 36.7234 | 16.21615 | -0.34683 | 0.476295 | 0.522186 |
| 9 | 75160000 | 75163599 | 19.88652 | 33.50389 | 56.94389 | 24.89228 | -0.38135 | 1.101448 | 0.522192 |
| Y | 8469600 | 8471599 | 15.5861 | 28.07477 | 18.03759 | 7.864821 | 1.039182 | 0.363035 | 0.522194 |
| 10 | 17753000 | 17754199 | 9.171525 | 17.08323 | 30.53265 | 12.73264 | -0.3821 | 0.154002 | 0.522198 |
| 3 | 87964000 | 87966199 | 10.92445 | 19.47933 | 35.28578 | 15.52377 | -0.45517 | 0.435788 | 0.522218 |
| 19 | 13376800 | 13377999 | 14.31011 | 26.34831 | 26.89096 | 11.39908 | 0.403346 | 0.253402 | 0.52223 |
| 10 | 25226400 | 25227399 | 9.477677 | 17.59873 | 31.54908 | 13.22078 | -0.39205 | 0.177434 | 0.522232 |
| 1 | 48208400 | 48210799 | 32.7011 | 54.11031 | 57.42977 | 24.83616 | 0.295009 | 1.331528 | 0.522265 |
| 7 | 98096600 | 98097999 | 8.893703 | 16.7202 | 29.65302 | 12.17301 | -0.35233 | 0.092263 | 0.522301 |
| 16 | 42823000 | 42823799 | 8.893703 | 16.7202 | 29.65302 | 12.17301 | -0.35233 | 0.092263 | 0.522301 |
| 6 | 131364000 | 131366599 | 17.73776 | 30.16834 | 53.73884 | 23.55299 | -0.44801 | 1.006077 | 0.522307 |
| 13 | 49169800 | 49176599 | 47.75063 | 76.2986 | 115.6656 | 48.60572 | -0.25749 | 2.083418 | 0.522307 |
| 15 | 3050000 | 3062999 | 452.9418 | 625.1519 | 582.6149 | 242.0768 | 0.250866 | 4.781479 | 0.522329 |
| 17 | 47934600 | 47935999 | 16.08056 | 28.63092 | 29.39694 | 12.9491 | 0.357942 | 0.468018 | 0.52233 |
| 14 | 42566000 | 42577599 | 24.9986 | 42.8498 | 36.90395 | 16.18463 | 0.597989 | 0.943712 | 0.52235 |
| 14 | 8279400 | 8280199 | 9.89861 | 18.07495 | 31.85676 | 13.65534 | -0.39077 | 0.278784 | 0.522357 |
| 10 | 96147000 | 96149799 | 50.87977 | 81.9575 | 82.59513 | 35.12329 | 0.344194 | 1.826656 | 0.522364 |
| 6 | 76772600 | 76777799 | 47.54517 | 77.0643 | 72.1942 | 30.91229 | 0.458918 | 1.656752 | 0.522369 |
| 5 | 142514800 | 142516999 | 33.29343 | 53.99615 | 84.54184 | 36.21515 | -0.27949 | 1.626249 | 0.522381 |
| 13 | 69716400 | 69717799 | 9.387958 | 17.39905 | 30.45879 | 12.79693 | -0.36023 | 0.1875 | 0.522391 |

| | | | | | | | | | |
|---|---|---|---|---|---|---|---|---|---|
| 5 | 67671600 | 67675199 | 43.13906 | 69.15562 | 106.5087 | 44.97173 | -0.27489 | 1.974328 | 0.522396 |
| X | 44687800 | 44692799 | 51.67107 | 83.51791 | 75.66564 | 32.32159 | 0.503217 | 1.727826 | 0.522406 |
| 4 | 113781400 | 113783999 | 29.49458 | 50.8628 | 41.24925 | 18.09524 | 0.686066 | 0.916876 | 0.522412 |
| 17 | 85287000 | 85288199 | 19.05791 | 33.69833 | 32.65699 | 14.44448 | 0.43582 | 0.547375 | 0.522416 |
| 2 | 131890200 | 131891399 | 12.62582 | 23.39236 | 24.96984 | 10.49541 | 0.347751 | 0.200641 | 0.522425 |
| 12 | 113550200 | 113555399 | 70.00463 | 111.1171 | 115.7981 | 48.24906 | 0.27228 | 2.308125 | 0.522435 |
| 6 | 15553200 | 15556399 | 76.02937 | 119.7712 | 125.2369 | 51.96566 | 0.258402 | 2.496073 | 0.522436 |
| 8 | 42408200 | 42409599 | 14.31834 | 26.75268 | 24.84283 | 10.30566 | 0.561701 | 0.163194 | 0.522445 |
| 12 | 113761800 | 113764799 | 33.00306 | 55.62 | 48.54274 | 21.22191 | 0.576732 | 1.132833 | 0.522459 |
| 1 | 134134800 | 134137399 | 21.6542 | 35.98176 | 63.19568 | 27.40899 | -0.42874 | 1.259649 | 0.522463 |
| X | 97832000 | 97833999 | 24.71505 | 43.14293 | 37.43432 | 16.48428 | 0.591474 | 0.756199 | 0.52247 |
| 5 | 87426600 | 87427399 | 12.67639 | 22.3542 | 38.31053 | 16.95421 | -0.38375 | 0.548109 | 0.522474 |
| 4 | 106962600 | 106964399 | 16.26175 | 28.1346 | 47.41145 | 20.80054 | -0.36359 | 0.83928 | 0.522495 |
| 12 | 23520800 | 23521399 | 12.10012 | 22.83778 | 19.96565 | 8.139335 | 0.668639 | 0.069724 | 0.522523 |
| X | 63019600 | 63020399 | 13.00621 | 24.34973 | 24.60451 | 10.17227 | 0.445272 | 0.142545 | 0.522546 |
| 9 | 78474600 | 78478199 | 29.69845 | 50.36526 | 42.55218 | 18.63833 | 0.623681 | 1.063467 | 0.522552 |
| 10 | 124710000 | 124711799 | 12.64294 | 22.67584 | 39.93177 | 17.4813 | -0.41012 | 0.419226 | 0.522568 |
| 2 | 110083000 | 110086199 | 29.98261 | 50.42354 | 50.74484 | 22.17049 | 0.371873 | 1.143888 | 0.522586 |
| 4 | 141985800 | 141993999 | 66.16306 | 103.5854 | 163.0298 | 67.62788 | -0.33767 | 2.614273 | 0.522601 |
| 1 | 8279800 | 8280199 | 9.666077 | 18.53256 | 20.26334 | 7.971706 | 0.394349 | -0.06672 | 0.522607 |
| X | 125934400 | 125936799 | 27.98408 | 48.59871 | 39.03026 | 17.13157 | 0.701932 | 0.843601 | 0.522609 |
| 14 | 38674200 | 38675399 | 12.50933 | 24.06583 | 21.77036 | 8.481085 | 0.678115 | -0.06635 | 0.52261 |
| 12 | 31405200 | 31408199 | 41.54836 | 68.12157 | 55.42066 | 23.85703 | 0.673629 | 1.463423 | 0.52261 |
| 12 | 79608600 | 79610399 | 13.8062 | 25.12074 | 26.39776 | 11.36579 | 0.346108 | 0.31483 | 0.522615 |
| X | 140530400 | 140531599 | 8.772655 | 16.59044 | 31.52876 | 12.79936 | -0.43769 | 0.057388 | 0.522619 |
| 11 | 96105800 | 96111799 | 65.74677 | 103.2893 | 155.4829 | 64.49291 | -0.26964 | 2.550107 | 0.522625 |
| 18 | 25325200 | 25328399 | 76.61357 | 120.6337 | 124.5299 | 51.66909 | 0.276376 | 2.506786 | 0.522626 |
| 1 | 115640800 | 115642399 | 18.93403 | 34.12637 | 28.85292 | 12.57084 | 0.646956 | 0.410131 | 0.522633 |
| 12 | 17133400 | 17141399 | 70.05308 | 109.3376 | 167.9887 | 69.70705 | -0.30645 | 2.6723 | 0.522657 |
| 2 | 8219800 | 8226799 | 106.2543 | 164.1401 | 169.1554 | 70.26354 | 0.257555 | 2.86878 | 0.52267 |
| 16 | 13370400 | 13372799 | 39.88882 | 64.95301 | 68.01345 | 29.20188 | 0.304289 | 1.560094 | 0.522704 |
| 4 | 147166800 | 147173799 | 254.6382 | 370.8986 | 351.6953 | 144.9396 | 0.303538 | 3.975118 | 0.522711 |
| 4 | 88640000 | 88640999 | 8.489014 | 15.86282 | 29.54414 | 12.2536 | -0.43387 | 0.129656 | 0.522716 |
| 15 | 48315200 | 48316399 | 15.09743 | 27.5027 | 27.87535 | 11.98423 | 0.399444 | 0.322536 | 0.52273 |
| 11 | 30378400 | 30379999 | 17.63382 | 31.98398 | 28.55298 | 12.34242 | 0.57625 | 0.365358 | 0.522732 |
| 15 | 39729600 | 39733799 | 43.31519 | 69.40993 | 106.8652 | 45.09455 | -0.2751 | 1.987486 | 0.522734 |
| 7 | 28004200 | 28008399 | 22.18943 | 36.63073 | 67.48235 | 29.13632 | -0.4983 | 1.338918 | 0.522757 |
| 4 | 151606200 | 151607999 | 19.98748 | 33.94664 | 54.85223 | 24.03666 | -0.30821 | 1.029797 | 0.522758 |
| 4 | 80726200 | 80729999 | 76.43718 | 120.4277 | 126.2692 | 52.39483 | 0.254585 | 2.495112 | 0.522768 |
| 6 | 83703600 | 83707399 | 31.01176 | 50.42976 | 79.84771 | 34.26607 | -0.29211 | 1.568649 | 0.522769 |
| 5 | 74664800 | 74666599 | 23.5988 | 41.30192 | 36.24486 | 15.99216 | 0.575173 | 0.70877 | 0.52277 |
| 13 | 17303400 | 17304999 | 16.213 | 29.67453 | 27.76688 | 11.86625 | 0.519567 | 0.303421 | 0.522773 |
| 16 | 71110800 | 71112199 | 16.13291 | 28.62634 | 29.79403 | 13.16179 | 0.335448 | 0.488545 | 0.522787 |
| 7 | 23362000 | 23367399 | 94.19472 | 147.8135 | 139.7494 | 57.95822 | 0.399262 | 2.586646 | 0.52279 |
| 6 | 44116600 | 44119799 | 40.53337 | 65.18906 | 100.0374 | 42.4439 | -0.26401 | 1.868471 | 0.522799 |
| 4 | 126014000 | 126015799 | 20.84959 | 35.06578 | 58.05936 | 25.3666 | -0.34388 | 1.120779 | 0.52283 |
| 9 | 7841000 | 7841599 | 6.831565 | 13.20259 | 25.49573 | 9.779971 | -0.38533 | -0.16551 | 0.522831 |

| | | | | | | | | | |
|---|---|---|---|---|---|---|---|---|---|
| 10 | 110750400 | 110752599 | 18.07718 | 32.07107 | 31.52455 | 13.92817 | 0.417258 | 0.504787 | 0.522838 |
| 4 | 92896200 | 92898999 | 28.99365 | 49.0022 | 49.07003 | 21.47096 | 0.379255 | 1.097062 | 0.52286 |
| 8 | 4921000 | 4923599 | 28.99365 | 49.0022 | 49.07003 | 21.47096 | 0.379255 | 1.097062 | 0.52286 |
| 4 | 78463400 | 78464799 | 24.1245 | 41.73734 | 36.25119 | 15.90426 | 0.588376 | 0.857812 | 0.52286 |
| 7 | 143898000 | 143914199 | 217.9705 | 315.5358 | 434.8868 | 179.349 | -0.24443 | 4.065438 | 0.522862 |
| X | 29396400 | 29400999 | 58.71649 | 94.6291 | 78.94009 | 33.59361 | 0.617276 | 1.824057 | 0.522875 |
| 12 | 8548400 | 8550999 | 18.07978 | 30.66267 | 54.79675 | 24.00192 | -0.45304 | 1.033948 | 0.522891 |
| X | 88372200 | 88373799 | 17.47538 | 32.49543 | 24.55974 | 10.27115 | 0.847726 | 0.2195 | 0.522897 |
| 2 | 166903600 | 166905599 | 9.369564 | 16.87476 | 33.37111 | 14.55313 | -0.57192 | 0.358523 | 0.52293 |
| 5 | 130097600 | 130099199 | 23.5113 | 40.19955 | 41.65773 | 18.26576 | 0.331676 | 0.967065 | 0.522949 |
| 2 | 76728200 | 76732199 | 51.89017 | 82.7321 | 125.3725 | 52.52331 | -0.26043 | 2.158522 | 0.522955 |
| 10 | 116596600 | 116603799 | 62.62236 | 98.86171 | 148.6456 | 61.71366 | -0.26252 | 2.446081 | 0.522959 |
| 5 | 3540800 | 3544599 | 32.33509 | 52.39899 | 83.57338 | 35.7787 | -0.30719 | 1.640612 | 0.522975 |
| Y | 16319600 | 16319999 | 3.31331 | 6.464482 | 19.7012 | 7.032972 | -0.93558 | -0.47854 | 0.522989 |
| 5 | 123501000 | 123503799 | 13.97345 | 24.50102 | 40.96231 | 18.10177 | -0.35009 | 0.636857 | 0.522995 |
| 5 | 49945800 | 49946799 | 8.262131 | 15.41321 | 30.57163 | 12.71551 | -0.52829 | 0.135179 | 0.522999 |
| 2 | 19888800 | 19889799 | 14.50216 | 26.57829 | 26.74536 | 11.40917 | 0.416842 | 0.278712 | 0.523009 |
| 18 | 7068800 | 7072799 | 74.68003 | 117.9621 | 124.9399 | 51.88617 | 0.242918 | 2.4372 | 0.523021 |
| 17 | 9412800 | 9414199 | 8.299148 | 15.62285 | 28.58892 | 11.71022 | -0.39419 | 0.077429 | 0.523029 |
| 3 | 3120200 | 3124399 | 62.44115 | 100.071 | 82.72402 | 34.91439 | 0.620657 | 1.999082 | 0.52305 |
| 13 | 58559200 | 58562399 | 42.92167 | 69.66099 | 71.13789 | 30.48928 | 0.336494 | 1.621577 | 0.523054 |
| 14 | 13659600 | 13661199 | 17.9329 | 32.17285 | 29.99352 | 13.12739 | 0.502466 | 0.432785 | 0.523072 |
| 3 | 104971200 | 104971999 | 11.91629 | 22.35374 | 24.11964 | 9.93821 | 0.354948 | 0.118169 | 0.523073 |
| 5 | 13707400 | 13708799 | 10.10266 | 18.71429 | 34.2692 | 14.40761 | -0.42613 | 0.201532 | 0.523081 |
| 12 | 71285600 | 71294399 | 122.9213 | 185.0668 | 300.7914 | 124.2335 | -0.42909 | 3.392494 | 0.523095 |
| 4 | 78179200 | 78180599 | 12.49033 | 23.57631 | 23.48021 | 9.570349 | 0.481734 | 0.079624 | 0.523114 |
| 1 | 117916000 | 117928999 | 262.9171 | 381.7229 | 363.4065 | 149.8163 | 0.293275 | 4.024238 | 0.523116 |
| 15 | 6014600 | 6019199 | 52.21707 | 84.1656 | 81.22635 | 34.57245 | 0.407648 | 1.808963 | 0.523124 |
| Y | 35905600 | 35907999 | 12.89789 | 23.88233 | 17.07814 | 7.184107 | 0.920418 | 0.184014 | 0.523148 |
| 5 | 145744800 | 145750399 | 135.3654 | 205.4557 | 212.0583 | 87.80039 | 0.235468 | 3.2279 | 0.523158 |
| 19 | 18204000 | 18204399 | 5.965642 | 11.55311 | 23.24613 | 8.830846 | -0.42955 | -0.21212 | 0.523176 |
| 10 | 103576800 | 103578199 | 13.93956 | 24.54573 | 40.30789 | 17.84336 | -0.32345 | 0.575093 | 0.523182 |
| 17 | 79644200 | 79645799 | 12.46266 | 22.11683 | 37.6894 | 16.64775 | -0.3719 | 0.479195 | 0.523188 |
| 8 | 26815800 | 26817799 | 17.45174 | 30.13016 | 49.53776 | 21.73202 | -0.32909 | 0.863138 | 0.523204 |
| 7 | 106697000 | 106697599 | 8.204623 | 15.6815 | 30.29784 | 12.02669 | -0.43322 | -0.02721 | 0.52321 |
| 8 | 47935000 | 47935999 | 8.671446 | 16.30869 | 30.22914 | 12.40136 | -0.41533 | 0.086054 | 0.523214 |
| 8 | 70114000 | 70114799 | 9.938762 | 19.08564 | 20.10771 | 7.874515 | 0.452567 | -0.08243 | 0.523215 |
| 13 | 29855200 | 29855599 | 6.110447 | 11.84354 | 23.84071 | 9.02102 | -0.42438 | -0.22019 | 0.523232 |
| 5 | 93913600 | 93922199 | 115.9914 | 180.6192 | 146.7584 | 60.91641 | 0.609003 | 2.732701 | 0.523246 |
| 10 | 59222800 | 59231599 | 94.25717 | 146.6384 | 153.7123 | 63.81174 | 0.240869 | 2.737823 | 0.523286 |
| 6 | 64636200 | 64640399 | 33.4566 | 54.17067 | 86.08896 | 36.83273 | -0.3032 | 1.660181 | 0.523315 |
| 7 | 23295000 | 23304199 | 105.1594 | 162.9244 | 147.189 | 61.13825 | 0.450531 | 2.813377 | 0.523328 |
| 7 | 6286400 | 6287999 | 8.905933 | 16.60014 | 29.48816 | 12.28251 | -0.37153 | 0.146819 | 0.523358 |
| 1 | 60388400 | 60390999 | 55.00294 | 87.43511 | 130.9894 | 54.66003 | -0.24847 | 2.256296 | 0.523359 |
| 2 | 15279800 | 15282799 | 26.93189 | 44.57315 | 72.86201 | 31.51446 | -0.32629 | 1.324953 | 0.523361 |
| 4 | 112610400 | 112616199 | 70.73099 | 113.0808 | 95.62862 | 40.21733 | 0.58402 | 2.087251 | 0.523389 |
| 15 | 49830600 | 49834399 | 73.32174 | 114.4664 | 169.5153 | 70.33883 | -0.25318 | 2.670204 | 0.5234 |

| | | | | | | | | | |
|---|---|---|---|---|---|---|---|---|---|
| 10 | 71463800 | 71466399 | 28.47195 | 48.06003 | 48.53829 | 21.23019 | 0.367031 | 1.107454 | 0.523406 |
| 1 | 28946200 | 28947399 | 19.75603 | 33.74708 | 54.931 | 24.08391 | -0.3176 | 0.975344 | 0.52341 |
| 7 | 140005400 | 140011199 | 61.07028 | 98.18136 | 84.29256 | 35.74424 | 0.571539 | 1.902868 | 0.523415 |
| 12 | 21611400 | 21616799 | 59.94769 | 96.09959 | 92.36304 | 38.99787 | 0.404069 | 1.990571 | 0.523429 |
| 6 | 6351200 | 6352399 | 8.676653 | 16.39199 | 29.8047 | 12.12586 | -0.37758 | 0.056351 | 0.523443 |
| 6 | 60333800 | 60337399 | 51.59399 | 83.34391 | 76.84636 | 32.80087 | 0.476937 | 1.744067 | 0.52345 |
| 7 | 13620600 | 13632999 | 173.384 | 260.5121 | 247.6678 | 102.2366 | 0.341454 | 3.439116 | 0.523496 |
| 3 | 50620600 | 50621999 | 7.727502 | 14.72045 | 29.54719 | 11.81202 | -0.49707 | -0.0126 | 0.523498 |
| 15 | 73279800 | 73280799 | 6.259076 | 12.14292 | 24.4843 | 9.224355 | -0.42095 | -0.2313 | 0.523509 |
| 5 | 65234600 | 65237999 | 25.81037 | 42.66408 | 68.5845 | 29.63897 | -0.30229 | 1.333598 | 0.523517 |
| 4 | 109457600 | 109461599 | 33.13632 | 53.79273 | 84.05746 | 36.03304 | -0.27536 | 1.606917 | 0.523535 |
| 9 | 63816800 | 63819799 | 33.13632 | 53.79273 | 84.05746 | 36.03304 | -0.27536 | 1.606917 | 0.523535 |
| 13 | 68243200 | 68259399 | 542.3279 | 734.9579 | 666.2237 | 277.6099 | 0.264218 | 5.067331 | 0.523535 |
| 12 | 4176200 | 4179799 | 19.48276 | 32.6455 | 58.99145 | 25.7235 | -0.46955 | 1.155189 | 0.523558 |
| 13 | 33756800 | 33758599 | 26.10402 | 44.38645 | 45.82935 | 20.08596 | 0.335761 | 1.03295 | 0.523574 |
| 17 | 88636200 | 88639599 | 37.2228 | 59.99884 | 94.42829 | 40.19301 | -0.29637 | 1.790501 | 0.52358 |
| 10 | 11132000 | 11133399 | 12.15096 | 21.76581 | 36.2353 | 15.88005 | -0.33039 | 0.417249 | 0.523582 |
| 12 | 15678000 | 15678799 | 10.70488 | 20.27069 | 22.14066 | 8.963709 | 0.358499 | 0.03978 | 0.523646 |
| 1 | 29446000 | 29448799 | 35.60154 | 59.97687 | 47.41801 | 20.72786 | 0.718892 | 1.144781 | 0.523655 |
| 6 | 122939200 | 122940799 | 23.96737 | 40.92002 | 43.18611 | 18.93371 | 0.305061 | 0.982952 | 0.523668 |
| 1 | 94871200 | 94871599 | 5.534666 | 10.75321 | 22.87282 | 8.531884 | -0.48462 | -0.28423 | 0.52367 |
| 2 | 49476200 | 49478199 | 15.35246 | 26.90088 | 45.09441 | 19.91649 | -0.35427 | 0.659424 | 0.523677 |
| 1 | 149267000 | 149270799 | 53.02389 | 85.03013 | 90.22112 | 38.11381 | 0.262225 | 1.970613 | 0.523681 |
| 19 | 39619600 | 39626599 | 99.03861 | 154.3904 | 152.3759 | 63.23774 | 0.329966 | 2.706263 | 0.523686 |
| 2 | 98887800 | 98888999 | 19.98076 | 35.03058 | 29.91679 | 13.21946 | 0.614099 | 0.651458 | 0.523693 |
| 9 | 77787600 | 77789599 | 18.68786 | 34.31453 | 26.56741 | 11.30404 | 0.795789 | 0.302697 | 0.523699 |
| 18 | 20869000 | 20872799 | 33.60591 | 54.55628 | 86.03929 | 36.88321 | -0.28861 | 1.607676 | 0.5237 |
| 16 | 31192000 | 31194599 | 30.91774 | 50.46454 | 82.37588 | 35.39906 | -0.33226 | 1.514003 | 0.523723 |
| X | 63806600 | 63808199 | 10.06958 | 18.72424 | 35.49543 | 14.84472 | -0.46979 | 0.185904 | 0.523741 |
| X | 3755800 | 3766199 | 79.58039 | 126.4167 | 99.42219 | 41.45866 | 0.678856 | 2.29164 | 0.523742 |
| 9 | 5358600 | 5360999 | 23.52591 | 41.23206 | 35.05383 | 15.48496 | 0.62059 | 0.683984 | 0.523743 |
| 13 | 88646800 | 88648399 | 15.34187 | 28.3391 | 26.23839 | 11.07385 | 0.547278 | 0.241288 | 0.52375 |
| 14 | 67731400 | 67733599 | 15.34187 | 28.3391 | 26.23839 | 11.07385 | 0.547278 | 0.241288 | 0.52375 |
| 4 | 66669400 | 66676199 | 84.93174 | 133.998 | 129.7128 | 53.83629 | 0.371202 | 2.472526 | 0.523783 |
| 8 | 110049600 | 110051399 | 14.99792 | 27.20957 | 28.30998 | 12.23369 | 0.356983 | 0.344726 | 0.5238 |
| 3 | 4806400 | 4808399 | 20.64158 | 36.48649 | 32.56224 | 14.40352 | 0.553978 | 0.561125 | 0.523801 |
| 16 | 10602200 | 10604999 | 22.79777 | 37.73642 | 65.38212 | 28.28202 | -0.4097 | 1.308832 | 0.523803 |
| 7 | 27640800 | 27643799 | 34.32206 | 56.82417 | 58.13424 | 25.1505 | 0.347903 | 1.329604 | 0.523812 |
| 9 | 100417400 | 100419799 | 15.93848 | 27.68698 | 50.6358 | 22.2293 | -0.48049 | 0.795368 | 0.52382 |
| 19 | 9581800 | 9588399 | 78.11791 | 124.6002 | 97.68232 | 40.94621 | 0.689514 | 2.164733 | 0.523828 |
| 1 | 89385400 | 89387399 | 25.87142 | 43.89472 | 46.37375 | 20.31499 | 0.302485 | 1.050746 | 0.523847 |
| 12 | 56359600 | 56360599 | 10.14911 | 18.76718 | 32.55217 | 13.71864 | -0.35061 | 0.209944 | 0.5239 |
| 10 | 98759600 | 98761999 | 24.81063 | 41.35746 | 66.49219 | 28.89934 | -0.30168 | 1.234385 | 0.523908 |
| 1 | 140737400 | 140738199 | 8.575497 | 16.03135 | 29.00265 | 12.02022 | -0.39197 | 0.122992 | 0.52391 |
| 8 | 81863000 | 81867999 | 53.23327 | 85.77364 | 81.85546 | 34.82352 | 0.423195 | 1.821436 | 0.523927 |
| 3 | 33226800 | 33227799 | 11.6057 | 22.01937 | 19.4894 | 7.857867 | 0.664786 | 0.028132 | 0.523931 |
| 10 | 29752400 | 29755399 | 32.64766 | 54.55107 | 47.57623 | 20.70906 | 0.577784 | 1.213955 | 0.523933 |

| | | | | | | | | | |
|---|---|---|---|---|---|---|---|---|---|
| 1 | 109798800 | 109799999 | 21.21756 | 35.53057 | 59.63723 | 25.99505 | -0.36352 | 1.167945 | 0.523938 |
| 7 | 143940000 | 143942199 | 17.94267 | 31.51156 | 32.96209 | 14.58629 | 0.322905 | 0.6178 | 0.523961 |
| 5 | 141006400 | 141012999 | 93.20152 | 145.3105 | 151.1628 | 62.73303 | 0.254284 | 2.700333 | 0.523965 |
| X | 87402200 | 87403999 | 28.41374 | 48.76045 | 43.02584 | 18.87123 | 0.563545 | 0.95818 | 0.523973 |
| 14 | 41806800 | 41808799 | 29.49084 | 49.23752 | 52.41637 | 22.80454 | 0.291169 | 1.217824 | 0.523987 |
| 12 | 43255200 | 43255799 | 5.38306 | 10.47439 | 21.83169 | 8.045274 | -0.43891 | -0.33124 | 0.524009 |
| 1 | 172798000 | 172798999 | 15.60241 | 27.15399 | 45.26835 | 19.89636 | -0.34669 | 0.765509 | 0.524016 |
| 15 | 87025600 | 87029199 | 44.62572 | 72.47504 | 70.78632 | 30.35355 | 0.401527 | 1.612342 | 0.524017 |
| 3 | 28529000 | 28531199 | 14.71928 | 25.79485 | 42.47272 | 18.76069 | -0.32841 | 0.650383 | 0.524018 |
| 13 | 120485600 | 120488599 | 18.09918 | 31.29582 | 51.72067 | 22.69071 | -0.33612 | 0.854999 | 0.52403 |
| 10 | 83558400 | 83561199 | 22.33791 | 37.37345 | 60.79846 | 26.48635 | -0.31857 | 1.181995 | 0.524031 |
| 10 | 42723000 | 42725399 | 16.50724 | 28.53254 | 47.93715 | 21.03058 | -0.35973 | 0.846917 | 0.524036 |
| 3 | 106342600 | 106344199 | 28.48043 | 47.59955 | 50.23127 | 21.86809 | 0.303727 | 1.201682 | 0.524049 |
| 5 | 3112800 | 3115399 | 54.45516 | 87.3729 | 77.49299 | 32.76394 | 0.52126 | 1.951405 | 0.524059 |
| 1 | 50606800 | 50607999 | 14.78548 | 26.95248 | 27.80225 | 11.9428 | 0.374762 | 0.314471 | 0.524067 |
| 17 | 53518200 | 53520599 | 25.86504 | 45.07608 | 36.71893 | 16.15253 | 0.681725 | 0.780261 | 0.524084 |
| 3 | 42182200 | 42187199 | 71.27981 | 112.876 | 119.0153 | 49.48897 | 0.252304 | 2.373657 | 0.524094 |
| 15 | 19954000 | 19956199 | 21.25306 | 37.44085 | 33.47243 | 14.81612 | 0.549467 | 0.599959 | 0.524099 |
| 13 | 38220200 | 38221999 | 12.7565 | 22.24716 | 43.20696 | 19.01513 | -0.56548 | 0.717054 | 0.524118 |
| 17 | 64518200 | 64522399 | 36.60467 | 58.96013 | 93.82394 | 39.89542 | -0.3135 | 1.812153 | 0.524126 |
| 11 | 55572000 | 55573999 | 22.70245 | 37.87798 | 61.83228 | 26.88955 | -0.32347 | 1.214666 | 0.524162 |
| 13 | 67606400 | 67608799 | 19.87186 | 34.78863 | 35.97809 | 15.87811 | 0.339276 | 0.683549 | 0.524189 |
| 1 | 142475400 | 142476599 | 13.46207 | 25.12335 | 25.49575 | 10.59752 | 0.432459 | 0.16594 | 0.524189 |
| X | 73524000 | 73525599 | 10.28211 | 18.89602 | 34.59465 | 14.70248 | -0.43643 | 0.253566 | 0.524192 |
| X | 54410400 | 54413199 | 25.212 | 44.36763 | 32.20771 | 14.25936 | 0.848059 | 0.634282 | 0.524202 |
| 9 | 123680200 | 123681199 | 8.635591 | 16.10019 | 29.50918 | 12.28616 | -0.41551 | 0.144764 | 0.524205 |
| 10 | 67192400 | 67193799 | 15.69264 | 27.38163 | 44.67447 | 19.66724 | -0.3155 | 0.726438 | 0.524216 |
| 10 | 41443800 | 41446999 | 21.59428 | 37.48189 | 39.34926 | 17.26582 | 0.316903 | 0.81964 | 0.524219 |
| 1 | 11120800 | 11123599 | 26.10549 | 45.69587 | 34.40432 | 15.18198 | 0.795154 | 0.716821 | 0.524224 |
| 5 | 134446400 | 134447399 | 13.43568 | 23.62865 | 40.32199 | 17.85411 | -0.37939 | 0.584739 | 0.524233 |
| 1 | 75613000 | 75614399 | 12.07933 | 21.41399 | 36.78727 | 16.26294 | -0.38391 | 0.48559 | 0.524275 |
| 11 | 67508000 | 67510399 | 11.6974 | 20.7925 | 36.23255 | 15.98235 | -0.40237 | 0.462062 | 0.524289 |
| 9 | 113129800 | 113132199 | 15.73606 | 27.17058 | 47.9383 | 21.0304 | -0.4303 | 0.853082 | 0.524307 |
| 2 | 4522400 | 4523599 | 8.7399 | 16.40299 | 29.07119 | 11.96876 | -0.35466 | 0.100499 | 0.524327 |
| 1 | 177988000 | 177993399 | 113.0214 | 173.9328 | 177.0196 | 73.51889 | 0.271404 | 2.942126 | 0.524331 |
| 14 | 83908400 | 83908999 | 6.686359 | 12.95318 | 25.06644 | 9.507504 | -0.37258 | -0.20578 | 0.524343 |
| 3 | 3889400 | 3895199 | 73.30543 | 116.2231 | 118.3747 | 49.26608 | 0.303711 | 2.346149 | 0.524345 |
| 10 | 82647800 | 82652599 | 32.37603 | 52.33816 | 85.89055 | 36.69613 | -0.35163 | 1.695236 | 0.52435 |
| 13 | 35188200 | 35199799 | 325.4405 | 452.9941 | 621.3224 | 257.8157 | -0.29416 | 4.649394 | 0.524361 |
| 8 | 122360400 | 122361399 | 11.70511 | 20.84257 | 35.7477 | 15.74527 | -0.37802 | 0.45102 | 0.524402 |
| 9 | 22166600 | 22168599 | 36.36605 | 59.51804 | 63.49645 | 27.31298 | 0.282546 | 1.477026 | 0.524406 |
| 7 | 31736200 | 31736999 | 21.0124 | 36.40721 | 37.44673 | 16.42952 | 0.345928 | 0.831835 | 0.524412 |
| 15 | 49285400 | 49285799 | 6.966769 | 13.59467 | 16.37679 | 5.830977 | 0.388952 | -0.44963 | 0.524431 |
| 1 | 118892000 | 118892999 | 10.74245 | 19.26558 | 34.28105 | 15.0086 | -0.42465 | 0.396292 | 0.524442 |
| 3 | 53770200 | 53771799 | 10.74245 | 19.26558 | 34.28105 | 15.0086 | -0.42465 | 0.396292 | 0.524442 |
| 12 | 96013400 | 96015199 | 23.52106 | 40.59661 | 41.42082 | 18.17099 | 0.356222 | 0.880521 | 0.524486 |
| 4 | 16057200 | 16058999 | 29.93739 | 49.85122 | 51.84728 | 22.51939 | 0.324832 | 1.248747 | 0.524494 |

| | | | | | | | | | |
|---|---|---|---|---|---|---|---|---|---|
| 3 | 6181000 | 6181399 | 8.103031 | 15.83718 | 15.1732 | 5.240822 | 0.752442 | -0.54565 | 0.524495 |
| 6 | 37656800 | 37661599 | 52.99678 | 85.1524 | 87.55542 | 37.09668 | 0.310801 | 1.909312 | 0.524506 |
| 17 | 17493400 | 17496399 | 48.248 | 77.76551 | 81.07724 | 34.50751 | 0.296375 | 1.8036 | 0.524525 |
| 4 | 43717800 | 43718999 | 13.88546 | 25.2196 | 25.8281 | 11.14554 | 0.380794 | 0.323366 | 0.524584 |
| 8 | 9850000 | 9850599 | 5.671938 | 11.00006 | 22.73301 | 8.578314 | -0.45815 | -0.23919 | 0.524587 |
| 10 | 26042400 | 26045199 | 19.04107 | 33.35375 | 34.99043 | 15.44988 | 0.31875 | 0.668086 | 0.524588 |
| X | 148675800 | 148677599 | 22.62101 | 39.61597 | 35.67444 | 15.74899 | 0.537962 | 0.691282 | 0.524597 |
| 10 | 124515800 | 124517999 | 33.94265 | 56.45077 | 49.80252 | 21.61335 | 0.561366 | 1.271096 | 0.52461 |
| 8 | 101244800 | 101245799 | 21.92192 | 36.6979 | 60.68432 | 26.44593 | -0.342 | 1.177108 | 0.524641 |
| 16 | 89817200 | 89818799 | 21.32761 | 35.85253 | 58.62928 | 25.61188 | -0.32608 | 1.126968 | 0.52466 |
| 4 | 83324400 | 83332199 | 103.6991 | 158.5153 | 228.601 | 94.83215 | -0.23846 | 3.076894 | 0.52467 |
| 4 | 55468000 | 55469799 | 10.24381 | 18.49606 | 33.29918 | 14.47585 | -0.43459 | 0.349203 | 0.52467 |
| 2 | 20187400 | 20191799 | 62.82852 | 99.13545 | 147.8768 | 61.37829 | -0.25166 | 2.459031 | 0.524678 |
| 1 | 53379600 | 53383999 | 89.47518 | 139.8726 | 146.1197 | 60.62263 | 0.251193 | 2.651229 | 0.52469 |
| 19 | 3075600 | 3077399 | 31.34984 | 52.53626 | 43.85906 | 19.12637 | 0.640664 | 1.17864 | 0.524699 |
| 2 | 132200200 | 132205199 | 64.57886 | 102.9966 | 104.3858 | 43.75123 | 0.31951 | 2.154313 | 0.52471 |
| 7 | 129168000 | 129170399 | 18.77734 | 31.86746 | 54.23587 | 23.76117 | -0.38267 | 1.033449 | 0.524733 |
| 3 | 32254400 | 32260999 | 59.9212 | 96.25316 | 87.36541 | 37.00223 | 0.489941 | 1.928307 | 0.524739 |
| 9 | 12325000 | 12328399 | 55.3927 | 88.93414 | 80.50869 | 34.07216 | 0.492859 | 1.933116 | 0.524746 |
| 3 | 84563800 | 84566599 | 31.83791 | 51.66383 | 81.97846 | 35.13126 | -0.29794 | 1.611408 | 0.524775 |
| 11 | 94100800 | 94102799 | 12.38103 | 21.77844 | 38.87985 | 17.21483 | -0.44377 | 0.575172 | 0.524776 |
| 5 | 3306200 | 3308999 | 34.01087 | 56.5954 | 48.70705 | 21.14598 | 0.597121 | 1.265198 | 0.524779 |
| 11 | 52601800 | 52604599 | 17.61902 | 30.49316 | 49.5444 | 21.73655 | -0.31134 | 0.842492 | 0.524788 |
| 9 | 105847400 | 105849999 | 17.69579 | 30.27795 | 51.10932 | 22.41133 | -0.3693 | 0.948583 | 0.524822 |
| 6 | 13531200 | 13534399 | 23.35419 | 40.62114 | 34.98989 | 15.37311 | 0.601335 | 0.792993 | 0.524827 |
| 1 | 123139600 | 123139999 | 4.4771 | 8.753428 | 19.01442 | 6.508982 | -0.39965 | -0.61192 | 0.524837 |
| 3 | 148042800 | 148043199 | 4.4771 | 8.753428 | 19.01442 | 6.508982 | -0.39965 | -0.61192 | 0.524837 |
| 8 | 46348600 | 46348999 | 4.4771 | 8.753428 | 19.01442 | 6.508982 | -0.39965 | -0.61192 | 0.524837 |
| 17 | 20005600 | 20011399 | 74.51303 | 118.7804 | 103.1738 | 43.21291 | 0.540945 | 2.18136 | 0.524845 |
| 11 | 58213600 | 58214799 | 7.872867 | 14.64057 | 28.98007 | 12.11839 | -0.53073 | 0.1523 | 0.524856 |
| 1 | 47112600 | 47113199 | 12.8667 | 23.82032 | 25.51181 | 10.73443 | 0.342033 | 0.209119 | 0.524878 |
| 10 | 66264600 | 66267199 | 30.66477 | 51.66503 | 50.19286 | 21.94147 | 0.422666 | 1.130433 | 0.524895 |
| 18 | 19095200 | 19099799 | 48.95285 | 79.15936 | 67.33987 | 28.78155 | 0.594519 | 1.704267 | 0.524927 |
| 5 | 15748600 | 15749799 | 15.0281 | 26.9554 | 27.90295 | 12.21471 | 0.352296 | 0.41123 | 0.52494 |
| 18 | 56477200 | 56483599 | 72.00619 | 114.1748 | 118.7119 | 49.41143 | 0.274126 | 2.340789 | 0.524943 |
| 11 | 113408000 | 113410199 | 21.55243 | 35.99329 | 61.26706 | 26.65967 | -0.38368 | 1.199213 | 0.524957 |
| 14 | 39779200 | 39782999 | 48.58122 | 78.33296 | 80.66026 | 34.34722 | 0.314905 | 1.792837 | 0.524963 |
| X | 89608400 | 89611399 | 38.18611 | 63.11121 | 58.91186 | 25.45619 | 0.479584 | 1.360875 | 0.525003 |
| 7 | 47672400 | 47674799 | 16.05435 | 27.91611 | 45.79687 | 20.1176 | -0.32383 | 0.780011 | 0.525022 |
| 11 | 15978000 | 15979999 | 19.60992 | 34.62571 | 33.07032 | 14.6314 | 0.45586 | 0.565154 | 0.52503 |
| 18 | 83575000 | 83575799 | 6.702035 | 12.87052 | 26.35239 | 10.31901 | -0.49737 | -0.09769 | 0.525032 |
| 2 | 39696200 | 39700399 | 94.19701 | 146.1922 | 151.1819 | 62.78384 | 0.257709 | 2.778842 | 0.525035 |
| 14 | 15960600 | 15962399 | 15.25418 | 28.45273 | 25.34342 | 10.54367 | 0.618282 | 0.183633 | 0.52504 |
| 16 | 95969600 | 95971399 | 11.13981 | 19.84443 | 35.85524 | 15.78697 | -0.45197 | 0.447714 | 0.525044 |
| 1 | 114010600 | 114012799 | 25.68884 | 43.76858 | 45.80244 | 20.07742 | 0.316824 | 1.011975 | 0.525087 |
| 7 | 8189200 | 8189999 | 7.440052 | 14.32131 | 27.06523 | 10.5271 | -0.37353 | -0.10836 | 0.525105 |
| 15 | 59640400 | 59644999 | 84.96798 | 131.5619 | 191.487 | 79.53866 | -0.23745 | 2.821129 | 0.52511 |

| | | | | | | | | | |
|---|---|---|---|---|---|---|---|---|---|
| 13 | 54026000 | 54033199 | 71.22399 | 111.5689 | 164.7222 | 68.32614 | -0.24499 | 2.608349 | 0.525124 |
| 8 | 107355000 | 107355799 | 5.821344 | 11.24621 | 24.22669 | 9.308768 | -0.54187 | -0.16857 | 0.525159 |
| 13 | 11940800 | 11943799 | 20.14075 | 35.19371 | 36.55185 | 16.10597 | 0.333195 | 0.71491 | 0.52521 |
| 13 | 53195800 | 53196799 | 7.786818 | 14.87364 | 27.28778 | 10.84653 | -0.3601 | -0.03162 | 0.525231 |
| 9 | 117458800 | 117465999 | 51.18143 | 81.5314 | 124.1311 | 51.94092 | -0.26849 | 2.18419 | 0.525242 |
| 5 | 102570200 | 102570999 | 8.971283 | 16.92343 | 32.66458 | 13.33123 | -0.46748 | 0.074662 | 0.525244 |
| 1 | 18587000 | 18590999 | 61.57976 | 97.99898 | 101.8682 | 42.56711 | 0.279677 | 2.220235 | 0.525258 |
| 15 | 8692400 | 8693599 | 9.064344 | 16.81441 | 29.9461 | 12.56613 | -0.38361 | 0.177986 | 0.52526 |
| 13 | 7980000 | 7981799 | 16.27797 | 29.65758 | 28.18445 | 12.11492 | 0.491721 | 0.329354 | 0.525269 |
| 7 | 74060600 | 74063799 | 29.55152 | 49.3598 | 52.39842 | 22.80285 | 0.29525 | 1.21449 | 0.525305 |
| 16 | 15658200 | 15662199 | 35.15788 | 56.7606 | 89.41175 | 38.13586 | -0.29493 | 1.740813 | 0.525323 |
| 8 | 100889400 | 100892799 | 35.48515 | 58.09462 | 61.79344 | 26.58372 | 0.287067 | 1.468809 | 0.525334 |
| 11 | 96724600 | 96727199 | 16.70529 | 28.84129 | 47.93956 | 21.03089 | -0.34462 | 0.859506 | 0.525337 |
| 16 | 16252400 | 16253999 | 17.52092 | 31.16804 | 27.10058 | 11.94641 | 0.595468 | 0.476426 | 0.525343 |
| 8 | 21490000 | 21499199 | 173.9819 | 260.8952 | 231.9316 | 95.69021 | 0.435693 | 3.475714 | 0.525352 |
| 14 | 87518600 | 87519199 | 6.388699 | 12.38096 | 24.48898 | 9.272979 | -0.40162 | -0.21734 | 0.525389 |
| 6 | 140920000 | 140921599 | 11.77926 | 21.34473 | 37.48665 | 16.22343 | -0.3953 | 0.347011 | 0.52539 |
| 2 | 9733600 | 9735999 | 21.97523 | 36.97024 | 59.21507 | 25.87394 | -0.29621 | 1.123245 | 0.525402 |
| X | 148680600 | 148691799 | 165.3359 | 249.1131 | 238.0312 | 98.33198 | 0.337718 | 3.383675 | 0.525416 |
| 5 | 7892600 | 7894999 | 22.69274 | 39.68677 | 36.8552 | 16.25128 | 0.493889 | 0.715879 | 0.525427 |
| 13 | 26981600 | 26983199 | 20.30324 | 35.9487 | 31.24541 | 13.8164 | 0.592466 | 0.535585 | 0.525447 |
| 9 | 20380000 | 20385599 | 65.41382 | 104.5254 | 99.49866 | 41.81399 | 0.412739 | 2.096465 | 0.525453 |
| 7 | 87290200 | 87292199 | 12.01864 | 21.64127 | 36.82305 | 16.06273 | -0.35705 | 0.386072 | 0.525456 |
| 15 | 55267800 | 55268999 | 14.54067 | 26.52564 | 27.25596 | 11.6974 | 0.381047 | 0.306402 | 0.525458 |
| 14 | 11022600 | 11023599 | 9.709852 | 17.81782 | 31.37718 | 13.3621 | -0.38301 | 0.247416 | 0.525472 |
| 13 | 66954400 | 66958799 | 92.59908 | 144.3495 | 150.2338 | 62.34889 | 0.253442 | 2.703189 | 0.525472 |
| 8 | 64254400 | 64254799 | 5.949764 | 11.57284 | 23.01467 | 8.506398 | -0.37619 | -0.30999 | 0.525476 |
| 5 | 90127600 | 90128999 | 16.33226 | 28.39476 | 46.31813 | 20.34451 | -0.3158 | 0.783114 | 0.525502 |
| 19 | 11396400 | 11403799 | 95.85592 | 148.7965 | 155.853 | 64.72177 | 0.239528 | 2.776971 | 0.525537 |
| 18 | 46327200 | 46329399 | 22.51364 | 38.92284 | 40.97441 | 17.97606 | 0.31189 | 0.862599 | 0.525545 |
| 8 | 5789000 | 5789799 | 7.925854 | 14.96956 | 28.11182 | 11.4436 | -0.4243 | 0.047211 | 0.525551 |
| 7 | 84907600 | 84910799 | 37.47101 | 61.82794 | 60.01347 | 25.90363 | 0.423145 | 1.377663 | 0.525558 |
| 12 | 110027600 | 110029799 | 19.20549 | 33.82207 | 33.78012 | 14.95334 | 0.39035 | 0.593594 | 0.52557 |
| 9 | 35917200 | 35926999 | 101.8784 | 158.8164 | 152.0018 | 63.08256 | 0.374237 | 2.708018 | 0.525574 |
| 19 | 56085400 | 56091199 | 49.89948 | 79.72129 | 123.2477 | 51.78192 | -0.28598 | 2.092116 | 0.525576 |
| 7 | 13721200 | 13739399 | 259.6416 | 379.1947 | 338.9604 | 139.6526 | 0.39234 | 3.936027 | 0.525603 |
| 5 | 141884000 | 141885999 | 31.56221 | 52.22266 | 55.13099 | 23.84104 | 0.302716 | 1.327848 | 0.525604 |
| 1 | 146084200 | 146085599 | 11.83478 | 22.21945 | 22.92975 | 9.434525 | 0.4198 | 0.106198 | 0.52561 |
| 5 | 149279400 | 149287799 | 85.64188 | 132.0855 | 196.6856 | 81.69968 | -0.27468 | 2.893167 | 0.525614 |
| 4 | 121216800 | 121225599 | 171.5238 | 256.8006 | 258.4662 | 106.6041 | 0.254642 | 3.505662 | 0.525616 |
| 5 | 150374600 | 150380399 | 68.98433 | 108.4771 | 160.6588 | 66.64794 | -0.2452 | 2.533861 | 0.525648 |
| 1 | 79008400 | 79013199 | 45.09082 | 72.93694 | 75.03483 | 32.08023 | 0.321419 | 1.694315 | 0.525649 |
| 9 | 76335000 | 76338599 | 101.663 | 156.979 | 163.1507 | 67.76949 | 0.244875 | 2.874435 | 0.525663 |
| 4 | 145558800 | 145559399 | 12.65011 | 24.58669 | 18.07768 | 6.723352 | 1.036566 | -0.2246 | 0.525671 |
| X | 107245600 | 107250199 | 72.82973 | 115.6694 | 104.0981 | 43.39962 | 0.484378 | 2.29158 | 0.525679 |
| 1 | 82977000 | 82979999 | 23.30975 | 38.80369 | 64.71745 | 28.10426 | -0.35442 | 1.240865 | 0.52568 |
| 18 | 5234000 | 5235999 | 26.53338 | 45.60788 | 43.54767 | 19.10162 | 0.450467 | 0.934417 | 0.525686 |

| | | | | | | | | | |
|---|---|---|---|---|---|---|---|---|---|
| 1 | 182137800 | 182142399 | 65.27506 | 103.7582 | 110.01 | 45.90303 | 0.249444 | 2.262158 | 0.525692 |
| 7 | 23760000 | 23760999 | 9.174627 | 17.00673 | 31.51682 | 13.23821 | -0.44208 | 0.184059 | 0.525697 |
| 10 | 46417400 | 46417999 | 10.53732 | 20.13557 | 22.23665 | 8.83118 | 0.366828 | -0.0237 | 0.525699 |
| 16 | 91485200 | 91485999 | 7.488473 | 14.15588 | 27.58909 | 11.21001 | -0.47491 | 0.035922 | 0.525709 |
| 1 | 141183000 | 141184799 | 24.52385 | 41.74451 | 43.75122 | 19.17683 | 0.314458 | 1.01384 | 0.525713 |
| 12 | 83884400 | 83885599 | 15.28903 | 27.91782 | 27.13882 | 11.63305 | 0.46157 | 0.305676 | 0.525714 |
| 12 | 10161200 | 10162399 | 13.35294 | 25.27297 | 24.13238 | 9.779935 | 0.549679 | 0.072694 | 0.525725 |
| 2 | 109847800 | 109848599 | 6.305385 | 12.05281 | 26.17778 | 10.38794 | -0.59941 | -0.05746 | 0.525752 |
| 2 | 34999200 | 35000199 | 5.988504 | 11.5365 | 24.17694 | 9.382015 | -0.51725 | -0.13815 | 0.525766 |
| 12 | 22720200 | 22725599 | 73.67696 | 116.8883 | 105.9379 | 44.11881 | 0.472737 | 2.324201 | 0.525767 |
| 13 | 3510600 | 3516599 | 59.86694 | 96.08275 | 88.75708 | 37.54223 | 0.463135 | 1.952667 | 0.525771 |
| 4 | 64909000 | 64911199 | 20.68549 | 36.08123 | 37.66223 | 16.57666 | 0.325809 | 0.744879 | 0.525778 |
| 4 | 147108800 | 147118199 | 163.7193 | 247.9291 | 219.8518 | 90.96396 | 0.451629 | 3.281563 | 0.525799 |
| 5 | 140139800 | 140143399 | 24.19611 | 40.28768 | 64.64361 | 28.07592 | -0.29872 | 1.244233 | 0.525806 |
| 13 | 90524800 | 90526999 | 19.13358 | 32.71342 | 52.73521 | 23.12287 | -0.30338 | 0.963138 | 0.525824 |
| 3 | 42457800 | 42460399 | 40.34685 | 65.33918 | 65.737 | 28.12914 | 0.355085 | 1.658822 | 0.525832 |
| X | 29416800 | 29423999 | 75.76729 | 120.9566 | 97.21415 | 40.80006 | 0.655053 | 2.136893 | 0.525844 |
| 5 | 46441600 | 46446199 | 93.81678 | 145.6039 | 149.1276 | 61.93061 | 0.271623 | 2.777275 | 0.525862 |
| 3 | 138533400 | 138535199 | 21.79387 | 37.74806 | 39.91447 | 17.51204 | 0.306009 | 0.840872 | 0.525868 |
| X | 43750200 | 43751199 | 18.79477 | 33.35808 | 31.62766 | 13.96881 | 0.469629 | 0.509539 | 0.52587 |
| X | 63764400 | 63769199 | 63.89593 | 102.1731 | 97.81126 | 41.14659 | 0.405727 | 2.074236 | 0.525891 |
| 17 | 94608600 | 94609999 | 37.21553 | 60.59927 | 62.61053 | 26.882 | 0.323562 | 1.551224 | 0.525894 |
| 19 | 44977200 | 44981399 | 22.28603 | 36.92529 | 65.48065 | 28.34369 | -0.44307 | 1.294252 | 0.525915 |
| X | 132958600 | 132960599 | 20.96806 | 36.84255 | 34.27895 | 15.17519 | 0.491185 | 0.631903 | 0.525915 |
| 4 | 113461600 | 113463799 | 59.32566 | 94.62337 | 100.2365 | 42.01523 | 0.255896 | 2.146681 | 0.525917 |
| 17 | 31367400 | 31370799 | 36.53169 | 60.12742 | 61.50546 | 26.51954 | 0.346301 | 1.407253 | 0.525926 |
| X | 9899000 | 9900399 | 15.41717 | 28.32446 | 27.36863 | 11.63625 | 0.478869 | 0.276253 | 0.525942 |
| 10 | 6699000 | 6699599 | 8.307165 | 15.46994 | 28.97136 | 12.08593 | -0.4488 | 0.146515 | 0.525952 |
| 17 | 72659200 | 72662799 | 27.46427 | 45.34899 | 72.72515 | 31.40496 | -0.299 | 1.353948 | 0.525958 |
| 13 | 7878000 | 7881199 | 63.94109 | 101.7426 | 107.9655 | 45.10621 | 0.249727 | 2.228805 | 0.525968 |
| 3 | 99271400 | 99272199 | 24.76262 | 42.01135 | 41.23053 | 18.06161 | 0.408413 | 1.04039 | 0.525969 |
| 14 | 97371000 | 97374799 | 45.28058 | 73.99885 | 64.34579 | 27.66346 | 0.575297 | 1.517952 | 0.526015 |
| 12 | 20841200 | 20867799 | 1003.29 | 1293.146 | 1113.118 | 466.6952 | 0.266488 | 5.875454 | 0.526017 |
| X | 16261200 | 16265599 | 50.78625 | 82.43293 | 69.47886 | 29.78033 | 0.612944 | 1.637236 | 0.526032 |
| 15 | 33839600 | 33841399 | 20.58763 | 36.26244 | 32.90711 | 14.56638 | 0.527763 | 0.595272 | 0.526076 |
| 18 | 42057200 | 42061399 | 43.16955 | 70.0648 | 71.61964 | 30.69624 | 0.335151 | 1.62315 | 0.526078 |
| 10 | 67256000 | 67256399 | 4.746694 | 9.27702 | 20.7966 | 7.188801 | -0.45752 | -0.56627 | 0.526082 |
| 5 | 109286800 | 109296199 | 144.6063 | 219.3799 | 218.1561 | 90.31258 | 0.288591 | 3.240346 | 0.526082 |
| 4 | 35308200 | 35310599 | 18.13281 | 31.17142 | 50.59946 | 22.19487 | -0.31202 | 0.90685 | 0.52609 |
| 10 | 14938000 | 14940599 | 26.4929 | 43.25627 | 81.81123 | 35.16058 | -0.54362 | 1.503049 | 0.526092 |
| X | 147517800 | 147520799 | 41.49434 | 67.69756 | 67.28538 | 28.90956 | 0.381259 | 1.535175 | 0.526106 |
| 8 | 78732200 | 78734199 | 18.30983 | 32.27278 | 33.28054 | 14.72964 | 0.344836 | 0.577208 | 0.526109 |
| 4 | 48525800 | 48531599 | 52.64967 | 83.72464 | 129.6918 | 54.13825 | -0.29609 | 2.242109 | 0.526112 |
| 9 | 76622600 | 76623999 | 16.04915 | 27.78079 | 46.88452 | 20.56968 | -0.36565 | 0.831346 | 0.526133 |
| 4 | 80846400 | 80849399 | 43.85506 | 70.15449 | 109.397 | 46.04627 | -0.29628 | 2.042886 | 0.526135 |
| 18 | 63881800 | 63883199 | 7.626661 | 14.57507 | 26.7385 | 10.61686 | -0.35863 | -0.03835 | 0.52614 |
| 10 | 23990400 | 23993399 | 44.87392 | 72.78972 | 72.27681 | 30.96496 | 0.376177 | 1.638748 | 0.52617 |

| | | | | | | | | | |
|---|---|---|---|---|---|---|---|---|---|
| 4 | 82646200 | 82648599 | 29.9533 | 49.96545 | 52.98957 | 23.04107 | 0.296637 | 1.228524 | 0.526174 |
| 5 | 107256800 | 107259199 | 12.21448 | 21.54978 | 37.83249 | 16.74108 | -0.41797 | 0.539478 | 0.526182 |
| 19 | 26777400 | 26779199 | 17.24794 | 30.80458 | 30.57242 | 13.4318 | 0.408947 | 0.456635 | 0.526187 |
| X | 26101400 | 26106199 | 58.89377 | 94.8962 | 79.49681 | 33.82107 | 0.610876 | 1.830567 | 0.526214 |
| 12 | 50385800 | 50387399 | 23.83265 | 41.1035 | 36.28976 | 15.91966 | 0.563896 | 0.882845 | 0.526229 |
| 7 | 139635600 | 139639799 | 36.76944 | 59.22358 | 93.82587 | 39.89434 | -0.30717 | 1.813552 | 0.52625 |
| 18 | 53900200 | 53903199 | 37.44787 | 61.61503 | 61.64912 | 26.57776 | 0.377893 | 1.412496 | 0.526266 |
| 10 | 103909200 | 103911999 | 36.08419 | 59.52768 | 53.70537 | 23.17889 | 0.528238 | 1.371915 | 0.526276 |
| 8 | 111297600 | 111298799 | 12.44846 | 22.20622 | 37.20919 | 16.3641 | -0.34334 | 0.446025 | 0.526284 |
| 16 | 27030800 | 27031199 | 8.717721 | 16.86367 | 19.31894 | 7.377953 | 0.367639 | -0.17714 | 0.52629 |
| 17 | 39283600 | 39300399 | 239.5526 | 353.0153 | 306.9216 | 126.3935 | 0.444674 | 3.792797 | 0.526292 |
| 1 | 117656800 | 117666199 | 34.73522 | 57.64858 | 50.93095 | 22.07127 | 0.55912 | 1.297326 | 0.526341 |
| 17 | 34391200 | 34398399 | 67.99424 | 106.8204 | 159.7086 | 66.24573 | -0.25986 | 2.551115 | 0.526351 |
| 15 | 8751200 | 8757199 | 60.45211 | 95.62231 | 143.6643 | 59.70461 | -0.25914 | 2.397466 | 0.526355 |
| 2 | 173055800 | 173057199 | 25.38339 | 43.13985 | 45.27658 | 19.84291 | 0.312014 | 1.030882 | 0.526378 |
| 5 | 8601400 | 8601999 | 5.811538 | 11.28173 | 22.78009 | 8.554719 | -0.4184 | -0.25361 | 0.526387 |
| 6 | 3116000 | 3122399 | 77.28281 | 122.0679 | 126.4612 | 52.51676 | 0.274733 | 2.44165 | 0.526416 |
| 6 | 93470400 | 93471599 | 13.86921 | 26.04321 | 25.05653 | 10.30681 | 0.520995 | 0.130635 | 0.526421 |
| 8 | 5252800 | 5254199 | 18.99005 | 33.36348 | 34.36578 | 15.21225 | 0.345025 | 0.620792 | 0.526428 |
| 1 | 24684600 | 24687799 | 36.2789 | 58.48021 | 92.22874 | 39.25923 | -0.29923 | 1.787726 | 0.526437 |
| 7 | 70933400 | 70933999 | 9.15184 | 17.7574 | 19.91434 | 7.497946 | 0.416998 | -0.2189 | 0.526448 |
| 9 | 4234800 | 4241799 | 115.5299 | 177.2065 | 184.2681 | 76.49463 | 0.23682 | 3.008784 | 0.52645 |
| 10 | 18239600 | 18240799 | 13.45097 | 24.50702 | 24.95285 | 10.72579 | 0.392912 | 0.301151 | 0.526466 |
| 9 | 110614000 | 110614399 | 5.052693 | 9.857055 | 20.93985 | 7.487124 | -0.42504 | -0.44196 | 0.526468 |
| 10 | 38273000 | 38276399 | 41.14371 | 67.26613 | 65.6835 | 28.24284 | 0.408833 | 1.503836 | 0.526471 |
| 9 | 120761400 | 120763799 | 16.94386 | 28.98503 | 50.56352 | 22.1716 | -0.41649 | 0.948311 | 0.526501 |
| 16 | 93501800 | 93507199 | 10.41859 | 18.37081 | 41.50618 | 18.36881 | -0.7798 | 0.544399 | 0.526511 |
| 5 | 72342600 | 72343799 | 7.157328 | 13.66049 | 26.70966 | 10.63442 | -0.45369 | -0.03672 | 0.526555 |
| 2 | 172906200 | 172917399 | 135.2571 | 203.1962 | 287.9686 | 118.8681 | -0.23433 | 3.43627 | 0.526568 |
| 1 | 170400000 | 170402199 | 20.27034 | 34.67829 | 60.5454 | 26.54873 | -0.41812 | 0.969594 | 0.526572 |
| 15 | 56749000 | 56749999 | 13.5353 | 23.65373 | 42.13555 | 18.56783 | -0.44103 | 0.691679 | 0.526594 |
| 6 | 146422800 | 146431599 | 111.8615 | 171.7058 | 178.5794 | 74.14056 | 0.23737 | 2.992378 | 0.52661 |
| 7 | 83428000 | 83430799 | 25.52922 | 43.35023 | 45.28044 | 19.84105 | 0.31877 | 1.038722 | 0.526611 |
| 2 | 37088200 | 37090399 | 30.5008 | 51.16299 | 51.86658 | 22.63218 | 0.361364 | 1.171567 | 0.526645 |
| 8 | 19436800 | 19443399 | 154.8342 | 233.3486 | 236.6712 | 97.77675 | 0.252046 | 3.376013 | 0.526669 |
| 3 | 129450000 | 129453599 | 30.5176 | 49.60739 | 79.7912 | 34.23709 | -0.31514 | 1.572394 | 0.52667 |
| 7 | 42153800 | 42155999 | 20.29653 | 34.34724 | 56.49052 | 24.72747 | -0.33393 | 1.062669 | 0.526682 |
| 8 | 18282200 | 18283399 | 20.36002 | 34.44805 | 55.90205 | 24.46845 | -0.31458 | 1.06611 | 0.52669 |
| Y | 22077800 | 22078799 | 9.932646 | 19.08145 | 14.09229 | 5.512504 | 0.962167 | -0.1051 | 0.526704 |
| 1 | 116314000 | 116316399 | 26.2709 | 43.47223 | 69.88215 | 30.22226 | -0.30218 | 1.322727 | 0.526715 |
| 13 | 61193000 | 61196399 | 43.75632 | 71.44071 | 65.91628 | 28.32895 | 0.489018 | 1.528199 | 0.526716 |
| 4 | 70589200 | 70590199 | 7.202522 | 13.65374 | 26.57399 | 10.73341 | -0.4648 | 0.012981 | 0.526736 |
| 4 | 25357400 | 25358199 | 10.49366 | 19.13577 | 32.83604 | 14.0988 | -0.35428 | 0.292055 | 0.526742 |
| 7 | 32332400 | 32335399 | 42.28634 | 68.82014 | 68.9795 | 29.61088 | 0.366651 | 1.571891 | 0.526753 |
| 14 | 16360600 | 16362199 | 11.507 | 20.23179 | 39.0133 | 17.27548 | -0.5542 | 0.578697 | 0.526761 |
| 8 | 60333600 | 60334399 | 15.09406 | 26.25001 | 44.76432 | 19.66577 | -0.37937 | 0.771713 | 0.526761 |
| 8 | 128594800 | 128597999 | 39.99714 | 65.97459 | 59.59628 | 25.72001 | 0.526401 | 1.38823 | 0.526763 |

| | | | | | | | | | |
|---|---|---|---|---|---|---|---|---|---|
| 5 | 45008800 | 45010199 | 14.10739 | 24.74947 | 40.91919 | 18.09112 | -0.33418 | 0.627819 | 0.526776 |
| 12 | 43881000 | 43884399 | 29.80717 | 48.72728 | 76.88097 | 33.05642 | -0.28167 | 1.487804 | 0.526784 |
| X | 5059400 | 5060799 | 27.56018 | 47.83855 | 39.65204 | 17.399 | 0.656571 | 0.850085 | 0.526785 |
| 7 | 119038800 | 119039399 | 8.332096 | 15.44285 | 29.45771 | 12.3756 | -0.4828 | 0.17659 | 0.526791 |
| 4 | 106635400 | 106640799 | 24.93067 | 40.99578 | 69.76983 | 30.07126 | -0.38647 | 1.396436 | 0.526801 |
| 11 | 105602000 | 105602999 | 18.48048 | 31.2313 | 60.68473 | 26.55238 | -0.57355 | 1.074433 | 0.526803 |
| 17 | 50413800 | 50415199 | 9.078916 | 16.98407 | 30.08981 | 12.45636 | -0.36051 | 0.127349 | 0.526833 |
| 8 | 76329400 | 76329799 | 8.427026 | 16.32627 | 18.31149 | 6.943457 | 0.407926 | -0.20688 | 0.526838 |
| 7 | 25878800 | 25898999 | 310.6904 | 445.3049 | 410.2767 | 169.4407 | 0.322144 | 4.217227 | 0.526849 |
| 5 | 38073600 | 38081799 | 81.6441 | 126.5016 | 186.8865 | 77.62747 | -0.25818 | 2.805668 | 0.526863 |
| X | 4841200 | 4844799 | 33.92878 | 57.10499 | 42.63412 | 18.63135 | 0.801271 | 1.145182 | 0.526897 |
| 1 | 166448600 | 166455399 | 83.21255 | 131.1474 | 131.337 | 54.5001 | 0.321116 | 2.492006 | 0.5269 |
| 11 | 31928000 | 31932999 | 82.62147 | 130.0355 | 134.0368 | 55.61041 | 0.278014 | 2.521372 | 0.526905 |
| 16 | 58770800 | 58773599 | 24.73977 | 42.82336 | 40.21328 | 17.6428 | 0.476211 | 0.858111 | 0.526929 |
| 7 | 14343000 | 14361599 | 241.9148 | 354.0689 | 334.8676 | 137.9455 | 0.313967 | 3.900623 | 0.526941 |
| 6 | 104641000 | 104642599 | 15.796 | 29.1458 | 26.79005 | 11.32365 | 0.55633 | 0.253113 | 0.526968 |
| 11 | 31544200 | 31547799 | 41.34534 | 66.43458 | 101.6616 | 43.07629 | -0.26159 | 1.898452 | 0.52698 |
| 10 | 116101800 | 116102799 | 7.159635 | 13.82938 | 26.62546 | 10.2383 | -0.38405 | -0.14833 | 0.526996 |
| 13 | 80367400 | 80369199 | 12.79481 | 22.76775 | 37.70915 | 16.61855 | -0.3287 | 0.465448 | 0.526996 |
| 17 | 91677600 | 91678399 | 8.057418 | 15.27943 | 28.22422 | 11.39194 | -0.38995 | 0.020593 | 0.527002 |
| 5 | 36393800 | 36396999 | 47.04532 | 74.80532 | 124.2695 | 51.86982 | -0.39687 | 2.240584 | 0.527015 |
| X | 60779800 | 60782799 | 27.99403 | 48.74469 | 37.3586 | 16.4251 | 0.769498 | 0.806834 | 0.527016 |
| X | 20044400 | 20046999 | 38.48986 | 64.58962 | 40.93842 | 17.86809 | 1.036628 | 1.18605 | 0.527029 |
| 1 | 10535000 | 10539799 | 107.4303 | 165.3653 | 172.3385 | 71.57695 | 0.237477 | 2.93642 | 0.527044 |
| 16 | 93583000 | 93586999 | 52.6083 | 84.52893 | 87.02454 | 36.8722 | 0.308981 | 1.907564 | 0.527045 |
| 7 | 40713000 | 40722399 | 134.6577 | 206.0773 | 191.774 | 79.57627 | 0.394303 | 3.061246 | 0.52706 |
| 14 | 113340000 | 113340999 | 17.68171 | 31.27413 | 32.24562 | 14.26168 | 0.346985 | 0.534461 | 0.527082 |
| 19 | 12941400 | 12948599 | 103.0628 | 160.5568 | 153.1713 | 63.57415 | 0.378163 | 2.719959 | 0.527087 |
| 15 | 38252000 | 38254599 | 17.91305 | 30.75532 | 54.41348 | 23.86681 | -0.43633 | 0.918731 | 0.527088 |
| 10 | 108317600 | 108319999 | 15.36034 | 26.80998 | 44.14611 | 19.43836 | -0.32867 | 0.719288 | 0.527099 |
| 11 | 39959200 | 39961999 | 34.74501 | 57.11441 | 60.68609 | 26.15224 | 0.29094 | 1.418251 | 0.527102 |
| 17 | 36805000 | 36807199 | 22.29564 | 37.41907 | 60.32279 | 26.33081 | -0.30561 | 1.147467 | 0.527113 |
| 11 | 59533200 | 59534399 | 6.950536 | 13.06376 | 27.53281 | 11.30744 | -0.59927 | 0.071011 | 0.527123 |
| 3 | 106768200 | 106774999 | 75.51986 | 119.9529 | 113.4249 | 47.29241 | 0.413274 | 2.291745 | 0.527144 |
| 17 | 55302600 | 55306999 | 79.09591 | 124.9694 | 126.7722 | 52.65393 | 0.305385 | 2.435851 | 0.527155 |
| 18 | 28565400 | 28568999 | 39.74004 | 65.04689 | 64.54593 | 27.76548 | 0.386843 | 1.482532 | 0.527184 |
| 15 | 94440800 | 94443799 | 19.70768 | 32.86725 | 68.83901 | 29.92853 | -0.68187 | 1.215112 | 0.527189 |
| 13 | 27970800 | 27972999 | 17.44673 | 31.65441 | 28.56148 | 12.34133 | 0.561299 | 0.362112 | 0.527211 |
| 14 | 61358600 | 61360399 | 27.25043 | 44.54073 | 74.07818 | 31.8495 | -0.35755 | 1.483379 | 0.527225 |
| 1 | 165786600 | 165787799 | 11.08105 | 20.07822 | 34.87182 | 15.09302 | -0.37883 | 0.34071 | 0.527244 |
| 16 | 91963400 | 91973199 | 84.56229 | 129.7931 | 224.1544 | 93.061 | -0.49366 | 2.989253 | 0.527245 |
| 10 | 41951200 | 41951799 | 4.910029 | 9.593602 | 20.72387 | 7.214708 | -0.41458 | -0.54083 | 0.527255 |
| 7 | 33882600 | 33886799 | 26.54369 | 43.48855 | 74.42434 | 32.02769 | -0.3967 | 1.447241 | 0.527269 |
| 4 | 20802600 | 20805799 | 33.6639 | 56.75514 | 47.97818 | 20.9775 | 0.622613 | 1.131379 | 0.527276 |
| 13 | 63860600 | 63861599 | 16.6204 | 30.40228 | 27.66895 | 11.83389 | 0.558416 | 0.308628 | 0.52728 |
| 5 | 100241200 | 100244199 | 19.89924 | 34.97321 | 33.54783 | 14.85463 | 0.446983 | 0.616097 | 0.527285 |
| 7 | 13651000 | 13652399 | 27.41139 | 46.95009 | 44.65016 | 19.58063 | 0.455428 | 0.97113 | 0.527293 |

| | | | | | | | | | |
|---|---|---|---|---|---|---|---|---|---|
| 13 | 12764000 | 12764399 | 7.356145 | 14.33729 | 13.74803 | 4.976416 | 0.693439 | -0.39789 | 0.527301 |
| 19 | 27789000 | 27790399 | 17.75107 | 32.19104 | 28.20974 | 12.19671 | 0.60301 | 0.369193 | 0.527323 |
| 6 | 68706600 | 68708199 | 18.56871 | 32.88522 | 32.01647 | 14.1567 | 0.429869 | 0.524433 | 0.527325 |
| 9 | 13664600 | 13666999 | 44.62198 | 71.98065 | 76.5043 | 32.59597 | 0.270044 | 1.772087 | 0.527332 |
| 17 | 70955600 | 70959399 | 42.69713 | 69.46761 | 68.54087 | 29.41933 | 0.388979 | 1.575555 | 0.527335 |
| 6 | 17890000 | 17891199 | 13.98467 | 26.02495 | 25.34535 | 10.58558 | 0.485712 | 0.186386 | 0.527363 |
| 6 | 124572400 | 124574199 | 18.51088 | 33.45105 | 28.97966 | 12.58484 | 0.615099 | 0.393415 | 0.52737 |
| 3 | 21456400 | 21457399 | 8.213015 | 15.66882 | 29.01252 | 11.56153 | -0.37711 | -0.01693 | 0.527373 |
| 17 | 47650600 | 47669199 | 255.147 | 364.9564 | 493.821 | 204.0084 | -0.23502 | 4.244393 | 0.527386 |
| 3 | 94582000 | 94588599 | 63.83243 | 100.6463 | 149.5665 | 62.06816 | -0.24706 | 2.475413 | 0.52739 |
| 14 | 4321200 | 4323599 | 30.33318 | 51.42686 | 47.94411 | 20.99822 | 0.482207 | 1.074401 | 0.527399 |
| 5 | 6319800 | 6336599 | 241.2253 | 352.6798 | 339.5967 | 139.903 | 0.286597 | 3.91706 | 0.527401 |
| 18 | 76720200 | 76721599 | 8.09183 | 15.24562 | 28.061 | 11.47734 | -0.40128 | 0.065842 | 0.527435 |
| X | 125399000 | 125400599 | 30.07782 | 50.96801 | 40.07025 | 17.54698 | 0.727274 | 1.071828 | 0.527438 |
| 12 | 79001800 | 79007399 | 67.02736 | 106.6203 | 110.8263 | 46.28151 | 0.279023 | 2.241578 | 0.52744 |
| 9 | 114853000 | 114855599 | 36.13966 | 58.30546 | 91.15014 | 38.84281 | -0.28516 | 1.763251 | 0.527508 |
| 7 | 26392800 | 26394799 | 34.13489 | 56.2451 | 59.72106 | 25.76228 | 0.293233 | 1.385188 | 0.527538 |
| 17 | 53466000 | 53466799 | 18.66255 | 32.69038 | 32.21276 | 14.22329 | 0.408897 | 0.663917 | 0.527542 |
| 15 | 86983000 | 86985599 | 21.3795 | 37.34668 | 37.55608 | 16.54524 | 0.379566 | 0.728094 | 0.527554 |
| 17 | 43841200 | 43843999 | 21.3795 | 37.34668 | 37.55608 | 16.54524 | 0.379566 | 0.728094 | 0.527554 |
| 11 | 12587800 | 12588399 | 9.751724 | 17.77457 | 31.85386 | 13.68501 | -0.41676 | 0.288067 | 0.527565 |
| 7 | 85653800 | 85665999 | 348.0638 | 489.8809 | 465.0271 | 192.6289 | 0.252725 | 4.48404 | 0.527582 |
| 11 | 108845600 | 108848199 | 17.11681 | 29.71938 | 48.46903 | 21.27155 | -0.31582 | 0.807856 | 0.527588 |
| 2 | 77654200 | 77655399 | 14.77237 | 27.24132 | 26.85092 | 11.35831 | 0.455359 | 0.248118 | 0.52759 |
| 10 | 126008200 | 126010199 | 20.83197 | 36.6796 | 33.46865 | 14.81602 | 0.519778 | 0.602264 | 0.52761 |
| 1 | 35610400 | 35613199 | 25.87629 | 44.35523 | 44.1006 | 19.34084 | 0.391564 | 0.955038 | 0.527619 |
| X | 24121200 | 24123799 | 35.01329 | 57.9964 | 56.92346 | 24.63484 | 0.407679 | 1.328059 | 0.527627 |
| 3 | 138336000 | 138337599 | 8.555728 | 16.13821 | 28.62722 | 11.6855 | -0.34606 | 0.064794 | 0.527632 |
| 12 | 11931200 | 11932799 | 16.99649 | 28.90235 | 52.63445 | 23.06861 | -0.47963 | 1.005891 | 0.527633 |
| 2 | 49487800 | 49498799 | 82.94792 | 127.7222 | 222.727 | 92.50887 | -0.50413 | 2.92521 | 0.527638 |
| 1 | 142469400 | 142470799 | 16.62966 | 29.42264 | 30.80887 | 13.62537 | 0.324944 | 0.521078 | 0.527641 |
| 17 | 50553000 | 50562399 | 150.1522 | 223.8461 | 316.7091 | 130.518 | -0.24196 | 3.583997 | 0.527644 |
| 12 | 114130600 | 114133199 | 53.05642 | 85.39307 | 84.46885 | 35.87708 | 0.369561 | 1.856478 | 0.527653 |
| 7 | 24282000 | 24289199 | 99.68437 | 150.8913 | 272.5541 | 112.754 | -0.57454 | 3.278507 | 0.527654 |
| 1 | 137538600 | 137538999 | 5.210324 | 10.16435 | 20.9705 | 7.500233 | -0.38432 | -0.44042 | 0.527654 |
| 10 | 30448600 | 30451799 | 48.99056 | 78.80971 | 83.1014 | 35.27069 | 0.276641 | 1.864107 | 0.527669 |
| 12 | 52249600 | 52252399 | 23.19934 | 41.01826 | 31.1187 | 13.76428 | 0.787086 | 0.570003 | 0.52771 |
| 4 | 147804000 | 147805199 | 13.50497 | 25.73902 | 23.16364 | 9.250334 | 0.652915 | 0.025019 | 0.527713 |
| 18 | 3537200 | 3547399 | 146.2736 | 220.988 | 227.2977 | 93.97474 | 0.234872 | 3.324495 | 0.527738 |
| 15 | 72459000 | 72467999 | 83.3798 | 128.53 | 196.2488 | 81.51848 | -0.31137 | 2.901889 | 0.527745 |
| 2 | 150863200 | 150864199 | 7.046863 | 13.4502 | 26.14737 | 10.40965 | -0.44431 | -0.03528 | 0.527769 |
| 19 | 5364000 | 5372399 | 181.7613 | 265.1482 | 391.9681 | 161.5113 | -0.33468 | 3.946487 | 0.52777 |
| 4 | 21134600 | 21135799 | 11.5091 | 20.7618 | 34.77374 | 15.13404 | -0.3322 | 0.367416 | 0.527771 |
| 17 | 39541800 | 39542399 | 21.66204 | 37.38945 | 36.53139 | 16.02605 | 0.418664 | 0.867908 | 0.527774 |
| 7 | 47773600 | 47782199 | 80.98045 | 127.3224 | 133.6707 | 55.45113 | 0.250429 | 2.538624 | 0.527782 |
| 17 | 12343200 | 12350799 | 66.06524 | 103.7626 | 156.2671 | 64.81602 | -0.27059 | 2.555249 | 0.52779 |
| 18 | 77834800 | 77838599 | 36.88455 | 59.57136 | 93.30146 | 39.81485 | -0.28609 | 1.732043 | 0.527805 |

| | | | | | | | | | |
|---|---|---|---|---|---|---|---|---|---|
| 7 | 138296200 | 138301199 | 39.17726 | 62.66969 | 117.9373 | 49.63897 | -0.56702 | 2.041043 | 0.527806 |
| 15 | 17358600 | 17359199 | 11.80871 | 22.12621 | 22.17347 | 9.154303 | 0.458919 | 0.120961 | 0.527814 |
| 4 | 144463400 | 144465599 | 26.70671 | 45.32346 | 46.8703 | 20.5341 | 0.333126 | 1.049953 | 0.527833 |
| 7 | 40758800 | 40762199 | 54.76922 | 87.81913 | 91.41643 | 38.61218 | 0.289542 | 1.975754 | 0.527848 |
| X | 166295000 | 166296399 | 15.74024 | 29.8344 | 22.89368 | 9.249658 | 0.866608 | 0.081874 | 0.527858 |
| 10 | 93676600 | 93684799 | 100.0704 | 153.0054 | 222.2379 | 92.1977 | -0.24848 | 3.0705 | 0.527884 |
| 9 | 114062600 | 114064599 | 23.58174 | 38.61861 | 78.8178 | 33.912 | -0.65018 | 1.451857 | 0.527895 |
| 1 | 85465400 | 85466199 | 27.0646 | 45.65647 | 48.00526 | 20.99315 | 0.308982 | 1.10365 | 0.527914 |
| 1 | 35173400 | 35175399 | 26.84383 | 44.42275 | 72.08852 | 31.17765 | -0.31575 | 1.32755 | 0.527923 |
| X | 164417400 | 164417999 | 10.14704 | 19.3643 | 21.24276 | 8.459961 | 0.373359 | -0.01963 | 0.527944 |
| 12 | 114861400 | 114865199 | 59.89344 | 96.36055 | 72.94515 | 30.96551 | 0.753999 | 1.873061 | 0.527946 |
| 14 | 56040000 | 56042599 | 19.65654 | 33.52908 | 53.79808 | 23.58447 | -0.29717 | 0.988453 | 0.527947 |
| X | 8128000 | 8131199 | 35.35303 | 58.91331 | 54.20032 | 23.55253 | 0.501196 | 1.259761 | 0.527978 |
| 10 | 117859800 | 117861799 | 14.3589 | 25.18067 | 43.4869 | 19.21988 | -0.39704 | 0.63704 | 0.527978 |
| 13 | 36095800 | 36098599 | 36.89155 | 60.52911 | 53.08569 | 22.85872 | 0.565883 | 1.43969 | 0.527987 |
| X | 130028200 | 130030199 | 20.33273 | 35.91409 | 33.02656 | 14.61106 | 0.510432 | 0.566411 | 0.527991 |
| 5 | 18114600 | 18115199 | 6.403017 | 12.36712 | 24.27186 | 9.336079 | -0.41179 | -0.16347 | 0.527991 |
| 4 | 117179000 | 117179999 | 8.23645 | 15.54777 | 28.09872 | 11.45137 | -0.37073 | 0.054663 | 0.527992 |
| 16 | 56026200 | 56028799 | 21.11437 | 37.247 | 32.30365 | 14.2949 | 0.59362 | 0.581429 | 0.527994 |
| 2 | 166221600 | 166222799 | 13.14048 | 23.16172 | 38.79331 | 17.16948 | -0.35113 | 0.55671 | 0.528006 |
| 15 | 37278800 | 37279999 | 13.14048 | 23.16172 | 38.79331 | 17.16948 | -0.35113 | 0.55671 | 0.528006 |
| 5 | 57145000 | 57145599 | 7.795842 | 14.85707 | 27.22413 | 10.87338 | -0.36438 | -0.01571 | 0.528011 |
| 4 | 59735200 | 59736399 | 14.69128 | 26.96396 | 27.38472 | 11.65889 | 0.40597 | 0.275589 | 0.528029 |
| 8 | 43366600 | 43367799 | 14.69128 | 26.96396 | 27.38472 | 11.65889 | 0.40597 | 0.275589 | 0.528029 |
| 7 | 61139800 | 61141599 | 26.58752 | 45.69608 | 43.56136 | 19.10756 | 0.452792 | 0.936205 | 0.52807 |
| 7 | 16777400 | 16778999 | 14.43213 | 25.25338 | 42.03429 | 18.54331 | -0.34385 | 0.673678 | 0.528092 |
| 1 | 55104600 | 55106199 | 7.242597 | 13.63293 | 27.5476 | 11.2851 | -0.53574 | 0.06624 | 0.528101 |
| 2 | 164047600 | 164048399 | 7.433057 | 14.33145 | 27.76977 | 10.75014 | -0.40302 | -0.12247 | 0.528101 |
| X | 17654600 | 17656599 | 24.26702 | 42.07757 | 40.87653 | 17.93473 | 0.428183 | 0.845527 | 0.528107 |
| 4 | 142968200 | 142972799 | 50.73355 | 80.74907 | 123.5778 | 51.64841 | -0.27725 | 2.211733 | 0.528113 |
| 11 | 7634800 | 7638399 | 44.7201 | 72.71946 | 69.62571 | 29.87906 | 0.431729 | 1.590803 | 0.52813 |
| 6 | 11925000 | 11927999 | 19.08968 | 32.69179 | 53.30194 | 23.37445 | -0.3194 | 0.947581 | 0.528135 |
| 18 | 53190400 | 53192799 | 13.29654 | 23.54064 | 38.72091 | 17.1223 | -0.32269 | 0.509213 | 0.528147 |
| 12 | 103950600 | 103952799 | 15.06283 | 26.28038 | 43.64826 | 19.2144 | -0.34101 | 0.723246 | 0.528151 |
| 17 | 22236600 | 22242599 | 83.45098 | 131.7344 | 128.9143 | 53.51983 | 0.356184 | 2.45863 | 0.528155 |
| 3 | 37754000 | 37759799 | 55.31651 | 87.95586 | 132.0918 | 55.13402 | -0.25165 | 2.249205 | 0.528159 |
| 2 | 120838400 | 120839399 | 6.741852 | 12.87804 | 25.62326 | 10.18423 | -0.47545 | -0.04589 | 0.528183 |
| 5 | 129577800 | 129579399 | 7.583748 | 14.07475 | 29.98663 | 12.5748 | -0.63951 | 0.161616 | 0.528206 |
| 12 | 115125400 | 115137599 | 161.2462 | 244.7701 | 210.8972 | 87.32316 | 0.495959 | 3.232614 | 0.528209 |
| 16 | 31220200 | 31223799 | 31.14133 | 50.7431 | 79.60442 | 34.18688 | -0.27676 | 1.539169 | 0.528212 |
| 15 | 76790400 | 76797999 | 92.78012 | 141.9286 | 216.5894 | 89.86456 | -0.31911 | 3.058516 | 0.528253 |
| 1 | 16356600 | 16358199 | 16.71926 | 30.95392 | 26.2608 | 11.04895 | 0.676047 | 0.24145 | 0.528261 |
| 1 | 21134600 | 21138199 | 39.34892 | 63.38463 | 98.26974 | 41.78992 | -0.27592 | 1.81548 | 0.528266 |
| 7 | 103838400 | 103839199 | 10.20374 | 19.62345 | 20.07746 | 7.829386 | 0.499931 | -0.09395 | 0.528269 |
| 13 | 10919000 | 10920999 | 18.55416 | 33.75282 | 28.00934 | 12.06279 | 0.684498 | 0.355005 | 0.52827 |
| 3 | 5438200 | 5439199 | 13.62922 | 24.81226 | 25.9135 | 11.14967 | 0.355566 | 0.309181 | 0.528311 |
| 17 | 8572600 | 8577799 | 46.64859 | 74.40883 | 116.3714 | 48.783 | -0.30507 | 2.136379 | 0.528334 |

| | | | | | | | | | |
|---|---|---|---|---|---|---|---|---|---|
| 1 | 162317000 | 162317999 | 11.41293 | 20.36589 | 35.29284 | 15.51688 | -0.39085 | 0.435973 | 0.528338 |
| 2 | 98265600 | 98268799 | 26.43611 | 43.69696 | 70.43807 | 30.43929 | -0.3063 | 1.336222 | 0.52835 |
| 6 | 94353800 | 94356999 | 30.63846 | 51.31709 | 51.36824 | 22.39358 | 0.379527 | 1.184629 | 0.528389 |
| 5 | 44013600 | 44018599 | 64.64562 | 102.7754 | 108.6484 | 45.34272 | 0.253899 | 2.254777 | 0.528392 |
| 6 | 60449800 | 60450999 | 22.64989 | 39.19173 | 39.98209 | 17.54117 | 0.35737 | 0.856358 | 0.528398 |
| 19 | 39243200 | 39252799 | 162.6389 | 244.648 | 243.7514 | 100.6509 | 0.275309 | 3.413379 | 0.528409 |
| 3 | 114980400 | 114988999 | 62.58235 | 100.1037 | 88.00306 | 37.03682 | 0.528898 | 2.059358 | 0.528452 |
| 19 | 50651200 | 50652999 | 10.67034 | 19.64633 | 34.64255 | 14.68528 | -0.38035 | 0.24341 | 0.528459 |
| 5 | 82022600 | 82024399 | 22.89337 | 39.96032 | 37.40839 | 16.47217 | 0.482174 | 0.744729 | 0.528465 |
| 14 | 113057400 | 113060599 | 30.03047 | 51.1469 | 46.26503 | 20.27959 | 0.52623 | 1.032353 | 0.528496 |
| 6 | 51254800 | 51255799 | 12.93622 | 23.8941 | 25.4325 | 10.73455 | 0.347496 | 0.220993 | 0.528506 |
| 1 | 66449800 | 66450399 | 4.807285 | 9.371145 | 20.26123 | 7.317253 | -0.46373 | -0.41487 | 0.528514 |
| 4 | 124923600 | 124924199 | 4.807285 | 9.371145 | 20.26123 | 7.317253 | -0.46373 | -0.41487 | 0.528514 |
| 12 | 78529200 | 78530399 | 16.70731 | 29.86984 | 30.07965 | 13.2038 | 0.388844 | 0.445196 | 0.528523 |
| 4 | 79852600 | 79860199 | 83.54967 | 131.8068 | 130.5513 | 54.1821 | 0.338077 | 2.47458 | 0.52854 |
| 10 | 71394600 | 71396799 | 22.44611 | 39.05699 | 39.19855 | 17.22647 | 0.381859 | 0.789027 | 0.528548 |
| 17 | 21991200 | 21992599 | 19.91886 | 34.88906 | 35.97052 | 15.88176 | 0.343739 | 0.676066 | 0.528578 |
| 3 | 90943000 | 90946199 | 40.5894 | 66.4943 | 64.07071 | 27.571 | 0.429661 | 1.474036 | 0.528581 |
| 18 | 7686400 | 7687399 | 10.49889 | 19.34669 | 33.56517 | 14.21253 | -0.35586 | 0.235067 | 0.528601 |
| 9 | 71677800 | 71689399 | 108.5612 | 165.3764 | 239.5576 | 99.29484 | -0.24881 | 3.14824 | 0.528626 |
| 5 | 145696200 | 145705199 | 366.5438 | 515.2117 | 490.7292 | 203.3176 | 0.245942 | 4.504322 | 0.528632 |
| 6 | 8301800 | 8303599 | 18.17818 | 31.49283 | 53.92155 | 23.65765 | -0.38637 | 0.842403 | 0.528647 |
| 1 | 65105400 | 65106799 | 15.3844 | 27.5676 | 28.86514 | 12.6463 | 0.335076 | 0.421933 | 0.528652 |
| 5 | 59570400 | 59572599 | 18.11475 | 32.03913 | 31.49904 | 13.93156 | 0.414999 | 0.533346 | 0.528674 |
| 4 | 143042600 | 143044799 | 26.17794 | 44.16546 | 46.43235 | 20.30596 | 0.309234 | 1.099771 | 0.528682 |
| 12 | 10881400 | 10884599 | 35.63858 | 59.18445 | 56.55457 | 24.51877 | 0.446378 | 1.2975 | 0.528688 |
| 4 | 122519800 | 122527399 | 109.2833 | 170.4389 | 147.7115 | 61.29777 | 0.517801 | 2.702985 | 0.528697 |
| 1 | 34121800 | 34122599 | 6.708898 | 12.96672 | 24.87074 | 9.536128 | -0.37423 | -0.16696 | 0.528713 |
| 18 | 28846200 | 28846999 | 6.995196 | 13.43019 | 25.77567 | 10.10034 | -0.40555 | -0.09262 | 0.528714 |
| 17 | 50439800 | 50442199 | 18.22833 | 30.93844 | 54.26998 | 23.77675 | -0.42624 | 1.027346 | 0.528731 |
| 1 | 171000400 | 171002399 | 13.20369 | 23.22442 | 39.38437 | 17.43838 | -0.36995 | 0.583391 | 0.52874 |
| X | 4396800 | 4397999 | 23.48195 | 41.07185 | 36.85105 | 16.25108 | 0.5434 | 0.719315 | 0.528743 |
| 11 | 73023000 | 73024799 | 15.88205 | 27.63844 | 45.25846 | 19.89101 | -0.32109 | 0.766824 | 0.528751 |
| 8 | 27371000 | 27372799 | 12.14593 | 21.6082 | 36.69978 | 16.1768 | -0.36494 | 0.460046 | 0.528771 |
| 2 | 173376400 | 173379199 | 28.33234 | 46.41245 | 73.95841 | 31.82527 | -0.29412 | 1.454439 | 0.528785 |
| 7 | 108315800 | 108317999 | 19.96581 | 34.14508 | 55.47667 | 24.32543 | -0.31466 | 0.967467 | 0.528794 |
| 7 | 27143200 | 27145199 | 15.81558 | 28.12735 | 29.33184 | 12.93272 | 0.334567 | 0.471603 | 0.528842 |
| 16 | 3252000 | 3254999 | 36.79997 | 60.62382 | 54.23512 | 23.39371 | 0.539607 | 1.391849 | 0.528849 |
| 12 | 102267400 | 102269399 | 17.40654 | 29.78993 | 50.57135 | 22.17584 | -0.3773 | 0.945804 | 0.52887 |
| 6 | 6358400 | 6359199 | 5.390712 | 10.4676 | 22.23256 | 8.331045 | -0.4879 | -0.26969 | 0.528885 |
| 6 | 130684200 | 130686999 | 38.78389 | 64.15072 | 56.80014 | 24.55733 | 0.555598 | 1.349187 | 0.528889 |
| 2 | 28729400 | 28731199 | 9.396117 | 16.85747 | 33.8925 | 14.83373 | -0.59911 | 0.384284 | 0.528898 |
| 1 | 35726400 | 35727999 | 11.76273 | 20.89864 | 36.30693 | 16.02161 | -0.39809 | 0.468686 | 0.528916 |
| 2 | 153669000 | 153670199 | 13.876 | 24.25469 | 42.64016 | 18.79385 | -0.42206 | 0.691593 | 0.528921 |
| 15 | 94035200 | 94036199 | 7.323665 | 13.97507 | 26.14935 | 10.41589 | -0.39087 | -0.03295 | 0.528925 |
| 2 | 72461400 | 72465599 | 52.22252 | 83.04022 | 126.3374 | 52.73472 | -0.27022 | 2.244137 | 0.528926 |
| 9 | 8767800 | 8769999 | 25.61313 | 43.75622 | 45.23149 | 19.83141 | 0.334963 | 0.986113 | 0.528941 |

| | | | | | | | | | |
|---|---|---|---|---|---|---|---|---|---|
| 10 | 25658400 | 25659199 | 9.580917 | 18.46769 | 20.33775 | 7.879175 | 0.404967 | -0.11551 | 0.528941 |
| 15 | 43004200 | 43005399 | 14.68968 | 25.4735 | 50.56183 | 22.18425 | -0.59799 | 0.815411 | 0.528942 |
| 4 | 78749400 | 78753599 | 51.85794 | 82.61475 | 124.2392 | 51.99106 | -0.25063 | 2.183149 | 0.528947 |
| 13 | 67090600 | 67094799 | 53.48635 | 85.78807 | 90.53515 | 38.25754 | 0.270316 | 1.964985 | 0.528955 |
| 10 | 115948800 | 115950599 | 30.40175 | 48.90162 | 104.2614 | 44.2453 | -0.73694 | 1.853779 | 0.528978 |
| 15 | 25343000 | 25345199 | 24.80342 | 42.7282 | 37.74366 | 16.55688 | 0.562794 | 0.898822 | 0.528979 |
| 4 | 6365400 | 6365999 | 5.098396 | 9.922122 | 21.24986 | 7.820467 | -0.47535 | -0.3404 | 0.528979 |
| X | 26828800 | 26839399 | 140.4663 | 215.0825 | 185.3707 | 76.9288 | 0.505525 | 3.053367 | 0.52898 |
| 2 | 73427000 | 73430199 | 44.63411 | 71.41243 | 109.9831 | 46.30371 | -0.27815 | 2.038081 | 0.528985 |
| 14 | 15922600 | 15923599 | 6.775352 | 12.82489 | 26.56288 | 10.7627 | -0.55757 | 0.019633 | 0.528987 |
| 16 | 36986200 | 36987599 | 13.00223 | 24.22971 | 25.52551 | 10.6362 | 0.376117 | 0.171697 | 0.529006 |
| 11 | 88145800 | 88153999 | 75.62223 | 117.9761 | 173.6192 | 72.04684 | -0.24492 | 2.683581 | 0.529011 |
| 15 | 41131000 | 41131599 | 5.802639 | 11.29557 | 22.47661 | 8.254793 | -0.3681 | -0.33458 | 0.529011 |
| 1 | 7588600 | 7589399 | 8.883249 | 16.4826 | 30.46798 | 12.78077 | -0.43685 | 0.174512 | 0.529017 |
| 1 | 177855600 | 177856399 | 6.693458 | 12.87679 | 26.43137 | 10.29754 | -0.49415 | -0.11452 | 0.529028 |
| 5 | 10211800 | 10215199 | 59.79437 | 95.40506 | 99.49925 | 41.72689 | 0.2789 | 2.133292 | 0.529046 |
| 3 | 15857600 | 15859999 | 31.63221 | 52.87707 | 52.97784 | 23.06653 | 0.378432 | 1.214372 | 0.529057 |
| 14 | 124671800 | 124674599 | 27.28463 | 47.14323 | 38.96916 | 17.09602 | 0.65887 | 0.882948 | 0.529073 |
| 5 | 57479600 | 57482199 | 24.62595 | 42.76483 | 40.23522 | 17.65995 | 0.474426 | 0.828658 | 0.5291 |
| 7 | 17921000 | 17922599 | 41.07359 | 66.48586 | 67.91049 | 29.04945 | 0.332719 | 1.671563 | 0.529109 |
| 13 | 94397600 | 94401799 | 23.32254 | 38.87358 | 63.01801 | 27.38787 | -0.31353 | 1.227191 | 0.529113 |
| 6 | 71346000 | 71346999 | 7.892358 | 15.07228 | 27.88159 | 11.08715 | -0.37302 | -0.03034 | 0.529119 |
| 3 | 6675800 | 6676599 | 9.980027 | 18.29509 | 32.91157 | 14.03482 | -0.41538 | 0.256604 | 0.52912 |
| 4 | 51893000 | 51895199 | 22.64729 | 38.87975 | 38.97148 | 17.09351 | 0.380195 | 0.920865 | 0.529158 |
| 3 | 62449800 | 62450799 | 15.24952 | 27.82511 | 24.24319 | 10.40144 | 0.618041 | 0.305394 | 0.52918 |
| 7 | 117048600 | 117049399 | 10.58953 | 19.03145 | 33.79828 | 14.77094 | -0.41996 | 0.381537 | 0.529188 |
| 10 | 18786200 | 18786999 | 5.373388 | 10.44803 | 21.75398 | 8.063458 | -0.44444 | -0.30849 | 0.529193 |
| 12 | 117617400 | 117618199 | 5.373388 | 10.44803 | 21.75398 | 8.063458 | -0.44444 | -0.30849 | 0.529193 |
| 8 | 89647800 | 89656399 | 60.15232 | 95.1192 | 142.3889 | 59.16546 | -0.25424 | 2.404599 | 0.529195 |
| 6 | 127862200 | 127862599 | 8.925609 | 17.30018 | 19.91494 | 7.535222 | 0.373552 | -0.20313 | 0.5292 |
| 14 | 41386800 | 41394599 | 71.51023 | 114.1221 | 100.4824 | 42.15165 | 0.523102 | 2.141014 | 0.529228 |
| 7 | 35038400 | 35039199 | 5.50162 | 10.71034 | 22.4206 | 8.229639 | -0.43877 | -0.33745 | 0.529234 |
| 17 | 18344600 | 18351599 | 111.4304 | 172.401 | 165.3197 | 68.66974 | 0.363087 | 2.844512 | 0.529267 |
| 10 | 24885000 | 24886599 | 16.525 | 29.33925 | 30.78052 | 13.59087 | 0.324551 | 0.490136 | 0.529277 |
| 16 | 71808000 | 71810999 | 30.74098 | 51.84321 | 49.64232 | 21.70713 | 0.443478 | 1.120969 | 0.529303 |
| 18 | 63071800 | 63074199 | 12.6637 | 22.31102 | 38.81415 | 17.18029 | -0.40588 | 0.558365 | 0.52931 |
| 18 | 82123800 | 82129599 | 56.71535 | 91.07629 | 89.13451 | 37.73442 | 0.380949 | 1.933104 | 0.529319 |
| 9 | 75394400 | 75398799 | 41.68839 | 66.99503 | 102.3525 | 43.37771 | -0.259 | 1.895247 | 0.529322 |
| 9 | 11206400 | 11209199 | 22.94397 | 38.54928 | 62.04998 | 27.10184 | -0.30346 | 1.139993 | 0.529325 |
| 1 | 69230600 | 69231599 | 13.40348 | 23.34633 | 47.87816 | 21.05437 | -0.64395 | 0.746081 | 0.529344 |
| 16 | 64484600 | 64485999 | 14.1096 | 26.74136 | 23.53149 | 9.509087 | 0.670112 | 0.068398 | 0.529352 |
| 4 | 111128400 | 111132199 | 55.02303 | 87.96477 | 90.72918 | 38.155 | 0.29797 | 2.068152 | 0.529357 |
| 6 | 111911400 | 111912999 | 24.4745 | 41.69308 | 44.24354 | 19.39384 | 0.296721 | 1.006281 | 0.529367 |
| 4 | 124152800 | 124156999 | 33.4473 | 55.41118 | 57.55547 | 24.91105 | 0.326122 | 1.318848 | 0.529369 |
| 7 | 61554200 | 61559599 | 62.99102 | 101.0481 | 89.70724 | 37.91641 | 0.519593 | 1.974826 | 0.529442 |
| 13 | 61951600 | 61976799 | 498.2425 | 686.11 | 596.2873 | 247.8476 | 0.348288 | 4.817612 | 0.529472 |
| X | 139796200 | 139802999 | 67.2427 | 107.168 | 105.1976 | 44.04756 | 0.36445 | 2.181905 | 0.529479 |

| | | | | | | | | | |
|---|---|---|---|---|---|---|---|---|---|
| 13 | 59020000 | 59023399 | 26.64569 | 44.01393 | 69.74091 | 30.12411 | -0.28162 | 1.345491 | 0.529489 |
| 3 | 93678800 | 93682999 | 43.80228 | 70.25851 | 106.9436 | 45.19419 | -0.25701 | 1.958113 | 0.529502 |
| 9 | 48595200 | 48597599 | 16.99848 | 29.25886 | 49.52824 | 21.72584 | -0.37184 | 0.888512 | 0.529521 |
| 19 | 10554200 | 10555199 | 6.130323 | 11.80725 | 24.17318 | 9.386169 | -0.48459 | -0.1341 | 0.529543 |
| 8 | 39683600 | 39684799 | 14.10826 | 26.36064 | 25.46453 | 10.56323 | 0.505383 | 0.163721 | 0.529547 |
| 5 | 22276000 | 22277199 | 17.3943 | 30.12912 | 49.5314 | 21.73139 | -0.32782 | 0.836549 | 0.529585 |
| 13 | 5693800 | 5695399 | 14.92499 | 27.65152 | 26.35528 | 11.07807 | 0.510006 | 0.222546 | 0.529595 |
| 6 | 140613400 | 140617799 | 56.517 | 89.69566 | 133.7137 | 55.70594 | -0.24365 | 2.3061 | 0.529597 |
| 10 | 91651200 | 91652999 | 35.75569 | 58.53702 | 62.24981 | 26.77995 | 0.287435 | 1.471748 | 0.529624 |
| 12 | 99371200 | 99376799 | 50.28977 | 80.01276 | 139.1476 | 58.12557 | -0.46199 | 2.227261 | 0.529636 |
| 12 | 115708400 | 115711399 | 23.77294 | 41.61065 | 31.49847 | 13.89905 | 0.78727 | 0.700358 | 0.529654 |
| 5 | 26882200 | 26885199 | 31.64769 | 52.65713 | 55.24467 | 23.98242 | 0.312194 | 1.263277 | 0.529656 |
| 18 | 8774000 | 8776799 | 69.26115 | 109.6539 | 109.7122 | 45.61517 | 0.327594 | 2.377366 | 0.529689 |
| 17 | 57964800 | 57968399 | 53.68934 | 86.3293 | 77.50988 | 32.87584 | 0.507357 | 1.88451 | 0.529714 |
| 6 | 69292800 | 69294399 | 22.13729 | 38.4515 | 39.38803 | 17.29049 | 0.352432 | 0.815072 | 0.529721 |
| 3 | 103969200 | 103969999 | 8.566735 | 16.12247 | 28.58123 | 11.71212 | -0.34963 | 0.079586 | 0.529729 |
| 3 | 124667400 | 124670199 | 28.00423 | 47.41001 | 48.51382 | 21.23846 | 0.34824 | 1.078764 | 0.529745 |
| 10 | 126102200 | 126102599 | 10.40378 | 19.75593 | 21.6756 | 8.723457 | 0.359646 | 0.016798 | 0.52976 |
| 7 | 51774400 | 51775599 | 13.29731 | 25.05511 | 24.56914 | 10.04778 | 0.50024 | 0.102444 | 0.529767 |
| 9 | 60958400 | 60960999 | 29.53693 | 48.15166 | 76.97812 | 33.06472 | -0.30337 | 1.528221 | 0.529793 |
| 13 | 23441600 | 23448399 | 77.02582 | 122.0084 | 121.9148 | 50.70161 | 0.330034 | 2.373553 | 0.529816 |
| 8 | 118528200 | 118529399 | 8.919109 | 16.51157 | 29.93065 | 12.59425 | -0.41176 | 0.188073 | 0.529826 |
| 8 | 25867600 | 25871999 | 32.09176 | 52.3352 | 85.36703 | 36.67279 | -0.33211 | 1.531479 | 0.529833 |
| 2 | 121597800 | 121598799 | 17.33227 | 31.12347 | 25.68674 | 11.23384 | 0.678192 | 0.416214 | 0.529841 |
| 5 | 99002000 | 99003599 | 13.46814 | 23.53676 | 43.62553 | 19.2247 | -0.4984 | 0.689413 | 0.529848 |
| 13 | 98088600 | 98090399 | 12.99393 | 22.92835 | 38.28861 | 16.94242 | -0.34618 | 0.543517 | 0.529879 |
| 2 | 35724400 | 35730399 | 70.15464 | 111.1252 | 118.0287 | 49.08564 | 0.242099 | 2.36642 | 0.52988 |
| 1 | 103245600 | 103254999 | 270.8596 | 390.2089 | 382.6465 | 157.9058 | 0.23957 | 4.139051 | 0.529881 |
| 3 | 36907200 | 36908399 | 11.52107 | 20.74207 | 34.75832 | 15.16503 | -0.33522 | 0.379255 | 0.529889 |
| 2 | 22309400 | 22309999 | 12.46619 | 23.05572 | 24.48315 | 10.31568 | 0.35274 | 0.20811 | 0.529889 |
| 4 | 21628400 | 21630599 | 15.45805 | 26.67345 | 47.42105 | 20.80308 | -0.4413 | 0.858396 | 0.529893 |
| 17 | 54052600 | 54054399 | 27.84502 | 45.21265 | 79.41909 | 34.05008 | -0.44325 | 1.59161 | 0.529923 |
| 8 | 77586400 | 77587599 | 7.902509 | 14.98978 | 28.23183 | 11.38841 | -0.41705 | 0.016804 | 0.529935 |
| 5 | 17261200 | 17264999 | 45.80854 | 74.06512 | 76.12103 | 32.52917 | 0.322165 | 1.708097 | 0.529946 |
| 1 | 19504400 | 19506999 | 23.18774 | 40.49751 | 36.80575 | 16.21269 | 0.524715 | 0.737067 | 0.52998 |
| 12 | 7207000 | 7208199 | 21.58792 | 37.85028 | 31.2568 | 13.81219 | 0.662406 | 0.662803 | 0.529983 |
| 8 | 68696800 | 68700199 | 47.80317 | 76.89916 | 81.28034 | 34.4975 | 0.273236 | 1.863911 | 0.529991 |
| 4 | 132340400 | 132343399 | 57.25642 | 91.67677 | 94.95752 | 40.02123 | 0.29431 | 2.023843 | 0.53 |
| 9 | 54038800 | 54039199 | 6.397257 | 12.39799 | 23.87886 | 9.040441 | -0.36282 | -0.21754 | 0.530002 |
| 10 | 47206600 | 47207999 | 16.37988 | 28.50814 | 46.30376 | 20.35193 | -0.3094 | 0.769347 | 0.530007 |
| 15 | 31798400 | 31799999 | 13.79496 | 24.31744 | 39.79611 | 17.613 | -0.31785 | 0.562039 | 0.530028 |
| X | 88114600 | 88116799 | 27.00051 | 47.23385 | 35.02689 | 15.44878 | 0.817129 | 0.733497 | 0.530047 |
| 3 | 157145800 | 157148199 | 12.68006 | 22.49417 | 37.22125 | 16.44618 | -0.32979 | 0.484453 | 0.530062 |
| 7 | 103904800 | 103907599 | 28.04688 | 45.45331 | 92.22362 | 39.48411 | -0.65336 | 1.639774 | 0.530074 |
| X | 29406800 | 29415799 | 103.7287 | 163.0322 | 130.6977 | 54.21537 | 0.638723 | 2.562224 | 0.530123 |
| 7 | 130982600 | 130983199 | 9.624966 | 18.60133 | 20.88847 | 8.012147 | 0.389864 | -0.14881 | 0.530124 |
| 9 | 39543200 | 39543799 | 9.624966 | 18.60133 | 20.88847 | 8.012147 | 0.389864 | -0.14881 | 0.530124 |

| | | | | | | | | | |
|---|---|---|---|---|---|---|---|---|---|
| 15 | 33194600 | 33195999 | 18.57541 | 32.67358 | 33.77434 | 14.95434 | 0.340222 | 0.601055 | 0.530132 |
| 6 | 71520800 | 71524199 | 24.08572 | 40.20611 | 65.43737 | 28.46636 | -0.31935 | 1.214229 | 0.530162 |
| 13 | 113134400 | 113136599 | 17.20688 | 29.61025 | 48.99665 | 21.49249 | -0.33915 | 0.89284 | 0.530164 |
| 15 | 99275400 | 99276999 | 18.08706 | 31.70972 | 32.58792 | 14.39946 | 0.348375 | 0.645953 | 0.530217 |
| 8 | 107279200 | 107286399 | 70.99992 | 111.3786 | 164.8135 | 68.35385 | -0.24663 | 2.582568 | 0.530226 |
| X | 33205400 | 33208399 | 51.85015 | 84.46578 | 63.95488 | 27.46353 | 0.771324 | 1.57868 | 0.53023 |
| 2 | 15369000 | 15370599 | 17.6275 | 31.18113 | 32.1997 | 14.24111 | 0.344736 | 0.532299 | 0.530245 |
| X | 108499400 | 108501199 | 22.75453 | 39.76615 | 37.02955 | 16.31822 | 0.490191 | 0.73041 | 0.530274 |
| 12 | 17675400 | 17679199 | 30.03413 | 48.8332 | 79.14675 | 33.96354 | -0.32585 | 1.567673 | 0.530281 |
| 14 | 43900600 | 43901199 | 12.38673 | 23.9474 | 20.80561 | 7.96689 | 0.75911 | -0.12493 | 0.530299 |
| 18 | 46604200 | 46606599 | 23.94886 | 41.07464 | 42.00294 | 18.42172 | 0.351225 | 0.938251 | 0.530301 |
| X | 90488000 | 90489799 | 23.69245 | 41.21535 | 39.24888 | 17.24587 | 0.457429 | 0.801405 | 0.530308 |
| 16 | 53501800 | 53503799 | 17.10491 | 31.26458 | 27.60504 | 11.81824 | 0.600859 | 0.316225 | 0.530319 |
| 12 | 52218200 | 52224199 | 76.82833 | 121.4986 | 110.8336 | 46.05544 | 0.459428 | 2.409123 | 0.530324 |
| 1 | 111810200 | 111813599 | 40.63913 | 66.0171 | 69.74173 | 29.91115 | 0.28893 | 1.598479 | 0.530348 |
| 4 | 117048400 | 117049399 | 8.092757 | 15.31428 | 27.61101 | 11.19401 | -0.36049 | 0.035776 | 0.53035 |
| 14 | 91171200 | 91171799 | 8.092757 | 15.31428 | 27.61101 | 11.19401 | -0.36049 | 0.035776 | 0.53035 |
| X | 127588400 | 127593999 | 50.35202 | 81.77547 | 68.81624 | 29.50892 | 0.615991 | 1.625534 | 0.530351 |
| 17 | 49800400 | 49801599 | 8.774779 | 16.16983 | 30.91994 | 13.09221 | -0.49509 | 0.21388 | 0.53036 |
| 7 | 42857400 | 42870599 | 246.1794 | 358.3921 | 352.8467 | 145.4219 | 0.248637 | 3.981657 | 0.530361 |
| 4 | 118712600 | 118714199 | 23.77898 | 40.9781 | 42.05413 | 18.44794 | 0.347595 | 0.899468 | 0.53037 |
| 2 | 87994800 | 87995999 | 16.71183 | 30.48937 | 28.21767 | 12.10864 | 0.531389 | 0.325933 | 0.530382 |
| 6 | 145977200 | 145981799 | 44.95482 | 71.98885 | 109.3807 | 46.10878 | -0.25724 | 2.011309 | 0.53039 |
| 4 | 77008200 | 77010799 | 31.34195 | 52.12855 | 55.21493 | 23.9632 | 0.298432 | 1.267051 | 0.530429 |
| 7 | 15707800 | 15711399 | 22.50349 | 37.64582 | 60.80051 | 26.48517 | -0.30818 | 1.184033 | 0.530433 |
| 11 | 28246600 | 28252399 | 59.25659 | 94.99952 | 83.53963 | 35.27682 | 0.532205 | 1.9845 | 0.530433 |
| 4 | 107121600 | 107123399 | 26.59397 | 45.55038 | 44.63989 | 19.57613 | 0.412178 | 0.966015 | 0.530438 |
| 17 | 54109800 | 54113799 | 60.4108 | 96.58295 | 88.51322 | 37.22572 | 0.468245 | 2.071601 | 0.530447 |
| X | 73811000 | 73812999 | 27.40374 | 46.81749 | 44.19032 | 19.37499 | 0.465812 | 0.995348 | 0.530459 |
| 1 | 58374800 | 58377799 | 31.01293 | 51.85883 | 52.48971 | 22.85872 | 0.36364 | 1.204563 | 0.530468 |
| 15 | 18518600 | 18521999 | 38.82137 | 63.12989 | 67.31721 | 28.88878 | 0.276745 | 1.575179 | 0.530513 |
| 1 | 103366800 | 103370399 | 41.44031 | 67.49411 | 68.38427 | 29.36343 | 0.351878 | 1.558866 | 0.530556 |
| 11 | 101206600 | 101207799 | 7.081428 | 13.36582 | 27.05199 | 11.02763 | -0.5322 | 0.044965 | 0.530582 |
| 15 | 102828800 | 102831399 | 23.32333 | 38.92023 | 62.47315 | 27.17104 | -0.29932 | 1.214623 | 0.530585 |
| X | 58721200 | 58724599 | 45.79006 | 74.00537 | 75.07917 | 32.06725 | 0.340219 | 1.721524 | 0.530631 |
| 17 | 94933000 | 94933799 | 13.4931 | 24.61967 | 25.94128 | 11.13033 | 0.345507 | 0.295763 | 0.530642 |
| 12 | 106599000 | 106600599 | 14.16418 | 24.82958 | 41.49326 | 18.33276 | -0.3494 | 0.643614 | 0.530655 |
| 10 | 117081200 | 117083199 | 14.85126 | 26.03638 | 43.49169 | 19.21718 | -0.34918 | 0.645331 | 0.530655 |
| X | 24525600 | 24527399 | 32.69764 | 54.90807 | 51.33643 | 22.41631 | 0.477702 | 1.1673 | 0.530667 |
| 13 | 63309200 | 63320999 | 117.1489 | 176.5649 | 267.9492 | 110.7006 | -0.32881 | 3.368133 | 0.530672 |
| 3 | 120814200 | 120815199 | 13.52063 | 24.76699 | 26.44566 | 11.28904 | 0.331434 | 0.275222 | 0.530696 |
| 10 | 67391600 | 67394599 | 38.55099 | 62.16169 | 95.94172 | 40.8558 | -0.2677 | 1.783809 | 0.530711 |
| 13 | 106106400 | 106108799 | 23.71849 | 41.5098 | 36.20904 | 15.97584 | 0.58377 | 0.709466 | 0.530729 |
| 3 | 101364200 | 101367199 | 39.71147 | 63.88051 | 98.1162 | 41.64134 | -0.2649 | 1.859964 | 0.530744 |
| 11 | 21210200 | 21213399 | 33.51137 | 55.81456 | 55.3162 | 24.02934 | 0.394121 | 1.258499 | 0.530749 |
| 8 | 98446400 | 98448799 | 27.31397 | 47.00964 | 42.94924 | 18.83986 | 0.514188 | 0.925184 | 0.530768 |
| 17 | 9970600 | 9971399 | 8.44399 | 15.85791 | 28.54943 | 11.74103 | -0.37563 | 0.093139 | 0.530772 |

| | | | | | | | | | |
|---|---|---|---|---|---|---|---|---|---|
| 12 | 87824200 | 87834399 | 230.9669 | 331.2155 | 467.7105 | 193.1442 | -0.29288 | 4.204845 | 0.530778 |
| 7 | 106969000 | 106971599 | 19.32893 | 32.74167 | 55.92416 | 24.48679 | -0.38819 | 1.049657 | 0.530778 |
| X | 118768400 | 118770599 | 24.27464 | 42.44244 | 36.78703 | 16.21772 | 0.593025 | 0.728734 | 0.530778 |
| 11 | 91163400 | 91164599 | 11.86785 | 21.30603 | 35.76965 | 15.6466 | -0.34041 | 0.401772 | 0.53078 |
| 7 | 31501200 | 31503199 | 44.27406 | 71.4475 | 76.02305 | 32.40735 | 0.269069 | 1.759612 | 0.530791 |
| 4 | 123901600 | 123902999 | 16.15535 | 28.75355 | 30.2661 | 13.33607 | 0.321783 | 0.470951 | 0.5308 |
| 6 | 97722200 | 97723599 | 14.9279 | 26.92633 | 28.25085 | 12.29679 | 0.338172 | 0.376106 | 0.530854 |
| X | 115874000 | 115884199 | 44.82237 | 73.08945 | 59.40269 | 25.51791 | 0.670324 | 1.54597 | 0.530882 |
| 9 | 78278800 | 78280799 | 16.83679 | 29.09255 | 47.93793 | 21.03071 | -0.33191 | 0.851839 | 0.530886 |
| 5 | 150301600 | 150302399 | 6.987667 | 13.48781 | 25.40532 | 9.800985 | -0.35737 | -0.14386 | 0.53089 |
| 14 | 68669400 | 68671199 | 12.28792 | 22.11051 | 37.84076 | 16.52077 | -0.36642 | 0.39234 | 0.53091 |
| 6 | 58136600 | 58149399 | 164.7073 | 241.0885 | 425.8912 | 175.4408 | -0.58682 | 3.892725 | 0.530914 |
| 1 | 13236200 | 13237399 | 6.883999 | 13.11745 | 26.12197 | 10.43629 | -0.4839 | -0.02866 | 0.530925 |
| 9 | 115229400 | 115231999 | 22.52826 | 37.61725 | 61.29795 | 26.67056 | -0.32094 | 1.204864 | 0.530952 |
| 14 | 68992000 | 68992599 | 5.961383 | 11.55728 | 23.28166 | 8.800004 | -0.4239 | -0.22756 | 0.530955 |
| 16 | 70178600 | 70181799 | 37.96498 | 61.83278 | 65.66932 | 28.19749 | 0.284088 | 1.549972 | 0.530972 |
| X | 18231000 | 18237399 | 20.83632 | 36.68754 | 29.16666 | 12.91158 | 0.717812 | 0.597622 | 0.530973 |
| 2 | 110833400 | 110833999 | 10.91486 | 20.70852 | 22.0657 | 8.896709 | 0.398728 | 0.026738 | 0.530974 |
| 14 | 91318400 | 91322399 | 69.92249 | 110.9541 | 116.4145 | 48.49161 | 0.262113 | 2.31758 | 0.531001 |
| 6 | 134015800 | 134019399 | 19.52795 | 33.07456 | 54.81919 | 24.00204 | -0.34488 | 1.052116 | 0.531038 |
| 7 | 109584200 | 109585799 | 9.564444 | 17.51766 | 31.36596 | 13.39191 | -0.40953 | 0.256943 | 0.53104 |
| 10 | 66904000 | 66909599 | 57.28268 | 90.92919 | 136.6062 | 56.92246 | -0.25454 | 2.300481 | 0.53104 |
| 4 | 43840800 | 43845199 | 51.32036 | 82.46253 | 86.18056 | 36.51654 | 0.287482 | 1.907087 | 0.531067 |
| 11 | 117778400 | 117798599 | 181.6169 | 263.0531 | 461.3178 | 190.2356 | -0.59105 | 4.055098 | 0.531075 |
| X | 3227400 | 3234399 | 109.2114 | 170.1319 | 133.8279 | 55.54578 | 0.65616 | 2.721456 | 0.531106 |
| 7 | 85033800 | 85037399 | 52.5897 | 84.54094 | 86.83559 | 36.81859 | 0.313222 | 1.893996 | 0.531107 |
| 14 | 123912200 | 123914799 | 22.14336 | 38.76298 | 36.28174 | 16.01134 | 0.482501 | 0.696516 | 0.531123 |
| 10 | 124597200 | 124599199 | 21.54796 | 38.10074 | 32.37475 | 14.31968 | 0.624472 | 0.56178 | 0.531128 |
| 15 | 47462600 | 47465199 | 31.5851 | 52.51905 | 55.1825 | 23.94475 | 0.310021 | 1.271988 | 0.531134 |
| 4 | 54072200 | 54074999 | 16.18458 | 28.091 | 46.34839 | 20.33643 | -0.3324 | 0.806575 | 0.531137 |
| 4 | 126711000 | 126712599 | 11.75593 | 20.78954 | 36.83802 | 16.29337 | -0.42978 | 0.509753 | 0.531139 |
| 9 | 120927200 | 120929399 | 22.5668 | 37.50659 | 63.39167 | 27.49826 | -0.3736 | 1.25771 | 0.531145 |
| 2 | 77435200 | 77437399 | 18.1998 | 30.72926 | 61.14012 | 26.74433 | -0.60754 | 1.083575 | 0.531159 |
| X | 31327200 | 31332599 | 69.96747 | 112.1657 | 87.25725 | 36.84402 | 0.708674 | 2.002521 | 0.531183 |
| 16 | 61189000 | 61190799 | 19.47035 | 33.27473 | 53.27538 | 23.3589 | -0.29367 | 0.969376 | 0.531184 |
| 13 | 8524800 | 8525399 | 10.18734 | 19.8165 | 19.50721 | 7.206545 | 0.628613 | -0.27208 | 0.531186 |
| X | 138026600 | 138029199 | 25.37727 | 44.48854 | 34.42928 | 15.21246 | 0.755657 | 0.690362 | 0.531206 |
| 12 | 10439800 | 10441399 | 12.03857 | 21.28436 | 37.28402 | 16.49141 | -0.41364 | 0.513428 | 0.531207 |
| 4 | 73639400 | 73640599 | 21.43847 | 37.3058 | 38.72554 | 17.0192 | 0.333403 | 0.782501 | 0.531213 |
| 1 | 143719600 | 143727599 | 77.14077 | 122.127 | 110.6527 | 46.00584 | 0.470454 | 2.381907 | 0.531235 |
| 3 | 10076600 | 10078599 | 14.60689 | 25.52608 | 42.5954 | 18.77029 | -0.34744 | 0.697041 | 0.531265 |
| 3 | 129035000 | 129037399 | 19.51217 | 34.18824 | 35.41848 | 15.64247 | 0.336727 | 0.666635 | 0.531305 |
| 12 | 114509000 | 114520799 | 192.5458 | 288.1996 | 257.9969 | 106.4052 | 0.423246 | 3.514621 | 0.531311 |
| 12 | 24076000 | 24122199 | 1409.458 | 1768.166 | 1501.933 | 631.4289 | 0.246334 | 6.31467 | 0.531329 |
| 14 | 7294800 | 7295799 | 9.990984 | 18.36199 | 31.89999 | 13.55628 | -0.36158 | 0.241211 | 0.531332 |
| 1 | 18733800 | 18736599 | 24.17155 | 42.12426 | 38.51585 | 16.94286 | 0.515919 | 0.775753 | 0.531335 |
| 4 | 137711000 | 137714999 | 19.55459 | 33.29632 | 54.29449 | 23.79871 | -0.3207 | 1.006174 | 0.531393 |

| | | | | | | | | | |
|---|---|---|---|---|---|---|---|---|---|
| 2 | 97827600 | 97828599 | 11.94981 | 22.46798 | 24.06171 | 9.875987 | 0.369792 | 0.102165 | 0.531414 |
| 6 | 27112200 | 27114999 | 40.89761 | 66.47081 | 69.67896 | 29.89545 | 0.300792 | 1.590456 | 0.531423 |
| X | 118478200 | 118479399 | 21.51887 | 37.42035 | 37.86861 | 16.63551 | 0.370104 | 0.794659 | 0.531424 |
| 7 | 130347400 | 130349999 | 12.41456 | 21.73745 | 41.06402 | 18.12544 | -0.52538 | 0.650025 | 0.531434 |
| X | 8517000 | 8537399 | 367.3141 | 520.7161 | 445.5058 | 184.3229 | 0.413079 | 4.38058 | 0.531463 |
| 6 | 74168000 | 74170599 | 39.36122 | 63.98149 | 68.42534 | 29.35994 | 0.27213 | 1.583517 | 0.531465 |
| 8 | 129493600 | 129495599 | 15.46992 | 27.06986 | 46.73312 | 20.61654 | -0.39665 | 0.682938 | 0.531474 |
| 6 | 96273800 | 96274599 | 12.33301 | 22.86631 | 24.51683 | 10.29491 | 0.342514 | 0.19364 | 0.531482 |
| 9 | 52072000 | 52072999 | 14.8647 | 27.21849 | 27.24964 | 11.63838 | 0.422988 | 0.28784 | 0.531488 |
| 9 | 15280000 | 15285199 | 70.80147 | 112.0427 | 116.6989 | 48.50914 | 0.26919 | 2.390723 | 0.53149 |
| 4 | 86493600 | 86494799 | 5.753188 | 10.98424 | 25.61728 | 10.19224 | -0.70387 | -0.05432 | 0.531491 |
| 8 | 52331200 | 52333999 | 29.95695 | 48.5248 | 90.61465 | 38.78112 | -0.53472 | 1.648997 | 0.531502 |
| 8 | 12073400 | 12075799 | 23.35585 | 40.36475 | 33.68213 | 14.77668 | 0.645437 | 0.85749 | 0.531505 |
| 4 | 39828200 | 39830599 | 22.29551 | 38.90047 | 38.03282 | 16.74273 | 0.419789 | 0.749042 | 0.531541 |
| 7 | 140194000 | 140195199 | 22.93026 | 38.15059 | 63.34232 | 27.49672 | -0.34781 | 1.249009 | 0.53156 |
| 2 | 111219800 | 111222999 | 55.16986 | 88.49242 | 91.47271 | 38.65646 | 0.300257 | 1.96644 | 0.531571 |
| X | 71347200 | 71348599 | 22.11427 | 38.53271 | 38.70938 | 17.02625 | 0.380779 | 0.769467 | 0.531574 |
| 12 | 40449000 | 40453599 | 52.09567 | 82.89151 | 125.2614 | 52.32976 | -0.2595 | 2.224373 | 0.53158 |
| 6 | 131695800 | 131703199 | 154.8232 | 233.4477 | 236.5983 | 97.75968 | 0.253734 | 3.366225 | 0.531591 |
| 15 | 23989400 | 23991399 | 19.2333 | 34.43138 | 30.46973 | 13.36014 | 0.575574 | 0.456777 | 0.531599 |
| 7 | 47424000 | 47432799 | 119.1812 | 183.3846 | 179.6242 | 74.59906 | 0.326466 | 2.948926 | 0.5316 |
| 9 | 4773400 | 4776399 | 24.63824 | 41.16428 | 67.12149 | 29.21504 | -0.32202 | 1.209819 | 0.5316 |
| 2 | 67346200 | 67347599 | 10.84904 | 19.68477 | 33.30608 | 14.39068 | -0.33989 | 0.325778 | 0.531615 |
| 1 | 25948000 | 25949999 | 18.96073 | 32.60846 | 52.77215 | 23.14819 | -0.3076 | 0.909506 | 0.531618 |
| 13 | 19506600 | 19507599 | 9.685331 | 17.77164 | 31.88327 | 13.57884 | -0.40997 | 0.247422 | 0.531668 |
| 1 | 83966600 | 83967399 | 5.804882 | 11.29208 | 22.39768 | 8.272134 | -0.37089 | -0.31467 | 0.531718 |
| 3 | 137033800 | 137034599 | 5.804882 | 11.29208 | 22.39768 | 8.272134 | -0.37089 | -0.31467 | 0.531718 |
| 6 | 13330400 | 13334599 | 73.1054 | 115.4646 | 121.2251 | 50.34116 | 0.255482 | 2.438433 | 0.531735 |
| X | 70688800 | 70689399 | 12.07139 | 21.6481 | 37.32585 | 16.34217 | -0.37954 | 0.413896 | 0.531738 |
| 13 | 47960200 | 47961199 | 7.299134 | 14.07945 | 26.55334 | 10.26622 | -0.36219 | -0.12969 | 0.531746 |
| 17 | 15911400 | 15913399 | 20.55778 | 35.68106 | 36.34896 | 15.94884 | 0.360108 | 0.811036 | 0.531752 |
| 3 | 23739400 | 23742399 | 74.69632 | 117.7129 | 119.1983 | 49.46228 | 0.305021 | 2.487214 | 0.531762 |
| 17 | 5595800 | 5597599 | 24.2769 | 41.56204 | 43.18147 | 18.93585 | 0.328262 | 0.961644 | 0.53177 |
| 1 | 84672200 | 84675999 | 26.15868 | 42.3489 | 94.54244 | 40.44566 | -0.79238 | 1.654349 | 0.531774 |
| 4 | 27256400 | 27258199 | 18.76132 | 33.7496 | 29.37574 | 12.82793 | 0.603151 | 0.420883 | 0.531775 |
| 4 | 96213000 | 96215999 | 31.67368 | 53.38721 | 42.08221 | 18.39829 | 0.723153 | 1.121897 | 0.53178 |
| 6 | 142170400 | 142171199 | 13.22667 | 24.46454 | 26.01192 | 10.95837 | 0.35122 | 0.218561 | 0.531793 |
| 15 | 46764400 | 46771599 | 129.3188 | 198.0435 | 174.1721 | 72.28393 | 0.476526 | 3.043491 | 0.531803 |
| 7 | 126871600 | 126873199 | 13.79491 | 24.12053 | 42.07287 | 18.54874 | -0.41106 | 0.681725 | 0.531812 |
| 7 | 12387800 | 12401599 | 189.7869 | 284.7997 | 247.5506 | 102.1539 | 0.469052 | 3.466043 | 0.531837 |
| 12 | 21244600 | 21248799 | 25.97 | 42.85959 | 69.71472 | 30.09426 | -0.31943 | 1.352619 | 0.531851 |
| 14 | 112498400 | 112499199 | 10.31957 | 19.87077 | 21.28327 | 8.270186 | 0.439592 | -0.09933 | 0.531852 |
| 1 | 26459400 | 26460999 | 18.20204 | 31.94295 | 33.42163 | 14.78022 | 0.322483 | 0.631103 | 0.531852 |
| X | 53715000 | 53715399 | 13.48628 | 25.74989 | 18.90625 | 7.52135 | 0.949559 | 0.004083 | 0.53186 |
| 15 | 7917800 | 7919799 | 10.54019 | 18.92773 | 36.79201 | 16.09007 | -0.55077 | 0.388194 | 0.531868 |
| 8 | 59611200 | 59613399 | 24.74026 | 42.67209 | 42.51316 | 18.6498 | 0.390413 | 0.895826 | 0.531874 |
| 11 | 3640800 | 3642599 | 15.83883 | 28.25577 | 29.79991 | 13.10496 | 0.320951 | 0.453772 | 0.531887 |

| | | | | | | | | | |
|---|---|---|---|---|---|---|---|---|---|
| X | 127042000 | 127048199 | 71.5841 | 114.7222 | 87.20328 | 36.80404 | 0.741515 | 2.014793 | 0.531897 |
| 6 | 70552600 | 70553399 | 12.5828 | 23.2687 | 24.08435 | 10.14924 | 0.389632 | 0.210572 | 0.531911 |
| 6 | 35676600 | 35681799 | 47.12188 | 75.33491 | 113.7546 | 47.8395 | -0.25093 | 2.066214 | 0.531912 |
| 9 | 91868200 | 91869199 | 11.8251 | 22.54222 | 23.23738 | 9.275553 | 0.458998 | 0.007527 | 0.53192 |
| 15 | 93981200 | 93981999 | 11.8251 | 22.54222 | 23.23738 | 9.275553 | 0.458998 | 0.007527 | 0.53192 |
| 13 | 65696400 | 65699399 | 40.37721 | 67.7119 | 47.47898 | 20.71414 | 0.891569 | 1.205634 | 0.531924 |
| 2 | 84152400 | 84153799 | 13.78019 | 24.24963 | 39.81051 | 17.62543 | -0.32338 | 0.580383 | 0.531935 |
| 2 | 74199800 | 74202999 | 35.95902 | 58.8425 | 61.62309 | 26.50561 | 0.3091 | 1.478922 | 0.531961 |
| 7 | 33670600 | 33676599 | 85.66211 | 134.7894 | 134.6671 | 55.8702 | 0.322735 | 2.528572 | 0.531961 |
| 7 | 41747800 | 41758199 | 157.0472 | 238.7185 | 185.845 | 76.98129 | 0.643736 | 3.202483 | 0.531986 |
| 3 | 69588400 | 69589199 | 11.92893 | 22.32787 | 24.11738 | 9.974672 | 0.349153 | 0.132475 | 0.532002 |
| 5 | 75282200 | 75285999 | 34.66574 | 56.10442 | 87.66602 | 37.49563 | -0.27939 | 1.674217 | 0.532015 |
| 7 | 88286000 | 88286799 | 11.04912 | 21.16252 | 18.58123 | 7.331511 | 0.704222 | -0.04808 | 0.532038 |
| 1 | 91494600 | 91495399 | 5.975984 | 11.54702 | 23.71631 | 9.105077 | -0.47386 | -0.17602 | 0.532057 |
| X | 148723600 | 148725799 | 27.60444 | 46.94492 | 46.86549 | 20.54033 | 0.384194 | 1.038161 | 0.532064 |
| 18 | 46982800 | 46983999 | 13.28888 | 23.34075 | 40.36998 | 17.86593 | -0.39896 | 0.601978 | 0.532072 |
| 4 | 148843200 | 148845599 | 30.2139 | 50.21753 | 53.49567 | 23.20594 | 0.290171 | 1.270835 | 0.532086 |
| 1 | 152641600 | 152642999 | 9.65628 | 17.37897 | 34.31207 | 14.97659 | -0.57076 | 0.36482 | 0.532108 |
| X | 60809200 | 60813399 | 33.79009 | 56.92024 | 49.65424 | 21.70481 | 0.577507 | 1.143372 | 0.532131 |
| X | 24314600 | 24316399 | 28.70196 | 48.43635 | 49.62159 | 21.7024 | 0.346435 | 1.109842 | 0.532163 |
| 3 | 105834200 | 105836199 | 16.49056 | 28.61221 | 46.87386 | 20.56648 | -0.32234 | 0.811501 | 0.532166 |
| 17 | 39536200 | 39543199 | 90.92504 | 143.126 | 133.9339 | 55.56881 | 0.417499 | 2.523816 | 0.53218 |
| 4 | 116096600 | 116097999 | 12.01055 | 22.53759 | 23.44674 | 9.655852 | 0.407228 | 0.112483 | 0.532186 |
| 15 | 18326000 | 18326999 | 12.01055 | 22.53759 | 23.44674 | 9.655852 | 0.407228 | 0.112483 | 0.532186 |
| 9 | 111530200 | 111532599 | 35.08141 | 57.4129 | 58.26774 | 25.06342 | 0.354381 | 1.472458 | 0.532197 |
| 8 | 75027800 | 75028999 | 13.94691 | 25.96971 | 19.49395 | 8.133734 | 0.860503 | 0.171042 | 0.532216 |
| 17 | 87335200 | 87337599 | 23.66218 | 40.9503 | 40.83499 | 17.91546 | 0.389904 | 0.856444 | 0.532249 |
| 4 | 154729400 | 154734399 | 70.53709 | 110.6737 | 162.2984 | 67.30943 | -0.2334 | 2.579278 | 0.532257 |
| 9 | 110903400 | 110904999 | 24.06375 | 41.5716 | 41.98586 | 18.41938 | 0.371256 | 0.878022 | 0.532264 |
| 5 | 37888200 | 37896799 | 72.37535 | 112.8406 | 170.8586 | 70.90684 | -0.28664 | 2.692418 | 0.532289 |
| 8 | 6185600 | 6186599 | 11.29214 | 20.54436 | 34.89199 | 15.02488 | -0.34237 | 0.315639 | 0.532292 |
| 16 | 60652600 | 60653399 | 18.12867 | 31.79701 | 33.06943 | 14.61779 | 0.331278 | 0.640238 | 0.532305 |
| 5 | 17518400 | 17523199 | 62.67966 | 100.026 | 102.6533 | 43.06014 | 0.302355 | 2.132818 | 0.532353 |
| 7 | 37909000 | 37915599 | 46.66874 | 74.32306 | 120.1404 | 50.25262 | -0.35511 | 2.18824 | 0.532409 |
| 17 | 37016600 | 37019199 | 29.7316 | 48.40677 | 77.9149 | 33.45142 | -0.31446 | 1.548584 | 0.532414 |
| 5 | 143210200 | 143211599 | 11.41615 | 20.21559 | 36.78521 | 16.26568 | -0.4673 | 0.490939 | 0.532415 |
| 16 | 57696200 | 57698999 | 16.67759 | 29.05274 | 47.3496 | 20.82353 | -0.31427 | 0.758464 | 0.53242 |
| 14 | 90699000 | 90702199 | 41.71259 | 68.11745 | 66.7341 | 28.6825 | 0.402955 | 1.522921 | 0.532421 |
| 4 | 16246000 | 16249399 | 29.72636 | 48.38443 | 77.91826 | 33.44938 | -0.31549 | 1.552318 | 0.532423 |
| 3 | 8679400 | 8681599 | 14.31197 | 25.16338 | 41.37085 | 18.31946 | -0.32593 | 0.597063 | 0.53247 |
| 11 | 22931000 | 22932399 | 9.372858 | 17.20482 | 32.41778 | 13.79936 | -0.47982 | 0.243089 | 0.532503 |
| 17 | 50325000 | 50328199 | 18.59946 | 31.84482 | 52.75055 | 23.13221 | -0.34222 | 0.946704 | 0.532511 |
| 14 | 123007400 | 123008999 | 16.71696 | 31.04298 | 25.12053 | 10.52599 | 0.747719 | 0.221517 | 0.532524 |
| 15 | 56775600 | 56778199 | 18.07538 | 30.88453 | 62.5959 | 27.4451 | -0.63269 | 0.97537 | 0.532529 |
| 16 | 17377200 | 17380999 | 24.58908 | 40.46253 | 69.8548 | 30.11685 | -0.40668 | 1.385844 | 0.532533 |
| 8 | 101303200 | 101304999 | 20.43552 | 35.87131 | 35.27754 | 15.6042 | 0.411459 | 0.643753 | 0.532536 |
| 14 | 59252600 | 59256399 | 48.51814 | 78.17782 | 81.34992 | 34.60945 | 0.29867 | 1.814566 | 0.532537 |

| | | | | | | | | | |
|---|---|---|---|---|---|---|---|---|---|
| 7 | 38137200 | 38152799 | 283.6896 | 408.298 | 394.9099 | 162.9882 | 0.258054 | 4.154559 | 0.532566 |
| 11 | 111381800 | 111382999 | 13.57078 | 24.0119 | 40.77048 | 18.03075 | -0.36882 | 0.519225 | 0.532576 |
| 10 | 93629000 | 93631399 | 13.98131 | 24.57373 | 40.36007 | 17.87185 | -0.32457 | 0.597425 | 0.532582 |
| 1 | 192023000 | 192026799 | 34.06616 | 56.24936 | 58.70654 | 25.35108 | 0.31887 | 1.358357 | 0.532587 |
| 15 | 60803600 | 60805199 | 17.64853 | 31.93593 | 29.06395 | 12.60007 | 0.545981 | 0.379691 | 0.532587 |
| 6 | 121794600 | 121800199 | 74.56592 | 118.6697 | 108.0351 | 45.14629 | 0.470743 | 2.231778 | 0.53259 |
| 7 | 33573800 | 33591799 | 253.2322 | 370.0701 | 340.328 | 140.2082 | 0.352294 | 3.925296 | 0.532593 |
| 13 | 111797800 | 111799199 | 16.14044 | 28.77686 | 30.23773 | 13.30411 | 0.325697 | 0.459844 | 0.532607 |
| 2 | 177144000 | 177148999 | 76.12746 | 120.3083 | 125.1624 | 51.99166 | 0.269311 | 2.427971 | 0.532619 |
| 7 | 8575400 | 8577399 | 27.36387 | 46.57733 | 41.52785 | 18.20224 | 0.546786 | 1.021365 | 0.532626 |
| 13 | 15441200 | 15442399 | 17.12739 | 30.59539 | 22.88767 | 10.05386 | 0.814684 | 0.439547 | 0.53264 |
| 15 | 97361800 | 97363799 | 25.26676 | 41.53749 | 69.72649 | 30.04923 | -0.36683 | 1.401405 | 0.53264 |
| 8 | 40760600 | 40761799 | 11.27198 | 20.11653 | 35.31989 | 15.52737 | -0.40942 | 0.435138 | 0.532642 |
| 1 | 122376800 | 122378599 | 16.23899 | 29.45939 | 28.64812 | 12.38075 | 0.453682 | 0.353848 | 0.532643 |
| 7 | 10087800 | 10100599 | 195.3329 | 292.382 | 256.7944 | 105.91 | 0.450805 | 3.515112 | 0.53265 |
| 5 | 129095800 | 129098199 | 14.15568 | 24.64954 | 43.7278 | 19.22583 | -0.43576 | 0.747566 | 0.532669 |
| 16 | 77982400 | 77984799 | 26.32231 | 44.49689 | 47.46162 | 20.76853 | 0.288527 | 1.083928 | 0.532669 |
| 17 | 55738200 | 55740399 | 20.12784 | 35.66982 | 31.8733 | 14.09155 | 0.553366 | 0.527618 | 0.5327 |
| 15 | 94589400 | 94590999 | 18.82752 | 33.49021 | 30.86085 | 13.60474 | 0.512004 | 0.491495 | 0.532701 |
| 17 | 92077800 | 92079999 | 18.82752 | 33.49021 | 30.86085 | 13.60474 | 0.512004 | 0.491495 | 0.532701 |
| 10 | 81571800 | 81574399 | 51.42659 | 83.33513 | 63.00512 | 26.97281 | 0.767136 | 1.665871 | 0.532701 |
| 4 | 67074800 | 67077799 | 29.6748 | 49.50496 | 52.96242 | 23.03046 | 0.284085 | 1.227584 | 0.532701 |
| 1 | 158665800 | 158668199 | 19.97653 | 33.48408 | 64.49185 | 28.12792 | -0.56134 | 1.158975 | 0.532702 |
| 5 | 7229200 | 7230799 | 25.40813 | 41.86178 | 68.75857 | 29.65966 | -0.33427 | 1.373984 | 0.532715 |
| 7 | 25839000 | 25841399 | 22.94607 | 40.08622 | 36.2385 | 15.96552 | 0.53233 | 0.730566 | 0.532731 |
| 6 | 99668800 | 99672599 | 24.71889 | 40.3962 | 81.15556 | 34.89026 | -0.62952 | 1.484533 | 0.532732 |
| 10 | 105995000 | 105997399 | 29.82513 | 48.70448 | 76.72273 | 32.97551 | -0.28045 | 1.503445 | 0.532735 |
| 15 | 98354200 | 98355199 | 9.15743 | 16.70967 | 31.40249 | 13.47259 | -0.48328 | 0.277196 | 0.532736 |
| 14 | 120591800 | 120600999 | 85.15847 | 131.3254 | 196.3937 | 81.57849 | -0.28098 | 2.894903 | 0.532746 |
| 4 | 131880600 | 131881799 | 21.57318 | 37.40593 | 39.44431 | 17.30633 | 0.310359 | 0.831279 | 0.532749 |
| 3 | 4702600 | 4707399 | 62.32581 | 99.62875 | 99.33805 | 41.76887 | 0.346561 | 2.082236 | 0.532769 |
| 10 | 66191400 | 66192599 | 11.94079 | 21.43697 | 36.22929 | 15.84766 | -0.35027 | 0.400313 | 0.532786 |
| 6 | 53101600 | 53102599 | 8.750285 | 16.38632 | 29.03372 | 11.99803 | -0.35855 | 0.1149 | 0.532805 |
| 12 | 70924400 | 70925599 | 16.2135 | 28.0606 | 46.88509 | 20.56983 | -0.35132 | 0.833828 | 0.532806 |
| 12 | 104052600 | 104054199 | 10.71118 | 19.36146 | 33.2909 | 14.45215 | -0.36734 | 0.346825 | 0.532809 |
| X | 70925600 | 70928399 | 32.09722 | 53.40927 | 55.20564 | 23.96677 | 0.333671 | 1.265492 | 0.532834 |
| 14 | 113874200 | 113874999 | 16.4725 | 29.25074 | 30.73227 | 13.56773 | 0.322521 | 0.487935 | 0.532854 |
| 18 | 37422800 | 37425199 | 47.23865 | 77.31288 | 62.15926 | 26.73782 | 0.689593 | 1.501999 | 0.532855 |
| X | 152957800 | 152963599 | 41.83901 | 68.5542 | 56.57449 | 24.34713 | 0.652719 | 1.471971 | 0.532867 |
| 13 | 61194400 | 61196199 | 29.9435 | 50.69844 | 48.5071 | 21.2362 | 0.444828 | 1.084128 | 0.53287 |
| 18 | 81179000 | 81183599 | 42.62086 | 68.48728 | 106.6029 | 45.17307 | -0.28607 | 1.899552 | 0.53287 |
| 16 | 89930400 | 89935599 | 43.52495 | 69.59914 | 109.7367 | 46.16364 | -0.31281 | 2.054346 | 0.532883 |
| 1 | 24875600 | 24877799 | 22.20797 | 38.59542 | 39.8322 | 17.49149 | 0.341668 | 0.806969 | 0.532885 |
| 8 | 110101400 | 110103799 | 22.20797 | 38.59542 | 39.8322 | 17.49149 | 0.341668 | 0.806969 | 0.532885 |
| 17 | 26899400 | 26901199 | 15.8787 | 28.3721 | 29.81806 | 13.09533 | 0.327348 | 0.444772 | 0.532888 |
| 15 | 95805200 | 95805999 | 7.003254 | 13.42232 | 25.71537 | 10.1271 | -0.40971 | -0.07605 | 0.532894 |
| X | 19307800 | 19309399 | 24.52932 | 42.46889 | 35.24939 | 15.46509 | 0.653866 | 0.852784 | 0.53293 |

| | | | | | | | | | |
|---|---|---|---|---|---|---|---|---|---|
| 3 | 92070200 | 92071599 | 13.32218 | 25.10894 | 23.95622 | 9.792518 | 0.53975 | 0.098226 | 0.532933 |
| 8 | 46905200 | 46907799 | 48.28278 | 77.9636 | 78.57776 | 33.52245 | 0.34813 | 1.750737 | 0.532937 |
| 3 | 103322600 | 103324999 | 14.91892 | 26.12541 | 42.51122 | 18.76599 | -0.31134 | 0.664746 | 0.53295 |
| X | 122909200 | 122910999 | 18.91691 | 33.82434 | 30.39759 | 13.342 | 0.552311 | 0.460457 | 0.532962 |
| 18 | 57806600 | 57808399 | 11.88109 | 21.18326 | 35.76086 | 15.73393 | -0.35412 | 0.445127 | 0.533023 |
| 8 | 57793000 | 57798399 | 28.16236 | 45.71085 | 88.40767 | 37.89341 | -0.58242 | 1.60268 | 0.53303 |
| 5 | 130033000 | 130041599 | 110.852 | 168.6822 | 241.0407 | 99.87697 | -0.23046 | 3.17206 | 0.533037 |
| 1 | 158416000 | 158416599 | 8.633239 | 16.02836 | 29.46194 | 12.34864 | -0.42727 | 0.169938 | 0.533038 |
| X | 125500000 | 125501799 | 24.27452 | 42.45026 | 31.6733 | 13.96554 | 0.808222 | 0.719098 | 0.533076 |
| 1 | 49388400 | 49390199 | 24.48927 | 42.33306 | 41.93472 | 18.39729 | 0.399215 | 0.873451 | 0.533092 |
| 14 | 34250200 | 34253199 | 37.727 | 61.67387 | 65.22204 | 28.04338 | 0.294001 | 1.498204 | 0.533108 |
| 6 | 62491800 | 62492599 | 10.78877 | 20.7224 | 21.76184 | 8.517024 | 0.458182 | -0.07034 | 0.533124 |
| 9 | 18992600 | 19000599 | 95.23741 | 149.791 | 134.3038 | 55.71631 | 0.478205 | 2.542529 | 0.533125 |
| 17 | 93142200 | 93145199 | 28.81328 | 49.39003 | 44.03476 | 19.31198 | 0.548299 | 0.968005 | 0.533126 |
| 12 | 87920000 | 87920999 | 14.78445 | 26.57798 | 27.81236 | 12.15203 | 0.33843 | 0.394199 | 0.53314 |
| 19 | 36779400 | 36781199 | 21.92893 | 38.29903 | 33.39106 | 14.70855 | 0.584471 | 0.726986 | 0.533158 |
| X | 81229000 | 81230999 | 19.6107 | 35.70025 | 27.39586 | 11.78807 | 0.797545 | 0.357715 | 0.533164 |
| 10 | 30226200 | 30229799 | 40.75291 | 66.56945 | 66.2358 | 28.47132 | 0.380911 | 1.514577 | 0.533198 |
| X | 24567200 | 24591599 | 340.4005 | 487.7549 | 395.4841 | 163.3384 | 0.506026 | 4.224104 | 0.5332 |
| X | 87633000 | 87638399 | 87.20268 | 137.5639 | 116.4506 | 48.3289 | 0.564301 | 2.476118 | 0.533217 |
| 4 | 122433000 | 122438399 | 69.86268 | 111.3585 | 105.5499 | 44.20329 | 0.415071 | 2.178268 | 0.533223 |
| 17 | 91595200 | 91595799 | 6.556007 | 12.67409 | 24.30887 | 9.310799 | -0.37268 | -0.17415 | 0.533227 |
| 13 | 13391000 | 13392399 | 20.79768 | 36.16296 | 38.29636 | 16.82246 | 0.304653 | 0.789888 | 0.533228 |
| 3 | 55728000 | 55729399 | 16.33628 | 30.26493 | 26.28001 | 11.04707 | 0.643596 | 0.234502 | 0.533228 |
| 13 | 56794400 | 56796399 | 22.34653 | 38.60531 | 40.99423 | 17.98435 | 0.299216 | 0.868421 | 0.533258 |
| 7 | 61073600 | 61081599 | 99.50011 | 156.038 | 139.7365 | 57.95546 | 0.476713 | 2.602458 | 0.533286 |
| 12 | 115913000 | 115915199 | 40.9065 | 67.00513 | 64.11282 | 27.58761 | 0.439651 | 1.47616 | 0.533287 |
| 10 | 5689800 | 5691599 | 14.90977 | 26.13977 | 42.46559 | 18.76427 | -0.30905 | 0.645505 | 0.533294 |
| 7 | 48076000 | 48076999 | 16.80536 | 29.72547 | 31.29946 | 13.84309 | 0.316824 | 0.526128 | 0.533295 |
| 18 | 63845200 | 63846399 | 9.88715 | 18.1748 | 31.40429 | 13.34198 | -0.3532 | 0.240242 | 0.533296 |
| 6 | 35648200 | 35649999 | 18.35528 | 33.18992 | 28.37004 | 12.31079 | 0.63486 | 0.386469 | 0.533298 |
| 10 | 72450200 | 72451999 | 18.45086 | 32.45637 | 33.81749 | 14.97327 | 0.328892 | 0.600308 | 0.533303 |
| 18 | 21072400 | 21073599 | 13.3502 | 24.81546 | 25.38985 | 10.62457 | 0.412845 | 0.188501 | 0.533304 |
| 9 | 114973200 | 114989599 | 349.0324 | 482.9386 | 659.2334 | 273.803 | -0.29596 | 4.740653 | 0.533309 |
| 13 | 66208400 | 66213999 | 232.7448 | 338.8396 | 316.0477 | 130.2554 | 0.326582 | 3.979519 | 0.533371 |
| 4 | 138077800 | 138081399 | 26.38468 | 42.90991 | 84.77507 | 36.38199 | -0.61044 | 1.561411 | 0.533375 |
| 1 | 174134200 | 174134599 | 6.717946 | 12.8815 | 25.16931 | 9.898802 | -0.43572 | -0.0841 | 0.533392 |
| 8 | 27784800 | 27787599 | 22.98583 | 38.29933 | 63.58126 | 27.62674 | -0.34775 | 1.229397 | 0.53342 |
| 12 | 44777000 | 44779399 | 16.07147 | 28.10696 | 47.2867 | 20.85103 | -0.3594 | 0.698946 | 0.533422 |
| 4 | 66089600 | 66091199 | 14.1834 | 25.89617 | 26.758 | 11.47134 | 0.373995 | 0.297386 | 0.533431 |
| 4 | 112930400 | 112935399 | 68.47006 | 109.6092 | 92.90392 | 39.14381 | 0.582766 | 2.043963 | 0.533438 |
| 19 | 16864600 | 16866399 | 19.13666 | 34.13196 | 31.05935 | 13.66118 | 0.53259 | 0.480543 | 0.533443 |
| 3 | 59499800 | 59500999 | 8.245685 | 15.72871 | 28.98554 | 11.55456 | -0.37048 | -0.01421 | 0.533444 |
| 3 | 9645600 | 9646599 | 9.773798 | 17.81165 | 31.87275 | 13.69616 | -0.41467 | 0.290449 | 0.533454 |
| X | 108620000 | 108621399 | 15.6273 | 28.56461 | 27.85675 | 11.92557 | 0.458657 | 0.307493 | 0.533468 |
| 3 | 154405600 | 154408399 | 15.4897 | 27.13412 | 46.70849 | 20.62471 | -0.39247 | 0.665948 | 0.533506 |
| 10 | 33098400 | 33099799 | 10.17337 | 18.73018 | 31.93381 | 13.53812 | -0.33201 | 0.233981 | 0.533516 |

| | | | | | | | | | |
|---|---|---|---|---|---|---|---|---|---|
| 15 | 70072000 | 70073199 | 9.463696 | 17.61224 | 31.60336 | 13.19841 | -0.38969 | 0.163682 | 0.53353 |
| X | 103961000 | 103965999 | 57.43258 | 92.22551 | 89.13623 | 37.73357 | 0.39894 | 1.935534 | 0.533564 |
| X | 4383200 | 4392399 | 122.7161 | 190.2071 | 156.8324 | 65.143 | 0.582715 | 2.816204 | 0.533565 |
| 5 | 131535600 | 131543399 | 81.90673 | 125.6745 | 224.2072 | 93.07859 | -0.54083 | 2.995819 | 0.533576 |
| 7 | 42582200 | 42587199 | 10.02997 | 19.34244 | 17.58436 | 6.803198 | 0.680478 | -0.12543 | 0.53364 |
| 6 | 138014200 | 138015999 | 20.49434 | 35.64374 | 37.80991 | 16.6112 | 0.302305 | 0.784877 | 0.533646 |
| 2 | 70039600 | 70041399 | 28.44563 | 48.03281 | 42.66404 | 18.66288 | 0.551516 | 1.094521 | 0.533648 |
| 8 | 87863600 | 87864999 | 9.400202 | 17.38108 | 30.43475 | 12.82755 | -0.36385 | 0.200993 | 0.533677 |
| 12 | 9873600 | 9874399 | 9.400202 | 17.38108 | 30.43475 | 12.82755 | -0.36385 | 0.200993 | 0.533677 |
| 17 | 71904600 | 71908399 | 27.9929 | 45.88802 | 73.42615 | 31.60545 | -0.29968 | 1.444362 | 0.533683 |
| 1 | 93048800 | 93049999 | 10.186 | 18.71103 | 31.90323 | 13.56688 | -0.33527 | 0.247042 | 0.533686 |
| 1 | 111361400 | 111368599 | 78.47855 | 124.3344 | 121.442 | 50.51012 | 0.363067 | 2.369584 | 0.533691 |
| 12 | 99444400 | 99445999 | 18.69944 | 32.88985 | 33.73075 | 14.93521 | 0.351469 | 0.601802 | 0.533725 |
| 15 | 49296200 | 49299199 | 30.14609 | 49.11687 | 78.96092 | 33.90939 | -0.31205 | 1.536928 | 0.533741 |
| X | 23847000 | 23850799 | 40.19472 | 66.94118 | 52.15644 | 22.65573 | 0.740171 | 1.276563 | 0.533744 |
| 11 | 21938600 | 21939999 | 16.25838 | 29.18654 | 29.77847 | 13.02653 | 0.373762 | 0.419018 | 0.533747 |
| 10 | 118767400 | 118769999 | 29.90636 | 50.05546 | 52.39739 | 22.83204 | 0.315352 | 1.194223 | 0.533762 |
| 4 | 121273200 | 121286999 | 211.624 | 313.5165 | 295.6546 | 121.7621 | 0.334655 | 3.701738 | 0.533776 |
| 8 | 61676400 | 61678199 | 12.06552 | 21.22965 | 38.87899 | 17.21344 | -0.48022 | 0.569469 | 0.533777 |
| 15 | 17429000 | 17429399 | 8.006463 | 15.56011 | 16.82474 | 6.26126 | 0.485915 | -0.27965 | 0.533777 |
| 16 | 29974200 | 29978799 | 49.89382 | 79.41702 | 123.5735 | 51.65066 | -0.30107 | 2.208527 | 0.533782 |
| 2 | 151284000 | 151284799 | 12.11284 | 23.08132 | 23.20206 | 9.269453 | 0.493829 | 0.012531 | 0.533802 |
| 7 | 37989000 | 37989999 | 12.11284 | 23.08132 | 23.20206 | 9.269453 | 0.493829 | 0.012531 | 0.533802 |
| 10 | 129502000 | 129504199 | 23.2689 | 40.54818 | 38.54649 | 16.95555 | 0.459945 | 0.771663 | 0.533895 |
| 13 | 72086600 | 72093199 | 59.58826 | 94.397 | 142.1816 | 59.13317 | -0.26105 | 2.360982 | 0.5339 |
| 16 | 79169000 | 79173799 | 78.31897 | 123.1677 | 127.8659 | 53.04476 | 0.266885 | 2.533038 | 0.533905 |
| 6 | 137921800 | 137923999 | 19.0786 | 32.25964 | 64.14069 | 28.06916 | -0.60676 | 1.073121 | 0.533917 |
| 4 | 46574200 | 46579599 | 86.0187 | 134.6438 | 142.1099 | 58.95114 | 0.237844 | 2.626724 | 0.533917 |
| 7 | 46957800 | 46977999 | 308.1948 | 440.3231 | 423.4116 | 174.9526 | 0.255884 | 4.263194 | 0.533923 |
| 11 | 4613800 | 4619799 | 87.55186 | 134.7378 | 200.4141 | 83.23507 | -0.27549 | 2.936115 | 0.533945 |
| 1 | 166952600 | 166954999 | 20.80502 | 35.01654 | 59.21143 | 25.87571 | -0.3742 | 1.113666 | 0.533947 |
| 1 | 16137200 | 16138199 | 17.12905 | 30.16857 | 29.82831 | 13.20373 | 0.404667 | 0.570063 | 0.533971 |
| 8 | 13576400 | 13581399 | 36.10148 | 57.81861 | 114.5715 | 48.30602 | -0.63932 | 1.999841 | 0.533972 |
| 4 | 22542000 | 22542999 | 20.52376 | 35.84898 | 36.68679 | 16.16146 | 0.354559 | 0.725486 | 0.533976 |
| X | 108985600 | 108987199 | 20.52376 | 35.84898 | 36.68679 | 16.16146 | 0.354559 | 0.725486 | 0.533976 |
| 14 | 4209000 | 4215799 | 90.16691 | 140.6213 | 148.0372 | 61.43315 | 0.237453 | 2.692599 | 0.533997 |
| 16 | 54041400 | 54042599 | 16.88694 | 30.18078 | 26.18973 | 11.49956 | 0.602224 | 0.441973 | 0.533999 |
| 4 | 16055800 | 16058799 | 41.44618 | 67.21709 | 70.49262 | 30.19643 | 0.297293 | 1.63588 | 0.534009 |
| 13 | 75790200 | 75793399 | 43.75159 | 71.32363 | 60.51543 | 25.99329 | 0.608128 | 1.547138 | 0.534015 |
| Y | 84475400 | 84477799 | 13.9709 | 25.46254 | 19.12053 | 8.215486 | 0.829425 | 0.287444 | 0.534025 |
| 18 | 13080000 | 13081199 | 9.386935 | 17.51107 | 30.57098 | 12.71585 | -0.34512 | 0.148674 | 0.534043 |
| 3 | 83291400 | 83293799 | 19.45772 | 34.47513 | 31.30409 | 13.84047 | 0.530046 | 0.522918 | 0.534044 |
| 3 | 97300000 | 97302599 | 17.60242 | 30.35769 | 50.07488 | 21.96693 | -0.33413 | 0.873676 | 0.534097 |
| 4 | 65915400 | 65916399 | 11.16184 | 21.65265 | 19.81563 | 7.468481 | 0.7059 | -0.19357 | 0.534104 |
| 12 | 22868400 | 22871799 | 48.91941 | 79.6618 | 57.97074 | 24.89043 | 0.827952 | 1.572263 | 0.53412 |
| 19 | 11416600 | 11418599 | 12.51313 | 22.13419 | 37.2595 | 16.47896 | -0.35623 | 0.511634 | 0.534139 |
| 15 | 47325600 | 47326199 | 8.923653 | 16.66609 | 29.52392 | 12.2563 | -0.36354 | 0.13532 | 0.534141 |

| | | | | | | | | | |
|---|---|---|---|---|---|---|---|---|---|
| 8 | 32497800 | 32499999 | 16.26777 | 27.57066 | 58.94349 | 25.8129 | -0.71049 | 1.040391 | 0.534142 |
| 12 | 49288800 | 49292599 | 33.73715 | 57.0108 | 47.39523 | 20.73721 | 0.646776 | 1.112394 | 0.534152 |
| X | 152941400 | 152944799 | 59.9661 | 96.21412 | 90.43286 | 38.23427 | 0.437718 | 1.965027 | 0.534156 |
| X | 156309800 | 156314999 | 78.59161 | 124.5944 | 119.5815 | 49.76444 | 0.38919 | 2.352586 | 0.534159 |
| 16 | 95738400 | 95747399 | 93.45614 | 142.9634 | 238.1799 | 98.82275 | -0.44562 | 3.059744 | 0.534173 |
| 1 | 48810200 | 48812799 | 26.64561 | 44.99252 | 48.00168 | 20.99765 | 0.288121 | 1.095133 | 0.534174 |
| 13 | 96017200 | 96018799 | 11.58629 | 20.97204 | 36.37305 | 15.76309 | -0.37863 | 0.348903 | 0.53418 |
| 12 | 37276400 | 37277799 | 15.94181 | 28.93279 | 28.7023 | 12.39744 | 0.425651 | 0.349698 | 0.534187 |
| 18 | 37118000 | 37122799 | 112.577 | 172.6215 | 168.1481 | 69.7997 | 0.330636 | 3.01356 | 0.534217 |
| 18 | 42067600 | 42070199 | 42.47196 | 69.72696 | 54.52154 | 23.48297 | 0.731308 | 1.446215 | 0.534223 |
| 7 | 10151400 | 10155599 | 26.22666 | 45.43268 | 35.36039 | 15.51406 | 0.746146 | 0.852666 | 0.534249 |
| 2 | 138746400 | 138748199 | 17.89721 | 31.7603 | 31.52959 | 13.92725 | 0.403257 | 0.501787 | 0.534251 |
| 3 | 91871000 | 91872199 | 17.38399 | 30.97625 | 31.18536 | 13.72768 | 0.386711 | 0.474442 | 0.534277 |
| 5 | 127315600 | 127323399 | 74.88877 | 116.5999 | 173.8235 | 72.15198 | -0.26574 | 2.717672 | 0.534299 |
| X | 73876400 | 73876599 | 6.693479 | 13.07729 | 15.36181 | 5.356076 | 0.452703 | -0.52803 | 0.534316 |
| X | 125227400 | 125230399 | 44.46021 | 72.48616 | 67.38381 | 28.94455 | 0.476856 | 1.553282 | 0.534316 |
| 3 | 91429400 | 91435199 | 100.8324 | 157.3822 | 148.7623 | 61.72885 | 0.393592 | 2.683692 | 0.534332 |
| X | 148801200 | 148804199 | 72.88272 | 115.2395 | 114.9104 | 47.74541 | 0.331019 | 2.412744 | 0.534333 |
| 9 | 6626600 | 6629199 | 34.26111 | 56.70124 | 51.45831 | 22.25634 | 0.520208 | 1.328794 | 0.53435 |
| 15 | 33444200 | 33445399 | 13.72325 | 25.71957 | 24.97037 | 10.3049 | 0.504003 | 0.139781 | 0.534361 |
| 13 | 38883200 | 38883599 | 4.348586 | 8.500225 | 18.32522 | 6.310042 | -0.39633 | -0.59611 | 0.534365 |
| 18 | 84066200 | 84067399 | 7.329119 | 13.92233 | 26.60526 | 10.6994 | -0.43341 | -0.00135 | 0.534374 |
| 13 | 94161200 | 94161999 | 9.372119 | 17.33235 | 30.42502 | 12.82023 | -0.36733 | 0.198515 | 0.534383 |
| 5 | 129031600 | 129044799 | 35.62904 | 58.90807 | 52.58198 | 22.72714 | 0.543977 | 1.344938 | 0.534383 |
| 9 | 63747800 | 63749199 | 12.59121 | 22.0562 | 40.5742 | 17.91572 | -0.48707 | 0.645266 | 0.534413 |
| 7 | 31573000 | 31587399 | 188.4209 | 281.3816 | 268.2439 | 110.5823 | 0.329584 | 3.55708 | 0.534421 |
| 1 | 159202400 | 159204799 | 18.25621 | 32.18122 | 33.23751 | 14.71032 | 0.3426 | 0.575089 | 0.534425 |
| 7 | 20774800 | 20777799 | 26.46395 | 45.84992 | 40.72845 | 17.86931 | 0.556439 | 0.86078 | 0.534433 |
| 10 | 9984400 | 9985599 | 10.28733 | 18.88864 | 32.93759 | 14.01534 | -0.36845 | 0.250442 | 0.534452 |
| 12 | 42063200 | 42064799 | 15.47968 | 27.82993 | 29.24807 | 12.77851 | 0.332331 | 0.401911 | 0.534466 |
| 17 | 19755400 | 19769599 | 44.24855 | 71.98631 | 62.15071 | 26.67579 | 0.581018 | 1.577887 | 0.53448 |
| 9 | 4400000 | 4408799 | 117.5897 | 181.4842 | 172.0887 | 71.4823 | 0.37655 | 2.891434 | 0.534485 |
| 11 | 71885400 | 71886999 | 24.82641 | 42.14883 | 44.28822 | 19.40516 | 0.310311 | 1.038514 | 0.534487 |
| 9 | 41221800 | 41223799 | 21.7242 | 38.01086 | 36.91617 | 16.28467 | 0.429554 | 0.703724 | 0.534489 |
| 14 | 44693200 | 44701399 | 100.587 | 154.1457 | 222.0306 | 92.15937 | -0.23385 | 3.024876 | 0.53449 |
| 1 | 58978200 | 58983399 | 85.53721 | 134.2272 | 140.0389 | 58.07825 | 0.257335 | 2.581084 | 0.534518 |
| 8 | 95093600 | 95095599 | 15.72741 | 27.33935 | 45.28316 | 19.88816 | -0.33764 | 0.781849 | 0.534523 |
| X | 24016000 | 24018799 | 63.00541 | 100.7565 | 96.63526 | 40.65606 | 0.403165 | 2.071906 | 0.53454 |
| 8 | 52386000 | 52386999 | 18.60142 | 32.63023 | 34.46808 | 15.23763 | 0.308864 | 0.642649 | 0.534585 |
| 17 | 7850200 | 7851999 | 18.60142 | 32.63023 | 34.46808 | 15.23763 | 0.308864 | 0.642649 | 0.534585 |
| 4 | 57428400 | 57431399 | 17.9692 | 31.15549 | 51.73192 | 22.69752 | -0.34204 | 0.831028 | 0.534586 |
| 5 | 38630800 | 38635599 | 45.1434 | 72.08607 | 113.8339 | 47.79226 | -0.31729 | 2.099619 | 0.534586 |
| 13 | 42276800 | 42282199 | 60.73694 | 97.73435 | 82.12272 | 34.87007 | 0.604122 | 1.873747 | 0.534593 |
| 5 | 62679000 | 62680799 | 19.85929 | 33.48729 | 56.97346 | 24.91182 | -0.3828 | 1.092889 | 0.534618 |
| 13 | 6827800 | 6828999 | 14.12849 | 26.02231 | 26.89814 | 11.39712 | 0.385538 | 0.249987 | 0.534625 |
| 6 | 118612000 | 118615799 | 45.60325 | 73.00162 | 110.6457 | 46.61758 | -0.25428 | 2.023155 | 0.534625 |
| 19 | 8140600 | 8145599 | 53.66216 | 86.21873 | 86.27198 | 36.55119 | 0.349898 | 1.908784 | 0.53465 |

| | | | | | | | | | |
|---|---|---|---|---|---|---|---|---|---|
| 19 | 40098000 | 40099399 | 19.48735 | 33.24961 | 53.25878 | 23.34856 | -0.29465 | 0.984571 | 0.534662 |
| 4 | 96441400 | 96441999 | 6.117897 | 11.81976 | 23.71223 | 9.108794 | -0.44096 | -0.17189 | 0.534679 |
| 3 | 12727600 | 12729399 | 32.14991 | 53.05157 | 56.2725 | 24.28961 | 0.295544 | 1.35985 | 0.534683 |
| 7 | 31864000 | 31868199 | 58.69507 | 93.98156 | 95.5731 | 40.2815 | 0.320801 | 2.026724 | 0.534703 |
| 9 | 72848600 | 72853199 | 43.48195 | 69.6428 | 107.2254 | 45.21261 | -0.27597 | 2.003164 | 0.534719 |
| 1 | 153058600 | 153059999 | 13.25968 | 23.49732 | 39.20358 | 17.33249 | -0.34271 | 0.49839 | 0.534728 |
| 14 | 21774800 | 21776999 | 13.25968 | 23.49732 | 39.20358 | 17.33249 | -0.34271 | 0.49839 | 0.534728 |
| 5 | 132478600 | 132479199 | 5.953511 | 11.57181 | 22.90203 | 8.514015 | -0.37641 | -0.28746 | 0.534729 |
| 1 | 24197400 | 24199999 | 33.12442 | 55.03206 | 56.43578 | 24.47429 | 0.344801 | 1.285164 | 0.534749 |
| 7 | 48140600 | 48143199 | 31.37467 | 52.06916 | 55.73117 | 24.15082 | 0.283198 | 1.29286 | 0.534751 |
| 1 | 8795200 | 8797799 | 29.93024 | 51.21863 | 44.00877 | 19.29797 | 0.601031 | 0.986706 | 0.534774 |
| 7 | 58632000 | 58636799 | 77.01267 | 121.8811 | 123.9294 | 51.51668 | 0.303916 | 2.394049 | 0.534782 |
| 8 | 115559000 | 115563599 | 34.66565 | 55.89039 | 90.72016 | 38.62658 | -0.34034 | 1.779109 | 0.534803 |
| X | 32546800 | 32553799 | 169.3763 | 256.6842 | 212.7491 | 88.04561 | 0.549907 | 3.269458 | 0.53482 |
| 7 | 85598000 | 85598999 | 12.59499 | 23.48889 | 25.08563 | 10.4393 | 0.35825 | 0.163528 | 0.534857 |
| 4 | 107189400 | 107190999 | 26.66314 | 45.08923 | 47.98957 | 21.00184 | 0.291651 | 1.082725 | 0.534862 |
| 2 | 12941200 | 12945599 | 32.19563 | 53.82102 | 53.07574 | 23.10974 | 0.401103 | 1.212128 | 0.534874 |
| 1 | 14103800 | 14104799 | 16.88725 | 29.1736 | 47.93853 | 21.03084 | -0.3279 | 0.855519 | 0.534876 |
| 5 | 44257800 | 44259399 | 13.8632 | 24.08381 | 44.80711 | 19.67141 | -0.50446 | 0.781338 | 0.534882 |
| 8 | 112133200 | 112135999 | 29.51205 | 48.15018 | 76.5625 | 32.89593 | -0.29481 | 1.515555 | 0.534883 |
| 1 | 12034800 | 12036199 | 11.64373 | 21.11691 | 35.89678 | 15.51902 | -0.34737 | 0.33914 | 0.534897 |
| X | 34917400 | 34921599 | 51.71469 | 83.77955 | 71.55831 | 30.62842 | 0.59131 | 1.678635 | 0.534902 |
| 14 | 56846200 | 56847999 | 18.1993 | 32.02126 | 33.404 | 14.78951 | 0.327409 | 0.596795 | 0.53491 |
| 15 | 96154800 | 96169399 | 234.2055 | 337.991 | 463.8197 | 191.3593 | -0.24257 | 4.112805 | 0.534949 |
| 16 | 12983800 | 12986799 | 44.76156 | 72.88806 | 68.52726 | 29.42257 | 0.459356 | 1.571323 | 0.53495 |
| X | 25377400 | 25381999 | 53.98622 | 87.81986 | 63.49981 | 27.2539 | 0.836202 | 1.60226 | 0.534967 |
| 1 | 54657600 | 54658599 | 9.396761 | 17.49206 | 30.52813 | 12.74396 | -0.34881 | 0.162598 | 0.534968 |
| 14 | 69398200 | 69400999 | 68.59947 | 109.2206 | 108.5348 | 45.38174 | 0.34522 | 2.212739 | 0.534995 |
| 3 | 111881400 | 111882399 | 18.84585 | 33.07623 | 34.33882 | 15.18716 | 0.333627 | 0.634325 | 0.534998 |
| 4 | 32036000 | 32037599 | 22.0222 | 37.96513 | 40.03162 | 17.56092 | 0.30876 | 0.884724 | 0.535019 |
| 9 | 25702200 | 25703799 | 21.8138 | 37.87567 | 39.86561 | 17.49608 | 0.313265 | 0.81725 | 0.535029 |
| 2 | 136968000 | 136971199 | 40.64131 | 67.16143 | 57.9557 | 25.04109 | 0.592583 | 1.367413 | 0.535051 |
| 4 | 126949600 | 126951799 | 15.04688 | 26.37713 | 42.97775 | 18.98875 | -0.31336 | 0.648176 | 0.535082 |
| 7 | 3239400 | 3244199 | 56.87589 | 90.78024 | 93.14138 | 39.07995 | 0.302998 | 2.118489 | 0.535092 |
| 10 | 88344800 | 88346999 | 12.17331 | 21.58402 | 36.70863 | 16.22676 | -0.36966 | 0.482015 | 0.535095 |
| 11 | 82095800 | 82097199 | 12.17331 | 21.58402 | 36.70863 | 16.22676 | -0.36966 | 0.482015 | 0.535095 |
| 10 | 120179600 | 120181799 | 19.55031 | 34.13162 | 36.1867 | 15.93616 | 0.303586 | 0.724756 | 0.535102 |
| 14 | 63176400 | 63178599 | 19.55031 | 34.13162 | 36.1867 | 15.93616 | 0.303586 | 0.724756 | 0.535102 |
| 7 | 116615200 | 116616199 | 15.10495 | 26.3011 | 44.2259 | 19.44417 | -0.35897 | 0.754102 | 0.535113 |
| 17 | 64537600 | 64539199 | 13.28344 | 24.35082 | 25.90787 | 11.04815 | 0.337518 | 0.267022 | 0.535128 |
| 10 | 54581000 | 54581999 | 15.95082 | 29.20951 | 27.74054 | 11.8468 | 0.498858 | 0.297659 | 0.53513 |
| 10 | 18890200 | 18891999 | 13.02992 | 22.74851 | 43.66569 | 19.23003 | -0.54868 | 0.70194 | 0.53513 |
| 2 | 57314400 | 57316399 | 21.28561 | 35.95325 | 57.59758 | 25.19766 | -0.29626 | 1.084157 | 0.535146 |
| 7 | 38266400 | 38275399 | 149.2202 | 225.4058 | 230.2725 | 95.20011 | 0.24448 | 3.328561 | 0.535147 |
| 2 | 8790400 | 8792999 | 27.76794 | 47.44453 | 40.41598 | 17.7203 | 0.61312 | 0.988733 | 0.535152 |
| 15 | 64533000 | 64533999 | 9.837083 | 17.99893 | 32.88341 | 14.05899 | -0.44022 | 0.266069 | 0.535154 |
| 11 | 65316800 | 65319399 | 19.62325 | 33.58205 | 53.83613 | 23.60737 | -0.29524 | 0.957958 | 0.535155 |

| | | | | | | | | | |
|---|---|---|---|---|---|---|---|---|---|
| 9 | 96245600 | 96247799 | 14.74922 | 25.65752 | 44.75363 | 19.66455 | -0.41185 | 0.764659 | 0.535171 |
| 16 | 78270400 | 78271799 | 11.4848 | 20.80205 | 34.81906 | 15.0772 | -0.32662 | 0.343401 | 0.535178 |
| 18 | 68633400 | 68636399 | 35.10909 | 56.83411 | 88.01874 | 37.65243 | -0.26632 | 1.669442 | 0.535194 |
| 11 | 6706600 | 6706999 | 7.808781 | 14.29104 | 34.02265 | 14.54149 | -0.81816 | 0.256019 | 0.535204 |
| 3 | 42770200 | 42776399 | 62.23498 | 99.38589 | 101.5369 | 42.63409 | 0.309963 | 2.111401 | 0.535223 |
| X | 116595000 | 116595799 | 17.88469 | 31.7443 | 30.33796 | 13.39867 | 0.458423 | 0.502558 | 0.535231 |
| 1 | 20842400 | 20844999 | 39.67138 | 65.00648 | 64.48773 | 27.75336 | 0.387927 | 1.470189 | 0.535252 |
| X | 66474400 | 66475599 | 26.87957 | 45.82514 | 44.75441 | 19.61898 | 0.416342 | 1.01264 | 0.535264 |
| 5 | 120191800 | 120194599 | 29.50604 | 48.19148 | 76.08483 | 32.70334 | -0.28349 | 1.500206 | 0.535264 |
| 7 | 14202200 | 14207799 | 65.57288 | 105.3517 | 83.95003 | 35.56553 | 0.677781 | 1.930473 | 0.535269 |
| 7 | 7164800 | 7168999 | 45.69853 | 73.83653 | 76.97765 | 32.86611 | 0.300491 | 1.728822 | 0.535298 |
| 16 | 63217600 | 63218799 | 8.733753 | 16.32252 | 28.99747 | 12.02391 | -0.36646 | 0.126587 | 0.535315 |
| 13 | 23579400 | 23586599 | 64.29327 | 101.4536 | 149.9595 | 62.24488 | -0.23844 | 2.458606 | 0.535319 |
| 17 | 17878000 | 17879599 | 10.60483 | 19.4934 | 34.04685 | 14.46533 | -0.36923 | 0.249121 | 0.535376 |
| 17 | 38766200 | 38801399 | 509.049 | 702.7834 | 568.4707 | 236.1895 | 0.455544 | 4.779232 | 0.535383 |
| 7 | 143186400 | 143187999 | 21.18951 | 37.06493 | 36.95406 | 16.29753 | 0.391873 | 0.704996 | 0.535384 |
| 5 | 83262400 | 83264599 | 18.5742 | 32.01367 | 52.80306 | 23.16329 | -0.33431 | 0.886765 | 0.535397 |
| 12 | 115346000 | 115349199 | 63.92907 | 101.9117 | 105.7053 | 44.27502 | 0.285433 | 2.169218 | 0.535415 |
| 2 | 143946000 | 143947599 | 15.88567 | 27.45227 | 47.41244 | 20.80025 | -0.39937 | 0.84551 | 0.535446 |
| 5 | 113997400 | 114000799 | 22.69363 | 37.84446 | 61.8399 | 26.88444 | -0.32493 | 1.219507 | 0.535456 |
| 2 | 3553800 | 3554799 | 8.683153 | 16.29805 | 29.61764 | 12.19175 | -0.39082 | 0.097971 | 0.535474 |
| 3 | 31641600 | 31642599 | 9.968595 | 18.48755 | 32.06952 | 13.46138 | -0.34658 | 0.190473 | 0.535476 |
| 12 | 44528400 | 44531399 | 27.34505 | 44.99813 | 71.23171 | 30.71063 | -0.28195 | 1.395164 | 0.535481 |
| 7 | 9481800 | 9482399 | 20.26818 | 35.57263 | 26.0575 | 11.52458 | 0.834363 | 0.629501 | 0.535489 |
| 1 | 100869600 | 100872199 | 23.13417 | 40.36979 | 37.97109 | 16.71713 | 0.475397 | 0.751183 | 0.535491 |
| 9 | 99869800 | 99871599 | 25.2754 | 42.11209 | 68.29093 | 29.67218 | -0.31411 | 1.241424 | 0.535492 |
| 2 | 122340400 | 122341199 | 12.25972 | 22.78958 | 24.09372 | 10.08152 | 0.366558 | 0.178974 | 0.535496 |
| 3 | 3488600 | 3504799 | 58.50218 | 93.84617 | 82.05445 | 34.6812 | 0.541438 | 1.963662 | 0.535511 |
| 7 | 123155600 | 123160599 | 55.07366 | 87.48291 | 131.1115 | 54.67056 | -0.25005 | 2.27908 | 0.535517 |
| 7 | 28822200 | 28825999 | 23.73275 | 39.32648 | 64.99894 | 28.13723 | -0.34182 | 1.298069 | 0.535519 |
| 14 | 4137600 | 4138799 | 7.211097 | 13.72593 | 26.63837 | 10.6669 | -0.44925 | -0.01276 | 0.535528 |
| 12 | 54872000 | 54873999 | 13.62082 | 24.03894 | 39.26589 | 17.37422 | -0.31447 | 0.546724 | 0.535547 |
| 7 | 36319000 | 36322399 | 32.03017 | 53.05075 | 56.85465 | 24.60366 | 0.281138 | 1.318209 | 0.535555 |
| 18 | 7069600 | 7072599 | 68.02738 | 107.9014 | 115.1357 | 47.93869 | 0.237239 | 2.327168 | 0.535585 |
| 10 | 79826800 | 79829199 | 23.12353 | 38.26732 | 65.38555 | 28.27926 | -0.38971 | 1.31261 | 0.535589 |
| 8 | 11611200 | 11612199 | 11.30302 | 20.01516 | 36.37138 | 16.08271 | -0.46501 | 0.492374 | 0.535597 |
| 11 | 17860000 | 17863999 | 38.6331 | 62.81463 | 66.74581 | 28.64206 | 0.281654 | 1.576114 | 0.535603 |
| 4 | 122470400 | 122470999 | 12.85381 | 23.8082 | 24.87716 | 10.46037 | 0.37772 | 0.202627 | 0.535636 |
| 11 | 27338800 | 27341199 | 28.86524 | 48.78141 | 43.70631 | 19.12357 | 0.539135 | 1.091108 | 0.535662 |
| 1 | 10536200 | 10539599 | 98.27899 | 152.087 | 158.5049 | 65.83897 | 0.243303 | 2.83274 | 0.535664 |
| 3 | 62291800 | 62294199 | 18.60707 | 32.62084 | 34.51612 | 15.25124 | 0.306486 | 0.652356 | 0.535673 |
| 12 | 39904400 | 39905199 | 7.744686 | 14.84791 | 27.46817 | 10.80782 | -0.35876 | -0.06681 | 0.535699 |
| 17 | 39257800 | 39262799 | 66.13657 | 106.106 | 88.02982 | 37.21377 | 0.617297 | 1.974677 | 0.535707 |
| 10 | 32898200 | 32898799 | 6.096494 | 11.84118 | 23.46351 | 8.773697 | -0.38683 | -0.26665 | 0.535719 |
| 6 | 11638000 | 11642799 | 46.78254 | 76.31691 | 65.55217 | 28.16376 | 0.591169 | 1.54684 | 0.535721 |
| 6 | 141155200 | 141157399 | 20.83584 | 35.12816 | 56.99301 | 24.91912 | -0.31452 | 1.098787 | 0.535724 |
| 1 | 149140600 | 149140999 | 9.746398 | 18.62559 | 19.69984 | 7.822471 | 0.429137 | -0.03863 | 0.535726 |

| | | | | | | | | | |
|---|---|---|---|---|---|---|---|---|---|
| 7 | 73485800 | 73486799 | 11.49993 | 21.59396 | 23.57998 | 9.69965 | 0.339521 | 0.105654 | 0.535737 |
| 7 | 3234600 | 3238599 | 34.02513 | 55.96122 | 60.15007 | 25.92715 | 0.27464 | 1.408482 | 0.535753 |
| 16 | 35323200 | 35325199 | 24.78757 | 42.08734 | 44.2757 | 19.4003 | 0.308594 | 1.036833 | 0.535755 |
| 8 | 48201600 | 48203799 | 16.62167 | 28.85659 | 47.39894 | 20.80052 | -0.32606 | 0.80533 | 0.535793 |
| 8 | 31493800 | 31495199 | 14.33441 | 25.10903 | 42.03182 | 18.55886 | -0.35184 | 0.657947 | 0.535797 |
| 5 | 142040800 | 142058199 | 267.1604 | 380.257 | 512.4496 | 211.8773 | -0.23636 | 4.316854 | 0.535799 |
| 7 | 80926600 | 80927199 | 6.847018 | 13.14843 | 25.78196 | 10.09679 | -0.43549 | -0.09661 | 0.535809 |
| 14 | 51525600 | 51526199 | 6.847018 | 13.14843 | 25.78196 | 10.09679 | -0.43549 | -0.09661 | 0.535809 |
| 7 | 23337600 | 23351799 | 202.0321 | 302.1023 | 258.1861 | 106.463 | 0.48884 | 3.535007 | 0.535815 |
| 4 | 5258200 | 5266199 | 112.3854 | 172.9324 | 178.6648 | 74.20091 | 0.249623 | 2.945078 | 0.535822 |
| 12 | 111925000 | 111926999 | 21.84205 | 36.56081 | 60.24093 | 26.25116 | -0.33694 | 1.174445 | 0.535823 |
| 2 | 110399000 | 110399599 | 8.50754 | 16.46367 | 18.24023 | 6.952994 | 0.417754 | -0.19034 | 0.535849 |
| 3 | 92476000 | 92477199 | 11.62904 | 21.78514 | 23.53758 | 9.720811 | 0.350069 | 0.121106 | 0.535856 |
| 7 | 41795400 | 41803799 | 110.0355 | 171.4076 | 151.2463 | 62.77584 | 0.490562 | 2.724377 | 0.53588 |
| 5 | 114360200 | 114368599 | 115.2943 | 174.2543 | 257.2297 | 106.3633 | -0.28549 | 3.311827 | 0.535897 |
| 12 | 24739200 | 24744399 | 51.71089 | 82.29315 | 124.4461 | 52.00128 | -0.26028 | 2.217284 | 0.535915 |
| 6 | 97911000 | 97911599 | 8.394375 | 16.37467 | 17.39732 | 6.22802 | 0.559627 | -0.41142 | 0.535933 |
| 9 | 19898400 | 19902199 | 53.45726 | 86.15069 | 81.84146 | 34.82701 | 0.430099 | 1.816723 | 0.535961 |
| 15 | 30316600 | 30319199 | 22.03713 | 38.95123 | 32.32126 | 14.29708 | 0.658322 | 0.568412 | 0.535978 |
| 17 | 53500200 | 53501799 | 12.81522 | 22.83972 | 37.73398 | 16.60761 | -0.32375 | 0.456559 | 0.535981 |
| 2 | 42496400 | 42497199 | 14.76429 | 28.01605 | 23.63066 | 9.524734 | 0.73458 | 0.068835 | 0.535982 |
| 4 | 146411000 | 146414599 | 19.24819 | 33.99746 | 29.11432 | 12.88035 | 0.612636 | 0.553186 | 0.535983 |
| 10 | 41665800 | 41679799 | 280.1542 | 396.6144 | 532.9861 | 220.5665 | -0.24009 | 4.396395 | 0.53599 |
| 5 | 65920600 | 65922399 | 11.27386 | 19.96715 | 36.31547 | 16.0574 | -0.46637 | 0.488018 | 0.536006 |
| 8 | 34211400 | 34215399 | 35.10128 | 58.06092 | 58.77837 | 25.41305 | 0.36291 | 1.341851 | 0.536014 |
| 1 | 7824800 | 7829399 | 51.82995 | 84.04962 | 70.26071 | 30.09503 | 0.62355 | 1.65614 | 0.536031 |
| 7 | 116492600 | 116494199 | 19.53127 | 34.15985 | 36.06179 | 15.90225 | 0.309779 | 0.696972 | 0.536037 |
| 17 | 56288200 | 56289399 | 15.67581 | 27.92978 | 29.34961 | 12.92064 | 0.325133 | 0.4597 | 0.536053 |
| 18 | 74157000 | 74158199 | 15.67581 | 27.92978 | 29.34961 | 12.92064 | 0.325133 | 0.4597 | 0.536053 |
| 7 | 33278600 | 33280599 | 36.88661 | 61.41754 | 54.88289 | 23.83655 | 0.543049 | 1.271754 | 0.536055 |
| 7 | 75600200 | 75601599 | 19.88871 | 33.64042 | 55.33657 | 24.21871 | -0.33404 | 1.06681 | 0.536081 |
| 3 | 67586800 | 67588199 | 13.45085 | 23.853 | 39.73373 | 17.56162 | -0.33991 | 0.494869 | 0.536083 |
| 7 | 120312200 | 120314199 | 16.76867 | 29.24283 | 51.12502 | 22.49909 | -0.41495 | 0.75024 | 0.536086 |
| 5 | 146060400 | 146060999 | 17.45998 | 30.69683 | 31.97352 | 14.15834 | 0.329137 | 0.597443 | 0.536088 |
| 5 | 109845600 | 109853199 | 134.5509 | 204.59 | 210.4513 | 87.18234 | 0.242482 | 3.191061 | 0.536089 |
| 8 | 105523200 | 105525199 | 16.62467 | 28.51984 | 49.51625 | 21.71735 | -0.40896 | 0.920454 | 0.536114 |
| 18 | 77702800 | 77710799 | 101.4546 | 155.36 | 223.4329 | 92.72635 | -0.23246 | 3.040057 | 0.536137 |
| 7 | 10349800 | 10360999 | 157.6536 | 239.7817 | 205.2851 | 85.04888 | 0.507437 | 3.192512 | 0.53616 |
| 7 | 116304200 | 116306199 | 25.96871 | 43.04787 | 68.1245 | 29.49795 | -0.2794 | 1.301491 | 0.536182 |
| 13 | 61258600 | 61266799 | 138.1179 | 210.2371 | 189.2095 | 78.40755 | 0.436355 | 3.16917 | 0.536188 |
| 3 | 31235000 | 31237999 | 81.39204 | 125.9124 | 187.2923 | 77.79663 | -0.26976 | 2.835736 | 0.536195 |
| 15 | 46546600 | 46547799 | 21.88684 | 37.74827 | 38.54295 | 16.9081 | 0.355165 | 0.879009 | 0.5362 |
| 4 | 101687600 | 101687999 | 5.22492 | 10.19306 | 21.00694 | 7.510999 | -0.38307 | -0.44356 | 0.536225 |
| 12 | 72369200 | 72372399 | 19.07455 | 32.63821 | 53.84794 | 23.61227 | -0.33663 | 0.955195 | 0.536238 |
| 8 | 116904200 | 116906199 | 16.40887 | 29.34752 | 30.13498 | 13.22382 | 0.361254 | 0.441268 | 0.536243 |
| 15 | 39854800 | 39858199 | 46.61067 | 75.28977 | 77.834 | 33.22005 | 0.312151 | 1.737344 | 0.536257 |
| 7 | 38194600 | 38224199 | 764.1689 | 1011.218 | 896.9008 | 374.9966 | 0.261209 | 5.448457 | 0.536264 |

| | | | | | | | | | |
|---|---|---|---|---|---|---|---|---|---|
| X | 77373600 | 77376999 | 50.42141 | 81.52198 | 75.70806 | 32.35197 | 0.468053 | 1.716213 | 0.536279 |
| 3 | 3974000 | 3979399 | 112.0286 | 171.9588 | 179.3234 | 74.44925 | 0.233481 | 2.993526 | 0.536305 |
| 18 | 37893400 | 37897999 | 202.6141 | 298.7887 | 299.2771 | 123.2463 | 0.241496 | 3.776285 | 0.536309 |
| 7 | 45025600 | 45028399 | 15.35445 | 26.74932 | 44.7088 | 19.66286 | -0.35042 | 0.746384 | 0.536326 |
| 7 | 56013000 | 56013599 | 10.04147 | 19.37434 | 21.40896 | 8.270952 | 0.40356 | -0.11731 | 0.536334 |
| 16 | 16175600 | 16176999 | 13.86067 | 24.08631 | 50.56382 | 22.20275 | -0.67813 | 0.783635 | 0.536347 |
| 4 | 32805800 | 32812399 | 59.32951 | 93.46266 | 160.3751 | 66.54581 | -0.45526 | 2.492835 | 0.536353 |
| 12 | 13902600 | 13906199 | 31.5542 | 52.9714 | 51.34413 | 22.41526 | 0.425799 | 1.165007 | 0.536355 |
| 17 | 25893400 | 25895799 | 27.72646 | 46.63326 | 49.10265 | 21.45357 | 0.30677 | 1.132492 | 0.536359 |
| 15 | 23551200 | 23551599 | 8.55256 | 16.54797 | 18.79687 | 7.170891 | 0.381259 | -0.18438 | 0.536365 |
| 4 | 35824800 | 35828399 | 27.72563 | 47.80225 | 41.79471 | 18.33441 | 0.577818 | 0.909352 | 0.536371 |
| 5 | 150780400 | 150782199 | 16.37127 | 28.42682 | 46.34319 | 20.33944 | -0.31506 | 0.802381 | 0.536403 |
| 2 | 119587600 | 119591799 | 33.69111 | 54.30612 | 91.76105 | 39.05767 | -0.39856 | 1.7841 | 0.536428 |
| 6 | 116485000 | 116485799 | 21.85034 | 37.66535 | 37.55105 | 16.47271 | 0.389305 | 0.881942 | 0.536446 |
| 4 | 156431400 | 156431999 | 7.615824 | 14.69598 | 30.30236 | 11.70274 | -0.48874 | -0.12082 | 0.536447 |
| X | 73471600 | 73473599 | 13.85954 | 24.55749 | 45.07656 | 19.92949 | -0.47982 | 0.515684 | 0.536454 |
| 19 | 39179600 | 39183199 | 68.70616 | 109.039 | 115.7719 | 48.23062 | 0.245193 | 2.311797 | 0.536476 |
| 3 | 67411800 | 67413599 | 15.37595 | 26.8713 | 43.59744 | 19.2126 | -0.30722 | 0.700929 | 0.536479 |
| 3 | 24021000 | 24026599 | 46.15397 | 74.68189 | 74.24277 | 31.74951 | 0.371154 | 1.68784 | 0.536491 |
| 7 | 17614200 | 17662399 | 740.5182 | 991.1277 | 761.0543 | 317.7201 | 0.48555 | 5.267819 | 0.536503 |
| X | 61881400 | 61886399 | 43.19645 | 70.87488 | 62.37372 | 26.85834 | 0.561037 | 1.463992 | 0.536556 |
| X | 32159800 | 32165599 | 61.62323 | 99.05588 | 74.62065 | 31.63618 | 0.759486 | 1.904058 | 0.536567 |
| 1 | 109182000 | 109188999 | 68.66877 | 109.7112 | 98.56839 | 41.41485 | 0.495785 | 2.103384 | 0.536575 |
| 16 | 19194400 | 19195799 | 23.28824 | 39.96744 | 39.63191 | 17.38274 | 0.395897 | 0.93182 | 0.536582 |
| 1 | 187664600 | 187665999 | 12.7825 | 22.78506 | 37.71701 | 16.59788 | -0.32649 | 0.45433 | 0.536595 |
| 17 | 53503000 | 53505999 | 50.59833 | 80.54056 | 122.7173 | 51.29469 | -0.27077 | 2.209374 | 0.536604 |
| X | 11266800 | 11268199 | 18.96986 | 33.7179 | 31.6045 | 13.94146 | 0.487276 | 0.501676 | 0.53661 |
| 7 | 11438200 | 11440199 | 28.16135 | 47.9536 | 46.83677 | 20.52987 | 0.415843 | 1.027332 | 0.536639 |
| 7 | 123156200 | 123160399 | 50.77585 | 80.8585 | 122.3966 | 51.19165 | -0.26065 | 2.195764 | 0.536646 |
| 12 | 103686400 | 103712999 | 346.4317 | 482.5066 | 640.8259 | 265.8834 | -0.24682 | 4.640807 | 0.536647 |
| 12 | 118799600 | 118802199 | 22.43344 | 37.66831 | 60.91886 | 26.599 | -0.31025 | 1.142698 | 0.536683 |
| 6 | 41405800 | 41414599 | 93.44087 | 146.9808 | 134.4304 | 55.76959 | 0.449664 | 2.538235 | 0.536684 |
| 8 | 57842800 | 57845799 | 24.85726 | 43.22281 | 39.05009 | 17.15383 | 0.532764 | 0.809134 | 0.5367 |
| X | 147814200 | 147819399 | 36.6985 | 61.43512 | 51.44685 | 22.42047 | 0.636243 | 1.212258 | 0.536705 |
| 14 | 16469000 | 16471199 | 15.09926 | 27.44744 | 28.26157 | 12.18272 | 0.374088 | 0.333401 | 0.536708 |
| 10 | 79478200 | 79480599 | 17.36715 | 29.77847 | 50.0523 | 21.95156 | -0.36267 | 0.926802 | 0.536723 |
| 13 | 64276400 | 64278599 | 11.64002 | 20.83833 | 35.7115 | 15.65825 | -0.37255 | 0.414892 | 0.53673 |
| 17 | 54886000 | 54886399 | 4.780579 | 9.318872 | 20.24244 | 7.312653 | -0.46999 | -0.41177 | 0.536733 |
| 5 | 63198200 | 63199999 | 15.6649 | 28.32737 | 24.67527 | 10.70646 | 0.607893 | 0.359503 | 0.536802 |
| 8 | 67770800 | 67772399 | 13.13202 | 24.18578 | 25.93902 | 10.99146 | 0.332618 | 0.240109 | 0.536807 |
| 8 | 21127400 | 21134199 | 222.7545 | 320.931 | 450.0038 | 185.7001 | -0.27601 | 4.135828 | 0.536828 |
| 11 | 70300400 | 70301199 | 6.585509 | 12.60711 | 25.12936 | 9.929178 | -0.47015 | -0.06886 | 0.536855 |
| 7 | 101258800 | 101259399 | 4.961235 | 9.67069 | 20.85741 | 7.53827 | -0.46198 | -0.40995 | 0.536899 |
| 16 | 21980000 | 21982799 | 19.30178 | 32.0518 | 69.62204 | 30.18252 | -0.7344 | 1.264924 | 0.536923 |
| 18 | 37221600 | 37225599 | 46.94565 | 74.91568 | 115.7378 | 48.54732 | -0.28676 | 2.122071 | 0.536926 |
| 5 | 113571200 | 113572399 | 14.13916 | 26.10316 | 26.49674 | 11.19123 | 0.415163 | 0.238323 | 0.536931 |
| 15 | 87996200 | 87998199 | 25.51143 | 43.20204 | 45.87369 | 20.08439 | 0.295106 | 1.064691 | 0.536934 |

| | | | | | | | | | |
|---|---|---|---|---|---|---|---|---|---|
| 19 | 41841200 | 41841599 | 5.512155 | 10.72441 | 21.80454 | 8.042363 | -0.40391 | -0.32379 | 0.536955 |
| 2 | 89795600 | 89799999 | 39.82239 | 64.13936 | 100.2825 | 42.63818 | -0.28831 | 1.820352 | 0.536968 |
| 4 | 58706200 | 58710999 | 57.07325 | 91.71797 | 87.63326 | 37.13888 | 0.416794 | 1.91091 | 0.536981 |
| 1 | 175318600 | 175321399 | 20.0243 | 33.39324 | 67.71668 | 29.43944 | -0.63538 | 1.216332 | 0.536985 |
| 3 | 87051800 | 87054799 | 23.33696 | 39.90778 | 41.65736 | 18.26584 | 0.321245 | 0.96475 | 0.537002 |
| 2 | 80538000 | 80539599 | 14.26362 | 24.91784 | 42.59921 | 18.76768 | -0.38229 | 0.699003 | 0.537008 |
| 13 | 69923000 | 69924599 | 13.12255 | 23.26126 | 38.1883 | 16.88261 | -0.31926 | 0.493332 | 0.537012 |
| 9 | 112522400 | 112524199 | 21.23173 | 37.51825 | 32.32194 | 14.2981 | 0.604238 | 0.564999 | 0.537019 |
| 2 | 170658400 | 170663199 | 69.83994 | 110.9034 | 115.4698 | 48.13302 | 0.274197 | 2.295471 | 0.537029 |
| 4 | 50250400 | 50251399 | 15.54258 | 28.47873 | 27.83389 | 11.87713 | 0.458673 | 0.292427 | 0.537086 |
| 9 | 23892600 | 23893999 | 22.10179 | 37.01009 | 61.22746 | 26.68766 | -0.34263 | 1.175575 | 0.537086 |
| 12 | 49041000 | 49041799 | 12.19603 | 22.91495 | 24.61492 | 10.11498 | 0.36428 | 0.111078 | 0.537099 |
| 2 | 46631800 | 46632799 | 13.41897 | 24.54146 | 25.87077 | 11.06771 | 0.347388 | 0.280735 | 0.53713 |
| 3 | 15996200 | 15998599 | 22.44571 | 39.28045 | 36.87874 | 16.27066 | 0.478227 | 0.70515 | 0.537131 |
| 8 | 21433000 | 21434799 | 28.94445 | 48.64855 | 50.73964 | 22.1642 | 0.320441 | 1.144818 | 0.537138 |
| 2 | 138308800 | 138310599 | 22.71084 | 39.18135 | 41.5332 | 18.22005 | 0.301359 | 0.88323 | 0.53714 |
| 2 | 160254600 | 160258799 | 36.26941 | 58.55324 | 90.38403 | 38.54898 | -0.26583 | 1.743118 | 0.537142 |
| 16 | 3312200 | 3314599 | 22.01895 | 39.42315 | 29.16078 | 12.78485 | 0.832966 | 0.472545 | 0.537144 |
| 16 | 15003200 | 15004999 | 14.06226 | 24.79986 | 41.3087 | 18.28078 | -0.34315 | 0.559403 | 0.537145 |
| 12 | 55631200 | 55632399 | 10.96327 | 20.83869 | 22.65091 | 9.097164 | 0.375242 | 0.016751 | 0.537147 |
| 18 | 4317400 | 4321999 | 67.1669 | 106.9706 | 108.0948 | 45.21583 | 0.321555 | 2.204397 | 0.537148 |
| 1 | 109501000 | 109503399 | 28.49038 | 48.09757 | 43.20273 | 18.89724 | 0.535432 | 1.097655 | 0.53715 |
| 13 | 95043800 | 95045799 | 14.58688 | 25.64086 | 41.85815 | 18.53608 | -0.31603 | 0.600562 | 0.537152 |
| 19 | 26894200 | 26895799 | 20.26888 | 35.80464 | 32.98282 | 14.59144 | 0.507941 | 0.564268 | 0.537163 |
| 11 | 27706200 | 27708199 | 19.26185 | 33.75221 | 35.49153 | 15.67574 | 0.315417 | 0.665211 | 0.537177 |
| 17 | 37414600 | 37418399 | 34.85451 | 57.90993 | 50.38933 | 21.85307 | 0.581086 | 1.284281 | 0.537202 |
| 2 | 83994000 | 83994999 | 6.838631 | 13.15555 | 25.85194 | 10.07294 | -0.4318 | -0.11344 | 0.537203 |
| X | 109124600 | 109126799 | 24.44689 | 42.85014 | 35.0077 | 15.46579 | 0.677809 | 0.688137 | 0.53724 |
| 9 | 117695000 | 117697399 | 17.80433 | 30.67445 | 49.53313 | 21.7286 | -0.30382 | 0.88437 | 0.537253 |
| 8 | 85845800 | 85847199 | 8.577652 | 16.1069 | 28.5426 | 11.74073 | -0.35342 | 0.094205 | 0.537267 |
| 15 | 93251000 | 93253399 | 24.42625 | 42.50771 | 39.06696 | 17.16996 | 0.508314 | 0.796555 | 0.537276 |
| 5 | 94487400 | 94488599 | 20.4737 | 35.83634 | 31.43843 | 13.8727 | 0.575588 | 0.682582 | 0.537282 |
| 17 | 24231600 | 24235799 | 31.83271 | 51.66962 | 81.45124 | 34.9124 | -0.28812 | 1.605516 | 0.537324 |
| 1 | 134096000 | 134098999 | 29.74886 | 49.72297 | 52.44285 | 22.83191 | 0.30448 | 1.20593 | 0.537336 |
| 11 | 70066600 | 70069999 | 28.47534 | 46.80065 | 73.65155 | 31.73744 | -0.27427 | 1.4141 | 0.53734 |
| 10 | 125346800 | 125349799 | 14.81114 | 25.82727 | 51.0965 | 22.48554 | -0.59246 | 0.740745 | 0.537355 |
| 4 | 115240800 | 115242999 | 29.79528 | 50.75963 | 46.25799 | 20.27694 | 0.515545 | 1.028394 | 0.537363 |
| 1 | 40565200 | 40566599 | 14.90838 | 27.54645 | 26.93368 | 11.36275 | 0.469895 | 0.239867 | 0.537363 |
| 14 | 5363400 | 5364399 | 6.401483 | 12.39204 | 23.81101 | 9.062902 | -0.36667 | -0.19932 | 0.537377 |
| 4 | 60414000 | 60416999 | 40.72002 | 66.26769 | 60.20128 | 25.84218 | 0.508077 | 1.563932 | 0.537402 |
| 1 | 30409000 | 30412199 | 24.57468 | 40.13088 | 80.91774 | 34.77865 | -0.63548 | 1.495899 | 0.537403 |
| 15 | 73882600 | 73883799 | 13.24749 | 24.39515 | 25.89335 | 10.97412 | 0.347207 | 0.241076 | 0.53741 |
| 4 | 134113600 | 134114799 | 6.864257 | 13.26397 | 25.43824 | 9.764094 | -0.37558 | -0.15979 | 0.537412 |
| 5 | 84811000 | 84815799 | 57.75615 | 92.79438 | 87.42973 | 37.04046 | 0.43652 | 1.917112 | 0.537427 |
| 8 | 72267200 | 72267799 | 21.47236 | 35.65442 | 63.82199 | 27.66714 | -0.45624 | 1.265401 | 0.537458 |
| 1 | 161270200 | 161272799 | 18.28113 | 31.47591 | 50.60823 | 22.19978 | -0.29785 | 0.893181 | 0.537504 |
| 3 | 64294200 | 64296999 | 27.12061 | 46.0942 | 46.29759 | 20.29044 | 0.375132 | 1.037163 | 0.537525 |

| | | | | | | | | | |
|---|---|---|---|---|---|---|---|---|---|
| 3 | 75767400 | 75769999 | 17.30199 | 30.0846 | 49.53528 | 21.75937 | -0.32933 | 0.789864 | 0.537537 |
| 9 | 55729200 | 55731399 | 41.71704 | 67.75026 | 70.86213 | 30.38576 | 0.302963 | 1.607903 | 0.537539 |
| 17 | 35413000 | 35416399 | 24.43589 | 40.23273 | 68.67591 | 29.61553 | -0.39002 | 1.37868 | 0.537569 |
| 5 | 26579400 | 26585999 | 88.93597 | 137.1092 | 198.7332 | 82.55039 | -0.23627 | 2.902623 | 0.537584 |
| 2 | 5648800 | 5649599 | 5.219294 | 10.14875 | 21.73958 | 8.055931 | -0.48408 | -0.30989 | 0.537588 |
| 8 | 54316400 | 54316799 | 4.148825 | 8.101936 | 20.54361 | 7.220444 | -0.65618 | -0.52348 | 0.53759 |
| 9 | 50919400 | 50920999 | 11.17522 | 20.10066 | 34.23396 | 14.95092 | -0.35949 | 0.380121 | 0.537631 |
| 16 | 9992000 | 10007199 | 367.9724 | 516.6208 | 495.4728 | 205.32 | 0.234326 | 4.52154 | 0.537639 |
| 9 | 36408600 | 36417399 | 154.7032 | 233.6414 | 233.0038 | 96.32323 | 0.278968 | 3.333788 | 0.537642 |
| X | 5477400 | 5499599 | 337.2893 | 481.7115 | 415.3617 | 171.6371 | 0.412588 | 4.271593 | 0.53768 |
| 17 | 41914000 | 41915799 | 27.49507 | 46.12313 | 49.12087 | 21.43468 | 0.290466 | 1.158426 | 0.537728 |
| 13 | 99446000 | 99447799 | 18.52306 | 32.4421 | 34.03053 | 15.02437 | 0.318899 | 0.66558 | 0.53775 |
| 3 | 27398200 | 27398999 | 6.549829 | 12.64911 | 24.26681 | 9.339561 | -0.37998 | -0.15936 | 0.537765 |
| 6 | 52496600 | 52498199 | 10.64206 | 19.35858 | 33.31431 | 14.34835 | -0.36141 | 0.307821 | 0.537765 |
| 2 | 8508400 | 8511199 | 25.12885 | 41.64613 | 66.95231 | 28.98581 | -0.30208 | 1.299865 | 0.537794 |
| 16 | 40155200 | 40156399 | 13.38979 | 25.16116 | 25.10439 | 10.31384 | 0.470342 | 0.122269 | 0.537819 |
| 5 | 107915000 | 107916599 | 22.94118 | 37.5875 | 78.57758 | 33.81545 | -0.68434 | 1.446239 | 0.537837 |
| 17 | 23667200 | 23667999 | 7.49982 | 14.30185 | 26.67876 | 10.64148 | -0.38796 | -0.02355 | 0.537856 |
| 3 | 16400200 | 16409199 | 92.80381 | 145.8673 | 136.8267 | 56.75808 | 0.412407 | 2.553227 | 0.537859 |
| 7 | 94762000 | 94762399 | 6.712775 | 12.91757 | 24.70895 | 9.618757 | -0.39113 | -0.11777 | 0.537869 |
| Y | 8437400 | 8443399 | 43.99314 | 71.1543 | 48.35698 | 20.67292 | 0.917817 | 1.676258 | 0.537876 |
| 8 | 26862200 | 26862999 | 10.1606 | 18.74962 | 31.97046 | 13.51144 | -0.32895 | 0.220784 | 0.537905 |
| 3 | 11111800 | 11115799 | 57.10644 | 91.37954 | 95.17239 | 40.07573 | 0.28541 | 2.045004 | 0.537905 |
| 1 | 8227400 | 8229799 | 31.37561 | 51.87573 | 54.57705 | 23.58951 | 0.307572 | 1.334996 | 0.537912 |
| 11 | 79867800 | 79868799 | 9.357779 | 17.5678 | 31.29847 | 12.87931 | -0.3619 | 0.108594 | 0.537915 |
| 14 | 90377800 | 90380599 | 52.3576 | 83.86129 | 87.60469 | 36.94124 | 0.282649 | 2.006005 | 0.537944 |
| X | 54563800 | 54565999 | 63.61729 | 101.2642 | 101.048 | 42.23748 | 0.338905 | 2.217474 | 0.537947 |
| 2 | 54362200 | 54366399 | 51.42048 | 82.82719 | 83.19998 | 35.3797 | 0.348867 | 1.828761 | 0.53799 |
| X | 73787800 | 73789999 | 23.81013 | 41.39589 | 38.29698 | 16.82143 | 0.499009 | 0.810329 | 0.537995 |
| 16 | 90820400 | 90822799 | 26.34428 | 44.46339 | 47.47802 | 20.76571 | 0.286887 | 1.097946 | 0.537995 |
| 9 | 88488800 | 88490599 | 15.13902 | 27.18874 | 28.78593 | 12.5878 | 0.32087 | 0.404995 | 0.53801 |
| 15 | 50443400 | 50444599 | 16.80992 | 30.524 | 28.75618 | 12.41274 | 0.500604 | 0.355268 | 0.538018 |
| Y | 87097600 | 87099599 | 16.47009 | 29.23061 | 21.66352 | 9.568417 | 0.822533 | 0.470632 | 0.538029 |
| 6 | 49548000 | 49549999 | 16.63704 | 28.918 | 46.84918 | 20.5748 | -0.30603 | 0.788115 | 0.538045 |
| 6 | 141273400 | 141274399 | 7.890603 | 15.00016 | 28.30489 | 11.36489 | -0.41392 | 0.001321 | 0.538054 |
| 7 | 3205600 | 3223999 | 51.61814 | 82.97805 | 74.85411 | 31.73781 | 0.500088 | 1.88737 | 0.538055 |
| 5 | 46013000 | 46015799 | 24.35052 | 39.57229 | 89.12497 | 38.23542 | -0.79979 | 1.570265 | 0.538081 |
| 7 | 133129600 | 133133999 | 75.51269 | 117.6791 | 172.8563 | 71.74071 | -0.2434 | 2.702351 | 0.538096 |
| 18 | 37421200 | 37426199 | 66.00814 | 106.0348 | 84.11407 | 35.62712 | 0.683998 | 1.935164 | 0.538098 |
| 9 | 49237000 | 49237799 | 6.712591 | 12.95964 | 24.80524 | 9.559862 | -0.3782 | -0.14939 | 0.53812 |
| 2 | 8243400 | 8244599 | 20.79943 | 37.71377 | 27.22353 | 11.77241 | 0.880706 | 0.38791 | 0.53813 |
| 6 | 32893800 | 32894799 | 13.8491 | 25.46082 | 26.93686 | 11.44214 | 0.349717 | 0.258647 | 0.538154 |
| 1 | 56537000 | 56538799 | 19.2172 | 33.65637 | 35.49889 | 15.67207 | 0.311013 | 0.672749 | 0.538177 |
| 2 | 51775600 | 51777599 | 52.53092 | 84.12664 | 87.5337 | 36.90313 | 0.288121 | 2.010858 | 0.538186 |
| 6 | 94141600 | 94143599 | 10.06004 | 18.36412 | 31.85282 | 13.65886 | -0.36818 | 0.282111 | 0.53823 |
| 10 | 30642400 | 30642799 | 8.194484 | 16.02711 | 14.73282 | 4.979312 | 0.840021 | -0.61672 | 0.53825 |
| 6 | 147343400 | 147346999 | 32.59686 | 54.60633 | 52.49429 | 22.88688 | 0.437742 | 1.191904 | 0.538255 |

| | | | | | | | | | |
|---|---|---|---|---|---|---|---|---|---|
| 11 | 39954000 | 39956799 | 39.20861 | 63.83113 | 66.46466 | 28.53458 | 0.312078 | 1.557874 | 0.538278 |
| 9 | 115539200 | 115540599 | 10.19849 | 18.69216 | 31.87851 | 13.59766 | -0.33873 | 0.259969 | 0.538284 |
| 3 | 122449400 | 122452199 | 24.57699 | 40.98305 | 66.59712 | 28.95165 | -0.31709 | 1.227837 | 0.538292 |
| X | 133954600 | 133955399 | 13.56068 | 23.9838 | 41.36568 | 18.29554 | -0.3912 | 0.52919 | 0.538298 |
| 1 | 44452400 | 44453799 | 14.73953 | 27.3169 | 26.36393 | 11.07675 | 0.492511 | 0.219063 | 0.538337 |
| 14 | 116009200 | 116010599 | 11.81625 | 21.11103 | 35.70896 | 15.68388 | -0.35545 | 0.429381 | 0.538382 |
| 6 | 140358600 | 140360399 | 19.18788 | 32.64454 | 53.22594 | 23.32876 | -0.32076 | 1.009945 | 0.538384 |
| 7 | 129933800 | 129935399 | 9.911336 | 18.06003 | 31.85029 | 13.68866 | -0.39409 | 0.291374 | 0.53839 |
| 18 | 46975000 | 46976999 | 33.62707 | 54.56725 | 84.66072 | 36.28078 | -0.26563 | 1.616329 | 0.538398 |
| 15 | 35713600 | 35715599 | 25.40191 | 44.17442 | 39.63388 | 17.4114 | 0.542877 | 0.813064 | 0.538401 |
| 18 | 43458800 | 43460199 | 7.956949 | 14.99691 | 27.53657 | 11.25596 | -0.39684 | 0.062072 | 0.538404 |
| 17 | 18489200 | 18492999 | 50.01839 | 81.7654 | 61.65487 | 26.50941 | 0.780732 | 1.525425 | 0.538428 |
| 1 | 117747000 | 117755399 | 155.4915 | 235.3655 | 225.5755 | 93.31918 | 0.339063 | 3.288504 | 0.538434 |
| 16 | 76329000 | 76329799 | 7.746422 | 14.76742 | 27.24933 | 10.87636 | -0.3741 | -0.02113 | 0.538436 |
| 4 | 80217400 | 80221199 | 44.54768 | 72.65866 | 66.56428 | 28.59681 | 0.498235 | 1.544026 | 0.538461 |
| 14 | 8535600 | 8537399 | 37.78243 | 60.78285 | 95.71389 | 40.62703 | -0.30056 | 1.854201 | 0.538467 |
| 10 | 66251200 | 66251799 | 11.02449 | 19.87501 | 33.75955 | 14.7043 | -0.35304 | 0.365163 | 0.53848 |
| 13 | 58705600 | 58707199 | 11.02449 | 19.87501 | 33.75955 | 14.7043 | -0.35304 | 0.365163 | 0.53848 |
| 17 | 66089600 | 66091199 | 12.06321 | 21.26666 | 37.8562 | 16.75413 | -0.43827 | 0.546969 | 0.538484 |
| 4 | 19420200 | 19421799 | 18.45676 | 32.65345 | 32.61522 | 14.42441 | 0.392633 | 0.536384 | 0.538486 |
| X | 79743600 | 79750399 | 73.1736 | 116.7445 | 100.9928 | 42.34923 | 0.548131 | 2.154219 | 0.538492 |
| 1 | 193492000 | 193494399 | 29.02483 | 48.92614 | 50.17464 | 21.93679 | 0.344904 | 1.120892 | 0.538494 |
| X | 147501200 | 147503999 | 31.31436 | 52.62942 | 45.34825 | 19.80885 | 0.595098 | 1.149637 | 0.538506 |
| 7 | 11864600 | 11871199 | 108.2138 | 168.9122 | 130.6455 | 54.211 | 0.682669 | 2.685611 | 0.538511 |
| 9 | 21131200 | 21136599 | 41.20519 | 66.10361 | 102.5099 | 43.33126 | -0.28349 | 1.948619 | 0.53855 |
| 13 | 75124000 | 75128799 | 43.83457 | 71.66457 | 65.35212 | 28.10023 | 0.507258 | 1.51028 | 0.538581 |
| 2 | 84508000 | 84509199 | 6.415109 | 12.37292 | 24.20707 | 9.361956 | -0.41424 | -0.14319 | 0.538601 |
| 15 | 80239400 | 80240199 | 6.415109 | 12.37292 | 24.20707 | 9.361956 | -0.41424 | -0.14319 | 0.538601 |
| 15 | 3927600 | 3928599 | 14.56619 | 26.50306 | 27.89778 | 12.01192 | 0.343764 | 0.325041 | 0.538629 |
| X | 154655400 | 154657399 | 23.16284 | 40.64957 | 35.07598 | 15.5121 | 0.599288 | 0.662576 | 0.538667 |
| 15 | 6536000 | 6536799 | 11.83417 | 22.35394 | 23.6947 | 9.643003 | 0.395222 | 0.073079 | 0.538686 |
| 6 | 67878200 | 67878599 | 7.970024 | 15.47866 | 17.30875 | 6.477213 | 0.430136 | -0.26028 | 0.538703 |
| 13 | 3258200 | 3290799 | 419.8415 | 583.427 | 549.7406 | 228.1718 | 0.244925 | 4.676821 | 0.538714 |
| 5 | 46283400 | 46285199 | 15.21505 | 27.70665 | 28.34024 | 12.18977 | 0.385794 | 0.325813 | 0.538736 |
| 4 | 112223200 | 112230599 | 185.9696 | 276.1981 | 280.1051 | 115.3875 | 0.233125 | 3.653411 | 0.538755 |
| X | 57595200 | 57600599 | 82.37275 | 130.1422 | 125.9661 | 52.31821 | 0.372983 | 2.440026 | 0.53876 |
| 6 | 25621400 | 25623599 | 50.38441 | 80.87905 | 86.46018 | 36.58114 | 0.253186 | 1.934761 | 0.538772 |
| 8 | 35961800 | 35963799 | 13.12894 | 23.06212 | 40.3683 | 17.86669 | -0.41615 | 0.599164 | 0.538794 |
| 7 | 34210600 | 34212399 | 14.05066 | 24.66377 | 41.89601 | 18.53177 | -0.37323 | 0.617821 | 0.538817 |
| 7 | 31827800 | 31829199 | 40.38022 | 65.41713 | 69.17233 | 29.60711 | 0.283972 | 1.653635 | 0.538817 |
| 1 | 138253600 | 138254799 | 7.592575 | 14.47815 | 27.25652 | 10.87285 | -0.40211 | -0.02499 | 0.538818 |
| 1 | 15339000 | 15340199 | 13.36862 | 23.43501 | 40.51149 | 17.899 | -0.3979 | 0.63765 | 0.538845 |
| 12 | 85871000 | 85871999 | 8.316506 | 15.41679 | 29.44481 | 12.36709 | -0.48456 | 0.174098 | 0.53885 |
| 5 | 24729000 | 24732399 | 26.90714 | 43.97327 | 75.33309 | 32.38734 | -0.40037 | 1.483607 | 0.538857 |
| 13 | 62013000 | 62029799 | 321.1989 | 460.4119 | 408.0423 | 168.5153 | 0.378276 | 4.216414 | 0.538863 |
| 7 | 125642400 | 125645799 | 33.45603 | 54.12116 | 87.15855 | 37.26568 | -0.32351 | 1.678281 | 0.538865 |
| 7 | 41647200 | 41651599 | 77.7848 | 123.3784 | 118.3724 | 49.28649 | 0.390417 | 2.33717 | 0.538877 |

| | | | | | | | | | |
|---|---|---|---|---|---|---|---|---|---|
| 17 | 50952400 | 50958199 | 72.38077 | 114.7198 | 120.1953 | 50.00748 | 0.262505 | 2.354164 | 0.538885 |
| 3 | 21552800 | 21554199 | 9.62604 | 17.51109 | 31.87557 | 13.72798 | -0.44126 | 0.299638 | 0.538903 |
| 15 | 92424600 | 92425199 | 11.96029 | 22.68922 | 23.67169 | 9.546684 | 0.428779 | 0.042431 | 0.538907 |
| 11 | 27634200 | 27634799 | 6.389832 | 12.40845 | 24.04442 | 9.005429 | -0.35699 | -0.25482 | 0.538919 |
| 11 | 73773800 | 73775599 | 18.5378 | 33.64084 | 28.61062 | 12.35941 | 0.646649 | 0.370959 | 0.538922 |
| 12 | 41420200 | 41424199 | 29.97376 | 49.21587 | 82.05922 | 35.34598 | -0.35804 | 1.434876 | 0.538935 |
| 6 | 9445800 | 9447999 | 19.24638 | 33.98991 | 33.20358 | 14.68983 | 0.423722 | 0.562841 | 0.538945 |
| X | 25882800 | 25904799 | 319.1602 | 458.8207 | 388.3882 | 160.3254 | 0.448658 | 4.17525 | 0.538948 |
| 11 | 62856800 | 62858599 | 13.9086 | 24.37385 | 41.96704 | 18.53713 | -0.39258 | 0.644341 | 0.538948 |
| 18 | 23338600 | 23340799 | 15.24381 | 27.42386 | 28.70517 | 12.53438 | 0.33856 | 0.394204 | 0.538949 |
| 3 | 27685000 | 27686599 | 11.354 | 20.45214 | 34.2557 | 14.93583 | -0.33388 | 0.373807 | 0.538961 |
| X | 30760000 | 30760599 | 11.18397 | 21.83061 | 16.16007 | 5.723668 | 1.089822 | -0.41124 | 0.538964 |
| X | 88924200 | 88929799 | 53.71721 | 86.74313 | 77.47127 | 33.05257 | 0.522275 | 1.75944 | 0.538988 |
| 5 | 58733800 | 58742399 | 168.0672 | 251.9008 | 255.8848 | 105.5619 | 0.242768 | 3.48488 | 0.538991 |
| 17 | 76315800 | 76317599 | 15.57464 | 28.69376 | 26.12408 | 11.06499 | 0.567543 | 0.256368 | 0.538993 |
| 8 | 46922400 | 46924799 | 19.1556 | 32.56679 | 53.77244 | 23.56696 | -0.3391 | 1.015504 | 0.538996 |
| 16 | 36314800 | 36316599 | 28.09154 | 45.81052 | 75.59976 | 32.47658 | -0.3487 | 1.519898 | 0.538999 |
| 4 | 67436800 | 67437599 | 7.469054 | 14.27947 | 26.19354 | 10.3894 | -0.35702 | -0.0451 | 0.539006 |
| 10 | 51077800 | 51078599 | 7.469054 | 14.27947 | 26.19354 | 10.3894 | -0.35702 | -0.0451 | 0.539006 |
| 16 | 15058000 | 15060799 | 20.95784 | 35.52563 | 56.5104 | 24.7487 | -0.28584 | 1.051621 | 0.539013 |
| 2 | 31066000 | 31066799 | 13.72803 | 25.03009 | 26.47477 | 11.36958 | 0.339307 | 0.303842 | 0.539014 |
| 7 | 29677600 | 29681599 | 36.28556 | 59.47297 | 63.09797 | 27.15806 | 0.291379 | 1.457137 | 0.539015 |
| 3 | 36394600 | 36396999 | 16.51781 | 28.74703 | 46.30131 | 20.35033 | -0.29739 | 0.770005 | 0.539019 |
| 10 | 9144800 | 9145399 | 5.368851 | 10.45426 | 21.29665 | 7.802651 | -0.39784 | -0.3516 | 0.539019 |
| 3 | 81552800 | 81554799 | 19.61774 | 34.60286 | 33.62005 | 14.87777 | 0.430685 | 0.577548 | 0.539032 |
| 3 | 28171800 | 28172399 | 6.253098 | 12.15849 | 24.7779 | 9.184675 | -0.41469 | -0.28839 | 0.53906 |
| 17 | 92960200 | 92962199 | 21.85784 | 37.97226 | 39.85927 | 17.49913 | 0.31725 | 0.810536 | 0.53907 |
| 11 | 103478200 | 103479399 | 11.95885 | 21.50948 | 35.73584 | 15.60725 | -0.32409 | 0.388689 | 0.539075 |
| 14 | 17468000 | 17468799 | 9.297151 | 16.96248 | 31.3797 | 13.46494 | -0.46102 | 0.278128 | 0.539082 |
| 2 | 68398200 | 68399399 | 14.82418 | 26.68905 | 28.32092 | 12.35628 | 0.319858 | 0.387039 | 0.539084 |
| 12 | 115082800 | 115084199 | 23.44756 | 40.53087 | 41.49603 | 18.20482 | 0.351899 | 0.870173 | 0.539092 |
| 10 | 122332800 | 122334399 | 12.99375 | 22.86521 | 38.83554 | 17.19385 | -0.37208 | 0.574324 | 0.539099 |
| 17 | 30311600 | 30315199 | 26.48203 | 44.00882 | 76.44427 | 33.15766 | -0.41321 | 1.280348 | 0.539106 |
| 3 | 144101800 | 144105599 | 42.12671 | 67.55626 | 106.0586 | 44.80541 | -0.30189 | 1.961115 | 0.539118 |
| 17 | 45842600 | 45843599 | 10.72659 | 20.43991 | 22.74175 | 9.08544 | 0.348774 | -0.00115 | 0.539118 |
| 9 | 62929400 | 62932799 | 24.27268 | 40.65729 | 67.78038 | 29.54943 | -0.35397 | 1.181164 | 0.539121 |
| 4 | 123248200 | 123250199 | 13.75785 | 24.27653 | 39.76163 | 17.59419 | -0.31855 | 0.54956 | 0.539133 |
| X | 148992400 | 148994999 | 42.58329 | 69.53542 | 66.77881 | 28.70113 | 0.431595 | 1.526184 | 0.539142 |
| 13 | 50270000 | 50274599 | 68.36785 | 107.443 | 157.4781 | 65.32336 | -0.23084 | 2.545485 | 0.539151 |
| 6 | 109677600 | 109680399 | 23.82206 | 39.96783 | 65.99263 | 28.79976 | -0.3401 | 1.162159 | 0.539153 |
| 11 | 62080800 | 62082999 | 29.11691 | 49.24749 | 49.06532 | 21.47378 | 0.386585 | 1.090555 | 0.539158 |
| 7 | 14002200 | 14006399 | 51.62653 | 84.18361 | 62.8733 | 27.00868 | 0.79201 | 1.563484 | 0.539182 |
| 1 | 101479000 | 101479399 | 8.80643 | 17.07199 | 19.39163 | 7.331586 | 0.394039 | -0.20757 | 0.53919 |
| 8 | 124785000 | 124785999 | 14.27774 | 25.94736 | 26.97512 | 11.63195 | 0.360371 | 0.328476 | 0.539205 |
| 14 | 69973200 | 69974799 | 12.61426 | 22.49594 | 37.20488 | 16.36591 | -0.32479 | 0.449089 | 0.53921 |
| 18 | 34113200 | 34113999 | 12.24774 | 23.2252 | 23.63309 | 9.539222 | 0.463388 | 0.047336 | 0.539263 |
| 8 | 48584800 | 48587199 | 20.03331 | 33.74939 | 56.96225 | 24.89991 | -0.37135 | 1.10196 | 0.539263 |

| | | | | | | | | | |
|---|---|---|---|---|---|---|---|---|---|
| 4 | 145262200 | 145264999 | 49.49567 | 80.79278 | 63.88845 | 27.45515 | 0.710848 | 1.545553 | 0.539275 |
| 1 | 17743000 | 17745399 | 39.17612 | 64.0991 | 65.50423 | 28.17373 | 0.344157 | 1.488625 | 0.539275 |
| X | 156824600 | 156827199 | 42.76759 | 70.4536 | 51.08902 | 22.03652 | 0.841759 | 1.406983 | 0.539277 |
| 5 | 57125600 | 57127599 | 55.86165 | 89.27245 | 94.3724 | 39.66633 | 0.26194 | 2.081484 | 0.539281 |
| 7 | 25485400 | 25491199 | 31.73715 | 51.37324 | 83.58544 | 35.75473 | -0.33729 | 1.663563 | 0.539282 |
| 17 | 23545400 | 23556599 | 211.8355 | 313.2482 | 301.9219 | 124.3392 | 0.300673 | 3.732703 | 0.539284 |
| 12 | 22371400 | 22374599 | 44.99757 | 72.60561 | 77.72772 | 33.12846 | 0.260092 | 1.766327 | 0.539291 |
| 6 | 47953400 | 47955799 | 18.1257 | 31.05274 | 51.6644 | 22.65703 | -0.34834 | 0.938177 | 0.539311 |
| 2 | 61358600 | 61359999 | 15.85329 | 28.23247 | 29.83228 | 13.13842 | 0.316801 | 0.464921 | 0.539318 |
| 19 | 3788600 | 3790399 | 9.600382 | 17.54217 | 31.37461 | 13.43736 | -0.41085 | 0.272123 | 0.539323 |
| 9 | 118232200 | 118234799 | 16.52295 | 28.73855 | 46.31247 | 20.34748 | -0.29823 | 0.778819 | 0.539328 |
| 16 | 54787400 | 54789399 | 30.32196 | 50.50022 | 54.08594 | 23.49426 | 0.282553 | 1.247843 | 0.539331 |
| 17 | 15731600 | 15733199 | 16.24695 | 29.73826 | 27.68739 | 11.83143 | 0.52664 | 0.301943 | 0.53935 |
| X | 3213800 | 3226599 | 154.3474 | 235.5413 | 193.0506 | 80.05351 | 0.574177 | 3.12448 | 0.539356 |
| 16 | 17404400 | 17408799 | 65.14853 | 103.708 | 108.5561 | 45.37964 | 0.270067 | 2.215206 | 0.539359 |
| 7 | 80872000 | 80874399 | 25.2379 | 42.88427 | 45.32331 | 19.86308 | 0.302077 | 1.034862 | 0.539375 |
| 7 | 13220600 | 13221399 | 22.63717 | 38.86663 | 41.07971 | 18.01837 | 0.304132 | 0.924356 | 0.539398 |
| 9 | 79629200 | 79630399 | 13.0606 | 24.28747 | 25.43823 | 10.63757 | 0.380133 | 0.183971 | 0.539452 |
| 5 | 87897400 | 87899599 | 17.0148 | 29.64207 | 49.51005 | 21.77453 | -0.34957 | 0.762059 | 0.53946 |
| 1 | 12524800 | 12525399 | 7.137416 | 13.7888 | 26.06916 | 10.01643 | -0.35722 | -0.15421 | 0.539472 |
| 11 | 114113000 | 114114599 | 20.78249 | 36.04747 | 37.88966 | 16.62439 | 0.314997 | 0.820732 | 0.539491 |
| 5 | 29613800 | 29614799 | 12.72105 | 22.5131 | 37.78919 | 16.71157 | -0.35171 | 0.510353 | 0.539491 |
| 7 | 110827800 | 110830199 | 22.93274 | 37.59989 | 77.59714 | 33.40338 | -0.66544 | 1.435719 | 0.539502 |
| 9 | 31989400 | 31992599 | 30.31887 | 50.319 | 53.51869 | 23.1927 | 0.292273 | 1.286651 | 0.539536 |
| 5 | 138008000 | 138012399 | 32.59619 | 52.96905 | 83.76862 | 35.93582 | -0.29129 | 1.583966 | 0.539571 |
| 14 | 106725200 | 106726599 | 16.4638 | 29.10944 | 30.37304 | 13.43445 | 0.329684 | 0.526646 | 0.539592 |
| 5 | 115698800 | 115701399 | 10.14596 | 18.14756 | 34.81111 | 15.27483 | -0.53467 | 0.409078 | 0.539601 |
| X | 121376200 | 121378399 | 25.94332 | 45.19323 | 38.45996 | 16.91387 | 0.619035 | 0.790882 | 0.53961 |
| 6 | 124557200 | 124557999 | 6.570804 | 12.62462 | 24.66807 | 9.646046 | -0.42763 | -0.10489 | 0.539625 |
| 2 | 150350600 | 150352799 | 25.33849 | 43.42414 | 37.26946 | 16.34468 | 0.602784 | 0.945325 | 0.539629 |
| 7 | 80097000 | 80098799 | 11.40976 | 20.25874 | 35.77608 | 15.79575 | -0.42228 | 0.467086 | 0.539641 |
| 9 | 119150800 | 119152599 | 11.40976 | 20.25874 | 35.77608 | 15.79575 | -0.42228 | 0.467086 | 0.539641 |
| 3 | 59690800 | 59691999 | 17.82016 | 31.28366 | 32.08727 | 14.19428 | 0.351212 | 0.622983 | 0.539643 |
| 8 | 73651400 | 73659399 | 162.34 | 243.5165 | 248.8124 | 102.6638 | 0.235434 | 3.468792 | 0.539649 |
| 2 | 138446000 | 138447599 | 12.20078 | 22.67836 | 24.55105 | 10.2739 | 0.332305 | 0.179002 | 0.539649 |
| 11 | 20333000 | 20335199 | 15.80191 | 27.5684 | 45.72244 | 20.1268 | -0.3392 | 0.728875 | 0.539658 |
| 4 | 8133200 | 8133799 | 6.84726 | 13.23337 | 24.88866 | 9.545572 | -0.34684 | -0.16568 | 0.539674 |
| 17 | 91788600 | 91790999 | 43.81461 | 71.0722 | 64.0283 | 27.43134 | 0.516222 | 1.63198 | 0.539677 |
| 10 | 107812000 | 107813799 | 15.4268 | 26.74688 | 45.84255 | 20.11353 | -0.38726 | 0.813111 | 0.539687 |
| 8 | 94265600 | 94268199 | 45.65282 | 73.67378 | 78.50031 | 33.46413 | 0.267118 | 1.763359 | 0.539705 |
| 15 | 4128600 | 4130999 | 19.43424 | 33.24374 | 53.29382 | 23.36873 | -0.29535 | 0.959964 | 0.539719 |
| 7 | 86744000 | 86750599 | 83.25309 | 131.9184 | 120.0217 | 49.92316 | 0.465501 | 2.37069 | 0.539726 |
| 6 | 132537000 | 132538199 | 12.46412 | 23.2525 | 25.02105 | 10.4066 | 0.347784 | 0.158448 | 0.539729 |
| 7 | 61028200 | 61030999 | 23.02104 | 41.00412 | 29.56634 | 13.01946 | 0.865334 | 0.510814 | 0.53974 |
| 19 | 13580400 | 13585399 | 71.55839 | 114.172 | 90.34479 | 37.88628 | 0.676501 | 2.144924 | 0.539744 |
| 1 | 33853000 | 33856399 | 30.78789 | 50.16353 | 79.66664 | 34.2127 | -0.29451 | 1.538348 | 0.539768 |
| 1 | 99127800 | 99131999 | 48.01325 | 77.32916 | 81.69718 | 34.73433 | 0.276018 | 1.827889 | 0.539776 |

| | | | | | | | | | |
|---|---|---|---|---|---|---|---|---|---|
| X | 81978400 | 81979799 | 16.2471 | 29.21982 | 29.75686 | 12.99697 | 0.377926 | 0.407424 | 0.539777 |
| 6 | 69779400 | 69781199 | 31.65139 | 52.4432 | 56.31522 | 24.37658 | 0.278301 | 1.311649 | 0.539779 |
| 13 | 24901600 | 24904199 | 20.78587 | 35.42295 | 57.67164 | 25.28073 | -0.31846 | 1.004574 | 0.539791 |
| 17 | 48284600 | 48286199 | 13.15617 | 23.05393 | 40.98735 | 18.10351 | -0.43844 | 0.640426 | 0.539793 |
| 13 | 67645200 | 67650199 | 51.06517 | 81.46592 | 122.3801 | 51.3125 | -0.2469 | 2.137683 | 0.539814 |
| 15 | 41760400 | 41767999 | 76.79185 | 121.7995 | 118.7213 | 49.43018 | 0.367562 | 2.335872 | 0.539823 |
| 13 | 3166800 | 3180799 | 179.9295 | 269.0016 | 265.6448 | 109.5346 | 0.280207 | 3.533721 | 0.539828 |
| 19 | 20437000 | 20439399 | 14.55398 | 25.12477 | 51.6036 | 22.63823 | -0.64847 | 0.856537 | 0.539851 |
| X | 125370000 | 125371999 | 17.41229 | 32.11518 | 26.70006 | 11.29178 | 0.699802 | 0.268428 | 0.539858 |
| 4 | 69638400 | 69640399 | 23.32133 | 38.54143 | 73.20027 | 31.62897 | -0.54207 | 1.335553 | 0.539881 |
| 5 | 13882400 | 13882599 | 3.341438 | 6.531157 | 15.20017 | 4.890551 | -0.41392 | -0.86196 | 0.539886 |
| 18 | 35245600 | 35247999 | 23.74373 | 39.78418 | 64.92321 | 28.30961 | -0.32325 | 1.172881 | 0.539915 |
| 15 | 73751600 | 73754599 | 25.87825 | 42.59109 | 69.38073 | 29.91458 | -0.32303 | 1.387799 | 0.539943 |
| 4 | 83576400 | 83576999 | 4.428282 | 8.640442 | 21.61063 | 7.705712 | -0.65603 | -0.46508 | 0.539945 |
| 16 | 13758000 | 13759799 | 12.84738 | 22.69472 | 37.78581 | 16.71626 | -0.34121 | 0.530197 | 0.539948 |
| 13 | 78345400 | 78347799 | 15.44034 | 26.29976 | 57.38222 | 25.15283 | -0.73897 | 0.989454 | 0.539955 |
| 16 | 21872000 | 21876599 | 40.25623 | 64.52954 | 114.8902 | 48.50534 | -0.48379 | 1.976155 | 0.539962 |
| 7 | 120825800 | 120827799 | 21.46351 | 37.125 | 39.44645 | 17.30597 | 0.298663 | 0.85094 | 0.539972 |
| 10 | 39444200 | 39445999 | 12.51303 | 22.20831 | 36.72039 | 16.21843 | -0.32829 | 0.479306 | 0.539974 |
| 11 | 94985200 | 94986599 | 11.47857 | 20.55483 | 35.71508 | 15.65625 | -0.39214 | 0.411783 | 0.539974 |
| 5 | 135375000 | 135382799 | 64.38317 | 100.4213 | 184.2083 | 76.44265 | -0.56259 | 2.683259 | 0.539975 |
| 15 | 32832800 | 32834799 | 30.55529 | 49.70805 | 79.07998 | 33.94172 | -0.29848 | 1.562743 | 0.539981 |
| 7 | 85738600 | 85741999 | 54.62024 | 87.51512 | 92.80209 | 39.15313 | 0.261625 | 1.999105 | 0.539983 |
| 10 | 92321000 | 92322999 | 23.53927 | 41.17311 | 36.78321 | 16.2215 | 0.549501 | 0.71801 | 0.539992 |
| 4 | 61823000 | 61825599 | 46.4755 | 75.11246 | 75.8512 | 32.3966 | 0.346771 | 1.718643 | 0.540003 |
| 17 | 76373200 | 76373799 | 4.481053 | 8.746941 | 19.24678 | 6.815208 | -0.46167 | -0.49036 | 0.54002 |
| 19 | 33545000 | 33546399 | 12.63753 | 22.32881 | 38.24536 | 16.91883 | -0.38198 | 0.525054 | 0.540026 |
| 13 | 106954600 | 106956799 | 26.28604 | 44.68962 | 46.86646 | 20.54027 | 0.313245 | 1.033776 | 0.540039 |
| 6 | 125342200 | 125344999 | 18.66133 | 31.84696 | 52.17823 | 22.87524 | -0.32707 | 0.977491 | 0.540051 |
| 8 | 52483600 | 52484799 | 12.30973 | 23.39135 | 23.70209 | 9.525084 | 0.475095 | 0.034689 | 0.540058 |
| 14 | 65411400 | 65416399 | 64.48431 | 101.1965 | 155.6868 | 64.56843 | -0.30221 | 2.571325 | 0.540098 |
| 3 | 75068000 | 75069799 | 14.07315 | 26.24905 | 24.78911 | 10.31357 | 0.534056 | 0.171028 | 0.5401 |
| 3 | 75622200 | 75622799 | 9.506691 | 17.65843 | 30.47834 | 12.76635 | -0.33699 | 0.17487 | 0.540112 |
| 14 | 42927400 | 42930199 | 19.78447 | 34.96178 | 33.031 | 14.61168 | 0.471863 | 0.557653 | 0.540118 |
| X | 127276200 | 127277799 | 21.6514 | 38.62909 | 30.24326 | 13.29878 | 0.74861 | 0.493091 | 0.540127 |
| 11 | 109302400 | 109304199 | 16.95742 | 29.90855 | 31.39175 | 13.89187 | 0.319861 | 0.558336 | 0.540142 |
| 2 | 68424800 | 68425199 | 4.778094 | 9.322669 | 19.77847 | 7.056668 | -0.42019 | -0.45833 | 0.540186 |
| 5 | 116380800 | 116385599 | 32.13758 | 52.19763 | 81.68945 | 35.03092 | -0.27686 | 1.592959 | 0.540194 |
| 14 | 47024200 | 47025999 | 11.4931 | 20.54471 | 35.71558 | 15.67969 | -0.39431 | 0.423204 | 0.540228 |
| 7 | 47953000 | 47953999 | 16.32336 | 29.54121 | 24.75298 | 10.72967 | 0.664732 | 0.361818 | 0.540236 |
| 6 | 67639000 | 67643199 | 111.3498 | 170.9757 | 179.0891 | 74.3555 | 0.227477 | 2.9863 | 0.540257 |
| 14 | 124757200 | 124758199 | 13.26136 | 24.94101 | 23.89328 | 9.802163 | 0.529637 | 0.11023 | 0.54026 |
| 1 | 141513800 | 141516399 | 23.40109 | 39.30579 | 66.10233 | 28.86813 | -0.3666 | 1.146538 | 0.540275 |
| 2 | 3264600 | 3265199 | 5.657138 | 11.01934 | 21.96996 | 8.014632 | -0.36213 | -0.36268 | 0.540279 |
| 14 | 92241400 | 92241999 | 5.657138 | 11.01934 | 21.96996 | 8.014632 | -0.36213 | -0.36268 | 0.540279 |
| 11 | 104375400 | 104377999 | 29.70094 | 49.36808 | 51.861 | 22.49781 | 0.31028 | 1.267108 | 0.540343 |
| 15 | 83075000 | 83079799 | 59.06571 | 93.30327 | 141.9212 | 58.93993 | -0.27846 | 2.429203 | 0.540355 |

| | | | | | | | | | |
|---|---|---|---|---|---|---|---|---|---|
| 15 | 43746200 | 43748999 | 42.97365 | 69.44066 | 74.38777 | 31.76435 | 0.261544 | 1.718762 | 0.540362 |
| 6 | 110312600 | 110315599 | 26.24799 | 43.04995 | 81.97724 | 35.29391 | -0.54977 | 1.439156 | 0.540371 |
| 3 | 138780200 | 138781999 | 13.50193 | 23.23645 | 54.76576 | 24.02318 | -0.84716 | 0.88247 | 0.540378 |
| 2 | 108615400 | 108616399 | 8.912696 | 16.46044 | 30.41777 | 12.84078 | -0.44292 | 0.201539 | 0.540382 |
| 15 | 82559000 | 82560599 | 22.41809 | 39.1146 | 33.90345 | 14.92497 | 0.592744 | 0.743939 | 0.540415 |
| 3 | 35642800 | 35645599 | 42.23882 | 68.46712 | 72.34727 | 30.97906 | 0.285728 | 1.647966 | 0.540417 |
| 10 | 23374400 | 23377799 | 30.69653 | 50.92223 | 54.6003 | 23.65387 | 0.280606 | 1.294625 | 0.540418 |
| 8 | 7951400 | 7951799 | 7.225394 | 14.09847 | 16.89155 | 6.020726 | 0.394328 | -0.44301 | 0.540423 |
| 11 | 50787600 | 50803199 | 334.9164 | 476.7402 | 438.6005 | 181.3409 | 0.31431 | 4.318809 | 0.540441 |
| 4 | 121289800 | 121301199 | 246.65 | 358.9161 | 355.2012 | 146.3991 | 0.240527 | 3.988572 | 0.540465 |
| 15 | 4662400 | 4665599 | 42.70392 | 69.54898 | 68.9694 | 29.61419 | 0.382762 | 1.564034 | 0.540473 |
| 1 | 12657400 | 12663399 | 86.03906 | 135.0819 | 139.8991 | 58.02404 | 0.268471 | 2.571722 | 0.540502 |
| 9 | 19297800 | 19301199 | 55.89075 | 89.50928 | 94.32783 | 39.76879 | 0.269818 | 2.016925 | 0.540521 |
| 15 | 44742400 | 44743199 | 12.90847 | 23.74426 | 24.56812 | 10.42833 | 0.382625 | 0.243901 | 0.540525 |
| 5 | 19996200 | 19997399 | 12.82261 | 24.17828 | 24.61664 | 10.05479 | 0.4479 | 0.093967 | 0.540532 |
| 8 | 113262000 | 113264799 | 24.84209 | 42.96794 | 41.91791 | 18.39024 | 0.421356 | 0.87009 | 0.540534 |
| 14 | 88828400 | 88829999 | 22.31344 | 38.86038 | 34.0947 | 14.9918 | 0.575174 | 0.770203 | 0.540542 |
| 7 | 18379400 | 18383599 | 53.50937 | 85.82113 | 91.06517 | 38.47888 | 0.262393 | 1.966862 | 0.540543 |
| 6 | 142174800 | 142176799 | 41.35562 | 66.4593 | 101.0435 | 42.82205 | -0.25204 | 1.894678 | 0.54055 |
| X | 60478000 | 60484799 | 63.21319 | 101.4937 | 87.56125 | 37.05887 | 0.562285 | 1.948287 | 0.540581 |
| 19 | 29831600 | 29832599 | 9.406969 | 17.47237 | 30.4931 | 12.77336 | -0.35259 | 0.17637 | 0.540586 |
| 12 | 113920600 | 113929999 | 153.184 | 232.558 | 214.7884 | 88.93733 | 0.395948 | 3.227484 | 0.540598 |
| 17 | 63346200 | 63348999 | 46.07041 | 74.42942 | 77.29226 | 32.99588 | 0.305924 | 1.730751 | 0.540602 |
| 11 | 42166000 | 42166999 | 11.91314 | 21.02488 | 37.3594 | 16.53068 | -0.43501 | 0.533729 | 0.540641 |
| 1 | 110721200 | 110722999 | 23.73944 | 40.9941 | 41.95161 | 18.40412 | 0.352102 | 0.878299 | 0.540643 |
| X | 120065000 | 120068599 | 42.18221 | 70.72722 | 45.77245 | 19.96754 | 1.006808 | 1.211109 | 0.540671 |
| 7 | 62317000 | 62323399 | 20.5576 | 36.08855 | 27.03617 | 11.95965 | 0.802221 | 0.630535 | 0.540677 |
| 11 | 29892000 | 29894199 | 15.87089 | 27.75317 | 45.15029 | 19.90717 | -0.31117 | 0.696012 | 0.540679 |
| 18 | 4706200 | 4708399 | 15.87089 | 27.75317 | 45.15029 | 19.90717 | -0.31117 | 0.696012 | 0.540679 |
| X | 94657400 | 94662599 | 52.75271 | 85.22659 | 76.77463 | 32.77567 | 0.510681 | 1.744321 | 0.54068 |
| 2 | 58968800 | 58969599 | 14.66981 | 26.91774 | 23.70651 | 10.09639 | 0.609722 | 0.267554 | 0.54072 |
| 7 | 4277000 | 4279799 | 30.50868 | 50.57641 | 53.50431 | 23.16825 | 0.299897 | 1.300646 | 0.540731 |
| X | 33736600 | 33739999 | 47.18721 | 77.07601 | 64.43318 | 27.69597 | 0.631492 | 1.53004 | 0.540732 |
| 11 | 43524000 | 43525999 | 16.94082 | 28.88491 | 52.14148 | 22.85748 | -0.46649 | 0.978475 | 0.54076 |
| 7 | 127053400 | 127056399 | 22.28208 | 38.93222 | 33.34081 | 14.69031 | 0.61024 | 0.723107 | 0.540761 |
| 1 | 163975800 | 163977199 | 12.18297 | 21.63843 | 36.22424 | 15.98938 | -0.34545 | 0.471131 | 0.540764 |
| 9 | 102400200 | 102407199 | 48.87321 | 77.61505 | 138.4819 | 57.73 | -0.50155 | 2.282059 | 0.540779 |
| 12 | 113523400 | 113528199 | 49.05434 | 79.47339 | 72.91722 | 31.21114 | 0.488269 | 1.669319 | 0.54078 |
| 7 | 41482000 | 41494999 | 202.2018 | 300.4502 | 284.4348 | 117.1768 | 0.333005 | 3.649078 | 0.540785 |
| 9 | 115885200 | 115887199 | 21.26285 | 37.17282 | 36.93006 | 16.2794 | 0.396926 | 0.713011 | 0.54079 |
| 13 | 65868800 | 65875199 | 96.55056 | 150.0512 | 153.6707 | 63.80429 | 0.273271 | 2.758208 | 0.540795 |
| 12 | 11026000 | 11029999 | 29.9131 | 51.00536 | 45.69226 | 20.03037 | 0.540334 | 1.019943 | 0.540808 |
| 9 | 64584600 | 64587799 | 19.62956 | 33.14195 | 56.34145 | 24.64496 | -0.38138 | 1.084545 | 0.540809 |
| 5 | 82327800 | 82328199 | 5.818337 | 11.28332 | 22.7443 | 8.584938 | -0.42336 | -0.23786 | 0.540821 |
| 1 | 60413400 | 60414999 | 18.61268 | 32.61154 | 34.56376 | 15.26473 | 0.304131 | 0.661989 | 0.540832 |
| 8 | 118563600 | 118569399 | 51.20647 | 81.64424 | 122.0941 | 51.15178 | -0.24128 | 2.15575 | 0.540851 |
| 17 | 16979800 | 16984199 | 68.53205 | 108.6589 | 116.1982 | 48.36192 | 0.233533 | 2.339373 | 0.540852 |

| | | | | | | | | | |
|---|---|---|---|---|---|---|---|---|---|
| X | 146962600 | 146964599 | 24.73007 | 43.02208 | 39.65469 | 17.42461 | 0.504192 | 0.804753 | 0.540863 |
| 10 | 20555400 | 20560399 | 41.21973 | 66.38187 | 107.6833 | 45.77681 | -0.34159 | 1.830806 | 0.540866 |
| 1 | 18586800 | 18590999 | 62.58202 | 99.52528 | 104.6951 | 43.70661 | 0.261448 | 2.244052 | 0.540877 |
| 15 | 89497600 | 89500199 | 63.42971 | 100.8152 | 105.6541 | 44.08122 | 0.266059 | 2.262109 | 0.54089 |
| 4 | 24057600 | 24060199 | 22.60417 | 39.66369 | 34.47077 | 15.24269 | 0.588852 | 0.65862 | 0.5409 |
| 1 | 93038400 | 93043399 | 54.287 | 87.17046 | 80.22196 | 33.95764 | 0.469388 | 1.928079 | 0.540901 |
| 13 | 51422600 | 51431399 | 105.4139 | 160.9311 | 229.9026 | 95.34398 | -0.22629 | 3.103368 | 0.540914 |
| 1 | 65225400 | 65226399 | 9.224687 | 17.10634 | 29.94141 | 12.56967 | -0.35915 | 0.181486 | 0.540943 |
| 3 | 143552200 | 143552999 | 8.47427 | 15.80753 | 28.99471 | 12.06148 | -0.41554 | 0.138622 | 0.540972 |
| 17 | 18951000 | 18954599 | 50.189 | 81.15592 | 75.21903 | 32.14795 | 0.471085 | 1.711347 | 0.54098 |
| 14 | 120788000 | 120790599 | 16.31831 | 28.18079 | 62.13833 | 27.26006 | -0.75079 | 0.873008 | 0.540982 |
| X | 126071400 | 126073599 | 31.60915 | 53.48273 | 41.057 | 17.97092 | 0.761474 | 1.091201 | 0.540983 |
| 18 | 11409400 | 11415799 | 64.62206 | 102.0198 | 150.4717 | 62.47219 | -0.2348 | 2.448078 | 0.54101 |
| 11 | 28090400 | 28096599 | 97.46912 | 151.2242 | 158.8708 | 65.97997 | 0.23464 | 2.785651 | 0.541027 |
| 18 | 22214400 | 22214999 | 5.658847 | 11.01711 | 21.89132 | 8.03129 | -0.36461 | -0.34236 | 0.541033 |
| 14 | 42545400 | 42546399 | 13.16254 | 24.80348 | 24.65422 | 10.08099 | 0.48142 | 0.10291 | 0.54105 |
| 9 | 61471800 | 61474599 | 26.02848 | 44.13405 | 46.3527 | 20.30198 | 0.310812 | 1.054012 | 0.541064 |
| 11 | 44456600 | 44458599 | 21.86379 | 36.50731 | 61.29783 | 26.67031 | -0.36403 | 1.201704 | 0.541091 |
| 15 | 6137000 | 6138399 | 14.57737 | 26.75851 | 27.42587 | 11.67427 | 0.393128 | 0.27464 | 0.541095 |
| 12 | 32077400 | 32078399 | 8.380548 | 15.77927 | 28.58898 | 11.70624 | -0.3802 | 0.073294 | 0.541104 |
| 13 | 78411400 | 78416199 | 42.98475 | 70.48911 | 63.11796 | 27.17254 | 0.535755 | 1.467864 | 0.541108 |
| 5 | 3645800 | 3646999 | 12.69526 | 22.29438 | 39.41817 | 17.44223 | -0.43036 | 0.600851 | 0.541108 |
| 11 | 61512200 | 61515999 | 28.1128 | 46.29973 | 73.24984 | 31.59176 | -0.28072 | 1.386895 | 0.541119 |
| 7 | 104488200 | 104492999 | 64.63452 | 102.9 | 107.9177 | 45.119 | 0.267448 | 2.211182 | 0.541135 |
| 1 | 118040400 | 118047399 | 127.5778 | 195.1148 | 195.8649 | 81.26148 | 0.284337 | 3.071584 | 0.541146 |
| 17 | 49876800 | 49878599 | 19.54779 | 34.21639 | 34.88353 | 15.39319 | 0.35968 | 0.680426 | 0.541148 |
| 12 | 51356600 | 51366199 | 273.9867 | 387.352 | 532.3663 | 220.3599 | -0.27448 | 4.415866 | 0.541199 |
| 10 | 37063400 | 37065399 | 19.88396 | 35.18632 | 31.24197 | 13.81645 | 0.561465 | 0.538116 | 0.54121 |
| 6 | 105938600 | 105939999 | 11.6086 | 22.27228 | 22.27446 | 8.74555 | 0.52385 | -0.05029 | 0.541224 |
| 4 | 63641800 | 63642599 | 6.408576 | 12.36173 | 24.21828 | 9.363227 | -0.41617 | -0.14602 | 0.541225 |
| 13 | 57710200 | 57712799 | 18.25411 | 31.50894 | 51.16687 | 22.44656 | -0.31141 | 0.870435 | 0.541231 |
| 7 | 26847400 | 26849999 | 20.49256 | 34.64642 | 55.92287 | 24.47202 | -0.30695 | 1.07208 | 0.541274 |
| 9 | 36527800 | 36528199 | 14.81298 | 26.54489 | 26.94155 | 11.80309 | 0.379766 | 0.412891 | 0.541286 |
| 1 | 55521400 | 55521999 | 9.263542 | 17.24618 | 30.53549 | 12.74439 | -0.36903 | 0.16158 | 0.541303 |
| 5 | 127180400 | 127181599 | 7.587103 | 14.55127 | 26.90038 | 10.57299 | -0.35627 | -0.07367 | 0.541313 |
| 6 | 83981600 | 83989999 | 86.5677 | 133.2342 | 199.4638 | 82.84133 | -0.28472 | 2.933964 | 0.54132 |
| 9 | 19698400 | 19711199 | 180.3198 | 270.0915 | 259.9272 | 107.2161 | 0.319863 | 3.499816 | 0.541324 |
| 14 | 53782200 | 53783199 | 6.866269 | 13.1632 | 25.16444 | 9.902744 | -0.40523 | -0.08014 | 0.541324 |
| 18 | 75645400 | 75646999 | 6.866269 | 13.1632 | 25.16444 | 9.902744 | -0.40523 | -0.08014 | 0.541324 |
| 18 | 19574400 | 19575399 | 16.00767 | 28.40713 | 29.84955 | 13.18514 | 0.321897 | 0.487737 | 0.541344 |
| 7 | 14103000 | 14104799 | 36.09878 | 59.00154 | 62.71976 | 26.96571 | 0.286398 | 1.496118 | 0.541358 |
| 5 | 90116000 | 90116599 | 9.675272 | 18.07219 | 31.70793 | 13.16005 | -0.34998 | 0.144411 | 0.54137 |
| 9 | 123195600 | 123196399 | 6.264156 | 12.08569 | 24.22302 | 9.359418 | -0.448 | -0.1501 | 0.541376 |
| 12 | 9714400 | 9716799 | 28.00706 | 47.10835 | 50.17648 | 21.92319 | 0.290302 | 1.135543 | 0.541382 |
| 6 | 146579600 | 146599199 | 251.4058 | 360.2259 | 486.1698 | 200.7932 | -0.22883 | 4.218825 | 0.541384 |
| 10 | 127824200 | 127825199 | 11.80477 | 22.10277 | 24.0471 | 9.940041 | 0.339117 | 0.127341 | 0.541385 |
| 11 | 73251600 | 73255399 | 48.22949 | 78.23384 | 72.1131 | 30.89513 | 0.483285 | 1.645216 | 0.541387 |

| | | | | | | | | | |
|---|---|---|---|---|---|---|---|---|---|
| X | 34772400 | 34784999 | 189.1937 | 284.5862 | 237.1128 | 97.90904 | 0.533117 | 3.421072 | 0.541413 |
| X | 146431200 | 146444999 | 173.7883 | 262.8904 | 219.6648 | 90.85532 | 0.536 | 3.306808 | 0.541419 |
| 5 | 58390400 | 58394999 | 123.9191 | 189.1756 | 196.3365 | 81.42297 | 0.23425 | 3.106028 | 0.541419 |
| 17 | 48959800 | 48961599 | 22.36888 | 39.00479 | 38.01105 | 16.72676 | 0.424317 | 0.756784 | 0.54142 |
| 16 | 44167600 | 44168799 | 10.58595 | 19.09914 | 33.31421 | 14.49631 | -0.38995 | 0.357924 | 0.541423 |
| 1 | 164971000 | 164973999 | 30.60165 | 49.95135 | 79.29876 | 34.07742 | -0.29209 | 1.511302 | 0.541435 |
| 4 | 73305400 | 73313599 | 143.4692 | 218.1316 | 212.5985 | 88.06956 | 0.320138 | 3.195422 | 0.541476 |
| 16 | 4104400 | 4107199 | 17.63222 | 30.03617 | 52.12218 | 22.84733 | -0.40985 | 0.992728 | 0.541542 |
| 4 | 108016200 | 108019599 | 29.10797 | 47.83503 | 76.14379 | 32.80982 | -0.29081 | 1.417878 | 0.541555 |
| 2 | 176879400 | 176880399 | 9.023886 | 16.96279 | 29.65306 | 12.17565 | -0.33213 | 0.093333 | 0.541558 |
| 1 | 21134800 | 21138199 | 39.34537 | 63.38878 | 97.73508 | 41.57193 | -0.26769 | 1.810368 | 0.541561 |
| 18 | 86411200 | 86414599 | 26.49383 | 43.56988 | 71.23997 | 30.70599 | -0.32892 | 1.398791 | 0.541574 |
| 16 | 82341000 | 82345199 | 47.9325 | 77.64007 | 66.08142 | 28.28093 | 0.59609 | 1.669045 | 0.541579 |
| 5 | 150954000 | 150963799 | 118.9664 | 180.1539 | 255.974 | 105.9089 | -0.22805 | 3.273184 | 0.541604 |
| 4 | 92372400 | 92373199 | 9.026369 | 16.8586 | 29.51255 | 12.25075 | -0.34683 | 0.133809 | 0.54161 |
| 4 | 7648200 | 7652599 | 113.7813 | 174.3276 | 180.4032 | 74.87774 | 0.242523 | 3.030031 | 0.541626 |
| 18 | 8227600 | 8228199 | 12.47309 | 22.17844 | 36.69275 | 16.18096 | -0.32771 | 0.466113 | 0.541626 |
| 1 | 5619000 | 5620399 | 14.65455 | 26.77483 | 27.84832 | 11.92849 | 0.365593 | 0.301023 | 0.54163 |
| 11 | 81520200 | 81521199 | 6.730796 | 12.92569 | 25.19671 | 9.866607 | -0.42581 | -0.09519 | 0.541644 |
| 13 | 62356800 | 62382999 | 443.1549 | 615.1438 | 557.3001 | 231.3492 | 0.299971 | 4.694442 | 0.541649 |
| 11 | 50473200 | 50479799 | 55.03382 | 87.3934 | 130.4934 | 54.39649 | -0.24515 | 2.286924 | 0.541653 |
| 2 | 39743200 | 39743999 | 19.83375 | 34.80372 | 35.39683 | 15.65269 | 0.363309 | 0.646509 | 0.541671 |
| 6 | 106821400 | 106824599 | 26.66555 | 45.38841 | 46.85932 | 20.53909 | 0.336042 | 1.025251 | 0.541683 |
| 3 | 126236600 | 126237999 | 11.14899 | 20.21558 | 33.79386 | 14.61355 | -0.32348 | 0.332267 | 0.541706 |
| 14 | 113182600 | 113183999 | 20.52396 | 35.93632 | 36.4207 | 16.07575 | 0.368302 | 0.684854 | 0.541715 |
| 7 | 96764200 | 96766799 | 38.49572 | 62.77653 | 66.93253 | 28.75314 | 0.279753 | 1.534515 | 0.541715 |
| 2 | 94576000 | 94577999 | 18.93821 | 32.54894 | 52.22903 | 22.90951 | -0.29556 | 0.91402 | 0.541726 |
| 18 | 7867800 | 7870199 | 16.32933 | 27.93863 | 55.33176 | 24.26277 | -0.59886 | 0.949438 | 0.54173 |
| 7 | 116715200 | 116715599 | 8.593413 | 16.64306 | 19.35917 | 7.352206 | 0.353609 | -0.19646 | 0.541735 |
| 19 | 21041800 | 21044199 | 24.85621 | 42.2246 | 44.80334 | 19.63442 | 0.296312 | 1.034465 | 0.541758 |
| 7 | 121235000 | 121236599 | 10.79718 | 19.757 | 33.40512 | 14.28068 | -0.329 | 0.275817 | 0.54178 |
| 7 | 9511000 | 9511599 | 15.72809 | 28.43658 | 24.74121 | 10.73747 | 0.6095 | 0.361833 | 0.541798 |
| 4 | 96291400 | 96293199 | 27.63956 | 46.69397 | 49.07147 | 21.46884 | 0.30981 | 1.095442 | 0.541819 |
| 4 | 13790800 | 13792199 | 14.81579 | 27.07398 | 27.20182 | 11.64912 | 0.415126 | 0.298094 | 0.541825 |
| 6 | 68186000 | 68187199 | 14.87912 | 26.78324 | 28.3693 | 12.38013 | 0.322358 | 0.389339 | 0.541848 |
| 2 | 39782200 | 39783599 | 19.65194 | 33.62768 | 53.82501 | 23.6023 | -0.29298 | 0.959709 | 0.54187 |
| 10 | 42241600 | 42243199 | 7.508642 | 14.37156 | 27.3333 | 10.80598 | -0.40394 | -0.05009 | 0.541871 |
| 15 | 100870600 | 100873199 | 11.89735 | 21.03838 | 36.81115 | 16.28168 | -0.41179 | 0.510543 | 0.54188 |
| 12 | 55867000 | 55868199 | 8.71762 | 15.88542 | 31.43527 | 13.50972 | -0.5587 | 0.282234 | 0.541884 |
| 12 | 114709800 | 114711199 | 22.78085 | 39.67601 | 39.2115 | 17.24214 | 0.404203 | 0.779736 | 0.541918 |
| 3 | 35923600 | 35931399 | 170.9196 | 257.2643 | 221.0912 | 91.30914 | 0.489267 | 3.402491 | 0.541931 |
| 13 | 88695400 | 88696999 | 15.25577 | 27.25935 | 28.29608 | 12.42679 | 0.344955 | 0.435729 | 0.54194 |
| 6 | 54984400 | 54985799 | 19.32767 | 33.75188 | 35.64562 | 15.70049 | 0.309133 | 0.718261 | 0.541975 |
| 8 | 71752200 | 71756999 | 54.16485 | 86.7474 | 92.9323 | 39.18191 | 0.24618 | 2.01267 | 0.541992 |
| 17 | 15794800 | 15797999 | 23.48135 | 39.06721 | 64.13335 | 27.83997 | -0.33167 | 1.247709 | 0.54201 |
| 3 | 27476600 | 27477199 | 3.814389 | 7.454806 | 17.68536 | 6.11546 | -0.53737 | -0.59216 | 0.542016 |
| 5 | 149120600 | 149121999 | 8.506458 | 15.60736 | 31.92312 | 13.5972 | -0.59783 | 0.238186 | 0.542022 |

| | | | | | | | | | |
|---|---|---|---|---|---|---|---|---|---|
| 1 | 73797400 | 73798399 | 9.520071 | 17.64148 | 30.44631 | 12.79407 | -0.34016 | 0.188498 | 0.542032 |
| 8 | 6332200 | 6334799 | 23.13488 | 38.72864 | 61.42343 | 26.76812 | -0.28211 | 1.17898 | 0.542054 |
| 9 | 97319400 | 97321799 | 15.46319 | 26.63513 | 47.94591 | 21.03219 | -0.45975 | 0.874661 | 0.542072 |
| 2 | 22685200 | 22690399 | 54.35356 | 87.59003 | 81.84488 | 34.82552 | 0.453803 | 1.820691 | 0.542074 |
| 10 | 15443400 | 15444199 | 12.01193 | 23.2288 | 20.82589 | 7.965038 | 0.71564 | -0.13325 | 0.542085 |
| 10 | 83590800 | 83593599 | 17.41088 | 29.69975 | 51.62045 | 22.6299 | -0.41196 | 0.976175 | 0.542108 |
| X | 110803800 | 110806399 | 34.83585 | 58.59896 | 41.5895 | 18.17177 | 0.874071 | 1.152776 | 0.542114 |
| 7 | 64725200 | 64728399 | 27.4069 | 44.8115 | 73.55724 | 31.6294 | -0.33833 | 1.478653 | 0.542118 |
| 2 | 20995800 | 20997599 | 15.5081 | 27.11444 | 44.09556 | 19.43951 | -0.31068 | 0.694194 | 0.542122 |
| 8 | 65758000 | 65758599 | 13.61265 | 25.16852 | 21.30164 | 8.978837 | 0.677272 | 0.210937 | 0.542143 |
| X | 38754600 | 38756599 | 24.03882 | 42.00234 | 30.43839 | 13.41161 | 0.850267 | 0.730554 | 0.542156 |
| 14 | 45918000 | 45921399 | 21.68671 | 36.28369 | 60.22252 | 26.23534 | -0.34743 | 1.178953 | 0.542158 |
| 13 | 57094400 | 57097199 | 21.97969 | 36.85654 | 59.73759 | 26.06098 | -0.31334 | 1.155235 | 0.542166 |
| 7 | 86061800 | 86065399 | 35.52686 | 59.27787 | 54.26819 | 23.60083 | 0.508169 | 1.245093 | 0.542168 |
| 13 | 29442800 | 29443799 | 9.528207 | 17.73458 | 30.52039 | 12.74336 | -0.32911 | 0.163616 | 0.542172 |
| 12 | 102310800 | 102312999 | 19.76689 | 33.66432 | 54.35382 | 23.82508 | -0.30659 | 1.002153 | 0.542178 |
| 12 | 67602400 | 67602999 | 15.94555 | 29.20919 | 22.83953 | 9.749555 | 0.777438 | 0.283332 | 0.542187 |
| 17 | 39124600 | 39132399 | 89.8974 | 142.2449 | 120.5422 | 50.10368 | 0.566244 | 2.409811 | 0.542205 |
| 1 | 54625800 | 54626599 | 6.681795 | 12.95934 | 25.16904 | 9.494902 | -0.37038 | -0.22415 | 0.542225 |
| 6 | 76136400 | 76136999 | 6.681795 | 12.95934 | 25.16904 | 9.494902 | -0.37038 | -0.22415 | 0.542225 |
| 6 | 25529200 | 25529599 | 16.04835 | 28.57832 | 29.67375 | 13.0692 | 0.341656 | 0.464343 | 0.542248 |
| 5 | 147062000 | 147066599 | 52.32335 | 83.19501 | 128.827 | 53.7703 | -0.29577 | 2.245341 | 0.542258 |
| 5 | 112933000 | 112936599 | 25.45953 | 41.68534 | 72.9712 | 31.39376 | -0.42956 | 1.452807 | 0.54227 |
| 12 | 12381200 | 12384599 | 43.51212 | 69.70411 | 106.5028 | 44.92064 | -0.26463 | 1.998007 | 0.542273 |
| 7 | 84428400 | 84428999 | 5.094538 | 9.925713 | 20.78985 | 7.561767 | -0.42822 | -0.38447 | 0.542278 |
| 14 | 17667400 | 17669999 | 25.49242 | 43.80299 | 38.80303 | 17.02026 | 0.558013 | 0.925955 | 0.542312 |
| 5 | 52720600 | 52727399 | 77.50165 | 120.6693 | 176.5387 | 73.27893 | -0.23864 | 2.718855 | 0.542333 |
| 16 | 77631000 | 77631999 | 9.014337 | 16.87269 | 29.55473 | 12.22328 | -0.34347 | 0.119615 | 0.542346 |
| 10 | 44648800 | 44650199 | 18.14912 | 32.74196 | 29.50606 | 12.84053 | 0.556571 | 0.401125 | 0.542347 |
| 3 | 151299400 | 151301599 | 25.03511 | 43.54616 | 39.63205 | 17.41301 | 0.52237 | 0.807958 | 0.542388 |
| 14 | 30469600 | 30472599 | 36.76414 | 60.89714 | 58.22722 | 25.19924 | 0.445274 | 1.332346 | 0.542396 |
| 10 | 28885800 | 28890799 | 64.40404 | 102.7063 | 94.00287 | 39.39331 | 0.466149 | 2.153563 | 0.542397 |
| 16 | 15977800 | 15978399 | 8.394629 | 15.84207 | 28.09217 | 11.45526 | -0.34423 | 0.058384 | 0.542405 |
| 13 | 102103200 | 102108399 | 39.07147 | 63.80334 | 66.06622 | 28.39526 | 0.323209 | 1.514516 | 0.542405 |
| 6 | 10185000 | 10186599 | 21.2893 | 37.30297 | 36.37419 | 16.06489 | 0.423788 | 0.678761 | 0.542405 |
| 13 | 111210600 | 111211199 | 6.851402 | 13.22688 | 24.81892 | 9.567685 | -0.3505 | -0.14813 | 0.542416 |
| X | 60783600 | 60789599 | 32.5984 | 55.25198 | 41.39789 | 18.12963 | 0.79632 | 1.078436 | 0.542421 |
| 7 | 34164400 | 34165199 | 7.012499 | 13.51067 | 25.31121 | 9.81904 | -0.35673 | -0.12378 | 0.542439 |
| 7 | 13439800 | 13443599 | 70.63577 | 112.1269 | 116.0933 | 48.37552 | 0.281755 | 2.308061 | 0.542448 |
| 12 | 62471400 | 62473799 | 25.20823 | 43.2498 | 38.88952 | 17.05672 | 0.536233 | 0.938773 | 0.54245 |
| 11 | 35612600 | 35614399 | 15.01567 | 27.36508 | 27.70927 | 11.90667 | 0.400891 | 0.31631 | 0.542462 |
| 11 | 103938200 | 103941399 | 28.15767 | 46.19344 | 72.89274 | 31.38601 | -0.2792 | 1.435411 | 0.542472 |
| 14 | 86825400 | 86828399 | 32.09729 | 53.26381 | 56.28599 | 24.38979 | 0.301592 | 1.297291 | 0.54248 |
| X | 147747000 | 147750599 | 32.31892 | 55.23632 | 43.99323 | 19.28919 | 0.709919 | 1.009053 | 0.542525 |
| 4 | 141523600 | 141531799 | 98.38713 | 150.8832 | 216.4498 | 89.85177 | -0.22719 | 3.010311 | 0.542525 |
| 3 | 123584000 | 123586199 | 19.2755 | 34.18941 | 31.93565 | 14.11246 | 0.490308 | 0.514879 | 0.542529 |
| 8 | 110184400 | 110185599 | 12.18153 | 21.57712 | 36.73264 | 16.2415 | -0.37162 | 0.49286 | 0.542536 |

| | | | | | | | | | |
|---|---|---|---|---|---|---|---|---|---|
| 3 | 113136400 | 113137399 | 16.707 | 29.85644 | 30.72365 | 13.49138 | 0.357668 | 0.451057 | 0.542547 |
| 18 | 22190000 | 22199599 | 123.2327 | 188.6391 | 192.9617 | 80.07363 | 0.25823 | 3.052124 | 0.542561 |
| 2 | 19918800 | 19920999 | 21.46271 | 36.1052 | 58.66641 | 25.6336 | -0.31689 | 1.120658 | 0.542575 |
| 10 | 25011600 | 25016599 | 53.76511 | 86.56751 | 84.05192 | 35.72022 | 0.397053 | 1.845876 | 0.542583 |
| 16 | 27528600 | 27529799 | 15.61301 | 28.47625 | 28.25209 | 12.12789 | 0.431214 | 0.31853 | 0.542598 |
| 7 | 6485600 | 6486199 | 12.5844 | 23.26571 | 24.57308 | 10.3588 | 0.360259 | 0.213067 | 0.542598 |
| 17 | 92265000 | 92266599 | 17.91192 | 31.95392 | 31.01751 | 13.64047 | 0.439826 | 0.468852 | 0.5426 |
| 14 | 78512800 | 78513799 | 12.1872 | 21.52336 | 37.78489 | 16.71665 | -0.41747 | 0.52687 | 0.542622 |
| 5 | 36940200 | 36945399 | 64.14909 | 101.2916 | 148.9016 | 61.82601 | -0.22979 | 2.443586 | 0.542624 |
| 2 | 109853600 | 109855799 | 30.07125 | 51.13519 | 46.80927 | 20.51553 | 0.508704 | 1.045749 | 0.542626 |
| 9 | 26159800 | 26160599 | 11.39815 | 21.36295 | 23.12022 | 9.540618 | 0.348925 | 0.11638 | 0.542635 |
| 8 | 86488200 | 86489399 | 10.50974 | 20.17739 | 18.57036 | 7.277354 | 0.645711 | -0.07642 | 0.54264 |
| X | 38988600 | 38989399 | 22.41086 | 38.99463 | 38.31555 | 16.83814 | 0.412534 | 0.791774 | 0.542647 |
| 6 | 24823000 | 24824199 | 11.04303 | 20.17834 | 34.99956 | 14.98979 | -0.36724 | 0.291165 | 0.542675 |
| 7 | 12118600 | 12131599 | 189.6371 | 281.875 | 284.0401 | 117.0182 | 0.243432 | 3.64059 | 0.542697 |
| Y | 35125800 | 35126799 | 8.686896 | 16.82375 | 14.16867 | 5.381497 | 0.814461 | -0.21528 | 0.542711 |
| Y | 5335800 | 5336799 | 10.4372 | 19.87652 | 15.63581 | 6.254526 | 0.842993 | -0.02239 | 0.542712 |
| X | 145289000 | 145292199 | 37.311 | 61.95454 | 56.55589 | 24.51736 | 0.51219 | 1.305648 | 0.542727 |
| 13 | 10938800 | 10946999 | 92.83099 | 144.3039 | 151.9825 | 63.10117 | 0.23316 | 2.751079 | 0.542729 |
| X | 59138000 | 59139399 | 13.64505 | 25.27708 | 26.03798 | 10.94653 | 0.398923 | 0.214593 | 0.542731 |
| 5 | 75243800 | 75246399 | 21.0752 | 35.18871 | 61.24186 | 26.64508 | -0.41559 | 1.199606 | 0.542736 |
| 15 | 76942200 | 76944799 | 35.02149 | 57.51005 | 61.86152 | 26.64725 | 0.272664 | 1.431158 | 0.542743 |
| 15 | 7597400 | 7600399 | 19.72629 | 34.4321 | 36.69414 | 16.15764 | 0.296151 | 0.729269 | 0.542745 |
| 18 | 7395800 | 7397599 | 19.83167 | 35.35582 | 30.23034 | 13.30154 | 0.62126 | 0.4833 | 0.542754 |
| 13 | 20958200 | 20962399 | 49.60926 | 80.35255 | 73.46853 | 31.44138 | 0.492948 | 1.676187 | 0.542756 |
| 13 | 68397000 | 68435199 | 918.7403 | 1196.14 | 1058.824 | 443.4946 | 0.24066 | 5.711779 | 0.542767 |
| 8 | 16953400 | 16955199 | 15.02891 | 27.57251 | 27.25644 | 11.61065 | 0.443873 | 0.278432 | 0.542774 |
| 6 | 129784600 | 129789999 | 91.55098 | 140.2558 | 231.7466 | 96.18634 | -0.43203 | 3.02912 | 0.542799 |
| 2 | 159274800 | 159276799 | 26.60274 | 46.00681 | 36.54589 | 16.03336 | 0.71651 | 0.872504 | 0.542803 |
| 8 | 17736400 | 17739199 | 27.53567 | 47.151 | 43.47807 | 19.06624 | 0.499542 | 0.96852 | 0.542815 |
| 7 | 16861000 | 16862599 | 25.32717 | 42.0287 | 66.35995 | 28.75482 | -0.27595 | 1.286753 | 0.542864 |
| 8 | 71894600 | 71896199 | 11.50749 | 20.5347 | 35.71908 | 15.70423 | -0.39657 | 0.434525 | 0.542865 |
| 11 | 31110000 | 31113599 | 39.86151 | 64.74908 | 69.54771 | 29.82631 | 0.264946 | 1.599061 | 0.542888 |
| 2 | 56100000 | 56100999 | 13.18059 | 24.11784 | 25.48475 | 10.89497 | 0.344925 | 0.276454 | 0.542933 |
| 7 | 53271600 | 53272599 | 13.18059 | 24.11784 | 25.48475 | 10.89497 | 0.344925 | 0.276454 | 0.542933 |
| 7 | 113928000 | 113930599 | 28.3209 | 47.68656 | 49.09958 | 21.45954 | 0.338992 | 1.124082 | 0.542954 |
| 6 | 48079800 | 48080999 | 17.98422 | 31.79863 | 32.70253 | 14.46473 | 0.350436 | 0.540278 | 0.542959 |
| 14 | 112525800 | 112532199 | 66.19222 | 105.119 | 112.4647 | 46.88392 | 0.235125 | 2.28937 | 0.542972 |
| 12 | 42283400 | 42286599 | 32.81348 | 55.72496 | 46.23208 | 20.25866 | 0.649932 | 1.067443 | 0.542982 |
| 4 | 115642000 | 115642599 | 6.274774 | 12.08295 | 24.16941 | 9.390328 | -0.4521 | -0.13005 | 0.542982 |
| 1 | 108779400 | 108783599 | 75.58056 | 119.3217 | 127.0367 | 52.74289 | 0.234851 | 2.451278 | 0.542996 |
| 14 | 100485400 | 100488399 | 53.43965 | 84.84969 | 129.4304 | 53.94566 | -0.27609 | 2.290341 | 0.542999 |
| 3 | 12775000 | 12776599 | 18.97014 | 33.29307 | 34.2979 | 15.16857 | 0.344677 | 0.635053 | 0.543009 |
| 1 | 23680600 | 23681799 | 14.61497 | 26.78148 | 26.69747 | 11.39081 | 0.430052 | 0.279655 | 0.543035 |
| X | 73676200 | 73677199 | 15.62585 | 27.95018 | 27.99535 | 12.28376 | 0.397448 | 0.434279 | 0.543045 |
| 10 | 18634800 | 18636399 | 11.45029 | 20.85966 | 35.99809 | 15.47556 | -0.36382 | 0.31124 | 0.543066 |
| 15 | 13402400 | 13406199 | 38.81614 | 63.83952 | 61.28623 | 26.41662 | 0.437139 | 1.419096 | 0.543069 |

| | | | | | | | | | |
|---|---|---|---|---|---|---|---|---|---|
| 12 | 91613400 | 91621799 | 147.08 | 218.8382 | 316.3862 | 130.3575 | -0.2757 | 3.617886 | 0.54307 |
| 18 | 7746200 | 7749799 | 33.24227 | 55.03468 | 58.00369 | 25.09346 | 0.30514 | 1.328115 | 0.543072 |
| X | 81079600 | 81080599 | 19.13759 | 33.70532 | 33.36359 | 14.76871 | 0.40336 | 0.592919 | 0.543112 |
| 12 | 115482400 | 115484999 | 62.43397 | 99.60388 | 103.9648 | 43.59176 | 0.277521 | 2.143096 | 0.543113 |
| 6 | 84329200 | 84331399 | 17.58857 | 30.44369 | 49.54864 | 21.73847 | -0.3138 | 0.840636 | 0.543126 |
| 8 | 21409400 | 21424199 | 226.3497 | 332.8664 | 321.7814 | 132.5153 | 0.288932 | 3.824524 | 0.54315 |
| 2 | 18272600 | 18275199 | 34.35233 | 57.27374 | 54.28753 | 23.59771 | 0.458147 | 1.247527 | 0.543179 |
| 11 | 58852400 | 58856399 | 41.07918 | 65.84935 | 103.6448 | 43.75832 | -0.30623 | 1.97433 | 0.543189 |
| 7 | 131215800 | 131216999 | 7.195716 | 13.70179 | 26.11003 | 10.44704 | -0.42233 | -0.01825 | 0.543192 |
| 6 | 10992800 | 10993399 | 9.621838 | 18.41799 | 20.23253 | 7.996558 | 0.381228 | -0.05312 | 0.543197 |
| 3 | 77580400 | 77580999 | 9.494244 | 17.67425 | 30.51904 | 12.73928 | -0.33379 | 0.161093 | 0.5432 |
| 8 | 3890200 | 3893399 | 16.82277 | 30.28522 | 25.99864 | 11.34405 | 0.623626 | 0.395936 | 0.543227 |
| X | 116028800 | 116029399 | 11.99113 | 22.4458 | 23.64829 | 9.779547 | 0.385182 | 0.13243 | 0.543245 |
| 3 | 40582600 | 40584599 | 17.66876 | 31.10528 | 32.90954 | 14.56895 | 0.30743 | 0.586216 | 0.543254 |
| 8 | 41326000 | 41331799 | 73.22679 | 115.9485 | 121.1066 | 50.35622 | 0.26579 | 2.377336 | 0.543262 |
| 18 | 42083800 | 42084799 | 7.214411 | 13.6742 | 26.57552 | 10.73748 | -0.46283 | 0.01564 | 0.543264 |
| 5 | 110186800 | 110196599 | 211.559 | 314.7427 | 277.0885 | 114.1648 | 0.439386 | 3.62911 | 0.543276 |
| 2 | 101471800 | 101474999 | 33.48056 | 54.14902 | 85.77147 | 36.66672 | -0.29994 | 1.683121 | 0.543279 |
| 9 | 38528200 | 38532599 | 92.85089 | 145.0106 | 149.6331 | 62.08641 | 0.26787 | 2.670723 | 0.543281 |
| X | 117320000 | 117324999 | 57.01024 | 91.2919 | 92.8563 | 39.1417 | 0.32069 | 2.022968 | 0.543309 |
| X | 21063800 | 21064399 | 15.56734 | 28.59695 | 26.92089 | 11.44773 | 0.51632 | 0.278694 | 0.543312 |
| 9 | 22218600 | 22224599 | 77.46273 | 122.1427 | 129.3561 | 53.68137 | 0.241044 | 2.47893 | 0.543314 |
| 13 | 58278600 | 58285199 | 61.31491 | 96.9354 | 143.1687 | 59.4826 | -0.23512 | 2.411233 | 0.543339 |
| 8 | 127950600 | 127951199 | 5.807082 | 11.28866 | 22.32934 | 8.292418 | -0.37418 | -0.29507 | 0.54334 |
| X | 78801200 | 78802799 | 22.7017 | 39.61902 | 38.11479 | 16.78143 | 0.443194 | 0.75045 | 0.54334 |
| 10 | 48081200 | 48081799 | 16.24072 | 28.78098 | 28.99814 | 12.82043 | 0.38133 | 0.499514 | 0.543343 |
| 5 | 125992400 | 125999599 | 81.20038 | 126.192 | 182.6504 | 75.83697 | -0.2253 | 2.753842 | 0.543385 |
| 2 | 98426800 | 98428399 | 11.61576 | 21.1688 | 39.33836 | 16.90384 | -0.46974 | 0.316298 | 0.543391 |
| 10 | 104388200 | 104392399 | 49.13761 | 78.93879 | 83.20225 | 35.24392 | 0.274835 | 1.906602 | 0.543432 |
| 14 | 52910400 | 52913999 | 38.1512 | 63.44827 | 54.96695 | 23.85413 | 0.587545 | 1.287549 | 0.543455 |
| 7 | 95898400 | 95899799 | 17.85875 | 30.99154 | 51.72309 | 22.69426 | -0.34904 | 0.823763 | 0.543479 |
| X | 116207600 | 116208599 | 18.18401 | 32.30526 | 31.23581 | 13.78451 | 0.442377 | 0.499774 | 0.543496 |
| 12 | 5670800 | 5673999 | 36.99946 | 61.49626 | 56.04863 | 24.31357 | 0.514515 | 1.29187 | 0.543498 |
| X | 88276200 | 88277799 | 19.14196 | 33.6822 | 33.40416 | 14.78938 | 0.40015 | 0.603038 | 0.543611 |
| 3 | 51233800 | 51235599 | 20.74861 | 36.14049 | 38.27689 | 16.83232 | 0.3048 | 0.766615 | 0.543612 |
| 18 | 61661800 | 61663599 | 26.44431 | 43.73607 | 69.27628 | 29.94932 | -0.281 | 1.329202 | 0.543622 |
| 12 | 30736600 | 30738999 | 27.87039 | 46.75685 | 50.18859 | 21.90181 | 0.279182 | 1.160373 | 0.543623 |
| 2 | 21701000 | 21701399 | 11.06753 | 20.89172 | 23.15453 | 9.436605 | 0.329299 | 0.067753 | 0.543633 |
| 5 | 109330800 | 109337799 | 96.66344 | 152.3409 | 128.0023 | 53.11607 | 0.57399 | 2.502816 | 0.543663 |
| 1 | 151178400 | 151179199 | 11.50711 | 21.79619 | 23.67852 | 9.580364 | 0.366891 | 0.048985 | 0.543675 |
| X | 8505200 | 8507599 | 33.4663 | 56.72915 | 46.83777 | 20.51263 | 0.656954 | 1.088882 | 0.543696 |
| 11 | 38547400 | 38548199 | 9.828929 | 18.87331 | 20.68614 | 8.102778 | 0.395861 | -0.08073 | 0.543698 |
| 8 | 120578800 | 120583199 | 26.35693 | 42.93024 | 83.91773 | 36.03409 | -0.5936 | 1.537878 | 0.543719 |
| 6 | 101815600 | 101816799 | 17.66301 | 31.88844 | 29.57278 | 12.85773 | 0.516308 | 0.393873 | 0.543722 |
| 1 | 93911800 | 93916399 | 37.87279 | 61.03936 | 94.5163 | 40.22436 | -0.27316 | 1.795062 | 0.54375 |
| 4 | 112775600 | 112786999 | 219.8889 | 325.0139 | 303.7491 | 125.0911 | 0.344546 | 3.740579 | 0.543757 |
| 11 | 21211400 | 21213399 | 24.14924 | 41.67805 | 42.4987 | 18.6438 | 0.357146 | 0.886715 | 0.543761 |

| | | | | | | | | | |
|---|---|---|---|---|---|---|---|---|---|
| 9 | 104446600 | 104447799 | 8.190997 | 15.15096 | 29.45417 | 12.40789 | -0.51279 | 0.186658 | 0.543812 |
| 11 | 49553200 | 49555999 | 17.19335 | 29.56449 | 48.99821 | 21.49268 | -0.34168 | 0.899078 | 0.543818 |
| 6 | 86181200 | 86182399 | 12.03103 | 22.56083 | 24.6041 | 10.14403 | 0.338802 | 0.121854 | 0.543822 |
| 15 | 98597200 | 98597999 | 5.541251 | 10.7487 | 22.75252 | 8.577895 | -0.4918 | -0.24638 | 0.543824 |
| 4 | 143864400 | 143875799 | 246.0612 | 359.0482 | 347.7462 | 143.2854 | 0.275496 | 3.947423 | 0.543831 |
| 15 | 34524800 | 34525399 | 9.587843 | 18.45062 | 19.81767 | 7.714087 | 0.434513 | -0.10144 | 0.543848 |
| 1 | 44399800 | 44401999 | 42.08951 | 68.12147 | 72.38769 | 30.96165 | 0.275565 | 1.675479 | 0.543851 |
| 2 | 128264600 | 128267799 | 24.30883 | 40.33351 | 65.79883 | 28.50908 | -0.323 | 1.283075 | 0.543853 |
| 3 | 102399400 | 102401199 | 11.8292 | 21.17236 | 35.22104 | 15.44605 | -0.32993 | 0.41804 | 0.543885 |
| 9 | 73064400 | 73066599 | 15.87043 | 27.65435 | 44.70629 | 19.66449 | -0.30245 | 0.747165 | 0.543888 |
| 10 | 83614200 | 83617799 | 54.61329 | 87.40281 | 93.41793 | 39.34506 | 0.248331 | 2.034769 | 0.543889 |
| 11 | 28119200 | 28119599 | 17.36481 | 30.58357 | 31.97527 | 14.15411 | 0.324795 | 0.578537 | 0.543906 |
| 16 | 3374000 | 3379399 | 52.62441 | 85.07509 | 75.35155 | 32.19536 | 0.536104 | 1.723344 | 0.543909 |
| 7 | 72969800 | 72979799 | 93.66057 | 143.6939 | 212.8283 | 88.35344 | -0.27278 | 2.998941 | 0.543915 |
| 3 | 142417000 | 142420999 | 67.63616 | 105.8802 | 162.1423 | 67.26049 | -0.29836 | 2.617031 | 0.543929 |
| 12 | 71747600 | 71749999 | 20.9739 | 36.46239 | 38.81357 | 17.04761 | 0.297207 | 0.794241 | 0.543961 |
| 10 | 115327600 | 115329399 | 26.14292 | 44.10569 | 46.95272 | 20.53344 | 0.291233 | 1.099637 | 0.54397 |
| 1 | 3321000 | 3321999 | 11.52736 | 20.6443 | 34.74568 | 15.23 | -0.34607 | 0.411862 | 0.543983 |
| 13 | 98775600 | 98779599 | 37.04026 | 59.77927 | 93.10934 | 39.69553 | -0.27932 | 1.753054 | 0.543995 |
| 9 | 11342000 | 11342399 | 6.119063 | 11.82122 | 23.70107 | 9.107363 | -0.44007 | -0.16904 | 0.544024 |
| 17 | 92817000 | 92819999 | 27.71375 | 46.64783 | 49.09834 | 21.45671 | 0.307375 | 1.125675 | 0.544037 |
| 5 | 95646800 | 95653399 | 93.92125 | 147.4651 | 139.7432 | 57.95978 | 0.396534 | 2.576324 | 0.544037 |
| X | 93164800 | 93169199 | 29.97221 | 51.31587 | 37.23584 | 16.32867 | 0.844136 | 0.977973 | 0.544048 |
| 12 | 10935000 | 10936199 | 15.06813 | 27.50262 | 24.24724 | 10.39917 | 0.601413 | 0.302076 | 0.544052 |
| 9 | 7608600 | 7610399 | 33.51733 | 55.53617 | 50.37477 | 21.80555 | 0.521035 | 1.311227 | 0.544058 |
| 3 | 40008600 | 40012399 | 27.97239 | 47.79338 | 40.34397 | 17.68871 | 0.626127 | 0.988124 | 0.544062 |
| 4 | 112285600 | 112290199 | 39.08074 | 64.18808 | 62.90518 | 27.09899 | 0.406733 | 1.438319 | 0.544066 |
| 2 | 17758600 | 17760599 | 18.6535 | 31.99253 | 51.12704 | 22.42339 | -0.29014 | 0.931466 | 0.54407 |
| 10 | 107997200 | 107998599 | 13.28293 | 23.30051 | 41.43424 | 18.31714 | -0.43887 | 0.623638 | 0.544077 |
| 18 | 37143600 | 37151399 | 144.7351 | 218.9989 | 223.4995 | 92.44908 | 0.248014 | 3.293993 | 0.544079 |
| 4 | 143903200 | 143908799 | 84.71308 | 133.8257 | 126.9805 | 52.73674 | 0.401499 | 2.441461 | 0.544102 |
| 6 | 137356000 | 137357799 | 15.23304 | 26.63947 | 43.0651 | 18.98881 | -0.30197 | 0.689003 | 0.544105 |
| 8 | 48871800 | 48876199 | 41.06582 | 66.02463 | 101.0113 | 42.83761 | -0.26017 | 1.880457 | 0.544124 |
| 13 | 90773000 | 90775199 | 22.78753 | 38.3267 | 60.92459 | 26.61879 | -0.28538 | 1.129783 | 0.544132 |
| 7 | 60928800 | 60932399 | 29.33634 | 51.01642 | 37.23643 | 16.35817 | 0.839389 | 0.829001 | 0.544132 |
| 15 | 14799200 | 14802599 | 40.49378 | 65.94671 | 68.77942 | 29.53216 | 0.310133 | 1.558968 | 0.544134 |
| 10 | 125390800 | 125393599 | 27.80229 | 47.35713 | 46.83663 | 20.53033 | 0.397907 | 1.022823 | 0.544134 |
| 4 | 17313200 | 17313799 | 12.76358 | 23.58331 | 25.06275 | 10.57354 | 0.350596 | 0.219017 | 0.544153 |
| 5 | 40621200 | 40624599 | 30.04953 | 50.79172 | 49.07249 | 21.47267 | 0.430705 | 1.098612 | 0.544159 |
| 1 | 110165400 | 110166799 | 22.07677 | 38.1835 | 40.49649 | 17.76662 | 0.301371 | 0.857171 | 0.544166 |
| 5 | 79177000 | 79177599 | 10.24835 | 19.63112 | 20.79599 | 8.202599 | 0.436483 | -0.04921 | 0.544194 |
| 6 | 108041800 | 108043199 | 10.35088 | 18.76266 | 33.28994 | 14.40119 | -0.40928 | 0.3259 | 0.544195 |
| 9 | 4331000 | 4332599 | 26.97443 | 46.09849 | 45.74209 | 20.05587 | 0.393869 | 0.989195 | 0.544208 |
| 4 | 151591800 | 151593799 | 14.46616 | 25.37035 | 41.45472 | 18.32268 | -0.31721 | 0.637383 | 0.544211 |
| 16 | 63788800 | 63789599 | 8.259549 | 15.52219 | 28.02235 | 11.51088 | -0.37845 | 0.084237 | 0.544221 |
| 19 | 61290000 | 61290999 | 16.01882 | 28.39129 | 29.41516 | 13.00447 | 0.341228 | 0.496466 | 0.544221 |
| 10 | 88022000 | 88023999 | 16.22119 | 28.2032 | 45.79839 | 20.11691 | -0.30919 | 0.782571 | 0.544235 |

| | | | | | | | | | |
|---|---|---|---|---|---|---|---|---|---|
| 4 | 144467000 | 144469199 | 29.09204 | 49.62124 | 46.27798 | 20.28766 | 0.48263 | 1.016493 | 0.544235 |
| 4 | 114036000 | 114046399 | 106.675 | 165.8401 | 157.6318 | 65.44574 | 0.381137 | 2.756167 | 0.544239 |
| 1 | 86199000 | 86201399 | 26.52402 | 44.63047 | 46.98091 | 20.52929 | 0.307285 | 1.124573 | 0.544242 |
| 8 | 54251200 | 54253199 | 31.31165 | 50.96686 | 79.49668 | 34.12749 | -0.26955 | 1.554614 | 0.544245 |
| X | 162718800 | 162719999 | 9.273998 | 17.36713 | 31.76294 | 13.12577 | -0.40402 | 0.127114 | 0.544249 |
| 8 | 39987800 | 39990399 | 27.18917 | 44.67021 | 71.10582 | 30.63565 | -0.29084 | 1.414253 | 0.54426 |
| 10 | 27671400 | 27672399 | 20.21163 | 34.28228 | 55.36035 | 24.24966 | -0.30733 | 1.044193 | 0.544266 |
| 8 | 56332400 | 56333599 | 12.59816 | 23.92974 | 23.66529 | 9.518449 | 0.508735 | 0.039626 | 0.544289 |
| 9 | 119703800 | 119707399 | 23.41056 | 40.51824 | 33.81457 | 14.83544 | 0.645864 | 0.84759 | 0.54429 |
| 11 | 28532200 | 28534599 | 17.73512 | 30.24347 | 51.62141 | 22.62975 | -0.38598 | 0.980745 | 0.544313 |
| 14 | 39842800 | 39843999 | 16.17161 | 28.10396 | 45.80015 | 20.11183 | -0.31447 | 0.787494 | 0.544332 |
| 18 | 53346800 | 53347999 | 10.41903 | 18.9953 | 32.88174 | 14.12248 | -0.3667 | 0.295899 | 0.544334 |
| 2 | 11265000 | 11267199 | 23.50445 | 40.51519 | 42.54575 | 18.66374 | 0.314558 | 0.895116 | 0.544343 |
| 2 | 86389600 | 86390599 | 11.22545 | 21.63212 | 21.25012 | 8.238637 | 0.566172 | -0.09763 | 0.544351 |
| 13 | 98557000 | 98560999 | 25.07617 | 41.81287 | 68.35274 | 29.71371 | -0.32571 | 1.23162 | 0.544368 |
| 4 | 16361800 | 16365399 | 63.09063 | 100.2779 | 103.7008 | 43.26694 | 0.285265 | 2.260849 | 0.544374 |
| 14 | 7307200 | 7307999 | 6.117157 | 11.81793 | 24.28195 | 9.329125 | -0.4759 | -0.17172 | 0.54438 |
| 5 | 109252200 | 109270799 | 353.2362 | 503.0084 | 429.8601 | 177.7158 | 0.421243 | 4.314666 | 0.544392 |
| 13 | 33483400 | 33490199 | 83.31734 | 131.8782 | 122.953 | 51.11455 | 0.42906 | 2.395308 | 0.544394 |
| 1 | 117948400 | 117952199 | 79.36486 | 125.2464 | 130.1973 | 54.04425 | 0.268804 | 2.463487 | 0.544406 |
| 9 | 24476000 | 24476799 | 7.275727 | 14.03592 | 26.5862 | 10.27552 | -0.36827 | -0.13252 | 0.544408 |
| 13 | 107530800 | 107531399 | 7.275727 | 14.03592 | 26.5862 | 10.27552 | -0.36827 | -0.13252 | 0.544408 |
| 7 | 95714000 | 95716399 | 20.95132 | 36.88759 | 33.42223 | 14.79564 | 0.529811 | 0.603014 | 0.544418 |
| 7 | 111592200 | 111595999 | 51.54783 | 82.05527 | 129.1851 | 54.00123 | -0.31785 | 2.212927 | 0.544422 |
| 17 | 21430800 | 21432199 | 23.376 | 40.40519 | 35.78712 | 15.70022 | 0.559974 | 0.86101 | 0.544427 |
| 11 | 64237000 | 64238199 | 17.35049 | 30.61101 | 32.37525 | 14.32624 | 0.309122 | 0.560021 | 0.544434 |
| 12 | 106157600 | 106160399 | 26.67094 | 43.85707 | 70.62158 | 30.43825 | -0.30695 | 1.400729 | 0.544449 |
| 3 | 21290000 | 21290599 | 8.541176 | 16.56216 | 19.35482 | 7.308613 | 0.35431 | -0.21767 | 0.54446 |
| X | 5794000 | 5797199 | 21.65807 | 36.80972 | 70.77531 | 31.01805 | -0.55842 | 1.044542 | 0.544482 |
| 14 | 5296400 | 5297999 | 12.06251 | 21.20272 | 39.88706 | 17.66336 | -0.51948 | 0.581544 | 0.544489 |
| 9 | 26282200 | 26282799 | 12.44997 | 23.08648 | 23.62413 | 9.918117 | 0.409988 | 0.193594 | 0.54449 |
| 13 | 41737200 | 41742799 | 49.81455 | 80.28534 | 81.5618 | 34.71089 | 0.333582 | 1.806903 | 0.544521 |
| 4 | 60183200 | 60185199 | 20.48535 | 34.71812 | 55.39449 | 24.25859 | -0.29018 | 1.050806 | 0.544528 |
| 3 | 43045000 | 43045799 | 11.9794 | 21.39677 | 35.70564 | 15.68598 | -0.33624 | 0.432464 | 0.544532 |
| 13 | 95531800 | 95534799 | 51.90686 | 84.01806 | 64.70272 | 27.67641 | 0.739121 | 1.689675 | 0.544536 |
| 2 | 144963800 | 144967399 | 23.66163 | 39.36307 | 63.54315 | 27.58179 | -0.3075 | 1.249811 | 0.544541 |
| 9 | 4136200 | 4144399 | 204.0255 | 301.9733 | 263.6583 | 108.5823 | 0.44443 | 3.715107 | 0.544585 |
| 16 | 93451200 | 93451999 | 9.615947 | 17.68097 | 31.43581 | 13.35053 | -0.39345 | 0.238317 | 0.544604 |
| 16 | 65950200 | 65950799 | 9.717581 | 17.99374 | 30.97365 | 13.02919 | -0.33737 | 0.196985 | 0.544619 |
| 2 | 102770200 | 102771799 | 12.47328 | 22.10777 | 37.18748 | 16.44055 | -0.35418 | 0.487949 | 0.544629 |
| 17 | 10689000 | 10690599 | 15.77695 | 27.53809 | 45.20816 | 19.90644 | -0.32419 | 0.725056 | 0.544632 |
| 2 | 35252400 | 35254199 | 21.35656 | 37.4307 | 35.1417 | 15.52402 | 0.477959 | 0.670217 | 0.544664 |
| X | 32554800 | 32568999 | 233.5207 | 347.0092 | 268.7002 | 110.6955 | 0.622883 | 3.652749 | 0.544678 |
| 9 | 24212600 | 24213999 | 13.60664 | 23.97213 | 39.28084 | 17.38688 | -0.32003 | 0.565279 | 0.54468 |
| 6 | 143443600 | 143446399 | 23.80226 | 39.83873 | 63.12831 | 27.50799 | -0.28086 | 1.184871 | 0.544682 |
| 8 | 109402600 | 109404999 | 16.46539 | 28.72403 | 47.32737 | 20.83242 | -0.32981 | 0.735887 | 0.544682 |
| 9 | 94524200 | 94524799 | 8.644711 | 16.08316 | 29.4876 | 12.3212 | -0.42008 | 0.158679 | 0.544683 |

| | | | | | | | | | |
|---|---|---|---|---|---|---|---|---|---|
| 4 | 143878200 | 143888599 | 153.3208 | 232.7261 | 216.4021 | 89.60094 | 0.386003 | 3.231708 | 0.544713 |
| 12 | 114146600 | 114160599 | 224.179 | 331.2827 | 303.4465 | 124.9662 | 0.373236 | 3.744749 | 0.544714 |
| 10 | 119078000 | 119080399 | 25.44457 | 43.12999 | 43.71446 | 19.14458 | 0.361912 | 1.050775 | 0.54472 |
| 2 | 47504200 | 47508799 | 36.55387 | 60.04829 | 54.75588 | 23.59008 | 0.510519 | 1.424769 | 0.544741 |
| 10 | 4275600 | 4277999 | 25.48006 | 43.2494 | 46.36457 | 20.31363 | 0.28146 | 1.044642 | 0.544792 |
| 9 | 14557600 | 14558199 | 9.494804 | 18.41615 | 20.48075 | 7.724337 | 0.426777 | -0.20722 | 0.544796 |
| 5 | 144611000 | 144613199 | 14.08804 | 24.35208 | 51.06292 | 22.40192 | -0.67769 | 0.842819 | 0.544801 |
| 1 | 180620600 | 180627999 | 59.53842 | 94.12161 | 140.8654 | 58.524 | -0.25424 | 2.411304 | 0.544801 |
| 7 | 126722400 | 126726399 | 37.93149 | 61.18797 | 93.94361 | 40.02619 | -0.25941 | 1.771092 | 0.544812 |
| 9 | 4079000 | 4080399 | 28.4213 | 48.39087 | 46.82597 | 20.52495 | 0.429164 | 1.028428 | 0.544812 |
| 14 | 11603000 | 11605399 | 28.4213 | 48.39087 | 46.82597 | 20.52495 | 0.429164 | 1.028428 | 0.544812 |
| X | 72682400 | 72685199 | 30.18585 | 50.78432 | 50.17812 | 21.92537 | 0.39839 | 1.143654 | 0.54482 |
| 8 | 14645800 | 14647799 | 18.28366 | 31.56061 | 50.61845 | 22.20599 | -0.29352 | 0.870552 | 0.544823 |
| 11 | 53718600 | 53721199 | 29.09863 | 48.63195 | 51.87637 | 22.58404 | 0.288204 | 1.204438 | 0.544836 |
| 7 | 38531600 | 38533999 | 40.82214 | 66.64286 | 67.27742 | 28.91278 | 0.359464 | 1.524659 | 0.544848 |
| X | 46311000 | 46312399 | 19.96534 | 36.58315 | 26.45597 | 11.28891 | 0.890745 | 0.323813 | 0.54486 |
| 12 | 21963000 | 22023599 | 2087.907 | 2535.502 | 2094.736 | 884.1349 | 0.2388 | 6.833413 | 0.544876 |
| 14 | 124318400 | 124319799 | 15.75957 | 28.61084 | 28.70965 | 12.39585 | 0.409565 | 0.346442 | 0.54488 |
| 3 | 98154400 | 98157999 | 36.75888 | 59.18488 | 94.36416 | 40.10191 | -0.31701 | 1.824699 | 0.544904 |
| 6 | 25250400 | 25253399 | 28.80707 | 48.87967 | 47.38641 | 20.75914 | 0.425939 | 1.059092 | 0.544904 |
| 5 | 89674200 | 89675599 | 13.79116 | 25.1404 | 26.87348 | 11.54356 | 0.323723 | 0.304754 | 0.544934 |
| 7 | 25398200 | 25402199 | 51.22678 | 82.1846 | 88.14012 | 37.25975 | 0.247573 | 1.954208 | 0.544957 |
| 17 | 18129800 | 18133399 | 52.95549 | 83.87933 | 134.4532 | 55.91546 | -0.35079 | 2.358269 | 0.544975 |
| 18 | 46433800 | 46436399 | 25.86421 | 44.07149 | 45.77061 | 20.06362 | 0.327659 | 1.009658 | 0.545001 |
| 13 | 105627600 | 105631399 | 33.34236 | 55.23065 | 58.07032 | 25.13179 | 0.308629 | 1.321173 | 0.545003 |
| Y | 86582400 | 86585599 | 26.45121 | 44.77003 | 32.16887 | 14.08191 | 0.856131 | 1.052342 | 0.545004 |
| 13 | 103849000 | 103859799 | 205.1313 | 297.6845 | 414.8028 | 171.013 | -0.25672 | 4.027562 | 0.545017 |
| 1 | 113075200 | 113076999 | 15.65385 | 28.82686 | 26.697 | 11.31451 | 0.542583 | 0.261276 | 0.545027 |
| 16 | 36308200 | 36309999 | 25.1451 | 41.77799 | 65.84668 | 28.55668 | -0.27326 | 1.27128 | 0.545044 |
| 16 | 90622000 | 90623999 | 16.11891 | 28.64414 | 30.24718 | 13.34521 | 0.315869 | 0.479755 | 0.54508 |
| 18 | 17355800 | 17359399 | 27.33905 | 47.2305 | 41.82052 | 18.34685 | 0.560266 | 0.891266 | 0.545085 |
| 12 | 114321000 | 114321799 | 20.98125 | 36.82123 | 35.16295 | 15.55065 | 0.45356 | 0.648713 | 0.545099 |
| 5 | 91029400 | 91030399 | 13.25852 | 24.5829 | 25.94481 | 10.89416 | 0.364955 | 0.203622 | 0.545102 |
| 18 | 30809000 | 30809799 | 7.557656 | 14.15421 | 28.02059 | 11.57758 | -0.51653 | 0.102749 | 0.54511 |
| 2 | 148693400 | 148696399 | 25.45808 | 43.42182 | 45.78518 | 20.07159 | 0.306115 | 1.000516 | 0.545122 |
| 2 | 173218200 | 173222599 | 38.24625 | 61.52731 | 95.96511 | 40.73201 | -0.28688 | 1.855084 | 0.545146 |
| 18 | 70975200 | 70978399 | 18.61253 | 32.79562 | 33.74192 | 14.93481 | 0.347923 | 0.582883 | 0.545149 |
| X | 140618600 | 140621399 | 29.23243 | 50.52426 | 40.64654 | 17.83207 | 0.698192 | 0.894652 | 0.54516 |
| 19 | 8271600 | 8285199 | 169.2858 | 254.2968 | 226.3985 | 93.45102 | 0.435794 | 3.440286 | 0.545164 |
| 4 | 62075200 | 62078999 | 45.02036 | 73.02299 | 72.92287 | 31.2404 | 0.367891 | 1.640172 | 0.545169 |
| X | 150260200 | 150261999 | 18.43629 | 33.6156 | 28.07433 | 12.05606 | 0.678028 | 0.341802 | 0.545193 |
| 12 | 63769600 | 63771199 | 21.94466 | 38.19066 | 39.29436 | 17.27033 | 0.346276 | 0.786416 | 0.545199 |
| 6 | 148356400 | 148357999 | 19.25649 | 33.76271 | 35.44643 | 15.66362 | 0.317656 | 0.65558 | 0.545224 |
| 4 | 22883000 | 22885999 | 28.31906 | 46.62496 | 73.72993 | 31.79464 | -0.2802 | 1.392749 | 0.545234 |
| 2 | 95643800 | 95644799 | 11.13573 | 20.23334 | 33.81645 | 14.58555 | -0.3208 | 0.320008 | 0.545244 |
| 8 | 128329400 | 128329999 | 8.440457 | 15.78447 | 28.48938 | 11.80044 | -0.38768 | 0.119345 | 0.545255 |
| 19 | 43068800 | 43070599 | 18.47137 | 33.60836 | 27.9686 | 12.04248 | 0.680546 | 0.352635 | 0.54527 |

| | | | | | | | | | |
|---|---|---|---|---|---|---|---|---|---|
| 7 | 33122000 | 33125599 | 60.0348 | 96.3161 | 91.30751 | 38.59904 | 0.425225 | 1.966891 | 0.545294 |
| 16 | 33435600 | 33437199 | 19.09674 | 33.42731 | 35.57924 | 15.70036 | 0.298034 | 0.681544 | 0.545298 |
| 4 | 91685200 | 91685799 | 8.85987 | 16.66325 | 29.11331 | 11.94337 | -0.33028 | 0.087005 | 0.545305 |
| 15 | 23419400 | 23419799 | 7.953281 | 15.5011 | 17.93258 | 6.491226 | 0.424191 | -0.37636 | 0.545308 |
| 10 | 85664200 | 85665999 | 17.53044 | 30.18347 | 48.99732 | 21.49314 | -0.31146 | 0.889577 | 0.545313 |
| 5 | 112105000 | 112106999 | 15.92528 | 27.51842 | 46.89551 | 20.57341 | -0.38007 | 0.847467 | 0.545347 |
| 17 | 56251400 | 56252999 | 13.49747 | 23.56104 | 42.64403 | 18.77822 | -0.46425 | 0.706155 | 0.54535 |
| 15 | 61785400 | 61787199 | 13.47646 | 23.81049 | 38.75639 | 17.14498 | -0.30874 | 0.533402 | 0.545353 |
| 17 | 20613000 | 20616999 | 44.31559 | 72.69806 | 62.08614 | 26.73247 | 0.604074 | 1.465966 | 0.545354 |
| 4 | 152834800 | 152850399 | 194.665 | 284.4911 | 391.7008 | 161.3723 | -0.22962 | 3.920044 | 0.545364 |
| 5 | 129651400 | 129653399 | 16.90535 | 29.22807 | 47.93696 | 21.03066 | -0.3249 | 0.849049 | 0.54537 |
| 10 | 83566800 | 83570799 | 27.37947 | 44.83996 | 72.60585 | 31.23984 | -0.31726 | 1.455025 | 0.54537 |
| 15 | 37245800 | 37247999 | 21.94365 | 36.84068 | 60.34159 | 26.3446 | -0.3285 | 1.141709 | 0.545374 |
| 2 | 177189200 | 177191999 | 21.59686 | 36.40925 | 58.71247 | 25.6705 | -0.30586 | 1.102141 | 0.545377 |
| 3 | 22313800 | 22314599 | 16.99007 | 30.01571 | 31.83804 | 14.08471 | 0.305647 | 0.541704 | 0.545378 |
| 17 | 26902200 | 26902599 | 3.983391 | 7.784858 | 18.27228 | 6.323494 | -0.52444 | -0.58677 | 0.545387 |
| 18 | 6894200 | 6898799 | 77.94871 | 122.6929 | 130.6205 | 54.1937 | 0.231571 | 2.515543 | 0.545387 |
| 8 | 10711400 | 10713799 | 19.50596 | 34.1975 | 35.37277 | 15.63006 | 0.338936 | 0.657015 | 0.545409 |
| 9 | 39612400 | 39613799 | 19.86638 | 34.85229 | 30.2145 | 13.35821 | 0.592359 | 0.639608 | 0.54541 |
| 14 | 37594400 | 37597199 | 26.8604 | 46.06811 | 44.63813 | 19.57747 | 0.428808 | 0.95637 | 0.545413 |
| 7 | 6564000 | 6582799 | 274.6046 | 395.1721 | 389.9689 | 160.9521 | 0.228905 | 4.155511 | 0.54542 |
| 7 | 5839200 | 5840199 | 46.80358 | 75.30079 | 75.82474 | 32.18796 | 0.343267 | 1.85646 | 0.545429 |
| 6 | 108855200 | 108856999 | 24.82428 | 43.02275 | 41.35059 | 18.14245 | 0.44332 | 0.851649 | 0.545439 |
| 2 | 47435000 | 47437799 | 37.84442 | 61.50563 | 64.27288 | 27.57505 | 0.305221 | 1.580838 | 0.545446 |
| 12 | 24036600 | 24037399 | 16.48892 | 29.82449 | 25.52208 | 11.07063 | 0.633588 | 0.365872 | 0.545473 |
| 12 | 101496800 | 101498999 | 36.41867 | 59.51066 | 63.887 | 27.46525 | 0.271985 | 1.500545 | 0.545485 |
| 9 | 7480400 | 7485399 | 43.54491 | 71.05259 | 67.46493 | 28.98813 | 0.447185 | 1.538612 | 0.545487 |
| 1 | 38872600 | 38875999 | 35.24563 | 56.93451 | 90.05766 | 38.43943 | -0.3 | 1.722201 | 0.545502 |
| 9 | 39354400 | 39355399 | 6.152278 | 11.77258 | 27.83461 | 11.01195 | -0.71592 | -0.06083 | 0.545527 |
| 8 | 21393600 | 21396799 | 55.83024 | 89.47251 | 93.65594 | 39.5259 | 0.280651 | 1.995129 | 0.545537 |
| 15 | 4033200 | 4035999 | 51.82847 | 83.19555 | 88.73689 | 37.54334 | 0.256377 | 1.937425 | 0.545554 |
| 5 | 147889800 | 147891599 | 22.00015 | 38.16076 | 40.41332 | 17.73169 | 0.304223 | 0.830072 | 0.545561 |
| 3 | 63202800 | 63203799 | 8.289054 | 15.43871 | 28.96131 | 12.07842 | -0.45111 | 0.14399 | 0.545566 |
| 3 | 121811800 | 121815799 | 28.31794 | 46.56738 | 73.06558 | 31.49221 | -0.26964 | 1.406905 | 0.545573 |
| 7 | 30994000 | 31009999 | 35.99964 | 59.34599 | 53.73194 | 23.18339 | 0.522693 | 1.379104 | 0.54559 |
| 16 | 96054600 | 96055799 | 9.038255 | 16.84472 | 29.47701 | 12.28041 | -0.35044 | 0.147845 | 0.545601 |
| 4 | 71656200 | 71660999 | 104.8847 | 161.6454 | 169.8966 | 70.57229 | 0.226598 | 2.91108 | 0.545612 |
| Y | 6173600 | 6176799 | 20.93738 | 36.45457 | 24.89241 | 10.94369 | 0.934636 | 0.747768 | 0.545621 |
| 6 | 19755000 | 19756799 | 17.66288 | 30.43635 | 48.99921 | 21.49451 | -0.29934 | 0.882069 | 0.545649 |
| 5 | 109024800 | 109027199 | 36.34112 | 59.65539 | 63.11443 | 27.18277 | 0.296298 | 1.439539 | 0.54565 |
| 14 | 42594600 | 42598799 | 41.06294 | 66.74446 | 57.89567 | 24.84123 | 0.573368 | 1.581216 | 0.545673 |
| 11 | 29363000 | 29364799 | 19.43704 | 34.28933 | 33.62175 | 14.87807 | 0.417612 | 0.574657 | 0.545682 |
| 5 | 102798600 | 102799999 | 23.82215 | 38.77238 | 84.17414 | 36.13428 | -0.74512 | 1.546669 | 0.545704 |
| 3 | 72089600 | 72090999 | 16.77475 | 29.77014 | 31.17264 | 13.7689 | 0.326828 | 0.493892 | 0.545707 |
| 1 | 29835800 | 29836799 | 12.80413 | 22.85973 | 37.74349 | 16.58742 | -0.32142 | 0.445356 | 0.545727 |
| 4 | 72056400 | 72058199 | 12.80413 | 22.85973 | 37.74349 | 16.58742 | -0.32142 | 0.445356 | 0.545727 |
| 14 | 18037000 | 18037999 | 16.31694 | 29.55786 | 22.88809 | 9.909428 | 0.779042 | 0.352566 | 0.54574 |

| | | | | | | | | | |
|---|---|---|---|---|---|---|---|---|---|
| 4 | 105351000 | 105352599 | 22.37882 | 39.2116 | 36.33707 | 16.04836 | 0.496977 | 0.685637 | 0.54574 |
| 4 | 61194800 | 61195999 | 8.538121 | 16.07126 | 28.5829 | 11.70954 | -0.35417 | 0.076981 | 0.545749 |
| 6 | 107880400 | 107880999 | 8.538121 | 16.07126 | 28.5829 | 11.70954 | -0.35417 | 0.076981 | 0.545749 |
| 13 | 84064600 | 84068399 | 38.14054 | 63.07248 | 58.9542 | 25.4847 | 0.477711 | 1.353629 | 0.545757 |
| 18 | 12182200 | 12188399 | 85.21281 | 134.2381 | 134.0934 | 55.64059 | 0.324256 | 2.504397 | 0.545786 |
| 1 | 129934200 | 129934999 | 12.0174 | 21.64411 | 36.30301 | 15.83123 | -0.33615 | 0.383803 | 0.545798 |
| 5 | 109904800 | 109909399 | 91.09861 | 141.8507 | 150.4642 | 62.455 | 0.224808 | 2.720871 | 0.545809 |
| X | 151268000 | 151268599 | 5.354457 | 10.43517 | 21.40185 | 7.754541 | -0.39257 | -0.3897 | 0.545816 |
| 3 | 59337400 | 59342999 | 61.61799 | 97.62329 | 146.5848 | 60.96789 | -0.25645 | 2.361684 | 0.545822 |
| 14 | 41309600 | 41314999 | 80.11275 | 126.3013 | 131.5119 | 54.57488 | 0.265295 | 2.484235 | 0.545851 |
| 12 | 76463200 | 76465599 | 20.28846 | 34.4781 | 54.87529 | 24.04948 | -0.28644 | 1.025253 | 0.545852 |
| 13 | 16716400 | 16716999 | 10.73726 | 19.40518 | 33.31183 | 14.46441 | -0.36508 | 0.349151 | 0.545873 |
| 16 | 72538400 | 72543199 | 57.57998 | 92.04575 | 97.88015 | 41.15794 | 0.253881 | 2.07404 | 0.545878 |
| 14 | 85465800 | 85467199 | 13.27047 | 25.12081 | 24.11278 | 9.768761 | 0.542433 | 0.070038 | 0.545904 |
| 9 | 6906600 | 6908999 | 33.91869 | 56.84773 | 51.96097 | 22.66116 | 0.510258 | 1.190635 | 0.545909 |
| 13 | 43547200 | 43550799 | 54.35249 | 86.38787 | 128.6141 | 53.6606 | -0.23967 | 2.259796 | 0.545922 |
| 4 | 112820400 | 112823799 | 52.26332 | 84.2726 | 81.27057 | 34.60847 | 0.409336 | 1.798572 | 0.545925 |
| 19 | 29228400 | 29229799 | 8.932439 | 16.49804 | 29.91582 | 12.6277 | -0.41546 | 0.201519 | 0.545939 |
| 7 | 48328800 | 48329599 | 18.70426 | 32.78972 | 34.10158 | 15.06749 | 0.331379 | 0.654483 | 0.545946 |
| 17 | 54154200 | 54156599 | 19.84855 | 35.43357 | 30.88525 | 13.5743 | 0.594956 | 0.478017 | 0.545971 |
| 4 | 87235200 | 87236999 | 19.01791 | 33.56719 | 33.06874 | 14.63186 | 0.411052 | 0.564672 | 0.54598 |
| X | 77627000 | 77628399 | 17.50353 | 31.18572 | 30.77555 | 13.54857 | 0.415434 | 0.476723 | 0.545986 |
| 10 | 57726200 | 57733399 | 74.19652 | 116.1403 | 170.1397 | 70.57862 | -0.23459 | 2.623474 | 0.546006 |
| 16 | 76325200 | 76326199 | 7.735007 | 14.77923 | 27.3216 | 10.85288 | -0.37072 | -0.03707 | 0.546008 |
| 7 | 84630000 | 84633799 | 57.05482 | 91.24458 | 96.87079 | 40.75778 | 0.2569 | 2.062134 | 0.546014 |
| 7 | 128951600 | 128953399 | 16.8887 | 30.67036 | 28.58703 | 12.33806 | 0.515714 | 0.352351 | 0.546017 |
| 6 | 70660400 | 70664599 | 79.56651 | 125.232 | 133.4118 | 55.35136 | 0.23055 | 2.517746 | 0.546039 |
| 14 | 68356600 | 68359399 | 28.79392 | 47.31464 | 74.28724 | 32.00861 | -0.27105 | 1.417522 | 0.546045 |
| 4 | 38479800 | 38480999 | 13.19125 | 24.68922 | 24.96406 | 10.32584 | 0.443086 | 0.143145 | 0.546045 |
| 9 | 36334000 | 36336599 | 42.19083 | 68.58008 | 70.50674 | 30.25287 | 0.328882 | 1.590917 | 0.546052 |
| 3 | 78289200 | 78289599 | 4.804317 | 9.373995 | 19.80046 | 7.062389 | -0.41435 | -0.46148 | 0.546054 |
| 8 | 54139600 | 54140199 | 4.804317 | 9.373995 | 19.80046 | 7.062389 | -0.41435 | -0.46148 | 0.546054 |
| 4 | 147232200 | 147239199 | 67.75181 | 108.5768 | 89.63345 | 37.82773 | 0.622552 | 2.008823 | 0.546063 |
| 19 | 39209200 | 39211999 | 45.43984 | 73.47652 | 76.17555 | 32.55677 | 0.309808 | 1.703921 | 0.546076 |
| 1 | 103223400 | 103225199 | 22.00743 | 38.23746 | 39.24132 | 17.22935 | 0.349571 | 0.806246 | 0.546093 |
| 6 | 121911000 | 121912399 | 20.09011 | 35.79316 | 30.79142 | 13.55588 | 0.612078 | 0.490739 | 0.546094 |
| 8 | 119308200 | 119311199 | 26.94087 | 45.65981 | 47.96517 | 21.00525 | 0.310531 | 1.064103 | 0.546095 |
| 9 | 3594000 | 3594799 | 10.58691 | 19.29084 | 33.39111 | 14.35105 | -0.36727 | 0.30158 | 0.546107 |
| 2 | 14928800 | 14930799 | 14.4433 | 25.41562 | 41.34211 | 18.30366 | -0.31022 | 0.587721 | 0.546139 |
| 13 | 61429600 | 61433399 | 47.16578 | 76.77066 | 68.3228 | 29.33027 | 0.537904 | 1.580854 | 0.546144 |
| 10 | 103097000 | 103102599 | 118.3571 | 180.9693 | 189.1885 | 78.4872 | 0.225571 | 3.072869 | 0.546168 |
| 9 | 3256400 | 3283799 | 546.6152 | 741.1512 | 690.236 | 287.5928 | 0.226002 | 5.059263 | 0.546189 |
| 3 | 95287600 | 95288799 | 15.32251 | 27.32646 | 28.39319 | 12.4895 | 0.34212 | 0.449227 | 0.546205 |
| 5 | 90727800 | 90730799 | 27.43489 | 46.00607 | 48.59491 | 21.20048 | 0.302267 | 1.160487 | 0.546217 |
| X | 55049000 | 55063599 | 104.3153 | 162.7361 | 134.8921 | 55.97634 | 0.58228 | 2.694417 | 0.546219 |
| 4 | 93777400 | 93777999 | 8.882597 | 16.63202 | 29.0259 | 11.99734 | -0.33727 | 0.115951 | 0.546239 |
| 12 | 57465800 | 57466999 | 8.882597 | 16.63202 | 29.0259 | 11.99734 | -0.33727 | 0.115951 | 0.546239 |

| | | | | | | | | | |
|---|---|---|---|---|---|---|---|---|---|
| 9 | 24684000 | 24690399 | 53.1314 | 84.14863 | 148.907 | 61.92291 | -0.49344 | 2.363626 | 0.54625 |
| 14 | 92035400 | 92039599 | 44.68206 | 72.90636 | 66.05624 | 28.3826 | 0.514542 | 1.538168 | 0.546257 |
| 6 | 133846800 | 133849999 | 24.12071 | 41.25828 | 42.56712 | 18.66512 | 0.337901 | 0.963734 | 0.546274 |
| 1 | 42413400 | 42415599 | 13.12134 | 23.36667 | 38.74631 | 17.06461 | -0.32978 | 0.464781 | 0.54629 |
| 5 | 111121400 | 111122799 | 12.12495 | 21.2577 | 44.10689 | 19.48973 | -0.65999 | 0.633322 | 0.546296 |
| 8 | 44958200 | 44960799 | 32.85483 | 54.55497 | 56.94229 | 24.68482 | 0.319395 | 1.291691 | 0.5463 |
| 4 | 148440600 | 148443799 | 31.98581 | 51.84267 | 83.70166 | 35.83855 | -0.32453 | 1.635077 | 0.546312 |
| 5 | 69658800 | 69659399 | 7.041984 | 13.38512 | 26.08212 | 10.47539 | -0.45936 | -0.00908 | 0.546315 |
| 4 | 66580000 | 66581399 | 15.28368 | 27.23162 | 26.96927 | 11.87255 | 0.410172 | 0.451636 | 0.546325 |
| 3 | 36061400 | 36062599 | 14.80258 | 27.49497 | 26.42002 | 11.0667 | 0.501996 | 0.208064 | 0.546347 |
| 1 | 9615200 | 9616999 | 30.07608 | 48.76492 | 81.54948 | 34.92632 | -0.37457 | 1.623903 | 0.546349 |
| 6 | 8954600 | 8959199 | 38.06861 | 61.18091 | 98.69245 | 41.82953 | -0.33716 | 1.885918 | 0.546366 |
| 4 | 20858400 | 20861399 | 51.85649 | 83.29001 | 88.32603 | 37.40329 | 0.265706 | 1.919436 | 0.546369 |
| 7 | 124771600 | 124772599 | 13.46992 | 23.82133 | 38.73805 | 17.13365 | -0.3069 | 0.522835 | 0.546384 |
| 6 | 99424400 | 99425399 | 10.91453 | 19.53438 | 34.29191 | 15.03914 | -0.40689 | 0.410931 | 0.546411 |
| X | 110372000 | 110374199 | 14.21909 | 25.09389 | 44.52639 | 19.702 | -0.43333 | 0.561692 | 0.546418 |
| 5 | 110080800 | 110087199 | 249.824 | 361.6482 | 354.7335 | 146.2934 | 0.246051 | 4.066326 | 0.546448 |
| 14 | 4821600 | 4824199 | 12.40317 | 21.85226 | 42.96515 | 19.0176 | -0.58151 | 0.565019 | 0.546464 |
| 4 | 81183200 | 81185199 | 16.00316 | 28.58527 | 30.26201 | 13.29391 | 0.316537 | 0.447804 | 0.54648 |
| 19 | 48145800 | 48146399 | 5.668924 | 11.02129 | 22.35789 | 8.29573 | -0.40971 | -0.30244 | 0.546481 |
| 2 | 166350000 | 166352199 | 31.5365 | 51.13095 | 81.67881 | 34.98077 | -0.30872 | 1.628529 | 0.546484 |
| 4 | 107523800 | 107527199 | 31.35315 | 50.8879 | 80.74373 | 34.60749 | -0.29758 | 1.605707 | 0.54649 |
| 4 | 122479400 | 122488199 | 133.4822 | 203.2421 | 207.8681 | 86.14737 | 0.252361 | 3.162691 | 0.546494 |
| 1 | 157029800 | 157031199 | 14.58667 | 27.10159 | 26.49091 | 11.09213 | 0.47809 | 0.206064 | 0.546504 |
| 3 | 126321200 | 126322399 | 10.14769 | 18.76936 | 32.0132 | 13.4868 | -0.32611 | 0.207447 | 0.546514 |
| 15 | 34344800 | 34346199 | 12.34036 | 22.02148 | 36.22886 | 15.92803 | -0.31648 | 0.443052 | 0.546517 |
| 17 | 56235600 | 56247799 | 166.1254 | 246.6078 | 347.4713 | 143.1316 | -0.24165 | 3.662446 | 0.54652 |
| 11 | 44182200 | 44184599 | 26.87186 | 44.11915 | 71.01643 | 30.58841 | -0.30727 | 1.421716 | 0.546523 |
| 17 | 6763800 | 6764799 | 16.57057 | 29.34889 | 30.70826 | 13.57803 | 0.326504 | 0.508213 | 0.54656 |
| 8 | 118513200 | 118517799 | 47.92324 | 76.55683 | 114.8491 | 48.24694 | -0.24278 | 2.090619 | 0.546562 |
| X | 96602800 | 96605399 | 28.79885 | 49.83616 | 40.67468 | 17.84508 | 0.677873 | 0.882738 | 0.546581 |
| 3 | 120878000 | 120880599 | 15.20405 | 26.59038 | 43.05261 | 18.98427 | -0.30426 | 0.687006 | 0.546609 |
| 14 | 88850600 | 88851999 | 11.16996 | 19.68598 | 41.96362 | 18.57308 | -0.69708 | 0.552412 | 0.546621 |
| 3 | 139820600 | 139824199 | 37.37655 | 60.17141 | 97.11068 | 41.26119 | -0.3348 | 1.829692 | 0.546623 |
| 13 | 13322600 | 13324999 | 35.08812 | 58.27409 | 57.02905 | 24.72558 | 0.412111 | 1.29704 | 0.546631 |
| 10 | 118797600 | 118799399 | 10.51014 | 19.20788 | 32.35113 | 13.85257 | -0.32485 | 0.279444 | 0.546636 |
| 4 | 61936400 | 61938799 | 37.77763 | 61.66083 | 65.60266 | 28.19108 | 0.283779 | 1.521402 | 0.546663 |
| 13 | 75363800 | 75364199 | 9.615991 | 18.61653 | 17.63953 | 6.710857 | 0.644198 | -0.18498 | 0.5467 |
| 13 | 87275200 | 87276799 | 10.16884 | 18.55921 | 32.32897 | 13.86633 | -0.37484 | 0.283038 | 0.546708 |
| 2 | 150482800 | 150486399 | 31.79502 | 51.6491 | 81.74917 | 35.06046 | -0.29292 | 1.588509 | 0.546726 |
| 5 | 70352400 | 70355199 | 34.7372 | 58.42474 | 50.23204 | 21.94703 | 0.598298 | 1.159438 | 0.546727 |
| 5 | 101019400 | 101021199 | 13.12417 | 23.17823 | 38.2442 | 16.91984 | -0.32857 | 0.533835 | 0.546727 |
| 6 | 13526800 | 13530399 | 97.06057 | 150.3995 | 158.9271 | 66.01386 | 0.224839 | 2.807963 | 0.546729 |
| 13 | 76151400 | 76155399 | 36.02064 | 59.86422 | 56.55719 | 24.53272 | 0.462831 | 1.29067 | 0.546731 |
| 15 | 30072000 | 30075799 | 47.93845 | 77.1619 | 82.85314 | 35.19485 | 0.251578 | 1.845899 | 0.546734 |
| 9 | 112812200 | 112814399 | 17.05912 | 29.26015 | 49.52322 | 21.72011 | -0.3725 | 0.922209 | 0.546736 |
| 10 | 91617400 | 91618199 | 10.40241 | 18.38188 | 40.4507 | 17.89399 | -0.74086 | 0.517159 | 0.546768 |

| | | | | | | | | | |
|---|---|---|---|---|---|---|---|---|---|
| 4 | 69625400 | 69626799 | 24.05282 | 41.49853 | 37.29777 | 16.36204 | 0.538408 | 0.882342 | 0.546798 |
| 2 | 54813000 | 54819599 | 119.389 | 183.4537 | 166.9082 | 69.30506 | 0.431333 | 2.973777 | 0.546802 |
| X | 23323400 | 23325999 | 22.85334 | 40.44863 | 31.7592 | 14.04466 | 0.738467 | 0.559268 | 0.54681 |
| 15 | 92598400 | 92599799 | 12.49818 | 22.33139 | 36.73693 | 16.13415 | -0.31539 | 0.437047 | 0.546818 |
| 2 | 141817000 | 141817799 | 6.879085 | 13.3038 | 26.66431 | 10.19458 | -0.4331 | -0.16855 | 0.546836 |
| 7 | 59303400 | 59304399 | 38.17083 | 62.19087 | 63.68579 | 27.34945 | 0.336997 | 1.548015 | 0.546837 |
| 18 | 59307800 | 59312599 | 38.40982 | 61.74062 | 97.47692 | 41.32536 | -0.30586 | 1.880991 | 0.546845 |
| 5 | 100197400 | 100204999 | 81.22747 | 126.1059 | 183.3779 | 76.1506 | -0.23316 | 2.77081 | 0.546847 |
| 7 | 48041000 | 48046599 | 105.4182 | 163.1196 | 166.948 | 69.34602 | 0.269337 | 2.839037 | 0.546883 |
| 9 | 123721000 | 123728399 | 192.6604 | 277.6463 | 479.9339 | 198.0443 | -0.57741 | 4.130907 | 0.546884 |
| 12 | 88472400 | 88475199 | 32.2943 | 53.6509 | 56.39344 | 24.45529 | 0.309316 | 1.283081 | 0.546886 |
| 14 | 117980000 | 117980599 | 8.664518 | 16.78092 | 19.30513 | 7.331367 | 0.36884 | -0.19805 | 0.54689 |
| 13 | 69560000 | 69561999 | 16.87552 | 30.20323 | 30.7007 | 13.46414 | 0.376554 | 0.44284 | 0.546891 |
| 16 | 47963200 | 47964199 | 9.082838 | 16.73113 | 30.39651 | 12.87726 | -0.42215 | 0.218266 | 0.546904 |
| 11 | 110330400 | 110333199 | 29.78132 | 48.7602 | 76.50627 | 32.91426 | -0.27221 | 1.465467 | 0.546923 |
| 15 | 49383000 | 49384799 | 16.06116 | 29.40762 | 27.69662 | 11.8302 | 0.510553 | 0.298595 | 0.546943 |
| 3 | 131121200 | 131124199 | 32.78103 | 53.12891 | 85.08479 | 36.42943 | -0.31294 | 1.637878 | 0.546953 |
| 4 | 122559400 | 122565199 | 94.314 | 147.8489 | 143.0257 | 59.32196 | 0.365077 | 2.606053 | 0.54696 |
| 5 | 117110800 | 117114799 | 48.62864 | 78.51409 | 79.18254 | 33.77604 | 0.347057 | 1.75466 | 0.546967 |
| 16 | 21170800 | 21175799 | 35.60544 | 57.49286 | 89.18308 | 38.0466 | -0.27254 | 1.73525 | 0.54697 |
| 2 | 151012000 | 151015999 | 63.67838 | 100.5219 | 147.6072 | 61.28044 | -0.22849 | 2.449555 | 0.546996 |
| 4 | 96042800 | 96044999 | 29.67952 | 49.3093 | 48.67934 | 21.11073 | 0.399594 | 1.269831 | 0.546997 |
| 5 | 12068600 | 12069399 | 7.298583 | 14.00685 | 25.7631 | 10.10746 | -0.34619 | -0.08465 | 0.54704 |
| 5 | 99791000 | 99792399 | 11.33788 | 20.38768 | 34.23098 | 14.95306 | -0.33924 | 0.383282 | 0.547041 |
| 2 | 87164200 | 87171799 | 106.2749 | 164.608 | 165.1096 | 68.58205 | 0.299469 | 2.819589 | 0.547052 |
| 9 | 42093600 | 42096799 | 23.80399 | 40.73907 | 43.67525 | 19.15187 | 0.283001 | 0.961073 | 0.547057 |
| 18 | 37367200 | 37380199 | 296.3511 | 424.8173 | 411.3804 | 169.8924 | 0.250563 | 4.214196 | 0.547067 |
| 17 | 14694400 | 14697399 | 18.60839 | 31.97279 | 51.14164 | 22.43234 | -0.29105 | 0.913879 | 0.547078 |
| 1 | 85009200 | 85018399 | 194.4977 | 288.0464 | 293.2477 | 120.7689 | 0.22381 | 3.704761 | 0.54708 |
| 13 | 3300600 | 3302799 | 25.04846 | 43.02579 | 42.44398 | 18.61711 | 0.403213 | 0.929277 | 0.547101 |
| 2 | 59918000 | 59920399 | 21.12735 | 35.20181 | 69.54631 | 30.21813 | -0.59796 | 1.229838 | 0.547113 |
| 2 | 36387800 | 36396399 | 109.2486 | 169.8818 | 156.7238 | 65.06642 | 0.424441 | 2.754298 | 0.54713 |
| 8 | 35501400 | 35505799 | 49.37583 | 78.86335 | 118.5254 | 49.77881 | -0.24571 | 2.098098 | 0.547142 |
| 1 | 33316000 | 33316799 | 17.59445 | 30.90861 | 32.46777 | 14.37093 | 0.316836 | 0.610188 | 0.547154 |
| 14 | 114935000 | 114935599 | 6.244579 | 12.11897 | 23.38039 | 8.794273 | -0.35632 | -0.24347 | 0.547162 |
| 8 | 21966600 | 21968599 | 24.99828 | 43.00272 | 42.99752 | 18.86075 | 0.384313 | 0.918049 | 0.547169 |
| 4 | 124375600 | 124384799 | 101.8393 | 155.7944 | 222.5856 | 92.35349 | -0.22408 | 3.060212 | 0.547176 |
| 2 | 37050800 | 37062999 | 182.8527 | 273.4761 | 265.6224 | 109.5334 | 0.304613 | 3.527766 | 0.54719 |
| 9 | 114029800 | 114031799 | 35.64403 | 58.75388 | 60.98097 | 26.30996 | 0.326194 | 1.38568 | 0.547197 |
| 4 | 112787800 | 112796399 | 137.0216 | 209.4967 | 194.9172 | 80.86286 | 0.393524 | 3.08161 | 0.547217 |
| 13 | 6485600 | 6486599 | 15.73292 | 27.33266 | 45.29598 | 19.88636 | -0.33847 | 0.790563 | 0.547219 |
| 7 | 28337600 | 28339999 | 21.34571 | 35.93886 | 57.52757 | 25.14247 | -0.29521 | 1.113569 | 0.54722 |
| 7 | 48252200 | 48254199 | 30.8724 | 52.08349 | 50.1848 | 21.94673 | 0.434572 | 1.120423 | 0.547234 |
| 6 | 69232800 | 69234799 | 14.82111 | 27.1304 | 27.84963 | 11.89957 | 0.386871 | 0.291721 | 0.547239 |
| 16 | 17562600 | 17564199 | 14.82111 | 27.1304 | 27.84963 | 11.89957 | 0.386871 | 0.291721 | 0.547239 |
| 3 | 17789400 | 17792199 | 17.94243 | 31.88888 | 27.63781 | 12.19257 | 0.599486 | 0.486844 | 0.547253 |
| 5 | 13274800 | 13277599 | 41.13771 | 67.28155 | 58.74237 | 25.26172 | 0.570247 | 1.49497 | 0.547258 |

| | | | | | | | | | |
|---|---|---|---|---|---|---|---|---|---|
| 1 | 102050400 | 102051399 | 17.68681 | 31.07187 | 31.59805 | 13.98636 | 0.36354 | 0.610351 | 0.547267 |
| 10 | 125595600 | 125601999 | 64.45084 | 102.7735 | 104.7031 | 43.87303 | 0.311717 | 2.158697 | 0.547277 |
| 14 | 83642600 | 83643999 | 17.7416 | 31.41123 | 32.11043 | 14.19907 | 0.359689 | 0.522454 | 0.547283 |
| 6 | 68513200 | 68516999 | 41.69927 | 68.31709 | 64.65242 | 27.82211 | 0.455642 | 1.477637 | 0.547292 |
| 3 | 144498200 | 144498599 | 7.270558 | 14.04234 | 27.29636 | 10.50079 | -0.39907 | -0.1469 | 0.547298 |
| 2 | 112894800 | 112896799 | 21.62108 | 38.07314 | 33.9882 | 15.04569 | 0.55136 | 0.607704 | 0.547304 |
| 18 | 37617600 | 37620599 | 104.2679 | 160.6158 | 162.8033 | 67.62355 | 0.278321 | 2.920415 | 0.547314 |
| 1 | 118115400 | 118118799 | 42.77217 | 70.32431 | 60.91091 | 26.25145 | 0.585049 | 1.434045 | 0.547337 |
| 1 | 129065800 | 129066399 | 4.962823 | 9.669515 | 20.79377 | 7.558847 | -0.46519 | -0.38901 | 0.547354 |
| 6 | 4174000 | 4182199 | 99.08229 | 153.9894 | 158.8286 | 65.94562 | 0.263047 | 2.757859 | 0.547354 |
| 12 | 72887400 | 72889599 | 32.37421 | 53.33978 | 56.19332 | 24.23968 | 0.304531 | 1.380234 | 0.547355 |
| 14 | 63074600 | 63075799 | 11.69097 | 20.85757 | 35.23763 | 15.49541 | -0.35475 | 0.437641 | 0.547357 |
| 5 | 150953800 | 150961799 | 105.5208 | 160.8539 | 230.9339 | 95.73864 | -0.23513 | 3.133511 | 0.547365 |
| 10 | 68196400 | 68201199 | 67.01156 | 106.3945 | 113.8488 | 47.44947 | 0.234541 | 2.297273 | 0.547372 |
| 3 | 95735400 | 95737199 | 15.67493 | 28.33607 | 29.26137 | 12.70163 | 0.363611 | 0.372003 | 0.547376 |
| X | 63483400 | 63485199 | 29.57136 | 50.20577 | 40.54603 | 17.76563 | 0.688714 | 1.054046 | 0.547395 |
| 4 | 142274000 | 142276799 | 20.20729 | 34.01111 | 56.95081 | 24.88795 | -0.35999 | 1.110976 | 0.547415 |
| 6 | 52225000 | 52227199 | 13.33192 | 23.41709 | 39.39496 | 17.4349 | -0.35897 | 0.602137 | 0.547427 |
| 15 | 47224200 | 47227999 | 39.9756 | 65.31167 | 66.6846 | 28.66614 | 0.343943 | 1.511216 | 0.547449 |
| 7 | 31015400 | 31019599 | 65.73277 | 104.8435 | 94.51703 | 39.61829 | 0.488244 | 2.15137 | 0.547461 |
| 12 | 23574200 | 23575399 | 18.48168 | 32.63954 | 28.62921 | 12.66605 | 0.578066 | 0.547743 | 0.54747 |
| 7 | 62265800 | 62269999 | 60.79483 | 97.53108 | 91.02662 | 38.47767 | 0.447629 | 1.968389 | 0.547472 |
| 6 | 61826400 | 61827399 | 12.39943 | 23.50027 | 24.22494 | 9.789197 | 0.443754 | 0.055415 | 0.547473 |
| 6 | 53305200 | 53306799 | 17.24754 | 30.54759 | 31.79828 | 14.06002 | 0.33412 | 0.51717 | 0.547486 |
| 6 | 132329000 | 132329799 | 17.24754 | 30.54759 | 31.79828 | 14.06002 | 0.33412 | 0.51717 | 0.547486 |
| 5 | 77372600 | 77373999 | 9.647592 | 17.98875 | 31.10414 | 12.94805 | -0.33261 | 0.153305 | 0.54749 |
| 18 | 36862400 | 36864799 | 13.48294 | 23.79977 | 38.77531 | 17.15655 | -0.31059 | 0.543883 | 0.547495 |
| 16 | 31999600 | 32000599 | 7.759568 | 14.82722 | 26.74499 | 10.62224 | -0.335 | -0.03718 | 0.547504 |
| 7 | 13188600 | 13189799 | 18.17158 | 31.88678 | 33.55388 | 14.83764 | 0.314461 | 0.634488 | 0.547509 |
| 7 | 59726000 | 59727199 | 13.60298 | 25.07916 | 26.51042 | 11.21775 | 0.355095 | 0.241198 | 0.547512 |
| 4 | 92739000 | 92739999 | 19.00764 | 32.45078 | 52.17752 | 22.87566 | -0.29994 | 0.976223 | 0.547521 |
| 3 | 12057400 | 12062399 | 39.62356 | 64.99361 | 64.01122 | 27.56003 | 0.398867 | 1.456894 | 0.54754 |
| 4 | 38739400 | 38741799 | 18.74728 | 33.46314 | 30.93249 | 13.59658 | 0.510455 | 0.470778 | 0.547541 |
| 12 | 82734000 | 82735399 | 12.45365 | 22.29477 | 36.72555 | 16.10296 | -0.31581 | 0.423427 | 0.547559 |
| 2 | 40096400 | 40098999 | 21.99923 | 38.36175 | 38.62162 | 16.99586 | 0.37765 | 0.756623 | 0.547571 |
| 2 | 156665600 | 156669199 | 62.10858 | 98.89401 | 106.1474 | 44.38841 | 0.234333 | 2.204714 | 0.547582 |
| 9 | 47366400 | 47367599 | 13.16583 | 24.13842 | 25.95719 | 11.06719 | 0.322518 | 0.266074 | 0.547585 |
| 7 | 15232200 | 15235999 | 48.90067 | 79.12605 | 75.37444 | 32.23337 | 0.432644 | 1.697012 | 0.547589 |
| 8 | 110686600 | 110690999 | 31.26083 | 50.49237 | 87.25761 | 37.2393 | -0.42745 | 1.717967 | 0.547589 |
| 2 | 47977400 | 47977799 | 4.052225 | 7.925953 | 17.31742 | 5.824488 | -0.38389 | -0.68632 | 0.547594 |
| 10 | 63027600 | 63029199 | 19.96238 | 34.75684 | 36.7529 | 16.158 | 0.30695 | 0.764699 | 0.547595 |
| 5 | 57121400 | 57122599 | 26.61984 | 45.31522 | 46.85399 | 20.53692 | 0.333881 | 1.023549 | 0.547608 |
| 1 | 188623000 | 188624999 | 13.72721 | 24.15523 | 40.80549 | 18.06612 | -0.36463 | 0.580232 | 0.547636 |
| 12 | 19751800 | 19776999 | 1152.022 | 1467.783 | 1257.813 | 527.9543 | 0.256182 | 6.063879 | 0.547638 |
| 14 | 54535800 | 54540399 | 52.08977 | 83.99583 | 81.26838 | 34.60916 | 0.404697 | 1.797149 | 0.547647 |
| 17 | 20708200 | 20713399 | 49.89066 | 80.9782 | 69.68263 | 29.86739 | 0.583034 | 1.633092 | 0.547664 |
| 4 | 27615200 | 27615999 | 13.6988 | 24.88835 | 25.51948 | 11.00807 | 0.379771 | 0.321565 | 0.547679 |

| | | | | | | | | | |
|---|---|---|---|---|---|---|---|---|---|
| X | 61025400 | 61027999 | 35.75218 | 59.19042 | 58.72804 | 25.40695 | 0.392006 | 1.335609 | 0.547684 |
| 1 | 8658200 | 8660199 | 26.92008 | 46.02787 | 40.43887 | 17.73133 | 0.568875 | 0.978954 | 0.547686 |
| 18 | 79268600 | 79269599 | 7.1613 | 13.69899 | 25.65848 | 10.16037 | -0.38466 | -0.05578 | 0.54769 |
| 7 | 47238000 | 47240199 | 54.04363 | 86.67634 | 91.67614 | 38.73591 | 0.267016 | 1.968841 | 0.547692 |
| 5 | 78284000 | 78285599 | 17.13506 | 31.16871 | 28.63108 | 12.33181 | 0.538606 | 0.345763 | 0.547693 |
| 9 | 13667800 | 13672199 | 84.09763 | 130.3493 | 187.8958 | 78.0457 | -0.22237 | 2.80106 | 0.547701 |
| 13 | 13311800 | 13318799 | 135.7842 | 206.2275 | 213.7801 | 88.53079 | 0.229995 | 3.214996 | 0.547776 |
| 11 | 51676400 | 51676999 | 7.645302 | 14.54484 | 26.63372 | 10.67678 | -0.36796 | -0.00663 | 0.547779 |
| 2 | 169457600 | 169460799 | 20.17278 | 34.19994 | 54.83908 | 24.01782 | -0.29726 | 1.046531 | 0.547781 |
| 7 | 68552400 | 68552799 | 7.645583 | 14.87069 | 17.29689 | 6.395406 | 0.389251 | -0.30766 | 0.547802 |
| 9 | 12561000 | 12561599 | 9.112353 | 17.03988 | 30.6239 | 12.68595 | -0.38164 | 0.132513 | 0.547809 |
| X | 167942400 | 167943199 | 9.112353 | 17.03988 | 30.6239 | 12.68595 | -0.38164 | 0.132513 | 0.547809 |
| 14 | 101324800 | 101328599 | 69.70258 | 110.2528 | 113.3567 | 47.10886 | 0.287315 | 2.39888 | 0.547827 |
| 14 | 85416800 | 85419599 | 16.32548 | 28.40242 | 45.77748 | 20.1157 | -0.29845 | 0.772526 | 0.547832 |
| 7 | 113981000 | 113982999 | 23.14709 | 40.75475 | 33.32076 | 14.75063 | 0.677479 | 0.617062 | 0.547874 |
| 6 | 9887800 | 9890599 | 41.5376 | 67.3169 | 72.29922 | 30.95389 | 0.262019 | 1.649912 | 0.547877 |
| 7 | 40862600 | 40885199 | 400.457 | 559.2549 | 526.7558 | 218.4635 | 0.252702 | 4.60155 | 0.547877 |
| 17 | 92706200 | 92708599 | 21.95211 | 38.47407 | 36.29915 | 16.03518 | 0.471124 | 0.678131 | 0.547909 |
| 6 | 65533400 | 65533999 | 9.351821 | 17.45242 | 30.03482 | 12.48452 | -0.32394 | 0.143553 | 0.547923 |
| 18 | 27201000 | 27202199 | 11.84328 | 21.08637 | 35.7199 | 15.73444 | -0.3604 | 0.451846 | 0.547944 |
| 5 | 39387800 | 39389399 | 23.12892 | 40.37252 | 32.00937 | 14.09503 | 0.721017 | 0.741209 | 0.547964 |
| X | 128010800 | 128012399 | 18.05387 | 32.93904 | 28.09254 | 12.05438 | 0.648613 | 0.335195 | 0.547966 |
| 17 | 50464400 | 50478599 | 127.0312 | 191.4825 | 273.4302 | 112.9687 | -0.24093 | 3.367878 | 0.547969 |
| 8 | 30645400 | 30655399 | 20.96623 | 36.64255 | 32.48584 | 14.31644 | 0.560542 | 0.711199 | 0.547973 |
| 15 | 67738000 | 67739399 | 15.686 | 27.38965 | 44.13237 | 19.43733 | -0.29744 | 0.715409 | 0.547976 |
| 5 | 143186200 | 143194999 | 103.8415 | 158.1825 | 231.1617 | 95.81402 | -0.26151 | 3.14827 | 0.547993 |
| 4 | 60335400 | 60341599 | 114.1538 | 174.7697 | 176.6815 | 73.32057 | 0.275387 | 3.044738 | 0.548003 |
| 6 | 131091600 | 131100799 | 118.6689 | 182.7886 | 177.3886 | 73.68126 | 0.340981 | 2.928871 | 0.548019 |
| 5 | 132649800 | 132650199 | 9.493433 | 18.21453 | 18.7738 | 7.370611 | 0.481764 | -0.07938 | 0.54803 |
| 9 | 122941800 | 122942999 | 11.16461 | 20.04436 | 34.72246 | 15.18799 | -0.38551 | 0.391779 | 0.54805 |
| 6 | 6674000 | 6674399 | 4.181271 | 8.165435 | 18.71663 | 6.576424 | -0.509 | -0.52306 | 0.548057 |
| 15 | 100989000 | 100991599 | 17.15389 | 29.14582 | 52.63665 | 23.06803 | -0.46785 | 1.013813 | 0.548065 |
| 5 | 55092200 | 55103199 | 170.0699 | 255.2486 | 253.7187 | 104.7033 | 0.275803 | 3.459232 | 0.548074 |
| 15 | 71520600 | 71522999 | 36.15196 | 58.20241 | 93.28834 | 39.63951 | -0.32472 | 1.827739 | 0.5481 |
| 8 | 110129000 | 110131199 | 19.80625 | 34.8935 | 34.16698 | 15.12343 | 0.418902 | 0.592698 | 0.548138 |
| 9 | 106684400 | 106686599 | 42.19348 | 69.11102 | 64.69697 | 27.8386 | 0.471148 | 1.481489 | 0.548151 |
| 15 | 88649400 | 88651199 | 48.68294 | 78.48758 | 64.7811 | 27.5812 | 0.633143 | 1.789972 | 0.548175 |
| 5 | 150338000 | 150340999 | 40.86024 | 66.45636 | 69.81582 | 29.96285 | 0.298376 | 1.579379 | 0.548182 |
| 3 | 138003200 | 138006799 | 50.37595 | 81.36877 | 78.21473 | 33.38095 | 0.416871 | 1.746607 | 0.54819 |
| 16 | 89578000 | 89580599 | 41.34736 | 67.16018 | 70.90904 | 30.40932 | 0.289612 | 1.60347 | 0.548221 |
| 8 | 29004400 | 29004999 | 10.28724 | 19.54136 | 21.21277 | 8.530974 | 0.376067 | 0.012898 | 0.548224 |
| 2 | 84303600 | 84305199 | 9.203518 | 16.98556 | 30.91787 | 13.06443 | -0.42245 | 0.208455 | 0.548229 |
| 1 | 127491600 | 127491999 | 5.343172 | 10.4076 | 22.46732 | 8.197893 | -0.47439 | -0.35919 | 0.548237 |
| 11 | 28119000 | 28125599 | 87.50075 | 137.8143 | 134.3284 | 55.73663 | 0.35939 | 2.509608 | 0.548238 |
| 2 | 15491400 | 15492799 | 12.31808 | 22.05917 | 36.23144 | 15.88199 | -0.31124 | 0.420361 | 0.548269 |
| 16 | 92886800 | 92888399 | 12.31808 | 22.05917 | 36.23144 | 15.88199 | -0.31124 | 0.420361 | 0.548269 |
| 4 | 62356000 | 62357399 | 11.72539 | 22.01173 | 23.49642 | 9.669437 | 0.371283 | 0.107778 | 0.548274 |

| | | | | | | | | | |
|---|---|---|---|---|---|---|---|---|---|
| 7 | 114158200 | 114158599 | 4.934104 | 9.633897 | 19.94169 | 7.028393 | -0.37007 | -0.50143 | 0.548287 |
| 8 | 8801000 | 8801999 | 8.503922 | 16.10826 | 29.33442 | 11.86787 | -0.37275 | 0.03476 | 0.548287 |
| 5 | 4545200 | 4546599 | 7.021345 | 13.37456 | 26.66171 | 10.65968 | -0.48667 | -0.02469 | 0.548292 |
| 14 | 52963800 | 52968599 | 57.50691 | 92.12024 | 95.00788 | 40.0705 | 0.301289 | 2.010997 | 0.548301 |
| 5 | 135516200 | 135520599 | 27.90135 | 45.51163 | 75.7634 | 32.54981 | -0.36102 | 1.514341 | 0.548318 |
| 4 | 5818800 | 5820799 | 15.6736 | 28.11354 | 29.76791 | 13.03098 | 0.319731 | 0.418875 | 0.548318 |
| 16 | 15221400 | 15222999 | 15.71587 | 27.43993 | 44.14185 | 19.44067 | -0.29507 | 0.717381 | 0.548327 |
| 12 | 8192400 | 8193999 | 13.24896 | 23.50985 | 39.20361 | 17.31807 | -0.34097 | 0.487537 | 0.548341 |
| 6 | 50381000 | 50382399 | 9.980624 | 18.47102 | 31.48401 | 13.25343 | -0.32435 | 0.201475 | 0.548359 |
| 13 | 73694600 | 73695599 | 12.53176 | 23.63961 | 24.6549 | 10.06255 | 0.414429 | 0.089178 | 0.548363 |
| 18 | 24413200 | 24414199 | 16.24165 | 28.77918 | 29.47149 | 13.03001 | 0.357922 | 0.501639 | 0.548365 |
| 14 | 84352200 | 84353599 | 14.26626 | 25.97491 | 27.44922 | 11.8088 | 0.339008 | 0.318481 | 0.548375 |
| 4 | 134717800 | 134719799 | 28.17655 | 47.07598 | 49.69133 | 21.62848 | 0.303396 | 1.204232 | 0.548378 |
| 7 | 5229600 | 5233199 | 34.37395 | 57.83231 | 50.22939 | 21.948 | 0.583721 | 1.155241 | 0.548397 |
| 13 | 94124800 | 94129399 | 43.65175 | 69.847 | 108.5474 | 45.70509 | -0.29099 | 2.033604 | 0.548418 |
| 2 | 74991200 | 74991799 | 5.690433 | 11.02252 | 23.83652 | 9.045979 | -0.532 | -0.22034 | 0.548424 |
| 1 | 41356800 | 41359599 | 28.67058 | 48.34439 | 50.17225 | 21.93787 | 0.327778 | 1.116628 | 0.548428 |
| 10 | 122115400 | 122118799 | 71.53254 | 113.0856 | 120.6268 | 50.11631 | 0.233677 | 2.414385 | 0.548432 |
| 3 | 128239600 | 128240199 | 7.084734 | 13.34492 | 27.02983 | 11.06014 | -0.53807 | 0.057266 | 0.548438 |
| 1 | 13221400 | 13222199 | 13.01572 | 23.97483 | 25.98381 | 11.00842 | 0.31788 | 0.239142 | 0.548445 |
| 11 | 111743000 | 111743799 | 7.29085 | 14.0167 | 25.83151 | 10.08311 | -0.3422 | -0.10138 | 0.548447 |
| 7 | 106160000 | 106169999 | 182.7718 | 271.8019 | 278.2146 | 114.6239 | 0.221482 | 3.630697 | 0.548487 |
| 5 | 86360000 | 86369399 | 261.3696 | 377.3176 | 374.741 | 154.6011 | 0.224169 | 4.108134 | 0.548493 |
| 4 | 9857400 | 9857999 | 6.547104 | 12.62557 | 24.78378 | 9.589193 | -0.42044 | -0.14178 | 0.548505 |
| 12 | 69126000 | 69129799 | 29.12194 | 47.76406 | 74.78862 | 32.19911 | -0.26821 | 1.441446 | 0.54852 |
| 19 | 38303200 | 38306399 | 26.11733 | 43.35271 | 68.79064 | 29.81416 | -0.28312 | 1.286613 | 0.548543 |
| 19 | 36224800 | 36225599 | 12.34244 | 23.6561 | 22.81528 | 8.984141 | 0.572086 | -0.03154 | 0.54855 |
| 9 | 58261000 | 58263799 | 18.13143 | 31.1729 | 50.06528 | 21.96064 | -0.29665 | 0.905176 | 0.548578 |
| 1 | 40559600 | 40559999 | 8.776461 | 17.15139 | 15.79242 | 5.471128 | 0.805832 | -0.52207 | 0.548582 |
| 8 | 123289200 | 123292399 | 21.53762 | 36.03908 | 59.64827 | 25.98737 | -0.34335 | 1.177049 | 0.548601 |
| 14 | 88157800 | 88162999 | 74.97661 | 119.1109 | 114.864 | 47.90087 | 0.385234 | 2.289106 | 0.548602 |
| 18 | 24586400 | 24590599 | 27.456 | 44.79937 | 74.91368 | 32.18847 | -0.36708 | 1.510808 | 0.548602 |
| 2 | 90107400 | 90109199 | 22.05515 | 38.75638 | 34.49452 | 15.27185 | 0.554657 | 0.634028 | 0.548604 |
| 11 | 99863400 | 99866999 | 41.45195 | 68.09241 | 62.50503 | 26.92824 | 0.500973 | 1.442724 | 0.548631 |
| 7 | 113081000 | 113082199 | 13.17921 | 24.89099 | 24.64547 | 10.03304 | 0.492039 | 0.086604 | 0.548654 |
| 8 | 18629600 | 18630199 | 11.21017 | 21.06362 | 23.11518 | 9.497935 | 0.33383 | 0.098285 | 0.548658 |
| X | 149034200 | 149039199 | 78.56932 | 124.5592 | 120.3429 | 50.08136 | 0.379638 | 2.352468 | 0.548662 |
| 15 | 90512800 | 90516199 | 18.7686 | 32.18751 | 51.6762 | 22.6641 | -0.29686 | 0.931926 | 0.548664 |
| 17 | 17108600 | 17113599 | 66.55575 | 105.9479 | 109.9243 | 45.95151 | 0.282813 | 2.217799 | 0.548684 |
| 16 | 32367600 | 32370399 | 27.93724 | 46.03939 | 72.71188 | 31.36804 | -0.27787 | 1.378156 | 0.548686 |
| 4 | 140369400 | 140370599 | 15.93475 | 27.35874 | 50.04504 | 21.95072 | -0.48394 | 0.9073 | 0.548706 |
| 7 | 53526400 | 53528999 | 26.60747 | 45.18462 | 47.40798 | 20.77378 | 0.312384 | 1.045393 | 0.548736 |
| 4 | 144910800 | 144912399 | 11.72136 | 22.33167 | 19.55248 | 7.814129 | 0.691705 | 0.00245 | 0.548752 |
| 3 | 38115800 | 38117999 | 27.31268 | 45.85118 | 49.11267 | 21.44114 | 0.28217 | 1.149659 | 0.548767 |
| 8 | 19486600 | 19491199 | 70.48229 | 111.8548 | 117.1967 | 48.82288 | 0.264259 | 2.313886 | 0.548773 |
| X | 104300400 | 104300999 | 13.99043 | 25.6593 | 26.50881 | 11.29696 | 0.380763 | 0.273988 | 0.548773 |
| 7 | 144606400 | 144608599 | 15.5249 | 28.38925 | 27.70284 | 11.85351 | 0.457846 | 0.301211 | 0.548778 |

| | | | | | | | | | |
|---|---|---|---|---|---|---|---|---|---|
| 19 | 39007400 | 39008999 | 14.91572 | 26.24315 | 44.48694 | 19.69655 | -0.36973 | 0.596176 | 0.548779 |
| 3 | 73363600 | 73364399 | 12.71225 | 22.37497 | 38.37989 | 16.99135 | -0.38595 | 0.570785 | 0.548789 |
| 15 | 83938000 | 83939599 | 12.71225 | 22.37497 | 38.37989 | 16.99135 | -0.38595 | 0.570785 | 0.548789 |
| 5 | 125372800 | 125374999 | 24.56705 | 40.58192 | 66.60688 | 28.77113 | -0.33215 | 1.338074 | 0.548796 |
| 6 | 3193000 | 3194999 | 27.79785 | 46.8788 | 49.62495 | 21.6994 | 0.299274 | 1.111375 | 0.548822 |
| 17 | 54628400 | 54629799 | 16.65101 | 29.55368 | 31.2232 | 13.79 | 0.314176 | 0.493085 | 0.548829 |
| 9 | 96280200 | 96282399 | 19.26553 | 33.04367 | 52.76551 | 23.14207 | -0.28911 | 0.934444 | 0.548851 |
| 8 | 78715200 | 78720399 | 41.38519 | 67.2679 | 70.35724 | 30.18799 | 0.304092 | 1.590254 | 0.548859 |
| 11 | 111503000 | 111503999 | 11.58635 | 21.66182 | 23.13132 | 9.585439 | 0.363397 | 0.134251 | 0.548875 |
| 9 | 64468600 | 64471199 | 34.11249 | 55.23525 | 85.56576 | 36.61015 | -0.26635 | 1.660645 | 0.548876 |
| 5 | 61258600 | 61260999 | 25.3863 | 43.72843 | 42.98256 | 18.85496 | 0.409277 | 0.908614 | 0.54888 |
| 3 | 59495200 | 59499199 | 39.71139 | 65.78898 | 57.30642 | 24.80333 | 0.579353 | 1.337257 | 0.548893 |
| 3 | 98085800 | 98088599 | 36.52504 | 59.81749 | 64.13637 | 27.59564 | 0.275775 | 1.46931 | 0.548915 |
| 14 | 79790400 | 79793599 | 40.28087 | 65.56734 | 68.7861 | 29.52967 | 0.301173 | 1.56477 | 0.548916 |
| 1 | 3836000 | 3837599 | 17.41177 | 31.58157 | 29.16947 | 12.60873 | 0.5276 | 0.365891 | 0.548942 |
| 6 | 99735000 | 99736199 | 13.32742 | 24.47576 | 26.444 | 11.249 | 0.318104 | 0.258988 | 0.548971 |
| 7 | 33655400 | 33679999 | 460.009 | 634.9456 | 590.1897 | 245.2216 | 0.254747 | 4.78073 | 0.548972 |
| 2 | 3835400 | 3838399 | 43.63671 | 70.42753 | 75.31394 | 32.1101 | 0.26207 | 1.755082 | 0.548985 |
| 18 | 42198400 | 42200199 | 14.42623 | 25.25936 | 41.49595 | 18.31581 | -0.325 | 0.66225 | 0.548985 |
| 10 | 83504800 | 83509799 | 41.17374 | 67.0789 | 68.86976 | 29.57494 | 0.33312 | 1.555522 | 0.548992 |
| 7 | 31010800 | 31011999 | 14.44963 | 26.65064 | 23.1655 | 9.797079 | 0.635595 | 0.235641 | 0.549009 |
| 13 | 34186000 | 34186799 | 7.032874 | 13.42856 | 25.61761 | 10.19062 | -0.41657 | -0.04083 | 0.549013 |
| 17 | 40200000 | 40200999 | 23.72234 | 40.56866 | 43.19789 | 18.94139 | 0.292686 | 0.968315 | 0.549014 |
| 5 | 113735800 | 113738599 | 30.02804 | 48.95741 | 77.03451 | 33.0902 | -0.28044 | 1.528124 | 0.549023 |
| 6 | 17389800 | 17393399 | 43.15055 | 69.24997 | 104.5365 | 44.21269 | -0.24412 | 1.94017 | 0.549027 |
| 12 | 102388200 | 102389999 | 17.61689 | 30.4141 | 49.00304 | 21.49741 | -0.30001 | 0.863867 | 0.54903 |
| 4 | 143340600 | 143344999 | 77.41447 | 122.5545 | 123.4619 | 51.33031 | 0.317643 | 2.386248 | 0.549041 |
| 12 | 24263400 | 24304799 | 1022.147 | 1319.404 | 1156.695 | 484.8964 | 0.242233 | 5.851831 | 0.549042 |
| X | 77188400 | 77190599 | 28.98125 | 49.46744 | 40.393 | 17.70878 | 0.673763 | 1.003038 | 0.549053 |
| X | 117051600 | 117056199 | 85.97701 | 135.4973 | 132.7879 | 55.10179 | 0.352263 | 2.497695 | 0.549075 |
| Y | 3940800 | 3943199 | 15.95588 | 28.43793 | 21.14942 | 9.308127 | 0.820942 | 0.439412 | 0.549083 |
| 12 | 99447800 | 99450399 | 28.20575 | 46.13441 | 73.71188 | 31.70009 | -0.29915 | 1.475348 | 0.549109 |
| 3 | 11011000 | 11013799 | 32.01989 | 54.06286 | 49.63495 | 21.71156 | 0.504071 | 1.116817 | 0.549114 |
| 10 | 21638600 | 21640999 | 15.828 | 27.54812 | 45.24553 | 19.88703 | -0.32547 | 0.762979 | 0.549128 |
| X | 94847800 | 94850399 | 23.09696 | 40.5365 | 35.0347 | 15.49467 | 0.596925 | 0.660521 | 0.549143 |
| 7 | 38784600 | 38791199 | 118.0369 | 181.5249 | 164.2861 | 68.22398 | 0.439821 | 2.957546 | 0.549151 |
| 12 | 21663800 | 21689399 | 559.1286 | 756.5581 | 703.9135 | 293.3843 | 0.224386 | 5.091948 | 0.549154 |
| 17 | 39084600 | 39086399 | 32.21273 | 54.08754 | 51.35352 | 22.42215 | 0.455513 | 1.165062 | 0.549157 |
| 13 | 64960600 | 64961999 | 15.25097 | 27.59793 | 28.7253 | 12.45297 | 0.353324 | 0.360933 | 0.549164 |
| 9 | 87025800 | 87029199 | 24.33676 | 40.25077 | 67.96123 | 29.38098 | -0.37285 | 1.321322 | 0.549165 |
| 1 | 8925000 | 8927399 | 23.97982 | 40.90317 | 43.72073 | 19.16666 | 0.286593 | 0.992103 | 0.549172 |
| 4 | 143220200 | 143233999 | 192.5374 | 286.4932 | 282.3594 | 116.3385 | 0.276844 | 3.621773 | 0.549179 |
| 12 | 102914800 | 102916399 | 14.34055 | 24.90517 | 44.81208 | 19.66967 | -0.45649 | 0.788959 | 0.549188 |
| 4 | 113817200 | 113822799 | 80.04398 | 126.638 | 125.1841 | 52.02982 | 0.344203 | 2.404417 | 0.549193 |
| 14 | 33421600 | 33423999 | 14.14405 | 24.71039 | 42.11332 | 18.55429 | -0.37767 | 0.698409 | 0.549199 |
| 18 | 76234000 | 76235799 | 22.15998 | 38.32425 | 40.97453 | 17.9763 | 0.289697 | 0.857674 | 0.549207 |
| 5 | 112388600 | 112391199 | 21.69515 | 36.40852 | 58.58548 | 25.57127 | -0.30286 | 1.146253 | 0.549212 |

| | | | | | | | | | |
|---|---|---|---|---|---|---|---|---|---|
| 4 | 150336400 | 150338199 | 10.3298 | 18.84724 | 32.32399 | 13.86948 | -0.35287 | 0.286358 | 0.549226 |
| 12 | 118558200 | 118559199 | 10.3298 | 18.84724 | 32.32399 | 13.86948 | -0.35287 | 0.286358 | 0.549226 |
| 2 | 97874400 | 97876199 | 17.09062 | 30.95487 | 29.08932 | 12.59655 | 0.500966 | 0.370017 | 0.549242 |
| 10 | 54131600 | 54134199 | 28.05531 | 48.58288 | 40.67949 | 17.84756 | 0.641262 | 0.872869 | 0.549246 |
| 3 | 76959400 | 76962799 | 28.27064 | 48.32597 | 44.58452 | 19.54872 | 0.498487 | 0.988667 | 0.549247 |
| 13 | 62791000 | 62798199 | 146.537 | 222.9928 | 209.4975 | 86.80787 | 0.374135 | 3.178719 | 0.549268 |
| 14 | 101478000 | 101479599 | 25.83948 | 44.3178 | 44.66041 | 19.58726 | 0.372414 | 0.951849 | 0.549273 |
| 1 | 65546200 | 65548399 | 17.57432 | 31.80741 | 29.0241 | 12.57995 | 0.542282 | 0.377356 | 0.549276 |
| 16 | 25668400 | 25669599 | 8.444592 | 15.96194 | 28.67671 | 11.65401 | -0.35827 | 0.051365 | 0.549283 |
| 7 | 13156600 | 13157799 | 27.62348 | 46.36034 | 48.05625 | 20.97634 | 0.32943 | 1.155187 | 0.549286 |
| 7 | 7797200 | 7797599 | 7.634178 | 14.49388 | 26.59477 | 10.70757 | -0.37662 | 0.006265 | 0.54929 |
| 15 | 5154000 | 5157999 | 30.5723 | 50.87002 | 54.60163 | 23.70397 | 0.279424 | 1.26041 | 0.549322 |
| 4 | 16265200 | 16265999 | 9.928939 | 18.92604 | 19.7203 | 7.872992 | 0.443914 | -0.01881 | 0.549323 |
| 14 | 110172200 | 110173799 | 17.10221 | 30.10719 | 31.4392 | 13.9181 | 0.32616 | 0.579618 | 0.549328 |
| 4 | 23420000 | 23424799 | 74.38009 | 117.4717 | 125.8652 | 52.26668 | 0.226125 | 2.440757 | 0.549331 |
| 13 | 33271600 | 33273999 | 23.04425 | 37.90986 | 75.13973 | 32.39154 | -0.60542 | 1.391385 | 0.549339 |
| 17 | 22322000 | 22335799 | 200.5757 | 297.8964 | 289.0808 | 119.0852 | 0.296721 | 3.655961 | 0.549343 |
| 1 | 191200400 | 191202999 | 14.78365 | 25.99784 | 42.90947 | 19.00002 | -0.33153 | 0.599459 | 0.549351 |
| 10 | 129282200 | 129283999 | 15.99852 | 28.7987 | 29.07342 | 12.68714 | 0.390931 | 0.395981 | 0.549355 |
| 10 | 47265000 | 47265799 | 11.08087 | 21.09841 | 23.26982 | 9.311871 | 0.359279 | 0.009354 | 0.549358 |
| 6 | 35262200 | 35268399 | 49.25764 | 78.42019 | 121.4515 | 50.77786 | -0.29399 | 2.201385 | 0.549375 |
| 11 | 51456200 | 51458999 | 36.56583 | 59.74596 | 64.49413 | 27.72538 | 0.263954 | 1.502097 | 0.549382 |
| 2 | 20062600 | 20064599 | 23.82314 | 41.20497 | 41.95355 | 18.40587 | 0.359922 | 0.865458 | 0.549388 |
| 3 | 4836200 | 4838399 | 31.15469 | 51.66421 | 55.74541 | 24.14413 | 0.271494 | 1.300082 | 0.549391 |
| 6 | 107384200 | 107384799 | 6.536271 | 12.70672 | 26.12852 | 9.69991 | -0.43047 | -0.27913 | 0.549404 |
| 7 | 38132000 | 38136399 | 56.01199 | 90.46192 | 76.14522 | 32.49291 | 0.607794 | 1.758123 | 0.54944 |
| 13 | 29019800 | 29022199 | 18.96941 | 32.30984 | 53.25247 | 23.34254 | -0.33612 | 0.996839 | 0.54945 |
| 5 | 130426000 | 130445999 | 235.6267 | 339.4163 | 458.7371 | 189.3085 | -0.22323 | 4.139898 | 0.549462 |
| 7 | 46915200 | 46915799 | 9.757003 | 18.79987 | 20.86321 | 8.091687 | 0.392477 | -0.10847 | 0.549464 |
| Y | 10417400 | 10422799 | 29.09891 | 48.7712 | 33.28814 | 14.51786 | 0.929845 | 1.152259 | 0.549475 |
| 5 | 131123400 | 131130599 | 86.42061 | 135.321 | 143.2759 | 59.43269 | 0.233688 | 2.619868 | 0.549477 |
| 7 | 51745400 | 51746999 | 15.9707 | 29.52842 | 26.80984 | 11.30056 | 0.576831 | 0.243367 | 0.549507 |
| 2 | 111271400 | 111275799 | 89.96682 | 140.8234 | 145.8787 | 60.51459 | 0.264782 | 2.629218 | 0.549511 |
| 10 | 4296800 | 4298599 | 10.38737 | 18.79175 | 32.79568 | 14.22237 | -0.3877 | 0.338073 | 0.549536 |
| 14 | 79668800 | 79669599 | 7.948596 | 15.07924 | 27.12087 | 10.93714 | -0.35004 | 0.016626 | 0.549544 |
| 1 | 30761600 | 30761999 | 7.280699 | 13.91846 | 27.91461 | 11.07412 | -0.48589 | -0.04593 | 0.54955 |
| 15 | 38998400 | 39001799 | 48.96974 | 78.97054 | 81.25133 | 34.6052 | 0.316308 | 1.790429 | 0.549552 |
| 8 | 113119400 | 113120799 | 15.19195 | 28.12419 | 26.91318 | 11.32458 | 0.503333 | 0.231015 | 0.549567 |
| 12 | 32701400 | 32702799 | 15.92766 | 29.09355 | 28.32591 | 12.13689 | 0.460155 | 0.314137 | 0.549579 |
| 12 | 42912800 | 42913799 | 6.8571 | 13.27998 | 25.6172 | 9.729982 | -0.36959 | -0.19556 | 0.549625 |
| 11 | 5641000 | 5646999 | 59.73946 | 94.71163 | 142.3808 | 59.25519 | -0.25723 | 2.339465 | 0.549627 |
| 10 | 59703400 | 59707999 | 47.9321 | 76.34991 | 120.3873 | 50.3695 | -0.31903 | 2.182441 | 0.549634 |
| 11 | 20326200 | 20329599 | 51.42073 | 81.87985 | 122.6265 | 51.28294 | -0.24538 | 2.198427 | 0.549641 |
| 12 | 80073200 | 80075399 | 20.47601 | 35.53323 | 36.91315 | 16.19618 | 0.33205 | 0.814051 | 0.549642 |
| 1 | 64670800 | 64673199 | 21.31812 | 37.15367 | 37.55221 | 16.51956 | 0.371814 | 0.757229 | 0.549644 |
| 1 | 144659200 | 144664199 | 29.19835 | 49.57727 | 42.54415 | 18.64167 | 0.601334 | 1.051299 | 0.549662 |
| 5 | 88565200 | 88569599 | 39.85374 | 65.56541 | 61.35018 | 26.4475 | 0.474202 | 1.418736 | 0.549667 |

| | | | | | | | | | |
|---|---|---|---|---|---|---|---|---|---|
| 14 | 18093200 | 18098599 | 47.32461 | 76.79085 | 72.15702 | 30.92132 | 0.456025 | 1.63615 | 0.549696 |
| 12 | 43708800 | 43711799 | 25.19709 | 43.60753 | 41.90119 | 18.38324 | 0.443337 | 0.866714 | 0.549724 |
| 17 | 81556400 | 81559399 | 24.31828 | 41.80787 | 43.60339 | 19.12617 | 0.323631 | 0.922935 | 0.549758 |
| 8 | 23853200 | 23855999 | 49.27057 | 79.42876 | 81.87168 | 34.85417 | 0.313122 | 1.800466 | 0.549779 |
| 11 | 65349200 | 65351799 | 31.49842 | 52.54805 | 47.60127 | 20.69939 | 0.523293 | 1.227001 | 0.549811 |
| 10 | 39265400 | 39268199 | 48.69077 | 78.22308 | 82.98472 | 35.15316 | 0.265551 | 1.905751 | 0.549815 |
| 12 | 3136000 | 3137799 | 21.48927 | 37.22905 | 39.93663 | 17.52188 | 0.285408 | 0.837758 | 0.549816 |
| 4 | 53143200 | 53147799 | 23.22466 | 38.43809 | 67.29805 | 29.1082 | -0.42491 | 1.309726 | 0.549839 |
| 7 | 34245200 | 34246799 | 17.91626 | 30.54456 | 51.61583 | 22.62684 | -0.3716 | 0.984747 | 0.549842 |
| 9 | 87469000 | 87471999 | 39.84285 | 64.57716 | 68.68721 | 29.40974 | 0.276118 | 1.642031 | 0.549849 |
| 19 | 54441000 | 54443599 | 23.86198 | 40.93312 | 43.65836 | 19.14804 | 0.290911 | 0.941288 | 0.549854 |
| 1 | 181148000 | 181151399 | 34.98152 | 57.91936 | 58.75436 | 25.4198 | 0.360103 | 1.331087 | 0.549873 |
| 1 | 95157400 | 95158799 | 18.4881 | 32.82639 | 32.0509 | 14.15116 | 0.427645 | 0.504275 | 0.549873 |
| 7 | 11253200 | 11254799 | 22.94657 | 40.08542 | 32.64364 | 14.38136 | 0.682323 | 0.726626 | 0.549898 |
| 18 | 13429800 | 13433799 | 42.85774 | 68.63261 | 106.1417 | 44.74534 | -0.28262 | 2.00751 | 0.549915 |
| 13 | 28065400 | 28067399 | 38.75862 | 62.36641 | 96.03492 | 40.77571 | -0.268 | 1.849229 | 0.549926 |
| 1 | 40638800 | 40639999 | 13.12465 | 24.01989 | 25.4355 | 10.8712 | 0.341988 | 0.274044 | 0.549929 |
| 17 | 30361000 | 30361599 | 10.81316 | 20.50969 | 22.72835 | 9.169435 | 0.342502 | 0.032589 | 0.549933 |
| 1 | 38416600 | 38418399 | 13.49455 | 23.64262 | 40.50146 | 17.88617 | -0.385 | 0.646008 | 0.549943 |
| 17 | 73407600 | 73410599 | 27.9629 | 47.18686 | 49.62235 | 21.70241 | 0.308795 | 1.106583 | 0.549962 |
| 5 | 50510000 | 50511399 | 15.77819 | 28.63794 | 28.83087 | 12.45184 | 0.404879 | 0.350253 | 0.549988 |
| 1 | 112221000 | 112223399 | 31.97859 | 52.77442 | 56.27096 | 24.29044 | 0.288075 | 1.358007 | 0.549996 |
| 7 | 52203800 | 52206199 | 24.72105 | 43.0638 | 39.05358 | 17.17487 | 0.527673 | 0.785363 | 0.550013 |
| 15 | 26040400 | 26041999 | 15.42431 | 27.95432 | 28.8441 | 12.48058 | 0.367843 | 0.355892 | 0.550026 |
| 15 | 82430000 | 82430999 | 7.296004 | 13.95307 | 26.20958 | 10.38627 | -0.39012 | -0.05174 | 0.550032 |
| 15 | 50694200 | 50695399 | 8.411509 | 15.69454 | 29.49922 | 12.26595 | -0.45074 | 0.133588 | 0.550038 |
| 4 | 146231400 | 146232799 | 23.59639 | 40.80689 | 36.28691 | 15.91973 | 0.554439 | 0.858738 | 0.550043 |
| 11 | 53041400 | 53044999 | 28.93326 | 48.66722 | 50.73869 | 22.16897 | 0.321051 | 1.138045 | 0.550056 |
| 5 | 69578400 | 69580999 | 48.10919 | 77.36499 | 83.07476 | 35.24163 | 0.249875 | 1.875257 | 0.550063 |
| 1 | 6714600 | 6715799 | 13.04712 | 24.30692 | 21.38995 | 8.916959 | 0.633395 | 0.163546 | 0.550064 |
| 7 | 41033600 | 41037399 | 51.22086 | 82.57783 | 80.2763 | 34.17767 | 0.397491 | 1.798602 | 0.5501 |
| 4 | 28723000 | 28724999 | 18.2594 | 32.37887 | 32.02701 | 14.15587 | 0.407891 | 0.509913 | 0.550103 |
| 10 | 23895000 | 23899999 | 66.17034 | 105.2711 | 110.8152 | 46.28538 | 0.261023 | 2.236195 | 0.550147 |
| 8 | 21203800 | 21205399 | 20.24951 | 34.04928 | 56.90348 | 24.85998 | -0.35717 | 1.121035 | 0.550175 |
| 16 | 26031800 | 26034399 | 35.7261 | 59.27963 | 57.64168 | 24.97549 | 0.421248 | 1.309147 | 0.550202 |
| 10 | 69071000 | 69078199 | 70.71457 | 109.4927 | 198.208 | 82.33039 | -0.55175 | 2.815855 | 0.550217 |
| 17 | 20597200 | 20611399 | 207.4733 | 308.1558 | 287.9169 | 118.6064 | 0.351426 | 3.656514 | 0.550217 |
| 2 | 92768600 | 92770199 | 16.72315 | 28.82116 | 47.94448 | 21.03192 | -0.34627 | 0.875918 | 0.550236 |
| 10 | 121415400 | 121416999 | 13.98275 | 24.48892 | 41.0142 | 18.10696 | -0.35257 | 0.656167 | 0.550236 |
| 15 | 97009800 | 97010799 | 7.287681 | 13.96094 | 26.27227 | 10.36026 | -0.38617 | -0.06802 | 0.550273 |
| 16 | 75882200 | 75882799 | 7.287681 | 13.96094 | 26.27227 | 10.36026 | -0.38617 | -0.06802 | 0.550273 |
| 13 | 69491200 | 69495799 | 59.09211 | 94.3273 | 101.0396 | 42.40006 | 0.241138 | 2.120572 | 0.550278 |
| 12 | 13479200 | 13480399 | 10.30062 | 18.87562 | 32.37179 | 13.81178 | -0.34734 | 0.260919 | 0.550301 |
| 2 | 110853400 | 110856999 | 48.30786 | 78.38772 | 71.72064 | 30.73472 | 0.494402 | 1.63731 | 0.550304 |
| 18 | 74540000 | 74544999 | 89.48707 | 139.7436 | 148.0553 | 61.43236 | 0.229718 | 2.669661 | 0.550307 |
| X | 148983400 | 148986999 | 49.73812 | 80.50192 | 74.94433 | 32.05554 | 0.466042 | 1.69327 | 0.550318 |
| 5 | 58390800 | 58394999 | 121.0364 | 184.9947 | 192.7807 | 79.97035 | 0.229729 | 3.08096 | 0.550348 |

| | | | | | | | | | |
|---|---|---|---|---|---|---|---|---|---|
| 5 | 133318200 | 133325999 | 120.5726 | 182.4726 | 259.8546 | 107.4934 | -0.23208 | 3.285781 | 0.550359 |
| 15 | 62944000 | 62945399 | 15.05124 | 26.37138 | 42.476 | 18.75902 | -0.29677 | 0.655876 | 0.550364 |
| X | 153805600 | 153807999 | 30.1507 | 51.39079 | 46.25008 | 20.27428 | 0.533668 | 1.025501 | 0.550364 |
| 2 | 8271600 | 8272999 | 13.39084 | 25.35775 | 22.8916 | 9.266996 | 0.630788 | 0.064134 | 0.550393 |
| 3 | 15194200 | 15194999 | 11.38564 | 21.38206 | 23.62035 | 9.714165 | 0.323333 | 0.10459 | 0.550397 |
| 5 | 91012800 | 91013399 | 11.38564 | 21.38206 | 23.62035 | 9.714165 | 0.323333 | 0.10459 | 0.550397 |
| 4 | 108157600 | 108158599 | 7.091768 | 13.32754 | 27.52202 | 11.30541 | -0.57061 | 0.072086 | 0.550416 |
| 13 | 15636600 | 15639199 | 20.62233 | 34.40256 | 61.16915 | 26.59881 | -0.44631 | 1.207937 | 0.550419 |
| 6 | 89071000 | 89073599 | 29.0954 | 47.2815 | 78.22068 | 33.55646 | -0.35595 | 1.575392 | 0.550443 |
| X | 5007000 | 5008399 | 15.43813 | 28.71022 | 26.45286 | 11.06088 | 0.564295 | 0.207532 | 0.550459 |
| 7 | 67498800 | 67501999 | 34.31383 | 55.43204 | 87.23884 | 37.23863 | -0.29258 | 1.719465 | 0.550463 |
| 11 | 31928800 | 31932999 | 76.94492 | 121.5228 | 127.6274 | 52.99854 | 0.254952 | 2.443694 | 0.550466 |
| 3 | 4782200 | 4782799 | 15.57626 | 27.84821 | 29.38323 | 12.89791 | 0.322241 | 0.438647 | 0.550469 |
| 3 | 28453000 | 28454599 | 14.68701 | 26.49831 | 28.35431 | 12.33804 | 0.310385 | 0.37435 | 0.550472 |
| 8 | 4620400 | 4625399 | 56.97455 | 91.33601 | 83.54777 | 35.27734 | 0.47535 | 1.983691 | 0.550473 |
| 8 | 91062600 | 91064999 | 23.34645 | 40.45067 | 40.83758 | 17.91722 | 0.372513 | 0.844957 | 0.550475 |
| 17 | 84039000 | 84041199 | 17.48024 | 30.10331 | 48.99998 | 21.49444 | -0.31538 | 0.885958 | 0.550491 |
| 13 | 65153600 | 65158199 | 37.21929 | 61.11255 | 62.7609 | 27.03343 | 0.339206 | 1.43664 | 0.550497 |
| 9 | 115562000 | 115564199 | 14.07579 | 24.78419 | 40.28918 | 17.83529 | -0.309 | 0.57585 | 0.550522 |
| 7 | 27414200 | 27423799 | 104.8224 | 161.7734 | 170.6278 | 70.87551 | 0.223078 | 2.885983 | 0.550527 |
| 11 | 21094000 | 21094999 | 6.835887 | 13.19781 | 24.83849 | 9.572211 | -0.35471 | -0.15097 | 0.550545 |
| 6 | 34231200 | 34234199 | 24.42528 | 40.64075 | 66.79066 | 28.995 | -0.33317 | 1.256166 | 0.550558 |
| 9 | 120183000 | 120183999 | 12.45146 | 23.08329 | 24.1103 | 10.12578 | 0.380195 | 0.196113 | 0.550575 |
| 11 | 3552000 | 3552999 | 12.45146 | 23.08329 | 24.1103 | 10.12578 | 0.380195 | 0.196113 | 0.550575 |
| 10 | 116167000 | 116170599 | 25.60215 | 42.43776 | 67.59845 | 29.26899 | -0.28882 | 1.299437 | 0.550578 |
| 4 | 72874200 | 72875999 | 19.90401 | 35.47134 | 30.79796 | 13.55563 | 0.599018 | 0.487696 | 0.550588 |
| 3 | 116344800 | 116346399 | 26.03174 | 43.95054 | 46.95226 | 20.53783 | 0.286253 | 1.093778 | 0.550605 |
| 9 | 43417000 | 43419999 | 32.56792 | 53.86038 | 57.94663 | 25.05019 | 0.275417 | 1.338269 | 0.550627 |
| 1 | 39336400 | 39341399 | 45.34883 | 72.35915 | 114.2835 | 47.92947 | -0.3187 | 2.124272 | 0.550638 |
| 4 | 142250000 | 142250599 | 5.661203 | 11.01402 | 21.82275 | 8.051295 | -0.3679 | -0.32237 | 0.550655 |
| 5 | 106513400 | 106514799 | 8.11573 | 15.22259 | 27.99983 | 11.53917 | -0.40897 | 0.095129 | 0.55066 |
| 11 | 33652600 | 33654999 | 15.11619 | 26.46465 | 43.04655 | 18.99854 | -0.31073 | 0.671418 | 0.550666 |
| 6 | 123195400 | 123196599 | 13.76069 | 24.19587 | 39.80826 | 17.62727 | -0.32699 | 0.588405 | 0.550679 |
| 2 | 97506600 | 97512199 | 96.94803 | 150.7023 | 157.8829 | 65.55103 | 0.240779 | 2.753665 | 0.550696 |
| 15 | 75155800 | 75162799 | 96.94803 | 150.7023 | 157.8829 | 65.55103 | 0.240779 | 2.753665 | 0.550696 |
| 1 | 61536800 | 61538999 | 38.75997 | 61.84396 | 124.2992 | 52.12768 | -0.66581 | 2.122367 | 0.550699 |
| 7 | 36007400 | 36010799 | 26.53449 | 43.53883 | 71.5929 | 30.82879 | -0.33832 | 1.427628 | 0.550708 |
| 7 | 17052000 | 17059199 | 92.103 | 144.6487 | 139.8767 | 58.01716 | 0.367666 | 2.570118 | 0.55071 |
| 14 | 120606600 | 120609199 | 15.19339 | 26.60204 | 43.01012 | 18.98381 | -0.30226 | 0.668091 | 0.550712 |
| 18 | 59607000 | 59609799 | 28.09449 | 48.14214 | 44.59377 | 19.55662 | 0.493295 | 0.968034 | 0.55072 |
| 1 | 142536000 | 142543799 | 139.6786 | 211.5039 | 220.2856 | 91.14085 | 0.219593 | 3.276881 | 0.550755 |
| 5 | 121163600 | 121171799 | 68.26902 | 106.9057 | 162.1786 | 67.27328 | -0.28443 | 2.611458 | 0.55076 |
| 6 | 48205400 | 48206199 | 9.737058 | 17.78965 | 31.35256 | 13.43005 | -0.39005 | 0.27306 | 0.550796 |
| 9 | 116816400 | 116821199 | 67.9282 | 105.621 | 188.0525 | 78.07387 | -0.52312 | 2.739955 | 0.550806 |
| 17 | 60646000 | 60649199 | 27.63432 | 47.24018 | 45.72512 | 20.0489 | 0.429625 | 0.989066 | 0.550809 |
| 10 | 121578200 | 121586999 | 111.5076 | 168.4756 | 278.9656 | 115.339 | -0.45146 | 3.318812 | 0.550819 |
| 19 | 61283600 | 61286599 | 30.77566 | 52.65827 | 44.56908 | 19.54345 | 0.622745 | 0.993429 | 0.550828 |

| | | | | | | | | | |
|---|---|---|---|---|---|---|---|---|---|
| 16 | 22426200 | 22427599 | 14.43204 | 25.19338 | 42.62088 | 18.76848 | -0.36732 | 0.710898 | 0.550839 |
| X | 15980400 | 15981799 | 13.68641 | 26.07058 | 23.76542 | 9.501619 | 0.633441 | 0.033243 | 0.550844 |
| 12 | 30610200 | 30611599 | 13.05556 | 22.86191 | 45.16096 | 19.93535 | -0.58963 | 0.659277 | 0.55085 |
| 1 | 103462400 | 103463399 | 11.86939 | 22.76125 | 22.85459 | 8.986202 | 0.51648 | -0.04065 | 0.550852 |
| 11 | 109892000 | 109893199 | 18.67026 | 32.93277 | 33.10903 | 14.65153 | 0.381557 | 0.569215 | 0.550857 |
| 2 | 6453400 | 6455199 | 12.97684 | 22.87932 | 38.28679 | 16.9444 | -0.34971 | 0.551771 | 0.550876 |
| 4 | 94792400 | 94793999 | 20.63535 | 35.88463 | 37.39305 | 16.4268 | 0.328039 | 0.788806 | 0.550899 |
| X | 128215200 | 128217799 | 20.91245 | 37.20328 | 31.33066 | 13.81047 | 0.641553 | 0.507243 | 0.55091 |
| 16 | 19843200 | 19844199 | 15.53122 | 27.02147 | 44.73944 | 19.65995 | -0.33702 | 0.766777 | 0.550911 |
| 2 | 37248400 | 37251599 | 21.26432 | 35.76764 | 60.92423 | 26.61927 | -0.38479 | 1.121217 | 0.550941 |
| 17 | 34569400 | 34571799 | 22.95061 | 38.475 | 60.89678 | 26.56249 | -0.27921 | 1.162299 | 0.550983 |
| 5 | 135617800 | 135622599 | 66.13598 | 105.3335 | 108.6175 | 45.43733 | 0.292457 | 2.201683 | 0.550986 |
| 1 | 40352400 | 40353599 | 12.64345 | 22.38455 | 37.21007 | 16.45418 | -0.33778 | 0.501727 | 0.550989 |
| 18 | 73013800 | 73016999 | 33.28582 | 55.84084 | 51.93098 | 22.66186 | 0.485369 | 1.178944 | 0.550995 |
| 7 | 31459600 | 31463199 | 64.70663 | 102.7054 | 103.8701 | 43.2781 | 0.315455 | 2.301855 | 0.551018 |
| 17 | 27455800 | 27501799 | 773.3611 | 1018.21 | 905.8636 | 378.9716 | 0.24952 | 5.529335 | 0.55103 |
| 19 | 57933800 | 57936799 | 33.30126 | 54.89954 | 59.121 | 25.50894 | 0.273213 | 1.376215 | 0.551034 |
| 6 | 80991400 | 80993399 | 22.50653 | 39.38783 | 36.8118 | 16.24146 | 0.484673 | 0.703842 | 0.55104 |
| 14 | 44276000 | 44282599 | 84.74408 | 132.9551 | 140.6933 | 58.34968 | 0.236667 | 2.584809 | 0.551043 |
| X | 41498800 | 41506399 | 95.7603 | 149.4056 | 151.3168 | 62.79145 | 0.293666 | 2.690642 | 0.551044 |
| 4 | 147733800 | 147740999 | 254.2791 | 369.7171 | 322.7935 | 133.0533 | 0.420104 | 4.000557 | 0.551072 |
| 12 | 59517600 | 59518399 | 9.482235 | 17.68956 | 30.56789 | 12.71369 | -0.3307 | 0.147164 | 0.551072 |
| 8 | 57003400 | 57005399 | 12.19536 | 21.62412 | 36.23429 | 16.01645 | -0.34817 | 0.482071 | 0.551073 |
| 18 | 17145800 | 17148199 | 19.79238 | 34.73972 | 30.75407 | 13.60247 | 0.562375 | 0.633118 | 0.551086 |
| 13 | 62721600 | 62723199 | 19.68542 | 34.3414 | 36.70013 | 16.15455 | 0.292022 | 0.736417 | 0.551105 |
| 5 | 122318000 | 122320599 | 18.35179 | 31.27829 | 52.16576 | 22.8674 | -0.3529 | 0.988013 | 0.551111 |
| 13 | 85086800 | 85088999 | 20.10945 | 34.28843 | 54.36353 | 23.83162 | -0.28016 | 0.993444 | 0.551115 |
| 17 | 47719400 | 47723199 | 24.03166 | 39.7354 | 67.31664 | 29.09687 | -0.37768 | 1.323603 | 0.551128 |
| 12 | 55558200 | 55560799 | 18.32066 | 31.30443 | 51.64968 | 22.64579 | -0.33687 | 0.963671 | 0.551135 |
| 7 | 123105800 | 123112599 | 85.10499 | 131.8284 | 189.8692 | 78.86647 | -0.22186 | 2.812834 | 0.551135 |
| 14 | 111422600 | 111425199 | 25.01479 | 43.05257 | 43.57632 | 19.11493 | 0.367015 | 0.916536 | 0.551148 |
| 7 | 34855000 | 34857399 | 13.84182 | 24.24299 | 41.03038 | 18.11467 | -0.36756 | 0.655455 | 0.551149 |
| 9 | 90297200 | 90299199 | 13.89954 | 24.60086 | 42.3559 | 18.7308 | -0.38869 | 0.520795 | 0.55115 |
| 4 | 78129200 | 78131399 | 15.95302 | 27.41093 | 57.05131 | 25.02452 | -0.66939 | 0.908675 | 0.551157 |
| 4 | 106165400 | 106166999 | 14.69926 | 27.11158 | 26.81834 | 11.34163 | 0.450395 | 0.245656 | 0.551173 |
| 7 | 23407000 | 23420599 | 205.9232 | 307.7491 | 260.0832 | 107.2336 | 0.504259 | 3.545845 | 0.551181 |
| 5 | 109692600 | 109711199 | 364.5587 | 496.449 | 845.8613 | 352.1996 | -0.63894 | 4.988654 | 0.551187 |
| 9 | 39905200 | 39905999 | 7.177617 | 13.72865 | 25.65222 | 10.16119 | -0.38132 | -0.05304 | 0.551194 |
| 6 | 13224600 | 13226599 | 14.42816 | 25.33731 | 40.86848 | 18.08384 | -0.29872 | 0.613694 | 0.551197 |
| 4 | 145570000 | 145575799 | 260.0367 | 375.3781 | 371.9277 | 153.4413 | 0.227547 | 4.108697 | 0.551201 |
| 12 | 88013600 | 88028799 | 279.2781 | 402.1815 | 393.1853 | 162.2631 | 0.243425 | 4.145559 | 0.551204 |
| 7 | 10458200 | 10470799 | 154.8622 | 236.2177 | 195.7588 | 81.16785 | 0.557705 | 3.133668 | 0.55121 |
| 2 | 155847000 | 155848799 | 15.79819 | 27.38063 | 52.73972 | 23.13942 | -0.55521 | 0.827726 | 0.551221 |
| 5 | 76966600 | 76968199 | 14.06127 | 24.80101 | 40.7841 | 18.04821 | -0.32465 | 0.557348 | 0.551228 |
| 9 | 49559200 | 49560999 | 18.98804 | 33.93859 | 30.95996 | 13.5932 | 0.53061 | 0.464903 | 0.551242 |
| 6 | 131617200 | 131624799 | 44.06015 | 71.36842 | 65.72881 | 28.13022 | 0.482558 | 1.6604 | 0.551252 |
| 9 | 98222400 | 98230599 | 84.23678 | 130.1512 | 190.34 | 79.06308 | -0.24669 | 2.859513 | 0.551264 |

| | | | | | | | | | |
|---|---|---|---|---|---|---|---|---|---|
| X | 105262800 | 105264799 | 34.99546 | 57.86851 | 58.5893 | 25.32605 | 0.362827 | 1.348169 | 0.551291 |
| 17 | 51383000 | 51384599 | 12.92963 | 24.3769 | 24.58596 | 10.04448 | 0.461026 | 0.095047 | 0.551318 |
| 6 | 10575400 | 10578799 | 36.73658 | 60.22851 | 63.20817 | 27.20861 | 0.307167 | 1.454293 | 0.551362 |
| 7 | 84628400 | 84632399 | 58.88098 | 94.123 | 98.9325 | 41.5989 | 0.270598 | 2.078381 | 0.551377 |
| 17 | 47204000 | 47206999 | 21.19921 | 36.84453 | 39.35772 | 17.2838 | 0.292197 | 0.800486 | 0.551381 |
| 7 | 20748600 | 20750999 | 50.52895 | 81.33603 | 69.13392 | 29.36946 | 0.588141 | 1.842926 | 0.551398 |
| 14 | 124030000 | 124031199 | 17.39885 | 29.85537 | 49.52253 | 21.72054 | -0.34341 | 0.920824 | 0.551407 |
| 14 | 99705000 | 99705999 | 7.831383 | 14.72894 | 27.52201 | 11.29052 | -0.42578 | 0.07576 | 0.551416 |
| 3 | 68566600 | 68568999 | 14.44238 | 25.2429 | 42.07998 | 18.54583 | -0.34596 | 0.692471 | 0.551422 |
| 13 | 61605400 | 61615599 | 149.1327 | 226.3816 | 216.8002 | 89.76728 | 0.343574 | 3.230282 | 0.551424 |
| 16 | 20868600 | 20877999 | 145.1483 | 220.6049 | 214.2677 | 88.75141 | 0.324693 | 3.203215 | 0.551472 |
| 12 | 113162200 | 113170999 | 114.4191 | 173.4792 | 248.0645 | 102.6757 | -0.23573 | 3.247191 | 0.551476 |
| 5 | 104325200 | 104330399 | 84.70167 | 132.736 | 140.3094 | 58.19732 | 0.236952 | 2.605889 | 0.551482 |
| 12 | 3106400 | 3109599 | 49.83588 | 80.12862 | 84.94412 | 36.02611 | 0.268092 | 1.88462 | 0.551487 |
| 2 | 18419600 | 18421799 | 20.89441 | 36.32318 | 38.86417 | 17.06953 | 0.289915 | 0.795498 | 0.551487 |
| 11 | 100835600 | 100836799 | 20.89441 | 36.32318 | 38.86417 | 17.06953 | 0.289915 | 0.795498 | 0.551487 |
| 4 | 61580800 | 61581799 | 11.24755 | 21.45875 | 23.30271 | 9.285586 | 0.386834 | -0.00251 | 0.551518 |
| X | 164650000 | 164654799 | 58.78715 | 94.27548 | 92.43753 | 39.05218 | 0.375986 | 1.979208 | 0.551519 |
| 3 | 96421800 | 96423399 | 16.99924 | 28.51529 | 63.67968 | 27.78702 | -0.77381 | 1.140963 | 0.551526 |
| 11 | 10988800 | 10993999 | 81.24755 | 127.7635 | 135.8753 | 56.3669 | 0.232034 | 2.536964 | 0.551571 |
| 10 | 35886800 | 35887399 | 7.312489 | 14.0567 | 25.81294 | 10.07917 | -0.33721 | -0.09858 | 0.551598 |
| 11 | 56478200 | 56478799 | 7.312489 | 14.0567 | 25.81294 | 10.07917 | -0.33721 | -0.09858 | 0.551598 |
| 3 | 134983200 | 134985199 | 20.3126 | 35.353 | 37.77354 | 16.6027 | 0.29199 | 0.773506 | 0.551623 |
| 10 | 81981000 | 82002199 | 462.8917 | 640.7965 | 569.4283 | 236.4851 | 0.323891 | 4.736696 | 0.551625 |
| 2 | 178844200 | 178846199 | 20.53588 | 35.73231 | 38.31558 | 16.83819 | 0.286868 | 0.779815 | 0.55163 |
| 3 | 130370000 | 130370999 | 9.656243 | 18.10779 | 32.99266 | 13.60229 | -0.39652 | 0.121198 | 0.551632 |
| 2 | 57616000 | 57617399 | 14.05452 | 24.80916 | 41.29927 | 18.27323 | -0.34179 | 0.549025 | 0.551634 |
| 5 | 148025000 | 148027799 | 30.45615 | 49.63394 | 79.07855 | 33.96287 | -0.2989 | 1.533895 | 0.551635 |
| 2 | 57525000 | 57525799 | 10.14823 | 19.3622 | 21.75827 | 8.669396 | 0.33822 | -0.01681 | 0.551638 |
| 5 | 145355800 | 145362199 | 121.8704 | 186.6947 | 191.5404 | 79.49773 | 0.254807 | 3.036347 | 0.551638 |
| 7 | 139732600 | 139736199 | 26.64844 | 43.44356 | 75.43655 | 32.40277 | -0.42223 | 1.520204 | 0.551646 |
| 7 | 35453400 | 35456999 | 27.39959 | 44.75393 | 74.71128 | 32.1137 | -0.36366 | 1.492761 | 0.551652 |
| 1 | 150213600 | 150214199 | 8.01869 | 14.97149 | 28.50066 | 11.83749 | -0.46672 | 0.127852 | 0.55166 |
| 10 | 86725200 | 86726999 | 13.30798 | 23.33294 | 40.45315 | 17.87597 | -0.40233 | 0.631528 | 0.551666 |
| 12 | 100768400 | 100769999 | 20.01905 | 34.25338 | 56.00954 | 24.56008 | -0.32369 | 0.964905 | 0.551683 |
| 7 | 65936200 | 65937599 | 7.478311 | 14.26621 | 26.14405 | 10.41988 | -0.36176 | -0.02908 | 0.551711 |
| 13 | 101868400 | 101869999 | 7.478311 | 14.26621 | 26.14405 | 10.41988 | -0.36176 | -0.02908 | 0.551711 |
| 9 | 55868000 | 55874399 | 176.8692 | 263.7807 | 270.3487 | 111.4212 | 0.223447 | 3.579184 | 0.551725 |
| 11 | 118044800 | 118048399 | 41.86849 | 67.19977 | 102.235 | 43.24646 | -0.25513 | 1.935013 | 0.551746 |
| 6 | 95400200 | 95406799 | 78.48381 | 124.3219 | 122.6226 | 50.99683 | 0.348792 | 2.373723 | 0.551746 |
| X | 160080600 | 160082999 | 37.28034 | 60.88612 | 65.02673 | 27.94978 | 0.278888 | 1.511341 | 0.551752 |
| 17 | 10257600 | 10259799 | 18.75357 | 32.97825 | 34.27981 | 15.17895 | 0.332083 | 0.605904 | 0.551775 |
| Y | 5879200 | 5879799 | 9.363273 | 18.063 | 14.12428 | 5.459089 | 0.896617 | -0.14979 | 0.551782 |
| X | 25355800 | 25364799 | 155.9937 | 238.2586 | 188.3894 | 78.13615 | 0.62692 | 3.108608 | 0.551796 |
| 17 | 39757400 | 39769799 | 175.9476 | 265.0501 | 239.1881 | 98.80475 | 0.419886 | 3.389638 | 0.551806 |
| 2 | 81541000 | 81542199 | 18.04908 | 31.92184 | 27.74069 | 12.26939 | 0.592278 | 0.528558 | 0.551829 |
| 12 | 55906400 | 55908599 | 30.47487 | 49.64751 | 78.3797 | 33.65854 | -0.28605 | 1.539097 | 0.551833 |

| | | | | | | | | | |
|---|---|---|---|---|---|---|---|---|---|
| 12 | 23042400 | 23062399 | 473.9956 | 651.5939 | 611.4615 | 254.2243 | 0.235077 | 4.843522 | 0.551834 |
| 4 | 7648000 | 7652599 | 114.6357 | 175.5456 | 182.7435 | 75.84003 | 0.233392 | 3.040904 | 0.551836 |
| 1 | 171303600 | 171322199 | 507.2229 | 677.3261 | 941.4531 | 393.1567 | -0.3737 | 5.300636 | 0.551864 |
| 11 | 90848600 | 90853199 | 21.00444 | 36.77008 | 32.3669 | 14.28397 | 0.570734 | 0.685101 | 0.551867 |
| 18 | 13105000 | 13109199 | 106.2638 | 160.9222 | 249.4152 | 103.1961 | -0.35331 | 3.270531 | 0.55187 |
| 16 | 70071800 | 70072999 | 19.46935 | 33.99224 | 35.75453 | 15.74641 | 0.315076 | 0.724174 | 0.55188 |
| 6 | 149222800 | 149226999 | 28.10565 | 47.53974 | 49.06408 | 21.47361 | 0.335875 | 1.086232 | 0.551891 |
| 1 | 62105400 | 62107199 | 29.48828 | 50.13248 | 46.81016 | 20.51555 | 0.48012 | 1.044665 | 0.551892 |
| 4 | 147784400 | 147787599 | 34.10393 | 56.50238 | 58.17104 | 25.17852 | 0.338873 | 1.320332 | 0.551903 |
| 1 | 135104400 | 135105999 | 15.86761 | 28.20946 | 29.86713 | 13.17284 | 0.312563 | 0.475972 | 0.551903 |
| 13 | 10408600 | 10411399 | 19.59218 | 32.58516 | 68.64378 | 29.79072 | -0.69017 | 1.246103 | 0.551928 |
| 19 | 57046200 | 57048399 | 25.56091 | 42.07738 | 68.76828 | 29.65308 | -0.32759 | 1.384325 | 0.551929 |
| 7 | 12155200 | 12156799 | 12.6106 | 22.41591 | 37.17649 | 16.39861 | -0.33159 | 0.469088 | 0.551951 |
| 4 | 89917200 | 89921399 | 70.13255 | 111.3712 | 115.6576 | 48.21264 | 0.277971 | 2.296267 | 0.551964 |
| 10 | 42480800 | 42485199 | 65.91848 | 104.9182 | 110.1447 | 46.03437 | 0.265639 | 2.222133 | 0.551977 |
| 10 | 118259000 | 118263399 | 44.68942 | 72.19284 | 75.98809 | 32.43594 | 0.286359 | 1.728999 | 0.55198 |
| 11 | 59528600 | 59532399 | 30.66324 | 49.86252 | 79.3269 | 34.04244 | -0.29899 | 1.566705 | 0.551981 |
| 9 | 121031200 | 121036399 | 60.53011 | 95.54285 | 143.9956 | 59.77811 | -0.26609 | 2.448374 | 0.551982 |
| 10 | 17148200 | 17149599 | 17.2828 | 30.55163 | 32.2436 | 14.26237 | 0.313144 | 0.536981 | 0.551982 |
| 10 | 93440600 | 93448999 | 52.29973 | 82.91087 | 145.5093 | 60.55299 | -0.4803 | 2.337174 | 0.551988 |
| 10 | 77944200 | 77951999 | 80.2642 | 124.5475 | 182.514 | 75.79758 | -0.24483 | 2.778804 | 0.551999 |
| 7 | 29677400 | 29679799 | 25.11778 | 42.68807 | 45.82234 | 20.08208 | 0.279694 | 1.031354 | 0.552012 |
| 6 | 141101200 | 141103599 | 31.21273 | 52.50941 | 51.30375 | 22.41689 | 0.414449 | 1.146525 | 0.552036 |
| 3 | 99441000 | 99444199 | 22.25577 | 37.46256 | 59.83129 | 26.14751 | -0.29211 | 1.118892 | 0.552044 |
| 12 | 95554000 | 95554999 | 11.3439 | 21.44626 | 23.6556 | 9.609625 | 0.340043 | 0.060325 | 0.552075 |
| 5 | 140094800 | 140097399 | 21.85404 | 36.69305 | 58.59985 | 25.58541 | -0.29203 | 1.14162 | 0.552078 |
| 3 | 94040400 | 94046999 | 95.70614 | 149.3336 | 150.5675 | 62.47997 | 0.300203 | 2.686557 | 0.552089 |
| 1 | 34692000 | 34697799 | 44.76257 | 71.79393 | 109.1922 | 46.13967 | -0.25598 | 1.961653 | 0.552091 |
| 8 | 65914200 | 65916799 | 21.81137 | 38.36256 | 34.54027 | 15.29371 | 0.538487 | 0.622607 | 0.55212 |
| 18 | 76249400 | 76250199 | 7.306198 | 13.99716 | 25.70381 | 10.13454 | -0.35055 | -0.06815 | 0.552126 |
| 10 | 15359000 | 15359399 | 5.195949 | 10.13103 | 21.4496 | 7.724025 | -0.42945 | -0.41231 | 0.552148 |
| 6 | 58377800 | 58378199 | 6.701758 | 12.89784 | 24.72 | 9.619914 | -0.3939 | -0.12057 | 0.552154 |
| 10 | 55769800 | 55770999 | 7.751687 | 14.83817 | 26.81408 | 10.59782 | -0.33113 | -0.05334 | 0.552231 |
| 7 | 5597000 | 5597999 | 41.74375 | 67.5246 | 71.91932 | 30.74881 | 0.27156 | 1.688348 | 0.552248 |
| X | 7491400 | 7492999 | 26.8804 | 45.82107 | 45.28088 | 19.84958 | 0.399374 | 1.014224 | 0.552251 |
| 15 | 49830000 | 49834399 | 76.34167 | 118.963 | 172.9958 | 71.79931 | -0.229 | 2.704135 | 0.552269 |
| 10 | 61558000 | 61560399 | 12.40132 | 21.85431 | 37.88108 | 16.76643 | -0.4002 | 0.554863 | 0.552286 |
| 8 | 3070600 | 3075199 | 64.10692 | 102.2536 | 92.75307 | 38.88067 | 0.479408 | 2.148577 | 0.552287 |
| X | 5502400 | 5504999 | 38.68794 | 63.972 | 58.84091 | 25.43416 | 0.500909 | 1.358696 | 0.552295 |
| 18 | 39898200 | 39900999 | 30.11034 | 50.61655 | 51.27817 | 22.40058 | 0.36237 | 1.150144 | 0.552328 |
| 17 | 66842200 | 66847999 | 63.37505 | 100.8949 | 107.9227 | 45.12109 | 0.239046 | 2.210634 | 0.552331 |
| 13 | 67014800 | 67029999 | 210.0845 | 311.1875 | 298.3957 | 122.8893 | 0.31039 | 3.704113 | 0.552356 |
| 5 | 18712400 | 18712799 | 4.349276 | 8.499323 | 18.26182 | 6.331325 | -0.40033 | -0.57197 | 0.552377 |
| 1 | 141112800 | 141118999 | 89.42308 | 137.8617 | 198.4397 | 82.42849 | -0.22624 | 2.903151 | 0.552396 |
| 14 | 42302200 | 42305599 | 46.08818 | 74.47893 | 77.33837 | 33.02746 | 0.306487 | 1.723045 | 0.552398 |
| 7 | 60907000 | 60910599 | 55.87477 | 89.85995 | 86.58145 | 36.73338 | 0.406045 | 1.885053 | 0.552402 |
| 8 | 126930400 | 126931599 | 12.23934 | 22.79969 | 24.46523 | 10.20073 | 0.348812 | 0.163703 | 0.552412 |

| | | | | | | | | | |
|---|---|---|---|---|---|---|---|---|---|
| 1 | 184397400 | 184399799 | 22.0228 | 36.85917 | 59.6698 | 26.00042 | -0.31152 | 1.176628 | 0.552422 |
| 4 | 111995000 | 111998799 | 59.1322 | 95.04201 | 77.18828 | 32.71993 | 0.651048 | 1.904543 | 0.55246 |
| 1 | 44994600 | 44994999 | 4.774156 | 9.328707 | 19.47544 | 6.769128 | -0.36354 | -0.55309 | 0.55248 |
| 8 | 37822800 | 37823199 | 4.774156 | 9.328707 | 19.47544 | 6.769128 | -0.36354 | -0.55309 | 0.55248 |
| 12 | 17531800 | 17534799 | 27.36206 | 44.89124 | 71.59483 | 30.82803 | -0.29441 | 1.432344 | 0.552495 |
| X | 148849800 | 148850399 | 16.8731 | 30.20319 | 26.61714 | 11.67184 | 0.581302 | 0.432835 | 0.552496 |
| 2 | 136553600 | 136556399 | 19.59023 | 33.28494 | 55.97766 | 24.53221 | -0.36579 | 1.02339 | 0.5525 |
| 18 | 51559000 | 51560599 | 13.64829 | 24.00018 | 39.3415 | 17.4203 | -0.32151 | 0.587734 | 0.552503 |
| 13 | 20200600 | 20200999 | 6.842769 | 13.19019 | 24.77226 | 9.596304 | -0.35869 | -0.13363 | 0.552512 |
| 4 | 71626200 | 71628199 | 23.5258 | 40.90194 | 40.26567 | 17.68626 | 0.409532 | 0.807981 | 0.552533 |
| 10 | 113954000 | 113955599 | 16.79949 | 29.91176 | 31.17387 | 13.7315 | 0.336316 | 0.474159 | 0.552541 |
| 2 | 19400400 | 19402999 | 31.62002 | 52.39512 | 56.33628 | 24.38694 | 0.276436 | 1.310192 | 0.55255 |
| 2 | 56797800 | 56800399 | 22.438 | 37.78864 | 59.82663 | 26.14953 | -0.27949 | 1.115764 | 0.55258 |
| X | 152757400 | 152759599 | 20.37636 | 35.57002 | 37.65373 | 16.58112 | 0.30575 | 0.732477 | 0.552647 |
| 5 | 149316000 | 149317399 | 15.9435 | 28.69129 | 29.67449 | 12.95416 | 0.356031 | 0.399674 | 0.55265 |
| 11 | 99152200 | 99153199 | 6.43166 | 12.3383 | 24.63444 | 9.675499 | -0.46445 | -0.09217 | 0.552683 |
| 12 | 87565400 | 87578199 | 216.9095 | 320.508 | 305.9596 | 126.001 | 0.313528 | 3.74499 | 0.552719 |
| 3 | 15295600 | 15296199 | 4.910884 | 9.591997 | 19.99338 | 7.00321 | -0.3713 | -0.52115 | 0.552725 |
| 14 | 117660000 | 117661199 | 11.77746 | 21.23575 | 35.24575 | 15.34865 | -0.31994 | 0.371456 | 0.552734 |
| 9 | 89502600 | 89506199 | 30.43173 | 49.23022 | 84.69614 | 36.21362 | -0.41847 | 1.671658 | 0.552735 |
| X | 26728400 | 26741999 | 315.8886 | 452.2244 | 358.0215 | 147.8869 | 0.539157 | 4.234567 | 0.552737 |
| 8 | 86159000 | 86159799 | 6.122319 | 11.81451 | 24.22776 | 9.355613 | -0.48 | -0.15419 | 0.552737 |
| 12 | 36278200 | 36282199 | 38.23372 | 62.73611 | 57.0426 | 24.56338 | 0.51402 | 1.444589 | 0.552755 |
| 13 | 64583400 | 64587599 | 30.55558 | 49.5938 | 79.97696 | 34.27904 | -0.32088 | 1.603957 | 0.552756 |
| 3 | 84694600 | 84701799 | 96.84709 | 148.3644 | 215.1017 | 89.27717 | -0.24362 | 3.029585 | 0.552773 |
| 3 | 155015600 | 155017599 | 14.87305 | 26.0185 | 42.52795 | 18.75719 | -0.31788 | 0.6797 | 0.552783 |
| 18 | 63767200 | 63768999 | 14.32755 | 25.03284 | 48.37667 | 21.31479 | -0.55836 | 0.707137 | 0.552799 |
| 7 | 15034800 | 15087199 | 1738.193 | 2154.66 | 1747.811 | 735.7904 | 0.295345 | 6.506953 | 0.552803 |
| X | 153786800 | 153788399 | 28.7372 | 48.73378 | 48.49783 | 21.24242 | 0.388414 | 1.067783 | 0.55281 |
| 13 | 22588200 | 22597199 | 182.7418 | 272.8148 | 271.1646 | 111.7795 | 0.268979 | 3.561822 | 0.552824 |
| 4 | 23839800 | 23840399 | 9.844687 | 18.84482 | 19.28156 | 7.620407 | 0.483454 | -0.05217 | 0.552829 |
| 5 | 132900800 | 132903599 | 16.75243 | 29.99693 | 26.20631 | 11.48855 | 0.594142 | 0.429775 | 0.552834 |
| 1 | 100880400 | 100881599 | 11.53317 | 21.71123 | 22.98478 | 9.414394 | 0.38846 | 0.086885 | 0.552848 |
| 3 | 129017600 | 129022199 | 28.18306 | 45.67839 | 94.24392 | 40.35193 | -0.67747 | 1.633632 | 0.55288 |
| 2 | 129569200 | 129571599 | 23.6963 | 41.24812 | 39.67321 | 17.43925 | 0.443079 | 0.791266 | 0.552905 |
| X | 26033200 | 26035199 | 21.31029 | 38.43021 | 28.17441 | 12.26687 | 0.852249 | 0.423439 | 0.552923 |
| 8 | 21694600 | 21695599 | 12.86525 | 24.83108 | 21.92603 | 8.46162 | 0.725769 | -0.09047 | 0.552929 |
| 1 | 162646000 | 162647999 | 16.37447 | 28.51855 | 45.76311 | 20.12309 | -0.2919 | 0.758639 | 0.552941 |
| 1 | 190262200 | 190263199 | 6.455534 | 12.31202 | 25.60167 | 10.20909 | -0.54313 | -0.04035 | 0.552964 |
| 8 | 97791800 | 97793999 | 22.21831 | 38.81464 | 38.0183 | 16.74959 | 0.417376 | 0.732099 | 0.552966 |
| 6 | 70629600 | 70630199 | 10.31174 | 19.70909 | 22.29441 | 8.849404 | 0.333511 | -0.02604 | 0.552993 |
| 11 | 103037200 | 103039399 | 12.98582 | 22.93791 | 37.76116 | 16.70555 | -0.32509 | 0.530978 | 0.552999 |
| 19 | 58379200 | 58380999 | 12.98582 | 22.93791 | 37.76116 | 16.70555 | -0.32509 | 0.530978 | 0.552999 |
| 12 | 54423400 | 54425999 | 24.97386 | 43.44264 | 39.60742 | 17.40295 | 0.519813 | 0.806048 | 0.553005 |
| 17 | 48450200 | 48452399 | 27.62892 | 46.7134 | 49.06614 | 21.47172 | 0.310596 | 1.08842 | 0.553007 |
| 10 | 121878600 | 121879599 | 7.439896 | 14.2496 | 26.26553 | 10.36374 | -0.35726 | -0.06408 | 0.553009 |
| 16 | 77687600 | 77688399 | 7.439896 | 14.2496 | 26.26553 | 10.36374 | -0.35726 | -0.06408 | 0.553009 |

| | | | | | | | | | |
|---|---|---|---|---|---|---|---|---|---|
| 7 | 26610200 | 26614999 | 61.56571 | 98.8284 | 79.08369 | 33.46051 | 0.670188 | 1.946677 | 0.553032 |
| 17 | 53597600 | 53598999 | 20.4716 | 34.67047 | 55.39962 | 24.25567 | -0.29235 | 1.056319 | 0.553034 |
| X | 20628200 | 20629399 | 18.42772 | 33.44537 | 28.65797 | 12.37788 | 0.636135 | 0.370065 | 0.553037 |
| X | 127126200 | 127127199 | 18.18434 | 31.25736 | 50.05854 | 21.95754 | -0.29255 | 0.908764 | 0.553053 |
| 12 | 46097200 | 46099399 | 21.28917 | 37.58511 | 32.26244 | 14.2744 | 0.608848 | 0.574014 | 0.553067 |
| X | 43775000 | 43777399 | 18.40064 | 33.24798 | 29.55105 | 12.83483 | 0.578288 | 0.394827 | 0.553073 |
| 17 | 26367600 | 26373199 | 67.90061 | 108.1931 | 107.8269 | 45.13494 | 0.342255 | 2.188196 | 0.553078 |
| 1 | 127345200 | 127349199 | 27.968 | 46.04886 | 71.99591 | 31.04741 | -0.26382 | 1.389615 | 0.553081 |
| 9 | 71548000 | 71551399 | 32.57427 | 53.65639 | 56.71923 | 24.46403 | 0.299523 | 1.384709 | 0.553157 |
| 18 | 63246600 | 63247399 | 12.94357 | 23.90204 | 25.55667 | 10.79042 | 0.34105 | 0.224956 | 0.553163 |
| 13 | 57525200 | 57526999 | 16.94832 | 30.83862 | 28.63968 | 12.33072 | 0.523239 | 0.342485 | 0.553171 |
| 3 | 144102000 | 144105599 | 39.51149 | 63.5002 | 99.73395 | 42.27135 | -0.29871 | 1.887851 | 0.553177 |
| 6 | 123981400 | 123982799 | 11.80435 | 21.43595 | 37.00381 | 15.97172 | -0.36799 | 0.33485 | 0.553185 |
| 1 | 85913200 | 85914999 | 17.90867 | 30.87508 | 50.6188 | 22.2053 | -0.32552 | 0.878416 | 0.553201 |
| 11 | 62168800 | 62170399 | 14.47075 | 25.48172 | 41.86754 | 18.5337 | -0.3242 | 0.581695 | 0.553209 |
| 3 | 157263400 | 157264799 | 13.66074 | 25.31335 | 25.8565 | 10.86601 | 0.41079 | 0.209098 | 0.55323 |
| 5 | 24086800 | 24089599 | 25.94834 | 44.58437 | 44.08655 | 19.33776 | 0.400009 | 0.935079 | 0.553247 |
| 5 | 118463200 | 118464399 | 12.87822 | 22.59867 | 39.47234 | 17.45511 | -0.41275 | 0.615617 | 0.553249 |
| 4 | 68883000 | 68884799 | 21.41789 | 37.33693 | 31.63915 | 13.92056 | 0.625174 | 0.748014 | 0.55326 |
| 14 | 4163600 | 4166399 | 18.43222 | 32.31752 | 34.51477 | 15.25193 | 0.293112 | 0.649597 | 0.553284 |
| 6 | 102763200 | 102768599 | 55.17571 | 88.89003 | 82.28917 | 35.00037 | 0.466697 | 1.828608 | 0.553284 |
| 17 | 76875400 | 76878199 | 45.10914 | 72.76531 | 78.39384 | 33.40033 | 0.250494 | 1.773953 | 0.553287 |
| 5 | 143589200 | 143593599 | 34.31758 | 55.45488 | 86.70724 | 37.02617 | -0.28268 | 1.710712 | 0.553325 |
| 9 | 88203400 | 88207599 | 59.22057 | 93.6656 | 139.6961 | 58.05276 | -0.24865 | 2.399497 | 0.553334 |
| 3 | 135597000 | 135600599 | 31.85128 | 53.00965 | 55.79614 | 24.22616 | 0.307545 | 1.261892 | 0.553356 |
| 16 | 45859200 | 45860999 | 16.28117 | 28.81825 | 30.32182 | 13.40884 | 0.318233 | 0.513055 | 0.553368 |
| 2 | 177543000 | 177545999 | 25.18138 | 43.47688 | 37.68091 | 16.53048 | 0.590799 | 0.88107 | 0.553371 |
| 8 | 28066600 | 28067399 | 10.15386 | 18.36302 | 33.38741 | 14.48518 | -0.44633 | 0.34326 | 0.553382 |
| 12 | 100044600 | 100048399 | 37.12264 | 60.81064 | 64.2923 | 27.66551 | 0.296084 | 1.466885 | 0.553394 |
| 4 | 111946400 | 111949999 | 37.00953 | 61.26721 | 58.86912 | 25.46659 | 0.438129 | 1.338981 | 0.553402 |
| 5 | 72015400 | 72016199 | 7.305527 | 14.06852 | 25.88629 | 10.0559 | -0.33317 | -0.11552 | 0.553404 |
| 3 | 16499400 | 16500199 | 13.1404 | 24.00247 | 24.97895 | 10.70273 | 0.364568 | 0.284342 | 0.553405 |
| 1 | 47240800 | 47241399 | 8.4173 | 16.34066 | 19.40525 | 7.285699 | 0.339303 | -0.2376 | 0.553422 |
| 13 | 37464800 | 37465399 | 8.4173 | 16.34066 | 19.40525 | 7.285699 | 0.339303 | -0.2376 | 0.553422 |
| 2 | 153042200 | 153044599 | 15.7117 | 27.18395 | 46.88507 | 20.56963 | -0.39706 | 0.832941 | 0.553441 |
| 12 | 101579200 | 101580799 | 15.17297 | 27.81994 | 23.33367 | 9.947833 | 0.679012 | 0.276729 | 0.553453 |
| 3 | 38179000 | 38183999 | 49.63381 | 79.11083 | 119.5353 | 50.05706 | -0.25664 | 2.165373 | 0.553552 |
| 12 | 113576800 | 113578199 | 21.27672 | 37.06806 | 38.76345 | 17.04851 | 0.323094 | 0.767324 | 0.553568 |
| 7 | 74149200 | 74151399 | 13.09866 | 23.13258 | 38.73311 | 17.13622 | -0.34975 | 0.533769 | 0.55357 |
| 13 | 88178800 | 88181399 | 20.76518 | 36.96139 | 30.71739 | 13.53393 | 0.660858 | 0.50066 | 0.553586 |
| 8 | 16539000 | 16544999 | 100.5819 | 156.1196 | 144.3648 | 59.95176 | 0.41901 | 2.7785 | 0.5536 |
| X | 79568600 | 79574199 | 76.78678 | 122.098 | 111.0928 | 46.37054 | 0.470288 | 2.265025 | 0.553639 |
| 1 | 8153400 | 8154599 | 10.8246 | 19.99094 | 37.57976 | 15.86474 | -0.46801 | 0.232845 | 0.553643 |
| 19 | 42338400 | 42340799 | 21.94129 | 36.89254 | 59.24222 | 25.88142 | -0.29997 | 1.126852 | 0.553651 |
| 2 | 91085800 | 91088799 | 18.54519 | 31.41444 | 55.35463 | 24.2375 | -0.43292 | 1.045986 | 0.553657 |
| 11 | 29369800 | 29370199 | 3.932452 | 7.685368 | 17.68279 | 6.118255 | -0.49438 | -0.58722 | 0.553662 |
| 13 | 33413400 | 33417199 | 30.63585 | 52.08465 | 46.23779 | 20.26473 | 0.55272 | 1.048047 | 0.553666 |

| | | | | | | | | | |
|---|---|---|---|---|---|---|---|---|---|
| 18 | 84128200 | 84131199 | 22.59305 | 38.01394 | 60.39017 | 26.38831 | -0.28449 | 1.124598 | 0.553677 |
| 2 | 4840600 | 4842199 | 17.39438 | 30.63591 | 32.3234 | 14.30819 | 0.311507 | 0.577109 | 0.553744 |
| X | 77245200 | 77248199 | 34.34779 | 58.18343 | 47.3754 | 20.74376 | 0.67686 | 1.096874 | 0.553748 |
| 10 | 96376400 | 96376999 | 7.852555 | 14.70047 | 28.00513 | 11.57925 | -0.46253 | 0.10742 | 0.553765 |
| 14 | 119713200 | 119714199 | 11.15527 | 20.46112 | 36.19451 | 15.42288 | -0.39046 | 0.271446 | 0.553771 |
| 10 | 71676400 | 71677399 | 14.20201 | 25.71817 | 26.77002 | 11.59492 | 0.354017 | 0.344376 | 0.553791 |
| 5 | 78387200 | 78389199 | 21.18438 | 37.07738 | 36.91286 | 16.28731 | 0.393925 | 0.695614 | 0.553837 |
| 3 | 107538000 | 107539199 | 10.19483 | 18.60782 | 31.83148 | 13.65187 | -0.3486 | 0.283043 | 0.553837 |
| 11 | 3152200 | 3153399 | 10.19483 | 18.60782 | 31.83148 | 13.65187 | -0.3486 | 0.283043 | 0.553837 |
| 17 | 33064800 | 33068999 | 32.09991 | 52.27924 | 81.56104 | 35.02087 | -0.26933 | 1.548642 | 0.553851 |
| X | 21995000 | 21997199 | 20.7601 | 36.86966 | 31.49188 | 13.9014 | 0.620053 | 0.518708 | 0.553852 |
| 10 | 116857000 | 116863399 | 46.72712 | 74.26429 | 138.1686 | 57.64399 | -0.56076 | 2.256544 | 0.553862 |
| 14 | 19973800 | 19975399 | 18.48815 | 32.47708 | 33.77881 | 14.95063 | 0.33092 | 0.616878 | 0.553885 |
| 8 | 21521800 | 21522599 | 13.45642 | 24.87624 | 26.44975 | 11.15091 | 0.350241 | 0.223021 | 0.553886 |
| 9 | 101415000 | 101415999 | 15.69788 | 28.55762 | 28.79842 | 12.40218 | 0.404993 | 0.335652 | 0.553889 |
| 4 | 52292800 | 52294199 | 11.06117 | 20.13046 | 34.40207 | 14.80809 | -0.35054 | 0.313822 | 0.55389 |
| 2 | 150705800 | 150707799 | 20.37058 | 36.6263 | 29.25983 | 12.78267 | 0.725856 | 0.435361 | 0.553895 |
| 1 | 16239600 | 16243199 | 34.56385 | 55.62214 | 102.314 | 43.44801 | -0.52373 | 1.842599 | 0.553899 |
| 7 | 80711800 | 80715199 | 23.87471 | 39.58357 | 64.50702 | 27.93493 | -0.32147 | 1.291176 | 0.5539 |
| 5 | 73168800 | 73171399 | 23.99742 | 39.78498 | 65.7379 | 28.46695 | -0.34144 | 1.291339 | 0.553903 |
| 3 | 103172400 | 103173799 | 12.54329 | 22.50562 | 38.88596 | 17.01804 | -0.38231 | 0.416181 | 0.55391 |
| 18 | 71361800 | 71364799 | 25.48171 | 44.17139 | 41.36807 | 18.15023 | 0.480721 | 0.854437 | 0.553939 |
| 2 | 36906000 | 36908199 | 24.63869 | 42.76542 | 40.1983 | 17.63817 | 0.475607 | 0.83406 | 0.553941 |
| X | 153191000 | 153192999 | 27.03396 | 46.15163 | 46.29844 | 20.29808 | 0.377959 | 1.000092 | 0.553949 |
| 13 | 75681200 | 75682999 | 16.19744 | 29.86848 | 26.69562 | 11.293 | 0.595512 | 0.258478 | 0.553955 |
| X | 15147400 | 15148799 | 20.59048 | 36.28343 | 34.16685 | 15.12268 | 0.47521 | 0.596108 | 0.553979 |
| 14 | 18193400 | 18198199 | 45.58154 | 74.00771 | 71.54317 | 30.67209 | 0.415978 | 1.619266 | 0.553981 |
| 1 | 166587400 | 166592999 | 59.87937 | 95.65139 | 100.6028 | 42.25892 | 0.268554 | 2.099431 | 0.553992 |
| 9 | 40179400 | 40181399 | 16.44862 | 28.57843 | 46.85503 | 20.57191 | -0.32329 | 0.792369 | 0.553996 |
| 13 | 43230200 | 43233599 | 37.39218 | 60.29016 | 92.68862 | 39.46797 | -0.26211 | 1.781658 | 0.553998 |
| 12 | 34506000 | 34506599 | 5.39476 | 10.49632 | 23.03867 | 8.495564 | -0.5145 | -0.32769 | 0.554007 |
| X | 5564400 | 5565399 | 19.47643 | 34.18362 | 35.39442 | 15.65429 | 0.337561 | 0.640945 | 0.554047 |
| 2 | 46697800 | 46700399 | 27.17863 | 46.5581 | 45.18021 | 19.8133 | 0.426408 | 0.968673 | 0.554059 |
| 3 | 145737400 | 145741799 | 35.41083 | 57.22679 | 88.98908 | 38.0051 | -0.2747 | 1.709469 | 0.554068 |
| 14 | 39923800 | 39926199 | 32.88416 | 55.43207 | 50.20499 | 21.95011 | 0.523555 | 1.134247 | 0.554088 |
| 17 | 39989400 | 39989999 | 15.9164 | 28.29757 | 29.43845 | 12.98325 | 0.337924 | 0.476027 | 0.554091 |
| 12 | 23032200 | 23072399 | 866.8079 | 1134.651 | 1005.062 | 420.7406 | 0.247488 | 5.624068 | 0.554121 |
| 15 | 83075200 | 83079799 | 58.06617 | 91.78765 | 139.5972 | 57.995 | -0.27746 | 2.41165 | 0.554123 |
| 11 | 21545200 | 21547799 | 41.54978 | 67.4635 | 71.39987 | 30.61129 | 0.285678 | 1.612228 | 0.554144 |
| 2 | 87432800 | 87435199 | 33.25478 | 55.27577 | 56.46803 | 24.49657 | 0.35037 | 1.279498 | 0.554146 |
| 14 | 33149200 | 33150599 | 20.2052 | 35.69548 | 32.93856 | 14.57158 | 0.505473 | 0.562123 | 0.554146 |
| 1 | 173760400 | 173761999 | 11.06033 | 20.31651 | 37.91583 | 16.12441 | -0.46509 | 0.267334 | 0.554167 |
| 4 | 112388600 | 112395799 | 113.7674 | 174.9334 | 180.5444 | 74.97415 | 0.25034 | 2.96086 | 0.55417 |
| 5 | 19686400 | 19688999 | 30.09372 | 50.19301 | 53.55528 | 23.2851 | 0.287887 | 1.230377 | 0.554175 |
| 5 | 117857000 | 117859799 | 19.41026 | 32.72115 | 55.84821 | 24.41777 | -0.38732 | 1.093684 | 0.554175 |
| 2 | 95188400 | 95189199 | 12.26127 | 22.78655 | 24.59102 | 10.29325 | 0.336688 | 0.181514 | 0.554176 |
| 1 | 64050600 | 64052399 | 16.2607 | 28.13637 | 46.89007 | 20.57188 | -0.34756 | 0.837539 | 0.554189 |

| | | | | | | | | | |
|---|---|---|---|---|---|---|---|---|---|
| 10 | 14653600 | 14654799 | 8.427719 | 15.99949 | 28.80852 | 11.60036 | -0.34991 | 0.020892 | 0.554189 |
| 3 | 26151400 | 26152399 | 13.56461 | 25.5407 | 25.13366 | 10.29096 | 0.49401 | 0.111419 | 0.554199 |
| 3 | 154207800 | 154208599 | 12.7081 | 22.54031 | 37.24409 | 16.45849 | -0.32777 | 0.486648 | 0.554212 |
| 15 | 28582400 | 28584199 | 20.33379 | 35.65157 | 35.81209 | 15.82521 | 0.380996 | 0.661053 | 0.554217 |
| 18 | 77280200 | 77283399 | 25.77663 | 42.70059 | 67.5708 | 29.24454 | -0.27941 | 1.30733 | 0.554219 |
| 5 | 79536400 | 79537599 | 12.44922 | 23.27533 | 25.03659 | 10.37474 | 0.352496 | 0.144409 | 0.554236 |
| 8 | 27012600 | 27017399 | 46.90797 | 75.0545 | 112.6474 | 47.42897 | -0.24093 | 2.039652 | 0.554244 |
| 8 | 116567800 | 116570399 | 16.77659 | 29.23924 | 48.42356 | 21.30226 | -0.33736 | 0.751905 | 0.554245 |
| X | 38232000 | 38233999 | 35.74938 | 58.88409 | 61.4266 | 26.49396 | 0.318527 | 1.396788 | 0.55425 |
| 9 | 27077800 | 27079199 | 14.22938 | 26.02536 | 27.3086 | 11.68175 | 0.354042 | 0.289555 | 0.554255 |
| 18 | 47381000 | 47382799 | 18.42177 | 31.18642 | 54.25018 | 23.74953 | -0.41435 | 1.051229 | 0.554274 |
| 14 | 14810400 | 14811999 | 11.4498 | 20.19768 | 37.38445 | 16.54331 | -0.49385 | 0.535448 | 0.554281 |
| 15 | 23155000 | 23156799 | 14.78021 | 25.88543 | 42.00631 | 18.54488 | -0.30736 | 0.662105 | 0.554291 |
| 9 | 19257000 | 19267399 | 168.5968 | 253.9523 | 242.3645 | 100.1143 | 0.339074 | 3.389555 | 0.554328 |
| 5 | 102733400 | 102734599 | 14.49414 | 26.67171 | 27.40668 | 11.62685 | 0.392804 | 0.259265 | 0.554355 |
| 2 | 31738000 | 31740399 | 12.55659 | 22.01256 | 40.00631 | 17.67649 | -0.46984 | 0.629568 | 0.554367 |
| 18 | 7151000 | 7151599 | 6.555454 | 12.6189 | 24.72535 | 9.616234 | -0.42474 | -0.12461 | 0.554384 |
| 13 | 111093800 | 111096399 | 27.78479 | 47.23863 | 47.40315 | 20.77551 | 0.376707 | 1.041955 | 0.554384 |
| X | 146237200 | 146239999 | 21.92382 | 38.5981 | 33.9412 | 15.02551 | 0.572866 | 0.611305 | 0.554395 |
| 13 | 45623000 | 45623799 | 5.692044 | 11.00148 | 23.16916 | 8.883299 | -0.50741 | -0.18448 | 0.554398 |
| 14 | 75068600 | 75069599 | 5.692044 | 11.00148 | 23.16916 | 8.883299 | -0.50741 | -0.18448 | 0.554398 |
| 5 | 40801400 | 40803399 | 15.67279 | 28.78776 | 27.2071 | 11.57103 | 0.509939 | 0.276669 | 0.554405 |
| 6 | 44717800 | 44719199 | 12.47287 | 23.86643 | 22.10234 | 8.744991 | 0.623304 | -0.01865 | 0.55442 |
| 2 | 25610200 | 25619999 | 274.1143 | 386.6547 | 539.9425 | 223.5796 | -0.30075 | 4.449301 | 0.554426 |
| 17 | 19302600 | 19306399 | 47.99856 | 78.13743 | 68.22417 | 29.28948 | 0.565564 | 1.582529 | 0.554432 |
| 6 | 149429400 | 149431999 | 21.575 | 37.8955 | 35.10895 | 15.53794 | 0.497097 | 0.639543 | 0.554434 |
| 5 | 86464600 | 86466599 | 21.94234 | 37.08883 | 58.73177 | 25.69937 | -0.27958 | 1.081757 | 0.554445 |
| 12 | 69527600 | 69530199 | 6.429913 | 12.33989 | 29.192 | 11.45237 | -0.70636 | -0.08555 | 0.554448 |
| 6 | 37806600 | 37808599 | 27.34958 | 46.77633 | 45.16277 | 19.80311 | 0.433268 | 0.981665 | 0.554461 |
| 10 | 87007800 | 87008599 | 10.92854 | 20.7214 | 23.21936 | 9.37435 | 0.325477 | 0.036459 | 0.554496 |
| 2 | 112568400 | 112569999 | 13.58716 | 25.29295 | 25.96818 | 10.84007 | 0.410799 | 0.183377 | 0.554499 |
| 6 | 149074400 | 149077799 | 29.60668 | 49.39942 | 52.98548 | 23.04287 | 0.280347 | 1.224532 | 0.554503 |
| 16 | 89450800 | 89455799 | 55.50528 | 89.05713 | 91.69907 | 38.7698 | 0.306367 | 1.957772 | 0.554506 |
| 15 | 12510400 | 12512799 | 18.01728 | 30.89942 | 50.0555 | 21.95332 | -0.30961 | 0.928368 | 0.554508 |
| 16 | 82435200 | 82437999 | 15.99949 | 28.79719 | 25.16162 | 10.98164 | 0.598106 | 0.389599 | 0.554514 |
| X | 33586400 | 33589199 | 22.05962 | 39.02391 | 32.4249 | 14.34051 | 0.657012 | 0.561482 | 0.554515 |
| 4 | 82704400 | 82706399 | 16.44599 | 29.89753 | 28.03881 | 12.08568 | 0.50793 | 0.34112 | 0.554534 |
| 17 | 34228200 | 34242399 | 200.8215 | 292.8744 | 401.0901 | 165.2746 | -0.22482 | 3.952308 | 0.554538 |
| 6 | 136409800 | 136432999 | 303.05 | 426.9996 | 570.322 | 236.2021 | -0.23816 | 4.466498 | 0.55456 |
| 13 | 83960600 | 83961199 | 6.251447 | 12.13148 | 23.35588 | 8.787844 | -0.35332 | -0.24053 | 0.554568 |
| 4 | 112891600 | 112892999 | 27.41927 | 46.28522 | 49.07356 | 21.46461 | 0.297136 | 1.103025 | 0.554606 |
| 1 | 178407600 | 178409399 | 20.33273 | 35.47377 | 37.66004 | 16.578 | 0.301511 | 0.73962 | 0.554619 |
| 19 | 39097800 | 39101399 | 45.72458 | 73.88591 | 67.81535 | 28.95789 | 0.484006 | 1.720395 | 0.554624 |
| 18 | 79791600 | 79794399 | 26.38882 | 44.96136 | 40.53742 | 17.76952 | 0.530953 | 1.008079 | 0.554642 |
| 3 | 105969800 | 105970799 | 15.71501 | 28.03792 | 29.85177 | 13.12657 | 0.307507 | 0.452942 | 0.554657 |
| 9 | 5064800 | 5066999 | 38.01396 | 61.83835 | 66.65216 | 28.607 | 0.261646 | 1.568458 | 0.554661 |
| Y | 42282800 | 42285399 | 13.91058 | 25.36315 | 18.59868 | 7.987107 | 0.8638 | 0.282685 | 0.554671 |

| | | | | | | | | | |
|---|---|---|---|---|---|---|---|---|---|
| 5 | 101798600 | 101802199 | 33.05092 | 54.66822 | 58.55639 | 25.31669 | 0.281781 | 1.337876 | 0.554674 |
| 2 | 111113000 | 111114599 | 30.28137 | 50.64977 | 43.86847 | 19.10764 | 0.587704 | 1.191794 | 0.554718 |
| 14 | 49087000 | 49089999 | 25.99556 | 44.29286 | 45.76263 | 20.06003 | 0.335079 | 1.01021 | 0.554719 |
| 13 | 54021800 | 54023199 | 14.45401 | 26.09654 | 27.81967 | 12.0946 | 0.316491 | 0.366548 | 0.554725 |
| 18 | 8702400 | 8704799 | 22.14538 | 38.73475 | 37.47517 | 16.52655 | 0.435142 | 0.713014 | 0.554745 |
| 10 | 113365200 | 113366599 | 14.14953 | 26.61916 | 25.05662 | 10.27436 | 0.555836 | 0.120912 | 0.554748 |
| 12 | 21632600 | 21639599 | 91.48043 | 143.0296 | 148.2151 | 61.49098 | 0.263015 | 2.650149 | 0.554754 |
| 13 | 93298000 | 93299599 | 14.46292 | 26.74589 | 26.92011 | 11.34402 | 0.429337 | 0.230477 | 0.55477 |
| 8 | 65458800 | 65461199 | 13.15876 | 24.61331 | 21.44443 | 8.879238 | 0.655714 | 0.140601 | 0.554795 |
| 10 | 22846000 | 22849399 | 36.48873 | 58.93838 | 91.54961 | 39.07267 | -0.27394 | 1.730403 | 0.554803 |
| 12 | 13907600 | 13911399 | 35.2277 | 58.58707 | 56.01367 | 24.30861 | 0.445725 | 1.278585 | 0.554809 |
| 5 | 68767000 | 68768799 | 16.99351 | 29.56241 | 47.36959 | 20.81433 | -0.2901 | 0.779433 | 0.554812 |
| 9 | 84769600 | 84772999 | 18.87758 | 33.1944 | 34.23789 | 15.16056 | 0.343142 | 0.606649 | 0.554832 |
| 1 | 69414600 | 69419199 | 87.4552 | 134.8108 | 196.467 | 81.60912 | -0.24423 | 2.904744 | 0.554833 |
| 18 | 46185400 | 46189799 | 48.50883 | 78.29567 | 79.33359 | 33.82655 | 0.339751 | 1.761984 | 0.554834 |
| 13 | 74831600 | 74833999 | 20.55298 | 36.09807 | 35.19173 | 15.57374 | 0.423929 | 0.634658 | 0.554838 |
| 4 | 21104600 | 21106399 | 22.90133 | 39.74269 | 35.70929 | 15.66798 | 0.540473 | 0.82362 | 0.554846 |
| 14 | 83932200 | 83934599 | 28.28346 | 48.30184 | 45.14379 | 19.79242 | 0.479662 | 0.997846 | 0.554849 |
| 2 | 155708400 | 155711799 | 20.57011 | 34.60393 | 56.91868 | 24.87002 | -0.3343 | 1.11843 | 0.554851 |
| 1 | 48353000 | 48353999 | 13.30149 | 24.67016 | 25.34245 | 10.63678 | 0.403898 | 0.199603 | 0.554854 |
| 5 | 3486200 | 3489399 | 17.16854 | 28.75375 | 64.12358 | 27.95279 | -0.77172 | 1.160934 | 0.55488 |
| X | 84851800 | 84855599 | 29.90753 | 50.63387 | 41.11786 | 18.00082 | 0.680682 | 1.079811 | 0.554891 |
| 5 | 40373400 | 40376599 | 46.03421 | 74.2707 | 78.77106 | 33.56873 | 0.273336 | 1.76952 | 0.554904 |
| 5 | 144607800 | 144607999 | 5.932093 | 11.5948 | 14.29186 | 4.929185 | 0.397635 | -0.58391 | 0.554906 |
| 6 | 123677800 | 123678199 | 8.983338 | 17.4327 | 19.92121 | 7.496009 | 0.390857 | -0.22318 | 0.554912 |
| 2 | 48379200 | 48382999 | 34.17535 | 56.10732 | 60.66819 | 26.1306 | 0.265026 | 1.430823 | 0.554923 |
| 14 | 120292600 | 120294599 | 20.84796 | 36.63032 | 35.22414 | 15.59318 | 0.443718 | 0.631731 | 0.554932 |
| 18 | 44512800 | 44514199 | 13.06019 | 22.81873 | 41.64687 | 18.34974 | -0.47587 | 0.691075 | 0.554934 |
| 2 | 78287200 | 78287999 | 12.01107 | 22.37999 | 24.55177 | 10.23254 | 0.317856 | 0.161661 | 0.554937 |
| 1 | 144984600 | 144988799 | 66.86808 | 106.1441 | 114.1622 | 47.56995 | 0.22689 | 2.303631 | 0.554946 |
| 2 | 47651000 | 47652799 | 35.13474 | 58.12552 | 52.07877 | 22.51893 | 0.538558 | 1.33415 | 0.55495 |
| 16 | 75788200 | 75789799 | 15.78204 | 29.18921 | 26.81939 | 11.29952 | 0.560187 | 0.239909 | 0.554954 |
| 2 | 111691200 | 111692999 | 36.91514 | 60.13025 | 63.12833 | 27.10763 | 0.300878 | 1.546595 | 0.554955 |
| 3 | 48911600 | 48915999 | 106.8419 | 164.2265 | 167.4509 | 69.53349 | 0.267428 | 2.962771 | 0.554974 |
| 13 | 33527800 | 33527999 | 6.22883 | 12.17124 | 13.79668 | 4.794336 | 0.508113 | -0.55376 | 0.554989 |
| X | 33871400 | 33874999 | 33.2566 | 56.68411 | 45.11933 | 19.77859 | 0.710274 | 1.038531 | 0.555008 |
| 2 | 60639200 | 60640599 | 10.43625 | 19.19932 | 32.97579 | 13.99459 | -0.34399 | 0.240823 | 0.555009 |
| 8 | 77973000 | 77975399 | 24.53761 | 40.51909 | 67.25692 | 29.04489 | -0.34844 | 1.340898 | 0.555013 |
| 10 | 129356000 | 129356799 | 9.23736 | 17.0898 | 29.92202 | 12.60159 | -0.36288 | 0.195024 | 0.555034 |
| 17 | 7108800 | 7109599 | 9.23736 | 17.0898 | 29.92202 | 12.60159 | -0.36288 | 0.195024 | 0.555034 |
| 6 | 130468400 | 130473799 | 51.75388 | 84.1847 | 66.31631 | 28.46227 | 0.712861 | 1.599118 | 0.555041 |
| 7 | 57001400 | 57003599 | 17.33887 | 30.84062 | 31.66416 | 13.95948 | 0.357009 | 0.485423 | 0.555046 |
| 4 | 52337400 | 52339799 | 15.11987 | 26.60159 | 47.73158 | 21.13323 | -0.45141 | 0.603134 | 0.555062 |
| 4 | 35724800 | 35726799 | 19.39526 | 34.15143 | 34.16383 | 15.12363 | 0.387814 | 0.595021 | 0.555064 |
| X | 158293400 | 158295199 | 19.76038 | 34.69787 | 34.91783 | 15.44948 | 0.378567 | 0.63607 | 0.55507 |
| 7 | 83471800 | 83473199 | 13.73356 | 25.2518 | 26.97812 | 11.45756 | 0.335989 | 0.25769 | 0.555081 |
| 13 | 76868400 | 76869199 | 10.04011 | 18.492 | 31.4227 | 13.31443 | -0.3265 | 0.230551 | 0.555082 |

| | | | | | | | | | |
|---|---|---|---|---|---|---|---|---|---|
| 10 | 6516400 | 6523599 | 113.743 | 174.3871 | 183.6407 | 76.2295 | 0.218124 | 3.016749 | 0.555082 |
| 9 | 100409800 | 100412399 | 25.73641 | 44.35636 | 42.96787 | 18.84883 | 0.430427 | 0.905369 | 0.555084 |
| 5 | 14071600 | 14073999 | 22.05155 | 38.85186 | 33.99216 | 15.04584 | 0.580622 | 0.605412 | 0.555091 |
| 1 | 120483400 | 120484999 | 12.25434 | 21.94746 | 36.71338 | 16.09174 | -0.33761 | 0.418056 | 0.555122 |
| 5 | 5960800 | 5961799 | 11.12819 | 21.00302 | 23.18666 | 9.452765 | 0.334709 | 0.070383 | 0.555126 |
| X | 4539200 | 4539799 | 11.93614 | 22.80402 | 22.80511 | 9.060731 | 0.509016 | -0.00408 | 0.555133 |
| 7 | 13254800 | 13269999 | 193.881 | 289.1551 | 274.9608 | 113.3223 | 0.331501 | 3.582296 | 0.555136 |
| X | 104549200 | 104550599 | 17.48736 | 31.71291 | 29.20627 | 12.62751 | 0.531625 | 0.368241 | 0.55514 |
| X | 153060400 | 153082999 | 413.8305 | 580.6711 | 493.407 | 204.4822 | 0.408087 | 4.533031 | 0.555142 |
| 19 | 39909400 | 39910199 | 19.79886 | 34.73034 | 30.81708 | 13.62346 | 0.559097 | 0.642897 | 0.555147 |
| 2 | 164384200 | 164385999 | 16.77444 | 29.6177 | 31.34436 | 13.86791 | 0.30855 | 0.545034 | 0.555158 |
| 7 | 108518200 | 108519599 | 9.196922 | 17.058 | 29.93211 | 12.56261 | -0.36269 | 0.178989 | 0.555159 |
| 16 | 8945800 | 8946799 | 9.196922 | 17.058 | 29.93211 | 12.56261 | -0.36269 | 0.178989 | 0.555159 |
| 18 | 55968000 | 55968999 | 10.04731 | 18.30549 | 31.82738 | 13.68096 | -0.37399 | 0.292299 | 0.555184 |
| 7 | 40749600 | 40756799 | 105.1142 | 163.6486 | 155.1726 | 64.41093 | 0.386228 | 2.730593 | 0.555208 |
| 9 | 20768200 | 20769799 | 11.35043 | 20.3735 | 34.23728 | 14.97854 | -0.34198 | 0.394954 | 0.555215 |
| X | 152548600 | 152551799 | 35.9124 | 59.35348 | 59.84953 | 25.8614 | 0.368572 | 1.356192 | 0.555225 |
| 13 | 28149800 | 28151199 | 18.62905 | 32.76117 | 34.32132 | 15.19716 | 0.320941 | 0.605159 | 0.555232 |
| 16 | 95987200 | 95988599 | 23.202 | 39.68397 | 41.16891 | 18.05192 | 0.330186 | 0.962559 | 0.555245 |
| X | 163351600 | 163353199 | 11.57954 | 21.02419 | 37.55396 | 16.21258 | -0.4169 | 0.337737 | 0.555282 |
| 12 | 114673800 | 114676799 | 45.70243 | 74.68511 | 65.48495 | 28.15278 | 0.563312 | 1.521105 | 0.555313 |
| 12 | 6725400 | 6726399 | 15.74291 | 28.18256 | 29.83954 | 13.08395 | 0.318297 | 0.432594 | 0.555326 |
| X | 108538400 | 108540199 | 11.47066 | 20.96752 | 37.75628 | 16.16293 | -0.42089 | 0.2957 | 0.555341 |
| 16 | 28671200 | 28675799 | 53.40239 | 84.67927 | 131.9435 | 54.92084 | -0.30855 | 2.328025 | 0.555388 |
| 1 | 121898000 | 121901399 | 36.11279 | 58.49601 | 93.7781 | 40.13493 | -0.31526 | 1.661605 | 0.555396 |
| Y | 4838600 | 4840399 | 13.26206 | 24.36599 | 18.09969 | 7.695011 | 0.855176 | 0.23325 | 0.5554 |
| 6 | 130044200 | 130059999 | 260.6381 | 380.3036 | 346.665 | 142.8405 | 0.36287 | 3.949838 | 0.555408 |
| 15 | 75409200 | 75410799 | 22.73239 | 39.3211 | 41.49649 | 18.20539 | 0.308502 | 0.860329 | 0.555414 |
| 1 | 66071400 | 66072399 | 12.14357 | 21.68417 | 35.70233 | 15.68808 | -0.31717 | 0.435544 | 0.555419 |
| 7 | 32798200 | 32801799 | 106.3141 | 163.5316 | 168.092 | 69.80651 | 0.256593 | 2.949513 | 0.55544 |
| 11 | 66842600 | 66844199 | 19.47348 | 32.29058 | 69.98715 | 30.31079 | -0.73132 | 1.284794 | 0.555442 |
| 12 | 82153000 | 82155799 | 34.94318 | 56.49479 | 87.11086 | 37.22316 | -0.2618 | 1.69642 | 0.55545 |
| 17 | 39010800 | 39011599 | 19.86192 | 34.70874 | 36.2105 | 15.95641 | 0.326969 | 0.71493 | 0.555473 |
| X | 37927000 | 37928999 | 25.96232 | 44.68112 | 43.57567 | 19.11462 | 0.420444 | 0.922311 | 0.555502 |
| 14 | 5434400 | 5434799 | 9.792882 | 18.73527 | 21.25559 | 8.413955 | 0.332916 | -0.04347 | 0.555519 |
| X | 43336200 | 43340399 | 71.04285 | 113.1472 | 97.37548 | 40.73381 | 0.552875 | 2.203868 | 0.55554 |
| 5 | 140894800 | 140899199 | 60.2171 | 95.22077 | 141.1104 | 58.63394 | -0.2397 | 2.405841 | 0.555552 |
| 3 | 121052600 | 121054199 | 12.85301 | 22.62142 | 38.35616 | 16.98104 | -0.36945 | 0.571541 | 0.555566 |
| 11 | 35589000 | 35600399 | 94.73065 | 144.2842 | 249.9906 | 103.6142 | -0.50711 | 3.148997 | 0.555567 |
| 18 | 58624600 | 58627799 | 24.48546 | 40.72874 | 64.13787 | 27.83766 | -0.27187 | 1.254752 | 0.555582 |
| 7 | 23284800 | 23290999 | 113.8108 | 175.6056 | 173.8005 | 72.19348 | 0.314532 | 2.895306 | 0.555589 |
| 19 | 9677200 | 9684999 | 145.0431 | 219.3088 | 227.5479 | 94.10253 | 0.223296 | 3.307816 | 0.555594 |
| 19 | 39494000 | 39498399 | 42.68843 | 69.25444 | 72.64599 | 31.12637 | 0.297364 | 1.631343 | 0.555601 |
| X | 7707600 | 7710999 | 46.48303 | 75.94795 | 64.06222 | 27.53944 | 0.618948 | 1.527833 | 0.555603 |
| 4 | 136898400 | 136900799 | 21.11727 | 35.81927 | 56.52146 | 24.75845 | -0.27425 | 1.046615 | 0.555616 |
| 12 | 79946400 | 79946999 | 10.62585 | 20.20481 | 22.74831 | 9.129133 | 0.326349 | 0.013347 | 0.55565 |
| 1 | 179494000 | 179496399 | 12.54272 | 22.10222 | 37.85636 | 16.75568 | -0.38319 | 0.555628 | 0.555652 |

| | | | | | | | | | |
|---|---|---|---|---|---|---|---|---|---|
| 8 | 92269400 | 92270999 | 12.54272 | 22.10222 | 37.85636 | 16.75568 | -0.38319 | 0.555628 | 0.555652 |
| 9 | 118609000 | 118610599 | 13.43215 | 23.77931 | 38.69947 | 17.11302 | -0.30752 | 0.510008 | 0.555667 |
| 18 | 31516400 | 31518399 | 13.43215 | 23.77931 | 38.69947 | 17.11302 | -0.30752 | 0.510008 | 0.555667 |
| 8 | 84548400 | 84554199 | 63.78222 | 100.5943 | 149.5763 | 62.07438 | -0.2476 | 2.469243 | 0.555672 |
| 7 | 6007000 | 6015999 | 148.7939 | 226.1928 | 192.5034 | 79.74144 | 0.515399 | 3.198325 | 0.555691 |
| 12 | 55566800 | 55568799 | 17.38438 | 30.6463 | 32.27965 | 14.28621 | 0.314415 | 0.56686 | 0.555698 |
| 18 | 32877600 | 32878599 | 10.40104 | 19.76315 | 20.52406 | 8.249027 | 0.439429 | 0.00703 | 0.555711 |
| Y | 8798400 | 8798999 | 8.741975 | 16.90685 | 15.22984 | 5.822325 | 0.709826 | -0.18819 | 0.555748 |
| X | 19589800 | 19595399 | 64.23124 | 103.1672 | 86.68457 | 36.70892 | 0.600966 | 1.938044 | 0.555751 |
| 17 | 64554400 | 64557199 | 25.09567 | 43.00121 | 44.67274 | 19.59117 | 0.328429 | 0.95627 | 0.55577 |
| 3 | 101084400 | 101086799 | 24.07007 | 41.86769 | 39.63464 | 17.41418 | 0.465724 | 0.801976 | 0.555779 |
| 13 | 65345400 | 65350199 | 57.18636 | 91.81236 | 90.46338 | 38.28411 | 0.370853 | 1.940566 | 0.555785 |
| 16 | 21606200 | 21608199 | 15.42676 | 26.98725 | 43.58041 | 19.22145 | -0.3004 | 0.686211 | 0.555805 |
| 12 | 115647000 | 115649399 | 58.07126 | 92.73526 | 98.59789 | 41.39914 | 0.252513 | 2.108273 | 0.555823 |
| 7 | 32157200 | 32159799 | 41.0214 | 67.26768 | 56.53963 | 24.34027 | 0.626797 | 1.460595 | 0.555871 |
| 17 | 25068600 | 25069799 | 8.815944 | 16.28208 | 29.94726 | 12.64181 | -0.43554 | 0.203003 | 0.555871 |
| 10 | 126351800 | 126354399 | 28.62164 | 48.15323 | 50.73177 | 22.16736 | 0.306001 | 1.135435 | 0.55589 |
| X | 37080800 | 37082599 | 23.84681 | 41.68283 | 32.68534 | 14.40609 | 0.736815 | 0.724797 | 0.555895 |
| 3 | 41644600 | 41645999 | 10.23569 | 18.56163 | 32.32106 | 13.97472 | -0.38155 | 0.322684 | 0.55591 |
| 5 | 40529000 | 40529599 | 6.100312 | 11.84884 | 22.85315 | 8.544053 | -0.34742 | -0.26688 | 0.555934 |
| 10 | 37612000 | 37612399 | 6.100312 | 11.84884 | 22.85315 | 8.544053 | -0.34742 | -0.26688 | 0.555934 |
| 7 | 51271400 | 51272799 | 15.65953 | 28.1394 | 29.7455 | 13.00123 | 0.323604 | 0.407312 | 0.555949 |
| 4 | 119949400 | 119957999 | 98.3002 | 150.1743 | 220.3162 | 91.38293 | -0.26382 | 3.086844 | 0.555952 |
| 3 | 62582800 | 62583399 | 11.26888 | 21.17044 | 23.15342 | 9.516309 | 0.338576 | 0.100878 | 0.555978 |
| 4 | 48712600 | 48716599 | 21.05681 | 35.7448 | 57.66673 | 25.26614 | -0.30613 | 1.039105 | 0.555983 |
| X | 30919000 | 30922599 | 50.32728 | 81.40351 | 66.65291 | 28.49568 | 0.649966 | 1.702769 | 0.555991 |
| 5 | 117161200 | 117167199 | 74.51485 | 116.1573 | 170.7629 | 70.86886 | -0.24427 | 2.696581 | 0.556018 |
| 7 | 30860200 | 30862399 | 12.82316 | 22.59121 | 38.30883 | 16.95673 | -0.36914 | 0.559199 | 0.55602 |
| 13 | 94843400 | 94844999 | 12.82316 | 22.59121 | 38.30883 | 16.95673 | -0.36914 | 0.559199 | 0.55602 |
| 7 | 13047200 | 13048799 | 13.92341 | 25.18494 | 25.89389 | 11.23144 | 0.370515 | 0.347712 | 0.556022 |
| 15 | 48080200 | 48081799 | 19.12844 | 33.35812 | 33.75736 | 14.85596 | 0.370301 | 0.733599 | 0.556025 |
| 2 | 120309400 | 120310599 | 14.99291 | 26.84287 | 28.37027 | 12.43855 | 0.320865 | 0.422672 | 0.556038 |
| 1 | 137644000 | 137644399 | 3.931039 | 7.686894 | 17.20225 | 5.866952 | -0.43608 | -0.64019 | 0.556038 |
| 11 | 114518400 | 114522799 | 47.11647 | 75.12318 | 117.0195 | 49.02584 | -0.29998 | 2.150058 | 0.556099 |
| 19 | 36323800 | 36327199 | 28.36221 | 46.32255 | 74.82156 | 32.16016 | -0.31631 | 1.496415 | 0.556108 |
| 19 | 39455800 | 39464599 | 125.028 | 191.4591 | 175.1113 | 72.67254 | 0.419956 | 3.043306 | 0.556113 |
| 10 | 35435000 | 35437399 | 29.24534 | 48.76464 | 51.89244 | 22.55821 | 0.291733 | 1.228944 | 0.556116 |
| 5 | 127753400 | 127763599 | 104.8992 | 160.2227 | 227.8452 | 94.50131 | -0.21914 | 3.093199 | 0.556125 |
| 6 | 79800600 | 79803799 | 65.52318 | 104.4802 | 95.67827 | 40.08966 | 0.465227 | 2.159758 | 0.556126 |
| 7 | 96763000 | 96766999 | 45.43752 | 73.40092 | 77.70416 | 33.1682 | 0.278113 | 1.732576 | 0.556128 |
| 17 | 92484600 | 92488799 | 41.78153 | 68.0584 | 69.01602 | 29.63605 | 0.350755 | 1.558275 | 0.556132 |
| 10 | 40516600 | 40520999 | 35.91739 | 58.04625 | 88.9883 | 38.00546 | -0.25419 | 1.71064 | 0.556145 |
| 7 | 116609800 | 116611199 | 13.17155 | 24.49078 | 25.39727 | 10.62265 | 0.394039 | 0.184979 | 0.556163 |
| 2 | 122798000 | 122798799 | 17.32649 | 30.48479 | 29.17985 | 12.91935 | 0.450987 | 0.584601 | 0.556165 |
| 15 | 87122800 | 87125999 | 30.29747 | 49.50162 | 76.88828 | 33.05295 | -0.25968 | 1.496821 | 0.556183 |
| 5 | 21587600 | 21589399 | 9.387025 | 17.19697 | 31.87057 | 13.60278 | -0.45878 | 0.253583 | 0.556216 |
| Y | 29065800 | 29067599 | 11.98796 | 22.37483 | 17.11388 | 7.112171 | 0.837356 | 0.128944 | 0.556245 |

| | | | | | | | | | |
|---|---|---|---|---|---|---|---|---|---|
| 7 | 5459400 | 5465599 | 120.2557 | 184.6734 | 167.3962 | 69.50194 | 0.436013 | 2.986339 | 0.556266 |
| 5 | 88299400 | 88301799 | 33.87903 | 54.72571 | 87.39925 | 37.30365 | -0.31375 | 1.722971 | 0.556293 |
| X | 11865000 | 11867799 | 23.95159 | 41.30036 | 42.56131 | 18.67076 | 0.34179 | 0.895475 | 0.556303 |
| 19 | 52765400 | 52767599 | 38.24907 | 62.52892 | 55.2222 | 23.74399 | 0.553287 | 1.491669 | 0.556311 |
| 9 | 114020600 | 114023199 | 38.68181 | 62.76181 | 64.70508 | 27.72637 | 0.322459 | 1.616485 | 0.556322 |
| 7 | 43211400 | 43212399 | 9.828099 | 18.07288 | 31.38226 | 13.32597 | -0.36006 | 0.235362 | 0.556333 |
| 5 | 129513000 | 129514999 | 18.86438 | 31.66797 | 62.75053 | 27.39468 | -0.60206 | 1.138157 | 0.556336 |
| 12 | 55929600 | 55931399 | 13.27795 | 23.40268 | 38.77095 | 17.15978 | -0.33555 | 0.557475 | 0.556348 |
| 8 | 114736800 | 114737999 | 15.99006 | 28.82782 | 29.71611 | 12.94265 | 0.363014 | 0.390235 | 0.556366 |
| 17 | 63866000 | 63866399 | 3.636446 | 7.113549 | 16.18191 | 5.387473 | -0.42685 | -0.73301 | 0.556367 |
| 9 | 43674800 | 43677199 | 25.97684 | 42.9811 | 67.47107 | 29.17741 | -0.26797 | 1.322552 | 0.556367 |
| 8 | 59506400 | 59507199 | 10.28563 | 18.89027 | 32.40657 | 13.78468 | -0.34464 | 0.248005 | 0.556382 |
| 19 | 40200400 | 40201999 | 21.54334 | 36.23021 | 58.04002 | 25.35765 | -0.29639 | 1.125609 | 0.556387 |
| 5 | 26814000 | 26815599 | 14.1067 | 24.75015 | 40.40651 | 17.86561 | -0.31601 | 0.62585 | 0.556391 |
| 11 | 91391200 | 91391799 | 7.282996 | 14.02674 | 25.90953 | 10.06162 | -0.33859 | -0.11833 | 0.556391 |
| 18 | 17062800 | 17063399 | 7.282996 | 14.02674 | 25.90953 | 10.06162 | -0.33859 | -0.11833 | 0.556391 |
| 2 | 177701800 | 177703399 | 20.23293 | 35.38797 | 37.05263 | 16.33956 | 0.321596 | 0.702187 | 0.55641 |
| 13 | 84724400 | 84727199 | 24.87415 | 43.14513 | 41.35226 | 18.14365 | 0.447608 | 0.845082 | 0.556425 |
| X | 109661400 | 109662399 | 19.41126 | 34.0304 | 35.05561 | 15.48951 | 0.345094 | 0.659749 | 0.556447 |
| 2 | 77165200 | 77166199 | 12.62546 | 22.48178 | 36.68835 | 16.15935 | -0.30685 | 0.458071 | 0.55645 |
| 3 | 38254800 | 38256199 | 10.71021 | 19.71609 | 34.63605 | 14.68623 | -0.37512 | 0.24586 | 0.556453 |
| 12 | 31314800 | 31317599 | 15.75205 | 27.20215 | 46.8969 | 20.57363 | -0.39694 | 0.851474 | 0.556454 |
| 2 | 39275600 | 39276999 | 19.94893 | 33.99675 | 53.81099 | 23.58839 | -0.27785 | 0.997079 | 0.556461 |
| 18 | 64420200 | 64423399 | 23.27804 | 39.1962 | 64.36663 | 28.13235 | -0.33215 | 1.124473 | 0.556487 |
| 2 | 114385400 | 114387999 | 26.76068 | 44.24936 | 69.90075 | 30.21459 | -0.27717 | 1.332806 | 0.556502 |
| 5 | 91736200 | 91737399 | 14.68352 | 27.7993 | 24.06703 | 9.747543 | 0.690751 | 0.082428 | 0.556503 |
| 2 | 33829000 | 33830599 | 19.47819 | 33.9492 | 35.73685 | 15.72161 | 0.313599 | 0.746227 | 0.556503 |
| 18 | 68307200 | 68308999 | 20.79835 | 36.22462 | 38.30462 | 16.84379 | 0.307136 | 0.768548 | 0.556505 |
| 10 | 115449800 | 115451799 | 16.83438 | 29.37027 | 48.95091 | 21.54824 | -0.34629 | 0.739597 | 0.556509 |
| 13 | 106754800 | 106758599 | 37.42294 | 62.12188 | 56.67737 | 24.56496 | 0.512881 | 1.306691 | 0.556522 |
| 4 | 71921800 | 71925399 | 44.12491 | 71.50679 | 73.96679 | 31.66679 | 0.316025 | 1.655587 | 0.556537 |
| 13 | 94462400 | 94463799 | 11.67624 | 21.05702 | 34.76815 | 15.13726 | -0.31205 | 0.370608 | 0.556545 |
| 10 | 108181200 | 108183199 | 17.25219 | 30.06536 | 53.90726 | 23.71325 | -0.45143 | 0.765337 | 0.556551 |
| 9 | 117955800 | 117959399 | 55.03971 | 87.99984 | 93.82651 | 39.46287 | 0.250348 | 2.066247 | 0.556562 |
| 10 | 82606600 | 82609799 | 22.75655 | 37.89331 | 62.26113 | 27.04233 | -0.33273 | 1.238062 | 0.556562 |
| 7 | 97345200 | 97346199 | 5.393972 | 10.46471 | 22.19046 | 8.359648 | -0.49267 | -0.25086 | 0.556568 |
| 16 | 57412200 | 57413599 | 15.84377 | 26.74656 | 60.45186 | 26.44186 | -0.7906 | 1.075676 | 0.556577 |
| 1 | 85302600 | 85307999 | 156.7585 | 236.8185 | 211.868 | 87.59426 | 0.4359 | 3.326342 | 0.556593 |
| 11 | 22997200 | 22999399 | 16.00919 | 28.35978 | 28.28199 | 12.50477 | 0.395464 | 0.497272 | 0.556601 |
| 9 | 47706200 | 47709999 | 21.21988 | 36.66508 | 37.92382 | 16.63743 | 0.336963 | 0.856237 | 0.556601 |
| 7 | 6199000 | 6201799 | 22.16181 | 37.11248 | 59.7104 | 26.02724 | -0.30269 | 1.170544 | 0.55661 |
| 6 | 95782600 | 95788599 | 94.0801 | 146.3649 | 155.1486 | 64.40784 | 0.224791 | 2.737468 | 0.556615 |
| 1 | 105766800 | 105768399 | 11.8566 | 21.07425 | 35.72962 | 15.76128 | -0.36301 | 0.462932 | 0.556622 |
| 8 | 39250800 | 39251799 | 10.76783 | 19.24334 | 34.30918 | 15.0655 | -0.43036 | 0.41925 | 0.556648 |
| 5 | 105010800 | 105036799 | 636.6415 | 854.7009 | 775.2146 | 323.4853 | 0.249763 | 5.21768 | 0.556651 |
| 1 | 61709800 | 61710799 | 18.2728 | 31.99498 | 32.69709 | 14.43231 | 0.356527 | 0.667917 | 0.556656 |
| 12 | 10204000 | 10210799 | 30.90946 | 50.11237 | 90.43633 | 38.73026 | -0.48437 | 1.632587 | 0.556662 |

| | | | | | | | | | |
|---|---|---|---|---|---|---|---|---|---|
| X | 25884800 | 25885999 | 24.72166 | 42.60068 | 42.05387 | 18.4478 | 0.403457 | 0.905264 | 0.55667 |
| 9 | 7763600 | 7764799 | 13.4868 | 23.60172 | 41.04176 | 18.10707 | -0.40657 | 0.665057 | 0.556691 |
| 1 | 171338000 | 171339399 | 26.02793 | 42.91347 | 68.36462 | 29.4986 | -0.28993 | 1.365814 | 0.556697 |
| 6 | 133542400 | 133544199 | 40.61598 | 65.96487 | 70.03051 | 30.0301 | 0.281597 | 1.605433 | 0.556712 |
| 16 | 92323800 | 92329599 | 54.92456 | 87.34514 | 132.5645 | 55.33926 | -0.26664 | 2.243674 | 0.556714 |
| 4 | 44551200 | 44552199 | 9.930941 | 17.98316 | 32.32882 | 14.0032 | -0.42897 | 0.328395 | 0.556721 |
| 1 | 20353400 | 20356199 | 23.44068 | 40.86128 | 39.12488 | 17.21361 | 0.449756 | 0.770391 | 0.556747 |
| 10 | 78317800 | 78319999 | 17.49008 | 30.09409 | 48.99985 | 21.49382 | -0.31607 | 0.894036 | 0.556748 |
| 2 | 18698800 | 18700599 | 11.72985 | 20.76727 | 36.28753 | 16.0462 | -0.40917 | 0.494771 | 0.55675 |
| 12 | 18027800 | 18029199 | 16.63625 | 29.57797 | 31.19932 | 13.75961 | 0.317842 | 0.482149 | 0.55676 |
| 14 | 89512600 | 89515399 | 24.55475 | 42.69963 | 39.61285 | 17.40176 | 0.494741 | 0.807733 | 0.556773 |
| 1 | 58035000 | 58035799 | 6.272106 | 12.07918 | 24.17672 | 9.390106 | -0.45297 | -0.13286 | 0.556777 |
| 12 | 81021600 | 81023599 | 22.4873 | 37.86385 | 59.78176 | 26.12828 | -0.27551 | 1.118992 | 0.556803 |
| 7 | 42590200 | 42604399 | 289.331 | 416.6684 | 396.6832 | 163.7061 | 0.28172 | 4.14666 | 0.556819 |
| 1 | 94574200 | 94575999 | 17.78386 | 31.74262 | 30.41362 | 13.36881 | 0.458806 | 0.462197 | 0.556837 |
| 3 | 45050800 | 45061199 | 183.012 | 272.8657 | 275.6424 | 113.5968 | 0.243944 | 3.585055 | 0.556849 |
| 10 | 70174800 | 70181199 | 63.94299 | 101.0129 | 148.2441 | 61.56743 | -0.22685 | 2.43187 | 0.556852 |
| 1 | 82774000 | 82774799 | 7.779916 | 14.79913 | 26.63307 | 10.67928 | -0.34369 | -0.00549 | 0.556856 |
| 3 | 10606800 | 10607399 | 6.685949 | 12.95373 | 24.47556 | 9.278894 | -0.33765 | -0.20888 | 0.556867 |
| 2 | 98449200 | 98450599 | 15.26313 | 27.74999 | 27.6087 | 11.89878 | 0.423262 | 0.330506 | 0.556876 |
| 11 | 43666200 | 43667399 | 7.233207 | 13.62064 | 27.02691 | 11.06437 | -0.50916 | 0.060959 | 0.556877 |
| 6 | 10026800 | 10027599 | 11.99623 | 21.31578 | 36.20197 | 15.97384 | -0.36588 | 0.465905 | 0.556894 |
| 9 | 54984600 | 54986999 | 27.34181 | 44.44399 | 77.35159 | 33.19001 | -0.42851 | 1.566428 | 0.556902 |
| 8 | 98948600 | 98951599 | 39.45854 | 64.02594 | 68.32657 | 29.27949 | 0.272827 | 1.61765 | 0.556923 |
| 6 | 140613600 | 140618799 | 61.47816 | 97.20363 | 142.649 | 59.26982 | -0.22578 | 2.408833 | 0.556963 |
| 2 | 44394000 | 44395599 | 9.526446 | 17.87502 | 31.84999 | 13.11796 | -0.36314 | 0.114878 | 0.556976 |
| 13 | 13711600 | 13720399 | 103.1924 | 157.2528 | 257.2815 | 106.6524 | -0.4239 | 3.141603 | 0.556999 |
| 7 | 26452200 | 26452999 | 20.76735 | 35.9442 | 35.96297 | 15.77801 | 0.385275 | 0.837243 | 0.557002 |
| 18 | 73569000 | 73572999 | 36.33038 | 58.35087 | 115.3268 | 48.83514 | -0.63079 | 1.908671 | 0.557006 |
| 8 | 25286800 | 25289799 | 33.7845 | 56.43665 | 54.22745 | 23.60052 | 0.43854 | 1.226634 | 0.557061 |
| 14 | 99255600 | 99258799 | 25.89376 | 44.2846 | 44.09072 | 19.33297 | 0.389033 | 0.973486 | 0.557061 |
| 13 | 76617200 | 76618999 | 18.30198 | 31.38031 | 50.5942 | 22.18916 | -0.30294 | 0.931183 | 0.557071 |
| 10 | 77787800 | 77788199 | 5.964889 | 11.55087 | 23.23116 | 8.827942 | -0.42883 | -0.20922 | 0.557071 |
| 1 | 37267600 | 37272399 | 53.39025 | 84.88868 | 126.4592 | 52.78034 | -0.24004 | 2.248571 | 0.557077 |
| 6 | 5704800 | 5706399 | 36.54433 | 59.55976 | 63.1978 | 27.14302 | 0.286125 | 1.536959 | 0.55708 |
| 6 | 14315400 | 14317199 | 24.36316 | 41.56357 | 44.76383 | 19.62419 | 0.275781 | 0.994053 | 0.557091 |
| 8 | 56426000 | 56427399 | 14.04556 | 25.74873 | 27.43822 | 11.70025 | 0.335449 | 0.277352 | 0.557107 |
| 12 | 71303800 | 71305199 | 14.04556 | 25.74873 | 27.43822 | 11.70025 | 0.335449 | 0.277352 | 0.557107 |
| 10 | 86742800 | 86747199 | 42.65478 | 68.29854 | 106.9546 | 45.07893 | -0.30088 | 2.010415 | 0.557112 |
| 1 | 35341200 | 35344399 | 31.43914 | 51.44179 | 85.64926 | 36.83869 | -0.35825 | 1.483281 | 0.557117 |
| 13 | 66323800 | 66332599 | 257.2906 | 373.4585 | 312.0914 | 128.6647 | 0.480861 | 4.026342 | 0.557127 |
| 11 | 73537600 | 73543799 | 63.19059 | 101.1183 | 97.58711 | 41.09433 | 0.39521 | 2.05113 | 0.557158 |
| 3 | 19389600 | 19390399 | 7.968096 | 14.98418 | 27.51151 | 11.28863 | -0.40111 | 0.076836 | 0.557158 |
| 4 | 44072000 | 44072999 | 7.968096 | 14.98418 | 27.51151 | 11.28863 | -0.40111 | 0.076836 | 0.557158 |
| 2 | 116317400 | 116320599 | 31.41161 | 52.23208 | 55.73479 | 24.18488 | 0.287813 | 1.27139 | 0.557196 |
| 9 | 13635200 | 13638399 | 29.6725 | 49.37179 | 50.84027 | 22.07075 | 0.33897 | 1.251187 | 0.557209 |
| 2 | 76588400 | 76590799 | 56.25107 | 89.93005 | 95.74249 | 40.26457 | 0.252374 | 2.071093 | 0.557246 |

| | | | | | | | | | |
|---|---|---|---|---|---|---|---|---|---|
| 14 | 116364400 | 116365399 | 10.43801 | 20.14725 | 18.6352 | 7.19002 | 0.659676 | -0.12569 | 0.557253 |
| 1 | 109221400 | 109222799 | 13.3705 | 24.78708 | 25.90474 | 10.8794 | 0.378726 | 0.204617 | 0.55726 |
| 1 | 115359800 | 115361999 | 20.07025 | 35.53194 | 32.38641 | 14.32128 | 0.524295 | 0.538216 | 0.557262 |
| X | 26375600 | 26382399 | 187.9662 | 283.3201 | 199.2585 | 82.32332 | 0.780033 | 3.379553 | 0.55729 |
| X | 51986800 | 51987799 | 14.78637 | 27.00589 | 27.92193 | 11.96544 | 0.373929 | 0.30582 | 0.557295 |
| 2 | 173565800 | 173566599 | 5.100751 | 9.919907 | 21.20448 | 7.847058 | -0.47986 | -0.32054 | 0.557331 |
| 16 | 67786800 | 67787399 | 7.137725 | 13.6765 | 26.279 | 10.35678 | -0.41525 | -0.07197 | 0.557342 |
| 3 | 93704000 | 93704599 | 5.668214 | 11.00491 | 22.22562 | 8.337889 | -0.41726 | -0.26108 | 0.557396 |
| 17 | 46694200 | 46695599 | 11.62445 | 20.92936 | 34.71542 | 15.14527 | -0.3207 | 0.377826 | 0.5574 |
| 15 | 16946600 | 16948599 | 24.77644 | 42.47501 | 37.36995 | 16.38919 | 0.567309 | 0.941242 | 0.557412 |
| 12 | 88042800 | 88047599 | 59.65154 | 95.63364 | 93.50897 | 39.48723 | 0.379474 | 1.988078 | 0.557417 |
| 12 | 114465000 | 114477799 | 197.0816 | 294.6359 | 264.227 | 108.9494 | 0.419128 | 3.538367 | 0.557434 |
| 5 | 125520400 | 125520999 | 5.056959 | 9.847439 | 21.26157 | 7.785518 | -0.47894 | -0.35449 | 0.557449 |
| 6 | 82656600 | 82659399 | 19.46534 | 33.13916 | 53.25358 | 23.34215 | -0.29981 | 1.003626 | 0.557471 |
| 9 | 11084000 | 11090999 | 112.856 | 173.712 | 159.0497 | 66.05749 | 0.424068 | 2.93728 | 0.557487 |
| 3 | 15408200 | 15409999 | 15.73324 | 28.96014 | 27.26689 | 11.56305 | 0.518409 | 0.266178 | 0.557522 |
| 3 | 120069400 | 120070999 | 15.99928 | 27.82641 | 45.26036 | 19.88465 | -0.31159 | 0.774383 | 0.557527 |
| 8 | 3826000 | 3836599 | 158.5782 | 239.0773 | 239.5231 | 98.96448 | 0.270201 | 3.36909 | 0.55755 |
| 12 | 78466200 | 78468999 | 17.63534 | 31.97894 | 29.07802 | 12.57321 | 0.549609 | 0.367678 | 0.557557 |
| Y | 87270000 | 87271799 | 13.68088 | 25.07156 | 17.56617 | 7.494105 | 0.935245 | 0.249079 | 0.55756 |
| 4 | 127994600 | 127995999 | 10.83669 | 19.54566 | 33.743 | 14.68841 | -0.37589 | 0.359661 | 0.557584 |
| 6 | 132435600 | 132437599 | 22.71129 | 39.15726 | 41.47743 | 18.19523 | 0.302095 | 0.886526 | 0.557622 |
| 13 | 108139200 | 108140999 | 23.69623 | 39.29488 | 64.54001 | 27.95277 | -0.33274 | 1.287795 | 0.557641 |
| 10 | 69970200 | 69971799 | 16.98388 | 29.57939 | 47.89021 | 21.05818 | -0.30486 | 0.763553 | 0.557654 |
| 5 | 150185800 | 150188799 | 14.71401 | 25.62901 | 44.71748 | 19.66446 | -0.41219 | 0.744939 | 0.557666 |
| 8 | 23043200 | 23043599 | 10.20436 | 19.47506 | 20.7728 | 8.271559 | 0.414048 | -0.01974 | 0.557671 |
| 12 | 59643000 | 59645799 | 32.31184 | 54.40658 | 50.76898 | 22.18918 | 0.480564 | 1.141417 | 0.557681 |
| 10 | 23334200 | 23337399 | 39.61168 | 64.25829 | 68.33242 | 29.27658 | 0.277579 | 1.622771 | 0.557691 |
| 7 | 21148400 | 21151799 | 31.47973 | 54.14368 | 41.72736 | 18.30351 | 0.758585 | 0.948067 | 0.557703 |
| 7 | 36654800 | 36656799 | 20.45619 | 34.7578 | 54.87563 | 24.04933 | -0.27486 | 1.027486 | 0.557719 |
| 18 | 79013400 | 79014399 | 5.377079 | 10.47974 | 21.46586 | 7.773061 | -0.39125 | -0.39588 | 0.557736 |
| 10 | 117309400 | 117312999 | 44.5515 | 72.28615 | 72.85825 | 31.22024 | 0.355012 | 1.633166 | 0.55774 |
| 11 | 120230800 | 120233999 | 26.60758 | 43.91753 | 69.80478 | 30.13585 | -0.28622 | 1.35281 | 0.557744 |
| 5 | 123260200 | 123262599 | 17.184 | 29.47213 | 50.05496 | 21.95319 | -0.37759 | 0.922673 | 0.55775 |
| 13 | 66420200 | 66423799 | 39.83337 | 66.66818 | 50.33149 | 21.93131 | 0.785422 | 1.226093 | 0.557753 |
| 16 | 17632800 | 17636999 | 34.3127 | 55.491 | 85.77433 | 36.66529 | -0.26485 | 1.687322 | 0.557753 |
| 18 | 90527800 | 90528599 | 6.872122 | 13.12726 | 25.62525 | 10.18439 | -0.44871 | -0.04744 | 0.557795 |
| 1 | 143130000 | 143131399 | 11.98191 | 21.47796 | 35.2151 | 15.42388 | -0.30789 | 0.409677 | 0.557811 |
| 14 | 107764400 | 107766799 | 17.04162 | 29.71846 | 59.02422 | 25.97516 | -0.59848 | 0.759407 | 0.557812 |
| Y | 89250600 | 89252399 | 13.79917 | 25.15718 | 19.12135 | 8.212728 | 0.812379 | 0.284135 | 0.557814 |
| 11 | 101292400 | 101294199 | 16.03238 | 28.7351 | 30.26079 | 13.25528 | 0.32694 | 0.427346 | 0.557819 |
| 3 | 37033000 | 37033799 | 14.1411 | 26.28049 | 22.83755 | 9.559064 | 0.646981 | 0.192404 | 0.557831 |
| 14 | 62864800 | 62866799 | 19.34284 | 33.7357 | 35.72998 | 15.72516 | 0.304923 | 0.736523 | 0.557838 |
| 6 | 89746800 | 89760199 | 397.3912 | 553.0792 | 531.1093 | 220.3862 | 0.219844 | 4.652728 | 0.557856 |
| 6 | 59776600 | 59777799 | 18.96309 | 33.30076 | 34.24805 | 15.15424 | 0.34706 | 0.625214 | 0.557858 |
| 1 | 62760800 | 62764399 | 29.97578 | 48.91747 | 76.7314 | 32.97112 | -0.27504 | 1.512851 | 0.557863 |
| 14 | 91232600 | 91239199 | 99.4979 | 154.3157 | 162.268 | 67.39449 | 0.23291 | 2.793708 | 0.557881 |

| | | | | | | | | | |
|---|---|---|---|---|---|---|---|---|---|
| 13 | 97685600 | 97687999 | 21.90028 | 36.87519 | 60.40816 | 26.402 | -0.32865 | 1.114515 | 0.557882 |
| 11 | 91988400 | 91991999 | 56.94044 | 90.86011 | 95.22705 | 39.94064 | 0.271981 | 2.127697 | 0.557884 |
| 13 | 91163200 | 91165199 | 11.46215 | 20.72733 | 34.26248 | 14.86636 | -0.31062 | 0.350833 | 0.557915 |
| 12 | 19287000 | 19309799 | 805.7399 | 1056.107 | 914.618 | 382.8252 | 0.281932 | 5.60182 | 0.557923 |
| 9 | 22183000 | 22197799 | 320.1809 | 458.2645 | 418.3728 | 172.8222 | 0.333317 | 4.238612 | 0.557968 |
| 17 | 55320000 | 55323199 | 41.69074 | 67.54567 | 72.88882 | 31.1997 | 0.254823 | 1.655988 | 0.557975 |
| X | 13760600 | 13763199 | 29.61449 | 50.31219 | 47.93171 | 21.00592 | 0.451219 | 1.055659 | 0.558003 |
| 9 | 104015400 | 104018999 | 26.42543 | 43.70807 | 69.31958 | 29.96956 | -0.28283 | 1.327762 | 0.558004 |
| X | 110093200 | 110094999 | 19.41289 | 34.42164 | 31.6598 | 13.99449 | 0.51253 | 0.520718 | 0.558015 |
| 19 | 13614800 | 13618599 | 33.74902 | 56.97082 | 48.525 | 21.22474 | 0.611865 | 1.122328 | 0.558025 |
| 4 | 98277600 | 98279599 | 28.16006 | 47.4925 | 49.08793 | 21.46497 | 0.33355 | 1.109801 | 0.558032 |
| 7 | 11314000 | 11334599 | 503.2889 | 689.6767 | 566.8338 | 235.7874 | 0.421697 | 4.892486 | 0.558036 |
| 4 | 5143000 | 5150199 | 33.58785 | 55.64016 | 50.90031 | 22.02953 | 0.50881 | 1.315453 | 0.558037 |
| 7 | 45984000 | 45987999 | 41.80757 | 67.83395 | 72.05657 | 30.87669 | 0.279414 | 1.625925 | 0.558057 |
| 10 | 43983000 | 43987599 | 33.08617 | 53.70705 | 84.26134 | 36.11831 | -0.28128 | 1.606792 | 0.558061 |
| 6 | 88334600 | 88336199 | 11.68775 | 21.03593 | 34.75311 | 15.16838 | -0.31516 | 0.382426 | 0.558076 |
| 7 | 128388200 | 128389599 | 20.4789 | 35.82759 | 31.01455 | 13.67991 | 0.594779 | 0.690102 | 0.558081 |
| 1 | 78223000 | 78225799 | 19.59368 | 33.45969 | 53.287 | 23.36255 | -0.28624 | 0.975827 | 0.558096 |
| 4 | 144363400 | 144372999 | 103.3801 | 161.2369 | 151.5276 | 62.88121 | 0.401081 | 2.698895 | 0.558117 |
| 8 | 28951000 | 28953599 | 34.05615 | 56.47072 | 58.09887 | 25.1619 | 0.339985 | 1.312323 | 0.55812 |
| 10 | 34971400 | 34972199 | 8.375221 | 15.97381 | 29.00316 | 11.56464 | -0.34975 | -0.01306 | 0.558129 |
| 11 | 48515400 | 48516399 | 9.169402 | 16.69652 | 31.40732 | 13.51 | -0.48703 | 0.289766 | 0.558131 |
| 1 | 69259800 | 69262199 | 58.64174 | 93.48979 | 99.17752 | 41.54505 | 0.253132 | 2.16002 | 0.558145 |
| 10 | 41806200 | 41808399 | 14.41872 | 25.20307 | 42.06502 | 18.54018 | -0.34778 | 0.690482 | 0.558146 |
| 2 | 173885600 | 173887599 | 24.22982 | 41.26292 | 43.71015 | 19.15953 | 0.299126 | 1.006465 | 0.558156 |
| 14 | 14776200 | 14781199 | 39.34459 | 64.80665 | 60.78203 | 26.21608 | 0.47152 | 1.402305 | 0.558161 |
| 4 | 142427800 | 142433999 | 62.59506 | 98.94969 | 145.8863 | 60.60871 | -0.23271 | 2.414342 | 0.558163 |
| 13 | 103759600 | 103761599 | 15.49071 | 27.0535 | 44.12149 | 19.43473 | -0.31486 | 0.710779 | 0.558167 |
| 5 | 149634200 | 149635399 | 12.11772 | 21.71439 | 35.70222 | 15.64149 | -0.31228 | 0.412758 | 0.558189 |
| 7 | 12206800 | 12237999 | 519.1986 | 712.5654 | 619.8977 | 257.7984 | 0.341945 | 4.870407 | 0.558248 |
| 4 | 61327000 | 61328199 | 50.44097 | 80.91777 | 79.35383 | 33.54163 | 0.376313 | 1.953562 | 0.558254 |
| 2 | 65617600 | 65618199 | 8.488873 | 16.48064 | 18.80166 | 7.05551 | 0.396746 | -0.24237 | 0.558293 |
| 9 | 54013600 | 54017399 | 45.82528 | 74.62462 | 68.81413 | 29.54637 | 0.487267 | 1.571449 | 0.558318 |
| 8 | 8642400 | 8644399 | 13.89429 | 24.37913 | 41.39857 | 18.3053 | -0.37272 | 0.6229 | 0.558319 |
| 17 | 83270800 | 83273799 | 22.47706 | 37.67639 | 59.74388 | 26.05754 | -0.28184 | 1.16144 | 0.55832 |
| 4 | 147158600 | 147173799 | 394.151 | 551.1146 | 522.354 | 216.5972 | 0.245412 | 4.583505 | 0.558322 |
| 1 | 144924200 | 144926399 | 16.82006 | 30.83686 | 27.06661 | 11.54086 | 0.613286 | 0.295428 | 0.558332 |
| 1 | 114016200 | 114017399 | 13.64645 | 25.45409 | 26.0248 | 10.82703 | 0.42077 | 0.172081 | 0.55838 |
| 2 | 65778200 | 65779999 | 10.63951 | 19.56342 | 32.97053 | 14.00194 | -0.31738 | 0.246671 | 0.558382 |
| 12 | 57731200 | 57732399 | 7.458169 | 14.22997 | 26.14982 | 10.4188 | -0.36561 | -0.03179 | 0.558393 |
| 17 | 19464400 | 19467599 | 65.8706 | 105.0687 | 105.0042 | 44.01729 | 0.33988 | 2.153587 | 0.558398 |
| 17 | 94786000 | 94786799 | 13.09263 | 23.21036 | 38.1663 | 16.87258 | -0.32161 | 0.491143 | 0.558413 |
| 13 | 86331600 | 86346799 | 357.7078 | 496.0229 | 648.5504 | 269.2726 | -0.23066 | 4.70768 | 0.558413 |
| X | 125561800 | 125564399 | 31.6925 | 53.90223 | 46.7958 | 20.50994 | 0.584968 | 1.051242 | 0.558419 |
| 4 | 150603200 | 150604199 | 8.079022 | 15.25803 | 27.56663 | 11.22294 | -0.36896 | 0.048233 | 0.558422 |
| 6 | 88302400 | 88304599 | 23.99389 | 41.53158 | 41.93758 | 18.39929 | 0.372029 | 0.859601 | 0.55843 |
| 14 | 43258000 | 43259799 | 20.26875 | 35.59734 | 29.46793 | 13.04026 | 0.658447 | 0.622229 | 0.558438 |

| | | | | | | | | | |
|---|---|---|---|---|---|---|---|---|---|
| 1 | 96072400 | 96073399 | 13.53696 | 25.25303 | 26.05791 | 10.83851 | 0.407934 | 0.171058 | 0.55847 |
| 7 | 30369400 | 30371199 | 10.69675 | 19.37915 | 32.79168 | 14.19522 | -0.34172 | 0.332408 | 0.558475 |
| 10 | 48227800 | 48229799 | 25.21416 | 43.87996 | 39.60105 | 17.40496 | 0.534529 | 0.801791 | 0.558508 |
| 19 | 59860400 | 59864199 | 46.94561 | 75.55125 | 81.39578 | 34.56745 | 0.246466 | 1.849023 | 0.558565 |
| 1 | 78749400 | 78750999 | 15.49424 | 28.01793 | 29.26791 | 12.69965 | 0.347422 | 0.368789 | 0.558566 |
| 11 | 76821800 | 76824999 | 42.90247 | 68.60554 | 119.5371 | 50.28827 | -0.4567 | 2.05599 | 0.558576 |
| 5 | 60314000 | 60314599 | 6.12237 | 11.81528 | 23.67106 | 9.139151 | -0.44589 | -0.15436 | 0.558587 |
| 7 | 62193600 | 62223599 | 563.156 | 766.2531 | 678.8347 | 282.6897 | 0.303155 | 5.005431 | 0.558606 |
| 6 | 51463800 | 51466799 | 39.00487 | 63.45537 | 68.50905 | 29.40483 | 0.259323 | 1.570414 | 0.558607 |
| 12 | 35778600 | 35782199 | 31.30896 | 52.31957 | 54.13108 | 23.56378 | 0.332152 | 1.215423 | 0.558614 |
| 12 | 21486800 | 21489199 | 20.12031 | 34.3591 | 54.37272 | 23.83856 | -0.2771 | 0.980068 | 0.558616 |
| 6 | 149092600 | 149095199 | 34.62573 | 57.03204 | 60.89194 | 26.26311 | 0.285066 | 1.391747 | 0.55862 |
| 13 | 11268600 | 11271199 | 52.97252 | 84.04162 | 130.2946 | 54.26295 | -0.30056 | 2.313565 | 0.558639 |
| 17 | 83560200 | 83565799 | 80.95005 | 125.0997 | 187.5973 | 77.92374 | -0.28258 | 2.854102 | 0.558656 |
| 16 | 86995200 | 87001399 | 53.61672 | 86.29358 | 84.84015 | 36.03402 | 0.378306 | 1.856587 | 0.558662 |
| 1 | 82130600 | 82133599 | 20.68635 | 36.07965 | 38.18014 | 16.80372 | 0.306111 | 0.74673 | 0.558662 |
| 19 | 40844200 | 40846799 | 20.68635 | 36.07965 | 38.18014 | 16.80372 | 0.306111 | 0.74673 | 0.558662 |
| 10 | 72719200 | 72720999 | 16.5566 | 29.71036 | 30.0942 | 13.16896 | 0.383538 | 0.421537 | 0.558674 |
| 1 | 182738800 | 182740599 | 22.88019 | 39.49741 | 42.03724 | 18.4415 | 0.295736 | 0.879187 | 0.558678 |
| 16 | 71691600 | 71692399 | 8.69751 | 16.3665 | 28.57683 | 11.71284 | -0.32833 | 0.080663 | 0.558679 |
| 6 | 38342600 | 38345199 | 25.78529 | 42.29811 | 70.68722 | 30.43533 | -0.36165 | 1.431214 | 0.558682 |
| 3 | 45053400 | 45057799 | 79.28795 | 125.0339 | 131.3853 | 54.52422 | 0.252444 | 2.478515 | 0.558688 |
| 4 | 70852000 | 70852999 | 10.93793 | 20.95972 | 22.86982 | 9.011084 | 0.394845 | -0.04254 | 0.558705 |
| 5 | 12767800 | 12773599 | 102.2255 | 158.3123 | 165.5004 | 68.74443 | 0.2397 | 2.821809 | 0.558706 |
| 2 | 18416600 | 18418799 | 23.80667 | 40.87738 | 43.63044 | 19.13728 | 0.290032 | 0.931651 | 0.55871 |
| 3 | 21334800 | 21335199 | 9.546375 | 17.60833 | 30.3992 | 12.85518 | -0.34713 | 0.215321 | 0.558714 |
| 17 | 81639000 | 81642399 | 37.64067 | 62.29362 | 58.93347 | 25.48923 | 0.460417 | 1.343606 | 0.558718 |
| 4 | 4554800 | 4556599 | 11.93875 | 21.36051 | 36.19693 | 15.87899 | -0.35701 | 0.42106 | 0.558719 |
| 10 | 119771000 | 119771599 | 6.393113 | 12.37668 | 23.83112 | 9.067691 | -0.36964 | -0.20221 | 0.558724 |
| 5 | 22811600 | 22812599 | 10.00443 | 18.55534 | 31.52831 | 13.23291 | -0.31664 | 0.190449 | 0.558726 |
| 4 | 96827400 | 96832399 | 87.77884 | 137.1934 | 145.7257 | 60.4604 | 0.226986 | 2.653087 | 0.558763 |
| 6 | 136415000 | 136433199 | 261.2063 | 372.8611 | 502.4912 | 207.6588 | -0.23217 | 4.274101 | 0.558772 |
| X | 21908000 | 21911199 | 27.0006 | 46.90731 | 40.15747 | 17.63025 | 0.610196 | 0.835792 | 0.558806 |
| 5 | 145858600 | 145860199 | 42.6493 | 69.15169 | 71.70614 | 30.71082 | 0.313254 | 1.644252 | 0.55882 |
| 3 | 103716200 | 103717399 | 9.520833 | 17.14612 | 32.87825 | 14.33932 | -0.52773 | 0.359426 | 0.558832 |
| 11 | 15755200 | 15758399 | 29.49456 | 49.61523 | 51.28595 | 22.40861 | 0.333398 | 1.140642 | 0.558833 |
| 13 | 22557400 | 22558799 | 14.63936 | 26.80625 | 27.85714 | 11.89761 | 0.369671 | 0.288369 | 0.558842 |
| 3 | 46063600 | 46065199 | 18.73242 | 33.84288 | 28.87646 | 12.54397 | 0.636496 | 0.395169 | 0.558845 |
| 15 | 65470200 | 65470999 | 9.187681 | 17.15496 | 29.50656 | 12.25422 | -0.32211 | 0.137391 | 0.558848 |
| 8 | 20173000 | 20182199 | 208.2706 | 302.9695 | 413.1887 | 170.3049 | -0.22235 | 3.991296 | 0.558858 |
| X | 125925600 | 125932999 | 210.442 | 309.5043 | 304.6531 | 125.4608 | 0.263014 | 3.821415 | 0.55886 |
| 4 | 113403600 | 113410799 | 109.7157 | 171.4808 | 144.1725 | 59.81594 | 0.563932 | 2.666906 | 0.558865 |
| 8 | 46753800 | 46757799 | 30.83455 | 50.12032 | 79.27158 | 34.01357 | -0.29099 | 1.573262 | 0.55887 |
| 14 | 17397800 | 17399399 | 14.21408 | 24.60199 | 45.87153 | 20.12519 | -0.50855 | 0.824193 | 0.5589 |
| 2 | 177001200 | 177002799 | 16.92794 | 29.80823 | 30.95572 | 13.70334 | 0.334222 | 0.574725 | 0.558944 |
| 6 | 82116200 | 82118999 | 36.42951 | 59.6306 | 63.64983 | 27.38041 | 0.281898 | 1.47342 | 0.558948 |
| 12 | 46201400 | 46203199 | 14.98236 | 26.37036 | 46.10892 | 20.41314 | -0.41394 | 0.59633 | 0.558984 |

| | | | | | | | | | |
|---|---|---|---|---|---|---|---|---|---|
| 10 | 62566200 | 62569199 | 42.97069 | 69.44458 | 75.02386 | 32.04115 | 0.249573 | 1.715329 | 0.558985 |
| 2 | 57102600 | 57104599 | 21.79997 | 38.09916 | 33.38425 | 14.7122 | 0.577328 | 0.717087 | 0.559001 |
| X | 43101200 | 43108999 | 137.511 | 210.2946 | 194.5398 | 80.71067 | 0.402065 | 3.077298 | 0.559044 |
| 4 | 83680200 | 83680999 | 10.77169 | 19.35045 | 33.82159 | 14.78652 | -0.39732 | 0.386976 | 0.559046 |
| 9 | 117923800 | 117925199 | 9.842337 | 18.14143 | 30.89652 | 13.07804 | -0.3288 | 0.222205 | 0.559051 |
| 9 | 97946400 | 97947599 | 14.08714 | 25.65665 | 27.45371 | 11.80619 | 0.321425 | 0.315186 | 0.559059 |
| 13 | 56558000 | 56562799 | 63.90449 | 100.557 | 150.1902 | 62.3106 | -0.25656 | 2.516124 | 0.55907 |
| 13 | 82271000 | 82273999 | 27.91013 | 48.10609 | 42.33954 | 18.57324 | 0.56824 | 0.913456 | 0.559081 |
| 11 | 29323200 | 29324199 | 9.842996 | 18.05976 | 31.35877 | 13.35665 | -0.36316 | 0.248371 | 0.559085 |
| 18 | 38349400 | 38350599 | 9.842996 | 18.05976 | 31.35877 | 13.35665 | -0.36316 | 0.248371 | 0.559085 |
| 6 | 68019200 | 68020599 | 19.76792 | 34.74139 | 29.82922 | 13.20774 | 0.606383 | 0.612756 | 0.559091 |
| X | 8396200 | 8419999 | 326.8754 | 470.9315 | 377.038 | 155.5902 | 0.532108 | 4.144174 | 0.5591 |
| 19 | 21754800 | 21756199 | 14.27945 | 24.86575 | 43.7089 | 19.21785 | -0.42272 | 0.746298 | 0.559117 |
| 2 | 58119000 | 58122199 | 28.93961 | 47.51417 | 74.30038 | 32.00292 | -0.26575 | 1.427562 | 0.559126 |
| 17 | 23593800 | 23595999 | 34.63618 | 56.11629 | 86.20971 | 36.90171 | -0.25355 | 1.651344 | 0.559141 |
| 7 | 42422000 | 42425799 | 77.79635 | 122.3532 | 129.8948 | 53.88687 | 0.234705 | 2.531459 | 0.559155 |
| 12 | 114161400 | 114164399 | 28.95917 | 49.80432 | 43.47533 | 19.0702 | 0.579545 | 0.940375 | 0.559157 |
| 7 | 56259000 | 56261399 | 26.39188 | 44.63547 | 47.99242 | 21.0038 | 0.277068 | 1.081678 | 0.559177 |
| 4 | 74592600 | 74594799 | 30.96558 | 50.41736 | 78.97021 | 33.90487 | -0.27532 | 1.548929 | 0.559204 |
| 2 | 51761400 | 51762199 | 6.416263 | 12.35423 | 24.1725 | 9.394087 | -0.42126 | -0.12881 | 0.559211 |
| 17 | 88046200 | 88059399 | 134.7132 | 201.825 | 290.7366 | 119.9298 | -0.26141 | 3.488405 | 0.559212 |
| 14 | 51141400 | 51142999 | 9.723249 | 18.11972 | 31.63775 | 13.18261 | -0.34739 | 0.160942 | 0.55922 |
| 19 | 8110000 | 8120399 | 181.8498 | 270.4186 | 277.8728 | 114.4825 | 0.215838 | 3.631191 | 0.55924 |
| 6 | 60575000 | 60576599 | 27.86515 | 47.07917 | 41.6805 | 18.23576 | 0.556304 | 1.087797 | 0.559244 |
| 1 | 117828200 | 117831399 | 62.46859 | 99.61036 | 105.3812 | 44.15579 | 0.257375 | 2.159393 | 0.559261 |
| 8 | 84226600 | 84244399 | 282.4432 | 408.1166 | 384.4675 | 158.5944 | 0.301686 | 4.096618 | 0.559302 |
| 5 | 103511600 | 103514199 | 18.21243 | 30.91469 | 54.2811 | 23.78237 | -0.42767 | 1.025664 | 0.559306 |
| 13 | 113450400 | 113451199 | 15.94837 | 28.49204 | 30.21588 | 13.27183 | 0.314088 | 0.44556 | 0.559337 |
| 2 | 127879400 | 127882399 | 32.33736 | 54.17272 | 46.98953 | 20.48713 | 0.585538 | 1.185603 | 0.559342 |
| 7 | 7022000 | 7029799 | 161.3031 | 241.7474 | 249.7068 | 103.0119 | 0.218604 | 3.484647 | 0.559371 |
| 14 | 97960800 | 97962799 | 18.06059 | 31.83721 | 33.36744 | 14.76775 | 0.321748 | 0.575686 | 0.559375 |
| 10 | 38716200 | 38718399 | 26.53028 | 45.69619 | 42.9352 | 18.83415 | 0.474088 | 0.91379 | 0.559381 |
| 3 | 63836400 | 63837599 | 9.233828 | 16.96453 | 30.87676 | 13.12766 | -0.42865 | 0.234812 | 0.559388 |
| 8 | 92454400 | 92474999 | 268.8269 | 381.8083 | 512.4012 | 211.9338 | -0.23347 | 4.348213 | 0.559413 |
| 14 | 3716000 | 3728799 | 164.7398 | 249.2064 | 226.8054 | 93.80921 | 0.41283 | 3.303994 | 0.559414 |
| 13 | 70638200 | 70638799 | 6.409595 | 12.45271 | 25.26927 | 9.428149 | -0.41724 | -0.26151 | 0.559427 |
| X | 94742600 | 94744599 | 19.36872 | 34.19985 | 33.67099 | 14.89716 | 0.412344 | 0.565679 | 0.559432 |
| 8 | 3355000 | 3358799 | 73.47688 | 116.4558 | 108.3855 | 45.09351 | 0.433116 | 2.352793 | 0.559436 |
| 4 | 146354800 | 146368999 | 264.2467 | 385.0602 | 352.496 | 145.263 | 0.354898 | 3.969781 | 0.559464 |
| 13 | 13483200 | 13485599 | 28.35207 | 48.01656 | 49.05362 | 21.47717 | 0.350628 | 1.077547 | 0.559492 |
| 8 | 43164800 | 43168799 | 117.8506 | 180.2438 | 189.6187 | 78.67062 | 0.216813 | 3.067077 | 0.5595 |
| 5 | 137935400 | 137936799 | 11.8539 | 21.14259 | 35.23507 | 15.49777 | -0.33539 | 0.440709 | 0.559521 |
| 3 | 98308800 | 98316199 | 70.24725 | 109.9438 | 163.8877 | 67.98602 | -0.25978 | 2.622346 | 0.559532 |
| 14 | 24578200 | 24579999 | 17.78723 | 31.33695 | 33.32217 | 14.74945 | 0.300517 | 0.578817 | 0.559533 |
| 8 | 13963600 | 13965599 | 12.52761 | 22.11851 | 37.30847 | 16.50677 | -0.36013 | 0.532778 | 0.55954 |
| 10 | 92637200 | 92640199 | 22.09223 | 38.34661 | 40.43277 | 17.74136 | 0.310764 | 0.825307 | 0.559548 |
| 1 | 178385000 | 178388599 | 43.8494 | 70.91121 | 75.81327 | 32.40574 | 0.265684 | 1.699251 | 0.559604 |

| | | | | | | | | | |
|---|---|---|---|---|---|---|---|---|---|
| 7 | 60763600 | 60767799 | 48.44651 | 78.37323 | 76.10029 | 32.53828 | 0.404755 | 1.699379 | 0.559645 |
| X | 21756200 | 21757999 | 27.00676 | 46.39053 | 44.11007 | 19.34792 | 0.456361 | 0.94439 | 0.559646 |
| 9 | 26556000 | 26558399 | 17.31869 | 30.67091 | 32.20497 | 14.2401 | 0.321478 | 0.517901 | 0.559647 |
| 3 | 16488600 | 16489199 | 15.67762 | 27.92713 | 29.83509 | 13.13668 | 0.301267 | 0.461874 | 0.55966 |
| 3 | 31765000 | 31766199 | 15.67762 | 27.92713 | 29.83509 | 13.13668 | 0.301267 | 0.461874 | 0.55966 |
| 7 | 85671600 | 85672399 | 14.27921 | 25.94378 | 27.46536 | 11.84691 | 0.334042 | 0.33082 | 0.559661 |
| 12 | 106993000 | 106994799 | 13.95613 | 24.35534 | 42.6359 | 18.77158 | -0.41645 | 0.712135 | 0.559699 |
| 10 | 74589800 | 74591999 | 22.53916 | 39.2868 | 39.1527 | 17.22503 | 0.392236 | 0.7663 | 0.559716 |
| 19 | 23848200 | 23850799 | 24.57426 | 40.98599 | 65.45643 | 28.45904 | -0.29213 | 1.225071 | 0.559717 |
| 13 | 31510400 | 31513199 | 24.86005 | 41.16566 | 65.64506 | 28.40319 | -0.29042 | 1.309765 | 0.559743 |
| 8 | 56289000 | 56294599 | 54.23626 | 87.07993 | 90.33882 | 38.23403 | 0.296714 | 1.935589 | 0.559754 |
| X | 116058800 | 116060199 | 21.09057 | 37.05047 | 35.26466 | 15.60894 | 0.458515 | 0.636614 | 0.559763 |
| 3 | 18045400 | 18045999 | 6.388254 | 12.38138 | 23.90398 | 9.047081 | -0.36623 | -0.22045 | 0.559765 |
| 4 | 51165600 | 51166999 | 17.1747 | 30.30542 | 32.37567 | 14.32599 | 0.294768 | 0.55712 | 0.559765 |
| 3 | 81755600 | 81758199 | 25.86106 | 44.77264 | 41.86606 | 18.36803 | 0.482526 | 0.866417 | 0.559773 |
| 8 | 66041000 | 66043999 | 26.96053 | 44.59063 | 69.84879 | 30.19719 | -0.26499 | 1.331683 | 0.559792 |
| 3 | 38681000 | 38681399 | 5.116919 | 9.957957 | 21.84062 | 8.039265 | -0.5104 | -0.34013 | 0.559793 |
| 12 | 18658000 | 18703199 | 1189.618 | 1517.831 | 1311.163 | 550.2701 | 0.246682 | 6.041351 | 0.559804 |
| 4 | 58685600 | 58688599 | 45.9396 | 74.66059 | 62.7765 | 26.93266 | 0.618021 | 1.597559 | 0.559804 |
| 7 | 3222200 | 3223999 | 29.16767 | 48.54941 | 51.86345 | 22.52053 | 0.286277 | 1.248844 | 0.559833 |
| 10 | 48223600 | 48223999 | 9.00794 | 17.39894 | 20.34417 | 7.824179 | 0.328567 | -0.14795 | 0.559902 |
| 8 | 98722200 | 98722599 | 5.508493 | 10.73042 | 21.35528 | 7.784547 | -0.35793 | -0.36749 | 0.559908 |
| 4 | 70965800 | 70974199 | 106.1431 | 164.3166 | 167.0524 | 69.38925 | 0.279471 | 2.829535 | 0.559921 |
| X | 48070000 | 48074799 | 44.03934 | 72.73968 | 57.46633 | 24.8209 | 0.719581 | 1.383956 | 0.559929 |
| 14 | 121258800 | 121259599 | 9.976942 | 19.05282 | 20.61722 | 8.198879 | 0.394059 | -0.03062 | 0.559931 |
| 1 | 80020600 | 80021999 | 13.62296 | 25.00088 | 26.90253 | 11.45505 | 0.322813 | 0.265756 | 0.559933 |
| 12 | 114135400 | 114137399 | 28.40005 | 47.89348 | 50.16408 | 21.93502 | 0.314576 | 1.115604 | 0.559942 |
| 3 | 108769200 | 108770799 | 13.95265 | 25.47519 | 27.36186 | 11.73004 | 0.318673 | 0.297913 | 0.559945 |
| 2 | 92550000 | 92552999 | 34.00465 | 56.4531 | 57.58081 | 24.95822 | 0.352523 | 1.297478 | 0.559947 |
| X | 65242200 | 65245599 | 61.26667 | 98.03791 | 97.47951 | 41.04801 | 0.352225 | 2.050168 | 0.559953 |
| 2 | 29646800 | 29647799 | 11.44584 | 21.49157 | 23.65527 | 9.731252 | 0.328394 | 0.10718 | 0.559966 |
| 9 | 122258200 | 122259399 | 10.03917 | 18.25147 | 32.30873 | 13.92518 | -0.4024 | 0.304745 | 0.55997 |
| 9 | 110081600 | 110086799 | 58.18475 | 92.83714 | 99.50533 | 41.72963 | 0.239552 | 2.134029 | 0.560014 |
| 1 | 35036000 | 35038599 | 19.03175 | 32.48865 | 52.17015 | 22.87224 | -0.29805 | 0.977954 | 0.560018 |
| 8 | 95346800 | 95347599 | 7.182291 | 13.78339 | 25.74089 | 10.09932 | -0.36743 | -0.08309 | 0.560024 |
| 18 | 23371800 | 23374799 | 40.237 | 65.23848 | 70.39831 | 30.15004 | 0.255844 | 1.637883 | 0.560042 |
| 2 | 85939000 | 85940799 | 23.60372 | 40.79285 | 35.32935 | 15.49932 | 0.592208 | 0.863583 | 0.560053 |
| 5 | 7551200 | 7552199 | 7.867851 | 15.10143 | 29.37979 | 11.5218 | -0.42681 | -0.07342 | 0.560054 |
| 4 | 61514000 | 61515599 | 11.84002 | 22.83754 | 21.89266 | 8.465251 | 0.605082 | -0.09665 | 0.560057 |
| 13 | 61980600 | 62011399 | 756.9858 | 1003.783 | 878.0801 | 367.0342 | 0.284124 | 5.415728 | 0.560061 |
| 1 | 48722400 | 48730399 | 86.08044 | 135.9989 | 127.912 | 53.12618 | 0.414304 | 2.442112 | 0.560069 |
| 14 | 40989400 | 40990799 | 13.15754 | 24.74308 | 24.48228 | 10.04564 | 0.483252 | 0.111818 | 0.560098 |
| 7 | 42358400 | 42359199 | 13.3453 | 25.78122 | 21.36208 | 8.208778 | 0.822468 | -0.10121 | 0.560125 |
| Y | 1746400 | 1748599 | 17.25319 | 30.4417 | 22.7075 | 10.04805 | 0.809867 | 0.536594 | 0.560127 |
| 5 | 128583200 | 128587799 | 53.85262 | 85.66401 | 126.6083 | 52.87275 | -0.22795 | 2.234077 | 0.560142 |
| 15 | 22368600 | 22369799 | 22.03079 | 38.10316 | 40.99392 | 17.98484 | 0.280751 | 0.857055 | 0.560145 |
| 1 | 79081200 | 79082999 | 14.95826 | 27.56804 | 27.37342 | 11.58852 | 0.444026 | 0.254008 | 0.560149 |

| | | | | | | | | | |
|---|---|---|---|---|---|---|---|---|---|
| 6 | 149135800 | 149137999 | 27.66021 | 46.70942 | 48.53351 | 21.2308 | 0.326036 | 1.096543 | 0.560165 |
| 2 | 98529200 | 98529999 | 8.031721 | 15.68016 | 17.56356 | 6.202994 | 0.502475 | -0.46001 | 0.560181 |
| 9 | 26527400 | 26528799 | 28.85263 | 48.54126 | 43.7858 | 19.13224 | 0.529229 | 1.129365 | 0.560188 |
| 15 | 73373600 | 73375599 | 19.92292 | 34.797 | 36.55698 | 16.10355 | 0.316606 | 0.719492 | 0.560207 |
| 6 | 43168400 | 43170599 | 13.35858 | 25.34646 | 23.57507 | 9.504066 | 0.593538 | 0.053265 | 0.56022 |
| 6 | 39862800 | 39863399 | 11.95772 | 22.88375 | 23.37418 | 9.244691 | 0.484327 | -0.0188 | 0.56023 |
| 2 | 166249400 | 166251799 | 24.49252 | 40.71908 | 64.11342 | 27.81695 | -0.27171 | 1.260943 | 0.560235 |
| 12 | 20304400 | 20358599 | 1987.135 | 2423.434 | 2031.525 | 856.9657 | 0.224045 | 6.765958 | 0.560236 |
| 3 | 103611000 | 103615199 | 24.08205 | 39.80488 | 74.65299 | 32.26018 | -0.52404 | 1.334447 | 0.560241 |
| 10 | 23825800 | 23826999 | 9.563203 | 17.67681 | 30.43007 | 12.83116 | -0.33988 | 0.204457 | 0.560267 |
| X | 26283200 | 26287799 | 30.90557 | 52.49423 | 40.96781 | 17.95286 | 0.737905 | 1.055486 | 0.560313 |
| 17 | 53678600 | 53680599 | 22.96881 | 39.35132 | 42.12991 | 18.47581 | 0.285201 | 0.946273 | 0.560316 |
| 5 | 122252400 | 122255599 | 30.79952 | 49.74897 | 85.73709 | 36.592 | -0.42338 | 1.717273 | 0.560349 |
| 10 | 63224600 | 63235399 | 91.83199 | 141.7597 | 205.0027 | 85.15405 | -0.23153 | 2.880482 | 0.560355 |
| 10 | 57633800 | 57636599 | 21.90543 | 38.32121 | 37.4425 | 16.51439 | 0.420915 | 0.708324 | 0.56037 |
| 9 | 3299600 | 3299999 | 10.4624 | 19.85978 | 22.22234 | 8.950611 | 0.330655 | 0.022239 | 0.560374 |
| 13 | 34667200 | 34668999 | 19.35038 | 34.24541 | 32.43208 | 14.34249 | 0.469097 | 0.537001 | 0.560374 |
| 5 | 139882000 | 139884599 | 20.35608 | 34.34006 | 55.89123 | 24.44187 | -0.319 | 1.090997 | 0.560377 |
| 8 | 61450000 | 61452199 | 29.03266 | 49.30325 | 41.57368 | 18.21726 | 0.626647 | 1.050489 | 0.56038 |
| 4 | 24474000 | 24481799 | 77.11003 | 122.4048 | 116.9976 | 48.75262 | 0.396919 | 2.31199 | 0.560382 |
| 6 | 118806000 | 118807999 | 18.00221 | 31.54639 | 32.54649 | 14.37553 | 0.342694 | 0.651545 | 0.560384 |
| 6 | 31683800 | 31684999 | 8.709334 | 16.35211 | 28.53538 | 11.74027 | -0.33185 | 0.095271 | 0.560385 |
| 8 | 108596200 | 108596999 | 8.709334 | 16.35211 | 28.53538 | 11.74027 | -0.33185 | 0.095271 | 0.560385 |
| 9 | 15281600 | 15284399 | 58.70971 | 93.62082 | 98.45957 | 41.25812 | 0.265997 | 2.153085 | 0.560401 |
| 15 | 97230200 | 97235399 | 61.67822 | 97.26739 | 146.4592 | 60.78008 | -0.2658 | 2.469403 | 0.560405 |
| X | 15719200 | 15722599 | 55.78931 | 89.7716 | 84.33575 | 35.80899 | 0.443476 | 1.870253 | 0.56041 |
| 8 | 11323600 | 11324599 | 9.068335 | 17.00555 | 29.59701 | 12.20204 | -0.33026 | 0.110418 | 0.560421 |
| 12 | 118308200 | 118312199 | 39.21327 | 63.24463 | 97.38953 | 41.48507 | -0.26401 | 1.778734 | 0.560432 |
| 7 | 17210400 | 17213399 | 34.92018 | 57.38824 | 61.81656 | 26.63663 | 0.271126 | 1.422495 | 0.560455 |
| 1 | 128921800 | 128922799 | 8.22135 | 15.65626 | 27.78663 | 11.11562 | -0.32113 | -0.00676 | 0.56048 |
| 6 | 133015800 | 133016799 | 8.22135 | 15.65626 | 27.78663 | 11.11562 | -0.32113 | -0.00676 | 0.56048 |
| 8 | 42786200 | 42788199 | 14.2562 | 24.98036 | 40.95545 | 18.08869 | -0.32213 | 0.647981 | 0.560497 |
| 5 | 89720400 | 89721799 | 12.61276 | 23.8987 | 24.17167 | 9.772754 | 0.47006 | 0.057635 | 0.560521 |
| 8 | 19344600 | 19345599 | 11.68628 | 22.41486 | 22.864 | 8.984435 | 0.4947 | -0.04462 | 0.560553 |
| 2 | 121217000 | 121218799 | 22.00453 | 37.99287 | 41.01417 | 17.99279 | 0.27537 | 0.871769 | 0.560585 |
| 1 | 48742400 | 48745199 | 29.71014 | 50.16861 | 50.17061 | 21.94661 | 0.381165 | 1.108682 | 0.560592 |
| 13 | 104083200 | 104088199 | 83.14592 | 130.2594 | 139.408 | 57.82519 | 0.218763 | 2.609112 | 0.560604 |
| Y | 9748200 | 9750199 | 16.09957 | 28.64216 | 21.14713 | 9.321325 | 0.829843 | 0.451396 | 0.560613 |
| 1 | 111362800 | 111363799 | 20.18492 | 35.13222 | 37.8016 | 16.61546 | 0.281966 | 0.772849 | 0.560624 |
| 3 | 78924000 | 78926799 | 29.98477 | 49.92214 | 54.07561 | 23.48488 | 0.266276 | 1.250153 | 0.560629 |
| 11 | 92216400 | 92217199 | 13.29428 | 24.52706 | 26.44669 | 11.1802 | 0.327322 | 0.232841 | 0.560646 |
| 4 | 110624000 | 110625199 | 19.96691 | 33.97333 | 53.79248 | 23.57726 | -0.27868 | 1.011965 | 0.560652 |
| 12 | 19715800 | 19745999 | 1225.324 | 1553.582 | 1344.261 | 564.5137 | 0.235162 | 6.143115 | 0.560653 |
| 13 | 49958200 | 49961199 | 29.55631 | 48.00303 | 80.14635 | 34.36808 | -0.37 | 1.587067 | 0.560654 |
| 11 | 88766400 | 88769599 | 46.0166 | 73.66279 | 111.4834 | 46.97008 | -0.25215 | 2.025966 | 0.56066 |
| 5 | 115226000 | 115228599 | 21.50099 | 36.07119 | 58.6118 | 25.57761 | -0.31693 | 1.147704 | 0.560696 |
| 10 | 74479200 | 74480199 | 9.624104 | 17.90973 | 31.05137 | 12.96916 | -0.34047 | 0.164666 | 0.560699 |

| | | | | | | | | | |
|---|---|---|---|---|---|---|---|---|---|
| 18 | 68212000 | 68215199 | 46.23325 | 74.71822 | 69.05915 | 29.49428 | 0.474199 | 1.717995 | 0.560707 |
| 5 | 122668800 | 122670399 | 21.13684 | 36.63554 | 39.42107 | 17.29593 | 0.28111 | 0.831017 | 0.560745 |
| 7 | 85997200 | 85998799 | 32.59618 | 54.04066 | 57.41629 | 24.86337 | 0.293717 | 1.310227 | 0.560769 |
| 4 | 21450000 | 21451599 | 12.07515 | 21.67114 | 36.73062 | 16.07135 | -0.35478 | 0.403413 | 0.560783 |
| 3 | 76033000 | 76037999 | 42.90307 | 70.40081 | 62.63422 | 26.97169 | 0.545362 | 1.458568 | 0.560795 |
| 10 | 3791800 | 3792799 | 9.077707 | 16.87984 | 29.45679 | 12.31931 | -0.35069 | 0.164242 | 0.560806 |
| 7 | 66795600 | 66805199 | 75.82298 | 117.3326 | 204.293 | 84.85821 | -0.49637 | 2.828506 | 0.560826 |
| 10 | 98801400 | 98801999 | 6.244496 | 12.14366 | 23.60356 | 8.738352 | -0.34562 | -0.2979 | 0.560836 |
| 5 | 34326600 | 34327199 | 5.545912 | 10.74541 | 22.70518 | 8.606035 | -0.49653 | -0.22787 | 0.560846 |
| 18 | 65869200 | 65871399 | 18.68234 | 31.88337 | 51.63943 | 22.63906 | -0.31046 | 0.977618 | 0.560875 |
| 9 | 117070200 | 117071599 | 29.59673 | 49.41178 | 52.98828 | 23.05261 | 0.280608 | 1.218157 | 0.560875 |
| 10 | 100151600 | 100153599 | 19.47541 | 34.4561 | 33.03905 | 14.61181 | 0.45125 | 0.543428 | 0.560883 |
| 8 | 48957800 | 48959199 | 19.66126 | 33.32933 | 54.30255 | 23.78211 | -0.32022 | 1.043919 | 0.560889 |
| 6 | 146612000 | 146613999 | 26.04242 | 43.93397 | 46.96124 | 20.53685 | 0.285406 | 1.10075 | 0.560917 |
| 5 | 109012400 | 109013799 | 20.08104 | 33.87624 | 55.8442 | 24.42133 | -0.33725 | 1.091593 | 0.560957 |
| 17 | 29165400 | 29167599 | 19.16899 | 32.17355 | 63.3618 | 27.65841 | -0.5933 | 1.140607 | 0.560968 |
| 1 | 33654000 | 33656999 | 21.52967 | 37.16832 | 39.52112 | 17.33772 | 0.297139 | 0.869355 | 0.560982 |
| 17 | 67445000 | 67446599 | 9.866453 | 17.98031 | 32.32354 | 13.89005 | -0.4218 | 0.288986 | 0.560987 |
| 6 | 39433800 | 39438599 | 64.59768 | 101.584 | 151.6003 | 62.89051 | -0.25608 | 2.529737 | 0.561013 |
| 15 | 34305200 | 34307399 | 14.35359 | 25.12915 | 43.53697 | 19.21474 | -0.40164 | 0.663908 | 0.561047 |
| 18 | 28928600 | 28929399 | 11.56228 | 21.70237 | 24.13031 | 9.932749 | 0.313011 | 0.110883 | 0.561072 |
| 4 | 4901600 | 4903199 | 28.26169 | 47.72319 | 42.21421 | 18.46622 | 0.557561 | 1.095254 | 0.561079 |
| 13 | 33433400 | 33435199 | 19.46034 | 34.22767 | 34.75034 | 15.38671 | 0.366037 | 0.609223 | 0.561081 |
| 3 | 51957000 | 51959999 | 27.74657 | 46.93527 | 49.06071 | 21.47254 | 0.317654 | 1.08631 | 0.561087 |
| 15 | 95679600 | 95686199 | 61.4018 | 96.87745 | 165.7227 | 68.78303 | -0.44912 | 2.463042 | 0.561103 |
| 7 | 15964000 | 15969799 | 58.11274 | 92.89338 | 99.11987 | 41.67669 | 0.248893 | 2.076737 | 0.561105 |
| 6 | 146199000 | 146200399 | 13.18378 | 23.50114 | 39.83448 | 17.52921 | -0.36037 | 0.462461 | 0.561123 |
| 18 | 63602600 | 63603999 | 16.90084 | 30.81695 | 28.73564 | 12.33955 | 0.520004 | 0.331609 | 0.561143 |
| X | 125950800 | 125953599 | 25.03068 | 43.56548 | 39.60016 | 17.40579 | 0.524231 | 0.799223 | 0.56115 |
| 6 | 10312800 | 10314399 | 17.1408 | 29.78158 | 47.38751 | 20.80593 | -0.28023 | 0.797546 | 0.561152 |
| 13 | 68739200 | 68741399 | 17.1408 | 29.78158 | 47.38751 | 20.80593 | -0.28023 | 0.797546 | 0.561152 |
| 11 | 14938400 | 14939199 | 11.38356 | 21.6125 | 23.74347 | 9.559376 | 0.357154 | 0.032595 | 0.561157 |
| 1 | 89330800 | 89336999 | 66.44136 | 104.7438 | 153.3636 | 63.64261 | -0.22587 | 2.480497 | 0.561159 |
| 11 | 31039000 | 31040999 | 12.28027 | 21.92129 | 36.18605 | 15.90489 | -0.32125 | 0.438575 | 0.561167 |
| 16 | 68695400 | 68696199 | 6.713887 | 13.01844 | 25.68748 | 9.701406 | -0.39387 | -0.21517 | 0.561178 |
| 9 | 32262600 | 32269999 | 58.29667 | 91.09582 | 177.9449 | 73.82942 | -0.65105 | 2.647442 | 0.561183 |
| 3 | 16142600 | 16143199 | 6.912067 | 13.11113 | 26.06252 | 10.51324 | -0.49306 | 0.005313 | 0.561186 |
| 16 | 21912600 | 21922599 | 93.72561 | 143.9645 | 206.9534 | 85.92852 | -0.22841 | 2.976006 | 0.561194 |
| 18 | 18435800 | 18444599 | 159.7135 | 240.1886 | 246.7366 | 101.8768 | 0.230928 | 3.418378 | 0.561207 |
| 6 | 52133600 | 52138199 | 28.04364 | 45.41933 | 90.17697 | 38.58988 | -0.62306 | 1.648042 | 0.561221 |
| 11 | 15229200 | 15229799 | 12.30826 | 22.72388 | 24.01577 | 10.14285 | 0.357179 | 0.215516 | 0.56124 |
| 14 | 43822200 | 43823799 | 29.00051 | 48.75003 | 43.79067 | 19.1296 | 0.535192 | 1.136665 | 0.561248 |
| 2 | 47687600 | 47688999 | 18.64656 | 32.79982 | 33.68808 | 14.91604 | 0.349339 | 0.599709 | 0.561249 |
| X | 29286800 | 29288399 | 16.47003 | 30.34761 | 27.25896 | 11.54349 | 0.587454 | 0.266819 | 0.561299 |
| 4 | 83874600 | 83877999 | 45.10339 | 71.80512 | 130.298 | 54.47514 | -0.52219 | 2.201704 | 0.561308 |
| X | 126551000 | 126552599 | 23.71889 | 41.60935 | 35.60628 | 15.74149 | 0.611309 | 0.672842 | 0.561316 |
| X | 32346000 | 32364199 | 220.641 | 328.2074 | 274.3549 | 113.0368 | 0.513896 | 3.633371 | 0.561343 |

| | | | | | | | | | |
|---|---|---|---|---|---|---|---|---|---|
| 7 | 129358600 | 129360199 | 17.6948 | 31.15498 | 32.7736 | 14.50841 | 0.315549 | 0.582761 | 0.561344 |
| 14 | 74994200 | 74996399 | 11.02623 | 19.95937 | 33.28951 | 14.42581 | -0.322 | 0.341201 | 0.561347 |
| 3 | 21024400 | 21024999 | 10.15896 | 19.34315 | 21.24643 | 8.502084 | 0.365584 | -0.00384 | 0.561353 |
| 7 | 98911000 | 98911599 | 9.002086 | 17.40695 | 20.36981 | 7.79349 | 0.33442 | -0.16558 | 0.56136 |
| 10 | 121882600 | 121884599 | 18.22568 | 31.06556 | 51.62283 | 22.62953 | -0.34757 | 0.987582 | 0.561403 |
| 3 | 32664800 | 32667599 | 24.30747 | 40.33456 | 65.23479 | 28.2663 | -0.31056 | 1.281742 | 0.561412 |
| 3 | 19410000 | 19410799 | 11.92256 | 21.06144 | 36.86484 | 16.30876 | -0.41276 | 0.523292 | 0.561426 |
| 17 | 53746200 | 53748599 | 22.4 | 38.95231 | 39.73463 | 17.45517 | 0.358377 | 0.797667 | 0.561426 |
| 8 | 25749800 | 25752399 | 46.54732 | 74.30272 | 115.0862 | 48.29488 | -0.28994 | 2.114414 | 0.561459 |
| 7 | 88648600 | 88649199 | 8.990222 | 16.62979 | 30.47653 | 12.83804 | -0.42805 | 0.198025 | 0.561482 |
| 11 | 5411800 | 5415799 | 62.11447 | 98.23301 | 143.9288 | 59.80872 | -0.22321 | 2.403655 | 0.561487 |
| 17 | 90244800 | 90245599 | 13.41996 | 24.93879 | 20.94697 | 8.768447 | 0.695651 | 0.181852 | 0.561507 |
| 5 | 149732200 | 149734999 | 25.29401 | 42.9759 | 46.35199 | 20.31375 | 0.272778 | 1.035142 | 0.561523 |
| 4 | 118212000 | 118214199 | 19.4997 | 34.20685 | 35.32667 | 15.61753 | 0.341166 | 0.647322 | 0.561535 |
| 6 | 13527000 | 13530399 | 95.71121 | 148.4166 | 157.769 | 65.52766 | 0.217063 | 2.794541 | 0.561541 |
| 8 | 101200400 | 101201799 | 14.81841 | 27.45072 | 26.93052 | 11.32174 | 0.468525 | 0.224064 | 0.561561 |
| 7 | 46935800 | 46945399 | 312.3963 | 443.9051 | 435.7875 | 180.2309 | 0.218128 | 4.342491 | 0.561579 |
| 13 | 40947600 | 40951399 | 30.29441 | 50.54225 | 54.10207 | 23.5271 | 0.283257 | 1.228936 | 0.561609 |
| 19 | 7844800 | 7846199 | 20.08209 | 34.24525 | 54.37555 | 23.83708 | -0.2823 | 0.991724 | 0.561643 |
| 2 | 35132800 | 35134599 | 16.53879 | 29.31381 | 31.30544 | 13.84008 | 0.297719 | 0.503128 | 0.561647 |
| 12 | 92809600 | 92811799 | 19.93286 | 35.3837 | 31.85574 | 14.06765 | 0.543826 | 0.513862 | 0.561655 |
| 10 | 91649800 | 91653199 | 44.50464 | 71.82689 | 77.71671 | 33.13367 | 0.245156 | 1.758801 | 0.56167 |
| 11 | 113425200 | 113427399 | 17.44252 | 29.92055 | 50.04168 | 21.94766 | -0.35527 | 0.926033 | 0.561695 |
| 10 | 118479600 | 118480799 | 9.740145 | 17.95218 | 30.9229 | 13.09009 | -0.34481 | 0.223688 | 0.561698 |
| 5 | 100091000 | 100096599 | 65.16559 | 102.223 | 170.9034 | 70.8798 | -0.42253 | 2.581686 | 0.5617 |
| 3 | 9373800 | 9375599 | 17.99667 | 31.58303 | 33.55244 | 14.83826 | 0.300764 | 0.631703 | 0.561719 |
| 10 | 16843200 | 16844999 | 18.27691 | 32.89349 | 29.96266 | 13.07842 | 0.538354 | 0.415963 | 0.561721 |
| 14 | 95922000 | 95923399 | 14.77769 | 26.4446 | 26.46299 | 11.60712 | 0.398847 | 0.419867 | 0.561721 |
| 7 | 24097000 | 24099399 | 17.18928 | 29.42818 | 51.11339 | 22.41415 | -0.41025 | 0.939837 | 0.561726 |
| X | 55256800 | 55259199 | 23.89942 | 40.0761 | 67.60331 | 29.49268 | -0.37084 | 1.171209 | 0.561727 |
| 12 | 77204000 | 77206999 | 16.52343 | 29.56599 | 24.7911 | 10.87461 | 0.652394 | 0.429167 | 0.561736 |
| 3 | 5275400 | 5276799 | 13.14308 | 23.2339 | 37.73847 | 16.69289 | -0.30539 | 0.52336 | 0.561776 |
| 1 | 132146200 | 132147599 | 13.93108 | 24.47554 | 39.83536 | 17.63348 | -0.31163 | 0.60127 | 0.561828 |
| 17 | 91117800 | 91119199 | 18.38012 | 32.82578 | 30.94452 | 13.59557 | 0.482714 | 0.464641 | 0.561829 |
| X | 52943000 | 52947199 | 68.2884 | 108.9317 | 104.1406 | 43.65883 | 0.403884 | 2.154905 | 0.56184 |
| 3 | 12002800 | 12005599 | 35.27632 | 58.74455 | 55.92029 | 24.28926 | 0.452129 | 1.267072 | 0.561847 |
| 1 | 16210400 | 16211399 | 12.03537 | 23.1622 | 22.40089 | 8.717872 | 0.583904 | -0.07269 | 0.561849 |
| 18 | 21197800 | 21200799 | 29.97658 | 50.61291 | 50.17852 | 21.94934 | 0.393579 | 1.109714 | 0.561849 |
| 4 | 15735400 | 15737199 | 28.54589 | 47.94562 | 51.26716 | 22.38885 | 0.284669 | 1.151514 | 0.561856 |
| X | 10807000 | 10808599 | 16.37678 | 30.10187 | 27.80571 | 11.81438 | 0.543996 | 0.281264 | 0.56186 |
| 8 | 107249400 | 107250599 | 12.64495 | 23.36682 | 25.48276 | 10.74905 | 0.313402 | 0.216558 | 0.561863 |
| 9 | 62310400 | 62311599 | 12.64495 | 23.36682 | 25.48276 | 10.74905 | 0.313402 | 0.216558 | 0.561863 |
| 10 | 26965600 | 26968999 | 21.91884 | 36.64807 | 68.54073 | 29.84676 | -0.51916 | 1.195379 | 0.561865 |
| 18 | 33169800 | 33170999 | 12.33483 | 22.8638 | 25.0196 | 10.50971 | 0.312871 | 0.196157 | 0.561876 |
| 3 | 35807800 | 35808999 | 11.69135 | 20.93233 | 34.74309 | 15.23233 | -0.3263 | 0.414974 | 0.561887 |
| 9 | 114828400 | 114830799 | 24.63079 | 41.03368 | 64.21126 | 27.89768 | -0.26275 | 1.237984 | 0.561901 |
| 15 | 22381200 | 22384599 | 39.87378 | 65.35085 | 65.06561 | 28.00436 | 0.382812 | 1.469116 | 0.561902 |

| | | | | | | | | | |
|---|---|---|---|---|---|---|---|---|---|
| 9 | 61673600 | 61675999 | 22.06758 | 38.03459 | 40.56377 | 17.79424 | 0.292376 | 0.888241 | 0.561916 |
| 10 | 50553600 | 50555799 | 15.96973 | 29.15884 | 24.68334 | 10.58139 | 0.660201 | 0.308517 | 0.561919 |
| 13 | 53175000 | 53176799 | 16.83222 | 29.03599 | 47.94095 | 21.03096 | -0.33532 | 0.866521 | 0.561921 |
| 8 | 8138600 | 8140199 | 18.90137 | 34.04924 | 29.98958 | 13.07484 | 0.587818 | 0.4161 | 0.561931 |
| 4 | 61445400 | 61454199 | 70.99779 | 113.0627 | 109.1479 | 45.65137 | 0.387239 | 2.209287 | 0.561934 |
| 7 | 104282000 | 104282599 | 8.7076 | 16.45514 | 28.68022 | 11.66134 | -0.3159 | 0.053567 | 0.561939 |
| X | 76237800 | 76240199 | 33.09853 | 55.33561 | 54.15 | 23.57888 | 0.412291 | 1.217849 | 0.56194 |
| 3 | 7277800 | 7283199 | 78.68985 | 124.1049 | 131.5348 | 54.58708 | 0.240191 | 2.47654 | 0.561943 |
| 1 | 38947200 | 38948199 | 9.63739 | 17.80903 | 31.45343 | 13.26764 | -0.37757 | 0.20548 | 0.561947 |
| 11 | 50447200 | 50448799 | 28.05587 | 47.29087 | 50.16413 | 21.93193 | 0.296342 | 1.118238 | 0.561969 |
| 11 | 56982600 | 56984799 | 21.11647 | 36.66185 | 39.37003 | 17.27769 | 0.28441 | 0.813964 | 0.561997 |
| 9 | 29960000 | 29961999 | 23.21831 | 40.43405 | 39.08698 | 17.18636 | 0.435782 | 0.780438 | 0.562015 |
| 17 | 33401000 | 33401399 | 7.680521 | 14.9372 | 17.86185 | 6.609461 | 0.349008 | -0.30139 | 0.562016 |
| 16 | 87436800 | 87437599 | 9.144511 | 17.11026 | 30.09207 | 12.453 | -0.35016 | 0.123272 | 0.56202 |
| 10 | 114489400 | 114491399 | 18.13831 | 32.18129 | 31.427 | 13.88443 | 0.426546 | 0.503396 | 0.562031 |
| 8 | 47518200 | 47519999 | 14.85111 | 25.91519 | 44.14887 | 19.43676 | -0.37761 | 0.718546 | 0.562036 |
| 10 | 125069800 | 125071999 | 18.60041 | 31.90804 | 51.1381 | 22.42861 | -0.29428 | 0.927918 | 0.562042 |
| 12 | 52208200 | 52211199 | 11.5809 | 20.64389 | 41.45776 | 18.24353 | -0.60364 | 0.454049 | 0.562046 |
| 11 | 4936600 | 4937999 | 7.858893 | 14.74212 | 28.028 | 11.54908 | -0.45547 | 0.095584 | 0.562059 |
| 10 | 4786200 | 4788599 | 19.26961 | 32.54199 | 61.28973 | 26.81138 | -0.52883 | 1.083476 | 0.562104 |
| 1 | 47194200 | 47195799 | 26.86084 | 45.47928 | 48.51541 | 21.23986 | 0.288437 | 1.073901 | 0.562104 |
| 10 | 106540800 | 106541399 | 5.3803 | 10.47689 | 21.3186 | 7.808337 | -0.39644 | -0.35464 | 0.562168 |
| 5 | 111816000 | 111819799 | 39.87021 | 64.12057 | 97.44117 | 41.34056 | -0.24998 | 1.867332 | 0.562193 |
| 7 | 22875000 | 22875999 | 17.55304 | 31.1194 | 27.20403 | 12.02624 | 0.585045 | 0.498282 | 0.562207 |
| 13 | 120693200 | 120700199 | 76.87052 | 119.6029 | 175.6836 | 72.9316 | -0.24522 | 2.730261 | 0.562217 |
| 9 | 30744000 | 30746599 | 17.74646 | 30.65952 | 49.54473 | 21.73552 | -0.30411 | 0.857936 | 0.562223 |
| 8 | 10591000 | 10595399 | 42.79167 | 68.79379 | 106.2265 | 45.04395 | -0.27371 | 1.885319 | 0.562235 |
| 13 | 35923600 | 35926399 | 25.38666 | 41.85428 | 67.71392 | 29.21727 | -0.31213 | 1.364348 | 0.562238 |
| 14 | 38038400 | 38039999 | 21.36286 | 36.87687 | 37.92533 | 16.63761 | 0.344889 | 0.865116 | 0.562239 |
| 15 | 12526800 | 12528999 | 34.83597 | 57.23124 | 60.0049 | 25.85244 | 0.309843 | 1.426658 | 0.562243 |
| 19 | 21687600 | 21689199 | 14.56734 | 25.67311 | 42.35984 | 18.74851 | -0.32994 | 0.572157 | 0.562244 |
| 7 | 38064600 | 38079799 | 211.4527 | 312.4617 | 307.2295 | 126.524 | 0.270967 | 3.743996 | 0.562244 |
| 11 | 43686200 | 43687399 | 10.0517 | 18.47142 | 31.39944 | 13.34567 | -0.33021 | 0.243641 | 0.562246 |
| 2 | 41126400 | 41126999 | 8.432942 | 15.87641 | 28.05195 | 11.4882 | -0.34392 | 0.075836 | 0.562258 |
| 2 | 80840600 | 80841199 | 8.432942 | 15.87641 | 28.05195 | 11.4882 | -0.34392 | 0.075836 | 0.562258 |
| X | 154405000 | 154409199 | 45.54391 | 73.7777 | 72.41734 | 30.99456 | 0.391052 | 1.669251 | 0.562261 |
| 1 | 158876400 | 158877399 | 8.474308 | 15.74238 | 28.95437 | 12.12636 | -0.42731 | 0.163887 | 0.562266 |
| 4 | 54824600 | 54827399 | 24.1502 | 42.20258 | 37.2954 | 16.4345 | 0.565114 | 0.735864 | 0.562272 |
| 17 | 84481800 | 84485599 | 42.39431 | 67.76602 | 109.1617 | 45.89758 | -0.34425 | 2.064259 | 0.562302 |
| 7 | 63039800 | 63042199 | 22.73964 | 39.91093 | 35.67871 | 15.78002 | 0.548549 | 0.660319 | 0.562305 |
| 5 | 116965400 | 116977799 | 135.2207 | 203.2946 | 284.208 | 117.3394 | -0.21372 | 3.421328 | 0.562348 |
| 8 | 123888200 | 123892599 | 32.68222 | 52.86053 | 84.38951 | 36.0773 | -0.31113 | 1.68038 | 0.56236 |
| 17 | 28744600 | 28745799 | 12.60244 | 23.24457 | 24.94723 | 10.55008 | 0.333616 | 0.224892 | 0.56237 |
| 8 | 117429000 | 117430799 | 20.87216 | 36.51169 | 37.08098 | 16.35421 | 0.365507 | 0.704912 | 0.562372 |
| 7 | 85641800 | 85647199 | 93.28077 | 145.7156 | 148.4535 | 61.59549 | 0.28653 | 2.667995 | 0.562387 |
| 14 | 51399600 | 51404799 | 59.03492 | 94.26781 | 100.8246 | 42.33007 | 0.243842 | 2.111528 | 0.562407 |
| 12 | 16552400 | 16553199 | 9.329148 | 17.48617 | 30.13617 | 12.43452 | -0.31753 | 0.115158 | 0.562425 |

| | | | | | | | | | |
|---|---|---|---|---|---|---|---|---|---|
| 3 | 65141800 | 65144199 | 21.84527 | 38.65566 | 32.28367 | 14.27723 | 0.64963 | 0.554995 | 0.562425 |
| 14 | 39387400 | 39389399 | 19.69581 | 34.91837 | 31.82792 | 14.07034 | 0.524966 | 0.519467 | 0.562447 |
| 14 | 117333600 | 117335599 | 17.53449 | 30.34103 | 51.74014 | 22.69975 | -0.38106 | 0.84389 | 0.562464 |
| 6 | 3250800 | 3252199 | 25.92225 | 43.93833 | 39.8137 | 17.43609 | 0.523259 | 1.05403 | 0.562484 |
| 8 | 117410200 | 117418199 | 88.7153 | 136.9283 | 198.0062 | 82.24794 | -0.23161 | 2.880399 | 0.562487 |
| 4 | 28614200 | 28615999 | 22.62737 | 39.59782 | 36.77848 | 16.22624 | 0.493576 | 0.704539 | 0.562492 |
| X | 84557400 | 84558399 | 12.20383 | 22.07166 | 39.61966 | 17.18778 | -0.42877 | 0.367389 | 0.562496 |
| 16 | 83869600 | 83873799 | 89.59595 | 139.5313 | 148.1342 | 61.48652 | 0.223703 | 2.717158 | 0.562498 |
| 3 | 97228000 | 97230599 | 12.01044 | 21.25653 | 37.23462 | 16.46617 | -0.41315 | 0.50059 | 0.562507 |
| 16 | 59155600 | 59160999 | 83.70405 | 130.1526 | 197.7167 | 82.086 | -0.29441 | 2.748773 | 0.562513 |
| 18 | 19449800 | 19452599 | 25.33123 | 43.79078 | 42.43622 | 18.61734 | 0.430794 | 0.877345 | 0.562521 |
| 3 | 57928200 | 57928799 | 5.809237 | 11.28531 | 22.27105 | 8.315533 | -0.37795 | -0.27578 | 0.562539 |
| 4 | 113367400 | 113373599 | 73.14333 | 116.7973 | 99.63535 | 41.83125 | 0.569606 | 2.125997 | 0.562543 |
| 7 | 25997000 | 25998399 | 19.94874 | 35.04672 | 35.26629 | 15.61047 | 0.378495 | 0.627579 | 0.562552 |
| 1 | 7443000 | 7444799 | 15.50201 | 28.60923 | 27.37492 | 11.56762 | 0.499072 | 0.251179 | 0.562557 |
| X | 61375200 | 61375799 | 13.67895 | 25.16291 | 26.04924 | 11.05618 | 0.382082 | 0.254281 | 0.562587 |
| 7 | 60069600 | 60071399 | 21.1554 | 37.13951 | 35.75444 | 15.81679 | 0.442097 | 0.647086 | 0.562607 |
| 5 | 131743600 | 131747799 | 37.71629 | 60.76622 | 93.69317 | 39.85462 | -0.26762 | 1.805244 | 0.562632 |
| 9 | 70218000 | 70219999 | 21.98011 | 36.90621 | 58.64231 | 25.6046 | -0.2848 | 1.140528 | 0.562638 |
| 14 | 120739000 | 120739999 | 11.95739 | 21.51149 | 35.21866 | 15.37858 | -0.30284 | 0.386428 | 0.562652 |
| 17 | 39305400 | 39319199 | 164.6999 | 249.3076 | 224.4993 | 92.87433 | 0.428943 | 3.290992 | 0.56267 |
| 13 | 33492200 | 33493799 | 20.26903 | 34.68074 | 56.05667 | 24.58072 | -0.30717 | 0.964297 | 0.56267 |
| X | 150172000 | 150173599 | 21.76408 | 38.19402 | 35.79525 | 15.82992 | 0.480765 | 0.657492 | 0.562684 |
| 12 | 14489200 | 14493999 | 118.5168 | 181.2401 | 191.0844 | 79.2764 | 0.213518 | 3.069928 | 0.562687 |
| 1 | 90462400 | 90465999 | 23.85112 | 39.99624 | 64.80586 | 28.27276 | -0.31292 | 1.167506 | 0.562688 |
| 5 | 18739000 | 18740999 | 24.90158 | 42.33639 | 45.28537 | 19.84734 | 0.284795 | 1.025867 | 0.562704 |
| 4 | 53870600 | 53873399 | 43.02788 | 70.52782 | 64.05928 | 27.57249 | 0.515046 | 1.476592 | 0.562709 |
| 17 | 5061200 | 5066999 | 25.45191 | 41.77729 | 77.69486 | 33.46139 | -0.5152 | 1.425912 | 0.562733 |
| 1 | 54477600 | 54478799 | 9.897152 | 18.07664 | 31.34898 | 13.43364 | -0.36728 | 0.276393 | 0.562749 |
| 3 | 138475600 | 138478399 | 16.86412 | 29.01134 | 48.47198 | 21.26214 | -0.35316 | 0.892606 | 0.562781 |
| 8 | 57765400 | 57767199 | 18.31154 | 31.37049 | 50.59144 | 22.18646 | -0.30348 | 0.939002 | 0.562782 |
| 2 | 162271000 | 162272799 | 14.00703 | 25.58204 | 26.76389 | 11.46944 | 0.356506 | 0.294061 | 0.562787 |
| 2 | 82337400 | 82337999 | 15.59709 | 27.80557 | 28.03211 | 12.33463 | 0.385331 | 0.454569 | 0.562788 |
| 17 | 57051600 | 57054399 | 31.21645 | 50.52559 | 83.78734 | 35.83858 | -0.36488 | 1.662226 | 0.562793 |
| 19 | 49552800 | 49555999 | 30.93084 | 51.55232 | 54.69139 | 23.76832 | 0.296148 | 1.241547 | 0.56282 |
| 8 | 34436000 | 34438799 | 16.91923 | 29.31223 | 48.47264 | 21.267 | -0.3364 | 0.827193 | 0.56284 |
| 15 | 29907600 | 29911799 | 43.5344 | 70.57097 | 73.78938 | 31.59782 | 0.300939 | 1.649533 | 0.562849 |
| 13 | 63272600 | 63277999 | 54.85928 | 88.14473 | 89.09149 | 37.74736 | 0.335491 | 1.91299 | 0.562865 |
| 7 | 3545400 | 3547599 | 22.37715 | 38.54981 | 41.57109 | 18.23586 | 0.276287 | 0.89465 | 0.562869 |
| 13 | 120614400 | 120618599 | 25.19172 | 42.9882 | 45.23325 | 19.8304 | 0.309162 | 0.993195 | 0.562875 |
| 16 | 52444400 | 52445399 | 15.76263 | 28.60485 | 25.20834 | 10.8896 | 0.595426 | 0.342306 | 0.562876 |
| 3 | 145683600 | 145685399 | 28.73774 | 46.80592 | 76.1862 | 32.71363 | -0.33019 | 1.539052 | 0.562909 |
| X | 33104600 | 33112999 | 150.3305 | 228.0696 | 192.6053 | 79.73561 | 0.52419 | 3.240787 | 0.562914 |
| 1 | 89833200 | 89836599 | 42.52571 | 68.95912 | 73.13705 | 31.32628 | 0.280911 | 1.64032 | 0.562927 |
| 1 | 14228200 | 14228599 | 5.5142 | 10.72107 | 21.75044 | 8.066782 | -0.40804 | -0.30409 | 0.562928 |
| 5 | 114358000 | 114367199 | 122.5484 | 184.8857 | 263.9834 | 109.0941 | -0.23969 | 3.349818 | 0.562931 |
| 3 | 119345200 | 119346999 | 19.44586 | 34.49705 | 31.27069 | 13.81676 | 0.533316 | 0.512158 | 0.562967 |

| | | | | | | | | | |
|---|---|---|---|---|---|---|---|---|---|
| 1 | 18286000 | 18287199 | 24.78349 | 42.32776 | 44.76195 | 19.62513 | 0.302318 | 0.98698 | 0.562973 |
| 4 | 135713600 | 135715599 | 10.47876 | 19.07186 | 32.8022 | 14.11821 | -0.35994 | 0.302173 | 0.562986 |
| 14 | 48541400 | 48541999 | 5.096032 | 9.924335 | 20.73676 | 7.58525 | -0.43201 | -0.36391 | 0.562987 |
| 16 | 68758800 | 68759999 | 8.688299 | 16.48649 | 29.39353 | 11.84761 | -0.33746 | 0.025904 | 0.562996 |
| 5 | 54705800 | 54707799 | 15.33054 | 27.31645 | 26.57235 | 11.69734 | 0.436067 | 0.451686 | 0.563006 |
| X | 66089000 | 66092599 | 34.51886 | 57.72715 | 53.61174 | 23.34883 | 0.487531 | 1.219734 | 0.563008 |
| 7 | 14385200 | 14433599 | 874.0597 | 1148.438 | 962.3331 | 402.6878 | 0.332963 | 5.563921 | 0.563069 |
| 1 | 59480400 | 59482399 | 23.63988 | 39.48853 | 61.96001 | 26.96513 | -0.26663 | 1.204424 | 0.563077 |
| 1 | 138150800 | 138156799 | 105.1481 | 162.3089 | 171.561 | 71.26307 | 0.220205 | 2.882629 | 0.563079 |
| 10 | 118840600 | 118844599 | 26.83064 | 44.03783 | 72.31171 | 31.14224 | -0.33617 | 1.424184 | 0.56308 |
| 2 | 19971000 | 19972399 | 13.15443 | 23.09528 | 39.40093 | 17.43112 | -0.37912 | 0.6072 | 0.563081 |
| 6 | 147697400 | 147698999 | 21.08439 | 36.86578 | 32.87868 | 14.49512 | 0.551916 | 0.704486 | 0.563092 |
| 9 | 84097400 | 84103599 | 60.27457 | 96.24541 | 102.1176 | 42.87172 | 0.255347 | 2.112761 | 0.563096 |
| X | 164651000 | 164654799 | 55.15248 | 88.63807 | 88.30202 | 37.42647 | 0.35681 | 1.906944 | 0.563102 |
| 2 | 123070400 | 123071399 | 21.8246 | 37.57309 | 39.55075 | 17.34926 | 0.310803 | 0.895014 | 0.563102 |
| 10 | 43282800 | 43284799 | 11.47678 | 20.55599 | 35.20491 | 15.42986 | -0.37121 | 0.409559 | 0.563103 |
| 3 | 113202400 | 113202999 | 11.30545 | 21.50603 | 19.52913 | 7.830332 | 0.635115 | 0.007714 | 0.563108 |
| 10 | 8269200 | 8271799 | 24.05814 | 40.23496 | 64.93313 | 28.28023 | -0.30724 | 1.193492 | 0.56311 |
| 14 | 6226200 | 6227799 | 10.85158 | 19.53386 | 33.73634 | 14.72186 | -0.37875 | 0.37152 | 0.563119 |
| 7 | 18338400 | 18348999 | 178.2356 | 265.314 | 273.0983 | 112.5271 | 0.214701 | 3.61281 | 0.563153 |
| 3 | 63322600 | 63325599 | 27.9393 | 47.38267 | 48.497 | 21.24233 | 0.347954 | 1.062819 | 0.563192 |
| 15 | 91494800 | 91497199 | 16.49804 | 28.68226 | 45.79115 | 20.11273 | -0.28488 | 0.783871 | 0.563206 |
| 4 | 94990000 | 94991799 | 18.6078 | 32.61961 | 35.01842 | 15.47218 | 0.285678 | 0.654306 | 0.563222 |
| 2 | 13972800 | 13974799 | 22.4336 | 38.8961 | 40.89828 | 17.94423 | 0.314329 | 0.835593 | 0.563241 |
| 16 | 72248200 | 72250999 | 22.41172 | 37.42192 | 71.42261 | 31.07539 | -0.54838 | 1.213349 | 0.563251 |
| 3 | 64077400 | 64082399 | 59.85026 | 93.71942 | 172.2703 | 71.45965 | -0.56109 | 2.610983 | 0.56326 |
| 1 | 22070200 | 22073199 | 52.21142 | 83.61639 | 88.05071 | 37.12205 | 0.270911 | 2.0102 | 0.563261 |
| 13 | 9307200 | 9307999 | 14.0969 | 25.84419 | 27.36051 | 11.66629 | 0.344686 | 0.275875 | 0.563262 |
| 6 | 134024200 | 134026199 | 18.20682 | 31.08694 | 51.10347 | 22.40497 | -0.33167 | 0.970859 | 0.563266 |
| 13 | 73630000 | 73634799 | 37.37948 | 60.27121 | 92.76646 | 39.50241 | -0.26374 | 1.780541 | 0.563277 |
| 7 | 6272200 | 6277999 | 39.99187 | 64.2755 | 99.90666 | 42.34767 | -0.28365 | 1.886881 | 0.563288 |
| 12 | 41857000 | 41859799 | 24.57895 | 42.96531 | 37.28855 | 16.43584 | 0.591152 | 0.733977 | 0.56329 |
| 12 | 119630600 | 119631599 | 27.6801 | 46.29589 | 48.62052 | 21.17665 | 0.310673 | 1.190412 | 0.563329 |
| 5 | 65179400 | 65180399 | 12.89451 | 23.76641 | 25.39645 | 10.75043 | 0.338763 | 0.231768 | 0.563347 |
| 6 | 104759800 | 104761799 | 23.99464 | 39.95158 | 63.00411 | 27.36358 | -0.27384 | 1.241402 | 0.563364 |
| 15 | 46517000 | 46518599 | 13.2141 | 25.25916 | 22.64231 | 8.98472 | 0.66628 | -0.00016 | 0.563365 |
| 6 | 40893200 | 40895799 | 20.6815 | 36.24803 | 36.43956 | 16.09753 | 0.380054 | 0.671858 | 0.563374 |
| 17 | 14385200 | 14388199 | 16.51806 | 28.39269 | 50.58926 | 22.19032 | -0.44596 | 0.898966 | 0.563381 |
| 8 | 60073000 | 60076399 | 28.26715 | 46.59136 | 73.34079 | 31.64174 | -0.27302 | 1.377379 | 0.563409 |
| 10 | 58457000 | 58459599 | 22.27331 | 37.38824 | 59.79162 | 26.10188 | -0.2941 | 1.144431 | 0.563433 |
| 13 | 53477600 | 53481199 | 39.1655 | 64.24795 | 64.51137 | 27.77642 | 0.371126 | 1.455555 | 0.563443 |
| 14 | 72768000 | 72771599 | 27.81254 | 45.60306 | 72.90002 | 31.38209 | -0.29817 | 1.440051 | 0.563443 |
| 8 | 43479000 | 43482599 | 31.95215 | 53.81815 | 50.76635 | 22.19019 | 0.465005 | 1.137187 | 0.563445 |
| 14 | 119054600 | 119055999 | 33.03886 | 53.40175 | 85.24246 | 36.41228 | -0.312 | 1.701376 | 0.563481 |
| 4 | 139773000 | 139784599 | 123.1794 | 185.4292 | 270.8286 | 111.8539 | -0.27515 | 3.393082 | 0.563498 |
| 6 | 60338200 | 60341799 | 35.45361 | 58.50318 | 53.6656 | 23.16437 | 0.504496 | 1.367393 | 0.56351 |
| 9 | 31858400 | 31862199 | 32.99433 | 53.45769 | 82.9927 | 35.5254 | -0.2686 | 1.645755 | 0.563518 |

| | | | | | | | | | |
|---|---|---|---|---|---|---|---|---|---|
| 18 | 6793600 | 6806599 | 135.2374 | 205.6611 | 211.8266 | 87.75556 | 0.240773 | 3.188065 | 0.563538 |
| 13 | 93967800 | 93971599 | 43.30908 | 70.41786 | 70.75607 | 30.36302 | 0.362137 | 1.588654 | 0.563546 |
| 7 | 105233400 | 105235399 | 11.78614 | 21.3337 | 35.31726 | 15.30562 | -0.31168 | 0.349844 | 0.563546 |
| X | 9461600 | 9475999 | 193.435 | 288.7898 | 270.301 | 111.4243 | 0.35568 | 3.563484 | 0.563548 |
| 3 | 158944200 | 158944999 | 14.75828 | 25.30987 | 54.22799 | 23.78326 | -0.7112 | 0.920223 | 0.563549 |
| X | 34771400 | 34774199 | 31.68073 | 53.4441 | 50.18095 | 21.94466 | 0.471819 | 1.125311 | 0.563556 |
| 13 | 62160000 | 62162399 | 21.46119 | 37.27145 | 39.86331 | 17.49744 | 0.290252 | 0.812198 | 0.563569 |
| 5 | 26631000 | 26631599 | 8.419089 | 16.01873 | 28.88785 | 11.57767 | -0.34621 | 0.005372 | 0.563572 |
| 18 | 9471000 | 9474399 | 66.91598 | 106.3101 | 114.032 | 47.55595 | 0.231972 | 2.279643 | 0.563577 |
| 15 | 56046400 | 56046999 | 8.463543 | 16.38132 | 17.68096 | 6.734439 | 0.455904 | -0.1963 | 0.563596 |
| 1 | 61331200 | 61332999 | 21.39205 | 37.31434 | 38.67161 | 17.02113 | 0.336111 | 0.750041 | 0.563608 |
| 13 | 119424400 | 119428399 | 39.2321 | 64.52122 | 62.41411 | 26.90236 | 0.426191 | 1.422356 | 0.563618 |
| 1 | 156591200 | 156595799 | 84.10988 | 130.3383 | 186.8052 | 77.59362 | -0.21435 | 2.805262 | 0.563649 |
| 9 | 117416000 | 117418399 | 40.05732 | 64.98109 | 70.48645 | 30.19941 | 0.249043 | 1.626845 | 0.5637 |
| 8 | 32171400 | 32173399 | 12.94385 | 22.90887 | 37.6929 | 16.66878 | -0.32333 | 0.5076 | 0.563701 |
| 6 | 75273600 | 75275199 | 20.66186 | 34.98344 | 55.38723 | 24.2483 | -0.27909 | 1.060171 | 0.563721 |
| 16 | 14775600 | 14778199 | 30.65317 | 50.0509 | 77.36279 | 33.24921 | -0.25329 | 1.507873 | 0.563724 |
| 7 | 87201600 | 87204399 | 35.84979 | 59.76204 | 48.15604 | 20.93063 | 0.69174 | 1.251289 | 0.563742 |
| 17 | 38596600 | 38608999 | 177.3754 | 267.3793 | 236.8686 | 97.8641 | 0.447401 | 3.37714 | 0.563744 |
| 4 | 60271800 | 60274999 | 117.8962 | 180.1733 | 188.184 | 78.06133 | 0.226316 | 3.082838 | 0.563772 |
| 11 | 64733600 | 64734599 | 8.943003 | 16.63138 | 29.47041 | 12.32247 | -0.37219 | 0.163228 | 0.563776 |
| 6 | 33209800 | 33213999 | 38.84072 | 63.17189 | 68.46529 | 29.38326 | 0.253537 | 1.573735 | 0.563782 |
| 14 | 113704600 | 113705999 | 14.29545 | 26.69631 | 26.00138 | 10.79557 | 0.492621 | 0.169887 | 0.563792 |
| 11 | 40427200 | 40427599 | 5.516367 | 10.746 | 21.3819 | 7.791914 | -0.35786 | -0.37056 | 0.563816 |
| 7 | 143537800 | 143543799 | 45.42232 | 72.75236 | 109.1413 | 46.02215 | -0.23855 | 2.005012 | 0.563824 |
| 1 | 100199600 | 100200799 | 14.8509 | 26.83385 | 28.8323 | 12.52273 | 0.306149 | 0.36793 | 0.563826 |
| 5 | 104730000 | 104752999 | 740.3758 | 982.4832 | 874.7879 | 365.6247 | 0.259666 | 5.403643 | 0.563831 |
| 9 | 115380000 | 115381999 | 21.38714 | 35.83531 | 58.54186 | 25.52692 | -0.32455 | 1.162102 | 0.563831 |
| 18 | 17706600 | 17709399 | 21.98049 | 38.34968 | 38.66808 | 17.02206 | 0.375648 | 0.750836 | 0.563838 |
| 7 | 23525600 | 23535599 | 139.2009 | 213.6049 | 182.956 | 75.95436 | 0.516871 | 3.009712 | 0.563844 |
| 19 | 50388600 | 50391999 | 50.13416 | 79.86861 | 120.4893 | 50.42184 | -0.25513 | 2.182191 | 0.563876 |
| 13 | 55862400 | 55863199 | 6.253468 | 12.0943 | 23.77336 | 9.087851 | -0.40555 | -0.18838 | 0.563882 |
| 14 | 27130200 | 27133999 | 43.64946 | 70.53789 | 76.25842 | 32.56624 | 0.248481 | 1.718869 | 0.563892 |
| 10 | 108793000 | 108794199 | 15.41096 | 27.51749 | 28.93882 | 12.71652 | 0.32594 | 0.444659 | 0.563898 |
| 1 | 100116000 | 100116599 | 11.61168 | 21.84064 | 24.1684 | 9.913314 | 0.324043 | 0.098957 | 0.563903 |
| 15 | 75079400 | 75080999 | 44.33216 | 71.90304 | 73.46094 | 31.46972 | 0.334988 | 1.63971 | 0.563912 |
| 14 | 105758200 | 105759199 | 9.356989 | 17.3493 | 29.92707 | 12.56599 | -0.33861 | 0.182487 | 0.563916 |
| 2 | 74693800 | 74695599 | 23.72061 | 41.46582 | 36.7239 | 16.18726 | 0.56191 | 0.726667 | 0.563925 |
| 4 | 117670600 | 117682999 | 118.5356 | 179.6567 | 252.8834 | 104.6587 | -0.21346 | 3.255781 | 0.563941 |
| 10 | 116431200 | 116432999 | 21.72346 | 37.58503 | 40.49618 | 17.76666 | 0.278764 | 0.852233 | 0.563942 |
| 6 | 27908400 | 27910599 | 30.42303 | 52.4605 | 41.77168 | 18.32439 | 0.712448 | 0.922573 | 0.563943 |
| 19 | 8741000 | 8744199 | 43.98559 | 70.5018 | 105.8106 | 44.66648 | -0.23791 | 1.980786 | 0.563953 |
| 1 | 178038800 | 178040599 | 18.56974 | 32.88309 | 32.52947 | 14.38388 | 0.406892 | 0.526543 | 0.563964 |
| 7 | 40861200 | 40873199 | 209.5675 | 309.5288 | 308.5689 | 127.075 | 0.250453 | 3.75125 | 0.563968 |
| 15 | 94669600 | 94672199 | 26.30177 | 44.52831 | 47.97967 | 21.00469 | 0.273985 | 1.071282 | 0.563993 |
| 14 | 77105200 | 77106999 | 15.39118 | 26.94408 | 43.02451 | 18.98761 | -0.28432 | 0.67283 | 0.564014 |
| 11 | 23844600 | 23845399 | 8.89645 | 16.47265 | 29.91942 | 12.58638 | -0.41454 | 0.185588 | 0.564017 |

| | | | | | | | | | |
|---|---|---|---|---|---|---|---|---|---|
| Y | 6730600 | 6733999 | 22.66362 | 39.05387 | 28.5671 | 12.53154 | 0.833694 | 0.871543 | 0.564022 |
| 4 | 62077600 | 62078599 | 15.19533 | 27.27954 | 29.3274 | 12.82868 | 0.298684 | 0.40952 | 0.564028 |
| 7 | 113266200 | 113267999 | 17.99849 | 30.92334 | 49.5293 | 21.72509 | -0.29288 | 0.910891 | 0.564048 |
| 9 | 18152800 | 18157199 | 51.05982 | 82.54369 | 76.44073 | 32.65971 | 0.471907 | 1.720885 | 0.564048 |
| 9 | 35761000 | 35764799 | 37.71954 | 62.06175 | 62.72927 | 27.04338 | 0.363328 | 1.415353 | 0.564051 |
| X | 126004800 | 126006599 | 21.85632 | 38.6006 | 28.36795 | 12.55038 | 0.832167 | 0.569738 | 0.564067 |
| 4 | 123663400 | 123667599 | 33.41522 | 54.36689 | 86.46528 | 37.11311 | -0.29808 | 1.568222 | 0.564073 |
| 15 | 18448400 | 18450599 | 23.44572 | 40.45445 | 42.51508 | 18.65085 | 0.313682 | 0.885156 | 0.564074 |
| 9 | 56684000 | 56685799 | 11.55498 | 20.54998 | 35.26441 | 15.5485 | -0.37976 | 0.456921 | 0.564103 |
| 6 | 59827200 | 59838599 | 245.2806 | 357.7041 | 351.6923 | 144.92 | 0.253016 | 3.955449 | 0.564107 |
| 4 | 90487200 | 90488199 | 12.00172 | 22.74876 | 20.36936 | 8.229431 | 0.645265 | 0.039659 | 0.564115 |
| 13 | 112979000 | 112983799 | 62.42155 | 99.41624 | 107.1629 | 44.828 | 0.228569 | 2.19734 | 0.564123 |
| 14 | 83749600 | 83750199 | 15.83301 | 28.10464 | 29.85208 | 13.18331 | 0.306615 | 0.484729 | 0.564126 |
| 7 | 51655400 | 51656399 | 9.621994 | 17.39393 | 33.31455 | 14.46108 | -0.52204 | 0.336353 | 0.56413 |
| 11 | 20131200 | 20131999 | 9.424391 | 18.13613 | 20.77924 | 8.088318 | 0.341129 | -0.10396 | 0.564134 |
| 12 | 70472600 | 70473799 | 9.424391 | 18.13613 | 20.77924 | 8.088318 | 0.341129 | -0.10396 | 0.564134 |
| 15 | 81380400 | 81382399 | 19.88742 | 33.64226 | 54.8009 | 23.98514 | -0.31995 | 1.06529 | 0.56415 |
| 15 | 22455800 | 22458599 | 47.39312 | 76.47048 | 80.15615 | 34.16322 | 0.290427 | 1.770157 | 0.564155 |
| 17 | 18595200 | 18601999 | 100.2748 | 156.4329 | 152.1327 | 63.13067 | 0.35204 | 2.695261 | 0.564189 |
| 1 | 150712800 | 150714599 | 16.96376 | 30.79358 | 29.14973 | 12.58761 | 0.493112 | 0.35704 | 0.564203 |
| 8 | 5459600 | 5459999 | 5.504468 | 10.73705 | 21.60907 | 7.739168 | -0.35127 | -0.43073 | 0.564214 |
| 2 | 18995800 | 18997399 | 16.02469 | 28.07682 | 51.09165 | 22.56391 | -0.47232 | 0.672532 | 0.56422 |
| 9 | 39917400 | 39918399 | 9.853164 | 18.35955 | 31.63643 | 13.18482 | -0.32887 | 0.161964 | 0.564241 |
| 3 | 16312800 | 16315199 | 50.08968 | 80.53515 | 86.44449 | 36.66142 | 0.250115 | 1.884924 | 0.564246 |
| 14 | 32313800 | 32315799 | 15.21411 | 26.51033 | 43.674 | 19.21025 | -0.32956 | 0.74204 | 0.564253 |
| 18 | 13101000 | 13109199 | 154.2507 | 228.8382 | 328.0369 | 135.1176 | -0.2673 | 3.671523 | 0.564262 |
| 6 | 16515200 | 16518199 | 42.20695 | 68.64922 | 70.54858 | 30.27773 | 0.330113 | 1.582352 | 0.564287 |
| 8 | 45313400 | 45316199 | 20.31406 | 35.92434 | 32.33015 | 14.29953 | 0.541888 | 0.550351 | 0.56429 |
| 6 | 68771600 | 68780199 | 168.4378 | 252.8314 | 229.0737 | 94.53611 | 0.409563 | 3.455374 | 0.564292 |
| 16 | 50153600 | 50157599 | 61.71492 | 98.15736 | 102.837 | 42.93759 | 0.267477 | 2.238712 | 0.564297 |
| 11 | 50464800 | 50467599 | 14.56327 | 25.31673 | 46.85107 | 20.57731 | -0.49713 | 0.774297 | 0.564328 |
| 3 | 17053600 | 17054999 | 13.73852 | 25.2096 | 26.85844 | 11.43835 | 0.336801 | 0.266706 | 0.564329 |
| 11 | 103393600 | 103395799 | 15.04332 | 26.299 | 42.55165 | 18.75717 | -0.30325 | 0.691782 | 0.564337 |
| X | 146438400 | 146439799 | 28.22318 | 47.60279 | 50.1629 | 21.93557 | 0.305859 | 1.113468 | 0.564339 |
| 7 | 41729400 | 41746999 | 260.5687 | 380.1353 | 349.4583 | 143.9941 | 0.350412 | 3.952148 | 0.564347 |
| 15 | 5016800 | 5018599 | 21.75551 | 36.58532 | 59.8164 | 26.13339 | -0.32589 | 1.124609 | 0.564351 |
| 19 | 16784200 | 16786599 | 14.58167 | 25.48544 | 42.06677 | 18.53944 | -0.33184 | 0.693163 | 0.564364 |
| 7 | 96663600 | 96665399 | 18.71302 | 33.72024 | 29.99918 | 13.07403 | 0.57376 | 0.412931 | 0.56437 |
| X | 148423200 | 148436999 | 141.5564 | 216.1303 | 200.3934 | 83.10803 | 0.39695 | 3.110481 | 0.564385 |
| 8 | 8212200 | 8213799 | 13.01107 | 24.35912 | 24.97228 | 10.32412 | 0.423857 | 0.139536 | 0.56439 |
| 5 | 109378600 | 109386599 | 110.011 | 170.9206 | 159.1029 | 66.06261 | 0.41055 | 2.768379 | 0.564395 |
| 16 | 13001600 | 13007799 | 71.9929 | 113.8052 | 122.2473 | 50.78772 | 0.223503 | 2.416272 | 0.564432 |
| 14 | 11560400 | 11565599 | 57.85062 | 92.88054 | 90.62593 | 38.35398 | 0.384956 | 1.940941 | 0.564438 |
| 17 | 82941200 | 82942999 | 16.12637 | 29.25169 | 29.21303 | 12.62671 | 0.415459 | 0.355276 | 0.564468 |
| X | 44710400 | 44711399 | 18.08333 | 31.77194 | 33.04774 | 14.62939 | 0.331151 | 0.614091 | 0.564478 |
| 12 | 17069000 | 17074999 | 58.12705 | 92.05138 | 137.9609 | 57.36805 | -0.25426 | 2.36708 | 0.564482 |
| 14 | 3797600 | 3801199 | 24.40401 | 40.84108 | 66.08866 | 28.79581 | -0.31112 | 1.188922 | 0.56449 |

| | | | | | | | | | |
|---|---|---|---|---|---|---|---|---|---|
| 3 | 64455600 | 64456199 | 15.39933 | 27.53936 | 29.38582 | 12.89609 | 0.30632 | 0.435555 | 0.564491 |
| 14 | 3264200 | 3265999 | 23.40571 | 40.72719 | 39.6552 | 17.42727 | 0.425345 | 0.793638 | 0.564498 |
| 7 | 5869600 | 5885399 | 254.2269 | 372.839 | 327.3548 | 134.8337 | 0.423942 | 3.870919 | 0.564502 |
| 14 | 110592600 | 110594399 | 24.62917 | 42.58906 | 37.13353 | 16.29112 | 0.582571 | 0.863746 | 0.564527 |
| 1 | 152268000 | 152273199 | 59.041 | 94.52553 | 96.8189 | 40.80006 | 0.310285 | 2.030863 | 0.564531 |
| 5 | 30488200 | 30492999 | 50.31923 | 80.84565 | 86.55936 | 36.67162 | 0.252473 | 1.908861 | 0.56454 |
| 10 | 86984400 | 86986999 | 22.96379 | 38.521 | 60.33179 | 26.32636 | -0.26408 | 1.155782 | 0.564559 |
| 10 | 15877400 | 15877999 | 6.253489 | 12.10832 | 23.25267 | 8.840937 | -0.36435 | -0.2066 | 0.564563 |
| 10 | 92598600 | 92599399 | 6.253489 | 12.10832 | 23.25267 | 8.840937 | -0.36435 | -0.2066 | 0.564563 |
| 6 | 49260400 | 49261599 | 9.657968 | 17.9739 | 30.51224 | 12.7426 | -0.30991 | 0.164633 | 0.564581 |
| 7 | 6491200 | 6494199 | 43.8495 | 70.83317 | 76.68393 | 32.73127 | 0.245843 | 1.731387 | 0.56459 |
| 15 | 100738600 | 100740999 | 16.76014 | 28.90429 | 47.94353 | 21.03193 | -0.34182 | 0.871387 | 0.564592 |
| 13 | 67086200 | 67094799 | 105.4706 | 163.4619 | 165.4932 | 68.74118 | 0.28673 | 2.808905 | 0.564598 |
| 2 | 6844800 | 6845599 | 11.84104 | 20.70035 | 45.7373 | 20.16114 | -0.75024 | 0.681403 | 0.564609 |
| 5 | 134598800 | 134600399 | 18.26092 | 31.50054 | 50.07913 | 21.96895 | -0.28107 | 0.875218 | 0.564634 |
| 8 | 117825800 | 117829199 | 45.47663 | 73.33633 | 79.46047 | 33.84182 | 0.24181 | 1.784164 | 0.564635 |
| 8 | 120916800 | 120919999 | 32.21142 | 52.06093 | 85.24455 | 36.4101 | -0.34876 | 1.700532 | 0.56464 |
| 18 | 41290800 | 41293199 | 22.12125 | 38.82267 | 35.03915 | 15.49617 | 0.534604 | 0.654309 | 0.564643 |
| 1 | 75358000 | 75363199 | 42.36649 | 67.40744 | 132.5853 | 55.38535 | -0.63918 | 2.219122 | 0.564654 |
| 7 | 27417200 | 27422399 | 71.68337 | 113.2388 | 120.1865 | 49.9144 | 0.240029 | 2.431869 | 0.564662 |
| 2 | 169855800 | 169859599 | 36.60473 | 58.91588 | 94.13981 | 39.98611 | -0.32077 | 1.835589 | 0.564674 |
| 7 | 54148400 | 54151199 | 35.45197 | 58.45585 | 60.84397 | 26.25438 | 0.322346 | 1.383374 | 0.564692 |
| 3 | 88276200 | 88281599 | 95.5223 | 146.2393 | 213.718 | 88.68944 | -0.25585 | 3.043185 | 0.564695 |
| 10 | 79331200 | 79333599 | 21.02307 | 36.54236 | 39.35654 | 17.28445 | 0.280414 | 0.797939 | 0.564696 |
| 4 | 156004400 | 156011199 | 120.7826 | 184.3248 | 192.5928 | 79.86415 | 0.224177 | 3.112491 | 0.564697 |
| 12 | 93481000 | 93484999 | 42.71528 | 69.43923 | 63.15496 | 27.09934 | 0.505335 | 1.585228 | 0.564739 |
| 8 | 52799000 | 52799799 | 13.52239 | 24.764 | 26.94755 | 11.50721 | 0.303931 | 0.277634 | 0.56474 |
| 9 | 36394600 | 36398999 | 44.85622 | 72.72072 | 74.01028 | 31.69467 | 0.339955 | 1.650175 | 0.56476 |
| 13 | 49894200 | 49904199 | 130.7161 | 196.9415 | 275.4778 | 113.7999 | -0.21178 | 3.378802 | 0.56483 |
| 7 | 48385000 | 48396599 | 189.0427 | 280.8628 | 285.3912 | 117.5694 | 0.230795 | 3.648859 | 0.564836 |
| 6 | 139628200 | 139629799 | 14.42525 | 25.45061 | 41.83822 | 18.51358 | -0.32397 | 0.559031 | 0.564853 |
| 13 | 60521000 | 60522999 | 14.70983 | 25.89608 | 42.88201 | 18.98375 | -0.33578 | 0.585157 | 0.564866 |
| 7 | 39011800 | 39019599 | 217.9706 | 319.7471 | 325.0117 | 133.8665 | 0.213124 | 3.862141 | 0.564873 |
| 16 | 15871600 | 15878199 | 75.31958 | 119.6119 | 116.6255 | 48.61757 | 0.368832 | 2.298856 | 0.564887 |
| 11 | 50425800 | 50431799 | 57.83756 | 91.65703 | 135.376 | 56.31577 | -0.23235 | 2.350282 | 0.564888 |
| 2 | 39149600 | 39154999 | 62.64355 | 100.52 | 90.26081 | 38.16731 | 0.503659 | 1.963754 | 0.564893 |
| 19 | 37737200 | 37742999 | 61.93669 | 98.09629 | 145.1676 | 60.36852 | -0.23587 | 2.36915 | 0.564902 |
| X | 125804400 | 125806399 | 28.8002 | 48.74927 | 49.05654 | 21.47494 | 0.372348 | 1.085221 | 0.56492 |
| 16 | 78240200 | 78241599 | 18.60215 | 32.629 | 34.97207 | 15.45942 | 0.287959 | 0.644611 | 0.564941 |
| 10 | 70764000 | 70774199 | 121.4788 | 183.603 | 260.1385 | 107.5663 | -0.22636 | 3.312634 | 0.564941 |
| 6 | 146589600 | 146592199 | 54.09516 | 86.57078 | 92.13029 | 38.80095 | 0.254522 | 2.036759 | 0.565005 |
| 5 | 69853200 | 69858399 | 74.89439 | 118.0675 | 126.5039 | 52.49627 | 0.224048 | 2.481729 | 0.56501 |
| 11 | 21362400 | 21363599 | 13.91752 | 24.42214 | 40.38487 | 17.85846 | -0.33448 | 0.62101 | 0.565017 |
| 5 | 132715600 | 132717199 | 31.60816 | 51.14795 | 93.46697 | 39.97218 | -0.50492 | 1.67543 | 0.565018 |
| 4 | 147051200 | 147062399 | 270.9419 | 395.2024 | 346.0216 | 142.5804 | 0.420487 | 3.955787 | 0.565019 |
| 3 | 76038200 | 76039599 | 11.83457 | 21.37407 | 35.29925 | 15.34016 | -0.31068 | 0.364184 | 0.565032 |
| 2 | 11879400 | 11881199 | 24.37545 | 41.98725 | 43.01422 | 18.86884 | 0.349818 | 0.904037 | 0.565035 |

| | | | | | | | | | |
|---|---|---|---|---|---|---|---|---|---|
| 14 | 7934600 | 7936599 | 18.81666 | 32.49885 | 53.35323 | 23.40612 | -0.3269 | 0.871833 | 0.565057 |
| 1 | 181806000 | 181807999 | 16.2584 | 29.18657 | 30.14729 | 13.18788 | 0.355853 | 0.417551 | 0.565077 |
| X | 79704800 | 79706399 | 18.78647 | 33.39391 | 32.11685 | 14.16676 | 0.450365 | 0.500806 | 0.565078 |
| X | 51084600 | 51089199 | 34.79462 | 57.88841 | 56.44498 | 24.5021 | 0.417557 | 1.276436 | 0.565092 |
| 7 | 43490200 | 43494999 | 43.52012 | 69.69289 | 106.2382 | 44.78554 | -0.26186 | 2.008865 | 0.565093 |
| 5 | 107332200 | 107334799 | 16.84183 | 29.15824 | 46.88088 | 20.5682 | -0.296 | 0.833485 | 0.565102 |
| 7 | 18435800 | 18438599 | 27.57957 | 46.41898 | 49.63765 | 21.69197 | 0.284608 | 1.126641 | 0.565114 |
| 1 | 79638400 | 79641599 | 23.77043 | 39.81646 | 62.61282 | 27.29678 | -0.26991 | 1.175324 | 0.565123 |
| 19 | 9649000 | 9651999 | 46.40733 | 74.98942 | 77.62541 | 33.14704 | 0.310909 | 1.728062 | 0.565132 |
| 11 | 113804000 | 113804599 | 12.37363 | 21.82983 | 37.33365 | 16.52045 | -0.38037 | 0.540319 | 0.565135 |
| 11 | 117385600 | 117391999 | 39.04779 | 62.37308 | 121.1975 | 50.90765 | -0.61539 | 2.086135 | 0.565147 |
| 11 | 19842600 | 19844599 | 17.28079 | 30.74231 | 31.62108 | 13.93858 | 0.35444 | 0.483209 | 0.565229 |
| 9 | 59133000 | 59135599 | 18.62004 | 31.8883 | 51.13 | 22.42203 | -0.29529 | 0.943557 | 0.565271 |
| 6 | 26991200 | 26993799 | 11.75318 | 20.8462 | 40.86145 | 18.05729 | -0.57306 | 0.485794 | 0.565273 |
| 10 | 67524200 | 67526399 | 19.77097 | 33.80902 | 53.2944 | 23.36839 | -0.2712 | 0.964599 | 0.565299 |
| 4 | 6847400 | 6850999 | 37.31681 | 62.64281 | 50.2696 | 21.94559 | 0.697463 | 1.18592 | 0.565308 |
| 17 | 77933800 | 77938399 | 62.92552 | 100.2404 | 107.4548 | 44.96342 | 0.236873 | 2.191159 | 0.56532 |
| 8 | 52575600 | 52576799 | 16.81518 | 29.71123 | 31.83848 | 14.08448 | 0.29106 | 0.538777 | 0.56532 |
| 3 | 67558400 | 67559399 | 7.611751 | 14.59933 | 26.31157 | 10.33995 | -0.31929 | -0.0738 | 0.565331 |
| 3 | 78887400 | 78889199 | 15.67563 | 27.32204 | 44.17257 | 19.43291 | -0.30257 | 0.74077 | 0.565333 |
| 3 | 143700400 | 143701399 | 8.317515 | 15.38184 | 29.9316 | 12.61251 | -0.51492 | 0.187675 | 0.565362 |
| 18 | 68020400 | 68022599 | 22.60666 | 38.06671 | 59.82802 | 26.14889 | -0.26898 | 1.117887 | 0.565373 |
| 14 | 82724400 | 82725999 | 16.22509 | 29.24433 | 29.621 | 12.9052 | 0.387769 | 0.391979 | 0.565388 |
| 13 | 4508000 | 4511399 | 27.61821 | 46.73296 | 49.06056 | 21.47453 | 0.311393 | 1.08136 | 0.565402 |
| Y | 88335000 | 88337199 | 14.79051 | 26.64551 | 20.12645 | 8.774038 | 0.807734 | 0.36227 | 0.565403 |
| 19 | 15689000 | 15689599 | 5.514807 | 10.74769 | 21.46044 | 7.775608 | -0.35548 | -0.3913 | 0.565406 |
| 4 | 147281000 | 147282399 | 47.85352 | 77.08461 | 82.17296 | 34.94496 | 0.263351 | 1.820238 | 0.565412 |
| 6 | 41990000 | 41990999 | 11.88177 | 22.18955 | 24.59332 | 10.21039 | 0.307706 | 0.146658 | 0.565471 |
| 1 | 170303000 | 170307199 | 21.82203 | 36.578 | 59.17156 | 25.80774 | -0.31054 | 1.158379 | 0.565474 |
| 17 | 62884000 | 62884599 | 11.6348 | 21.79149 | 23.65658 | 9.773337 | 0.343358 | 0.125199 | 0.565475 |
| 7 | 12429000 | 12440599 | 164.7485 | 249.0286 | 229.666 | 94.9694 | 0.392794 | 3.319753 | 0.565477 |
| 8 | 76294600 | 76296399 | 20.82582 | 36.72808 | 34.04886 | 15.068 | 0.498005 | 0.587413 | 0.565479 |
| 19 | 3198200 | 3207199 | 153.4712 | 230.9402 | 240.3553 | 99.25804 | 0.212863 | 3.404784 | 0.565522 |
| 18 | 46251800 | 46252599 | 13.65385 | 25.16776 | 26.95206 | 11.40783 | 0.33593 | 0.242175 | 0.565526 |
| X | 24015400 | 24041399 | 441.1618 | 615.311 | 524.9851 | 217.7726 | 0.393466 | 4.623358 | 0.565544 |
| 4 | 122160000 | 122160399 | 8.476718 | 16.49144 | 18.32365 | 6.761146 | 0.456391 | -0.30331 | 0.565555 |
| 9 | 68943600 | 68948999 | 43.92326 | 70.46515 | 106.6948 | 45.10127 | -0.24914 | 1.95174 | 0.565575 |
| 4 | 55415600 | 55419599 | 38.89284 | 62.64283 | 94.83325 | 40.32188 | -0.24178 | 1.818069 | 0.56558 |
| 8 | 39644200 | 39647399 | 33.93734 | 56.03866 | 59.65815 | 25.76265 | 0.290425 | 1.361394 | 0.565598 |
| 15 | 56260200 | 56261599 | 17.59562 | 32.22699 | 26.39623 | 11.2701 | 0.710992 | 0.304965 | 0.565614 |
| X | 44393800 | 44394599 | 15.85026 | 29.23133 | 27.42354 | 11.59929 | 0.526637 | 0.259542 | 0.565635 |
| 5 | 125450200 | 125452199 | 35.23384 | 57.71986 | 62.84233 | 27.04223 | 0.253927 | 1.462411 | 0.565645 |
| 7 | 26730200 | 26731599 | 22.95857 | 39.49332 | 42.57765 | 18.67657 | 0.276153 | 0.910262 | 0.565647 |
| 7 | 61812400 | 61814199 | 22.43326 | 39.08309 | 39.13401 | 17.21152 | 0.385319 | 0.771346 | 0.565647 |
| 14 | 81092400 | 81094399 | 19.63723 | 34.9252 | 32.03616 | 14.12468 | 0.51877 | 0.502774 | 0.565655 |
| 2 | 166337200 | 166338399 | 14.93654 | 25.96546 | 44.26122 | 19.43975 | -0.37882 | 0.776126 | 0.565657 |
| 9 | 19440800 | 19442999 | 38.1297 | 62.04965 | 66.78004 | 28.66576 | 0.26415 | 1.562594 | 0.565671 |

| | | | | | | | | | |
|---|---|---|---|---|---|---|---|---|---|
| 14 | 52291200 | 52292399 | 17.64699 | 30.98035 | 32.57042 | 14.40829 | 0.31574 | 0.622168 | 0.565698 |
| 5 | 91980000 | 91980199 | 4.389256 | 8.571604 | 18.74878 | 6.587676 | -0.44409 | -0.52465 | 0.565716 |
| 2 | 87207600 | 87209599 | 29.48315 | 49.05877 | 52.96272 | 22.99281 | 0.271076 | 1.253993 | 0.565743 |
| 5 | 9360000 | 9360999 | 10.27047 | 18.90513 | 32.44752 | 13.7597 | -0.34216 | 0.234958 | 0.565744 |
| X | 126994600 | 126997399 | 34.56322 | 57.58555 | 55.90986 | 24.29231 | 0.423704 | 1.259203 | 0.565772 |
| 14 | 110200000 | 110202999 | 33.34482 | 54.84379 | 59.55822 | 25.67234 | 0.259858 | 1.405638 | 0.565787 |
| X | 158314400 | 158316999 | 15.37938 | 26.88832 | 46.7683 | 20.61709 | -0.40694 | 0.702917 | 0.565801 |
| 4 | 81322400 | 81324399 | 42.00491 | 68.14484 | 72.59261 | 31.10326 | 0.275154 | 1.629805 | 0.565805 |
| 3 | 15509200 | 15510799 | 11.11296 | 19.73154 | 40.87936 | 18.04911 | -0.65168 | 0.473763 | 0.565817 |
| 10 | 15748000 | 15753599 | 110.7299 | 170.932 | 157.551 | 65.44356 | 0.417631 | 2.882378 | 0.565829 |
| 10 | 111949800 | 111951399 | 16.07647 | 29.71983 | 26.76992 | 11.28587 | 0.587944 | 0.244344 | 0.565831 |
| 7 | 38406800 | 38426999 | 376.8049 | 528.8527 | 504.4528 | 209.0516 | 0.241709 | 4.527014 | 0.565859 |
| 13 | 100379800 | 100383999 | 25.81651 | 42.61792 | 67.95652 | 29.34082 | -0.29076 | 1.348355 | 0.565869 |
| 15 | 49329200 | 49330799 | 17.1162 | 30.23318 | 32.28692 | 14.28379 | 0.29575 | 0.544558 | 0.565871 |
| 9 | 66065200 | 66069399 | 30.17212 | 49.09511 | 78.15756 | 33.54841 | -0.29928 | 1.555872 | 0.565888 |
| 9 | 13690400 | 13693599 | 34.18233 | 57.74845 | 49.07489 | 21.47047 | 0.615295 | 1.120254 | 0.565894 |
| 14 | 9559800 | 9560599 | 10.55218 | 20.11319 | 22.74849 | 9.082755 | 0.325972 | -0.00501 | 0.565927 |
| 1 | 91158000 | 91158999 | 7.60221 | 14.53271 | 26.19714 | 10.39357 | -0.33267 | -0.04393 | 0.565929 |
| 5 | 27454800 | 27456399 | 16.43466 | 29.00969 | 29.94413 | 13.24913 | 0.344254 | 0.543441 | 0.565931 |
| 6 | 41240800 | 41245799 | 42.19026 | 68.9591 | 66.90073 | 28.76348 | 0.417931 | 1.511161 | 0.565939 |
| 8 | 128388200 | 128388999 | 14.2283 | 25.09184 | 40.78324 | 18.04837 | -0.30796 | 0.560242 | 0.565943 |
| X | 116572800 | 116575199 | 25.47591 | 44.2032 | 41.35164 | 18.14355 | 0.482588 | 0.845998 | 0.565957 |
| 7 | 85514000 | 85515599 | 17.89752 | 31.98497 | 31.00533 | 13.61472 | 0.443223 | 0.457658 | 0.565958 |
| 1 | 166373000 | 166377399 | 60.97742 | 97.15519 | 104.9597 | 43.93073 | 0.22598 | 2.183138 | 0.565963 |
| 12 | 60551000 | 60553199 | 19.86829 | 35.00397 | 34.08415 | 15.08667 | 0.426839 | 0.591342 | 0.565964 |
| X | 147559600 | 147561999 | 29.90828 | 51.29447 | 38.06788 | 16.69585 | 0.811967 | 0.961079 | 0.565977 |
| 3 | 133658400 | 133660799 | 22.50755 | 39.1307 | 40.30994 | 17.70544 | 0.344339 | 0.803275 | 0.565989 |
| 9 | 35988200 | 35989399 | 27.27942 | 45.91237 | 49.08945 | 21.4522 | 0.284921 | 1.128475 | 0.565997 |
| 5 | 68649200 | 68652199 | 29.79532 | 49.87739 | 52.97666 | 23.08673 | 0.294351 | 1.19097 | 0.566004 |
| 5 | 121995200 | 122000399 | 73.33973 | 113.9915 | 174.3595 | 72.3931 | -0.30469 | 2.747931 | 0.566005 |
| 2 | 27938800 | 27941799 | 20.01491 | 33.80558 | 55.36037 | 24.21864 | -0.32777 | 1.07818 | 0.566017 |
| 12 | 65211200 | 65215599 | 53.92309 | 87.06526 | 78.62381 | 33.53916 | 0.506165 | 1.764037 | 0.566048 |
| 9 | 39806600 | 39807399 | 25.57486 | 44.01232 | 38.80266 | 17.0209 | 0.565366 | 0.91361 | 0.566085 |
| 2 | 145332200 | 145333999 | 13.03668 | 22.94366 | 45.5911 | 20.18427 | -0.59749 | 0.584534 | 0.566089 |
| 1 | 88828600 | 88832399 | 37.73532 | 60.88169 | 92.29007 | 39.33013 | -0.2408 | 1.764436 | 0.566111 |
| 3 | 103821200 | 103822799 | 12.07945 | 21.29581 | 37.38522 | 16.5456 | -0.41819 | 0.547038 | 0.566112 |
| 14 | 123908400 | 123909399 | 15.70151 | 27.29839 | 44.75525 | 19.65815 | -0.32296 | 0.778142 | 0.566114 |
| 3 | 16892400 | 16894799 | 19.73107 | 34.8334 | 33.54201 | 14.84063 | 0.443969 | 0.567988 | 0.566135 |
| 11 | 19813800 | 19820399 | 54.50015 | 86.71795 | 127.8039 | 53.39241 | -0.22346 | 2.225946 | 0.566139 |
| 18 | 43483800 | 43486999 | 30.86182 | 50.17611 | 78.56878 | 33.71482 | -0.2763 | 1.570991 | 0.566148 |
| 1 | 48854000 | 48854999 | 17.4589 | 30.12328 | 48.46985 | 21.26311 | -0.29825 | 0.86794 | 0.566163 |
| X | 42855000 | 42856199 | 19.21345 | 34.19558 | 31.62677 | 13.9359 | 0.507601 | 0.49597 | 0.566164 |
| 5 | 144350600 | 144353399 | 26.27978 | 44.86037 | 46.29905 | 20.29821 | 0.337084 | 0.996755 | 0.566184 |
| 7 | 30349000 | 30353999 | 26.69309 | 43.836 | 70.4937 | 30.3663 | -0.30577 | 1.41735 | 0.566186 |
| 7 | 52183800 | 52186199 | 26.69309 | 43.836 | 70.4937 | 30.3663 | -0.30577 | 1.41735 | 0.566186 |
| 1 | 65578200 | 65581599 | 75.6463 | 119.1513 | 124.9242 | 51.83498 | 0.254387 | 2.499773 | 0.566226 |
| 6 | 54799400 | 54802799 | 44.07788 | 71.41961 | 74.02826 | 31.68952 | 0.312856 | 1.65784 | 0.566232 |

| | | | | | | | | | |
|---|---|---|---|---|---|---|---|---|---|
| 1 | 130307600 | 130315199 | 94.10366 | 146.9185 | 151.2174 | 62.74589 | 0.271153 | 2.676424 | 0.566252 |
| 7 | 18184400 | 18187999 | 39.0614 | 63.51499 | 68.94989 | 29.58855 | 0.250949 | 1.578991 | 0.566259 |
| 6 | 39735400 | 39736399 | 8.081403 | 15.32933 | 27.11906 | 10.93922 | -0.32694 | 0.017751 | 0.566276 |
| 18 | 32728800 | 32730199 | 10.17196 | 18.73232 | 31.40825 | 13.31074 | -0.30766 | 0.231521 | 0.566285 |
| 10 | 120705200 | 120705799 | 4.182942 | 8.16251 | 19.19174 | 6.832041 | -0.56222 | -0.47436 | 0.566302 |
| X | 148450800 | 148452599 | 27.1926 | 46.29541 | 47.39363 | 20.77364 | 0.348241 | 1.026189 | 0.566312 |
| 12 | 93920400 | 93920799 | 5.242373 | 10.21765 | 21.47127 | 7.770516 | -0.42715 | -0.40046 | 0.566318 |
| 1 | 9817600 | 9818799 | 9.585227 | 17.0991 | 34.44228 | 15.1469 | -0.60649 | 0.423998 | 0.566336 |
| 11 | 116803400 | 116804799 | 8.110208 | 15.17805 | 28.49152 | 11.78662 | -0.44276 | 0.109575 | 0.56634 |
| 16 | 89432600 | 89434399 | 46.66399 | 74.4183 | 115.4455 | 48.3813 | -0.29369 | 2.143069 | 0.566343 |
| 7 | 34615200 | 34616999 | 15.85503 | 28.22965 | 30.32083 | 13.35595 | 0.293161 | 0.46709 | 0.566349 |
| X | 87798800 | 87801399 | 20.01033 | 35.56933 | 32.02483 | 14.12619 | 0.545089 | 0.508811 | 0.566353 |
| 3 | 65946200 | 65946799 | 5.94955 | 11.5731 | 22.42101 | 8.283029 | -0.33806 | -0.31325 | 0.566356 |
| 13 | 3732400 | 3734999 | 24.95155 | 42.50915 | 45.80272 | 20.07742 | 0.274784 | 1.008656 | 0.566361 |
| 4 | 5899800 | 5901799 | 19.73078 | 34.60391 | 35.88127 | 15.8595 | 0.33544 | 0.654096 | 0.566376 |
| 1 | 74644400 | 74646799 | 21.06968 | 35.2486 | 59.05875 | 25.72709 | -0.36091 | 1.177606 | 0.566433 |
| 6 | 106555800 | 106556799 | 10.96897 | 20.88794 | 23.29536 | 9.319634 | 0.343845 | 0.008211 | 0.566434 |
| 18 | 79113000 | 79114999 | 10.69226 | 19.25274 | 33.74082 | 14.71839 | -0.39941 | 0.368345 | 0.566459 |
| 6 | 147360200 | 147363199 | 30.7144 | 51.73115 | 45.30829 | 19.80419 | 0.57165 | 1.128121 | 0.566462 |
| 8 | 4821000 | 4826399 | 62.14607 | 99.31195 | 100.368 | 42.18419 | 0.326631 | 2.089256 | 0.566467 |
| 2 | 71967200 | 71968599 | 9.960058 | 18.39352 | 31.44042 | 13.27371 | -0.33154 | 0.212389 | 0.566473 |
| 8 | 113342800 | 113345199 | 24.32373 | 41.63327 | 44.71673 | 19.60875 | 0.280362 | 0.963592 | 0.566498 |
| 9 | 117546600 | 117553999 | 70.93349 | 110.9679 | 163.4723 | 67.81686 | -0.24326 | 2.631012 | 0.566498 |
| 4 | 8460800 | 8463999 | 63.6661 | 101.4331 | 107.4446 | 44.96716 | 0.254255 | 2.188704 | 0.566509 |
| 10 | 7320200 | 7323199 | 26.79543 | 45.90668 | 39.81739 | 17.46165 | 0.587667 | 0.958448 | 0.566523 |
| 19 | 9251000 | 9255199 | 53.19297 | 85.35622 | 89.24281 | 37.73746 | 0.284499 | 1.948512 | 0.566537 |
| 5 | 137216000 | 137220199 | 37.12345 | 59.7807 | 93.00083 | 39.53101 | -0.28135 | 1.821705 | 0.566543 |
| 13 | 83875200 | 83875999 | 5.654123 | 11.0075 | 21.86396 | 8.023701 | -0.3639 | -0.33932 | 0.566553 |
| 11 | 99540400 | 99541999 | 17.18665 | 29.63457 | 47.94266 | 21.03143 | -0.30612 | 0.873211 | 0.566557 |
| 1 | 110264000 | 110265399 | 16.3873 | 28.93209 | 29.41061 | 13.01253 | 0.36626 | 0.539232 | 0.566566 |
| 1 | 151465800 | 151467799 | 16.64373 | 29.00061 | 46.27573 | 20.35428 | -0.28388 | 0.752858 | 0.566596 |
| 7 | 89497600 | 89499999 | 16.64373 | 29.00061 | 46.27573 | 20.35428 | -0.28388 | 0.752858 | 0.566596 |
| 7 | 139589600 | 139597399 | 136.5308 | 207.0498 | 216.5409 | 89.63372 | 0.215465 | 3.246128 | 0.566601 |
| 12 | 5858000 | 5859399 | 8.06034 | 15.35734 | 27.23496 | 10.88339 | -0.31866 | -0.01341 | 0.566604 |
| 13 | 58874200 | 58877399 | 25.7918 | 42.85172 | 67.05705 | 29.08125 | -0.26299 | 1.274109 | 0.566607 |
| 8 | 86091200 | 86091999 | 6.592427 | 12.59892 | 25.09965 | 9.965572 | -0.47587 | -0.05263 | 0.566608 |
| 7 | 29796600 | 29805599 | 137.7277 | 209.343 | 214.523 | 88.85926 | 0.247529 | 3.199396 | 0.566634 |
| 5 | 7688400 | 7689199 | 11.33033 | 21.46727 | 23.68621 | 9.577867 | 0.345339 | 0.045213 | 0.566647 |
| 12 | 115703800 | 115707999 | 54.44216 | 87.56052 | 86.57874 | 36.735 | 0.368839 | 1.88008 | 0.566674 |
| 18 | 6373000 | 6373799 | 7.644325 | 14.48164 | 26.55852 | 10.74224 | -0.38164 | 0.021659 | 0.566697 |
| 4 | 34455400 | 34456399 | 10.30346 | 19.09128 | 32.59851 | 13.7003 | -0.32525 | 0.199973 | 0.566699 |
| 7 | 15703400 | 15706799 | 15.73005 | 28.43336 | 25.2179 | 10.9476 | 0.58168 | 0.364133 | 0.566702 |
| 1 | 143170200 | 143172199 | 15.1393 | 28.04073 | 26.27479 | 11.04844 | 0.533841 | 0.224521 | 0.566715 |
| 17 | 50608600 | 50609199 | 6.547657 | 12.68786 | 23.87321 | 9.043663 | -0.33018 | -0.21331 | 0.566725 |
| X | 8366000 | 8383999 | 263.4693 | 385.8015 | 328.5358 | 135.3331 | 0.465928 | 3.896245 | 0.566744 |
| 10 | 122903200 | 122907599 | 59.71464 | 94.79728 | 146.6835 | 61.12651 | -0.29696 | 2.303994 | 0.56675 |
| 18 | 47786600 | 47788199 | 19.00627 | 33.25392 | 35.54935 | 15.68118 | 0.291693 | 0.686971 | 0.566752 |

| | | | | | | | | | |
|---|---|---|---|---|---|---|---|---|---|
| X | 34494200 | 34497599 | 56.36769 | 90.14266 | 89.94979 | 37.84401 | 0.346138 | 2.062487 | 0.566752 |
| 2 | 49003200 | 49005199 | 30.36037 | 51.70546 | 46.23651 | 20.26699 | 0.542638 | 1.032291 | 0.566769 |
| 7 | 62507800 | 62510199 | 33.40751 | 56.30865 | 49.65906 | 21.71077 | 0.561774 | 1.1348 | 0.566772 |
| 10 | 9867400 | 9868399 | 12.79483 | 23.95209 | 20.96575 | 8.669066 | 0.650577 | 0.130779 | 0.566773 |
| 18 | 58697600 | 58698399 | 10.10429 | 19.23773 | 21.72141 | 8.693434 | 0.325624 | -0.00374 | 0.566831 |
| 1 | 162802000 | 162805599 | 57.58803 | 92.0983 | 98.12408 | 41.28571 | 0.251794 | 2.061118 | 0.566845 |
| 14 | 14655800 | 14659399 | 30.73737 | 51.10216 | 54.68745 | 23.7276 | 0.283543 | 1.267709 | 0.566849 |
| 14 | 16841200 | 16841399 | 4.003263 | 7.823328 | 18.22876 | 6.316051 | -0.5131 | -0.57531 | 0.566866 |
| 16 | 29955800 | 29957399 | 16.34443 | 28.87973 | 29.52607 | 13.0615 | 0.358673 | 0.532247 | 0.566867 |
| 4 | 113496000 | 113497399 | 28.59803 | 49.14978 | 44.03418 | 19.31494 | 0.541973 | 0.945188 | 0.566891 |
| 16 | 80513200 | 80514999 | 18.51752 | 33.25677 | 29.86904 | 13.06128 | 0.556736 | 0.429255 | 0.566891 |
| 7 | 41183400 | 41218799 | 560.9092 | 764.7834 | 664.829 | 276.7682 | 0.33348 | 4.972861 | 0.566924 |
| 2 | 75534000 | 75536999 | 20.00845 | 33.76772 | 55.90389 | 24.4505 | -0.3435 | 1.085065 | 0.566929 |
| 5 | 53904200 | 53905799 | 15.18723 | 26.53206 | 43.08119 | 18.98034 | -0.30842 | 0.703668 | 0.566942 |
| 3 | 16378400 | 16381399 | 46.34803 | 75.32028 | 71.02267 | 30.46925 | 0.453018 | 1.604114 | 0.566943 |
| 3 | 130257000 | 130259399 | 21.81468 | 37.87408 | 40.38949 | 17.72509 | 0.294431 | 0.819023 | 0.566946 |
| 12 | 113451400 | 113455999 | 37.6182 | 61.68393 | 64.39778 | 27.7226 | 0.314806 | 1.456005 | 0.566974 |
| 5 | 44032400 | 44033799 | 16.126 | 29.37572 | 28.82831 | 12.3943 | 0.445678 | 0.332266 | 0.566995 |
| 8 | 68479600 | 68481599 | 14.61793 | 25.69607 | 41.35359 | 18.31256 | -0.29543 | 0.600641 | 0.567011 |
| 14 | 18053200 | 18055199 | 14.95688 | 25.53213 | 56.32012 | 24.69162 | -0.75413 | 0.966034 | 0.567012 |
| 10 | 58634800 | 58637399 | 24.50126 | 40.54542 | 66.25692 | 28.65519 | -0.32573 | 1.314827 | 0.567019 |
| 19 | 15096200 | 15097799 | 18.29385 | 32.19269 | 33.18291 | 14.69116 | 0.344405 | 0.592001 | 0.567045 |
| 19 | 32488200 | 32491999 | 41.35853 | 66.98419 | 72.78931 | 31.14913 | 0.244311 | 1.662196 | 0.567057 |
| X | 66826800 | 66829999 | 41.98217 | 68.98244 | 62.97097 | 27.13214 | 0.509184 | 1.443377 | 0.567094 |
| 14 | 44478400 | 44479599 | 12.09923 | 23.10875 | 23.25978 | 9.247178 | 0.498063 | -0.00323 | 0.567095 |
| 2 | 19449000 | 19451799 | 27.63343 | 44.96617 | 76.48532 | 32.83113 | -0.39443 | 1.550021 | 0.567111 |
| 5 | 41864400 | 41866199 | 15.03662 | 25.87618 | 52.66546 | 23.10184 | -0.63646 | 0.888082 | 0.567112 |
| 19 | 21020000 | 21023599 | 50.88433 | 81.58544 | 87.18528 | 36.82195 | 0.251688 | 1.972165 | 0.567132 |
| 11 | 95674400 | 95679599 | 40.64589 | 65.31031 | 100.5322 | 42.59738 | -0.27009 | 1.895998 | 0.567135 |
| X | 116038400 | 116044799 | 118.1548 | 182.7551 | 149.7963 | 62.22176 | 0.588948 | 2.849766 | 0.567139 |
| 16 | 4614600 | 4631799 | 259.7508 | 377.9877 | 359.729 | 148.265 | 0.29695 | 3.989354 | 0.56714 |
| 1 | 40187000 | 40187399 | 6.994429 | 13.43097 | 25.21847 | 9.877232 | -0.37346 | -0.09554 | 0.567145 |
| 15 | 35552400 | 35553199 | 6.994429 | 13.43097 | 25.21847 | 9.877232 | -0.37346 | -0.09554 | 0.567145 |
| 16 | 22651200 | 22651999 | 9.495612 | 17.78887 | 30.67828 | 12.66942 | -0.31953 | 0.121406 | 0.567155 |
| 8 | 80256400 | 80262799 | 58.5994 | 92.96554 | 137.1658 | 57.12258 | -0.22934 | 2.318596 | 0.567201 |
| 8 | 74269800 | 74271999 | 40.02201 | 64.89856 | 70.49581 | 30.19477 | 0.246492 | 1.633886 | 0.567214 |
| 5 | 91777000 | 91778999 | 15.50187 | 27.12093 | 43.54837 | 19.20868 | -0.29241 | 0.682904 | 0.567214 |
| 8 | 79316800 | 79319799 | 20.74846 | 34.7408 | 58.5063 | 25.50014 | -0.36825 | 1.167086 | 0.567216 |
| 5 | 77742000 | 77742999 | 8.872577 | 16.75866 | 29.2244 | 11.89497 | -0.31799 | 0.06001 | 0.567231 |
| 6 | 59507800 | 59508199 | 4.100951 | 8.016708 | 17.74256 | 6.098882 | -0.43159 | -0.60985 | 0.567232 |
| 7 | 55723600 | 55724799 | 25.72849 | 44.00387 | 39.97101 | 17.52662 | 0.521024 | 0.970549 | 0.567241 |
| 17 | 13538200 | 13542799 | 102.4553 | 157.2705 | 238.8239 | 99.15246 | -0.30821 | 2.994943 | 0.567247 |
| 6 | 39817600 | 39821399 | 37.65372 | 60.78705 | 95.18233 | 40.59458 | -0.28643 | 1.749943 | 0.56727 |
| 7 | 101448600 | 101457799 | 118.287 | 181.5864 | 187.0002 | 77.63767 | 0.251708 | 2.993547 | 0.567276 |
| X | 152973800 | 152978799 | 63.95445 | 102.9958 | 78.91778 | 33.54891 | 0.738403 | 1.854536 | 0.567281 |
| 6 | 117676400 | 117677199 | 6.864399 | 13.20659 | 24.70403 | 9.622609 | -0.35995 | -0.11374 | 0.567291 |
| 10 | 5320800 | 5321399 | 6.864399 | 13.20659 | 24.70403 | 9.622609 | -0.35995 | -0.11374 | 0.567291 |

| | | | | | | | | | |
|---|---|---|---|---|---|---|---|---|---|
| 12 | 88204600 | 88216999 | 168.8052 | 254.3324 | 242.3048 | 100.0966 | 0.341917 | 3.384176 | 0.567358 |
| 14 | 61975200 | 61978999 | 36.05438 | 58.31033 | 89.99035 | 38.46958 | -0.26258 | 1.689487 | 0.567361 |
| 7 | 11438200 | 11470399 | 496.6503 | 686.2483 | 575.0411 | 238.8871 | 0.405866 | 4.766173 | 0.567391 |
| X | 53185000 | 53191199 | 87.84 | 139.1622 | 119.7843 | 49.81997 | 0.545179 | 2.378968 | 0.567417 |
| 5 | 121027800 | 121032599 | 52.04099 | 82.82514 | 123.2935 | 51.52539 | -0.23746 | 2.214872 | 0.56742 |
| 1 | 61456200 | 61458799 | 16.09745 | 28.02492 | 46.82232 | 20.5837 | -0.35014 | 0.760884 | 0.567427 |
| 4 | 81690200 | 81692799 | 19.34811 | 34.50906 | 30.81761 | 13.55487 | 0.559253 | 0.478547 | 0.567446 |
| 12 | 49980600 | 49982199 | 16.85583 | 30.74972 | 28.09059 | 12.05528 | 0.549712 | 0.325382 | 0.56745 |
| 4 | 146026400 | 146042999 | 49.27144 | 79.90081 | 64.02862 | 27.42566 | 0.684341 | 1.647975 | 0.567451 |
| 3 | 54433000 | 54435799 | 34.14275 | 56.32721 | 60.33434 | 26.04036 | 0.281328 | 1.370768 | 0.567469 |
| 14 | 44994800 | 45003399 | 241.7743 | 351.7306 | 354.6798 | 146.1881 | 0.213122 | 3.992093 | 0.56748 |
| Y | 53878000 | 53879599 | 12.3138 | 22.86941 | 17.6168 | 7.380467 | 0.81816 | 0.163901 | 0.567481 |
| 15 | 96155800 | 96170999 | 246.6479 | 354.5336 | 477.9596 | 197.3041 | -0.22266 | 4.172147 | 0.567487 |
| 8 | 80625000 | 80625599 | 8.823109 | 17.10385 | 19.88456 | 7.518914 | 0.359607 | -0.20897 | 0.56751 |
| 4 | 143466400 | 143481399 | 393.2809 | 549.4502 | 526.4139 | 218.3083 | 0.228702 | 4.595481 | 0.56753 |
| 5 | 106668200 | 106670599 | 18.34477 | 31.55529 | 50.06117 | 21.95917 | -0.2788 | 0.903153 | 0.567543 |
| 2 | 137392400 | 137393399 | 9.631175 | 18.59084 | 21.40966 | 8.258574 | 0.34612 | -0.12834 | 0.567544 |
| 12 | 47425400 | 47426999 | 20.06232 | 34.90186 | 37.35868 | 16.41576 | 0.289445 | 0.7792 | 0.567546 |
| 12 | 114272800 | 114284599 | 234.4164 | 345.0576 | 319.593 | 131.6101 | 0.351941 | 3.810387 | 0.567565 |
| 11 | 21041000 | 21041999 | 9.750222 | 17.77609 | 31.35 | 13.46451 | -0.39346 | 0.28569 | 0.567565 |
| 17 | 74314000 | 74316199 | 19.93231 | 33.94476 | 53.81242 | 23.5877 | -0.28027 | 1.002834 | 0.567572 |
| 1 | 119051200 | 119053999 | 17.37507 | 30.20206 | 52.3119 | 22.97446 | -0.40233 | 0.795685 | 0.567582 |
| 7 | 10132000 | 10133999 | 29.48634 | 49.80397 | 50.17205 | 21.94902 | 0.37062 | 1.104898 | 0.567626 |
| 1 | 141690800 | 141691599 | 20.67115 | 35.9414 | 38.83715 | 17.05956 | 0.275703 | 0.791039 | 0.567643 |
| 9 | 36103600 | 36106399 | 29.39235 | 49.90031 | 48.48447 | 21.24374 | 0.422859 | 1.060545 | 0.567654 |
| 6 | 29334000 | 29336799 | 16.50373 | 28.67502 | 45.80315 | 20.11044 | -0.28569 | 0.792585 | 0.567683 |
| 2 | 24081000 | 24088399 | 52.07707 | 83.71054 | 87.97731 | 37.2986 | 0.280063 | 1.895741 | 0.567705 |
| 16 | 73950600 | 73954199 | 42.01504 | 69.01756 | 63.0898 | 27.1802 | 0.507014 | 1.44515 | 0.567748 |
| 13 | 95567000 | 95567799 | 7.167558 | 13.65234 | 26.11507 | 10.44231 | -0.4276 | -0.02365 | 0.567804 |
| 12 | 114478800 | 114492599 | 231.8109 | 340.7282 | 325.8065 | 134.1771 | 0.303987 | 3.833019 | 0.567816 |
| 12 | 22666000 | 22668199 | 47.0258 | 75.88604 | 70.02167 | 29.848 | 0.47415 | 1.762216 | 0.567819 |
| 4 | 69353400 | 69354799 | 19.75098 | 34.39206 | 36.34808 | 15.98099 | 0.307855 | 0.763483 | 0.56782 |
| 11 | 73813600 | 73813999 | 6.093245 | 11.8593 | 23.10313 | 8.496294 | -0.34006 | -0.32538 | 0.567842 |
| 4 | 145350800 | 145356999 | 157.6444 | 239.7652 | 208.3521 | 86.31926 | 0.48595 | 3.193206 | 0.567853 |
| X | 45035800 | 45037599 | 14.1706 | 25.08494 | 43.98652 | 19.45122 | -0.41471 | 0.525923 | 0.567858 |
| 3 | 9270000 | 9273399 | 26.15643 | 44.08876 | 47.48913 | 20.76235 | 0.274309 | 1.108062 | 0.567866 |
| 6 | 134768600 | 134771399 | 15.55717 | 26.86891 | 47.93747 | 21.03021 | -0.44629 | 0.848793 | 0.567884 |
| 7 | 49319400 | 49321999 | 21.54834 | 36.52647 | 58.78907 | 25.74662 | -0.30284 | 1.054375 | 0.567888 |
| X | 4377200 | 4378199 | 21.38016 | 37.24628 | 38.30212 | 16.84505 | 0.347271 | 0.769366 | 0.567897 |
| 15 | 103918800 | 103920999 | 21.18137 | 37.35288 | 33.39376 | 14.77827 | 0.549946 | 0.587649 | 0.567921 |
| 3 | 145464200 | 145465399 | 9.562843 | 17.51907 | 30.85662 | 13.17 | -0.38555 | 0.254525 | 0.567978 |
| 19 | 29091200 | 29094199 | 26.59495 | 44.08831 | 69.39699 | 30.05003 | -0.27171 | 1.30283 | 0.567995 |
| 7 | 12911800 | 12913999 | 21.20128 | 37.03632 | 38.07788 | 16.77538 | 0.347749 | 0.727311 | 0.568011 |
| 7 | 26647200 | 26650599 | 48.4637 | 78.01168 | 82.74772 | 35.1529 | 0.269247 | 1.84291 | 0.568043 |
| 5 | 109305800 | 109308999 | 57.61345 | 91.99939 | 99.58891 | 41.81217 | 0.226533 | 2.107925 | 0.568057 |
| 18 | 26215400 | 26217799 | 22.09971 | 38.42756 | 33.70237 | 14.80461 | 0.575517 | 0.790329 | 0.568057 |
| X | 78955800 | 78957999 | 29.37517 | 50.12714 | 46.28398 | 20.29104 | 0.497172 | 1.014732 | 0.568062 |

| | | | | | | | | | |
|---|---|---|---|---|---|---|---|---|---|
| 3 | 27763400 | 27765399 | 16.57129 | 28.44041 | 48.9986 | 21.49105 | -0.3979 | 0.913461 | 0.568064 |
| 1 | 143161800 | 143164399 | 27.72174 | 47.21207 | 47.38897 | 20.77216 | 0.376651 | 1.025555 | 0.568075 |
| X | 55034000 | 55045199 | 162.9658 | 246.8269 | 222.4785 | 92.05532 | 0.42828 | 3.279398 | 0.568077 |
| 5 | 124005600 | 124006599 | 8.298199 | 15.62472 | 28.05826 | 11.4883 | -0.36666 | 0.074756 | 0.56808 |
| 7 | 85581600 | 85593199 | 183.8108 | 275.2231 | 263.1695 | 108.5461 | 0.328664 | 3.507116 | 0.568081 |
| 2 | 36269600 | 36274199 | 72.65679 | 115.8954 | 91.43348 | 38.32951 | 0.680461 | 2.155134 | 0.568091 |
| 2 | 67026200 | 67027799 | 15.96003 | 27.8611 | 45.17716 | 19.89433 | -0.3067 | 0.720516 | 0.568099 |
| 1 | 101143600 | 101147799 | 43.24471 | 70.69597 | 65.98638 | 28.37232 | 0.473692 | 1.508931 | 0.568129 |
| 9 | 118311200 | 118314399 | 21.4421 | 36.07423 | 58.68701 | 25.6434 | -0.31865 | 1.119051 | 0.56813 |
| 8 | 23218000 | 23221599 | 17.95958 | 31.1742 | 50.64536 | 22.22289 | -0.31034 | 0.81898 | 0.568131 |
| 8 | 46009000 | 46010399 | 14.49456 | 26.21202 | 28.34417 | 12.29778 | 0.297955 | 0.359203 | 0.568151 |
| 9 | 113012200 | 113013599 | 16.03983 | 28.97119 | 29.75774 | 12.93077 | 0.370331 | 0.380711 | 0.568167 |
| X | 68963400 | 68965199 | 23.30958 | 40.66335 | 38.14704 | 16.79134 | 0.479375 | 0.760294 | 0.568171 |
| 10 | 119978000 | 119990799 | 167.8045 | 248.7545 | 344.1738 | 141.7533 | -0.21725 | 3.68621 | 0.568183 |
| 15 | 51987000 | 51989399 | 35.64365 | 57.50766 | 89.29209 | 38.05307 | -0.27526 | 1.760252 | 0.568192 |
| 1 | 87430600 | 87433999 | 29.17893 | 47.65385 | 74.87718 | 32.18343 | -0.27667 | 1.499683 | 0.568195 |
| 14 | 82047800 | 82050399 | 22.19925 | 38.8782 | 36.80137 | 16.24679 | 0.466362 | 0.690998 | 0.568197 |
| 11 | 6863800 | 6868999 | 57.9841 | 93.27789 | 85.40127 | 36.24712 | 0.48004 | 1.878642 | 0.568228 |
| 16 | 87413600 | 87415399 | 17.55092 | 30.08935 | 49.52463 | 21.71989 | -0.33248 | 0.929277 | 0.568242 |
| 3 | 27526400 | 27528999 | 32.33801 | 52.39598 | 82.34191 | 35.24765 | -0.2861 | 1.643166 | 0.568267 |
| 12 | 79774400 | 79776399 | 27.96893 | 47.92384 | 45.15062 | 19.80072 | 0.468933 | 0.969092 | 0.568281 |
| 3 | 52324000 | 52327799 | 31.27954 | 51.15413 | 80.69548 | 34.70146 | -0.28106 | 1.487922 | 0.568288 |
| 14 | 40964800 | 40965399 | 5.663513 | 11.011 | 21.76432 | 8.074125 | -0.37169 | -0.30269 | 0.568293 |
| 1 | 162402400 | 162402999 | 9.420511 | 18.37929 | 18.05199 | 6.448695 | 0.675024 | -0.40134 | 0.568294 |
| 10 | 105203600 | 105209599 | 53.61415 | 86.05682 | 90.44279 | 38.26162 | 0.27754 | 1.941799 | 0.568309 |
| 10 | 74481200 | 74481799 | 7.025544 | 13.48765 | 25.1974 | 9.875422 | -0.36642 | -0.08998 | 0.568313 |
| 18 | 27519800 | 27521799 | 16.41115 | 29.52588 | 28.96124 | 12.64549 | 0.431591 | 0.400876 | 0.568313 |
| 18 | 37433800 | 37436999 | 70.16013 | 111.4145 | 94.39597 | 39.34941 | 0.571127 | 2.29117 | 0.568343 |
| 17 | 23585800 | 23591399 | 56.34601 | 90.71577 | 84.10591 | 35.73902 | 0.46337 | 1.85129 | 0.568346 |
| 11 | 28116000 | 28118199 | 29.74659 | 50.12405 | 44.22911 | 19.33516 | 0.56088 | 1.118177 | 0.568374 |
| 7 | 34545400 | 34551799 | 58.84601 | 93.10344 | 138.2693 | 57.46928 | -0.24216 | 2.390519 | 0.568375 |
| 14 | 31117000 | 31118799 | 20.71715 | 34.90343 | 56.40574 | 24.65707 | -0.30882 | 1.105331 | 0.568463 |
| 18 | 13235000 | 13237799 | 29.52008 | 49.45044 | 51.27302 | 22.35394 | 0.329101 | 1.184945 | 0.568464 |
| 17 | 17515800 | 17518599 | 29.83141 | 51.22966 | 43.42921 | 19.04912 | 0.621148 | 0.955324 | 0.568466 |
| 8 | 85338000 | 85338999 | 8.29344 | 15.54889 | 28.00269 | 11.54918 | -0.37935 | 0.101343 | 0.568478 |
| 4 | 145116400 | 145137999 | 141.806 | 215.6247 | 192.9156 | 79.91824 | 0.443639 | 3.191939 | 0.568481 |
| 15 | 72136800 | 72139399 | 24.16545 | 41.86033 | 41.92187 | 18.39282 | 0.384114 | 0.853712 | 0.568499 |
| 4 | 145447600 | 145449199 | 23.04224 | 40.06627 | 33.70322 | 14.80495 | 0.635554 | 0.796289 | 0.568509 |
| X | 126513800 | 126515599 | 33.2837 | 55.85502 | 52.45565 | 22.8954 | 0.471332 | 1.177993 | 0.56852 |
| 11 | 3587000 | 3588999 | 27.07952 | 46.8749 | 41.82732 | 18.35094 | 0.549769 | 0.873537 | 0.568541 |
| 10 | 48440600 | 48441199 | 13.19679 | 24.09533 | 25.51289 | 10.93727 | 0.339396 | 0.289128 | 0.568547 |
| 2 | 53880400 | 53882599 | 23.75505 | 39.77 | 62.62341 | 27.29244 | -0.27183 | 1.180331 | 0.568569 |
| 14 | 11017600 | 11019199 | 9.992279 | 18.57699 | 31.57586 | 13.20973 | -0.31377 | 0.176792 | 0.568576 |
| 19 | 35876400 | 35877199 | 9.992279 | 18.57699 | 31.57586 | 13.20973 | -0.31377 | 0.176792 | 0.568576 |
| 1 | 35528400 | 35531599 | 34.63541 | 57.56912 | 56.0293 | 24.30581 | 0.420018 | 1.282053 | 0.568577 |
| 1 | 21204400 | 21206599 | 27.80536 | 47.47064 | 40.89389 | 17.92869 | 0.596772 | 0.995001 | 0.568587 |
| 1 | 34940200 | 34943399 | 32.48765 | 53.90073 | 57.49832 | 24.91165 | 0.287962 | 1.299817 | 0.568593 |

| | | | | | | | | | |
|---|---|---|---|---|---|---|---|---|---|
| 12 | 93989400 | 93990799 | 9.048812 | 17.04538 | 29.70317 | 12.15296 | -0.32332 | 0.081272 | 0.568595 |
| 7 | 84863000 | 84866799 | 42.02059 | 69.79836 | 54.49437 | 23.62919 | 0.73711 | 1.313581 | 0.568609 |
| 7 | 144448800 | 144451599 | 20.40982 | 34.24771 | 57.44227 | 25.07053 | -0.36244 | 1.141698 | 0.568613 |
| 4 | 106713800 | 106716399 | 21.32446 | 35.85558 | 57.53736 | 25.13665 | -0.2989 | 1.124035 | 0.568638 |
| 9 | 68840400 | 68844199 | 28.51483 | 46.57063 | 74.87657 | 32.18351 | -0.30972 | 1.496798 | 0.568672 |
| 7 | 75890600 | 75896599 | 56.41012 | 89.24303 | 153.2281 | 63.6843 | -0.45127 | 2.392869 | 0.568685 |
| 2 | 54199000 | 54199999 | 10.63057 | 19.09903 | 33.84875 | 14.79702 | -0.41702 | 0.386112 | 0.568698 |
| 2 | 174602200 | 174604199 | 15.19777 | 26.41495 | 45.27892 | 19.88454 | -0.38705 | 0.779119 | 0.568717 |
| 4 | 40461600 | 40465599 | 46.73519 | 75.40687 | 80.02483 | 34.10607 | 0.272581 | 1.769808 | 0.568735 |
| 15 | 45445600 | 45448399 | 22.84943 | 40.19246 | 34.45299 | 15.25475 | 0.609097 | 0.627522 | 0.568735 |
| 13 | 53187600 | 53191799 | 43.76056 | 70.23176 | 106.235 | 44.93379 | -0.24702 | 1.939254 | 0.568749 |
| 10 | 3466200 | 3469799 | 28.95592 | 47.54121 | 73.6672 | 31.73026 | -0.25259 | 1.427705 | 0.568762 |
| 10 | 85752200 | 85753399 | 10.05391 | 18.24067 | 32.30316 | 13.95867 | -0.40537 | 0.317079 | 0.568767 |
| 5 | 10285400 | 10287199 | 24.19407 | 40.29157 | 63.52459 | 27.59306 | -0.27346 | 1.241496 | 0.568775 |
| 7 | 7533000 | 7534999 | 22.39853 | 40.15431 | 29.1337 | 12.75915 | 0.861737 | 0.467459 | 0.568783 |
| 8 | 65672200 | 65676399 | 44.62842 | 71.56554 | 106.9638 | 45.18535 | -0.23117 | 1.967227 | 0.568802 |
| 18 | 32815800 | 32816799 | 18.45303 | 32.28647 | 33.23475 | 14.66039 | 0.346172 | 0.682036 | 0.568803 |
| 4 | 16788000 | 16789799 | 17.2618 | 31.32094 | 29.09584 | 12.57119 | 0.519573 | 0.361188 | 0.568823 |
| 4 | 90038400 | 90039999 | 21.63061 | 37.43236 | 40.45467 | 17.74857 | 0.274371 | 0.848528 | 0.568834 |
| 1 | 29960600 | 29961599 | 16.66344 | 29.72458 | 31.20277 | 13.72277 | 0.327546 | 0.462223 | 0.568834 |
| 4 | 155515000 | 155516999 | 16.66344 | 29.72458 | 31.20277 | 13.72277 | 0.327546 | 0.462223 | 0.568834 |
| 1 | 117606800 | 117608599 | 35.14048 | 57.66475 | 61.69532 | 26.56773 | 0.280054 | 1.442066 | 0.568838 |
| 12 | 27956400 | 27957199 | 6.250347 | 12.08538 | 25.52446 | 9.768112 | -0.51049 | -0.18209 | 0.568841 |
| 3 | 128742200 | 128744599 | 18.03744 | 32.53673 | 30.07501 | 13.09002 | 0.520081 | 0.402529 | 0.568865 |
| 3 | 145784000 | 145792599 | 100.9644 | 154.2359 | 222.1539 | 92.144 | -0.23739 | 3.088403 | 0.568893 |
| 16 | 56228200 | 56228999 | 15.82425 | 28.48024 | 29.35698 | 12.81355 | 0.361349 | 0.400278 | 0.56892 |
| 18 | 13609000 | 13613999 | 31.0997 | 50.75493 | 78.71476 | 33.82417 | -0.25867 | 1.515186 | 0.568921 |
| 13 | 114196600 | 114197999 | 12.95413 | 22.97553 | 37.16849 | 16.42578 | -0.2977 | 0.486068 | 0.568927 |
| X | 31874600 | 31887799 | 222.6059 | 328.738 | 272.5005 | 112.2211 | 0.516305 | 3.755238 | 0.568933 |
| 10 | 32279000 | 32279799 | 12.66207 | 23.3443 | 25.49066 | 10.78634 | 0.308364 | 0.229824 | 0.568936 |
| 6 | 23520600 | 23522799 | 23.70874 | 41.24827 | 39.63283 | 17.41579 | 0.444382 | 0.796854 | 0.568956 |
| 12 | 24049200 | 24050599 | 20.21454 | 35.4293 | 30.93802 | 13.66708 | 0.582234 | 0.660145 | 0.568979 |
| 3 | 71344600 | 71346399 | 16.75723 | 29.11664 | 47.388 | 20.80682 | -0.3125 | 0.797511 | 0.568987 |
| 19 | 41074000 | 41076199 | 18.50643 | 31.61744 | 51.11299 | 22.41023 | -0.30753 | 0.966103 | 0.568995 |
| 5 | 104335200 | 104340399 | 167.105 | 249.7349 | 257.0893 | 105.9992 | 0.219766 | 3.538707 | 0.56901 |
| 19 | 21719400 | 21719999 | 6.704799 | 12.9746 | 24.36996 | 9.291046 | -0.33642 | -0.18786 | 0.569066 |
| 5 | 148214400 | 148230399 | 171.3806 | 253.0411 | 352.6687 | 145.2356 | -0.23317 | 3.753149 | 0.569076 |
| 13 | 65725600 | 65729999 | 48.86514 | 79.10773 | 75.53506 | 32.31385 | 0.429898 | 1.686701 | 0.56909 |
| 18 | 81420000 | 81421399 | 14.70398 | 26.86655 | 27.76748 | 11.89309 | 0.374529 | 0.299553 | 0.569103 |
| 12 | 119407800 | 119408199 | 7.584131 | 14.77005 | 17.39313 | 6.36087 | 0.387052 | -0.34297 | 0.56911 |
| 14 | 92778800 | 92781599 | 31.70128 | 52.95259 | 54.68085 | 23.7967 | 0.334953 | 1.222382 | 0.569114 |
| 7 | 12073400 | 12076399 | 51.2043 | 82.29186 | 87.83521 | 37.22802 | 0.257431 | 1.899781 | 0.569115 |
| 18 | 50281400 | 50283599 | 20.46705 | 35.02683 | 57.71998 | 25.3105 | -0.33489 | 0.965548 | 0.569128 |
| 16 | 59711600 | 59713199 | 17.3474 | 30.11492 | 47.93224 | 21.03385 | -0.28074 | 0.812376 | 0.569133 |
| X | 87615400 | 87616799 | 17.3474 | 30.11492 | 47.93224 | 21.03385 | -0.28074 | 0.812376 | 0.569133 |
| 9 | 66057800 | 66058399 | 5.674169 | 11.01688 | 22.24019 | 8.340843 | -0.41679 | -0.26403 | 0.569154 |
| 1 | 117488200 | 117489799 | 26.79885 | 45.60739 | 47.39442 | 20.77334 | 0.326579 | 1.027368 | 0.569164 |

| | | | | | | | | | |
|---|---|---|---|---|---|---|---|---|---|
| 10 | 26658200 | 26660199 | 15.69287 | 28.49488 | 28.67127 | 12.37649 | 0.405766 | 0.344085 | 0.569165 |
| 7 | 59442600 | 59449399 | 95.70545 | 149.0592 | 135.1491 | 56.096 | 0.451193 | 2.717537 | 0.569175 |
| 11 | 15222400 | 15224799 | 29.11099 | 48.95267 | 51.28923 | 22.40764 | 0.313932 | 1.141614 | 0.569217 |
| 6 | 129884200 | 129887199 | 25.57142 | 44.70095 | 37.28474 | 16.43439 | 0.648273 | 0.740094 | 0.569224 |
| 12 | 68758200 | 68758799 | 6.560401 | 12.71037 | 24.42364 | 9.259599 | -0.36102 | -0.20729 | 0.569243 |
| 10 | 91166600 | 91168999 | 27.00223 | 44.47253 | 69.55706 | 29.99963 | -0.26412 | 1.382161 | 0.56925 |
| 9 | 90010600 | 90012799 | 24.65559 | 41.91477 | 43.69952 | 19.15218 | 0.321587 | 1.022899 | 0.56928 |
| 7 | 5650800 | 5653199 | 32.28697 | 54.5816 | 49.62308 | 21.71456 | 0.51824 | 1.108311 | 0.569287 |
| 10 | 125211600 | 125215399 | 26.12168 | 44.37673 | 47.40907 | 20.77669 | 0.28644 | 1.040408 | 0.569324 |
| 17 | 85405400 | 85407599 | 34.2349 | 56.8701 | 57.09414 | 24.75773 | 0.375342 | 1.289489 | 0.569332 |
| 1 | 101637000 | 101640199 | 36.05326 | 59.36755 | 54.79378 | 23.63028 | 0.494458 | 1.394335 | 0.569342 |
| X | 15719400 | 15722599 | 55.29241 | 88.99035 | 84.32145 | 35.81361 | 0.431499 | 1.863204 | 0.56935 |
| 4 | 101784400 | 101787799 | 31.56928 | 51.19602 | 92.96529 | 39.82129 | -0.49293 | 1.630179 | 0.569352 |
| 12 | 64964400 | 64965199 | 9.503847 | 17.57357 | 30.41019 | 12.81635 | -0.34717 | 0.199504 | 0.569358 |
| 17 | 83905200 | 83906799 | 9.503847 | 17.57357 | 30.41019 | 12.81635 | -0.34717 | 0.199504 | 0.569358 |
| 9 | 70514200 | 70514999 | 7.137006 | 13.78942 | 25.48136 | 9.785917 | -0.3238 | -0.15724 | 0.569384 |
| 13 | 86404000 | 86408199 | 46.06558 | 74.70646 | 73.25913 | 31.38077 | 0.393835 | 1.643818 | 0.569392 |
| 2 | 163628800 | 163632999 | 24.14042 | 40.14909 | 63.58175 | 27.59356 | -0.27999 | 1.254187 | 0.569392 |
| X | 137254400 | 137257599 | 39.04457 | 63.95908 | 65.52954 | 28.19793 | 0.341324 | 1.473664 | 0.569396 |
| 15 | 82607600 | 82610399 | 27.86823 | 47.52846 | 41.46367 | 18.17697 | 0.578461 | 1.005836 | 0.569396 |
| 11 | 73174000 | 73175399 | 10.93605 | 19.47076 | 34.80661 | 15.33235 | -0.43738 | 0.444806 | 0.5694 |
| 4 | 92514000 | 92520599 | 82.14105 | 129.5377 | 132.907 | 55.15597 | 0.286881 | 2.480429 | 0.569421 |
| 3 | 136741200 | 136742399 | 8.61089 | 15.99391 | 28.93743 | 12.12153 | -0.40415 | 0.164898 | 0.569427 |
| 7 | 56901400 | 56901799 | 8.61089 | 15.99391 | 28.93743 | 12.12153 | -0.40415 | 0.164898 | 0.569427 |
| 10 | 50022200 | 50024799 | 23.78699 | 39.94851 | 64.33564 | 28.09417 | -0.30424 | 1.149538 | 0.569432 |
| 4 | 94316000 | 94319199 | 36.54855 | 59.11925 | 91.94132 | 39.30835 | -0.27342 | 1.688056 | 0.569459 |
| 14 | 61812600 | 61814799 | 11.8025 | 21.20468 | 34.70734 | 15.17228 | -0.30353 | 0.392711 | 0.569464 |
| 16 | 26681000 | 26681999 | 11.8025 | 21.20468 | 34.70734 | 15.17228 | -0.30353 | 0.392711 | 0.569464 |
| 5 | 104621800 | 104624399 | 30.12724 | 49.91089 | 50.2837 | 21.76368 | 0.370065 | 1.30399 | 0.569474 |
| 12 | 84039600 | 84043599 | 41.1441 | 66.02094 | 103.0441 | 43.57246 | -0.29241 | 1.939772 | 0.569488 |
| X | 91860600 | 91863999 | 60.8756 | 97.56018 | 93.55354 | 39.48397 | 0.406918 | 2.003427 | 0.569492 |
| 3 | 9959000 | 9961199 | 18.08326 | 31.71444 | 34.01274 | 15.03359 | 0.287079 | 0.642112 | 0.569497 |
| 6 | 143192800 | 143194999 | 18.77682 | 32.1785 | 51.13434 | 22.42515 | -0.28225 | 0.938108 | 0.569499 |
| 10 | 4883800 | 4888599 | 56.77078 | 89.73374 | 152.9793 | 63.5523 | -0.44205 | 2.414508 | 0.569511 |
| 12 | 21001800 | 21009799 | 180.0269 | 267.3803 | 268.7124 | 110.6947 | 0.246253 | 3.652044 | 0.569527 |
| 17 | 78782400 | 78786399 | 55.55094 | 88.24078 | 130.0795 | 54.24009 | -0.22625 | 2.279138 | 0.569533 |
| 7 | 58575200 | 58577199 | 13.64525 | 24.22334 | 40.28072 | 17.7875 | -0.3365 | 0.491327 | 0.569541 |
| 8 | 91542000 | 91543999 | 12.28196 | 21.84788 | 36.67392 | 16.16685 | -0.34827 | 0.460867 | 0.569562 |
| 13 | 109699200 | 109700399 | 8.224161 | 15.77153 | 30.5223 | 11.99993 | -0.42243 | -0.0569 | 0.569567 |
| 14 | 26110800 | 26120399 | 101.3299 | 155.0461 | 219.7695 | 91.18928 | -0.21244 | 3.055921 | 0.569568 |
| X | 20945400 | 20946399 | 23.04654 | 40.41612 | 36.21565 | 16.0063 | 0.54518 | 0.675435 | 0.569588 |
| 4 | 153044000 | 153053599 | 78.77062 | 121.8712 | 184.1145 | 76.47661 | -0.29198 | 2.831501 | 0.569595 |
| X | 81129000 | 81132199 | 45.27903 | 73.46027 | 73.13511 | 31.33689 | 0.372604 | 1.634911 | 0.569603 |
| 8 | 57477400 | 57483599 | 76.03236 | 119.0052 | 174.6586 | 72.4537 | -0.23733 | 2.626788 | 0.569621 |
| 7 | 41805000 | 41807399 | 43.97946 | 71.29286 | 73.03675 | 31.27567 | 0.330385 | 1.651729 | 0.56964 |
| 2 | 170531600 | 170532199 | 5.831582 | 11.28298 | 23.21544 | 8.857617 | -0.46793 | -0.20108 | 0.569643 |
| 2 | 39176600 | 39179799 | 27.63009 | 47.00731 | 46.82762 | 20.52439 | 0.387184 | 1.030776 | 0.569644 |

| | | | | | | | | | |
|---|---|---|---|---|---|---|---|---|---|
| 4 | 153712800 | 153719599 | 68.04761 | 106.8513 | 156.0252 | 64.71371 | -0.22636 | 2.561322 | 0.569646 |
| 15 | 24155800 | 24158399 | 24.54487 | 41.87745 | 44.72149 | 19.60578 | 0.287867 | 0.991723 | 0.569649 |
| 12 | 115744200 | 115749799 | 70.0721 | 112.1611 | 94.26074 | 39.7091 | 0.594869 | 2.050506 | 0.569652 |
| 7 | 26297000 | 26299599 | 42.07413 | 68.01622 | 72.98536 | 31.18272 | 0.259846 | 1.70581 | 0.569665 |
| 9 | 61493400 | 61495199 | 19.26274 | 33.91582 | 34.32702 | 15.19607 | 0.371205 | 0.597727 | 0.569666 |
| 17 | 3287200 | 3290599 | 22.03459 | 37.17774 | 61.02166 | 26.68678 | -0.33134 | 1.097389 | 0.569676 |
| 3 | 4632600 | 4635999 | 36.31857 | 59.80938 | 62.14116 | 26.79984 | 0.324166 | 1.3997 | 0.569681 |
| 1 | 21447400 | 21449799 | 38.64675 | 62.92967 | 68.01393 | 29.20195 | 0.258693 | 1.556061 | 0.569688 |
| 4 | 112811800 | 112828199 | 255.7371 | 373.1457 | 347.8962 | 143.3481 | 0.330333 | 3.949913 | 0.569702 |
| 15 | 49939600 | 49940599 | 8.992362 | 17.01195 | 30.44243 | 12.34912 | -0.35104 | 0.051507 | 0.569704 |
| 11 | 111244200 | 111251599 | 98.37527 | 153.1827 | 156.7996 | 65.08462 | 0.276139 | 2.725162 | 0.569715 |
| 15 | 3050000 | 3070999 | 610.9328 | 822.8307 | 755.1711 | 314.9712 | 0.237393 | 5.166294 | 0.569723 |
| 4 | 153364200 | 153373399 | 111.0734 | 168.6004 | 241.8242 | 100.1272 | -0.23876 | 3.222332 | 0.569733 |
| 2 | 30104800 | 30110999 | 46.95011 | 75.1186 | 111.7234 | 47.03701 | -0.22789 | 2.041703 | 0.569749 |
| 6 | 52228200 | 52231399 | 44.36165 | 70.8776 | 110.5134 | 46.43517 | -0.29808 | 2.081718 | 0.569788 |
| 8 | 104390400 | 104392799 | 25.91475 | 43.13092 | 69.50856 | 30.18062 | -0.30517 | 1.255276 | 0.569805 |
| X | 118207000 | 118212199 | 69.84241 | 111.2936 | 108.0288 | 45.22308 | 0.380359 | 2.189459 | 0.569808 |
| 12 | 119748200 | 119750599 | 24.49842 | 42.56729 | 40.76473 | 17.89869 | 0.449077 | 0.821216 | 0.569812 |
| 4 | 67773600 | 67775799 | 32.66836 | 54.43625 | 48.68396 | 21.15461 | 0.541846 | 1.246234 | 0.569815 |
| 1 | 182578600 | 182579399 | 8.275469 | 15.44963 | 28.46349 | 11.82372 | -0.42064 | 0.127382 | 0.569818 |
| 4 | 8757200 | 8758399 | 12.90479 | 23.0358 | 38.23082 | 16.80383 | -0.32914 | 0.446141 | 0.569821 |
| X | 45631000 | 45633199 | 29.90932 | 51.24453 | 38.19662 | 16.75095 | 0.805625 | 0.971775 | 0.569835 |
| 16 | 76162200 | 76164799 | 18.75752 | 32.19961 | 51.1426 | 22.43182 | -0.28119 | 0.922397 | 0.56986 |
| 13 | 37396800 | 37397199 | 6.832121 | 13.33703 | 16.41724 | 5.806961 | 0.366933 | -0.4734 | 0.569903 |
| 2 | 13954000 | 13955599 | 18.37747 | 31.70849 | 51.13905 | 22.43407 | -0.30153 | 0.877749 | 0.569906 |
| 3 | 14475400 | 14477799 | 28.04279 | 47.65882 | 42.57518 | 18.65897 | 0.543662 | 1.039104 | 0.56991 |
| 14 | 110728200 | 110733999 | 70.30242 | 111.5371 | 118.265 | 49.25368 | 0.246684 | 2.322139 | 0.569951 |
| 5 | 86988200 | 86992399 | 70.36611 | 112.0178 | 99.42378 | 41.56419 | 0.507738 | 2.218138 | 0.569951 |
| 1 | 35826600 | 35829199 | 18.60689 | 31.60034 | 59.11558 | 25.90451 | -0.5187 | 1.030621 | 0.569973 |
| 6 | 68164600 | 68167999 | 59.42884 | 94.94254 | 101.0255 | 42.44336 | 0.251987 | 2.096096 | 0.569974 |
| 1 | 165268200 | 165269599 | 11.93642 | 21.16606 | 36.31834 | 16.05292 | -0.3819 | 0.482648 | 0.569978 |
| 13 | 62119200 | 62134999 | 223.0553 | 328.6057 | 318.2756 | 131.0688 | 0.288567 | 3.79504 | 0.56998 |
| 15 | 6849600 | 6851599 | 12.47148 | 22.18019 | 36.18647 | 15.9549 | -0.30746 | 0.463952 | 0.57001 |
| 12 | 109986600 | 109992199 | 47.44021 | 75.70954 | 114.597 | 48.07298 | -0.25721 | 2.120639 | 0.570035 |
| 1 | 30080200 | 30082799 | 22.19582 | 37.12211 | 59.65568 | 25.9825 | -0.30094 | 1.185204 | 0.570038 |
| 15 | 59547800 | 59551399 | 23.11237 | 38.69776 | 60.89291 | 26.53986 | -0.2708 | 1.175995 | 0.570056 |
| 10 | 110060600 | 110061399 | 13.3961 | 23.39588 | 46.26778 | 20.3804 | -0.59129 | 0.706074 | 0.570058 |
| 10 | 38381400 | 38381799 | 5.797991 | 11.30282 | 22.78094 | 8.223177 | -0.36412 | -0.39631 | 0.570083 |
| 9 | 24239400 | 24239999 | 5.524984 | 10.70342 | 22.68313 | 8.603086 | -0.50047 | -0.22207 | 0.570093 |
| 2 | 125089000 | 125090599 | 10.623 | 18.95676 | 34.32863 | 15.09264 | -0.45391 | 0.427504 | 0.570106 |
| 6 | 42426600 | 42427999 | 13.70155 | 24.08677 | 39.89004 | 17.66423 | -0.33625 | 0.593926 | 0.570122 |
| 3 | 25207800 | 25212599 | 114.5898 | 175.6068 | 185.6989 | 77.07913 | 0.211616 | 3.026299 | 0.570137 |
| 8 | 35127600 | 35129999 | 32.81014 | 53.21543 | 82.05603 | 35.15181 | -0.25739 | 1.623191 | 0.570153 |
| 15 | 95889800 | 95893199 | 45.72625 | 73.86408 | 78.04829 | 33.3132 | 0.280765 | 1.736417 | 0.57021 |
| 19 | 36462600 | 36465599 | 33.44992 | 55.52378 | 50.88858 | 22.0552 | 0.50634 | 1.292431 | 0.570212 |
| 17 | 15783600 | 15784999 | 12.668 | 22.42862 | 36.7321 | 16.24274 | -0.31628 | 0.501787 | 0.570232 |
| 1 | 88476800 | 88479999 | 28.13445 | 46.22213 | 71.8766 | 30.96741 | -0.25728 | 1.416524 | 0.570239 |

| | | | | | | | | | |
|---|---|---|---|---|---|---|---|---|---|
| 7 | 143342400 | 143346399 | 34.946 | 56.60345 | 86.28769 | 36.92797 | -0.24278 | 1.656686 | 0.570242 |
| 15 | 93958200 | 93960599 | 14.05849 | 26.27053 | 20.60574 | 8.546114 | 0.803654 | 0.150379 | 0.570251 |
| 14 | 103081200 | 103082799 | 11.17368 | 20.1022 | 33.72889 | 14.72768 | -0.33785 | 0.377858 | 0.570257 |
| 3 | 21971400 | 21972999 | 12.44494 | 22.13192 | 36.67016 | 16.16882 | -0.3298 | 0.4639 | 0.570263 |
| 13 | 54568800 | 54574999 | 70.22998 | 110.0501 | 160.5399 | 66.58897 | -0.22728 | 2.601169 | 0.570272 |
| 15 | 10163400 | 10170399 | 136.4185 | 207.8093 | 189.1124 | 78.3853 | 0.421251 | 3.153112 | 0.570275 |
| 16 | 8879400 | 8882199 | 36.52977 | 59.64753 | 65.07013 | 27.96625 | 0.248103 | 1.511015 | 0.5703 |
| 8 | 113199800 | 113201399 | 12.6202 | 22.72395 | 39.47159 | 17.21854 | -0.38699 | 0.393738 | 0.570341 |
| 12 | 96974600 | 96975599 | 11.29529 | 20.43655 | 33.75331 | 14.63562 | -0.30877 | 0.345302 | 0.570348 |
| 14 | 52354600 | 52355799 | 11.29529 | 20.43655 | 33.75331 | 14.63562 | -0.30877 | 0.345302 | 0.570348 |
| 12 | 22383200 | 22389799 | 104.6338 | 162.2471 | 148.489 | 61.67363 | 0.432793 | 2.798475 | 0.570352 |
| 6 | 122601200 | 122606599 | 51.63971 | 81.9518 | 140.9383 | 58.71335 | -0.44956 | 2.304848 | 0.570378 |
| 5 | 100361800 | 100364599 | 20.71253 | 34.77101 | 57.47564 | 25.09209 | -0.34151 | 1.137558 | 0.570383 |
| 4 | 146208600 | 146209399 | 40.04839 | 65.68428 | 64.13257 | 27.61118 | 0.411277 | 1.458088 | 0.570407 |
| 11 | 27465600 | 27466999 | 23.02027 | 39.69616 | 42.5619 | 18.67105 | 0.284789 | 0.88967 | 0.57041 |
| X | 84837200 | 84837999 | 14.02531 | 25.89655 | 26.53604 | 11.20574 | 0.401936 | 0.237348 | 0.570439 |
| 12 | 91607200 | 91609599 | 13.44595 | 23.76154 | 38.22292 | 16.90828 | -0.29174 | 0.529169 | 0.570453 |
| 6 | 59787600 | 59789799 | 18.40833 | 32.35331 | 34.32036 | 15.19685 | 0.302669 | 0.610256 | 0.570456 |
| 11 | 91045000 | 91046799 | 18.40833 | 32.35331 | 34.32036 | 15.19685 | 0.302669 | 0.610256 | 0.570456 |
| X | 107114000 | 107115199 | 14.11002 | 25.8621 | 27.47355 | 11.71827 | 0.339758 | 0.279773 | 0.570462 |
| 2 | 143275000 | 143276199 | 13.37368 | 23.42787 | 45.14874 | 19.93635 | -0.55414 | 0.655164 | 0.570469 |
| X | 132122600 | 132125399 | 38.05956 | 62.22498 | 63.31705 | 27.22549 | 0.349647 | 1.499067 | 0.570478 |
| 7 | 23511800 | 23515599 | 53.03808 | 85.52551 | 81.80353 | 34.83733 | 0.421263 | 1.800134 | 0.570489 |
| 3 | 152419600 | 152420399 | 7.603003 | 14.4688 | 26.64438 | 10.67414 | -0.37579 | -0.01201 | 0.570491 |
| 19 | 29491400 | 29494199 | 20.94787 | 35.62438 | 55.99081 | 24.53979 | -0.26822 | 1.022639 | 0.570502 |
| 11 | 107353000 | 107353599 | 12.90301 | 24.10633 | 24.63693 | 10.22161 | 0.425103 | 0.155628 | 0.570517 |
| 18 | 77634400 | 77637999 | 30.47254 | 50.53782 | 54.62291 | 23.65951 | 0.269052 | 1.295808 | 0.570521 |
| 7 | 42007600 | 42023399 | 237.0379 | 350.0015 | 304.2228 | 125.2848 | 0.44772 | 3.760374 | 0.570522 |
| 8 | 30477800 | 30479399 | 13.42106 | 24.53925 | 26.36916 | 11.28451 | 0.319573 | 0.283134 | 0.570533 |
| 8 | 54430000 | 54431599 | 17.84998 | 32.08838 | 29.93853 | 13.08118 | 0.502869 | 0.418976 | 0.57054 |
| X | 146679600 | 146683399 | 29.65368 | 50.33819 | 48.48287 | 21.2423 | 0.435429 | 1.061615 | 0.570553 |
| 9 | 76443000 | 76443599 | 5.091326 | 9.928671 | 20.33508 | 7.306943 | -0.38047 | -0.43007 | 0.570554 |
| 6 | 74503000 | 74503599 | 9.603827 | 18.44144 | 21.29095 | 8.339205 | 0.32194 | -0.08023 | 0.570578 |
| 12 | 18008800 | 18012799 | 135.765 | 205.4925 | 208.4121 | 86.21704 | 0.257392 | 3.283667 | 0.570597 |
| 10 | 110835800 | 110839799 | 62.9844 | 100.1998 | 107.9817 | 45.10019 | 0.227191 | 2.234874 | 0.570605 |
| X | 73797400 | 73807199 | 125.8101 | 194.2222 | 169.8388 | 70.54764 | 0.493587 | 2.891256 | 0.570611 |
| 18 | 52856000 | 52857799 | 17.60637 | 32.45394 | 27.23267 | 11.5263 | 0.685735 | 0.274311 | 0.570629 |
| Y | 89943600 | 89944799 | 13.23843 | 24.2494 | 19.65926 | 8.392393 | 0.72598 | 0.254961 | 0.570631 |
| 5 | 66067200 | 66069799 | 23.63834 | 39.00743 | 66.37725 | 28.65157 | -0.38414 | 1.34864 | 0.570632 |
| 4 | 117092600 | 117094799 | 22.33539 | 37.33154 | 60.23802 | 26.2255 | -0.30685 | 1.192045 | 0.570653 |
| 1 | 79985400 | 79985999 | 8.917647 | 16.51307 | 29.41634 | 12.37356 | -0.38638 | 0.185563 | 0.570672 |
| 3 | 82879200 | 82880999 | 27.37464 | 45.86713 | 49.12913 | 21.42238 | 0.282287 | 1.171203 | 0.570674 |
| 14 | 111781600 | 111783199 | 13.6341 | 23.94455 | 39.37561 | 17.4243 | -0.32635 | 0.605688 | 0.570685 |
| 17 | 29299800 | 29306199 | 50.76912 | 80.26453 | 150.4679 | 62.51577 | -0.57856 | 2.405125 | 0.570689 |
| 7 | 66834400 | 66836399 | 13.71674 | 24.16134 | 40.27282 | 17.82667 | -0.34484 | 0.568006 | 0.570693 |
| 18 | 7170000 | 7170599 | 5.517974 | 10.74426 | 21.31416 | 7.811241 | -0.36081 | -0.35016 | 0.57075 |
| 15 | 19550200 | 19551399 | 14.10059 | 25.62707 | 27.46948 | 11.84417 | 0.316542 | 0.327549 | 0.570751 |

| | | | | | | | | | |
|---|---|---|---|---|---|---|---|---|---|
| Y | 39773000 | 39775599 | 15.78419 | 28.13941 | 21.14948 | 9.306068 | 0.805991 | 0.436352 | 0.570769 |
| 4 | 5354400 | 5357999 | 28.16534 | 46.54541 | 76.89157 | 33.22328 | -0.34162 | 1.349995 | 0.570777 |
| 1 | 43548000 | 43548799 | 14.2599 | 25.76607 | 26.53394 | 11.52443 | 0.367324 | 0.360218 | 0.570779 |
| 16 | 9283400 | 9283999 | 5.235434 | 10.18111 | 21.20141 | 7.850426 | -0.4432 | -0.31614 | 0.570785 |
| 8 | 69775000 | 69779599 | 50.48566 | 81.32998 | 83.33658 | 35.44321 | 0.320407 | 1.822629 | 0.570786 |
| 10 | 95572200 | 95573999 | 19.45606 | 33.38995 | 52.76436 | 23.14266 | -0.27394 | 0.930718 | 0.570788 |
| 3 | 21556800 | 21570199 | 223.6609 | 328.6573 | 326.6752 | 134.5372 | 0.247751 | 3.836149 | 0.570798 |
| 19 | 37772000 | 37774199 | 20.26187 | 34.32381 | 55.97031 | 24.50739 | -0.32159 | 1.05233 | 0.570834 |
| 19 | 20640400 | 20641799 | 16.77819 | 30.46613 | 29.15827 | 12.58643 | 0.47769 | 0.353784 | 0.570855 |
| X | 119010200 | 119012199 | 19.21418 | 35.07724 | 27.4328 | 11.76239 | 0.773529 | 0.338795 | 0.570873 |
| 16 | 51071200 | 51073599 | 14.51433 | 25.36 | 51.0734 | 22.50344 | -0.61766 | 0.71195 | 0.570878 |
| 14 | 17742200 | 17743599 | 12.86043 | 22.61436 | 38.38109 | 16.9951 | -0.3713 | 0.581689 | 0.570921 |
| 18 | 48634600 | 48637199 | 28.57365 | 48.55814 | 47.92682 | 21.00427 | 0.400314 | 1.046757 | 0.570938 |
| 7 | 90060000 | 90063599 | 85.59476 | 133.9482 | 143.2714 | 59.43444 | 0.218374 | 2.629549 | 0.570959 |
| X | 25393000 | 25397799 | 45.54169 | 75.04475 | 58.68506 | 25.31598 | 0.733391 | 1.415693 | 0.570992 |
| 4 | 53364000 | 53364999 | 8.565595 | 16.1241 | 28.04526 | 11.48794 | -0.32185 | 0.076916 | 0.571012 |
| 6 | 82194000 | 82198199 | 46.02277 | 74.44008 | 76.7559 | 32.81464 | 0.318086 | 1.69945 | 0.571037 |
| 5 | 34122000 | 34122999 | 14.41376 | 26.71737 | 27.00811 | 11.34496 | 0.426595 | 0.218602 | 0.571057 |
| 12 | 35243600 | 35245599 | 26.63629 | 44.70448 | 46.48315 | 20.28996 | 0.324898 | 1.147492 | 0.571076 |
| 14 | 51120400 | 51124199 | 41.31805 | 67.04839 | 72.00001 | 30.8554 | 0.263954 | 1.62242 | 0.571085 |
| 4 | 39774200 | 39775599 | 9.210571 | 17.12483 | 29.44273 | 12.31595 | -0.32977 | 0.165256 | 0.571094 |
| 3 | 146105400 | 146110999 | 47.16827 | 75.14674 | 130.2174 | 54.49591 | -0.4547 | 2.179205 | 0.571115 |
| X | 69115400 | 69117199 | 24.30366 | 42.28256 | 33.11744 | 14.55258 | 0.738186 | 0.794931 | 0.571123 |
| 5 | 95631200 | 95636799 | 50.72779 | 81.83353 | 81.24289 | 34.61681 | 0.368276 | 1.782586 | 0.571136 |
| 17 | 59836600 | 59837999 | 17.82282 | 31.67422 | 32.11405 | 14.16808 | 0.3743 | 0.494433 | 0.571148 |
| 8 | 20522200 | 20524399 | 27.57121 | 47.70668 | 41.80669 | 18.34166 | 0.575632 | 0.879084 | 0.571176 |
| 4 | 130738800 | 130740799 | 30.16551 | 49.26992 | 76.14105 | 32.72783 | -0.25268 | 1.501783 | 0.571211 |
| 13 | 23140000 | 23142399 | 35.22052 | 58.34225 | 50.36065 | 21.79531 | 0.592409 | 1.322195 | 0.571231 |
| 1 | 101787000 | 101789199 | 25.6199 | 44.31364 | 42.45247 | 18.62477 | 0.447521 | 0.875221 | 0.571271 |
| 1 | 153607600 | 153609799 | 10.70968 | 19.36328 | 32.78797 | 14.22963 | -0.34506 | 0.34452 | 0.571278 |
| 7 | 129361800 | 129362599 | 10.70968 | 19.36328 | 32.78797 | 14.22963 | -0.34506 | 0.34452 | 0.571278 |
| 1 | 147971600 | 147973799 | 44.17873 | 71.4032 | 76.90064 | 32.84644 | 0.254198 | 1.716607 | 0.571291 |
| 10 | 21824200 | 21827399 | 29.09987 | 47.79344 | 74.34889 | 32.02839 | -0.25806 | 1.423845 | 0.571322 |
| 4 | 147940000 | 147942199 | 30.89644 | 51.36568 | 55.71864 | 24.1747 | 0.264145 | 1.271706 | 0.571325 |
| 4 | 127799800 | 127802999 | 27.86817 | 45.61187 | 72.61372 | 31.23521 | -0.29327 | 1.464225 | 0.571329 |
| 1 | 104091200 | 104092999 | 21.28659 | 37.68468 | 32.28862 | 14.27809 | 0.613094 | 0.546142 | 0.571349 |
| 7 | 103177200 | 103178199 | 10.61903 | 19.34126 | 33.91797 | 14.58513 | -0.38637 | 0.306328 | 0.571355 |
| 14 | 112146200 | 112147199 | 11.13906 | 21.60364 | 21.05994 | 7.945721 | 0.614708 | -0.18569 | 0.57136 |
| 8 | 3722000 | 3723199 | 8.31721 | 15.68964 | 28.62476 | 11.6824 | -0.38543 | 0.065237 | 0.571381 |
| X | 23950000 | 23979399 | 423.9504 | 595.078 | 488.0857 | 202.2659 | 0.459578 | 4.528518 | 0.571385 |
| 16 | 29562600 | 29565799 | 24.73591 | 40.84428 | 75.83753 | 32.74918 | -0.50975 | 1.348857 | 0.571387 |
| 17 | 89578400 | 89579799 | 14.9523 | 27.30014 | 28.31675 | 12.13953 | 0.368674 | 0.307662 | 0.571404 |
| 12 | 107735200 | 107736799 | 10.51949 | 19.10742 | 32.29605 | 13.93546 | -0.33712 | 0.31457 | 0.571411 |
| 17 | 44091800 | 44094199 | 28.07706 | 47.59387 | 43.16268 | 18.90313 | 0.521856 | 1.062358 | 0.571415 |
| 7 | 41967400 | 41970799 | 59.02806 | 94.9508 | 85.48187 | 36.27773 | 0.504165 | 1.882781 | 0.571419 |
| 8 | 79229400 | 79233799 | 41.15344 | 66.20061 | 99.85142 | 42.37827 | -0.23879 | 1.861778 | 0.57143 |
| 5 | 66150800 | 66152799 | 16.07941 | 27.8086 | 46.37216 | 20.34435 | -0.34861 | 0.839855 | 0.571431 |

| | | | | | | | | | |
|---|---|---|---|---|---|---|---|---|---|
| 5 | 122336400 | 122340199 | 53.93146 | 85.55743 | 130.1432 | 54.19634 | -0.27324 | 2.315757 | 0.57145 |
| 13 | 37592200 | 37596999 | 59.36398 | 94.9316 | 99.36828 | 41.8049 | 0.277188 | 2.067191 | 0.571458 |
| 14 | 112521200 | 112524999 | 28.05071 | 48.02535 | 45.14167 | 19.79552 | 0.47205 | 0.975612 | 0.571477 |
| 6 | 91121600 | 91124599 | 15.94912 | 27.54667 | 52.18173 | 22.89219 | -0.53232 | 0.858259 | 0.571483 |
| 12 | 19289200 | 19309599 | 786.1374 | 1032.212 | 898.1244 | 375.8484 | 0.277675 | 5.573567 | 0.571518 |
| 9 | 93398600 | 93399399 | 9.566659 | 18.48991 | 20.87505 | 8.004446 | 0.382326 | -0.15168 | 0.57154 |
| 12 | 101683800 | 101686199 | 13.71792 | 23.90549 | 47.34757 | 20.82732 | -0.59387 | 0.740473 | 0.571541 |
| 19 | 48458600 | 48460599 | 11.14881 | 20.0564 | 34.21121 | 14.93894 | -0.36168 | 0.377827 | 0.571546 |
| 4 | 118546800 | 118548799 | 11.31016 | 20.34112 | 34.20801 | 14.94098 | -0.34154 | 0.38099 | 0.57155 |
| 10 | 82187800 | 82225199 | 779.7746 | 1032.284 | 900.8207 | 376.6168 | 0.285236 | 5.442376 | 0.571563 |
| 13 | 45641400 | 45643399 | 13.27142 | 23.34852 | 39.29833 | 17.39954 | -0.35935 | 0.577855 | 0.571588 |
| 1 | 78268200 | 78269999 | 15.692 | 27.47863 | 44.04992 | 19.44489 | -0.29003 | 0.670595 | 0.571591 |
| 13 | 61213800 | 61218199 | 42.91411 | 70.38648 | 64.20936 | 27.64459 | 0.509123 | 1.467195 | 0.571594 |
| 14 | 114713800 | 114715799 | 21.0786 | 35.27454 | 67.42793 | 29.37865 | -0.55057 | 1.180822 | 0.571601 |
| 6 | 129476600 | 129482599 | 64.03141 | 102.608 | 94.15986 | 39.73411 | 0.470084 | 2.008492 | 0.571621 |
| 11 | 110658800 | 110660199 | 21.6741 | 36.33499 | 58.60078 | 25.56099 | -0.30615 | 1.156449 | 0.571627 |
| 17 | 18629800 | 18634599 | 52.8616 | 85.17325 | 74.52646 | 31.70474 | 0.548082 | 1.818476 | 0.57163 |
| 4 | 9577000 | 9578599 | 15.6846 | 28.5946 | 28.809 | 12.37356 | 0.408768 | 0.323286 | 0.571638 |
| 5 | 96297600 | 96298999 | 15.6846 | 28.5946 | 28.809 | 12.37356 | 0.408768 | 0.323286 | 0.571638 |
| 4 | 34429600 | 34430799 | 15.03653 | 26.2385 | 43.11785 | 18.98273 | -0.32577 | 0.719422 | 0.57164 |
| 5 | 21897400 | 21898199 | 11.44039 | 21.76582 | 23.80187 | 9.540634 | 0.369469 | 0.019822 | 0.57164 |
| 5 | 82725000 | 82725599 | 6.535617 | 12.66538 | 23.89783 | 9.050117 | -0.33418 | -0.21622 | 0.571642 |
| 5 | 8702800 | 8703599 | 6.562944 | 12.63402 | 24.16821 | 9.398048 | -0.38972 | -0.12476 | 0.57165 |
| 8 | 121659800 | 121663199 | 29.54129 | 48.17855 | 75.822 | 32.573 | -0.28035 | 1.522117 | 0.57168 |
| 12 | 30388000 | 30389799 | 15.39713 | 27.97001 | 28.72436 | 12.39269 | 0.376999 | 0.339919 | 0.571686 |
| 18 | 53311400 | 53313599 | 17.19079 | 29.8001 | 47.40484 | 20.79888 | -0.2803 | 0.825172 | 0.571688 |
| 13 | 73464200 | 73472199 | 71.2375 | 111.5142 | 162.8616 | 67.55911 | -0.23008 | 2.619928 | 0.571688 |
| 6 | 131177400 | 131181999 | 101.2027 | 156.8993 | 164.0898 | 68.15506 | 0.240321 | 2.80188 | 0.571695 |
| 7 | 118709400 | 118712199 | 26.82369 | 45.27514 | 49.07543 | 21.46475 | 0.265296 | 1.100282 | 0.571699 |
| 9 | 98851600 | 98854199 | 30.00784 | 50.09771 | 54.07481 | 23.52518 | 0.27128 | 1.221628 | 0.571708 |
| X | 9246800 | 9255999 | 137.7543 | 210.2091 | 199.5434 | 82.74499 | 0.362411 | 3.121554 | 0.571733 |
| 10 | 43708800 | 43713199 | 51.06869 | 81.27344 | 122.2159 | 51.07137 | -0.25213 | 2.215547 | 0.571734 |
| 3 | 6434800 | 6436599 | 32.74042 | 54.76197 | 45.97221 | 20.02376 | 0.63275 | 1.20563 | 0.571739 |
| 4 | 130390200 | 130391399 | 7.103738 | 13.31377 | 29.05708 | 11.99108 | -0.65561 | 0.091987 | 0.571747 |
| 3 | 129050000 | 129050599 | 7.612425 | 14.52026 | 26.14409 | 10.42263 | -0.33709 | -0.02792 | 0.571749 |
| 10 | 19467400 | 19468199 | 7.612425 | 14.52026 | 26.14409 | 10.42263 | -0.33709 | -0.02792 | 0.571749 |
| 7 | 106766000 | 106766799 | 6.99423 | 13.47932 | 24.76649 | 9.599864 | -0.32809 | -0.12957 | 0.571762 |
| 12 | 52482800 | 52484599 | 16.96199 | 29.41846 | 46.87297 | 20.56584 | -0.28254 | 0.81716 | 0.571775 |
| 5 | 150212600 | 150215399 | 19.13095 | 32.71675 | 51.66141 | 22.65248 | -0.27359 | 0.959897 | 0.571793 |
| 5 | 105113000 | 105114199 | 14.72698 | 25.87206 | 41.34558 | 18.30092 | -0.28549 | 0.609297 | 0.571827 |
| 8 | 48622200 | 48623599 | 14.72698 | 25.87206 | 41.34558 | 18.30092 | -0.28549 | 0.609297 | 0.571827 |
| 17 | 84181200 | 84186999 | 62.80662 | 99.07984 | 146.5922 | 60.83855 | -0.24014 | 2.463381 | 0.571833 |
| 6 | 62824000 | 62827399 | 34.46688 | 57.17514 | 58.19262 | 25.20962 | 0.355506 | 1.307651 | 0.571845 |
| 2 | 156665000 | 156679399 | 277.8759 | 393.8951 | 522.0697 | 216.0031 | -0.2183 | 4.37716 | 0.57186 |
| 7 | 38437600 | 38446399 | 115.8567 | 179.869 | 158.6071 | 65.86408 | 0.487766 | 2.783844 | 0.571878 |
| 5 | 109413600 | 109423599 | 137.4609 | 210.4586 | 192.7206 | 79.97708 | 0.418027 | 3.053889 | 0.571908 |
| 15 | 87999200 | 88008999 | 91.17528 | 140.2418 | 201.6655 | 83.74888 | -0.22733 | 2.948073 | 0.57191 |

| | | | | | | | | | |
|---|---|---|---|---|---|---|---|---|---|
| 6 | 43475000 | 43476799 | 21.7152 | 38.03954 | 36.83237 | 16.26375 | 0.433837 | 0.684842 | 0.571918 |
| 11 | 35597800 | 35599999 | 21.16952 | 35.5736 | 57.52243 | 25.12553 | -0.30991 | 1.128728 | 0.57192 |
| 15 | 93702000 | 93703199 | 8.977317 | 16.91631 | 29.72401 | 12.15466 | -0.33496 | 0.076044 | 0.571924 |
| 3 | 100130200 | 100131999 | 19.74951 | 32.8459 | 68.08549 | 29.54785 | -0.66696 | 1.246703 | 0.571938 |
| 4 | 148224400 | 148225599 | 8.305983 | 15.47124 | 28.45897 | 11.86763 | -0.42261 | 0.143951 | 0.571942 |
| 9 | 47314600 | 47315599 | 8.305983 | 15.47124 | 28.45897 | 11.86763 | -0.42261 | 0.143951 | 0.571942 |
| 6 | 132141600 | 132143199 | 38.39943 | 62.72172 | 66.95698 | 28.78075 | 0.279639 | 1.51238 | 0.571945 |
| 17 | 4733000 | 4733999 | 12.93148 | 23.92961 | 26.05705 | 10.97048 | 0.317771 | 0.214107 | 0.571965 |
| 2 | 59858800 | 59860799 | 18.25176 | 32.02328 | 34.46547 | 15.23905 | 0.281981 | 0.637097 | 0.57197 |
| 15 | 20359000 | 20360799 | 21.812 | 37.95547 | 30.70159 | 13.49274 | 0.691705 | 0.773092 | 0.57198 |
| 2 | 36251800 | 36257999 | 174.7256 | 261.4605 | 264.1055 | 108.9192 | 0.248786 | 3.51697 | 0.571986 |
| 8 | 82230800 | 82232199 | 15.44731 | 28.11555 | 28.77007 | 12.38239 | 0.384472 | 0.329997 | 0.571991 |
| 10 | 58183000 | 58185199 | 13.60562 | 23.9731 | 38.7702 | 17.16046 | -0.3011 | 0.563237 | 0.571992 |
| 7 | 26538000 | 26540999 | 38.00791 | 61.84593 | 67.2649 | 28.87295 | 0.248932 | 1.564618 | 0.572002 |
| 5 | 93627000 | 93628799 | 19.54769 | 34.90244 | 31.46067 | 13.8252 | 0.546971 | 0.47631 | 0.572003 |
| 5 | 90288000 | 90289799 | 18.53295 | 31.97433 | 52.82506 | 23.17364 | -0.33642 | 0.876786 | 0.572017 |
| X | 45777400 | 45781399 | 60.82087 | 97.32705 | 97.20459 | 40.93373 | 0.345859 | 2.049398 | 0.572059 |
| 4 | 72855200 | 72856799 | 18.49097 | 32.3507 | 34.08632 | 15.03518 | 0.312507 | 0.682551 | 0.572062 |
| 11 | 12510400 | 12514399 | 50.87851 | 80.97608 | 122.4368 | 51.16829 | -0.25992 | 2.21273 | 0.572075 |
| 4 | 113248000 | 113253799 | 82.68411 | 130.8342 | 124.8909 | 51.91183 | 0.394727 | 2.404238 | 0.572079 |
| 8 | 64338000 | 64339599 | 19.56455 | 34.11054 | 36.7703 | 16.17948 | 0.279569 | 0.744797 | 0.57208 |
| 2 | 100825600 | 100827799 | 36.69307 | 60.01365 | 65.21641 | 28.04607 | 0.255347 | 1.487963 | 0.572112 |
| 6 | 140026200 | 140027799 | 14.24891 | 24.87814 | 42.60599 | 18.76403 | -0.38494 | 0.706274 | 0.572119 |
| 9 | 117439000 | 117441599 | 32.6068 | 53.96187 | 58.06467 | 25.11317 | 0.27521 | 1.328497 | 0.572126 |
| 1 | 138828800 | 138830599 | 21.6875 | 37.65476 | 40.40939 | 17.73426 | 0.285416 | 0.818387 | 0.572129 |
| 2 | 132640800 | 132642199 | 15.86968 | 27.65536 | 44.18338 | 19.4355 | -0.28547 | 0.745326 | 0.572136 |
| 7 | 71772400 | 71775199 | 47.94977 | 76.48903 | 115.5511 | 48.44326 | -0.25509 | 2.135587 | 0.572147 |
| 15 | 73003800 | 73005399 | 16.4868 | 28.79843 | 46.76868 | 20.60316 | -0.30881 | 0.717691 | 0.57215 |
| 16 | 91614600 | 91616799 | 18.35051 | 31.33056 | 51.10899 | 22.40729 | -0.32066 | 0.971424 | 0.572169 |
| X | 153630400 | 153632799 | 26.46853 | 45.44312 | 44.65871 | 19.58813 | 0.408788 | 0.946813 | 0.572169 |
| 11 | 14422400 | 14426399 | 30.14204 | 50.75657 | 51.2961 | 22.42003 | 0.365856 | 1.133865 | 0.572215 |
| 5 | 138361600 | 138364599 | 43.60817 | 69.53161 | 125.5718 | 52.61109 | -0.5129 | 2.149108 | 0.572222 |
| 15 | 22347600 | 22351399 | 37.05959 | 61.1129 | 61.61544 | 26.58843 | 0.368204 | 1.3859 | 0.572242 |
| 7 | 109889600 | 109896599 | 86.83538 | 133.4582 | 202.2399 | 83.97811 | -0.30378 | 2.961325 | 0.572258 |
| 6 | 65007000 | 65008999 | 19.85117 | 34.86747 | 30.08202 | 13.31314 | 0.59919 | 0.61994 | 0.572264 |
| 10 | 58683400 | 58685799 | 17.71641 | 30.68727 | 50.1008 | 21.98121 | -0.31812 | 0.834591 | 0.572274 |
| 2 | 60103600 | 60105999 | 27.40625 | 44.81251 | 83.604 | 35.95007 | -0.52259 | 1.482917 | 0.572309 |
| 15 | 35371000 | 35372199 | 9.39571 | 17.49409 | 29.99848 | 12.51803 | -0.32306 | 0.16004 | 0.572313 |
| X | 13192200 | 13194599 | 20.4768 | 34.89621 | 54.89557 | 24.06382 | -0.26899 | 1.001296 | 0.572328 |
| 16 | 15964200 | 15964799 | 6.106564 | 11.82491 | 23.84229 | 9.061278 | -0.43407 | -0.21061 | 0.572332 |
| 9 | 103903000 | 103905599 | 28.47002 | 47.82645 | 51.27787 | 22.39581 | 0.280745 | 1.148331 | 0.572333 |
| 16 | 60921600 | 60922399 | 12.55755 | 22.46028 | 37.80732 | 16.59003 | -0.34738 | 0.434624 | 0.572358 |
| 2 | 88034800 | 88038799 | 52.74253 | 85.10551 | 80.42055 | 34.27776 | 0.439794 | 1.778524 | 0.572365 |
| 13 | 67213000 | 67219599 | 113.4965 | 175.9518 | 164.1651 | 68.18674 | 0.404795 | 2.808577 | 0.572381 |
| 2 | 173302200 | 173303999 | 16.2408 | 29.45553 | 29.16849 | 12.60945 | 0.427391 | 0.356173 | 0.572382 |
| 12 | 105492800 | 105496199 | 25.32293 | 41.4928 | 78.00918 | 33.57186 | -0.53182 | 1.448445 | 0.572422 |
| 18 | 87733600 | 87734599 | 11.23629 | 21.4839 | 23.35165 | 9.25952 | 0.391733 | -0.01842 | 0.572445 |

| | | | | | | | | | |
|---|---|---|---|---|---|---|---|---|---|
| Y | 10165400 | 10166199 | 7.323755 | 13.53897 | 29.48344 | 12.43002 | -0.67607 | 0.181812 | 0.572453 |
| 13 | 3192000 | 3200999 | 119.4136 | 183.4038 | 186.3105 | 77.35638 | 0.271914 | 2.984429 | 0.572455 |
| 8 | 34607200 | 34610399 | 27.78734 | 45.62386 | 72.46023 | 31.21077 | -0.28804 | 1.42247 | 0.572472 |
| 2 | 51962600 | 51963599 | 9.477861 | 17.17909 | 31.86528 | 13.78597 | -0.47259 | 0.318682 | 0.57248 |
| 3 | 20345600 | 20347999 | 22.39146 | 38.96678 | 35.06419 | 15.41076 | 0.538654 | 0.781419 | 0.572495 |
| 7 | 112087800 | 112089599 | 16.03699 | 27.79213 | 45.83611 | 20.11043 | -0.33221 | 0.819382 | 0.572511 |
| 17 | 20830800 | 20833799 | 58.52238 | 93.8133 | 83.69968 | 35.33917 | 0.511207 | 1.9877 | 0.572514 |
| 15 | 88751800 | 88753599 | 17.82166 | 30.65464 | 49.00066 | 21.49396 | -0.28966 | 0.898833 | 0.57252 |
| 7 | 54031200 | 54035599 | 62.2604 | 99.26306 | 106.0197 | 44.4138 | 0.243402 | 2.163729 | 0.572538 |
| 5 | 127318600 | 127324999 | 59.82567 | 94.6154 | 139.6883 | 58.04725 | -0.23414 | 2.400816 | 0.572541 |
| 7 | 3111400 | 3112799 | 33.85117 | 55.60661 | 60.66145 | 26.13397 | 0.252612 | 1.423158 | 0.572559 |
| 6 | 90054000 | 90061399 | 91.23761 | 142.4407 | 150.3919 | 62.40351 | 0.23442 | 2.676504 | 0.572597 |
| 16 | 81091600 | 81093399 | 16.0926 | 29.67156 | 27.27879 | 11.54161 | 0.554969 | 0.259976 | 0.572599 |
| 10 | 71013600 | 71016399 | 27.86351 | 46.65943 | 50.74024 | 22.11687 | 0.26051 | 1.180079 | 0.572604 |
| Y | 752400 | 752799 | 3.025183 | 5.914773 | 17.15595 | 5.871042 | -0.8082 | -0.64382 | 0.572618 |
| 3 | 69759000 | 69760599 | 13.57041 | 23.93228 | 39.24252 | 17.36644 | -0.32055 | 0.55285 | 0.572625 |
| 11 | 61139600 | 61141599 | 12.37974 | 21.88157 | 36.8113 | 16.28321 | -0.35563 | 0.519376 | 0.572626 |
| 3 | 27940800 | 27941599 | 6.998792 | 13.39016 | 26.22202 | 10.37898 | -0.44826 | -0.05957 | 0.572639 |
| 13 | 67021400 | 67023999 | 40.66135 | 66.23328 | 69.4082 | 29.80439 | 0.303484 | 1.556734 | 0.572704 |
| 3 | 157358000 | 157358799 | 9.646342 | 17.99054 | 30.56062 | 12.71685 | -0.30671 | 0.150731 | 0.572787 |
| 4 | 82656200 | 82657999 | 24.16155 | 42.16059 | 38.44463 | 16.9251 | 0.519951 | 0.757813 | 0.572792 |
| 15 | 70525400 | 70528199 | 31.00912 | 52.81093 | 46.7953 | 20.51194 | 0.55585 | 1.036646 | 0.572792 |
| 4 | 154893200 | 154895799 | 31.55426 | 51.13434 | 81.49865 | 34.89083 | -0.3061 | 1.639168 | 0.572831 |
| 2 | 76135200 | 76137399 | 27.0155 | 46.52522 | 38.1336 | 16.72778 | 0.670293 | 0.912067 | 0.572835 |
| 3 | 125930400 | 125932399 | 19.77969 | 34.70831 | 35.87334 | 15.86319 | 0.340106 | 0.646421 | 0.572846 |
| 8 | 65363800 | 65364799 | 15.48141 | 27.82701 | 29.74944 | 12.99993 | 0.307605 | 0.404163 | 0.572852 |
| X | 153669000 | 153670799 | 19.53164 | 34.58594 | 33.04468 | 14.61173 | 0.456926 | 0.534993 | 0.572873 |
| 4 | 148006800 | 148008999 | 19.33674 | 32.73655 | 54.83521 | 24.0059 | -0.36015 | 1.053765 | 0.572898 |
| 2 | 166601400 | 166603199 | 16.46247 | 28.6396 | 45.77378 | 20.11356 | -0.28643 | 0.773181 | 0.572918 |
| 1 | 108324600 | 108326999 | 22.23314 | 39.20257 | 33.95692 | 15.02788 | 0.595437 | 0.598114 | 0.572933 |
| 2 | 66919800 | 66921999 | 26.05303 | 43.9175 | 46.96998 | 20.5358 | 0.284571 | 1.107684 | 0.572943 |
| 1 | 17775000 | 17777399 | 18.10327 | 30.79098 | 58.03779 | 25.43722 | -0.52926 | 1.013444 | 0.572951 |
| 6 | 131925600 | 131926799 | 20.80529 | 36.3086 | 38.11778 | 16.78251 | 0.317573 | 0.738335 | 0.572952 |
| 15 | 24759200 | 24760999 | 23.52605 | 40.17735 | 43.21492 | 18.94675 | 0.277878 | 0.979531 | 0.57297 |
| 1 | 20110000 | 20113599 | 37.87151 | 62.1041 | 63.93722 | 27.52523 | 0.334919 | 1.454244 | 0.573 |
| 5 | 145202200 | 145206599 | 59.02434 | 94.08448 | 100.2752 | 41.99542 | 0.246143 | 2.164392 | 0.573039 |
| 9 | 100877600 | 100882399 | 49.60393 | 79.67086 | 86.42642 | 36.59805 | 0.233037 | 1.916996 | 0.573051 |
| 7 | 110801200 | 110806599 | 61.878 | 98.65481 | 105.4932 | 44.19413 | 0.241735 | 2.162155 | 0.573072 |
| 7 | 25773000 | 25801199 | 397.1847 | 555.997 | 519.8689 | 215.5281 | 0.266683 | 4.566618 | 0.573088 |
| 3 | 59967800 | 59969199 | 8.877678 | 16.55394 | 28.9567 | 12.05317 | -0.34885 | 0.141712 | 0.573105 |
| 14 | 47059200 | 47059999 | 6.008362 | 11.53338 | 24.64644 | 9.663067 | -0.55945 | -0.10411 | 0.573105 |
| 17 | 36755600 | 36756799 | 13.79263 | 24.23512 | 39.34642 | 17.41875 | -0.30799 | 0.598528 | 0.573133 |
| 3 | 58504200 | 58514799 | 168.1319 | 253.5695 | 239.3472 | 98.90221 | 0.356547 | 3.365281 | 0.573148 |
| 9 | 27590200 | 27591999 | 20.86201 | 36.97984 | 32.34719 | 14.30037 | 0.584136 | 0.530942 | 0.573155 |
| 9 | 114600000 | 114602199 | 27.76252 | 45.28665 | 74.45603 | 31.98852 | -0.34302 | 1.514249 | 0.573168 |
| 18 | 9152000 | 9153799 | 20.99303 | 35.47564 | 55.92643 | 24.47004 | -0.27305 | 1.078589 | 0.573172 |
| 7 | 107594800 | 107595399 | 8.229835 | 15.76021 | 29.19458 | 11.52442 | -0.3651 | -0.04632 | 0.573196 |

| | | | | | | | | | |
|---|---|---|---|---|---|---|---|---|---|
| 5 | 106170800 | 106171599 | 11.52749 | 20.57571 | 34.74057 | 15.27069 | -0.35348 | 0.432377 | 0.5732 |
| 8 | 96090400 | 96091599 | 15.4886 | 28.06579 | 28.88336 | 12.50038 | 0.371489 | 0.358237 | 0.573213 |
| 5 | 126424200 | 126425799 | 26.09058 | 44.29187 | 47.41455 | 20.77453 | 0.283469 | 1.045994 | 0.573234 |
| 19 | 48528200 | 48529999 | 14.70333 | 25.90524 | 43.41169 | 19.21518 | -0.35248 | 0.576997 | 0.57324 |
| 7 | 131357200 | 131360399 | 38.591 | 62.18246 | 94.20628 | 40.07938 | -0.2421 | 1.805131 | 0.573241 |
| 13 | 85213200 | 85213999 | 12.44692 | 23.37003 | 25.16555 | 10.35348 | 0.359573 | 0.119969 | 0.573248 |
| 3 | 85535400 | 85542199 | 59.25045 | 94.49582 | 102.3829 | 42.91012 | 0.223301 | 2.148671 | 0.573249 |
| 3 | 98154800 | 98162399 | 82.19395 | 127.6507 | 183.5857 | 76.23297 | -0.21686 | 2.76524 | 0.573257 |
| 12 | 104165200 | 104168599 | 41.003 | 65.95241 | 99.07425 | 42.04269 | -0.23311 | 1.864562 | 0.57326 |
| 13 | 23461200 | 23464199 | 39.54889 | 64.22801 | 70.03203 | 30.02944 | 0.24306 | 1.603051 | 0.573301 |
| 3 | 45165600 | 45166599 | 11.52052 | 22.1494 | 21.7393 | 8.484502 | 0.558458 | -0.07256 | 0.573336 |
| 9 | 92907000 | 92910799 | 31.72437 | 53.27845 | 51.90053 | 22.66391 | 0.4186 | 1.161851 | 0.573343 |
| 15 | 68505000 | 68506799 | 16.09394 | 27.79164 | 52.73116 | 23.13309 | -0.53476 | 0.860594 | 0.573343 |
| 16 | 30333400 | 30336999 | 36.92593 | 60.11363 | 61.94903 | 26.59572 | 0.327056 | 1.554363 | 0.573361 |
| 8 | 74729400 | 74732399 | 40.85627 | 66.56481 | 69.40522 | 29.80539 | 0.310952 | 1.554638 | 0.573374 |
| 10 | 19172600 | 19174399 | 18.7248 | 33.66727 | 30.5133 | 13.33023 | 0.544881 | 0.426815 | 0.573377 |
| 10 | 116308400 | 116310599 | 17.00664 | 29.30628 | 47.94367 | 21.03149 | -0.32238 | 0.877129 | 0.57339 |
| 13 | 54339200 | 54340999 | 14.59038 | 25.63508 | 40.84258 | 18.08006 | -0.28105 | 0.60656 | 0.573392 |
| 2 | 107749800 | 107752399 | 39.58547 | 63.6291 | 97.47864 | 41.32488 | -0.26253 | 1.884591 | 0.573393 |
| 14 | 42597600 | 42598799 | 10.24537 | 19.42211 | 21.1837 | 8.55616 | 0.363665 | 0.025632 | 0.573406 |
| 1 | 184903200 | 184904599 | 17.25225 | 29.9558 | 47.93268 | 21.03672 | -0.28847 | 0.806718 | 0.573413 |
| 8 | 91025000 | 91027199 | 11.91094 | 21.46878 | 35.74512 | 15.57305 | -0.325 | 0.374612 | 0.573415 |
| 17 | 17402800 | 17411999 | 110.9978 | 173.4899 | 124.8042 | 51.78005 | 0.788597 | 2.66317 | 0.573418 |
| X | 80591200 | 80592799 | 24.59911 | 42.87608 | 39.12368 | 17.21194 | 0.519021 | 0.779035 | 0.573465 |
| 7 | 59729800 | 59732199 | 25.31204 | 44.31347 | 36.08715 | 15.92655 | 0.68251 | 0.713043 | 0.573474 |
| 14 | 124784000 | 124787799 | 38.54998 | 63.24677 | 64.00866 | 27.56154 | 0.359822 | 1.450436 | 0.573476 |
| 5 | 105515800 | 105519399 | 37.55821 | 62.03542 | 60.11358 | 25.96447 | 0.425694 | 1.365176 | 0.573477 |
| 14 | 45979400 | 45980199 | 10.90781 | 19.49139 | 34.27974 | 15.05393 | -0.41096 | 0.420092 | 0.573497 |
| 10 | 76828400 | 76832599 | 35.91641 | 58.12725 | 88.76865 | 37.96661 | -0.24649 | 1.67575 | 0.573521 |
| 8 | 48075400 | 48089599 | 155.8047 | 232.351 | 322.8084 | 133.0403 | -0.21526 | 3.579005 | 0.573533 |
| 8 | 97298000 | 97298999 | 9.130688 | 17.23074 | 30.92584 | 12.61137 | -0.36214 | 0.070298 | 0.573551 |
| 2 | 168833800 | 168836599 | 23.54278 | 39.12616 | 62.87356 | 27.27228 | -0.301 | 1.261875 | 0.573558 |
| 6 | 72454800 | 72458399 | 37.2991 | 60.78552 | 65.66236 | 28.2008 | 0.260274 | 1.539608 | 0.573564 |
| 4 | 150960800 | 150969199 | 137.4122 | 204.2557 | 342.4042 | 141.0635 | -0.49036 | 3.635025 | 0.573571 |
| 2 | 37150400 | 37153199 | 33.59542 | 56.06833 | 48.14059 | 20.93908 | 0.600295 | 1.229178 | 0.573576 |
| 6 | 130896800 | 130904799 | 142.2794 | 217.0246 | 183.5199 | 76.09321 | 0.528591 | 3.128003 | 0.57358 |
| 9 | 4181800 | 4182799 | 8.787477 | 17.12185 | 19.1068 | 6.943395 | 0.470999 | -0.34672 | 0.573582 |
| 2 | 108365400 | 108366599 | 8.594309 | 16.30762 | 29.36139 | 11.83546 | -0.35118 | 0.027453 | 0.573585 |
| 9 | 61444200 | 61448999 | 46.97213 | 74.97395 | 113.7582 | 47.73137 | -0.26044 | 2.115791 | 0.57359 |
| 13 | 64861400 | 64862799 | 23.06018 | 39.90477 | 35.78555 | 15.70008 | 0.542464 | 0.849595 | 0.573593 |
| 7 | 142398200 | 142398799 | 16.69651 | 29.47915 | 31.44328 | 13.91177 | 0.297411 | 0.5464 | 0.5736 |
| 9 | 19690000 | 19690599 | 16.69651 | 29.47915 | 31.44328 | 13.91177 | 0.297411 | 0.5464 | 0.5736 |
| 9 | 113225000 | 113227399 | 19.22793 | 32.93067 | 52.7683 | 23.14055 | -0.29448 | 0.946522 | 0.573604 |
| 3 | 95157400 | 95158799 | 7.030563 | 13.39601 | 25.58492 | 10.22288 | -0.4241 | -0.02761 | 0.573624 |
| 19 | 21225000 | 21226399 | 18.02652 | 31.78402 | 33.19559 | 14.69106 | 0.326634 | 0.570084 | 0.573628 |
| 8 | 44236400 | 44240399 | 45.27472 | 73.43705 | 73.40267 | 31.44642 | 0.366639 | 1.641743 | 0.573634 |
| 5 | 98940400 | 98941399 | 10.13701 | 18.58627 | 31.85469 | 13.58073 | -0.34559 | 0.255128 | 0.573634 |

| | | | | | | | | | |
|---|---|---|---|---|---|---|---|---|---|
| 14 | 96858400 | 96859599 | 10.13701 | 18.58627 | 31.85469 | 13.58073 | -0.34559 | 0.255128 | 0.573634 |
| 8 | 75314400 | 75316999 | 33.50126 | 55.42643 | 59.199 | 25.59877 | 0.285895 | 1.336042 | 0.573651 |
| 4 | 89917400 | 89921399 | 68.65542 | 109.1047 | 114.4546 | 47.74586 | 0.264428 | 2.273894 | 0.573677 |
| 16 | 63648200 | 63649399 | 11.28122 | 20.45281 | 33.7729 | 14.60676 | -0.30611 | 0.333168 | 0.573684 |
| 19 | 10433600 | 10435599 | 16.13964 | 27.66091 | 48.99787 | 21.48831 | -0.438 | 0.92569 | 0.573729 |
| 5 | 96811600 | 96813399 | 13.68384 | 25.69664 | 25.61469 | 10.53534 | 0.4703 | 0.129624 | 0.573737 |
| X | 155409800 | 155413199 | 65.77084 | 105.2106 | 98.42705 | 41.4276 | 0.439473 | 2.065656 | 0.573739 |
| 7 | 74868000 | 74870199 | 25.71224 | 44.6724 | 40.7223 | 17.87906 | 0.519938 | 0.828447 | 0.573742 |
| 13 | 22995000 | 23002799 | 133.0213 | 202.917 | 205.0433 | 85.01888 | 0.271925 | 3.125134 | 0.573755 |
| 10 | 24859200 | 24861999 | 33.25639 | 55.15568 | 50.89226 | 22.04386 | 0.496574 | 1.301128 | 0.573758 |
| 12 | 56182600 | 56185599 | 44.89563 | 72.72522 | 74.99037 | 32.09506 | 0.32003 | 1.667494 | 0.573774 |
| 7 | 111138400 | 111139399 | 14.33838 | 26.58267 | 26.98202 | 11.33108 | 0.420875 | 0.216106 | 0.573781 |
| 6 | 122688400 | 122690999 | 20.13948 | 35.03029 | 35.68655 | 15.67938 | 0.360242 | 0.776216 | 0.5738 |
| 9 | 40753600 | 40754799 | 13.66363 | 24.77166 | 25.97667 | 11.23514 | 0.345001 | 0.333797 | 0.57381 |
| 17 | 86309600 | 86311199 | 14.20946 | 25.11713 | 42.88296 | 18.96874 | -0.3774 | 0.537221 | 0.573833 |
| 16 | 17765000 | 17765599 | 4.940558 | 9.626263 | 20.21031 | 7.345458 | -0.42984 | -0.38931 | 0.573861 |
| 1 | 155121600 | 155122999 | 13.93772 | 24.54814 | 39.28013 | 17.3876 | -0.28607 | 0.571024 | 0.573872 |
| 13 | 98667000 | 98670199 | 28.45233 | 46.73493 | 72.50556 | 31.23573 | -0.25407 | 1.419937 | 0.573876 |
| 4 | 48690400 | 48690999 | 6.705005 | 12.92808 | 24.2088 | 9.370856 | -0.35304 | -0.13786 | 0.573886 |
| 3 | 39476200 | 39477599 | 14.5466 | 25.59823 | 41.31026 | 18.28943 | -0.29895 | 0.586374 | 0.573896 |
| 6 | 120476000 | 120480399 | 37.6017 | 60.4602 | 95.57595 | 40.53748 | -0.3071 | 1.869655 | 0.57395 |
| 2 | 166367800 | 166379199 | 137.1831 | 204.2562 | 336.216 | 138.5376 | -0.46184 | 3.605276 | 0.573959 |
| 1 | 185814800 | 185816199 | 28.11643 | 46.35135 | 72.12744 | 31.12065 | -0.25632 | 1.374433 | 0.57399 |
| 16 | 23025400 | 23028799 | 32.64815 | 54.33767 | 56.41658 | 24.49612 | 0.327216 | 1.263843 | 0.573991 |
| 14 | 110526800 | 110528799 | 23.33407 | 41.07284 | 33.84383 | 14.983 | 0.666155 | 0.621925 | 0.574025 |
| 7 | 12992800 | 12995999 | 34.6631 | 55.87318 | 90.06286 | 38.33513 | -0.3307 | 1.785486 | 0.574039 |
| 4 | 96803600 | 96805599 | 27.82664 | 47.61417 | 45.14249 | 19.79495 | 0.459516 | 0.979213 | 0.574054 |
| 2 | 58377800 | 58381399 | 33.26405 | 53.99508 | 83.01267 | 35.58273 | -0.25209 | 1.607431 | 0.574085 |
| X | 61306000 | 61306799 | 17.43378 | 31.1278 | 29.86004 | 13.1219 | 0.458006 | 0.462634 | 0.574101 |
| 10 | 52016400 | 52017399 | 10.50494 | 18.71393 | 38.37148 | 16.89453 | -0.63358 | 0.446389 | 0.574108 |
| 10 | 3906600 | 3910199 | 27.68615 | 45.39769 | 71.60198 | 30.82385 | -0.27874 | 1.439938 | 0.574112 |
| 3 | 149170200 | 149171399 | 12.3662 | 22.82082 | 24.55235 | 10.37567 | 0.331111 | 0.220463 | 0.574118 |
| 4 | 138678600 | 138681599 | 41.02003 | 66.5046 | 72.34222 | 30.98139 | 0.244213 | 1.640549 | 0.574118 |
| X | 66193200 | 66196399 | 36.78178 | 61.90149 | 49.66915 | 21.70513 | 0.697718 | 1.162431 | 0.574135 |
| 3 | 70840600 | 70841199 | 9.26759 | 17.80928 | 20.23802 | 7.90957 | 0.347512 | -0.09407 | 0.574188 |
| 14 | 6556400 | 6559399 | 35.26294 | 56.9216 | 89.12433 | 38.00585 | -0.28651 | 1.743148 | 0.574193 |
| 4 | 113639600 | 113649199 | 152.0295 | 230.6464 | 221.0875 | 91.52672 | 0.341573 | 3.242121 | 0.574196 |
| 3 | 91298200 | 91301199 | 25.24485 | 43.77984 | 41.29571 | 18.11873 | 0.470425 | 0.846966 | 0.574199 |
| 19 | 11399400 | 11403599 | 77.48991 | 121.8615 | 128.8955 | 53.47174 | 0.239944 | 2.532986 | 0.574228 |
| 13 | 50571600 | 50572999 | 9.872157 | 18.03419 | 31.33005 | 13.42254 | -0.36974 | 0.273996 | 0.574238 |
| 7 | 26569800 | 26576999 | 126.7815 | 193.7832 | 199.0504 | 82.57191 | 0.250548 | 3.083581 | 0.57424 |
| 4 | 112724800 | 112727799 | 41.0929 | 67.36092 | 65.14496 | 28.04063 | 0.424767 | 1.470575 | 0.574241 |
| 5 | 22950400 | 22952199 | 14.59255 | 25.42308 | 43.1808 | 18.99135 | -0.37334 | 0.740701 | 0.574277 |
| 8 | 67839600 | 67840999 | 14.34346 | 26.27632 | 27.90539 | 11.91088 | 0.339253 | 0.284073 | 0.574278 |
| 15 | 87537800 | 87541399 | 27.22475 | 44.54128 | 72.51508 | 31.18831 | -0.32588 | 1.467956 | 0.574304 |
| 9 | 71382200 | 71384399 | 17.62211 | 30.58691 | 48.47347 | 21.26921 | -0.27464 | 0.815422 | 0.574304 |
| 10 | 61863000 | 61872399 | 77.4511 | 119.8517 | 201.2915 | 83.61145 | -0.44442 | 2.829349 | 0.574321 |

| | | | | | | | | | |
|---|---|---|---|---|---|---|---|---|---|
| 1 | 68916800 | 68920199 | 29.99436 | 50.76668 | 49.61081 | 21.71708 | 0.41442 | 1.088817 | 0.574343 |
| 19 | 7783600 | 7785799 | 28.53576 | 48.85579 | 39.76536 | 17.4379 | 0.678931 | 0.972266 | 0.574368 |
| 1 | 12145600 | 12146199 | 12.35151 | 22.84089 | 25.02945 | 10.54757 | 0.307612 | 0.20962 | 0.574373 |
| 9 | 103539200 | 103542799 | 36.02784 | 59.04176 | 64.12754 | 27.59946 | 0.257561 | 1.460093 | 0.574405 |
| 5 | 42027400 | 42028399 | 13.4625 | 23.83366 | 38.20606 | 16.89474 | -0.28571 | 0.51007 | 0.574411 |
| 6 | 130856400 | 130858399 | 56.60646 | 90.61547 | 95.93514 | 40.41964 | 0.262442 | 2.032504 | 0.574412 |
| 18 | 26674400 | 26674999 | 8.570962 | 16.60715 | 17.61916 | 6.677142 | 0.486912 | -0.21316 | 0.574417 |
| 18 | 64836000 | 64836799 | 9.787469 | 18.20854 | 31.04372 | 12.97224 | -0.31694 | 0.168201 | 0.574443 |
| 12 | 104251800 | 104255199 | 38.07885 | 63.25907 | 56.14669 | 24.34781 | 0.552578 | 1.299467 | 0.574459 |
| 14 | 3406000 | 3407799 | 43.50344 | 70.16732 | 73.25683 | 31.20717 | 0.295568 | 1.773305 | 0.574462 |
| 2 | 173162600 | 173166999 | 29.24619 | 47.55537 | 77.18047 | 33.1206 | -0.32767 | 1.564926 | 0.574474 |
| 5 | 45007000 | 45009399 | 27.29017 | 46.21903 | 48.51397 | 21.24102 | 0.311742 | 1.07447 | 0.574488 |
| 15 | 12062000 | 12067599 | 44.27536 | 70.94176 | 106.5461 | 44.95333 | -0.23953 | 1.991196 | 0.574495 |
| 14 | 41912800 | 41920999 | 79.08373 | 125.1972 | 112.6695 | 46.84326 | 0.480162 | 2.38597 | 0.574522 |
| 3 | 126187800 | 126188399 | 15.51596 | 27.60481 | 29.38988 | 12.95423 | 0.305181 | 0.467752 | 0.574535 |
| 8 | 81712600 | 81713599 | 9.492963 | 17.67588 | 29.98444 | 12.5113 | -0.30784 | 0.158535 | 0.57454 |
| 5 | 101378600 | 101382199 | 31.19134 | 52.14088 | 53.60964 | 23.34184 | 0.341091 | 1.209397 | 0.57455 |
| 4 | 24887800 | 24888999 | 17.67752 | 31.08902 | 33.44618 | 14.80953 | 0.28291 | 0.598346 | 0.574567 |
| 12 | 59363800 | 59365199 | 24.1614 | 40.22437 | 63.00619 | 27.36239 | -0.26409 | 1.243373 | 0.574567 |
| 14 | 27145800 | 27147199 | 18.94923 | 33.24838 | 35.43637 | 15.66889 | 0.296014 | 0.642376 | 0.574575 |
| 14 | 37612800 | 37619799 | 39.07784 | 63.87975 | 59.19053 | 25.44968 | 0.484117 | 1.496626 | 0.574588 |
| 3 | 126013400 | 126014599 | 13.61576 | 24.82827 | 26.87829 | 11.54119 | 0.305879 | 0.301446 | 0.574608 |
| 16 | 84606600 | 84609399 | 19.02943 | 33.36999 | 35.40736 | 15.64849 | 0.302366 | 0.650778 | 0.574659 |
| 8 | 84095000 | 84096399 | 12.4181 | 22.34716 | 37.2961 | 16.28127 | -0.3303 | 0.390945 | 0.574683 |
| 2 | 5287400 | 5290799 | 26.80868 | 45.43277 | 48.50691 | 21.24207 | 0.28723 | 1.065104 | 0.574717 |
| 7 | 106080200 | 106083199 | 26.88643 | 45.60979 | 47.95339 | 21.006 | 0.30932 | 1.055248 | 0.57472 |
| 16 | 79080800 | 79083199 | 18.77495 | 32.4597 | 53.37849 | 23.41792 | -0.32905 | 0.861725 | 0.574726 |
| 6 | 12216000 | 12224599 | 174.8314 | 261.4808 | 266.1684 | 109.7587 | 0.23699 | 3.526619 | 0.574745 |
| X | 167640200 | 167642199 | 22.92082 | 40.2925 | 29.03199 | 12.85612 | 0.85818 | 0.627515 | 0.574765 |
| 14 | 48798600 | 48800799 | 28.11096 | 47.0362 | 50.19904 | 21.86994 | 0.287576 | 1.190361 | 0.574774 |
| 12 | 20984400 | 20989399 | 63.03804 | 100.2942 | 108.5997 | 45.36398 | 0.22045 | 2.230817 | 0.57479 |
| 4 | 103135600 | 103135999 | 14.37299 | 26.39809 | 27.3239 | 11.62203 | 0.379458 | 0.267343 | 0.574798 |
| 1 | 179129800 | 179132199 | 17.06751 | 29.30298 | 55.972 | 24.55036 | -0.54626 | 0.920188 | 0.574816 |
| 15 | 8080000 | 8080999 | 12.18144 | 21.64017 | 35.72456 | 15.76607 | -0.3252 | 0.46898 | 0.574841 |
| 19 | 39709800 | 39712199 | 39.10551 | 63.68882 | 68.44877 | 29.39069 | 0.267016 | 1.556834 | 0.574856 |
| 14 | 10542000 | 10545599 | 32.33946 | 53.57365 | 58.01685 | 25.11003 | 0.266142 | 1.317214 | 0.574863 |
| 7 | 61117000 | 61118999 | 20.08715 | 35.48485 | 33.50426 | 14.82202 | 0.472548 | 0.563421 | 0.574867 |
| 6 | 146071800 | 146075199 | 21.48411 | 35.75138 | 67.71575 | 29.39908 | -0.53724 | 1.24524 | 0.574877 |
| 7 | 10046600 | 10050399 | 86.49783 | 135.4423 | 144.7035 | 60.02484 | 0.220701 | 2.620941 | 0.574888 |
| 18 | 33548000 | 33551399 | 16.6926 | 30.18389 | 25.22264 | 10.94483 | 0.667701 | 0.370563 | 0.5749 |
| 5 | 140900000 | 140915599 | 182.1535 | 267.5879 | 370.0788 | 152.4152 | -0.22894 | 3.836929 | 0.574911 |
| 10 | 67747200 | 67749199 | 12.82026 | 22.5452 | 38.83478 | 17.19576 | -0.39273 | 0.579521 | 0.574912 |
| 6 | 54800000 | 54802799 | 40.7651 | 66.36209 | 69.46529 | 29.82232 | 0.304467 | 1.564552 | 0.574914 |
| 8 | 104418400 | 104421599 | 28.79433 | 47.26342 | 73.0408 | 31.45695 | -0.24888 | 1.430191 | 0.574929 |
| 4 | 80224800 | 80227799 | 30.51791 | 51.47448 | 50.1984 | 21.95126 | 0.417178 | 1.118698 | 0.574936 |
| 14 | 38264400 | 38266399 | 19.1198 | 33.80476 | 29.11743 | 12.87913 | 0.6049 | 0.54192 | 0.574948 |
| 17 | 40477400 | 40477999 | 11.16291 | 20.04558 | 34.2158 | 14.96369 | -0.36413 | 0.389532 | 0.574951 |

| | | | | | | | | | |
|---|---|---|---|---|---|---|---|---|---|
| 8 | 38884600 | 38887799 | 30.86109 | 52.37667 | 48.48123 | 21.24021 | 0.492557 | 1.068199 | 0.574956 |
| 15 | 31755000 | 31758999 | 45.61841 | 72.81919 | 113.8565 | 47.77819 | -0.30356 | 2.11044 | 0.574961 |
| 6 | 83112600 | 83115599 | 15.84125 | 27.08755 | 50.03514 | 21.93918 | -0.49876 | 0.945868 | 0.574962 |
| 12 | 98328000 | 98328799 | 12.7254 | 23.43866 | 22.67869 | 9.608936 | 0.480751 | 0.230658 | 0.574965 |
| 13 | 109557800 | 109559199 | 16.40223 | 30.52589 | 26.38459 | 11.01842 | 0.657254 | 0.210014 | 0.57499 |
| 1 | 106657200 | 106660599 | 38.0695 | 61.82833 | 65.24764 | 27.97887 | 0.290187 | 1.596903 | 0.575013 |
| 3 | 90372000 | 90373799 | 10.36838 | 18.38103 | 35.39127 | 15.64577 | -0.54736 | 0.470188 | 0.575018 |
| 1 | 30725800 | 30730599 | 80.2971 | 126.3586 | 135.0975 | 56.0494 | 0.225164 | 2.522807 | 0.575019 |
| 1 | 27107800 | 27109599 | 19.77924 | 34.75877 | 30.62649 | 13.55993 | 0.569033 | 0.613333 | 0.575028 |
| 14 | 63090400 | 63092799 | 19.45984 | 34.01225 | 36.67912 | 16.16538 | 0.279301 | 0.709487 | 0.575044 |
| 2 | 3995200 | 3997199 | 16.20943 | 28.75773 | 30.17371 | 13.33124 | 0.323562 | 0.489294 | 0.575063 |
| 15 | 37162800 | 37166999 | 41.50068 | 66.76025 | 100.4607 | 42.63781 | -0.2354 | 1.862485 | 0.575091 |
| 4 | 143317400 | 143325199 | 96.18408 | 150.8233 | 143.4251 | 59.48588 | 0.390071 | 2.601198 | 0.575094 |
| 9 | 4395200 | 4397399 | 28.53213 | 48.30669 | 43.68107 | 19.12313 | 0.525984 | 1.074279 | 0.575099 |
| 17 | 5727400 | 5728199 | 14.88377 | 27.63324 | 26.98742 | 11.31134 | 0.478213 | 0.213098 | 0.575142 |
| 12 | 115393400 | 115417599 | 451.1825 | 624.8899 | 575.3039 | 238.9068 | 0.273536 | 4.728684 | 0.575147 |
| 4 | 91262400 | 91263599 | 12.69835 | 22.26022 | 39.98753 | 17.66763 | -0.45319 | 0.63029 | 0.575149 |
| 13 | 66922000 | 66952599 | 646.468 | 868.4002 | 777.6423 | 324.4703 | 0.269035 | 5.208615 | 0.575158 |
| 15 | 31561600 | 31564399 | 24.09863 | 39.81612 | 73.40762 | 31.71313 | -0.49944 | 1.339143 | 0.575164 |
| 4 | 14047600 | 14049799 | 24.3915 | 39.88176 | 81.85476 | 35.19749 | -0.6598 | 1.475717 | 0.57517 |
| 3 | 7715400 | 7717799 | 37.27483 | 61.00626 | 65.20837 | 28.0495 | 0.27984 | 1.481263 | 0.575189 |
| 1 | 55735000 | 55737199 | 18.26691 | 31.34977 | 50.60146 | 22.19403 | -0.30439 | 0.921539 | 0.5752 |
| 8 | 110793600 | 110796199 | 33.81228 | 55.44558 | 58.94403 | 25.37527 | 0.288759 | 1.44272 | 0.575233 |
| X | 96353400 | 96357399 | 49.88586 | 81.05874 | 69.98799 | 30.02239 | 0.57969 | 1.615783 | 0.575234 |
| 18 | 9088800 | 9089399 | 6.563753 | 12.7024 | 24.35185 | 9.281611 | -0.36494 | -0.18916 | 0.575235 |
| 5 | 146043200 | 146046999 | 51.2688 | 82.60366 | 83.85925 | 35.67306 | 0.33406 | 1.821141 | 0.575236 |
| 12 | 17746400 | 17749999 | 19.39942 | 32.81025 | 54.25648 | 23.74543 | -0.3416 | 1.064339 | 0.575253 |
| 5 | 8943400 | 8944799 | 10.94576 | 19.55296 | 34.33551 | 15.08249 | -0.40879 | 0.424584 | 0.575261 |
| 13 | 62076800 | 62087599 | 35.45041 | 58.76894 | 52.03988 | 22.5342 | 0.555735 | 1.313635 | 0.575281 |
| 6 | 8384000 | 8385199 | 15.27001 | 28.05743 | 27.83228 | 11.83147 | 0.441133 | 0.273813 | 0.575296 |
| 1 | 66950400 | 66952799 | 28.88425 | 49.14181 | 47.92771 | 21.00658 | 0.41777 | 1.03945 | 0.575298 |
| 6 | 130486200 | 130497999 | 168.1017 | 254.2539 | 227.5389 | 94.10813 | 0.4369 | 3.307368 | 0.575347 |
| 6 | 10371200 | 10373199 | 20.54797 | 36.11146 | 35.14368 | 15.56056 | 0.426388 | 0.624759 | 0.575364 |
| 8 | 43164600 | 43168799 | 118.2076 | 180.7127 | 191.166 | 79.30475 | 0.20836 | 3.075886 | 0.575384 |
| 1 | 137605200 | 137606999 | 17.57815 | 30.37761 | 49.00734 | 21.49996 | -0.30162 | 0.853754 | 0.575406 |
| 1 | 140887000 | 140887999 | 13.75354 | 24.98334 | 25.57483 | 11.0344 | 0.382044 | 0.323926 | 0.575409 |
| 8 | 34920200 | 34922799 | 24.48281 | 40.55881 | 65.75271 | 28.45841 | -0.31415 | 1.301603 | 0.575419 |
| 4 | 118050800 | 118054999 | 32.49962 | 52.80084 | 81.03288 | 34.75674 | -0.24829 | 1.589065 | 0.575424 |
| 15 | 15086400 | 15092399 | 59.26936 | 94.72229 | 100.565 | 42.2717 | 0.255795 | 2.084804 | 0.575434 |
| 6 | 141546400 | 141548199 | 19.06761 | 32.79785 | 52.24561 | 22.91735 | -0.28487 | 0.906626 | 0.575448 |
| 13 | 102574200 | 102575999 | 12.02709 | 21.77941 | 37.47922 | 16.23362 | -0.36665 | 0.354931 | 0.575453 |
| 7 | 7165600 | 7168599 | 39.72982 | 64.56213 | 70.11514 | 30.07892 | 0.249657 | 1.591318 | 0.575495 |
| 6 | 52228000 | 52231399 | 45.34179 | 72.3913 | 112.1966 | 47.09421 | -0.29057 | 2.105337 | 0.575531 |
| 10 | 31627800 | 31628999 | 12.53259 | 23.63749 | 20.01866 | 8.173032 | 0.712588 | 0.079781 | 0.575531 |
| 9 | 44132000 | 44132799 | 9.030231 | 17.5486 | 20.03503 | 7.459917 | 0.406288 | -0.25795 | 0.575538 |
| 17 | 33213600 | 33227799 | 218.8996 | 323.0956 | 312.463 | 128.6777 | 0.293321 | 3.763497 | 0.575543 |
| 4 | 120463600 | 120468599 | 32.4243 | 52.34105 | 95.34074 | 40.65835 | -0.50422 | 1.742558 | 0.575543 |

| | | | | | | | | | |
|---|---|---|---|---|---|---|---|---|---|
| 2 | 18681000 | 18683199 | 15.72332 | 26.48972 | 64.85655 | 28.35114 | -0.90593 | 1.094569 | 0.575548 |
| X | 125888000 | 125889599 | 19.14477 | 35.57701 | 25.09543 | 10.50516 | 0.946018 | 0.239132 | 0.57555 |
| 17 | 68328800 | 68329999 | 19.12509 | 33.47977 | 35.46868 | 15.65269 | 0.304601 | 0.678223 | 0.575551 |
| 2 | 95090000 | 95091999 | 127.0391 | 194.4396 | 153.7506 | 63.80017 | 0.629146 | 3.053809 | 0.575556 |
| 16 | 93876200 | 93876799 | 6.402581 | 12.36753 | 23.71497 | 9.117566 | -0.3778 | -0.16649 | 0.575556 |
| X | 52076600 | 52078599 | 27.19684 | 47.20964 | 40.25674 | 17.66476 | 0.615895 | 0.849958 | 0.575568 |
| 18 | 68622400 | 68625799 | 31.03293 | 50.3523 | 80.04625 | 34.30057 | -0.30073 | 1.611048 | 0.575595 |
| 13 | 13064200 | 13064599 | 15.36528 | 27.81059 | 23.80716 | 10.31817 | 0.633989 | 0.350774 | 0.575609 |
| 19 | 39123200 | 39125399 | 37.49098 | 61.64459 | 55.87626 | 24.08217 | 0.519685 | 1.418174 | 0.575613 |
| 7 | 11617800 | 11636799 | 271.2343 | 395.797 | 344.1056 | 141.7847 | 0.431255 | 3.950048 | 0.575623 |
| 3 | 33519000 | 33520399 | 18.21581 | 31.97911 | 34.04117 | 15.05897 | 0.297927 | 0.628701 | 0.575631 |
| 15 | 46187000 | 46192199 | 50.12326 | 80.60141 | 85.68016 | 36.34529 | 0.26433 | 1.878091 | 0.575634 |
| 1 | 140914400 | 140915199 | 8.065392 | 15.45372 | 28.03106 | 11.04782 | -0.33279 | -0.05602 | 0.575639 |
| 5 | 143988000 | 143993399 | 52.38272 | 83.28051 | 124.9695 | 52.15469 | -0.25062 | 2.248993 | 0.575643 |
| 6 | 123660800 | 123664999 | 69.54895 | 110.2764 | 119.072 | 49.56039 | 0.219692 | 2.339597 | 0.575647 |
| 6 | 7152000 | 7154999 | 37.75753 | 61.66937 | 65.93475 | 28.34071 | 0.277315 | 1.508801 | 0.575652 |
| 5 | 17779000 | 17779999 | 9.650561 | 17.88084 | 30.43342 | 12.79104 | -0.32058 | 0.189495 | 0.575662 |
| 8 | 106586800 | 106587999 | 11.72099 | 20.77379 | 35.76345 | 15.81108 | -0.38724 | 0.481829 | 0.575664 |
| 17 | 4896200 | 4898199 | 11.72099 | 20.77379 | 35.76345 | 15.81108 | -0.38724 | 0.481829 | 0.575664 |
| 4 | 33637000 | 33639799 | 18.21482 | 31.39194 | 51.16654 | 22.44534 | -0.31727 | 0.883144 | 0.575664 |
| 5 | 118169600 | 118175199 | 49.90868 | 79.48353 | 119.9961 | 50.19286 | -0.2567 | 2.190937 | 0.575683 |
| 7 | 89762600 | 89766599 | 37.74387 | 62.29914 | 60.67987 | 26.19661 | 0.418245 | 1.374051 | 0.575688 |
| 18 | 7312000 | 7313799 | 17.33854 | 31.60644 | 28.02511 | 12.03853 | 0.591576 | 0.332907 | 0.575707 |
| 18 | 44077200 | 44077799 | 10.96371 | 20.65735 | 22.68724 | 9.278716 | 0.338353 | 0.078814 | 0.57572 |
| 10 | 120429600 | 120431999 | 32.70868 | 54.66414 | 54.71765 | 23.82064 | 0.379722 | 1.2201 | 0.575721 |
| 12 | 106098600 | 106101599 | 30.31813 | 49.34821 | 78.27223 | 33.60148 | -0.29369 | 1.551247 | 0.575723 |
| 14 | 123873200 | 123876199 | 31.3779 | 52.19827 | 55.28669 | 23.9975 | 0.298443 | 1.26302 | 0.575733 |
| 11 | 15895000 | 15895399 | 9.065121 | 17.4846 | 20.35138 | 7.870054 | 0.328019 | -0.12773 | 0.575741 |
| 4 | 20461600 | 20464599 | 37.19322 | 61.15048 | 55.94842 | 24.11252 | 0.506154 | 1.416417 | 0.575766 |
| 16 | 68850600 | 68852199 | 15.4561 | 27.68569 | 29.75024 | 13.03821 | 0.297374 | 0.424961 | 0.575773 |
| 10 | 121066200 | 121070599 | 47.22686 | 75.52901 | 112.4796 | 47.3266 | -0.23045 | 2.05421 | 0.57578 |
| 10 | 115318800 | 115319999 | 8.179992 | 15.61175 | 28.4606 | 11.33247 | -0.354 | -0.02247 | 0.575781 |
| 18 | 53707800 | 53711199 | 33.28003 | 53.94555 | 83.31134 | 35.67397 | -0.26045 | 1.635849 | 0.575789 |
| 9 | 65815800 | 65816999 | 7.948802 | 14.95034 | 27.50499 | 11.28283 | -0.40394 | 0.074238 | 0.575829 |
| 7 | 131471200 | 131471799 | 9.657625 | 18.77597 | 20.07189 | 7.447496 | 0.504819 | -0.26265 | 0.575832 |
| 13 | 70517400 | 70517999 | 9.657625 | 18.77597 | 20.07189 | 7.447496 | 0.504819 | -0.26265 | 0.575832 |
| 2 | 133684600 | 133687399 | 23.89886 | 40.77337 | 43.69366 | 19.15511 | 0.282894 | 0.988656 | 0.575841 |
| X | 19736800 | 19737799 | 12.00546 | 21.67109 | 35.81228 | 15.57354 | -0.31223 | 0.369678 | 0.575856 |
| 14 | 8010800 | 8014399 | 63.25283 | 100.8084 | 107.304 | 44.92909 | 0.247745 | 2.176671 | 0.57591 |
| 13 | 92562200 | 92565799 | 25.11565 | 40.85344 | 85.87211 | 36.85522 | -0.69943 | 1.555582 | 0.57592 |
| 6 | 122809400 | 122810799 | 10.98863 | 19.77363 | 34.2144 | 14.9369 | -0.38197 | 0.37466 | 0.575921 |
| 10 | 106304600 | 106306599 | 15.87225 | 27.46065 | 46.3638 | 20.34091 | -0.36644 | 0.833677 | 0.575924 |
| 5 | 8028600 | 8031999 | 28.44514 | 46.98458 | 77.01971 | 33.26729 | -0.33059 | 1.355232 | 0.575926 |
| 18 | 74475200 | 74479599 | 67.98782 | 107.7489 | 115.8861 | 48.21092 | 0.224678 | 2.351011 | 0.57593 |
| 8 | 92383000 | 92385799 | 19.58741 | 33.39836 | 53.28001 | 23.35663 | -0.28904 | 0.989222 | 0.575935 |
| X | 73954600 | 73956999 | 26.68543 | 45.52821 | 46.84566 | 20.53696 | 0.341379 | 1.00464 | 0.575991 |
| 8 | 101905000 | 101905999 | 11.14023 | 20.97882 | 23.18721 | 9.491716 | 0.328209 | 0.085184 | 0.575991 |

| | | | | | | | | | |
|---|---|---|---|---|---|---|---|---|---|
| 18 | 8513800 | 8515199 | 20.34479 | 34.35191 | 55.36266 | 24.21732 | -0.30479 | 1.0825 | 0.575996 |
| 5 | 99924600 | 99927399 | 19.5731 | 34.19656 | 36.0417 | 15.87965 | 0.312014 | 0.712289 | 0.576014 |
| 4 | 113219800 | 113221999 | 31.87605 | 52.95752 | 56.88584 | 24.66929 | 0.278137 | 1.283066 | 0.576017 |
| 5 | 85653400 | 85655799 | 23.23531 | 39.91478 | 43.14697 | 18.92551 | 0.272006 | 0.92586 | 0.576019 |
| 2 | 150891600 | 150894199 | 28.95619 | 47.72659 | 76.90322 | 33.17855 | -0.30665 | 1.382277 | 0.576034 |
| 5 | 58673600 | 58674399 | 9.932943 | 18.92098 | 21.21385 | 8.481297 | 0.337043 | -0.01039 | 0.576037 |
| 6 | 136154600 | 136156199 | 11.77582 | 21.23777 | 34.7287 | 15.11905 | -0.2983 | 0.369174 | 0.57604 |
| 1 | 138790200 | 138791599 | 13.87367 | 25.6801 | 27.00751 | 11.36645 | 0.367832 | 0.221577 | 0.576055 |
| 17 | 40196800 | 40201799 | 59.68918 | 95.52915 | 98.08831 | 41.31388 | 0.306168 | 2.042603 | 0.576058 |
| 18 | 77392400 | 77397599 | 47.94721 | 76.46174 | 115.979 | 48.60225 | -0.2614 | 2.145303 | 0.576079 |
| 6 | 10188000 | 10188999 | 11.83961 | 22.2584 | 24.62813 | 10.11031 | 0.322931 | 0.103752 | 0.576132 |
| 12 | 14089800 | 14090799 | 11.83961 | 22.2584 | 24.62813 | 10.11031 | 0.322931 | 0.103752 | 0.576132 |
| 16 | 83869800 | 83873799 | 89.09324 | 138.7782 | 148.1361 | 61.48536 | 0.216125 | 2.713232 | 0.576152 |
| 7 | 56792000 | 56793999 | 47.50935 | 76.60793 | 69.32087 | 29.52024 | 0.501007 | 1.784682 | 0.576169 |
| 15 | 72655000 | 72657399 | 20.09965 | 34.22443 | 53.81275 | 23.58754 | -0.26851 | 1.005095 | 0.576178 |
| 18 | 21234800 | 21236599 | 17.93113 | 31.52779 | 33.85825 | 14.99268 | 0.285216 | 0.601849 | 0.576183 |
| 15 | 64969000 | 64970599 | 25.29366 | 43.1081 | 46.32766 | 20.30814 | 0.278595 | 1.007315 | 0.576193 |
| 1 | 50476200 | 50478199 | 25.15735 | 43.75329 | 40.14847 | 17.63851 | 0.510546 | 0.810401 | 0.576235 |
| 14 | 106957400 | 106960399 | 36.76112 | 58.83301 | 115.4352 | 48.61897 | -0.62628 | 2.023445 | 0.576252 |
| 13 | 104404800 | 104405399 | 5.80073 | 11.29854 | 21.96454 | 8.017278 | -0.3267 | -0.35816 | 0.576263 |
| 17 | 6396800 | 6397199 | 5.80073 | 11.29854 | 21.96454 | 8.017278 | -0.3267 | -0.35816 | 0.576263 |
| 14 | 37348000 | 37351199 | 39.87237 | 64.80833 | 70.20839 | 30.12391 | 0.253512 | 1.586786 | 0.576264 |
| 1 | 184347000 | 184347599 | 5.230818 | 10.18638 | 20.73331 | 7.588319 | -0.3952 | -0.35942 | 0.576285 |
| 10 | 49629200 | 49629599 | 5.230818 | 10.18638 | 20.73331 | 7.588319 | -0.3952 | -0.35942 | 0.576285 |
| 1 | 44398600 | 44402399 | 50.95168 | 81.86736 | 87.32225 | 36.99849 | 0.258 | 1.905077 | 0.576297 |
| 14 | 120792600 | 120794999 | 23.87565 | 41.58926 | 39.03085 | 17.1644 | 0.478337 | 0.780002 | 0.576309 |
| 4 | 143466600 | 143483799 | 416.2147 | 578.9872 | 551.1228 | 228.7086 | 0.231794 | 4.66128 | 0.576339 |
| 2 | 47563400 | 47564599 | 9.317594 | 17.50345 | 30.19774 | 12.41303 | -0.31472 | 0.100715 | 0.57634 |
| 7 | 113410600 | 113411999 | 9.317594 | 17.50345 | 30.19774 | 12.41303 | -0.31472 | 0.100715 | 0.57634 |
| 2 | 139950400 | 139952599 | 15.20035 | 26.52145 | 43.11937 | 18.98209 | -0.31042 | 0.722062 | 0.576356 |
| 13 | 41228000 | 41230399 | 13.51839 | 23.7408 | 39.4195 | 17.44339 | -0.34007 | 0.607021 | 0.57636 |
| X | 72679000 | 72681199 | 13.99295 | 24.66053 | 39.81475 | 17.62209 | -0.29847 | 0.567041 | 0.576404 |
| 16 | 97829600 | 97842199 | 270.1495 | 391.8776 | 371.7102 | 153.2536 | 0.297327 | 4.03874 | 0.576419 |
| X | 167711200 | 167712199 | 8.634873 | 16.20525 | 29.07528 | 11.97122 | -0.3719 | 0.102021 | 0.576426 |
| 4 | 146257600 | 146270599 | 269.373 | 389.0824 | 383.5822 | 158.2372 | 0.235592 | 4.101636 | 0.576441 |
| 18 | 38913800 | 38915999 | 16.8723 | 29.78156 | 31.93082 | 14.12823 | 0.28983 | 0.55138 | 0.576442 |
| 12 | 114523400 | 114533199 | 95.94088 | 150.18 | 131.9589 | 54.74003 | 0.502031 | 2.630477 | 0.576444 |
| X | 3262000 | 3268599 | 78.57597 | 124.1788 | 106.574 | 44.27035 | 0.546561 | 2.426397 | 0.57645 |
| 13 | 3558200 | 3560199 | 24.2574 | 41.45939 | 44.71894 | 19.60742 | 0.273925 | 0.975434 | 0.57646 |
| 6 | 26939600 | 26941599 | 17.18913 | 29.89446 | 54.41233 | 23.90478 | -0.47318 | 0.796844 | 0.576502 |
| 6 | 147444800 | 147447999 | 31.31028 | 52.27967 | 53.63354 | 23.33534 | 0.344286 | 1.221169 | 0.576509 |
| 3 | 20780200 | 20780799 | 6.544892 | 12.65475 | 23.76326 | 9.094614 | -0.34163 | -0.18007 | 0.576516 |
| 6 | 99897400 | 99902399 | 47.88682 | 76.56492 | 113.7459 | 47.84211 | -0.22736 | 2.063193 | 0.57654 |
| X | 80858400 | 80861399 | 33.69888 | 56.84748 | 50.19692 | 21.95151 | 0.560129 | 1.132264 | 0.576543 |
| 18 | 39057800 | 39061199 | 31.06901 | 50.35861 | 90.44412 | 38.72654 | -0.47778 | 1.637647 | 0.576549 |
| 8 | 48531200 | 48532999 | 15.04555 | 26.37882 | 41.93027 | 18.52834 | -0.2778 | 0.644272 | 0.57655 |
| 5 | 67891800 | 67894199 | 20.76324 | 36.32285 | 37.53612 | 16.55559 | 0.340435 | 0.702815 | 0.57656 |

| | | | | | | | | | |
|---|---|---|---|---|---|---|---|---|---|
| 6 | 132049200 | 132053199 | 49.12149 | 79.68829 | 72.43567 | 31.03549 | 0.503479 | 1.643794 | 0.576584 |
| 7 | 25980600 | 25984399 | 67.65489 | 107.342 | 115.3067 | 48.024 | 0.228039 | 2.319732 | 0.576585 |
| 11 | 85001800 | 85002599 | 9.936391 | 18.03077 | 31.8509 | 13.76008 | -0.40121 | 0.316186 | 0.576586 |
| 4 | 21683800 | 21690399 | 62.38941 | 98.24059 | 147.8727 | 61.35464 | -0.26696 | 2.500498 | 0.576593 |
| 7 | 6000800 | 6001599 | 30.62073 | 50.77211 | 50.24839 | 21.76133 | 0.395872 | 1.298569 | 0.576611 |
| 7 | 9344000 | 9345599 | 36.69702 | 59.73179 | 62.57739 | 26.86398 | 0.303173 | 1.555036 | 0.576628 |
| 15 | 31240200 | 31243399 | 28.15591 | 46.15737 | 72.27736 | 31.11157 | -0.26853 | 1.443697 | 0.576645 |
| 7 | 117066600 | 117067199 | 5.802447 | 11.29587 | 21.88633 | 8.034148 | -0.32928 | -0.33788 | 0.576656 |
| 1 | 26038000 | 26040399 | 25.82802 | 44.27167 | 44.62886 | 19.57248 | 0.371704 | 0.954913 | 0.576657 |
| 6 | 22476000 | 22476799 | 10.19709 | 18.69426 | 31.36273 | 13.37311 | -0.31479 | 0.257546 | 0.576658 |
| 17 | 60740800 | 60741999 | 10.19709 | 18.69426 | 31.36273 | 13.37311 | -0.31479 | 0.257546 | 0.576658 |
| 3 | 26528800 | 26530999 | 23.72379 | 40.48583 | 43.17203 | 18.92685 | 0.290041 | 0.984772 | 0.576667 |
| 3 | 93089000 | 93091199 | 12.68465 | 22.72557 | 37.80216 | 16.5644 | -0.32905 | 0.424065 | 0.576714 |
| 1 | 179463400 | 179464199 | 11.0666 | 19.76987 | 34.2779 | 15.05652 | -0.39074 | 0.423179 | 0.57672 |
| 6 | 13366200 | 13368599 | 27.3706 | 46.56544 | 47.38285 | 20.76773 | 0.356698 | 1.031251 | 0.576722 |
| 14 | 59607200 | 59610799 | 62.80112 | 99.13861 | 145.1533 | 60.26214 | -0.22429 | 2.447231 | 0.576731 |
| 2 | 43000200 | 43004799 | 37.10617 | 60.83592 | 64.35993 | 27.70469 | 0.295649 | 1.456154 | 0.576737 |
| 7 | 58992800 | 59003599 | 143.671 | 218.4666 | 213.468 | 88.43314 | 0.316616 | 3.192023 | 0.576747 |
| 2 | 19135400 | 19136799 | 15.88277 | 28.58708 | 29.63256 | 12.93297 | 0.352955 | 0.397376 | 0.576748 |
| 10 | 11215200 | 11220399 | 57.63616 | 91.47235 | 133.6842 | 55.69397 | -0.21508 | 2.306416 | 0.576752 |
| 2 | 140073200 | 140073799 | 7.451553 | 14.18449 | 26.65054 | 10.67039 | -0.40385 | -0.01585 | 0.576761 |
| 3 | 68374400 | 68375599 | 7.451553 | 14.18449 | 26.65054 | 10.67039 | -0.40385 | -0.01585 | 0.576761 |
| 2 | 83078800 | 83080999 | 15.33483 | 26.83332 | 43.02053 | 18.97863 | -0.2902 | 0.678285 | 0.576764 |
| 5 | 94497800 | 94503199 | 140.2152 | 213.4892 | 210.6788 | 87.31166 | 0.303886 | 3.166109 | 0.576768 |
| X | 118171400 | 118175599 | 70.37561 | 111.9282 | 112.8494 | 47.11578 | 0.322718 | 2.252259 | 0.576786 |
| 16 | 13896200 | 13898799 | 27.10704 | 46.73198 | 43.50733 | 19.08565 | 0.487662 | 0.911431 | 0.576807 |
| 17 | 30106000 | 30109399 | 28.59968 | 46.4652 | 78.85998 | 33.82496 | -0.39308 | 1.577714 | 0.57681 |
| 13 | 120426000 | 120428599 | 14.5797 | 25.04621 | 55.33654 | 24.27222 | -0.75503 | 0.903544 | 0.576818 |
| 6 | 131438000 | 131439399 | 13.98745 | 25.60212 | 26.76171 | 11.43609 | 0.360458 | 0.281397 | 0.576826 |
| 3 | 144930000 | 144933599 | 37.94418 | 62.17909 | 64.98647 | 27.96853 | 0.312856 | 1.464721 | 0.576839 |
| 5 | 20181800 | 20184999 | 34.56748 | 56.59017 | 61.0598 | 26.2677 | 0.266518 | 1.467915 | 0.576861 |
| 4 | 100120200 | 100122599 | 21.01958 | 35.67726 | 56.48355 | 24.74683 | -0.27889 | 1.042188 | 0.576868 |
| 8 | 13670200 | 13671799 | 8.749197 | 16.38806 | 28.5058 | 11.77527 | -0.3316 | 0.112281 | 0.576877 |
| 13 | 62652400 | 62678399 | 601.8018 | 815.1693 | 713.7444 | 297.433 | 0.313559 | 5.076434 | 0.576903 |
| 1 | 81421000 | 81423399 | 16.08542 | 27.80185 | 52.74127 | 23.13816 | -0.53425 | 0.852305 | 0.576905 |
| X | 77770800 | 77774199 | 42.44079 | 69.67927 | 63.17567 | 27.21097 | 0.518463 | 1.451803 | 0.576906 |
| 13 | 15269800 | 15276599 | 30.03986 | 50.5034 | 45.87798 | 20.04218 | 0.519068 | 1.141572 | 0.576935 |
| 5 | 146571800 | 146576199 | 40.51208 | 65.17892 | 97.8424 | 41.53486 | -0.23163 | 1.855671 | 0.576951 |
| 5 | 125402000 | 125403199 | 14.19572 | 25.68862 | 27.90397 | 12.09676 | 0.292321 | 0.352784 | 0.576963 |
| 6 | 131744600 | 131753399 | 139.5933 | 213.0579 | 203.1542 | 84.24616 | 0.356185 | 3.116451 | 0.576969 |
| 5 | 148618400 | 148622599 | 41.69397 | 66.8765 | 101.9132 | 43.06804 | -0.25863 | 1.954341 | 0.576984 |
| 3 | 9989200 | 9990599 | 8.262028 | 15.69624 | 27.70644 | 11.1396 | -0.31943 | 0.011546 | 0.576987 |
| 7 | 23643400 | 23649599 | 61.94923 | 99.55395 | 87.33989 | 37.01464 | 0.539688 | 1.917966 | 0.576993 |
| 1 | 77643400 | 77648799 | 51.09519 | 81.91551 | 87.63218 | 37.00518 | 0.249983 | 1.976297 | 0.576999 |
| 11 | 66036000 | 66041799 | 70.79453 | 112.1208 | 120.6336 | 50.16489 | 0.22333 | 2.372666 | 0.577002 |
| 2 | 100760000 | 100763399 | 38.09649 | 62.4033 | 65.36269 | 28.12561 | 0.309551 | 1.472622 | 0.57701 |
| 4 | 56844000 | 56846199 | 22.64035 | 38.86259 | 42.1091 | 18.46952 | 0.26835 | 0.927673 | 0.577019 |

| | | | | | | | | | |
|---|---|---|---|---|---|---|---|---|---|
| 6 | 7554200 | 7557599 | 32.31197 | 52.36976 | 82.52995 | 35.33608 | -0.28971 | 1.636053 | 0.577022 |
| 7 | 43443000 | 43445199 | 13.75959 | 24.1967 | 39.29769 | 17.40079 | -0.30832 | 0.586391 | 0.577022 |
| 18 | 25389600 | 25394599 | 70.08247 | 110.0135 | 159.9265 | 66.32923 | -0.22026 | 2.57012 | 0.577029 |
| 12 | 64421200 | 64422399 | 13.43122 | 23.78048 | 38.18362 | 16.88451 | -0.2881 | 0.507896 | 0.577037 |
| 4 | 71054000 | 71058199 | 47.8568 | 77.83614 | 68.52692 | 29.40974 | 0.552651 | 1.591267 | 0.577055 |
| 9 | 109627000 | 109634599 | 96.78705 | 147.9746 | 244.1382 | 101.2815 | -0.43224 | 3.072416 | 0.577081 |
| 7 | 51184000 | 51185399 | 11.60879 | 21.18191 | 36.5885 | 15.69827 | -0.36336 | 0.304433 | 0.577092 |
| 2 | 28881800 | 28886199 | 40.99765 | 65.74728 | 103.2333 | 43.61348 | -0.30203 | 1.958158 | 0.577103 |
| 13 | 19147800 | 19150599 | 26.61954 | 44.18713 | 69.98185 | 30.3308 | -0.2804 | 1.288982 | 0.577107 |
| 2 | 55893400 | 55896199 | 26.34339 | 45.63195 | 41.83155 | 18.35318 | 0.511165 | 0.863642 | 0.577117 |
| 3 | 32974600 | 32979199 | 51.77947 | 83.3393 | 85.15093 | 36.1677 | 0.322938 | 1.852681 | 0.577132 |
| 3 | 35917000 | 35919199 | 23.95701 | 39.89536 | 63.06593 | 27.39322 | -0.27732 | 1.238383 | 0.577141 |
| 18 | 64558000 | 64561799 | 32.32778 | 52.46653 | 80.87128 | 34.66057 | -0.25598 | 1.609186 | 0.577149 |
| 13 | 13585800 | 13590199 | 49.15977 | 79.02399 | 85.25404 | 36.14593 | 0.242428 | 1.887954 | 0.577151 |
| 1 | 64803200 | 64805399 | 21.02275 | 36.72858 | 38.07696 | 16.77616 | 0.335802 | 0.724651 | 0.577152 |
| X | 12258600 | 12262199 | 33.19532 | 55.70438 | 52.43729 | 22.88679 | 0.467984 | 1.17908 | 0.577171 |
| 1 | 28839200 | 28841999 | 39.52569 | 64.19285 | 70.08055 | 30.05112 | 0.241319 | 1.60181 | 0.577185 |
| 7 | 41322600 | 41324199 | 15.56956 | 28.79672 | 27.43411 | 11.55824 | 0.508491 | 0.240441 | 0.577209 |
| 2 | 6733400 | 6734199 | 7.328044 | 13.92336 | 26.07201 | 10.48007 | -0.40365 | -0.00413 | 0.577215 |
| 9 | 107983400 | 107984599 | 13.19084 | 23.17197 | 38.91383 | 17.22365 | -0.35634 | 0.5994 | 0.577238 |
| 8 | 65787600 | 65789399 | 17.63498 | 30.39524 | 48.47058 | 21.26276 | -0.28565 | 0.878553 | 0.577248 |
| 3 | 18748400 | 18748999 | 6.242213 | 12.14763 | 23.70863 | 8.728267 | -0.34414 | -0.31763 | 0.577248 |
| 1 | 82078800 | 82081799 | 34.15734 | 56.75466 | 56.56503 | 24.53228 | 0.385806 | 1.285185 | 0.577275 |
| 1 | 91374600 | 91376999 | 23.86509 | 40.64913 | 43.2492 | 18.95776 | 0.292973 | 1.004936 | 0.577277 |
| 1 | 98329400 | 98330999 | 24.12521 | 41.23577 | 44.72937 | 19.61208 | 0.265852 | 0.974868 | 0.577278 |
| 1 | 171804200 | 171805799 | 15.11574 | 26.46586 | 42.527 | 18.77044 | -0.29321 | 0.669492 | 0.577329 |
| 15 | 70168400 | 70171199 | 38.53316 | 62.71751 | 68.45644 | 29.3873 | 0.244081 | 1.563085 | 0.577355 |
| 13 | 19175000 | 19175999 | 8.058763 | 15.46547 | 28.72829 | 11.27131 | -0.36084 | -0.06957 | 0.57736 |
| 6 | 134421800 | 134423399 | 13.2181 | 23.19874 | 44.57859 | 19.71577 | -0.54996 | 0.623321 | 0.577367 |
| 17 | 95139800 | 95142599 | 32.94011 | 53.52641 | 81.93237 | 35.14735 | -0.24434 | 1.585115 | 0.577377 |
| X | 26209200 | 26210199 | 18.44398 | 32.36258 | 32.67148 | 14.4466 | 0.374156 | 0.637442 | 0.577382 |
| 13 | 13414600 | 13416999 | 27.88241 | 47.15669 | 49.60462 | 21.70942 | 0.308539 | 1.088346 | 0.577393 |
| 3 | 16103000 | 16110599 | 108.354 | 167.0514 | 175.8742 | 73.05516 | 0.224568 | 2.907271 | 0.577398 |
| 4 | 113296600 | 113298599 | 56.94669 | 91.22501 | 85.09118 | 35.88933 | 0.446007 | 2.006203 | 0.577403 |
| 8 | 35608600 | 35610599 | 16.19591 | 28.69052 | 30.86289 | 13.64591 | 0.287017 | 0.505821 | 0.577405 |
| 4 | 43137400 | 43139599 | 19.52603 | 34.16764 | 36.5319 | 16.11632 | 0.291469 | 0.689487 | 0.577422 |
| 2 | 133813600 | 133813999 | 5.951254 | 11.57598 | 22.39269 | 8.27509 | -0.33578 | -0.31024 | 0.577437 |
| 7 | 26474000 | 26476599 | 34.3608 | 56.44359 | 61.82227 | 26.63403 | 0.246833 | 1.425436 | 0.577476 |
| 5 | 144633800 | 144635799 | 14.71415 | 25.73044 | 42.03051 | 18.53165 | -0.31694 | 0.684503 | 0.57748 |
| 17 | 65448200 | 65449399 | 14.66727 | 25.50936 | 48.95822 | 21.50906 | -0.5491 | 0.775417 | 0.57749 |
| 11 | 14791000 | 14791999 | 19.3014 | 32.92208 | 52.18719 | 22.8782 | -0.27974 | 0.984914 | 0.577497 |
| 19 | 53965200 | 53969399 | 57.57447 | 92.18902 | 96.36965 | 40.61839 | 0.281128 | 2.023437 | 0.57752 |
| 7 | 6564800 | 6568199 | 49.6618 | 79.73467 | 86.00148 | 36.39788 | 0.240645 | 1.926604 | 0.577528 |
| 2 | 39801200 | 39803199 | 30.95668 | 52.38205 | 48.50768 | 21.23254 | 0.491686 | 1.091102 | 0.577537 |
| 8 | 3591200 | 3594599 | 25.73046 | 42.37141 | 68.20186 | 29.41322 | -0.30546 | 1.380633 | 0.577556 |
| Y | 5512600 | 5513199 | 8.130963 | 15.7865 | 14.20034 | 5.326856 | 0.737268 | -0.26359 | 0.577557 |
| 18 | 90228000 | 90228799 | 17.03749 | 29.27967 | 48.47468 | 21.26266 | -0.3403 | 0.903018 | 0.577568 |

| | | | | | | | | | |
|---|---|---|---|---|---|---|---|---|---|
| 6 | 142254800 | 142255999 | 14.28858 | 25.21687 | 40.80311 | 18.05448 | -0.30099 | 0.554139 | 0.577569 |
| 16 | 78223800 | 78227199 | 19.21575 | 32.76216 | 56.03251 | 24.56311 | -0.38923 | 0.98919 | 0.577581 |
| 3 | 151761000 | 151763399 | 19.93852 | 34.01034 | 53.27923 | 23.35703 | -0.2628 | 0.987976 | 0.577585 |
| 16 | 37613600 | 37621599 | 83.31217 | 128.1482 | 216.6342 | 89.96528 | -0.46049 | 2.946016 | 0.577612 |
| 5 | 54299400 | 54301799 | 59.78699 | 95.2126 | 101.2136 | 42.33459 | 0.248497 | 2.196126 | 0.577614 |
| X | 155650600 | 155651999 | 9.843133 | 18.38042 | 31.69968 | 13.16307 | -0.32592 | 0.147993 | 0.577622 |
| 18 | 80050800 | 80054399 | 32.4137 | 52.89515 | 84.29573 | 36.2213 | -0.29795 | 1.522298 | 0.577647 |
| 15 | 91367800 | 91368999 | 13.66903 | 25.19622 | 26.54543 | 11.23542 | 0.359743 | 0.243658 | 0.577657 |
| X | 8582000 | 8601799 | 267.2462 | 390.4 | 340.852 | 140.4283 | 0.426667 | 3.933485 | 0.57766 |
| X | 75313000 | 75323199 | 167.348 | 250.7774 | 253.584 | 104.609 | 0.249113 | 3.489536 | 0.577761 |
| 7 | 40364200 | 40365599 | 14.5801 | 25.55774 | 40.93242 | 18.0844 | -0.28852 | 0.643821 | 0.577765 |
| 9 | 5061600 | 5068799 | 99.27555 | 153.5553 | 164.1009 | 68.16356 | 0.206613 | 2.842137 | 0.577766 |
| 13 | 36809600 | 36810599 | 16.6812 | 29.50415 | 31.84138 | 14.08293 | 0.281473 | 0.527438 | 0.577781 |
| 6 | 31083200 | 31087199 | 35.80099 | 57.75248 | 89.29914 | 38.0484 | -0.26952 | 1.764866 | 0.577806 |
| 1 | 84579600 | 84582199 | 21.96253 | 38.05916 | 40.93657 | 17.96074 | 0.281602 | 0.83843 | 0.577809 |
| 6 | 9643000 | 9646199 | 57.45542 | 92.1443 | 83.8957 | 35.44613 | 0.482736 | 1.971695 | 0.577811 |
| 11 | 74159600 | 74161199 | 26.36434 | 44.55253 | 48.53097 | 21.23424 | 0.258289 | 1.08859 | 0.577814 |
| 4 | 113471600 | 113481399 | 132.6961 | 204.0237 | 180.5693 | 74.98373 | 0.471799 | 2.970128 | 0.577834 |
| X | 123022800 | 123025799 | 42.23061 | 69.35483 | 63.64114 | 27.41491 | 0.501386 | 1.449497 | 0.577838 |
| 8 | 115385400 | 115386999 | 16.71579 | 29.61942 | 31.79986 | 14.05943 | 0.289969 | 0.508237 | 0.57784 |
| 1 | 39239600 | 39241399 | 21.53026 | 36.1886 | 57.50166 | 25.11832 | -0.28468 | 1.129343 | 0.577845 |
| X | 153694800 | 153697199 | 31.74462 | 53.18672 | 53.54892 | 23.34891 | 0.371309 | 1.190275 | 0.577845 |
| 6 | 88516600 | 88518799 | 15.93758 | 27.80394 | 45.7185 | 20.12454 | -0.32691 | 0.729552 | 0.577877 |
| 1 | 96382200 | 96383799 | 27.02156 | 45.52493 | 49.62248 | 21.69186 | 0.257195 | 1.117754 | 0.577878 |
| 7 | 22560400 | 22569399 | 137.6528 | 211.7377 | 176.0909 | 73.1284 | 0.561973 | 2.962866 | 0.577882 |
| 5 | 113043000 | 113055399 | 203.4795 | 295.4413 | 408.5731 | 168.4356 | -0.24507 | 4.019751 | 0.577887 |
| 13 | 67295600 | 67314399 | 223.269 | 330.142 | 304.4715 | 125.3891 | 0.364236 | 3.735158 | 0.577891 |
| 7 | 51893800 | 51895199 | 12.11371 | 22.81603 | 24.61476 | 10.0731 | 0.362634 | 0.093887 | 0.577896 |
| 9 | 18287000 | 18288599 | 12.11371 | 22.81603 | 24.61476 | 10.0731 | 0.362634 | 0.093887 | 0.577896 |
| 2 | 39458200 | 39458999 | 10.00575 | 18.43676 | 30.89126 | 13.08147 | -0.30583 | 0.225633 | 0.5779 |
| 11 | 10492800 | 10493199 | 4.477711 | 8.752421 | 18.32123 | 6.312513 | -0.35499 | -0.59113 | 0.577906 |
| 10 | 98797000 | 98799799 | 23.73157 | 39.79932 | 63.23889 | 27.5907 | -0.2849 | 1.161952 | 0.577907 |
| 7 | 90059800 | 90063599 | 86.10595 | 134.6933 | 144.4349 | 59.9196 | 0.214258 | 2.637146 | 0.577938 |
| 9 | 19773800 | 19785799 | 168.2284 | 252.4463 | 228.4679 | 94.27904 | 0.410825 | 3.461068 | 0.577939 |
| 15 | 16435400 | 16436199 | 9.00087 | 16.88851 | 29.06073 | 11.96956 | -0.31321 | 0.102622 | 0.577945 |
| 13 | 35403200 | 35405599 | 42.23173 | 68.58515 | 72.08283 | 30.90887 | 0.295992 | 1.610048 | 0.577958 |
| 19 | 25437000 | 25439199 | 19.98528 | 34.73353 | 36.86707 | 16.18969 | 0.301346 | 0.791082 | 0.577958 |
| 3 | 31410200 | 31413599 | 35.52965 | 58.2678 | 63.54683 | 27.35798 | 0.252078 | 1.449612 | 0.577959 |
| 12 | 118640600 | 118642399 | 17.55954 | 31.84727 | 29.04042 | 12.5541 | 0.545663 | 0.365329 | 0.577983 |
| 3 | 100385600 | 100390199 | 31.48063 | 51.08585 | 81.52121 | 34.9362 | -0.30606 | 1.608257 | 0.577998 |
| 6 | 52401000 | 52403999 | 23.22526 | 38.97234 | 62.10243 | 27.10459 | -0.28902 | 1.152844 | 0.578013 |
| 11 | 114451600 | 114458199 | 74.96588 | 116.5364 | 173.6675 | 72.10419 | -0.26699 | 2.745493 | 0.578017 |
| 7 | 122498400 | 122499399 | 10.58886 | 20.06156 | 21.54879 | 8.714365 | 0.383448 | 0.03173 | 0.578026 |
| 1 | 117374600 | 117376799 | 30.44373 | 50.74044 | 54.59062 | 23.72453 | 0.276057 | 1.242716 | 0.578032 |
| 1 | 111265600 | 111267399 | 16.62 | 29.81278 | 30.64843 | 13.41579 | 0.362057 | 0.426044 | 0.578052 |
| 3 | 56056000 | 56057599 | 16.62 | 29.81278 | 30.64843 | 13.41579 | 0.362057 | 0.426044 | 0.578052 |
| 13 | 66854000 | 66864599 | 316.4663 | 448.4595 | 434.3127 | 179.7033 | 0.233766 | 4.382321 | 0.578054 |

| | | | | | | | | | |
|---|---|---|---|---|---|---|---|---|---|
| 13 | 66854200 | 66864599 | 316.4663 | 448.4595 | 434.3127 | 179.7033 | 0.233766 | 4.382321 | 0.578054 |
| 3 | 158905400 | 158906399 | 20.34846 | 35.62387 | 30.52722 | 13.47277 | 0.60934 | 0.678038 | 0.578056 |
| 7 | 10790000 | 10790399 | 13.40357 | 24.88041 | 18.93211 | 7.937126 | 0.835295 | 0.181378 | 0.578057 |
| 5 | 35304200 | 35309399 | 61.86673 | 97.62758 | 144.0039 | 59.7738 | -0.23537 | 2.455868 | 0.578067 |
| 17 | 89475800 | 89477399 | 12.40652 | 22.25372 | 36.72074 | 16.07442 | -0.31676 | 0.409677 | 0.578068 |
| 1 | 23204400 | 23205399 | 14.7038 | 27.62934 | 25.63367 | 10.53195 | 0.57466 | 0.136096 | 0.578072 |
| 12 | 56733600 | 56745799 | 109.1236 | 164.8159 | 279.3322 | 115.4778 | -0.48544 | 3.325153 | 0.578082 |
| X | 78961200 | 78962799 | 23.88462 | 41.45902 | 40.91531 | 17.95344 | 0.405907 | 0.835158 | 0.578086 |
| 18 | 70421800 | 70422599 | 8.576982 | 15.96853 | 28.9421 | 12.07869 | -0.40276 | 0.148561 | 0.578098 |
| 1 | 30928000 | 30930599 | 17.77012 | 31.36477 | 33.24496 | 14.70983 | 0.306091 | 0.558213 | 0.578103 |
| X | 91289200 | 91294599 | 72.31228 | 114.9815 | 114.1628 | 47.65129 | 0.344475 | 2.260706 | 0.578114 |
| 11 | 9925200 | 9927999 | 37.28511 | 60.86598 | 65.81262 | 28.28284 | 0.260596 | 1.515379 | 0.578126 |
| 13 | 23197400 | 23198199 | 9.908532 | 18.9516 | 21.74887 | 8.615643 | 0.315257 | -0.03944 | 0.578136 |
| 4 | 16562200 | 16563999 | 17.35352 | 30.80716 | 32.18567 | 14.2125 | 0.330424 | 0.498504 | 0.578144 |
| 13 | 61303200 | 61307399 | 90.97639 | 141.5743 | 151.6097 | 62.93619 | 0.210391 | 2.731612 | 0.578169 |
| 2 | 156765200 | 156768599 | 45.18656 | 72.20079 | 110.711 | 46.52295 | -0.2739 | 2.080797 | 0.578171 |
| 5 | 144918400 | 144923799 | 65.8284 | 103.7124 | 150.221 | 62.33342 | -0.21098 | 2.492574 | 0.578177 |
| 7 | 33920000 | 33922599 | 20.98184 | 35.32154 | 57.00406 | 24.91267 | -0.30702 | 1.11149 | 0.578193 |
| 4 | 14873600 | 14874799 | 10.09632 | 18.72185 | 32.06452 | 13.46181 | -0.32863 | 0.191472 | 0.578205 |
| 2 | 169910600 | 169912999 | 24.92485 | 41.51644 | 64.84718 | 28.17092 | -0.26014 | 1.240286 | 0.578209 |
| 2 | 149141600 | 149149199 | 101.055 | 157.7228 | 152.5576 | 63.30401 | 0.360368 | 2.687245 | 0.578244 |
| 13 | 58057400 | 58059999 | 24.21145 | 40.11276 | 64.4829 | 27.9105 | -0.30191 | 1.301001 | 0.578253 |
| 3 | 52987000 | 52989799 | 20.24438 | 34.62174 | 54.95572 | 24.09702 | -0.28121 | 0.966986 | 0.578262 |
| 9 | 23915800 | 23916799 | 18.71567 | 32.86626 | 34.83947 | 15.41593 | 0.303703 | 0.625731 | 0.578288 |
| 17 | 81715200 | 81715799 | 5.411821 | 10.48692 | 22.7088 | 8.602358 | -0.53085 | -0.2321 | 0.578297 |
| X | 119118400 | 119120599 | 25.76698 | 44.43179 | 43.55014 | 19.10456 | 0.413782 | 0.903819 | 0.578299 |
| X | 33597400 | 33600999 | 48.86513 | 79.4648 | 60.10123 | 25.79234 | 0.771134 | 1.593718 | 0.578309 |
| 6 | 136096600 | 136099599 | 23.93099 | 41.27028 | 43.03124 | 18.87699 | 0.324869 | 0.89232 | 0.57831 |
| 3 | 12644200 | 12646399 | 31.61277 | 52.71952 | 55.2725 | 24.02968 | 0.313137 | 1.238255 | 0.578313 |
| 12 | 21027400 | 21036599 | 102.3357 | 157.9195 | 167.3251 | 69.50368 | 0.216383 | 2.88587 | 0.578329 |
| 3 | 61083800 | 61084999 | 14.93605 | 27.99563 | 25.47485 | 10.51113 | 0.59714 | 0.150957 | 0.578334 |
| 15 | 6305000 | 6306199 | 19.16775 | 32.82729 | 51.65844 | 22.65383 | -0.26829 | 0.947953 | 0.578346 |
| 18 | 49818600 | 49824399 | 75.94302 | 120.2752 | 123.8553 | 51.50473 | 0.286492 | 2.375639 | 0.578351 |
| 2 | 86329000 | 86332999 | 72.16916 | 114.2699 | 122.8491 | 51.07989 | 0.224169 | 2.378597 | 0.578384 |
| 7 | 64280600 | 64283799 | 27.30939 | 44.81351 | 72.36589 | 31.16259 | -0.31227 | 1.42812 | 0.578399 |
| 18 | 41558000 | 41559199 | 14.07003 | 26.86256 | 22.56744 | 8.979202 | 0.755763 | 0.016552 | 0.578407 |
| 1 | 3900800 | 3902599 | 16.81787 | 29.28278 | 46.28885 | 20.35077 | -0.27046 | 0.764381 | 0.578417 |
| 4 | 64372800 | 64373999 | 14.00823 | 23.98765 | 54.67443 | 23.97616 | -0.80008 | 0.934629 | 0.57842 |
| 6 | 10918600 | 10930399 | 332.1549 | 462.5895 | 611.9919 | 253.9227 | -0.24132 | 4.641372 | 0.578431 |
| 13 | 25177600 | 25179999 | 17.37559 | 30.05646 | 47.93664 | 21.03084 | -0.28464 | 0.846258 | 0.578439 |
| 19 | 23707000 | 23711199 | 66.49343 | 105.937 | 108.903 | 45.5771 | 0.297401 | 2.192106 | 0.578488 |
| 10 | 32208400 | 32209799 | 18.33109 | 32.48431 | 30.79043 | 13.61291 | 0.468342 | 0.511481 | 0.578501 |
| 4 | 134772200 | 134774199 | 16.18059 | 28.17328 | 44.71621 | 19.65941 | -0.27621 | 0.759371 | 0.578509 |
| 16 | 96933800 | 96936199 | 20.02456 | 35.40702 | 33.55341 | 14.84102 | 0.467883 | 0.55434 | 0.578521 |
| 14 | 121782800 | 121787799 | 37.7214 | 60.7348 | 94.00887 | 39.9511 | -0.27437 | 1.825796 | 0.578534 |
| 1 | 31787800 | 31790399 | 33.74744 | 56.15221 | 49.27822 | 21.39268 | 0.56891 | 1.26226 | 0.578537 |
| 1 | 12527800 | 12528599 | 13.48025 | 24.63703 | 26.91385 | 11.52371 | 0.295574 | 0.287933 | 0.578546 |

| | | | | | | | | | |
|---|---|---|---|---|---|---|---|---|---|
| 16 | 87176000 | 87178999 | 22.8053 | 39.16807 | 42.61053 | 18.6901 | 0.262719 | 0.923799 | 0.578609 |
| 10 | 91263800 | 91265399 | 12.36529 | 21.89988 | 36.26008 | 16.03296 | -0.33167 | 0.495898 | 0.57861 |
| 15 | 76793600 | 76797599 | 74.52915 | 115.7035 | 175.8777 | 73.03632 | -0.29703 | 2.768712 | 0.578629 |
| 5 | 138824400 | 138830599 | 58.41072 | 92.29605 | 155.3397 | 64.52231 | -0.42394 | 2.423734 | 0.578644 |
| 4 | 92970800 | 92972799 | 13.41667 | 23.90993 | 40.3244 | 17.74878 | -0.35363 | 0.464058 | 0.578651 |
| X | 24209600 | 24246399 | 607.7735 | 825.2208 | 677.8431 | 282.3688 | 0.409135 | 5.039859 | 0.578668 |
| 7 | 40736800 | 40748799 | 176.5128 | 264.2375 | 265.0887 | 109.3329 | 0.259168 | 3.509539 | 0.57867 |
| 9 | 122105400 | 122115599 | 144.8803 | 216.3668 | 302.1618 | 124.5655 | -0.22085 | 3.550345 | 0.578679 |
| 14 | 41955000 | 41960199 | 99.27096 | 154.3082 | 132.4951 | 55.01058 | 0.527452 | 2.752958 | 0.578765 |
| 13 | 66402000 | 66402799 | 13.00143 | 24.84822 | 23.8821 | 9.481411 | 0.566454 | 0.004765 | 0.578775 |
| 1 | 20098800 | 20099399 | 6.863935 | 13.2647 | 24.86511 | 9.539556 | -0.34216 | -0.16282 | 0.578784 |
| 1 | 57858000 | 57858799 | 6.863935 | 13.2647 | 24.86511 | 9.539556 | -0.34216 | -0.16282 | 0.578784 |
| 6 | 33476200 | 33476999 | 6.863935 | 13.2647 | 24.86511 | 9.539556 | -0.34216 | -0.16282 | 0.578784 |
| 3 | 91054600 | 91069399 | 234.5042 | 344.6232 | 327.2098 | 134.7566 | 0.313927 | 3.836399 | 0.578785 |
| 5 | 117652000 | 117655799 | 27.65118 | 45.3658 | 71.74463 | 30.89259 | -0.28233 | 1.431789 | 0.5788 |
| 14 | 83658600 | 83665199 | 98.30727 | 153.3197 | 154.3718 | 64.06173 | 0.301686 | 2.697794 | 0.578823 |
| 12 | 114801600 | 114803599 | 31.94569 | 53.21472 | 55.75909 | 24.22399 | 0.314082 | 1.253949 | 0.578828 |
| 3 | 65482600 | 65486599 | 27.94195 | 45.53253 | 83.71924 | 35.9548 | -0.5051 | 1.534272 | 0.578884 |
| 14 | 14875600 | 14877199 | 27.74407 | 46.97008 | 39.6447 | 17.35578 | 0.625087 | 1.068714 | 0.578892 |
| 7 | 103640800 | 103641399 | 9.382166 | 18.02775 | 20.19983 | 7.896792 | 0.367072 | -0.09286 | 0.578906 |
| 6 | 100789000 | 100789599 | 6.396873 | 12.39853 | 23.30643 | 8.819451 | -0.32729 | -0.22066 | 0.578914 |
| 7 | 55143600 | 55146599 | 55.96294 | 89.73964 | 93.50739 | 39.49967 | 0.288254 | 1.976132 | 0.578926 |
| 5 | 105061800 | 105065799 | 65.95671 | 104.9933 | 110.7647 | 46.30218 | 0.258798 | 2.218376 | 0.57893 |
| 6 | 84697400 | 84700799 | 34.99446 | 59.20341 | 40.80828 | 17.86138 | 0.91617 | 1.102461 | 0.578931 |
| 1 | 49971400 | 49975399 | 44.84737 | 72.44882 | 78.04526 | 33.31454 | 0.25306 | 1.732294 | 0.578943 |
| 10 | 16312600 | 16312999 | 9.896745 | 18.96231 | 21.32041 | 8.403458 | 0.352015 | -0.05423 | 0.578965 |
| 7 | 42416800 | 42417599 | 12.92035 | 23.91465 | 25.94827 | 10.92118 | 0.322818 | 0.210126 | 0.578972 |
| 4 | 118791000 | 118791999 | 7.928252 | 15.11092 | 26.67853 | 10.65321 | -0.31104 | -0.01737 | 0.578982 |
| 2 | 70740400 | 70744399 | 29.22875 | 47.65322 | 87.00061 | 37.37085 | -0.49447 | 1.531685 | 0.578983 |
| 5 | 58739200 | 58742399 | 74.3555 | 117.4068 | 127.152 | 52.79513 | 0.210417 | 2.445832 | 0.579 |
| 4 | 109512800 | 109515399 | 20.10806 | 33.72317 | 65.13613 | 28.41883 | -0.56548 | 1.152675 | 0.579006 |
| 14 | 42621000 | 42623399 | 31.63503 | 53.29575 | 44.73132 | 19.55355 | 0.632883 | 1.127681 | 0.579008 |
| 8 | 69965400 | 69966999 | 17.56639 | 31.29291 | 31.55368 | 13.89294 | 0.384039 | 0.476029 | 0.579018 |
| 11 | 5393200 | 5396399 | 21.29118 | 35.40279 | 62.21824 | 26.99783 | -0.42966 | 1.247067 | 0.579072 |
| 3 | 124039600 | 124040199 | 13.47525 | 24.90517 | 26.55282 | 11.19796 | 0.346361 | 0.226994 | 0.579074 |
| 9 | 12269600 | 12277999 | 168.1608 | 251.5991 | 260.9002 | 107.5942 | 0.210761 | 3.517315 | 0.579078 |
| 9 | 115385000 | 115388799 | 37.39603 | 60.18249 | 94.99079 | 40.34076 | -0.30334 | 1.840177 | 0.579091 |
| 10 | 75920600 | 75921399 | 13.46318 | 24.66011 | 26.90817 | 11.48751 | 0.300136 | 0.275219 | 0.579095 |
| 1 | 107748400 | 107751799 | 44.6998 | 72.05512 | 77.27069 | 32.89243 | 0.256038 | 1.79438 | 0.579112 |
| X | 19346600 | 19348199 | 17.55682 | 31.33064 | 30.79635 | 13.5395 | 0.422489 | 0.467835 | 0.579123 |
| 12 | 5884000 | 5885599 | 19.34548 | 33.09494 | 52.18871 | 22.88432 | -0.27154 | 0.959643 | 0.579127 |
| 2 | 71824000 | 71825999 | 19.80591 | 34.56908 | 37.14155 | 16.35405 | 0.284343 | 0.729848 | 0.579147 |
| 16 | 13980800 | 13981799 | 7.322683 | 13.97632 | 25.61076 | 10.19654 | -0.36027 | -0.03578 | 0.579157 |
| 15 | 14494000 | 14495599 | 16.68005 | 30.22084 | 29.70989 | 12.86016 | 0.43674 | 0.367318 | 0.579164 |
| 3 | 77159400 | 77162399 | 22.55913 | 39.52388 | 36.7604 | 16.23405 | 0.491617 | 0.686957 | 0.579168 |
| 6 | 15684400 | 15685199 | 9.74524 | 17.40258 | 38.4432 | 16.89164 | -0.73792 | 0.428744 | 0.579171 |
| 12 | 17285400 | 17302799 | 196.6763 | 287.4715 | 389.9021 | 160.6299 | -0.20776 | 3.917931 | 0.579173 |

| | | | | | | | | | |
|---|---|---|---|---|---|---|---|---|---|
| 13 | 33690600 | 33692399 | 37.06469 | 60.8072 | 56.40898 | 24.28872 | 0.485059 | 1.443293 | 0.579182 |
| 1 | 18222600 | 18223199 | 16.51393 | 29.18852 | 31.39632 | 13.88797 | 0.285891 | 0.53298 | 0.579183 |
| 7 | 103302200 | 103305399 | 41.57876 | 67.73875 | 62.02563 | 26.63585 | 0.497812 | 1.551793 | 0.57919 |
| 14 | 50549600 | 50552199 | 35.52261 | 58.99282 | 50.95879 | 22.09302 | 0.591527 | 1.291865 | 0.579227 |
| 11 | 10038000 | 10040399 | 24.2569 | 41.93757 | 42.45435 | 18.62535 | 0.368027 | 0.870877 | 0.579261 |
| 3 | 66986000 | 66989999 | 34.35212 | 56.88932 | 59.26193 | 25.64316 | 0.32195 | 1.327746 | 0.57927 |
| 16 | 88359200 | 88359999 | 8.271692 | 15.58102 | 27.52768 | 11.26345 | -0.34273 | 0.069469 | 0.579284 |
| X | 129948200 | 129949999 | 21.21868 | 37.22278 | 36.35669 | 16.07306 | 0.421412 | 0.660846 | 0.579311 |
| X | 91945600 | 91946799 | 8.921625 | 16.96835 | 33.65481 | 13.49916 | -0.4839 | 0.023587 | 0.579315 |
| 1 | 15356000 | 15357599 | 17.03561 | 29.40558 | 47.41647 | 20.80128 | -0.30105 | 0.86268 | 0.579331 |
| 1 | 8738600 | 8740399 | 21.81469 | 38.31347 | 35.02913 | 15.50159 | 0.516039 | 0.641005 | 0.579368 |
| 1 | 162927400 | 162928799 | 10.92687 | 19.99169 | 33.39126 | 14.2773 | -0.31177 | 0.276757 | 0.579375 |
| 4 | 54601600 | 54603399 | 12.59496 | 22.43106 | 36.66694 | 16.14773 | -0.30921 | 0.455849 | 0.57943 |
| 11 | 87146000 | 87148399 | 12.82522 | 22.72524 | 36.7067 | 16.2289 | -0.29599 | 0.493985 | 0.579435 |
| 12 | 109369200 | 109371199 | 16.29699 | 27.97335 | 48.47797 | 21.26293 | -0.40616 | 0.912404 | 0.579441 |
| 16 | 19704000 | 19705399 | 19.61683 | 34.36212 | 36.51661 | 16.12369 | 0.300266 | 0.674557 | 0.57945 |
| 5 | 15099200 | 15112599 | 258.4081 | 375.7891 | 363.9093 | 149.9981 | 0.270968 | 3.999184 | 0.579474 |
| 7 | 64268800 | 64278399 | 86.64165 | 133.5995 | 195.9553 | 81.39641 | -0.25313 | 2.89689 | 0.579507 |
| 12 | 20667200 | 20684199 | 326.4458 | 464.5683 | 400.5502 | 165.6157 | 0.407524 | 4.320903 | 0.579508 |
| 4 | 71187600 | 71188999 | 12.77949 | 22.78927 | 36.68641 | 16.13864 | -0.28607 | 0.449968 | 0.579519 |
| 11 | 109625000 | 109627199 | 17.66997 | 30.74539 | 51.16947 | 22.48685 | -0.34466 | 0.784791 | 0.579552 |
| 5 | 92293800 | 92294599 | 8.382855 | 15.85703 | 27.60178 | 11.19955 | -0.31136 | 0.04064 | 0.579559 |
| 12 | 82686400 | 82689799 | 25.28313 | 41.54363 | 76.57948 | 32.9956 | -0.50185 | 1.412046 | 0.579561 |
| 16 | 53782000 | 53782999 | 9.518623 | 17.64332 | 29.92204 | 12.56938 | -0.31471 | 0.185981 | 0.579567 |
| 7 | 60622800 | 60629199 | 132.4949 | 202.2361 | 184.5734 | 76.54194 | 0.419366 | 3.111302 | 0.579586 |
| 13 | 22084200 | 22086999 | 28.43664 | 48.65521 | 39.7398 | 17.4258 | 0.673754 | 0.976442 | 0.579592 |
| 7 | 112243200 | 112244999 | 16.82262 | 29.18231 | 46.34721 | 20.33528 | -0.2778 | 0.81476 | 0.579593 |
| 13 | 22415200 | 22417599 | 45.15868 | 72.27825 | 108.2101 | 45.58095 | -0.23679 | 2.027963 | 0.579601 |
| 12 | 78962800 | 78964199 | 13.26342 | 24.21236 | 25.89595 | 11.10414 | 0.324499 | 0.290051 | 0.579614 |
| 18 | 88305000 | 88306599 | 16.57181 | 29.3476 | 31.20372 | 13.79743 | 0.303395 | 0.510329 | 0.579616 |
| 9 | 90336800 | 90338399 | 20.86416 | 36.38266 | 38.70175 | 17.03119 | 0.298613 | 0.7512 | 0.579619 |
| 17 | 61086200 | 61087799 | 17.86656 | 32.05215 | 30.46828 | 13.33577 | 0.474414 | 0.432739 | 0.579625 |
| 3 | 28566600 | 28568199 | 15.08138 | 26.33194 | 42.60712 | 18.76468 | -0.30334 | 0.712312 | 0.579625 |
| X | 127625400 | 127628999 | 39.47635 | 64.27877 | 68.40472 | 29.36944 | 0.281005 | 1.561391 | 0.579626 |
| 18 | 30400000 | 30404599 | 104.5701 | 158.8107 | 236.6514 | 98.00058 | -0.2932 | 3.210333 | 0.579645 |
| 18 | 30400200 | 30404599 | 104.5701 | 158.8107 | 236.6514 | 98.00058 | -0.2932 | 3.210333 | 0.579645 |
| 3 | 66236600 | 66238999 | 19.99628 | 34.85068 | 37.72969 | 16.59786 | 0.273277 | 0.752434 | 0.579655 |
| 12 | 92850400 | 92858199 | 213.0238 | 315.2389 | 274.5503 | 113.0678 | 0.447832 | 3.719508 | 0.579658 |
| 1 | 8072200 | 8072799 | 11.1061 | 21.04757 | 23.74435 | 9.596426 | 0.3145 | 0.042986 | 0.579679 |
| 19 | 10916400 | 10918399 | 17.08052 | 29.19977 | 50.04612 | 21.94357 | -0.39131 | 0.953219 | 0.579682 |
| 9 | 30554000 | 30554599 | 19.33456 | 32.46151 | 56.80734 | 24.80213 | -0.42332 | 1.133577 | 0.579694 |
| 10 | 125660200 | 125662599 | 29.66002 | 48.62241 | 75.26684 | 32.39818 | -0.25202 | 1.450611 | 0.57971 |
| 18 | 12732200 | 12733199 | 11.27911 | 19.99957 | 35.33226 | 15.61768 | -0.42382 | 0.475082 | 0.579728 |
| X | 89136600 | 89138799 | 28.6738 | 49.66313 | 41.22326 | 18.08624 | 0.653894 | 0.875555 | 0.579753 |
| 19 | 8085200 | 8086599 | 25.35756 | 43.0033 | 46.91044 | 20.54726 | 0.256412 | 1.052879 | 0.579776 |
| 1 | 102321400 | 102324999 | 29.50558 | 49.42025 | 51.89514 | 22.62347 | 0.310697 | 1.181398 | 0.579783 |
| 3 | 133156600 | 133158599 | 26.84281 | 44.35369 | 69.11767 | 29.86148 | -0.25758 | 1.3431 | 0.579789 |

| | | | | | | | | | |
|---|---|---|---|---|---|---|---|---|---|
| 14 | 49303200 | 49304399 | 9.374044 | 17.25979 | 30.3736 | 12.87543 | -0.37725 | 0.222659 | 0.579794 |
| 10 | 12742200 | 12745399 | 19.93027 | 33.41468 | 57.97094 | 25.28774 | -0.41103 | 1.148712 | 0.579795 |
| 13 | 67412800 | 67415999 | 40.10037 | 65.9843 | 62.91203 | 27.12309 | 0.447382 | 1.421013 | 0.579803 |
| 1 | 12935000 | 12936399 | 8.794557 | 16.24967 | 29.4261 | 12.41444 | -0.41277 | 0.198042 | 0.579812 |
| 14 | 44298400 | 44300799 | 19.46724 | 34.20911 | 35.30185 | 15.62942 | 0.342323 | 0.621214 | 0.579828 |
| 9 | 69213600 | 69214599 | 8.270923 | 15.67866 | 27.65004 | 11.16869 | -0.32395 | 0.026956 | 0.579844 |
| 17 | 47095200 | 47096199 | 8.270923 | 15.67866 | 27.65004 | 11.16869 | -0.32395 | 0.026956 | 0.579844 |
| 12 | 88566000 | 88568599 | 28.0752 | 47.44929 | 49.61673 | 21.71008 | 0.316976 | 1.093135 | 0.579868 |
| 9 | 27707800 | 27709199 | 18.47317 | 31.53769 | 51.65157 | 22.64504 | -0.32646 | 0.971863 | 0.579898 |
| 7 | 13294000 | 13314399 | 266.51 | 387.0152 | 368.3155 | 151.8369 | 0.294057 | 4.022209 | 0.579935 |
| 1 | 46126200 | 46129599 | 47.27175 | 76.11678 | 82.90209 | 35.23444 | 0.231745 | 1.833294 | 0.579938 |
| 4 | 101352600 | 101355199 | 18.04543 | 30.36129 | 62.23494 | 27.19412 | -0.6507 | 1.112592 | 0.57995 |
| 11 | 71545600 | 71546199 | 11.00713 | 21.24823 | 21.94712 | 8.46325 | 0.501919 | -0.11426 | 0.579952 |
| 7 | 62672000 | 62674999 | 26.87452 | 43.67335 | 84.72136 | 36.34578 | -0.58501 | 1.574887 | 0.579957 |
| 14 | 50730600 | 50734999 | 55.62832 | 89.66608 | 82.31964 | 35.03885 | 0.479602 | 1.815037 | 0.57996 |
| 10 | 34949800 | 34951599 | 20.06097 | 35.13963 | 35.82688 | 15.81896 | 0.359575 | 0.673436 | 0.579972 |
| 3 | 26342400 | 26350599 | 79.8437 | 126.7557 | 117.0289 | 48.77008 | 0.446987 | 2.312352 | 0.57999 |
| 12 | 115771800 | 115788999 | 217.4964 | 323.4164 | 281.0904 | 115.8078 | 0.457254 | 3.637586 | 0.579993 |
| 14 | 64055800 | 64057399 | 15.62718 | 26.94056 | 46.9105 | 20.57853 | -0.41167 | 0.867336 | 0.580004 |
| 12 | 62121200 | 62125199 | 68.38911 | 108.4056 | 117.7716 | 49.00473 | 0.210454 | 2.346906 | 0.580017 |
| 14 | 86190800 | 86191399 | 8.432961 | 16.31752 | 18.83378 | 7.183017 | 0.359048 | -0.18568 | 0.58003 |
| 2 | 100926000 | 100926599 | 10.48507 | 20.34449 | 21.09592 | 7.942453 | 0.529274 | -0.19983 | 0.580035 |
| 3 | 115633400 | 115634799 | 13.35219 | 24.81267 | 25.92142 | 10.85087 | 0.38261 | 0.191038 | 0.580043 |
| 11 | 108860200 | 108863199 | 32.15128 | 53.21769 | 58.00816 | 25.09188 | 0.256672 | 1.325741 | 0.580063 |
| 1 | 150043000 | 150045599 | 29.6776 | 49.50053 | 54.06072 | 23.50532 | 0.254436 | 1.230352 | 0.580069 |
| 5 | 128564800 | 128572599 | 78.14298 | 121.6818 | 175.7953 | 72.96909 | -0.22039 | 2.71687 | 0.580076 |
| X | 8538200 | 8540999 | 33.90381 | 57.48832 | 47.92585 | 20.99109 | 0.643038 | 1.088065 | 0.580086 |
| 16 | 57314200 | 57315799 | 16.1536 | 29.49129 | 28.76882 | 12.33449 | 0.456631 | 0.318367 | 0.580099 |
| 2 | 149243600 | 149244599 | 13.7204 | 25.23358 | 26.86616 | 11.40634 | 0.340905 | 0.253764 | 0.580107 |
| 14 | 17897200 | 17898199 | 19.94114 | 34.91979 | 30.46702 | 13.44914 | 0.583407 | 0.670574 | 0.580122 |
| 4 | 79691000 | 79693999 | 32.92916 | 55.25139 | 52.48909 | 22.90774 | 0.454721 | 1.176246 | 0.580141 |
| 12 | 4315400 | 4318399 | 26.4615 | 43.27957 | 79.7716 | 34.30554 | -0.50477 | 1.47612 | 0.580143 |
| 2 | 86041000 | 86044799 | 56.53397 | 90.52637 | 96.35782 | 40.61574 | 0.255176 | 2.022449 | 0.580146 |
| 8 | 22947000 | 22948599 | 23.53532 | 40.45123 | 43.64339 | 19.14356 | 0.274867 | 0.922559 | 0.580222 |
| 3 | 4710600 | 4715399 | 32.56526 | 54.23647 | 49.77959 | 21.62366 | 0.504513 | 1.250705 | 0.580263 |
| 8 | 54425200 | 54425999 | 15.41254 | 27.51449 | 29.42045 | 12.93047 | 0.301906 | 0.446852 | 0.580294 |
| 7 | 61214800 | 61219999 | 69.80357 | 111.1852 | 109.765 | 45.92331 | 0.355343 | 2.202654 | 0.580298 |
| 2 | 8362800 | 8363199 | 5.224795 | 10.19319 | 20.40315 | 7.291563 | -0.34063 | -0.44705 | 0.58031 |
| 7 | 75667200 | 75667399 | 5.224795 | 10.19319 | 20.40315 | 7.291563 | -0.34063 | -0.44705 | 0.58031 |
| X | 34798400 | 34800399 | 27.54416 | 47.37357 | 44.08553 | 19.3379 | 0.487669 | 0.935712 | 0.580313 |
| 10 | 86785600 | 86789199 | 20.89488 | 35.33985 | 57.6435 | 25.22784 | -0.32215 | 1.069472 | 0.580317 |
| 17 | 83588400 | 83590599 | 16.6486 | 28.99364 | 45.75537 | 20.11866 | -0.26803 | 0.759964 | 0.580361 |
| 17 | 87430600 | 87433199 | 27.46006 | 45.15837 | 70.7656 | 30.50138 | -0.2678 | 1.401359 | 0.580372 |
| 2 | 81406000 | 81407799 | 14.06134 | 24.90442 | 42.88909 | 18.96397 | -0.38868 | 0.515181 | 0.580374 |
| 5 | 145785200 | 145793799 | 166.7788 | 250.711 | 251.2623 | 103.7341 | 0.266017 | 3.428241 | 0.580375 |
| 4 | 71841400 | 71842399 | 18.56939 | 32.59075 | 34.54121 | 15.27658 | 0.304212 | 0.636258 | 0.580379 |
| 12 | 55920600 | 55926199 | 61.88526 | 98.85307 | 102.3351 | 42.98503 | 0.291347 | 2.103106 | 0.580383 |

| | | | | | | | | | |
|---|---|---|---|---|---|---|---|---|---|
| 7 | 34238000 | 34244199 | 60.77235 | 96.26588 | 140.2468 | 58.32639 | -0.21315 | 2.3639 | 0.580389 |
| 7 | 41465200 | 41467599 | 51.13548 | 82.13192 | 88.45965 | 37.45986 | 0.243389 | 1.919959 | 0.580418 |
| 12 | 117003000 | 117005199 | 19.65742 | 33.61734 | 52.73475 | 23.12313 | -0.2641 | 0.964154 | 0.580425 |
| 17 | 70749800 | 70751199 | 17.48784 | 30.88412 | 32.14554 | 14.22175 | 0.332368 | 0.546887 | 0.580429 |
| 9 | 78760800 | 78761799 | 20.89402 | 36.16648 | 38.00624 | 16.6745 | 0.314848 | 0.841384 | 0.580436 |
| 12 | 11612800 | 11613399 | 7.792881 | 15.18916 | 17.93862 | 6.489602 | 0.395319 | -0.38095 | 0.580437 |
| 3 | 69394400 | 69397799 | 21.78448 | 36.96954 | 58.75703 | 25.74162 | -0.2846 | 1.04717 | 0.580454 |
| 7 | 48164400 | 48175399 | 187.8687 | 279.958 | 280.6176 | 115.6382 | 0.25445 | 3.595916 | 0.580456 |
| 11 | 27558600 | 27568599 | 111.4844 | 171.2699 | 182.1891 | 75.64787 | 0.205778 | 2.976366 | 0.580459 |
| 6 | 26207400 | 26208199 | 12.31023 | 22.72173 | 24.50561 | 10.35331 | 0.327714 | 0.217998 | 0.580473 |
| 15 | 41961600 | 41963199 | 17.06842 | 30.67271 | 29.40817 | 12.85322 | 0.463337 | 0.413447 | 0.580473 |
| 12 | 87948000 | 87955399 | 92.07258 | 144.0148 | 148.6271 | 61.65932 | 0.269361 | 2.642692 | 0.580476 |
| 11 | 15813800 | 15814399 | 8.412288 | 15.34481 | 35.00474 | 15.02442 | -0.76171 | 0.279677 | 0.580489 |
| 17 | 20866000 | 20874999 | 99.93637 | 156.6616 | 143.4217 | 59.48624 | 0.444519 | 2.609228 | 0.58049 |
| 16 | 32479400 | 32481399 | 13.28708 | 23.3412 | 39.34928 | 17.41648 | -0.36213 | 0.597987 | 0.580494 |
| 2 | 126430400 | 126432399 | 18.26903 | 31.49059 | 49.53876 | 21.73139 | -0.26614 | 0.881681 | 0.580511 |
| 12 | 114889400 | 114892999 | 70.40122 | 111.8441 | 116.4143 | 48.54817 | 0.275251 | 2.284021 | 0.580514 |
| 9 | 71783800 | 71785799 | 17.91009 | 31.99359 | 31.64885 | 13.90237 | 0.413979 | 0.463511 | 0.580567 |
| 3 | 56968000 | 56969599 | 27.36244 | 44.46402 | 85.23888 | 36.56645 | -0.56801 | 1.578437 | 0.580574 |
| 5 | 89976600 | 89979999 | 36.95162 | 59.52431 | 91.95057 | 39.10392 | -0.27057 | 1.810122 | 0.580583 |
| 1 | 123805800 | 123807999 | 23.04497 | 40.3654 | 36.72853 | 16.21669 | 0.523118 | 0.693118 | 0.580585 |
| 13 | 50903000 | 50906599 | 25.27119 | 41.99257 | 65.35465 | 28.34567 | -0.25512 | 1.268902 | 0.580605 |
| 9 | 124079200 | 124093399 | 258.3108 | 378.2224 | 333.6348 | 137.4341 | 0.415016 | 3.896706 | 0.580647 |
| 2 | 108222000 | 108222599 | 10.74081 | 20.41632 | 23.25099 | 9.337722 | 0.30883 | 0.017261 | 0.580649 |
| 16 | 94077600 | 94078799 | 36.17464 | 58.31523 | 91.24286 | 38.84221 | -0.28769 | 1.787782 | 0.580658 |
| 12 | 24641200 | 24643599 | 13.81347 | 24.15513 | 41.07044 | 18.10815 | -0.37426 | 0.679914 | 0.580685 |
| 17 | 56161600 | 56174799 | 215.5129 | 319.9195 | 290.537 | 119.6789 | 0.391651 | 3.667987 | 0.580701 |
| 17 | 38438600 | 38442199 | 53.06126 | 85.38101 | 77.00645 | 32.69651 | 0.502118 | 1.861396 | 0.580703 |
| 5 | 103377400 | 103378999 | 25.71904 | 43.56138 | 47.45188 | 20.77647 | 0.258405 | 1.065983 | 0.580707 |
| 17 | 12139600 | 12143199 | 32.6798 | 53.86486 | 59.03463 | 25.46964 | 0.247858 | 1.377415 | 0.580708 |
| 1 | 110540800 | 110541199 | 7.683039 | 14.93369 | 16.82937 | 6.25577 | 0.428139 | -0.28839 | 0.580725 |
| 12 | 10975400 | 10977599 | 18.56786 | 32.68615 | 34.24588 | 15.16073 | 0.321276 | 0.592977 | 0.580729 |
| 19 | 55001800 | 55005599 | 39.2199 | 63.07639 | 96.06487 | 40.75821 | -0.25308 | 1.865116 | 0.580731 |
| 1 | 121454000 | 121457399 | 35.00594 | 56.40376 | 90.44259 | 38.47615 | -0.32382 | 1.798799 | 0.580732 |
| 3 | 60220400 | 60221799 | 14.69641 | 27.29442 | 26.99577 | 11.30975 | 0.460518 | 0.209593 | 0.580737 |
| 8 | 48630000 | 48634599 | 53.1385 | 84.55313 | 125.7803 | 52.54855 | -0.23685 | 2.222631 | 0.580741 |
| 4 | 115243800 | 115245799 | 25.52432 | 43.44986 | 46.87501 | 20.54612 | 0.272851 | 1.020291 | 0.580744 |
| 5 | 54930400 | 54931199 | 11.66658 | 20.96728 | 34.22503 | 14.95734 | -0.29921 | 0.389595 | 0.580768 |
| X | 150927000 | 150928199 | 14.77748 | 27.18669 | 26.51454 | 11.25198 | 0.467846 | 0.264157 | 0.58077 |
| 7 | 18226000 | 18237399 | 59.76426 | 95.74661 | 84.9552 | 35.83655 | 0.518087 | 2.005261 | 0.580794 |
| 5 | 111180800 | 111183799 | 21.45835 | 36.16245 | 57.03244 | 24.93317 | -0.27386 | 1.104087 | 0.580827 |
| 13 | 18108400 | 18111799 | 40.07402 | 64.88664 | 67.85809 | 29.03323 | 0.299172 | 1.663561 | 0.580837 |
| 3 | 104602400 | 104605399 | 28.99716 | 48.4514 | 52.94922 | 23.04781 | 0.253452 | 1.208307 | 0.580851 |
| 7 | 62373000 | 62387799 | 216.0329 | 319.4482 | 304.7054 | 125.4855 | 0.315674 | 3.734131 | 0.580851 |
| 14 | 120753200 | 120754799 | 13.63445 | 25.31151 | 26.56744 | 11.13667 | 0.374962 | 0.203605 | 0.580858 |
| 15 | 82372200 | 82374199 | 20.08761 | 35.21991 | 36.40131 | 16.08557 | 0.340144 | 0.661569 | 0.5809 |
| 19 | 13333800 | 13337999 | 67.74478 | 107.7934 | 112.391 | 46.95416 | 0.27505 | 2.233694 | 0.580903 |

| | | | | | | | | | |
|---|---|---|---|---|---|---|---|---|---|
| 7 | 130434800 | 130437399 | 73.1992 | 113.6915 | 174.3551 | 72.39902 | -0.30927 | 2.759197 | 0.580904 |
| 2 | 104634400 | 104635599 | 12.27106 | 22.0226 | 35.70271 | 15.62168 | -0.29082 | 0.404346 | 0.580905 |
| 3 | 46421200 | 46423199 | 20.95201 | 36.90959 | 34.55974 | 15.29748 | 0.483071 | 0.600402 | 0.580914 |
| 11 | 7099400 | 7104799 | 34.96886 | 56.41504 | 88.70364 | 37.79893 | -0.29348 | 1.759833 | 0.580916 |
| 3 | 106634200 | 106636399 | 39.14408 | 64.61793 | 60.74113 | 26.22519 | 0.469405 | 1.377297 | 0.580956 |
| 1 | 96923800 | 96925399 | 13.58002 | 24.87083 | 26.86253 | 11.47009 | 0.314488 | 0.276162 | 0.580983 |
| 17 | 33490400 | 33493199 | 34.48177 | 57.28788 | 50.90241 | 22.07485 | 0.551035 | 1.28575 | 0.580994 |
| 14 | 45998400 | 45999199 | 6.433725 | 12.36101 | 24.65855 | 9.642859 | -0.45657 | -0.10333 | 0.580997 |
| 6 | 16426600 | 16428999 | 30.72469 | 52.18911 | 41.48965 | 18.18176 | 0.711326 | 1.054816 | 0.581033 |
| 6 | 67632200 | 67633599 | 26.62871 | 45.43315 | 40.96171 | 17.95751 | 0.531205 | 0.995802 | 0.581034 |
| X | 153854400 | 153859799 | 84.8474 | 133.8425 | 132.2537 | 54.88684 | 0.341404 | 2.478852 | 0.58104 |
| 4 | 129818000 | 129819599 | 12.67509 | 22.41994 | 36.75777 | 16.25716 | -0.31835 | 0.512487 | 0.581046 |
| 8 | 70426200 | 70427599 | 10.53919 | 18.62587 | 36.42674 | 16.11353 | -0.57161 | 0.507524 | 0.581048 |
| 1 | 29933000 | 29934799 | 18.28305 | 31.1561 | 58.59932 | 25.68728 | -0.5257 | 0.998392 | 0.58106 |
| 14 | 53909400 | 53910799 | 13.19218 | 24.46447 | 25.91173 | 10.8775 | 0.359984 | 0.201125 | 0.581083 |
| 13 | 73486600 | 73487799 | 13.80282 | 25.3165 | 27.40958 | 11.68002 | 0.313236 | 0.271558 | 0.581096 |
| 8 | 10302600 | 10303399 | 6.884889 | 13.17484 | 25.12136 | 9.937701 | -0.40817 | -0.061 | 0.581108 |
| 8 | 71636000 | 71638399 | 26.45276 | 45.59563 | 43.49423 | 19.07979 | 0.452585 | 0.908636 | 0.581109 |
| 6 | 41469400 | 41472199 | 27.34089 | 45.84059 | 47.53415 | 20.73547 | 0.32877 | 1.159529 | 0.581112 |
| 17 | 39611400 | 39614999 | 30.03734 | 51.00787 | 42.50762 | 18.62634 | 0.643465 | 1.053461 | 0.581147 |
| 8 | 33858600 | 33860799 | 34.69594 | 57.63766 | 50.35029 | 21.83384 | 0.575488 | 1.287826 | 0.581172 |
| 15 | 90713400 | 90716199 | 29.78817 | 50.34331 | 50.17365 | 21.95362 | 0.386074 | 1.100583 | 0.581225 |
| 12 | 115277200 | 115279799 | 38.54476 | 63.94795 | 56.76197 | 24.59203 | 0.552342 | 1.31568 | 0.581237 |
| X | 5559800 | 5561599 | 27.18996 | 46.69163 | 44.64732 | 19.5834 | 0.448182 | 0.948409 | 0.581247 |
| X | 79008800 | 79014199 | 61.59222 | 98.67661 | 95.79225 | 40.40886 | 0.388684 | 2.013584 | 0.581265 |
| 1 | 117837400 | 117844799 | 96.1643 | 151.2704 | 135.8573 | 56.36182 | 0.475951 | 2.539949 | 0.581272 |
| X | 122457800 | 122464199 | 63.48008 | 101.6086 | 86.53137 | 36.45248 | 0.57574 | 2.040232 | 0.581277 |
| 4 | 87413400 | 87416599 | 28.49858 | 46.76401 | 72.31545 | 31.14062 | -0.24994 | 1.433628 | 0.581289 |
| 14 | 121136200 | 121139599 | 32.81923 | 53.41835 | 82.55715 | 35.44075 | -0.25637 | 1.561285 | 0.581289 |
| 17 | 64039400 | 64045999 | 57.59099 | 92.31667 | 94.41571 | 39.86127 | 0.314483 | 1.989359 | 0.581315 |
| 13 | 113910000 | 113912799 | 22.08611 | 37.41142 | 59.33063 | 25.97819 | -0.28162 | 1.064162 | 0.581317 |
| X | 8575000 | 8578199 | 47.58569 | 77.66206 | 66.73196 | 28.67535 | 0.591133 | 1.54442 | 0.581319 |
| 1 | 143453000 | 143455999 | 30.80123 | 51.42446 | 54.67468 | 23.7868 | 0.292983 | 1.223595 | 0.581351 |
| 14 | 44281000 | 44283199 | 42.55298 | 68.8219 | 74.98891 | 32.05718 | 0.238547 | 1.694201 | 0.581355 |
| 9 | 11845000 | 11847799 | 38.91117 | 62.65456 | 94.60539 | 40.20844 | -0.23855 | 1.827404 | 0.581394 |
| 17 | 65676600 | 65677799 | 12.91783 | 22.86222 | 38.18212 | 16.88521 | -0.34489 | 0.507539 | 0.581396 |
| 2 | 168462000 | 168467999 | 75.63955 | 117.956 | 170.4384 | 70.73012 | -0.21896 | 2.690318 | 0.581418 |
| 9 | 60637000 | 60641799 | 56.50168 | 89.68034 | 131.2782 | 54.69678 | -0.21727 | 2.302487 | 0.581452 |
| 5 | 115908000 | 115912599 | 56.14532 | 89.71663 | 97.69812 | 41.05826 | 0.219036 | 2.085081 | 0.581458 |
| 12 | 45131400 | 45133799 | 16.83443 | 30.50574 | 29.05945 | 12.57594 | 0.48192 | 0.364443 | 0.581489 |
| 14 | 56861400 | 56864799 | 31.43582 | 51.20861 | 79.14507 | 33.98615 | -0.2556 | 1.54254 | 0.581493 |
| 7 | 121018600 | 121022599 | 48.34996 | 77.89341 | 83.08191 | 35.33748 | 0.262783 | 1.820195 | 0.581511 |
| 8 | 117120400 | 117123799 | 36.40479 | 59.99368 | 62.09502 | 26.78793 | 0.33009 | 1.392615 | 0.581557 |
| 3 | 27363000 | 27364799 | 14.85002 | 25.91547 | 43.62548 | 19.20734 | -0.36044 | 0.716681 | 0.581577 |
| 5 | 121005800 | 121007799 | 21.44692 | 36.3116 | 56.52002 | 24.74401 | -0.2547 | 1.06371 | 0.581586 |
| 10 | 125860000 | 125860399 | 5.947342 | 11.58318 | 22.56026 | 8.243396 | -0.3308 | -0.3503 | 0.581595 |
| 13 | 50685400 | 50688799 | 28.89225 | 49.739 | 43.44779 | 19.05871 | 0.578818 | 0.930634 | 0.58162 |

| | | | | | | | | | |
|---|---|---|---|---|---|---|---|---|---|
| 1 | 117504600 | 117508799 | 58.00189 | 92.81019 | 98.26643 | 41.37718 | 0.261652 | 2.046334 | 0.581654 |
| 7 | 17987800 | 17991599 | 71.25995 | 113.0333 | 119.7358 | 49.85601 | 0.247801 | 2.330466 | 0.581656 |
| 3 | 150041200 | 150044999 | 46.82308 | 75.33373 | 81.13556 | 34.44347 | 0.24646 | 1.857875 | 0.58166 |
| 15 | 100163000 | 100176399 | 148.8923 | 222.5202 | 307.8659 | 126.935 | -0.20644 | 3.537316 | 0.581672 |
| 4 | 117011200 | 117013399 | 29.42987 | 47.79694 | 78.11528 | 33.49674 | -0.33919 | 1.588381 | 0.581697 |
| 1 | 33675000 | 33677399 | 19.99701 | 33.41499 | 66.11463 | 28.77953 | -0.60004 | 1.192897 | 0.581718 |
| 9 | 84876000 | 84878999 | 27.34827 | 46.98096 | 44.02274 | 19.30966 | 0.477233 | 0.941382 | 0.581718 |
| 3 | 33649600 | 33654199 | 46.03241 | 74.41412 | 77.7893 | 33.23462 | 0.297451 | 1.715013 | 0.58173 |
| 2 | 23195000 | 23200199 | 67.26396 | 107.4823 | 101.2371 | 42.54488 | 0.427983 | 2.098976 | 0.581738 |
| 2 | 87999800 | 88004799 | 65.39277 | 104.1189 | 109.0648 | 45.60568 | 0.269383 | 2.211468 | 0.581753 |
| 14 | 6452400 | 6458199 | 83.43054 | 128.8498 | 210.0304 | 87.24228 | -0.40355 | 2.86962 | 0.581757 |
| 17 | 45453800 | 45455799 | 21.36164 | 35.86463 | 57.49397 | 25.10183 | -0.29745 | 1.139178 | 0.581759 |
| 11 | 117417400 | 117419399 | 19.31254 | 32.75213 | 54.31156 | 23.78912 | -0.34558 | 1.03783 | 0.581767 |
| 15 | 15931000 | 15933199 | 18.74889 | 33.45882 | 31.45099 | 13.82699 | 0.486238 | 0.47296 | 0.581778 |
| X | 31842000 | 31846799 | 59.38817 | 95.97484 | 73.47977 | 31.38091 | 0.74531 | 1.747101 | 0.581786 |
| 3 | 87184400 | 87188599 | 47.01923 | 76.10152 | 76.44079 | 32.69567 | 0.356653 | 1.688131 | 0.58181 |
| 10 | 82599200 | 82600599 | 10.12947 | 18.22429 | 32.84236 | 14.34012 | -0.43989 | 0.369823 | 0.581814 |
| 8 | 72777600 | 72790199 | 130.1435 | 198.3257 | 206.139 | 85.45235 | 0.230093 | 3.144141 | 0.581818 |
| 5 | 140907000 | 140914799 | 114.9758 | 174.3902 | 247.3012 | 102.3715 | -0.22329 | 3.238781 | 0.581825 |
| 1 | 81668600 | 81671799 | 74.90802 | 117.9655 | 124.4109 | 51.62071 | 0.245685 | 2.502066 | 0.581828 |
| 18 | 61592000 | 61593599 | 13.00361 | 23.09484 | 37.21194 | 16.42604 | -0.29065 | 0.479543 | 0.581829 |
| 6 | 112983800 | 112987199 | 32.97345 | 54.49318 | 59.16577 | 25.56496 | 0.262148 | 1.346558 | 0.581869 |
| 10 | 34983000 | 34983599 | 6.717221 | 12.88226 | 24.62945 | 9.681847 | -0.40387 | -0.087 | 0.581882 |
| 1 | 79975200 | 79977199 | 24.32116 | 40.50854 | 63.0308 | 27.38084 | -0.25454 | 1.239079 | 0.581882 |
| 5 | 128137400 | 128139199 | 11.90629 | 21.58253 | 36.40324 | 15.74759 | -0.33696 | 0.343264 | 0.58194 |
| 1 | 108959600 | 108960399 | 11.95159 | 22.27282 | 24.51456 | 10.21419 | 0.313307 | 0.15913 | 0.581976 |
| 15 | 28136600 | 28138599 | 12.86461 | 22.61142 | 44.52985 | 19.71859 | -0.58533 | 0.595751 | 0.581986 |
| 1 | 164920800 | 164921599 | 9.168812 | 17.08595 | 29.46545 | 12.27814 | -0.32991 | 0.148871 | 0.581991 |
| 17 | 52021800 | 52024399 | 25.35543 | 42.28061 | 68.19034 | 29.64403 | -0.3062 | 1.234889 | 0.581995 |
| 7 | 41489000 | 41494999 | 128.1863 | 195.592 | 202.5361 | 83.98429 | 0.236958 | 3.119377 | 0.582006 |
| 9 | 42969800 | 42973199 | 31.88806 | 53.44104 | 53.04583 | 23.13338 | 0.391657 | 1.185125 | 0.582013 |
| Y | 86806200 | 86807399 | 10.77247 | 20.36462 | 16.652 | 6.765471 | 0.768201 | 0.03439 | 0.582037 |
| 5 | 84980800 | 84983399 | 9.623695 | 17.51405 | 35.47663 | 15.27098 | -0.59432 | 0.303723 | 0.582087 |
| X | 24133800 | 24137199 | 73.01168 | 115.9225 | 117.5052 | 48.97403 | 0.312594 | 2.304823 | 0.582111 |
| 4 | 70616000 | 70619199 | 32.13513 | 54.13976 | 51.3025 | 22.42622 | 0.458551 | 1.138279 | 0.582112 |
| 3 | 100749200 | 100751999 | 17.67219 | 30.29863 | 49.52874 | 21.72177 | -0.32271 | 0.92809 | 0.582114 |
| 18 | 33931000 | 33937799 | 56.22049 | 89.22102 | 131.3212 | 54.70672 | -0.22533 | 2.30732 | 0.582142 |
| 15 | 41894200 | 41898199 | 50.67482 | 81.43035 | 87.76643 | 37.19187 | 0.243168 | 1.902559 | 0.582186 |
| 7 | 104361000 | 104362599 | 14.26132 | 25.80873 | 27.80716 | 12.05398 | 0.303867 | 0.35134 | 0.582191 |
| 9 | 8779600 | 8781799 | 36.61057 | 60.82296 | 50.40684 | 21.85951 | 0.651204 | 1.291415 | 0.582198 |
| 6 | 146046400 | 146051399 | 83.37328 | 130.4337 | 138.7762 | 57.57141 | 0.225681 | 2.633875 | 0.582206 |
| 7 | 17702600 | 17705199 | 43.09705 | 70.63466 | 64.34425 | 27.69406 | 0.510716 | 1.474672 | 0.58222 |
| 15 | 18403800 | 18406799 | 30.24457 | 51.45706 | 47.35572 | 20.75591 | 0.501152 | 1.042232 | 0.582244 |
| 4 | 107817000 | 107818199 | 9.076007 | 16.80093 | 29.41239 | 12.37728 | -0.36185 | 0.189043 | 0.582267 |
| 13 | 106110800 | 106115799 | 74.89615 | 118.7571 | 109.54 | 45.59377 | 0.446643 | 2.34079 | 0.582271 |
| 5 | 134727600 | 134729599 | 18.40169 | 31.65733 | 50.61372 | 22.20163 | -0.29006 | 0.900029 | 0.582296 |
| 17 | 43816800 | 43817399 | 6.856774 | 13.28072 | 25.02525 | 9.500538 | -0.33531 | -0.19865 | 0.582306 |

| | | | | | | | | | |
|---|---|---|---|---|---|---|---|---|---|
| 17 | 20131600 | 20134599 | 79.21544 | 124.4429 | 130.2035 | 54.00691 | 0.254431 | 2.555956 | 0.582311 |
| 11 | 19862600 | 19868599 | 49.21422 | 78.48714 | 117.3488 | 49.18051 | -0.24058 | 2.144139 | 0.582313 |
| 2 | 122881400 | 122882599 | 9.824863 | 18.41859 | 32.98241 | 13.60481 | -0.37223 | 0.12483 | 0.582313 |
| 14 | 125037600 | 125039399 | 40.48606 | 65.71694 | 71.35646 | 30.58602 | 0.248385 | 1.614654 | 0.582323 |
| 4 | 47055400 | 47058199 | 22.02436 | 36.77764 | 60.19919 | 26.1933 | -0.32739 | 1.20092 | 0.58233 |
| 2 | 8200600 | 8203999 | 33.01284 | 54.35343 | 59.57694 | 25.69154 | 0.247063 | 1.392222 | 0.582337 |
| 3 | 151810400 | 151812799 | 23.07148 | 38.23719 | 63.88647 | 27.65888 | -0.35737 | 1.292349 | 0.582345 |
| 12 | 114624400 | 114653199 | 410.9901 | 576.1175 | 509.5957 | 211.2236 | 0.348744 | 4.547177 | 0.582362 |
| 14 | 46345400 | 46347799 | 33.94809 | 55.00152 | 83.84685 | 35.89032 | -0.24243 | 1.647825 | 0.582371 |
| 2 | 98034200 | 98038999 | 48.91968 | 79.04495 | 78.60086 | 33.56129 | 0.368668 | 1.734549 | 0.582374 |
| X | 81054400 | 81056399 | 21.12811 | 37.35972 | 33.50951 | 14.82149 | 0.546923 | 0.561457 | 0.582405 |
| 17 | 92645000 | 92645799 | 13.66721 | 25.30459 | 22.78425 | 9.585605 | 0.590565 | 0.207507 | 0.582405 |
| 3 | 67145200 | 67148199 | 27.80671 | 47.82674 | 44.02324 | 19.31059 | 0.50327 | 0.933893 | 0.582416 |
| X | 87919600 | 87920599 | 15.65618 | 29.59132 | 24.62606 | 10.00925 | 0.743399 | 0.105431 | 0.582431 |
| X | 152961200 | 152963799 | 33.29078 | 56.1314 | 50.75972 | 22.19431 | 0.525821 | 1.134738 | 0.582464 |
| 7 | 59821800 | 59823999 | 32.58803 | 54.53015 | 52.97116 | 23.07839 | 0.422899 | 1.208213 | 0.582476 |
| 5 | 18600200 | 18601599 | 13.55172 | 25.1083 | 26.427 | 11.10758 | 0.367772 | 0.210616 | 0.582489 |
| 12 | 55318600 | 55319999 | 10.72053 | 19.23342 | 33.75649 | 14.77431 | -0.40471 | 0.391779 | 0.582536 |
| X | 122559000 | 122562799 | 39.64505 | 66.44441 | 50.30943 | 21.94075 | 0.781158 | 1.21109 | 0.582554 |
| 1 | 8190400 | 8200999 | 259.7605 | 375.1859 | 378.2235 | 156.0273 | 0.203384 | 4.100581 | 0.582566 |
| 18 | 4292200 | 4294999 | 25.50638 | 42.03679 | 68.41358 | 29.51474 | -0.32093 | 1.367524 | 0.582568 |
| 1 | 10986400 | 10988799 | 19.98738 | 34.33604 | 57.19422 | 25.08706 | -0.34946 | 0.929193 | 0.582569 |
| 2 | 142553600 | 142554999 | 10.50872 | 19.20993 | 31.83497 | 13.62742 | -0.30132 | 0.277046 | 0.582588 |
| 4 | 90133000 | 90134599 | 14.74225 | 25.9568 | 41.83822 | 18.52115 | -0.29686 | 0.583206 | 0.58261 |
| 12 | 81064000 | 81068199 | 31.65971 | 51.38582 | 80.28702 | 34.41191 | -0.27541 | 1.605323 | 0.582613 |
| 7 | 11775400 | 11787799 | 177.3648 | 267.0451 | 242.666 | 100.2274 | 0.40922 | 3.400819 | 0.582627 |
| 1 | 19328200 | 19329799 | 14.68882 | 26.49533 | 28.84848 | 12.55678 | 0.285148 | 0.376633 | 0.582633 |
| 6 | 25617400 | 25624999 | 107.5855 | 165.6298 | 176.8202 | 73.4399 | 0.202947 | 2.933707 | 0.582641 |
| 3 | 124869400 | 124871399 | 23.64709 | 40.89544 | 35.8237 | 15.71652 | 0.576067 | 0.858854 | 0.582655 |
| 15 | 25411400 | 25415399 | 25.45127 | 41.35488 | 86.92379 | 37.28694 | -0.70062 | 1.574619 | 0.582661 |
| 3 | 46151400 | 46153399 | 19.77849 | 33.79814 | 52.74527 | 23.1263 | -0.2569 | 0.970636 | 0.582671 |
| 4 | 61668000 | 61671799 | 55.40766 | 88.79166 | 94.54098 | 39.89695 | 0.256048 | 1.997029 | 0.582678 |
| 9 | 36254200 | 36256399 | 28.49865 | 48.45489 | 47.93934 | 21.01061 | 0.397109 | 1.045137 | 0.582736 |
| 15 | 90854800 | 90856599 | 17.76347 | 30.72116 | 48.47138 | 21.26532 | -0.26938 | 0.848006 | 0.582742 |
| 16 | 82367600 | 82370199 | 24.74475 | 41.20123 | 64.85485 | 28.16768 | -0.27128 | 1.24307 | 0.582753 |
| 6 | 86310800 | 86313799 | 29.47684 | 48.15501 | 75.03824 | 32.25623 | -0.26443 | 1.495248 | 0.582763 |
| 12 | 54248800 | 54250799 | 13.50513 | 23.63414 | 40.5547 | 17.89241 | -0.38736 | 0.665147 | 0.582771 |
| X | 90928800 | 90931999 | 38.81481 | 64.00102 | 61.68499 | 26.61733 | 0.432988 | 1.392789 | 0.582772 |
| 19 | 61186800 | 61188599 | 16.45222 | 28.30825 | 47.95244 | 21.03437 | -0.37267 | 0.89093 | 0.582784 |
| 13 | 53896400 | 53896999 | 5.965267 | 11.55357 | 22.69432 | 8.617087 | -0.39436 | -0.21522 | 0.582793 |
| 9 | 119706800 | 119707399 | 13.38946 | 24.57895 | 26.98826 | 11.48737 | 0.294415 | 0.263853 | 0.582802 |
| 1 | 13733400 | 13736399 | 31.6123 | 52.40479 | 56.90808 | 24.64176 | 0.262213 | 1.305523 | 0.582807 |
| 11 | 43788200 | 43788599 | 3.754668 | 7.343442 | 16.66598 | 5.638654 | -0.44435 | -0.67121 | 0.58281 |
| 10 | 38513800 | 38514999 | 9.862007 | 18.34121 | 31.04225 | 12.97881 | -0.30678 | 0.173236 | 0.582815 |
| 6 | 89854200 | 89856999 | 41.02331 | 67.10806 | 66.84528 | 28.74844 | 0.380811 | 1.496653 | 0.582817 |
| 16 | 71484800 | 71486199 | 22.75553 | 37.89503 | 61.71165 | 26.80519 | -0.31991 | 1.236691 | 0.582824 |
| 7 | 26655600 | 26656599 | 16.75669 | 29.6386 | 31.74616 | 14.04073 | 0.292184 | 0.526063 | 0.582829 |

| | | | | | | | | | |
|---|---|---|---|---|---|---|---|---|---|
| 1 | 23140000 | 23142199 | 20.39993 | 34.16813 | 59.0893 | 25.75943 | -0.40649 | 1.161274 | 0.582834 |
| 7 | 17663200 | 17665599 | 26.06785 | 45.49119 | 38.40886 | 16.91034 | 0.630653 | 0.767998 | 0.582852 |
| 6 | 130790000 | 130793799 | 51.36965 | 83.29696 | 72.56028 | 31.0783 | 0.564163 | 1.658169 | 0.582853 |
| 10 | 116697000 | 116698399 | 10.77583 | 19.92514 | 35.34649 | 14.89722 | -0.38317 | 0.220423 | 0.582854 |
| 1 | 43947600 | 43949599 | 16.32187 | 30.01589 | 27.16208 | 11.53309 | 0.573862 | 0.274895 | 0.582868 |
| X | 8555600 | 8558999 | 50.56505 | 81.94095 | 74.10696 | 31.72611 | 0.509353 | 1.670021 | 0.582907 |
| 17 | 11059000 | 11059999 | 10.53299 | 19.09268 | 32.29156 | 13.96936 | -0.34039 | 0.326838 | 0.582913 |
| 2 | 154309400 | 154310799 | 10.98793 | 20.01879 | 32.82001 | 14.10804 | -0.2899 | 0.301976 | 0.582921 |
| 14 | 80054400 | 80057799 | 28.63022 | 48.2756 | 50.7235 | 22.1788 | 0.309995 | 1.116492 | 0.582934 |
| 4 | 140992400 | 140993999 | 16.63532 | 28.91978 | 45.7875 | 20.11062 | -0.27296 | 0.78452 | 0.582947 |
| 1 | 136627800 | 136630599 | 17.58504 | 30.30794 | 49.53887 | 21.73136 | -0.32124 | 0.878318 | 0.582953 |
| 4 | 101863600 | 101864999 | 13.57299 | 24.10785 | 39.73209 | 17.53739 | -0.32336 | 0.48478 | 0.582954 |
| 16 | 43537400 | 43538999 | 17.3154 | 31.35408 | 28.99342 | 12.55898 | 0.523653 | 0.371795 | 0.582977 |
| 9 | 120616800 | 120622399 | 85.00443 | 131.4818 | 188.1355 | 78.14697 | -0.21403 | 2.839187 | 0.582993 |
| 5 | 53740400 | 53741999 | 13.04378 | 22.87354 | 39.50382 | 17.45893 | -0.39651 | 0.628204 | 0.582993 |
| 15 | 8085400 | 8088199 | 42.40036 | 68.76883 | 63.42436 | 27.16988 | 0.482237 | 1.630131 | 0.583018 |
| 15 | 98688800 | 98691599 | 22.60405 | 37.43408 | 64.37916 | 27.85759 | -0.39903 | 1.299954 | 0.58302 |
| 4 | 150440000 | 150440999 | 11.75315 | 22.00015 | 24.63559 | 10.18799 | 0.297548 | 0.131474 | 0.583025 |
| 5 | 7861800 | 7862599 | 11.75315 | 22.00015 | 24.63559 | 10.18799 | 0.297548 | 0.131474 | 0.583025 |
| 1 | 166373600 | 166376999 | 54.52626 | 87.19479 | 94.50985 | 39.76054 | 0.226943 | 2.059987 | 0.583027 |
| 7 | 33620000 | 33620599 | 13.06328 | 24.10747 | 26.52705 | 11.21002 | 0.298823 | 0.230892 | 0.583037 |
| 6 | 92924600 | 92926999 | 37.05911 | 60.6171 | 65.789 | 28.29312 | 0.257255 | 1.48863 | 0.583053 |
| 10 | 81833000 | 81847799 | 474.5694 | 651.077 | 622.6465 | 258.9565 | 0.204867 | 4.874413 | 0.583073 |
| 7 | 114936200 | 114938199 | 26.90982 | 45.71214 | 41.612 | 18.23574 | 0.516563 | 1.038362 | 0.583077 |
| 4 | 52194600 | 52195399 | 9.505454 | 17.77243 | 30.07507 | 12.46162 | -0.29616 | 0.133035 | 0.583077 |
| 12 | 103275200 | 103277199 | 19.57113 | 34.1008 | 36.80853 | 16.18997 | 0.277602 | 0.753789 | 0.583086 |
| 14 | 36762400 | 36763399 | 8.393538 | 15.84345 | 27.55454 | 11.23087 | -0.31579 | 0.055685 | 0.583091 |
| 6 | 90041800 | 90042799 | 23.24736 | 40.19041 | 35.32845 | 15.49909 | 0.570943 | 0.858666 | 0.583149 |
| 9 | 5925000 | 5926799 | 67.95109 | 107.6538 | 114.8977 | 47.78778 | 0.23533 | 2.361206 | 0.583158 |
| 16 | 45973600 | 45976199 | 14.03557 | 24.67079 | 40.79996 | 18.06763 | -0.33477 | 0.59392 | 0.583159 |
| 18 | 37150200 | 37151599 | 23.86623 | 41.58733 | 39.68315 | 17.45508 | 0.454715 | 0.779367 | 0.583163 |
| 1 | 81426200 | 81430199 | 39.59354 | 65.06382 | 63.55574 | 27.38521 | 0.411722 | 1.433894 | 0.583167 |
| 6 | 88972800 | 88976199 | 24.42905 | 40.38701 | 74.01904 | 31.99092 | -0.49083 | 1.333904 | 0.583199 |
| 17 | 22515400 | 22528199 | 200.099 | 298.3508 | 277.3793 | 114.3133 | 0.363675 | 3.588428 | 0.583236 |
| 7 | 46930800 | 46947199 | 391.2237 | 545.8663 | 529.6794 | 219.7069 | 0.208456 | 4.615065 | 0.583285 |
| 5 | 75146800 | 75158199 | 190.3497 | 278.8616 | 382.1871 | 157.429 | -0.21964 | 3.880932 | 0.583294 |
| 9 | 71423200 | 71425399 | 14.08455 | 24.86862 | 39.74524 | 17.58472 | -0.28302 | 0.544956 | 0.583299 |
| 4 | 153102400 | 153108199 | 46.25653 | 74.13257 | 110.9279 | 46.81784 | -0.23386 | 1.988469 | 0.583317 |
| 8 | 61788600 | 61791399 | 26.24392 | 44.61728 | 47.40005 | 20.7741 | 0.294609 | 1.033645 | 0.583328 |
| 3 | 49905600 | 49908999 | 34.07118 | 56.69553 | 56.52696 | 24.54089 | 0.385456 | 1.26927 | 0.583335 |
| X | 146516200 | 146517199 | 15.15451 | 27.65428 | 28.42113 | 12.19258 | 0.381567 | 0.314842 | 0.583345 |
| 12 | 23872600 | 23892399 | 571.1246 | 769.9716 | 710.9974 | 296.5027 | 0.229987 | 5.149558 | 0.583347 |
| 15 | 77752200 | 77754399 | 13.27691 | 23.40354 | 38.2627 | 16.93444 | -0.31647 | 0.555426 | 0.583355 |
| 16 | 29826800 | 29833999 | 52.2002 | 82.60512 | 147.3758 | 61.2604 | -0.50615 | 2.383084 | 0.583357 |
| 8 | 31914800 | 31917399 | 18.74567 | 31.85976 | 59.15359 | 25.9247 | -0.50781 | 1.023844 | 0.583357 |
| 9 | 78427400 | 78429199 | 11.58741 | 20.56972 | 35.30965 | 15.59266 | -0.38163 | 0.470153 | 0.583358 |
| 13 | 91468400 | 91477999 | 92.72301 | 144.4501 | 154.3981 | 64.08325 | 0.214323 | 2.713729 | 0.583377 |

| | | | | | | | | | |
|---|---|---|---|---|---|---|---|---|---|
| 3 | 11792000 | 11794199 | 31.84875 | 53.11794 | 54.76254 | 23.80935 | 0.337254 | 1.236289 | 0.583387 |
| 4 | 80164600 | 80165999 | 16.4478 | 28.31476 | 53.24189 | 23.35494 | -0.52272 | 0.891658 | 0.583425 |
| 7 | 4200800 | 4203799 | 23.55699 | 39.00914 | 64.45392 | 27.88842 | -0.34146 | 1.303907 | 0.58343 |
| 6 | 48649200 | 48650399 | 10.46962 | 19.26689 | 31.91439 | 13.53823 | -0.29138 | 0.238362 | 0.583432 |
| X | 148929400 | 148930799 | 22.70575 | 39.59396 | 39.19682 | 17.24904 | 0.401974 | 0.763173 | 0.583443 |
| 8 | 117048000 | 117050199 | 18.43883 | 32.30766 | 35.06302 | 15.48551 | 0.270071 | 0.661186 | 0.583451 |
| 10 | 82494800 | 82495399 | 9.370883 | 18.00061 | 20.82119 | 8.146802 | 0.321119 | -0.08557 | 0.583459 |
| 11 | 85125200 | 85126399 | 6.167389 | 11.76796 | 25.08949 | 9.99498 | -0.57683 | -0.04551 | 0.583465 |
| X | 54954000 | 54954999 | 18.58678 | 32.80714 | 33.79353 | 14.95247 | 0.347314 | 0.565792 | 0.583479 |
| 5 | 21849000 | 21851399 | 15.84968 | 27.47501 | 45.83096 | 20.10833 | -0.34851 | 0.81501 | 0.583498 |
| 17 | 80647000 | 80652999 | 52.13538 | 82.92145 | 127.2532 | 53.1324 | -0.28236 | 2.235126 | 0.583501 |
| X | 78954400 | 78966199 | 163.8282 | 247.7497 | 227.6491 | 94.14885 | 0.398643 | 3.311081 | 0.583507 |
| 13 | 61524800 | 61545799 | 297.4046 | 428.8823 | 392.9984 | 162.156 | 0.338791 | 4.127379 | 0.583511 |
| 18 | 37719800 | 37722599 | 41.93067 | 68.13434 | 71.48167 | 30.66375 | 0.299309 | 1.598541 | 0.583513 |
| X | 90917000 | 90918599 | 26.953 | 45.77675 | 47.95379 | 21.0137 | 0.314746 | 1.048629 | 0.583525 |
| 11 | 50717200 | 50720399 | 19.58508 | 33.34066 | 53.81319 | 23.58731 | -0.30622 | 1.00405 | 0.583533 |
| 16 | 73453600 | 73454399 | 11.83087 | 20.721 | 44.14768 | 19.49097 | -0.6979 | 0.649189 | 0.583533 |
| 7 | 17428000 | 17429399 | 52.03682 | 83.35307 | 89.9989 | 37.95469 | 0.23518 | 2.00427 | 0.583543 |
| 7 | 40801200 | 40809399 | 140.1776 | 213.1294 | 194.4609 | 80.55628 | 0.415251 | 3.193089 | 0.583586 |
| X | 114694000 | 114694599 | 8.681842 | 16.85131 | 19.4336 | 7.305087 | 0.380414 | -0.22741 | 0.583587 |
| 6 | 5688000 | 5689799 | 12.52201 | 22.19534 | 36.23286 | 16.01834 | -0.31092 | 0.488064 | 0.583594 |
| 6 | 69295200 | 69299799 | 55.28665 | 88.92463 | 87.56576 | 37.15736 | 0.374924 | 1.881765 | 0.583597 |
| 1 | 17308600 | 17311199 | 22.63962 | 39.33161 | 40.84879 | 17.9339 | 0.332503 | 0.814361 | 0.5836 |
| 12 | 114671800 | 114672999 | 15.917 | 27.92715 | 49.40078 | 21.84295 | -0.43155 | 0.648705 | 0.5836 |
| 2 | 150946600 | 150953799 | 84.66805 | 133.1319 | 138.8306 | 57.59228 | 0.260247 | 2.540518 | 0.583633 |
| 3 | 142459000 | 142461399 | 20.44396 | 35.71189 | 37.55787 | 16.54586 | 0.315115 | 0.722019 | 0.583656 |
| 2 | 139907600 | 139908799 | 9.223302 | 17.10815 | 29.42281 | 12.34765 | -0.33355 | 0.178961 | 0.58366 |
| X | 95900000 | 95903399 | 39.50418 | 64.19814 | 68.1523 | 29.23778 | 0.282486 | 1.591029 | 0.583672 |
| 3 | 27038000 | 27042999 | 50.35617 | 80.33359 | 118.6022 | 49.72746 | -0.22192 | 2.135589 | 0.583686 |
| 5 | 52523400 | 52528399 | 57.57675 | 90.95345 | 139.6258 | 57.98734 | -0.29168 | 2.426735 | 0.583689 |
| 11 | 62809000 | 62810199 | 15.37586 | 27.40565 | 29.40623 | 12.94146 | 0.295547 | 0.45582 | 0.583701 |
| 7 | 72681800 | 72682999 | 7.803849 | 14.84238 | 26.62842 | 10.68088 | -0.33939 | -0.0028 | 0.583702 |
| 4 | 23079400 | 23080599 | 11.18202 | 20.92622 | 22.60067 | 9.350124 | 0.348824 | 0.121879 | 0.583722 |
| 8 | 70386400 | 70389599 | 15.37349 | 26.73603 | 44.23285 | 19.43231 | -0.33595 | 0.771212 | 0.583735 |
| 17 | 21314000 | 21314999 | 16.16915 | 28.58101 | 28.58878 | 12.6459 | 0.390278 | 0.525139 | 0.583739 |
| 6 | 131913800 | 131921599 | 86.73669 | 137.4938 | 119.0289 | 49.52968 | 0.537665 | 2.361646 | 0.583758 |
| 5 | 142030400 | 142039999 | 184.5108 | 275.3312 | 276.1342 | 113.818 | 0.255441 | 3.569941 | 0.583765 |
| 15 | 49818800 | 49819399 | 11.80852 | 21.92647 | 23.53755 | 9.863712 | 0.342827 | 0.17808 | 0.583779 |
| 11 | 86765600 | 86769599 | 24.309 | 40.46846 | 63.03924 | 27.37605 | -0.25617 | 1.243836 | 0.583802 |
| 1 | 126873400 | 126878399 | 41.2914 | 66.94491 | 72.50301 | 31.05037 | 0.250502 | 1.640027 | 0.583806 |
| 7 | 27435200 | 27437599 | 13.01985 | 22.80623 | 40.01918 | 17.67011 | -0.41949 | 0.64547 | 0.583814 |
| 16 | 65067600 | 65072399 | 54.31546 | 87.59898 | 80.72751 | 34.38601 | 0.475033 | 1.794653 | 0.583822 |
| 13 | 14117200 | 14121599 | 67.16645 | 107.2019 | 104.6907 | 43.92389 | 0.374095 | 2.135433 | 0.583822 |
| 5 | 111338400 | 111340399 | 12.82898 | 22.64591 | 37.28221 | 16.49584 | -0.32549 | 0.536472 | 0.583833 |
| 1 | 43724800 | 43730799 | 104.7318 | 159.6197 | 228.3205 | 94.65077 | -0.23005 | 3.136929 | 0.583848 |
| X | 129221000 | 129221799 | 8.134155 | 15.57403 | 29.71874 | 11.73772 | -0.40791 | -0.04188 | 0.583853 |
| 16 | 17805800 | 17813599 | 237.4947 | 345.2143 | 347.7719 | 143.3534 | 0.213344 | 4.004065 | 0.583898 |

| | | | | | | | | | |
|---|---|---|---|---|---|---|---|---|---|
| 7 | 32080600 | 32085399 | 48.49943 | 78.95375 | 68.51155 | 29.41298 | 0.574448 | 1.581455 | 0.583906 |
| 6 | 146505200 | 146507799 | 22.13405 | 37.14032 | 58.65276 | 25.59856 | -0.27596 | 1.147743 | 0.583917 |
| 8 | 45665200 | 45665799 | 9.243564 | 17.79256 | 20.3392 | 7.911401 | 0.346152 | -0.10657 | 0.58392 |
| X | 80927400 | 80929199 | 19.19011 | 33.15549 | 57.82003 | 25.36598 | -0.41366 | 0.87583 | 0.583922 |
| 5 | 98789000 | 98789799 | 13.93501 | 25.28715 | 27.00684 | 11.66714 | 0.320006 | 0.334206 | 0.583945 |
| 3 | 65977800 | 65979399 | 11.32608 | 20.61235 | 33.84848 | 14.57001 | -0.29356 | 0.313296 | 0.583973 |
| 17 | 38170400 | 38173399 | 52.19352 | 83.58409 | 89.93119 | 37.91235 | 0.239842 | 2.011758 | 0.58398 |
| 6 | 7019000 | 7021599 | 40.48282 | 66.06821 | 68.40461 | 29.39384 | 0.322771 | 1.528743 | 0.583988 |
| Y | 1888200 | 1891199 | 48.0696 | 77.27825 | 62.82062 | 26.63797 | 0.649985 | 1.87233 | 0.583996 |
| 15 | 95826800 | 95829599 | 21.8985 | 36.9975 | 58.20418 | 25.46492 | -0.27021 | 1.084002 | 0.583998 |
| X | 5351000 | 5358399 | 80.81898 | 128.5665 | 110.0087 | 45.94393 | 0.559432 | 2.253614 | 0.584001 |
| 1 | 188207400 | 188209399 | 25.15271 | 43.17977 | 44.08726 | 19.33733 | 0.35367 | 0.937813 | 0.584051 |
| 18 | 83093600 | 83095799 | 12.61819 | 22.34188 | 37.18453 | 16.44259 | -0.33958 | 0.499553 | 0.584053 |
| 17 | 9302600 | 9306199 | 55.93246 | 89.44531 | 97.05514 | 40.83253 | 0.225383 | 2.060612 | 0.584062 |
| 15 | 9268000 | 9269199 | 28.98963 | 48.19781 | 50.2567 | 21.80561 | 0.320978 | 1.259937 | 0.584069 |
| 16 | 77820800 | 77823999 | 41.58787 | 67.29835 | 73.11821 | 31.27026 | 0.243444 | 1.680976 | 0.584077 |
| 2 | 32350000 | 32351399 | 11.81189 | 21.42051 | 35.36126 | 15.28873 | -0.30537 | 0.340062 | 0.584092 |
| 8 | 35580200 | 35581799 | 11.81189 | 21.42051 | 35.36126 | 15.28873 | -0.30537 | 0.340062 | 0.584092 |
| 9 | 121308600 | 121310199 | 17.19408 | 29.70072 | 47.41362 | 20.80049 | -0.28641 | 0.856865 | 0.584099 |
| 2 | 148573600 | 148574599 | 10.98126 | 19.8448 | 33.23995 | 14.43444 | -0.33028 | 0.348603 | 0.584103 |
| 12 | 46677000 | 46678199 | 10.98126 | 19.8448 | 33.23995 | 14.43444 | -0.33028 | 0.348603 | 0.584103 |
| 4 | 14220200 | 14223999 | 35.72271 | 59.13047 | 58.62223 | 25.35787 | 0.393173 | 1.339509 | 0.584104 |
| X | 8627800 | 8648599 | 327.0063 | 467.3016 | 424.735 | 175.4934 | 0.337462 | 4.262352 | 0.58411 |
| 1 | 56787200 | 56789399 | 14.39383 | 25.22058 | 40.93883 | 18.08078 | -0.30787 | 0.648697 | 0.584124 |
| 12 | 108469400 | 108471199 | 17.53942 | 29.99766 | 50.05162 | 21.94678 | -0.35267 | 0.950727 | 0.584166 |
| 13 | 105116400 | 105117999 | 17.622 | 32.02257 | 29.09751 | 12.54832 | 0.553024 | 0.355549 | 0.584166 |
| 15 | 51853800 | 51855799 | 15.28647 | 26.49907 | 50.58364 | 22.19359 | -0.54218 | 0.819815 | 0.584172 |
| 6 | 59674600 | 59675199 | 11.2819 | 21.14773 | 23.16487 | 9.557006 | 0.332045 | 0.115326 | 0.584181 |
| 11 | 63758400 | 63759799 | 14.60325 | 26.14504 | 26.00028 | 11.39954 | 0.408205 | 0.414542 | 0.584187 |
| 3 | 134044800 | 134045399 | 10.89098 | 20.79019 | 23.3183 | 9.280719 | 0.341959 | -0.01026 | 0.584218 |
| 3 | 155459800 | 155462599 | 38.33115 | 62.28793 | 67.76576 | 29.0706 | 0.247048 | 1.586176 | 0.584218 |
| 10 | 21824000 | 21827399 | 29.75758 | 48.79057 | 75.48585 | 32.49477 | -0.25118 | 1.446959 | 0.58422 |
| 13 | 75974400 | 75978599 | 35.8861 | 58.04588 | 88.78745 | 37.95908 | -0.24957 | 1.685898 | 0.584221 |
| 9 | 39790800 | 39793399 | 19.43343 | 34.57974 | 31.91028 | 14.06335 | 0.510351 | 0.496134 | 0.584226 |
| 15 | 94706800 | 94708799 | 17.11428 | 29.72226 | 46.86088 | 20.56895 | -0.26712 | 0.802518 | 0.58425 |
| X | 158813600 | 158815199 | 14.15733 | 24.9665 | 40.31361 | 17.84056 | -0.29831 | 0.561674 | 0.584257 |
| 7 | 85916600 | 85917599 | 16.18456 | 28.2664 | 44.64767 | 19.667 | -0.26884 | 0.716051 | 0.584264 |
| 14 | 46835200 | 46837399 | 17.02415 | 30.34509 | 31.69916 | 13.94996 | 0.333947 | 0.470515 | 0.584265 |
| 3 | 95045000 | 95045999 | 7.610012 | 14.06183 | 29.4626 | 12.42883 | -0.62161 | 0.186265 | 0.584274 |
| 4 | 38577200 | 38578799 | 19.3295 | 34.33142 | 32.48338 | 14.3382 | 0.47301 | 0.508383 | 0.584283 |
| 15 | 73951400 | 73953799 | 20.87062 | 35.26706 | 55.89558 | 24.45618 | -0.28069 | 1.07971 | 0.584288 |
| 15 | 25043400 | 25045199 | 24.62738 | 42.16852 | 44.73692 | 19.61818 | 0.298241 | 0.963468 | 0.584294 |
| 11 | 110685200 | 110687799 | 29.0408 | 48.60334 | 52.93406 | 23.06488 | 0.258343 | 1.191966 | 0.584306 |
| 18 | 21673200 | 21673799 | 10.8063 | 21.08573 | 17.41002 | 6.205164 | 0.925995 | -0.39189 | 0.584312 |
| 2 | 28250800 | 28252599 | 12.1884 | 21.57134 | 36.25903 | 16.03476 | -0.3538 | 0.501505 | 0.584322 |
| 14 | 11179600 | 11182399 | 24.4439 | 42.22543 | 37.77497 | 16.57201 | 0.545472 | 0.874351 | 0.584343 |
| 8 | 121549800 | 121551599 | 19.49648 | 33.16653 | 52.70229 | 23.09907 | -0.28378 | 1.011172 | 0.584373 |

| | | | | | | | | | |
|---|---|---|---|---|---|---|---|---|---|
| 4 | 112194200 | 112202399 | 130.6216 | 201.1589 | 176.9283 | 73.488 | 0.482543 | 2.937369 | 0.58438 |
| X | 164511400 | 164512599 | 18.20903 | 31.55921 | 51.71264 | 22.6888 | -0.32303 | 0.837264 | 0.584392 |
| X | 54452000 | 54457799 | 68.90102 | 110.3153 | 94.95649 | 40.0173 | 0.560815 | 2.043021 | 0.584393 |
| 1 | 34220400 | 34222599 | 16.87621 | 28.95747 | 49.00027 | 21.49141 | -0.37214 | 0.916415 | 0.584409 |
| 9 | 55758400 | 55759199 | 5.799316 | 11.30075 | 22.05375 | 8.003698 | -0.32483 | -0.37877 | 0.584437 |
| 10 | 124856200 | 124860399 | 39.54842 | 63.84054 | 102.8267 | 43.85021 | -0.32732 | 1.756269 | 0.584439 |
| 6 | 78914800 | 78917199 | 27.75412 | 46.95297 | 42.70173 | 18.68998 | 0.51773 | 1.077839 | 0.584456 |
| X | 49234800 | 49237599 | 16.69321 | 29.16552 | 54.97458 | 24.22155 | -0.52278 | 0.730103 | 0.584463 |
| 13 | 21703200 | 21706599 | 53.01891 | 85.49087 | 73.00249 | 31.08752 | 0.584436 | 1.796043 | 0.584472 |
| 13 | 58788600 | 58791799 | 20.43657 | 34.64691 | 54.87865 | 24.0351 | -0.27971 | 1.045931 | 0.584487 |
| 1 | 89090200 | 89092399 | 18.45735 | 31.67214 | 50.06445 | 21.95825 | -0.27433 | 0.924025 | 0.584506 |
| 17 | 18963000 | 18966799 | 48.68425 | 78.96818 | 64.10474 | 27.46329 | 0.666037 | 1.64085 | 0.584507 |
| 4 | 112117800 | 112119799 | 23.0948 | 40.48404 | 35.54304 | 15.70355 | 0.574315 | 0.678404 | 0.584526 |
| 11 | 31638600 | 31639599 | 7.625672 | 14.4485 | 26.55654 | 10.73799 | -0.38469 | 0.019003 | 0.584528 |
| 9 | 38996000 | 38999399 | 63.87554 | 101.9927 | 88.09356 | 36.98269 | 0.55162 | 2.115645 | 0.584565 |
| 3 | 79694600 | 79696399 | 15.03151 | 26.31008 | 41.98702 | 18.52716 | -0.28348 | 0.671033 | 0.584586 |
| 8 | 72704400 | 72705999 | 14.07465 | 24.65194 | 40.96168 | 18.08506 | -0.34141 | 0.652832 | 0.584596 |
| 1 | 111200400 | 111202799 | 29.23141 | 50.11575 | 45.12923 | 19.79225 | 0.534055 | 0.969971 | 0.584614 |
| 1 | 80723200 | 80724199 | 11.76692 | 22.3787 | 23.72971 | 9.519228 | 0.412141 | 0.023227 | 0.584632 |
| 6 | 53815800 | 53817999 | 16.48608 | 28.79964 | 46.23012 | 20.36701 | -0.29209 | 0.715812 | 0.584636 |
| 1 | 91099200 | 91100399 | 14.28068 | 25.94021 | 27.95949 | 12.06368 | 0.30797 | 0.333163 | 0.584643 |
| 9 | 37941600 | 37944199 | 31.19691 | 51.69537 | 56.8222 | 24.59774 | 0.244779 | 1.310182 | 0.584643 |
| 5 | 31543400 | 31543999 | 7.597166 | 14.62316 | 26.47296 | 10.29791 | -0.31205 | -0.10728 | 0.584646 |
| 8 | 81754600 | 81762399 | 82.76568 | 130.3577 | 122.3891 | 50.7827 | 0.413358 | 2.503745 | 0.584653 |
| 11 | 67314400 | 67316799 | 17.06081 | 30.3118 | 31.59909 | 13.94406 | 0.334062 | 0.488945 | 0.584662 |
| 1 | 13527200 | 13533599 | 88.68212 | 138.8007 | 128.8355 | 53.44505 | 0.422854 | 2.626533 | 0.584685 |
| 2 | 26723600 | 26735399 | 179.8911 | 264.7594 | 362.3108 | 149.2013 | -0.21111 | 3.807279 | 0.58469 |
| X | 8142800 | 8146999 | 58.23865 | 93.61959 | 84.71389 | 35.91767 | 0.495724 | 1.905903 | 0.584712 |
| X | 42298000 | 42298799 | 8.435361 | 15.98247 | 28.18875 | 11.39821 | -0.32546 | 0.03348 | 0.584725 |
| 14 | 3534200 | 3535199 | 14.40215 | 26.15125 | 28.41289 | 12.26493 | 0.295991 | 0.336407 | 0.584726 |
| 6 | 39814200 | 39816599 | 31.14966 | 52.49112 | 51.30793 | 22.4301 | 0.413879 | 1.131289 | 0.584736 |
| 14 | 94971800 | 94974199 | 24.97455 | 42.4755 | 46.35037 | 20.31468 | 0.25621 | 1.025034 | 0.584747 |
| X | 108919000 | 108920599 | 19.03727 | 34.14483 | 30.9882 | 13.56586 | 0.540886 | 0.444524 | 0.584788 |
| 13 | 85746200 | 85749599 | 38.55652 | 62.55206 | 65.74571 | 28.17013 | 0.294555 | 1.619529 | 0.584791 |
| 7 | 97408200 | 97409999 | 15.27502 | 26.66301 | 43.13676 | 18.99479 | -0.30313 | 0.718804 | 0.584791 |
| 18 | 37013200 | 37016399 | 56.65214 | 90.59785 | 98.28551 | 41.35123 | 0.225715 | 2.061999 | 0.584794 |
| 6 | 144201800 | 144202799 | 8.988525 | 16.90307 | 29.11081 | 11.94492 | -0.31011 | 0.088078 | 0.584802 |
| 12 | 101992000 | 101992999 | 8.988525 | 16.90307 | 29.11081 | 11.94492 | -0.31011 | 0.088078 | 0.584802 |
| 10 | 15566600 | 15568199 | 16.38126 | 29.39109 | 30.73959 | 13.45316 | 0.337667 | 0.424342 | 0.584815 |
| 13 | 64811800 | 64812999 | 22.26479 | 38.45745 | 41.5508 | 18.22842 | 0.274335 | 0.871364 | 0.58483 |
| 8 | 110786200 | 110789399 | 45.26992 | 72.47048 | 108.0715 | 45.5352 | -0.23083 | 2.020914 | 0.584843 |
| 11 | 85671400 | 85674399 | 24.97889 | 41.32537 | 66.30193 | 28.66966 | -0.29935 | 1.319032 | 0.584847 |
| 6 | 81522800 | 81523199 | 4.347197 | 8.502036 | 17.87009 | 6.063485 | -0.34105 | -0.64973 | 0.584862 |
| 7 | 92565800 | 92566599 | 8.163009 | 15.17742 | 28.45336 | 11.90172 | -0.45333 | 0.154297 | 0.584868 |
| 7 | 115060600 | 115065399 | 57.26604 | 91.42286 | 96.84446 | 40.64621 | 0.257382 | 2.116133 | 0.584885 |
| X | 54437000 | 54448999 | 137.0031 | 210.7558 | 176.9966 | 73.50536 | 0.547971 | 2.961405 | 0.584907 |
| 10 | 52576400 | 52577399 | 11.84011 | 21.15916 | 34.72444 | 15.24813 | -0.31169 | 0.427222 | 0.584915 |

| | | | | | | | | | |
|---|---|---|---|---|---|---|---|---|---|
| 1 | 93439400 | 93440999 | 13.32658 | 23.32534 | 40.53375 | 17.88554 | -0.40558 | 0.660418 | 0.584923 |
| 2 | 82876000 | 82878399 | 22.86202 | 40.04897 | 36.72762 | 16.21767 | 0.511834 | 0.690392 | 0.584929 |
| Y | 2069600 | 2074599 | 29.22889 | 48.81357 | 36.38204 | 15.83131 | 0.803586 | 1.195077 | 0.58494 |
| 8 | 29030400 | 29031999 | 18.7246 | 34.20106 | 28.04028 | 12.01511 | 0.706471 | 0.333622 | 0.584952 |
| 10 | 20631600 | 20632999 | 18.83771 | 32.99549 | 35.60618 | 15.72094 | 0.278326 | 0.670624 | 0.584957 |
| 11 | 104807200 | 104809599 | 17.57602 | 31.80278 | 29.54978 | 12.81167 | 0.516047 | 0.379658 | 0.584983 |
| 16 | 47377200 | 47378599 | 17.83828 | 30.28879 | 58.49167 | 25.62952 | -0.56426 | 1.030289 | 0.584986 |
| 7 | 40613200 | 40614399 | 19.83418 | 34.67972 | 37.05783 | 16.33753 | 0.292309 | 0.704151 | 0.584991 |
| 12 | 109154200 | 109156599 | 21.56313 | 36.58649 | 57.67107 | 25.26431 | -0.27279 | 1.045753 | 0.584992 |
| 7 | 73884800 | 73885599 | 8.609588 | 16.06163 | 28.46546 | 11.83921 | -0.36624 | 0.137073 | 0.585001 |
| 16 | 92807200 | 92808199 | 8.609588 | 16.06163 | 28.46546 | 11.83921 | -0.36624 | 0.137073 | 0.585001 |
| 19 | 24046600 | 24047799 | 15.03096 | 27.14455 | 29.32822 | 12.74672 | 0.297582 | 0.373429 | 0.585004 |
| 17 | 18713400 | 18714399 | 17.57947 | 31.24847 | 32.20657 | 14.20632 | 0.351036 | 0.492811 | 0.585013 |
| 16 | 86737600 | 86739199 | 14.95019 | 27.95265 | 25.98415 | 10.76644 | 0.561786 | 0.167677 | 0.585021 |
| 16 | 5451000 | 5456599 | 60.43562 | 96.27505 | 104.8715 | 43.88315 | 0.213808 | 2.187595 | 0.585061 |
| 4 | 37364400 | 37368199 | 77.82001 | 122.6046 | 132.0911 | 54.81049 | 0.215423 | 2.495982 | 0.585061 |
| 6 | 142017000 | 142020999 | 44.12411 | 70.76821 | 105.1043 | 44.41065 | -0.22177 | 1.960785 | 0.585078 |
| 17 | 29152400 | 29162599 | 112.3533 | 170.3119 | 244.9414 | 101.3766 | -0.24427 | 3.250362 | 0.585086 |
| 1 | 24755200 | 24756399 | 20.2152 | 35.42774 | 31.42333 | 13.88057 | 0.559818 | 0.662089 | 0.585106 |
| X | 78219800 | 78225199 | 117.8905 | 181.3777 | 182.1643 | 75.65281 | 0.290243 | 2.951706 | 0.585117 |
| X | 61239600 | 61241599 | 37.16726 | 61.07429 | 63.0623 | 27.17214 | 0.331946 | 1.431546 | 0.58512 |
| 7 | 22249200 | 22249599 | 9.30479 | 18.16794 | 17.58218 | 6.195468 | 0.71384 | -0.45043 | 0.585126 |
| 19 | 26307200 | 26308799 | 14.75002 | 27.12178 | 27.95413 | 11.87146 | 0.387491 | 0.267558 | 0.585133 |
| 17 | 38715800 | 38720599 | 24.45858 | 42.70927 | 33.15461 | 14.60272 | 0.751175 | 0.742533 | 0.585149 |
| 1 | 106017800 | 106018999 | 11.24966 | 20.39437 | 34.27919 | 14.82685 | -0.33177 | 0.333149 | 0.585153 |
| 1 | 130527600 | 130529399 | 36.01029 | 57.93652 | 93.43261 | 39.66329 | -0.33469 | 1.848721 | 0.585171 |
| 8 | 34310000 | 34312599 | 19.3704 | 33.32371 | 54.46441 | 23.89072 | -0.32182 | 0.908317 | 0.585219 |
| 16 | 28926400 | 28927199 | 13.01749 | 23.972 | 26.49069 | 11.22705 | 0.289633 | 0.241602 | 0.585227 |
| 14 | 118932400 | 118934599 | 19.52186 | 32.99235 | 54.82004 | 23.98657 | -0.34864 | 1.070172 | 0.585267 |
| 7 | 42588000 | 42604599 | 320.7729 | 458.1755 | 431.6508 | 178.3647 | 0.285035 | 4.268323 | 0.585274 |
| 10 | 45147200 | 45149399 | 18.12533 | 31.9783 | 33.75065 | 14.93484 | 0.312023 | 0.566076 | 0.585289 |
| 14 | 64405400 | 64407599 | 18.12533 | 31.9783 | 33.75065 | 14.93484 | 0.312023 | 0.566076 | 0.585289 |
| 10 | 55608400 | 55610399 | 22.72757 | 38.16176 | 60.79131 | 26.54312 | -0.28839 | 1.147599 | 0.5853 |
| 15 | 16110800 | 16113999 | 31.89293 | 53.59125 | 51.89998 | 22.6694 | 0.42704 | 1.157231 | 0.585308 |
| 5 | 54292200 | 54294999 | 28.7562 | 46.88378 | 84.43119 | 36.26744 | -0.47468 | 1.529777 | 0.585326 |
| 16 | 46054600 | 46055999 | 14.04663 | 26.42268 | 25.64335 | 10.51689 | 0.511956 | 0.122502 | 0.585332 |
| 8 | 6801400 | 6801799 | 5.520437 | 10.74156 | 21.25591 | 7.833897 | -0.36464 | -0.33008 | 0.585343 |
| 7 | 58782600 | 58788199 | 76.47958 | 120.4532 | 129.7328 | 53.82961 | 0.215525 | 2.502809 | 0.585366 |
| 7 | 131774200 | 131776799 | 20.93931 | 35.47781 | 55.42524 | 24.27005 | -0.25993 | 1.054072 | 0.585369 |
| 13 | 27862000 | 27865199 | 98.56978 | 152.6861 | 163.7519 | 68.01806 | 0.20311 | 2.815524 | 0.585369 |
| 18 | 18017600 | 18018999 | 9.192345 | 17.40913 | 32.24982 | 13.05235 | -0.39765 | 0.05098 | 0.58537 |
| 3 | 41089600 | 41093399 | 34.33338 | 56.80009 | 59.82636 | 25.86909 | 0.305941 | 1.339397 | 0.585391 |
| 17 | 8965000 | 8967799 | 19.02068 | 32.26856 | 53.21871 | 23.31294 | -0.33764 | 1.033663 | 0.585409 |
| 8 | 14065400 | 14070199 | 41.1591 | 66.23508 | 99.38809 | 42.20504 | -0.23066 | 1.849831 | 0.585409 |
| 5 | 105053800 | 105055199 | 17.7427 | 31.40943 | 32.61927 | 14.42442 | 0.336969 | 0.524561 | 0.58541 |
| 14 | 14880200 | 14882799 | 19.53476 | 34.33184 | 30.73879 | 13.61055 | 0.54614 | 0.611851 | 0.585431 |
| 8 | 37849400 | 37852599 | 57.74531 | 92.15117 | 99.96586 | 41.93254 | 0.222461 | 2.127175 | 0.585434 |

| | | | | | | | | | |
|---|---|---|---|---|---|---|---|---|---|
| 7 | 25968800 | 25970999 | 33.5606 | 55.84534 | 56.98398 | 24.73913 | 0.352198 | 1.270293 | 0.585444 |
| 17 | 94648400 | 94653399 | 155.0034 | 232.9525 | 244.1778 | 100.8026 | 0.201018 | 3.430007 | 0.585451 |
| 16 | 3460000 | 3480999 | 373.887 | 522.4655 | 501.3794 | 207.9195 | 0.226649 | 4.59091 | 0.585452 |
| 19 | 15989400 | 15991399 | 29.64165 | 48.59902 | 74.73955 | 32.1731 | -0.24252 | 1.448055 | 0.585454 |
| 16 | 25956600 | 25958999 | 30.19676 | 51.22835 | 49.03941 | 21.48211 | 0.444211 | 1.069774 | 0.58546 |
| 5 | 74062800 | 74065599 | 32.86748 | 53.20078 | 83.20385 | 35.59044 | -0.28046 | 1.666236 | 0.585477 |
| 7 | 19074800 | 19077599 | 16.66682 | 28.75897 | 47.41512 | 20.80047 | -0.33315 | 0.862371 | 0.585479 |
| 6 | 71885600 | 71886799 | 6.881375 | 13.14587 | 25.09159 | 9.970812 | -0.41594 | -0.04756 | 0.585493 |
| 13 | 25341000 | 25341799 | 6.881375 | 13.14587 | 25.09159 | 9.970812 | -0.41594 | -0.04756 | 0.585493 |
| 13 | 15445000 | 15448799 | 34.31685 | 56.83517 | 59.28883 | 25.65612 | 0.319921 | 1.326296 | 0.5855 |
| 2 | 3712800 | 3715399 | 15.56114 | 27.05765 | 44.24339 | 19.43481 | -0.31908 | 0.775687 | 0.585512 |
| 8 | 12488200 | 12490399 | 16.56683 | 28.48514 | 47.95238 | 21.03388 | -0.36406 | 0.89597 | 0.585519 |
| 3 | 68553400 | 68558399 | 43.59213 | 69.96684 | 104.9951 | 44.41458 | -0.23542 | 1.936186 | 0.585535 |
| 17 | 9823800 | 9824399 | 5.804692 | 11.29238 | 21.81828 | 8.054311 | -0.33264 | -0.31794 | 0.585536 |
| 12 | 113855200 | 113858999 | 48.21689 | 78.28835 | 72.29644 | 30.99556 | 0.481903 | 1.626191 | 0.585543 |
| 1 | 97214400 | 97215199 | 12.01645 | 23.1213 | 23.00772 | 8.959019 | 0.542758 | -0.06815 | 0.585546 |
| 3 | 158339400 | 158340999 | 20.32788 | 35.66335 | 31.74332 | 14.0346 | 0.554648 | 0.645401 | 0.585563 |
| X | 148698600 | 148710599 | 117.3215 | 181.7713 | 165.2702 | 68.64866 | 0.441351 | 2.820921 | 0.585567 |
| 7 | 83531200 | 83533999 | 23.17913 | 37.78946 | 82.16867 | 35.29505 | -0.74567 | 1.517761 | 0.585569 |
| 19 | 32390200 | 32391799 | 11.5405 | 20.56349 | 34.7543 | 15.29893 | -0.35631 | 0.443616 | 0.58557 |
| 4 | 102914200 | 102916999 | 20.27688 | 34.57259 | 53.82102 | 23.59372 | -0.25387 | 0.994129 | 0.585591 |
| 13 | 45470800 | 45471799 | 13.12035 | 23.26418 | 37.16476 | 16.42779 | -0.27985 | 0.489068 | 0.585612 |
| 3 | 142392400 | 142396199 | 69.99871 | 109.4048 | 162.6116 | 67.4662 | -0.25712 | 2.647008 | 0.585633 |
| X | 80406600 | 80407999 | 19.77397 | 35.04069 | 33.0491 | 14.61155 | 0.475944 | 0.529469 | 0.585659 |
| 7 | 28404400 | 28408399 | 43.96413 | 70.4556 | 105.2092 | 44.4013 | -0.23094 | 1.985289 | 0.585686 |
| Y | 40489000 | 40490799 | 11.90221 | 22.16046 | 18.15437 | 7.575922 | 0.734489 | 0.147065 | 0.585686 |
| 13 | 79457400 | 79458199 | 8.549051 | 16.05956 | 28.00808 | 11.51313 | -0.32994 | 0.088968 | 0.585698 |
| 2 | 67774600 | 67777999 | 24.28208 | 42.36912 | 38.41669 | 16.91235 | 0.528046 | 0.758483 | 0.5857 |
| X | 153506400 | 153508999 | 36.96084 | 61.33907 | 58.25715 | 25.2452 | 0.455135 | 1.31183 | 0.585709 |
| 13 | 68230200 | 68241999 | 289.5851 | 414.9624 | 414.0724 | 171.0143 | 0.206766 | 4.218957 | 0.585717 |
| 5 | 109754200 | 109755999 | 20.37719 | 35.56842 | 38.17149 | 16.80822 | 0.286051 | 0.734341 | 0.585725 |
| 6 | 131208800 | 131209399 | 11.34976 | 21.50006 | 23.75958 | 9.611345 | 0.343131 | 0.049423 | 0.585734 |
| 8 | 74117000 | 74119199 | 23.30149 | 39.24558 | 62.64171 | 27.38054 | -0.29122 | 1.121655 | 0.585755 |
| 9 | 25898000 | 25904599 | 99.20924 | 154.4323 | 158.2451 | 65.68777 | 0.274268 | 2.731063 | 0.58576 |
| 13 | 41373400 | 41378999 | 92.31893 | 142.0602 | 202.2624 | 84.00155 | -0.21257 | 2.939557 | 0.585765 |
| 14 | 81413400 | 81417599 | 43.64862 | 71.41476 | 66.01795 | 28.39438 | 0.488404 | 1.499294 | 0.585793 |
| 7 | 16612400 | 16623999 | 141.0542 | 208.5815 | 364.4307 | 150.0819 | -0.557 | 3.725535 | 0.5858 |
| 11 | 97962000 | 97964599 | 56.3302 | 89.42137 | 130.7531 | 54.48603 | -0.21545 | 2.297823 | 0.585801 |
| 16 | 78710000 | 78712399 | 22.3556 | 39.37455 | 34.47001 | 15.25835 | 0.579563 | 0.611049 | 0.585816 |
| 3 | 68605200 | 68606799 | 13.4245 | 23.7037 | 38.2202 | 16.90992 | -0.29567 | 0.537533 | 0.585818 |
| 4 | 8865600 | 8866599 | 11.44516 | 20.31933 | 35.34022 | 15.60478 | -0.40025 | 0.46934 | 0.585848 |
| X | 151671400 | 151674599 | 53.1452 | 85.60987 | 73.65237 | 31.32211 | 0.572035 | 1.827445 | 0.585868 |
| 17 | 89786800 | 89788999 | 33.65489 | 54.56387 | 83.17349 | 35.62008 | -0.24137 | 1.6344 | 0.585871 |
| 2 | 176740200 | 176741399 | 16.00804 | 28.98706 | 29.12731 | 12.61682 | 0.404483 | 0.36267 | 0.585884 |
| 1 | 45825000 | 45825599 | 5.811123 | 11.29671 | 22.96696 | 8.528059 | -0.41438 | -0.29781 | 0.585895 |
| X | 82912200 | 82920599 | 115.5649 | 178.8314 | 168.8633 | 70.14158 | 0.38551 | 2.843046 | 0.585901 |
| 9 | 23442600 | 23442999 | 5.944975 | 11.57884 | 22.59975 | 8.255342 | -0.33397 | -0.35336 | 0.585976 |

| | | | | | | | | | |
|---|---|---|---|---|---|---|---|---|---|
| 6 | 123199800 | 123204599 | 43.34097 | 70.56501 | 69.31472 | 29.75916 | 0.396088 | 1.567044 | 0.585976 |
| 10 | 81802600 | 81806999 | 74.25278 | 116.963 | 120.3976 | 49.95712 | 0.280915 | 2.496189 | 0.586004 |
| 10 | 45413000 | 45413799 | 13.30866 | 23.37947 | 38.8671 | 17.2033 | -0.34197 | 0.598066 | 0.586006 |
| 8 | 86163800 | 86164799 | 8.566937 | 16.2298 | 28.19143 | 11.40222 | -0.30414 | 0.034596 | 0.586016 |
| 17 | 94649000 | 94653199 | 149.6975 | 225.448 | 236.6679 | 97.75693 | 0.201508 | 3.388164 | 0.586023 |
| 2 | 178421200 | 178425799 | 46.91946 | 74.94008 | 112.685 | 47.32597 | -0.24639 | 2.093812 | 0.586029 |
| 4 | 15254600 | 15256799 | 18.78425 | 33.11996 | 34.2162 | 15.14275 | 0.342361 | 0.577563 | 0.586032 |
| 9 | 103862800 | 103865199 | 21.88758 | 36.59844 | 69.18694 | 30.12952 | -0.53466 | 1.193692 | 0.586049 |
| 15 | 79921800 | 79923399 | 16.27828 | 28.456 | 46.75178 | 20.60742 | -0.3255 | 0.703779 | 0.586051 |
| X | 55109800 | 55111799 | 33.77759 | 56.40276 | 55.29674 | 24.05988 | 0.409664 | 1.233542 | 0.586054 |
| 6 | 85802200 | 85804599 | 21.53624 | 37.48753 | 39.79148 | 17.49107 | 0.301479 | 0.781157 | 0.586093 |
| 13 | 59345400 | 59346399 | 11.10724 | 20.27166 | 33.34923 | 14.30414 | -0.29297 | 0.292746 | 0.586113 |
| 7 | 15522800 | 15571199 | 1090.104 | 1397.932 | 1246.216 | 522.7552 | 0.208618 | 5.958412 | 0.58612 |
| 12 | 19336600 | 19367199 | 768.4376 | 1013.06 | 928.6599 | 388.4514 | 0.208247 | 5.508682 | 0.586128 |
| 4 | 96249800 | 96251599 | 22.4076 | 37.83052 | 59.31797 | 25.94658 | -0.2655 | 1.093482 | 0.586157 |
| 10 | 83601600 | 83604199 | 32.20019 | 53.23687 | 58.51066 | 25.28898 | 0.244683 | 1.340244 | 0.58616 |
| 7 | 107516000 | 107522199 | 84.27399 | 132.9433 | 133.1544 | 55.26114 | 0.321952 | 2.477079 | 0.58616 |
| 1 | 186716800 | 186718999 | 18.99939 | 31.91548 | 62.82568 | 27.43881 | -0.59267 | 1.130312 | 0.586192 |
| 13 | 98784200 | 98785599 | 15.65533 | 27.34032 | 43.597 | 19.2034 | -0.28252 | 0.711561 | 0.586215 |
| 4 | 99032400 | 99034199 | 14.73676 | 27.04197 | 27.834 | 11.85343 | 0.386283 | 0.276534 | 0.586236 |
| 11 | 18174600 | 18176599 | 17.72812 | 30.41744 | 59.86261 | 26.25548 | -0.58943 | 0.933354 | 0.586251 |
| 19 | 39274800 | 39279399 | 45.33223 | 73.55519 | 73.66987 | 31.56875 | 0.364145 | 1.633688 | 0.586264 |
| 16 | 16793800 | 16797999 | 51.37604 | 81.86807 | 120.3237 | 50.37045 | -0.21717 | 2.173825 | 0.586284 |
| X | 77047800 | 77049599 | 27.06736 | 46.44355 | 44.66978 | 19.59205 | 0.439656 | 0.955654 | 0.586328 |
| X | 153167800 | 153168799 | 17.07054 | 30.23512 | 32.29673 | 14.28032 | 0.297031 | 0.516311 | 0.586341 |
| 4 | 103455000 | 103455799 | 11.47406 | 20.71197 | 33.73334 | 14.66939 | -0.29137 | 0.360523 | 0.586345 |
| 5 | 74345200 | 74346599 | 11.47406 | 20.71197 | 33.73334 | 14.66939 | -0.29137 | 0.360523 | 0.586345 |
| 10 | 116630600 | 116632599 | 13.2768 | 23.57015 | 37.67187 | 16.63494 | -0.27957 | 0.483311 | 0.586355 |
| 5 | 39171400 | 39172199 | 14.64072 | 25.64403 | 41.50653 | 18.32599 | -0.30355 | 0.65946 | 0.586367 |
| 5 | 151531200 | 151532799 | 16.52067 | 28.85806 | 46.23436 | 20.36798 | -0.28927 | 0.717789 | 0.586369 |
| 4 | 13672800 | 13677399 | 59.46715 | 95.38911 | 93.18432 | 39.3822 | 0.3817 | 1.971124 | 0.58637 |
| 5 | 111435400 | 111436399 | 7.516854 | 14.21139 | 26.53949 | 10.78055 | -0.4127 | 0.03576 | 0.586392 |
| 3 | 14358400 | 14364199 | 36.71717 | 60.5465 | 54.86218 | 23.67396 | 0.521744 | 1.378957 | 0.586407 |
| 7 | 7471800 | 7473599 | 26.5097 | 46.02182 | 35.43975 | 15.5521 | 0.762184 | 0.836658 | 0.586424 |
| 17 | 29672800 | 29674799 | 14.66026 | 25.59984 | 42.64255 | 18.78175 | -0.34483 | 0.710259 | 0.586438 |
| 1 | 171544800 | 171557599 | 233.3114 | 341.3706 | 341.3799 | 140.6309 | 0.233099 | 3.904598 | 0.586441 |
| 6 | 14543800 | 14544399 | 7.151618 | 13.84635 | 26.2426 | 9.981039 | -0.34656 | -0.18695 | 0.586447 |
| Y | 9624200 | 9633199 | 80.22772 | 126.3335 | 84.91043 | 35.23078 | 0.894559 | 2.495099 | 0.586472 |
| 12 | 59380200 | 59381199 | 8.687259 | 16.48818 | 28.82666 | 11.61367 | -0.30878 | 0.023145 | 0.586496 |
| 7 | 22973000 | 22974999 | 17.0266 | 30.36083 | 27.12531 | 11.93336 | 0.558461 | 0.45842 | 0.5865 |
| 3 | 69172600 | 69173399 | 14.98857 | 27.01821 | 29.29306 | 12.7608 | 0.290411 | 0.382975 | 0.586527 |
| 10 | 106679000 | 106683199 | 36.87265 | 60.9917 | 59.52557 | 25.73633 | 0.415565 | 1.346257 | 0.586528 |
| 1 | 35340600 | 35344399 | 33.41991 | 54.46821 | 89.12216 | 38.27752 | -0.33722 | 1.545261 | 0.58655 |
| 1 | 107099800 | 107104399 | 62.23511 | 99.76742 | 93.67466 | 39.55224 | 0.437719 | 1.993104 | 0.586553 |
| 15 | 32860200 | 32863199 | 26.53171 | 45.69285 | 43.47961 | 19.07289 | 0.45584 | 0.915474 | 0.586559 |
| 7 | 26830200 | 26832799 | 26.16325 | 43.10217 | 68.41867 | 29.51196 | -0.28526 | 1.374822 | 0.586566 |
| 6 | 39232000 | 39234599 | 16.44697 | 28.51627 | 46.87312 | 20.56571 | -0.32744 | 0.81629 | 0.58657 |

| | | | | | | | | | |
|---|---|---|---|---|---|---|---|---|---|
| X | 112239200 | 112241999 | 34.39196 | 57.75565 | 44.776 | 19.55218 | 0.747008 | 1.167573 | 0.586574 |
| 13 | 106896600 | 106897799 | 8.404202 | 15.75715 | 27.98371 | 11.5406 | -0.35968 | 0.099826 | 0.586589 |
| 6 | 84259400 | 84264199 | 64.57736 | 102.6463 | 111.8947 | 46.68812 | 0.20941 | 2.262252 | 0.5866 |
| 13 | 51169800 | 51174399 | 34.43234 | 55.6356 | 87.58946 | 37.3988 | -0.29275 | 1.711872 | 0.586609 |
| 1 | 65868000 | 65868599 | 9.204451 | 17.87279 | 20.55892 | 7.704835 | 0.38711 | -0.23145 | 0.586611 |
| 18 | 41732200 | 41732799 | 5.663297 | 11.02619 | 21.9197 | 8.039253 | -0.36544 | -0.3454 | 0.586615 |
| 1 | 7431200 | 7433799 | 54.05918 | 86.50134 | 94.03997 | 39.59753 | 0.223597 | 2.040864 | 0.586683 |
| 13 | 77778200 | 77792599 | 403.6303 | 553.392 | 716.4367 | 297.9942 | -0.23293 | 4.883158 | 0.586694 |
| 1 | 119695200 | 119696599 | 14.6488 | 27.07376 | 27.44634 | 11.575 | 0.418249 | 0.236423 | 0.586707 |
| 6 | 131239200 | 131243799 | 48.17958 | 77.72376 | 72.46835 | 30.87857 | 0.458643 | 1.774017 | 0.586715 |
| 12 | 110165600 | 110166599 | 6.106664 | 11.83946 | 22.73607 | 8.591995 | -0.35554 | -0.22935 | 0.586724 |
| 3 | 144483800 | 144485999 | 17.75376 | 30.73232 | 48.47261 | 21.26644 | -0.26864 | 0.839631 | 0.586764 |
| 5 | 150356000 | 150357999 | 33.68209 | 55.19451 | 57.76199 | 24.85925 | 0.311016 | 1.45071 | 0.586765 |
| 13 | 60134000 | 60136799 | 22.40188 | 37.92422 | 61.03714 | 26.72067 | -0.3029 | 1.071456 | 0.586774 |
| 6 | 8693200 | 8696599 | 27.40653 | 46.03472 | 49.64786 | 21.68251 | 0.272263 | 1.143295 | 0.58678 |
| 9 | 116039400 | 116040399 | 14.85194 | 26.8006 | 27.05301 | 11.7692 | 0.39462 | 0.374442 | 0.586792 |
| 8 | 78320800 | 78321999 | 14.10667 | 25.63788 | 27.83369 | 12.00134 | 0.297919 | 0.326076 | 0.586809 |
| 16 | 76247600 | 76249799 | 19.06564 | 32.71742 | 51.68897 | 22.67084 | -0.27365 | 0.927005 | 0.586837 |
| 6 | 127529600 | 127530399 | 11.62228 | 20.85243 | 34.69023 | 15.18225 | -0.32809 | 0.398913 | 0.586845 |
| 15 | 4144000 | 4145599 | 17.1696 | 31.23322 | 28.54782 | 12.29513 | 0.545639 | 0.3443 | 0.586858 |
| Y | 2260600 | 2262399 | 13.40078 | 24.56306 | 18.09166 | 7.716233 | 0.864013 | 0.247182 | 0.58687 |
| 12 | 23440800 | 23443199 | 22.82523 | 39.86434 | 33.84924 | 14.91036 | 0.622471 | 0.732958 | 0.586882 |
| 14 | 54566600 | 54568399 | 11.61321 | 20.79368 | 35.69088 | 15.64708 | -0.37477 | 0.412632 | 0.586933 |
| 11 | 66588000 | 66590399 | 29.30415 | 49.71361 | 49.04006 | 21.48233 | 0.401014 | 1.06594 | 0.586962 |
| 4 | 102729200 | 102732999 | 28.31346 | 48.11554 | 48.47888 | 21.2462 | 0.370717 | 1.04723 | 0.586964 |
| 10 | 33756600 | 33763199 | 105.4548 | 163.5537 | 149.013 | 61.88936 | 0.439505 | 2.795433 | 0.586974 |
| 12 | 19449200 | 19454399 | 56.08186 | 90.64008 | 64.93256 | 27.73399 | 0.841035 | 1.732555 | 0.586976 |
| 5 | 4257200 | 4259399 | 46.94677 | 75.54956 | 82.58333 | 35.06938 | 0.225494 | 1.850898 | 0.587023 |
| 16 | 83307400 | 83311599 | 49.61412 | 80.20631 | 77.94017 | 33.30022 | 0.402595 | 1.719646 | 0.587027 |
| 3 | 77026800 | 77030799 | 39.20912 | 64.04552 | 66.73261 | 28.68452 | 0.314482 | 1.510889 | 0.58703 |
| 3 | 26622800 | 26628599 | 71.85769 | 113.9312 | 121.1601 | 50.42684 | 0.241542 | 2.343315 | 0.587046 |
| 14 | 51120600 | 51125799 | 54.64415 | 87.54569 | 94.56096 | 39.88983 | 0.234942 | 2.002645 | 0.587051 |
| 7 | 83872600 | 83875999 | 28.46034 | 46.25357 | 85.77457 | 36.79945 | -0.52019 | 1.576057 | 0.587069 |
| 6 | 72923200 | 72927999 | 67.16201 | 105.6128 | 153.329 | 63.60736 | -0.21644 | 2.53119 | 0.587079 |
| 4 | 151506800 | 151513199 | 67.70513 | 106.6133 | 154.636 | 64.16102 | -0.21355 | 2.505798 | 0.587091 |
| 8 | 126206000 | 126209399 | 28.38785 | 46.37862 | 73.56659 | 31.62427 | -0.28987 | 1.492805 | 0.587125 |
| 2 | 87237000 | 87241599 | 43.1699 | 70.94317 | 62.6457 | 26.99346 | 0.556996 | 1.442645 | 0.587132 |
| 12 | 70963600 | 70966799 | 26.34066 | 44.64273 | 47.42344 | 20.76804 | 0.294416 | 1.059817 | 0.587134 |
| 5 | 133704800 | 133707799 | 27.9756 | 45.47161 | 86.35904 | 37.05371 | -0.55433 | 1.572319 | 0.587135 |
| 4 | 91788600 | 91789599 | 8.243852 | 15.73231 | 27.84546 | 11.0895 | -0.31137 | -0.01986 | 0.587148 |
| 1 | 17593200 | 17597999 | 106.8212 | 164.2486 | 173.473 | 72.03737 | 0.217055 | 2.956666 | 0.587148 |
| 12 | 18658200 | 18689599 | 860.1356 | 1124.959 | 1020.971 | 427.438 | 0.211235 | 5.637614 | 0.587163 |
| 5 | 98929400 | 98931199 | 15.05946 | 26.46275 | 42.37208 | 18.75924 | -0.28833 | 0.608959 | 0.587186 |
| 5 | 87194400 | 87197199 | 26.301 | 45.20934 | 44.61629 | 19.57037 | 0.402977 | 0.933582 | 0.587198 |
| 4 | 131938200 | 131939799 | 12.65031 | 22.37815 | 36.73045 | 16.24478 | -0.31999 | 0.510323 | 0.587211 |
| 17 | 4778400 | 4779399 | 11.58917 | 21.65708 | 24.12986 | 10.00704 | 0.30155 | 0.139446 | 0.587211 |
| 12 | 63004600 | 63007999 | 18.87107 | 33.40522 | 28.99948 | 12.82387 | 0.594185 | 0.526212 | 0.587211 |

| | | | | | | | | | |
|---|---|---|---|---|---|---|---|---|---|
| 1 | 11959000 | 11959599 | 15.06999 | 27.49931 | 24.73477 | 10.61149 | 0.572402 | 0.304456 | 0.587214 |
| X | 155408000 | 155413399 | 90.0259 | 142.2481 | 127.5829 | 52.99249 | 0.482883 | 2.44324 | 0.587216 |
| 1 | 29655800 | 29656399 | 9.070165 | 17.69701 | 18.06896 | 6.446954 | 0.621007 | -0.41075 | 0.587216 |
| 5 | 124001800 | 124004399 | 21.59996 | 35.59542 | 73.30837 | 31.62522 | -0.65942 | 1.365569 | 0.587224 |
| 3 | 62158800 | 62159599 | 8.27138 | 15.81076 | 29.05394 | 11.52943 | -0.36043 | -0.02743 | 0.587232 |
| 18 | 37118400 | 37122599 | 106.3419 | 163.4943 | 162.3641 | 67.42325 | 0.305699 | 2.959035 | 0.587234 |
| 13 | 66353400 | 66365199 | 369.7174 | 519.9563 | 440.1089 | 182.3292 | 0.416965 | 4.495957 | 0.587279 |
| 8 | 117432200 | 117435799 | 46.5178 | 75.33704 | 75.88314 | 32.47212 | 0.353492 | 1.674507 | 0.587297 |
| 10 | 68931600 | 68934399 | 37.89967 | 61.04368 | 92.87087 | 39.48796 | -0.24888 | 1.815094 | 0.58732 |
| 13 | 111634800 | 111638999 | 38.32143 | 61.49031 | 109.066 | 46.11736 | -0.47654 | 1.94132 | 0.587336 |
| 7 | 117861600 | 117863399 | 13.08392 | 23.31359 | 37.69058 | 16.59151 | -0.29296 | 0.458211 | 0.587339 |
| 19 | 7721200 | 7728399 | 78.50567 | 121.7258 | 203.0911 | 84.35283 | -0.43269 | 2.793606 | 0.587351 |
| 13 | 60726600 | 60730599 | 24.67329 | 41.99715 | 44.73576 | 19.6083 | 0.290987 | 1.012978 | 0.587351 |
| X | 148137200 | 148144199 | 73.37353 | 116.6976 | 101.3694 | 42.32343 | 0.537366 | 2.250928 | 0.587354 |
| 17 | 28703800 | 28705999 | 19.09525 | 31.8283 | 67.01315 | 29.12457 | -0.68928 | 1.218738 | 0.587378 |
| 12 | 38797400 | 38799799 | 32.522 | 52.59538 | 83.74102 | 35.79642 | -0.30742 | 1.682377 | 0.587396 |
| 7 | 85656600 | 85666199 | 311.3169 | 441.8251 | 430.8206 | 178.2137 | 0.226093 | 4.360381 | 0.587399 |
| 4 | 80145600 | 80151599 | 96.08931 | 149.7609 | 154.3423 | 64.05583 | 0.267367 | 2.706823 | 0.587405 |
| 9 | 9312600 | 9313599 | 12.75783 | 22.29288 | 45.73839 | 20.15386 | -0.64402 | 0.693516 | 0.587408 |
| 6 | 147377200 | 147378399 | 12.33608 | 21.80836 | 36.75663 | 16.25844 | -0.35839 | 0.515066 | 0.587427 |
| 3 | 139336400 | 139337599 | 14.45605 | 26.09407 | 28.31307 | 12.31274 | 0.290849 | 0.368837 | 0.587453 |
| 8 | 100421400 | 100423999 | 46.23084 | 74.39851 | 80.21665 | 34.06508 | 0.24527 | 1.847609 | 0.587458 |
| 17 | 73774800 | 73775799 | 13.2809 | 24.61218 | 26.56655 | 11.16276 | 0.332377 | 0.210134 | 0.587462 |
| 17 | 86211200 | 86214399 | 26.91508 | 44.48459 | 69.33767 | 29.96187 | -0.25796 | 1.337803 | 0.587468 |
| 1 | 101849000 | 101850799 | 19.39599 | 34.3922 | 32.40131 | 14.32207 | 0.477707 | 0.517806 | 0.587472 |
| 10 | 10397600 | 10400799 | 63.98044 | 100.9952 | 145.9304 | 60.58322 | -0.20531 | 2.449917 | 0.587477 |
| 1 | 85246200 | 85266599 | 1035.692 | 1332.811 | 1189.765 | 498.9067 | 0.211737 | 5.901156 | 0.587484 |
| 7 | 26733600 | 26737999 | 41.16156 | 66.82728 | 72.039 | 30.8833 | 0.259078 | 1.612483 | 0.587504 |
| 10 | 71768400 | 71773399 | 43.96489 | 71.29849 | 74.05705 | 31.72218 | 0.311026 | 1.641161 | 0.587512 |
| 1 | 178959000 | 178961799 | 47.26976 | 76.05461 | 83.18833 | 35.31639 | 0.224231 | 1.857532 | 0.587518 |
| 15 | 35758600 | 35758999 | 4.390412 | 8.5707 | 18.69987 | 6.6124 | -0.44864 | -0.50185 | 0.587529 |
| 11 | 23117000 | 23118999 | 23.04383 | 39.64181 | 42.54531 | 18.66241 | 0.282589 | 0.909058 | 0.587539 |
| X | 105389600 | 105391799 | 16.76684 | 29.25209 | 48.40196 | 21.30643 | -0.33573 | 0.740982 | 0.587541 |
| 18 | 85638600 | 85639999 | 19.31704 | 33.05365 | 51.66029 | 22.65302 | -0.25868 | 0.956267 | 0.587542 |
| 7 | 38840400 | 38849399 | 154.1798 | 232.8444 | 236.5742 | 97.79834 | 0.252085 | 3.334621 | 0.587565 |
| 4 | 112459800 | 112461599 | 22.02684 | 39.00918 | 32.24163 | 14.25629 | 0.665011 | 0.547377 | 0.587566 |
| 9 | 74670800 | 74671399 | 6.103517 | 11.83417 | 22.75081 | 8.59485 | -0.35715 | -0.23227 | 0.587572 |
| 15 | 66326800 | 66329399 | 37.16833 | 61.14539 | 63.30921 | 27.29162 | 0.328579 | 1.415391 | 0.587585 |
| 2 | 88364000 | 88366999 | 29.87342 | 51.05081 | 46.24085 | 20.27449 | 0.524974 | 1.001923 | 0.587589 |
| 1 | 105707800 | 105711599 | 25.31989 | 41.98357 | 65.21198 | 28.24204 | -0.25248 | 1.29472 | 0.5876 |
| 2 | 42118600 | 42121399 | 27.13998 | 46.98173 | 41.79165 | 18.33532 | 0.554221 | 0.872303 | 0.587606 |
| 3 | 52513800 | 52516399 | 22.0163 | 37.97438 | 41.5517 | 18.22803 | 0.255671 | 0.881677 | 0.587657 |
| 15 | 50346200 | 50347399 | 28.41063 | 45.88296 | 94.29089 | 40.23439 | -0.6769 | 1.724002 | 0.587674 |
| 17 | 11999600 | 12001999 | 19.7133 | 34.30496 | 36.8534 | 16.19678 | 0.284255 | 0.772242 | 0.587681 |
| 5 | 110019600 | 110021599 | 38.92682 | 63.75589 | 65.65983 | 28.25206 | 0.333758 | 1.473641 | 0.587717 |
| 9 | 72081800 | 72082799 | 12.9826 | 23.13666 | 37.21516 | 16.38007 | -0.28536 | 0.457436 | 0.587722 |
| 4 | 60788000 | 60789399 | 10.98106 | 19.92658 | 32.7718 | 14.15782 | -0.29913 | 0.324348 | 0.587726 |

| | | | | | | | | | |
|---|---|---|---|---|---|---|---|---|---|
| 2 | 10641400 | 10642599 | 12.34871 | 23.19973 | 24.51886 | 10.07704 | 0.386976 | 0.11061 | 0.587767 |
| 2 | 179688600 | 179690399 | 20.60982 | 35.14928 | 54.92053 | 24.07622 | -0.25913 | 0.994356 | 0.58777 |
| 15 | 88287600 | 88299199 | 128.3211 | 193.437 | 270.5479 | 111.7794 | -0.21095 | 3.367099 | 0.587787 |
| 10 | 34959000 | 34960599 | 14.37048 | 25.3179 | 40.77746 | 18.04933 | -0.29547 | 0.571293 | 0.587802 |
| 11 | 66105200 | 66111199 | 58.83579 | 93.0597 | 138.1532 | 57.41241 | -0.24197 | 2.397191 | 0.587818 |
| 14 | 60769200 | 60773999 | 40.64683 | 65.37176 | 98.47443 | 41.78096 | -0.23731 | 1.866789 | 0.587838 |
| 13 | 63631800 | 63635999 | 25.35665 | 41.85245 | 66.70335 | 28.79675 | -0.29002 | 1.348879 | 0.587853 |
| 3 | 38353600 | 38355199 | 10.47656 | 19.15456 | 31.81945 | 13.61773 | -0.30469 | 0.274643 | 0.587857 |
| 4 | 12097400 | 12099599 | 34.66667 | 57.06132 | 61.97141 | 26.72114 | 0.260158 | 1.40126 | 0.587862 |
| 15 | 44511600 | 44514999 | 23.45128 | 40.85448 | 39.61664 | 17.4232 | 0.431393 | 0.777917 | 0.587871 |
| 15 | 89718800 | 89719399 | 6.871122 | 13.19615 | 24.65917 | 9.654058 | -0.36528 | -0.09689 | 0.587875 |
| 2 | 17756000 | 17757599 | 15.06114 | 26.28187 | 42.61386 | 18.76103 | -0.30643 | 0.71952 | 0.587878 |
| 7 | 67281800 | 67282599 | 9.94152 | 18.52495 | 31.64135 | 13.18595 | -0.31647 | 0.160455 | 0.58788 |
| 5 | 51987000 | 51988199 | 7.890173 | 15.0749 | 26.75119 | 10.62751 | -0.31219 | -0.03601 | 0.587883 |
| 1 | 106179400 | 106180999 | 19.84882 | 34.83044 | 35.80086 | 15.83145 | 0.347958 | 0.645092 | 0.587888 |
| 16 | 58960200 | 58961399 | 15.2375 | 28.13559 | 27.35616 | 11.55159 | 0.476696 | 0.245271 | 0.587902 |
| 1 | 67107000 | 67109199 | 19.12202 | 32.81141 | 51.13761 | 22.42887 | -0.25405 | 0.928904 | 0.587918 |
| 11 | 107240800 | 107243799 | 23.97456 | 39.92496 | 62.47717 | 27.13765 | -0.26273 | 1.238523 | 0.587919 |
| X | 125746200 | 125752599 | 67.12734 | 107.8262 | 85.98154 | 36.41495 | 0.676421 | 1.940102 | 0.587926 |
| 11 | 113284400 | 113293599 | 102.216 | 154.968 | 263.3537 | 109.002 | -0.48467 | 3.246557 | 0.58793 |
| 14 | 18195000 | 18197199 | 30.65475 | 51.50844 | 52.41538 | 22.89422 | 0.355863 | 1.155053 | 0.587956 |
| 3 | 3505800 | 3510199 | 40.54255 | 67.01298 | 59.66416 | 25.78319 | 0.547651 | 1.366471 | 0.587972 |
| Y | 37992200 | 37994199 | 12.7557 | 23.56603 | 17.59549 | 7.424406 | 0.855463 | 0.196514 | 0.587972 |
| 5 | 38073600 | 38076999 | 30.31419 | 49.37855 | 77.16145 | 33.13363 | -0.27146 | 1.540215 | 0.587982 |
| 8 | 17348400 | 17349399 | 12.09763 | 22.67664 | 25.15748 | 10.37921 | 0.313624 | 0.127059 | 0.588003 |
| 4 | 38332800 | 38333399 | 6.693717 | 12.99597 | 24.63058 | 9.239881 | -0.32739 | -0.24302 | 0.588006 |
| 14 | 61817400 | 61818399 | 10.4622 | 19.17044 | 31.84269 | 13.58725 | -0.30152 | 0.261865 | 0.588016 |
| 3 | 108913200 | 108914799 | 11.60794 | 20.86419 | 34.68929 | 15.15883 | -0.32577 | 0.387273 | 0.588067 |
| 5 | 75531400 | 75533799 | 34.8028 | 56.3854 | 85.61806 | 36.64806 | -0.23677 | 1.651829 | 0.588069 |
| 8 | 72741000 | 72743999 | 16.52266 | 28.12143 | 57.43783 | 25.17563 | -0.64443 | 1.000167 | 0.588076 |
| 11 | 52023400 | 52028199 | 78.55582 | 124.151 | 129.0646 | 53.61368 | 0.270418 | 2.428814 | 0.588109 |
| 18 | 60607800 | 60608999 | 9.484394 | 17.25968 | 31.40284 | 13.51826 | -0.43995 | 0.296357 | 0.588118 |
| 4 | 68610400 | 68610999 | 8.41253 | 15.91073 | 27.59873 | 11.20192 | -0.30653 | 0.043285 | 0.588125 |
| 7 | 3647600 | 3649599 | 11.94215 | 21.15427 | 36.34449 | 16.0697 | -0.38434 | 0.493497 | 0.588134 |
| 14 | 37479800 | 37480399 | 15.19953 | 27.10434 | 28.92793 | 12.72582 | 0.30361 | 0.450581 | 0.588135 |
| 12 | 43614600 | 43617199 | 37.37532 | 60.76095 | 65.35375 | 28.04325 | 0.264038 | 1.575808 | 0.588145 |
| 14 | 94142400 | 94143599 | 14.55249 | 26.12504 | 27.48619 | 12.02345 | 0.329775 | 0.401867 | 0.588157 |
| 1 | 170880600 | 170882399 | 15.1148 | 26.44614 | 45.12255 | 19.90456 | -0.37977 | 0.67922 | 0.588161 |
| 8 | 105018400 | 105020599 | 19.18455 | 32.54135 | 53.21979 | 23.31228 | -0.32557 | 1.035878 | 0.588193 |
| 1 | 154670200 | 154672799 | 26.4795 | 43.69379 | 67.96477 | 29.33624 | -0.25514 | 1.355744 | 0.588196 |
| X | 125433600 | 125435799 | 19.35667 | 35.07844 | 28.92554 | 12.51703 | 0.689157 | 0.383318 | 0.588207 |
| 16 | 55294000 | 55294999 | 9.393388 | 18.0176 | 20.19514 | 7.934931 | 0.359903 | -0.07622 | 0.588224 |
| 5 | 85529400 | 85531799 | 27.46795 | 47.38119 | 43.49358 | 19.07995 | 0.508116 | 0.90818 | 0.588225 |
| 5 | 47823600 | 47824199 | 6.409458 | 12.45318 | 24.66237 | 9.197356 | -0.3817 | -0.26471 | 0.588232 |
| 18 | 74294800 | 74298599 | 26.85065 | 44.22756 | 68.94225 | 29.7358 | -0.25921 | 1.380488 | 0.588247 |
| 11 | 87958400 | 87960199 | 8.168158 | 15.11043 | 29.93554 | 12.60873 | -0.5402 | 0.184205 | 0.588247 |
| Y | 4877800 | 4878599 | 4.866105 | 9.400608 | 23.14425 | 8.893668 | -0.73297 | -0.18583 | 0.588282 |

| | | | | | | | | | |
|---|---|---|---|---|---|---|---|---|---|
| 13 | 60329400 | 60332399 | 28.25254 | 46.61333 | 72.84644 | 31.44161 | -0.26205 | 1.364681 | 0.588292 |
| 13 | 99714400 | 99717599 | 29.53997 | 49.16371 | 54.03371 | 23.46118 | 0.245401 | 1.253498 | 0.588313 |
| 8 | 119644800 | 119649199 | 81.11015 | 127.3973 | 138.1024 | 57.28198 | 0.20317 | 2.560506 | 0.588321 |
| 2 | 89256400 | 89258199 | 17.20965 | 31.0457 | 28.88335 | 12.57025 | 0.51046 | 0.390953 | 0.58833 |
| 8 | 3418600 | 3419799 | 16.94149 | 29.92911 | 32.28751 | 14.28362 | 0.281281 | 0.541635 | 0.588341 |
| 4 | 19233800 | 19236399 | 28.23029 | 47.30626 | 51.81567 | 22.59547 | 0.250137 | 1.175444 | 0.588356 |
| 5 | 115042200 | 115044999 | 48.4288 | 77.24203 | 115.6667 | 48.48205 | -0.24265 | 2.140495 | 0.588364 |
| X | 102010400 | 102012799 | 22.18136 | 39.14706 | 34.00944 | 15.04837 | 0.591763 | 0.58924 | 0.588378 |
| 16 | 36370200 | 36371999 | 15.79977 | 27.64998 | 44.584 | 19.67022 | -0.29851 | 0.680699 | 0.588386 |
| 2 | 43986800 | 43988599 | 21.576 | 37.91438 | 35.7017 | 15.80631 | 0.473886 | 0.635071 | 0.588387 |
| 10 | 88485200 | 88488399 | 52.57131 | 83.54874 | 125.1573 | 52.21329 | -0.24868 | 2.260692 | 0.588394 |
| 16 | 37454600 | 37455199 | 5.398206 | 10.46091 | 22.15621 | 8.391647 | -0.49826 | -0.23231 | 0.588403 |
| 8 | 3846000 | 3864599 | 386.8463 | 540.0734 | 524.6585 | 217.6046 | 0.207652 | 4.606223 | 0.588405 |
| 4 | 91641400 | 91646399 | 112.0311 | 171.9533 | 183.6862 | 76.25997 | 0.198726 | 2.994592 | 0.588421 |
| 2 | 86447400 | 86456999 | 164.2272 | 246.9115 | 251.0904 | 103.6672 | 0.245173 | 3.424393 | 0.588445 |
| 16 | 14383000 | 14384999 | 20.68663 | 36.28175 | 36.29198 | 16.04158 | 0.387042 | 0.65886 | 0.588452 |
| 16 | 19119800 | 19124399 | 52.45768 | 84.61578 | 81.99261 | 34.9318 | 0.403037 | 1.789384 | 0.588472 |
| 7 | 61443400 | 61452399 | 122.9849 | 189.775 | 154.7772 | 64.2915 | 0.593519 | 2.894351 | 0.58849 |
| 11 | 20732600 | 20736599 | 61.54114 | 97.99903 | 105.6353 | 44.17959 | 0.228352 | 2.202399 | 0.588497 |
| 3 | 96272400 | 96280599 | 256.6807 | 364.2482 | 504.8627 | 208.8456 | -0.28122 | 4.35995 | 0.5885 |
| 3 | 39706800 | 39708999 | 27.60599 | 48.00776 | 39.50423 | 17.35453 | 0.667144 | 0.823153 | 0.588509 |
| 8 | 126395000 | 126399999 | 105.4169 | 160.4634 | 230.5161 | 95.52527 | -0.23772 | 3.163299 | 0.58851 |
| 3 | 55689600 | 55691799 | 28.65869 | 46.489 | 87.03684 | 37.28961 | -0.5364 | 1.616069 | 0.58852 |
| 2 | 5301400 | 5303199 | 16.38643 | 28.9683 | 30.96037 | 13.69464 | 0.295261 | 0.530109 | 0.588551 |
| 17 | 39736200 | 39737999 | 29.66621 | 50.09364 | 50.72522 | 22.18912 | 0.363184 | 1.108546 | 0.588553 |
| 15 | 37527000 | 37528799 | 11.12948 | 20.14734 | 33.24613 | 14.40604 | -0.30685 | 0.339759 | 0.588564 |
| 10 | 11458800 | 11463199 | 34.53122 | 57.94482 | 52.50474 | 22.9159 | 0.522779 | 1.181604 | 0.588571 |
| 1 | 66518000 | 66519199 | 12.14498 | 22.76628 | 25.09938 | 10.35455 | 0.32238 | 0.125533 | 0.588574 |
| 4 | 33301600 | 33303999 | 29.92927 | 49.80981 | 54.09866 | 23.48879 | 0.262388 | 1.253377 | 0.588584 |
| 19 | 18072400 | 18075799 | 19.49072 | 34.40912 | 34.14435 | 15.10713 | 0.400913 | 0.568513 | 0.588598 |
| 4 | 121477800 | 121490599 | 274.3474 | 396.0915 | 390.1826 | 160.9695 | 0.236104 | 4.10787 | 0.588623 |
| 18 | 52579600 | 52581799 | 17.93025 | 30.57833 | 56.97871 | 24.97842 | -0.51213 | 0.987351 | 0.588657 |
| X | 113909600 | 113911799 | 24.79168 | 43.11897 | 40.7511 | 17.9037 | 0.468247 | 0.81026 | 0.588667 |
| 4 | 57977000 | 57981999 | 62.12533 | 98.32946 | 142.4753 | 59.22891 | -0.20624 | 2.384802 | 0.588668 |
| 18 | 8166000 | 8172799 | 73.47934 | 117.0524 | 108.2528 | 45.29725 | 0.449554 | 2.20117 | 0.588672 |
| 13 | 102961800 | 102964199 | 47.32926 | 76.234 | 82.88284 | 35.24275 | 0.234881 | 1.82422 | 0.588682 |
| 19 | 39631400 | 39638599 | 110.4159 | 170.5754 | 173.8472 | 72.2124 | 0.273598 | 2.87088 | 0.588718 |
| 19 | 39699400 | 39701199 | 30.7358 | 51.4122 | 47.56387 | 20.71783 | 0.492946 | 1.1974 | 0.588724 |
| 7 | 9469200 | 9469799 | 9.625495 | 18.60043 | 21.45769 | 8.234448 | 0.350631 | -0.14578 | 0.588728 |
| 1 | 33282000 | 33284199 | 23.65329 | 39.60215 | 61.42088 | 26.76943 | -0.24996 | 1.180122 | 0.588736 |
| 8 | 38294600 | 38296999 | 24.39101 | 42.66577 | 37.2456 | 16.42634 | 0.58271 | 0.722044 | 0.588737 |
| 9 | 94220600 | 94221199 | 6.827398 | 13.13099 | 27.03116 | 10.53921 | -0.49967 | -0.11035 | 0.588739 |
| 11 | 30185400 | 30185999 | 7.758582 | 14.82859 | 26.19092 | 10.39723 | -0.30423 | -0.04002 | 0.588741 |
| 12 | 97525200 | 97528599 | 31.49579 | 51.52265 | 82.40421 | 35.43999 | -0.30058 | 1.486622 | 0.588763 |
| 9 | 19433600 | 19434399 | 12.12734 | 22.58867 | 25.02377 | 10.43548 | 0.30303 | 0.165245 | 0.588764 |
| 7 | 22082600 | 22082999 | 11.30684 | 21.50386 | 20.02918 | 8.034635 | 0.598135 | 0.010499 | 0.588768 |
| 10 | 48597600 | 48600399 | 46.9707 | 75.57662 | 82.45122 | 35.00561 | 0.228053 | 1.856009 | 0.588772 |

| | | | | | | | | | |
|---|---|---|---|---|---|---|---|---|---|
| 2 | 177246200 | 177247399 | 10.13566 | 19.55679 | 21.34174 | 8.244123 | 0.420645 | -0.12049 | 0.58878 |
| 7 | 85965400 | 85966599 | 13.82688 | 25.09387 | 27.40197 | 11.8361 | 0.288058 | 0.331832 | 0.588781 |
| 7 | 94044000 | 94044599 | 8.200586 | 15.14189 | 33.0755 | 13.96698 | -0.68269 | 0.204121 | 0.588816 |
| 4 | 90980400 | 90980799 | 4.674257 | 9.124077 | 19.28338 | 6.828095 | -0.40577 | -0.49195 | 0.588818 |
| Y | 35685200 | 35687999 | 15.2718 | 27.34382 | 20.63464 | 9.046515 | 0.803666 | 0.404416 | 0.588826 |
| 7 | 112575800 | 112576599 | 6.549411 | 12.64958 | 23.71043 | 9.121117 | -0.346 | -0.16237 | 0.588828 |
| 15 | 73889800 | 73890599 | 6.549411 | 12.64958 | 23.71043 | 9.121117 | -0.346 | -0.16237 | 0.588828 |
| 1 | 105244600 | 105250399 | 46.54037 | 75.09722 | 80.08779 | 34.13565 | 0.26562 | 1.767235 | 0.588829 |
| Y | 9859600 | 9862199 | 16.89673 | 29.85251 | 22.18672 | 9.814985 | 0.815732 | 0.518181 | 0.588848 |
| 14 | 101325200 | 101328599 | 66.71265 | 105.7074 | 109.9046 | 45.71451 | 0.273214 | 2.354902 | 0.588892 |
| 13 | 73086400 | 73088999 | 18.68676 | 32.02329 | 50.58651 | 22.18485 | -0.2736 | 0.939417 | 0.588894 |
| 8 | 27194400 | 27195799 | 11.70971 | 21.11366 | 34.78513 | 15.14799 | -0.30899 | 0.372919 | 0.588906 |
| 9 | 20862600 | 20864999 | 17.80329 | 30.25467 | 52.11393 | 22.83928 | -0.39975 | 1.015536 | 0.588906 |
| 16 | 61413400 | 61417599 | 69.71206 | 110.4399 | 120.3356 | 50.04862 | 0.20547 | 2.363685 | 0.588932 |
| 7 | 100727200 | 100728799 | 17.17428 | 30.49715 | 32.19021 | 14.21129 | 0.315912 | 0.495496 | 0.588935 |
| 5 | 92008600 | 92011399 | 30.52696 | 50.63818 | 55.69853 | 24.12874 | 0.2439 | 1.295364 | 0.588945 |
| X | 30998400 | 30999999 | 27.69363 | 47.11913 | 47.40594 | 20.778 | 0.373177 | 1.03578 | 0.588946 |
| 3 | 104544600 | 104547199 | 17.33689 | 29.70788 | 50.05693 | 21.95248 | -0.36642 | 0.931103 | 0.588981 |
| 8 | 28740000 | 28745599 | 47.7836 | 77.09815 | 81.28936 | 34.64482 | 0.281947 | 1.772982 | 0.588985 |
| 1 | 33106800 | 33107399 | 10.52071 | 19.96739 | 22.25452 | 8.966492 | 0.33613 | 0.024914 | 0.588992 |
| X | 145994200 | 145995799 | 15.35903 | 28.63798 | 25.85685 | 10.76385 | 0.598041 | 0.187217 | 0.589 |
| 2 | 168077400 | 168082599 | 68.68439 | 107.8343 | 156.4313 | 64.88083 | -0.21709 | 2.564454 | 0.58902 |
| 12 | 78130800 | 78131799 | 14.38546 | 25.30149 | 40.28421 | 17.83702 | -0.27982 | 0.589602 | 0.589025 |
| 17 | 76074400 | 76076199 | 15.75677 | 28.15132 | 30.36003 | 13.33451 | 0.290259 | 0.446135 | 0.589028 |
| 4 | 23847000 | 23850399 | 36.90106 | 60.15197 | 66.05943 | 28.37394 | 0.236815 | 1.537137 | 0.589041 |
| 10 | 4122800 | 4123999 | 13.0334 | 23.94663 | 26.4936 | 11.26317 | 0.284791 | 0.254642 | 0.589043 |
| 13 | 98810800 | 98812199 | 13.0334 | 23.94663 | 26.4936 | 11.26317 | 0.284791 | 0.254642 | 0.589043 |
| 13 | 47312200 | 47315799 | 19.91147 | 33.96763 | 53.28903 | 23.36151 | -0.26488 | 0.986252 | 0.589054 |
| 12 | 85538600 | 85541199 | 13.83227 | 24.25809 | 39.96432 | 17.66359 | -0.32887 | 0.632214 | 0.58908 |
| X | 49898400 | 49899999 | 12.80146 | 22.86462 | 36.71021 | 16.12757 | -0.28088 | 0.440966 | 0.589083 |
| 17 | 19207200 | 19207999 | 15.34631 | 27.63995 | 29.78334 | 12.98794 | 0.298298 | 0.391603 | 0.589115 |
| 7 | 66655200 | 66659199 | 32.59368 | 54.57398 | 54.13285 | 23.59389 | 0.392765 | 1.20036 | 0.589153 |
| 1 | 175806200 | 175807199 | 15.24539 | 27.19318 | 28.03487 | 12.33025 | 0.353645 | 0.448445 | 0.589176 |
| 18 | 12290600 | 12292799 | 17.90758 | 31.46387 | 33.53718 | 14.84634 | 0.296028 | 0.61327 | 0.589192 |
| 1 | 111965600 | 111967799 | 18.90686 | 33.33417 | 34.17523 | 15.1248 | 0.353258 | 0.578325 | 0.589193 |
| 5 | 95418400 | 95427999 | 123.0723 | 190.278 | 148.1019 | 61.51811 | 0.662995 | 2.858684 | 0.589207 |
| 12 | 3369400 | 3371599 | 15.8466 | 27.68702 | 43.59117 | 19.20636 | -0.26412 | 0.706962 | 0.589211 |
| 9 | 16948000 | 16949999 | 19.99876 | 35.19528 | 34.5936 | 15.31551 | 0.412741 | 0.604248 | 0.589228 |
| 7 | 15475200 | 15520199 | 746.1322 | 992.6611 | 858.4368 | 358.6706 | 0.305438 | 5.362864 | 0.589246 |
| 17 | 70112800 | 70113399 | 11.10515 | 20.83216 | 22.18547 | 9.141236 | 0.373923 | 0.106429 | 0.589262 |
| 5 | 61005600 | 61006599 | 14.08867 | 25.65323 | 27.95289 | 12.0245 | 0.295091 | 0.317549 | 0.589266 |
| 11 | 24285400 | 24286799 | 13.16391 | 23.20627 | 37.80738 | 16.73265 | -0.31121 | 0.554731 | 0.589291 |
| 17 | 72758600 | 72758999 | 9.192758 | 17.69377 | 20.83815 | 8.106784 | 0.302929 | -0.10642 | 0.589297 |
| 18 | 74358000 | 74363599 | 73.35761 | 116.4554 | 118.8406 | 49.52365 | 0.302714 | 2.307917 | 0.589298 |
| 2 | 173842000 | 173848399 | 91.44143 | 140.8479 | 199.3959 | 82.8222 | -0.20327 | 2.920239 | 0.589305 |
| 2 | 40071400 | 40073199 | 20.29895 | 35.36989 | 38.21881 | 16.81055 | 0.275954 | 0.757544 | 0.589316 |
| 5 | 76716000 | 76722599 | 75.40663 | 117.8969 | 168.4912 | 69.90314 | -0.20023 | 2.644947 | 0.589319 |

| | | | | | | | | | |
|---|---|---|---|---|---|---|---|---|---|
| 12 | 46215800 | 46216399 | 5.098192 | 9.922309 | 20.69263 | 7.611775 | -0.43658 | -0.34369 | 0.589339 |
| 6 | 123114200 | 123117799 | 43.14442 | 69.30278 | 103.1807 | 43.6993 | -0.22268 | 1.911707 | 0.589347 |
| 6 | 131875200 | 131893199 | 326.637 | 466.9893 | 427.2207 | 176.5077 | 0.328746 | 4.255003 | 0.589355 |
| 12 | 107544400 | 107546199 | 20.42997 | 35.46995 | 37.46875 | 16.4433 | 0.308217 | 0.810376 | 0.589361 |
| 2 | 49371200 | 49374999 | 46.78126 | 74.85401 | 110.4779 | 46.51754 | -0.21671 | 2.038828 | 0.589373 |
| 6 | 130112400 | 130118399 | 65.67863 | 105.1387 | 97.21725 | 40.96008 | 0.457413 | 2.042713 | 0.589378 |
| 12 | 62928800 | 62930599 | 16.56203 | 29.27085 | 31.44506 | 13.90978 | 0.287731 | 0.535133 | 0.589391 |
| 4 | 130898800 | 130903399 | 41.51765 | 66.77909 | 99.68412 | 42.30047 | -0.22403 | 1.866626 | 0.589443 |
| 5 | 143155000 | 143160799 | 80.51964 | 125.2039 | 178.2053 | 73.9852 | -0.2005 | 2.743446 | 0.589447 |
| 19 | 13679600 | 13680599 | 10.40146 | 20.23159 | 19.48301 | 7.201666 | 0.658725 | -0.26928 | 0.589449 |
| 9 | 4245200 | 4245999 | 9.8116 | 18.93717 | 20.90399 | 8.067375 | 0.406971 | -0.12281 | 0.58945 |
| 11 | 24569200 | 24571199 | 14.86545 | 25.96135 | 42.5786 | 18.75503 | -0.32283 | 0.705707 | 0.589456 |
| 12 | 31038400 | 31042599 | 26.28378 | 43.63139 | 67.64666 | 29.31945 | -0.24976 | 1.28587 | 0.589457 |
| 6 | 139689600 | 139693199 | 33.27584 | 55.12416 | 58.6875 | 25.40007 | 0.290621 | 1.319577 | 0.589463 |
| 16 | 3459600 | 3480999 | 377.6344 | 527.2726 | 507.3076 | 210.4045 | 0.221723 | 4.603248 | 0.589469 |
| 4 | 112548000 | 112554399 | 104.4374 | 162.6874 | 140.3427 | 58.25018 | 0.523126 | 2.719183 | 0.589491 |
| 7 | 41294400 | 41307599 | 183.4766 | 275.7551 | 249.8715 | 103.1542 | 0.411034 | 3.43553 | 0.589522 |
| 11 | 73968400 | 73970399 | 25.306 | 43.82167 | 42.45536 | 18.6267 | 0.431722 | 0.863821 | 0.589524 |
| 17 | 71587600 | 71590999 | 28.76858 | 47.16624 | 73.63559 | 31.6975 | -0.26421 | 1.44365 | 0.58953 |
| 1 | 181621800 | 181622799 | 12.30571 | 21.89573 | 35.68569 | 15.72782 | -0.30558 | 0.458753 | 0.589536 |
| 15 | 27966600 | 27968599 | 12.30571 | 21.89573 | 35.68569 | 15.72782 | -0.30558 | 0.458753 | 0.589536 |
| 13 | 112538400 | 112540599 | 15.39506 | 26.93749 | 42.52467 | 18.75915 | -0.26785 | 0.680384 | 0.589547 |
| 17 | 22627000 | 22627399 | 6.916548 | 13.50208 | 16.34891 | 5.780972 | 0.389189 | -0.4782 | 0.589581 |
| 7 | 97386600 | 97388399 | 9.388831 | 17.24818 | 30.3613 | 12.90966 | -0.38077 | 0.235764 | 0.589584 |
| 13 | 25078800 | 25080599 | 12.26485 | 21.93675 | 35.67552 | 15.65459 | -0.29822 | 0.425045 | 0.589593 |
| 8 | 118637000 | 118637599 | 6.712252 | 12.96029 | 24.24712 | 9.340347 | -0.34483 | -0.1524 | 0.589603 |
| 8 | 118445000 | 118452799 | 76.8091 | 119.7167 | 171.7388 | 71.27537 | -0.20916 | 2.700152 | 0.589632 |
| 4 | 48527800 | 48530599 | 38.05011 | 61.29341 | 93.02359 | 39.55984 | -0.24516 | 1.811252 | 0.589637 |
| 3 | 16571000 | 16572599 | 20.56844 | 36.05808 | 31.85618 | 14.07549 | 0.565485 | 0.659905 | 0.589638 |
| 7 | 97633200 | 97634799 | 15.18028 | 26.53746 | 42.53953 | 18.75125 | -0.28993 | 0.692476 | 0.589638 |
| 5 | 52524000 | 52528399 | 54.76006 | 86.69397 | 133.9563 | 55.69457 | -0.2985 | 2.371708 | 0.589642 |
| 3 | 32577600 | 32583799 | 36.81533 | 60.47734 | 55.81588 | 24.04668 | 0.493213 | 1.427606 | 0.589652 |
| 1 | 128014400 | 128019799 | 102.0282 | 157.484 | 168.4558 | 69.97324 | 0.202976 | 2.88246 | 0.589653 |
| 5 | 40759800 | 40762799 | 63.72749 | 101.2006 | 107.7592 | 44.92044 | 0.24192 | 2.287609 | 0.589655 |
| 1 | 97726200 | 97728599 | 28.22802 | 48.06114 | 41.42401 | 18.15712 | 0.595426 | 1.020668 | 0.589661 |
| X | 153221400 | 153223399 | 32.60038 | 55.09257 | 50.18476 | 21.95806 | 0.515431 | 1.112486 | 0.589672 |
| 8 | 20018000 | 20018599 | 6.660018 | 12.93707 | 27.38263 | 10.23048 | -0.4811 | -0.25512 | 0.589685 |
| Y | 41642800 | 41643599 | 12.34074 | 22.85989 | 18.13174 | 7.623139 | 0.772725 | 0.180078 | 0.589686 |
| 18 | 75732800 | 75734399 | 24.27874 | 39.96605 | 67.51431 | 29.11216 | -0.37514 | 1.380011 | 0.589687 |
| 6 | 14662600 | 14663599 | 19.24236 | 33.61773 | 36.68453 | 16.16258 | 0.262268 | 0.714091 | 0.589689 |
| 10 | 11455000 | 11456199 | 21.923 | 38.02029 | 40.97442 | 17.97778 | 0.279127 | 0.833075 | 0.589751 |
| 10 | 66224400 | 66225999 | 16.92578 | 30.7155 | 29.75163 | 12.85231 | 0.459966 | 0.360825 | 0.589768 |
| 9 | 18130200 | 18131599 | 10.37744 | 19.25701 | 33.74213 | 14.15224 | -0.35999 | 0.196463 | 0.589769 |
| 13 | 114325600 | 114327999 | 18.56333 | 31.87467 | 51.14801 | 22.43467 | -0.2959 | 0.918246 | 0.589778 |
| 7 | 144458400 | 144459799 | 14.7313 | 25.86608 | 40.85287 | 18.07482 | -0.26857 | 0.617233 | 0.589803 |
| 16 | 32196600 | 32197399 | 7.79531 | 14.96779 | 27.50477 | 10.77489 | -0.34271 | -0.0778 | 0.589804 |
| 3 | 135183000 | 135185399 | 16.60502 | 28.95316 | 46.25974 | 20.35609 | -0.2857 | 0.741915 | 0.589826 |

| | | | | | | | | | |
|---|---|---|---|---|---|---|---|---|---|
| 5 | 8383200 | 8390999 | 71.58333 | 113.8001 | 115.3448 | 48.13521 | 0.314481 | 2.265282 | 0.589827 |
| 6 | 76158200 | 76159199 | 9.271648 | 17.22785 | 29.98041 | 12.55183 | -0.34771 | 0.172823 | 0.589829 |
| 13 | 68382400 | 68386199 | 115.389 | 176.4441 | 183.8141 | 76.25886 | 0.230693 | 3.07008 | 0.589837 |
| 4 | 120451600 | 120454599 | 17.56553 | 30.46768 | 48.4766 | 21.2695 | -0.28062 | 0.82021 | 0.589861 |
| 17 | 22276400 | 22280799 | 45.68889 | 73.22049 | 110.2348 | 46.52304 | -0.24269 | 1.990371 | 0.589863 |
| 4 | 141523600 | 141528199 | 60.65985 | 95.96283 | 139.4161 | 57.94274 | -0.21062 | 2.394848 | 0.589864 |
| 17 | 64523800 | 64524599 | 11.48743 | 20.6948 | 33.72362 | 14.70177 | -0.29444 | 0.372408 | 0.589881 |
| 16 | 36029600 | 36030599 | 8.745557 | 16.24181 | 28.92004 | 12.11651 | -0.38162 | 0.165908 | 0.589886 |
| 15 | 31782000 | 31786399 | 58.32009 | 92.6477 | 136.6495 | 56.98469 | -0.22693 | 2.279775 | 0.589895 |
| 1 | 122941000 | 122944799 | 38.54613 | 63.48041 | 62.86226 | 27.11131 | 0.393318 | 1.406206 | 0.589899 |
| 4 | 113650800 | 113657999 | 92.33094 | 145.2684 | 137.7419 | 57.14513 | 0.397982 | 2.535336 | 0.5899 |
| 14 | 116902000 | 116904399 | 29.40142 | 49.65167 | 44.76739 | 19.58356 | 0.530057 | 1.100587 | 0.589917 |
| X | 142801000 | 142804399 | 41.02452 | 67.87248 | 58.55817 | 25.32144 | 0.592987 | 1.35642 | 0.589967 |
| X | 145987800 | 145990199 | 29.52574 | 48.47414 | 76.46214 | 32.93336 | -0.27822 | 1.433624 | 0.589982 |
| 15 | 97213400 | 97226999 | 145.732 | 216.1703 | 354.3216 | 145.9421 | -0.46074 | 3.672594 | 0.589984 |
| 15 | 59098600 | 59105799 | 100.8099 | 155.851 | 166.9482 | 69.34646 | 0.202662 | 2.852874 | 0.58999 |
| 2 | 139029800 | 139030999 | 11.49471 | 21.09122 | 39.03842 | 16.62665 | -0.45509 | 0.277474 | 0.589992 |
| 3 | 102141800 | 102146599 | 103.6911 | 160.0671 | 170.1734 | 70.68665 | 0.211919 | 2.881948 | 0.589999 |
| 9 | 19052200 | 19055999 | 61.55492 | 98.21552 | 104.6791 | 43.90003 | 0.247655 | 2.137237 | 0.590007 |
| 1 | 23009600 | 23011799 | 17.89473 | 30.88617 | 49.54487 | 21.73495 | -0.29384 | 0.866805 | 0.590021 |
| 8 | 56057600 | 56058599 | 16.2574 | 29.18845 | 25.75281 | 11.26432 | 0.582195 | 0.408953 | 0.590024 |
| 2 | 168882400 | 168884999 | 37.89304 | 61.08398 | 92.02287 | 39.1731 | -0.23327 | 1.789187 | 0.590031 |
| 1 | 117579600 | 117587199 | 166.3976 | 249.5488 | 257.3793 | 106.1924 | 0.221627 | 3.47314 | 0.590085 |
| 4 | 66357200 | 66357999 | 7.158942 | 13.83086 | 25.45316 | 9.778184 | -0.318 | -0.15438 | 0.590102 |
| 1 | 36662600 | 36668199 | 39.89534 | 64.06123 | 116.0555 | 49.12578 | -0.50588 | 1.921892 | 0.590114 |
| 2 | 45820400 | 45820999 | 4.64368 | 9.067831 | 18.74184 | 6.593085 | -0.36455 | -0.51501 | 0.590135 |
| 5 | 109401200 | 109407199 | 84.36417 | 133.2734 | 130.1553 | 54.05505 | 0.35996 | 2.443185 | 0.59014 |
| 10 | 18073400 | 18077599 | 35.73116 | 58.5844 | 64.0815 | 27.58539 | 0.247697 | 1.453919 | 0.590143 |
| 18 | 61667600 | 61670399 | 16.64756 | 28.83342 | 46.35254 | 20.33665 | -0.29573 | 0.827387 | 0.590154 |
| 1 | 188507800 | 188509999 | 16.67619 | 28.95441 | 45.81646 | 20.10745 | -0.27226 | 0.803771 | 0.590156 |
| 2 | 41436200 | 41437599 | 14.54035 | 26.5631 | 28.40544 | 12.16889 | 0.325737 | 0.302482 | 0.59017 |
| 2 | 39451400 | 39452199 | 8.055109 | 15.28179 | 27.12345 | 10.93745 | -0.33146 | 0.015083 | 0.590184 |
| 12 | 114071800 | 114073799 | 28.06865 | 47.62028 | 49.03902 | 21.48224 | 0.339247 | 1.062178 | 0.590213 |
| 1 | 151998600 | 152001599 | 42.52566 | 68.87368 | 74.70922 | 31.97046 | 0.246859 | 1.665673 | 0.590215 |
| 11 | 19645200 | 19647599 | 25.35388 | 43.72133 | 37.80629 | 16.58487 | 0.59387 | 0.89471 | 0.590223 |
| 7 | 29089600 | 29091999 | 20.38528 | 34.23347 | 56.93191 | 24.86013 | -0.35025 | 1.131919 | 0.590258 |
| 2 | 36592200 | 36601599 | 136.9568 | 209.0003 | 205.1472 | 85.06947 | 0.314212 | 3.118953 | 0.590267 |
| 4 | 113969800 | 113981599 | 281.1958 | 403.1274 | 403.7515 | 166.7396 | 0.202105 | 4.211307 | 0.590277 |
| X | 134118400 | 134119599 | 18.01188 | 31.93982 | 32.25776 | 14.25791 | 0.378287 | 0.511893 | 0.590284 |
| 5 | 17200000 | 17201399 | 16.93542 | 30.30553 | 31.12179 | 13.65068 | 0.361576 | 0.443632 | 0.590298 |
| 7 | 132729800 | 132731799 | 33.59714 | 55.4571 | 60.19823 | 25.98946 | 0.26241 | 1.364744 | 0.590298 |
| X | 103826200 | 103826799 | 8.741573 | 16.51681 | 28.67152 | 11.66158 | -0.3103 | 0.056204 | 0.590306 |
| 15 | 15994000 | 15998799 | 97.40911 | 151.0118 | 161.6647 | 67.14819 | 0.206617 | 2.800703 | 0.590315 |
| 10 | 88295200 | 88296599 | 26.66296 | 43.47773 | 74.13854 | 31.84803 | -0.39581 | 1.517967 | 0.590316 |
| 5 | 144556400 | 144567799 | 133.222 | 200.0808 | 283.396 | 116.9715 | -0.2339 | 3.440484 | 0.590331 |
| 7 | 53419600 | 53422799 | 31.60171 | 53.6583 | 47.91284 | 20.99412 | 0.544246 | 1.065477 | 0.590341 |
| 17 | 44560400 | 44562599 | 27.28931 | 46.11149 | 49.61216 | 21.70671 | 0.276002 | 1.092628 | 0.590353 |

| | | | | | | | | | |
|---|---|---|---|---|---|---|---|---|---|
| 6 | 72523600 | 72525399 | 16.20421 | 28.6777 | 30.91156 | 13.67004 | 0.283616 | 0.516539 | 0.590358 |
| 11 | 46794200 | 46797199 | 30.10749 | 51.2682 | 47.35498 | 20.75704 | 0.496081 | 1.034339 | 0.590378 |
| 12 | 114595600 | 114599999 | 96.08991 | 148.8995 | 159.5537 | 66.27414 | 0.204756 | 2.807488 | 0.590379 |
| 12 | 85760600 | 85764199 | 36.15663 | 59.3021 | 64.23474 | 27.65533 | 0.261944 | 1.449713 | 0.590382 |
| 3 | 9232600 | 9234999 | 16.03665 | 27.94328 | 44.18487 | 19.43482 | -0.27066 | 0.747935 | 0.590405 |
| X | 116848400 | 116858199 | 140.646 | 215.2508 | 193.1052 | 80.12943 | 0.447169 | 3.06377 | 0.590413 |
| 2 | 15502800 | 15504599 | 16.61677 | 28.94017 | 45.75004 | 20.11718 | -0.27055 | 0.758031 | 0.59042 |
| 15 | 8810400 | 8812399 | 24.5204 | 40.68103 | 64.01129 | 27.73292 | -0.27097 | 1.28542 | 0.590421 |
| 14 | 94681800 | 94682799 | 7.42582 | 14.34188 | 26.68386 | 10.26629 | -0.33642 | -0.1458 | 0.590445 |
| 18 | 89946800 | 89947399 | 11.91906 | 22.1292 | 23.15152 | 9.703442 | 0.379929 | 0.180558 | 0.590455 |
| 9 | 62179800 | 62181999 | 14.50909 | 25.17367 | 49.50475 | 21.7208 | -0.58451 | 0.807487 | 0.590455 |
| 18 | 82450400 | 82452399 | 23.23247 | 39.9201 | 43.08836 | 18.89995 | 0.274096 | 0.921088 | 0.590473 |
| 9 | 42801200 | 42804599 | 31.24395 | 51.00705 | 78.83058 | 33.87799 | -0.25334 | 1.510413 | 0.590479 |
| 4 | 12151200 | 12152799 | 35.59859 | 58.15316 | 63.21178 | 27.172 | 0.253776 | 1.502615 | 0.590486 |
| 8 | 123767800 | 123769799 | 13.47325 | 23.60851 | 39.97579 | 17.65605 | -0.36837 | 0.642033 | 0.590508 |
| 8 | 112382000 | 112383799 | 11.01266 | 19.98036 | 32.79077 | 14.16915 | -0.29618 | 0.326701 | 0.590523 |
| 15 | 7863000 | 7866999 | 30.94378 | 50.47516 | 77.43434 | 33.26789 | -0.24352 | 1.521464 | 0.590543 |
| 9 | 105079800 | 105082199 | 24.82108 | 42.42303 | 45.78235 | 20.07369 | 0.273206 | 0.97969 | 0.590556 |
| 12 | 113744400 | 113748999 | 67.85099 | 108.1271 | 108.764 | 45.53483 | 0.329091 | 2.183995 | 0.590557 |
| 6 | 137745000 | 137746999 | 24.98801 | 42.73698 | 45.77361 | 20.07094 | 0.284219 | 0.974363 | 0.590557 |
| 5 | 27645600 | 27647199 | 13.58823 | 24.20402 | 40.84989 | 17.98737 | -0.35508 | 0.469279 | 0.590558 |
| 16 | 30530200 | 30532399 | 21.98558 | 36.85099 | 58.62549 | 25.56907 | -0.2865 | 1.159008 | 0.590596 |
| 1 | 69979400 | 69979999 | 7.497376 | 14.23902 | 26.06902 | 10.48832 | -0.37219 | 0.002363 | 0.590615 |
| 13 | 3310400 | 3312999 | 38.08038 | 61.49282 | 93.2324 | 39.77711 | -0.23958 | 1.74024 | 0.59064 |
| 1 | 180971200 | 180971999 | 6.14429 | 11.79251 | 24.11109 | 9.457297 | -0.49624 | -0.10027 | 0.590677 |
| 1 | 148207600 | 148214999 | 70.76965 | 112.6023 | 113.1557 | 47.27116 | 0.328101 | 2.238245 | 0.590677 |
| 2 | 94923800 | 94927799 | 50.72685 | 81.48639 | 88.68671 | 37.56343 | 0.228549 | 1.914219 | 0.590701 |
| 8 | 126349800 | 126357999 | 107.2676 | 163.1733 | 233.0343 | 96.54984 | -0.23002 | 3.177353 | 0.590722 |
| 4 | 67247400 | 67247999 | 7.439112 | 14.25048 | 25.70803 | 10.13891 | -0.32574 | -0.06696 | 0.590771 |
| 7 | 31976200 | 32037799 | 1329.011 | 1680.589 | 1448.226 | 608.339 | 0.237201 | 6.186436 | 0.590801 |
| 16 | 64913000 | 64915399 | 16.00473 | 27.8215 | 44.7506 | 19.6543 | -0.29561 | 0.781349 | 0.590804 |
| 5 | 33390200 | 33390799 | 9.62374 | 18.41548 | 21.26048 | 8.4109 | 0.308685 | -0.04739 | 0.59081 |
| 14 | 40525000 | 40526999 | 12.27089 | 22.13955 | 36.29989 | 15.7951 | -0.30171 | 0.373724 | 0.590813 |
| 16 | 90271800 | 90275199 | 35.5711 | 57.65355 | 88.48327 | 37.88581 | -0.25165 | 1.644982 | 0.590864 |
| 4 | 112355000 | 112356999 | 34.65045 | 57.59471 | 57.64766 | 25.00803 | 0.379767 | 1.285455 | 0.590867 |
| 9 | 109728000 | 109729999 | 9.692834 | 17.83546 | 30.36474 | 12.88276 | -0.33064 | 0.229491 | 0.590873 |
| 17 | 23341000 | 23343399 | 21.97567 | 36.55399 | 68.28545 | 29.63782 | -0.51742 | 1.252486 | 0.59088 |
| 1 | 43982600 | 43983999 | 12.40421 | 23.18486 | 25.62709 | 10.62433 | 0.313231 | 0.148525 | 0.590881 |
| 13 | 57927000 | 57931399 | 21.52588 | 35.71639 | 69.20531 | 29.98525 | -0.57024 | 1.281549 | 0.590882 |
| 16 | 28672200 | 28675799 | 48.4236 | 77.1628 | 117.571 | 49.21762 | -0.26907 | 2.170097 | 0.590884 |
| 12 | 59709800 | 59713199 | 32.25953 | 53.83915 | 55.30984 | 24.05748 | 0.342362 | 1.232034 | 0.590884 |
| 8 | 72853200 | 72861799 | 86.62741 | 134.0417 | 190.0922 | 78.95958 | -0.20075 | 2.8332 | 0.59089 |
| 4 | 147848000 | 147935799 | 2632.696 | 3116.069 | 3551.788 | 1504.633 | -0.26322 | 7.238338 | 0.590895 |
| 1 | 9136400 | 9137399 | 8.201567 | 15.68613 | 27.95312 | 11.07468 | -0.31465 | -0.0386 | 0.590896 |
| 17 | 26656200 | 26659199 | 30.44504 | 49.24298 | 82.75754 | 35.38008 | -0.38496 | 1.675468 | 0.590901 |
| 7 | 23453200 | 23455999 | 30.36616 | 51.87314 | 46.23365 | 20.27071 | 0.548086 | 1.007066 | 0.590904 |
| 8 | 115140400 | 115141399 | 14.92706 | 26.77139 | 28.77039 | 12.59554 | 0.298235 | 0.41115 | 0.590905 |

| | | | | | | | | | |
|---|---|---|---|---|---|---|---|---|---|
| 11 | 85635800 | 85638199 | 15.19629 | 26.46486 | 43.68058 | 19.20657 | -0.33236 | 0.749072 | 0.590929 |
| 1 | 94129200 | 94132399 | 34.19148 | 56.34362 | 61.3869 | 26.48193 | 0.25624 | 1.386097 | 0.590935 |
| Y | 28869600 | 28872199 | 16.77724 | 29.63971 | 22.70473 | 10.04426 | 0.7724 | 0.519482 | 0.590943 |
| 4 | 28744000 | 28744599 | 8.009902 | 15.55372 | 18.37886 | 6.88621 | 0.349846 | -0.25096 | 0.590989 |
| 7 | 128644400 | 128645399 | 9.550096 | 17.69838 | 29.93203 | 12.57675 | -0.31081 | 0.188474 | 0.590991 |
| 10 | 59377200 | 59386999 | 116.0797 | 175.5234 | 280.7446 | 116.1037 | -0.40058 | 3.30364 | 0.590996 |
| 7 | 106806200 | 106810599 | 110.5968 | 167.8514 | 242.4295 | 100.3733 | -0.24896 | 3.22654 | 0.591018 |
| 10 | 58561200 | 58561599 | 7.011874 | 13.51178 | 24.75204 | 9.597412 | -0.3239 | -0.12675 | 0.591019 |
| 15 | 65381200 | 65383199 | 34.09392 | 56.05496 | 60.8161 | 26.2104 | 0.260962 | 1.411546 | 0.591019 |
| 13 | 87379000 | 87383799 | 76.77768 | 121.3983 | 127.4736 | 52.96552 | 0.256519 | 2.414992 | 0.591055 |
| 5 | 88759000 | 88765199 | 51.80515 | 82.21367 | 140.2985 | 58.44633 | -0.43842 | 2.30518 | 0.591082 |
| Y | 41880600 | 41883799 | 22.74884 | 39.04258 | 31.09975 | 13.64052 | 0.710236 | 0.912099 | 0.591087 |
| 5 | 112059000 | 112062999 | 62.83703 | 99.12521 | 144.9148 | 60.14159 | -0.22292 | 2.464348 | 0.59109 |
| 13 | 58140200 | 58142599 | 17.14793 | 29.68143 | 46.87935 | 20.56739 | -0.27051 | 0.836593 | 0.591106 |
| 10 | 59405200 | 59409599 | 31.32937 | 51.32313 | 84.3602 | 36.30163 | -0.33885 | 1.468577 | 0.591119 |
| 6 | 52855600 | 52858199 | 31.0506 | 51.98178 | 54.09491 | 23.57492 | 0.323777 | 1.196596 | 0.591122 |
| 1 | 8191800 | 8200399 | 242.2214 | 351.6776 | 356.5185 | 146.9765 | 0.202658 | 4.021881 | 0.591134 |
| 1 | 25966200 | 25968599 | 41.81084 | 67.75458 | 73.63246 | 31.52293 | 0.244922 | 1.651522 | 0.591136 |
| 2 | 168942200 | 168947199 | 70.19996 | 109.1873 | 190.3701 | 79.03243 | -0.49259 | 2.734774 | 0.591166 |
| 8 | 88405200 | 88405999 | 11.35866 | 21.42348 | 24.16129 | 9.864223 | 0.301948 | 0.077945 | 0.591167 |
| 12 | 88079000 | 88079599 | 11.35866 | 21.42348 | 24.16129 | 9.864223 | 0.301948 | 0.077945 | 0.591167 |
| X | 8187000 | 8204399 | 276.6374 | 403.3334 | 347.034 | 143.0041 | 0.445032 | 3.963792 | 0.59118 |
| 5 | 109277600 | 109278999 | 29.52438 | 50.11535 | 49.03466 | 21.48362 | 0.412793 | 1.062622 | 0.591182 |
| 5 | 125852600 | 125868999 | 195.1732 | 285.7344 | 385.8096 | 158.9279 | -0.19907 | 3.892479 | 0.59119 |
| 10 | 23335200 | 23337199 | 34.62951 | 56.66442 | 60.60683 | 26.06807 | 0.278636 | 1.472411 | 0.591199 |
| 1 | 15297800 | 15300199 | 23.33466 | 40.62414 | 40.20609 | 17.675 | 0.402008 | 0.787926 | 0.591206 |
| 2 | 144450600 | 144452799 | 27.40861 | 47.28392 | 43.48038 | 19.07422 | 0.505599 | 0.90636 | 0.591228 |
| 8 | 10853800 | 10854599 | 7.29785 | 14.0078 | 25.2069 | 9.884474 | -0.31413 | -0.08756 | 0.591271 |
| 6 | 133237000 | 133240599 | 42.70324 | 68.41596 | 119.7196 | 50.50379 | -0.45981 | 1.997068 | 0.591273 |
| 3 | 18008200 | 18009799 | 22.0517 | 38.33308 | 32.24812 | 14.16325 | 0.635289 | 0.791755 | 0.591274 |
| 13 | 119240400 | 119241199 | 14.89389 | 26.07213 | 41.48428 | 18.30706 | -0.27916 | 0.668405 | 0.591302 |
| 5 | 26489000 | 26492799 | 46.33857 | 74.73432 | 79.98294 | 34.06905 | 0.259701 | 1.780663 | 0.591306 |
| 16 | 30852200 | 30854999 | 18.65852 | 31.84971 | 51.11495 | 22.40952 | -0.29728 | 0.974284 | 0.591311 |
| X | 23864800 | 23878599 | 160.246 | 243.9298 | 205.1213 | 85.00241 | 0.534309 | 3.177453 | 0.59132 |
| 7 | 35305400 | 35308399 | 23.62604 | 39.44913 | 61.41232 | 26.71982 | -0.2553 | 1.207915 | 0.591321 |
| 1 | 130350000 | 130352199 | 14.92761 | 26.03367 | 42.64573 | 18.76819 | -0.32114 | 0.727961 | 0.591338 |
| 14 | 105263000 | 105264199 | 10.60004 | 19.49936 | 32.43284 | 13.7654 | -0.29799 | 0.241779 | 0.591338 |
| X | 153789400 | 153793199 | 54.55463 | 87.72687 | 87.65111 | 37.18107 | 0.353599 | 1.889168 | 0.591342 |
| 2 | 150834200 | 150834799 | 5.11418 | 9.959546 | 21.32735 | 7.802625 | -0.4681 | -0.36363 | 0.591346 |
| 2 | 76304200 | 76305799 | 17.97167 | 31.15096 | 49.00934 | 21.5028 | -0.26457 | 0.830721 | 0.591361 |
| 1 | 122575800 | 122581599 | 54.59435 | 88.10415 | 80.13999 | 34.16345 | 0.494881 | 1.77828 | 0.591395 |
| 4 | 52246200 | 52247599 | 15.44004 | 28.42625 | 27.85277 | 11.80822 | 0.461589 | 0.26426 | 0.59141 |
| 7 | 98735000 | 98737399 | 12.97012 | 22.82829 | 38.30789 | 16.95955 | -0.35471 | 0.570195 | 0.59142 |
| 17 | 18990000 | 19004799 | 190.5321 | 286.0892 | 251.1078 | 103.6384 | 0.455762 | 3.455766 | 0.591432 |
| 5 | 59879200 | 59880599 | 17.32884 | 30.15507 | 47.90138 | 21.05208 | -0.2776 | 0.777582 | 0.591442 |
| 5 | 146483200 | 146485599 | 17.73278 | 31.42569 | 28.15636 | 12.44822 | 0.549515 | 0.505526 | 0.59146 |
| 2 | 181117600 | 181119599 | 17.51002 | 31.00886 | 32.0649 | 14.17824 | 0.343266 | 0.517312 | 0.59147 |

| | | | | | | | | | |
|---|---|---|---|---|---|---|---|---|---|
| 5 | 97629600 | 97631199 | 15.20328 | 27.73792 | 28.86387 | 12.3855 | 0.363311 | 0.31575 | 0.59147 |
| 13 | 37418200 | 37419999 | 25.93878 | 44.06166 | 47.42513 | 20.7831 | 0.275797 | 1.042696 | 0.591508 |
| 1 | 49868800 | 49871199 | 51.9939 | 83.4751 | 90.10035 | 38.12992 | 0.239533 | 1.932881 | 0.591518 |
| 14 | 55343800 | 55346199 | 16.21067 | 27.88068 | 48.99658 | 21.4921 | -0.42589 | 0.890942 | 0.59152 |
| 11 | 68304000 | 68306399 | 28.97872 | 48.47834 | 52.9533 | 23.06693 | 0.254096 | 1.195369 | 0.591537 |
| 6 | 27996600 | 27998599 | 15.1763 | 26.6207 | 42.41656 | 18.75074 | -0.28134 | 0.637225 | 0.591547 |
| 10 | 40456800 | 40458999 | 19.24326 | 32.9891 | 51.68384 | 22.66676 | -0.26183 | 0.937212 | 0.591557 |
| 3 | 136278600 | 136279199 | 6.241481 | 12.12264 | 22.87209 | 8.553432 | -0.3167 | -0.26551 | 0.591567 |
| 16 | 19399600 | 19400999 | 15.63592 | 29.13398 | 26.42679 | 11.01408 | 0.590226 | 0.195863 | 0.591572 |
| 1 | 71902200 | 71903399 | 17.59164 | 30.96848 | 32.99887 | 14.6086 | 0.29731 | 0.587567 | 0.591572 |
| 1 | 115315400 | 115325799 | 131.7432 | 199.1038 | 291.9442 | 120.7154 | -0.27591 | 3.318317 | 0.591582 |
| 6 | 55118800 | 55123399 | 44.64827 | 71.94544 | 77.87027 | 33.1312 | 0.242076 | 1.804614 | 0.591586 |
| 14 | 41438000 | 41443399 | 167.2578 | 251.6723 | 249.2752 | 102.9401 | 0.284204 | 3.409385 | 0.591607 |
| 8 | 34927600 | 34928999 | 20.59451 | 35.98741 | 38.13122 | 16.80218 | 0.304559 | 0.720617 | 0.591634 |
| 5 | 53047400 | 53052399 | 56.2043 | 89.04792 | 133.2526 | 55.42327 | -0.25145 | 2.354644 | 0.591647 |
| 15 | 32813600 | 32815999 | 54.18654 | 86.71441 | 94.90339 | 39.96714 | 0.214137 | 2.037455 | 0.591686 |
| 8 | 81010200 | 81011599 | 18.99118 | 33.36021 | 35.38301 | 15.66046 | 0.302989 | 0.624774 | 0.591689 |
| 13 | 10721400 | 10722399 | 19.61775 | 33.50537 | 52.19992 | 22.88619 | -0.25457 | 0.974345 | 0.591703 |
| 5 | 125077800 | 125080799 | 22.91741 | 37.96686 | 63.85698 | 27.63859 | -0.36695 | 1.29645 | 0.591728 |
| 5 | 40173200 | 40173799 | 7.282254 | 14.02771 | 25.33624 | 9.834165 | -0.30574 | -0.12129 | 0.59173 |
| 16 | 48458200 | 48458999 | 7.282254 | 14.02771 | 25.33624 | 9.834165 | -0.30574 | -0.12129 | 0.59173 |
| 4 | 47291200 | 47300799 | 107.7606 | 163.7574 | 236.5828 | 97.99032 | -0.24785 | 3.197592 | 0.591732 |
| 12 | 48215400 | 48216199 | 11.60961 | 22.27039 | 22.85496 | 8.977888 | 0.486234 | -0.04739 | 0.591752 |
| 11 | 87367400 | 87369799 | 26.17899 | 44.28182 | 48.52379 | 21.23728 | 0.249773 | 1.079371 | 0.591753 |
| 14 | 65411000 | 65415999 | 65.44307 | 102.7373 | 153.2629 | 63.56606 | -0.25744 | 2.564256 | 0.591758 |
| 9 | 98719600 | 98727599 | 83.82687 | 129.2869 | 210.9655 | 87.63131 | -0.40657 | 2.894282 | 0.591764 |
| 4 | 21190400 | 21191799 | 12.59825 | 22.51614 | 36.18056 | 15.88626 | -0.28161 | 0.433436 | 0.591771 |
| 1 | 96168000 | 96169999 | 17.34679 | 30.11631 | 47.39875 | 20.80081 | -0.26456 | 0.810601 | 0.591771 |
| 1 | 83708400 | 83708799 | 3.931729 | 7.686147 | 17.15994 | 5.893816 | -0.44173 | -0.61531 | 0.59182 |
| 16 | 36144600 | 36145199 | 3.931729 | 7.686147 | 17.15994 | 5.893816 | -0.44173 | -0.61531 | 0.59182 |
| 15 | 22756000 | 22756799 | 11.50695 | 21.60273 | 23.69002 | 9.748282 | 0.333538 | 0.10977 | 0.591825 |
| 8 | 127986400 | 127987799 | 15.06152 | 27.6117 | 28.41337 | 12.11579 | 0.385543 | 0.287176 | 0.591827 |
| 5 | 79326400 | 79327199 | 7.021728 | 13.56655 | 25.43207 | 9.767449 | -0.34373 | -0.15565 | 0.591832 |
| 9 | 31702000 | 31704399 | 16.40788 | 29.55786 | 30.127 | 13.1345 | 0.378162 | 0.397425 | 0.591843 |
| 9 | 7267200 | 7269799 | 19.22648 | 32.65319 | 60.83473 | 26.658 | -0.51291 | 1.033521 | 0.591867 |
| 1 | 149462200 | 149465799 | 30.26591 | 51.00242 | 51.30231 | 22.42772 | 0.372644 | 1.127527 | 0.591868 |
| 1 | 79387200 | 79388199 | 15.67475 | 27.32278 | 43.65051 | 19.20425 | -0.28543 | 0.738925 | 0.591873 |
| 6 | 9647000 | 9648399 | 20.73945 | 37.05545 | 30.17437 | 13.25237 | 0.693152 | 0.476363 | 0.591875 |
| 3 | 97056200 | 97058799 | 28.44632 | 46.84656 | 72.09304 | 31.09215 | -0.24094 | 1.388016 | 0.591894 |
| 9 | 54063800 | 54070799 | 73.16598 | 114.7786 | 165.3708 | 68.58489 | -0.20815 | 2.58418 | 0.591895 |
| X | 18368400 | 18372199 | 47.56419 | 76.85017 | 79.01059 | 33.73365 | 0.320543 | 1.733722 | 0.5919 |
| 2 | 112003000 | 112007199 | 63.14148 | 100.4831 | 108.477 | 45.32791 | 0.225155 | 2.222403 | 0.591903 |
| 7 | 18328400 | 18331999 | 69.08383 | 109.4539 | 119.3534 | 49.64226 | 0.204458 | 2.361408 | 0.591906 |
| 13 | 93790200 | 93795799 | 59.4532 | 94.27004 | 136.7042 | 56.90014 | -0.20515 | 2.335185 | 0.591913 |
| X | 8224400 | 8251599 | 352.3957 | 502.7444 | 424.9433 | 175.6283 | 0.439772 | 4.288365 | 0.591916 |
| 3 | 138738800 | 138739999 | 15.0868 | 26.32483 | 42.62731 | 18.76615 | -0.30448 | 0.721482 | 0.591942 |
| 9 | 109413200 | 109415399 | 34.26748 | 55.55106 | 84.03208 | 35.98493 | -0.23053 | 1.636859 | 0.591953 |

| | | | | | | | | | |
|---|---|---|---|---|---|---|---|---|---|
| 2 | 44172400 | 44173599 | 10.44765 | 19.18657 | 31.87211 | 13.55855 | -0.29852 | 0.248957 | 0.59197 |
| 17 | 72771200 | 72772399 | 10.44765 | 19.18657 | 31.87211 | 13.55855 | -0.29852 | 0.248957 | 0.59197 |
| 13 | 49967800 | 49968599 | 10.45628 | 19.28645 | 31.95309 | 13.51248 | -0.28846 | 0.225199 | 0.591986 |
| 15 | 28320800 | 28322599 | 10.45628 | 19.28645 | 31.95309 | 13.51248 | -0.28846 | 0.225199 | 0.591986 |
| 15 | 86986600 | 86990799 | 52.76491 | 84.62706 | 91.86796 | 38.81804 | 0.229591 | 1.967244 | 0.591994 |
| 4 | 62785400 | 62792599 | 75.9092 | 118.0823 | 172.6055 | 71.65607 | -0.23841 | 2.733593 | 0.591994 |
| 14 | 95047400 | 95048399 | 19.62109 | 34.17014 | 35.8855 | 15.77884 | 0.317021 | 0.760995 | 0.592006 |
| 10 | 41815000 | 41822199 | 70.00824 | 109.8552 | 158.5226 | 65.74371 | -0.21005 | 2.57592 | 0.592032 |
| 10 | 122529400 | 122532999 | 26.29684 | 43.44282 | 67.46114 | 29.14175 | -0.25257 | 1.34102 | 0.592048 |
| 16 | 9677800 | 9679799 | 24.25798 | 41.1416 | 41.25205 | 18.06923 | 0.377568 | 1.040995 | 0.592077 |
| 15 | 39730200 | 39733799 | 41.33264 | 66.41358 | 99.2282 | 42.04465 | -0.22716 | 1.897682 | 0.592086 |
| 9 | 96235000 | 96236599 | 10.82046 | 19.72444 | 32.30906 | 13.87893 | -0.28799 | 0.296299 | 0.592112 |
| 13 | 42628800 | 42630399 | 10.82046 | 19.72444 | 32.30906 | 13.87893 | -0.28799 | 0.296299 | 0.592112 |
| 19 | 52971400 | 52975599 | 45.80751 | 73.29253 | 108.8473 | 45.82795 | -0.22571 | 2.038845 | 0.592117 |
| 2 | 151123800 | 151126799 | 53.70685 | 86.2318 | 80.29633 | 33.98487 | 0.452309 | 1.928733 | 0.592119 |
| 13 | 34089200 | 34089799 | 6.413614 | 12.4008 | 23.73827 | 9.085467 | -0.36838 | -0.17848 | 0.592139 |
| 2 | 36409000 | 36410399 | 20.65433 | 35.16513 | 54.88443 | 24.05708 | -0.25787 | 1.011008 | 0.592153 |
| 8 | 61010600 | 61012199 | 14.46948 | 25.48538 | 40.82657 | 18.07205 | -0.28768 | 0.577629 | 0.592155 |
| 2 | 11794200 | 11795599 | 18.81084 | 32.21053 | 50.59259 | 22.18611 | -0.26563 | 0.946016 | 0.592156 |
| 17 | 82052000 | 82053999 | 19.51731 | 34.32074 | 35.30098 | 15.63117 | 0.347301 | 0.613322 | 0.592195 |
| 5 | 70651600 | 70653799 | 20.7713 | 36.36385 | 36.87742 | 16.27525 | 0.367315 | 0.68818 | 0.592229 |
| 13 | 105060400 | 105061399 | 9.781168 | 18.11572 | 30.95244 | 13.01613 | -0.32685 | 0.193004 | 0.592288 |
| 16 | 32842400 | 32844399 | 9.781168 | 18.11572 | 30.95244 | 13.01613 | -0.32685 | 0.193004 | 0.592288 |
| 5 | 109417600 | 109420599 | 53.00659 | 85.2045 | 88.50626 | 37.52274 | 0.296916 | 1.89863 | 0.5923 |
| 18 | 56105200 | 56106799 | 17.59785 | 30.36122 | 48.47083 | 21.26351 | -0.28703 | 0.86855 | 0.592304 |
| 8 | 5579400 | 5582199 | 43.04418 | 68.9688 | 104.4667 | 44.07579 | -0.25173 | 1.991007 | 0.592318 |
| 12 | 40450400 | 40453599 | 43.04418 | 68.9688 | 104.4667 | 44.07579 | -0.25173 | 1.991007 | 0.592318 |
| 6 | 131067800 | 131075599 | 108.099 | 168.4436 | 154.3109 | 64.04482 | 0.436859 | 2.717447 | 0.592351 |
| 7 | 75690800 | 75693999 | 30.51795 | 49.63337 | 77.46818 | 33.24653 | -0.27132 | 1.564247 | 0.592353 |
| 1 | 43725000 | 43730799 | 103.0313 | 157.1619 | 224.7817 | 93.2022 | -0.22893 | 3.118991 | 0.592354 |
| 7 | 17489000 | 17514799 | 435.0461 | 606.3978 | 540.0757 | 224.0544 | 0.330628 | 4.63237 | 0.592362 |
| 6 | 99551600 | 99552999 | 8.743928 | 16.31025 | 28.45043 | 11.83542 | -0.34391 | 0.138105 | 0.592364 |
| 9 | 21526400 | 21527999 | 10.26124 | 18.53294 | 32.32806 | 14.04857 | -0.38873 | 0.34693 | 0.592374 |
| 12 | 84039400 | 84043199 | 40.16262 | 64.50977 | 99.59485 | 42.17731 | -0.27502 | 1.906683 | 0.59238 |
| 3 | 14807800 | 14811399 | 36.53986 | 59.98919 | 64.26012 | 27.67768 | 0.278522 | 1.439645 | 0.592412 |
| 17 | 18646400 | 18651799 | 71.79324 | 114.3051 | 110.6626 | 46.27121 | 0.382798 | 2.216786 | 0.592425 |
| 3 | 137556200 | 137559399 | 43.04747 | 69.09728 | 103.6326 | 43.84267 | -0.23446 | 1.935493 | 0.592426 |
| 12 | 91244000 | 91244999 | 8.93235 | 16.65046 | 28.97371 | 12.0669 | -0.34159 | 0.14678 | 0.592434 |
| 1 | 103345600 | 103347799 | 31.55353 | 51.1354 | 91.11946 | 39.0109 | -0.4667 | 1.642174 | 0.592438 |
| 18 | 72654800 | 72655399 | 9.973103 | 18.37999 | 30.88042 | 13.07361 | -0.30965 | 0.223178 | 0.592439 |
| 5 | 39354600 | 39355199 | 14.03428 | 25.35853 | 27.44253 | 11.91899 | 0.295989 | 0.359279 | 0.592446 |
| 18 | 7959600 | 7961399 | 18.04165 | 32.1611 | 32.12666 | 14.13722 | 0.398198 | 0.477668 | 0.59245 |
| 6 | 59801000 | 59804599 | 41.60939 | 68.43619 | 62.5957 | 26.98151 | 0.506832 | 1.428008 | 0.592469 |
| X | 151675400 | 151677799 | 13.65864 | 24.34854 | 42.4986 | 18.70097 | -0.4026 | 0.469146 | 0.592482 |
| 5 | 104719200 | 104721799 | 25.45998 | 42.52767 | 68.92181 | 29.99439 | -0.31326 | 1.216123 | 0.592488 |
| 1 | 11799000 | 11801199 | 21.59013 | 36.63291 | 57.10119 | 25.01479 | -0.25665 | 1.045884 | 0.592491 |
| Y | 20316000 | 20316999 | 10.05541 | 19.1509 | 16.70011 | 6.679177 | 0.696029 | -0.02294 | 0.592517 |

| | | | | | | | | | |
|---|---|---|---|---|---|---|---|---|---|
| 15 | 73216800 | 73219999 | 23.70243 | 41.22631 | 40.77311 | 17.91387 | 0.402939 | 0.803632 | 0.592517 |
| X | 42805400 | 42808199 | 20.08335 | 35.54204 | 33.55928 | 14.84102 | 0.473615 | 0.545966 | 0.592519 |
| 9 | 12409000 | 12409799 | 13.28183 | 24.19459 | 25.92096 | 11.14439 | 0.319653 | 0.302587 | 0.592522 |
| 9 | 100399400 | 100401999 | 23.21919 | 40.43216 | 39.62297 | 17.4211 | 0.416139 | 0.782259 | 0.592524 |
| 17 | 16496800 | 16498399 | 21.47342 | 37.42169 | 33.14591 | 14.58054 | 0.56161 | 0.75548 | 0.592535 |
| 17 | 90761400 | 90764399 | 27.74293 | 44.89346 | 91.08288 | 38.9527 | -0.65537 | 1.665587 | 0.592539 |
| 4 | 103451600 | 103451999 | 13.20911 | 24.07825 | 24.59969 | 10.56746 | 0.38885 | 0.296933 | 0.592567 |
| 2 | 150352800 | 150357199 | 40.9598 | 65.9662 | 101.1651 | 43.00861 | -0.26064 | 1.824754 | 0.592567 |
| X | 119966800 | 119968199 | 26.03442 | 44.15797 | 45.87943 | 20.0966 | 0.326585 | 1.055898 | 0.592579 |
| 4 | 113317600 | 113320399 | 44.37924 | 72.13551 | 71.94174 | 30.86818 | 0.372608 | 1.596907 | 0.592586 |
| 2 | 116361400 | 116394599 | 691.9101 | 911.2607 | 1117.208 | 467.3717 | -0.21258 | 5.524081 | 0.592633 |
| 8 | 102898400 | 102899399 | 13.35007 | 24.26947 | 25.52496 | 11.00208 | 0.344004 | 0.315015 | 0.592647 |
| 15 | 21917600 | 21925199 | 171.5094 | 255.5646 | 260.8983 | 107.5115 | 0.22798 | 3.593438 | 0.59266 |
| 13 | 65379200 | 65383599 | 36.78601 | 61.30798 | 55.99944 | 24.33601 | 0.511505 | 1.263083 | 0.592662 |
| 1 | 11040200 | 11041399 | 13.43703 | 24.71783 | 27.02411 | 11.46998 | 0.303616 | 0.253327 | 0.592665 |
| X | 9250400 | 9255999 | 104.1059 | 161.5124 | 161.2362 | 66.96285 | 0.308154 | 2.792773 | 0.592685 |
| 10 | 56388800 | 56393399 | 41.53206 | 68.00572 | 66.37714 | 28.55766 | 0.410875 | 1.484797 | 0.592687 |
| 17 | 56065200 | 56066199 | 11.4967 | 20.91144 | 34.36577 | 14.80278 | -0.2953 | 0.318938 | 0.592698 |
| 3 | 44148800 | 44152999 | 83.94467 | 131.7374 | 141.7329 | 58.78134 | 0.213111 | 2.579612 | 0.592715 |
| 19 | 28639800 | 28640999 | 7.647086 | 14.39121 | 27.5138 | 11.27536 | -0.45793 | 0.066881 | 0.592717 |
| 8 | 18545800 | 18549399 | 31.52951 | 53.24214 | 50.7424 | 22.19691 | 0.450357 | 1.115 | 0.592723 |
| 4 | 23525000 | 23528199 | 53.69023 | 86.23495 | 90.55446 | 38.34674 | 0.279854 | 1.922963 | 0.592733 |
| 5 | 32693600 | 32700999 | 106.4582 | 164.433 | 173.1964 | 71.94213 | 0.225954 | 2.872405 | 0.592744 |
| 2 | 20544400 | 20547399 | 21.72428 | 36.79786 | 57.66779 | 25.24997 | -0.26453 | 1.06068 | 0.592762 |
| 2 | 30879800 | 30886799 | 66.92894 | 105.0616 | 154.3079 | 63.99883 | -0.23509 | 2.567134 | 0.592785 |
| 13 | 113507400 | 113511199 | 35.25089 | 56.93053 | 88.13411 | 37.60777 | -0.26937 | 1.727995 | 0.592792 |
| 17 | 88115800 | 88117199 | 10.93948 | 19.97645 | 32.83022 | 14.07186 | -0.29086 | 0.287036 | 0.592793 |
| 2 | 19034600 | 19035599 | 12.44525 | 23.53088 | 24.68463 | 10.02544 | 0.412054 | 0.071666 | 0.592794 |
| 5 | 94497600 | 94503199 | 141.2398 | 214.9215 | 213.5964 | 88.50506 | 0.293 | 3.178587 | 0.592805 |
| X | 95662800 | 95662999 | 8.339529 | 16.20034 | 19.46674 | 7.270028 | 0.330037 | -0.25489 | 0.592805 |
| 5 | 145907400 | 145915599 | 109.2658 | 169.359 | 166.6386 | 69.21621 | 0.327969 | 2.809077 | 0.592831 |
| 2 | 97929800 | 97930999 | 13.94883 | 26.31036 | 25.11667 | 10.25297 | 0.540773 | 0.102612 | 0.592832 |
| 5 | 65830000 | 65832799 | 29.2481 | 49.48168 | 50.15479 | 21.95225 | 0.361899 | 1.091474 | 0.592838 |
| 7 | 3528800 | 3535599 | 108.7796 | 167.3263 | 179.1722 | 74.40812 | 0.197658 | 2.950613 | 0.592866 |
| 2 | 87769200 | 87771799 | 28.00611 | 47.54768 | 41.47292 | 18.17168 | 0.577784 | 1.044362 | 0.592869 |
| 17 | 87725600 | 87729999 | 30.74718 | 49.41452 | 105.3248 | 44.64062 | -0.73813 | 1.88087 | 0.592889 |
| Y | 90181800 | 90182799 | 9.094552 | 17.54698 | 14.66426 | 5.665592 | 0.802202 | -0.15213 | 0.592893 |
| 12 | 14490200 | 14493999 | 109.1872 | 167.7152 | 178.9437 | 74.29875 | 0.201289 | 2.978189 | 0.592902 |
| 13 | 90432800 | 90435999 | 46.80011 | 75.61977 | 69.54477 | 29.6943 | 0.481081 | 1.725792 | 0.592919 |
| 11 | 6480000 | 6482199 | 18.24466 | 30.60317 | 62.67615 | 27.34842 | -0.64946 | 1.144838 | 0.592921 |
| 16 | 16216200 | 16216999 | 15.01029 | 26.82087 | 28.89105 | 12.68803 | 0.29197 | 0.436254 | 0.592926 |
| 3 | 98657600 | 98658199 | 9.799442 | 18.1938 | 30.46164 | 12.76738 | -0.29417 | 0.179389 | 0.592941 |
| 17 | 62755800 | 62757199 | 9.799442 | 18.1938 | 30.46164 | 12.76738 | -0.29417 | 0.179389 | 0.592941 |
| 13 | 69257800 | 69258999 | 13.74768 | 26.00059 | 24.62787 | 9.996567 | 0.559014 | 0.08136 | 0.592943 |
| 19 | 13996800 | 14000199 | 50.26661 | 80.65393 | 87.89751 | 37.16403 | 0.224684 | 1.949521 | 0.592952 |
| 5 | 14215800 | 14216599 | 10.25886 | 19.81361 | 21.9994 | 8.463724 | 0.40186 | -0.1291 | 0.592956 |
| 6 | 115274800 | 115275999 | 10.25886 | 19.81361 | 21.9994 | 8.463724 | 0.40186 | -0.1291 | 0.592956 |

| | | | | | | | | | |
|---|---|---|---|---|---|---|---|---|---|
| 3 | 32129800 | 32132599 | 40.15264 | 65.30398 | 70.39941 | 30.21305 | 0.261181 | 1.577678 | 0.592956 |
| 10 | 41524400 | 41525999 | 18.18225 | 31.86538 | 34.14587 | 15.08337 | 0.288447 | 0.655874 | 0.592967 |
| 4 | 72205600 | 72207399 | 18.02904 | 31.28988 | 50.63221 | 22.21569 | -0.30459 | 0.822742 | 0.592984 |
| 5 | 6364000 | 6365999 | 27.92986 | 47.0386 | 50.70299 | 22.16182 | 0.273267 | 1.126079 | 0.593009 |
| 2 | 107783000 | 107794599 | 155.1406 | 233.5375 | 243.2983 | 100.4832 | 0.211906 | 3.39787 | 0.593011 |
| 5 | 86014200 | 86018799 | 35.32715 | 58.66126 | 58.24373 | 25.24931 | 0.391268 | 1.299127 | 0.593021 |
| 6 | 100461200 | 100464999 | 48.37121 | 77.89492 | 83.92305 | 35.67353 | 0.247531 | 1.831401 | 0.593028 |
| 1 | 60768200 | 60772999 | 48.60198 | 78.0783 | 85.14513 | 36.0669 | 0.225844 | 1.908315 | 0.593048 |
| 2 | 53482400 | 53483199 | 6.835444 | 13.19829 | 24.2716 | 9.349324 | -0.32089 | -0.15398 | 0.593054 |
| 16 | 97651800 | 97653399 | 18.75415 | 31.71768 | 53.71479 | 23.50808 | -0.37584 | 1.061211 | 0.593064 |
| 6 | 97348000 | 97349599 | 22.46131 | 39.02608 | 40.84771 | 17.93465 | 0.32134 | 0.811826 | 0.593082 |
| 17 | 88556000 | 88557999 | 21.77273 | 36.73788 | 57.05897 | 24.95417 | -0.25178 | 1.094481 | 0.593087 |
| 1 | 192216800 | 192218199 | 13.36158 | 24.42327 | 26.96023 | 11.54171 | 0.280854 | 0.286999 | 0.593099 |
| 7 | 9799800 | 9804599 | 78.61668 | 124.6193 | 122.7012 | 51.05664 | 0.352212 | 2.356779 | 0.593145 |
| 17 | 87563000 | 87569599 | 74.5861 | 116.5826 | 166.6246 | 69.13066 | -0.20063 | 2.648865 | 0.593149 |
| 2 | 3862800 | 3863399 | 9.000781 | 16.99878 | 29.22784 | 11.89938 | -0.29828 | 0.061106 | 0.593175 |
| 17 | 22429000 | 22432399 | 77.22284 | 122.5295 | 119.4034 | 49.733 | 0.368467 | 2.327816 | 0.59318 |
| 1 | 178576000 | 178577799 | 15.86089 | 27.77163 | 44.05154 | 19.44399 | -0.27484 | 0.673321 | 0.59321 |
| 15 | 22194200 | 22195599 | 20.79967 | 36.16042 | 39.33109 | 17.27519 | 0.266218 | 0.793485 | 0.593213 |
| 5 | 145941600 | 145943599 | 38.58902 | 62.77632 | 68.89684 | 29.57084 | 0.235705 | 1.571703 | 0.593218 |
| 1 | 183412200 | 183414599 | 19.72738 | 33.87224 | 53.88773 | 23.6363 | -0.28359 | 0.927317 | 0.593218 |
| X | 64385400 | 64387199 | 23.74301 | 41.37435 | 39.70621 | 17.46568 | 0.446539 | 0.778706 | 0.593226 |
| 18 | 64899200 | 64900399 | 11.59404 | 20.96326 | 34.24043 | 14.85908 | -0.2937 | 0.351723 | 0.593239 |
| 11 | 9435600 | 9440599 | 47.32244 | 76.95211 | 70.67071 | 30.32765 | 0.491715 | 1.595523 | 0.593263 |
| 3 | 98500600 | 98504999 | 102.473 | 158.228 | 169.3197 | 70.33196 | 0.202791 | 2.87507 | 0.593266 |
| 7 | 48055000 | 48056999 | 71.66853 | 113.9251 | 103.8325 | 43.32693 | 0.46739 | 2.265113 | 0.593297 |
| 11 | 79488000 | 79488999 | 11.60097 | 21.63732 | 23.64203 | 9.836836 | 0.325835 | 0.150912 | 0.593322 |
| 12 | 118917800 | 118919399 | 21.37833 | 36.09915 | 56.98172 | 24.92599 | -0.27488 | 1.089205 | 0.593353 |
| 15 | 26266000 | 26266599 | 7.75138 | 14.93886 | 27.75051 | 10.75578 | -0.34418 | -0.11433 | 0.593359 |
| 13 | 106025000 | 106025799 | 7.269846 | 13.97792 | 25.8508 | 10.08734 | -0.34711 | -0.10417 | 0.593376 |
| 2 | 88899600 | 88899999 | 7.713814 | 15.01089 | 18.44206 | 6.791393 | 0.316918 | -0.31484 | 0.593388 |
| 18 | 42510200 | 42513399 | 24.57656 | 40.92985 | 64.28646 | 27.92455 | -0.26812 | 1.239728 | 0.593394 |
| 1 | 52803200 | 52803599 | 4.001782 | 7.827262 | 17.29227 | 5.81779 | -0.39882 | -0.68294 | 0.593403 |
| 13 | 35758400 | 35758799 | 5.528682 | 10.75738 | 21.85081 | 8.054537 | -0.40291 | -0.32982 | 0.59341 |
| 5 | 52728200 | 52733599 | 71.31756 | 113.1713 | 119.3488 | 49.71544 | 0.254789 | 2.317377 | 0.59342 |
| 6 | 65477800 | 65479399 | 7.661735 | 14.51836 | 30.99765 | 12.53093 | -0.59895 | 0.030972 | 0.593421 |
| 14 | 40417200 | 40419599 | 24.27079 | 42.68407 | 34.35769 | 15.21316 | 0.699372 | 0.638102 | 0.593428 |
| 7 | 23662000 | 23676599 | 193.0191 | 286.1178 | 295.3255 | 121.6337 | 0.205159 | 3.688647 | 0.593439 |
| 14 | 6521400 | 6522399 | 11.80184 | 22.59066 | 23.35652 | 9.231557 | 0.467432 | -0.0243 | 0.593463 |
| 7 | 23541000 | 23565999 | 328.5493 | 471.6136 | 404.7696 | 167.1267 | 0.426592 | 4.19734 | 0.593478 |
| 17 | 47586600 | 47590399 | 36.2515 | 58.47884 | 89.60067 | 38.17656 | -0.25642 | 1.764683 | 0.593492 |
| 5 | 32890800 | 32892199 | 14.65586 | 26.22391 | 26.99323 | 11.84072 | 0.35834 | 0.421236 | 0.5935 |
| 5 | 149323200 | 149324399 | 24.18868 | 41.14445 | 44.80042 | 19.63557 | 0.259415 | 1.019876 | 0.593522 |
| X | 113236800 | 113242199 | 82.01924 | 129.3382 | 132.9538 | 55.17498 | 0.284105 | 2.483496 | 0.593535 |
| 7 | 103531000 | 103532999 | 17.04344 | 29.57775 | 46.87544 | 20.56731 | -0.27455 | 0.814248 | 0.593535 |
| 6 | 131481000 | 131482799 | 19.77811 | 34.97991 | 33.51322 | 14.82251 | 0.452239 | 0.549245 | 0.59354 |
| 10 | 43740200 | 43744199 | 53.99703 | 86.53795 | 93.92237 | 39.64066 | 0.228599 | 1.991459 | 0.593558 |

| | | | | | | | | | |
|---|---|---|---|---|---|---|---|---|---|
| 13 | 13193600 | 13197599 | 46.01063 | 74.36735 | 78.39338 | 33.48598 | 0.285148 | 1.719339 | 0.593591 |
| 7 | 44978400 | 44983399 | 33.36188 | 53.84866 | 86.69998 | 36.96826 | -0.32675 | 1.742542 | 0.593594 |
| 1 | 166203400 | 166204399 | 9.30589 | 17.521 | 30.26701 | 12.39399 | -0.31218 | 0.086104 | 0.593617 |
| 7 | 41825000 | 41842399 | 267.8755 | 390.2366 | 357.0371 | 147.1388 | 0.355298 | 3.974672 | 0.593632 |
| 1 | 110956200 | 110956799 | 9.159516 | 17.57821 | 19.26963 | 7.559906 | 0.394322 | -0.08443 | 0.593668 |
| 2 | 140158400 | 140167599 | 57.36338 | 91.77303 | 83.5094 | 35.15503 | 0.479529 | 2.045244 | 0.593673 |
| 1 | 25182800 | 25183799 | 9.33595 | 17.61058 | 30.89435 | 12.60989 | -0.33036 | 0.076638 | 0.59368 |
| 6 | 143065000 | 143068399 | 28.92292 | 47.53525 | 73.23019 | 31.55558 | -0.24365 | 1.414093 | 0.59369 |
| 2 | 125108400 | 125108799 | 3.754296 | 7.344033 | 16.17895 | 5.38984 | -0.38182 | -0.72776 | 0.59369 |
| 5 | 31761200 | 31762199 | 9.31705 | 17.39436 | 29.49727 | 12.25293 | -0.3022 | 0.138426 | 0.593696 |
| 17 | 17456800 | 17495799 | 915.0921 | 1198.672 | 992.8313 | 415.5912 | 0.345309 | 5.613862 | 0.593704 |
| 18 | 4142200 | 4143799 | 18.95798 | 33.3862 | 34.36848 | 15.21379 | 0.347022 | 0.594115 | 0.593734 |
| 18 | 58969000 | 58971199 | 22.25533 | 36.96497 | 62.23468 | 26.9843 | -0.36807 | 1.26344 | 0.593754 |
| 12 | 87845000 | 87846399 | 21.98515 | 38.0234 | 41.49689 | 18.20549 | 0.260167 | 0.85694 | 0.593809 |
| 1 | 159209200 | 159210199 | 7.902906 | 15.15064 | 26.88477 | 10.57948 | -0.29918 | -0.06574 | 0.593815 |
| 18 | 15863000 | 15864399 | 29.8091 | 48.26048 | 81.45742 | 34.8488 | -0.38983 | 1.656011 | 0.593835 |
| 9 | 63807400 | 63812999 | 56.97607 | 90.40265 | 131.9356 | 54.95203 | -0.21339 | 2.312564 | 0.59386 |
| 1 | 46384200 | 46386599 | 22.17459 | 38.96161 | 34.98085 | 15.48561 | 0.542106 | 0.636753 | 0.593872 |
| 2 | 54030800 | 54033199 | 19.77796 | 35.30574 | 31.36373 | 13.78518 | 0.567597 | 0.477961 | 0.593877 |
| 10 | 11365000 | 11371199 | 140.3522 | 213.0163 | 217.1935 | 89.9255 | 0.253034 | 3.231811 | 0.593892 |
| 7 | 6758800 | 6763399 | 34.91771 | 57.25931 | 61.9151 | 26.65452 | 0.264292 | 1.446143 | 0.593894 |
| 3 | 133007200 | 133013599 | 95.9137 | 149.2179 | 158.4365 | 65.77276 | 0.222394 | 2.738926 | 0.593895 |
| 16 | 91303400 | 91304199 | 11.54811 | 20.50645 | 35.25147 | 15.56281 | -0.38363 | 0.46575 | 0.593898 |
| 12 | 106067800 | 106069799 | 27.12161 | 45.41385 | 48.08234 | 20.9574 | 0.298944 | 1.174772 | 0.593951 |
| 8 | 20585400 | 20587799 | 25.52163 | 42.12025 | 66.70531 | 28.79547 | -0.2809 | 1.350727 | 0.593961 |
| 15 | 27303800 | 27305599 | 18.94465 | 33.08716 | 35.75055 | 15.74795 | 0.276487 | 0.71621 | 0.593965 |
| 7 | 61102200 | 61103799 | 16.26948 | 29.90103 | 28.37941 | 12.06012 | 0.505019 | 0.282748 | 0.593971 |
| 2 | 88409400 | 88410799 | 28.84012 | 48.24851 | 52.94074 | 23.06207 | 0.247617 | 1.194888 | 0.594008 |
| 4 | 145596600 | 145604399 | 173.9435 | 261.9726 | 244.8151 | 101.1231 | 0.369238 | 3.39292 | 0.594013 |
| 5 | 112117400 | 112131599 | 126.5236 | 189.6515 | 308.679 | 127.3436 | -0.43678 | 3.478934 | 0.59404 |
| 2 | 132089400 | 132090199 | 5.385356 | 10.47231 | 21.21081 | 7.856042 | -0.40444 | -0.31473 | 0.594056 |
| 8 | 117788800 | 117792599 | 39.85035 | 64.07698 | 96.84857 | 41.07831 | -0.24251 | 1.872252 | 0.594108 |
| 5 | 104093600 | 104101599 | 119.79 | 181.1712 | 255.0621 | 105.4894 | -0.21635 | 3.29849 | 0.594116 |
| 4 | 88502400 | 88505999 | 37.47004 | 60.39783 | 92.15288 | 39.22487 | -0.25164 | 1.792565 | 0.594169 |
| 6 | 136264600 | 136266199 | 17.3742 | 30.56538 | 32.99771 | 14.61002 | 0.27813 | 0.59274 | 0.594174 |
| 12 | 103488200 | 103490199 | 18.61933 | 33.29075 | 31.4923 | 13.82301 | 0.478728 | 0.460936 | 0.594206 |
| 13 | 71241200 | 71243799 | 17.45155 | 30.29904 | 47.39788 | 20.8008 | -0.25593 | 0.809354 | 0.594211 |
| 17 | 27593800 | 27626799 | 459.5142 | 621.2421 | 828.3464 | 345.2916 | -0.29586 | 5.103698 | 0.594221 |
| 12 | 64769200 | 64773799 | 52.09007 | 83.59757 | 90.85593 | 38.42736 | 0.228989 | 1.946341 | 0.594233 |
| 7 | 33746800 | 33756799 | 138.253 | 210.1304 | 215.9697 | 89.45708 | 0.243188 | 3.20027 | 0.594256 |
| 1 | 5092800 | 5094199 | 13.78057 | 24.08526 | 41.56907 | 18.32008 | -0.39589 | 0.687199 | 0.594261 |
| 2 | 17626200 | 17627799 | 19.54771 | 34.30684 | 35.83556 | 15.84866 | 0.324841 | 0.641585 | 0.594263 |
| 8 | 27076600 | 27079199 | 31.13076 | 50.28276 | 93.44614 | 39.8809 | -0.53204 | 1.722446 | 0.594266 |
| 6 | 8701000 | 8706399 | 74.30943 | 117.8025 | 108.9194 | 45.32614 | 0.442907 | 2.345278 | 0.594275 |
| 13 | 43236400 | 43237599 | 19.53886 | 34.14862 | 37.12656 | 16.36187 | 0.267536 | 0.710078 | 0.594275 |
| 14 | 81116200 | 81117199 | 10.38598 | 19.06885 | 31.90761 | 13.57799 | -0.30885 | 0.252865 | 0.594285 |
| 1 | 48848200 | 48848999 | 9.958447 | 18.39522 | 30.91023 | 13.04536 | -0.30663 | 0.209902 | 0.594293 |

| | | | | | | | | | |
|---|---|---|---|---|---|---|---|---|---|
| 2 | 4686200 | 4690999 | 52.46541 | 84.35952 | 87.893 | 37.2787 | 0.293064 | 1.885703 | 0.594308 |
| 2 | 56112000 | 56112599 | 7.76881 | 14.81445 | 26.13836 | 10.42645 | -0.30876 | -0.02405 | 0.594342 |
| 12 | 3431000 | 3433599 | 17.4551 | 30.20239 | 47.40639 | 20.79839 | -0.26173 | 0.841438 | 0.594387 |
| 12 | 86976000 | 86979599 | 30.10696 | 48.8384 | 87.90567 | 37.66189 | -0.47978 | 1.619346 | 0.594397 |
| 10 | 68050800 | 68054399 | 41.40672 | 67.71574 | 67.38969 | 28.97953 | 0.381835 | 1.502566 | 0.594426 |
| 1 | 169837800 | 169840399 | 22.75418 | 38.99384 | 42.11184 | 18.46827 | 0.272715 | 0.942167 | 0.594445 |
| 12 | 93411800 | 93412399 | 9.12503 | 16.69709 | 30.38058 | 12.98787 | -0.43364 | 0.257282 | 0.594446 |
| 18 | 8013800 | 8014599 | 10.83951 | 20.64785 | 23.25891 | 9.298935 | 0.32994 | 0.002802 | 0.594454 |
| 10 | 58070800 | 58076799 | 113.1538 | 171.5013 | 244.7854 | 101.3076 | -0.23349 | 3.253592 | 0.594501 |
| 2 | 163347000 | 163349399 | 16.54335 | 28.70508 | 45.82987 | 20.10845 | -0.28518 | 0.811721 | 0.594524 |
| 14 | 7245200 | 7247999 | 43.96857 | 70.92103 | 77.55524 | 33.04038 | 0.228923 | 1.773707 | 0.594528 |
| 7 | 140151800 | 140152799 | 8.092731 | 15.52887 | 28.69058 | 11.26023 | -0.35329 | -0.06679 | 0.594529 |
| 1 | 175644200 | 175644799 | 5.658645 | 11.01737 | 21.3097 | 7.814159 | -0.32534 | -0.34567 | 0.59454 |
| X | 144936400 | 144938999 | 31.03702 | 52.4304 | 50.73452 | 22.19602 | 0.42851 | 1.110186 | 0.594557 |
| 3 | 42233200 | 42234399 | 9.95234 | 18.50997 | 31.03607 | 12.97532 | -0.29359 | 0.171732 | 0.594565 |
| 6 | 24322600 | 24323599 | 9.95234 | 18.50997 | 31.03607 | 12.97532 | -0.29359 | 0.171732 | 0.594565 |
| 11 | 65845200 | 65846399 | 9.95234 | 18.50997 | 31.03607 | 12.97532 | -0.29359 | 0.171732 | 0.594565 |
| 12 | 76792800 | 76793799 | 35.79849 | 58.45431 | 62.58284 | 26.89738 | 0.27518 | 1.509632 | 0.594569 |
| 12 | 86913000 | 86914599 | 16.94247 | 29.79399 | 29.18008 | 12.92076 | 0.417717 | 0.586928 | 0.59459 |
| 10 | 112893400 | 112895999 | 25.23601 | 43.80391 | 41.2786 | 18.11263 | 0.472044 | 0.838492 | 0.594594 |
| 1 | 117845600 | 117850799 | 58.22635 | 93.76265 | 83.8398 | 35.63673 | 0.515878 | 1.84815 | 0.594597 |
| X | 94490600 | 94491199 | 10.7231 | 20.51618 | 22.86798 | 9.050608 | 0.358747 | -0.02869 | 0.594622 |
| 5 | 138565200 | 138568599 | 30.64568 | 51.10628 | 55.21918 | 24.00636 | 0.26984 | 1.235685 | 0.594628 |
| 6 | 102630400 | 102631999 | 16.19422 | 27.61836 | 56.37071 | 24.71193 | -0.64291 | 0.979176 | 0.594637 |
| 7 | 117102800 | 117106399 | 25.02536 | 41.23783 | 68.3707 | 29.49437 | -0.34772 | 1.367746 | 0.594643 |
| 3 | 43838600 | 43839199 | 15.2613 | 26.47111 | 44.81038 | 19.66092 | -0.36927 | 0.807069 | 0.594644 |
| 15 | 83289600 | 83290799 | 11.5552 | 20.92212 | 34.84328 | 15.0944 | -0.31955 | 0.348081 | 0.594652 |
| 18 | 54717000 | 54717999 | 12.57802 | 23.45665 | 25.51009 | 10.61639 | 0.331743 | 0.162047 | 0.594669 |
| 18 | 37071000 | 37078199 | 131.1211 | 199.7964 | 179.5608 | 74.43259 | 0.439605 | 3.145235 | 0.594681 |
| 13 | 36986800 | 36987399 | 6.690029 | 12.94822 | 23.81994 | 9.074116 | -0.30589 | -0.19382 | 0.594722 |
| 17 | 92071400 | 92072799 | 12.97087 | 23.04516 | 36.68221 | 16.18719 | -0.27292 | 0.475188 | 0.59474 |
| 16 | 86976600 | 86979599 | 24.67728 | 42.40053 | 44.67031 | 19.59381 | 0.30896 | 0.931023 | 0.594743 |
| 8 | 80948000 | 80949199 | 12.80585 | 24.31819 | 23.60753 | 9.500209 | 0.534349 | 0.041872 | 0.594745 |
| 15 | 89858400 | 89859999 | 10.49203 | 19.06091 | 32.27601 | 13.92373 | -0.33965 | 0.312211 | 0.594769 |
| 4 | 40016600 | 40018399 | 11.94254 | 21.53203 | 34.72243 | 15.12196 | -0.27864 | 0.372363 | 0.594778 |
| 3 | 21835800 | 21838799 | 29.05125 | 49.2433 | 49.59467 | 21.71977 | 0.371178 | 1.073576 | 0.594785 |
| 9 | 4858000 | 4859999 | 44.37174 | 71.55876 | 77.96024 | 33.20539 | 0.234043 | 1.78092 | 0.594795 |
| 5 | 49708600 | 49710799 | 25.11013 | 41.26618 | 76.63312 | 33.02102 | -0.51242 | 1.408887 | 0.594795 |
| 9 | 3880800 | 3888999 | 101.7716 | 157.4468 | 168.1877 | 69.86104 | 0.207448 | 2.840711 | 0.5948 |
| 1 | 48861600 | 48862399 | 17.23868 | 30.35968 | 32.50951 | 14.39077 | 0.290469 | 0.579813 | 0.594823 |
| 13 | 13241200 | 13242399 | 16.49167 | 30.76314 | 25.84039 | 10.74913 | 0.702563 | 0.194749 | 0.594828 |
| X | 97783200 | 97788199 | 84.57168 | 130.6285 | 190.8623 | 79.28017 | -0.2457 | 2.866786 | 0.594829 |
| 8 | 39887400 | 39889399 | 16.46809 | 28.72043 | 45.19566 | 19.89047 | -0.26376 | 0.737542 | 0.594839 |
| 11 | 33053400 | 33055599 | 15.99297 | 27.76775 | 45.29306 | 19.87756 | -0.31591 | 0.798612 | 0.594859 |
| 19 | 20750400 | 20751799 | 15.12 | 27.05594 | 29.29029 | 12.84771 | 0.285873 | 0.427949 | 0.594865 |
| 15 | 72803800 | 72810599 | 64.20624 | 101.0029 | 149.8114 | 62.15115 | -0.24687 | 2.521252 | 0.594898 |
| 8 | 9996000 | 9998199 | 34.55151 | 56.54223 | 61.62209 | 26.50568 | 0.251762 | 1.473086 | 0.594904 |

| | | | | | | | | | |
|---|---|---|---|---|---|---|---|---|---|
| 9 | 72812000 | 72813799 | 10.24052 | 18.49875 | 32.30214 | 14.03423 | -0.39016 | 0.344607 | 0.594923 |
| 13 | 8004400 | 8005399 | 12.38077 | 22.80114 | 24.09593 | 10.2111 | 0.353988 | 0.231203 | 0.594933 |
| 2 | 80309400 | 80312199 | 28.47218 | 47.52057 | 51.85716 | 22.5564 | 0.255549 | 1.217181 | 0.594944 |
| 14 | 57114000 | 57116199 | 17.61408 | 30.4169 | 47.93901 | 21.03079 | -0.26823 | 0.86043 | 0.594955 |
| 16 | 87150200 | 87151399 | 7.923511 | 14.97163 | 27.03828 | 10.99649 | -0.36706 | 0.04179 | 0.594995 |
| 4 | 91585600 | 91590199 | 48.26527 | 77.90555 | 80.93557 | 34.51132 | 0.303861 | 1.762278 | 0.595003 |
| 8 | 55966200 | 55973799 | 66.657 | 103.9898 | 179.6946 | 74.56791 | -0.47629 | 2.680628 | 0.595015 |
| 17 | 22905600 | 22922799 | 311.2543 | 445.9034 | 417.8485 | 172.5805 | 0.297032 | 4.224972 | 0.595016 |
| 1 | 27524600 | 27529399 | 42.60648 | 69.23191 | 71.84768 | 30.82057 | 0.314902 | 1.597838 | 0.595064 |
| 6 | 143442600 | 143453799 | 111.6572 | 170.0657 | 237.5178 | 98.44469 | -0.19637 | 3.15265 | 0.595077 |
| 18 | 44507600 | 44508399 | 11.8879 | 21.04639 | 35.79279 | 15.82754 | -0.37029 | 0.495627 | 0.595089 |
| 7 | 48566800 | 48568399 | 12.65995 | 22.43848 | 36.20429 | 16.0059 | -0.29427 | 0.488868 | 0.595112 |
| 5 | 135972200 | 135974799 | 23.193 | 39.73017 | 42.59811 | 18.68094 | 0.282966 | 0.947303 | 0.595126 |
| 3 | 7811000 | 7813999 | 26.58899 | 44.16643 | 69.47608 | 30.12573 | -0.27058 | 1.280075 | 0.595139 |
| 5 | 8246400 | 8256999 | 128.7705 | 196.7977 | 201.1341 | 83.43377 | 0.257639 | 3.087008 | 0.595155 |
| 3 | 146057800 | 146059799 | 18.02468 | 30.88999 | 49.52781 | 21.72063 | -0.29495 | 0.934558 | 0.595155 |
| 14 | 105998600 | 106000599 | 16.02352 | 29.2012 | 28.68271 | 12.32528 | 0.444502 | 0.326039 | 0.595194 |
| 5 | 28105800 | 28113599 | 72.28493 | 113.5356 | 165.1639 | 68.50652 | -0.22067 | 2.559306 | 0.595201 |
| X | 43654200 | 43659399 | 27.17005 | 46.86638 | 37.64619 | 16.51478 | 0.699818 | 0.899658 | 0.595207 |
| 7 | 62748600 | 62749599 | 11.57966 | 20.52697 | 35.30074 | 15.60596 | -0.38548 | 0.47889 | 0.595219 |
| 10 | 37903400 | 37905399 | 26.5728 | 45.07029 | 48.50239 | 21.24542 | 0.275943 | 1.058528 | 0.595227 |
| 5 | 41516000 | 41519599 | 33.87473 | 56.20069 | 58.6783 | 25.42246 | 0.318863 | 1.305729 | 0.595235 |
| X | 97784000 | 97788199 | 81.71001 | 126.4373 | 185.7015 | 77.13578 | -0.25118 | 2.831714 | 0.595242 |
| 11 | 13808200 | 13808799 | 11.22607 | 21.04166 | 23.62788 | 9.749145 | 0.296131 | 0.115377 | 0.595282 |
| 3 | 10517000 | 10526199 | 209.8381 | 308.6053 | 318.0727 | 130.9875 | 0.196612 | 3.821078 | 0.595302 |
| 11 | 42866200 | 42867199 | 10.6413 | 19.02144 | 37.86459 | 16.62359 | -0.58828 | 0.424731 | 0.595304 |
| 12 | 119877800 | 119879999 | 18.16261 | 33.12486 | 28.58274 | 12.27055 | 0.631367 | 0.338479 | 0.595313 |
| 12 | 114138200 | 114140999 | 42.05679 | 68.42608 | 71.13102 | 30.53113 | 0.313909 | 1.578171 | 0.595332 |
| 5 | 35704400 | 35708199 | 30.68401 | 49.84472 | 78.52629 | 33.67843 | -0.28614 | 1.585243 | 0.595336 |
| 8 | 46248000 | 46249799 | 15.85929 | 27.58858 | 44.22189 | 19.43091 | -0.29046 | 0.77011 | 0.59534 |
| 9 | 35578600 | 35579999 | 19.13125 | 32.58808 | 52.1824 | 22.87363 | -0.29458 | 0.995937 | 0.59537 |
| 7 | 75722600 | 75725199 | 22.1219 | 38.82075 | 35.56908 | 15.72948 | 0.512984 | 0.656274 | 0.595375 |
| 13 | 60005400 | 60006599 | 14.7539 | 26.61418 | 28.7549 | 12.51519 | 0.296115 | 0.375197 | 0.59538 |
| 17 | 92654600 | 92656999 | 32.44571 | 54.10629 | 55.87203 | 24.28967 | 0.334966 | 1.241737 | 0.595381 |
| 11 | 48625200 | 48626399 | 13.09062 | 23.07802 | 38.74929 | 17.14938 | -0.35476 | 0.552468 | 0.595411 |
| 14 | 94819800 | 94826399 | 103.4222 | 160.3866 | 165.1215 | 68.58061 | 0.263286 | 2.796753 | 0.595412 |
| 6 | 133294800 | 133297199 | 30.82238 | 51.8477 | 51.33192 | 22.42854 | 0.39532 | 1.143189 | 0.595439 |
| 9 | 31385600 | 31387599 | 24.38023 | 42.15835 | 37.71354 | 16.54557 | 0.54576 | 0.864241 | 0.595442 |
| 15 | 15135200 | 15136199 | 20.21539 | 35.37315 | 27.92168 | 12.31763 | 0.726994 | 0.678038 | 0.59545 |
| 5 | 142248800 | 142252999 | 42.6042 | 69.17656 | 72.62116 | 31.13523 | 0.297391 | 1.615561 | 0.595454 |
| 13 | 48528800 | 48530199 | 10.1511 | 18.9026 | 32.17548 | 13.42402 | -0.31273 | 0.169094 | 0.595463 |
| 17 | 4944000 | 4945399 | 9.341691 | 17.36651 | 29.42821 | 12.31241 | -0.30938 | 0.166269 | 0.595466 |
| 11 | 51185600 | 51189799 | 43.16795 | 69.18876 | 103.6952 | 43.77109 | -0.23595 | 1.980245 | 0.595472 |
| 12 | 115227800 | 115245799 | 292.0933 | 422.1827 | 384.1516 | 158.458 | 0.352151 | 4.094024 | 0.595475 |
| 15 | 60906800 | 60907399 | 5.227809 | 10.18978 | 20.26914 | 7.33064 | -0.34697 | -0.40418 | 0.595493 |
| X | 123071000 | 123071799 | 17.88743 | 31.72919 | 31.80821 | 14.05546 | 0.389261 | 0.508962 | 0.595525 |
| X | 80276600 | 80279399 | 30.7269 | 51.87576 | 44.73745 | 19.56882 | 0.593999 | 1.107552 | 0.595533 |

| | | | | | | | | | |
|---|---|---|---|---|---|---|---|---|---|
| 3 | 93928600 | 93933399 | 36.16899 | 59.45392 | 62.20503 | 26.80658 | 0.313021 | 1.419266 | 0.595543 |
| 11 | 32910800 | 32914399 | 32.11053 | 52.1621 | 80.63984 | 34.58492 | -0.25902 | 1.588265 | 0.59555 |
| 6 | 79297000 | 79299199 | 39.53388 | 64.11524 | 70.44233 | 30.17471 | 0.230281 | 1.630405 | 0.595566 |
| 12 | 60757200 | 60760999 | 88.81012 | 138.2381 | 146.4509 | 60.79238 | 0.226196 | 2.725459 | 0.595571 |
| 1 | 186409000 | 186410399 | 8.261724 | 15.46071 | 27.96481 | 11.56985 | -0.38962 | 0.110572 | 0.595578 |
| 8 | 64210800 | 64214599 | 28.90753 | 48.85512 | 50.17098 | 21.95247 | 0.342982 | 1.096791 | 0.595588 |
| 17 | 40832600 | 40833599 | 10.50676 | 19.33167 | 31.92161 | 13.54469 | -0.28701 | 0.240808 | 0.595591 |
| 16 | 71189000 | 71192199 | 23.93013 | 41.52039 | 40.76017 | 17.88403 | 0.413087 | 0.834292 | 0.595608 |
| 12 | 103853400 | 103887799 | 494.4737 | 668.8754 | 837.5525 | 349.0995 | -0.20445 | 5.095986 | 0.595615 |
| 2 | 55525400 | 55526999 | 21.08729 | 35.96231 | 57.15249 | 25.0546 | -0.2836 | 0.998328 | 0.595617 |
| 5 | 78584400 | 78584799 | 6.238896 | 12.1257 | 22.94914 | 8.536169 | -0.31408 | -0.28468 | 0.595635 |
| 10 | 41319600 | 41319999 | 6.742685 | 12.90391 | 25.12265 | 9.935587 | -0.43729 | -0.06219 | 0.595651 |
| X | 90914600 | 90920199 | 67.54115 | 108.0168 | 99.07476 | 41.68619 | 0.467579 | 2.075972 | 0.595676 |
| 6 | 109343800 | 109344999 | 12.17001 | 21.8448 | 35.23217 | 15.41369 | -0.28306 | 0.403527 | 0.595706 |
| 10 | 29834600 | 29836999 | 23.32491 | 38.5667 | 72.61948 | 31.38889 | -0.52959 | 1.329834 | 0.595711 |
| 4 | 122307600 | 122309399 | 19.69764 | 34.67318 | 30.63013 | 13.56017 | 0.566169 | 0.594586 | 0.595715 |
| 4 | 147300400 | 147300999 | 12.28799 | 23.75292 | 22.03816 | 8.444814 | 0.664369 | -0.12013 | 0.59573 |
| 2 | 108934200 | 108935399 | 9.04874 | 16.83252 | 28.92795 | 12.09018 | -0.32814 | 0.159177 | 0.595744 |
| 3 | 56803000 | 56804199 | 9.04874 | 16.83252 | 28.92795 | 12.09018 | -0.32814 | 0.159177 | 0.595744 |
| 11 | 12720000 | 12720799 | 9.04874 | 16.83252 | 28.92795 | 12.09018 | -0.32814 | 0.159177 | 0.595744 |
| 7 | 112225800 | 112226199 | 4.38707 | 8.57328 | 18.28242 | 6.336551 | -0.39016 | -0.57523 | 0.595774 |
| 16 | 76717000 | 76717799 | 9.286584 | 17.78953 | 19.24234 | 7.588913 | 0.406258 | -0.06674 | 0.595786 |
| 10 | 43437200 | 43440399 | 25.38242 | 43.26138 | 46.31467 | 20.30252 | 0.284061 | 1.006155 | 0.595794 |
| 19 | 37211200 | 37213599 | 23.72808 | 39.26857 | 64.42688 | 27.86502 | -0.33138 | 1.311751 | 0.595804 |
| 1 | 94188000 | 94189599 | 18.77226 | 31.87603 | 59.11691 | 25.90383 | -0.50626 | 1.032846 | 0.595807 |
| 2 | 30654200 | 30657799 | 28.46401 | 46.32048 | 76.37297 | 32.78362 | -0.34951 | 1.5533 | 0.595833 |
| X | 80671400 | 80674799 | 37.48789 | 61.98442 | 60.50263 | 26.1515 | 0.415421 | 1.356777 | 0.595845 |
| X | 17842600 | 17844999 | 41.63198 | 67.81467 | 69.89364 | 30.01289 | 0.327457 | 1.561168 | 0.59585 |
| 11 | 49341800 | 49344399 | 22.14085 | 38.35673 | 41.46206 | 18.19113 | 0.274383 | 0.84267 | 0.595855 |
| 8 | 68696200 | 68700199 | 50.43157 | 80.93383 | 88.64337 | 37.49011 | 0.217822 | 1.944141 | 0.595868 |
| 1 | 189550400 | 189552599 | 12.6438 | 22.32439 | 37.25611 | 16.4835 | -0.34497 | 0.531415 | 0.595882 |
| 7 | 124615600 | 124616799 | 8.716539 | 16.44066 | 28.07908 | 11.46305 | -0.29183 | 0.065814 | 0.595926 |
| 3 | 19166400 | 19170999 | 71.12335 | 112.5921 | 121.9813 | 50.71405 | 0.212845 | 2.381561 | 0.595937 |
| 7 | 29235800 | 29237399 | 12.40488 | 22.25554 | 36.19902 | 15.84318 | -0.29591 | 0.407441 | 0.59594 |
| 7 | 51987600 | 51988199 | 11.39723 | 21.5859 | 24.24855 | 9.813082 | 0.318783 | 0.05061 | 0.59595 |
| 18 | 66794400 | 66795199 | 8.696276 | 16.36801 | 28.03817 | 11.48751 | -0.30042 | 0.077995 | 0.595967 |
| 16 | 85124600 | 85125599 | 8.684394 | 16.38252 | 28.08108 | 11.46017 | -0.29682 | 0.063192 | 0.59597 |
| 7 | 19694200 | 19713599 | 239.1792 | 343.8015 | 459.4763 | 189.6738 | -0.21009 | 4.17089 | 0.595971 |
| 15 | 97103800 | 97104799 | 13.59786 | 24.07185 | 38.183 | 16.88475 | -0.27067 | 0.510864 | 0.595974 |
| X | 61084000 | 61088999 | 47.21244 | 76.76629 | 71.16552 | 30.53898 | 0.478116 | 1.597526 | 0.595982 |
| 9 | 114851000 | 114855599 | 55.02691 | 87.52245 | 127.3684 | 53.18219 | -0.20583 | 2.23927 | 0.595988 |
| 10 | 100793800 | 100794599 | 8.633951 | 15.92443 | 29.40794 | 12.43699 | -0.44357 | 0.205482 | 0.595997 |
| 11 | 45719000 | 45720799 | 27.13869 | 44.30138 | 81.295 | 34.93604 | -0.50039 | 1.507327 | 0.595999 |
| X | 62408200 | 62411199 | 55.26453 | 88.73741 | 89.4013 | 37.84148 | 0.339044 | 1.936946 | 0.596009 |
| 10 | 108257200 | 108260799 | 33.43503 | 54.43997 | 84.66066 | 36.34846 | -0.265 | 1.557424 | 0.596013 |
| 11 | 110794600 | 110795199 | 6.685539 | 12.95428 | 23.89168 | 9.053155 | -0.3023 | -0.21199 | 0.596014 |
| 7 | 72697400 | 72699199 | 20.36156 | 35.42989 | 38.82864 | 17.06393 | 0.255537 | 0.779054 | 0.596037 |

| | | | | | | | | | |
|---|---|---|---|---|---|---|---|---|---|
| 9 | 4977400 | 4979599 | 42.23532 | 67.5623 | 106.367 | 44.76987 | -0.31004 | 2.040869 | 0.596088 |
| 2 | 53301800 | 53303599 | 20.65761 | 34.81796 | 55.91331 | 24.44497 | -0.29979 | 1.098981 | 0.596103 |
| 2 | 73262400 | 73265399 | 20.65761 | 34.81796 | 55.91331 | 24.44497 | -0.29979 | 1.098981 | 0.596103 |
| 4 | 143531400 | 143539999 | 133.8161 | 205.9053 | 179.0919 | 74.37809 | 0.497722 | 2.955127 | 0.596106 |
| X | 30624200 | 30624599 | 5.804963 | 11.28412 | 22.30663 | 8.286498 | -0.37315 | -0.29208 | 0.59611 |
| 1 | 181594600 | 181598399 | 29.41611 | 48.36352 | 75.07742 | 32.35658 | -0.25445 | 1.412069 | 0.596116 |
| 10 | 115641000 | 115642999 | 19.92014 | 34.88595 | 37.00214 | 16.33512 | 0.302996 | 0.679922 | 0.596121 |
| 12 | 58839000 | 58840599 | 19.32934 | 34.06987 | 34.09005 | 15.08791 | 0.38796 | 0.582725 | 0.596129 |
| 16 | 55686200 | 55688999 | 23.70792 | 39.3178 | 63.39307 | 27.45772 | -0.30606 | 1.286608 | 0.596132 |
| 13 | 92615400 | 92616199 | 16.32519 | 28.99206 | 30.44561 | 13.44004 | 0.323602 | 0.488575 | 0.596151 |
| X | 71556400 | 71560999 | 39.19108 | 64.31844 | 57.01403 | 24.55292 | 0.550727 | 1.446587 | 0.596157 |
| 7 | 3897000 | 3901199 | 44.14618 | 71.243 | 78.30882 | 33.38287 | 0.222346 | 1.760082 | 0.596185 |
| 18 | 3727400 | 3732599 | 57.22946 | 91.72112 | 94.96352 | 40.08191 | 0.29653 | 1.99404 | 0.596187 |
| 11 | 84296200 | 84298799 | 29.26242 | 48.02061 | 73.72776 | 31.74968 | -0.23964 | 1.43398 | 0.596196 |
| 4 | 111694400 | 111696799 | 17.44681 | 31.08171 | 27.66254 | 12.17908 | 0.563366 | 0.468931 | 0.59621 |
| 8 | 26812000 | 26814999 | 20.76019 | 35.21282 | 54.88821 | 24.04286 | -0.25665 | 1.043122 | 0.596226 |
| 1 | 95778200 | 95779799 | 17.25455 | 31.03556 | 30.56746 | 13.34961 | 0.425726 | 0.413205 | 0.596235 |
| 4 | 23525200 | 23528199 | 52.6968 | 84.68715 | 89.3697 | 37.87623 | 0.273735 | 1.903669 | 0.596247 |
| 6 | 143580800 | 143583199 | 14.71154 | 25.80228 | 40.88868 | 18.07344 | -0.27339 | 0.634822 | 0.596253 |
| 1 | 14451200 | 14452999 | 26.45663 | 44.85072 | 48.50451 | 21.24311 | 0.268768 | 1.060709 | 0.59626 |
| 14 | 24560200 | 24561999 | 21.38156 | 36.97215 | 40.51726 | 17.7756 | 0.254223 | 0.855634 | 0.596292 |
| 6 | 143417400 | 143418999 | 17.5843 | 30.44796 | 47.93735 | 21.0318 | -0.26595 | 0.835402 | 0.596298 |
| 7 | 9451600 | 9452399 | 18.09042 | 32.04582 | 28.63752 | 12.66203 | 0.55304 | 0.513608 | 0.596304 |
| 18 | 17142000 | 17144999 | 39.73058 | 64.56067 | 70.6954 | 30.32691 | 0.237705 | 1.592423 | 0.596308 |
| 4 | 20971000 | 20974399 | 42.514 | 69.31406 | 69.68861 | 29.93472 | 0.364037 | 1.548442 | 0.596338 |
| 10 | 108199000 | 108206399 | 83.03537 | 128.4465 | 215.1077 | 89.35051 | -0.44072 | 2.839845 | 0.596361 |
| 4 | 61437400 | 61440399 | 22.19951 | 38.6828 | 39.1074 | 17.20176 | 0.371476 | 0.766811 | 0.596368 |
| 8 | 38301600 | 38302399 | 8.42201 | 15.89477 | 27.55533 | 11.23481 | -0.31137 | 0.058312 | 0.596378 |
| 11 | 50757200 | 50772199 | 242.989 | 354.5844 | 353.2357 | 145.5465 | 0.234945 | 3.946202 | 0.596381 |
| 17 | 74645400 | 74646799 | 11.79496 | 20.72312 | 43.00881 | 19.03866 | -0.66038 | 0.594171 | 0.596386 |
| X | 128161200 | 128162799 | 29.23119 | 49.03553 | 50.71576 | 22.13065 | 0.332683 | 1.169325 | 0.596386 |
| 7 | 58853400 | 58854399 | 25.42674 | 43.47685 | 38.50368 | 16.88288 | 0.557449 | 0.970472 | 0.596398 |
| 4 | 133339200 | 133342399 | 35.80679 | 57.74509 | 89.1622 | 37.97586 | -0.26797 | 1.773422 | 0.596408 |
| X | 5051800 | 5062199 | 173.3833 | 262.6843 | 218.878 | 90.57416 | 0.541941 | 3.276001 | 0.596414 |
| 4 | 115193400 | 115195399 | 20.09216 | 35.01003 | 38.29976 | 16.8463 | 0.258346 | 0.758154 | 0.596425 |
| 12 | 24219800 | 24259799 | 1077.45 | 1380.748 | 1236.183 | 518.5815 | 0.201431 | 5.969617 | 0.596431 |
| 4 | 127092400 | 127095799 | 24.14698 | 40.1423 | 62.99953 | 27.33257 | -0.26703 | 1.259022 | 0.596441 |
| 2 | 20150400 | 20152199 | 12.18279 | 21.71759 | 35.22995 | 15.50251 | -0.29711 | 0.446835 | 0.596453 |
| 11 | 26337000 | 26337599 | 6.554681 | 12.61955 | 24.17524 | 9.397715 | -0.39173 | -0.12757 | 0.596477 |
| 4 | 88648400 | 88651999 | 41.99063 | 67.94655 | 74.56291 | 31.88689 | 0.228973 | 1.681699 | 0.596479 |
| 14 | 52764800 | 52769399 | 43.95762 | 71.52681 | 70.94979 | 30.4546 | 0.381455 | 1.579105 | 0.596483 |
| 10 | 81012600 | 81014599 | 147.0842 | 221.526 | 210.4082 | 86.91163 | 0.345837 | 3.386016 | 0.596514 |
| 18 | 4711800 | 4719799 | 77.26807 | 122.1786 | 127.7154 | 53.06706 | 0.263066 | 2.414404 | 0.596515 |
| 4 | 12103400 | 12107599 | 80.01648 | 126.5973 | 126.5208 | 52.58599 | 0.328471 | 2.405362 | 0.596524 |
| 7 | 19793200 | 19804599 | 153.2228 | 232.0752 | 205.8502 | 85.17557 | 0.451435 | 3.273656 | 0.596539 |
| 2 | 157197000 | 157200399 | 34.13159 | 56.04257 | 61.09393 | 26.31539 | 0.25343 | 1.431624 | 0.596546 |
| 17 | 53489200 | 53490799 | 33.965 | 55.75426 | 60.50631 | 26.05935 | 0.259769 | 1.434106 | 0.596561 |

| | | | | | | | | | |
|---|---|---|---|---|---|---|---|---|---|
| 7 | 39177800 | 39180999 | 25.50494 | 43.86794 | 44.63856 | 19.5805 | 0.359169 | 0.92526 | 0.596576 |
| 7 | 19219800 | 19226199 | 50.80266 | 80.65552 | 126.0973 | 52.55169 | -0.31164 | 2.289429 | 0.596618 |
| 8 | 93606200 | 93606999 | 11.75717 | 21.16576 | 34.69071 | 15.13677 | -0.3038 | 0.378698 | 0.596627 |
| Y | 8735400 | 8737199 | 16.0451 | 28.55567 | 20.62772 | 9.08958 | 0.861351 | 0.447012 | 0.596629 |
| 2 | 126845200 | 126846599 | 20.08622 | 35.02154 | 38.26592 | 16.83802 | 0.260156 | 0.749172 | 0.596629 |
| 13 | 62586200 | 62604599 | 463.1219 | 638.6935 | 602.9165 | 250.5439 | 0.231212 | 4.794123 | 0.596629 |
| 15 | 89503800 | 89506999 | 34.34952 | 55.53758 | 85.50684 | 36.53995 | -0.2595 | 1.695378 | 0.596634 |
| 18 | 19538200 | 19539399 | 15.08671 | 27.09338 | 29.23379 | 12.78422 | 0.293528 | 0.404957 | 0.596636 |
| 3 | 74783000 | 74786799 | 20.89002 | 36.77935 | 29.83943 | 13.2096 | 0.688291 | 0.596862 | 0.59664 |
| 2 | 25742800 | 25744399 | 15.42615 | 26.808 | 44.26598 | 19.43793 | -0.33314 | 0.783786 | 0.596664 |
| 15 | 92948200 | 92950799 | 22.95791 | 38.00463 | 71.0791 | 30.74808 | -0.51977 | 1.311922 | 0.59667 |
| 2 | 74200000 | 74202999 | 34.6418 | 56.78941 | 61.60288 | 26.51666 | 0.259639 | 1.453051 | 0.596697 |
| 11 | 8272000 | 8276799 | 55.32699 | 89.25274 | 81.25024 | 34.61986 | 0.493113 | 1.791396 | 0.596712 |
| X | 59591000 | 59592599 | 23.9309 | 41.19544 | 41.60985 | 18.25245 | 0.37027 | 0.911265 | 0.596716 |
| 18 | 7968000 | 7971799 | 80.30469 | 126.2378 | 136.9311 | 56.80234 | 0.203199 | 2.543702 | 0.596726 |
| 7 | 36024400 | 36024799 | 7.115758 | 13.7493 | 25.51052 | 9.794013 | -0.32951 | -0.1601 | 0.596729 |
| 17 | 92224800 | 92225399 | 7.115758 | 13.7493 | 25.51052 | 9.794013 | -0.32951 | -0.1601 | 0.596729 |
| 2 | 116671600 | 116672399 | 8.530526 | 16.29916 | 29.68413 | 11.79145 | -0.34939 | -0.02225 | 0.596757 |
| 6 | 3584200 | 3585199 | 10.2344 | 19.43534 | 22.19528 | 8.931587 | 0.302338 | 0.015758 | 0.596759 |
| 13 | 12816000 | 12884199 | 3053.27 | 3569.603 | 4130.732 | 1753.216 | -0.30317 | 7.432127 | 0.596759 |
| 4 | 113017000 | 113028999 | 162.0591 | 245.3801 | 227.7137 | 94.21262 | 0.385905 | 3.284539 | 0.596774 |
| 7 | 8069400 | 8071999 | 34.86267 | 57.64027 | 51.43643 | 22.23182 | 0.5444 | 1.34317 | 0.596794 |
| 14 | 91312200 | 91313799 | 17.33569 | 31.11214 | 31.13244 | 13.62191 | 0.401249 | 0.429294 | 0.596802 |
| 14 | 50429200 | 50434999 | 81.55484 | 128.1114 | 138.7385 | 57.54675 | 0.204734 | 2.557891 | 0.596806 |
| 4 | 132851000 | 132852599 | 22.43284 | 39.08442 | 35.01299 | 15.39956 | 0.545242 | 0.765617 | 0.596815 |
| 19 | 20251800 | 20253399 | 18.53749 | 32.68874 | 34.2969 | 15.17811 | 0.320322 | 0.576038 | 0.596819 |
| 13 | 56034400 | 56043399 | 72.16588 | 111.8158 | 196.173 | 81.48482 | -0.50588 | 2.80312 | 0.596828 |
| 16 | 95113000 | 95113799 | 10.23295 | 18.87615 | 31.97855 | 13.52167 | -0.3199 | 0.225711 | 0.596837 |
| 19 | 42992800 | 42994399 | 10.23295 | 18.87615 | 31.97855 | 13.52167 | -0.3199 | 0.225711 | 0.596837 |
| 11 | 10673000 | 10674199 | 14.00201 | 25.84631 | 27.47938 | 11.60849 | 0.348303 | 0.23834 | 0.596856 |
| 17 | 87232400 | 87235199 | 27.84099 | 45.76653 | 70.626 | 30.43607 | -0.2459 | 1.409084 | 0.596866 |
| 5 | 28093400 | 28099199 | 58.15717 | 92.32442 | 133.9107 | 55.80357 | -0.20377 | 2.299442 | 0.596872 |
| 19 | 28315600 | 28317399 | 18.60562 | 32.80995 | 34.2177 | 15.14295 | 0.328865 | 0.574677 | 0.59689 |
| 10 | 45628000 | 45629399 | 15.62619 | 27.70835 | 29.36004 | 12.97752 | 0.309157 | 0.488288 | 0.596901 |
| 1 | 118029800 | 118035599 | 99.28089 | 154.1392 | 162.7225 | 67.57277 | 0.228389 | 2.775778 | 0.596903 |
| 13 | 67470000 | 67471599 | 12.22449 | 21.97762 | 36.21617 | 15.82288 | -0.31292 | 0.392678 | 0.596908 |
| 14 | 37988600 | 37989199 | 9.711304 | 18.93565 | 18.59519 | 6.69338 | 0.665419 | -0.37038 | 0.596917 |
| 9 | 119519600 | 119521999 | 16.05805 | 27.91043 | 44.76601 | 19.6594 | -0.29144 | 0.785158 | 0.596919 |
| 2 | 68007200 | 68008399 | 8.706574 | 16.4568 | 28.13219 | 11.43322 | -0.28748 | 0.050853 | 0.596943 |
| 4 | 20825800 | 20828999 | 62.76318 | 98.84551 | 146.8392 | 60.92715 | -0.24782 | 2.498426 | 0.596952 |
| 1 | 47103600 | 47107199 | 62.57424 | 99.71597 | 107.7357 | 45.10298 | 0.226097 | 2.179367 | 0.596957 |
| 1 | 186094200 | 186096799 | 24.32516 | 41.36952 | 45.31242 | 19.8597 | 0.250856 | 1.02199 | 0.597005 |
| 3 | 153464400 | 153470199 | 37.19435 | 60.02325 | 91.31029 | 38.9246 | -0.2455 | 1.754716 | 0.597012 |
| 6 | 70487000 | 70487599 | 9.659623 | 18.54331 | 21.85185 | 8.565888 | 0.291722 | -0.07454 | 0.597012 |
| 5 | 80078600 | 80079799 | 11.7718 | 21.35672 | 34.8258 | 15.04936 | -0.28736 | 0.335381 | 0.597019 |
| 5 | 141041400 | 141046599 | 49.48127 | 79.04662 | 118.4197 | 49.74766 | -0.2408 | 2.091362 | 0.597024 |
| 6 | 72267400 | 72270399 | 23.40106 | 39.19296 | 62.43934 | 27.21905 | -0.28849 | 1.178557 | 0.597024 |

| | | | | | | | | | |
|---|---|---|---|---|---|---|---|---|---|
| 15 | 98985000 | 98987199 | 17.27189 | 29.44768 | 50.56354 | 22.16565 | -0.39451 | 0.979973 | 0.597031 |
| 4 | 52034800 | 52035399 | 7.328758 | 14.11164 | 25.86383 | 10.05049 | -0.32766 | -0.1127 | 0.597037 |
| 8 | 94310000 | 94312999 | 23.8646 | 39.63454 | 62.88446 | 27.26446 | -0.28277 | 1.270415 | 0.597054 |
| 15 | 46569400 | 46576199 | 178.9187 | 267.228 | 242.822 | 100.1048 | 0.3989 | 3.550841 | 0.597068 |
| 12 | 14643400 | 14646799 | 38.04133 | 61.80428 | 67.63357 | 29.0095 | 0.238384 | 1.59034 | 0.597107 |
| 4 | 155332400 | 155335399 | 18.04999 | 30.80277 | 50.57419 | 22.17197 | -0.33 | 0.974555 | 0.597111 |
| 1 | 183253400 | 183258599 | 45.44393 | 72.7635 | 107.5386 | 45.32412 | -0.21756 | 2.015113 | 0.597111 |
| 6 | 37871200 | 37873399 | 24.94524 | 41.61955 | 64.28351 | 27.95536 | -0.24402 | 1.222988 | 0.597169 |
| 19 | 56645000 | 56647799 | 27.17739 | 45.61735 | 50.18122 | 21.9056 | 0.243882 | 1.15206 | 0.597185 |
| 1 | 112773400 | 112774399 | 11.6458 | 22.20164 | 23.80966 | 9.502888 | 0.402439 | 0.006474 | 0.597206 |
| 13 | 8316600 | 8320199 | 54.49441 | 87.38064 | 93.69813 | 39.57739 | 0.246881 | 1.975974 | 0.597208 |
| 12 | 113579800 | 113583799 | 49.12138 | 79.19642 | 83.06965 | 35.36887 | 0.288163 | 1.797889 | 0.597227 |
| 5 | 18125200 | 18127599 | 24.36138 | 42.01075 | 38.28805 | 16.79622 | 0.518219 | 0.888784 | 0.597229 |
| 4 | 48512000 | 48515399 | 49.02302 | 78.83094 | 85.90903 | 36.44155 | 0.228494 | 1.877775 | 0.597231 |
| 2 | 6836000 | 6837799 | 14.9908 | 27.48719 | 28.38399 | 12.10018 | 0.380701 | 0.284753 | 0.597234 |
| 15 | 51988400 | 51990199 | 26.92838 | 44.10855 | 71.26791 | 30.66634 | -0.31399 | 1.452783 | 0.597237 |
| 9 | 15217000 | 15218599 | 10.91003 | 19.60098 | 33.29834 | 14.5565 | -0.3565 | 0.385588 | 0.597243 |
| X | 63833600 | 63834599 | 16.05036 | 28.99035 | 29.37452 | 12.76417 | 0.390142 | 0.382187 | 0.59725 |
| 3 | 12430800 | 12440199 | 93.66814 | 146.7471 | 146.9517 | 60.95413 | 0.314595 | 2.616786 | 0.597253 |
| 7 | 28404600 | 28408399 | 43.63666 | 69.95304 | 104.0562 | 43.93592 | -0.22483 | 1.974673 | 0.597266 |
| 3 | 28205400 | 28207199 | 35.47056 | 57.28585 | 87.25095 | 37.23138 | -0.24586 | 1.729806 | 0.597274 |
| X | 3155800 | 3159999 | 38.68154 | 65.29813 | 46.81881 | 20.47854 | 0.859615 | 1.141075 | 0.597283 |
| X | 90193000 | 90197599 | 63.43029 | 101.6863 | 93.59723 | 39.52125 | 0.466438 | 1.997503 | 0.597287 |
| 10 | 84042600 | 84044399 | 35.28049 | 58.21905 | 54.21682 | 23.40256 | 0.482803 | 1.366825 | 0.5973 |
| 12 | 110086200 | 110094799 | 71.81983 | 112.1172 | 166.122 | 68.93109 | -0.25396 | 2.668789 | 0.5973 |
| 10 | 96646800 | 96649799 | 36.79157 | 59.50075 | 90.9662 | 38.88537 | -0.24904 | 1.694425 | 0.59731 |
| X | 15309400 | 15310799 | 16.19794 | 30.1462 | 27.00335 | 11.27648 | 0.606307 | 0.210548 | 0.597338 |
| 9 | 57586000 | 57587599 | 12.79025 | 22.88492 | 36.7168 | 16.1065 | -0.27841 | 0.429632 | 0.597368 |
| 18 | 60125600 | 60126399 | 12.79025 | 22.88492 | 36.7168 | 16.1065 | -0.27841 | 0.429632 | 0.597368 |
| 2 | 26549200 | 26549999 | 10.60441 | 20.03522 | 22.69094 | 9.230392 | 0.300781 | 0.056497 | 0.597423 |
| X | 80789600 | 80804199 | 87.96119 | 139.5627 | 98.7857 | 41.14391 | 0.828968 | 2.333366 | 0.597424 |
| 7 | 5641000 | 5643199 | 44.43553 | 71.8412 | 76.82355 | 32.82703 | 0.264996 | 1.709339 | 0.59743 |
| X | 79590600 | 79591599 | 16.25595 | 29.73844 | 28.39736 | 12.14386 | 0.490035 | 0.309708 | 0.597435 |
| 1 | 35754600 | 35756399 | 18.50091 | 31.69046 | 50.0583 | 21.95241 | -0.27365 | 0.941466 | 0.597437 |
| 6 | 41226600 | 41231599 | 41.61865 | 67.66982 | 71.19198 | 30.55015 | 0.296002 | 1.584675 | 0.597452 |
| 6 | 143092600 | 143093599 | 11.32611 | 20.49353 | 33.26053 | 14.42088 | -0.28346 | 0.345324 | 0.597453 |
| 14 | 93254800 | 93257599 | 21.50316 | 37.85881 | 35.07627 | 15.52789 | 0.49796 | 0.610998 | 0.59747 |
| 18 | 25325800 | 25328399 | 72.12503 | 113.8566 | 121.0616 | 50.25997 | 0.236625 | 2.450054 | 0.597476 |
| 11 | 31395600 | 31399999 | 76.32086 | 120.1189 | 128.3085 | 53.23378 | 0.226634 | 2.514352 | 0.597481 |
| 5 | 67564600 | 67567199 | 19.48037 | 33.9473 | 36.74352 | 16.16277 | 0.273578 | 0.749908 | 0.597493 |
| X | 91932200 | 91935999 | 41.10695 | 67.50296 | 64.65511 | 27.85056 | 0.439656 | 1.449118 | 0.597502 |
| 7 | 37351800 | 37353199 | 13.46031 | 23.6151 | 39.41211 | 17.42535 | -0.34761 | 0.620594 | 0.597502 |
| 6 | 142947200 | 142948199 | 6.579078 | 12.59831 | 24.60025 | 9.712359 | -0.43991 | -0.07447 | 0.597525 |
| 1 | 29360000 | 29361399 | 19.00718 | 33.253 | 36.05791 | 15.90449 | 0.271249 | 0.68888 | 0.597538 |
| 17 | 36255200 | 36256599 | 20.59974 | 35.64642 | 37.44874 | 16.42975 | 0.315063 | 0.83999 | 0.597555 |
| 10 | 54757600 | 54758999 | 15.82697 | 27.71394 | 44.04143 | 19.44053 | -0.27756 | 0.671303 | 0.597575 |
| 12 | 73333200 | 73333999 | 8.282097 | 15.56563 | 27.50345 | 11.29642 | -0.34724 | 0.084175 | 0.597598 |

| | | | | | | | | | |
|---|---|---|---|---|---|---|---|---|---|
| 19 | 42430400 | 42431599 | 8.282097 | 15.56563 | 27.50345 | 11.29642 | -0.34724 | 0.084175 | 0.597598 |
| 6 | 131029600 | 131036399 | 135.4462 | 206.3118 | 190.2909 | 78.87188 | 0.401776 | 3.154197 | 0.597602 |
| 3 | 3216600 | 3217999 | 21.08832 | 37.0665 | 35.73658 | 15.82504 | 0.440012 | 0.628702 | 0.597609 |
| 8 | 52469400 | 52472999 | 32.0447 | 53.7467 | 53.56381 | 23.3711 | 0.385923 | 1.179647 | 0.597618 |
| 4 | 112126200 | 112134199 | 27.02344 | 46.21923 | 39.79337 | 17.44874 | 0.597874 | 0.973242 | 0.597626 |
| 3 | 87548200 | 87549799 | 13.66343 | 24.18062 | 38.74056 | 17.13234 | -0.28505 | 0.517313 | 0.597629 |
| 2 | 52015800 | 52018999 | 54.93993 | 88.13448 | 93.04895 | 39.32997 | 0.270042 | 1.964063 | 0.597633 |
| X | 148962200 | 148966199 | 55.54092 | 89.32234 | 87.95351 | 37.31507 | 0.374778 | 1.887214 | 0.597644 |
| 6 | 131977600 | 131979599 | 28.23936 | 47.94188 | 49.0335 | 21.48417 | 0.349116 | 1.057176 | 0.597649 |
| 12 | 96612000 | 96613999 | 37.82675 | 62.9581 | 56.60355 | 24.57769 | 0.534325 | 1.280922 | 0.597654 |
| 6 | 134105400 | 134106799 | 14.44164 | 25.31791 | 40.4378 | 17.86759 | -0.28452 | 0.641148 | 0.597669 |
| 6 | 141968200 | 141968599 | 7.881486 | 15.08523 | 26.82416 | 10.60463 | -0.30862 | -0.05215 | 0.597684 |
| 16 | 95518200 | 95524599 | 75.24437 | 117.6087 | 167.5556 | 69.51709 | -0.19606 | 2.649776 | 0.597691 |
| 13 | 33137600 | 33140399 | 38.15944 | 62.40417 | 56.97421 | 24.50042 | 0.505779 | 1.485989 | 0.5977 |
| X | 136018000 | 136023199 | 63.35717 | 101.2764 | 100.2079 | 42.13405 | 0.357636 | 2.086075 | 0.59771 |
| 15 | 26680800 | 26682599 | 37.2144 | 59.66089 | 110.5057 | 46.66295 | -0.54041 | 1.968819 | 0.597736 |
| 16 | 27484400 | 27486199 | 15.9705 | 29.04769 | 28.62967 | 12.33263 | 0.437265 | 0.33601 | 0.597736 |
| 11 | 15996600 | 16002599 | 92.06145 | 143.3365 | 154.5069 | 64.13395 | 0.201506 | 2.722923 | 0.597794 |
| 1 | 54330000 | 54331799 | 33.21403 | 54.51831 | 58.87426 | 25.35581 | 0.26679 | 1.430228 | 0.597795 |
| X | 76979000 | 76988399 | 164.1946 | 246.7495 | 250.8447 | 103.553 | 0.245057 | 3.434364 | 0.597796 |
| 1 | 177751400 | 177754599 | 77.98755 | 122.4606 | 126.0507 | 52.2814 | 0.277512 | 2.562301 | 0.597802 |
| 4 | 14582400 | 14585799 | 30.39171 | 50.86926 | 54.07068 | 23.5616 | 0.293318 | 1.196514 | 0.597807 |
| 4 | 30228600 | 30231999 | 34.08976 | 56.31078 | 51.9919 | 22.4577 | 0.49524 | 1.350198 | 0.597819 |
| 15 | 13225800 | 13228799 | 27.38199 | 46.21603 | 50.15419 | 21.93637 | 0.26363 | 1.104386 | 0.597824 |
| 2 | 35275400 | 35277799 | 28.61526 | 47.06022 | 72.01284 | 31.03956 | -0.23358 | 1.40525 | 0.59783 |
| 11 | 82452400 | 82454399 | 17.9621 | 31.4846 | 34.10259 | 15.06629 | 0.27293 | 0.651087 | 0.59785 |
| 16 | 30748400 | 30753399 | 55.47272 | 88.15989 | 128.457 | 53.58993 | -0.20881 | 2.263128 | 0.597863 |
| X | 67440200 | 67447599 | 83.2674 | 131.996 | 105.7674 | 44.01192 | 0.649047 | 2.35662 | 0.597881 |
| 12 | 109486400 | 109518399 | 489.8397 | 655.0166 | 1006.992 | 420.447 | -0.5171 | 5.278655 | 0.597894 |
| Y | 8434600 | 8435999 | 14.24704 | 25.7223 | 21.17365 | 9.205496 | 0.687148 | 0.349081 | 0.597908 |
| 12 | 113650400 | 113652799 | 34.43147 | 56.88185 | 53.09691 | 22.93685 | 0.479515 | 1.350672 | 0.59791 |
| 14 | 41418800 | 41419999 | 16.98293 | 30.44447 | 26.69648 | 11.69215 | 0.589817 | 0.425937 | 0.597912 |
| 3 | 77859000 | 77861599 | 33.19506 | 54.89572 | 59.71588 | 25.81445 | 0.259512 | 1.340899 | 0.597922 |
| 7 | 47646600 | 47647399 | 19.02784 | 33.23052 | 36.18308 | 15.9379 | 0.265311 | 0.716796 | 0.59794 |
| 1 | 191520400 | 191527999 | 82.5268 | 128.2618 | 182.2804 | 75.68263 | -0.19886 | 2.753739 | 0.597955 |
| X | 34751000 | 34758799 | 115.9787 | 179.9738 | 162.671 | 67.55644 | 0.451627 | 2.793403 | 0.597959 |
| 14 | 114021200 | 114025999 | 90.89072 | 141.9488 | 125.962 | 52.26472 | 0.485247 | 2.665406 | 0.59796 |
| 7 | 15032200 | 15032999 | 17.88358 | 31.52623 | 33.86951 | 14.98986 | 0.28621 | 0.574849 | 0.598002 |
| 3 | 16006400 | 16013799 | 91.81823 | 144.3775 | 139.452 | 57.85021 | 0.370682 | 2.545031 | 0.598006 |
| 13 | 34035000 | 34037999 | 35.08521 | 58.0981 | 59.83282 | 25.88856 | 0.338418 | 1.332129 | 0.598032 |
| X | 109028000 | 109031599 | 41.56816 | 68.60883 | 60.74286 | 26.22312 | 0.555644 | 1.388252 | 0.598038 |
| 6 | 95450800 | 95454599 | 41.51113 | 67.28671 | 73.56956 | 31.50215 | 0.236542 | 1.64615 | 0.598073 |
| 1 | 156887200 | 156888599 | 15.34544 | 26.82358 | 42.5412 | 18.75045 | -0.27458 | 0.695159 | 0.5981 |
| 4 | 79948600 | 79949999 | 14.09066 | 24.85924 | 39.24395 | 17.36575 | -0.26575 | 0.553304 | 0.59812 |
| 16 | 7629800 | 7633199 | 43.31755 | 69.56146 | 103.0191 | 43.61238 | -0.21554 | 1.920739 | 0.598131 |
| 18 | 10373800 | 10376599 | 26.55754 | 44.88345 | 48.52877 | 21.23391 | 0.268913 | 1.086348 | 0.598138 |
| 16 | 12150600 | 12151799 | 17.29618 | 30.65701 | 32.29928 | 14.27938 | 0.317268 | 0.510724 | 0.598153 |

| | | | | | | | | | |
|---|---|---|---|---|---|---|---|---|---|
| 11 | 40979200 | 40983399 | 175.3391 | 262.3502 | 212.0009 | 87.42904 | 0.570361 | 3.516308 | 0.598174 |
| 4 | 8044400 | 8045999 | 13.86703 | 25.59959 | 23.29997 | 9.841655 | 0.57108 | 0.226813 | 0.598192 |
| 6 | 105996600 | 105998199 | 15.64928 | 29.08836 | 26.93168 | 11.27011 | 0.556251 | 0.21207 | 0.598201 |
| 9 | 24727000 | 24727599 | 6.244302 | 12.144 | 23.00266 | 8.511832 | -0.30798 | -0.30116 | 0.598235 |
| 11 | 31631200 | 31632399 | 13.13413 | 23.47246 | 38.26901 | 16.80412 | -0.30249 | 0.442435 | 0.59824 |
| 3 | 63604800 | 63606799 | 15.70471 | 27.45798 | 43.05763 | 18.98181 | -0.2583 | 0.695078 | 0.598274 |
| 1 | 124393200 | 124396199 | 24.1482 | 41.20261 | 45.27957 | 19.85051 | 0.246765 | 0.991642 | 0.598274 |
| 14 | 116713800 | 116715199 | 12.7599 | 24.21842 | 24.82109 | 9.999492 | 0.455919 | 0.050509 | 0.598276 |
| 16 | 12160400 | 12165199 | 56.89829 | 91.11068 | 96.97345 | 40.87587 | 0.255292 | 2.023148 | 0.598282 |
| 12 | 21672000 | 21675399 | 63.85887 | 101.464 | 110.3396 | 46.02099 | 0.212323 | 2.272105 | 0.598293 |
| 6 | 60495800 | 60498999 | 42.00197 | 67.51917 | 100.1582 | 42.46615 | -0.21603 | 1.88536 | 0.598297 |
| 8 | 29897000 | 29898399 | 14.86772 | 26.80364 | 29.34065 | 12.77957 | 0.276861 | 0.3821 | 0.598308 |
| X | 117826800 | 117834399 | 67.4414 | 108.0075 | 95.7394 | 40.36382 | 0.518984 | 2.032082 | 0.598329 |
| 13 | 64698000 | 64699799 | 29.61256 | 49.73093 | 52.40543 | 22.88412 | 0.305611 | 1.155955 | 0.598334 |
| 14 | 102525200 | 102527399 | 22.19312 | 38.2682 | 42.07132 | 18.45582 | 0.248805 | 0.885808 | 0.598336 |
| 8 | 105260600 | 105262199 | 9.076951 | 16.68572 | 30.36935 | 12.90203 | -0.42784 | 0.228967 | 0.598342 |
| X | 111905400 | 111907399 | 21.07158 | 37.33615 | 33.00274 | 14.59136 | 0.569074 | 0.539607 | 0.598349 |
| 16 | 85516400 | 85517399 | 7.633155 | 14.49514 | 26.06245 | 10.48836 | -0.34689 | 0.003497 | 0.598372 |
| 19 | 40989400 | 40992599 | 43.71781 | 70.63892 | 77.25878 | 32.98777 | 0.231578 | 1.724385 | 0.598378 |
| 7 | 11975200 | 11993199 | 250.5863 | 367.1661 | 336.871 | 138.7612 | 0.359263 | 3.884927 | 0.598399 |
| 10 | 49189000 | 49191199 | 23.55294 | 40.93557 | 39.62271 | 17.40022 | 0.43359 | 0.80854 | 0.598409 |
| 14 | 114174000 | 114176799 | 24.21994 | 42.33603 | 38.44909 | 16.9468 | 0.526038 | 0.735688 | 0.59841 |
| 16 | 95935400 | 95937599 | 16.68272 | 28.94512 | 45.82597 | 20.1067 | -0.27316 | 0.812357 | 0.598412 |
| 18 | 9430200 | 9433599 | 47.16103 | 76.23418 | 78.64078 | 33.59589 | 0.31646 | 1.718526 | 0.598419 |
| 15 | 60813400 | 60815199 | 12.93352 | 23.23522 | 39.41213 | 17.22989 | -0.35505 | 0.409425 | 0.598419 |
| X | 53016200 | 53024999 | 89.94862 | 141.7097 | 118.6734 | 49.24274 | 0.578403 | 2.504683 | 0.598464 |
| 3 | 96268200 | 96280999 | 337.8469 | 469.4578 | 620.7617 | 257.6517 | -0.24386 | 4.675528 | 0.598498 |
| 1 | 88995600 | 88998399 | 18.18542 | 31.0068 | 51.10661 | 22.40381 | -0.33584 | 0.98255 | 0.598514 |
| 2 | 86849200 | 86850799 | 15.57987 | 28.28643 | 29.23625 | 12.6222 | 0.367182 | 0.345517 | 0.598549 |
| 14 | 70191000 | 70192399 | 14.98102 | 26.85806 | 29.31184 | 12.83666 | 0.275893 | 0.415653 | 0.598551 |
| 5 | 22904600 | 22906199 | 14.78815 | 27.38614 | 27.52661 | 11.57723 | 0.43353 | 0.228072 | 0.598566 |
| 11 | 31655200 | 31660799 | 53.77407 | 86.21136 | 93.0123 | 39.27785 | 0.237771 | 1.979134 | 0.598572 |
| 4 | 112355400 | 112366999 | 177.1805 | 266.3019 | 252.8522 | 104.3857 | 0.343689 | 3.433769 | 0.598575 |
| 10 | 95568800 | 95570999 | 23.40921 | 40.35448 | 43.04583 | 18.88316 | 0.292005 | 0.89318 | 0.598575 |
| X | 149166200 | 149168199 | 38.14817 | 62.3789 | 66.32409 | 28.51998 | 0.28654 | 1.497455 | 0.598588 |
| 13 | 22529400 | 22532199 | 26.81793 | 46.20367 | 44.03836 | 19.31822 | 0.453641 | 0.915144 | 0.598608 |
| X | 161513800 | 161514999 | 7.721438 | 14.80381 | 32.11959 | 12.63683 | -0.58601 | -0.05023 | 0.598614 |
| 2 | 108924400 | 108925999 | 10.43578 | 19.03241 | 33.33415 | 14.3106 | -0.3838 | 0.289661 | 0.598616 |
| 7 | 4814200 | 4817199 | 20.94448 | 34.97895 | 58.98492 | 25.66707 | -0.37007 | 1.196605 | 0.598619 |
| 7 | 24100600 | 24108199 | 65.51886 | 102.6245 | 156.5433 | 64.93404 | -0.29206 | 2.605791 | 0.598632 |
| 6 | 41303000 | 41308799 | 70.5069 | 111.8142 | 120.3894 | 50.1172 | 0.224022 | 2.335355 | 0.598656 |
| X | 23720000 | 23722399 | 29.54751 | 50.09959 | 49.04315 | 21.48019 | 0.41213 | 1.074282 | 0.59867 |
| 5 | 77972800 | 77975799 | 28.12187 | 46.28684 | 70.85071 | 30.54927 | -0.23352 | 1.393157 | 0.598693 |
| 2 | 4349000 | 4350399 | 13.59489 | 23.91329 | 38.78957 | 17.1749 | -0.3064 | 0.581531 | 0.598693 |
| 19 | 30006800 | 30008799 | 13.84055 | 24.13477 | 47.33805 | 20.83159 | -0.57993 | 0.730157 | 0.598702 |
| 10 | 22682200 | 22684199 | 18.98154 | 32.80787 | 53.36565 | 23.41209 | -0.31342 | 0.866026 | 0.598708 |
| 4 | 33233600 | 33236799 | 28.67715 | 48.25405 | 50.74658 | 22.17487 | 0.308537 | 1.132729 | 0.59872 |

| | | | | | | | | | |
|---|---|---|---|---|---|---|---|---|---|
| 2 | 169809200 | 169817199 | 97.30558 | 149.5722 | 212.3855 | 88.19274 | -0.20989 | 2.96298 | 0.598733 |
| 1 | 87135000 | 87136199 | 13.6096 | 24.8113 | 27.01449 | 11.60303 | 0.297714 | 0.305293 | 0.598745 |
| 14 | 91713200 | 91715399 | 13.44503 | 23.7627 | 37.71329 | 16.68245 | -0.27232 | 0.527082 | 0.598751 |
| 16 | 11269800 | 11273599 | 46.06452 | 74.17773 | 81.04496 | 34.44838 | 0.227318 | 1.827852 | 0.598756 |
| 7 | 34958800 | 34959999 | 14.56919 | 25.57069 | 40.36317 | 17.85182 | -0.26771 | 0.622326 | 0.598757 |
| 6 | 130731600 | 130733199 | 17.04353 | 29.71615 | 47.87365 | 21.06555 | -0.2975 | 0.749517 | 0.598777 |
| 4 | 8618400 | 8619799 | 15.49596 | 28.01435 | 29.77849 | 12.92506 | 0.322143 | 0.371089 | 0.598786 |
| 12 | 88317000 | 88321599 | 72.60188 | 113.522 | 163.4692 | 67.819 | -0.211 | 2.640979 | 0.598786 |
| 14 | 18314800 | 18318199 | 24.93386 | 42.41057 | 46.34376 | 20.31195 | 0.254209 | 1.023338 | 0.598806 |
| 17 | 43473400 | 43475199 | 13.85347 | 25.1081 | 26.28913 | 11.37467 | 0.346516 | 0.335577 | 0.598807 |
| 18 | 11284400 | 11296599 | 214.6359 | 311.5227 | 416.5485 | 171.7301 | -0.19705 | 4.026365 | 0.598811 |
| X | 61188400 | 61189199 | 14.89145 | 26.9705 | 27.5214 | 11.91825 | 0.383315 | 0.355109 | 0.598816 |
| 9 | 78995600 | 79006399 | 115.9312 | 176.1555 | 245.0808 | 101.507 | -0.19365 | 3.202713 | 0.598821 |
| 8 | 36140200 | 36148199 | 74.16755 | 115.2247 | 192.9125 | 80.10175 | -0.43547 | 2.756992 | 0.598828 |
| 5 | 37770000 | 37778199 | 65.63662 | 103.5515 | 148.6698 | 61.70297 | -0.19674 | 2.464801 | 0.598831 |
| 6 | 59021000 | 59022199 | 11.83303 | 21.5346 | 36.51924 | 15.71995 | -0.33986 | 0.322564 | 0.598838 |
| 15 | 36466800 | 36471799 | 55.16122 | 87.71314 | 127.3879 | 53.17321 | -0.20331 | 2.246953 | 0.598839 |
| 18 | 85628600 | 85630199 | 14.74144 | 25.95837 | 41.31398 | 18.28865 | -0.27859 | 0.581176 | 0.598847 |
| 13 | 113791800 | 113794399 | 19.21941 | 33.10738 | 52.81207 | 23.16686 | -0.28646 | 0.894615 | 0.598855 |
| 1 | 192954600 | 192955999 | 8.205999 | 15.11245 | 29.43527 | 12.47395 | -0.52185 | 0.210907 | 0.598878 |
| 14 | 62260600 | 62263799 | 28.64407 | 47.19667 | 73.24233 | 31.59474 | -0.25268 | 1.38383 | 0.598894 |
| 13 | 14571400 | 14573999 | 23.42706 | 40.67976 | 41.35057 | 18.14863 | 0.36329 | 0.824997 | 0.598914 |
| 2 | 23180400 | 23186999 | 161.2305 | 241.2923 | 248.8054 | 102.6115 | 0.219228 | 3.511383 | 0.598918 |
| 12 | 22978200 | 23031399 | 1735.307 | 2138.122 | 1830.742 | 771.2621 | 0.208507 | 6.602943 | 0.598919 |
| 17 | 42590800 | 42592399 | 19.1108 | 33.5896 | 35.29961 | 15.63112 | 0.316141 | 0.615572 | 0.598935 |
| 10 | 102151400 | 102151799 | 5.527781 | 10.73362 | 21.67867 | 8.124325 | -0.41589 | -0.26856 | 0.598949 |
| 15 | 88262400 | 88272599 | 109.0667 | 166.0271 | 233.0144 | 96.56195 | -0.20407 | 3.163076 | 0.598964 |
| 10 | 126578400 | 126581199 | 29.34999 | 47.88148 | 74.78066 | 32.12929 | -0.26905 | 1.515549 | 0.598969 |
| Y | 41493600 | 41494999 | 11.60167 | 21.65548 | 18.16736 | 7.549014 | 0.705371 | 0.128468 | 0.598981 |
| 15 | 95777000 | 95778199 | 12.2434 | 22.9909 | 20.94459 | 8.615013 | 0.599164 | 0.105518 | 0.598995 |
| 15 | 33828800 | 33833599 | 92.13656 | 143.2571 | 154.9727 | 64.34048 | 0.194841 | 2.746691 | 0.599019 |
| 12 | 48928000 | 48929399 | 23.06966 | 39.88986 | 35.328 | 15.49897 | 0.560215 | 0.856203 | 0.599039 |
| 3 | 4799400 | 4800399 | 10.50428 | 19.12406 | 31.79553 | 13.67978 | -0.31077 | 0.29982 | 0.59905 |
| 3 | 51515400 | 51526399 | 106.5696 | 162.3912 | 229.9241 | 95.31039 | -0.21554 | 3.141019 | 0.599059 |
| 13 | 68620800 | 68624599 | 82.72761 | 130.1951 | 137.7858 | 57.16538 | 0.239966 | 2.522437 | 0.599072 |
| 7 | 55057000 | 55061999 | 81.13186 | 127.3199 | 138.6373 | 57.49731 | 0.195757 | 2.577958 | 0.599079 |
| 12 | 115668600 | 115678199 | 124.3653 | 192.3665 | 167.5755 | 69.60681 | 0.501227 | 2.854302 | 0.59909 |
| 17 | 7205000 | 7205999 | 14.16482 | 24.74708 | 41.62095 | 18.33755 | -0.35856 | 0.698504 | 0.599104 |
| 6 | 132179600 | 132182399 | 35.6131 | 58.81435 | 61.07814 | 26.37852 | 0.326083 | 1.362783 | 0.599114 |
| X | 33407200 | 33424199 | 398.0819 | 562.9846 | 455.1774 | 188.3989 | 0.491197 | 4.417593 | 0.599133 |
| 16 | 52063800 | 52064599 | 6.287277 | 12.06581 | 24.10768 | 9.458486 | -0.46399 | -0.09906 | 0.599146 |
| 14 | 82415600 | 82416999 | 17.96881 | 30.96047 | 48.47124 | 21.26284 | -0.25929 | 0.883395 | 0.599153 |
| 2 | 139741800 | 139746599 | 52.91743 | 85.56948 | 77.08691 | 32.9471 | 0.512092 | 1.719162 | 0.599166 |
| 10 | 11731600 | 11732599 | 19.61163 | 34.18738 | 37.30688 | 16.414 | 0.262013 | 0.748589 | 0.599169 |
| 8 | 67866600 | 67867999 | 26.75685 | 44.9301 | 48.57475 | 21.20987 | 0.268897 | 1.146216 | 0.599175 |
| 1 | 88038800 | 88042399 | 50.24916 | 80.91757 | 85.35026 | 36.2792 | 0.27804 | 1.836256 | 0.599183 |
| 7 | 135732800 | 135734999 | 17.56223 | 31.10557 | 27.2709 | 12.05813 | 0.580398 | 0.50907 | 0.599196 |

| | | | | | | | | | |
|---|---|---|---|---|---|---|---|---|---|
| 2 | 73698000 | 73698799 | 6.842025 | 13.19101 | 24.2145 | 9.375644 | -0.32527 | -0.13661 | 0.5992 |
| 5 | 92615200 | 92627199 | 122.5244 | 185.0304 | 259.5273 | 107.2856 | -0.21282 | 3.329562 | 0.599267 |
| 17 | 21864000 | 21865799 | 17.81285 | 30.96283 | 49.53974 | 21.75689 | -0.28816 | 0.797522 | 0.599272 |
| 2 | 139801600 | 139802999 | 19.73199 | 34.83089 | 34.06173 | 15.07094 | 0.421727 | 0.57005 | 0.599286 |
| 16 | 79583000 | 79585199 | 19.81915 | 34.88442 | 34.59566 | 15.31597 | 0.399985 | 0.601399 | 0.599287 |
| 3 | 64146600 | 64148199 | 21.23326 | 37.51196 | 33.37004 | 14.76245 | 0.558069 | 0.569143 | 0.599334 |
| 8 | 60238800 | 60241199 | 21.23326 | 37.51196 | 33.37004 | 14.76245 | 0.558069 | 0.569143 | 0.599334 |
| 11 | 28961000 | 28962599 | 17.99271 | 31.59011 | 34.49642 | 15.26145 | 0.261193 | 0.625821 | 0.599339 |
| 5 | 33654000 | 33657799 | 28.19958 | 46.08277 | 73.09814 | 31.42582 | -0.28965 | 1.487301 | 0.599361 |
| 6 | 10919200 | 10930399 | 329.1583 | 458.7785 | 603.6767 | 250.4419 | -0.23237 | 4.629295 | 0.599368 |
| 17 | 92345600 | 92349199 | 61.83331 | 98.36793 | 106.3285 | 44.40929 | 0.222832 | 2.233723 | 0.599373 |
| 10 | 70083000 | 70087199 | 38.63057 | 62.8815 | 68.5965 | 29.44835 | 0.244959 | 1.560913 | 0.599406 |
| 9 | 9549800 | 9550999 | 12.88439 | 22.59397 | 44.07859 | 19.48073 | -0.57172 | 0.633329 | 0.599415 |
| 13 | 50489000 | 50491199 | 16.49431 | 28.4403 | 47.94142 | 21.03088 | -0.36529 | 0.868031 | 0.599421 |
| 6 | 136602200 | 136603399 | 10.39829 | 18.93012 | 31.83957 | 13.69962 | -0.32716 | 0.301275 | 0.599423 |
| 3 | 91222400 | 91228999 | 83.92218 | 131.9694 | 124.3664 | 51.59251 | 0.406216 | 2.535793 | 0.59944 |
| 7 | 16630400 | 16631799 | 9.235982 | 17.0916 | 29.40844 | 12.38101 | -0.33751 | 0.19252 | 0.599455 |
| 4 | 143909600 | 143926199 | 214.1468 | 318.8976 | 279.1599 | 115.0294 | 0.448828 | 3.612516 | 0.599463 |
| 15 | 95996400 | 96002599 | 41.09548 | 65.70677 | 120.4742 | 50.67243 | -0.53033 | 2.057376 | 0.599469 |
| 4 | 121421200 | 121431599 | 148.0389 | 225.2035 | 214.3789 | 88.82173 | 0.354764 | 3.186249 | 0.599472 |
| 4 | 31706600 | 31708599 | 11.437 | 20.52015 | 35.69883 | 15.62357 | -0.39242 | 0.397983 | 0.599484 |
| 14 | 90346200 | 90347199 | 9.49785 | 18.39223 | 17.01496 | 6.465978 | 0.679233 | -0.19386 | 0.599497 |
| 4 | 53510200 | 53512599 | 13.96211 | 24.51572 | 39.37907 | 17.4229 | -0.29262 | 0.611301 | 0.599502 |
| 13 | 43079200 | 43086399 | 93.20025 | 143.2313 | 203.1398 | 84.35102 | -0.20839 | 2.965623 | 0.599504 |
| 5 | 139497800 | 139499199 | 10.83806 | 19.9589 | 33.54944 | 14.21826 | -0.3114 | 0.241911 | 0.599524 |
| 3 | 105623400 | 105624599 | 7.771153 | 14.68772 | 27.04389 | 10.99248 | -0.39409 | 0.038052 | 0.599531 |
| 14 | 46908600 | 46911199 | 17.11953 | 29.63541 | 46.87704 | 20.56644 | -0.27268 | 0.834736 | 0.599538 |
| 17 | 23549800 | 23556799 | 158.8416 | 239.1787 | 245.9711 | 101.595 | 0.230943 | 3.394307 | 0.599539 |
| 4 | 43358600 | 43360599 | 25.86589 | 43.90535 | 47.96068 | 21.01308 | 0.254461 | 1.048145 | 0.599555 |
| 5 | 79167600 | 79172599 | 72.64808 | 115.2798 | 106.6592 | 44.4284 | 0.443267 | 2.315486 | 0.599558 |
| 15 | 91783400 | 91784199 | 14.55264 | 26.30613 | 28.88475 | 12.53884 | 0.2756 | 0.363834 | 0.599559 |
| 16 | 64577000 | 64578599 | 14.85983 | 26.02904 | 41.44226 | 18.29834 | -0.28017 | 0.656834 | 0.599597 |
| 11 | 10044400 | 10045199 | 17.0985 | 30.43768 | 26.76604 | 11.79205 | 0.579974 | 0.46959 | 0.5996 |
| 16 | 6534000 | 6535399 | 8.577724 | 16.34924 | 29.54351 | 11.79656 | -0.34448 | -0.00367 | 0.599611 |
| 15 | 31687000 | 31691199 | 36.20966 | 58.45722 | 90.09024 | 38.42404 | -0.26352 | 1.740769 | 0.59962 |
| 9 | 19456400 | 19460199 | 56.35674 | 90.26623 | 96.51535 | 40.69836 | 0.249122 | 2.01402 | 0.599621 |
| 14 | 94168200 | 94176199 | 117.8041 | 180.7694 | 166.0659 | 68.94231 | 0.415866 | 2.998884 | 0.599631 |
| 9 | 123158600 | 123159999 | 14.84899 | 27.24425 | 27.79343 | 11.83832 | 0.398785 | 0.277481 | 0.599652 |
| 1 | 93613000 | 93619999 | 35.5271 | 56.74922 | 121.8654 | 51.18811 | -0.75931 | 2.08274 | 0.599653 |
| 14 | 98466800 | 98471199 | 43.62481 | 71.18058 | 68.78723 | 29.55593 | 0.421787 | 1.538698 | 0.599659 |
| 5 | 134153800 | 134157399 | 41.95659 | 67.35125 | 101.5501 | 42.96651 | -0.24198 | 1.9284 | 0.599667 |
| 18 | 37580800 | 37583199 | 73.26793 | 115.5918 | 124.9869 | 51.87397 | 0.211752 | 2.465278 | 0.599673 |
| 13 | 87088000 | 87088599 | 12.4292 | 22.93239 | 24.95188 | 10.54745 | 0.314383 | 0.221454 | 0.599677 |
| 2 | 92922000 | 92923999 | 22.47239 | 37.62982 | 59.18756 | 25.79814 | -0.27015 | 1.171657 | 0.599688 |
| 17 | 18295600 | 18299199 | 63.18247 | 100.8056 | 105.8385 | 44.38204 | 0.269193 | 2.142858 | 0.599689 |
| 7 | 104482200 | 104483599 | 24.82185 | 42.15514 | 46.36738 | 20.31822 | 0.244562 | 1.037442 | 0.599702 |
| X | 18175800 | 18181999 | 73.9865 | 117.5437 | 117.2004 | 48.87756 | 0.337214 | 2.287329 | 0.599708 |

| | | | | | | | | | |
|---|---|---|---|---|---|---|---|---|---|
| 17 | 34994800 | 35004599 | 125.1065 | 189.0328 | 262.1386 | 108.3838 | -0.19574 | 3.319273 | 0.599731 |
| 5 | 26603400 | 26606199 | 36.2176 | 59.28787 | 65.20613 | 28.05069 | 0.238932 | 1.474895 | 0.599735 |
| 1 | 20775600 | 20776199 | 5.094411 | 9.925824 | 20.22124 | 7.351462 | -0.38782 | -0.38784 | 0.599738 |
| 14 | 28105400 | 28105999 | 5.094411 | 9.925824 | 20.22124 | 7.351462 | -0.38782 | -0.38784 | 0.599738 |
| 19 | 27259200 | 27260399 | 15.21092 | 27.25251 | 29.84888 | 13.079 | 0.270321 | 0.423266 | 0.599757 |
| 5 | 13777200 | 13781199 | 34.54835 | 56.99456 | 61.43961 | 26.51705 | 0.272163 | 1.374077 | 0.599768 |
| 7 | 81310000 | 81311999 | 22.51862 | 38.62309 | 42.14534 | 18.48423 | 0.258103 | 0.934954 | 0.599774 |
| 8 | 15623400 | 15625799 | 45.66844 | 73.50746 | 78.12285 | 33.18508 | 0.266373 | 1.841696 | 0.599776 |
| 15 | 26306400 | 26309999 | 27.1419 | 44.64749 | 69.57278 | 29.99076 | -0.25953 | 1.396668 | 0.599781 |
| 3 | 46400400 | 46402599 | 41.16479 | 66.0319 | 101.4832 | 42.89049 | -0.27075 | 1.950336 | 0.599787 |
| 13 | 68158000 | 68178199 | 403.8266 | 564.5654 | 530.6687 | 220.0496 | 0.257214 | 4.585926 | 0.599788 |
| 2 | 151171800 | 151174399 | 30.05575 | 50.96319 | 49.04509 | 21.48128 | 0.436499 | 1.071936 | 0.599851 |
| 11 | 67871400 | 67872399 | 7.030608 | 13.54401 | 25.29296 | 9.815187 | -0.35225 | -0.12096 | 0.599894 |
| 9 | 20403600 | 20404399 | 6.387809 | 12.3818 | 23.32628 | 8.824168 | -0.33044 | -0.22357 | 0.599905 |
| 2 | 122722600 | 122724999 | 21.53764 | 37.73844 | 36.19832 | 15.98734 | 0.447202 | 0.676834 | 0.599909 |
| 3 | 50209400 | 50210399 | 9.342834 | 17.28577 | 29.89345 | 12.58873 | -0.34544 | 0.193535 | 0.599913 |
| 13 | 48088400 | 48091199 | 29.07339 | 49.5494 | 47.35937 | 20.76038 | 0.447087 | 1.022514 | 0.599914 |
| 12 | 27905400 | 27906199 | 12.67012 | 23.77739 | 25.64536 | 10.55968 | 0.356049 | 0.124721 | 0.599946 |
| 17 | 4826200 | 4827999 | 20.69273 | 35.90971 | 39.44332 | 17.30634 | 0.251946 | 0.818738 | 0.599953 |
| 2 | 86076000 | 86077999 | 25.95164 | 44.38324 | 46.29038 | 20.29747 | 0.322376 | 0.978715 | 0.599959 |
| 16 | 62412200 | 62415199 | 26.27704 | 45.14591 | 44.03677 | 19.31588 | 0.419503 | 0.935054 | 0.599968 |
| 14 | 106687800 | 106693799 | 150.4077 | 226.0922 | 232.5106 | 95.9914 | 0.228772 | 3.424483 | 0.600007 |
| 13 | 120655600 | 120658399 | 27.71423 | 44.91472 | 88.46854 | 37.87676 | -0.61072 | 1.633588 | 0.600008 |
| 7 | 63447600 | 63455799 | 69.30752 | 108.9864 | 156.0867 | 64.75133 | -0.19683 | 2.533205 | 0.600012 |
| X | 74966200 | 74968599 | 28.22374 | 48.24808 | 46.8161 | 20.52727 | 0.426118 | 0.993332 | 0.600015 |
| 18 | 3224800 | 3227999 | 63.14014 | 100.4316 | 109.8657 | 45.87698 | 0.205357 | 2.240358 | 0.600075 |
| 1 | 80028000 | 80030799 | 27.34201 | 46.15338 | 50.157 | 21.9383 | 0.261581 | 1.102754 | 0.600079 |
| 15 | 96361600 | 96364799 | 28.01947 | 45.86111 | 72.09416 | 31.01211 | -0.27507 | 1.463426 | 0.600086 |
| 17 | 55047800 | 55052599 | 57.26602 | 91.90541 | 92.43343 | 39.09529 | 0.340634 | 1.951815 | 0.600094 |
| 7 | 66511800 | 66515199 | 23.62933 | 39.56903 | 60.89035 | 26.54116 | -0.2387 | 1.177136 | 0.600102 |
| 3 | 108149200 | 108151599 | 41.69693 | 67.61291 | 73.61435 | 31.52993 | 0.243126 | 1.639513 | 0.600112 |
| 10 | 108309600 | 108313999 | 58.99081 | 94.20779 | 102.3434 | 42.97443 | 0.221556 | 2.107905 | 0.600133 |
| 1 | 13674400 | 13678799 | 68.69244 | 109 | 118.3233 | 49.2857 | 0.21305 | 2.316671 | 0.600139 |
| 10 | 77439400 | 77445799 | 53.1729 | 84.44262 | 127.8356 | 53.29134 | -0.26491 | 2.281992 | 0.600149 |
| 11 | 12744400 | 12748999 | 85.39343 | 133.2604 | 141.0808 | 58.54251 | 0.23004 | 2.678651 | 0.600151 |
| 3 | 123338600 | 123339999 | 14.92713 | 26.21943 | 41.36474 | 18.30689 | -0.26693 | 0.614184 | 0.600157 |
| 16 | 8059600 | 8060799 | 10.54548 | 19.39443 | 32.45475 | 13.7791 | -0.30662 | 0.245703 | 0.600163 |
| 9 | 35930800 | 35931599 | 13.9047 | 25.15124 | 25.59985 | 11.10382 | 0.385337 | 0.349177 | 0.600163 |
| 6 | 58299400 | 58311199 | 140.2093 | 213.6136 | 189.8392 | 78.68983 | 0.455595 | 3.15139 | 0.600163 |
| 9 | 15141800 | 15147799 | 81.44872 | 126.3064 | 180.488 | 74.96321 | -0.20924 | 2.790527 | 0.600172 |
| 9 | 116135000 | 116138599 | 33.00818 | 55.22201 | 54.74238 | 23.84703 | 0.39363 | 1.20971 | 0.600185 |
| 8 | 86727800 | 86730399 | 28.17371 | 48.43569 | 44.57063 | 19.55043 | 0.50358 | 0.940317 | 0.600209 |
| 2 | 36854400 | 36854999 | 16.67058 | 29.36859 | 29.64006 | 13.11972 | 0.375652 | 0.567097 | 0.600212 |
| 13 | 120118600 | 120123799 | 50.56586 | 81.19834 | 89.0791 | 37.70898 | 0.216474 | 1.925363 | 0.600229 |
| 2 | 157196800 | 157200199 | 34.47725 | 56.59487 | 61.65677 | 26.55479 | 0.254195 | 1.436377 | 0.60025 |
| 2 | 144429600 | 144430999 | 19.41815 | 33.91818 | 37.19472 | 16.38532 | 0.255187 | 0.718633 | 0.600253 |
| 10 | 81730800 | 81732999 | 46.7348 | 75.6046 | 63.843 | 27.29754 | 0.605823 | 1.692932 | 0.600257 |

| | | | | | | | | | |
|---|---|---|---|---|---|---|---|---|---|
| 8 | 60882000 | 60884399 | 18.92532 | 32.56715 | 50.61345 | 22.2017 | -0.24921 | 0.90099 | 0.60029 |
| 6 | 100526800 | 100529599 | 28.66806 | 46.87294 | 73.24931 | 31.49683 | -0.26765 | 1.480606 | 0.600311 |
| 12 | 22203600 | 22208599 | 106.0403 | 162.9391 | 167.4192 | 69.51708 | 0.255985 | 2.969011 | 0.60037 |
| 1 | 183611600 | 183614199 | 20.22651 | 34.64887 | 54.98012 | 24.11095 | -0.28038 | 0.951556 | 0.600372 |
| 13 | 50077000 | 50080399 | 48.41491 | 78.0481 | 83.00676 | 35.33666 | 0.26802 | 1.799523 | 0.600397 |
| 2 | 69043000 | 69045399 | 17.82032 | 30.65613 | 48.47322 | 21.26268 | -0.27399 | 0.897152 | 0.600398 |
| 11 | 84640400 | 84643399 | 17.82032 | 30.65613 | 48.47322 | 21.26268 | -0.27399 | 0.897152 | 0.600398 |
| 9 | 23327200 | 23328399 | 17.85184 | 32.20766 | 26.03639 | 11.32971 | 0.712582 | 0.392913 | 0.600425 |
| 8 | 127537600 | 127541599 | 39.00173 | 62.83178 | 94.62571 | 40.24623 | -0.23393 | 1.809979 | 0.600429 |
| 12 | 21604200 | 21607999 | 37.97668 | 62.45198 | 56.48124 | 24.34436 | 0.522984 | 1.41785 | 0.60043 |
| 5 | 131491800 | 131495799 | 30.95158 | 50.50577 | 76.84758 | 33.02007 | -0.23133 | 1.51648 | 0.600434 |
| 11 | 19535200 | 19535999 | 7.140992 | 13.78344 | 24.82637 | 9.578678 | -0.29338 | -0.14277 | 0.600434 |
| 18 | 37665400 | 37698599 | 604.3108 | 817.4432 | 723.0515 | 301.3749 | 0.296895 | 5.098524 | 0.600436 |
| 7 | 108690600 | 108693799 | 24.71495 | 41.29182 | 64.95596 | 28.27221 | -0.27045 | 1.206509 | 0.600438 |
| 13 | 23208400 | 23209599 | 12.06289 | 21.7798 | 35.77155 | 15.55133 | -0.30344 | 0.365922 | 0.600454 |
| 10 | 14706400 | 14709999 | 39.65898 | 64.69681 | 68.85278 | 29.58194 | 0.282768 | 1.533052 | 0.600459 |
| 5 | 145336400 | 145343199 | 154.3909 | 232.2239 | 243.9042 | 100.712 | 0.199171 | 3.413234 | 0.600474 |
| 5 | 46603000 | 46603799 | 8.349984 | 15.89926 | 27.7943 | 11.12159 | -0.30001 | -0.00561 | 0.600477 |
| 5 | 149847600 | 149848799 | 8.349984 | 15.89926 | 27.7943 | 11.12159 | -0.30001 | -0.00561 | 0.600477 |
| 1 | 154874800 | 154876599 | 14.28494 | 25.94772 | 28.33341 | 12.22515 | 0.289006 | 0.331689 | 0.600497 |
| 1 | 63984200 | 63985199 | 5.808822 | 11.29313 | 21.7781 | 8.081436 | -0.33727 | -0.3013 | 0.600503 |
| 10 | 112079800 | 112080399 | 5.808822 | 11.29313 | 21.7781 | 8.081436 | -0.33727 | -0.3013 | 0.600503 |
| 10 | 14003200 | 14008399 | 83.53125 | 128.77 | 191.2402 | 79.43798 | -0.27136 | 2.900982 | 0.600528 |
| 3 | 9099600 | 9103799 | 28.39737 | 46.62669 | 74.40758 | 32.0499 | -0.29496 | 1.423637 | 0.600548 |
| 9 | 114819200 | 114822399 | 42.65735 | 68.47952 | 101.4564 | 42.93005 | -0.21663 | 1.929052 | 0.600558 |
| 17 | 8288800 | 8290199 | 14.06601 | 24.79523 | 39.24075 | 17.36721 | -0.26985 | 0.561519 | 0.600576 |
| 5 | 105562200 | 105568799 | 61.29156 | 96.90115 | 140.721 | 58.46645 | -0.21075 | 2.409803 | 0.600585 |
| 3 | 155085000 | 155088399 | 26.78068 | 46.05643 | 44.59456 | 19.56113 | 0.430485 | 0.931 | 0.600621 |
| 3 | 135261800 | 135263999 | 20.26786 | 35.40824 | 37.55695 | 16.54662 | 0.302888 | 0.719358 | 0.600627 |
| 8 | 69828200 | 69847199 | 458.9485 | 633.6449 | 597.2835 | 248.1589 | 0.234932 | 4.777019 | 0.600635 |
| 10 | 30343400 | 30344999 | 13.08664 | 24.77418 | 24.68807 | 10.00067 | 0.488857 | 0.069005 | 0.600639 |
| 18 | 34699000 | 34699599 | 6.097798 | 11.85256 | 22.34313 | 8.304573 | -0.30689 | -0.2893 | 0.600645 |
| X | 45305800 | 45310399 | 43.44694 | 71.51138 | 62.06574 | 26.76145 | 0.582771 | 1.425259 | 0.600646 |
| 6 | 14414800 | 14415199 | 5.498044 | 10.71934 | 21.47599 | 7.740244 | -0.35228 | -0.40617 | 0.600647 |
| 17 | 62894200 | 62896799 | 31.81393 | 53.43107 | 51.91626 | 22.67137 | 0.422244 | 1.160759 | 0.600684 |
| 3 | 74527800 | 74530599 | 75.98023 | 119.5954 | 128.4948 | 53.31179 | 0.218383 | 2.513944 | 0.600723 |
| 15 | 39300600 | 39301399 | 10.95346 | 20.52197 | 21.097 | 8.711139 | 0.421626 | 0.107845 | 0.600737 |
| X | 74982800 | 74984799 | 17.5854 | 32.57991 | 26.76464 | 11.2504 | 0.72317 | 0.245509 | 0.600753 |
| 7 | 62262200 | 62264999 | 48.27583 | 77.71445 | 84.78463 | 36.0217 | 0.228886 | 1.843343 | 0.600763 |
| 4 | 123062400 | 123063599 | 13.74808 | 25.51219 | 27.06281 | 11.35052 | 0.359117 | 0.207127 | 0.600773 |
| 17 | 48810400 | 48811399 | 12.48763 | 23.03007 | 25.49534 | 10.78362 | 0.289093 | 0.226391 | 0.600803 |
| 1 | 34953000 | 34954799 | 22.37799 | 38.71616 | 42.02137 | 18.43578 | 0.2682 | 0.857126 | 0.60081 |
| 10 | 124534600 | 124535399 | 11.28767 | 20.55106 | 33.3146 | 14.33269 | -0.27463 | 0.308561 | 0.600826 |
| 7 | 107669000 | 107670999 | 31.86306 | 53.09783 | 49.22901 | 21.39234 | 0.489957 | 1.241433 | 0.600832 |
| 1 | 98346600 | 98347199 | 9.965482 | 19.05516 | 22.30613 | 8.843534 | 0.285951 | -0.03385 | 0.600842 |
| 18 | 25885000 | 25888799 | 29.79382 | 48.66762 | 75.0948 | 32.27927 | -0.25039 | 1.497327 | 0.600843 |
| 12 | 35329200 | 35330199 | 8.708107 | 16.3536 | 28.00288 | 11.51665 | -0.30422 | 0.092625 | 0.600849 |

| | | | | | | | | | |
|---|---|---|---|---|---|---|---|---|---|
| X | 6907200 | 6908399 | 13.53827 | 25.2496 | 26.59604 | 11.06669 | 0.377974 | 0.17363 | 0.600856 |
| 18 | 8735600 | 8736799 | 16.51557 | 29.56887 | 31.21743 | 13.6869 | 0.322375 | 0.43885 | 0.600888 |
| 4 | 81373200 | 81379799 | 69.73004 | 109.3149 | 157.9453 | 65.50978 | -0.21296 | 2.592921 | 0.600901 |
| 1 | 81811800 | 81813199 | 13.46168 | 23.83503 | 37.69428 | 16.66804 | -0.26618 | 0.507958 | 0.60091 |
| X | 115107600 | 115108199 | 12.21079 | 22.99031 | 24.71601 | 10.12091 | 0.367568 | 0.100651 | 0.600911 |
| 9 | 17971600 | 17971999 | 5.801671 | 11.30064 | 21.99792 | 8.027051 | -0.32876 | -0.36122 | 0.600914 |
| 10 | 94375200 | 94378999 | 47.39828 | 75.82839 | 111.4139 | 46.90031 | -0.21051 | 2.044209 | 0.600929 |
| 18 | 56777200 | 56783199 | 69.32405 | 109.0579 | 156.2178 | 64.80926 | -0.19663 | 2.525374 | 0.600945 |
| 12 | 52215200 | 52216399 | 19.21731 | 33.83924 | 34.68093 | 15.35244 | 0.353061 | 0.594189 | 0.600953 |
| 7 | 41175800 | 41176199 | 13.8435 | 25.07232 | 27.42209 | 11.87536 | 0.283424 | 0.344007 | 0.600988 |
| 1 | 89768000 | 89769999 | 18.29735 | 31.54004 | 49.00256 | 21.49619 | -0.24819 | 0.881799 | 0.601003 |
| 17 | 84182200 | 84185199 | 44.27104 | 70.94622 | 105.4004 | 44.48082 | -0.2236 | 1.985824 | 0.601011 |
| 12 | 51875400 | 51876799 | 12.01933 | 21.29805 | 35.73352 | 15.79852 | -0.35047 | 0.485625 | 0.601011 |
| 16 | 88570600 | 88570999 | 5.699875 | 11.01735 | 23.17699 | 8.883524 | -0.50591 | -0.18733 | 0.601015 |
| 7 | 131125400 | 131126999 | 15.82776 | 28.27364 | 30.76428 | 13.51388 | 0.277206 | 0.446921 | 0.601031 |
| 9 | 114058400 | 114061399 | 41.71617 | 67.50207 | 74.39239 | 31.81383 | 0.222833 | 1.682214 | 0.601035 |
| 2 | 177151400 | 177155199 | 34.84807 | 57.42819 | 62.02518 | 26.75779 | 0.268809 | 1.387567 | 0.601044 |
| 18 | 9650200 | 9652799 | 20.94344 | 36.68299 | 36.83669 | 16.26393 | 0.381435 | 0.681417 | 0.601049 |
| 6 | 42053200 | 42054599 | 11.73209 | 22.30672 | 24.32604 | 9.763759 | 0.371618 | 0.027566 | 0.60108 |
| 6 | 61456600 | 61460199 | 45.11941 | 72.95848 | 76.93629 | 32.88196 | 0.285366 | 1.70635 | 0.601087 |
| 5 | 146476400 | 146481999 | 55.51318 | 88.63079 | 92.52043 | 38.8353 | 0.278599 | 2.109605 | 0.601087 |
| 4 | 26292600 | 26295799 | 27.23338 | 46.63556 | 46.27485 | 20.29277 | 0.394339 | 0.973686 | 0.601088 |
| 8 | 110951800 | 110955799 | 32.77673 | 53.25561 | 80.69675 | 34.61474 | -0.22989 | 1.586181 | 0.601103 |
| 6 | 14463000 | 14464199 | 11.42613 | 21.69402 | 23.69761 | 9.540266 | 0.365053 | 0.031041 | 0.601111 |
| 5 | 110104600 | 110106599 | 17.00222 | 30.86506 | 29.06832 | 12.5516 | 0.500346 | 0.355582 | 0.60113 |
| 5 | 23390000 | 23394999 | 78.68901 | 122.418 | 174.6568 | 72.50662 | -0.20338 | 2.733497 | 0.601144 |
| X | 121590000 | 121600199 | 78.12206 | 123.8216 | 110.5232 | 45.98425 | 0.493375 | 2.35612 | 0.601152 |
| 3 | 25785000 | 25786399 | 14.31215 | 26.10272 | 28.3718 | 12.17981 | 0.300337 | 0.309307 | 0.601174 |
| 4 | 16639400 | 16641799 | 24.34981 | 42.47277 | 39.59821 | 17.42875 | 0.488156 | 0.767372 | 0.60118 |
| 8 | 55901000 | 55904199 | 53.3463 | 85.68808 | 90.54948 | 38.34839 | 0.27088 | 1.920333 | 0.601205 |
| 6 | 134216000 | 134220599 | 47.688 | 76.19477 | 114.1116 | 47.94932 | -0.24006 | 2.083164 | 0.601206 |
| 6 | 136072000 | 136073799 | 10.9495 | 20.07901 | 32.9084 | 14.02708 | -0.28127 | 0.263961 | 0.601218 |
| 14 | 75316600 | 75318199 | 8.92736 | 16.57106 | 28.93018 | 12.12911 | -0.35385 | 0.171923 | 0.601225 |
| 4 | 143648400 | 143653799 | 139.1592 | 211.477 | 216.951 | 89.85964 | 0.245697 | 3.204961 | 0.601255 |
| X | 5276000 | 5286199 | 194.4108 | 289.9315 | 242.6422 | 100.0016 | 0.515582 | 3.582609 | 0.601256 |
| 7 | 133130200 | 133134999 | 78.32615 | 121.9853 | 173.4104 | 71.97755 | -0.19694 | 2.714866 | 0.601256 |
| 3 | 47957600 | 47959199 | 18.1297 | 31.4687 | 50.10031 | 21.98338 | -0.28127 | 0.819727 | 0.601273 |
| 15 | 49296000 | 49299199 | 31.61441 | 51.42806 | 78.98216 | 33.89903 | -0.24789 | 1.562914 | 0.60128 |
| 11 | 120474800 | 120476399 | 15.62661 | 28.37967 | 28.63285 | 12.35712 | 0.401985 | 0.341727 | 0.601291 |
| 12 | 48222600 | 48226399 | 36.66914 | 60.1467 | 64.84938 | 27.92074 | 0.268631 | 1.450768 | 0.601296 |
| 10 | 122115000 | 122118999 | 74.72305 | 117.8504 | 128.7769 | 53.44265 | 0.196233 | 2.473068 | 0.601299 |
| 13 | 90590200 | 90592199 | 26.75688 | 44.2538 | 68.1768 | 29.47435 | -0.24105 | 1.328916 | 0.601301 |
| 15 | 24323000 | 24324199 | 17.19347 | 30.51042 | 26.27361 | 11.60591 | 0.607366 | 0.488012 | 0.601301 |
| 12 | 119623800 | 119627399 | 25.50596 | 44.12037 | 42.9629 | 18.84876 | 0.424023 | 0.873257 | 0.601318 |
| 2 | 15486200 | 15487399 | 12.39152 | 23.21234 | 25.65058 | 10.59438 | 0.317878 | 0.134353 | 0.601327 |
| X | 161521200 | 161523799 | 14.91732 | 26.22581 | 46.64948 | 20.65719 | -0.43904 | 0.611843 | 0.601373 |
| 14 | 56639800 | 56640599 | 10.22146 | 18.6576 | 31.34181 | 13.44081 | -0.32228 | 0.283048 | 0.601375 |

| | | | | | | | | | |
|---|---|---|---|---|---|---|---|---|---|
| 18 | 56030000 | 56030599 | 7.274201 | 14.03781 | 25.41554 | 9.813359 | -0.30216 | -0.13851 | 0.601375 |
| 15 | 97297000 | 97299199 | 23.58677 | 39.38639 | 62.02525 | 26.98772 | -0.27192 | 1.20624 | 0.601382 |
| 13 | 61851000 | 61868799 | 49.2195 | 79.27264 | 69.68248 | 29.62655 | 0.540763 | 1.82375 | 0.601386 |
| 12 | 112330800 | 112336399 | 53.67431 | 85.5422 | 125.7527 | 52.65321 | -0.21731 | 2.17291 | 0.601393 |
| 1 | 88722200 | 88725399 | 44.85848 | 71.73193 | 108.6096 | 45.68981 | -0.25446 | 2.056317 | 0.601395 |
| 5 | 89662000 | 89666399 | 81.34527 | 127.6828 | 138.5083 | 57.44546 | 0.201446 | 2.574358 | 0.601406 |
| 5 | 77415800 | 77418199 | 17.84584 | 30.51524 | 50.05541 | 21.94806 | -0.32828 | 0.953619 | 0.601414 |
| 13 | 66733200 | 66735399 | 82.73267 | 129.4742 | 137.8867 | 57.20043 | 0.224681 | 2.628004 | 0.601466 |
| 9 | 84580400 | 84581799 | 11.96287 | 21.41381 | 34.68809 | 15.21101 | -0.29177 | 0.416669 | 0.601481 |
| 10 | 126154400 | 126156399 | 19.5048 | 34.04871 | 37.17079 | 16.3685 | 0.261475 | 0.726524 | 0.601485 |
| 18 | 19761600 | 19765399 | 35.27256 | 57.79687 | 63.02634 | 27.12411 | 0.251715 | 1.457434 | 0.601491 |
| 3 | 136990000 | 136992399 | 18.07155 | 31.88634 | 33.71012 | 14.91663 | 0.3096 | 0.563945 | 0.601504 |
| 6 | 107563200 | 107566199 | 15.17914 | 27.65576 | 23.53225 | 10.11511 | 0.649808 | 0.309277 | 0.601505 |
| X | 15932400 | 15934999 | 26.76843 | 46.91176 | 35.49102 | 15.67718 | 0.788365 | 0.705902 | 0.601505 |
| 17 | 35041600 | 35051399 | 151.3532 | 225.0729 | 314.3673 | 129.5184 | -0.2267 | 3.628207 | 0.601507 |
| 7 | 80702200 | 80703799 | 12.62097 | 22.40573 | 36.16404 | 15.9692 | -0.29361 | 0.475773 | 0.601513 |
| 1 | 16517200 | 16520599 | 54.3528 | 87.27663 | 90.69634 | 38.39176 | 0.294481 | 1.933968 | 0.601522 |
| 12 | 19584200 | 19585799 | 34.71009 | 57.34602 | 45.62402 | 19.70962 | 0.709393 | 1.345164 | 0.601551 |
| X | 6244600 | 6247999 | 29.78358 | 51.03098 | 45.67945 | 20.03257 | 0.542546 | 0.978602 | 0.601564 |
| 19 | 39812800 | 39815799 | 19.23282 | 33.80978 | 30.30843 | 13.42133 | 0.544482 | 0.606279 | 0.601572 |
| 1 | 74323400 | 74325199 | 10.65124 | 18.90991 | 34.84992 | 15.38823 | -0.48345 | 0.460902 | 0.601579 |
| 6 | 105403600 | 105404599 | 14.04825 | 25.33873 | 26.99315 | 11.74988 | 0.316554 | 0.368919 | 0.601584 |
| 7 | 6202600 | 6207999 | 69.37196 | 109.0777 | 156.7135 | 65.01057 | -0.20149 | 2.536188 | 0.601592 |
| 16 | 20811600 | 20815399 | 54.71988 | 87.55405 | 96.04638 | 40.43943 | 0.210529 | 2.043732 | 0.601604 |
| 2 | 95553400 | 95555199 | 17.90336 | 31.47506 | 33.98378 | 15.04864 | 0.277605 | 0.605281 | 0.601608 |
| 11 | 113468000 | 113470399 | 19.25541 | 33.59774 | 36.76192 | 16.18372 | 0.258255 | 0.732432 | 0.601616 |
| 5 | 52670600 | 52680399 | 107.5335 | 163.8575 | 230.0804 | 95.37475 | -0.20358 | 3.141861 | 0.601629 |
| 15 | 27397200 | 27398599 | 16.99558 | 30.18794 | 32.19474 | 14.21007 | 0.301289 | 0.492485 | 0.601631 |
| 12 | 106809800 | 106814599 | 38.83208 | 62.60365 | 94.30147 | 40.148 | -0.23299 | 1.787741 | 0.601653 |
| 17 | 9302400 | 9306199 | 56.58905 | 90.4417 | 98.7615 | 41.51555 | 0.215305 | 2.079782 | 0.601659 |
| 14 | 15813400 | 15816599 | 28.24751 | 47.61297 | 50.72685 | 22.17782 | 0.289983 | 1.117504 | 0.60167 |
| 10 | 29960000 | 29961399 | 14.75077 | 25.83935 | 40.96057 | 18.08645 | -0.27374 | 0.656211 | 0.60167 |
| 4 | 147269000 | 147270799 | 17.93377 | 32.10663 | 26.84605 | 11.77073 | 0.656636 | 0.438607 | 0.601741 |
| 13 | 62112200 | 62125999 | 228.5836 | 335.5601 | 333.5824 | 137.3903 | 0.246201 | 3.852159 | 0.601741 |
| 11 | 17123000 | 17124799 | 34.75543 | 57.90177 | 49.80414 | 21.63849 | 0.597852 | 1.254563 | 0.601764 |
| 9 | 7374800 | 7377999 | 39.63594 | 64.74105 | 60.26197 | 25.90292 | 0.476808 | 1.510567 | 0.601765 |
| 3 | 24882200 | 24883999 | 14.2264 | 25.09434 | 39.74072 | 17.58619 | -0.27048 | 0.556132 | 0.601771 |
| 16 | 66861400 | 66862999 | 10.92522 | 19.99366 | 32.86172 | 14.04612 | -0.28833 | 0.274351 | 0.601799 |
| 4 | 74169600 | 74170999 | 14.33414 | 24.87374 | 50.04096 | 21.95617 | -0.61724 | 0.804861 | 0.601801 |
| 3 | 50107400 | 50108199 | 12.9832 | 23.67788 | 24.54391 | 10.53713 | 0.368459 | 0.291236 | 0.60181 |
| 5 | 89153600 | 89156799 | 52.64005 | 83.68689 | 125.5387 | 52.3905 | -0.25018 | 2.252197 | 0.60181 |
| 12 | 114673600 | 114684399 | 174.3637 | 262.3821 | 248.8658 | 102.7729 | 0.346746 | 3.410418 | 0.601815 |
| 19 | 5233800 | 5235799 | 49.2823 | 79.23435 | 86.82117 | 36.81937 | 0.220352 | 1.885186 | 0.601867 |
| X | 146457400 | 146479799 | 432.7824 | 604.1452 | 530.3413 | 219.9649 | 0.353667 | 4.610142 | 0.601875 |
| 18 | 5307600 | 5314399 | 104.3158 | 162.9348 | 151.5627 | 62.88627 | 0.417491 | 2.675326 | 0.601884 |
| 6 | 125382400 | 125385999 | 29.47573 | 48.1561 | 74.46091 | 32.00875 | -0.25321 | 1.494078 | 0.601898 |
| 14 | 42431600 | 42432399 | 19.41212 | 33.92837 | 37.15677 | 16.37569 | 0.257113 | 0.709389 | 0.601902 |

| | | | | | | | | | |
|---|---|---|---|---|---|---|---|---|---|
| 5 | 106011400 | 106013399 | 11.39882 | 20.70122 | 34.30338 | 14.80463 | -0.30932 | 0.32417 | 0.601904 |
| 16 | 5150800 | 5153599 | 32.04022 | 52.85282 | 58.43927 | 25.22129 | 0.235619 | 1.36593 | 0.601922 |
| 12 | 51214000 | 51215399 | 11.30252 | 19.97856 | 40.38779 | 17.86507 | -0.61928 | 0.517739 | 0.60193 |
| 11 | 73413000 | 73413999 | 15.79901 | 29.13623 | 27.86895 | 11.788 | 0.498523 | 0.258047 | 0.601936 |
| 12 | 103824600 | 103837599 | 146.5945 | 218.6605 | 304.2058 | 125.3785 | -0.21696 | 3.573078 | 0.601951 |
| 1 | 122738000 | 122740599 | 19.58577 | 34.31538 | 35.86041 | 15.83694 | 0.324151 | 0.667256 | 0.601965 |
| 17 | 57951800 | 57953599 | 16.15871 | 29.17532 | 30.22728 | 13.14077 | 0.357448 | 0.383878 | 0.601984 |
| 13 | 29859200 | 29861799 | 24.01283 | 41.73086 | 41.32042 | 18.14468 | 0.401141 | 0.817287 | 0.601987 |
| 14 | 61087600 | 61089599 | 13.14921 | 23.22574 | 37.25719 | 16.4827 | -0.28789 | 0.531824 | 0.602007 |
| 1 | 6196000 | 6197199 | 11.57946 | 20.9796 | 34.26577 | 14.83222 | -0.29129 | 0.339636 | 0.602016 |
| 4 | 9156200 | 9156599 | 5.806894 | 11.28896 | 21.76034 | 8.077294 | -0.33649 | -0.29831 | 0.602017 |
| 3 | 21377600 | 21379999 | 18.24115 | 31.04829 | 58.07418 | 25.45468 | -0.51808 | 1.006588 | 0.602019 |
| 4 | 22769400 | 22770999 | 20.43962 | 35.62684 | 38.78386 | 17.06265 | 0.26543 | 0.756243 | 0.602027 |
| 3 | 98500800 | 98505399 | 104.1706 | 160.6922 | 172.3138 | 71.57601 | 0.198723 | 2.893902 | 0.602027 |
| 14 | 20278600 | 20280599 | 13.94705 | 24.35888 | 41.59369 | 18.32188 | -0.38056 | 0.699171 | 0.602031 |
| 19 | 56931000 | 56934199 | 32.99437 | 53.40154 | 82.8211 | 35.42445 | -0.26855 | 1.666553 | 0.602058 |
| 4 | 146258400 | 146266999 | 220.4032 | 323.6538 | 326.2906 | 134.3844 | 0.226411 | 3.84703 | 0.602101 |
| 2 | 29484600 | 29489599 | 42.35609 | 67.90595 | 102.368 | 43.22871 | -0.24387 | 1.970489 | 0.602105 |
| 13 | 60173000 | 60181999 | 134.0307 | 201.7891 | 279.5639 | 115.4655 | -0.19889 | 3.393183 | 0.602111 |
| Y | 2286000 | 2286999 | 12.03573 | 22.3575 | 18.14333 | 7.598956 | 0.743914 | 0.161913 | 0.602118 |
| 6 | 124380200 | 124382199 | 16.28294 | 28.09922 | 46.90569 | 20.57705 | -0.35069 | 0.862482 | 0.602152 |
| 15 | 87270200 | 87276399 | 68.99895 | 108.4286 | 155.283 | 64.41234 | -0.19752 | 2.546272 | 0.602162 |
| 12 | 84920600 | 84924199 | 30.75689 | 49.74578 | 82.88979 | 35.43578 | -0.37271 | 1.676138 | 0.602188 |
| 8 | 100886200 | 100886999 | 12.23828 | 22.83158 | 25.61622 | 10.65382 | 0.288164 | 0.158971 | 0.602206 |
| 11 | 90717600 | 90722399 | 41.56307 | 67.59092 | 70.70375 | 30.34252 | 0.304382 | 1.581048 | 0.602215 |
| 4 | 39218600 | 39219399 | 15.98518 | 29.11723 | 25.20299 | 10.83656 | 0.625554 | 0.323358 | 0.602222 |
| 1 | 7351000 | 7354399 | 37.25026 | 61.06926 | 65.35048 | 28.13052 | 0.279203 | 1.459919 | 0.602229 |
| 13 | 32960000 | 32960599 | 6.384637 | 12.41577 | 23.63707 | 8.753164 | -0.31672 | -0.2965 | 0.602241 |
| 11 | 30972000 | 30973999 | 16.8384 | 29.25024 | 45.80615 | 20.10905 | -0.25727 | 0.797666 | 0.602263 |
| 1 | 81030800 | 81032599 | 15.10337 | 27.68987 | 28.34586 | 12.08614 | 0.392892 | 0.285696 | 0.602263 |
| 12 | 51748200 | 51749799 | 21.1895 | 36.98513 | 38.71622 | 17.04774 | 0.321909 | 0.741585 | 0.602265 |
| 6 | 133904600 | 133906399 | 13.78792 | 24.44361 | 39.20534 | 17.33183 | -0.28579 | 0.498728 | 0.602272 |
| 13 | 10606400 | 10606999 | 9.209444 | 17.04562 | 29.39398 | 12.37182 | -0.34059 | 0.190037 | 0.60228 |
| 4 | 16775000 | 16775799 | 15.54046 | 27.57191 | 28.06572 | 12.39951 | 0.36745 | 0.483263 | 0.602281 |
| 3 | 57086800 | 57087999 | 11.34528 | 21.44413 | 24.1918 | 9.832034 | 0.30718 | 0.063038 | 0.602319 |
| 3 | 17274200 | 17276799 | 19.1237 | 33.62405 | 30.19309 | 13.36977 | 0.542004 | 0.602104 | 0.602324 |
| 13 | 81210000 | 81216799 | 61.06404 | 95.88951 | 165.0807 | 68.46999 | -0.46354 | 2.558669 | 0.602337 |
| 4 | 5787600 | 5788799 | 8.223199 | 15.56266 | 27.06799 | 10.97156 | -0.30889 | 0.034157 | 0.602363 |
| 8 | 4718800 | 4722999 | 45.69596 | 74.73236 | 66.56853 | 28.6267 | 0.541422 | 1.512733 | 0.602371 |
| 5 | 68610400 | 68612199 | 22.11647 | 38.59095 | 34.45105 | 15.16667 | 0.550408 | 0.742488 | 0.602382 |
| X | 24065600 | 24081199 | 179.898 | 271.5238 | 233.7576 | 96.61596 | 0.490197 | 3.353068 | 0.602405 |
| 2 | 69461400 | 69462399 | 11.77418 | 21.2398 | 34.21417 | 14.89068 | -0.27645 | 0.36689 | 0.602412 |
| 15 | 57655200 | 57657599 | 17.62703 | 29.99882 | 56.9323 | 24.9539 | -0.53889 | 1.005248 | 0.602422 |
| 11 | 55024800 | 55026799 | 16.46211 | 28.72806 | 45.18125 | 19.89199 | -0.26284 | 0.72845 | 0.602444 |
| 2 | 125782800 | 125783999 | 11.30275 | 20.65769 | 33.91398 | 14.52066 | -0.28827 | 0.288244 | 0.602446 |
| 17 | 44729600 | 44731399 | 12.00454 | 21.44724 | 34.72187 | 15.25045 | -0.2924 | 0.430308 | 0.602468 |
| X | 90200000 | 90201999 | 32.17308 | 53.70886 | 55.19846 | 24.01299 | 0.341872 | 1.232378 | 0.602514 |

| | | | | | | | | | |
|---|---|---|---|---|---|---|---|---|---|
| 8 | 47817400 | 47822799 | 46.55976 | 74.46756 | 110.5091 | 46.50234 | -0.22526 | 2.050411 | 0.602517 |
| Y | 10758400 | 10760199 | 13.20647 | 24.14253 | 19.65209 | 8.411877 | 0.717358 | 0.26535 | 0.602544 |
| 16 | 66235400 | 66237199 | 18.76778 | 33.01562 | 29.22669 | 12.94048 | 0.56275 | 0.5924 | 0.602551 |
| 12 | 114560200 | 114563199 | 51.4499 | 82.49918 | 90.18013 | 38.09167 | 0.219243 | 1.97219 | 0.602554 |
| 3 | 130892600 | 130893599 | 14.69796 | 26.38327 | 27.74156 | 12.13629 | 0.330143 | 0.401267 | 0.602555 |
| 11 | 63170800 | 63171599 | 12.2174 | 23.15654 | 24.78898 | 10.01596 | 0.389765 | 0.054378 | 0.602557 |
| 6 | 100421800 | 100427199 | 52.37164 | 83.35812 | 122.1419 | 51.04984 | -0.21458 | 2.212456 | 0.602568 |
| 10 | 106354400 | 106355399 | 8.908654 | 16.82423 | 29.21297 | 11.89417 | -0.31203 | 0.062644 | 0.602573 |
| 6 | 119867000 | 119871199 | 23.13547 | 38.58009 | 61.37615 | 26.68231 | -0.28647 | 1.219533 | 0.602615 |
| 11 | 108714800 | 108718999 | 60.85546 | 96.92717 | 105.7225 | 44.22869 | 0.211637 | 2.194829 | 0.602631 |
| 7 | 18197400 | 18199799 | 108.7485 | 166.8759 | 171.2357 | 71.08872 | 0.256439 | 2.997469 | 0.602633 |
| 2 | 122376800 | 122378199 | 27.81975 | 47.13226 | 49.5963 | 21.71718 | 0.308083 | 1.07251 | 0.602636 |
| 2 | 73309000 | 73312399 | 28.68842 | 46.8533 | 73.67628 | 31.66739 | -0.27776 | 1.497467 | 0.602654 |
| 5 | 82211200 | 82212799 | 20.27033 | 35.77355 | 33.40636 | 14.78159 | 0.487716 | 0.573194 | 0.602656 |
| 8 | 16531200 | 16532599 | 11.10825 | 20.39924 | 34.02089 | 14.47312 | -0.30431 | 0.259299 | 0.602662 |
| 17 | 35163800 | 35164999 | 10.05042 | 18.4737 | 30.88407 | 13.12214 | -0.30591 | 0.241196 | 0.602669 |
| 4 | 99425600 | 99427999 | 21.39559 | 37.57501 | 36.26963 | 16.04969 | 0.43833 | 0.644113 | 0.60267 |
| 13 | 45863600 | 45865199 | 10.17055 | 18.73446 | 30.886 | 13.08491 | -0.28306 | 0.229058 | 0.602672 |
| 17 | 33555800 | 33566999 | 236.9474 | 348.4367 | 325.8877 | 134.2067 | 0.336515 | 3.827326 | 0.602681 |
| 10 | 126190800 | 126193199 | 26.3659 | 44.5498 | 49.06682 | 21.46791 | 0.242404 | 1.090096 | 0.602684 |
| 4 | 127716600 | 127717999 | 18.90604 | 33.00102 | 35.75638 | 15.74491 | 0.272399 | 0.723422 | 0.60269 |
| 19 | 22774200 | 22774799 | 5.368709 | 10.45445 | 20.72986 | 7.591391 | -0.35851 | -0.35492 | 0.602691 |
| 10 | 4432000 | 4435799 | 29.62338 | 48.48068 | 74.07912 | 31.86461 | -0.2344 | 1.469619 | 0.602698 |
| 10 | 70801600 | 70803799 | 19.10362 | 33.41513 | 36.63598 | 16.15678 | 0.255496 | 0.694789 | 0.602726 |
| 1 | 53691400 | 53691799 | 8.896773 | 17.16354 | 20.35584 | 7.866984 | 0.301945 | -0.13182 | 0.602738 |
| 2 | 16024400 | 16028399 | 35.79661 | 57.79092 | 87.78575 | 37.44161 | -0.24264 | 1.741372 | 0.602752 |
| 10 | 92470400 | 92473999 | 45.91338 | 74.62986 | 72.09939 | 30.93604 | 0.418407 | 1.600378 | 0.602752 |
| 11 | 44033800 | 44034399 | 6.38286 | 12.3866 | 23.39969 | 8.803985 | -0.32702 | -0.24212 | 0.602798 |
| 17 | 20348200 | 20352999 | 41.72451 | 68.84361 | 60.88499 | 26.27937 | 0.557048 | 1.390562 | 0.60282 |
| 10 | 89092600 | 89093999 | 11.78772 | 21.06786 | 35.17828 | 15.44597 | -0.33667 | 0.42493 | 0.602831 |
| 10 | 55093800 | 55097799 | 47.33121 | 76.63597 | 76.60871 | 32.77679 | 0.364077 | 1.68083 | 0.602832 |
| 5 | 117528200 | 117534199 | 111.2115 | 170.6817 | 182.687 | 75.8443 | 0.195804 | 2.996277 | 0.602833 |
| 7 | 14240800 | 14276999 | 651.9661 | 875.223 | 790.6687 | 329.937 | 0.255431 | 5.21883 | 0.602839 |
| 9 | 119159200 | 119161599 | 14.94301 | 26.18514 | 41.46808 | 18.31588 | -0.27231 | 0.653308 | 0.602851 |
| 1 | 102756000 | 102758199 | 22.2839 | 38.92694 | 38.49223 | 16.95776 | 0.403644 | 0.73268 | 0.602852 |
| 6 | 80355200 | 80356399 | 21.42885 | 37.3203 | 33.64265 | 14.79315 | 0.53623 | 0.76431 | 0.602858 |
| 11 | 15989800 | 15994199 | 38.50389 | 62.14085 | 94.03291 | 40.08881 | -0.23779 | 1.756192 | 0.602878 |
| 3 | 145686200 | 145687399 | 9.958758 | 18.31071 | 31.36571 | 13.32135 | -0.34088 | 0.236326 | 0.602898 |
| 19 | 3795200 | 3799799 | 66.37342 | 105.372 | 115.4187 | 48.09946 | 0.200778 | 2.299821 | 0.602907 |
| 5 | 109320400 | 109325599 | 85.61311 | 134.6312 | 139.8761 | 58.02675 | 0.265689 | 2.540504 | 0.602923 |
| 7 | 11802000 | 11806399 | 56.03448 | 90.47425 | 77.57475 | 33.0916 | 0.580758 | 1.764574 | 0.602935 |
| 6 | 71748000 | 71748999 | 12.2511 | 22.80645 | 25.60496 | 10.68794 | 0.283069 | 0.172855 | 0.602943 |
| 14 | 123801600 | 123802799 | 12.2511 | 22.80645 | 25.60496 | 10.68794 | 0.283069 | 0.172855 | 0.602943 |
| 7 | 81058200 | 81060599 | 24.23418 | 42.21961 | 40.15098 | 17.65852 | 0.45944 | 0.783172 | 0.602946 |
| 10 | 15220600 | 15224199 | 41.68259 | 67.64007 | 73.17133 | 31.35741 | 0.253383 | 1.6242 | 0.602958 |
| 4 | 73752600 | 73753399 | 23.0236 | 39.39874 | 43.19659 | 18.9418 | 0.25078 | 0.959038 | 0.602967 |
| 1 | 178582800 | 178584999 | 31.99549 | 51.9373 | 79.70288 | 34.16469 | -0.24935 | 1.603829 | 0.602973 |

| | | | | | | | | | |
|---|---|---|---|---|---|---|---|---|---|
| 6 | 36200600 | 36201999 | 14.375 | 25.23392 | 40.34131 | 17.84457 | -0.28605 | 0.617471 | 0.602982 |
| 7 | 64154600 | 64156399 | 14.375 | 25.23392 | 40.34131 | 17.84457 | -0.28605 | 0.617471 | 0.602982 |
| 3 | 66018400 | 66021199 | 38.71706 | 62.77715 | 66.83837 | 28.62642 | 0.275246 | 1.631535 | 0.602984 |
| 5 | 98032600 | 98034199 | 11.88094 | 21.50135 | 35.81883 | 15.52656 | -0.32112 | 0.350712 | 0.602995 |
| 4 | 80152400 | 80154799 | 36.96536 | 60.59851 | 55.21803 | 23.76833 | 0.510424 | 1.45265 | 0.603003 |
| 14 | 37380600 | 37381399 | 15.24865 | 27.18756 | 28.98109 | 12.75094 | 0.305331 | 0.452813 | 0.603025 |
| 2 | 138463400 | 138466399 | 86.25473 | 134.5303 | 143.3862 | 59.50252 | 0.219737 | 2.689333 | 0.603036 |
| 12 | 74011000 | 74011799 | 7.457108 | 14.23113 | 25.60794 | 10.19808 | -0.33485 | -0.03462 | 0.603037 |
| 4 | 147806000 | 147808399 | 45.84907 | 74.15376 | 68.69449 | 29.36666 | 0.472122 | 1.694823 | 0.603038 |
| 5 | 23389200 | 23394999 | 81.56454 | 126.6172 | 179.8127 | 74.67117 | -0.19913 | 2.772477 | 0.603039 |
| 6 | 11003400 | 11005399 | 23.34068 | 40.60603 | 40.7711 | 17.91546 | 0.381238 | 0.798513 | 0.603057 |
| 17 | 58534600 | 58537799 | 30.39483 | 51.61357 | 48.47393 | 21.24053 | 0.471639 | 1.060427 | 0.603066 |
| 6 | 131780600 | 131782999 | 36.31882 | 59.80914 | 55.30406 | 23.85107 | 0.491866 | 1.395949 | 0.603076 |
| 2 | 161987800 | 161988799 | 12.61157 | 23.21451 | 24.14494 | 10.23888 | 0.376301 | 0.237071 | 0.603081 |
| 7 | 135606200 | 135608199 | 16.9556 | 30.0705 | 32.16953 | 14.21742 | 0.295493 | 0.501163 | 0.603088 |
| 12 | 30933200 | 30936199 | 21.99351 | 36.45094 | 69.87643 | 30.25234 | -0.55507 | 1.294835 | 0.603091 |
| 18 | 46973800 | 46976799 | 39.87439 | 64.28628 | 96.72314 | 41.18086 | -0.23124 | 1.790837 | 0.603109 |
| 2 | 150202000 | 150204999 | 26.09854 | 43.31671 | 66.51278 | 28.82471 | -0.23588 | 1.285876 | 0.603111 |
| 12 | 115738400 | 115743399 | 60.36007 | 97.11237 | 85.22549 | 36.17915 | 0.541249 | 1.878607 | 0.603127 |
| 3 | 47744200 | 47749399 | 141.531 | 213.6614 | 222.2367 | 91.86039 | 0.217801 | 3.338577 | 0.603144 |
| X | 29316000 | 29321199 | 25.93955 | 45.22554 | 33.21539 | 14.61532 | 0.830707 | 0.772587 | 0.603165 |
| 3 | 64205800 | 64206599 | 10.84243 | 19.80301 | 32.34109 | 13.8589 | -0.28134 | 0.28611 | 0.603175 |
| 3 | 49331400 | 49334399 | 25.35914 | 43.00072 | 47.4409 | 20.77883 | 0.24015 | 1.054417 | 0.603178 |
| 6 | 124639600 | 124640599 | 7.789453 | 14.72327 | 26.51946 | 10.77783 | -0.36208 | 0.037972 | 0.603189 |
| 17 | 49059800 | 49069999 | 106.3852 | 163.886 | 175.689 | 72.97639 | 0.197637 | 2.920174 | 0.60319 |
| 4 | 38750200 | 38750599 | 8.222924 | 15.93231 | 18.80689 | 7.128264 | 0.334863 | -0.2109 | 0.60321 |
| 19 | 51847200 | 51847599 | 7.970853 | 15.47758 | 18.88822 | 7.078224 | 0.303036 | -0.25074 | 0.603226 |
| 17 | 86129400 | 86130599 | 13.9606 | 25.15116 | 25.9741 | 11.32287 | 0.3596 | 0.371601 | 0.603242 |
| 2 | 107715200 | 107716799 | 12.33232 | 21.86918 | 35.70673 | 15.78191 | -0.31093 | 0.480742 | 0.603249 |
| 13 | 3397400 | 3404199 | 161.6396 | 243.1973 | 220.2903 | 90.96844 | 0.412911 | 3.408836 | 0.603261 |
| 14 | 6510400 | 6513999 | 27.77029 | 47.93542 | 43.46752 | 19.06891 | 0.525864 | 0.90309 | 0.60327 |
| 17 | 91666200 | 91667399 | 10.14485 | 18.77371 | 30.94504 | 13.02772 | -0.27633 | 0.202446 | 0.603284 |
| 7 | 15681800 | 15684799 | 16.27957 | 27.94969 | 47.96075 | 21.03645 | -0.39194 | 0.908959 | 0.60329 |
| 12 | 98916200 | 98916799 | 5.215228 | 10.15579 | 20.71805 | 7.584994 | -0.39815 | -0.35637 | 0.603296 |
| 14 | 35968800 | 35969199 | 5.215228 | 10.15579 | 20.71805 | 7.584994 | -0.39815 | -0.35637 | 0.603296 |
| 8 | 75083000 | 75084399 | 15.49768 | 27.79962 | 30.27251 | 13.25121 | 0.27952 | 0.41798 | 0.603305 |
| 6 | 136125800 | 136130199 | 34.24113 | 55.5862 | 84.39357 | 36.17709 | -0.23393 | 1.608286 | 0.603311 |
| 5 | 86695800 | 86698799 | 23.00545 | 39.56462 | 42.14202 | 18.48535 | 0.29363 | 0.915181 | 0.603328 |
| 13 | 51871800 | 51876199 | 38.40739 | 61.86197 | 92.87057 | 39.48832 | -0.22969 | 1.816242 | 0.603334 |
| 7 | 59507400 | 59508999 | 28.36293 | 47.74619 | 42.76117 | 18.68794 | 0.539569 | 1.121685 | 0.603337 |
| 4 | 12369400 | 12371599 | 18.28594 | 31.33362 | 50.06394 | 21.95552 | -0.29009 | 0.935596 | 0.603348 |
| 18 | 9913600 | 9918399 | 49.95884 | 80.52442 | 82.90728 | 35.28719 | 0.314436 | 1.804955 | 0.603371 |
| 1 | 181215200 | 181217799 | 41.35159 | 66.42624 | 98.98267 | 41.92434 | -0.22379 | 1.906508 | 0.603376 |
| 7 | 34859000 | 34859199 | 4.511027 | 8.820041 | 18.42924 | 6.303502 | -0.34396 | -0.61902 | 0.60339 |
| 19 | 9648800 | 9651999 | 47.90217 | 77.30235 | 81.07625 | 34.56143 | 0.289835 | 1.770034 | 0.603408 |
| 9 | 4238200 | 4241799 | 73.06168 | 115.2883 | 125.3376 | 52.02449 | 0.204142 | 2.460247 | 0.603408 |
| X | 43292200 | 43293599 | 21.91155 | 37.70551 | 40.60938 | 17.81338 | 0.277831 | 0.901993 | 0.603411 |

| | | | | | | | | | |
|---|---|---|---|---|---|---|---|---|---|
| 2 | 39525200 | 39527399 | 28.31487 | 48.91573 | 42.31067 | 18.56188 | 0.593884 | 0.895769 | 0.603411 |
| 4 | 16319000 | 16320199 | 10.81664 | 20.68136 | 19.55802 | 7.755426 | 0.590957 | -0.03285 | 0.603414 |
| 12 | 24069000 | 24071399 | 47.22936 | 76.18429 | 67.54382 | 28.77704 | 0.531023 | 1.773718 | 0.603466 |
| 3 | 106320200 | 106322399 | 39.097 | 63.52793 | 69.96969 | 30.01448 | 0.229294 | 1.592264 | 0.603467 |
| 5 | 85902000 | 85902399 | 4.910813 | 9.592137 | 19.38691 | 6.787374 | -0.3265 | -0.52477 | 0.603469 |
| 14 | 113262200 | 113265199 | 25.82118 | 44.16398 | 39.00622 | 17.10362 | 0.561444 | 0.970802 | 0.603497 |
| 12 | 101750600 | 101752399 | 21.41282 | 37.51214 | 37.39065 | 16.51112 | 0.3922 | 0.683334 | 0.603511 |
| 16 | 28900200 | 28901599 | 19.6703 | 34.5241 | 31.2987 | 13.8429 | 0.528282 | 0.634363 | 0.603516 |
| 6 | 85829400 | 85830199 | 9.459457 | 17.61637 | 29.98076 | 12.50631 | -0.31235 | 0.15601 | 0.603526 |
| 4 | 48124600 | 48125999 | 11.33867 | 20.47486 | 33.25206 | 14.45366 | -0.28673 | 0.357346 | 0.603526 |
| 12 | 38873600 | 38875799 | 15.11902 | 27.41264 | 29.28488 | 12.66459 | 0.318282 | 0.350278 | 0.603528 |
| 7 | 126168200 | 126168999 | 11.72075 | 21.84802 | 24.59856 | 10.24491 | 0.281545 | 0.157083 | 0.603542 |
| 8 | 110833600 | 110843399 | 127.2469 | 191.6812 | 267.915 | 110.6705 | -0.2109 | 3.381289 | 0.603556 |
| 7 | 17706000 | 17710599 | 76.88339 | 122.2632 | 114.1144 | 47.63626 | 0.433693 | 2.260873 | 0.603561 |
| 11 | 100077800 | 100080999 | 28.11801 | 46.24251 | 70.81359 | 30.52266 | -0.23462 | 1.402962 | 0.603562 |
| 10 | 41980400 | 41981799 | 27.84709 | 45.75649 | 70.58345 | 30.41209 | -0.24559 | 1.414623 | 0.603594 |
| 7 | 31121600 | 31124199 | 49.14866 | 79.92848 | 69.72484 | 29.92265 | 0.565812 | 1.597504 | 0.603597 |
| 5 | 18800600 | 18802399 | 14.52627 | 26.59601 | 28.41734 | 12.13884 | 0.329733 | 0.289837 | 0.603603 |
| 18 | 53959600 | 53961399 | 20.45595 | 35.58927 | 38.88008 | 17.08507 | 0.260154 | 0.782889 | 0.603611 |
| 18 | 70914000 | 70916399 | 18.88084 | 32.38147 | 51.69009 | 22.67027 | -0.28867 | 0.930737 | 0.603615 |
| X | 85626400 | 85627999 | 17.09805 | 30.1692 | 32.46656 | 14.3663 | 0.284382 | 0.558483 | 0.603625 |
| 18 | 85463200 | 85464999 | 17.02138 | 30.62666 | 26.18547 | 11.43258 | 0.629183 | 0.402832 | 0.603633 |
| 14 | 5528800 | 5532799 | 91.86748 | 142.5953 | 150.6616 | 62.56653 | 0.226922 | 2.775081 | 0.603635 |
| X | 12330600 | 12338199 | 46.99778 | 75.35725 | 117.4443 | 49.6049 | -0.29165 | 1.977652 | 0.603662 |
| 9 | 23996200 | 23997199 | 12.4111 | 22.94096 | 25.56979 | 10.78175 | 0.282759 | 0.214592 | 0.60373 |
| 8 | 34181400 | 34182999 | 10.21063 | 18.52383 | 31.79736 | 13.7412 | -0.36067 | 0.318 | 0.60374 |
| 19 | 5822000 | 5824599 | 21.52919 | 37.31917 | 40.97462 | 17.9776 | 0.252192 | 0.834679 | 0.603743 |
| 18 | 74539000 | 74544999 | 96.79696 | 150.3318 | 161.2513 | 66.95859 | 0.205798 | 2.768975 | 0.603747 |
| 3 | 135396400 | 135397599 | 8.870438 | 16.09727 | 31.87543 | 13.76982 | -0.56499 | 0.305661 | 0.603748 |
| 1 | 34641000 | 34645399 | 158.7088 | 235.6535 | 323.2823 | 133.1708 | -0.2028 | 3.656227 | 0.60376 |
| 1 | 43761200 | 43763199 | 19.49782 | 34.38526 | 34.69293 | 15.3531 | 0.376471 | 0.580883 | 0.603762 |
| 11 | 48429200 | 48430599 | 13.455 | 23.84614 | 37.67336 | 16.65559 | -0.2642 | 0.497185 | 0.603766 |
| 19 | 32487800 | 32492199 | 44.98766 | 72.55194 | 79.98325 | 34.06697 | 0.216983 | 1.781972 | 0.603766 |
| 3 | 13415200 | 13418399 | 36.23764 | 59.34196 | 65.20116 | 28.05281 | 0.240565 | 1.470911 | 0.603767 |
| 9 | 64388600 | 64392799 | 26.01775 | 42.79037 | 68.86935 | 29.6852 | -0.30611 | 1.395406 | 0.603773 |
| 10 | 53710200 | 53712599 | 13.50528 | 24.96063 | 22.33064 | 9.417385 | 0.597342 | 0.214938 | 0.603774 |
| 5 | 30993000 | 30995599 | 30.1717 | 50.11587 | 52.89231 | 22.93425 | 0.303584 | 1.279624 | 0.603808 |
| 3 | 127725600 | 127727599 | 15.80958 | 27.63992 | 43.56049 | 19.20266 | -0.26561 | 0.695677 | 0.603821 |
| X | 13635600 | 13637999 | 20.71227 | 36.48726 | 35.16172 | 15.56394 | 0.441836 | 0.600926 | 0.603824 |
| 1 | 39110200 | 39113399 | 28.95819 | 47.72244 | 74.50788 | 32.143 | -0.26131 | 1.382915 | 0.603846 |
| 6 | 11179400 | 11180999 | 17.77174 | 30.71161 | 47.9382 | 21.0309 | -0.25414 | 0.854601 | 0.603854 |
| 4 | 87295600 | 87298199 | 14.84491 | 25.97278 | 41.99362 | 18.52363 | -0.30229 | 0.675785 | 0.603866 |
| 8 | 10583200 | 10585599 | 14.84491 | 25.97278 | 41.99362 | 18.52363 | -0.30229 | 0.675785 | 0.603866 |
| 16 | 75793800 | 75794599 | 6.834002 | 13.25423 | 24.56071 | 9.266423 | -0.30322 | -0.22302 | 0.60387 |
| 7 | 29462600 | 29465599 | 28.96039 | 47.53479 | 72.46485 | 31.20856 | -0.22926 | 1.430738 | 0.603878 |
| 1 | 120374200 | 120375799 | 19.45553 | 34.02142 | 37.14933 | 16.37941 | 0.261357 | 0.702073 | 0.603883 |
| 1 | 65578400 | 65581999 | 75.99895 | 119.6095 | 127.6205 | 52.94818 | 0.228252 | 2.516573 | 0.603893 |

| | | | | | | | | | |
|---|---|---|---|---|---|---|---|---|---|
| 3 | 78543400 | 78546399 | 52.85241 | 85.19621 | 74.08819 | 31.53736 | 0.557711 | 1.804987 | 0.603897 |
| 12 | 73085600 | 73088399 | 23.315 | 38.98605 | 60.28355 | 26.25262 | -0.24559 | 1.191238 | 0.603905 |
| 10 | 50910600 | 50911399 | 9.738924 | 17.95447 | 30.40769 | 12.86756 | -0.32015 | 0.221217 | 0.603908 |
| 2 | 153375800 | 153376599 | 9.37486 | 17.53449 | 29.54233 | 12.23351 | -0.28896 | 0.12939 | 0.603919 |
| 2 | 171229200 | 171231199 | 19.28205 | 33.91523 | 35.2656 | 15.61475 | 0.331738 | 0.608396 | 0.60393 |
| 13 | 69538000 | 69544399 | 56.42715 | 89.6983 | 131.1904 | 54.74345 | -0.2139 | 2.257054 | 0.603953 |
| 18 | 82400800 | 82401799 | 13.25052 | 25.07095 | 25.27864 | 10.25153 | 0.470948 | 0.077037 | 0.603969 |
| Y | 32950800 | 32951399 | 3.125569 | 6.113271 | 16.11588 | 5.43474 | -0.6524 | -0.69741 | 0.603975 |
| 5 | 58933400 | 58935799 | 18.3895 | 31.66787 | 50.07957 | 21.96799 | -0.27404 | 0.890283 | 0.603978 |
| 10 | 105676400 | 105678799 | 18.3895 | 31.66787 | 50.07957 | 21.96799 | -0.27404 | 0.890283 | 0.603978 |
| X | 130708000 | 130708599 | 7.914197 | 15.42179 | 14.57327 | 5.28928 | 0.709862 | -0.37899 | 0.604029 |
| 1 | 44242000 | 44242599 | 5.955368 | 11.56837 | 22.2671 | 8.31881 | -0.34298 | -0.27144 | 0.60403 |
| 1 | 133498600 | 133506599 | 125.2443 | 191.089 | 203.2663 | 84.28566 | 0.19811 | 3.119283 | 0.604054 |
| 5 | 44013200 | 44018799 | 67.31719 | 106.766 | 116.4049 | 48.46309 | 0.206085 | 2.32862 | 0.604065 |
| 14 | 105197800 | 105199199 | 17.36007 | 30.59253 | 33.40906 | 14.78712 | 0.262497 | 0.574424 | 0.604085 |
| 2 | 114009800 | 114010799 | 10.02208 | 18.66894 | 31.62867 | 13.188 | -0.30512 | 0.165516 | 0.604088 |
| 11 | 90261800 | 90264199 | 16.96006 | 29.28439 | 47.41123 | 20.79918 | -0.30687 | 0.85718 | 0.604098 |
| 8 | 75709000 | 75711599 | 27.34253 | 45.91272 | 50.71799 | 22.14545 | 0.237846 | 1.148886 | 0.604147 |
| 2 | 23805600 | 23810999 | 73.89795 | 116.5841 | 127.3737 | 52.86419 | 0.196792 | 2.466052 | 0.604155 |
| 11 | 44460200 | 44464399 | 51.08816 | 81.99525 | 90.08289 | 38.10437 | 0.213542 | 1.942494 | 0.604193 |
| 8 | 90818400 | 90823799 | 70.8879 | 112.2841 | 122.4849 | 50.9383 | 0.203633 | 2.369053 | 0.604194 |
| 7 | 82514600 | 82519599 | 36.61217 | 58.54746 | 119.5887 | 50.31093 | -0.68551 | 2.047929 | 0.604209 |
| 14 | 112334000 | 112337599 | 35.37301 | 58.89715 | 57.10224 | 24.8004 | 0.425755 | 1.26875 | 0.604215 |
| X | 62213600 | 62217399 | 43.70123 | 72.43422 | 55.73499 | 24.13132 | 0.757858 | 1.342837 | 0.604223 |
| 9 | 76233600 | 76234399 | 10.18207 | 19.49918 | 22.28411 | 8.796007 | 0.325823 | -0.04767 | 0.604237 |
| X | 146445200 | 146448199 | 50.7116 | 81.81751 | 82.16792 | 35.01654 | 0.351927 | 1.781319 | 0.604239 |
| 1 | 85785000 | 85815399 | 600.9319 | 799.3988 | 985.5965 | 411.8063 | -0.2063 | 5.362738 | 0.604254 |
| 8 | 73374200 | 73375999 | 17.90406 | 31.56627 | 33.74679 | 14.93517 | 0.293135 | 0.571369 | 0.604276 |
| 1 | 58679000 | 58692399 | 243.3262 | 348.2444 | 468.5781 | 193.5629 | -0.22613 | 4.234793 | 0.60428 |
| 13 | 68530200 | 68533799 | 59.73108 | 95.55751 | 99.5536 | 41.90701 | 0.284589 | 2.053808 | 0.60428 |
| 15 | 31763400 | 31764599 | 12.41669 | 23.58633 | 24.24164 | 9.749133 | 0.453664 | 0.038583 | 0.60429 |
| 13 | 72539600 | 72544399 | 53.74861 | 85.25987 | 128.8853 | 53.66779 | -0.26439 | 2.317516 | 0.604291 |
| 5 | 117785600 | 117787799 | 19.68009 | 33.31175 | 53.75467 | 23.53156 | -0.30645 | 1.056747 | 0.604293 |
| 16 | 78576200 | 78577999 | 15.66261 | 28.42488 | 29.38393 | 12.69253 | 0.36684 | 0.351684 | 0.604311 |
| 1 | 17593400 | 17597999 | 105.6381 | 162.5171 | 172.9543 | 71.82729 | 0.206688 | 2.945703 | 0.604319 |
| 7 | 27132400 | 27134999 | 22.6253 | 37.97352 | 58.67096 | 25.6103 | -0.24451 | 1.147285 | 0.604339 |
| 8 | 71511200 | 71511999 | 8.453788 | 15.71118 | 28.43139 | 11.9002 | -0.40358 | 0.158837 | 0.60435 |
| 8 | 91113800 | 91116999 | 24.69608 | 42.03988 | 46.35527 | 20.31831 | 0.241467 | 1.016558 | 0.604356 |
| 4 | 28623200 | 28624799 | 19.47792 | 33.35131 | 51.66566 | 22.6566 | -0.24581 | 0.950863 | 0.604359 |
| 7 | 14371200 | 14384399 | 187.7962 | 281.9026 | 255.2615 | 105.3449 | 0.410497 | 3.460482 | 0.604371 |
| 7 | 41528200 | 41554799 | 62.24387 | 99.63771 | 87.74877 | 36.96943 | 0.527525 | 2.035915 | 0.604374 |
| 10 | 112980200 | 112981999 | 26.84422 | 45.67237 | 42.06874 | 18.43866 | 0.499876 | 1.023572 | 0.604375 |
| 17 | 23049200 | 23051799 | 33.34599 | 55.66692 | 55.88557 | 24.31191 | 0.375512 | 1.234231 | 0.604388 |
| 9 | 37618000 | 37619199 | 11.44213 | 20.65796 | 33.71334 | 14.65747 | -0.29418 | 0.358204 | 0.604395 |
| 9 | 113382400 | 113383599 | 11.44213 | 20.65796 | 33.71334 | 14.65747 | -0.29418 | 0.358204 | 0.604395 |
| 12 | 12772600 | 12776199 | 34.7044 | 55.91229 | 99.31663 | 42.2452 | -0.47138 | 1.805389 | 0.604414 |
| 9 | 92234800 | 92236199 | 9.609454 | 18.01635 | 32.93527 | 13.58334 | -0.40118 | 0.125304 | 0.604421 |

| | | | | | | | | | |
|---|---|---|---|---|---|---|---|---|---|
| 3 | 106644200 | 106647599 | 76.0725 | 120.8305 | 118.0138 | 49.20567 | 0.366658 | 2.295097 | 0.604434 |
| X | 24057200 | 24058599 | 25.53692 | 44.32554 | 36.07021 | 15.82708 | 0.682923 | 0.835623 | 0.604474 |
| 10 | 128907200 | 128908999 | 17.74865 | 31.6023 | 32.06146 | 14.12227 | 0.374864 | 0.481237 | 0.604489 |
| 15 | 31022600 | 31024399 | 17.63012 | 31.00491 | 33.90132 | 15.01114 | 0.259528 | 0.598267 | 0.604497 |
| 12 | 62593600 | 62596799 | 64.64206 | 101.4317 | 153.1137 | 63.50024 | -0.27498 | 2.573291 | 0.604502 |
| 14 | 41305200 | 41325199 | 231.8949 | 340.5659 | 332.8269 | 137.0755 | 0.271406 | 3.846029 | 0.604518 |
| 13 | 104811400 | 104817199 | 151.4622 | 225.9823 | 309.0464 | 127.3803 | -0.19189 | 3.569081 | 0.604522 |
| 17 | 66201800 | 66205999 | 38.03313 | 62.44383 | 64.99773 | 27.99596 | 0.319828 | 1.443109 | 0.604529 |
| 9 | 12321600 | 12322199 | 10.55193 | 20.15515 | 22.84312 | 9.079961 | 0.329182 | -0.01657 | 0.604531 |
| 6 | 54929400 | 54932199 | 24.90787 | 42.57334 | 45.76682 | 20.06696 | 0.278736 | 0.978517 | 0.604534 |
| Y | 8218200 | 8219599 | 11.41174 | 21.35777 | 17.65484 | 7.30303 | 0.73159 | 0.108067 | 0.604534 |
| 3 | 22329600 | 22334599 | 57.24677 | 91.62135 | 97.90531 | 41.23752 | 0.248707 | 2.037741 | 0.60459 |
| 4 | 20477600 | 20478199 | 6.837057 | 13.15698 | 24.72586 | 9.624817 | -0.36644 | -0.11934 | 0.604593 |
| 6 | 10333200 | 10345599 | 133.8167 | 203.7293 | 188.2551 | 78.0166 | 0.398543 | 3.163934 | 0.604595 |
| 9 | 19037600 | 19041199 | 27.17755 | 46.336 | 41.04984 | 17.9951 | 0.556392 | 1.007235 | 0.6046 |
| 19 | 61167000 | 61173999 | 101.9192 | 159.8486 | 143.0319 | 59.32157 | 0.47802 | 2.603164 | 0.604623 |
| 19 | 13111400 | 13124399 | 178.393 | 267.205 | 265.7344 | 109.6115 | 0.272511 | 3.499507 | 0.604639 |
| 1 | 30447200 | 30448199 | 9.151011 | 17.20359 | 29.10388 | 11.94799 | -0.28506 | 0.091747 | 0.60464 |
| 14 | 36143000 | 36144599 | 19.78622 | 34.95652 | 34.06955 | 15.07181 | 0.427075 | 0.561802 | 0.604642 |
| 12 | 24490400 | 24500199 | 75.64484 | 119.7239 | 113.1394 | 47.03284 | 0.409491 | 2.389418 | 0.604658 |
| 16 | 62407600 | 62408599 | 15.3776 | 27.40301 | 29.89006 | 13.15672 | 0.2718 | 0.457998 | 0.604697 |
| 5 | 78530800 | 78531399 | 11.54569 | 21.53281 | 24.09884 | 10.02801 | 0.291137 | 0.150949 | 0.604707 |
| 2 | 43315800 | 43316399 | 9.13214 | 17.61115 | 20.83824 | 8.061892 | 0.30332 | -0.12633 | 0.604719 |
| 10 | 21327400 | 21328799 | 14.16604 | 25.73499 | 28.37869 | 12.24276 | 0.275137 | 0.330782 | 0.604719 |
| 15 | 96174600 | 96179399 | 43.6459 | 69.90936 | 104.771 | 44.18152 | -0.23701 | 2.001014 | 0.604724 |
| 1 | 50050200 | 50052599 | 22.8868 | 39.68663 | 41.97418 | 18.41635 | 0.306038 | 0.839379 | 0.604733 |
| 4 | 37516000 | 37519399 | 30.26594 | 50.70711 | 53.55091 | 23.34916 | 0.302533 | 1.183869 | 0.604757 |
| 13 | 82473600 | 82474399 | 15.93558 | 27.565 | 45.85946 | 20.11953 | -0.34516 | 0.837495 | 0.604777 |
| 17 | 33095400 | 33099599 | 42.98842 | 68.90618 | 103.1735 | 43.55584 | -0.23445 | 1.977125 | 0.604797 |
| X | 167921400 | 167924199 | 30.39328 | 50.59692 | 55.71303 | 24.19439 | 0.242713 | 1.254953 | 0.604809 |
| 15 | 101686200 | 101686999 | 7.19057 | 13.66682 | 25.55301 | 10.26404 | -0.40023 | -0.00804 | 0.604811 |
| 16 | 77820200 | 77827799 | 92.64269 | 144.1334 | 156.0305 | 64.77363 | 0.194522 | 2.73703 | 0.604812 |
| 4 | 90140200 | 90151799 | 143.447 | 218.7075 | 208.1063 | 86.27856 | 0.357996 | 3.139692 | 0.604832 |
| 8 | 65428200 | 65430799 | 31.81363 | 53.05544 | 48.68426 | 21.16569 | 0.504761 | 1.232229 | 0.604856 |
| 1 | 16809400 | 16811399 | 19.7101 | 34.52859 | 28.97772 | 12.79591 | 0.639111 | 0.66002 | 0.604867 |
| 12 | 119555800 | 119557999 | 27.05524 | 45.36288 | 49.65052 | 21.65905 | 0.251193 | 1.163463 | 0.604875 |
| 1 | 78539800 | 78543399 | 42.58341 | 69.0273 | 73.90289 | 31.64572 | 0.266817 | 1.646293 | 0.604876 |
| 4 | 11175600 | 11184799 | 250.4329 | 358.127 | 475.2259 | 196.3338 | -0.20725 | 4.247142 | 0.604878 |
| 8 | 121086400 | 121088399 | 15.90198 | 27.48794 | 45.85654 | 20.11783 | -0.34938 | 0.842162 | 0.604901 |
| 7 | 77009400 | 77009999 | 12.35688 | 21.84569 | 36.28691 | 16.05058 | -0.33734 | 0.515128 | 0.604938 |
| 12 | 114669400 | 114672799 | 55.58823 | 89.36757 | 89.36498 | 37.89495 | 0.351952 | 1.896961 | 0.604944 |
| 12 | 48298600 | 48299399 | 15.13725 | 27.13071 | 28.96949 | 12.68966 | 0.30731 | 0.420234 | 0.60497 |
| X | 31871000 | 31873599 | 47.28434 | 77.32519 | 65.64563 | 28.22918 | 0.610512 | 1.515601 | 0.604977 |
| 15 | 44482200 | 44483199 | 11.17214 | 20.10374 | 33.22675 | 14.5058 | -0.31602 | 0.375593 | 0.604979 |
| 9 | 80804000 | 80805999 | 16.8024 | 30.12513 | 31.15677 | 13.64397 | 0.353015 | 0.431394 | 0.604981 |
| 3 | 51046200 | 51049999 | 23.90569 | 40.06939 | 62.09719 | 27.08323 | -0.24894 | 1.16778 | 0.604991 |
| 5 | 15757400 | 15758799 | 26.69238 | 45.10886 | 48.5288 | 21.23357 | 0.276093 | 1.08687 | 0.604994 |

| | | | | | | | | | |
|---|---|---|---|---|---|---|---|---|---|
| 1 | 153802000 | 153806399 | 40.79094 | 66.11394 | 72.40363 | 31.00104 | 0.234101 | 1.644755 | 0.604999 |
| 3 | 42676600 | 42677399 | 13.61887 | 25.23339 | 21.46243 | 9.020595 | 0.673565 | 0.201478 | 0.605024 |
| 5 | 89696600 | 89697199 | 6.741511 | 12.9448 | 25.18675 | 9.865926 | -0.42318 | -0.09241 | 0.605035 |
| 6 | 36480600 | 36483399 | 18.16124 | 31.98655 | 33.69762 | 14.9165 | 0.313574 | 0.583123 | 0.605051 |
| 10 | 107942000 | 107944999 | 19.72607 | 33.87415 | 53.33872 | 23.39565 | -0.26874 | 0.925657 | 0.605061 |
| 7 | 42051000 | 42063999 | 191.2553 | 287.1031 | 253.5125 | 104.6239 | 0.446802 | 3.460611 | 0.605065 |
| 4 | 20041600 | 20042599 | 6.856927 | 13.21823 | 24.20406 | 9.374676 | -0.32177 | -0.13379 | 0.605073 |
| 8 | 98478600 | 98478999 | 6.856927 | 13.21823 | 24.20406 | 9.374676 | -0.32177 | -0.13379 | 0.605073 |
| 19 | 6871200 | 6871799 | 6.856927 | 13.21823 | 24.20406 | 9.374676 | -0.32177 | -0.13379 | 0.605073 |
| 14 | 8046000 | 8052599 | 40.0924 | 65.31612 | 61.31617 | 26.33102 | 0.461943 | 1.548434 | 0.605098 |
| X | 126159200 | 126160399 | 15.31761 | 28.6869 | 26.01556 | 10.74939 | 0.600457 | 0.160805 | 0.605102 |
| 8 | 81642400 | 81645999 | 32.62673 | 53.0914 | 80.20873 | 34.42938 | -0.22393 | 1.562573 | 0.605108 |
| 12 | 115283000 | 115299599 | 253.7909 | 370.5201 | 353.0248 | 145.4525 | 0.299854 | 3.940603 | 0.605115 |
| X | 77316200 | 77319399 | 35.06706 | 59.57017 | 46.76939 | 20.49599 | 0.729358 | 1.075587 | 0.605133 |
| 15 | 45269600 | 45271399 | 12.56567 | 22.68078 | 38.99697 | 16.9599 | -0.36961 | 0.377347 | 0.605141 |
| 3 | 31241600 | 31245599 | 40.60879 | 65.7973 | 71.7769 | 30.72531 | 0.239266 | 1.650291 | 0.605148 |
| 9 | 105506600 | 105507799 | 7.683707 | 14.64292 | 27.20673 | 10.86712 | -0.38378 | -0.01416 | 0.605158 |
| 4 | 96192800 | 96196999 | 76.60834 | 121.0454 | 129.2637 | 53.69018 | 0.231437 | 2.433181 | 0.605158 |
| 1 | 170281800 | 170285599 | 36.23519 | 58.56018 | 87.87004 | 37.52815 | -0.22329 | 1.710414 | 0.605161 |
| 4 | 145590200 | 145610399 | 552.9249 | 751.2496 | 701.5516 | 292.2149 | 0.225059 | 5.02747 | 0.605163 |
| 3 | 144643600 | 144645199 | 21.50695 | 36.37641 | 56.47813 | 24.71809 | -0.25106 | 1.074129 | 0.605169 |
| 8 | 73278000 | 73282199 | 52.87029 | 84.66781 | 93.11897 | 39.25617 | 0.208227 | 2.013734 | 0.605195 |
| 4 | 100298000 | 100300599 | 40.21835 | 64.66823 | 96.67977 | 41.00628 | -0.22677 | 1.874153 | 0.605209 |
| 1 | 85277200 | 85278799 | 52.58523 | 84.16375 | 89.47316 | 37.68784 | 0.256237 | 2.030478 | 0.605212 |
| 3 | 127491800 | 127493199 | 10.84611 | 19.68881 | 32.28759 | 13.94232 | -0.29443 | 0.321104 | 0.605214 |
| 15 | 96785400 | 96795399 | 144.386 | 216.1861 | 296.5698 | 122.3216 | -0.19175 | 3.501819 | 0.605224 |
| 7 | 27448000 | 27462999 | 179.6679 | 268.8019 | 269.545 | 111.1582 | 0.259055 | 3.521276 | 0.605241 |
| 13 | 81974400 | 81975999 | 15.64501 | 27.81983 | 30.30614 | 13.364 | 0.27173 | 0.47289 | 0.605252 |
| X | 51251600 | 51255199 | 35.07517 | 58.17065 | 58.04305 | 25.14078 | 0.384117 | 1.31572 | 0.605272 |
| 2 | 93674200 | 93674999 | 6.55954 | 12.66829 | 23.69999 | 9.119965 | -0.34328 | -0.15953 | 0.605276 |
| 14 | 76953400 | 76956599 | 21.79519 | 37.90875 | 40.30563 | 17.7084 | 0.298931 | 0.793054 | 0.605311 |
| 5 | 127960000 | 127967199 | 87.53822 | 137.0601 | 146.0132 | 60.56862 | 0.224726 | 2.621761 | 0.605327 |
| 13 | 68958400 | 68959799 | 18.56021 | 32.53567 | 35.48643 | 15.67876 | 0.262849 | 0.654217 | 0.605338 |
| 18 | 60012400 | 60013599 | 13.98343 | 25.63312 | 27.91959 | 11.90661 | 0.30383 | 0.277348 | 0.605341 |
| 5 | 110166000 | 110174999 | 163.2122 | 245.247 | 252.8364 | 104.3723 | 0.224709 | 3.434787 | 0.605344 |
| 2 | 172000200 | 172000999 | 7.579184 | 14.49033 | 26.20708 | 10.39414 | -0.33724 | -0.04666 | 0.605351 |
| 7 | 86245000 | 86245799 | 7.136596 | 13.79003 | 24.89985 | 9.558034 | -0.28997 | -0.16027 | 0.605357 |
| 12 | 91042000 | 91046799 | 55.87717 | 88.57231 | 131.5699 | 54.74606 | -0.24021 | 2.340876 | 0.605358 |
| 9 | 22105600 | 22108199 | 20.96214 | 35.6004 | 54.87855 | 24.04787 | -0.24055 | 1.034172 | 0.60536 |
| 9 | 3237200 | 3241599 | 83.98176 | 132.3398 | 111.4776 | 46.25828 | 0.570213 | 2.495336 | 0.605378 |
| 13 | 12952200 | 12952999 | 15.89884 | 29.02754 | 25.12396 | 10.77114 | 0.628153 | 0.308505 | 0.605378 |
| 17 | 73242800 | 73250599 | 51.85284 | 82.07444 | 147.7567 | 61.42591 | -0.51889 | 2.376798 | 0.605388 |
| 13 | 64976000 | 64982999 | 192.793 | 286.2583 | 253.4576 | 104.4075 | 0.428501 | 3.657706 | 0.605391 |
| 3 | 95861000 | 95864199 | 27.61118 | 45.48699 | 69.74005 | 30.08177 | -0.23535 | 1.37957 | 0.605413 |
| 7 | 80522000 | 80523799 | 12.36953 | 22.07461 | 35.74557 | 15.71489 | -0.29359 | 0.443099 | 0.605437 |
| 1 | 96542000 | 96542799 | 14.26478 | 25.75665 | 27.97181 | 12.15971 | 0.290213 | 0.367103 | 0.605438 |
| 1 | 115689400 | 115689999 | 14.26478 | 25.75665 | 27.97181 | 12.15971 | 0.290213 | 0.367103 | 0.605438 |

| | | | | | | | | | |
|---|---|---|---|---|---|---|---|---|---|
| 12 | 76781200 | 76796599 | 411.2333 | 561.9371 | 731.8077 | 304.548 | -0.24631 | 4.935233 | 0.605441 |
| 2 | 28670200 | 28671799 | 19.52423 | 34.05421 | 33.92478 | 14.93126 | 0.393152 | 0.736927 | 0.605442 |
| X | 133986000 | 133988799 | 20.9595 | 35.70148 | 55.45547 | 24.30831 | -0.25092 | 1.007964 | 0.605447 |
| 1 | 134991400 | 134993399 | 24.36601 | 41.55853 | 45.81641 | 20.08525 | 0.242137 | 0.99724 | 0.605468 |
| 6 | 62424000 | 62426199 | 25.66872 | 44.29514 | 38.16805 | 16.74393 | 0.598974 | 0.888259 | 0.605475 |
| X | 44303600 | 44310999 | 82.31452 | 130.7434 | 115.8063 | 48.28466 | 0.507457 | 2.30245 | 0.605477 |
| 17 | 94899800 | 94901599 | 21.43009 | 36.42694 | 56.55582 | 24.78646 | -0.25076 | 1.029224 | 0.605479 |
| 19 | 53571400 | 53578599 | 88.37564 | 138.4285 | 131.003 | 54.33984 | 0.395817 | 2.613554 | 0.605491 |
| 18 | 9280000 | 9280999 | 7.851508 | 14.70188 | 27.49515 | 11.36397 | -0.43555 | 0.104802 | 0.605497 |
| 15 | 46984200 | 46984999 | 18.54152 | 32.45101 | 34.24515 | 15.1099 | 0.310528 | 0.680432 | 0.60551 |
| 5 | 128180400 | 128184999 | 50.19021 | 80.70804 | 87.345 | 37.05083 | 0.238514 | 1.880061 | 0.605511 |
| 12 | 5555800 | 5558599 | 19.44001 | 34.12472 | 35.24125 | 15.58848 | 0.341098 | 0.635494 | 0.605518 |
| 3 | 98198200 | 98199999 | 26.95005 | 45.43297 | 49.61309 | 21.69189 | 0.254647 | 1.113589 | 0.605522 |
| 7 | 56425600 | 56428599 | 25.14605 | 43.07254 | 45.75607 | 20.0657 | 0.296322 | 0.961298 | 0.605525 |
| 3 | 19019600 | 19022199 | 21.21729 | 37.2461 | 36.20618 | 16.01588 | 0.428094 | 0.647724 | 0.605536 |
| 5 | 9379000 | 9380999 | 18.59754 | 31.91087 | 50.06798 | 21.95969 | -0.26368 | 0.924611 | 0.605548 |
| 8 | 121843000 | 121843999 | 11.42888 | 20.24343 | 35.33173 | 15.62216 | -0.40713 | 0.486754 | 0.605554 |
| 8 | 7735400 | 7736399 | 7.966946 | 14.98549 | 26.99233 | 11.07124 | -0.37317 | 0.074172 | 0.605566 |
| Y | 5073200 | 5076999 | 19.83129 | 34.50599 | 28.4431 | 12.49955 | 0.664438 | 0.75836 | 0.605571 |
| 17 | 86282200 | 86283799 | 10.89919 | 19.54968 | 33.28671 | 14.57138 | -0.36122 | 0.394948 | 0.605605 |
| 2 | 91805200 | 91806999 | 14.27595 | 24.83139 | 43.73159 | 19.21409 | -0.42565 | 0.76215 | 0.605614 |
| 3 | 44712800 | 44715399 | 68.88991 | 109.4937 | 102.1084 | 42.60185 | 0.434179 | 2.2665 | 0.605618 |
| 9 | 112899600 | 112901799 | 31.04719 | 51.57855 | 56.84178 | 24.64954 | 0.241299 | 1.280984 | 0.605621 |
| 2 | 108110000 | 108110399 | 5.507071 | 10.73276 | 20.83974 | 7.549365 | -0.31456 | -0.39161 | 0.605622 |
| 10 | 8514800 | 8516199 | 18.55708 | 32.62974 | 34.90628 | 15.45663 | 0.290919 | 0.608513 | 0.605623 |
| 12 | 8584400 | 8586799 | 16.59917 | 28.80565 | 46.34105 | 20.33282 | -0.29626 | 0.808487 | 0.605628 |
| 1 | 82910200 | 82925399 | 211.5667 | 306.854 | 414.4687 | 170.8851 | -0.21258 | 4.036156 | 0.60565 |
| 12 | 48566800 | 48567599 | 7.778124 | 14.88956 | 26.23892 | 10.36891 | -0.29458 | -0.05346 | 0.605654 |
| 13 | 65075600 | 65087999 | 161.2832 | 242.9276 | 246.4091 | 101.7841 | 0.25118 | 3.389147 | 0.605669 |
| 5 | 63909600 | 63911999 | 19.8636 | 33.92916 | 53.25652 | 23.34966 | -0.26521 | 0.977459 | 0.605687 |
| 3 | 78759400 | 78760799 | 21.82 | 37.70977 | 41.51481 | 18.21294 | 0.247435 | 0.862783 | 0.605705 |
| X | 77074000 | 77076199 | 29.3402 | 49.62695 | 50.14994 | 21.94871 | 0.366285 | 1.094769 | 0.605714 |
| 10 | 51346000 | 51348399 | 21.57121 | 37.90924 | 35.58731 | 15.75682 | 0.478153 | 0.631649 | 0.605725 |
| 2 | 85005800 | 85006599 | 11.11035 | 21.20361 | 23.27233 | 9.267359 | 0.371983 | -0.00795 | 0.605729 |
| 7 | 106132200 | 106133599 | 18.22075 | 31.96615 | 34.58589 | 15.2912 | 0.274576 | 0.640477 | 0.605753 |
| 5 | 41739000 | 41740999 | 30.88267 | 51.39304 | 56.2944 | 24.44101 | 0.250214 | 1.2607 | 0.605765 |
| 6 | 30732000 | 30740999 | 208.4901 | 299.2908 | 491.9605 | 203.1256 | -0.51045 | 4.189188 | 0.605776 |
| 5 | 146480400 | 146481399 | 23.07358 | 39.44825 | 39.71407 | 17.41342 | 0.373311 | 0.964804 | 0.605796 |
| 13 | 62806800 | 62808399 | 25.05739 | 42.60278 | 46.89235 | 20.55169 | 0.243782 | 1.029986 | 0.605799 |
| 11 | 26505400 | 26507199 | 15.33032 | 27.91471 | 23.86773 | 10.2669 | 0.642843 | 0.317874 | 0.605802 |
| 7 | 42685600 | 42689399 | 77.92714 | 122.9524 | 111.6845 | 46.35487 | 0.462901 | 2.463416 | 0.605829 |
| 2 | 13348200 | 13349399 | 22.3028 | 38.26767 | 41.61299 | 18.25132 | 0.26313 | 0.929171 | 0.605835 |
| 3 | 105602600 | 105604599 | 14.25512 | 24.87548 | 42.11351 | 18.54081 | -0.36844 | 0.713594 | 0.60585 |
| 18 | 3497800 | 3499799 | 13.29397 | 23.54446 | 37.18544 | 16.44214 | -0.26412 | 0.502869 | 0.605855 |
| 12 | 64201200 | 64204199 | 62.95018 | 100.0761 | 109.4751 | 45.68596 | 0.204606 | 2.255867 | 0.605855 |
| 10 | 33608000 | 33617399 | 125.3743 | 191.937 | 197.097 | 81.7916 | 0.252763 | 3.050911 | 0.605858 |
| 19 | 35910600 | 35914599 | 30.06738 | 51.00543 | 49.59164 | 21.72357 | 0.421854 | 1.070778 | 0.60586 |

| | | | | | | | | | |
|---|---|---|---|---|---|---|---|---|---|
| 16 | 58947800 | 58951199 | 43.66473 | 70.51407 | 77.81254 | 33.20047 | 0.217821 | 1.739579 | 0.605861 |
| 16 | 68716400 | 68720199 | 38.34674 | 63.24635 | 61.34392 | 26.47319 | 0.423958 | 1.384135 | 0.605869 |
| 16 | 15361400 | 15365999 | 60.49621 | 97.0243 | 82.97157 | 35.05751 | 0.573036 | 1.973121 | 0.60587 |
| 7 | 66009400 | 66015399 | 103.5603 | 159.6784 | 171.9528 | 71.42621 | 0.192131 | 2.902886 | 0.605897 |
| 6 | 111565200 | 111565599 | 4.804223 | 9.374078 | 19.22473 | 6.853699 | -0.37141 | -0.46498 | 0.605913 |
| 10 | 18425000 | 18426399 | 10.35938 | 18.70574 | 33.25112 | 14.45399 | -0.41654 | 0.34777 | 0.605929 |
| 15 | 16488200 | 16490399 | 26.20645 | 45.48374 | 41.82108 | 18.34968 | 0.507417 | 0.846182 | 0.605932 |
| 13 | 43137600 | 43138999 | 13.60564 | 24.17205 | 38.70245 | 17.07922 | -0.28153 | 0.482689 | 0.605938 |
| 16 | 77774200 | 77775399 | 13.18592 | 24.55357 | 26.57911 | 11.08932 | 0.335709 | 0.180363 | 0.605944 |
| 16 | 80736200 | 80737199 | 13.06274 | 24.28441 | 21.4275 | 8.964784 | 0.625497 | 0.177349 | 0.605946 |
| Y | 2282000 | 2284999 | 24.05603 | 41.04371 | 31.68431 | 13.89032 | 0.754125 | 0.971836 | 0.605948 |
| 11 | 92140400 | 92141599 | 25.75185 | 42.31299 | 68.60886 | 29.56108 | -0.31727 | 1.408473 | 0.605967 |
| 8 | 31886600 | 31887999 | 10.66771 | 19.3345 | 32.26513 | 13.96008 | -0.32143 | 0.327739 | 0.605968 |
| 8 | 103550600 | 103551799 | 11.33481 | 21.70787 | 23.39341 | 9.232467 | 0.410146 | -0.03335 | 0.605986 |
| 3 | 16886600 | 16888799 | 27.34154 | 46.30191 | 49.59644 | 21.71437 | 0.282515 | 1.074731 | 0.606006 |
| 8 | 62189200 | 62196399 | 69.91773 | 109.0604 | 182.8628 | 75.88381 | -0.43307 | 2.684697 | 0.606007 |
| X | 21887600 | 21888599 | 8.114198 | 15.63672 | 32.78975 | 12.71214 | -0.51819 | -0.09667 | 0.606042 |
| 11 | 20342600 | 20344799 | 38.61972 | 63.13102 | 67.21595 | 28.90033 | 0.284195 | 1.500126 | 0.606053 |
| X | 7184800 | 7185799 | 8.617484 | 16.41506 | 30.6718 | 12.2628 | -0.3939 | 0.00465 | 0.606055 |
| 9 | 114050200 | 114053399 | 19.57785 | 33.65483 | 52.79121 | 23.15624 | -0.26291 | 0.915466 | 0.606066 |
| 14 | 74066600 | 74068599 | 22.05885 | 37.34248 | 60.33375 | 26.41253 | -0.30831 | 1.073319 | 0.606081 |
| 3 | 13964400 | 13967999 | 24.39172 | 40.93792 | 66.72874 | 29.12744 | -0.32165 | 1.158908 | 0.606138 |
| 14 | 82550000 | 82552199 | 21.88598 | 38.4004 | 36.80662 | 16.27448 | 0.448456 | 0.662137 | 0.606199 |
| 15 | 11881800 | 11883799 | 15.00304 | 26.3371 | 41.86453 | 18.51878 | -0.27792 | 0.622663 | 0.606213 |
| 9 | 48419400 | 48421399 | 22.49897 | 38.65049 | 42.62639 | 18.69718 | 0.243151 | 0.920837 | 0.606221 |
| 15 | 52774200 | 52784199 | 129.915 | 198.6154 | 181.5864 | 75.33108 | 0.418652 | 3.078432 | 0.606229 |
| 5 | 37676600 | 37680599 | 33.58754 | 54.46741 | 82.9092 | 35.51302 | -0.23907 | 1.627236 | 0.606243 |
| 17 | 9539000 | 9540199 | 11.32226 | 20.33054 | 33.70925 | 14.74288 | -0.32249 | 0.390429 | 0.606291 |
| 18 | 66672600 | 66674799 | 23.13193 | 38.8732 | 59.79873 | 26.12021 | -0.23822 | 1.137848 | 0.606291 |
| 5 | 50087800 | 50089599 | 18.97767 | 33.97402 | 31.47581 | 13.80118 | 0.509684 | 0.455837 | 0.606295 |
| 11 | 3709000 | 3712799 | 23.89304 | 39.45867 | 66.09195 | 28.54377 | -0.36143 | 1.336718 | 0.606296 |
| 13 | 92760800 | 92766999 | 87.18927 | 134.7091 | 190.6167 | 79.17805 | -0.19922 | 2.860958 | 0.606308 |
| 9 | 42276600 | 42278799 | 20.23834 | 34.40798 | 54.823 | 24.02741 | -0.28775 | 1.024135 | 0.606315 |
| 6 | 136535600 | 136541599 | 68.48063 | 107.3368 | 177.5164 | 73.62854 | -0.40782 | 2.598927 | 0.606318 |
| 1 | 177913600 | 177916199 | 24.65976 | 42.24097 | 44.6628 | 19.58631 | 0.302978 | 0.955831 | 0.606321 |
| 17 | 92995600 | 92996599 | 11.11925 | 20.25546 | 32.80001 | 14.10167 | -0.27243 | 0.302898 | 0.606334 |
| 16 | 86990000 | 86990599 | 9.384822 | 17.51515 | 29.50424 | 12.26262 | -0.29289 | 0.143513 | 0.606337 |
| 18 | 16081600 | 16083399 | 16.69898 | 30.68153 | 27.68352 | 11.76902 | 0.57684 | 0.284113 | 0.606337 |
| 10 | 33480000 | 33483999 | 44.55857 | 72.39806 | 63.05027 | 27.04512 | 0.567179 | 1.601435 | 0.606344 |
| 16 | 85236800 | 85237999 | 12.2246 | 22.81602 | 24.99551 | 10.38799 | 0.32278 | 0.152336 | 0.606348 |
| 3 | 75093400 | 75096199 | 43.84806 | 71.00004 | 76.06907 | 32.54719 | 0.264369 | 1.673053 | 0.606354 |
| 3 | 31453000 | 31456399 | 47.6779 | 76.95707 | 71.96941 | 30.6879 | 0.455225 | 1.756899 | 0.606354 |
| X | 78942400 | 78942999 | 12.01821 | 22.68532 | 24.74714 | 10.0866 | 0.352103 | 0.082205 | 0.60637 |
| 9 | 90978800 | 90981199 | 20.07013 | 33.35684 | 68.09703 | 29.53922 | -0.6451 | 1.255312 | 0.606385 |
| 1 | 127148800 | 127150199 | 14.75521 | 27.00796 | 28.33909 | 12.1098 | 0.355187 | 0.291713 | 0.606391 |
| 4 | 112308400 | 112328399 | 382.3567 | 536.6992 | 511.8117 | 212.0979 | 0.242211 | 4.525752 | 0.606404 |
| 17 | 81160000 | 81161599 | 9.980023 | 19.2133 | 17.55384 | 6.824746 | 0.666606 | -0.11114 | 0.606408 |

| | | | | | | | | | |
|---|---|---|---|---|---|---|---|---|---|
| 9 | 118793400 | 118797199 | 31.93448 | 51.90603 | 80.16304 | 34.39505 | -0.2568 | 1.575756 | 0.606421 |
| X | 95984200 | 95986999 | 40.71863 | 66.31123 | 70.48176 | 30.26283 | 0.282869 | 1.561544 | 0.606468 |
| 18 | 51992600 | 51994599 | 16.31043 | 28.5169 | 45.14091 | 19.90015 | -0.27197 | 0.700087 | 0.606472 |
| 7 | 103980400 | 103982999 | 20.65837 | 36.0399 | 37.54196 | 16.52545 | 0.328599 | 0.739685 | 0.606478 |
| 17 | 19260800 | 19268799 | 112.7223 | 175.9603 | 128.2163 | 53.20267 | 0.768693 | 2.686871 | 0.60649 |
| 5 | 54065800 | 54070199 | 40.88885 | 67.54648 | 60.71781 | 26.22808 | 0.533869 | 1.376577 | 0.606495 |
| X | 21226400 | 21229399 | 31.72825 | 52.91964 | 54.59465 | 23.73712 | 0.336457 | 1.240737 | 0.606505 |
| 16 | 3830600 | 3837199 | 49.49666 | 79.10627 | 117.3353 | 49.32308 | -0.22576 | 2.077943 | 0.606506 |
| 6 | 140089400 | 140090799 | 10.64938 | 19.54726 | 31.8758 | 13.56936 | -0.27253 | 0.254774 | 0.606508 |
| 8 | 94153200 | 94156599 | 22.06446 | 36.77785 | 60.11785 | 26.128 | -0.32536 | 1.221602 | 0.606515 |
| 3 | 91467800 | 91475799 | 204.88 | 304.8692 | 256.8809 | 105.8412 | 0.502876 | 3.622356 | 0.606526 |
| 14 | 104855000 | 104855599 | 11.69151 | 21.7122 | 23.58897 | 9.883402 | 0.325952 | 0.177074 | 0.606542 |
| 14 | 84522200 | 84523999 | 15.88439 | 27.85569 | 45.0713 | 19.92003 | -0.3034 | 0.650417 | 0.606546 |
| 19 | 34367200 | 34368999 | 20.55488 | 35.87153 | 38.69295 | 17.03573 | 0.27873 | 0.738843 | 0.606553 |
| X | 13252400 | 13256199 | 29.68186 | 48.7864 | 77.19768 | 33.26631 | -0.28212 | 1.420133 | 0.60656 |
| 11 | 35542200 | 35544199 | 11.51238 | 20.66299 | 33.73311 | 14.7587 | -0.30047 | 0.39585 | 0.60657 |
| 16 | 8250800 | 8254799 | 50.26951 | 81.18256 | 71.099 | 30.33717 | 0.550569 | 1.747467 | 0.606576 |
| 18 | 54781200 | 54781799 | 8.532813 | 16.1712 | 27.65018 | 11.17446 | -0.28073 | 0.029193 | 0.606606 |
| 11 | 33703800 | 33709799 | 43.31007 | 69.26185 | 108.2861 | 45.55929 | -0.30051 | 2.049413 | 0.60665 |
| 9 | 93266000 | 93266799 | 10.29535 | 20.02468 | 20.12263 | 7.439628 | 0.597871 | -0.26742 | 0.606661 |
| 7 | 68182600 | 68187599 | 89.32973 | 139.1505 | 149.4192 | 62.01763 | 0.207596 | 2.714148 | 0.606663 |
| 5 | 95136400 | 95136999 | 10.33312 | 19.92391 | 22.51821 | 8.715997 | 0.368521 | -0.10601 | 0.606705 |
| 11 | 115735800 | 115737199 | 18.35197 | 31.3884 | 50.05519 | 21.9484 | -0.28763 | 0.954624 | 0.606728 |
| 2 | 18323600 | 18331199 | 153.9192 | 233.0108 | 207.9316 | 86.02295 | 0.442106 | 3.28422 | 0.606729 |
| 4 | 143345800 | 143350999 | 82.21227 | 129.5878 | 135.5349 | 56.24304 | 0.2587 | 2.490897 | 0.606737 |
| 11 | 52615800 | 52618399 | 19.77234 | 33.65871 | 52.72667 | 23.11102 | -0.2632 | 1.004858 | 0.606744 |
| 3 | 74148200 | 74149999 | 24.73506 | 42.58183 | 44.08845 | 19.33973 | 0.334422 | 0.912214 | 0.606744 |
| 10 | 68353200 | 68355399 | 16.37332 | 28.65171 | 46.19468 | 20.37963 | -0.2983 | 0.689141 | 0.606747 |
| X | 148506000 | 148513799 | 81.50864 | 128.2344 | 137.0375 | 56.85247 | 0.225591 | 2.529109 | 0.606805 |
| 7 | 41767600 | 41775799 | 149.3203 | 225.2196 | 237.5325 | 98.15519 | 0.1967 | 3.359097 | 0.606811 |
| 19 | 4294600 | 4295999 | 14.84304 | 26.66172 | 29.33367 | 12.82548 | 0.265925 | 0.403238 | 0.606821 |
| 11 | 121815600 | 121816799 | 14.92674 | 26.11319 | 41.52572 | 18.3156 | -0.27825 | 0.679885 | 0.606845 |
| X | 17485600 | 17487199 | 24.55876 | 42.59591 | 41.95602 | 18.40852 | 0.40847 | 0.846826 | 0.606882 |
| 1 | 179240200 | 179243199 | 40.03288 | 65.10338 | 70.883 | 30.4196 | 0.246824 | 1.57969 | 0.606884 |
| 8 | 129933800 | 129936399 | 46.18856 | 74.73484 | 77.46185 | 33.12514 | 0.311069 | 1.692924 | 0.606894 |
| X | 59089800 | 59091599 | 15.79259 | 27.64761 | 44.60758 | 19.68675 | -0.29914 | 0.678571 | 0.606894 |
| X | 14742000 | 14743599 | 18.11992 | 32.62493 | 25.16652 | 10.97906 | 0.777628 | 0.405609 | 0.6069 |
| 3 | 43100400 | 43104599 | 50.19892 | 80.51388 | 87.9204 | 37.15163 | 0.22113 | 1.960491 | 0.606915 |
| X | 153467000 | 153474199 | 120.5363 | 185.5665 | 181.7476 | 75.48635 | 0.327153 | 2.940782 | 0.606938 |
| 9 | 38481600 | 38485999 | 35.68876 | 59.21486 | 51.47369 | 22.30224 | 0.582455 | 1.305143 | 0.606986 |
| 10 | 115150800 | 115151999 | 9.711994 | 17.75129 | 30.82698 | 13.19801 | -0.36854 | 0.268263 | 0.607005 |
| 3 | 153885200 | 153890199 | 36.19318 | 58.61054 | 90.39262 | 38.67839 | -0.25989 | 1.663515 | 0.60702 |
| 4 | 125069200 | 125071799 | 16.39677 | 28.17608 | 47.95373 | 21.0341 | -0.37999 | 0.899795 | 0.607023 |
| 13 | 9487800 | 9490799 | 31.79123 | 52.86646 | 56.844 | 24.66721 | 0.276852 | 1.27332 | 0.607036 |
| 1 | 60605200 | 60606399 | 11.67516 | 20.87437 | 34.22732 | 15.02376 | -0.31003 | 0.421924 | 0.607042 |
| 12 | 113261800 | 113264399 | 33.02372 | 54.50949 | 60.23021 | 26.00291 | 0.236817 | 1.362133 | 0.60706 |
| X | 81128600 | 81137199 | 95.09905 | 149.6039 | 138.3794 | 57.4086 | 0.433558 | 2.539219 | 0.607064 |

| | | | | | | | | | |
|---|---|---|---|---|---|---|---|---|---|
| 15 | 86595400 | 86598199 | 22.73349 | 38.25718 | 59.30245 | 25.91444 | -0.24918 | 1.12216 | 0.607074 |
| 14 | 84357600 | 84364399 | 49.08717 | 79.08875 | 73.23615 | 31.15413 | 0.466469 | 1.813714 | 0.60708 |
| 1 | 29674400 | 29675999 | 19.48589 | 33.33727 | 51.66065 | 22.65289 | -0.24644 | 0.95859 | 0.607091 |
| 4 | 140934600 | 140935599 | 11.10796 | 20.82773 | 23.16863 | 9.55385 | 0.310498 | 0.111694 | 0.6071 |
| 5 | 66110800 | 66113399 | 17.78142 | 30.56542 | 49.00285 | 21.49449 | -0.29418 | 0.903252 | 0.607117 |
| 1 | 107108400 | 107110599 | 35.6005 | 58.22975 | 64.44262 | 27.71384 | 0.229187 | 1.485357 | 0.607127 |
| 2 | 111570000 | 111572799 | 24.44329 | 42.03607 | 38.40707 | 16.84702 | 0.513921 | 0.914777 | 0.60715 |
| 9 | 25865400 | 25867999 | 22.06184 | 38.97583 | 34.02055 | 15.05022 | 0.58558 | 0.578148 | 0.607172 |
| 7 | 4610400 | 4615799 | 40.84166 | 65.55063 | 99.80771 | 42.21864 | -0.25633 | 1.932017 | 0.607174 |
| 13 | 29623800 | 29625399 | 13.29716 | 24.75764 | 26.54732 | 11.07839 | 0.34894 | 0.181378 | 0.607179 |
| 7 | 85034200 | 85037399 | 50.28816 | 80.92 | 86.76912 | 36.84247 | 0.253029 | 1.860474 | 0.607223 |
| 5 | 149331000 | 149333399 | 24.45177 | 40.77502 | 62.54712 | 27.19133 | -0.23408 | 1.225551 | 0.607249 |
| 6 | 144491000 | 144493599 | 33.42007 | 55.30931 | 59.699 | 25.82052 | 0.270818 | 1.333905 | 0.607251 |
| 17 | 75987200 | 75989399 | 22.29778 | 38.69426 | 41.42856 | 18.17738 | 0.288609 | 0.828228 | 0.607251 |
| 2 | 144190000 | 144191199 | 11.62761 | 21.14688 | 34.3499 | 14.79828 | -0.27883 | 0.319851 | 0.607266 |
| 2 | 67138600 | 67139799 | 10.60055 | 19.62762 | 33.13518 | 13.9395 | -0.31009 | 0.206957 | 0.607276 |
| X | 112109800 | 112111799 | 20.96646 | 36.85399 | 35.7735 | 15.84169 | 0.430322 | 0.627964 | 0.607288 |
| 7 | 12993000 | 12996199 | 34.98446 | 56.38204 | 89.47978 | 38.07898 | -0.30857 | 1.790457 | 0.607294 |
| 7 | 23457200 | 23476399 | 250.0058 | 367.6317 | 322.851 | 132.9563 | 0.427315 | 3.829088 | 0.607295 |
| 7 | 37882400 | 37884399 | 11.44484 | 20.23127 | 35.88215 | 15.87187 | -0.43128 | 0.510305 | 0.607304 |
| 1 | 189274600 | 189277399 | 29.69561 | 50.19752 | 50.17891 | 21.95734 | 0.38172 | 1.097302 | 0.607317 |
| 7 | 21994600 | 21994999 | 17.32304 | 30.87762 | 26.77666 | 11.78377 | 0.601146 | 0.463759 | 0.607319 |
| 3 | 43850600 | 43853999 | 26.91084 | 46.4863 | 43.48569 | 19.07731 | 0.481349 | 0.892745 | 0.607323 |
| 2 | 57852800 | 57857199 | 82.95266 | 129.9587 | 141.5986 | 58.73376 | 0.19298 | 2.608624 | 0.607336 |
| 1 | 94467400 | 94469799 | 23.31657 | 40.49679 | 40.77327 | 17.89772 | 0.376966 | 0.819447 | 0.607344 |
| 17 | 48920800 | 48924199 | 41.01783 | 65.89575 | 98.37219 | 41.67095 | -0.22627 | 1.902958 | 0.607358 |
| 19 | 13671400 | 13673599 | 22.5596 | 39.52163 | 37.29247 | 16.46794 | 0.4709 | 0.688886 | 0.607377 |
| 4 | 12138600 | 12140999 | 23.07903 | 38.62518 | 60.92766 | 26.54774 | -0.27435 | 1.179433 | 0.607404 |
| 6 | 145657200 | 145662799 | 47.89482 | 76.55524 | 112.2135 | 47.17795 | -0.20847 | 2.071944 | 0.607406 |
| 14 | 113913800 | 113914199 | 7.069009 | 13.79883 | 16.93458 | 5.994637 | 0.368614 | -0.46988 | 0.607447 |
| 6 | 130184800 | 130189799 | 54.23858 | 87.29196 | 87.1378 | 37.00838 | 0.356365 | 1.860659 | 0.607453 |
| 8 | 42502600 | 42504999 | 20.48992 | 35.9808 | 36.30855 | 16.06552 | 0.374488 | 0.639958 | 0.607456 |
| 6 | 18389000 | 18389999 | 8.373219 | 15.9772 | 27.85681 | 11.09695 | -0.29038 | -0.0187 | 0.607475 |
| X | 136312400 | 136313199 | 8.373219 | 15.9772 | 27.85681 | 11.09695 | -0.29038 | -0.0187 | 0.607475 |
| 3 | 4618200 | 4621799 | 28.7466 | 48.53321 | 44.78975 | 19.59179 | 0.49658 | 1.100593 | 0.607479 |
| 14 | 92069200 | 92069999 | 9.447897 | 17.31189 | 30.39439 | 12.96925 | -0.38032 | 0.253572 | 0.60748 |
| 9 | 91065400 | 91065999 | 8.553617 | 16.14124 | 27.54849 | 11.23483 | -0.2895 | 0.059405 | 0.607509 |
| 7 | 59606000 | 59606599 | 11.70049 | 22.09849 | 24.71692 | 10.06184 | 0.317481 | 0.073213 | 0.607511 |
| 2 | 6427200 | 6429199 | 17.89316 | 30.88735 | 49.00767 | 21.49936 | -0.27807 | 0.865091 | 0.607518 |
| 2 | 166703200 | 166704599 | 11.02383 | 19.80644 | 33.2412 | 14.53089 | -0.33928 | 0.384167 | 0.607519 |
| 13 | 57077400 | 57078599 | 19.53994 | 34.00696 | 36.8521 | 16.19732 | 0.271777 | 0.769664 | 0.607528 |
| 3 | 54062000 | 54064399 | 15.92508 | 27.427 | 46.91126 | 20.57824 | -0.38637 | 0.876706 | 0.607544 |
| 9 | 28801800 | 28803399 | 15.48913 | 27.641 | 30.29853 | 13.32054 | 0.265797 | 0.449826 | 0.60755 |
| Y | 20645400 | 20646199 | 10.79297 | 20.35603 | 17.16625 | 7.008761 | 0.717948 | 0.052116 | 0.607551 |
| 4 | 83134000 | 83134999 | 9.80499 | 18.71778 | 21.76811 | 8.663067 | 0.290499 | -0.02457 | 0.607553 |
| 14 | 86850600 | 86852199 | 23.00106 | 39.93605 | 35.74185 | 15.686 | 0.546298 | 0.818831 | 0.607564 |
| 15 | 46589000 | 46589999 | 7.908791 | 15.2592 | 29.7146 | 11.4803 | -0.40756 | -0.11847 | 0.60757 |

| | | | | | | | | | |
|---|---|---|---|---|---|---|---|---|---|
| 4 | 10318200 | 10320599 | 18.79171 | 32.23557 | 50.06518 | 21.958 | -0.24907 | 0.928747 | 0.607572 |
| 14 | 17518800 | 17521799 | 18.24639 | 32.03053 | 34.92281 | 15.44807 | 0.263336 | 0.629263 | 0.60758 |
| 1 | 162714800 | 162715399 | 8.718758 | 16.86221 | 20.39799 | 7.79754 | 0.288317 | -0.17103 | 0.607606 |
| 17 | 22565000 | 22568399 | 34.3571 | 57.6003 | 52.54754 | 22.92144 | 0.512997 | 1.188517 | 0.607628 |
| 14 | 120593600 | 120601999 | 83.07708 | 128.4 | 186.5583 | 77.49213 | -0.23694 | 2.852365 | 0.607633 |
| 6 | 123056200 | 123056599 | 12.05779 | 22.31369 | 24.58421 | 10.34987 | 0.300991 | 0.202663 | 0.607668 |
| 7 | 19632600 | 19636399 | 65.99318 | 104.8894 | 113.9223 | 47.53081 | 0.214571 | 2.265162 | 0.607689 |
| 9 | 13660200 | 13660999 | 6.969389 | 13.45479 | 24.8523 | 9.580088 | -0.32858 | -0.1497 | 0.607694 |
| 10 | 33895000 | 33895999 | 11.13082 | 19.66332 | 36.37132 | 16.09034 | -0.49207 | 0.514967 | 0.607697 |
| 4 | 26666200 | 26667399 | 13.26733 | 24.81584 | 26.07774 | 10.7981 | 0.38677 | 0.150915 | 0.6077 |
| 15 | 48069400 | 48070399 | 7.293286 | 14.0896 | 25.47418 | 9.788894 | -0.29333 | -0.15311 | 0.607719 |
| 2 | 55638600 | 55641799 | 29.97391 | 50.04056 | 54.6434 | 23.7724 | 0.254574 | 1.221488 | 0.607721 |
| 14 | 12787600 | 12789999 | 26.84957 | 45.86715 | 40.51016 | 17.76134 | 0.561205 | 0.987736 | 0.607729 |
| 6 | 109420600 | 109421199 | 9.142365 | 17.5988 | 20.81456 | 8.09494 | 0.297001 | -0.10923 | 0.60773 |
| 13 | 87368400 | 87368999 | 7.863896 | 15.02362 | 27.33134 | 10.85962 | -0.3483 | -0.0359 | 0.60774 |
| 7 | 12023000 | 12024599 | 34.74732 | 56.78282 | 60.38982 | 25.96234 | 0.285545 | 1.493491 | 0.607761 |
| 13 | 66979000 | 66983599 | 68.77558 | 109.3179 | 113.4807 | 47.34922 | 0.279846 | 2.269199 | 0.607761 |
| 5 | 53780000 | 53780799 | 12.32505 | 23.3577 | 24.76702 | 10.0097 | 0.402962 | 0.055489 | 0.607766 |
| 1 | 124835600 | 124839399 | 33.51345 | 54.68489 | 95.16821 | 40.89178 | -0.42479 | 1.533973 | 0.607775 |
| 9 | 111576800 | 111582199 | 61.61824 | 98.1136 | 107.916 | 45.12824 | 0.199123 | 2.204527 | 0.607778 |
| 16 | 62131800 | 62133199 | 20.40128 | 35.71738 | 37.49473 | 16.5482 | 0.317818 | 0.687992 | 0.607779 |
| 12 | 105896000 | 105902199 | 68.29748 | 107.0024 | 157.1058 | 65.16578 | -0.23677 | 2.603458 | 0.607789 |
| 15 | 100163800 | 100179799 | 187.154 | 275.1078 | 371.5123 | 152.9995 | -0.19369 | 3.827863 | 0.607819 |
| 13 | 45287600 | 45290999 | 28.88943 | 47.63706 | 75.29962 | 32.49262 | -0.27881 | 1.37427 | 0.607849 |
| 2 | 99399000 | 99402399 | 58.36324 | 93.2332 | 101.6047 | 42.68218 | 0.217483 | 2.098434 | 0.607849 |
| 3 | 71690000 | 71690999 | 11.94405 | 22.29705 | 25.13912 | 10.44399 | 0.282534 | 0.151791 | 0.607876 |
| 13 | 9524600 | 9528599 | 50.86931 | 82.07947 | 72.85439 | 31.05116 | 0.529831 | 1.773386 | 0.607915 |
| 8 | 112244200 | 112245999 | 20.64696 | 35.83412 | 39.42021 | 17.29625 | 0.249748 | 0.816858 | 0.607918 |
| 4 | 100361000 | 100362399 | 28.70628 | 46.92746 | 73.0945 | 31.42832 | -0.26306 | 1.484571 | 0.607941 |
| 6 | 109481800 | 109484399 | 13.0458 | 22.92974 | 43.48948 | 19.25779 | -0.5312 | 0.596653 | 0.607941 |
| 2 | 150103400 | 150107399 | 48.62318 | 78.581 | 78.02482 | 33.32362 | 0.371037 | 1.72421 | 0.607965 |
| 9 | 42258800 | 42259799 | 12.39307 | 21.86469 | 36.86519 | 16.31308 | -0.35983 | 0.540387 | 0.607983 |
| 13 | 9294000 | 9295599 | 14.42692 | 25.33907 | 39.84033 | 17.63121 | -0.262 | 0.609715 | 0.607985 |
| 9 | 25934400 | 25940399 | 130.3784 | 198.8879 | 184.2503 | 76.39753 | 0.397158 | 3.122875 | 0.607987 |
| 5 | 123467800 | 123468999 | 12.99264 | 23.82246 | 26.46074 | 11.28087 | 0.276121 | 0.265126 | 0.608009 |
| 9 | 102941800 | 102945399 | 30.85311 | 50.12862 | 77.94468 | 33.43352 | -0.26701 | 1.582109 | 0.608036 |
| X | 162214000 | 162218399 | 22.92138 | 38.77268 | 67.72857 | 29.64284 | -0.42068 | 1.089773 | 0.608041 |
| 3 | 3271600 | 3280399 | 54.38481 | 87.65501 | 73.61842 | 31.33214 | 0.607623 | 1.811442 | 0.608043 |
| 15 | 87067800 | 87075999 | 82.98254 | 128.4948 | 184.1216 | 76.47925 | -0.21486 | 2.81742 | 0.608049 |
| 12 | 3439800 | 3441799 | 16.01998 | 27.96902 | 43.61077 | 19.20596 | -0.25029 | 0.718843 | 0.608076 |
| 3 | 41364000 | 41366199 | 21.75153 | 37.53657 | 40.9947 | 17.98396 | 0.258466 | 0.873884 | 0.60808 |
| 13 | 38841000 | 38847399 | 66.31195 | 103.9487 | 172.49 | 71.54293 | -0.41252 | 2.594292 | 0.608102 |
| 9 | 11114000 | 11116399 | 25.40654 | 42.45332 | 67.84327 | 29.53156 | -0.29312 | 1.210249 | 0.60811 |
| 12 | 20823800 | 20880799 | 1958.143 | 2388.685 | 2037.941 | 859.6441 | 0.199036 | 6.761899 | 0.608119 |
| 4 | 143487200 | 143494799 | 124.201 | 190.8372 | 187.8242 | 77.9894 | 0.317986 | 2.978657 | 0.608123 |
| 17 | 7252800 | 7255999 | 27.46535 | 45.49146 | 71.10189 | 30.76971 | -0.26165 | 1.317736 | 0.608123 |
| 7 | 13230800 | 13238399 | 144.2337 | 218.2585 | 228.347 | 94.45667 | 0.2123 | 3.292439 | 0.608125 |

| | | | | | | | | | |
|---|---|---|---|---|---|---|---|---|---|
| 2 | 68799400 | 68801599 | 24.65897 | 42.97425 | 40.1453 | 17.65979 | 0.485187 | 0.781418 | 0.608128 |
| 8 | 55974800 | 55982399 | 88.94372 | 139.299 | 147.4549 | 61.16495 | 0.23423 | 2.618975 | 0.608135 |
| X | 151070600 | 151074999 | 45.42969 | 73.7932 | 73.09485 | 31.34736 | 0.381538 | 1.613578 | 0.608142 |
| 13 | 82332000 | 82333799 | 13.38819 | 23.83677 | 38.70738 | 17.05055 | -0.30008 | 0.466425 | 0.608156 |
| 13 | 22932200 | 22938199 | 79.17252 | 124.4364 | 134.732 | 55.88884 | 0.205612 | 2.544449 | 0.608168 |
| 3 | 130615400 | 130616799 | 12.96868 | 24.05043 | 26.51459 | 11.1303 | 0.302773 | 0.20165 | 0.60817 |
| 3 | 12285400 | 12289399 | 48.66373 | 78.3282 | 85.24686 | 36.21093 | 0.232154 | 1.849196 | 0.608174 |
| 14 | 87291600 | 87294199 | 20.38102 | 36.0963 | 33.4801 | 14.80375 | 0.499585 | 0.539105 | 0.608182 |
| 3 | 70621400 | 70621999 | 6.251104 | 12.13208 | 22.78613 | 8.56929 | -0.31716 | -0.24368 | 0.608191 |
| 2 | 168963000 | 168969199 | 93.30164 | 143.1389 | 205.5442 | 85.32914 | -0.22819 | 2.999675 | 0.608218 |
| 13 | 96763600 | 96766199 | 17.58489 | 31.07172 | 33.20744 | 14.69002 | 0.294797 | 0.544859 | 0.608219 |
| 5 | 150027400 | 150028599 | 14.74711 | 25.94786 | 40.8045 | 18.06596 | -0.26176 | 0.589301 | 0.608243 |
| 8 | 25400000 | 25401799 | 13.39979 | 23.82148 | 38.17157 | 16.83591 | -0.28221 | 0.475307 | 0.608262 |
| 8 | 70432800 | 70434999 | 13.39979 | 23.82148 | 38.17157 | 16.83591 | -0.28221 | 0.475307 | 0.608262 |
| 10 | 40380200 | 40382799 | 15.713 | 27.56215 | 44.00043 | 19.45092 | -0.28406 | 0.643733 | 0.608268 |
| 13 | 14713400 | 14714399 | 16.39431 | 28.95503 | 31.00966 | 13.71902 | 0.291832 | 0.540632 | 0.608276 |
| 4 | 40299000 | 40302199 | 22.36374 | 37.67525 | 59.89445 | 26.18164 | -0.28549 | 1.110916 | 0.60829 |
| 1 | 101214000 | 101215599 | 21.76368 | 37.52412 | 41.01498 | 17.99237 | 0.257035 | 0.882063 | 0.608297 |
| 18 | 6138400 | 6140199 | 17.0384 | 30.31124 | 32.22028 | 14.20266 | 0.307342 | 0.483736 | 0.608313 |
| 19 | 9979600 | 9980599 | 13.30507 | 24.32435 | 26.9187 | 11.52134 | 0.27733 | 0.284596 | 0.608314 |
| X | 16561000 | 16563799 | 40.80612 | 67.41993 | 60.77904 | 26.25725 | 0.529714 | 1.373734 | 0.608323 |
| 8 | 59691600 | 59692599 | 15.18988 | 27.77375 | 28.87873 | 12.35825 | 0.366907 | 0.303205 | 0.608327 |
| 18 | 35468800 | 35471399 | 28.9013 | 48.25627 | 53.46763 | 23.26283 | 0.233693 | 1.217183 | 0.608334 |
| 12 | 18095200 | 18125799 | 865.0163 | 1128.399 | 1033.671 | 432.8717 | 0.194057 | 5.679671 | 0.608338 |
| 17 | 57317400 | 57320399 | 15.84639 | 27.60196 | 43.66939 | 19.20381 | -0.27153 | 0.750521 | 0.608343 |
| 1 | 35491200 | 35495799 | 41.11335 | 66.06778 | 100.1825 | 42.45511 | -0.24831 | 1.89308 | 0.608364 |
| 9 | 76693200 | 76693999 | 10.75958 | 20.55042 | 22.7294 | 9.036031 | 0.362994 | -0.01854 | 0.608367 |
| X | 121771600 | 121772599 | 19.36316 | 34.21826 | 33.75816 | 14.93326 | 0.410005 | 0.558776 | 0.608367 |
| 5 | 138718600 | 138719999 | 10.29725 | 18.87884 | 31.32872 | 13.35768 | -0.29936 | 0.256075 | 0.608388 |
| 1 | 139869400 | 139872799 | 46.34748 | 74.66583 | 82.13832 | 34.93511 | 0.218266 | 1.817066 | 0.608407 |
| X | 53762200 | 53764199 | 39.82366 | 65.50775 | 52.03896 | 22.43233 | 0.709783 | 1.418903 | 0.608421 |
| 12 | 108743600 | 108753199 | 117.6396 | 180.5626 | 189.5883 | 78.71038 | 0.223565 | 2.997047 | 0.608457 |
| 9 | 11943000 | 11948999 | 77.3777 | 122.1677 | 131.2263 | 54.48432 | 0.222142 | 2.451246 | 0.608458 |
| 14 | 34109400 | 34113399 | 65.75954 | 104.3136 | 113.62 | 47.31186 | 0.207749 | 2.322849 | 0.608484 |
| 14 | 120256000 | 120257599 | 11.76018 | 21.25728 | 34.23413 | 14.86191 | -0.27379 | 0.354939 | 0.608498 |
| 4 | 7100200 | 7103399 | 28.99602 | 47.64258 | 73.02611 | 31.46406 | -0.23649 | 1.419589 | 0.608524 |
| 1 | 182360200 | 182361599 | 16.26375 | 28.83752 | 31.20954 | 13.79557 | 0.278685 | 0.495757 | 0.60853 |
| X | 19791400 | 19791999 | 14.06963 | 26.16007 | 26.59847 | 11.12547 | 0.422983 | 0.199538 | 0.608539 |
| 1 | 31525600 | 31529799 | 46.73649 | 76.29426 | 67.87402 | 29.16863 | 0.541377 | 1.539662 | 0.608579 |
| 14 | 114201800 | 114205999 | 53.81188 | 86.52397 | 88.74102 | 37.63803 | 0.315709 | 1.888413 | 0.608602 |
| 7 | 33285800 | 33365199 | 1693.446 | 2102.959 | 1746.268 | 734.9836 | 0.264207 | 6.478371 | 0.60862 |
| 18 | 36598600 | 36600199 | 16.72587 | 29.84218 | 26.71795 | 11.74831 | 0.555999 | 0.452345 | 0.608624 |
| 19 | 8740800 | 8743999 | 44.6408 | 71.54635 | 105.2978 | 44.44468 | -0.20987 | 1.983979 | 0.608626 |
| 12 | 103686600 | 103707199 | 295.2225 | 417.3437 | 545.7014 | 225.8824 | -0.20272 | 4.417738 | 0.608639 |
| X | 164715600 | 164718199 | 23.77914 | 41.82971 | 34.93629 | 15.4686 | 0.646511 | 0.63371 | 0.608649 |
| 16 | 69206000 | 69206399 | 27.93621 | 48.36063 | 42.33658 | 18.5744 | 0.577241 | 0.877453 | 0.608681 |
| 1 | 44598400 | 44599999 | 15.70881 | 27.88133 | 30.38881 | 13.42149 | 0.26956 | 0.486062 | 0.608682 |

| | | | | | | | | | |
|---|---|---|---|---|---|---|---|---|---|
| 2 | 40438600 | 40439799 | 9.291796 | 17.42308 | 29.59329 | 12.20231 | -0.29603 | 0.109953 | 0.608714 |
| 1 | 17534600 | 17538199 | 38.98855 | 62.92202 | 96.01849 | 40.93447 | -0.24993 | 1.75807 | 0.608733 |
| 5 | 64272400 | 64275799 | 48.27101 | 77.80368 | 83.47688 | 35.52946 | 0.255113 | 1.804384 | 0.608747 |
| 18 | 7073600 | 7074999 | 12.51407 | 23.17655 | 26.03399 | 10.94832 | 0.27418 | 0.204668 | 0.608748 |
| 6 | 86337800 | 86338799 | 11.3214 | 20.4065 | 33.22413 | 14.47443 | -0.29253 | 0.366947 | 0.60875 |
| 2 | 36833800 | 36838399 | 60.79184 | 97.649 | 90.62063 | 38.37939 | 0.457956 | 1.931663 | 0.608781 |
| X | 97466800 | 97470199 | 28.73233 | 49.85407 | 41.21021 | 18.08151 | 0.660391 | 0.860671 | 0.608818 |
| 1 | 118711200 | 118712799 | 12.59487 | 22.35833 | 36.64471 | 16.1821 | -0.31575 | 0.475718 | 0.608844 |
| 8 | 117515200 | 117521399 | 56.84574 | 90.27543 | 130.4524 | 54.38225 | -0.19789 | 2.287144 | 0.608846 |
| 5 | 66509400 | 66511199 | 17.41531 | 30.16424 | 47.40388 | 20.79793 | -0.26322 | 0.831166 | 0.608849 |
| 7 | 12297600 | 12298199 | 12.7665 | 23.84257 | 26.14295 | 10.85317 | 0.32321 | 0.158287 | 0.608855 |
| 4 | 62024400 | 62028199 | 53.00297 | 84.4311 | 122.5769 | 51.28888 | -0.20004 | 2.186903 | 0.608863 |
| 8 | 15966400 | 15972199 | 52.98873 | 84.45025 | 123.0994 | 51.54283 | -0.20508 | 2.170918 | 0.608876 |
| 13 | 120750600 | 120752599 | 54.73877 | 86.81618 | 129.6434 | 53.97453 | -0.24702 | 2.323508 | 0.608903 |
| 4 | 22514800 | 22516199 | 12.81755 | 23.84626 | 25.41257 | 10.6187 | 0.355999 | 0.177924 | 0.608913 |
| 17 | 94813000 | 94815399 | 28.10588 | 48.37336 | 43.99831 | 19.30005 | 0.520587 | 0.928917 | 0.608935 |
| 5 | 95671200 | 95672999 | 16.83103 | 30.06193 | 31.6904 | 13.92221 | 0.322531 | 0.456281 | 0.608947 |
| 7 | 11744200 | 11770999 | 2010.652 | 2414.012 | 2787.529 | 1178.402 | -0.26097 | 7.013294 | 0.608952 |
| 1 | 3723200 | 3724999 | 21.46664 | 37.63618 | 31.80315 | 14.05316 | 0.629426 | 0.664036 | 0.608953 |
| 13 | 7133200 | 7135399 | 21.29214 | 36.82535 | 40.47569 | 17.75748 | 0.249962 | 0.851939 | 0.608965 |
| 9 | 115617200 | 115621399 | 39.74225 | 64.5681 | 69.78807 | 29.93374 | 0.256171 | 1.592837 | 0.608966 |
| 12 | 114204400 | 114217999 | 248.8005 | 362.4269 | 360.1805 | 148.4351 | 0.235961 | 3.973617 | 0.608985 |
| 8 | 14329800 | 14340999 | 157.3243 | 233.1326 | 326.5575 | 134.4927 | -0.23549 | 3.690746 | 0.60899 |
| 2 | 36509400 | 36510599 | 19.80816 | 34.71715 | 36.37346 | 16.06826 | 0.320504 | 0.664374 | 0.60901 |
| 17 | 24534800 | 24536999 | 21.97433 | 37.03945 | 57.03102 | 24.93407 | -0.23934 | 1.105195 | 0.60902 |
| 7 | 41320800 | 41321799 | 8.164465 | 15.22099 | 31.6405 | 13.17475 | -0.59626 | 0.144373 | 0.609023 |
| 2 | 36807000 | 36810199 | 35.62852 | 59.64855 | 54.23283 | 23.63575 | 0.517996 | 1.211297 | 0.609024 |
| 1 | 20114600 | 20115999 | 20.59096 | 35.79799 | 39.40108 | 17.30606 | 0.249307 | 0.794095 | 0.609033 |
| 10 | 64814600 | 64815399 | 6.405256 | 12.38674 | 23.19913 | 8.872095 | -0.3364 | -0.18441 | 0.609061 |
| 13 | 69348600 | 69350199 | 17.65498 | 31.13078 | 33.83004 | 14.97167 | 0.269818 | 0.569842 | 0.60907 |
| 10 | 19971800 | 19973199 | 17.17532 | 31.44202 | 28.73788 | 12.27795 | 0.553439 | 0.310837 | 0.609086 |
| 8 | 115220000 | 115223599 | 26.12298 | 43.22836 | 66.3779 | 28.70718 | -0.23621 | 1.320382 | 0.609088 |
| 8 | 59153000 | 59153999 | 10.6743 | 19.50663 | 31.83032 | 13.63065 | -0.27947 | 0.280384 | 0.609093 |
| 2 | 144542600 | 144543999 | 8.813532 | 16.6204 | 28.62353 | 11.6895 | -0.30464 | 0.066972 | 0.609095 |
| X | 36861000 | 36865399 | 67.1726 | 106.7015 | 108.8849 | 45.40251 | 0.303622 | 2.28619 | 0.609099 |
| 12 | 34408600 | 34409199 | 8.048521 | 15.37314 | 26.74399 | 10.63093 | -0.28446 | -0.03212 | 0.609105 |
| 17 | 41346800 | 41348599 | 28.35641 | 46.55258 | 71.68649 | 30.87591 | -0.24359 | 1.427903 | 0.609109 |
| 3 | 22723800 | 22725999 | 21.51931 | 37.77096 | 36.17168 | 15.99871 | 0.449537 | 0.651234 | 0.609117 |
| 17 | 39448000 | 39463999 | 226.926 | 336.4136 | 295.4455 | 121.6844 | 0.438219 | 3.693077 | 0.609122 |
| 9 | 67698200 | 67700199 | 19.06789 | 33.31663 | 36.68334 | 16.16318 | 0.249397 | 0.711425 | 0.609127 |
| X | 10300400 | 10302399 | 24.14708 | 42.29203 | 37.22578 | 16.43499 | 0.570837 | 0.70191 | 0.609129 |
| 8 | 10208200 | 10209399 | 12.72864 | 22.76011 | 37.8032 | 16.59202 | -0.32842 | 0.437717 | 0.609143 |
| 2 | 124574800 | 124576999 | 16.98989 | 30.89181 | 29.80225 | 12.84293 | 0.46805 | 0.351009 | 0.609194 |
| 1 | 32104200 | 32105999 | 33.7683 | 55.91281 | 46.64006 | 20.17915 | 0.641545 | 1.318757 | 0.609194 |
| 17 | 28508400 | 28513199 | 45.48952 | 72.64692 | 110.6407 | 46.45904 | -0.26495 | 2.096448 | 0.609196 |
| 17 | 33456000 | 33461799 | 91.30438 | 143.6377 | 139.4542 | 57.85467 | 0.363792 | 2.536772 | 0.609204 |
| 10 | 55540400 | 55542799 | 20.33398 | 35.87829 | 33.97242 | 15.03269 | 0.467741 | 0.577381 | 0.609204 |

| | | | | | | | | | |
|---|---|---|---|---|---|---|---|---|---|
| 16 | 11371000 | 11372599 | 10.66537 | 19.65717 | 32.48329 | 13.7509 | -0.28571 | 0.233562 | 0.609208 |
| 6 | 139022600 | 139023399 | 9.032622 | 16.94924 | 28.52413 | 11.74719 | -0.28098 | 0.102567 | 0.609226 |
| 14 | 80386000 | 80389799 | 52.65911 | 84.33774 | 92.67551 | 39.07494 | 0.209632 | 2.009694 | 0.609248 |
| 1 | 34562000 | 34565199 | 20.26608 | 34.50651 | 53.27305 | 23.35091 | -0.24222 | 1.005776 | 0.609255 |
| 8 | 76697600 | 76698599 | 15.4747 | 27.80894 | 29.88337 | 13.06083 | 0.3003 | 0.407879 | 0.609263 |
| 13 | 85798600 | 85799399 | 11.06068 | 19.72835 | 34.26589 | 15.07115 | -0.39465 | 0.432245 | 0.609296 |
| 12 | 88625600 | 88629199 | 36.85292 | 61.09963 | 59.46031 | 25.74902 | 0.419968 | 1.322064 | 0.609303 |
| 5 | 111802400 | 111804999 | 28.38955 | 47.6209 | 52.36095 | 22.84771 | 0.244577 | 1.165697 | 0.609325 |
| 7 | 39308400 | 39309399 | 9.649006 | 17.88254 | 29.90692 | 12.56537 | -0.29503 | 0.18698 | 0.609328 |
| 2 | 124668600 | 124669599 | 10.51792 | 19.10911 | 31.79067 | 13.71332 | -0.31406 | 0.312224 | 0.609348 |
| 10 | 93645000 | 93646599 | 14.40375 | 25.94618 | 28.42965 | 12.39532 | 0.274631 | 0.382123 | 0.609361 |
| 7 | 27693400 | 27703599 | 158.5004 | 240.0829 | 227.6497 | 94.1974 | 0.355314 | 3.276819 | 0.609389 |
| 15 | 29740600 | 29743399 | 26.32772 | 45.71585 | 41.8313 | 18.35943 | 0.514503 | 0.841446 | 0.609405 |
| 3 | 27967400 | 27969799 | 24.8005 | 41.14579 | 64.1174 | 27.77893 | -0.25707 | 1.284845 | 0.609407 |
| 4 | 13916600 | 13917199 | 10.10713 | 19.49537 | 21.97629 | 8.497207 | 0.374135 | -0.11148 | 0.609415 |
| 9 | 118536800 | 118538199 | 14.48026 | 26.23367 | 28.83726 | 12.48339 | 0.276452 | 0.349456 | 0.609425 |
| 5 | 139925400 | 139927799 | 31.63324 | 52.28933 | 57.27304 | 24.75074 | 0.249641 | 1.34178 | 0.609431 |
| 1 | 185034000 | 185041399 | 73.29368 | 114.0486 | 186.7908 | 77.54143 | -0.40198 | 2.729318 | 0.609433 |
| 7 | 14049600 | 14107599 | 945.3302 | 1232.042 | 1070.251 | 448.2328 | 0.269336 | 5.695303 | 0.609444 |
| 1 | 85467000 | 85484599 | 464.6789 | 639.09 | 614.1953 | 255.3418 | 0.201388 | 4.835465 | 0.609448 |
| 9 | 76473400 | 76474999 | 12.62216 | 22.2856 | 37.73511 | 16.69551 | -0.36593 | 0.531349 | 0.609467 |
| 11 | 12095200 | 12101199 | 66.93965 | 105.2934 | 153.5534 | 63.70284 | -0.22257 | 2.527419 | 0.609471 |
| 2 | 107517400 | 107519199 | 18.2708 | 31.99781 | 35.10717 | 15.49869 | 0.254445 | 0.668023 | 0.609575 |
| 1 | 102160000 | 102162199 | 22.85998 | 39.81544 | 39.62117 | 17.42273 | 0.394129 | 0.777085 | 0.609582 |
| X | 112286800 | 112288599 | 13.69791 | 24.41457 | 42.49964 | 18.70399 | -0.39881 | 0.471375 | 0.609611 |
| 4 | 71656400 | 71663999 | 132.8559 | 201.7454 | 215.2214 | 89.12307 | 0.18843 | 3.217469 | 0.609611 |
| 6 | 65927600 | 65930599 | 34.35809 | 56.29628 | 62.27561 | 26.80059 | 0.231122 | 1.456538 | 0.609623 |
| 15 | 14112800 | 14114799 | 27.00948 | 46.53897 | 43.45149 | 19.06084 | 0.483269 | 0.912842 | 0.60963 |
| 1 | 117828200 | 117836199 | 122.0631 | 187.6051 | 187.0262 | 77.66106 | 0.299812 | 2.973231 | 0.609632 |
| X | 96493600 | 96495199 | 11.70097 | 21.30215 | 35.99147 | 15.48561 | -0.33399 | 0.319291 | 0.609643 |
| 1 | 21467000 | 21468199 | 8.353847 | 15.80457 | 27.60532 | 11.19737 | -0.31612 | 0.037995 | 0.609652 |
| 2 | 51836200 | 51837999 | 120.1309 | 180.5874 | 271.2644 | 111.992 | -0.31739 | 3.421198 | 0.609677 |
| 3 | 47744000 | 47748599 | 137.1142 | 207.3458 | 215.153 | 88.98006 | 0.223332 | 3.302965 | 0.609681 |
| 14 | 86845400 | 86849799 | 43.50301 | 71.20041 | 66.60496 | 28.65036 | 0.471683 | 1.494828 | 0.609685 |
| 6 | 17839800 | 17841999 | 20.34343 | 34.40853 | 54.29328 | 23.76189 | -0.27425 | 1.067003 | 0.609687 |
| X | 16068000 | 16072999 | 77.44764 | 123.354 | 93.7754 | 39.23055 | 0.731623 | 2.208443 | 0.609709 |
| 3 | 151718400 | 151723199 | 86.54784 | 135.479 | 146.1261 | 60.61688 | 0.206654 | 2.626997 | 0.609733 |
| X | 156312200 | 156314999 | 45.75261 | 74.14119 | 75.74561 | 32.42726 | 0.333866 | 1.662335 | 0.609765 |
| 15 | 12456400 | 12458599 | 17.29012 | 29.84446 | 47.41197 | 20.7993 | -0.27976 | 0.862072 | 0.609774 |
| 15 | 9913600 | 9914799 | 14.60327 | 25.61377 | 40.4116 | 17.86337 | -0.26693 | 0.634183 | 0.609782 |
| 7 | 62109200 | 62114799 | 111.7738 | 171.3354 | 182.728 | 75.84866 | 0.199652 | 3.020113 | 0.609824 |
| 16 | 61182400 | 61182999 | 8.368866 | 16.25802 | 19.42537 | 7.252106 | 0.337568 | -0.25946 | 0.609839 |
| 9 | 19126000 | 19127599 | 29.66846 | 49.32307 | 54.59956 | 23.68881 | 0.234934 | 1.266205 | 0.60985 |
| 5 | 17672800 | 17674399 | 19.92143 | 34.12163 | 52.76797 | 23.14058 | -0.24335 | 0.949862 | 0.609855 |
| 12 | 3112000 | 3113399 | 24.93361 | 42.21865 | 44.35995 | 19.42096 | 0.310327 | 1.063472 | 0.609861 |
| 1 | 136492800 | 136495399 | 14.39741 | 25.28878 | 39.81851 | 17.62247 | -0.26414 | 0.607642 | 0.609877 |
| 4 | 21515800 | 21520599 | 32.62353 | 54.97885 | 51.87027 | 22.67645 | 0.464846 | 1.137665 | 0.609879 |

| | | | | | | | | | |
|---|---|---|---|---|---|---|---|---|---|
| 18 | 19686600 | 19687799 | 19.99459 | 35.00859 | 31.00443 | 13.68478 | 0.56201 | 0.674519 | 0.609879 |
| 9 | 117543600 | 117545799 | 19.0191 | 32.26979 | 52.68745 | 23.08107 | -0.32315 | 1.032115 | 0.6099 |
| 10 | 124441200 | 124442999 | 27.52467 | 46.60913 | 43.18419 | 18.9066 | 0.491008 | 1.068418 | 0.609905 |
| 14 | 121860000 | 121861799 | 22.00397 | 36.79345 | 59.1371 | 25.75319 | -0.3012 | 1.185166 | 0.609945 |
| 3 | 41169800 | 41170999 | 14.26922 | 25.9677 | 28.45062 | 12.24702 | 0.286619 | 0.323198 | 0.609948 |
| 4 | 46399600 | 46400599 | 12.26438 | 22.78049 | 25.60075 | 10.72347 | 0.27784 | 0.186587 | 0.609949 |
| 4 | 136211600 | 136212599 | 12.26438 | 22.78049 | 25.60075 | 10.72347 | 0.27784 | 0.186587 | 0.609949 |
| 1 | 139903200 | 139903999 | 17.64344 | 30.98553 | 33.99459 | 15.04298 | 0.254467 | 0.618234 | 0.609959 |
| 15 | 13343800 | 13345599 | 17.94542 | 31.88193 | 32.57895 | 14.37704 | 0.362732 | 0.497388 | 0.609962 |
| 5 | 118554000 | 118556199 | 21.0288 | 35.4252 | 55.88537 | 24.42891 | -0.27418 | 1.106489 | 0.609982 |
| 15 | 62251600 | 62253599 | 21.00472 | 37.06847 | 34.53852 | 15.28264 | 0.491114 | 0.58208 | 0.610017 |
| 1 | 48400000 | 48404199 | 78.02427 | 123.183 | 114.2872 | 47.45023 | 0.433139 | 2.450166 | 0.610021 |
| 17 | 88385800 | 88388599 | 26.47834 | 43.69517 | 67.39748 | 29.09243 | -0.24299 | 1.354468 | 0.610036 |
| 5 | 135640400 | 135642199 | 15.65442 | 27.34106 | 43.07331 | 18.97374 | -0.2651 | 0.709683 | 0.610038 |
| 7 | 85412400 | 85420999 | 88.4456 | 139.5275 | 134.0151 | 55.61905 | 0.38235 | 2.479604 | 0.610078 |
| X | 5155800 | 5158199 | 15.49793 | 29.00789 | 26.61272 | 11.00687 | 0.583031 | 0.168551 | 0.610081 |
| 16 | 9915800 | 9918599 | 34.46265 | 55.86005 | 83.90144 | 35.92548 | -0.22047 | 1.640844 | 0.61009 |
| 8 | 5833800 | 5836999 | 40.19977 | 65.16015 | 72.1883 | 30.91037 | 0.217588 | 1.643376 | 0.610108 |
| 16 | 23647200 | 23649399 | 24.81757 | 42.59691 | 45.20734 | 19.82833 | 0.298137 | 0.941147 | 0.610139 |
| 15 | 54444800 | 54445799 | 14.67272 | 26.73731 | 28.87434 | 12.4089 | 0.308645 | 0.318287 | 0.610152 |
| X | 8736200 | 8741399 | 79.07859 | 125.6624 | 116.078 | 48.42056 | 0.447621 | 2.285803 | 0.610169 |
| 15 | 52130200 | 52131399 | 10.94979 | 19.87334 | 32.75305 | 14.14658 | -0.30207 | 0.321994 | 0.61017 |
| 12 | 53578200 | 53579799 | 16.36435 | 29.00562 | 31.80091 | 14.05903 | 0.260002 | 0.502266 | 0.610176 |
| 15 | 33895200 | 33897199 | 16.13142 | 28.14421 | 44.13442 | 19.42817 | -0.25867 | 0.728569 | 0.610178 |
| 4 | 144271000 | 144275999 | 109.2637 | 165.3788 | 245.7018 | 101.6429 | -0.29296 | 3.28069 | 0.610191 |
| 16 | 42752000 | 42754799 | 40.16767 | 66.09707 | 63.60972 | 27.42423 | 0.433959 | 1.419895 | 0.610202 |
| 11 | 72821800 | 72824199 | 25.29775 | 43.25212 | 46.30393 | 20.30293 | 0.284794 | 0.978856 | 0.610217 |
| 1 | 75768800 | 75770399 | 21.80925 | 37.72551 | 41.49701 | 18.20564 | 0.248902 | 0.854472 | 0.610244 |
| 19 | 8292400 | 8303599 | 253.4018 | 367.1173 | 374.2352 | 154.3199 | 0.191642 | 4.055287 | 0.610249 |
| 17 | 10179400 | 10181599 | 11.58633 | 20.48261 | 43.48051 | 19.23264 | -0.68972 | 0.521263 | 0.610265 |
| 14 | 31927200 | 31927799 | 7.785022 | 14.87691 | 26.18198 | 10.39707 | -0.29921 | -0.03729 | 0.610267 |
| X | 67560200 | 67560999 | 12.09782 | 22.98417 | 19.56869 | 7.86726 | 0.724469 | 0.025696 | 0.610267 |
| X | 154812800 | 154813599 | 8.491158 | 16.14 | 30.54438 | 12.26701 | -0.41759 | 0.019105 | 0.610296 |
| 12 | 88030000 | 88039999 | 130.5353 | 199.6976 | 199.4282 | 82.74558 | 0.292154 | 3.066561 | 0.610317 |
| 10 | 110518800 | 110520199 | 10.41807 | 19.21953 | 31.94925 | 13.50745 | -0.29315 | 0.222737 | 0.61032 |
| 14 | 117003800 | 117005999 | 16.67654 | 29.91577 | 30.56027 | 13.37661 | 0.371035 | 0.424621 | 0.610321 |
| 19 | 56625000 | 56638799 | 148.2649 | 221.477 | 305.6424 | 126.0065 | -0.20339 | 3.545599 | 0.610323 |
| 12 | 115367800 | 115382799 | 189.5646 | 284.6147 | 254.2023 | 104.9134 | 0.430556 | 3.457072 | 0.610323 |
| 7 | 84927800 | 84937399 | 197.8766 | 292.1372 | 303.212 | 124.8683 | 0.191748 | 3.75692 | 0.610325 |
| 1 | 110604800 | 110618199 | 147.8521 | 225.0727 | 212.6802 | 88.13592 | 0.366195 | 3.171858 | 0.610325 |
| 7 | 48065600 | 48071199 | 62.96693 | 100.7613 | 99.65206 | 41.96444 | 0.359968 | 2.05048 | 0.610353 |
| 17 | 44051400 | 44053399 | 23.61562 | 41.21071 | 39.59452 | 17.43206 | 0.444971 | 0.756875 | 0.610382 |
| 13 | 82174800 | 82176999 | 30.43266 | 51.01121 | 53.55347 | 23.35744 | 0.311049 | 1.17934 | 0.610391 |
| 11 | 48914200 | 48914599 | 6.11063 | 11.83362 | 22.69069 | 8.620778 | -0.36064 | -0.21101 | 0.61042 |
| 5 | 90300600 | 90302199 | 14.65806 | 25.76917 | 41.81345 | 18.51583 | -0.30744 | 0.597133 | 0.610421 |
| 11 | 65975600 | 65976599 | 9.516707 | 17.56127 | 29.87007 | 12.62393 | -0.3253 | 0.210381 | 0.610443 |
| 18 | 30301800 | 30304399 | 26.56607 | 45.61395 | 45.1502 | 19.80382 | 0.398333 | 0.944297 | 0.61047 |

| | | | | | | | | | |
|---|---|---|---|---|---|---|---|---|---|
| 10 | 66752000 | 66754399 | 17.40873 | 30.09744 | 47.93789 | 21.03105 | -0.28308 | 0.847833 | 0.610475 |
| 13 | 62035800 | 62073599 | 592.7652 | 802.9681 | 723.6237 | 301.5476 | 0.27205 | 5.075498 | 0.610475 |
| 1 | 186144800 | 186147799 | 24.5565 | 40.59417 | 64.96231 | 28.07493 | -0.29565 | 1.32835 | 0.610512 |
| 3 | 119777200 | 119778599 | 15.9211 | 28.2882 | 30.88773 | 13.62962 | 0.267917 | 0.482486 | 0.610515 |
| 6 | 18988400 | 18992999 | 55.82742 | 88.88124 | 131.3675 | 54.92509 | -0.2267 | 2.209616 | 0.610515 |
| 6 | 88666800 | 88668999 | 23.99416 | 41.00915 | 45.26783 | 19.84807 | 0.240737 | 0.975915 | 0.610517 |
| 14 | 37350600 | 37351199 | 19.30134 | 33.67554 | 36.79436 | 16.1973 | 0.260352 | 0.734392 | 0.610517 |
| 6 | 37323000 | 37325199 | 12.02973 | 21.34483 | 35.24532 | 15.57045 | -0.32651 | 0.474788 | 0.61053 |
| 6 | 130847600 | 130861199 | 229.42 | 338.3239 | 317.5158 | 130.7573 | 0.335408 | 3.780413 | 0.610541 |
| 6 | 87980800 | 87982199 | 10.3121 | 18.86436 | 31.30947 | 13.38964 | -0.30266 | 0.268896 | 0.610548 |
| 16 | 84887400 | 84888599 | 10.3121 | 18.86436 | 31.30947 | 13.38964 | -0.30266 | 0.268896 | 0.610548 |
| X | 126252000 | 126253599 | 25.90565 | 44.14645 | 47.40415 | 20.77985 | 0.279825 | 1.013251 | 0.610569 |
| 1 | 103244400 | 103253799 | 270.2397 | 389.1147 | 393.5733 | 162.426 | 0.194185 | 4.146811 | 0.61057 |
| 7 | 110110200 | 110112599 | 30.32736 | 51.71365 | 47.34921 | 20.75681 | 0.509 | 1.023497 | 0.610573 |
| 13 | 51447400 | 51456199 | 79.73594 | 123.8242 | 177.7131 | 73.79517 | -0.21393 | 2.764632 | 0.610617 |
| 17 | 38836800 | 38847399 | 102.7338 | 161.179 | 142.3746 | 59.04694 | 0.496968 | 2.597773 | 0.610618 |
| 17 | 12536800 | 12537999 | 10.37038 | 18.80801 | 31.79384 | 13.74483 | -0.33901 | 0.321256 | 0.610629 |
| 17 | 71508600 | 71510599 | 13.71723 | 24.23524 | 38.69085 | 17.11608 | -0.28116 | 0.532815 | 0.610631 |
| 10 | 84143200 | 84145199 | 19.96426 | 33.97676 | 52.71863 | 23.10703 | -0.24948 | 1.008818 | 0.610632 |
| 2 | 138674600 | 138681599 | 30.34393 | 50.68075 | 44.95263 | 19.56193 | 0.553566 | 1.209076 | 0.610641 |
| 1 | 26543200 | 26546199 | 28.40273 | 47.88651 | 51.2548 | 22.41033 | 0.283421 | 1.118576 | 0.610647 |
| 13 | 59869600 | 59871399 | 21.24147 | 36.76569 | 40.51622 | 17.77567 | 0.246514 | 0.846662 | 0.610669 |
| X | 9506600 | 9508999 | 30.79668 | 51.6568 | 53.5022 | 23.34517 | 0.330579 | 1.176044 | 0.610683 |
| 15 | 68857800 | 68860599 | 34.1001 | 55.2491 | 83.32449 | 35.66734 | -0.22695 | 1.647104 | 0.610687 |
| 2 | 77805600 | 77813199 | 95.47756 | 148.7925 | 156.2917 | 64.86589 | 0.239819 | 2.708978 | 0.610688 |
| 18 | 6323400 | 6335199 | 163.4897 | 246.1336 | 248.4056 | 102.5956 | 0.257837 | 3.397865 | 0.6107 |
| 8 | 59203200 | 59205799 | 46.37655 | 75.04222 | 68.78031 | 29.41356 | 0.488103 | 1.686909 | 0.610711 |
| 5 | 81942000 | 81943399 | 22.70705 | 37.4039 | 73.51786 | 31.70988 | -0.59259 | 1.372723 | 0.610722 |
| 12 | 9741200 | 9743799 | 22.41829 | 38.81747 | 42.02157 | 18.43633 | 0.27221 | 0.85062 | 0.610754 |
| 3 | 144128000 | 144130199 | 25.67646 | 43.38849 | 47.99985 | 21.00161 | 0.236037 | 1.085459 | 0.610779 |
| 17 | 44983200 | 44985199 | 12.84569 | 22.69702 | 36.78391 | 16.27226 | -0.30232 | 0.526032 | 0.610819 |
| 13 | 44501200 | 44507799 | 79.64786 | 123.0371 | 202.9603 | 84.30539 | -0.42044 | 2.863117 | 0.610827 |
| 4 | 71025400 | 71028199 | 37.73181 | 61.86797 | 65.06025 | 28.00736 | 0.304327 | 1.456258 | 0.61085 |
| 6 | 19994200 | 19995999 | 16.27782 | 29.34101 | 29.53133 | 12.86524 | 0.396757 | 0.390543 | 0.610871 |
| 9 | 114710400 | 114723799 | 131.4339 | 197.7827 | 274.009 | 113.1564 | -0.19948 | 3.401997 | 0.610873 |
| 6 | 132087000 | 132090399 | 43.3694 | 70.49564 | 72.42721 | 31.07688 | 0.329853 | 1.594488 | 0.610912 |
| 8 | 25824600 | 25827199 | 28.5956 | 47.85784 | 52.40958 | 22.83623 | 0.250375 | 1.188326 | 0.610919 |
| 2 | 48193600 | 48195999 | 25.12522 | 43.87927 | 37.77443 | 16.6389 | 0.602668 | 0.749709 | 0.610926 |
| 7 | 123239200 | 123240999 | 13.26354 | 23.49273 | 37.15982 | 16.43051 | -0.26632 | 0.50069 | 0.610935 |
| 12 | 21541600 | 21547399 | 58.7182 | 93.61117 | 102.1167 | 42.77595 | 0.212909 | 2.15975 | 0.610944 |
| 2 | 99853400 | 99855999 | 40.07597 | 65.42737 | 60.90024 | 26.17254 | 0.476302 | 1.518034 | 0.610945 |
| 16 | 97645800 | 97646799 | 11.2934 | 21.12772 | 22.69343 | 9.393718 | 0.356421 | 0.127003 | 0.61095 |
| 4 | 70340000 | 70341599 | 19.41416 | 33.35827 | 51.69451 | 22.67487 | -0.24557 | 0.917703 | 0.61095 |
| 6 | 84260600 | 84264399 | 59.4559 | 94.77319 | 104.408 | 43.72676 | 0.198499 | 2.166768 | 0.610958 |
| 14 | 34326400 | 34330199 | 35.94751 | 57.90859 | 90.76086 | 38.59968 | -0.29139 | 1.805906 | 0.610977 |
| 6 | 60401200 | 60410199 | 121.8415 | 186.7463 | 193.8871 | 80.47938 | 0.238226 | 3.026192 | 0.610983 |
| X | 153137600 | 153140799 | 33.55328 | 56.74306 | 50.17988 | 21.96064 | 0.558087 | 1.110944 | 0.610984 |

| | | | | | | | | | |
|---|---|---|---|---|---|---|---|---|---|
| 12 | 109033400 | 109036199 | 21.66309 | 36.3454 | 57.52334 | 25.10361 | -0.27906 | 1.14689 | 0.611007 |
| 2 | 103349800 | 103350599 | 5.514676 | 10.74783 | 20.86496 | 7.55623 | -0.31451 | -0.3947 | 0.611018 |
| 18 | 13012600 | 13013199 | 5.514676 | 10.74783 | 20.86496 | 7.55623 | -0.31451 | -0.3947 | 0.611018 |
| 14 | 101652200 | 101652799 | 8.964065 | 16.93203 | 29.24004 | 11.90049 | -0.3043 | 0.058473 | 0.611025 |
| 15 | 96842800 | 96843799 | 7.625186 | 14.6754 | 26.44628 | 10.29092 | -0.30565 | -0.10447 | 0.611037 |
| 14 | 96974400 | 96974999 | 6.379578 | 12.38973 | 23.48168 | 8.787876 | -0.3246 | -0.26096 | 0.611065 |
| 4 | 123701000 | 123704799 | 33.31794 | 54.07856 | 81.58002 | 34.96673 | -0.22486 | 1.610184 | 0.611066 |
| X | 169076200 | 169119799 | 3207.838 | 3660.375 | 5316.416 | 2265.363 | -0.66672 | 7.813127 | 0.611068 |
| 8 | 4137000 | 4139599 | 47.56466 | 76.42571 | 82.76219 | 35.06671 | 0.236229 | 1.899593 | 0.611073 |
| X | 147565000 | 147569599 | 29.70603 | 50.40909 | 49.58885 | 21.72452 | 0.405029 | 1.06637 | 0.611073 |
| 5 | 127973000 | 127984599 | 120.5686 | 184.6438 | 194.9407 | 80.90327 | 0.213172 | 3.041986 | 0.611099 |
| 19 | 39414600 | 39415799 | 33.52408 | 55.19111 | 61.2978 | 26.43281 | 0.228055 | 1.395771 | 0.611121 |
| 3 | 73054000 | 73054599 | 12.53026 | 23.15211 | 26.03296 | 10.98292 | 0.269467 | 0.218065 | 0.611144 |
| 12 | 71311000 | 71318599 | 66.18644 | 104.2024 | 149.5673 | 62.05644 | -0.19869 | 2.507111 | 0.611169 |
| 5 | 59073400 | 59074399 | 11.81128 | 22.30506 | 24.68972 | 10.05333 | 0.331929 | 0.074305 | 0.611172 |
| 3 | 12579600 | 12580999 | 21.26947 | 36.87832 | 40.49448 | 17.76708 | 0.252198 | 0.831664 | 0.611179 |
| 12 | 42822800 | 42823999 | 16.18335 | 29.40452 | 29.7844 | 12.84659 | 0.397158 | 0.347793 | 0.61118 |
| X | 85396200 | 85400199 | 56.56588 | 90.53797 | 97.88017 | 41.23103 | 0.232078 | 2.03663 | 0.611198 |
| 9 | 24842200 | 24845399 | 35.83659 | 59.06504 | 61.95275 | 26.72803 | 0.311056 | 1.390944 | 0.611207 |
| 12 | 96445000 | 96449199 | 34.74551 | 57.50483 | 59.93705 | 25.92409 | 0.32103 | 1.33482 | 0.611242 |
| 1 | 73458600 | 73463399 | 41.71461 | 69.19731 | 57.36512 | 24.85096 | 0.650716 | 1.32985 | 0.611243 |
| 10 | 48611800 | 48617399 | 62.02685 | 99.27816 | 99.35205 | 41.85075 | 0.343294 | 2.043483 | 0.611269 |
| 2 | 89735600 | 89736399 | 10.70817 | 19.36511 | 32.28801 | 14.00854 | -0.32258 | 0.342212 | 0.611289 |
| 16 | 48245600 | 48246599 | 10.63668 | 19.56808 | 31.90789 | 13.54134 | -0.26928 | 0.241772 | 0.61129 |
| 13 | 18087200 | 18092999 | 99.68757 | 152.8832 | 215.2744 | 89.36432 | -0.20032 | 3.010387 | 0.611313 |
| 9 | 86633600 | 86639599 | 63.49862 | 100.1803 | 144.5263 | 59.98382 | -0.20367 | 2.462222 | 0.611338 |
| 10 | 14023400 | 14025399 | 13.755 | 24.27998 | 38.21928 | 16.91053 | -0.2613 | 0.543363 | 0.611358 |
| 7 | 106260600 | 106263999 | 57.78895 | 91.56656 | 136.1855 | 56.64564 | -0.24253 | 2.356433 | 0.611359 |
| 14 | 124785000 | 124787799 | 33.39578 | 55.4987 | 57.62947 | 24.99686 | 0.326841 | 1.282639 | 0.611373 |
| 6 | 58617200 | 58617799 | 5.259186 | 10.20556 | 21.65165 | 8.147918 | -0.49186 | -0.25828 | 0.611377 |
| 17 | 88485800 | 88487199 | 7.497436 | 14.14907 | 26.50336 | 10.80713 | -0.42241 | 0.045555 | 0.611421 |
| 4 | 67108800 | 67110999 | 17.07755 | 30.98864 | 29.63404 | 12.80268 | 0.478035 | 0.360264 | 0.611428 |
| 3 | 12002800 | 12009799 | 105.007 | 162.0914 | 173.6251 | 72.12045 | 0.201039 | 2.883373 | 0.611445 |
| 4 | 5090800 | 5098199 | 88.4534 | 138.9124 | 143.1576 | 59.37774 | 0.275911 | 2.568952 | 0.611468 |
| 9 | 111138000 | 111142199 | 46.54298 | 74.51926 | 109.991 | 46.35394 | -0.21582 | 2.017827 | 0.61147 |
| 17 | 92345800 | 92349199 | 61.50856 | 97.86392 | 106.3205 | 44.4139 | 0.21574 | 2.229338 | 0.611517 |
| 10 | 119102200 | 119104799 | 35.19195 | 57.62612 | 64.02009 | 27.54396 | 0.224601 | 1.467466 | 0.611521 |
| 9 | 19070600 | 19072199 | 36.51994 | 60.15671 | 48.77744 | 21.03875 | 0.681041 | 1.389768 | 0.611539 |
| 2 | 57168800 | 57170599 | 16.29312 | 28.80523 | 31.34761 | 13.86575 | 0.269187 | 0.527887 | 0.611541 |
| 15 | 45841400 | 45843399 | 19.75059 | 33.83851 | 52.22534 | 22.90314 | -0.2404 | 0.945887 | 0.611562 |
| 1 | 136685000 | 136688399 | 35.17524 | 56.68282 | 99.64462 | 42.39769 | -0.45607 | 1.799053 | 0.611564 |
| 5 | 60681400 | 60683399 | 33.3002 | 55.45572 | 50.32463 | 21.85924 | 0.520859 | 1.255212 | 0.611566 |
| 12 | 24481200 | 24485199 | 74.47195 | 117.9949 | 124.248 | 51.67902 | 0.254838 | 2.364953 | 0.611566 |
| 1 | 88295200 | 88296399 | 10.96474 | 19.85975 | 32.74107 | 14.17768 | -0.30488 | 0.334206 | 0.611569 |
| 12 | 17689600 | 17697399 | 62.92428 | 99.41902 | 143.5756 | 59.63324 | -0.20348 | 2.426245 | 0.611579 |
| X | 118084600 | 118086599 | 25.58932 | 44.47977 | 41.3138 | 18.14413 | 0.493185 | 0.824139 | 0.611586 |
| 16 | 28294200 | 28297999 | 34.53539 | 56.71651 | 62.95919 | 27.12103 | 0.227414 | 1.429657 | 0.611591 |

| | | | | | | | | | |
|---|---|---|---|---|---|---|---|---|---|
| 13 | 4615600 | 4623399 | 114.7966 | 175.8451 | 187.2301 | 77.7074 | 0.201226 | 3.0341 | 0.611602 |
| 6 | 60072000 | 60072399 | 9.201874 | 17.67966 | 20.82572 | 8.143677 | 0.295812 | -0.08958 | 0.611604 |
| 4 | 60080800 | 60083199 | 34.69289 | 55.95138 | 97.95455 | 41.72324 | -0.44874 | 1.774066 | 0.611609 |
| 12 | 115940200 | 115941799 | 24.88618 | 42.75892 | 44.63872 | 19.57995 | 0.321993 | 0.930412 | 0.611648 |
| 17 | 93164200 | 93165399 | 15.77991 | 28.57006 | 25.72562 | 11.14525 | 0.561913 | 0.356755 | 0.611652 |
| 9 | 23842000 | 23842999 | 8.189314 | 15.60115 | 27.23992 | 10.88822 | -0.29691 | -0.01226 | 0.611659 |
| 4 | 112464400 | 112467799 | 57.25267 | 91.45396 | 99.8074 | 41.92382 | 0.21536 | 2.099102 | 0.611661 |
| 5 | 76634600 | 76635999 | 13.00235 | 23.09739 | 36.70566 | 16.19972 | -0.27063 | 0.477394 | 0.611663 |
| 2 | 4811400 | 4814999 | 52.20347 | 83.68831 | 92.34057 | 38.99062 | 0.205351 | 1.979835 | 0.611693 |
| 9 | 12268600 | 12277999 | 173.8564 | 259.4972 | 271.856 | 112.06 | 0.192881 | 3.563184 | 0.611704 |
| 3 | 114980200 | 114983199 | 43.20327 | 69.90247 | 76.38555 | 32.66417 | 0.234952 | 1.687521 | 0.611707 |
| 16 | 65368400 | 65370799 | 19.88872 | 33.70301 | 53.73698 | 23.53189 | -0.28905 | 1.049591 | 0.611707 |
| 4 | 12421400 | 12422399 | 7.620834 | 14.40751 | 27.0495 | 10.98848 | -0.4213 | 0.03431 | 0.611716 |
| 16 | 86326400 | 86327199 | 11.1407 | 21.1606 | 23.73297 | 9.546392 | 0.328434 | 0.026109 | 0.611736 |
| 1 | 88656000 | 88658999 | 24.9459 | 41.49036 | 63.64868 | 27.62337 | -0.23425 | 1.25627 | 0.611752 |
| X | 133451800 | 133455399 | 17.41676 | 30.40406 | 54.47538 | 23.9889 | -0.45019 | 0.742749 | 0.611759 |
| 6 | 16359000 | 16360399 | 17.40279 | 30.69328 | 33.41171 | 14.78584 | 0.267603 | 0.566261 | 0.611808 |
| X | 54905200 | 54906599 | 13.50728 | 25.73159 | 24.3218 | 9.722282 | 0.581422 | 0.030617 | 0.611822 |
| 13 | 46155800 | 46159999 | 19.0798 | 32.22174 | 63.00614 | 27.5623 | -0.58285 | 1.082556 | 0.611844 |
| 15 | 83124800 | 83129999 | 44.62959 | 71.56372 | 105.026 | 44.36326 | -0.20496 | 1.968338 | 0.611845 |
| 4 | 40873600 | 40877599 | 21.44026 | 35.65233 | 61.70076 | 26.77414 | -0.40755 | 1.246169 | 0.611846 |
| 19 | 56340000 | 56341599 | 22.55116 | 38.70299 | 42.6594 | 18.71063 | 0.243798 | 0.930522 | 0.611893 |
| 17 | 53310200 | 53314999 | 53.64122 | 86.13438 | 91.3063 | 38.6502 | 0.265857 | 1.932432 | 0.611908 |
| 10 | 64867800 | 64871399 | 39.72089 | 65.00605 | 67.2902 | 28.94443 | 0.325919 | 1.488102 | 0.611926 |
| 6 | 132434000 | 132437599 | 34.066 | 56.08404 | 60.89851 | 26.26078 | 0.260505 | 1.394789 | 0.611931 |
| 10 | 90469200 | 90471799 | 33.86809 | 56.32579 | 50.91232 | 22.09472 | 0.526443 | 1.270812 | 0.61197 |
| X | 13891400 | 13893799 | 24.16374 | 41.7628 | 43.06145 | 18.8915 | 0.341679 | 0.876896 | 0.611973 |
| 19 | 9251600 | 9255599 | 51.74442 | 83.05613 | 89.58879 | 37.90065 | 0.240095 | 1.938028 | 0.612011 |
| 11 | 104938400 | 104939799 | 17.33036 | 31.78646 | 28.83174 | 12.28609 | 0.567242 | 0.303005 | 0.612023 |
| 2 | 30758400 | 30760399 | 19.50796 | 33.92098 | 35.40939 | 15.55447 | 0.325406 | 0.779929 | 0.612029 |
| 3 | 8160800 | 8164199 | 27.2656 | 47.17659 | 42.91572 | 18.82821 | 0.521962 | 0.879462 | 0.612049 |
| 12 | 21160400 | 21163199 | 26.85708 | 43.67752 | 84.39053 | 36.21677 | -0.57844 | 1.561386 | 0.612058 |
| 11 | 36361400 | 36363599 | 11.74719 | 21.39643 | 35.44198 | 15.24014 | -0.30532 | 0.313091 | 0.612072 |
| 11 | 121668400 | 121670799 | 20.3414 | 35.46054 | 38.73144 | 17.04107 | 0.260579 | 0.752354 | 0.612098 |
| 7 | 58990400 | 58991999 | 16.19111 | 29.56131 | 28.69528 | 12.30215 | 0.463587 | 0.316896 | 0.61213 |
| 13 | 100035400 | 100037999 | 22.85583 | 38.07661 | 60.72627 | 26.38354 | -0.28997 | 1.229872 | 0.612136 |
| 5 | 127274200 | 127303199 | 341.1023 | 476.3716 | 615.4405 | 255.2434 | -0.20304 | 4.599668 | 0.612148 |
| 15 | 57543400 | 57544799 | 20.86685 | 36.59128 | 36.81874 | 16.27195 | 0.378564 | 0.663573 | 0.612173 |
| 5 | 145410400 | 145414199 | 64.27068 | 102.6388 | 104.5375 | 43.89524 | 0.31444 | 2.114211 | 0.612187 |
| 13 | 66993200 | 66999599 | 118.8816 | 185.1861 | 128.5926 | 53.37332 | 0.835787 | 2.72688 | 0.612209 |
| X | 106574600 | 106576599 | 22.2234 | 38.80024 | 33.98434 | 14.96671 | 0.577928 | 0.735448 | 0.61222 |
| 6 | 9886800 | 9894199 | 98.67539 | 153.095 | 164.1007 | 68.15199 | 0.205682 | 2.787661 | 0.61223 |
| 19 | 3276400 | 3278399 | 14.43025 | 25.19451 | 41.59863 | 18.32027 | -0.33232 | 0.707151 | 0.612244 |
| X | 73783200 | 73785599 | 18.84237 | 32.57765 | 52.25713 | 22.926 | -0.29322 | 0.861957 | 0.612285 |
| 2 | 170056600 | 170059799 | 34.46843 | 55.77026 | 84.13377 | 35.97781 | -0.22915 | 1.676602 | 0.612303 |
| 16 | 5039400 | 5041199 | 13.76245 | 24.27075 | 38.24026 | 16.92293 | -0.26313 | 0.553759 | 0.612307 |
| X | 13820200 | 13825799 | 5.256286 | 10.15757 | 25.34553 | 9.725268 | -0.75051 | -0.17903 | 0.612337 |

| | | | | | | | | | |
|---|---|---|---|---|---|---|---|---|---|
| 9 | 38967200 | 38969599 | 23.06302 | 40.28962 | 39.02282 | 17.19196 | 0.433485 | 0.737984 | 0.612338 |
| 4 | 146286000 | 146299399 | 313.7248 | 449.7319 | 420.972 | 173.853 | 0.299548 | 4.21526 | 0.612341 |
| 6 | 88736200 | 88738599 | 18.90175 | 32.04319 | 52.66664 | 23.06595 | -0.33268 | 1.0405 | 0.612367 |
| 17 | 60576800 | 60579999 | 29.05307 | 49.60234 | 46.79537 | 20.51606 | 0.466201 | 1.005983 | 0.612381 |
| 17 | 48352800 | 48354599 | 16.20274 | 28.14407 | 44.77396 | 19.65506 | -0.27984 | 0.794499 | 0.612382 |
| 18 | 38067800 | 38068799 | 10.16443 | 18.64243 | 30.83358 | 13.13975 | -0.29388 | 0.252698 | 0.612393 |
| 8 | 9365600 | 9366399 | 9.268595 | 17.8082 | 20.7664 | 8.120026 | 0.309831 | -0.09115 | 0.612396 |
| 3 | 107444800 | 107446999 | 20.08207 | 34.17484 | 53.27414 | 23.35038 | -0.25628 | 1.009239 | 0.612405 |
| 9 | 109467800 | 109469599 | 28.71156 | 48.11879 | 52.92193 | 23.07971 | 0.244225 | 1.176311 | 0.612407 |
| 6 | 130577400 | 130580799 | 91.50951 | 142.4453 | 153.5548 | 63.74088 | 0.201186 | 2.727879 | 0.612425 |
| 13 | 14419200 | 14420399 | 10.76677 | 19.79976 | 32.4255 | 13.76825 | -0.27618 | 0.245184 | 0.61244 |
| 10 | 3821800 | 3824199 | 29.04695 | 47.48802 | 73.66479 | 31.67441 | -0.25708 | 1.485234 | 0.612456 |
| 5 | 60087600 | 60088599 | 12.52039 | 22.99148 | 25.03707 | 10.65189 | 0.30677 | 0.250125 | 0.61247 |
| X | 162186800 | 162189799 | 34.49364 | 57.10189 | 59.72174 | 25.83531 | 0.316209 | 1.334842 | 0.612476 |
| 6 | 127874400 | 127875799 | 17.26623 | 30.57719 | 32.66623 | 14.44413 | 0.29658 | 0.517836 | 0.612479 |
| 11 | 73802600 | 73808799 | 71.55026 | 114.1729 | 105.6211 | 44.30005 | 0.451748 | 2.147557 | 0.612509 |
| 4 | 59166400 | 59168399 | 22.35854 | 38.6975 | 41.42552 | 18.17458 | 0.288003 | 0.850453 | 0.612511 |
| 5 | 55475600 | 55476799 | 12.36715 | 23.17688 | 25.03192 | 10.33116 | 0.351107 | 0.127635 | 0.612516 |
| 2 | 90364200 | 90365399 | 11.35109 | 20.45644 | 33.24811 | 14.48805 | -0.29015 | 0.369255 | 0.61252 |
| 6 | 128075200 | 128077199 | 13.15615 | 23.21653 | 37.28201 | 16.49673 | -0.28991 | 0.542294 | 0.612534 |
| 7 | 41708600 | 41710399 | 18.26759 | 33.79442 | 27.3115 | 11.50309 | 0.744817 | 0.260888 | 0.61254 |
| X | 147778400 | 147782199 | 31.74446 | 53.67757 | 50.7339 | 22.20228 | 0.462389 | 1.104841 | 0.612543 |
| 8 | 36455400 | 36456999 | 11.07939 | 20.30941 | 33.42878 | 14.25633 | -0.28809 | 0.267331 | 0.61255 |
| 16 | 88957000 | 88961199 | 44.4765 | 71.73729 | 79.38577 | 33.81838 | 0.211742 | 1.776693 | 0.612551 |
| 7 | 29380600 | 29383799 | 44.49114 | 71.78007 | 79.43778 | 33.8522 | 0.212101 | 1.769259 | 0.612551 |
| X | 156900800 | 156904399 | 35.73143 | 60.21921 | 50.78098 | 22.20055 | 0.626314 | 1.147536 | 0.612565 |
| 6 | 8606200 | 8608799 | 82.04151 | 128.4907 | 139.0342 | 57.67193 | 0.202603 | 2.614199 | 0.612567 |
| 13 | 61661000 | 61672999 | 171.8997 | 259.1239 | 243.7826 | 100.7205 | 0.360699 | 3.376452 | 0.612576 |
| 2 | 19959000 | 19960799 | 18.7449 | 32.2134 | 50.07289 | 21.96368 | -0.24988 | 0.911133 | 0.612578 |
| 7 | 43387000 | 43387799 | 8.706274 | 16.59218 | 29.56316 | 11.80752 | -0.32484 | -0.00252 | 0.612591 |
| 5 | 48575000 | 48575799 | 5.364427 | 10.46032 | 20.32626 | 7.312305 | -0.30667 | -0.42079 | 0.612593 |
| 17 | 65680200 | 65681799 | 17.72707 | 31.16976 | 33.98409 | 15.04833 | 0.263666 | 0.602448 | 0.612594 |
| 9 | 22054400 | 22061599 | 80.00212 | 125.7731 | 136.5241 | 56.63416 | 0.202258 | 2.540191 | 0.6126 |
| 13 | 49957800 | 49961199 | 31.50056 | 51.03572 | 82.40843 | 35.2744 | -0.32523 | 1.64115 | 0.612604 |
| 5 | 24389200 | 24393999 | 52.70254 | 83.85629 | 122.5696 | 51.20452 | -0.21156 | 2.223827 | 0.612609 |
| 3 | 77468200 | 77472399 | 53.238 | 85.29522 | 93.95355 | 39.63524 | 0.206781 | 2.000942 | 0.612613 |
| 13 | 89205400 | 89206599 | 17.81224 | 30.96433 | 48.99589 | 21.51916 | -0.2722 | 0.795728 | 0.612616 |
| X | 140655800 | 140659799 | 24.99908 | 41.76496 | 65.38387 | 28.45778 | -0.26342 | 1.211191 | 0.612624 |
| 12 | 17934000 | 17970199 | 1218.457 | 1547.445 | 1380.038 | 579.4425 | 0.193977 | 6.116337 | 0.612629 |
| X | 7401200 | 7402599 | 18.12493 | 31.88567 | 33.98551 | 15.04742 | 0.296557 | 0.60016 | 0.61263 |
| 14 | 73811600 | 73814799 | 35.92319 | 58.67457 | 63.48027 | 27.28591 | 0.260289 | 1.502406 | 0.612682 |
| 7 | 137476000 | 137478399 | 22.98413 | 38.55918 | 59.20421 | 25.83605 | -0.23548 | 1.154475 | 0.612686 |
| 12 | 71603800 | 71605199 | 16.04852 | 28.5056 | 31.32901 | 13.82801 | 0.258208 | 0.485446 | 0.612691 |
| 13 | 8704200 | 8704799 | 6.700629 | 12.98263 | 23.86756 | 9.046889 | -0.29776 | -0.20908 | 0.612693 |
| 19 | 8305600 | 8306999 | 24.19418 | 41.78261 | 35.86353 | 15.73332 | 0.604951 | 0.874494 | 0.612695 |
| 4 | 122778800 | 122780599 | 13.77322 | 24.13088 | 39.95839 | 17.64635 | -0.33638 | 0.645497 | 0.612703 |
| 4 | 34850200 | 34856199 | 64.95571 | 103.4654 | 110.3967 | 46.18883 | 0.243446 | 2.197075 | 0.612715 |

| | | | | | | | | | |
|---|---|---|---|---|---|---|---|---|---|
| 10 | 97157600 | 97161399 | 33.57327 | 55.78014 | 51.44962 | 22.31261 | 0.497263 | 1.282042 | 0.612717 |
| 13 | 33821600 | 33825999 | 58.57597 | 94.1831 | 77.36198 | 32.81128 | 0.635388 | 1.891626 | 0.612725 |
| 7 | 42858600 | 42865799 | 154.5915 | 232.2563 | 245.7191 | 101.4299 | 0.187207 | 3.436223 | 0.612729 |
| Y | 58474000 | 58474399 | 2.739473 | 5.358199 | 15.09466 | 4.96299 | -0.71229 | -0.79886 | 0.612734 |
| 19 | 29829400 | 29830399 | 13.687 | 24.02022 | 39.4516 | 17.44735 | -0.32441 | 0.619715 | 0.61274 |
| 19 | 23250600 | 23251399 | 6.698777 | 12.93646 | 23.70589 | 9.12467 | -0.31439 | -0.15826 | 0.612753 |
| 3 | 23604800 | 23607199 | 25.30678 | 43.81729 | 42.99346 | 18.86271 | 0.41344 | 0.865559 | 0.612757 |
| 13 | 97124200 | 97126999 | 25.47198 | 42.27794 | 64.70535 | 28.04195 | -0.2311 | 1.283202 | 0.612757 |
| 15 | 68589800 | 68591599 | 15.70463 | 28.05675 | 30.81144 | 13.53335 | 0.264114 | 0.446086 | 0.612762 |
| 4 | 84801200 | 84803399 | 17.59396 | 29.99315 | 56.97953 | 24.97799 | -0.54004 | 0.988597 | 0.612804 |
| 7 | 129091800 | 129093599 | 16.21079 | 30.17141 | 26.95075 | 11.25371 | 0.610169 | 0.209027 | 0.612831 |
| 7 | 48185600 | 48191599 | 65.96799 | 105.7784 | 93.45478 | 39.47469 | 0.525822 | 1.992331 | 0.612832 |
| 19 | 19928600 | 19929799 | 11.95354 | 21.63532 | 34.78722 | 15.07712 | -0.26989 | 0.350761 | 0.612856 |
| 12 | 54279800 | 54283599 | 40.2447 | 65.48001 | 70.87635 | 30.42219 | 0.255772 | 1.573958 | 0.612879 |
| 10 | 78739800 | 78741599 | 26.8329 | 45.38901 | 49.60176 | 21.70923 | 0.253673 | 1.082377 | 0.612887 |
| 5 | 133503400 | 133504999 | 23.03391 | 39.38382 | 43.21259 | 18.94755 | 0.249539 | 0.966725 | 0.612932 |
| 3 | 19990800 | 19992999 | 22.22492 | 38.8317 | 33.80168 | 14.89322 | 0.586753 | 0.723008 | 0.612939 |
| 18 | 21669400 | 21669799 | 15.43144 | 26.80135 | 44.28096 | 19.43728 | -0.33405 | 0.792478 | 0.61295 |
| 3 | 72964000 | 72965199 | 9.01343 | 16.39025 | 30.90262 | 13.31548 | -0.49205 | 0.294214 | 0.612976 |
| 6 | 57667400 | 57669199 | 12.51245 | 22.076 | 36.80979 | 16.28846 | -0.34402 | 0.539028 | 0.612979 |
| 3 | 23378000 | 23379399 | 12.50135 | 22.45823 | 36.25807 | 15.85138 | -0.28387 | 0.402692 | 0.613001 |
| 13 | 40779400 | 40780599 | 11.98936 | 21.38637 | 34.70602 | 15.26376 | -0.29721 | 0.439354 | 0.613004 |
| 1 | 184530800 | 184534999 | 45.49037 | 73.38487 | 80.76006 | 34.41126 | 0.22 | 1.774461 | 0.613013 |
| 3 | 8610000 | 8611399 | 8.378131 | 15.66481 | 30.07529 | 12.4615 | -0.4774 | 0.119492 | 0.613018 |
| 16 | 55857800 | 55862399 | 48.72277 | 77.30773 | 139.5758 | 58.13454 | -0.51984 | 2.307922 | 0.613054 |
| 1 | 85308800 | 85324399 | 661.0125 | 882.3215 | 829.2487 | 346.3192 | 0.190122 | 5.309041 | 0.613088 |
| 10 | 121171800 | 121173999 | 34.27134 | 55.63319 | 83.66787 | 35.86503 | -0.22031 | 1.609664 | 0.613096 |
| X | 112841000 | 112846799 | 82.70716 | 130.8563 | 112.7974 | 46.88245 | 0.541559 | 2.403713 | 0.613107 |
| 17 | 74728800 | 74730399 | 9.800611 | 18.31248 | 30.60829 | 12.69641 | -0.27988 | 0.140264 | 0.613108 |
| 13 | 66892000 | 66892999 | 15.71829 | 28.03191 | 30.83514 | 13.56381 | 0.260313 | 0.457311 | 0.613114 |
| 15 | 45630000 | 45631199 | 15.71829 | 28.03191 | 30.83514 | 13.56381 | 0.260313 | 0.457311 | 0.613114 |
| 14 | 29592000 | 29592999 | 10.7931 | 20.346 | 22.57044 | 9.223187 | 0.324353 | 0.070965 | 0.613147 |
| 9 | 82583600 | 82585599 | 37.11937 | 59.73084 | 92.71894 | 39.36982 | -0.27947 | 1.844053 | 0.613159 |
| 18 | 7823200 | 7832399 | 116.2398 | 176.4254 | 245.8523 | 101.7921 | -0.1973 | 3.225323 | 0.613163 |
| X | 153218800 | 153233399 | 229.433 | 338.2213 | 317.9848 | 130.9499 | 0.332356 | 3.786594 | 0.613169 |
| 6 | 62163200 | 62165399 | 23.61008 | 41.24392 | 33.34158 | 14.68872 | 0.693147 | 0.734513 | 0.613178 |
| 18 | 89384800 | 89386599 | 19.05719 | 33.49751 | 35.26103 | 15.61479 | 0.313713 | 0.613489 | 0.613192 |
| 4 | 46847600 | 46851199 | 42.63456 | 68.95785 | 75.80716 | 32.40837 | 0.225815 | 1.692036 | 0.613193 |
| 1 | 130446800 | 130447999 | 10.24783 | 19.042 | 32.11472 | 13.44512 | -0.30367 | 0.18135 | 0.61322 |
| 7 | 55316200 | 55317799 | 29.20889 | 47.61934 | 74.62504 | 32.05459 | -0.27461 | 1.525398 | 0.613221 |
| 5 | 147731200 | 147759599 | 466.1159 | 633.5331 | 791.1994 | 329.5371 | -0.19368 | 5.020311 | 0.613266 |
| X | 89496800 | 89499999 | 32.82252 | 56.00101 | 46.20984 | 20.25828 | 0.658688 | 1.028038 | 0.613269 |
| 6 | 81513200 | 81515799 | 24.59659 | 42.86855 | 40.12233 | 17.65046 | 0.482442 | 0.77948 | 0.613273 |
| 4 | 89724800 | 89726999 | 16.53741 | 28.96871 | 50.54084 | 22.31646 | -0.4118 | 0.678801 | 0.613333 |
| 5 | 38469800 | 38471199 | 15.56307 | 27.22879 | 42.52635 | 18.75834 | -0.25246 | 0.68309 | 0.613351 |
| 18 | 41077000 | 41086999 | 154.6515 | 233.1352 | 212.6614 | 87.86373 | 0.405275 | 3.368794 | 0.613383 |
| 1 | 42844000 | 42852199 | 77.43324 | 122.0023 | 131.6937 | 54.64598 | 0.212727 | 2.493908 | 0.613393 |

| | | | | | | | | | |
|---|---|---|---|---|---|---|---|---|---|
| 3 | 40800800 | 40801799 | 10.06679 | 18.23121 | 31.80385 | 13.77388 | -0.38581 | 0.326974 | 0.613393 |
| 1 | 31578200 | 31579599 | 16.70722 | 29.55382 | 31.69831 | 14.0193 | 0.290214 | 0.523901 | 0.613408 |
| 17 | 65602400 | 65603799 | 15.57178 | 28.38563 | 29.37129 | 12.61691 | 0.370456 | 0.324736 | 0.613415 |
| 2 | 56907800 | 56912399 | 46.33355 | 74.24775 | 110.4395 | 46.60386 | -0.22544 | 1.993418 | 0.613461 |
| 13 | 70627000 | 70627799 | 5.970524 | 11.55154 | 22.6636 | 8.648002 | -0.39985 | -0.19997 | 0.613469 |
| 1 | 5907200 | 5909799 | 24.38751 | 42.6846 | 37.20315 | 16.41573 | 0.58495 | 0.712713 | 0.613489 |
| 5 | 92652400 | 92654599 | 16.21241 | 27.91514 | 47.42486 | 20.80339 | -0.37685 | 0.879607 | 0.613495 |
| 16 | 7704600 | 7707399 | 29.24533 | 49.66056 | 49.0317 | 21.4848 | 0.399739 | 1.057056 | 0.613506 |
| 3 | 138349800 | 138353599 | 32.26737 | 52.35194 | 82.30679 | 35.26759 | -0.28494 | 1.613564 | 0.613512 |
| 11 | 73303200 | 73303799 | 9.621352 | 17.91275 | 29.97399 | 12.50953 | -0.28865 | 0.159555 | 0.613522 |
| 3 | 105530800 | 105532999 | 12.11457 | 21.718 | 34.68017 | 15.18826 | -0.26996 | 0.408299 | 0.613541 |
| X | 147635200 | 147661799 | 338.4605 | 482.2822 | 443.4217 | 183.3025 | 0.316709 | 4.304509 | 0.613542 |
| 3 | 132305000 | 132305399 | 4.513544 | 8.817597 | 18.22147 | 6.361497 | -0.35451 | -0.54664 | 0.613547 |
| 5 | 143635200 | 143636999 | 24.05343 | 40.91662 | 44.81072 | 19.64017 | 0.251138 | 1.019329 | 0.613554 |
| 18 | 3351200 | 3361599 | 116.0136 | 178.5485 | 183.8298 | 76.34789 | 0.254817 | 2.943444 | 0.613576 |
| 14 | 103878800 | 103880799 | 18.75941 | 32.8033 | 36.16726 | 15.94606 | 0.24746 | 0.696865 | 0.613592 |
| 6 | 29020200 | 29026399 | 31.70161 | 50.95701 | 106.6527 | 45.2145 | -0.71169 | 1.875627 | 0.613626 |
| 6 | 137848600 | 137851399 | 27.29867 | 46.23422 | 49.59751 | 21.71554 | 0.280363 | 1.073073 | 0.613693 |
| 14 | 55691200 | 55693999 | 29.90393 | 49.14704 | 76.31123 | 32.88308 | -0.25501 | 1.418514 | 0.613702 |
| 12 | 88030000 | 88033399 | 53.09831 | 85.38745 | 88.94804 | 37.73277 | 0.293559 | 1.884986 | 0.613703 |
| 6 | 145486600 | 145494399 | 60.2392 | 95.04621 | 157.9759 | 65.56877 | -0.40758 | 2.46004 | 0.613704 |
| 15 | 81376000 | 81379599 | 46.98158 | 75.06583 | 111.22 | 46.7341 | -0.22455 | 2.084177 | 0.613738 |
| 8 | 21127600 | 21131399 | 123.4238 | 186.197 | 264.6207 | 109.3557 | -0.23305 | 3.350034 | 0.613751 |
| 4 | 60587600 | 60595799 | 147.241 | 222.3633 | 234.8941 | 97.10354 | 0.195949 | 3.333211 | 0.613754 |
| 17 | 88922600 | 88933799 | 135.633 | 203.5919 | 282.0574 | 116.4022 | -0.20268 | 3.451802 | 0.613762 |
| 2 | 128089400 | 128091599 | 18.13651 | 31.24432 | 48.47157 | 21.26287 | -0.24623 | 0.885812 | 0.613769 |
| 2 | 60123800 | 60128799 | 71.69452 | 113.4131 | 124.4924 | 51.73884 | 0.193119 | 2.400355 | 0.613786 |
| 18 | 8139400 | 8143199 | 42.42987 | 68.60858 | 75.40766 | 32.2281 | 0.225743 | 1.700601 | 0.613796 |
| Y | 6739800 | 6743399 | 22.34128 | 38.47269 | 29.08412 | 12.75809 | 0.786201 | 0.876545 | 0.61383 |
| 4 | 143713800 | 143734999 | 423.236 | 592.0277 | 471.1377 | 195.3474 | 0.498126 | 4.577854 | 0.613895 |
| 6 | 6177600 | 6180399 | 26.10074 | 44.07687 | 48.53664 | 21.23231 | 0.242702 | 1.09456 | 0.613896 |
| 13 | 27471800 | 27474999 | 73.91971 | 116.5988 | 127.7741 | 53.02741 | 0.192262 | 2.470025 | 0.613903 |
| 3 | 11196200 | 11199799 | 37.60106 | 61.30149 | 67.81563 | 29.12973 | 0.226444 | 1.53835 | 0.613906 |
| 17 | 16618000 | 16619399 | 16.39036 | 29.14092 | 31.794 | 14.02183 | 0.269494 | 0.482675 | 0.613924 |
| 10 | 82493400 | 82493999 | 6.708819 | 12.89075 | 24.12918 | 9.433317 | -0.36613 | -0.1066 | 0.613933 |
| 16 | 48640000 | 48640799 | 6.708819 | 12.89075 | 24.12918 | 9.433317 | -0.36613 | -0.1066 | 0.613933 |
| 2 | 109110800 | 109111599 | 7.312976 | 13.98858 | 25.11284 | 9.945429 | -0.32398 | -0.05472 | 0.613942 |
| 4 | 147123400 | 147126799 | 83.39542 | 132.5437 | 113.7559 | 47.45967 | 0.553764 | 2.28426 | 0.613976 |
| X | 10275000 | 10276599 | 12.64478 | 22.67238 | 36.22913 | 15.86435 | -0.27077 | 0.415116 | 0.613997 |
| 9 | 120472800 | 120475399 | 23.82842 | 39.73274 | 61.34864 | 26.66912 | -0.24344 | 1.224204 | 0.614018 |
| 4 | 87390200 | 87393999 | 27.01403 | 44.69523 | 69.17797 | 29.91462 | -0.24765 | 1.328822 | 0.614055 |
| 1 | 26579800 | 26580599 | 8.522202 | 16.18657 | 27.71368 | 11.14827 | -0.27683 | 0.01381 | 0.614077 |
| 15 | 13398200 | 13399799 | 18.94365 | 33.45444 | 34.26648 | 15.16016 | 0.355698 | 0.563292 | 0.614079 |
| 2 | 131836800 | 131840799 | 26.67465 | 43.85347 | 68.87428 | 29.68238 | -0.27113 | 1.402584 | 0.614119 |
| 4 | 47167600 | 47169799 | 14.0604 | 24.72066 | 39.27514 | 17.39151 | -0.27691 | 0.589979 | 0.614126 |
| 5 | 133255400 | 133256799 | 10.88421 | 20.0435 | 34.09518 | 14.44991 | -0.3289 | 0.240433 | 0.614149 |
| 14 | 6819200 | 6823599 | 67.40199 | 107.1725 | 114.972 | 47.98843 | 0.232958 | 2.255749 | 0.614154 |

| | | | | | | | | | |
|---|---|---|---|---|---|---|---|---|---|
| X | 51060000 | 51063399 | 18.17765 | 31.49525 | 49.54392 | 21.73708 | -0.26446 | 0.836836 | 0.614158 |
| 10 | 69818800 | 69831999 | 325.9659 | 451.4982 | 695.8682 | 288.9702 | -0.46952 | 4.724325 | 0.614168 |
| 17 | 38036200 | 38044399 | 91.95349 | 144.0568 | 147.9988 | 61.38863 | 0.277665 | 2.615838 | 0.614174 |
| 1 | 194982400 | 194983399 | 20.21202 | 35.34572 | 37.66044 | 16.60534 | 0.296644 | 0.707385 | 0.61419 |
| 18 | 82436600 | 82437599 | 9.51517 | 17.75623 | 29.49458 | 12.26117 | -0.27324 | 0.144545 | 0.6142 |
| X | 107144200 | 107146399 | 23.61726 | 41.18309 | 39.69806 | 17.46952 | 0.440283 | 0.769114 | 0.6142 |
| 4 | 8398000 | 8400199 | 42.36775 | 67.78719 | 105.5239 | 44.42712 | -0.2935 | 2.034023 | 0.6142 |
| 13 | 93710000 | 93714999 | 37.76701 | 62.38983 | 61.26931 | 26.46646 | 0.406524 | 1.365232 | 0.614203 |
| 12 | 82134600 | 82136399 | 16.30112 | 28.34954 | 44.73138 | 19.65144 | -0.26798 | 0.775795 | 0.61421 |
| 19 | 41450600 | 41452999 | 17.22028 | 29.83079 | 47.93796 | 21.03235 | -0.29531 | 0.828562 | 0.614219 |
| 1 | 113511600 | 113513999 | 18.71648 | 32.86511 | 35.35156 | 15.64149 | 0.282697 | 0.627714 | 0.61422 |
| 13 | 104156200 | 104158399 | 18.71648 | 32.86511 | 35.35156 | 15.64149 | 0.282697 | 0.627714 | 0.61422 |
| 4 | 102554200 | 102556399 | 16.76708 | 29.25085 | 46.24894 | 20.35795 | -0.2706 | 0.735481 | 0.61423 |
| 13 | 17777200 | 17778799 | 18.63689 | 32.7445 | 35.3806 | 15.66207 | 0.276307 | 0.619151 | 0.614233 |
| 4 | 90243800 | 90245199 | 19.53486 | 34.25816 | 36.49973 | 16.132 | 0.296609 | 0.656656 | 0.614282 |
| 3 | 78252600 | 78255799 | 40.5286 | 65.28853 | 97.13426 | 41.31038 | -0.21647 | 1.816167 | 0.614283 |
| 12 | 105937800 | 105940799 | 32.93947 | 53.45564 | 81.12637 | 34.76815 | -0.23376 | 1.610597 | 0.614327 |
| 15 | 43154000 | 43157399 | 36.40701 | 58.90299 | 88.1813 | 37.70679 | -0.21826 | 1.68266 | 0.614329 |
| 4 | 48849400 | 48850599 | 28.88817 | 48.07959 | 52.91836 | 22.97712 | 0.243348 | 1.250649 | 0.614334 |
| 5 | 80201600 | 80202999 | 16.5734 | 29.8611 | 29.47239 | 12.84639 | 0.424294 | 0.394591 | 0.614337 |
| 10 | 122138800 | 122141999 | 33.99266 | 55.92417 | 61.9079 | 26.68815 | 0.232387 | 1.405244 | 0.614359 |
| 12 | 52017600 | 52018399 | 9.649395 | 17.79923 | 30.36932 | 12.84168 | -0.3305 | 0.213838 | 0.614371 |
| 7 | 63676000 | 63678399 | 24.69331 | 40.82189 | 65.03793 | 28.10834 | -0.28931 | 1.327347 | 0.614372 |
| 12 | 22078600 | 22082799 | 128.0661 | 194.4584 | 201.9337 | 83.61888 | 0.227183 | 3.217154 | 0.61439 |
| 11 | 43530200 | 43534599 | 45.4667 | 73.28467 | 81.12835 | 34.5305 | 0.210082 | 1.798644 | 0.614408 |
| 15 | 37007200 | 37007999 | 6.419077 | 12.33913 | 24.13799 | 9.42554 | -0.42774 | -0.11462 | 0.614421 |
| 16 | 16763400 | 16766199 | 22.74026 | 38.1879 | 59.2787 | 25.88474 | -0.25131 | 1.140924 | 0.614485 |
| 7 | 12464600 | 12469399 | 56.76039 | 90.60315 | 99.23907 | 41.64313 | 0.208984 | 2.11795 | 0.614503 |
| 1 | 49685800 | 49687599 | 17.27384 | 31.05667 | 31.2068 | 13.63386 | 0.396859 | 0.419181 | 0.614509 |
| 6 | 144882200 | 144886999 | 56.32386 | 89.36185 | 130.3053 | 54.26938 | -0.21225 | 2.313699 | 0.614514 |
| 3 | 111309000 | 111309999 | 10.93624 | 20.09993 | 32.95247 | 14.00288 | -0.27854 | 0.25103 | 0.614519 |
| 6 | 16261400 | 16262599 | 13.68512 | 25.69294 | 26.16901 | 10.76767 | 0.438943 | 0.132262 | 0.614529 |
| 6 | 133625000 | 133628199 | 40.2884 | 66.5711 | 60.29015 | 26.04781 | 0.523061 | 1.368831 | 0.614541 |
| 1 | 151068200 | 151069599 | 12.78733 | 22.61919 | 36.70279 | 16.23274 | -0.30365 | 0.511114 | 0.614573 |
| 18 | 90513000 | 90516599 | 87.95169 | 136.8472 | 140.4436 | 58.30223 | 0.271529 | 2.730314 | 0.614574 |
| 3 | 153417200 | 153421599 | 36.79486 | 59.22395 | 103.8907 | 44.13004 | -0.45514 | 1.838994 | 0.614592 |
| 14 | 71021000 | 71025199 | 53.19695 | 84.66051 | 122.8945 | 51.35497 | -0.20131 | 2.217782 | 0.614607 |
| 11 | 91988800 | 91991999 | 53.49853 | 85.53747 | 91.84135 | 38.62904 | 0.2403 | 2.062465 | 0.61461 |
| 6 | 8107400 | 8107999 | 11.5038 | 21.41741 | 23.57173 | 9.837599 | 0.311655 | 0.159772 | 0.614629 |
| 14 | 32538800 | 32541599 | 19.93907 | 33.9373 | 52.72703 | 23.11089 | -0.25139 | 1.007114 | 0.614647 |
| 1 | 30798200 | 30800199 | 20.07925 | 34.11592 | 53.80632 | 23.58062 | -0.27342 | 1.023851 | 0.614654 |
| X | 135114800 | 135116999 | 21.79059 | 37.9708 | 38.81209 | 17.07201 | 0.356019 | 0.769417 | 0.614682 |
| 12 | 94090200 | 94090799 | 5.949335 | 11.57337 | 21.83603 | 8.06308 | -0.2995 | -0.31652 | 0.614685 |
| 16 | 58720000 | 58722199 | 25.71058 | 43.43332 | 46.93155 | 20.53236 | 0.269785 | 1.085071 | 0.614685 |
| X | 8248800 | 8249399 | 12.92116 | 24.03067 | 25.14626 | 10.51362 | 0.382449 | 0.185953 | 0.614696 |
| 7 | 56307400 | 56311199 | 45.08281 | 72.16124 | 107.4644 | 45.27081 | -0.22909 | 2.024049 | 0.614706 |
| 7 | 143202800 | 143208199 | 56.50397 | 89.71435 | 129.5895 | 54.01148 | -0.19759 | 2.292394 | 0.614722 |

| | | | | | | | | | |
|---|---|---|---|---|---|---|---|---|---|
| 19 | 50851800 | 50853599 | 19.03845 | 34.13948 | 31.5132 | 13.79824 | 0.516375 | 0.446747 | 0.614729 |
| 18 | 10029400 | 10029999 | 5.245579 | 10.21536 | 20.74932 | 7.591918 | -0.3925 | -0.36246 | 0.614756 |
| 7 | 105128600 | 105131999 | 21.70641 | 36.34579 | 58.00952 | 25.28406 | -0.29107 | 1.169821 | 0.614775 |
| 9 | 6386000 | 6388599 | 23.93161 | 41.4614 | 42.44416 | 18.62206 | 0.352559 | 0.851112 | 0.614776 |
| 12 | 19249800 | 19251599 | 29.88267 | 51.01153 | 47.35268 | 20.76017 | 0.489496 | 1.012648 | 0.61478 |
| 2 | 3487200 | 3489599 | 24.59654 | 41.74863 | 45.3559 | 19.87154 | 0.262405 | 1.042215 | 0.614807 |
| 8 | 21789800 | 21797199 | 73.37128 | 116.8127 | 112.1551 | 46.89256 | 0.394721 | 2.222204 | 0.614832 |
| 13 | 98215000 | 98220799 | 67.5566 | 106.0612 | 153.6826 | 63.74058 | -0.21545 | 2.56423 | 0.614838 |
| 1 | 94027000 | 94027599 | 9.043532 | 16.93237 | 28.4903 | 11.77711 | -0.28498 | 0.116951 | 0.614855 |
| X | 113991000 | 113993599 | 23.29245 | 40.91572 | 36.11603 | 15.98484 | 0.566745 | 0.65111 | 0.614862 |
| 8 | 84574000 | 84575199 | 17.26511 | 30.7547 | 32.24154 | 14.19639 | 0.328388 | 0.477961 | 0.614862 |
| 9 | 12938000 | 12938599 | 6.982227 | 13.49481 | 24.33097 | 9.329747 | -0.28638 | -0.16744 | 0.614864 |
| 5 | 65329800 | 65333199 | 33.08389 | 54.50031 | 60.6603 | 26.1648 | 0.225357 | 1.387634 | 0.614923 |
| 3 | 48399000 | 48401199 | 23.14843 | 40.3296 | 40.17543 | 17.6697 | 0.392722 | 0.77649 | 0.614949 |
| 3 | 124075800 | 124076599 | 6.986816 | 13.48889 | 24.26607 | 9.352632 | -0.29015 | -0.14987 | 0.614952 |
| 9 | 10540800 | 10541399 | 5.946819 | 11.57652 | 21.90878 | 8.044628 | -0.2965 | -0.33645 | 0.614974 |
| 6 | 100984200 | 100985199 | 17.41044 | 29.90477 | 48.47683 | 21.26326 | -0.31007 | 0.91136 | 0.614981 |
| 12 | 75769800 | 75770999 | 9.860943 | 18.34341 | 30.50758 | 12.75035 | -0.28121 | 0.170688 | 0.614997 |
| X | 32835600 | 32836999 | 20.17318 | 35.31244 | 31.97631 | 14.11077 | 0.530107 | 0.681083 | 0.614998 |
| 15 | 19736800 | 19739999 | 32.69981 | 54.59491 | 55.35475 | 24.08286 | 0.361209 | 1.228615 | 0.615031 |
| 12 | 6255400 | 6256399 | 9.533137 | 17.86038 | 30.12346 | 12.43887 | -0.28727 | 0.121358 | 0.615035 |
| 2 | 116318400 | 116320599 | 26.22949 | 44.32175 | 49.06552 | 21.4677 | 0.235085 | 1.089576 | 0.615043 |
| 9 | 55856600 | 55859799 | 60.72114 | 96.77464 | 106.1667 | 44.45356 | 0.204285 | 2.172834 | 0.615064 |
| 4 | 10008200 | 10010399 | 21.41593 | 37.25361 | 40.36637 | 17.73644 | 0.271833 | 0.791104 | 0.615092 |
| 1 | 47495400 | 47496599 | 11.91529 | 21.56993 | 34.77479 | 15.06839 | -0.27362 | 0.348425 | 0.615093 |
| 15 | 68986800 | 68988599 | 10.02082 | 18.26484 | 31.30078 | 13.4476 | -0.35288 | 0.287544 | 0.615097 |
| 10 | 81864200 | 81870599 | 120.9847 | 185.5477 | 192.695 | 79.99134 | 0.238513 | 3.014103 | 0.615108 |
| 14 | 46439200 | 46440599 | 15.43396 | 27.90726 | 29.74172 | 12.90621 | 0.318534 | 0.36876 | 0.615115 |
| 13 | 11974000 | 11975199 | 14.25511 | 25.05705 | 39.29704 | 17.40203 | -0.25836 | 0.594901 | 0.615131 |
| 10 | 12743200 | 12745399 | 17.08442 | 29.1622 | 50.04665 | 21.94113 | -0.39353 | 0.966781 | 0.615141 |
| 3 | 111701400 | 111701599 | 4.679995 | 9.119708 | 19.66331 | 7.135068 | -0.46578 | -0.4018 | 0.615148 |
| 8 | 124876200 | 124880799 | 33.87763 | 55.04384 | 84.63549 | 36.30401 | -0.25102 | 1.58931 | 0.615163 |
| 7 | 62110000 | 62114399 | 104.311 | 160.5296 | 170.6814 | 70.88663 | 0.208396 | 2.939237 | 0.615174 |
| 3 | 62630800 | 62632599 | 16.91525 | 29.13007 | 53.80478 | 23.60212 | -0.497 | 0.890682 | 0.615211 |
| 6 | 111570800 | 111571599 | 10.4944 | 19.23072 | 31.33797 | 13.37068 | -0.27384 | 0.261864 | 0.615238 |
| 17 | 67586400 | 67587399 | 10.4944 | 19.23072 | 31.33797 | 13.37068 | -0.27384 | 0.261864 | 0.615238 |
| 6 | 69067000 | 69068399 | 14.24773 | 26.48366 | 26.98918 | 11.2939 | 0.418646 | 0.200055 | 0.615238 |
| 12 | 10671600 | 10674599 | 28.74285 | 48.53982 | 50.72512 | 22.18981 | 0.317842 | 1.103129 | 0.615251 |
| 16 | 15103200 | 15104999 | 13.891 | 24.51576 | 38.70856 | 17.12761 | -0.26587 | 0.546204 | 0.615261 |
| 3 | 24136800 | 24137999 | 18.21406 | 32.22027 | 33.72367 | 14.91498 | 0.325513 | 0.539004 | 0.615265 |
| 2 | 89303200 | 89306399 | 33.10391 | 54.94028 | 58.59114 | 25.39059 | 0.288433 | 1.300559 | 0.615276 |
| 14 | 118714000 | 118716799 | 26.25151 | 44.47858 | 48.50888 | 21.24149 | 0.25661 | 1.064013 | 0.615318 |
| 10 | 114673400 | 114674799 | 8.783578 | 16.55884 | 29.74356 | 12.15319 | -0.36584 | 0.069728 | 0.615325 |
| 13 | 26867000 | 26872599 | 79.85586 | 125.2568 | 134.8135 | 55.91248 | 0.211985 | 2.585894 | 0.615374 |
| X | 111387400 | 111391599 | 42.56682 | 69.43481 | 60.30874 | 25.91031 | 0.575094 | 1.53779 | 0.615406 |
| 7 | 86490000 | 86493599 | 71.29462 | 113.0195 | 121.8128 | 50.68999 | 0.221988 | 2.347115 | 0.615412 |
| 5 | 57832800 | 57834599 | 20.451 | 35.82315 | 37.48648 | 16.55187 | 0.322402 | 0.680519 | 0.615419 |

| | | | | | | | | | |
|---|---|---|---|---|---|---|---|---|---|
| 6 | 132256000 | 132259199 | 49.26179 | 79.93024 | 73.14108 | 31.34187 | 0.494128 | 1.640485 | 0.615434 |
| 3 | 90849000 | 90849799 | 15.58608 | 28.05175 | 30.32006 | 13.23381 | 0.29316 | 0.399337 | 0.615436 |
| 1 | 151387600 | 151389799 | 10.69686 | 20.48947 | 19.59919 | 7.731392 | 0.581714 | -0.05021 | 0.615466 |
| 14 | 38396200 | 38398199 | 16.44266 | 29.73624 | 30.18993 | 13.0979 | 0.388349 | 0.376059 | 0.615516 |
| 2 | 54780000 | 54783399 | 30.42572 | 52.02743 | 45.65499 | 20.01863 | 0.570617 | 0.996727 | 0.615531 |
| 8 | 31465400 | 31468199 | 21.47706 | 36.14254 | 56.46491 | 24.67442 | -0.26033 | 1.116374 | 0.615584 |
| 6 | 132471600 | 132476799 | 106.0041 | 164.1276 | 153.0159 | 63.5588 | 0.404477 | 2.825879 | 0.61559 |
| 15 | 5740800 | 5748799 | 121.0254 | 185.4616 | 173.7395 | 72.11465 | 0.386219 | 3.027589 | 0.61565 |
| 6 | 73225200 | 73226599 | 11.59135 | 20.8783 | 34.18414 | 14.90586 | -0.30168 | 0.373253 | 0.615653 |
| 9 | 92629400 | 92632799 | 39.43161 | 64.09262 | 70.638 | 30.30848 | 0.228783 | 1.586814 | 0.615671 |
| X | 17485000 | 17495599 | 134.2512 | 205.0232 | 203.3077 | 84.32112 | 0.300343 | 3.104638 | 0.615699 |
| 13 | 33920400 | 33921599 | 19.60644 | 34.19687 | 37.77867 | 16.62724 | 0.244407 | 0.741408 | 0.615713 |
| X | 85235400 | 85236599 | 11.38328 | 20.62687 | 33.81963 | 14.63671 | -0.2962 | 0.340205 | 0.615725 |
| 2 | 127634000 | 127635599 | 10.54583 | 19.15642 | 31.813 | 13.72599 | -0.31159 | 0.314585 | 0.615728 |
| 4 | 55254200 | 55256599 | 32.72183 | 53.80936 | 59.46862 | 25.63175 | 0.234547 | 1.406798 | 0.615745 |
| 4 | 86945000 | 86947199 | 19.46179 | 33.03476 | 53.24457 | 23.32809 | -0.30464 | 1.029687 | 0.615756 |
| 7 | 6543600 | 6548799 | 77.14103 | 121.8398 | 130.5369 | 54.20839 | 0.226285 | 2.441475 | 0.615756 |
| 12 | 60894400 | 60895799 | 18.44483 | 32.46769 | 34.79689 | 15.40484 | 0.288739 | 0.594249 | 0.615776 |
| 2 | 109291800 | 109294999 | 31.42103 | 53.35369 | 48.47048 | 21.23872 | 0.519423 | 1.064808 | 0.615785 |
| 4 | 119499800 | 119500799 | 12.48957 | 23.44548 | 25.6533 | 10.55767 | 0.336 | 0.121076 | 0.615804 |
| 6 | 78192200 | 78195199 | 52.87161 | 84.15528 | 122.2536 | 51.09767 | -0.20216 | 2.212612 | 0.615815 |
| 1 | 168940400 | 168940999 | 9.759889 | 17.34793 | 38.40318 | 16.9403 | -0.74521 | 0.455636 | 0.615829 |
| 14 | 53749400 | 53751999 | 20.76999 | 36.59017 | 35.08567 | 15.53017 | 0.448911 | 0.599569 | 0.615854 |
| 3 | 104300000 | 104300799 | 9.557935 | 17.594 | 29.87481 | 12.66827 | -0.32613 | 0.226064 | 0.615856 |
| 1 | 48673600 | 48676599 | 32.57398 | 54.34988 | 55.89833 | 24.30949 | 0.340719 | 1.235883 | 0.615872 |
| Y | 6178000 | 6180999 | 30.23092 | 50.25715 | 37.94392 | 16.46229 | 0.785077 | 1.249301 | 0.615879 |
| X | 34995200 | 35000599 | 54.06387 | 87.42366 | 77.0232 | 32.92003 | 0.544176 | 1.722695 | 0.615886 |
| X | 95729800 | 95732199 | 33.19492 | 55.40552 | 55.79492 | 24.26993 | 0.371152 | 1.237873 | 0.615897 |
| 18 | 9246600 | 9250799 | 55.31159 | 87.96242 | 126.9796 | 53.00915 | -0.19444 | 2.243399 | 0.615903 |
| 10 | 54407400 | 54411599 | 121.3388 | 185.4092 | 197.4816 | 81.91553 | 0.197943 | 3.088175 | 0.615922 |
| 16 | 85240200 | 85241199 | 13.40759 | 24.96013 | 26.5148 | 11.06714 | 0.362027 | 0.182391 | 0.615947 |
| 1 | 4513200 | 4516799 | 33.67939 | 56.31585 | 55.3354 | 24.09751 | 0.406361 | 1.217666 | 0.615966 |
| 1 | 14516600 | 14519399 | 35.79005 | 58.90714 | 63.07718 | 27.19729 | 0.280529 | 1.407533 | 0.615983 |
| 16 | 67100600 | 67100999 | 5.088399 | 9.931345 | 19.88651 | 7.055513 | -0.33181 | -0.47733 | 0.615983 |
| 4 | 58911000 | 58912799 | 16.24847 | 27.97489 | 46.91301 | 20.57881 | -0.35809 | 0.881534 | 0.615988 |
| 2 | 12470800 | 12472399 | 28.50944 | 48.51129 | 42.52105 | 18.63708 | 0.571203 | 1.029461 | 0.615994 |
| 1 | 171307000 | 171323199 | 500.376 | 669.8219 | 899.1569 | 375.3654 | -0.31993 | 5.261457 | 0.616003 |
| 2 | 160666600 | 160667599 | 17.38603 | 29.39244 | 58.96038 | 25.80076 | -0.61916 | 1.066705 | 0.616017 |
| 17 | 80330800 | 80333399 | 35.70454 | 58.54044 | 64.69321 | 27.84865 | 0.232931 | 1.453775 | 0.616021 |
| 7 | 142218400 | 142226999 | 140.0084 | 209.7946 | 288.5465 | 119.0304 | -0.19445 | 3.48639 | 0.616061 |
| 1 | 95622800 | 95624199 | 23.18388 | 40.29806 | 36.05735 | 15.83547 | 0.546619 | 0.802747 | 0.616061 |
| 7 | 92333400 | 92334999 | 13.54076 | 24.95631 | 27.51033 | 11.64602 | 0.294351 | 0.243671 | 0.616069 |
| 10 | 54872800 | 54873199 | 5.089645 | 9.930214 | 19.81462 | 7.073624 | -0.33472 | -0.45523 | 0.616098 |
| 7 | 42463600 | 42508999 | 737.1024 | 983.7495 | 831.1786 | 347.1318 | 0.343528 | 5.312786 | 0.616115 |
| 13 | 28488600 | 28490999 | 30.44343 | 49.79878 | 75.29843 | 32.38341 | -0.2198 | 1.479263 | 0.616142 |
| 7 | 13870200 | 13875399 | 50.3864 | 81.37642 | 80.36024 | 34.29161 | 0.377832 | 1.74859 | 0.616153 |
| X | 18383200 | 18384199 | 21.48629 | 37.27875 | 40.49402 | 17.76723 | 0.267957 | 0.827544 | 0.616161 |

| | | | | | | | | | |
|---|---|---|---|---|---|---|---|---|---|
| 4 | 5797800 | 5800999 | 24.02949 | 39.4239 | 76.20935 | 32.81507 | -0.57103 | 1.428075 | 0.616161 |
| 14 | 14739200 | 14744599 | 66.04897 | 105.5962 | 88.11689 | 37.05861 | 0.603208 | 2.079892 | 0.616189 |
| 7 | 42717000 | 42720999 | 73.14514 | 115.9201 | 123.1682 | 51.23948 | 0.242174 | 2.359079 | 0.616191 |
| 1 | 60822400 | 60824199 | 16.15512 | 29.48677 | 29.29957 | 12.56593 | 0.429904 | 0.320745 | 0.616193 |
| 3 | 106654400 | 106655399 | 19.70996 | 34.58244 | 36.4588 | 16.12069 | 0.311732 | 0.649716 | 0.616194 |
| 9 | 10221800 | 10224999 | 25.62971 | 42.80984 | 68.34822 | 29.74418 | -0.29172 | 1.216738 | 0.616195 |
| 13 | 98218400 | 98222599 | 51.92411 | 82.50431 | 124.0787 | 51.75335 | -0.25459 | 2.265607 | 0.6162 |
| 19 | 38473800 | 38480199 | 57.30821 | 90.89965 | 131.7132 | 54.8413 | -0.20352 | 2.322478 | 0.6162 |
| 19 | 39166600 | 39174199 | 88.46406 | 139.3545 | 137.0593 | 56.87095 | 0.346586 | 2.507107 | 0.616214 |
| 9 | 76722800 | 76723199 | 7.201613 | 13.74484 | 25.61125 | 10.19737 | -0.38367 | -0.03423 | 0.616254 |
| X | 89295400 | 89300399 | 61.37114 | 98.89575 | 68.34657 | 29.08062 | 0.88797 | 1.827889 | 0.616258 |
| 9 | 3315400 | 3316199 | 18.64023 | 32.56602 | 34.79606 | 15.33062 | 0.29265 | 0.709101 | 0.61627 |
| 7 | 133715800 | 133719599 | 27.06517 | 44.77596 | 68.8302 | 29.76246 | -0.23789 | 1.326638 | 0.616271 |
| 1 | 116977600 | 116979999 | 25.31483 | 43.46002 | 45.22478 | 19.83633 | 0.326556 | 0.943347 | 0.616294 |
| 11 | 52555400 | 52556599 | 10.85873 | 19.67135 | 32.28383 | 13.97649 | -0.29789 | 0.333327 | 0.616305 |
| 14 | 58121600 | 58123399 | 10.85873 | 19.67135 | 32.28383 | 13.97649 | -0.29789 | 0.333327 | 0.616305 |
| 3 | 131247000 | 131251399 | 60.61761 | 96.06087 | 137.7792 | 57.31444 | -0.19014 | 2.353351 | 0.616339 |
| 4 | 102737400 | 102741399 | 38.23959 | 61.50677 | 94.09453 | 39.92881 | -0.2593 | 1.859255 | 0.616344 |
| 7 | 12699200 | 12700399 | 10.33526 | 18.42237 | 33.89594 | 14.91667 | -0.47751 | 0.432679 | 0.616349 |
| 3 | 14497600 | 14500999 | 33.05179 | 55.2851 | 55.29369 | 24.08448 | 0.380882 | 1.212666 | 0.616419 |
| 3 | 19981400 | 19983799 | 20.88407 | 36.34337 | 39.83941 | 17.50943 | 0.255205 | 0.78159 | 0.616433 |
| 4 | 37628800 | 37630599 | 22.40636 | 38.65503 | 42.59292 | 18.68491 | 0.24572 | 0.883604 | 0.616435 |
| X | 132383200 | 132385999 | 32.1204 | 54.21316 | 43.1741 | 18.88338 | 0.708599 | 1.116031 | 0.61651 |
| 14 | 113182200 | 113190399 | 91.06519 | 142.6492 | 147.9976 | 61.38883 | 0.263398 | 2.616624 | 0.616516 |
| 15 | 75076400 | 75079999 | 61.76902 | 98.48195 | 107.1702 | 44.89722 | 0.216512 | 2.16277 | 0.616521 |
| 9 | 103729000 | 103732999 | 65.53913 | 104.0123 | 114.254 | 47.59799 | 0.196207 | 2.308594 | 0.616526 |
| 10 | 6163600 | 6169399 | 66.49135 | 106.3004 | 101.8215 | 42.82053 | 0.404548 | 2.08391 | 0.616532 |
| 13 | 3762200 | 3763199 | 7.94416 | 15.01163 | 26.51473 | 10.78207 | -0.33474 | 0.041712 | 0.616567 |
| 3 | 49472200 | 49473399 | 12.04584 | 22.69198 | 24.59101 | 10.06054 | 0.356357 | 0.091273 | 0.616577 |
| 5 | 134906800 | 134911799 | 52.30921 | 83.29102 | 122.2721 | 51.13168 | -0.21669 | 2.198836 | 0.616598 |
| 11 | 10200400 | 10201199 | 11.59058 | 21.65472 | 24.63724 | 10.22146 | 0.27109 | 0.142039 | 0.61661 |
| 12 | 63006400 | 63007999 | 49.39712 | 79.21843 | 80.97439 | 34.21033 | 0.316099 | 1.961451 | 0.616615 |
| 5 | 128228600 | 128230999 | 20.75685 | 35.47701 | 55.50679 | 24.33748 | -0.26061 | 0.975534 | 0.616616 |
| 16 | 17814600 | 17819599 | 46.66796 | 74.71714 | 112.165 | 47.268 | -0.24026 | 2.018665 | 0.616639 |
| 8 | 16100200 | 16100799 | 11.22007 | 21.25624 | 24.25632 | 9.810484 | 0.296976 | 0.04684 | 0.616641 |
| 10 | 45192400 | 45193999 | 12.52108 | 23.66531 | 25.25651 | 10.26591 | 0.386555 | 0.077021 | 0.616641 |
| 13 | 3208800 | 3217799 | 103.5301 | 160.303 | 149.2498 | 61.99443 | 0.406451 | 2.823885 | 0.616657 |
| 7 | 45576400 | 45578799 | 19.55971 | 34.11773 | 37.74943 | 16.61509 | 0.242151 | 0.739453 | 0.616666 |
| 7 | 131147400 | 131151799 | 34.23499 | 55.4793 | 84.10987 | 36.00911 | -0.23421 | 1.642625 | 0.616667 |
| 6 | 137399200 | 137400999 | 12.06989 | 21.67563 | 35.17114 | 15.38068 | -0.29163 | 0.396684 | 0.616698 |
| 11 | 73836400 | 73843599 | 86.55573 | 135.5593 | 145.7427 | 60.4547 | 0.211856 | 2.61843 | 0.616707 |
| 13 | 51406200 | 51407399 | 14.58447 | 25.55249 | 40.44735 | 17.86246 | -0.27166 | 0.65153 | 0.616715 |
| 6 | 137711800 | 137713399 | 12.39318 | 21.77189 | 37.94255 | 16.7945 | -0.40949 | 0.591308 | 0.616728 |
| 5 | 126392400 | 126395399 | 34.52766 | 56.72874 | 62.99055 | 27.13958 | 0.227253 | 1.424162 | 0.616731 |
| 5 | 130716400 | 130717599 | 11.42703 | 21.31902 | 23.64576 | 9.833601 | 0.304899 | 0.147344 | 0.616744 |
| 14 | 3114800 | 3125799 | 188.8469 | 281.4789 | 283.0131 | 116.628 | 0.250306 | 3.592651 | 0.616749 |
| 5 | 151453800 | 151455799 | 38.55589 | 62.5529 | 68.61821 | 29.40156 | 0.233055 | 1.620176 | 0.616778 |

| | | | | | | | | | |
|---|---|---|---|---|---|---|---|---|---|
| 11 | 98946600 | 98948799 | 23.29583 | 39.73399 | 43.21714 | 18.9458 | 0.261537 | 0.988912 | 0.61678 |
| 17 | 22532400 | 22534799 | 16.26525 | 28.06137 | 52.18311 | 22.89201 | -0.50612 | 0.869556 | 0.616783 |
| 12 | 62434000 | 62434799 | 10.65054 | 20.15823 | 23.24114 | 9.418676 | 0.280081 | 0.046788 | 0.616793 |
| 8 | 76246000 | 76247999 | 22.99738 | 40.20483 | 38.42309 | 16.93741 | 0.452669 | 0.72492 | 0.616794 |
| 10 | 38536800 | 38538399 | 12.87612 | 23.07102 | 37.74239 | 16.53675 | -0.30538 | 0.42123 | 0.616796 |
| 18 | 4049400 | 4052799 | 46.66881 | 75.35833 | 70.78457 | 30.1982 | 0.44955 | 1.742926 | 0.616807 |
| 2 | 148832200 | 148836599 | 40.955 | 67.30682 | 64.25792 | 27.68876 | 0.44474 | 1.436967 | 0.616821 |
| 7 | 7349400 | 7354199 | 83.2801 | 130.9667 | 139.7538 | 57.97623 | 0.227221 | 2.538231 | 0.616835 |
| X | 59932400 | 59940599 | 106.9612 | 165.9626 | 162.1933 | 67.35913 | 0.338936 | 2.792553 | 0.616849 |
| 5 | 76119800 | 76121599 | 14.56275 | 25.44088 | 41.56127 | 18.31061 | -0.31705 | 0.698591 | 0.61685 |
| 3 | 24676800 | 24677399 | 8.252096 | 15.71591 | 27.21625 | 10.88585 | -0.28549 | -0.00686 | 0.616852 |
| 3 | 86861000 | 86862399 | 20.00495 | 34.99216 | 37.53929 | 16.55533 | 0.286632 | 0.699398 | 0.616858 |
| 5 | 62186200 | 62187399 | 15.98403 | 28.29495 | 30.86125 | 13.64713 | 0.266708 | 0.511405 | 0.616897 |
| 7 | 63351200 | 63355199 | 50.35285 | 80.7486 | 88.52944 | 37.40043 | 0.215122 | 1.965365 | 0.616899 |
| 19 | 17383000 | 17384999 | 23.63724 | 39.72014 | 61.48982 | 26.85842 | -0.24736 | 1.141758 | 0.6169 |
| Y | 6986000 | 6986599 | 3.702853 | 7.229678 | 18.14068 | 6.396479 | -0.64272 | -0.52294 | 0.616912 |
| 4 | 115165000 | 115166399 | 24.8822 | 42.3115 | 46.89209 | 20.55184 | 0.233955 | 1.027748 | 0.616933 |
| 3 | 52411800 | 52417799 | 69.54183 | 110.1193 | 121.2738 | 50.42704 | 0.189561 | 2.374427 | 0.616946 |
| X | 105562400 | 105564399 | 36.53084 | 59.88188 | 63.9672 | 27.53353 | 0.281709 | 1.460908 | 0.616953 |
| 15 | 5173000 | 5173999 | 9.867868 | 18.11112 | 30.34924 | 12.92111 | -0.31149 | 0.245933 | 0.616994 |
| 12 | 20601600 | 20604399 | 23.22835 | 40.41404 | 33.51861 | 14.72938 | 0.655777 | 0.786926 | 0.616994 |
| 18 | 8140400 | 8142999 | 35.47821 | 58.03149 | 63.79879 | 27.43726 | 0.238781 | 1.485113 | 0.617006 |
| 4 | 5998200 | 6002199 | 58.3022 | 93.06824 | 102.6094 | 43.06003 | 0.199614 | 2.119891 | 0.617006 |
| 7 | 81429800 | 81431599 | 35.72074 | 58.28872 | 64.34845 | 27.64955 | 0.230349 | 1.518204 | 0.617034 |
| 13 | 86574800 | 86577399 | 19.03543 | 34.01779 | 31.39127 | 13.78445 | 0.513839 | 0.465681 | 0.617045 |
| 4 | 52580000 | 52580799 | 8.523547 | 16.08694 | 27.5477 | 11.2309 | -0.2941 | 0.05678 | 0.617046 |
| 18 | 34364400 | 34367799 | 23.70424 | 39.83357 | 62.7224 | 27.39698 | -0.2719 | 1.14039 | 0.617047 |
| X | 31981800 | 31983199 | 13.61861 | 25.03746 | 27.52959 | 11.69367 | 0.29466 | 0.259176 | 0.617049 |
| 12 | 24426400 | 24447999 | 614.7291 | 824.2737 | 778.856 | 325.0638 | 0.188945 | 5.236277 | 0.617051 |
| 3 | 90819200 | 90821399 | 32.47267 | 53.92759 | 58.08974 | 25.18456 | 0.274048 | 1.289029 | 0.617074 |
| 18 | 41468200 | 41471599 | 62.67025 | 99.61403 | 107.718 | 44.945 | 0.221016 | 2.260875 | 0.617082 |
| 11 | 15272600 | 15273599 | 14.83532 | 27.4024 | 27.97537 | 11.80753 | 0.407374 | 0.242357 | 0.617102 |
| 18 | 52393600 | 52394399 | 9.833718 | 19.11578 | 20.68003 | 7.681374 | 0.48644 | -0.25503 | 0.61714 |
| 4 | 55332000 | 55334399 | 21.77653 | 36.88756 | 56.53264 | 24.75318 | -0.23237 | 1.060944 | 0.617146 |
| 7 | 106436400 | 106439599 | 30.32296 | 51.04091 | 52.40835 | 22.9034 | 0.343027 | 1.138921 | 0.617153 |
| 1 | 182860200 | 182864799 | 39.34506 | 63.38907 | 94.01809 | 39.99161 | -0.21182 | 1.808765 | 0.6172 |
| 16 | 23006800 | 23007999 | 17.81818 | 31.28683 | 34.49506 | 15.26214 | 0.247383 | 0.623022 | 0.617211 |
| 2 | 20504400 | 20507599 | 53.14494 | 84.69015 | 123.1088 | 51.53913 | -0.20127 | 2.174363 | 0.617228 |
| 15 | 12529800 | 12533799 | 71.02107 | 112.2905 | 123.28 | 51.22174 | 0.192301 | 2.411396 | 0.61724 |
| 9 | 23827000 | 23827799 | 9.467487 | 17.70844 | 29.53492 | 12.22962 | -0.27472 | 0.127874 | 0.617259 |
| 8 | 54136600 | 54137599 | 13.4945 | 25.05458 | 27.04569 | 11.33536 | 0.334645 | 0.201119 | 0.617296 |
| 8 | 75302400 | 75303399 | 13.4945 | 25.05458 | 27.04569 | 11.33536 | 0.334645 | 0.201119 | 0.617296 |
| 6 | 104859800 | 104861399 | 14.588 | 25.7609 | 41.30112 | 18.27263 | -0.28751 | 0.549473 | 0.617305 |
| 18 | 62956200 | 62958199 | 14.2329 | 25.08567 | 39.24015 | 17.36746 | -0.25319 | 0.564403 | 0.617313 |
| 1 | 20389800 | 20394199 | 43.68553 | 71.01635 | 63.64902 | 27.31131 | 0.526485 | 1.58962 | 0.617314 |
| 16 | 73228200 | 73229399 | 15.27443 | 26.66423 | 42.62223 | 18.76922 | -0.2858 | 0.716934 | 0.617315 |
| 9 | 23518600 | 23520399 | 24.79044 | 41.25859 | 63.07681 | 27.38746 | -0.22923 | 1.248233 | 0.617315 |

| | | | | | | | | | |
|---|---|---|---|---|---|---|---|---|---|
| 6 | 13677000 | 13678399 | 14.65036 | 25.00741 | 58.49479 | 25.64491 | -0.83855 | 0.966137 | 0.617319 |
| X | 32420800 | 32423799 | 59.60918 | 95.54894 | 82.42945 | 34.79924 | 0.559525 | 1.991861 | 0.617323 |
| 12 | 36853800 | 36855399 | 21.76272 | 38.1889 | 36.7381 | 16.2459 | 0.443099 | 0.658034 | 0.617376 |
| 6 | 52364400 | 52367999 | 45.01032 | 72.74068 | 78.80638 | 33.65651 | 0.245522 | 1.721498 | 0.617377 |
| 8 | 5822800 | 5827399 | 73.33814 | 115.81 | 127.0291 | 52.74628 | 0.192174 | 2.442822 | 0.617381 |
| 13 | 14911200 | 14911999 | 11.32994 | 21.10934 | 22.59823 | 9.419493 | 0.352812 | 0.1511 | 0.617381 |
| 11 | 59092000 | 59093799 | 9.631883 | 17.45716 | 31.37075 | 13.57309 | -0.42715 | 0.319602 | 0.617405 |
| 5 | 131375800 | 131377799 | 16.91443 | 29.31516 | 46.87193 | 20.56494 | -0.28785 | 0.821927 | 0.617408 |
| 2 | 94633600 | 94635799 | 38.56687 | 62.959 | 54.64561 | 23.48392 | 0.576826 | 1.51206 | 0.617414 |
| 2 | 39405600 | 39405999 | 7.785388 | 15.17383 | 17.98836 | 6.511685 | 0.390883 | -0.37313 | 0.617415 |
| 13 | 38555400 | 38557799 | 21.91564 | 38.28519 | 39.10199 | 17.22683 | 0.357292 | 0.732509 | 0.617424 |
| 3 | 91247800 | 91249399 | 17.92266 | 32.19958 | 31.02692 | 13.56365 | 0.456186 | 0.425751 | 0.617477 |
| 5 | 124041400 | 124043399 | 20.77794 | 35.9351 | 38.96104 | 17.09296 | 0.2696 | 0.847676 | 0.617478 |
| 15 | 33062000 | 33075199 | 186.555 | 277.1862 | 289.3254 | 119.191 | 0.192146 | 3.646688 | 0.617485 |
| 17 | 35598400 | 35599999 | 12.1874 | 21.57211 | 35.76275 | 15.81492 | -0.33388 | 0.499394 | 0.617496 |
| 4 | 86942800 | 86943999 | 10.1957 | 18.69636 | 30.85025 | 13.15013 | -0.29062 | 0.255122 | 0.617506 |
| X | 153052400 | 153055199 | 40.26847 | 66.30962 | 63.55848 | 27.41031 | 0.440163 | 1.412903 | 0.617507 |
| 16 | 9986600 | 9988399 | 24.30942 | 41.2811 | 45.32605 | 19.86136 | 0.247093 | 1.03495 | 0.617561 |
| 1 | 51773800 | 51775199 | 15.00805 | 26.33225 | 41.36656 | 18.29049 | -0.26097 | 0.630516 | 0.617568 |
| 17 | 54291200 | 54293599 | 25.84088 | 43.68891 | 48.52766 | 21.23588 | 0.230252 | 1.082092 | 0.617571 |
| 17 | 30936800 | 30938399 | 17.66147 | 30.52353 | 47.41035 | 20.79938 | -0.24694 | 0.854099 | 0.61758 |
| 7 | 12822600 | 12823399 | 6.704455 | 12.97527 | 23.8008 | 9.069694 | -0.30179 | -0.19093 | 0.617585 |
| 4 | 71352400 | 71353199 | 14.60003 | 25.53407 | 41.03645 | 18.09528 | -0.29341 | 0.681994 | 0.617588 |
| 8 | 51815000 | 51817799 | 29.19907 | 48.79856 | 54.0417 | 23.52632 | 0.234355 | 1.20823 | 0.617592 |
| 1 | 48336000 | 48336999 | 13.87516 | 25.67619 | 27.53783 | 11.59373 | 0.339357 | 0.224076 | 0.617612 |
| 2 | 173210800 | 173216599 | 92.00187 | 140.8274 | 210.1874 | 87.22111 | -0.28637 | 3.043903 | 0.617616 |
| 17 | 20622600 | 20623999 | 31.7036 | 52.94658 | 55.80259 | 24.28201 | 0.305586 | 1.225169 | 0.617635 |
| 6 | 142152200 | 142155799 | 40.23643 | 66.06379 | 64.78195 | 27.90347 | 0.405685 | 1.446151 | 0.617637 |
| 17 | 48864800 | 48866799 | 22.03348 | 38.09806 | 42.03771 | 18.44263 | 0.244344 | 0.860517 | 0.617641 |
| X | 126697000 | 126698399 | 24.14815 | 41.54276 | 44.17101 | 19.37554 | 0.296191 | 0.919046 | 0.617645 |
| 6 | 15451400 | 15453599 | 18.49115 | 32.81797 | 33.07602 | 14.60896 | 0.381734 | 0.508548 | 0.617652 |
| 1 | 133403400 | 133405399 | 19.63515 | 33.5631 | 51.66248 | 22.65209 | -0.237 | 0.966839 | 0.617668 |
| 11 | 85329600 | 85334199 | 71.89467 | 112.6469 | 160.1619 | 66.43287 | -0.19039 | 2.604374 | 0.617674 |
| 12 | 48237000 | 48238799 | 22.06041 | 38.4958 | 34.40866 | 15.14867 | 0.548587 | 0.740529 | 0.617679 |
| 14 | 98975400 | 98976599 | 17.3111 | 29.44706 | 56.91645 | 24.94599 | -0.56527 | 1.008156 | 0.617733 |
| 2 | 136874400 | 136877399 | 18.27637 | 31.10563 | 57.49203 | 25.19934 | -0.50088 | 1.008421 | 0.617754 |
| 4 | 111400200 | 111402199 | 19.72935 | 34.6076 | 30.92779 | 13.67213 | 0.548985 | 0.644223 | 0.617761 |
| 7 | 47161200 | 47162999 | 45.81224 | 73.69803 | 76.22095 | 32.35117 | 0.304602 | 1.858323 | 0.617768 |
| 2 | 172736400 | 172744199 | 75.66027 | 117.0477 | 198.3595 | 82.39368 | -0.45769 | 2.833883 | 0.617802 |
| 6 | 23087200 | 23087799 | 10.16189 | 19.52435 | 17.58369 | 6.875715 | 0.679658 | -0.09025 | 0.617803 |
| 2 | 86139200 | 86140599 | 17.01884 | 30.14273 | 32.75479 | 14.48294 | 0.272328 | 0.516248 | 0.617806 |
| 13 | 10881000 | 10881799 | 9.141771 | 17.11303 | 29.00906 | 11.99545 | -0.29642 | 0.11805 | 0.617811 |
| 13 | 3371800 | 3385199 | 350.7321 | 495.9971 | 429.9498 | 177.9585 | 0.39065 | 4.413873 | 0.617821 |
| 4 | 112223200 | 112237599 | 297.3264 | 425.6949 | 424.2267 | 175.2313 | 0.207433 | 4.231693 | 0.617832 |
| X | 81214200 | 81216799 | 34.00497 | 56.72629 | 56.4068 | 24.52755 | 0.389353 | 1.246167 | 0.617836 |
| 6 | 57417200 | 57418399 | 13.84663 | 26.11495 | 25.70774 | 10.4965 | 0.496612 | 0.104216 | 0.617855 |
| 5 | 141915200 | 141915999 | 8.633627 | 16.10385 | 28.4764 | 11.84703 | -0.36313 | 0.139611 | 0.61788 |

| | | | | | | | | | |
|---|---|---|---|---|---|---|---|---|---|
| 5 | 14945200 | 14948599 | 32.26699 | 52.59297 | 79.25848 | 34.04202 | -0.21868 | 1.536936 | 0.617891 |
| 12 | 54643800 | 54647999 | 42.33042 | 68.01908 | 100.01 | 42.37794 | -0.20402 | 1.898755 | 0.617891 |
| 12 | 17290000 | 17294199 | 47.9372 | 76.57556 | 112.1596 | 47.11398 | -0.20836 | 2.091744 | 0.617895 |
| 5 | 6241200 | 6242599 | 15.4025 | 27.53332 | 30.36621 | 13.33112 | 0.258531 | 0.439965 | 0.617896 |
| 3 | 72254200 | 72256999 | 49.56835 | 79.80898 | 85.81849 | 36.47011 | 0.250025 | 1.839849 | 0.617907 |
| 15 | 20912800 | 20915999 | 26.53551 | 45.48244 | 45.70735 | 20.0454 | 0.376178 | 0.959783 | 0.617915 |
| 5 | 71019600 | 71021399 | 18.9113 | 32.35668 | 51.13756 | 22.42365 | -0.27481 | 0.952454 | 0.61793 |
| 2 | 110166800 | 110167799 | 13.00976 | 23.80088 | 26.47166 | 11.31971 | 0.271143 | 0.277915 | 0.617935 |
| 4 | 20569800 | 20572199 | 19.37084 | 33.42295 | 54.99276 | 24.12465 | -0.33039 | 0.886116 | 0.617969 |
| 2 | 59394800 | 59396199 | 24.26549 | 41.73714 | 38.40692 | 16.84704 | 0.50371 | 0.912388 | 0.617975 |
| 4 | 35317600 | 35318199 | 10.11862 | 19.21781 | 22.23211 | 8.944093 | 0.28435 | 0.014628 | 0.617994 |
| 4 | 143120000 | 143125799 | 65.45441 | 103.0981 | 149.0605 | 61.8499 | -0.20864 | 2.496088 | 0.618002 |
| 6 | 117695800 | 117705999 | 183.7475 | 270.2072 | 363.138 | 149.547 | -0.18618 | 3.821385 | 0.618011 |
| 12 | 51745000 | 51749199 | 37.0468 | 60.54904 | 66.90558 | 28.76502 | 0.230615 | 1.500138 | 0.618027 |
| 14 | 103296600 | 103299199 | 40.85187 | 66.09965 | 73.17573 | 31.29218 | 0.216262 | 1.680476 | 0.618041 |
| 4 | 38921000 | 38922999 | 13.56227 | 24.18058 | 41.97349 | 18.46743 | -0.39439 | 0.468376 | 0.618058 |
| 8 | 3566400 | 3567999 | 13.422 | 23.63652 | 38.26051 | 16.93673 | -0.30276 | 0.566467 | 0.618064 |
| 8 | 46215600 | 46217199 | 14.60769 | 25.60649 | 40.43952 | 17.86686 | -0.2683 | 0.643907 | 0.618066 |
| 19 | 9546800 | 9547999 | 19.67306 | 34.3576 | 37.64948 | 16.58389 | 0.256113 | 0.721876 | 0.618086 |
| 2 | 40025200 | 40028199 | 26.02603 | 43.44726 | 72.94219 | 31.73209 | -0.36404 | 1.228568 | 0.618093 |
| 10 | 83622000 | 83624799 | 22.90296 | 38.10358 | 61.17787 | 26.55637 | -0.29959 | 1.246787 | 0.618105 |
| 12 | 87057400 | 87058599 | 13.34298 | 23.68005 | 37.71304 | 16.65761 | -0.27456 | 0.48771 | 0.618107 |
| 6 | 25495600 | 25496799 | 10.4169 | 19.34482 | 32.65779 | 13.68338 | -0.30593 | 0.187399 | 0.618109 |
| 5 | 150139200 | 150141999 | 22.88199 | 38.6383 | 60.47786 | 26.45536 | -0.26299 | 1.094318 | 0.618141 |
| X | 99786200 | 99788999 | 31.87311 | 54.06954 | 49.59278 | 21.72418 | 0.50575 | 1.078451 | 0.618154 |
| 10 | 70194600 | 70196799 | 17.35579 | 30.67801 | 32.65668 | 14.44534 | 0.300619 | 0.537695 | 0.618155 |
| 2 | 36281000 | 36286599 | 74.49853 | 118.2458 | 120.4882 | 50.20139 | 0.304611 | 2.314067 | 0.618161 |
| 7 | 10282800 | 10288199 | 48.17601 | 78.74497 | 66.24736 | 28.48288 | 0.623092 | 1.525244 | 0.618163 |
| 2 | 79962000 | 79962399 | 7.778289 | 15.09777 | 17.28748 | 6.497875 | 0.390058 | -0.24867 | 0.618171 |
| 12 | 98904600 | 98906199 | 14.09317 | 24.97343 | 40.2349 | 17.78859 | -0.29256 | 0.50677 | 0.618181 |
| 14 | 39943400 | 39944599 | 15.19739 | 27.48635 | 29.82839 | 12.93979 | 0.292992 | 0.366988 | 0.618189 |
| 7 | 45424400 | 45425799 | 15.40853 | 26.83017 | 43.1886 | 18.98833 | -0.29628 | 0.753702 | 0.61819 |
| 18 | 36413600 | 36415599 | 15.40853 | 26.83017 | 43.1886 | 18.98833 | -0.29628 | 0.753702 | 0.61819 |
| 14 | 48207800 | 48211999 | 47.70999 | 76.35036 | 111.7962 | 47.08159 | -0.20509 | 2.035092 | 0.618193 |
| 16 | 43896600 | 43901599 | 54.25491 | 86.86451 | 95.57117 | 40.27538 | 0.207283 | 2.024368 | 0.618209 |
| Y | 5296600 | 5297399 | 8.590854 | 16.5967 | 15.75269 | 6.052598 | 0.628197 | -0.17153 | 0.618253 |
| 7 | 11999600 | 12008999 | 100.3211 | 157.4295 | 143.4347 | 59.48677 | 0.452606 | 2.591674 | 0.618266 |
| 10 | 117278000 | 117279999 | 28.92155 | 48.61774 | 52.3577 | 22.8721 | 0.274462 | 1.14673 | 0.618273 |
| 2 | 82032400 | 82034399 | 24.45489 | 40.71337 | 62.5135 | 27.14859 | -0.23545 | 1.242932 | 0.618279 |
| 7 | 85138200 | 85140799 | 25.26071 | 44.27197 | 36.60548 | 16.17023 | 0.660705 | 0.697393 | 0.618313 |
| 6 | 72951400 | 72954199 | 23.1035 | 39.69589 | 43.63047 | 19.1377 | 0.248036 | 0.922164 | 0.618317 |
| 7 | 29534600 | 29537799 | 33.28553 | 53.97135 | 81.54282 | 34.92474 | -0.22845 | 1.629123 | 0.61832 |
| 1 | 112844000 | 112850799 | 65.43831 | 104.4588 | 105.8419 | 44.41603 | 0.321225 | 2.128778 | 0.618361 |
| 2 | 88269000 | 88269999 | 16.59102 | 30.33283 | 24.20742 | 10.36043 | 0.746392 | 0.305947 | 0.618361 |
| 8 | 16057800 | 16059399 | 15.62515 | 27.36698 | 43.49753 | 19.20426 | -0.27785 | 0.664623 | 0.618374 |
| 17 | 10075000 | 10078199 | 36.39507 | 60.30524 | 59.50675 | 25.75872 | 0.399918 | 1.325811 | 0.618379 |
| 5 | 19257600 | 19261199 | 41.46518 | 67.38137 | 71.96044 | 30.87028 | 0.273784 | 1.597323 | 0.618379 |

| | | | | | | | | | |
|---|---|---|---|---|---|---|---|---|---|
| 16 | 68795200 | 68799799 | 37.25253 | 61.37408 | 62.89433 | 27.12964 | 0.344213 | 1.396371 | 0.618386 |
| 13 | 6344400 | 6345599 | 7.452993 | 14.30261 | 25.20111 | 9.888099 | -0.28475 | -0.08358 | 0.618406 |
| X | 25625800 | 25629799 | 41.96395 | 69.99224 | 52.71348 | 22.9193 | 0.789028 | 1.269048 | 0.618414 |
| 5 | 99798600 | 99803399 | 43.31636 | 69.47837 | 102.3705 | 43.2609 | -0.21015 | 1.956644 | 0.618422 |
| 2 | 171310000 | 171312799 | 30.59782 | 51.43847 | 52.94584 | 23.12942 | 0.339498 | 1.152465 | 0.618446 |
| 9 | 88026000 | 88028799 | 20.08415 | 34.41951 | 53.32107 | 23.38419 | -0.24573 | 0.946064 | 0.618451 |
| 7 | 63013200 | 63013599 | 6.090973 | 11.86267 | 22.59328 | 8.257747 | -0.29962 | -0.34885 | 0.618454 |
| 10 | 119097400 | 119101199 | 41.61946 | 66.92463 | 99.26914 | 42.10775 | -0.2154 | 1.873705 | 0.618462 |
| 4 | 128299400 | 128302399 | 22.65835 | 38.23581 | 59.25558 | 25.91558 | -0.24861 | 1.100529 | 0.618472 |
| X | 125921400 | 125933599 | 258.3398 | 374.0222 | 371.3277 | 153.1344 | 0.228817 | 4.066633 | 0.618494 |
| 4 | 144013400 | 144025799 | 152.96 | 230.5766 | 242.3605 | 100.1333 | 0.20086 | 3.370715 | 0.618532 |
| 4 | 122806400 | 122809199 | 30.05459 | 49.91554 | 55.6771 | 24.1398 | 0.223919 | 1.280398 | 0.618547 |
| 12 | 59684200 | 59686399 | 48.41207 | 78.05371 | 73.3782 | 31.24296 | 0.445849 | 1.7924 | 0.618582 |
| X | 33418000 | 33423799 | 151.8639 | 231.7143 | 203.145 | 84.2363 | 0.476915 | 3.127863 | 0.618601 |
| 5 | 99937600 | 99939399 | 15.55027 | 27.25355 | 42.45775 | 18.7555 | -0.2489 | 0.654489 | 0.618608 |
| 8 | 27702000 | 27704999 | 23.46896 | 40.80835 | 36.08894 | 15.85295 | 0.56357 | 0.800411 | 0.618611 |
| 7 | 13434000 | 13445199 | 149.149 | 226.3804 | 223.1789 | 92.40525 | 0.301629 | 3.232779 | 0.618635 |
| 9 | 89389000 | 89391599 | 23.9998 | 39.94477 | 61.87044 | 26.86461 | -0.24794 | 1.244922 | 0.618637 |
| 9 | 19806200 | 19818799 | 229.5717 | 335.1861 | 344.8267 | 142.0737 | 0.189315 | 3.936893 | 0.618642 |
| 11 | 22689600 | 22698199 | 99.49346 | 152.363 | 213.7289 | 88.69993 | -0.19648 | 3.037723 | 0.618643 |
| 6 | 19798400 | 19799399 | 9.671917 | 18.07844 | 30.06819 | 12.46487 | -0.27191 | 0.13663 | 0.618648 |
| 12 | 67722400 | 67724399 | 21.76208 | 37.89412 | 40.31236 | 17.72399 | 0.298374 | 0.778644 | 0.618652 |
| 12 | 24306200 | 24315199 | 157.0283 | 237.0732 | 215.4451 | 89.05504 | 0.412523 | 3.339161 | 0.618656 |
| 4 | 49618800 | 49622199 | 42.34999 | 68.13671 | 101.8506 | 43.23711 | -0.22553 | 1.860178 | 0.61866 |
| 10 | 95328400 | 95331799 | 41.98852 | 68.01763 | 74.76068 | 31.9973 | 0.228125 | 1.659852 | 0.618671 |
| 13 | 58563400 | 58566599 | 29.6023 | 48.6078 | 73.77148 | 31.77666 | -0.22265 | 1.427939 | 0.618679 |
| 12 | 62366800 | 62369799 | 31.27614 | 52.51075 | 53.51868 | 23.36633 | 0.353675 | 1.167384 | 0.618681 |
| 17 | 68749200 | 68750399 | 13.65247 | 24.78487 | 27.40639 | 11.83352 | 0.270383 | 0.328572 | 0.61869 |
| 17 | 18602800 | 18618999 | 404.8214 | 561.4591 | 542.8004 | 225.3595 | 0.205058 | 4.705748 | 0.618695 |
| 9 | 122766600 | 122770599 | 62.77714 | 100.4236 | 100.7513 | 42.40695 | 0.338799 | 2.060488 | 0.618695 |
| 11 | 37523600 | 37525199 | 11.49544 | 20.91419 | 33.84239 | 14.57295 | -0.27279 | 0.316583 | 0.618698 |
| 4 | 132857200 | 132858599 | 9.128587 | 16.74142 | 29.90125 | 12.74367 | -0.40365 | 0.244402 | 0.618707 |
| 12 | 8668600 | 8672399 | 57.87562 | 92.65539 | 87.62447 | 36.92284 | 0.425101 | 2.026513 | 0.618716 |
| 2 | 33655400 | 33657199 | 16.7391 | 29.57235 | 32.42425 | 14.34404 | 0.258199 | 0.542226 | 0.618717 |
| 15 | 53715000 | 53719199 | 72.92276 | 115.2759 | 126.5958 | 52.59453 | 0.191658 | 2.41864 | 0.618734 |
| 1 | 28971600 | 28974199 | 18.40108 | 32.36429 | 34.79163 | 15.40502 | 0.283888 | 0.602241 | 0.618749 |
| 5 | 151516200 | 151518199 | 25.83867 | 43.09521 | 68.83792 | 29.92941 | -0.29238 | 1.234392 | 0.61875 |
| 7 | 18368000 | 18369199 | 24.71444 | 40.9121 | 64.46002 | 27.8846 | -0.2731 | 1.312654 | 0.618782 |
| 13 | 51210600 | 51213199 | 26.77873 | 44.10426 | 69.12604 | 29.81368 | -0.26719 | 1.379155 | 0.618801 |
| 4 | 115111200 | 115113799 | 35.76101 | 57.8859 | 86.30774 | 36.91885 | -0.21175 | 1.67122 | 0.618805 |
| 5 | 101123000 | 101124199 | 18.73565 | 32.83744 | 35.99761 | 15.90224 | 0.255578 | 0.658915 | 0.618822 |
| 7 | 122651000 | 122652399 | 18.73565 | 32.83744 | 35.99761 | 15.90224 | 0.255578 | 0.658915 | 0.618822 |
| 19 | 39467000 | 39478999 | 189.634 | 282.2919 | 257.6293 | 106.1537 | 0.388257 | 3.613346 | 0.618835 |
| 15 | 13143800 | 13145799 | 21.78355 | 38.1036 | 38.52727 | 16.99119 | 0.371748 | 0.711432 | 0.61884 |
| 1 | 11815200 | 11818999 | 51.64264 | 83.01286 | 88.82156 | 37.65694 | 0.254253 | 1.89371 | 0.618873 |
| X | 96270200 | 96270799 | 11.82628 | 22.37765 | 24.79253 | 10.053 | 0.336353 | 0.063517 | 0.618874 |
| 14 | 77988200 | 77991599 | 41.92699 | 68.94421 | 63.22585 | 27.25073 | 0.502942 | 1.431431 | 0.618874 |

| | | | | | | | | | |
|---|---|---|---|---|---|---|---|---|---|
| 5 | 14886000 | 14888399 | 28.92734 | 48.67718 | 45.36533 | 19.82368 | 0.482344 | 1.13017 | 0.618909 |
| 8 | 116930400 | 116933799 | 37.87449 | 61.14652 | 90.8403 | 38.74524 | -0.21067 | 1.744444 | 0.61891 |
| 6 | 59491600 | 59493799 | 20.12083 | 35.69255 | 32.88935 | 14.53261 | 0.50992 | 0.52112 | 0.618944 |
| Y | 10153600 | 10154999 | 12.25626 | 22.6425 | 19.16868 | 8.088605 | 0.675779 | 0.197572 | 0.618947 |
| 5 | 15038200 | 15097199 | 1380.826 | 1726.416 | 2104.469 | 885.0643 | -0.27965 | 6.369264 | 0.618947 |
| 4 | 37301200 | 37303199 | 38.71455 | 62.90348 | 70.02004 | 30.03502 | 0.213959 | 1.591431 | 0.618967 |
| 3 | 4596400 | 4598599 | 26.26966 | 44.57762 | 48.49591 | 21.24569 | 0.260276 | 1.0513 | 0.618982 |
| 10 | 67483800 | 67486199 | 19.56634 | 34.10791 | 37.78475 | 16.62409 | 0.240327 | 0.748487 | 0.618994 |
| X | 24499600 | 24503999 | 51.27409 | 83.74214 | 64.6377 | 27.78274 | 0.746252 | 1.543009 | 0.619033 |
| 6 | 128662600 | 128670799 | 120.2357 | 182.1984 | 251.0127 | 103.8782 | -0.18274 | 3.25968 | 0.619059 |
| 14 | 53803400 | 53804399 | 30.88673 | 51.50808 | 44.9834 | 19.55635 | 0.575951 | 1.227843 | 0.619063 |
| 4 | 83901600 | 83904599 | 21.07703 | 35.40928 | 64.64049 | 28.23252 | -0.48436 | 1.138463 | 0.619072 |
| 17 | 38287800 | 38290199 | 21.16366 | 37.10051 | 37.41156 | 16.52971 | 0.37559 | 0.672351 | 0.619073 |
| 10 | 35381000 | 35382399 | 15.22141 | 28.02417 | 28.46022 | 12.06549 | 0.410411 | 0.264792 | 0.619074 |
| 8 | 28596000 | 28598999 | 30.84006 | 51.06721 | 56.81198 | 24.58107 | 0.227364 | 1.316855 | 0.619086 |
| X | 107635400 | 107636999 | 18.74873 | 34.07085 | 29.59967 | 12.7645 | 0.617567 | 0.365291 | 0.619089 |
| 3 | 147069000 | 147070399 | 24.38493 | 41.97161 | 38.85919 | 17.04578 | 0.495088 | 0.906649 | 0.61909 |
| 17 | 55743800 | 55744399 | 9.392011 | 18.17183 | 21.53622 | 8.21669 | 0.320359 | -0.16605 | 0.619095 |
| 1 | 145984800 | 145989199 | 45.99578 | 73.84689 | 116.1394 | 49.15117 | -0.30249 | 1.935504 | 0.619098 |
| 7 | 13265800 | 13267599 | 46.25142 | 74.46481 | 79.86172 | 33.93626 | 0.253778 | 1.836681 | 0.619104 |
| 11 | 15232400 | 15233399 | 30.22959 | 50.01982 | 53.44844 | 23.1141 | 0.285323 | 1.322148 | 0.619111 |
| 13 | 81318000 | 81319999 | 21.95504 | 38.02575 | 40.3502 | 17.70319 | 0.300781 | 0.840148 | 0.619121 |
| 7 | 42006200 | 42012199 | 105.4143 | 164.0837 | 160.164 | 66.48475 | 0.344219 | 2.735941 | 0.619122 |
| 1 | 87442600 | 87443799 | 11.95438 | 21.51563 | 34.19347 | 14.92428 | -0.2597 | 0.3819 | 0.619135 |
| 12 | 104836400 | 104838399 | 11.59234 | 20.96513 | 33.72403 | 14.63066 | -0.27147 | 0.349417 | 0.619135 |
| Y | 5823800 | 5825599 | 13.88 | 25.13079 | 20.66304 | 8.951701 | 0.691949 | 0.328262 | 0.619141 |
| Y | 82718600 | 82720799 | 13.88 | 25.13079 | 20.66304 | 8.951701 | 0.691949 | 0.328262 | 0.619141 |
| 5 | 93260400 | 93261199 | 8.512023 | 16.09943 | 27.59829 | 11.20097 | -0.28997 | 0.041747 | 0.619165 |
| 1 | 49737400 | 49738599 | 18.21157 | 31.93333 | 34.97679 | 15.45719 | 0.256763 | 0.646748 | 0.619166 |
| 2 | 122731200 | 122734599 | 46.97201 | 76.13193 | 66.24452 | 28.36382 | 0.565102 | 1.653055 | 0.619175 |
| 8 | 64075800 | 64084399 | 170.3788 | 254.2647 | 267.7878 | 110.3796 | 0.185035 | 3.565633 | 0.619218 |
| 8 | 72643600 | 72644799 | 26.94596 | 46.72867 | 42.36219 | 18.58664 | 0.527549 | 0.857934 | 0.61922 |
| 2 | 36840400 | 36844999 | 59.88505 | 96.0914 | 93.86061 | 39.68809 | 0.382426 | 1.962625 | 0.619229 |
| 8 | 96723400 | 96726599 | 50.59908 | 80.57277 | 119.1595 | 49.8341 | -0.22721 | 2.196353 | 0.619249 |
| 17 | 9886000 | 9888799 | 34.64332 | 55.99013 | 85.00679 | 36.30746 | -0.24007 | 1.706157 | 0.619264 |
| 7 | 43733600 | 43748799 | 219.6383 | 322.3936 | 330.6649 | 136.1896 | 0.200998 | 3.85226 | 0.619268 |
| 13 | 82447800 | 82449399 | 13.16519 | 24.34283 | 27.01714 | 11.38695 | 0.28908 | 0.22107 | 0.61927 |
| 7 | 55308200 | 55309999 | 16.5428 | 29.7335 | 30.59749 | 13.371 | 0.362159 | 0.412218 | 0.619308 |
| 17 | 40922400 | 40924399 | 34.3591 | 57.26033 | 49.8045 | 21.64314 | 0.581781 | 1.249102 | 0.619321 |
| X | 8440400 | 8442199 | 36.02225 | 59.30646 | 62.46614 | 26.93712 | 0.304443 | 1.405272 | 0.619328 |
| 1 | 147045000 | 147048399 | 46.85386 | 75.68857 | 80.15241 | 34.21305 | 0.277648 | 1.737508 | 0.619368 |
| 7 | 53192400 | 53193199 | 8.370952 | 15.87229 | 27.11026 | 10.94481 | -0.27786 | 0.022663 | 0.619373 |
| 11 | 40532600 | 40537199 | 38.85561 | 63.5933 | 67.24491 | 28.9255 | 0.295312 | 1.484792 | 0.619388 |
| 11 | 70951200 | 70967199 | 142.1038 | 215.7202 | 199.4956 | 82.6032 | 0.393985 | 3.225614 | 0.619399 |
| 9 | 25744800 | 25746199 | 14.5119 | 26.62974 | 28.4374 | 12.11015 | 0.333683 | 0.277064 | 0.619402 |
| 10 | 3120400 | 3122799 | 16.63075 | 29.01914 | 45.18261 | 19.89121 | -0.24842 | 0.731089 | 0.619418 |
| 4 | 107370200 | 107373399 | 27.75125 | 45.45052 | 71.88518 | 30.93025 | -0.28336 | 1.457307 | 0.61944 |

| | | | | | | | | | |
|---|---|---|---|---|---|---|---|---|---|
| X | 17697200 | 17698199 | 17.88043 | 31.76453 | 32.28461 | 14.24789 | 0.370735 | 0.500241 | 0.619463 |
| 7 | 79664000 | 79666199 | 12.15458 | 21.60036 | 35.19881 | 15.52926 | -0.30634 | 0.464619 | 0.61948 |
| 1 | 99000600 | 99004199 | 37.85175 | 62.17968 | 65.03042 | 28.01646 | 0.313305 | 1.435875 | 0.619518 |
| 8 | 116343000 | 116345399 | 16.77189 | 29.01997 | 46.87745 | 20.56632 | -0.30303 | 0.836325 | 0.619542 |
| 11 | 13604200 | 13604999 | 7.561629 | 14.5065 | 26.34832 | 10.34762 | -0.33013 | -0.07934 | 0.619593 |
| 6 | 68510000 | 68511199 | 18.84098 | 33.08244 | 35.31442 | 15.62454 | 0.293628 | 0.628446 | 0.6196 |
| X | 76247200 | 76249199 | 25.55814 | 43.94927 | 45.18858 | 19.82163 | 0.344191 | 0.928723 | 0.619619 |
| 11 | 11074200 | 11076999 | 47.90271 | 77.22258 | 72.10785 | 30.697 | 0.455367 | 1.79463 | 0.619624 |
| 6 | 133003000 | 133003599 | 9.614817 | 18.42708 | 21.80568 | 8.588099 | 0.279232 | -0.06082 | 0.61963 |
| 8 | 109896000 | 109896399 | 9.614817 | 18.42708 | 21.80568 | 8.588099 | 0.279232 | -0.06082 | 0.61963 |
| 15 | 58267000 | 58269399 | 22.31307 | 37.59415 | 57.57973 | 25.17067 | -0.23176 | 1.110945 | 0.619642 |
| 15 | 35737600 | 35740399 | 33.78266 | 56.27564 | 50.33443 | 21.86753 | 0.541668 | 1.253369 | 0.619666 |
| 15 | 51596800 | 51598599 | 20.56111 | 35.13114 | 54.4087 | 23.85539 | -0.24605 | 0.975891 | 0.619674 |
| X | 113176600 | 113177799 | 9.25171 | 17.77639 | 20.33863 | 7.951947 | 0.33806 | -0.08971 | 0.619691 |
| 5 | 24161200 | 24162199 | 14.15408 | 25.43326 | 26.95879 | 11.78357 | 0.319706 | 0.391045 | 0.619696 |
| 3 | 137142000 | 137145399 | 30.29804 | 51.74167 | 47.35173 | 20.76043 | 0.510073 | 1.011432 | 0.619722 |
| 12 | 87568200 | 87586199 | 281.4503 | 406.9071 | 390.2455 | 160.9661 | 0.276681 | 4.088672 | 0.619736 |
| 1 | 127994000 | 127994799 | 13.13582 | 24.18047 | 26.96211 | 11.4329 | 0.276089 | 0.245024 | 0.619739 |
| 11 | 10819800 | 10820999 | 13.18618 | 24.1101 | 26.96474 | 11.53919 | 0.262439 | 0.28366 | 0.619742 |
| 13 | 79309800 | 79311999 | 26.34515 | 44.25776 | 48.5703 | 21.21339 | 0.247283 | 1.138847 | 0.619745 |
| 2 | 163492000 | 163498999 | 92.25594 | 142.1276 | 198.9575 | 82.64212 | -0.18687 | 2.916757 | 0.6198 |
| 12 | 78382200 | 78386199 | 61.6175 | 98.686 | 97.96815 | 41.30693 | 0.355923 | 2.022309 | 0.619835 |
| 2 | 22303200 | 22305599 | 22.17439 | 38.6808 | 39.65018 | 17.45235 | 0.35183 | 0.753926 | 0.619838 |
| 3 | 37968000 | 37969599 | 14.80814 | 25.83295 | 48.39653 | 21.30305 | -0.51402 | 0.731477 | 0.61985 |
| 5 | 93789600 | 93792399 | 60.96361 | 97.24894 | 105.9762 | 44.42931 | 0.215314 | 2.145198 | 0.619856 |
| 3 | 144596200 | 144597799 | 17.89759 | 31.42698 | 34.41459 | 15.22684 | 0.257045 | 0.621688 | 0.619857 |
| 3 | 62599400 | 62606799 | 177.2448 | 265.0103 | 243.2748 | 100.3123 | 0.385591 | 3.531052 | 0.61987 |
| 15 | 19432400 | 19434599 | 22.40597 | 39.10267 | 39.64388 | 17.4546 | 0.367741 | 0.749465 | 0.619888 |
| 7 | 55494600 | 55496999 | 16.8592 | 29.4567 | 47.3033 | 20.84214 | -0.29261 | 0.717652 | 0.619891 |
| 11 | 24473400 | 24473999 | 6.860062 | 13.27338 | 24.36432 | 9.294374 | -0.30428 | -0.18367 | 0.619898 |
| 13 | 22352200 | 22359399 | 125.1908 | 191.1638 | 203.1174 | 84.23986 | 0.200658 | 3.10291 | 0.619898 |
| 9 | 105582600 | 105584599 | 32.79429 | 53.27151 | 79.99355 | 34.30712 | -0.21716 | 1.5912 | 0.619946 |
| 2 | 57443000 | 57443599 | 10.29319 | 18.62147 | 32.37069 | 14.03797 | -0.38159 | 0.339532 | 0.619947 |
| 12 | 88282000 | 88285399 | 32.07052 | 52.31645 | 79.42957 | 34.1259 | -0.22854 | 1.523798 | 0.61996 |
| 2 | 139615400 | 139617199 | 16.57459 | 29.21789 | 31.39048 | 13.89277 | 0.286014 | 0.560758 | 0.619994 |
| 11 | 6441800 | 6443199 | 16.57459 | 29.21789 | 31.39048 | 13.89277 | 0.286014 | 0.560758 | 0.619994 |
| 18 | 45122600 | 45126599 | 40.91994 | 67.26268 | 64.71999 | 27.89023 | 0.433636 | 1.436406 | 0.620008 |
| 2 | 36638000 | 36645799 | 101.9544 | 158.557 | 161.906 | 67.21679 | 0.278254 | 2.749488 | 0.620019 |
| 2 | 19661000 | 19664399 | 22.82036 | 37.99826 | 61.8935 | 26.88207 | -0.32038 | 1.234371 | 0.620022 |
| 12 | 32472200 | 32474199 | 31.41343 | 52.60957 | 48.0812 | 20.95923 | 0.510458 | 1.187406 | 0.620026 |
| X | 9583200 | 9585199 | 21.52378 | 37.70154 | 37.02876 | 16.34949 | 0.413636 | 0.688791 | 0.620034 |
| 4 | 30183400 | 30185199 | 19.39746 | 34.38855 | 32.92133 | 14.55379 | 0.454565 | 0.51993 | 0.620041 |
| 5 | 102096200 | 102097799 | 15.21706 | 26.66335 | 41.43722 | 18.30109 | -0.24536 | 0.654738 | 0.620044 |
| 15 | 47231200 | 47232599 | 13.46512 | 24.65749 | 27.41686 | 11.70867 | 0.272758 | 0.277631 | 0.620062 |
| X | 92571000 | 92573999 | 22.93137 | 38.56076 | 62.46088 | 27.28832 | -0.31234 | 1.137 | 0.620066 |
| 1 | 79468000 | 79473799 | 61.14781 | 98.06583 | 94.70896 | 40.01467 | 0.397857 | 1.979885 | 0.620097 |
| 12 | 116142400 | 116144399 | 18.33764 | 32.15315 | 34.93886 | 15.43993 | 0.268127 | 0.647468 | 0.620097 |

| | | | | | | | | | |
|---|---|---|---|---|---|---|---|---|---|
| 7 | 42362400 | 42371399 | 140.0641 | 213.8583 | 205.3074 | 85.14661 | 0.34683 | 3.10974 | 0.620124 |
| 5 | 72720200 | 72730599 | 177.5039 | 261.6414 | 353.2921 | 145.4898 | -0.1898 | 3.781913 | 0.620124 |
| 2 | 148004200 | 148008199 | 39.11324 | 64.38065 | 63.99435 | 27.59345 | 0.387622 | 1.41549 | 0.620125 |
| 8 | 17387400 | 17391199 | 64.25126 | 102.0877 | 112.7414 | 47.02284 | 0.190141 | 2.273585 | 0.620138 |
| 11 | 93429600 | 93431599 | 15.81849 | 27.55525 | 43.65874 | 19.19994 | -0.27366 | 0.748585 | 0.620145 |
| 18 | 52279000 | 52282599 | 36.98727 | 60.65627 | 65.47154 | 28.1862 | 0.26691 | 1.454125 | 0.620157 |
| 6 | 93570400 | 93572999 | 25.81406 | 44.38406 | 45.17021 | 19.8135 | 0.358845 | 0.929902 | 0.620165 |
| 4 | 55125200 | 55128399 | 23.44693 | 39.13881 | 60.86356 | 26.47626 | -0.25375 | 1.209387 | 0.620167 |
| 19 | 27469200 | 27471599 | 41.12542 | 66.6137 | 73.14941 | 31.30568 | 0.229399 | 1.661533 | 0.62017 |
| 10 | 96403800 | 96405399 | 15.79818 | 28.32165 | 30.72726 | 13.45709 | 0.284265 | 0.424185 | 0.620177 |
| 19 | 51427800 | 51429399 | 15.48924 | 27.13409 | 42.45226 | 18.74578 | -0.25512 | 0.660043 | 0.620214 |
| 8 | 48492400 | 48496199 | 24.61545 | 40.7801 | 65.82974 | 28.49249 | -0.30804 | 1.300588 | 0.620215 |
| 5 | 30428000 | 30428799 | 13.84869 | 24.31602 | 39.4231 | 17.44204 | -0.30582 | 0.612633 | 0.62026 |
| 5 | 80990400 | 80993799 | 20.72317 | 36.43976 | 35.62966 | 15.77684 | 0.419866 | 0.619621 | 0.62026 |
| 4 | 66598600 | 66602199 | 39.75393 | 64.51616 | 71.63095 | 30.69933 | 0.216095 | 1.618772 | 0.620277 |
| 15 | 60079800 | 60083599 | 41.38798 | 67.00602 | 74.43529 | 31.84444 | 0.212085 | 1.671175 | 0.620283 |
| 13 | 87518200 | 87520399 | 23.07418 | 40.05755 | 36.68603 | 16.09899 | 0.513099 | 0.821128 | 0.620298 |
| 6 | 10249400 | 10251599 | 20.53515 | 36.14307 | 35.71322 | 15.81086 | 0.405395 | 0.610199 | 0.620337 |
| 2 | 39556800 | 39558399 | 19.5766 | 33.65685 | 52.24545 | 22.91693 | -0.24784 | 0.913794 | 0.620352 |
| 4 | 52012200 | 52013599 | 16.00657 | 28.57921 | 31.26194 | 13.7381 | 0.269211 | 0.452191 | 0.620355 |
| 12 | 82898600 | 82900399 | 26.18671 | 43.52661 | 68.14431 | 29.56157 | -0.26357 | 1.27287 | 0.620358 |
| 15 | 53888000 | 53889599 | 23.34977 | 39.06098 | 60.2509 | 26.24552 | -0.24202 | 1.187835 | 0.620359 |
| 16 | 37474400 | 37474999 | 4.104745 | 8.013673 | 18.14096 | 6.405978 | -0.49851 | -0.51178 | 0.620375 |
| 17 | 50479400 | 50480399 | 18.46505 | 32.26864 | 33.8194 | 14.90312 | 0.320398 | 0.702659 | 0.620385 |
| 15 | 22456200 | 22463999 | 107.9767 | 167.2896 | 169.4905 | 70.40165 | 0.285323 | 2.816698 | 0.620407 |
| 17 | 64562600 | 64565399 | 20.63643 | 36.17156 | 36.82624 | 16.26932 | 0.361668 | 0.668411 | 0.620433 |
| X | 146547600 | 146551799 | 44.49235 | 72.33156 | 63.76654 | 27.36224 | 0.550281 | 1.590282 | 0.620436 |
| 16 | 93070800 | 93072399 | 13.55979 | 23.94116 | 38.19639 | 16.89959 | -0.28056 | 0.53831 | 0.62044 |
| 2 | 59379600 | 59381799 | 5.387327 | 10.47063 | 26.56251 | 9.880069 | -0.73293 | -0.28559 | 0.620448 |
| 17 | 30958000 | 30959799 | 13.93451 | 24.6574 | 38.69739 | 17.11366 | -0.25562 | 0.518895 | 0.620453 |
| 11 | 8515200 | 8517599 | 18.80438 | 31.77618 | 59.59274 | 26.07397 | -0.52256 | 1.075745 | 0.62046 |
| 12 | 39191200 | 39193399 | 20.8323 | 35.99279 | 39.0135 | 17.11545 | 0.269738 | 0.857839 | 0.620471 |
| 19 | 27720600 | 27722599 | 16.34166 | 28.4705 | 44.17844 | 19.43017 | -0.24366 | 0.751207 | 0.620471 |
| 13 | 33100400 | 33102199 | 21.82185 | 38.05016 | 39.67923 | 17.46062 | 0.327102 | 0.757695 | 0.620471 |
| 1 | 144128000 | 144130799 | 56.30222 | 89.84089 | 97.47976 | 40.88496 | 0.222047 | 2.128045 | 0.620476 |
| 13 | 97182000 | 97185199 | 29.40015 | 48.92872 | 54.56414 | 23.69072 | 0.224466 | 1.254399 | 0.62049 |
| 14 | 53823000 | 53824199 | 12.31334 | 22.91855 | 25.63569 | 10.70375 | 0.288205 | 0.17538 | 0.620496 |
| X | 142746800 | 142747999 | 12.31334 | 22.91855 | 25.63569 | 10.70375 | 0.288205 | 0.17538 | 0.620496 |
| 1 | 30565000 | 30567199 | 20.83044 | 36.34401 | 32.50212 | 14.30795 | 0.547786 | 0.732835 | 0.620496 |
| 16 | 86937000 | 86937799 | 12.91883 | 22.83602 | 37.83549 | 16.73602 | -0.33358 | 0.526101 | 0.620514 |
| 15 | 64479600 | 64480799 | 11.40515 | 20.8013 | 33.85516 | 14.53343 | -0.27848 | 0.29937 | 0.620551 |
| 1 | 111428600 | 111431199 | 20.83353 | 36.44391 | 38.00152 | 16.76012 | 0.327429 | 0.703413 | 0.620557 |
| 2 | 20544600 | 20547599 | 22.04483 | 37.30268 | 57.67702 | 25.24618 | -0.24523 | 1.070525 | 0.620567 |
| 3 | 132797000 | 132797999 | 11.84632 | 21.02825 | 35.21792 | 15.55282 | -0.3466 | 0.469579 | 0.620573 |
| 3 | 45845400 | 45848799 | 26.04826 | 44.34597 | 47.94361 | 21.01466 | 0.269816 | 1.022798 | 0.620573 |
| 7 | 68304000 | 68305199 | 17.4433 | 32.08172 | 27.67894 | 11.7508 | 0.642578 | 0.284805 | 0.620587 |
| 19 | 7026800 | 7029599 | 26.71721 | 44.78855 | 48.59353 | 21.19569 | 0.263827 | 1.163094 | 0.620588 |

| | | | | | | | | | |
|---|---|---|---|---|---|---|---|---|---|
| 3 | 12920000 | 12922599 | 22.60344 | 40.32658 | 31.24876 | 13.74124 | 0.763462 | 0.501648 | 0.620596 |
| 8 | 60790400 | 60793199 | 25.7504 | 44.32403 | 44.6071 | 19.56716 | 0.375214 | 0.918464 | 0.6206 |
| 16 | 94082600 | 94088999 | 63.84513 | 100.5792 | 146.208 | 60.66756 | -0.21625 | 2.492038 | 0.620614 |
| 2 | 46274600 | 46279799 | 113.8328 | 174.7737 | 186.8905 | 77.59363 | 0.197597 | 2.990123 | 0.620636 |
| 9 | 18412800 | 18414599 | 15.12061 | 27.65216 | 28.8488 | 12.34256 | 0.36222 | 0.300799 | 0.620686 |
| 17 | 21419200 | 21425199 | 65.15842 | 102.6962 | 146.9852 | 60.9938 | -0.19336 | 2.48398 | 0.620695 |
| 1 | 101970800 | 101972599 | 20.28527 | 35.74921 | 35.12931 | 15.54836 | 0.413953 | 0.593079 | 0.620738 |
| 15 | 58077000 | 58079599 | 15.90787 | 27.06775 | 57.40269 | 25.15968 | -0.69844 | 1.000281 | 0.620757 |
| 16 | 38223000 | 38224799 | 13.59697 | 24.07313 | 37.66966 | 16.65735 | -0.25108 | 0.508753 | 0.620763 |
| 13 | 104151000 | 104154199 | 27.10159 | 44.68773 | 68.63159 | 29.61494 | -0.23729 | 1.365871 | 0.620772 |
| 7 | 141750800 | 141752399 | 14.41192 | 26.33801 | 28.33805 | 12.13453 | 0.31714 | 0.297682 | 0.62078 |
| X | 32536200 | 32538999 | 67.67862 | 108.0822 | 104.003 | 43.67132 | 0.396295 | 2.120035 | 0.620803 |
| 19 | 49336000 | 49336999 | 10.4709 | 19.40023 | 32.04733 | 13.46988 | -0.27783 | 0.200947 | 0.620836 |
| 10 | 56596400 | 56599199 | 33.42535 | 55.3399 | 51.9834 | 22.48956 | 0.470685 | 1.322123 | 0.62084 |
| 1 | 47911800 | 47913399 | 22.27455 | 38.73799 | 35.65189 | 15.66262 | 0.50629 | 0.79135 | 0.620852 |
| X | 81244000 | 81248199 | 36.5562 | 60.95533 | 56.51697 | 24.56863 | 0.489994 | 1.258783 | 0.620855 |
| 15 | 92711200 | 92712399 | 21.85407 | 36.5915 | 58.03145 | 25.29295 | -0.28195 | 1.170309 | 0.620874 |
| 13 | 69421400 | 69423399 | 20.26224 | 35.54097 | 36.8814 | 16.30349 | 0.33434 | 0.656612 | 0.620883 |
| 4 | 58405400 | 58408199 | 29.93485 | 49.15852 | 74.98737 | 32.30169 | -0.22992 | 1.428442 | 0.6209 |
| 11 | 91960000 | 91962399 | 31.97315 | 53.75729 | 45.83465 | 20.02365 | 0.610168 | 1.146846 | 0.620906 |
| 6 | 132212800 | 132219599 | 85.51398 | 135.3533 | 126.2876 | 52.5005 | 0.42797 | 2.398086 | 0.620914 |
| 13 | 100004600 | 100005599 | 17.80966 | 30.66823 | 47.945 | 21.03161 | -0.25738 | 0.887398 | 0.620925 |
| 7 | 99167800 | 99170799 | 21.0311 | 36.52833 | 40.42553 | 17.74539 | 0.24126 | 0.810168 | 0.620942 |
| 5 | 144465800 | 144467199 | 19.62536 | 33.49516 | 51.65907 | 22.6478 | -0.24017 | 0.980331 | 0.62096 |
| 4 | 91864800 | 91866599 | 15.98296 | 28.44243 | 31.25976 | 13.77566 | 0.259647 | 0.472241 | 0.620967 |
| 10 | 40185600 | 40188599 | 20.56728 | 35.03578 | 53.83389 | 23.59763 | -0.23533 | 1.002723 | 0.620972 |
| 18 | 30307200 | 30307999 | 9.338156 | 17.47274 | 29.00567 | 12.00325 | -0.26711 | 0.124223 | 0.621003 |
| 6 | 62122200 | 62122799 | 10.17828 | 18.62661 | 30.82132 | 13.1741 | -0.29762 | 0.265546 | 0.621017 |
| 18 | 18058600 | 18058999 | 6.3894 | 12.40906 | 22.8667 | 8.55631 | -0.28367 | -0.26118 | 0.621038 |
| 3 | 152037800 | 152039599 | 12.36048 | 21.79125 | 36.83702 | 16.30299 | -0.36413 | 0.546521 | 0.62106 |
| 8 | 19059200 | 19059999 | 18.78518 | 33.11771 | 34.73047 | 15.37071 | 0.320777 | 0.579608 | 0.621063 |
| 19 | 33650000 | 33650999 | 6.993514 | 13.48024 | 24.20927 | 9.379279 | -0.29469 | -0.13255 | 0.621064 |
| 15 | 71519600 | 71522999 | 43.17022 | 69.18584 | 102.9992 | 43.47103 | -0.22647 | 1.983061 | 0.621069 |
| 1 | 38258200 | 38267799 | 94.74864 | 147.302 | 159.0569 | 66.03721 | 0.197351 | 2.749801 | 0.621109 |
| 1 | 38811200 | 38814999 | 33.83624 | 55.90279 | 60.90417 | 26.31082 | 0.257205 | 1.353894 | 0.621136 |
| 4 | 91891200 | 91893999 | 33.83624 | 55.90279 | 60.90417 | 26.31082 | 0.257205 | 1.353894 | 0.621136 |
| 11 | 99070400 | 99071999 | 15.13897 | 26.49679 | 42.48085 | 18.74424 | -0.29025 | 0.671647 | 0.621179 |
| 12 | 56654800 | 56662599 | 52.02561 | 82.26598 | 155.2718 | 64.51763 | -0.58819 | 2.40104 | 0.621186 |
| 4 | 36700600 | 36702199 | 14.48111 | 26.84483 | 26.88588 | 11.29211 | 0.439614 | 0.215592 | 0.621186 |
| 15 | 49593200 | 49601399 | 123.703 | 188.9325 | 201.7821 | 83.69022 | 0.193484 | 3.09771 | 0.621198 |
| 2 | 94272400 | 94274599 | 21.83484 | 38.02189 | 40.27016 | 17.70575 | 0.304645 | 0.777374 | 0.621212 |
| 14 | 58672400 | 58673999 | 16.72977 | 29.06072 | 45.81485 | 20.11252 | -0.26679 | 0.798918 | 0.621215 |
| X | 8757800 | 8760399 | 37.34103 | 61.97728 | 57.58405 | 24.95553 | 0.486822 | 1.313745 | 0.621219 |
| 6 | 37721600 | 37726199 | 66.92672 | 106.1674 | 117.2098 | 48.80774 | 0.188342 | 2.323965 | 0.621237 |
| 18 | 7843000 | 7846599 | 64.59153 | 102.6837 | 113.2542 | 47.26222 | 0.192739 | 2.258493 | 0.621242 |
| 16 | 93941800 | 93942399 | 5.668018 | 11.00516 | 21.67567 | 8.127803 | -0.38067 | -0.26425 | 0.621253 |
| 9 | 120544000 | 120548799 | 39.31056 | 63.3517 | 94.92934 | 40.39606 | -0.22606 | 1.799167 | 0.621262 |

| | | | | | | | | | |
|---|---|---|---|---|---|---|---|---|---|
| X | 9784200 | 9786999 | 21.93006 | 38.6672 | 35.0522 | 15.51268 | 0.530134 | 0.598559 | 0.621273 |
| X | 32013000 | 32015799 | 32.64701 | 54.21211 | 47.6554 | 20.65948 | 0.566436 | 1.28382 | 0.621278 |
| X | 12269200 | 12271199 | 21.77334 | 38.10751 | 38.04821 | 16.78801 | 0.389966 | 0.703476 | 0.621305 |
| 8 | 113130800 | 113131399 | 6.977558 | 13.50085 | 24.40588 | 9.309814 | -0.28306 | -0.18526 | 0.621332 |
| 12 | 24125600 | 24160399 | 915.1561 | 1188.723 | 1091.23 | 457.1721 | 0.184867 | 5.749279 | 0.621337 |
| 4 | 143796800 | 143797999 | 13.3902 | 25.27772 | 25.15088 | 10.24887 | 0.483553 | 0.091475 | 0.621344 |
| 7 | 108277200 | 108278399 | 23.55271 | 40.19541 | 43.69314 | 19.15537 | 0.262443 | 0.984083 | 0.621353 |
| 18 | 73984800 | 73986799 | 17.02814 | 30.35877 | 31.56987 | 13.89053 | 0.340427 | 0.46688 | 0.621354 |
| 4 | 72413800 | 72418399 | 85.76278 | 134.1773 | 145.8649 | 60.51188 | 0.194702 | 2.634291 | 0.621388 |
| 17 | 48579200 | 48580599 | 11.83861 | 21.16093 | 34.22499 | 15.02614 | -0.29058 | 0.425016 | 0.621427 |
| 10 | 129259800 | 129262599 | 22.27298 | 38.92522 | 39.07046 | 17.21785 | 0.382305 | 0.728637 | 0.621437 |
| 3 | 94949600 | 94950199 | 4.272393 | 8.330333 | 19.14835 | 6.894852 | -0.54656 | -0.43706 | 0.621445 |
| 4 | 147523400 | 147597199 | 2961.88 | 3471.637 | 3815.47 | 1618.76 | -0.22502 | 7.391785 | 0.621454 |
| 16 | 47904000 | 47906199 | 30.3581 | 51.13405 | 45.34237 | 19.81943 | 0.55396 | 1.128211 | 0.621454 |
| X | 89114000 | 89115799 | 24.49351 | 42.44632 | 41.4779 | 18.19826 | 0.419713 | 0.854743 | 0.621465 |
| 13 | 103555000 | 103556599 | 14.13389 | 25.78557 | 28.37812 | 12.17782 | 0.282818 | 0.305994 | 0.621485 |
| 15 | 19283800 | 19293799 | 168.1093 | 251.7403 | 264.2591 | 108.9978 | 0.194262 | 3.501784 | 0.621488 |
| 7 | 106026000 | 106027399 | 17.3603 | 29.90654 | 47.42332 | 20.80303 | -0.27759 | 0.882085 | 0.621497 |
| 3 | 109247200 | 109249599 | 16.16288 | 28.11256 | 44.21576 | 19.42641 | -0.2633 | 0.773334 | 0.621506 |
| 19 | 11570200 | 11571999 | 16.16288 | 28.11256 | 44.21576 | 19.42641 | -0.2633 | 0.773334 | 0.621506 |
| 2 | 151055200 | 151057999 | 35.24104 | 57.11014 | 85.36319 | 36.54591 | -0.21378 | 1.645914 | 0.621516 |
| 7 | 130604800 | 130611199 | 71.5153 | 113.4615 | 121.0562 | 50.41616 | 0.237727 | 2.324188 | 0.621527 |
| 16 | 48373600 | 48375599 | 17.02319 | 30.61964 | 31.17561 | 13.61512 | 0.378229 | 0.413736 | 0.621537 |
| 8 | 23023400 | 23025599 | 30.14299 | 49.0924 | 76.58457 | 32.88413 | -0.26918 | 1.541189 | 0.621572 |
| 6 | 96897800 | 96899599 | 17.88888 | 30.63649 | 56.50826 | 24.78059 | -0.49658 | 0.947504 | 0.621581 |
| 19 | 22843600 | 22845599 | 16.77859 | 29.76453 | 32.17404 | 14.21621 | 0.280817 | 0.498166 | 0.62159 |
| 6 | 130252800 | 130255999 | 25.95195 | 45.13242 | 41.23137 | 18.11356 | 0.516958 | 0.817421 | 0.621608 |
| 19 | 3261800 | 3263599 | 17.66847 | 31.30359 | 33.17882 | 14.66947 | 0.308251 | 0.517359 | 0.621624 |
| 2 | 149526800 | 149529599 | 29.70542 | 50.02912 | 45.85895 | 20.04456 | 0.506174 | 1.125336 | 0.621629 |
| 16 | 46288800 | 46294999 | 104.6994 | 163.4724 | 153.8008 | 63.81739 | 0.400696 | 2.682704 | 0.621636 |
| 2 | 88197400 | 88200999 | 28.3208 | 47.21879 | 51.85119 | 22.53901 | 0.24659 | 1.227899 | 0.621653 |
| 15 | 76841600 | 76847399 | 61.70974 | 98.53136 | 104.6198 | 43.91861 | 0.25421 | 2.117946 | 0.621657 |
| 3 | 100216600 | 100222399 | 108.0991 | 166.3599 | 179.9648 | 74.74222 | 0.18348 | 2.941252 | 0.621658 |
| 16 | 37623200 | 37626199 | 32.78113 | 54.05829 | 60.08708 | 25.92906 | 0.227912 | 1.374171 | 0.621676 |
| 10 | 23897000 | 23899999 | 44.27371 | 71.44798 | 79.02793 | 33.68892 | 0.21329 | 1.760258 | 0.621684 |
| 5 | 4651200 | 4651799 | 7.490089 | 14.32044 | 25.64248 | 10.1693 | -0.3219 | -0.04522 | 0.621697 |
| X | 19229200 | 19230599 | 18.62795 | 32.97102 | 33.72629 | 14.91446 | 0.358847 | 0.536479 | 0.621708 |
| 11 | 23058400 | 23062399 | 79.10496 | 125.1153 | 129.0011 | 53.60822 | 0.283163 | 2.411732 | 0.62174 |
| 15 | 61642600 | 61643999 | 9.951042 | 18.51177 | 30.4965 | 12.74482 | -0.26783 | 0.169184 | 0.621743 |
| 8 | 68863400 | 68864399 | 9.229039 | 17.19649 | 28.95554 | 12.06767 | -0.29541 | 0.151363 | 0.621761 |
| 18 | 82794000 | 82795999 | 21.08643 | 35.86647 | 54.91256 | 24.06756 | -0.2305 | 1.020052 | 0.621767 |
| 1 | 184482600 | 184484199 | 14.2991 | 25.76676 | 28.33497 | 12.34858 | 0.269668 | 0.377512 | 0.621779 |
| 7 | 111059000 | 111060599 | 17.69188 | 31.25131 | 33.79775 | 14.95206 | 0.277734 | 0.551229 | 0.621786 |
| 11 | 21827000 | 21829999 | 32.21854 | 53.68616 | 56.9327 | 24.73878 | 0.296768 | 1.251356 | 0.621789 |
| 12 | 114262400 | 114264399 | 29.99032 | 50.57744 | 51.84341 | 22.66963 | 0.345586 | 1.121111 | 0.621792 |
| 17 | 65567800 | 65568399 | 9.786811 | 18.95276 | 21.57617 | 8.19736 | 0.383191 | -0.1772 | 0.621793 |
| 3 | 23517400 | 23519999 | 30.00988 | 51.22578 | 47.3486 | 20.75828 | 0.495616 | 1.013204 | 0.621793 |

| | | | | | | | | | |
|---|---|---|---|---|---|---|---|---|---|
| 3 | 101585800 | 101588399 | 26.1023 | 44.409 | 47.93975 | 21.01187 | 0.271771 | 1.027321 | 0.621806 |
| 2 | 18612400 | 18614799 | 17.51761 | 29.60485 | 61.78283 | 27.03307 | -0.67627 | 1.069864 | 0.621814 |
| X | 24082400 | 24087199 | 56.614 | 91.01231 | 89.04899 | 37.75832 | 0.383257 | 1.902092 | 0.621832 |
| 7 | 84750000 | 84758599 | 118.6947 | 181.9624 | 192.9288 | 80.08401 | 0.208199 | 3.021299 | 0.621833 |
| 1 | 14152600 | 14155999 | 54.66547 | 86.87327 | 126.0871 | 52.59899 | -0.20321 | 2.263385 | 0.621835 |
| Y | 41496400 | 41499599 | 22.89236 | 39.25855 | 30.64414 | 13.43985 | 0.739199 | 0.918927 | 0.62184 |
| X | 40399800 | 40400399 | 7.882535 | 15.31709 | 18.4108 | 6.85937 | 0.333119 | -0.27137 | 0.621841 |
| 17 | 39575800 | 39579599 | 47.16283 | 76.46371 | 75.30912 | 32.25199 | 0.387252 | 1.656202 | 0.621843 |
| 8 | 100103800 | 100104799 | 10.1826 | 18.84008 | 30.95102 | 13.03352 | -0.27166 | 0.204929 | 0.621847 |
| 13 | 120748000 | 120752599 | 70.25007 | 110.1102 | 157.6738 | 65.3984 | -0.20023 | 2.596854 | 0.621886 |
| 3 | 45051000 | 45052399 | 44.31699 | 71.4515 | 79.20959 | 33.72598 | 0.208539 | 1.78801 | 0.621891 |
| X | 153350600 | 153353599 | 40.99989 | 67.61684 | 62.47449 | 26.96018 | 0.493873 | 1.396525 | 0.621903 |
| 17 | 72153600 | 72154599 | 10.83688 | 19.9618 | 33.01141 | 13.98541 | -0.28761 | 0.239451 | 0.621951 |
| 11 | 85329800 | 85334399 | 72.90262 | 114.1237 | 161.8342 | 67.13272 | -0.1876 | 2.620872 | 0.621952 |
| 2 | 17055200 | 17057799 | 27.71368 | 47.36654 | 46.8065 | 20.52273 | 0.399778 | 0.99111 | 0.621955 |
| 10 | 90272600 | 90274599 | 20.55278 | 37.00656 | 29.78419 | 12.993 | 0.716567 | 0.429193 | 0.62196 |
| X | 43292400 | 43295599 | 35.63404 | 58.57313 | 63.55396 | 27.38759 | 0.260773 | 1.423645 | 0.621964 |
| 9 | 81495200 | 81496999 | 28.40103 | 47.81101 | 42.265 | 18.47118 | 0.55833 | 1.121821 | 0.621967 |
| 17 | 68544000 | 68544799 | 8.09068 | 15.31705 | 26.54096 | 10.75023 | -0.30233 | 0.030313 | 0.621971 |
| 4 | 81731200 | 81733599 | 22.36315 | 39.25846 | 36.09344 | 15.9663 | 0.508112 | 0.653331 | 0.621976 |
| 13 | 21391600 | 21397799 | 121.045 | 185.4306 | 195.3757 | 81.08957 | 0.216439 | 3.036242 | 0.621982 |
| 3 | 106980000 | 106981399 | 10.53142 | 19.09436 | 31.79031 | 13.74845 | -0.31752 | 0.324509 | 0.622008 |
| 7 | 40984000 | 40989799 | 119.2874 | 182.0992 | 195.3665 | 81.02014 | 0.186404 | 3.105812 | 0.622019 |
| 5 | 77691000 | 77692399 | 15.4422 | 27.70463 | 30.22971 | 13.23066 | 0.276685 | 0.415706 | 0.622045 |
| 6 | 20305800 | 20306799 | 15.4422 | 27.70463 | 30.22971 | 13.23066 | 0.276685 | 0.415706 | 0.622045 |
| 12 | 21460400 | 21464599 | 36.85955 | 60.39221 | 65.93661 | 28.37557 | 0.249948 | 1.46568 | 0.622047 |
| 10 | 61847000 | 61850799 | 24.46839 | 40.64941 | 63.01167 | 27.32521 | -0.24934 | 1.267582 | 0.62206 |
| 6 | 80182200 | 80182999 | 12.83415 | 23.49288 | 25.98219 | 11.10188 | 0.280018 | 0.272158 | 0.622083 |
| 17 | 39416600 | 39428599 | 265.3649 | 384.264 | 341.5758 | 140.8614 | 0.388465 | 4.062797 | 0.622087 |
| 5 | 109849600 | 109853399 | 52.90646 | 84.88208 | 92.78778 | 39.22638 | 0.220131 | 1.957168 | 0.622107 |
| 13 | 78988600 | 78990399 | 18.84438 | 33.48449 | 33.05869 | 14.58675 | 0.412338 | 0.503698 | 0.622116 |
| 1 | 31113600 | 31115199 | 13.73299 | 25.02093 | 27.96259 | 12.01945 | 0.259466 | 0.310955 | 0.622133 |
| 8 | 35519400 | 35521599 | 24.21348 | 41.36822 | 45.80602 | 20.08343 | 0.23623 | 0.981566 | 0.622137 |
| X | 91291600 | 91292799 | 24.21348 | 41.36822 | 45.80602 | 20.08343 | 0.23623 | 0.981566 | 0.622137 |
| 5 | 66405000 | 66406599 | 11.94095 | 21.53422 | 34.20825 | 14.89366 | -0.25679 | 0.370082 | 0.622151 |
| 10 | 8561600 | 8563199 | 16.97002 | 29.89296 | 32.91101 | 14.56787 | 0.250564 | 0.574759 | 0.622169 |
| 6 | 12493600 | 12494599 | 11.50307 | 21.58974 | 24.62255 | 10.13655 | 0.276284 | 0.110935 | 0.622171 |
| 13 | 66720600 | 66761999 | 995.6469 | 1290.175 | 1129.852 | 473.4648 | 0.249374 | 5.789225 | 0.622183 |
| 14 | 44695000 | 44695999 | 18.42194 | 32.33293 | 35.43264 | 15.67123 | 0.256027 | 0.634028 | 0.622199 |
| 3 | 91446800 | 91449199 | 29.24427 | 49.10998 | 52.91818 | 23.1073 | 0.273603 | 1.157504 | 0.62223 |
| X | 153334600 | 153340599 | 111.8022 | 173.4511 | 166.448 | 69.12748 | 0.36507 | 2.795543 | 0.622243 |
| 7 | 78064800 | 78066599 | 16.47424 | 28.33272 | 53.22217 | 23.3456 | -0.52151 | 0.901508 | 0.622252 |
| 10 | 68244200 | 68253399 | 135.4635 | 203.6086 | 279.129 | 115.2347 | -0.18581 | 3.425844 | 0.622255 |
| 11 | 7651000 | 7651599 | 5.951803 | 11.57028 | 21.77368 | 8.084446 | -0.30301 | -0.29691 | 0.622264 |
| 13 | 33981000 | 33983999 | 38.62042 | 62.40733 | 97.75369 | 41.74272 | -0.28538 | 1.724335 | 0.622267 |
| 1 | 30766800 | 30770399 | 50.77194 | 81.55136 | 89.74515 | 38.0066 | 0.212461 | 1.91809 | 0.622272 |
| 1 | 127440400 | 127445199 | 57.58665 | 91.53821 | 133.4806 | 55.69709 | -0.20964 | 2.259758 | 0.622281 |

| | | | | | | | | | |
|---|---|---|---|---|---|---|---|---|---|
| 1 | 147079800 | 147082799 | 61.78595 | 98.67889 | 92.41372 | 38.80898 | 0.435444 | 2.106269 | 0.622289 |
| 4 | 7094600 | 7099399 | 51.80479 | 83.16392 | 90.71671 | 38.38544 | 0.224178 | 1.935258 | 0.622295 |
| 7 | 108086800 | 108089599 | 25.15311 | 43.76933 | 40.66197 | 17.87012 | 0.492873 | 0.803485 | 0.622308 |
| 10 | 81963000 | 81973799 | 275.7287 | 398.4049 | 390.5731 | 161.1135 | 0.244199 | 4.097132 | 0.622314 |
| 16 | 62033800 | 62034799 | 16.69391 | 29.66323 | 32.24494 | 14.22772 | 0.274239 | 0.48866 | 0.622322 |
| 10 | 6198200 | 6200799 | 28.28721 | 47.72379 | 43.73026 | 19.12424 | 0.506714 | 1.102338 | 0.622328 |
| 15 | 94538000 | 94540999 | 16.40431 | 28.61071 | 45.71718 | 20.12066 | -0.28585 | 0.735488 | 0.622332 |
| 3 | 134761000 | 134762799 | 30.29778 | 49.46117 | 75.04715 | 32.25174 | -0.22681 | 1.507582 | 0.622339 |
| 16 | 7842600 | 7845599 | 24.44284 | 41.02857 | 64.94625 | 28.35149 | -0.27948 | 1.157737 | 0.622351 |
| 5 | 21390400 | 21391799 | 10.56246 | 19.4338 | 32.42838 | 13.76005 | -0.3025 | 0.239335 | 0.622389 |
| 19 | 20716000 | 20716399 | 9.999056 | 19.0015 | 21.26007 | 8.542798 | 0.333965 | 0.00795 | 0.622392 |
| 7 | 55078000 | 55078599 | 7.766578 | 14.74672 | 26.05543 | 10.48823 | -0.32243 | 0.004629 | 0.622421 |
| X | 126557200 | 126559799 | 24.53438 | 42.78621 | 40.13676 | 17.66358 | 0.479297 | 0.77183 | 0.622438 |
| 7 | 83595800 | 83599399 | 22.09409 | 37.24315 | 57.06693 | 24.95014 | -0.2324 | 1.104085 | 0.622438 |
| 1 | 100769200 | 100770799 | 17.15667 | 29.8662 | 46.29174 | 20.34926 | -0.24223 | 0.769553 | 0.622438 |
| 5 | 60273600 | 60275999 | 28.44942 | 47.85971 | 51.8283 | 22.64595 | 0.266446 | 1.136261 | 0.622438 |
| 13 | 67632000 | 67633599 | 17.40309 | 30.50948 | 31.67308 | 13.99478 | 0.333852 | 0.640703 | 0.622439 |
| 16 | 26627200 | 26627799 | 7.326968 | 13.92438 | 25.54418 | 10.26312 | -0.3736 | -0.0069 | 0.622459 |
| 16 | 8920600 | 8927999 | 104.2194 | 160.6876 | 174.331 | 72.41409 | 0.181364 | 2.904061 | 0.622468 |
| Y | 40968200 | 40968799 | 4.558081 | 8.873848 | 20.1488 | 7.393367 | -0.55427 | -0.36182 | 0.622472 |
| 18 | 75164200 | 75165799 | 13.10785 | 23.06938 | 38.28361 | 16.949 | -0.33882 | 0.570951 | 0.622483 |
| 14 | 100987200 | 100988599 | 8.878413 | 16.48628 | 28.90219 | 12.11129 | -0.35969 | 0.166917 | 0.622489 |
| 14 | 83622000 | 83622999 | 11.91045 | 22.14309 | 25.10244 | 10.50161 | 0.266609 | 0.177011 | 0.622497 |
| 3 | 138964400 | 138968599 | 60.74255 | 97.10848 | 89.70715 | 37.72517 | 0.45666 | 2.071907 | 0.622501 |
| 1 | 97739200 | 97740399 | 11.39705 | 20.70296 | 33.7781 | 14.57358 | -0.28676 | 0.321827 | 0.622518 |
| 3 | 29051000 | 29052199 | 14.70522 | 25.80844 | 40.34452 | 17.84299 | -0.25382 | 0.623056 | 0.622518 |
| 7 | 107486000 | 107488799 | 27.35029 | 47.18232 | 44.01485 | 19.30867 | 0.484914 | 0.906238 | 0.622521 |
| 6 | 89572000 | 89572799 | 7.486962 | 14.25386 | 25.57181 | 10.2355 | -0.3366 | -0.01606 | 0.622527 |
| 8 | 31200400 | 31202399 | 17.8118 | 30.20777 | 57.97776 | 25.39515 | -0.55558 | 1.038242 | 0.622531 |
| 2 | 98264200 | 98268799 | 28.23777 | 46.52099 | 71.03191 | 30.6395 | -0.22944 | 1.380892 | 0.622533 |
| 3 | 12108800 | 12114599 | 88.49244 | 138.1858 | 150.3939 | 62.40292 | 0.190941 | 2.669879 | 0.622539 |
| 2 | 42752000 | 42752599 | 7.289017 | 14.09702 | 25.56693 | 9.773609 | -0.29071 | -0.17079 | 0.622542 |
| 7 | 12455600 | 12457199 | 27.82772 | 47.29367 | 42.62675 | 18.68158 | 0.530942 | 1.039709 | 0.622568 |
| 6 | 146216800 | 146218199 | 15.37638 | 26.96939 | 41.90879 | 18.52496 | -0.24527 | 0.640084 | 0.622577 |
| 18 | 26919800 | 26920599 | 11.11901 | 21.02153 | 24.2489 | 9.850211 | 0.276227 | 0.06084 | 0.622577 |
| 8 | 72882000 | 72884399 | 23.98803 | 40.78141 | 44.80135 | 19.63514 | 0.246505 | 1.0233 | 0.622579 |
| 3 | 39897200 | 39898799 | 22.05489 | 38.18072 | 41.44399 | 18.18281 | 0.268131 | 0.847354 | 0.622593 |
| 7 | 15627200 | 15630399 | 31.4891 | 52.06602 | 57.95047 | 25.04846 | 0.22652 | 1.335896 | 0.6226 |
| 8 | 69871600 | 69872999 | 15.97061 | 27.85162 | 43.62496 | 19.19869 | -0.25704 | 0.733171 | 0.622608 |
| 12 | 113157200 | 113170799 | 181.4135 | 266.5517 | 360.8881 | 148.6282 | -0.19801 | 3.83411 | 0.622623 |
| 13 | 42138600 | 42140199 | 14.0183 | 25.80964 | 27.98471 | 11.86358 | 0.31653 | 0.253935 | 0.622632 |
| 6 | 88586200 | 88587199 | 11.3584 | 21.23545 | 24.09793 | 9.986135 | 0.27595 | 0.133298 | 0.622654 |
| 7 | 47233200 | 47236799 | 49.25304 | 79.57277 | 80.20398 | 34.23979 | 0.348925 | 1.737344 | 0.622673 |
| 13 | 61065400 | 61068999 | 40.84382 | 66.49021 | 71.12379 | 30.53433 | 0.273306 | 1.566756 | 0.622687 |
| 13 | 27030000 | 27030399 | 7.162384 | 13.75138 | 24.65594 | 9.661152 | -0.30755 | -0.09169 | 0.622693 |
| 5 | 56433400 | 56434799 | 9.608715 | 18.01919 | 32.37427 | 13.34668 | -0.37583 | 0.122676 | 0.622703 |
| 5 | 128276800 | 128277999 | 13.61113 | 24.80808 | 27.50822 | 11.81868 | 0.271249 | 0.307665 | 0.622705 |

| | | | | | | | | | |
|---|---|---|---|---|---|---|---|---|---|
| 13 | 98695400 | 98699399 | 37.84831 | 61.06231 | 91.32554 | 38.91773 | -0.22147 | 1.763687 | 0.622712 |
| 4 | 26981200 | 26981799 | 7.583802 | 14.64516 | 26.67498 | 10.2689 | -0.30675 | -0.14168 | 0.622727 |
| 16 | 68700400 | 68704799 | 38.65335 | 63.18364 | 66.2283 | 28.47529 | 0.306614 | 1.498121 | 0.622738 |
| 18 | 3976000 | 3976599 | 12.83125 | 22.7169 | 36.23124 | 16.02137 | -0.27823 | 0.502644 | 0.622774 |
| 1 | 28209600 | 28210999 | 28.31558 | 46.51878 | 70.68216 | 30.45254 | -0.22395 | 1.417219 | 0.622777 |
| 9 | 116185200 | 116185599 | 7.505745 | 14.64966 | 17.51586 | 6.212773 | 0.403047 | -0.45524 | 0.622794 |
| 4 | 129299800 | 129300799 | 7.277621 | 13.97056 | 25.21743 | 9.885375 | -0.31842 | -0.09034 | 0.622803 |
| 17 | 58957200 | 58958199 | 10.19406 | 18.81746 | 30.92123 | 13.06226 | -0.27525 | 0.218289 | 0.622811 |
| 7 | 34271600 | 34292199 | 261.1026 | 372.6546 | 487.1241 | 201.3135 | -0.18841 | 4.276371 | 0.622823 |
| 8 | 7387200 | 7388999 | 17.05583 | 31.29903 | 28.81636 | 12.27095 | 0.546443 | 0.297209 | 0.622843 |
| 5 | 99959400 | 99962799 | 70.86465 | 112.0086 | 122.9371 | 51.07131 | 0.192338 | 2.418546 | 0.622846 |
| 11 | 96013600 | 96015599 | 15.01012 | 26.19641 | 42.58693 | 18.75079 | -0.3103 | 0.715592 | 0.622851 |
| 8 | 7975800 | 7977199 | 14.01766 | 26.11909 | 27.09289 | 11.29142 | 0.398088 | 0.184323 | 0.622868 |
| 2 | 58264600 | 58265799 | 13.85242 | 25.45397 | 27.97259 | 11.89032 | 0.294486 | 0.263528 | 0.622869 |
| 17 | 30620200 | 30621199 | 9.599631 | 18.0579 | 31.39577 | 12.87569 | -0.32335 | 0.096191 | 0.622872 |
| 2 | 21880000 | 21883599 | 44.63292 | 71.96602 | 79.92673 | 34.03447 | 0.206005 | 1.786487 | 0.622873 |
| 6 | 68068800 | 68069599 | 12.21325 | 23.03335 | 21.40391 | 8.739921 | 0.578922 | 0.078888 | 0.622873 |
| 4 | 35578200 | 35580399 | 40.21264 | 65.45397 | 70.97258 | 30.46793 | 0.253639 | 1.56773 | 0.622874 |
| 4 | 39752800 | 39753799 | 12.42606 | 22.91539 | 25.58132 | 10.82071 | 0.277296 | 0.227878 | 0.622912 |
| 9 | 72657400 | 72659999 | 18.20506 | 31.82122 | 33.73209 | 14.86737 | 0.303922 | 0.696009 | 0.622936 |
| 5 | 28742000 | 28744999 | 43.85358 | 70.69358 | 77.20668 | 32.86671 | 0.229823 | 1.791496 | 0.622939 |
| 15 | 98083400 | 98087199 | 22.21644 | 37.6653 | 58.80797 | 25.75621 | -0.25915 | 1.055991 | 0.622965 |
| 3 | 97456200 | 97457399 | 10.09315 | 18.72563 | 30.98067 | 12.99765 | -0.27825 | 0.186427 | 0.622988 |
| 17 | 33811800 | 33813599 | 12.44928 | 22.30117 | 35.18268 | 15.41816 | -0.25321 | 0.416765 | 0.622991 |
| 7 | 42363000 | 42378199 | 232.0238 | 341.5651 | 326.2787 | 134.3634 | 0.307535 | 3.809184 | 0.623047 |
| 1 | 190357200 | 190357999 | 6.539587 | 12.63145 | 23.72152 | 9.122516 | -0.34869 | -0.16522 | 0.623057 |
| 6 | 107266400 | 107268199 | 17.92203 | 31.12134 | 48.47139 | 21.27426 | -0.24967 | 0.809998 | 0.623063 |
| 9 | 14362000 | 14364199 | 20.97854 | 35.32425 | 55.92031 | 24.44077 | -0.27928 | 1.108532 | 0.623075 |
| 5 | 28874000 | 28875599 | 15.22359 | 26.76754 | 42.35511 | 18.75511 | -0.27114 | 0.601777 | 0.623084 |
| 13 | 25348800 | 25350199 | 11.22626 | 20.19459 | 33.27999 | 14.53301 | -0.31187 | 0.380185 | 0.623109 |
| 19 | 8065600 | 8075399 | 115.3608 | 178.0383 | 177.3004 | 73.64715 | 0.305874 | 2.890116 | 0.623121 |
| 6 | 130559200 | 130561799 | 23.03947 | 40.47809 | 36.07777 | 15.97015 | 0.552756 | 0.646237 | 0.623133 |
| 1 | 165275200 | 165278999 | 46.0861 | 73.94948 | 117.8908 | 49.84887 | -0.32301 | 1.955505 | 0.623145 |
| 1 | 7633600 | 7635199 | 18.45009 | 32.669 | 29.12208 | 12.87737 | 0.5564 | 0.521665 | 0.623178 |
| 3 | 117978600 | 117979199 | 6.863756 | 13.20759 | 24.15931 | 9.405855 | -0.32719 | -0.11669 | 0.623184 |
| 13 | 120477000 | 120477799 | 6.863756 | 13.20759 | 24.15931 | 9.405855 | -0.32719 | -0.11669 | 0.623184 |
| 12 | 103722200 | 103743599 | 275.1444 | 391.4007 | 508.117 | 210.1058 | -0.18327 | 4.32546 | 0.623191 |
| 6 | 110340000 | 110341599 | 11.52162 | 22.14769 | 22.32974 | 8.719266 | 0.519337 | -0.06963 | 0.623206 |
| 15 | 21956400 | 21959399 | 32.83982 | 54.34757 | 59.72945 | 25.83322 | 0.244814 | 1.331272 | 0.623222 |
| 6 | 58036800 | 58043399 | 111.6073 | 172.0986 | 161.4386 | 67.05876 | 0.391103 | 2.902963 | 0.623222 |
| 10 | 11128800 | 11131199 | 27.35286 | 46.27786 | 50.14565 | 21.94862 | 0.265893 | 1.083344 | 0.623263 |
| 9 | 86265000 | 86268399 | 29.78676 | 50.54861 | 49.59145 | 21.72599 | 0.408892 | 1.065231 | 0.623268 |
| 6 | 147073800 | 147077999 | 60.2577 | 95.9119 | 104.8432 | 43.82089 | 0.207475 | 2.213834 | 0.623275 |
| 6 | 52437200 | 52441799 | 4.946882 | 9.618742 | 23.45316 | 8.693234 | -0.67003 | -0.30894 | 0.623287 |
| 19 | 18006800 | 18008199 | 14.04782 | 26.41859 | 26.20413 | 10.75125 | 0.480264 | 0.125156 | 0.623294 |
| 9 | 44772400 | 44774599 | 21.1185 | 36.65935 | 40.41224 | 17.73324 | 0.24674 | 0.81751 | 0.623316 |
| 7 | 84538800 | 84541999 | 85.76005 | 133.7168 | 141.7224 | 58.81468 | 0.227464 | 2.694079 | 0.623332 |

| | | | | | | | | | |
|---|---|---|---|---|---|---|---|---|---|
| 4 | 54820400 | 54821599 | 14.71079 | 25.80294 | 40.37305 | 17.84668 | -0.25512 | 0.632859 | 0.623336 |
| 2 | 172820600 | 172822399 | 23.30277 | 38.46846 | 72.70228 | 31.38997 | -0.53515 | 1.346018 | 0.62334 |
| 5 | 71599200 | 71599799 | 6.116224 | 11.81851 | 23.17329 | 8.894781 | -0.40749 | -0.17778 | 0.623346 |
| 5 | 36403400 | 36407799 | 36.55025 | 58.97827 | 89.29268 | 38.0599 | -0.23875 | 1.755713 | 0.623356 |
| 14 | 84026400 | 84028999 | 29.63578 | 49.45921 | 54.63353 | 23.763 | 0.238034 | 1.223859 | 0.623362 |
| 15 | 80228200 | 80230599 | 17.34224 | 29.80052 | 47.95251 | 21.03356 | -0.29945 | 0.904324 | 0.623379 |
| 16 | 39748200 | 39750199 | 16.22995 | 28.87576 | 31.16251 | 13.73529 | 0.285517 | 0.473878 | 0.623396 |
| 19 | 15862800 | 15865999 | 28.19361 | 47.8634 | 49.58148 | 21.72415 | 0.330757 | 1.057043 | 0.623397 |
| 2 | 95686200 | 95687799 | 12.6901 | 23.66081 | 20.91641 | 8.707687 | 0.628379 | 0.153867 | 0.623423 |
| 5 | 121367400 | 121369199 | 16.25281 | 28.75767 | 31.44705 | 13.90744 | 0.262907 | 0.520811 | 0.623425 |
| 7 | 107503600 | 107505399 | 22.03771 | 38.27109 | 41.43896 | 18.18858 | 0.272618 | 0.818405 | 0.623445 |
| 12 | 25802400 | 25803999 | 17.11347 | 31.06307 | 29.02218 | 12.53368 | 0.511579 | 0.356481 | 0.62348 |
| 1 | 47881200 | 47882199 | 12.96615 | 24.30251 | 22.42607 | 9.253703 | 0.577184 | 0.127979 | 0.623498 |
| 1 | 67873000 | 67873799 | 15.06961 | 26.3473 | 41.53518 | 18.31048 | -0.26579 | 0.689978 | 0.62352 |
| 8 | 36892400 | 36894399 | 29.01827 | 48.42067 | 54.03871 | 23.50183 | 0.22329 | 1.223896 | 0.62354 |
| 6 | 102554400 | 102559199 | 55.87788 | 89.63237 | 82.83875 | 35.0103 | 0.461548 | 1.964082 | 0.62354 |
| 16 | 8098000 | 8101799 | 30.16335 | 51.3457 | 48.46739 | 21.24405 | 0.464827 | 1.039177 | 0.623548 |
| 8 | 123160600 | 123163199 | 24.54822 | 40.60382 | 64.43579 | 27.85844 | -0.28354 | 1.321167 | 0.623555 |
| 17 | 93578000 | 93578599 | 8.137683 | 15.78365 | 19.40894 | 7.320587 | 0.283536 | -0.22459 | 0.623565 |
| 17 | 47471000 | 47472599 | 12.93376 | 23.58625 | 24.96201 | 10.71746 | 0.338443 | 0.291229 | 0.623566 |
| 18 | 36997800 | 37051999 | 1083.289 | 1394.355 | 1218.392 | 510.9004 | 0.242862 | 5.898555 | 0.623566 |
| 4 | 61388400 | 61388799 | 8.414414 | 16.34493 | 20.01061 | 7.476424 | 0.302034 | -0.2533 | 0.623589 |
| 9 | 114533000 | 114535799 | 25.55101 | 43.55584 | 47.4038 | 20.78022 | 0.260529 | 1.008717 | 0.623595 |
| 19 | 12343400 | 12344399 | 14.46504 | 26.01945 | 27.08099 | 11.82646 | 0.34708 | 0.38938 | 0.623595 |
| 8 | 58516400 | 58517199 | 8.270024 | 15.68042 | 27.10943 | 10.94522 | -0.29486 | 0.024206 | 0.623609 |
| 18 | 19069800 | 19071799 | 34.35872 | 56.87422 | 51.97327 | 22.48218 | 0.510331 | 1.328528 | 0.623624 |
| 8 | 23151200 | 23152199 | 7.127426 | 13.68507 | 25.22366 | 9.881928 | -0.34757 | -0.09433 | 0.623633 |
| 19 | 44983400 | 44986999 | 18.09024 | 30.72119 | 51.60791 | 22.61426 | -0.36391 | 1.020715 | 0.623634 |
| 10 | 77639400 | 77649199 | 127.2293 | 191.9509 | 264.4277 | 109.2782 | -0.18804 | 3.350128 | 0.623646 |
| 9 | 45487600 | 45489399 | 11.09866 | 20.0951 | 33.22706 | 14.3946 | -0.30967 | 0.337422 | 0.623659 |
| 9 | 24776200 | 24777399 | 12.58366 | 22.53463 | 35.66997 | 15.63596 | -0.25838 | 0.419835 | 0.62367 |
| 4 | 5998400 | 6002799 | 59.93546 | 95.56827 | 105.5064 | 44.20639 | 0.195948 | 2.157072 | 0.623678 |
| 14 | 77445000 | 77446199 | 14.7576 | 25.9283 | 40.32761 | 17.85175 | -0.24648 | 0.607382 | 0.623735 |
| 4 | 17080800 | 17081799 | 15.90268 | 28.15338 | 30.48011 | 13.47836 | 0.277549 | 0.510674 | 0.623737 |
| 15 | 9243400 | 9246199 | 26.20066 | 44.52667 | 48.48732 | 21.24996 | 0.259066 | 1.039598 | 0.623755 |
| 17 | 37861800 | 37864199 | 28.93697 | 48.96004 | 50.6988 | 22.19101 | 0.331162 | 1.090293 | 0.623779 |
| 7 | 73479800 | 73482199 | 19.04352 | 33.35041 | 36.52052 | 16.12237 | 0.257128 | 0.673878 | 0.623781 |
| 3 | 17043400 | 17047599 | 70.51219 | 111.6363 | 122.9037 | 51.10024 | 0.189835 | 2.379318 | 0.623789 |
| 2 | 107786000 | 107792799 | 104.8741 | 161.6091 | 173.2047 | 71.94646 | 0.198341 | 2.912834 | 0.623798 |
| 1 | 157078400 | 157078999 | 5.948946 | 11.58023 | 21.88135 | 8.037009 | -0.29413 | -0.33341 | 0.623833 |
| 17 | 23048000 | 23065799 | 231.7694 | 341.1958 | 326.4578 | 134.4372 | 0.305203 | 3.808803 | 0.623833 |
| 3 | 9011800 | 9013199 | 12.16997 | 21.73696 | 34.7193 | 15.25277 | -0.27325 | 0.433392 | 0.623834 |
| 3 | 146892400 | 146892999 | 7.900186 | 15.06311 | 26.13759 | 10.42887 | -0.28545 | -0.02289 | 0.623853 |
| X | 77553200 | 77555599 | 46.66469 | 75.4995 | 77.21866 | 33.01929 | 0.330182 | 1.697316 | 0.623861 |
| 5 | 55225200 | 55226599 | 19.52075 | 34.17551 | 37.00763 | 16.3331 | 0.273183 | 0.681954 | 0.623866 |
| 13 | 61730200 | 61746999 | 282.4318 | 408.2179 | 390.669 | 161.1464 | 0.279379 | 4.093101 | 0.623873 |
| 5 | 74136200 | 74140199 | 28.11564 | 45.78316 | 83.01345 | 35.64249 | -0.48572 | 1.544876 | 0.62388 |

| | | | | | | | | | |
|---|---|---|---|---|---|---|---|---|---|
| 6 | 49798200 | 49800999 | 15.54083 | 26.58864 | 55.31733 | 24.25637 | -0.66966 | 0.945539 | 0.623889 |
| 10 | 69095000 | 69097399 | 19.5287 | 32.64814 | 64.37275 | 28.02979 | -0.59494 | 1.185683 | 0.623899 |
| 6 | 121207200 | 121208799 | 9.326735 | 17.48978 | 29.04781 | 11.97602 | -0.26353 | 0.109901 | 0.623899 |
| 3 | 159207000 | 159208999 | 14.97796 | 26.36203 | 43.39161 | 19.21023 | -0.32739 | 0.588688 | 0.6239 |
| 2 | 179180800 | 179189999 | 114.3078 | 173.6805 | 240.1041 | 99.44427 | -0.18453 | 3.202858 | 0.623908 |
| 1 | 34622800 | 34624199 | 11.59566 | 20.62004 | 34.82986 | 15.35834 | -0.35693 | 0.459185 | 0.623908 |
| 3 | 7845800 | 7846199 | 5.807945 | 11.29889 | 22.4563 | 8.288738 | -0.37401 | -0.32069 | 0.623924 |
| 1 | 63361400 | 63363599 | 37.60904 | 61.13233 | 67.53252 | 28.97667 | 0.22549 | 1.580725 | 0.623924 |
| 7 | 84928200 | 84930599 | 56.21647 | 89.72612 | 97.28347 | 40.81684 | 0.223509 | 2.121487 | 0.623932 |
| X | 36989600 | 36991399 | 35.09581 | 57.81767 | 60.06773 | 25.90978 | 0.324593 | 1.394995 | 0.62394 |
| 1 | 119205400 | 119206799 | 12.41331 | 22.35426 | 35.72305 | 15.58086 | -0.26713 | 0.384148 | 0.623954 |
| 9 | 38800000 | 38802599 | 25.77188 | 43.791 | 47.94672 | 21.01277 | 0.251148 | 1.037508 | 0.623965 |
| 6 | 123333800 | 123337999 | 53.08018 | 84.57581 | 122.9618 | 51.46822 | -0.20168 | 2.180199 | 0.623971 |
| 16 | 37486400 | 37488199 | 12.25663 | 22.16393 | 35.81134 | 15.53848 | -0.27806 | 0.35949 | 0.623975 |
| 18 | 32788600 | 32789999 | 16.24503 | 29.20969 | 30.648 | 13.38867 | 0.334656 | 0.408208 | 0.623993 |
| 2 | 51337000 | 51338399 | 19.23131 | 33.63468 | 37.11641 | 16.3671 | 0.246144 | 0.697387 | 0.624014 |
| 16 | 76681800 | 76683999 | 35.80824 | 59.29878 | 52.65761 | 22.78489 | 0.551506 | 1.325268 | 0.624019 |
| 9 | 8114400 | 8117199 | 57.14538 | 90.59986 | 131.892 | 54.89055 | -0.21088 | 2.335696 | 0.624026 |
| 17 | 65814000 | 65815399 | 17.44883 | 31.42279 | 31.20432 | 13.60975 | 0.415391 | 0.410702 | 0.624043 |
| 1 | 156554400 | 156554999 | 13.63177 | 24.76525 | 25.63199 | 11.05733 | 0.366512 | 0.323017 | 0.624045 |
| 6 | 123355600 | 123356199 | 10.50093 | 20.01037 | 23.33286 | 9.321126 | 0.281981 | -0.00064 | 0.624126 |
| 1 | 45595200 | 45598999 | 82.26569 | 129.5944 | 123.0272 | 51.04871 | 0.39761 | 2.499813 | 0.624129 |
| X | 149312000 | 149313399 | 10.26263 | 18.96653 | 32.54886 | 13.72765 | -0.33565 | 0.217375 | 0.624135 |
| 6 | 127892600 | 127897999 | 37.32514 | 59.91424 | 108.0068 | 45.69568 | -0.49923 | 1.92694 | 0.624149 |
| 5 | 55156800 | 55157799 | 13.91216 | 24.69706 | 39.72051 | 17.54018 | -0.28871 | 0.490802 | 0.62415 |
| 1 | 14787600 | 14792599 | 54.0119 | 86.93411 | 87.28586 | 37.07556 | 0.348138 | 1.85714 | 0.624151 |
| 7 | 12123600 | 12125799 | 36.63914 | 59.97554 | 65.69562 | 28.26079 | 0.244774 | 1.478225 | 0.624152 |
| 3 | 49931800 | 49932599 | 8.564613 | 16.53301 | 19.32283 | 7.451633 | 0.32586 | -0.146 | 0.624158 |
| 16 | 36905400 | 36906799 | 18.66264 | 32.68958 | 36.10949 | 15.94347 | 0.244771 | 0.669823 | 0.624163 |
| 13 | 67253600 | 67256799 | 37.68982 | 60.70542 | 93.1111 | 39.58986 | -0.26064 | 1.813459 | 0.624171 |
| 9 | 55676000 | 55678599 | 25.02737 | 42.81069 | 45.75023 | 20.06093 | 0.287382 | 0.97144 | 0.62422 |
| 4 | 61920200 | 61923799 | 55.24522 | 88.62322 | 83.72943 | 35.39017 | 0.429927 | 1.959812 | 0.624221 |
| 15 | 85898200 | 85899799 | 19.90683 | 34.57562 | 36.96673 | 16.22709 | 0.290944 | 0.801008 | 0.62423 |
| 15 | 3185800 | 3186799 | 15.79325 | 28.06988 | 30.45255 | 13.43402 | 0.277504 | 0.479521 | 0.624255 |
| 15 | 97042400 | 97042799 | 5.091197 | 9.928784 | 19.75397 | 7.09471 | -0.33828 | -0.43352 | 0.62428 |
| 1 | 88384600 | 88388999 | 37.53562 | 60.58013 | 90.1064 | 38.41643 | -0.21291 | 1.752436 | 0.62428 |
| 15 | 57390000 | 57391599 | 15.56429 | 28.93571 | 26.90836 | 11.25735 | 0.550109 | 0.209555 | 0.624284 |
| 2 | 74193600 | 74197399 | 39.28559 | 64.10185 | 68.89792 | 29.60372 | 0.268738 | 1.528416 | 0.62432 |
| 7 | 75465000 | 75470599 | 61.96193 | 98.78284 | 107.7007 | 45.11527 | 0.213692 | 2.165559 | 0.624341 |
| 3 | 69995600 | 69996599 | 8.565988 | 16.23158 | 27.64364 | 11.17545 | -0.27523 | 0.031854 | 0.624352 |
| 10 | 61027000 | 61027999 | 8.565988 | 16.23158 | 27.64364 | 11.17545 | -0.27523 | 0.031854 | 0.624352 |
| 18 | 42515400 | 42516399 | 6.708107 | 12.86624 | 24.63291 | 9.680066 | -0.40578 | -0.08976 | 0.624356 |
| 10 | 78933000 | 78933599 | 6.254943 | 12.12541 | 22.73195 | 8.595529 | -0.32191 | -0.2251 | 0.624364 |
| 1 | 117854600 | 117860599 | 72.20822 | 114.8704 | 115.0069 | 48.03248 | 0.333171 | 2.245128 | 0.624365 |
| 15 | 30002400 | 30004999 | 31.64485 | 51.4291 | 78.17508 | 33.53912 | -0.23406 | 1.577866 | 0.624377 |
| X | 63131600 | 63133999 | 26.16492 | 45.26344 | 43.51741 | 19.09207 | 0.442452 | 0.877778 | 0.624384 |
| 7 | 108125600 | 108131399 | 101.8068 | 155.8238 | 216.3911 | 89.79387 | -0.18255 | 3.049593 | 0.624385 |

| | | | | | | | | | |
|---|---|---|---|---|---|---|---|---|---|
| 1 | 191894200 | 191899799 | 69.84719 | 110.6504 | 121.8425 | 50.67457 | 0.190244 | 2.364443 | 0.624388 |
| 6 | 148541200 | 148542799 | 9.088092 | 16.86398 | 28.92656 | 12.1329 | -0.329 | 0.17543 | 0.624388 |
| 10 | 31365600 | 31366999 | 18.76552 | 33.41512 | 32.49724 | 14.31341 | 0.435611 | 0.488381 | 0.624405 |
| X | 91490000 | 91492199 | 25.30072 | 43.84793 | 42.98191 | 18.85818 | 0.415092 | 0.857191 | 0.624411 |
| 2 | 56910400 | 56911199 | 17.2692 | 29.84407 | 47.93843 | 21.03102 | -0.29506 | 0.855818 | 0.624413 |
| 14 | 37109800 | 37110799 | 10.3008 | 18.97184 | 30.8691 | 13.08014 | -0.26455 | 0.230028 | 0.624434 |
| 7 | 74677000 | 74678199 | 15.11074 | 27.22866 | 29.7493 | 12.96535 | 0.278871 | 0.386122 | 0.624448 |
| 6 | 14664600 | 14665799 | 13.98397 | 25.26615 | 27.88117 | 12.11039 | 0.267813 | 0.359252 | 0.624454 |
| 10 | 53332600 | 53335999 | 30.77202 | 51.09845 | 56.78318 | 24.6165 | 0.229315 | 1.286182 | 0.624459 |
| 13 | 20542800 | 20546599 | 55.09781 | 88.13206 | 97.0383 | 40.83955 | 0.204712 | 2.05204 | 0.624477 |
| 10 | 3131600 | 3133999 | 21.7574 | 36.6342 | 57.05755 | 24.9375 | -0.25595 | 1.112005 | 0.624521 |
| 4 | 135529000 | 135532399 | 37.29904 | 60.06295 | 91.73358 | 38.99186 | -0.25481 | 1.822282 | 0.624529 |
| 5 | 11507400 | 11509799 | 28.49648 | 46.76732 | 71.16916 | 30.64887 | -0.22675 | 1.431194 | 0.624529 |
| 4 | 77897600 | 77899599 | 21.01643 | 36.7433 | 33.4703 | 14.75471 | 0.521626 | 0.70768 | 0.624531 |
| 11 | 3559600 | 3561799 | 19.06276 | 32.72045 | 50.61049 | 22.19826 | -0.24314 | 0.923692 | 0.624548 |
| 13 | 51181600 | 51194199 | 153.0585 | 228.1149 | 310.0578 | 127.7718 | -0.18449 | 3.589495 | 0.624561 |
| 1 | 183295600 | 183300199 | 102.817 | 158.2656 | 164.3123 | 68.24351 | 0.242979 | 2.933765 | 0.624562 |
| 19 | 26093400 | 26094399 | 9.054305 | 16.91576 | 28.46292 | 11.80915 | -0.28922 | 0.131172 | 0.624568 |
| 14 | 13163600 | 13165599 | 20.09295 | 35.00845 | 38.81379 | 17.07149 | 0.239112 | 0.759988 | 0.624585 |
| 19 | 49859200 | 49859599 | 7.260598 | 14.1469 | 17.96133 | 6.526641 | 0.287403 | -0.36917 | 0.624588 |
| 18 | 29431600 | 29433399 | 11.94115 | 21.65739 | 34.81867 | 15.05198 | -0.26735 | 0.338634 | 0.624593 |
| 2 | 108821000 | 108821599 | 5.365846 | 10.45837 | 20.26522 | 7.333473 | -0.31019 | -0.39959 | 0.6246 |
| 8 | 34974400 | 34975799 | 13.83172 | 24.25898 | 39.46091 | 17.44222 | -0.3106 | 0.630251 | 0.624607 |
| 8 | 111782800 | 111783999 | 9.52212 | 17.49074 | 30.33977 | 12.90282 | -0.36007 | 0.236725 | 0.624648 |
| 17 | 63233600 | 63234799 | 9.52212 | 17.49074 | 30.33977 | 12.90282 | -0.36007 | 0.236725 | 0.624648 |
| 6 | 132020400 | 132023999 | 68.79425 | 109.3152 | 116.6248 | 48.64668 | 0.239996 | 2.275057 | 0.62466 |
| 3 | 15399800 | 15400399 | 13.64382 | 24.97088 | 27.52656 | 11.76438 | 0.28472 | 0.284869 | 0.62468 |
| 14 | 93239000 | 93240399 | 13.86466 | 26.15568 | 25.10007 | 10.24281 | 0.533528 | 0.099985 | 0.624681 |
| 2 | 171741000 | 171741799 | 5.809051 | 11.2856 | 21.71202 | 8.103009 | -0.34083 | -0.27898 | 0.624683 |
| 4 | 91464400 | 91465199 | 14.12352 | 25.46393 | 27.85 | 12.12915 | 0.278085 | 0.37205 | 0.624685 |
| 11 | 65414400 | 65417599 | 33.91574 | 56.62048 | 56.45003 | 24.55813 | 0.38551 | 1.236789 | 0.624706 |
| 12 | 114396200 | 114397599 | 32.27616 | 53.47236 | 58.63147 | 25.37721 | 0.248236 | 1.315616 | 0.624715 |
| 1 | 75866600 | 75871599 | 68.36006 | 108.5089 | 118.5297 | 49.38821 | 0.204502 | 2.306693 | 0.624715 |
| 9 | 33304400 | 33305199 | 19.86619 | 33.67242 | 53.21502 | 23.30491 | -0.27633 | 1.046386 | 0.624726 |
| 1 | 151507800 | 151509199 | 14.38977 | 26.17743 | 28.91617 | 12.45318 | 0.274349 | 0.326465 | 0.624732 |
| X | 104658600 | 104658999 | 8.518281 | 16.50082 | 18.38204 | 6.974645 | 0.417501 | -0.20106 | 0.624765 |
| 14 | 74641800 | 74642999 | 10.171 | 18.86303 | 30.98663 | 13.00673 | -0.26826 | 0.191424 | 0.624776 |
| 15 | 65342000 | 65344199 | 22.58356 | 38.10065 | 58.19631 | 25.45049 | -0.22783 | 1.099609 | 0.624784 |
| 1 | 161436000 | 161441999 | 84.36935 | 132.5816 | 125.9716 | 52.25373 | 0.393659 | 2.550904 | 0.624818 |
| 17 | 22514600 | 22521599 | 112.0259 | 173.7836 | 166.5508 | 69.17102 | 0.36685 | 2.797759 | 0.624832 |
| 1 | 100977200 | 100979199 | 20.88985 | 35.62292 | 54.94743 | 24.08779 | -0.24067 | 0.995471 | 0.624833 |
| 3 | 138432200 | 138435799 | 27.23298 | 45.04866 | 68.83286 | 29.76136 | -0.22923 | 1.328516 | 0.624858 |
| 1 | 160247600 | 160249199 | 17.07063 | 30.04096 | 32.55476 | 14.41298 | 0.272916 | 0.58712 | 0.624859 |
| X | 132397400 | 132398599 | 12.86163 | 24.84951 | 22.62103 | 8.688174 | 0.688645 | -0.10469 | 0.624893 |
| 15 | 31374600 | 31378399 | 35.06958 | 56.67853 | 86.6082 | 36.99101 | -0.24942 | 1.705696 | 0.624896 |
| 1 | 166192600 | 166193399 | 7.645749 | 14.39203 | 26.99072 | 11.05666 | -0.42984 | 0.064209 | 0.624898 |
| 16 | 95717800 | 95721799 | 35.79035 | 57.64229 | 90.61766 | 38.52606 | -0.29615 | 1.812807 | 0.624907 |

| | | | | | | | | | |
|---|---|---|---|---|---|---|---|---|---|
| Y | 43182000 | 43182599 | 9.473117 | 18.12965 | 16.73499 | 6.618054 | 0.629207 | -0.06513 | 0.624932 |
| 3 | 112765400 | 112765799 | 11.13055 | 20.99828 | 24.22432 | 9.884956 | 0.270424 | 0.075771 | 0.625023 |
| 3 | 65538600 | 65539599 | 8.835032 | 16.69761 | 28.13062 | 11.43548 | -0.2671 | 0.051954 | 0.625035 |
| 13 | 106515000 | 106517199 | 16.83932 | 29.16131 | 45.83424 | 20.10916 | -0.26331 | 0.829989 | 0.625051 |
| 7 | 10126600 | 10136999 | 132.4158 | 203.0846 | 193.1226 | 80.16917 | 0.365532 | 3.018273 | 0.625052 |
| 17 | 20892200 | 20896799 | 55.47491 | 89.33481 | 75.46018 | 32.07638 | 0.597859 | 1.839313 | 0.625056 |
| 13 | 67224200 | 67227399 | 49.98881 | 80.82847 | 77.70086 | 33.20668 | 0.418455 | 1.71279 | 0.625063 |
| 1 | 39831600 | 39837999 | 81.59899 | 128.2234 | 140.0315 | 58.08559 | 0.192996 | 2.552088 | 0.625083 |
| 5 | 133550400 | 133551399 | 16.31414 | 28.33835 | 44.76015 | 19.64935 | -0.26961 | 0.793249 | 0.625094 |
| 1 | 128516600 | 128517399 | 7.927277 | 15.11244 | 26.13263 | 10.43028 | -0.28063 | -0.02018 | 0.625109 |
| 4 | 75534600 | 75536399 | 16.43194 | 29.26063 | 31.81574 | 14.01291 | 0.275745 | 0.473866 | 0.625119 |
| 15 | 7806000 | 7811799 | 62.46473 | 99.61857 | 107.454 | 45.03327 | 0.229658 | 2.155195 | 0.625126 |
| 7 | 87347200 | 87348399 | 17.06435 | 30.24995 | 32.75854 | 14.48205 | 0.277791 | 0.507674 | 0.625134 |
| 1 | 170093400 | 170099199 | 47.86765 | 77.36804 | 70.52507 | 30.12459 | 0.494301 | 1.718073 | 0.625139 |
| 14 | 80421200 | 80423199 | 15.87652 | 27.15541 | 54.22313 | 23.77604 | -0.61057 | 0.950435 | 0.625151 |
| 14 | 85561400 | 85563199 | 14.13829 | 26.65349 | 25.6739 | 10.49039 | 0.526826 | 0.108995 | 0.625169 |
| 14 | 86906400 | 86913399 | 62.94371 | 101.115 | 89.94108 | 38.09678 | 0.519189 | 1.931276 | 0.625174 |
| 5 | 121225800 | 121236799 | 107.1617 | 163.7586 | 226.8716 | 94.10814 | -0.18095 | 3.084107 | 0.625177 |
| 16 | 9992200 | 9996999 | 65.67215 | 104.6401 | 110.0526 | 46.06526 | 0.264686 | 2.18631 | 0.625203 |
| 10 | 19420600 | 19423999 | 42.34657 | 68.46864 | 76.17271 | 32.55186 | 0.208119 | 1.701449 | 0.625212 |
| 18 | 90455000 | 90456399 | 19.38163 | 33.23316 | 52.19182 | 22.88993 | -0.26509 | 0.93988 | 0.625215 |
| 3 | 106648600 | 106650199 | 33.42318 | 55.87014 | 55.36069 | 24.10382 | 0.394251 | 1.218995 | 0.625238 |
| 10 | 99309800 | 99311199 | 16.54852 | 29.29574 | 32.34017 | 14.30097 | 0.249501 | 0.518106 | 0.625268 |
| 5 | 110134000 | 110137799 | 28.5601 | 48.12385 | 51.28517 | 22.41966 | 0.289561 | 1.121625 | 0.625272 |
| 11 | 46618000 | 46619199 | 28.54341 | 46.57115 | 73.52844 | 31.59265 | -0.28453 | 1.511291 | 0.625287 |
| 14 | 113740800 | 113743799 | 20.2084 | 34.67645 | 55.00499 | 24.12514 | -0.27953 | 0.93594 | 0.62529 |
| X | 61836800 | 61839599 | 43.56 | 70.72377 | 74.01993 | 31.73337 | 0.301666 | 1.618761 | 0.62532 |
| 4 | 113203200 | 113208599 | 69.81414 | 111.7013 | 98.89805 | 41.63789 | 0.519146 | 2.064597 | 0.625325 |
| 18 | 67255800 | 67257399 | 9.811878 | 18.17854 | 29.90154 | 12.56862 | -0.27167 | 0.190468 | 0.625348 |
| 12 | 109230600 | 109231999 | 24.15145 | 41.1412 | 44.30269 | 19.41971 | 0.275772 | 1.008043 | 0.625354 |
| 17 | 7707800 | 7711399 | 39.47474 | 64.12784 | 70.80375 | 30.36743 | 0.225432 | 1.595572 | 0.625388 |
| 3 | 35785600 | 35787799 | 17.72379 | 30.76677 | 47.93723 | 21.03537 | -0.25029 | 0.812404 | 0.62542 |
| 5 | 103227400 | 103228399 | 7.757205 | 14.92806 | 26.46506 | 10.30107 | -0.28287 | -0.10324 | 0.625428 |
| 8 | 28515600 | 28518799 | 17.73855 | 31.87771 | 31.03339 | 13.56328 | 0.441682 | 0.42261 | 0.625433 |
| 1 | 41050000 | 41054799 | 47.05831 | 75.77734 | 83.63873 | 35.55041 | 0.212659 | 1.830681 | 0.625452 |
| 7 | 86404400 | 86405999 | 13.35418 | 24.80988 | 26.45732 | 11.07914 | 0.352742 | 0.193572 | 0.625455 |
| 8 | 113145000 | 113147599 | 29.92759 | 49.11067 | 74.36237 | 32.02256 | -0.2197 | 1.436793 | 0.625459 |
| 6 | 117550200 | 117553199 | 22.14288 | 37.31768 | 57.03226 | 24.93343 | -0.22861 | 1.107329 | 0.625473 |
| 11 | 20913000 | 20915399 | 28.11402 | 47.38489 | 51.27026 | 22.41496 | 0.267768 | 1.118477 | 0.625473 |
| 5 | 133715000 | 133716799 | 19.47203 | 33.84331 | 34.92544 | 15.33756 | 0.3418 | 0.784968 | 0.625476 |
| 5 | 71350000 | 71352799 | 28.96955 | 48.6067 | 52.94522 | 23.10653 | 0.258028 | 1.164035 | 0.625477 |
| 2 | 11742800 | 11744999 | 19.26826 | 33.57811 | 36.83818 | 16.20454 | 0.254309 | 0.750516 | 0.625479 |
| 19 | 29406400 | 29410199 | 29.38927 | 48.29038 | 73.92066 | 31.84995 | -0.23464 | 1.417922 | 0.625498 |
| 8 | 76022200 | 76023599 | 13.18843 | 23.48845 | 37.74302 | 16.62137 | -0.28438 | 0.464886 | 0.625521 |
| 6 | 29048200 | 29049399 | 12.39761 | 22.96403 | 26.07257 | 10.96249 | 0.259158 | 0.203676 | 0.625539 |
| 3 | 7714400 | 7717799 | 40.44958 | 65.77806 | 71.36409 | 30.62556 | 0.251908 | 1.582894 | 0.625544 |
| X | 151692000 | 151693399 | 11.71863 | 21.09271 | 34.26702 | 14.95515 | -0.29088 | 0.382458 | 0.625552 |

| | | | | | | | | | |
|---|---|---|---|---|---|---|---|---|---|
| 5 | 98135800 | 98136599 | 12.59535 | 23.43327 | 26.01837 | 10.87152 | 0.297438 | 0.178448 | 0.625557 |
| 2 | 4606800 | 4609199 | 17.16994 | 29.13438 | 58.51595 | 25.63494 | -0.62085 | 1.030928 | 0.625558 |
| 2 | 59709800 | 59715199 | 38.95579 | 62.58231 | 107.4819 | 45.52911 | -0.4281 | 1.904645 | 0.625566 |
| 1 | 55895600 | 55897399 | 14.06504 | 24.79637 | 38.727 | 17.13944 | -0.25079 | 0.55947 | 0.625572 |
| 12 | 97586800 | 97589199 | 20.30077 | 35.71518 | 35.07285 | 15.5287 | 0.413837 | 0.609853 | 0.625581 |
| 16 | 90637800 | 90640799 | 18.77785 | 32.05486 | 51.12565 | 22.41429 | -0.28842 | 0.973117 | 0.625598 |
| 9 | 10060000 | 10061199 | 13.28231 | 24.60839 | 27.08834 | 11.38603 | 0.303886 | 0.212644 | 0.625611 |
| 11 | 119867000 | 119867999 | 11.48586 | 20.6968 | 33.21963 | 14.47784 | -0.27241 | 0.370132 | 0.625617 |
| 16 | 48572400 | 48574199 | 15.26754 | 27.79016 | 29.29986 | 12.60938 | 0.341647 | 0.329066 | 0.625647 |
| 1 | 45145000 | 45146999 | 15.82942 | 29.04224 | 28.87422 | 12.29916 | 0.435892 | 0.28881 | 0.625659 |
| 8 | 89989200 | 89992799 | 33.22353 | 55.59912 | 55.29935 | 24.09427 | 0.388867 | 1.208223 | 0.625673 |
| 9 | 32756000 | 32758599 | 8.964201 | 16.05735 | 38.45607 | 16.85336 | -0.85121 | 0.401828 | 0.625682 |
| 2 | 145113400 | 145114999 | 15.83326 | 27.61555 | 43.11323 | 18.97498 | -0.25224 | 0.730643 | 0.625709 |
| 12 | 62552400 | 62552999 | 10.07359 | 19.29062 | 22.36951 | 8.830777 | 0.305647 | -0.04453 | 0.625713 |
| 8 | 103680000 | 103680599 | 10.69883 | 20.2908 | 23.28062 | 9.39423 | 0.292603 | 0.034221 | 0.625748 |
| 2 | 18193800 | 18197399 | 40.61522 | 66.13474 | 70.63507 | 30.3273 | 0.275751 | 1.56145 | 0.625748 |
| 7 | 78070400 | 78072199 | 13.77693 | 25.79655 | 26.68887 | 11.03119 | 0.411502 | 0.151727 | 0.625786 |
| 13 | 43230600 | 43233799 | 36.40076 | 58.79157 | 87.68352 | 37.41877 | -0.21557 | 1.72849 | 0.625798 |
| 9 | 8990800 | 8991799 | 12.00296 | 21.67681 | 34.77601 | 15.11398 | -0.26912 | 0.365096 | 0.625815 |
| 4 | 66784200 | 66784799 | 4.389164 | 8.571669 | 18.17838 | 6.384096 | -0.39916 | -0.52825 | 0.625875 |
| 1 | 68057200 | 68058599 | 17.81015 | 31.70519 | 28.13895 | 12.39668 | 0.566775 | 0.477185 | 0.625904 |
| 8 | 19092600 | 19098199 | 44.7905 | 71.31623 | 128.7996 | 53.85833 | -0.51515 | 2.197481 | 0.625907 |
| 13 | 53178400 | 53181999 | 39.91385 | 64.06783 | 98.07027 | 41.48994 | -0.26378 | 1.928572 | 0.625915 |
| 14 | 39485200 | 39486799 | 25.76964 | 42.269 | 69.05082 | 29.72973 | -0.32899 | 1.430301 | 0.625939 |
| 14 | 123977200 | 123978199 | 15.599 | 28.25108 | 25.72997 | 11.1429 | 0.545892 | 0.353523 | 0.62594 |
| 3 | 58815800 | 58816599 | 6.570581 | 12.60573 | 24.09992 | 9.463556 | -0.40234 | -0.09387 | 0.625956 |
| 7 | 25672800 | 25675599 | 46.81355 | 75.34996 | 83.76131 | 35.57975 | 0.201647 | 1.845448 | 0.625967 |
| 9 | 22344000 | 22346399 | 18.10852 | 31.82836 | 34.86689 | 15.43966 | 0.256521 | 0.608691 | 0.625989 |
| 19 | 35736000 | 35737399 | 15.10752 | 26.48626 | 41.4444 | 18.31293 | -0.25508 | 0.6464 | 0.625997 |
| 19 | 13592200 | 13593199 | 10.12139 | 18.96208 | 33.55701 | 13.85716 | -0.35674 | 0.132127 | 0.625999 |
| 4 | 115233400 | 115237799 | 79.98438 | 125.6287 | 138.1124 | 57.28613 | 0.182922 | 2.560207 | 0.626001 |
| 12 | 57243400 | 57244199 | 7.269036 | 13.97873 | 25.28184 | 9.860521 | -0.31445 | -0.10711 | 0.626007 |
| 19 | 45995000 | 46000799 | 69.08263 | 108.6009 | 157.1491 | 65.18959 | -0.21205 | 2.537869 | 0.626021 |
| 4 | 146463400 | 146466799 | 54.17704 | 87.03008 | 91.40159 | 38.71422 | 0.279924 | 1.91906 | 0.626026 |
| 13 | 83307800 | 83309399 | 15.76937 | 27.52049 | 44.12219 | 19.42631 | -0.29049 | 0.721328 | 0.626036 |
| 3 | 108282200 | 108283199 | 8.441913 | 15.86138 | 27.49942 | 11.30032 | -0.32061 | 0.087838 | 0.626042 |
| 3 | 33675400 | 33677199 | 20.93727 | 35.09553 | 63.99924 | 27.91362 | -0.48275 | 1.16172 | 0.626061 |
| Y | 8048800 | 8049399 | 4.857173 | 9.405917 | 22.13538 | 8.411062 | -0.65292 | -0.22793 | 0.626066 |
| 9 | 120699800 | 120702799 | 23.29909 | 38.89718 | 60.28257 | 26.22571 | -0.24884 | 1.207518 | 0.626067 |
| 2 | 126840200 | 126843599 | 31.51607 | 51.21667 | 77.99905 | 33.46209 | -0.23682 | 1.580011 | 0.626071 |
| 8 | 99571400 | 99575599 | 58.01986 | 92.19655 | 133.9355 | 55.86854 | -0.20466 | 2.27135 | 0.626073 |
| 12 | 17877600 | 17883199 | 84.60348 | 132.5245 | 144.8371 | 60.07801 | 0.188393 | 2.612645 | 0.626103 |
| 5 | 127805000 | 127808199 | 38.5252 | 62.53235 | 69.32269 | 29.71366 | 0.218489 | 1.610458 | 0.626103 |
| 6 | 64353800 | 64356999 | 31.27151 | 52.55687 | 53.00023 | 23.15129 | 0.368872 | 1.154961 | 0.626107 |
| 7 | 17222800 | 17226199 | 83.83677 | 131.6794 | 124.3433 | 51.57497 | 0.402015 | 2.559669 | 0.626142 |
| 8 | 46392000 | 46393199 | 9.822016 | 18.27814 | 29.9717 | 12.51809 | -0.26028 | 0.165617 | 0.626148 |
| 2 | 6264800 | 6266799 | 22.75232 | 39.11011 | 43.10309 | 18.90663 | 0.244267 | 0.915737 | 0.626156 |

| | | | | | | | | | |
|---|---|---|---|---|---|---|---|---|---|
| 13 | 96664400 | 96667199 | 26.61548 | 44.1951 | 67.68045 | 29.34009 | -0.23198 | 1.28364 | 0.626157 |
| 15 | 94622200 | 94623199 | 8.665994 | 16.3137 | 28.0367 | 11.48371 | -0.30501 | 0.075389 | 0.626163 |
| 3 | 52327200 | 52327799 | 9.224818 | 17.83049 | 21.49242 | 8.238411 | 0.289767 | -0.15247 | 0.626221 |
| X | 97379200 | 97382199 | 18.51826 | 32.09188 | 51.17557 | 22.45309 | -0.28403 | 0.838633 | 0.62624 |
| X | 136018600 | 136022799 | 56.55919 | 90.74411 | 93.12239 | 39.36894 | 0.311168 | 1.963657 | 0.626246 |
| 5 | 125799000 | 125805599 | 99.71515 | 152.6141 | 215.4257 | 89.39213 | -0.20616 | 3.049196 | 0.626268 |
| 18 | 52388600 | 52389599 | 21.54964 | 36.10983 | 57.46046 | 25.05637 | -0.28675 | 1.161296 | 0.62627 |
| X | 116988800 | 116993999 | 91.5178 | 144.3175 | 117.7214 | 48.85368 | 0.617309 | 2.486478 | 0.626276 |
| 1 | 145130800 | 145132599 | 22.09068 | 38.62066 | 33.42517 | 14.73456 | 0.595185 | 0.714468 | 0.626282 |
| 1 | 121935000 | 121937399 | 21.85894 | 38.18881 | 39.07629 | 17.21627 | 0.354581 | 0.730533 | 0.626313 |
| 14 | 104990000 | 104990599 | 5.511452 | 10.75132 | 21.03274 | 7.52681 | -0.31024 | -0.43734 | 0.62632 |
| 16 | 80100200 | 80100599 | 5.511452 | 10.75132 | 21.03274 | 7.52681 | -0.31024 | -0.43734 | 0.62632 |
| 19 | 9908400 | 9910399 | 16.34121 | 28.88421 | 31.88678 | 14.10482 | 0.248588 | 0.532131 | 0.626324 |
| 12 | 115269600 | 115272199 | 42.52762 | 69.09806 | 71.96407 | 30.86861 | 0.309749 | 1.60333 | 0.626344 |
| X | 142718800 | 142720799 | 21.04108 | 37.61288 | 31.92557 | 14.01567 | 0.634328 | 0.480743 | 0.626346 |
| 4 | 60189600 | 60194599 | 113.8243 | 174.2631 | 187.4159 | 77.77497 | 0.186172 | 3.04564 | 0.62635 |
| 5 | 49600600 | 49602399 | 20.05853 | 34.90781 | 38.85195 | 17.07578 | 0.233394 | 0.775824 | 0.626361 |
| 14 | 16540200 | 16543199 | 19.53881 | 33.06372 | 62.51241 | 27.3631 | -0.53435 | 1.06635 | 0.626384 |
| 11 | 31396000 | 31399999 | 74.31664 | 117.1142 | 126.6263 | 52.5446 | 0.210563 | 2.487438 | 0.626386 |
| 7 | 81273400 | 81276599 | 24.61283 | 41.13135 | 62.59916 | 27.25065 | -0.22281 | 1.203506 | 0.626388 |
| 3 | 3310400 | 3313199 | 31.62994 | 52.96541 | 54.70705 | 23.84561 | 0.334506 | 1.194979 | 0.626398 |
| 7 | 11210600 | 11240999 | 866.095 | 1129.426 | 1041.556 | 436.1889 | 0.183922 | 5.685158 | 0.626426 |
| 19 | 9067600 | 9069199 | 12.15321 | 21.67286 | 34.70346 | 15.26608 | -0.27822 | 0.44242 | 0.62643 |
| 4 | 19133400 | 19133999 | 9.273343 | 17.04355 | 34.05734 | 14.47251 | -0.56194 | 0.24131 | 0.626445 |
| 11 | 117085400 | 117086599 | 6.279091 | 12.0732 | 23.60665 | 9.211131 | -0.42561 | -0.11878 | 0.62645 |
| 2 | 55815600 | 55816999 | 12.95962 | 24.20778 | 25.45613 | 10.56468 | 0.382891 | 0.153693 | 0.626454 |
| 7 | 35941400 | 35944399 | 30.2555 | 49.55704 | 74.86408 | 32.21339 | -0.21743 | 1.460398 | 0.626469 |
| 13 | 85816400 | 85817999 | 16.15653 | 28.94195 | 31.22681 | 13.68468 | 0.29161 | 0.432637 | 0.626503 |
| 7 | 143422200 | 143426599 | 42.02268 | 67.3221 | 102.5589 | 43.26107 | -0.26024 | 1.99256 | 0.62651 |
| 19 | 30940400 | 30943599 | 40.86783 | 65.68111 | 97.22625 | 41.20948 | -0.21337 | 1.890177 | 0.626511 |
| 4 | 149535600 | 149537999 | 18.40538 | 32.2113 | 35.61762 | 15.71489 | 0.243245 | 0.680154 | 0.626515 |
| 2 | 41793400 | 41793999 | 6.686053 | 12.91275 | 23.71652 | 9.125891 | -0.31762 | -0.1611 | 0.62652 |
| 12 | 21150600 | 21153999 | 20.20035 | 34.36969 | 54.38 | 23.83472 | -0.27776 | 1.011234 | 0.626538 |
| 5 | 83869600 | 83872199 | 31.25422 | 52.04966 | 48.15249 | 20.91603 | 0.493138 | 1.246357 | 0.626539 |
| 15 | 87331400 | 87332999 | 32.98595 | 54.23995 | 60.12088 | 25.91212 | 0.230242 | 1.407476 | 0.626554 |
| 1 | 18240800 | 18242599 | 23.68609 | 40.80605 | 44.13692 | 19.36142 | 0.271911 | 0.90261 | 0.626586 |
| 8 | 18287200 | 18289599 | 20.21561 | 35.66734 | 34.53912 | 15.28364 | 0.435538 | 0.578669 | 0.626595 |
| 5 | 70709200 | 70714799 | 77.16271 | 121.6524 | 132.979 | 55.18383 | 0.195296 | 2.482297 | 0.626607 |
| 6 | 39182200 | 39184399 | 13.90954 | 24.05555 | 50.54672 | 22.1758 | -0.68065 | 0.834919 | 0.626625 |
| 1 | 110114600 | 110117399 | 52.04455 | 83.46405 | 92.48282 | 39.07239 | 0.199891 | 1.968449 | 0.626626 |
| 4 | 32229400 | 32231999 | 15.22651 | 26.64919 | 41.48766 | 18.30553 | -0.24783 | 0.673841 | 0.62664 |
| 14 | 107721400 | 107725799 | 87.76561 | 136.813 | 149.4345 | 62.01729 | 0.18384 | 2.700135 | 0.626641 |
| 19 | 40134400 | 40135799 | 15.7075 | 27.36343 | 43.16141 | 18.9819 | -0.26711 | 0.748022 | 0.626647 |
| 17 | 70822600 | 70825999 | 56.02996 | 88.85164 | 130.1281 | 54.16878 | -0.21922 | 2.328402 | 0.62666 |
| 2 | 178059000 | 178060999 | 18.22812 | 32.18005 | 34.30416 | 15.17784 | 0.298144 | 0.562042 | 0.626692 |
| 18 | 87363000 | 87363799 | 10.43129 | 19.20475 | 31.37803 | 13.30279 | -0.2712 | 0.233457 | 0.626692 |
| 6 | 68755200 | 68757199 | 27.01692 | 46.52179 | 38.65742 | 16.9575 | 0.65055 | 0.913751 | 0.626704 |

| | | | | | | | | | |
|---|---|---|---|---|---|---|---|---|---|
| 18 | 45043800 | 45045999 | 27.12665 | 46.71895 | 44.57379 | 19.55288 | 0.452097 | 0.919827 | 0.626747 |
| 4 | 55504800 | 55505599 | 6.10238 | 11.84579 | 22.22845 | 8.350641 | -0.31437 | -0.25113 | 0.626749 |
| 12 | 112723000 | 112724599 | 12.95991 | 23.18219 | 36.71243 | 16.10826 | -0.25994 | 0.43273 | 0.626762 |
| 18 | 40840200 | 40844799 | 45.63565 | 73.79171 | 78.55878 | 33.5745 | 0.271596 | 1.706216 | 0.626774 |
| 13 | 27551400 | 27551999 | 20.08222 | 34.87081 | 37.96686 | 16.66328 | 0.26476 | 0.807101 | 0.626788 |
| 17 | 66210000 | 66212599 | 23.43611 | 39.32951 | 59.82041 | 26.11001 | -0.22199 | 1.152304 | 0.626804 |
| 4 | 118513600 | 118515799 | 15.66773 | 27.26237 | 43.69161 | 19.20101 | -0.29023 | 0.763804 | 0.626814 |
| 3 | 75130800 | 75131799 | 12.0972 | 21.84819 | 34.73942 | 15.09705 | -0.25645 | 0.363657 | 0.626814 |
| 5 | 138479000 | 138489799 | 156.2123 | 236.7904 | 227.8508 | 94.30241 | 0.335029 | 3.26078 | 0.626837 |
| X | 6051200 | 6052799 | 17.74239 | 31.63961 | 31.79255 | 13.98574 | 0.390298 | 0.475255 | 0.626844 |
| 3 | 38161600 | 38162399 | 6.85277 | 13.28972 | 24.52648 | 9.256462 | -0.29744 | -0.2201 | 0.626849 |
| 1 | 167023600 | 167025199 | 20.70177 | 34.4303 | 67.01052 | 29.08172 | -0.57631 | 1.249522 | 0.62687 |
| 15 | 15994400 | 15998999 | 95.39623 | 148.0161 | 161.0236 | 66.8735 | 0.184408 | 2.785297 | 0.626876 |
| 11 | 40695000 | 40696799 | 14.07682 | 26.24369 | 25.88443 | 10.77777 | 0.470904 | 0.176164 | 0.626882 |
| 5 | 78879800 | 78881799 | 19.10541 | 33.84076 | 33.52516 | 14.82335 | 0.404995 | 0.528992 | 0.626894 |
| 12 | 20178400 | 20179799 | 32.27337 | 53.68703 | 44.50675 | 19.31819 | 0.650906 | 1.259558 | 0.626908 |
| 7 | 14343800 | 14344599 | 13.23713 | 23.34765 | 38.35789 | 16.9745 | -0.32275 | 0.552812 | 0.626914 |
| 5 | 66714200 | 66716199 | 18.77228 | 32.26109 | 49.53605 | 21.72822 | -0.23221 | 0.911253 | 0.626921 |
| 1 | 94076600 | 94079399 | 24.7419 | 40.80551 | 74.59684 | 32.19187 | -0.48777 | 1.362187 | 0.626968 |
| 17 | 22091400 | 22092199 | 17.3465 | 30.45469 | 33.10104 | 14.64378 | 0.267992 | 0.620619 | 0.62697 |
| 14 | 88725600 | 88725999 | 6.997343 | 13.53821 | 24.38184 | 9.303592 | -0.27776 | -0.18238 | 0.626977 |
| 1 | 58845800 | 58848399 | 42.77039 | 69.35328 | 74.54022 | 31.9264 | 0.261684 | 1.640728 | 0.626978 |
| 15 | 39735400 | 39736199 | 6.434858 | 12.3451 | 24.10476 | 9.466 | -0.43185 | -0.09231 | 0.626979 |
| 9 | 3763600 | 3764399 | 11.10936 | 20.82552 | 23.66902 | 9.764037 | 0.279236 | 0.114322 | 0.626999 |
| 14 | 82280800 | 82281399 | 11.10936 | 20.82552 | 23.66902 | 9.764037 | 0.279236 | 0.114322 | 0.626999 |
| X | 62887400 | 62890799 | 35.15796 | 59.49723 | 49.05924 | 21.47463 | 0.658708 | 1.108977 | 0.627001 |
| 17 | 68218400 | 68221199 | 20.33273 | 34.42201 | 53.76507 | 23.53726 | -0.25958 | 1.058356 | 0.627003 |
| 4 | 79903000 | 79904999 | 44.51639 | 71.80653 | 79.90737 | 34.04359 | 0.203825 | 1.775651 | 0.627014 |
| 4 | 121432400 | 121447599 | 231.4161 | 341.18 | 320.8395 | 132.1257 | 0.332168 | 3.784702 | 0.627015 |
| X | 79180600 | 79183599 | 40.16999 | 65.51663 | 69.19803 | 29.72799 | 0.293534 | 1.540189 | 0.627036 |
| 1 | 110924400 | 110928799 | 52.31806 | 84.22061 | 87.2717 | 37.07772 | 0.303003 | 1.851586 | 0.627056 |
| 8 | 57668600 | 57668999 | 8.218014 | 15.9561 | 18.40342 | 6.901469 | 0.383756 | -0.24369 | 0.62707 |
| 16 | 49937400 | 49939399 | 15.50686 | 27.98658 | 25.19745 | 10.95991 | 0.558246 | 0.370543 | 0.627076 |
| 18 | 75471000 | 75475199 | 39.02673 | 62.95929 | 93.21374 | 39.71894 | -0.20701 | 1.769286 | 0.627114 |
| 6 | 109338600 | 109340799 | 22.04809 | 38.87299 | 35.0124 | 15.49525 | 0.539314 | 0.599316 | 0.627116 |
| 1 | 188578600 | 188579599 | 10.19317 | 19.68809 | 21.99629 | 8.459816 | 0.393139 | -0.13196 | 0.627131 |
| 14 | 44994600 | 45009199 | 334.3335 | 472.5517 | 473.4165 | 195.967 | 0.18114 | 4.421563 | 0.627136 |
| 15 | 38934800 | 38937399 | 43.89252 | 71.01743 | 66.71993 | 28.53086 | 0.452434 | 1.682797 | 0.627147 |
| 9 | 71316800 | 71318999 | 23.49623 | 40.80104 | 41.31432 | 18.13304 | 0.368765 | 0.823747 | 0.627166 |
| 9 | 106950000 | 106951199 | 17.44733 | 30.79767 | 33.41437 | 14.78453 | 0.272868 | 0.558036 | 0.627183 |
| X | 65241600 | 65250799 | 129.8349 | 198.8151 | 197.3157 | 81.88709 | 0.302205 | 3.048817 | 0.62721 |
| 13 | 33567800 | 33570999 | 42.35382 | 69.12842 | 69.77173 | 29.98236 | 0.359563 | 1.532811 | 0.627233 |
| 17 | 48523800 | 48530399 | 82.93323 | 129.0002 | 184.5984 | 76.63558 | -0.20789 | 2.738529 | 0.627239 |
| 2 | 19288600 | 19291999 | 30.63712 | 49.8852 | 76.58768 | 32.88259 | -0.24643 | 1.546131 | 0.62725 |
| 10 | 85140000 | 85141799 | 13.74079 | 24.21264 | 38.23788 | 16.92466 | -0.267 | 0.561956 | 0.627259 |
| 8 | 66809400 | 66812599 | 23.35507 | 39.33232 | 62.02695 | 27.1111 | -0.27382 | 1.123405 | 0.627262 |
| 4 | 5672200 | 5674599 | 22.58124 | 37.9059 | 58.71197 | 25.63089 | -0.24813 | 1.144112 | 0.627263 |

| | | | | | | | | | |
|---|---|---|---|---|---|---|---|---|---|
| 3 | 140089000 | 140091199 | 32.68454 | 54.76876 | 47.5253 | 20.72417 | 0.584967 | 1.184563 | 0.627263 |
| 3 | 21789800 | 21792799 | 25.81929 | 43.15281 | 70.67734 | 30.76962 | -0.32847 | 1.213042 | 0.627297 |
| 15 | 88510200 | 88515399 | 40.45232 | 65.13221 | 97.25614 | 41.33145 | -0.22267 | 1.830238 | 0.627326 |
| X | 66581200 | 66589199 | 79.4365 | 125.6939 | 112.2129 | 46.64185 | 0.49117 | 2.399338 | 0.627345 |
| 17 | 35581800 | 35588399 | 124.6132 | 188.539 | 258.4983 | 106.9237 | -0.1777 | 3.292434 | 0.627346 |
| 14 | 91319800 | 91322799 | 58.52005 | 93.4583 | 102.7694 | 43.15482 | 0.204112 | 2.107041 | 0.627367 |
| X | 88829800 | 88831799 | 24.63052 | 42.38811 | 44.68068 | 19.59932 | 0.308736 | 0.91811 | 0.62737 |
| 5 | 116253200 | 116255199 | 15.37918 | 26.86713 | 42.05995 | 18.5303 | -0.25585 | 0.704548 | 0.627385 |
| 6 | 10512000 | 10515799 | 35.83623 | 59.41377 | 59.97887 | 25.97351 | 0.367219 | 1.319405 | 0.627392 |
| X | 94612200 | 94616199 | 38.03511 | 62.74033 | 62.91028 | 27.15062 | 0.376254 | 1.385195 | 0.627398 |
| 19 | 21678800 | 21686399 | 85.65167 | 135.5191 | 127.491 | 52.99044 | 0.415628 | 2.407049 | 0.627401 |
| 14 | 26318200 | 26320399 | 20.10357 | 35.43439 | 35.13151 | 15.54886 | 0.401231 | 0.590206 | 0.627402 |
| 11 | 92661000 | 92661799 | 9.382652 | 17.3206 | 29.3784 | 12.4096 | -0.32055 | 0.206917 | 0.627402 |
| 14 | 6610400 | 6613599 | 28.31829 | 48.31982 | 47.91706 | 21.00686 | 0.394417 | 1.010443 | 0.62741 |
| 4 | 131330200 | 131332199 | 17.40034 | 30.26651 | 47.37 | 20.81293 | -0.25644 | 0.781391 | 0.627424 |
| 3 | 28216200 | 28216999 | 10.63699 | 19.36324 | 31.77102 | 13.67138 | -0.2921 | 0.300741 | 0.627438 |
| 2 | 164211800 | 164213599 | 12.16878 | 21.84727 | 34.72292 | 15.18815 | -0.26181 | 0.401284 | 0.627447 |
| 16 | 84148800 | 84150199 | 15.74876 | 29.18097 | 28.00995 | 11.7718 | 0.500251 | 0.233085 | 0.627456 |
| 10 | 78912200 | 78916599 | 79.22285 | 125.4419 | 126.0871 | 52.42693 | 0.321129 | 2.386422 | 0.627517 |
| 3 | 4612800 | 4613799 | 11.77087 | 21.07921 | 34.66668 | 15.1961 | -0.31324 | 0.411299 | 0.627517 |
| 9 | 92215800 | 92217199 | 11.77087 | 21.07921 | 34.66668 | 15.1961 | -0.31324 | 0.411299 | 0.627517 |
| 5 | 41131800 | 41132599 | 8.740675 | 16.51873 | 28.12675 | 11.43471 | -0.28201 | 0.053492 | 0.627525 |
| 8 | 47482800 | 47485799 | 25.77152 | 42.70607 | 65.29973 | 28.268 | -0.22998 | 1.302064 | 0.627546 |
| 2 | 15419000 | 15420799 | 19.3654 | 33.44375 | 55.00876 | 24.13234 | -0.32965 | 0.877922 | 0.62755 |
| 3 | 12942800 | 12943399 | 10.46485 | 19.8556 | 23.26206 | 9.378194 | 0.263594 | 0.027759 | 0.627564 |
| 7 | 104542000 | 104542799 | 10.46485 | 19.8556 | 23.26206 | 9.378194 | 0.263594 | 0.027759 | 0.627564 |
| 5 | 149208000 | 149217799 | 94.00574 | 144.5474 | 202.932 | 84.27098 | -0.19315 | 2.954651 | 0.627612 |
| 12 | 76516200 | 76517999 | 15.20313 | 27.09938 | 29.89262 | 13.15486 | 0.255894 | 0.454943 | 0.627681 |
| 2 | 126367600 | 126370599 | 30.42966 | 49.72816 | 75.25547 | 32.35353 | -0.22202 | 1.492377 | 0.627688 |
| 7 | 60010600 | 60011399 | 11.28462 | 21.14299 | 24.17049 | 9.979745 | 0.26985 | 0.120574 | 0.627705 |
| 14 | 51676400 | 51676999 | 11.47214 | 21.44105 | 24.17063 | 10.02184 | 0.285145 | 0.138408 | 0.627715 |
| 5 | 38284800 | 38288199 | 29.13637 | 47.63225 | 73.35629 | 31.54129 | -0.24679 | 1.48309 | 0.627736 |
| 19 | 9179600 | 9181399 | 15.89792 | 28.56486 | 25.28574 | 11.0529 | 0.578068 | 0.400421 | 0.627754 |
| 17 | 30867800 | 30870199 | 22.8472 | 38.5269 | 60.4917 | 26.45045 | -0.26758 | 1.104981 | 0.62779 |
| 3 | 30476200 | 30479599 | 28.37251 | 47.79728 | 51.80864 | 22.64699 | 0.265228 | 1.125317 | 0.627804 |
| 7 | 65613600 | 65614799 | 14.70354 | 26.67417 | 29.37648 | 12.69536 | 0.275405 | 0.345233 | 0.627809 |
| 16 | 15815800 | 15816999 | 14.70354 | 26.67417 | 29.37648 | 12.69536 | 0.275405 | 0.345233 | 0.627809 |
| 18 | 48350800 | 48352599 | 14.70354 | 26.67417 | 29.37648 | 12.69536 | 0.275405 | 0.345233 | 0.627809 |
| 9 | 114884800 | 114887799 | 29.5355 | 48.09697 | 75.15499 | 32.26882 | -0.27161 | 1.54247 | 0.627809 |
| 14 | 52829200 | 52833999 | 63.9274 | 102.0886 | 93.56001 | 39.28471 | 0.466453 | 2.111829 | 0.627811 |
| 1 | 17364600 | 17365199 | 10.10761 | 18.30134 | 35.91599 | 15.55882 | -0.55496 | 0.338057 | 0.627825 |
| 7 | 18488000 | 18490399 | 19.73535 | 34.41737 | 38.26355 | 16.83931 | 0.235257 | 0.743931 | 0.627828 |
| 2 | 150950600 | 150953599 | 54.03666 | 86.69061 | 93.74321 | 39.627 | 0.235638 | 1.960252 | 0.627831 |
| 6 | 30912800 | 30915199 | 20.48394 | 35.8189 | 38.0367 | 16.77031 | 0.301255 | 0.70737 | 0.627834 |
| 7 | 40757600 | 40779999 | 324.6426 | 462.3604 | 449.4795 | 185.8223 | 0.235572 | 4.31065 | 0.627866 |
| 3 | 137363800 | 137365599 | 15.41311 | 26.76095 | 50.59885 | 22.20608 | -0.52814 | 0.803331 | 0.627868 |
| 6 | 23811000 | 23812399 | 18.34778 | 31.33331 | 50.05554 | 21.94585 | -0.29052 | 0.968207 | 0.627893 |

| | | | | | | | | | |
|---|---|---|---|---|---|---|---|---|---|
| 18 | 37579800 | 37583199 | 80.49169 | 126.2922 | 138.7737 | 57.55315 | 0.182462 | 2.581279 | 0.627946 |
| 13 | 61730200 | 61734399 | 103.0084 | 159.0482 | 172.5512 | 71.67425 | 0.182957 | 2.877446 | 0.627966 |
| X | 115059400 | 115062999 | 45.104 | 73.30541 | 73.13089 | 31.37584 | 0.372033 | 1.601923 | 0.627967 |
| 7 | 24721000 | 24726199 | 28.27213 | 45.98585 | 75.40676 | 32.36313 | -0.34218 | 1.556757 | 0.62797 |
| 1 | 148448400 | 148450399 | 16.93715 | 30.30209 | 31.63632 | 13.87887 | 0.337701 | 0.44584 | 0.628045 |
| 2 | 147878200 | 147884199 | 53.06302 | 84.1638 | 128.2596 | 53.40184 | -0.27617 | 2.318984 | 0.628053 |
| 1 | 167458000 | 167461799 | 42.67626 | 68.41808 | 101.8534 | 43.01025 | -0.22583 | 1.971163 | 0.628084 |
| 14 | 54683000 | 54683999 | 7.911136 | 15.05024 | 26.08989 | 10.45928 | -0.29 | -0.00713 | 0.628116 |
| Y | 81285200 | 81285799 | 4.98524 | 9.652718 | 22.13353 | 8.415256 | -0.6165 | -0.22373 | 0.628125 |
| 15 | 39200800 | 39201399 | 10.05092 | 19.13478 | 22.21183 | 8.891018 | 0.285506 | -0.00364 | 0.628127 |
| 2 | 108082800 | 108084599 | 18.20734 | 32.23951 | 33.68853 | 14.89656 | 0.328375 | 0.528386 | 0.628146 |
| 15 | 45714200 | 45715599 | 18.20734 | 32.23951 | 33.68853 | 14.89656 | 0.328375 | 0.528386 | 0.628146 |
| 3 | 51677200 | 51680399 | 31.56929 | 51.60353 | 78.40439 | 33.7104 | -0.22752 | 1.493157 | 0.628151 |
| 1 | 99943000 | 99944399 | 15.67268 | 28.62782 | 29.35404 | 12.57741 | 0.385738 | 0.313182 | 0.628164 |
| 2 | 99341200 | 99344799 | 41.87861 | 68.16113 | 71.76592 | 30.80776 | 0.2959 | 1.57271 | 0.628172 |
| 6 | 139996200 | 139998199 | 18.34459 | 32.19643 | 35.50612 | 15.70342 | 0.24708 | 0.635358 | 0.62821 |
| X | 163392000 | 163394399 | 33.70117 | 56.83057 | 51.87784 | 22.68407 | 0.512325 | 1.13671 | 0.628211 |
| 7 | 107783600 | 107791199 | 98.84815 | 153.844 | 160.3547 | 66.56516 | 0.24949 | 2.733519 | 0.628216 |
| X | 106924800 | 106925999 | 16.05604 | 28.92785 | 30.77452 | 13.41459 | 0.317085 | 0.397086 | 0.628217 |
| 6 | 74318200 | 74319199 | 12.31743 | 21.88397 | 35.19628 | 15.53161 | -0.28772 | 0.467642 | 0.628229 |
| 3 | 100995400 | 100997199 | 18.43098 | 32.40989 | 34.94502 | 15.4736 | 0.279698 | 0.607769 | 0.628233 |
| 5 | 146195000 | 146199999 | 987.2989 | 1255.079 | 1541.269 | 647.0771 | -0.2661 | 6.09999 | 0.628276 |
| 4 | 16184200 | 16186199 | 24.68688 | 42.85342 | 42.41499 | 18.61199 | 0.401578 | 0.837615 | 0.628306 |
| 7 | 15127400 | 15166599 | 1110.118 | 1429.979 | 1201.132 | 503.6259 | 0.300939 | 5.886441 | 0.62831 |
| 4 | 51493600 | 51494999 | 13.72446 | 24.22827 | 38.19585 | 16.89985 | -0.2635 | 0.541225 | 0.628325 |
| 5 | 17138800 | 17140999 | 20.62954 | 36.18473 | 32.19229 | 14.23049 | 0.555294 | 0.650856 | 0.628344 |
| 4 | 115848000 | 115849599 | 28.78456 | 48.39005 | 52.34832 | 22.86838 | 0.267987 | 1.14623 | 0.628352 |
| 11 | 66518600 | 66519799 | 12.42866 | 22.22844 | 35.16123 | 15.42968 | -0.25844 | 0.425932 | 0.628353 |
| 14 | 119238000 | 119239599 | 16.09052 | 28.68038 | 31.19116 | 13.72662 | 0.276012 | 0.46197 | 0.628353 |
| 8 | 48867600 | 48870999 | 23.93653 | 39.61534 | 71.70112 | 31.01214 | -0.47264 | 1.318622 | 0.628364 |
| 2 | 30914400 | 30915799 | 13.9792 | 24.57766 | 38.83651 | 17.19731 | -0.26889 | 0.591398 | 0.628368 |
| 9 | 57173400 | 57174999 | 10.48891 | 19.14093 | 31.29431 | 13.42469 | -0.28398 | 0.284911 | 0.628369 |
| 10 | 102863800 | 102866199 | 21.17621 | 35.56746 | 56.42761 | 24.64244 | -0.28251 | 1.132581 | 0.628386 |
| 10 | 11628400 | 11629599 | 18.33554 | 31.68396 | 49.00236 | 21.49772 | -0.24086 | 0.860811 | 0.62839 |
| 15 | 12652600 | 12656599 | 37.96307 | 61.71685 | 67.84901 | 29.11403 | 0.232543 | 1.576788 | 0.628405 |
| 19 | 8253400 | 8255799 | 18.58853 | 32.84538 | 29.66274 | 13.12193 | 0.536586 | 0.545717 | 0.628408 |
| 5 | 117155800 | 117159799 | 49.78765 | 79.4955 | 115.9231 | 48.66369 | -0.20272 | 2.109422 | 0.628435 |
| X | 107448400 | 107450599 | 23.87178 | 41.30518 | 36.32168 | 15.93524 | 0.570735 | 0.856573 | 0.628441 |
| 16 | 94749400 | 94753199 | 39.87432 | 64.11556 | 95.47612 | 40.49622 | -0.22107 | 1.872525 | 0.628479 |
| 12 | 56182400 | 56185599 | 45.90322 | 74.23681 | 78.42234 | 33.52344 | 0.283056 | 1.703397 | 0.628508 |
| 12 | 80836600 | 80837999 | 11.34742 | 20.37677 | 33.23658 | 14.53553 | -0.29878 | 0.390463 | 0.628548 |
| 11 | 8037200 | 8039399 | 18.27414 | 31.97642 | 34.04266 | 15.01793 | 0.297559 | 0.677839 | 0.628549 |
| 14 | 67643600 | 67644199 | 10.31613 | 19.96803 | 22.14705 | 8.433573 | 0.41753 | -0.16151 | 0.628569 |
| 16 | 44411400 | 44412799 | 17.42103 | 30.92222 | 32.60549 | 14.39977 | 0.316914 | 0.49925 | 0.628602 |
| 4 | 20295000 | 20296999 | 33.6103 | 54.08064 | 101.1922 | 42.96359 | -0.5485 | 1.845834 | 0.628632 |
| 13 | 14108600 | 14110199 | 18.66119 | 33.16798 | 33.06485 | 14.58613 | 0.398647 | 0.500684 | 0.628649 |
| 5 | 97057200 | 97058399 | 10.81898 | 19.72649 | 31.79584 | 13.65435 | -0.26456 | 0.293924 | 0.628661 |

| | | | | | | | | | |
|---|---|---|---|---|---|---|---|---|---|
| X | 33233400 | 33238399 | 51.93446 | 84.44005 | 57.05535 | 24.4835 | 0.933068 | 1.598143 | 0.628683 |
| 13 | 15905800 | 15909199 | 49.91131 | 80.61209 | 81.09694 | 34.60761 | 0.351205 | 1.749526 | 0.628717 |
| 5 | 105987800 | 105988599 | 12.90202 | 23.56734 | 25.56941 | 10.95555 | 0.305034 | 0.284867 | 0.628725 |
| 12 | 21103000 | 21117399 | 291.8109 | 417.2446 | 423.2238 | 174.8533 | 0.180022 | 4.250605 | 0.628738 |
| 15 | 23669800 | 23670999 | 16.198 | 29.36209 | 30.28978 | 13.103 | 0.368253 | 0.362268 | 0.628739 |
| 11 | 12737800 | 12740799 | 38.15944 | 62.38323 | 67.70637 | 29.11198 | 0.256672 | 1.498827 | 0.628745 |
| 2 | 138463200 | 138466399 | 86.27059 | 134.5094 | 145.0522 | 60.1968 | 0.202485 | 2.695399 | 0.628811 |
| 11 | 28826400 | 28827799 | 21.45583 | 37.1559 | 41.03121 | 18.00189 | 0.243712 | 0.844351 | 0.628816 |
| 3 | 99461600 | 99463199 | 12.92486 | 23.01111 | 36.14864 | 15.92385 | -0.25237 | 0.459769 | 0.628827 |
| 4 | 99899000 | 99902999 | 25.59696 | 42.03835 | 76.6429 | 33.01597 | -0.48632 | 1.41838 | 0.628831 |
| 6 | 123223800 | 123226199 | 30.14343 | 50.59331 | 53.52611 | 23.365 | 0.299934 | 1.165093 | 0.628836 |
| 11 | 90978600 | 90979399 | 10.6609 | 20.54989 | 22.48387 | 8.710049 | 0.413276 | -0.10229 | 0.628867 |
| 5 | 66746200 | 66748199 | 19.21876 | 32.87177 | 51.68135 | 22.66149 | -0.26742 | 0.956973 | 0.628874 |
| Y | 9950800 | 9953999 | 17.99563 | 31.58538 | 23.77024 | 10.51334 | 0.795179 | 0.60744 | 0.62888 |
| 8 | 129355600 | 129356799 | 9.360574 | 17.43945 | 28.9413 | 12.06424 | -0.27501 | 0.152387 | 0.628883 |
| 12 | 15144400 | 15146199 | 18.28107 | 33.08895 | 30.67113 | 13.29261 | 0.520073 | 0.386643 | 0.628915 |
| 10 | 74645400 | 74645999 | 7.887548 | 15.17484 | 27.07079 | 10.54491 | -0.29313 | -0.09906 | 0.62892 |
| 11 | 6872000 | 6875399 | 32.9668 | 53.57225 | 80.04518 | 34.33895 | -0.20947 | 1.584294 | 0.628941 |
| 18 | 6331000 | 6334199 | 38.91625 | 63.58554 | 67.85165 | 29.16861 | 0.280481 | 1.50727 | 0.628941 |
| 9 | 107541400 | 107545799 | 27.21027 | 44.57977 | 70.32995 | 30.26546 | -0.27957 | 1.448485 | 0.628962 |
| 3 | 128422800 | 128423399 | 5.208123 | 10.16822 | 19.93172 | 7.033277 | -0.29365 | -0.49184 | 0.628962 |
| 3 | 3402800 | 3413599 | 223.3957 | 326.3016 | 336.3937 | 138.5933 | 0.186847 | 3.929233 | 0.628991 |
| 7 | 29653000 | 29653599 | 7.144629 | 13.6709 | 25.12048 | 9.939379 | -0.35643 | -0.06137 | 0.629006 |
| 4 | 55262400 | 55264599 | 16.96659 | 30.05298 | 32.71157 | 14.46356 | 0.269924 | 0.514069 | 0.629033 |
| 1 | 117595000 | 117596199 | 20.10868 | 35.16664 | 38.1194 | 16.80836 | 0.271846 | 0.705292 | 0.629033 |
| 19 | 47003000 | 47005199 | 20.77866 | 34.99212 | 57.03774 | 24.93012 | -0.32139 | 1.106137 | 0.629042 |
| 10 | 34685200 | 34686999 | 13.72228 | 24.1576 | 38.74512 | 17.15235 | -0.28978 | 0.572123 | 0.629051 |
| 1 | 117413200 | 117418799 | 85.72404 | 134.6989 | 115.9989 | 48.11648 | 0.53521 | 2.550797 | 0.629058 |
| 14 | 41101800 | 41112199 | 119.5478 | 183.8045 | 187.6855 | 77.9386 | 0.265626 | 2.964887 | 0.62906 |
| 1 | 133925600 | 133929399 | 52.3142 | 83.9189 | 92.27275 | 38.99948 | 0.211434 | 1.95965 | 0.629069 |
| 16 | 22164600 | 22167999 | 30.74495 | 50.28631 | 75.99422 | 32.6813 | -0.21916 | 1.481223 | 0.629085 |
| 5 | 103291800 | 103292599 | 14.61909 | 25.03359 | 54.20745 | 23.77138 | -0.72674 | 0.933544 | 0.629123 |
| 8 | 92160400 | 92165199 | 87.0197 | 134.0256 | 193.1269 | 80.21733 | -0.22887 | 2.91966 | 0.629136 |
| 8 | 121593400 | 121594799 | 11.50488 | 20.7688 | 33.24754 | 14.45708 | -0.26642 | 0.36055 | 0.629143 |
| 8 | 102938200 | 102939399 | 9.825674 | 18.16166 | 29.87847 | 12.59887 | -0.27516 | 0.203935 | 0.629153 |
| 13 | 92680200 | 92692399 | 190.4459 | 282.1835 | 296.3127 | 122.035 | 0.179819 | 3.69573 | 0.629159 |
| X | 75804200 | 75805399 | 14.31802 | 26.10735 | 28.48366 | 12.23139 | 0.294978 | 0.313158 | 0.629165 |
| 18 | 63136000 | 63142399 | 67.8786 | 107.8409 | 117.1259 | 48.84693 | 0.214018 | 2.280321 | 0.629182 |
| 12 | 21931400 | 21962199 | 895.8907 | 1170.737 | 935.2544 | 391.5862 | 0.39405 | 5.651436 | 0.629195 |
| 4 | 148057200 | 148059999 | 38.9582 | 63.37904 | 69.76401 | 29.9434 | 0.231439 | 1.570127 | 0.6292 |
| 1 | 181190000 | 181191599 | 15.2792 | 27.23642 | 29.79114 | 13.10962 | 0.267898 | 0.453541 | 0.629202 |
| 2 | 15436600 | 15440399 | 47.79917 | 77.02302 | 68.67234 | 29.21793 | 0.52118 | 1.80427 | 0.629204 |
| 5 | 71974400 | 71978599 | 35.89322 | 58.16275 | 87.29368 | 37.37036 | -0.2198 | 1.64943 | 0.629223 |
| 6 | 132166200 | 132175799 | 128.2406 | 197.8761 | 175.5404 | 72.91619 | 0.472313 | 2.900614 | 0.629223 |
| 1 | 26554000 | 26555799 | 19.54445 | 34.10253 | 36.60058 | 16.11283 | 0.28577 | 0.730541 | 0.629233 |
| 4 | 143358400 | 143360199 | 38.4758 | 63.16842 | 65.49489 | 28.20969 | 0.325478 | 1.446487 | 0.629237 |
| 5 | 93903800 | 93905199 | 20.31402 | 35.65105 | 36.87293 | 16.30718 | 0.339132 | 0.648937 | 0.629281 |

| | | | | | | | | | |
|---|---|---|---|---|---|---|---|---|---|
| 13 | 11933200 | 11934599 | 9.84644 | 18.23919 | 29.91197 | 12.57617 | -0.26749 | 0.192958 | 0.629316 |
| 10 | 121689400 | 121692199 | 26.60199 | 44.72616 | 49.62338 | 21.67876 | 0.23167 | 1.135121 | 0.629336 |
| 5 | 146195400 | 146199999 | 983.2834 | 1250.298 | 1536.54 | 645.075 | -0.26679 | 6.095777 | 0.629337 |
| 12 | 41719200 | 41720799 | 10.08352 | 18.33308 | 35.98033 | 15.50661 | -0.55029 | 0.313565 | 0.629345 |
| 2 | 20841000 | 20845399 | 49.75562 | 80.82658 | 72.54877 | 31.11505 | 0.52348 | 1.620035 | 0.62936 |
| 8 | 44187200 | 44191799 | 37.30876 | 61.63429 | 60.75423 | 26.24439 | 0.401101 | 1.361758 | 0.629374 |
| 9 | 24946400 | 24946799 | 5.209128 | 10.16645 | 19.85436 | 7.050417 | -0.29644 | -0.46941 | 0.629376 |
| 3 | 15574200 | 15576199 | 18.08986 | 31.45692 | 52.81087 | 23.19934 | -0.3572 | 0.799152 | 0.629382 |
| 19 | 8178400 | 8183199 | 59.47498 | 95.004 | 103.3351 | 43.40573 | 0.220166 | 2.101156 | 0.629388 |
| 3 | 87190000 | 87193799 | 31.81882 | 52.1169 | 83.83201 | 36.07199 | -0.30781 | 1.471242 | 0.629399 |
| 15 | 62089600 | 62093399 | 39.67512 | 63.45589 | 115.8861 | 48.76454 | -0.52404 | 2.04667 | 0.629407 |
| 14 | 12426800 | 12427999 | 14.05734 | 24.80561 | 38.70795 | 17.12786 | -0.24905 | 0.54911 | 0.629411 |
| Y | 8429400 | 8430599 | 13.03701 | 23.83999 | 19.65292 | 8.409029 | 0.699533 | 0.262007 | 0.629414 |
| 8 | 66811200 | 66812199 | 12.3488 | 22.12002 | 35.23062 | 15.43985 | -0.26673 | 0.41818 | 0.629422 |
| 1 | 48979200 | 48980399 | 13.77232 | 25.61836 | 27.03911 | 11.30149 | 0.369747 | 0.192004 | 0.629431 |
| 4 | 134168600 | 134170999 | 24.67634 | 41.02387 | 62.96631 | 27.31887 | -0.23499 | 1.261682 | 0.629466 |
| 9 | 20242600 | 20243599 | 14.59499 | 26.22765 | 28.94192 | 12.64953 | 0.262197 | 0.399267 | 0.629474 |
| 10 | 78727400 | 78733999 | 70.69815 | 112.5978 | 99.24889 | 41.51727 | 0.518411 | 2.203909 | 0.629479 |
| 10 | 116447600 | 116452199 | 37.50249 | 60.51033 | 91.39765 | 38.9534 | -0.23553 | 1.759667 | 0.629494 |
| 6 | 61464600 | 61465199 | 11.85453 | 22.48128 | 24.80776 | 10.01174 | 0.34769 | 0.046814 | 0.629535 |
| 13 | 14380800 | 14383799 | 40.19354 | 65.49489 | 70.52418 | 30.28758 | 0.264764 | 1.55159 | 0.629593 |
| 8 | 70180600 | 70181599 | 13.90062 | 25.55049 | 22.88047 | 9.721692 | 0.588426 | 0.250611 | 0.629597 |
| 10 | 72513400 | 72516799 | 28.2087 | 46.55468 | 71.77938 | 30.98574 | -0.24252 | 1.3593 | 0.629602 |
| X | 77125600 | 77127799 | 25.58733 | 44.11814 | 44.62973 | 19.5781 | 0.368372 | 0.904791 | 0.629606 |
| 19 | 21783200 | 21785199 | 15.35049 | 26.81887 | 42.04489 | 18.52447 | -0.25796 | 0.702568 | 0.629641 |
| 12 | 99457400 | 99461999 | 42.81895 | 69.27793 | 76.44665 | 32.68935 | 0.220814 | 1.686676 | 0.629656 |
| 8 | 108487800 | 108490399 | 34.65119 | 57.30812 | 61.00812 | 26.37445 | 0.29057 | 1.344292 | 0.629657 |
| 19 | 23707000 | 23714199 | 92.32461 | 144.5712 | 149.5631 | 62.04047 | 0.267134 | 2.624597 | 0.629661 |
| X | 31374400 | 31379799 | 130.1727 | 201.3744 | 165.8886 | 68.90608 | 0.581918 | 2.85609 | 0.629676 |
| 4 | 50721800 | 50725399 | 36.49636 | 60.18293 | 62.62342 | 27.02317 | 0.322818 | 1.386744 | 0.629701 |
| 6 | 41411800 | 41414399 | 27.39721 | 46.51895 | 49.03326 | 21.48501 | 0.305725 | 1.050516 | 0.62971 |
| 16 | 57209800 | 57213799 | 1772.083 | 2145.467 | 2550.907 | 1077.111 | -0.29083 | 6.88099 | 0.629711 |
| 15 | 43698600 | 43700999 | 26.41204 | 44.67036 | 49.06337 | 21.47261 | 0.246362 | 1.080286 | 0.629711 |
| 10 | 58282600 | 58283399 | 6.861874 | 13.13593 | 24.5916 | 9.717214 | -0.38101 | -0.06935 | 0.62975 |
| 13 | 100704600 | 100706399 | 15.61383 | 27.37704 | 43.98969 | 19.43948 | -0.29354 | 0.647312 | 0.629762 |
| 1 | 183163800 | 183165199 | 11.55422 | 21.69166 | 24.0283 | 9.887343 | 0.318277 | 0.10677 | 0.629802 |
| 18 | 60582600 | 60586399 | 82.00287 | 129.0107 | 138.5873 | 57.49537 | 0.218071 | 2.52979 | 0.629833 |
| X | 79572000 | 79574799 | 41.55066 | 67.82444 | 68.79573 | 29.56405 | 0.352588 | 1.531992 | 0.629899 |
| 7 | 95963000 | 95968799 | 55.50552 | 88.75707 | 98.53987 | 41.45353 | 0.192306 | 2.063257 | 0.629904 |
| 4 | 29246000 | 29247999 | 38.72822 | 62.7638 | 67.32207 | 28.82294 | 0.263898 | 1.642026 | 0.629921 |
| 16 | 9555000 | 9556399 | 12.96101 | 22.96721 | 36.1752 | 15.99372 | -0.25988 | 0.492661 | 0.629943 |
| 13 | 17681800 | 17682399 | 6.39564 | 12.35495 | 23.72652 | 9.119145 | -0.37993 | -0.16934 | 0.629952 |
| 7 | 142990000 | 142993199 | 18.8458 | 32.16492 | 50.58265 | 22.1759 | -0.26806 | 0.976703 | 0.629953 |
| 1 | 14153200 | 14156199 | 51.7009 | 82.304 | 120.2463 | 50.26896 | -0.21012 | 2.205854 | 0.629995 |
| 17 | 75386600 | 75388399 | 22.38681 | 39.17076 | 38.43849 | 16.9564 | 0.414671 | 0.708865 | 0.629997 |
| 13 | 67872000 | 67872599 | 7.294411 | 13.95465 | 25.11524 | 9.94315 | -0.32747 | -0.05745 | 0.630007 |
| 13 | 98066600 | 98067999 | 11.7986 | 20.99564 | 35.84882 | 15.79831 | -0.37168 | 0.459901 | 0.630011 |

| | | | | | | | | | |
|---|---|---|---|---|---|---|---|---|---|
| 14 | 65106400 | 65107599 | 9.784793 | 18.21184 | 29.96723 | 12.51276 | -0.26513 | 0.163096 | 0.630026 |
| 18 | 45992800 | 45996399 | 50.31034 | 80.78394 | 89.36868 | 37.82866 | 0.204363 | 1.927754 | 0.630054 |
| 16 | 48845200 | 48847599 | 42.98048 | 69.49563 | 76.94001 | 32.881 | 0.21516 | 1.701642 | 0.630056 |
| 16 | 24858600 | 24860799 | 13.78469 | 24.24418 | 38.30861 | 16.96236 | -0.26872 | 0.584452 | 0.630058 |
| X | 118053800 | 118055999 | 36.71872 | 60.99657 | 58.77702 | 25.48759 | 0.434426 | 1.301653 | 0.630062 |
| 6 | 149442000 | 149446399 | 58.43393 | 94.09708 | 85.93013 | 36.52533 | 0.485582 | 1.851355 | 0.630066 |
| 18 | 83788600 | 83792599 | 32.26425 | 52.47138 | 79.04634 | 33.92332 | -0.22043 | 1.568162 | 0.630071 |
| 1 | 18362200 | 18362799 | 6.098466 | 11.86972 | 22.46557 | 8.260384 | -0.29796 | -0.32564 | 0.630082 |
| 6 | 26496400 | 26496799 | 6.098466 | 11.86972 | 22.46557 | 8.260384 | -0.29796 | -0.32564 | 0.630082 |
| 1 | 73444000 | 73447799 | 32.72458 | 55.05173 | 53.00979 | 23.1622 | 0.435397 | 1.153098 | 0.630086 |
| 4 | 90157600 | 90166599 | 164.2392 | 247.2208 | 251.1519 | 103.7253 | 0.24813 | 3.401838 | 0.630093 |
| 6 | 131687400 | 131689599 | 36.70113 | 60.29668 | 64.90651 | 27.96448 | 0.271874 | 1.431688 | 0.630193 |
| 7 | 7519800 | 7531799 | 60.47686 | 96.69779 | 90.2466 | 37.95986 | 0.442186 | 2.069958 | 0.630205 |
| 14 | 120581000 | 120590999 | 105.7459 | 161.896 | 226.7841 | 94.11234 | -0.19477 | 3.045957 | 0.630206 |
| 4 | 120996400 | 120998599 | 21.81496 | 38.42594 | 35.64402 | 15.77774 | 0.496619 | 0.609991 | 0.630207 |
| 4 | 147598400 | 147662399 | 2194.769 | 2640.62 | 3148.843 | 1330.746 | -0.30427 | 6.98064 | 0.630228 |
| 6 | 146423000 | 146426999 | 75.92859 | 119.4611 | 130.5295 | 54.15284 | 0.19363 | 2.522227 | 0.630255 |
| 2 | 149396200 | 149397399 | 10.92284 | 20.12115 | 33.00301 | 13.98089 | -0.27603 | 0.237965 | 0.630256 |
| 7 | 12540400 | 12552999 | 199.1901 | 298.0236 | 234.7602 | 96.81408 | 0.607116 | 3.522482 | 0.630265 |
| 2 | 25632800 | 25635399 | 49.90184 | 80.15042 | 89.06366 | 37.71558 | 0.198408 | 1.917294 | 0.630279 |
| 17 | 85596000 | 85599199 | 36.67945 | 59.87956 | 66.83113 | 28.72094 | 0.215041 | 1.514876 | 0.630302 |
| 12 | 47932400 | 47932999 | 9.849744 | 18.83561 | 22.33287 | 8.851779 | 0.268193 | -0.03502 | 0.630305 |
| 8 | 15664000 | 15665999 | 18.42133 | 32.74874 | 33.1512 | 14.62155 | 0.376925 | 0.499065 | 0.630307 |
| 12 | 55421000 | 55423799 | 27.47214 | 46.15342 | 51.26422 | 22.39033 | 0.229928 | 1.144108 | 0.630316 |
| 9 | 49046000 | 49048599 | 24.20415 | 40.64595 | 65.4039 | 28.55927 | -0.30299 | 1.154669 | 0.630342 |
| 1 | 88827000 | 88832399 | 47.24098 | 75.633 | 109.8953 | 46.31004 | -0.19328 | 2.021123 | 0.630345 |
| 8 | 46910200 | 46912999 | 23.0766 | 40.05259 | 41.9115 | 18.38945 | 0.321534 | 0.830273 | 0.63035 |
| 12 | 22470600 | 22470999 | 10.21631 | 19.6517 | 19.12766 | 7.454556 | 0.573029 | -0.09545 | 0.630375 |
| 1 | 178837400 | 178838799 | 11.94781 | 21.42942 | 34.17622 | 14.96145 | -0.26773 | 0.402939 | 0.630381 |
| 14 | 80321600 | 80323999 | 21.64331 | 37.74303 | 39.67823 | 17.4614 | 0.3155 | 0.755078 | 0.630385 |
| 15 | 43334400 | 43335399 | 11.17024 | 21.11909 | 24.19949 | 9.829446 | 0.285535 | 0.059297 | 0.630389 |
| 11 | 12745000 | 12749199 | 82.03377 | 128.3054 | 136.9958 | 56.83427 | 0.220294 | 2.636973 | 0.630405 |
| 2 | 55986800 | 55989799 | 31.33909 | 53.24969 | 49.02448 | 21.48507 | 0.500368 | 1.060269 | 0.630419 |
| 8 | 11600200 | 11600999 | 6.545511 | 12.6268 | 23.67285 | 9.150394 | -0.35319 | -0.14776 | 0.63042 |
| 18 | 19113000 | 19113799 | 11.82769 | 22.53689 | 24.36897 | 9.736614 | 0.389327 | 0.013136 | 0.630428 |
| 5 | 77712200 | 77714799 | 21.91426 | 37.7008 | 41.62829 | 18.26019 | 0.241952 | 0.905355 | 0.630433 |
| 2 | 71206800 | 71208199 | 14.31626 | 25.74777 | 28.35922 | 12.38632 | 0.265357 | 0.389276 | 0.630437 |
| 14 | 21820200 | 21821599 | 13.20226 | 24.08779 | 26.9791 | 11.57635 | 0.257904 | 0.29628 | 0.630438 |
| 10 | 41669800 | 41680799 | 235.7928 | 338.6064 | 450.5 | 185.9965 | -0.20499 | 4.185154 | 0.630459 |
| 14 | 33013800 | 33015999 | 15.29566 | 26.72221 | 43.05825 | 18.97056 | -0.29751 | 0.702376 | 0.630462 |
| 18 | 4299600 | 4301599 | 12.33013 | 22.32396 | 36.92724 | 15.99832 | -0.31021 | 0.356741 | 0.630467 |
| 1 | 85968200 | 85970399 | 30.52644 | 50.84115 | 56.29966 | 24.45599 | 0.234541 | 1.250539 | 0.63048 |
| Y | 4687600 | 4698199 | 73.6332 | 116.2989 | 85.97099 | 35.7013 | 0.760819 | 2.426335 | 0.630483 |
| 3 | 10632600 | 10634599 | 29.87409 | 49.64728 | 55.68584 | 24.15418 | 0.216 | 1.272337 | 0.630494 |
| 18 | 82804400 | 82805999 | 16.40316 | 28.54029 | 45.7635 | 20.11076 | -0.29118 | 0.769345 | 0.630499 |
| 11 | 5405400 | 5416399 | 139.8751 | 210.0107 | 286.4675 | 118.2299 | -0.17999 | 3.448053 | 0.630501 |
| 8 | 99242000 | 99243999 | 23.87972 | 41.55225 | 41.28555 | 18.14383 | 0.396362 | 0.799262 | 0.630503 |

| | | | | | | | | | |
|---|---|---|---|---|---|---|---|---|---|
| 5 | 111778800 | 111783399 | 57.91557 | 91.46651 | 138.11 | 57.35082 | -0.26812 | 2.432687 | 0.630523 |
| 1 | 122947600 | 122949799 | 17.40313 | 31.61746 | 30.24532 | 13.04713 | 0.479132 | 0.358418 | 0.630546 |
| 4 | 34712600 | 34721999 | 98.60616 | 152.4737 | 164.752 | 68.4341 | 0.190381 | 2.846763 | 0.630557 |
| 7 | 34172800 | 34176599 | 25.91324 | 43.0659 | 65.42555 | 28.37936 | -0.22038 | 1.269039 | 0.630566 |
| 4 | 92881800 | 92883599 | 20.92497 | 36.68806 | 37.37879 | 16.51751 | 0.360728 | 0.667562 | 0.630567 |
| 13 | 86685600 | 86686999 | 15.28851 | 28.01319 | 28.86861 | 12.31867 | 0.382547 | 0.291472 | 0.630579 |
| X | 94779000 | 94780999 | 15.77141 | 27.73385 | 47.72341 | 21.13175 | -0.39162 | 0.612393 | 0.63058 |
| 3 | 43492600 | 43499999 | 106.1696 | 163.2508 | 175.4772 | 72.8699 | 0.19175 | 2.957241 | 0.630599 |
| 1 | 130356800 | 130358799 | 18.18403 | 32.53575 | 31.46541 | 13.80287 | 0.44751 | 0.452507 | 0.630627 |
| 4 | 118678000 | 118679199 | 11.23385 | 20.28449 | 33.30021 | 14.47557 | -0.30203 | 0.358782 | 0.630647 |
| X | 33357800 | 33371599 | 555.1136 | 761.6377 | 620.3241 | 257.9874 | 0.436596 | 4.876301 | 0.630657 |
| 2 | 36080600 | 36081599 | 5.370863 | 10.4515 | 20.68651 | 7.618325 | -0.36329 | -0.3348 | 0.630662 |
| 7 | 42295400 | 42327199 | 634.4775 | 854.8185 | 767.2815 | 320.0069 | 0.269926 | 5.161829 | 0.630673 |
| 12 | 16659600 | 16663799 | 29.78889 | 48.75048 | 73.50235 | 31.61627 | -0.21519 | 1.470154 | 0.630683 |
| 12 | 20691200 | 20723599 | 1279.692 | 1616.404 | 1443.314 | 606.3458 | 0.184291 | 6.204156 | 0.630683 |
| 9 | 121155400 | 121155799 | 9.035058 | 17.53968 | 17.20158 | 6.459056 | 0.612993 | -0.23601 | 0.63073 |
| 2 | 19710000 | 19712399 | 19.79174 | 34.58543 | 37.58421 | 16.56148 | 0.268116 | 0.71334 | 0.630746 |
| 12 | 21039000 | 21042399 | 40.35206 | 64.75675 | 99.03386 | 41.88336 | -0.26288 | 1.934838 | 0.630754 |
| 16 | 47625600 | 47627599 | 20.72962 | 36.42119 | 36.19464 | 16.02236 | 0.396361 | 0.631559 | 0.630759 |
| 4 | 104861200 | 104862199 | 13.11342 | 23.18903 | 36.70227 | 16.23349 | -0.26807 | 0.517015 | 0.630765 |
| 5 | 75614400 | 75618399 | 30.80226 | 50.15009 | 76.58871 | 32.88208 | -0.23891 | 1.547775 | 0.630807 |
| 14 | 34109200 | 34113399 | 65.76428 | 104.3068 | 114.7715 | 47.78477 | 0.192936 | 2.32711 | 0.630822 |
| 1 | 136404200 | 136406399 | 15.55732 | 27.13878 | 42.61857 | 18.75923 | -0.26054 | 0.727435 | 0.630844 |
| X | 156408600 | 156411999 | 40.84726 | 66.70082 | 60.86364 | 26.15885 | 0.505158 | 1.518369 | 0.630849 |
| X | 82530200 | 82531599 | 20.67169 | 35.99944 | 39.35831 | 17.30574 | 0.259198 | 0.771539 | 0.630852 |
| 5 | 107212000 | 107215199 | 22.91048 | 38.524 | 58.73463 | 25.66004 | -0.22538 | 1.12962 | 0.630858 |
| 8 | 109046200 | 109050599 | 42.45382 | 69.21962 | 70.37903 | 30.23187 | 0.347923 | 1.547012 | 0.630863 |
| 4 | 96326000 | 96326599 | 12.08766 | 22.45957 | 25.60559 | 10.72074 | 0.257662 | 0.183072 | 0.630868 |
| 16 | 68891600 | 68894199 | 33.7664 | 56.02645 | 50.94616 | 22.07399 | 0.517548 | 1.295411 | 0.630868 |
| 15 | 69513400 | 69515399 | 17.20467 | 30.88718 | 31.05513 | 13.5843 | 0.394516 | 0.424737 | 0.63087 |
| 14 | 115732600 | 115733999 | 13.49353 | 23.70226 | 38.39229 | 16.99194 | -0.30453 | 0.600961 | 0.63088 |
| X | 20939200 | 20951399 | 157.558 | 240.0201 | 207.3225 | 85.93412 | 0.496522 | 3.162101 | 0.630932 |
| 2 | 118772800 | 118775599 | 26.38331 | 44.40413 | 49.62185 | 21.68485 | 0.221339 | 1.124596 | 0.630938 |
| 6 | 40052600 | 40054999 | 15.4746 | 26.99775 | 43.09811 | 18.97175 | -0.28426 | 0.723418 | 0.630943 |
| 6 | 102156400 | 102158399 | 18.28127 | 32.57416 | 32.55454 | 14.33073 | 0.397058 | 0.481481 | 0.630949 |
| 10 | 76352400 | 76356799 | 46.15018 | 74.52898 | 80.63721 | 34.40637 | 0.246226 | 1.744656 | 0.630951 |
| 9 | 100939400 | 100942599 | 34.32329 | 55.47709 | 84.349 | 36.03002 | -0.242 | 1.703308 | 0.630951 |
| 6 | 68262000 | 68262199 | 3.523366 | 6.889417 | 15.15394 | 4.918095 | -0.346 | -0.83052 | 0.630951 |
| 2 | 31440600 | 31441999 | 14.91478 | 26.04833 | 41.58075 | 18.31587 | -0.2838 | 0.7059 | 0.630953 |
| 12 | 54450200 | 54452599 | 30.20808 | 49.20182 | 75.7258 | 32.51622 | -0.24961 | 1.542689 | 0.630955 |
| 2 | 22626600 | 22632599 | 59.24035 | 93.85252 | 136.2561 | 56.6708 | -0.20796 | 2.357758 | 0.630968 |
| 1 | 122637800 | 122639999 | 30.5876 | 50.05148 | 75.40533 | 32.43337 | -0.21423 | 1.474363 | 0.630969 |
| 17 | 21521800 | 21525999 | 45.87897 | 74.76887 | 70.63741 | 30.33763 | 0.45319 | 1.562389 | 0.630984 |
| 14 | 62792400 | 62794399 | 14.06649 | 24.65561 | 39.91193 | 17.63243 | -0.3039 | 0.639299 | 0.630999 |
| X | 22445800 | 22446999 | 15.65583 | 28.12059 | 30.37804 | 13.28255 | 0.292328 | 0.413271 | 0.631008 |
| 7 | 112720200 | 112721399 | 12.7641 | 23.37508 | 25.1118 | 10.72345 | 0.322592 | 0.268805 | 0.631009 |
| 7 | 59279600 | 59280199 | 7.348073 | 14.17059 | 26.49063 | 10.24875 | -0.3498 | -0.12403 | 0.631034 |

| | | | | | | | | | |
|---|---|---|---|---|---|---|---|---|---|
| 3 | 113091400 | 113093799 | 29.62345 | 49.58658 | 54.08403 | 23.56838 | 0.256184 | 1.191765 | 0.631039 |
| 1 | 18830800 | 18834999 | 71.47324 | 112.1944 | 158.8217 | 65.87086 | -0.18202 | 2.569677 | 0.63104 |
| 4 | 16140200 | 16146399 | 75.18636 | 119.3336 | 121.2529 | 50.51834 | 0.308645 | 2.315961 | 0.631041 |
| X | 64196600 | 64197599 | 31.73119 | 52.52758 | 55.58666 | 24.0461 | 0.299383 | 1.326011 | 0.631066 |
| 4 | 59548600 | 59549999 | 11.90228 | 20.99327 | 36.37266 | 16.09597 | -0.39911 | 0.537901 | 0.631082 |
| 13 | 87427800 | 87428799 | 9.821597 | 18.42541 | 31.28727 | 12.89024 | -0.29459 | 0.116942 | 0.631088 |
| 1 | 47885000 | 47887399 | 21.05761 | 36.87272 | 37.95613 | 16.75425 | 0.345951 | 0.689304 | 0.631107 |
| 5 | 132098800 | 132099999 | 11.73393 | 21.41793 | 35.49493 | 15.22223 | -0.30356 | 0.300599 | 0.631127 |
| 7 | 26551600 | 26576999 | 485.3446 | 666.5793 | 634.698 | 263.9237 | 0.212731 | 4.858264 | 0.63113 |
| 4 | 147497000 | 147500399 | 46.22327 | 74.87374 | 77.01056 | 32.95901 | 0.323635 | 1.671317 | 0.631135 |
| 2 | 18278000 | 18286999 | 152.7185 | 230.9336 | 235.7759 | 97.50779 | 0.246656 | 3.308915 | 0.631151 |
| 4 | 96148400 | 96149599 | 16.06087 | 28.00328 | 43.65746 | 19.21051 | -0.25018 | 0.739026 | 0.631156 |
| 9 | 41743800 | 41745399 | 22.71751 | 39.35788 | 42.51983 | 18.65525 | 0.275266 | 0.847075 | 0.631158 |
| 2 | 3075400 | 3077399 | 26.71864 | 44.78673 | 49.12134 | 21.42464 | 0.248232 | 1.16453 | 0.631166 |
| 12 | 63031600 | 63033599 | 28.19436 | 47.86216 | 42.64892 | 18.68635 | 0.547283 | 1.051423 | 0.631179 |
| 15 | 96785600 | 96795399 | 142.9907 | 214.2861 | 291.4498 | 120.2305 | -0.17821 | 3.484963 | 0.631194 |
| 7 | 104465800 | 104468399 | 33.80394 | 55.85354 | 60.93503 | 26.3255 | 0.255203 | 1.352467 | 0.631202 |
| 17 | 20889000 | 20891199 | 29.79755 | 50.51863 | 50.15402 | 21.96609 | 0.391782 | 1.073939 | 0.631204 |
| 4 | 37235400 | 37239199 | 31.3609 | 52.68986 | 53.5409 | 23.38514 | 0.357931 | 1.1596 | 0.631229 |
| 12 | 63346000 | 63348399 | 51.38826 | 82.34574 | 91.26306 | 38.51025 | 0.198235 | 1.992027 | 0.631229 |
| X | 26589800 | 26602599 | 153.9038 | 234.4812 | 208.9393 | 86.60737 | 0.451804 | 3.158479 | 0.631234 |
| 15 | 16291600 | 16296199 | 38.08578 | 62.16535 | 67.68104 | 29.08447 | 0.250569 | 1.519506 | 0.631256 |
| 5 | 62451800 | 62452799 | 13.14669 | 23.60779 | 39.40996 | 17.23534 | -0.33242 | 0.41485 | 0.631277 |
| 1 | 111398600 | 111401199 | 33.94554 | 56.44395 | 58.20902 | 25.25802 | 0.336818 | 1.279034 | 0.631282 |
| X | 110113200 | 110113999 | 14.5793 | 26.47143 | 27.55027 | 11.89341 | 0.357958 | 0.338807 | 0.631295 |
| X | 60612200 | 60613399 | 19.69069 | 34.57199 | 35.50035 | 15.70587 | 0.349682 | 0.639384 | 0.631298 |
| 10 | 119428200 | 119429799 | 11.78641 | 21.22456 | 33.69107 | 14.69526 | -0.25737 | 0.37646 | 0.631319 |
| 6 | 89892200 | 89895599 | 36.14665 | 60.32852 | 56.53558 | 24.59115 | 0.474599 | 1.247 | 0.631329 |
| 1 | 17534400 | 17538199 | 39.97276 | 64.46782 | 97.20123 | 41.40341 | -0.23372 | 1.779314 | 0.631335 |
| 7 | 76359000 | 76360599 | 19.39664 | 32.81338 | 53.19627 | 23.28312 | -0.31305 | 1.061297 | 0.631335 |
| 8 | 44860400 | 44862999 | 42.05518 | 68.1306 | 75.2311 | 32.20038 | 0.221602 | 1.659869 | 0.631374 |
| 7 | 23517400 | 23524599 | 148.0001 | 224.7282 | 198.7219 | 82.2891 | 0.458861 | 3.222449 | 0.631379 |
| 14 | 54198600 | 54201599 | 16.80551 | 29.30245 | 45.19848 | 19.88897 | -0.23505 | 0.742789 | 0.631387 |
| X | 144986400 | 144989599 | 30.74307 | 52.87022 | 44.53354 | 19.53436 | 0.630874 | 0.949874 | 0.631396 |
| 14 | 13813200 | 13814199 | 14.31274 | 26.9667 | 26.26584 | 10.74282 | 0.510068 | 0.116911 | 0.631403 |
| 18 | 8180200 | 8183199 | 58.35369 | 92.98749 | 101.0827 | 42.31598 | 0.217261 | 2.175499 | 0.631405 |
| 11 | 15605200 | 15605799 | 7.43009 | 14.26028 | 25.2112 | 9.888826 | -0.28943 | -0.08635 | 0.631426 |
| 18 | 25865400 | 25869199 | 33.1981 | 54.40972 | 58.28097 | 25.08423 | 0.277632 | 1.448027 | 0.631427 |
| 3 | 43676600 | 43677399 | 8.135633 | 15.33028 | 27.01465 | 11.04677 | -0.33803 | 0.065703 | 0.631431 |
| X | 33453000 | 33455799 | 20.90616 | 36.75732 | 27.48059 | 12.1687 | 0.805404 | 0.611795 | 0.631445 |
| 14 | 91676000 | 91678399 | 25.32519 | 43.80944 | 43.50695 | 19.08747 | 0.395779 | 0.872523 | 0.631445 |
| 3 | 129180800 | 129181599 | 10.49368 | 19.35678 | 31.42757 | 13.28899 | -0.25919 | 0.225187 | 0.631467 |
| 5 | 92538400 | 92541999 | 31.12017 | 50.69427 | 76.69696 | 32.93485 | -0.22488 | 1.541012 | 0.631483 |
| 15 | 33137400 | 33142199 | 96.72675 | 149.9412 | 163.7545 | 68.01707 | 0.177754 | 2.801531 | 0.631485 |
| 14 | 47472000 | 47475799 | 49.69658 | 79.41621 | 115.1438 | 48.39356 | -0.19315 | 2.082013 | 0.631493 |
| 11 | 96976600 | 96980599 | 53.68493 | 85.39942 | 123.3303 | 51.50563 | -0.19458 | 2.232416 | 0.6315 |
| X | 128328800 | 128330199 | 10.32099 | 19.21343 | 32.16685 | 13.42695 | -0.28951 | 0.172637 | 0.631507 |

| | | | | | | | | | |
|---|---|---|---|---|---|---|---|---|---|
| X | 115439000 | 115442399 | 36.23759 | 60.22492 | 58.84493 | 25.52511 | 0.41445 | 1.29335 | 0.63151 |
| 17 | 12484400 | 12486399 | 10.4202 | 18.67553 | 33.28049 | 14.57852 | -0.42738 | 0.39487 | 0.631511 |
| X | 148869000 | 148870199 | 10.15926 | 18.88634 | 31.02829 | 12.98194 | -0.26508 | 0.177774 | 0.631534 |
| 12 | 81504000 | 81517399 | 198.913 | 296.9546 | 278.1983 | 114.6787 | 0.354299 | 3.564917 | 0.631544 |
| 11 | 29861800 | 29863599 | 18.59732 | 32.48776 | 35.74789 | 15.74898 | 0.250344 | 0.710886 | 0.631568 |
| 3 | 53629200 | 53631399 | 24.60915 | 41.80516 | 46.3592 | 20.31629 | 0.23281 | 1.033531 | 0.63159 |
| 2 | 36706200 | 36728799 | 342.2215 | 485.7791 | 463.793 | 191.8469 | 0.256797 | 4.360097 | 0.63162 |
| X | 33214000 | 33217199 | 44.45507 | 71.89127 | 77.4605 | 33.11019 | 0.25454 | 1.702537 | 0.631632 |
| 2 | 10302600 | 10304199 | 12.28071 | 22.00986 | 34.67681 | 15.19027 | -0.25088 | 0.411421 | 0.631632 |
| 10 | 36132800 | 36133799 | 11.41762 | 20.78503 | 33.29559 | 14.32677 | -0.25782 | 0.309479 | 0.631634 |
| 4 | 60603000 | 60603599 | 9.785144 | 18.74646 | 22.34024 | 8.80815 | 0.267715 | -0.05398 | 0.63169 |
| 7 | 71967400 | 71969999 | 33.02505 | 54.50731 | 60.79747 | 26.24636 | 0.223263 | 1.363409 | 0.631706 |
| 4 | 151761800 | 151765799 | 36.07771 | 58.22221 | 87.95025 | 37.49306 | -0.23531 | 1.75164 | 0.63171 |
| 9 | 113241200 | 113241999 | 12.52232 | 22.98925 | 25.52547 | 10.86338 | 0.278552 | 0.25256 | 0.631755 |
| 7 | 17227800 | 17229999 | 35.1844 | 58.01378 | 53.05558 | 22.89352 | 0.508455 | 1.376724 | 0.631756 |
| 3 | 136430800 | 136433399 | 21.08313 | 36.08693 | 58.89889 | 25.82779 | -0.32113 | 0.96689 | 0.63176 |
| 15 | 97388200 | 97391199 | 54.84898 | 87.00086 | 128.9682 | 53.69953 | -0.23629 | 2.3191 | 0.631762 |
| 8 | 89226400 | 89228999 | 32.67841 | 53.72147 | 59.50241 | 25.64298 | 0.23118 | 1.409596 | 0.631765 |
| 3 | 80851200 | 80855599 | 33.32481 | 56.31765 | 49.07679 | 21.47338 | 0.579042 | 1.11095 | 0.631766 |
| 1 | 32444400 | 32446799 | 29.00004 | 48.44722 | 54.0485 | 23.52355 | 0.223775 | 1.211107 | 0.631767 |
| X | 29972200 | 29973399 | 16.01979 | 29.02507 | 23.42458 | 10.1393 | 0.720005 | 0.350751 | 0.631769 |
| 10 | 95576600 | 95578999 | 22.28022 | 38.41531 | 42.56234 | 18.67117 | 0.237549 | 0.886294 | 0.63178 |
| 8 | 51051200 | 51054399 | 40.83225 | 66.41998 | 62.31737 | 26.74608 | 0.461248 | 1.573474 | 0.631784 |
| 3 | 140569800 | 140570199 | 5.06657 | 9.902151 | 20.25004 | 7.000578 | -0.32787 | -0.56663 | 0.63179 |
| 2 | 28479800 | 28481399 | 14.42201 | 25.45681 | 39.74376 | 17.58513 | -0.24956 | 0.550782 | 0.6318 |
| 12 | 18798400 | 18917599 | 3961.782 | 4558.394 | 3705.593 | 1576.816 | 0.182765 | 7.682822 | 0.631827 |
| X | 42907600 | 42911199 | 22.68614 | 38.56518 | 69.07251 | 30.2723 | -0.45634 | 1.045957 | 0.631837 |
| 9 | 6474800 | 6475599 | 13.09871 | 24.61298 | 26.23482 | 10.77917 | 0.37562 | 0.122842 | 0.631837 |
| 15 | 12496000 | 12497599 | 14.3296 | 26.07264 | 28.88109 | 12.43528 | 0.270441 | 0.324094 | 0.631845 |
| 3 | 3767000 | 3767599 | 17.51592 | 30.76277 | 34.03547 | 15.06157 | 0.24243 | 0.617498 | 0.631861 |
| X | 19389600 | 19391399 | 24.52643 | 42.3699 | 43.59303 | 19.12444 | 0.344664 | 0.883511 | 0.631872 |
| 9 | 20016200 | 20018399 | 24.77802 | 42.34868 | 46.31102 | 20.30546 | 0.254145 | 0.979563 | 0.631873 |
| 4 | 40662600 | 40666599 | 44.19405 | 71.45525 | 78.18641 | 33.41208 | 0.232008 | 1.706386 | 0.631874 |
| 1 | 181147200 | 181150199 | 31.7154 | 52.77205 | 57.45816 | 24.94406 | 0.258848 | 1.265507 | 0.631884 |
| 1 | 25013000 | 25013799 | 8.030558 | 15.30637 | 26.68943 | 10.65688 | -0.29364 | -0.01892 | 0.631889 |
| 1 | 144340000 | 144344399 | 45.26993 | 73.81425 | 70.52768 | 30.29618 | 0.437463 | 1.554594 | 0.631913 |
| 13 | 48235000 | 48235999 | 9.471258 | 17.60495 | 29.41343 | 12.30855 | -0.28946 | 0.167281 | 0.631943 |
| 1 | 23868000 | 23870399 | 19.03066 | 32.6092 | 51.69173 | 22.66936 | -0.27886 | 0.939144 | 0.631964 |
| 3 | 6486400 | 6491999 | 60.83796 | 97.20112 | 103.7084 | 43.57518 | 0.248227 | 2.096994 | 0.631965 |
| 15 | 79147800 | 79149399 | 14.72178 | 25.7921 | 40.42919 | 17.85388 | -0.25767 | 0.652244 | 0.631971 |
| 18 | 10849400 | 10850199 | 8.880232 | 16.63526 | 27.96939 | 11.55162 | -0.28286 | 0.110712 | 0.631981 |
| 17 | 41326800 | 41327999 | 19.55779 | 33.21144 | 52.67449 | 23.08122 | -0.28131 | 1.029254 | 0.632015 |
| 6 | 18876000 | 18876999 | 12.06613 | 23.17556 | 22.89335 | 8.960221 | 0.54565 | -0.05469 | 0.632021 |
| 18 | 65532600 | 65535399 | 37.75951 | 60.81206 | 91.44647 | 38.87688 | -0.23225 | 1.81811 | 0.63203 |
| 12 | 55126600 | 55127599 | 8.149749 | 15.11911 | 28.92397 | 12.13582 | -0.48605 | 0.16558 | 0.632044 |
| 12 | 56957800 | 56960599 | 54.69209 | 87.41233 | 96.71082 | 40.65511 | 0.19652 | 2.077338 | 0.632079 |
| 13 | 17427200 | 17429199 | 23.37788 | 40.7736 | 40.13881 | 17.66564 | 0.409929 | 0.763161 | 0.632092 |

| | | | | | | | | | |
|---|---|---|---|---|---|---|---|---|---|
| 14 | 3981400 | 3982799 | 15.40598 | 27.3351 | 28.82837 | 12.73575 | 0.316347 | 0.480955 | 0.632094 |
| X | 14339200 | 14340399 | 17.0202 | 30.7379 | 30.79389 | 13.38321 | 0.406142 | 0.391742 | 0.632103 |
| 3 | 4235400 | 4236399 | 15.97135 | 28.37292 | 30.93458 | 13.65214 | 0.26998 | 0.484692 | 0.632134 |
| 10 | 47486200 | 47489599 | 54.04534 | 86.59561 | 95.51782 | 40.30004 | 0.20492 | 1.999652 | 0.632139 |
| 19 | 24201200 | 24203199 | 15.53307 | 26.96458 | 44.27098 | 19.4269 | -0.32523 | 0.797982 | 0.632171 |
| 17 | 4636600 | 4637999 | 14.8284 | 27.71426 | 27.17005 | 11.26516 | 0.485124 | 0.173355 | 0.632171 |
| 15 | 30349000 | 30350799 | 18.58237 | 33.09801 | 32.50374 | 14.31298 | 0.421847 | 0.485342 | 0.632193 |
| 1 | 142599200 | 142601399 | 21.98206 | 38.40229 | 39.05128 | 17.20479 | 0.36347 | 0.731215 | 0.632196 |
| 13 | 41373600 | 41378999 | 91.9389 | 141.613 | 198.0069 | 82.24519 | -0.1854 | 2.920023 | 0.632231 |
| 5 | 68719200 | 68722999 | 38.14013 | 61.52851 | 93.37421 | 39.78691 | -0.24262 | 1.766153 | 0.632254 |
| 11 | 78333800 | 78335199 | 17.10653 | 29.73139 | 45.78673 | 20.10706 | -0.23325 | 0.790272 | 0.632259 |
| 19 | 9229000 | 9229999 | 16.19797 | 28.68734 | 31.84519 | 14.0809 | 0.241732 | 0.510056 | 0.632266 |
| 18 | 58160800 | 58161599 | 8.065419 | 15.27155 | 26.53811 | 10.74562 | -0.30628 | 0.027662 | 0.632266 |
| 7 | 41074800 | 41075199 | 5.501064 | 10.74268 | 21.98044 | 7.724744 | -0.35058 | -0.49739 | 0.632271 |
| 18 | 18268200 | 18268599 | 5.501064 | 10.74268 | 21.98044 | 7.724744 | -0.35058 | -0.49739 | 0.632271 |
| 1 | 25059200 | 25060599 | 15.30575 | 27.5129 | 30.25829 | 13.22163 | 0.267003 | 0.403263 | 0.632279 |
| 1 | 109208400 | 109209799 | 15.30575 | 27.5129 | 30.25829 | 13.22163 | 0.267003 | 0.403263 | 0.632279 |
| 17 | 66697000 | 66698599 | 15.87283 | 28.20108 | 31.33228 | 13.82634 | 0.242849 | 0.48243 | 0.632304 |
| 8 | 86130600 | 86132999 | 13.1631 | 23.29462 | 36.73156 | 16.24397 | -0.26212 | 0.510686 | 0.632308 |
| X | 49335800 | 49338599 | 24.28659 | 42.41625 | 39.58218 | 17.43559 | 0.487033 | 0.750569 | 0.632313 |
| 9 | 94479400 | 94480999 | 13.72366 | 25.58195 | 27.1296 | 11.29874 | 0.367243 | 0.179755 | 0.632319 |
| 12 | 41316600 | 41324599 | 204.3201 | 296.3888 | 405.2186 | 167.0687 | -0.22985 | 4.033283 | 0.632322 |
| 19 | 13372600 | 13372999 | 4.609766 | 9.012776 | 18.39204 | 6.296563 | -0.31037 | -0.61066 | 0.632329 |
| 6 | 142112400 | 142116799 | 73.45547 | 114.749 | 163.565 | 67.8653 | -0.1974 | 2.658062 | 0.632333 |
| 1 | 18960800 | 18964399 | 37.34735 | 60.37688 | 90.76762 | 38.78167 | -0.22552 | 1.70356 | 0.632339 |
| 5 | 114697400 | 114703799 | 75.85575 | 118.19 | 168.5291 | 69.94689 | -0.20084 | 2.705761 | 0.63234 |
| 8 | 15668800 | 15675199 | 90.29315 | 140.927 | 152.0631 | 63.09877 | 0.202792 | 2.681566 | 0.63234 |
| 8 | 45401600 | 45407999 | 111.7531 | 167.7235 | 302.1072 | 124.6718 | -0.58124 | 3.451658 | 0.632362 |
| 13 | 25708800 | 25709999 | 20.83485 | 35.3326 | 54.31229 | 23.7892 | -0.23652 | 1.0466 | 0.632388 |
| 3 | 158579800 | 158581399 | 26.26246 | 44.35368 | 49.07235 | 21.4672 | 0.23581 | 1.092191 | 0.632399 |
| 9 | 114218400 | 114219799 | 18.51146 | 31.55663 | 50.58102 | 22.17306 | -0.29574 | 0.985514 | 0.632408 |
| 7 | 26834000 | 26846199 | 131.7119 | 198.102 | 274.1982 | 113.2194 | -0.19879 | 3.411532 | 0.632409 |
| 5 | 27170400 | 27173399 | 25.26766 | 43.15328 | 46.86492 | 20.54709 | 0.264044 | 0.990974 | 0.63242 |
| 6 | 114431800 | 114432999 | 19.72785 | 34.28604 | 37.42785 | 16.43592 | 0.261099 | 0.791582 | 0.632428 |
| 4 | 19150800 | 19151799 | 28.40659 | 47.28136 | 50.75638 | 22.03945 | 0.279302 | 1.247037 | 0.632433 |
| 12 | 104909400 | 104911599 | 21.01305 | 35.59586 | 54.84493 | 24.01453 | -0.23993 | 1.057532 | 0.632437 |
| 2 | 89560400 | 89561199 | 13.71617 | 24.23621 | 38.17567 | 16.88778 | -0.26178 | 0.530733 | 0.632453 |
| 13 | 53394400 | 53399999 | 63.12806 | 99.70162 | 142.3022 | 59.09254 | -0.187 | 2.436077 | 0.632456 |
| 7 | 18286400 | 18306599 | 303.1586 | 433.9817 | 428.7172 | 177.0902 | 0.219842 | 4.23432 | 0.632458 |
| 10 | 96315200 | 96317599 | 17.40348 | 30.17537 | 46.86995 | 20.56433 | -0.24611 | 0.820933 | 0.632467 |
| X | 103840200 | 103842399 | 29.9281 | 50.33923 | 52.93124 | 23.12675 | 0.308856 | 1.145667 | 0.63248 |
| 5 | 95115000 | 95116799 | 17.41841 | 31.32522 | 26.3469 | 11.50796 | 0.65187 | 0.406247 | 0.6325 |
| 12 | 16617800 | 16621799 | 27.11792 | 44.71494 | 68.01794 | 29.35022 | -0.22347 | 1.366013 | 0.63253 |
| 18 | 26827200 | 26830199 | 33.71313 | 54.43404 | 95.13125 | 40.58103 | -0.44426 | 1.73787 | 0.63253 |
| 16 | 48738800 | 48740799 | 30.27365 | 51.20127 | 51.2684 | 22.43762 | 0.379432 | 1.098822 | 0.632532 |
| 5 | 14468600 | 14469599 | 12.81041 | 23.92333 | 21.48358 | 8.919637 | 0.609237 | 0.147526 | 0.632549 |
| 13 | 40979400 | 40985399 | 68.77163 | 108.1863 | 153.8819 | 63.83998 | -0.18651 | 2.524195 | 0.632572 |

| | | | | | | | | | |
|---|---|---|---|---|---|---|---|---|---|
| 3 | 75276400 | 75280199 | 39.27294 | 63.87895 | 70.67904 | 30.33405 | 0.223851 | 1.576616 | 0.632595 |
| 1 | 71507000 | 71507599 | 5.525229 | 10.73634 | 21.16784 | 7.887213 | -0.37373 | -0.29089 | 0.632623 |
| 9 | 123023800 | 123027999 | 84.43429 | 131.857 | 142.1104 | 58.96546 | 0.205105 | 2.666573 | 0.63263 |
| 2 | 28836400 | 28839999 | 24.18809 | 40.18703 | 62.3585 | 27.04349 | -0.25073 | 1.266814 | 0.63264 |
| 12 | 50736600 | 50739599 | 19.6607 | 34.47484 | 36.98359 | 16.34425 | 0.286707 | 0.659327 | 0.63266 |
| 10 | 112872400 | 112872999 | 7.005308 | 13.52368 | 24.25167 | 9.350025 | -0.28568 | -0.14703 | 0.63266 |
| 7 | 134571800 | 134574199 | 24.48165 | 41.98732 | 45.25338 | 19.84727 | 0.275899 | 0.949005 | 0.632666 |
| 12 | 50456800 | 50457999 | 15.46601 | 28.07647 | 29.79913 | 12.86707 | 0.329074 | 0.346949 | 0.632666 |
| X | 81525200 | 81526399 | 21.3441 | 37.42569 | 31.36419 | 13.86055 | 0.641377 | 0.66137 | 0.632675 |
| 1 | 148512400 | 148514999 | 26.08306 | 44.60006 | 46.82802 | 20.53292 | 0.312722 | 0.980897 | 0.632698 |
| 13 | 62252000 | 62254199 | 24.13342 | 41.5229 | 44.66051 | 19.59033 | 0.279579 | 0.915921 | 0.6327 |
| 13 | 29515400 | 29515999 | 5.224677 | 10.19332 | 19.81022 | 7.076213 | -0.29772 | -0.45054 | 0.632711 |
| 12 | 21663600 | 21666999 | 33.75296 | 55.43591 | 60.76287 | 26.17579 | 0.245641 | 1.423181 | 0.632725 |
| 4 | 22370000 | 22373599 | 36.44001 | 60.20662 | 61.74432 | 26.67439 | 0.344181 | 1.36033 | 0.632739 |
| 4 | 12171200 | 12172199 | 10.11806 | 18.60239 | 30.83761 | 13.09823 | -0.29391 | 0.23728 | 0.632742 |
| 5 | 79389000 | 79390199 | 11.69385 | 21.88767 | 25.12647 | 10.39217 | 0.261605 | 0.131519 | 0.632775 |
| 14 | 89616600 | 89617199 | 11.69385 | 21.88767 | 25.12647 | 10.39217 | 0.261605 | 0.131519 | 0.632775 |
| 13 | 20114400 | 20116399 | 24.20325 | 41.59667 | 44.19456 | 19.38529 | 0.297046 | 0.928794 | 0.632786 |
| 1 | 6091000 | 6091799 | 12.67654 | 23.51961 | 26.55618 | 11.1407 | 0.269263 | 0.197155 | 0.632787 |
| 9 | 22878800 | 22879799 | 11.94935 | 22.55384 | 25.26897 | 10.30073 | 0.313506 | 0.082269 | 0.632789 |
| 8 | 98380200 | 98382199 | 18.42576 | 33.12809 | 31.61327 | 13.81131 | 0.470901 | 0.427304 | 0.632793 |
| 4 | 81931400 | 81931799 | 7.281512 | 14.02867 | 24.76914 | 9.609335 | -0.27251 | -0.12426 | 0.632796 |
| 6 | 69534000 | 69540599 | 70.236 | 111.8297 | 113.1031 | 47.294 | 0.319902 | 2.21563 | 0.632806 |
| 10 | 125590000 | 125591999 | 19.57756 | 34.25052 | 37.59628 | 16.58289 | 0.253746 | 0.695309 | 0.632815 |
| 4 | 143201800 | 143204399 | 28.4926 | 48.05673 | 42.76647 | 18.70115 | 0.548877 | 1.107489 | 0.632817 |
| 2 | 169055000 | 169060399 | 68.82951 | 108.0722 | 154.7527 | 64.18538 | -0.19826 | 2.562164 | 0.632824 |
| 5 | 92087800 | 92090799 | 22.40317 | 37.83661 | 57.6557 | 25.22222 | -0.22435 | 1.088976 | 0.632828 |
| 9 | 78521800 | 78524599 | 18.06418 | 31.09839 | 49.00536 | 21.49658 | -0.26902 | 0.890171 | 0.632833 |
| 14 | 7446000 | 7453999 | 107.6543 | 166.1217 | 177.3809 | 73.68067 | 0.205218 | 2.890691 | 0.63284 |
| 6 | 66222000 | 66224599 | 36.31704 | 59.43 | 65.86899 | 28.33216 | 0.227469 | 1.479111 | 0.632847 |
| 11 | 52046400 | 52050199 | 48.81613 | 78.05227 | 114.9959 | 48.36922 | -0.21539 | 2.063002 | 0.632868 |
| 3 | 33505600 | 33507199 | 15.03289 | 27.00112 | 29.18673 | 12.76188 | 0.290986 | 0.402674 | 0.632909 |
| 6 | 143516800 | 143520399 | 46.44578 | 74.67066 | 80.8982 | 34.30539 | 0.236609 | 1.87991 | 0.632928 |
| 3 | 21189800 | 21190799 | 13.59061 | 25.04944 | 27.4448 | 11.61748 | 0.302963 | 0.242182 | 0.632935 |
| 4 | 155396000 | 155397999 | 19.8889 | 33.99244 | 52.22201 | 22.89718 | -0.23435 | 0.967849 | 0.632947 |
| 5 | 130834200 | 130838799 | 49.03437 | 78.26763 | 114.1818 | 47.91091 | -0.20386 | 2.116064 | 0.632962 |
| 8 | 74223600 | 74225399 | 18.26043 | 31.1317 | 56.97945 | 24.97812 | -0.48643 | 0.991908 | 0.633005 |
| X | 123048200 | 123054399 | 79.34046 | 126.1505 | 115.6435 | 48.26689 | 0.459399 | 2.268969 | 0.633009 |
| 13 | 93942400 | 93943199 | 8.408569 | 16.04216 | 27.83767 | 11.09296 | -0.28376 | -0.01598 | 0.633037 |
| 12 | 12036000 | 12036999 | 9.325152 | 17.63184 | 30.42179 | 12.35509 | -0.30021 | 0.059004 | 0.63304 |
| 10 | 63369800 | 63370799 | 10.04418 | 18.30855 | 30.82009 | 13.24005 | -0.327 | 0.287551 | 0.633069 |
| 8 | 41270400 | 41271199 | 15.58084 | 27.83863 | 30.85798 | 13.55254 | 0.250905 | 0.445251 | 0.633094 |
| 4 | 103403400 | 103403999 | 5.665789 | 11.02401 | 21.26891 | 7.840845 | -0.32927 | -0.32866 | 0.633124 |
| 10 | 14003000 | 14008399 | 85.03783 | 131.0142 | 191.2417 | 79.43772 | -0.24713 | 2.912638 | 0.633124 |
| 13 | 65337200 | 65343399 | 68.74222 | 109.8565 | 102.445 | 43.05919 | 0.442542 | 2.097389 | 0.633165 |
| 19 | 24373800 | 24375799 | 19.49697 | 33.93167 | 37.32434 | 16.40472 | 0.250242 | 0.769558 | 0.633173 |
| 7 | 25536200 | 25538199 | 16.63571 | 29.01203 | 44.66678 | 19.6574 | -0.23233 | 0.738316 | 0.633196 |

| | | | | | | | | | |
|---|---|---|---|---|---|---|---|---|---|
| 6 | 48355200 | 48357199 | 15.94699 | 28.68499 | 30.69734 | 13.40582 | 0.306708 | 0.40419 | 0.633235 |
| 16 | 89396800 | 89397999 | 9.161728 | 17.18934 | 28.51572 | 11.74624 | -0.26082 | 0.103628 | 0.633254 |
| 3 | 69168600 | 69173399 | 50.54721 | 80.68921 | 116.3669 | 48.83364 | -0.18715 | 2.116425 | 0.633261 |
| 14 | 63300800 | 63301199 | 8.926485 | 17.29826 | 21.04022 | 7.968789 | 0.293313 | -0.1969 | 0.633292 |
| 15 | 75190800 | 75198799 | 98.55822 | 153.9767 | 153.6552 | 63.75303 | 0.316481 | 2.667437 | 0.633296 |
| 14 | 98733200 | 98733999 | 7.913806 | 15.24865 | 27.74087 | 10.7586 | -0.31508 | -0.11027 | 0.633309 |
| 7 | 42605400 | 42648999 | 663.9703 | 893.3574 | 693.7729 | 289.4064 | 0.476857 | 5.182824 | 0.633332 |
| 1 | 58837800 | 58838599 | 15.19865 | 27.27375 | 30.32249 | 13.26878 | 0.250121 | 0.413997 | 0.633347 |
| 15 | 12106200 | 12109599 | 34.88121 | 56.52693 | 84.83845 | 36.32121 | -0.2197 | 1.644009 | 0.633362 |
| 15 | 37335600 | 37336599 | 7.06075 | 13.39245 | 25.54396 | 10.30523 | -0.43392 | 0.006347 | 0.63337 |
| 15 | 76924800 | 76927999 | 26.09116 | 43.39547 | 66.01196 | 28.64916 | -0.22216 | 1.261495 | 0.633385 |
| 1 | 8942600 | 8945399 | 35.14888 | 57.53885 | 64.06335 | 27.55918 | 0.221264 | 1.47012 | 0.633395 |
| 2 | 23597600 | 23599599 | 30.67462 | 51.4604 | 46.98753 | 20.50465 | 0.511781 | 1.167327 | 0.633404 |
| 17 | 52785200 | 52785999 | 11.86614 | 21.20672 | 34.25287 | 15.04042 | -0.28867 | 0.427265 | 0.633405 |
| 14 | 92286400 | 92289999 | 59.44192 | 94.98402 | 89.06866 | 37.43147 | 0.434352 | 2.084896 | 0.633413 |
| 15 | 89724800 | 89728199 | 25.82839 | 42.79081 | 65.19043 | 28.21636 | -0.22469 | 1.30644 | 0.633423 |
| 6 | 132177800 | 132182999 | 57.89286 | 92.99774 | 92.06453 | 38.99415 | 0.365004 | 1.925628 | 0.633426 |
| 3 | 86804800 | 86806199 | 9.938909 | 18.52857 | 30.5472 | 12.72011 | -0.26477 | 0.155319 | 0.633447 |
| 10 | 126001400 | 126002599 | 9.938909 | 18.52857 | 30.5472 | 12.72011 | -0.26477 | 0.155319 | 0.633447 |
| 13 | 19957400 | 19958199 | 9.938909 | 18.52857 | 30.5472 | 12.72011 | -0.26477 | 0.155319 | 0.633447 |
| 13 | 54760800 | 54762999 | 11.0503 | 19.78357 | 33.27409 | 14.58892 | -0.34526 | 0.407339 | 0.633465 |
| 4 | 86087600 | 86090199 | 33.33597 | 54.93658 | 61.35253 | 26.46766 | 0.220679 | 1.382664 | 0.633478 |
| 4 | 18269400 | 18274999 | 58.49861 | 93.67246 | 98.85044 | 41.66652 | 0.267593 | 2.02514 | 0.633506 |
| 13 | 7020400 | 7021199 | 7.289376 | 14.01862 | 24.7085 | 9.635374 | -0.27692 | -0.10725 | 0.633532 |
| 9 | 121029800 | 121045399 | 192.0714 | 281.3076 | 375.7629 | 154.7854 | -0.18297 | 3.885513 | 0.633542 |
| X | 164228000 | 164231799 | 37.49479 | 61.73417 | 63.79676 | 27.51155 | 0.331853 | 1.408625 | 0.633557 |
| 7 | 133062200 | 133066999 | 40.52328 | 65.60212 | 73.21767 | 31.32213 | 0.205257 | 1.669436 | 0.633573 |
| 5 | 94875400 | 94875599 | 8.608473 | 16.0632 | 27.95275 | 11.62149 | -0.33955 | 0.134491 | 0.633583 |
| X | 8736000 | 8741199 | 76.25865 | 121.3162 | 114.8047 | 47.94926 | 0.414402 | 2.250237 | 0.633584 |
| 9 | 3335400 | 3335799 | 8.438796 | 16.30893 | 19.35379 | 7.423481 | 0.311576 | -0.16481 | 0.633611 |
| X | 4483800 | 4488999 | 61.20808 | 98.71942 | 80.03599 | 34.09331 | 0.659626 | 1.808238 | 0.633627 |
| 6 | 30707400 | 30708599 | 8.219297 | 15.65936 | 26.67392 | 10.66041 | -0.26101 | -0.01236 | 0.633632 |
| 10 | 109858000 | 109860999 | 25.86189 | 44.25936 | 46.83702 | 20.53827 | 0.301664 | 0.973855 | 0.633635 |
| 8 | 22422600 | 22424999 | 16.68303 | 29.17301 | 46.20817 | 20.3734 | -0.27284 | 0.701914 | 0.633644 |
| 1 | 31596000 | 31596999 | 12.18141 | 21.71967 | 34.7313 | 15.28038 | -0.27631 | 0.444652 | 0.633653 |
| 4 | 146834600 | 146836199 | 53.39798 | 85.75282 | 75.05358 | 31.77546 | 0.541886 | 1.923081 | 0.633662 |
| 12 | 105684600 | 105686199 | 10.25976 | 18.53458 | 31.83608 | 13.83073 | -0.36629 | 0.34463 | 0.633672 |
| 13 | 22881400 | 22885799 | 40.95737 | 67.31458 | 65.26771 | 28.12456 | 0.422549 | 1.439002 | 0.633693 |
| 4 | 14766000 | 14767399 | 10.13353 | 18.58916 | 30.81717 | 13.12949 | -0.29711 | 0.250274 | 0.633696 |
| 5 | 114188200 | 114191199 | 22.36918 | 37.5673 | 58.05227 | 25.3503 | -0.24465 | 1.14129 | 0.633696 |
| 9 | 73950600 | 73951199 | 8.2788 | 15.66311 | 27.06817 | 10.97871 | -0.30001 | 0.039452 | 0.633697 |
| 19 | 9875400 | 9877199 | 11.5436 | 20.46752 | 35.23867 | 15.57697 | -0.38718 | 0.474508 | 0.63371 |
| X | 95900200 | 95903399 | 37.68398 | 61.38725 | 67.03007 | 28.78385 | 0.244448 | 1.551157 | 0.633724 |
| 6 | 86911600 | 86914999 | 26.48212 | 45.10582 | 41.09427 | 18.01314 | 0.515949 | 1.013079 | 0.633735 |
| 2 | 168127200 | 168128599 | 28.53777 | 47.71763 | 53.47126 | 23.28546 | 0.217394 | 1.200277 | 0.63375 |
| 6 | 66463200 | 66467399 | 51.0201 | 82.04108 | 77.70873 | 32.96237 | 0.430391 | 1.878941 | 0.633755 |
| 14 | 111286200 | 111288599 | 22.87121 | 40.01635 | 33.22914 | 14.65778 | 0.654502 | 0.705271 | 0.633763 |

| | | | | | | | | | |
|---|---|---|---|---|---|---|---|---|---|
| 1 | 70134400 | 70136999 | 29.12654 | 49.89124 | 46.22609 | 20.27184 | 0.492854 | 0.977432 | 0.633767 |
| Y | 88724000 | 88726799 | 22.4491 | 38.53646 | 31.15508 | 13.66502 | 0.689012 | 0.90914 | 0.633791 |
| X | 121894200 | 121894999 | 17.77584 | 31.25268 | 34.02529 | 15.06685 | 0.265753 | 0.604536 | 0.6338 |
| 14 | 86862200 | 86863399 | 14.96712 | 26.87581 | 29.7825 | 13.02531 | 0.255256 | 0.406342 | 0.633801 |
| 12 | 115894600 | 115895999 | 11.48906 | 21.10768 | 36.25042 | 15.41254 | -0.34576 | 0.265323 | 0.633803 |
| 7 | 44438400 | 44446399 | 69.7255 | 108.4137 | 185.647 | 77.07537 | -0.46698 | 2.740627 | 0.633816 |
| 7 | 33592800 | 33618599 | 373.1088 | 526.9687 | 486.3511 | 201.3458 | 0.29839 | 4.434707 | 0.633825 |
| 10 | 113734800 | 113735999 | 14.40773 | 25.36697 | 39.25763 | 17.37887 | -0.23838 | 0.577514 | 0.633839 |
| 7 | 144468600 | 144469399 | 6.716447 | 12.86017 | 24.59596 | 9.713228 | -0.41089 | -0.07328 | 0.633846 |
| 3 | 133791600 | 133793799 | 37.65056 | 61.62766 | 57.96965 | 24.93671 | 0.463813 | 1.474933 | 0.633861 |
| X | 72162800 | 72164799 | 11.2024 | 20.51841 | 33.97231 | 14.50398 | -0.29733 | 0.277003 | 0.633865 |
| 18 | 6828800 | 6829399 | 9.173546 | 17.17366 | 28.47917 | 11.775 | -0.26456 | 0.118 | 0.633866 |
| 12 | 76783800 | 76791999 | 315.6181 | 440.4104 | 587.2127 | 243.5683 | -0.24975 | 4.611528 | 0.633868 |
| 8 | 93151400 | 93153599 | 19.51291 | 33.14625 | 52.15964 | 22.85792 | -0.27005 | 1.024326 | 0.633875 |
| 11 | 9054800 | 9056199 | 19.67801 | 34.42204 | 37.65636 | 16.60768 | 0.258729 | 0.699311 | 0.633895 |
| 10 | 126029600 | 126032599 | 30.46478 | 49.99973 | 76.69431 | 33.02886 | -0.23823 | 1.438764 | 0.633898 |
| 12 | 49048000 | 49051399 | 41.33914 | 67.27031 | 62.95303 | 27.02268 | 0.465336 | 1.569571 | 0.633899 |
| 18 | 37805200 | 37811399 | 167.6716 | 250.5081 | 264.5435 | 109.0663 | 0.182631 | 3.544872 | 0.633909 |
| 12 | 86245000 | 86247199 | 18.95577 | 33.23491 | 36.50377 | 16.13063 | 0.252837 | 0.656006 | 0.63394 |
| X | 164249200 | 164250199 | 26.81074 | 45.06545 | 49.62242 | 21.67663 | 0.242579 | 1.138822 | 0.633953 |
| 11 | 18592400 | 18594799 | 18.87199 | 33.40309 | 33.52031 | 14.82336 | 0.38606 | 0.534504 | 0.633965 |
| 5 | 9150400 | 9155999 | 36.53109 | 58.91297 | 91.03771 | 38.77451 | -0.2692 | 1.772464 | 0.633968 |
| 14 | 121071400 | 121073399 | 14.945 | 25.95896 | 49.51317 | 21.73528 | -0.54055 | 0.794061 | 0.634018 |
| 19 | 9674800 | 9675999 | 16.28457 | 28.96219 | 31.70873 | 13.98041 | 0.264633 | 0.478251 | 0.63402 |
| 12 | 78718200 | 78720999 | 29.35277 | 48.66051 | 51.31552 | 22.22057 | 0.30445 | 1.295488 | 0.634021 |
| 4 | 5018000 | 5020599 | 31.78284 | 53.31337 | 47.51394 | 20.73285 | 0.546595 | 1.171879 | 0.63403 |
| 4 | 121160200 | 121165999 | 98.07815 | 152.6935 | 160.3324 | 66.55275 | 0.239229 | 2.728665 | 0.634047 |
| 1 | 102192200 | 102193999 | 33.9282 | 55.82936 | 62.36614 | 26.88799 | 0.219491 | 1.404647 | 0.634073 |
| 4 | 11174800 | 11185399 | 262.3531 | 373.9964 | 487.7049 | 201.5936 | -0.18666 | 4.293712 | 0.634094 |
| 14 | 113203200 | 113204199 | 9.293137 | 17.54019 | 29.78249 | 12.14088 | -0.28211 | 0.068625 | 0.634098 |
| 1 | 86467000 | 86469999 | 16.82166 | 29.10732 | 45.83585 | 20.10937 | -0.26629 | 0.836551 | 0.634099 |
| 3 | 5144200 | 5147999 | 63.70311 | 101.5125 | 109.4159 | 45.8049 | 0.229504 | 2.18317 | 0.634108 |
| 13 | 82704000 | 82705999 | 18.33312 | 32.46823 | 33.00373 | 14.59321 | 0.367874 | 0.52351 | 0.634116 |
| 16 | 30549400 | 30553399 | 30.78121 | 50.45048 | 76.66017 | 32.99519 | -0.22546 | 1.455805 | 0.63415 |
| 9 | 87070400 | 87077199 | 97.27574 | 148.2539 | 243.0434 | 100.7537 | -0.42662 | 3.136724 | 0.634162 |
| 4 | 147677600 | 147681399 | 92.93493 | 146.062 | 142.8867 | 59.27127 | 0.351807 | 2.55674 | 0.63418 |
| 17 | 4254600 | 4256999 | 18.5718 | 32.87884 | 33.56404 | 14.84206 | 0.36163 | 0.530757 | 0.634218 |
| 1 | 83650400 | 83651199 | 8.870599 | 16.76226 | 28.12511 | 11.43694 | -0.26148 | 0.054592 | 0.634226 |
| 2 | 74716400 | 74718399 | 13.51056 | 23.8519 | 37.78514 | 16.71796 | -0.27011 | 0.541882 | 0.634244 |
| 1 | 37922200 | 37923799 | 9.988598 | 18.28854 | 30.80706 | 13.15914 | -0.32269 | 0.25978 | 0.634245 |
| 2 | 83110000 | 83111199 | 9.181283 | 17.43442 | 31.20101 | 12.55714 | -0.34104 | 0.030207 | 0.634249 |
| 4 | 9464000 | 9464599 | 7.009677 | 13.41613 | 24.58723 | 9.721203 | -0.35129 | -0.06542 | 0.634278 |
| 10 | 13995400 | 13996399 | 7.009677 | 13.41613 | 24.58723 | 9.721203 | -0.35129 | -0.06542 | 0.634278 |
| 7 | 108405800 | 108406199 | 9.583137 | 18.46242 | 21.95312 | 8.517642 | 0.292998 | -0.10659 | 0.6343 |
| 15 | 12867600 | 12868999 | 10.03754 | 18.49661 | 30.38562 | 12.86615 | -0.27723 | 0.225621 | 0.63431 |
| 15 | 49305800 | 49309599 | 39.8118 | 65.56366 | 64.06359 | 27.62922 | 0.412559 | 1.411519 | 0.634339 |
| 14 | 3364800 | 3367599 | 36.54576 | 59.97351 | 56.4057 | 24.29033 | 0.465436 | 1.438016 | 0.634342 |

| | | | | | | | | | |
|---|---|---|---|---|---|---|---|---|---|
| 12 | 20753400 | 20783599 | 1123.345 | 1433.724 | 1293.402 | 542.7995 | 0.184577 | 6.035724 | 0.63437 |
| 3 | 10039400 | 10040799 | 9.486048 | 17.59069 | 29.38919 | 12.33814 | -0.29275 | 0.180966 | 0.634393 |
| 4 | 29585600 | 29586399 | 9.486048 | 17.59069 | 29.38919 | 12.33814 | -0.29275 | 0.180966 | 0.634393 |
| 10 | 42007200 | 42012599 | 50.35542 | 80.20729 | 118.6596 | 49.64435 | -0.22729 | 2.185322 | 0.634421 |
| 18 | 32874600 | 32875599 | 8.705341 | 16.59415 | 28.98444 | 11.57086 | -0.2957 | -0.00533 | 0.634431 |
| 12 | 113865000 | 113866799 | 25.48983 | 44.17491 | 42.94047 | 18.83999 | 0.427067 | 0.856575 | 0.634435 |
| 13 | 5175600 | 5178399 | 24.25354 | 42.17153 | 41.25787 | 18.12325 | 0.418413 | 0.809919 | 0.63445 |
| 13 | 120629400 | 120632399 | 31.97584 | 51.89938 | 79.18641 | 33.9404 | -0.24122 | 1.605082 | 0.634458 |
| 3 | 49001400 | 49003399 | 19.70128 | 34.95793 | 33.50883 | 14.8029 | 0.453591 | 0.518655 | 0.634463 |
| 15 | 15066400 | 15067799 | 25.49081 | 41.74916 | 77.38127 | 33.29532 | -0.51158 | 1.454358 | 0.634506 |
| X | 144522800 | 144523399 | 12.97302 | 23.83448 | 26.57515 | 11.29862 | 0.273814 | 0.256122 | 0.634509 |
| 13 | 61481200 | 61497199 | 287.1664 | 414.0002 | 402.3336 | 166.0141 | 0.253601 | 4.13094 | 0.634515 |
| 2 | 82975600 | 82977199 | 17.05372 | 30.86394 | 30.74114 | 13.32484 | 0.416762 | 0.37839 | 0.634515 |
| 14 | 52694000 | 52695599 | 17.05372 | 30.86394 | 30.74114 | 13.32484 | 0.416762 | 0.37839 | 0.634515 |
| 6 | 122740400 | 122742599 | 15.20936 | 26.33396 | 46.3579 | 20.3388 | -0.42629 | 0.821886 | 0.63452 |
| 18 | 62174200 | 62177599 | 38.6806 | 62.88417 | 70.15593 | 30.10407 | 0.211419 | 1.580387 | 0.634529 |
| 1 | 163611000 | 163611599 | 10.27525 | 19.88869 | 22.14386 | 8.432733 | 0.412453 | -0.1599 | 0.634545 |
| 1 | 140682400 | 140683599 | 12.42597 | 23.24499 | 19.38113 | 8.023524 | 0.718358 | 0.125883 | 0.634548 |
| 11 | 13821800 | 13825799 | 52.2837 | 83.78421 | 92.45193 | 39.01447 | 0.204586 | 1.990827 | 0.634557 |
| 2 | 6250200 | 6251999 | 24.59218 | 41.93892 | 46.32695 | 20.30886 | 0.239199 | 0.998227 | 0.634557 |
| 3 | 56275800 | 56276799 | 11.72796 | 20.96206 | 34.28267 | 15.05213 | -0.30638 | 0.426424 | 0.63456 |
| 2 | 3078400 | 3084399 | 76.17371 | 121.1032 | 116.4962 | 48.61787 | 0.389836 | 2.267736 | 0.634561 |
| 13 | 78520400 | 78521999 | 16.90551 | 29.7126 | 31.97382 | 14.15804 | 0.282128 | 0.596901 | 0.634575 |
| 8 | 75695400 | 75699399 | 53.4876 | 85.50081 | 92.37035 | 38.8392 | 0.231057 | 2.069262 | 0.634587 |
| 15 | 25647200 | 25649599 | 27.14614 | 44.90031 | 68.08897 | 29.4376 | -0.21828 | 1.331106 | 0.634591 |
| 6 | 69269200 | 69271199 | 23.20326 | 40.09799 | 37.2967 | 16.36238 | 0.489563 | 0.863728 | 0.634615 |
| 16 | 22168800 | 22172199 | 22.61596 | 37.49063 | 70.00576 | 30.31284 | -0.51731 | 1.293525 | 0.634616 |
| 12 | 98056400 | 98061599 | 50.67733 | 81.07068 | 126.0557 | 53.06054 | -0.2922 | 2.052916 | 0.634632 |
| 4 | 23521200 | 23523999 | 57.72651 | 92.66248 | 82.68785 | 34.98369 | 0.513202 | 1.943312 | 0.634633 |
| 10 | 79179600 | 79180799 | 11.68252 | 21.36001 | 36.09658 | 15.4477 | -0.32933 | 0.294256 | 0.634638 |
| 7 | 64911400 | 64915199 | 40.00706 | 64.98915 | 71.97085 | 30.86678 | 0.221042 | 1.598966 | 0.634678 |
| 5 | 84871000 | 84874799 | 27.84524 | 47.69702 | 46.80401 | 20.5253 | 0.410373 | 0.97341 | 0.634705 |
| 6 | 146045600 | 146051399 | 87.45629 | 136.2576 | 148.1319 | 61.48164 | 0.189994 | 2.709012 | 0.634721 |
| 16 | 46016800 | 46019599 | 23.17817 | 39.09396 | 61.63863 | 26.95383 | -0.27358 | 1.105307 | 0.63473 |
| 4 | 42242600 | 42243199 | 7.777543 | 14.73547 | 26.02298 | 10.52343 | -0.3275 | 0.020034 | 0.634731 |
| 4 | 55480400 | 55481799 | 14.87066 | 26.02017 | 40.978 | 18.07701 | -0.26442 | 0.673962 | 0.63474 |
| 13 | 90741000 | 90741999 | 10.70844 | 20.10295 | 22.14479 | 9.112962 | 0.326678 | 0.096554 | 0.634762 |
| 11 | 108245800 | 108246599 | 9.405495 | 18.00663 | 20.71283 | 8.182784 | 0.315514 | -0.05693 | 0.634763 |
| 1 | 174115800 | 174118199 | 44.68386 | 71.95382 | 77.52263 | 32.95079 | 0.247494 | 1.826436 | 0.634764 |
| 8 | 43582800 | 43584399 | 16.14329 | 28.13223 | 43.64414 | 19.19833 | -0.24336 | 0.744822 | 0.634767 |
| 14 | 48176600 | 48178599 | 21.42338 | 37.04535 | 40.97484 | 17.97636 | 0.240817 | 0.85429 | 0.634777 |
| 10 | 111024000 | 111025199 | 12.62507 | 23.22134 | 25.96067 | 11.02048 | 0.271172 | 0.242959 | 0.634782 |
| 14 | 86125800 | 86129599 | 41.66101 | 67.27869 | 73.43573 | 31.33326 | 0.233564 | 1.733598 | 0.634784 |
| 12 | 62034000 | 62035599 | 20.64854 | 35.83223 | 39.93598 | 17.52249 | 0.230952 | 0.818627 | 0.634785 |
| X | 164426000 | 164432599 | 138.0665 | 210.1958 | 188.3264 | 78.04574 | 0.443025 | 3.166458 | 0.634794 |
| 12 | 3186200 | 3195799 | 67.68159 | 107.8486 | 98.92192 | 41.40983 | 0.461873 | 2.18429 | 0.634804 |
| 10 | 15331400 | 15333399 | 26.88442 | 45.27748 | 49.63308 | 21.69411 | 0.248948 | 1.118164 | 0.63481 |

| | | | | | | | | | |
|---|---|---|---|---|---|---|---|---|---|
| 2 | 105004000 | 105007399 | 30.74113 | 50.24669 | 75.94662 | 32.65293 | -0.22007 | 1.490409 | 0.634815 |
| 7 | 9303400 | 9305399 | 29.69029 | 49.86259 | 44.36345 | 19.37267 | 0.549048 | 1.150785 | 0.634816 |
| X | 153998400 | 153999799 | 17.315 | 30.89277 | 32.26772 | 14.18883 | 0.335022 | 0.469084 | 0.634834 |
| 8 | 76448400 | 76449799 | 14.85056 | 27.35919 | 28.47826 | 12.06246 | 0.375927 | 0.257962 | 0.63487 |
| 8 | 40583400 | 40585199 | 14.80657 | 26.91402 | 29.34199 | 12.64926 | 0.292104 | 0.333911 | 0.634871 |
| 6 | 133068200 | 133069799 | 23.89703 | 40.60243 | 43.80749 | 19.19863 | 0.272381 | 1.028658 | 0.634875 |
| 9 | 44179000 | 44180999 | 20.1105 | 34.97458 | 38.90906 | 17.0936 | 0.234 | 0.786532 | 0.63489 |
| 7 | 23583000 | 23585799 | 79.36815 | 124.5571 | 135.5771 | 56.22846 | 0.19636 | 2.576345 | 0.634898 |
| 11 | 3488200 | 3490799 | 18.08635 | 31.08275 | 49.00444 | 21.49504 | -0.2702 | 0.906232 | 0.634904 |
| 12 | 83643600 | 83644199 | 5.52844 | 10.75754 | 21.27332 | 7.837938 | -0.36386 | -0.3331 | 0.634907 |
| 7 | 60408600 | 60408999 | 7.421726 | 14.26938 | 25.27509 | 9.863806 | -0.28537 | -0.10309 | 0.634936 |
| 16 | 20634400 | 20637799 | 32.7344 | 53.19309 | 80.88235 | 34.69737 | -0.23476 | 1.582431 | 0.634963 |
| X | 106633200 | 106634799 | 14.31177 | 25.30595 | 40.27311 | 17.81318 | -0.27604 | 0.533248 | 0.634981 |
| 1 | 14830400 | 14831399 | 8.632425 | 15.92561 | 28.90655 | 12.22077 | -0.41841 | 0.203003 | 0.634986 |
| 1 | 94321600 | 94322599 | 13.30119 | 23.44962 | 37.2806 | 16.49932 | -0.27612 | 0.553446 | 0.634997 |
| 14 | 14980600 | 14981799 | 9.893368 | 17.96327 | 31.30499 | 13.51429 | -0.38138 | 0.309136 | 0.635014 |
| 7 | 62171400 | 62175599 | 41.95838 | 68.14375 | 64.61117 | 27.7058 | 0.444219 | 1.603742 | 0.635017 |
| X | 102172800 | 102173799 | 16.85635 | 29.94365 | 31.87235 | 14.06658 | 0.304427 | 0.493129 | 0.635034 |
| 5 | 41094600 | 41096399 | 15.66797 | 28.32334 | 29.65222 | 12.87137 | 0.343658 | 0.370529 | 0.635066 |
| 10 | 72509800 | 72511199 | 17.37954 | 30.80588 | 32.58208 | 14.4043 | 0.311399 | 0.507887 | 0.635075 |
| 13 | 91377600 | 91378999 | 12.13526 | 21.91305 | 34.75459 | 15.10699 | -0.25292 | 0.365979 | 0.635083 |
| 8 | 65263200 | 65266399 | 28.58224 | 46.95681 | 71.45202 | 30.78421 | -0.22611 | 1.416518 | 0.635118 |
| 16 | 20861400 | 20863999 | 39.3949 | 63.92812 | 71.63308 | 30.69833 | 0.202752 | 1.619164 | 0.635138 |
| 6 | 147960400 | 147965599 | 83.64643 | 131.1389 | 144.0645 | 59.75219 | 0.181928 | 2.597545 | 0.635155 |
| 2 | 143268800 | 143271199 | 26.26 | 45.09284 | 45.70583 | 20.04766 | 0.364276 | 0.943006 | 0.635162 |
| X | 74744800 | 74746999 | 32.6886 | 54.72476 | 55.29102 | 24.09642 | 0.366262 | 1.202118 | 0.635164 |
| 1 | 171270600 | 171272399 | 22.05892 | 37.21674 | 57.0962 | 24.96963 | -0.23415 | 1.095514 | 0.635181 |
| 17 | 72070800 | 72072599 | 29.82802 | 50.43377 | 51.27748 | 22.43973 | 0.357401 | 1.098227 | 0.635197 |
| 11 | 24025600 | 24032199 | 50.41246 | 79.39377 | 162.019 | 67.22553 | -0.70524 | 2.492081 | 0.635202 |
| 1 | 181684000 | 181685599 | 16.26558 | 28.92218 | 31.84005 | 14.04087 | 0.256752 | 0.48186 | 0.635242 |
| 14 | 104366200 | 104367599 | 12.52543 | 21.98997 | 38.42184 | 16.99734 | -0.41337 | 0.60194 | 0.635248 |
| 8 | 107955800 | 107958199 | 19.21774 | 33.51041 | 37.27366 | 16.40236 | 0.234532 | 0.741404 | 0.635274 |
| 8 | 92359200 | 92360999 | 13.7574 | 24.19836 | 38.28365 | 16.95101 | -0.27053 | 0.582358 | 0.635299 |
| 8 | 20009400 | 20010399 | 11.32494 | 21.72727 | 23.46531 | 9.214048 | 0.413788 | -0.04966 | 0.635317 |
| 6 | 24446400 | 24448599 | 21.46297 | 37.44093 | 34.53639 | 15.20176 | 0.503393 | 0.74299 | 0.635321 |
| 11 | 96575800 | 96576999 | 16.11098 | 28.51457 | 31.30125 | 13.8422 | 0.257315 | 0.514306 | 0.635343 |
| 4 | 28617400 | 28619799 | 25.04658 | 43.49456 | 42.39369 | 18.60715 | 0.423697 | 0.834122 | 0.635354 |
| 7 | 95041600 | 95044399 | 23.12411 | 40.39747 | 38.97063 | 17.16927 | 0.439176 | 0.73669 | 0.635375 |
| 7 | 81403600 | 81405199 | 16.13882 | 28.70589 | 31.39195 | 13.83956 | 0.266658 | 0.478888 | 0.635377 |
| 19 | 39947000 | 39957799 | 308.1786 | 431.7143 | 567.2746 | 235.1552 | -0.22301 | 4.55289 | 0.635392 |
| X | 64141200 | 64149599 | 122.5966 | 188.3665 | 188.0048 | 78.06406 | 0.297791 | 2.980361 | 0.635401 |
| 2 | 98467800 | 98470599 | 19.85228 | 35.41726 | 32.45511 | 14.27206 | 0.522621 | 0.484554 | 0.635408 |
| 4 | 143635600 | 143638399 | 43.15289 | 70.33208 | 70.97724 | 30.48452 | 0.358306 | 1.55477 | 0.635442 |
| 14 | 114474400 | 114476399 | 18.10011 | 31.84151 | 34.83221 | 15.42235 | 0.258979 | 0.598653 | 0.635442 |
| 13 | 65031200 | 65033599 | 27.86535 | 47.39604 | 42.10631 | 18.4547 | 0.551899 | 1.032519 | 0.635456 |
| 6 | 50263400 | 50269799 | 52.46009 | 83.72634 | 123.6853 | 51.89038 | -0.22213 | 2.125121 | 0.635473 |
| 15 | 41342800 | 41343199 | 8.007654 | 15.55787 | 19.47605 | 7.265102 | 0.272877 | -0.26358 | 0.63549 |

| | | | | | | | | | |
|---|---|---|---|---|---|---|---|---|---|
| 9 | 99472800 | 99473799 | 10.83188 | 19.70854 | 31.78705 | 13.68651 | -0.26789 | 0.306404 | 0.6355 |
| X | 102239000 | 102242799 | 43.04927 | 69.87017 | 74.51047 | 31.93546 | 0.274212 | 1.624678 | 0.635506 |
| 9 | 87360200 | 87364999 | 55.15897 | 88.81472 | 86.48292 | 36.75511 | 0.392945 | 1.846486 | 0.635534 |
| X | 38457600 | 38462399 | 55.87658 | 90.30711 | 78.46365 | 33.51282 | 0.56335 | 1.740918 | 0.635562 |
| 9 | 81944200 | 81945999 | 16.63961 | 29.27681 | 31.53903 | 13.96382 | 0.281211 | 0.583309 | 0.635572 |
| 10 | 100418000 | 100420399 | 22.05372 | 38.23467 | 42.00654 | 18.43044 | 0.251326 | 0.837219 | 0.635578 |
| 17 | 72680200 | 72681999 | 22.05372 | 38.23467 | 42.00654 | 18.43044 | 0.251326 | 0.837219 | 0.635578 |
| 4 | 100959200 | 100962799 | 54.84945 | 87.98268 | 94.5412 | 39.95611 | 0.244504 | 1.965517 | 0.635588 |
| 7 | 80156800 | 80158399 | 13.79032 | 25.12089 | 28.00138 | 12.03883 | 0.2631 | 0.313335 | 0.635602 |
| 4 | 20960000 | 20962199 | 22.90219 | 39.9005 | 40.75336 | 17.92385 | 0.356893 | 0.776997 | 0.635629 |
| 19 | 12721400 | 12723799 | 21.29126 | 36.09233 | 54.8795 | 24.03476 | -0.22095 | 1.051415 | 0.635636 |
| X | 56546000 | 56547999 | 21.97658 | 38.23987 | 39.39133 | 17.31115 | 0.344721 | 0.791518 | 0.635643 |
| 5 | 14993400 | 14995799 | 32.80022 | 54.1278 | 60.26706 | 26.01458 | 0.225777 | 1.363211 | 0.635649 |
| 3 | 126157200 | 126158599 | 10.01427 | 18.21374 | 31.78115 | 13.6907 | -0.38139 | 0.300023 | 0.635652 |
| 9 | 34593400 | 34594999 | 13.69233 | 24.35887 | 39.73603 | 17.51542 | -0.30738 | 0.474609 | 0.635677 |
| 12 | 24424600 | 24448199 | 634.8861 | 849.1509 | 807.0775 | 336.9674 | 0.176834 | 5.276712 | 0.635681 |
| 1 | 104820400 | 104823999 | 50.61075 | 81.75409 | 80.95371 | 34.55321 | 0.374225 | 1.744595 | 0.635681 |
| 19 | 7973000 | 7975399 | 34.23964 | 56.86118 | 52.01983 | 22.55275 | 0.508973 | 1.289629 | 0.63569 |
| 12 | 109152400 | 109153399 | 16.44061 | 29.85002 | 30.33179 | 13.09471 | 0.391861 | 0.355748 | 0.635709 |
| 19 | 39438600 | 39439799 | 14.12214 | 26.6156 | 26.27585 | 10.74143 | 0.491295 | 0.113218 | 0.635709 |
| 6 | 133709000 | 133723999 | 195.1956 | 286.617 | 385.434 | 158.7437 | -0.18914 | 3.846228 | 0.635709 |
| 14 | 87784600 | 87786199 | 22.45926 | 38.85354 | 42.51877 | 18.65396 | 0.256287 | 0.857629 | 0.635717 |
| 15 | 57634200 | 57637199 | 30.83875 | 50.99203 | 56.78964 | 24.54715 | 0.22566 | 1.333125 | 0.635724 |
| 15 | 72210600 | 72212799 | 23.22524 | 40.22752 | 42.46006 | 18.62889 | 0.30851 | 0.849572 | 0.635731 |
| 1 | 29314800 | 29315799 | 8.824069 | 16.71289 | 28.19576 | 11.4098 | -0.26338 | 0.036825 | 0.635744 |
| 16 | 3422000 | 3423999 | 18.61765 | 32.26976 | 50.64514 | 22.22048 | -0.26113 | 0.83564 | 0.635757 |
| 6 | 69508400 | 69519799 | 119.1306 | 183.5215 | 183.3237 | 76.1477 | 0.299441 | 2.926533 | 0.635757 |
| 6 | 14445800 | 14447799 | 26.98163 | 45.8922 | 42.12473 | 18.46275 | 0.504895 | 1.028663 | 0.635777 |
| 12 | 65481600 | 65483799 | 39.6464 | 64.8246 | 60.31698 | 25.93626 | 0.478383 | 1.49626 | 0.635797 |
| 5 | 25441400 | 25443799 | 30.41367 | 49.40159 | 77.59893 | 33.27873 | -0.28193 | 1.588407 | 0.635798 |
| 8 | 85432200 | 85434999 | 21.28702 | 35.40467 | 60.57824 | 26.29071 | -0.39111 | 1.24299 | 0.635812 |
| 7 | 108092400 | 108097199 | 56.83002 | 90.73038 | 100.9795 | 42.38411 | 0.186251 | 2.113957 | 0.635821 |
| 16 | 6166200 | 6168399 | 17.86378 | 30.42255 | 51.10383 | 22.40047 | -0.36338 | 0.991301 | 0.63583 |
| 11 | 20181200 | 20182199 | 8.722814 | 16.5568 | 28.25043 | 11.37987 | -0.27345 | 0.023058 | 0.635843 |
| 12 | 76228800 | 76229999 | 8.722814 | 16.5568 | 28.25043 | 11.37987 | -0.27345 | 0.023058 | 0.635843 |
| 3 | 21712600 | 21720399 | 93.17699 | 145.7149 | 152.1613 | 63.12671 | 0.252207 | 2.648118 | 0.63586 |
| 2 | 150079000 | 150083399 | 100.5891 | 157.2101 | 153.2524 | 63.58339 | 0.350657 | 2.663907 | 0.635865 |
| 3 | 129449800 | 129463199 | 162.1184 | 241.0701 | 326.9182 | 134.6887 | -0.18421 | 3.63052 | 0.635882 |
| 13 | 36184200 | 36189799 | 53.1061 | 84.6964 | 124.6906 | 52.25952 | -0.21833 | 2.148329 | 0.635891 |
| 11 | 66865000 | 66868399 | 53.71658 | 86.47096 | 77.30344 | 32.84213 | 0.515561 | 1.849628 | 0.635895 |
| 3 | 70134000 | 70135599 | 15.90179 | 27.44166 | 45.86678 | 20.12129 | -0.35265 | 0.856918 | 0.63597 |
| 10 | 51465200 | 51466799 | 14.20428 | 24.89429 | 40.40433 | 17.84825 | -0.30775 | 0.641968 | 0.635993 |
| 4 | 143052000 | 143056399 | 43.63926 | 69.88574 | 104.0781 | 43.87691 | -0.22826 | 2.006421 | 0.636005 |
| 7 | 46148800 | 46152199 | 20.37253 | 34.32273 | 54.79974 | 23.95501 | -0.2914 | 1.101902 | 0.636016 |
| 11 | 88984600 | 88986799 | 27.91568 | 45.92386 | 70.33244 | 30.31921 | -0.23461 | 1.397154 | 0.636019 |
| 10 | 93150400 | 93153599 | 26.22099 | 44.48263 | 49.03907 | 21.48182 | 0.241248 | 1.055655 | 0.636023 |
| 3 | 37755400 | 37759399 | 47.54091 | 76.08546 | 110.5452 | 46.55969 | -0.19377 | 2.032204 | 0.636042 |

| | | | | | | | | | |
|---|---|---|---|---|---|---|---|---|---|
| 14 | 57417200 | 57418399 | 12.47273 | 22.37958 | 35.20441 | 15.40652 | -0.24763 | 0.407519 | 0.636072 |
| 4 | 121193600 | 121198999 | 59.82154 | 95.91749 | 96.16324 | 40.61537 | 0.343635 | 1.985422 | 0.636076 |
| 4 | 33680600 | 33681399 | 7.16856 | 13.8108 | 24.74666 | 9.601126 | -0.29304 | -0.12269 | 0.636104 |
| 8 | 29681600 | 29684999 | 38.60778 | 62.24607 | 92.53437 | 39.39894 | -0.21383 | 1.787627 | 0.636109 |
| 2 | 35416600 | 35421399 | 52.172 | 83.00141 | 121.9499 | 50.9365 | -0.21922 | 2.226539 | 0.63614 |
| 3 | 158042400 | 158043999 | 20.10875 | 35.40208 | 35.78841 | 15.8433 | 0.372847 | 0.605853 | 0.636158 |
| 5 | 22562000 | 22564199 | 17.01877 | 29.51825 | 45.82693 | 20.10691 | -0.24511 | 0.817382 | 0.636159 |
| 3 | 19748000 | 19749799 | 15.32648 | 26.65788 | 44.2149 | 19.42607 | -0.33967 | 0.767385 | 0.636189 |
| X | 80718600 | 80719999 | 25.73802 | 43.82709 | 47.94578 | 21.01598 | 0.252891 | 1.019995 | 0.636191 |
| 7 | 41509800 | 41511799 | 24.83712 | 43.33532 | 40.08455 | 17.64602 | 0.499521 | 0.766135 | 0.636199 |
| 6 | 149425000 | 149425599 | 8.701373 | 16.88695 | 20.52156 | 7.718885 | 0.303582 | -0.22565 | 0.636205 |
| 12 | 113947000 | 113949199 | 27.04217 | 45.61548 | 50.69612 | 22.16943 | 0.229335 | 1.108537 | 0.636218 |
| 5 | 21445400 | 21448999 | 62.64459 | 99.54273 | 108.8685 | 45.41109 | 0.20425 | 2.269653 | 0.636226 |
| 13 | 57611800 | 57612999 | 12.67445 | 22.61026 | 35.7161 | 15.70692 | -0.2584 | 0.447001 | 0.636244 |
| 4 | 98162000 | 98163599 | 38.79893 | 62.97309 | 69.7024 | 29.87517 | 0.220702 | 1.614494 | 0.636256 |
| 3 | 106346000 | 106349799 | 99.4864 | 153.8145 | 167.8402 | 69.71693 | 0.176096 | 2.851203 | 0.636322 |
| 1 | 113516800 | 113517599 | 11.42272 | 22.17461 | 21.13691 | 7.937356 | 0.652943 | -0.1985 | 0.636331 |
| 16 | 39804600 | 39805599 | 9.719822 | 17.88634 | 29.87115 | 12.67218 | -0.30276 | 0.229479 | 0.636334 |
| 11 | 73872800 | 73876599 | 51.73661 | 82.53609 | 119.2303 | 49.99103 | -0.19056 | 2.138371 | 0.636342 |
| 7 | 33455200 | 33473799 | 303.6159 | 436.4957 | 412.1401 | 170.1288 | 0.291192 | 4.173243 | 0.636385 |
| 4 | 125269400 | 125273199 | 29.1155 | 47.61793 | 73.45706 | 31.58941 | -0.24879 | 1.476501 | 0.636396 |
| X | 129425200 | 129426999 | 13.7499 | 24.38943 | 38.15254 | 16.85919 | -0.2495 | 0.49227 | 0.63642 |
| 1 | 81168600 | 81169999 | 8.882916 | 16.88597 | 29.45379 | 11.82707 | -0.30111 | 0.01695 | 0.63644 |
| 15 | 91192800 | 91195399 | 19.84151 | 34.04493 | 53.35063 | 23.39943 | -0.26207 | 0.932348 | 0.636443 |
| 6 | 133504200 | 133506399 | 16.52883 | 30.14258 | 29.8152 | 12.80103 | 0.43547 | 0.329736 | 0.636446 |
| 3 | 10874600 | 10876999 | 27.31205 | 45.98182 | 50.69054 | 22.154 | 0.240962 | 1.127671 | 0.63646 |
| 19 | 14879000 | 14880799 | 15.5972 | 28.84175 | 27.92146 | 11.76695 | 0.484986 | 0.241427 | 0.63647 |
| 7 | 73552000 | 73554399 | 20.13427 | 35.33917 | 36.87181 | 16.30809 | 0.326542 | 0.646155 | 0.636471 |
| X | 153718800 | 153721999 | 38.81184 | 64.36073 | 51.01791 | 22.09624 | 0.715026 | 1.319294 | 0.636495 |
| 9 | 77582000 | 77582399 | 3.982444 | 7.785382 | 17.16901 | 5.895152 | -0.42497 | -0.6186 | 0.636497 |
| 6 | 126227400 | 126228999 | 18.00829 | 31.08083 | 47.94064 | 21.03116 | -0.23732 | 0.868985 | 0.636497 |
| 5 | 84933800 | 84937199 | 36.99706 | 61.5042 | 58.36481 | 25.32171 | 0.456476 | 1.29113 | 0.636499 |
| 5 | 7255400 | 7256999 | 15.91151 | 27.94753 | 46.1046 | 20.40404 | -0.33126 | 0.627041 | 0.636523 |
| 3 | 16061800 | 16066399 | 84.5481 | 132.2374 | 145.5932 | 60.40104 | 0.176077 | 2.64004 | 0.636538 |
| 1 | 32704800 | 32704999 | 4.799507 | 9.378537 | 18.89566 | 6.562302 | -0.31242 | -0.56002 | 0.636542 |
| 12 | 54842000 | 54842799 | 6.121163 | 11.81525 | 23.13652 | 8.925725 | -0.41248 | -0.16019 | 0.636553 |
| 18 | 75921400 | 75928399 | 40.38261 | 65.84179 | 60.1174 | 25.82379 | 0.502747 | 1.539068 | 0.636576 |
| 13 | 76642000 | 76644199 | 20.28086 | 36.09985 | 32.35504 | 14.2553 | 0.552515 | 0.500256 | 0.636586 |
| 7 | 14021000 | 14048799 | 435.3601 | 603.3953 | 581.9219 | 241.6283 | 0.207568 | 4.717437 | 0.636588 |
| 3 | 85364000 | 85365199 | 8.672059 | 16.39765 | 27.59125 | 11.20457 | -0.264 | 0.045495 | 0.636593 |
| 18 | 15741000 | 15741599 | 6.087798 | 11.84985 | 22.5333 | 8.280221 | -0.30485 | -0.33171 | 0.63663 |
| 1 | 66055800 | 66058399 | 43.34216 | 69.94435 | 78.11972 | 33.30156 | 0.199324 | 1.759418 | 0.636632 |
| 1 | 43893800 | 43894599 | 8.100863 | 15.3036 | 26.51 | 10.78632 | -0.30758 | 0.045448 | 0.636661 |
| 7 | 53297800 | 53298999 | 13.61423 | 24.15634 | 37.66337 | 16.63864 | -0.24441 | 0.489326 | 0.636678 |
| 3 | 142390800 | 142395999 | 80.75724 | 125.3408 | 177.0034 | 73.50658 | -0.19124 | 2.775249 | 0.636679 |
| 18 | 43820400 | 43820999 | 5.954235 | 11.56727 | 21.7212 | 8.108615 | -0.30701 | -0.2776 | 0.63669 |
| 19 | 59861000 | 59863799 | 40.90344 | 66.14783 | 73.5478 | 31.4378 | 0.209297 | 1.692977 | 0.636695 |

| | | | | | | | | | |
|---|---|---|---|---|---|---|---|---|---|
| 11 | 5766000 | 5769199 | 29.95305 | 49.07164 | 73.59349 | 31.6694 | -0.20681 | 1.457211 | 0.6367 |
| 1 | 118876400 | 118877599 | 15.02438 | 26.31668 | 40.92703 | 18.0705 | -0.24639 | 0.657635 | 0.636706 |
| 7 | 32919200 | 32990799 | 1538.831 | 1923.677 | 1650.121 | 693.9929 | 0.227059 | 6.371923 | 0.636726 |
| 1 | 71625200 | 71630999 | 45.55622 | 73.47637 | 80.9729 | 34.49304 | 0.217649 | 1.778823 | 0.636742 |
| 11 | 3494000 | 3498399 | 40.70331 | 65.80327 | 72.3531 | 30.91445 | 0.224822 | 1.700604 | 0.636746 |
| 10 | 82401600 | 82403399 | 28.99852 | 48.82524 | 52.37979 | 22.89287 | 0.28001 | 1.132047 | 0.636753 |
| 2 | 40453800 | 40454999 | 17.27621 | 30.5618 | 33.21348 | 14.68928 | 0.271434 | 0.530593 | 0.636758 |
| 3 | 84386600 | 84387999 | 13.99009 | 24.31083 | 43.26223 | 18.99641 | -0.44062 | 0.774704 | 0.636758 |
| 3 | 84989400 | 84991199 | 16.5603 | 29.70414 | 31.12311 | 13.62417 | 0.334635 | 0.426001 | 0.636763 |
| 12 | 5666000 | 5669999 | 47.53779 | 76.08928 | 110.6368 | 46.60588 | -0.19471 | 2.028649 | 0.636764 |
| X | 8453600 | 8455399 | 27.29451 | 46.68447 | 47.37601 | 20.77366 | 0.361819 | 0.986335 | 0.636791 |
| 8 | 86409400 | 86416399 | 126.401 | 193.3765 | 200.8873 | 83.35133 | 0.235305 | 3.065586 | 0.636795 |
| 19 | 7212000 | 7213199 | 8.78761 | 16.12803 | 30.36882 | 12.9299 | -0.47882 | 0.235224 | 0.636826 |
| Y | 39202200 | 39205799 | 17.76578 | 31.15891 | 25.32818 | 11.19541 | 0.68474 | 0.621738 | 0.636829 |
| 7 | 62964800 | 62965999 | 14.00122 | 25.24651 | 27.90349 | 12.15059 | 0.263256 | 0.371173 | 0.636831 |
| X | 94308600 | 94313999 | 40.73105 | 67.53191 | 58.5472 | 25.35428 | 0.586243 | 1.331979 | 0.636852 |
| 1 | 34050000 | 34051399 | 11.35299 | 19.91256 | 44.61711 | 19.72211 | -0.77039 | 0.621433 | 0.636878 |
| 10 | 120813600 | 120815599 | 14.76594 | 25.69332 | 42.70859 | 18.76905 | -0.34249 | 0.759317 | 0.636903 |
| 14 | 100434000 | 100437199 | 27.99848 | 47.26239 | 51.25726 | 22.41967 | 0.26451 | 1.105745 | 0.636913 |
| 14 | 121049200 | 121051399 | 17.13382 | 29.55704 | 46.89019 | 20.57003 | -0.27792 | 0.866147 | 0.636914 |
| 11 | 119608800 | 119610999 | 22.38691 | 38.39472 | 42.67642 | 18.71706 | 0.231591 | 0.936021 | 0.636925 |
| 1 | 7162600 | 7165199 | 21.76758 | 37.95817 | 39.65546 | 17.45092 | 0.324452 | 0.755746 | 0.636929 |
| 9 | 11744600 | 11746799 | 20.18231 | 35.29625 | 38.06326 | 16.78394 | 0.279162 | 0.70399 | 0.636932 |
| 6 | 37721800 | 37729399 | 82.30003 | 129.0246 | 142.379 | 59.05329 | 0.175434 | 2.596681 | 0.636948 |
| 6 | 28097600 | 28098799 | 12.70781 | 23.6396 | 25.98207 | 10.85862 | 0.311653 | 0.17946 | 0.63695 |
| 7 | 131324600 | 131326199 | 9.8544 | 18.04966 | 30.32266 | 12.94555 | -0.31811 | 0.256418 | 0.636953 |
| 6 | 3200400 | 3206999 | 358.5066 | 497.2709 | 632.0239 | 262.3994 | -0.1894 | 4.703525 | 0.636959 |
| 3 | 133276600 | 133277999 | 14.56888 | 26.276 | 29.3926 | 12.79543 | 0.246347 | 0.378038 | 0.636969 |
| 8 | 99508800 | 99511799 | 28.49226 | 47.79061 | 52.92953 | 23.09503 | 0.234125 | 1.166052 | 0.636978 |
| 12 | 67503000 | 67508999 | 51.81409 | 83.60599 | 83.18694 | 35.45612 | 0.365494 | 1.779319 | 0.636988 |
| 10 | 33909400 | 33910199 | 5.076906 | 9.887218 | 20.16949 | 7.376478 | -0.39666 | -0.3642 | 0.636992 |
| 13 | 111643200 | 111644599 | 12.30572 | 21.97705 | 34.68999 | 15.24129 | -0.25646 | 0.434229 | 0.636994 |
| 10 | 54002400 | 54003599 | 10.89225 | 20.78787 | 23.87515 | 9.506919 | 0.307372 | -0.00744 | 0.63704 |
| 6 | 58882600 | 58885999 | 56.06794 | 89.05614 | 128.6955 | 53.65944 | -0.19768 | 2.281688 | 0.637045 |
| 12 | 20753400 | 20801599 | 1589.788 | 1973.145 | 1736.437 | 730.894 | 0.179626 | 6.486686 | 0.637057 |
| 14 | 34084200 | 34088799 | 52.22346 | 82.95696 | 124.2063 | 51.7922 | -0.24859 | 2.273346 | 0.637071 |
| 2 | 101812600 | 101813999 | 7.921935 | 15.03758 | 26.05024 | 10.49221 | -0.29487 | 0.008433 | 0.637076 |
| 8 | 31676800 | 31677799 | 7.921935 | 15.03758 | 26.05024 | 10.49221 | -0.29487 | 0.008433 | 0.637076 |
| X | 49555000 | 49555799 | 16.70163 | 29.54439 | 30.58363 | 13.52627 | 0.341678 | 0.526848 | 0.637087 |
| 6 | 25156200 | 25158199 | 17.07442 | 30.71029 | 27.03679 | 11.80809 | 0.586688 | 0.405914 | 0.637102 |
| 9 | 113305800 | 113307599 | 33.39897 | 54.25847 | 81.47903 | 34.94643 | -0.21716 | 1.589969 | 0.637113 |
| 7 | 38490000 | 38492399 | 32.28806 | 54.48181 | 44.71919 | 19.55761 | 0.665057 | 1.118275 | 0.637117 |
| 2 | 143215200 | 143216799 | 22.14152 | 37.96499 | 41.63305 | 18.25912 | 0.250885 | 0.934667 | 0.637117 |
| 8 | 19000400 | 19001199 | 7.153994 | 13.66328 | 25.08132 | 9.971953 | -0.36134 | -0.04521 | 0.637125 |
| 3 | 6481400 | 6483599 | 21.27797 | 37.3535 | 36.13427 | 15.98438 | 0.434993 | 0.646391 | 0.637141 |
| 14 | 46694200 | 46694999 | 8.695043 | 16.36952 | 27.50401 | 11.26419 | -0.27223 | 0.075323 | 0.637143 |
| Y | 6525400 | 6526999 | 10.92456 | 20.5553 | 17.15431 | 7.032194 | 0.728179 | 0.06803 | 0.637157 |

| | | | | | | | | | |
|---|---|---|---|---|---|---|---|---|---|
| 2 | 98050200 | 98051199 | 17.25666 | 30.52341 | 33.33857 | 14.74497 | 0.264353 | 0.534132 | 0.637163 |
| 7 | 12147400 | 12156799 | 100.5599 | 155.6294 | 168.8801 | 70.14851 | 0.185194 | 2.832645 | 0.637185 |
| 10 | 82774200 | 82777999 | 47.74969 | 76.73073 | 85.20367 | 36.10644 | 0.200281 | 1.897788 | 0.637196 |
| 3 | 143821600 | 143823399 | 15.09372 | 27.25291 | 29.7504 | 12.93316 | 0.282402 | 0.374279 | 0.63721 |
| 3 | 38144400 | 38147399 | 33.34957 | 55.00546 | 61.37511 | 26.48687 | 0.222436 | 1.372679 | 0.637215 |
| X | 33891000 | 33895399 | 33.67646 | 56.59517 | 53.57085 | 23.40009 | 0.460015 | 1.166621 | 0.637216 |
| 6 | 104879600 | 104880799 | 15.87182 | 28.91275 | 29.85185 | 12.83433 | 0.372718 | 0.331314 | 0.637218 |
| 10 | 49389200 | 49390999 | 16.71581 | 29.20928 | 48.39344 | 21.3239 | -0.33765 | 0.713494 | 0.637256 |
| 1 | 21090000 | 21094199 | 38.5389 | 62.338 | 100.3547 | 42.90499 | -0.32322 | 1.695822 | 0.637262 |
| 12 | 87956200 | 87972999 | 201.8551 | 300.2262 | 292.5502 | 120.532 | 0.292655 | 3.631438 | 0.637269 |
| 10 | 67424600 | 67426199 | 12.78274 | 23.55161 | 26.46026 | 11.20643 | 0.266464 | 0.235728 | 0.637281 |
| X | 8064000 | 8072999 | 106.8407 | 166.159 | 156.2382 | 64.86356 | 0.397137 | 2.753751 | 0.637338 |
| 2 | 177457600 | 177460999 | 38.67009 | 64.44026 | 56.65663 | 24.61973 | 0.566416 | 1.270634 | 0.637341 |
| 4 | 12402400 | 12405599 | 51.41073 | 81.85745 | 119.3861 | 49.9223 | -0.20732 | 2.200114 | 0.637357 |
| 7 | 70639000 | 70640599 | 14.91293 | 27.68144 | 27.47923 | 11.52084 | 0.454302 | 0.214111 | 0.637367 |
| 13 | 86861200 | 86863199 | 17.6442 | 30.98446 | 34.4937 | 15.26284 | 0.233481 | 0.62022 | 0.637369 |
| 19 | 9364200 | 9368199 | 39.50932 | 63.58694 | 94.19723 | 40.00691 | -0.21195 | 1.84183 | 0.637389 |
| 12 | 23916000 | 23917999 | 27.52821 | 46.4823 | 42.21419 | 18.46593 | 0.519639 | 1.091995 | 0.637395 |
| 7 | 26144600 | 26171199 | 377.288 | 531.4079 | 500.6924 | 207.3771 | 0.264585 | 4.474849 | 0.637395 |
| 15 | 78259600 | 78263999 | 35.78504 | 57.84668 | 85.86536 | 36.68341 | -0.20727 | 1.703038 | 0.637395 |
| 6 | 25148200 | 25149599 | 21.78849 | 36.65896 | 56.48374 | 24.68108 | -0.24044 | 1.118997 | 0.637421 |
| 2 | 144744400 | 144745399 | 12.14652 | 22.76384 | 25.63533 | 10.57986 | 0.291474 | 0.128156 | 0.637424 |
| 8 | 11546800 | 11547999 | 10.14343 | 18.77589 | 30.41611 | 12.80057 | -0.25105 | 0.199941 | 0.637459 |
| 13 | 47054000 | 47056999 | 28.62975 | 48.01915 | 52.94125 | 23.09951 | 0.240653 | 1.166544 | 0.637496 |
| 13 | 62810800 | 62833999 | 487.9531 | 670.7799 | 630.9975 | 262.3468 | 0.23157 | 4.844207 | 0.637524 |
| 15 | 45511200 | 45516399 | 46.64092 | 75.41166 | 79.28936 | 33.88353 | 0.289412 | 1.710584 | 0.63755 |
| X | 5231000 | 5235199 | 95.85223 | 150.7569 | 140.7611 | 58.3951 | 0.419803 | 2.544833 | 0.637573 |
| 12 | 41020400 | 41021999 | 13.45121 | 25.14652 | 26.61753 | 11.03146 | 0.375378 | 0.157107 | 0.637626 |
| 10 | 121427000 | 121427799 | 6.114528 | 11.82788 | 22.6541 | 8.652142 | -0.36614 | -0.19292 | 0.637633 |
| 13 | 50751000 | 50753199 | 23.66336 | 41.07601 | 42.447 | 18.62583 | 0.339614 | 0.832936 | 0.637634 |
| 16 | 35331800 | 35334199 | 21.20417 | 36.54882 | 40.07686 | 17.58068 | 0.252471 | 0.882569 | 0.637644 |
| 8 | 94264600 | 94268799 | 56.20123 | 89.95989 | 98.58425 | 41.53127 | 0.212623 | 2.034049 | 0.637652 |
| 19 | 17527600 | 17529399 | 14.16582 | 24.74563 | 46.25665 | 20.37832 | -0.51051 | 0.706194 | 0.637672 |
| 12 | 106661600 | 106664999 | 25.39935 | 42.25539 | 64.30859 | 27.91471 | -0.22282 | 1.254299 | 0.637687 |
| 18 | 20134200 | 20135599 | 10.85738 | 19.59325 | 32.26239 | 14.03469 | -0.30721 | 0.35511 | 0.637689 |
| X | 48587800 | 48589799 | 18.80998 | 33.73002 | 32.12439 | 14.06577 | 0.47161 | 0.44728 | 0.637697 |
| 2 | 15580800 | 15582999 | 15.6412 | 27.47922 | 45.03644 | 19.93528 | -0.32173 | 0.621592 | 0.63771 |
| 18 | 61381600 | 61388799 | 84.49189 | 130.2712 | 209.1932 | 86.89525 | -0.38382 | 2.900551 | 0.637717 |
| Y | 4240600 | 4242399 | 11.75753 | 21.84516 | 18.66896 | 7.816954 | 0.670151 | 0.159924 | 0.637732 |
| 15 | 103563200 | 103564799 | 18.04669 | 30.92828 | 49.00299 | 21.4905 | -0.2778 | 0.934681 | 0.637752 |
| 12 | 23584600 | 23584999 | 12.80892 | 23.92608 | 21.00331 | 8.716827 | 0.642278 | 0.144937 | 0.637759 |
| 11 | 81258000 | 81259999 | 20.57792 | 35.78122 | 39.30561 | 17.26591 | 0.251973 | 0.789023 | 0.637762 |
| 13 | 96883800 | 96886999 | 30.13532 | 49.27056 | 73.98377 | 31.81232 | -0.21027 | 1.485334 | 0.637792 |
| 5 | 83706600 | 83709199 | 60.31111 | 95.99688 | 105.8795 | 44.25401 | 0.194589 | 2.214793 | 0.637807 |
| 3 | 14354200 | 14356599 | 30.50042 | 51.53492 | 44.75298 | 19.5804 | 0.584112 | 1.101177 | 0.637838 |
| 5 | 64842600 | 64846799 | 44.90326 | 72.47658 | 79.87308 | 34.05686 | 0.218831 | 1.756139 | 0.637856 |
| 18 | 61043400 | 61047199 | 24.70614 | 40.84337 | 64.44401 | 27.8518 | -0.27533 | 1.327489 | 0.637857 |

| | | | | | | | | | |
|---|---|---|---|---|---|---|---|---|---|
| 3 | 9984400 | 9985799 | 11.91369 | 21.57215 | 34.25235 | 14.8376 | -0.25145 | 0.34611 | 0.637876 |
| 19 | 20243800 | 20245399 | 11.91369 | 21.57215 | 34.25235 | 14.8376 | -0.25145 | 0.34611 | 0.637876 |
| 5 | 92336000 | 92336999 | 17.71415 | 31.19991 | 34.40944 | 15.23152 | 0.248077 | 0.584241 | 0.637879 |
| 4 | 24230000 | 24231599 | 19.81448 | 34.61662 | 38.21674 | 16.83744 | 0.24559 | 0.720476 | 0.637887 |
| 3 | 3154800 | 3155399 | 15.61716 | 27.76434 | 30.45486 | 13.43201 | 0.261895 | 0.476497 | 0.637916 |
| 10 | 24962000 | 24962999 | 15.71315 | 27.35654 | 43.17979 | 18.98268 | -0.26815 | 0.756956 | 0.637925 |
| 10 | 89687000 | 89689399 | 26.31376 | 44.3953 | 49.61476 | 21.69781 | 0.221332 | 1.102298 | 0.637926 |
| 10 | 46158200 | 46159999 | 18.57417 | 32.01341 | 49.00623 | 21.49772 | -0.227 | 0.883008 | 0.637932 |
| 15 | 94009200 | 94011799 | 33.11338 | 54.82413 | 59.80832 | 25.8751 | 0.255495 | 1.326238 | 0.637933 |
| X | 146481000 | 146488599 | 105.5899 | 164.5824 | 158.5062 | 65.7831 | 0.365121 | 2.713168 | 0.637943 |
| 3 | 61355200 | 61361199 | 45.62018 | 73.92083 | 76.58348 | 32.78407 | 0.31361 | 1.662031 | 0.637962 |
| 1 | 11375000 | 11376999 | 18.33748 | 31.82193 | 50.09513 | 21.97996 | -0.26509 | 0.824122 | 0.637978 |
| 6 | 8799000 | 8800799 | 22.25864 | 38.4719 | 42.57766 | 18.67928 | 0.239958 | 0.866546 | 0.637987 |
| 17 | 47167400 | 47169599 | 14.07287 | 24.65254 | 39.9408 | 17.63657 | -0.3051 | 0.648956 | 0.637995 |
| 1 | 113261800 | 113264399 | 29.68975 | 50.46477 | 49.58002 | 21.72914 | 0.40702 | 1.051905 | 0.63802 |
| 8 | 15682200 | 15688999 | 87.34553 | 137.1436 | 129.7907 | 53.83207 | 0.398446 | 2.569616 | 0.638029 |
| 5 | 10334800 | 10335199 | 4.642527 | 9.069261 | 18.21802 | 6.364109 | -0.31474 | -0.54177 | 0.638055 |
| 15 | 23970200 | 23973999 | 12.11938 | 22.80699 | 21.4746 | 8.800925 | 0.556092 | 0.092537 | 0.638074 |
| 6 | 68007600 | 68008199 | 6.849408 | 13.2973 | 24.62353 | 9.242537 | -0.29491 | -0.23872 | 0.638087 |
| 10 | 119893600 | 119894799 | 18.25197 | 31.36232 | 49.00469 | 21.49505 | -0.25739 | 0.908615 | 0.638121 |
| 13 | 27471600 | 27474999 | 74.59238 | 117.5749 | 130.092 | 53.98428 | 0.177572 | 2.485088 | 0.638147 |
| 2 | 70604200 | 70604999 | 11.31796 | 21.48649 | 24.27748 | 9.775798 | 0.316522 | 0.032678 | 0.638149 |
| 19 | 21090400 | 21092599 | 14.86235 | 26.21477 | 41.8095 | 18.50186 | -0.28077 | 0.563463 | 0.638191 |
| 16 | 77145800 | 77147599 | 22.42947 | 38.90306 | 35.78236 | 15.69978 | 0.506742 | 0.826489 | 0.638208 |
| 7 | 59266400 | 59270799 | 59.01504 | 94.87441 | 77.50463 | 32.86442 | 0.642982 | 1.896429 | 0.638223 |
| 10 | 24608400 | 24610199 | 10.12024 | 18.96438 | 32.98399 | 13.61514 | -0.33139 | 0.129508 | 0.638223 |
| 9 | 116691400 | 116695999 | 41.29467 | 66.45886 | 97.72644 | 41.5045 | -0.20138 | 1.846237 | 0.638228 |
| 2 | 18396800 | 18397799 | 8.559842 | 16.04804 | 27.45648 | 11.32428 | -0.30625 | 0.100829 | 0.638261 |
| 5 | 144320600 | 144322999 | 34.57794 | 56.82458 | 63.51921 | 27.37005 | 0.217807 | 1.422622 | 0.638298 |
| 6 | 69523000 | 69523599 | 10.52209 | 19.96485 | 23.20748 | 9.355558 | 0.274526 | 0.02621 | 0.638313 |
| 15 | 44977400 | 44978399 | 13.30055 | 23.38487 | 37.83236 | 16.75261 | -0.30274 | 0.584014 | 0.638334 |
| 10 | 82244800 | 82280199 | 674.4104 | 902.9588 | 821.5634 | 342.9634 | 0.241312 | 5.261063 | 0.638338 |
| 4 | 62693000 | 62696599 | 24.83202 | 41.11465 | 64.0101 | 27.69365 | -0.25595 | 1.308881 | 0.638338 |
| 17 | 18501600 | 18504199 | 28.00176 | 47.99104 | 46.81339 | 20.5303 | 0.419105 | 0.970969 | 0.638351 |
| 3 | 63979600 | 63980199 | 10.43841 | 19.46239 | 33.36253 | 13.88926 | -0.32054 | 0.164922 | 0.638366 |
| 8 | 116093400 | 116096199 | 22.23938 | 38.62087 | 41.387 | 18.16564 | 0.287394 | 0.817792 | 0.638368 |
| X | 109857600 | 109859399 | 16.27804 | 28.45613 | 45.16806 | 19.90967 | -0.27569 | 0.705367 | 0.638372 |
| 2 | 61451200 | 61452999 | 25.58186 | 44.48944 | 35.34393 | 15.52725 | 0.717546 | 0.806775 | 0.638375 |
| 6 | 8610200 | 8610999 | 15.08815 | 27.46713 | 25.25333 | 10.86619 | 0.538265 | 0.319287 | 0.638376 |
| 14 | 50253200 | 50258199 | 57.1413 | 91.60027 | 96.1827 | 40.61027 | 0.276673 | 1.99034 | 0.638381 |
| 16 | 20809400 | 20815199 | 71.56254 | 113.2496 | 125.2382 | 52.05935 | 0.182898 | 2.391395 | 0.638386 |
| 8 | 121589600 | 121591399 | 16.68421 | 29.04072 | 44.72064 | 19.65734 | -0.23282 | 0.767131 | 0.638388 |
| X | 144344200 | 144345799 | 19.22532 | 34.81732 | 30.65121 | 13.27478 | 0.594777 | 0.39081 | 0.63842 |
| 2 | 125275400 | 125276999 | 23.24243 | 39.01638 | 59.28554 | 25.88153 | -0.22059 | 1.147187 | 0.638425 |
| 1 | 16918400 | 16919999 | 20.95366 | 36.34857 | 40.43418 | 17.74088 | 0.233561 | 0.823538 | 0.63846 |
| X | 67049000 | 67051199 | 23.61449 | 41.22139 | 33.89794 | 14.92694 | 0.668525 | 0.745513 | 0.638551 |
| 1 | 117811000 | 117813999 | 75.39288 | 118.9763 | 131.4226 | 54.55242 | 0.181294 | 2.461167 | 0.638552 |

| | | | | | | | | | |
|---|---|---|---|---|---|---|---|---|---|
| 19 | 16901600 | 16903999 | 20.32862 | 35.66188 | 32.2412 | 14.25379 | 0.532243 | 0.647364 | 0.638555 |
| 7 | 47056000 | 47070999 | 178.202 | 268.3566 | 248.5255 | 102.6527 | 0.382125 | 3.396404 | 0.638585 |
| 2 | 139712200 | 139716199 | 48.61557 | 78.34928 | 84.69025 | 36.04011 | 0.244193 | 1.80914 | 0.638597 |
| 5 | 123408800 | 123410599 | 13.90585 | 24.49987 | 38.23756 | 16.92502 | -0.25013 | 0.564834 | 0.638599 |
| 9 | 42522600 | 42524199 | 8.678718 | 15.96505 | 29.44874 | 12.49879 | -0.44517 | 0.223604 | 0.638616 |
| 3 | 24343200 | 24344199 | 15.1518 | 27.58473 | 24.82531 | 10.68119 | 0.569115 | 0.319303 | 0.638641 |
| 2 | 50997200 | 50997799 | 6.242569 | 12.10306 | 22.76125 | 8.601164 | -0.32644 | -0.23094 | 0.63869 |
| 13 | 49969600 | 49971999 | 25.79631 | 44.04095 | 47.38688 | 20.77535 | 0.277231 | 0.994819 | 0.6387 |
| 1 | 50920000 | 50922599 | 23.74989 | 40.02135 | 62.72112 | 27.41952 | -0.26491 | 1.117927 | 0.638711 |
| 7 | 70402600 | 70403599 | 12.12517 | 21.62632 | 34.67571 | 15.25181 | -0.28013 | 0.440186 | 0.638734 |
| 2 | 13258400 | 13263799 | 61.74995 | 97.95064 | 143.3641 | 59.69584 | -0.21797 | 2.326622 | 0.638746 |
| 4 | 24944800 | 24945999 | 8.715519 | 16.4423 | 27.54244 | 11.23881 | -0.26341 | 0.063122 | 0.638754 |
| 1 | 111372000 | 111373799 | 16.72508 | 30.29674 | 30.15915 | 13.05769 | 0.418433 | 0.368179 | 0.638785 |
| 4 | 18703600 | 18705199 | 7.508653 | 14.17648 | 30.30254 | 12.3448 | -0.61053 | 0.052108 | 0.638793 |
| X | 55126600 | 55127599 | 9.420382 | 17.75995 | 31.41931 | 12.83717 | -0.34329 | 0.084571 | 0.638796 |
| 12 | 73280400 | 73282599 | 18.48232 | 32.3959 | 36.06568 | 15.93334 | 0.233509 | 0.657484 | 0.638798 |
| 4 | 135365000 | 135367999 | 24.97307 | 41.64597 | 63.10479 | 27.43445 | -0.21644 | 1.228093 | 0.638798 |
| 10 | 80259000 | 80266599 | 62.26177 | 97.98117 | 145.6872 | 60.44314 | -0.24997 | 2.512458 | 0.638804 |
| 16 | 8631200 | 8634399 | 20.30252 | 34.62099 | 52.73187 | 23.11652 | -0.2224 | 0.992664 | 0.638806 |
| 8 | 92920800 | 92923399 | 20.02018 | 33.90426 | 53.22026 | 23.30113 | -0.26667 | 1.054105 | 0.638809 |
| 4 | 125105800 | 125110399 | 44.31293 | 71.04512 | 103.5303 | 43.72131 | -0.19503 | 1.972481 | 0.638832 |
| 11 | 4513000 | 4514599 | 15.80865 | 27.23955 | 51.62469 | 22.64618 | -0.53366 | 0.878735 | 0.638848 |
| 2 | 171842800 | 171845999 | 34.41233 | 55.70262 | 83.87266 | 35.87712 | -0.2259 | 1.666205 | 0.638854 |
| 7 | 19607800 | 19611999 | 34.41233 | 55.70262 | 83.87266 | 35.87712 | -0.2259 | 1.666205 | 0.638854 |
| 14 | 124016200 | 124018599 | 24.96869 | 43.47243 | 41.23745 | 18.12933 | 0.463055 | 0.797145 | 0.638856 |
| 18 | 53907600 | 53909399 | 16.60363 | 29.31367 | 32.33144 | 14.30487 | 0.249149 | 0.546261 | 0.63886 |
| 10 | 38704400 | 38709599 | 75.2888 | 119.9417 | 95.26621 | 39.86626 | 0.668834 | 2.199835 | 0.638872 |
| 6 | 144811600 | 144814199 | 19.46029 | 33.85239 | 36.82554 | 16.18085 | 0.266125 | 0.774651 | 0.638874 |
| X | 26515600 | 26520199 | 49.39638 | 80.19163 | 73.98495 | 31.71603 | 0.483092 | 1.634306 | 0.638903 |
| 17 | 36325000 | 36329199 | 49.42866 | 79.2855 | 87.00656 | 36.77046 | 0.214203 | 1.957244 | 0.638909 |
| 4 | 153700600 | 153701999 | 13.14239 | 23.85958 | 25.03858 | 10.81074 | 0.345518 | 0.319361 | 0.638914 |
| 6 | 100459000 | 100464799 | 55.75822 | 88.75213 | 128.9461 | 53.89681 | -0.20233 | 2.214811 | 0.638915 |
| 10 | 81980600 | 82006199 | 505.739 | 694.0646 | 639.7666 | 266.0631 | 0.258438 | 4.870852 | 0.638917 |
| 18 | 69454000 | 69456399 | 18.89651 | 32.60789 | 50.08788 | 21.97301 | -0.23161 | 0.874937 | 0.638928 |
| 6 | 138313200 | 138322399 | 150.1098 | 227.8993 | 223.797 | 92.66857 | 0.307597 | 3.227953 | 0.638945 |
| 17 | 84331200 | 84332599 | 22.52165 | 38.61889 | 43.17844 | 18.93696 | 0.223077 | 0.93825 | 0.638948 |
| 12 | 88640200 | 88641199 | 14.0177 | 24.75801 | 39.18674 | 17.33658 | -0.26906 | 0.538598 | 0.638965 |
| 6 | 35341000 | 35343799 | 25.4456 | 42.15734 | 64.70392 | 28.00611 | -0.23541 | 1.302931 | 0.638977 |
| 10 | 100322600 | 100323599 | 8.327091 | 15.4084 | 28.43135 | 11.97098 | -0.43756 | 0.182642 | 0.63899 |
| 19 | 48534600 | 48536799 | 21.66832 | 37.90672 | 38.47139 | 16.96831 | 0.366299 | 0.707434 | 0.63899 |
| 2 | 31250800 | 31252199 | 15.06904 | 26.16772 | 43.76174 | 19.20743 | -0.35159 | 0.7905 | 0.639008 |
| 16 | 55462400 | 55465199 | 40.95167 | 66.53942 | 72.49512 | 31.09707 | 0.244877 | 1.59848 | 0.63902 |
| 4 | 58610400 | 58612399 | 20.9566 | 34.56964 | 72.21708 | 31.16758 | -0.67931 | 1.349953 | 0.639061 |
| 7 | 68146400 | 68148599 | 21.38093 | 36.24553 | 55.46432 | 24.29112 | -0.23019 | 1.050149 | 0.639072 |
| 2 | 162688400 | 162692599 | 47.89967 | 76.67647 | 111.2508 | 46.87143 | -0.19144 | 2.026751 | 0.639073 |
| 15 | 75151800 | 75154199 | 29.90352 | 50.06216 | 54.60611 | 23.79792 | 0.256143 | 1.193369 | 0.639076 |
| 10 | 71057200 | 71058999 | 20.90984 | 35.38269 | 54.89274 | 24.02757 | -0.25001 | 1.063523 | 0.639083 |

| | | | | | | | | | |
|---|---|---|---|---|---|---|---|---|---|
| 19 | 8528800 | 8531199 | 29.13296 | 48.50569 | 54.05913 | 23.47825 | 0.225301 | 1.246555 | 0.639097 |
| 1 | 12449000 | 12452399 | 38.63151 | 63.99888 | 60.6679 | 26.25793 | 0.457695 | 1.337683 | 0.639097 |
| 6 | 35433400 | 35434599 | 13.26258 | 23.49381 | 36.65143 | 16.20534 | -0.24639 | 0.49857 | 0.639107 |
| 18 | 59145800 | 59146799 | 13.26258 | 23.49381 | 36.65143 | 16.20534 | -0.24639 | 0.49857 | 0.639107 |
| 14 | 62485000 | 62486599 | 13.24978 | 23.36052 | 37.73172 | 16.69875 | -0.29899 | 0.551253 | 0.639113 |
| 3 | 64153000 | 64160199 | 68.8441 | 109.1265 | 120.7705 | 50.25244 | 0.183767 | 2.347404 | 0.639115 |
| 17 | 92905000 | 92906399 | 11.66274 | 22.29094 | 23.26687 | 9.23582 | 0.447655 | -0.01375 | 0.639117 |
| 16 | 97879800 | 97881399 | 23.46814 | 40.26482 | 44.70311 | 19.60654 | 0.233247 | 0.938103 | 0.639128 |
| 17 | 69316000 | 69318199 | 23.46814 | 40.26482 | 44.70311 | 19.60654 | 0.233247 | 0.938103 | 0.639128 |
| Y | 5934600 | 5937399 | 23.82112 | 40.69559 | 30.65378 | 13.43994 | 0.789582 | 0.956967 | 0.639137 |
| 12 | 72778000 | 72781599 | 46.10444 | 74.1871 | 82.44796 | 35.00716 | 0.201486 | 1.852134 | 0.639137 |
| 14 | 38272000 | 38273199 | 16.23 | 29.23588 | 26.62764 | 11.60966 | 0.539566 | 0.390122 | 0.639163 |
| 1 | 47728200 | 47729599 | 16.37777 | 29.88538 | 29.22185 | 12.53667 | 0.452526 | 0.322588 | 0.639187 |
| 6 | 24934600 | 24940799 | 62.23176 | 99.39211 | 104.8616 | 44.03722 | 0.263825 | 2.10864 | 0.639204 |
| 1 | 170793000 | 170796599 | 28.58462 | 46.81542 | 74.23051 | 31.93939 | -0.28717 | 1.455224 | 0.639212 |
| 6 | 129235600 | 129235999 | 4.609142 | 9.014067 | 18.48038 | 6.281537 | -0.30777 | -0.63568 | 0.639246 |
| 18 | 3506800 | 3508799 | 16.00672 | 27.00242 | 59.91813 | 26.20276 | -0.76424 | 1.081705 | 0.639262 |
| 2 | 177871800 | 177876599 | 52.33313 | 84.09727 | 80.10968 | 33.94851 | 0.421057 | 1.901963 | 0.639265 |
| 7 | 23775400 | 23777599 | 17.09428 | 29.74596 | 45.76176 | 20.11134 | -0.23161 | 0.772736 | 0.639291 |
| 13 | 62605800 | 62615999 | 216.0472 | 318.8383 | 319.6285 | 131.6284 | 0.241302 | 3.766096 | 0.639345 |
| X | 80476600 | 80485199 | 74.50457 | 118.7012 | 110.6972 | 46.32847 | 0.43777 | 2.201766 | 0.639347 |
| 1 | 33964400 | 33966799 | 21.96451 | 37.78823 | 42.0897 | 18.4626 | 0.229337 | 0.90403 | 0.639356 |
| 8 | 35508600 | 35509199 | 8.84198 | 16.81452 | 28.90451 | 11.59722 | -0.27937 | 0.011481 | 0.639359 |
| 13 | 6881400 | 6882199 | 10.1069 | 19.23408 | 22.76576 | 9.120038 | 0.256825 | 0.001851 | 0.639387 |
| 14 | 60948400 | 60950599 | 18.09702 | 31.21093 | 47.94064 | 21.03068 | -0.23165 | 0.874144 | 0.639393 |
| 5 | 132958400 | 132959399 | 8.81225 | 16.62174 | 28.0761 | 11.46101 | -0.2763 | 0.064282 | 0.639402 |
| 7 | 56266600 | 56267599 | 14.72721 | 25.78676 | 40.45682 | 17.85739 | -0.25892 | 0.661829 | 0.639433 |
| 1 | 182539400 | 182541999 | 21.31675 | 36.79678 | 41.03383 | 18.00128 | 0.228629 | 0.870225 | 0.639436 |
| 17 | 77063600 | 77066799 | 50.22209 | 80.6596 | 89.75272 | 38.00366 | 0.196316 | 1.919789 | 0.639441 |
| 5 | 127865800 | 127869599 | 41.68778 | 67.41373 | 75.52081 | 32.27961 | 0.198395 | 1.695569 | 0.639456 |
| 14 | 88275800 | 88278799 | 46.83386 | 75.31537 | 83.31021 | 35.34236 | 0.207185 | 1.873591 | 0.639466 |
| 6 | 146593000 | 146597199 | 89.49058 | 139.3357 | 152.6446 | 63.36081 | 0.178209 | 2.722245 | 0.639467 |
| 8 | 60977000 | 60977399 | 8.642757 | 16.0868 | 28.46549 | 11.88502 | -0.36821 | 0.153567 | 0.63948 |
| 4 | 73768400 | 73770399 | 16.59395 | 29.32401 | 32.28868 | 14.28328 | 0.252044 | 0.535779 | 0.639493 |
| 9 | 117721600 | 117723399 | 16.98804 | 29.87079 | 32.99663 | 14.61103 | 0.244824 | 0.595046 | 0.639493 |
| 17 | 53459000 | 53463799 | 101.9196 | 158.9727 | 156.9437 | 65.1282 | 0.330189 | 2.697541 | 0.639505 |
| 1 | 113619000 | 113620799 | 18.83889 | 32.9925 | 36.61843 | 16.16578 | 0.237924 | 0.674484 | 0.639535 |
| 12 | 11052200 | 11055999 | 35.98281 | 59.61877 | 60.57317 | 26.21932 | 0.357918 | 1.327115 | 0.639546 |
| 19 | 8289600 | 8304399 | 314.5072 | 447.2839 | 451.1127 | 186.5425 | 0.180451 | 4.330362 | 0.63955 |
| 4 | 112301800 | 112307599 | 57.35473 | 92.36552 | 86.542 | 36.78905 | 0.448732 | 1.847002 | 0.639553 |
| 18 | 32172200 | 32176599 | 63.86153 | 101.5641 | 112.402 | 46.9265 | 0.188389 | 2.245007 | 0.639572 |
| 16 | 10009600 | 10011599 | 20.3516 | 34.34468 | 54.28275 | 23.74101 | -0.27677 | 1.086502 | 0.639586 |
| 3 | 19711800 | 19713999 | 14.77745 | 25.68593 | 49.50797 | 21.74257 | -0.5555 | 0.782816 | 0.639602 |
| 1 | 73857200 | 73858599 | 12.24285 | 22.79578 | 25.50573 | 10.64282 | 0.287943 | 0.168822 | 0.639634 |
| 17 | 20597400 | 20600999 | 59.54783 | 95.27107 | 100.9491 | 42.4987 | 0.260346 | 2.053535 | 0.63964 |
| X | 156970200 | 156970599 | 10.17336 | 19.54528 | 22.97748 | 8.986292 | 0.298272 | -0.07334 | 0.639661 |
| 1 | 43032000 | 43035199 | 46.4658 | 74.96391 | 82.22145 | 35.03725 | 0.224943 | 1.774254 | 0.639664 |

| | | | | | | | | | |
|---|---|---|---|---|---|---|---|---|---|
| 13 | 60341200 | 60344599 | 40.69282 | 65.2746 | 99.3985 | 42.00919 | -0.25745 | 1.949629 | 0.639693 |
| X | 4093400 | 4096199 | 25.8239 | 45.24157 | 37.14723 | 16.4042 | 0.670779 | 0.707507 | 0.639694 |
| 3 | 3486400 | 3487399 | 18.95807 | 33.06593 | 36.33023 | 15.98973 | 0.252311 | 0.736356 | 0.639695 |
| X | 132939800 | 132940999 | 18.03315 | 32.25363 | 32.22978 | 14.14267 | 0.400642 | 0.458964 | 0.63971 |
| X | 126441200 | 126441599 | 10.20861 | 19.4656 | 22.83062 | 9.108336 | 0.275474 | -0.00846 | 0.639726 |
| 16 | 82007000 | 82009199 | 35.73317 | 58.68649 | 53.05834 | 22.8565 | 0.522703 | 1.424585 | 0.639731 |
| 14 | 10223400 | 10225399 | 22.96205 | 40.0346 | 40.14454 | 17.66435 | 0.383354 | 0.764961 | 0.639734 |
| X | 14938600 | 14939999 | 16.74132 | 30.0689 | 30.83967 | 13.48493 | 0.366829 | 0.422791 | 0.639741 |
| 5 | 109531800 | 109535399 | 64.32604 | 102.498 | 110.3666 | 46.19843 | 0.230848 | 2.185739 | 0.639762 |
| Y | 20311800 | 20313799 | 14.1368 | 25.54411 | 20.13589 | 8.745384 | 0.749919 | 0.339842 | 0.639779 |
| Y | 9634600 | 9642999 | 62.41563 | 99.28368 | 77.22481 | 32.24904 | 0.696557 | 2.224945 | 0.639782 |
| 15 | 76882800 | 76884599 | 14.95117 | 26.07516 | 42.14954 | 18.54999 | -0.30196 | 0.72806 | 0.639788 |
| 4 | 73340000 | 73341599 | 16.13887 | 28.48687 | 30.9579 | 13.69776 | 0.270363 | 0.543892 | 0.639792 |
| 10 | 78485800 | 78486999 | 13.88998 | 24.51682 | 38.19531 | 16.90012 | -0.24656 | 0.544137 | 0.639798 |
| 1 | 117615000 | 117620999 | 91.55194 | 144.2097 | 138.3643 | 57.41198 | 0.38222 | 2.511371 | 0.639799 |
| X | 156065800 | 156066799 | 17.37036 | 30.19366 | 46.85035 | 20.57575 | -0.24395 | 0.793411 | 0.639826 |
| 13 | 46138800 | 46140199 | 11.98122 | 21.72619 | 34.82894 | 15.05978 | -0.26332 | 0.340983 | 0.639867 |
| 14 | 110346200 | 110347199 | 11.98122 | 21.72619 | 34.82894 | 15.05978 | -0.26332 | 0.340983 | 0.639867 |
| 12 | 5495200 | 5496399 | 8.901221 | 16.52982 | 28.40484 | 11.90172 | -0.33069 | 0.166885 | 0.639882 |
| 2 | 148942200 | 148951999 | 162.915 | 244.4694 | 258.7362 | 106.7691 | 0.185149 | 3.461641 | 0.639902 |
| 4 | 19761000 | 19762599 | 18.00123 | 32.21756 | 31.47204 | 13.80257 | 0.433312 | 0.449413 | 0.639902 |
| 2 | 87807400 | 87807799 | 6.675577 | 13.03151 | 16.98548 | 6.007344 | 0.285159 | -0.47462 | 0.639918 |
| 2 | 18992400 | 18994999 | 23.1848 | 40.12463 | 42.45854 | 18.62776 | 0.304621 | 0.856088 | 0.639929 |
| 10 | 129443200 | 129444399 | 18.82387 | 33.10368 | 30.25163 | 13.39514 | 0.516984 | 0.598522 | 0.639941 |
| 12 | 46782800 | 46783799 | 14.74224 | 27.03196 | 24.75215 | 10.55163 | 0.553058 | 0.274827 | 0.639952 |
| 6 | 133556400 | 133558399 | 22.48904 | 39.62985 | 35.57248 | 15.74476 | 0.544016 | 0.609206 | 0.639962 |
| 8 | 67739600 | 67740199 | 8.758377 | 16.37337 | 27.96274 | 11.58906 | -0.30904 | 0.123941 | 0.639963 |
| 5 | 110022400 | 110029799 | 158.8851 | 240.1615 | 238.5453 | 98.64053 | 0.285812 | 3.318349 | 0.639964 |
| 13 | 105639400 | 105640999 | 17.19677 | 30.44954 | 33.26 | 14.70712 | 0.264751 | 0.52133 | 0.639973 |
| 12 | 18212400 | 18244199 | 1461.143 | 1824.112 | 1597.842 | 672.1083 | 0.194875 | 6.392838 | 0.639998 |
| 10 | 17692200 | 17694199 | 36.04839 | 58.92556 | 66.2422 | 28.48128 | 0.205988 | 1.495296 | 0.640006 |
| X | 97192200 | 97195799 | 36.91629 | 60.63029 | 63.48898 | 27.34995 | 0.31148 | 1.438812 | 0.640064 |
| 17 | 94246200 | 94247399 | 16.87963 | 30.00937 | 26.75556 | 11.80024 | 0.559063 | 0.475379 | 0.640069 |
| 1 | 177795200 | 177798399 | 64.98528 | 103.1299 | 113.4898 | 47.27806 | 0.193572 | 2.3102 | 0.640069 |
| 3 | 4246200 | 4248199 | 30.50135 | 51.01445 | 55.21289 | 24.04793 | 0.267343 | 1.204762 | 0.640088 |
| 12 | 22873000 | 22977399 | 2884.907 | 3410.75 | 2858.71 | 1211.231 | 0.178647 | 7.255934 | 0.64009 |
| 14 | 50207400 | 50207999 | 10.84644 | 19.9389 | 32.43645 | 13.77982 | -0.26687 | 0.250066 | 0.640095 |
| 13 | 32829800 | 32831799 | 17.75042 | 31.59875 | 32.5773 | 14.35205 | 0.351617 | 0.4834 | 0.640108 |
| 12 | 7467000 | 7468799 | 20.6118 | 35.05653 | 53.26562 | 23.34558 | -0.21951 | 1.017686 | 0.640112 |
| 1 | 192352000 | 192359599 | 65.73494 | 102.828 | 173.488 | 71.97241 | -0.4388 | 2.631305 | 0.640115 |
| 9 | 39176000 | 39177599 | 20.65775 | 36.04295 | 39.26542 | 17.28534 | 0.264535 | 0.744621 | 0.640116 |
| 10 | 42474600 | 42485599 | 112.3422 | 171.4409 | 239.7526 | 99.4199 | -0.19603 | 3.114009 | 0.64013 |
| 17 | 84842600 | 84844999 | 22.22322 | 37.05148 | 68.56281 | 29.80285 | -0.50389 | 1.225306 | 0.640136 |
| 12 | 103420800 | 103425399 | 28.06009 | 47.28105 | 51.79944 | 22.64443 | 0.249891 | 1.122684 | 0.640139 |
| 4 | 101504200 | 101505599 | 13.79387 | 24.08088 | 41.10834 | 18.10314 | -0.38022 | 0.703872 | 0.640144 |
| 2 | 88793800 | 88795799 | 22.37776 | 39.20398 | 37.826 | 16.70387 | 0.438896 | 0.688085 | 0.64018 |
| 7 | 29557600 | 29559799 | 5.099417 | 9.921189 | 24.17219 | 8.917129 | -0.66342 | -0.32398 | 0.6402 |

| | | | | | | | | | |
|---|---|---|---|---|---|---|---|---|---|
| 5 | 8176400 | 8177599 | 17.45615 | 30.29225 | 46.34131 | 20.33291 | -0.22392 | 0.814386 | 0.640202 |
| 10 | 55529200 | 55529999 | 14.05072 | 24.72852 | 38.74437 | 17.15303 | -0.25636 | 0.577841 | 0.640217 |
| 15 | 45905600 | 45909599 | 17.14849 | 30.56841 | 27.54219 | 12.12002 | 0.546013 | 0.461425 | 0.640219 |
| 8 | 65694600 | 65695399 | 12.35514 | 22.83591 | 26.03809 | 10.98027 | 0.249894 | 0.214608 | 0.640221 |
| 10 | 119731000 | 119735199 | 67.20301 | 106.8828 | 115.2853 | 48.13305 | 0.225498 | 2.247748 | 0.640254 |
| 4 | 68328600 | 68329999 | 13.74653 | 24.29047 | 37.68775 | 16.67185 | -0.23999 | 0.530799 | 0.640259 |
| 15 | 24441000 | 24441599 | 13.43426 | 24.49323 | 27.51238 | 11.81593 | 0.253048 | 0.304358 | 0.640261 |
| 7 | 73053600 | 73057399 | 39.06015 | 62.77743 | 95.08504 | 40.30336 | -0.24634 | 1.885577 | 0.640262 |
| 9 | 100974800 | 100975599 | 8.270186 | 15.81376 | 27.90543 | 11.06304 | -0.30098 | -0.03312 | 0.640285 |
| 12 | 61794200 | 61795199 | 8.270186 | 15.81376 | 27.90543 | 11.06304 | -0.30098 | -0.03312 | 0.640285 |
| 9 | 5657800 | 5660199 | 54.60321 | 87.53953 | 83.2476 | 35.15367 | 0.419487 | 1.978035 | 0.640304 |
| 6 | 108704600 | 108706799 | 21.158 | 35.75729 | 54.84774 | 23.99866 | -0.23366 | 1.077734 | 0.640319 |
| 12 | 41073800 | 41074599 | 8.596362 | 16.00925 | 27.92828 | 11.64986 | -0.34713 | 0.14596 | 0.64032 |
| X | 66737200 | 66739199 | 23.94951 | 41.60649 | 32.3062 | 14.18327 | 0.750536 | 0.812315 | 0.640326 |
| 10 | 75265400 | 75268999 | 42.67614 | 68.902 | 76.98885 | 32.83943 | 0.19948 | 1.744051 | 0.640333 |
| 5 | 55731800 | 55732599 | 10.12044 | 18.69322 | 30.37777 | 12.82179 | -0.25872 | 0.210868 | 0.640354 |
| 6 | 68792600 | 68796399 | 53.49059 | 85.54637 | 94.50433 | 39.76332 | 0.199675 | 2.055296 | 0.640367 |
| X | 29349400 | 29353999 | 49.03665 | 80.04891 | 68.01269 | 29.2282 | 0.607606 | 1.546083 | 0.640399 |
| 5 | 43216600 | 43217599 | 19.29463 | 34.13725 | 29.54998 | 13.06866 | 0.598138 | 0.536392 | 0.640403 |
| 7 | 32817800 | 32818399 | 11.92565 | 22.1186 | 25.11045 | 10.53927 | 0.261234 | 0.190667 | 0.640429 |
| 5 | 30258800 | 30260999 | 26.01261 | 44.78572 | 45.18859 | 19.82237 | 0.371699 | 0.921152 | 0.640443 |
| X | 90896800 | 90898599 | 26.01261 | 44.78572 | 45.18859 | 19.82237 | 0.371699 | 0.921152 | 0.640443 |
| 3 | 95575400 | 95577399 | 14.90344 | 26.06123 | 41.02195 | 18.08684 | -0.2636 | 0.685387 | 0.640448 |
| 17 | 53684400 | 53685999 | 15.93692 | 28.71678 | 30.81674 | 13.43093 | 0.30482 | 0.396217 | 0.640455 |
| 19 | 57963600 | 57966799 | 31.64491 | 52.55099 | 58.04058 | 25.16262 | 0.238067 | 1.286949 | 0.640473 |
| 6 | 142479800 | 142481999 | 25.36059 | 43.13329 | 47.9502 | 21.01591 | 0.229544 | 1.028554 | 0.640486 |
| 1 | 166595000 | 166599799 | 66.9892 | 106.1183 | 114.3491 | 47.55699 | 0.221409 | 2.363229 | 0.640493 |
| 9 | 8396600 | 8400399 | 57.61476 | 91.88838 | 101.8973 | 42.70739 | 0.189872 | 2.146528 | 0.640532 |
| 3 | 96957600 | 96960599 | 22.33136 | 37.33512 | 58.59466 | 25.51448 | -0.26692 | 1.187817 | 0.640541 |
| X | 5117400 | 5121599 | 27.83226 | 48.02479 | 44.57462 | 19.55437 | 0.492323 | 0.908315 | 0.640543 |
| 14 | 87478400 | 87480199 | 21.01007 | 35.79654 | 55.51268 | 24.33387 | -0.24864 | 1.003384 | 0.640549 |
| 6 | 16209000 | 16210599 | 21.19792 | 38.29349 | 29.30962 | 12.73366 | 0.792528 | 0.415398 | 0.640551 |
| 1 | 121281600 | 121283399 | 18.36774 | 32.14515 | 35.22808 | 15.5429 | 0.256255 | 0.681554 | 0.640562 |
| 2 | 39820600 | 39822399 | 27.57198 | 46.33678 | 51.78995 | 22.62226 | 0.221028 | 1.143601 | 0.640568 |
| 10 | 47609000 | 47611599 | 21.74104 | 37.82564 | 35.14433 | 15.44361 | 0.492673 | 0.781851 | 0.640592 |
| 6 | 129821000 | 129832599 | 172.2228 | 257.9572 | 267.4711 | 110.3273 | 0.212306 | 3.499112 | 0.64061 |
| 16 | 48713000 | 48717599 | 43.64331 | 70.79299 | 75.23675 | 32.23287 | 0.278339 | 1.636671 | 0.640645 |
| 19 | 57934800 | 57936999 | 29.12272 | 48.51644 | 54.06557 | 23.48954 | 0.225427 | 1.240289 | 0.640655 |
| 14 | 45588400 | 45589199 | 8.438141 | 15.78737 | 27.45656 | 11.36382 | -0.3335 | 0.11413 | 0.640661 |
| 1 | 113966800 | 113968599 | 15.8986 | 28.83187 | 30.33667 | 13.11617 | 0.340441 | 0.358131 | 0.640667 |
| 9 | 86273400 | 86274599 | 15.62938 | 27.70549 | 30.325 | 13.40629 | 0.262358 | 0.492553 | 0.64067 |
| 5 | 113272800 | 113274199 | 18.38181 | 31.4888 | 49.52699 | 21.71954 | -0.26738 | 0.941011 | 0.640671 |
| 5 | 116895000 | 116903599 | 105.1319 | 162.2948 | 175.4056 | 72.86002 | 0.188208 | 2.881361 | 0.640684 |
| 17 | 94490200 | 94492799 | 10.01471 | 19.36381 | 17.02426 | 6.515394 | 0.74286 | -0.16339 | 0.640694 |
| 1 | 102171400 | 102173399 | 22.36662 | 38.95597 | 40.77806 | 17.9314 | 0.321548 | 0.778098 | 0.640697 |
| 14 | 89769200 | 89771799 | 18.39967 | 32.78155 | 32.51023 | 14.31254 | 0.407979 | 0.4823 | 0.640712 |
| 4 | 48290600 | 48297399 | 80.60697 | 127.7146 | 126.3378 | 52.54806 | 0.344857 | 2.371757 | 0.640721 |

| | | | | | | | | | |
|---|---|---|---|---|---|---|---|---|---|
| 1 | 191764600 | 191767399 | 40.25868 | 65.13467 | 71.75656 | 30.6835 | 0.22321 | 1.681616 | 0.640723 |
| 19 | 53761400 | 53763399 | 16.79092 | 29.13789 | 45.29322 | 19.87305 | -0.24685 | 0.809344 | 0.640724 |
| 2 | 151086000 | 151089399 | 46.06927 | 74.22886 | 82.47677 | 35.08676 | 0.204134 | 1.810168 | 0.640738 |
| 17 | 55697000 | 55698799 | 25.15091 | 43.25159 | 39.40002 | 17.28255 | 0.51821 | 0.919697 | 0.640752 |
| 19 | 8347400 | 8349799 | 20.14331 | 35.30991 | 37.47576 | 16.55733 | 0.30208 | 0.6675 | 0.640753 |
| 6 | 75573200 | 75573799 | 8.210933 | 15.57657 | 26.57424 | 10.7172 | -0.27485 | 0.016119 | 0.640757 |
| 8 | 91111600 | 91116999 | 44.38743 | 71.86935 | 77.25638 | 33.05382 | 0.259526 | 1.676338 | 0.640772 |
| 13 | 118993400 | 118999799 | 70.26121 | 110.6035 | 157.5704 | 65.37587 | -0.18807 | 2.512828 | 0.640794 |
| 3 | 77538200 | 77538999 | 12.22487 | 23.01694 | 25.13198 | 10.29116 | 0.344543 | 0.097633 | 0.640808 |
| 4 | 17597800 | 17599399 | 15.46129 | 28.74087 | 27.49632 | 11.50533 | 0.509346 | 0.211087 | 0.640816 |
| 16 | 44262000 | 44262599 | 15.66292 | 27.79857 | 30.82622 | 13.61522 | 0.244649 | 0.486005 | 0.640817 |
| 4 | 120682200 | 120684399 | 13.29285 | 23.29785 | 39.44791 | 17.42635 | -0.36837 | 0.635298 | 0.640822 |
| 1 | 132345000 | 132345999 | 8.561461 | 15.98067 | 27.92605 | 11.60401 | -0.34537 | 0.12941 | 0.640867 |
| 7 | 72576800 | 72577799 | 8.561461 | 15.98067 | 27.92605 | 11.60401 | -0.34537 | 0.12941 | 0.640867 |
| 15 | 26907600 | 26912999 | 57.13352 | 90.31099 | 149.9951 | 62.31395 | -0.40437 | 2.414186 | 0.640874 |
| 3 | 9897800 | 9901999 | 10.42634 | 18.35429 | 41.99923 | 18.59312 | -0.79934 | 0.560705 | 0.640886 |
| 7 | 27982800 | 27990799 | 62.67102 | 97.96266 | 174.632 | 72.45255 | -0.51893 | 2.643053 | 0.640926 |
| 19 | 50742600 | 50744999 | 19.62504 | 34.42005 | 36.32933 | 16.05816 | 0.309857 | 0.651961 | 0.640942 |
| 12 | 32404600 | 32407199 | 30.51475 | 51.32891 | 52.44244 | 22.91359 | 0.350089 | 1.143511 | 0.640944 |
| 9 | 55853200 | 55859799 | 74.65495 | 116.9268 | 164.9379 | 68.41661 | -0.17945 | 2.612512 | 0.640948 |
| 17 | 76061200 | 76065999 | 60.38197 | 96.50954 | 103.0506 | 43.32232 | 0.247713 | 2.085196 | 0.640953 |
| 18 | 63545600 | 63547199 | 12.82509 | 22.94843 | 36.21985 | 15.88782 | -0.25473 | 0.429677 | 0.64097 |
| 8 | 31542400 | 31550399 | 98.40134 | 152.9345 | 162.907 | 67.63878 | 0.216622 | 2.757273 | 0.640986 |
| 7 | 62488800 | 62490599 | 34.44158 | 56.67833 | 62.96894 | 27.14873 | 0.22741 | 1.405699 | 0.641012 |
| 1 | 141623400 | 141625199 | 23.50987 | 39.43967 | 60.30569 | 26.31608 | -0.22952 | 1.158496 | 0.641016 |
| 13 | 88307600 | 88311399 | 89.41434 | 139.344 | 153.3331 | 63.63794 | 0.172824 | 2.706043 | 0.641016 |
| 11 | 20099600 | 20101599 | 23.82126 | 41.45341 | 41.26649 | 18.13625 | 0.393575 | 0.797346 | 0.641026 |
| 1 | 67647800 | 67648599 | 7.959903 | 15.28062 | 27.49682 | 10.77827 | -0.31347 | -0.07381 | 0.641047 |
| 15 | 12935800 | 12942599 | 64.71739 | 102.1753 | 146.4507 | 60.80428 | -0.19352 | 2.445345 | 0.64105 |
| 2 | 49537600 | 49539399 | 12.70278 | 22.3861 | 36.86533 | 16.31677 | -0.32669 | 0.554416 | 0.641072 |
| 11 | 11986400 | 11988199 | 15.37161 | 26.73715 | 43.20109 | 18.981 | -0.30193 | 0.767594 | 0.641074 |
| 2 | 156885200 | 156888999 | 37.36799 | 60.39343 | 89.35699 | 38.16525 | -0.20303 | 1.711639 | 0.64108 |
| 2 | 20503400 | 20507799 | 57.26522 | 91.02866 | 130.6717 | 54.52585 | -0.18701 | 2.258619 | 0.641081 |
| 6 | 71110600 | 71111399 | 6.103098 | 11.83458 | 22.19295 | 8.380046 | -0.32083 | -0.2354 | 0.641089 |
| 11 | 47877200 | 47880399 | 24.02256 | 39.69343 | 66.20402 | 28.60256 | -0.35532 | 1.32988 | 0.641115 |
| 1 | 95770800 | 95771199 | 10.47691 | 19.83506 | 23.24649 | 9.415375 | 0.257295 | 0.043026 | 0.641126 |
| 4 | 55358600 | 55359799 | 10.66516 | 20.14626 | 23.13459 | 9.414554 | 0.280062 | 0.057668 | 0.641155 |
| 13 | 41396000 | 41397199 | 14.21105 | 25.86024 | 28.9221 | 12.45095 | 0.256903 | 0.323181 | 0.641156 |
| Y | 6050600 | 6051399 | 8.261465 | 16.00314 | 14.71115 | 5.582136 | 0.690635 | -0.22193 | 0.64117 |
| X | 33620800 | 33626399 | 52.58642 | 85.31846 | 73.74256 | 31.59792 | 0.576215 | 1.652239 | 0.641172 |
| 18 | 68873400 | 68874199 | 11.04422 | 20.92889 | 24.26404 | 9.807865 | 0.274984 | 0.043065 | 0.641194 |
| 19 | 35288600 | 35289199 | 7.284674 | 14.10459 | 25.67099 | 9.76159 | -0.28855 | -0.18873 | 0.641225 |
| 1 | 49868200 | 49874399 | 101.9386 | 157.6158 | 171.4642 | 71.22222 | 0.180559 | 2.850935 | 0.641226 |
| 2 | 37145800 | 37149399 | 49.45144 | 79.68284 | 85.37966 | 36.32465 | 0.256522 | 1.816038 | 0.641229 |
| 7 | 85743200 | 85746199 | 48.62235 | 78.25142 | 86.53282 | 36.75024 | 0.208942 | 1.852972 | 0.641242 |
| 13 | 69047400 | 69048199 | 13.78295 | 24.17824 | 38.91182 | 17.20239 | -0.29539 | 0.624186 | 0.641246 |
| 3 | 69838800 | 69840599 | 14.88207 | 26.0855 | 40.40293 | 17.84955 | -0.24051 | 0.645345 | 0.641246 |

| | | | | | | | | | |
|---|---|---|---|---|---|---|---|---|---|
| 4 | 143763800 | 143775599 | 241.6662 | 351.6529 | 363.1559 | 149.6782 | 0.179064 | 3.988897 | 0.641265 |
| 4 | 123609400 | 123612399 | 14.46914 | 25.16546 | 42.7371 | 18.77623 | -0.37326 | 0.764928 | 0.641303 |
| Y | 5156800 | 5157599 | 9.07577 | 17.39415 | 18.70575 | 7.366992 | 0.416149 | -0.07632 | 0.641351 |
| 12 | 76503200 | 76509999 | 55.30953 | 87.89043 | 126.3245 | 52.69421 | -0.18926 | 2.265933 | 0.641351 |
| 17 | 48679400 | 48683399 | 39.80611 | 64.68724 | 71.48029 | 30.66496 | 0.224679 | 1.59014 | 0.641373 |
| 10 | 111236400 | 111246199 | 100.9756 | 155.0575 | 217.521 | 90.3128 | -0.19353 | 2.98435 | 0.641378 |
| 17 | 18530000 | 18533799 | 36.50669 | 60.49769 | 60.59858 | 26.23359 | 0.378395 | 1.326853 | 0.641393 |
| 13 | 112017000 | 112017999 | 14.77471 | 26.97227 | 24.80032 | 10.6338 | 0.541434 | 0.30021 | 0.641398 |
| 6 | 139621400 | 139622399 | 10.27635 | 19.13786 | 31.62416 | 13.19163 | -0.27011 | 0.167554 | 0.641431 |
| X | 7626200 | 7630599 | 50.23636 | 81.28572 | 77.70603 | 33.22997 | 0.427562 | 1.702869 | 0.641434 |
| 7 | 89892400 | 89893799 | 14.23337 | 26.01978 | 28.34516 | 12.13286 | 0.29968 | 0.294348 | 0.641436 |
| 18 | 21625000 | 21627999 | 39.68321 | 64.24094 | 70.8047 | 30.28946 | 0.223291 | 1.66679 | 0.641452 |
| 9 | 8544000 | 8545599 | 14.0498 | 24.61175 | 40.43034 | 17.85251 | -0.32506 | 0.648868 | 0.641455 |
| 7 | 23779000 | 23812799 | 982.1268 | 1256.177 | 1464.178 | 614.3409 | -0.18192 | 6.000612 | 0.641457 |
| 8 | 93925600 | 93926399 | 6.708697 | 13.03321 | 24.69266 | 9.216481 | -0.31984 | -0.25899 | 0.641478 |
| 15 | 96675800 | 96676799 | 13.1694 | 23.28376 | 36.75745 | 16.25849 | -0.2643 | 0.521332 | 0.641494 |
| 6 | 81956000 | 81958199 | 17.87329 | 31.45366 | 34.28566 | 15.17928 | 0.264216 | 0.591691 | 0.641496 |
| 4 | 112830400 | 112848999 | 257.0118 | 373.108 | 373.8456 | 154.1216 | 0.219365 | 4.026292 | 0.641511 |
| 3 | 152268800 | 152269999 | 8.884183 | 16.12415 | 30.94638 | 13.36648 | -0.51958 | 0.305684 | 0.641522 |
| 17 | 40361800 | 40363199 | 23.5864 | 41.03599 | 36.01107 | 15.82442 | 0.574683 | 0.792324 | 0.641525 |
| 1 | 128014800 | 128019799 | 99.17168 | 153.2721 | 167.2911 | 69.48899 | 0.175325 | 2.857681 | 0.641547 |
| 9 | 11613800 | 11615399 | 17.0735 | 30.71174 | 31.09623 | 13.57986 | 0.386096 | 0.412304 | 0.64155 |
| 7 | 25933800 | 25937599 | 70.87626 | 112.5847 | 119.2105 | 49.70802 | 0.250388 | 2.288334 | 0.64161 |
| X | 58973000 | 58976199 | 41.71476 | 67.95769 | 70.3763 | 30.22147 | 0.320728 | 1.561118 | 0.641624 |
| 3 | 64604400 | 64605799 | 11.58951 | 20.87984 | 33.67392 | 14.6791 | -0.27971 | 0.370978 | 0.641662 |
| 4 | 4932600 | 4935599 | 28.38271 | 47.63431 | 52.90843 | 23.09422 | 0.230001 | 1.160477 | 0.641663 |
| 4 | 35660800 | 35663999 | 33.45912 | 55.95499 | 48.64817 | 21.18714 | 0.582323 | 1.210142 | 0.641683 |
| 6 | 79264400 | 79264999 | 7.872295 | 15.09622 | 26.33285 | 10.35389 | -0.27376 | -0.0714 | 0.641713 |
| 19 | 34107800 | 34111799 | 55.50518 | 88.58957 | 96.0696 | 40.30673 | 0.223205 | 2.120117 | 0.641725 |
| 5 | 128310000 | 128311399 | 17.3456 | 30.62013 | 33.83542 | 14.97064 | 0.246487 | 0.555827 | 0.641758 |
| 13 | 20960000 | 20961799 | 27.058 | 46.22666 | 47.36662 | 20.76762 | 0.347579 | 0.992842 | 0.641771 |
| 4 | 112299400 | 112300999 | 21.79747 | 38.5936 | 33.91983 | 14.99906 | 0.576778 | 0.552109 | 0.641773 |
| 17 | 13362400 | 13364599 | 16.71454 | 29.44975 | 32.50937 | 14.38942 | 0.246997 | 0.571201 | 0.641774 |
| 10 | 21979400 | 21980999 | 12.64873 | 23.36184 | 26.50511 | 11.18809 | 0.255945 | 0.221532 | 0.641802 |
| 2 | 177634400 | 177637199 | 19.91 | 34.95973 | 35.68156 | 15.78706 | 0.357942 | 0.633958 | 0.641813 |
| 18 | 56497200 | 56500799 | 27.22196 | 45.96519 | 50.16552 | 21.94412 | 0.255409 | 1.097855 | 0.641818 |
| X | 64755000 | 64758399 | 25.53323 | 43.59669 | 47.39409 | 20.7787 | 0.262503 | 0.993698 | 0.641822 |
| 12 | 114844800 | 114849399 | 69.76435 | 111.3174 | 107.4397 | 45.05957 | 0.390584 | 2.146497 | 0.641825 |
| 2 | 40284400 | 40288799 | 74.01269 | 117.0689 | 126.9366 | 52.75208 | 0.210717 | 2.406562 | 0.641828 |
| 18 | 11508000 | 11512599 | 45.9948 | 74.5136 | 77.12798 | 33.01251 | 0.314664 | 1.667378 | 0.641829 |
| 18 | 37182600 | 37193999 | 148.7474 | 226.7864 | 209.8908 | 87.0182 | 0.397969 | 3.141651 | 0.641835 |
| 15 | 22261200 | 22262399 | 12.57971 | 23.45438 | 26.04298 | 10.84239 | 0.301509 | 0.164618 | 0.641836 |
| 19 | 39571400 | 39591399 | 294.0014 | 422.9295 | 412.228 | 170.149 | 0.246223 | 4.163214 | 0.641865 |
| 16 | 78027400 | 78029399 | 24.0242 | 41.26013 | 45.22515 | 19.83692 | 0.251912 | 0.932886 | 0.641867 |
| 6 | 68763000 | 68765799 | 28.73392 | 49.31227 | 46.23165 | 20.27738 | 0.476344 | 0.960219 | 0.641923 |
| 7 | 25827000 | 25828399 | 44.50942 | 71.69059 | 78.80292 | 33.50694 | 0.219065 | 1.818701 | 0.641923 |
| 3 | 17536600 | 17537199 | 7.318147 | 14.13667 | 25.44649 | 9.781355 | -0.2871 | -0.15025 | 0.641923 |

| | | | | | | | | | |
|---|---|---|---|---|---|---|---|---|---|
| 7 | 106378200 | 106378999 | 13.37049 | 24.24478 | 26.51402 | 11.46486 | 0.285067 | 0.331975 | 0.641939 |
| 4 | 156462600 | 156464799 | 22.72583 | 39.43459 | 36.72173 | 16.11216 | 0.489084 | 0.824641 | 0.641941 |
| 7 | 118845800 | 118846199 | 8.088928 | 15.71584 | 19.44864 | 7.25463 | 0.288692 | -0.26518 | 0.641941 |
| 14 | 54953600 | 54954199 | 8.088928 | 15.71584 | 19.44864 | 7.25463 | 0.288692 | -0.26518 | 0.641941 |
| 16 | 58878600 | 58879599 | 9.632059 | 18.01105 | 29.52808 | 12.23277 | -0.25064 | 0.131475 | 0.641957 |
| 7 | 85682200 | 85682799 | 10.35281 | 19.62843 | 20.30526 | 8.199021 | 0.440396 | 0.025461 | 0.641958 |
| 1 | 177571200 | 177579999 | 134.0977 | 204.3684 | 190.2002 | 78.84707 | 0.389455 | 3.142581 | 0.641996 |
| 19 | 7823000 | 7825999 | 57.71865 | 92.48761 | 97.63592 | 41.19799 | 0.268248 | 2.0031 | 0.642003 |
| 1 | 171417600 | 171419199 | 29.64527 | 48.02737 | 88.03343 | 37.6807 | -0.50769 | 1.645265 | 0.642034 |
| 10 | 24378200 | 24381399 | 30.24325 | 52.07825 | 44.53594 | 19.53619 | 0.609472 | 0.936591 | 0.642038 |
| 9 | 93755200 | 93755599 | 7.104693 | 13.83818 | 17.40273 | 6.353363 | 0.295118 | -0.35651 | 0.642054 |
| 8 | 89116400 | 89122599 | 49.24084 | 78.55653 | 114.2092 | 47.88743 | -0.1997 | 2.133106 | 0.642069 |
| 5 | 131712000 | 131712599 | 9.556611 | 18.53083 | 21.65973 | 8.179705 | 0.354046 | -0.19802 | 0.642092 |
| 8 | 110879000 | 110883999 | 42.78143 | 68.68434 | 100.5014 | 42.53128 | -0.19858 | 1.925497 | 0.64211 |
| 3 | 93887400 | 93888399 | 10.16914 | 18.7366 | 30.36716 | 12.86069 | -0.25821 | 0.226593 | 0.64211 |
| 9 | 77443800 | 77446999 | 35.76274 | 58.36425 | 65.43703 | 28.11852 | 0.208208 | 1.517976 | 0.642137 |
| 13 | 18123200 | 18125799 | 29.16557 | 47.22584 | 88.49098 | 37.86214 | -0.54007 | 1.654459 | 0.642166 |
| X | 46152400 | 46155199 | 50.28212 | 80.88935 | 86.18379 | 36.58022 | 0.261804 | 1.870605 | 0.642182 |
| 6 | 132068600 | 132073999 | 53.02942 | 85.87279 | 76.60518 | 32.77827 | 0.52817 | 1.691115 | 0.642243 |
| 7 | 34118400 | 34120799 | 27.96268 | 47.05824 | 51.81761 | 22.64382 | 0.242482 | 1.131537 | 0.64226 |
| 16 | 33834000 | 33842399 | 165.6698 | 244.8133 | 334.8642 | 137.9046 | -0.205 | 3.73872 | 0.642263 |
| 1 | 120245600 | 120252399 | 57.85874 | 91.45332 | 136.6626 | 56.77382 | -0.25219 | 2.411724 | 0.642287 |
| 8 | 23310400 | 23315599 | 65.23011 | 103.5918 | 115.2887 | 48.06151 | 0.17833 | 2.289134 | 0.642293 |
| 14 | 82502200 | 82505799 | 48.02133 | 76.87227 | 112.178 | 47.26299 | -0.19972 | 2.025911 | 0.642305 |
| 8 | 72690400 | 72691999 | 23.54789 | 40.60663 | 44.13798 | 19.36245 | 0.265127 | 0.893909 | 0.642326 |
| 14 | 118548400 | 118550799 | 20.65938 | 35.95875 | 39.81547 | 17.50086 | 0.240754 | 0.777092 | 0.642343 |
| 7 | 41026600 | 41032799 | 78.98983 | 125.6137 | 115.7432 | 48.31649 | 0.452241 | 2.263819 | 0.642347 |
| 18 | 65529000 | 65536399 | 74.75113 | 116.8875 | 164.1567 | 68.10395 | -0.17491 | 2.6419 | 0.642356 |
| 10 | 95794800 | 95798599 | 43.25739 | 70.78478 | 68.24729 | 29.35475 | 0.428034 | 1.498922 | 0.642407 |
| 5 | 148871200 | 148874399 | 42.11846 | 67.68833 | 99.04652 | 41.97854 | -0.19685 | 1.892468 | 0.642427 |
| 9 | 35585800 | 35587799 | 19.96389 | 34.39913 | 55.57553 | 24.37929 | -0.30466 | 0.900173 | 0.642433 |
| 8 | 80499400 | 80501399 | 22.53224 | 39.23626 | 40.83865 | 17.95569 | 0.329806 | 0.783885 | 0.642459 |
| 10 | 63498800 | 63499599 | 9.274076 | 17.11932 | 29.41631 | 12.42644 | -0.33887 | 0.208401 | 0.642464 |
| 18 | 85780600 | 85781999 | 31.58843 | 52.01727 | 56.10396 | 24.19566 | 0.270427 | 1.385266 | 0.642488 |
| 1 | 48187200 | 48191199 | 43.90418 | 71.00262 | 77.29614 | 33.04127 | 0.239686 | 1.696948 | 0.642489 |
| 6 | 146092800 | 146096399 | 37.73991 | 61.07962 | 93.96861 | 40.19124 | -0.25713 | 1.681372 | 0.642558 |
| 2 | 88001600 | 88004799 | 45.56288 | 73.39124 | 81.21274 | 34.53422 | 0.209553 | 1.820685 | 0.642566 |
| 9 | 9088600 | 9089799 | 11.03827 | 20.39616 | 35.30602 | 14.89432 | -0.34875 | 0.228835 | 0.642567 |
| 5 | 57806400 | 57807399 | 25.15502 | 42.53283 | 47.46619 | 20.77189 | 0.223492 | 1.077446 | 0.64257 |
| 7 | 121814000 | 121819999 | 48.34014 | 77.2668 | 112.6253 | 47.35088 | -0.20044 | 2.071756 | 0.642575 |
| 14 | 89858400 | 89858999 | 9.373798 | 17.53655 | 29.00906 | 12.00799 | -0.26219 | 0.126784 | 0.642586 |
| 14 | 27282400 | 27293599 | 143.7282 | 217.78 | 229.0936 | 94.80486 | 0.206043 | 3.265207 | 0.642591 |
| 14 | 6508600 | 6509199 | 13.95596 | 25.24546 | 25.19747 | 10.92848 | 0.413638 | 0.349201 | 0.642615 |
| 7 | 41952400 | 41965799 | 204.5318 | 304.0571 | 293.7302 | 121.0121 | 0.304479 | 3.641673 | 0.642656 |
| X | 76384400 | 76387599 | 41.49709 | 67.53767 | 71.50679 | 30.6961 | 0.287879 | 1.574146 | 0.642668 |
| X | 116125200 | 116127399 | 30.53848 | 51.64167 | 44.21676 | 19.35061 | 0.604471 | 1.0943 | 0.64267 |
| 12 | 106293800 | 106310799 | 270.672 | 390.1854 | 398.584 | 164.4689 | 0.181537 | 4.130385 | 0.642738 |

| | | | | | | | | | |
|---|---|---|---|---|---|---|---|---|---|
| 14 | 53710200 | 53714599 | 51.51524 | 82.81691 | 78.90193 | 33.45657 | 0.421567 | 1.887712 | 0.642769 |
| 8 | 87044800 | 87045199 | 5.070485 | 9.896489 | 19.27158 | 6.836045 | -0.29086 | -0.47769 | 0.642783 |
| 13 | 98463800 | 98465799 | 20.23617 | 35.06095 | 36.55059 | 16.03632 | 0.326668 | 0.82805 | 0.642792 |
| 19 | 33348600 | 33351199 | 23.9915 | 41.2813 | 38.3477 | 16.82125 | 0.490165 | 0.906396 | 0.642802 |
| 3 | 34005800 | 34008399 | 19.20221 | 32.6897 | 60.8993 | 26.69334 | -0.51238 | 1.011454 | 0.642818 |
| 7 | 29455200 | 29460599 | 63.45522 | 100.9525 | 112.1592 | 46.84739 | 0.183344 | 2.234639 | 0.642845 |
| 17 | 49852200 | 49854199 | 18.95265 | 33.07449 | 36.79172 | 16.19838 | 0.234596 | 0.729117 | 0.642849 |
| 2 | 97730600 | 97732199 | 25.2351 | 42.75271 | 47.98273 | 21.01054 | 0.215489 | 1.062598 | 0.642888 |
| 12 | 46345600 | 46346399 | 7.437365 | 14.19563 | 25.61039 | 10.19575 | -0.33845 | -0.03733 | 0.642894 |
| 1 | 84357000 | 84359999 | 19.21932 | 32.51139 | 61.93201 | 27.10609 | -0.54517 | 1.067576 | 0.642902 |
| 13 | 94344400 | 94345599 | 17.33166 | 30.14915 | 46.30469 | 20.34576 | -0.22918 | 0.780952 | 0.642907 |
| X | 42656000 | 42659799 | 57.01281 | 91.41695 | 95.54476 | 40.35612 | 0.283798 | 1.982836 | 0.642922 |
| 6 | 132594000 | 132597999 | 41.44778 | 68.13527 | 65.347 | 28.1613 | 0.438358 | 1.437467 | 0.642925 |
| 1 | 170743400 | 170744999 | 15.50633 | 27.61783 | 30.81919 | 13.57173 | 0.238545 | 0.46317 | 0.642926 |
| 7 | 64913400 | 64915199 | 27.18954 | 45.60106 | 51.26251 | 22.36626 | 0.212734 | 1.161625 | 0.642929 |
| 3 | 74844800 | 74845799 | 17.41029 | 30.67467 | 33.94876 | 15.02662 | 0.24328 | 0.578578 | 0.642932 |
| 6 | 3157200 | 3160399 | 21.90669 | 38.14809 | 40.22753 | 17.68732 | 0.310859 | 0.776104 | 0.642952 |
| 7 | 27136400 | 27142199 | 60.74477 | 96.22713 | 137.1511 | 57.0404 | -0.18151 | 2.361762 | 0.642976 |
| 15 | 62965400 | 62968599 | 52.38369 | 83.88289 | 93.55936 | 39.43778 | 0.187907 | 2.015436 | 0.642978 |
| X | 32178200 | 32180399 | 19.66319 | 36.10203 | 28.1738 | 11.98907 | 0.784434 | 0.31409 | 0.642995 |
| 4 | 52085800 | 52088599 | 44.13016 | 71.41255 | 62.07972 | 26.54952 | 0.564401 | 1.679525 | 0.643 |
| 1 | 54426000 | 54426999 | 8.980632 | 16.82011 | 28.47954 | 11.76613 | -0.29373 | 0.111817 | 0.643006 |
| 11 | 88823200 | 88825799 | 16.71933 | 28.82533 | 46.38628 | 20.34861 | -0.29843 | 0.870809 | 0.643006 |
| 1 | 28231200 | 28233599 | 23.3366 | 40.04145 | 44.71333 | 19.61112 | 0.22496 | 0.93752 | 0.643037 |
| 13 | 65463200 | 65500799 | 784.929 | 1037.306 | 936.2867 | 391.5282 | 0.234043 | 5.470371 | 0.643045 |
| 1 | 120982600 | 120983999 | 12.9209 | 22.9297 | 36.13128 | 15.95971 | -0.25951 | 0.479594 | 0.643047 |
| 1 | 85286800 | 85289199 | 33.84301 | 56.66919 | 47.53413 | 20.71861 | 0.633772 | 1.195773 | 0.643052 |
| 7 | 81860200 | 81863999 | 26.05161 | 42.7902 | 68.14126 | 29.35517 | -0.29154 | 1.412215 | 0.643085 |
| 3 | 128992200 | 128994399 | 20.01845 | 33.41523 | 64.91597 | 28.23878 | -0.57363 | 1.20452 | 0.643108 |
| 15 | 80356000 | 80357199 | 9.196809 | 17.25232 | 28.5185 | 11.75066 | -0.25585 | 0.10621 | 0.64311 |
| 5 | 65051400 | 65054799 | 50.21448 | 80.04016 | 116.4381 | 48.76326 | -0.20195 | 2.162278 | 0.64312 |
| 7 | 11744200 | 11759599 | 1746.152 | 2118.239 | 2486.477 | 1049.628 | -0.26946 | 6.849863 | 0.643131 |
| 7 | 13920000 | 13932199 | 166.6489 | 250.572 | 254.9988 | 105.2831 | 0.244223 | 3.424577 | 0.643135 |
| 14 | 37609200 | 37610599 | 14.58881 | 27.04072 | 26.8497 | 11.27906 | 0.451675 | 0.216583 | 0.643157 |
| 9 | 86479000 | 86479999 | 11.73689 | 21.04895 | 34.28216 | 15.00914 | -0.29755 | 0.405775 | 0.643162 |
| 18 | 10055600 | 10064199 | 135.3445 | 207.0295 | 202.0426 | 83.82796 | 0.325224 | 3.072074 | 0.643175 |
| 10 | 74710800 | 74712199 | 21.97605 | 37.77528 | 42.10767 | 18.47 | 0.227991 | 0.912029 | 0.643184 |
| X | 89211400 | 89214199 | 37.42882 | 61.83834 | 61.53972 | 26.5854 | 0.387529 | 1.367571 | 0.643186 |
| 19 | 14059600 | 14062799 | 38.5133 | 62.83082 | 69.29166 | 29.77102 | 0.231558 | 1.530206 | 0.643226 |
| 1 | 66920000 | 66922199 | 19.63921 | 34.4016 | 36.9358 | 16.3089 | 0.285374 | 0.67315 | 0.643232 |
| 5 | 8534400 | 8536199 | 24.26697 | 41.73448 | 38.91522 | 17.06993 | 0.484685 | 0.914063 | 0.643257 |
| 1 | 158189800 | 158190799 | 7.767805 | 14.81585 | 25.59743 | 10.20585 | -0.27802 | -0.02687 | 0.643267 |
| 16 | 21740000 | 21740799 | 7.767805 | 14.81585 | 25.59743 | 10.20585 | -0.27802 | -0.02687 | 0.643267 |
| 9 | 91618600 | 91623399 | 77.36285 | 122.862 | 105.2355 | 43.87136 | 0.555553 | 2.297818 | 0.643295 |
| 12 | 114110200 | 114115599 | 124.2498 | 190.6753 | 173.3864 | 71.98218 | 0.430682 | 3.001249 | 0.643295 |
| 3 | 32825800 | 32828599 | 37.1985 | 60.05251 | 88.972 | 37.9462 | -0.20681 | 1.743174 | 0.643322 |
| 12 | 47903600 | 47905399 | 22.21984 | 38.84101 | 38.42799 | 16.93791 | 0.402839 | 0.721488 | 0.643329 |

| | | | | | | | | | |
|---|---|---|---|---|---|---|---|---|---|
| 6 | 66226800 | 66229999 | 38.76946 | 63.03232 | 70.58455 | 30.28864 | 0.20613 | 1.581607 | 0.643333 |
| 5 | 114861000 | 114863999 | 33.365 | 53.86451 | 84.94135 | 36.22778 | -0.29648 | 1.736315 | 0.643334 |
| X | 90388800 | 90391199 | 29.49061 | 49.78006 | 51.80568 | 22.6594 | 0.3239 | 1.113768 | 0.643334 |
| 12 | 9305200 | 9306599 | 11.10539 | 20.40431 | 32.9447 | 14.00565 | -0.25708 | 0.254425 | 0.643335 |
| 8 | 37027400 | 37029199 | 12.93255 | 22.91887 | 36.14639 | 15.98069 | -0.2618 | 0.490476 | 0.643336 |
| 1 | 49003000 | 49003799 | 7.413232 | 14.27864 | 25.34847 | 9.841622 | -0.28169 | -0.12005 | 0.643379 |
| 2 | 83771000 | 83771799 | 7.413232 | 14.27864 | 25.34847 | 9.841622 | -0.28169 | -0.12005 | 0.643379 |
| 1 | 71681200 | 71684999 | 31.11296 | 50.66795 | 76.7006 | 32.93296 | -0.226 | 1.544769 | 0.643399 |
| 2 | 116756800 | 116758199 | 19.91571 | 34.89596 | 37.48293 | 16.55449 | 0.284834 | 0.672315 | 0.643416 |
| 4 | 134638000 | 134640999 | 24.00763 | 40.86564 | 45.83518 | 20.09025 | 0.216942 | 1.013378 | 0.643425 |
| X | 133375200 | 133378199 | 76.7671 | 121.1314 | 129.1641 | 53.61191 | 0.232025 | 2.466768 | 0.643434 |
| 7 | 62448000 | 62450999 | 53.39114 | 85.86566 | 89.79234 | 38.09563 | 0.288141 | 1.882521 | 0.643437 |
| 16 | 82972000 | 82972599 | 7.847514 | 15.23859 | 18.91323 | 7.085695 | 0.279563 | -0.2521 | 0.643465 |
| 2 | 162648800 | 162650999 | 15.56619 | 27.35729 | 43.41331 | 19.22266 | -0.27548 | 0.611444 | 0.643473 |
| 2 | 70314600 | 70319199 | 42.77367 | 68.731 | 100.3651 | 42.52816 | -0.19416 | 1.898568 | 0.64349 |
| 13 | 50538200 | 50542399 | 83.5319 | 130.5818 | 143.1382 | 59.38563 | 0.18186 | 2.647486 | 0.643495 |
| 15 | 27387600 | 27391399 | 31.28088 | 51.00504 | 76.21942 | 32.74125 | -0.20612 | 1.527609 | 0.643499 |
| 5 | 105137000 | 105138599 | 25.75582 | 43.317 | 46.45608 | 20.297 | 0.280389 | 1.123486 | 0.643501 |
| 10 | 59579600 | 59581199 | 13.39812 | 23.72967 | 37.13356 | 16.4191 | -0.251 | 0.501489 | 0.643503 |
| 6 | 14623800 | 14624399 | 7.456504 | 14.38491 | 25.38493 | 9.810743 | -0.26638 | -0.13159 | 0.643512 |
| 2 | 107907400 | 107908799 | 15.96042 | 27.9487 | 43.50065 | 19.20245 | -0.24773 | 0.670083 | 0.643549 |
| 11 | 97917800 | 97921399 | 50.4394 | 80.411 | 116.8362 | 48.94035 | -0.19996 | 2.158927 | 0.643558 |
| 4 | 96930600 | 96933199 | 19.30917 | 33.77166 | 37.0541 | 16.33995 | 0.254302 | 0.696085 | 0.643569 |
| 15 | 62999200 | 63001199 | 19.25688 | 32.55786 | 59.65586 | 26.10573 | -0.48919 | 1.071896 | 0.643607 |
| 7 | 13498800 | 13507199 | 115.1869 | 178.1299 | 154.3535 | 64.1148 | 0.5086 | 2.852554 | 0.643609 |
| 19 | 13364200 | 13367199 | 24.51321 | 42.51693 | 42.97248 | 18.8545 | 0.371273 | 0.845352 | 0.643624 |
| 12 | 74184000 | 74185799 | 18.45114 | 32.92598 | 32.54624 | 14.30904 | 0.414074 | 0.473435 | 0.643627 |
| 11 | 119021400 | 119022399 | 13.00108 | 23.09994 | 36.20179 | 15.97454 | -0.25044 | 0.475243 | 0.643629 |
| 1 | 143737800 | 143738599 | 17.55272 | 30.15111 | 47.95658 | 21.03511 | -0.28278 | 0.910284 | 0.643633 |
| 8 | 61189400 | 61192599 | 43.70623 | 70.58422 | 78.92626 | 33.67759 | 0.198877 | 1.739502 | 0.643672 |
| 3 | 47978200 | 47984399 | 65.738 | 104.7451 | 111.1101 | 46.50795 | 0.252363 | 2.187281 | 0.643683 |
| 3 | 74998600 | 74999799 | 17.76342 | 31.55361 | 33.21719 | 14.66065 | 0.320065 | 0.500119 | 0.64369 |
| Y | 1883800 | 1891199 | 112.7687 | 172.623 | 144.8457 | 60.10701 | 0.543734 | 3.041466 | 0.643693 |
| 4 | 23421200 | 23424599 | 59.438 | 94.6882 | 105.2899 | 44.05977 | 0.184172 | 2.185745 | 0.643702 |
| 5 | 107859200 | 107859799 | 11.41631 | 21.33529 | 24.64215 | 10.21849 | 0.250038 | 0.13845 | 0.643708 |
| 10 | 74638600 | 74639399 | 12.56133 | 23.30909 | 26.59038 | 11.15294 | 0.254834 | 0.196156 | 0.643736 |
| 8 | 108601400 | 108603399 | 18.33752 | 32.43282 | 34.19771 | 15.12518 | 0.31476 | 0.54188 | 0.643741 |
| 8 | 49550600 | 49552799 | 35.49007 | 58.33062 | 52.55876 | 22.64842 | 0.52794 | 1.413317 | 0.643784 |
| 12 | 115499600 | 115512199 | 195.4469 | 292.0264 | 276.9577 | 114.186 | 0.337716 | 3.549048 | 0.643807 |
| 13 | 61673800 | 61702399 | 418.7047 | 586.0153 | 529.5301 | 219.5396 | 0.315704 | 4.570762 | 0.64382 |
| 11 | 63120200 | 63122399 | 20.6205 | 35.04379 | 53.2576 | 23.34059 | -0.21998 | 1.02504 | 0.64382 |
| 19 | 11067400 | 11069999 | 18.41628 | 31.80369 | 48.47206 | 21.26467 | -0.22009 | 0.864298 | 0.643822 |
| 2 | 60776000 | 60778799 | 20.89472 | 35.83071 | 58.93853 | 25.84903 | -0.33195 | 0.949058 | 0.643845 |
| 6 | 32601600 | 32602599 | 21.27736 | 36.69976 | 40.09811 | 17.59037 | 0.257894 | 0.878051 | 0.643847 |
| 17 | 64996400 | 64998399 | 32.41363 | 54.81712 | 51.30212 | 22.45195 | 0.476587 | 1.106346 | 0.643855 |
| 7 | 26432400 | 26433599 | 24.84961 | 42.71633 | 38.43682 | 16.85984 | 0.53581 | 0.921341 | 0.643865 |
| 4 | 24282000 | 24286399 | 17.76539 | 31.56945 | 28.60569 | 12.62116 | 0.535489 | 0.488125 | 0.64392 |

| | | | | | | | | | |
|---|---|---|---|---|---|---|---|---|---|
| 6 | 29309200 | 29311399 | 20.8691 | 35.48559 | 54.38112 | 23.83403 | -0.23198 | 1.020216 | 0.643926 |
| 7 | 36701800 | 36705399 | 20.8691 | 35.48559 | 54.38112 | 23.83403 | -0.23198 | 1.020216 | 0.643926 |
| 6 | 70219800 | 70222199 | 17.38521 | 30.52287 | 33.45978 | 14.80257 | 0.255533 | 0.619204 | 0.643953 |
| 7 | 34298800 | 34302199 | 25.44686 | 42.20263 | 64.16016 | 27.7904 | -0.2216 | 1.291047 | 0.643972 |
| 4 | 19708600 | 19709599 | 8.421042 | 15.8964 | 27.02558 | 11.01376 | -0.28278 | 0.055612 | 0.644008 |
| 4 | 8171000 | 8173199 | 15.99016 | 28.01533 | 43.4912 | 19.20671 | -0.24397 | 0.662545 | 0.644038 |
| 4 | 43299000 | 43302799 | 46.14414 | 74.3113 | 83.27813 | 35.40129 | 0.190918 | 1.827787 | 0.644039 |
| 18 | 31493800 | 31495999 | 12.75769 | 23.77749 | 21.45888 | 8.939473 | 0.598136 | 0.158971 | 0.64404 |
| 6 | 144877800 | 144880999 | 33.22403 | 54.0395 | 81.05389 | 34.78657 | -0.21404 | 1.568852 | 0.644041 |
| 16 | 16750000 | 16753799 | 50.82874 | 81.59888 | 90.47075 | 38.28269 | 0.200773 | 1.933304 | 0.644154 |
| 10 | 16821200 | 16822999 | 18.41088 | 32.78737 | 33.14694 | 14.59817 | 0.380226 | 0.488094 | 0.64417 |
| 4 | 49522600 | 49523199 | 6.382411 | 12.38702 | 22.81954 | 8.581524 | -0.2903 | -0.24528 | 0.644182 |
| 3 | 134433600 | 134436999 | 28.82188 | 48.53554 | 52.37851 | 22.89341 | 0.271491 | 1.12993 | 0.644202 |
| 17 | 22415000 | 22417799 | 61.62288 | 98.2359 | 108.2311 | 45.33332 | 0.198518 | 2.167303 | 0.644204 |
| Y | 90515600 | 90516199 | 5.132318 | 9.94917 | 21.62816 | 8.178134 | -0.53323 | -0.24389 | 0.644229 |
| 17 | 68132200 | 68145399 | 271.4364 | 385.9526 | 499.5077 | 206.5615 | -0.17949 | 4.331675 | 0.644236 |
| 14 | 11970400 | 11972599 | 17.03993 | 30.30755 | 32.72756 | 14.42883 | 0.284565 | 0.485893 | 0.644256 |
| 2 | 87393800 | 87394399 | 9.202738 | 17.67832 | 21.3523 | 8.353659 | 0.259246 | -0.08665 | 0.644288 |
| 6 | 130597400 | 130602799 | 40.99815 | 66.1686 | 112.8277 | 48.10372 | -0.40976 | 1.766741 | 0.644311 |
| 1 | 115067800 | 115069999 | 24.10939 | 40.45228 | 61.51675 | 26.84743 | -0.22174 | 1.15827 | 0.644313 |
| 4 | 113110600 | 113112199 | 37.23682 | 60.63647 | 68.31102 | 29.32997 | 0.199073 | 1.553786 | 0.644318 |
| 2 | 51019000 | 51023799 | 42.55589 | 68.89597 | 76.36813 | 32.67104 | 0.215201 | 1.674642 | 0.644323 |
| 3 | 23652600 | 23653799 | 8.823561 | 16.84835 | 30.29135 | 12.04883 | -0.33283 | -0.01437 | 0.644338 |
| 6 | 133310200 | 133317399 | 46.86594 | 75.79876 | 67.28537 | 28.76437 | 0.533501 | 1.694043 | 0.644345 |
| 14 | 66355000 | 66359999 | 51.75797 | 83.03397 | 92.4576 | 39.08286 | 0.193456 | 1.957702 | 0.644345 |
| 10 | 97438200 | 97439599 | 19.08234 | 33.29659 | 37.26642 | 16.40607 | 0.225728 | 0.731655 | 0.644351 |
| 3 | 22888600 | 22889999 | 22.9397 | 39.54541 | 43.64317 | 19.14521 | 0.243233 | 0.894283 | 0.644352 |
| 2 | 94506200 | 94507999 | 15.77646 | 27.67367 | 44.50851 | 19.67565 | -0.29492 | 0.64232 | 0.644381 |
| 1 | 19706600 | 19709599 | 25.22967 | 43.06829 | 46.84582 | 20.53794 | 0.261573 | 0.992174 | 0.644388 |
| 7 | 43668800 | 43670999 | 24.87474 | 42.1954 | 47.44184 | 20.78165 | 0.212971 | 1.049468 | 0.644392 |
| 12 | 44198600 | 44203599 | 93.30828 | 145.0713 | 159.2797 | 66.12913 | 0.173413 | 2.748845 | 0.644392 |
| 17 | 27068400 | 27071799 | 28.48675 | 46.62247 | 71.62586 | 30.80983 | -0.2421 | 1.464747 | 0.644405 |
| X | 104076200 | 104077599 | 15.89403 | 27.8322 | 43.00783 | 18.98477 | -0.23719 | 0.671486 | 0.644406 |
| 9 | 99375800 | 99376799 | 10.92219 | 19.65977 | 32.81766 | 14.32206 | -0.32956 | 0.373879 | 0.644424 |
| 15 | 49857200 | 49857999 | 8.706881 | 16.3551 | 27.47487 | 11.29506 | -0.27631 | 0.089976 | 0.644448 |
| 5 | 107732600 | 107735399 | 29.75996 | 48.96292 | 74.53567 | 32.13256 | -0.2259 | 1.404666 | 0.644459 |
| 1 | 191861800 | 191864999 | 31.08049 | 51.99724 | 48.63551 | 21.18679 | 0.477175 | 1.197793 | 0.644459 |
| 10 | 54402600 | 54411799 | 229.9474 | 336.1343 | 347.9259 | 143.3359 | 0.182135 | 3.918794 | 0.644476 |
| 7 | 71772000 | 71774999 | 51.38882 | 81.85125 | 118.552 | 49.59757 | -0.19679 | 2.188261 | 0.644498 |
| 3 | 12006800 | 12011199 | 72.40229 | 114.338 | 127.066 | 52.76297 | 0.17339 | 2.441558 | 0.644501 |
| 12 | 78059800 | 78061399 | 11.81132 | 21.59271 | 36.09209 | 15.44828 | -0.31386 | 0.295189 | 0.644508 |
| 5 | 44101200 | 44101999 | 6.968904 | 13.45522 | 24.2812 | 9.355517 | -0.29455 | -0.15271 | 0.644525 |
| 6 | 147027400 | 147037799 | 127.2982 | 192.5234 | 262.877 | 108.721 | -0.17227 | 3.301279 | 0.644529 |
| 7 | 82038000 | 82040999 | 28.22149 | 46.53982 | 70.53697 | 30.43864 | -0.21824 | 1.36824 | 0.64455 |
| 12 | 58176800 | 58179399 | 26.80227 | 46.59718 | 40.60554 | 17.83525 | 0.584698 | 0.823292 | 0.644553 |
| X | 109286800 | 109288799 | 14.99957 | 26.5218 | 46.14509 | 20.41047 | -0.40428 | 0.546889 | 0.644559 |
| 8 | 21585000 | 21591399 | 95.26557 | 148.4055 | 158.8644 | 65.9374 | 0.212118 | 2.716968 | 0.644579 |

| | | | | | | | | | |
|---|---|---|---|---|---|---|---|---|---|
| 7 | 27472200 | 27489799 | 290.6517 | 417.1816 | 418.7406 | 172.8973 | 0.200678 | 4.195313 | 0.644584 |
| 3 | 36102400 | 36103199 | 10.89476 | 20.39707 | 22.65393 | 9.365596 | 0.309674 | 0.117219 | 0.644596 |
| 8 | 116017600 | 116022399 | 40.36265 | 65.06977 | 96.20247 | 40.95619 | -0.20613 | 1.793016 | 0.644609 |
| 17 | 48904400 | 48906199 | 13.807 | 24.21895 | 38.936 | 17.2122 | -0.2938 | 0.626237 | 0.644625 |
| 1 | 75972600 | 75975199 | 15.21235 | 27.46374 | 29.70503 | 12.91543 | 0.295431 | 0.375159 | 0.644634 |
| X | 127013600 | 127015599 | 25.48913 | 43.78465 | 45.75261 | 20.06839 | 0.320722 | 0.9395 | 0.644638 |
| 6 | 122080000 | 122083199 | 46.07231 | 73.80103 | 106.8563 | 45.06571 | -0.18729 | 2.001995 | 0.644646 |
| 7 | 41971600 | 41975199 | 57.51928 | 92.01618 | 98.60188 | 41.50452 | 0.244026 | 2.055263 | 0.644652 |
| 4 | 57815200 | 57820199 | 49.58823 | 79.77153 | 87.84669 | 37.28497 | 0.214174 | 1.867865 | 0.644656 |
| 16 | 51147400 | 51147799 | 5.806027 | 11.29606 | 21.26449 | 7.843755 | -0.29481 | -0.32423 | 0.644662 |
| 9 | 36395200 | 36398599 | 36.56569 | 60.04514 | 65.40951 | 28.17543 | 0.254487 | 1.437984 | 0.644666 |
| 18 | 38423800 | 38425399 | 14.52482 | 26.35798 | 29.38274 | 12.69322 | 0.258327 | 0.341979 | 0.644674 |
| 4 | 61305000 | 61307199 | 27.18251 | 46.01488 | 39.14534 | 17.13666 | 0.613732 | 1.0661 | 0.644679 |
| 10 | 120037200 | 120041199 | 26.75048 | 44.16595 | 67.55939 | 29.1722 | -0.23103 | 1.348119 | 0.644722 |
| 2 | 51833000 | 51838199 | 147.5138 | 219.5192 | 308.8436 | 127.2518 | -0.23621 | 3.616001 | 0.644755 |
| 2 | 21894200 | 21899999 | 67.4351 | 107.8325 | 102.529 | 43.13126 | 0.415537 | 2.079726 | 0.644768 |
| 12 | 114191000 | 114196999 | 105.3589 | 163.0073 | 172.67 | 71.72286 | 0.219629 | 2.840614 | 0.644779 |
| 10 | 25235200 | 25236599 | 14.69357 | 25.74424 | 40.40927 | 17.846 | -0.25969 | 0.650215 | 0.64478 |
| 3 | 53304400 | 53306199 | 21.38433 | 36.96752 | 41.55022 | 18.22864 | 0.21778 | 0.859093 | 0.644795 |
| 5 | 104708200 | 104710999 | 24.57236 | 41.79339 | 46.89675 | 20.55423 | 0.216162 | 1.024962 | 0.644801 |
| 11 | 48257600 | 48260399 | 34.13548 | 55.46445 | 83.11411 | 35.65221 | -0.21385 | 1.591317 | 0.644801 |
| X | 15464400 | 15466199 | 19.71033 | 34.92271 | 34.05079 | 15.05486 | 0.428095 | 0.531516 | 0.644803 |
| 3 | 135823400 | 135824599 | 24.95983 | 41.1279 | 74.92008 | 32.31906 | -0.48312 | 1.376622 | 0.644829 |
| 5 | 40316000 | 40317399 | 13.1047 | 23.28531 | 36.14814 | 15.95014 | -0.23696 | 0.473851 | 0.64487 |
| 19 | 39572000 | 39580999 | 123.2477 | 189.2346 | 192.3695 | 79.86876 | 0.270565 | 2.99296 | 0.644873 |
| 10 | 70133400 | 70134799 | 12.09986 | 21.73524 | 34.16809 | 14.93868 | -0.2458 | 0.394472 | 0.644888 |
| 4 | 112277600 | 112284799 | 105.8079 | 164.6049 | 162.7718 | 67.57285 | 0.324896 | 2.746321 | 0.644908 |
| 4 | 120891600 | 120893199 | 13.18103 | 23.36914 | 36.70834 | 16.22758 | -0.25555 | 0.491317 | 0.644914 |
| 4 | 68344200 | 68344999 | 13.82688 | 24.26665 | 38.92858 | 17.21661 | -0.29065 | 0.618469 | 0.644925 |
| 1 | 116789000 | 116790599 | 19.89204 | 34.82642 | 28.58181 | 12.61469 | 0.671272 | 0.670373 | 0.64493 |
| 18 | 39489200 | 39491599 | 14.2484 | 24.98695 | 39.398 | 17.41274 | -0.26605 | 0.632425 | 0.644936 |
| 11 | 95745400 | 95747999 | 23.66435 | 39.51389 | 60.25351 | 26.21597 | -0.22549 | 1.208194 | 0.644937 |
| 2 | 102111200 | 102113199 | 13.5587 | 23.94203 | 37.68548 | 16.67315 | -0.2611 | 0.536236 | 0.64494 |
| 4 | 151142200 | 151145599 | 28.25246 | 47.38848 | 52.89697 | 23.08085 | 0.222909 | 1.166644 | 0.644944 |
| 6 | 132348000 | 132350599 | 25.146 | 43.83548 | 40.63752 | 17.87951 | 0.496206 | 0.780211 | 0.644957 |
| 1 | 82981400 | 82982799 | 13.01205 | 22.85142 | 37.92308 | 16.78112 | -0.33933 | 0.600495 | 0.644962 |
| 17 | 54120800 | 54125199 | 62.07162 | 99.02346 | 107.2064 | 44.94949 | 0.224837 | 2.143791 | 0.644993 |
| 14 | 118651000 | 118659999 | 95.60722 | 146.7284 | 206.7517 | 85.83462 | -0.20055 | 2.992691 | 0.644995 |
| 7 | 47198200 | 47201399 | 53.19649 | 85.45485 | 91.76887 | 38.86974 | 0.247848 | 1.917987 | 0.645008 |
| 16 | 22327000 | 22328999 | 16.67499 | 29.51432 | 32.7957 | 14.50293 | 0.239928 | 0.521005 | 0.645024 |
| 10 | 8326200 | 8329599 | 30.2836 | 49.62114 | 74.22332 | 31.9426 | -0.20296 | 1.456523 | 0.645025 |
| 8 | 39472800 | 39473599 | 9.234605 | 17.09339 | 28.89881 | 12.16225 | -0.3119 | 0.190013 | 0.645027 |
| 12 | 69762400 | 69767199 | 139.6333 | 210.9962 | 225.9777 | 93.4346 | 0.176802 | 3.319305 | 0.645041 |
| 7 | 60924400 | 60927799 | 34.11317 | 57.08108 | 55.88498 | 24.34758 | 0.411572 | 1.213307 | 0.645053 |
| 17 | 48450400 | 48453599 | 31.09297 | 50.91489 | 76.67554 | 32.98904 | -0.2131 | 1.467308 | 0.645061 |
| 9 | 62517400 | 62524199 | 73.25621 | 114.5139 | 162.3045 | 67.33769 | -0.18847 | 2.645779 | 0.645061 |
| 13 | 29684800 | 29686799 | 17.60905 | 30.6082 | 46.8454 | 20.57343 | -0.22434 | 0.792801 | 0.645067 |

| | | | | | | | | | |
|---|---|---|---|---|---|---|---|---|---|
| 18 | 9353400 | 9367799 | 242.4129 | 354.0474 | 355.5404 | 146.483 | 0.224581 | 3.933759 | 0.645071 |
| 2 | 122372200 | 122372599 | 9.68785 | 18.49647 | 21.8013 | 8.673986 | 0.271817 | -0.02573 | 0.645076 |
| 19 | 39921800 | 39924399 | 49.15587 | 79.02632 | 87.87542 | 37.26321 | 0.199088 | 1.887785 | 0.645079 |
| 4 | 13914000 | 13917599 | 35.56474 | 58.77265 | 62.14342 | 26.85063 | 0.300293 | 1.357616 | 0.645093 |
| 1 | 23264600 | 23265199 | 6.680557 | 12.961 | 23.38737 | 8.809859 | -0.26293 | -0.23357 | 0.645107 |
| 14 | 69254800 | 69257199 | 18.05907 | 31.74364 | 34.82747 | 15.42275 | 0.254269 | 0.606606 | 0.645118 |
| 3 | 43133000 | 43133999 | 19.44685 | 33.30956 | 50.60096 | 22.19031 | -0.21762 | 0.945834 | 0.645142 |
| 13 | 68354000 | 68372999 | 394.2358 | 550.5065 | 540.6917 | 224.247 | 0.192142 | 4.603019 | 0.645153 |
| 18 | 15503800 | 15505799 | 17.42535 | 31.38792 | 31.09446 | 13.55784 | 0.418911 | 0.406962 | 0.645159 |
| 12 | 61846200 | 61849199 | 23.22515 | 40.79683 | 36.61283 | 16.20449 | 0.542901 | 0.651021 | 0.645169 |
| 5 | 28412000 | 28417199 | 35.15669 | 56.62039 | 103.3276 | 43.92815 | -0.51098 | 1.817062 | 0.64519 |
| 8 | 42940000 | 42940999 | 12.21023 | 21.1884 | 48.94237 | 21.47415 | -0.81509 | 0.796393 | 0.645196 |
| 13 | 3702800 | 3704199 | 12.113 | 21.7198 | 34.17311 | 14.96351 | -0.2485 | 0.406067 | 0.645219 |
| 13 | 101607000 | 101608999 | 13.24064 | 23.51948 | 36.62768 | 16.16601 | -0.24187 | 0.476815 | 0.645233 |
| X | 58721000 | 58724199 | 40.67026 | 66.02107 | 72.20318 | 30.95073 | 0.238215 | 1.617096 | 0.64524 |
| 13 | 25538200 | 25542199 | 58.65932 | 93.64722 | 103.9423 | 43.62526 | 0.190109 | 2.118025 | 0.645275 |
| 5 | 129519800 | 129524399 | 42.14956 | 68.38548 | 64.4696 | 27.62463 | 0.451149 | 1.624532 | 0.645284 |
| 5 | 110002400 | 110004399 | 20.53748 | 35.90761 | 38.59098 | 17.01272 | 0.284075 | 0.711256 | 0.645295 |
| 4 | 142546400 | 142547999 | 16.69955 | 29.47417 | 32.91635 | 14.56453 | 0.231216 | 0.552603 | 0.645309 |
| 4 | 69130200 | 69131199 | 9.619977 | 17.91427 | 29.44078 | 12.28227 | -0.26232 | 0.156995 | 0.645313 |
| 5 | 10258400 | 10261799 | 45.56623 | 73.46557 | 81.57152 | 34.73161 | 0.206273 | 1.792531 | 0.645315 |
| 3 | 50938600 | 50939399 | 8.188176 | 15.60247 | 26.68231 | 10.66029 | -0.26652 | -0.01507 | 0.645343 |
| 14 | 4211600 | 4215799 | 67.63759 | 107.1391 | 117.461 | 48.84976 | 0.196426 | 2.363536 | 0.645349 |
| 3 | 137147800 | 137148599 | 29.92736 | 48.5759 | 77.43854 | 33.19192 | -0.30419 | 1.601753 | 0.645352 |
| 12 | 98982000 | 98982999 | 8.22208 | 15.56395 | 26.53249 | 10.74953 | -0.27951 | 0.031426 | 0.645369 |
| 7 | 143463800 | 143465199 | 12.28559 | 22.25794 | 35.85959 | 15.52302 | -0.27166 | 0.34977 | 0.645375 |
| 9 | 82171200 | 82177999 | 60.03773 | 96.30541 | 95.6743 | 40.43521 | 0.357521 | 1.972999 | 0.645397 |
| 6 | 38240200 | 38242199 | 12.90664 | 22.94315 | 36.12675 | 15.93499 | -0.2571 | 0.468618 | 0.645433 |
| 1 | 28532400 | 28533599 | 12.59511 | 22.52006 | 35.15764 | 15.43162 | -0.2398 | 0.429026 | 0.645438 |
| 14 | 77293600 | 77295399 | 26.50121 | 45.05245 | 48.48901 | 21.25127 | 0.276036 | 1.039591 | 0.645454 |
| 3 | 21241600 | 21246999 | 68.5358 | 108.7696 | 119.702 | 49.86827 | 0.193546 | 2.312035 | 0.645469 |
| 5 | 70710600 | 70728599 | 80.78576 | 127.7336 | 115.4859 | 47.98419 | 0.472063 | 2.415405 | 0.645493 |
| 15 | 96874600 | 96877799 | 47.4175 | 75.73363 | 110.8303 | 46.54536 | -0.20735 | 2.094939 | 0.645494 |
| 14 | 6945800 | 6947199 | 10.65272 | 19.68443 | 31.99233 | 13.49575 | -0.25824 | 0.217833 | 0.645494 |
| 1 | 59205000 | 59206799 | 11.50519 | 20.89325 | 33.30159 | 14.37329 | -0.25322 | 0.326563 | 0.645505 |
| 3 | 33989600 | 33990399 | 11.50519 | 20.89325 | 33.30159 | 14.37329 | -0.25322 | 0.326563 | 0.645505 |
| X | 43036400 | 43037599 | 16.19142 | 29.10547 | 30.85155 | 13.48065 | 0.320189 | 0.413359 | 0.645517 |
| 3 | 149864200 | 149864799 | 11.99096 | 21.31963 | 34.70494 | 15.30529 | -0.30442 | 0.459433 | 0.645523 |
| X | 60906800 | 60913599 | 29.50168 | 50.19028 | 42.53842 | 18.64441 | 0.619573 | 1.036717 | 0.645529 |
| 6 | 101844400 | 101848199 | 48.03624 | 76.85348 | 111.0915 | 46.76848 | -0.18692 | 2.042804 | 0.645538 |
| 4 | 40664400 | 40667999 | 35.26448 | 57.22125 | 85.52889 | 36.65138 | -0.21205 | 1.621195 | 0.645544 |
| 8 | 98761200 | 98761799 | 13.41603 | 24.3404 | 25.15398 | 10.86948 | 0.367057 | 0.32733 | 0.645548 |
| 2 | 129262800 | 129264399 | 15.59929 | 28.01756 | 30.84011 | 13.48519 | 0.265258 | 0.41323 | 0.645561 |
| 1 | 8727800 | 8733199 | 66.31203 | 105.1013 | 114.661 | 47.70502 | 0.204278 | 2.348757 | 0.645567 |
| 15 | 19555200 | 19560399 | 58.1764 | 93.26495 | 96.88965 | 40.91234 | 0.292143 | 1.988541 | 0.64557 |
| 13 | 50022600 | 50026399 | 64.77209 | 103.1413 | 112.1084 | 46.88506 | 0.21629 | 2.206806 | 0.645588 |
| 2 | 36297600 | 36298999 | 45.66262 | 73.45705 | 79.69991 | 33.82758 | 0.235616 | 1.859599 | 0.645591 |

| | | | | | | | | | |
|---|---|---|---|---|---|---|---|---|---|
| 3 | 22702000 | 22705399 | 38.02708 | 62.60205 | 64.5754 | 27.84572 | 0.334364 | 1.410132 | 0.645592 |
| 13 | 68666800 | 68681399 | 186.857 | 278.3819 | 286.6154 | 118.1068 | 0.21549 | 3.599917 | 0.645594 |
| 2 | 129964600 | 129966399 | 15.40202 | 27.04497 | 41.8746 | 18.52757 | -0.24007 | 0.622519 | 0.645607 |
| 1 | 107750400 | 107751799 | 31.60891 | 52.06284 | 53.92707 | 23.25908 | 0.328777 | 1.381645 | 0.645615 |
| 5 | 52849200 | 52850199 | 12.26593 | 22.77746 | 26.11282 | 10.94183 | 0.248868 | 0.18912 | 0.645631 |
| 11 | 111105800 | 111108799 | 43.86224 | 71.13105 | 66.497 | 28.48356 | 0.462581 | 1.636492 | 0.64566 |
| 7 | 113539600 | 113543599 | 46.45237 | 74.77527 | 83.82779 | 35.61258 | 0.189632 | 1.841614 | 0.645671 |
| X | 107260000 | 107260599 | 14.28363 | 25.93306 | 28.9589 | 12.50242 | 0.256604 | 0.33784 | 0.64568 |
| 12 | 42188200 | 42189199 | 12.23966 | 22.82888 | 26.14012 | 10.87597 | 0.258502 | 0.161546 | 0.645682 |
| 6 | 26196600 | 26198199 | 13.06108 | 23.47193 | 38.32777 | 16.7513 | -0.30023 | 0.40579 | 0.645683 |
| 16 | 91919600 | 91928399 | 89.81468 | 138.7652 | 192.5809 | 79.99395 | -0.17125 | 2.862182 | 0.645692 |
| 18 | 83509600 | 83512399 | 19.92534 | 34.02753 | 51.67103 | 22.65414 | -0.21774 | 0.9756 | 0.645704 |
| 15 | 49157600 | 49158799 | 15.02246 | 27.73179 | 28.50704 | 12.04154 | 0.396709 | 0.248277 | 0.645716 |
| X | 29378000 | 29381599 | 62.61779 | 99.52882 | 110.5215 | 46.11389 | 0.182737 | 2.261228 | 0.645725 |
| 8 | 125351800 | 125353999 | 24.30653 | 41.66376 | 45.76323 | 20.06997 | 0.248444 | 0.951372 | 0.645736 |
| 17 | 18950600 | 18954599 | 51.20564 | 82.65671 | 82.2224 | 35.0628 | 0.366472 | 1.765303 | 0.645756 |
| 4 | 17010400 | 17013799 | 35.53639 | 58.8548 | 61.06664 | 26.42539 | 0.327667 | 1.331585 | 0.645789 |
| 14 | 77617400 | 77618999 | 12.6208 | 22.48757 | 35.16874 | 15.48185 | -0.24518 | 0.451548 | 0.645797 |
| 11 | 67405200 | 67407399 | 17.4736 | 30.26653 | 46.34975 | 20.33547 | -0.22592 | 0.831391 | 0.64583 |
| 4 | 146061600 | 146064199 | 35.46053 | 59.91476 | 50.75967 | 22.20835 | 0.619759 | 1.125891 | 0.645834 |
| 15 | 40964400 | 40966799 | 21.78226 | 36.33098 | 66.14762 | 28.76036 | -0.48043 | 1.219884 | 0.64584 |
| 7 | 50687000 | 50688599 | 16.62358 | 29.53666 | 32.33076 | 14.26625 | 0.264373 | 0.490055 | 0.64584 |
| X | 165997000 | 165997599 | 5.496858 | 10.72235 | 20.96051 | 7.505497 | -0.30869 | -0.43106 | 0.645846 |
| 9 | 18868000 | 18868999 | 13.39437 | 24.37326 | 27.48597 | 11.8332 | 0.245062 | 0.314427 | 0.645857 |
| 3 | 71326200 | 71326999 | 8.910923 | 17.00212 | 31.40151 | 12.51069 | -0.37309 | -0.00328 | 0.64586 |
| X | 5028200 | 5050999 | 421.6547 | 591.4774 | 510.73 | 211.6709 | 0.384494 | 4.538101 | 0.64586 |
| 1 | 20304400 | 20305399 | 9.874219 | 18.48004 | 31.20457 | 12.90816 | -0.29103 | 0.133821 | 0.645864 |
| 6 | 103941000 | 103943199 | 24.29567 | 41.40858 | 46.34062 | 20.31387 | 0.22038 | 1.002887 | 0.645869 |
| 9 | 10963400 | 10964799 | 24.08221 | 40.87199 | 44.33652 | 19.4244 | 0.264578 | 1.039725 | 0.645881 |
| 16 | 48389000 | 48390399 | 19.96796 | 34.88668 | 38.10645 | 16.78943 | 0.26074 | 0.717889 | 0.645882 |
| 2 | 138294000 | 138298599 | 37.91155 | 61.17766 | 89.88655 | 38.31526 | -0.19549 | 1.758742 | 0.645887 |
| 15 | 59740600 | 59743199 | 30.48862 | 49.66472 | 75.37288 | 32.36609 | -0.22933 | 1.539736 | 0.645888 |
| 5 | 66329000 | 66334799 | 59.40892 | 94.21298 | 134.4983 | 55.9898 | -0.18239 | 2.330027 | 0.645891 |
| 2 | 25631600 | 25635799 | 58.45697 | 93.3005 | 103.8636 | 43.57657 | 0.185454 | 2.125041 | 0.645892 |
| 15 | 6234600 | 6235999 | 17.35076 | 30.10801 | 46.34712 | 20.33544 | -0.23279 | 0.81563 | 0.645894 |
| 13 | 54659000 | 54663999 | 54.75243 | 88.14263 | 87.96158 | 37.37279 | 0.35726 | 1.853571 | 0.645913 |
| 13 | 9085800 | 9086599 | 10.52564 | 20.39699 | 21.619 | 8.188391 | 0.489308 | -0.18002 | 0.645927 |
| Y | 9852600 | 9856799 | 25.50536 | 43.19612 | 32.73139 | 14.33087 | 0.779839 | 1.044038 | 0.645966 |
| 13 | 14254000 | 14259199 | 54.35945 | 87.28168 | 93.03753 | 39.37891 | 0.257727 | 1.934584 | 0.645972 |
| 3 | 13443200 | 13444999 | 23.61862 | 39.06188 | 71.05177 | 30.71645 | -0.47985 | 1.325371 | 0.645982 |
| 16 | 69188200 | 69189399 | 12.60567 | 22.20172 | 42.98503 | 19.02765 | -0.55941 | 0.572116 | 0.645987 |
| 12 | 69141200 | 69142799 | 12.54887 | 22.47597 | 35.64947 | 15.62484 | -0.26126 | 0.417572 | 0.646016 |
| 10 | 87420400 | 87421399 | 8.657068 | 16.48417 | 31.21822 | 12.49134 | -0.41403 | 0.010154 | 0.646049 |
| 18 | 24799000 | 24806399 | 112.9332 | 170.008 | 287.7189 | 118.8303 | -0.48752 | 3.392266 | 0.646065 |
| 5 | 120047000 | 120047999 | 9.647452 | 17.88424 | 29.38419 | 12.34145 | -0.26922 | 0.184461 | 0.646071 |
| 13 | 27624000 | 27625399 | 15.16985 | 27.32388 | 30.29479 | 13.20992 | 0.257412 | 0.390698 | 0.646075 |
| 7 | 113941400 | 113942799 | 14.43071 | 26.30917 | 28.84782 | 12.38993 | 0.287247 | 0.312601 | 0.646081 |

| | | | | | | | | | |
|---|---|---|---|---|---|---|---|---|---|
| 19 | 14532400 | 14534799 | 18.99458 | 33.26674 | 36.48664 | 16.10824 | 0.254798 | 0.671858 | 0.646133 |
| 3 | 111674600 | 111675399 | 18.73887 | 33.00572 | 27.47395 | 12.16141 | 0.651984 | 0.577594 | 0.646133 |
| 17 | 33520400 | 33528399 | 138.3656 | 209.7846 | 224.8393 | 93.06236 | 0.179902 | 3.251577 | 0.646137 |
| 12 | 96565400 | 96568399 | 36.71931 | 60.37614 | 64.93447 | 27.98625 | 0.273619 | 1.422036 | 0.646139 |
| 7 | 10908800 | 10960799 | 1326.722 | 1671.71 | 1501.22 | 630.8319 | 0.172514 | 6.243863 | 0.646161 |
| 12 | 116816200 | 116820199 | 42.91894 | 69.27912 | 76.59663 | 32.66309 | 0.214309 | 1.74898 | 0.646177 |
| 17 | 88816200 | 88819999 | 28.19553 | 46.11492 | 71.43325 | 30.71947 | -0.25462 | 1.47457 | 0.646186 |
| 9 | 87883800 | 87884399 | 11.57865 | 21.67477 | 25.15983 | 10.40376 | 0.246073 | 0.130474 | 0.646194 |
| 9 | 110939600 | 110940199 | 8.726079 | 16.42524 | 27.50901 | 11.26939 | -0.26772 | 0.07793 | 0.646195 |
| X | 77307600 | 77310999 | 36.98583 | 60.99917 | 63.2698 | 27.30382 | 0.327504 | 1.386393 | 0.646212 |
| X | 95041800 | 95044799 | 26.21658 | 43.66687 | 67.2724 | 29.22399 | -0.24037 | 1.246002 | 0.646216 |
| X | 103024200 | 103026399 | 30.70219 | 51.76608 | 52.37458 | 22.9003 | 0.364407 | 1.128585 | 0.646235 |
| 3 | 93667400 | 93670399 | 44.50774 | 71.87446 | 78.31797 | 33.41553 | 0.235943 | 1.739215 | 0.646245 |
| 10 | 128341800 | 128343199 | 14.43105 | 25.3331 | 39.36114 | 17.41147 | -0.24492 | 0.617636 | 0.646249 |
| 2 | 83972400 | 83973599 | 10.4015 | 19.23847 | 31.45702 | 13.2528 | -0.26584 | 0.206924 | 0.646252 |
| 16 | 32010200 | 32011799 | 14.25416 | 25.05823 | 38.78916 | 17.17673 | -0.23954 | 0.592892 | 0.646311 |
| 6 | 78549200 | 78550199 | 11.16703 | 21.35035 | 23.91004 | 9.482391 | 0.34853 | -0.01831 | 0.646327 |
| 1 | 184930600 | 184933399 | 20.45583 | 34.68854 | 53.25137 | 23.32424 | -0.23464 | 1.04298 | 0.646332 |
| 19 | 47689200 | 47694199 | 42.33232 | 68.56146 | 75.94429 | 32.49931 | 0.216724 | 1.665048 | 0.646335 |
| 8 | 93964200 | 93965799 | 11.77565 | 21.49546 | 34.95625 | 14.99002 | -0.27624 | 0.300609 | 0.646336 |
| X | 122005800 | 122008599 | 34.02245 | 56.92691 | 55.84982 | 24.33163 | 0.408615 | 1.214378 | 0.646341 |
| 10 | 54577800 | 54578399 | 10.18347 | 19.29989 | 21.76541 | 8.795814 | 0.315513 | 0.029922 | 0.646364 |
| 14 | 17775000 | 17775399 | 5.390275 | 10.46796 | 21.13957 | 7.914133 | -0.41429 | -0.27605 | 0.646366 |
| 10 | 45459400 | 45461399 | 13.60308 | 24.3222 | 39.85136 | 17.49188 | -0.30934 | 0.444748 | 0.646369 |
| 2 | 73923000 | 73926199 | 36.50614 | 59.85064 | 65.97006 | 28.39748 | 0.236603 | 1.45683 | 0.646372 |
| 1 | 45135800 | 45137399 | 24.3496 | 42.1557 | 43.53586 | 19.10062 | 0.339775 | 0.861414 | 0.646383 |
| 5 | 141036800 | 141044199 | 82.13235 | 127.7679 | 178.3191 | 74.02758 | -0.17171 | 2.736327 | 0.646405 |
| 10 | 66803800 | 66806799 | 23.25489 | 38.48699 | 72.94366 | 31.54944 | -0.53899 | 1.315273 | 0.646409 |
| 5 | 135509400 | 135514799 | 58.71203 | 93.07274 | 133.566 | 55.58073 | -0.19041 | 2.342265 | 0.646416 |
| 11 | 74048600 | 74050599 | 20.04376 | 35.32362 | 35.72685 | 15.81283 | 0.372621 | 0.593602 | 0.646419 |
| 7 | 27009000 | 27016799 | 74.89208 | 117.0749 | 164.4302 | 68.21939 | -0.17532 | 2.646302 | 0.646423 |
| 9 | 79247800 | 79250799 | 48.27227 | 77.03101 | 112.7436 | 47.28991 | -0.20893 | 2.124057 | 0.646433 |
| 4 | 18473600 | 18483599 | 142.3172 | 214.773 | 229.8706 | 95.00531 | 0.176161 | 3.345558 | 0.646436 |
| 10 | 68243000 | 68251999 | 129.5918 | 195.3793 | 266.3126 | 110.0336 | -0.17362 | 3.364666 | 0.646446 |
| 8 | 60870600 | 60875999 | 36.19741 | 59.59415 | 55.9325 | 24.11952 | 0.470235 | 1.397248 | 0.646452 |
| 6 | 107225200 | 107226999 | 21.93649 | 38.10051 | 41.40293 | 18.17432 | 0.26739 | 0.814627 | 0.646469 |
| 4 | 98550000 | 98554799 | 50.86825 | 81.60714 | 91.24227 | 38.5695 | 0.187552 | 1.955011 | 0.646483 |
| 13 | 27861400 | 27864999 | 101.3025 | 156.5868 | 171.2837 | 71.14739 | 0.172317 | 2.856222 | 0.646498 |
| 5 | 139073800 | 139074399 | 9.559643 | 17.52189 | 29.84834 | 12.73107 | -0.33691 | 0.249682 | 0.646504 |
| 5 | 144314000 | 144319399 | 66.2106 | 103.9662 | 151.7446 | 62.93825 | -0.22571 | 2.558946 | 0.646518 |
| 4 | 30814000 | 30816599 | 29.55083 | 49.6429 | 53.48511 | 23.35994 | 0.273775 | 1.154828 | 0.646524 |
| 2 | 48940400 | 48943999 | 40.66168 | 66.72493 | 66.472 | 28.6247 | 0.38267 | 1.456024 | 0.64653 |
| X | 34975200 | 34994199 | 294.5475 | 424.0339 | 404.931 | 167.1175 | 0.276823 | 4.153243 | 0.646535 |
| 7 | 140296200 | 140297799 | 11.15374 | 20.31957 | 32.2991 | 13.88524 | -0.2456 | 0.302909 | 0.646555 |
| 18 | 7091600 | 7093599 | 20.0853 | 33.74256 | 55.29116 | 24.14895 | -0.32888 | 1.126536 | 0.646573 |
| 10 | 78658400 | 78659599 | 20.91648 | 36.54486 | 38.54795 | 16.98425 | 0.310852 | 0.722573 | 0.646644 |
| 13 | 113974800 | 113977199 | 33.77629 | 55.89947 | 53.0829 | 22.95916 | 0.455017 | 1.328501 | 0.646657 |

| | | | | | | | | | |
|---|---|---|---|---|---|---|---|---|---|
| 9 | 89041200 | 89042399 | 13.92188 | 24.48407 | 38.28366 | 16.95151 | -0.25375 | 0.585202 | 0.646666 |
| 3 | 85211000 | 85220199 | 97.46997 | 149.7936 | 206.5416 | 85.76364 | -0.16777 | 2.967399 | 0.646666 |
| 18 | 46710800 | 46715799 | 24.61695 | 42.1627 | 38.41214 | 16.84501 | 0.516984 | 0.949965 | 0.646674 |
| 9 | 78154600 | 78159999 | 41.62383 | 66.88481 | 98.27275 | 41.64283 | -0.20297 | 1.895392 | 0.646679 |
| 12 | 54827400 | 54827799 | 7.311167 | 14.05892 | 24.6986 | 9.634671 | -0.27235 | -0.10446 | 0.646691 |
| 13 | 37444600 | 37448799 | 31.52853 | 52.65587 | 56.36332 | 24.5265 | 0.28328 | 1.224556 | 0.646699 |
| 5 | 109193200 | 109199799 | 118.97 | 182.0301 | 197.241 | 81.84071 | 0.174683 | 3.060158 | 0.646717 |
| 12 | 77599600 | 77600599 | 8.058226 | 15.36018 | 26.13142 | 10.43253 | -0.25785 | -0.01903 | 0.646719 |
| Y | 89953400 | 89953999 | 9.365223 | 18.23523 | 20.25489 | 7.423576 | 0.466275 | -0.3078 | 0.64672 |
| 6 | 106369800 | 106371799 | 31.29353 | 52.21673 | 56.38298 | 24.52123 | 0.27073 | 1.232229 | 0.64674 |
| 4 | 96072200 | 96074599 | 27.45921 | 46.7275 | 49.02615 | 21.4884 | 0.312769 | 1.031991 | 0.646775 |
| 6 | 22435600 | 22437199 | 38.58638 | 62.63395 | 70.37145 | 30.164 | 0.199284 | 1.611534 | 0.646801 |
| 1 | 148569600 | 148571599 | 15.15584 | 27.67786 | 27.58351 | 11.82193 | 0.42579 | 0.303269 | 0.646804 |
| 16 | 87388600 | 87389599 | 11.56513 | 21.69755 | 25.17608 | 10.37144 | 0.250931 | 0.116156 | 0.646808 |
| X | 148878800 | 148880599 | 38.84923 | 63.02088 | 66.09503 | 28.3178 | 0.297602 | 1.624771 | 0.646809 |
| 17 | 89080200 | 89082799 | 18.66032 | 32.65256 | 36.05532 | 15.90599 | 0.245153 | 0.683472 | 0.64682 |
| 18 | 22772800 | 22773799 | 9.852907 | 18.12882 | 29.84918 | 12.66515 | -0.28274 | 0.230447 | 0.646829 |
| 6 | 69956400 | 69957999 | 18.55276 | 33.17003 | 31.9774 | 14.03655 | 0.451462 | 0.460884 | 0.646832 |
| 10 | 85608000 | 85609399 | 18.55473 | 32.5428 | 35.95304 | 15.89182 | 0.244366 | 0.646483 | 0.646856 |
| 18 | 69627800 | 69629999 | 28.05905 | 47.01636 | 52.90293 | 23.06868 | 0.211425 | 1.176247 | 0.646874 |
| 11 | 17839400 | 17841399 | 21.98974 | 37.42024 | 59.34114 | 26.00952 | -0.28096 | 1.025107 | 0.646878 |
| 1 | 100732000 | 100733399 | 19.43642 | 33.8702 | 37.22017 | 16.37228 | 0.251739 | 0.749805 | 0.646954 |
| X | 121352600 | 121356199 | 67.28237 | 107.1293 | 112.0079 | 46.83746 | 0.272116 | 2.213975 | 0.646964 |
| 5 | 93064400 | 93067599 | 24.61269 | 40.99821 | 61.97901 | 26.92564 | -0.21309 | 1.237266 | 0.646994 |
| 7 | 14142200 | 14153599 | 139.9706 | 214.2128 | 199.8899 | 82.94437 | 0.390401 | 3.063716 | 0.647001 |
| 7 | 56305000 | 56306199 | 15.45143 | 27.44173 | 29.53862 | 13.0391 | 0.288164 | 0.480151 | 0.647006 |
| 15 | 93952600 | 93956799 | 50.58077 | 80.83188 | 117.6506 | 49.44952 | -0.19867 | 2.083705 | 0.64702 |
| 12 | 21023200 | 21025799 | 23.16648 | 39.65095 | 43.12285 | 18.90976 | 0.262294 | 0.954927 | 0.64704 |
| 1 | 104326000 | 104329999 | 10.2254 | 19.63745 | 18.63064 | 7.294601 | 0.603605 | -0.08162 | 0.647042 |
| 13 | 113309600 | 113311999 | 18.43978 | 31.86957 | 48.47164 | 21.265 | -0.21684 | 0.85786 | 0.647048 |
| 17 | 38334600 | 38338199 | 43.20916 | 70.073 | 75.60506 | 32.38529 | 0.256312 | 1.641431 | 0.647051 |
| 11 | 73447800 | 73450199 | 41.31477 | 67.036 | 73.43175 | 31.46653 | 0.235254 | 1.627911 | 0.647054 |
| 3 | 26933200 | 26935799 | 25.26171 | 41.9033 | 64.19238 | 27.8079 | -0.23256 | 1.28766 | 0.647087 |
| 7 | 47117200 | 47119199 | 21.52537 | 37.75282 | 37.27778 | 16.47733 | 0.40563 | 0.664865 | 0.64709 |
| 1 | 98194800 | 98195399 | 7.140585 | 13.78405 | 24.26055 | 9.355943 | -0.2596 | -0.14577 | 0.647092 |
| 10 | 100517000 | 100517399 | 7.140585 | 13.78405 | 24.26055 | 9.355943 | -0.2596 | -0.14577 | 0.647092 |
| X | 18995200 | 18996599 | 19.93602 | 35.03739 | 36.36598 | 16.10229 | 0.334133 | 0.625062 | 0.647131 |
| 8 | 87230000 | 87232799 | 28.95638 | 48.62872 | 44.36343 | 19.37233 | 0.512984 | 1.147563 | 0.647152 |
| 1 | 9426800 | 9427399 | 7.624563 | 14.67672 | 25.87252 | 10.06272 | -0.27339 | -0.10742 | 0.647196 |
| 16 | 52387400 | 52387799 | 7.476577 | 14.34624 | 25.1916 | 9.887599 | -0.27997 | -0.08081 | 0.647211 |
| 13 | 85617200 | 85619799 | 44.80477 | 71.81855 | 104.157 | 43.97188 | -0.18849 | 1.979841 | 0.647236 |
| 12 | 114717200 | 114729599 | 212.5593 | 316.0233 | 293.4675 | 120.9051 | 0.361565 | 3.640837 | 0.647288 |
| 15 | 58070400 | 58071399 | 13.80324 | 25.09142 | 28.01367 | 12.07593 | 0.258377 | 0.325712 | 0.647289 |
| 12 | 103970600 | 103971999 | 15.4409 | 27.90662 | 30.3675 | 13.18555 | 0.288327 | 0.37483 | 0.647295 |
| 15 | 46023600 | 46024999 | 14.48175 | 26.41141 | 23.75183 | 10.1968 | 0.571892 | 0.299898 | 0.6473 |
| 7 | 13012200 | 13013599 | 20.56183 | 35.55228 | 36.59074 | 16.05294 | 0.344499 | 0.847961 | 0.647317 |
| 6 | 116983800 | 116985599 | 17.2664 | 30.75046 | 32.74991 | 14.42281 | 0.305553 | 0.480128 | 0.647318 |

| | | | | | | | | | |
|---|---|---|---|---|---|---|---|---|---|
| X | 40356800 | 40357199 | 9.708242 | 18.66455 | 22.44804 | 8.761869 | 0.26844 | -0.0855 | 0.647319 |
| 1 | 58651600 | 58653999 | 13.69937 | 24.25234 | 38.13707 | 16.86433 | -0.25835 | 0.509486 | 0.647322 |
| 7 | 54301800 | 54303199 | 13.60322 | 24.06413 | 37.18209 | 16.44449 | -0.23334 | 0.51733 | 0.647354 |
| X | 153019800 | 153034599 | 143.6601 | 219.6525 | 201.7892 | 83.71429 | 0.411855 | 3.084097 | 0.647439 |
| 7 | 26716000 | 26722599 | 122.7802 | 187.8162 | 172.7025 | 71.6552 | 0.411022 | 3.064253 | 0.647447 |
| 6 | 121672800 | 121678399 | 59.90142 | 95.75608 | 91.47984 | 38.46628 | 0.40818 | 2.075536 | 0.647448 |
| 14 | 56346800 | 56349999 | 31.87385 | 53.25103 | 56.37515 | 24.53701 | 0.299138 | 1.222177 | 0.647475 |
| 1 | 33484400 | 33485999 | 14.27505 | 25.03215 | 39.42085 | 17.42194 | -0.26422 | 0.63447 | 0.647478 |
| 12 | 109248600 | 109249199 | 7.136186 | 13.79065 | 24.3249 | 9.332883 | -0.25575 | -0.1633 | 0.647485 |
| 13 | 41271600 | 41272599 | 7.136186 | 13.79065 | 24.3249 | 9.332883 | -0.25575 | -0.1633 | 0.647485 |
| 8 | 96105400 | 96106199 | 12.33216 | 22.69858 | 25.51222 | 10.81972 | 0.264605 | 0.236132 | 0.647497 |
| 6 | 10649800 | 10650799 | 9.976224 | 18.47707 | 29.89616 | 12.57186 | -0.2485 | 0.193953 | 0.647498 |
| 4 | 26781400 | 26781999 | 7.184927 | 13.86368 | 25.3529 | 9.792908 | -0.31602 | -0.13409 | 0.647531 |
| 10 | 119344400 | 119359199 | 258.243 | 369.7576 | 480.2564 | 198.3732 | -0.17455 | 4.229361 | 0.647532 |
| 10 | 63390400 | 63392599 | 25.35099 | 43.15135 | 47.94654 | 21.01565 | 0.230418 | 1.021184 | 0.647536 |
| 11 | 45996800 | 45999199 | 25.35099 | 43.15135 | 47.94654 | 21.01565 | 0.230418 | 1.021184 | 0.647536 |
| 1 | 158848000 | 158850399 | 29.44134 | 47.82223 | 85.13488 | 36.51057 | -0.46213 | 1.589497 | 0.647546 |
| 14 | 50340000 | 50341799 | 19.09967 | 33.88546 | 27.68569 | 12.23198 | 0.682887 | 0.507103 | 0.647604 |
| 6 | 132337800 | 132338999 | 18.82962 | 31.74056 | 60.08694 | 26.27057 | -0.53617 | 1.098233 | 0.647607 |
| 18 | 31750400 | 31752399 | 27.06698 | 45.44962 | 51.24467 | 22.37529 | 0.208444 | 1.149342 | 0.647613 |
| 6 | 4753800 | 4757999 | 77.69533 | 119.9501 | 202.8675 | 84.26711 | -0.45704 | 2.873713 | 0.647614 |
| 15 | 7716000 | 7717799 | 14.67317 | 25.69332 | 40.93183 | 18.06775 | -0.28105 | 0.65972 | 0.647625 |
| 4 | 129413600 | 129415399 | 15.54852 | 27.56128 | 30.42965 | 13.45263 | 0.250056 | 0.493938 | 0.647643 |
| 3 | 91566800 | 91569399 | 31.30326 | 52.86137 | 51.86688 | 22.68897 | 0.408497 | 1.114211 | 0.647663 |
| 1 | 78944000 | 78945799 | 18.97459 | 32.36018 | 50.5885 | 22.17712 | -0.25975 | 0.983101 | 0.647698 |
| X | 5760000 | 5762999 | 34.33228 | 56.95425 | 59.75957 | 25.88918 | 0.311789 | 1.310587 | 0.64771 |
| 18 | 49516000 | 49516999 | 11.93971 | 21.65996 | 34.29401 | 14.82074 | -0.24509 | 0.336304 | 0.64771 |
| 5 | 100076000 | 100076599 | 6.106305 | 11.83999 | 22.18347 | 8.379113 | -0.31952 | -0.23248 | 0.647716 |
| 14 | 13302400 | 13303399 | 6.106305 | 11.83999 | 22.18347 | 8.379113 | -0.31952 | -0.23248 | 0.647716 |
| 18 | 60481800 | 60482599 | 12.58453 | 23.67442 | 25.66793 | 10.52543 | 0.353118 | 0.107655 | 0.647718 |
| 10 | 82304200 | 82305599 | 18.53355 | 32.43195 | 36.08911 | 15.9214 | 0.234114 | 0.682771 | 0.647719 |
| 8 | 10185800 | 10187599 | 11.88409 | 21.27275 | 34.26638 | 15.02611 | -0.28337 | 0.418123 | 0.647818 |
| 7 | 6679200 | 6681799 | 33.64751 | 54.5355 | 81.36138 | 34.83629 | -0.21081 | 1.639393 | 0.647825 |
| 8 | 98754600 | 98757999 | 23.7562 | 41.37234 | 40.67605 | 17.88549 | 0.411518 | 0.78481 | 0.647833 |
| 17 | 27983200 | 27985199 | 34.75884 | 56.76767 | 62.58041 | 26.89824 | 0.233279 | 1.503371 | 0.647849 |
| 13 | 29128600 | 29131199 | 23.29764 | 38.85285 | 60.27239 | 26.20346 | -0.25022 | 1.218758 | 0.647867 |
| 1 | 21171000 | 21171799 | 8.047458 | 15.37458 | 26.18705 | 10.40458 | -0.25353 | -0.03496 | 0.647875 |
| 9 | 3242600 | 3253199 | 140.4201 | 213.683 | 217.781 | 90.24012 | 0.256749 | 3.177543 | 0.647884 |
| 5 | 139455000 | 139458199 | 21.90861 | 37.05796 | 55.99792 | 24.50836 | -0.21219 | 1.072538 | 0.647892 |
| 13 | 60028400 | 60029599 | 10.67152 | 19.26693 | 32.23929 | 14.01637 | -0.32987 | 0.349591 | 0.647899 |
| 9 | 88448200 | 88457999 | 188.6028 | 280.1041 | 294.4593 | 121.3004 | 0.181295 | 3.655216 | 0.647924 |
| 3 | 103577600 | 103579399 | 14.94135 | 26.32809 | 41.31774 | 18.28786 | -0.25798 | 0.575944 | 0.647925 |
| 4 | 112558600 | 112567399 | 96.84232 | 151.997 | 146.2049 | 60.63704 | 0.374628 | 2.586679 | 0.647943 |
| 1 | 20501200 | 20503999 | 64.19529 | 100.2014 | 176.6108 | 73.28431 | -0.50421 | 2.670428 | 0.647985 |
| 5 | 76915000 | 76916599 | 14.95821 | 27.06682 | 29.79925 | 12.91935 | 0.272772 | 0.361435 | 0.647987 |
| 2 | 100639200 | 100641199 | 16.71112 | 29.2642 | 47.79797 | 21.09998 | -0.31699 | 0.681405 | 0.647995 |
| 13 | 28554800 | 28555599 | 9.962699 | 18.49568 | 29.92448 | 12.54337 | -0.24511 | 0.180367 | 0.648018 |

| | | | | | | | | | |
|---|---|---|---|---|---|---|---|---|---|
| 7 | 40944000 | 40946599 | 32.67102 | 54.59039 | 56.43055 | 24.56326 | 0.333435 | 1.222755 | 0.648021 |
| 9 | 36060800 | 36077999 | 217.8421 | 320.5958 | 292.2029 | 120.3305 | 0.374831 | 3.810558 | 0.648026 |
| 9 | 55763000 | 55765199 | 16.42685 | 29.52199 | 31.16036 | 13.61818 | 0.325733 | 0.413606 | 0.64803 |
| 6 | 58806800 | 58808999 | 18.77628 | 32.05598 | 50.59448 | 22.18163 | -0.27332 | 0.971511 | 0.648038 |
| 12 | 38981000 | 38981399 | 8.548274 | 16.55333 | 20.4354 | 7.765989 | 0.267126 | -0.19317 | 0.648056 |
| 4 | 52144400 | 52145199 | 11.98775 | 21.38798 | 34.20809 | 15.04212 | -0.27617 | 0.437165 | 0.648059 |
| 19 | 6702800 | 6703599 | 11.98775 | 21.38798 | 34.20809 | 15.04212 | -0.27617 | 0.437165 | 0.648059 |
| 15 | 93557600 | 93560799 | 30.30296 | 49.45517 | 74.41964 | 31.97866 | -0.21519 | 1.511581 | 0.648059 |
| 1 | 100361600 | 100362999 | 14.86955 | 27.46701 | 27.91878 | 11.78285 | 0.41352 | 0.240859 | 0.648061 |
| 19 | 12173600 | 12176199 | 28.77181 | 48.13772 | 53.5134 | 23.3126 | 0.228795 | 1.193196 | 0.648081 |
| 8 | 59388200 | 59389999 | 25.906 | 43.5935 | 48.55941 | 21.21949 | 0.225986 | 1.123748 | 0.648082 |
| X | 75670600 | 75673799 | 38.02389 | 62.5816 | 64.37988 | 27.75856 | 0.338171 | 1.415268 | 0.648126 |
| 5 | 47990600 | 47991999 | 19.29705 | 33.68366 | 37.77005 | 16.63164 | 0.223132 | 0.728999 | 0.648146 |
| 10 | 72030600 | 72032599 | 20.06765 | 35.27036 | 36.22673 | 16.04126 | 0.348972 | 0.622376 | 0.648151 |
| 9 | 77657600 | 77658599 | 7.971828 | 14.93117 | 26.95671 | 11.13638 | -0.38495 | 0.10067 | 0.648154 |
| 2 | 7751800 | 7759999 | 89.36495 | 140.221 | 146.3059 | 60.67664 | 0.25711 | 2.587848 | 0.648155 |
| 6 | 83873600 | 83874199 | 6.379325 | 12.38995 | 22.891 | 8.562805 | -0.28743 | -0.26414 | 0.648168 |
| 18 | 85855600 | 85857199 | 12.54683 | 24.03912 | 23.36603 | 9.210597 | 0.55922 | -0.0263 | 0.648173 |
| Y | 5263600 | 5265999 | 19.2167 | 33.4899 | 27.97702 | 12.30703 | 0.645459 | 0.732117 | 0.64818 |
| 10 | 98909600 | 98912599 | 29.99189 | 48.8198 | 75.90069 | 32.58404 | -0.26483 | 1.548454 | 0.648183 |
| Y | 86799800 | 86800999 | 10.09333 | 19.13054 | 17.72687 | 7.162596 | 0.596373 | 0.013753 | 0.648191 |
| 1 | 192501000 | 192503599 | 15.93343 | 27.8061 | 43.59614 | 19.19439 | -0.25841 | 0.722106 | 0.648196 |
| 14 | 26046000 | 26047199 | 15.93343 | 27.8061 | 43.59614 | 19.19439 | -0.25841 | 0.722106 | 0.648196 |
| 14 | 39739400 | 39744199 | 51.29911 | 82.6295 | 86.77003 | 36.8964 | 0.284894 | 1.831148 | 0.648209 |
| X | 77569000 | 77572199 | 35.76953 | 59.32232 | 60.48477 | 26.19842 | 0.352985 | 1.317223 | 0.648212 |
| 16 | 16452600 | 16456599 | 91.45496 | 142.3381 | 156.8201 | 65.09785 | 0.169595 | 2.729919 | 0.648219 |
| 12 | 110147600 | 110150399 | 16.59 | 28.88263 | 44.69668 | 19.64933 | -0.24001 | 0.761353 | 0.648223 |
| 1 | 13139200 | 13140999 | 13.12618 | 23.25634 | 36.17493 | 15.99409 | -0.24199 | 0.495647 | 0.648231 |
| 6 | 58112400 | 58113199 | 12.30577 | 22.90392 | 26.06078 | 10.88168 | 0.263234 | 0.1739 | 0.648238 |
| 14 | 41765200 | 41766599 | 27.6104 | 46.15222 | 51.26238 | 22.31904 | 0.230169 | 1.200077 | 0.648246 |
| 11 | 76276400 | 76278599 | 16.92969 | 29.84552 | 33.00043 | 14.60514 | 0.24499 | 0.567974 | 0.648271 |
| 6 | 145493400 | 145493799 | 10.93102 | 20.7169 | 24.28861 | 9.815264 | 0.259427 | 0.041951 | 0.64828 |
| 18 | 73652200 | 73654599 | 23.89188 | 40.73196 | 45.83241 | 20.09156 | 0.212684 | 0.999633 | 0.648294 |
| 8 | 61701400 | 61705999 | 56.31883 | 90.18266 | 98.3919 | 41.47368 | 0.219619 | 2.021792 | 0.648304 |
| 8 | 35155000 | 35157799 | 18.09592 | 31.01983 | 49.53563 | 21.7245 | -0.28925 | 0.929837 | 0.648308 |
| 6 | 139757800 | 139759399 | 22.61574 | 39.18998 | 42.4913 | 18.64286 | 0.270077 | 0.843348 | 0.648322 |
| 10 | 16696000 | 16697199 | 10.85149 | 19.17256 | 39.91079 | 17.65568 | -0.66143 | 0.519575 | 0.648325 |
| 8 | 70550000 | 70551799 | 15.96968 | 27.85235 | 43.10246 | 18.96975 | -0.23967 | 0.731318 | 0.648341 |
| 4 | 14245800 | 14246599 | 16.01164 | 28.48871 | 30.94987 | 13.64102 | 0.276456 | 0.475911 | 0.648354 |
| 3 | 3486600 | 3492999 | 119.4296 | 182.347 | 198.1491 | 82.1773 | 0.16817 | 3.102918 | 0.648387 |
| 11 | 20978800 | 20983799 | 59.61183 | 94.95137 | 105.8197 | 44.27258 | 0.180718 | 2.190785 | 0.648391 |
| 3 | 34688600 | 34691399 | 25.77024 | 42.70743 | 64.73551 | 28.02533 | -0.21744 | 1.300745 | 0.648408 |
| 9 | 58683200 | 58686199 | 20.05068 | 34.47574 | 53.91107 | 23.64762 | -0.25842 | 0.916201 | 0.64841 |
| 2 | 13156800 | 13165999 | 81.3911 | 128.6989 | 131.1751 | 54.50479 | 0.299483 | 2.41921 | 0.648443 |
| 14 | 60767800 | 60769999 | 19.24412 | 33.61504 | 37.7053 | 16.61052 | 0.222695 | 0.717851 | 0.648447 |
| 12 | 115911600 | 115919999 | 99.14544 | 155.1981 | 151.3238 | 62.77305 | 0.3521 | 2.634756 | 0.64845 |
| 15 | 7772600 | 7774599 | 17.73887 | 31.24733 | 34.29172 | 15.17903 | 0.255073 | 0.580776 | 0.64847 |

| | | | | | | | | | |
|---|---|---|---|---|---|---|---|---|---|
| X | 55285600 | 55286199 | 13.5013 | 24.82514 | 27.56687 | 11.70736 | 0.280794 | 0.258218 | 0.64848 |
| X | 126493400 | 126495799 | 22.38402 | 39.18187 | 31.43891 | 13.87345 | 0.703938 | 0.694839 | 0.64851 |
| 15 | 12997400 | 12998199 | 10.37933 | 19.61832 | 22.66139 | 9.210076 | 0.273409 | 0.050122 | 0.648545 |
| 6 | 13152800 | 13155399 | 20.92959 | 35.10729 | 63.46219 | 27.69165 | -0.47016 | 1.153624 | 0.648548 |
| 1 | 60830200 | 60832399 | 30.89316 | 51.16511 | 57.89307 | 25.0522 | 0.203044 | 1.316434 | 0.648582 |
| 1 | 143230400 | 143231799 | 17.41307 | 30.89492 | 33.23729 | 14.68407 | 0.287933 | 0.504975 | 0.648606 |
| 13 | 100836600 | 100838999 | 17.4777 | 30.3654 | 46.32464 | 20.33862 | -0.21972 | 0.799051 | 0.648607 |
| 17 | 29531800 | 29533399 | 15.73345 | 28.00441 | 31.35459 | 13.81495 | 0.233332 | 0.470602 | 0.64863 |
| 17 | 27024200 | 27026399 | 23.23054 | 39.18816 | 59.90468 | 26.19678 | -0.22901 | 1.104085 | 0.648637 |
| 1 | 69770600 | 69771399 | 9.515126 | 17.56278 | 29.3571 | 12.40287 | -0.29995 | 0.207899 | 0.648655 |
| 3 | 32948800 | 32951799 | 22.0623 | 38.43177 | 40.79619 | 17.94108 | 0.301487 | 0.774847 | 0.648662 |
| 13 | 80852800 | 80854799 | 15.9365 | 28.50918 | 31.20476 | 13.69063 | 0.26981 | 0.438655 | 0.648674 |
| 7 | 22509000 | 22537799 | 1007.217 | 1303.7 | 1156.665 | 484.7546 | 0.22896 | 5.807521 | 0.648729 |
| 5 | 129013600 | 129015199 | 10.51635 | 19.11081 | 31.28848 | 13.49268 | -0.29079 | 0.309876 | 0.64873 |
| 14 | 102687200 | 102690199 | 22.57508 | 39.09085 | 34.83229 | 15.28216 | 0.551859 | 0.840529 | 0.648735 |
| 2 | 97721200 | 97723399 | 15.10055 | 26.66322 | 43.95923 | 19.44905 | -0.32777 | 0.558138 | 0.648751 |
| 9 | 115780000 | 115781799 | 20.53004 | 35.59563 | 39.97951 | 17.54112 | 0.219665 | 0.826514 | 0.648781 |
| 1 | 10512400 | 10513199 | 9.661189 | 18.76781 | 21.21925 | 7.924468 | 0.416202 | -0.23711 | 0.648792 |
| 10 | 11351600 | 11358599 | 83.89676 | 130.1099 | 181.3954 | 75.33944 | -0.17365 | 2.79027 | 0.648817 |
| 8 | 65498800 | 65499599 | 7.770588 | 14.90339 | 25.74423 | 10.11815 | -0.25863 | -0.07273 | 0.648821 |
| X | 79743000 | 79744799 | 30.55912 | 51.29035 | 53.46656 | 23.33882 | 0.321323 | 1.169997 | 0.648834 |
| 17 | 53414000 | 53415199 | 19.21525 | 32.82789 | 51.10807 | 22.40804 | -0.25333 | 0.971164 | 0.648839 |
| 4 | 62793600 | 62796999 | 32.28936 | 52.47932 | 78.29292 | 33.59098 | -0.20708 | 1.578163 | 0.648841 |
| 2 | 100772800 | 100774799 | 16.5868 | 28.98981 | 45.69618 | 20.1389 | -0.26576 | 0.710624 | 0.648854 |
| 8 | 127714600 | 127719999 | 51.97599 | 82.79387 | 119.9446 | 50.18639 | -0.197 | 2.185003 | 0.64888 |
| 9 | 92310200 | 92312399 | 35.77624 | 58.69278 | 65.21713 | 28.08123 | 0.225426 | 1.448227 | 0.648887 |
| 4 | 134694200 | 134696199 | 17.66732 | 30.61792 | 46.87593 | 20.56665 | -0.22538 | 0.829044 | 0.648907 |
| 5 | 120967800 | 120970999 | 33.81992 | 55.57125 | 62.42087 | 26.89522 | 0.2107 | 1.419959 | 0.648913 |
| 5 | 126759200 | 126760399 | 10.11742 | 18.81591 | 30.4893 | 12.74798 | -0.24467 | 0.172714 | 0.648915 |
| 19 | 13730200 | 13731599 | 20.18087 | 35.13792 | 39.31377 | 17.28406 | 0.225865 | 0.769451 | 0.648945 |
| 7 | 61790200 | 61798399 | 169.7026 | 253.8692 | 268.8957 | 110.8884 | 0.180003 | 3.52149 | 0.648955 |
| 17 | 12274800 | 12281599 | 78.41661 | 123.2804 | 136.804 | 56.75026 | 0.170468 | 2.539955 | 0.648966 |
| 1 | 93883200 | 93891999 | 96.25472 | 149.0182 | 162.7999 | 67.62283 | 0.17586 | 2.823685 | 0.648999 |
| 5 | 10260600 | 10261999 | 23.3193 | 39.04384 | 59.18948 | 25.79727 | -0.21712 | 1.176896 | 0.649009 |
| 13 | 80274200 | 80274999 | 10.04413 | 19.11832 | 22.30137 | 8.930354 | 0.278666 | 0.000612 | 0.649037 |
| 9 | 38863600 | 38869799 | 63.63827 | 101.4226 | 109.1879 | 45.71779 | 0.23141 | 2.176232 | 0.649038 |
| 6 | 129923600 | 129925599 | 28.26052 | 46.60131 | 69.86718 | 30.14881 | -0.20261 | 1.369794 | 0.649046 |
| 18 | 54389800 | 54390999 | 10.62241 | 19.59158 | 31.41434 | 13.28584 | -0.24161 | 0.226162 | 0.649053 |
| 14 | 51964200 | 51967399 | 20.48651 | 35.2733 | 60.66172 | 26.60984 | -0.39477 | 0.914521 | 0.649062 |
| 4 | 105482200 | 105485199 | 32.20083 | 52.15315 | 91.13899 | 39.00203 | -0.43946 | 1.655294 | 0.649076 |
| X | 159084800 | 159086999 | 40.12396 | 65.08617 | 72.6956 | 31.14484 | 0.206905 | 1.630082 | 0.649077 |
| 6 | 35676000 | 35681199 | 48.913 | 78.13235 | 112.4702 | 47.24279 | -0.1834 | 2.09355 | 0.64909 |
| 8 | 41560000 | 41560999 | 9.009216 | 16.98561 | 28.06813 | 11.46702 | -0.24558 | 0.070611 | 0.649096 |
| 9 | 61520200 | 61522399 | 24.25191 | 41.29474 | 46.36745 | 20.32408 | 0.215477 | 1.013203 | 0.649103 |
| 7 | 125298000 | 125298999 | 16.39194 | 29.13761 | 32.29406 | 14.24489 | 0.246748 | 0.484828 | 0.649134 |
| 9 | 8254800 | 8256599 | 16.39194 | 29.13761 | 32.29406 | 14.24489 | 0.246748 | 0.484828 | 0.649134 |
| 13 | 120333000 | 120334799 | 26.06392 | 42.73651 | 69.19242 | 29.78622 | -0.31638 | 1.433831 | 0.649152 |

| | | | | | | | | | |
|---|---|---|---|---|---|---|---|---|---|
| X | 161479400 | 161482599 | 16.801 | 29.31466 | 47.32719 | 20.83464 | -0.30031 | 0.739227 | 0.649167 |
| 1 | 120433800 | 120435599 | 11.21372 | 20.21603 | 32.77812 | 14.27692 | -0.28613 | 0.366098 | 0.649169 |
| 12 | 91486600 | 91488199 | 21.79995 | 37.73917 | 42.00599 | 18.42938 | 0.232116 | 0.84785 | 0.649175 |
| 7 | 6186800 | 6193999 | 66.5128 | 104.5532 | 150.081 | 62.25695 | -0.20052 | 2.537636 | 0.649195 |
| 7 | 27577400 | 27593399 | 271.6438 | 390.5487 | 399.7476 | 165.0121 | 0.174881 | 4.169721 | 0.649208 |
| 17 | 30007600 | 30009599 | 19.99098 | 33.7807 | 53.76532 | 23.52413 | -0.28677 | 1.072009 | 0.649234 |
| 12 | 51027000 | 51028799 | 15.9843 | 28.81734 | 30.10569 | 13.11238 | 0.343548 | 0.388763 | 0.649238 |
| 6 | 83899600 | 83900399 | 13.66753 | 23.91346 | 45.17013 | 19.92526 | -0.52546 | 0.675635 | 0.649242 |
| 10 | 82614800 | 82616399 | 17.82446 | 31.27668 | 35.04315 | 15.49578 | 0.224308 | 0.634832 | 0.64925 |
| 1 | 53994600 | 53996599 | 15.18118 | 26.79115 | 45.58701 | 20.1714 | -0.37334 | 0.5686 | 0.64928 |
| 13 | 100259800 | 100260599 | 10.3806 | 19.78721 | 23.29886 | 9.301434 | 0.268404 | -0.00606 | 0.64929 |
| 4 | 10437600 | 10439999 | 22.25751 | 37.30672 | 58.69252 | 25.59827 | -0.27054 | 1.158419 | 0.649299 |
| X | 108806400 | 108809399 | 35.30516 | 59.18499 | 54.78375 | 23.89568 | 0.492201 | 1.197142 | 0.649339 |
| 14 | 48575000 | 48578399 | 26.95617 | 44.53828 | 66.93409 | 28.91687 | -0.20545 | 1.341348 | 0.649346 |
| X | 114733800 | 114735999 | 25.86351 | 44.3313 | 46.30635 | 20.30812 | 0.320799 | 0.961491 | 0.649354 |
| 5 | 98949600 | 98953399 | 32.00905 | 52.89562 | 59.64231 | 25.7696 | 0.207806 | 1.345366 | 0.649357 |
| 7 | 29498400 | 29501399 | 30.93459 | 50.57422 | 75.30568 | 32.38 | -0.19827 | 1.488336 | 0.649369 |
| 12 | 114954600 | 114956999 | 38.76657 | 63.22429 | 69.73162 | 29.95664 | 0.231148 | 1.536897 | 0.649413 |
| 11 | 40970600 | 40971999 | 21.62845 | 37.77356 | 34.44241 | 15.17149 | 0.520129 | 0.727395 | 0.649418 |
| 12 | 3669200 | 3674399 | 42.65013 | 70.09859 | 65.43297 | 28.19611 | 0.477247 | 1.443035 | 0.649425 |
| 16 | 3689800 | 3690999 | 10.9778 | 19.84786 | 32.23047 | 13.98786 | -0.28522 | 0.343996 | 0.649431 |
| 6 | 35560800 | 35562399 | 17.72933 | 31.66576 | 32.68478 | 14.3593 | 0.35282 | 0.464818 | 0.649467 |
| 19 | 28132600 | 28134799 | 34.08483 | 54.87203 | 99.79544 | 42.40174 | -0.50668 | 1.829782 | 0.649486 |
| 6 | 145877400 | 145884199 | 109.6379 | 166.707 | 231.8329 | 96.03964 | -0.19219 | 3.186383 | 0.649489 |
| 2 | 168864800 | 168867399 | 32.96991 | 53.45542 | 80.25125 | 34.36959 | -0.21937 | 1.62979 | 0.649501 |
| 12 | 114179800 | 114194999 | 190.8077 | 285.7416 | 269.6442 | 111.2198 | 0.347893 | 3.506272 | 0.649517 |
| 17 | 18907400 | 18909799 | 22.23431 | 39.19063 | 35.53968 | 15.72946 | 0.529406 | 0.604225 | 0.649527 |
| 8 | 20058800 | 20064399 | 252.8069 | 362.4963 | 479.8296 | 198.1522 | -0.19979 | 4.207548 | 0.649527 |
| 3 | 123087200 | 123089199 | 15.41296 | 26.90722 | 42.09672 | 18.53831 | -0.25498 | 0.715745 | 0.649536 |
| 15 | 83692800 | 83694799 | 25.77435 | 42.37566 | 76.04944 | 32.77538 | -0.46298 | 1.406458 | 0.649538 |
| 9 | 38845400 | 38846799 | 26.60214 | 45.19387 | 42.14501 | 18.46986 | 0.481895 | 1.037149 | 0.649539 |
| 4 | 113299400 | 113316399 | 232.0312 | 342.528 | 280.4126 | 115.479 | 0.533783 | 3.762041 | 0.649547 |
| 17 | 44496400 | 44497999 | 13.01168 | 23.79848 | 26.96952 | 11.53651 | 0.243908 | 0.280318 | 0.649552 |
| 2 | 97944200 | 97948599 | 39.39116 | 64.98818 | 54.43813 | 23.49591 | 0.635029 | 1.379934 | 0.649562 |
| 7 | 20756000 | 20758599 | 38.78687 | 62.85667 | 66.06911 | 28.28577 | 0.293052 | 1.643521 | 0.649569 |
| 4 | 47467400 | 47472599 | 102.359 | 158.0342 | 172.6481 | 71.71456 | 0.17285 | 2.878743 | 0.649583 |
| Y | 5884200 | 5885799 | 13.06373 | 23.83263 | 20.16685 | 8.6544 | 0.659426 | 0.277093 | 0.649614 |
| Y | 87466200 | 87468199 | 13.06373 | 23.83263 | 20.16685 | 8.6544 | 0.659426 | 0.277093 | 0.649614 |
| 17 | 87886800 | 87895399 | 101.3562 | 155.4752 | 214.2266 | 88.93194 | -0.16883 | 3.005574 | 0.649656 |
| 3 | 49083400 | 49095199 | 168.6649 | 254.088 | 250.3591 | 103.4203 | 0.293225 | 3.387925 | 0.649669 |
| X | 123022800 | 123038399 | 292.3425 | 421.6277 | 400.3901 | 165.2036 | 0.28747 | 4.126174 | 0.649676 |
| 5 | 93085200 | 93091999 | 197.3533 | 283.6569 | 484.2162 | 199.8891 | -0.56314 | 4.170619 | 0.649719 |
| 7 | 47620200 | 47621199 | 12.1052 | 23.37123 | 23.18124 | 8.933292 | 0.561109 | -0.10088 | 0.64973 |
| 18 | 85714200 | 85720599 | 78.06571 | 122.9977 | 135.7157 | 56.31485 | 0.18109 | 2.496577 | 0.649746 |
| 1 | 50408200 | 50413799 | 75.08838 | 117.687 | 168.9763 | 70.08677 | -0.20421 | 2.600476 | 0.649746 |
| 13 | 77913200 | 77919199 | 75.81945 | 120.277 | 121.8815 | 50.75427 | 0.311811 | 2.32951 | 0.649748 |
| 16 | 93201400 | 93211199 | 61.40611 | 96.31994 | 166.6196 | 69.10357 | -0.4717 | 2.57977 | 0.649763 |

| | | | | | | | | | |
|---|---|---|---|---|---|---|---|---|---|
| Y | 6853000 | 6855799 | 22.27868 | 38.25 | 31.15418 | 13.6648 | 0.678354 | 0.906767 | 0.649767 |
| 3 | 61680800 | 61681599 | 7.584791 | 14.55417 | 25.20497 | 9.892279 | -0.26064 | -0.08237 | 0.649771 |
| 4 | 142718000 | 142718799 | 7.584791 | 14.55417 | 25.20497 | 9.892279 | -0.26064 | -0.08237 | 0.649771 |
| X | 157740600 | 157744399 | 47.31016 | 76.5086 | 79.08437 | 33.8044 | 0.314286 | 1.708625 | 0.649778 |
| 7 | 28909000 | 28925799 | 226.0366 | 323.6953 | 499.5143 | 206.3227 | -0.42288 | 4.225931 | 0.649797 |
| 2 | 110296800 | 110300599 | 52.22477 | 84.04151 | 88.59749 | 37.62222 | 0.277602 | 1.861294 | 0.649809 |
| 7 | 61788800 | 61798599 | 182.8444 | 272.3048 | 285.5478 | 117.6625 | 0.188565 | 3.606247 | 0.649811 |
| X | 30856400 | 30876799 | 1003.054 | 1303.87 | 1089.741 | 456.5147 | 0.321286 | 5.738533 | 0.649838 |
| 4 | 46848200 | 46851399 | 40.16312 | 65.14602 | 72.96638 | 31.25956 | 0.202757 | 1.629068 | 0.649869 |
| 2 | 19249200 | 19252199 | 20.89545 | 35.53099 | 53.82647 | 23.59097 | -0.21536 | 1.020346 | 0.6499 |
| 10 | 39263600 | 39267999 | 55.65137 | 88.83581 | 98.16008 | 41.19228 | 0.196419 | 2.116049 | 0.649902 |
| 10 | 19519800 | 19522199 | 18.08218 | 31.43107 | 49.00385 | 21.52157 | -0.25102 | 0.79702 | 0.649908 |
| 9 | 13972800 | 13978599 | 89.71055 | 138.434 | 192.008 | 79.75646 | -0.17175 | 2.885059 | 0.649923 |
| 5 | 14625600 | 14626399 | 7.788363 | 14.72441 | 25.99785 | 10.56099 | -0.33287 | 0.035254 | 0.649987 |
| 10 | 24325600 | 24328199 | 21.5409 | 37.70713 | 38.46143 | 16.97278 | 0.359092 | 0.697353 | 0.649992 |
| 4 | 123730400 | 123749799 | 340.0727 | 470.1905 | 704.626 | 292.6909 | -0.43167 | 4.752283 | 0.649994 |
| 18 | 15947800 | 15950599 | 18.55388 | 31.95369 | 49.00669 | 21.49742 | -0.22995 | 0.889311 | 0.650011 |
| X | 149156200 | 149160799 | 71.33166 | 114.1291 | 99.27479 | 41.79641 | 0.544641 | 2.066574 | 0.650021 |
| 7 | 95965400 | 95967199 | 31.92645 | 52.65537 | 59.49647 | 25.67554 | 0.204357 | 1.369458 | 0.65003 |
| 2 | 163635200 | 163639199 | 46.25788 | 74.42206 | 83.04948 | 35.25437 | 0.195283 | 1.857394 | 0.650037 |
| 2 | 107631200 | 107635599 | 40.20153 | 66.36981 | 62.56029 | 27.01238 | 0.465466 | 1.380682 | 0.650066 |
| 13 | 88859200 | 88861799 | 19.44676 | 33.52452 | 51.70944 | 22.6837 | -0.23783 | 0.889161 | 0.650078 |
| 5 | 13787600 | 13787999 | 4.802967 | 9.375264 | 18.70304 | 6.623266 | -0.32322 | -0.49074 | 0.650092 |
| 11 | 73781400 | 73785399 | 45.91496 | 74.62293 | 73.89213 | 31.70228 | 0.382754 | 1.603692 | 0.650094 |
| 17 | 9496800 | 9497999 | 8.52113 | 16.18812 | 27.15981 | 10.92034 | -0.24713 | 0.011039 | 0.650097 |
| 16 | 21917200 | 21921799 | 66.20505 | 104.9526 | 116.0546 | 48.29448 | 0.185097 | 2.341805 | 0.650098 |
| 5 | 136436000 | 136437199 | 17.16855 | 30.16111 | 33.53835 | 14.84508 | 0.235176 | 0.609885 | 0.650102 |
| 13 | 26255000 | 26256599 | 18.75188 | 33.19227 | 29.55197 | 13.06834 | 0.557947 | 0.527526 | 0.650103 |
| 12 | 84921200 | 84924199 | 28.10209 | 45.71802 | 74.89076 | 32.14391 | -0.34051 | 1.552735 | 0.650107 |
| 18 | 31413400 | 31414599 | 18.26929 | 32.04686 | 35.12284 | 15.52678 | 0.256125 | 0.644491 | 0.650111 |
| 6 | 140746800 | 140748199 | 16.99988 | 30.40282 | 32.18997 | 14.12627 | 0.317324 | 0.450298 | 0.650118 |
| X | 158770200 | 158773599 | 31.34201 | 52.73149 | 51.81038 | 22.63886 | 0.406514 | 1.150324 | 0.650156 |
| X | 91038200 | 91041799 | 32.76341 | 55.48859 | 50.7349 | 22.21342 | 0.510147 | 1.0951 | 0.650179 |
| 5 | 144377600 | 144381999 | 45.27061 | 72.54477 | 104.8847 | 44.26012 | -0.18452 | 1.988849 | 0.650188 |
| 14 | 42972800 | 42975799 | 38.96941 | 63.58895 | 69.43398 | 29.83456 | 0.246075 | 1.527519 | 0.650204 |
| 9 | 5814800 | 5819799 | 47.21033 | 76.60287 | 75.69064 | 32.43478 | 0.383649 | 1.638756 | 0.650205 |
| 14 | 35124600 | 35126399 | 16.79939 | 28.904 | 53.77022 | 23.5863 | -0.50745 | 0.899953 | 0.65022 |
| 10 | 8576200 | 8577199 | 18.79918 | 31.88686 | 52.63843 | 23.05744 | -0.33877 | 1.037746 | 0.650228 |
| 1 | 168969600 | 168971799 | 22.73799 | 39.92302 | 37.22306 | 16.46827 | 0.488109 | 0.656476 | 0.650241 |
| 6 | 121695000 | 121698999 | 34.37774 | 57.13118 | 59.31317 | 25.72736 | 0.327174 | 1.288578 | 0.650257 |
| 9 | 55773600 | 55775199 | 16.73456 | 30.58408 | 29.23385 | 12.5174 | 0.486907 | 0.316728 | 0.650263 |
| 4 | 67777000 | 67779999 | 27.85542 | 47.37981 | 43.08442 | 18.88341 | 0.518287 | 1.031107 | 0.650273 |
| 1 | 35932000 | 35937599 | 94.11896 | 144.856 | 201.759 | 83.79454 | -0.1807 | 2.93625 | 0.650293 |
| 6 | 36364600 | 36367999 | 32.74373 | 52.80921 | 96.42052 | 41.0712 | -0.50914 | 1.768812 | 0.650296 |
| 16 | 93298600 | 93306999 | 114.3135 | 175.5453 | 189.3689 | 78.62469 | 0.185165 | 2.98619 | 0.6503 |
| 6 | 23187400 | 23188399 | 9.765502 | 18.775 | 22.42216 | 8.751278 | 0.278159 | -0.08708 | 0.650307 |
| X | 5379800 | 5381799 | 31.88141 | 54.01815 | 40.53391 | 17.74941 | 0.794329 | 1.080998 | 0.650308 |

| | | | | | | | | | |
|---|---|---|---|---|---|---|---|---|---|
| 17 | 13309000 | 13317399 | 80.17829 | 124.2122 | 178.5623 | 74.16987 | -0.21898 | 2.806917 | 0.650335 |
| 5 | 40209200 | 40210399 | 17.48205 | 30.66847 | 34.09139 | 15.07186 | 0.235673 | 0.635119 | 0.650405 |
| 3 | 103286800 | 103292999 | 63.48217 | 99.26905 | 172.6837 | 71.64134 | -0.48323 | 2.637343 | 0.650421 |
| 3 | 10512800 | 10526199 | 235.4815 | 343.0026 | 356.799 | 147.0426 | 0.170036 | 3.972545 | 0.650427 |
| 7 | 54175200 | 54177599 | 17.92422 | 32.19534 | 31.5536 | 13.79644 | 0.431665 | 0.427991 | 0.650431 |
| 9 | 77230000 | 77231399 | 16.42948 | 29.89338 | 30.35566 | 13.07017 | 0.395253 | 0.343528 | 0.650433 |
| 14 | 118211200 | 118212799 | 14.84638 | 26.03978 | 40.35445 | 17.83789 | -0.24139 | 0.633584 | 0.65044 |
| 5 | 34817200 | 34818399 | 11.69062 | 20.79762 | 34.24673 | 15.09588 | -0.32034 | 0.453414 | 0.650454 |
| 12 | 15539000 | 15539599 | 9.054623 | 17.50374 | 21.49877 | 8.235971 | 0.263628 | -0.15661 | 0.650481 |
| 14 | 117852800 | 117855399 | 19.32137 | 34.08354 | 34.57998 | 15.30226 | 0.368474 | 0.574506 | 0.65049 |
| 10 | 49821800 | 49822599 | 10.09157 | 18.72752 | 30.44383 | 12.76794 | -0.25265 | 0.183901 | 0.650494 |
| 13 | 30564400 | 30566199 | 22.78546 | 38.19768 | 58.1208 | 25.35145 | -0.22248 | 1.159687 | 0.650499 |
| 6 | 51640200 | 51643399 | 46.25432 | 74.48178 | 83.4255 | 35.45914 | 0.191491 | 1.830021 | 0.650504 |
| 2 | 178102000 | 178104999 | 28.74334 | 47.27948 | 70.95736 | 30.58698 | -0.20556 | 1.401798 | 0.650534 |
| 8 | 15136800 | 15140399 | 31.97257 | 52.94587 | 59.1504 | 25.59397 | 0.22129 | 1.320542 | 0.650542 |
| 11 | 31162800 | 31165599 | 62.80645 | 99.88152 | 111.5459 | 46.56614 | 0.175284 | 2.246509 | 0.650551 |
| 8 | 103565200 | 103566199 | 17.37634 | 30.99725 | 32.70033 | 14.381 | 0.320487 | 0.469856 | 0.650553 |
| 10 | 105011800 | 105013399 | 17.37634 | 30.99725 | 32.70033 | 14.381 | 0.320487 | 0.469856 | 0.650553 |
| 19 | 39708400 | 39712799 | 55.13004 | 88.46286 | 94.42627 | 39.92819 | 0.254688 | 1.956857 | 0.650555 |
| 7 | 23965200 | 23975999 | 117.7665 | 178.5092 | 244.8302 | 101.3291 | -0.17592 | 3.25271 | 0.650557 |
| 8 | 40738400 | 40740199 | 16.3596 | 28.1955 | 46.38213 | 20.34658 | -0.33024 | 0.870489 | 0.650561 |
| 7 | 121054000 | 121056199 | 20.00402 | 34.7146 | 38.44872 | 16.86994 | 0.240005 | 0.813692 | 0.650562 |
| 4 | 146451400 | 146454399 | 120.919 | 185.1167 | 168.4158 | 69.8895 | 0.427284 | 3.04724 | 0.650564 |
| X | 152854200 | 152857199 | 52.12958 | 84.08397 | 83.99944 | 35.78537 | 0.359126 | 1.792597 | 0.650589 |
| 19 | 9403400 | 9404999 | 17.84855 | 31.40259 | 34.90965 | 15.45628 | 0.236019 | 0.597156 | 0.650592 |
| 13 | 34654600 | 34655799 | 14.78516 | 26.50846 | 29.45888 | 12.90042 | 0.250205 | 0.416221 | 0.650611 |
| 7 | 38563000 | 38567799 | 133.412 | 202.2353 | 217.4072 | 89.98089 | 0.175297 | 3.254052 | 0.650625 |
| 5 | 8355200 | 8357799 | 24.49022 | 42.56413 | 42.41316 | 18.62552 | 0.392133 | 0.821101 | 0.650628 |
| 2 | 36697000 | 36698599 | 31.18734 | 52.05798 | 47.06699 | 20.47423 | 0.526141 | 1.22242 | 0.650634 |
| 1 | 82902400 | 82920999 | 251.8031 | 360.3213 | 469.2367 | 193.8396 | -0.17921 | 4.237563 | 0.650661 |
| 5 | 89392000 | 89393399 | 19.44627 | 34.26608 | 30.68994 | 13.58384 | 0.547056 | 0.582906 | 0.650697 |
| 10 | 62043000 | 62044799 | 15.8896 | 27.49819 | 44.81876 | 19.66323 | -0.3156 | 0.830303 | 0.650701 |
| 14 | 49598600 | 49600799 | 16.45779 | 29.4639 | 31.69055 | 13.89494 | 0.295558 | 0.438803 | 0.650709 |
| 10 | 70340600 | 70344999 | 39.48551 | 63.57615 | 93.89509 | 39.90364 | -0.2068 | 1.827551 | 0.650716 |
| X | 32016600 | 32017599 | 10.96624 | 21.21468 | 22.72224 | 8.674919 | 0.463955 | -0.1446 | 0.650716 |
| 10 | 105565200 | 105566999 | 15.552 | 27.39231 | 45.55738 | 20.16612 | -0.34182 | 0.589278 | 0.650725 |
| 15 | 10516600 | 10517999 | 8.254967 | 15.41736 | 27.9347 | 11.5964 | -0.39618 | 0.122248 | 0.65075 |
| 8 | 71652800 | 71655799 | 28.71801 | 48.3961 | 52.36315 | 22.89179 | 0.267841 | 1.124181 | 0.650756 |
| X | 103389200 | 103392799 | 30.64085 | 50.18629 | 75.33559 | 32.41575 | -0.2083 | 1.463737 | 0.65077 |
| 6 | 123335800 | 123337999 | 41.71826 | 67.03265 | 98.36184 | 41.67707 | -0.20117 | 1.899703 | 0.650787 |
| 17 | 38264800 | 38266199 | 16.96539 | 29.96133 | 32.96114 | 14.5826 | 0.253254 | 0.549368 | 0.650797 |
| 2 | 83300800 | 83303799 | 31.58545 | 53.06696 | 53.57634 | 23.4006 | 0.36719 | 1.159003 | 0.650799 |
| 17 | 90105600 | 90107599 | 19.34421 | 33.98889 | 36.37089 | 16.10111 | 0.290245 | 0.624465 | 0.650811 |
| 8 | 126609000 | 126610399 | 12.79457 | 22.61343 | 36.22849 | 16.0256 | -0.28576 | 0.519642 | 0.650815 |
| 14 | 18286000 | 18288599 | 21.86357 | 35.76247 | 78.3051 | 33.67595 | -0.7522 | 1.464848 | 0.650836 |
| 17 | 24034000 | 24036399 | 71.50178 | 112.8305 | 122.0494 | 50.66051 | 0.211428 | 2.45693 | 0.650859 |
| 6 | 89852400 | 89857999 | 67.85238 | 108.0395 | 113.3028 | 47.38054 | 0.267756 | 2.211292 | 0.650878 |

| | | | | | | | | | |
|---|---|---|---|---|---|---|---|---|---|
| 19 | 9379200 | 9387399 | 78.59784 | 124.6878 | 123.5882 | 51.46638 | 0.343726 | 2.333955 | 0.650886 |
| 1 | 120462600 | 120463599 | 10.69523 | 20.28326 | 23.73725 | 9.579217 | 0.263547 | 0.032755 | 0.650887 |
| 2 | 65570800 | 65572999 | 23.36986 | 40.30636 | 44.13836 | 19.36272 | 0.254489 | 0.891485 | 0.650892 |
| 4 | 144581000 | 144582999 | 38.49281 | 63.18824 | 58.13445 | 25.03807 | 0.497348 | 1.441974 | 0.650901 |
| 7 | 47519400 | 47522399 | 46.77208 | 75.28494 | 84.50076 | 35.89504 | 0.187778 | 1.844231 | 0.650907 |
| 11 | 96416000 | 96420399 | 34.92749 | 56.5125 | 83.74934 | 35.81317 | -0.20356 | 1.676565 | 0.650926 |
| 1 | 44463800 | 44465599 | 16.70388 | 30.51881 | 29.83782 | 12.78082 | 0.454334 | 0.320588 | 0.650927 |
| Y | 3931000 | 3934399 | 24.60391 | 41.74891 | 33.74977 | 14.7853 | 0.686976 | 1.026085 | 0.650932 |
| 15 | 93066400 | 93067999 | 28.14524 | 46.13313 | 70.56601 | 30.37305 | -0.23489 | 1.444131 | 0.650939 |
| 4 | 143445800 | 143465799 | 363.6803 | 513.9295 | 487.9123 | 201.9741 | 0.258416 | 4.426308 | 0.65097 |
| 15 | 25245600 | 25246399 | 13.46572 | 23.73616 | 37.2805 | 16.49976 | -0.25875 | 0.556334 | 0.650972 |
| 10 | 42206200 | 42208399 | 16.43378 | 28.6674 | 44.12766 | 19.42335 | -0.23205 | 0.731879 | 0.650973 |
| 10 | 43881600 | 43882799 | 9.663073 | 17.78842 | 29.83551 | 12.65075 | -0.30874 | 0.224586 | 0.650981 |
| 5 | 119080200 | 119082599 | 18.18254 | 31.4828 | 48.46907 | 21.26506 | -0.23367 | 0.841765 | 0.650987 |
| X | 41578200 | 41579799 | 24.39559 | 41.84294 | 44.74445 | 19.62418 | 0.28728 | 0.94855 | 0.650988 |
| 14 | 75020400 | 75021199 | 13.29624 | 23.65093 | 36.67322 | 16.16725 | -0.23436 | 0.470228 | 0.650991 |
| 11 | 14664400 | 14665999 | 16.02238 | 28.55136 | 31.78258 | 13.99002 | 0.242498 | 0.465548 | 0.651007 |
| 17 | 93067200 | 93068199 | 16.02238 | 28.55136 | 31.78258 | 13.99002 | 0.242498 | 0.465548 | 0.651007 |
| 4 | 155531800 | 155532799 | 9.140401 | 17.11441 | 28.47357 | 11.76942 | -0.26911 | 0.115435 | 0.651015 |
| 8 | 112891000 | 112899399 | 92.65897 | 144.5173 | 137.4531 | 57.04046 | 0.383808 | 2.691932 | 0.65103 |
| 3 | 77610600 | 77614199 | 24.73046 | 40.88903 | 72.96729 | 31.53833 | -0.45248 | 1.332182 | 0.651031 |
| 7 | 29569400 | 29587799 | 419.3375 | 581.4689 | 574.7225 | 238.6208 | 0.172838 | 4.709502 | 0.651044 |
| 7 | 39200600 | 39202399 | 18.77228 | 32.77432 | 35.64793 | 15.6992 | 0.266725 | 0.717518 | 0.651052 |
| 10 | 120600200 | 120602999 | 29.46124 | 48.10662 | 73.89186 | 31.75793 | -0.24421 | 1.500689 | 0.651065 |
| 13 | 13453800 | 13454599 | 11.44374 | 20.75107 | 32.73727 | 14.15577 | -0.24039 | 0.331761 | 0.651072 |
| 4 | 152247200 | 152248599 | 13.56873 | 23.81993 | 39.33763 | 17.40143 | -0.33266 | 0.609407 | 0.651112 |
| 1 | 14449600 | 14453399 | 40.31339 | 65.59723 | 71.66606 | 30.76209 | 0.24245 | 1.571651 | 0.65113 |
| 4 | 119275400 | 119277799 | 15.67161 | 27.51194 | 42.90303 | 18.97886 | -0.25046 | 0.627946 | 0.651144 |
| 2 | 97995600 | 97996799 | 14.87439 | 26.6166 | 29.88287 | 13.10757 | 0.233779 | 0.428474 | 0.651146 |
| 1 | 59361800 | 59363399 | 16.16809 | 28.58253 | 31.88713 | 14.10455 | 0.233569 | 0.529193 | 0.651148 |
| 11 | 26623600 | 26625999 | 18.11003 | 32.00774 | 34.19371 | 15.12555 | 0.295542 | 0.547312 | 0.651149 |
| 13 | 77951200 | 77957599 | 90.48606 | 142.0879 | 145.6542 | 60.41233 | 0.283439 | 2.5732 | 0.65116 |
| X | 8585800 | 8588599 | 34.15521 | 56.89177 | 58.1156 | 25.24677 | 0.350651 | 1.263281 | 0.651172 |
| 8 | 8370200 | 8374599 | 35.41522 | 58.45981 | 62.24479 | 26.87357 | 0.290101 | 1.367833 | 0.651183 |
| 7 | 65297200 | 65299599 | 20.38564 | 34.95381 | 53.88983 | 23.63458 | -0.23878 | 0.942831 | 0.651186 |
| 7 | 132230000 | 132230799 | 9.058031 | 16.69686 | 29.36049 | 12.42678 | -0.37451 | 0.210874 | 0.651211 |
| 6 | 96946200 | 96946799 | 8.767125 | 16.95285 | 20.96165 | 8.019415 | 0.25615 | -0.1651 | 0.65122 |
| 9 | 38238600 | 38239199 | 12.58986 | 23.06326 | 25.11456 | 10.72003 | 0.303597 | 0.265444 | 0.651244 |
| 4 | 24159400 | 24160599 | 11.63862 | 21.78343 | 25.18705 | 10.41793 | 0.251531 | 0.13304 | 0.651246 |
| 11 | 109804000 | 109804399 | 11.63862 | 21.78343 | 25.18705 | 10.41793 | 0.251531 | 0.13304 | 0.651246 |
| 15 | 35124400 | 35125599 | 13.93221 | 25.17708 | 27.83376 | 12.08712 | 0.265284 | 0.356927 | 0.65125 |
| 7 | 107475200 | 107478399 | 29.13358 | 48.16525 | 77.69154 | 33.57843 | -0.30728 | 1.347939 | 0.651255 |
| 15 | 61151800 | 61154399 | 14.54628 | 26.51791 | 24.29496 | 10.43546 | 0.544878 | 0.304658 | 0.651257 |
| 13 | 78092600 | 78102999 | 137.102 | 208.6304 | 217.1067 | 89.96024 | 0.226704 | 3.177192 | 0.651279 |
| 17 | 66870200 | 66875399 | 50.32037 | 79.92367 | 136.0271 | 56.71405 | -0.43339 | 2.278662 | 0.651292 |
| 12 | 114690000 | 114704399 | 212.3706 | 314.6918 | 307.3965 | 126.6007 | 0.284187 | 3.698439 | 0.651292 |
| 12 | 47877800 | 47880999 | 55.25704 | 89.27315 | 66.89111 | 28.55656 | 0.775822 | 1.744657 | 0.651295 |

| | | | | | | | | | |
|---|---|---|---|---|---|---|---|---|---|
| 12 | 92206200 | 92206999 | 7.777456 | 14.89077 | 25.68646 | 10.14572 | -0.26328 | -0.05633 | 0.651302 |
| X | 40814800 | 40819599 | 54.12733 | 86.9441 | 77.50301 | 32.82396 | 0.51591 | 1.918582 | 0.651309 |
| 13 | 14983200 | 14985999 | 40.22246 | 64.70229 | 95.0177 | 40.32612 | -0.20027 | 1.860464 | 0.651326 |
| 10 | 14434600 | 14438199 | 53.66808 | 85.31684 | 123.5438 | 51.55531 | -0.19942 | 2.251966 | 0.651329 |
| 5 | 110073400 | 110087399 | 355.84 | 499.381 | 501.0934 | 207.6624 | 0.168509 | 4.527336 | 0.651334 |
| 14 | 50836600 | 50845199 | 104.1436 | 162.39 | 158.4879 | 65.77177 | 0.346387 | 2.706599 | 0.651334 |
| 8 | 75976000 | 75983199 | 89.7918 | 140.3161 | 152.255 | 63.17045 | 0.196071 | 2.658185 | 0.651341 |
| 3 | 120352200 | 120353199 | 15.10427 | 27.43866 | 25.28387 | 10.90801 | 0.53266 | 0.331617 | 0.651343 |
| 1 | 130828800 | 130829399 | 7.592799 | 14.54431 | 25.15059 | 9.920184 | -0.26518 | -0.06589 | 0.651369 |
| 5 | 137511600 | 137513799 | 12.18641 | 21.57288 | 35.26933 | 15.59635 | -0.31379 | 0.497281 | 0.65137 |
| 1 | 130799800 | 130801999 | 13.92703 | 24.67073 | 38.16274 | 16.87358 | -0.23427 | 0.506075 | 0.651395 |
| 13 | 110534400 | 110536799 | 24.91049 | 41.6008 | 62.64296 | 27.25814 | -0.20751 | 1.210892 | 0.651407 |
| 1 | 176676800 | 176681399 | 42.57839 | 68.45494 | 100.7697 | 42.7337 | -0.20477 | 1.880457 | 0.651431 |
| 6 | 70985200 | 70986199 | 10.97251 | 20.04288 | 31.80445 | 13.62698 | -0.24003 | 0.284652 | 0.651451 |
| 1 | 108904600 | 108905999 | 16.66664 | 29.528 | 32.7563 | 14.48272 | 0.242824 | 0.510296 | 0.651458 |
| 5 | 91789000 | 91789599 | 5.811168 | 11.28231 | 21.67266 | 8.131285 | -0.3456 | -0.25995 | 0.651469 |
| 9 | 36313000 | 36316599 | 33.41136 | 55.65244 | 58.11645 | 25.24704 | 0.318923 | 1.260135 | 0.651476 |
| 3 | 148514400 | 148515199 | 7.304742 | 13.99905 | 24.61363 | 9.695407 | -0.28688 | -0.07392 | 0.651478 |
| 6 | 31513400 | 31514599 | 7.304742 | 13.99905 | 24.61363 | 9.695407 | -0.28688 | -0.07392 | 0.651478 |
| 5 | 127533000 | 127535399 | 28.83059 | 47.36499 | 71.25392 | 30.69891 | -0.2096 | 1.41964 | 0.651481 |
| 17 | 37643000 | 37644399 | 21.07027 | 36.32472 | 39.5957 | 17.36971 | 0.261085 | 0.880263 | 0.651486 |
| 16 | 38702600 | 38703199 | 9.516966 | 18.17939 | 21.29295 | 8.462693 | 0.282196 | -0.03246 | 0.651518 |
| 14 | 94288600 | 94290199 | 20.17651 | 35.30637 | 38.02607 | 16.77568 | 0.280948 | 0.69462 | 0.651542 |
| X | 20513200 | 20516799 | 45.56622 | 73.84731 | 75.44983 | 32.30326 | 0.333964 | 1.661812 | 0.651552 |
| 18 | 9196600 | 9198599 | 13.05485 | 23.35326 | 36.67252 | 16.09004 | -0.24795 | 0.431318 | 0.651568 |
| 1 | 146697800 | 146699199 | 21.08892 | 35.70918 | 54.34737 | 23.7936 | -0.22238 | 1.058706 | 0.651574 |
| 4 | 119778200 | 119788999 | 137.9005 | 206.6535 | 283.4533 | 116.9312 | -0.19041 | 3.484135 | 0.651583 |
| 13 | 5249400 | 5252199 | 30.85395 | 51.5864 | 55.77898 | 24.2892 | 0.268723 | 1.210219 | 0.651609 |
| 6 | 82317400 | 82318799 | 16.26599 | 28.00561 | 46.40345 | 20.3553 | -0.34075 | 0.881651 | 0.651614 |
| X | 35590000 | 35594399 | 49.13359 | 79.29818 | 82.40441 | 35.13253 | 0.303331 | 1.770358 | 0.651626 |
| 1 | 62681800 | 62685199 | 28.02436 | 46.96214 | 52.91272 | 23.07417 | 0.209477 | 1.174678 | 0.651631 |
| 4 | 78749600 | 78753999 | 54.81335 | 87.24001 | 124.3189 | 51.95601 | -0.17457 | 2.216628 | 0.651638 |
| 5 | 92190000 | 92190999 | 7.643465 | 14.54781 | 25.56745 | 10.23972 | -0.30784 | -0.01221 | 0.651652 |
| 18 | 46692200 | 46697199 | 51.76019 | 82.29697 | 136.3972 | 56.92817 | -0.3937 | 2.248986 | 0.651663 |
| 10 | 99241400 | 99241799 | 6.104329 | 11.85689 | 22.26314 | 8.322091 | -0.30822 | -0.26711 | 0.651687 |
| 12 | 119294200 | 119294999 | 9.370732 | 17.26235 | 29.35272 | 12.43418 | -0.32723 | 0.217731 | 0.651716 |
| 6 | 27507000 | 27510199 | 68.20979 | 107.9556 | 119.2151 | 49.55837 | 0.185061 | 2.384641 | 0.651718 |
| 15 | 12766000 | 12766799 | 7.763612 | 14.91623 | 25.81107 | 10.09331 | -0.25434 | -0.08935 | 0.65172 |
| 7 | 45306800 | 45308799 | 31.19047 | 51.80599 | 57.98662 | 25.14246 | 0.218906 | 1.284117 | 0.651727 |
| 15 | 38924600 | 38928999 | 63.30089 | 100.7875 | 111.4768 | 46.61384 | 0.19101 | 2.20895 | 0.651729 |
| 19 | 46954400 | 46956199 | 11.9528 | 21.5178 | 33.6859 | 14.69845 | -0.2378 | 0.379633 | 0.651736 |
| 1 | 82496600 | 82498199 | 12.04481 | 21.80169 | 34.75522 | 15.06224 | -0.25771 | 0.349318 | 0.65175 |
| 15 | 63569600 | 63570199 | 9.978272 | 19.03467 | 22.81581 | 9.094063 | 0.245234 | -0.01505 | 0.651768 |
| 2 | 58762800 | 58764799 | 22.49212 | 38.53979 | 43.1797 | 18.93639 | 0.219908 | 0.944291 | 0.651785 |
| 15 | 100106800 | 100108399 | 10.16286 | 18.6442 | 30.32218 | 12.91744 | -0.26937 | 0.250269 | 0.651801 |
| 16 | 8104800 | 8106999 | 25.12742 | 42.95173 | 46.86358 | 20.54796 | 0.257556 | 0.982819 | 0.651812 |
| 1 | 144723200 | 144724199 | 14.50845 | 26.5697 | 28.31638 | 12.0918 | 0.333447 | 0.285948 | 0.651842 |

| | | | | | | | | | |
|---|---|---|---|---|---|---|---|---|---|
| 14 | 6395200 | 6398599 | 62.98003 | 99.41572 | 141.6804 | 58.81874 | -0.18543 | 2.448379 | 0.651844 |
| 12 | 20990600 | 21000799 | 135.5508 | 206.2588 | 192.7464 | 79.86506 | 0.381712 | 3.174658 | 0.651851 |
| 3 | 108242800 | 108243399 | 6.248533 | 12.13658 | 22.27657 | 8.329348 | -0.27654 | -0.26574 | 0.651892 |
| 19 | 9661400 | 9663199 | 26.19078 | 44.32128 | 48.52298 | 21.23976 | 0.251209 | 1.078369 | 0.651895 |
| 2 | 52299600 | 52301999 | 42.78175 | 69.90414 | 69.80429 | 30.00845 | 0.376089 | 1.519506 | 0.651917 |
| 8 | 57530200 | 57531199 | 16.51037 | 29.0544 | 31.11095 | 13.77382 | 0.29005 | 0.580533 | 0.651921 |
| 13 | 30615200 | 30617799 | 31.242 | 52.41501 | 54.65036 | 23.84939 | 0.320963 | 1.17534 | 0.651927 |
| 19 | 54164800 | 54167599 | 20.94247 | 35.41026 | 54.33272 | 23.77681 | -0.23414 | 1.070757 | 0.651932 |
| 11 | 50717800 | 50725799 | 93.51586 | 143.7823 | 200.3642 | 83.20371 | -0.18253 | 2.956033 | 0.65195 |
| X | 153493600 | 153497599 | 67.57049 | 107.3249 | 116.5441 | 48.59227 | 0.213956 | 2.28917 | 0.651964 |
| 14 | 32235000 | 32235799 | 8.233733 | 15.4834 | 26.96463 | 11.06461 | -0.32581 | 0.076325 | 0.651974 |
| 3 | 29974000 | 29975599 | 17.49841 | 30.45193 | 46.82872 | 20.58141 | -0.23095 | 0.77653 | 0.651975 |
| 8 | 9740600 | 9742599 | 13.53986 | 23.70482 | 44.64327 | 19.7032 | -0.52106 | 0.66553 | 0.65198 |
| 13 | 92685800 | 92693199 | 129.038 | 196.0467 | 212.2853 | 87.92051 | 0.167519 | 3.20594 | 0.651981 |
| 2 | 98038400 | 98038999 | 15.11288 | 26.98405 | 29.95817 | 13.16495 | 0.248159 | 0.445137 | 0.652001 |
| 15 | 29713400 | 29715199 | 20.42222 | 35.48417 | 39.94458 | 17.53843 | 0.216895 | 0.802029 | 0.652004 |
| 12 | 65450400 | 65452199 | 14.90849 | 26.05549 | 41.04741 | 18.08952 | -0.26479 | 0.694737 | 0.652011 |
| 4 | 105299200 | 105302199 | 12.48968 | 23.36573 | 18.99532 | 7.862918 | 0.754974 | 0.125836 | 0.652017 |
| 13 | 85861600 | 85862399 | 9.031472 | 16.95103 | 27.99248 | 11.52371 | -0.25337 | 0.099928 | 0.652037 |
| X | 162824800 | 162826799 | 15.3188 | 27.01036 | 45.03705 | 19.93155 | -0.34473 | 0.577619 | 0.652059 |
| 7 | 10404200 | 10406799 | 52.06965 | 83.56099 | 92.76871 | 39.23364 | 0.198273 | 1.947971 | 0.652061 |
| 3 | 99066600 | 99070799 | 36.83751 | 60.43442 | 66.04508 | 28.43785 | 0.249314 | 1.449526 | 0.652092 |
| 7 | 10105800 | 10125799 | 262.5852 | 385.5248 | 330.0701 | 135.9426 | 0.461807 | 3.854871 | 0.652162 |
| 11 | 70934200 | 70938799 | 44.06914 | 71.35525 | 67.99102 | 29.09006 | 0.433074 | 1.669251 | 0.652168 |
| 5 | 118539200 | 118545999 | 63.48537 | 100.2995 | 141.7756 | 58.88382 | -0.17266 | 2.42893 | 0.652168 |
| 5 | 23547800 | 23548199 | 7.118649 | 13.81706 | 25.92554 | 9.745771 | -0.31655 | -0.2297 | 0.652191 |
| 1 | 95854800 | 95856199 | 12.78127 | 23.97725 | 26.16618 | 10.78077 | 0.338395 | 0.128407 | 0.652191 |
| 6 | 73275000 | 73277799 | 29.06244 | 48.90235 | 52.40364 | 22.89896 | 0.281546 | 1.136153 | 0.652191 |
| 4 | 53909000 | 53910599 | 17.64551 | 31.35433 | 33.14263 | 14.62374 | 0.31433 | 0.49571 | 0.652214 |
| 2 | 3693200 | 3694399 | 10.93627 | 19.98027 | 31.78813 | 13.61694 | -0.2437 | 0.282254 | 0.652218 |
| 14 | 37974600 | 37975199 | 11.2771 | 20.70795 | 33.47321 | 14.24141 | -0.25954 | 0.260252 | 0.65222 |
| 3 | 104347000 | 104349599 | 15.38309 | 26.63766 | 44.80868 | 19.65917 | -0.36084 | 0.820986 | 0.652226 |
| 3 | 139188200 | 139190199 | 19.6166 | 34.83573 | 32.29775 | 14.2582 | 0.501813 | 0.508597 | 0.652238 |
| 13 | 11778600 | 11783599 | 57.76351 | 92.6794 | 84.24146 | 35.61618 | 0.485895 | 1.958301 | 0.652244 |
| 3 | 106828600 | 106830599 | 24.4267 | 40.88169 | 61.51126 | 26.80246 | -0.20639 | 1.185666 | 0.652245 |
| 14 | 77582400 | 77586799 | 33.10909 | 53.93732 | 80.4412 | 34.54341 | -0.20421 | 1.547484 | 0.652252 |
| 4 | 122609000 | 122616399 | 93.96398 | 147.0548 | 150.5099 | 62.43683 | 0.281941 | 2.633485 | 0.652256 |
| 6 | 149083800 | 149084399 | 9.670799 | 18.08052 | 29.52896 | 12.23654 | -0.24531 | 0.134032 | 0.652257 |
| 5 | 18957600 | 18963199 | 59.89805 | 95.69014 | 103.8153 | 43.61413 | 0.224011 | 2.097576 | 0.652263 |
| 4 | 63363600 | 63365199 | 19.14942 | 33.45714 | 37.13108 | 16.35987 | 0.237909 | 0.712014 | 0.652266 |
| 5 | 56626600 | 56629599 | 32.53275 | 54.20256 | 49.75613 | 21.61856 | 0.504346 | 1.248512 | 0.652269 |
| 6 | 29963800 | 29964399 | 7.60312 | 14.61341 | 25.25434 | 9.864862 | -0.25099 | -0.09628 | 0.652276 |
| 2 | 170381200 | 170382399 | 12.15288 | 21.60179 | 34.70265 | 15.30769 | -0.28567 | 0.462462 | 0.652283 |
| 7 | 55310800 | 55312799 | 17.61645 | 31.4187 | 32.61616 | 14.34663 | 0.343278 | 0.471506 | 0.652287 |
| 2 | 15769200 | 15770999 | 19.57697 | 34.32367 | 31.31159 | 13.83613 | 0.519286 | 0.649756 | 0.652327 |
| 6 | 149127800 | 149130599 | 47.64595 | 77.00484 | 70.58094 | 30.146 | 0.486269 | 1.718716 | 0.652335 |
| 12 | 107826800 | 107831199 | 37.83335 | 61.51766 | 68.95008 | 29.58855 | 0.204926 | 1.574986 | 0.652349 |

| | | | | | | | | | |
|---|---|---|---|---|---|---|---|---|---|
| 8 | 48860400 | 48865599 | 57.30435 | 91.66757 | 100.5856 | 42.33639 | 0.209758 | 2.054369 | 0.652362 |
| 2 | 73078400 | 73079199 | 11.84185 | 22.50752 | 24.87053 | 9.989701 | 0.351655 | 0.031472 | 0.652372 |
| 3 | 48911800 | 48916399 | 105.5384 | 162.1915 | 174.5527 | 72.48056 | 0.189378 | 2.966813 | 0.652392 |
| 11 | 15880000 | 15880599 | 8.046198 | 15.64275 | 19.51772 | 7.243164 | 0.28471 | -0.27984 | 0.652401 |
| 4 | 122653800 | 122655599 | 55.11023 | 87.26466 | 131.3041 | 54.59706 | -0.26001 | 2.367792 | 0.652401 |
| 10 | 126682200 | 126687799 | 54.93474 | 87.44277 | 125.405 | 52.41785 | -0.18359 | 2.212748 | 0.652423 |
| 3 | 4412000 | 4415999 | 31.35247 | 52.36837 | 56.36074 | 24.52731 | 0.275464 | 1.222548 | 0.652483 |
| 8 | 34351000 | 34354999 | 41.88896 | 67.84677 | 75.76674 | 32.42452 | 0.205094 | 1.664028 | 0.652532 |
| 13 | 57180400 | 57183399 | 28.82779 | 48.26752 | 53.43646 | 23.29008 | 0.234835 | 1.189001 | 0.652534 |
| 2 | 161383800 | 161386599 | 23.0943 | 39.85124 | 43.58132 | 19.11865 | 0.256497 | 0.883704 | 0.652539 |
| 10 | 81416200 | 81417999 | 15.37818 | 26.7326 | 43.21892 | 18.9817 | -0.30284 | 0.776382 | 0.652544 |
| 14 | 121902800 | 121903999 | 14.42282 | 25.98046 | 28.77729 | 12.54694 | 0.258789 | 0.380659 | 0.652561 |
| 3 | 7728600 | 7733399 | 48.97542 | 78.95569 | 84.81426 | 36.1085 | 0.253746 | 1.800495 | 0.652565 |
| 10 | 74275000 | 74275599 | 7.451876 | 14.39417 | 25.46699 | 9.791874 | -0.26309 | -0.14898 | 0.652641 |
| 13 | 58559000 | 58561599 | 39.00453 | 63.61632 | 69.92701 | 30.04126 | 0.236047 | 1.535066 | 0.652644 |
| 6 | 137025200 | 137028399 | 42.50691 | 68.57594 | 75.55476 | 32.19573 | 0.218455 | 1.765454 | 0.652658 |
| 2 | 43018200 | 43019999 | 14.88462 | 26.98793 | 29.88205 | 12.92267 | 0.267078 | 0.350807 | 0.652663 |
| 16 | 62845400 | 62847799 | 18.71421 | 32.72378 | 36.64055 | 16.15454 | 0.225239 | 0.69677 | 0.652685 |
| 17 | 23628200 | 23629199 | 13.40662 | 24.55489 | 27.38028 | 11.69012 | 0.268831 | 0.275216 | 0.652704 |
| 14 | 99768800 | 99770599 | 26.30389 | 44.68268 | 41.07763 | 18.00154 | 0.502328 | 1.034052 | 0.65271 |
| 6 | 118544800 | 118550599 | 47.21255 | 75.72809 | 114.4598 | 48.37006 | -0.24693 | 1.964248 | 0.652714 |
| 10 | 89234000 | 89240399 | 76.04654 | 119.9795 | 132.8442 | 55.13591 | 0.177616 | 2.467282 | 0.652728 |
| 13 | 82275600 | 82276599 | 6.742109 | 12.90494 | 24.59815 | 9.723529 | -0.40624 | -0.06505 | 0.652734 |
| 11 | 86750800 | 86752799 | 16.73447 | 28.94946 | 53.85433 | 23.62712 | -0.50592 | 0.848964 | 0.652752 |
| 1 | 31011600 | 31013799 | 20.9831 | 36.78483 | 37.93754 | 16.76243 | 0.343251 | 0.671546 | 0.652752 |
| 6 | 109554800 | 109559799 | 63.22638 | 101.1181 | 102.9046 | 43.29865 | 0.317908 | 2.069981 | 0.652758 |
| 8 | 87141800 | 87143599 | 18.8458 | 33.47991 | 33.57799 | 14.81853 | 0.389593 | 0.505841 | 0.652758 |
| 16 | 71888600 | 71892599 | 36.9455 | 60.96523 | 63.3363 | 27.3387 | 0.325479 | 1.379278 | 0.652766 |
| 10 | 33776200 | 33778799 | 28.25009 | 48.15001 | 49.02354 | 21.49002 | 0.356349 | 1.022237 | 0.652782 |
| 13 | 7855000 | 7858799 | 29.55766 | 50.94109 | 43.95951 | 19.28383 | 0.596622 | 0.924104 | 0.652794 |
| 3 | 72686600 | 72687599 | 9.949743 | 18.51357 | 29.96047 | 12.51599 | -0.24179 | 0.166633 | 0.652816 |
| 11 | 7644400 | 7645399 | 13.57957 | 24.00186 | 37.17958 | 16.44618 | -0.2375 | 0.525778 | 0.652835 |
| 17 | 22854400 | 22855199 | 9.870494 | 18.32363 | 29.946 | 12.55321 | -0.25929 | 0.181866 | 0.652851 |
| Y | 9624400 | 9629599 | 53.71747 | 85.96333 | 66.87711 | 28.1641 | 0.705512 | 2.02317 | 0.652857 |
| 4 | 136868000 | 136869999 | 13.92454 | 24.36725 | 39.9959 | 17.64534 | -0.32375 | 0.665343 | 0.652875 |
| 13 | 52125200 | 52126999 | 16.57185 | 28.40367 | 47.96352 | 21.03481 | -0.36936 | 0.924143 | 0.65288 |
| 4 | 34581600 | 34583799 | 14.06406 | 24.79752 | 38.21568 | 16.91275 | -0.23155 | 0.557419 | 0.652883 |
| 6 | 84736200 | 84740199 | 64.31149 | 101.4306 | 144.0171 | 59.76741 | -0.18107 | 2.469226 | 0.652886 |
| 19 | 38083600 | 38087799 | 31.09202 | 50.65403 | 76.81076 | 32.98505 | -0.22807 | 1.538451 | 0.652892 |
| 8 | 124230000 | 124231199 | 13.77851 | 24.12811 | 39.48295 | 17.42905 | -0.31932 | 0.65317 | 0.652898 |
| 6 | 5854000 | 5855399 | 11.63318 | 20.91961 | 33.19194 | 14.49863 | -0.25807 | 0.382791 | 0.652904 |
| 14 | 36948800 | 36951199 | 17.84296 | 31.35197 | 34.32437 | 15.19549 | 0.257283 | 0.609689 | 0.652932 |
| 2 | 44140800 | 44141399 | 4.962557 | 9.669697 | 19.66602 | 7.14212 | -0.38398 | -0.39576 | 0.652935 |
| 14 | 22334600 | 22335199 | 4.962557 | 9.669697 | 19.66602 | 7.14212 | -0.38398 | -0.39576 | 0.652935 |
| 10 | 106174200 | 106183999 | 140.9927 | 214.4266 | 219.4664 | 90.92226 | 0.249939 | 3.189049 | 0.652936 |
| 4 | 35431400 | 35432199 | 7.9329 | 15.20896 | 26.29796 | 10.34695 | -0.26153 | -0.06587 | 0.652938 |
| 15 | 14118400 | 14120399 | 37.62081 | 61.67947 | 58.62542 | 25.23615 | 0.449757 | 1.454735 | 0.652938 |

| | | | | | | | | | |
|---|---|---|---|---|---|---|---|---|---|
| 17 | 83154200 | 83155799 | 20.37884 | 35.56522 | 39.21478 | 17.26576 | 0.247156 | 0.738066 | 0.652939 |
| 8 | 47226000 | 47235199 | 90.81427 | 140.275 | 193.8996 | 80.5418 | -0.16576 | 2.867457 | 0.652953 |
| 8 | 34710400 | 34715799 | 82.04291 | 128.2663 | 138.0197 | 57.26153 | 0.208659 | 2.644336 | 0.652957 |
| 9 | 88202200 | 88207199 | 66.02016 | 103.9395 | 147.5008 | 61.19893 | -0.18229 | 2.506902 | 0.652985 |
| 13 | 29078800 | 29084199 | 73.29 | 116.0862 | 125.7165 | 52.28157 | 0.214055 | 2.371228 | 0.653003 |
| 9 | 12127400 | 12144199 | 430.9262 | 589.0112 | 731.6215 | 304.4571 | -0.17767 | 4.931359 | 0.653015 |
| 5 | 150809000 | 150810799 | 20.37654 | 35.40721 | 39.92238 | 17.52943 | 0.21454 | 0.800132 | 0.653019 |
| 16 | 68786200 | 68788399 | 18.57006 | 32.68245 | 35.27182 | 15.61564 | 0.278595 | 0.597022 | 0.653022 |
| 14 | 55459400 | 55468999 | 94.14427 | 144.8161 | 200.5838 | 83.30028 | -0.17327 | 2.947203 | 0.653023 |
| 14 | 94877200 | 94887799 | 80.52045 | 126.8888 | 121.563 | 50.44342 | 0.384849 | 2.492427 | 0.653034 |
| 8 | 112903400 | 112907599 | 41.62965 | 66.9936 | 97.73109 | 41.50255 | -0.19002 | 1.848982 | 0.653039 |
| 15 | 68406200 | 68408399 | 14.81704 | 25.98792 | 40.8528 | 18.05793 | -0.26196 | 0.633499 | 0.653042 |
| 9 | 81061400 | 81065399 | 47.08831 | 75.4912 | 110.6726 | 46.73417 | -0.20381 | 1.979205 | 0.653052 |
| 10 | 117724400 | 117727199 | 24.07476 | 40.21738 | 60.93372 | 26.5195 | -0.21636 | 1.203053 | 0.653082 |
| 13 | 104812200 | 104816999 | 147.5045 | 220.4988 | 297.2667 | 122.5705 | -0.16877 | 3.532408 | 0.653086 |
| 5 | 131514200 | 131516599 | 29.35845 | 48.9959 | 55.15281 | 23.98871 | 0.210918 | 1.224305 | 0.653095 |
| 3 | 13188200 | 13190199 | 18.51611 | 33.42816 | 31.08938 | 13.51736 | 0.512199 | 0.402553 | 0.653104 |
| 15 | 16121400 | 16121999 | 7.314978 | 13.93549 | 25.04556 | 10.01109 | -0.3377 | -0.02541 | 0.653104 |
| 8 | 23156600 | 23159799 | 18.63301 | 31.8169 | 50.06198 | 21.94848 | -0.26879 | 0.969338 | 0.65311 |
| X | 136617800 | 136619199 | 9.820509 | 18.42769 | 30.72893 | 12.65519 | -0.26811 | 0.114308 | 0.653134 |
| 5 | 111712000 | 111715799 | 46.58256 | 74.54983 | 107.5735 | 45.30644 | -0.18391 | 2.031523 | 0.653142 |
| 2 | 33197400 | 33201999 | 26.47055 | 43.21941 | 80.83459 | 34.74073 | -0.52762 | 1.500675 | 0.653144 |
| 11 | 82671800 | 82672399 | 4.913575 | 9.586952 | 19.20702 | 6.853473 | -0.33829 | -0.45713 | 0.65315 |
| X | 33655600 | 33658799 | 45.76135 | 73.98101 | 79.44448 | 33.94477 | 0.258864 | 1.714122 | 0.653163 |
| 2 | 43773400 | 43773799 | 7.238619 | 14.08194 | 16.85788 | 6.223079 | 0.350992 | -0.31818 | 0.653182 |
| 5 | 148970800 | 148974799 | 35.23775 | 57.07248 | 84.05874 | 35.97242 | -0.19332 | 1.656579 | 0.653196 |
| 2 | 72688600 | 72689599 | 10.32489 | 18.8519 | 30.79075 | 13.19966 | -0.28189 | 0.279199 | 0.653201 |
| 6 | 120786400 | 120789999 | 41.79156 | 67.18551 | 97.89451 | 41.51046 | -0.19013 | 1.881139 | 0.653206 |
| 2 | 102088000 | 102089999 | 16.30267 | 28.28231 | 44.77793 | 19.64679 | -0.2733 | 0.808593 | 0.65323 |
| 4 | 87414200 | 87416799 | 24.99777 | 41.42661 | 63.47461 | 27.47916 | -0.23288 | 1.298965 | 0.653237 |
| 8 | 11640600 | 11644199 | 32.82353 | 53.17024 | 80.6022 | 34.497 | -0.23455 | 1.648089 | 0.653253 |
| 14 | 86759000 | 86760199 | 24.33037 | 41.16387 | 44.35235 | 19.41286 | 0.274078 | 1.066199 | 0.653292 |
| 12 | 52462400 | 52464999 | 38.42812 | 63.20758 | 57.11815 | 24.62106 | 0.524252 | 1.416145 | 0.653299 |
| 13 | 30629000 | 30630799 | 14.34144 | 26.57443 | 28.06259 | 11.79326 | 0.363384 | 0.221126 | 0.653304 |
| 15 | 12664600 | 12668399 | 27.77753 | 45.23952 | 84.3142 | 36.203 | -0.52517 | 1.541294 | 0.653306 |
| 17 | 40851800 | 40854399 | 47.54416 | 75.64059 | 130.58 | 54.54286 | -0.45141 | 2.225321 | 0.653312 |
| 2 | 105002400 | 105002799 | 5.504247 | 10.73742 | 20.39329 | 7.29662 | -0.26699 | -0.43768 | 0.653333 |
| 2 | 139149000 | 139149399 | 5.504247 | 10.73742 | 20.39329 | 7.29662 | -0.26699 | -0.43768 | 0.653333 |
| X | 32555600 | 32558399 | 52.1943 | 84.73995 | 73.28151 | 31.41614 | 0.576341 | 1.637305 | 0.653336 |
| 11 | 5485400 | 5485999 | 5.119144 | 9.956631 | 20.66472 | 7.641407 | -0.43723 | -0.3268 | 0.653348 |
| 18 | 7027000 | 7030199 | 29.08752 | 47.92982 | 72.23534 | 31.16104 | -0.21058 | 1.38257 | 0.65336 |
| 6 | 123768800 | 123771799 | 18.37745 | 31.30342 | 58.11095 | 25.47231 | -0.50711 | 0.999696 | 0.653366 |
| 18 | 49655000 | 49656999 | 23.52357 | 40.31662 | 45.25467 | 19.84674 | 0.217278 | 0.949332 | 0.653375 |
| 15 | 38546200 | 38548999 | 29.38392 | 49.06322 | 55.14367 | 23.99239 | 0.21313 | 1.219456 | 0.653376 |
| 1 | 117833600 | 117836399 | 41.40921 | 67.88036 | 67.57985 | 29.08886 | 0.383021 | 1.47215 | 0.653419 |
| 14 | 106476000 | 106477999 | 21.68054 | 36.85239 | 56.60506 | 24.80803 | -0.23539 | 1.028621 | 0.653435 |
| 3 | 71880200 | 71882799 | 41.44679 | 67.11041 | 74.71636 | 31.96792 | 0.209047 | 1.666736 | 0.653441 |

| | | | | | | | | | |
|---|---|---|---|---|---|---|---|---|---|
| 16 | 47222200 | 47224399 | 21.51081 | 37.54562 | 34.53073 | 15.20468 | 0.507743 | 0.735862 | 0.653463 |
| 4 | 69364400 | 69365199 | 11.84184 | 21.35932 | 33.7438 | 14.68814 | -0.24843 | 0.369233 | 0.653479 |
| 17 | 7940400 | 7941999 | 13.07923 | 23.31992 | 36.14455 | 15.9025 | -0.23187 | 0.451675 | 0.653494 |
| 17 | 18880400 | 18885399 | 66.23497 | 105.814 | 105.5332 | 44.33604 | 0.345251 | 2.107048 | 0.653531 |
| 6 | 145508800 | 145510799 | 18.05135 | 31.10736 | 48.47263 | 21.26354 | -0.2526 | 0.880368 | 0.653564 |
| 2 | 177063000 | 177066399 | 25.40127 | 43.05707 | 48.50957 | 21.24465 | 0.210006 | 1.059159 | 0.653567 |
| 2 | 176423000 | 176424999 | 16.76166 | 29.72189 | 32.80463 | 14.50003 | 0.25074 | 0.50388 | 0.653584 |
| 3 | 114776000 | 114776599 | 7.19392 | 13.70445 | 25.05943 | 10.01727 | -0.36191 | -0.02386 | 0.653586 |
| 18 | 59430000 | 59431599 | 16.94267 | 30.54078 | 31.15674 | 13.57258 | 0.37765 | 0.399746 | 0.6536 |
| 7 | 108148400 | 108150599 | 33.00671 | 54.2865 | 61.24237 | 26.39804 | 0.20502 | 1.405767 | 0.653607 |
| 11 | 104585800 | 104586399 | 11.01072 | 20.77858 | 24.1756 | 9.858674 | 0.258686 | 0.070518 | 0.653647 |
| 1 | 47017600 | 47020599 | 43.54574 | 69.6889 | 117.0989 | 49.31738 | -0.40321 | 2.032093 | 0.653647 |
| 19 | 58467000 | 58469399 | 33.99357 | 56.47993 | 59.31964 | 25.72614 | 0.310484 | 1.289329 | 0.653687 |
| 9 | 122767200 | 122770399 | 57.32264 | 91.97032 | 95.30346 | 40.2919 | 0.297147 | 1.961999 | 0.653692 |
| 10 | 22021200 | 22023199 | 19.94701 | 34.91303 | 37.46971 | 16.53354 | 0.285963 | 0.687992 | 0.653705 |
| 4 | 17458000 | 17460599 | 20.75984 | 36.33233 | 38.54789 | 17.00795 | 0.302574 | 0.697245 | 0.653707 |
| 5 | 86009800 | 86012999 | 38.65322 | 63.27427 | 67.64285 | 29.0988 | 0.279799 | 1.483899 | 0.653711 |
| 18 | 66445600 | 66446799 | 13.82216 | 25.51603 | 28.01836 | 11.83511 | 0.302067 | 0.237374 | 0.653713 |
| Y | 86533000 | 86534399 | 10.02132 | 19.04669 | 16.68689 | 6.703032 | 0.683704 | -0.00993 | 0.653737 |
| 8 | 74090200 | 74093399 | 55.81742 | 89.41887 | 84.78367 | 35.76138 | 0.422485 | 2.004049 | 0.65374 |
| 9 | 119335200 | 119339199 | 39.25754 | 63.25947 | 93.08387 | 39.60459 | -0.20003 | 1.804661 | 0.653741 |
| 9 | 89760600 | 89763999 | 60.90953 | 96.0351 | 141.461 | 58.70425 | -0.23435 | 2.47323 | 0.653778 |
| 8 | 14437800 | 14439199 | 9.010507 | 16.8772 | 27.95608 | 11.54856 | -0.26191 | 0.111765 | 0.653783 |
| 3 | 85745600 | 85747199 | 17.19358 | 30.2725 | 33.94667 | 15.02771 | 0.223995 | 0.583799 | 0.653799 |
| X | 76975000 | 76988599 | 216.7823 | 319.6758 | 321.0716 | 132.2216 | 0.237546 | 3.779656 | 0.653816 |
| 15 | 37752000 | 37753599 | 19.18058 | 33.4271 | 37.27958 | 16.39925 | 0.230621 | 0.748471 | 0.653821 |
| 19 | 35361400 | 35361999 | 6.829902 | 13.26071 | 24.05914 | 9.023699 | -0.26497 | -0.24484 | 0.653822 |
| X | 126182200 | 126188599 | 68.25231 | 108.9912 | 105.1717 | 44.15795 | 0.392168 | 2.123602 | 0.653837 |
| 6 | 124593600 | 124602999 | 219.5559 | 321.6041 | 337.1234 | 138.8677 | 0.166716 | 3.885014 | 0.653878 |
| X | 78212200 | 78226599 | 268.6094 | 388.9018 | 384.6007 | 158.6075 | 0.234471 | 4.067251 | 0.653889 |
| 3 | 14819000 | 14820599 | 20.36998 | 35.41713 | 39.89381 | 17.52351 | 0.216037 | 0.791432 | 0.65389 |
| 6 | 59215400 | 59219199 | 95.59239 | 148.0592 | 162.6644 | 67.56641 | 0.168252 | 2.815579 | 0.653935 |
| 6 | 138062400 | 138063399 | 9.502326 | 17.66398 | 28.9014 | 12.09309 | -0.25917 | 0.16725 | 0.653986 |
| 13 | 57184600 | 57188399 | 28.59733 | 46.90345 | 71.32597 | 30.70827 | -0.22616 | 1.437056 | 0.654029 |
| 17 | 65744600 | 65745799 | 10.20189 | 18.15649 | 37.87205 | 16.68823 | -0.65893 | 0.44927 | 0.654047 |
| 4 | 146271600 | 146272999 | 31.90048 | 52.94787 | 58.10324 | 25.18074 | 0.24725 | 1.292446 | 0.654071 |
| Y | 84950000 | 84950799 | 9.487874 | 18.12112 | 17.24907 | 6.857644 | 0.578281 | -0.0461 | 0.654075 |
| 1 | 31432000 | 31432799 | 9.514017 | 17.75815 | 28.9647 | 12.03622 | -0.24661 | 0.141962 | 0.654098 |
| 12 | 72176000 | 72177799 | 10.8797 | 19.57018 | 39.46971 | 17.23673 | -0.6022 | 0.386968 | 0.654103 |
| 14 | 34981400 | 34983199 | 20.46331 | 36.67793 | 31.95164 | 13.99555 | 0.599161 | 0.460218 | 0.654105 |
| 14 | 27253400 | 27254599 | 17.17774 | 30.30015 | 33.87256 | 14.98941 | 0.22941 | 0.563294 | 0.654134 |
| 7 | 107868000 | 107870199 | 36.78799 | 60.42325 | 57.51128 | 24.7755 | 0.448775 | 1.429911 | 0.654144 |
| 12 | 35846800 | 35847799 | 10.58128 | 20.0696 | 23.76459 | 9.587773 | 0.247216 | 0.031635 | 0.654167 |
| 19 | 12519600 | 12524399 | 60.0362 | 95.58471 | 105.7621 | 44.22122 | 0.190406 | 2.206694 | 0.654167 |
| 16 | 82780600 | 82785799 | 60.93206 | 97.57105 | 99.68285 | 42.01867 | 0.31427 | 2.027065 | 0.654181 |
| 12 | 21209400 | 21210599 | 8.11112 | 15.29018 | 26.48752 | 10.82391 | -0.31296 | 0.060403 | 0.654185 |
| 7 | 19718000 | 19718399 | 8.468941 | 16.41851 | 20.49852 | 7.748377 | 0.258838 | -0.20982 | 0.654209 |

| | | | | | | | | | |
|---|---|---|---|---|---|---|---|---|---|
| 1 | 103444400 | 103444999 | 10.16493 | 19.56439 | 23.03649 | 8.96276 | 0.302919 | -0.09011 | 0.654234 |
| 10 | 56720800 | 56723399 | 18.71666 | 33.2684 | 32.98298 | 14.54933 | 0.406472 | 0.499286 | 0.654246 |
| 11 | 53446200 | 53448199 | 19.18991 | 32.58538 | 51.62773 | 22.62226 | -0.27985 | 1.025626 | 0.654248 |
| 2 | 151300600 | 151302199 | 26.38218 | 44.21533 | 48.07177 | 20.96465 | 0.260839 | 1.16416 | 0.654255 |
| 5 | 150212800 | 150215999 | 21.7649 | 36.82134 | 55.43683 | 24.26408 | -0.20687 | 1.070493 | 0.654266 |
| X | 94042000 | 94042799 | 10.05518 | 18.76785 | 31.68239 | 13.16601 | -0.29595 | 0.154117 | 0.654275 |
| 5 | 40374600 | 40376799 | 38.31178 | 62.25113 | 69.54006 | 29.82987 | 0.208872 | 1.588246 | 0.654286 |
| 13 | 25538400 | 25542199 | 57.66512 | 92.10666 | 102.7583 | 43.15928 | 0.183515 | 2.101182 | 0.654324 |
| 12 | 110719800 | 110723599 | 35.72768 | 57.94038 | 86.21317 | 36.92936 | -0.20632 | 1.632831 | 0.654328 |
| 15 | 3762400 | 3763599 | 14.58161 | 26.74587 | 28.99089 | 12.35327 | 0.311756 | 0.281914 | 0.654339 |
| 3 | 91118200 | 91125599 | 186.7108 | 278.1244 | 256.6067 | 105.7395 | 0.373425 | 3.601436 | 0.654341 |
| 11 | 113814000 | 113816999 | 28.49841 | 46.5576 | 72.63464 | 31.22325 | -0.26607 | 1.49081 | 0.65435 |
| 4 | 99461000 | 99462599 | 14.10462 | 24.7522 | 38.88345 | 17.19554 | -0.26067 | 0.619932 | 0.654354 |
| 15 | 64397800 | 64401399 | 46.89305 | 75.58084 | 83.51129 | 35.543 | 0.212776 | 1.804567 | 0.654354 |
| 14 | 7354200 | 7358399 | 39.10241 | 64.23243 | 66.23742 | 28.53614 | 0.333587 | 1.441128 | 0.65436 |
| 5 | 94877200 | 94881199 | 42.45418 | 68.83953 | 65.6186 | 28.10487 | 0.434463 | 1.636871 | 0.654362 |
| 3 | 133081400 | 133090599 | 113.1866 | 172.6539 | 239.291 | 99.21846 | -0.18359 | 3.123138 | 0.654367 |
| 6 | 95276800 | 95278399 | 14.92243 | 27.35996 | 28.88721 | 12.31584 | 0.348637 | 0.284747 | 0.65437 |
| 16 | 83199600 | 83201999 | 23.15038 | 38.25924 | 70.92522 | 30.64613 | -0.5072 | 1.332937 | 0.654384 |
| 18 | 32793600 | 32795399 | 14.12135 | 26.43861 | 26.00443 | 10.75024 | 0.483198 | 0.150711 | 0.654387 |
| 14 | 119718800 | 119721599 | 35.95438 | 58.10935 | 85.80417 | 36.64827 | -0.20004 | 1.709016 | 0.654406 |
| 8 | 35214800 | 35217599 | 23.36544 | 39.43521 | 61.06867 | 26.70981 | -0.24766 | 1.100636 | 0.654434 |
| X | 161764600 | 161770599 | 54.26548 | 87.10208 | 93.87199 | 39.7122 | 0.241355 | 1.945739 | 0.654452 |
| 2 | 24006000 | 24006399 | 7.493768 | 14.56568 | 17.83279 | 6.629188 | 0.308849 | -0.28929 | 0.654462 |
| 8 | 86042800 | 86043799 | 12.3024 | 21.89894 | 34.68528 | 15.28155 | -0.26417 | 0.454418 | 0.654493 |
| 17 | 46641800 | 46644999 | 22.26722 | 37.58838 | 56.5271 | 24.72478 | -0.20537 | 1.092452 | 0.654501 |
| 8 | 78633000 | 78634599 | 30.03004 | 49.03859 | 74.25067 | 31.9129 | -0.22344 | 1.504401 | 0.654512 |
| 2 | 27867200 | 27868399 | 10.49294 | 19.23284 | 30.82398 | 13.14692 | -0.24958 | 0.259443 | 0.654517 |
| 3 | 28205800 | 28209399 | 51.12822 | 81.66169 | 119.6755 | 50.26132 | -0.2094 | 2.102529 | 0.654539 |
| 12 | 81436600 | 81438799 | 22.36857 | 39.23791 | 37.20775 | 16.44857 | 0.46374 | 0.66697 | 0.654554 |
| 2 | 101975200 | 101976599 | 11.66378 | 20.97125 | 33.21915 | 14.51249 | -0.25573 | 0.385083 | 0.65456 |
| 16 | 31342800 | 31344799 | 18.66816 | 32.67782 | 36.65755 | 16.17649 | 0.222644 | 0.68126 | 0.65456 |
| 10 | 126692600 | 126694199 | 11.39195 | 20.60537 | 33.30046 | 14.44458 | -0.27755 | 0.349991 | 0.654573 |
| 2 | 41939800 | 41942599 | 31.13757 | 50.79054 | 75.5795 | 32.47083 | -0.19964 | 1.522271 | 0.65458 |
| 5 | 21155800 | 21157199 | 9.912474 | 17.99905 | 30.83621 | 13.31092 | -0.35663 | 0.309149 | 0.65458 |
| 1 | 34328200 | 34330599 | 31.58445 | 52.02254 | 57.76529 | 24.91452 | 0.228705 | 1.383377 | 0.654585 |
| 17 | 70008600 | 70011799 | 22.23679 | 38.75574 | 40.76968 | 17.93525 | 0.314546 | 0.768537 | 0.654609 |
| 1 | 181997400 | 182004999 | 125.0366 | 190.4377 | 207.2411 | 85.89495 | 0.163252 | 3.154749 | 0.654649 |
| 16 | 48201400 | 48208599 | 59.66391 | 95.20021 | 104.4787 | 43.81727 | 0.205525 | 2.133602 | 0.654687 |
| 15 | 29353000 | 29353999 | 11.78357 | 20.76178 | 40.92429 | 18.11406 | -0.5849 | 0.55773 | 0.654697 |
| 17 | 66460400 | 66471399 | 145.5004 | 219.6767 | 236.7922 | 97.88005 | 0.166497 | 3.337857 | 0.654715 |
| 13 | 95648400 | 95652599 | 27.9204 | 45.55568 | 81.55205 | 35.0404 | -0.46531 | 1.514377 | 0.654734 |
| 1 | 60907600 | 60909999 | 29.94534 | 49.08344 | 73.06138 | 31.44704 | -0.19574 | 1.450631 | 0.654735 |
| 4 | 121124800 | 121129199 | 59.31857 | 94.63574 | 105.566 | 44.26447 | 0.181847 | 2.139761 | 0.654749 |
| 16 | 83753600 | 83754799 | 21.54119 | 37.40529 | 40.87565 | 17.94026 | 0.259272 | 0.814503 | 0.654764 |
| 13 | 31457400 | 31458799 | 13.00289 | 22.99553 | 36.25885 | 16.03709 | -0.26241 | 0.51629 | 0.654772 |
| 7 | 118261800 | 118266199 | 43.60927 | 70.02644 | 101.3683 | 42.91027 | -0.18277 | 1.921429 | 0.654777 |

| | | | | | | | | | |
|---|---|---|---|---|---|---|---|---|---|
| 15 | 13172000 | 13173999 | 14.36269 | 25.12597 | 40.98072 | 18.07494 | -0.3148 | 0.67329 | 0.654807 |
| 19 | 45179600 | 45187199 | 119.3762 | 180.3368 | 252.1706 | 104.2555 | -0.20799 | 3.321717 | 0.654813 |
| 9 | 10370600 | 10374199 | 28.55483 | 49.43411 | 43.43546 | 19.05655 | 0.57204 | 0.881871 | 0.654817 |
| 17 | 17141200 | 17142399 | 26.56272 | 44.48549 | 48.07956 | 20.95841 | 0.269399 | 1.172822 | 0.654823 |
| 18 | 9695000 | 9709599 | 150.8185 | 225.5006 | 303.6177 | 125.1944 | -0.16663 | 3.529129 | 0.654848 |
| X | 56624000 | 56625799 | 19.6088 | 34.63966 | 34.30606 | 15.17674 | 0.404328 | 0.565918 | 0.654849 |
| 2 | 100204200 | 100204799 | 11.62187 | 22.24708 | 23.94897 | 9.464621 | 0.40972 | -0.02537 | 0.654851 |
| 1 | 114984200 | 114985199 | 11.33577 | 21.70602 | 23.972 | 9.465436 | 0.374371 | -0.03048 | 0.65486 |
| 5 | 132740200 | 132741999 | 19.51877 | 34.31536 | 36.33227 | 16.08857 | 0.305591 | 0.617332 | 0.65486 |
| 2 | 72318400 | 72319999 | 20.84978 | 36.41175 | 33.57167 | 14.78746 | 0.504239 | 0.723501 | 0.654866 |
| 15 | 93109400 | 93112399 | 24.95602 | 41.42733 | 63.02579 | 27.31669 | -0.22249 | 1.278022 | 0.654867 |
| 7 | 108145600 | 108161799 | 265.8745 | 384.2949 | 392.1382 | 161.752 | 0.186906 | 4.090975 | 0.654874 |
| 5 | 17979400 | 17980599 | 14.05636 | 24.80676 | 38.19478 | 16.90039 | -0.22974 | 0.547046 | 0.654875 |
| 13 | 90442400 | 90445599 | 28.18085 | 47.46771 | 51.28868 | 22.41874 | 0.269679 | 1.122627 | 0.654899 |
| 7 | 4390200 | 4391999 | 16.2855 | 28.69138 | 31.00689 | 13.72462 | 0.27732 | 0.565355 | 0.654902 |
| 8 | 115814600 | 115816799 | 22.80521 | 39.46992 | 43.03788 | 18.88196 | 0.2616 | 0.855964 | 0.654906 |
| 10 | 38248000 | 38250599 | 19.47794 | 34.44815 | 34.6021 | 15.30418 | 0.384363 | 0.5497 | 0.654913 |
| 3 | 74772200 | 74773399 | 14.87761 | 27.28386 | 23.8824 | 10.179 | 0.618132 | 0.274833 | 0.654966 |
| X | 102448400 | 102449199 | 7.726541 | 14.8733 | 25.91009 | 10.07678 | -0.25691 | -0.109 | 0.654967 |
| 10 | 7293000 | 7295999 | 39.06425 | 63.36892 | 71.53817 | 30.64959 | 0.191539 | 1.625547 | 0.654972 |
| 2 | 77176200 | 77178799 | 24.28969 | 41.30369 | 46.35713 | 20.31736 | 0.215733 | 1.023426 | 0.654983 |
| 1 | 124848000 | 124851199 | 49.5913 | 79.69245 | 89.18806 | 37.79652 | 0.189054 | 1.901197 | 0.654984 |
| 6 | 38806000 | 38807999 | 16.32741 | 28.61067 | 45.0788 | 19.91046 | -0.2653 | 0.664251 | 0.654985 |
| 3 | 98264400 | 98268399 | 41.14429 | 66.52897 | 74.35782 | 31.77889 | 0.201558 | 1.695672 | 0.655 |
| 12 | 115977000 | 115978399 | 18.32109 | 32.24802 | 35.34662 | 15.64839 | 0.256493 | 0.595521 | 0.655006 |
| 7 | 9788000 | 9805399 | 238.492 | 351.505 | 323.5013 | 133.2214 | 0.362835 | 3.791193 | 0.655055 |
| 4 | 44478200 | 44480999 | 26.1331 | 43.5693 | 67.17121 | 29.19758 | -0.24139 | 1.237507 | 0.655056 |
| 17 | 19899600 | 19904399 | 54.85291 | 88.55735 | 82.01451 | 34.98086 | 0.469667 | 1.768868 | 0.655064 |
| 10 | 104425600 | 104426599 | 16.94227 | 30.09179 | 32.66108 | 14.41649 | 0.275951 | 0.492426 | 0.655096 |
| 17 | 63522200 | 63523599 | 15.10465 | 26.65177 | 42.88894 | 18.97818 | -0.29335 | 0.564391 | 0.655101 |
| 7 | 131600200 | 131602799 | 23.09057 | 40.18663 | 35.14118 | 15.44538 | 0.579956 | 0.786096 | 0.655105 |
| 14 | 42955400 | 42957199 | 15.41812 | 28.17678 | 29.33445 | 12.56124 | 0.364366 | 0.307456 | 0.655108 |
| 13 | 57117200 | 57119799 | 48.73001 | 77.88889 | 111.6957 | 46.95916 | -0.17696 | 2.073648 | 0.655113 |
| 10 | 35363400 | 35364999 | 17.90484 | 31.47088 | 34.98221 | 15.49094 | 0.235715 | 0.609291 | 0.65515 |
| 13 | 83345400 | 83348999 | 46.91198 | 75.60055 | 83.87508 | 35.69111 | 0.206619 | 1.806614 | 0.655165 |
| 17 | 78778000 | 78779599 | 16.05562 | 28.014 | 43.12012 | 18.98254 | -0.23174 | 0.728091 | 0.655167 |
| 12 | 59521400 | 59526599 | 54.01577 | 86.508 | 96.60435 | 40.72965 | 0.186378 | 2.013873 | 0.655174 |
| 12 | 3878000 | 3880999 | 30.56827 | 50.50235 | 56.15995 | 24.26089 | 0.227771 | 1.342663 | 0.655223 |
| 4 | 147784200 | 147787199 | 30.62019 | 51.14934 | 55.79834 | 24.28379 | 0.255982 | 1.21798 | 0.655232 |
| 8 | 5267200 | 5268999 | 32.74831 | 53.92996 | 61.19321 | 26.39103 | 0.19745 | 1.390905 | 0.655294 |
| 11 | 23723000 | 23724399 | 12.77174 | 24.2966 | 24.86924 | 9.970377 | 0.463521 | 0.03357 | 0.65532 |
| X | 79296200 | 79297599 | 15.54718 | 28.88736 | 28.07111 | 11.75313 | 0.486455 | 0.216132 | 0.655327 |
| 4 | 93842800 | 93845399 | 36.01812 | 58.76043 | 66.13586 | 28.41518 | 0.202264 | 1.52242 | 0.655327 |
| 4 | 70575800 | 70581199 | 48.51063 | 78.21486 | 83.886 | 35.7183 | 0.25617 | 1.794366 | 0.655354 |
| 3 | 91553400 | 91558799 | 67.39586 | 107.3902 | 111.6346 | 46.7296 | 0.28157 | 2.189718 | 0.655354 |
| 9 | 22859800 | 22860399 | 7.555671 | 14.15698 | 27.0095 | 11.15116 | -0.46257 | 0.097501 | 0.65536 |
| 4 | 124008000 | 124010799 | 22.74421 | 38.31435 | 57.63888 | 25.19529 | -0.20603 | 1.112478 | 0.65536 |

| | | | | | | | | | |
|---|---|---|---|---|---|---|---|---|---|
| X | 45226200 | 45233799 | 160.2389 | 242.5066 | 204.7518 | 84.69957 | 0.521554 | 3.292976 | 0.655367 |
| 8 | 31777600 | 31779999 | 22.36119 | 38.97094 | 40.75033 | 17.92627 | 0.323148 | 0.7692 | 0.655386 |
| 1 | 184348800 | 184351399 | 23.5267 | 39.64294 | 61.05983 | 26.69279 | -0.23999 | 1.115001 | 0.655425 |
| 2 | 51326200 | 51326799 | 10.45893 | 19.17401 | 30.80681 | 13.13633 | -0.25306 | 0.257023 | 0.655468 |
| 11 | 29079200 | 29081199 | 10.45893 | 19.17401 | 30.80681 | 13.13633 | -0.25306 | 0.257023 | 0.655468 |
| 9 | 81553000 | 81559799 | 87.6936 | 137.559 | 146.7009 | 60.84225 | 0.225273 | 2.589978 | 0.655478 |
| 16 | 95309000 | 95311999 | 18.53053 | 31.53915 | 50.57669 | 22.16833 | -0.29674 | 1.000475 | 0.655502 |
| 10 | 95021200 | 95023199 | 22.19575 | 37.69681 | 58.2738 | 25.53594 | -0.24474 | 1.039977 | 0.655522 |
| 15 | 96081600 | 96082999 | 13.56165 | 24.52805 | 27.02156 | 11.72208 | 0.271581 | 0.349654 | 0.655532 |
| 13 | 37329800 | 37332399 | 31.47866 | 51.23216 | 76.53632 | 32.8551 | -0.20757 | 1.555609 | 0.655542 |
| 5 | 87596800 | 87597399 | 8.08544 | 15.7197 | 19.49151 | 7.23099 | 0.293007 | -0.28445 | 0.655546 |
| 12 | 115676400 | 115677799 | 24.81095 | 42.81279 | 44.62317 | 19.57569 | 0.325582 | 0.893277 | 0.655576 |
| 2 | 143646200 | 143646799 | 14.09626 | 24.69388 | 39.45674 | 17.42305 | -0.28509 | 0.649043 | 0.655597 |
| 14 | 51535600 | 51545799 | 158.6018 | 239.0743 | 222.2183 | 91.81068 | 0.378085 | 3.37142 | 0.655646 |
| 5 | 98857000 | 98858399 | 12.02296 | 21.63174 | 34.23738 | 14.94877 | -0.25395 | 0.386495 | 0.655666 |
| 6 | 125085800 | 125087999 | 19.2636 | 33.58524 | 37.8113 | 16.63735 | 0.217185 | 0.745201 | 0.655667 |
| 8 | 19571000 | 19581399 | 153.2409 | 230.5225 | 247.9908 | 102.4028 | 0.164773 | 3.411499 | 0.655678 |
| 4 | 12690200 | 12691999 | 23.71688 | 40.32478 | 44.82191 | 19.64432 | 0.229731 | 1.022209 | 0.655696 |
| 13 | 73537000 | 73542399 | 57.97426 | 92.237 | 135.419 | 56.5663 | -0.21813 | 2.234053 | 0.655711 |
| 14 | 32074800 | 32076399 | 15.76855 | 27.94112 | 30.96353 | 13.68966 | 0.244619 | 0.501241 | 0.655729 |
| X | 94509400 | 94513199 | 51.02759 | 82.20274 | 86.6096 | 36.83505 | 0.280342 | 1.827511 | 0.655753 |
| 11 | 89800600 | 89802399 | 12.66328 | 22.78445 | 36.26431 | 15.83339 | -0.26215 | 0.39418 | 0.655761 |
| 14 | 45080800 | 45084399 | 25.01434 | 41.26936 | 65.58885 | 28.30874 | -0.28613 | 1.351345 | 0.655769 |
| 3 | 14381000 | 14381799 | 11.30275 | 21.13489 | 24.06134 | 9.968196 | 0.27147 | 0.130741 | 0.655775 |
| 3 | 79264200 | 79264599 | 6.992895 | 13.43254 | 24.12271 | 9.438949 | -0.30825 | -0.10138 | 0.655779 |
| 7 | 100490800 | 100493399 | 24.77238 | 41.28288 | 62.00159 | 26.94314 | -0.20367 | 1.232951 | 0.655795 |
| 15 | 79849200 | 79850999 | 24.77238 | 41.28288 | 62.00159 | 26.94314 | -0.20367 | 1.232951 | 0.655795 |
| 14 | 8315000 | 8319599 | 97.50406 | 150.702 | 164.4194 | 68.29612 | 0.175961 | 2.855265 | 0.655805 |
| 10 | 129648800 | 129658999 | 162.5183 | 243.8028 | 259.7717 | 107.188 | 0.175079 | 3.467007 | 0.655824 |
| X | 76553000 | 76555799 | 42.77812 | 69.52954 | 71.31974 | 30.60017 | 0.332146 | 1.597839 | 0.655842 |
| 3 | 144346400 | 144351199 | 35.78399 | 57.71526 | 104.8159 | 44.64997 | -0.50149 | 1.774398 | 0.655842 |
| 17 | 88198800 | 88211599 | 150.8587 | 224.9875 | 307.1843 | 126.6024 | -0.19009 | 3.575897 | 0.655858 |
| 19 | 55047000 | 55051799 | 36.40298 | 58.7272 | 87.89254 | 37.45207 | -0.2225 | 1.762034 | 0.655861 |
| X | 130919600 | 130920799 | 14.23228 | 26.12896 | 28.05555 | 11.93987 | 0.326897 | 0.274221 | 0.655863 |
| 3 | 61469200 | 61470399 | 8.18492 | 15.71141 | 27.56829 | 10.8086 | -0.27841 | -0.07418 | 0.655864 |
| 6 | 13034600 | 13035999 | 16.27252 | 30.49156 | 26.58925 | 10.97649 | 0.657827 | 0.167359 | 0.655923 |
| 13 | 69515400 | 69520599 | 55.53369 | 89.29116 | 90.82995 | 38.5232 | 0.327522 | 1.89275 | 0.655926 |
| 6 | 134264400 | 134265599 | 13.83074 | 25.08888 | 28.39624 | 12.27291 | 0.236063 | 0.336498 | 0.655945 |
| 14 | 54585400 | 54588599 | 58.34794 | 92.50985 | 131.8901 | 54.8918 | -0.18075 | 2.336599 | 0.655974 |
| 7 | 47381000 | 47393199 | 163.359 | 246.821 | 241.2297 | 99.73742 | 0.308602 | 3.327991 | 0.655994 |
| 3 | 19535600 | 19536599 | 13.19608 | 23.94141 | 26.03953 | 11.25209 | 0.293533 | 0.326389 | 0.656006 |
| X | 16653200 | 16653999 | 13.77629 | 25.43008 | 28.07393 | 11.85938 | 0.294529 | 0.238869 | 0.656007 |
| 6 | 38140400 | 38142199 | 20.21894 | 35.08695 | 38.37017 | 16.83547 | 0.258148 | 0.813084 | 0.656011 |
| 12 | 88004200 | 88012799 | 102.8079 | 159.2271 | 169.7929 | 70.52739 | 0.211181 | 2.819628 | 0.656012 |
| 14 | 72028000 | 72029999 | 12.74954 | 22.3341 | 43.60527 | 19.2545 | -0.57269 | 0.650023 | 0.656022 |
| 13 | 37871600 | 37875199 | 28.94571 | 47.46246 | 71.27986 | 30.68503 | -0.20831 | 1.441919 | 0.656029 |
| 9 | 117458800 | 117462199 | 27.28783 | 45.03204 | 67.4831 | 29.12984 | -0.20148 | 1.356507 | 0.656058 |

| | | | | | | | | | |
|---|---|---|---|---|---|---|---|---|---|
| 1 | 50369000 | 50369599 | 5.8086 | 11.29336 | 21.21218 | 7.86772 | -0.29886 | -0.30454 | 0.656062 |
| 10 | 68421200 | 68421999 | 5.8086 | 11.29336 | 21.21218 | 7.86772 | -0.29886 | -0.30454 | 0.656062 |
| 17 | 93263800 | 93266999 | 27.03874 | 46.13753 | 48.47445 | 21.25125 | 0.311387 | 1.006472 | 0.656064 |
| 4 | 83541400 | 83544599 | 29.43085 | 48.34156 | 72.03103 | 31.03112 | -0.19601 | 1.422493 | 0.656084 |
| 7 | 12486800 | 12487999 | 26.40603 | 43.32055 | 68.58037 | 29.5294 | -0.28372 | 1.429319 | 0.656084 |
| 3 | 93776000 | 93778799 | 34.27064 | 55.47645 | 82.56231 | 35.31812 | -0.20899 | 1.664957 | 0.656088 |
| 6 | 130936600 | 130940799 | 46.68345 | 75.51701 | 79.32882 | 33.91736 | 0.291442 | 1.699399 | 0.65612 |
| 10 | 37906400 | 37908199 | 21.67239 | 38.2586 | 34.99138 | 15.48207 | 0.517941 | 0.583289 | 0.656121 |
| 4 | 40038400 | 40039399 | 11.07653 | 21.04197 | 23.71488 | 9.535992 | 0.321684 | 0.023427 | 0.656131 |
| 3 | 32433800 | 32435399 | 13.15469 | 24.15403 | 27.46964 | 11.68826 | 0.244244 | 0.26047 | 0.656148 |
| 11 | 28052000 | 28052999 | 15.11656 | 27.41705 | 24.36811 | 10.53401 | 0.582669 | 0.339174 | 0.656157 |
| 7 | 99141600 | 99145199 | 39.73559 | 64.06825 | 94.13185 | 40.08223 | -0.19686 | 1.787742 | 0.656168 |
| 16 | 52381800 | 52383799 | 20.87317 | 36.57493 | 37.91656 | 16.74652 | 0.335758 | 0.677063 | 0.65617 |
| 12 | 114074600 | 114077799 | 60.09793 | 95.9866 | 104.2014 | 43.76158 | 0.222772 | 2.107523 | 0.65617 |
| 10 | 71404000 | 71410799 | 85.64725 | 132.0496 | 210.7537 | 87.54348 | -0.37503 | 2.901195 | 0.656177 |
| 1 | 85162200 | 85180399 | 589.6919 | 795.366 | 760.5408 | 317.1413 | 0.180294 | 5.142909 | 0.656185 |
| 12 | 72963800 | 72965199 | 16.52472 | 28.24298 | 54.75795 | 24.0091 | -0.56843 | 0.961324 | 0.656188 |
| 5 | 123356200 | 123358399 | 39.34004 | 63.87431 | 71.82724 | 30.79406 | 0.198358 | 1.607096 | 0.656209 |
| 2 | 149380200 | 149381199 | 17.22 | 30.40022 | 33.87577 | 14.98831 | 0.234513 | 0.555055 | 0.656227 |
| 15 | 94041200 | 94042999 | 14.07949 | 24.77975 | 38.26002 | 16.93851 | -0.23522 | 0.577916 | 0.656228 |
| 10 | 80338000 | 80340199 | 21.14107 | 35.65208 | 54.84874 | 23.98348 | -0.23807 | 1.095475 | 0.656248 |
| 16 | 9680600 | 9684999 | 38.41264 | 62.03909 | 91.49494 | 39.04374 | -0.1995 | 1.73577 | 0.656251 |
| 14 | 75204400 | 75209399 | 36.56983 | 59.28887 | 89.89456 | 38.49803 | -0.2338 | 1.642369 | 0.656254 |
| 13 | 67743800 | 67754799 | 150.9896 | 228.6127 | 234.1007 | 96.85423 | 0.243949 | 3.282651 | 0.656255 |
| 14 | 31558000 | 31561199 | 44.58607 | 72.18129 | 77.79344 | 33.28031 | 0.255616 | 1.680109 | 0.656259 |
| 15 | 91167000 | 91169399 | 23.48983 | 39.14294 | 60.22989 | 26.17215 | -0.23846 | 1.228603 | 0.656295 |
| 13 | 30210200 | 30211399 | 16.16691 | 28.19131 | 43.1162 | 18.97367 | -0.22272 | 0.735897 | 0.656299 |
| Y | 5017800 | 5020199 | 19.79694 | 34.43075 | 27.99665 | 12.29988 | 0.683943 | 0.763471 | 0.656299 |
| 10 | 27561000 | 27561999 | 12.85633 | 23.83921 | 27.0756 | 11.36767 | 0.26003 | 0.203172 | 0.656314 |
| 15 | 21921600 | 21925199 | 106.9475 | 164.0931 | 169.9044 | 70.53486 | 0.243322 | 2.998734 | 0.656317 |
| 5 | 87183600 | 87186599 | 37.33573 | 61.58701 | 55.39557 | 23.9075 | 0.532571 | 1.378531 | 0.656331 |
| 2 | 126812800 | 126815199 | 23.28205 | 39.1042 | 58.70238 | 25.63576 | -0.20311 | 1.142187 | 0.656335 |
| 15 | 6294200 | 6296799 | 20.89424 | 36.52142 | 39.12119 | 17.24249 | 0.288845 | 0.718499 | 0.656338 |
| 13 | 33075400 | 33078599 | 38.85672 | 63.08541 | 71.14516 | 30.50015 | 0.194123 | 1.607287 | 0.656339 |
| 3 | 71354400 | 71355199 | 17.4717 | 30.05073 | 47.42476 | 20.80265 | -0.2713 | 0.895233 | 0.656377 |
| 9 | 4846600 | 4860999 | 253.9008 | 362.6473 | 472.9247 | 195.4208 | -0.18391 | 4.265192 | 0.656382 |
| 1 | 9480000 | 9480799 | 7.886807 | 15.17601 | 26.48373 | 10.31117 | -0.26091 | -0.10201 | 0.656386 |
| 8 | 41410000 | 41411799 | 17.27679 | 29.8556 | 46.35679 | 20.33711 | -0.24653 | 0.85035 | 0.65639 |
| 14 | 94667600 | 94668599 | 17.23428 | 30.36911 | 33.94922 | 15.02637 | 0.228957 | 0.575704 | 0.656401 |
| 2 | 28244200 | 28244999 | 9.367669 | 17.33734 | 28.87785 | 12.15567 | -0.2909 | 0.191007 | 0.656415 |
| 16 | 11783800 | 11786999 | 28.71353 | 47.72239 | 52.91111 | 22.95189 | 0.232672 | 1.265779 | 0.656461 |
| 13 | 5738800 | 5742999 | 58.96878 | 94.61739 | 83.7814 | 35.42405 | 0.523663 | 1.958913 | 0.656477 |
| 1 | 26207000 | 26207399 | 7.59576 | 14.62545 | 25.3226 | 9.840761 | -0.24678 | -0.11319 | 0.656488 |
| 15 | 87250000 | 87250599 | 7.59576 | 14.62545 | 25.3226 | 9.840761 | -0.24678 | -0.11319 | 0.656488 |
| 10 | 70125600 | 70128199 | 17.56422 | 30.02225 | 55.93445 | 24.52513 | -0.51145 | 0.96262 | 0.656498 |
| 3 | 114475200 | 114476399 | 14.9458 | 27.094 | 29.91532 | 12.93996 | 0.270996 | 0.353155 | 0.656524 |
| 1 | 134312400 | 134313399 | 11.9258 | 21.55527 | 33.71215 | 14.63648 | -0.23181 | 0.355853 | 0.656555 |

| | | | | | | | | | |
|---|---|---|---|---|---|---|---|---|---|
| 6 | 56965600 | 56966799 | 18.31286 | 32.08326 | 36.10671 | 15.94491 | 0.217968 | 0.664351 | 0.656584 |
| 3 | 12928800 | 12929799 | 12.82641 | 23.67632 | 27.01633 | 11.41286 | 0.246923 | 0.227461 | 0.656591 |
| 15 | 70727800 | 70733599 | 80.27289 | 126.5833 | 121.7407 | 50.52156 | 0.380002 | 2.478389 | 0.656591 |
| X | 147791000 | 147796599 | 54.3941 | 87.77493 | 83.53543 | 35.60775 | 0.429662 | 1.782672 | 0.656606 |
| 15 | 38931200 | 38933999 | 43.96219 | 71.13219 | 77.87369 | 33.30115 | 0.232291 | 1.687902 | 0.656625 |
| 17 | 3893200 | 3895999 | 27.18777 | 46.30802 | 49.02487 | 21.48943 | 0.300067 | 1.023515 | 0.656626 |
| 15 | 4195400 | 4197399 | 21.53078 | 37.73679 | 37.8516 | 16.72141 | 0.383115 | 0.676429 | 0.656632 |
| 17 | 69363200 | 69367399 | 32.86656 | 53.67488 | 82.75589 | 35.56947 | -0.2496 | 1.510963 | 0.656642 |
| 14 | 73727400 | 73731199 | 42.23145 | 68.31385 | 76.74153 | 32.81065 | 0.194858 | 1.69093 | 0.656645 |
| 4 | 118843400 | 118845199 | 17.52663 | 30.74478 | 34.13308 | 15.0895 | 0.23755 | 0.63717 | 0.656647 |
| 14 | 114546000 | 114548799 | 22.58285 | 39.42892 | 40.14912 | 17.682 | 0.361539 | 0.744877 | 0.656665 |
| 1 | 65456800 | 65457599 | 14.34565 | 25.18588 | 39.80498 | 17.60939 | -0.26962 | 0.613425 | 0.656666 |
| 1 | 112667200 | 112667999 | 15.53658 | 27.57725 | 30.87696 | 13.63642 | 0.23097 | 0.485196 | 0.656691 |
| 1 | 75662400 | 75663999 | 12.98163 | 22.87488 | 36.80914 | 16.29228 | -0.29372 | 0.555946 | 0.656693 |
| 7 | 48243400 | 48249599 | 67.90988 | 108.3735 | 94.76707 | 39.75258 | 0.532956 | 2.137353 | 0.656695 |
| 4 | 136775600 | 136777999 | 40.65467 | 65.39873 | 95.43192 | 40.50298 | -0.19112 | 1.85921 | 0.656695 |
| 1 | 48538200 | 48542799 | 33.35665 | 55.6362 | 57.58674 | 25.0382 | 0.331615 | 1.244709 | 0.656697 |
| 12 | 82162000 | 82163799 | 11.47912 | 20.81153 | 32.75704 | 14.16746 | -0.23717 | 0.334109 | 0.656727 |
| 1 | 81972400 | 81973399 | 12.45221 | 22.87085 | 24.65758 | 10.48758 | 0.321696 | 0.249163 | 0.656748 |
| 1 | 127630600 | 127631599 | 13.42844 | 23.784 | 36.65101 | 16.20565 | -0.22883 | 0.501549 | 0.656755 |
| 6 | 120786600 | 120789999 | 41.29463 | 66.41957 | 96.81869 | 41.08382 | -0.18984 | 1.864372 | 0.656773 |
| 6 | 65127600 | 65129199 | 14.22189 | 24.94196 | 39.37508 | 17.4035 | -0.26787 | 0.630379 | 0.656791 |
| 11 | 49233000 | 49234399 | 11.07679 | 20.0896 | 32.82937 | 14.19213 | -0.2902 | 0.331403 | 0.656824 |
| 13 | 106744000 | 106745199 | 19.06424 | 33.19036 | 36.38198 | 15.99411 | 0.255381 | 0.762018 | 0.656842 |
| 17 | 66253200 | 66254599 | 14.73791 | 26.48151 | 29.2411 | 12.7815 | 0.260808 | 0.398667 | 0.65685 |
| 16 | 49791800 | 49794199 | 21.5755 | 37.50734 | 40.84233 | 17.93841 | 0.264523 | 0.799111 | 0.656852 |
| 2 | 66893200 | 66893799 | 8.559973 | 16.53879 | 20.37988 | 7.826621 | 0.255604 | -0.15747 | 0.656853 |
| 2 | 70710200 | 70713199 | 17.39174 | 30.1864 | 46.33515 | 20.33069 | -0.22879 | 0.810623 | 0.656863 |
| 9 | 55281000 | 55281599 | 10.29228 | 19.53357 | 23.26788 | 9.375063 | 0.24051 | 0.023962 | 0.656882 |
| 12 | 62084600 | 62084999 | 7.189572 | 13.66793 | 25.03392 | 10.05085 | -0.37006 | -0.01082 | 0.656897 |
| 6 | 96871200 | 96873199 | 20.34164 | 35.87452 | 35.66257 | 15.7821 | 0.397779 | 0.58702 | 0.656903 |
| 5 | 109121400 | 109121999 | 15.86191 | 27.66596 | 42.60807 | 18.75283 | -0.23262 | 0.730743 | 0.65692 |
| 19 | 55155800 | 55159999 | 48.26223 | 77.8213 | 84.26979 | 35.88584 | 0.242499 | 1.792723 | 0.656936 |
| 2 | 23861600 | 23864599 | 33.33115 | 55.13754 | 60.90628 | 26.33457 | 0.237454 | 1.338059 | 0.656962 |
| 2 | 128926600 | 128929199 | 34.73888 | 57.51659 | 53.69351 | 23.22996 | 0.479611 | 1.326391 | 0.656989 |
| X | 81686600 | 81692399 | 56.17124 | 90.40847 | 75.59038 | 32.10718 | 0.611687 | 1.859012 | 0.657003 |
| 2 | 176931800 | 176935199 | 55.47373 | 88.19586 | 125.4941 | 52.3743 | -0.17405 | 2.251302 | 0.657009 |
| 12 | 115427400 | 115430199 | 43.84715 | 71.3279 | 73.22492 | 31.4281 | 0.331762 | 1.584471 | 0.657031 |
| 16 | 15569200 | 15570199 | 9.702322 | 18.02192 | 29.42844 | 12.32648 | -0.25725 | 0.17582 | 0.657041 |
| 14 | 89273800 | 89278999 | 87.79667 | 137.2721 | 150.5513 | 62.46015 | 0.181281 | 2.647527 | 0.65705 |
| 10 | 68868800 | 68873199 | 52.22913 | 83.7121 | 94.0116 | 39.6837 | 0.179584 | 1.987094 | 0.657064 |
| 5 | 5372800 | 5373999 | 16.77004 | 29.15761 | 44.72792 | 19.64677 | -0.22756 | 0.781569 | 0.657065 |
| 11 | 50960400 | 50963599 | 37.78787 | 62.0786 | 65.69576 | 28.30385 | 0.296331 | 1.434363 | 0.657112 |
| X | 4784400 | 4785799 | 25.03484 | 43.00107 | 37.46579 | 16.43352 | 0.581935 | 0.928318 | 0.657115 |
| 4 | 119027000 | 119028199 | 14.01727 | 24.50803 | 40.60555 | 17.90103 | -0.33691 | 0.684695 | 0.657139 |
| 11 | 75506600 | 75508999 | 15.52113 | 27.02922 | 42.65145 | 18.75347 | -0.26777 | 0.750604 | 0.657165 |
| 10 | 69515400 | 69523199 | 74.52627 | 118.266 | 122.8536 | 51.17663 | 0.276569 | 2.320572 | 0.657198 |

| | | | | | | | | | |
|---|---|---|---|---|---|---|---|---|---|
| 3 | 33582400 | 33586999 | 38.54724 | 63.61589 | 62.53755 | 26.99588 | 0.404932 | 1.378493 | 0.657207 |
| Y | 1883800 | 1887399 | 59.17493 | 94.18777 | 84.59391 | 35.35703 | 0.490408 | 2.205369 | 0.657218 |
| 14 | 47069200 | 47070199 | 7.458922 | 14.13855 | 26.56058 | 10.73 | -0.41516 | 0.012567 | 0.657221 |
| 9 | 70260400 | 70266599 | 50.77612 | 80.73121 | 134.7548 | 56.24193 | -0.40393 | 2.248062 | 0.657235 |
| 4 | 46772000 | 46774599 | 27.39056 | 45.84465 | 51.80655 | 22.57458 | 0.205289 | 1.184899 | 0.657237 |
| 6 | 36682000 | 36683799 | 16.00914 | 28.78577 | 31.24979 | 13.65148 | 0.285908 | 0.408737 | 0.657249 |
| 1 | 65927600 | 65932399 | 55.47913 | 88.79833 | 98.58939 | 41.52964 | 0.193721 | 2.034324 | 0.657257 |
| 10 | 4535000 | 4538999 | 31.41039 | 51.3731 | 76.63588 | 32.95687 | -0.20053 | 1.483988 | 0.657268 |
| 1 | 89115400 | 89117999 | 17.24218 | 29.96594 | 45.783 | 20.10493 | -0.22189 | 0.790919 | 0.657275 |
| X | 22689000 | 22690599 | 23.02758 | 40.03215 | 41.93792 | 18.41974 | 0.320279 | 0.810779 | 0.657283 |
| 19 | 29016000 | 29020999 | 40.17952 | 65.12456 | 73.40589 | 31.43075 | 0.192659 | 1.644302 | 0.657292 |
| 10 | 122919200 | 122922199 | 25.78596 | 42.69037 | 64.68048 | 27.982 | -0.21689 | 1.312688 | 0.657317 |
| 3 | 148854000 | 148856799 | 22.35344 | 38.99406 | 35.00048 | 15.40597 | 0.542636 | 0.74893 | 0.657337 |
| 2 | 89594200 | 89594799 | 15.36132 | 27.27909 | 30.39388 | 13.41771 | 0.238389 | 0.48004 | 0.657345 |
| 5 | 58476200 | 58483399 | 184.013 | 269.0171 | 370.9138 | 152.8052 | -0.23115 | 3.913033 | 0.657351 |
| 8 | 64079600 | 64083999 | 131.858 | 199.8309 | 214.2934 | 88.68531 | 0.17855 | 3.257747 | 0.657364 |
| 6 | 130665200 | 130668199 | 30.2794 | 51.47258 | 50.13496 | 21.97254 | 0.419505 | 1.05455 | 0.65737 |
| 8 | 47115200 | 47116199 | 15.08947 | 26.85894 | 29.50061 | 12.99777 | 0.261 | 0.463072 | 0.657398 |
| 6 | 126447600 | 126450599 | 28.92996 | 47.57852 | 70.91086 | 30.56478 | -0.19562 | 1.404977 | 0.657418 |
| 11 | 65194600 | 65199199 | 43.67728 | 70.639 | 78.23405 | 33.44401 | 0.215034 | 1.697358 | 0.657428 |
| 14 | 120581200 | 120591199 | 106.9172 | 163.6401 | 226.9484 | 94.17398 | -0.18071 | 3.052326 | 0.657437 |
| 3 | 11074000 | 11075799 | 18.3551 | 31.78377 | 49.01846 | 21.50614 | -0.23638 | 0.839182 | 0.657445 |
| 16 | 44726600 | 44727799 | 10.62054 | 19.37868 | 31.26983 | 13.41632 | -0.26543 | 0.285841 | 0.657446 |
| 6 | 44560800 | 44562999 | 22.00821 | 38.34021 | 40.77599 | 17.93298 | 0.298741 | 0.772914 | 0.657455 |
| 15 | 12989200 | 12991599 | 31.50419 | 52.71504 | 55.8155 | 24.31718 | 0.298879 | 1.203839 | 0.657457 |
| 18 | 62425400 | 62426399 | 8.736617 | 16.40826 | 27.48297 | 11.30153 | -0.27219 | 0.092566 | 0.657463 |
| 5 | 102320400 | 102323599 | 27.47388 | 46.15113 | 51.80942 | 22.62753 | 0.214637 | 1.145564 | 0.657464 |
| 9 | 25351400 | 25352399 | 12.21945 | 21.98276 | 34.6549 | 15.13251 | -0.24875 | 0.385903 | 0.657474 |
| 3 | 84674600 | 84681599 | 63.65194 | 101.1881 | 112.2113 | 46.82613 | 0.184892 | 2.256598 | 0.657503 |
| 9 | 78522000 | 78524999 | 18.52134 | 31.84101 | 50.07999 | 21.96704 | -0.26676 | 0.905162 | 0.657513 |
| 12 | 40906000 | 40910399 | 58.47787 | 93.32991 | 104.3905 | 43.79501 | 0.178556 | 2.12674 | 0.657518 |
| X | 107065400 | 107067199 | 20.78509 | 36.26051 | 39.76554 | 17.5041 | 0.254941 | 0.747148 | 0.657523 |
| 2 | 17989600 | 17991799 | 36.34375 | 59.26104 | 66.71298 | 28.65774 | 0.201443 | 1.530719 | 0.657549 |
| X | 16171800 | 16173799 | 17.76896 | 31.87787 | 26.22342 | 11.47757 | 0.682254 | 0.426994 | 0.657553 |
| 9 | 27823400 | 27824199 | 12.72453 | 22.50592 | 36.31655 | 16.06223 | -0.29535 | 0.514712 | 0.657556 |
| 13 | 62882200 | 62884999 | 23.09385 | 39.70884 | 44.14829 | 19.36527 | 0.231749 | 0.91588 | 0.657558 |
| 7 | 47999800 | 48004999 | 88.08475 | 137.5181 | 151.9323 | 63.04272 | 0.16905 | 2.674853 | 0.657562 |
| 17 | 67191600 | 67192999 | 9.421644 | 17.34979 | 29.39213 | 12.45708 | -0.32207 | 0.222633 | 0.657564 |
| 17 | 22190600 | 22204799 | 261.1442 | 380.9935 | 361.7837 | 149.072 | 0.303751 | 3.951938 | 0.657566 |
| 18 | 65413400 | 65427399 | 134.8762 | 201.3326 | 322.3125 | 132.8606 | -0.41837 | 3.558102 | 0.65757 |
| 8 | 77588600 | 77589799 | 12.94429 | 23.08666 | 35.6617 | 15.68585 | -0.22663 | 0.448662 | 0.657579 |
| 3 | 152620000 | 152623799 | 47.28333 | 76.16817 | 84.88713 | 36.10059 | 0.19941 | 1.820764 | 0.657604 |
| 10 | 106223200 | 106227999 | 70.94466 | 112.302 | 124.992 | 51.96407 | 0.173939 | 2.385081 | 0.657607 |
| 13 | 8684800 | 8685199 | 7.653026 | 14.8634 | 18.35844 | 6.8686 | 0.287875 | -0.26256 | 0.657616 |
| 13 | 60003000 | 60006399 | 24.02393 | 40.15532 | 62.12533 | 27.04782 | -0.2465 | 1.196319 | 0.657617 |
| 15 | 75412600 | 75414999 | 43.30353 | 69.99986 | 78.13642 | 33.38116 | 0.202924 | 1.710022 | 0.657623 |
| 11 | 10181200 | 10182599 | 17.90911 | 31.45918 | 35.02849 | 15.50344 | 0.233317 | 0.619251 | 0.657641 |

| | | | | | | | | | |
|---|---|---|---|---|---|---|---|---|---|
| 8 | 51725600 | 51727199 | 29.76714 | 49.80508 | 45.9829 | 20.03254 | 0.495906 | 1.190564 | 0.657648 |
| 8 | 128453400 | 128454999 | 23.78409 | 40.96104 | 44.69118 | 19.60436 | 0.259386 | 0.907916 | 0.657667 |
| 3 | 99279800 | 99280399 | 6.090847 | 11.86285 | 21.98611 | 8.032005 | -0.25994 | -0.35219 | 0.65768 |
| 6 | 63599200 | 63599599 | 6.090847 | 11.86285 | 21.98611 | 8.032005 | -0.25994 | -0.35219 | 0.65768 |
| 13 | 49392400 | 49393599 | 13.0815 | 23.87049 | 27.0259 | 11.59458 | 0.242669 | 0.295353 | 0.657695 |
| 14 | 24815400 | 24816999 | 15.86899 | 27.8941 | 45.5389 | 20.1661 | -0.31632 | 0.613031 | 0.657695 |
| 3 | 124620600 | 124625999 | 95.5352 | 148.1721 | 163.3815 | 67.85679 | 0.164523 | 2.791638 | 0.657697 |
| 5 | 113186800 | 113195399 | 75.76171 | 117.7889 | 170.2375 | 70.67871 | -0.22269 | 2.742248 | 0.657709 |
| 18 | 35555200 | 35559199 | 55.42805 | 88.80866 | 96.84564 | 40.85856 | 0.221314 | 2.001225 | 0.657712 |
| 6 | 102950400 | 102950799 | 4.909887 | 9.593889 | 18.86208 | 6.556601 | -0.27773 | -0.55185 | 0.657714 |
| 5 | 30434200 | 30436199 | 36.75874 | 60.54994 | 53.61059 | 23.12336 | 0.554516 | 1.394371 | 0.657715 |
| 4 | 122045800 | 122047599 | 30.32767 | 50.72447 | 45.47136 | 19.80514 | 0.538348 | 1.196805 | 0.657721 |
| 3 | 54680000 | 54681599 | 19.1006 | 32.95112 | 50.08322 | 21.97078 | -0.21647 | 0.879198 | 0.657729 |
| 4 | 38971400 | 38972999 | 15.86131 | 28.83873 | 29.73041 | 12.81255 | 0.372454 | 0.339795 | 0.657739 |
| 7 | 18523800 | 18528999 | 39.28553 | 63.27414 | 94.80566 | 40.30906 | -0.22701 | 1.816689 | 0.657746 |
| 2 | 58754400 | 58755199 | 9.996155 | 18.2231 | 31.27967 | 13.43555 | -0.35514 | 0.285162 | 0.657755 |
| 7 | 31567800 | 31570199 | 22.78855 | 39.70804 | 40.71217 | 17.9073 | 0.351338 | 0.773126 | 0.65778 |
| 1 | 102729000 | 102738799 | 162.3782 | 243.3447 | 261.0323 | 107.6829 | 0.164074 | 3.486626 | 0.657789 |
| 6 | 84627800 | 84629599 | 17.04955 | 30.54156 | 32.2211 | 14.12045 | 0.323877 | 0.441216 | 0.657817 |
| 15 | 91537200 | 91538199 | 16.72241 | 29.4369 | 32.55438 | 14.41196 | 0.243908 | 0.581409 | 0.657831 |
| 17 | 92487200 | 92488999 | 21.9639 | 37.91232 | 42.56278 | 18.672 | 0.218928 | 0.875079 | 0.657848 |
| 6 | 7167200 | 7172799 | 75.90862 | 120.3375 | 124.8782 | 51.96718 | 0.276612 | 2.350328 | 0.657854 |
| X | 35675600 | 35680799 | 40.49666 | 65.35168 | 104.873 | 44.7056 | -0.32248 | 1.769981 | 0.657863 |
| 6 | 27506800 | 27510199 | 69.70797 | 110.2028 | 122.0634 | 50.71356 | 0.179435 | 2.412705 | 0.657885 |
| 17 | 92133200 | 92133999 | 10.80798 | 20.49459 | 23.70911 | 9.570331 | 0.279611 | 0.033875 | 0.657896 |
| 6 | 124537400 | 124538799 | 18.5119 | 32.52362 | 35.92825 | 15.90397 | 0.24457 | 0.620407 | 0.657897 |
| 13 | 93655800 | 93675799 | 223.4326 | 323.6459 | 424.5546 | 175.0734 | -0.17226 | 4.05616 | 0.657914 |
| 5 | 149055400 | 149057599 | 27.72217 | 46.33237 | 52.35632 | 22.79325 | 0.205362 | 1.201822 | 0.657944 |
| X | 153829000 | 153831599 | 35.53001 | 58.92498 | 60.39409 | 26.1591 | 0.345507 | 1.317801 | 0.657959 |
| 2 | 170241800 | 170243599 | 13.13045 | 23.48087 | 36.70806 | 16.11003 | -0.24162 | 0.435825 | 0.657972 |
| 12 | 58834000 | 58836999 | 39.49645 | 64.62665 | 68.27602 | 29.36647 | 0.296409 | 1.492345 | 0.657973 |
| 4 | 21776000 | 21776799 | 7.339949 | 14.01026 | 25.0797 | 9.983618 | -0.32622 | -0.03589 | 0.657979 |
| 10 | 104700800 | 104701199 | 7.318028 | 14.04729 | 24.65107 | 9.665004 | -0.27757 | -0.0877 | 0.657983 |
| 1 | 116048800 | 116051399 | 20.46675 | 35.54374 | 39.90752 | 17.51664 | 0.220465 | 0.807523 | 0.658 |
| 16 | 20638800 | 20641599 | 19.81108 | 33.56654 | 52.70504 | 23.07897 | -0.26711 | 1.047047 | 0.658001 |
| X | 18370000 | 18371999 | 34.2553 | 56.4284 | 63.02895 | 27.18614 | 0.220239 | 1.392662 | 0.65803 |
| 1 | 57657800 | 57658399 | 7.776776 | 15.01337 | 27.22685 | 10.49585 | -0.30214 | -0.13172 | 0.658046 |
| 12 | 13897600 | 13900799 | 43.74161 | 70.66975 | 78.96952 | 33.71352 | 0.200503 | 1.728589 | 0.658049 |
| 2 | 32008600 | 32011599 | 33.56067 | 54.93178 | 60.43916 | 25.99877 | 0.238285 | 1.467004 | 0.65805 |
| X | 67498800 | 67502999 | 50.48589 | 81.54926 | 81.6926 | 34.86687 | 0.357509 | 1.748472 | 0.658052 |
| 2 | 90572800 | 90576199 | 23.87675 | 40.9038 | 45.78676 | 20.07938 | 0.221173 | 0.955647 | 0.658063 |
| 1 | 133477800 | 133481199 | 27.66739 | 46.40842 | 52.36152 | 22.84762 | 0.207442 | 1.162487 | 0.658065 |
| 4 | 143373600 | 143373999 | 14.12065 | 24.7351 | 39.47926 | 17.4321 | -0.28344 | 0.65107 | 0.658089 |
| 6 | 35941600 | 35942399 | 8.494114 | 16.03383 | 27.54736 | 11.22715 | -0.29864 | 0.054153 | 0.658111 |
| X | 31671400 | 31672799 | 24.95016 | 43.59131 | 39.45276 | 17.38262 | 0.530923 | 0.746806 | 0.658125 |
| 10 | 72328600 | 72331999 | 30.09825 | 50.16316 | 56.24615 | 24.44355 | 0.216615 | 1.242877 | 0.658144 |
| 5 | 76269000 | 76274999 | 69.97397 | 111.3185 | 115.9556 | 48.43015 | 0.276114 | 2.241803 | 0.658149 |

| | | | | | | | | | |
|---|---|---|---|---|---|---|---|---|---|
| 13 | 33942200 | 33943599 | 14.51353 | 25.64013 | 40.21648 | 17.79126 | -0.25592 | 0.541035 | 0.658157 |
| 8 | 22433800 | 22434399 | 6.661677 | 12.93549 | 24.10193 | 9.031823 | -0.30233 | -0.2521 | 0.658167 |
| 13 | 21386400 | 21397599 | 197.7193 | 293.0489 | 304.6395 | 125.4682 | 0.194738 | 3.69189 | 0.6582 |
| 7 | 52947000 | 52948999 | 21.97443 | 38.20994 | 41.37071 | 18.17319 | 0.272795 | 0.799211 | 0.658203 |
| 17 | 70303600 | 70304999 | 16.26114 | 28.74175 | 31.98008 | 14.14619 | 0.237412 | 0.533502 | 0.658226 |
| 17 | 16762800 | 16764799 | 17.60498 | 31.04184 | 34.29786 | 15.17878 | 0.245909 | 0.569767 | 0.658229 |
| 13 | 7383000 | 7383599 | 15.40782 | 27.35729 | 30.44324 | 13.44132 | 0.24012 | 0.482242 | 0.65823 |
| 11 | 63404400 | 63405999 | 17.9813 | 32.55263 | 31.25146 | 13.54086 | 0.469998 | 0.384841 | 0.658255 |
| 8 | 34536800 | 34537999 | 12.39811 | 22.3769 | 35.23206 | 15.32376 | -0.24321 | 0.370032 | 0.658271 |
| 11 | 31105400 | 31105999 | 5.227687 | 10.18991 | 19.70026 | 7.121488 | -0.30552 | -0.40759 | 0.658289 |
| 13 | 93166800 | 93168399 | 14.89589 | 26.66646 | 29.24428 | 12.82285 | 0.267568 | 0.422587 | 0.658327 |
| 3 | 42137200 | 42144599 | 149.5627 | 225.167 | 242.918 | 100.3293 | 0.161689 | 3.395125 | 0.658334 |
| 8 | 18151200 | 18153799 | 28.20465 | 47.67911 | 44.77123 | 19.59128 | 0.471679 | 1.087111 | 0.658351 |
| 7 | 68184200 | 68187999 | 77.38 | 121.6277 | 133.3827 | 55.32967 | 0.18732 | 2.543979 | 0.658377 |
| 18 | 70010400 | 70012799 | 16.78886 | 29.32909 | 45.15459 | 19.8934 | -0.23212 | 0.715401 | 0.658383 |
| 3 | 16581600 | 16582599 | 14.40534 | 26.14445 | 29.42367 | 12.7089 | 0.244887 | 0.341078 | 0.658383 |
| X | 68046000 | 68049599 | 38.31658 | 62.43301 | 69.50099 | 29.84756 | 0.216883 | 1.549676 | 0.658384 |
| 1 | 18659200 | 18660599 | 11.09249 | 20.42712 | 32.99474 | 13.98349 | -0.25446 | 0.241385 | 0.658402 |
| 17 | 94948000 | 94948799 | 15.29024 | 27.28759 | 30.44241 | 13.38318 | 0.240801 | 0.4504 | 0.658413 |
| 7 | 52970600 | 52971799 | 16.96751 | 29.4122 | 45.30079 | 19.87571 | -0.23393 | 0.820391 | 0.658419 |
| 5 | 62886800 | 62887399 | 10.5597 | 20.48348 | 22.31156 | 8.411264 | 0.45732 | -0.1906 | 0.658419 |
| 9 | 58658800 | 58659799 | 13.67154 | 24.76246 | 27.91719 | 12.0888 | 0.239953 | 0.343089 | 0.658449 |
| 10 | 63205000 | 63207199 | 23.18047 | 38.95132 | 59.20268 | 25.85916 | -0.22088 | 1.139581 | 0.658463 |
| 14 | 82307600 | 82308399 | 9.126019 | 17.23701 | 28.6737 | 11.67133 | -0.25078 | 0.059497 | 0.658482 |
| 9 | 10993000 | 10993599 | 11.38812 | 21.1998 | 24.11404 | 10.06573 | 0.264318 | 0.161311 | 0.658513 |
| 4 | 60150400 | 60151199 | 7.48307 | 14.33382 | 25.14496 | 9.918552 | -0.28523 | -0.06433 | 0.658532 |
| 6 | 57029000 | 57030199 | 30.26105 | 50.1411 | 56.73063 | 24.55692 | 0.203214 | 1.309133 | 0.658535 |
| 8 | 62501800 | 62502199 | 6.107699 | 11.82995 | 22.15242 | 8.409587 | -0.32602 | -0.217 | 0.658553 |
| 14 | 3374400 | 3375399 | 23.76115 | 39.51744 | 62.24861 | 27.01459 | -0.27216 | 1.256086 | 0.658558 |
| 5 | 15864200 | 15864999 | 8.159044 | 15.54587 | 27.25268 | 10.88977 | -0.3025 | -0.01496 | 0.65856 |
| 14 | 103547800 | 103550199 | 38.85773 | 63.59961 | 67.59934 | 29.07843 | 0.287987 | 1.48695 | 0.658563 |
| 12 | 65473600 | 65475999 | 20.27737 | 35.14747 | 38.93295 | 17.08179 | 0.239423 | 0.825246 | 0.658571 |
| 17 | 45471600 | 45476799 | 48.76053 | 77.74012 | 114.1332 | 47.81287 | -0.21478 | 2.153387 | 0.658576 |
| 5 | 19426600 | 19428399 | 15.19701 | 26.69348 | 40.81074 | 18.06171 | -0.22194 | 0.613668 | 0.658576 |
| X | 96801400 | 96804999 | 54.05949 | 86.94004 | 90.08458 | 38.21931 | 0.301385 | 1.885905 | 0.658589 |
| 8 | 84441000 | 84442599 | 11.41792 | 20.25187 | 34.32706 | 15.16946 | -0.36421 | 0.471613 | 0.658612 |
| 4 | 128721600 | 128724199 | 34.08871 | 56.1342 | 62.99701 | 27.16832 | 0.213249 | 1.396484 | 0.658623 |
| 16 | 31558800 | 31559799 | 11.1478 | 20.47844 | 32.94846 | 14.01078 | -0.25218 | 0.256861 | 0.658639 |
| 1 | 98092800 | 98094399 | 12.94934 | 22.98125 | 35.65571 | 15.74902 | -0.23712 | 0.479649 | 0.658661 |
| 5 | 68540000 | 68542599 | 17.74018 | 31.87365 | 31.56035 | 13.79614 | 0.417159 | 0.424853 | 0.658664 |
| 7 | 9599000 | 9600399 | 20.57812 | 35.9159 | 31.14811 | 13.7148 | 0.592033 | 0.725893 | 0.658679 |
| X | 149828800 | 149831199 | 32.2494 | 54.16351 | 54.06742 | 23.61106 | 0.383609 | 1.169048 | 0.65871 |
| 17 | 7106800 | 7107999 | 13.27648 | 23.84336 | 38.8631 | 16.99467 | -0.29808 | 0.413465 | 0.658715 |
| 16 | 90437800 | 90439799 | 13.87598 | 24.44946 | 38.21348 | 16.91405 | -0.2522 | 0.562719 | 0.658733 |
| 8 | 17978400 | 17980199 | 13.89982 | 25.78467 | 27.51524 | 11.54582 | 0.349752 | 0.209138 | 0.658759 |
| 2 | 21825800 | 21827599 | 19.45311 | 35.1385 | 31.06813 | 13.49911 | 0.585474 | 0.406726 | 0.658776 |
| 7 | 42024800 | 42049599 | 663.7523 | 883.7612 | 839.632 | 350.7757 | 0.170911 | 5.349067 | 0.658783 |

| | | | | | | | | | |
|---|---|---|---|---|---|---|---|---|---|
| 3 | 107406200 | 107410199 | 57.14313 | 91.14008 | 101.7018 | 42.62801 | 0.18093 | 2.144913 | 0.658789 |
| 2 | 169299000 | 169303999 | 53.9439 | 86.42246 | 95.79166 | 40.40804 | 0.197689 | 2.002001 | 0.65883 |
| 1 | 168860800 | 168862799 | 13.38332 | 23.84312 | 37.14449 | 16.35341 | -0.23996 | 0.459912 | 0.658833 |
| 7 | 21021800 | 21025999 | 87.81817 | 136.7879 | 150.4468 | 62.44471 | 0.172974 | 2.714011 | 0.658861 |
| 4 | 96151400 | 96154799 | 43.10374 | 69.82984 | 76.46668 | 32.72942 | 0.233635 | 1.663155 | 0.658869 |
| 16 | 42503400 | 42504999 | 16.21842 | 29.65438 | 29.89078 | 12.79134 | 0.41148 | 0.313004 | 0.658883 |
| 5 | 142665200 | 142665799 | 15.15139 | 27.09613 | 29.97262 | 13.15354 | 0.254799 | 0.436082 | 0.658888 |
| 1 | 78618800 | 78620999 | 17.45684 | 30.70119 | 34.40334 | 15.23333 | 0.224285 | 0.59744 | 0.658913 |
| 12 | 95768400 | 95770199 | 12.08048 | 21.25609 | 42.4514 | 18.79508 | -0.60472 | 0.575574 | 0.658917 |
| 6 | 57896400 | 57896999 | 15.59991 | 27.62978 | 29.58215 | 13.0801 | 0.293855 | 0.502676 | 0.658927 |
| 1 | 61412400 | 61413999 | 13.59506 | 23.98468 | 37.22833 | 16.47385 | -0.24141 | 0.546744 | 0.658927 |
| 7 | 13595200 | 13619599 | 53.20366 | 85.99603 | 68.23981 | 29.14996 | 0.693973 | 1.725763 | 0.65893 |
| 4 | 115078000 | 115078799 | 8.078237 | 15.431 | 26.17845 | 10.40456 | -0.24796 | -0.03223 | 0.658941 |
| 6 | 31622200 | 31624399 | 15.04858 | 26.37475 | 40.40459 | 17.8488 | -0.22472 | 0.648099 | 0.658966 |
| 15 | 5757600 | 5759399 | 18.56314 | 32.88744 | 34.16922 | 15.10747 | 0.336905 | 0.525754 | 0.658977 |
| 7 | 59652200 | 59656799 | 102.7098 | 159.1788 | 141.2978 | 58.69111 | 0.476226 | 2.806719 | 0.659016 |
| 15 | 33150600 | 33151199 | 8.67478 | 16.50853 | 27.77794 | 11.12812 | -0.24673 | 0.002074 | 0.659024 |
| 6 | 69824400 | 69829799 | 78.12589 | 123.7315 | 113.8197 | 47.33129 | 0.449037 | 2.376204 | 0.659028 |
| 3 | 152338800 | 152342199 | 38.43391 | 62.87445 | 67.88163 | 29.19453 | 0.264902 | 1.48911 | 0.659031 |
| 18 | 30143000 | 30149399 | 88.59478 | 138.0994 | 153.0695 | 63.52629 | 0.162639 | 2.701769 | 0.659044 |
| 6 | 134946800 | 134948799 | 18.6732 | 32.28943 | 50.10396 | 21.98145 | -0.24562 | 0.854125 | 0.659064 |
| 16 | 12902200 | 12905199 | 23.54873 | 39.47514 | 59.86497 | 26.11136 | -0.21777 | 1.162967 | 0.659072 |
| 10 | 82307400 | 82314799 | 81.05324 | 128.3324 | 128.8505 | 53.57432 | 0.322636 | 2.388167 | 0.659076 |
| 7 | 6608200 | 6629199 | 655.514 | 873.4734 | 830.3471 | 346.8597 | 0.171186 | 5.336317 | 0.659087 |
| 14 | 36818600 | 36823199 | 29.76431 | 48.23895 | 92.83456 | 39.7471 | -0.57739 | 1.636448 | 0.659097 |
| 8 | 26351400 | 26355599 | 38.16031 | 61.48569 | 92.20783 | 39.22657 | -0.22757 | 1.80217 | 0.659105 |
| 2 | 176936000 | 176937199 | 9.878267 | 18.98998 | 22.40495 | 8.746967 | 0.294989 | -0.08586 | 0.659111 |
| 15 | 49596400 | 49601199 | 100.2755 | 155.0627 | 170.3911 | 70.77643 | 0.166199 | 2.847748 | 0.659116 |
| 7 | 86536400 | 86537599 | 16.77248 | 29.08208 | 45.29446 | 19.8731 | -0.24992 | 0.816014 | 0.659125 |
| 5 | 97143000 | 97146599 | 25.2003 | 42.48046 | 46.97348 | 20.53739 | 0.236615 | 1.103926 | 0.659137 |
| 18 | 47996600 | 47998399 | 17.31556 | 30.89388 | 32.66422 | 14.36313 | 0.317342 | 0.467621 | 0.659164 |
| 12 | 114384200 | 114402799 | 230.4498 | 340.2945 | 318.851 | 131.309 | 0.339527 | 3.758866 | 0.659169 |
| 4 | 47467200 | 47472599 | 103.0424 | 158.998 | 174.3313 | 72.41395 | 0.166997 | 2.889635 | 0.659173 |
| 1 | 195007800 | 195010799 | 39.53576 | 64.54206 | 60.90093 | 26.17234 | 0.456612 | 1.516836 | 0.659175 |
| 5 | 47341600 | 47342399 | 12.60374 | 22.42234 | 35.15064 | 15.49415 | -0.24998 | 0.460447 | 0.659181 |
| 6 | 68127400 | 68129599 | 16.49261 | 28.92028 | 47.76948 | 21.11129 | -0.3332 | 0.657465 | 0.659191 |
| 8 | 91539800 | 91541199 | 13.10276 | 23.12918 | 36.72665 | 16.25002 | -0.27376 | 0.535953 | 0.659194 |
| 5 | 48674000 | 48674599 | 4.933925 | 9.634102 | 18.75183 | 6.602475 | -0.28062 | -0.5086 | 0.659216 |
| 10 | 31872800 | 31873799 | 12.42473 | 23.89755 | 22.95968 | 8.950356 | 0.590842 | -0.06329 | 0.659218 |
| 13 | 23167800 | 23172799 | 77.72459 | 122.8247 | 116.0824 | 48.21839 | 0.407564 | 2.42863 | 0.659242 |
| 17 | 49550400 | 49550999 | 5.071531 | 9.894978 | 19.21782 | 6.859295 | -0.295 | -0.45558 | 0.659263 |
| 2 | 112020600 | 112021199 | 8.079314 | 15.33212 | 26.04504 | 10.49619 | -0.2675 | 0.012233 | 0.659266 |
| 4 | 16197400 | 16197999 | 8.079314 | 15.33212 | 26.04504 | 10.49619 | -0.2675 | 0.012233 | 0.659266 |
| 11 | 30704800 | 30705599 | 8.079314 | 15.33212 | 26.04504 | 10.49619 | -0.2675 | 0.012233 | 0.659266 |
| 14 | 75732000 | 75734399 | 22.82082 | 39.14692 | 44.19673 | 19.38416 | 0.209133 | 0.936647 | 0.659268 |
| 8 | 76838200 | 76840199 | 19.04151 | 33.75556 | 34.09513 | 15.07287 | 0.37772 | 0.521783 | 0.65927 |
| 6 | 112378200 | 112383399 | 51.77066 | 82.22798 | 137.4009 | 57.2874 | -0.40693 | 2.279518 | 0.659275 |

| | | | | | | | | | |
|---|---|---|---|---|---|---|---|---|---|
| 6 | 100545800 | 100548599 | 20.70136 | 34.74374 | 55.86631 | 24.38573 | -0.30179 | 1.138342 | 0.659279 |
| 7 | 94321600 | 94322599 | 14.68203 | 26.98815 | 24.34661 | 10.34393 | 0.578088 | 0.260983 | 0.659318 |
| 7 | 36920000 | 36922799 | 26.90378 | 45.71855 | 49.5712 | 21.72487 | 0.265254 | 1.044328 | 0.659332 |
| 10 | 118463600 | 118466199 | 20.45984 | 34.18138 | 66.19 | 28.80862 | -0.56912 | 1.195876 | 0.65934 |
| 2 | 39787200 | 39788999 | 19.24305 | 34.00849 | 34.59717 | 15.30418 | 0.36568 | 0.555108 | 0.659341 |
| 13 | 67474800 | 67477199 | 20.14509 | 35.67418 | 34.61064 | 15.304 | 0.435048 | 0.541739 | 0.659351 |
| 12 | 116493200 | 116494399 | 14.04859 | 24.81611 | 38.17443 | 16.88825 | -0.22793 | 0.536589 | 0.659407 |
| 17 | 74366600 | 74367399 | 14.04859 | 24.81611 | 38.17443 | 16.88825 | -0.22793 | 0.536589 | 0.659407 |
| 17 | 38761800 | 38764599 | 25.33217 | 44.32759 | 38.88073 | 17.15255 | 0.576036 | 0.725884 | 0.659407 |
| 6 | 121733000 | 121735799 | 39.69588 | 64.03714 | 93.8994 | 40.00977 | -0.19317 | 1.771747 | 0.659419 |
| 1 | 117810600 | 117820999 | 180.2359 | 269.8551 | 272.2461 | 112.2885 | 0.251302 | 3.507642 | 0.65942 |
| 9 | 96846400 | 96848399 | 16.77654 | 29.15148 | 44.74252 | 19.64581 | -0.2284 | 0.790295 | 0.659421 |
| X | 81479600 | 81486399 | 120.5748 | 185.4055 | 188.4297 | 78.24893 | 0.272558 | 2.964251 | 0.659443 |
| 14 | 59866800 | 59867999 | 7.031012 | 13.43101 | 24.56364 | 9.76212 | -0.35481 | -0.0465 | 0.659473 |
| 4 | 152900000 | 152901799 | 18.51003 | 31.5579 | 50.05648 | 21.94332 | -0.28066 | 0.983921 | 0.659478 |
| 15 | 10042600 | 10045199 | 20.37858 | 35.08522 | 58.43646 | 25.63366 | -0.34879 | 0.910404 | 0.659483 |
| 13 | 51499200 | 51500599 | 10.47333 | 19.15807 | 30.79278 | 13.16985 | -0.25669 | 0.269838 | 0.65949 |
| 17 | 12985200 | 12990399 | 61.44288 | 97.18186 | 137.8757 | 57.29666 | -0.17666 | 2.399336 | 0.659518 |
| 6 | 130475200 | 130478399 | 48.84331 | 79.00319 | 79.33072 | 33.91574 | 0.35628 | 1.70747 | 0.659518 |
| 16 | 48064000 | 48070199 | 57.8787 | 92.35756 | 102.6371 | 43.04876 | 0.187583 | 2.12964 | 0.659526 |
| 7 | 104364400 | 104365999 | 16.0619 | 29.1259 | 30.22736 | 13.07009 | 0.359863 | 0.357574 | 0.659531 |
| 6 | 34199200 | 34200399 | 11.62485 | 21.15205 | 33.30247 | 14.3384 | -0.23345 | 0.315142 | 0.659543 |
| 1 | 25971800 | 25976399 | 83.00941 | 130.1457 | 144.3228 | 59.85907 | 0.168441 | 2.596753 | 0.659544 |
| 5 | 38631000 | 38636599 | 49.54577 | 78.92976 | 116.5593 | 48.77595 | -0.22444 | 2.17926 | 0.659555 |
| 6 | 7151200 | 7154999 | 45.54341 | 73.55591 | 80.23226 | 34.23768 | 0.23468 | 1.739038 | 0.659564 |
| 13 | 63330600 | 63334399 | 23.7473 | 40.02651 | 61.59223 | 26.92795 | -0.23858 | 1.114961 | 0.65957 |
| X | 147391200 | 147391799 | 11.10379 | 21.13333 | 23.86733 | 9.559311 | 0.324765 | 0.016409 | 0.65957 |
| 1 | 116199200 | 116201599 | 34.27488 | 57.07399 | 50.89779 | 22.107 | 0.545929 | 1.259135 | 0.659585 |
| 16 | 16051000 | 16054999 | 32.04922 | 52.34349 | 78.41682 | 33.70529 | -0.20813 | 1.506036 | 0.659588 |
| 4 | 8034400 | 8034999 | 7.146973 | 13.85623 | 25.14531 | 9.502985 | -0.27561 | -0.21138 | 0.659593 |
| 15 | 95898400 | 95898799 | 6.538296 | 12.76411 | 16.44424 | 5.811314 | 0.303451 | -0.47976 | 0.659596 |
| 9 | 40047200 | 40048599 | 14.55932 | 26.11493 | 29.41472 | 12.87635 | 0.231147 | 0.410822 | 0.65961 |
| 11 | 120825400 | 120830799 | 31.09348 | 50.26849 | 92.90733 | 39.69774 | -0.52265 | 1.693418 | 0.659615 |
| 5 | 118971600 | 118972799 | 13.91336 | 24.58157 | 37.68728 | 16.67215 | -0.22295 | 0.533729 | 0.65962 |
| 14 | 96287000 | 96288399 | 15.32963 | 27.18693 | 29.89752 | 13.21328 | 0.25603 | 0.486618 | 0.659651 |
| 10 | 32396200 | 32396999 | 15.49243 | 26.77568 | 50.56267 | 22.18236 | -0.52747 | 0.849393 | 0.659667 |
| 14 | 17427000 | 17427799 | 8.60808 | 15.99688 | 27.92312 | 11.6885 | -0.35192 | 0.159816 | 0.659669 |
| 2 | 23692200 | 23692799 | 17.45472 | 30.57571 | 32.73612 | 14.45479 | 0.28965 | 0.656277 | 0.65967 |
| 1 | 3842000 | 3843599 | 13.86568 | 24.54302 | 38.13628 | 16.86449 | -0.24129 | 0.512451 | 0.659695 |
| 13 | 91343800 | 91346799 | 19.87578 | 33.65884 | 60.83715 | 26.63578 | -0.46955 | 1.060672 | 0.659708 |
| 3 | 120354200 | 120357599 | 30.41929 | 51.98299 | 47.90039 | 21.0021 | 0.500318 | 1.006059 | 0.659724 |
| 18 | 84152400 | 84155399 | 28.73499 | 47.77552 | 53.4449 | 23.18928 | 0.219882 | 1.262455 | 0.659739 |
| 2 | 174956000 | 174967799 | 96.60491 | 150.7978 | 156.7701 | 65.05109 | 0.256553 | 2.681502 | 0.659749 |
| 13 | 69390400 | 69391999 | 19.57747 | 33.55947 | 50.6084 | 22.19497 | -0.20695 | 0.938619 | 0.659751 |
| 12 | 103894200 | 103907199 | 272.6257 | 387.3458 | 499.3302 | 206.5148 | -0.1749 | 4.342965 | 0.659767 |
| 12 | 52125200 | 52128799 | 33.19404 | 54.70038 | 61.88615 | 26.69739 | 0.202005 | 1.383504 | 0.659778 |
| 8 | 117227200 | 117229199 | 16.26386 | 28.47186 | 44.5616 | 19.66599 | -0.25577 | 0.677361 | 0.659805 |

| | | | | | | | | | |
|---|---|---|---|---|---|---|---|---|---|
| 8 | 55806200 | 55808199 | 11.97754 | 21.48385 | 33.69882 | 14.75182 | -0.24374 | 0.402977 | 0.659849 |
| 10 | 106972800 | 106974399 | 11.97754 | 21.48385 | 33.69882 | 14.75182 | -0.24374 | 0.402977 | 0.659849 |
| 8 | 117115600 | 117119399 | 42.64348 | 69.46181 | 72.06964 | 30.9472 | 0.31782 | 1.562134 | 0.659869 |
| 18 | 73396800 | 73397999 | 14.87398 | 26.1963 | 40.25271 | 17.81823 | -0.22805 | 0.577867 | 0.659889 |
| X | 88891200 | 88897199 | 95.74708 | 150.1084 | 131.1488 | 54.39536 | 0.51194 | 2.603435 | 0.659904 |
| 9 | 72003400 | 72004599 | 7.756523 | 14.9293 | 25.88747 | 10.07131 | -0.25043 | -0.10619 | 0.659908 |
| 2 | 45486000 | 45486999 | 9.939078 | 18.32528 | 30.34571 | 12.8396 | -0.28839 | 0.218251 | 0.659929 |
| 3 | 23368600 | 23369799 | 17.58588 | 30.53605 | 46.32943 | 20.33349 | -0.21197 | 0.806439 | 0.659945 |
| X | 153815600 | 153817599 | 30.62346 | 52.0089 | 43.03885 | 18.85974 | 0.653597 | 1.056097 | 0.659961 |
| 15 | 12549000 | 12550999 | 11.85802 | 21.02036 | 34.74545 | 15.36048 | -0.3288 | 0.47837 | 0.659966 |
| 1 | 33577800 | 33578999 | 13.49423 | 24.81169 | 27.98693 | 11.88604 | 0.257955 | 0.256736 | 0.659967 |
| 2 | 57307400 | 57311599 | 56.81809 | 90.19732 | 128.3294 | 53.47695 | -0.17603 | 2.297299 | 0.659976 |
| 3 | 65020000 | 65020799 | 11.45823 | 21.65023 | 24.70714 | 10.04873 | 0.289613 | 0.066845 | 0.659981 |
| 7 | 75058200 | 75059599 | 17.06521 | 30.24782 | 33.26486 | 14.70624 | 0.255599 | 0.509799 | 0.659989 |
| 1 | 44430000 | 44431799 | 28.85618 | 48.12363 | 54.58064 | 23.72926 | 0.200102 | 1.229635 | 0.659995 |
| 7 | 123566000 | 123569599 | 46.63841 | 75.09716 | 82.58681 | 35.10056 | 0.217785 | 1.829866 | 0.659995 |
| 9 | 115302800 | 115306799 | 52.73026 | 84.35496 | 93.62161 | 39.40818 | 0.193465 | 2.04619 | 0.660024 |
| 7 | 40391200 | 40392399 | 9.795419 | 18.02513 | 30.3293 | 12.86685 | -0.31404 | 0.228013 | 0.660025 |
| 13 | 60932400 | 60932999 | 5.953263 | 11.56582 | 21.73455 | 8.111145 | -0.30817 | -0.28057 | 0.66003 |
| X | 150586000 | 150588599 | 27.63678 | 47.83863 | 42.44998 | 18.62411 | 0.55797 | 0.88054 | 0.660069 |
| X | 97829400 | 97830399 | 17.6869 | 31.15528 | 33.53479 | 14.84403 | 0.283245 | 0.583638 | 0.660073 |
| 2 | 150885600 | 150886999 | 12.46132 | 23.49982 | 25.74118 | 10.51201 | 0.343249 | 0.091924 | 0.660092 |
| 1 | 58856400 | 58860999 | 90.49823 | 140.8924 | 155.5026 | 64.54798 | 0.167391 | 2.72475 | 0.660112 |
| 2 | 150027600 | 150042999 | 235.4886 | 345.6432 | 341.2942 | 140.5679 | 0.255725 | 3.85595 | 0.660112 |
| 7 | 106009400 | 106012599 | 49.49379 | 79.97512 | 80.99911 | 34.58671 | 0.342307 | 1.736517 | 0.660117 |
| 15 | 58087400 | 58088799 | 12.41356 | 22.24553 | 34.64939 | 15.17966 | -0.23464 | 0.412273 | 0.660123 |
| 16 | 4258200 | 4260399 | 12.41356 | 22.24553 | 34.64939 | 15.17966 | -0.23464 | 0.412273 | 0.660123 |
| 12 | 7191400 | 7194799 | 66.21144 | 104.1657 | 148.1412 | 61.45889 | -0.1862 | 2.521606 | 0.660142 |
| 1 | 46542600 | 46545199 | 22.36375 | 37.8913 | 57.70285 | 25.2674 | -0.22331 | 1.060633 | 0.660143 |
| 8 | 129850000 | 129853399 | 41.38958 | 67.00324 | 75.60636 | 32.34354 | 0.189445 | 1.673276 | 0.660167 |
| 3 | 60921400 | 60922999 | 15.65313 | 27.96334 | 31.27015 | 13.73541 | 0.237985 | 0.446038 | 0.660203 |
| 6 | 78416600 | 78420399 | 36.42048 | 59.1075 | 91.89135 | 39.38302 | -0.26846 | 1.622765 | 0.660209 |
| 5 | 129202200 | 129209199 | 82.61951 | 128.0566 | 179.0361 | 74.36584 | -0.17833 | 2.799757 | 0.66021 |
| 17 | 36447800 | 36450799 | 25.44428 | 42.15858 | 64.14196 | 27.76444 | -0.22281 | 1.301614 | 0.660223 |
| 7 | 33746600 | 33754399 | 112.9409 | 173.5768 | 187.2105 | 77.7371 | 0.18631 | 2.969187 | 0.660224 |
| 13 | 83346200 | 83348799 | 42.1414 | 68.18127 | 76.38719 | 32.66378 | 0.198953 | 1.685162 | 0.660251 |
| 5 | 95458400 | 95459399 | 14.47218 | 27.06558 | 27.15245 | 11.24582 | 0.453047 | 0.164646 | 0.660273 |
| 2 | 143218000 | 143222399 | 61.37071 | 98.07211 | 104.5138 | 43.92617 | 0.250254 | 2.09205 | 0.66032 |
| 7 | 133432400 | 133434399 | 19.47266 | 33.2713 | 50.59308 | 22.18244 | -0.21955 | 0.968946 | 0.660342 |
| 6 | 129917200 | 129925599 | 161.6005 | 241.9011 | 259.2306 | 106.9157 | 0.163978 | 3.507385 | 0.660351 |
| 9 | 5792600 | 5794799 | 23.31578 | 40.49796 | 36.57977 | 16.05759 | 0.532953 | 0.813819 | 0.660355 |
| 1 | 69415600 | 69419199 | 83.69289 | 129.4624 | 182.7853 | 75.92458 | -0.19467 | 2.837679 | 0.660357 |
| 4 | 5747600 | 5749999 | 24.23989 | 41.47276 | 46.32828 | 20.31476 | 0.223873 | 0.969644 | 0.660366 |
| 11 | 84977600 | 84980599 | 22.19151 | 37.3919 | 56.99783 | 24.91681 | -0.22485 | 1.110569 | 0.660366 |
| 12 | 115549400 | 115557599 | 151.6306 | 229.0963 | 239.9617 | 99.21261 | 0.20874 | 3.325574 | 0.660376 |
| 13 | 77338000 | 77338999 | 14.03314 | 25.54432 | 28.92803 | 12.44873 | 0.239312 | 0.319893 | 0.660409 |
| 3 | 15370200 | 15372999 | 48.38014 | 77.70521 | 87.02529 | 36.85159 | 0.187106 | 1.914332 | 0.66041 |

| | | | | | | | | | |
|---|---|---|---|---|---|---|---|---|---|
| 15 | 13710200 | 13715599 | 94.37003 | 147.1647 | 139.5053 | 57.89322 | 0.388455 | 2.696217 | 0.660433 |
| 14 | 75169000 | 75170999 | 15.61304 | 27.60653 | 29.46911 | 13.0345 | 0.296962 | 0.51834 | 0.66044 |
| 6 | 70403400 | 70408399 | 75.52499 | 117.652 | 167.0913 | 69.35213 | -0.19528 | 2.708411 | 0.660441 |
| 4 | 21072600 | 21082599 | 214.3161 | 313.9908 | 329.2697 | 135.6309 | 0.166431 | 3.881497 | 0.660447 |
| 6 | 60570600 | 60572999 | 18.86195 | 33.21959 | 35.20272 | 15.58284 | 0.305155 | 0.590496 | 0.660463 |
| 14 | 117516400 | 117518799 | 23.81018 | 40.72996 | 45.78902 | 20.078 | 0.214627 | 0.967568 | 0.66051 |
| 18 | 54522400 | 54523399 | 9.957051 | 18.64013 | 30.66232 | 12.67801 | -0.25328 | 0.129689 | 0.660512 |
| 14 | 105610000 | 105610799 | 7.135504 | 13.72789 | 24.16176 | 9.41194 | -0.2734 | -0.11423 | 0.66053 |
| 15 | 103965800 | 103968599 | 11.99466 | 22.62657 | 21.51129 | 8.779417 | 0.547172 | 0.076716 | 0.660545 |
| 5 | 104514200 | 104517599 | 82.85829 | 129.8092 | 144.3144 | 59.86033 | 0.163937 | 2.609998 | 0.660559 |
| 6 | 55899400 | 55900199 | 9.436827 | 18.12149 | 21.84003 | 8.553291 | 0.260492 | -0.08131 | 0.66059 |
| 15 | 24675200 | 24677999 | 17.21028 | 29.67637 | 54.93575 | 24.09918 | -0.4999 | 0.880848 | 0.66059 |
| 2 | 48228800 | 48230599 | 25.59853 | 42.78118 | 65.51638 | 28.52165 | -0.23181 | 1.208202 | 0.660618 |
| 7 | 49306000 | 49307599 | 13.44616 | 23.68236 | 37.27933 | 16.50194 | -0.26248 | 0.564503 | 0.660618 |
| 6 | 144355000 | 144356399 | 14.82369 | 25.92453 | 41.4835 | 18.29198 | -0.28743 | 0.6813 | 0.660623 |
| 10 | 43713800 | 43715199 | 11.81267 | 22.30276 | 25.23882 | 10.28171 | 0.299663 | 0.077005 | 0.660638 |
| 7 | 31627800 | 31640799 | 391.566 | 543.9812 | 531.1523 | 220.4663 | 0.193159 | 4.680067 | 0.66064 |
| 13 | 36123800 | 36128999 | 41.32369 | 66.35634 | 97.71002 | 41.3617 | -0.20733 | 1.917062 | 0.660669 |
| 16 | 59649400 | 59650399 | 12.52231 | 23.66237 | 25.8163 | 10.49838 | 0.354363 | 0.079728 | 0.66067 |
| 2 | 98642200 | 98644399 | 25.92133 | 44.54785 | 46.2732 | 20.29703 | 0.329385 | 0.93681 | 0.660677 |
| 13 | 61336600 | 61346199 | 117.346 | 181.8416 | 170.7372 | 70.91945 | 0.395204 | 2.81856 | 0.660692 |
| 17 | 83973600 | 83977999 | 35.87865 | 58.51387 | 66.0539 | 28.3766 | 0.197681 | 1.527154 | 0.660696 |
| 6 | 50447200 | 50451399 | 41.32148 | 67.04293 | 74.35257 | 31.85972 | 0.217336 | 1.626754 | 0.660705 |
| 16 | 39385800 | 39388399 | 24.09023 | 41.75086 | 43.55053 | 19.10767 | 0.32577 | 0.85179 | 0.66071 |
| 18 | 68910600 | 68912399 | 28.27421 | 47.2248 | 53.43563 | 23.25375 | 0.203494 | 1.212215 | 0.660713 |
| 13 | 104714000 | 104717799 | 30.64757 | 49.90532 | 75.37726 | 32.36364 | -0.22284 | 1.545141 | 0.660749 |
| 5 | 69983400 | 69984199 | 11.01474 | 20.78228 | 24.2546 | 9.894808 | 0.254289 | 0.074683 | 0.660772 |
| 17 | 24461000 | 24463999 | 34.52244 | 56.44979 | 63.31492 | 27.22594 | 0.209588 | 1.484544 | 0.660784 |
| 12 | 3621800 | 3623999 | 17.27944 | 29.52171 | 54.81101 | 24.03164 | -0.50644 | 0.965905 | 0.660794 |
| 14 | 73421200 | 73423399 | 20.59608 | 36.24728 | 36.16676 | 16.01193 | 0.391269 | 0.610864 | 0.660795 |
| 6 | 131995800 | 132000399 | 97.17009 | 150.8631 | 164.7662 | 68.42139 | 0.179451 | 2.775428 | 0.660797 |
| 12 | 115431400 | 115434399 | 34.50749 | 57.7922 | 55.88057 | 24.35931 | 0.429485 | 1.205949 | 0.660811 |
| 1 | 99279200 | 99279799 | 6.533986 | 12.6358 | 23.21577 | 8.878055 | -0.30966 | -0.18599 | 0.660812 |
| 9 | 63774600 | 63778999 | 68.19635 | 108.0656 | 120.8477 | 50.26867 | 0.168334 | 2.356191 | 0.660836 |
| 7 | 9984400 | 9986799 | 98.5974 | 152.0839 | 158.2762 | 65.74507 | 0.241769 | 2.892614 | 0.660873 |
| 1 | 12339800 | 12340799 | 11.07656 | 21.0422 | 18.53104 | 7.451388 | 0.675297 | 0.011193 | 0.660879 |
| 9 | 115264000 | 115269599 | 68.3167 | 108.1996 | 120.46 | 50.09343 | 0.174154 | 2.370025 | 0.660912 |
| 11 | 74509200 | 74512799 | 22.95241 | 38.47273 | 58.12278 | 25.35031 | -0.2122 | 1.161753 | 0.660914 |
| 8 | 52203600 | 52204399 | 10.97122 | 20.88281 | 24.38865 | 9.766343 | 0.276539 | 0.013799 | 0.660922 |
| 7 | 43243000 | 43245599 | 24.20887 | 40.76935 | 64.48968 | 28.18734 | -0.27836 | 1.125506 | 0.660933 |
| X | 101869200 | 101872599 | 24.69381 | 41.31922 | 63.66297 | 27.736 | -0.24047 | 1.193071 | 0.660935 |
| 8 | 21592800 | 21613799 | 273.0118 | 396.8512 | 375.3002 | 154.7003 | 0.304549 | 4.007856 | 0.660964 |
| 17 | 95136000 | 95142599 | 87.87991 | 137.4917 | 150.2911 | 62.34799 | 0.186815 | 2.637389 | 0.660965 |
| Y | 34255800 | 34256399 | 4.156978 | 8.09824 | 19.63775 | 7.152187 | -0.63784 | -0.39609 | 0.660979 |
| 7 | 11784800 | 11785599 | 26.75333 | 44.83273 | 50.17953 | 21.88244 | 0.21908 | 1.168969 | 0.660981 |
| 2 | 102549000 | 102549799 | 6.981798 | 13.49536 | 23.76231 | 9.107351 | -0.25175 | -0.17048 | 0.661016 |
| X | 4423200 | 4467999 | 782.179 | 1039.925 | 864.5761 | 361.2686 | 0.361292 | 5.374638 | 0.661019 |

| | | | | | | | | | |
|---|---|---|---|---|---|---|---|---|---|
| 3 | 106345000 | 106350599 | 115.0849 | 176.3591 | 192.5751 | 79.93288 | 0.165329 | 3.027836 | 0.661023 |
| 11 | 96417400 | 96420199 | 24.759 | 41.24027 | 62.01025 | 26.93853 | -0.20536 | 1.237731 | 0.661035 |
| 17 | 71514200 | 71516799 | 19.25133 | 33.28287 | 51.7366 | 22.69759 | -0.24816 | 0.862079 | 0.661046 |
| 17 | 39514600 | 39521599 | 128.4551 | 196.2979 | 184.258 | 76.4317 | 0.380188 | 3.086968 | 0.661046 |
| 9 | 56181600 | 56185399 | 44.65522 | 72.51631 | 74.94764 | 32.13578 | 0.320011 | 1.616418 | 0.661097 |
| 5 | 82606800 | 82607199 | 6.852946 | 13.14352 | 24.09094 | 9.468151 | -0.34344 | -0.08869 | 0.661114 |
| 1 | 27049200 | 27049599 | 12.5916 | 23.0607 | 25.59509 | 10.92885 | 0.275881 | 0.26786 | 0.661131 |
| 12 | 3358600 | 3360799 | 23.35365 | 38.48583 | 72.94476 | 31.46851 | -0.54003 | 1.370186 | 0.661136 |
| 15 | 91603800 | 91607799 | 44.16903 | 71.20998 | 80.32398 | 34.19862 | 0.183485 | 1.789649 | 0.66114 |
| 18 | 62408400 | 62410599 | 37.40474 | 60.34462 | 88.55874 | 37.73701 | -0.19422 | 1.764214 | 0.661146 |
| 7 | 86320600 | 86327199 | 196.3481 | 291.4788 | 268.0389 | 110.4118 | 0.373651 | 3.663024 | 0.661151 |
| 18 | 45157000 | 45158399 | 10.19281 | 18.81993 | 30.3999 | 12.83758 | -0.25029 | 0.215806 | 0.66116 |
| 8 | 19444200 | 19448599 | 71.42257 | 112.9324 | 126.3487 | 52.49843 | 0.165168 | 2.411518 | 0.661197 |
| 17 | 47375000 | 47381199 | 59.4514 | 94.31329 | 133.1937 | 55.4661 | -0.16632 | 2.32036 | 0.6612 |
| 6 | 21352800 | 21353999 | 11.41752 | 21.34684 | 24.01896 | 9.952738 | 0.287915 | 0.131783 | 0.66121 |
| 4 | 33265200 | 33270999 | 50.1261 | 80.02495 | 116.2663 | 48.79828 | -0.19767 | 2.111146 | 0.661216 |
| 9 | 30268200 | 30268599 | 6.239764 | 12.15191 | 22.58682 | 8.260154 | -0.26545 | -0.34434 | 0.661232 |
| 11 | 23196400 | 23202399 | 44.32567 | 71.64306 | 79.39458 | 33.91311 | 0.213126 | 1.716561 | 0.661234 |
| 1 | 181846400 | 181848399 | 11.90075 | 20.93321 | 37.38549 | 16.55323 | -0.44436 | 0.568618 | 0.661237 |
| 17 | 38404800 | 38406599 | 25.8285 | 43.978 | 48.48236 | 21.25105 | 0.241768 | 1.020414 | 0.66124 |
| 1 | 172989200 | 172990199 | 8.368627 | 15.37641 | 33.52638 | 14.25237 | -0.68721 | 0.233958 | 0.66126 |
| 17 | 62418800 | 62419399 | 11.83892 | 22.05598 | 25.17353 | 10.49428 | 0.261207 | 0.164714 | 0.661261 |
| 8 | 88238200 | 88239599 | 14.01325 | 24.74546 | 38.77658 | 17.15578 | -0.25441 | 0.544319 | 0.661272 |
| 1 | 189450400 | 189453599 | 21.62058 | 36.44301 | 55.40482 | 24.22306 | -0.22108 | 1.10176 | 0.661272 |
| 11 | 73995800 | 74006199 | 162.8856 | 245.1446 | 253.9136 | 104.8614 | 0.220003 | 3.404975 | 0.66129 |
| 3 | 11111600 | 11124799 | 196.0219 | 290.4174 | 303.5261 | 125.0046 | 0.186484 | 3.699375 | 0.661293 |
| 13 | 93164000 | 93165599 | 19.2418 | 33.68681 | 37.163 | 16.40154 | 0.246704 | 0.683872 | 0.661337 |
| 17 | 27583600 | 27592999 | 115.7384 | 175.1224 | 247.5339 | 102.3904 | -0.22155 | 3.287582 | 0.661353 |
| X | 119068400 | 119071599 | 55.62882 | 89.23102 | 95.60167 | 40.40303 | 0.248711 | 1.968939 | 0.661375 |
| 4 | 119451400 | 119452799 | 6.856281 | 13.21924 | 23.65757 | 9.158519 | -0.28823 | -0.13677 | 0.661385 |
| 5 | 15740600 | 15740999 | 5.641444 | 11.0328 | 14.90874 | 5.069063 | 0.285699 | -0.63303 | 0.661405 |
| 5 | 65126000 | 65128199 | 15.83858 | 27.60998 | 42.61472 | 18.74924 | -0.23587 | 0.737851 | 0.661413 |
| 14 | 3183200 | 3214799 | 535.5343 | 727.9426 | 704.8712 | 293.5775 | 0.173435 | 5.020169 | 0.661434 |
| 1 | 38384800 | 38386199 | 14.99376 | 26.96258 | 29.93517 | 13.07614 | 0.25406 | 0.400713 | 0.661438 |
| 1 | 158028200 | 158030799 | 23.36089 | 40.32718 | 44.12838 | 19.35887 | 0.255807 | 0.88332 | 0.661441 |
| 5 | 119048000 | 119048599 | 6.53997 | 12.70058 | 22.86131 | 8.55919 | -0.25082 | -0.25686 | 0.661455 |
| 4 | 66822400 | 66824399 | 35.74096 | 57.79201 | 84.94965 | 36.30565 | -0.19274 | 1.692953 | 0.66147 |
| 14 | 41380800 | 41383599 | 31.83734 | 53.03927 | 49.75149 | 21.61534 | 0.473268 | 1.246879 | 0.661476 |
| 10 | 39870400 | 39878199 | 93.25605 | 145.0471 | 159.4456 | 66.19413 | 0.172107 | 2.741547 | 0.661477 |
| 10 | 5801800 | 5802599 | 10.01255 | 18.54091 | 29.90693 | 12.57956 | -0.24417 | 0.196441 | 0.661533 |
| 1 | 140172600 | 140174599 | 12.93442 | 22.52661 | 46.83329 | 20.59334 | -0.66355 | 0.743572 | 0.661558 |
| X | 4365400 | 4381799 | 185.0166 | 277.9578 | 257.6454 | 106.352 | 0.377825 | 3.445475 | 0.661563 |
| 7 | 23082200 | 23174999 | 2076.756 | 2529.619 | 2159.964 | 911.2948 | 0.195665 | 6.785714 | 0.661577 |
| 3 | 61078000 | 61079599 | 14.6535 | 25.61403 | 47.83488 | 21.08365 | -0.5092 | 0.701313 | 0.661602 |
| 10 | 124889000 | 124891599 | 44.88495 | 72.6433 | 78.46061 | 33.55829 | 0.252097 | 1.686408 | 0.661634 |
| X | 7476200 | 7479199 | 39.36496 | 64.52551 | 67.17513 | 28.91393 | 0.318729 | 1.469664 | 0.661645 |
| 2 | 166546400 | 166548599 | 23.53056 | 39.24641 | 60.19029 | 26.16992 | -0.23369 | 1.220495 | 0.661655 |

| | | | | | | | | | |
|---|---|---|---|---|---|---|---|---|---|
| 17 | 28529600 | 28530599 | 7.181092 | 13.78573 | 24.65027 | 9.662085 | -0.30375 | -0.08891 | 0.661666 |
| 3 | 147950200 | 147952199 | 21.64415 | 37.74128 | 40.21111 | 17.69495 | 0.296255 | 0.756922 | 0.661673 |
| 17 | 40217000 | 40218599 | 15.72113 | 28.99055 | 28.39107 | 12.01001 | 0.464512 | 0.258049 | 0.661683 |
| 4 | 25558000 | 25561799 | 36.58644 | 59.25475 | 87.76594 | 37.5579 | -0.2015 | 1.662529 | 0.661684 |
| 12 | 78447600 | 78448399 | 14.31931 | 26.10335 | 28.9921 | 12.45352 | 0.269081 | 0.31553 | 0.661693 |
| 13 | 3462200 | 3490599 | 614.0956 | 824.4171 | 792.3871 | 330.652 | 0.166097 | 5.217262 | 0.661721 |
| 4 | 129303800 | 129304599 | 16.23963 | 28.78285 | 32.344 | 14.2993 | 0.224647 | 0.503591 | 0.661732 |
| 11 | 121714200 | 121715799 | 19.35985 | 34.81589 | 32.11316 | 14.02685 | 0.519968 | 0.433727 | 0.661752 |
| 9 | 102360000 | 102364999 | 66.1052 | 104.2862 | 146.3425 | 60.73573 | -0.16388 | 2.465458 | 0.661762 |
| 6 | 73259400 | 73269599 | 124.7457 | 187.245 | 299.4862 | 123.5941 | -0.40982 | 3.45159 | 0.661781 |
| 13 | 51016800 | 51024599 | 127.9065 | 195.0448 | 209.5735 | 86.89007 | 0.182981 | 3.131656 | 0.661783 |
| 17 | 40084600 | 40087199 | 35.01182 | 58.33347 | 58.25496 | 25.31138 | 0.383145 | 1.261322 | 0.661798 |
| 12 | 41082200 | 41084599 | 21.41385 | 37.50815 | 38.45136 | 16.97729 | 0.351873 | 0.68719 | 0.661807 |
| 6 | 130585000 | 130591799 | 81.20673 | 129.0381 | 117.565 | 49.03858 | 0.467396 | 2.287513 | 0.661809 |
| 1 | 173620400 | 173623799 | 58.60327 | 93.42386 | 102.9969 | 43.14202 | 0.19763 | 2.164038 | 0.661835 |
| 14 | 15965200 | 15968599 | 43.16979 | 69.61959 | 77.89738 | 33.17814 | 0.195624 | 1.778292 | 0.661848 |
| 9 | 76353600 | 76355399 | 21.5547 | 37.28966 | 40.57812 | 17.80256 | 0.264639 | 0.854695 | 0.661869 |
| 1 | 47906800 | 47908799 | 20.60139 | 36.21473 | 36.81612 | 16.30318 | 0.363948 | 0.626316 | 0.661875 |
| 3 | 20430800 | 20435599 | 38.22207 | 61.88186 | 97.32352 | 41.63702 | -0.28839 | 1.676236 | 0.661891 |
| 2 | 111261200 | 111263999 | 32.67711 | 54.27628 | 59.21845 | 25.67691 | 0.255666 | 1.289025 | 0.661892 |
| X | 95436000 | 95436999 | 13.84815 | 25.2217 | 28.04012 | 12.05819 | 0.266763 | 0.315714 | 0.661902 |
| 6 | 47985400 | 47986999 | 14.19132 | 25.04057 | 38.68524 | 17.11801 | -0.23477 | 0.549882 | 0.661931 |
| 9 | 120246000 | 120246799 | 5.830692 | 11.28355 | 22.12675 | 8.434215 | -0.39765 | -0.20722 | 0.661946 |
| 9 | 19619000 | 19619799 | 14.49907 | 26.66039 | 24.35123 | 10.34137 | 0.56072 | 0.257584 | 0.661955 |
| Y | 38282400 | 38283799 | 10.22379 | 19.33404 | 17.71182 | 7.190828 | 0.606621 | 0.030126 | 0.662019 |
| 18 | 37770400 | 37786199 | 274.0366 | 394.2419 | 406.0953 | 167.6155 | 0.166659 | 4.160112 | 0.662038 |
| 15 | 97519600 | 97534599 | 248.3278 | 356.1661 | 460.2663 | 190.0655 | -0.16479 | 4.20397 | 0.66204 |
| 4 | 15860800 | 15862599 | 26.56182 | 43.62857 | 68.09737 | 29.33635 | -0.26266 | 1.414775 | 0.662063 |
| 16 | 5123600 | 5125999 | 18.00812 | 30.84652 | 48.48291 | 21.26155 | -0.26644 | 0.937317 | 0.662071 |
| 5 | 65043400 | 65056399 | 157.8994 | 234.9221 | 315.3735 | 129.9394 | -0.16891 | 3.620286 | 0.662087 |
| 15 | 76806400 | 76807599 | 10.50586 | 19.21406 | 30.81255 | 13.18157 | -0.25354 | 0.272244 | 0.662115 |
| 3 | 16129200 | 16129799 | 8.397005 | 16.22761 | 19.84946 | 7.614824 | 0.267611 | -0.16464 | 0.662124 |
| 7 | 103776600 | 103784399 | 88.83635 | 138.224 | 151.979 | 63.09088 | 0.172209 | 2.733009 | 0.662131 |
| 13 | 65318200 | 65320999 | 31.00554 | 52.1031 | 54.0996 | 23.63058 | 0.326897 | 1.156874 | 0.662147 |
| 8 | 10565400 | 10565799 | 4.773346 | 9.329957 | 18.33845 | 6.324234 | -0.26707 | -0.58446 | 0.662161 |
| 9 | 95978400 | 95986599 | 176.3146 | 263.0661 | 279.198 | 115.0779 | 0.173655 | 3.571043 | 0.662163 |
| 13 | 33756400 | 33769199 | 158.7263 | 239.801 | 242.0226 | 100.0626 | 0.262167 | 3.328732 | 0.662183 |
| 4 | 22116200 | 22116999 | 8.246489 | 15.47305 | 26.9466 | 11.0988 | -0.33009 | 0.090917 | 0.662184 |
| 14 | 83781400 | 83782599 | 18.15356 | 32.07768 | 34.34689 | 15.19397 | 0.292375 | 0.552985 | 0.662187 |
| 11 | 12049400 | 12051199 | 35.29805 | 57.8713 | 65.10312 | 28.02456 | 0.207311 | 1.455137 | 0.662205 |
| 9 | 42018400 | 42020199 | 14.40682 | 25.36829 | 38.74602 | 17.15199 | -0.21939 | 0.575482 | 0.662214 |
| 19 | 41800800 | 41802599 | 14.40682 | 25.36829 | 38.74602 | 17.15199 | -0.21939 | 0.575482 | 0.662214 |
| 7 | 52100200 | 52102599 | 25.40395 | 43.01273 | 47.9791 | 21.0051 | 0.224175 | 1.06582 | 0.662225 |
| 1 | 190879200 | 190879999 | 11.20014 | 21.0775 | 24.67894 | 10.11383 | 0.243864 | 0.091661 | 0.66223 |
| 16 | 11780200 | 11781199 | 11.20014 | 21.0775 | 24.67894 | 10.11383 | 0.243864 | 0.091661 | 0.66223 |
| 7 | 27589000 | 27589799 | 19.1162 | 33.39299 | 37.73341 | 16.62334 | 0.21215 | 0.717166 | 0.662234 |
| 2 | 95628800 | 95630399 | 31.85055 | 51.83212 | 77.57792 | 33.30099 | -0.21035 | 1.559954 | 0.662239 |

| | | | | | | | | | |
|---|---|---|---|---|---|---|---|---|---|
| 12 | 119533600 | 119539399 | 114.6542 | 176.0348 | 190.5496 | 79.11283 | 0.180009 | 2.990212 | 0.66225 |
| 1 | 191764000 | 191767399 | 42.55696 | 68.67799 | 76.70313 | 32.69816 | 0.199404 | 1.757902 | 0.662252 |
| 8 | 117758000 | 117761599 | 26.64815 | 43.91734 | 66.76438 | 28.80292 | -0.22286 | 1.370822 | 0.662257 |
| 13 | 67733600 | 67741799 | 115.7342 | 179.3597 | 170.9382 | 71.00288 | 0.373801 | 2.816903 | 0.662283 |
| 5 | 145328200 | 145333799 | 77.43728 | 122.9001 | 122.3148 | 50.95821 | 0.338435 | 2.318803 | 0.662324 |
| 14 | 106532200 | 106538599 | 59.61288 | 95.68513 | 95.07569 | 40.22134 | 0.358348 | 1.953549 | 0.662339 |
| 14 | 54645800 | 54648399 | 17.24914 | 29.75359 | 47.41129 | 20.79857 | -0.2844 | 0.866641 | 0.662348 |
| 1 | 190274000 | 190274399 | 8.473756 | 16.40888 | 20.46501 | 7.777577 | 0.253055 | -0.19158 | 0.662354 |
| 13 | 85033200 | 85034599 | 19.20968 | 33.02799 | 50.07973 | 21.967 | -0.21408 | 0.908516 | 0.662381 |
| 12 | 115902800 | 115907999 | 65.79744 | 105.3332 | 86.54839 | 36.46717 | 0.627527 | 2.038477 | 0.662405 |
| 8 | 124455800 | 124456599 | 5.373019 | 10.44854 | 20.65202 | 7.647786 | -0.36855 | -0.31499 | 0.662415 |
| 15 | 38480200 | 38484599 | 38.70988 | 62.36062 | 91.30434 | 38.83269 | -0.19332 | 1.809827 | 0.662428 |
| 11 | 27232800 | 27236999 | 77.58951 | 122.6178 | 113.9651 | 47.34017 | 0.43172 | 2.427259 | 0.66243 |
| 17 | 70127400 | 70127799 | 10.05928 | 19.32796 | 22.99017 | 8.988781 | 0.282035 | -0.07455 | 0.662442 |
| 15 | 56038600 | 56041199 | 40.2441 | 65.71089 | 59.60767 | 25.61833 | 0.513596 | 1.518382 | 0.662458 |
| 10 | 88880800 | 88882399 | 18.59206 | 31.99214 | 48.47351 | 21.26298 | -0.21269 | 0.897501 | 0.662464 |
| 1 | 84905600 | 84908599 | 25.90942 | 42.9584 | 64.80016 | 28.06248 | -0.21042 | 1.295178 | 0.662498 |
| 19 | 54632600 | 54635799 | 22.99426 | 39.74879 | 43.58372 | 19.12073 | 0.25327 | 0.868474 | 0.662503 |
| 4 | 79768200 | 79769399 | 7.280253 | 14.1123 | 25.78704 | 9.752991 | -0.28689 | -0.20694 | 0.662505 |
| 12 | 21365400 | 21366799 | 8.596833 | 16.07967 | 27.45296 | 11.36785 | -0.30757 | 0.117736 | 0.662512 |
| 4 | 72251000 | 72252399 | 19.76825 | 34.47808 | 38.18114 | 16.80407 | 0.240765 | 0.740697 | 0.662547 |
| 1 | 109696400 | 109697399 | 13.44223 | 23.86739 | 36.64761 | 16.18061 | -0.22184 | 0.482033 | 0.662549 |
| 19 | 40768400 | 40769799 | 13.44223 | 23.86739 | 36.64761 | 16.18061 | -0.22184 | 0.482033 | 0.662549 |
| 1 | 142636800 | 142637599 | 13.29994 | 24.30344 | 27.54336 | 11.7957 | 0.242966 | 0.290861 | 0.662552 |
| 5 | 59912800 | 59914399 | 16.50864 | 29.19645 | 32.8361 | 14.52313 | 0.222185 | 0.52868 | 0.662554 |
| 11 | 26875600 | 26876999 | 14.91401 | 26.78482 | 29.7382 | 13.00415 | 0.252568 | 0.404059 | 0.662561 |
| 7 | 35366200 | 35367999 | 15.98926 | 27.835 | 43.15887 | 18.97253 | -0.24266 | 0.758277 | 0.662568 |
| 5 | 17124600 | 17125199 | 8.831901 | 16.95652 | 19.27602 | 7.553399 | 0.343793 | -0.0924 | 0.662573 |
| 13 | 21668000 | 21673199 | 73.03812 | 115.5238 | 128.0628 | 53.21927 | 0.178799 | 2.405158 | 0.662589 |
| 10 | 129573200 | 129574399 | 10.71185 | 19.87816 | 33.20348 | 13.92575 | -0.29184 | 0.194435 | 0.6626 |
| 10 | 11137200 | 11140599 | 34.71608 | 57.07088 | 64.16287 | 27.65135 | 0.2097 | 1.418736 | 0.662622 |
| 5 | 25419200 | 25421599 | 18.07601 | 31.23432 | 47.40669 | 20.79766 | -0.21392 | 0.855949 | 0.66264 |
| 1 | 75537800 | 75540799 | 34.51146 | 56.95566 | 62.64789 | 27.04357 | 0.243321 | 1.369472 | 0.662643 |
| 5 | 103372400 | 103374399 | 27.1102 | 44.89663 | 67.05015 | 29.01336 | -0.19616 | 1.31522 | 0.662648 |
| 1 | 4870200 | 4870799 | 7.437159 | 14.32553 | 24.76284 | 9.612698 | -0.24294 | -0.12021 | 0.662653 |
| 12 | 118936800 | 118937199 | 7.5684 | 14.57444 | 25.34118 | 9.844741 | -0.2527 | -0.116 | 0.662657 |
| 6 | 111555400 | 111556599 | 22.90279 | 39.89815 | 41.29084 | 18.15927 | 0.337971 | 0.778823 | 0.662659 |
| 1 | 16911800 | 16912399 | 8.953752 | 17.24 | 20.37504 | 7.918365 | 0.299856 | -0.11185 | 0.662693 |
| 10 | 112341800 | 112342399 | 8.953752 | 17.24 | 20.37504 | 7.918365 | 0.299856 | -0.11185 | 0.662693 |
| 3 | 3999000 | 4003199 | 63.94485 | 101.879 | 94.36483 | 39.49398 | 0.447565 | 2.185156 | 0.662705 |
| 6 | 119405600 | 119407599 | 16.88944 | 29.04114 | 53.80584 | 23.60149 | -0.50195 | 0.903173 | 0.662718 |
| Y | 715000 | 715799 | 8.840433 | 17.02532 | 15.72331 | 6.106949 | 0.65245 | -0.13424 | 0.662742 |
| 11 | 63746200 | 63748599 | 24.40116 | 41.63816 | 46.8763 | 20.55068 | 0.212124 | 0.994631 | 0.662745 |
| 12 | 91838800 | 91842399 | 30.99485 | 50.31632 | 77.96394 | 33.42091 | -0.26316 | 1.598177 | 0.662751 |
| 4 | 114762600 | 114766999 | 36.92616 | 59.44538 | 90.63988 | 38.5109 | -0.25296 | 1.828599 | 0.662752 |
| 12 | 114655400 | 114657999 | 45.74219 | 74.04395 | 78.57952 | 33.61359 | 0.277653 | 1.684875 | 0.662757 |
| 8 | 87334400 | 87338599 | 45.28322 | 72.91353 | 81.95961 | 34.83294 | 0.186276 | 1.83018 | 0.662768 |

| | | | | | | | | | |
|---|---|---|---|---|---|---|---|---|---|
| 6 | 44667400 | 44670399 | 36.59497 | 59.13734 | 86.54258 | 36.95774 | -0.18736 | 1.713677 | 0.662785 |
| 9 | 67955000 | 67957399 | 28.77542 | 46.91267 | 83.71279 | 35.95817 | -0.46152 | 1.531209 | 0.662803 |
| 14 | 32500200 | 32502199 | 20.35672 | 35.43725 | 39.83592 | 17.5115 | 0.219074 | 0.773856 | 0.66281 |
| 10 | 31683000 | 31685199 | 17.80165 | 30.98882 | 47.89726 | 21.05496 | -0.23828 | 0.774563 | 0.662818 |
| 7 | 109913600 | 109915199 | 16.5171 | 29.18382 | 32.87624 | 14.54363 | 0.219314 | 0.53923 | 0.662821 |
| 17 | 95002800 | 95004399 | 24.51458 | 42.18699 | 39.34717 | 17.25974 | 0.484425 | 0.908926 | 0.662824 |
| 6 | 128307000 | 128308399 | 16.75455 | 29.54119 | 33.0005 | 14.60473 | 0.230344 | 0.565087 | 0.66285 |
| 7 | 47973800 | 47976599 | 39.56934 | 64.35032 | 71.94667 | 30.87621 | 0.208664 | 1.579573 | 0.662873 |
| 11 | 79037200 | 79039599 | 17.0343 | 29.94601 | 33.5314 | 14.84844 | 0.225294 | 0.599144 | 0.662892 |
| 9 | 33420200 | 33421599 | 17.37521 | 30.7571 | 33.76797 | 14.93256 | 0.257276 | 0.526401 | 0.662901 |
| 17 | 21009800 | 21011799 | 39.08878 | 63.86292 | 60.85828 | 26.16157 | 0.443213 | 1.505348 | 0.662907 |
| 1 | 169560800 | 169562399 | 14.10018 | 26.07858 | 28.01174 | 11.80188 | 0.336237 | 0.228537 | 0.662926 |
| 7 | 58678600 | 58685999 | 85.13872 | 134.4748 | 134.4873 | 55.84953 | 0.325535 | 2.44915 | 0.662928 |
| 2 | 126076400 | 126077399 | 15.1029 | 27.48584 | 29.92897 | 12.88284 | 0.295425 | 0.33197 | 0.662935 |
| 7 | 66676000 | 66677799 | 15.01796 | 26.32269 | 40.38446 | 17.84083 | -0.22691 | 0.646064 | 0.662936 |
| 14 | 15926600 | 15927199 | 6.977127 | 13.5014 | 23.82774 | 9.084913 | -0.24796 | -0.18833 | 0.663035 |
| 4 | 96150600 | 96153599 | 37.78318 | 61.64755 | 69.05058 | 29.66882 | 0.209379 | 1.528324 | 0.663055 |
| 1 | 145893000 | 145895399 | 19.46297 | 34.49306 | 34.01353 | 15.03763 | 0.411955 | 0.526358 | 0.663056 |
| 11 | 68290000 | 68291599 | 15.48812 | 26.9901 | 42.61829 | 18.7469 | -0.26871 | 0.739696 | 0.663057 |
| 9 | 120430600 | 120446799 | 88.44531 | 138.6349 | 131.6488 | 54.6038 | 0.391669 | 2.602364 | 0.66307 |
| 3 | 133082600 | 133089999 | 101.1248 | 155.306 | 214.3956 | 89.0167 | -0.17022 | 2.982288 | 0.66307 |
| 13 | 70777600 | 70780999 | 22.54096 | 38.26118 | 59.39667 | 26.02347 | -0.25085 | 1.047514 | 0.663076 |
| 9 | 99254600 | 99255799 | 10.41476 | 19.22321 | 30.88307 | 13.04766 | -0.24343 | 0.217781 | 0.663083 |
| 2 | 177109400 | 177111599 | 14.90811 | 27.14771 | 25.25773 | 10.86391 | 0.521568 | 0.315994 | 0.663114 |
| 15 | 14733000 | 14734599 | 33.59388 | 55.01694 | 61.5443 | 26.48033 | 0.2148 | 1.461397 | 0.663123 |
| X | 157902200 | 157905399 | 81.12188 | 127.9545 | 136.5762 | 56.68006 | 0.23014 | 2.47794 | 0.663171 |
| 13 | 86370400 | 86372799 | 22.0154 | 38.13191 | 42.53485 | 18.6617 | 0.229194 | 0.84554 | 0.663172 |
| 3 | 63584000 | 63584599 | 9.247147 | 17.07707 | 28.89447 | 12.19875 | -0.31628 | 0.203448 | 0.663175 |
| 5 | 145233400 | 145235199 | 16.46565 | 28.72345 | 43.61124 | 19.19623 | -0.21227 | 0.731979 | 0.663224 |
| 1 | 99186600 | 99188199 | 17.68621 | 31.07269 | 34.98835 | 15.48781 | 0.217138 | 0.614362 | 0.663224 |
| X | 79782400 | 79785399 | 34.11402 | 57.07708 | 56.45557 | 24.5947 | 0.396864 | 1.214716 | 0.663226 |
| 13 | 87235400 | 87235799 | 8.739438 | 16.83282 | 20.35156 | 7.901858 | 0.2682 | -0.11871 | 0.663228 |
| X | 89028600 | 89030599 | 24.44653 | 42.39377 | 43.5394 | 19.10314 | 0.348277 | 0.848349 | 0.663233 |
| X | 121283400 | 121285799 | 35.09182 | 58.37415 | 58.66462 | 25.463 | 0.374121 | 1.282114 | 0.663239 |
| 15 | 31936600 | 31941999 | 84.57748 | 133.3422 | 138.3866 | 57.42789 | 0.27007 | 2.489459 | 0.663275 |
| 13 | 88563400 | 88565999 | 29.51014 | 50.16142 | 50.13593 | 21.97285 | 0.382323 | 1.051233 | 0.6633 |
| 7 | 106576600 | 106580599 | 42.99473 | 70.28657 | 69.45039 | 29.86164 | 0.391719 | 1.511293 | 0.663322 |
| 19 | 42662800 | 42675399 | 122.6558 | 185.581 | 251.9722 | 104.2249 | -0.16361 | 3.291175 | 0.663329 |
| 7 | 80443000 | 80445399 | 30.61657 | 51.75599 | 50.7529 | 22.20872 | 0.409298 | 1.09984 | 0.663334 |
| 16 | 58658800 | 58663599 | 57.38913 | 92.09793 | 94.474 | 39.95502 | 0.312104 | 1.953718 | 0.663351 |
| 11 | 29230000 | 29236199 | 69.27532 | 109.9284 | 104.5815 | 43.56316 | 0.403151 | 2.313033 | 0.663359 |
| 3 | 86919400 | 86921399 | 12.91659 | 23.26975 | 37.3145 | 16.26695 | -0.27167 | 0.388708 | 0.663369 |
| X | 103681600 | 103683199 | 21.10399 | 35.48238 | 56.3522 | 24.61836 | -0.28384 | 1.126502 | 0.663371 |
| X | 87305600 | 87308799 | 48.19283 | 77.91416 | 80.62832 | 34.45188 | 0.312171 | 1.71715 | 0.663382 |
| 4 | 61365400 | 61367199 | 17.89018 | 31.58608 | 27.87062 | 12.33112 | 0.569511 | 0.546664 | 0.663405 |
| 7 | 86221000 | 86221999 | 8.563317 | 16.12736 | 26.98713 | 11.04575 | -0.26544 | 0.071568 | 0.663405 |
| 1 | 98521200 | 98522599 | 31.12579 | 50.72406 | 75.53509 | 32.44074 | -0.20164 | 1.534963 | 0.663459 |

| | | | | | | | | | |
|---|---|---|---|---|---|---|---|---|---|
| 13 | 110966200 | 110967799 | 11.12632 | 20.52771 | 33.59365 | 14.20117 | -0.2705 | 0.23315 | 0.663476 |
| 6 | 42759000 | 42759999 | 9.783456 | 18.2135 | 29.43446 | 12.28557 | -0.2388 | 0.160539 | 0.66349 |
| 6 | 42920000 | 42922999 | 27.61372 | 47.16916 | 48.47155 | 21.25182 | 0.34352 | 1.000002 | 0.663493 |
| 12 | 84298400 | 84301599 | 24.11064 | 40.36699 | 60.36614 | 26.30936 | -0.19756 | 1.180295 | 0.663495 |
| 11 | 109704000 | 109705399 | 17.61719 | 30.88378 | 34.59557 | 15.28566 | 0.224615 | 0.647525 | 0.663512 |
| 6 | 141687200 | 141688999 | 17.45303 | 31.63502 | 30.68092 | 13.27329 | 0.45662 | 0.371391 | 0.663523 |
| 10 | 61453200 | 61453999 | 14.00324 | 25.24423 | 28.3911 | 12.36655 | 0.237986 | 0.373453 | 0.663551 |
| 15 | 43940400 | 43942399 | 17.81994 | 32.15408 | 30.49343 | 13.26713 | 0.483383 | 0.397864 | 0.663583 |
| 10 | 59354800 | 59355999 | 15.39214 | 27.77335 | 30.21174 | 13.1446 | 0.28664 | 0.380639 | 0.663583 |
| 12 | 114816200 | 114835399 | 345.5199 | 489.0484 | 481.2266 | 199.1538 | 0.209067 | 4.402136 | 0.663607 |
| 7 | 41178200 | 41182599 | 60.08854 | 96.01641 | 103.9314 | 43.67704 | 0.227687 | 2.093152 | 0.663619 |
| 4 | 144093000 | 144105199 | 149.0672 | 225.272 | 238.7615 | 98.7231 | 0.191962 | 3.319903 | 0.663629 |
| 5 | 72020400 | 72024799 | 55.90966 | 89.6758 | 85.02049 | 35.92777 | 0.424645 | 1.965673 | 0.663635 |
| 3 | 4812200 | 4814199 | 21.30694 | 36.80823 | 41.53289 | 18.22065 | 0.211909 | 0.863697 | 0.663635 |
| 3 | 123910800 | 123913199 | 19.7494 | 34.565 | 38.08014 | 16.80201 | 0.248486 | 0.690514 | 0.663654 |
| 19 | 35230400 | 35231599 | 15.2313 | 26.75205 | 40.82991 | 18.06925 | -0.2194 | 0.615741 | 0.663654 |
| 10 | 60000800 | 60004999 | 51.6376 | 82.33614 | 118.1231 | 49.49184 | -0.18141 | 2.152528 | 0.663688 |
| 7 | 25710400 | 25714599 | 67.58805 | 107.1565 | 120.1822 | 50.01743 | 0.164839 | 2.341313 | 0.663715 |
| 13 | 81873800 | 81876999 | 31.96122 | 53.01788 | 59.0574 | 25.58399 | 0.225756 | 1.30348 | 0.66374 |
| 14 | 76838200 | 76839599 | 14.57642 | 26.68248 | 28.87519 | 12.33777 | 0.310984 | 0.290768 | 0.663742 |
| 14 | 107208400 | 107208799 | 6.812641 | 13.28809 | 17.47839 | 6.264037 | 0.25477 | -0.42556 | 0.663743 |
| 7 | 47275600 | 47279599 | 47.83611 | 77.52567 | 66.87695 | 28.63275 | 0.577432 | 1.657201 | 0.66375 |
| 13 | 57108000 | 57122199 | 201.0715 | 294.5824 | 389.9335 | 160.6194 | -0.16943 | 3.882011 | 0.663785 |
| X | 114753400 | 114772199 | 319.5836 | 457.4333 | 431.9601 | 178.4335 | 0.284683 | 4.238645 | 0.663788 |
| 9 | 106265000 | 106268799 | 33.05056 | 54.4657 | 61.851 | 26.68259 | 0.196665 | 1.383085 | 0.663789 |
| 11 | 115267800 | 115269799 | 24.26012 | 41.15219 | 45.88573 | 20.09991 | 0.224986 | 1.046575 | 0.663815 |
| 3 | 5924800 | 5925999 | 1963.322 | 2361.207 | 2699.36 | 1140.852 | -0.24401 | 6.986353 | 0.663844 |
| 16 | 53495600 | 53498999 | 37.49272 | 61.41862 | 58.5469 | 25.19352 | 0.445139 | 1.465077 | 0.663846 |
| 18 | 12428000 | 12430199 | 19.033 | 32.84144 | 50.09237 | 21.97494 | -0.22153 | 0.875546 | 0.66389 |
| 6 | 57202600 | 57203199 | 9.810367 | 18.18039 | 29.37918 | 12.34475 | -0.24586 | 0.187953 | 0.663902 |
| 4 | 16599600 | 16605199 | 65.64527 | 104.5442 | 112.9463 | 47.24336 | 0.225214 | 2.20449 | 0.663925 |
| 2 | 89377200 | 89380599 | 31.02303 | 52.47903 | 51.84963 | 22.69335 | 0.398637 | 1.0992 | 0.663929 |
| 10 | 8390400 | 8391399 | 11.25581 | 21.0008 | 24.14622 | 10.04074 | 0.253376 | 0.146353 | 0.663949 |
| 4 | 24235600 | 24239399 | 45.06611 | 72.28385 | 104.3395 | 44.09196 | -0.18064 | 1.958952 | 0.663955 |
| 17 | 21375200 | 21378799 | 32.83498 | 53.94559 | 60.62672 | 26.12069 | 0.209878 | 1.418002 | 0.664003 |
| 10 | 68036800 | 68038399 | 14.53227 | 25.52666 | 39.28479 | 17.38683 | -0.23136 | 0.606387 | 0.664035 |
| 4 | 146148800 | 146152399 | 43.02892 | 69.66311 | 65.0405 | 27.82536 | 0.462222 | 1.667435 | 0.66405 |
| 8 | 58339400 | 58342999 | 36.90383 | 60.93111 | 63.39563 | 27.3748 | 0.323389 | 1.372128 | 0.664072 |
| 10 | 112439600 | 112441599 | 15.79641 | 27.17013 | 52.6956 | 23.11471 | -0.56747 | 0.894678 | 0.664073 |
| 10 | 25848200 | 25850799 | 29.94606 | 50.51414 | 45.2802 | 19.80108 | 0.538323 | 1.111097 | 0.664081 |
| 14 | 82743200 | 82745599 | 26.22387 | 45.17845 | 39.23937 | 17.21303 | 0.587145 | 0.907027 | 0.664102 |
| 7 | 114003600 | 114006599 | 32.31235 | 52.04476 | 98.82948 | 42.02196 | -0.56786 | 1.811822 | 0.664103 |
| 6 | 138724200 | 138725599 | 17.86789 | 30.17217 | 58.96816 | 25.79512 | -0.58181 | 1.078622 | 0.664104 |
| 18 | 85904800 | 85907799 | 30.19215 | 51.24416 | 50.7031 | 22.21401 | 0.396704 | 1.067202 | 0.664107 |
| 4 | 135461000 | 135464999 | 40.50947 | 65.22433 | 94.4984 | 40.15958 | -0.17918 | 1.83079 | 0.664139 |
| 13 | 69758000 | 69760399 | 16.00408 | 27.37434 | 54.79478 | 24.02675 | -0.61425 | 0.949135 | 0.664153 |
| 11 | 113843800 | 113865399 | 378.3547 | 523.4792 | 653.1137 | 271.2638 | -0.16634 | 4.742423 | 0.664161 |

| | | | | | | | | | |
|---|---|---|---|---|---|---|---|---|---|
| 12 | 93474600 | 93478999 | 36.64346 | 60.09581 | 53.01964 | 22.82607 | 0.557136 | 1.445672 | 0.664163 |
| 5 | 147920200 | 147929799 | 106.1442 | 161.9489 | 222.7963 | 92.38411 | -0.17262 | 3.114794 | 0.664219 |
| 2 | 16773400 | 16774999 | 26.65661 | 43.47162 | 80.60293 | 34.6262 | -0.5163 | 1.5191 | 0.66423 |
| 11 | 113105200 | 113106599 | 12.58679 | 23.50833 | 26.68398 | 11.07668 | 0.27368 | 0.157292 | 0.664231 |
| 5 | 64575600 | 64578999 | 32.14682 | 53.22418 | 59.71969 | 25.83687 | 0.215034 | 1.323738 | 0.664233 |
| 8 | 117108400 | 117113399 | 41.09308 | 66.14974 | 97.35683 | 41.3614 | -0.20223 | 1.837693 | 0.664234 |
| 11 | 11269400 | 11270799 | 13.16692 | 24.33975 | 27.54329 | 11.61278 | 0.260868 | 0.223562 | 0.664241 |
| 1 | 100763800 | 100765199 | 16.9733 | 30.18741 | 32.80836 | 14.46512 | 0.275407 | 0.487292 | 0.664243 |
| 1 | 148146000 | 148147199 | 19.82353 | 34.42713 | 38.41571 | 16.86198 | 0.229389 | 0.802572 | 0.664247 |
| 10 | 96025400 | 96030399 | 49.57962 | 79.64415 | 89.65888 | 37.97564 | 0.179939 | 1.912728 | 0.664256 |
| 7 | 64284800 | 64286399 | 12.26649 | 22.02855 | 34.16471 | 14.94064 | -0.22664 | 0.397617 | 0.66426 |
| 1 | 57557000 | 57559999 | 25.80372 | 42.59271 | 65.70207 | 28.36556 | -0.24307 | 1.347603 | 0.664278 |
| 2 | 40466000 | 40467599 | 14.13574 | 25.78265 | 28.89444 | 12.40317 | 0.256501 | 0.308371 | 0.66429 |
| 14 | 118412400 | 118413599 | 12.27919 | 22.01182 | 34.17001 | 14.96558 | -0.22943 | 0.409192 | 0.664309 |
| 14 | 91233400 | 91235999 | 56.52295 | 90.2594 | 101.5817 | 42.64974 | 0.17053 | 2.107543 | 0.664311 |
| Y | 4422800 | 4431999 | 83.58607 | 130.7887 | 101.1577 | 41.9646 | 0.685299 | 2.620733 | 0.664321 |
| 5 | 22029800 | 22032799 | 26.42975 | 44.3734 | 49.11118 | 21.44184 | 0.235063 | 1.144358 | 0.664337 |
| 8 | 127367000 | 127369799 | 36.94154 | 60.15864 | 68.21798 | 29.29056 | 0.189715 | 1.553048 | 0.664342 |
| 9 | 103474200 | 103475399 | 10.75267 | 19.61257 | 31.7594 | 13.63263 | -0.27108 | 0.289152 | 0.664346 |
| 14 | 120433400 | 120434799 | 13.76356 | 24.49311 | 38.19011 | 16.82962 | -0.24207 | 0.472557 | 0.664351 |
| 9 | 17197800 | 17198999 | 9.332943 | 17.61648 | 29.21062 | 11.9061 | -0.2477 | 0.068556 | 0.664358 |
| X | 139498600 | 139501999 | 50.64417 | 81.93939 | 65.35536 | 27.94682 | 0.68812 | 1.69904 | 0.664363 |
| 3 | 95618400 | 95621799 | 38.48797 | 62.63986 | 70.29088 | 30.17407 | 0.204152 | 1.563922 | 0.664389 |
| 3 | 46059200 | 46059999 | 12.78062 | 23.27065 | 24.53039 | 10.55271 | 0.342381 | 0.298067 | 0.664395 |
| 17 | 22458800 | 22459999 | 14.11546 | 24.92968 | 38.22107 | 16.9095 | -0.22309 | 0.540879 | 0.664461 |
| 7 | 90513400 | 90516599 | 26.25928 | 43.67384 | 65.44532 | 28.40267 | -0.20055 | 1.262093 | 0.664479 |
| 12 | 12845200 | 12846999 | 15.47332 | 27.51612 | 30.80015 | 13.58117 | 0.232791 | 0.472019 | 0.664492 |
| 7 | 127723600 | 127727399 | 25.8666 | 42.57147 | 66.56978 | 28.70248 | -0.26418 | 1.385789 | 0.664506 |
| 9 | 50290200 | 50292199 | 18.24192 | 31.62363 | 47.93814 | 21.03289 | -0.21119 | 0.828467 | 0.664515 |
| 15 | 38930000 | 38933599 | 49.87863 | 80.25775 | 88.1391 | 37.42196 | 0.218552 | 1.860423 | 0.664518 |
| 4 | 91489400 | 91493399 | 43.59395 | 70.09408 | 102.7861 | 43.59456 | -0.19907 | 1.880465 | 0.66452 |
| 17 | 21590200 | 21594999 | 44.59341 | 71.4899 | 103.2024 | 43.57828 | -0.1816 | 1.973859 | 0.664529 |
| 6 | 81954200 | 81954599 | 5.245438 | 10.21545 | 20.18559 | 7.382185 | -0.35238 | -0.36579 | 0.66453 |
| 14 | 78687800 | 78689999 | 28.45868 | 48.5051 | 49.02415 | 21.49027 | 0.366835 | 1.021628 | 0.664531 |
| 5 | 87317000 | 87320399 | 38.43384 | 63.32806 | 64.17433 | 27.68321 | 0.360394 | 1.398899 | 0.664532 |
| 9 | 21623200 | 21628399 | 53.84921 | 85.56554 | 123.7385 | 51.61287 | -0.19814 | 2.265808 | 0.664536 |
| 2 | 23129000 | 23130399 | 10.36727 | 18.88602 | 30.81367 | 13.24733 | -0.28285 | 0.294163 | 0.664554 |
| 6 | 6327200 | 6333599 | 74.48694 | 118.2728 | 121.5715 | 50.6722 | 0.292579 | 2.303063 | 0.664565 |
| 18 | 50186000 | 50188199 | 13.8671 | 24.38935 | 38.74237 | 17.15439 | -0.27655 | 0.583033 | 0.664597 |
| 8 | 64307800 | 64309199 | 14.68193 | 26.98781 | 28.98843 | 12.31615 | 0.327704 | 0.269997 | 0.664625 |
| 19 | 47948400 | 47952999 | 50.75047 | 80.89919 | 116.327 | 48.7206 | -0.1851 | 2.161869 | 0.664631 |
| 1 | 190731600 | 190733799 | 23.4358 | 40.33885 | 44.70142 | 19.60852 | 0.236893 | 0.911147 | 0.664633 |
| 7 | 19697400 | 19713799 | 206.3603 | 300.7065 | 395.6437 | 163.0442 | -0.16819 | 3.964774 | 0.66468 |
| 7 | 3773800 | 3774399 | 10.26302 | 19.4008 | 22.70055 | 9.223656 | 0.255454 | 0.04902 | 0.664684 |
| 7 | 66693000 | 66696199 | 25.8018 | 42.63149 | 65.16313 | 28.15201 | -0.22979 | 1.336115 | 0.664703 |
| 4 | 56901200 | 56902799 | 19.1223 | 33.38233 | 37.7688 | 16.63225 | 0.210276 | 0.726352 | 0.664708 |
| 13 | 5795400 | 5796199 | 10.10066 | 18.61728 | 30.34006 | 12.84278 | -0.26588 | 0.221683 | 0.664714 |

| | | | | | | | | | |
|---|---|---|---|---|---|---|---|---|---|
| 16 | 27239800 | 27242199 | 29.01168 | 48.9579 | 52.37689 | 22.90729 | 0.284105 | 1.113024 | 0.664766 |
| 1 | 167757200 | 167759199 | 22.21471 | 37.50754 | 56.55958 | 24.74055 | -0.20933 | 1.089182 | 0.664771 |
| X | 78815000 | 78815799 | 12.67409 | 23.6846 | 24.71171 | 10.24827 | 0.396203 | 0.153561 | 0.664777 |
| 13 | 11435800 | 11439599 | 42.25897 | 68.02926 | 100.6742 | 42.77323 | -0.21008 | 1.839838 | 0.664783 |
| X | 101951800 | 101955199 | 20.2178 | 34.6011 | 53.81345 | 23.59752 | -0.25156 | 0.962477 | 0.664788 |
| 11 | 73877400 | 73890799 | 153.2973 | 232.211 | 233.7787 | 96.73469 | 0.268997 | 3.273977 | 0.664796 |
| 7 | 47302800 | 47315599 | 185.0997 | 277.2618 | 271.3588 | 111.9323 | 0.295637 | 3.500404 | 0.664808 |
| 18 | 9594400 | 9597399 | 36.92521 | 60.11473 | 68.22128 | 29.28889 | 0.188297 | 1.55678 | 0.664822 |
| 3 | 70694600 | 70695399 | 11.70886 | 21.86849 | 24.70363 | 10.25472 | 0.280996 | 0.147141 | 0.664839 |
| 12 | 74330600 | 74332799 | 17.2609 | 29.74772 | 47.41431 | 20.79934 | -0.28502 | 0.874822 | 0.664849 |
| 5 | 21186000 | 21186399 | 5.361703 | 10.46405 | 19.87665 | 7.060383 | -0.25782 | -0.46785 | 0.664853 |
| 14 | 62817800 | 62820799 | 14.62932 | 25.40107 | 49.51342 | 21.73025 | -0.57181 | 0.797641 | 0.664865 |
| 17 | 77194600 | 77195799 | 12.54604 | 22.71326 | 36.89988 | 15.98788 | -0.2847 | 0.356187 | 0.664883 |
| 15 | 84777600 | 84779599 | 47.80327 | 76.3371 | 111.0514 | 46.62724 | -0.19904 | 2.101788 | 0.66489 |
| 8 | 129816600 | 129818399 | 18.81577 | 32.6567 | 51.75093 | 22.70657 | -0.27464 | 0.82655 | 0.664902 |
| 4 | 71153000 | 71154199 | 11.79641 | 21.4525 | 33.81839 | 14.57048 | -0.23592 | 0.32078 | 0.664911 |
| 6 | 68375000 | 68382999 | 99.24359 | 154.5212 | 162.2028 | 67.32827 | 0.239758 | 2.728092 | 0.664917 |
| 11 | 64459800 | 64460999 | 13.60763 | 25.25448 | 27.55385 | 11.55465 | 0.318679 | 0.204643 | 0.66493 |
| 5 | 100133600 | 100138599 | 45.61642 | 73.20984 | 106.4252 | 45.01386 | -0.1898 | 1.943154 | 0.664934 |
| 7 | 24425000 | 24428399 | 19.34903 | 32.92898 | 51.10646 | 22.40036 | -0.24976 | 1.005902 | 0.66494 |
| 15 | 35866400 | 35867599 | 8.316273 | 15.84105 | 27.24456 | 10.89292 | -0.27584 | -0.01111 | 0.664942 |
| 7 | 34634400 | 34637599 | 36.26636 | 58.99785 | 65.22271 | 27.99389 | 0.225254 | 1.564537 | 0.664952 |
| 14 | 40863200 | 40863999 | 10.87919 | 20.04956 | 32.46135 | 13.74321 | -0.25689 | 0.233047 | 0.664961 |
| 7 | 51808400 | 51810599 | 18.43807 | 32.9615 | 32.54754 | 14.28808 | 0.417019 | 0.462247 | 0.664965 |
| 19 | 26102600 | 26103399 | 10.54326 | 19.40015 | 31.40685 | 13.32503 | -0.25835 | 0.2408 | 0.664968 |
| 1 | 145650800 | 145651799 | 12.25365 | 22.16906 | 34.76063 | 15.07362 | -0.2344 | 0.354879 | 0.664976 |
| 8 | 92148600 | 92152399 | 67.85838 | 106.5955 | 151.1132 | 62.67955 | -0.18328 | 2.55257 | 0.664988 |
| 8 | 121458400 | 121459599 | 7.466812 | 14.22083 | 25.04123 | 10.01519 | -0.30913 | -0.02155 | 0.664997 |
| 13 | 50537400 | 50542399 | 88.62903 | 137.998 | 152.7771 | 63.41557 | 0.163083 | 2.720757 | 0.665007 |
| 7 | 40061000 | 40062199 | 13.46665 | 23.82678 | 36.7037 | 16.23233 | -0.22879 | 0.514438 | 0.665014 |
| 2 | 90091400 | 90092199 | 7.554582 | 14.59122 | 26.1116 | 10.04403 | -0.28064 | -0.14755 | 0.66503 |
| 7 | 41695000 | 41702199 | 170.6468 | 254.5076 | 271.1999 | 111.7726 | 0.167352 | 3.578028 | 0.665054 |
| 15 | 94015400 | 94017399 | 22.01743 | 38.5484 | 39.01503 | 17.21976 | 0.370351 | 0.699289 | 0.66508 |
| 13 | 35672600 | 35674599 | 17.4229 | 30.91906 | 33.11746 | 14.62846 | 0.294227 | 0.501387 | 0.665084 |
| 2 | 85792400 | 85800199 | 93.36627 | 145.6157 | 157.5948 | 65.39911 | 0.197629 | 2.695567 | 0.6651 |
| 15 | 19454200 | 19456999 | 35.6288 | 57.4676 | 86.87222 | 37.00844 | -0.23717 | 1.76693 | 0.665124 |
| 18 | 59398000 | 59401399 | 35.43476 | 57.29976 | 84.22045 | 35.99632 | -0.19255 | 1.691254 | 0.665131 |
| 15 | 94383200 | 94385999 | 34.50349 | 56.86332 | 63.17059 | 27.25253 | 0.228281 | 1.386442 | 0.665143 |
| 11 | 10249000 | 10252199 | 56.50505 | 90.50536 | 85.9919 | 36.26182 | 0.419222 | 2.010725 | 0.665185 |
| 4 | 19064400 | 19073399 | 95.23128 | 148.3611 | 159.9322 | 66.37998 | 0.2021 | 2.715596 | 0.665186 |
| 6 | 57979400 | 57980199 | 17.1305 | 30.078 | 33.54526 | 14.84135 | 0.230889 | 0.617736 | 0.665187 |
| X | 61563800 | 61566399 | 21.97757 | 38.03536 | 42.53431 | 18.661 | 0.225307 | 0.852066 | 0.665198 |
| 17 | 74829200 | 74832799 | 25.33161 | 43.1879 | 47.93941 | 21.01526 | 0.232176 | 1.006318 | 0.6652 |
| 15 | 28973600 | 28975999 | 12.60507 | 23.63958 | 22.43644 | 9.248801 | 0.537955 | 0.120716 | 0.665209 |
| 14 | 123094600 | 123096599 | 22.7739 | 39.52979 | 42.46407 | 18.63266 | 0.283861 | 0.828894 | 0.66523 |
| 1 | 12119400 | 12121399 | 14.5401 | 26.74395 | 28.40378 | 12.05741 | 0.344576 | 0.262672 | 0.665257 |
| 5 | 127805400 | 127808199 | 34.90256 | 57.01089 | 64.3377 | 27.65506 | 0.199711 | 1.50206 | 0.66526 |

| | | | | | | | | | |
|---|---|---|---|---|---|---|---|---|---|
| 7 | 14850600 | 14852599 | 28.42897 | 48.58657 | 47.90797 | 21.00559 | 0.402912 | 0.995936 | 0.665276 |
| 7 | 41125600 | 41142399 | 329.7255 | 470.6747 | 444.0271 | 183.4977 | 0.282196 | 4.278474 | 0.665283 |
| 4 | 79385200 | 79385999 | 12.24245 | 22.9922 | 25.64176 | 10.54468 | 0.309372 | 0.114837 | 0.665297 |
| 5 | 62571000 | 62572599 | 12.24245 | 22.9922 | 25.64176 | 10.54468 | 0.309372 | 0.114837 | 0.665297 |
| X | 91059800 | 91085999 | 368.1756 | 519.7637 | 492.3355 | 203.8386 | 0.260235 | 4.44167 | 0.665304 |
| 18 | 48143400 | 48143799 | 20.23255 | 36.1069 | 26.51483 | 11.65678 | 0.840215 | 0.472766 | 0.665332 |
| 1 | 115907800 | 115909999 | 19.48867 | 34.22338 | 36.27645 | 16.05164 | 0.303693 | 0.631679 | 0.665341 |
| 6 | 43782200 | 43782799 | 14.91421 | 26.55963 | 29.02677 | 12.78386 | 0.268552 | 0.457858 | 0.66536 |
| 18 | 3469800 | 3474799 | 63.26532 | 100.5944 | 113.0266 | 47.17619 | 0.16629 | 2.251666 | 0.665394 |
| 17 | 84850000 | 84850599 | 9.348241 | 17.45774 | 28.44404 | 11.80953 | -0.24403 | 0.135795 | 0.6654 |
| 4 | 147133600 | 147136399 | 43.31752 | 70.29656 | 66.34634 | 28.4336 | 0.449822 | 1.623288 | 0.66543 |
| 13 | 82296000 | 82296799 | 11.78478 | 21.22657 | 33.18585 | 14.47067 | -0.23526 | 0.374187 | 0.665439 |
| 1 | 18873800 | 18875199 | 16.31469 | 29.56273 | 30.77678 | 13.31981 | 0.354642 | 0.365478 | 0.665505 |
| 7 | 118958000 | 118960799 | 26.23247 | 44.31611 | 50.14051 | 21.93644 | 0.20372 | 1.092587 | 0.665522 |
| 16 | 16281000 | 16286399 | 47.50268 | 76.79811 | 80.56286 | 34.42376 | 0.292559 | 1.715643 | 0.665528 |
| 18 | 37153000 | 37156399 | 89.06716 | 138.3699 | 145.1612 | 60.27445 | 0.238157 | 2.758905 | 0.665551 |
| 3 | 117368200 | 117370399 | 20.25306 | 34.38603 | 52.72676 | 23.10288 | -0.23297 | 1.030278 | 0.66556 |
| 4 | 52598200 | 52601599 | 27.96204 | 45.96011 | 69.01448 | 29.74011 | -0.20671 | 1.408583 | 0.66559 |
| 18 | 37827400 | 37830799 | 152.1448 | 228.3102 | 238.853 | 98.56443 | 0.201853 | 3.458865 | 0.66559 |
| 5 | 141253400 | 141254799 | 14.44042 | 25.31963 | 39.42259 | 17.42126 | -0.24788 | 0.637237 | 0.665601 |
| 3 | 44221200 | 44221799 | 9.982907 | 18.76027 | 31.92602 | 13.11515 | -0.29443 | 0.108782 | 0.665624 |
| 6 | 57319800 | 57323599 | 66.60125 | 105.7241 | 118.3673 | 49.3238 | 0.169121 | 2.303889 | 0.665647 |
| 8 | 49008000 | 49009199 | 11.43865 | 20.66137 | 32.69623 | 14.20692 | -0.24939 | 0.353614 | 0.665658 |
| 9 | 3243400 | 3253399 | 134.8352 | 205.6336 | 212.418 | 88.07172 | 0.239886 | 3.131669 | 0.665658 |
| 2 | 100067000 | 100068799 | 17.2578 | 31.02779 | 31.61542 | 13.81239 | 0.376636 | 0.417707 | 0.665676 |
| 3 | 91396400 | 91401799 | 56.58729 | 90.77939 | 96.0292 | 40.59116 | 0.267258 | 1.966126 | 0.665686 |
| 18 | 83683800 | 83689599 | 96.99526 | 149.9716 | 164.5551 | 68.35235 | 0.168181 | 2.848305 | 0.665701 |
| 11 | 113007200 | 113009799 | 21.23365 | 36.96017 | 40.86462 | 17.96259 | 0.242904 | 0.781554 | 0.665716 |
| 7 | 56915000 | 56917199 | 17.4837 | 31.24076 | 32.65555 | 14.34116 | 0.334986 | 0.459498 | 0.665718 |
| 7 | 47144400 | 47156199 | 25.61047 | 44.14168 | 39.29402 | 17.23725 | 0.551951 | 0.901004 | 0.665725 |
| 7 | 87233400 | 87233999 | 6.675597 | 12.92249 | 23.26784 | 8.85674 | -0.27441 | -0.19979 | 0.665732 |
| 2 | 82167200 | 82174799 | 51.50247 | 83.01156 | 75.69557 | 32.21727 | 0.48924 | 1.806069 | 0.665749 |
| 9 | 30783600 | 30783999 | 5.506962 | 10.73294 | 20.26131 | 7.336309 | -0.27357 | -0.39501 | 0.665803 |
| 14 | 99391000 | 99391599 | 6.537625 | 12.70448 | 22.93725 | 8.54159 | -0.24807 | -0.27595 | 0.665845 |
| 14 | 86632600 | 86636399 | 45.0347 | 72.10871 | 106.6945 | 44.9688 | -0.21924 | 2.012294 | 0.665851 |
| 13 | 68182600 | 68213799 | 755.7476 | 997.3209 | 944.1709 | 394.8945 | 0.163195 | 5.492428 | 0.665866 |
| X | 39515000 | 39516199 | 15.4923 | 28.05224 | 30.40796 | 13.17209 | 0.296127 | 0.36518 | 0.665871 |
| 5 | 109815000 | 109833599 | 292.6503 | 424.2658 | 382.0872 | 157.5436 | 0.371305 | 4.0502 | 0.665871 |
| 3 | 104884000 | 104886599 | 25.77291 | 43.46332 | 49.07485 | 21.45878 | 0.206626 | 1.105689 | 0.665884 |
| 5 | 136772000 | 136784399 | 183.8219 | 269.3198 | 364.3697 | 150.0889 | -0.20089 | 3.878889 | 0.665891 |
| 15 | 56811600 | 56812399 | 9.218833 | 17.65349 | 20.32 | 8.022088 | 0.316271 | -0.05942 | 0.665895 |
| 19 | 25932400 | 25935799 | 33.81382 | 56.06495 | 60.43234 | 26.17123 | 0.272922 | 1.31277 | 0.665911 |
| 1 | 156708600 | 156710399 | 15.27878 | 26.65476 | 50.56052 | 22.25599 | -0.53199 | 0.735755 | 0.665915 |
| 9 | 100692600 | 100694999 | 26.92034 | 46.23572 | 46.24036 | 20.28223 | 0.38359 | 0.944474 | 0.665922 |
| 7 | 86672200 | 86675999 | 49.76885 | 80.1996 | 86.12964 | 36.64723 | 0.25335 | 1.814547 | 0.665939 |
| 14 | 22285600 | 22290199 | 26.18976 | 43.56442 | 66.12316 | 28.69968 | -0.21903 | 1.258959 | 0.665945 |
| 13 | 116100400 | 116104199 | 45.12664 | 72.24803 | 104.8016 | 44.16364 | -0.19083 | 2.017426 | 0.665961 |

| | | | | | | | | | |
|---|---|---|---|---|---|---|---|---|---|
| 9 | 54493400 | 54494399 | 6.277209 | 12.07094 | 23.60633 | 9.208018 | -0.42583 | -0.12157 | 0.665972 |
| 4 | 8953000 | 8954999 | 20.18391 | 34.12523 | 53.22013 | 23.28942 | -0.25743 | 1.068885 | 0.665983 |
| 6 | 118177000 | 118178799 | 11.03299 | 19.8649 | 32.26634 | 14.07641 | -0.29024 | 0.370182 | 0.665991 |
| 5 | 142613800 | 142617599 | 22.11026 | 37.16426 | 56.49096 | 24.67697 | -0.22101 | 1.128428 | 0.665996 |
| 10 | 58568400 | 58569999 | 12.10939 | 21.64026 | 34.16385 | 15.00203 | -0.25624 | 0.426699 | 0.666005 |
| 17 | 58145600 | 58146999 | 13.58877 | 25.05214 | 22.43265 | 9.492682 | 0.592774 | 0.230676 | 0.666008 |
| 18 | 76317400 | 76321199 | 30.03419 | 48.91422 | 74.58277 | 32.02436 | -0.23624 | 1.542526 | 0.66601 |
| 11 | 73322800 | 73328999 | 94.5705 | 147.6919 | 139.6131 | 57.92904 | 0.394112 | 2.669927 | 0.666015 |
| 16 | 49960800 | 49964999 | 80.60961 | 125.0509 | 175.9268 | 73.06475 | -0.18633 | 2.783194 | 0.666023 |
| 1 | 72750000 | 72751799 | 11.04741 | 19.92677 | 32.28627 | 14.05337 | -0.28447 | 0.360622 | 0.666042 |
| 1 | 89356400 | 89360199 | 34.77232 | 56.17841 | 84.50462 | 36.07623 | -0.22733 | 1.715358 | 0.66606 |
| 4 | 147340200 | 147341199 | 14.26302 | 27.18999 | 25.55588 | 10.20073 | 0.591764 | 0.037632 | 0.666082 |
| 7 | 18530000 | 18534399 | 31.31924 | 50.99433 | 75.9543 | 32.6103 | -0.20284 | 1.549109 | 0.666097 |
| 18 | 40877200 | 40880799 | 39.7018 | 63.97654 | 92.80159 | 39.48555 | -0.17942 | 1.803511 | 0.666131 |
| 17 | 54121600 | 54125199 | 57.12988 | 91.40178 | 100.668 | 42.37983 | 0.204623 | 2.049649 | 0.666133 |
| 4 | 33577000 | 33579399 | 29.55174 | 49.70141 | 52.97405 | 23.14478 | 0.289255 | 1.142334 | 0.666144 |
| 7 | 38863600 | 38875199 | 206.9118 | 305.6888 | 316.6043 | 130.3845 | 0.19575 | 3.747201 | 0.666148 |
| 14 | 39793800 | 39794399 | 9.91984 | 18.12359 | 30.36715 | 13.00796 | -0.31734 | 0.274001 | 0.666196 |
| 6 | 95380400 | 95382599 | 15.55053 | 28.34123 | 25.65496 | 11.02325 | 0.561785 | 0.316745 | 0.666198 |
| 19 | 50101400 | 50104999 | 42.80267 | 68.73532 | 99.26733 | 42.02471 | -0.17929 | 1.91809 | 0.666214 |
| 10 | 28781600 | 28785399 | 37.98579 | 61.35792 | 90.6679 | 38.69751 | -0.20213 | 1.728299 | 0.666232 |
| 2 | 79672200 | 79673599 | 12.40497 | 23.34355 | 25.67681 | 10.52378 | 0.332994 | 0.103979 | 0.666237 |
| 9 | 7907000 | 7910399 | 45.19706 | 73.01544 | 69.50207 | 29.66868 | 0.431168 | 1.726707 | 0.666241 |
| 9 | 15753600 | 15758199 | 41.9414 | 67.56349 | 100.8822 | 42.90348 | -0.22174 | 1.818685 | 0.666264 |
| X | 153408400 | 153411999 | 62.62433 | 100.0024 | 106.5673 | 44.74294 | 0.249147 | 2.115206 | 0.666285 |
| 1 | 114214400 | 114217199 | 32.29 | 52.44481 | 78.19058 | 33.53032 | -0.20705 | 1.591571 | 0.6663 |
| X | 61368400 | 61371599 | 29.6739 | 50.51926 | 49.58105 | 21.73242 | 0.408872 | 1.037276 | 0.666302 |
| 18 | 38313200 | 38314999 | 22.4018 | 38.50555 | 43.68242 | 19.16072 | 0.202786 | 0.917425 | 0.666304 |
| 3 | 12732800 | 12734799 | 18.782 | 33.37202 | 33.54227 | 14.80073 | 0.386558 | 0.503637 | 0.666363 |
| 13 | 91022400 | 91024399 | 22.05495 | 38.23183 | 42.53538 | 18.6624 | 0.2332 | 0.838974 | 0.666391 |
| 18 | 6187600 | 6191599 | 35.43485 | 57.26478 | 84.63056 | 36.143 | -0.20144 | 1.709104 | 0.666408 |
| 15 | 49618400 | 49619399 | 11.31048 | 20.51685 | 32.25319 | 13.94014 | -0.23475 | 0.328527 | 0.666409 |
| 9 | 29740600 | 29741599 | 8.555643 | 16.25119 | 27.15275 | 10.92105 | -0.24136 | 0.013719 | 0.666438 |
| 13 | 112144600 | 112146799 | 25.87145 | 43.7605 | 49.59609 | 21.70642 | 0.201325 | 1.079704 | 0.666455 |
| 16 | 4065600 | 4071999 | 101.0924 | 157.0976 | 147.739 | 61.34216 | 0.396041 | 2.759783 | 0.666462 |
| 13 | 65889400 | 65896999 | 238.0732 | 347.34 | 306.2642 | 126.1951 | 0.410642 | 3.947256 | 0.66647 |
| 4 | 10587400 | 10588199 | 12.65798 | 23.80294 | 26.23841 | 10.76675 | 0.328734 | 0.112915 | 0.666511 |
| 19 | 55044800 | 55049199 | 49.85248 | 79.93288 | 88.98065 | 37.58154 | 0.192897 | 1.969847 | 0.666514 |
| 18 | 47276200 | 47287599 | 122.9795 | 185.3949 | 292.0856 | 120.6797 | -0.38253 | 3.366191 | 0.666519 |
| 13 | 68694600 | 68698999 | 47.09138 | 75.73583 | 85.45261 | 36.25562 | 0.178766 | 1.870841 | 0.66654 |
| 7 | 138895000 | 138901999 | 50.00843 | 79.85503 | 114.1575 | 47.92846 | -0.174 | 2.104563 | 0.666553 |
| 1 | 64567600 | 64569199 | 18.31357 | 32.26429 | 35.31654 | 15.63221 | 0.258925 | 0.585364 | 0.666563 |
| 4 | 85832200 | 85833599 | 18.31357 | 32.26429 | 35.31654 | 15.63221 | 0.258925 | 0.585364 | 0.666563 |
| 1 | 3499800 | 3502199 | 19.61321 | 35.05384 | 33.07249 | 14.52148 | 0.482344 | 0.473947 | 0.666565 |
| 7 | 52190400 | 52194399 | 41.19806 | 67.27604 | 70.00745 | 30.0892 | 0.316177 | 1.525815 | 0.666569 |
| 18 | 3942200 | 3944199 | 26.23904 | 45.03883 | 46.81834 | 20.535 | 0.327973 | 0.949283 | 0.666578 |
| 9 | 93167200 | 93169599 | 18.13142 | 30.71841 | 57.98517 | 25.39052 | -0.53175 | 1.048199 | 0.666579 |

| | | | | | | | | | |
|---|---|---|---|---|---|---|---|---|---|
| 18 | 35817200 | 35820199 | 33.60535 | 55.2313 | 62.83878 | 27.07791 | 0.192452 | 1.418973 | 0.666582 |
| 1 | 148338200 | 148339999 | 26.83267 | 43.8016 | 81.10869 | 34.8559 | -0.51337 | 1.50787 | 0.666594 |
| 3 | 97698000 | 97702199 | 39.17468 | 63.20991 | 92.15429 | 39.27706 | -0.18452 | 1.763327 | 0.666611 |
| 15 | 34867400 | 34870199 | 38.33068 | 62.18117 | 69.28728 | 29.68601 | 0.210346 | 1.620774 | 0.666647 |
| 2 | 21078200 | 21081199 | 37.89464 | 62.07448 | 56.27682 | 24.21626 | 0.517332 | 1.466442 | 0.666663 |
| 11 | 61470600 | 61472199 | 15.19205 | 26.70097 | 40.78248 | 18.05847 | -0.22056 | 0.603697 | 0.666682 |
| 2 | 85308200 | 85314799 | 128.7037 | 196.4579 | 185.9078 | 77.09612 | 0.367244 | 3.109123 | 0.666684 |
| 8 | 42532200 | 42536799 | 44.16284 | 71.01397 | 105.7746 | 44.86683 | -0.22143 | 1.881445 | 0.666751 |
| 2 | 22633600 | 22635199 | 115.2355 | 177.287 | 155.9212 | 64.75376 | 0.481529 | 2.949098 | 0.666763 |
| 12 | 6749200 | 6756199 | 58.44961 | 93.52066 | 89.67969 | 37.75815 | 0.404217 | 2.046081 | 0.666774 |
| 5 | 4429000 | 4434399 | 70.61829 | 110.9403 | 155.3097 | 64.4209 | -0.16511 | 2.552718 | 0.666806 |
| 14 | 41797000 | 41799799 | 37.98671 | 62.82485 | 53.29332 | 23.03932 | 0.617215 | 1.346656 | 0.666829 |
| 2 | 150239000 | 150250399 | 303.1586 | 432.968 | 439.5344 | 181.6258 | 0.177161 | 4.268773 | 0.666831 |
| 16 | 17001600 | 17002999 | 15.92423 | 28.28195 | 31.86979 | 14.06792 | 0.222306 | 0.486782 | 0.666847 |
| 6 | 92154800 | 92156599 | 22.49964 | 38.95003 | 43.05177 | 18.88819 | 0.242169 | 0.852868 | 0.666863 |
| 6 | 107114600 | 107115799 | 12.29819 | 22.73487 | 26.00269 | 10.96257 | 0.245606 | 0.212132 | 0.666865 |
| 6 | 127193600 | 127195999 | 17.09737 | 29.47069 | 47.41394 | 20.79928 | -0.29841 | 0.872398 | 0.666871 |
| X | 62404400 | 62414399 | 127.2558 | 195.6142 | 188.711 | 78.36219 | 0.347311 | 2.973979 | 0.666875 |
| 7 | 139635200 | 139639599 | 38.37368 | 61.8003 | 91.16481 | 38.75632 | -0.20465 | 1.818366 | 0.666888 |
| 8 | 67523400 | 67523999 | 10.65842 | 20.3879 | 23.96191 | 9.489615 | 0.281706 | -0.02573 | 0.666889 |
| 12 | 6656800 | 6659199 | 19.23089 | 34.0484 | 34.64802 | 15.32115 | 0.366371 | 0.537557 | 0.666915 |
| 8 | 15622600 | 15628999 | 75.13602 | 118.3148 | 131.9227 | 54.7369 | 0.165387 | 2.505618 | 0.666918 |
| 11 | 31453400 | 31456199 | 21.97769 | 37.77343 | 42.62603 | 18.6973 | 0.210295 | 0.913699 | 0.666921 |
| 6 | 16601400 | 16605799 | 69.24306 | 109.86 | 101.3923 | 42.22809 | 0.446645 | 2.316643 | 0.66695 |
| 13 | 119063200 | 119063799 | 12.44247 | 22.88739 | 26.09078 | 11.07445 | 0.243708 | 0.243477 | 0.666956 |
| 10 | 86477000 | 86479599 | 15.10632 | 26.23851 | 48.9803 | 21.50103 | -0.5096 | 0.794812 | 0.666962 |
| 14 | 92982400 | 92983199 | 13.43995 | 24.71064 | 28.05087 | 11.91402 | 0.248972 | 0.25822 | 0.666968 |
| 10 | 20921800 | 20922999 | 15.6856 | 27.31268 | 42.65297 | 18.75286 | -0.25284 | 0.753199 | 0.66698 |
| 10 | 84490800 | 84491599 | 5.535016 | 10.72578 | 21.6214 | 8.18997 | -0.42752 | -0.23119 | 0.666991 |
| 2 | 98556000 | 98563199 | 56.55718 | 91.00713 | 89.47065 | 37.98924 | 0.37802 | 1.870242 | 0.666991 |
| 1 | 146053800 | 146054399 | 8.347767 | 15.90215 | 26.67518 | 10.6637 | -0.23959 | -0.01121 | 0.667002 |
| 16 | 84723600 | 84724199 | 5.646868 | 11.00407 | 21.3929 | 7.760308 | -0.31754 | -0.38055 | 0.667011 |
| 18 | 40286600 | 40290199 | 30.30522 | 51.0852 | 52.9715 | 23.15971 | 0.328964 | 1.126718 | 0.667013 |
| 3 | 5649200 | 5652599 | 37.96458 | 62.63697 | 63.99486 | 27.62112 | 0.349433 | 1.384423 | 0.667025 |
| 3 | 9153400 | 9158999 | 59.25141 | 94.09429 | 133.704 | 55.73686 | -0.17372 | 2.289394 | 0.667049 |
| 7 | 119351000 | 119352799 | 18.02728 | 31.28902 | 47.94031 | 21.03479 | -0.22625 | 0.81556 | 0.66705 |
| 1 | 3472000 | 3473599 | 14.18748 | 26.16666 | 28.5361 | 12.06869 | 0.310684 | 0.246721 | 0.667065 |
| 12 | 62944000 | 62944999 | 15.12377 | 26.92014 | 29.82697 | 13.1415 | 0.248219 | 0.461623 | 0.667077 |
| 1 | 100487200 | 100487599 | 8.275819 | 15.99596 | 19.35714 | 7.420418 | 0.284343 | -0.16895 | 0.667098 |
| 8 | 31995800 | 31997799 | 16.41166 | 29.55065 | 31.16314 | 13.5943 | 0.32874 | 0.401956 | 0.66711 |
| 2 | 86417600 | 86421399 | 117.3962 | 180.4339 | 170.4181 | 70.76478 | 0.377673 | 2.968769 | 0.66711 |
| 17 | 43946600 | 43949799 | 30.04384 | 49.84144 | 56.7782 | 24.59804 | 0.193483 | 1.293739 | 0.667129 |
| 3 | 16434000 | 16437399 | 80.51132 | 126.1529 | 139.8051 | 57.98893 | 0.168694 | 2.604787 | 0.667155 |
| 19 | 56640800 | 56644199 | 29.29542 | 47.84263 | 73.31578 | 31.51196 | -0.24072 | 1.497832 | 0.667177 |
| 17 | 72176000 | 72180199 | 41.7624 | 67.1084 | 98.44592 | 41.71704 | -0.20069 | 1.894633 | 0.667177 |
| 7 | 60499600 | 60502399 | 27.83485 | 48.03668 | 44.54362 | 19.54086 | 0.493643 | 0.905274 | 0.667182 |
| 4 | 33836600 | 33838199 | 13.27232 | 23.57687 | 36.14462 | 15.95217 | -0.21918 | 0.47687 | 0.667183 |

| | | | | | | | | | |
|---|---|---|---|---|---|---|---|---|---|
| 14 | 62198800 | 62200199 | 13.08747 | 24.48086 | 26.63602 | 11.02816 | 0.337006 | 0.149916 | 0.667188 |
| 18 | 52671000 | 52672999 | 15.50982 | 27.11233 | 41.50687 | 18.29537 | -0.22381 | 0.694052 | 0.667209 |
| 16 | 95992600 | 95993799 | 9.469751 | 17.60642 | 28.88912 | 12.08471 | -0.26312 | 0.164737 | 0.667215 |
| X | 9484800 | 9485599 | 20.82551 | 36.15015 | 40.51171 | 17.77669 | 0.223291 | 0.819367 | 0.667216 |
| 3 | 32180800 | 32183199 | 20.89762 | 36.31671 | 40.9409 | 17.97791 | 0.214799 | 0.802668 | 0.667232 |
| X | 119876400 | 119876999 | 9.923852 | 19.16797 | 22.60437 | 8.693563 | 0.316661 | -0.12988 | 0.667233 |
| 12 | 79103600 | 79106399 | 35.11219 | 56.75626 | 84.03326 | 35.89846 | -0.20369 | 1.701776 | 0.66724 |
| 7 | 124574000 | 124575599 | 11.80557 | 21.30265 | 33.21564 | 14.45081 | -0.22922 | 0.364632 | 0.667245 |
| 16 | 65604400 | 65605599 | 11.80557 | 21.30265 | 33.21564 | 14.45081 | -0.22922 | 0.364632 | 0.667245 |
| 2 | 117266600 | 117269199 | 15.20644 | 26.58754 | 41.00729 | 18.07834 | -0.23443 | 0.688774 | 0.667247 |
| 5 | 83598000 | 83600399 | 39.98789 | 64.37041 | 93.82797 | 39.86197 | -0.1883 | 1.835797 | 0.667265 |
| 7 | 41100600 | 41101799 | 19.95141 | 34.54831 | 36.97973 | 16.22433 | 0.28843 | 0.830172 | 0.667281 |
| X | 17820000 | 17822999 | 29.81313 | 50.26146 | 52.92042 | 23.13818 | 0.307056 | 1.126299 | 0.667298 |
| 10 | 83618200 | 83619599 | 11.50347 | 20.77113 | 32.74436 | 14.23411 | -0.24408 | 0.358257 | 0.66732 |
| 14 | 66693000 | 66703399 | 89.4601 | 139.3875 | 154.747 | 64.22665 | 0.159826 | 2.710204 | 0.667321 |
| 5 | 76826800 | 76829799 | 30.57169 | 50.07215 | 74.33809 | 31.98631 | -0.19244 | 1.461331 | 0.667325 |
| 1 | 172492200 | 172494599 | 21.888 | 36.94297 | 55.99929 | 24.49286 | -0.21684 | 1.09037 | 0.667328 |
| 17 | 43845400 | 43846999 | 10.73679 | 19.62534 | 31.77776 | 13.60424 | -0.26853 | 0.276519 | 0.667353 |
| 3 | 133077400 | 133079999 | 26.34642 | 44.53309 | 49.06895 | 21.47121 | 0.241704 | 1.084049 | 0.667353 |
| 5 | 147118000 | 147129599 | 111.971 | 168.8872 | 283.6015 | 117.1926 | -0.47381 | 3.352978 | 0.667355 |
| 1 | 106046600 | 106048799 | 27.98217 | 45.83443 | 70.52589 | 30.34699 | -0.24382 | 1.451812 | 0.667362 |
| 16 | 56628400 | 56628799 | 5.652457 | 11.01077 | 20.77415 | 7.573476 | -0.28159 | -0.36632 | 0.667388 |
| 1 | 153743600 | 153745599 | 14.03858 | 24.66878 | 38.76907 | 17.16406 | -0.26148 | 0.595932 | 0.667399 |
| 8 | 74485200 | 74487799 | 25.9126 | 43.69133 | 49.60942 | 21.69139 | 0.198538 | 1.107692 | 0.667403 |
| 7 | 5617800 | 5620599 | 47.15494 | 76.49709 | 66.13037 | 28.3337 | 0.575712 | 1.63802 | 0.667423 |
| 12 | 68531800 | 68533199 | 12.05706 | 21.91043 | 34.84011 | 15.02445 | -0.24941 | 0.327298 | 0.667424 |
| 3 | 144604200 | 144604799 | 8.270786 | 16.00244 | 19.88941 | 7.588442 | 0.252407 | -0.18372 | 0.667428 |
| 19 | 39373400 | 39383599 | 107.3432 | 167.0007 | 143.9792 | 59.77095 | 0.522554 | 2.743118 | 0.667432 |
| 2 | 115897200 | 115898599 | 7.717658 | 14.33225 | 28.44228 | 11.91702 | -0.53693 | 0.15502 | 0.667434 |
| 4 | 21071800 | 21076799 | 159.857 | 239.1566 | 252.3804 | 104.0791 | 0.185405 | 3.517777 | 0.667434 |
| 9 | 37916400 | 37918199 | 18.14897 | 32.14271 | 29.18245 | 12.90352 | 0.530279 | 0.517902 | 0.667457 |
| 5 | 116696800 | 116698999 | 13.16232 | 23.29599 | 36.23102 | 16.02224 | -0.24225 | 0.508579 | 0.667479 |
| 3 | 138534200 | 138538399 | 39.68188 | 63.88053 | 93.06243 | 39.53918 | -0.18741 | 1.834255 | 0.667491 |
| 7 | 136344000 | 136345399 | 14.25271 | 25.99268 | 28.85489 | 12.38818 | 0.26987 | 0.309294 | 0.667491 |
| 6 | 82550000 | 82550799 | 8.029392 | 15.30757 | 26.13645 | 10.43115 | -0.26289 | -0.02174 | 0.667501 |
| 2 | 12669600 | 12673199 | 28.65743 | 48.22295 | 52.92825 | 23.1285 | 0.247149 | 1.136109 | 0.667535 |
| 16 | 89371400 | 89373199 | 14.04958 | 24.72935 | 38.2353 | 16.92726 | -0.23724 | 0.575814 | 0.667543 |
| 19 | 3805200 | 3806599 | 14.04958 | 24.72935 | 38.2353 | 16.92726 | -0.23724 | 0.575814 | 0.667543 |
| X | 37935000 | 37937399 | 23.92839 | 41.7026 | 34.96909 | 15.38315 | 0.640324 | 0.770518 | 0.667547 |
| 10 | 57362000 | 57363199 | 11.29404 | 20.28746 | 33.18058 | 14.50709 | -0.30263 | 0.385897 | 0.667551 |
| 16 | 60767800 | 60768999 | 10.8175 | 19.72854 | 31.28579 | 13.43127 | -0.24091 | 0.291546 | 0.667557 |
| 1 | 135448200 | 135449799 | 16.14871 | 28.12671 | 43.14176 | 18.97099 | -0.22706 | 0.751953 | 0.667562 |
| 10 | 19897000 | 19899599 | 16.14871 | 28.12671 | 43.14176 | 18.97099 | -0.22706 | 0.751953 | 0.667562 |
| 8 | 66020800 | 66024599 | 32.46481 | 53.44645 | 60.53854 | 26.10525 | 0.199821 | 1.394453 | 0.667589 |
| 1 | 4624600 | 4629999 | 43.77572 | 70.34188 | 101.7681 | 43.12329 | -0.18078 | 1.90055 | 0.667605 |
| 9 | 122187800 | 122191599 | 50.55168 | 80.68227 | 114.9175 | 48.21347 | -0.16949 | 2.121522 | 0.667605 |
| X | 79269000 | 79274799 | 64.23013 | 102.5058 | 108.1086 | 45.35196 | 0.263158 | 2.136495 | 0.667616 |

| | | | | | | | | | |
|---|---|---|---|---|---|---|---|---|---|
| 2 | 85526400 | 85528799 | 21.03372 | 36.7055 | 33.62315 | 14.80318 | 0.513518 | 0.735293 | 0.667628 |
| 10 | 129189000 | 129204399 | 292.6012 | 418.6643 | 429.2524 | 177.3248 | 0.16552 | 4.240839 | 0.667633 |
| 5 | 110172800 | 110175199 | 22.71352 | 39.45457 | 41.88537 | 18.3874 | 0.300893 | 0.816669 | 0.66764 |
| 18 | 6260200 | 6263799 | 117.0957 | 178.8225 | 193.528 | 80.26485 | 0.1743 | 3.097875 | 0.667647 |
| 4 | 22632400 | 22633599 | 19.83571 | 34.56309 | 38.86277 | 17.09391 | 0.218993 | 0.758659 | 0.66765 |
| 17 | 95156400 | 95158199 | 16.46104 | 29.6943 | 31.21669 | 13.58627 | 0.335431 | 0.39251 | 0.66765 |
| 1 | 60460800 | 60461999 | 15.23558 | 27.85187 | 29.34318 | 12.56007 | 0.347641 | 0.304124 | 0.667652 |
| 1 | 7418200 | 7442199 | 472.301 | 643.2507 | 790.2579 | 329.0411 | -0.16704 | 4.988641 | 0.667654 |
| 1 | 141696200 | 141698399 | 25.03783 | 43.53249 | 42.34611 | 18.60053 | 0.426703 | 0.816929 | 0.667655 |
| 12 | 70228800 | 70232399 | 72.3825 | 114.2175 | 124.1573 | 51.53482 | 0.204355 | 2.46115 | 0.667658 |
| 16 | 51944000 | 51945599 | 17.00141 | 29.8548 | 33.09803 | 14.64492 | 0.239528 | 0.615016 | 0.667673 |
| 10 | 66655600 | 66658199 | 33.5592 | 55.13858 | 62.87938 | 27.0919 | 0.188911 | 1.421746 | 0.66768 |
| X | 123053200 | 123054799 | 22.65411 | 39.46821 | 41.32629 | 18.17581 | 0.321241 | 0.777504 | 0.667737 |
| 9 | 113463600 | 113467399 | 42.30549 | 67.97002 | 98.19569 | 41.60118 | -0.17893 | 1.901749 | 0.667766 |
| 4 | 108557200 | 108558599 | 11.95963 | 21.41707 | 33.68262 | 14.76483 | -0.24894 | 0.412228 | 0.667767 |
| 5 | 17981400 | 17982199 | 11.61271 | 21.17469 | 33.32981 | 14.31226 | -0.23065 | 0.302692 | 0.667771 |
| 7 | 25987800 | 26063399 | 1797.584 | 2213.026 | 1940.657 | 817.6515 | 0.172859 | 6.616324 | 0.66783 |
| 11 | 28078600 | 28079199 | 10.49944 | 19.79678 | 22.73592 | 9.287345 | 0.27581 | 0.070325 | 0.667836 |
| 5 | 60618000 | 60620399 | 12.71164 | 22.37606 | 41.92192 | 18.5591 | -0.51273 | 0.5748 | 0.66784 |
| 7 | 43063200 | 43069799 | 53.49937 | 85.28597 | 121.4033 | 50.85075 | -0.17051 | 2.163213 | 0.667862 |
| 11 | 65527200 | 65529599 | 17.6391 | 30.99185 | 34.94941 | 15.47138 | 0.214935 | 0.612285 | 0.667866 |
| 10 | 9721600 | 9726199 | 53.08718 | 84.86681 | 91.94301 | 38.66344 | 0.22715 | 2.067881 | 0.667868 |
| 14 | 107162000 | 107164399 | 18.01715 | 30.49205 | 58.45516 | 25.58799 | -0.55394 | 1.059682 | 0.667908 |
| 5 | 91645000 | 91646999 | 21.88932 | 38.37407 | 38.43528 | 16.98257 | 0.385319 | 0.677635 | 0.667914 |
| 3 | 11432400 | 11432799 | 9.299815 | 17.77569 | 20.75458 | 8.23724 | 0.288232 | -0.04192 | 0.66792 |
| 12 | 87605000 | 87608799 | 74.62374 | 117.4477 | 129.5253 | 53.73861 | 0.180644 | 2.51479 | 0.667945 |
| 7 | 92717800 | 92719599 | 13.40528 | 23.72254 | 36.64845 | 16.20731 | -0.23299 | 0.510103 | 0.667977 |
| 2 | 120643200 | 120647599 | 78.68882 | 122.4069 | 171.2548 | 71.09526 | -0.17525 | 2.734546 | 0.667986 |
| 4 | 17176200 | 17177999 | 20.41453 | 35.93319 | 36.16917 | 16.01251 | 0.378746 | 0.608018 | 0.667989 |
| 19 | 52919800 | 52925399 | 63.62282 | 100.5004 | 141.2653 | 58.66705 | -0.16477 | 2.432309 | 0.668045 |
| 18 | 17724200 | 17727999 | 44.89706 | 72.87729 | 75.07888 | 32.18172 | 0.324038 | 1.624252 | 0.668061 |
| 15 | 72026600 | 72030799 | 35.18411 | 57.01589 | 84.97707 | 36.37966 | -0.20971 | 1.644698 | 0.668061 |
| 6 | 97783800 | 97785599 | 12.9649 | 22.88981 | 36.25669 | 16.04153 | -0.27001 | 0.533104 | 0.668061 |
| 6 | 47922400 | 47924199 | 25.34551 | 41.94803 | 63.98098 | 27.67339 | -0.22645 | 1.316001 | 0.668066 |
| 1 | 129722400 | 129725999 | 36.24731 | 59.4217 | 66.41993 | 28.5902 | 0.216456 | 1.458587 | 0.668071 |
| 6 | 89887200 | 89890399 | 39.57016 | 64.44029 | 71.49944 | 30.69968 | 0.22104 | 1.55986 | 0.668097 |
| 1 | 166001200 | 166003199 | 12.96978 | 24.29512 | 19.08394 | 7.882426 | 0.807111 | 0.123933 | 0.668098 |
| 11 | 52023000 | 52028199 | 79.75647 | 125.8063 | 136.3216 | 56.57472 | 0.208479 | 2.475722 | 0.66812 |
| 14 | 82716800 | 82718199 | 17.11252 | 30.13227 | 33.53375 | 14.84467 | 0.235074 | 0.583121 | 0.668128 |
| 7 | 6227600 | 6237999 | 169.6987 | 253.0517 | 270.0161 | 111.2811 | 0.16517 | 3.579987 | 0.668199 |
| 5 | 6474400 | 6475599 | 9.441372 | 17.74202 | 29.09327 | 11.95212 | -0.24134 | 0.096482 | 0.668221 |
| 10 | 51905000 | 51905999 | 9.441372 | 17.74202 | 29.09327 | 11.95212 | -0.24134 | 0.096482 | 0.668221 |
| 3 | 35683800 | 35687199 | 47.1629 | 76.13564 | 81.66739 | 34.82894 | 0.257997 | 1.755471 | 0.668222 |
| 1 | 117509600 | 117511799 | 27.81276 | 47.1475 | 50.69166 | 22.20068 | 0.277157 | 1.068423 | 0.668226 |
| 2 | 146481600 | 146483199 | 17.96442 | 31.48145 | 35.59917 | 15.72442 | 0.211086 | 0.656924 | 0.668232 |
| 3 | 107106800 | 107108399 | 15.21422 | 27.24677 | 30.84336 | 13.51964 | 0.22261 | 0.427708 | 0.668235 |
| 7 | 87314600 | 87315999 | 17.62955 | 30.4793 | 46.35633 | 20.3372 | -0.21663 | 0.848795 | 0.668242 |

| | | | | | | | | | |
|---|---|---|---|---|---|---|---|---|---|
| 3 | 137305400 | 137306799 | 17.82944 | 30.18907 | 57.95102 | 25.37106 | -0.55588 | 1.052648 | 0.668243 |
| 5 | 65057600 | 65067199 | 105.8107 | 161.8787 | 222.6909 | 92.39835 | -0.16949 | 3.059692 | 0.668259 |
| 3 | 133044800 | 133048199 | 20.74382 | 34.64234 | 58.51568 | 25.46218 | -0.37259 | 1.193478 | 0.668292 |
| 15 | 84521400 | 84522999 | 15.52974 | 27.74593 | 31.31515 | 13.75391 | 0.224877 | 0.445203 | 0.668354 |
| 6 | 138549600 | 138549999 | 11.62929 | 21.59006 | 24.17087 | 10.13142 | 0.282773 | 0.181106 | 0.668357 |
| 17 | 83993400 | 83998199 | 56.6087 | 89.94734 | 127.4334 | 53.15229 | -0.16865 | 2.269573 | 0.668364 |
| 3 | 113176800 | 113179199 | 67.17716 | 107.0077 | 112.841 | 47.21384 | 0.260436 | 2.19756 | 0.668371 |
| 3 | 12110200 | 12114799 | 80.56949 | 126.5283 | 140.8194 | 58.40775 | 0.165057 | 2.563022 | 0.668375 |
| 1 | 61149400 | 61149999 | 7.429395 | 14.33673 | 24.83247 | 9.589319 | -0.23883 | -0.13741 | 0.668375 |
| 5 | 25208600 | 25209399 | 7.429395 | 14.33673 | 24.83247 | 9.589319 | -0.23883 | -0.13741 | 0.668375 |
| 15 | 83156800 | 83161999 | 45.67282 | 72.89121 | 122.9031 | 51.55958 | -0.41259 | 2.120172 | 0.668389 |
| 9 | 115142800 | 115151199 | 81.08631 | 126.2702 | 175.8992 | 73.01177 | -0.16786 | 2.717503 | 0.668391 |
| 1 | 7067400 | 7068799 | 16.1157 | 28.81572 | 31.71006 | 13.91803 | 0.261714 | 0.443952 | 0.668403 |
| 1 | 124848400 | 124851199 | 46.46456 | 74.82053 | 84.54851 | 35.93661 | 0.178787 | 1.831378 | 0.668424 |
| 5 | 122845600 | 122846199 | 6.404877 | 12.38727 | 22.65283 | 8.65905 | -0.30149 | -0.18746 | 0.668433 |
| 8 | 27338400 | 27339399 | 7.765477 | 14.74787 | 25.52541 | 10.27009 | -0.29223 | 0.00186 | 0.668437 |
| 14 | 102656000 | 102658599 | 28.47387 | 48.6991 | 47.90716 | 21.00647 | 0.406435 | 0.99008 | 0.66844 |
| 16 | 59613800 | 59614599 | 15.55774 | 27.23917 | 41.46592 | 18.30177 | -0.21564 | 0.669779 | 0.668468 |
| 2 | 100136600 | 100137599 | 15.88873 | 28.17564 | 31.85099 | 14.07782 | 0.216423 | 0.495477 | 0.668471 |
| 18 | 50829200 | 50829999 | 20.77774 | 35.9164 | 40.07083 | 17.57957 | 0.228493 | 0.856059 | 0.668474 |
| Y | 10361800 | 10369799 | 66.90017 | 106.0665 | 83.29443 | 34.66568 | 0.678178 | 2.328097 | 0.668485 |
| 9 | 20257800 | 20268199 | 141.0729 | 214.0464 | 226.1861 | 93.64042 | 0.20115 | 3.237988 | 0.668493 |
| 7 | 15506400 | 15508999 | 37.69782 | 62.1251 | 64.59789 | 27.86778 | 0.323452 | 1.396672 | 0.668494 |
| 4 | 124857800 | 124859399 | 12.77682 | 22.62733 | 35.67222 | 15.77322 | -0.26157 | 0.496176 | 0.668503 |
| 4 | 71333200 | 71334199 | 10.76952 | 19.68641 | 31.27673 | 13.38868 | -0.24104 | 0.276522 | 0.668515 |
| 5 | 59986800 | 59987399 | 9.457259 | 18.09763 | 21.79099 | 8.619791 | 0.248523 | -0.04845 | 0.668535 |
| 12 | 115952200 | 115954399 | 51.91601 | 82.57484 | 120.4405 | 50.28978 | -0.20904 | 2.234934 | 0.668536 |
| 6 | 73180000 | 73184599 | 34.89733 | 56.63223 | 83.21842 | 35.66442 | -0.18737 | 1.616016 | 0.668542 |
| X | 75076400 | 75077199 | 15.27615 | 27.51863 | 29.01565 | 12.65027 | 0.33012 | 0.392237 | 0.668553 |
| 8 | 83544400 | 83545999 | 13.56651 | 23.61728 | 46.83989 | 20.59066 | -0.59591 | 0.753951 | 0.668553 |
| 2 | 5684200 | 5689399 | 43.57332 | 69.9139 | 101.7656 | 43.02714 | -0.19207 | 1.944884 | 0.668568 |
| 15 | 68513000 | 68514399 | 14.92534 | 26.37033 | 41.82042 | 18.50045 | -0.27161 | 0.546929 | 0.66857 |
| 17 | 50285400 | 50291399 | 70.52881 | 111.4321 | 123.8673 | 51.44706 | 0.173573 | 2.427915 | 0.668595 |
| 12 | 12235000 | 12236399 | 14.61136 | 26.62381 | 29.362 | 12.61948 | 0.278335 | 0.318272 | 0.668603 |
| 17 | 15017400 | 15020399 | 34.56217 | 56.6742 | 64.57117 | 27.79753 | 0.189107 | 1.449634 | 0.668605 |
| 5 | 33340600 | 33347799 | 86.00844 | 132.4915 | 210.7299 | 87.53106 | -0.371 | 2.917972 | 0.668607 |
| 3 | 84800000 | 84801799 | 15.63796 | 27.98583 | 31.25402 | 13.70781 | 0.241322 | 0.434716 | 0.668612 |
| 6 | 104362200 | 104363399 | 10.15179 | 19.3559 | 23.34561 | 9.315181 | 0.234983 | -0.00836 | 0.668615 |
| Y | 6481200 | 6483799 | 18.54544 | 32.39774 | 26.92154 | 11.86061 | 0.653283 | 0.692301 | 0.668631 |
| 9 | 26364600 | 26366399 | 16.56448 | 28.76434 | 44.78474 | 19.65633 | -0.24885 | 0.80146 | 0.668643 |
| 9 | 15103400 | 15109999 | 81.1719 | 127.338 | 142.1255 | 58.94423 | 0.159767 | 2.585242 | 0.668663 |
| 12 | 19683000 | 19749599 | 2632.027 | 3129.514 | 2655.702 | 1124.258 | 0.169129 | 7.166447 | 0.668665 |
| 16 | 52438600 | 52441199 | 24.23492 | 42.22144 | 40.62853 | 17.86869 | 0.442413 | 0.78179 | 0.668674 |
| 1 | 137231400 | 137232199 | 7.879879 | 15.08718 | 25.6986 | 10.14991 | -0.24565 | -0.05789 | 0.668731 |
| 3 | 159359000 | 159360399 | 16.56021 | 28.18063 | 56.29778 | 24.6756 | -0.61242 | 1.003832 | 0.668754 |
| 7 | 57157800 | 57160799 | 29.65814 | 48.40464 | 73.16541 | 31.44014 | -0.22156 | 1.508994 | 0.668758 |
| 11 | 27139000 | 27151999 | 153.5269 | 232.4181 | 235.879 | 97.58516 | 0.256646 | 3.28598 | 0.66876 |

| | | | | | | | | | |
|---|---|---|---|---|---|---|---|---|---|
| 1 | 166389000 | 166392799 | 37.4119 | 62.08771 | 60.64719 | 26.28092 | 0.414732 | 1.313916 | 0.668813 |
| 7 | 7082600 | 7093799 | 326.3648 | 460.6847 | 462.6167 | 191.538 | 0.175839 | 4.440241 | 0.668816 |
| 6 | 133190800 | 133194799 | 35.8489 | 59.55524 | 60.00927 | 26.02302 | 0.370121 | 1.296771 | 0.668817 |
| 7 | 4883800 | 4886799 | 37.40231 | 61.17549 | 68.14314 | 29.30502 | 0.219805 | 1.491512 | 0.668838 |
| 9 | 119994000 | 119995199 | 11.37493 | 21.21558 | 24.60831 | 10.239 | 0.239933 | 0.149959 | 0.668845 |
| 12 | 41940000 | 41940599 | 9.260502 | 17.759 | 20.85035 | 8.197285 | 0.293646 | -0.07019 | 0.668857 |
| 11 | 29058200 | 29060599 | 23.35596 | 40.28187 | 37.83708 | 16.59845 | 0.474897 | 0.882494 | 0.66887 |
| 8 | 28510600 | 28512799 | 20.40441 | 34.82864 | 52.21666 | 22.89246 | -0.19953 | 0.982358 | 0.668876 |
| 18 | 54838200 | 54841399 | 34.68127 | 56.48868 | 90.7262 | 38.95757 | -0.31109 | 1.560296 | 0.668878 |
| 4 | 35781400 | 35781799 | 10.92435 | 20.52558 | 24.18178 | 9.93701 | 0.232071 | 0.098858 | 0.66888 |
| 10 | 75799400 | 75800399 | 10.92435 | 20.52558 | 24.18178 | 9.93701 | 0.232071 | 0.098858 | 0.66888 |
| 9 | 121406000 | 121407599 | 20.67269 | 35.68647 | 39.54442 | 17.34791 | 0.237786 | 0.865372 | 0.6689 |
| X | 135397800 | 135398999 | 22.27683 | 38.26463 | 43.19004 | 18.94435 | 0.209905 | 0.923109 | 0.668904 |
| 13 | 107976600 | 107977199 | 5.522645 | 10.73914 | 20.65563 | 7.651203 | -0.33049 | -0.31358 | 0.66891 |
| 5 | 148081200 | 148094199 | 157.4426 | 232.7823 | 362.8817 | 149.4438 | -0.39279 | 3.72958 | 0.668912 |
| 11 | 15746400 | 15747399 | 12.00619 | 22.60568 | 25.73428 | 10.53742 | 0.285057 | 0.098153 | 0.668916 |
| 1 | 93772800 | 93773599 | 8.938171 | 16.55645 | 28.41889 | 11.94871 | -0.33243 | 0.183024 | 0.668936 |
| 7 | 84906800 | 84908399 | 18.14406 | 31.94119 | 35.34788 | 15.64848 | 0.242785 | 0.592667 | 0.668955 |
| 4 | 87628800 | 87631199 | 21.4952 | 37.22016 | 42.02224 | 18.43664 | 0.21175 | 0.844746 | 0.668989 |
| 14 | 45986800 | 45990599 | 32.27363 | 53.41388 | 60.17301 | 26.02617 | 0.209289 | 1.33083 | 0.66899 |
| 3 | 31662400 | 31662999 | 6.670468 | 12.92729 | 23.3348 | 8.835057 | -0.27077 | -0.21801 | 0.669019 |
| 6 | 10678000 | 10678599 | 10.99149 | 19.9141 | 31.75446 | 13.74537 | -0.25654 | 0.331903 | 0.669043 |
| 5 | 143137200 | 143141599 | 53.65469 | 85.40021 | 121.4464 | 50.7591 | -0.17151 | 2.213127 | 0.669046 |
| 1 | 33749800 | 33751399 | 13.96932 | 25.14341 | 27.87889 | 12.1674 | 0.256483 | 0.380684 | 0.669054 |
| 13 | 108590800 | 108595999 | 68.37892 | 108.421 | 121.3898 | 50.52497 | 0.167468 | 2.338926 | 0.669075 |
| 2 | 136071000 | 136075199 | 42.16517 | 67.79408 | 97.80049 | 41.47765 | -0.17554 | 1.87908 | 0.669097 |
| 16 | 22257000 | 22259999 | 16.91597 | 28.56554 | 60.04982 | 26.26851 | -0.68665 | 1.074065 | 0.669103 |
| 6 | 68101000 | 68109199 | 111.2213 | 171.3036 | 183.6444 | 76.27888 | 0.197407 | 2.927924 | 0.669124 |
| 6 | 136455000 | 136461799 | 69.73607 | 111.1693 | 111.3857 | 46.65454 | 0.335286 | 2.176138 | 0.669144 |
| 15 | 38606000 | 38618399 | 271.1194 | 386.0056 | 493.9682 | 204.2259 | -0.16132 | 4.312624 | 0.669147 |
| 1 | 119666400 | 119672799 | 85.36513 | 132.7148 | 190.5598 | 79.11649 | -0.21335 | 2.751669 | 0.669148 |
| 6 | 51491800 | 51495199 | 45.13141 | 72.79847 | 81.95656 | 34.91655 | 0.187102 | 1.776705 | 0.669177 |
| 12 | 7186600 | 7187999 | 11.57791 | 21.23998 | 34.53794 | 14.71356 | -0.26957 | 0.269443 | 0.669177 |
| 1 | 23640000 | 23641199 | 10.55891 | 19.4371 | 31.36224 | 13.29852 | -0.25355 | 0.234425 | 0.669184 |
| 1 | 46297800 | 46300399 | 40.98435 | 66.48777 | 74.30008 | 31.83435 | 0.206218 | 1.628363 | 0.669234 |
| X | 32761200 | 32763199 | 42.60697 | 69.29896 | 61.19483 | 26.26321 | 0.548396 | 1.580125 | 0.669241 |
| 1 | 3985400 | 3986799 | 14.54396 | 25.51677 | 39.34753 | 17.39706 | -0.23416 | 0.626148 | 0.669256 |
| 5 | 5852400 | 5858399 | 68.69351 | 109.0605 | 120.1367 | 50.06806 | 0.19269 | 2.301369 | 0.669261 |
| 3 | 101237800 | 101239999 | 16.59951 | 29.06387 | 46.1684 | 20.37773 | -0.27712 | 0.679077 | 0.669288 |
| 7 | 5832600 | 5863999 | 559.35 | 757.9448 | 727.5582 | 303.1724 | 0.181459 | 5.069985 | 0.669303 |
| 10 | 61291400 | 61292399 | 10.81372 | 20.01726 | 32.57508 | 13.70859 | -0.25764 | 0.210415 | 0.669313 |
| 13 | 3102800 | 3134599 | 426.6521 | 594.1574 | 563.6705 | 233.8759 | 0.238223 | 4.645444 | 0.669317 |
| 8 | 129941000 | 129953799 | 187.6251 | 281.0648 | 271.3528 | 111.9314 | 0.315407 | 3.499989 | 0.669326 |
| 4 | 113455400 | 113466199 | 157.6936 | 238.329 | 241.3239 | 99.78537 | 0.257912 | 3.321211 | 0.669326 |
| 10 | 57013000 | 57015399 | 34.4168 | 56.28902 | 63.79968 | 27.43651 | 0.194751 | 1.48272 | 0.669326 |
| 19 | 26045800 | 26048599 | 19.36325 | 34.21885 | 34.55817 | 15.28712 | 0.376074 | 0.555884 | 0.669341 |
| 4 | 143411200 | 143415999 | 105.6904 | 163.3492 | 154.6857 | 64.25297 | 0.380005 | 2.857346 | 0.669355 |

| | | | | | | | | | |
|---|---|---|---|---|---|---|---|---|---|
| 1 | 59931000 | 59932199 | 9.770925 | 18.22899 | 29.47399 | 12.25773 | -0.23537 | 0.146606 | 0.669361 |
| 4 | 113549400 | 113552399 | 39.1691 | 64.11498 | 68.506 | 29.4694 | 0.280459 | 1.483984 | 0.669365 |
| 13 | 14937400 | 14938599 | 13.04429 | 23.26137 | 36.12225 | 15.89053 | -0.23455 | 0.449443 | 0.669409 |
| 5 | 147738400 | 147761199 | 366.2644 | 508.1831 | 635.1862 | 263.7001 | -0.16486 | 4.699004 | 0.669418 |
| 6 | 90208800 | 90210799 | 70.54401 | 110.1097 | 160.7409 | 66.69921 | -0.23269 | 2.670957 | 0.669459 |
| 19 | 18512600 | 18514799 | 15.33211 | 27.31486 | 30.82248 | 13.57011 | 0.222767 | 0.460122 | 0.66949 |
| 18 | 78221200 | 78229399 | 61.95875 | 97.43384 | 162.3428 | 67.34694 | -0.41484 | 2.528916 | 0.669511 |
| 13 | 28300800 | 28301599 | 12.26641 | 22.57642 | 25.60143 | 10.8583 | 0.252049 | 0.237613 | 0.669526 |
| 2 | 144471400 | 144476399 | 60.94556 | 96.56221 | 135.7362 | 56.45698 | -0.16135 | 2.358163 | 0.669551 |
| 2 | 141061600 | 141063799 | 28.57778 | 48.60758 | 50.12771 | 21.97037 | 0.337484 | 1.042916 | 0.669555 |
| 4 | 43300800 | 43302599 | 30.77363 | 50.89204 | 57.81643 | 24.99359 | 0.197124 | 1.333728 | 0.669567 |
| 6 | 142010800 | 142013999 | 44.04859 | 71.128 | 80.11545 | 34.17936 | 0.188185 | 1.744447 | 0.669579 |
| 13 | 65386600 | 65391999 | 98.04181 | 150.1492 | 210.2704 | 87.26583 | -0.19399 | 3.035209 | 0.669609 |
| 17 | 36500000 | 36500999 | 9.82417 | 18.1635 | 29.36124 | 12.37648 | -0.24958 | 0.20144 | 0.669612 |
| 4 | 60548800 | 60549599 | 12.88112 | 23.4397 | 25.11559 | 10.81249 | 0.318507 | 0.305176 | 0.669622 |
| 5 | 98007600 | 98008199 | 7.14348 | 13.71895 | 24.11789 | 9.442684 | -0.27852 | -0.09738 | 0.669639 |
| 10 | 98565200 | 98565999 | 7.14348 | 13.71895 | 24.11789 | 9.442684 | -0.27852 | -0.09738 | 0.669639 |
| 9 | 115389600 | 115392399 | 24.43301 | 40.87158 | 60.93643 | 26.54352 | -0.19322 | 1.190772 | 0.669648 |
| 2 | 90129600 | 90135199 | 105.6391 | 162.5188 | 178.3222 | 74.05655 | 0.162651 | 2.946201 | 0.669672 |
| 1 | 5390200 | 5396399 | 60.90534 | 97.3602 | 103.8182 | 43.65401 | 0.249873 | 2.079942 | 0.669674 |
| 2 | 18276600 | 18286999 | 164.274 | 247.3065 | 253.884 | 104.8574 | 0.233204 | 3.400028 | 0.669674 |
| 1 | 30773200 | 30774399 | 13.50672 | 25.65319 | 26.00051 | 10.4597 | 0.473766 | 0.055948 | 0.669679 |
| X | 153854600 | 153860199 | 84.02555 | 132.4623 | 138.1412 | 57.32547 | 0.263015 | 2.491536 | 0.669682 |
| 1 | 97300200 | 97301799 | 20.62338 | 36.012 | 39.13496 | 17.23663 | 0.268005 | 0.730293 | 0.669748 |
| 4 | 16630200 | 16632399 | 23.26577 | 39.76554 | 44.73996 | 19.61667 | 0.213297 | 0.970968 | 0.66976 |
| 7 | 110076400 | 110080399 | 34.46771 | 57.68944 | 56.46546 | 24.60455 | 0.411956 | 1.212303 | 0.66978 |
| 15 | 57568800 | 57569799 | 8.40162 | 15.75958 | 26.94263 | 11.10258 | -0.30413 | 0.094559 | 0.669781 |
| 13 | 69004200 | 69005599 | 11.69895 | 20.84894 | 33.76773 | 14.86275 | -0.29501 | 0.442312 | 0.669795 |
| X | 152919400 | 152929199 | 199.9895 | 297.7793 | 290.7806 | 119.8163 | 0.291069 | 3.613491 | 0.669795 |
| 4 | 73464600 | 73465599 | 16.96294 | 30.22149 | 32.7963 | 14.43861 | 0.278974 | 0.476264 | 0.669823 |
| 12 | 32428400 | 32429999 | 18.28264 | 31.0954 | 56.93395 | 24.95338 | -0.48746 | 1.01425 | 0.669839 |
| 3 | 129065200 | 129066399 | 11.58766 | 20.88138 | 33.16639 | 14.45365 | -0.25752 | 0.368702 | 0.669863 |
| 1 | 4592000 | 4597799 | 48.38453 | 77.16811 | 113.3969 | 47.52783 | -0.21555 | 2.140341 | 0.669865 |
| 14 | 50831400 | 50835599 | 99.27886 | 154.1161 | 147.7218 | 61.34469 | 0.36741 | 2.77645 | 0.669874 |
| 7 | 55264000 | 55264999 | 10.7397 | 19.49926 | 31.83229 | 13.74253 | -0.28743 | 0.320212 | 0.669877 |
| 4 | 112540200 | 112543199 | 36.41124 | 60.79571 | 57.16127 | 24.86991 | 0.46983 | 1.24299 | 0.669923 |
| 13 | 11012600 | 11015199 | 30.93276 | 50.26024 | 93.12435 | 39.94596 | -0.51933 | 1.591548 | 0.669923 |
| 14 | 86833200 | 86837599 | 56.49058 | 90.31195 | 101.2126 | 42.56435 | 0.178417 | 2.071819 | 0.669933 |
| 14 | 83763200 | 83763599 | 11.76545 | 21.78268 | 24.63461 | 10.36461 | 0.264272 | 0.198196 | 0.669936 |
| X | 116849200 | 116851599 | 35.80598 | 59.64653 | 58.80745 | 25.5487 | 0.401633 | 1.267344 | 0.66994 |
| 7 | 7159600 | 7162399 | 35.24982 | 58.16367 | 55.3482 | 23.89009 | 0.451662 | 1.367954 | 0.669943 |
| 3 | 90948000 | 90951799 | 58.29348 | 92.59794 | 130.758 | 54.52345 | -0.16432 | 2.280649 | 0.669957 |
| 1 | 38749400 | 38750799 | 12.73066 | 22.99347 | 36.83926 | 16.00853 | -0.26745 | 0.371392 | 0.669961 |
| 1 | 7129800 | 7137599 | 183.0638 | 272.5308 | 288.6356 | 118.9304 | 0.173762 | 3.612366 | 0.669996 |
| 8 | 34827800 | 34831199 | 26.49864 | 44.08381 | 67.91826 | 29.48139 | -0.24053 | 1.259247 | 0.670018 |
| 14 | 47472200 | 47475799 | 49.68898 | 79.42692 | 113.4523 | 47.70211 | -0.17118 | 2.072707 | 0.67007 |
| 4 | 153188600 | 153197599 | 93.7384 | 144.3954 | 197.9339 | 82.2157 | -0.15674 | 2.919614 | 0.67009 |

| | | | | | | | | | |
|---|---|---|---|---|---|---|---|---|---|
| 1 | 160640400 | 160642999 | 35.60848 | 58.51375 | 51.95774 | 22.38758 | 0.54895 | 1.417585 | 0.670099 |
| 11 | 82875600 | 82876799 | 9.154772 | 17.32591 | 28.73661 | 11.6476 | -0.24104 | 0.047085 | 0.670132 |
| 2 | 20257400 | 20265199 | 117.5153 | 180.2371 | 194.3955 | 80.69788 | 0.184054 | 3.012118 | 0.670181 |
| 1 | 133869400 | 133871999 | 32.28142 | 52.57133 | 77.33914 | 33.20725 | -0.18484 | 1.548656 | 0.670187 |
| 7 | 131459200 | 131459999 | 10.30473 | 19.51444 | 23.25185 | 9.412077 | 0.234314 | 0.03926 | 0.670213 |
| 6 | 59791400 | 59792199 | 10.74207 | 20.65884 | 23.58324 | 9.196377 | 0.343919 | -0.07666 | 0.670229 |
| 1 | 48360200 | 48369199 | 137.5777 | 209.5525 | 193.5324 | 80.21474 | 0.399865 | 3.155976 | 0.670233 |
| 12 | 3424600 | 3426999 | 20.59393 | 35.63696 | 40.04689 | 17.56964 | 0.218419 | 0.845248 | 0.670243 |
| 13 | 76110400 | 76111799 | 10.784 | 19.67094 | 31.26524 | 13.41946 | -0.24409 | 0.289159 | 0.670255 |
| X | 118482000 | 118487399 | 51.28918 | 83.18009 | 76.12225 | 32.60787 | 0.493402 | 1.657282 | 0.670324 |
| X | 120011600 | 120013999 | 36.99655 | 61.16766 | 62.77675 | 27.13312 | 0.343247 | 1.357774 | 0.67033 |
| 9 | 57297200 | 57301799 | 29.80294 | 50.13843 | 53.51785 | 23.38443 | 0.287209 | 1.142326 | 0.670333 |
| 10 | 70731200 | 70732599 | 16.95811 | 29.78059 | 33.05255 | 14.62556 | 0.237861 | 0.612947 | 0.670392 |
| 14 | 80818000 | 80819799 | 16.2549 | 28.94708 | 32.32118 | 14.23482 | 0.237684 | 0.472991 | 0.670394 |
| 3 | 22807600 | 22809999 | 42.51904 | 68.88622 | 76.58454 | 32.78113 | 0.21188 | 1.659566 | 0.670401 |
| 6 | 17391000 | 17393399 | 36.8855 | 59.65128 | 87.00339 | 37.19028 | -0.18133 | 1.69186 | 0.670422 |
| 2 | 61641400 | 61643799 | 24.01045 | 39.84383 | 61.79438 | 26.77655 | -0.25015 | 1.278918 | 0.670451 |
| 2 | 14343000 | 14343799 | 14.54346 | 26.55574 | 29.44464 | 12.62184 | 0.273213 | 0.307253 | 0.670453 |
| 4 | 145560400 | 145568999 | 406.8142 | 565.1098 | 563.0042 | 233.6906 | 0.163982 | 4.682532 | 0.670482 |
| 7 | 26679600 | 26680199 | 12.98888 | 23.0172 | 35.70412 | 15.78536 | -0.23782 | 0.49272 | 0.670493 |
| 11 | 17456600 | 17459399 | 24.90702 | 43.36439 | 41.76426 | 18.36075 | 0.441205 | 0.797035 | 0.670498 |
| X | 49065200 | 49065999 | 11.05674 | 20.94868 | 24.32282 | 9.83552 | 0.272904 | 0.047277 | 0.670503 |
| 9 | 67422400 | 67424999 | 24.9689 | 42.95648 | 45.72351 | 20.05656 | 0.294522 | 0.921443 | 0.670562 |
| 10 | 57325800 | 57327199 | 10.59622 | 19.63478 | 31.49887 | 13.23719 | -0.23577 | 0.199428 | 0.670584 |
| 15 | 47257200 | 47258599 | 14.24677 | 26.21644 | 28.44727 | 12.06834 | 0.314518 | 0.258309 | 0.670597 |
| 5 | 130979800 | 130982399 | 25.68292 | 42.74301 | 64.97253 | 28.21014 | -0.22109 | 1.25036 | 0.670602 |
| 7 | 41555600 | 41567999 | 179.3947 | 269.5834 | 260.5486 | 107.5581 | 0.317899 | 3.43841 | 0.670606 |
| 5 | 53831000 | 53831599 | 7.919395 | 15.23697 | 26.45716 | 10.30423 | -0.2539 | -0.09921 | 0.670611 |
| 12 | 64134000 | 64134799 | 7.919395 | 15.23697 | 26.45716 | 10.30423 | -0.2539 | -0.09921 | 0.670611 |
| 4 | 70654400 | 70657599 | 30.30898 | 50.76054 | 55.21811 | 24.0705 | 0.25998 | 1.189704 | 0.670626 |
| 3 | 104883600 | 104886799 | 28.44847 | 47.50559 | 53.98221 | 23.4884 | 0.197389 | 1.215622 | 0.670629 |
| 7 | 23578200 | 23603399 | 434.1426 | 602.0594 | 584.2584 | 242.5762 | 0.199442 | 4.708863 | 0.670629 |
| 10 | 66722000 | 66725199 | 31.89228 | 52.06075 | 77.13218 | 33.14699 | -0.19266 | 1.512364 | 0.670691 |
| 15 | 8946600 | 8951599 | 154.2614 | 231.7025 | 250.4661 | 103.3825 | 0.155861 | 3.441295 | 0.670693 |
| 9 | 110943800 | 110944999 | 16.57129 | 29.44233 | 32.79266 | 14.47065 | 0.239298 | 0.489997 | 0.670711 |
| 12 | 51759600 | 51761199 | 12.58122 | 22.65519 | 35.1962 | 15.3523 | -0.22666 | 0.385052 | 0.670727 |
| Y | 74674200 | 74676799 | 12.88676 | 23.49466 | 21.72528 | 9.330689 | 0.53182 | 0.284761 | 0.670798 |
| 8 | 69663800 | 69675799 | 273.1701 | 393.9219 | 403.0393 | 166.2998 | 0.179739 | 4.125738 | 0.670839 |
| 7 | 11211600 | 11217399 | 176.3384 | 261.9926 | 276.6646 | 113.9746 | 0.175313 | 3.645737 | 0.670843 |
| 11 | 3965000 | 3967799 | 36.74921 | 59.27137 | 87.75635 | 37.38232 | -0.20738 | 1.770326 | 0.670853 |
| 3 | 66587600 | 66590399 | 23.03726 | 39.04217 | 61.09993 | 26.75696 | -0.26257 | 1.064279 | 0.670868 |
| 15 | 41862800 | 41863799 | 9.372914 | 18.03606 | 21.3051 | 8.294344 | 0.296902 | -0.10382 | 0.670879 |
| 8 | 4075200 | 4079799 | 80.53303 | 126.5933 | 139.9062 | 58.03627 | 0.176254 | 2.545468 | 0.670886 |
| 15 | 31653000 | 31656799 | 27.08134 | 44.66404 | 67.53444 | 29.14418 | -0.21475 | 1.361921 | 0.670887 |
| 2 | 94434000 | 94439799 | 82.35983 | 129.7142 | 124.3541 | 51.59783 | 0.38325 | 2.505287 | 0.670889 |
| 13 | 9578800 | 9580599 | 33.25342 | 54.8884 | 61.89646 | 26.7222 | 0.207543 | 1.364882 | 0.670911 |
| 2 | 104700800 | 104702999 | 23.97695 | 40.90712 | 46.33904 | 20.31485 | 0.20314 | 0.992551 | 0.670914 |

| | | | | | | | | | |
|---|---|---|---|---|---|---|---|---|---|
| 8 | 114410200 | 114412799 | 17.60254 | 31.44964 | 32.61429 | 14.32437 | 0.346218 | 0.46033 | 0.67092 |
| 16 | 74208200 | 74210199 | 14.53513 | 25.45638 | 39.9247 | 17.62599 | -0.25849 | 0.655145 | 0.670954 |
| 3 | 47610200 | 47612599 | 30.57075 | 50.87304 | 57.3871 | 24.91511 | 0.207937 | 1.260999 | 0.670955 |
| 15 | 5596800 | 5597599 | 8.721894 | 16.55873 | 27.69589 | 11.15125 | -0.24424 | 0.020295 | 0.670968 |
| 3 | 5098000 | 5103199 | 66.18284 | 105.727 | 105.6073 | 44.36494 | 0.342978 | 2.105841 | 0.671016 |
| 18 | 83559200 | 83561399 | 17.95788 | 30.8986 | 47.42209 | 20.80172 | -0.23112 | 0.894384 | 0.671018 |
| 9 | 31488000 | 31490799 | 20.93093 | 35.56399 | 53.26774 | 23.34465 | -0.19914 | 1.027817 | 0.671027 |
| X | 113373800 | 113377999 | 49.64426 | 80.1883 | 82.16159 | 35.06619 | 0.325006 | 1.747663 | 0.671028 |
| 1 | 154781400 | 154782199 | 10.9803 | 20.6273 | 24.21183 | 9.952148 | 0.237235 | 0.101452 | 0.671031 |
| 7 | 19747400 | 19747999 | 10.9803 | 20.6273 | 24.21183 | 9.952148 | 0.237235 | 0.101452 | 0.671031 |
| 4 | 57420200 | 57421199 | 9.053167 | 16.91751 | 27.94301 | 11.58894 | -0.26215 | 0.128576 | 0.671035 |
| 6 | 123323800 | 123326199 | 23.34802 | 40.58368 | 42.41445 | 18.62744 | 0.323565 | 0.812598 | 0.671037 |
| 1 | 102994800 | 102995799 | 15.20031 | 27.27085 | 30.82492 | 13.49108 | 0.226198 | 0.416233 | 0.671042 |
| 14 | 3826400 | 3831999 | 49.22381 | 78.75681 | 113.1636 | 47.64363 | -0.17824 | 2.044186 | 0.67105 |
| X | 7529400 | 7533999 | 47.99288 | 77.58564 | 76.88793 | 32.85077 | 0.374427 | 1.721541 | 0.671062 |
| 14 | 44746400 | 44748799 | 19.31218 | 34.09936 | 34.55128 | 15.28669 | 0.370828 | 0.564163 | 0.671072 |
| 2 | 171801400 | 171805799 | 50.55444 | 80.67799 | 114.8198 | 48.16512 | -0.16851 | 2.124848 | 0.671075 |
| 10 | 63303000 | 63304599 | 12.74067 | 23.41347 | 27.04521 | 11.49454 | 0.223296 | 0.252731 | 0.671106 |
| 12 | 106777600 | 106779999 | 22.47137 | 38.56458 | 43.67097 | 19.15425 | 0.204878 | 0.930277 | 0.67112 |
| 17 | 48382000 | 48383399 | 20.50936 | 35.65217 | 39.9606 | 17.55059 | 0.223239 | 0.797148 | 0.67113 |
| 4 | 136984400 | 136986399 | 19.52621 | 33.904 | 36.96932 | 16.22506 | 0.262573 | 0.802686 | 0.671135 |
| 4 | 71800200 | 71802599 | 26.24877 | 44.6098 | 49.02825 | 21.48708 | 0.245733 | 1.03832 | 0.671137 |
| 1 | 117794200 | 117805199 | 46.2109 | 74.75549 | 69.90339 | 29.88834 | 0.458948 | 1.692026 | 0.671142 |
| 19 | 13924600 | 13927399 | 46.75839 | 75.18795 | 84.71797 | 35.93537 | 0.180454 | 1.875174 | 0.671146 |
| 12 | 81751200 | 81752799 | 23.95387 | 41.07251 | 45.76298 | 20.07039 | 0.227963 | 0.946682 | 0.671152 |
| X | 81188800 | 81190999 | 28.45495 | 48.65342 | 48.46659 | 21.25131 | 0.388406 | 0.994608 | 0.671166 |
| 7 | 7005000 | 7008999 | 40.82747 | 65.65801 | 95.28276 | 40.423 | -0.18368 | 1.868568 | 0.671194 |
| X | 148657400 | 148671399 | 210.4076 | 311.777 | 309.0578 | 127.2847 | 0.262972 | 3.699154 | 0.671196 |
| 2 | 83408600 | 83413599 | 60.61377 | 97.05653 | 88.7071 | 37.3893 | 0.474566 | 2.025607 | 0.671198 |
| 1 | 46184400 | 46190999 | 64.72696 | 103.5559 | 102.9172 | 43.3265 | 0.352618 | 2.060671 | 0.671224 |
| 9 | 122991800 | 122992599 | 8.546526 | 16.06228 | 26.95589 | 11.0719 | -0.27383 | 0.083662 | 0.671243 |
| 6 | 66603200 | 66604199 | 13.66185 | 24.94298 | 28.43119 | 12.18923 | 0.233392 | 0.298431 | 0.671262 |
| 17 | 6851600 | 6852599 | 12.83723 | 22.70864 | 35.76289 | 15.81689 | -0.2605 | 0.511239 | 0.671269 |
| 18 | 17484600 | 17488599 | 51.56441 | 82.10468 | 118.2447 | 49.44713 | -0.18909 | 2.198847 | 0.671288 |
| 18 | 17484800 | 17488599 | 51.56441 | 82.10468 | 118.2447 | 49.44713 | -0.18909 | 2.198847 | 0.671288 |
| 19 | 39155800 | 39156599 | 16.82643 | 29.77411 | 32.92358 | 14.56029 | 0.246979 | 0.52751 | 0.671303 |
| 14 | 122491200 | 122493399 | 21.17629 | 36.58474 | 41.55062 | 18.22846 | 0.202593 | 0.863083 | 0.671308 |
| 15 | 63444600 | 63445799 | 10.99805 | 20.29902 | 33.0549 | 13.96597 | -0.26268 | 0.229699 | 0.67132 |
| 2 | 55561200 | 55561799 | 6.546926 | 12.68907 | 22.73707 | 8.604932 | -0.25879 | -0.21954 | 0.671337 |
| 16 | 37091000 | 37091799 | 6.546926 | 12.68907 | 22.73707 | 8.604932 | -0.25879 | -0.21954 | 0.671337 |
| 17 | 42376800 | 42379599 | 23.64539 | 41.06209 | 43.00733 | 18.8767 | 0.320422 | 0.8293 | 0.671342 |
| 14 | 102652200 | 102654999 | 43.83801 | 71.03087 | 76.46148 | 32.73109 | 0.258513 | 1.662439 | 0.671348 |
| 17 | 35188800 | 35190199 | 13.98551 | 24.48526 | 39.52937 | 17.44633 | -0.29975 | 0.659938 | 0.671361 |
| 8 | 23034000 | 23043599 | 91.30884 | 140.5545 | 194.947 | 80.96725 | -0.17444 | 2.932281 | 0.671374 |
| 16 | 44183800 | 44184599 | 12.72767 | 24.32457 | 25.0602 | 9.949503 | 0.466707 | 0.003799 | 0.671381 |
| 3 | 74143200 | 74143599 | 6.386716 | 12.41283 | 22.35593 | 8.316383 | -0.24314 | -0.28354 | 0.671392 |
| 16 | 89520400 | 89527799 | 95.37573 | 146.0371 | 231.8262 | 96.20746 | -0.37492 | 3.040813 | 0.671446 |

| | | | | | | | | | |
|---|---|---|---|---|---|---|---|---|---|
| 3 | 147341000 | 147341599 | 10.36834 | 19.5982 | 22.26937 | 9.050122 | 0.297666 | 0.051584 | 0.671481 |
| 10 | 119033600 | 119034599 | 16.04567 | 27.92941 | 43.18286 | 18.98145 | -0.23851 | 0.76213 | 0.671489 |
| 5 | 41510400 | 41512399 | 21.01806 | 35.87289 | 54.42304 | 23.85959 | -0.21667 | 0.986901 | 0.671544 |
| 3 | 4403400 | 4405599 | 21.16802 | 36.73509 | 41.48093 | 18.20038 | 0.212177 | 0.822929 | 0.671558 |
| X | 17695600 | 17714399 | 223.9719 | 331.1537 | 312.8746 | 128.8496 | 0.329282 | 3.737921 | 0.67158 |
| 4 | 114607400 | 114608799 | 16.48902 | 30.28062 | 29.39487 | 12.50507 | 0.471432 | 0.289505 | 0.671595 |
| 18 | 8031400 | 8035799 | 48.08727 | 77.55333 | 84.33032 | 35.92006 | 0.236798 | 1.787058 | 0.671602 |
| 18 | 52900200 | 52902199 | 14.39138 | 25.22202 | 39.40733 | 17.40773 | -0.25298 | 0.642859 | 0.671608 |
| 17 | 30010400 | 30011199 | 10.91589 | 19.90421 | 31.75409 | 13.63554 | -0.25 | 0.292465 | 0.671614 |
| Y | 35115200 | 35117799 | 17.10057 | 30.06124 | 24.26871 | 10.74681 | 0.69446 | 0.57878 | 0.671616 |
| 11 | 59831200 | 59832599 | 16.42475 | 28.90563 | 31.48861 | 13.94084 | 0.265234 | 0.578357 | 0.671625 |
| 19 | 29981000 | 29981999 | 9.034465 | 16.84916 | 27.91308 | 11.61543 | -0.27046 | 0.140125 | 0.671649 |
| 7 | 118856200 | 118857199 | 10.23214 | 19.06303 | 31.07981 | 12.95593 | -0.25019 | 0.162464 | 0.671656 |
| 17 | 32173400 | 32176599 | 42.14641 | 67.73329 | 97.6579 | 41.39004 | -0.17554 | 1.89274 | 0.671697 |
| 5 | 94504000 | 94508999 | 191.7735 | 286.8091 | 243.9228 | 100.5847 | 0.496078 | 3.529141 | 0.671709 |
| 8 | 75330400 | 75331999 | 15.87326 | 28.90828 | 30.38169 | 13.06614 | 0.34697 | 0.333673 | 0.671723 |
| 9 | 103081200 | 103081799 | 7.781755 | 15.00084 | 26.52931 | 10.27569 | -0.27253 | -0.11746 | 0.671745 |
| 9 | 20220600 | 20222999 | 26.0605 | 43.53205 | 67.31582 | 29.29602 | -0.2458 | 1.215368 | 0.671755 |
| 4 | 36448200 | 36448599 | 4.930914 | 9.637633 | 18.97931 | 6.550581 | -0.27173 | -0.57904 | 0.671755 |
| 2 | 89718200 | 89723199 | 59.09126 | 94.53781 | 90.46138 | 38.08186 | 0.407136 | 2.049848 | 0.671771 |
| 3 | 18501800 | 18503599 | 27.2438 | 46.16048 | 50.67879 | 22.19177 | 0.247131 | 1.071787 | 0.67179 |
| 1 | 104330800 | 104338399 | 139.4846 | 211.5281 | 226.9602 | 93.94648 | 0.178559 | 3.247681 | 0.671815 |
| 17 | 23703200 | 23703799 | 5.673586 | 11.03141 | 21.80154 | 8.083038 | -0.37178 | -0.30868 | 0.671837 |
| 14 | 53012400 | 53013199 | 13.44488 | 23.35719 | 48.42532 | 21.25996 | -0.65981 | 0.78703 | 0.67185 |
| 10 | 127861000 | 127862199 | 13.5635 | 24.5256 | 27.50222 | 11.93411 | 0.245852 | 0.35196 | 0.671855 |
| 17 | 49500600 | 49503599 | 37.55379 | 60.4992 | 89.95706 | 38.2606 | -0.21551 | 1.806402 | 0.671868 |
| 8 | 15795600 | 15816999 | 283.2108 | 407.8268 | 410.863 | 169.5595 | 0.200043 | 4.147433 | 0.671881 |
| 3 | 78782600 | 78783999 | 11.39351 | 20.70644 | 32.73567 | 14.11538 | -0.24094 | 0.317136 | 0.671892 |
| 1 | 111629200 | 111633599 | 64.61394 | 102.5363 | 114.855 | 47.84479 | 0.167967 | 2.310818 | 0.671901 |
| 4 | 91381600 | 91384199 | 20.23694 | 34.86543 | 55.60646 | 24.39277 | -0.2862 | 0.901373 | 0.671922 |
| 13 | 67194400 | 67197399 | 37.74588 | 62.72088 | 60.00009 | 26.02312 | 0.444935 | 1.302271 | 0.671934 |
| 7 | 22453200 | 22455399 | 21.76362 | 38.18447 | 37.80364 | 16.71487 | 0.401878 | 0.661955 | 0.67195 |
| 2 | 81687000 | 81688599 | 12.34509 | 22.3251 | 35.7853 | 15.52613 | -0.26674 | 0.358048 | 0.671956 |
| 6 | 41971000 | 41973799 | 22.75165 | 39.79509 | 39.51496 | 17.42606 | 0.397745 | 0.718049 | 0.671964 |
| 8 | 117454000 | 117456799 | 38.49345 | 62.86551 | 69.42265 | 29.83891 | 0.230705 | 1.514798 | 0.671979 |
| 1 | 24678600 | 24680799 | 22.45984 | 39.24829 | 39.53261 | 17.4202 | 0.377182 | 0.729904 | 0.671979 |
| 4 | 60339400 | 60341599 | 64.50966 | 102.325 | 111.5093 | 46.43026 | 0.206969 | 2.323948 | 0.671983 |
| 11 | 117594800 | 117595399 | 9.850588 | 18.83407 | 22.86683 | 9.067769 | 0.23355 | -0.03217 | 0.671997 |
| 2 | 150159000 | 150167199 | 164.9507 | 248.7519 | 249.1101 | 102.9338 | 0.271157 | 3.366915 | 0.672021 |
| 4 | 4456000 | 4460999 | 76.77979 | 120.8134 | 135.3831 | 56.16724 | 0.157363 | 2.521649 | 0.672023 |
| 12 | 35397200 | 35399599 | 23.86011 | 41.62662 | 40.63135 | 17.88565 | 0.422172 | 0.761918 | 0.672038 |
| 7 | 31631400 | 31644399 | 409.7197 | 566.9957 | 556.4248 | 231.0973 | 0.18023 | 4.739513 | 0.672045 |
| 7 | 134053800 | 134056199 | 23.82267 | 40.8499 | 45.77045 | 20.07376 | 0.219953 | 0.946103 | 0.672071 |
| 9 | 18399400 | 18401599 | 18.44088 | 31.45285 | 57.52332 | 25.2174 | -0.48523 | 0.989981 | 0.672078 |
| 6 | 139775400 | 139776199 | 5.660803 | 11.01454 | 20.69053 | 7.626311 | -0.29018 | -0.32893 | 0.672096 |
| 3 | 159420000 | 159422199 | 20.17932 | 34.61765 | 52.80235 | 23.15859 | -0.22327 | 0.935412 | 0.672118 |
| 17 | 92742600 | 92745999 | 34.50175 | 57.11155 | 60.56974 | 26.20095 | 0.296039 | 1.331351 | 0.672131 |

| | | | | | | | | | |
|---|---|---|---|---|---|---|---|---|---|
| 14 | 43949800 | 43952399 | 28.05475 | 46.17505 | 69.84705 | 30.11616 | -0.21653 | 1.390715 | 0.672139 |
| 3 | 63960400 | 63961399 | 13.28461 | 24.78209 | 27.12972 | 11.28445 | 0.322969 | 0.170097 | 0.672145 |
| 1 | 88454800 | 88457199 | 18.3602 | 32.37339 | 35.32232 | 15.63217 | 0.264035 | 0.577247 | 0.672152 |
| 3 | 142673600 | 142675599 | 16.6785 | 29.35887 | 32.998 | 14.60842 | 0.220571 | 0.581333 | 0.672153 |
| 10 | 43306400 | 43309599 | 41.96991 | 67.47851 | 97.28136 | 41.25604 | -0.17465 | 1.878663 | 0.67217 |
| X | 55037400 | 55042399 | 79.69192 | 126.2141 | 128.6312 | 53.49108 | 0.301455 | 2.38148 | 0.672171 |
| 12 | 99391000 | 99393799 | 21.97598 | 36.96918 | 55.92734 | 24.43691 | -0.21406 | 1.121247 | 0.672194 |
| 1 | 78590400 | 78592199 | 15.40732 | 26.91673 | 41.56252 | 18.31134 | -0.23603 | 0.704644 | 0.672199 |
| 1 | 31974800 | 31977199 | 42.8477 | 69.23346 | 78.4367 | 33.49099 | 0.180862 | 1.722279 | 0.672214 |
| 7 | 112789800 | 112790799 | 15.30754 | 27.51016 | 30.76639 | 13.4461 | 0.242778 | 0.405514 | 0.672234 |
| 5 | 147979200 | 147986999 | 111.8191 | 171.7788 | 188.1216 | 78.11079 | 0.163835 | 2.977902 | 0.672245 |
| 6 | 61471200 | 61478599 | 87.24427 | 137.2881 | 142.6101 | 59.16607 | 0.266694 | 2.52857 | 0.672281 |
| 6 | 32806800 | 32812399 | 76.0409 | 120.6321 | 124.2246 | 51.73162 | 0.2887 | 2.332786 | 0.672306 |
| 3 | 12603200 | 12605999 | 37.24972 | 60.23859 | 88.12547 | 37.66854 | -0.18572 | 1.694774 | 0.672311 |
| 13 | 99292800 | 99294199 | 21.73859 | 36.95224 | 55.47371 | 24.31226 | -0.2024 | 1.028881 | 0.672312 |
| 5 | 66379000 | 66381799 | 22.43879 | 37.7875 | 56.51155 | 24.70006 | -0.19751 | 1.113761 | 0.672337 |
| 2 | 22244400 | 22245599 | 14.30052 | 25.33765 | 39.72513 | 17.56365 | -0.25345 | 0.509814 | 0.672361 |
| 5 | 147976200 | 147978199 | 17.65255 | 30.64768 | 46.33705 | 20.33511 | -0.20689 | 0.810213 | 0.672377 |
| 7 | 86555400 | 86558599 | 33.06784 | 54.89451 | 59.31555 | 25.70907 | 0.269501 | 1.294971 | 0.672377 |
| 2 | 174991200 | 175009599 | 330.6959 | 469.8754 | 465.9708 | 192.7162 | 0.203446 | 4.345021 | 0.672382 |
| 8 | 3826200 | 3828599 | 35.87465 | 58.58317 | 66.73966 | 28.68474 | 0.185825 | 1.50906 | 0.672385 |
| 4 | 140947800 | 140948599 | 10.63214 | 19.9943 | 23.20657 | 9.521943 | 0.255401 | 0.088812 | 0.672401 |
| 11 | 68216800 | 68220999 | 29.71815 | 48.78887 | 72.7194 | 31.32099 | -0.19677 | 1.428649 | 0.672405 |
| 11 | 26168800 | 26178799 | 94.51865 | 147.7589 | 155.06 | 64.33172 | 0.244677 | 2.655641 | 0.672421 |
| 12 | 21038800 | 21042399 | 42.14312 | 67.57064 | 100.2083 | 42.32251 | -0.22015 | 1.965458 | 0.67243 |
| 6 | 63023600 | 63024599 | 14.66578 | 26.2139 | 29.38049 | 12.8994 | 0.235239 | 0.432283 | 0.672473 |
| 10 | 67751400 | 67752599 | 14.22914 | 25.84478 | 28.80495 | 12.42956 | 0.25944 | 0.33167 | 0.672475 |
| X | 149556800 | 149561999 | 41.93234 | 69.13048 | 64.23919 | 27.71761 | 0.48548 | 1.405143 | 0.672482 |
| 1 | 72160600 | 72161799 | 15.66564 | 27.33078 | 42.07646 | 18.52191 | -0.2321 | 0.724294 | 0.672504 |
| X | 145288600 | 145291999 | 37.69291 | 62.39176 | 62.26879 | 26.93527 | 0.383569 | 1.346423 | 0.67251 |
| 6 | 12214200 | 12224999 | 189.5562 | 281.6484 | 296.1822 | 122.0158 | 0.181417 | 3.647781 | 0.672523 |
| 1 | 119666400 | 119670599 | 48.46663 | 78.40243 | 79.99293 | 34.20159 | 0.33346 | 1.704311 | 0.672531 |
| 7 | 41280800 | 41283399 | 25.09995 | 41.33205 | 73.83315 | 31.84146 | -0.45537 | 1.383059 | 0.672532 |
| 9 | 99828400 | 99829999 | 24.88716 | 41.27764 | 63.39231 | 27.4593 | -0.23604 | 1.29087 | 0.672561 |
| 10 | 43856000 | 43857799 | 12.74159 | 22.73967 | 35.13293 | 15.44494 | -0.22648 | 0.441171 | 0.672572 |
| 19 | 12961200 | 12976799 | 153.3167 | 231.8108 | 214.4067 | 88.66702 | 0.388935 | 3.309639 | 0.672578 |
| 2 | 160248600 | 160251399 | 24.59047 | 40.62208 | 72.26502 | 31.21597 | -0.44805 | 1.34215 | 0.67258 |
| 18 | 17680000 | 17681999 | 12.44048 | 21.91907 | 41.95089 | 18.56848 | -0.54258 | 0.565165 | 0.672582 |
| 2 | 111670000 | 111671399 | 13.83439 | 25.77913 | 27.6509 | 11.52282 | 0.35016 | 0.183334 | 0.672582 |
| 1 | 23491600 | 23494399 | 28.32657 | 48.43669 | 48.46762 | 21.25186 | 0.381987 | 0.994044 | 0.672595 |
| 19 | 38981600 | 38982599 | 14.35681 | 26.53266 | 28.56441 | 12.04742 | 0.332023 | 0.236962 | 0.672601 |
| 16 | 70069800 | 70072999 | 23.92339 | 41.72438 | 34.79896 | 15.31535 | 0.648012 | 0.758327 | 0.672634 |
| 6 | 25781200 | 25782399 | 17.47535 | 30.70993 | 33.62023 | 14.88536 | 0.25774 | 0.608969 | 0.672635 |
| 6 | 26063200 | 26069799 | 101.153 | 156.6692 | 148.0904 | 61.51275 | 0.384967 | 2.816244 | 0.672658 |
| 15 | 12776000 | 12777799 | 21.94142 | 37.94212 | 42.53378 | 18.66031 | 0.221537 | 0.858555 | 0.67266 |
| 11 | 55462600 | 55466199 | 33.02604 | 53.8515 | 80.8038 | 34.71076 | -0.21209 | 1.532226 | 0.672663 |
| 3 | 7156000 | 7157199 | 12.81399 | 22.97443 | 35.7098 | 15.63772 | -0.23108 | 0.416009 | 0.672673 |

| | | | | | | | | | |
|---|---|---|---|---|---|---|---|---|---|
| 7 | 81906000 | 81911199 | 43.28723 | 69.40642 | 101.0853 | 42.69419 | -0.19407 | 1.966803 | 0.672706 |
| 9 | 36129000 | 36131199 | 43.07042 | 69.55364 | 78.73815 | 33.59498 | 0.181058 | 1.740306 | 0.672709 |
| 13 | 111861400 | 111872599 | 284.2704 | 400.5263 | 529.6165 | 219.3449 | -0.22371 | 4.465888 | 0.672729 |
| 5 | 14746800 | 14749199 | 26.46342 | 44.98167 | 40.12888 | 17.58707 | 0.54574 | 1.030377 | 0.67274 |
| 18 | 4539600 | 4540599 | 16.88778 | 29.93795 | 33.26569 | 14.70615 | 0.240853 | 0.506811 | 0.672742 |
| 2 | 142940000 | 142942199 | 19.08945 | 33.5135 | 35.75972 | 15.81948 | 0.294139 | 0.631888 | 0.67278 |
| 9 | 95510000 | 95512999 | 43.27487 | 70.19686 | 66.93238 | 28.67557 | 0.434532 | 1.631441 | 0.672781 |
| 9 | 4032200 | 4033599 | 21.31082 | 36.97098 | 41.53122 | 18.22226 | 0.219658 | 0.828562 | 0.67279 |
| 8 | 15773000 | 15775199 | 19.69134 | 34.68526 | 35.65202 | 15.78128 | 0.34901 | 0.594571 | 0.672816 |
| 7 | 126212000 | 126213599 | 15.51545 | 27.10638 | 41.53103 | 18.29778 | -0.22497 | 0.703352 | 0.672833 |
| 13 | 69218600 | 69220199 | 12.1643 | 21.65993 | 34.2222 | 15.07354 | -0.26025 | 0.451422 | 0.672835 |
| 4 | 21678200 | 21680399 | 25.79841 | 41.84568 | 86.48545 | 37.05109 | -0.6787 | 1.611686 | 0.672836 |
| 1 | 19868200 | 19870199 | 29.83458 | 48.24896 | 89.92306 | 38.43876 | -0.53401 | 1.683265 | 0.672839 |
| Y | 20756400 | 20759799 | 22.84211 | 39.02296 | 33.21864 | 14.56448 | 0.613957 | 0.957585 | 0.67284 |
| 10 | 85933800 | 85936599 | 37.3943 | 61.00838 | 69.09931 | 29.68892 | 0.193267 | 1.527503 | 0.672847 |
| 6 | 68793800 | 68796199 | 38.38353 | 62.23066 | 69.67721 | 29.84026 | 0.202659 | 1.633778 | 0.672857 |
| 17 | 69643200 | 69647599 | 44.50728 | 71.89698 | 80.42198 | 34.32757 | 0.198805 | 1.733389 | 0.672857 |
| 1 | 121708800 | 121711799 | 25.43405 | 42.40607 | 63.17282 | 27.46049 | -0.19201 | 1.23102 | 0.672871 |
| 18 | 37863800 | 37892599 | 1388.466 | 1722.843 | 1965.952 | 827.5254 | -0.19551 | 6.490388 | 0.672894 |
| 4 | 125190200 | 125192799 | 27.31916 | 45.03946 | 67.36669 | 29.06721 | -0.19928 | 1.369397 | 0.672904 |
| 11 | 8841000 | 8841399 | 4.935056 | 9.632768 | 18.69961 | 6.625969 | -0.28493 | -0.48599 | 0.672908 |
| X | 81222000 | 81223399 | 23.10239 | 40.00065 | 42.57721 | 18.68009 | 0.296583 | 0.856996 | 0.672941 |
| 17 | 38699400 | 38705799 | 69.34407 | 110.8163 | 104.8956 | 44.08805 | 0.421008 | 2.100134 | 0.67295 |
| 15 | 84914200 | 84917199 | 21.4414 | 36.31981 | 54.34468 | 23.79516 | -0.19789 | 1.057595 | 0.672951 |
| 8 | 21217800 | 21220599 | 43.66739 | 69.42881 | 132.5073 | 55.30394 | -0.59685 | 2.241897 | 0.672952 |
| 8 | 96245000 | 96249199 | 55.69168 | 88.8361 | 96.25374 | 40.35136 | 0.223558 | 2.138601 | 0.672983 |
| 10 | 28798000 | 28800199 | 30.41004 | 49.79465 | 73.7077 | 31.71163 | -0.18832 | 1.463746 | 0.672994 |
| 17 | 54859000 | 54860199 | 24.96192 | 42.37498 | 47.96017 | 21.01347 | 0.20343 | 1.042633 | 0.673005 |
| 6 | 74634600 | 74636199 | 17.71998 | 31.27179 | 34.22468 | 15.14376 | 0.259979 | 0.560202 | 0.673005 |
| 10 | 39635600 | 39638399 | 61.65988 | 97.99275 | 108.0336 | 45.06961 | 0.192946 | 2.26428 | 0.673009 |
| 17 | 19001400 | 19004399 | 46.77394 | 75.74091 | 78.75528 | 33.69758 | 0.307536 | 1.680274 | 0.673033 |
| 1 | 82940000 | 82941599 | 15.01966 | 27.16666 | 30.34735 | 13.16386 | 0.251507 | 0.366073 | 0.673043 |
| 10 | 118270000 | 118271799 | 16.29691 | 28.43462 | 43.08606 | 18.96738 | -0.20935 | 0.727485 | 0.673047 |
| 14 | 45717000 | 45719199 | 16.29691 | 28.43462 | 43.08606 | 18.96738 | -0.20935 | 0.727485 | 0.673047 |
| 18 | 20926800 | 20927799 | 10.29794 | 19.10074 | 30.44311 | 12.7765 | -0.22487 | 0.189902 | 0.673065 |
| 3 | 77351800 | 77354599 | 21.50058 | 37.56796 | 39.60806 | 17.45208 | 0.311614 | 0.727088 | 0.673067 |
| 4 | 101943600 | 101945999 | 22.79145 | 37.72013 | 70.51587 | 30.49945 | -0.51917 | 1.313215 | 0.673068 |
| 2 | 163151400 | 163153599 | 17.64685 | 30.959 | 33.92476 | 14.99877 | 0.256021 | 0.632468 | 0.67312 |
| 1 | 78760800 | 78763999 | 40.99658 | 65.85298 | 96.56945 | 40.89954 | -0.20081 | 1.905938 | 0.673122 |
| 3 | 58628200 | 58629199 | 9.045738 | 17.05165 | 28.06747 | 11.46996 | -0.24011 | 0.073229 | 0.673164 |
| 11 | 24002800 | 24003999 | 9.642595 | 17.99592 | 28.9522 | 12.03354 | -0.22733 | 0.142994 | 0.673172 |
| 3 | 35466600 | 35468999 | 27.39175 | 46.52857 | 43.65969 | 19.13282 | 0.472929 | 1.042281 | 0.673181 |
| 13 | 60173200 | 60183799 | 153.908 | 229.7553 | 307.9156 | 126.9275 | -0.16203 | 3.55891 | 0.673198 |
| 8 | 99872000 | 99874599 | 34.78208 | 56.89517 | 63.464 | 27.29369 | 0.217847 | 1.478727 | 0.673214 |
| 11 | 38533800 | 38534599 | 12.56182 | 22.44735 | 35.76006 | 15.7038 | -0.26897 | 0.437139 | 0.673244 |
| 1 | 53608400 | 53609199 | 9.359388 | 17.4412 | 28.42076 | 11.84268 | -0.24836 | 0.14982 | 0.673264 |
| 6 | 15527200 | 15527799 | 9.359388 | 17.4412 | 28.42076 | 11.84268 | -0.24836 | 0.14982 | 0.673264 |

| | | | | | | | | | |
|---|---|---|---|---|---|---|---|---|---|
| 8 | 67812200 | 67813799 | 15.57236 | 27.34134 | 41.87618 | 18.52669 | -0.22446 | 0.625319 | 0.673276 |
| 17 | 19463000 | 19486799 | 312.9672 | 449.9971 | 416.0876 | 171.7549 | 0.321561 | 4.172419 | 0.673282 |
| 4 | 60752600 | 60754399 | 18.07004 | 32.3261 | 27.95907 | 12.26657 | 0.607699 | 0.447541 | 0.673311 |
| 12 | 101540200 | 101541599 | 14.61347 | 26.86029 | 24.79236 | 10.53429 | 0.545023 | 0.260985 | 0.673316 |
| X | 24133600 | 24177399 | 640.004 | 864.6684 | 744.5224 | 310.3823 | 0.333927 | 5.118767 | 0.673319 |
| 2 | 170450800 | 170456199 | 54.48281 | 86.78776 | 123.0299 | 51.47791 | -0.16572 | 2.188779 | 0.673319 |
| 14 | 72864200 | 72865199 | 9.622853 | 18.60491 | 18.69614 | 7.157445 | 0.551584 | -0.16097 | 0.673322 |
| 10 | 93482200 | 93485799 | 24.91093 | 41.42359 | 62.56764 | 27.1503 | -0.21197 | 1.25676 | 0.673337 |
| 17 | 10037200 | 10038399 | 14.33188 | 26.30115 | 28.97394 | 12.33752 | 0.289131 | 0.276113 | 0.673348 |
| 4 | 52379800 | 52382399 | 40.65319 | 65.92706 | 74.19826 | 31.78238 | 0.195527 | 1.634707 | 0.673349 |
| 16 | 17403400 | 17408199 | 65.51579 | 104.1504 | 115.8146 | 48.32947 | 0.18092 | 2.25979 | 0.673349 |
| 2 | 86590600 | 86591999 | 12.2399 | 22.06406 | 34.17691 | 14.88784 | -0.22117 | 0.374147 | 0.67335 |
| 13 | 100798800 | 100800599 | 15.76682 | 27.44626 | 43.20363 | 18.99155 | -0.26415 | 0.76081 | 0.673354 |
| X | 41848800 | 41850599 | 23.17245 | 39.89534 | 44.19387 | 19.38602 | 0.237545 | 0.908272 | 0.673382 |
| 16 | 71511200 | 71512799 | 14.21086 | 26.27964 | 27.98059 | 11.79097 | 0.348559 | 0.229519 | 0.673382 |
| 15 | 47165000 | 47165399 | 5.968977 | 11.54895 | 22.12216 | 8.440032 | -0.36477 | -0.20016 | 0.67339 |
| 11 | 20302800 | 20304799 | 14.66149 | 25.76621 | 39.74714 | 17.5987 | -0.23485 | 0.59914 | 0.673392 |
| 14 | 50298200 | 50300399 | 14.22337 | 24.57082 | 50.54995 | 22.17646 | -0.65068 | 0.846351 | 0.673402 |
| 1 | 129968200 | 129969199 | 15.93706 | 28.2568 | 31.89508 | 14.09921 | 0.21851 | 0.497669 | 0.673411 |
| 5 | 98598400 | 98598799 | 5.364311 | 10.46048 | 19.74616 | 7.100186 | -0.26452 | -0.42424 | 0.673438 |
| 1 | 87549400 | 87550999 | 16.33404 | 28.39252 | 43.71452 | 19.19865 | -0.23274 | 0.78286 | 0.67344 |
| X | 29378200 | 29381599 | 61.29775 | 97.4905 | 109.3477 | 45.65382 | 0.169185 | 2.241642 | 0.673452 |
| X | 152864600 | 152886799 | 446.2683 | 620.0886 | 572.6574 | 237.6872 | 0.273775 | 4.680337 | 0.673456 |
| 18 | 88286800 | 88287399 | 8.400944 | 15.93023 | 26.56813 | 10.72442 | -0.24323 | 0.022576 | 0.67349 |
| 10 | 87781800 | 87782999 | 13.27437 | 25.11128 | 25.82008 | 10.47533 | 0.442097 | 0.078201 | 0.673512 |
| 17 | 13732200 | 13733799 | 16.43286 | 28.66821 | 43.60027 | 19.19223 | -0.21472 | 0.730021 | 0.673516 |
| 11 | 15749200 | 15750999 | 29.93965 | 50.35933 | 46.94816 | 20.51271 | 0.48186 | 1.138819 | 0.67352 |
| 15 | 43353400 | 43354599 | 18.46394 | 31.82994 | 47.94202 | 21.03101 | -0.20355 | 0.880821 | 0.67354 |
| X | 139712200 | 139714799 | 18.46394 | 31.82994 | 47.94202 | 21.03101 | -0.20355 | 0.880821 | 0.67354 |
| 14 | 8199200 | 8199999 | 11.89824 | 21.59374 | 33.75787 | 14.58143 | -0.22648 | 0.331618 | 0.673556 |
| 7 | 12881800 | 12883399 | 39.6587 | 64.1231 | 68.92421 | 29.45449 | 0.257688 | 1.692068 | 0.673561 |
| 10 | 104034400 | 104035999 | 14.2887 | 26.30167 | 27.84874 | 11.80949 | 0.349894 | 0.254372 | 0.67357 |
| 12 | 18082400 | 18168599 | 2633.679 | 3134.945 | 2681.699 | 1135.079 | 0.159204 | 7.149077 | 0.673585 |
| 17 | 93698600 | 93702199 | 32.37721 | 52.95296 | 80.41378 | 34.5812 | -0.22626 | 1.488167 | 0.673593 |
| 7 | 85064200 | 85075199 | 130.702 | 200.8452 | 191.888 | 79.67784 | 0.360919 | 2.980605 | 0.673652 |
| 5 | 104661000 | 104671399 | 181.9474 | 273.565 | 258.2608 | 106.6288 | 0.352434 | 3.428894 | 0.673668 |
| 3 | 141316200 | 141316599 | 5.947122 | 11.58359 | 21.36724 | 7.80005 | -0.25164 | -0.35699 | 0.673668 |
| 17 | 70795800 | 70796199 | 5.947122 | 11.58359 | 21.36724 | 7.80005 | -0.25164 | -0.35699 | 0.673668 |
| 7 | 25050400 | 25052799 | 41.2963 | 66.3822 | 96.51801 | 40.92043 | -0.18725 | 1.883176 | 0.673706 |
| 2 | 21760600 | 21761799 | 11.78154 | 21.94767 | 25.65165 | 10.69478 | 0.22687 | 0.164747 | 0.673712 |
| 3 | 28821200 | 28823199 | 18.64328 | 32.7307 | 36.45138 | 16.12566 | 0.232911 | 0.633008 | 0.673726 |
| 15 | 25331400 | 25333799 | 19.42005 | 34.00337 | 36.94275 | 16.30615 | 0.268344 | 0.677923 | 0.673731 |
| 12 | 51145800 | 51147599 | 15.02549 | 26.52319 | 41.80118 | 18.4953 | -0.26326 | 0.556 | 0.673732 |
| 9 | 35921600 | 35923399 | 30.92693 | 51.42236 | 57.93965 | 25.14035 | 0.209522 | 1.27244 | 0.673769 |
| 18 | 44374400 | 44375799 | 11.93828 | 21.66254 | 33.77194 | 14.59075 | -0.22261 | 0.333972 | 0.673804 |
| 7 | 104506200 | 104506999 | 10.78103 | 20.34999 | 24.23683 | 9.878643 | 0.22614 | 0.068337 | 0.673849 |
| 1 | 5723800 | 5725799 | 19.31342 | 33.4358 | 52.83888 | 23.18214 | -0.27144 | 0.852577 | 0.673849 |

| | | | | | | | | | |
|---|---|---|---|---|---|---|---|---|---|
| 4 | 135100400 | 135104399 | 39.10787 | 63.43215 | 72.0138 | 30.85066 | 0.183294 | 1.629072 | 0.67387 |
| 6 | 124521200 | 124523399 | 18.0464 | 31.62584 | 35.52705 | 15.69288 | 0.220465 | 0.655607 | 0.673876 |
| 16 | 38296600 | 38298999 | 13.41965 | 23.6378 | 37.25208 | 16.48957 | -0.26416 | 0.562391 | 0.673892 |
| 18 | 67094200 | 67098199 | 31.1782 | 50.60927 | 77.27254 | 33.12227 | -0.24208 | 1.601147 | 0.673893 |
| 3 | 28650800 | 28652599 | 16.75829 | 29.26151 | 44.62832 | 19.65556 | -0.21868 | 0.720756 | 0.673901 |
| 4 | 112809800 | 112815799 | 112.8241 | 173.3054 | 188.0586 | 78.0836 | 0.176967 | 2.981205 | 0.673912 |
| 10 | 43716600 | 43725799 | 93.76153 | 145.7486 | 160.6097 | 66.68319 | 0.167926 | 2.751607 | 0.673913 |
| X | 96085800 | 96087999 | 27.01876 | 46.01583 | 49.02416 | 21.48896 | 0.291053 | 1.025829 | 0.673919 |
| 3 | 33600000 | 33600599 | 12.06903 | 21.89182 | 34.27489 | 14.81479 | -0.22926 | 0.337206 | 0.673924 |
| 17 | 56230200 | 56252599 | 276.7581 | 394.846 | 512.7234 | 211.9067 | -0.17942 | 4.2839 | 0.673964 |
| 19 | 61193400 | 61196799 | 31.22196 | 51.13157 | 76.20228 | 32.78691 | -0.19804 | 1.465169 | 0.673971 |
| X | 153394000 | 153400199 | 99.3018 | 155.72 | 149.1059 | 61.84167 | 0.380215 | 2.605374 | 0.673992 |
| 2 | 170513200 | 170518399 | 56.06491 | 89.12396 | 127.3576 | 53.14496 | -0.18041 | 2.254968 | 0.674015 |
| 18 | 75486600 | 75488199 | 13.86077 | 24.28806 | 45.13793 | 19.93637 | -0.50203 | 0.653777 | 0.674016 |
| 12 | 105726000 | 105727799 | 15.47739 | 27.39094 | 29.94662 | 13.24336 | 0.263146 | 0.508996 | 0.674027 |
| 2 | 76535400 | 76537599 | 23.30099 | 39.72869 | 44.77514 | 19.62824 | 0.210347 | 0.993692 | 0.674045 |
| 13 | 56369400 | 56374599 | 43.80706 | 70.17959 | 102.4149 | 43.19927 | -0.1984 | 1.995228 | 0.674053 |
| X | 34909000 | 34910399 | 19.613 | 34.60625 | 35.77379 | 15.82979 | 0.341992 | 0.578954 | 0.674055 |
| 3 | 77275000 | 77275799 | 9.148507 | 17.20694 | 28.0186 | 11.49334 | -0.22933 | 0.086429 | 0.674066 |
| 9 | 39831400 | 39831999 | 12.33936 | 22.85503 | 26.14298 | 10.99282 | 0.248659 | 0.205173 | 0.674081 |
| 1 | 27631600 | 27632999 | 10.06434 | 18.76009 | 30.54066 | 12.72004 | -0.24705 | 0.156344 | 0.67409 |
| X | 57440800 | 57445199 | 48.49466 | 78.44474 | 79.9293 | 34.17341 | 0.335337 | 1.705488 | 0.674112 |
| 2 | 150258600 | 150268399 | 283.9961 | 407.5551 | 418.8111 | 172.9313 | 0.166509 | 4.197512 | 0.674117 |
| 10 | 10500800 | 10502399 | 12.82676 | 24.16872 | 26.28361 | 10.74875 | 0.352021 | 0.102002 | 0.674128 |
| 2 | 5379800 | 5384599 | 285.7971 | 402.372 | 532.3326 | 220.4974 | -0.22556 | 4.477312 | 0.674136 |
| 2 | 39256800 | 39258599 | 21.47039 | 37.42715 | 35.08615 | 15.43682 | 0.480105 | 0.753862 | 0.674144 |
| 16 | 88460000 | 88461799 | 17.28078 | 30.12075 | 45.72955 | 20.11829 | -0.21236 | 0.750536 | 0.674157 |
| 12 | 23238400 | 23241599 | 69.10229 | 109.2266 | 120.7422 | 50.16033 | 0.182107 | 2.416679 | 0.674163 |
| 7 | 41414800 | 41415799 | 21.43015 | 37.03807 | 42.08642 | 18.46374 | 0.202059 | 0.862662 | 0.674172 |
| 8 | 19629400 | 19641599 | 201.348 | 298.0489 | 310.8039 | 127.9984 | 0.188547 | 3.714235 | 0.674196 |
| 2 | 109719200 | 109721799 | 19.19338 | 32.63774 | 59.21556 | 25.95295 | -0.47459 | 1.019823 | 0.674201 |
| 5 | 92982400 | 92984799 | 16.29067 | 28.44167 | 43.0666 | 18.96634 | -0.20827 | 0.718336 | 0.674221 |
| 8 | 72914400 | 72917199 | 16.77036 | 28.76794 | 53.22659 | 23.34455 | -0.50052 | 0.924739 | 0.674255 |
| 15 | 62983600 | 62983999 | 7.851266 | 15.23257 | 18.90494 | 7.120768 | 0.272525 | -0.23334 | 0.674263 |
| 18 | 8056000 | 8056999 | 22.6539 | 39.30181 | 33.85199 | 14.85293 | 0.601213 | 0.823819 | 0.674308 |
| 4 | 20072600 | 20074599 | 24.79378 | 42.47336 | 39.50895 | 17.32625 | 0.487241 | 0.952516 | 0.674315 |
| 6 | 111916400 | 111917799 | 15.34774 | 27.29572 | 30.84983 | 13.60186 | 0.219091 | 0.471202 | 0.674332 |
| 18 | 61389600 | 61391599 | 14.06549 | 24.71647 | 38.28712 | 16.95092 | -0.24066 | 0.59601 | 0.674334 |
| X | 19504400 | 19509999 | 52.02058 | 84.0976 | 80.96239 | 34.592 | 0.416098 | 1.72767 | 0.674334 |
| 4 | 89275400 | 89276199 | 6.718417 | 12.90989 | 23.59599 | 9.223417 | -0.33134 | -0.10674 | 0.674361 |
| X | 120785200 | 120789799 | 51.72041 | 83.47184 | 84.32003 | 35.94852 | 0.344053 | 1.774655 | 0.674369 |
| 7 | 41015200 | 41025399 | 152.7399 | 232.7762 | 213.3857 | 88.4582 | 0.411271 | 3.151812 | 0.67437 |
| 4 | 112159000 | 112165599 | 65.60746 | 104.742 | 107.9358 | 45.3028 | 0.297106 | 2.125634 | 0.674392 |
| 10 | 60682200 | 60688399 | 61.38797 | 97.28195 | 137.1182 | 57.03965 | -0.16501 | 2.353019 | 0.674407 |
| 4 | 22600200 | 22602999 | 25.09701 | 43.17431 | 45.71588 | 20.05318 | 0.301991 | 0.922035 | 0.674415 |
| 7 | 43645000 | 43652399 | 54.18372 | 86.00418 | 125.501 | 52.2915 | -0.21266 | 2.297903 | 0.674418 |
| 15 | 21888000 | 21891999 | 47.80866 | 76.7635 | 85.24754 | 36.08176 | 0.198771 | 1.923993 | 0.674422 |

| | | | | | | | | | |
|---|---|---|---|---|---|---|---|---|---|
| 6 | 132765200 | 132774799 | 172.4027 | 258.4294 | 268.6206 | 110.8184 | 0.209785 | 3.484365 | 0.674438 |
| 1 | 91405600 | 91407999 | 31.26431 | 51.12326 | 75.35877 | 32.40513 | -0.18355 | 1.486502 | 0.674448 |
| 4 | 37834000 | 37834999 | 16.72974 | 29.83647 | 27.6738 | 12.17294 | 0.504922 | 0.456713 | 0.674479 |
| 14 | 66438600 | 66441999 | 29.856 | 50.17765 | 54.06075 | 23.61451 | 0.273793 | 1.152145 | 0.674489 |
| 16 | 89502600 | 89504999 | 18.66842 | 32.6457 | 36.60743 | 16.14076 | 0.223052 | 0.694775 | 0.674503 |
| 8 | 22689000 | 22690399 | 14.42687 | 25.44773 | 38.72889 | 17.13846 | -0.21327 | 0.557043 | 0.674504 |
| 9 | 39307200 | 39307999 | 12.7852 | 22.62073 | 35.70142 | 15.78913 | -0.26367 | 0.506908 | 0.674506 |
| 2 | 166182800 | 166184199 | 13.22186 | 24.22488 | 26.62416 | 11.36194 | 0.290779 | 0.274808 | 0.674525 |
| 3 | 64101000 | 64103599 | 18.47565 | 32.64807 | 34.07317 | 15.07278 | 0.328806 | 0.549646 | 0.674528 |
| 1 | 15264400 | 15264999 | 7.293615 | 13.95543 | 24.57692 | 9.72541 | -0.29566 | -0.06031 | 0.674531 |
| X | 105770200 | 105775199 | 13.55303 | 23.75386 | 44.61743 | 19.71002 | -0.51722 | 0.648366 | 0.674532 |
| 10 | 45685200 | 45686399 | 9.966983 | 18.62226 | 30.05561 | 12.46838 | -0.22977 | 0.141259 | 0.67455 |
| 14 | 109984200 | 109985599 | 9.966983 | 18.62226 | 30.05561 | 12.46838 | -0.22977 | 0.141259 | 0.67455 |
| 13 | 86472800 | 86473399 | 9.69964 | 18.477 | 21.79879 | 8.714724 | 0.264495 | -0.00988 | 0.674576 |
| 1 | 54003600 | 54004599 | 10.89249 | 20.03332 | 31.87915 | 13.53408 | -0.23491 | 0.243698 | 0.674587 |
| 1 | 20771800 | 20773799 | 24.66928 | 42.08595 | 47.40291 | 20.78116 | 0.211344 | 0.997343 | 0.674597 |
| 7 | 9965400 | 9970599 | 94.76229 | 148.853 | 126.8273 | 52.60591 | 0.550268 | 2.566479 | 0.674623 |
| 14 | 29640200 | 29642399 | 32.61304 | 53.06644 | 77.87322 | 33.42625 | -0.18212 | 1.561786 | 0.674631 |
| 11 | 87384600 | 87385599 | 10.98218 | 20.81453 | 24.24687 | 9.797726 | 0.268368 | 0.040403 | 0.674658 |
| 6 | 119849600 | 119856199 | 68.5056 | 108.7566 | 120.1403 | 50.06697 | 0.188553 | 2.302448 | 0.674676 |
| 13 | 85746800 | 85749599 | 35.76977 | 58.22731 | 64.09725 | 27.51727 | 0.232038 | 1.55367 | 0.674708 |
| 17 | 72885600 | 72888199 | 27.69643 | 46.36805 | 52.90589 | 23.05721 | 0.191384 | 1.183723 | 0.67471 |
| 1 | 184429000 | 184430399 | 21.18719 | 36.57072 | 41.56909 | 18.23606 | 0.201156 | 0.871328 | 0.674717 |
| 1 | 93765000 | 93767399 | 19.10782 | 33.2612 | 36.9119 | 16.22553 | 0.237693 | 0.765766 | 0.674722 |
| 2 | 81580800 | 81581999 | 16.81493 | 29.28746 | 44.17336 | 19.42478 | -0.20293 | 0.757072 | 0.674743 |
| 14 | 82543400 | 82544799 | 16.93699 | 30.10024 | 27.29138 | 12.04031 | 0.534727 | 0.479743 | 0.674747 |
| 4 | 35391800 | 35393599 | 17.64295 | 30.66756 | 46.31494 | 20.34065 | -0.20513 | 0.792909 | 0.674794 |
| 1 | 95775200 | 95776999 | 20.37301 | 35.80788 | 35.55251 | 15.74303 | 0.39749 | 0.620513 | 0.674799 |
| 18 | 19670400 | 19672199 | 17.91688 | 32.36652 | 31.72287 | 13.78203 | 0.43775 | 0.396698 | 0.674806 |
| 14 | 46828600 | 46830599 | 16.58787 | 29.208 | 32.41631 | 14.35005 | 0.238743 | 0.575093 | 0.674823 |
| 4 | 62049000 | 62063199 | 408.814 | 567.6327 | 567.3918 | 235.5283 | 0.158556 | 4.688888 | 0.674835 |
| 16 | 83195000 | 83198399 | 49.43342 | 79.46919 | 89.32026 | 37.87358 | 0.183564 | 1.889141 | 0.674843 |
| 18 | 46264000 | 46265599 | 13.89713 | 25.36601 | 28.34961 | 12.15816 | 0.261161 | 0.300287 | 0.674844 |
| 12 | 41315200 | 41324399 | 225.1033 | 324.3481 | 429.4822 | 177.2292 | -0.19328 | 4.134512 | 0.674864 |
| 18 | 75885800 | 75889199 | 12.0439 | 21.39442 | 40.33724 | 17.80276 | -0.51608 | 0.476539 | 0.674885 |
| 9 | 89760200 | 89763999 | 63.40511 | 99.77787 | 144.7949 | 60.07274 | -0.21497 | 2.513589 | 0.67492 |
| 7 | 16369200 | 16370599 | 10.90044 | 19.91837 | 31.77205 | 13.60701 | -0.24735 | 0.279855 | 0.674935 |
| 6 | 31780200 | 31781799 | 17.08393 | 30.01409 | 33.13651 | 14.66789 | 0.245597 | 0.609226 | 0.674981 |
| 9 | 4419800 | 4420399 | 9.526906 | 18.35304 | 22.50186 | 8.731975 | 0.248625 | -0.10644 | 0.674985 |
| 11 | 7983000 | 7985799 | 27.58929 | 45.56765 | 67.5945 | 29.19479 | -0.18677 | 1.347211 | 0.67501 |
| 4 | 112640600 | 112643199 | 31.63553 | 53.0851 | 55.20626 | 24.09481 | 0.324693 | 1.17586 | 0.675015 |
| 14 | 29710800 | 29711799 | 7.910056 | 15.05152 | 25.55313 | 10.23928 | -0.25945 | -0.00992 | 0.67503 |
| 2 | 87761600 | 87766999 | 71.14353 | 112.5135 | 126.1038 | 52.40193 | 0.162841 | 2.40723 | 0.675053 |
| 8 | 115054600 | 115058199 | 55.7308 | 89.0629 | 100.4861 | 42.23574 | 0.168142 | 2.083393 | 0.675067 |
| 10 | 27970400 | 27970999 | 7.341438 | 13.95282 | 25.03043 | 10.05527 | -0.34102 | -0.007 | 0.675132 |
| 3 | 120534400 | 120538199 | 52.36932 | 82.87573 | 144.6836 | 60.14241 | -0.47483 | 2.382191 | 0.675166 |
| X | 81120600 | 81122799 | 23.6022 | 40.92163 | 43.56382 | 19.11374 | 0.296631 | 0.84618 | 0.675174 |

| | | | | | | | | | |
|---|---|---|---|---|---|---|---|---|---|
| 13 | 72100200 | 72105599 | 44.25038 | 70.77153 | 106.9308 | 44.99439 | -0.2512 | 2.046881 | 0.67522 |
| X | 36914200 | 36917599 | 39.38752 | 63.96568 | 70.26142 | 30.12771 | 0.23252 | 1.60604 | 0.675241 |
| 8 | 111818800 | 111819799 | 9.910513 | 18.94905 | 22.82756 | 9.051638 | 0.244388 | -0.03374 | 0.675246 |
| 8 | 10777800 | 10783999 | 31.59026 | 52.63789 | 50.30142 | 21.85696 | 0.446483 | 1.243249 | 0.675248 |
| 14 | 88356800 | 88358799 | 16.63259 | 29.13697 | 46.15749 | 20.38188 | -0.27315 | 0.671574 | 0.675281 |
| 8 | 16072800 | 16073999 | 14.14431 | 26.27271 | 23.84647 | 9.988475 | 0.583835 | 0.197472 | 0.675283 |
| X | 16165200 | 16165799 | 9.891252 | 18.97535 | 22.95064 | 9.013787 | 0.252097 | -0.06197 | 0.6753 |
| 3 | 54497200 | 54499599 | 26.0779 | 43.99233 | 50.14373 | 21.92838 | 0.193024 | 1.104389 | 0.675307 |
| X | 112871400 | 112884399 | 266.2586 | 384.5752 | 392.2506 | 161.8136 | 0.18653 | 4.102163 | 0.675308 |
| 17 | 19886200 | 19887399 | 14.78084 | 27.88513 | 26.89126 | 10.97287 | 0.527276 | 0.114977 | 0.675325 |
| 15 | 4720200 | 4721399 | 9.469644 | 17.83248 | 29.14602 | 11.93137 | -0.23239 | 0.084448 | 0.675358 |
| 18 | 74232600 | 74233599 | 9.469644 | 17.83248 | 29.14602 | 11.93137 | -0.23239 | 0.084448 | 0.675358 |
| 10 | 22633200 | 22635599 | 24.7631 | 41.36399 | 61.49131 | 26.76083 | -0.18901 | 1.207621 | 0.67537 |
| 3 | 145324000 | 145326199 | 24.73928 | 41.3276 | 61.51849 | 26.77405 | -0.19093 | 1.206082 | 0.675379 |
| 15 | 59218600 | 59222199 | 24.73928 | 41.3276 | 61.51849 | 26.77405 | -0.19093 | 1.206082 | 0.675379 |
| 12 | 21591400 | 21594399 | 31.2601 | 51.03091 | 75.21367 | 32.32302 | -0.18509 | 1.509917 | 0.675391 |
| 2 | 85857000 | 85857999 | 13.00497 | 23.99203 | 27.53074 | 11.63933 | 0.238069 | 0.233378 | 0.675411 |
| 18 | 9685200 | 9687199 | 19.45583 | 34.23896 | 36.27318 | 16.05919 | 0.305207 | 0.605209 | 0.675412 |
| 16 | 17769400 | 17776199 | 77.74395 | 122.1836 | 136.4284 | 56.59211 | 0.161195 | 2.545585 | 0.675425 |
| 7 | 56334800 | 56347199 | 245.3518 | 352.0224 | 454.5268 | 187.685 | -0.16306 | 4.198497 | 0.675426 |
| 11 | 108713200 | 108718999 | 69.97112 | 110.7259 | 124.292 | 51.6648 | 0.161309 | 2.3918 | 0.675451 |
| 1 | 46110200 | 46111999 | 18.1514 | 32.3966 | 33.1183 | 14.55847 | 0.365977 | 0.471729 | 0.675464 |
| 3 | 93356400 | 93357799 | 14.42506 | 26.13209 | 29.3072 | 12.68778 | 0.247515 | 0.349432 | 0.675465 |
| 13 | 22767400 | 22768199 | 31.21409 | 51.37591 | 55.5659 | 23.95869 | 0.266146 | 1.388878 | 0.675469 |
| 19 | 15410400 | 15411199 | 11.70965 | 21.86435 | 25.63462 | 10.64582 | 0.226692 | 0.148251 | 0.67548 |
| 9 | 101636000 | 101636799 | 12.0384 | 22.32517 | 24.69493 | 10.36648 | 0.298655 | 0.193141 | 0.675491 |
| 1 | 53413200 | 53414599 | 15.32799 | 26.45141 | 50.55984 | 22.18009 | -0.54538 | 0.861663 | 0.675519 |
| 1 | 130665000 | 130666399 | 11.12959 | 20.24153 | 31.75727 | 13.68102 | -0.22887 | 0.310606 | 0.675525 |
| 4 | 45385800 | 45387599 | 25.92591 | 44.09436 | 49.02641 | 21.48758 | 0.229339 | 1.032753 | 0.675545 |
| 8 | 37943600 | 37948799 | 48.77241 | 78.34849 | 88.81724 | 37.61988 | 0.169933 | 1.911173 | 0.675553 |
| 17 | 39650000 | 39655999 | 64.75409 | 103.7714 | 98.73294 | 41.66716 | 0.418121 | 2.009538 | 0.675561 |
| 18 | 42094800 | 42097999 | 24.64164 | 42.48401 | 45.17435 | 19.81696 | 0.296609 | 0.900679 | 0.675588 |
| 3 | 99215400 | 99216799 | 13.59913 | 24.85684 | 26.73256 | 11.44462 | 0.317684 | 0.285 | 0.675589 |
| 4 | 142418000 | 142419999 | 19.29109 | 32.81441 | 51.11769 | 22.4044 | -0.25529 | 1.008208 | 0.675594 |
| Y | 56466400 | 56468999 | 16.41132 | 28.9051 | 25.29223 | 11.19575 | 0.580085 | 0.556588 | 0.675612 |
| 10 | 29177600 | 29179399 | 16.24438 | 28.91372 | 30.51511 | 13.44512 | 0.31766 | 0.4681 | 0.675626 |
| 19 | 23640200 | 23640799 | 9.148169 | 17.76314 | 21.16101 | 7.931801 | 0.336449 | -0.23124 | 0.675632 |
| 5 | 101757200 | 101767799 | 116.3155 | 176.8298 | 242.6007 | 100.4951 | -0.17288 | 3.191362 | 0.675645 |
| 3 | 63355000 | 63356799 | 25.2604 | 42.21718 | 63.22017 | 27.52249 | -0.19956 | 1.204997 | 0.675664 |
| 10 | 8262600 | 8274399 | 113.7798 | 173.3866 | 236.3604 | 97.97987 | -0.16086 | 3.142629 | 0.675688 |
| 2 | 39824200 | 39827599 | 19.5914 | 34.30889 | 31.43199 | 13.87564 | 0.513248 | 0.668975 | 0.675698 |
| 1 | 75691200 | 75692199 | 12.87038 | 22.60523 | 37.45559 | 16.57587 | -0.33698 | 0.597762 | 0.6757 |
| 15 | 17691600 | 17695199 | 44.46574 | 71.38549 | 102.519 | 43.38238 | -0.17173 | 1.930779 | 0.675718 |
| 12 | 114956400 | 114967799 | 126.6007 | 194.2495 | 196.5361 | 81.59099 | 0.276631 | 3.008258 | 0.675724 |
| 3 | 59425400 | 59426599 | 13.22891 | 23.12567 | 44.68884 | 19.6983 | -0.55799 | 0.688441 | 0.675756 |
| 5 | 86334400 | 86335599 | 12.88317 | 23.0622 | 35.64072 | 15.62815 | -0.22438 | 0.42379 | 0.675757 |
| 1 | 17404800 | 17405399 | 10.46823 | 19.4053 | 30.97032 | 13.00806 | -0.22765 | 0.195917 | 0.67577 |

| | | | | | | | | | |
|---|---|---|---|---|---|---|---|---|---|
| 6 | 57031000 | 57033599 | 26.50182 | 43.88364 | 65.76202 | 28.45352 | -0.20104 | 1.315336 | 0.6758 |
| 16 | 27478600 | 27478999 | 5.801 | 11.2933 | 21.28461 | 7.811899 | -0.28884 | -0.33816 | 0.675809 |
| 4 | 48823800 | 48826199 | 19.53415 | 34.02737 | 37.9047 | 16.66934 | 0.232435 | 0.760666 | 0.675834 |
| 18 | 10598400 | 10599799 | 14.91509 | 26.39626 | 42.88171 | 18.96362 | -0.30531 | 0.530002 | 0.675844 |
| 18 | 8745800 | 8747799 | 24.47485 | 41.97602 | 46.29403 | 20.30369 | 0.242715 | 0.947565 | 0.675861 |
| 15 | 51933800 | 51935199 | 22.19277 | 38.52837 | 36.21125 | 15.89089 | 0.475865 | 0.815321 | 0.675864 |
| 5 | 146861800 | 146864799 | 29.7333 | 49.74329 | 47.03311 | 20.48872 | 0.461624 | 1.19184 | 0.675868 |
| 4 | 143541000 | 143712799 | 3447.248 | 3995.925 | 4325.594 | 1838.417 | -0.21948 | 7.566288 | 0.675909 |
| 12 | 48042600 | 48044399 | 19.33264 | 34.02415 | 36.30775 | 16.07431 | 0.294876 | 0.604461 | 0.675911 |
| 16 | 25382400 | 25385199 | 27.68838 | 44.99945 | 84.65388 | 36.31893 | -0.54069 | 1.576901 | 0.675929 |
| 12 | 83276600 | 83278399 | 16.96615 | 29.95915 | 33.45674 | 14.80221 | 0.231648 | 0.55144 | 0.675931 |
| 10 | 81833000 | 81870599 | 141.4527 | 217.6295 | 151.9345 | 63.0984 | 0.814412 | 2.958076 | 0.675931 |
| X | 24479000 | 24498199 | 228.805 | 338.8727 | 307.6915 | 126.7174 | 0.388881 | 3.710896 | 0.675945 |
| 3 | 3246400 | 3246799 | 9.568118 | 18.27428 | 21.32507 | 8.478225 | 0.287334 | -0.02973 | 0.675953 |
| 5 | 84843200 | 84843599 | 9.568118 | 18.27428 | 21.32507 | 8.478225 | 0.287334 | -0.02973 | 0.675953 |
| 10 | 35751600 | 35753999 | 39.90484 | 64.8293 | 73.03043 | 31.32341 | 0.196608 | 1.598442 | 0.675971 |
| 19 | 28389200 | 28390599 | 15.76897 | 28.36032 | 31.33134 | 13.6845 | 0.26112 | 0.407013 | 0.675989 |
| 14 | 68026600 | 68028799 | 25.28784 | 43.11768 | 47.93701 | 21.01438 | 0.229904 | 1.004601 | 0.675992 |
| 3 | 120618800 | 120620599 | 26.41513 | 43.37016 | 68.01319 | 29.2951 | -0.26964 | 1.419875 | 0.675993 |
| 14 | 8243600 | 8244399 | 17.0402 | 29.93678 | 33.58059 | 14.86426 | 0.222702 | 0.609145 | 0.675994 |
| X | 25698000 | 25751199 | 729.2134 | 974.5324 | 827.6253 | 345.5842 | 0.338072 | 5.291917 | 0.676009 |
| 5 | 135895400 | 135900199 | 36.85665 | 59.10713 | 109.8446 | 46.40742 | -0.54465 | 1.955795 | 0.67601 |
| 17 | 84553400 | 84555399 | 14.80835 | 26.91076 | 29.86114 | 12.87696 | 0.266483 | 0.336255 | 0.676014 |
| 7 | 137187000 | 137188399 | 16.04561 | 28.03446 | 42.56157 | 18.7526 | -0.21178 | 0.707844 | 0.676015 |
| 1 | 26063000 | 26063599 | 10.77367 | 20.18221 | 21.74212 | 8.979716 | 0.354757 | 0.110931 | 0.67602 |
| 14 | 37825200 | 37841399 | 285.6515 | 404.4896 | 515.3197 | 213.2475 | -0.16279 | 4.393005 | 0.676065 |
| 15 | 3694600 | 3696799 | 19.86913 | 34.79343 | 38.01534 | 16.78111 | 0.260327 | 0.681754 | 0.676087 |
| 13 | 94895400 | 94896599 | 9.599271 | 17.84043 | 29.39823 | 12.3045 | -0.27002 | 0.168293 | 0.676099 |
| 17 | 20208800 | 20212399 | 36.09675 | 60.015 | 59.98538 | 26.02704 | 0.381863 | 1.289474 | 0.676123 |
| 5 | 76849400 | 76850999 | 13.55673 | 23.87514 | 37.72909 | 16.7013 | -0.26827 | 0.56519 | 0.676151 |
| 14 | 85225000 | 85225799 | 7.250793 | 13.99552 | 24.85889 | 9.590756 | -0.27396 | -0.14434 | 0.676153 |
| 4 | 36137600 | 36139199 | 23.0515 | 39.91166 | 43.09042 | 18.90524 | 0.276118 | 0.856879 | 0.676168 |
| 3 | 10504400 | 10505399 | 10.81454 | 19.5631 | 31.73405 | 13.76324 | -0.28305 | 0.338465 | 0.676178 |
| 5 | 14684600 | 14686799 | 20.02896 | 35.13478 | 37.37344 | 16.52218 | 0.298732 | 0.653754 | 0.676187 |
| 9 | 41925800 | 41927799 | 25.2637 | 42.13105 | 62.60845 | 27.21883 | -0.18846 | 1.227649 | 0.676213 |
| X | 23516800 | 23522399 | 90.29949 | 141.35 | 150.1759 | 62.29692 | 0.228384 | 2.631005 | 0.676238 |
| 17 | 22352600 | 22365999 | 246.1779 | 360.0098 | 355.9767 | 146.6488 | 0.248675 | 3.914522 | 0.676277 |
| 11 | 79971200 | 79972399 | 7.486015 | 14.25522 | 25.04664 | 10.02059 | -0.30609 | -0.01886 | 0.676311 |
| 3 | 16527600 | 16528199 | 8.053139 | 15.21806 | 26.4975 | 10.77432 | -0.31463 | 0.04018 | 0.67632 |
| 15 | 92370600 | 92373199 | 47.90948 | 76.57502 | 109.5351 | 46.04877 | -0.17336 | 2.073086 | 0.676324 |
| 13 | 21570400 | 21577799 | 86.60181 | 135.83 | 147.7359 | 61.27229 | 0.196784 | 2.592217 | 0.676326 |
| 1 | 38511200 | 38511599 | 4.554548 | 8.875964 | 19.12318 | 6.931817 | -0.46326 | -0.40975 | 0.676331 |
| 7 | 5287400 | 5290199 | 50.15073 | 80.86514 | 85.70123 | 36.49465 | 0.273472 | 1.797655 | 0.676351 |
| 11 | 12612800 | 12616799 | 57.71495 | 91.46845 | 131.0668 | 54.52659 | -0.1886 | 2.348396 | 0.67641 |
| 7 | 132493800 | 132501799 | 75.74527 | 118.2035 | 166.3356 | 69.02221 | -0.18004 | 2.676839 | 0.67641 |
| 13 | 97676200 | 97677399 | 15.76947 | 27.93943 | 31.44926 | 13.90474 | 0.22211 | 0.503364 | 0.676441 |
| 6 | 113404200 | 113405999 | 18.24457 | 31.52035 | 47.40685 | 20.79763 | -0.20087 | 0.858404 | 0.676465 |

| | | | | | | | | | |
|---|---|---|---|---|---|---|---|---|---|
| X | 106226800 | 106227799 | 10.77187 | 19.96176 | 34.81748 | 14.63032 | -0.35528 | 0.210986 | 0.676469 |
| 4 | 29236000 | 29236599 | 10.74007 | 20.85209 | 22.40816 | 8.406004 | 0.483328 | -0.2048 | 0.676478 |
| 3 | 15513800 | 15516799 | 30.32294 | 51.46055 | 43.60805 | 19.10489 | 0.619465 | 1.062115 | 0.676483 |
| 19 | 39034600 | 39036199 | 23.75326 | 40.01482 | 60.45186 | 26.42503 | -0.21214 | 1.118881 | 0.676503 |
| 6 | 86484600 | 86487399 | 26.39991 | 43.73461 | 65.38224 | 28.29802 | -0.19763 | 1.306751 | 0.676507 |
| 1 | 76207400 | 76210999 | 46.6471 | 75.41802 | 80.10139 | 34.22841 | 0.274824 | 1.714839 | 0.676512 |
| 5 | 50421000 | 50431399 | 128.9239 | 196.5775 | 211.3117 | 87.60883 | 0.182249 | 3.133413 | 0.676521 |
| 18 | 81388400 | 81390599 | 17.70775 | 31.46665 | 33.06187 | 14.58743 | 0.322888 | 0.494314 | 0.676539 |
| 4 | 141670200 | 141671999 | 26.85818 | 44.13499 | 67.66396 | 29.15514 | -0.23665 | 1.408833 | 0.676551 |
| 7 | 14037000 | 14049999 | 231.5289 | 338.2353 | 353.641 | 145.6967 | 0.166766 | 3.928234 | 0.676566 |
| 6 | 147050800 | 147052799 | 22.21078 | 37.44612 | 56.55433 | 24.72729 | -0.21165 | 1.101486 | 0.676576 |
| 5 | 32282600 | 32284599 | 12.81168 | 22.74405 | 35.18869 | 15.53865 | -0.23273 | 0.476694 | 0.676586 |
| 15 | 93938800 | 93940399 | 25.13883 | 43.27996 | 45.71629 | 20.05383 | 0.305738 | 0.915841 | 0.676588 |
| 1 | 13674800 | 13679399 | 67.86747 | 107.6739 | 120.5392 | 50.20076 | 0.168465 | 2.320939 | 0.676642 |
| 6 | 128593600 | 128594799 | 23.44162 | 39.25389 | 58.68561 | 25.57982 | -0.19715 | 1.174395 | 0.676654 |
| 6 | 92217000 | 92217799 | 8.891644 | 17.17179 | 21.46289 | 8.260087 | 0.232357 | -0.14318 | 0.676684 |
| 5 | 76650400 | 76652399 | 14.92973 | 26.0317 | 47.87135 | 21.06508 | -0.4875 | 0.737415 | 0.676699 |
| 12 | 8626200 | 8628399 | 33.67345 | 56.33566 | 48.64133 | 21.18943 | 0.592282 | 1.207249 | 0.676709 |
| 2 | 95111600 | 95112799 | 9.938021 | 19.09938 | 22.98006 | 8.978873 | 0.266044 | -0.08014 | 0.676729 |
| 13 | 59498200 | 59499599 | 13.13953 | 23.46074 | 36.18942 | 15.90318 | -0.22368 | 0.444926 | 0.676731 |
| 1 | 130176800 | 130178599 | 28.1232 | 48.23859 | 39.34829 | 17.25757 | 0.676343 | 0.956372 | 0.676736 |
| 9 | 53923400 | 53928599 | 74.83925 | 117.808 | 130.6752 | 54.21696 | 0.172523 | 2.512229 | 0.676743 |
| 12 | 65298200 | 65299799 | 17.00024 | 29.55068 | 44.74513 | 19.65133 | -0.20881 | 0.787937 | 0.676758 |
| 7 | 18107800 | 18123799 | 492.4088 | 671.2326 | 653.3315 | 271.9914 | 0.170114 | 4.974529 | 0.676766 |
| 1 | 10503200 | 10504999 | 18.3384 | 32.4305 | 34.71558 | 15.35459 | 0.293011 | 0.54397 | 0.676791 |
| 5 | 139861400 | 139861799 | 8.574273 | 16.521 | 19.83975 | 7.69471 | 0.279378 | -0.12574 | 0.676791 |
| 18 | 86166400 | 86167399 | 11.14493 | 21.15471 | 20.0522 | 8.077049 | 0.567913 | 0.022172 | 0.676791 |
| 1 | 102189000 | 102193999 | 72.61555 | 115.24 | 123.1983 | 51.32294 | 0.235272 | 2.31827 | 0.676795 |
| 6 | 71433600 | 71437799 | 26.3683 | 44.68234 | 50.12997 | 21.95222 | 0.215983 | 1.06644 | 0.676797 |
| 4 | 155766800 | 155767399 | 7.714499 | 14.39864 | 30.58788 | 12.71172 | -0.62481 | 0.131045 | 0.676818 |
| 8 | 46071600 | 46073199 | 14.52508 | 26.01119 | 29.39879 | 12.88703 | 0.224804 | 0.420046 | 0.676825 |
| X | 122418800 | 122419399 | 13.80874 | 25.54391 | 23.88445 | 10.06063 | 0.535496 | 0.217479 | 0.676855 |
| 13 | 82527000 | 82527799 | 9.339067 | 17.36947 | 28.38866 | 11.86838 | -0.25661 | 0.161175 | 0.676859 |
| 3 | 62586400 | 62587399 | 17.06611 | 30.04981 | 33.9874 | 15.04557 | 0.211757 | 0.583044 | 0.676875 |
| 10 | 26409000 | 26409999 | 22.29067 | 38.91207 | 34.91384 | 15.37467 | 0.543192 | 0.737889 | 0.676881 |
| 14 | 6620400 | 6620999 | 8.979227 | 16.82136 | 27.94784 | 11.54196 | -0.26612 | 0.109196 | 0.676903 |
| 15 | 84163800 | 84164799 | 11.88714 | 21.10271 | 34.2989 | 15.14576 | -0.30324 | 0.469686 | 0.676908 |
| 11 | 44462400 | 44464399 | 15.03478 | 26.30677 | 40.46789 | 17.85164 | -0.23067 | 0.674756 | 0.676951 |
| 11 | 19518400 | 19521399 | 32.70756 | 53.6911 | 60.53813 | 26.07325 | 0.204723 | 1.428587 | 0.67697 |
| 6 | 66791200 | 66792999 | 23.39342 | 39.24778 | 58.74361 | 25.63585 | -0.19888 | 1.152934 | 0.676973 |
| 4 | 106458200 | 106460399 | 17.34306 | 30.45988 | 34.53136 | 15.2813 | 0.20743 | 0.61668 | 0.676991 |
| 12 | 19577400 | 19578399 | 21.71145 | 37.71162 | 34.74043 | 15.25023 | 0.504745 | 0.804292 | 0.677038 |
| X | 20884200 | 20914999 | 520.4361 | 715.725 | 624.1954 | 259.502 | 0.341341 | 4.839783 | 0.677057 |
| 6 | 131723000 | 131724799 | 21.64641 | 37.95444 | 38.40575 | 16.97187 | 0.370569 | 0.672847 | 0.677061 |
| 7 | 43346200 | 43350399 | 37.53486 | 60.7092 | 89.01017 | 38.05265 | -0.18869 | 1.690765 | 0.677062 |
| X | 36977600 | 36980799 | 24.56448 | 41.14698 | 64.20206 | 27.98982 | -0.25861 | 1.182483 | 0.677067 |
| 5 | 11187000 | 11188399 | 16.51264 | 28.66363 | 44.26518 | 19.42374 | -0.23725 | 0.804522 | 0.677078 |

| | | | | | | | | | |
|---|---|---|---|---|---|---|---|---|---|
| 4 | 54821000 | 54823599 | 25.92064 | 43.94124 | 49.58136 | 21.7146 | 0.207813 | 1.061648 | 0.677098 |
| 16 | 10041600 | 10045399 | 41.3181 | 66.51533 | 95.88422 | 40.73877 | -0.17223 | 1.837433 | 0.677107 |
| 1 | 114301200 | 114302199 | 15.5082 | 27.61535 | 31.31456 | 13.79232 | 0.215346 | 0.465341 | 0.677112 |
| 17 | 50185200 | 50187799 | 19.29225 | 32.86879 | 50.59087 | 22.17635 | -0.23765 | 0.993449 | 0.677116 |
| 1 | 126906800 | 126910999 | 44.12914 | 70.90203 | 102.2995 | 43.34158 | -0.17704 | 1.905153 | 0.677117 |
| 3 | 159170200 | 159171799 | 11.94185 | 22.47951 | 21.479 | 8.798342 | 0.535609 | 0.088854 | 0.677128 |
| 9 | 117945000 | 117948399 | 35.00076 | 57.01604 | 92.08344 | 39.54213 | -0.31898 | 1.559562 | 0.677134 |
| X | 122012800 | 122015199 | 33.80985 | 56.42431 | 56.89874 | 24.74798 | 0.369222 | 1.24334 | 0.67714 |
| 16 | 68379000 | 68380599 | 16.68633 | 29.90742 | 32.23216 | 14.11872 | 0.293697 | 0.434994 | 0.677155 |
| 3 | 110494200 | 110495999 | 16.9046 | 29.83179 | 33.28738 | 14.72912 | 0.232172 | 0.554074 | 0.677164 |
| 2 | 14988200 | 14990999 | 23.79711 | 39.71764 | 59.73736 | 25.984 | -0.20574 | 1.212146 | 0.677173 |
| 3 | 102091000 | 102091799 | 7.741681 | 14.85214 | 25.19874 | 9.895735 | -0.23203 | -0.0784 | 0.677175 |
| 17 | 56789400 | 56813399 | 407.2337 | 558.8571 | 696.1196 | 289.5006 | -0.17589 | 4.868566 | 0.677176 |
| 8 | 58152200 | 58156199 | 29.30109 | 49.80054 | 50.68149 | 22.21175 | 0.356411 | 1.053393 | 0.677187 |
| 1 | 179234800 | 179238599 | 36.61049 | 59.972 | 66.60597 | 28.66117 | 0.22522 | 1.466467 | 0.677196 |
| 3 | 34445400 | 34452199 | 72.73223 | 114.9342 | 129.0506 | 53.60479 | 0.159302 | 2.426821 | 0.677198 |
| 8 | 45241800 | 45245599 | 31.90391 | 52.89902 | 59.65775 | 25.83604 | 0.207904 | 1.307883 | 0.677198 |
| 2 | 122297400 | 122298599 | 10.2073 | 18.52655 | 30.80319 | 13.3037 | -0.31428 | 0.313325 | 0.6772 |
| 3 | 66808000 | 66810199 | 25.96361 | 42.46784 | 79.25014 | 34.08206 | -0.52253 | 1.472417 | 0.6772 |
| 1 | 165894600 | 165898399 | 67.76527 | 107.3072 | 120.4649 | 50.09095 | 0.161954 | 2.373069 | 0.677203 |
| 15 | 35384800 | 35388199 | 57.45578 | 92.01473 | 99.52882 | 41.96087 | 0.232268 | 2.021407 | 0.677213 |
| 8 | 19733600 | 19735999 | 43.46707 | 70.35582 | 77.86231 | 33.30492 | 0.217174 | 1.679952 | 0.677232 |
| 14 | 57994600 | 57996199 | 9.433522 | 17.3326 | 29.39449 | 12.49629 | -0.32675 | 0.235746 | 0.677253 |
| 18 | 5677400 | 5679999 | 40.16318 | 64.75128 | 93.85696 | 39.96008 | -0.17758 | 1.789698 | 0.677273 |
| 4 | 144046000 | 144053799 | 84.27263 | 132.5338 | 141.9625 | 58.89287 | 0.221806 | 2.536532 | 0.67728 |
| 18 | 8162200 | 8164999 | 49.42448 | 79.65099 | 75.7424 | 32.23129 | 0.42864 | 1.80787 | 0.677284 |
| 6 | 66306600 | 66308799 | 20.08765 | 35.22089 | 32.39634 | 14.3161 | 0.50762 | 0.655624 | 0.677331 |
| 15 | 94435800 | 94439599 | 39.68354 | 63.95908 | 92.3484 | 39.3028 | -0.17245 | 1.797248 | 0.677343 |
| 1 | 29843000 | 29846199 | 34.45992 | 56.75161 | 63.56566 | 27.41481 | 0.216132 | 1.397121 | 0.677381 |
| 6 | 123274200 | 123277799 | 38.76203 | 63.23541 | 70.35894 | 30.22937 | 0.218804 | 1.532952 | 0.677402 |
| 10 | 45751200 | 45753599 | 20.41389 | 34.7369 | 52.2 | 22.87944 | -0.20354 | 1.010596 | 0.677412 |
| 10 | 30955800 | 30958199 | 14.33418 | 24.91136 | 47.90889 | 21.03813 | -0.55216 | 0.782308 | 0.677413 |
| 18 | 10041200 | 10046399 | 66.11269 | 104.8823 | 118.2457 | 49.24199 | 0.158205 | 2.321871 | 0.677424 |
| 11 | 7274200 | 7275799 | 25.58451 | 43.91446 | 39.81583 | 17.4636 | 0.524312 | 0.935973 | 0.677438 |
| 5 | 116947400 | 116948399 | 10.29567 | 18.80189 | 30.76895 | 13.18739 | -0.28462 | 0.276805 | 0.677455 |
| 14 | 13488200 | 13488799 | 6.3842 | 12.41638 | 22.43252 | 8.299185 | -0.24042 | -0.30301 | 0.67749 |
| 1 | 3921400 | 3923399 | 35.58407 | 58.03637 | 65.96825 | 28.34033 | 0.187826 | 1.526403 | 0.677511 |
| 17 | 84178000 | 84181399 | 20.43022 | 34.65296 | 52.7247 | 23.09514 | -0.22184 | 1.039763 | 0.677547 |
| 3 | 155108800 | 155109799 | 10.29387 | 18.88206 | 30.29862 | 12.90972 | -0.2504 | 0.251223 | 0.67755 |
| 5 | 18386600 | 18387199 | 9.194498 | 17.69107 | 21.91687 | 8.534817 | 0.229042 | -0.10052 | 0.677613 |
| 2 | 179562000 | 179564799 | 18.28258 | 32.2026 | 29.22915 | 12.93831 | 0.527514 | 0.575777 | 0.677614 |
| 12 | 116007400 | 116009799 | 18.28258 | 32.2026 | 29.22915 | 12.93831 | 0.527514 | 0.575777 | 0.677614 |
| 4 | 58950000 | 58951399 | 13.095 | 23.90064 | 26.89271 | 11.53411 | 0.251487 | 0.291499 | 0.677627 |
| 12 | 59492200 | 59492999 | 13.19865 | 24.45628 | 22.85416 | 9.604738 | 0.538459 | 0.199551 | 0.677646 |
| 9 | 119936200 | 119940199 | 39.67129 | 63.8241 | 93.45554 | 39.67072 | -0.19586 | 1.853325 | 0.677656 |
| 3 | 84603800 | 84606799 | 30.74409 | 50.13587 | 75.75104 | 32.54149 | -0.22189 | 1.522685 | 0.677678 |
| 19 | 40881800 | 40883999 | 26.62465 | 43.50364 | 79.14391 | 34.02342 | -0.48699 | 1.489768 | 0.677698 |

| | | | | | | | | | |
|---|---|---|---|---|---|---|---|---|---|
| 12 | 106653200 | 106655199 | 12.28931 | 21.84201 | 34.66983 | 15.29458 | -0.26862 | 0.46328 | 0.677705 |
| 4 | 147509000 | 147519399 | 266.0872 | 387.0769 | 375.4316 | 154.7424 | 0.269132 | 3.995718 | 0.677705 |
| 18 | 44413400 | 44413799 | 5.799081 | 11.30112 | 20.85519 | 7.561465 | -0.24346 | -0.38555 | 0.677723 |
| 2 | 5134400 | 5137599 | 24.29078 | 40.38996 | 61.36455 | 26.62714 | -0.22034 | 1.255418 | 0.677735 |
| 13 | 60707200 | 60709999 | 30.43118 | 50.5837 | 56.89787 | 24.68348 | 0.211865 | 1.270704 | 0.677755 |
| 13 | 51368000 | 51372399 | 31.56054 | 51.47928 | 76.44323 | 32.84163 | -0.19669 | 1.521963 | 0.677756 |
| 4 | 57807200 | 57809199 | 22.95576 | 38.3212 | 59.18585 | 25.7474 | -0.24389 | 1.205587 | 0.677757 |
| 2 | 7757200 | 7757999 | 18.51736 | 32.32935 | 35.36691 | 15.57546 | 0.258767 | 0.71951 | 0.67778 |
| 1 | 110622200 | 110623199 | 17.8617 | 31.37539 | 35.49453 | 15.70949 | 0.21044 | 0.619167 | 0.677781 |
| 12 | 65947000 | 65949199 | 20.6711 | 36.31778 | 36.68916 | 16.24104 | 0.37274 | 0.631473 | 0.677785 |
| 11 | 91887400 | 91888799 | 10.15494 | 18.75826 | 29.86849 | 12.60563 | -0.22921 | 0.21085 | 0.677792 |
| 1 | 37723400 | 37725999 | 28.75039 | 47.83998 | 54.53257 | 23.67449 | 0.192927 | 1.254273 | 0.677793 |
| 1 | 55578600 | 55580199 | 30.7481 | 50.28248 | 74.197 | 31.90633 | -0.18499 | 1.482948 | 0.677832 |
| 18 | 27943200 | 27943999 | 12.95734 | 23.18746 | 35.6829 | 15.65092 | -0.2184 | 0.428307 | 0.677851 |
| 3 | 3333600 | 3334999 | 16.78239 | 29.90796 | 32.8009 | 14.43733 | 0.264014 | 0.473216 | 0.677861 |
| 5 | 75463800 | 75468599 | 68.73984 | 107.6554 | 155.9719 | 64.6979 | -0.218 | 2.608605 | 0.677872 |
| 2 | 146533400 | 146535199 | 18.41068 | 32.34982 | 35.85607 | 15.87369 | 0.239648 | 0.616252 | 0.677905 |
| 18 | 56354800 | 56357399 | 38.6005 | 62.00379 | 94.53316 | 40.03679 | -0.2568 | 1.903215 | 0.677917 |
| 5 | 25378200 | 25381799 | 41.97582 | 68.19928 | 65.21464 | 27.97281 | 0.432547 | 1.59638 | 0.677921 |
| 5 | 78113400 | 78116799 | 49.43692 | 78.94155 | 112.8059 | 47.36023 | -0.1734 | 2.105558 | 0.677933 |
| 16 | 79274000 | 79274999 | 14.91595 | 26.55717 | 29.50245 | 12.99565 | 0.244902 | 0.460029 | 0.677981 |
| 19 | 52920200 | 52925199 | 61.12668 | 96.73248 | 136.0623 | 56.55811 | -0.16366 | 2.386764 | 0.677997 |
| 12 | 3877800 | 3880999 | 30.91115 | 50.98045 | 57.22936 | 24.69709 | 0.213744 | 1.364829 | 0.678004 |
| 19 | 8121400 | 8124599 | 34.8841 | 57.7273 | 62.14134 | 26.87527 | 0.27466 | 1.339905 | 0.678031 |
| 11 | 9314800 | 9315999 | 16.18316 | 27.76698 | 53.18556 | 23.32698 | -0.55007 | 0.922398 | 0.678035 |
| 17 | 93009200 | 93010599 | 15.74383 | 27.25901 | 50.59195 | 22.19641 | -0.50224 | 0.832035 | 0.678038 |
| X | 54529000 | 54537999 | 19.17714 | 34.25015 | 27.93419 | 12.27269 | 0.690702 | 0.466036 | 0.678073 |
| 2 | 145447400 | 145449599 | 19.23577 | 32.80145 | 51.13341 | 22.41576 | -0.25582 | 0.984094 | 0.678076 |
| 11 | 81117800 | 81120399 | 22.24937 | 37.53715 | 55.98654 | 24.48417 | -0.19356 | 1.096299 | 0.678116 |
| X | 107254800 | 107258399 | 56.7538 | 90.85967 | 99.90106 | 42.09607 | 0.208117 | 2.030658 | 0.678148 |
| 17 | 21249400 | 21258599 | 221.5066 | 324.3777 | 341.7245 | 140.7652 | 0.159199 | 3.889934 | 0.678149 |
| 11 | 12597600 | 12598599 | 13.89426 | 25.00693 | 27.98369 | 12.21399 | 0.243419 | 0.382112 | 0.678176 |
| X | 74748000 | 74752599 | 86.41682 | 136.4273 | 135.9647 | 56.44918 | 0.330015 | 2.462528 | 0.67819 |
| 8 | 127288200 | 127289399 | 11.08875 | 20.29665 | 31.80492 | 13.59102 | -0.21954 | 0.272887 | 0.678199 |
| 6 | 54951600 | 54957799 | 60.47021 | 95.69515 | 135.1748 | 56.18962 | -0.16974 | 2.385726 | 0.678206 |
| 1 | 67945200 | 67947799 | 41.2659 | 67.13147 | 64.17345 | 27.54361 | 0.434409 | 1.572975 | 0.678207 |
| 12 | 19601600 | 19617199 | 452.7372 | 624.9623 | 527.5587 | 219.1957 | 0.393814 | 4.778732 | 0.678213 |
| 11 | 17509600 | 17513199 | 84.82547 | 132.7837 | 148.0761 | 61.42573 | 0.158595 | 2.625275 | 0.678234 |
| 11 | 5738000 | 5739999 | 13.60419 | 23.81305 | 39.4884 | 17.42539 | -0.33845 | 0.657984 | 0.678244 |
| 12 | 114568200 | 114583599 | 226.0709 | 334.0674 | 319.5188 | 131.5851 | 0.310845 | 3.746665 | 0.678268 |
| 2 | 122856400 | 122857199 | 8.924242 | 16.78664 | 28.03738 | 11.49929 | -0.26497 | 0.086882 | 0.678268 |
| 17 | 22929200 | 22936799 | 144.6294 | 220.2785 | 216.7586 | 89.83965 | 0.308376 | 3.162667 | 0.678278 |
| 6 | 57310000 | 57316399 | 114.5131 | 176.4018 | 185.6617 | 77.1185 | 0.224104 | 2.925982 | 0.67828 |
| 6 | 59262400 | 59265199 | 22.17587 | 38.67675 | 40.72132 | 17.92189 | 0.31336 | 0.757622 | 0.678285 |
| 8 | 58478000 | 58479199 | 11.3731 | 20.42395 | 32.77073 | 14.33116 | -0.27502 | 0.390498 | 0.6783 |
| 8 | 69677800 | 69689199 | 243.1133 | 355.6449 | 353.1147 | 145.4668 | 0.243175 | 3.909447 | 0.678335 |
| 6 | 68374800 | 68377199 | 33.76837 | 54.87677 | 80.23038 | 34.41929 | -0.17814 | 1.584953 | 0.678335 |

| | | | | | | | | | |
|---|---|---|---|---|---|---|---|---|---|
| 2 | 176454800 | 176456799 | 31.06804 | 51.27355 | 58.37741 | 25.20035 | 0.193737 | 1.357717 | 0.67835 |
| 5 | 37157000 | 37158399 | 14.20116 | 25.68016 | 29.39572 | 12.75481 | 0.216265 | 0.359703 | 0.678354 |
| 16 | 20402400 | 20403199 | 10.23716 | 19.43195 | 23.24435 | 9.362533 | 0.234685 | 0.021288 | 0.678356 |
| 5 | 72742200 | 72744999 | 25.56441 | 42.10573 | 74.56354 | 32.15931 | -0.44278 | 1.381309 | 0.678362 |
| 2 | 97571800 | 97572399 | 10.47813 | 19.83299 | 23.76795 | 9.631069 | 0.224739 | 0.045757 | 0.678387 |
| 6 | 102905200 | 102906999 | 21.54148 | 37.70524 | 38.99559 | 17.20739 | 0.33921 | 0.699266 | 0.678441 |
| 13 | 46595400 | 46595999 | 10.36639 | 19.63183 | 23.19206 | 9.387232 | 0.246303 | 0.037717 | 0.678445 |
| 18 | 76040800 | 76042399 | 10.66436 | 19.33756 | 31.26236 | 13.51822 | -0.2753 | 0.323082 | 0.678459 |
| 5 | 147731600 | 147736599 | 73.21285 | 114.7043 | 159.2694 | 66.06315 | -0.15634 | 2.606303 | 0.678471 |
| 5 | 59601000 | 59601999 | 15.85858 | 27.77604 | 41.94702 | 18.51983 | -0.20425 | 0.665604 | 0.678472 |
| 13 | 22141400 | 22147199 | 80.39434 | 126.7146 | 138.1545 | 57.32962 | 0.198743 | 2.491239 | 0.678499 |
| 15 | 83401000 | 83402999 | 18.52176 | 32.49907 | 36.52583 | 16.15092 | 0.219856 | 0.642081 | 0.678515 |
| 7 | 26521800 | 26525399 | 17.39907 | 30.82511 | 27.06381 | 11.96591 | 0.578214 | 0.502529 | 0.678524 |
| 10 | 53114000 | 53114999 | 12.58051 | 22.53858 | 34.64579 | 15.18156 | -0.21593 | 0.41539 | 0.678525 |
| 12 | 37418800 | 37420599 | 12.58051 | 22.53858 | 34.64579 | 15.18156 | -0.21593 | 0.41539 | 0.678525 |
| 15 | 88300000 | 88301399 | 12.58051 | 22.53858 | 34.64579 | 15.18156 | -0.21593 | 0.41539 | 0.678525 |
| 10 | 11956200 | 11957799 | 13.15098 | 24.3679 | 27.56824 | 11.58593 | 0.264519 | 0.210169 | 0.678527 |
| 9 | 123126600 | 123127599 | 8.87905 | 16.63689 | 27.44775 | 11.33175 | -0.25524 | 0.108089 | 0.678539 |
| 3 | 51070800 | 51072999 | 21.03931 | 36.51402 | 41.4978 | 18.20781 | 0.202969 | 0.822295 | 0.678571 |
| 6 | 138678200 | 138678799 | 10.97456 | 20.17682 | 31.89489 | 13.54757 | -0.22564 | 0.248581 | 0.678573 |
| 3 | 24745600 | 24747399 | 16.41408 | 28.69721 | 44.10488 | 19.4429 | -0.22938 | 0.702271 | 0.678575 |
| 4 | 135874800 | 135876399 | 10.69222 | 19.38467 | 31.28907 | 13.53275 | -0.27311 | 0.325429 | 0.678583 |
| 10 | 3555400 | 3559199 | 54.41373 | 86.95356 | 97.57812 | 41.01043 | 0.175842 | 2.082826 | 0.678586 |
| 15 | 93558000 | 93560799 | 28.77857 | 47.2373 | 70.1711 | 30.22096 | -0.19199 | 1.4282 | 0.678593 |
| 13 | 113141600 | 113145199 | 36.94455 | 60.41298 | 68.00785 | 29.24441 | 0.204401 | 1.494278 | 0.678603 |
| 2 | 173516600 | 173517799 | 9.666006 | 17.96243 | 28.89674 | 12.09647 | -0.23536 | 0.170772 | 0.678617 |
| X | 111294800 | 111298399 | 50.18018 | 80.58006 | 90.49133 | 38.30757 | 0.182862 | 1.926455 | 0.678622 |
| 4 | 87187800 | 87188599 | 10.53116 | 20.14492 | 19.11396 | 7.56919 | 0.58822 | -0.04078 | 0.678656 |
| 9 | 56607200 | 56608599 | 17.82518 | 31.27552 | 35.54651 | 15.71733 | 0.203767 | 0.636801 | 0.678707 |
| 18 | 37800000 | 37804399 | 145.7523 | 219.444 | 231.0099 | 95.41256 | 0.197118 | 3.393536 | 0.67873 |
| 1 | 147046000 | 147048399 | 40.40677 | 65.68187 | 73.1763 | 31.39862 | 0.213283 | 1.588628 | 0.678736 |
| 3 | 51209200 | 51210599 | 18.23338 | 31.911 | 36.12126 | 15.9374 | 0.209608 | 0.679359 | 0.678781 |
| 10 | 28796800 | 28800199 | 37.68323 | 60.91093 | 88.8984 | 37.97583 | -0.18312 | 1.707962 | 0.678783 |
| 7 | 59330400 | 59334599 | 40.23958 | 65.40309 | 61.76332 | 26.49981 | 0.451026 | 1.58217 | 0.67882 |
| 13 | 64644600 | 64648999 | 73.15179 | 115.3838 | 129.0835 | 53.57059 | 0.162452 | 2.470969 | 0.678825 |
| 1 | 102641600 | 102644199 | 31.96918 | 53.15702 | 58.65201 | 25.45002 | 0.239612 | 1.273805 | 0.678836 |
| 8 | 111519200 | 111525799 | 73.9485 | 115.53 | 161.7555 | 67.11388 | -0.17149 | 2.656589 | 0.678872 |
| 11 | 9988200 | 9988799 | 8.594842 | 16.64065 | 21.00523 | 7.988958 | 0.234478 | -0.18719 | 0.678876 |
| 4 | 66481600 | 66482599 | 12.78476 | 23.54933 | 26.97475 | 11.42828 | 0.238341 | 0.238191 | 0.678906 |
| 2 | 74265200 | 74269399 | 64.75656 | 102.7906 | 115.9121 | 48.29826 | 0.15871 | 2.302969 | 0.678907 |
| 7 | 14616000 | 14620599 | 68.31645 | 108.8457 | 113.4572 | 47.48568 | 0.277468 | 2.193496 | 0.678966 |
| 6 | 31085400 | 31088599 | 30.46291 | 49.8189 | 73.45799 | 31.58919 | -0.18382 | 1.482316 | 0.678977 |
| 17 | 37934800 | 37936399 | 13.89852 | 25.95098 | 27.71833 | 11.51149 | 0.360227 | 0.172 | 0.678988 |
| 7 | 118251400 | 118252599 | 8.992518 | 16.80967 | 27.92029 | 11.57312 | -0.26992 | 0.123472 | 0.678994 |
| 6 | 114944400 | 114948199 | 31.29943 | 50.88419 | 76.49246 | 32.82305 | -0.21769 | 1.570036 | 0.679001 |
| 14 | 80280600 | 80281999 | 17.0328 | 31.40308 | 28.91631 | 12.23551 | 0.552745 | 0.271421 | 0.679005 |
| 11 | 14793800 | 14796599 | 54.87467 | 87.72154 | 99.32897 | 41.76745 | 0.163445 | 2.07238 | 0.679019 |

| | | | | | | | | | |
|---|---|---|---|---|---|---|---|---|---|
| 7 | 31184000 | 31191999 | 126.5368 | 195.0474 | 158.7232 | 65.93098 | 0.595563 | 2.915965 | 0.679025 |
| 2 | 170811600 | 170813799 | 19.27065 | 33.13001 | 49.53809 | 21.72936 | -0.19396 | 0.910431 | 0.679036 |
| 6 | 36598400 | 36599799 | 13.45269 | 25.14338 | 27.17022 | 11.26499 | 0.345279 | 0.159699 | 0.679041 |
| 7 | 115666800 | 115667799 | 11.50394 | 20.89597 | 32.7881 | 14.14742 | -0.23043 | 0.324221 | 0.679078 |
| 7 | 108125400 | 108129999 | 93.48197 | 143.7934 | 197.7275 | 82.11374 | -0.16284 | 2.947702 | 0.679113 |
| X | 124903600 | 124904199 | 16.40507 | 29.61611 | 25.33296 | 11.01492 | 0.632448 | 0.378399 | 0.679121 |
| 4 | 92246200 | 92247199 | 13.00903 | 23.65251 | 26.49095 | 11.41663 | 0.253853 | 0.31316 | 0.679148 |
| 15 | 27974600 | 27979599 | 39.77663 | 63.92729 | 109.1546 | 46.26523 | -0.419 | 1.893586 | 0.67916 |
| X | 42656400 | 42659599 | 52.21562 | 83.94357 | 90.77258 | 38.49053 | 0.239267 | 1.894819 | 0.679163 |
| 17 | 14304000 | 14304999 | 8.207852 | 15.67663 | 26.18037 | 10.40807 | -0.22601 | -0.03106 | 0.679174 |
| 12 | 3148800 | 3150599 | 15.52338 | 27.59495 | 31.33883 | 13.823 | 0.211778 | 0.476386 | 0.679199 |
| 19 | 7795600 | 7798199 | 21.13686 | 36.96385 | 39.03759 | 17.21287 | 0.309121 | 0.710594 | 0.6792 |
| 10 | 108210600 | 108213799 | 30.40648 | 49.75314 | 73.66416 | 31.684 | -0.18926 | 1.47305 | 0.679201 |
| 12 | 35830400 | 35831999 | 16.08334 | 28.25092 | 45.02514 | 19.92725 | -0.28169 | 0.625873 | 0.679203 |
| 18 | 35224000 | 35226599 | 25.3744 | 42.97584 | 49.04997 | 21.47591 | 0.191301 | 1.066186 | 0.679241 |
| 17 | 29153600 | 29163199 | 111.3923 | 169.1765 | 233.5839 | 96.73113 | -0.18322 | 3.210613 | 0.679268 |
| 6 | 68686600 | 68703599 | 202.0002 | 299.2717 | 309.4987 | 127.4642 | 0.201677 | 3.699835 | 0.679268 |
| 18 | 18173400 | 18180799 | 148.3482 | 224.4045 | 211.6561 | 87.54256 | 0.361315 | 3.298709 | 0.679291 |
| 1 | 39212800 | 39220599 | 84.15848 | 129.3687 | 213.9072 | 88.82513 | -0.42926 | 2.958565 | 0.67933 |
| 5 | 151475000 | 151477399 | 22.66579 | 39.4924 | 41.27096 | 18.15259 | 0.323953 | 0.774294 | 0.679339 |
| 17 | 71234800 | 71236799 | 14.65322 | 25.77243 | 39.72613 | 17.58992 | -0.23341 | 0.589085 | 0.679343 |
| 1 | 171720400 | 171723399 | 49.68313 | 79.3506 | 112.8953 | 47.41122 | -0.16681 | 2.098175 | 0.679371 |
| 8 | 19471000 | 19472599 | 18.28534 | 32.13083 | 35.89105 | 15.88971 | 0.228538 | 0.615513 | 0.679371 |
| 1 | 54360000 | 54367199 | 104.5968 | 161.0915 | 178.2184 | 74.02458 | 0.151978 | 2.924916 | 0.679374 |
| 3 | 159487800 | 159488599 | 6.819853 | 13.17041 | 23.72232 | 9.130557 | -0.29038 | -0.15983 | 0.679379 |
| 8 | 109210200 | 109212999 | 19.906 | 34.72713 | 39.26748 | 17.28493 | 0.210951 | 0.741207 | 0.67938 |
| 12 | 116069600 | 116073999 | 50.81534 | 81.82217 | 87.56065 | 37.21406 | 0.257059 | 1.839211 | 0.679429 |
| 5 | 61445000 | 61447799 | 26.21968 | 43.53025 | 64.88278 | 28.12388 | -0.1931 | 1.280843 | 0.67944 |
| 1 | 110271000 | 110273999 | 46.92377 | 75.81246 | 71.97928 | 30.73008 | 0.435049 | 1.729023 | 0.679458 |
| 1 | 55078000 | 55081999 | 69.24028 | 109.6332 | 123.5891 | 51.38776 | 0.155855 | 2.376695 | 0.67948 |
| 2 | 131579200 | 131581799 | 19.15841 | 33.4486 | 37.68213 | 16.59509 | 0.216297 | 0.723096 | 0.679496 |
| 15 | 87210600 | 87214199 | 29.93037 | 49.0064 | 72.44828 | 31.16903 | -0.18649 | 1.462839 | 0.679497 |
| 4 | 122410400 | 122414599 | 34.58718 | 56.80639 | 64.62934 | 27.84149 | 0.192084 | 1.430575 | 0.679522 |
| 1 | 111402200 | 111403799 | 15.1832 | 28.15359 | 28.5932 | 12.00499 | 0.420253 | 0.225141 | 0.679526 |
| 17 | 7577400 | 7579799 | 22.45017 | 38.86857 | 43.03892 | 18.88261 | 0.239585 | 0.851012 | 0.67956 |
| X | 14899000 | 14899599 | 9.450702 | 18.28223 | 22.11027 | 8.442512 | 0.29052 | -0.16009 | 0.679573 |
| 14 | 97575400 | 97576799 | 18.97747 | 33.19068 | 37.68823 | 16.61923 | 0.205173 | 0.697898 | 0.679598 |
| 2 | 23152200 | 23152799 | 8.179587 | 15.86218 | 19.9676 | 7.537386 | 0.249226 | -0.21853 | 0.679607 |
| 16 | 70548800 | 70550999 | 13.45844 | 24.87283 | 22.45679 | 9.471219 | 0.584441 | 0.21643 | 0.679608 |
| 5 | 76242800 | 76245999 | 20.69985 | 36.25071 | 38.48388 | 16.98896 | 0.301654 | 0.685784 | 0.679612 |
| 2 | 75300400 | 75302199 | 11.31052 | 20.64264 | 32.311 | 13.85884 | -0.22141 | 0.293692 | 0.679613 |
| 11 | 7962200 | 7962799 | 11.31052 | 20.64264 | 32.311 | 13.85884 | -0.22141 | 0.293692 | 0.679613 |
| 3 | 71878200 | 71882999 | 56.24754 | 90.02036 | 99.58509 | 41.94388 | 0.198786 | 2.037414 | 0.679644 |
| 10 | 25293000 | 25295399 | 19.42156 | 32.88402 | 59.11898 | 25.88219 | -0.46176 | 1.059986 | 0.679651 |
| 19 | 9674800 | 9685599 | 174.171 | 259.9926 | 278.4977 | 114.8 | 0.16096 | 3.562088 | 0.679684 |
| 10 | 75939600 | 75943399 | 41.82609 | 67.2648 | 96.65831 | 41.00729 | -0.16949 | 1.870563 | 0.679689 |
| 18 | 7054000 | 7063399 | 170.6472 | 254.5781 | 274.4195 | 113.1056 | 0.151094 | 3.572513 | 0.679702 |

| | | | | | | | | | |
|---|---|---|---|---|---|---|---|---|---|
| 14 | 104047800 | 104049199 | 13.83904 | 25.48131 | 28.52149 | 12.09004 | 0.270983 | 0.252976 | 0.679704 |
| 13 | 61483200 | 61484599 | 46.68764 | 75.068 | 84.35842 | 35.77859 | 0.184159 | 1.879115 | 0.679725 |
| 6 | 90882200 | 90882999 | 9.935645 | 18.18853 | 30.38569 | 12.98211 | -0.31039 | 0.263658 | 0.679778 |
| Y | 8385200 | 8387199 | 15.52471 | 27.53461 | 22.69747 | 10.03067 | 0.669348 | 0.470018 | 0.679838 |
| 1 | 54793400 | 54793799 | 3.930525 | 7.687405 | 16.10649 | 5.448257 | -0.33108 | -0.67329 | 0.679848 |
| 10 | 67654400 | 67657599 | 30.52497 | 51.10669 | 55.7676 | 24.30535 | 0.255542 | 1.194727 | 0.679856 |
| 1 | 164051800 | 164057199 | 61.36599 | 96.33814 | 147.5869 | 61.21203 | -0.29568 | 2.560808 | 0.679859 |
| 14 | 4198200 | 4200599 | 38.82899 | 62.1329 | 115.8578 | 48.78634 | -0.55328 | 2.028874 | 0.679869 |
| 15 | 102111400 | 102112999 | 13.31992 | 23.29236 | 39.56493 | 17.44455 | -0.37287 | 0.673524 | 0.679873 |
| 2 | 50356000 | 50358999 | 24.13047 | 41.25016 | 45.22218 | 19.82837 | 0.250513 | 0.971827 | 0.679935 |
| 7 | 125035800 | 125039199 | 29.57686 | 48.05674 | 85.39647 | 36.63084 | -0.45915 | 1.582481 | 0.679936 |
| 7 | 71986000 | 71988199 | 22.7991 | 39.97619 | 38.31668 | 16.93275 | 0.448422 | 0.678426 | 0.679958 |
| X | 105256400 | 105258799 | 22.7991 | 39.97619 | 38.31668 | 16.93275 | 0.448422 | 0.678426 | 0.679958 |
| 6 | 36893800 | 36895599 | 21.5255 | 37.57491 | 34.40586 | 15.15062 | 0.513931 | 0.73261 | 0.679987 |
| 12 | 52932400 | 52935399 | 27.44972 | 44.7675 | 80.78768 | 34.70626 | -0.47731 | 1.519751 | 0.680016 |
| 3 | 135259400 | 135260799 | 16.15455 | 28.20759 | 42.55618 | 18.74304 | -0.20293 | 0.715781 | 0.68002 |
| 11 | 34579400 | 34581599 | 19.66008 | 34.26081 | 38.8615 | 17.09453 | 0.206427 | 0.756054 | 0.680025 |
| 14 | 118437200 | 118438599 | 13.88678 | 25.18003 | 28.93701 | 12.51316 | 0.213811 | 0.341176 | 0.680029 |
| 15 | 3389400 | 3393399 | 88.56484 | 137.6373 | 147.2029 | 61.11906 | 0.210767 | 2.753021 | 0.68003 |
| 6 | 39435000 | 39439199 | 62.18823 | 98.1984 | 139.3685 | 57.86863 | -0.17908 | 2.440223 | 0.680075 |
| 7 | 112234600 | 112237999 | 30.06587 | 49.73419 | 56.16359 | 24.28333 | 0.205756 | 1.326775 | 0.680085 |
| 15 | 34972200 | 34972799 | 6.237135 | 12.10743 | 22.25227 | 8.359286 | -0.28549 | -0.25272 | 0.680088 |
| 4 | 71379400 | 71380799 | 17.92359 | 31.7134 | 34.1618 | 15.10809 | 0.284193 | 0.533823 | 0.680118 |
| 13 | 29530400 | 29533399 | 39.87156 | 64.29182 | 93.08252 | 39.63886 | -0.17561 | 1.784984 | 0.680136 |
| 13 | 111712400 | 111719599 | 87.1025 | 134.8137 | 185.409 | 77.01415 | -0.15622 | 2.827665 | 0.680141 |
| 5 | 109497600 | 109507999 | 119.1887 | 183.7374 | 183.9289 | 76.40097 | 0.29715 | 2.915562 | 0.680144 |
| 5 | 42769000 | 42769799 | 13.37432 | 24.2402 | 27.48078 | 11.88996 | 0.232834 | 0.336628 | 0.680158 |
| 16 | 84342800 | 84343799 | 13.37432 | 24.2402 | 27.48078 | 11.88996 | 0.232834 | 0.336628 | 0.680158 |
| 5 | 105540800 | 105543999 | 41.84771 | 67.92797 | 75.04199 | 32.16605 | 0.223549 | 1.619476 | 0.680165 |
| 19 | 6851400 | 6861999 | 101.9888 | 157.5357 | 174.2816 | 72.39291 | 0.155233 | 2.869635 | 0.680167 |
| 18 | 75945800 | 75947999 | 27.15762 | 46.02425 | 50.6858 | 22.19627 | 0.242648 | 1.068462 | 0.680194 |
| 3 | 110946600 | 110947599 | 15.67596 | 28.33405 | 25.34633 | 11.0041 | 0.569519 | 0.365575 | 0.680195 |
| 6 | 114903800 | 114905199 | 21.36266 | 36.86283 | 42.08753 | 18.46334 | 0.194693 | 0.875478 | 0.680202 |
| 14 | 83685000 | 83693999 | 129.6614 | 197.9985 | 186.7701 | 77.46082 | 0.372286 | 3.101678 | 0.68024 |
| 10 | 77018000 | 77021599 | 33.21406 | 54.51837 | 62.25413 | 26.81145 | 0.186458 | 1.432076 | 0.680245 |
| 16 | 89790000 | 89799199 | 131.9566 | 198.5777 | 268.9346 | 111.0621 | -0.16669 | 3.400932 | 0.680255 |
| 17 | 38217000 | 38217999 | 16.83523 | 29.84776 | 33.22509 | 14.68778 | 0.238263 | 0.504621 | 0.680273 |
| 4 | 23437800 | 23441399 | 64.14769 | 101.2585 | 141.936 | 58.92471 | -0.16157 | 2.44968 | 0.680274 |
| 14 | 78149400 | 78151199 | 17.72962 | 32.03809 | 31.73346 | 13.78133 | 0.422974 | 0.393498 | 0.680294 |
| 15 | 82376600 | 82394199 | 236.5121 | 341.8932 | 444.278 | 183.2563 | -0.16156 | 4.088213 | 0.680302 |
| 4 | 85331400 | 85333799 | 19.26214 | 33.14416 | 49.53985 | 21.73071 | -0.19315 | 0.902423 | 0.680302 |
| 10 | 57013800 | 57016199 | 31.11489 | 51.28451 | 57.18454 | 24.67124 | 0.223097 | 1.373782 | 0.68031 |
| 17 | 18916600 | 18918399 | 18.11561 | 32.64044 | 32.2086 | 14.03868 | 0.425062 | 0.413916 | 0.680314 |
| 10 | 98675800 | 98684999 | 115.2892 | 177.99 | 181.3294 | 75.32059 | 0.273492 | 2.886041 | 0.680343 |
| 5 | 109867600 | 109874399 | 199.9981 | 295.6522 | 312.22 | 128.5798 | 0.168532 | 3.736479 | 0.680351 |
| 7 | 41154400 | 41157999 | 55.19919 | 88.22512 | 99.95524 | 42.02055 | 0.162364 | 2.078081 | 0.680377 |
| 6 | 132486000 | 132486999 | 12.59877 | 22.76888 | 35.79568 | 15.54341 | -0.23948 | 0.365919 | 0.680408 |

| | | | | | | | | | |
|---|---|---|---|---|---|---|---|---|---|
| 8 | 19337600 | 19340799 | 32.46251 | 53.73459 | 60.32215 | 26.0934 | 0.214291 | 1.327366 | 0.680413 |
| 5 | 18717000 | 18718199 | 9.618602 | 17.91578 | 28.91142 | 12.05678 | -0.23571 | 0.154433 | 0.680478 |
| 11 | 43554200 | 43554999 | 7.407839 | 14.2192 | 25.22188 | 9.889782 | -0.29404 | -0.08913 | 0.680509 |
| 2 | 144471600 | 144476399 | 60.61458 | 96.06538 | 134.5631 | 55.98201 | -0.1559 | 2.349963 | 0.680543 |
| 3 | 8868200 | 8872599 | 47.07359 | 75.36522 | 107.3199 | 45.22498 | -0.16422 | 2.02011 | 0.680574 |
| 4 | 8296800 | 8299799 | 29.2919 | 48.73848 | 55.62555 | 24.14812 | 0.191156 | 1.258161 | 0.680621 |
| X | 117631800 | 117633399 | 19.20895 | 33.80845 | 36.34199 | 16.08927 | 0.284464 | 0.603713 | 0.680634 |
| 10 | 48728600 | 48729599 | 12.40421 | 22.76972 | 26.06485 | 11.09472 | 0.23473 | 0.254039 | 0.680637 |
| 1 | 126981000 | 126982399 | 12.42817 | 22.63506 | 37.63865 | 16.18624 | -0.31101 | 0.323312 | 0.680643 |
| 4 | 8545200 | 8550799 | 57.28452 | 91.9923 | 94.77067 | 40.12178 | 0.307115 | 1.935631 | 0.680656 |
| 12 | 102370600 | 102373599 | 31.07584 | 51.42758 | 58.99498 | 25.51531 | 0.183202 | 1.326862 | 0.680672 |
| 5 | 48043600 | 48046199 | 30.04841 | 50.82156 | 52.38872 | 22.92809 | 0.33762 | 1.098029 | 0.680676 |
| 1 | 127777400 | 127781199 | 33.95411 | 56.17542 | 61.57321 | 26.62534 | 0.24861 | 1.338099 | 0.680687 |
| 13 | 7384400 | 7385599 | 20.61817 | 36.17408 | 37.89655 | 16.75568 | 0.3207 | 0.656364 | 0.680719 |
| 1 | 5513000 | 5513999 | 13.12321 | 23.98789 | 27.54755 | 11.79295 | 0.224342 | 0.287526 | 0.680724 |
| 10 | 81803600 | 81806999 | 69.61664 | 109.9279 | 118.7137 | 49.29228 | 0.214574 | 2.440808 | 0.680785 |
| X | 8659200 | 8671799 | 165.7994 | 251.7198 | 223.7172 | 92.63553 | 0.451438 | 3.233225 | 0.680798 |
| 18 | 18866000 | 18869799 | 38.50274 | 62.76543 | 70.41911 | 30.24696 | 0.206051 | 1.541377 | 0.680804 |
| 2 | 7612800 | 7619199 | 94.0644 | 146.8327 | 157.493 | 65.351 | 0.211534 | 2.681404 | 0.680818 |
| 11 | 66083600 | 66086999 | 28.15653 | 48.20771 | 47.9065 | 21.00799 | 0.392106 | 0.979556 | 0.680826 |
| 2 | 88402400 | 88410799 | 127.6875 | 194.9642 | 207.9031 | 86.22343 | 0.195393 | 3.104502 | 0.680853 |
| 8 | 126549200 | 126550599 | 13.38881 | 24.22271 | 27.02463 | 11.71901 | 0.253776 | 0.34643 | 0.680864 |
| 3 | 29008000 | 29008599 | 9.607989 | 18.6902 | 21.38979 | 7.899754 | 0.413934 | -0.27394 | 0.680875 |
| 5 | 100376000 | 100378199 | 25.09555 | 42.49976 | 47.97951 | 21.00665 | 0.206939 | 1.063102 | 0.680899 |
| 6 | 40881000 | 40881399 | 4.640835 | 9.071345 | 17.74529 | 6.112831 | -0.25874 | -0.59317 | 0.680916 |
| 10 | 130425000 | 130428999 | 75.04466 | 118.1521 | 131.776 | 54.67491 | 0.16482 | 2.507543 | 0.680938 |
| 7 | 64177800 | 64181799 | 35.74294 | 57.96759 | 84.95099 | 36.38984 | -0.18434 | 1.631897 | 0.68096 |
| 15 | 7395400 | 7397199 | 15.1029 | 26.49787 | 40.3884 | 17.85754 | -0.21738 | 0.632739 | 0.68097 |
| 6 | 13979000 | 13982999 | 98.89925 | 152.6173 | 167.223 | 69.46125 | 0.168053 | 2.885072 | 0.680972 |
| 3 | 43297000 | 43298799 | 25.09041 | 42.53944 | 48.50887 | 21.24575 | 0.192664 | 1.056424 | 0.68099 |
| 16 | 97153200 | 97154399 | 9.631508 | 17.90164 | 28.88393 | 12.08805 | -0.2395 | 0.168262 | 0.681005 |
| 8 | 120333800 | 120339399 | 61.07417 | 96.69264 | 136.1537 | 56.6089 | -0.16471 | 2.375456 | 0.681005 |
| 17 | 81303600 | 81304599 | 14.85619 | 26.8243 | 30.34723 | 13.19246 | 0.231311 | 0.374807 | 0.681027 |
| 4 | 79471400 | 79473799 | 36.95943 | 60.51942 | 57.96541 | 24.93884 | 0.438029 | 1.468024 | 0.68103 |
| 17 | 54847000 | 54849199 | 22.11094 | 38.40808 | 36.23978 | 15.90908 | 0.470379 | 0.807968 | 0.681031 |
| 7 | 92904800 | 92905799 | 11.7819 | 22.18914 | 25.78022 | 10.5516 | 0.256611 | 0.096003 | 0.681039 |
| X | 126959800 | 126962799 | 29.63215 | 50.18437 | 44.72653 | 19.58263 | 0.546865 | 1.077866 | 0.681046 |
| 8 | 3360800 | 3363199 | 17.07468 | 29.76933 | 45.18754 | 19.88312 | -0.21209 | 0.744131 | 0.681051 |
| X | 49459600 | 49465599 | 110.6236 | 170.2129 | 181.5993 | 75.4191 | 0.203252 | 2.947992 | 0.681107 |
| 3 | 55308600 | 55309999 | 12.87017 | 23.45212 | 26.51635 | 11.39585 | 0.242909 | 0.299811 | 0.681111 |
| 7 | 129835600 | 129840799 | 29.89081 | 48.65 | 84.45637 | 36.25555 | -0.42367 | 1.556369 | 0.681143 |
| 12 | 8687000 | 8689599 | 18.85273 | 32.4042 | 49.54296 | 21.73135 | -0.22611 | 0.908262 | 0.681145 |
| 13 | 45559800 | 45562399 | 19.41355 | 33.12661 | 50.60002 | 22.18313 | -0.22639 | 0.978953 | 0.681147 |
| 11 | 21118000 | 21121399 | 43.33975 | 69.41383 | 117.2451 | 49.43619 | -0.40942 | 2.00659 | 0.681147 |
| 2 | 3099800 | 3101599 | 21.69117 | 38.11918 | 31.45328 | 13.92614 | 0.663339 | 0.626355 | 0.681149 |
| 8 | 33641000 | 33642399 | 28.20503 | 45.69747 | 87.60821 | 37.50075 | -0.57211 | 1.642022 | 0.681152 |
| 16 | 88495200 | 88496199 | 12.31161 | 23.11278 | 26.20842 | 10.78506 | 0.284905 | 0.120067 | 0.681204 |

| | | | | | | | | | |
|---|---|---|---|---|---|---|---|---|---|
| 14 | 68790400 | 68793199 | 20.4649 | 35.0321 | 52.75984 | 23.13594 | -0.20536 | 0.958614 | 0.681208 |
| 3 | 20526000 | 20528799 | 27.93607 | 48.42531 | 44.00736 | 19.30821 | 0.524004 | 0.869092 | 0.681278 |
| 3 | 90884400 | 90886399 | 26.30655 | 44.80617 | 49.0253 | 21.48958 | 0.252629 | 1.019643 | 0.681287 |
| 19 | 57819200 | 57819799 | 7.317822 | 14.13748 | 24.8703 | 9.555313 | -0.25351 | -0.15328 | 0.681288 |
| 5 | 37919400 | 37921599 | 21.3294 | 36.38786 | 55.53939 | 24.34813 | -0.2255 | 0.992942 | 0.681289 |
| 19 | 9487000 | 9489399 | 39.99383 | 64.7581 | 73.26744 | 31.3481 | 0.185898 | 1.663641 | 0.681307 |
| 1 | 143387400 | 143388399 | 10.4124 | 19.35154 | 31.01344 | 12.98065 | -0.23017 | 0.179792 | 0.681337 |
| 9 | 37859000 | 37860199 | 21.21009 | 37.01514 | 40.2227 | 17.7093 | 0.268083 | 0.743915 | 0.681351 |
| X | 67929600 | 67932399 | 19.3827 | 33.59353 | 55.06107 | 24.15792 | -0.32353 | 0.846482 | 0.681394 |
| 17 | 62690400 | 62692199 | 18.0794 | 31.59566 | 35.24661 | 15.53244 | 0.230649 | 0.700991 | 0.681403 |
| X | 34808200 | 34808999 | 13.50258 | 24.82116 | 28.07941 | 11.92916 | 0.253768 | 0.260661 | 0.681428 |
| 16 | 5455800 | 5463399 | 91.12344 | 141.717 | 157.7098 | 65.47442 | 0.154308 | 2.741982 | 0.681429 |
| 8 | 19431800 | 19435799 | 61.24399 | 97.69085 | 108.1695 | 45.34543 | 0.192246 | 2.1471 | 0.681432 |
| 15 | 97519800 | 97529399 | 166.3392 | 246.5047 | 327.0598 | 134.7003 | -0.15722 | 3.691377 | 0.681433 |
| 2 | 111266000 | 111275199 | 146.9487 | 222.7503 | 232.1755 | 96.09257 | 0.219813 | 3.258575 | 0.681447 |
| 11 | 118286400 | 118288399 | 23.91391 | 39.65274 | 62.19573 | 26.93598 | -0.26638 | 1.290274 | 0.681448 |
| 19 | 36617600 | 36621199 | 36.86821 | 60.72442 | 65.06962 | 28.06474 | 0.27986 | 1.401526 | 0.681453 |
| Y | 41248000 | 41249799 | 14.75037 | 26.31631 | 22.70453 | 9.983923 | 0.6089 | 0.42815 | 0.681478 |
| 10 | 72005000 | 72006799 | 19.72971 | 34.84306 | 35.71367 | 15.80038 | 0.354627 | 0.569381 | 0.681525 |
| 5 | 122185600 | 122188999 | 24.46866 | 40.17764 | 66.81234 | 28.77887 | -0.35395 | 1.411273 | 0.681529 |
| 10 | 86833800 | 86834599 | 10.08999 | 18.72941 | 29.91055 | 12.53989 | -0.22678 | 0.181372 | 0.681532 |
| 12 | 88250800 | 88253999 | 70.05055 | 108.5146 | 193.2682 | 80.27832 | -0.52779 | 2.806253 | 0.681541 |
| 8 | 10396400 | 10398199 | 22.227 | 38.21133 | 43.68231 | 19.16077 | 0.191807 | 0.915041 | 0.681542 |
| 14 | 50486400 | 50487199 | 12.50411 | 23.41761 | 21.03234 | 8.690466 | 0.614628 | 0.126116 | 0.68156 |
| 9 | 98851800 | 98855999 | 43.79697 | 70.26581 | 100.7993 | 42.61221 | -0.17125 | 1.94935 | 0.681593 |
| 17 | 86730200 | 86735199 | 55.6666 | 88.50717 | 124.7422 | 52.06323 | -0.16024 | 2.2503 | 0.681619 |
| 3 | 106499200 | 106505999 | 68.31404 | 108.8022 | 113.8228 | 47.61468 | 0.271681 | 2.203417 | 0.68163 |
| 5 | 18652600 | 18653799 | 8.410577 | 15.914 | 26.53058 | 10.75842 | -0.24841 | 0.037834 | 0.681653 |
| 5 | 73341800 | 73342999 | 16.87533 | 29.96498 | 33.25166 | 14.68044 | 0.244067 | 0.495963 | 0.681668 |
| 10 | 58845600 | 58847199 | 14.98113 | 26.27513 | 40.33573 | 17.82904 | -0.22786 | 0.634309 | 0.681685 |
| X | 61592600 | 61599399 | 81.28057 | 128.1798 | 137.8813 | 57.22026 | 0.218755 | 2.481965 | 0.681689 |
| 4 | 36587400 | 36588999 | 22.89079 | 39.94349 | 34.95733 | 15.38991 | 0.579016 | 0.747765 | 0.681692 |
| 18 | 16219400 | 16221999 | 31.85931 | 52.67197 | 60.19601 | 26.01697 | 0.188469 | 1.340386 | 0.681701 |
| 8 | 7036000 | 7036399 | 6.383602 | 12.46255 | 16.4475 | 5.808984 | 0.269705 | -0.48455 | 0.681708 |
| 12 | 11753000 | 11753999 | 11.57433 | 20.98527 | 32.7121 | 14.14709 | -0.22344 | 0.332663 | 0.681717 |
| 10 | 88483400 | 88488399 | 70.06144 | 109.882 | 154.1977 | 63.95256 | -0.17041 | 2.585372 | 0.681719 |
| 2 | 23693600 | 23695399 | 25.40558 | 42.92516 | 49.06432 | 21.46651 | 0.189089 | 1.087432 | 0.681723 |
| 11 | 34518400 | 34521599 | 19.3841 | 34.93937 | 31.53563 | 13.73878 | 0.553376 | 0.41842 | 0.681745 |
| 9 | 49141400 | 49142599 | 7.776448 | 14.73662 | 25.49992 | 10.30715 | -0.29766 | 0.01729 | 0.681762 |
| 16 | 64824600 | 64825799 | 15.12823 | 27.40974 | 30.42124 | 13.16779 | 0.262932 | 0.358714 | 0.681786 |
| 7 | 49408600 | 49410999 | 22.76925 | 37.76598 | 68.90595 | 29.84821 | -0.48393 | 1.28675 | 0.681789 |
| 1 | 142360800 | 142362199 | 29.34366 | 49.27183 | 43.34817 | 18.92722 | 0.565209 | 1.150424 | 0.681792 |
| 14 | 124619600 | 124623199 | 33.91621 | 56.8639 | 55.8891 | 24.37979 | 0.405911 | 1.1917 | 0.681794 |
| 5 | 53301000 | 53302999 | 17.48103 | 30.2556 | 45.83125 | 20.10751 | -0.21048 | 0.838068 | 0.681795 |
| 13 | 107059200 | 107064199 | 52.96252 | 85.11503 | 92.26649 | 39.10451 | 0.235091 | 1.905909 | 0.681798 |
| 7 | 130724800 | 130726599 | 17.2392 | 30.12805 | 47.83887 | 21.08203 | -0.27636 | 0.717613 | 0.68181 |
| 16 | 32395000 | 32401199 | 52.64136 | 83.60727 | 137.7963 | 57.44974 | -0.38716 | 2.282831 | 0.681828 |

| | | | | | | | | | |
|---|---|---|---|---|---|---|---|---|---|
| 13 | 26181600 | 26183199 | 18.81008 | 33.0605 | 36.36741 | 16.10373 | 0.25057 | 0.616009 | 0.681829 |
| 2 | 21015000 | 21015999 | 6.842213 | 13.12223 | 24.6316 | 9.682131 | -0.37816 | -0.08856 | 0.681845 |
| 8 | 45931000 | 45933399 | 17.02665 | 29.76807 | 46.21148 | 20.37149 | -0.24393 | 0.707283 | 0.681848 |
| 13 | 94439400 | 94440199 | 9.352405 | 17.57828 | 28.55253 | 11.72608 | -0.22696 | 0.095355 | 0.681879 |
| 5 | 138500800 | 138504999 | 52.72364 | 83.8301 | 139.5803 | 58.2576 | -0.40034 | 2.249665 | 0.681905 |
| 3 | 138355800 | 138363399 | 115.1964 | 175.3695 | 239.2793 | 99.16008 | -0.16334 | 3.163186 | 0.681909 |
| Y | 5892800 | 5893399 | 6.219453 | 11.76541 | 25.54265 | 10.36279 | -0.62653 | 0.014393 | 0.68192 |
| 3 | 153075600 | 153079999 | 47.64 | 76.76449 | 85.61654 | 36.42577 | 0.198826 | 1.813169 | 0.681951 |
| 14 | 73153400 | 73155199 | 22.26788 | 37.00719 | 67.82136 | 29.41837 | -0.49006 | 1.261432 | 0.68197 |
| 10 | 39162400 | 39166199 | 41.41055 | 66.73286 | 99.10444 | 42.16986 | -0.21324 | 1.808564 | 0.681972 |
| 4 | 24892400 | 24894399 | 21.53855 | 37.4821 | 41.37849 | 18.18579 | 0.245039 | 0.786596 | 0.681976 |
| 18 | 47054400 | 47055199 | 7.477429 | 14.20956 | 25.01809 | 10.05288 | -0.3148 | -0.00586 | 0.681996 |
| 4 | 72923200 | 72923799 | 15.28654 | 27.51778 | 30.87405 | 13.47247 | 0.239862 | 0.397567 | 0.682005 |
| 10 | 12461800 | 12462799 | 10.37742 | 20.00622 | 23.09426 | 8.942958 | 0.337208 | -0.10462 | 0.682039 |
| 19 | 54994600 | 54996799 | 25.08594 | 41.86795 | 62.06483 | 26.99606 | -0.18494 | 1.217905 | 0.682041 |
| 1 | 140935400 | 140940999 | 111.8588 | 172.8431 | 159.5505 | 66.27378 | 0.416556 | 2.865543 | 0.682055 |
| 17 | 20932200 | 20943599 | 175.2045 | 262.5642 | 270.3396 | 111.5222 | 0.223152 | 3.490259 | 0.682082 |
| X | 26114200 | 26115599 | 42.16668 | 68.32672 | 75.63765 | 32.3798 | 0.218316 | 1.657637 | 0.682096 |
| 1 | 18830000 | 18834799 | 74.49071 | 116.7147 | 161.6686 | 67.05784 | -0.15278 | 2.605448 | 0.682098 |
| 17 | 25925400 | 25929199 | 23.5856 | 39.15252 | 69.54122 | 30.14066 | -0.44518 | 1.280959 | 0.682102 |
| 7 | 105966400 | 105986999 | 370.5048 | 514.2846 | 637.4882 | 264.6383 | -0.15224 | 4.693197 | 0.682103 |
| 7 | 31258200 | 31259199 | 22.67945 | 38.81281 | 44.23543 | 19.39745 | 0.195079 | 0.957643 | 0.682117 |
| 11 | 19631400 | 19635399 | 40.42899 | 66.07103 | 69.80436 | 30.01034 | 0.295048 | 1.509127 | 0.682122 |
| 16 | 26771800 | 26773399 | 14.08102 | 24.87404 | 37.6868 | 16.67244 | -0.20603 | 0.536656 | 0.682152 |
| 18 | 19943000 | 19945799 | 37.10254 | 60.62213 | 68.5495 | 29.46861 | 0.197136 | 1.506272 | 0.682203 |
| 7 | 5081600 | 5083799 | 16.33752 | 28.31732 | 44.27813 | 19.42696 | -0.25558 | 0.817238 | 0.682207 |
| 19 | 39316600 | 39317799 | 17.29684 | 30.96418 | 32.76753 | 14.36787 | 0.319006 | 0.448795 | 0.682212 |
| 1 | 194149800 | 194152799 | 43.12404 | 69.20636 | 99.65295 | 42.14624 | -0.17615 | 1.938642 | 0.682219 |
| 4 | 100419400 | 100420599 | 12.22582 | 22.08299 | 34.19799 | 14.85965 | -0.21851 | 0.362247 | 0.68223 |
| 4 | 91724600 | 91726399 | 29.4735 | 49.83672 | 45.29016 | 19.81988 | 0.518805 | 1.091355 | 0.68227 |
| 6 | 50417200 | 50418599 | 12.60409 | 22.61761 | 34.66748 | 15.17022 | -0.21038 | 0.406137 | 0.68229 |
| 7 | 38419400 | 38421399 | 71.12701 | 112.1982 | 122.1575 | 50.69672 | 0.201653 | 2.466565 | 0.682368 |
| 3 | 92571400 | 92571999 | 8.143417 | 15.77484 | 19.93024 | 7.561683 | 0.236794 | -0.20311 | 0.682379 |
| 18 | 74545800 | 74557199 | 124.7172 | 188.7257 | 255.0824 | 105.5159 | -0.15691 | 3.288411 | 0.6824 |
| 7 | 43792800 | 43793199 | 8.997668 | 17.26248 | 20.79501 | 8.165299 | 0.257933 | -0.0797 | 0.682427 |
| X | 152948800 | 152949799 | 19.28293 | 33.81632 | 37.07638 | 16.38686 | 0.255529 | 0.657173 | 0.682446 |
| 14 | 45606600 | 45607599 | 13.02268 | 24.75099 | 26.03311 | 10.45749 | 0.422569 | 0.047204 | 0.682451 |
| 17 | 88130200 | 88130799 | 8.008238 | 15.32897 | 26.26494 | 10.38133 | -0.25539 | -0.05384 | 0.682451 |
| 13 | 102555400 | 102558599 | 29.77383 | 48.87401 | 71.98958 | 31.00497 | -0.17978 | 1.43152 | 0.682472 |
| 5 | 40411800 | 40412799 | 13.94133 | 25.27443 | 28.97517 | 12.53246 | 0.217186 | 0.343524 | 0.682479 |
| 17 | 45226000 | 45226799 | 13.85604 | 25.2297 | 28.93396 | 12.4465 | 0.221575 | 0.316602 | 0.682492 |
| 7 | 62553400 | 62555599 | 30.53882 | 50.93171 | 56.21412 | 24.44001 | 0.239333 | 1.238794 | 0.682493 |
| 15 | 35049600 | 35050799 | 17.45115 | 31.41198 | 32.24682 | 14.07284 | 0.36718 | 0.415221 | 0.682497 |
| 12 | 68033200 | 68033599 | 4.777943 | 9.322901 | 18.65326 | 6.648731 | -0.3349 | -0.46534 | 0.682511 |
| 4 | 136586800 | 136587599 | 15.24698 | 27.03888 | 30.00425 | 13.26105 | 0.243188 | 0.488005 | 0.682514 |
| 14 | 19855000 | 19856999 | 17.55309 | 30.86132 | 34.98274 | 15.49079 | 0.207725 | 0.603631 | 0.682515 |
| 3 | 22122000 | 22124599 | 29.32585 | 49.14139 | 53.56062 | 23.35618 | 0.257105 | 1.177047 | 0.682518 |

| | | | | | | | | | |
|---|---|---|---|---|---|---|---|---|---|
| 7 | 39012000 | 39016199 | 173.8437 | 258.6905 | 276.9982 | 114.1302 | 0.157365 | 3.61801 | 0.682544 |
| 1 | 147968400 | 147970599 | 48.24888 | 77.68756 | 86.80184 | 36.89009 | 0.194901 | 1.836929 | 0.68258 |
| 17 | 88050000 | 88059399 | 108.146 | 164.565 | 227.431 | 94.2378 | -0.18229 | 3.169646 | 0.682605 |
| 9 | 36449000 | 36449999 | 14.05538 | 24.80792 | 37.68407 | 16.67402 | -0.21027 | 0.54498 | 0.682606 |
| 4 | 150008800 | 150013199 | 49.88414 | 79.69924 | 112.9728 | 47.46703 | -0.16093 | 2.088552 | 0.682632 |
| 8 | 91527200 | 91531799 | 55.55418 | 89.23335 | 94.04552 | 39.82821 | 0.274693 | 1.925468 | 0.682634 |
| 19 | 22990400 | 22993999 | 28.33255 | 47.3799 | 54.03104 | 23.53093 | 0.192172 | 1.19811 | 0.682636 |
| 9 | 117753400 | 117754399 | 7.151336 | 13.71017 | 24.08222 | 9.475929 | -0.284 | -0.08076 | 0.682647 |
| 17 | 36468600 | 36469199 | 7.151336 | 13.71017 | 24.08222 | 9.475929 | -0.284 | -0.08076 | 0.682647 |
| 6 | 12369000 | 12370799 | 28.27594 | 46.03007 | 82.43361 | 35.38957 | -0.46821 | 1.54913 | 0.682647 |
| 19 | 21327000 | 21327999 | 10.4566 | 19.08996 | 30.76471 | 13.19059 | -0.26295 | 0.280132 | 0.682647 |
| 1 | 46711600 | 46712799 | 15.00007 | 26.26257 | 40.42048 | 17.84023 | -0.23149 | 0.663246 | 0.682651 |
| 4 | 68913800 | 68914399 | 16.43051 | 29.05734 | 32.92197 | 14.5612 | 0.21169 | 0.530055 | 0.682653 |
| 6 | 78246000 | 78248999 | 33.25206 | 54.14097 | 79.27349 | 34.03527 | -0.17829 | 1.55435 | 0.682682 |
| 5 | 120600800 | 120604399 | 32.60131 | 52.99946 | 77.82588 | 33.39494 | -0.18399 | 1.574111 | 0.682689 |
| Y | 9103000 | 9104999 | 13.76401 | 24.81719 | 21.68783 | 9.444737 | 0.599784 | 0.35373 | 0.682695 |
| 5 | 143578600 | 143585599 | 61.42059 | 97.82658 | 110.191 | 46.09763 | 0.165282 | 2.195338 | 0.68271 |
| 2 | 111074200 | 111081599 | 157.2696 | 237.6515 | 215.2753 | 89.01018 | 0.418277 | 3.321157 | 0.68271 |
| 13 | 85186600 | 85188799 | 24.07742 | 40.34284 | 59.84872 | 26.09671 | -0.18608 | 1.170722 | 0.682716 |
| 8 | 40876200 | 40877799 | 12.15549 | 21.45644 | 36.75408 | 16.26225 | -0.38264 | 0.528938 | 0.682717 |
| 4 | 90255200 | 90257999 | 35.39315 | 57.79079 | 66.0833 | 28.40155 | 0.180354 | 1.509314 | 0.68275 |
| 7 | 59301200 | 59314999 | 198.7338 | 296.0098 | 292.1589 | 120.3884 | 0.276105 | 3.607224 | 0.682768 |
| 13 | 50655400 | 50660999 | 80.51928 | 126.2488 | 141.4787 | 58.67912 | 0.153383 | 2.593753 | 0.682777 |
| 1 | 128562200 | 128565199 | 20.77944 | 35.34033 | 52.72884 | 23.11025 | -0.19336 | 1.018362 | 0.682792 |
| 18 | 37645600 | 37656799 | 326.336 | 460.8232 | 464.8689 | 192.4581 | 0.169833 | 4.434945 | 0.682813 |
| 1 | 84918400 | 84920199 | 14.80273 | 25.9957 | 40.27617 | 17.8221 | -0.24115 | 0.611845 | 0.682844 |
| 9 | 7870400 | 7873999 | 23.96938 | 39.93076 | 60.26142 | 26.18118 | -0.21057 | 1.233034 | 0.682857 |
| 19 | 39398000 | 39404399 | 82.58926 | 130.2549 | 121.1013 | 50.25717 | 0.428959 | 2.478666 | 0.682878 |
| 6 | 8618800 | 8623399 | 38.1983 | 62.7504 | 66.82254 | 28.78882 | 0.287349 | 1.437607 | 0.682891 |
| 1 | 137595800 | 137597799 | 13.79494 | 24.97694 | 28.36453 | 12.28806 | 0.228999 | 0.346301 | 0.682898 |
| 5 | 46711400 | 46712199 | 10.79387 | 20.32894 | 24.22231 | 9.913471 | 0.220633 | 0.083151 | 0.6829 |
| 2 | 180201200 | 180203599 | 16.03805 | 28.52387 | 32.30214 | 14.242 | 0.21625 | 0.478777 | 0.682914 |
| 19 | 9354000 | 9362399 | 114.2759 | 176.358 | 182.3702 | 75.75307 | 0.25154 | 2.892088 | 0.682926 |
| 2 | 65327600 | 65328199 | 6.824918 | 13.20988 | 23.2623 | 8.859846 | -0.24335 | -0.19559 | 0.682933 |
| 17 | 37581000 | 37583399 | 19.27886 | 32.49259 | 60.15878 | 26.30069 | -0.50433 | 1.099775 | 0.683002 |
| 7 | 35716000 | 35716999 | 9.183985 | 17.15985 | 27.92679 | 11.58464 | -0.24136 | 0.129616 | 0.683005 |
| 4 | 10339400 | 10341199 | 16.97237 | 29.89019 | 33.90674 | 15.00932 | 0.207491 | 0.578829 | 0.683023 |
| 5 | 72036000 | 72037999 | 20.28751 | 35.47051 | 39.16685 | 17.26658 | 0.245274 | 0.711768 | 0.683025 |
| 19 | 46082600 | 46085399 | 25.6221 | 42.73177 | 63.72876 | 27.70713 | -0.1936 | 1.229583 | 0.683026 |
| 5 | 55780200 | 55782399 | 19.72193 | 34.53401 | 38.12785 | 16.8301 | 0.245509 | 0.68438 | 0.683045 |
| 1 | 48139800 | 48140399 | 7.288639 | 14.01958 | 24.15646 | 9.415522 | -0.24376 | -0.1102 | 0.683053 |
| X | 126505400 | 126508599 | 42.11762 | 69.23404 | 66.61426 | 28.70701 | 0.433723 | 1.44016 | 0.683059 |
| 18 | 82745800 | 82748799 | 30.76738 | 49.78887 | 89.38558 | 38.22755 | -0.47938 | 1.669485 | 0.683075 |
| 5 | 110043400 | 110051399 | 132.4603 | 201.1495 | 219.6309 | 90.94969 | 0.15496 | 3.217857 | 0.68312 |
| X | 153467200 | 153471599 | 91.2327 | 142.7568 | 152.8073 | 63.39094 | 0.217195 | 2.63767 | 0.683124 |
| 4 | 81826400 | 81827399 | 11.61772 | 20.93676 | 32.6879 | 14.2441 | -0.23266 | 0.368731 | 0.683174 |
| 4 | 23656600 | 23659199 | 21.04029 | 35.69259 | 53.82857 | 23.57965 | -0.2092 | 1.041109 | 0.683179 |

| | | | | | | | | | |
|---|---|---|---|---|---|---|---|---|---|
| 1 | 190472200 | 190474399 | 22.51595 | 38.03352 | 57.14063 | 24.99818 | -0.20402 | 1.086353 | 0.6832 |
| 3 | 7015600 | 7016199 | 11.85941 | 22.47106 | 21.55091 | 8.713333 | 0.546506 | 0.045577 | 0.683208 |
| 10 | 30191600 | 30192999 | 26.28983 | 44.68306 | 41.1405 | 18.03031 | 0.500348 | 1.031262 | 0.683209 |
| 16 | 47410000 | 47412799 | 25.80902 | 44.01602 | 48.47805 | 21.25193 | 0.243468 | 1.005525 | 0.683211 |
| 1 | 100920600 | 100922799 | 32.02828 | 53.29093 | 49.75683 | 21.60023 | 0.479931 | 1.261314 | 0.683213 |
| 1 | 24368600 | 24370199 | 31.45266 | 51.84274 | 58.85986 | 25.39434 | 0.197136 | 1.375528 | 0.683217 |
| 12 | 82566200 | 82569399 | 28.04712 | 46.33093 | 68.867 | 29.74755 | -0.18973 | 1.346684 | 0.683225 |
| 12 | 15100200 | 15103199 | 22.90699 | 38.19216 | 67.95584 | 29.53934 | -0.44744 | 1.228593 | 0.683241 |
| 4 | 61114400 | 61115199 | 8.558586 | 16.04939 | 26.93867 | 11.10637 | -0.27835 | 0.098197 | 0.683246 |
| 18 | 5901800 | 5902599 | 11.13574 | 20.78412 | 23.69442 | 9.846983 | 0.26645 | 0.14274 | 0.683274 |
| 1 | 164747600 | 164749999 | 16.97358 | 30.46591 | 32.18579 | 14.08138 | 0.323436 | 0.427502 | 0.683293 |
| 5 | 141758200 | 141761799 | 37.7584 | 61.60689 | 69.67763 | 29.9382 | 0.195405 | 1.52818 | 0.683293 |
| 2 | 136711600 | 136714799 | 46.96587 | 75.52166 | 85.83673 | 36.40999 | 0.167951 | 1.876512 | 0.683341 |
| 1 | 97251800 | 97252799 | 14.24355 | 25.17802 | 38.17714 | 16.88709 | -0.2068 | 0.531141 | 0.683351 |
| 4 | 135519200 | 135520799 | 17.2753 | 29.85679 | 45.83363 | 20.10767 | -0.23012 | 0.848622 | 0.683351 |
| 7 | 20809800 | 20811199 | 59.07547 | 94.04632 | 102.1011 | 42.68495 | 0.217647 | 2.208401 | 0.683358 |
| X | 77927400 | 77938399 | 207.4724 | 305.5442 | 320.9732 | 132.1792 | 0.171013 | 3.799945 | 0.683378 |
| 7 | 60631000 | 60634999 | 38.82101 | 64.55027 | 59.56932 | 25.84813 | 0.496698 | 1.296352 | 0.683403 |
| 8 | 65887200 | 65889599 | 16.76975 | 29.70162 | 33.34389 | 14.74341 | 0.225682 | 0.516772 | 0.683451 |
| 4 | 128320200 | 128321399 | 12.43666 | 22.4478 | 34.72608 | 15.10258 | -0.21773 | 0.370061 | 0.683458 |
| 4 | 51104000 | 51106999 | 20.60255 | 36.04287 | 38.50144 | 16.98181 | 0.292701 | 0.700621 | 0.683473 |
| 3 | 139341400 | 139342799 | 14.28884 | 25.94232 | 29.35434 | 12.67327 | 0.237351 | 0.336366 | 0.683484 |
| 11 | 48717400 | 48719999 | 19.73067 | 34.28247 | 38.94352 | 17.09886 | 0.203858 | 0.796956 | 0.683487 |
| 13 | 95058200 | 95059199 | 14.17628 | 26.96007 | 26.05071 | 10.45099 | 0.545633 | 0.053183 | 0.68351 |
| 2 | 47582000 | 47585999 | 39.65304 | 64.27218 | 72.82329 | 31.18163 | 0.185175 | 1.642049 | 0.683514 |
| 1 | 89992000 | 89993399 | 13.61419 | 24.80163 | 28.50736 | 12.25538 | 0.219103 | 0.312402 | 0.683521 |
| 9 | 19915800 | 19920999 | 67.83077 | 108.1485 | 111.8999 | 46.87982 | 0.289208 | 2.170399 | 0.683546 |
| 19 | 49434200 | 49434599 | 13.76408 | 24.79297 | 26.62128 | 11.60742 | 0.303801 | 0.37443 | 0.683597 |
| 13 | 13469000 | 13470799 | 21.88086 | 38.13162 | 35.12769 | 15.45228 | 0.505228 | 0.761023 | 0.6836 |
| 12 | 96882400 | 96884999 | 20.67204 | 35.51062 | 56.16321 | 24.63479 | -0.2751 | 0.928805 | 0.683608 |
| 9 | 98013000 | 98020199 | 78.43923 | 123.576 | 137.5326 | 57.06812 | 0.168599 | 2.499155 | 0.683618 |
| 1 | 144826200 | 144830799 | 62.08714 | 98.80843 | 111.6114 | 46.63975 | 0.159974 | 2.222151 | 0.683618 |
| 1 | 18841800 | 18842999 | 16.78718 | 29.93405 | 27.73408 | 12.20113 | 0.506444 | 0.458946 | 0.683673 |
| 8 | 85842000 | 85842599 | 8.869255 | 17.04772 | 20.83478 | 8.137437 | 0.244597 | -0.09759 | 0.683686 |
| 7 | 23896400 | 23908599 | 185.2647 | 272.2125 | 358.4269 | 147.6146 | -0.15783 | 3.834284 | 0.683712 |
| 5 | 5110400 | 5113999 | 34.88414 | 58.0785 | 59.37168 | 25.78475 | 0.349564 | 1.269808 | 0.683714 |
| 1 | 166014400 | 166018599 | 37.04703 | 60.39146 | 69.01449 | 29.64347 | 0.179561 | 1.538153 | 0.683728 |
| 11 | 45258600 | 45259599 | 13.5168 | 23.74352 | 37.92136 | 16.78374 | -0.28422 | 0.601041 | 0.683737 |
| 6 | 143473600 | 143476399 | 31.00311 | 51.4465 | 58.5425 | 25.36705 | 0.195056 | 1.294199 | 0.683765 |
| 14 | 44460800 | 44461599 | 12.37349 | 22.81105 | 26.54743 | 11.23432 | 0.216959 | 0.23036 | 0.683766 |
| 17 | 55837800 | 55840799 | 28.20218 | 47.13578 | 54.01168 | 23.51428 | 0.185294 | 1.2041 | 0.683771 |
| 14 | 119340000 | 119340399 | 4.669597 | 9.127494 | 18.36979 | 6.329257 | -0.30062 | -0.59272 | 0.683795 |
| 18 | 26790400 | 26792999 | 19.87439 | 34.19525 | 51.71575 | 22.68512 | -0.21018 | 0.907106 | 0.683798 |
| 10 | 59076600 | 59080399 | 41.84008 | 67.20242 | 97.17747 | 41.15197 | -0.18078 | 1.910602 | 0.683807 |
| 16 | 82303800 | 82304999 | 13.67298 | 23.96648 | 44.07769 | 19.47322 | -0.48707 | 0.645099 | 0.683841 |
| 2 | 78145400 | 78150599 | 71.42902 | 112.9762 | 127.2188 | 52.86793 | 0.15618 | 2.404785 | 0.683885 |
| 19 | 33710600 | 33711599 | 17.75708 | 31.38603 | 27.40469 | 12.12228 | 0.585234 | 0.533314 | 0.68389 |

| | | | | | | | | | |
|---|---|---|---|---|---|---|---|---|---|
| 14 | 50288000 | 50289399 | 15.68418 | 27.49141 | 41.39902 | 18.28959 | -0.20018 | 0.651311 | 0.683915 |
| 2 | 169319800 | 169321399 | 22.5887 | 38.09308 | 56.53504 | 24.72079 | -0.18646 | 1.102079 | 0.683928 |
| 12 | 90560800 | 90562599 | 24.2068 | 41.381 | 46.86379 | 20.54817 | 0.203964 | 0.977172 | 0.683932 |
| 3 | 52297400 | 52298799 | 16.01076 | 27.88989 | 42.63621 | 18.75014 | -0.22219 | 0.749455 | 0.683976 |
| 7 | 70923200 | 70923799 | 6.254602 | 12.126 | 22.17985 | 8.382708 | -0.28591 | -0.22823 | 0.683983 |
| X | 149073800 | 149076199 | 38.65914 | 63.25235 | 68.76613 | 29.57655 | 0.255116 | 1.491547 | 0.684001 |
| 8 | 97351800 | 97358599 | 140.3677 | 212.3979 | 231.5831 | 95.79381 | 0.152711 | 3.291931 | 0.684024 |
| 8 | 71980400 | 71983199 | 14.64506 | 25.43082 | 48.44712 | 21.26355 | -0.53875 | 0.795968 | 0.68406 |
| 6 | 32950400 | 32950799 | 5.193937 | 10.13662 | 19.82824 | 7.043273 | -0.29841 | -0.46624 | 0.684077 |
| 13 | 39221000 | 39223399 | 48.4469 | 77.25203 | 112.7828 | 47.25727 | -0.20646 | 2.148115 | 0.684097 |
| 8 | 3974200 | 3977399 | 34.40447 | 55.85973 | 81.43207 | 34.91144 | -0.17523 | 1.60441 | 0.684107 |
| X | 161134400 | 161136399 | 15.30633 | 26.84632 | 41.43771 | 18.31661 | -0.23542 | 0.641495 | 0.68411 |
| 18 | 8085800 | 8092199 | 140.6223 | 213.5279 | 201.7817 | 83.55639 | 0.363145 | 3.219656 | 0.684143 |
| 5 | 139572600 | 139574399 | 26.9207 | 45.23131 | 51.77631 | 22.61465 | 0.186683 | 1.143565 | 0.684167 |
| 8 | 112981400 | 112981999 | 8.6421 | 16.731 | 21.01185 | 7.994026 | 0.241657 | -0.18429 | 0.684167 |
| 6 | 104930000 | 104933799 | 37.75624 | 62.14245 | 65.24612 | 28.13232 | 0.308737 | 1.410999 | 0.684169 |
| 12 | 78382800 | 78385999 | 53.2077 | 85.61739 | 90.06809 | 38.24289 | 0.280523 | 1.865186 | 0.684181 |
| 3 | 144500400 | 144501799 | 17.78261 | 32.06953 | 31.63169 | 13.7716 | 0.426554 | 0.403856 | 0.684181 |
| 6 | 149194200 | 149196199 | 35.57062 | 58.285 | 66.34064 | 28.5504 | 0.190113 | 1.462362 | 0.684185 |
| 13 | 71772200 | 71773399 | 11.68892 | 20.7989 | 33.7571 | 14.87745 | -0.29938 | 0.451247 | 0.6842 |
| 2 | 20184600 | 20186199 | 31.18062 | 51.62902 | 59.06821 | 25.55657 | 0.187066 | 1.319931 | 0.684202 |
| 4 | 63839600 | 63841199 | 19.13986 | 33.60391 | 27.55964 | 12.19249 | 0.672178 | 0.620005 | 0.684215 |
| 13 | 23429400 | 23429799 | 5.653789 | 11.00816 | 20.72285 | 7.597632 | -0.28567 | -0.34595 | 0.684218 |
| 17 | 67305600 | 67308799 | 65.3981 | 103.7583 | 117.089 | 48.76489 | 0.156972 | 2.317672 | 0.684222 |
| 9 | 120857600 | 120859199 | 15.10117 | 26.3881 | 41.06711 | 18.0958 | -0.24719 | 0.699398 | 0.684237 |
| 15 | 94019600 | 94024799 | 70.94792 | 112.7562 | 120.0449 | 50.08162 | 0.243253 | 2.274486 | 0.684261 |
| 17 | 39827800 | 39830999 | 35.56212 | 59.14648 | 59.97597 | 26.02914 | 0.361143 | 1.283647 | 0.684289 |
| 17 | 61028400 | 61030799 | 17.65742 | 30.77977 | 47.33731 | 20.82669 | -0.23089 | 0.754241 | 0.684294 |
| 8 | 118191200 | 118192199 | 13.23208 | 24.24288 | 27.38691 | 11.68791 | 0.250564 | 0.271856 | 0.684318 |
| 9 | 83859200 | 83862199 | 38.70361 | 62.79335 | 71.5863 | 30.67362 | 0.177628 | 1.621173 | 0.684356 |
| 10 | 47809600 | 47810599 | 12.9071 | 24.15056 | 26.64527 | 11.02652 | 0.317575 | 0.146314 | 0.68436 |
| 18 | 4895000 | 4895999 | 10.16456 | 19.05318 | 31.83051 | 13.13228 | -0.27285 | 0.126839 | 0.684361 |
| 19 | 9363800 | 9367599 | 40.64739 | 65.40671 | 94.37266 | 40.07089 | -0.17434 | 1.849286 | 0.68441 |
| 17 | 33370800 | 33371199 | 8.209808 | 15.57787 | 26.03672 | 10.49553 | -0.24482 | 0.013361 | 0.68441 |
| 16 | 65724400 | 65726399 | 17.58504 | 30.5371 | 45.79813 | 20.10134 | -0.19533 | 0.80466 | 0.684438 |
| 1 | 108899600 | 108900999 | 15.25346 | 27.81727 | 29.84942 | 12.81543 | 0.318554 | 0.319029 | 0.684445 |
| 12 | 44308200 | 44309799 | 32.42791 | 53.34133 | 61.10649 | 26.34089 | 0.183061 | 1.404121 | 0.684446 |
| 15 | 22711000 | 22712999 | 15.52709 | 27.2987 | 41.81248 | 18.51517 | -0.22449 | 0.603347 | 0.684495 |
| 3 | 77097200 | 77097799 | 12.17683 | 21.6454 | 34.24126 | 15.10356 | -0.2634 | 0.462514 | 0.684504 |
| 4 | 46184000 | 46186799 | 19.36279 | 33.71451 | 38.33441 | 16.8538 | 0.202731 | 0.763673 | 0.684505 |
| 3 | 98148200 | 98149199 | 6.715773 | 12.88378 | 23.56906 | 9.256141 | -0.33925 | -0.0928 | 0.68451 |
| 14 | 97162600 | 97164799 | 22.18571 | 38.02301 | 43.19368 | 18.94285 | 0.200108 | 0.941369 | 0.684529 |
| 5 | 62489400 | 62489799 | 5.665556 | 11.02421 | 20.70131 | 7.627971 | -0.28984 | -0.33194 | 0.684533 |
| 18 | 47330000 | 47330599 | 5.665556 | 11.02421 | 20.70131 | 7.627971 | -0.28984 | -0.33194 | 0.684533 |
| 11 | 60875800 | 60877999 | 19.5669 | 33.57372 | 50.07486 | 21.96269 | -0.1909 | 0.929126 | 0.684554 |
| 2 | 22135400 | 22136799 | 13.76222 | 24.49588 | 37.67131 | 16.59806 | -0.22208 | 0.470393 | 0.68456 |
| 4 | 8489800 | 8491799 | 11.73769 | 21.82317 | 25.61544 | 10.71518 | 0.216806 | 0.176029 | 0.684565 |

| | | | | | | | | | |
|---|---|---|---|---|---|---|---|---|---|
| 8 | 47644200 | 47648599 | 61.53957 | 97.15193 | 138.8723 | 57.65594 | -0.18964 | 2.445146 | 0.68457 |
| 1 | 160067200 | 160069199 | 18.72872 | 32.70754 | 37.22707 | 16.39757 | 0.201683 | 0.717206 | 0.684576 |
| 8 | 47610400 | 47611399 | 12.94943 | 23.88837 | 27.59256 | 11.66625 | 0.228802 | 0.234868 | 0.684577 |
| 8 | 65842600 | 65843599 | 11.72115 | 21.20048 | 33.18451 | 14.39456 | -0.23226 | 0.348001 | 0.684601 |
| 5 | 100444800 | 100449399 | 43.31671 | 69.73223 | 103.7545 | 44.08218 | -0.21787 | 1.84385 | 0.684613 |
| 1 | 103508600 | 103517399 | 80.01936 | 127.0207 | 123.0095 | 51.25679 | 0.378044 | 2.316745 | 0.684641 |
| 19 | 3141400 | 3155799 | 114.6042 | 177.5951 | 150.4593 | 62.49654 | 0.543362 | 2.814242 | 0.684648 |
| 7 | 113588200 | 113592399 | 38.54534 | 62.26712 | 90.50974 | 38.63429 | -0.17827 | 1.728513 | 0.684648 |
| 17 | 33180600 | 33181999 | 18.59293 | 32.64432 | 36.4172 | 16.11139 | 0.230407 | 0.630944 | 0.684656 |
| 10 | 87320600 | 87322599 | 18.79577 | 32.04156 | 50.06271 | 21.94587 | -0.25907 | 0.985038 | 0.684659 |
| 10 | 95996600 | 95999199 | 33.70629 | 55.54465 | 63.06085 | 27.20418 | 0.19699 | 1.385812 | 0.684669 |
| 13 | 76847200 | 76849199 | 24.57449 | 42.32303 | 45.73897 | 20.064 | 0.273052 | 0.911652 | 0.684677 |
| 4 | 89143000 | 89144999 | 24.06941 | 40.28422 | 59.84401 | 26.07619 | -0.18804 | 1.182306 | 0.684677 |
| 9 | 52429400 | 52431799 | 27.05519 | 45.59075 | 51.78426 | 22.63824 | 0.197968 | 1.118416 | 0.684721 |
| 6 | 101612000 | 101613199 | 13.6507 | 25.44426 | 27.66059 | 11.52117 | 0.331453 | 0.179783 | 0.68473 |
| 8 | 7084200 | 7085799 | 16.3005 | 29.59916 | 30.80491 | 13.29698 | 0.357507 | 0.353365 | 0.684731 |
| 8 | 11688800 | 11692799 | 36.99678 | 60.2605 | 68.44784 | 29.39137 | 0.187461 | 1.548312 | 0.684743 |
| 13 | 77918400 | 77918999 | 13.03636 | 23.95167 | 26.88507 | 11.42985 | 0.26366 | 0.253166 | 0.68476 |
| 5 | 115551800 | 115553999 | 27.66131 | 46.27621 | 52.32156 | 22.79228 | 0.204612 | 1.19302 | 0.684768 |
| 1 | 9316000 | 9317399 | 14.13267 | 26.12566 | 28.61987 | 12.06626 | 0.307643 | 0.235006 | 0.684787 |
| X | 12370800 | 12372799 | 14.47331 | 25.63679 | 40.24872 | 17.79612 | -0.25556 | 0.51491 | 0.684799 |
| 2 | 28319200 | 28320999 | 29.22514 | 48.56152 | 55.64864 | 24.13554 | 0.185149 | 1.272197 | 0.684799 |
| 14 | 44554800 | 44555399 | 9.174257 | 17.29169 | 28.06204 | 11.47024 | -0.22023 | 0.074314 | 0.684821 |
| 6 | 14453200 | 14454399 | 17.44885 | 30.79313 | 34.41784 | 15.22925 | 0.230027 | 0.562162 | 0.684823 |
| 1 | 8538400 | 8541799 | 36.72237 | 59.26729 | 86.90873 | 37.05287 | -0.19242 | 1.749992 | 0.684837 |
| 10 | 47583800 | 47585199 | 10.66587 | 19.83947 | 32.70024 | 13.66821 | -0.2687 | 0.182257 | 0.684843 |
| 14 | 55683600 | 55686799 | 66.70685 | 104.8512 | 148.3994 | 61.55884 | -0.18025 | 2.539449 | 0.684885 |
| 6 | 137201000 | 137201599 | 9.063814 | 16.90114 | 27.92741 | 11.62459 | -0.2669 | 0.142661 | 0.684895 |
| 17 | 37527400 | 37530199 | 31.5996 | 53.23107 | 54.09943 | 23.64799 | 0.357852 | 1.139295 | 0.684897 |
| 17 | 21228800 | 21229799 | 8.075589 | 15.26149 | 25.97954 | 10.56224 | -0.28179 | 0.040101 | 0.684928 |
| X | 8616800 | 8623399 | 92.07974 | 145.0262 | 141.5399 | 58.72887 | 0.357557 | 2.515771 | 0.684937 |
| 1 | 147819400 | 147820999 | 28.91535 | 47.66158 | 70.49592 | 30.41485 | -0.18339 | 1.376966 | 0.684948 |
| 3 | 155989400 | 155990799 | 11.49479 | 21.09173 | 34.00363 | 14.48198 | -0.25672 | 0.268495 | 0.684954 |
| 8 | 35613800 | 35616399 | 24.06507 | 40.44208 | 60.45044 | 26.40502 | -0.19698 | 1.140342 | 0.684962 |
| 10 | 41282000 | 41288599 | 81.82306 | 129.1515 | 121.3786 | 50.38872 | 0.41432 | 2.459054 | 0.684997 |
| 12 | 55574000 | 55575599 | 11.90022 | 21.3905 | 33.63806 | 14.6989 | -0.24587 | 0.386782 | 0.684999 |
| 6 | 59048600 | 59053399 | 100.5998 | 155.0925 | 170.5705 | 70.85211 | 0.161614 | 2.904013 | 0.685015 |
| 14 | 57085600 | 57095399 | 109.7207 | 169.7299 | 176.9711 | 73.50948 | 0.242254 | 2.84663 | 0.68504 |
| 15 | 25661000 | 25662199 | 12.65063 | 23.35936 | 27.0227 | 11.41054 | 0.227687 | 0.224016 | 0.685048 |
| 10 | 8580800 | 8584199 | 39.56771 | 64.54952 | 71.09461 | 30.5454 | 0.233421 | 1.534758 | 0.685084 |
| 2 | 20193200 | 20194599 | 32.76054 | 53.82991 | 61.64502 | 26.5608 | 0.182936 | 1.418652 | 0.685103 |
| 11 | 113630400 | 113634599 | 47.23293 | 75.45577 | 109.1666 | 45.8612 | -0.19051 | 2.087028 | 0.685106 |
| X | 147732200 | 147744999 | 155.5963 | 236.0808 | 230.1527 | 95.28236 | 0.317337 | 3.242539 | 0.685119 |
| 17 | 26611800 | 26615599 | 29.91347 | 48.98195 | 72.49902 | 31.19172 | -0.18819 | 1.461501 | 0.685145 |
| 9 | 40294400 | 40296799 | 35.78503 | 57.94037 | 84.45475 | 36.13338 | -0.17877 | 1.665843 | 0.685185 |
| 10 | 119696200 | 119710599 | 156.3713 | 232.1495 | 356.1966 | 146.7271 | -0.3644 | 3.659753 | 0.685192 |
| 1 | 111678600 | 111685399 | 66.70979 | 105.3599 | 148.0051 | 61.46114 | -0.16409 | 2.438752 | 0.685196 |

| | | | | | | | | | |
|---|---|---|---|---|---|---|---|---|---|
| 11 | 110458000 | 110458799 | 11.41015 | 21.51637 | 24.64471 | 10.06513 | 0.279004 | 0.079095 | 0.685198 |
| 14 | 40267400 | 40270399 | 32.57753 | 52.61694 | 92.57106 | 39.50485 | -0.4533 | 1.725837 | 0.685208 |
| 3 | 91607800 | 91609999 | 26.5982 | 45.3581 | 42.58771 | 18.66958 | 0.472854 | 1.003464 | 0.685214 |
| 18 | 84994000 | 84996399 | 15.87112 | 27.88929 | 43.40417 | 19.21632 | -0.24756 | 0.615006 | 0.685218 |
| 15 | 81081600 | 81083599 | 18.84153 | 32.57923 | 49.55958 | 21.7426 | -0.21708 | 0.854859 | 0.685243 |
| 6 | 135399200 | 135401199 | 21.90693 | 37.8521 | 42.53326 | 18.65962 | 0.217884 | 0.865005 | 0.685256 |
| 3 | 30730600 | 30733399 | 32.04176 | 53.17598 | 58.69904 | 25.43684 | 0.238733 | 1.294225 | 0.685257 |
| 14 | 100009400 | 100012199 | 19.91266 | 34.68236 | 38.74209 | 17.03626 | 0.228108 | 0.761173 | 0.68526 |
| 12 | 73685000 | 73687399 | 17.09257 | 29.74792 | 44.70092 | 19.64718 | -0.19777 | 0.769104 | 0.685278 |
| 7 | 64671800 | 64672399 | 10.42887 | 19.91725 | 23.90865 | 9.506381 | 0.246004 | -0.01637 | 0.685283 |
| 1 | 47823800 | 47830399 | 120.6356 | 185.0558 | 175.507 | 72.85862 | 0.369609 | 3.007626 | 0.685286 |
| 5 | 145659200 | 145665799 | 114.7286 | 177.1302 | 181.5292 | 75.40356 | 0.264955 | 2.884117 | 0.6853 |
| 15 | 31555600 | 31560399 | 50.45758 | 80.59506 | 115.3939 | 48.46698 | -0.1758 | 2.095957 | 0.685305 |
| 6 | 101305800 | 101308599 | 18.09399 | 31.07691 | 47.94966 | 21.03199 | -0.23935 | 0.910859 | 0.685344 |
| 14 | 6870800 | 6872199 | 17.46424 | 31.04597 | 28.67015 | 12.6456 | 0.50862 | 0.484291 | 0.685345 |
| 8 | 75443800 | 75445599 | 14.0476 | 24.81727 | 37.66184 | 16.66109 | -0.20838 | 0.534509 | 0.685364 |
| 3 | 112342400 | 112345199 | 37.71008 | 62.11217 | 56.58669 | 24.40621 | 0.513301 | 1.396512 | 0.685369 |
| 6 | 114971200 | 114971999 | 10.65104 | 19.43556 | 30.78294 | 13.20638 | -0.23848 | 0.285847 | 0.685374 |
| 6 | 118550000 | 118553799 | 34.21399 | 55.67325 | 82.42364 | 35.37989 | -0.19487 | 1.565992 | 0.685375 |
| 9 | 98790400 | 98794599 | 33.83395 | 56.30698 | 59.32982 | 25.7597 | 0.305995 | 1.270446 | 0.685387 |
| 1 | 112403200 | 112404199 | 13.90192 | 25.15571 | 28.94536 | 12.54899 | 0.209661 | 0.353286 | 0.685393 |
| 6 | 110311200 | 110311599 | 6.088966 | 11.83628 | 22.39128 | 8.317416 | -0.31209 | -0.29527 | 0.685394 |
| 19 | 8371800 | 8372799 | 14.24958 | 25.67605 | 28.78265 | 12.54459 | 0.241937 | 0.37748 | 0.685399 |
| 1 | 172090600 | 172094599 | 43.82997 | 70.61315 | 79.44093 | 33.78997 | 0.186073 | 1.809909 | 0.685403 |
| 19 | 38979200 | 38980599 | 18.8284 | 32.01759 | 50.58289 | 22.16946 | -0.27553 | 1.008989 | 0.685423 |
| 18 | 33176600 | 33177999 | 19.6798 | 32.90241 | 63.85235 | 27.80408 | -0.57214 | 1.184753 | 0.685452 |
| 2 | 111499200 | 111500599 | 15.28686 | 27.75304 | 30.33056 | 13.09547 | 0.287142 | 0.346024 | 0.685461 |
| 8 | 12588800 | 12590999 | 25.87141 | 43.12212 | 64.4069 | 27.99154 | -0.19581 | 1.23518 | 0.685476 |
| 3 | 101146200 | 101147199 | 9.961225 | 18.49772 | 29.39725 | 12.31805 | -0.21907 | 0.177834 | 0.68548 |
| 7 | 27429600 | 27431199 | 15.95715 | 27.73054 | 43.70744 | 19.19256 | -0.26654 | 0.782681 | 0.685486 |
| 8 | 44877200 | 44882199 | 66.17234 | 104.223 | 145.9559 | 60.56126 | -0.16273 | 2.498354 | 0.685491 |
| 2 | 141260600 | 141261999 | 10.88169 | 20.18675 | 33.19399 | 13.92795 | -0.26979 | 0.197935 | 0.685498 |
| 17 | 44810200 | 44811599 | 16.662 | 29.37995 | 33.41102 | 14.78622 | 0.204626 | 0.562891 | 0.685507 |
| 19 | 37765400 | 37767799 | 23.17394 | 39.60202 | 45.28598 | 19.85582 | 0.189955 | 0.973755 | 0.68551 |
| 6 | 144815000 | 144816999 | 22.14354 | 37.24526 | 55.92851 | 24.43626 | -0.2034 | 1.123359 | 0.685511 |
| 16 | 70355000 | 70355999 | 9.931167 | 18.10595 | 30.37556 | 13.04602 | -0.32156 | 0.28663 | 0.685517 |
| X | 71448800 | 71450199 | 13.97165 | 25.43105 | 29.00204 | 12.4814 | 0.229402 | 0.321353 | 0.685535 |
| 10 | 43731000 | 43734799 | 36.24548 | 59.08863 | 67.72526 | 29.0904 | 0.175395 | 1.535212 | 0.685536 |
| 3 | 139632000 | 139634799 | 35.6705 | 58.22334 | 66.17295 | 28.43646 | 0.188757 | 1.513931 | 0.685543 |
| 5 | 26532400 | 26534799 | 53.40671 | 85.43689 | 96.83358 | 40.76004 | 0.16323 | 2.048143 | 0.685561 |
| 6 | 4695400 | 4696999 | 14.595 | 26.65123 | 29.3803 | 12.59343 | 0.281376 | 0.305784 | 0.685589 |
| 16 | 82641400 | 82642599 | 12.03147 | 22.71038 | 25.18913 | 10.2667 | 0.327531 | 0.079184 | 0.685622 |
| 1 | 133318600 | 133320999 | 22.21977 | 37.19192 | 57.45426 | 25.03633 | -0.2441 | 1.176105 | 0.685627 |
| 3 | 90689000 | 90689399 | 7.920861 | 15.03885 | 25.52068 | 10.27413 | -0.26468 | 0.005667 | 0.685648 |
| 4 | 90226000 | 90228799 | 28.47963 | 48.0832 | 52.37293 | 22.9089 | 0.258299 | 1.106589 | 0.685651 |
| 19 | 7388800 | 7392599 | 37.70368 | 61.3185 | 70.05272 | 30.06381 | 0.17763 | 1.573179 | 0.685656 |
| 16 | 64985400 | 64986599 | 15.14637 | 27.17852 | 30.78499 | 13.47176 | 0.223238 | 0.413957 | 0.685672 |

| | | | | | | | | | |
|---|---|---|---|---|---|---|---|---|---|
| 10 | 49049200 | 49050799 | 19.30788 | 32.62999 | 60.13782 | 26.31319 | -0.49752 | 1.078348 | 0.685726 |
| X | 15774800 | 15777199 | 34.11167 | 57.09026 | 47.55076 | 20.71926 | 0.644057 | 1.205108 | 0.685728 |
| 7 | 130292200 | 130295199 | 21.62935 | 36.37762 | 55.3901 | 24.20046 | -0.2234 | 1.120774 | 0.685738 |
| 5 | 137192600 | 137197199 | 92.52945 | 142.2083 | 197.6586 | 82.07566 | -0.17923 | 2.964309 | 0.685749 |
| 6 | 131174600 | 131181999 | 125.261 | 191.5625 | 204.365 | 84.78714 | 0.196564 | 3.072988 | 0.68575 |
| 3 | 144684200 | 144686399 | 21.25828 | 37.12047 | 40.21535 | 17.71243 | 0.272532 | 0.73677 | 0.685766 |
| 4 | 93670800 | 93675799 | 43.94238 | 71.56365 | 72.88309 | 31.29428 | 0.344368 | 1.567575 | 0.68579 |
| 18 | 85301800 | 85305399 | 39.83582 | 64.20384 | 92.42835 | 39.33629 | -0.16822 | 1.797555 | 0.685802 |
| 11 | 102625600 | 102626199 | 7.455875 | 14.38617 | 24.81394 | 9.585327 | -0.23295 | -0.13458 | 0.685804 |
| X | 103671600 | 103672599 | 11.88177 | 22.4248 | 25.80374 | 10.52019 | 0.274915 | 0.082332 | 0.685804 |
| 16 | 18800200 | 18805799 | 59.88864 | 95.57479 | 106.3445 | 44.61034 | 0.185956 | 2.129899 | 0.685809 |
| 11 | 12510800 | 12514799 | 52.97982 | 84.28188 | 120.9223 | 50.50343 | -0.18507 | 2.22876 | 0.685814 |
| 13 | 64645200 | 64648999 | 69.3155 | 109.6168 | 122.8555 | 51.05078 | 0.162807 | 2.406406 | 0.685825 |
| 11 | 119969600 | 119971199 | 14.60655 | 25.6084 | 39.42434 | 17.42058 | -0.23165 | 0.640001 | 0.685826 |
| 13 | 56918600 | 56919399 | 11.82935 | 21.38484 | 33.24584 | 14.43074 | -0.22277 | 0.35499 | 0.685834 |
| 1 | 119221200 | 119223599 | 17.31669 | 29.76698 | 46.90861 | 20.57589 | -0.26935 | 0.899184 | 0.685843 |
| 17 | 58075600 | 58076799 | 11.44215 | 20.75312 | 32.22676 | 13.93088 | -0.2174 | 0.329431 | 0.68585 |
| 4 | 71838000 | 71842599 | 47.15038 | 76.06234 | 84.09913 | 35.83374 | 0.213249 | 1.778684 | 0.68586 |
| 6 | 140259600 | 140266199 | 119.7882 | 182.8341 | 201.4614 | 83.5446 | 0.147743 | 3.109692 | 0.685871 |
| 16 | 75488400 | 75488999 | 7.040445 | 13.41875 | 24.54602 | 9.801734 | -0.36105 | -0.03053 | 0.685877 |
| 13 | 102957800 | 102960999 | 36.44007 | 59.05942 | 87.21611 | 37.34207 | -0.19624 | 1.648351 | 0.68589 |
| 5 | 10011800 | 10014599 | 30.74527 | 50.60264 | 78.28413 | 33.7538 | -0.24893 | 1.404766 | 0.685892 |
| 13 | 33990400 | 33993399 | 39.58179 | 64.84461 | 68.13741 | 29.32117 | 0.30496 | 1.473343 | 0.685919 |
| X | 85980200 | 85981399 | 16.80012 | 30.15636 | 32.29671 | 14.1292 | 0.304087 | 0.428077 | 0.685923 |
| 15 | 49790200 | 49795999 | 53.25585 | 85.53405 | 92.65009 | 39.23191 | 0.235075 | 1.922746 | 0.685937 |
| 6 | 114418200 | 114420599 | 25.67032 | 43.39704 | 49.60007 | 21.70464 | 0.18917 | 1.082947 | 0.685949 |
| X | 33062600 | 33064199 | 22.18761 | 38.95483 | 31.62673 | 13.99178 | 0.686795 | 0.64597 | 0.685951 |
| 1 | 20158800 | 20161999 | 47.83742 | 77.0661 | 86.08639 | 36.61428 | 0.196218 | 1.820783 | 0.685956 |
| 1 | 177383200 | 177383799 | 6.835484 | 13.15841 | 23.62552 | 9.187563 | -0.29967 | -0.12525 | 0.685965 |
| 2 | 144422800 | 144424799 | 13.36877 | 24.43611 | 27.33926 | 11.70416 | 0.261217 | 0.285526 | 0.685967 |
| 8 | 44653400 | 44653799 | 6.409157 | 12.34612 | 23.09889 | 8.965219 | -0.35566 | -0.13755 | 0.685968 |
| 8 | 21464800 | 21469399 | 63.3939 | 101.4157 | 103.799 | 43.69315 | 0.310136 | 2.06155 | 0.68599 |
| 7 | 121487600 | 121493399 | 50.5123 | 80.65249 | 114.1644 | 47.92534 | -0.15991 | 2.108066 | 0.685996 |
| 19 | 3376000 | 3377999 | 16.65114 | 28.98996 | 43.66605 | 19.19661 | -0.20104 | 0.761561 | 0.686014 |
| 19 | 59510600 | 59512199 | 12.00047 | 21.68254 | 33.74993 | 14.65937 | -0.22517 | 0.360504 | 0.68602 |
| 1 | 85568400 | 85582199 | 269.7141 | 388.6902 | 401.5182 | 165.6861 | 0.165004 | 4.134226 | 0.686035 |
| 11 | 12752200 | 12752599 | 5.207016 | 10.17017 | 19.4079 | 6.800843 | -0.24627 | -0.5183 | 0.686035 |
| 6 | 38277800 | 38282199 | 51.91535 | 83.33257 | 93.4464 | 39.53411 | 0.184242 | 1.941001 | 0.686039 |
| 3 | 11393600 | 11394799 | 13.16776 | 23.97251 | 27.4477 | 11.80958 | 0.223702 | 0.308763 | 0.686058 |
| 7 | 47870000 | 47870399 | 5.34994 | 10.44474 | 19.92673 | 7.035724 | -0.25567 | -0.48705 | 0.686067 |
| 1 | 100842000 | 100844199 | 24.54559 | 42.71427 | 42.33506 | 18.60598 | 0.399925 | 0.802471 | 0.686089 |
| 15 | 88349200 | 88350999 | 13.94523 | 24.46236 | 38.41245 | 16.9813 | -0.26005 | 0.624997 | 0.686095 |
| 4 | 145555200 | 145557999 | 88.7717 | 139.7213 | 142.2144 | 59.00353 | 0.296261 | 2.527164 | 0.686124 |
| 6 | 99845400 | 99852599 | 69.29822 | 108.003 | 181.0519 | 75.14122 | -0.4337 | 2.698693 | 0.68613 |
| 2 | 111509000 | 111510799 | 30.76586 | 51.25558 | 47.61681 | 20.68824 | 0.487114 | 1.240573 | 0.686132 |
| 5 | 37633600 | 37635399 | 23.7506 | 41.36784 | 35.52371 | 15.62122 | 0.606074 | 0.778647 | 0.686163 |
| 5 | 10076000 | 10076599 | 8.915183 | 17.13404 | 20.86073 | 8.150247 | 0.249893 | -0.0948 | 0.686175 |

| | | | | | | | | | |
|---|---|---|---|---|---|---|---|---|---|
| 1 | 100604200 | 100610399 | 67.28728 | 107.1633 | 113.6082 | 47.52268 | 0.252479 | 2.203107 | 0.686178 |
| 8 | 44485200 | 44487399 | 19.63365 | 34.32177 | 38.69864 | 17.05784 | 0.21529 | 0.710439 | 0.686183 |
| 4 | 130720400 | 130724799 | 78.28545 | 121.5641 | 171.9921 | 71.42024 | -0.19338 | 2.765619 | 0.68619 |
| X | 45361400 | 45362199 | 20.14757 | 35.66501 | 28.15384 | 12.4499 | 0.73093 | 0.535639 | 0.686196 |
| 8 | 67125000 | 67127399 | 20.21192 | 35.43469 | 32.84494 | 14.51292 | 0.496514 | 0.658292 | 0.686214 |
| 6 | 117701400 | 117706399 | 82.27504 | 128.0758 | 176.2678 | 73.16865 | -0.15082 | 2.724819 | 0.686222 |
| 19 | 38477200 | 38480199 | 41.41324 | 66.89776 | 75.62208 | 32.27807 | 0.183654 | 1.727229 | 0.686288 |
| 2 | 80431400 | 80435999 | 39.77325 | 64.15077 | 94.23415 | 40.14338 | -0.19606 | 1.774992 | 0.686309 |
| 3 | 149585200 | 149585599 | 5.223392 | 10.19476 | 19.28706 | 6.843524 | -0.25061 | -0.47612 | 0.686312 |
| 11 | 19518200 | 19521399 | 33.55664 | 54.93609 | 62.14174 | 26.7333 | 0.198555 | 1.465254 | 0.686361 |
| 17 | 20965400 | 20979399 | 341.273 | 475.9905 | 594.919 | 246.7824 | -0.15717 | 4.619442 | 0.686411 |
| 7 | 17683800 | 17690399 | 102.303 | 159.0967 | 166.0064 | 68.91927 | 0.247091 | 2.750525 | 0.686419 |
| 16 | 58126600 | 58127599 | 13.88529 | 25.38709 | 28.97992 | 12.40235 | 0.233023 | 0.293943 | 0.68643 |
| 12 | 115311000 | 115315199 | 50.75776 | 81.98391 | 82.85689 | 35.36128 | 0.344656 | 1.747938 | 0.686431 |
| 4 | 127762000 | 127775399 | 214.7863 | 314.8163 | 334.3285 | 137.711 | 0.148828 | 3.874665 | 0.686443 |
| 17 | 17633400 | 17634799 | 9.394532 | 17.30737 | 28.87004 | 12.22858 | -0.29941 | 0.217716 | 0.686488 |
| 10 | 79296200 | 79297999 | 18.92436 | 32.79802 | 50.10545 | 21.98363 | -0.22245 | 0.83701 | 0.686488 |
| 11 | 105855000 | 105856199 | 11.34502 | 20.57585 | 32.27491 | 13.95262 | -0.23168 | 0.330878 | 0.68653 |
| 2 | 15024000 | 15026599 | 31.35231 | 52.36926 | 49.18639 | 21.40544 | 0.471336 | 1.217214 | 0.686543 |
| 9 | 120360600 | 120364799 | 38.988 | 62.84453 | 91.2203 | 38.82785 | -0.1799 | 1.791628 | 0.68657 |
| 1 | 106366800 | 106367599 | 9.710242 | 18.15128 | 29.53025 | 12.24048 | -0.23992 | 0.136588 | 0.686577 |
| 5 | 127260400 | 127262999 | 38.71221 | 62.90695 | 71.80513 | 30.80336 | 0.177954 | 1.589867 | 0.686607 |
| 12 | 6854200 | 6855399 | 26.91303 | 44.95364 | 48.62358 | 21.16037 | 0.268359 | 1.202796 | 0.686607 |
| 12 | 75664800 | 75665399 | 10.67493 | 20.11226 | 23.74093 | 9.710464 | 0.234944 | 0.079393 | 0.686618 |
| 18 | 17151200 | 17152199 | 13.88372 | 24.44381 | 37.7288 | 16.70216 | -0.23461 | 0.570923 | 0.686618 |
| 14 | 106170400 | 106171599 | 29.61483 | 49.13918 | 56.15862 | 24.33285 | 0.188918 | 1.291068 | 0.686631 |
| 10 | 89233400 | 89237999 | 62.11315 | 98.82703 | 111.9374 | 46.76103 | 0.155693 | 2.229936 | 0.686678 |
| 2 | 46467800 | 46469799 | 15.51986 | 27.19228 | 40.9051 | 18.06515 | -0.19862 | 0.656255 | 0.686684 |
| 3 | 114649200 | 114650799 | 22.12448 | 38.57392 | 35.5146 | 15.627 | 0.506018 | 0.755224 | 0.686684 |
| 1 | 47796200 | 47799799 | 30.93523 | 51.78696 | 55.8235 | 24.32776 | 0.273083 | 1.196177 | 0.686722 |
| 7 | 38551200 | 38553199 | 29.45903 | 49.69337 | 52.95541 | 23.15756 | 0.289706 | 1.117679 | 0.68674 |
| 8 | 107180600 | 107183799 | 37.18671 | 60.63883 | 69.10582 | 29.68621 | 0.183858 | 1.53345 | 0.686741 |
| 5 | 4846200 | 4849599 | 37.83199 | 60.70017 | 107.0478 | 45.25859 | -0.46839 | 1.942359 | 0.686744 |
| X | 80292000 | 80294799 | 30.71982 | 51.93358 | 44.74921 | 19.58194 | 0.59536 | 1.097972 | 0.686776 |
| 10 | 125046600 | 125047799 | 15.65035 | 28.15625 | 31.25988 | 13.6495 | 0.254139 | 0.402425 | 0.686779 |
| 10 | 47454200 | 47457599 | 36.06162 | 59.07197 | 66.96487 | 28.81541 | 0.195761 | 1.467427 | 0.68678 |
| Y | 85099800 | 85100999 | 29.05137 | 48.79718 | 40.27853 | 17.59059 | 0.656819 | 1.141566 | 0.686783 |
| 1 | 132004400 | 132005399 | 12.93596 | 23.91938 | 27.60741 | 11.63546 | 0.233144 | 0.221597 | 0.686795 |
| 17 | 30314400 | 30315199 | 12.08811 | 23.0125 | 25.50995 | 10.2118 | 0.351518 | 0.025282 | 0.686809 |
| 6 | 10732000 | 10737599 | 53.5619 | 86.38487 | 85.42023 | 36.38554 | 0.373629 | 1.795563 | 0.686828 |
| 6 | 86720800 | 86721399 | 9.067741 | 17.16063 | 28.72076 | 11.64199 | -0.25368 | 0.048629 | 0.686855 |
| 7 | 143212600 | 143213599 | 11.11217 | 20.8211 | 24.68649 | 10.19179 | 0.217642 | 0.119572 | 0.686856 |
| 3 | 124556600 | 124560599 | 28.30262 | 47.65171 | 44.85623 | 19.60442 | 0.468012 | 1.123453 | 0.68686 |
| 7 | 102536400 | 102537199 | 7.398639 | 14.22661 | 25.29021 | 9.866473 | -0.29039 | -0.10588 | 0.686889 |
| X | 15357800 | 15360999 | 31.11762 | 53.47583 | 46.20168 | 20.26563 | 0.59421 | 0.959587 | 0.68691 |
| 9 | 95951600 | 95952999 | 19.25687 | 32.97327 | 49.53822 | 21.72369 | -0.20177 | 0.946234 | 0.686932 |
| 9 | 46515800 | 46520199 | 43.84022 | 70.32887 | 101.143 | 42.75164 | -0.175 | 1.953263 | 0.68694 |

| | | | | | | | | | |
|---|---|---|---|---|---|---|---|---|---|
| 1 | 118169800 | 118173599 | 48.61453 | 78.79412 | 77.72508 | 33.27943 | 0.38458 | 1.66283 | 0.686942 |
| 17 | 33178600 | 33181799 | 29.08116 | 48.75666 | 54.08029 | 23.59051 | 0.231941 | 1.172466 | 0.686951 |
| 17 | 70276000 | 70277799 | 24.04843 | 39.40465 | 76.05912 | 32.7327 | -0.56961 | 1.444351 | 0.686951 |
| 15 | 37135200 | 37136799 | 9.867289 | 18.03833 | 29.82062 | 12.76432 | -0.29765 | 0.266802 | 0.686977 |
| 8 | 43717000 | 43723199 | 62.76994 | 100.1621 | 108.754 | 45.61409 | 0.221046 | 2.137664 | 0.686992 |
| 2 | 20672800 | 20675199 | 25.53898 | 42.44821 | 63.76525 | 27.66111 | -0.20419 | 1.264279 | 0.686999 |
| 17 | 88274400 | 88277399 | 25.7425 | 42.93432 | 63.79416 | 27.73628 | -0.18831 | 1.228529 | 0.687024 |
| 13 | 35789200 | 35791399 | 15.02749 | 26.40247 | 39.77241 | 17.60599 | -0.20065 | 0.606865 | 0.687036 |
| 8 | 82945600 | 82945999 | 4.30777 | 8.422966 | 17.19323 | 5.874664 | -0.30697 | -0.62502 | 0.687038 |
| 5 | 18939600 | 18940399 | 10.99285 | 19.53428 | 38.36389 | 16.92677 | -0.5738 | 0.466601 | 0.687039 |
| 15 | 82411600 | 82414399 | 59.95833 | 95.37842 | 106.0271 | 44.27915 | 0.182352 | 2.23331 | 0.687047 |
| 12 | 51254400 | 51257399 | 41.52289 | 67.13553 | 76.64061 | 32.75097 | 0.170968 | 1.700643 | 0.687049 |
| 13 | 89110200 | 89111199 | 14.01031 | 26.46108 | 26.94031 | 10.96688 | 0.4524 | 0.100113 | 0.687066 |
| 6 | 60053200 | 60055999 | 29.28295 | 49.57056 | 51.8304 | 22.68981 | 0.31719 | 1.086179 | 0.687091 |
| 19 | 18096600 | 18097199 | 8.444533 | 16.30049 | 19.87168 | 7.664872 | 0.265423 | -0.14426 | 0.687106 |
| 9 | 31374600 | 31376999 | 20.60674 | 34.86202 | 53.25695 | 23.30998 | -0.22778 | 1.063364 | 0.687113 |
| 16 | 20725400 | 20726399 | 17.87634 | 31.34476 | 35.62697 | 15.7442 | 0.203788 | 0.648585 | 0.687127 |
| Y | 1120000 | 1120799 | 6.485381 | 12.29063 | 25.03935 | 10.11989 | -0.53056 | 0.003867 | 0.687132 |
| 4 | 156239200 | 156255399 | 294.2616 | 415.3905 | 528.3714 | 218.7607 | -0.16501 | 4.438534 | 0.687142 |
| 7 | 73021200 | 73023399 | 16.77562 | 29.15228 | 44.21908 | 19.41697 | -0.21143 | 0.788502 | 0.68715 |
| 1 | 156766600 | 156767799 | 8.146456 | 15.12068 | 27.91229 | 11.70325 | -0.43405 | 0.160518 | 0.68716 |
| 12 | 119630600 | 119637799 | 89.46903 | 139.9666 | 152.805 | 63.39141 | 0.188522 | 2.6392 | 0.68717 |
| 15 | 82492600 | 82494799 | 41.65801 | 66.99821 | 96.04978 | 40.7524 | -0.16601 | 1.868285 | 0.687201 |
| 6 | 102035400 | 102037399 | 14.01915 | 24.97001 | 39.74881 | 17.50347 | -0.27128 | 0.469629 | 0.687207 |
| 14 | 119482200 | 119484399 | 22.35195 | 37.82334 | 56.59176 | 24.77134 | -0.198 | 1.070191 | 0.687226 |
| 9 | 38854800 | 38856399 | 16.76299 | 29.71855 | 33.30702 | 14.72424 | 0.228602 | 0.50601 | 0.687256 |
| 11 | 112682800 | 112684399 | 14.80033 | 26.11129 | 40.27982 | 17.82413 | -0.23257 | 0.560785 | 0.687268 |
| 17 | 72204800 | 72206599 | 18.40344 | 31.32504 | 50.04904 | 21.93718 | -0.29135 | 1.00005 | 0.687279 |
| 17 | 76361000 | 76361799 | 9.974758 | 18.47909 | 29.37418 | 12.34806 | -0.22269 | 0.191441 | 0.687287 |
| 4 | 143943800 | 143957199 | 241.0288 | 351.7466 | 362.5452 | 149.3813 | 0.185936 | 3.945205 | 0.687288 |
| 12 | 91207000 | 91207799 | 13.75426 | 24.80725 | 28.00805 | 12.19241 | 0.233167 | 0.369416 | 0.68729 |
| 3 | 82496800 | 82498999 | 23.41147 | 40.40503 | 44.67406 | 19.59812 | 0.240878 | 0.886856 | 0.687293 |
| 18 | 6402000 | 6403199 | 20.79245 | 36.03685 | 40.97494 | 17.97766 | 0.201838 | 0.831289 | 0.687295 |
| 13 | 88471200 | 88472599 | 11.55635 | 21.95904 | 21.08755 | 8.474972 | 0.552262 | 0.022566 | 0.687299 |
| 4 | 113187600 | 113191599 | 58.03546 | 93.00937 | 99.14003 | 41.8423 | 0.254587 | 1.999036 | 0.687313 |
| 11 | 58786000 | 58787399 | 12.84031 | 23.65886 | 26.38162 | 11.17237 | 0.277076 | 0.234243 | 0.687339 |
| 8 | 18092600 | 18093999 | 16.02523 | 28.38236 | 32.3417 | 14.30034 | 0.204176 | 0.509197 | 0.687361 |
| 15 | 24160400 | 24162399 | 21.7929 | 37.49446 | 42.60959 | 18.69068 | 0.200476 | 0.903314 | 0.687366 |
| 12 | 24507400 | 24512399 | 48.61964 | 78.18162 | 88.61925 | 37.59073 | 0.171863 | 1.879658 | 0.687386 |
| 17 | 93340000 | 93342999 | 28.41281 | 48.22319 | 50.6968 | 22.21111 | 0.309471 | 1.059524 | 0.687404 |
| 2 | 150311400 | 150313799 | 50.94338 | 81.37033 | 116.532 | 48.94439 | -0.17613 | 2.09982 | 0.687412 |
| 3 | 111133600 | 111134599 | 13.95305 | 25.15819 | 28.34434 | 12.34346 | 0.235552 | 0.371143 | 0.687416 |
| 15 | 37877200 | 37883999 | 140.416 | 212.3227 | 232.1904 | 96.02387 | 0.147568 | 3.306052 | 0.687453 |
| 17 | 28767400 | 28768799 | 10.609 | 19.01072 | 32.36685 | 14.1803 | -0.36172 | 0.398065 | 0.687456 |
| 6 | 64360800 | 64361399 | 6.809 | 13.17861 | 23.85413 | 9.086741 | -0.28336 | -0.1954 | 0.687506 |
| 8 | 64897200 | 64898999 | 16.01659 | 28.39453 | 32.30033 | 14.27928 | 0.207136 | 0.498412 | 0.687506 |
| 1 | 48406800 | 48407999 | 20.35834 | 34.5221 | 52.69174 | 23.07889 | -0.22627 | 1.044247 | 0.68751 |

| | | | | | | | | | |
|---|---|---|---|---|---|---|---|---|---|
| 12 | 107683600 | 107686799 | 19.19814 | 33.04095 | 49.54495 | 21.73308 | -0.19779 | 0.898818 | 0.687526 |
| 10 | 69432600 | 69436799 | 33.54117 | 55.25303 | 63.02894 | 27.18635 | 0.189945 | 1.389652 | 0.687529 |
| 17 | 55103000 | 55105599 | 19.96074 | 35.34893 | 35.13876 | 15.53742 | 0.400162 | 0.540881 | 0.687539 |
| 6 | 132892800 | 132899199 | 64.02856 | 102.3343 | 106.0239 | 44.57001 | 0.291091 | 2.090973 | 0.687546 |
| 14 | 97635600 | 97637799 | 10.20662 | 18.12138 | 40.38483 | 17.83151 | -0.75629 | 0.466704 | 0.68756 |
| 16 | 96009800 | 96012599 | 26.79359 | 44.25975 | 65.7687 | 28.409 | -0.18921 | 1.342739 | 0.687577 |
| 12 | 99071200 | 99075999 | 54.17989 | 86.66443 | 98.42683 | 41.42435 | 0.160089 | 2.052026 | 0.687578 |
| 13 | 65227800 | 65242799 | 24.79731 | 42.84957 | 38.77637 | 17.01149 | 0.529053 | 0.874444 | 0.687612 |
| 1 | 48100200 | 48103799 | 35.84794 | 59.22677 | 63.37753 | 27.37943 | 0.283032 | 1.361039 | 0.687616 |
| 1 | 159042800 | 159043799 | 9.524173 | 18.07142 | 31.76484 | 12.80519 | -0.31738 | 0.040567 | 0.687627 |
| 16 | 15946800 | 15955399 | 101.844 | 158.2212 | 167.3749 | 69.49803 | 0.226164 | 2.7679 | 0.687632 |
| X | 133375400 | 133378399 | 77.27962 | 121.852 | 132.6518 | 55.04894 | 0.201371 | 2.482951 | 0.687632 |
| 13 | 25787200 | 25789199 | 18.6238 | 33.27716 | 33.06861 | 14.52284 | 0.407516 | 0.46752 | 0.68764 |
| 14 | 42026600 | 42029999 | 40.80846 | 66.46466 | 63.58902 | 27.30435 | 0.434395 | 1.555811 | 0.687646 |
| 14 | 99282800 | 99284399 | 14.74424 | 25.70893 | 41.12483 | 18.09666 | -0.28703 | 0.726902 | 0.687656 |
| 2 | 139363000 | 139364399 | 11.563 | 20.5423 | 33.81636 | 14.92347 | -0.32091 | 0.461497 | 0.687658 |
| 5 | 126604600 | 126607599 | 30.21173 | 50.5908 | 55.74662 | 24.29881 | 0.241496 | 1.192201 | 0.687665 |
| 3 | 99789600 | 99794799 | 48.82651 | 78.04223 | 111.4838 | 46.86931 | -0.17143 | 2.072768 | 0.687683 |
| 4 | 72006400 | 72008599 | 26.54388 | 45.4175 | 47.92896 | 21.01675 | 0.305709 | 0.981628 | 0.687694 |
| 12 | 26642200 | 26644199 | 15.03075 | 26.51401 | 40.75128 | 18.03327 | -0.22758 | 0.562248 | 0.687694 |
| 7 | 121830400 | 121834399 | 35.52646 | 57.54037 | 84.26256 | 36.05978 | -0.18507 | 1.656061 | 0.687694 |
| 14 | 40203200 | 40203799 | 7.445665 | 14.24344 | 24.57215 | 9.729242 | -0.26689 | -0.05639 | 0.687697 |
| 2 | 35691400 | 35694799 | 26.78909 | 44.17142 | 66.27896 | 28.59948 | -0.2038 | 1.364268 | 0.687746 |
| 6 | 6981200 | 6982399 | 24.33474 | 41.77386 | 46.29292 | 20.3047 | 0.236019 | 0.939149 | 0.687751 |
| 7 | 88919000 | 88928799 | 137.5733 | 207.1091 | 277.1843 | 114.4806 | -0.14913 | 3.395449 | 0.687779 |
| 9 | 92524600 | 92528199 | 75.13261 | 118.1588 | 131.0748 | 54.3761 | 0.171365 | 2.530382 | 0.687787 |
| 1 | 95164800 | 95167199 | 30.11837 | 48.76016 | 88.78306 | 37.98224 | -0.49908 | 1.658895 | 0.687794 |
| 3 | 48688400 | 48690999 | 30.81368 | 50.24131 | 74.32262 | 31.926 | -0.19156 | 1.526846 | 0.687797 |
| 17 | 93936800 | 93937599 | 7.630367 | 14.6641 | 25.24098 | 9.862881 | -0.24541 | -0.09349 | 0.687798 |
| 7 | 101877600 | 101879799 | 15.1492 | 26.65075 | 40.74977 | 18.04236 | -0.22172 | 0.591552 | 0.687803 |
| X | 85658400 | 85659999 | 14.24344 | 25.31482 | 39.72908 | 17.52663 | -0.25258 | 0.4859 | 0.687809 |
| 10 | 5825000 | 5825999 | 8.64814 | 16.4274 | 27.15983 | 10.92317 | -0.22664 | 0.012171 | 0.687816 |
| 19 | 14148400 | 14150999 | 21.12561 | 36.41193 | 41.05481 | 18.00966 | 0.212328 | 0.882333 | 0.687819 |
| 7 | 134886400 | 134887599 | 12.79384 | 23.02176 | 35.72346 | 15.59587 | -0.2257 | 0.392786 | 0.687819 |
| 5 | 146603200 | 146612599 | 132.2055 | 200.776 | 219.5351 | 90.91185 | 0.152985 | 3.215487 | 0.687823 |
| 7 | 121865800 | 121868199 | 28.62232 | 47.67844 | 54.51637 | 23.68403 | 0.18852 | 1.242826 | 0.687844 |
| 12 | 43933200 | 43933599 | 4.423716 | 8.642311 | 18.83292 | 6.572323 | -0.43096 | -0.5511 | 0.68787 |
| 3 | 31283600 | 31285199 | 10.36715 | 18.81105 | 30.80034 | 13.30746 | -0.29263 | 0.316586 | 0.687891 |
| 2 | 97143800 | 97145399 | 18.79678 | 32.41769 | 48.47147 | 21.26355 | -0.19292 | 0.879217 | 0.687894 |
| 13 | 67188000 | 67190799 | 53.85201 | 87.12935 | 79.68227 | 34.07315 | 0.49124 | 1.710644 | 0.687894 |
| X | 77061600 | 77063399 | 24.94436 | 43.18764 | 44.62162 | 19.57701 | 0.339189 | 0.865822 | 0.687914 |
| 7 | 26208600 | 26214599 | 66.5094 | 105.8895 | 114.771 | 47.98721 | 0.220099 | 2.215354 | 0.687946 |
| 10 | 103539800 | 103541599 | 13.82466 | 24.70642 | 40.89738 | 17.95834 | -0.32467 | 0.448573 | 0.68795 |
| 7 | 78734000 | 78736399 | 17.12524 | 29.62967 | 45.3105 | 19.87883 | -0.22423 | 0.83791 | 0.687953 |
| 15 | 54955600 | 54957199 | 16.28509 | 28.3631 | 43.13108 | 18.96582 | -0.21473 | 0.752617 | 0.687957 |
| 5 | 150261400 | 150263199 | 17.9321 | 31.39033 | 35.64978 | 15.73218 | 0.204889 | 0.67402 | 0.687969 |
| 3 | 90876400 | 90880999 | 71.46677 | 113.1487 | 126.505 | 52.59785 | 0.167633 | 2.381831 | 0.68797 |

| | | | | | | | | | |
|---|---|---|---|---|---|---|---|---|---|
| 18 | 57088600 | 57091199 | 25.19519 | 43.52348 | 45.18553 | 19.82305 | 0.331673 | 0.888386 | 0.688005 |
| 5 | 132755400 | 132758999 | 27.83965 | 46.10464 | 69.60321 | 30.11809 | -0.21182 | 1.316712 | 0.688025 |
| 15 | 3388600 | 3393799 | 94.8528 | 146.705 | 158.9554 | 66.02625 | 0.18669 | 2.841508 | 0.688085 |
| 3 | 91866000 | 91872799 | 22.57936 | 39.25571 | 36.63741 | 16.09239 | 0.48612 | 0.798151 | 0.688101 |
| 4 | 6539000 | 6540999 | 22.27873 | 37.49936 | 55.95575 | 24.45345 | -0.19432 | 1.117027 | 0.68811 |
| 1 | 182121200 | 182124399 | 31.0184 | 51.52168 | 58.54745 | 25.38613 | 0.197142 | 1.283523 | 0.688125 |
| 3 | 11998000 | 11998999 | 25.89073 | 43.4308 | 48.06182 | 20.97223 | 0.235372 | 1.152877 | 0.688217 |
| 3 | 145783400 | 145791199 | 100.362 | 153.5851 | 211.0059 | 87.55561 | -0.16728 | 3.051925 | 0.688227 |
| 18 | 28589800 | 28592399 | 19.83667 | 34.67621 | 38.62135 | 17.02358 | 0.232798 | 0.70983 | 0.688238 |
| X | 137547400 | 137549199 | 22.98892 | 40.35048 | 38.32648 | 16.95179 | 0.46152 | 0.665253 | 0.688288 |
| 3 | 90009600 | 90012399 | 28.9455 | 47.50666 | 70.17243 | 30.22028 | -0.18389 | 1.429966 | 0.688302 |
| 13 | 104656600 | 104657999 | 13.05681 | 23.47951 | 36.7329 | 16.04546 | -0.23818 | 0.399034 | 0.688309 |
| 18 | 52167400 | 52168199 | 8.489363 | 16.23441 | 27.38199 | 10.85254 | -0.23653 | -0.03636 | 0.688318 |
| 6 | 86488200 | 86490999 | 20.23496 | 34.46911 | 51.66967 | 22.64894 | -0.19975 | 0.999323 | 0.688323 |
| 12 | 85057600 | 85058999 | 18.31892 | 32.07281 | 36.65472 | 16.17793 | 0.195891 | 0.675823 | 0.688327 |
| 10 | 87143400 | 87150199 | 74.64063 | 117.9872 | 130.8153 | 54.34648 | 0.177875 | 2.422097 | 0.68833 |
| 4 | 128516600 | 128518799 | 33.57964 | 55.18873 | 63.44245 | 27.33797 | 0.177568 | 1.41883 | 0.688334 |
| 2 | 55947000 | 55949199 | 24.42123 | 42.27614 | 44.05709 | 19.32925 | 0.326689 | 0.861917 | 0.688339 |
| 3 | 43094600 | 43104599 | 94.56329 | 147.5946 | 158.2864 | 65.68099 | 0.211623 | 2.683733 | 0.688342 |
| 18 | 89995600 | 89996399 | 14.56103 | 26.1124 | 29.9058 | 13.09369 | 0.207058 | 0.413055 | 0.688362 |
| 3 | 31938800 | 31941399 | 27.58338 | 46.31452 | 52.87953 | 23.08843 | 0.190387 | 1.153241 | 0.688372 |
| 10 | 39206600 | 39209399 | 37.83801 | 62.08839 | 67.33841 | 28.99722 | 0.260374 | 1.450284 | 0.6884 |
| 2 | 174992000 | 174996199 | 71.01783 | 112.2495 | 126.3606 | 52.49322 | 0.155846 | 2.419288 | 0.688432 |
| 18 | 15112400 | 15113199 | 10.92028 | 19.99956 | 31.29066 | 13.361 | -0.2165 | 0.267119 | 0.688484 |
| 6 | 132139600 | 132157399 | 242.4801 | 355.6322 | 347.3943 | 143.0872 | 0.27021 | 3.869185 | 0.688522 |
| 18 | 46906200 | 46907999 | 18.19139 | 32.28452 | 34.1579 | 15.08737 | 0.312002 | 0.509013 | 0.688524 |
| 5 | 58908800 | 58909799 | 9.848886 | 18.36842 | 29.48261 | 12.26812 | -0.22516 | 0.151683 | 0.688538 |
| 17 | 74594600 | 74596199 | 21.83429 | 36.78986 | 55.37773 | 24.20855 | -0.20672 | 1.107136 | 0.688563 |
| 3 | 16674600 | 16676399 | 21.03157 | 36.9345 | 37.85024 | 16.74866 | 0.352352 | 0.644457 | 0.688567 |
| 12 | 8547800 | 8551199 | 22.14039 | 37.00807 | 58.0762 | 25.28538 | -0.26682 | 1.187776 | 0.688575 |
| 9 | 109285400 | 109294999 | 137.9239 | 209.1581 | 226.7627 | 93.86427 | 0.16355 | 3.246666 | 0.688587 |
| 6 | 64042600 | 64043199 | 13.45664 | 24.87589 | 21.98217 | 9.267812 | 0.615623 | 0.213939 | 0.688587 |
| 8 | 34739400 | 34740399 | 9.125934 | 17.12898 | 27.97592 | 11.51614 | -0.23786 | 0.098406 | 0.688593 |
| 18 | 53994600 | 53996199 | 14.64624 | 25.38937 | 49.51119 | 21.72305 | -0.57281 | 0.814791 | 0.688623 |
| 13 | 99108200 | 99112999 | 50.00153 | 80.24649 | 90.80938 | 38.40822 | 0.170847 | 1.944076 | 0.688625 |
| 1 | 65658200 | 65661599 | 40.65927 | 65.91889 | 74.73691 | 32.00665 | 0.184573 | 1.6405 | 0.688634 |
| 16 | 81336000 | 81336799 | 9.17004 | 17.46011 | 30.14648 | 12.0637 | -0.2824 | 0.009132 | 0.688651 |
| 12 | 26884000 | 26886999 | 38.20141 | 61.64893 | 88.91682 | 37.9047 | -0.16875 | 1.756255 | 0.688656 |
| 18 | 78170200 | 78174199 | 33.65479 | 54.42883 | 81.5447 | 34.85878 | -0.21979 | 1.683092 | 0.688656 |
| 7 | 42300200 | 42301999 | 26.02682 | 44.22773 | 49.57023 | 21.72442 | 0.217597 | 1.040481 | 0.688658 |
| 4 | 22414600 | 22416399 | 17.44252 | 31.34428 | 32.13675 | 14.04605 | 0.367338 | 0.423098 | 0.68866 |
| 18 | 18486600 | 18487199 | 8.347737 | 16.02059 | 27.55886 | 10.8114 | -0.25079 | -0.0702 | 0.68866 |
| 12 | 9438400 | 9440599 | 15.78146 | 27.58311 | 41.99622 | 18.50876 | -0.21603 | 0.69733 | 0.68875 |
| 12 | 23952200 | 23953399 | 33.57658 | 55.55415 | 46.12815 | 19.94745 | 0.648083 | 1.325958 | 0.688771 |
| 3 | 28819000 | 28820399 | 12.63352 | 23.17323 | 26.59517 | 11.33204 | 0.230108 | 0.262277 | 0.688776 |
| 15 | 82316600 | 82320599 | 64.63681 | 102.6672 | 116.2094 | 48.45363 | 0.154202 | 2.284592 | 0.688777 |
| 1 | 21021400 | 21022799 | 12.48897 | 22.04165 | 35.79166 | 15.83691 | -0.30586 | 0.53276 | 0.688777 |

| | | | | | | | | | |
|---|---|---|---|---|---|---|---|---|---|
| 10 | 83058000 | 83060199 | 15.36842 | 27.41698 | 31.3387 | 13.7815 | 0.205462 | 0.453368 | 0.688792 |
| 4 | 3355800 | 3368999 | 279.1681 | 399.6619 | 413.5365 | 170.8179 | 0.153109 | 4.231861 | 0.688795 |
| 1 | 173064600 | 173066799 | 18.75369 | 33.18774 | 34.60797 | 15.30489 | 0.330883 | 0.537916 | 0.688805 |
| 18 | 37143000 | 37146599 | 88.20464 | 137.2729 | 151.4557 | 62.87147 | 0.167493 | 2.729324 | 0.68883 |
| 3 | 23015400 | 23016399 | 17.17287 | 29.7328 | 45.31527 | 19.88134 | -0.2191 | 0.833215 | 0.688833 |
| X | 64151000 | 64158999 | 67.31749 | 107.4293 | 95.02284 | 39.8605 | 0.516498 | 2.137173 | 0.688859 |
| 9 | 100897200 | 100901399 | 35.35157 | 56.87533 | 101.3852 | 43.03775 | -0.47901 | 1.850295 | 0.688868 |
| 11 | 14447600 | 14449199 | 18.56388 | 32.88477 | 34.68956 | 15.3379 | 0.315022 | 0.527868 | 0.688877 |
| 6 | 57720000 | 57721199 | 10.42813 | 19.03678 | 31.25476 | 13.40176 | -0.28986 | 0.280128 | 0.688892 |
| 12 | 69591800 | 69592999 | 9.542074 | 17.61374 | 28.86793 | 12.19506 | -0.27139 | 0.207881 | 0.688898 |
| 2 | 156615200 | 156618199 | 21.93587 | 36.76147 | 57.57995 | 25.11039 | -0.26417 | 1.158054 | 0.688911 |
| 15 | 46435800 | 46437399 | 17.53259 | 30.73477 | 34.67363 | 15.31984 | 0.214544 | 0.648845 | 0.688929 |
| 1 | 136927600 | 136931799 | 47.76191 | 76.89924 | 86.65513 | 36.82481 | 0.182557 | 1.837952 | 0.688942 |
| 3 | 23557800 | 23559799 | 18.42379 | 32.73923 | 34.18547 | 15.08311 | 0.332051 | 0.503358 | 0.688949 |
| 3 | 4220000 | 4221799 | 21.96548 | 37.91026 | 43.09009 | 18.90325 | 0.201114 | 0.876791 | 0.688964 |
| 7 | 55136000 | 55146999 | 186.9582 | 277.6642 | 296.5138 | 122.1469 | 0.158667 | 3.654514 | 0.688967 |
| 13 | 112397800 | 112398999 | 11.52406 | 20.93607 | 33.23394 | 14.33676 | -0.24734 | 0.320392 | 0.688976 |
| 1 | 15911000 | 15911999 | 10.97108 | 20.04513 | 31.29035 | 13.40264 | -0.21618 | 0.282259 | 0.689 |
| X | 117573800 | 117575999 | 25.25493 | 43.58217 | 45.20006 | 19.82881 | 0.33289 | 0.898385 | 0.689004 |
| 4 | 95872400 | 95878399 | 67.61071 | 107.4389 | 118.9118 | 49.60262 | 0.18715 | 2.274734 | 0.689028 |
| 7 | 43832800 | 43834799 | 24.62561 | 42.32685 | 39.93995 | 17.51912 | 0.467432 | 0.922175 | 0.689038 |
| 3 | 114381400 | 114384399 | 42.03015 | 68.04422 | 75.37309 | 32.24514 | 0.216208 | 1.675817 | 0.68905 |
| 12 | 87727800 | 87729599 | 19.85249 | 33.88592 | 51.68532 | 22.65948 | -0.22434 | 0.977997 | 0.689052 |
| 15 | 20977800 | 20979599 | 17.04649 | 29.70375 | 45.19522 | 19.87959 | -0.21562 | 0.751183 | 0.689055 |
| 7 | 89464600 | 89465599 | 14.08371 | 26.72024 | 26.54592 | 10.7035 | 0.499641 | 0.070093 | 0.689077 |
| 1 | 172958000 | 172959999 | 14.56735 | 25.80418 | 40.21567 | 17.78139 | -0.24515 | 0.513526 | 0.689103 |
| 9 | 75326200 | 75328399 | 21.73146 | 36.71841 | 54.90394 | 24.02165 | -0.19711 | 1.079825 | 0.689118 |
| 2 | 83836200 | 83836999 | 11.26333 | 20.73508 | 32.98649 | 13.98609 | -0.23308 | 0.244801 | 0.689134 |
| 4 | 45291400 | 45292799 | 12.93474 | 22.99898 | 35.14458 | 15.49843 | -0.21371 | 0.466513 | 0.689143 |
| 2 | 123500600 | 123501199 | 5.974705 | 11.54804 | 22.09942 | 8.472326 | -0.37047 | -0.18513 | 0.689157 |
| 17 | 95056400 | 95058199 | 15.12912 | 26.13464 | 51.64416 | 22.65647 | -0.59303 | 0.850717 | 0.689165 |
| X | 76837200 | 76842999 | 60.9175 | 97.4805 | 101.3669 | 42.68578 | 0.287732 | 2.051987 | 0.689178 |
| 11 | 94653400 | 94655599 | 22.56019 | 38.13513 | 56.56692 | 24.75248 | -0.18558 | 1.081093 | 0.689198 |
| 14 | 3100800 | 3135199 | 480.8252 | 663.1106 | 622.0853 | 258.5094 | 0.240188 | 4.795016 | 0.689209 |
| 17 | 66618400 | 66619199 | 7.469342 | 14.36008 | 24.69325 | 9.638369 | -0.24248 | -0.10044 | 0.689258 |
| 10 | 110879400 | 110881399 | 30.47488 | 50.93433 | 56.34781 | 24.53135 | 0.235822 | 1.212482 | 0.689275 |
| 7 | 59173400 | 59173799 | 10.05183 | 19.31144 | 18.13721 | 7.093681 | 0.61952 | -0.08854 | 0.689281 |
| 9 | 79161600 | 79162799 | 9.249312 | 17.00684 | 28.87712 | 12.26388 | -0.32759 | 0.227459 | 0.689292 |
| 9 | 37798400 | 37799399 | 11.959 | 22.26673 | 26.1616 | 10.91694 | 0.218332 | 0.170853 | 0.6893 |
| 6 | 5337200 | 5341399 | 36.86508 | 59.63495 | 86.54015 | 37.00084 | -0.17366 | 1.685099 | 0.689343 |
| 14 | 106576400 | 106577199 | 6.846104 | 13.23519 | 23.19601 | 8.883678 | -0.24366 | -0.17481 | 0.689353 |
| 6 | 148047400 | 148048999 | 19.97444 | 34.97086 | 38.59633 | 17.0347 | 0.245945 | 0.687708 | 0.689367 |
| 2 | 108466200 | 108467199 | 11.41441 | 20.78918 | 32.25435 | 13.87038 | -0.21135 | 0.304751 | 0.689393 |
| 12 | 31337200 | 31340599 | 24.56194 | 41.1349 | 60.99557 | 26.58872 | -0.18543 | 1.178175 | 0.689398 |
| 18 | 67786400 | 67791199 | 46.41999 | 74.27811 | 106.1583 | 44.70055 | -0.17033 | 2.035724 | 0.689413 |
| 8 | 12526800 | 12528199 | 23.47083 | 39.16348 | 59.16787 | 25.73231 | -0.21215 | 1.213137 | 0.689415 |
| 3 | 40560600 | 40560999 | 6.238437 | 12.12622 | 21.78205 | 8.094408 | -0.23794 | -0.29114 | 0.689436 |

| | | | | | | | | | |
|---|---|---|---|---|---|---|---|---|---|
| 1 | 38220200 | 38221799 | 9.369118 | 17.26371 | 28.84813 | 12.21626 | -0.30186 | 0.215263 | 0.689451 |
| 1 | 107046800 | 107049199 | 18.8793 | 32.1216 | 57.54688 | 25.22274 | -0.45623 | 1.012322 | 0.689497 |
| 10 | 18143600 | 18147799 | 61.10111 | 98.03355 | 97.16894 | 41.08116 | 0.361207 | 1.968496 | 0.689504 |
| 5 | 8548000 | 8548799 | 8.978863 | 16.7471 | 28.40797 | 11.81941 | -0.3046 | 0.137631 | 0.68952 |
| 4 | 74375400 | 74377799 | 38.95602 | 63.27419 | 72.18871 | 30.95747 | 0.178081 | 1.601176 | 0.689531 |
| 5 | 109619800 | 109631199 | 60.25997 | 96.53735 | 86.00038 | 36.27549 | 0.512317 | 2.010952 | 0.689539 |
| X | 147804200 | 147805799 | 36.54741 | 59.58388 | 66.36817 | 28.50801 | 0.216653 | 1.537235 | 0.689563 |
| 1 | 100682000 | 100683199 | 16.0887 | 28.31337 | 29.17555 | 12.91686 | 0.345091 | 0.572685 | 0.689566 |
| 12 | 106580800 | 106587599 | 75.47523 | 117.726 | 164.3068 | 68.18371 | -0.16872 | 2.685918 | 0.689577 |
| 2 | 170635600 | 170645599 | 111.5331 | 170.252 | 232.0073 | 96.21409 | -0.15842 | 3.107361 | 0.689598 |
| 3 | 3299600 | 3301599 | 32.95672 | 54.23166 | 62.25933 | 26.84212 | 0.180207 | 1.402213 | 0.689599 |
| 13 | 59058200 | 59060799 | 24.78714 | 41.11577 | 62.45006 | 27.05298 | -0.22024 | 1.285441 | 0.689599 |
| 1 | 144246000 | 144247599 | 15.88744 | 28.86538 | 30.88425 | 13.32183 | 0.318571 | 0.348301 | 0.689626 |
| 11 | 28746000 | 28746399 | 6.995159 | 13.63691 | 17.46748 | 6.303455 | 0.284526 | -0.39645 | 0.689636 |
| 7 | 41343600 | 41357599 | 269.6095 | 389.4547 | 398.8454 | 164.532 | 0.180813 | 4.100688 | 0.689647 |
| 1 | 79721800 | 79723199 | 13.27276 | 23.69605 | 36.16421 | 15.89355 | -0.20852 | 0.44576 | 0.68965 |
| 12 | 77821600 | 77824599 | 25.75551 | 42.33792 | 74.95354 | 32.30089 | -0.44348 | 1.406784 | 0.689674 |
| 11 | 23054800 | 23062599 | 121.4128 | 186.5632 | 193.2473 | 80.2418 | 0.244371 | 2.977954 | 0.689686 |
| 15 | 23245200 | 23247399 | 12.30816 | 23.25033 | 17.45129 | 7.101455 | 0.889099 | 0.0584 | 0.689686 |
| 7 | 42731400 | 42755599 | 358.4791 | 508.2004 | 479.8724 | 198.5411 | 0.270848 | 4.379804 | 0.689689 |
| 2 | 65594000 | 65594999 | 10.47841 | 19.25401 | 30.32541 | 12.89059 | -0.22111 | 0.244071 | 0.689704 |
| 2 | 115671200 | 115671999 | 13.95306 | 25.30156 | 28.85477 | 12.47676 | 0.224673 | 0.339727 | 0.689718 |
| 7 | 62535200 | 62536199 | 21.05556 | 36.3457 | 41.58678 | 18.24386 | 0.191724 | 0.870717 | 0.689739 |
| 7 | 84931600 | 84935999 | 108.8981 | 167.1552 | 185.0231 | 76.81566 | 0.147553 | 2.992401 | 0.689739 |
| 6 | 142175000 | 142181799 | 110.5169 | 168.7366 | 228.6986 | 94.84661 | -0.15036 | 3.10285 | 0.689748 |
| 8 | 115690000 | 115693799 | 41.86926 | 67.04912 | 114.8696 | 48.4241 | -0.43008 | 2.008296 | 0.689782 |
| 1 | 179555800 | 179557599 | 22.57766 | 37.56047 | 59.51997 | 25.83579 | -0.28069 | 1.245006 | 0.689829 |
| 2 | 176774800 | 176775599 | 8.646655 | 16.55466 | 28.64149 | 11.31347 | -0.26849 | -0.04289 | 0.68983 |
| 14 | 121201400 | 121202399 | 8.984654 | 16.90765 | 27.48814 | 11.26569 | -0.22603 | 0.080086 | 0.689838 |
| 12 | 66229000 | 66229599 | 11.20389 | 21.08027 | 24.75733 | 10.14946 | 0.239607 | 0.095799 | 0.68985 |
| 6 | 106862600 | 106864199 | 20.47404 | 35.54765 | 40.50909 | 17.77794 | 0.199249 | 0.814325 | 0.689875 |
| 3 | 111051800 | 111056599 | 55.06524 | 88.54958 | 91.97339 | 39.0156 | 0.297736 | 1.888714 | 0.689876 |
| 16 | 5458600 | 5463399 | 74.5769 | 117.6448 | 132.7482 | 55.09231 | 0.150191 | 2.469586 | 0.689877 |
| 7 | 25905800 | 25935599 | 462.4122 | 639.9972 | 601.6134 | 249.8601 | 0.242482 | 4.740018 | 0.689888 |
| 3 | 22619200 | 22621599 | 32.76114 | 53.54643 | 80.2368 | 34.49701 | -0.20768 | 1.499455 | 0.689915 |
| 8 | 18332600 | 18334199 | 15.91688 | 28.37254 | 31.70079 | 13.95021 | 0.237289 | 0.461093 | 0.689989 |
| 6 | 59880800 | 59891599 | 155.5553 | 234.3793 | 250.5037 | 103.4848 | 0.176204 | 3.381134 | 0.690007 |
| 18 | 32653200 | 32657399 | 39.961 | 64.39752 | 93.19874 | 39.65748 | -0.17606 | 1.800811 | 0.690009 |
| 5 | 90086600 | 90089199 | 27.56932 | 47.34636 | 47.34987 | 20.76884 | 0.383673 | 0.950576 | 0.69004 |
| X | 121149600 | 121167799 | 235.3161 | 345.9643 | 337.0464 | 138.8039 | 0.277407 | 3.831117 | 0.690043 |
| 1 | 136730600 | 136741799 | 125.3938 | 189.7526 | 256.4099 | 106.0656 | -0.15655 | 3.288171 | 0.690048 |
| 17 | 69589000 | 69599199 | 153.5352 | 228.6307 | 305.5959 | 125.9222 | -0.16146 | 3.604197 | 0.690051 |
| 14 | 50718800 | 50720999 | 17.74035 | 30.9708 | 50.05853 | 22.04472 | -0.30226 | 0.735055 | 0.690062 |
| 12 | 72467000 | 72469799 | 31.73058 | 52.50982 | 60.1753 | 26.02536 | 0.184647 | 1.329643 | 0.690072 |
| 18 | 59872400 | 59874199 | 14.89317 | 27.31978 | 28.25735 | 12.03924 | 0.378534 | 0.278424 | 0.690106 |
| 14 | 90488000 | 90488999 | 10.10442 | 18.83603 | 29.99814 | 12.49271 | -0.21524 | 0.156308 | 0.690126 |
| 7 | 29488200 | 29503199 | 187.0981 | 275.3309 | 360.4259 | 148.4249 | -0.14731 | 3.810271 | 0.690149 |

| | | | | | | | | | |
|---|---|---|---|---|---|---|---|---|---|
| 8 | 25891200 | 25891999 | 11.93959 | 22.57925 | 25.87217 | 10.50336 | 0.286441 | 0.070057 | 0.690168 |
| 1 | 127655800 | 127656999 | 18.20349 | 32.01016 | 35.92425 | 15.90718 | 0.222114 | 0.606862 | 0.690179 |
| 17 | 18347800 | 18351599 | 65.95689 | 104.978 | 113.8169 | 47.56833 | 0.219217 | 2.225523 | 0.690179 |
| 1 | 97150000 | 97150799 | 8.32675 | 15.9294 | 26.8176 | 10.61745 | -0.23241 | -0.04313 | 0.690185 |
| 3 | 131678200 | 131678999 | 8.32675 | 15.9294 | 26.8176 | 10.61745 | -0.23241 | -0.04313 | 0.690185 |
| 6 | 38988800 | 38990399 | 20.12214 | 34.1978 | 51.64924 | 22.63576 | -0.21099 | 1.020988 | 0.690186 |
| 4 | 10066800 | 10068199 | 19.80123 | 34.65094 | 38.6304 | 17.04281 | 0.231493 | 0.694434 | 0.690192 |
| 1 | 98249200 | 98249999 | 9.892778 | 18.08171 | 29.84626 | 12.77814 | -0.29548 | 0.269204 | 0.690192 |
| 5 | 151545200 | 151546199 | 17.48922 | 30.80758 | 34.87841 | 15.43872 | 0.210458 | 0.581297 | 0.690214 |
| 9 | 88740600 | 88741599 | 10.75814 | 19.30452 | 36.8316 | 16.11765 | -0.52429 | 0.398172 | 0.690241 |
| 12 | 73494000 | 73498599 | 31.32947 | 50.9447 | 75.80098 | 32.52905 | -0.20265 | 1.567758 | 0.690265 |
| 1 | 7936400 | 7940399 | 35.53751 | 58.56658 | 55.25196 | 23.83619 | 0.463464 | 1.385419 | 0.690302 |
| 3 | 13582400 | 13583199 | 8.705654 | 16.3566 | 26.95152 | 11.07555 | -0.24812 | 0.087325 | 0.690302 |
| 17 | 48495800 | 48516799 | 301.1608 | 425.4633 | 538.3073 | 222.8455 | -0.15619 | 4.427362 | 0.690313 |
| X | 96720800 | 96726399 | 78.40504 | 124.2801 | 127.4077 | 53.01345 | 0.294024 | 2.359323 | 0.690343 |
| 16 | 56184000 | 56187799 | 40.17013 | 65.54513 | 71.29675 | 30.63443 | 0.251541 | 1.532452 | 0.690347 |
| 8 | 55526400 | 55526999 | 10.44272 | 19.19196 | 30.3085 | 12.88019 | -0.22482 | 0.241634 | 0.690353 |
| 9 | 20485200 | 20487599 | 24.29765 | 41.85112 | 39.39939 | 17.28312 | 0.471373 | 0.901493 | 0.690366 |
| 7 | 139297600 | 139300399 | 24.04421 | 40.31504 | 59.88512 | 26.12397 | -0.18799 | 1.162528 | 0.690369 |
| 7 | 28582200 | 28583999 | 18.89111 | 32.23955 | 50.05164 | 21.94296 | -0.24949 | 0.978558 | 0.690371 |
| 11 | 109692800 | 109695599 | 16.21939 | 28.26519 | 44.14083 | 19.41696 | -0.253 | 0.743446 | 0.690417 |
| X | 79605600 | 79608199 | 37.50735 | 61.6806 | 66.0493 | 28.46844 | 0.279997 | 1.424711 | 0.690423 |
| 1 | 35284400 | 35289799 | 40.29274 | 64.93485 | 93.88115 | 39.95029 | -0.17453 | 1.800413 | 0.690439 |
| 4 | 40059600 | 40065599 | 94.09191 | 145.897 | 162.6695 | 67.56364 | 0.148241 | 2.795806 | 0.690444 |
| 1 | 142428000 | 142428599 | 8.819679 | 17.10794 | 16.66259 | 6.282152 | 0.617122 | -0.22775 | 0.690452 |
| 14 | 82502600 | 82505799 | 45.54774 | 73.06865 | 104.6079 | 44.21671 | -0.16851 | 1.954296 | 0.690457 |
| 19 | 61098400 | 61112599 | 215.8084 | 320.8082 | 298.8339 | 123.1142 | 0.356919 | 3.64451 | 0.690461 |
| 10 | 68755600 | 68757399 | 11.18198 | 20.0937 | 32.25004 | 14.09578 | -0.27499 | 0.382812 | 0.690467 |
| X | 72317600 | 72318799 | 8.899597 | 16.7095 | 27.47856 | 11.30527 | -0.24644 | 0.096221 | 0.690474 |
| 1 | 161357800 | 161365399 | 114.2489 | 175.6669 | 167.0731 | 69.37959 | 0.368092 | 2.960039 | 0.690493 |
| 13 | 22428600 | 22433199 | 43.21946 | 70.7443 | 69.52868 | 29.90923 | 0.40074 | 1.494695 | 0.690512 |
| 2 | 125993800 | 125994399 | 5.530968 | 10.75573 | 20.66241 | 7.651401 | -0.32894 | -0.31658 | 0.690513 |
| 3 | 51897800 | 51899599 | 9.489119 | 17.68079 | 28.40366 | 11.83801 | -0.22835 | 0.150845 | 0.690518 |
| 17 | 16357200 | 16357799 | 12.36272 | 21.90308 | 34.77927 | 15.3771 | -0.27134 | 0.48953 | 0.690533 |
| 15 | 59390800 | 59391599 | 12.17548 | 22.95581 | 22.01343 | 8.992992 | 0.533674 | 0.083112 | 0.690547 |
| 1 | 177810600 | 177815999 | 50.03523 | 80.29126 | 91.22481 | 38.57712 | 0.164852 | 1.945814 | 0.690548 |
| 2 | 87111200 | 87113399 | 25.19794 | 42.98021 | 40.04905 | 17.55701 | 0.483819 | 0.992298 | 0.690589 |
| 1 | 139874000 | 139878799 | 49.99977 | 80.91819 | 80.02281 | 34.22548 | 0.378953 | 1.696532 | 0.690599 |
| 17 | 52992400 | 52997599 | 43.85844 | 71.177 | 76.46982 | 32.77304 | 0.263394 | 1.628597 | 0.690606 |
| 1 | 162796200 | 162801199 | 96.49479 | 150.3413 | 160.8365 | 66.75454 | 0.213177 | 2.716358 | 0.690616 |
| 4 | 108011800 | 108015199 | 51.85227 | 83.45965 | 78.83823 | 33.48825 | 0.436001 | 1.850576 | 0.690621 |
| 9 | 112611400 | 112611799 | 6.235807 | 12.12937 | 21.84835 | 8.074529 | -0.23471 | -0.31069 | 0.69063 |
| 10 | 51458200 | 51458799 | 6.235807 | 12.12937 | 21.84835 | 8.074529 | -0.23471 | -0.31069 | 0.69063 |
| 18 | 37274600 | 37279599 | 51.99226 | 82.74912 | 119.1197 | 49.78239 | -0.18915 | 2.212463 | 0.690636 |
| Y | 87750000 | 87751199 | 9.942595 | 17.58645 | 35.00563 | 15.48227 | -0.59623 | 0.49159 | 0.69064 |
| 11 | 63356800 | 63359599 | 31.47927 | 51.97224 | 59.45169 | 25.67092 | 0.18705 | 1.356682 | 0.690682 |
| 6 | 84736800 | 84742599 | 100.0017 | 153.0137 | 210.4663 | 87.32907 | -0.1691 | 3.055173 | 0.690688 |

| | | | | | | | | | |
|---|---|---|---|---|---|---|---|---|---|
| 10 | 15784400 | 15784999 | 9.074887 | 16.99222 | 27.97344 | 11.56215 | -0.2533 | 0.116901 | 0.690695 |
| 5 | 6799000 | 6799599 | 6.381963 | 12.38744 | 22.24737 | 8.362274 | -0.25318 | -0.24843 | 0.690702 |
| 10 | 96468200 | 96468799 | 6.381963 | 12.38744 | 22.24737 | 8.362274 | -0.25318 | -0.24843 | 0.690702 |
| 16 | 83990800 | 83991399 | 6.381963 | 12.38744 | 22.24737 | 8.362274 | -0.25318 | -0.24843 | 0.690702 |
| X | 20291400 | 20293399 | 31.94215 | 54.63635 | 37.68057 | 16.52242 | 0.917032 | 0.995656 | 0.690707 |
| 5 | 40760000 | 40763199 | 63.25297 | 100.4072 | 112.3345 | 46.80911 | 0.170127 | 2.299785 | 0.690733 |
| 17 | 79610200 | 79613399 | 25.4169 | 42.18465 | 63.09661 | 27.34392 | -0.1981 | 1.28085 | 0.690751 |
| 4 | 116743200 | 116743799 | 8.520059 | 16.18968 | 26.61084 | 10.69458 | -0.2171 | 0.008265 | 0.690751 |
| 12 | 13134400 | 13136399 | 22.61672 | 38.05192 | 56.50135 | 24.68867 | -0.18726 | 1.122734 | 0.690758 |
| 13 | 91747800 | 91750199 | 31.71006 | 52.28314 | 60.08901 | 25.92803 | 0.179698 | 1.371789 | 0.690769 |
| 7 | 26771000 | 26773799 | 22.76003 | 39.37926 | 37.32738 | 16.37645 | 0.462778 | 0.851713 | 0.690796 |
| X | 102735800 | 102739199 | 21.33929 | 36.44328 | 56.02842 | 24.56461 | -0.23555 | 0.987516 | 0.690812 |
| 2 | 53345400 | 53348199 | 19.02124 | 33.56143 | 30.58619 | 13.53435 | 0.522676 | 0.56473 | 0.690814 |
| 4 | 39390800 | 39391399 | 11.27435 | 21.16089 | 25.21212 | 10.37891 | 0.213912 | 0.111454 | 0.690832 |
| 6 | 137902800 | 137905399 | 23.0919 | 39.55436 | 45.28202 | 19.85783 | 0.188907 | 0.953608 | 0.690876 |
| 14 | 27409400 | 27412799 | 65.44724 | 103.4431 | 144.3375 | 59.9608 | -0.15352 | 2.420603 | 0.690885 |
| 17 | 54425800 | 54429199 | 38.60462 | 63.25467 | 59.82342 | 25.74534 | 0.456594 | 1.464898 | 0.690924 |
| 1 | 103165200 | 103166199 | 14.61816 | 26.26361 | 29.78981 | 13.02248 | 0.222257 | 0.400055 | 0.690927 |
| 11 | 107268600 | 107271199 | 22.20978 | 36.63678 | 71.89753 | 31.02963 | -0.5895 | 1.351988 | 0.69093 |
| 8 | 53317600 | 53321199 | 42.0496 | 68.44534 | 73.59159 | 31.59248 | 0.265819 | 1.572745 | 0.690962 |
| 14 | 12214200 | 12214999 | 9.20923 | 17.56413 | 31.42094 | 12.52924 | -0.32865 | 0.001748 | 0.690966 |
| 17 | 36339000 | 36343599 | 37.0502 | 59.68882 | 89.36248 | 38.00855 | -0.22535 | 1.804292 | 0.690967 |
| 7 | 11990800 | 11992599 | 35.55707 | 58.25353 | 64.95012 | 27.95009 | 0.219795 | 1.466283 | 0.690971 |
| 7 | 19770600 | 19780599 | 136.6327 | 208.4847 | 188.7644 | 78.2741 | 0.430505 | 3.119859 | 0.690974 |
| 11 | 82226600 | 82228199 | 24.51592 | 40.56612 | 62.84115 | 27.17628 | -0.2488 | 1.314368 | 0.690982 |
| 3 | 36309800 | 36314399 | 46.0696 | 73.32038 | 128.5483 | 53.72033 | -0.47313 | 2.211954 | 0.690984 |
| 6 | 130187200 | 130189799 | 29.52123 | 49.55647 | 54.61064 | 23.84069 | 0.24132 | 1.162839 | 0.691018 |
| 4 | 113693200 | 113695599 | 55.34046 | 88.72141 | 96.96657 | 40.94683 | 0.219155 | 1.985826 | 0.691068 |
| 1 | 17612200 | 17613799 | 16.64524 | 30.0103 | 31.73152 | 13.8197 | 0.326538 | 0.397969 | 0.691088 |
| 2 | 37367600 | 37367999 | 7.451505 | 14.39492 | 24.88661 | 9.564156 | -0.2293 | -0.15201 | 0.691103 |
| 1 | 110084200 | 110086599 | 25.11341 | 42.96923 | 47.38089 | 20.7764 | 0.242451 | 0.972128 | 0.691118 |
| 9 | 116764600 | 116767599 | 27.25102 | 45.83683 | 52.3266 | 22.86264 | 0.19065 | 1.135684 | 0.691122 |
| 10 | 9211800 | 9214599 | 22.76565 | 39.22214 | 44.1381 | 19.36197 | 0.214929 | 0.896822 | 0.691127 |
| 5 | 73763400 | 73766199 | 37.97887 | 61.69613 | 70.30375 | 30.15239 | 0.180024 | 1.595613 | 0.691133 |
| 4 | 37507400 | 37508599 | 23.40615 | 40.35909 | 38.37554 | 16.83455 | 0.457315 | 0.886018 | 0.691143 |
| 4 | 125236600 | 125240999 | 40.1533 | 64.60299 | 93.6153 | 39.74159 | -0.18077 | 1.852322 | 0.691154 |
| 5 | 144823200 | 144827199 | 40.35545 | 64.93818 | 93.52501 | 39.71207 | -0.17164 | 1.848831 | 0.691158 |
| 6 | 136392200 | 136393599 | 14.14543 | 25.44202 | 28.85225 | 12.60104 | 0.223504 | 0.388375 | 0.691171 |
| 1 | 177407600 | 177413199 | 65.42294 | 104.0034 | 116.7507 | 48.72043 | 0.167305 | 2.260517 | 0.691173 |
| 17 | 48239800 | 48240799 | 12.61204 | 23.44873 | 27.15735 | 11.36022 | 0.236024 | 0.187563 | 0.691177 |
| 14 | 8160000 | 8161399 | 21.36428 | 35.81667 | 55.8632 | 24.36735 | -0.25794 | 1.153345 | 0.691179 |
| 3 | 87381000 | 87383799 | 37.24688 | 61.99254 | 59.51354 | 25.84111 | 0.43996 | 1.280335 | 0.691179 |
| X | 8602600 | 8623599 | 319.0937 | 458.5454 | 420.3099 | 173.5133 | 0.333322 | 4.181343 | 0.691201 |
| 16 | 4551600 | 4553799 | 15.10852 | 26.93375 | 30.28587 | 13.32667 | 0.228245 | 0.452664 | 0.69122 |
| 4 | 120984800 | 120985799 | 20.79921 | 35.39836 | 52.72739 | 23.1109 | -0.19079 | 1.012672 | 0.691245 |
| 7 | 58686800 | 58690999 | 54.89601 | 88.23961 | 80.92566 | 34.30777 | 0.476157 | 1.896387 | 0.691278 |
| 9 | 15710000 | 15711599 | 14.48403 | 26.22796 | 29.85467 | 12.93155 | 0.225823 | 0.354088 | 0.691295 |

| | | | | | | | | | |
|---|---|---|---|---|---|---|---|---|---|
| 7 | 16121000 | 16121999 | 8.046395 | 15.37601 | 25.63556 | 10.18061 | -0.22225 | -0.0378 | 0.691316 |
| 14 | 84547400 | 84547999 | 8.046395 | 15.37601 | 25.63556 | 10.18061 | -0.22225 | -0.0378 | 0.691316 |
| 12 | 9583400 | 9584599 | 16.8527 | 29.33325 | 44.19909 | 19.42778 | -0.20156 | 0.767894 | 0.691332 |
| 4 | 121472200 | 121476799 | 67.40887 | 107.3169 | 115.4354 | 48.2621 | 0.230996 | 2.216714 | 0.691341 |
| 11 | 73744800 | 73749399 | 53.28725 | 85.42299 | 96.00591 | 40.53631 | 0.178794 | 1.984527 | 0.691369 |
| 13 | 22755400 | 22769799 | 167.9356 | 252.3464 | 262.1728 | 108.2266 | 0.213585 | 3.436997 | 0.691373 |
| 13 | 107158000 | 107161999 | 34.73536 | 56.31365 | 81.80534 | 35.03326 | -0.17215 | 1.634824 | 0.691392 |
| 4 | 3248600 | 3261599 | 407.6408 | 570.3397 | 483.5988 | 200.5073 | 0.406972 | 4.574282 | 0.691416 |
| 3 | 12286400 | 12288799 | 36.95999 | 60.28317 | 68.91289 | 29.60585 | 0.179693 | 1.53148 | 0.691418 |
| 2 | 134977000 | 134978199 | 12.19766 | 22.70083 | 26.18302 | 10.93425 | 0.244125 | 0.176918 | 0.691419 |
| 2 | 111215000 | 111222999 | 132.6515 | 202.4631 | 211.8178 | 87.8389 | 0.22246 | 3.114753 | 0.69142 |
| 11 | 63787000 | 63789999 | 22.40847 | 37.82926 | 55.99767 | 24.49369 | -0.18267 | 1.09147 | 0.691437 |
| 17 | 55298600 | 55308799 | 160.9716 | 242.3211 | 254.7808 | 105.2262 | 0.198687 | 3.400484 | 0.69145 |
| 3 | 142736000 | 142737399 | 17.62827 | 30.69808 | 46.28017 | 20.34872 | -0.2024 | 0.766511 | 0.691459 |
| 13 | 72244400 | 72246199 | 12.46621 | 22.11339 | 34.68401 | 15.32593 | -0.25307 | 0.477273 | 0.691496 |
| Y | 67981800 | 67982799 | 13.14263 | 24.33095 | 20.44733 | 8.603589 | 0.689355 | 0.198007 | 0.691505 |
| 12 | 106915200 | 106919399 | 37.87384 | 61.73182 | 70.27164 | 30.18212 | 0.185046 | 1.542378 | 0.691526 |
| 8 | 8548400 | 8556199 | 63.92908 | 100.8755 | 142.076 | 58.9716 | -0.16889 | 2.457404 | 0.691545 |
| 1 | 18751800 | 18753799 | 20.23883 | 35.44329 | 32.81135 | 14.48477 | 0.498244 | 0.674121 | 0.691546 |
| 12 | 98146000 | 98146999 | 9.018621 | 16.7868 | 27.88365 | 11.64161 | -0.27824 | 0.151549 | 0.691549 |
| 1 | 27981400 | 27982799 | 19.3819 | 33.86754 | 38.2685 | 16.86233 | 0.212317 | 0.71644 | 0.691553 |
| 1 | 9128200 | 9129399 | 21.66345 | 36.59407 | 54.85123 | 23.99671 | -0.20051 | 1.084227 | 0.691557 |
| 5 | 127054400 | 127057599 | 40.2156 | 64.87457 | 94.50539 | 40.2672 | -0.18374 | 1.773893 | 0.691601 |
| 15 | 86350400 | 86355599 | 53.57068 | 85.85555 | 96.72765 | 40.82529 | 0.174842 | 1.992273 | 0.691614 |
| 9 | 68439000 | 68442399 | 28.04355 | 47.65713 | 50.13508 | 21.97205 | 0.308605 | 1.046293 | 0.691626 |
| 19 | 43335400 | 43339599 | 21.69055 | 37.8519 | 35.03622 | 15.42535 | 0.498522 | 0.740362 | 0.691628 |
| 3 | 8193400 | 8195199 | 20.03486 | 34.30131 | 51.15327 | 22.43171 | -0.19125 | 0.951531 | 0.691631 |
| 16 | 15018800 | 15019799 | 8.99702 | 16.89306 | 27.45675 | 11.29557 | -0.22997 | 0.094699 | 0.691634 |
| 15 | 95344200 | 95346599 | 22.01141 | 38.58358 | 38.94313 | 17.20526 | 0.374255 | 0.680287 | 0.691645 |
| 14 | 114357800 | 114358399 | 10.72414 | 20.24571 | 24.21333 | 9.865866 | 0.220388 | 0.065707 | 0.691751 |
| 16 | 97810200 | 97815799 | 119.6721 | 184.8915 | 179.4425 | 74.53654 | 0.344211 | 2.875793 | 0.691758 |
| X | 32865000 | 32866999 | 26.14558 | 45.39137 | 36.06747 | 15.82793 | 0.717253 | 0.836527 | 0.691764 |
| 4 | 146048200 | 146051199 | 35.25872 | 59.45336 | 43.59433 | 19.06167 | 0.827242 | 1.134542 | 0.691786 |
| 7 | 42087800 | 42107599 | 424.8627 | 587.6943 | 589.7954 | 244.974 | 0.147303 | 4.746884 | 0.691817 |
| 3 | 53083000 | 53083799 | 13.15681 | 24.43149 | 27.66697 | 11.59325 | 0.266492 | 0.200646 | 0.691823 |
| 2 | 110851000 | 110860599 | 130.7835 | 199.7647 | 209.6276 | 86.94601 | 0.218969 | 3.099107 | 0.69184 |
| 9 | 77922400 | 77924599 | 12.95591 | 22.89782 | 35.73107 | 15.80567 | -0.24796 | 0.520491 | 0.691869 |
| 11 | 15023600 | 15024599 | 15.02164 | 26.41025 | 39.74517 | 17.5997 | -0.19913 | 0.596853 | 0.691917 |
| 16 | 94073200 | 94078799 | 83.34334 | 128.0352 | 214.7468 | 89.16612 | -0.45053 | 2.971233 | 0.691929 |
| X | 146529400 | 146546799 | 230.6473 | 341.7343 | 308.6103 | 127.0991 | 0.397228 | 3.704484 | 0.691943 |
| 14 | 116720800 | 116721799 | 11.77016 | 22.21683 | 25.82389 | 10.52572 | 0.260898 | 0.081245 | 0.691962 |
| 3 | 96295800 | 96298799 | 34.66762 | 55.90974 | 96.91687 | 41.2804 | -0.43452 | 1.773004 | 0.691985 |
| 8 | 4502400 | 4503199 | 12.20766 | 22.66759 | 26.60229 | 11.14803 | 0.215046 | 0.189145 | 0.692002 |
| X | 89132400 | 89134599 | 27.0137 | 46.20994 | 48.46859 | 21.25295 | 0.314324 | 0.983769 | 0.692003 |
| 7 | 18431200 | 18431999 | 13.85737 | 25.41195 | 23.33536 | 9.946322 | 0.549045 | 0.263548 | 0.692038 |
| 4 | 145260000 | 145341199 | 1756.952 | 2155.556 | 2463.342 | 1038.319 | -0.2142 | 6.671342 | 0.692056 |
| 10 | 3270600 | 3273799 | 38.49758 | 62.58625 | 71.0121 | 30.47139 | 0.187174 | 1.583252 | 0.692056 |

| | | | | | | | | | |
|---|---|---|---|---|---|---|---|---|---|
| 18 | 73553200 | 73556399 | 31.26073 | 51.77173 | 58.58523 | 25.351 | 0.202828 | 1.316158 | 0.692065 |
| 13 | 103708400 | 103711599 | 25.83345 | 42.17557 | 81.35545 | 34.96348 | -0.57213 | 1.500266 | 0.692068 |
| 1 | 103673000 | 103675599 | 50.75656 | 81.02802 | 114.9002 | 48.22203 | -0.16274 | 2.115416 | 0.692073 |
| 4 | 143495800 | 143498799 | 53.34255 | 85.69535 | 93.16052 | 39.46311 | 0.230339 | 1.916203 | 0.692075 |
| 7 | 25968800 | 25984399 | 223.4961 | 329.0162 | 333.8755 | 137.4919 | 0.220391 | 3.808253 | 0.692085 |
| 11 | 74046000 | 74046999 | 16.52167 | 29.04172 | 32.11797 | 14.22284 | 0.24307 | 0.594666 | 0.692103 |
| 6 | 118257000 | 118262599 | 52.47996 | 83.58491 | 118.5883 | 49.60924 | -0.16714 | 2.189997 | 0.692106 |
| 7 | 123287800 | 123289799 | 13.6856 | 24.36735 | 37.63521 | 16.57764 | -0.22815 | 0.465953 | 0.692119 |
| 17 | 80324400 | 80324999 | 6.680145 | 12.96155 | 22.80834 | 8.587551 | -0.22626 | -0.23672 | 0.69212 |
| 5 | 149611000 | 149613399 | 16.29174 | 28.35738 | 43.14978 | 18.96677 | -0.21572 | 0.761523 | 0.692133 |
| 18 | 71088000 | 71089999 | 16.54652 | 29.72724 | 31.63988 | 13.8316 | 0.313424 | 0.416709 | 0.692139 |
| 10 | 19606800 | 19612199 | 55.69209 | 89.21317 | 98.97903 | 41.74542 | 0.196139 | 2.009804 | 0.692176 |
| 15 | 93670400 | 93671999 | 13.33223 | 23.79727 | 37.13674 | 16.32383 | -0.24094 | 0.446497 | 0.692208 |
| 13 | 82677000 | 82677599 | 7.63669 | 14.65042 | 25.18625 | 9.891401 | -0.25041 | -0.07683 | 0.692214 |
| 4 | 113956600 | 113958399 | 13.74121 | 24.53872 | 38.22118 | 16.7939 | -0.2377 | 0.450266 | 0.692251 |
| 2 | 149062200 | 149077399 | 345.8452 | 488.3017 | 493.5961 | 204.3562 | 0.166678 | 4.438038 | 0.692261 |
| X | 71064400 | 71066599 | 25.75823 | 44.18698 | 47.37356 | 20.77752 | 0.283466 | 0.95346 | 0.692278 |
| 14 | 24896600 | 24897399 | 10.56635 | 19.54208 | 30.91339 | 13.02494 | -0.21844 | 0.207893 | 0.692292 |
| 15 | 74699600 | 74702999 | 25.03468 | 42.03728 | 65.0679 | 28.41111 | -0.2473 | 1.154612 | 0.692312 |
| 7 | 24275200 | 24280999 | 46.46873 | 74.69553 | 84.37243 | 35.77031 | 0.176257 | 1.886464 | 0.692345 |
| 14 | 86195800 | 86198199 | 22.60676 | 39.20596 | 43.01559 | 18.87335 | 0.253261 | 0.83665 | 0.692376 |
| 17 | 64071800 | 64074199 | 29.95791 | 50.18579 | 48.07748 | 20.96126 | 0.442761 | 1.180213 | 0.69238 |
| 2 | 181557000 | 181566199 | 219.7193 | 318.0321 | 413.8284 | 170.667 | -0.16164 | 4.066813 | 0.692386 |
| 17 | 23067400 | 23081599 | 206.2381 | 307.1492 | 293.4162 | 120.905 | 0.323016 | 3.610879 | 0.692395 |
| 1 | 86387000 | 86389399 | 22.11733 | 37.06135 | 57.01836 | 24.8637 | -0.23836 | 1.160254 | 0.692409 |
| 18 | 49494800 | 49496199 | 12.39421 | 23.20651 | 26.72827 | 11.04826 | 0.257583 | 0.139581 | 0.692417 |
| 9 | 9400400 | 9406199 | 62.77623 | 99.9996 | 111.863 | 46.80607 | 0.17545 | 2.193028 | 0.692417 |
| 8 | 86704200 | 86709799 | 65.76403 | 104.9217 | 110.1397 | 46.18407 | 0.269427 | 2.148253 | 0.692459 |
| 4 | 9190200 | 9191399 | 13.88181 | 24.38297 | 38.29327 | 16.94733 | -0.26045 | 0.601107 | 0.692479 |
| 7 | 7014800 | 7035599 | 352.8165 | 496.0648 | 505.6947 | 209.5084 | 0.14843 | 4.499445 | 0.692484 |
| 13 | 67215400 | 67227599 | 52.27509 | 84.23665 | 76.11701 | 32.38608 | 0.501955 | 1.815982 | 0.692491 |
| X | 54929600 | 54930999 | 23.51674 | 40.5416 | 44.70174 | 19.60959 | 0.244589 | 0.89864 | 0.692492 |
| 9 | 35153400 | 35154599 | 11.0191 | 21.22385 | 23.65651 | 9.184792 | 0.383799 | -0.08823 | 0.692501 |
| 15 | 4405000 | 4407199 | 15.20709 | 26.45816 | 42.17733 | 18.53436 | -0.28248 | 0.761395 | 0.692527 |
| 6 | 68605600 | 68605999 | 8.010612 | 15.55238 | 20.0009 | 7.504518 | 0.22665 | -0.24144 | 0.692532 |
| 7 | 92704000 | 92704399 | 8.010612 | 15.55238 | 20.0009 | 7.504518 | 0.22665 | -0.24144 | 0.692532 |
| 6 | 130159800 | 130162199 | 23.66276 | 41.60776 | 37.10685 | 16.43101 | 0.552148 | 0.640973 | 0.692552 |
| 14 | 45662200 | 45666199 | 39.78138 | 65.10536 | 69.58704 | 29.93245 | 0.279835 | 1.490216 | 0.692557 |
| 5 | 142049600 | 142058799 | 160.4786 | 238.3587 | 316.5088 | 130.3849 | -0.15538 | 3.650285 | 0.692565 |
| 2 | 39694800 | 39699999 | 8.688989 | 16.90456 | 17.20217 | 6.346894 | 0.582726 | -0.30241 | 0.6926 |
| 1 | 79357400 | 79359199 | 26.51943 | 43.84686 | 65.67281 | 28.38546 | -0.20044 | 1.333054 | 0.692626 |
| 11 | 109873000 | 109874599 | 17.27152 | 30.75972 | 33.14966 | 14.5988 | 0.288375 | 0.478669 | 0.692639 |
| 6 | 138029600 | 138030199 | 6.248875 | 12.16938 | 22.54832 | 8.248562 | -0.26086 | -0.34129 | 0.692679 |
| 8 | 80360600 | 80363199 | 36.17889 | 59.2662 | 67.0531 | 28.85382 | 0.198591 | 1.466481 | 0.692684 |
| 13 | 97639000 | 97642199 | 31.5609 | 52.13434 | 60.12117 | 25.96939 | 0.175427 | 1.34975 | 0.692717 |
| 1 | 44477800 | 44479599 | 24.96408 | 42.78384 | 40.50504 | 17.7637 | 0.462003 | 0.95036 | 0.69272 |
| 6 | 74100000 | 74100999 | 10.49001 | 19.81278 | 23.7522 | 9.668603 | 0.21854 | 0.060815 | 0.692725 |

| | | | | | | | | | |
|---|---|---|---|---|---|---|---|---|---|
| 7 | 19218200 | 19218999 | 9.660123 | 17.87046 | 28.85215 | 12.15239 | -0.24699 | 0.195473 | 0.692736 |
| 16 | 10619800 | 10623599 | 28.91361 | 47.37885 | 71.33854 | 30.7018 | -0.21244 | 1.448673 | 0.692756 |
| 13 | 18110000 | 18111799 | 31.30021 | 51.53836 | 57.14308 | 24.64292 | 0.23069 | 1.386938 | 0.692767 |
| X | 88998400 | 88999999 | 19.23063 | 34.07706 | 34.72726 | 15.35356 | 0.364913 | 0.530458 | 0.692784 |
| 14 | 55683800 | 55686799 | 66.70133 | 104.8581 | 147.8859 | 61.347 | -0.175 | 2.536375 | 0.69279 |
| 10 | 36976200 | 36980199 | 32.13753 | 53.58141 | 58.11943 | 25.26344 | 0.264232 | 1.244793 | 0.692795 |
| 12 | 114836400 | 114839599 | 41.40684 | 67.56419 | 71.93849 | 30.91036 | 0.282345 | 1.53638 | 0.692797 |
| 19 | 45686400 | 45690399 | 39.53732 | 63.55983 | 93.52641 | 39.65671 | -0.20428 | 1.87791 | 0.692803 |
| 18 | 79973600 | 79978999 | 61.78033 | 98.5322 | 109.3388 | 45.82646 | 0.188894 | 2.152967 | 0.692819 |
| 4 | 121464000 | 121468799 | 64.23038 | 102.8566 | 101.5618 | 42.8132 | 0.363455 | 2.033538 | 0.692836 |
| 3 | 128048000 | 128051599 | 40.43051 | 65.82965 | 72.72658 | 31.22455 | 0.227074 | 1.565321 | 0.692868 |
| 5 | 4252400 | 4259599 | 82.48052 | 129.35 | 145.2351 | 60.23588 | 0.15081 | 2.591693 | 0.692868 |
| 5 | 118990000 | 118995799 | 54.83636 | 87.06848 | 124.6897 | 51.97043 | -0.18515 | 2.288875 | 0.692871 |
| 10 | 17152600 | 17156799 | 29.57184 | 50.13714 | 51.27449 | 22.45699 | 0.34909 | 1.07314 | 0.692874 |
| 2 | 152219800 | 152220999 | 12.4441 | 22.88461 | 26.58809 | 11.28942 | 0.216079 | 0.245926 | 0.692891 |
| 9 | 99624200 | 99624599 | 13.22306 | 24.22099 | 27.11635 | 11.57593 | 0.263899 | 0.277223 | 0.692908 |
| 1 | 167830200 | 167834399 | 64.20116 | 101.4957 | 142.1276 | 59.04924 | -0.15842 | 2.412866 | 0.69292 |
| 13 | 59021400 | 59024599 | 32.93628 | 54.27137 | 62.34859 | 26.89599 | 0.179953 | 1.385326 | 0.692925 |
| 4 | 152902600 | 152919599 | 246.1509 | 349.5391 | 538.474 | 222.7245 | -0.43265 | 4.350076 | 0.692928 |
| 1 | 60151200 | 60154599 | 38.74194 | 63.18009 | 70.94733 | 30.4782 | 0.205191 | 1.537899 | 0.692936 |
| 3 | 97725400 | 97727399 | 10.45729 | 19.17579 | 30.29385 | 12.91321 | -0.22847 | 0.254598 | 0.692957 |
| 10 | 114256200 | 114256999 | 7.71897 | 14.72722 | 25.60107 | 10.20068 | -0.28656 | -0.03229 | 0.692975 |
| 11 | 34731200 | 34733199 | 10.35297 | 18.90435 | 30.31057 | 12.99279 | -0.25511 | 0.279203 | 0.69299 |
| 18 | 60147600 | 60148599 | 10.35297 | 18.90435 | 30.31057 | 12.99279 | -0.25511 | 0.279203 | 0.69299 |
| 8 | 52315000 | 52318199 | 36.56942 | 59.52796 | 68.22268 | 29.28815 | 0.174013 | 1.557226 | 0.693 |
| 1 | 165087600 | 165089799 | 30.99251 | 51.812 | 49.17759 | 21.41302 | 0.456167 | 1.206747 | 0.693003 |
| 13 | 54039400 | 54041399 | 23.16742 | 38.68641 | 58.58828 | 25.49238 | -0.21559 | 1.204387 | 0.693004 |
| 8 | 92148800 | 92152399 | 67.34755 | 105.8515 | 148.7929 | 61.72155 | -0.17046 | 2.540869 | 0.693009 |
| X | 47607200 | 47608999 | 12.36433 | 22.22892 | 34.7391 | 15.17761 | -0.2364 | 0.395075 | 0.69301 |
| 7 | 24656400 | 24660799 | 36.30411 | 59.29639 | 67.79391 | 29.14 | 0.180923 | 1.509641 | 0.693012 |
| 13 | 33695200 | 33706399 | 135.9339 | 206.6843 | 220.1385 | 91.19426 | 0.192237 | 3.191854 | 0.693014 |
| 15 | 4309600 | 4311199 | 23.05427 | 40.09541 | 36.56604 | 16.06458 | 0.51934 | 0.795189 | 0.69302 |
| 17 | 40708000 | 40711799 | 40.72883 | 66.09878 | 74.76374 | 32.04213 | 0.189357 | 1.621664 | 0.693054 |
| 9 | 86655400 | 86657199 | 13.18984 | 24.10504 | 27.9725 | 11.97775 | 0.208916 | 0.288453 | 0.693076 |
| 17 | 55723600 | 55724399 | 13.18984 | 24.10504 | 27.9725 | 11.97775 | 0.208916 | 0.288453 | 0.693076 |
| 4 | 37365000 | 37368799 | 76.0058 | 119.8007 | 134.4191 | 55.77977 | 0.157419 | 2.486185 | 0.693078 |
| 9 | 19668400 | 19671999 | 52.65899 | 84.79108 | 89.31852 | 37.96135 | 0.279796 | 1.844958 | 0.693078 |
| 17 | 79337400 | 79339399 | 17.28857 | 30.95612 | 32.66226 | 14.31908 | 0.323229 | 0.445115 | 0.69313 |
| 18 | 9226200 | 9227799 | 11.47532 | 21.14712 | 34.67166 | 14.67999 | -0.27483 | 0.244949 | 0.693133 |
| 4 | 61339200 | 61340199 | 15.4045 | 27.75609 | 24.84012 | 10.82677 | 0.56512 | 0.37924 | 0.693155 |
| 6 | 121952200 | 121952999 | 15.25091 | 27.13788 | 30.26355 | 13.3374 | 0.238585 | 0.46461 | 0.693156 |
| 17 | 21852000 | 21853199 | 14.65614 | 27.91685 | 26.1241 | 10.44537 | 0.595984 | 0.048131 | 0.693159 |
| 5 | 54870000 | 54871399 | 15.50361 | 27.11878 | 40.96638 | 18.06687 | -0.20465 | 0.682777 | 0.693163 |
| 17 | 7189000 | 7195799 | 161.1207 | 240.9857 | 260.1251 | 107.2676 | 0.152477 | 3.522746 | 0.693164 |
| 9 | 109422800 | 109424799 | 21.94234 | 38.56184 | 37.77506 | 16.72561 | 0.417143 | 0.638767 | 0.693193 |
| 1 | 150672000 | 150674599 | 30.40929 | 51.4089 | 52.95228 | 23.17159 | 0.338746 | 1.103862 | 0.693202 |
| 19 | 3255000 | 3257599 | 26.47642 | 43.74558 | 65.16366 | 28.15199 | -0.19268 | 1.33916 | 0.693207 |

| | | | | | | | | | |
|---|---|---|---|---|---|---|---|---|---|
| 17 | 36673200 | 36673799 | 6.10226 | 11.86148 | 21.75244 | 8.083663 | -0.26667 | -0.28954 | 0.693214 |
| 12 | 105703800 | 105706599 | 30.96567 | 50.48787 | 74.37712 | 31.94912 | -0.18561 | 1.527221 | 0.69324 |
| 7 | 62398200 | 62420599 | 331.3071 | 473.2961 | 447.2939 | 184.8238 | 0.28075 | 4.267284 | 0.693253 |
| 12 | 40358200 | 40362599 | 51.62462 | 83.01778 | 91.07427 | 38.63486 | 0.219024 | 1.883678 | 0.693268 |
| 2 | 90253200 | 90255199 | 12.64446 | 22.45776 | 34.69577 | 15.3149 | -0.23025 | 0.471531 | 0.693277 |
| 10 | 98010200 | 98011999 | 13.1667 | 23.20288 | 43.46246 | 19.23683 | -0.51198 | 0.56695 | 0.693304 |
| 6 | 39261800 | 39265599 | 29.61077 | 48.6539 | 71.44984 | 30.78524 | -0.17488 | 1.418904 | 0.693313 |
| 13 | 55062000 | 55065399 | 35.07444 | 56.84171 | 82.33909 | 35.25193 | -0.16859 | 1.643713 | 0.693324 |
| 18 | 76855400 | 76858199 | 27.73087 | 46.41751 | 53.44116 | 23.28789 | 0.178461 | 1.186703 | 0.693339 |
| 3 | 13585600 | 13586199 | 9.30096 | 17.77447 | 21.26508 | 8.44387 | 0.252665 | -0.03908 | 0.693358 |
| 2 | 87762800 | 87766999 | 64.1861 | 101.935 | 115.0359 | 47.95666 | 0.158297 | 2.289167 | 0.693361 |
| 5 | 137739600 | 137741799 | 19.42008 | 33.77084 | 38.42712 | 16.88093 | 0.201567 | 0.783243 | 0.693375 |
| 11 | 106817600 | 106820399 | 30.49339 | 50.59563 | 57.95867 | 25.11234 | 0.18547 | 1.293163 | 0.693397 |
| 9 | 62317400 | 62319599 | 21.11166 | 35.91839 | 53.27473 | 23.35019 | -0.18477 | 1.017043 | 0.693399 |
| X | 6629800 | 6630399 | 11.16536 | 21.2958 | 23.9163 | 9.535902 | 0.338618 | 0.003441 | 0.693407 |
| 14 | 42988000 | 42989399 | 16.14116 | 28.97672 | 26.58384 | 11.62884 | 0.525874 | 0.408875 | 0.693416 |
| 11 | 23593000 | 23595799 | 31.93796 | 53.91919 | 53.54799 | 23.42253 | 0.391159 | 1.121449 | 0.693417 |
| 7 | 100409600 | 100412199 | 25.57821 | 41.63179 | 81.6763 | 35.0624 | -0.59945 | 1.547704 | 0.69342 |
| 16 | 27736600 | 27739799 | 44.13504 | 71.19649 | 81.22153 | 34.60665 | 0.168159 | 1.772497 | 0.693442 |
| Y | 5510400 | 5511399 | 10.74606 | 20.1171 | 19.23234 | 7.952104 | 0.524084 | 0.105533 | 0.693454 |
| 15 | 40061000 | 40064599 | 52.30417 | 84.14456 | 90.75295 | 38.52111 | 0.244245 | 1.872238 | 0.693504 |
| 12 | 62411800 | 62413199 | 16.05965 | 28.00586 | 42.62327 | 18.75743 | -0.21556 | 0.735325 | 0.693506 |
| 5 | 116943200 | 116945399 | 21.39349 | 36.30585 | 53.82565 | 23.58114 | -0.18458 | 1.039965 | 0.693517 |
| 6 | 19784000 | 19790599 | 127.5924 | 195.0958 | 184.018 | 76.34265 | 0.373889 | 3.074801 | 0.693519 |
| 11 | 19008200 | 19009399 | 11.68479 | 20.86398 | 33.25682 | 14.61405 | -0.27049 | 0.428866 | 0.693523 |
| 12 | 103856000 | 103864999 | 136.1367 | 205.0131 | 274.1152 | 113.2229 | -0.14732 | 3.388364 | 0.693524 |
| 15 | 33357800 | 33360999 | 26.45051 | 44.86065 | 50.13188 | 21.95885 | 0.221647 | 1.05813 | 0.693526 |
| 1 | 58672800 | 58675799 | 19.9534 | 34.42925 | 53.37412 | 23.4146 | -0.24487 | 0.885249 | 0.693546 |
| 10 | 63596400 | 63597199 | 8.541162 | 16.15915 | 26.52126 | 10.75736 | -0.22654 | 0.038943 | 0.693617 |
| 16 | 89352800 | 89353599 | 8.541162 | 16.15915 | 26.52126 | 10.75736 | -0.22654 | 0.038943 | 0.693617 |
| 3 | 93666200 | 93670599 | 52.25197 | 83.87376 | 92.80996 | 39.26545 | 0.203394 | 1.938754 | 0.693623 |
| 8 | 8569600 | 8572599 | 33.11693 | 53.822 | 78.35388 | 33.61605 | -0.1719 | 1.581328 | 0.693654 |
| 4 | 96201800 | 96202799 | 7.419784 | 14.35057 | 24.99774 | 9.554032 | -0.23291 | -0.17256 | 0.693657 |
| 15 | 43204800 | 43205399 | 7.419784 | 14.35057 | 24.99774 | 9.554032 | -0.23291 | -0.17256 | 0.693657 |
| 1 | 34465600 | 34474999 | 201.4908 | 294.2046 | 383.1798 | 157.8747 | -0.15073 | 3.933861 | 0.693662 |
| 9 | 122188000 | 122191599 | 49.89512 | 79.68167 | 112.5912 | 47.27723 | -0.15706 | 2.102126 | 0.693678 |
| 5 | 84970200 | 84974999 | 69.17086 | 109.666 | 122.8434 | 51.12499 | 0.166643 | 2.344114 | 0.693695 |
| 8 | 84203800 | 84207399 | 44.58465 | 71.42893 | 102.8756 | 43.39695 | -0.17939 | 1.993899 | 0.693701 |
| 4 | 4290600 | 4292599 | 22.48207 | 38.67414 | 44.17155 | 19.37574 | 0.193214 | 0.909915 | 0.693721 |
| X | 140927400 | 140927799 | 6.243915 | 12.14468 | 21.82617 | 8.068789 | -0.23137 | -0.30768 | 0.693725 |
| 7 | 86940000 | 86943199 | 32.01222 | 54.22334 | 52.40447 | 22.9453 | 0.430464 | 1.093724 | 0.693735 |
| 7 | 135379600 | 135382799 | 31.342 | 52.0199 | 59.12733 | 25.62421 | 0.196723 | 1.293316 | 0.693735 |
| 5 | 21913000 | 21915399 | 20.12133 | 34.35743 | 51.13089 | 22.41707 | -0.18875 | 0.978024 | 0.693746 |
| 13 | 88087600 | 88089199 | 17.74791 | 30.65774 | 46.35379 | 20.33555 | -0.20849 | 0.85588 | 0.69385 |
| 15 | 96616400 | 96618599 | 17.74791 | 30.65774 | 46.35379 | 20.33555 | -0.20849 | 0.85588 | 0.69385 |
| 17 | 17333600 | 17333999 | 6.397607 | 12.43353 | 22.33328 | 8.310486 | -0.23926 | -0.28056 | 0.693894 |
| 9 | 97231200 | 97232199 | 10.71047 | 20.26338 | 24.24243 | 9.832944 | 0.225595 | 0.050697 | 0.693928 |

| | | | | | | | | | |
|---|---|---|---|---|---|---|---|---|---|
| 17 | 5569000 | 5570799 | 15.78427 | 27.66537 | 41.89041 | 18.50585 | -0.20819 | 0.65194 | 0.693932 |
| 16 | 87232200 | 87234399 | 13.09494 | 23.07529 | 36.75212 | 16.26694 | -0.27911 | 0.554594 | 0.693953 |
| 7 | 121414400 | 121415399 | 19.15828 | 33.48214 | 38.23942 | 16.85175 | 0.19691 | 0.711787 | 0.69396 |
| 11 | 108899400 | 108904199 | 1995.868 | 2396.367 | 2717.944 | 1148.979 | -0.23502 | 7.012535 | 0.693967 |
| 5 | 52633400 | 52642199 | 78.96566 | 123.4565 | 172.8592 | 71.71564 | -0.17077 | 2.647041 | 0.693971 |
| 3 | 46925200 | 46925599 | 5.799729 | 11.30264 | 20.88048 | 7.56836 | -0.24517 | -0.38864 | 0.693979 |
| 7 | 54894000 | 54895999 | 18.80298 | 33.07672 | 36.33879 | 16.08953 | 0.252822 | 0.605997 | 0.693989 |
| 18 | 65620800 | 65628199 | 58.75265 | 92.83253 | 150.5353 | 62.52544 | -0.37036 | 2.424566 | 0.693995 |
| 6 | 102288200 | 102289599 | 15.36728 | 28.02123 | 29.81321 | 12.80194 | 0.33055 | 0.319949 | 0.693996 |
| X | 60388800 | 60390399 | 27.47479 | 46.99242 | 39.486 | 17.314 | 0.633168 | 0.979678 | 0.69404 |
| 13 | 33293000 | 33294199 | 17.41567 | 30.24868 | 45.2675 | 19.8708 | -0.19218 | 0.800346 | 0.694042 |
| 17 | 94626400 | 94627999 | 19.03998 | 32.83253 | 49.00963 | 21.49954 | -0.19068 | 0.880312 | 0.694046 |
| 14 | 39483600 | 39486799 | 31.68925 | 51.43827 | 77.21493 | 33.09732 | -0.21762 | 1.603568 | 0.694076 |
| 15 | 12832600 | 12834799 | 25.72378 | 43.14613 | 45.99801 | 20.07031 | 0.289056 | 1.151704 | 0.69408 |
| 17 | 6759800 | 6761799 | 19.34778 | 32.93068 | 50.5812 | 22.17034 | -0.2348 | 1.004325 | 0.6941 |
| 3 | 23304400 | 23307199 | 25.52985 | 43.43874 | 49.02615 | 21.48818 | 0.207863 | 1.026572 | 0.694108 |
| 3 | 110080000 | 110082799 | 25.57915 | 43.34098 | 49.58332 | 21.71222 | 0.187955 | 1.064421 | 0.694109 |
| 7 | 75690600 | 75692599 | 25.60347 | 42.48643 | 63.06635 | 27.32731 | -0.18716 | 1.28425 | 0.69411 |
| 2 | 149126400 | 149131799 | 80.22353 | 126.1059 | 140.2966 | 58.19818 | 0.166673 | 2.54571 | 0.694159 |
| 10 | 8272600 | 8273599 | 32.50278 | 53.37493 | 59.27206 | 25.53201 | 0.226926 | 1.425314 | 0.694199 |
| 7 | 35436800 | 35437999 | 11.60204 | 21.37226 | 35.22364 | 14.92166 | -0.28298 | 0.248377 | 0.69422 |
| 6 | 38104800 | 38105999 | 13.81127 | 24.21412 | 38.46913 | 16.99851 | -0.27672 | 0.634034 | 0.694225 |
| 6 | 58544400 | 58545999 | 16.50948 | 28.88328 | 43.53772 | 19.20371 | -0.20162 | 0.689704 | 0.694238 |
| 18 | 81037200 | 81039199 | 17.99309 | 31.01067 | 46.89622 | 20.57209 | -0.20933 | 0.880116 | 0.694307 |
| 10 | 13659200 | 13661199 | 15.20251 | 26.80982 | 41.27715 | 18.26695 | -0.23022 | 0.567195 | 0.694338 |
| 12 | 21370200 | 21373199 | 22.78825 | 38.09465 | 58.09736 | 25.29613 | -0.22573 | 1.189398 | 0.694349 |
| 1 | 5747000 | 5747399 | 7.231765 | 14.09053 | 18.51978 | 6.730765 | 0.237147 | -0.36887 | 0.694358 |
| 12 | 101466600 | 101467799 | 15.19665 | 26.82127 | 41.79995 | 18.49531 | -0.24725 | 0.558906 | 0.694367 |
| 4 | 15853400 | 15854799 | 10.62794 | 19.73686 | 31.54883 | 13.21863 | -0.2274 | 0.188353 | 0.694376 |
| 18 | 68604800 | 68605799 | 6.879461 | 13.14808 | 24.04896 | 9.547469 | -0.35358 | -0.05324 | 0.694383 |
| 3 | 3330400 | 3332399 | 25.56152 | 43.93848 | 46.81909 | 20.5366 | 0.292775 | 0.933724 | 0.694425 |
| 8 | 70137000 | 70140799 | 57.36038 | 91.87856 | 87.35095 | 36.83687 | 0.418388 | 2.011408 | 0.694469 |
| 2 | 15400600 | 15402799 | 18.26755 | 31.69826 | 47.40196 | 20.79825 | -0.19132 | 0.820066 | 0.694479 |
| 17 | 11189600 | 11191599 | 14.31386 | 26.64748 | 28.15989 | 11.75365 | 0.36963 | 0.193987 | 0.69449 |
| 11 | 20971800 | 20972799 | 11.26369 | 21.19426 | 24.26771 | 9.947558 | 0.276307 | 0.095741 | 0.694494 |
| 16 | 21475200 | 21481199 | 70.96798 | 111.4436 | 154.2592 | 63.98172 | -0.14927 | 2.561208 | 0.694519 |
| 3 | 72424600 | 72425799 | 17.52114 | 31.14863 | 28.24745 | 12.45865 | 0.534807 | 0.484348 | 0.694519 |
| 17 | 21035000 | 21039799 | 56.25039 | 90.20579 | 97.656 | 41.25575 | 0.233313 | 1.977355 | 0.69453 |
| 8 | 17095400 | 17095999 | 8.678896 | 16.76588 | 20.90287 | 8.033462 | 0.237533 | -0.15317 | 0.694536 |
| 3 | 4389800 | 4390999 | 13.73752 | 25.42444 | 28.02995 | 11.79906 | 0.299743 | 0.221605 | 0.694542 |
| 12 | 105363600 | 105366999 | 47.92742 | 76.55125 | 109.5938 | 46.02877 | -0.17566 | 2.094403 | 0.69456 |
| 16 | 83074000 | 83074799 | 9.471208 | 18.24785 | 22.49712 | 8.727238 | 0.240883 | -0.10926 | 0.694577 |
| 9 | 49958600 | 49959999 | 15.36688 | 27.82845 | 30.86354 | 13.36745 | 0.26325 | 0.362812 | 0.694586 |
| 14 | 57262800 | 57263599 | 7.7668 | 14.81724 | 25.06219 | 9.987724 | -0.24696 | -0.02969 | 0.694595 |
| 16 | 7908800 | 7909599 | 7.7668 | 14.81724 | 25.06219 | 9.987724 | -0.24696 | -0.02969 | 0.694595 |
| 3 | 32912800 | 32915999 | 36.85689 | 59.52952 | 86.71234 | 37.00532 | -0.18156 | 1.727439 | 0.694613 |
| 2 | 164508200 | 164511599 | 41.12984 | 66.27686 | 95.79099 | 40.75671 | -0.17427 | 1.806179 | 0.694615 |

| | | | | | | | | | |
|---|---|---|---|---|---|---|---|---|---|
| 10 | 33621400 | 33630999 | 179.5906 | 266.7621 | 287.0547 | 118.2521 | 0.147858 | 3.650415 | 0.694616 |
| 11 | 117070400 | 117075199 | 76.4065 | 119.3038 | 164.6602 | 68.32298 | -0.15146 | 2.667549 | 0.694623 |
| 7 | 47651600 | 47656399 | 58.54727 | 93.55599 | 104.3583 | 43.85803 | 0.18443 | 2.09014 | 0.694634 |
| 4 | 29566800 | 29569799 | 30.86395 | 50.74267 | 77.12202 | 33.2374 | -0.22407 | 1.416634 | 0.694634 |
| 3 | 72252600 | 72256999 | 59.66303 | 95.40451 | 104.0265 | 43.76119 | 0.218268 | 2.069945 | 0.694636 |
| 6 | 99814400 | 99815399 | 9.606772 | 18.04755 | 29.08593 | 11.95503 | -0.21705 | 0.100142 | 0.69464 |
| 16 | 36248600 | 36252599 | 36.83279 | 60.8179 | 64.05223 | 27.6596 | 0.305872 | 1.370565 | 0.694674 |
| 17 | 22578800 | 22595599 | 189.2012 | 283.0512 | 277.9335 | 114.6165 | 0.289285 | 3.525475 | 0.694678 |
| 5 | 75926600 | 75927799 | 17.23159 | 29.89388 | 45.28282 | 19.86831 | -0.20988 | 0.813131 | 0.694679 |
| 9 | 103157800 | 103158399 | 8.907297 | 17.14666 | 21.40214 | 8.322955 | 0.220392 | -0.10874 | 0.694736 |
| 14 | 50404000 | 50406199 | 30.33062 | 50.48707 | 57.42746 | 24.93718 | 0.195955 | 1.256033 | 0.694775 |
| 3 | 16340800 | 16345199 | 40.19066 | 66.41328 | 63.19206 | 27.3026 | 0.452044 | 1.370453 | 0.694777 |
| 14 | 53651600 | 53654199 | 42.39049 | 68.79194 | 62.13827 | 26.63011 | 0.513018 | 1.620683 | 0.694817 |
| 7 | 40474400 | 40479199 | 45.37184 | 72.6639 | 104.928 | 44.23848 | -0.18379 | 2.006009 | 0.694827 |
| 11 | 59248000 | 59249799 | 17.42186 | 30.24156 | 45.28079 | 19.86925 | -0.19301 | 0.808961 | 0.694828 |
| 2 | 161968400 | 161969999 | 17.06606 | 30.24568 | 33.77408 | 14.93172 | 0.233616 | 0.511923 | 0.694832 |
| 8 | 79287400 | 79288599 | 17.06606 | 30.24568 | 33.77408 | 14.93172 | 0.233616 | 0.511923 | 0.694832 |
| 11 | 45205600 | 45207599 | 17.06606 | 30.24568 | 33.77408 | 14.93172 | 0.233616 | 0.511923 | 0.694832 |
| 15 | 16613600 | 16614799 | 15.33916 | 27.46161 | 31.30217 | 13.7245 | 0.212344 | 0.43077 | 0.694832 |
| X | 75793200 | 75794999 | 18.36145 | 32.63378 | 34.15259 | 15.06653 | 0.328873 | 0.501152 | 0.694836 |
| 11 | 74548600 | 74549599 | 12.89432 | 22.69781 | 36.92338 | 16.34619 | -0.30962 | 0.569659 | 0.694837 |
| 17 | 42903400 | 42905399 | 18.38782 | 32.23173 | 36.50777 | 16.12927 | 0.208598 | 0.655359 | 0.694842 |
| 11 | 115499000 | 115500599 | 9.61893 | 17.58565 | 29.90173 | 12.79782 | -0.33743 | 0.267324 | 0.694847 |
| 7 | 30679000 | 30682599 | 38.65816 | 62.31003 | 90.54577 | 38.53848 | -0.18158 | 1.792032 | 0.694859 |
| 10 | 11212400 | 11219599 | 75.31698 | 118.2068 | 165.8311 | 68.77638 | -0.16924 | 2.577305 | 0.694924 |
| X | 152916200 | 152929599 | 239.5798 | 351.7513 | 342.5322 | 141.0752 | 0.276168 | 3.852905 | 0.694924 |
| 4 | 54624000 | 54625599 | 19.71117 | 34.3079 | 39.36476 | 17.30261 | 0.189645 | 0.772484 | 0.694927 |
| X | 76376000 | 76379399 | 45.47517 | 73.46865 | 80.87877 | 34.52748 | 0.221996 | 1.734315 | 0.694943 |
| 13 | 72449600 | 72452599 | 20.54465 | 34.98521 | 52.21366 | 22.88678 | -0.19355 | 1.003721 | 0.694968 |
| X | 120605000 | 120606399 | 19.87202 | 34.87859 | 37.03216 | 16.3789 | 0.301691 | 0.648094 | 0.694973 |
| 1 | 170368600 | 170370199 | 25.18842 | 43.1613 | 47.37888 | 20.77799 | 0.249276 | 0.960217 | 0.694977 |
| X | 34418200 | 34418799 | 5.496771 | 10.72259 | 20.3652 | 7.28877 | -0.26673 | -0.43454 | 0.694978 |
| 15 | 54539400 | 54540999 | 15.26856 | 27.15343 | 30.94478 | 13.64434 | 0.207286 | 0.472597 | 0.69498 |
| 4 | 109420800 | 109423799 | 25.39726 | 42.46941 | 63.27124 | 27.55434 | -0.19219 | 1.19901 | 0.694996 |
| 7 | 26789400 | 26793999 | 44.27014 | 71.39467 | 81.23599 | 34.60047 | 0.17144 | 1.780222 | 0.694999 |
| 11 | 15209800 | 15211199 | 18.26719 | 32.00275 | 36.5811 | 16.15419 | 0.195551 | 0.66426 | 0.695014 |
| 12 | 47498200 | 47498999 | 16.65968 | 28.89262 | 44.26885 | 19.42324 | -0.22622 | 0.813727 | 0.695015 |
| 6 | 78013000 | 78014399 | 14.85924 | 26.11153 | 39.25607 | 17.38004 | -0.19779 | 0.602045 | 0.695027 |
| 9 | 97217200 | 97228799 | 114.9109 | 175.1139 | 237.3465 | 98.38922 | -0.15254 | 3.142854 | 0.695036 |
| X | 76388800 | 76392199 | 29.54551 | 50.61155 | 47.89707 | 21.00472 | 0.462466 | 0.981028 | 0.695053 |
| 12 | 114919600 | 114922199 | 37.3002 | 61.17477 | 67.30242 | 28.97548 | 0.23951 | 1.454494 | 0.695058 |
| 7 | 141772400 | 141777599 | 89.16668 | 139.1749 | 153.3483 | 63.63178 | 0.172686 | 2.680779 | 0.695063 |
| 13 | 28777800 | 28779799 | 20.22716 | 35.63302 | 36.77354 | 16.27736 | 0.343476 | 0.600535 | 0.695066 |
| 7 | 139655600 | 139656799 | 9.938834 | 18.41594 | 29.361 | 12.33941 | -0.22686 | 0.18895 | 0.695077 |
| 5 | 16386000 | 16386599 | 7.28823 | 14.0984 | 24.3926 | 9.31573 | -0.22183 | -0.17691 | 0.69508 |
| 3 | 13270800 | 13271599 | 13.97149 | 25.14148 | 28.36458 | 12.38294 | 0.231312 | 0.38295 | 0.695081 |
| 7 | 102908400 | 102909599 | 13.91689 | 25.13171 | 28.95803 | 12.58642 | 0.205341 | 0.365281 | 0.695094 |

| | | | | | | | | | |
|---|---|---|---|---|---|---|---|---|---|
| 3 | 22338800 | 22339399 | 10.96898 | 20.6478 | 24.73866 | 10.1339 | 0.211675 | 0.08951 | 0.695099 |
| Y | 5872800 | 5873599 | 9.525503 | 18.09994 | 18.7344 | 7.529381 | 0.444729 | -0.00605 | 0.695128 |
| 1 | 104097000 | 104099199 | 23.68714 | 40.63512 | 45.20879 | 19.82812 | 0.230125 | 0.939144 | 0.695129 |
| 7 | 104643200 | 104644599 | 16.351 | 28.45666 | 43.17388 | 18.97574 | -0.21141 | 0.765371 | 0.695145 |
| 1 | 27308800 | 27311599 | 18.18747 | 32.11147 | 34.66782 | 15.33844 | 0.279759 | 0.558464 | 0.695153 |
| 1 | 46374800 | 46375799 | 9.884357 | 18.47396 | 30.0593 | 12.46297 | -0.2411 | 0.136154 | 0.695162 |
| 2 | 56785200 | 56786199 | 9.884357 | 18.47396 | 30.0593 | 12.46297 | -0.2411 | 0.136154 | 0.695162 |
| X | 60120200 | 60122799 | 23.92615 | 41.38566 | 44.64886 | 19.5884 | 0.276793 | 0.869497 | 0.695198 |
| 9 | 6757200 | 6757599 | 7.135776 | 13.79126 | 23.75677 | 9.110563 | -0.22114 | -0.16634 | 0.695223 |
| 17 | 81597000 | 81601599 | 65.30641 | 103.5848 | 117.0618 | 48.74033 | 0.154527 | 2.326362 | 0.695225 |
| 4 | 10773000 | 10775199 | 14.59776 | 26.89315 | 24.78026 | 10.49627 | 0.550653 | 0.247954 | 0.695234 |
| X | 22372600 | 22373799 | 18.78597 | 33.39879 | 27.70581 | 12.21927 | 0.662825 | 0.492394 | 0.695241 |
| 12 | 35850800 | 35853599 | 17.11519 | 29.24916 | 54.27354 | 23.79651 | -0.50553 | 0.961982 | 0.695242 |
| 17 | 33843800 | 33847199 | 28.42577 | 47.82763 | 53.44892 | 23.35521 | 0.221277 | 1.138173 | 0.695247 |
| 14 | 48081600 | 48082599 | 9.231988 | 17.84575 | 21.46605 | 8.225295 | 0.292433 | -0.15693 | 0.695266 |
| 17 | 88589200 | 88592799 | 31.10317 | 51.8395 | 57.5145 | 24.99521 | 0.231739 | 1.24411 | 0.695338 |
| 3 | 68872200 | 68874199 | 22.74597 | 39.10862 | 44.72196 | 19.61688 | 0.191395 | 0.91757 | 0.695351 |
| 12 | 98431000 | 98432999 | 25.53179 | 43.59122 | 41.54367 | 18.21364 | 0.451627 | 0.986573 | 0.695355 |
| 1 | 101376400 | 101377799 | 13.82383 | 25.51255 | 28.55331 | 12.0653 | 0.27438 | 0.23985 | 0.695371 |
| 2 | 66483600 | 66484199 | 11.29847 | 21.34551 | 25.31566 | 10.29913 | 0.234524 | 0.07 | 0.695377 |
| 2 | 23145200 | 23146599 | 12.3699 | 21.9717 | 43.5128 | 19.20562 | -0.58685 | 0.484504 | 0.695385 |
| 3 | 73370600 | 73371799 | 14.35314 | 26.77333 | 28.26345 | 11.75756 | 0.375271 | 0.184257 | 0.695406 |
| X | 64144400 | 64149999 | 98.8871 | 153.8739 | 162.8953 | 67.62177 | 0.226908 | 2.739724 | 0.695407 |
| 5 | 37797400 | 37802599 | 58.85844 | 93.74374 | 105.8401 | 44.27615 | 0.161891 | 2.193555 | 0.695424 |
| 4 | 45681800 | 45683399 | 20.68522 | 35.78742 | 41.03183 | 18.00165 | 0.189409 | 0.847493 | 0.695425 |
| 5 | 38070000 | 38072199 | 23.26791 | 40.01626 | 45.24271 | 19.84545 | 0.207809 | 0.91843 | 0.695431 |
| 4 | 4661400 | 4663399 | 35.78228 | 57.81167 | 84.50387 | 36.07717 | -0.186 | 1.717696 | 0.695445 |
| 4 | 152827000 | 152831399 | 45.75629 | 73.37133 | 104.1724 | 44.00355 | -0.15728 | 1.968792 | 0.695486 |
| 7 | 102735000 | 102735599 | 7.707222 | 14.73621 | 25.65396 | 10.17261 | -0.2825 | -0.04833 | 0.695498 |
| 2 | 51888400 | 51889999 | 13.95488 | 24.74761 | 37.65487 | 16.64234 | -0.2098 | 0.495327 | 0.695527 |
| 14 | 115865200 | 115867199 | 18.59716 | 33.09594 | 34.18164 | 15.06274 | 0.348982 | 0.495447 | 0.695534 |
| 12 | 23209800 | 23273999 | 1546.363 | 1924.254 | 1735.754 | 730.3878 | 0.14781 | 6.444999 | 0.695536 |
| 16 | 5173600 | 5178399 | 55.47578 | 88.12168 | 124.7314 | 52.01023 | -0.16771 | 2.276686 | 0.695552 |
| 10 | 108535600 | 108536199 | 8.286552 | 15.64778 | 26.51258 | 10.79702 | -0.27672 | 0.051812 | 0.69557 |
| 17 | 7752600 | 7754799 | 22.68981 | 38.91966 | 44.7343 | 19.61983 | 0.183396 | 0.937714 | 0.695599 |
| X | 18448200 | 18452199 | 44.95815 | 72.91129 | 63.34865 | 27.13581 | 0.568108 | 1.640453 | 0.695601 |
| 10 | 77569400 | 77572799 | 45.7855 | 73.37166 | 104.2694 | 44.00211 | -0.15968 | 1.989092 | 0.695607 |
| 6 | 37767600 | 37773199 | 52.8333 | 84.4133 | 123.7209 | 51.98495 | -0.20906 | 2.093339 | 0.695635 |
| 18 | 5105200 | 5117799 | 195.5292 | 289.9739 | 269.5809 | 111.0357 | 0.356591 | 3.679932 | 0.695636 |
| 13 | 30067800 | 30075799 | 91.33935 | 141.9125 | 158.7075 | 65.89833 | 0.146103 | 2.759239 | 0.695672 |
| 1 | 81669400 | 81671199 | 65.91598 | 104.3662 | 114.1412 | 47.45856 | 0.199467 | 2.372911 | 0.695713 |
| 9 | 19955600 | 19962399 | 98.968 | 154.5074 | 140.3784 | 58.2486 | 0.450877 | 2.67918 | 0.695716 |
| 9 | 22561200 | 22563399 | 19.15352 | 33.49279 | 38.20394 | 16.84417 | 0.198672 | 0.702488 | 0.695785 |
| 4 | 58391400 | 58394199 | 25.74794 | 42.78788 | 63.13945 | 27.38626 | -0.17851 | 1.267854 | 0.695792 |
| 4 | 34342000 | 34355399 | 291.7398 | 411.2134 | 528.1432 | 218.7204 | -0.18114 | 4.460432 | 0.695807 |
| X | 75694200 | 75696799 | 27.01841 | 46.3299 | 47.91704 | 21.0152 | 0.335092 | 0.963739 | 0.695821 |
| 14 | 90410000 | 90411599 | 19.66172 | 34.47129 | 38.02498 | 16.80228 | 0.24667 | 0.663227 | 0.695824 |

| | | | | | | | | | |
|---|---|---|---|---|---|---|---|---|---|
| 12 | 113722600 | 113728199 | 86.22475 | 135.1738 | 148.3642 | 61.53589 | 0.183162 | 2.601623 | 0.695835 |
| 17 | 94265000 | 94268799 | 41.79682 | 67.93547 | 74.15308 | 31.81614 | 0.242615 | 1.591839 | 0.695847 |
| 12 | 81579000 | 81579799 | 7.168714 | 13.74193 | 24.0841 | 9.479752 | -0.2809 | -0.078 | 0.695851 |
| 17 | 22135000 | 22177399 | 861.8147 | 1130.445 | 1020.964 | 427.3024 | 0.222455 | 5.591253 | 0.69587 |
| 7 | 10044600 | 10050399 | 100.5669 | 155.8975 | 170.2373 | 70.70951 | 0.177969 | 2.804277 | 0.695887 |
| 11 | 70184400 | 70186599 | 19.10645 | 32.94021 | 49.00602 | 21.49781 | -0.18585 | 0.883953 | 0.695889 |
| 4 | 88031200 | 88032599 | 18.79267 | 33.00685 | 36.51128 | 16.1579 | 0.242753 | 0.629352 | 0.695892 |
| 1 | 8437400 | 8437999 | 10.31512 | 19.70249 | 23.92482 | 9.510278 | 0.230015 | -0.01753 | 0.695907 |
| 12 | 47326200 | 47327399 | 14.86906 | 26.02128 | 39.9569 | 17.62879 | -0.22808 | 0.670128 | 0.695925 |
| 2 | 109011000 | 109011799 | 14.72618 | 26.87444 | 29.96173 | 12.85243 | 0.264721 | 0.312953 | 0.695936 |
| 14 | 45425800 | 45427799 | 23.65628 | 41.09781 | 37.15843 | 16.31371 | 0.531683 | 0.815196 | 0.695956 |
| 5 | 64990600 | 64994399 | 42.72366 | 69.37423 | 75.7382 | 32.4729 | 0.24106 | 1.615329 | 0.69596 |
| 2 | 58146800 | 58147199 | 4.432511 | 8.638516 | 19.12532 | 6.92862 | -0.50152 | -0.41432 | 0.695973 |
| 3 | 109176000 | 109177799 | 16.32728 | 28.40014 | 43.18025 | 18.97214 | -0.21459 | 0.772284 | 0.695975 |
| 18 | 58450200 | 58453399 | 50.1994 | 80.5134 | 91.6185 | 38.7134 | 0.161759 | 1.961991 | 0.695976 |
| 2 | 108873800 | 108874599 | 10.35019 | 19.14889 | 30.41923 | 12.81073 | -0.22354 | 0.205899 | 0.695986 |
| 8 | 20597600 | 20606799 | 114.2561 | 175.6539 | 190.2939 | 79.02086 | 0.180262 | 2.964432 | 0.695993 |
| 2 | 70373200 | 70376199 | 33.52817 | 55.61964 | 60.96094 | 26.40942 | 0.24899 | 1.307202 | 0.695996 |
| 15 | 38606800 | 38623999 | 340 | 474.7634 | 590.8216 | 245.0387 | -0.14925 | 4.601835 | 0.696045 |
| 1 | 60418200 | 60421799 | 26.3167 | 44.29074 | 50.70341 | 22.1573 | 0.186603 | 1.124324 | 0.696049 |
| 6 | 44636800 | 44638799 | 21.66961 | 37.90317 | 39.54252 | 17.43849 | 0.326721 | 0.711235 | 0.696065 |
| 3 | 129229800 | 129232399 | 20.18625 | 34.6087 | 51.70442 | 22.67595 | -0.19355 | 0.939949 | 0.696075 |
| 11 | 91307800 | 91309399 | 18.2355 | 32.04526 | 35.85492 | 15.87454 | 0.226096 | 0.613433 | 0.696076 |
| 3 | 28744200 | 28745999 | 17.18101 | 30.14451 | 34.13007 | 15.09071 | 0.209336 | 0.631614 | 0.696081 |
| X | 110710600 | 110712799 | 26.07961 | 44.79998 | 46.8325 | 20.54208 | 0.320181 | 0.943823 | 0.696087 |
| 7 | 79981800 | 79983399 | 20.72225 | 34.80085 | 54.77909 | 23.92072 | -0.27112 | 1.13187 | 0.696092 |
| 3 | 15836000 | 15839199 | 28.9298 | 49.15805 | 50.69445 | 22.21701 | 0.337237 | 1.051696 | 0.696096 |
| 18 | 67364000 | 67364599 | 7.872569 | 15.01611 | 25.59496 | 10.20422 | -0.25909 | -0.02842 | 0.696099 |
| 1 | 147647000 | 147648799 | 16.25479 | 29.00374 | 32.18859 | 14.15298 | 0.24778 | 0.458228 | 0.696101 |
| 1 | 48802800 | 48804799 | 17.05648 | 30.85399 | 31.79972 | 13.79202 | 0.36716 | 0.382992 | 0.696104 |
| 15 | 16386800 | 16389799 | 26.33306 | 45.3631 | 46.2439 | 20.28565 | 0.356991 | 0.916479 | 0.696121 |
| 6 | 130301400 | 130304399 | 21.36517 | 37.75605 | 35.54127 | 15.72205 | 0.477158 | 0.571315 | 0.696135 |
| 1 | 103527000 | 103527599 | 7.575884 | 14.56519 | 24.70651 | 9.64301 | -0.22347 | -0.10201 | 0.696146 |
| 8 | 23046600 | 23052199 | 56.61244 | 89.81151 | 128.0106 | 53.30882 | -0.17955 | 2.315519 | 0.69615 |
| 7 | 63351800 | 63355199 | 46.4268 | 74.64083 | 84.58195 | 35.86798 | 0.171922 | 1.880409 | 0.696156 |
| 6 | 28954000 | 28956199 | 15.67459 | 27.5071 | 41.34632 | 18.28457 | -0.19757 | 0.631847 | 0.696166 |
| 13 | 54988800 | 54991199 | 28.07802 | 46.19648 | 68.6155 | 29.58057 | -0.19045 | 1.395223 | 0.696173 |
| 5 | 106184200 | 106187199 | 28.70357 | 47.32727 | 70.01007 | 30.20923 | -0.18341 | 1.37106 | 0.696186 |
| 13 | 60549600 | 60553399 | 26.90455 | 44.68297 | 66.04647 | 28.63523 | -0.18103 | 1.280526 | 0.696193 |
| 8 | 61587600 | 61589999 | 38.42903 | 62.65447 | 70.84961 | 30.43351 | 0.194912 | 1.540939 | 0.696199 |
| Y | 4948600 | 4952999 | 38.52888 | 62.38683 | 57.53179 | 24.61569 | 0.480191 | 1.650455 | 0.696206 |
| 10 | 124429600 | 124430199 | 12.8801 | 24.2027 | 26.82609 | 11.02172 | 0.319839 | 0.123366 | 0.69621 |
| 4 | 142398400 | 142400399 | 14.52204 | 25.35934 | 41.0399 | 18.07625 | -0.30368 | 0.702014 | 0.696223 |
| 3 | 61942800 | 61945999 | 27.77499 | 46.12047 | 70.1061 | 30.39163 | -0.22133 | 1.288836 | 0.696232 |
| 11 | 26417200 | 26417999 | 17.3664 | 29.8984 | 46.39173 | 20.35012 | -0.24652 | 0.886807 | 0.696241 |
| 13 | 26599800 | 26600999 | 11.61132 | 21.1773 | 32.80584 | 14.08299 | -0.20744 | 0.300316 | 0.696266 |
| 6 | 66811000 | 66812799 | 12.17308 | 23.26696 | 21.05095 | 8.355584 | 0.652991 | -0.01184 | 0.69627 |

| | | | | | | | | | |
|---|---|---|---|---|---|---|---|---|---|
| 5 | 76370600 | 76372999 | 36.51758 | 58.75629 | 101.3804 | 43.04105 | -0.43194 | 1.850976 | 0.696285 |
| 5 | 8173600 | 8174799 | 17.886 | 31.51944 | 35.39045 | 15.66409 | 0.222616 | 0.580976 | 0.696299 |
| 10 | 129894400 | 129896799 | 22.11523 | 38.40046 | 42.47459 | 18.64154 | 0.241906 | 0.820705 | 0.696314 |
| 14 | 75324800 | 75327399 | 27.29139 | 46.59618 | 49.02306 | 21.49279 | 0.309509 | 1.001667 | 0.696321 |
| X | 90652200 | 90652799 | 10.93179 | 20.10698 | 31.3586 | 13.31185 | -0.20578 | 0.243691 | 0.696337 |
| X | 21829600 | 21830199 | 13.01335 | 24.6219 | 25.90072 | 10.50408 | 0.410695 | 0.079115 | 0.696355 |
| 5 | 148947400 | 148949799 | 30.652 | 49.93273 | 74.22489 | 31.87356 | -0.19945 | 1.539114 | 0.6964 |
| 14 | 53043400 | 53045599 | 22.42784 | 38.60885 | 44.19411 | 19.38606 | 0.190347 | 0.904899 | 0.696406 |
| 4 | 13943000 | 13943599 | 9.953616 | 18.40033 | 29.34087 | 12.37018 | -0.23035 | 0.202427 | 0.696427 |
| 19 | 8493800 | 8496599 | 40.44828 | 65.43706 | 74.63832 | 31.91419 | 0.173109 | 1.684966 | 0.696459 |
| 2 | 68742600 | 68744399 | 15.35944 | 27.13954 | 44.48831 | 19.68035 | -0.31926 | 0.551322 | 0.696476 |
| 17 | 54217200 | 54219399 | 25.56588 | 43.12037 | 49.60723 | 21.69251 | 0.179695 | 1.103414 | 0.696478 |
| X | 96950600 | 96952399 | 12.30441 | 22.21707 | 34.2327 | 14.88133 | -0.21149 | 0.36689 | 0.696486 |
| 7 | 4882200 | 4886999 | 48.44029 | 77.99967 | 87.40369 | 37.14858 | 0.190819 | 1.836253 | 0.69649 |
| 12 | 57273400 | 57274999 | 13.39333 | 23.82988 | 36.11305 | 15.91677 | -0.20137 | 0.46667 | 0.696494 |
| 3 | 112291000 | 112292399 | 15.14236 | 27.03106 | 30.80136 | 13.53786 | 0.210381 | 0.445874 | 0.696504 |
| 15 | 5909400 | 5911999 | 52.49952 | 84.05622 | 95.56942 | 40.27622 | 0.160009 | 2.019864 | 0.696521 |
| 8 | 115321400 | 115323599 | 35.76304 | 57.8012 | 84.10713 | 35.92431 | -0.17891 | 1.706594 | 0.696558 |
| 17 | 39020600 | 39022199 | 24.71962 | 42.30267 | 47.39124 | 20.78123 | 0.219764 | 0.970418 | 0.696561 |
| 1 | 180356200 | 180361799 | 56.45757 | 89.73907 | 125.6241 | 52.41629 | -0.15089 | 2.25902 | 0.696571 |
| 7 | 58706600 | 58708799 | 36.85476 | 60.33888 | 51.43991 | 22.12991 | 0.605447 | 1.464352 | 0.696619 |
| 16 | 12788600 | 12790199 | 16.96784 | 29.49878 | 44.21394 | 19.41988 | -0.19419 | 0.784233 | 0.696621 |
| 7 | 27920800 | 27936399 | 229.9827 | 333.0248 | 428.649 | 176.7695 | -0.14537 | 4.061992 | 0.696623 |
| 3 | 4788000 | 4790799 | 40.71018 | 65.74455 | 72.6392 | 30.99423 | 0.216112 | 1.731605 | 0.696626 |
| 1 | 149368200 | 149370599 | 20.2165 | 35.68538 | 36.20154 | 16.01778 | 0.369161 | 0.578022 | 0.696679 |
| 1 | 184042400 | 184043399 | 11.49316 | 21.42854 | 25.08055 | 10.44173 | 0.226157 | 0.15356 | 0.696681 |
| 13 | 114403200 | 114404999 | 20.69771 | 36.17563 | 39.71975 | 17.50538 | 0.253479 | 0.720996 | 0.696689 |
| 4 | 111912800 | 111916399 | 49.72365 | 79.57499 | 114.0793 | 48.04497 | -0.1746 | 2.038075 | 0.696707 |
| 3 | 103133600 | 103138199 | 84.05971 | 131.7024 | 147.1699 | 61.04421 | 0.156663 | 2.610158 | 0.696729 |
| X | 74915600 | 74918999 | 22.18438 | 37.80245 | 59.35988 | 26.02057 | -0.26649 | 1.014253 | 0.696737 |
| 15 | 33845200 | 33846999 | 24.25409 | 40.60926 | 59.83703 | 26.07961 | -0.17631 | 1.179358 | 0.69674 |
| 3 | 130580200 | 130581799 | 15.39713 | 27.37345 | 31.38633 | 13.84271 | 0.19819 | 0.475571 | 0.696773 |
| 17 | 48630200 | 48631999 | 18.9933 | 32.55337 | 49.00576 | 21.49187 | -0.20434 | 0.937414 | 0.696774 |
| 4 | 145358000 | 145368199 | 397.6074 | 552.3056 | 557.3622 | 231.3498 | 0.145414 | 4.682299 | 0.696778 |
| 10 | 25776200 | 25778199 | 28.74616 | 47.7611 | 54.52153 | 23.64515 | 0.190687 | 1.271308 | 0.696778 |
| 6 | 11388000 | 11389799 | 24.44385 | 41.61561 | 47.40983 | 20.78074 | 0.194504 | 1.014045 | 0.69679 |
| 5 | 57006600 | 57008799 | 19.51772 | 34.59119 | 33.93356 | 15.00218 | 0.419321 | 0.524968 | 0.696806 |
| 2 | 174648800 | 174649599 | 9.512864 | 17.76007 | 28.4389 | 11.81314 | -0.21971 | 0.139376 | 0.696843 |
| 6 | 99024000 | 99024599 | 6.676856 | 12.96598 | 22.87925 | 8.56821 | -0.22309 | -0.2555 | 0.696845 |
| 13 | 4460200 | 4461399 | 12.23999 | 22.35816 | 36.62109 | 15.69068 | -0.28591 | 0.302773 | 0.696845 |
| 18 | 80052800 | 80054599 | 18.78071 | 32.58571 | 49.56191 | 21.7459 | -0.21574 | 0.826135 | 0.696873 |
| X | 34797000 | 34820999 | 298.1351 | 431.4405 | 393.528 | 162.2996 | 0.350388 | 4.077472 | 0.696887 |
| 17 | 77607600 | 77608599 | 30.14752 | 49.17691 | 73.1717 | 31.43669 | -0.19946 | 1.517868 | 0.696926 |
| 7 | 89515000 | 89517399 | 23.72619 | 39.63212 | 59.82703 | 26.03662 | -0.21108 | 1.20112 | 0.69694 |
| 6 | 41855200 | 41859799 | 46.62713 | 74.56679 | 106.8391 | 44.95072 | -0.17485 | 2.054559 | 0.696942 |
| 15 | 27025400 | 27026199 | 13.12106 | 22.81674 | 47.37495 | 20.8115 | -0.66175 | 0.769578 | 0.696961 |
| 11 | 31129600 | 31132599 | 38.80024 | 63.85993 | 57.13611 | 24.63537 | 0.538939 | 1.409981 | 0.696964 |

| | | | | | | | | | |
|---|---|---|---|---|---|---|---|---|---|
| 3 | 46399400 | 46402599 | 47.06103 | 75.26209 | 107.3272 | 45.15725 | -0.16804 | 2.053419 | 0.696974 |
| 7 | 96469600 | 96470999 | 9.009231 | 16.8787 | 27.43184 | 11.32757 | -0.23416 | 0.109143 | 0.697005 |
| 13 | 119320800 | 119322999 | 33.96246 | 54.92518 | 81.67521 | 34.91356 | -0.20908 | 1.683759 | 0.697009 |
| 9 | 88434400 | 88435999 | 22.71029 | 38.88928 | 44.75735 | 19.62728 | 0.181195 | 0.953298 | 0.697017 |
| 14 | 78058600 | 78059599 | 12.19268 | 22.69104 | 26.62057 | 11.11741 | 0.219274 | 0.175438 | 0.697034 |
| 16 | 84660800 | 84664199 | 30.89176 | 51.08109 | 58.98252 | 25.4955 | 0.173686 | 1.335314 | 0.697061 |
| 5 | 6718600 | 6720399 | 17.30225 | 30.7018 | 33.67386 | 14.87551 | 0.260153 | 0.502718 | 0.697064 |
| 1 | 20294800 | 20296399 | 12.87216 | 22.60446 | 42.47719 | 18.79542 | -0.5179 | 0.607728 | 0.697077 |
| 3 | 127085000 | 127088599 | 30.92423 | 50.49694 | 74.05316 | 31.82758 | -0.17755 | 1.50422 | 0.697094 |
| 6 | 58502000 | 58505199 | 25.07451 | 41.60731 | 63.10067 | 27.34156 | -0.21806 | 1.281604 | 0.697113 |
| 2 | 26405600 | 26407799 | 12.89287 | 22.54972 | 39.13144 | 17.25609 | -0.40333 | 0.669515 | 0.69712 |
| 4 | 130121600 | 130125199 | 34.73292 | 57.0398 | 65.24397 | 28.10502 | 0.184294 | 1.432203 | 0.697125 |
| 8 | 35107200 | 35108399 | 10.09891 | 18.61874 | 29.81977 | 12.6182 | -0.24058 | 0.219212 | 0.697146 |
| 2 | 88431600 | 88434999 | 55.38622 | 88.59772 | 100.4567 | 42.28116 | 0.162533 | 2.05341 | 0.697173 |
| 6 | 71625600 | 71628599 | 24.25459 | 41.28957 | 47.42451 | 20.78702 | 0.182832 | 1.016354 | 0.697174 |
| 3 | 78515600 | 78517399 | 18.45805 | 32.97574 | 33.70534 | 14.80446 | 0.367121 | 0.470295 | 0.697177 |
| 1 | 31329400 | 31330799 | 21.48421 | 37.55494 | 40.17862 | 17.70947 | 0.290673 | 0.723025 | 0.697178 |
| 4 | 146976200 | 146979999 | 206.5246 | 309.2137 | 270.1419 | 111.4205 | 0.458795 | 3.514358 | 0.697192 |
| 15 | 87715400 | 87717199 | 14.63144 | 25.66554 | 39.41674 | 17.42497 | -0.22813 | 0.632304 | 0.697197 |
| 17 | 29503800 | 29506199 | 19.08708 | 31.85796 | 64.19926 | 27.92557 | -0.62614 | 1.203622 | 0.697221 |
| 2 | 17963000 | 17967399 | 54.65253 | 86.92495 | 122.3091 | 51.06804 | -0.15734 | 2.237921 | 0.697223 |
| 15 | 4829600 | 4831799 | 21.59577 | 37.55292 | 41.95235 | 18.42913 | 0.227907 | 0.799067 | 0.69723 |
| 12 | 54520800 | 54522799 | 22.28602 | 38.71607 | 42.45344 | 18.63801 | 0.254446 | 0.814639 | 0.697245 |
| 14 | 96485400 | 96486999 | 16.05792 | 28.33646 | 32.02372 | 14.17009 | 0.214138 | 0.549386 | 0.697262 |
| Y | 89015400 | 89015799 | 3.986362 | 7.783043 | 17.6039 | 6.209625 | -0.49581 | -0.52013 | 0.697279 |
| 13 | 85181000 | 85182399 | 17.3633 | 30.21165 | 45.75397 | 20.10691 | -0.20914 | 0.774047 | 0.697293 |
| X | 96538800 | 96541599 | 24.96258 | 43.5972 | 41.1563 | 18.12906 | 0.470454 | 0.755674 | 0.69738 |
| X | 18277400 | 18280199 | 32.52173 | 54.0918 | 59.10517 | 25.65189 | 0.253733 | 1.275691 | 0.697384 |
| 12 | 47660400 | 47661799 | 18.11064 | 31.93713 | 35.39413 | 15.66358 | 0.241797 | 0.575722 | 0.697386 |
| 14 | 22837400 | 22839599 | 18.11064 | 31.93713 | 35.39413 | 15.66358 | 0.241797 | 0.575722 | 0.697386 |
| 7 | 5214400 | 5214999 | 14.82159 | 26.29518 | 27.14374 | 11.99268 | 0.347214 | 0.475684 | 0.697395 |
| 2 | 110039200 | 110040799 | 15.73074 | 27.5236 | 41.49272 | 18.30323 | -0.20162 | 0.681972 | 0.697436 |
| 10 | 114677600 | 114678399 | 9.030323 | 16.95281 | 27.46531 | 11.30224 | -0.22547 | 0.097285 | 0.697461 |
| 12 | 32565600 | 32567199 | 13.01622 | 24.35164 | 26.61892 | 11.01797 | 0.330522 | 0.147353 | 0.697495 |
| 2 | 40704200 | 40705399 | 13.40425 | 23.72351 | 36.14306 | 15.98343 | -0.21291 | 0.507996 | 0.697505 |
| 17 | 37893600 | 37895799 | 23.57132 | 41.04042 | 42.36081 | 18.62346 | 0.341697 | 0.790878 | 0.697512 |
| X | 56096200 | 56099799 | 31.6318 | 53.0346 | 55.14587 | 24.05579 | 0.324989 | 1.186706 | 0.697513 |
| 14 | 48117600 | 48119599 | 15.18417 | 26.71292 | 40.24626 | 17.81933 | -0.20043 | 0.59163 | 0.697543 |
| 4 | 145218600 | 145222799 | 64.71223 | 103.3555 | 107.5656 | 45.17386 | 0.283502 | 2.111393 | 0.697555 |
| 9 | 25773000 | 25775999 | 49.42498 | 79.23661 | 89.18862 | 37.66155 | 0.176722 | 1.974605 | 0.697574 |
| 10 | 81915000 | 81921599 | 108.703 | 166.9455 | 185.5905 | 77.05694 | 0.14192 | 2.981985 | 0.697612 |
| 2 | 154080600 | 154088199 | 91.83888 | 141.7383 | 192.6571 | 80.02602 | -0.14246 | 2.883193 | 0.69766 |
| 7 | 103054800 | 103057799 | 26.45684 | 44.72521 | 50.69367 | 22.18297 | 0.201119 | 1.08647 | 0.697664 |
| 16 | 85063600 | 85064599 | 18.19664 | 32.0271 | 35.89513 | 15.8914 | 0.22452 | 0.596792 | 0.69767 |
| 17 | 54187200 | 54198399 | 301.692 | 429.9493 | 445.2718 | 184.0629 | 0.145252 | 4.299507 | 0.697677 |
| 1 | 130126600 | 130127799 | 8.498032 | 16.11485 | 26.56203 | 10.72123 | -0.22676 | 0.021033 | 0.6977 |
| 12 | 88139600 | 88145799 | 38.42611 | 63.0218 | 57.44693 | 24.73251 | 0.510204 | 1.453925 | 0.697703 |

| | | | | | | | | | |
|---|---|---|---|---|---|---|---|---|---|
| 17 | 23517600 | 23517999 | 7.131298 | 13.79799 | 23.82166 | 9.087949 | -0.21726 | -0.18415 | 0.697709 |
| 12 | 24677000 | 24678199 | 7.495491 | 14.24171 | 25.02695 | 10.05968 | -0.31218 | -0.00318 | 0.697718 |
| 6 | 38188600 | 38193799 | 70.34771 | 110.4734 | 152.9405 | 63.43508 | -0.14945 | 2.560614 | 0.697736 |
| 17 | 30992400 | 30994599 | 20.73594 | 35.76921 | 39.4975 | 17.32694 | 0.242502 | 0.870543 | 0.697748 |
| 19 | 35905000 | 35906999 | 16.21859 | 28.33537 | 43.55134 | 19.18812 | -0.22976 | 0.707045 | 0.697762 |
| 5 | 65011200 | 65014999 | 45.64237 | 73.90872 | 79.10164 | 33.84582 | 0.265979 | 1.678788 | 0.69777 |
| X | 32530600 | 32535399 | 88.82063 | 140.8839 | 123.4954 | 51.41987 | 0.520566 | 2.348295 | 0.697773 |
| 1 | 178033200 | 178035799 | 29.90283 | 49.14873 | 72.1735 | 31.10104 | -0.17467 | 1.415424 | 0.697788 |
| 4 | 142632800 | 142635599 | 22.3713 | 37.8003 | 56.0225 | 24.5112 | -0.18439 | 1.082818 | 0.697799 |
| 10 | 93568800 | 93571199 | 22.35075 | 37.90941 | 56.60444 | 24.79438 | -0.19492 | 1.050371 | 0.697824 |
| 4 | 151999200 | 152001399 | 26.66977 | 43.93349 | 66.12521 | 28.52182 | -0.20872 | 1.376651 | 0.697831 |
| 8 | 41658600 | 41659599 | 13.48594 | 24.6294 | 28.43715 | 12.18708 | 0.215279 | 0.295106 | 0.697838 |
| 12 | 91210200 | 91213599 | 35.9382 | 59.73773 | 60.07943 | 26.06373 | 0.372875 | 1.289644 | 0.697839 |
| 6 | 130581800 | 130583799 | 26.06154 | 44.83247 | 46.243 | 20.28438 | 0.339653 | 0.926474 | 0.697845 |
| 10 | 62943600 | 62949199 | 61.75615 | 98.3732 | 111.0211 | 46.45285 | 0.162687 | 2.191969 | 0.697852 |
| X | 126038200 | 126038999 | 22.19051 | 38.27834 | 43.15811 | 18.93279 | 0.212758 | 0.88592 | 0.697866 |
| 2 | 144604800 | 144609199 | 37.33728 | 60.38882 | 87.88473 | 37.57114 | -0.17801 | 1.689359 | 0.697883 |
| 16 | 19466000 | 19473599 | 121.6432 | 186.0508 | 175.3129 | 72.73589 | 0.375625 | 3.066333 | 0.697888 |
| 6 | 62497600 | 62499599 | 22.36307 | 38.83976 | 36.68172 | 16.10145 | 0.468926 | 0.81103 | 0.69789 |
| 1 | 5318200 | 5319799 | 14.02296 | 24.75295 | 37.65881 | 16.66255 | -0.21251 | 0.542843 | 0.697913 |
| 19 | 12829600 | 12830399 | 9.691804 | 17.92576 | 28.87001 | 12.16296 | -0.24354 | 0.197952 | 0.697923 |
| 19 | 13324200 | 13338999 | 201.4572 | 299.6706 | 299.4278 | 123.3671 | 0.256641 | 3.630021 | 0.697941 |
| 12 | 74356000 | 74357999 | 14.71153 | 26.85524 | 25.23636 | 10.82163 | 0.509992 | 0.3002 | 0.697942 |
| X | 156579000 | 156582999 | 49.15167 | 79.38305 | 83.23918 | 35.52056 | 0.291518 | 1.750796 | 0.697976 |
| 3 | 82571000 | 82571999 | 11.41811 | 21.16408 | 23.67051 | 9.943326 | 0.281839 | 0.186198 | 0.698044 |
| 17 | 58569400 | 58570599 | 17.95338 | 31.49681 | 36.01894 | 15.92467 | 0.19488 | 0.639492 | 0.69807 |
| 6 | 118034000 | 118038199 | 34.56325 | 56.14107 | 81.5477 | 34.97182 | -0.16949 | 1.596325 | 0.698083 |
| 4 | 114266200 | 114268799 | 18.02954 | 30.88378 | 47.96468 | 21.03432 | -0.2492 | 0.93744 | 0.698087 |
| 2 | 11743600 | 11746399 | 23.30267 | 39.00311 | 58.1017 | 25.31789 | -0.19194 | 1.179049 | 0.698093 |
| 17 | 8125600 | 8132599 | 120.651 | 181.8993 | 255.3692 | 105.512 | -0.21618 | 3.363391 | 0.698102 |
| 11 | 41002000 | 41003199 | 17.66907 | 30.35979 | 46.9143 | 20.57812 | -0.24127 | 0.905748 | 0.698127 |
| X | 148904000 | 148905799 | 18.27636 | 33.69372 | 29.47866 | 12.47451 | 0.626245 | 0.283854 | 0.698132 |
| 5 | 126072600 | 126074599 | 14.19351 | 24.95894 | 38.23305 | 16.92948 | -0.22446 | 0.586701 | 0.698161 |
| 6 | 131658000 | 131674799 | 209.9287 | 311.6727 | 306.2158 | 126.1392 | 0.278391 | 3.66479 | 0.698182 |
| 3 | 42223200 | 42224399 | 10.08333 | 18.63284 | 29.84172 | 12.58801 | -0.23733 | 0.205907 | 0.698183 |
| 11 | 114217400 | 114221799 | 53.66289 | 85.39031 | 120.5004 | 50.34497 | -0.16082 | 2.221713 | 0.698189 |
| 17 | 92651400 | 92653399 | 26.48017 | 44.11556 | 66.57986 | 28.92714 | -0.21072 | 1.246384 | 0.698209 |
| 6 | 114943600 | 114948399 | 34.25617 | 55.45955 | 81.4542 | 34.84711 | -0.18982 | 1.661658 | 0.698221 |
| 14 | 37779600 | 37782199 | 32.28261 | 53.10401 | 61.07285 | 26.32674 | 0.177467 | 1.403709 | 0.69823 |
| 1 | 185752800 | 185757799 | 62.93767 | 100.0953 | 113.7515 | 47.49024 | 0.150229 | 2.24501 | 0.698235 |
| 1 | 35941800 | 35945999 | 35.40849 | 57.90962 | 66.76578 | 28.71183 | 0.170165 | 1.487042 | 0.69824 |
| X | 147794200 | 147796599 | 30.61572 | 51.87772 | 52.38596 | 22.93986 | 0.367382 | 1.085009 | 0.698243 |
| 3 | 26910800 | 26913199 | 27.569 | 45.07786 | 79.43562 | 34.15744 | -0.44047 | 1.481633 | 0.698251 |
| 4 | 74369400 | 74371399 | 38.14588 | 61.9762 | 70.92436 | 30.42174 | 0.17408 | 1.593453 | 0.698252 |
| 10 | 12261600 | 12262799 | 23.90159 | 39.13212 | 76.52851 | 32.92334 | -0.58891 | 1.454555 | 0.698262 |
| 3 | 103145200 | 103148999 | 59.61674 | 94.99875 | 108.0664 | 45.2386 | 0.151814 | 2.178002 | 0.698264 |
| 15 | 14108400 | 14110199 | 16.53315 | 29.52858 | 32.76228 | 14.39365 | 0.249093 | 0.456758 | 0.698274 |

| | | | | | | | | | |
|---|---|---|---|---|---|---|---|---|---|
| 19 | 58710000 | 58711199 | 16.53315 | 29.52858 | 32.76228 | 14.39365 | 0.249093 | 0.456758 | 0.698274 |
| 17 | 51909800 | 51910399 | 12.29583 | 22.71935 | 26.61858 | 11.22973 | 0.210745 | 0.21859 | 0.698285 |
| 2 | 35422800 | 35423999 | 11.78081 | 21.76037 | 24.65693 | 10.40694 | 0.258276 | 0.211607 | 0.698319 |
| 13 | 54440000 | 54441799 | 12.98395 | 22.82065 | 36.87 | 16.32635 | -0.30091 | 0.584401 | 0.698326 |
| 7 | 22992600 | 23023399 | 930.7262 | 1205.203 | 1131.295 | 474.1025 | 0.14824 | 5.799799 | 0.698338 |
| 7 | 6157400 | 6160399 | 22.12666 | 37.05475 | 57.00725 | 24.84945 | -0.23831 | 1.166866 | 0.698345 |
| 9 | 6709400 | 6711599 | 23.75618 | 40.2709 | 43.85676 | 19.20731 | 0.25888 | 1.048228 | 0.698346 |
| 19 | 53048200 | 53049399 | 12.25218 | 22.04756 | 33.65196 | 14.69124 | -0.20202 | 0.383673 | 0.698355 |
| 2 | 90129400 | 90135199 | 105.9919 | 162.9957 | 180.6655 | 75.02558 | 0.147614 | 2.95439 | 0.698356 |
| 15 | 54641200 | 54643999 | 11.22625 | 21.39734 | 20.48883 | 8.181088 | 0.563983 | -0.00507 | 0.698373 |
| 6 | 131938000 | 131939199 | 42.67202 | 68.85087 | 78.58751 | 33.49343 | 0.167755 | 1.763243 | 0.698386 |
| 12 | 80166600 | 80167599 | 10.44471 | 19.45027 | 31.06818 | 12.96099 | -0.22154 | 0.16854 | 0.698387 |
| 10 | 66709800 | 66711599 | 14.34844 | 26.26553 | 29.47421 | 12.59348 | 0.259122 | 0.291298 | 0.69839 |
| 8 | 39185200 | 39185799 | 7.140178 | 13.78466 | 23.70151 | 9.136032 | -0.22545 | -0.14877 | 0.69841 |
| 3 | 57988000 | 57990399 | 42.71745 | 69.50963 | 74.24575 | 31.8695 | 0.274959 | 1.579119 | 0.698435 |
| 11 | 96099800 | 96104999 | 70.71512 | 111.8571 | 126.6957 | 52.65294 | 0.147873 | 2.401439 | 0.698484 |
| 5 | 6272000 | 6273399 | 30.09559 | 48.79568 | 86.74121 | 37.14936 | -0.46253 | 1.627772 | 0.698492 |
| 5 | 127018000 | 127018999 | 18.87673 | 32.90537 | 35.94326 | 15.81377 | 0.260752 | 0.744828 | 0.698503 |
| 9 | 79804800 | 79811399 | 101.8933 | 154.4505 | 259.1971 | 107.2756 | -0.46678 | 3.250019 | 0.698513 |
| 5 | 40975000 | 40975399 | 7.215108 | 14.02634 | 17.34694 | 6.440074 | 0.296512 | -0.2983 | 0.698514 |
| 13 | 65347000 | 65350199 | 40.22851 | 65.62896 | 71.78516 | 30.84233 | 0.243395 | 1.536189 | 0.698518 |
| 17 | 6696800 | 6698399 | 9.220306 | 17.03492 | 28.37185 | 11.971 | -0.29318 | 0.198488 | 0.698564 |
| 3 | 114144800 | 114146199 | 13.69137 | 24.01131 | 44.06873 | 19.4779 | -0.48404 | 0.63743 | 0.698566 |
| 5 | 116907400 | 116909799 | 21.62575 | 37.43909 | 42.53493 | 18.66146 | 0.202662 | 0.847113 | 0.698567 |
| 13 | 81014800 | 81015799 | 9.585238 | 18.46576 | 22.48477 | 8.724875 | 0.258082 | -0.10802 | 0.698582 |
| 3 | 36345200 | 36347999 | 36.89425 | 59.56014 | 86.37735 | 36.83864 | -0.17606 | 1.742891 | 0.698596 |
| 2 | 66753600 | 66755999 | 30.19075 | 50.49395 | 56.31397 | 24.52719 | 0.224199 | 1.205072 | 0.698599 |
| 19 | 39949800 | 39964399 | 357.3083 | 495.4142 | 621.1085 | 257.8834 | -0.17024 | 4.709318 | 0.6986 |
| 14 | 41341400 | 41356599 | 169.7963 | 256.0752 | 219.435 | 90.67365 | 0.495916 | 3.363895 | 0.698606 |
| 2 | 174882000 | 174886999 | 106.8972 | 164.1474 | 179.6727 | 74.59832 | 0.164108 | 2.984582 | 0.698615 |
| 3 | 25238200 | 25239399 | 12.75168 | 22.95529 | 35.18922 | 15.35498 | -0.20783 | 0.388223 | 0.698622 |
| 19 | 8149000 | 8152599 | 58.82904 | 93.91968 | 105.9109 | 44.45251 | 0.167293 | 2.118212 | 0.69863 |
| 2 | 4530200 | 4532399 | 18.85014 | 33.1869 | 36.34527 | 16.08976 | 0.257831 | 0.598009 | 0.698644 |
| 3 | 146640200 | 146641799 | 18.85014 | 33.1869 | 36.34527 | 16.08976 | 0.257831 | 0.598009 | 0.698644 |
| 6 | 132622600 | 132623999 | 21.18712 | 36.94565 | 34.49932 | 15.18181 | 0.485662 | 0.745505 | 0.698662 |
| 7 | 19453000 | 19457999 | 79.71176 | 123.9166 | 171.3438 | 71.1392 | -0.15907 | 2.746796 | 0.698683 |
| 6 | 67975400 | 67976599 | 21.40098 | 37.20314 | 34.23827 | 15.03762 | 0.506278 | 0.790637 | 0.698702 |
| 8 | 62459400 | 62465599 | 49.56646 | 79.40733 | 117.3262 | 49.48998 | -0.21631 | 2.006322 | 0.698706 |
| 18 | 36926800 | 36927399 | 7.125784 | 13.68652 | 24.12025 | 9.440532 | -0.28195 | -0.10016 | 0.698714 |
| 16 | 34409800 | 34412399 | 26.62395 | 45.15639 | 42.19016 | 18.4804 | 0.47911 | 1.051658 | 0.698727 |
| 17 | 21093200 | 21093799 | 13.69041 | 24.74035 | 28.42682 | 12.34452 | 0.210106 | 0.357453 | 0.698753 |
| 17 | 28769800 | 28772199 | 20.4616 | 34.62253 | 52.71356 | 23.07337 | -0.2229 | 1.061314 | 0.698754 |
| 11 | 20725200 | 20726199 | 10.84228 | 20.6438 | 24.37006 | 9.752479 | 0.261577 | 0.008398 | 0.698777 |
| 12 | 87763200 | 87765199 | 31.88049 | 52.8326 | 60.23437 | 26.07636 | 0.192191 | 1.313727 | 0.698821 |
| 14 | 13588200 | 13591799 | 53.41666 | 85.17195 | 121.0515 | 50.71807 | -0.16792 | 2.157723 | 0.698828 |
| 8 | 21121200 | 21124199 | 39.324 | 63.33923 | 91.54985 | 38.92754 | -0.1751 | 1.815192 | 0.69883 |
| 6 | 120109200 | 120112399 | 34.67117 | 56.09332 | 82.76541 | 35.39034 | -0.19731 | 1.679591 | 0.698848 |

| | | | | | | | | | |
|---|---|---|---|---|---|---|---|---|---|
| 5 | 133678600 | 133679799 | 12.06398 | 22.05433 | 35.53527 | 15.2102 | -0.2614 | 0.29466 | 0.698852 |
| 4 | 90957000 | 90957599 | 10.88447 | 20.4085 | 24.14686 | 9.957399 | 0.221792 | 0.110741 | 0.69887 |
| Y | 20603000 | 20604199 | 7.891203 | 14.58583 | 27.95129 | 11.78642 | -0.49331 | 0.183228 | 0.698878 |
| 13 | 85134200 | 85135999 | 25.51199 | 43.20121 | 49.58973 | 21.71107 | 0.183053 | 1.068241 | 0.698893 |
| 18 | 37403800 | 37404999 | 11.60313 | 21.85893 | 25.7874 | 10.54924 | 0.235273 | 0.092311 | 0.698905 |
| X | 8542000 | 8574199 | 498.6788 | 688.3091 | 614.3477 | 255.2599 | 0.313257 | 4.783211 | 0.698907 |
| 2 | 47507600 | 47509399 | 23.72957 | 39.80171 | 58.74804 | 25.63371 | -0.17882 | 1.157084 | 0.698907 |
| 11 | 116677800 | 116680199 | 21.49443 | 36.402 | 54.32181 | 23.78363 | -0.19399 | 1.060919 | 0.698916 |
| 17 | 22388000 | 22396199 | 89.29252 | 140.1459 | 149.0734 | 61.82662 | 0.229635 | 2.582028 | 0.698968 |
| 14 | 72258000 | 72258799 | 12.27138 | 22.2891 | 34.84032 | 15.03365 | -0.22528 | 0.332928 | 0.698996 |
| 11 | 21447000 | 21448999 | 17.73409 | 31.15244 | 35.53015 | 15.72577 | 0.198807 | 0.618431 | 0.699037 |
| 12 | 49362800 | 49363399 | 11.60658 | 21.62794 | 25.65698 | 10.69179 | 0.206071 | 0.161195 | 0.699065 |
| 3 | 3756200 | 3759799 | 124.1687 | 188.7488 | 204.3964 | 84.66666 | 0.168053 | 3.194058 | 0.69907 |
| 3 | 5593400 | 5594999 | 23.93244 | 41.10636 | 38.46644 | 16.87242 | 0.479209 | 0.923538 | 0.699074 |
| 16 | 77913600 | 77915999 | 32.29855 | 52.43605 | 77.50369 | 33.22518 | -0.19516 | 1.600163 | 0.699074 |
| 6 | 52325200 | 52331799 | 91.58118 | 142.6342 | 159.0821 | 66.02993 | 0.152731 | 2.718098 | 0.699087 |
| 7 | 12521200 | 12522399 | 12.27767 | 22.01379 | 33.66597 | 14.74217 | -0.20777 | 0.40696 | 0.69909 |
| Y | 43727800 | 43728399 | 6.778149 | 12.81241 | 25.50682 | 10.36516 | -0.50506 | 0.02411 | 0.699113 |
| 8 | 30288000 | 30288599 | 10.05605 | 19.09414 | 22.80871 | 9.180988 | 0.238111 | 0.018944 | 0.699114 |
| 2 | 107535400 | 107537399 | 27.48319 | 46.28216 | 52.31718 | 22.86675 | 0.204904 | 1.127372 | 0.699126 |
| 3 | 64923000 | 64923399 | 9.397919 | 17.95423 | 21.32876 | 8.474683 | 0.262488 | -0.03362 | 0.699138 |
| 6 | 64788000 | 64790199 | 19.75914 | 34.76143 | 36.76115 | 16.27628 | 0.307557 | 0.610813 | 0.699139 |
| 3 | 88578400 | 88581199 | 20.64518 | 35.85936 | 41.00214 | 17.99891 | 0.194452 | 0.810029 | 0.699143 |
| 6 | 26151000 | 26156999 | 112.8462 | 173.7364 | 166.7482 | 69.25704 | 0.356393 | 2.93355 | 0.69919 |
| X | 74765600 | 74768199 | 22.05368 | 37.36101 | 57.02708 | 24.97049 | -0.22646 | 1.064162 | 0.699207 |
| 1 | 65565400 | 65565999 | 11.98574 | 21.57834 | 33.20614 | 14.48808 | -0.21299 | 0.379665 | 0.699214 |
| 12 | 84092600 | 84098599 | 67.7309 | 107.6669 | 118.8174 | 49.58035 | 0.191823 | 2.264383 | 0.699281 |
| 9 | 22964400 | 22967399 | 29.24627 | 48.7167 | 56.21578 | 24.42196 | 0.175315 | 1.245545 | 0.699284 |
| 19 | 12480400 | 12480799 | 13.87111 | 25.59354 | 20.0477 | 8.474043 | 0.785715 | 0.222719 | 0.699297 |
| 13 | 36926800 | 36928799 | 16.94059 | 30.29523 | 32.67361 | 14.33913 | 0.290714 | 0.450249 | 0.699317 |
| 19 | 8396800 | 8398199 | 16.94059 | 30.29523 | 32.67361 | 14.33913 | 0.290714 | 0.450249 | 0.699317 |
| 15 | 13108000 | 13113599 | 81.69997 | 127.5894 | 179.6928 | 74.55926 | -0.18044 | 2.669884 | 0.699324 |
| 15 | 30935400 | 30936799 | 19.20426 | 33.40418 | 37.8922 | 16.64864 | 0.206009 | 0.776962 | 0.699325 |
| 17 | 26708600 | 26716799 | 109.0427 | 166.1002 | 225.4334 | 93.4388 | -0.15498 | 3.148948 | 0.699368 |
| 2 | 98421200 | 98424999 | 17.93322 | 31.69633 | 34.71256 | 15.35497 | 0.259885 | 0.546463 | 0.699401 |
| 16 | 9938400 | 9945999 | 98.54021 | 152.7347 | 169.2994 | 70.32139 | 0.156234 | 2.80472 | 0.699407 |
| 4 | 69703800 | 69705799 | 25.10443 | 41.7192 | 61.95915 | 26.87463 | -0.18772 | 1.265023 | 0.699412 |
| 5 | 115893800 | 115897199 | 36.04161 | 58.28744 | 84.55782 | 36.14562 | -0.17355 | 1.688052 | 0.699416 |
| 16 | 7383800 | 7384999 | 16.77945 | 30.14266 | 26.95927 | 11.78645 | 0.562844 | 0.40981 | 0.699443 |
| 17 | 64244400 | 64245199 | 11.31731 | 21.08632 | 22.28217 | 9.287374 | 0.372035 | 0.152568 | 0.699448 |
| 5 | 106065800 | 106068599 | 25.24757 | 42.1572 | 62.08589 | 27.01311 | -0.17554 | 1.213517 | 0.699456 |
| 13 | 26724800 | 26727599 | 38.55906 | 63.03108 | 59.00767 | 25.37096 | 0.469375 | 1.49772 | 0.69947 |
| 2 | 61484800 | 61487799 | 19.05347 | 31.84634 | 64.92891 | 28.26788 | -0.6431 | 1.185954 | 0.69947 |
| 12 | 27209000 | 27210599 | 10.97632 | 19.93218 | 31.25261 | 13.48947 | -0.22973 | 0.31733 | 0.69948 |
| 1 | 89270400 | 89272799 | 19.87458 | 34.00817 | 50.61181 | 22.19329 | -0.18839 | 0.955304 | 0.699507 |
| 6 | 102479800 | 102485999 | 48.07763 | 77.57201 | 84.51778 | 36.02042 | 0.234674 | 1.774258 | 0.69952 |
| 10 | 22838200 | 22841799 | 39.55828 | 64.37251 | 72.62852 | 31.17592 | 0.196209 | 1.570804 | 0.699527 |

| | | | | | | | | | |
|---|---|---|---|---|---|---|---|---|---|
| 14 | 89526400 | 89530599 | 58.80144 | 93.9002 | 105.5633 | 44.32304 | 0.172137 | 2.109788 | 0.69954 |
| 4 | 95975600 | 95977199 | 13.6626 | 24.28859 | 37.60525 | 16.58619 | -0.23301 | 0.474783 | 0.699547 |
| 2 | 156651400 | 156653799 | 15.94783 | 27.8712 | 41.99783 | 18.50795 | -0.20116 | 0.700007 | 0.699578 |
| 10 | 22639800 | 22641799 | 20.26863 | 35.52272 | 38.49931 | 17.00677 | 0.271969 | 0.672097 | 0.699588 |
| 1 | 44540800 | 44544199 | 33.25328 | 54.96132 | 61.53254 | 26.58952 | 0.218007 | 1.346744 | 0.6996 |
| 15 | 16612000 | 16612799 | 15.3746 | 26.9725 | 40.35094 | 17.83983 | -0.19067 | 0.634192 | 0.699635 |
| 4 | 113313800 | 113335199 | 491.9842 | 674.3467 | 659.9352 | 274.5047 | 0.170283 | 4.889396 | 0.699652 |
| 1 | 164051200 | 164056399 | 61.84997 | 97.10671 | 146.5721 | 60.79186 | -0.27419 | 2.559117 | 0.699657 |
| 14 | 105650800 | 105653399 | 31.80326 | 52.97714 | 50.84045 | 22.08706 | 0.440424 | 1.248108 | 0.699665 |
| 9 | 38984000 | 38987599 | 62.26267 | 98.39115 | 138.1684 | 57.39255 | -0.16293 | 2.420704 | 0.699668 |
| 1 | 42553400 | 42555199 | 18.55931 | 32.62237 | 36.4339 | 16.13418 | 0.228877 | 0.614527 | 0.699669 |
| 7 | 6439400 | 6445599 | 61.42305 | 97.20449 | 135.7939 | 56.44743 | -0.15377 | 2.385722 | 0.699672 |
| 4 | 70004800 | 70006799 | 34.33516 | 56.24965 | 64.34481 | 27.68919 | 0.182863 | 1.462293 | 0.699732 |
| 3 | 131192800 | 131194399 | 10.27 | 19.01244 | 29.87416 | 12.57046 | -0.20726 | 0.198427 | 0.699742 |
| 9 | 85825200 | 85826399 | 10.27 | 19.01244 | 29.87416 | 12.57046 | -0.20726 | 0.198427 | 0.699742 |
| 5 | 83611400 | 83611999 | 7.454987 | 14.23344 | 24.54149 | 9.764142 | -0.27238 | -0.04027 | 0.699753 |
| 7 | 121830600 | 121834399 | 34.52372 | 56.01705 | 82.10618 | 35.18363 | -0.18395 | 1.616564 | 0.699757 |
| 3 | 87374200 | 87378199 | 55.35453 | 88.34828 | 99.72417 | 41.83946 | 0.165487 | 2.121265 | 0.699769 |
| 1 | 35172400 | 35176199 | 39.75537 | 64.22498 | 96.10867 | 41.02681 | -0.22005 | 1.732616 | 0.699781 |
| 4 | 31126200 | 31129199 | 40.41983 | 65.35769 | 74.59693 | 31.88119 | 0.171411 | 1.696221 | 0.699796 |
| 9 | 92151800 | 92153199 | 15.37919 | 26.96464 | 40.37958 | 17.84359 | -0.19208 | 0.643932 | 0.699801 |
| X | 118753000 | 118753799 | 7.842946 | 15.15112 | 29.08119 | 11.17613 | -0.37879 | -0.13438 | 0.699807 |
| 4 | 24443600 | 24444599 | 14.79226 | 26.94018 | 25.31758 | 10.88793 | 0.507412 | 0.315077 | 0.699819 |
| 9 | 51209000 | 51210199 | 16.25556 | 28.75232 | 32.92239 | 14.56094 | 0.196595 | 0.527108 | 0.699826 |
| 12 | 22320000 | 22321399 | 28.81854 | 48.40395 | 43.85757 | 19.15311 | 0.52283 | 1.14561 | 0.699889 |
| 1 | 191398200 | 191398999 | 7.592034 | 14.54527 | 24.60538 | 9.700487 | -0.23299 | -0.06877 | 0.69989 |
| 3 | 146691200 | 146692199 | 7.592034 | 14.54527 | 24.60538 | 9.700487 | -0.23299 | -0.06877 | 0.69989 |
| 12 | 69759200 | 69767199 | 168.6614 | 251.7345 | 273.0481 | 112.5507 | 0.14274 | 3.563719 | 0.69991 |
| 11 | 57319400 | 57321399 | 11.57015 | 20.89634 | 32.66724 | 14.19649 | -0.23214 | 0.354501 | 0.699911 |
| 13 | 29776400 | 29779399 | 39.21673 | 64.88105 | 63.55502 | 27.48199 | 0.410393 | 1.357125 | 0.699919 |
| 10 | 69854000 | 69857799 | 31.8374 | 52.49908 | 59.96374 | 25.87523 | 0.188762 | 1.373123 | 0.699926 |
| X | 126100600 | 126103599 | 39.08304 | 63.85647 | 57.87519 | 24.87962 | 0.515503 | 1.505685 | 0.699967 |
| 11 | 95817200 | 95820599 | 46.9586 | 75.66658 | 85.5433 | 36.39464 | 0.179316 | 1.81171 | 0.699978 |
| 8 | 127583200 | 127587599 | 36.39842 | 58.79411 | 85.31084 | 36.41128 | -0.17583 | 1.724443 | 0.70002 |
| 2 | 6361200 | 6362599 | 11.26039 | 20.47686 | 31.73126 | 13.67195 | -0.21138 | 0.311521 | 0.700024 |
| 14 | 69388800 | 69397399 | 83.67406 | 131.9632 | 123.2074 | 51.13107 | 0.422864 | 2.479372 | 0.700029 |
| 15 | 7791800 | 7793999 | 21.66486 | 36.51442 | 54.83918 | 23.97512 | -0.20345 | 1.103516 | 0.700033 |
| 16 | 92966800 | 92969199 | 21.66486 | 36.51442 | 54.83918 | 23.97512 | -0.20345 | 1.103516 | 0.700033 |
| 1 | 51979000 | 51980999 | 31.25659 | 51.79169 | 59.58606 | 25.7933 | 0.179143 | 1.313701 | 0.700036 |
| X | 14987600 | 14989199 | 22.20893 | 38.80286 | 40.71789 | 17.94028 | 0.318506 | 0.735568 | 0.700055 |
| 12 | 62594200 | 62596799 | 63.10735 | 99.20314 | 145.0515 | 60.17083 | -0.22703 | 2.534767 | 0.700055 |
| 7 | 41814800 | 41822199 | 142.2701 | 215.0227 | 235.7274 | 97.4718 | 0.143398 | 3.31631 | 0.700058 |
| 9 | 41083000 | 41083999 | 10.512 | 19.77552 | 22.75114 | 9.32936 | 0.268948 | 0.085075 | 0.700066 |
| 5 | 131572600 | 131575799 | 27.82548 | 46.46184 | 52.86169 | 22.99978 | 0.195648 | 1.21427 | 0.700091 |
| 1 | 5280000 | 5281799 | 16.18016 | 28.93252 | 31.62739 | 13.88113 | 0.271167 | 0.442539 | 0.700095 |
| 15 | 10567600 | 10568999 | 12.15169 | 22.03389 | 34.2999 | 14.83222 | -0.22123 | 0.341905 | 0.700113 |
| 7 | 12503000 | 12509599 | 74.62462 | 116.423 | 163.284 | 67.75783 | -0.17553 | 2.681086 | 0.700138 |

| | | | | | | | | | |
|---|---|---|---|---|---|---|---|---|---|
| 3 | 137883200 | 137885199 | 15.94772 | 27.96336 | 42.39503 | 18.73529 | -0.21002 | 0.646973 | 0.700162 |
| 5 | 128465600 | 128484999 | 228.8166 | 331.4227 | 427.6906 | 176.3682 | -0.14873 | 4.057685 | 0.700172 |
| 3 | 31707000 | 31710599 | 40.33352 | 65.01592 | 93.1816 | 39.66468 | -0.16156 | 1.794365 | 0.700184 |
| 6 | 130598600 | 130601799 | 27.99422 | 46.551 | 75.71289 | 32.85464 | -0.31861 | 1.274384 | 0.700191 |
| 15 | 10778000 | 10779399 | 13.34498 | 23.40387 | 37.99489 | 16.79335 | -0.30769 | 0.625813 | 0.700208 |
| 9 | 25678200 | 25681199 | 27.39898 | 46.33409 | 51.78403 | 22.6625 | 0.221403 | 1.089533 | 0.700218 |
| 17 | 36592000 | 36594399 | 27.39898 | 46.33409 | 51.78403 | 22.6625 | 0.221403 | 1.089533 | 0.700218 |
| 4 | 44350800 | 44351799 | 13.7915 | 24.91421 | 28.51491 | 12.38825 | 0.215514 | 0.362102 | 0.70022 |
| 10 | 61168200 | 61169199 | 13.7915 | 24.91421 | 28.51491 | 12.38825 | 0.215514 | 0.362102 | 0.70022 |
| 7 | 70401800 | 70409399 | 70.08677 | 111.0981 | 124.7228 | 51.89782 | 0.1632 | 2.349553 | 0.700239 |
| 6 | 55258000 | 55261999 | 51.66307 | 82.34529 | 116.3556 | 48.72553 | -0.16012 | 2.164851 | 0.700245 |
| 10 | 34614600 | 34619199 | 81.8293 | 127.2889 | 174.191 | 72.31447 | -0.14343 | 2.737418 | 0.700252 |
| 6 | 105677000 | 105677999 | 17.46248 | 30.2829 | 45.29478 | 19.87333 | -0.1917 | 0.819393 | 0.700254 |
| 3 | 70355000 | 70357399 | 23.02073 | 39.53715 | 45.24314 | 19.84481 | 0.190036 | 0.92835 | 0.700267 |
| 2 | 111347600 | 111350999 | 59.52028 | 94.98738 | 106.9915 | 44.88203 | 0.168364 | 2.131074 | 0.700271 |
| 2 | 149160600 | 149162799 | 24.64949 | 41.98354 | 47.94579 | 21.01639 | 0.191109 | 1.012164 | 0.700282 |
| 12 | 62325600 | 62329799 | 40.368 | 66.14082 | 69.26874 | 29.80958 | 0.30985 | 1.476013 | 0.700321 |
| 13 | 62681400 | 62707799 | 732.7251 | 969.5566 | 929.8902 | 388.7921 | 0.148341 | 5.449087 | 0.700327 |
| 1 | 17469000 | 17472199 | 46.84095 | 75.44416 | 85.80552 | 36.48318 | 0.169909 | 1.825898 | 0.70034 |
| 17 | 84191800 | 84194399 | 16.38574 | 28.61993 | 43.08438 | 18.97929 | -0.19968 | 0.715054 | 0.700347 |
| 7 | 107819600 | 107828199 | 104.8254 | 161.7937 | 178.791 | 74.2663 | 0.156011 | 2.885093 | 0.700357 |
| 11 | 31444600 | 31445599 | 11.08452 | 21.09108 | 24.92988 | 9.990475 | 0.258203 | 0.017727 | 0.70039 |
| X | 153861000 | 153863999 | 34.68011 | 58.13823 | 45.90607 | 20.02363 | 0.720791 | 1.188932 | 0.700394 |
| 11 | 82218600 | 82225599 | 55.64096 | 88.35492 | 125.1483 | 52.16683 | -0.16917 | 2.286514 | 0.70041 |
| 9 | 4672200 | 4672999 | 9.782119 | 18.21515 | 28.90552 | 12.06015 | -0.21219 | 0.15798 | 0.700418 |
| 8 | 44141200 | 44143399 | 25.23556 | 43.56679 | 45.14301 | 19.80403 | 0.334121 | 0.890466 | 0.700421 |
| 7 | 84481000 | 84482199 | 23.33684 | 40.38441 | 37.83477 | 16.59914 | 0.479696 | 0.854366 | 0.700442 |
| 17 | 14712800 | 14716399 | 24.61203 | 40.85409 | 61.91593 | 26.83465 | -0.21699 | 1.276107 | 0.700446 |
| 3 | 20477800 | 20478799 | 12.29713 | 23.13812 | 26.25628 | 10.76325 | 0.288248 | 0.105571 | 0.700466 |
| 15 | 96467800 | 96469599 | 20.69623 | 35.77399 | 41.05158 | 18.00981 | 0.187928 | 0.855829 | 0.700493 |
| 2 | 95146800 | 95151399 | 44.63287 | 72.09899 | 81.15242 | 34.63882 | 0.189796 | 1.733571 | 0.7005 |
| 3 | 91836600 | 91837399 | 24.79674 | 41.9875 | 47.99197 | 21.01116 | 0.189266 | 1.066492 | 0.700546 |
| 14 | 106027800 | 106028999 | 11.76097 | 21.52609 | 33.93125 | 14.50054 | -0.22815 | 0.28318 | 0.700549 |
| 3 | 138936200 | 138936799 | 9.97679 | 18.60467 | 29.47174 | 12.26616 | -0.20671 | 0.15271 | 0.70058 |
| 1 | 183976000 | 183979199 | 23.45411 | 40.03878 | 45.80744 | 20.08276 | 0.188996 | 0.984091 | 0.700587 |
| 1 | 124862000 | 124863199 | 11.04219 | 20.51639 | 34.39085 | 14.39963 | -0.29521 | 0.192899 | 0.700593 |
| 11 | 64350400 | 64351999 | 11.39175 | 20.70818 | 32.21872 | 13.88832 | -0.21769 | 0.314787 | 0.700606 |
| 9 | 106681000 | 106682799 | 16.62362 | 28.52129 | 52.68217 | 23.1061 | -0.49802 | 0.922499 | 0.700609 |
| 3 | 106757400 | 106759599 | 35.15743 | 57.7569 | 65.40399 | 28.17806 | 0.198912 | 1.42793 | 0.700637 |
| 15 | 97447400 | 97450199 | 18.20227 | 32.27434 | 24.81504 | 10.96912 | 0.769356 | 0.497413 | 0.700652 |
| 1 | 67283800 | 67287199 | 37.50541 | 61.49669 | 67.84539 | 29.2063 | 0.235377 | 1.458797 | 0.700653 |
| 12 | 35647200 | 35648599 | 19.67488 | 34.43112 | 38.65586 | 17.05459 | 0.221442 | 0.693735 | 0.700664 |
| 16 | 72549000 | 72550599 | 17.2595 | 30.58761 | 33.65442 | 14.8789 | 0.254658 | 0.511335 | 0.7007 |
| 8 | 81402400 | 81407799 | 70.08887 | 110.279 | 152.4636 | 63.25311 | -0.14536 | 2.521321 | 0.700717 |
| 1 | 137219400 | 137221199 | 16.89598 | 29.91979 | 33.80961 | 14.94987 | 0.216113 | 0.519679 | 0.700719 |
| Y | 40301400 | 40301799 | 4.72032 | 9.170292 | 20.61647 | 7.69525 | -0.5633 | -0.29793 | 0.700749 |
| 15 | 46808000 | 46809399 | 13.93404 | 24.55297 | 37.25312 | 16.48881 | -0.20938 | 0.562864 | 0.700763 |

| | | | | | | | | | |
|---|---|---|---|---|---|---|---|---|---|
| 4 | 113391000 | 113391799 | 19.93739 | 34.67153 | 39.9346 | 17.54351 | 0.184089 | 0.787576 | 0.700771 |
| 15 | 36324800 | 36327799 | 26.71247 | 44.89308 | 51.78216 | 22.61898 | 0.175714 | 1.139873 | 0.700781 |
| 1 | 154944000 | 154947199 | 24.03364 | 40.6382 | 44.37709 | 19.42026 | 0.2548 | 1.070592 | 0.700829 |
| X | 64547400 | 64554999 | 149.8712 | 227.8569 | 193.1561 | 80.01461 | 0.52123 | 3.197927 | 0.700837 |
| 7 | 39311000 | 39312399 | 19.96604 | 34.7829 | 39.87095 | 17.53505 | 0.191227 | 0.762989 | 0.700839 |
| 18 | 69373600 | 69375399 | 16.42526 | 28.67546 | 43.05554 | 18.96097 | -0.19619 | 0.719018 | 0.700839 |
| 14 | 14193800 | 14198999 | 36.14593 | 58.20906 | 99.60171 | 42.33938 | -0.41838 | 1.820796 | 0.700859 |
| 6 | 74865000 | 74865999 | 13.09676 | 24.04037 | 28.05268 | 11.94129 | 0.207135 | 0.264381 | 0.700888 |
| 15 | 28392400 | 28394999 | 21.77962 | 37.7691 | 42.50789 | 18.65056 | 0.21666 | 0.832735 | 0.700914 |
| 13 | 104999600 | 105000399 | 10.29354 | 19.53163 | 23.79745 | 9.592945 | 0.207502 | 0.026721 | 0.70094 |
| 1 | 78491600 | 78492999 | 15.86275 | 27.58483 | 42.69342 | 18.75513 | -0.24012 | 0.773487 | 0.700959 |
| 3 | 152522000 | 152524599 | 17.8846 | 30.97275 | 46.33997 | 20.33107 | -0.19265 | 0.833272 | 0.700967 |
| 7 | 7351400 | 7353399 | 46.11471 | 74.2914 | 84.73717 | 36.04086 | 0.166172 | 1.816207 | 0.700968 |
| X | 88369600 | 88370199 | 8.863075 | 17.15544 | 21.01387 | 8.002113 | 0.276479 | -0.17719 | 0.700979 |
| 15 | 20645200 | 20646999 | 17.68649 | 31.04031 | 34.92652 | 15.44662 | 0.218017 | 0.628662 | 0.700987 |
| 16 | 55160600 | 55162399 | 21.31562 | 37.55592 | 36.64166 | 16.21852 | 0.424107 | 0.604105 | 0.700988 |
| 3 | 158460600 | 158462999 | 19.01343 | 33.57501 | 35.1181 | 15.53718 | 0.325299 | 0.562547 | 0.700988 |
| 10 | 19420400 | 19423999 | 43.1849 | 69.71674 | 79.99705 | 34.11838 | 0.161062 | 1.749081 | 0.700997 |
| 16 | 94632000 | 94634399 | 21.75266 | 36.8391 | 54.3586 | 23.79969 | -0.17788 | 1.060319 | 0.70101 |
| 17 | 80115400 | 80119999 | 32.06474 | 52.20153 | 76.87802 | 33.00536 | -0.18676 | 1.550811 | 0.701019 |
| 15 | 24047600 | 24048999 | 15.10667 | 27.43434 | 30.33873 | 13.09397 | 0.270521 | 0.342764 | 0.701021 |
| 4 | 73374600 | 73379799 | 53.26761 | 85.9303 | 73.63059 | 31.37308 | 0.580235 | 1.784831 | 0.701024 |
| 8 | 72637600 | 72640999 | 62.56671 | 100.0278 | 105.6958 | 44.4509 | 0.263195 | 2.080592 | 0.701029 |
| 6 | 13198800 | 13199199 | 9.925489 | 18.89482 | 22.33324 | 8.94078 | 0.2603 | -0.00053 | 0.701046 |
| 18 | 6611800 | 6612599 | 8.231966 | 15.55281 | 25.97485 | 10.56638 | -0.25513 | 0.043838 | 0.70105 |
| 4 | 147183000 | 147191999 | 128.0174 | 194.945 | 214.0959 | 88.73163 | 0.149865 | 3.15933 | 0.701053 |
| 3 | 42183400 | 42187199 | 60.4081 | 96.2031 | 109.2583 | 45.7003 | 0.153247 | 2.197441 | 0.701054 |
| 3 | 90875200 | 90898599 | 344.0308 | 487.1824 | 485.1257 | 200.751 | 0.192636 | 4.394895 | 0.701056 |
| 6 | 32890800 | 32892999 | 22.96907 | 40.62613 | 34.35797 | 15.19594 | 0.631478 | 0.570483 | 0.70106 |
| 8 | 21461400 | 21463799 | 28.36561 | 48.14823 | 50.70007 | 22.21326 | 0.307125 | 1.057848 | 0.701062 |
| 2 | 40868400 | 40869799 | 12.41505 | 23.45732 | 26.39608 | 10.73644 | 0.309911 | 0.081341 | 0.701088 |
| 15 | 32359200 | 32360999 | 20.62804 | 36.00496 | 34.07255 | 15.00297 | 0.466713 | 0.729911 | 0.701093 |
| 8 | 82213600 | 82215399 | 11.53894 | 20.63008 | 38.84732 | 17.05218 | -0.50879 | 0.429582 | 0.7011 |
| 5 | 10116000 | 10122799 | 82.04842 | 128.6861 | 143.9399 | 59.69802 | 0.156416 | 2.588205 | 0.701115 |
| 3 | 17345400 | 17348799 | 44.50545 | 71.82544 | 81.92845 | 34.92756 | 0.169076 | 1.761435 | 0.701133 |
| 1 | 99130600 | 99131399 | 18.13272 | 31.24946 | 46.89312 | 20.57069 | -0.19827 | 0.880721 | 0.701139 |
| 13 | 106515200 | 106518599 | 31.41547 | 50.8839 | 88.3963 | 37.82982 | -0.43081 | 1.652798 | 0.70114 |
| 13 | 42077200 | 42077799 | 8.335552 | 16.16492 | 19.90343 | 7.512555 | 0.279704 | -0.22172 | 0.701146 |
| 17 | 12237400 | 12241199 | 42.04621 | 67.70984 | 98.37604 | 41.81751 | -0.18295 | 1.830629 | 0.701163 |
| 8 | 62043000 | 62044799 | 14.70599 | 25.67815 | 40.55139 | 17.86003 | -0.26847 | 0.704769 | 0.701205 |
| 13 | 98657800 | 98662799 | 60.46036 | 96.01433 | 135.7574 | 56.59297 | -0.16657 | 2.291583 | 0.701227 |
| 13 | 83980200 | 83981599 | 17.61276 | 30.34859 | 46.37099 | 20.34155 | -0.22429 | 0.878084 | 0.701265 |
| 13 | 66793400 | 66798399 | 47.07681 | 77.00436 | 69.73283 | 29.98941 | 0.517888 | 1.51525 | 0.701272 |
| 7 | 133078200 | 133079999 | 18.18136 | 31.84021 | 36.04409 | 15.91191 | 0.2094 | 0.667841 | 0.701283 |
| 16 | 80064800 | 80067599 | 36.93626 | 59.57804 | 87.08941 | 37.10223 | -0.18881 | 1.768749 | 0.701358 |
| 7 | 6672600 | 6674199 | 12.54011 | 22.59044 | 34.66104 | 15.11109 | -0.20838 | 0.380478 | 0.701362 |
| 1 | 46539400 | 46541799 | 24.94253 | 41.70447 | 61.50157 | 26.78182 | -0.17747 | 1.198276 | 0.701382 |

| | | | | | | | | | |
|---|---|---|---|---|---|---|---|---|---|
| 7 | 27102200 | 27113799 | 116.4257 | 176.8536 | 237.4371 | 98.33223 | -0.14262 | 3.208778 | 0.701398 |
| 8 | 33622400 | 33623399 | 11.79147 | 21.32517 | 32.71666 | 14.1937 | -0.20337 | 0.350356 | 0.701421 |
| 10 | 41712600 | 41716999 | 48.01562 | 77.19143 | 87.50557 | 37.10516 | 0.171264 | 1.884337 | 0.701431 |
| 1 | 3654200 | 3655199 | 10.80298 | 19.74883 | 30.78379 | 13.17662 | -0.21348 | 0.276525 | 0.701453 |
| 8 | 62667800 | 62668399 | 10.80298 | 19.74883 | 30.78379 | 13.17662 | -0.21348 | 0.276525 | 0.701453 |
| 19 | 39385800 | 39387399 | 26.85841 | 45.64892 | 42.16256 | 18.47823 | 0.495844 | 1.03545 | 0.701454 |
| 7 | 43381800 | 43383799 | 16.378 | 28.64017 | 43.04309 | 18.97821 | -0.19726 | 0.696482 | 0.701468 |
| 10 | 108258400 | 108260799 | 25.2318 | 42.27083 | 63.29235 | 27.59571 | -0.19945 | 1.178935 | 0.701468 |
| 3 | 139521400 | 139523199 | 21.91746 | 38.27841 | 40.704 | 17.92968 | 0.299316 | 0.738125 | 0.701479 |
| X | 104691000 | 104692399 | 25.29455 | 42.76955 | 49.05842 | 21.4688 | 0.184114 | 1.081511 | 0.701496 |
| 2 | 177073800 | 177074599 | 14.03486 | 25.54126 | 29.44065 | 12.67316 | 0.21365 | 0.32225 | 0.701504 |
| X | 51976200 | 51978399 | 28.37881 | 47.89808 | 52.32426 | 22.88546 | 0.254187 | 1.112082 | 0.701507 |
| X | 122956000 | 122956599 | 17.23996 | 30.83638 | 26.80463 | 11.76161 | 0.600685 | 0.442704 | 0.701508 |
| X | 153787000 | 153793199 | 90.7508 | 142.3632 | 149.7184 | 62.09334 | 0.245484 | 2.593541 | 0.70151 |
| 14 | 107016000 | 107016399 | 6.092628 | 11.86021 | 21.31688 | 7.828099 | -0.2227 | -0.33529 | 0.701515 |
| 4 | 59026600 | 59036799 | 120.708 | 183.1205 | 245.812 | 101.761 | -0.14394 | 3.236153 | 0.701523 |
| 7 | 52260600 | 52262799 | 19.38038 | 33.82428 | 38.74437 | 17.05914 | 0.192409 | 0.731441 | 0.701553 |
| 8 | 4107200 | 4118799 | 161.0992 | 241.8534 | 261.2578 | 107.8226 | 0.156229 | 3.452479 | 0.701584 |
| 3 | 40547800 | 40549999 | 23.01609 | 39.70439 | 44.6741 | 19.59786 | 0.215566 | 0.888333 | 0.70162 |
| 16 | 59319000 | 59319599 | 7.56762 | 14.57543 | 24.77056 | 9.618342 | -0.21928 | -0.11897 | 0.701628 |
| 1 | 5554400 | 5554799 | 12.88686 | 23.43332 | 26.54303 | 11.43728 | 0.237928 | 0.312246 | 0.701641 |
| 15 | 29130800 | 29134799 | 53.66445 | 85.55875 | 120.2415 | 50.37192 | -0.15189 | 2.159568 | 0.701647 |
| 6 | 30419000 | 30419799 | 8.413607 | 15.81814 | 26.45452 | 10.85937 | -0.26791 | 0.079932 | 0.70165 |
| 4 | 38855600 | 38864799 | 212.7362 | 307.7606 | 408.4344 | 168.4543 | -0.19082 | 4.074205 | 0.701704 |
| 4 | 112238400 | 112246799 | 97.79128 | 152.5922 | 160.2101 | 66.48107 | 0.241901 | 2.689245 | 0.701725 |
| 2 | 109747400 | 109749799 | 15.72431 | 26.8362 | 54.74798 | 24.00194 | -0.64197 | 0.969232 | 0.701729 |
| 3 | 91476600 | 91480599 | 48.44779 | 78.70902 | 75.99556 | 32.59545 | 0.418584 | 1.617572 | 0.701759 |
| X | 21842800 | 21843999 | 14.15652 | 26.04104 | 29.08879 | 12.34651 | 0.272792 | 0.263731 | 0.701764 |
| 16 | 47831600 | 47837799 | 73.62361 | 117.0051 | 120.9052 | 50.43898 | 0.286198 | 2.276491 | 0.701772 |
| 10 | 42032000 | 42038399 | 102.1251 | 157.8556 | 175.1254 | 72.74313 | 0.151959 | 2.857147 | 0.701794 |
| 16 | 55725600 | 55729199 | 27.97578 | 45.81092 | 69.96137 | 30.10045 | -0.23316 | 1.454634 | 0.701794 |
| 11 | 46801200 | 46802199 | 15.2829 | 26.50306 | 48.98489 | 21.49238 | -0.49566 | 0.814528 | 0.701809 |
| 3 | 32730400 | 32732199 | 18.39374 | 31.33693 | 49.53085 | 21.71164 | -0.27565 | 0.991001 | 0.701829 |
| 3 | 127222600 | 127225399 | 23.25255 | 39.15222 | 57.6428 | 25.1934 | -0.17504 | 1.118851 | 0.701836 |
| 13 | 62509600 | 62533999 | 509.7457 | 699.3873 | 658.0823 | 273.691 | 0.228378 | 4.874767 | 0.701841 |
| 12 | 109899200 | 109902599 | 58.838 | 93.14279 | 132.1288 | 54.93262 | -0.17545 | 2.376386 | 0.701858 |
| 7 | 26616000 | 26618999 | 39.22645 | 63.88711 | 72.08348 | 30.95161 | 0.197022 | 1.557705 | 0.701873 |
| 7 | 16503600 | 16510199 | 78.49662 | 122.1366 | 169.8588 | 70.51326 | -0.16639 | 2.729673 | 0.701883 |
| 7 | 45234200 | 45235799 | 16.05608 | 27.83647 | 43.78285 | 19.2098 | -0.26384 | 0.814212 | 0.701919 |
| 12 | 103034200 | 103038799 | 37.57348 | 60.99847 | 69.74556 | 29.89886 | 0.174247 | 1.605723 | 0.701922 |
| 10 | 124521600 | 124523399 | 21.09638 | 36.05887 | 53.8608 | 23.61586 | -0.19391 | 0.975307 | 0.701967 |
| 16 | 8173200 | 8175599 | 26.69686 | 43.83277 | 79.26755 | 34.14267 | -0.47488 | 1.424724 | 0.70204 |
| 7 | 47161400 | 47190599 | 458.34 | 633.4732 | 611.6747 | 254.0954 | 0.20174 | 4.761148 | 0.702052 |
| 4 | 101841000 | 101843399 | 13.8555 | 24.16527 | 45.76441 | 20.14141 | -0.5294 | 0.726548 | 0.702053 |
| 16 | 10157000 | 10158399 | 18.7497 | 32.29031 | 47.9425 | 21.03058 | -0.18328 | 0.89015 | 0.702061 |
| 14 | 81323600 | 81326799 | 34.80781 | 56.90217 | 65.41862 | 28.12798 | 0.173834 | 1.492315 | 0.702062 |
| 17 | 24037200 | 24043599 | 212.6783 | 310.885 | 316.9198 | 130.5624 | 0.204104 | 3.916167 | 0.70207 |

| | | | | | | | | | |
|---|---|---|---|---|---|---|---|---|---|
| 4 | 66902200 | 66902999 | 16.72777 | 30.04174 | 27.57417 | 12.05801 | 0.525632 | 0.41348 | 0.702071 |
| 5 | 14970200 | 14972199 | 32.29514 | 53.21281 | 60.75551 | 26.20833 | 0.18877 | 1.380423 | 0.702104 |
| 10 | 40416600 | 40421999 | 44.78062 | 72.33687 | 81.22005 | 34.66719 | 0.193315 | 1.733894 | 0.702107 |
| 5 | 22110400 | 22113199 | 13.67184 | 23.90929 | 44.66213 | 19.69302 | -0.50943 | 0.68315 | 0.702116 |
| 5 | 93863400 | 93910199 | 798.7962 | 1055.812 | 954.3845 | 399.0879 | 0.232179 | 5.467899 | 0.70212 |
| 17 | 22289600 | 22292199 | 30.70749 | 51.38026 | 48.09369 | 20.95258 | 0.476186 | 1.196488 | 0.702122 |
| 14 | 65417400 | 65418399 | 16.5791 | 29.85377 | 26.43449 | 11.53073 | 0.57934 | 0.392731 | 0.702135 |
| 5 | 34093000 | 34097799 | 55.27203 | 87.71668 | 126.8423 | 52.84137 | -0.19984 | 2.303013 | 0.702146 |
| 3 | 57188800 | 57191199 | 24.27276 | 41.25502 | 47.43469 | 20.78883 | 0.180998 | 1.031083 | 0.702148 |
| 2 | 104750200 | 104752399 | 29.76053 | 49.84581 | 55.70772 | 24.2851 | 0.221181 | 1.189186 | 0.702152 |
| 19 | 49971000 | 49973399 | 19.62411 | 34.34438 | 38.62911 | 17.04367 | 0.218764 | 0.691727 | 0.702168 |
| 17 | 57799200 | 57801599 | 29.63046 | 49.20838 | 56.71434 | 24.58856 | 0.176874 | 1.281722 | 0.702187 |
| 9 | 124054600 | 124060599 | 71.30514 | 114.028 | 104.3799 | 43.91374 | 0.470352 | 2.084427 | 0.702195 |
| 7 | 47563800 | 47566799 | 37.35733 | 61.27369 | 58.56171 | 25.21328 | 0.442093 | 1.449889 | 0.702203 |
| 4 | 61794200 | 61802199 | 82.32658 | 129.8412 | 139.038 | 57.70097 | 0.225367 | 2.479948 | 0.70222 |
| 2 | 169270000 | 169271799 | 16.93574 | 29.80141 | 33.90374 | 15.01043 | 0.202856 | 0.586888 | 0.702238 |
| 15 | 24357000 | 24359799 | 46.09838 | 73.64234 | 107.5399 | 45.177 | -0.20393 | 2.08584 | 0.702245 |
| 11 | 79636000 | 79637999 | 28.4471 | 46.37707 | 81.89458 | 35.17707 | -0.44643 | 1.529532 | 0.702247 |
| 17 | 54449200 | 54449999 | 8.03002 | 15.39807 | 25.75727 | 10.12873 | -0.21382 | -0.07033 | 0.702262 |
| 8 | 73926000 | 73928199 | 14.12708 | 25.04319 | 38.16578 | 16.87249 | -0.21241 | 0.500453 | 0.702268 |
| 1 | 67928200 | 67928999 | 11.23239 | 21.00791 | 23.26887 | 9.636328 | 0.312148 | 0.131126 | 0.702284 |
| 6 | 136516400 | 136518199 | 23.21305 | 39.96124 | 45.22543 | 19.83847 | 0.206626 | 0.908627 | 0.702311 |
| 7 | 29152600 | 29154999 | 19.22879 | 33.09435 | 49.00702 | 21.49713 | -0.17973 | 0.898938 | 0.702321 |
| 18 | 24480200 | 24482199 | 20.22967 | 35.21456 | 40.37919 | 17.74983 | 0.19058 | 0.775017 | 0.702332 |
| 12 | 19215000 | 19286199 | 2157.22 | 2609.283 | 2291.921 | 967.8825 | 0.144562 | 6.896332 | 0.702337 |
| 16 | 83745400 | 83747199 | 15.03608 | 27.13447 | 30.85156 | 13.42075 | 0.223541 | 0.380291 | 0.70234 |
| 1 | 59388000 | 59390999 | 54.96753 | 87.36982 | 123.265 | 51.43147 | -0.16213 | 2.257216 | 0.70234 |
| 14 | 3205000 | 3207599 | 50.80294 | 81.51588 | 92.17016 | 38.97156 | 0.171637 | 1.949089 | 0.702375 |
| 7 | 26481400 | 26483199 | 18.34471 | 32.02873 | 35.8574 | 15.79173 | 0.225548 | 0.71873 | 0.702403 |
| 7 | 5277200 | 5322199 | 775.709 | 1025.446 | 953.588 | 398.7476 | 0.191498 | 5.465141 | 0.702407 |
| 3 | 76408400 | 76409399 | 13.79794 | 24.84232 | 27.88288 | 12.16463 | 0.239316 | 0.377513 | 0.702417 |
| 3 | 124928000 | 124929399 | 15.31925 | 26.84712 | 40.36747 | 17.83126 | -0.19802 | 0.649533 | 0.702445 |
| 12 | 68159600 | 68162799 | 24.97474 | 43.16125 | 45.15507 | 19.80995 | 0.320633 | 0.881044 | 0.702448 |
| 1 | 60008600 | 60009999 | 11.42743 | 20.89818 | 32.31461 | 13.82535 | -0.2017 | 0.281987 | 0.702451 |
| 19 | 15981600 | 15983599 | 26.15296 | 43.83571 | 49.12425 | 21.42504 | 0.217233 | 1.161822 | 0.702487 |
| 16 | 14086400 | 14087999 | 12.62956 | 22.39725 | 34.68173 | 15.32832 | -0.23488 | 0.480276 | 0.70249 |
| 1 | 88648400 | 88651399 | 29.41293 | 48.14276 | 71.87774 | 30.91923 | -0.20094 | 1.464448 | 0.702496 |
| 5 | 104935200 | 104943999 | 153.4348 | 233.2019 | 195.7937 | 81.09961 | 0.534733 | 3.20375 | 0.702508 |
| 12 | 60768200 | 60768599 | 8.242102 | 15.9144 | 19.32946 | 7.445525 | 0.272262 | -0.15421 | 0.702509 |
| 9 | 39794200 | 39797999 | 49.36826 | 79.47137 | 88.40521 | 37.55881 | 0.200842 | 1.848297 | 0.702526 |
| 6 | 108249000 | 108251399 | 17.908 | 30.55429 | 55.36675 | 24.27266 | -0.47191 | 0.980834 | 0.702533 |
| 3 | 59336400 | 59340799 | 52.79192 | 84.17912 | 118.4486 | 49.6302 | -0.15345 | 2.153056 | 0.702565 |
| 13 | 8613000 | 8614199 | 8.486225 | 16.12787 | 26.61476 | 10.69269 | -0.22262 | 0.005582 | 0.702567 |
| 9 | 115021000 | 115022599 | 13.72229 | 24.43387 | 37.13485 | 16.35657 | -0.20489 | 0.466006 | 0.702588 |
| 1 | 17397600 | 17398799 | 10.88018 | 20.18918 | 32.63842 | 13.69 | -0.24502 | 0.195413 | 0.7026 |
| 1 | 178899400 | 178900199 | 10.34868 | 19.8308 | 19.60869 | 7.725895 | 0.53574 | -0.05811 | 0.702601 |
| 2 | 45780000 | 45781399 | 11.55646 | 21.12626 | 32.82352 | 14.04971 | -0.20912 | 0.285322 | 0.702603 |

| | | | | | | | | | |
|---|---|---|---|---|---|---|---|---|---|
| 5 | 91364800 | 91366399 | 10.06164 | 18.74814 | 30.54515 | 12.72987 | -0.24827 | 0.162919 | 0.702608 |
| 18 | 28445800 | 28448599 | 23.54225 | 39.92442 | 63.41551 | 27.77658 | -0.28392 | 1.062667 | 0.702609 |
| 6 | 146384600 | 146386199 | 12.04184 | 21.93421 | 34.34946 | 14.76942 | -0.22477 | 0.312578 | 0.70262 |
| 5 | 148631200 | 148633199 | 23.11882 | 39.50195 | 45.3127 | 19.8673 | 0.185526 | 0.976666 | 0.702634 |
| 9 | 46342000 | 46349399 | 76.02384 | 118.8902 | 163.3023 | 67.74876 | -0.14279 | 2.639906 | 0.702646 |
| 3 | 112954400 | 112956999 | 25.94597 | 45.17071 | 42.27709 | 18.58555 | 0.482315 | 0.80359 | 0.702648 |
| 16 | 87027600 | 87034399 | 84.83779 | 131.6762 | 180.2043 | 74.83568 | -0.146 | 2.776379 | 0.702672 |
| 5 | 10307000 | 10307999 | 13.76967 | 24.35777 | 36.67544 | 16.22052 | -0.19623 | 0.518183 | 0.702693 |
| X | 20725000 | 20731599 | 70.65305 | 112.615 | 111.2876 | 46.60421 | 0.354983 | 2.185596 | 0.702725 |
| 10 | 59142200 | 59145199 | 42.40696 | 68.15541 | 98.08721 | 41.575 | -0.17285 | 1.89017 | 0.702734 |
| 4 | 3354400 | 3369999 | 298.5281 | 424.9524 | 440.2161 | 182.0082 | 0.143195 | 4.315957 | 0.702739 |
| 7 | 6489600 | 6492399 | 36.92745 | 60.11128 | 69.36859 | 29.78018 | 0.16409 | 1.559035 | 0.702748 |
| 17 | 60319000 | 60319599 | 10.46143 | 19.67902 | 23.21036 | 9.518785 | 0.232929 | 0.085137 | 0.702766 |
| 14 | 85764600 | 85770799 | 71.32155 | 113.3107 | 121.6532 | 50.73498 | 0.230665 | 2.285533 | 0.702823 |
| 9 | 9449000 | 9449999 | 14.94375 | 27.09008 | 30.32789 | 13.11854 | 0.250857 | 0.351679 | 0.702867 |
| X | 73949400 | 73952399 | 28.88886 | 48.41609 | 53.96608 | 23.53513 | 0.225069 | 1.180422 | 0.702872 |
| 8 | 100859200 | 100861799 | 32.16202 | 54.01026 | 55.78276 | 24.35829 | 0.334583 | 1.170216 | 0.702879 |
| 9 | 4848400 | 4855599 | 162.4907 | 240.0849 | 327.2497 | 134.7686 | -0.20012 | 3.740316 | 0.702892 |
| 1 | 35114800 | 35116999 | 19.12804 | 32.65075 | 49.53715 | 21.71777 | -0.21651 | 0.974621 | 0.702892 |
| 5 | 105247000 | 105248599 | 13.58862 | 24.20354 | 36.64465 | 16.13634 | -0.19905 | 0.463013 | 0.702904 |
| 10 | 4429000 | 4430799 | 19.35588 | 33.60909 | 37.46602 | 16.44331 | 0.230833 | 0.801936 | 0.702917 |
| 5 | 79008800 | 79011399 | 33.52459 | 55.38243 | 62.58697 | 27.03596 | 0.204531 | 1.35546 | 0.702931 |
| 12 | 49208000 | 49209199 | 16.30225 | 28.42854 | 42.58516 | 18.7405 | -0.19291 | 0.734719 | 0.702943 |
| 11 | 92801200 | 92802199 | 15.66474 | 27.33151 | 41.56238 | 18.29658 | -0.21438 | 0.722432 | 0.702955 |
| 9 | 23652000 | 23653599 | 14.95709 | 26.00002 | 47.90864 | 21.04113 | -0.49074 | 0.78188 | 0.70296 |
| 10 | 10340000 | 10343399 | 25.63796 | 44.07194 | 46.80971 | 20.53252 | 0.297386 | 0.932523 | 0.702975 |
| 1 | 147050800 | 147053799 | 28.08281 | 47.22892 | 53.44176 | 23.34883 | 0.203329 | 1.14075 | 0.702978 |
| 18 | 8306600 | 8308799 | 34.94323 | 56.43702 | 83.79477 | 35.75888 | -0.20908 | 1.724681 | 0.702985 |
| 5 | 93570800 | 93588999 | 221.0124 | 326.8582 | 281.2548 | 115.8281 | 0.4644 | 3.730608 | 0.702985 |
| 3 | 46824000 | 46826399 | 21.51624 | 37.82005 | 37.83375 | 16.75406 | 0.387077 | 0.63448 | 0.702987 |
| 8 | 126131000 | 126133799 | 41.95228 | 67.42126 | 96.53255 | 40.91305 | -0.16551 | 1.89141 | 0.702992 |
| 3 | 137720000 | 137720799 | 12.77121 | 23.56721 | 27.59856 | 11.66367 | 0.209512 | 0.231429 | 0.703002 |
| X | 107790200 | 107791799 | 14.61434 | 25.70444 | 38.79043 | 17.17556 | -0.20268 | 0.590584 | 0.703006 |
| 11 | 109169800 | 109173799 | 56.59673 | 89.96148 | 125.8565 | 52.51393 | -0.14997 | 2.258397 | 0.703018 |
| 4 | 145150400 | 145152599 | 47.80609 | 76.89384 | 87.92155 | 37.30914 | 0.159807 | 1.870117 | 0.703051 |
| 7 | 29319800 | 29324999 | 54.66096 | 87.66207 | 85.09864 | 35.954 | 0.390422 | 1.968806 | 0.703062 |
| 18 | 84129400 | 84133199 | 35.95862 | 58.94783 | 66.90954 | 28.80083 | 0.194376 | 1.458997 | 0.703064 |
| 17 | 78476800 | 78477999 | 12.72778 | 23.82201 | 26.65452 | 11.02487 | 0.297982 | 0.142709 | 0.703071 |
| 10 | 95053800 | 95055799 | 16.12663 | 27.70875 | 52.15502 | 22.87685 | -0.52473 | 0.905789 | 0.703078 |
| 12 | 61172400 | 61173599 | 18.43568 | 32.48553 | 30.31258 | 13.4168 | 0.488228 | 0.575964 | 0.703108 |
| 11 | 22915800 | 22925799 | 177.8093 | 262.5391 | 343.1434 | 141.313 | -0.14052 | 3.753789 | 0.703115 |
| 13 | 7528400 | 7530599 | 21.01401 | 36.74759 | 39.59577 | 17.45842 | 0.280332 | 0.711798 | 0.703141 |
| 7 | 66952000 | 66954999 | 38.34084 | 62.74401 | 60.24234 | 25.91231 | 0.4341 | 1.481272 | 0.703147 |
| 12 | 28577600 | 28578399 | 15.72371 | 28.22563 | 31.81658 | 13.91855 | 0.230607 | 0.418567 | 0.703151 |
| 13 | 107238200 | 107240199 | 14.23749 | 25.07962 | 37.73012 | 16.70101 | -0.19744 | 0.568542 | 0.703159 |
| 8 | 100390000 | 100391999 | 28.06246 | 46.72541 | 53.4383 | 23.20927 | 0.188191 | 1.24368 | 0.703175 |
| 7 | 76386800 | 76389199 | 31.37167 | 52.32394 | 58.04012 | 25.23488 | 0.232055 | 1.239422 | 0.703185 |

| | | | | | | | | | |
|---|---|---|---|---|---|---|---|---|---|
| 16 | 75749200 | 75749599 | 5.944488 | 11.57311 | 21.34376 | 7.831007 | -0.25894 | -0.34279 | 0.703208 |
| 14 | 28407200 | 28408199 | 9.177371 | 17.07744 | 27.87936 | 11.64541 | -0.25395 | 0.155089 | 0.703209 |
| 8 | 69025600 | 69028799 | 29.52217 | 50.3878 | 49.58167 | 21.7371 | 0.40564 | 1.014459 | 0.703211 |
| 4 | 34717400 | 34721599 | 64.86118 | 102.8548 | 115.4092 | 48.0411 | 0.164487 | 2.33099 | 0.703219 |
| X | 118701600 | 118711399 | 137.5898 | 210.7764 | 174.1157 | 72.26581 | 0.567173 | 3.040722 | 0.703244 |
| X | 161957400 | 161958399 | 17.28124 | 31.03022 | 25.82133 | 11.29319 | 0.666533 | 0.41764 | 0.70325 |
| 11 | 91994800 | 91999999 | 41.86946 | 67.97399 | 75.62896 | 32.42142 | 0.213524 | 1.617002 | 0.703272 |
| 2 | 173623400 | 173626599 | 21.56226 | 36.44266 | 54.36098 | 23.7861 | -0.19364 | 1.076148 | 0.703275 |
| 5 | 148401000 | 148404199 | 21.56226 | 36.44266 | 54.36098 | 23.7861 | -0.19364 | 1.076148 | 0.703275 |
| 16 | 62822800 | 62823999 | 11.68236 | 21.04585 | 32.74088 | 14.27345 | -0.22771 | 0.373343 | 0.703318 |
| 3 | 99106200 | 99108399 | 22.00631 | 38.00109 | 43.14387 | 18.92716 | 0.203078 | 0.875291 | 0.703327 |
| 17 | 74171800 | 74173999 | 15.4453 | 27.18119 | 42.30752 | 18.73075 | -0.24719 | 0.591099 | 0.70334 |
| 17 | 61318800 | 61321799 | 40.97908 | 66.57073 | 64.52375 | 27.67366 | 0.41291 | 1.596631 | 0.703341 |
| 14 | 3902200 | 3906999 | 40.46251 | 65.294 | 95.66303 | 40.77746 | -0.19149 | 1.763537 | 0.703343 |
| 2 | 30470000 | 30473399 | 30.05938 | 49.44127 | 72.84513 | 31.39993 | -0.17907 | 1.407344 | 0.70336 |
| 15 | 57276000 | 57277399 | 16.8142 | 29.28863 | 43.64991 | 19.19561 | -0.18572 | 0.755241 | 0.703373 |
| 18 | 6846000 | 6848799 | 33.98131 | 55.60546 | 62.81825 | 27.01955 | 0.199949 | 1.471604 | 0.703375 |
| 15 | 12525800 | 12533999 | 122.0362 | 185.9459 | 205.661 | 85.25077 | 0.140373 | 3.145388 | 0.703403 |
| 1 | 29830400 | 29830799 | 6.840538 | 13.19264 | 23.11938 | 8.942858 | -0.25737 | -0.14258 | 0.703444 |
| X | 161172800 | 161174199 | 12.62175 | 22.89497 | 36.422 | 15.74235 | -0.2521 | 0.346447 | 0.703448 |
| 1 | 41413200 | 41414199 | 13.2218 | 24.0608 | 27.98634 | 12.04765 | 0.200713 | 0.313497 | 0.703498 |
| 6 | 49253000 | 49257199 | 23.44624 | 40.54461 | 37.71422 | 16.54589 | 0.489587 | 0.85839 | 0.703512 |
| 6 | 136449400 | 136451799 | 18.7559 | 32.80839 | 37.65725 | 16.6078 | 0.18965 | 0.693205 | 0.703564 |
| 15 | 64285800 | 64286999 | 18.7559 | 32.80839 | 37.65725 | 16.6078 | 0.18965 | 0.693205 | 0.703564 |
| 18 | 58450400 | 58453399 | 49.05229 | 78.73847 | 89.91301 | 38.0404 | 0.158122 | 1.934512 | 0.703617 |
| 13 | 97477000 | 97480199 | 26.84288 | 45.24487 | 51.78975 | 22.64245 | 0.186851 | 1.114668 | 0.703663 |
| 5 | 104887200 | 104925199 | 884.9025 | 1154.434 | 1083.149 | 453.5877 | 0.159633 | 5.678897 | 0.703667 |
| 3 | 78459400 | 78462999 | 40.48663 | 66.06899 | 61.58021 | 26.46057 | 0.473897 | 1.524162 | 0.703669 |
| 17 | 89038800 | 89040999 | 29.21449 | 48.66773 | 56.23453 | 24.43137 | 0.17337 | 1.244034 | 0.703674 |
| 5 | 119097400 | 119103199 | 50.12334 | 80.40236 | 91.96673 | 38.86859 | 0.154558 | 1.959143 | 0.703678 |
| 12 | 21043200 | 21063799 | 654.3222 | 871.996 | 843.7011 | 352.432 | 0.145914 | 5.33459 | 0.703693 |
| 11 | 99603200 | 99605199 | 15.34122 | 27.88917 | 22.98947 | 9.909238 | 0.693513 | 0.325526 | 0.703698 |
| 5 | 60344000 | 60349199 | 98.13209 | 152.8336 | 145.3025 | 60.31065 | 0.382802 | 2.71971 | 0.703701 |
| 4 | 127777600 | 127783599 | 65.46497 | 104.4301 | 97.44603 | 40.8535 | 0.438789 | 2.146446 | 0.703716 |
| 8 | 46604400 | 46605999 | 13.26523 | 23.35092 | 36.78022 | 16.28278 | -0.26372 | 0.567722 | 0.703719 |
| 17 | 94534600 | 94539199 | 93.30308 | 144.5152 | 160.0215 | 66.46858 | 0.156963 | 2.814474 | 0.703731 |
| 6 | 107506000 | 107507999 | 16.64185 | 28.83822 | 44.27074 | 19.42358 | -0.22927 | 0.82037 | 0.703769 |
| 10 | 10393000 | 10400799 | 118.6127 | 180.1162 | 241.0792 | 99.8307 | -0.13861 | 3.216043 | 0.703779 |
| X | 110267000 | 110268599 | 23.37845 | 39.90801 | 44.30971 | 19.42606 | 0.232292 | 0.986808 | 0.703788 |
| 18 | 57209200 | 57209999 | 10.58259 | 20.06763 | 24.30312 | 9.80965 | 0.214354 | 0.034386 | 0.703806 |
| 18 | 41723400 | 41725199 | 14.41576 | 26.33211 | 29.38819 | 12.59193 | 0.264048 | 0.30246 | 0.703807 |
| 8 | 41019400 | 41020799 | 15.97137 | 28.45906 | 32.24775 | 14.19537 | 0.21687 | 0.465496 | 0.703815 |
| 1 | 36027600 | 36037199 | 121.5444 | 184.337 | 247.0058 | 102.2465 | -0.14171 | 3.242277 | 0.703815 |
| 17 | 43744800 | 43746199 | 21.67551 | 37.14875 | 41.67125 | 18.27519 | 0.218261 | 0.937211 | 0.703881 |
| 2 | 177169600 | 177182799 | 182.3353 | 272.4655 | 282.7669 | 116.5836 | 0.20796 | 3.544633 | 0.70389 |
| 17 | 9205800 | 9207399 | 14.03269 | 24.61595 | 38.84212 | 17.17136 | -0.26726 | 0.62359 | 0.703903 |
| 1 | 109131000 | 109133999 | 24.84041 | 43.27871 | 41.72012 | 18.34942 | 0.439927 | 0.786282 | 0.703907 |

| | | | | | | | | | |
|---|---|---|---|---|---|---|---|---|---|
| 7 | 78492800 | 78494599 | 17.32052 | 29.95941 | 45.31761 | 19.88178 | -0.20858 | 0.842237 | 0.703916 |
| 10 | 122565800 | 122567199 | 21.4868 | 36.25251 | 54.30996 | 23.75134 | -0.1998 | 1.092935 | 0.703944 |
| 8 | 78450200 | 78452199 | 13.45314 | 23.6761 | 36.80997 | 16.29672 | -0.24507 | 0.572684 | 0.703948 |
| 10 | 81926800 | 81928399 | 32.57826 | 54.41784 | 58.09744 | 25.28368 | 0.287044 | 1.228001 | 0.703956 |
| 4 | 105401000 | 105402999 | 20.11445 | 35.36733 | 37.37491 | 16.54982 | 0.308458 | 0.621926 | 0.703976 |
| 5 | 115869000 | 115871799 | 24.41958 | 40.64904 | 60.88107 | 26.43738 | -0.19968 | 1.242073 | 0.703986 |
| 9 | 112810000 | 112810999 | 12.04142 | 21.28197 | 34.84609 | 15.41415 | -0.31679 | 0.513578 | 0.703998 |
| 10 | 83219600 | 83230799 | 99.84179 | 153.2648 | 207.2194 | 86.03171 | -0.14072 | 2.989947 | 0.704004 |
| 2 | 84030000 | 84031399 | 19.90188 | 35.20933 | 35.66079 | 15.77141 | 0.372719 | 0.551393 | 0.704007 |
| 15 | 33685800 | 33688199 | 21.26095 | 36.14191 | 53.27648 | 23.34937 | -0.17611 | 1.024954 | 0.704039 |
| 1 | 113732000 | 113733999 | 21.07476 | 37.10614 | 36.63399 | 16.21734 | 0.406682 | 0.609254 | 0.704046 |
| 18 | 67205000 | 67216199 | 253.1739 | 362.7741 | 461.4794 | 190.5946 | -0.14344 | 4.217882 | 0.704069 |
| 10 | 43445400 | 43450799 | 60.00476 | 95.92259 | 92.34919 | 38.83259 | 0.397073 | 2.074715 | 0.704088 |
| Y | 76197600 | 76199399 | 11.94704 | 22.02929 | 19.68333 | 8.327302 | 0.595765 | 0.206505 | 0.704096 |
| 10 | 56342000 | 56344199 | 17.25509 | 30.16477 | 46.74461 | 20.60497 | -0.24153 | 0.708567 | 0.704096 |
| 18 | 76657200 | 76661599 | 45.69646 | 73.3154 | 104.787 | 44.29994 | -0.16593 | 1.950812 | 0.704099 |
| 5 | 56998200 | 57000599 | 22.44339 | 39.29061 | 38.88553 | 17.1613 | 0.402348 | 0.700077 | 0.704115 |
| 10 | 125395400 | 125398399 | 34.55685 | 57.36732 | 60.58038 | 26.25759 | 0.302533 | 1.299871 | 0.704116 |
| 5 | 42227200 | 42227999 | 10.5517 | 19.56098 | 30.95627 | 13.0013 | -0.21575 | 0.194419 | 0.704127 |
| 5 | 119482200 | 119486399 | 38.50869 | 62.06253 | 89.79989 | 38.21578 | -0.17557 | 1.79496 | 0.704142 |
| 3 | 101639600 | 101642999 | 40.96394 | 66.51767 | 74.4964 | 31.94077 | 0.204325 | 1.609212 | 0.704147 |
| 8 | 71558200 | 71559199 | 10.50441 | 19.21617 | 30.30668 | 12.96079 | -0.22926 | 0.269839 | 0.704176 |
| 4 | 33077000 | 33079599 | 23.99222 | 40.01191 | 59.69744 | 25.9538 | -0.19411 | 1.222046 | 0.704219 |
| 10 | 40197600 | 40202999 | 96.86349 | 149.6364 | 165.1734 | 68.60948 | 0.158581 | 2.86403 | 0.704265 |
| 3 | 55314000 | 55315199 | 18.18044 | 31.86869 | 36.60536 | 16.17259 | 0.188639 | 0.655962 | 0.704271 |
| 15 | 36481400 | 36482799 | 18.4153 | 32.46375 | 35.87128 | 15.87578 | 0.24583 | 0.581413 | 0.704281 |
| 6 | 111437600 | 111440599 | 22.56157 | 38.22557 | 56.58069 | 24.77584 | -0.1824 | 1.061429 | 0.70429 |
| 13 | 22845000 | 22846999 | 23.1339 | 39.06394 | 58.18842 | 25.45392 | -0.19164 | 1.095375 | 0.704294 |
| 4 | 145554400 | 145557799 | 92.12675 | 144.6671 | 148.1523 | 61.44862 | 0.284889 | 2.575444 | 0.704301 |
| 11 | 3959200 | 3960799 | 10.93132 | 19.81206 | 31.70436 | 13.71763 | -0.26149 | 0.327216 | 0.704316 |
| 17 | 69589000 | 69602199 | 179.8036 | 264.9333 | 346.9315 | 142.8698 | -0.14607 | 3.788193 | 0.704317 |
| 2 | 22769400 | 22770999 | 16.68781 | 29.03416 | 43.69411 | 19.20078 | -0.19981 | 0.77234 | 0.704329 |
| 11 | 116674400 | 116679199 | 50.87626 | 81.10711 | 114.7732 | 48.07593 | -0.1619 | 2.158017 | 0.704341 |
| 8 | 73962800 | 73983399 | 518.1257 | 697.8314 | 839.2788 | 350.0516 | -0.1526 | 5.16505 | 0.704345 |
| 17 | 20824400 | 20830199 | 99.35261 | 154.4367 | 148.1342 | 61.5037 | 0.367874 | 2.753221 | 0.704351 |
| 14 | 51256400 | 51258799 | 29.1007 | 49.08594 | 53.47057 | 23.38241 | 0.258158 | 1.119255 | 0.704381 |
| 8 | 46017600 | 46020799 | 40.68599 | 66.39002 | 71.93871 | 30.91081 | 0.257125 | 1.533508 | 0.704388 |
| 13 | 64270400 | 64273799 | 29.72275 | 49.78669 | 55.72334 | 24.29319 | 0.219051 | 1.187624 | 0.704394 |
| 6 | 42151000 | 42151599 | 14.74249 | 25.77168 | 40.03282 | 17.64604 | -0.24455 | 0.688265 | 0.704399 |
| 14 | 102307000 | 102308199 | 14.74249 | 25.77168 | 40.03282 | 17.64604 | -0.24455 | 0.688265 | 0.704399 |
| 6 | 82573600 | 82579399 | 52.55811 | 84.43045 | 93.44431 | 39.58014 | 0.204454 | 1.917486 | 0.704427 |
| 7 | 26546200 | 26547199 | 10.2559 | 19.03122 | 29.9044 | 12.54284 | -0.204 | 0.184877 | 0.704443 |
| 7 | 18670800 | 18677199 | 55.05215 | 87.54542 | 124.9085 | 52.14168 | -0.17768 | 2.242632 | 0.704452 |
| 11 | 9393800 | 9401599 | 161.5459 | 241.255 | 255.8082 | 105.4563 | 0.17623 | 3.550595 | 0.704468 |
| 5 | 78142000 | 78147199 | 69.44926 | 110.3118 | 107.3392 | 44.75548 | 0.372031 | 2.285508 | 0.704472 |
| 11 | 107122200 | 107123399 | 14.23241 | 26.05661 | 29.50722 | 12.6053 | 0.246333 | 0.29036 | 0.704474 |
| 3 | 18104600 | 18110399 | 47.98926 | 76.55738 | 114.0102 | 47.80188 | -0.23439 | 2.130235 | 0.70449 |

| | | | | | | | | | |
|---|---|---|---|---|---|---|---|---|---|
| 1 | 102389200 | 102390199 | 11.90832 | 22.78458 | 24.51631 | 9.702055 | 0.408445 | -0.01742 | 0.704524 |
| 18 | 30110200 | 30110799 | 7.642547 | 14.54929 | 25.04272 | 10.02486 | -0.27733 | -0.01501 | 0.704552 |
| 2 | 82941000 | 82942199 | 10.37172 | 20.0161 | 19.66841 | 7.59355 | 0.571861 | -0.12254 | 0.704563 |
| 11 | 88246400 | 88248199 | 16.79323 | 29.21838 | 43.67418 | 19.19236 | -0.19016 | 0.771078 | 0.704601 |
| 3 | 31597400 | 31599399 | 32.59886 | 54.3027 | 58.73511 | 25.51688 | 0.268317 | 1.25534 | 0.704627 |
| 7 | 108098400 | 108101599 | 25.01024 | 42.86374 | 40.01669 | 17.54956 | 0.48216 | 0.950485 | 0.704637 |
| 2 | 80655000 | 80657999 | 25.63515 | 44.3181 | 45.13823 | 19.80277 | 0.359277 | 0.880769 | 0.70465 |
| 12 | 105500600 | 105501799 | 14.3143 | 25.05602 | 39.53307 | 17.44509 | -0.26679 | 0.665385 | 0.704683 |
| 5 | 31477400 | 31480399 | 11.43271 | 20.40765 | 38.34629 | 16.85486 | -0.50679 | 0.4401 | 0.704711 |
| 6 | 74910600 | 74912399 | 18.92358 | 32.97912 | 37.84353 | 16.64731 | 0.189745 | 0.748976 | 0.704722 |
| 17 | 16018000 | 16018799 | 18.01787 | 31.79477 | 29.71705 | 13.14944 | 0.486508 | 0.5557 | 0.704722 |
| 5 | 30031800 | 30032799 | 9.366226 | 17.71459 | 29.2737 | 11.8816 | -0.23734 | 0.056212 | 0.704725 |
| 6 | 102962600 | 102963799 | 9.366226 | 17.71459 | 29.2737 | 11.8816 | -0.23734 | 0.056212 | 0.704725 |
| 7 | 98400800 | 98404399 | 24.45845 | 40.54123 | 62.42908 | 27.03039 | -0.24008 | 1.292223 | 0.704729 |
| 8 | 29118600 | 29120399 | 23.26847 | 39.84559 | 45.8098 | 20.08882 | 0.182735 | 0.957575 | 0.704772 |
| 8 | 48241800 | 48249399 | 112.1548 | 170.8936 | 229.9111 | 95.30359 | -0.14196 | 3.143612 | 0.704774 |
| 7 | 103359000 | 103361999 | 25.0685 | 42.75479 | 47.93453 | 21.01373 | 0.21785 | 1.000608 | 0.704798 |
| 2 | 167873400 | 167876199 | 18.69052 | 31.9389 | 49.00937 | 21.48828 | -0.23245 | 0.963692 | 0.704804 |
| 1 | 13861400 | 13863999 | 36.24398 | 59.5422 | 66.5608 | 28.67652 | 0.217519 | 1.432777 | 0.704804 |
| 16 | 8380000 | 8381799 | 23.87666 | 39.97362 | 59.31005 | 25.84842 | -0.18631 | 1.177279 | 0.704829 |
| 7 | 38051800 | 38056799 | 67.31624 | 106.8682 | 120.3833 | 50.16817 | 0.160422 | 2.302971 | 0.704833 |
| 5 | 81725200 | 81725799 | 6.973296 | 13.58254 | 17.42997 | 6.361548 | 0.267005 | -0.35797 | 0.704877 |
| 7 | 61874800 | 61875199 | 8.048727 | 15.63616 | 20.04206 | 7.482068 | 0.238355 | -0.2574 | 0.704904 |
| 5 | 45552200 | 45555799 | 49.41298 | 78.89285 | 111.5749 | 46.83462 | -0.1587 | 2.108853 | 0.704916 |
| 12 | 98935600 | 98939999 | 37.86586 | 61.0793 | 88.12112 | 37.5432 | -0.16993 | 1.76846 | 0.704939 |
| 3 | 21207600 | 21214999 | 172.1802 | 256.7352 | 279.0248 | 114.9924 | 0.138601 | 3.582318 | 0.704953 |
| 16 | 37791200 | 37793799 | 17.74686 | 30.74002 | 45.81589 | 20.10125 | -0.18704 | 0.830895 | 0.704961 |
| 5 | 102556400 | 102558199 | 18.53391 | 31.22133 | 60.02165 | 26.23674 | -0.55828 | 1.104128 | 0.704968 |
| 16 | 16234000 | 16234599 | 15.50567 | 27.4813 | 31.35386 | 13.86162 | 0.202754 | 0.496008 | 0.704971 |
| 14 | 57783000 | 57787599 | 29.79904 | 48.25944 | 87.52605 | 37.45378 | -0.49293 | 1.651463 | 0.704987 |
| X | 113159800 | 113161599 | 25.38976 | 42.56393 | 67.14936 | 29.28974 | -0.27455 | 1.178579 | 0.705022 |
| 18 | 37062200 | 37066799 | 113.7707 | 172.8547 | 233.4994 | 96.70717 | -0.15125 | 3.203934 | 0.705046 |
| 6 | 137529200 | 137531399 | 17.79567 | 31.00277 | 46.80731 | 20.586 | -0.20444 | 0.762026 | 0.705049 |
| 6 | 135192600 | 135195999 | 24.87582 | 41.10579 | 63.30806 | 27.35254 | -0.24057 | 1.332633 | 0.705058 |
| 4 | 115410000 | 115410599 | 8.317176 | 16.07317 | 19.90305 | 7.635742 | 0.250614 | -0.16305 | 0.705059 |
| 2 | 158022800 | 158024199 | 9.881805 | 18.18322 | 29.36247 | 12.45749 | -0.25457 | 0.230436 | 0.705098 |
| 2 | 46892200 | 46895599 | 35.06843 | 57.987 | 63.28072 | 27.35328 | 0.2549 | 1.35022 | 0.705099 |
| 4 | 112482600 | 112492599 | 130.8717 | 201.1166 | 194.7195 | 80.85415 | 0.341812 | 2.979971 | 0.705107 |
| 10 | 121003200 | 121011799 | 76.59703 | 119.4259 | 166.5655 | 69.12492 | -0.16821 | 2.692856 | 0.705125 |
| 6 | 20693400 | 20694399 | 10.16574 | 19.56259 | 23.61244 | 9.191398 | 0.266737 | -0.08716 | 0.705127 |
| 6 | 142111800 | 142116599 | 78.50042 | 122.2166 | 168.4248 | 69.91162 | -0.15254 | 2.720016 | 0.705145 |
| 17 | 95102200 | 95118999 | 267.6384 | 388.1429 | 389.8095 | 160.7202 | 0.214623 | 4.042416 | 0.705145 |
| 1 | 12680800 | 12683999 | 27.19008 | 45.04099 | 66.63784 | 28.84029 | -0.1827 | 1.309985 | 0.705159 |
| 4 | 119354200 | 119355599 | 11.99698 | 21.55856 | 33.2141 | 14.51697 | -0.21632 | 0.391393 | 0.705186 |
| 5 | 43163400 | 43165399 | 24.93861 | 42.09534 | 48.01444 | 21.00099 | 0.192113 | 1.093602 | 0.705186 |
| 6 | 126379800 | 126382599 | 23.96766 | 40.91971 | 46.86587 | 20.54696 | 0.187467 | 0.986601 | 0.705202 |
| 15 | 10889600 | 10893399 | 30.91357 | 50.60451 | 73.71204 | 31.70961 | -0.16535 | 1.468935 | 0.705214 |

| | | | | | | | | | |
|---|---|---|---|---|---|---|---|---|---|
| 16 | 69092200 | 69094399 | 28.88229 | 48.18246 | 55.67619 | 24.2104 | 0.173274 | 1.227584 | 0.70525 |
| 12 | 20175800 | 20176399 | 19.06424 | 33.48534 | 28.54413 | 12.63255 | 0.616785 | 0.615409 | 0.705259 |
| 12 | 60248800 | 60249399 | 8.424238 | 16.00711 | 26.61017 | 10.69503 | -0.23293 | 0.009813 | 0.705267 |
| 15 | 94388200 | 94389999 | 14.06468 | 24.71697 | 37.78292 | 16.72879 | -0.22158 | 0.594007 | 0.705272 |
| 11 | 29464800 | 29466599 | 15.47892 | 27.06198 | 40.9742 | 18.06264 | -0.20797 | 0.690164 | 0.705278 |
| 4 | 15461800 | 15464799 | 25.70593 | 44.70127 | 42.8618 | 18.82895 | 0.447396 | 0.818344 | 0.705287 |
| 4 | 34368600 | 34371599 | 30.53041 | 50.54946 | 58.45523 | 25.29034 | 0.171678 | 1.318452 | 0.70529 |
| 12 | 19222800 | 19227799 | 197.504 | 290.5628 | 303.1936 | 124.8574 | 0.179286 | 3.814212 | 0.705291 |
| 12 | 62215200 | 62216999 | 21.73226 | 37.55656 | 42.53356 | 18.65985 | 0.20665 | 0.862551 | 0.705305 |
| 2 | 96806000 | 96807599 | 21.05401 | 36.88239 | 38.9831 | 17.21386 | 0.307954 | 0.683661 | 0.705306 |
| 3 | 28595400 | 28596999 | 12.11123 | 22.1367 | 35.53319 | 15.21276 | -0.25607 | 0.297063 | 0.705311 |
| 13 | 23388800 | 23391999 | 30.1975 | 50.06057 | 57.90049 | 25.07187 | 0.171548 | 1.302858 | 0.705344 |
| 13 | 31796200 | 31800199 | 43.74225 | 70.17747 | 100.461 | 42.46867 | -0.16825 | 1.948184 | 0.705358 |
| 3 | 106000600 | 106001399 | 15.55924 | 27.54721 | 31.44785 | 13.9061 | 0.201423 | 0.50896 | 0.705362 |
| 9 | 54822200 | 54824999 | 33.56359 | 54.87341 | 61.49218 | 26.44065 | 0.210736 | 1.483908 | 0.705388 |
| 4 | 120781400 | 120792399 | 120.6036 | 184.0635 | 204.1123 | 84.64234 | 0.138467 | 3.11207 | 0.705389 |
| 7 | 33590800 | 33593999 | 46.15794 | 74.43019 | 84.36901 | 35.92933 | 0.176716 | 1.789211 | 0.705391 |
| 15 | 19943000 | 19944199 | 18.65432 | 32.92374 | 30.2607 | 13.38953 | 0.51083 | 0.560363 | 0.705395 |
| 1 | 40448200 | 40449399 | 15.96396 | 28.39056 | 32.38561 | 14.27945 | 0.205914 | 0.480174 | 0.705407 |
| 19 | 7601600 | 7602799 | 12.76712 | 22.71169 | 34.64769 | 15.2732 | -0.21091 | 0.461319 | 0.70541 |
| 3 | 28193200 | 28194599 | 13.13831 | 23.46352 | 35.67924 | 15.67619 | -0.20293 | 0.442731 | 0.705413 |
| 12 | 113572400 | 113573799 | 15.23793 | 27.96838 | 30.03519 | 12.78662 | 0.325987 | 0.285017 | 0.70542 |
| 4 | 7622800 | 7625799 | 27.14654 | 44.97665 | 66.43284 | 28.75499 | -0.18021 | 1.310857 | 0.705427 |
| 1 | 8903000 | 8903799 | 17.29549 | 30.09577 | 44.7104 | 19.64916 | -0.18135 | 0.773603 | 0.705435 |
| 15 | 33404200 | 33406599 | 22.54337 | 37.9402 | 56.55105 | 24.7127 | -0.19281 | 1.117908 | 0.705457 |
| 5 | 20850800 | 20851999 | 25.37938 | 42.75539 | 49.08144 | 21.45492 | 0.182727 | 1.112 | 0.705462 |
| 11 | 14682600 | 14685599 | 32.5544 | 53.27742 | 79.87152 | 34.35616 | -0.20706 | 1.479475 | 0.705483 |
| 5 | 118684400 | 118685799 | 17.42337 | 30.86569 | 34.28419 | 15.15855 | 0.241009 | 0.520002 | 0.705505 |
| 8 | 49435400 | 49440199 | 48.46529 | 77.81809 | 89.0547 | 37.69338 | 0.1555 | 1.92589 | 0.705508 |
| 6 | 64147400 | 64151599 | 83.53664 | 129.6436 | 177.2444 | 73.60756 | -0.14466 | 2.777475 | 0.705541 |
| 1 | 170421400 | 170425399 | 82.67502 | 128.4154 | 175.855 | 73.02134 | -0.14609 | 2.762221 | 0.705543 |
| 12 | 42135000 | 42135599 | 9.963532 | 18.88281 | 22.74574 | 9.192244 | 0.22072 | 0.028972 | 0.705564 |
| 10 | 65831000 | 65832999 | 19.00262 | 33.32964 | 37.52861 | 16.58879 | 0.217175 | 0.652274 | 0.705607 |
| 8 | 118433000 | 118435399 | 19.91982 | 33.78376 | 52.19198 | 22.86108 | -0.24378 | 1.037152 | 0.705609 |
| 10 | 47577800 | 47579599 | 18.3238 | 32.46757 | 34.65499 | 15.32185 | 0.298184 | 0.522713 | 0.705614 |
| 17 | 46652400 | 46677199 | 539.9148 | 725.8384 | 866.0578 | 361.3194 | -0.14381 | 5.194465 | 0.705618 |
| 5 | 75873400 | 75875199 | 19.56482 | 34.45202 | 36.83488 | 16.30586 | 0.292476 | 0.601336 | 0.705643 |
| 4 | 52163600 | 52165399 | 17.01197 | 29.98111 | 34.41237 | 15.23099 | 0.191023 | 0.572752 | 0.705645 |
| 6 | 69000200 | 69005599 | 63.63204 | 101.4299 | 111.9164 | 46.87132 | 0.19624 | 2.173167 | 0.705648 |
| 16 | 72320200 | 72320999 | 10.31144 | 18.95797 | 29.81787 | 12.66506 | -0.2183 | 0.238226 | 0.705671 |
| 3 | 8743600 | 8744799 | 8.077288 | 15.43266 | 25.63088 | 10.18207 | -0.21687 | -0.03507 | 0.705681 |
| 1 | 45806800 | 45810399 | 33.89819 | 54.95285 | 80.56976 | 34.50247 | -0.18554 | 1.633089 | 0.705716 |
| 6 | 88137000 | 88139399 | 22.89822 | 39.29533 | 45.26156 | 19.85182 | 0.180483 | 0.935647 | 0.705723 |
| 7 | 48391400 | 48393399 | 72.66633 | 114.466 | 126.6232 | 52.54047 | 0.177206 | 2.496395 | 0.705724 |
| 16 | 47051200 | 47052399 | 13.7115 | 24.86974 | 28.94216 | 12.51064 | 0.196101 | 0.337926 | 0.705725 |
| 18 | 46706600 | 46708799 | 14.19454 | 24.89741 | 38.84377 | 17.17066 | -0.25099 | 0.626367 | 0.705738 |
| 13 | 67357400 | 67360399 | 29.27097 | 49.42344 | 52.95624 | 23.16465 | 0.281887 | 1.108603 | 0.70574 |

| | | | | | | | | | |
|---|---|---|---|---|---|---|---|---|---|
| 6 | 145301400 | 145302599 | 18.29093 | 31.97106 | 36.74161 | 16.19371 | 0.187963 | 0.702027 | 0.705744 |
| 8 | 126337800 | 126342199 | 47.87451 | 76.62142 | 108.0614 | 45.51525 | -0.15089 | 2.031317 | 0.705768 |
| 3 | 25329600 | 25334799 | 70.01732 | 111.0953 | 108.2239 | 45.0761 | 0.36892 | 2.317932 | 0.70577 |
| 18 | 51249600 | 51252399 | 24.80599 | 41.00204 | 64.03831 | 27.67348 | -0.26077 | 1.326752 | 0.705794 |
| 19 | 39565400 | 39570199 | 41.40019 | 67.91511 | 68.84404 | 29.64216 | 0.357496 | 1.463812 | 0.705795 |
| 15 | 88916200 | 88917799 | 21.7734 | 36.67459 | 54.87107 | 23.98455 | -0.1981 | 1.107697 | 0.705816 |
| 7 | 59006400 | 59016199 | 99.81626 | 154.7241 | 169.897 | 70.56885 | 0.169885 | 2.80334 | 0.705852 |
| 9 | 8410800 | 8411399 | 9.076117 | 17.46497 | 21.93571 | 8.539806 | 0.209992 | -0.10175 | 0.705865 |
| X | 121577000 | 121580999 | 37.91159 | 63.14081 | 49.27798 | 21.40581 | 0.737765 | 1.26842 | 0.705876 |
| 1 | 15599000 | 15600599 | 26.31925 | 43.51242 | 64.5935 | 27.91742 | -0.18764 | 1.331586 | 0.705893 |
| 11 | 49694200 | 49695799 | 15.34965 | 27.29342 | 31.3421 | 13.82133 | 0.196064 | 0.47336 | 0.705909 |
| 1 | 126355200 | 126359799 | 15.40419 | 27.95798 | 26.19826 | 11.31541 | 0.507457 | 0.340459 | 0.705919 |
| 3 | 115212800 | 115216399 | 35.27731 | 58.03733 | 65.34063 | 28.16792 | 0.208136 | 1.412685 | 0.705921 |
| 1 | 112881800 | 112883999 | 18.2767 | 31.8218 | 48.98411 | 21.53528 | -0.23244 | 0.775174 | 0.70594 |
| 3 | 21287800 | 21288399 | 10.18697 | 19.29375 | 23.28354 | 9.422496 | 0.216566 | 0.038147 | 0.705998 |
| 8 | 6531200 | 6532799 | 19.64918 | 34.38818 | 31.54437 | 13.91776 | 0.511549 | 0.680488 | 0.705998 |
| 19 | 40496000 | 40496999 | 11.98821 | 21.46933 | 33.2145 | 14.55901 | -0.22519 | 0.412268 | 0.706004 |
| 4 | 118890800 | 118891999 | 12.35696 | 21.99362 | 34.25124 | 15.09174 | -0.23978 | 0.456687 | 0.706016 |
| 12 | 50294600 | 50295199 | 7.811405 | 14.92839 | 25.63275 | 10.17743 | -0.26336 | -0.03741 | 0.706017 |
| 11 | 39973400 | 39975599 | 18.06739 | 31.73371 | 35.89786 | 15.88674 | 0.210366 | 0.620556 | 0.706039 |
| X | 117046400 | 117056999 | 158.1253 | 239.0505 | 243.6705 | 100.7649 | 0.248676 | 3.316028 | 0.70604 |
| Y | 90212400 | 90213799 | 11.5431 | 21.82746 | 17.84418 | 7.24562 | 0.768974 | 0.043519 | 0.706073 |
| 1 | 60718400 | 60719599 | 12.36941 | 22.65941 | 26.03932 | 11.11477 | 0.226198 | 0.264498 | 0.706087 |
| 7 | 26484000 | 26486199 | 17.04484 | 30.09199 | 34.3816 | 15.21201 | 0.198601 | 0.554211 | 0.706095 |
| 13 | 72795000 | 72798199 | 28.56134 | 46.97744 | 69.23648 | 29.84445 | -0.17948 | 1.400476 | 0.706113 |
| 1 | 78107800 | 78111599 | 28.01523 | 45.86583 | 83.97742 | 36.12754 | -0.49441 | 1.465959 | 0.706133 |
| 11 | 22915600 | 22924599 | 172.9376 | 255.8791 | 334.8107 | 137.8845 | -0.13931 | 3.718644 | 0.706134 |
| 3 | 142696400 | 142696999 | 7.146676 | 13.77492 | 23.65406 | 9.165216 | -0.23056 | -0.13146 | 0.706137 |
| 7 | 69255800 | 69256599 | 7.146676 | 13.77492 | 23.65406 | 9.165216 | -0.23056 | -0.13146 | 0.706137 |
| 18 | 18376800 | 18377999 | 16.91119 | 29.71129 | 33.62248 | 14.88329 | 0.210108 | 0.608405 | 0.70614 |
| 8 | 93365400 | 93369399 | 35.93963 | 58.09177 | 84.10322 | 35.92677 | -0.17147 | 1.704775 | 0.706155 |
| 14 | 113608000 | 113608599 | 9.007678 | 17.25178 | 20.79332 | 8.205339 | 0.250551 | -0.06324 | 0.706158 |
| 7 | 126884400 | 126887999 | 51.56565 | 82.66501 | 94.60746 | 39.94722 | 0.152694 | 1.980218 | 0.706163 |
| X | 88901600 | 88905599 | 51.03744 | 82.16527 | 89.39529 | 37.98407 | 0.232952 | 1.848654 | 0.706176 |
| 3 | 3985400 | 3985799 | 9.154319 | 17.58434 | 21.86956 | 8.557379 | 0.216814 | -0.08652 | 0.706179 |
| 19 | 60012600 | 60013399 | 8.785896 | 16.91747 | 20.89755 | 8.120504 | 0.236617 | -0.11293 | 0.706224 |
| 2 | 20187400 | 20195799 | 116.382 | 176.968 | 236.9444 | 98.15819 | -0.13749 | 3.187652 | 0.706234 |
| 11 | 13341400 | 13347599 | 97.44929 | 151.8712 | 144.2706 | 59.87639 | 0.384712 | 2.707979 | 0.706264 |
| 2 | 39633000 | 39635999 | 29.29553 | 49.82517 | 51.24096 | 22.45818 | 0.341476 | 1.047663 | 0.706284 |
| 9 | 4711400 | 4712199 | 13.69463 | 25.34285 | 23.44521 | 9.870304 | 0.551469 | 0.213993 | 0.706287 |
| 14 | 66590000 | 66594599 | 52.77903 | 84.45725 | 95.51733 | 40.22251 | 0.166764 | 2.036253 | 0.706289 |
| 7 | 9915800 | 9919199 | 41.76268 | 68.66389 | 56.0669 | 24.16367 | 0.669924 | 1.431013 | 0.70629 |
| 9 | 53318200 | 53319999 | 15.56298 | 28.11033 | 31.35803 | 13.6259 | 0.251923 | 0.380297 | 0.706314 |
| 3 | 157824000 | 157824999 | 16.91577 | 29.82047 | 33.82964 | 14.97218 | 0.207872 | 0.566469 | 0.70633 |
| 6 | 144506000 | 144508399 | 33.62899 | 55.38221 | 63.49079 | 27.38232 | 0.182678 | 1.395094 | 0.706355 |
| 12 | 72966000 | 72969399 | 40.24324 | 65.22742 | 75.04525 | 32.13253 | 0.163147 | 1.646446 | 0.706363 |
| 3 | 100039600 | 100042599 | 35.1952 | 57.78541 | 65.90614 | 28.38744 | 0.188301 | 1.435999 | 0.706378 |

| | | | | | | | | | |
|---|---|---|---|---|---|---|---|---|---|
| 18 | 52191200 | 52192199 | 13.55129 | 24.54187 | 28.45863 | 12.32501 | 0.199502 | 0.344511 | 0.70638 |
| 3 | 116993400 | 117000199 | 133.7611 | 201.1612 | 269.6518 | 111.3387 | -0.15274 | 3.415042 | 0.706382 |
| 2 | 30066000 | 30067199 | 12.01216 | 21.36275 | 33.75392 | 14.88269 | -0.26129 | 0.457322 | 0.706394 |
| 1 | 36167200 | 36170999 | 30.92875 | 50.4186 | 74.48804 | 31.99467 | -0.18994 | 1.528443 | 0.706395 |
| 12 | 22428800 | 22429399 | 13.78812 | 25.34069 | 23.42317 | 9.953889 | 0.542929 | 0.252095 | 0.706399 |
| 8 | 35576400 | 35579399 | 32.97729 | 53.44713 | 78.97687 | 33.81436 | -0.19705 | 1.637077 | 0.706407 |
| X | 37871000 | 37873799 | 27.09881 | 46.77778 | 38.19696 | 16.75677 | 0.676494 | 0.894439 | 0.706465 |
| 2 | 29527600 | 29529599 | 16.36913 | 28.84643 | 32.99939 | 14.60578 | 0.195864 | 0.567489 | 0.70649 |
| 1 | 37824600 | 37827199 | 31.75064 | 53.16642 | 56.92669 | 24.81278 | 0.282796 | 1.198803 | 0.706547 |
| 4 | 61886000 | 61887399 | 12.4119 | 22.49471 | 34.78622 | 15.05172 | -0.21251 | 0.346026 | 0.706553 |
| 1 | 132017600 | 132018799 | 9.313072 | 17.39889 | 27.91008 | 11.58013 | -0.22109 | 0.130655 | 0.706568 |
| 4 | 57330200 | 57332599 | 15.65994 | 27.42082 | 40.91493 | 18.06009 | -0.18698 | 0.666541 | 0.706569 |
| 1 | 69198600 | 69199199 | 7.587585 | 14.63889 | 24.82562 | 9.592511 | -0.20936 | -0.13333 | 0.706569 |
| 15 | 37857000 | 37862799 | 85.19009 | 133.6245 | 147.9819 | 61.37391 | 0.170875 | 2.591142 | 0.706586 |
| 4 | 112438400 | 112441599 | 59.24684 | 95.50363 | 70.39111 | 29.96354 | 0.795787 | 1.812853 | 0.706615 |
| 17 | 67739200 | 67741599 | 24.58711 | 41.37153 | 62.73639 | 27.41402 | -0.21763 | 1.133757 | 0.706626 |
| 9 | 35559600 | 35561399 | 22.26482 | 37.67919 | 55.44897 | 24.27176 | -0.17406 | 1.069929 | 0.706657 |
| 18 | 5799200 | 5801799 | 19.87883 | 34.37579 | 55.04032 | 24.14715 | -0.29059 | 0.868769 | 0.706662 |
| X | 65010400 | 65010999 | 12.62343 | 23.47583 | 26.20024 | 10.95517 | 0.290046 | 0.186504 | 0.706662 |
| 15 | 62966200 | 62968599 | 45.95738 | 73.89158 | 83.83985 | 35.55708 | 0.170224 | 1.877566 | 0.706667 |
| 10 | 114756400 | 114757999 | 10.7378 | 19.72045 | 30.80018 | 13.10162 | -0.21007 | 0.248428 | 0.706691 |
| 7 | 133930000 | 133931799 | 21.76731 | 37.66725 | 43.10413 | 18.91093 | 0.192266 | 0.854415 | 0.706693 |
| 2 | 49219600 | 49222199 | 28.99941 | 48.35166 | 55.7046 | 24.21449 | 0.17757 | 1.232865 | 0.706758 |
| 12 | 107898600 | 107901399 | 20.28536 | 34.30608 | 52.71672 | 23.07103 | -0.23624 | 1.064593 | 0.706762 |
| 18 | 60528800 | 60531199 | 18.79551 | 33.09375 | 36.31313 | 16.07524 | 0.25506 | 0.595905 | 0.706772 |
| 8 | 48612000 | 48613599 | 19.3576 | 33.33912 | 49.54755 | 21.7347 | -0.18475 | 0.893156 | 0.706798 |
| 18 | 71248000 | 71249599 | 22.39955 | 39.15273 | 34.26352 | 15.10155 | 0.579085 | 0.718324 | 0.706819 |
| 14 | 43263800 | 43265399 | 33.95287 | 55.53776 | 63.72107 | 27.40406 | 0.177345 | 1.480228 | 0.706841 |
| 18 | 77676000 | 77681599 | 55.79091 | 88.79536 | 124.0295 | 51.83467 | -0.14577 | 2.217586 | 0.706862 |
| 7 | 138249400 | 138252999 | 30.77235 | 50.25323 | 73.42233 | 31.55744 | -0.1721 | 1.502674 | 0.706901 |
| 2 | 87889200 | 87890399 | 15.14813 | 26.65193 | 40.2274 | 17.81066 | -0.20305 | 0.589534 | 0.706922 |
| 13 | 12689400 | 12693399 | 38.49914 | 62.14642 | 89.20229 | 38.03983 | -0.16136 | 1.748217 | 0.706934 |
| X | 64788000 | 64791999 | 42.99831 | 70.4327 | 70.01147 | 30.12599 | 0.38487 | 1.487188 | 0.70696 |
| 8 | 22335800 | 22336199 | 8.158866 | 15.90617 | 19.83201 | 7.152953 | 0.322137 | -0.37936 | 0.706978 |
| 13 | 65196400 | 65202399 | 76.81802 | 121.2196 | 135.2969 | 56.15929 | 0.166298 | 2.464927 | 0.706981 |
| 19 | 42625200 | 42627999 | 15.02499 | 26.24598 | 40.52889 | 17.85474 | -0.2362 | 0.700847 | 0.706991 |
| 1 | 3560400 | 3561599 | 12.23016 | 21.97157 | 33.63127 | 14.7027 | -0.20755 | 0.393085 | 0.706999 |
| 10 | 51691000 | 51693599 | 15.794 | 27.7594 | 42.84367 | 18.97143 | -0.2357 | 0.608843 | 0.707004 |
| 17 | 74194600 | 74196999 | 52.42285 | 83.82652 | 93.81763 | 39.46623 | 0.180723 | 2.05954 | 0.707048 |
| 7 | 69842800 | 69843799 | 12.44203 | 21.77479 | 43.63895 | 19.25396 | -0.61018 | 0.663689 | 0.707053 |
| 15 | 45889600 | 45890199 | 9.965809 | 18.62437 | 29.5144 | 12.23908 | -0.20308 | 0.138665 | 0.707077 |
| 6 | 140014200 | 140017599 | 30.97496 | 51.59329 | 58.04638 | 25.21632 | 0.211682 | 1.251286 | 0.707078 |
| 9 | 22218400 | 22220199 | 17.4617 | 30.1203 | 45.84865 | 20.11307 | -0.21851 | 0.867598 | 0.707085 |
| 1 | 156230400 | 156234399 | 52.2973 | 83.72127 | 94.96796 | 40.01478 | 0.163123 | 2.023006 | 0.707109 |
| 15 | 54174400 | 54180399 | 48.30432 | 77.33968 | 110.6932 | 46.6502 | -0.1715 | 2.019369 | 0.707111 |
| 4 | 62134000 | 62136799 | 33.86899 | 55.20027 | 81.40402 | 34.96265 | -0.18761 | 1.54167 | 0.707121 |
| 12 | 13899600 | 13901799 | 26.8038 | 44.36177 | 65.24798 | 28.22149 | -0.17423 | 1.320839 | 0.707136 |

| | | | | | | | | | |
|---|---|---|---|---|---|---|---|---|---|
| 3 | 5283200 | 5284999 | 16.09955 | 27.9148 | 44.77186 | 19.6438 | -0.29215 | 0.810879 | 0.70715 |
| 3 | 113090800 | 113093999 | 33.47495 | 55.36439 | 62.08875 | 26.84216 | 0.215664 | 1.341083 | 0.707156 |
| X | 149029000 | 149040399 | 145.4948 | 221.5184 | 221.7602 | 91.90295 | 0.283165 | 3.169603 | 0.707173 |
| 12 | 81659800 | 81661399 | 16.13398 | 28.8263 | 25.71377 | 11.29347 | 0.562269 | 0.436492 | 0.70718 |
| 1 | 118205800 | 118207399 | 14.82777 | 27.11519 | 29.95387 | 12.81625 | 0.280151 | 0.301319 | 0.707181 |
| 6 | 22534000 | 22534599 | 11.34495 | 21.24265 | 25.22427 | 10.42596 | 0.214306 | 0.128381 | 0.707186 |
| 18 | 3729200 | 3731999 | 37.18179 | 60.89485 | 68.31043 | 29.39196 | 0.210735 | 1.472589 | 0.707189 |
| 18 | 66135000 | 66135999 | 13.76344 | 24.19302 | 37.30771 | 16.52106 | -0.23408 | 0.588423 | 0.707189 |
| 4 | 122555200 | 122564399 | 150.6541 | 227.5945 | 243.5385 | 100.6878 | 0.177784 | 3.327796 | 0.707194 |
| 4 | 25451000 | 25452999 | 17.29014 | 30.10485 | 44.69519 | 19.65003 | -0.18035 | 0.764747 | 0.707196 |
| 17 | 87627800 | 87630199 | 37.39394 | 60.28823 | 87.77417 | 37.37153 | -0.18383 | 1.782325 | 0.707198 |
| 1 | 143470000 | 143472399 | 18.95136 | 33.46034 | 35.71401 | 15.8013 | 0.296247 | 0.565981 | 0.707242 |
| 6 | 145986400 | 145992599 | 76.9584 | 120.7188 | 170.355 | 70.65251 | -0.17832 | 2.588388 | 0.707248 |
| 12 | 24057000 | 24057599 | 16.04613 | 28.73681 | 24.41163 | 10.70034 | 0.634484 | 0.419893 | 0.707248 |
| 14 | 107515400 | 107521999 | 163.4178 | 245.5449 | 231.7332 | 95.65962 | 0.352052 | 3.434709 | 0.707267 |
| 5 | 12678000 | 12679399 | 14.85355 | 27.35059 | 29.55886 | 12.52896 | 0.321332 | 0.262872 | 0.707268 |
| 18 | 60598600 | 60606799 | 100.5194 | 154.2663 | 208.8484 | 86.70502 | -0.1429 | 2.99494 | 0.707274 |
| 4 | 32469400 | 32469999 | 14.2495 | 25.16787 | 37.68633 | 16.67273 | -0.18922 | 0.53958 | 0.707286 |
| 7 | 26391800 | 26399999 | 93.55034 | 146.2124 | 157.0111 | 65.14271 | 0.211244 | 2.659465 | 0.707308 |
| 5 | 8308600 | 8309399 | 16.43115 | 28.33889 | 45.37091 | 19.90341 | -0.29085 | 0.866178 | 0.707308 |
| 8 | 46373000 | 46377799 | 41.66931 | 66.9415 | 96.04572 | 40.6847 | -0.16916 | 1.903598 | 0.70731 |
| 15 | 56451600 | 56455799 | 41.45271 | 67.67721 | 71.59874 | 30.77071 | 0.292101 | 1.526594 | 0.707327 |
| 8 | 57124200 | 57125799 | 16.66075 | 29.97132 | 32.23314 | 14.07555 | 0.299706 | 0.411933 | 0.707328 |
| 12 | 62122000 | 62125199 | 63.7614 | 101.2564 | 115.3684 | 48.09333 | 0.144449 | 2.288865 | 0.707372 |
| 5 | 45399400 | 45402799 | 36.57889 | 59.21395 | 85.25843 | 36.47189 | -0.1614 | 1.670976 | 0.707401 |
| 2 | 166534400 | 166537599 | 34.44176 | 55.70509 | 81.803 | 34.97092 | -0.19092 | 1.682573 | 0.707447 |
| 1 | 8480600 | 8481799 | 18.51679 | 32.74866 | 29.72302 | 13.14618 | 0.530044 | 0.536561 | 0.70748 |
| 2 | 53651800 | 53653799 | 23.12857 | 38.28945 | 69.33372 | 29.9943 | -0.47322 | 1.311336 | 0.7075 |
| 1 | 25948800 | 25951999 | 34.09383 | 56.23718 | 63.65664 | 27.47234 | 0.201849 | 1.37647 | 0.707506 |
| 6 | 18955200 | 18957399 | 23.3242 | 39.03593 | 58.08125 | 25.30766 | -0.1902 | 1.180608 | 0.707523 |
| 15 | 61786800 | 61790399 | 38.1523 | 62.22614 | 70.46239 | 30.27123 | 0.193261 | 1.532974 | 0.707563 |
| 7 | 36587400 | 36589199 | 15.40479 | 27.2011 | 42.87336 | 18.96853 | -0.26319 | 0.557687 | 0.707568 |
| 1 | 56175600 | 56176799 | 18.16703 | 31.9793 | 28.40389 | 12.57456 | 0.558411 | 0.581149 | 0.70757 |
| 11 | 73754400 | 73760199 | 59.06853 | 94.50147 | 91.15641 | 38.37446 | 0.395557 | 2.049717 | 0.707586 |
| 12 | 67696000 | 67699999 | 28.83889 | 47.52696 | 70.64353 | 30.47621 | -0.19062 | 1.377036 | 0.707586 |
| 2 | 148944000 | 148944799 | 30.65229 | 49.96782 | 73.69826 | 31.65541 | -0.18748 | 1.529126 | 0.707616 |
| 12 | 20267600 | 20290999 | 1435.163 | 1792.427 | 1610.873 | 677.5575 | 0.158491 | 6.386135 | 0.707622 |
| 5 | 70276600 | 70279199 | 28.42728 | 47.77369 | 53.49412 | 23.36659 | 0.218297 | 1.145822 | 0.70764 |
| 18 | 4517000 | 4518399 | 14.40067 | 25.28522 | 38.32454 | 16.95602 | -0.20936 | 0.611603 | 0.707656 |
| 2 | 18196200 | 18197199 | 12.36885 | 22.4207 | 34.77549 | 15.04372 | -0.21669 | 0.34368 | 0.707666 |
| 3 | 130899400 | 130901599 | 17.21321 | 29.08754 | 59.54755 | 26.05424 | -0.64854 | 1.068188 | 0.70768 |
| 5 | 146050200 | 146053999 | 43.54739 | 71.0431 | 72.85499 | 31.30193 | 0.336183 | 1.543849 | 0.707685 |
| 11 | 68525000 | 68526799 | 9.982833 | 18.37004 | 29.31718 | 12.43743 | -0.23794 | 0.228958 | 0.707689 |
| X | 125014200 | 125036799 | 303.6493 | 437.8614 | 410.6066 | 169.4228 | 0.305381 | 4.129918 | 0.707696 |
| 16 | 46930800 | 46931799 | 16.00406 | 27.89774 | 42.10112 | 18.52297 | -0.20354 | 0.738623 | 0.707704 |
| 10 | 43965400 | 43967799 | 29.13166 | 49.39849 | 51.82032 | 22.69711 | 0.312556 | 1.071414 | 0.707718 |
| 10 | 120321800 | 120325999 | 56.40974 | 90.15142 | 102.5249 | 43.09504 | 0.156769 | 2.083708 | 0.707747 |

| | | | | | | | | | |
|---|---|---|---|---|---|---|---|---|---|
| 9 | 96785400 | 96786399 | 8.040018 | 15.29695 | 25.54528 | 10.23879 | -0.2364 | -0.00878 | 0.707759 |
| 13 | 50180800 | 50181799 | 14.57739 | 25.56079 | 38.89618 | 17.18919 | -0.21501 | 0.636002 | 0.707761 |
| 17 | 66017600 | 66021399 | 40.81243 | 65.63838 | 94.15742 | 39.94959 | -0.16688 | 1.865654 | 0.707767 |
| 11 | 20732200 | 20736599 | 63.52874 | 100.9414 | 114.6007 | 47.79915 | 0.150346 | 2.274035 | 0.707784 |
| 7 | 18328400 | 18358399 | 439.9802 | 607.4011 | 609.214 | 253.1195 | 0.145264 | 4.777591 | 0.707785 |
| 11 | 29626600 | 29633599 | 74.68762 | 118.1184 | 131.2611 | 54.54501 | 0.175119 | 2.40969 | 0.707791 |
| 6 | 127932200 | 127933799 | 14.92035 | 26.2343 | 39.27569 | 17.39153 | -0.19149 | 0.596185 | 0.707803 |
| 17 | 23499400 | 23502599 | 45.8974 | 73.6825 | 105.6361 | 44.70013 | -0.16932 | 1.933312 | 0.707809 |
| 3 | 91314400 | 91314999 | 15.78439 | 27.91248 | 31.54181 | 13.95098 | 0.21551 | 0.524745 | 0.707822 |
| 5 | 33221000 | 33222799 | 16.92049 | 30.3624 | 32.77918 | 14.34426 | 0.29224 | 0.431161 | 0.707825 |
| 9 | 95156000 | 95157199 | 12.04122 | 22.93277 | 24.91525 | 9.964969 | 0.380953 | 0.01822 | 0.707829 |
| 9 | 55682400 | 55684799 | 23.26545 | 39.45128 | 60.50592 | 26.50139 | -0.23345 | 1.061105 | 0.707841 |
| 6 | 29620200 | 29622599 | 25.69864 | 43.46992 | 50.13449 | 21.94232 | 0.176215 | 1.079046 | 0.707848 |
| 19 | 17379200 | 17381799 | 25.69864 | 43.46992 | 50.13449 | 21.94232 | 0.176215 | 1.079046 | 0.707848 |
| 6 | 74406200 | 74409999 | 31.81559 | 53.08461 | 58.10173 | 25.26775 | 0.251255 | 1.236023 | 0.707848 |
| 10 | 18301400 | 18305799 | 47.69528 | 76.46502 | 109.8216 | 46.37553 | -0.17424 | 1.978555 | 0.707852 |
| 3 | 20500200 | 20502199 | 20.36497 | 34.71642 | 51.6812 | 22.65535 | -0.18967 | 0.992392 | 0.707862 |
| 17 | 38120800 | 38122599 | 14.43332 | 25.25273 | 46.74681 | 20.62034 | -0.49666 | 0.681452 | 0.707869 |
| 6 | 68338400 | 68342799 | 69.73812 | 109.8073 | 151.6889 | 62.9377 | -0.14338 | 2.50642 | 0.707883 |
| 7 | 29290800 | 29297199 | 84.46277 | 132.2969 | 148.6106 | 61.6436 | 0.148793 | 2.61423 | 0.707903 |
| 5 | 104791600 | 104806199 | 334.0985 | 476.8185 | 399.5534 | 165.1243 | 0.452798 | 4.280265 | 0.707909 |
| 4 | 22853800 | 22854799 | 9.56682 | 18.50385 | 22.68976 | 8.670383 | 0.269437 | -0.15575 | 0.707927 |
| X | 112159800 | 112160399 | 8.092351 | 15.71172 | 18.9519 | 7.108581 | 0.31964 | -0.24187 | 0.707946 |
| 7 | 72572000 | 72573399 | 12.7961 | 22.8744 | 34.67268 | 15.22157 | -0.19752 | 0.432162 | 0.707955 |
| 14 | 83697400 | 83698999 | 29.92342 | 49.49631 | 56.68845 | 24.50949 | 0.185484 | 1.327639 | 0.707969 |
| 3 | 127274600 | 127275199 | 5.520018 | 10.74201 | 20.14185 | 7.416256 | -0.28612 | -0.33666 | 0.707995 |
| 9 | 62472400 | 62473999 | 24.66299 | 41.74493 | 47.9888 | 21.00732 | 0.180915 | 1.066989 | 0.707999 |
| 10 | 38435000 | 38435599 | 8.062554 | 15.45869 | 25.74415 | 10.12689 | -0.20764 | -0.06756 | 0.708017 |
| 11 | 16307600 | 16310199 | 33.09906 | 53.89955 | 78.04047 | 33.50763 | -0.162 | 1.55172 | 0.70802 |
| X | 49460400 | 49470199 | 145.6884 | 220.7298 | 231.2288 | 95.6857 | 0.211959 | 3.269595 | 0.708042 |
| 8 | 26760600 | 26765399 | 40.86153 | 66.08746 | 76.04077 | 32.50727 | 0.160147 | 1.690232 | 0.708056 |
| 13 | 14393600 | 14400599 | 92.84127 | 144.312 | 161.9968 | 67.25954 | 0.141218 | 2.753126 | 0.708092 |
| 4 | 92425400 | 92428799 | 38.6767 | 63.44685 | 59.24205 | 25.50775 | 0.475693 | 1.451589 | 0.708094 |
| 7 | 54768800 | 54770999 | 15.35288 | 27.86113 | 30.88413 | 13.34153 | 0.266368 | 0.35071 | 0.708108 |
| 14 | 92173800 | 92178799 | 45.738 | 74.21492 | 76.95576 | 32.97725 | 0.314189 | 1.634163 | 0.708118 |
| 14 | 45961200 | 45963999 | 33.39674 | 55.52093 | 60.3916 | 26.20166 | 0.260169 | 1.282606 | 0.708156 |
| 17 | 58868600 | 58869799 | 16.61997 | 29.60491 | 26.83775 | 11.81722 | 0.536897 | 0.462679 | 0.708165 |
| 1 | 44942200 | 44942799 | 9.382081 | 17.98138 | 22.39146 | 8.81832 | 0.206661 | -0.06032 | 0.708165 |
| 14 | 4532000 | 4532599 | 10.91995 | 20.73712 | 24.88077 | 10.01235 | 0.231547 | 0.029387 | 0.708196 |
| 11 | 54715000 | 54717999 | 36.58659 | 59.19979 | 85.13347 | 36.40636 | -0.16023 | 1.68015 | 0.70821 |
| 12 | 17892000 | 17901999 | 139.0753 | 210.1754 | 230.5853 | 95.34291 | 0.142212 | 3.316505 | 0.70822 |
| 14 | 79422600 | 79428799 | 80.09474 | 124.8775 | 170.5258 | 70.76857 | -0.13777 | 2.696598 | 0.708251 |
| 15 | 95924800 | 95925399 | 10.9258 | 20.52351 | 24.70012 | 10.15404 | 0.201047 | 0.101507 | 0.708269 |
| 13 | 84390000 | 84391199 | 18.87963 | 33.19111 | 31.28145 | 13.85202 | 0.472596 | 0.60463 | 0.708289 |
| 12 | 88080400 | 88089399 | 230.4832 | 336.2986 | 357.554 | 147.3274 | 0.140921 | 3.946683 | 0.708303 |
| 13 | 26860600 | 26861599 | 9.863435 | 18.5095 | 30.16619 | 12.42199 | -0.23493 | 0.114257 | 0.708341 |
| 18 | 40960000 | 40961999 | 18.47844 | 32.40323 | 37.04712 | 16.37296 | 0.195233 | 0.65172 | 0.708342 |

| | | | | | | | | | |
|---|---|---|---|---|---|---|---|---|---|
| 4 | 11092600 | 11094199 | 14.49641 | 26.41129 | 29.92226 | 12.86213 | 0.239663 | 0.319719 | 0.708362 |
| 11 | 94593800 | 94596999 | 22.59804 | 38.76047 | 44.75455 | 19.62864 | 0.176904 | 0.938904 | 0.708365 |
| 7 | 29550600 | 29553199 | 22.10646 | 37.06899 | 56.48561 | 24.64223 | -0.22458 | 1.152169 | 0.708366 |
| 18 | 57253000 | 57253999 | 15.17497 | 27.26907 | 30.92831 | 13.51805 | 0.222843 | 0.408368 | 0.7084 |
| 7 | 47651800 | 47656399 | 56.55824 | 90.47192 | 101.9785 | 42.92124 | 0.171016 | 2.055564 | 0.708412 |
| 2 | 8195400 | 8199399 | 61.66499 | 98.3812 | 95.89288 | 40.20961 | 0.37619 | 2.137934 | 0.708444 |
| 5 | 54111600 | 54114599 | 28.00695 | 46.79144 | 54.01811 | 23.51136 | 0.174571 | 1.206984 | 0.708445 |
| 8 | 129937200 | 129938599 | 38.73914 | 62.70026 | 69.98596 | 29.93485 | 0.204853 | 1.669344 | 0.708469 |
| 8 | 43792800 | 43795999 | 33.0987 | 54.84678 | 60.99055 | 26.40218 | 0.22801 | 1.316975 | 0.708502 |
| 2 | 89706600 | 89707399 | 26.93904 | 45.00328 | 50.72245 | 22.07569 | 0.209162 | 1.203499 | 0.708507 |
| 1 | 34302400 | 34303999 | 10.86689 | 19.47174 | 32.26424 | 14.13603 | -0.3232 | 0.397507 | 0.708516 |
| 1 | 111313600 | 111314399 | 14.70212 | 26.30988 | 30.37858 | 13.32435 | 0.193556 | 0.427586 | 0.708518 |
| 7 | 103443600 | 103445599 | 17.46704 | 30.52816 | 47.29389 | 20.84296 | -0.24107 | 0.715118 | 0.708521 |
| 1 | 30081200 | 30083399 | 25.63773 | 42.49665 | 62.98433 | 27.27111 | -0.18503 | 1.297801 | 0.708526 |
| 16 | 61158200 | 61160599 | 25.98802 | 44.48952 | 41.51975 | 18.20709 | 0.482403 | 0.965869 | 0.708532 |
| 4 | 103693200 | 103694199 | 18.76832 | 32.26622 | 47.94731 | 21.03158 | -0.18499 | 0.906236 | 0.708548 |
| 6 | 129829800 | 129831599 | 33.43432 | 55.04605 | 63.49465 | 27.38071 | 0.173659 | 1.397516 | 0.708564 |
| 18 | 6134800 | 6136799 | 12.82502 | 22.22666 | 49.48145 | 21.70983 | -0.76285 | 0.809271 | 0.708585 |
| 7 | 42145000 | 42152799 | 106.4418 | 164.3634 | 179.6984 | 74.64304 | 0.171925 | 2.877194 | 0.708589 |
| 4 | 76031000 | 76033599 | 24.59466 | 40.70387 | 72.32132 | 31.28002 | -0.44606 | 1.321545 | 0.708593 |
| 16 | 84798000 | 84802199 | 42.35062 | 68.19874 | 98.54908 | 41.88989 | -0.17517 | 1.83123 | 0.70861 |
| 13 | 113983200 | 113989799 | 50.81578 | 81.23102 | 115.226 | 48.45033 | -0.16114 | 2.072959 | 0.708651 |
| 2 | 176450400 | 176456799 | 77.89722 | 122.6334 | 138.4534 | 57.44482 | 0.147146 | 2.512525 | 0.708652 |
| 18 | 80564400 | 80566599 | 22.00974 | 36.92509 | 55.88857 | 24.38938 | -0.2148 | 1.148291 | 0.708662 |
| 2 | 40287200 | 40288799 | 46.21445 | 74.32027 | 84.27702 | 35.7532 | 0.171456 | 1.873214 | 0.708677 |
| 2 | 37036200 | 37036999 | 13.12179 | 24.41961 | 23.41892 | 9.780681 | 0.508136 | 0.176302 | 0.708698 |
| 1 | 104954400 | 104955199 | 18.20031 | 31.09423 | 48.49062 | 21.26044 | -0.25571 | 0.962804 | 0.708708 |
| 3 | 129353800 | 129354799 | 10.32168 | 19.06015 | 29.86349 | 12.60902 | -0.20651 | 0.214303 | 0.708729 |
| 6 | 130437200 | 130452799 | 221.4131 | 326.6455 | 329.5522 | 135.7146 | 0.231636 | 3.772799 | 0.708733 |
| X | 74755600 | 74756999 | 19.16089 | 33.56392 | 37.18539 | 16.41917 | 0.240653 | 0.675678 | 0.708736 |
| 3 | 103751400 | 103754999 | 36.26664 | 59.05827 | 68.17212 | 29.27063 | 0.16409 | 1.551485 | 0.70874 |
| 5 | 58474000 | 58484399 | 228.1179 | 328.3603 | 429.5555 | 177.2835 | -0.17724 | 4.1496 | 0.708746 |
| 7 | 41804600 | 41822599 | 309.8545 | 442.4174 | 448.3153 | 185.2622 | 0.179398 | 4.27335 | 0.70875 |
| 5 | 140962600 | 140965999 | 53.00091 | 84.52202 | 118.3299 | 49.5883 | -0.14597 | 2.150367 | 0.708756 |
| 12 | 27480400 | 27483199 | 25.0239 | 42.66729 | 48.48163 | 21.25311 | 0.198604 | 1.005079 | 0.708771 |
| 13 | 9094600 | 9097799 | 42.24028 | 68.56267 | 76.16939 | 32.64841 | 0.215437 | 1.622491 | 0.708773 |
| 15 | 46749200 | 46750399 | 15.73863 | 27.50635 | 41.5421 | 18.30754 | -0.20418 | 0.700708 | 0.708775 |
| 3 | 5623000 | 5624399 | 27.60146 | 46.73979 | 51.77916 | 22.66968 | 0.234154 | 1.079212 | 0.708791 |
| X | 126764400 | 126768199 | 35.75818 | 59.97637 | 56.48328 | 24.64527 | 0.467387 | 1.195534 | 0.708791 |
| 7 | 97414000 | 97416399 | 19.12437 | 32.72507 | 49.00925 | 21.4906 | -0.19729 | 0.951771 | 0.708811 |
| 4 | 101722400 | 101723999 | 20.0302 | 34.68697 | 39.5147 | 17.33655 | 0.198975 | 0.834202 | 0.708811 |
| 8 | 76932600 | 76933799 | 9.798394 | 18.47421 | 30.31769 | 12.38526 | -0.23588 | 0.082452 | 0.708817 |
| 15 | 57118600 | 57119599 | 9.798394 | 18.47421 | 30.31769 | 12.38526 | -0.23588 | 0.082452 | 0.708817 |
| X | 80906200 | 80906799 | 11.9162 | 22.43456 | 25.81089 | 10.57014 | 0.270339 | 0.101228 | 0.708828 |
| 3 | 18485400 | 18486599 | 9.483534 | 17.96942 | 29.93778 | 12.10404 | -0.24435 | 0.044914 | 0.708846 |
| 10 | 71362600 | 71363999 | 13.88006 | 24.62792 | 37.09861 | 16.3892 | -0.1951 | 0.488802 | 0.708847 |
| 6 | 20139000 | 20143999 | 81.17938 | 127.4353 | 142.5096 | 59.10574 | 0.157711 | 2.57079 | 0.70886 |

| | | | | | | | | | |
|---|---|---|---|---|---|---|---|---|---|
| 13 | 93786400 | 93793599 | 77.1974 | 121.8029 | 136.0591 | 56.47206 | 0.164966 | 2.467001 | 0.708865 |
| X | 98808200 | 98810599 | 20.41721 | 35.918 | 30.31035 | 13.42023 | 0.6318 | 0.607405 | 0.708868 |
| 10 | 99575000 | 99577999 | 23.5414 | 39.08789 | 69.66689 | 30.1998 | -0.45019 | 1.276522 | 0.708881 |
| 4 | 22907400 | 22907799 | 7.393433 | 14.38866 | 19.00913 | 6.995396 | 0.213769 | -0.32046 | 0.708882 |
| 4 | 76646800 | 76647199 | 6.394657 | 12.35554 | 22.6248 | 8.687503 | -0.31037 | -0.1754 | 0.708889 |
| 5 | 55919400 | 55924599 | 43.63557 | 71.45797 | 70.11587 | 30.1676 | 0.40337 | 1.491417 | 0.708904 |
| 1 | 92478600 | 92482799 | 41.49262 | 67.48898 | 74.14137 | 31.81995 | 0.234094 | 1.581298 | 0.708904 |
| X | 114165200 | 114166799 | 24.26054 | 41.60382 | 46.84954 | 20.54714 | 0.212815 | 0.94982 | 0.708914 |
| 2 | 42704800 | 42705199 | 10.06564 | 19.07469 | 22.30277 | 9.014868 | 0.263633 | 0.03155 | 0.708954 |
| X | 20077600 | 20078199 | 11.25527 | 21.22195 | 24.79369 | 10.12789 | 0.251369 | 0.083708 | 0.708963 |
| 2 | 21417000 | 21418999 | 21.68474 | 37.66004 | 42.50078 | 18.65523 | 0.213119 | 0.816905 | 0.708967 |
| 9 | 68140200 | 68142599 | 17.06745 | 29.77792 | 44.63857 | 19.65036 | -0.19402 | 0.733218 | 0.708981 |
| 2 | 177534400 | 177536999 | 22.26751 | 37.51378 | 55.42379 | 24.2275 | -0.17998 | 1.108671 | 0.708988 |
| X | 8573800 | 8578199 | 58.37703 | 93.98149 | 90.36518 | 38.3963 | 0.410896 | 1.859852 | 0.70899 |
| 1 | 91591000 | 91595599 | 41.01093 | 66.38496 | 76.25404 | 32.61868 | 0.163699 | 1.673482 | 0.709021 |
| 17 | 27861400 | 27863399 | 16.51185 | 28.66459 | 43.74931 | 19.19835 | -0.22034 | 0.802746 | 0.70908 |
| 2 | 177046400 | 177056399 | 109.5261 | 169.6025 | 176.6443 | 73.37336 | 0.244964 | 2.828 | 0.709117 |
| 13 | 74691400 | 74697999 | 87.85357 | 137.4114 | 153.0361 | 63.48863 | 0.159568 | 2.642254 | 0.709183 |
| 2 | 68384800 | 68385599 | 14.56921 | 25.95912 | 29.02998 | 12.77952 | 0.235986 | 0.451758 | 0.709186 |
| 3 | 154360400 | 154361599 | 11.09311 | 20.09928 | 31.69985 | 13.72082 | -0.24097 | 0.33046 | 0.709198 |
| 17 | 31625400 | 31628999 | 25.77093 | 42.54812 | 65.21584 | 28.16 | -0.2339 | 1.343475 | 0.709229 |
| 6 | 126394800 | 126395599 | 8.078269 | 15.33352 | 25.51594 | 10.27818 | -0.23733 | 0.009471 | 0.709231 |
| 3 | 115702200 | 115703199 | 8.582717 | 16.33649 | 27.20279 | 10.89326 | -0.23076 | 0.000835 | 0.709316 |
| 13 | 93421200 | 93424199 | 32.86473 | 53.97908 | 62.28222 | 26.83072 | 0.171855 | 1.423054 | 0.70935 |
| 13 | 49996600 | 49999999 | 38.17474 | 61.56569 | 88.80067 | 37.82333 | -0.16991 | 1.774344 | 0.709358 |
| 5 | 31353600 | 31358199 | 48.47671 | 77.31201 | 112.0538 | 46.96227 | -0.19574 | 2.143505 | 0.70937 |
| 13 | 74024000 | 74029199 | 79.21784 | 123.1157 | 170.5528 | 70.81364 | -0.16205 | 2.750618 | 0.709382 |
| 1 | 40817000 | 40820799 | 46.76289 | 74.93286 | 105.7889 | 44.63875 | -0.15033 | 1.992704 | 0.709382 |
| 3 | 58185000 | 58185599 | 7.9542 | 15.16483 | 25.59145 | 10.21278 | -0.24518 | -0.02029 | 0.709386 |
| 18 | 29616200 | 29616799 | 10.54531 | 19.91417 | 23.77873 | 9.68254 | 0.224041 | 0.063449 | 0.709398 |
| 14 | 39312400 | 39313199 | 16.82659 | 29.17719 | 44.2695 | 19.42342 | -0.21221 | 0.816236 | 0.709405 |
| X | 115821600 | 115822999 | 15.87092 | 28.64755 | 31.41055 | 13.66003 | 0.276101 | 0.388136 | 0.709434 |
| 5 | 89473800 | 89476999 | 54.51781 | 86.69266 | 121.8689 | 50.87106 | -0.15631 | 2.244662 | 0.709438 |
| 1 | 111852200 | 111853799 | 21.34287 | 36.89035 | 42.57807 | 18.67948 | 0.179569 | 0.860704 | 0.709481 |
| 15 | 68266800 | 68268999 | 19.46946 | 34.20222 | 37.38292 | 16.54637 | 0.259773 | 0.629182 | 0.709484 |
| 1 | 107311400 | 107312999 | 13.351 | 24.43949 | 28.48332 | 12.17023 | 0.20484 | 0.281507 | 0.709488 |
| 2 | 61830200 | 61832199 | 27.36439 | 45.98906 | 52.88854 | 23.10231 | 0.179981 | 1.142826 | 0.709497 |
| 13 | 61294800 | 61308399 | 220.3973 | 324.2685 | 336.7052 | 138.657 | 0.186486 | 3.817097 | 0.709508 |
| 13 | 67284000 | 67285999 | 17.569 | 31.09554 | 34.16425 | 15.10825 | 0.255983 | 0.527914 | 0.709537 |
| 8 | 81890400 | 81891199 | 11.34987 | 20.71075 | 32.32605 | 13.8684 | -0.21743 | 0.296083 | 0.709539 |
| 1 | 4987600 | 4994999 | 75.50571 | 119.1774 | 133.9738 | 55.61683 | 0.156242 | 2.456517 | 0.70954 |
| 1 | 58855600 | 58860999 | 92.21971 | 143.2664 | 160.9024 | 66.81069 | 0.13989 | 2.762507 | 0.709541 |
| 2 | 40314400 | 40314599 | 3.930036 | 7.687827 | 15.60561 | 5.197775 | -0.26588 | -0.72988 | 0.709552 |
| 15 | 65587600 | 65588399 | 11.8166 | 21.2851 | 32.71016 | 14.26222 | -0.21039 | 0.374218 | 0.709583 |
| 19 | 8135600 | 8135999 | 5.504136 | 10.7376 | 19.80141 | 7.081395 | -0.22413 | -0.44117 | 0.709609 |
| 10 | 62938800 | 62941799 | 29.45201 | 48.26147 | 70.6674 | 30.41307 | -0.17223 | 1.451356 | 0.709669 |
| X | 80442200 | 80445399 | 40.23975 | 66.64241 | 63.07947 | 27.29582 | 0.459846 | 1.348051 | 0.709735 |

| | | | | | | | | | |
|---|---|---|---|---|---|---|---|---|---|
| 6 | 27112800 | 27115399 | 35.9305 | 58.72029 | 67.878 | 29.18232 | 0.16566 | 1.499744 | 0.709743 |
| 8 | 81806000 | 81808399 | 19.25242 | 33.60235 | 38.76918 | 17.07051 | 0.182077 | 0.730763 | 0.709757 |
| 16 | 31286400 | 31292399 | 55.11096 | 87.66054 | 122.759 | 51.26111 | -0.15039 | 2.235522 | 0.709766 |
| 15 | 100177200 | 100178999 | 30.78375 | 50.87523 | 58.90754 | 25.45367 | 0.169682 | 1.342167 | 0.709804 |
| 10 | 73690400 | 73693399 | 46.42956 | 74.95047 | 72.85373 | 31.06944 | 0.399816 | 1.752935 | 0.709804 |
| 7 | 42510000 | 42521199 | 155.7783 | 235.32 | 246.1535 | 101.7667 | 0.21043 | 3.33042 | 0.709808 |
| 11 | 18182000 | 18182799 | 10.91118 | 19.53807 | 37.32127 | 16.36088 | -0.52786 | 0.412789 | 0.709812 |
| 3 | 20473000 | 20474999 | 24.44065 | 41.74089 | 47.39435 | 20.77921 | 0.199957 | 0.985779 | 0.709816 |
| 5 | 49351200 | 49354599 | 45.08815 | 72.7305 | 82.69114 | 35.23064 | 0.172939 | 1.775442 | 0.709822 |
| 12 | 64508000 | 64509999 | 29.48221 | 49.6422 | 54.06836 | 23.63124 | 0.258235 | 1.134375 | 0.709845 |
| 13 | 48629600 | 48630799 | 13.39789 | 24.36793 | 28.48154 | 12.2692 | 0.193118 | 0.319137 | 0.709876 |
| 13 | 29300000 | 29300799 | 8.720973 | 16.56066 | 27.14609 | 10.92475 | -0.2147 | 0.017529 | 0.709883 |
| X | 67237000 | 67237599 | 10.61817 | 20.2693 | 23.9358 | 9.526599 | 0.268852 | -0.00819 | 0.709892 |
| 14 | 114379800 | 114380999 | 14.32032 | 25.61567 | 28.90038 | 12.68057 | 0.226355 | 0.423888 | 0.709901 |
| 2 | 41404200 | 41404599 | 7.531115 | 14.61924 | 17.37679 | 6.527915 | 0.338046 | -0.25137 | 0.70992 |
| 8 | 117525200 | 117533399 | 93.36936 | 142.6766 | 232.181 | 96.30971 | -0.41297 | 3.081059 | 0.70995 |
| 4 | 81696400 | 81699199 | 43.80207 | 70.25874 | 100.0259 | 42.27127 | -0.16066 | 1.955983 | 0.709987 |
| 10 | 52642600 | 52643999 | 17.92005 | 31.40322 | 36.06908 | 15.93178 | 0.188614 | 0.656835 | 0.710001 |
| 16 | 30972400 | 30978199 | 111.4004 | 169.3494 | 230.1963 | 95.35532 | -0.15957 | 3.191827 | 0.710007 |
| 19 | 39263600 | 39266799 | 39.62762 | 64.90944 | 69.62137 | 29.95775 | 0.275364 | 1.479231 | 0.710009 |
| 10 | 59885000 | 59896999 | 117.9161 | 179.5571 | 243.7449 | 101.0187 | -0.15567 | 3.15958 | 0.710038 |
| 9 | 35485000 | 35485999 | 10.81276 | 19.56447 | 31.2366 | 13.54352 | -0.25998 | 0.336153 | 0.710047 |
| 17 | 60503200 | 60504799 | 17.41975 | 30.15976 | 45.29724 | 19.87342 | -0.19818 | 0.832608 | 0.710067 |
| 1 | 59557400 | 59559999 | 21.73888 | 37.43817 | 42.57804 | 18.6774 | 0.199617 | 0.893455 | 0.710071 |
| 15 | 70674400 | 70675799 | 11.50042 | 21.06892 | 33.39499 | 14.2532 | -0.23499 | 0.272586 | 0.710085 |
| 17 | 54263400 | 54263999 | 11.20166 | 21.07539 | 25.21214 | 10.33647 | 0.212608 | 0.094325 | 0.710118 |
| 8 | 74819000 | 74819799 | 17.82589 | 31.27436 | 36.05273 | 15.94012 | 0.1834 | 0.638768 | 0.710135 |
| 18 | 77663800 | 77665999 | 20.39376 | 34.57035 | 52.73752 | 23.09655 | -0.2257 | 1.043648 | 0.710171 |
| 3 | 71484400 | 71486999 | 17.0269 | 30.56321 | 32.11228 | 14.0486 | 0.331201 | 0.426076 | 0.710214 |
| 15 | 82040000 | 82041999 | 15.82442 | 27.62445 | 41.54073 | 18.29301 | -0.19825 | 0.715921 | 0.710243 |
| 17 | 52635200 | 52636199 | 20.61851 | 35.74142 | 40.975 | 17.97778 | 0.190049 | 0.828791 | 0.710245 |
| 1 | 147967800 | 147973999 | 104.9551 | 162.1431 | 178.4682 | 74.13184 | 0.162741 | 2.868429 | 0.710275 |
| 18 | 24655000 | 24656199 | 18.50038 | 32.55118 | 36.37436 | 16.10461 | 0.228565 | 0.602408 | 0.710303 |
| 6 | 133708200 | 133717399 | 122.2225 | 185.6445 | 250.6864 | 103.8168 | -0.15107 | 3.212292 | 0.71031 |
| 9 | 34438000 | 34440999 | 33.04044 | 54.39543 | 62.96359 | 27.15125 | 0.16859 | 1.395487 | 0.71031 |
| 12 | 74618000 | 74621399 | 40.35494 | 64.93889 | 92.92415 | 39.45829 | -0.1623 | 1.847894 | 0.710312 |
| 9 | 116527600 | 116532999 | 62.35468 | 99.52462 | 108.5101 | 45.52809 | 0.21547 | 2.128529 | 0.710318 |
| 1 | 165860800 | 165862999 | 28.55744 | 48.27839 | 44.19527 | 19.3396 | 0.508166 | 1.085523 | 0.71035 |
| X | 162691400 | 162693399 | 36.45295 | 61.58044 | 40.88663 | 17.88772 | 0.970083 | 1.122475 | 0.71037 |
| 18 | 41683000 | 41684799 | 22.2472 | 37.71032 | 55.46504 | 24.29071 | -0.17322 | 1.055633 | 0.710384 |
| X | 64816600 | 64818399 | 19.09066 | 33.56773 | 36.91871 | 16.34776 | 0.251114 | 0.615023 | 0.710386 |
| 18 | 42767200 | 42769799 | 34.64821 | 57.44822 | 53.66575 | 23.24079 | 0.478817 | 1.310993 | 0.710391 |
| 16 | 26906200 | 26906799 | 12.29147 | 22.53795 | 24.22632 | 10.32759 | 0.32368 | 0.256285 | 0.710403 |
| 13 | 113431600 | 113436799 | 53.82267 | 86.27193 | 97.16111 | 41.01729 | 0.175602 | 1.987157 | 0.710404 |
| 17 | 21037200 | 21039799 | 34.42921 | 56.83245 | 63.71781 | 27.50962 | 0.215893 | 1.368682 | 0.71043 |
| 11 | 83189400 | 83191199 | 9.482856 | 17.59378 | 28.35481 | 11.8956 | -0.24034 | 0.175916 | 0.710435 |
| 5 | 40665400 | 40667199 | 17.73193 | 31.65679 | 33.72597 | 14.8221 | 0.307042 | 0.469193 | 0.710458 |

| | | | | | | | | | |
|---|---|---|---|---|---|---|---|---|---|
| 2 | 17676000 | 17676599 | 5.36712 | 10.45662 | 19.65709 | 7.151195 | -0.27353 | -0.38212 | 0.710469 |
| 7 | 85027200 | 85029799 | 59.3527 | 94.02059 | 131.9604 | 54.88123 | -0.15933 | 2.361509 | 0.710498 |
| 14 | 44835800 | 44837599 | 17.10011 | 30.10994 | 34.3694 | 15.21465 | 0.198394 | 0.581617 | 0.710502 |
| 2 | 57444400 | 57444999 | 10.62467 | 20.6891 | 21.59427 | 7.909139 | 0.556586 | -0.28966 | 0.710513 |
| 10 | 25463000 | 25463599 | 11.5958 | 21.06897 | 32.23284 | 13.90276 | -0.19397 | 0.320425 | 0.71052 |
| 13 | 18871400 | 18872799 | 15.68356 | 27.49243 | 40.87954 | 18.06126 | -0.18197 | 0.649362 | 0.710525 |
| 1 | 163283600 | 163285799 | 19.59676 | 34.21462 | 39.25859 | 17.28959 | 0.190036 | 0.72878 | 0.710528 |
| 14 | 78652400 | 78654399 | 17.22729 | 29.98118 | 44.69427 | 19.64376 | -0.18638 | 0.769761 | 0.710541 |
| 19 | 8220400 | 8221399 | 20.25829 | 34.70014 | 51.14669 | 22.42968 | -0.17428 | 0.949628 | 0.710592 |
| 3 | 144450800 | 144451599 | 7.436436 | 14.32658 | 24.20269 | 9.390648 | -0.20936 | -0.12317 | 0.710596 |
| Y | 4426800 | 4430599 | 42.53316 | 68.59756 | 59.0663 | 25.15978 | 0.572728 | 1.762491 | 0.710605 |
| Y | 4670200 | 4671799 | 12.27987 | 22.52611 | 20.18535 | 8.600037 | 0.584404 | 0.23958 | 0.710624 |
| 5 | 126462200 | 126463799 | 17.72628 | 31.12447 | 35.42768 | 15.67435 | 0.20147 | 0.622855 | 0.710655 |
| 6 | 59907400 | 59910799 | 29.94389 | 50.69698 | 52.40402 | 22.94196 | 0.333676 | 1.087678 | 0.710662 |
| 10 | 7102400 | 7105399 | 23.40274 | 39.12847 | 58.71007 | 25.56562 | -0.20237 | 1.189335 | 0.710663 |
| 1 | 44420000 | 44420999 | 19.65271 | 34.13133 | 38.92419 | 17.08544 | 0.198086 | 0.801822 | 0.710671 |
| 18 | 39995800 | 39996999 | 13.90683 | 25.59478 | 29.07521 | 12.33219 | 0.249259 | 0.257878 | 0.710707 |
| 2 | 4396400 | 4399199 | 20.69854 | 35.90119 | 41.51469 | 18.21489 | 0.177888 | 0.825799 | 0.710714 |
| 18 | 79611000 | 79613599 | 40.77474 | 66.51946 | 72.37483 | 31.09558 | 0.251021 | 1.538538 | 0.710734 |
| 13 | 26232600 | 26233399 | 11.18192 | 20.17165 | 31.76149 | 13.82301 | -0.24339 | 0.359217 | 0.71075 |
| 16 | 57872000 | 57873599 | 14.99563 | 27.47509 | 29.9825 | 12.79491 | 0.3004 | 0.291995 | 0.710758 |
| X | 143783400 | 143784599 | 16.28202 | 29.42658 | 31.8682 | 13.83728 | 0.295156 | 0.383511 | 0.710766 |
| 9 | 77577800 | 77580999 | 46.20992 | 74.60396 | 82.66241 | 35.25738 | 0.211461 | 1.752139 | 0.710787 |
| 16 | 92569600 | 92572199 | 35.58069 | 57.68789 | 83.04749 | 35.56693 | -0.15905 | 1.636561 | 0.710813 |
| 15 | 17570400 | 17575199 | 96.99345 | 149.8867 | 165.4767 | 68.73535 | 0.158707 | 2.857431 | 0.710823 |
| 14 | 104755600 | 104756799 | 18.22724 | 31.94877 | 36.63842 | 16.18635 | 0.191003 | 0.657999 | 0.710839 |
| 11 | 15234400 | 15238199 | 54.58144 | 86.79907 | 122.7828 | 51.25563 | -0.16521 | 2.244259 | 0.710874 |
| 7 | 139901600 | 139903399 | 18.59222 | 32.08163 | 47.40973 | 20.79811 | -0.17594 | 0.871543 | 0.71088 |
| 8 | 75665600 | 75670599 | 44.3236 | 71.64682 | 80.64728 | 34.45261 | 0.190826 | 1.714139 | 0.710888 |
| 2 | 98498800 | 98501999 | 38.79892 | 63.60336 | 68.62213 | 29.53787 | 0.267327 | 1.464532 | 0.710889 |
| 2 | 110858000 | 110858599 | 14.26042 | 25.5886 | 29.46597 | 12.89462 | 0.199814 | 0.406836 | 0.710932 |
| 1 | 16607600 | 16608199 | 6.852113 | 13.29123 | 23.35776 | 8.806769 | -0.22593 | -0.22637 | 0.710941 |
| X | 80323800 | 80330399 | 85.71603 | 134.9539 | 142.9989 | 59.33143 | 0.238649 | 2.518647 | 0.710973 |
| 16 | 9906400 | 9908199 | 16.09217 | 28.18264 | 42.42963 | 18.73148 | -0.19998 | 0.666933 | 0.710988 |
| 1 | 17276600 | 17278399 | 18.07706 | 31.32624 | 46.33951 | 20.33126 | -0.1761 | 0.829176 | 0.711006 |
| 7 | 3225600 | 3251999 | 294.895 | 420.3534 | 434.6391 | 179.6612 | 0.147868 | 4.295378 | 0.711008 |
| 19 | 9520400 | 9537599 | 426.3441 | 581.9156 | 721.6133 | 300.359 | -0.17736 | 4.953509 | 0.71101 |
| 5 | 70400600 | 70402399 | 25.7165 | 43.56594 | 50.12609 | 21.94871 | 0.179712 | 1.066481 | 0.711033 |
| 17 | 65979600 | 65980999 | 16.38343 | 28.97332 | 33.37933 | 14.76356 | 0.187632 | 0.529989 | 0.711052 |
| 6 | 67995600 | 67996799 | 16.55835 | 29.12286 | 33.08533 | 14.64954 | 0.204805 | 0.59071 | 0.711067 |
| 10 | 74573400 | 74575799 | 39.26727 | 63.99095 | 72.07529 | 30.95468 | 0.200125 | 1.550214 | 0.711089 |
| 14 | 60698600 | 60699799 | 13.7271 | 24.84688 | 28.95014 | 12.54633 | 0.192034 | 0.350058 | 0.711117 |
| 16 | 46895600 | 46896599 | 8.545152 | 16.27114 | 26.66058 | 10.6676 | -0.20684 | -0.00466 | 0.711132 |
| 15 | 33069000 | 33074999 | 76.74636 | 120.7764 | 137.0479 | 56.85891 | 0.13945 | 2.518919 | 0.711143 |
| 4 | 84953200 | 84954399 | 11.17509 | 19.77407 | 39.4073 | 17.4277 | -0.59782 | 0.506426 | 0.71115 |
| 5 | 23289400 | 23290399 | 7.879075 | 15.08816 | 25.14426 | 9.926214 | -0.21363 | -0.06076 | 0.711157 |
| 10 | 116788200 | 116792199 | 43.34729 | 69.43695 | 115.3583 | 48.65241 | -0.3853 | 2.001405 | 0.711161 |

| | | | | | | | | | |
|---|---|---|---|---|---|---|---|---|---|
| 9 | 9595400 | 9602399 | 92.60243 | 143.0944 | 194.7028 | 80.87525 | -0.1426 | 2.860764 | 0.71117 |
| 8 | 115970400 | 115973799 | 27.82515 | 46.71588 | 53.46222 | 23.34141 | 0.186984 | 1.154097 | 0.711216 |
| 1 | 19386200 | 19387599 | 13.8309 | 24.81609 | 27.48188 | 12.02709 | 0.255657 | 0.398637 | 0.711222 |
| 14 | 35717800 | 35718399 | 8.829782 | 16.83686 | 27.85044 | 11.11048 | -0.21697 | -0.0098 | 0.711223 |
| 15 | 9139600 | 9141399 | 13.72777 | 24.15391 | 37.25144 | 16.49292 | -0.23379 | 0.576219 | 0.711224 |
| 4 | 33191400 | 33192199 | 8.2264 | 15.7669 | 26.30052 | 10.35694 | -0.21147 | -0.06071 | 0.711255 |
| 17 | 43364000 | 43366599 | 17.31129 | 30.069 | 44.75474 | 19.64638 | -0.18427 | 0.799813 | 0.711261 |
| 3 | 74177000 | 74181199 | 54.40313 | 87.27171 | 96.83066 | 40.92724 | 0.198464 | 1.964455 | 0.71127 |
| 12 | 25326000 | 25327799 | 18.35071 | 31.78839 | 47.40153 | 20.79698 | -0.18782 | 0.833937 | 0.711273 |
| 2 | 174846200 | 174846999 | 7.887348 | 15.07831 | 25.0954 | 9.955482 | -0.21833 | -0.04454 | 0.711282 |
| 16 | 27554400 | 27555399 | 15.21518 | 26.66615 | 39.89442 | 17.6231 | -0.19067 | 0.648904 | 0.711305 |
| 16 | 22187400 | 22190199 | 17.24931 | 30.05433 | 44.67155 | 19.64809 | -0.18197 | 0.753891 | 0.711323 |
| 10 | 48679000 | 48682799 | 38.89382 | 63.16289 | 72.87991 | 31.25014 | 0.161589 | 1.603392 | 0.711325 |
| 15 | 75835200 | 75836599 | 12.33184 | 21.81105 | 34.77872 | 15.38212 | -0.27846 | 0.506677 | 0.711367 |
| 3 | 98006800 | 98014799 | 55.67131 | 87.83577 | 153.5034 | 63.7113 | -0.47987 | 2.456797 | 0.711394 |
| 2 | 65136600 | 65137199 | 4.649557 | 9.060517 | 19.11935 | 6.936463 | -0.43378 | -0.40211 | 0.711398 |
| 7 | 36854400 | 36866399 | 10.39731 | 18.93171 | 36.00663 | 15.48912 | -0.50353 | 0.307737 | 0.711437 |
| 2 | 97936800 | 97937799 | 16.02293 | 28.25129 | 32.02248 | 14.17189 | 0.209368 | 0.557624 | 0.711444 |
| 18 | 70980000 | 70982399 | 26.40732 | 44.79549 | 42.68632 | 18.69856 | 0.450735 | 1.049318 | 0.711456 |
| 10 | 7040000 | 7043799 | 72.31469 | 114.1893 | 129.781 | 53.88791 | 0.140956 | 2.444919 | 0.711461 |
| 1 | 13579400 | 13587199 | 110.0865 | 167.9779 | 225.7786 | 93.62253 | -0.13901 | 3.114358 | 0.711468 |
| 17 | 22680800 | 22691199 | 173.6105 | 259.8043 | 275.0119 | 113.4182 | 0.181345 | 3.516674 | 0.71149 |
| 2 | 13165000 | 13165999 | 9.637438 | 18.14311 | 29.13861 | 11.93429 | -0.20784 | 0.088135 | 0.711491 |
| 2 | 84488200 | 84490999 | 25.01005 | 41.37029 | 62.84631 | 27.1724 | -0.22072 | 1.320015 | 0.711494 |
| 14 | 73147600 | 73148199 | 14.41534 | 26.56565 | 24.7861 | 10.49414 | 0.533171 | 0.244532 | 0.711503 |
| 1 | 120454800 | 120456599 | 12.81266 | 22.85699 | 36.60148 | 16.09619 | -0.27852 | 0.444098 | 0.711522 |
| 13 | 42186400 | 42187399 | 11.45131 | 22.0814 | 24.33309 | 9.416704 | 0.404794 | -0.09129 | 0.711524 |
| 17 | 83992200 | 83995799 | 44.41129 | 71.24034 | 101.0817 | 42.72146 | -0.15572 | 1.954204 | 0.711545 |
| 1 | 29546200 | 29546799 | 11.61959 | 21.60622 | 25.65252 | 10.72847 | 0.200776 | 0.175018 | 0.71155 |
| X | 38770000 | 38771199 | 15.78417 | 28.74297 | 30.50049 | 13.11897 | 0.333345 | 0.336604 | 0.711553 |
| 17 | 36407400 | 36418399 | 140.7288 | 210.7954 | 281.771 | 116.2269 | -0.15382 | 3.493133 | 0.711579 |
| 8 | 104528400 | 104529799 | 14.23948 | 24.93143 | 38.96319 | 17.19733 | -0.25329 | 0.65741 | 0.711589 |
| 10 | 122633200 | 122635599 | 21.22656 | 35.89097 | 53.81866 | 23.55193 | -0.20113 | 1.070292 | 0.7116 |
| 11 | 17510200 | 17513199 | 78.92835 | 124.0248 | 140.5566 | 58.30328 | 0.139614 | 2.550416 | 0.7116 |
| 2 | 45650600 | 45651199 | 7.284477 | 13.96412 | 24.07593 | 9.475985 | -0.258 | -0.07956 | 0.71161 |
| 15 | 34164600 | 34167199 | 41.7769 | 67.53283 | 77.57933 | 33.14364 | 0.161637 | 1.70888 | 0.711612 |
| 7 | 32286600 | 32288399 | 62.51151 | 99.21994 | 109.0049 | 45.41698 | 0.196209 | 2.302602 | 0.711617 |
| X | 8875600 | 8876999 | 11.56829 | 21.34367 | 36.38839 | 15.38209 | -0.32907 | 0.244128 | 0.711646 |
| 4 | 63873000 | 63874599 | 13.65601 | 23.92429 | 44.08657 | 19.46859 | -0.48994 | 0.652713 | 0.711652 |
| 5 | 106681000 | 106682199 | 12.79877 | 23.50996 | 27.58516 | 11.73101 | 0.200334 | 0.257607 | 0.711655 |
| 14 | 114286000 | 114286999 | 12.79877 | 23.50996 | 27.58516 | 11.73101 | 0.200334 | 0.257607 | 0.711655 |
| 18 | 85714400 | 85717199 | 39.53321 | 64.05665 | 73.83093 | 31.60534 | 0.160131 | 1.649732 | 0.711678 |
| 17 | 72039000 | 72042799 | 35.39248 | 57.2772 | 82.76824 | 35.39925 | -0.16697 | 1.672088 | 0.711687 |
| 17 | 40346600 | 40348399 | 20.21138 | 35.71038 | 36.18 | 16.0052 | 0.371529 | 0.567655 | 0.711716 |
| 1 | 139935800 | 139940199 | 47.21003 | 76.45462 | 80.21209 | 34.32341 | 0.294769 | 1.678787 | 0.711717 |
| X | 114180400 | 114182199 | 30.17196 | 51.52501 | 40.39199 | 17.70903 | 0.732564 | 1.005153 | 0.711747 |
| 2 | 34474000 | 34476199 | 19.49762 | 34.12429 | 38.65454 | 17.05543 | 0.208622 | 0.691027 | 0.71175 |

| | | | | | | | | | |
|---|---|---|---|---|---|---|---|---|---|
| 5 | 67974400 | 67976199 | 16.31544 | 28.80468 | 32.55648 | 14.40445 | 0.214361 | 0.548216 | 0.71175 |
| 5 | 111136600 | 111138199 | 14.25729 | 25.77095 | 29.93589 | 12.99595 | 0.194807 | 0.364334 | 0.711759 |
| 9 | 91125600 | 91126199 | 6.855796 | 13.28296 | 23.2911 | 8.829346 | -0.23009 | -0.20796 | 0.711776 |
| 11 | 13954400 | 13955599 | 15.02196 | 27.51674 | 30.05452 | 12.82982 | 0.299069 | 0.295892 | 0.711812 |
| 15 | 93091000 | 93095399 | 52.18384 | 83.29167 | 116.8053 | 49.00917 | -0.14704 | 2.12193 | 0.711817 |
| 14 | 116913200 | 116913799 | 9.458232 | 18.09647 | 22.32369 | 8.834747 | 0.21316 | -0.04559 | 0.711818 |
| 14 | 116971200 | 116973599 | 27.22554 | 45.11966 | 66.0182 | 28.58083 | -0.16667 | 1.305607 | 0.711819 |
| 14 | 13938200 | 13941399 | 39.13415 | 63.96213 | 70.53268 | 30.3247 | 0.233598 | 1.508016 | 0.711847 |
| 5 | 47299200 | 47302199 | 39.42465 | 63.9484 | 73.85854 | 31.64184 | 0.158592 | 1.629482 | 0.711857 |
| 18 | 29814800 | 29815199 | 8.464475 | 16.42745 | 21.11497 | 7.941529 | 0.223923 | -0.22518 | 0.71186 |
| 4 | 89265400 | 89267599 | 15.6334 | 27.45297 | 41.30163 | 18.27502 | -0.19893 | 0.619935 | 0.711867 |
| 5 | 33111000 | 33112799 | 15.6334 | 27.45297 | 41.30163 | 18.27502 | -0.19893 | 0.619935 | 0.711867 |
| 5 | 62035600 | 62038399 | 49.78996 | 79.79274 | 90.16442 | 38.05257 | 0.170513 | 1.987212 | 0.711885 |
| 2 | 4934000 | 4936199 | 14.26949 | 25.13372 | 37.7561 | 16.71282 | -0.19531 | 0.570654 | 0.711888 |
| 4 | 74708000 | 74708799 | 9.325488 | 17.38482 | 27.8896 | 11.61367 | -0.22535 | 0.144717 | 0.711899 |
| 17 | 22500800 | 22514399 | 223.4794 | 331.7919 | 304.932 | 125.6104 | 0.374658 | 3.66814 | 0.71191 |
| 9 | 91522000 | 91523799 | 12.40638 | 21.8482 | 40.90796 | 18.10864 | -0.51148 | 0.567113 | 0.711915 |
| 5 | 148160600 | 148162599 | 16.61242 | 28.94469 | 43.11501 | 18.96349 | -0.18498 | 0.748866 | 0.711923 |
| 7 | 12311600 | 12316399 | 78.10081 | 123.7066 | 114.6785 | 47.69131 | 0.438054 | 2.373298 | 0.711955 |
| 13 | 35035200 | 35041999 | 125.639 | 191.5005 | 211.5483 | 87.69966 | 0.142513 | 3.139881 | 0.711998 |
| 4 | 69805600 | 69806199 | 9.346988 | 18.0419 | 22.56591 | 8.705445 | 0.227893 | -0.12769 | 0.71203 |
| 17 | 70820200 | 70825999 | 70.18648 | 110.15 | 152.4131 | 63.21105 | -0.1494 | 2.572905 | 0.71205 |
| 13 | 51220800 | 51223399 | 21.92631 | 37.71707 | 43.1176 | 18.91341 | 0.191864 | 0.905589 | 0.712058 |
| 11 | 57629800 | 57632799 | 20.70729 | 35.65172 | 39.62258 | 17.3808 | 0.232826 | 0.889855 | 0.712061 |
| 1 | 58687400 | 58691799 | 57.77252 | 91.78016 | 127.5622 | 53.19651 | -0.1413 | 2.276097 | 0.712078 |
| 2 | 100017800 | 100019799 | 25.39074 | 42.7415 | 49.08506 | 21.45147 | 0.182124 | 1.118871 | 0.712087 |
| X | 148595600 | 148602199 | 74.05047 | 117.7249 | 105.0704 | 43.84847 | 0.497745 | 2.26475 | 0.712089 |
| 10 | 129292400 | 129296799 | 11.88266 | 22.42289 | 21.06621 | 8.591611 | 0.565268 | 0.072935 | 0.712095 |
| 3 | 91460800 | 91462199 | 15.85272 | 28.63116 | 26.61255 | 11.56513 | 0.513336 | 0.371904 | 0.712118 |
| X | 153224400 | 153225399 | 30.73716 | 51.13052 | 57.28091 | 24.86247 | 0.217903 | 1.268489 | 0.712136 |
| 17 | 31451400 | 31452399 | 13.77913 | 24.93662 | 28.98992 | 12.56636 | 0.195134 | 0.352394 | 0.712167 |
| 9 | 21388000 | 21390199 | 20.59546 | 34.93185 | 52.19185 | 22.86146 | -0.1957 | 1.040431 | 0.712191 |
| 14 | 50179400 | 50182399 | 47.65388 | 76.65947 | 87.91282 | 37.31282 | 0.1558 | 1.865825 | 0.712196 |
| 1 | 57624000 | 57626599 | 58.47586 | 93.21533 | 106.6902 | 44.68544 | 0.143553 | 2.164855 | 0.71221 |
| 8 | 129913400 | 129914199 | 10.89949 | 20.3918 | 24.14789 | 9.994939 | 0.216329 | 0.125053 | 0.712226 |
| 17 | 23092200 | 23093399 | 14.61383 | 26.3932 | 30.42916 | 13.22399 | 0.204633 | 0.373046 | 0.712228 |
| 17 | 35511000 | 35512599 | 12.85947 | 22.98774 | 35.10197 | 15.41 | -0.20838 | 0.430708 | 0.712275 |
| 8 | 111214400 | 111214999 | 8.05165 | 15.28532 | 25.50965 | 10.27232 | -0.24134 | 0.006798 | 0.712276 |
| 5 | 90335800 | 90337599 | 18.8705 | 33.04562 | 37.55307 | 16.57744 | 0.203878 | 0.675006 | 0.71231 |
| 15 | 3097800 | 3099199 | 23.23307 | 40.02714 | 43.54036 | 19.09975 | 0.263585 | 0.895721 | 0.712335 |
| 2 | 132623200 | 132623999 | 8.404505 | 15.69931 | 26.90633 | 11.16629 | -0.31599 | 0.120647 | 0.712336 |
| 1 | 170081400 | 170084999 | 52.77328 | 84.40309 | 95.79526 | 40.30913 | 0.160853 | 2.054899 | 0.712351 |
| 17 | 85911800 | 85913199 | 12.06887 | 21.77246 | 33.15946 | 14.43077 | -0.19597 | 0.366393 | 0.712368 |
| 12 | 104926600 | 104927799 | 14.05439 | 24.80907 | 37.17589 | 16.44879 | -0.19062 | 0.542912 | 0.712374 |
| 5 | 49863400 | 49863799 | 5.798365 | 11.29946 | 20.8309 | 7.554931 | -0.24181 | -0.38246 | 0.712376 |
| 8 | 66070000 | 66071599 | 15.14186 | 26.49474 | 40.42995 | 17.83526 | -0.21922 | 0.673452 | 0.712377 |
| 4 | 4659200 | 4663399 | 57.27972 | 91.05158 | 126.606 | 52.82926 | -0.14111 | 2.258162 | 0.712411 |

| | | | | | | | | | |
|---|---|---|---|---|---|---|---|---|---|
| 5 | 72840400 | 72842199 | 18.20426 | 31.56986 | 46.86623 | 20.56282 | -0.18105 | 0.823075 | 0.71245 |
| 18 | 7297400 | 7298199 | 8.756203 | 16.89817 | 20.28209 | 7.83073 | 0.285237 | -0.14032 | 0.712468 |
| 11 | 20424800 | 20429199 | 30.03759 | 49.20777 | 72.64802 | 31.2619 | -0.18429 | 1.4552 | 0.71247 |
| 7 | 39057000 | 39059399 | 40.01599 | 64.34214 | 93.16258 | 39.51385 | -0.18071 | 1.872043 | 0.71247 |
| 8 | 119602600 | 119606999 | 42.67517 | 68.90333 | 79.39958 | 33.86956 | 0.155184 | 1.743694 | 0.712481 |
| 11 | 35631800 | 35638599 | 45.65562 | 72.85902 | 123.0192 | 51.60353 | -0.4147 | 2.121672 | 0.712482 |
| 3 | 131457600 | 131460599 | 47.05739 | 75.77853 | 86.11451 | 36.60459 | 0.17074 | 1.830373 | 0.712491 |
| 5 | 107096200 | 107096799 | 7.90823 | 15.14227 | 25.13913 | 9.927428 | -0.20837 | -0.05801 | 0.712499 |
| 5 | 135691600 | 135692399 | 7.90823 | 15.14227 | 25.13913 | 9.927428 | -0.20837 | -0.05801 | 0.712499 |
| 7 | 16883800 | 16897399 | 236.0836 | 343.0258 | 361.8462 | 149.1612 | 0.146406 | 4.009704 | 0.712537 |
| 1 | 147977000 | 147979999 | 32.20944 | 53.5617 | 59.75795 | 25.93163 | 0.223687 | 1.27528 | 0.712545 |
| 14 | 117673000 | 117673999 | 16.39135 | 28.95995 | 33.41726 | 14.78313 | 0.184842 | 0.540531 | 0.712565 |
| 8 | 126902400 | 126904599 | 19.54193 | 33.26311 | 51.6711 | 22.64817 | -0.25122 | 1.001762 | 0.712567 |
| 2 | 94637200 | 94642199 | 51.3363 | 81.87992 | 114.8605 | 48.14457 | -0.1486 | 2.142551 | 0.712568 |
| 3 | 23828400 | 23828799 | 5.20529 | 10.17323 | 19.59951 | 6.776368 | -0.24261 | -0.56539 | 0.712588 |
| 10 | 129138000 | 129141399 | 54.26462 | 86.31265 | 122.0152 | 50.94913 | -0.16399 | 2.234696 | 0.712614 |
| 12 | 21275600 | 21278399 | 21.8956 | 36.93567 | 54.90221 | 24.00906 | -0.18868 | 1.094352 | 0.71262 |
| 1 | 120502400 | 120508399 | 74.40263 | 117.4357 | 133.2255 | 55.30605 | 0.1431 | 2.456323 | 0.712625 |
| X | 73668200 | 73670999 | 17.63205 | 30.76137 | 48.94376 | 21.54484 | -0.27946 | 0.746925 | 0.712626 |
| 11 | 29529800 | 29531199 | 27.55419 | 46.56727 | 44.79349 | 19.59946 | 0.43704 | 1.08713 | 0.712664 |
| 16 | 26948800 | 26954799 | 76.11145 | 119.8304 | 136.273 | 56.54069 | 0.136784 | 2.510022 | 0.712676 |
| 1 | 109252200 | 109257799 | 60.39994 | 96.45376 | 106.8923 | 44.88166 | 0.192767 | 2.110915 | 0.712689 |
| X | 144852400 | 144856399 | 44.22163 | 71.57701 | 79.58764 | 34.04287 | 0.210415 | 1.683593 | 0.712692 |
| 11 | 21425200 | 21428399 | 36.47188 | 59.81569 | 67.21328 | 28.93698 | 0.209083 | 1.452497 | 0.712693 |
| 7 | 138257200 | 138260399 | 21.79491 | 36.60119 | 55.387 | 24.18573 | -0.21458 | 1.134856 | 0.7127 |
| 4 | 16248000 | 16249399 | 15.41784 | 26.81835 | 42.20792 | 18.54498 | -0.26401 | 0.767803 | 0.712704 |
| 17 | 55551800 | 55552399 | 9.231709 | 17.7052 | 21.25677 | 8.355145 | 0.260878 | -0.07439 | 0.712708 |
| 13 | 120339800 | 120341599 | 16.12135 | 28.15395 | 42.0218 | 18.50947 | -0.18754 | 0.711933 | 0.712713 |
| X | 3350400 | 3352199 | 22.1764 | 39.22792 | 28.37515 | 12.54959 | 0.855902 | 0.556068 | 0.712726 |
| 11 | 112965000 | 112967599 | 22.92788 | 38.44201 | 57.04876 | 24.88613 | -0.18654 | 1.157303 | 0.712743 |
| 7 | 103698800 | 103700199 | 15.61864 | 27.85144 | 31.75038 | 13.96758 | 0.208801 | 0.457209 | 0.712746 |
| 16 | 35190600 | 35192799 | 23.51697 | 40.0724 | 46.35986 | 20.32194 | 0.172623 | 1.002444 | 0.71277 |
| 13 | 113872200 | 113874199 | 26.74722 | 44.38792 | 64.85485 | 28.10384 | -0.16443 | 1.288097 | 0.712774 |
| 2 | 78147000 | 78149999 | 58.33693 | 92.95741 | 106.2122 | 44.4611 | 0.145432 | 2.177487 | 0.712792 |
| 2 | 5747200 | 5748399 | 16.08438 | 28.85089 | 32.32083 | 14.14831 | 0.238871 | 0.427059 | 0.7128 |
| 2 | 36296400 | 36299999 | 62.86851 | 100.0239 | 113.6127 | 47.45762 | 0.151568 | 2.233721 | 0.712808 |
| 6 | 73316800 | 73318799 | 25.53753 | 42.63393 | 62.71147 | 27.28229 | -0.17383 | 1.215858 | 0.712826 |
| 9 | 100883400 | 100887599 | 23.78678 | 41.29465 | 37.68295 | 16.53647 | 0.518299 | 0.82619 | 0.712858 |
| 15 | 13107800 | 13113599 | 82.20097 | 128.3442 | 179.69 | 74.55976 | -0.17216 | 2.673932 | 0.71288 |
| X | 7480000 | 7483799 | 44.31045 | 71.79393 | 75.63986 | 32.37857 | 0.289437 | 1.665833 | 0.712897 |
| 5 | 109679400 | 109681399 | 26.11711 | 44.18446 | 50.66914 | 22.17734 | 0.184458 | 1.081136 | 0.712901 |
| 7 | 67925200 | 67926399 | 10.79668 | 19.65739 | 30.75392 | 13.23081 | -0.22353 | 0.299307 | 0.712905 |
| 8 | 25834400 | 25837199 | 27.63035 | 45.69702 | 66.99684 | 28.96359 | -0.16973 | 1.332694 | 0.712918 |
| 3 | 129694800 | 129695599 | 8.258515 | 15.6006 | 25.98501 | 10.57391 | -0.25146 | 0.046472 | 0.71292 |
| 6 | 96681000 | 96683799 | 8.721187 | 15.90724 | 33.45976 | 14.36246 | -0.64511 | 0.281551 | 0.712923 |
| 17 | 47737600 | 47740399 | 15.99586 | 27.7655 | 43.23715 | 18.97218 | -0.24926 | 0.800118 | 0.712925 |
| 2 | 48245800 | 48247799 | 15.58545 | 28.27603 | 30.81249 | 13.31479 | 0.290474 | 0.352508 | 0.712926 |

| | | | | | | | | | |
|---|---|---|---|---|---|---|---|---|---|
| 7 | 23862800 | 23864399 | 22.63086 | 38.84579 | 44.75327 | 19.62926 | 0.180287 | 0.932825 | 0.712949 |
| 9 | 39951800 | 39953199 | 15.52523 | 27.59249 | 31.83389 | 14.04384 | 0.18897 | 0.47854 | 0.71295 |
| 15 | 3136600 | 3139399 | 30.31021 | 50.63052 | 56.9021 | 24.76384 | 0.213152 | 1.218288 | 0.712973 |
| X | 102492200 | 102494799 | 33.41654 | 55.24753 | 61.22796 | 26.46332 | 0.232785 | 1.348976 | 0.713007 |
| 16 | 35988000 | 35988799 | 14.05751 | 25.32571 | 29.41889 | 12.82458 | 0.190911 | 0.380312 | 0.713016 |
| 17 | 42296000 | 42298799 | 21.94655 | 38.51552 | 38.92272 | 17.21348 | 0.372494 | 0.662328 | 0.713022 |
| 17 | 87884400 | 87893999 | 111.3942 | 169.9386 | 227.7861 | 94.45056 | -0.13528 | 3.119298 | 0.713028 |
| 12 | 8262200 | 8263999 | 16.33465 | 30.13259 | 30.13135 | 12.74004 | 0.435446 | 0.266604 | 0.713043 |
| 14 | 11950200 | 11952799 | 33.15662 | 54.96991 | 53.05294 | 22.9739 | 0.431903 | 1.307437 | 0.713043 |
| X | 87720400 | 87722399 | 18.81536 | 34.06914 | 32.34248 | 14.01021 | 0.486451 | 0.392731 | 0.713049 |
| 18 | 90134200 | 90137999 | 35.98194 | 59.12062 | 57.45327 | 24.75428 | 0.419084 | 1.423773 | 0.713068 |
| 10 | 40197400 | 40202999 | 97.20773 | 150.1231 | 166.3345 | 69.09189 | 0.152834 | 2.869776 | 0.713097 |
| 7 | 87714200 | 87714799 | 10.88466 | 20.80176 | 20.63106 | 8.19147 | 0.521259 | -0.02442 | 0.713103 |
| 2 | 92656600 | 92657999 | 17.03613 | 29.94315 | 34.52129 | 15.28649 | 0.183283 | 0.603157 | 0.713106 |
| 3 | 88648800 | 88659999 | 142.6838 | 213.4152 | 285.8554 | 117.8786 | -0.15865 | 3.520476 | 0.713107 |
| 8 | 54529000 | 54531399 | 18.52181 | 31.54759 | 58.13037 | 25.48071 | -0.49647 | 1.00025 | 0.713116 |
| 19 | 42603200 | 42605399 | 32.78505 | 53.43094 | 77.4463 | 33.26221 | -0.16274 | 1.53977 | 0.713121 |
| 7 | 33165400 | 33168199 | 41.28168 | 67.59025 | 70.41653 | 30.2945 | 0.317018 | 1.488366 | 0.713124 |
| 2 | 105708000 | 105720599 | 194.7228 | 290.3084 | 293.3567 | 120.8957 | 0.243407 | 3.589556 | 0.713131 |
| 1 | 58845800 | 58855399 | 101.2296 | 157.502 | 166.715 | 69.20856 | 0.22696 | 2.741215 | 0.713158 |
| 5 | 136725800 | 136729399 | 38.97732 | 62.78794 | 91.02264 | 38.70972 | -0.17917 | 1.810338 | 0.713182 |
| 2 | 112541800 | 112543599 | 13.20104 | 23.39187 | 36.08748 | 15.95503 | -0.23036 | 0.492082 | 0.713183 |
| 2 | 31758800 | 31762599 | 33.67092 | 54.41187 | 81.82842 | 34.9444 | -0.22654 | 1.706665 | 0.713196 |
| 3 | 18090600 | 18092799 | 17.84994 | 31.60881 | 34.72468 | 15.35452 | 0.256525 | 0.526671 | 0.713213 |
| 14 | 71696400 | 71697599 | 16.87818 | 29.62184 | 33.19319 | 14.68009 | 0.224248 | 0.624156 | 0.713227 |
| 5 | 150003200 | 150004999 | 11.62964 | 20.8462 | 32.69276 | 14.3193 | -0.24408 | 0.401538 | 0.713229 |
| 6 | 68387600 | 68390399 | 38.56676 | 63.01211 | 70.53814 | 30.32308 | 0.211568 | 1.510697 | 0.713259 |
| 5 | 52546400 | 52549999 | 39.79276 | 64.13182 | 91.98742 | 39.14658 | -0.16303 | 1.797464 | 0.713266 |
| 13 | 114510000 | 114516199 | 82.6234 | 129.9097 | 144.0582 | 59.76048 | 0.171626 | 2.542067 | 0.713268 |
| 7 | 6200400 | 6244199 | 541.3979 | 727.6719 | 868.2924 | 362.2635 | -0.14421 | 5.197855 | 0.71328 |
| 8 | 15180200 | 15181599 | 15.19368 | 27.11256 | 31.34228 | 13.77993 | 0.189482 | 0.450302 | 0.713341 |
| 3 | 125011000 | 125012599 | 20.33354 | 34.74717 | 51.69656 | 22.66651 | -0.18833 | 0.969677 | 0.713402 |
| 6 | 7258800 | 7262199 | 44.25305 | 71.61089 | 79.2537 | 33.8941 | 0.216798 | 1.686095 | 0.713403 |
| 14 | 70137400 | 70138599 | 10.334 | 19.0391 | 29.84744 | 12.64171 | -0.21054 | 0.227553 | 0.71341 |
| 1 | 104158800 | 104159799 | 14.1081 | 25.41237 | 29.46084 | 12.84571 | 0.193661 | 0.382617 | 0.713421 |
| 15 | 42791600 | 42795799 | 35.23484 | 58.18783 | 63.42874 | 27.39311 | 0.256322 | 1.362778 | 0.713426 |
| X | 76383400 | 76387799 | 48.9871 | 79.05411 | 84.74987 | 36.12799 | 0.258329 | 1.774004 | 0.713466 |
| 3 | 16067400 | 16070399 | 50.91091 | 81.60625 | 93.74941 | 39.57816 | 0.147065 | 1.982664 | 0.713489 |
| 11 | 81965400 | 81968199 | 28.7895 | 47.96996 | 55.6218 | 24.16841 | 0.168398 | 1.241268 | 0.713494 |
| 5 | 119508200 | 119509399 | 13.72951 | 24.22328 | 36.69741 | 16.23873 | -0.20659 | 0.545453 | 0.713519 |
| 10 | 25643800 | 25645599 | 14.33573 | 25.10637 | 39.52597 | 17.4495 | -0.2636 | 0.657856 | 0.713525 |
| 13 | 67459400 | 67472799 | 159.8279 | 242.1643 | 237.8651 | 98.43344 | 0.304837 | 3.271599 | 0.71353 |
| X | 103660400 | 103662799 | 18.27462 | 31.68511 | 46.86988 | 20.56429 | -0.17591 | 0.82682 | 0.713573 |
| 1 | 147467400 | 147467999 | 14.09786 | 25.26929 | 28.53876 | 12.50037 | 0.226821 | 0.41078 | 0.713578 |
| 5 | 86886600 | 86886999 | 4.931669 | 9.636752 | 18.28642 | 6.351355 | -0.22674 | -0.55879 | 0.713597 |
| 19 | 13896600 | 13910999 | 217.6406 | 315.853 | 405.7875 | 167.2954 | -0.13954 | 4.027652 | 0.713617 |
| X | 38145800 | 38146799 | 14.40081 | 26.53831 | 29.13127 | 12.33418 | 0.30047 | 0.256566 | 0.713653 |

| | | | | | | | | | |
|---|---|---|---|---|---|---|---|---|---|
| 10 | 103098000 | 103102399 | 107.9029 | 165.6492 | 184.735 | 76.69745 | 0.136897 | 2.989499 | 0.713671 |
| 3 | 41607600 | 41610399 | 31.22432 | 52.69974 | 54.10481 | 23.66409 | 0.343357 | 1.121297 | 0.713671 |
| 6 | 122998200 | 123000399 | 21.78105 | 38.11255 | 40.09476 | 17.6877 | 0.314775 | 0.707793 | 0.713678 |
| 18 | 36405800 | 36408599 | 16.22787 | 28.1916 | 51.14662 | 22.4545 | -0.46873 | 0.800506 | 0.713686 |
| 14 | 5527200 | 5531799 | 98.10436 | 151.308 | 165.6237 | 68.79714 | 0.168883 | 2.89458 | 0.713687 |
| 6 | 111173000 | 111174199 | 13.31756 | 25.00584 | 27.31122 | 11.23539 | 0.33902 | 0.130241 | 0.71369 |
| 16 | 30972800 | 30978199 | 109.3518 | 166.3984 | 226.6555 | 93.91622 | -0.16141 | 3.171167 | 0.713724 |
| 4 | 124050600 | 124055199 | 36.95708 | 59.49014 | 100.8414 | 42.84007 | -0.40562 | 1.835658 | 0.713725 |
| 7 | 132936600 | 132938999 | 42.02497 | 68.10742 | 77.16691 | 33.03815 | 0.185033 | 1.652527 | 0.713727 |
| 7 | 34153200 | 34154599 | 14.08715 | 24.86462 | 37.20272 | 16.46096 | -0.18843 | 0.545049 | 0.713729 |
| 14 | 91410600 | 91414799 | 39.22951 | 63.52819 | 73.04595 | 31.25612 | 0.162842 | 1.660744 | 0.713734 |
| 11 | 77985400 | 77985999 | 5.670864 | 11.0196 | 20.6255 | 7.683607 | -0.2995 | -0.29272 | 0.71375 |
| 2 | 152223400 | 152224199 | 10.04975 | 18.77765 | 30.05118 | 12.4657 | -0.21788 | 0.139744 | 0.713751 |
| 2 | 47441200 | 47442999 | 18.20006 | 32.26156 | 29.18482 | 12.90229 | 0.535986 | 0.509335 | 0.713755 |
| 3 | 64727000 | 64731399 | 45.39617 | 73.55549 | 68.84209 | 29.46942 | 0.459812 | 1.658248 | 0.713758 |
| 8 | 87276800 | 87277999 | 11.66172 | 21.5931 | 25.04702 | 10.53649 | 0.22776 | 0.195731 | 0.71376 |
| 5 | 125686200 | 125697799 | 174.2463 | 257.8843 | 336.4178 | 138.5468 | -0.13459 | 3.714436 | 0.713778 |
| 2 | 103817000 | 103818599 | 12.72284 | 22.86953 | 34.64247 | 15.13758 | -0.19214 | 0.395377 | 0.713782 |
| 14 | 45370200 | 45372399 | 12.72284 | 22.86953 | 34.64247 | 15.13758 | -0.19214 | 0.395377 | 0.713782 |
| 1 | 17491400 | 17493199 | 19.19783 | 33.41048 | 38.36623 | 16.86246 | 0.188466 | 0.769956 | 0.713786 |
| 13 | 56558200 | 56565399 | 81.72155 | 126.8742 | 173.9353 | 72.22931 | -0.14819 | 2.770102 | 0.713798 |
| Y | 10229200 | 10231599 | 12.88232 | 23.38786 | 22.22962 | 9.597707 | 0.487463 | 0.309644 | 0.713819 |
| 14 | 92005400 | 92007799 | 24.68234 | 41.93456 | 48.4916 | 21.25135 | 0.172644 | 1.035825 | 0.713834 |
| 9 | 18156400 | 18157599 | 18.63366 | 32.97338 | 35.17058 | 15.55388 | 0.298442 | 0.539228 | 0.713869 |
| 7 | 35779000 | 35781199 | 26.31559 | 43.46666 | 64.57129 | 27.89023 | -0.18877 | 1.34184 | 0.713897 |
| 13 | 42723000 | 42724399 | 14.99141 | 26.20454 | 40.48302 | 17.84317 | -0.23688 | 0.68956 | 0.713898 |
| 8 | 73972600 | 73981999 | 398.7861 | 547.0794 | 683.6198 | 284.3174 | -0.18097 | 4.87371 | 0.713916 |
| 5 | 7672200 | 7674199 | 31.932 | 53.00181 | 47.13411 | 20.42738 | 0.549719 | 1.285466 | 0.713919 |
| 13 | 84080600 | 84081199 | 10.82897 | 20.47675 | 24.81437 | 10.07618 | 0.206134 | 0.058711 | 0.713932 |
| X | 84710800 | 84713599 | 28.859 | 48.76233 | 52.89092 | 23.14105 | 0.264446 | 1.104469 | 0.713933 |
| 6 | 79283800 | 79284799 | 19.94996 | 33.99534 | 50.59486 | 22.17842 | -0.18928 | 0.995042 | 0.713946 |
| 6 | 12277200 | 12279599 | 27.72318 | 45.77171 | 67.07609 | 28.96343 | -0.16933 | 1.350731 | 0.713949 |
| 6 | 130225400 | 130238599 | 148.594 | 225.7328 | 229.2906 | 94.96027 | 0.259569 | 3.215696 | 0.71395 |
| 10 | 45361600 | 45363199 | 17.64633 | 30.45755 | 45.84752 | 20.11295 | -0.20225 | 0.863644 | 0.71395 |
| 7 | 110607000 | 110609599 | 35.11613 | 56.866 | 82.03291 | 35.10116 | -0.16361 | 1.660104 | 0.713962 |
| 9 | 55545200 | 55551999 | 79.27386 | 125.22 | 135.9794 | 56.47113 | 0.206974 | 2.443119 | 0.713967 |
| 18 | 28535400 | 28536199 | 10.62143 | 20.01006 | 24.22731 | 9.910579 | 0.198223 | 0.079457 | 0.713985 |
| 4 | 38855400 | 38864799 | 214.9864 | 310.8449 | 409.6156 | 168.9538 | -0.18138 | 4.082513 | 0.713995 |
| 12 | 62585800 | 62588599 | 27.45199 | 46.71294 | 43.06824 | 18.87695 | 0.498493 | 1.024923 | 0.714014 |
| 16 | 3050000 | 3078399 | 1967.55 | 2401.222 | 2109.164 | 889.6363 | 0.157681 | 6.755201 | 0.714018 |
| 10 | 54883000 | 54885599 | 26.56372 | 45.42615 | 48.46866 | 21.25246 | 0.289607 | 0.983294 | 0.714025 |
| 12 | 12170200 | 12170599 | 8.175575 | 15.87061 | 20.55572 | 7.720744 | 0.215095 | -0.23397 | 0.71406 |
| 12 | 19593200 | 19594599 | 37.13115 | 60.71512 | 56.79258 | 24.42123 | 0.47083 | 1.487211 | 0.714077 |
| X | 16555600 | 16566599 | 143.9602 | 219.3231 | 220.0637 | 91.21749 | 0.280637 | 3.156601 | 0.714084 |
| 6 | 35452000 | 35455399 | 59.22755 | 94.23491 | 105.0829 | 43.89704 | 0.178164 | 2.224952 | 0.714102 |
| 2 | 3141800 | 3144999 | 24.52699 | 41.03768 | 60.45635 | 26.338 | -0.17607 | 1.186732 | 0.714105 |
| 5 | 55131200 | 55132999 | 17.63135 | 30.03745 | 55.31421 | 24.24671 | -0.49529 | 0.994761 | 0.714107 |

| | | | | | | | | | |
|---|---|---|---|---|---|---|---|---|---|
| 1 | 54304200 | 54305399 | 11.54868 | 20.69668 | 32.78957 | 14.36427 | -0.25849 | 0.405221 | 0.714108 |
| X | 55088600 | 55093599 | 81.61451 | 128.9341 | 135.3217 | 56.2017 | 0.256237 | 2.443361 | 0.714119 |
| 18 | 39731600 | 39732199 | 9.688141 | 17.84026 | 28.84134 | 12.2239 | -0.25528 | 0.222112 | 0.714131 |
| 12 | 116470800 | 116471599 | 12.47153 | 22.83811 | 26.1278 | 11.1581 | 0.23235 | 0.269329 | 0.714154 |
| 12 | 109873800 | 109882399 | 118.2239 | 179.2468 | 240.459 | 99.5275 | -0.1437 | 3.247726 | 0.714165 |
| 14 | 89101400 | 89106999 | 62.45369 | 99.7265 | 108.4665 | 45.53814 | 0.219669 | 2.116419 | 0.714167 |
| 7 | 29330400 | 29331199 | 12.17313 | 22.96054 | 26.33048 | 10.74895 | 0.278041 | 0.08982 | 0.71418 |
| 2 | 83940200 | 83942199 | 18.39004 | 32.54624 | 35.24181 | 15.585 | 0.276977 | 0.537656 | 0.714186 |
| 15 | 95446600 | 95449799 | 34.70599 | 57.3137 | 63.82 | 27.56183 | 0.225742 | 1.363772 | 0.714196 |
| 13 | 3218800 | 3243999 | 458.709 | 629.4195 | 630.7672 | 262.3272 | 0.137619 | 4.871371 | 0.71421 |
| 14 | 8134200 | 8135799 | 54.47444 | 86.30964 | 128.029 | 53.2514 | -0.23877 | 2.353191 | 0.714217 |
| 1 | 149888600 | 149889199 | 8.380275 | 15.96519 | 26.11908 | 10.43985 | -0.20327 | -0.01131 | 0.714242 |
| 13 | 60583600 | 60585199 | 15.78059 | 27.58365 | 41.47648 | 18.28084 | -0.1981 | 0.695437 | 0.714263 |
| 10 | 68197000 | 68201599 | 61.94038 | 98.56235 | 112.6656 | 47.07197 | 0.142648 | 2.226498 | 0.714286 |
| 14 | 42116600 | 42120599 | 23.73045 | 41.00999 | 38.80435 | 17.02382 | 0.464951 | 0.867988 | 0.714294 |
| 12 | 8657600 | 8666599 | 116.7193 | 178.9152 | 197.2043 | 81.85757 | 0.152146 | 3.023138 | 0.714331 |
| 1 | 31960400 | 31962999 | 28.39721 | 47.74194 | 54.00778 | 23.59377 | 0.203684 | 1.144816 | 0.714335 |
| 12 | 7191800 | 7194799 | 64.85773 | 102.1853 | 142.9974 | 59.33618 | -0.16131 | 2.490774 | 0.714341 |
| 13 | 103004400 | 103006999 | 26.34601 | 45.58763 | 45.69799 | 20.04889 | 0.38251 | 0.87887 | 0.714348 |
| 15 | 97266000 | 97267599 | 15.63301 | 27.35964 | 40.92306 | 18.05586 | -0.19051 | 0.674034 | 0.714368 |
| 8 | 51212200 | 51213799 | 16.43396 | 29.30925 | 32.69707 | 14.38242 | 0.239936 | 0.463472 | 0.71437 |
| 7 | 103816000 | 103817599 | 14.70854 | 25.80494 | 38.84104 | 17.1728 | -0.19941 | 0.62696 | 0.714386 |
| 10 | 117798800 | 117806799 | 100.6629 | 155.4237 | 173.3457 | 72.00429 | 0.143083 | 2.875634 | 0.714393 |
| 1 | 136900200 | 136902399 | 17.71495 | 30.688 | 45.8111 | 20.09922 | -0.18933 | 0.829031 | 0.714394 |
| 9 | 98332200 | 98334599 | 29.07783 | 47.36786 | 82.03202 | 35.22595 | -0.41927 | 1.541153 | 0.714424 |
| 4 | 12145400 | 12149199 | 71.20366 | 112.9331 | 125.228 | 52.13785 | 0.18177 | 2.333548 | 0.714426 |
| 3 | 35098600 | 35099999 | 10.12749 | 18.80038 | 29.39185 | 12.32122 | -0.19598 | 0.181345 | 0.714428 |
| 10 | 71951000 | 71952999 | 19.02546 | 33.27008 | 38.23035 | 16.85634 | 0.188136 | 0.701794 | 0.714432 |
| 15 | 80867200 | 80868999 | 13.65026 | 23.99765 | 44.51388 | 19.7116 | -0.49936 | 0.597782 | 0.714436 |
| 12 | 54524000 | 54524799 | 12.23248 | 23.11877 | 26.40659 | 10.73442 | 0.289214 | 0.077607 | 0.714442 |
| 4 | 151977400 | 151981999 | 61.98547 | 97.76601 | 139.5328 | 57.90675 | -0.18846 | 2.46548 | 0.714443 |
| 4 | 61645800 | 61647199 | 11.3348 | 21.91741 | 23.93442 | 9.157728 | 0.433327 | -0.13016 | 0.714461 |
| 13 | 91371800 | 91372399 | 5.808379 | 11.29359 | 20.65551 | 7.657635 | -0.26007 | -0.30778 | 0.714468 |
| 12 | 9963200 | 9966199 | 43.20008 | 69.17933 | 114.3522 | 48.20498 | -0.37856 | 2.011138 | 0.714472 |
| 16 | 18952400 | 18953199 | 16.06571 | 28.47556 | 32.844 | 14.52025 | 0.187214 | 0.502729 | 0.714472 |
| 2 | 128131800 | 128134599 | 36.55425 | 59.76617 | 68.63502 | 29.51255 | 0.175533 | 1.494641 | 0.714486 |
| 8 | 15821400 | 15822999 | 11.10159 | 20.58411 | 33.17577 | 13.93004 | -0.24163 | 0.203929 | 0.714486 |
| 8 | 18640800 | 18642999 | 23.08803 | 39.59106 | 45.23636 | 19.83956 | 0.191793 | 0.941105 | 0.714493 |
| 1 | 85824600 | 85828999 | 27.02383 | 44.25936 | 68.68934 | 29.55522 | -0.2562 | 1.449861 | 0.714494 |
| X | 81312000 | 81312999 | 15.72361 | 28.48701 | 24.42455 | 10.5728 | 0.633193 | 0.351255 | 0.714503 |
| 6 | 13423800 | 13426799 | 36.62354 | 60.14528 | 67.10231 | 28.90566 | 0.220182 | 1.438906 | 0.714508 |
| 18 | 88220200 | 88221799 | 17.47218 | 31.26948 | 33.18131 | 14.55267 | 0.314665 | 0.450424 | 0.714537 |
| 13 | 87654800 | 87662999 | 125.1017 | 190.4088 | 211.3048 | 87.5616 | 0.134249 | 3.168376 | 0.714558 |
| 8 | 103713000 | 103715399 | 25.54059 | 43.41735 | 49.56838 | 21.72425 | 0.19115 | 1.035477 | 0.714583 |
| 8 | 60174600 | 60176399 | 19.24615 | 33.66988 | 38.22407 | 16.85944 | 0.205563 | 0.697126 | 0.7146 |
| 10 | 61053800 | 61055599 | 27.14273 | 45.42567 | 52.31633 | 22.79539 | 0.178057 | 1.186907 | 0.714661 |
| 7 | 23274600 | 23292799 | 250.4567 | 366.693 | 356.1636 | 146.7151 | 0.27604 | 3.897694 | 0.714695 |

| | | | | | | | | | |
|---|---|---|---|---|---|---|---|---|---|
| 11 | 29911000 | 29917399 | 64.323 | 102.5057 | 113.1012 | 47.3511 | 0.195885 | 2.18238 | 0.714708 |
| 6 | 35450400 | 35455599 | 73.88384 | 116.4729 | 132.2588 | 54.88436 | 0.140339 | 2.481808 | 0.714715 |
| 1 | 129481000 | 129482199 | 20.14117 | 35.19294 | 33.07363 | 14.57296 | 0.476831 | 0.710974 | 0.714722 |
| 7 | 12568400 | 12575799 | 143.2149 | 217.3025 | 230.8176 | 95.55863 | 0.193728 | 3.238257 | 0.714737 |
| 10 | 122534800 | 122536799 | 19.68868 | 34.20903 | 39.43068 | 17.3126 | 0.182861 | 0.796847 | 0.714753 |
| 15 | 35733600 | 35735799 | 26.99223 | 44.5985 | 65.72459 | 28.39462 | -0.17722 | 1.341633 | 0.714754 |
| 15 | 22395200 | 22397199 | 23.45544 | 40.61937 | 38.28859 | 16.79862 | 0.471058 | 0.848355 | 0.71478 |
| X | 102091600 | 102098999 | 90.89516 | 142.3452 | 153.3573 | 63.61375 | 0.208774 | 2.623377 | 0.714784 |
| 17 | 13245600 | 13247599 | 16.51876 | 28.74552 | 43.17539 | 18.97512 | -0.19696 | 0.76795 | 0.714822 |
| 13 | 74024200 | 74029199 | 77.6938 | 120.8844 | 167.7301 | 69.62983 | -0.16308 | 2.730002 | 0.714843 |
| 6 | 37792200 | 37793999 | 17.91013 | 31.45605 | 36.03743 | 15.94788 | 0.192381 | 0.623229 | 0.714863 |
| 14 | 48004400 | 48006199 | 19.71777 | 33.9962 | 50.63429 | 22.21214 | -0.18756 | 0.88697 | 0.714865 |
| 7 | 52736000 | 52737799 | 22.30658 | 39.259 | 37.72648 | 16.70233 | 0.445416 | 0.624512 | 0.714873 |
| 5 | 127798800 | 127799999 | 18.13558 | 31.65195 | 35.36967 | 15.57332 | 0.22814 | 0.721413 | 0.714887 |
| 5 | 128042200 | 128047399 | 61.19463 | 97.6362 | 108.56 | 45.52465 | 0.186623 | 2.138716 | 0.714887 |
| 1 | 31468800 | 31469999 | 20.38033 | 35.76905 | 32.73532 | 14.47787 | 0.514734 | 0.641625 | 0.71489 |
| 11 | 62843600 | 62846999 | 39.65852 | 63.90748 | 91.15815 | 38.78688 | -0.15524 | 1.801487 | 0.714902 |
| 7 | 123103400 | 123110199 | 99.09497 | 153.4359 | 171.4656 | 71.22229 | 0.143371 | 2.824319 | 0.714912 |
| 3 | 53139600 | 53144199 | 40.94435 | 65.9849 | 94.65046 | 40.2776 | -0.1632 | 1.801734 | 0.714931 |
| 13 | 94150600 | 94151599 | 8.86655 | 16.65406 | 26.94715 | 11.07919 | -0.22261 | 0.090985 | 0.714942 |
| 15 | 7597200 | 7598199 | 8.86655 | 16.65406 | 26.94715 | 11.07919 | -0.22261 | 0.090985 | 0.714942 |
| 9 | 69395800 | 69401999 | 56.4147 | 89.23824 | 147.0147 | 61.09775 | -0.39185 | 2.395164 | 0.714944 |
| 17 | 93596200 | 93597399 | 13.46112 | 23.66907 | 36.84031 | 16.31305 | -0.24715 | 0.582825 | 0.714949 |
| 7 | 31022400 | 31024599 | 37.61186 | 61.27946 | 70.65808 | 30.34353 | 0.166176 | 1.549101 | 0.714956 |
| 8 | 9134600 | 9138399 | 66.51514 | 105.6817 | 118.9224 | 49.59944 | 0.163041 | 2.277044 | 0.714958 |
| 1 | 17511000 | 17512199 | 13.066 | 23.46337 | 35.66977 | 15.59936 | -0.19812 | 0.406169 | 0.714967 |
| 10 | 119123000 | 119125799 | 22.10338 | 37.30173 | 54.88539 | 24.00474 | -0.17399 | 1.092788 | 0.714969 |
| 4 | 113895800 | 113902399 | 187.5306 | 277.364 | 297.8807 | 122.6752 | 0.144946 | 3.725851 | 0.714977 |
| X | 158036800 | 158040399 | 43.09338 | 69.86562 | 75.68026 | 32.41079 | 0.2503 | 1.650312 | 0.715003 |
| Y | 40104000 | 40107399 | 22.69516 | 38.67908 | 36.29466 | 15.91007 | 0.47362 | 0.984707 | 0.715028 |
| 8 | 86552200 | 86552799 | 10.28599 | 19.52234 | 22.8313 | 9.19845 | 0.267561 | 0.025424 | 0.715043 |
| X | 58689800 | 58690399 | 11.4719 | 21.81183 | 24.9402 | 10.01045 | 0.304337 | 0.029642 | 0.715072 |
| 3 | 84298200 | 84300999 | 28.944 | 48.09707 | 54.60732 | 23.68483 | 0.198439 | 1.267756 | 0.715077 |
| 12 | 28432200 | 28433199 | 7.622307 | 14.4516 | 24.98622 | 10.08903 | -0.29514 | 0.010759 | 0.715086 |
| 2 | 137909000 | 137909399 | 4.932676 | 9.635571 | 18.22524 | 6.372354 | -0.23064 | -0.53527 | 0.715097 |
| 1 | 7149200 | 7150999 | 20.44163 | 35.08523 | 52.23808 | 22.91203 | -0.18834 | 0.933586 | 0.715102 |
| 4 | 25558800 | 25561399 | 32.26923 | 52.6991 | 76.48412 | 32.87373 | -0.1625 | 1.508157 | 0.715131 |
| X | 17228200 | 17228799 | 10.59766 | 20.34251 | 24.09604 | 9.447634 | 0.284292 | -0.05673 | 0.71514 |
| 14 | 62248000 | 62249999 | 19.02498 | 33.116 | 37.85658 | 16.64047 | 0.194983 | 0.765557 | 0.715144 |
| 10 | 110700600 | 110700999 | 8.916009 | 17.13273 | 21.38334 | 8.3585 | 0.213644 | -0.09187 | 0.715149 |
| X | 63582800 | 63584399 | 10.38806 | 20.09546 | 18.71633 | 7.146731 | 0.664279 | -0.15918 | 0.715185 |
| 16 | 30303200 | 30304399 | 8.810716 | 16.73161 | 27.15226 | 10.9265 | -0.20066 | 0.01598 | 0.71519 |
| 12 | 56197200 | 56198199 | 10.39126 | 19.14423 | 30.3149 | 12.84003 | -0.22542 | 0.226082 | 0.715209 |
| 6 | 128435800 | 128437999 | 22.43768 | 38.31794 | 43.27216 | 18.97182 | 0.207931 | 0.976247 | 0.715218 |
| 19 | 18894600 | 18902999 | 65.83233 | 104.8586 | 101.3904 | 42.41894 | 0.385238 | 2.195801 | 0.715223 |
| X | 144689400 | 144695999 | 13.87841 | 24.34418 | 45.64038 | 20.17595 | -0.5146 | 0.638388 | 0.715239 |
| 5 | 5854200 | 5857999 | 52.687 | 84.4165 | 96.57144 | 40.74261 | 0.15236 | 1.99863 | 0.715244 |

| | | | | | | | | | |
|---|---|---|---|---|---|---|---|---|---|
| 1 | 12286400 | 12288999 | 27.14603 | 45.47172 | 52.36415 | 22.82907 | 0.17807 | 1.174588 | 0.715251 |
| 5 | 20296600 | 20297399 | 10.51697 | 19.19797 | 30.30735 | 12.99637 | -0.2332 | 0.28253 | 0.715254 |
| 14 | 46633800 | 46636599 | 19.7624 | 33.73918 | 50.06892 | 21.95119 | -0.18473 | 0.976082 | 0.715255 |
| 17 | 83942200 | 83945399 | 27.16631 | 45.45103 | 52.36213 | 22.81089 | 0.177516 | 1.187694 | 0.715262 |
| 7 | 84753800 | 84758599 | 70.23917 | 111.3367 | 125.4853 | 52.21374 | 0.15747 | 2.350438 | 0.715264 |
| 4 | 96542600 | 96543999 | 14.42545 | 25.59308 | 39.18056 | 17.31039 | -0.21836 | 0.497677 | 0.715266 |
| 7 | 41077000 | 41079199 | 16.81447 | 28.95217 | 53.76924 | 23.58642 | -0.50477 | 0.8937 | 0.715274 |
| 12 | 108530600 | 108534399 | 34.69742 | 56.36024 | 81.67564 | 35.02726 | -0.16612 | 1.595439 | 0.715291 |
| 6 | 133529600 | 133549399 | 335.8119 | 475.7801 | 482.8927 | 199.8105 | 0.165856 | 4.387088 | 0.715308 |
| 16 | 41774200 | 41777799 | 56.47327 | 89.71814 | 125.0721 | 52.15924 | -0.14559 | 2.274032 | 0.715309 |
| 13 | 52520600 | 52521399 | 11.53905 | 22.20283 | 24.81485 | 9.66257 | 0.376517 | -0.06861 | 0.715331 |
| 10 | 19884800 | 19894999 | 266.0102 | 379.2953 | 479.5373 | 198.21 | -0.14169 | 4.290851 | 0.715342 |
| 15 | 87312600 | 87318799 | 66.59484 | 104.9028 | 145.5752 | 60.40439 | -0.14946 | 2.494765 | 0.715342 |
| 6 | 145325600 | 145327599 | 20.38994 | 35.25704 | 40.55764 | 17.79331 | 0.184543 | 0.851017 | 0.71536 |
| 7 | 32301000 | 32301599 | 15.11848 | 26.81381 | 30.05776 | 13.28353 | 0.228791 | 0.487198 | 0.715379 |
| 4 | 96188600 | 96198199 | 133.3584 | 203.3133 | 216.4104 | 89.72009 | 0.196305 | 3.138425 | 0.715416 |
| X | 163487000 | 163490199 | 37.66865 | 61.42971 | 70.06772 | 30.10031 | 0.182747 | 1.536637 | 0.71543 |
| 17 | 70105400 | 70106599 | 16.07 | 28.11326 | 42.46193 | 18.72854 | -0.20462 | 0.683886 | 0.715459 |
| 1 | 101867400 | 101868599 | 24.09864 | 40.96542 | 47.43438 | 20.78895 | 0.170908 | 1.028848 | 0.715484 |
| Y | 43797200 | 43798199 | 6.637555 | 12.55083 | 25.02147 | 10.16076 | -0.50581 | 0.02028 | 0.715508 |
| 9 | 71579600 | 71580599 | 13.9152 | 25.32672 | 29.47844 | 12.68742 | 0.199941 | 0.321336 | 0.715511 |
| 15 | 65916000 | 65918599 | 18.64956 | 32.93448 | 35.75052 | 15.81678 | 0.272183 | 0.5623 | 0.715536 |
| 7 | 43466200 | 43468399 | 20.41464 | 35.47389 | 40.3803 | 17.73065 | 0.200733 | 0.798769 | 0.715564 |
| 13 | 67252600 | 67257199 | 47.32356 | 75.65847 | 108.8808 | 45.79078 | -0.18168 | 2.06221 | 0.715582 |
| 7 | 3299600 | 3307399 | 77.12444 | 119.7637 | 169.5409 | 70.40213 | -0.19417 | 2.763554 | 0.715583 |
| 2 | 22071400 | 22075799 | 55.64487 | 88.29104 | 125.7523 | 52.37701 | -0.17827 | 2.309735 | 0.715601 |
| 13 | 43178400 | 43179999 | 19.84707 | 34.13056 | 50.62803 | 22.20764 | -0.18251 | 0.909959 | 0.715618 |
| 4 | 128772800 | 128775199 | 15.03869 | 27.68241 | 24.81686 | 10.52369 | 0.589031 | 0.257238 | 0.715624 |
| X | 9225600 | 9227199 | 24.82043 | 42.80596 | 45.225 | 19.83938 | 0.30614 | 0.902878 | 0.715702 |
| 7 | 27579800 | 27584799 | 122.5937 | 186.4623 | 199.6919 | 82.73224 | 0.184732 | 3.182267 | 0.715706 |
| 17 | 54247200 | 54251199 | 59.4913 | 94.67006 | 107.3445 | 44.85196 | 0.154392 | 2.221757 | 0.715719 |
| X | 90354200 | 90375599 | 262.3465 | 383.5747 | 310.2819 | 127.8259 | 0.537921 | 3.918523 | 0.71574 |
| 3 | 98900600 | 98903399 | 30.43003 | 49.93133 | 72.62375 | 31.27283 | -0.16184 | 1.438429 | 0.715743 |
| 8 | 78104400 | 78106399 | 21.38234 | 36.1774 | 53.82559 | 23.55951 | -0.18986 | 1.065371 | 0.715765 |
| 1 | 115536400 | 115536999 | 10.57924 | 19.5255 | 30.34886 | 12.82312 | -0.19608 | 0.218745 | 0.715783 |
| 19 | 57591800 | 57593999 | 17.50647 | 30.77862 | 35.45027 | 15.6979 | 0.184726 | 0.60355 | 0.715784 |
| 2 | 177020000 | 177022399 | 36.46414 | 59.70318 | 68.03732 | 29.27106 | 0.187947 | 1.475949 | 0.715791 |
| 17 | 13043400 | 13054999 | 146.1467 | 220.6538 | 241.993 | 100.03 | 0.141581 | 3.3382 | 0.715793 |
| 4 | 5053200 | 5053799 | 13.81055 | 25.83944 | 23.38242 | 9.677254 | 0.60176 | 0.148712 | 0.715824 |
| 13 | 86334400 | 86345799 | 333.3081 | 465.2349 | 581.9452 | 241.3721 | -0.15723 | 4.607361 | 0.715843 |
| 19 | 30940600 | 30943599 | 40.85413 | 65.70093 | 93.91525 | 39.84288 | -0.1619 | 1.868842 | 0.71585 |
| 4 | 52679800 | 52681399 | 12.03283 | 21.34214 | 40.31027 | 17.81377 | -0.51998 | 0.485271 | 0.715855 |
| 2 | 93207200 | 93208399 | 10.45459 | 19.43136 | 30.47527 | 12.75413 | -0.19889 | 0.179764 | 0.71587 |
| X | 88691800 | 88693599 | 20.73668 | 36.34998 | 39.04889 | 17.25233 | 0.284768 | 0.674006 | 0.715875 |
| 7 | 32065000 | 32073799 | 88.48635 | 140.0744 | 109.3585 | 45.4654 | 0.684827 | 2.393519 | 0.71591 |
| X | 88804800 | 88806599 | 24.3432 | 42.52937 | 33.79953 | 14.89152 | 0.717679 | 0.737845 | 0.715936 |
| 11 | 88765800 | 88769399 | 49.44572 | 79.02705 | 111.4969 | 46.8707 | -0.1536 | 2.07779 | 0.715962 |

| | | | | | | | | | |
|---|---|---|---|---|---|---|---|---|---|
| 8 | 25308400 | 25311199 | 29.61568 | 49.63569 | 55.6945 | 24.28937 | 0.215438 | 1.182131 | 0.716005 |
| 2 | 87721600 | 87722599 | 25.47916 | 43.03532 | 49.59534 | 21.69667 | 0.177324 | 1.092215 | 0.716006 |
| 17 | 92556200 | 92558399 | 61.00825 | 96.93648 | 107.6291 | 44.8913 | 0.182444 | 2.269902 | 0.716063 |
| 2 | 17987800 | 17992199 | 58.60687 | 93.48209 | 106.7289 | 44.73998 | 0.148095 | 2.144791 | 0.716069 |
| 15 | 86599000 | 86603999 | 58.60562 | 92.85929 | 130.652 | 54.34364 | -0.16262 | 2.353566 | 0.716071 |
| 7 | 88637200 | 88637999 | 8.390188 | 15.9484 | 26.07287 | 10.47075 | -0.20816 | 0.004352 | 0.716089 |
| 9 | 34280400 | 34281399 | 8.390188 | 15.9484 | 26.07287 | 10.47075 | -0.20816 | 0.004352 | 0.716089 |
| 2 | 39420400 | 39423999 | 52.48763 | 83.69001 | 116.8687 | 48.96508 | -0.14257 | 2.154297 | 0.716101 |
| 10 | 104390400 | 104394399 | 45.10963 | 72.59713 | 82.96188 | 35.23296 | 0.161609 | 1.845921 | 0.716104 |
| 5 | 76435800 | 76437399 | 16.00995 | 27.8908 | 42.12243 | 18.52517 | -0.2047 | 0.747621 | 0.716113 |
| 12 | 11160200 | 11161199 | 12.30009 | 22.58689 | 25.98033 | 11.05321 | 0.22807 | 0.249178 | 0.71612 |
| 13 | 10999800 | 11001999 | 23.64826 | 40.5524 | 46.30689 | 20.30909 | 0.192696 | 0.945398 | 0.71613 |
| 1 | 5412600 | 5421799 | 259.7218 | 376.5111 | 343.986 | 141.8344 | 0.350513 | 4.046146 | 0.71619 |
| 2 | 15312800 | 15316399 | 33.76815 | 55.13482 | 82.62854 | 35.51181 | -0.20896 | 1.518141 | 0.716241 |
| 12 | 116244200 | 116245999 | 16.67812 | 28.87169 | 44.29067 | 19.43173 | -0.22848 | 0.830693 | 0.716246 |
| 13 | 86865800 | 86866799 | 11.74923 | 21.9871 | 25.598 | 10.59066 | 0.240769 | 0.132608 | 0.716249 |
| 16 | 48698800 | 48699799 | 13.1116 | 23.3837 | 35.15179 | 15.46233 | -0.18764 | 0.449541 | 0.716288 |
| 10 | 89305800 | 89307999 | 23.29596 | 39.96607 | 45.77001 | 20.07435 | 0.18861 | 0.939047 | 0.716302 |
| 3 | 66016800 | 66021199 | 48.09296 | 77.21151 | 88.46163 | 37.4359 | 0.153653 | 1.927987 | 0.716307 |
| 6 | 143910000 | 143910999 | 13.33474 | 24.46448 | 28.49816 | 12.1395 | 0.208681 | 0.26871 | 0.716314 |
| 12 | 72942000 | 72943999 | 14.97818 | 27.03481 | 30.81849 | 13.40337 | 0.219917 | 0.377974 | 0.716329 |
| 5 | 109277400 | 109285999 | 107.5593 | 166.6652 | 175.2695 | 72.80203 | 0.231765 | 2.81584 | 0.716386 |
| 18 | 37493000 | 37498399 | 56.4869 | 91.1467 | 71.09317 | 30.30182 | 0.715981 | 1.782491 | 0.716392 |
| 9 | 122104600 | 122113199 | 138.6798 | 208.1025 | 276.6493 | 114.1611 | -0.14356 | 3.458436 | 0.716435 |
| 5 | 16605200 | 16605799 | 6.664773 | 12.93261 | 22.82764 | 8.592083 | -0.23064 | -0.23965 | 0.71646 |
| X | 94811200 | 94816799 | 64.73951 | 103.5004 | 106.9298 | 44.96919 | 0.295615 | 2.083747 | 0.716466 |
| 6 | 20117000 | 20121799 | 77.76396 | 121.3998 | 165.8245 | 68.80746 | -0.13673 | 2.672614 | 0.716469 |
| 4 | 18922000 | 18922999 | 10.08049 | 18.87322 | 30.10818 | 12.44664 | -0.20923 | 0.128123 | 0.716478 |
| 14 | 46863000 | 46865599 | 18.10251 | 31.39605 | 46.33891 | 20.33151 | -0.17261 | 0.822555 | 0.716481 |
| 2 | 12643000 | 12644199 | 12.38807 | 22.54013 | 35.40461 | 15.24403 | -0.23038 | 0.323879 | 0.71651 |
| 16 | 61972600 | 61973599 | 17.08565 | 30.27615 | 26.87295 | 11.88102 | 0.563016 | 0.50019 | 0.716512 |
| 11 | 52053000 | 52054999 | 30.72111 | 49.90747 | 75.42659 | 32.35021 | -0.22624 | 1.584709 | 0.716513 |
| 10 | 35846800 | 35848999 | 30.76059 | 51.49692 | 49.16527 | 21.42672 | 0.447674 | 1.190173 | 0.716515 |
| 7 | 63492200 | 63493999 | 27.89304 | 47.89331 | 40.26614 | 17.6616 | 0.632947 | 0.94475 | 0.716518 |
| 9 | 87573200 | 87574399 | 11.2974 | 20.66825 | 31.81271 | 13.60357 | -0.19444 | 0.278646 | 0.716532 |
| 3 | 135748200 | 135749999 | 18.90059 | 32.71257 | 48.47534 | 21.26754 | -0.17906 | 0.846738 | 0.716536 |
| 17 | 84547400 | 84549999 | 22.343 | 38.71539 | 36.75532 | 16.1262 | 0.460889 | 0.834743 | 0.716544 |
| 6 | 48729400 | 48730199 | 10.63242 | 19.57509 | 30.33774 | 12.86171 | -0.19524 | 0.234406 | 0.716546 |
| 6 | 139498400 | 139499999 | 16.4579 | 29.44504 | 32.83004 | 14.40241 | 0.243605 | 0.446898 | 0.716573 |
| 9 | 4265400 | 4268599 | 46.51988 | 75.05685 | 72.87706 | 31.05834 | 0.400513 | 1.767135 | 0.716587 |
| 4 | 126954600 | 126955199 | 5.523527 | 10.76098 | 20.23843 | 7.366013 | -0.27606 | -0.38035 | 0.716596 |
| 12 | 21413200 | 21415399 | 15.64372 | 27.44042 | 40.83297 | 18.0506 | -0.18314 | 0.637607 | 0.716607 |
| 9 | 5075200 | 5076599 | 25.47973 | 42.91113 | 49.0796 | 21.4521 | 0.188068 | 1.116848 | 0.716611 |
| 13 | 32770400 | 32774999 | 51.06948 | 81.6828 | 117.1677 | 49.30631 | -0.17632 | 2.057993 | 0.716612 |
| 6 | 149140400 | 149149999 | 132.0775 | 200.8336 | 220.5298 | 91.35812 | 0.148359 | 3.190459 | 0.716624 |
| 9 | 110128800 | 110132399 | 58.60781 | 93.42685 | 106.892 | 44.77072 | 0.14411 | 2.164198 | 0.716674 |
| X | 131555400 | 131556599 | 19.16579 | 33.53359 | 37.26249 | 16.4371 | 0.236439 | 0.694617 | 0.716677 |

| | | | | | | | | | |
|---|---|---|---|---|---|---|---|---|---|
| X | 94504600 | 94508599 | 36.10142 | 59.77695 | 63.38581 | 27.42464 | 0.296402 | 1.337996 | 0.716684 |
| 3 | 108840400 | 108846399 | 82.95431 | 128.9898 | 175.8278 | 72.99813 | -0.13821 | 2.743585 | 0.716704 |
| 8 | 122562000 | 122566799 | 93.39385 | 143.487 | 197.0814 | 81.83207 | -0.1625 | 2.970493 | 0.716706 |
| 15 | 20013400 | 20014199 | 24.06058 | 40.69724 | 45.91542 | 20.09549 | 0.207911 | 1.069705 | 0.716709 |
| 3 | 103639000 | 103640399 | 13.96499 | 24.51186 | 37.9068 | 16.76635 | -0.23805 | 0.615244 | 0.716732 |
| 8 | 44177400 | 44178599 | 8.370218 | 15.98228 | 26.17369 | 10.41156 | -0.19871 | -0.02718 | 0.716778 |
| 11 | 19022200 | 19024399 | 22.52349 | 38.02531 | 56.02943 | 24.50795 | -0.1761 | 1.090354 | 0.71678 |
| 11 | 55728000 | 55728399 | 10.82894 | 19.71261 | 30.7789 | 13.2445 | -0.22075 | 0.301682 | 0.716802 |
| 7 | 32086600 | 32092599 | 62.77415 | 100.8058 | 80.0133 | 33.87303 | 0.682526 | 1.937637 | 0.716807 |
| 17 | 88032000 | 88039799 | 77.7394 | 121.2937 | 165.9585 | 68.867 | -0.13978 | 2.68153 | 0.716817 |
| 2 | 142145200 | 142155799 | 163.9467 | 243.9411 | 319.7375 | 131.7386 | -0.13429 | 3.620185 | 0.716838 |
| 18 | 53571000 | 53572399 | 19.47919 | 34.17513 | 37.98607 | 16.79499 | 0.235706 | 0.650779 | 0.716838 |
| 10 | 57642800 | 57643399 | 5.245297 | 10.21554 | 19.6321 | 7.176392 | -0.31189 | -0.36913 | 0.71684 |
| 15 | 77730600 | 77734599 | 26.21452 | 43.11676 | 65.93697 | 28.42202 | -0.23246 | 1.395148 | 0.716844 |
| 2 | 34715800 | 34718199 | 21.09023 | 35.7785 | 52.73248 | 23.09973 | -0.17609 | 1.04137 | 0.716903 |
| 5 | 107397400 | 107399999 | 21.09023 | 35.7785 | 52.73248 | 23.09973 | -0.17609 | 1.04137 | 0.716903 |
| 13 | 77933600 | 77947999 | 176.3758 | 264.5001 | 271.8428 | 112.1585 | 0.226638 | 3.478194 | 0.716949 |
| 6 | 88140600 | 88147199 | 57.48958 | 91.03062 | 130.2208 | 54.1461 | -0.18726 | 2.368498 | 0.716958 |
| 11 | 109403800 | 109404799 | 16.47196 | 28.97616 | 32.99685 | 14.60984 | 0.201361 | 0.586521 | 0.716968 |
| X | 5992200 | 5996199 | 94.79718 | 147.0519 | 163.0526 | 67.71854 | 0.156686 | 2.79028 | 0.717006 |
| 18 | 75115600 | 75118199 | 23.21949 | 39.81421 | 45.22753 | 19.83559 | 0.200116 | 0.941686 | 0.717014 |
| 14 | 119393000 | 119394599 | 12.35493 | 22.31345 | 34.17401 | 14.85151 | -0.20284 | 0.363131 | 0.717019 |
| 14 | 66146800 | 66148999 | 28.79684 | 48.35507 | 54.54858 | 23.81813 | 0.207707 | 1.158775 | 0.717025 |
| 1 | 51754800 | 51757199 | 11.46801 | 20.21408 | 40.41522 | 17.88726 | -0.60506 | 0.549916 | 0.717038 |
| X | 143068200 | 143071599 | 26.17694 | 43.55539 | 64.76283 | 28.11473 | -0.18927 | 1.258575 | 0.71704 |
| 4 | 153433200 | 153434399 | 14.09409 | 24.85397 | 37.22758 | 16.47499 | -0.19049 | 0.555437 | 0.717042 |
| 18 | 82282200 | 82285199 | 31.67549 | 52.50263 | 60.26019 | 26.09089 | 0.182564 | 1.310366 | 0.717045 |
| 3 | 8453400 | 8456399 | 78.52571 | 123.0671 | 137.2538 | 56.9295 | 0.15978 | 2.599621 | 0.717052 |
| 18 | 6720600 | 6722599 | 17.73529 | 30.34257 | 54.3076 | 23.81358 | -0.4535 | 0.954129 | 0.717054 |
| 16 | 20770200 | 20775199 | 49.26607 | 79.34745 | 88.30054 | 37.54216 | 0.201189 | 1.830741 | 0.717076 |
| X | 13122200 | 13124399 | 22.55152 | 38.84886 | 44.70223 | 19.60937 | 0.182984 | 0.899124 | 0.717085 |
| 11 | 17879200 | 17883399 | 62.15838 | 98.68485 | 141.8524 | 59.11815 | -0.1907 | 2.302399 | 0.717099 |
| 6 | 11666600 | 11667399 | 10.81985 | 20.28655 | 24.21639 | 9.986723 | 0.209339 | 0.112263 | 0.717102 |
| 1 | 33717200 | 33720599 | 38.33815 | 61.87216 | 88.4675 | 37.71518 | -0.1562 | 1.754444 | 0.717108 |
| 3 | 11437000 | 11440199 | 31.98005 | 53.138 | 59.77825 | 25.92621 | 0.211666 | 1.282631 | 0.717118 |
| 11 | 39934200 | 39935399 | 18.20961 | 31.99451 | 36.47349 | 16.14618 | 0.199413 | 0.618848 | 0.717123 |
| 3 | 126801000 | 126802999 | 15.09772 | 27.24722 | 30.77788 | 13.38777 | 0.232824 | 0.378852 | 0.717147 |
| 2 | 116371800 | 116393999 | 408.7874 | 563.1023 | 688.5616 | 286.1872 | -0.14375 | 4.810431 | 0.717156 |
| 3 | 140533400 | 140534799 | 17.75394 | 31.31758 | 35.39638 | 15.66363 | 0.213722 | 0.569941 | 0.71719 |
| 13 | 29476600 | 29477599 | 9.135117 | 17.22483 | 27.52025 | 11.24115 | -0.19723 | 0.069011 | 0.717203 |
| 5 | 146884600 | 146885999 | 9.774526 | 18.12196 | 28.84519 | 12.1131 | -0.22369 | 0.182941 | 0.717219 |
| 4 | 91458800 | 91460599 | 19.05534 | 33.71025 | 35.09638 | 15.52192 | 0.332978 | 0.543745 | 0.71722 |
| 12 | 38396000 | 38398799 | 19.05534 | 33.71025 | 35.09638 | 15.52192 | 0.332978 | 0.543745 | 0.71722 |
| 3 | 16103400 | 16103999 | 10.50316 | 20.00559 | 24.42057 | 9.764968 | 0.215105 | 0.004967 | 0.717222 |
| 10 | 41509800 | 41510599 | 10.50316 | 20.00559 | 24.42057 | 9.764968 | 0.215105 | 0.004967 | 0.717222 |
| 19 | 8382800 | 8392799 | 121.8697 | 184.8244 | 246.3349 | 101.9678 | -0.13402 | 3.243022 | 0.717227 |
| 7 | 110834800 | 110838799 | 35.18609 | 57.89912 | 65.46125 | 28.22236 | 0.202133 | 1.408532 | 0.717235 |

| | | | | | | | | | |
|---|---|---|---|---|---|---|---|---|---|
| 13 | 43896200 | 43897999 | 28.75809 | 48.02969 | 55.65615 | 24.21901 | 0.169217 | 1.215886 | 0.717239 |
| 3 | 77140800 | 77141599 | 7.915632 | 15.13066 | 25.09426 | 9.958271 | -0.21345 | -0.04181 | 0.717249 |
| Y | 44010600 | 44013199 | 14.92197 | 26.52384 | 23.21705 | 10.24129 | 0.585089 | 0.453503 | 0.717268 |
| 15 | 84416200 | 84427999 | 116.7451 | 176.6786 | 275.7653 | 114.0748 | -0.36428 | 3.28613 | 0.717278 |
| 1 | 4025600 | 4026999 | 15.78414 | 28.56132 | 31.3055 | 13.57238 | 0.2791 | 0.370124 | 0.71728 |
| 7 | 76481800 | 76483399 | 26.8299 | 43.74842 | 79.90443 | 34.32447 | -0.49474 | 1.522205 | 0.717287 |
| 9 | 122707000 | 122708999 | 16.14496 | 28.33613 | 42.89412 | 18.97108 | -0.2078 | 0.634256 | 0.717291 |
| 14 | 120198600 | 120201799 | 41.82746 | 67.21976 | 95.67572 | 40.54912 | -0.15699 | 1.892307 | 0.717297 |
| 7 | 127819000 | 127819999 | 12.16975 | 22.15903 | 34.87105 | 15.00062 | -0.23231 | 0.315859 | 0.717305 |
| X | 77089200 | 77092399 | 46.37797 | 75.18498 | 78.71451 | 33.70832 | 0.299211 | 1.658463 | 0.717305 |
| 14 | 13448800 | 13452399 | 35.36386 | 57.34852 | 82.64861 | 35.40164 | -0.16036 | 1.630269 | 0.717327 |
| 7 | 114161800 | 114162599 | 7.920008 | 14.97476 | 25.46496 | 10.34237 | -0.27897 | 0.03364 | 0.717348 |
| 14 | 95857000 | 95857599 | 8.86326 | 16.93307 | 28.4951 | 11.32173 | -0.23633 | -0.02014 | 0.71738 |
| 12 | 109899400 | 109902799 | 60.32111 | 95.40953 | 133.9372 | 55.66052 | -0.16139 | 2.397818 | 0.717412 |
| 5 | 100317000 | 100319199 | 7.942212 | 15.01378 | 30.93848 | 12.57065 | -0.55501 | 0.048361 | 0.717418 |
| 15 | 92617200 | 92624999 | 76.61675 | 121.6222 | 108.8598 | 45.36178 | 0.491535 | 2.3101 | 0.717427 |
| 15 | 72697800 | 72699399 | 21.55013 | 37.1378 | 43.14508 | 18.92652 | 0.169336 | 0.889469 | 0.71746 |
| 6 | 85864200 | 85872599 | 97.35739 | 150.9306 | 168.6457 | 70.04785 | 0.144898 | 2.800431 | 0.717488 |
| 16 | 44714200 | 44714799 | 8.868623 | 16.76587 | 27.04364 | 10.98698 | -0.20373 | 0.049163 | 0.717511 |
| 7 | 116604800 | 116606399 | 12.59203 | 23.25548 | 26.99275 | 11.39501 | 0.223014 | 0.221546 | 0.717523 |
| 17 | 71731200 | 71732199 | 11.5947 | 21.46798 | 25.13079 | 10.57242 | 0.214856 | 0.197224 | 0.717525 |
| 2 | 128445800 | 128449599 | 42.8216 | 69.91832 | 72.67993 | 31.23638 | 0.317557 | 1.531759 | 0.717527 |
| 12 | 115350000 | 115352599 | 28.17208 | 47.47795 | 45.33901 | 19.82125 | 0.447324 | 1.113214 | 0.717528 |
| 12 | 107739400 | 107742599 | 29.92213 | 49.18073 | 71.54494 | 30.83025 | -0.16113 | 1.415567 | 0.71754 |
| 5 | 19028000 | 19029599 | 27.637 | 46.01708 | 50.72704 | 22.03174 | 0.241188 | 1.24171 | 0.717544 |
| 3 | 13858800 | 13861399 | 34.63564 | 57.76077 | 49.22164 | 21.3991 | 0.611324 | 1.245766 | 0.717593 |
| 17 | 89282800 | 89284199 | 18.63881 | 32.74016 | 36.92722 | 16.34347 | 0.214665 | 0.625124 | 0.717598 |
| X | 151305400 | 151307599 | 23.49975 | 40.36388 | 45.79053 | 20.08497 | 0.202616 | 0.932201 | 0.717611 |
| X | 77124600 | 77127999 | 35.69141 | 59.3009 | 61.61594 | 26.72202 | 0.326021 | 1.298242 | 0.717632 |
| 10 | 69192200 | 69194599 | 32.55044 | 53.01581 | 76.98826 | 33.05805 | -0.1661 | 1.545708 | 0.717636 |
| 9 | 90735800 | 90737399 | 15.37407 | 27.10829 | 41.27565 | 18.26684 | -0.21434 | 0.570083 | 0.717636 |
| 7 | 41391200 | 41407199 | 44.09276 | 71.32069 | 69.55305 | 29.73599 | 0.398332 | 1.690697 | 0.717651 |
| 18 | 48841200 | 48843199 | 17.48289 | 31.0075 | 34.18323 | 15.10662 | 0.252439 | 0.507848 | 0.717653 |
| 7 | 61184800 | 61185399 | 10.88563 | 19.84572 | 31.32748 | 13.45517 | -0.23447 | 0.296318 | 0.717666 |
| X | 15524000 | 15527399 | 44.15971 | 71.69609 | 76.9319 | 32.98131 | 0.265775 | 1.623072 | 0.717674 |
| 19 | 8149200 | 8152599 | 58.33388 | 93.14564 | 105.8966 | 44.45764 | 0.155846 | 2.112288 | 0.71769 |
| 12 | 24508400 | 24515799 | 62.09211 | 98.26161 | 135.9189 | 56.49886 | -0.13953 | 2.386876 | 0.717699 |
| 7 | 42156800 | 42162199 | 65.61069 | 104.2941 | 116.7907 | 48.73346 | 0.170714 | 2.261223 | 0.71772 |
| 4 | 27003600 | 27006199 | 22.70656 | 39.30254 | 43.55091 | 19.10709 | 0.238416 | 0.853995 | 0.717735 |
| 10 | 50403800 | 50404399 | 15.16541 | 27.06202 | 30.97827 | 13.61994 | 0.203816 | 0.451757 | 0.717747 |
| 5 | 145848000 | 145857799 | 192.9698 | 286.072 | 306.9796 | 126.4348 | 0.149251 | 3.687684 | 0.717768 |
| 8 | 91107400 | 91109399 | 29.94328 | 49.54628 | 57.23135 | 24.75012 | 0.173268 | 1.324485 | 0.717769 |
| 8 | 126650200 | 126651799 | 17.19556 | 30.26899 | 34.94989 | 15.47254 | 0.181865 | 0.587854 | 0.71777 |
| 7 | 17169600 | 17172599 | 47.73463 | 77.00899 | 85.22995 | 36.31809 | 0.211932 | 1.77979 | 0.717777 |
| 5 | 6471200 | 6471999 | 13.30105 | 24.10611 | 27.57644 | 11.93214 | 0.220039 | 0.338074 | 0.717782 |
| 13 | 64198800 | 64201199 | 29.55375 | 48.83723 | 73.69466 | 31.84077 | -0.21141 | 1.350434 | 0.717796 |
| 6 | 130247200 | 130248799 | 14.01116 | 26.45675 | 27.51651 | 11.20676 | 0.421235 | 0.102808 | 0.717799 |

| | | | | | | | | | |
|---|---|---|---|---|---|---|---|---|---|
| 17 | 20291600 | 20299999 | 102.6403 | 159.8425 | 165.8273 | 68.83067 | 0.256939 | 2.727679 | 0.717801 |
| 14 | 14188400 | 14192799 | 44.11071 | 70.64355 | 101.8265 | 42.93027 | -0.18116 | 2.004804 | 0.717826 |
| 19 | 35331400 | 35332399 | 10.53689 | 19.58017 | 31.00547 | 12.97977 | -0.21328 | 0.1808 | 0.717854 |
| 13 | 50400200 | 50401599 | 11.09348 | 20.18428 | 31.2268 | 13.44548 | -0.20839 | 0.30588 | 0.717864 |
| 3 | 57262400 | 57264599 | 25.19317 | 43.35795 | 46.81554 | 20.53581 | 0.274097 | 0.921 | 0.717906 |
| 8 | 74344000 | 74347199 | 42.63317 | 69.10176 | 77.40529 | 33.14328 | 0.201608 | 1.650431 | 0.717916 |
| 9 | 58226600 | 58230199 | 31.26384 | 51.18262 | 74.29071 | 31.9598 | -0.1602 | 1.469595 | 0.717927 |
| 19 | 27215400 | 27215999 | 6.092835 | 11.84474 | 21.21597 | 7.875795 | -0.23245 | -0.29875 | 0.717928 |
| 9 | 105480600 | 105492399 | 154.7325 | 230.7179 | 303.2697 | 124.9841 | -0.1356 | 3.580803 | 0.717934 |
| 6 | 66853800 | 66854199 | 4.90984 | 9.593985 | 18.26665 | 6.346416 | -0.23111 | -0.55553 | 0.717937 |
| 8 | 75506000 | 75524399 | 312.4395 | 444.2026 | 460.8131 | 190.5639 | 0.139339 | 4.33501 | 0.71794 |
| 9 | 100512200 | 100515399 | 29.23872 | 48.84263 | 56.24291 | 24.47757 | 0.178252 | 1.215369 | 0.717941 |
| 9 | 101663800 | 101665399 | 13.62663 | 24.26734 | 36.66612 | 16.14798 | -0.19617 | 0.465236 | 0.717949 |
| 17 | 15357600 | 15358399 | 6.848374 | 13.18395 | 23.08779 | 8.977018 | -0.26328 | -0.1254 | 0.717962 |
| 12 | 119956600 | 119958799 | 22.04752 | 36.98732 | 65.82448 | 28.72478 | -0.44781 | 1.157415 | 0.717968 |
| 3 | 69057000 | 69058999 | 15.02066 | 26.41154 | 39.23053 | 17.37142 | -0.18027 | 0.594842 | 0.717969 |
| 16 | 83547400 | 83549199 | 15.88441 | 29.25778 | 30.08034 | 12.74465 | 0.393374 | 0.270339 | 0.718001 |
| 14 | 83663800 | 83665799 | 36.78518 | 60.10647 | 69.0801 | 29.69722 | 0.173799 | 1.505362 | 0.718005 |
| 15 | 27505200 | 27507399 | 15.49181 | 27.13114 | 40.39738 | 17.83426 | -0.18394 | 0.661881 | 0.718034 |
| 5 | 101947800 | 101950399 | 33.43157 | 54.87459 | 63.91147 | 27.52506 | 0.158033 | 1.434759 | 0.718079 |
| 11 | 57857600 | 57860399 | 27.05564 | 44.84672 | 65.44646 | 28.33707 | -0.16286 | 1.302383 | 0.718135 |
| 13 | 7227200 | 7229399 | 21.04083 | 36.61209 | 40.81236 | 17.93579 | 0.230951 | 0.782683 | 0.718147 |
| 14 | 88122800 | 88123799 | 16.8823 | 29.61608 | 32.28563 | 14.27342 | 0.263839 | 0.630025 | 0.718158 |
| 7 | 108728800 | 108729999 | 9.05203 | 16.91926 | 27.42754 | 11.37075 | -0.23483 | 0.125978 | 0.718175 |
| 9 | 89927400 | 89931399 | 52.8337 | 84.60184 | 96.96238 | 40.87102 | 0.148698 | 2.016843 | 0.71818 |
| 12 | 14628000 | 14633799 | 54.54653 | 87.62119 | 93.76168 | 39.71226 | 0.252891 | 1.919499 | 0.718181 |
| 17 | 33954200 | 33962399 | 84.08373 | 130.8736 | 179.0959 | 74.34411 | -0.14277 | 2.728197 | 0.718185 |
| 16 | 29331200 | 29336399 | 51.67162 | 82.30201 | 116.38 | 48.68915 | -0.16224 | 2.187127 | 0.718191 |
| 7 | 14432000 | 14433599 | 22.75955 | 39.30188 | 43.66743 | 19.15684 | 0.234155 | 0.880393 | 0.718211 |
| 10 | 54993000 | 54994199 | 10.70716 | 19.88493 | 31.54445 | 13.21593 | -0.21668 | 0.18685 | 0.718211 |
| 6 | 131685000 | 131686599 | 36.97985 | 60.32979 | 69.48588 | 29.85458 | 0.169132 | 1.528819 | 0.718231 |
| 8 | 79372600 | 79373999 | 19.71365 | 34.18243 | 39.0057 | 17.1139 | 0.196942 | 0.82085 | 0.718245 |
| 14 | 45988600 | 45990999 | 23.10779 | 39.39098 | 45.33471 | 19.87321 | 0.180284 | 0.997605 | 0.718248 |
| 15 | 9498800 | 9504599 | 115.0332 | 176.9334 | 169.5151 | 70.39696 | 0.357903 | 2.952896 | 0.718256 |
| 10 | 81527000 | 81529199 | 41.28749 | 67.18186 | 64.7436 | 27.79062 | 0.422995 | 1.570405 | 0.718256 |
| 18 | 49645200 | 49645999 | 10.29675 | 19.73867 | 24.04003 | 9.461699 | 0.238913 | -0.04982 | 0.718294 |
| 14 | 97398600 | 97399199 | 6.087317 | 11.85033 | 21.33836 | 7.833593 | -0.22543 | -0.33832 | 0.718307 |
| 11 | 65547600 | 65551999 | 84.55398 | 132.16 | 149.8601 | 62.17539 | 0.132874 | 2.65181 | 0.718324 |
| 14 | 105992400 | 105996999 | 54.27909 | 86.85609 | 98.95038 | 41.6671 | 0.156213 | 2.039811 | 0.718345 |
| 3 | 99273000 | 99273999 | 16.46402 | 28.72545 | 42.56597 | 18.73813 | -0.17727 | 0.728262 | 0.718371 |
| 10 | 78909800 | 78916599 | 102.3874 | 159.6014 | 163.0559 | 67.67066 | 0.280127 | 2.712187 | 0.718396 |
| 1 | 4074400 | 4075399 | 14.74493 | 26.25422 | 29.98367 | 13.20698 | 0.205234 | 0.459155 | 0.718403 |
| 18 | 5803600 | 5807399 | 38.00064 | 61.34265 | 87.93187 | 37.4988 | -0.15942 | 1.746194 | 0.71841 |
| 19 | 56986600 | 56987999 | 14.87384 | 26.47538 | 30.41666 | 13.40126 | 0.196347 | 0.462147 | 0.718456 |
| 11 | 13348800 | 13349999 | 16.99153 | 29.86668 | 34.48103 | 15.26938 | 0.181226 | 0.601072 | 0.718463 |
| 2 | 98895000 | 98895399 | 7.68734 | 14.92775 | 19.45867 | 7.289279 | 0.209412 | -0.25323 | 0.718482 |
| 15 | 41048400 | 41048999 | 9.101772 | 17.26977 | 28.26606 | 11.395 | -0.21539 | 0.026432 | 0.718496 |

| | | | | | | | | | |
|---|---|---|---|---|---|---|---|---|---|
| 8 | 93581800 | 93583399 | 15.00494 | 26.3349 | 39.29771 | 17.38026 | -0.18709 | 0.622625 | 0.718505 |
| 1 | 151023600 | 151024399 | 11.94074 | 22.28515 | 26.09821 | 10.84715 | 0.227125 | 0.152895 | 0.718536 |
| 15 | 94906000 | 94907599 | 12.02802 | 21.95586 | 34.39518 | 14.74965 | -0.22281 | 0.300081 | 0.718539 |
| 18 | 74464200 | 74466399 | 32.34391 | 53.10811 | 60.56773 | 26.0889 | 0.188441 | 1.423682 | 0.718552 |
| 16 | 44159000 | 44160999 | 17.31455 | 30.67862 | 34.20977 | 15.12508 | 0.235374 | 0.515639 | 0.718566 |
| 13 | 35322200 | 35322999 | 9.203844 | 17.38249 | 28.11096 | 11.44439 | -0.21012 | 0.062059 | 0.718572 |
| 10 | 121050000 | 121050599 | 9.851431 | 18.83254 | 23.40738 | 9.286578 | 0.199286 | -0.02933 | 0.718608 |
| 16 | 11100000 | 11102599 | 19.54484 | 34.00137 | 39.39359 | 17.30984 | 0.175639 | 0.778729 | 0.71863 |
| 18 | 44395200 | 44398599 | 117.8127 | 179.824 | 199.9069 | 82.90052 | 0.135011 | 3.108386 | 0.718634 |
| 4 | 113062400 | 113074799 | 169.3337 | 254.9554 | 259.2713 | 107.086 | 0.247008 | 3.3992 | 0.718641 |
| 1 | 38621600 | 38626599 | 48.0231 | 76.79013 | 108.2499 | 45.53663 | -0.15165 | 2.058614 | 0.718649 |
| 13 | 13300600 | 13319799 | 64.24399 | 102.6086 | 95.05321 | 39.91962 | 0.451184 | 2.108096 | 0.718654 |
| X | 163677400 | 163677999 | 11.18385 | 21.13152 | 25.3368 | 10.3051 | 0.219356 | 0.068905 | 0.718669 |
| 10 | 3985000 | 3987599 | 19.90279 | 34.05706 | 50.07357 | 21.9573 | -0.17091 | 0.955428 | 0.718674 |
| 17 | 18729800 | 18754999 | 402.1147 | 562.4229 | 548.4419 | 227.4038 | 0.204865 | 4.580077 | 0.718712 |
| 1 | 90422800 | 90428599 | 62.26899 | 99.1182 | 113.0694 | 47.26245 | 0.146104 | 2.216252 | 0.718713 |
| 5 | 96192400 | 96196599 | 37.42179 | 62.30634 | 60.11225 | 26.10715 | 0.432823 | 1.277468 | 0.71872 |
| 2 | 103899800 | 103903399 | 45.086 | 72.34509 | 101.9441 | 43.10633 | -0.14527 | 1.946192 | 0.718729 |
| 13 | 77135000 | 77146799 | 208.4368 | 302.7925 | 394.9889 | 162.8268 | -0.16016 | 4.011616 | 0.718739 |
| 12 | 83100000 | 83102799 | 19.0228 | 33.37901 | 31.40781 | 13.88812 | 0.474897 | 0.635082 | 0.718742 |
| 12 | 94061400 | 94062399 | 12.55247 | 22.57435 | 34.13191 | 14.90833 | -0.18906 | 0.389965 | 0.718773 |
| 6 | 130390400 | 130396599 | 85.39299 | 134.4842 | 143.3897 | 59.49579 | 0.230003 | 2.511357 | 0.718787 |
| 4 | 120270800 | 120272599 | 18.09468 | 31.40947 | 46.33375 | 20.33129 | -0.17162 | 0.81401 | 0.718791 |
| 19 | 21212600 | 21215599 | 31.91876 | 52.07373 | 75.22169 | 32.31913 | -0.15676 | 1.520476 | 0.718798 |
| 12 | 14647600 | 14651999 | 94.32972 | 146.0098 | 162.8522 | 67.64465 | 0.145798 | 2.827477 | 0.718802 |
| 4 | 61935400 | 61939599 | 48.51996 | 78.07633 | 88.72747 | 37.6751 | 0.169431 | 1.858567 | 0.718828 |
| 13 | 117452800 | 117456199 | 39.89142 | 64.85591 | 73.76131 | 31.65173 | 0.183785 | 1.584894 | 0.718845 |
| 10 | 109311200 | 109312599 | 16.55127 | 29.13324 | 33.53272 | 14.84531 | 0.186428 | 0.582606 | 0.718852 |
| 12 | 29968600 | 29969199 | 9.842802 | 18.84841 | 23.44921 | 9.257379 | 0.204527 | -0.04548 | 0.718866 |
| 2 | 99697400 | 99697999 | 6.542451 | 12.69647 | 22.22891 | 8.362974 | -0.2175 | -0.24121 | 0.718916 |
| 17 | 48274600 | 48277999 | 30.19921 | 49.18043 | 84.64779 | 36.34539 | -0.41076 | 1.548367 | 0.718925 |
| 2 | 150339600 | 150342999 | 38.99243 | 62.91717 | 89.74865 | 38.2528 | -0.15308 | 1.761095 | 0.718962 |
| 10 | 40897200 | 40899599 | 22.87691 | 39.07076 | 45.30469 | 19.86305 | 0.169848 | 0.978502 | 0.719011 |
| 3 | 35425000 | 35425799 | 9.289746 | 17.54531 | 28.11261 | 11.44424 | -0.19714 | 0.060521 | 0.719019 |
| 9 | 11838400 | 11840799 | 19.95148 | 35.03771 | 37.88314 | 16.76321 | 0.275333 | 0.637506 | 0.719025 |
| 8 | 109641800 | 109644399 | 20.41088 | 34.81974 | 51.13819 | 22.41861 | -0.17 | 0.986716 | 0.719033 |
| 1 | 27715600 | 27716799 | 14.09522 | 25.65246 | 29.3701 | 12.64196 | 0.223197 | 0.320792 | 0.719066 |
| X | 120963400 | 120965799 | 26.94868 | 46.73974 | 45.13761 | 19.80445 | 0.436819 | 0.861386 | 0.719105 |
| 3 | 17189800 | 17190799 | 18.34037 | 32.03609 | 36.81245 | 16.21709 | 0.188143 | 0.713275 | 0.71915 |
| 13 | 15446600 | 15448799 | 21.44059 | 37.02748 | 42.56322 | 18.67248 | 0.185098 | 0.867749 | 0.719164 |
| 6 | 136549800 | 136555799 | 55.0899 | 88.47485 | 95.65946 | 40.50282 | 0.237656 | 1.930125 | 0.719171 |
| 13 | 10916400 | 10916799 | 7.396367 | 14.3844 | 18.97869 | 7.023413 | 0.208348 | -0.3007 | 0.719172 |
| 1 | 38303400 | 38305599 | 18.41233 | 31.80892 | 46.87741 | 20.56533 | -0.17163 | 0.859135 | 0.719195 |
| 15 | 84028600 | 84035399 | 56.34383 | 89.51852 | 124.8366 | 52.06455 | -0.14599 | 2.272496 | 0.719204 |
| 5 | 37835200 | 37847999 | 160.249 | 240.7305 | 261.2313 | 107.8301 | 0.15048 | 3.440519 | 0.719217 |
| 17 | 66784200 | 66784999 | 7.42408 | 14.34438 | 24.33117 | 9.343278 | -0.2012 | -0.15789 | 0.719251 |
| 8 | 76442600 | 76444599 | 29.94036 | 49.73329 | 57.86221 | 25.0898 | 0.163313 | 1.280682 | 0.719305 |

| | | | | | | | | | |
|---|---|---|---|---|---|---|---|---|---|
| 6 | 24474600 | 24476999 | 21.27756 | 36.46748 | 55.54584 | 24.36002 | -0.22143 | 0.953453 | 0.719306 |
| 17 | 11757400 | 11758999 | 14.27365 | 25.9569 | 29.97961 | 12.91694 | 0.210044 | 0.330256 | 0.719317 |
| 1 | 179290400 | 179294599 | 45.91306 | 73.65729 | 103.8948 | 43.91765 | -0.14704 | 1.954257 | 0.719325 |
| 5 | 86840200 | 86840599 | 4.937473 | 9.629912 | 18.6253 | 6.681006 | -0.29532 | -0.44197 | 0.719362 |
| X | 149595200 | 149597399 | 15.14139 | 26.53256 | 40.46466 | 17.87258 | -0.2181 | 0.6542 | 0.719365 |
| 12 | 119789000 | 119791399 | 44.24834 | 71.05009 | 100.3279 | 42.46755 | -0.14709 | 1.922821 | 0.719377 |
| 18 | 66685400 | 66687199 | 14.99912 | 26.34042 | 39.26594 | 17.37498 | -0.18565 | 0.612744 | 0.71941 |
| 19 | 54877600 | 54879199 | 14.99912 | 26.34042 | 39.26594 | 17.37498 | -0.18565 | 0.612744 | 0.71941 |
| 9 | 87367000 | 87370199 | 36.58715 | 59.31536 | 86.15368 | 36.89523 | -0.17198 | 1.641863 | 0.719428 |
| 2 | 23875200 | 23876399 | 13.86692 | 24.64642 | 37.10279 | 16.3676 | -0.19278 | 0.477862 | 0.719434 |
| 11 | 9324200 | 9326599 | 20.522 | 35.94007 | 39.01511 | 17.22381 | 0.269588 | 0.684985 | 0.719521 |
| 12 | 113955400 | 113956599 | 15.49546 | 27.35886 | 43.39319 | 19.19884 | -0.27236 | 0.558387 | 0.719522 |
| 1 | 15099000 | 15101399 | 19.46467 | 34.48768 | 35.07271 | 15.50666 | 0.36758 | 0.530589 | 0.719537 |
| 4 | 31020800 | 31021399 | 7.435038 | 14.19753 | 24.53618 | 9.758868 | -0.27558 | -0.04299 | 0.719549 |
| 6 | 49510400 | 49511199 | 12.22849 | 21.73033 | 34.30552 | 15.13597 | -0.26054 | 0.466932 | 0.719559 |
| 14 | 3698600 | 3701799 | 46.07556 | 74.51399 | 81.48135 | 34.83016 | 0.233299 | 1.705554 | 0.71957 |
| 5 | 41799000 | 41801599 | 41.95235 | 68.32445 | 74.12095 | 31.82608 | 0.253529 | 1.566086 | 0.719578 |
| 4 | 71921600 | 71923199 | 22.47416 | 38.82892 | 44.13067 | 19.36037 | 0.201529 | 0.871101 | 0.71959 |
| 13 | 120718600 | 120721399 | 36.34766 | 59.25682 | 68.43693 | 29.39638 | 0.164439 | 1.534154 | 0.719595 |
| 7 | 42278400 | 42292799 | 250.3901 | 364.1288 | 378.7278 | 156.106 | 0.167957 | 3.998943 | 0.719619 |
| 16 | 79795000 | 79798399 | 44.2186 | 70.96227 | 100.8013 | 42.63132 | -0.15662 | 1.93971 | 0.719619 |
| 5 | 104946400 | 104966399 | 546.5935 | 740.2955 | 730.256 | 304.3195 | 0.141422 | 5.07745 | 0.719647 |
| 5 | 20197200 | 20200799 | 30.33967 | 50.99898 | 55.207 | 24.11642 | 0.266965 | 1.153009 | 0.719674 |
| 18 | 88368200 | 88372399 | 42.55616 | 69.19705 | 75.33446 | 32.32829 | 0.246789 | 1.589577 | 0.719677 |
| 8 | 105507600 | 105511599 | 52.75513 | 84.63801 | 81.49301 | 34.45068 | 0.402844 | 1.952205 | 0.719678 |
| 1 | 25981000 | 25983799 | 28.94948 | 48.54436 | 55.1133 | 24.04377 | 0.198543 | 1.174029 | 0.71969 |
| 4 | 20407800 | 20411799 | 64.14258 | 101.1422 | 141.1941 | 58.59424 | -0.15689 | 2.472843 | 0.719697 |
| 10 | 69427800 | 69428599 | 11.72526 | 21.84139 | 26.14099 | 10.89999 | 0.19252 | 0.16478 | 0.719717 |
| 4 | 20826600 | 20828999 | 60.40066 | 95.36451 | 136.4624 | 56.65858 | -0.19103 | 2.441539 | 0.719725 |
| 11 | 13876600 | 13879799 | 61.95358 | 98.39212 | 110.9575 | 46.25772 | 0.159372 | 2.284524 | 0.719726 |
| 8 | 47492400 | 47493999 | 19.82793 | 34.82209 | 37.91178 | 16.77644 | 0.26542 | 0.636775 | 0.719751 |
| 3 | 122905400 | 122908199 | 42.25628 | 67.50202 | 118.03 | 49.58119 | -0.4634 | 2.086437 | 0.719752 |
| 1 | 93032200 | 93037599 | 49.6735 | 79.77629 | 91.05 | 38.55131 | 0.159736 | 1.919428 | 0.719791 |
| 17 | 22094400 | 22111599 | 215.8923 | 320.9261 | 303.6916 | 125.1152 | 0.33417 | 3.644836 | 0.71983 |
| 5 | 63805600 | 63808399 | 20.235 | 33.59744 | 68.11408 | 29.52635 | -0.63528 | 1.266582 | 0.719838 |
| 9 | 80199200 | 80199999 | 8.730066 | 16.71052 | 28.57762 | 11.29569 | -0.25231 | -0.03738 | 0.719848 |
| 4 | 113162200 | 113170399 | 118.2879 | 183.2988 | 148.893 | 61.84597 | 0.603977 | 2.816603 | 0.719867 |
| 13 | 72607400 | 72614199 | 52.12965 | 83.87537 | 91.43653 | 38.81901 | 0.229069 | 1.867802 | 0.71987 |
| 5 | 55903200 | 55904599 | 12.63367 | 23.59135 | 27.1823 | 11.2872 | 0.25157 | 0.158357 | 0.719878 |
| X | 19326800 | 19328799 | 20.21608 | 35.74861 | 36.25229 | 16.0345 | 0.37079 | 0.560737 | 0.719885 |
| 7 | 117652000 | 117660199 | 92.45163 | 142.7846 | 194.7022 | 80.87525 | -0.14631 | 2.869763 | 0.719893 |
| 4 | 121135200 | 121137199 | 18.80368 | 33.53681 | 32.94451 | 14.4908 | 0.422397 | 0.478086 | 0.719904 |
| 7 | 11926200 | 11932599 | 54.16109 | 87.39733 | 86.14319 | 36.71846 | 0.379158 | 1.782923 | 0.719908 |
| 4 | 11321600 | 11322399 | 8.063125 | 15.27389 | 25.48158 | 10.30828 | -0.24659 | 0.022184 | 0.719913 |
| 6 | 111788600 | 111789999 | 18.7379 | 33.61081 | 27.49714 | 12.03668 | 0.689902 | 0.43641 | 0.71992 |
| 8 | 69956200 | 69963399 | 147.8186 | 222.5505 | 244.8178 | 101.115 | 0.133661 | 3.394506 | 0.719935 |
| 14 | 92642600 | 92650999 | 87.73081 | 137.102 | 154.0476 | 63.91388 | 0.14587 | 2.657513 | 0.719938 |

| | | | | | | | | | |
|---|---|---|---|---|---|---|---|---|---|
| 7 | 59008400 | 59014399 | 69.48093 | 110.2055 | 123.7008 | 51.50427 | 0.164283 | 2.329836 | 0.719954 |
| 7 | 7355800 | 7362599 | 87.83067 | 137.691 | 133.6817 | 55.44616 | 0.359892 | 2.598549 | 0.719975 |
| 1 | 171813400 | 171817999 | 44.02528 | 71.59492 | 75.6202 | 32.4524 | 0.290493 | 1.590544 | 0.719978 |
| 3 | 34532600 | 34533999 | 12.72152 | 22.67299 | 34.60731 | 15.23133 | -0.21024 | 0.44797 | 0.719981 |
| 16 | 20727400 | 20729199 | 12.72152 | 22.67299 | 34.60731 | 15.23133 | -0.21024 | 0.44797 | 0.719981 |
| 3 | 74859200 | 74860199 | 11.42556 | 21.53081 | 25.79458 | 10.54688 | 0.213745 | 0.088615 | 0.719989 |
| 5 | 63134000 | 63138799 | 100.9914 | 154.5507 | 209.3852 | 86.88342 | -0.14711 | 3.052216 | 0.720002 |
| 3 | 138148200 | 138149199 | 9.337755 | 17.37096 | 27.87507 | 11.64921 | -0.22984 | 0.158625 | 0.720007 |
| 1 | 8547000 | 8547999 | 10.8711 | 20.60193 | 24.81891 | 10.031 | 0.220117 | 0.04203 | 0.720036 |
| 17 | 91857800 | 91860399 | 21.27423 | 36.86748 | 42.578 | 18.68093 | 0.179507 | 0.838684 | 0.720046 |
| 13 | 52640200 | 52644199 | 48.63844 | 78.14787 | 90.01137 | 38.136 | 0.147347 | 1.905969 | 0.72006 |
| 7 | 31513200 | 31514399 | 26.02512 | 44.4015 | 38.59548 | 16.92011 | 0.583899 | 0.994282 | 0.720061 |
| 18 | 89882800 | 89884799 | 22.55731 | 38.83939 | 44.13778 | 19.36147 | 0.200637 | 0.900678 | 0.720061 |
| 14 | 27138000 | 27141599 | 31.67482 | 51.96547 | 77.69562 | 33.45458 | -0.2017 | 1.444545 | 0.720062 |
| 2 | 150226200 | 150235799 | 124.2136 | 190.1769 | 203.9199 | 84.62475 | 0.190529 | 3.049003 | 0.720074 |
| 12 | 80570200 | 80573599 | 25.27437 | 41.89192 | 62.46327 | 27.04441 | -0.19373 | 1.295749 | 0.720076 |
| 11 | 7982600 | 7984799 | 23.75309 | 39.8416 | 58.17588 | 25.38423 | -0.16328 | 1.157221 | 0.720089 |
| 8 | 101637000 | 101638599 | 19.80247 | 34.57327 | 39.18693 | 17.25787 | 0.207518 | 0.728171 | 0.720094 |
| 2 | 92705400 | 92706999 | 15.49736 | 27.12529 | 40.4254 | 17.83798 | -0.18522 | 0.67141 | 0.72014 |
| 5 | 12040000 | 12044199 | 48.88113 | 78.42047 | 89.62695 | 37.8876 | 0.155948 | 1.951122 | 0.720143 |
| X | 158920600 | 158921599 | 13.5854 | 24.85643 | 28.10348 | 12.0159 | 0.248144 | 0.288746 | 0.720156 |
| 17 | 41741200 | 41741799 | 9.084539 | 17.44995 | 21.90854 | 8.573118 | 0.203696 | -0.08494 | 0.720157 |
| 4 | 8733000 | 8735999 | 25.86363 | 43.00429 | 63.22825 | 27.43542 | -0.17326 | 1.260654 | 0.720158 |
| 2 | 105708600 | 105712999 | 82.39228 | 129.8354 | 141.4156 | 58.68122 | 0.199942 | 2.497382 | 0.720167 |
| 14 | 55531000 | 55533199 | 25.71024 | 43.74976 | 49.02707 | 21.48875 | 0.218023 | 1.024268 | 0.72019 |
| 1 | 124013200 | 124013599 | 8.096253 | 15.39922 | 25.55095 | 10.2477 | -0.22747 | -0.0034 | 0.720191 |
| 19 | 25372600 | 25374799 | 21.72428 | 37.3055 | 43.19014 | 18.94424 | 0.173298 | 0.922127 | 0.720229 |
| 14 | 47673800 | 47679199 | 63.44433 | 100.9084 | 115.2274 | 48.11127 | 0.143425 | 2.243571 | 0.720243 |
| 14 | 55444600 | 55446599 | 21.59458 | 36.72233 | 53.82632 | 23.59103 | -0.1679 | 1.023647 | 0.720247 |
| 3 | 132227600 | 132229399 | 19.60218 | 33.5262 | 49.53872 | 21.72191 | -0.17812 | 0.958614 | 0.720254 |
| 4 | 98042000 | 98047999 | 43.03301 | 68.76736 | 119.4822 | 50.21662 | -0.45374 | 2.075741 | 0.720306 |
| 6 | 132683200 | 132685199 | 21.68141 | 37.87308 | 40.6825 | 17.92227 | 0.284755 | 0.733499 | 0.720317 |
| 16 | 17528200 | 17529799 | 14.52437 | 25.53325 | 38.23253 | 16.93025 | -0.19193 | 0.592376 | 0.720393 |
| 2 | 152255600 | 152266599 | 89.76074 | 138.8115 | 188.7605 | 78.40666 | -0.14086 | 2.844072 | 0.720393 |
| 5 | 19262200 | 19264199 | 17.45052 | 30.87265 | 28.34358 | 12.53528 | 0.51371 | 0.523181 | 0.720428 |
| 1 | 49455000 | 49459999 | 80.92318 | 127.1169 | 143.6737 | 59.59366 | 0.143119 | 2.558781 | 0.72043 |
| 18 | 31347000 | 31347999 | 10.72872 | 20.08275 | 23.64838 | 9.779581 | 0.22541 | 0.118834 | 0.720432 |
| 15 | 18459200 | 18459599 | 4.930851 | 9.637702 | 18.35989 | 6.333563 | -0.22356 | -0.58277 | 0.720432 |
| 19 | 13585800 | 13592799 | 88.69338 | 138.9888 | 134.8446 | 55.93075 | 0.360493 | 2.606331 | 0.720435 |
| 12 | 83052800 | 83053999 | 13.34332 | 23.78567 | 36.09095 | 15.88179 | -0.20169 | 0.453331 | 0.720441 |
| 6 | 68631400 | 68640999 | 163.7571 | 245.8806 | 264.1139 | 109.0057 | 0.164554 | 3.450033 | 0.72045 |
| 1 | 165743600 | 165745199 | 14.39854 | 26.35862 | 29.41546 | 12.56748 | 0.266945 | 0.289823 | 0.720451 |
| 3 | 36007200 | 36011599 | 29.24445 | 48.0965 | 69.81049 | 30.09068 | -0.15753 | 1.404703 | 0.720453 |
| 16 | 86908800 | 86909599 | 9.231603 | 17.31876 | 27.99346 | 11.53299 | -0.22333 | 0.106156 | 0.720477 |
| 1 | 11063400 | 11065999 | 19.32757 | 33.0673 | 50.61658 | 22.19511 | -0.22896 | 0.952546 | 0.720479 |
| 7 | 86303600 | 86304999 | 14.00447 | 25.59539 | 29.4661 | 12.61633 | 0.220349 | 0.29727 | 0.720489 |
| 1 | 118129400 | 118132999 | 34.99648 | 57.64931 | 64.94689 | 28.0134 | 0.207866 | 1.39426 | 0.7205 |

| | | | | | | | | | |
|---|---|---|---|---|---|---|---|---|---|
| 4 | 134912400 | 134914199 | 20.76776 | 36.31125 | 33.5583 | 14.79443 | 0.500953 | 0.706298 | 0.720516 |
| 5 | 94005600 | 94006199 | 14.24044 | 25.47145 | 29.004 | 12.7266 | 0.21325 | 0.425299 | 0.720519 |
| 14 | 38016000 | 38018599 | 29.68961 | 49.69416 | 55.24809 | 24.07468 | 0.228624 | 1.19197 | 0.720524 |
| 15 | 10395800 | 10397799 | 18.24238 | 32.78589 | 33.21203 | 14.51444 | 0.38487 | 0.430891 | 0.720531 |
| 17 | 77798800 | 77801599 | 24.8479 | 42.37389 | 48.48148 | 21.25331 | 0.188717 | 1.002805 | 0.720535 |
| 15 | 87773600 | 87781999 | 109.8847 | 168.6732 | 188.887 | 78.42006 | 0.130817 | 2.992662 | 0.720535 |
| 5 | 57743200 | 57744799 | 17.44204 | 30.31297 | 44.73369 | 19.644 | -0.17196 | 0.791781 | 0.720546 |
| 17 | 67343800 | 67345799 | 17.77989 | 31.19557 | 36.01763 | 15.92543 | 0.181118 | 0.636715 | 0.720564 |
| 7 | 18189200 | 18206999 | 311.0139 | 442.8111 | 454.8906 | 188.0693 | 0.155554 | 4.31483 | 0.720566 |
| 1 | 178050600 | 178059599 | 292.8822 | 413.9078 | 518.9148 | 214.8055 | -0.14249 | 4.422119 | 0.720594 |
| 10 | 68148600 | 68149399 | 9.73138 | 18.64245 | 22.87515 | 9.020444 | 0.225383 | -0.05386 | 0.720595 |
| 7 | 39145200 | 39149399 | 54.42055 | 87.54058 | 92.56473 | 39.28446 | 0.27257 | 1.878415 | 0.720612 |
| 4 | 125515400 | 125518999 | 31.92486 | 52.06545 | 75.17057 | 32.29305 | -0.15637 | 1.52559 | 0.720705 |
| 2 | 46703000 | 46704999 | 20.14156 | 35.69443 | 35.64095 | 15.75724 | 0.394139 | 0.535304 | 0.720708 |
| 12 | 86436000 | 86437599 | 17.0416 | 29.93459 | 34.56862 | 15.29959 | 0.180939 | 0.613133 | 0.72071 |
| 9 | 8687400 | 8688199 | 12.14256 | 21.77857 | 33.19666 | 14.53271 | -0.20266 | 0.403871 | 0.720712 |
| 4 | 143123000 | 143124199 | 9.629992 | 17.90331 | 28.36394 | 11.86618 | -0.2129 | 0.165718 | 0.720726 |
| 1 | 60669400 | 60671599 | 21.49757 | 37.50098 | 41.32975 | 18.19188 | 0.247839 | 0.754169 | 0.720735 |
| 5 | 107168200 | 107169599 | 14.87099 | 26.36032 | 29.05461 | 12.84474 | 0.252495 | 0.486423 | 0.720764 |
| 14 | 76503000 | 76504799 | 17.7668 | 31.08625 | 35.69139 | 15.74424 | 0.189253 | 0.680774 | 0.720769 |
| 16 | 94827800 | 94831399 | 38.51595 | 62.67085 | 71.99206 | 30.90173 | 0.170217 | 1.568558 | 0.72077 |
| X | 88932400 | 88934799 | 22.94966 | 39.9663 | 42.4333 | 18.65775 | 0.301301 | 0.787602 | 0.720789 |
| 7 | 107515400 | 107517199 | 30.4608 | 51.09763 | 55.79373 | 24.34633 | 0.25453 | 1.173422 | 0.720809 |
| Y | 21039400 | 21040399 | 9.65584 | 18.30948 | 18.25751 | 7.370063 | 0.49255 | 0.007813 | 0.720811 |
| 10 | 96034400 | 96038199 | 34.91657 | 56.6044 | 81.15533 | 34.75352 | -0.15332 | 1.636528 | 0.720823 |
| 4 | 59625800 | 59626799 | 7.504838 | 14.22843 | 25.01413 | 10.10098 | -0.31856 | 0.012296 | 0.720837 |
| 3 | 66955000 | 66966199 | 166.1051 | 249.9981 | 260.0132 | 107.3814 | 0.214086 | 3.405704 | 0.72085 |
| 1 | 65164200 | 65165799 | 10.81087 | 19.64231 | 30.75611 | 13.26622 | -0.22712 | 0.311714 | 0.720854 |
| 12 | 17151800 | 17153999 | 17.62519 | 30.87509 | 35.63985 | 15.73727 | 0.181521 | 0.661045 | 0.720856 |
| 5 | 73738800 | 73742799 | 40.16454 | 64.74922 | 91.97098 | 39.1538 | -0.1485 | 1.791001 | 0.72086 |
| 11 | 52481600 | 52489799 | 90.9624 | 141.5193 | 159.8052 | 66.34065 | 0.133693 | 2.736058 | 0.720873 |
| 15 | 38951600 | 38952999 | 19.81935 | 34.76192 | 38.07433 | 16.82992 | 0.256988 | 0.659693 | 0.720875 |
| 8 | 37011600 | 37013399 | 20.91145 | 36.28965 | 42.0384 | 18.44497 | 0.175522 | 0.823428 | 0.720878 |
| X | 90545000 | 90548599 | 35.99972 | 58.84459 | 67.78483 | 29.14428 | 0.170899 | 1.498473 | 0.720901 |
| 5 | 143157000 | 143160799 | 68.65795 | 108.0026 | 148.5459 | 61.62588 | -0.13814 | 2.525139 | 0.72091 |
| 10 | 22260600 | 22261999 | 8.680967 | 16.38673 | 26.47536 | 10.78998 | -0.2102 | 0.055101 | 0.720919 |
| 13 | 20541800 | 20546599 | 62.37631 | 99.19919 | 113.2325 | 47.27138 | 0.143804 | 2.244626 | 0.720924 |
| 14 | 17700200 | 17702999 | 21.93075 | 36.80478 | 55.95071 | 24.42161 | -0.2212 | 1.141953 | 0.720924 |
| 12 | 24462800 | 24487199 | 425.0894 | 589.8468 | 587.3677 | 243.8451 | 0.163073 | 4.699969 | 0.72094 |
| 2 | 32005400 | 32011399 | 81.39283 | 127.2545 | 142.3706 | 59.06638 | 0.152563 | 2.643965 | 0.720952 |
| 10 | 126029000 | 126032599 | 33.95556 | 55.36119 | 81.79938 | 35.13705 | -0.18999 | 1.53911 | 0.720963 |
| 9 | 25599600 | 25600399 | 10.67366 | 19.81807 | 31.54755 | 13.22145 | -0.22157 | 0.19086 | 0.720967 |
| 13 | 113886600 | 113888399 | 14.83339 | 26.67573 | 30.81786 | 13.46106 | 0.196766 | 0.398346 | 0.720968 |
| 6 | 141649000 | 141651399 | 26.96528 | 44.58289 | 65.22916 | 28.19321 | -0.16679 | 1.33306 | 0.720981 |
| 2 | 53220200 | 53220999 | 6.679464 | 12.88617 | 23.12584 | 8.937043 | -0.29059 | -0.14949 | 0.720983 |
| 13 | 40520000 | 40522399 | 17.15403 | 29.87157 | 44.17624 | 19.42339 | -0.17468 | 0.762261 | 0.720996 |
| 15 | 3679600 | 3680799 | 16.18885 | 29.10431 | 26.78499 | 11.70258 | 0.522901 | 0.40326 | 0.72102 |

| | | | | | | | | | |
|---|---|---|---|---|---|---|---|---|---|
| 2 | 60001600 | 60006599 | 49.50686 | 79.63852 | 90.04021 | 38.21451 | 0.176226 | 1.87048 | 0.721024 |
| 14 | 121089000 | 121099399 | 124.7116 | 188.7018 | 251.2304 | 103.9199 | -0.13525 | 3.289946 | 0.721031 |
| 7 | 94433600 | 94434399 | 12.03245 | 22.34956 | 26.60825 | 11.14558 | 0.194899 | 0.185633 | 0.721049 |
| 16 | 29836000 | 29837199 | 12.92002 | 23.01692 | 34.63283 | 15.24806 | -0.18994 | 0.453251 | 0.721057 |
| X | 22576000 | 22578799 | 34.75377 | 57.67755 | 61.45786 | 26.6326 | 0.289696 | 1.309289 | 0.721108 |
| 13 | 92749600 | 92754999 | 46.22229 | 74.09034 | 104.2017 | 43.99046 | -0.1443 | 1.9807 | 0.721156 |
| 19 | 45981800 | 45984599 | 37.58154 | 61.25481 | 70.75309 | 30.38881 | 0.16406 | 1.542773 | 0.721172 |
| 10 | 93524200 | 93524999 | 7.428514 | 14.26205 | 24.10977 | 9.447677 | -0.22387 | -0.09218 | 0.721182 |
| 13 | 14982600 | 14985999 | 42.34762 | 68.0362 | 96.72914 | 40.97855 | -0.15588 | 1.903062 | 0.721184 |
| 13 | 41604400 | 41606199 | 12.90677 | 23.03313 | 34.62356 | 15.22176 | -0.18713 | 0.442051 | 0.721186 |
| 5 | 41578800 | 41580199 | 11.10818 | 20.16912 | 31.22329 | 13.47891 | -0.21169 | 0.318239 | 0.721195 |
| 8 | 78433800 | 78435399 | 11.10818 | 20.16912 | 31.22329 | 13.47891 | -0.21169 | 0.318239 | 0.721195 |
| 7 | 87648200 | 87650799 | 31.0764 | 50.49942 | 88.41893 | 37.93077 | -0.43769 | 1.586693 | 0.721208 |
| 8 | 14849800 | 14855599 | 62.35315 | 99.45796 | 110.59 | 46.35872 | 0.186171 | 2.150826 | 0.721215 |
| 15 | 49742800 | 49743999 | 13.51324 | 24.19529 | 37.17842 | 16.29947 | -0.21611 | 0.429188 | 0.721216 |
| 15 | 89085400 | 89089599 | 50.3759 | 80.52427 | 112.8282 | 47.43914 | -0.14348 | 2.07416 | 0.721245 |
| 6 | 27124200 | 27125999 | 35.87014 | 58.9304 | 54.62872 | 23.53619 | 0.486949 | 1.424109 | 0.721249 |
| 17 | 42521800 | 42523599 | 17.67887 | 31.52318 | 28.64695 | 12.60303 | 0.53422 | 0.467444 | 0.721264 |
| 15 | 35396000 | 35406199 | 131.9894 | 201.4024 | 214.4692 | 88.93327 | 0.196674 | 3.120354 | 0.721274 |
| 17 | 88060600 | 88069199 | 79.68298 | 123.0599 | 203.0408 | 84.33892 | -0.42103 | 2.866685 | 0.721319 |
| 15 | 4502200 | 4503199 | 12.43264 | 22.32561 | 33.6543 | 14.71816 | -0.18584 | 0.398514 | 0.721337 |
| 4 | 64386600 | 64388199 | 16.03341 | 28.21419 | 43.9236 | 19.44651 | -0.24742 | 0.599798 | 0.721355 |
| 13 | 13618200 | 13620799 | 26.37417 | 44.84289 | 43.1477 | 18.91056 | 0.437005 | 1.029028 | 0.72137 |
| 2 | 101626600 | 101629999 | 21.83123 | 36.56987 | 55.8899 | 24.36654 | -0.22873 | 1.163028 | 0.721379 |
| 5 | 109644400 | 109652199 | 83.20077 | 130.6943 | 145.5071 | 60.35412 | 0.164832 | 2.560113 | 0.721392 |
| 8 | 122561800 | 122569599 | 120.0279 | 181.7199 | 243.9058 | 100.9085 | -0.14623 | 3.277059 | 0.721396 |
| 4 | 53801200 | 53808799 | 263.3633 | 379.6215 | 400.7805 | 165.3769 | 0.133925 | 4.1308 | 0.721424 |
| 12 | 106174800 | 106176599 | 20.03392 | 34.22266 | 50.61073 | 22.18978 | -0.1797 | 0.971212 | 0.721467 |
| 12 | 23680000 | 23681399 | 19.20313 | 33.83417 | 28.00412 | 12.39531 | 0.660452 | 0.57938 | 0.721467 |
| 7 | 80624400 | 80624799 | 8.803564 | 17.07932 | 21.68292 | 8.169018 | 0.239369 | -0.21348 | 0.72148 |
| 18 | 4698200 | 4700399 | 22.96662 | 39.68477 | 44.68504 | 19.60365 | 0.215058 | 0.874985 | 0.721503 |
| 6 | 18358600 | 18359399 | 10.30706 | 18.86916 | 29.7839 | 12.72435 | -0.22951 | 0.261668 | 0.72153 |
| 17 | 22879600 | 22879999 | 13.28961 | 24.32584 | 28.55077 | 12.19992 | 0.194897 | 0.28298 | 0.721541 |
| 12 | 114937800 | 114939999 | 22.68436 | 39.79093 | 39.46414 | 17.44572 | 0.399499 | 0.675358 | 0.721584 |
| 8 | 76480800 | 76485199 | 49.96481 | 80.6278 | 86.01679 | 36.66565 | 0.265223 | 1.774937 | 0.721596 |
| 2 | 84350000 | 84351799 | 15.47928 | 27.65438 | 31.77953 | 13.95846 | 0.198875 | 0.445155 | 0.721599 |
| 1 | 129058000 | 129059599 | 16.0263 | 28.38087 | 32.8409 | 14.52142 | 0.182038 | 0.511312 | 0.721607 |
| 3 | 59683800 | 59687399 | 42.4872 | 68.65603 | 79.08731 | 33.77201 | 0.157025 | 1.719168 | 0.721633 |
| 2 | 3118800 | 3120399 | 15.20632 | 26.95994 | 30.91462 | 13.66633 | 0.195768 | 0.490081 | 0.721635 |
| 2 | 97668200 | 97669399 | 15.20632 | 26.95994 | 30.91462 | 13.66633 | 0.195768 | 0.490081 | 0.721635 |
| 4 | 84737400 | 84741799 | 57.38071 | 91.82848 | 102.5135 | 43.17368 | 0.185611 | 2.043092 | 0.721664 |
| 3 | 158692600 | 158694799 | 26.6887 | 45.33687 | 41.67127 | 18.26175 | 0.502747 | 1.038966 | 0.721677 |
| 13 | 98970400 | 98972199 | 14.07748 | 24.78189 | 37.2524 | 16.49166 | -0.19662 | 0.57386 | 0.721682 |
| 5 | 7601400 | 7612399 | 256.9576 | 370.9215 | 393.0197 | 162.1436 | 0.130734 | 4.109326 | 0.721682 |
| 10 | 93614600 | 93617599 | 22.06309 | 37.21235 | 54.90341 | 24.00844 | -0.17798 | 1.096496 | 0.721683 |
| 10 | 127628000 | 127634999 | 153.7762 | 229.7392 | 301.4068 | 124.2634 | -0.13029 | 3.54463 | 0.721684 |
| 9 | 120606200 | 120608599 | 16.71772 | 27.98556 | 63.04056 | 27.47254 | -0.78618 | 1.161985 | 0.721694 |

| | | | | | | | | | |
|---|---|---|---|---|---|---|---|---|---|
| 4 | 135678600 | 135679999 | 15.49855 | 27.04679 | 41.04919 | 18.07287 | -0.21154 | 0.717927 | 0.721695 |
| 2 | 97068200 | 97069399 | 11.2014 | 20.5458 | 31.81744 | 13.5577 | -0.19979 | 0.26101 | 0.7217 |
| 5 | 130922200 | 130925399 | 33.08839 | 54.3259 | 63.34237 | 27.28307 | 0.156594 | 1.430006 | 0.72171 |
| 17 | 20103800 | 20119599 | 52.49767 | 83.30318 | 138.0158 | 57.48884 | -0.39597 | 2.309649 | 0.721717 |
| 6 | 68207200 | 68209599 | 30.45774 | 51.79071 | 51.81774 | 22.71058 | 0.38096 | 1.054164 | 0.721723 |
| 2 | 176912600 | 176916399 | 44.85086 | 72.35979 | 81.69089 | 34.81196 | 0.183361 | 1.768655 | 0.721724 |
| 8 | 113265600 | 113268199 | 26.61595 | 45.33796 | 50.11586 | 21.96781 | 0.23805 | 1.022762 | 0.721746 |
| 5 | 64404800 | 64405999 | 9.432202 | 17.3345 | 28.89929 | 12.2816 | -0.30184 | 0.2333 | 0.721769 |
| 13 | 14289200 | 14291599 | 21.26985 | 37.0869 | 41.33612 | 18.18949 | 0.231569 | 0.758606 | 0.721779 |
| X | 67825000 | 67827399 | 23.62273 | 41.11954 | 42.96155 | 18.88456 | 0.32433 | 0.797805 | 0.72179 |
| 4 | 74858000 | 74861199 | 31.42416 | 51.3526 | 74.15168 | 31.87882 | -0.1545 | 1.495014 | 0.721804 |
| 12 | 103838800 | 103847599 | 91.80552 | 143.6336 | 155.7297 | 64.6042 | 0.198561 | 2.639666 | 0.721811 |
| 1 | 187555800 | 187557799 | 22.15784 | 37.57074 | 55.50599 | 24.31098 | -0.17967 | 1.050625 | 0.721821 |
| X | 96298400 | 96299999 | 15.65788 | 27.6747 | 45.59137 | 20.16717 | -0.3261 | 0.554383 | 0.721847 |
| 2 | 45010200 | 45013399 | 27.07088 | 45.44316 | 52.87316 | 23.08219 | 0.163248 | 1.153693 | 0.721853 |
| 1 | 117577800 | 117588799 | 200.2442 | 296.5082 | 314.2643 | 129.424 | 0.165534 | 3.708872 | 0.721856 |
| 5 | 81439200 | 81440599 | 14.7409 | 26.83705 | 30.46564 | 13.10772 | 0.236046 | 0.327752 | 0.721858 |
| 17 | 40359200 | 40360199 | 14.7409 | 26.83705 | 30.46564 | 13.10772 | 0.236046 | 0.327752 | 0.721858 |
| 5 | 87352000 | 87357999 | 52.69838 | 84.77612 | 91.90088 | 39.00661 | 0.236867 | 1.874941 | 0.721863 |
| 5 | 12767600 | 12770399 | 57.22457 | 91.36734 | 104.6858 | 43.94453 | 0.144537 | 2.113736 | 0.721895 |
| 13 | 107891000 | 107892799 | 17.31132 | 30.36731 | 35.08452 | 15.51021 | 0.180236 | 0.636273 | 0.721915 |
| 1 | 183195200 | 183196799 | 25.27617 | 42.11088 | 61.45049 | 26.69903 | -0.16225 | 1.23752 | 0.721921 |
| 14 | 80096200 | 80101799 | 73.90011 | 116.628 | 132.9813 | 55.20121 | 0.135654 | 2.458766 | 0.721928 |
| 17 | 19116400 | 19129399 | 215.3024 | 313.4403 | 402.7501 | 165.9891 | -0.13538 | 3.980739 | 0.721944 |
| 18 | 54480800 | 54482199 | 13.54475 | 24.72633 | 28.98344 | 12.42779 | 0.193227 | 0.299887 | 0.721948 |
| 4 | 74423400 | 74424399 | 18.07697 | 30.88209 | 48.49184 | 21.2609 | -0.26552 | 0.963975 | 0.721983 |
| 4 | 102738200 | 102742599 | 43.26647 | 69.36295 | 99.87972 | 42.17558 | -0.17795 | 1.970487 | 0.721985 |
| X | 147517000 | 147520799 | 44.87472 | 72.67062 | 79.82625 | 34.15759 | 0.2286 | 1.674734 | 0.721992 |
| 5 | 103141400 | 103142399 | 8.847771 | 16.58482 | 26.91652 | 11.10501 | -0.23112 | 0.10289 | 0.721992 |
| 9 | 12759600 | 12761199 | 15.76335 | 27.95046 | 32.39063 | 14.31756 | 0.180756 | 0.496805 | 0.722034 |
| 3 | 44334000 | 44334799 | 10.14669 | 19.179 | 23.24781 | 9.447999 | 0.204721 | 0.050633 | 0.722035 |
| 18 | 74759400 | 74764799 | 44.14777 | 71.12464 | 81.92953 | 34.84813 | 0.152031 | 1.811299 | 0.722041 |
| 1 | 90914000 | 90917999 | 27.7174 | 46.339 | 53.9926 | 23.51028 | 0.161284 | 1.199574 | 0.722048 |
| 14 | 115623600 | 115624599 | 10.28807 | 19.28678 | 31.27405 | 12.90036 | -0.23 | 0.125266 | 0.722065 |
| X | 108431200 | 108433599 | 23.75621 | 40.82999 | 46.30489 | 20.31098 | 0.203183 | 0.927124 | 0.722092 |
| 6 | 72044600 | 72047799 | 34.34958 | 56.37966 | 65.25188 | 28.10197 | 0.167041 | 1.436875 | 0.722109 |
| 8 | 71614200 | 71614599 | 6.793236 | 13.19051 | 24.13086 | 9.045092 | -0.27679 | -0.25074 | 0.72211 |
| 9 | 61446800 | 61451399 | 51.53934 | 82.13883 | 115.3098 | 48.28007 | -0.15089 | 2.168854 | 0.722118 |
| 6 | 90243200 | 90245999 | 19.23547 | 33.70997 | 38.11567 | 16.83644 | 0.211266 | 0.668666 | 0.722121 |
| 17 | 74224800 | 74227399 | 15.7222 | 27.54241 | 40.92547 | 18.07171 | -0.1809 | 0.661024 | 0.722125 |
| 4 | 153362600 | 153371799 | 113.8209 | 172.9466 | 232.2835 | 96.2062 | -0.14285 | 3.201538 | 0.722127 |
| 12 | 60088800 | 60089399 | 8.29287 | 16.08492 | 19.31965 | 7.286327 | 0.315964 | -0.22778 | 0.72214 |
| 14 | 62528200 | 62531199 | 17.58721 | 30.44462 | 45.29763 | 19.87349 | -0.18473 | 0.835092 | 0.722169 |
| 7 | 40937200 | 40939199 | 65.30357 | 103.4413 | 115.2681 | 47.93762 | 0.172993 | 2.363059 | 0.722217 |
| 13 | 33808200 | 33809799 | 25.53124 | 43.43596 | 49.56865 | 21.72575 | 0.191922 | 1.028139 | 0.72223 |
| 6 | 8635800 | 8638999 | 27.50676 | 46.10785 | 53.45488 | 23.31476 | 0.168393 | 1.169931 | 0.722251 |
| 12 | 35895400 | 35896199 | 10.59558 | 19.87972 | 23.68146 | 9.754757 | 0.213478 | 0.103412 | 0.722262 |

| | | | | | | | | | |
|---|---|---|---|---|---|---|---|---|---|
| 2 | 39486800 | 39488599 | 18.8436 | 33.07929 | 36.87257 | 16.3107 | 0.231485 | 0.634382 | 0.722293 |
| 10 | 94999800 | 95001799 | 20.95808 | 36.36667 | 42.05348 | 18.45153 | 0.178061 | 0.825307 | 0.722308 |
| 3 | 23228800 | 23229399 | 29.30472 | 48.5306 | 50.21532 | 21.72864 | 0.33177 | 1.308769 | 0.722313 |
| 13 | 37219800 | 37222799 | 44.88046 | 72.24155 | 82.67194 | 35.11916 | 0.159937 | 1.841514 | 0.72234 |
| 2 | 156050400 | 156053799 | 47.04679 | 75.64135 | 86.73844 | 36.78511 | 0.154957 | 1.88258 | 0.72237 |
| 11 | 17108400 | 17110599 | 21.61985 | 36.29254 | 55.39214 | 24.18181 | -0.2269 | 1.137917 | 0.722376 |
| 1 | 66053800 | 66058199 | 57.39031 | 91.72255 | 104.1497 | 43.78073 | 0.15907 | 2.083714 | 0.722409 |
| 2 | 72019800 | 72022399 | 31.04436 | 50.82042 | 73.81881 | 31.75603 | -0.16131 | 1.467992 | 0.722416 |
| 3 | 93908800 | 93910799 | 18.23462 | 32.21836 | 35.19565 | 15.56969 | 0.263278 | 0.552247 | 0.722422 |
| 10 | 81962800 | 81979799 | 389.6958 | 545.6079 | 541.1601 | 224.349 | 0.18184 | 4.56449 | 0.722429 |
| 14 | 52948200 | 52948999 | 14.28517 | 25.7176 | 29.9495 | 13.06749 | 0.18657 | 0.388059 | 0.722463 |
| 9 | 120520400 | 120525999 | 42.94305 | 68.85683 | 99.26236 | 41.92634 | -0.17926 | 1.964366 | 0.722487 |
| 9 | 23890600 | 23891799 | 11.38852 | 20.62092 | 32.17252 | 13.93689 | -0.22633 | 0.336918 | 0.722492 |
| 13 | 43679800 | 43681999 | 30.58523 | 50.55635 | 58.93248 | 25.46765 | 0.160029 | 1.338869 | 0.722494 |
| 8 | 43812200 | 43813399 | 11.31842 | 20.7905 | 32.41 | 13.78305 | -0.20691 | 0.257814 | 0.722535 |
| 13 | 93934400 | 93935999 | 15.52905 | 27.17911 | 40.44539 | 17.84589 | -0.18302 | 0.673421 | 0.722543 |
| 10 | 13509800 | 13511999 | 26.76572 | 44.91508 | 52.31685 | 22.83425 | 0.161683 | 1.156361 | 0.72255 |
| 8 | 36289600 | 36294199 | 37.52073 | 60.46915 | 88.45916 | 37.64431 | -0.19138 | 1.792933 | 0.722631 |
| 5 | 110070400 | 110071999 | 17.38518 | 30.04805 | 45.82333 | 20.10318 | -0.22069 | 0.847396 | 0.722644 |
| 4 | 112038600 | 112044399 | 66.65855 | 106.705 | 104.1761 | 43.89065 | 0.379126 | 2.047671 | 0.722671 |
| 17 | 94608000 | 94614399 | 97.77451 | 151.4078 | 170.2533 | 70.71868 | 0.134542 | 2.821719 | 0.72268 |
| 13 | 38448800 | 38451999 | 38.03616 | 61.94442 | 71.39666 | 30.65595 | 0.166284 | 1.555088 | 0.722686 |
| 5 | 45507800 | 45509399 | 15.36485 | 28.12634 | 30.50106 | 13.03208 | 0.308185 | 0.301106 | 0.722687 |
| 8 | 5618800 | 5621799 | 53.61666 | 86.20632 | 81.30011 | 34.47993 | 0.436427 | 1.886855 | 0.722701 |
| 1 | 194998400 | 195005999 | 99.65165 | 154.0312 | 173.3464 | 72.00419 | 0.131321 | 2.856061 | 0.722701 |
| 3 | 34431200 | 34432199 | 10.56475 | 19.54414 | 30.38044 | 12.79593 | -0.19296 | 0.205394 | 0.722708 |
| 12 | 61400200 | 61403799 | 38.9377 | 63.84296 | 59.87873 | 25.77641 | 0.468958 | 1.45764 | 0.722719 |
| 10 | 66327600 | 66329199 | 12.42027 | 22.3439 | 33.64853 | 14.69314 | -0.18294 | 0.386838 | 0.722745 |
| 2 | 171116800 | 171117999 | 8.350249 | 15.80813 | 25.99011 | 10.52759 | -0.22752 | 0.029806 | 0.722758 |
| 10 | 34614800 | 34619199 | 80.97477 | 126.0729 | 171.3529 | 71.1267 | -0.13258 | 2.721295 | 0.722761 |
| 18 | 81871000 | 81873599 | 35.48698 | 58.02538 | 66.90063 | 28.76754 | 0.169878 | 1.488716 | 0.72279 |
| 7 | 113893600 | 113894999 | 12.4103 | 22.24915 | 33.6353 | 14.73011 | -0.19139 | 0.407814 | 0.722796 |
| 11 | 78136000 | 78137199 | 9.004257 | 16.79937 | 27.38337 | 11.38747 | -0.24663 | 0.135008 | 0.722866 |
| X | 127796400 | 127801199 | 63.37854 | 101.3346 | 106.1179 | 44.63397 | 0.276301 | 2.080547 | 0.722869 |
| 6 | 73372200 | 73373799 | 19.20641 | 33.52437 | 38.74325 | 17.05984 | 0.179662 | 0.728798 | 0.722883 |
| 5 | 37655400 | 37657599 | 26.04061 | 43.7366 | 50.69807 | 22.13996 | 0.168598 | 1.142082 | 0.722893 |
| 11 | 102972600 | 102974799 | 22.32809 | 37.70533 | 56.04347 | 24.51604 | -0.18862 | 1.086562 | 0.722903 |
| 4 | 9858800 | 9861399 | 24.90415 | 41.26884 | 63.65109 | 27.55436 | -0.24248 | 1.2962 | 0.722924 |
| 4 | 40059000 | 40061799 | 68.33895 | 108.0539 | 121.6486 | 50.54489 | 0.156149 | 2.409651 | 0.72296 |
| 11 | 82417400 | 82419799 | 14.93712 | 26.19815 | 39.39717 | 17.4143 | -0.19798 | 0.635803 | 0.72297 |
| 15 | 80180600 | 80181799 | 12.94139 | 23.0912 | 34.6511 | 15.23593 | -0.1847 | 0.444288 | 0.722982 |
| 4 | 58656200 | 58658799 | 16.19143 | 28.25458 | 42.07432 | 18.52364 | -0.18419 | 0.725045 | 0.722984 |
| 12 | 59692000 | 59693199 | 16.19143 | 28.25458 | 42.07432 | 18.52364 | -0.18419 | 0.725045 | 0.722984 |
| 9 | 4309000 | 4310399 | 13.38941 | 23.36036 | 45.24501 | 19.91711 | -0.56152 | 0.718741 | 0.722994 |
| 1 | 23518600 | 23519799 | 13.13826 | 23.95816 | 28.05682 | 12.04686 | 0.19354 | 0.302499 | 0.722998 |
| 5 | 42528600 | 42529399 | 9.638348 | 18.39651 | 22.82756 | 9.087994 | 0.197061 | -0.0228 | 0.723 |
| 12 | 24328600 | 24375399 | 1206.709 | 1530.15 | 1416.073 | 594.6644 | 0.138325 | 6.141159 | 0.723002 |

| | | | | | | | | | |
|---|---|---|---|---|---|---|---|---|---|
| 15 | 74732600 | 74735799 | 73.05781 | 115.2325 | 129.8733 | 53.89817 | 0.151749 | 2.472394 | 0.723012 |
| 6 | 112218200 | 112220199 | 24.01683 | 40.65509 | 46.41753 | 20.32007 | 0.19078 | 1.062458 | 0.723012 |
| 14 | 113321400 | 113324599 | 33.71129 | 54.97039 | 80.21078 | 34.45636 | -0.17185 | 1.53394 | 0.72302 |
| 5 | 130107400 | 130110799 | 27.5768 | 45.43288 | 66.93769 | 28.87368 | -0.17803 | 1.37615 | 0.72302 |
| X | 164613000 | 164621799 | 134.1466 | 205.4168 | 206.272 | 85.6039 | 0.285303 | 3.051029 | 0.72305 |
| 14 | 109431600 | 109433199 | 24.86889 | 42.20462 | 49.03587 | 21.48286 | 0.165773 | 1.046867 | 0.723056 |
| 10 | 48478600 | 48479999 | 18.12537 | 31.97861 | 29.83162 | 13.20062 | 0.489312 | 0.559964 | 0.723077 |
| 3 | 51741800 | 51742799 | 12.59682 | 23.053 | 27.06316 | 11.56747 | 0.194292 | 0.275092 | 0.723079 |
| 1 | 88221600 | 88239399 | 1664.802 | 2062.189 | 1853.908 | 780.5225 | 0.145991 | 6.518207 | 0.723121 |
| 4 | 70277000 | 70278999 | 16.57328 | 28.89637 | 43.08347 | 18.95781 | -0.1863 | 0.737914 | 0.723125 |
| 3 | 119181400 | 119182199 | 9.361567 | 17.56037 | 27.98209 | 11.53077 | -0.20332 | 0.107215 | 0.723168 |
| 10 | 55279400 | 55280399 | 11.7596 | 21.52899 | 33.39625 | 14.26704 | -0.20479 | 0.280773 | 0.723169 |
| 4 | 92774600 | 92783199 | 219.443 | 322.9001 | 299.5744 | 123.3659 | 0.349018 | 3.813427 | 0.723173 |
| 11 | 95816000 | 95820799 | 58.48102 | 93.39008 | 106.0879 | 44.54436 | 0.157178 | 2.109169 | 0.723185 |
| 3 | 66834600 | 66836799 | 21.59151 | 37.83203 | 39.48825 | 17.44006 | 0.32599 | 0.684266 | 0.723186 |
| 1 | 181566600 | 181569399 | 29.33811 | 48.67488 | 56.2248 | 24.35975 | 0.173492 | 1.289316 | 0.723205 |
| 19 | 46017000 | 46019199 | 29.22341 | 48.12629 | 69.93327 | 30.16069 | -0.15842 | 1.38788 | 0.723205 |
| 3 | 123740600 | 123741399 | 11.07279 | 20.88336 | 25.26697 | 10.31663 | 0.201183 | 0.078526 | 0.723206 |
| 13 | 43083200 | 43086399 | 32.47141 | 52.72628 | 76.92266 | 32.98052 | -0.17613 | 1.597065 | 0.723212 |
| 12 | 7915000 | 7916399 | 20.73799 | 35.72321 | 40.62706 | 17.82171 | 0.199889 | 0.886952 | 0.723215 |
| 17 | 58519600 | 58521999 | 31.75113 | 52.38375 | 61.20415 | 26.41623 | 0.156379 | 1.3657 | 0.723219 |
| 2 | 106645400 | 106647599 | 25.65303 | 43.69702 | 48.47796 | 21.24982 | 0.232668 | 1.013588 | 0.723248 |
| 8 | 82938200 | 82941399 | 26.5997 | 44.14838 | 64.25246 | 27.8451 | -0.15876 | 1.286315 | 0.723252 |
| 14 | 100112000 | 100113799 | 17.24074 | 29.88599 | 44.76624 | 19.64118 | -0.19399 | 0.819895 | 0.72326 |
| 4 | 79873200 | 79875799 | 20.91873 | 35.07724 | 62.36091 | 27.20531 | -0.44621 | 1.155829 | 0.723265 |
| 11 | 34180600 | 34181799 | 10.47005 | 19.07882 | 30.25972 | 13.00445 | -0.24216 | 0.290336 | 0.723278 |
| 15 | 48957400 | 48958399 | 19.78603 | 33.64472 | 50.59143 | 22.17313 | -0.20455 | 1.013378 | 0.723334 |
| 1 | 117606800 | 117614199 | 135.9676 | 206.1587 | 225.9858 | 93.53826 | 0.147494 | 3.250906 | 0.723339 |
| 12 | 12878000 | 12878599 | 5.387193 | 10.50046 | 20.30359 | 7.342955 | -0.30764 | -0.40578 | 0.723351 |
| 13 | 45169400 | 45170599 | 22.01296 | 38.32994 | 36.1454 | 15.89165 | 0.47152 | 0.774248 | 0.723358 |
| 7 | 35760600 | 35762799 | 12.80069 | 22.6085 | 35.26865 | 15.60319 | -0.24785 | 0.526029 | 0.723372 |
| 11 | 28481800 | 28484999 | 46.69226 | 75.49195 | 82.23565 | 35.14096 | 0.23837 | 1.713574 | 0.723381 |
| 9 | 6787000 | 6788399 | 14.49197 | 26.45385 | 30.02771 | 12.87657 | 0.239396 | 0.311104 | 0.723394 |
| 13 | 57901400 | 57903999 | 35.93873 | 58.62908 | 68.2897 | 29.34024 | 0.152922 | 1.525419 | 0.723411 |
| 11 | 22736200 | 22739999 | 32.92959 | 53.46531 | 77.67865 | 33.30238 | -0.17033 | 1.599229 | 0.723412 |
| 11 | 99646800 | 99647199 | 6.967096 | 13.51438 | 23.39891 | 8.821933 | -0.20531 | -0.22796 | 0.723418 |
| 1 | 98715000 | 98717399 | 17.4594 | 30.54736 | 47.81589 | 21.09201 | -0.25598 | 0.698373 | 0.723454 |
| 4 | 115594600 | 115596199 | 13.10257 | 23.19994 | 35.1735 | 15.55328 | -0.20556 | 0.500009 | 0.723485 |
| 7 | 59006400 | 59007599 | 12.66962 | 23.33448 | 27.52962 | 11.66526 | 0.195922 | 0.239684 | 0.7235 |
| 13 | 43108400 | 43112199 | 31.39741 | 51.52131 | 75.75106 | 32.62021 | -0.17742 | 1.441462 | 0.723526 |
| 1 | 44935600 | 44936599 | 12.85717 | 23.46687 | 27.48514 | 11.78801 | 0.194162 | 0.292024 | 0.723527 |
| 6 | 128722600 | 128723799 | 12.85717 | 23.46687 | 27.48514 | 11.78801 | 0.194162 | 0.292024 | 0.723527 |
| 1 | 183302600 | 183303999 | 36.31867 | 59.04274 | 67.35195 | 28.9006 | 0.179446 | 1.57647 | 0.723533 |
| 8 | 4734800 | 4739399 | 48.25366 | 77.75503 | 87.12381 | 37.06822 | 0.192185 | 1.813248 | 0.723557 |
| 1 | 120488800 | 120490399 | 17.52868 | 30.74129 | 35.62407 | 15.74557 | 0.175963 | 0.643055 | 0.723597 |
| 10 | 34164800 | 34165999 | 13.32641 | 24.15191 | 27.92601 | 12.0835 | 0.204344 | 0.336608 | 0.723667 |
| 18 | 19117000 | 19122399 | 43.97045 | 71.28705 | 78.2396 | 33.50668 | 0.231327 | 1.648662 | 0.723685 |

| | | | | | | | | | |
|---|---|---|---|---|---|---|---|---|---|
| 1 | 119289200 | 119290399 | 17.46628 | 30.68891 | 35.45476 | 15.69539 | 0.180195 | 0.611464 | 0.723711 |
| 14 | 42453000 | 42455399 | 35.40047 | 59.07901 | 48.70137 | 21.18259 | 0.658851 | 1.235249 | 0.72374 |
| 19 | 9774400 | 9775399 | 15.06947 | 26.4449 | 39.34458 | 17.39918 | -0.18269 | 0.626742 | 0.723774 |
| 14 | 106274200 | 106275599 | 11.34508 | 20.7552 | 32.29101 | 13.80836 | -0.21022 | 0.277174 | 0.723777 |
| 6 | 146606200 | 146610199 | 44.91971 | 72.17459 | 102.6612 | 43.49688 | -0.15647 | 1.904766 | 0.723778 |
| 13 | 68024000 | 68037599 | 198.6531 | 292.6744 | 315.9554 | 130.1134 | 0.132225 | 3.791034 | 0.723798 |
| 12 | 103633800 | 103636999 | 32.12416 | 52.88839 | 61.65634 | 26.58785 | 0.158415 | 1.393336 | 0.723819 |
| 14 | 99161200 | 99162799 | 18.43083 | 31.78415 | 46.88565 | 20.56783 | -0.17351 | 0.875593 | 0.723826 |
| 9 | 76398200 | 76399999 | 42.38337 | 68.4133 | 79.15562 | 33.75348 | 0.148898 | 1.751767 | 0.723829 |
| 8 | 3691000 | 3693199 | 13.2553 | 23.42208 | 35.69967 | 15.79244 | -0.21439 | 0.52421 | 0.723835 |
| 11 | 105069400 | 105073199 | 28.54418 | 46.99446 | 68.74312 | 29.64387 | -0.16813 | 1.388013 | 0.723838 |
| 5 | 31275600 | 31278199 | 17.35049 | 30.13954 | 45.25457 | 19.86681 | -0.19704 | 0.796559 | 0.723844 |
| 12 | 31951400 | 31952999 | 13.85782 | 25.22682 | 29.44696 | 12.67102 | 0.195908 | 0.318962 | 0.723861 |
| 8 | 124819000 | 124827399 | 90.66219 | 139.9847 | 189.9465 | 78.89991 | -0.13949 | 2.87353 | 0.723891 |
| 17 | 37781600 | 37783999 | 47.32388 | 76.33308 | 72.6079 | 30.93267 | 0.4297 | 1.778518 | 0.723915 |
| 4 | 27212800 | 27215799 | 33.37565 | 55.76912 | 49.74242 | 21.65236 | 0.545616 | 1.220448 | 0.723976 |
| 4 | 106738200 | 106742599 | 38.86643 | 63.136 | 72.9797 | 31.29936 | 0.159369 | 1.597303 | 0.723978 |
| 3 | 31494600 | 31509599 | 188.9597 | 278.1653 | 360.714 | 148.5441 | -0.13323 | 3.804759 | 0.723984 |
| X | 117616000 | 117616799 | 15.50553 | 28.00501 | 31.4245 | 13.65573 | 0.243588 | 0.381739 | 0.723992 |
| 2 | 168669000 | 168671399 | 20.88295 | 35.29691 | 53.82119 | 23.55039 | -0.22524 | 1.071377 | 0.724012 |
| 4 | 85809600 | 85810399 | 13.74581 | 24.20793 | 36.75199 | 16.2688 | -0.21061 | 0.566096 | 0.724037 |
| 14 | 71362600 | 71366399 | 26.766 | 44.46178 | 68.0022 | 29.48724 | -0.23024 | 1.275655 | 0.724043 |
| 12 | 114172800 | 114178799 | 67.20967 | 107.3046 | 110.6868 | 46.46032 | 0.29575 | 2.127711 | 0.724044 |
| 2 | 40364600 | 40366199 | 16.4814 | 28.80758 | 42.51637 | 18.7386 | -0.17128 | 0.702616 | 0.724046 |
| 2 | 5551800 | 5552199 | 7.842255 | 15.24704 | 20.0891 | 7.452143 | 0.206826 | -0.28399 | 0.724048 |
| 13 | 50655200 | 50660999 | 81.37534 | 127.4659 | 144.957 | 60.12784 | 0.131109 | 2.611933 | 0.724058 |
| 18 | 6323600 | 6328399 | 79.53767 | 125.3329 | 139.8553 | 58.03353 | 0.165034 | 2.495474 | 0.72406 |
| 19 | 55962200 | 55965599 | 33.92266 | 55.11787 | 79.06431 | 33.91467 | -0.151 | 1.587988 | 0.724081 |
| 9 | 18084600 | 18085999 | 20.9483 | 36.21957 | 42.10193 | 18.47077 | 0.16955 | 0.857125 | 0.724082 |
| 1 | 164014200 | 164016999 | 26.97436 | 45.60364 | 51.79391 | 22.66496 | 0.19822 | 1.088949 | 0.724098 |
| 2 | 171689200 | 171692599 | 26.97436 | 45.60364 | 51.79391 | 22.66496 | 0.19822 | 1.088949 | 0.724098 |
| 6 | 12360000 | 12365599 | 54.74875 | 88.02694 | 93.67822 | 39.72983 | 0.26249 | 1.893783 | 0.724113 |
| 4 | 99313000 | 99316399 | 27.18085 | 44.89839 | 66.25136 | 28.61703 | -0.17908 | 1.346208 | 0.724118 |
| 9 | 75796200 | 75796799 | 9.926493 | 18.8925 | 22.84699 | 9.150784 | 0.226901 | 0.002262 | 0.724125 |
| 4 | 41739800 | 41741399 | 13.07058 | 23.03444 | 36.72345 | 16.25396 | -0.28057 | 0.552476 | 0.72414 |
| 6 | 69773400 | 69775199 | 22.46131 | 39.24431 | 40.61645 | 17.89573 | 0.33817 | 0.733659 | 0.724161 |
| 3 | 23222000 | 23230199 | 94.5719 | 145.645 | 195.96 | 81.39317 | -0.13014 | 2.924338 | 0.724168 |
| 10 | 75694600 | 75699999 | 60.00942 | 94.7813 | 135.2894 | 56.18192 | -0.18702 | 2.431299 | 0.724192 |
| 4 | 82710200 | 82712599 | 18.66747 | 32.88986 | 29.88769 | 13.22907 | 0.526277 | 0.578914 | 0.724194 |
| 19 | 39202200 | 39214399 | 153.5067 | 232.3525 | 241.5086 | 99.9094 | 0.222052 | 3.289566 | 0.724209 |
| 14 | 95661800 | 95662999 | 12.03966 | 21.8079 | 33.19072 | 14.3714 | -0.19033 | 0.342371 | 0.724225 |
| 17 | 36005600 | 36006799 | 12.03966 | 21.8079 | 33.19072 | 14.3714 | -0.19033 | 0.342371 | 0.724225 |
| 18 | 66653800 | 66655799 | 16.22246 | 29.04907 | 32.17667 | 14.10543 | 0.253446 | 0.435653 | 0.724255 |
| 7 | 109486000 | 109488199 | 21.89894 | 37.59476 | 43.70912 | 19.1715 | 0.167165 | 0.926156 | 0.724268 |
| 3 | 96224200 | 96230399 | 76.61842 | 121.0938 | 134.1531 | 55.72857 | 0.178829 | 2.427737 | 0.724271 |
| 19 | 51766600 | 51766999 | 6.996606 | 13.53957 | 23.2429 | 8.860478 | -0.20769 | -0.18852 | 0.724311 |
| 8 | 126727000 | 126729999 | 27.06048 | 44.6821 | 65.89746 | 28.45648 | -0.17847 | 1.346555 | 0.724315 |

| | | | | | | | | | |
|---|---|---|---|---|---|---|---|---|---|
| 14 | 80323600 | 80328399 | 74.05034 | 116.0673 | 159.0302 | 65.96098 | -0.13666 | 2.599518 | 0.724317 |
| 18 | 46214400 | 46215599 | 16.13163 | 28.72459 | 32.86664 | 14.4763 | 0.202554 | 0.474338 | 0.724344 |
| 9 | 9425000 | 9426999 | 21.36188 | 37.17561 | 41.91102 | 18.42013 | 0.21486 | 0.785835 | 0.724371 |
| 3 | 17777600 | 17779799 | 20.1297 | 35.35612 | 32.27587 | 14.28386 | 0.518408 | 0.628352 | 0.724377 |
| 6 | 75303600 | 75304999 | 16.68076 | 29.3561 | 33.98598 | 15.04649 | 0.177931 | 0.585384 | 0.72439 |
| 14 | 3462200 | 3466199 | 34.79983 | 56.77425 | 65.27515 | 28.04562 | 0.171713 | 1.520779 | 0.724395 |
| X | 61789400 | 61792199 | 18.17673 | 32.32663 | 29.10646 | 12.83333 | 0.54535 | 0.487569 | 0.724395 |
| 14 | 86435000 | 86437999 | 60.00891 | 96.63044 | 70.57097 | 29.9955 | 0.807137 | 1.84337 | 0.724447 |
| 2 | 71542200 | 71543199 | 12.50161 | 23.71226 | 22.02018 | 8.883713 | 0.595272 | 0.045547 | 0.724458 |
| 13 | 66383600 | 66383999 | 11.33264 | 21.0599 | 23.26052 | 9.736868 | 0.303731 | 0.17151 | 0.724499 |
| 2 | 65601200 | 65605599 | 68.42438 | 108.6566 | 121.9274 | 50.82532 | 0.166337 | 2.295872 | 0.724516 |
| 4 | 15236000 | 15239799 | 52.85235 | 83.93483 | 121.8257 | 50.78675 | -0.20419 | 2.280295 | 0.724516 |
| 3 | 16506800 | 16511199 | 79.44512 | 124.8442 | 141.9809 | 58.89451 | 0.13464 | 2.55004 | 0.724543 |
| X | 152373800 | 152374799 | 15.07971 | 27.26402 | 30.48653 | 13.23105 | 0.249943 | 0.372179 | 0.724559 |
| 7 | 27964800 | 27966199 | 10.60214 | 19.39596 | 30.25802 | 12.93907 | -0.21379 | 0.268374 | 0.724578 |
| 14 | 92021000 | 92021799 | 9.700021 | 18.1762 | 29.03585 | 11.98623 | -0.20901 | 0.119739 | 0.724586 |
| 1 | 58790400 | 58792199 | 10.21653 | 18.38802 | 32.72793 | 14.28443 | -0.42221 | 0.363806 | 0.72459 |
| 16 | 97816800 | 97817799 | 35.11331 | 57.29282 | 66.48777 | 28.56792 | 0.158419 | 1.520012 | 0.724608 |
| 1 | 147555400 | 147556399 | 9.951561 | 18.31585 | 29.29647 | 12.42555 | -0.24107 | 0.226512 | 0.724645 |
| X | 52511200 | 52512199 | 14.5659 | 26.21205 | 29.53719 | 12.89433 | 0.233519 | 0.393561 | 0.724663 |
| 5 | 151597200 | 151597999 | 19.15889 | 33.48069 | 38.75485 | 17.07781 | 0.177609 | 0.713673 | 0.724664 |
| 5 | 77206600 | 77209599 | 18.12254 | 31.26342 | 46.36473 | 20.33953 | -0.18104 | 0.870845 | 0.724665 |
| 14 | 50800200 | 50804399 | 53.0747 | 85.24819 | 94.6873 | 40.09819 | 0.199059 | 1.923233 | 0.724695 |
| 4 | 143879400 | 143899399 | 329.6806 | 469.5641 | 465.6794 | 192.5177 | 0.206894 | 4.310228 | 0.7247 |
| 11 | 16967800 | 16975199 | 88.47528 | 136.7715 | 185.0623 | 76.87066 | -0.13409 | 2.850829 | 0.724706 |
| 14 | 113085200 | 113085999 | 12.82284 | 23.90047 | 21.04515 | 8.76615 | 0.63377 | 0.158936 | 0.724713 |
| 5 | 47645000 | 47646199 | 22.42514 | 39.03968 | 35.67803 | 15.68416 | 0.516619 | 0.779523 | 0.724715 |
| 2 | 62691400 | 62691999 | 7.753718 | 15.07574 | 19.53344 | 7.242787 | 0.232163 | -0.28563 | 0.724733 |
| 9 | 53260400 | 53262599 | 33.62396 | 55.81956 | 61.48244 | 26.64887 | 0.242006 | 1.300875 | 0.724757 |
| X | 164910000 | 164912399 | 18.94184 | 32.64112 | 47.94188 | 21.0307 | -0.16752 | 0.886269 | 0.724768 |
| X | 135827200 | 135832199 | 71.53914 | 113.3432 | 126.7215 | 52.70665 | 0.168459 | 2.366242 | 0.72478 |
| 14 | 113273000 | 113275399 | 22.26215 | 38.46322 | 44.1622 | 19.37421 | 0.186997 | 0.871712 | 0.724785 |
| 16 | 18889400 | 18891199 | 31.74339 | 52.39587 | 61.22945 | 26.43421 | 0.156268 | 1.359947 | 0.724822 |
| 2 | 176995200 | 176999799 | 58.18208 | 93.1145 | 90.08051 | 37.93955 | 0.391877 | 2.037485 | 0.724866 |
| Y | 3825400 | 3826399 | 11.01454 | 20.52669 | 19.20837 | 8.003229 | 0.545783 | 0.135944 | 0.724897 |
| 12 | 105428600 | 105431199 | 20.63568 | 35.32309 | 52.21897 | 22.89868 | -0.17866 | 0.959313 | 0.724899 |
| 6 | 3120000 | 3123999 | 56.97115 | 91.09604 | 102.7733 | 43.23129 | 0.169121 | 2.069428 | 0.724902 |
| 6 | 53587200 | 53587599 | 6.661142 | 12.93595 | 22.90351 | 8.57469 | -0.22793 | -0.25846 | 0.724939 |
| 14 | 80035200 | 80038999 | 46.25359 | 74.13723 | 104.0168 | 43.90942 | -0.14089 | 1.982688 | 0.724946 |
| 15 | 81344400 | 81348599 | 63.55847 | 100.5874 | 138.5973 | 57.61376 | -0.13389 | 2.387056 | 0.724974 |
| 17 | 38738400 | 38741999 | 33.03133 | 55.59222 | 46.37415 | 20.26551 | 0.641751 | 1.144909 | 0.724981 |
| 10 | 56385800 | 56387999 | 37.31093 | 60.65349 | 70.33801 | 30.18148 | 0.155691 | 1.580648 | 0.72499 |
| 6 | 142091800 | 142096799 | 62.82839 | 99.90407 | 114.5868 | 47.82949 | 0.13669 | 2.250984 | 0.724997 |
| X | 155088400 | 155091399 | 22.93135 | 40.14261 | 40.59979 | 17.917 | 0.371409 | 0.709171 | 0.725004 |
| 6 | 130337400 | 130338199 | 8.512533 | 16.33344 | 27.54942 | 10.81421 | -0.22338 | -0.06622 | 0.72501 |
| 18 | 88328800 | 88329599 | 8.512533 | 16.33344 | 27.54942 | 10.81421 | -0.22338 | -0.06622 | 0.72501 |
| 15 | 94593000 | 94595599 | 26.31729 | 44.06476 | 68.44042 | 29.82979 | -0.25215 | 1.194044 | 0.725017 |

| | | | | | | | | | |
|---|---|---|---|---|---|---|---|---|---|
| 17 | 93477200 | 93479399 | 20.65779 | 36.1318 | 39.59345 | 17.46031 | 0.256115 | 0.706424 | 0.725018 |
| 10 | 24092200 | 24093199 | 13.48534 | 23.79585 | 36.28636 | 16.05638 | -0.21566 | 0.543977 | 0.725026 |
| 8 | 125698800 | 125701999 | 36.07493 | 58.75407 | 68.21657 | 29.29117 | 0.155845 | 1.548563 | 0.725027 |
| 13 | 16420400 | 16421399 | 14.96525 | 26.62955 | 30.51141 | 13.44665 | 0.200122 | 0.466583 | 0.725037 |
| 19 | 25557000 | 25557999 | 13.61641 | 24.29131 | 36.66964 | 16.12589 | -0.19346 | 0.454088 | 0.725047 |
| 6 | 137737800 | 137740999 | 18.73884 | 31.69611 | 60.79898 | 26.60956 | -0.55511 | 1.065104 | 0.725054 |
| X | 146703400 | 146704599 | 15.58936 | 28.53059 | 25.67746 | 10.97464 | 0.575113 | 0.294025 | 0.725058 |
| 10 | 47486400 | 47489799 | 54.05727 | 86.57306 | 98.97823 | 41.72948 | 0.152476 | 2.015604 | 0.725088 |
| 13 | 91960200 | 91963799 | 51.45676 | 82.48565 | 94.96251 | 40.09358 | 0.144084 | 1.982803 | 0.725093 |
| 19 | 9523200 | 9533999 | 341.9374 | 475.049 | 600.1297 | 249.0958 | -0.17832 | 4.67854 | 0.725105 |
| 11 | 73854600 | 73855999 | 14.88314 | 26.31063 | 39.69404 | 17.55781 | -0.19963 | 0.536447 | 0.725119 |
| 10 | 122137600 | 122141999 | 40.99555 | 66.40531 | 76.41215 | 32.70275 | 0.162063 | 1.659621 | 0.725129 |
| 12 | 79091200 | 79094399 | 32.1537 | 52.33207 | 75.9594 | 32.60773 | -0.16607 | 1.557311 | 0.725165 |
| 12 | 39415400 | 39415999 | 6.383982 | 12.41669 | 21.84295 | 8.077216 | -0.20158 | -0.30627 | 0.725208 |
| 11 | 99361400 | 99362999 | 13.67549 | 24.75786 | 28.91031 | 12.52628 | 0.18896 | 0.347721 | 0.725231 |
| 1 | 149298400 | 149300399 | 17.64018 | 30.55777 | 45.30263 | 19.87611 | -0.17927 | 0.83038 | 0.72524 |
| 8 | 78224600 | 78226399 | 20.69779 | 35.77213 | 41.56782 | 18.23622 | 0.169852 | 0.857556 | 0.725268 |
| 6 | 122892600 | 122893999 | 19.34136 | 32.82193 | 50.58368 | 22.16055 | -0.24012 | 1.030356 | 0.725272 |
| 1 | 12582800 | 12587599 | 60.5417 | 96.62776 | 108.5417 | 45.53922 | 0.172474 | 2.12961 | 0.725291 |
| 8 | 19339200 | 19340799 | 15.11225 | 26.59121 | 40.20862 | 17.80203 | -0.20567 | 0.587436 | 0.725293 |
| 13 | 62249400 | 62251199 | 15.11225 | 26.59121 | 40.20862 | 17.80203 | -0.20567 | 0.587436 | 0.725293 |
| X | 18633200 | 18634199 | 9.161748 | 17.70658 | 22.06837 | 8.463654 | 0.241685 | -0.14784 | 0.725302 |
| 6 | 7394000 | 7394999 | 16.67858 | 28.95097 | 43.75085 | 19.19775 | -0.20612 | 0.80527 | 0.725346 |
| 5 | 109690200 | 109694199 | 40.59446 | 65.46513 | 94.32554 | 40.17425 | -0.16852 | 1.781036 | 0.72535 |
| 6 | 39040600 | 39041199 | 11.34372 | 20.57838 | 31.77127 | 13.73081 | -0.20864 | 0.328545 | 0.725353 |
| 6 | 90271600 | 90275199 | 67.84481 | 106.8934 | 147.2496 | 61.10077 | -0.13862 | 2.493506 | 0.725368 |
| 10 | 58063200 | 58064599 | 27.74036 | 46.22303 | 53.42818 | 23.21563 | 0.172903 | 1.23496 | 0.725379 |
| 13 | 111864000 | 111872599 | 263.1943 | 373.773 | 484.5175 | 200.4041 | -0.18358 | 4.348859 | 0.725379 |
| 10 | 3363200 | 3366999 | 20.91048 | 35.51268 | 52.19832 | 22.87339 | -0.17211 | 1.030328 | 0.725389 |
| 1 | 141067000 | 141072399 | 101.6795 | 156.9186 | 175.8283 | 73.0357 | 0.135859 | 2.885131 | 0.725393 |
| 12 | 111961200 | 111963799 | 32.16528 | 52.26841 | 76.33023 | 32.74463 | -0.17652 | 1.581882 | 0.725405 |
| 8 | 26724000 | 26725999 | 8.793401 | 17.10639 | 17.27092 | 6.376187 | 0.594045 | -0.29636 | 0.725407 |
| 6 | 91745800 | 91749199 | 42.6476 | 68.58513 | 96.82907 | 41.07902 | -0.14407 | 1.872281 | 0.72542 |
| 9 | 45737600 | 45741999 | 34.43092 | 55.91619 | 80.18685 | 34.38372 | -0.15125 | 1.599862 | 0.725431 |
| 6 | 73665800 | 73666999 | 23.54883 | 40.03124 | 46.38296 | 20.32817 | 0.169949 | 1.024661 | 0.725446 |
| 1 | 66917400 | 66919199 | 29.59325 | 49.53706 | 56.25876 | 24.5164 | 0.198102 | 1.195098 | 0.725473 |
| X | 149146000 | 149168799 | 414.386 | 578.9086 | 550.2625 | 228.2002 | 0.240057 | 4.598522 | 0.725521 |
| 15 | 8924800 | 8928999 | 39.39842 | 63.35892 | 106.5926 | 45.22017 | -0.39655 | 1.871294 | 0.725536 |
| 13 | 59177200 | 59179599 | 17.64647 | 30.66037 | 45.2686 | 19.87569 | -0.17266 | 0.798016 | 0.725559 |
| 14 | 106856400 | 106857799 | 10.14569 | 18.66383 | 29.31316 | 12.44108 | -0.21543 | 0.232369 | 0.725568 |
| 7 | 40095000 | 40096399 | 17.42489 | 30.33826 | 44.68845 | 19.64656 | -0.16909 | 0.765407 | 0.725588 |
| 18 | 37634400 | 37657799 | 571.2212 | 769.7631 | 757.6154 | 315.9649 | 0.137356 | 5.157157 | 0.725592 |
| 4 | 31417200 | 31421199 | 30.8706 | 50.36428 | 75.33315 | 32.36715 | -0.20696 | 1.515496 | 0.725613 |
| 10 | 129064200 | 129066799 | 30.54179 | 50.05853 | 72.69256 | 31.28761 | -0.16022 | 1.451017 | 0.725684 |
| 5 | 53936800 | 53938599 | 12.56253 | 22.845 | 35.93454 | 15.48282 | -0.23307 | 0.329511 | 0.725717 |
| 6 | 63644400 | 63644999 | 9.124558 | 17.13037 | 27.44293 | 11.2923 | -0.20968 | 0.095764 | 0.725717 |
| 4 | 70679400 | 70680599 | 14.88541 | 26.17822 | 38.7425 | 17.15473 | -0.17487 | 0.592082 | 0.725746 |

| | | | | | | | | | |
|---|---|---|---|---|---|---|---|---|---|
| 17 | 11998600 | 12001399 | 18.30964 | 31.5216 | 46.89893 | 20.57259 | -0.18628 | 0.891245 | 0.725749 |
| 14 | 121458200 | 121458999 | 8.683102 | 16.49494 | 26.60423 | 10.69817 | -0.19069 | 0.01208 | 0.725758 |
| 18 | 19379600 | 19381399 | 15.9089 | 27.95833 | 42.35515 | 18.7534 | -0.2088 | 0.61309 | 0.725783 |
| 6 | 84689600 | 84691999 | 22.40224 | 36.89064 | 72.18762 | 31.13213 | -0.58628 | 1.376442 | 0.725798 |
| 10 | 32887000 | 32888999 | 29.42666 | 49.22391 | 55.76974 | 24.29261 | 0.20148 | 1.198928 | 0.725807 |
| 18 | 50555000 | 50555799 | 9.073885 | 17.18189 | 28.19868 | 11.41682 | -0.22488 | 0.03905 | 0.725821 |
| 5 | 50647600 | 50648399 | 6.09554 | 11.84208 | 21.16955 | 7.901106 | -0.23672 | -0.27943 | 0.725845 |
| 18 | 69298600 | 69299999 | 18.08113 | 31.71726 | 36.49744 | 16.13464 | 0.185895 | 0.64218 | 0.725871 |
| X | 126647600 | 126658399 | 119.2285 | 183.2859 | 170.5548 | 70.8234 | 0.39934 | 2.966599 | 0.72588 |
| 2 | 5509600 | 5512599 | 40.91924 | 66.33367 | 76.32962 | 32.68631 | 0.16314 | 1.645419 | 0.725903 |
| 5 | 101254000 | 101255799 | 16.09625 | 28.09427 | 42.02866 | 18.50594 | -0.19094 | 0.719152 | 0.725919 |
| 4 | 24066200 | 24069599 | 29.58375 | 49.93471 | 53.53083 | 23.4136 | 0.281142 | 1.112758 | 0.725924 |
| 8 | 16486000 | 16488799 | 26.31184 | 43.94368 | 66.06495 | 28.74825 | -0.20521 | 1.219195 | 0.725929 |
| 8 | 16729800 | 16730599 | 7.733037 | 14.86422 | 24.70028 | 9.646367 | -0.19478 | -0.098 | 0.725944 |
| 14 | 6351400 | 6351799 | 7.733037 | 14.86422 | 24.70028 | 9.646367 | -0.19478 | -0.098 | 0.725944 |
| 7 | 12492400 | 12510799 | 233.0351 | 339.7343 | 362.5402 | 149.3951 | 0.134425 | 3.959502 | 0.725961 |
| 1 | 125725800 | 125729799 | 42.44162 | 68.27811 | 96.53754 | 40.97641 | -0.14557 | 1.859869 | 0.725969 |
| 8 | 121455800 | 121457599 | 11.91525 | 21.46397 | 32.65202 | 14.2359 | -0.19598 | 0.372788 | 0.725969 |
| 6 | 123106600 | 123110999 | 43.70443 | 71.5754 | 58.5145 | 25.17677 | 0.665944 | 1.481626 | 0.72597 |
| X | 167577800 | 167579999 | 22.59199 | 38.94889 | 43.6928 | 19.16702 | 0.219895 | 0.894292 | 0.725974 |
| 7 | 94020800 | 94023599 | 36.14283 | 58.78137 | 67.39198 | 28.92217 | 0.172621 | 1.569914 | 0.726008 |
| 7 | 49769400 | 49772399 | 27.90621 | 46.02568 | 67.00902 | 28.91777 | -0.16028 | 1.364098 | 0.726017 |
| 6 | 133734400 | 133740199 | 92.73672 | 143.8196 | 162.1568 | 67.34902 | 0.132305 | 2.791603 | 0.726046 |
| 6 | 143425400 | 143429599 | 37.34817 | 60.33608 | 86.25074 | 36.81862 | -0.15415 | 1.722019 | 0.726058 |
| 7 | 143417200 | 143421399 | 35.3734 | 56.87215 | 101.5003 | 43.04461 | -0.48192 | 1.872291 | 0.726069 |
| 3 | 15794200 | 15798799 | 23.39337 | 40.75515 | 35.92417 | 15.79964 | 0.568411 | 0.772081 | 0.726083 |
| 18 | 89960000 | 89962799 | 25.97962 | 43.21432 | 64.45306 | 27.97455 | -0.19385 | 1.256136 | 0.726088 |
| 2 | 3053000 | 3053799 | 17.98824 | 31.44854 | 36.26962 | 15.9885 | 0.182906 | 0.696759 | 0.726088 |
| 3 | 105496600 | 105498599 | 15.30664 | 26.71664 | 41.02975 | 18.06633 | -0.22854 | 0.713315 | 0.726089 |
| X | 77242600 | 77244199 | 24.76439 | 42.87598 | 37.77452 | 16.57295 | 0.568191 | 0.857451 | 0.726094 |
| 16 | 75479800 | 75480599 | 9.064283 | 17.01377 | 27.47416 | 11.30902 | -0.2209 | 0.099871 | 0.726114 |
| 3 | 90913000 | 90914599 | 29.42541 | 49.23252 | 56.24986 | 24.50511 | 0.189474 | 1.19957 | 0.726141 |
| 3 | 102141200 | 102146399 | 103.9172 | 160.1883 | 179.3834 | 74.51285 | 0.135474 | 2.908924 | 0.726169 |
| X | 18183200 | 18192999 | 149.6801 | 226.3209 | 241.6295 | 99.92611 | 0.182115 | 3.310656 | 0.726183 |
| 12 | 115435200 | 115438599 | 37.86019 | 62.63061 | 64.1844 | 27.75226 | 0.345363 | 1.352713 | 0.726185 |
| 6 | 133876600 | 133878399 | 16.70738 | 28.73219 | 54.36165 | 23.84539 | -0.5321 | 0.900662 | 0.726191 |
| 9 | 19583800 | 19584399 | 15.47424 | 27.3991 | 31.06118 | 13.73488 | 0.211651 | 0.508227 | 0.7262 |
| 4 | 93293200 | 93295999 | 32.23107 | 53.53367 | 51.96816 | 22.5327 | 0.423677 | 1.281607 | 0.726216 |
| 19 | 7810800 | 7821799 | 25.81141 | 44.01161 | 42.12049 | 18.4646 | 0.445411 | 1.001431 | 0.726241 |
| 9 | 98759400 | 98771199 | 164.9634 | 247.7176 | 265.6408 | 109.639 | 0.167108 | 3.448755 | 0.726244 |
| 6 | 108931000 | 108932599 | 16.40262 | 28.38073 | 50.59275 | 22.19631 | -0.44451 | 0.841958 | 0.726283 |
| 6 | 70734600 | 70738199 | 47.87936 | 76.50113 | 108.2232 | 45.47558 | -0.15793 | 2.082386 | 0.726298 |
| X | 135998800 | 136002199 | 25.49291 | 42.66237 | 63.75485 | 27.77851 | -0.19658 | 1.193632 | 0.726319 |
| 5 | 32403400 | 32406799 | 28.62197 | 47.05035 | 68.49101 | 29.516 | -0.16195 | 1.408949 | 0.726325 |
| 6 | 3290800 | 3293599 | 40.2375 | 64.63952 | 94.26427 | 39.92847 | -0.1926 | 1.902848 | 0.72636 |
| 5 | 12896800 | 12903199 | 99.39442 | 153.896 | 172.1645 | 71.5126 | 0.141795 | 2.824262 | 0.726363 |
| 11 | 88778200 | 88780599 | 17.83709 | 31.24583 | 35.53302 | 15.68991 | 0.202823 | 0.660442 | 0.726379 |

| | | | | | | | | | |
|---|---|---|---|---|---|---|---|---|---|
| 1 | 144097800 | 144102199 | 75.80554 | 119.9462 | 116.9509 | 48.61075 | 0.364122 | 2.396331 | 0.726381 |
| 17 | 73626400 | 73628199 | 14.65338 | 26.22691 | 30.33474 | 13.3033 | 0.19114 | 0.425328 | 0.726383 |
| 7 | 40353400 | 40355799 | 28.63791 | 47.90227 | 55.16172 | 24.02706 | 0.178127 | 1.196953 | 0.726395 |
| X | 57075400 | 57079199 | 58.07273 | 92.94523 | 101.4089 | 42.71452 | 0.218806 | 2.043012 | 0.726398 |
| 3 | 145237800 | 145239399 | 23.6392 | 41.14693 | 36.52222 | 16.05372 | 0.558402 | 0.787268 | 0.726399 |
| 16 | 52018400 | 52020199 | 12.89986 | 22.6249 | 42.53572 | 18.80152 | -0.51845 | 0.629489 | 0.72642 |
| 19 | 50100600 | 50103999 | 40.69533 | 65.42309 | 93.68885 | 39.72706 | -0.16514 | 1.879406 | 0.72644 |
| 9 | 35506400 | 35506999 | 9.592247 | 18.44016 | 23.01536 | 8.985249 | 0.215261 | -0.08379 | 0.72645 |
| 1 | 83531600 | 83532799 | 16.04331 | 28.35695 | 32.92038 | 14.5621 | 0.176358 | 0.532585 | 0.726453 |
| 1 | 23998000 | 23999599 | 15.19579 | 26.52468 | 40.55621 | 17.85846 | -0.22208 | 0.712731 | 0.726485 |
| 5 | 44303000 | 44308799 | 60.56714 | 96.43315 | 110.4258 | 46.17328 | 0.141003 | 2.205461 | 0.726498 |
| 3 | 25002400 | 25004199 | 16.47266 | 29.95884 | 31.36802 | 13.51345 | 0.350665 | 0.346759 | 0.726586 |
| 3 | 12792400 | 12792999 | 15.61143 | 27.6108 | 31.54185 | 13.95058 | 0.19998 | 0.521798 | 0.72659 |
| 11 | 45473400 | 45474199 | 10.94359 | 20.08833 | 30.80956 | 13.11453 | -0.18456 | 0.254246 | 0.726604 |
| 6 | 42137000 | 42140599 | 69.94819 | 110.4243 | 124.4977 | 51.68747 | 0.15224 | 2.447678 | 0.72661 |
| 5 | 109710600 | 109711399 | 8.526617 | 16.30652 | 26.79326 | 10.61759 | -0.19856 | -0.03647 | 0.726637 |
| 8 | 57964000 | 57967199 | 47.62261 | 76.46194 | 87.70357 | 37.1192 | 0.152131 | 1.925229 | 0.726639 |
| 2 | 133774000 | 133774999 | 15.57418 | 27.20755 | 41.0541 | 18.08705 | -0.2029 | 0.705428 | 0.726647 |
| 2 | 50126000 | 50129199 | 29.4504 | 50.25071 | 50.69714 | 22.22562 | 0.369715 | 1.018804 | 0.726669 |
| 11 | 54739000 | 54739799 | 9.911504 | 18.94778 | 23.37679 | 9.273866 | 0.209567 | -0.03089 | 0.726681 |
| Y | 41927200 | 41928999 | 9.937617 | 19.21384 | 12.51954 | 4.792434 | 1.170026 | -0.18037 | 0.726685 |
| 4 | 14153000 | 14153599 | 10.22128 | 19.43704 | 23.86684 | 9.576451 | 0.202544 | 0.012812 | 0.726692 |
| 12 | 14647800 | 14651999 | 93.31286 | 144.5285 | 161.8272 | 67.21863 | 0.140911 | 2.815129 | 0.726695 |
| 18 | 29628800 | 29630199 | 15.05431 | 26.91406 | 31.36676 | 13.76901 | 0.179405 | 0.438195 | 0.726707 |
| 10 | 129873000 | 129878199 | 46.26512 | 74.8313 | 81.27273 | 34.74541 | 0.243279 | 1.700481 | 0.726711 |
| 1 | 117877800 | 117893999 | 222.9926 | 329.2041 | 329.6031 | 135.7366 | 0.243604 | 3.762165 | 0.726712 |
| 17 | 85245600 | 85246399 | 13.91864 | 25.12893 | 29.4527 | 12.8052 | 0.180591 | 0.367572 | 0.726714 |
| 14 | 113793600 | 113798199 | 46.47337 | 75.3401 | 79.22029 | 33.92506 | 0.292915 | 1.655363 | 0.726717 |
| 13 | 86491000 | 86491999 | 10.90398 | 20.0193 | 30.79418 | 13.10471 | -0.18865 | 0.251816 | 0.726726 |
| 17 | 61966200 | 61970799 | 47.66138 | 77.23655 | 80.18555 | 34.32876 | 0.310798 | 1.668154 | 0.726729 |
| X | 38038000 | 38040199 | 33.19939 | 54.13844 | 80.88985 | 34.74872 | -0.2058 | 1.536096 | 0.726751 |
| 17 | 8357000 | 8359199 | 25.48517 | 43.92278 | 39.84088 | 17.4771 | 0.524955 | 0.902096 | 0.726751 |
| 2 | 96024400 | 96027399 | 40.87898 | 66.41785 | 61.59387 | 26.41992 | 0.476707 | 1.593251 | 0.726775 |
| X | 7503800 | 7506399 | 28.81571 | 48.64843 | 52.31349 | 22.88259 | 0.276891 | 1.112689 | 0.726784 |
| 15 | 52338200 | 52342199 | 61.63462 | 98.03048 | 112.6838 | 47.0495 | 0.133912 | 2.241312 | 0.726787 |
| 3 | 131054000 | 131056199 | 33.1176 | 54.53182 | 63.45538 | 27.36531 | 0.161129 | 1.395293 | 0.726788 |
| 4 | 102602000 | 102602799 | 9.485981 | 18.17941 | 21.9009 | 8.626621 | 0.25439 | -0.05768 | 0.726826 |
| 2 | 120643600 | 120647599 | 77.64925 | 120.9467 | 166.105 | 68.944 | -0.14709 | 2.711114 | 0.726832 |
| 15 | 49991600 | 49993199 | 24.37222 | 40.43255 | 61.18874 | 26.50886 | -0.21486 | 1.284379 | 0.726859 |
| 9 | 71597200 | 71599599 | 18.80842 | 32.87711 | 38.23658 | 16.85327 | 0.170801 | 0.706436 | 0.726861 |
| 18 | 13037000 | 13037799 | 14.25351 | 25.45555 | 28.56945 | 12.55223 | 0.232791 | 0.434439 | 0.726931 |
| 8 | 69635000 | 69637199 | 16.29081 | 28.28995 | 43.21079 | 18.96587 | -0.22149 | 0.79466 | 0.726957 |
| 6 | 123739000 | 123739999 | 17.59717 | 30.43423 | 45.30542 | 19.87636 | -0.18572 | 0.84348 | 0.726959 |
| 10 | 68316000 | 68319799 | 36.58329 | 59.6398 | 69.4948 | 29.85056 | 0.151641 | 1.536889 | 0.726983 |
| 7 | 132517400 | 132520999 | 50.2594 | 80.81072 | 91.45374 | 38.78774 | 0.173976 | 1.885707 | 0.726983 |
| 2 | 47559600 | 47560199 | 8.584477 | 15.59664 | 33.9403 | 14.64031 | -0.69859 | 0.303223 | 0.726996 |
| 11 | 82963400 | 82965399 | 19.01896 | 32.85914 | 48.47501 | 21.26625 | -0.1732 | 0.862163 | 0.727007 |

| | | | | | | | | | |
|---|---|---|---|---|---|---|---|---|---|
| 18 | 83217800 | 83221199 | 40.33704 | 65.41379 | 75.31713 | 32.26143 | 0.162724 | 1.634945 | 0.727016 |
| 2 | 123720400 | 123722799 | 18.55409 | 31.68432 | 55.93651 | 24.52416 | -0.43426 | 0.976446 | 0.727031 |
| 5 | 142204200 | 142209799 | 66.14525 | 104.1708 | 144.1563 | 59.81384 | -0.14568 | 2.499421 | 0.727065 |
| 18 | 27731600 | 27732999 | 10.42634 | 19.21018 | 29.80933 | 12.62471 | -0.1961 | 0.22607 | 0.727094 |
| 4 | 21078800 | 21082399 | 42.13263 | 68.33391 | 76.8492 | 32.92061 | 0.196751 | 1.63648 | 0.727105 |
| 12 | 69595600 | 69598199 | 42.95688 | 68.53509 | 120.6532 | 50.59317 | -0.47517 | 2.127903 | 0.727133 |
| 2 | 36649400 | 36653399 | 56.61609 | 90.68847 | 100.7533 | 42.49026 | 0.194093 | 2.014275 | 0.727145 |
| 6 | 68023200 | 68025399 | 10.72218 | 19.42929 | 36.43934 | 15.77013 | -0.48917 | 0.338849 | 0.727171 |
| 16 | 76357600 | 76359199 | 29.37299 | 47.43334 | 91.73532 | 39.13985 | -0.5897 | 1.725298 | 0.727255 |
| 1 | 67536000 | 67536599 | 10.43686 | 19.31514 | 29.86843 | 12.57355 | -0.18477 | 0.201903 | 0.727271 |
| 7 | 6570800 | 6571599 | 10.43686 | 19.31514 | 29.86843 | 12.57355 | -0.18477 | 0.201903 | 0.727271 |
| X | 5453600 | 5459399 | 82.47312 | 130.6598 | 131.4386 | 54.66721 | 0.320591 | 2.377785 | 0.727274 |
| 3 | 128528000 | 128529799 | 24.15601 | 40.97356 | 47.44222 | 20.78141 | 0.170672 | 1.046181 | 0.727299 |
| 10 | 86962200 | 86965199 | 33.39692 | 54.77759 | 63.92073 | 27.52052 | 0.154846 | 1.442925 | 0.727311 |
| 16 | 7926200 | 7930799 | 29.55703 | 48.56115 | 70.39892 | 30.33126 | -0.15638 | 1.417846 | 0.727316 |
| 15 | 19393800 | 19395799 | 17.61764 | 30.72035 | 45.71887 | 20.11872 | -0.18365 | 0.746761 | 0.727341 |
| 10 | 9550400 | 9552399 | 16.97433 | 29.40456 | 44.27203 | 19.42395 | -0.20145 | 0.825363 | 0.727365 |
| 1 | 96459600 | 96461599 | 24.23807 | 40.4964 | 59.27163 | 25.79862 | -0.16662 | 1.200866 | 0.727367 |
| 1 | 173193200 | 173196199 | 27.35845 | 44.78047 | 68.65044 | 29.53107 | -0.2389 | 1.458682 | 0.727372 |
| X | 115144600 | 115145599 | 14.82641 | 27.22924 | 30.09979 | 12.80377 | 0.285461 | 0.279741 | 0.727381 |
| 3 | 65110400 | 65111399 | 10.51292 | 19.81686 | 23.6498 | 9.665058 | 0.219637 | 0.071546 | 0.727384 |
| 8 | 44883600 | 44886399 | 22.19629 | 37.60598 | 54.93724 | 24.05606 | -0.16352 | 1.057945 | 0.727393 |
| 5 | 21369400 | 21371399 | 15.98757 | 28.01936 | 41.3847 | 18.28115 | -0.1724 | 0.654779 | 0.727399 |
| 18 | 38776600 | 38780799 | 30.71517 | 50.43627 | 73.40158 | 31.61762 | -0.16231 | 1.42953 | 0.727417 |
| 19 | 53072200 | 53073199 | 10.55537 | 19.4404 | 30.30886 | 12.84303 | -0.20355 | 0.229505 | 0.727421 |
| 12 | 115171600 | 115174199 | 37.48699 | 61.58259 | 67.768 | 29.19629 | 0.240143 | 1.438031 | 0.727429 |
| 6 | 129046400 | 129051199 | 52.36887 | 83.70288 | 119.7607 | 50.34788 | -0.17377 | 2.080912 | 0.727432 |
| 19 | 49554200 | 49554799 | 12.97738 | 23.82441 | 28.08851 | 11.95436 | 0.19265 | 0.263429 | 0.727467 |
| 13 | 102431600 | 102432199 | 9.341033 | 17.87389 | 22.34917 | 8.842474 | 0.194366 | -0.04677 | 0.727468 |
| 17 | 56437400 | 56438599 | 13.96656 | 25.0527 | 27.18239 | 11.89874 | 0.285108 | 0.403226 | 0.72748 |
| X | 23858200 | 23861799 | 42.34456 | 69.50716 | 69.47353 | 29.921 | 0.378118 | 1.460282 | 0.727517 |
| X | 34584200 | 34586799 | 53.13355 | 85.39471 | 79.96607 | 33.89409 | 0.445944 | 1.900787 | 0.72752 |
| 6 | 99897000 | 99899999 | 29.57568 | 48.64959 | 70.43008 | 30.35995 | -0.1537 | 1.403958 | 0.727545 |
| 3 | 29375400 | 29375799 | 6.096496 | 11.87402 | 21.36235 | 7.802767 | -0.21661 | -0.35247 | 0.72756 |
| 18 | 35181600 | 35183599 | 16.01709 | 27.97349 | 41.53555 | 18.29596 | -0.17993 | 0.711345 | 0.727566 |
| 4 | 10871400 | 10874799 | 25.71878 | 42.66068 | 63.14191 | 27.35249 | -0.18309 | 1.287815 | 0.727577 |
| Y | 42483000 | 42485399 | 13.16793 | 23.80395 | 22.21491 | 9.643016 | 0.508631 | 0.335844 | 0.727579 |
| 7 | 132913600 | 132916999 | 33.30745 | 54.01366 | 78.50697 | 33.62759 | -0.1725 | 1.626177 | 0.727579 |
| 17 | 94758000 | 94761399 | 38.08904 | 62.73376 | 66.90849 | 28.85756 | 0.286603 | 1.406696 | 0.727579 |
| 7 | 28429800 | 28432599 | 38.83536 | 62.97258 | 72.38575 | 31.00364 | 0.165064 | 1.635407 | 0.727596 |
| 10 | 75592200 | 75594999 | 20.68506 | 36.08225 | 33.67058 | 14.82023 | 0.486842 | 0.739113 | 0.7276 |
| 3 | 53162800 | 53165799 | 20.23128 | 34.55086 | 50.60698 | 22.18767 | -0.1659 | 0.975248 | 0.727605 |
| 8 | 74686600 | 74689199 | 23.82095 | 41.08314 | 45.75551 | 20.07212 | 0.230256 | 0.896944 | 0.727619 |
| 10 | 79514400 | 79515599 | 9.371673 | 17.54067 | 27.95507 | 11.56265 | -0.20781 | 0.121565 | 0.727639 |
| 10 | 43323600 | 43326199 | 20.15285 | 34.47631 | 51.16039 | 22.43342 | -0.18435 | 0.958074 | 0.727665 |
| 8 | 33469800 | 33471999 | 27.76411 | 46.8874 | 52.88681 | 23.13552 | 0.208115 | 1.103871 | 0.727669 |
| 8 | 53777600 | 53778399 | 7.586878 | 14.48346 | 24.53144 | 9.7627 | -0.24745 | -0.03911 | 0.727681 |

| | | | | | | | | | |
|---|---|---|---|---|---|---|---|---|---|
| 17 | 61627800 | 61629399 | 16.83783 | 29.67865 | 34.4131 | 15.23086 | 0.176505 | 0.569872 | 0.727686 |
| 9 | 6489400 | 6492599 | 42.86023 | 69.43884 | 78.42661 | 33.56955 | 0.189164 | 1.661184 | 0.727734 |
| 2 | 176507400 | 176508799 | 12.383 | 22.28047 | 33.62353 | 14.68021 | -0.18591 | 0.384542 | 0.727747 |
| 18 | 29842600 | 29843599 | 8.819459 | 16.85584 | 27.94091 | 11.09255 | -0.2139 | -0.02571 | 0.727755 |
| 1 | 147544400 | 147546399 | 17.74712 | 31.40221 | 34.68308 | 15.33852 | 0.248296 | 0.532969 | 0.727763 |
| 7 | 55595000 | 55598999 | 121.7957 | 184.2297 | 246.8491 | 102.097 | -0.14486 | 3.296546 | 0.727766 |
| 15 | 10625200 | 10629999 | 36.77318 | 59.22041 | 101.1608 | 43.00344 | -0.41594 | 1.820877 | 0.727782 |
| 11 | 35937600 | 35938399 | 10.52936 | 19.18009 | 30.31294 | 13.0331 | -0.23722 | 0.295095 | 0.727811 |
| 1 | 155626400 | 155627999 | 17.10376 | 29.51932 | 45.32218 | 19.88245 | -0.23075 | 0.859178 | 0.727812 |
| 8 | 126452400 | 126455399 | 24.54901 | 41.48798 | 48.00375 | 21.00404 | 0.171553 | 1.08063 | 0.727816 |
| 4 | 98107200 | 98108999 | 15.82288 | 27.71943 | 40.88951 | 18.05619 | -0.17054 | 0.659709 | 0.727827 |
| 4 | 140615200 | 140619999 | 87.59082 | 135.1319 | 185.6211 | 77.10358 | -0.15804 | 2.887531 | 0.727838 |
| 4 | 138905200 | 138907199 | 34.99729 | 57.06857 | 65.1984 | 28.00767 | 0.180371 | 1.528852 | 0.727844 |
| 10 | 18114600 | 18117599 | 39.76288 | 64.42736 | 74.88335 | 32.05533 | 0.148055 | 1.652356 | 0.727846 |
| 5 | 41670600 | 41672399 | 23.64874 | 40.75637 | 45.76179 | 20.07442 | 0.218372 | 0.90263 | 0.72785 |
| 3 | 129227000 | 129230199 | 31.23027 | 51.36097 | 58.27461 | 25.11802 | 0.196783 | 1.400702 | 0.727874 |
| 6 | 49283200 | 49287399 | 35.08319 | 57.03891 | 82.89987 | 35.57325 | -0.16914 | 1.581436 | 0.72791 |
| 4 | 54282800 | 54285999 | 32.55067 | 53.10561 | 76.55521 | 32.8923 | -0.15383 | 1.521761 | 0.727911 |
| 3 | 129548000 | 129550199 | 18.00948 | 31.54231 | 36.04279 | 15.91266 | 0.195938 | 0.665111 | 0.727944 |
| 4 | 112405800 | 112406999 | 18.73232 | 32.9475 | 36.92161 | 16.34807 | 0.224363 | 0.609356 | 0.72796 |
| 17 | 18938600 | 18945599 | 129.5845 | 198.6592 | 176.9387 | 73.44667 | 0.459458 | 3.022671 | 0.727971 |
| 2 | 94184000 | 94184799 | 9.302223 | 17.92608 | 22.50458 | 8.724812 | 0.215706 | -0.11332 | 0.727984 |
| 4 | 86889800 | 86890799 | 11.34477 | 20.31166 | 32.26015 | 14.14384 | -0.26307 | 0.406855 | 0.727996 |
| 14 | 6399400 | 6403199 | 58.379 | 93.06354 | 107.0066 | 44.8197 | 0.136993 | 2.16483 | 0.728013 |
| 3 | 50315600 | 50317199 | 15.35563 | 27.43647 | 31.82239 | 13.976 | 0.185743 | 0.444319 | 0.728014 |
| 7 | 101240400 | 101241599 | 17.22154 | 30.47499 | 28.28074 | 12.50688 | 0.498319 | 0.518059 | 0.728015 |
| 4 | 8573400 | 8573999 | 15.45207 | 26.84191 | 42.76562 | 18.77421 | -0.28172 | 0.789012 | 0.728055 |
| 6 | 108050000 | 108053599 | 29.20498 | 48.03806 | 69.9034 | 30.13245 | -0.16115 | 1.401936 | 0.72806 |
| 11 | 8298600 | 8301599 | 34.75667 | 56.99602 | 66.28662 | 28.53663 | 0.159559 | 1.450954 | 0.728074 |
| 11 | 116339000 | 116342199 | 41.01736 | 66.26659 | 76.37617 | 32.60522 | 0.155906 | 1.721818 | 0.728076 |
| 18 | 85225000 | 85226799 | 17.22552 | 30.08604 | 45.14446 | 19.8865 | -0.1953 | 0.719431 | 0.728082 |
| 17 | 20821600 | 20823599 | 24.53583 | 42.77026 | 42.85745 | 18.85579 | 0.384477 | 0.781013 | 0.728093 |
| 4 | 130778200 | 130782799 | 49.22322 | 79.00674 | 91.12708 | 38.54952 | 0.143574 | 1.937585 | 0.7281 |
| 2 | 5073400 | 5076199 | 38.87684 | 63.43952 | 71.67843 | 30.79925 | 0.196949 | 1.529226 | 0.728104 |
| 11 | 5643400 | 5646999 | 47.49993 | 76.1357 | 107.874 | 45.5384 | -0.15504 | 1.983769 | 0.72811 |
| 7 | 29449800 | 29454199 | 17.12005 | 31.55756 | 24.11489 | 10.20668 | 0.819944 | 0.26479 | 0.72812 |
| 4 | 65908400 | 65909999 | 10.74515 | 19.32009 | 36.32334 | 15.86826 | -0.50151 | 0.384265 | 0.728123 |
| 7 | 33366000 | 33452199 | 1641.136 | 2041.505 | 1789.089 | 752.8595 | 0.188988 | 6.450812 | 0.728176 |
| 5 | 17471400 | 17473399 | 15.38053 | 27.57473 | 31.83457 | 13.94126 | 0.195261 | 0.42382 | 0.728214 |
| X | 77001000 | 77002999 | 36.18278 | 59.20009 | 67.86417 | 29.18851 | 0.178847 | 1.485757 | 0.728219 |
| 9 | 105483600 | 105486399 | 49.93382 | 79.75753 | 111.8546 | 46.97962 | -0.14595 | 2.097342 | 0.728225 |
| 2 | 17042400 | 17045599 | 34.37183 | 56.60972 | 64.82925 | 27.96044 | 0.184202 | 1.395439 | 0.728229 |
| 4 | 8142800 | 8144399 | 21.79395 | 38.06102 | 41.26045 | 18.17429 | 0.271651 | 0.739315 | 0.728235 |
| 4 | 147848200 | 147920199 | 2255.773 | 2711.53 | 2978.631 | 1258.975 | -0.18722 | 6.995032 | 0.728241 |
| 13 | 9891000 | 9894999 | 35.146 | 57.54211 | 66.39604 | 28.56435 | 0.16995 | 1.469564 | 0.728262 |
| 1 | 161141200 | 161144399 | 43.92225 | 70.76708 | 80.99709 | 34.45558 | 0.161434 | 1.809965 | 0.728274 |
| 7 | 65465800 | 65468199 | 49.96543 | 80.11152 | 92.26884 | 38.96849 | 0.143824 | 1.973634 | 0.728292 |

| | | | | | | | | | |
|---|---|---|---|---|---|---|---|---|---|
| 2 | 46526800 | 46527799 | 9.511965 | 17.56582 | 28.34294 | 11.96617 | -0.24851 | 0.202928 | 0.728295 |
| 14 | 65513800 | 65516199 | 19.21751 | 33.21895 | 49.01411 | 21.50309 | -0.17327 | 0.859898 | 0.728304 |
| 18 | 88233400 | 88237799 | 47.38071 | 75.73802 | 107.161 | 45.05755 | -0.15746 | 2.067822 | 0.728333 |
| 12 | 114654000 | 114668199 | 204.4636 | 303.9294 | 305.6394 | 125.9173 | 0.246453 | 3.643304 | 0.728334 |
| 16 | 90994200 | 90995399 | 16.60777 | 29.35941 | 33.92035 | 15.00394 | 0.183426 | 0.537177 | 0.72835 |
| 4 | 125165400 | 125166799 | 14.08515 | 24.77322 | 37.27998 | 16.50683 | -0.19866 | 0.584012 | 0.728364 |
| 14 | 27282800 | 27285199 | 30.05496 | 49.27873 | 71.48047 | 30.77071 | -0.15839 | 1.44471 | 0.728394 |
| 2 | 100767000 | 100768199 | 13.56252 | 25.33951 | 27.70538 | 11.49389 | 0.327651 | 0.163319 | 0.728414 |
| 16 | 48037200 | 48038199 | 11.45927 | 21.27032 | 25.1598 | 10.54869 | 0.203539 | 0.182659 | 0.728427 |
| 13 | 41429600 | 41432999 | 36.42367 | 59.69203 | 68.16112 | 29.33578 | 0.18557 | 1.463933 | 0.728438 |
| 6 | 140267200 | 140274199 | 88.05617 | 137.5332 | 155.2145 | 64.40174 | 0.138891 | 2.667373 | 0.728444 |
| 4 | 8336800 | 8337599 | 9.49486 | 18.05429 | 21.77275 | 8.73761 | 0.228035 | -0.00074 | 0.72846 |
| 6 | 124265600 | 124267399 | 20.35854 | 35.4345 | 40.88529 | 17.9709 | 0.181573 | 0.77749 | 0.728467 |
| 19 | 25386600 | 25389199 | 34.97915 | 56.02656 | 110.9164 | 46.77183 | -0.63774 | 1.995407 | 0.728477 |
| 16 | 23461800 | 23462599 | 14.20032 | 24.95438 | 37.75654 | 16.71656 | -0.20692 | 0.594751 | 0.728477 |
| 5 | 86761600 | 86763799 | 27.59437 | 46.4246 | 53.42753 | 23.34523 | 0.179037 | 1.136035 | 0.728487 |
| 12 | 56318200 | 56319199 | 9.837755 | 18.39159 | 28.98766 | 12.01342 | -0.1944 | 0.135028 | 0.728514 |
| 1 | 160970200 | 160970599 | 5.527958 | 10.75787 | 20.14686 | 7.415852 | -0.28456 | -0.33968 | 0.72853 |
| 15 | 99056800 | 99062399 | 39.18543 | 63.24165 | 89.73688 | 38.25797 | -0.14513 | 1.756173 | 0.728538 |
| X | 125728600 | 125729799 | 13.99772 | 25.36605 | 29.51211 | 12.77137 | 0.195636 | 0.348193 | 0.728546 |
| 2 | 135968000 | 135969799 | 16.5514 | 29.29036 | 33.84616 | 14.968 | 0.183675 | 0.524423 | 0.728549 |
| 3 | 9154800 | 9158999 | 51.16755 | 81.7446 | 114.1238 | 47.94639 | -0.13915 | 2.091662 | 0.728557 |
| 8 | 110308400 | 110311799 | 39.39169 | 63.49335 | 90.31364 | 38.44046 | -0.15074 | 1.794832 | 0.72856 |
| 13 | 18026800 | 18030599 | 42.62926 | 68.61691 | 97.36252 | 41.35885 | -0.14973 | 1.844115 | 0.728561 |
| 13 | 22382000 | 22386999 | 66.39813 | 105.5117 | 119.437 | 49.82154 | 0.154716 | 2.27323 | 0.728568 |
| 7 | 137048600 | 137051399 | 19.93825 | 34.84429 | 39.72291 | 17.5051 | 0.199436 | 0.717575 | 0.728574 |
| 15 | 73031000 | 73032799 | 18.68943 | 32.633 | 37.82274 | 16.65804 | 0.175715 | 0.722341 | 0.728583 |
| X | 148511400 | 148513199 | 32.66311 | 53.1242 | 77.06516 | 33.07402 | -0.16596 | 1.569418 | 0.728586 |
| 9 | 86175600 | 86177199 | 16.95028 | 30.07899 | 26.8792 | 11.87338 | 0.554372 | 0.488536 | 0.728613 |
| 6 | 3131800 | 3136799 | 188.2645 | 277.8115 | 286.4124 | 117.9493 | 0.200781 | 3.763435 | 0.728616 |
| 8 | 112877400 | 112879199 | 17.94417 | 31.58642 | 35.9644 | 15.9217 | 0.202013 | 0.59529 | 0.728621 |
| 14 | 50700800 | 50704799 | 50.1815 | 80.57662 | 92.31198 | 39.0742 | 0.153995 | 1.929135 | 0.728627 |
| X | 156580400 | 156583199 | 38.83972 | 63.50738 | 70.47706 | 30.3054 | 0.224898 | 1.502034 | 0.728647 |
| 3 | 8828200 | 8829599 | 8.268444 | 15.58584 | 25.96955 | 10.61329 | -0.25727 | 0.061421 | 0.728647 |
| 7 | 60445200 | 60449599 | 54.60426 | 87.78225 | 94.14125 | 39.91806 | 0.251134 | 1.898345 | 0.728659 |
| 2 | 50880400 | 50882999 | 24.75718 | 41.82376 | 48.53447 | 21.23379 | 0.167341 | 1.085944 | 0.72867 |
| 4 | 73629600 | 73632799 | 35.98253 | 58.40533 | 86.11323 | 36.91062 | -0.19192 | 1.617943 | 0.728687 |
| 13 | 53974400 | 53976399 | 23.59793 | 40.3242 | 46.87888 | 20.55414 | 0.166274 | 0.979053 | 0.72869 |
| 10 | 67752000 | 67754599 | 19.85539 | 34.11953 | 50.08346 | 21.96827 | -0.16763 | 0.91626 | 0.728701 |
| 11 | 53116600 | 53118999 | 27.39748 | 45.26994 | 65.92365 | 28.48143 | -0.16015 | 1.340064 | 0.728707 |
| 14 | 40843000 | 40844599 | 13.31403 | 23.43528 | 36.34599 | 16.09076 | -0.24134 | 0.569899 | 0.728713 |
| 3 | 8092200 | 8092999 | 12.65253 | 23.35689 | 27.54437 | 11.63487 | 0.199743 | 0.226496 | 0.728721 |
| 5 | 38764400 | 38766599 | 19.89427 | 34.74801 | 39.73017 | 17.50169 | 0.195112 | 0.724814 | 0.728758 |
| 6 | 74993800 | 74994199 | 6.546561 | 12.68967 | 22.18073 | 8.390323 | -0.22253 | -0.22265 | 0.728767 |
| 17 | 75861400 | 75863599 | 19.84285 | 34.04329 | 50.08186 | 21.96556 | -0.17123 | 0.930295 | 0.728774 |
| 17 | 81925600 | 81927599 | 19.77244 | 34.72682 | 37.88206 | 16.76418 | 0.26256 | 0.634706 | 0.728794 |
| 13 | 86275600 | 86275999 | 12.32333 | 22.66555 | 26.62908 | 11.30477 | 0.200378 | 0.244964 | 0.728846 |

| | | | | | | | | | |
|---|---|---|---|---|---|---|---|---|---|
| 14 | 44230000 | 44231599 | 35.28077 | 56.98656 | 83.20789 | 35.51195 | -0.18488 | 1.723359 | 0.728848 |
| 6 | 93307800 | 93310199 | 30.12117 | 50.39696 | 56.82658 | 24.75646 | 0.208431 | 1.20262 | 0.728879 |
| 13 | 68039000 | 68042999 | 130.9823 | 198.1545 | 207.9182 | 85.99914 | 0.207784 | 3.292945 | 0.72889 |
| 4 | 35622200 | 35623399 | 11.88545 | 21.49629 | 32.66633 | 14.17091 | -0.18984 | 0.348916 | 0.728899 |
| 11 | 9714200 | 9719599 | 68.26938 | 108.3547 | 122.6621 | 51.10521 | 0.152955 | 2.310902 | 0.72891 |
| 11 | 9714400 | 9719799 | 68.26938 | 108.3547 | 122.6621 | 51.10521 | 0.152955 | 2.310902 | 0.72891 |
| 5 | 129748400 | 129752599 | 58.2568 | 93.14382 | 104.5079 | 43.95566 | 0.176888 | 2.073395 | 0.728942 |
| 3 | 44838200 | 44840599 | 12.86632 | 22.60951 | 41.93636 | 18.56674 | -0.49918 | 0.595756 | 0.728943 |
| 7 | 31659400 | 31663599 | 60.72675 | 96.64034 | 110.5543 | 46.19593 | 0.141696 | 2.223227 | 0.728945 |
| 17 | 74600200 | 74604399 | 13.07862 | 22.8378 | 44.7174 | 19.69551 | -0.57696 | 0.704353 | 0.728956 |
| 3 | 8403400 | 8403799 | 8.01959 | 15.53589 | 19.95116 | 7.567533 | 0.214127 | -0.20442 | 0.72897 |
| 7 | 44321600 | 44324399 | 21.44898 | 36.17226 | 53.7904 | 23.52088 | -0.18923 | 1.095147 | 0.728979 |
| 12 | 60620800 | 60621199 | 7.267412 | 14.13492 | 18.45334 | 6.823867 | 0.22492 | -0.30536 | 0.728996 |
| 2 | 48120800 | 48123399 | 24.49159 | 42.58965 | 43.44343 | 19.08447 | 0.358475 | 0.812869 | 0.728999 |
| 3 | 69984600 | 69985199 | 5.662635 | 11.02675 | 20.18437 | 7.392191 | -0.24525 | -0.35536 | 0.729014 |
| 17 | 75208400 | 75215599 | 126.5925 | 189.271 | 310.9971 | 128.2362 | -0.4538 | 3.526598 | 0.729016 |
| 4 | 8506600 | 8509199 | 20.8513 | 36.09245 | 42.05404 | 18.45039 | 0.16637 | 0.845074 | 0.729023 |
| 6 | 133770200 | 133771199 | 15.53682 | 28.4966 | 30.53944 | 13.01394 | 0.32788 | 0.291754 | 0.729036 |
| 14 | 124987000 | 125001399 | 172.6911 | 259.1337 | 271.1228 | 111.8798 | 0.201691 | 3.464901 | 0.729051 |
| 16 | 88511600 | 88511999 | 5.947013 | 11.5838 | 20.78497 | 7.583889 | -0.21139 | -0.36033 | 0.729058 |
| 15 | 9385400 | 9389399 | 36.42467 | 60.38709 | 62.42927 | 27.03393 | 0.332962 | 1.321923 | 0.729071 |
| 19 | 13214000 | 13233799 | 255.8919 | 373.1823 | 373.7148 | 153.9945 | 0.226533 | 3.95778 | 0.729121 |
| 7 | 26468000 | 26469799 | 15.46911 | 27.15509 | 40.28221 | 17.81875 | -0.17867 | 0.623046 | 0.729129 |
| 10 | 38675400 | 38680999 | 53.97665 | 85.9479 | 119.3449 | 49.90906 | -0.13652 | 2.199808 | 0.729181 |
| 16 | 97776600 | 97780399 | 37.20298 | 60.21558 | 86.06725 | 36.81398 | -0.15106 | 1.674566 | 0.729182 |
| 13 | 13511800 | 13515199 | 65.5594 | 103.1014 | 144.9283 | 60.12307 | -0.16981 | 2.5281 | 0.729189 |
| X | 78569200 | 78571199 | 16.85643 | 30.30162 | 32.84782 | 14.35263 | 0.287955 | 0.421059 | 0.729239 |
| 16 | 64358400 | 64359799 | 14.48159 | 25.61142 | 38.71661 | 17.12404 | -0.20206 | 0.529914 | 0.729252 |
| 2 | 125454600 | 125455399 | 7.280006 | 14.03056 | 23.6544 | 9.167873 | -0.20494 | -0.13021 | 0.72927 |
| 3 | 5247600 | 5248199 | 7.280006 | 14.03056 | 23.6544 | 9.167873 | -0.20494 | -0.13021 | 0.72927 |
| 17 | 27505000 | 27522199 | 389.1115 | 540.2391 | 540.7876 | 224.4871 | 0.15638 | 4.688895 | 0.729325 |
| 14 | 121208200 | 121209399 | 16.48171 | 28.70287 | 42.62712 | 18.74311 | -0.18068 | 0.755317 | 0.729338 |
| 5 | 64330400 | 64331399 | 10.12129 | 19.05923 | 22.22486 | 9.098392 | 0.251306 | 0.072432 | 0.729354 |
| 17 | 23539200 | 23542999 | 35.45349 | 59.32337 | 58.25597 | 25.38044 | 0.40723 | 1.219135 | 0.729374 |
| 14 | 8633800 | 8634399 | 12.33437 | 22.31247 | 34.26949 | 14.86284 | -0.20465 | 0.357238 | 0.729374 |
| 2 | 144665000 | 144666399 | 20.76839 | 35.79884 | 41.13261 | 18.04383 | 0.185364 | 0.882337 | 0.729408 |
| 7 | 15463600 | 15466399 | 58.47881 | 93.18897 | 104.9908 | 43.9534 | 0.165789 | 2.177403 | 0.729452 |
| 3 | 17739200 | 17739999 | 15.22715 | 27.44473 | 24.84309 | 10.82402 | 0.549088 | 0.376058 | 0.729501 |
| 1 | 5945000 | 5945599 | 14.05338 | 25.18826 | 28.96879 | 12.68927 | 0.200593 | 0.410741 | 0.729565 |
| 9 | 121425200 | 121426799 | 21.80316 | 37.4332 | 43.21883 | 18.95657 | 0.177276 | 0.925692 | 0.729628 |
| 6 | 19499800 | 19503999 | 50.58187 | 81.48809 | 76.78997 | 32.66073 | 0.441045 | 1.818496 | 0.729638 |
| 13 | 69269000 | 69270599 | 16.75153 | 29.09749 | 43.70435 | 19.18559 | -0.1974 | 0.793501 | 0.729656 |
| 17 | 30718000 | 30724599 | 66.96638 | 106.5858 | 117.4622 | 49.09248 | 0.195695 | 2.225225 | 0.729662 |
| 8 | 10664600 | 10665599 | 9.934994 | 18.53401 | 28.9329 | 12.03389 | -0.18508 | 0.147596 | 0.729692 |
| 8 | 21391600 | 21396799 | 75.53686 | 119.5358 | 131.9809 | 54.8614 | 0.18514 | 2.396186 | 0.729698 |
| Y | 38979800 | 38980399 | 5.698507 | 11.01804 | 21.58045 | 8.259896 | -0.40147 | -0.19647 | 0.729724 |
| 6 | 147347800 | 147350399 | 26.0744 | 44.80383 | 47.35243 | 20.77032 | 0.304338 | 0.937735 | 0.729752 |

| | | | | | | | | | |
|---|---|---|---|---|---|---|---|---|---|
| X | 71616200 | 71616999 | 6.420838 | 12.47628 | 23.43879 | 8.735443 | -0.30477 | -0.26816 | 0.72976 |
| 7 | 35108800 | 35113399 | 40.20264 | 64.94008 | 73.2587 | 31.26799 | 0.186543 | 1.723781 | 0.729775 |
| 9 | 58599600 | 58603799 | 42.67927 | 68.89783 | 79.93371 | 34.08967 | 0.145127 | 1.749056 | 0.729795 |
| 14 | 106372800 | 106374399 | 32.86139 | 54.34725 | 61.81877 | 26.72463 | 0.195344 | 1.343231 | 0.729799 |
| X | 4432600 | 4435599 | 56.67984 | 91.62605 | 82.18395 | 35.11204 | 0.517934 | 1.738711 | 0.729815 |
| 8 | 35999200 | 36000399 | 17.9718 | 31.47128 | 36.65189 | 16.17938 | 0.168785 | 0.670374 | 0.729829 |
| 16 | 12158600 | 12159199 | 8.05494 | 14.88625 | 31.51172 | 13.29048 | -0.63606 | 0.195663 | 0.729829 |
| 8 | 49236200 | 49239399 | 28.36784 | 46.90081 | 68.36032 | 29.54612 | -0.16143 | 1.337452 | 0.729831 |
| 6 | 86730800 | 86732199 | 14.97009 | 26.21364 | 39.88687 | 17.60673 | -0.21513 | 0.659298 | 0.729835 |
| 1 | 28039200 | 28040599 | 15.44642 | 27.45093 | 31.92175 | 14.08324 | 0.177729 | 0.479934 | 0.729845 |
| 5 | 51406400 | 51408599 | 20.92873 | 36.59167 | 40.23987 | 17.73963 | 0.251243 | 0.718285 | 0.72986 |
| 17 | 22209600 | 22210999 | 23.30479 | 40.7093 | 41.77396 | 18.40325 | 0.350638 | 0.746088 | 0.729867 |
| 9 | 53259000 | 53263399 | 45.70451 | 73.92362 | 81.54247 | 34.8604 | 0.220947 | 1.701364 | 0.729876 |
| 18 | 53531400 | 53533999 | 19.29241 | 33.55566 | 38.34635 | 16.8476 | 0.195291 | 0.777508 | 0.729891 |
| 8 | 75479000 | 75481399 | 17.89366 | 32.05187 | 33.73406 | 14.78417 | 0.327229 | 0.449759 | 0.729966 |
| 7 | 102377400 | 102378599 | 9.845258 | 18.55947 | 30.30227 | 12.38246 | -0.2287 | 0.085073 | 0.729969 |
| 4 | 8881000 | 8889199 | 151.715 | 227.9433 | 251.4933 | 103.8144 | 0.126744 | 3.435469 | 0.729971 |
| 10 | 98373200 | 98373599 | 5.089445 | 9.930383 | 18.66184 | 6.655607 | -0.24754 | -0.46223 | 0.729993 |
| 15 | 93880000 | 93881599 | 14.88782 | 26.98075 | 30.91511 | 13.37747 | 0.217359 | 0.35545 | 0.729995 |
| 16 | 19853000 | 19857599 | 49.44714 | 79.44437 | 91.46662 | 38.75053 | 0.147873 | 1.908433 | 0.730007 |
| 3 | 91769200 | 91771199 | 27.2763 | 46.44134 | 43.64145 | 19.1291 | 0.471321 | 1.021416 | 0.730028 |
| 2 | 141924000 | 141926199 | 24.55978 | 42.36828 | 46.27278 | 20.29919 | 0.258375 | 0.897176 | 0.730031 |
| 10 | 103712800 | 103713999 | 8.077059 | 15.56528 | 26.54359 | 10.29035 | -0.22184 | -0.11215 | 0.730033 |
| 3 | 138364200 | 138366599 | 45.91633 | 73.83649 | 84.64279 | 35.90531 | 0.155676 | 1.8724 | 0.730036 |
| 12 | 23842600 | 23895199 | 1395.002 | 1746.911 | 1597.26 | 671.6349 | 0.137515 | 6.343324 | 0.730045 |
| 11 | 53067600 | 53069399 | 18.58788 | 32.65273 | 36.89461 | 16.32989 | 0.212038 | 0.623051 | 0.73006 |
| 14 | 15965000 | 15966999 | 29.9964 | 49.54408 | 55.50989 | 23.97557 | 0.216997 | 1.345476 | 0.730079 |
| 5 | 17696800 | 17698999 | 17.96039 | 31.6479 | 35.8352 | 15.8612 | 0.210366 | 0.581658 | 0.730093 |
| 13 | 14388600 | 14389999 | 37.45732 | 61.29424 | 51.93474 | 22.33841 | 0.613904 | 1.474722 | 0.73015 |
| 5 | 126027200 | 126031799 | 48.59712 | 78.30132 | 87.73852 | 37.32482 | 0.191982 | 1.817018 | 0.730161 |
| 5 | 59381200 | 59382199 | 16.31808 | 28.91587 | 33.26414 | 14.7067 | 0.190736 | 0.506454 | 0.730169 |
| 3 | 85104400 | 85107399 | 19.73266 | 33.77535 | 49.54108 | 21.72429 | -0.16742 | 0.951473 | 0.73017 |
| 5 | 77501400 | 77504399 | 19.73266 | 33.77535 | 49.54108 | 21.72429 | -0.16742 | 0.951473 | 0.73017 |
| 16 | 88303000 | 88303999 | 12.64555 | 23.57139 | 21.52255 | 8.964123 | 0.581802 | 0.157947 | 0.730214 |
| 13 | 10732200 | 10733599 | 11.35644 | 20.86525 | 32.3835 | 13.76736 | -0.20062 | 0.253899 | 0.730222 |
| 6 | 79007200 | 79010399 | 31.06806 | 50.95774 | 73.89559 | 31.8151 | -0.1578 | 1.44517 | 0.730241 |
| 1 | 52438200 | 52439199 | 28.11933 | 47.37616 | 45.89145 | 20.06111 | 0.426881 | 1.117423 | 0.730262 |
| 1 | 43059600 | 43063999 | 55.20391 | 88.42316 | 100.0434 | 42.18851 | 0.167772 | 2.012362 | 0.730297 |
| 1 | 43059800 | 43063999 | 55.20391 | 88.42316 | 100.0434 | 42.18851 | 0.167772 | 2.012362 | 0.730297 |
| 10 | 10487400 | 10489799 | 18.45995 | 31.66911 | 47.43356 | 20.80395 | -0.19687 | 0.923632 | 0.73033 |
| 14 | 40170600 | 40173999 | 30.47678 | 50.61642 | 58.55015 | 25.38536 | 0.171544 | 1.282346 | 0.730344 |
| 10 | 70977400 | 70979399 | 19.20397 | 33.38119 | 52.89948 | 23.23254 | -0.27458 | 0.806222 | 0.73035 |
| 12 | 74395400 | 74395799 | 6.099708 | 11.84973 | 21.16241 | 7.900831 | -0.23521 | -0.27646 | 0.73036 |
| 12 | 80672200 | 80672799 | 6.099708 | 11.84973 | 21.16241 | 7.900831 | -0.23521 | -0.27646 | 0.73036 |
| 5 | 69891000 | 69891199 | 4.512524 | 8.818583 | 17.12496 | 5.932985 | -0.25555 | -0.57769 | 0.730361 |
| X | 125273000 | 125275199 | 30.94905 | 52.404 | 53.4855 | 23.41612 | 0.352054 | 1.095199 | 0.730374 |
| 6 | 145020800 | 145021799 | 11.55418 | 20.82955 | 32.63553 | 14.21503 | -0.23755 | 0.364115 | 0.73039 |

| | | | | | | | | | |
|---|---|---|---|---|---|---|---|---|---|
| 3 | 7838200 | 7844999 | 130.9242 | 199.2209 | 184.7006 | 76.53149 | 0.393141 | 3.172956 | 0.730422 |
| 3 | 15871800 | 15874399 | 38.13665 | 62.17721 | 71.33058 | 30.64 | 0.174174 | 1.541335 | 0.730458 |
| 10 | 118158400 | 118160199 | 14.54632 | 25.70781 | 38.15256 | 16.87694 | -0.17616 | 0.534859 | 0.730459 |
| 7 | 47997800 | 48001599 | 52.00284 | 83.0449 | 115.9341 | 48.67862 | -0.13974 | 2.105778 | 0.730497 |
| 8 | 66023400 | 66025199 | 18.36778 | 31.61117 | 47.42709 | 20.80406 | -0.19832 | 0.896544 | 0.730499 |
| 11 | 116401200 | 116403599 | 23.57467 | 39.6557 | 57.65098 | 25.18948 | -0.15693 | 1.12831 | 0.730515 |
| 12 | 113722400 | 113726599 | 75.17408 | 118.6683 | 134.2619 | 55.73986 | 0.147131 | 2.455304 | 0.730519 |
| 13 | 67080800 | 67082599 | 21.97772 | 36.81371 | 55.90765 | 24.37363 | -0.21965 | 1.163517 | 0.730524 |
| 8 | 73985200 | 73988999 | 58.347 | 92.88951 | 133.2554 | 55.71189 | -0.18378 | 2.214145 | 0.730533 |
| 1 | 5710400 | 5710999 | 9.330939 | 17.62041 | 28.10798 | 11.44607 | -0.19099 | 0.063155 | 0.730536 |
| 13 | 86497400 | 86498399 | 9.351028 | 17.35602 | 27.86786 | 11.68506 | -0.23425 | 0.172381 | 0.730556 |
| 17 | 33336600 | 33337399 | 7.578025 | 14.65477 | 24.99001 | 9.556717 | -0.20321 | -0.1684 | 0.730567 |
| 3 | 25452000 | 25453199 | 13.42822 | 24.32201 | 28.50457 | 12.34299 | 0.184513 | 0.343615 | 0.730573 |
| 1 | 99110800 | 99112199 | 21.74049 | 37.96068 | 35.03042 | 15.42832 | 0.502985 | 0.733208 | 0.73058 |
| 16 | 15343600 | 15356799 | 201.2123 | 298.1125 | 313.5706 | 129.1416 | 0.177272 | 3.700224 | 0.730602 |
| 12 | 51255200 | 51258399 | 36.21659 | 58.66982 | 83.66388 | 35.80883 | -0.14654 | 1.654276 | 0.730613 |
| 10 | 103354400 | 103355999 | 16.61462 | 29.34382 | 33.95629 | 15.02267 | 0.180644 | 0.547677 | 0.730626 |
| 7 | 102297000 | 102297599 | 8.741145 | 16.83036 | 21.40719 | 8.319884 | 0.194165 | -0.11278 | 0.730638 |
| 8 | 29026400 | 29029399 | 43.21634 | 69.731 | 80.94119 | 34.49775 | 0.143609 | 1.765191 | 0.730654 |
| 11 | 73382400 | 73385399 | 25.22777 | 43.38576 | 46.80591 | 20.53112 | 0.275032 | 0.925971 | 0.730659 |
| 1 | 58841000 | 58842599 | 14.95813 | 26.51248 | 29.41449 | 13.00406 | 0.242863 | 0.487171 | 0.73066 |
| 3 | 69952600 | 69955599 | 31.4668 | 52.83618 | 47.48832 | 20.73478 | 0.534332 | 1.158392 | 0.73067 |
| 15 | 47076800 | 47079799 | 30.85852 | 50.21478 | 86.43684 | 37.10297 | -0.41172 | 1.562037 | 0.730677 |
| 3 | 122470000 | 122472799 | 13.02779 | 24.46306 | 22.97709 | 9.451547 | 0.555517 | 0.118826 | 0.730735 |
| 11 | 29626800 | 29628999 | 35.63143 | 58.2556 | 67.82756 | 29.16503 | 0.155728 | 1.493748 | 0.730741 |
| 2 | 58048600 | 58050199 | 9.577985 | 17.96058 | 28.49082 | 11.74677 | -0.19803 | 0.109384 | 0.730745 |
| 5 | 126781600 | 126787599 | 59.97667 | 95.1685 | 131.3822 | 54.72433 | -0.13326 | 2.31428 | 0.730752 |
| 5 | 126114800 | 126119999 | 48.78566 | 78.46224 | 89.97573 | 38.17576 | 0.155455 | 1.874904 | 0.730785 |
| 5 | 125655200 | 125656999 | 25.4446 | 42.97628 | 50.13157 | 21.93116 | 0.159858 | 1.092081 | 0.730801 |
| 19 | 15247000 | 15247799 | 10.19857 | 18.96494 | 29.9646 | 12.53071 | -0.20831 | 0.175201 | 0.730816 |
| X | 141748600 | 141750399 | 23.46131 | 40.63025 | 44.15218 | 19.3712 | 0.266872 | 0.858808 | 0.730846 |
| 13 | 38490400 | 38492799 | 24.64204 | 41.59089 | 47.48315 | 20.76803 | 0.190677 | 1.090404 | 0.730847 |
| 12 | 114897600 | 114899399 | 35.57301 | 58.28054 | 67.49572 | 29.04573 | 0.165079 | 1.464759 | 0.730848 |
| 4 | 78641400 | 78643799 | 34.97756 | 58.07676 | 53.13008 | 23.03139 | 0.509036 | 1.295408 | 0.730863 |
| 13 | 9886400 | 9890199 | 58.38447 | 93.18797 | 106.678 | 44.75952 | 0.145271 | 2.125732 | 0.730878 |
| 7 | 28758600 | 28761399 | 30.09482 | 48.79614 | 76.14829 | 32.61349 | -0.27492 | 1.622608 | 0.730901 |
| 12 | 19043000 | 19214199 | 5658.203 | 6311.114 | 5153.327 | 2204.292 | 0.130628 | 8.172164 | 0.73091 |
| 5 | 77925600 | 77927199 | 10.79992 | 20.04969 | 31.54141 | 13.22139 | -0.20496 | 0.191862 | 0.730912 |
| 2 | 35709200 | 35711199 | 17.72579 | 30.76429 | 45.27699 | 19.86601 | -0.16833 | 0.812117 | 0.730914 |
| 2 | 126033600 | 126035799 | 19.20916 | 33.6688 | 38.00214 | 16.78837 | 0.213603 | 0.66328 | 0.730927 |
| 3 | 126410000 | 126411399 | 17.2264 | 30.46925 | 34.19935 | 15.12574 | 0.224932 | 0.532622 | 0.73093 |
| 19 | 39967000 | 39967799 | 13.83753 | 25.7711 | 23.8978 | 9.967905 | 0.557864 | 0.179228 | 0.730934 |
| 16 | 79170400 | 79173799 | 66.67817 | 105.5381 | 118.7065 | 49.34851 | 0.158683 | 2.383306 | 0.730953 |
| 9 | 36034800 | 36037799 | 46.45523 | 74.34993 | 104.823 | 44.15127 | -0.15037 | 2.029859 | 0.730967 |
| 8 | 27866800 | 27867599 | 6.837196 | 13.2492 | 22.74102 | 8.613907 | -0.19875 | -0.21398 | 0.730985 |
| 4 | 145342200 | 145349799 | 115.7278 | 180.0976 | 167.2316 | 69.4208 | 0.415967 | 2.747027 | 0.731002 |
| 12 | 116770600 | 116773599 | 24.00074 | 40.92478 | 46.32106 | 20.30606 | 0.204097 | 0.995463 | 0.731009 |

| | | | | | | | | | |
|---|---|---|---|---|---|---|---|---|---|
| 17 | 28884400 | 28886799 | 14.14941 | 24.86128 | 37.86636 | 16.76312 | -0.21617 | 0.600251 | 0.731011 |
| 5 | 30425200 | 30428599 | 32.45819 | 52.82504 | 76.07306 | 32.65594 | -0.1548 | 1.558045 | 0.731013 |
| 6 | 8244800 | 8250799 | 185.4511 | 274.4351 | 297.0527 | 122.3349 | 0.134373 | 3.716271 | 0.731044 |
| 7 | 41977200 | 41986799 | 38.92344 | 64.28578 | 54.98219 | 23.74693 | 0.605315 | 1.367038 | 0.731053 |
| 1 | 71194400 | 71195199 | 10.47694 | 19.25615 | 29.81345 | 12.66864 | -0.19614 | 0.241626 | 0.731082 |
| 11 | 10166600 | 10167199 | 10.47694 | 19.25615 | 29.81345 | 12.66864 | -0.19614 | 0.241626 | 0.731082 |
| 12 | 13438600 | 13439999 | 10.47694 | 19.25615 | 29.81345 | 12.66864 | -0.19614 | 0.241626 | 0.731082 |
| 10 | 48685600 | 48687399 | 23.33281 | 38.96138 | 58.04609 | 25.25597 | -0.19203 | 1.205002 | 0.73109 |
| 1 | 132325000 | 132327199 | 17.25451 | 30.04738 | 44.1586 | 19.41597 | -0.16576 | 0.760992 | 0.731106 |
| 12 | 74649800 | 74651799 | 17.25451 | 30.04738 | 44.1586 | 19.41597 | -0.16576 | 0.760992 | 0.731106 |
| 15 | 95834200 | 95838399 | 57.06826 | 90.64426 | 126.1845 | 52.61205 | -0.14387 | 2.279902 | 0.731107 |
| 12 | 34987000 | 34990399 | 65.82795 | 104.4237 | 119.807 | 49.88892 | 0.132909 | 2.324336 | 0.731125 |
| 4 | 94364800 | 94365599 | 8.466115 | 16.05678 | 26.56247 | 10.71794 | -0.23178 | 0.018368 | 0.731147 |
| 10 | 125515400 | 125516999 | 20.28708 | 35.54361 | 33.30717 | 14.70918 | 0.480787 | 0.668529 | 0.731148 |
| X | 75629600 | 75632199 | 29.23658 | 49.25469 | 53.98868 | 23.6002 | 0.249226 | 1.132467 | 0.731162 |
| 4 | 39767600 | 39768399 | 10.76467 | 19.94561 | 30.95288 | 13.00891 | -0.18823 | 0.200402 | 0.731174 |
| 13 | 63667400 | 63669199 | 8.87358 | 16.48997 | 27.37327 | 11.45862 | -0.28025 | 0.159299 | 0.731176 |
| 13 | 83283200 | 83286799 | 51.20633 | 81.60602 | 114.5825 | 47.97386 | -0.15119 | 2.169065 | 0.731187 |
| 13 | 10661200 | 10662399 | 15.6639 | 27.64278 | 43.96061 | 19.45179 | -0.27625 | 0.568993 | 0.731191 |
| 3 | 131152600 | 131154399 | 14.59582 | 25.74489 | 38.21697 | 16.91204 | -0.17749 | 0.557883 | 0.731197 |
| 11 | 28453800 | 28455199 | 13.3494 | 24.65635 | 28.5989 | 12.07164 | 0.22429 | 0.231987 | 0.731242 |
| 8 | 18879800 | 18886799 | 101.0028 | 155.9958 | 175.8289 | 73.03566 | 0.128215 | 2.871279 | 0.731249 |
| 17 | 8888200 | 8891399 | 38.05229 | 61.29114 | 88.70508 | 37.71816 | -0.17692 | 1.812843 | 0.731276 |
| 10 | 7359600 | 7362799 | 53.82456 | 86.26857 | 97.77348 | 41.27116 | 0.166399 | 1.991884 | 0.73129 |
| 9 | 69513200 | 69513999 | 12.04234 | 22.3152 | 26.18383 | 11.00308 | 0.212898 | 0.200695 | 0.731293 |
| 10 | 74355600 | 74360399 | 59.30028 | 94.43076 | 108.6414 | 45.43692 | 0.134495 | 2.199525 | 0.731322 |
| 12 | 36155800 | 36158799 | 17.08874 | 30.00188 | 34.49685 | 15.26116 | 0.187004 | 0.619637 | 0.731335 |
| 5 | 118525800 | 118529999 | 75.5242 | 118.723 | 134.411 | 55.75697 | 0.141537 | 2.540376 | 0.731346 |
| 5 | 129294400 | 129296999 | 24.74899 | 41.10452 | 61.41469 | 26.62774 | -0.19646 | 1.269106 | 0.731349 |
| 1 | 97350400 | 97353199 | 30.66316 | 51.3317 | 49.16026 | 21.42399 | 0.443237 | 1.191247 | 0.731382 |
| 17 | 40078800 | 40080399 | 57.1588 | 91.49953 | 77.38845 | 32.60544 | 0.58579 | 2.027055 | 0.731398 |
| 3 | 73001000 | 73002199 | 17.65228 | 30.97285 | 36.05137 | 15.94087 | 0.16953 | 0.635989 | 0.731434 |
| 17 | 31684200 | 31686399 | 17.65228 | 30.97285 | 36.05137 | 15.94087 | 0.16953 | 0.635989 | 0.731434 |
| 15 | 80231800 | 80233199 | 14.87467 | 26.09381 | 38.84269 | 17.17209 | -0.18347 | 0.629739 | 0.731465 |
| 7 | 143823400 | 143825799 | 15.65697 | 27.26921 | 41.61424 | 18.29886 | -0.21971 | 0.747606 | 0.731465 |
| 3 | 131067200 | 131076599 | 94.84727 | 145.0085 | 233.6833 | 96.94422 | -0.39833 | 3.070891 | 0.731469 |
| 7 | 19297600 | 19300399 | 24.44025 | 40.73067 | 59.76004 | 25.96958 | -0.17005 | 1.229785 | 0.731473 |
| 6 | 5637400 | 5639199 | 8.740307 | 16.31463 | 26.91857 | 11.18523 | -0.26272 | 0.130356 | 0.73151 |
| 10 | 41824800 | 41826199 | 18.03353 | 31.68834 | 36.47221 | 16.14699 | 0.185642 | 0.616031 | 0.731514 |
| Y | 41881800 | 41883799 | 16.91685 | 29.63431 | 28.3663 | 12.52556 | 0.450267 | 0.642445 | 0.731519 |
| 7 | 117099600 | 117101999 | 35.9563 | 58.75553 | 68.40866 | 29.4092 | 0.155206 | 1.502229 | 0.731531 |
| 5 | 87787800 | 87789399 | 13.99052 | 25.12469 | 28.88601 | 12.63217 | 0.202558 | 0.396895 | 0.73158 |
| 14 | 118480000 | 118484799 | 45.14313 | 73.30564 | 77.76451 | 33.34258 | 0.282467 | 1.621617 | 0.731616 |
| 13 | 17652600 | 17656399 | 34.443 | 56.93613 | 63.75528 | 27.55283 | 0.217822 | 1.352749 | 0.731633 |
| 4 | 70328200 | 70328999 | 10.30712 | 18.44471 | 36.8349 | 16.15629 | -0.59204 | 0.411856 | 0.731643 |
| 3 | 92455400 | 92455799 | 8.565789 | 16.68981 | 20.38362 | 7.40796 | 0.341398 | -0.34762 | 0.731647 |
| 2 | 68684000 | 68685999 | 19.70574 | 34.46563 | 39.13787 | 17.259 | 0.204971 | 0.701715 | 0.731679 |

| | | | | | | | | | |
|---|---|---|---|---|---|---|---|---|---|
| 4 | 147663400 | 147672399 | 272.4266 | 397.5884 | 370.2362 | 152.5494 | 0.332406 | 3.949003 | 0.731736 |
| 4 | 100687000 | 100691399 | 28.75004 | 47.61599 | 70.71244 | 30.5997 | -0.18812 | 1.32126 | 0.731761 |
| 1 | 113532600 | 113534199 | 13.85948 | 24.79107 | 38.22394 | 16.7716 | -0.22182 | 0.439811 | 0.731783 |
| 10 | 70784200 | 70785999 | 11.33023 | 20.39633 | 31.73851 | 13.84932 | -0.22899 | 0.371959 | 0.731797 |
| 12 | 80364200 | 80366599 | 23.21665 | 39.66144 | 46.34931 | 20.32148 | 0.158696 | 0.978714 | 0.731801 |
| 14 | 112911800 | 112914599 | 27.85984 | 47.54533 | 49.58241 | 21.73724 | 0.322047 | 1.007219 | 0.731808 |
| 7 | 41852600 | 41866799 | 204.0395 | 303.2226 | 267.3372 | 110.1348 | 0.435815 | 3.645996 | 0.731858 |
| 5 | 10876600 | 10879599 | 31.05331 | 51.56714 | 59.65212 | 25.86078 | 0.171508 | 1.28768 | 0.731875 |
| 7 | 61290000 | 61292999 | 46.52405 | 75.59701 | 65.05454 | 27.90634 | 0.584454 | 1.604416 | 0.73189 |
| 7 | 133894400 | 133896799 | 20.69301 | 35.37571 | 51.69574 | 22.66689 | -0.16242 | 0.968389 | 0.731891 |
| 18 | 6253800 | 6267199 | 206.1437 | 303.1375 | 327.9085 | 135.039 | 0.126767 | 3.823177 | 0.731894 |
| 13 | 82350600 | 82360199 | 138.721 | 208.9336 | 276.3033 | 114.1309 | -0.1313 | 3.386497 | 0.731907 |
| 17 | 23044200 | 23046599 | 27.84106 | 45.88293 | 67.32846 | 29.04617 | -0.17196 | 1.376286 | 0.731917 |
| 3 | 27338600 | 27341199 | 21.25892 | 36.05884 | 52.73363 | 23.09911 | -0.1649 | 1.043582 | 0.731924 |
| 7 | 23836400 | 23842999 | 96.12399 | 147.8143 | 198.4641 | 82.41637 | -0.12877 | 2.954523 | 0.731931 |
| 16 | 54652600 | 54653999 | 10.36913 | 18.88378 | 36.00543 | 15.48514 | -0.50704 | 0.305361 | 0.731966 |
| 6 | 132097000 | 132101399 | 37.61133 | 62.28266 | 64.11269 | 27.74085 | 0.339085 | 1.341552 | 0.731985 |
| 4 | 11665000 | 11665599 | 9.796599 | 18.72986 | 23.39563 | 9.278872 | 0.192388 | -0.03206 | 0.732012 |
| 18 | 15363600 | 15366999 | 67.19712 | 106.7594 | 104.0066 | 43.37604 | 0.370643 | 2.276802 | 0.732045 |
| 7 | 53252000 | 53252999 | 12.47314 | 22.83537 | 26.61492 | 11.36999 | 0.205309 | 0.271741 | 0.73206 |
| 6 | 103564600 | 103566399 | 19.43193 | 34.28385 | 36.82255 | 16.29414 | 0.286997 | 0.580166 | 0.732071 |
| 18 | 26065000 | 26067399 | 19.43193 | 34.28385 | 36.82255 | 16.29414 | 0.286997 | 0.580166 | 0.732071 |
| Y | 41810000 | 41810399 | 5.289389 | 10.23013 | 21.59217 | 8.252625 | -0.50609 | -0.20608 | 0.732121 |
| 18 | 9701600 | 9709199 | 101.6192 | 155.6427 | 208.8259 | 86.66827 | -0.13215 | 3.034893 | 0.732148 |
| 7 | 70172200 | 70178999 | 65.38179 | 104.0768 | 116.1727 | 48.56197 | 0.177467 | 2.218435 | 0.732161 |
| 7 | 3996000 | 3999599 | 45.81255 | 73.75589 | 85.53815 | 36.34673 | 0.141045 | 1.837008 | 0.732162 |
| 2 | 177294200 | 177297999 | 44.49217 | 71.92572 | 81.30679 | 34.73835 | 0.184841 | 1.712183 | 0.732162 |
| 2 | 111537800 | 111539799 | 21.91352 | 38.05904 | 35.12349 | 15.41748 | 0.502021 | 0.805465 | 0.732166 |
| 3 | 51536000 | 51541999 | 67.88677 | 107.3594 | 152.1599 | 63.22839 | -0.17539 | 2.409997 | 0.732169 |
| 4 | 105569800 | 105572399 | 32.89891 | 54.58749 | 61.46495 | 26.63172 | 0.210234 | 1.30381 | 0.73217 |
| X | 125382800 | 125385199 | 36.20218 | 58.87503 | 66.13808 | 28.38354 | 0.201929 | 1.571461 | 0.732182 |
| 14 | 122470600 | 122472799 | 15.42667 | 26.69788 | 50.05138 | 21.95895 | -0.5168 | 0.833742 | 0.732198 |
| 16 | 64156200 | 64159399 | 27.60268 | 46.73614 | 52.33064 | 22.91028 | 0.2188 | 1.080734 | 0.732252 |
| 3 | 57433000 | 57436799 | 34.92899 | 56.73781 | 81.48213 | 34.94492 | -0.15303 | 1.597158 | 0.732262 |
| 12 | 22302600 | 22304399 | 26.33197 | 44.75541 | 40.65608 | 17.81804 | 0.519738 | 1.031393 | 0.732276 |
| 17 | 22343200 | 22351599 | 107.3885 | 166.8874 | 171.0433 | 71.01788 | 0.272121 | 2.764511 | 0.732279 |
| X | 25366400 | 25368799 | 40.18203 | 66.64385 | 63.03034 | 27.30201 | 0.461171 | 1.332759 | 0.73228 |
| 14 | 109271800 | 109273599 | 18.64153 | 32.21767 | 47.40805 | 20.79838 | -0.16928 | 0.858708 | 0.732283 |
| 18 | 3343200 | 3348399 | 89.85642 | 140.8329 | 135.3493 | 56.13919 | 0.374276 | 2.604514 | 0.732286 |
| 3 | 91802400 | 91807799 | 47.74216 | 77.59642 | 76.64387 | 32.88408 | 0.386436 | 1.607427 | 0.732295 |
| 1 | 18315600 | 18316999 | 20.05552 | 35.08233 | 33.05962 | 14.58034 | 0.472952 | 0.693635 | 0.732355 |
| 6 | 23674200 | 23675199 | 13.87483 | 24.10682 | 47.3867 | 20.80544 | -0.58352 | 0.783488 | 0.732356 |
| 8 | 22265600 | 22268999 | 34.16339 | 56.20379 | 64.8061 | 27.93729 | 0.173713 | 1.407667 | 0.732373 |
| 1 | 165894800 | 165900399 | 78.62729 | 123.4486 | 140.9581 | 58.46346 | 0.127871 | 2.566941 | 0.732376 |
| 1 | 3889200 | 3889799 | 12.32944 | 22.87921 | 27.17466 | 11.39772 | 0.197214 | 0.196681 | 0.732386 |
| 1 | 27474800 | 27475599 | 8.559902 | 16.39374 | 27.42642 | 10.8237 | -0.21864 | -0.04706 | 0.732388 |
| 3 | 21712600 | 21714399 | 31.41653 | 51.99345 | 60.70674 | 26.25636 | 0.157794 | 1.328628 | 0.732411 |

| | | | | | | | | | |
|---|---|---|---|---|---|---|---|---|---|
| 3 | 84936200 | 84940599 | 31.41653 | 51.99345 | 60.70674 | 26.25636 | 0.157794 | 1.328628 | 0.732411 |
| 10 | 32765600 | 32766999 | 15.0133 | 26.19438 | 40.56097 | 17.85552 | -0.24036 | 0.717286 | 0.732416 |
| 17 | 54208400 | 54210599 | 32.38461 | 53.40653 | 62.40885 | 26.93142 | 0.155896 | 1.372754 | 0.732432 |
| 6 | 131774400 | 131777199 | 27.26458 | 46.66291 | 41.96698 | 18.40282 | 0.535457 | 0.973128 | 0.732449 |
| 2 | 87252800 | 87253399 | 10.77204 | 20.59564 | 19.65837 | 7.7957 | 0.578077 | -0.03128 | 0.732486 |
| 17 | 61713600 | 61715199 | 14.76607 | 26.21156 | 40.76644 | 18.00285 | -0.24064 | 0.499227 | 0.73249 |
| 9 | 71467000 | 71468399 | 20.1761 | 35.20586 | 40.32981 | 17.75494 | 0.192372 | 0.742394 | 0.732513 |
| 7 | 112869800 | 112872799 | 21.22587 | 35.83699 | 53.81738 | 23.54085 | -0.20335 | 1.082747 | 0.732518 |
| 1 | 116349000 | 116349599 | 6.819518 | 13.21584 | 22.75486 | 8.616416 | -0.20319 | -0.21689 | 0.732543 |
| 2 | 15727200 | 15727999 | 6.819518 | 13.21584 | 22.75486 | 8.616416 | -0.20319 | -0.21689 | 0.732543 |
| 10 | 126063600 | 126064599 | 11.09504 | 21.02121 | 24.77579 | 10.01874 | 0.250527 | 0.044257 | 0.732547 |
| 2 | 31113400 | 31119799 | 63.72629 | 100.9393 | 139.326 | 57.94177 | -0.13543 | 2.367388 | 0.73257 |
| 14 | 73746600 | 73748799 | 16.91705 | 29.70572 | 33.99618 | 15.04192 | 0.193744 | 0.614852 | 0.732574 |
| 6 | 69332400 | 69346599 | 153.8343 | 232.9516 | 240.9068 | 99.6744 | 0.229905 | 3.28047 | 0.732574 |
| 2 | 151032000 | 151036399 | 44.18759 | 71.03308 | 101.8437 | 43.18122 | -0.16705 | 1.886635 | 0.732576 |
| 8 | 82411200 | 82412599 | 18.15245 | 32.13371 | 35.20909 | 15.56976 | 0.260058 | 0.53253 | 0.732581 |
| 5 | 57131400 | 57135799 | 53.35613 | 85.84929 | 92.008 | 39.062 | 0.253587 | 1.868274 | 0.732588 |
| 4 | 50630800 | 50631199 | 4.774 | 9.328951 | 17.68796 | 6.138543 | -0.22362 | -0.56414 | 0.732595 |
| 5 | 79258800 | 79259599 | 10.70014 | 19.65489 | 30.78549 | 13.0921 | -0.21404 | 0.245995 | 0.732596 |
| 3 | 34171200 | 34172399 | 13.68223 | 24.3716 | 36.59735 | 16.11478 | -0.18738 | 0.461612 | 0.732607 |
| 14 | 79953600 | 79955599 | 15.54147 | 27.44384 | 31.05905 | 13.74137 | 0.212462 | 0.53651 | 0.732613 |
| 2 | 150911800 | 150917599 | 68.34268 | 108.4837 | 122.4212 | 51.01083 | 0.157672 | 2.308336 | 0.73262 |
| 3 | 40012000 | 40012599 | 8.980921 | 17.18457 | 29.21386 | 11.55898 | -0.24559 | -0.03217 | 0.732628 |
| 5 | 149030600 | 149031799 | 13.18428 | 23.5008 | 35.7069 | 15.71356 | -0.20324 | 0.456186 | 0.732635 |
| 8 | 22295600 | 22298399 | 31.93852 | 52.63554 | 61.71272 | 26.62346 | 0.150764 | 1.380216 | 0.732643 |
| 10 | 51489600 | 51492199 | 18.26193 | 32.00982 | 37.05374 | 16.36994 | 0.177395 | 0.656589 | 0.732656 |
| 7 | 23970800 | 23976199 | 75.40831 | 117.8502 | 160.7637 | 66.70006 | -0.1336 | 2.651233 | 0.732669 |
| 13 | 5860000 | 5862399 | 19.34984 | 33.97 | 37.9761 | 16.7997 | 0.227446 | 0.640284 | 0.732672 |
| 7 | 41676400 | 41701999 | 396.5068 | 553.8948 | 552.6362 | 229.1869 | 0.170048 | 4.598203 | 0.732697 |
| 11 | 32086800 | 32089799 | 19.10207 | 32.61357 | 49.01338 | 21.48852 | -0.20285 | 0.971283 | 0.732742 |
| 18 | 48759800 | 48760999 | 9.819689 | 18.08008 | 28.81507 | 12.21498 | -0.23517 | 0.223083 | 0.732742 |
| 7 | 40688800 | 40703199 | 221.3929 | 324.2652 | 347.9807 | 143.3409 | 0.132765 | 3.887212 | 0.732779 |
| 7 | 38602200 | 38603599 | 13.49979 | 25.04337 | 23.92539 | 10.03904 | 0.508688 | 0.19954 | 0.732794 |
| 9 | 6351200 | 6351999 | 7.458957 | 14.29402 | 24.0735 | 9.483722 | -0.2253 | -0.07285 | 0.732798 |
| 4 | 37298400 | 37299599 | 15.60469 | 27.62187 | 31.98241 | 14.14308 | 0.181062 | 0.513243 | 0.732803 |
| 18 | 85628800 | 85632599 | 44.55772 | 72.04833 | 81.22578 | 34.71408 | 0.189125 | 1.706711 | 0.732805 |
| X | 103749200 | 103751999 | 19.6502 | 34.07065 | 52.86375 | 23.19413 | -0.24464 | 0.840725 | 0.732809 |
| X | 166909600 | 166912399 | 20.99381 | 36.23544 | 59.59981 | 26.14598 | -0.32997 | 0.894116 | 0.732835 |
| 6 | 149371200 | 149375599 | 48.76663 | 77.7335 | 127.5322 | 53.41047 | -0.37515 | 2.163023 | 0.73285 |
| 1 | 148437000 | 148438199 | 14.20671 | 24.9506 | 37.79222 | 16.72386 | -0.20846 | 0.604738 | 0.732861 |
| 5 | 45955400 | 45967799 | 182.8799 | 273.8442 | 280.1281 | 115.542 | 0.231458 | 3.506917 | 0.732862 |
| 2 | 174005800 | 174007999 | 17.10236 | 29.65173 | 44.25326 | 19.41623 | -0.18862 | 0.817502 | 0.732863 |
| 4 | 67785600 | 67786399 | 13.02149 | 24.13585 | 22.85844 | 9.602297 | 0.519706 | 0.196063 | 0.732881 |
| 17 | 68349400 | 68350599 | 14.48278 | 25.2012 | 46.84238 | 20.58603 | -0.50301 | 0.763681 | 0.732893 |
| 10 | 34839800 | 34841999 | 17.95351 | 31.18669 | 45.79395 | 20.10345 | -0.16487 | 0.802984 | 0.732899 |
| 2 | 120335400 | 120337599 | 20.85887 | 36.09111 | 40.95596 | 17.96841 | 0.20409 | 0.845032 | 0.732908 |
| 7 | 64476600 | 64477999 | 16.42438 | 29.27962 | 33.32043 | 14.66191 | 0.211221 | 0.469287 | 0.732911 |

| | | | | | | | | | |
|---|---|---|---|---|---|---|---|---|---|
| 1 | 16940400 | 16942199 | 33.15603 | 54.93818 | 61.94068 | 26.81227 | 0.208244 | 1.321939 | 0.732933 |
| 17 | 42003600 | 42005599 | 15.24512 | 26.62153 | 48.92979 | 21.55114 | -0.48657 | 0.719057 | 0.732939 |
| 7 | 47191800 | 47202199 | 129.9547 | 199.1939 | 204.3778 | 84.83465 | 0.255359 | 3.02949 | 0.732945 |
| 8 | 74237000 | 74238999 | 22.78357 | 38.41621 | 55.97503 | 24.47485 | -0.15998 | 1.104353 | 0.732949 |
| Y | 6552600 | 6555199 | 26.37466 | 44.18088 | 38.40546 | 16.74384 | 0.582345 | 1.155321 | 0.732951 |
| 14 | 17524400 | 17525999 | 31.6608 | 52.63391 | 47.11627 | 20.4417 | 0.540431 | 1.267687 | 0.732962 |
| 5 | 7795000 | 7797199 | 27.74976 | 45.79354 | 66.43444 | 28.67683 | -0.15484 | 1.356651 | 0.733023 |
| 3 | 17930400 | 17931799 | 15.0523 | 27.10268 | 31.35437 | 13.67792 | 0.196067 | 0.394362 | 0.733027 |
| 13 | 40209000 | 40209599 | 7.161032 | 13.75341 | 23.58262 | 9.231669 | -0.24217 | -0.09752 | 0.733048 |
| 3 | 27268200 | 27269999 | 13.82913 | 24.58319 | 37.08153 | 16.356 | -0.19559 | 0.475653 | 0.733056 |
| 5 | 21388000 | 21389199 | 10.86077 | 19.66863 | 37.48733 | 16.23584 | -0.51316 | 0.344386 | 0.733059 |
| 11 | 61542400 | 61544399 | 17.74549 | 30.74156 | 45.29057 | 19.87085 | -0.17035 | 0.829143 | 0.733059 |
| 15 | 32124000 | 32125399 | 14.38087 | 25.98495 | 30.40091 | 13.20378 | 0.184046 | 0.367516 | 0.733073 |
| 17 | 57750000 | 57751199 | 14.38087 | 25.98495 | 30.40091 | 13.20378 | 0.184046 | 0.367516 | 0.733073 |
| 8 | 77173000 | 77175799 | 16.32769 | 28.49119 | 42.06059 | 18.51712 | -0.1718 | 0.725723 | 0.733078 |
| 10 | 78707000 | 78708199 | 22.08282 | 37.75468 | 43.25543 | 18.96621 | 0.18742 | 0.964 | 0.733082 |
| 17 | 48168400 | 48170399 | 22.08282 | 37.75468 | 43.25543 | 18.96621 | 0.18742 | 0.964 | 0.733082 |
| 8 | 119615800 | 119618399 | 25.2909 | 43.11262 | 49.02445 | 21.49067 | 0.197432 | 1.007775 | 0.733103 |
| 4 | 95935400 | 95940199 | 93.27499 | 143.6023 | 193.6286 | 80.42132 | -0.13359 | 2.929736 | 0.733132 |
| 14 | 46687000 | 46689999 | 25.39086 | 42.39862 | 61.55516 | 26.78305 | -0.155 | 1.212596 | 0.733135 |
| 7 | 109889200 | 109892599 | 48.84936 | 77.92808 | 111.3201 | 46.67282 | -0.17445 | 2.132273 | 0.733141 |
| 9 | 57302600 | 57306799 | 44.53871 | 71.59858 | 102.4592 | 43.44312 | -0.16428 | 1.887342 | 0.733148 |
| 2 | 103894200 | 103896599 | 22.98459 | 38.84143 | 57.17332 | 25.01566 | -0.17462 | 1.08417 | 0.733148 |
| 8 | 21849000 | 21850599 | 27.66168 | 46.16271 | 53.39381 | 23.22317 | 0.171967 | 1.220048 | 0.733149 |
| 1 | 75752800 | 75753799 | 11.47261 | 21.25028 | 24.6835 | 10.3787 | 0.226706 | 0.19372 | 0.733156 |
| 6 | 84004000 | 84007399 | 21.80069 | 36.95541 | 54.95999 | 24.06991 | -0.18924 | 1.050214 | 0.733188 |
| 13 | 66474600 | 66475199 | 33.69269 | 55.0691 | 60.74194 | 26.1154 | 0.233321 | 1.49065 | 0.73319 |
| 12 | 22608000 | 22609199 | 33.77098 | 55.80506 | 47.17931 | 20.38418 | 0.622048 | 1.342152 | 0.733191 |
| 4 | 115233600 | 115239999 | 93.51129 | 145.4059 | 163.7204 | 67.97148 | 0.137252 | 2.747843 | 0.733192 |
| 2 | 17496000 | 17497399 | 16.25043 | 29.2003 | 32.21034 | 14.07894 | 0.262299 | 0.414953 | 0.733209 |
| 1 | 150959000 | 150960399 | 20.47568 | 34.7312 | 51.64726 | 22.62341 | -0.18883 | 1.040024 | 0.733211 |
| 3 | 88175000 | 88176999 | 18.67743 | 32.63816 | 37.67244 | 16.60033 | 0.181551 | 0.707873 | 0.733226 |
| 10 | 11014200 | 11018199 | 39.76637 | 65.15812 | 70.30377 | 30.2556 | 0.267007 | 1.47549 | 0.733228 |
| 6 | 130732400 | 130734999 | 28.71542 | 48.32403 | 53.97405 | 23.58601 | 0.222129 | 1.139765 | 0.733229 |
| 5 | 25160800 | 25162399 | 22.46899 | 37.89832 | 55.41522 | 24.23251 | -0.16499 | 1.100206 | 0.733231 |
| 19 | 47953800 | 47955199 | 13.60075 | 23.97831 | 36.25545 | 16.04554 | -0.20409 | 0.552998 | 0.73325 |
| 17 | 67781800 | 67793999 | 150.8764 | 225.16 | 296.0895 | 122.0453 | -0.13509 | 3.56404 | 0.733257 |
| 16 | 86893200 | 86893799 | 9.186123 | 17.70406 | 22.51586 | 8.726756 | 0.197737 | -0.11456 | 0.733262 |
| 9 | 9403000 | 9405399 | 38.66685 | 62.62781 | 71.41881 | 30.56375 | 0.174929 | 1.653991 | 0.733295 |
| 1 | 113653800 | 113658599 | 48.44964 | 78.36464 | 82.21575 | 35.14856 | 0.293097 | 1.705207 | 0.733305 |
| 4 | 60587800 | 60589799 | 74.69332 | 117.5491 | 133.6205 | 55.43717 | 0.136966 | 2.517611 | 0.733316 |
| 5 | 7623400 | 7624799 | 19.14196 | 33.34817 | 38.88923 | 17.10359 | 0.166568 | 0.757194 | 0.733337 |
| 5 | 100235400 | 100235799 | 9.183032 | 17.64109 | 20.90885 | 8.179336 | 0.287255 | -0.08521 | 0.733363 |
| 14 | 13588600 | 13591599 | 50.43138 | 80.57897 | 112.8642 | 47.42592 | -0.14362 | 2.088074 | 0.733373 |
| 17 | 71640200 | 71643799 | 29.40591 | 48.37871 | 69.85293 | 30.11337 | -0.1498 | 1.400919 | 0.733376 |
| 16 | 70696400 | 70697399 | 15.34093 | 27.45891 | 31.80906 | 13.94926 | 0.188963 | 0.432985 | 0.73339 |
| 3 | 44192000 | 44192399 | 8.642509 | 16.73005 | 21.57198 | 8.211115 | 0.203209 | -0.18121 | 0.733393 |

| | | | | | | | | | |
|---|---|---|---|---|---|---|---|---|---|
| 3 | 80283400 | 80284399 | 15.27637 | 27.0162 | 29.94564 | 13.24479 | 0.242965 | 0.514542 | 0.7334 |
| X | 145020200 | 145022999 | 23.55279 | 41.10862 | 42.31088 | 18.63014 | 0.346159 | 0.75839 | 0.733408 |
| 3 | 111093200 | 111095199 | 19.3438 | 33.52892 | 51.19082 | 22.45991 | -0.22149 | 0.84079 | 0.733417 |
| X | 75465800 | 75466599 | 12.55158 | 23.1202 | 27.56525 | 11.67821 | 0.181132 | 0.238716 | 0.733463 |
| 2 | 72438000 | 72439799 | 22.14371 | 38.20183 | 44.15089 | 19.36837 | 0.177051 | 0.882641 | 0.73347 |
| 1 | 58657000 | 58659199 | 21.80373 | 37.59895 | 43.63158 | 19.14031 | 0.171069 | 0.884242 | 0.733478 |
| X | 14368600 | 14369999 | 19.49144 | 34.38565 | 36.85183 | 16.30739 | 0.290116 | 0.582294 | 0.733489 |
| 3 | 9074800 | 9077599 | 42.08222 | 68.15801 | 78.1269 | 33.43391 | 0.167494 | 1.666358 | 0.73349 |
| 14 | 72763200 | 72764999 | 23.95397 | 40.00521 | 58.65705 | 25.52447 | -0.16912 | 1.204951 | 0.733505 |
| 12 | 105887800 | 105889599 | 26.72297 | 44.6961 | 51.79764 | 22.56077 | 0.169066 | 1.189793 | 0.733507 |
| 7 | 73249400 | 73250199 | 14.98946 | 26.75527 | 30.98086 | 13.61797 | 0.187547 | 0.448687 | 0.733522 |
| 1 | 58902600 | 58915999 | 215.3705 | 314.5767 | 410.7557 | 169.2304 | -0.15393 | 3.930951 | 0.733546 |
| 10 | 75964600 | 75965799 | 16.5436 | 29.30478 | 33.81123 | 14.94967 | 0.186372 | 0.513735 | 0.733555 |
| 5 | 75464000 | 75468599 | 68.36714 | 107.1869 | 151.4823 | 62.82865 | -0.18099 | 2.58877 | 0.733566 |
| 18 | 74775200 | 74777999 | 29.62329 | 48.4807 | 70.58572 | 30.36199 | -0.16486 | 1.467795 | 0.733579 |
| 19 | 39619200 | 39621399 | 46.21746 | 74.46428 | 85.71131 | 36.46039 | 0.153259 | 1.81531 | 0.733625 |
| 17 | 54204000 | 54210999 | 78.03018 | 123.2342 | 136.5118 | 56.68726 | 0.178107 | 2.445499 | 0.733627 |
| 1 | 176918000 | 176920399 | 23.69931 | 39.92011 | 58.25098 | 25.46223 | -0.16223 | 1.116603 | 0.733631 |
| 4 | 75614200 | 75615199 | 12.15494 | 22.17928 | 34.92359 | 14.98323 | -0.2307 | 0.303389 | 0.733633 |
| 7 | 38010600 | 38012799 | 28.77425 | 47.19943 | 68.99375 | 29.70571 | -0.16917 | 1.435318 | 0.733656 |
| 5 | 46964400 | 46965599 | 12.66106 | 23.12188 | 26.63375 | 11.41485 | 0.219076 | 0.287783 | 0.733663 |
| 6 | 123996800 | 123997199 | 9.651398 | 18.3797 | 22.80599 | 9.123043 | 0.191128 | -0.00698 | 0.733667 |
| X | 9886400 | 9888199 | 14.95335 | 26.47418 | 40.75762 | 18.02289 | -0.2278 | 0.523755 | 0.733694 |
| 19 | 53708600 | 53710799 | 17.28941 | 30.1061 | 44.16814 | 19.41933 | -0.16323 | 0.762922 | 0.7337 |
| 4 | 92360200 | 92361399 | 13.20773 | 24.08026 | 28.48191 | 12.2323 | 0.178813 | 0.303412 | 0.73371 |
| 2 | 107647400 | 107648799 | 19.41518 | 33.92314 | 39.22637 | 17.28358 | 0.179002 | 0.716929 | 0.733726 |
| 4 | 123798400 | 123799199 | 6.697256 | 12.87431 | 23.06118 | 9.004163 | -0.30163 | -0.11525 | 0.733738 |
| 15 | 99690000 | 99690799 | 11.53375 | 21.53771 | 25.72543 | 10.682 | 0.20063 | 0.148734 | 0.733748 |
| 13 | 30281000 | 30282999 | 18.69699 | 32.26388 | 47.40617 | 20.79658 | -0.16775 | 0.870324 | 0.733764 |
| 1 | 161141400 | 161144399 | 42.94705 | 69.25935 | 79.33184 | 33.78846 | 0.161893 | 1.780831 | 0.733794 |
| 14 | 52051000 | 52053599 | 25.42292 | 43.09449 | 49.5875 | 21.71683 | 0.179555 | 1.057759 | 0.73381 |
| 8 | 67049000 | 67051799 | 21.73581 | 37.83338 | 36.11446 | 15.87436 | 0.453882 | 0.776663 | 0.733811 |
| 11 | 58532600 | 58534999 | 16.41138 | 28.68866 | 42.48799 | 18.728 | -0.17636 | 0.698639 | 0.733816 |
| 4 | 81695600 | 81697599 | 37.65203 | 61.12586 | 70.74156 | 30.32578 | 0.156869 | 1.608034 | 0.733832 |
| 13 | 96443400 | 96445999 | 41.00428 | 66.79362 | 64.15844 | 27.55036 | 0.428832 | 1.554849 | 0.73384 |
| 7 | 71577400 | 71578199 | 9.542344 | 18.14179 | 21.80651 | 8.753997 | 0.232574 | 0.001949 | 0.733864 |
| 5 | 109927600 | 109946399 | 336.3308 | 478.5532 | 471.8632 | 195.1072 | 0.213754 | 4.325527 | 0.733873 |
| 14 | 37830800 | 37840799 | 224.6071 | 324.1354 | 418.1873 | 172.5329 | -0.15359 | 4.111642 | 0.733903 |
| 2 | 36635200 | 36636399 | 25.15827 | 42.8889 | 49.02405 | 21.49059 | 0.189996 | 1.007221 | 0.733912 |
| 12 | 21065200 | 21096999 | 926.3539 | 1200.779 | 1145.334 | 479.9372 | 0.126633 | 5.783017 | 0.733926 |
| 17 | 39388800 | 39395599 | 40.03205 | 65.98714 | 55.60806 | 23.99189 | 0.625886 | 1.392639 | 0.733928 |
| 5 | 5833000 | 5836799 | 91.45808 | 141.7454 | 158.0845 | 65.66381 | 0.147308 | 2.801882 | 0.73399 |
| 4 | 34875400 | 34880599 | 59.43956 | 94.42467 | 130.5947 | 54.45893 | -0.13429 | 2.279653 | 0.734017 |
| 15 | 45528800 | 45532599 | 40.7917 | 66.17612 | 75.53685 | 32.36457 | 0.175699 | 1.627728 | 0.734017 |
| 12 | 40359400 | 40362199 | 39.25989 | 63.90831 | 73.15676 | 31.4065 | 0.17571 | 1.566226 | 0.734019 |
| X | 9468000 | 9470399 | 41.04159 | 66.73944 | 74.23512 | 31.85741 | 0.216021 | 1.587252 | 0.73402 |
| 12 | 4903200 | 4904599 | 15.21552 | 26.95005 | 30.96258 | 13.69036 | 0.192505 | 0.500898 | 0.734046 |

| | | | | | | | | | |
|---|---|---|---|---|---|---|---|---|---|
| 1 | 39465200 | 39468399 | 29.73491 | 48.72529 | 70.81445 | 30.47642 | -0.1615 | 1.450803 | 0.734075 |
| 14 | 122494600 | 122495199 | 6.957417 | 13.46555 | 23.27136 | 8.865711 | -0.21717 | -0.19429 | 0.734076 |
| 1 | 170901400 | 170906999 | 115.5135 | 173.492 | 293.5056 | 121.1455 | -0.48993 | 3.437787 | 0.734093 |
| 4 | 8647000 | 8648999 | 14.00085 | 24.37048 | 46.83762 | 20.58811 | -0.55097 | 0.755934 | 0.734098 |
| 17 | 47732600 | 47735799 | 25.90117 | 43.5043 | 50.69204 | 22.13792 | 0.161129 | 1.141582 | 0.734125 |
| 3 | 14967800 | 14968999 | 8.635353 | 16.4434 | 26.6685 | 10.67007 | -0.19254 | -0.00621 | 0.734138 |
| 4 | 23477600 | 23478399 | 15.17661 | 26.99294 | 31.34538 | 13.81967 | 0.180231 | 0.47033 | 0.734192 |
| 10 | 90941600 | 90942999 | 11.13713 | 20.13938 | 31.22963 | 13.55059 | -0.21878 | 0.342603 | 0.734203 |
| 3 | 121896800 | 121899399 | 17.3047 | 29.83181 | 45.33921 | 19.88923 | -0.21643 | 0.87165 | 0.734211 |
| 14 | 122070000 | 122079799 | 175.2014 | 260.5735 | 284.9937 | 117.4182 | 0.126093 | 3.627729 | 0.734257 |
| 7 | 95942800 | 95944399 | 12.13992 | 22.06373 | 33.80552 | 14.57605 | -0.19576 | 0.327334 | 0.734258 |
| 2 | 65139400 | 65139999 | 7.46199 | 14.37417 | 24.19343 | 9.39014 | -0.20418 | -0.12036 | 0.734273 |
| 10 | 73129000 | 73130799 | 30.99826 | 51.66824 | 58.49719 | 25.42339 | 0.202703 | 1.247217 | 0.734292 |
| 4 | 61867800 | 61875199 | 147.1044 | 222.3591 | 212.4905 | 87.86724 | 0.341539 | 3.313148 | 0.734326 |
| 16 | 56813200 | 56816799 | 29.95146 | 49.87216 | 57.44406 | 24.94951 | 0.177881 | 1.250596 | 0.734334 |
| 3 | 86029000 | 86030599 | 11.59765 | 21.20292 | 32.3084 | 13.82811 | -0.18102 | 0.285328 | 0.734365 |
| 12 | 113529400 | 113532799 | 55.43366 | 89.03023 | 96.36342 | 40.80318 | 0.236173 | 1.929802 | 0.734386 |
| 6 | 136112600 | 136115399 | 19.76738 | 34.52783 | 39.7521 | 17.51181 | 0.185229 | 0.724131 | 0.734394 |
| 4 | 22807800 | 22808599 | 12.97529 | 23.62386 | 27.58149 | 11.86605 | 0.196113 | 0.309302 | 0.7344 |
| 7 | 59879200 | 59879999 | 9.614538 | 18.27556 | 22.24676 | 8.934336 | 0.213475 | 0.003182 | 0.734417 |
| 19 | 13264800 | 13265599 | 18.54099 | 32.45149 | 37.70585 | 16.63748 | 0.172193 | 0.684393 | 0.734423 |
| 5 | 93767200 | 93804199 | 559.7581 | 761.6357 | 722.6656 | 300.942 | 0.204202 | 5.005495 | 0.734424 |
| 16 | 91850400 | 91852199 | 41.68058 | 66.92673 | 95.73581 | 40.52351 | -0.16566 | 1.919061 | 0.734442 |
| 6 | 59652200 | 59652599 | 9.558737 | 17.8047 | 28.42866 | 11.85498 | -0.21987 | 0.155903 | 0.73445 |
| 12 | 105057200 | 105059199 | 17.30007 | 30.088 | 44.20084 | 19.41884 | -0.1653 | 0.780594 | 0.734453 |
| 11 | 42079600 | 42080199 | 7.629748 | 14.6654 | 24.6879 | 9.64207 | -0.21283 | -0.09642 | 0.734485 |
| 8 | 76286600 | 76287599 | 16.21753 | 28.65718 | 33.41774 | 14.7829 | 0.169805 | 0.537603 | 0.734486 |
| 3 | 29030000 | 29030999 | 10.15591 | 18.893 | 29.42983 | 12.29892 | -0.1873 | 0.170136 | 0.734543 |
| 1 | 73325800 | 73339799 | 181.7665 | 268.853 | 351.3585 | 144.6988 | -0.138 | 3.725874 | 0.734545 |
| 15 | 7324800 | 7325199 | 7.451134 | 14.39567 | 24.31276 | 9.339163 | -0.19511 | -0.15503 | 0.734556 |
| 4 | 21648600 | 21651799 | 43.06795 | 69.42526 | 79.93441 | 34.02673 | 0.153737 | 1.792589 | 0.734561 |
| 3 | 47985200 | 47987799 | 28.51128 | 48.36194 | 52.36122 | 22.93624 | 0.267167 | 1.067485 | 0.734614 |
| X | 144825000 | 144825999 | 15.74231 | 28.4774 | 31.46024 | 13.64441 | 0.267857 | 0.375353 | 0.734638 |
| X | 79350800 | 79351599 | 11.83925 | 22.33885 | 26.38262 | 10.762 | 0.237211 | 0.086568 | 0.734658 |
| 4 | 113275600 | 113283599 | 88.18853 | 138.7773 | 146.4263 | 60.74953 | 0.244222 | 2.530253 | 0.734662 |
| 13 | 79337800 | 79338399 | 9.868082 | 18.32861 | 28.90544 | 12.10851 | -0.2074 | 0.176799 | 0.734677 |
| 16 | 90931800 | 90937399 | 60.8146 | 96.70327 | 110.9129 | 46.29663 | 0.136748 | 2.247562 | 0.734692 |
| 2 | 135399800 | 135402199 | 33.88473 | 55.37901 | 63.7683 | 27.41584 | 0.171262 | 1.490703 | 0.734713 |
| 14 | 8051400 | 8052999 | 27.46524 | 46.1742 | 53.41101 | 23.33293 | 0.171691 | 1.142316 | 0.734719 |
| 19 | 39512000 | 39512799 | 19.83361 | 33.71889 | 50.58418 | 22.1696 | -0.20116 | 1.016769 | 0.734729 |
| 2 | 28036000 | 28036599 | 9.509805 | 18.19003 | 22.87868 | 9.061693 | 0.184497 | -0.04 | 0.734738 |
| 13 | 88862600 | 88862999 | 5.798963 | 11.3013 | 20.27088 | 7.34609 | -0.20209 | -0.38894 | 0.734748 |
| 5 | 106849200 | 106851799 | 25.25743 | 42.31821 | 62.13373 | 27.09235 | -0.17124 | 1.177646 | 0.734749 |
| 12 | 11492600 | 11494199 | 17.65618 | 31.58324 | 33.70229 | 14.7873 | 0.306243 | 0.455721 | 0.734762 |
| X | 35723600 | 35724199 | 9.788385 | 18.71474 | 22.90487 | 9.083413 | 0.222411 | -0.03061 | 0.73479 |
| 7 | 137002000 | 137004399 | 20.52813 | 35.9271 | 39.5832 | 17.46486 | 0.248333 | 0.696354 | 0.734798 |
| 1 | 45649800 | 45652999 | 62.93757 | 100.0397 | 115.2385 | 48.08328 | 0.129938 | 2.261508 | 0.734827 |

| | | | | | | | | | |
|---|---|---|---|---|---|---|---|---|---|
| 10 | 22563800 | 22564399 | 7.991244 | 15.45032 | 18.88528 | 7.229905 | 0.272129 | -0.17743 | 0.734843 |
| 12 | 80460600 | 80465599 | 68.26637 | 108.3607 | 122.772 | 51.15611 | 0.151885 | 2.307983 | 0.734862 |
| 8 | 76577400 | 76579799 | 29.69738 | 49.8459 | 47.5178 | 20.74261 | 0.449804 | 1.159481 | 0.734873 |
| 5 | 150191800 | 150193799 | 21.63587 | 36.86884 | 54.37671 | 23.83624 | -0.17632 | 1.006404 | 0.734896 |
| 16 | 46576800 | 46577799 | 12.35239 | 22.46997 | 35.42485 | 15.25718 | -0.23565 | 0.327708 | 0.734904 |
| 1 | 47103000 | 47112999 | 156.4051 | 235.4741 | 255.9996 | 105.7329 | 0.150629 | 3.397019 | 0.734908 |
| 12 | 31609000 | 31610199 | 12.59445 | 22.05359 | 43.10828 | 19.02861 | -0.57434 | 0.654947 | 0.734911 |
| 13 | 5029800 | 5033999 | 91.99106 | 141.1185 | 197.1609 | 81.84813 | -0.18873 | 3.000082 | 0.734913 |
| 3 | 119505000 | 119506199 | 13.2447 | 23.61852 | 35.12225 | 15.45074 | -0.17226 | 0.450371 | 0.734919 |
| 10 | 40339600 | 40342799 | 16.59984 | 29.82823 | 26.9632 | 11.78543 | 0.547795 | 0.406666 | 0.734979 |
| 17 | 42391600 | 42392799 | 17.47983 | 31.01417 | 29.19404 | 12.8975 | 0.479603 | 0.497355 | 0.734981 |
| 1 | 117318800 | 117322799 | 32.9349 | 54.90792 | 59.33844 | 25.79246 | 0.269538 | 1.24813 | 0.73499 |
| 17 | 39976000 | 39978199 | 23.92478 | 41.71913 | 35.43524 | 15.59358 | 0.621922 | 0.763403 | 0.734993 |
| 19 | 3222600 | 3223199 | 17.5437 | 30.7162 | 35.25781 | 15.56167 | 0.189738 | 0.669978 | 0.735034 |
| 11 | 62870600 | 62873199 | 22.63258 | 38.29397 | 56.62682 | 24.78586 | -0.18116 | 1.071244 | 0.735036 |
| 1 | 95709800 | 95711599 | 19.55489 | 33.58993 | 49.0072 | 21.49593 | -0.15896 | 0.917904 | 0.735046 |
| 4 | 62108400 | 62122399 | 248.0555 | 362.698 | 365.4194 | 150.5403 | 0.221432 | 3.917161 | 0.735092 |
| 9 | 39850200 | 39850799 | 8.054687 | 15.62091 | 19.98479 | 7.541772 | 0.22675 | -0.21986 | 0.7351 |
| 2 | 76594800 | 76595199 | 7.871377 | 15.0172 | 25.05244 | 9.983211 | -0.22765 | -0.03124 | 0.735102 |
| 14 | 84467400 | 84472599 | 80.7134 | 127.1306 | 123.431 | 51.21533 | 0.365066 | 2.500969 | 0.735108 |
| 6 | 92710200 | 92713399 | 49.45028 | 79.5941 | 89.86397 | 38.17209 | 0.179272 | 1.851299 | 0.735125 |
| 7 | 75465400 | 75470599 | 58.67989 | 93.67837 | 107.0642 | 44.9344 | 0.147936 | 2.119548 | 0.735141 |
| 10 | 119900400 | 119901999 | 18.45415 | 32.28491 | 37.68405 | 16.62152 | 0.165555 | 0.689809 | 0.735158 |
| 1 | 85106600 | 85112199 | 55.83039 | 89.75262 | 81.52043 | 34.56611 | 0.490331 | 1.895349 | 0.735185 |
| 1 | 85314800 | 85324199 | 454.7934 | 624.856 | 632.6227 | 263.045 | 0.124777 | 4.851445 | 0.735203 |
| 6 | 116475600 | 116489199 | 243.7402 | 353.8816 | 377.5144 | 155.6322 | 0.129146 | 4.023032 | 0.735213 |
| 14 | 42651800 | 42652399 | 8.995738 | 16.89458 | 26.93109 | 11.07496 | -0.20164 | 0.092052 | 0.735222 |
| 1 | 11941200 | 11941999 | 9.952006 | 18.40203 | 28.82527 | 12.1486 | -0.2044 | 0.199931 | 0.73523 |
| 4 | 6158800 | 6159999 | 28.31495 | 46.57254 | 67.87305 | 29.25684 | -0.16324 | 1.401323 | 0.735287 |
| 14 | 118744400 | 118746199 | 14.7528 | 26.39068 | 30.91629 | 13.56448 | 0.172542 | 0.432054 | 0.7353 |
| 5 | 133011600 | 133015999 | 31.35754 | 51.014 | 90.09122 | 38.66807 | -0.44888 | 1.568697 | 0.735309 |
| 9 | 115417200 | 115418599 | 13.60412 | 24.62732 | 28.99437 | 12.56355 | 0.177341 | 0.349165 | 0.735311 |
| 5 | 65058400 | 65066599 | 100.324 | 154.0575 | 207.0875 | 85.98095 | -0.132 | 2.982963 | 0.735324 |
| 5 | 46924400 | 46925599 | 19.56577 | 33.97661 | 39.48225 | 17.32919 | 0.171166 | 0.804852 | 0.735328 |
| 12 | 28890200 | 28895999 | 57.2279 | 91.36075 | 105.2179 | 44.15975 | 0.136926 | 2.117877 | 0.73534 |
| X | 141962600 | 141965799 | 46.99139 | 75.95062 | 71.96118 | 30.73744 | 0.438674 | 1.719175 | 0.735355 |
| 17 | 64773400 | 64774999 | 12.27616 | 22.01576 | 33.16477 | 14.5201 | -0.18592 | 0.404726 | 0.73537 |
| 13 | 80119400 | 80123599 | 49.72615 | 80.0281 | 90.03215 | 38.23658 | 0.184176 | 1.854864 | 0.735371 |
| X | 162543400 | 162545599 | 17.1846 | 30.38774 | 34.42553 | 15.22654 | 0.211849 | 0.539692 | 0.735375 |
| 2 | 23480200 | 23482599 | 40.1178 | 64.87906 | 75.11249 | 32.10774 | 0.151173 | 1.692066 | 0.735376 |
| 1 | 141067400 | 141072399 | 99.80974 | 154.2151 | 173.6259 | 72.12043 | 0.130281 | 2.862596 | 0.735378 |
| 7 | 106553800 | 106555999 | 20.10573 | 35.17363 | 39.6601 | 17.49282 | 0.215182 | 0.701647 | 0.735383 |
| 16 | 58797000 | 58798599 | 31.77044 | 52.2771 | 61.06937 | 26.32857 | 0.155132 | 1.398313 | 0.735387 |
| 15 | 68515200 | 68516999 | 21.58214 | 36.42324 | 53.80881 | 23.53411 | -0.17978 | 1.088691 | 0.7354 |
| 13 | 58600200 | 58601799 | 11.52012 | 20.6532 | 32.2685 | 14.13156 | -0.2383 | 0.40072 | 0.735404 |
| 4 | 48675200 | 48680399 | 42.60474 | 68.36239 | 97.75634 | 41.33438 | -0.16642 | 1.941434 | 0.735406 |
| 17 | 95190800 | 95193399 | 33.24228 | 54.81022 | 62.62717 | 27.02339 | 0.188026 | 1.375284 | 0.735434 |

| | | | | | | | | | |
|---|---|---|---|---|---|---|---|---|---|
| 16 | 59197800 | 59200199 | 24.32293 | 41.76423 | 46.26831 | 20.29432 | 0.236394 | 0.934395 | 0.735443 |
| 9 | 38713400 | 38714999 | 16.35947 | 29.4937 | 32.39272 | 14.10849 | 0.272233 | 0.39994 | 0.735447 |
| 6 | 123845400 | 123847599 | 20.43079 | 35.72941 | 33.92304 | 14.95782 | 0.462037 | 0.699527 | 0.73546 |
| X | 26372800 | 26374799 | 33.8375 | 56.38401 | 60.388 | 26.24024 | 0.28257 | 1.260076 | 0.735462 |
| 4 | 65221600 | 65223599 | 16.94814 | 29.64336 | 44.05185 | 19.42641 | -0.18126 | 0.695702 | 0.735474 |
| 4 | 112086400 | 112088399 | 27.21369 | 45.97544 | 44.28222 | 19.37366 | 0.435138 | 1.088325 | 0.735498 |
| 6 | 143554600 | 143555999 | 14.59652 | 25.62499 | 38.35252 | 16.96609 | -0.19116 | 0.61647 | 0.735503 |
| 6 | 129973000 | 129982599 | 91.95788 | 142.854 | 210.3161 | 87.32897 | -0.25032 | 2.768966 | 0.735532 |
| 5 | 110160200 | 110165199 | 106.4814 | 164.3407 | 182.1457 | 75.65981 | 0.15167 | 2.887134 | 0.735553 |
| 6 | 145578200 | 145580199 | 19.93958 | 34.00888 | 50.06805 | 21.94911 | -0.17346 | 0.985937 | 0.735555 |
| 7 | 10163000 | 10163799 | 23.92234 | 40.46564 | 45.92199 | 20.0987 | 0.199527 | 1.069178 | 0.735571 |
| 1 | 66316000 | 66317199 | 11.97463 | 21.48778 | 32.70217 | 14.31039 | -0.19999 | 0.398499 | 0.735586 |
| 13 | 71637800 | 71641199 | 26.95803 | 45.27999 | 52.85345 | 23.08203 | 0.158628 | 1.14807 | 0.735617 |
| 7 | 103136600 | 103138399 | 24.88891 | 41.64481 | 62.51541 | 27.23568 | -0.20299 | 1.193831 | 0.73562 |
| X | 7520600 | 7525599 | 48.44074 | 78.23714 | 82.82184 | 35.34381 | 0.277895 | 1.750338 | 0.735621 |
| 13 | 78271600 | 78273599 | 17.6872 | 31.26154 | 35.30378 | 15.61648 | 0.215901 | 0.546987 | 0.735628 |
| 1 | 142237600 | 142246399 | 137.615 | 209.0295 | 227.3597 | 94.15728 | 0.160789 | 3.213793 | 0.735682 |
| 4 | 126562400 | 126563599 | 21.87062 | 38.15862 | 35.54468 | 15.64727 | 0.489402 | 0.744853 | 0.735686 |
| 1 | 104200400 | 104201599 | 19.78508 | 34.69517 | 32.44323 | 14.3385 | 0.483978 | 0.652145 | 0.735728 |
| 13 | 29127000 | 29131199 | 30.75361 | 50.16051 | 73.38407 | 31.52677 | -0.17528 | 1.519377 | 0.735746 |
| 2 | 97557400 | 97557999 | 11.3714 | 21.0706 | 24.59907 | 10.33799 | 0.219645 | 0.18874 | 0.735766 |
| 7 | 61382200 | 61385399 | 37.02192 | 61.22955 | 54.84877 | 23.71185 | 0.538913 | 1.347509 | 0.735768 |
| 14 | 92043200 | 92045399 | 26.9399 | 44.06432 | 77.51369 | 33.33517 | -0.43754 | 1.475209 | 0.735773 |
| 3 | 44125800 | 44126399 | 12.6657 | 23.32176 | 27.62407 | 11.70903 | 0.190255 | 0.243621 | 0.735799 |
| 14 | 63981600 | 63982999 | 12.72442 | 22.75842 | 34.11422 | 14.96966 | -0.18124 | 0.425451 | 0.735819 |
| 3 | 43846800 | 43848199 | 22.88314 | 39.77462 | 37.12188 | 16.30281 | 0.486058 | 0.803139 | 0.735822 |
| 7 | 41335600 | 41363399 | 436.8825 | 606.5021 | 592.8667 | 246.1096 | 0.190494 | 4.693293 | 0.735842 |
| 5 | 61933200 | 61936199 | 51.83911 | 82.6006 | 115.4298 | 48.31739 | -0.14462 | 2.174854 | 0.735847 |
| 17 | 7822000 | 7823999 | 20.78176 | 36.06684 | 42.0535 | 18.45163 | 0.166207 | 0.822794 | 0.735848 |
| 2 | 52303400 | 52307999 | 54.98147 | 88.50095 | 92.2156 | 39.17355 | 0.294897 | 1.85962 | 0.735875 |
| 13 | 81045600 | 81047599 | 15.49385 | 27.22972 | 40.24368 | 17.81895 | -0.17334 | 0.605262 | 0.735877 |
| 5 | 150806200 | 150807599 | 12.01644 | 21.72541 | 33.15197 | 14.38805 | -0.19633 | 0.352111 | 0.735888 |
| X | 130369400 | 130370999 | 16.12853 | 28.28626 | 42.96918 | 18.99234 | -0.21261 | 0.649446 | 0.735898 |
| 5 | 28791200 | 28792599 | 16.25624 | 28.2485 | 43.18099 | 18.96062 | -0.22258 | 0.784083 | 0.735902 |
| 12 | 102378400 | 102380399 | 40.71812 | 65.35888 | 95.24839 | 40.29617 | -0.19301 | 1.928591 | 0.735953 |
| 2 | 170960600 | 170966799 | 78.84465 | 123.929 | 141.4624 | 58.68121 | 0.12957 | 2.545817 | 0.735958 |
| 15 | 99626200 | 99627399 | 14.09482 | 25.63956 | 29.98594 | 12.91471 | 0.192433 | 0.326976 | 0.73597 |
| 19 | 18269200 | 18269799 | 8.165231 | 15.62885 | 25.69416 | 10.15688 | -0.19624 | -0.05277 | 0.735972 |
| 6 | 89877000 | 89879199 | 27.63579 | 46.33259 | 53.96028 | 23.53791 | 0.161918 | 1.171036 | 0.735973 |
| X | 39371000 | 39372999 | 23.82798 | 41.36433 | 44.63615 | 19.58721 | 0.277494 | 0.837124 | 0.735991 |
| 9 | 57749400 | 57754999 | 69.14256 | 109.5254 | 125.4907 | 52.18972 | 0.132936 | 2.3677 | 0.735996 |
| 11 | 78694000 | 78695599 | 18.75815 | 32.67419 | 37.8767 | 16.65657 | 0.175082 | 0.755332 | 0.735997 |
| 7 | 13902200 | 13905399 | 73.86154 | 116.4906 | 132.9203 | 55.16208 | 0.133886 | 2.474718 | 0.736007 |
| X | 146557000 | 146560599 | 42.23714 | 69.12995 | 59.93304 | 25.78059 | 0.581131 | 1.489211 | 0.736019 |
| 8 | 34711400 | 34714999 | 71.63489 | 112.8356 | 125.2428 | 51.96734 | 0.172272 | 2.49589 | 0.73605 |
| 6 | 30407600 | 30408199 | 6.829465 | 13.2614 | 22.87337 | 8.570917 | -0.1912 | -0.25118 | 0.736058 |
| 8 | 116058200 | 116060199 | 20.1974 | 35.11504 | 40.93606 | 17.98067 | 0.166648 | 0.792474 | 0.736066 |

| | | | | | | | | | |
|---|---|---|---|---|---|---|---|---|---|
| 16 | 52958600 | 52960399 | 15.79022 | 28.8174 | 30.92996 | 13.26709 | 0.319105 | 0.321245 | 0.736082 |
| 15 | 49707600 | 49709199 | 18.70067 | 32.35412 | 47.4023 | 20.79653 | -0.16285 | 0.847303 | 0.736093 |
| 8 | 13432400 | 13434999 | 21.89393 | 36.93722 | 54.36124 | 23.77334 | -0.17437 | 1.092856 | 0.736104 |
| 3 | 141457200 | 141458399 | 13.76882 | 24.35914 | 36.17186 | 15.99743 | -0.17622 | 0.516082 | 0.736106 |
| 10 | 74509400 | 74511999 | 15.8139 | 27.15701 | 52.14771 | 22.87327 | -0.55358 | 0.908993 | 0.736142 |
| 3 | 139082400 | 139085199 | 22.7076 | 39.63475 | 41.75327 | 18.38499 | 0.312665 | 0.751134 | 0.736165 |
| 17 | 76591400 | 76591999 | 6.841495 | 13.24243 | 22.68907 | 8.640102 | -0.20343 | -0.19578 | 0.736166 |
| 15 | 51520600 | 51523199 | 28.57758 | 48.18273 | 54.01875 | 23.61903 | 0.216752 | 1.121238 | 0.736178 |
| 3 | 128563800 | 128564399 | 9.137083 | 17.46533 | 21.26954 | 8.440664 | 0.227965 | -0.04298 | 0.736196 |
| 15 | 12028600 | 12029399 | 10.77439 | 19.78898 | 30.2981 | 12.88687 | -0.18126 | 0.248423 | 0.736217 |
| 8 | 89604400 | 89608199 | 30.4031 | 49.75674 | 71.93045 | 30.94038 | -0.15468 | 1.469486 | 0.736222 |
| 4 | 52240800 | 52242999 | 20.26825 | 34.02792 | 61.2399 | 26.7368 | -0.46379 | 1.139502 | 0.73623 |
| 7 | 59028600 | 59044599 | 291.9897 | 417.7561 | 437.718 | 180.8242 | 0.134109 | 4.241977 | 0.73623 |
| 2 | 173271800 | 173273399 | 18.44131 | 32.16979 | 37.32831 | 16.43066 | 0.173889 | 0.732175 | 0.736235 |
| 14 | 107063200 | 107068199 | 68.28374 | 108.2014 | 124.3212 | 51.71224 | 0.129288 | 2.358575 | 0.736239 |
| 6 | 44666600 | 44670399 | 41.67139 | 67.07263 | 94.52007 | 40.14926 | -0.13988 | 1.84346 | 0.736243 |
| 1 | 26271000 | 26272199 | 11.56608 | 20.54 | 38.79096 | 17.12426 | -0.51863 | 0.472014 | 0.73626 |
| 4 | 5006000 | 5009199 | 41.27845 | 67.03311 | 65.71358 | 28.17655 | 0.396087 | 1.604389 | 0.736323 |
| 3 | 19996400 | 19999199 | 31.99115 | 53.23726 | 59.7952 | 25.96078 | 0.214104 | 1.265657 | 0.73634 |
| 10 | 130419400 | 130424199 | 48.05729 | 77.61099 | 84.27498 | 35.95985 | 0.241062 | 1.747278 | 0.736361 |
| 1 | 8208200 | 8211999 | 54.88325 | 87.91298 | 99.94761 | 42.15065 | 0.160887 | 2.010856 | 0.736396 |
| 3 | 46688600 | 46690599 | 20.92706 | 35.31653 | 53.26271 | 23.29503 | -0.20947 | 1.085576 | 0.736404 |
| 13 | 61978600 | 62034999 | 1249.699 | 1588.701 | 1436.22 | 602.9723 | 0.175813 | 6.100634 | 0.736416 |
| 13 | 8252600 | 8253999 | 14.05646 | 24.65991 | 37.86092 | 16.73889 | -0.22785 | 0.621704 | 0.736439 |
| 1 | 89356200 | 89359399 | 33.77007 | 54.68592 | 79.63735 | 34.07598 | -0.17722 | 1.655077 | 0.736448 |
| 5 | 128133400 | 128135399 | 18.38493 | 31.96053 | 47.37713 | 20.80846 | -0.1785 | 0.794833 | 0.736484 |
| 18 | 41195000 | 41195999 | 12.14527 | 22.84903 | 26.85858 | 11.01288 | 0.237819 | 0.108699 | 0.736486 |
| 8 | 44879000 | 44882199 | 58.84593 | 93.23396 | 130.0521 | 54.09229 | -0.15024 | 2.356503 | 0.736495 |
| 17 | 10358600 | 10362199 | 36.35751 | 59.30752 | 69.05385 | 29.66769 | 0.153348 | 1.526405 | 0.736527 |
| 4 | 6478400 | 6480599 | 22.97828 | 39.71498 | 37.77531 | 16.57243 | 0.457438 | 0.861155 | 0.736535 |
| 4 | 112009200 | 112011799 | 26.94646 | 45.96941 | 43.09632 | 18.89313 | 0.475114 | 1.00211 | 0.736578 |
| 2 | 153866800 | 153868399 | 19.57348 | 33.56483 | 49.00822 | 21.49382 | -0.16044 | 0.93368 | 0.736594 |
| 9 | 32232000 | 32234199 | 19.57348 | 33.56483 | 49.00822 | 21.49382 | -0.16044 | 0.93368 | 0.736594 |
| 5 | 15648200 | 15651799 | 64.57034 | 102.7849 | 115.8411 | 48.42322 | 0.163591 | 2.217825 | 0.736601 |
| 9 | 111716800 | 111719599 | 31.99588 | 52.91376 | 61.38554 | 26.53662 | 0.166919 | 1.337549 | 0.736611 |
| 8 | 19499400 | 19501199 | 13.96456 | 24.44562 | 44.62383 | 19.69276 | -0.47644 | 0.667652 | 0.736614 |
| 16 | 21505600 | 21507199 | 13.64376 | 24.42106 | 37.68722 | 16.52705 | -0.22264 | 0.432247 | 0.736618 |
| 3 | 152767000 | 152768199 | 13.91239 | 24.84293 | 37.69086 | 16.56171 | -0.19999 | 0.451148 | 0.736653 |
| 10 | 78581000 | 78585199 | 34.25443 | 55.46113 | 80.28548 | 34.34912 | -0.16886 | 1.65955 | 0.736677 |
| 9 | 28175800 | 28176999 | 12.82573 | 23.45439 | 27.59318 | 11.80602 | 0.190298 | 0.28325 | 0.736681 |
| 1 | 4752000 | 4752599 | 6.829333 | 13.205 | 22.65107 | 8.668112 | -0.21213 | -0.18072 | 0.736682 |
| 14 | 103700200 | 103702399 | 11.43544 | 20.28027 | 38.30168 | 16.92776 | -0.51967 | 0.480001 | 0.736693 |
| 13 | 100061400 | 100062399 | 8.098798 | 15.30635 | 25.47382 | 10.35521 | -0.24893 | 0.040021 | 0.7367 |
| 9 | 35889200 | 35896799 | 67.80318 | 107.8344 | 119.05 | 49.70297 | 0.191927 | 2.249215 | 0.736738 |
| 12 | 71965000 | 71965799 | 8.529618 | 16.17584 | 26.02548 | 10.50317 | -0.19165 | 0.020946 | 0.736739 |
| 14 | 114036600 | 114038199 | 16.21898 | 28.12278 | 50.06658 | 21.96689 | -0.44199 | 0.820825 | 0.736751 |
| 18 | 58813600 | 58817399 | 42.90169 | 69.09577 | 99.84487 | 42.44925 | -0.17495 | 1.826003 | 0.736763 |

| | | | | | | | | | |
|---|---|---|---|---|---|---|---|---|---|
| 8 | 3917800 | 3918599 | 9.92269 | 18.55111 | 28.97281 | 12.00645 | -0.18154 | 0.133512 | 0.736777 |
| X | 47373200 | 47376199 | 23.17466 | 39.92928 | 43.706 | 19.17246 | 0.255063 | 0.903355 | 0.736785 |
| 9 | 40641600 | 40643399 | 11.98434 | 21.58084 | 32.706 | 14.2657 | -0.19075 | 0.377395 | 0.736808 |
| 15 | 103896400 | 103897999 | 14.46553 | 26.46966 | 29.97239 | 12.81306 | 0.245554 | 0.294637 | 0.736812 |
| 9 | 47911000 | 47913799 | 22.53263 | 38.01542 | 55.47314 | 24.25973 | -0.16215 | 1.095836 | 0.736828 |
| 16 | 78468000 | 78468999 | 11.61683 | 21.82971 | 26.29248 | 10.80182 | 0.200668 | 0.109576 | 0.736871 |
| 12 | 6699400 | 6700199 | 14.52883 | 26.00684 | 30.38168 | 13.32265 | 0.176994 | 0.424483 | 0.736908 |
| 8 | 72661600 | 72662999 | 18.22035 | 31.46155 | 46.35045 | 20.33385 | -0.17134 | 0.861396 | 0.73691 |
| 13 | 67634600 | 67642599 | 114.7651 | 177.0701 | 186.8018 | 77.59396 | 0.222474 | 2.896715 | 0.736921 |
| 7 | 41871000 | 41872399 | 20.07399 | 35.17939 | 39.10444 | 17.27316 | 0.235737 | 0.674645 | 0.736928 |
| 12 | 88655400 | 88657599 | 15.85044 | 28.66923 | 31.85959 | 13.81987 | 0.258931 | 0.374759 | 0.736966 |
| 5 | 7154200 | 7155799 | 14.99153 | 27.22209 | 30.90051 | 13.33836 | 0.23299 | 0.344197 | 0.736996 |
| 9 | 65251400 | 65254399 | 29.06117 | 47.36247 | 81.56792 | 35.03252 | -0.41081 | 1.533675 | 0.737026 |
| 14 | 52817800 | 52819399 | 18.44843 | 32.29342 | 37.64683 | 16.61315 | 0.167324 | 0.680371 | 0.73704 |
| 6 | 123960400 | 123960999 | 8.303223 | 16.09761 | 21.06705 | 7.962406 | 0.19149 | -0.21095 | 0.737084 |
| 6 | 70592400 | 70594199 | 40.84118 | 65.64217 | 93.70158 | 39.71895 | -0.16095 | 1.886531 | 0.737085 |
| 1 | 108207600 | 108208799 | 10.21661 | 19.08392 | 30.04306 | 12.46843 | -0.19484 | 0.14333 | 0.737095 |
| 5 | 39403600 | 39404399 | 12.04085 | 22.03913 | 24.62958 | 10.52497 | 0.265113 | 0.263351 | 0.737121 |
| 9 | 88440200 | 88457999 | 290.9158 | 416.6414 | 435.7729 | 179.9921 | 0.138138 | 4.227483 | 0.737129 |
| 6 | 89281600 | 89284399 | 31.9854 | 52.72958 | 61.67478 | 26.61067 | 0.15442 | 1.377338 | 0.737141 |
| 12 | 36350800 | 36353599 | 16.56586 | 28.44459 | 54.33707 | 23.83345 | -0.54636 | 0.914228 | 0.737143 |
| 9 | 73089000 | 73089599 | 7.78679 | 14.98839 | 25.29838 | 9.840541 | -0.21124 | -0.10634 | 0.737155 |
| 7 | 56335000 | 56342199 | 191.4462 | 279.7238 | 367.6311 | 151.4583 | -0.16284 | 3.922578 | 0.737157 |
| 18 | 41706600 | 41708399 | 16.81886 | 29.28122 | 43.14762 | 18.9684 | -0.16951 | 0.762326 | 0.737157 |
| Y | 58440400 | 58440999 | 4.840093 | 9.415963 | 19.59554 | 7.233115 | -0.43794 | -0.33917 | 0.737159 |
| 13 | 60166200 | 60171399 | 28.58865 | 47.04379 | 68.04443 | 29.33627 | -0.15217 | 1.395168 | 0.737195 |
| 12 | 32481200 | 32482199 | 9.006922 | 17.13109 | 27.76159 | 11.13465 | -0.19429 | 0.009742 | 0.737212 |
| 9 | 30929800 | 30933799 | 30.54666 | 50.38765 | 76.10221 | 32.84389 | -0.21311 | 1.375309 | 0.737237 |
| 16 | 36214000 | 36215399 | 12.52381 | 22.61168 | 34.16701 | 14.85404 | -0.18383 | 0.366334 | 0.737237 |
| X | 38051800 | 38053599 | 30.15362 | 51.12649 | 44.69592 | 19.57607 | 0.574643 | 1.070169 | 0.737249 |
| 9 | 93208200 | 93210199 | 36.32329 | 59.10006 | 68.56661 | 29.43085 | 0.155966 | 1.565134 | 0.737274 |
| 14 | 15896800 | 15898999 | 21.0354 | 36.61693 | 34.63862 | 15.22635 | 0.466865 | 0.769732 | 0.73733 |
| 10 | 14433800 | 14438199 | 57.93286 | 91.87169 | 128.0313 | 53.29739 | -0.14764 | 2.327813 | 0.737343 |
| 4 | 92891600 | 92894799 | 37.3338 | 61.02679 | 58.42295 | 25.11935 | 0.437161 | 1.493766 | 0.737356 |
| 5 | 109937400 | 109946199 | 130.8948 | 199.5755 | 216.7696 | 89.87089 | 0.167268 | 3.135334 | 0.737372 |
| 16 | 95457400 | 95458799 | 34.47324 | 56.4194 | 66.05728 | 28.41437 | 0.148738 | 1.475586 | 0.737406 |
| 5 | 68663400 | 68664399 | 12.60662 | 23.63695 | 22.94848 | 9.463625 | 0.504989 | 0.123344 | 0.737452 |
| 12 | 23686600 | 23688799 | 56.06587 | 89.70291 | 83.20619 | 35.03054 | 0.452017 | 2.04192 | 0.737457 |
| 10 | 42269600 | 42271399 | 17.06158 | 29.61153 | 44.23483 | 19.40875 | -0.18973 | 0.807074 | 0.737463 |
| 11 | 68588600 | 68593999 | 56.61153 | 90.02397 | 124.1593 | 51.82822 | -0.1288 | 2.245783 | 0.737465 |
| 13 | 94127600 | 94130199 | 35.43629 | 57.68016 | 66.34147 | 28.47995 | 0.168838 | 1.554904 | 0.737493 |
| 1 | 189235600 | 189243999 | 98.53115 | 151.7009 | 204.9458 | 85.12227 | -0.13636 | 2.931335 | 0.737495 |
| X | 74994600 | 74996599 | 25.79421 | 43.86242 | 50.1078 | 21.96122 | 0.190363 | 1.035333 | 0.73753 |
| 12 | 5097600 | 5114999 | 667.8301 | 879.033 | 1041.572 | 435.7588 | -0.16424 | 5.533308 | 0.73754 |
| 10 | 117409200 | 117410799 | 10.76061 | 19.80844 | 30.31752 | 12.85549 | -0.17761 | 0.235373 | 0.737544 |
| 13 | 15017200 | 15018999 | 13.57563 | 24.2276 | 36.13429 | 15.88547 | -0.17588 | 0.449664 | 0.73755 |
| 4 | 74584000 | 74585799 | 23.19867 | 40.45164 | 42.31073 | 18.61881 | 0.322791 | 0.767916 | 0.73757 |

| | | | | | | | | | |
|---|---|---|---|---|---|---|---|---|---|
| 6 | 132517600 | 132519799 | 29.67222 | 49.90342 | 55.18703 | 24.11156 | 0.236281 | 1.146214 | 0.737572 |
| 5 | 39558800 | 39559599 | 6.391408 | 12.40623 | 21.67519 | 8.145816 | -0.21297 | -0.24865 | 0.737579 |
| 12 | 26806400 | 26807199 | 6.391408 | 12.40623 | 21.67519 | 8.145816 | -0.21297 | -0.24865 | 0.737579 |
| 8 | 52598600 | 52599799 | 17.70791 | 31.03191 | 36.17019 | 15.97663 | 0.167662 | 0.657438 | 0.737609 |
| 17 | 27742200 | 27745999 | 47.22841 | 75.56515 | 105.743 | 44.52024 | -0.14009 | 2.039065 | 0.737616 |
| 11 | 18586200 | 18586799 | 11.0954 | 19.79329 | 37.31072 | 16.40824 | -0.51193 | 0.438595 | 0.73766 |
| 5 | 145334600 | 145354799 | 387.415 | 541.3175 | 550.9866 | 228.4947 | 0.141562 | 4.593748 | 0.737661 |
| 3 | 34759800 | 34762199 | 21.32448 | 36.23045 | 53.3069 | 23.36169 | -0.17363 | 1.027758 | 0.737671 |
| 3 | 13127600 | 13132199 | 77.89999 | 122.5421 | 139.8931 | 58.03626 | 0.130303 | 2.527974 | 0.737675 |
| 1 | 85044200 | 85072599 | 1317.616 | 1664.839 | 1527.226 | 641.577 | 0.145826 | 6.199223 | 0.73768 |
| Y | 41649200 | 41653399 | 23.76348 | 40.22298 | 36.82646 | 16.12102 | 0.508047 | 1.048767 | 0.737708 |
| 6 | 50423200 | 50423599 | 8.730275 | 16.96476 | 21.21864 | 7.904107 | 0.276287 | -0.25265 | 0.737732 |
| X | 60353400 | 60355399 | 21.78129 | 38.03518 | 41.31572 | 18.19751 | 0.268849 | 0.742577 | 0.737795 |
| 1 | 183592600 | 183593599 | 6.093957 | 11.8796 | 21.52671 | 7.772024 | -0.21199 | -0.39394 | 0.7378 |
| 3 | 134388400 | 134390999 | 22.05809 | 36.86264 | 64.61105 | 28.12675 | -0.42581 | 1.197411 | 0.737807 |
| 19 | 61291800 | 61293999 | 21.03771 | 36.37119 | 42.59266 | 18.68601 | 0.1588 | 0.857627 | 0.737813 |
| 13 | 119362400 | 119364399 | 17.59167 | 31.52322 | 33.77559 | 14.79751 | 0.301952 | 0.445811 | 0.737813 |
| 12 | 5789800 | 5791199 | 19.00318 | 33.32803 | 38.04885 | 16.81765 | 0.197329 | 0.65423 | 0.737818 |
| 17 | 84343200 | 84345399 | 19.71361 | 33.71832 | 49.54251 | 21.72365 | -0.17007 | 0.957438 | 0.737828 |
| 15 | 86201200 | 86202799 | 22.77996 | 38.90338 | 45.32232 | 19.87071 | 0.163166 | 0.979672 | 0.73785 |
| 6 | 68338000 | 68342799 | 71.74543 | 112.8332 | 154.0404 | 63.90341 | -0.12771 | 2.531828 | 0.737859 |
| 3 | 98214400 | 98216199 | 23.29966 | 39.83786 | 46.32959 | 20.31426 | 0.165809 | 0.96991 | 0.737867 |
| 2 | 113502200 | 113503599 | 12.46033 | 22.85705 | 27.09548 | 11.54419 | 0.183727 | 0.261328 | 0.737876 |
| 13 | 8234200 | 8234799 | 9.757929 | 18.78612 | 18.17149 | 7.064457 | 0.585499 | -0.11065 | 0.737878 |
| 10 | 35395800 | 35400599 | 46.69178 | 75.61354 | 69.29138 | 29.64983 | 0.489482 | 1.673686 | 0.737878 |
| 8 | 16068000 | 16071599 | 23.71285 | 40.94243 | 45.73765 | 20.06496 | 0.22614 | 0.885136 | 0.737879 |
| 13 | 15989400 | 15989999 | 6.252292 | 12.10973 | 21.60897 | 8.20417 | -0.25704 | -0.21588 | 0.737905 |
| 10 | 129726800 | 129729399 | 28.41413 | 47.14314 | 70.17428 | 30.404 | -0.19129 | 1.299634 | 0.737916 |
| 5 | 41023400 | 41024399 | 15.66454 | 27.41524 | 40.42703 | 17.83723 | -0.17 | 0.674125 | 0.737932 |
| 18 | 12900200 | 12901799 | 15.66454 | 27.41524 | 40.42703 | 17.83723 | -0.17 | 0.674125 | 0.737932 |
| 11 | 20345800 | 20347199 | 14.41633 | 25.76819 | 29.96751 | 13.15706 | 0.182384 | 0.432807 | 0.737976 |
| 9 | 74527400 | 74527999 | 10.14547 | 18.91379 | 29.46535 | 12.26948 | -0.18332 | 0.156272 | 0.737985 |
| 4 | 62101000 | 62107599 | 104.9905 | 162.6677 | 175.9329 | 73.07764 | 0.191206 | 2.815292 | 0.737994 |
| X | 88143600 | 88147199 | 42.60137 | 69.18498 | 76.86393 | 32.95883 | 0.216081 | 1.613582 | 0.737995 |
| 6 | 11414800 | 11421999 | 74.71833 | 118.2545 | 131.7285 | 54.75975 | 0.172508 | 2.392862 | 0.737998 |
| X | 93808000 | 93809599 | 14.91154 | 26.40577 | 40.21338 | 17.78147 | -0.21214 | 0.519471 | 0.737998 |
| 5 | 88298400 | 88301399 | 38.27719 | 61.70056 | 88.74306 | 37.77482 | -0.16659 | 1.791612 | 0.738002 |
| 7 | 7490600 | 7496599 | 69.27615 | 110.1886 | 104.5407 | 43.65176 | 0.410435 | 2.24604 | 0.738003 |
| 5 | 140768800 | 140771399 | 17.06776 | 29.54019 | 44.77589 | 19.64449 | -0.2115 | 0.832458 | 0.738014 |
| 10 | 113323800 | 113325199 | 13.22362 | 24.05831 | 28.48638 | 12.2666 | 0.174865 | 0.315854 | 0.738021 |
| 18 | 34155800 | 34156799 | 13.22362 | 24.05831 | 28.48638 | 12.2666 | 0.174865 | 0.315854 | 0.738021 |
| 13 | 62781000 | 62801799 | 343.4661 | 488.1057 | 478.4039 | 197.8525 | 0.220622 | 4.343794 | 0.738031 |
| 10 | 119305200 | 119307999 | 27.55733 | 45.55165 | 65.95578 | 28.50291 | -0.15189 | 1.336085 | 0.738033 |
| 16 | 23849800 | 23852399 | 27.55733 | 45.55165 | 65.95578 | 28.50291 | -0.15189 | 1.336085 | 0.738033 |
| 12 | 78477600 | 78480399 | 29.57983 | 49.65631 | 55.21459 | 24.1048 | 0.228345 | 1.161611 | 0.73806 |
| 3 | 90489000 | 90490199 | 11.88102 | 21.6179 | 32.73948 | 14.09621 | -0.17793 | 0.314615 | 0.738066 |
| 6 | 68969400 | 68973599 | 91.33197 | 142.0596 | 161.3523 | 66.98534 | 0.125029 | 2.741073 | 0.738067 |

| | | | | | | | | | |
|---|---|---|---|---|---|---|---|---|---|
| 14 | 31556400 | 31562199 | 61.85214 | 98.78889 | 108.8713 | 45.72312 | 0.20107 | 2.109506 | 0.738075 |
| 19 | 50387800 | 50391399 | 53.23051 | 84.77771 | 118.1516 | 49.42448 | -0.14144 | 2.193349 | 0.738104 |
| 11 | 70254000 | 70254799 | 26.85507 | 44.04669 | 67.03741 | 28.86211 | -0.22724 | 1.430971 | 0.738105 |
| 4 | 146714800 | 146715999 | 12.7082 | 24.11395 | 25.94692 | 10.45886 | 0.385008 | 0.05178 | 0.738121 |
| 2 | 12501600 | 12502599 | 8.219842 | 15.56652 | 25.47689 | 10.31231 | -0.21984 | 0.025955 | 0.738142 |
| 4 | 34532200 | 34533199 | 8.237657 | 15.63177 | 25.51121 | 10.28222 | -0.21017 | 0.01327 | 0.73815 |
| 3 | 82388600 | 82390199 | 9.986474 | 18.58734 | 28.91005 | 12.07121 | -0.18411 | 0.164047 | 0.738161 |
| 3 | 146037000 | 146039999 | 19.28512 | 33.10614 | 48.47704 | 21.26272 | -0.16425 | 0.922985 | 0.738163 |
| 17 | 56185400 | 56186599 | 10.50731 | 18.98936 | 30.7609 | 13.35697 | -0.28211 | 0.339481 | 0.738167 |
| 9 | 55860600 | 55886599 | 495.4121 | 680.7309 | 655.4673 | 272.5396 | 0.197287 | 4.851692 | 0.738179 |
| 18 | 85890600 | 85892399 | 23.79173 | 41.01141 | 38.80376 | 17.02224 | 0.464226 | 0.88953 | 0.738192 |
| 17 | 82467400 | 82469799 | 33.327 | 55.14498 | 62.65124 | 27.09391 | 0.19703 | 1.336075 | 0.738195 |
| 1 | 118137600 | 118159599 | 238.0366 | 349.0775 | 354.8039 | 146.1401 | 0.212801 | 3.869954 | 0.738225 |
| 19 | 14205600 | 14206599 | 11.07021 | 19.38856 | 43.64263 | 19.26783 | -0.77677 | 0.643175 | 0.738238 |
| 13 | 68584200 | 68637799 | 1292.391 | 1621.144 | 1784.884 | 750.3928 | -0.12803 | 6.31612 | 0.738239 |
| 5 | 105355600 | 105357799 | 16.62684 | 29.02465 | 42.54656 | 18.73571 | -0.16167 | 0.721766 | 0.73826 |
| 14 | 7922200 | 7924799 | 29.59521 | 49.69372 | 55.70562 | 24.32274 | 0.216767 | 1.162219 | 0.738277 |
| 14 | 117783200 | 117786399 | 51.819 | 82.11771 | 140.922 | 58.62182 | -0.44851 | 2.347125 | 0.738278 |
| 9 | 17718600 | 17719999 | 13.99545 | 26.18047 | 28.32487 | 11.72651 | 0.345303 | 0.161612 | 0.738288 |
| 3 | 97487200 | 97490199 | 28.8998 | 47.07069 | 82.09771 | 35.25221 | -0.42965 | 1.541983 | 0.738294 |
| X | 156273400 | 156278199 | 52.4701 | 84.96838 | 65.72683 | 28.1239 | 0.733368 | 1.684821 | 0.73831 |
| 5 | 141197400 | 141199399 | 21.18414 | 36.1636 | 52.78303 | 23.14097 | -0.16101 | 0.984387 | 0.738317 |
| Y | 6183400 | 6185999 | 22.61685 | 38.50083 | 35.78611 | 15.68568 | 0.486937 | 0.99296 | 0.738399 |
| 12 | 52912800 | 52914999 | 21.7815 | 38.1104 | 40.634 | 17.92441 | 0.295504 | 0.709697 | 0.738414 |
| 13 | 67663800 | 67667799 | 41.8977 | 67.91264 | 77.69355 | 33.26778 | 0.171356 | 1.649777 | 0.738425 |
| 5 | 105168200 | 105170399 | 21.92797 | 38.11409 | 36.70632 | 16.12023 | 0.441012 | 0.798555 | 0.738431 |
| X | 37568400 | 37570399 | 17.49265 | 30.56128 | 48.38325 | 21.31702 | -0.27205 | 0.726122 | 0.738433 |
| 16 | 4456800 | 4459399 | 14.54817 | 25.51304 | 38.36752 | 16.95753 | -0.19809 | 0.631997 | 0.738439 |
| 4 | 71208400 | 71214599 | 117.0148 | 179.4783 | 197.7927 | 82.10868 | 0.153051 | 3.01053 | 0.738441 |
| 2 | 58010400 | 58012799 | 19.36134 | 33.71143 | 56.24722 | 24.72895 | -0.34832 | 0.789723 | 0.738446 |
| 10 | 41127800 | 41130799 | 22.20021 | 37.1171 | 65.23001 | 28.40439 | -0.42968 | 1.192822 | 0.738457 |
| 10 | 53632600 | 53633999 | 17.02121 | 29.96656 | 34.95059 | 15.47239 | 0.167488 | 0.584996 | 0.738459 |
| 12 | 115187800 | 115208199 | 295.9177 | 426.5443 | 419.9608 | 173.2915 | 0.234571 | 4.134572 | 0.738469 |
| 8 | 5623200 | 5626799 | 42.03151 | 68.17896 | 77.31902 | 33.12511 | 0.184905 | 1.634913 | 0.738489 |
| 3 | 98648600 | 98650399 | 17.66518 | 30.73236 | 45.75141 | 20.1039 | -0.18456 | 0.777273 | 0.738495 |
| 5 | 120397400 | 120399399 | 22.039 | 36.84124 | 56.37695 | 24.54666 | -0.23049 | 1.187709 | 0.738506 |
| 2 | 110790200 | 110792599 | 28.32745 | 47.91672 | 45.27899 | 19.81723 | 0.462612 | 1.082094 | 0.738555 |
| 16 | 19852600 | 19854399 | 19.67245 | 33.76047 | 49.54449 | 21.73037 | -0.16764 | 0.926279 | 0.73856 |
| 7 | 36992800 | 36997199 | 31.39643 | 51.1743 | 74.59917 | 32.04088 | -0.17083 | 1.530854 | 0.738563 |
| X | 102208800 | 102210399 | 19.73206 | 34.60167 | 32.88913 | 14.53538 | 0.460436 | 0.652056 | 0.738667 |
| 12 | 20685200 | 20752199 | 2343.254 | 2813.889 | 2485.879 | 1050.853 | 0.125656 | 7.01046 | 0.738695 |
| 5 | 86847400 | 86848599 | 13.44021 | 25.17005 | 27.79137 | 11.4838 | 0.318237 | 0.148191 | 0.738703 |
| 3 | 36194200 | 36196799 | 22.81104 | 39.90703 | 41.19099 | 18.1683 | 0.34224 | 0.719732 | 0.738734 |
| 16 | 60718800 | 60720799 | 25.09102 | 43.06064 | 47.92184 | 21.01871 | 0.229899 | 0.947469 | 0.73875 |
| 4 | 44603200 | 44604599 | 11.27262 | 20.97782 | 25.15617 | 10.50666 | 0.188009 | 0.165404 | 0.738776 |
| 9 | 120022800 | 120026599 | 40.50851 | 65.14562 | 92.71714 | 39.33504 | -0.15562 | 1.867211 | 0.738776 |
| 10 | 113633000 | 113633399 | 6.837608 | 13.32438 | 17.45949 | 6.334009 | 0.245265 | -0.38007 | 0.738807 |

| | | | | | | | | | |
|---|---|---|---|---|---|---|---|---|---|
| 17 | 83436400 | 83440399 | 28.27111 | 46.53079 | 67.43051 | 29.07411 | -0.15477 | 1.391703 | 0.738822 |
| X | 62676600 | 62678999 | 23.12031 | 40.42818 | 31.99113 | 14.1046 | 0.724007 | 0.71689 | 0.738827 |
| 6 | 37839600 | 37841599 | 18.20193 | 31.68442 | 47.35655 | 20.81677 | -0.19018 | 0.774716 | 0.738879 |
| 1 | 14172600 | 14173999 | 20.21259 | 35.43313 | 33.34536 | 14.73308 | 0.474736 | 0.66024 | 0.738881 |
| 11 | 15132600 | 15138999 | 14.89264 | 27.13239 | 24.58683 | 10.56908 | 0.559563 | 0.309792 | 0.738883 |
| 6 | 123660800 | 123667399 | 88.86591 | 139.0415 | 154.8911 | 64.25594 | 0.159578 | 2.637496 | 0.738885 |
| 17 | 63831200 | 63832799 | 13.47872 | 24.40913 | 28.54601 | 12.36368 | 0.18746 | 0.345955 | 0.738893 |
| 15 | 96143800 | 96145799 | 22.28069 | 38.25708 | 44.7527 | 19.62952 | 0.158414 | 0.928093 | 0.738901 |
| X | 27146400 | 27147599 | 18.80305 | 33.32392 | 27.81001 | 12.29499 | 0.651788 | 0.514108 | 0.738926 |
| 11 | 73800200 | 73801799 | 12.0413 | 22.8043 | 22.06357 | 8.93003 | 0.532605 | 0.052086 | 0.738932 |
| 13 | 20524200 | 20526399 | 13.92397 | 25.51658 | 24.3608 | 10.39309 | 0.492422 | 0.270811 | 0.738944 |
| 16 | 50377200 | 50378999 | 12.35515 | 22.10629 | 33.73553 | 14.79879 | -0.20624 | 0.422961 | 0.738956 |
| 6 | 55003200 | 55004999 | 21.64502 | 37.11381 | 42.23122 | 18.52146 | 0.197944 | 0.935758 | 0.738972 |
| 5 | 16496800 | 16497799 | 8.574294 | 16.6754 | 14.70547 | 5.441563 | 0.784514 | -0.30049 | 0.738977 |
| 17 | 69910200 | 69918199 | 73.71416 | 116.6043 | 131.6535 | 54.71412 | 0.152482 | 2.404128 | 0.738993 |
| 2 | 18530600 | 18532399 | 23.96911 | 39.48579 | 72.01287 | 31.06173 | -0.48492 | 1.37363 | 0.739014 |
| 3 | 106016600 | 106018599 | 15.68295 | 27.49345 | 40.36244 | 17.83395 | -0.16363 | 0.647412 | 0.739017 |
| 7 | 57186200 | 57186999 | 12.11038 | 22.24035 | 26.114 | 11.10884 | 0.199021 | 0.249697 | 0.739068 |
| 13 | 36711800 | 36713799 | 17.35014 | 30.44928 | 35.57895 | 15.73485 | 0.164031 | 0.630507 | 0.73907 |
| 9 | 23595400 | 23596599 | 26.19592 | 43.22534 | 64.43546 | 27.81271 | -0.19385 | 1.354423 | 0.739108 |
| 13 | 51487000 | 51490999 | 31.90308 | 51.96659 | 75.13033 | 32.26143 | -0.15959 | 1.542018 | 0.739154 |
| 14 | 41835000 | 41835599 | 8.412795 | 16.2105 | 20.35976 | 7.895498 | 0.215218 | -0.12682 | 0.739166 |
| X | 26518800 | 26520799 | 23.06055 | 40.23382 | 41.85784 | 18.42614 | 0.330795 | 0.762885 | 0.739167 |
| 13 | 111140800 | 111141999 | 13.99599 | 24.77806 | 37.07889 | 16.39632 | -0.18718 | 0.509026 | 0.739189 |
| 18 | 74698000 | 74699799 | 15.57502 | 27.67076 | 32.37234 | 14.28583 | 0.168721 | 0.482898 | 0.739195 |
| 5 | 60883000 | 60884199 | 17.09702 | 29.74254 | 43.66631 | 19.18712 | -0.16441 | 0.774305 | 0.739205 |
| 11 | 15324600 | 15327399 | 24.96422 | 42.78324 | 47.9218 | 21.01633 | 0.220246 | 0.957753 | 0.739212 |
| 18 | 11554800 | 11555999 | 20.56974 | 35.03264 | 51.1363 | 22.41479 | -0.16155 | 1.002269 | 0.739218 |
| 18 | 36949600 | 36992199 | 756.5682 | 998.4673 | 966.9679 | 404.4261 | 0.130529 | 5.491537 | 0.73922 |
| 12 | 75418200 | 75423199 | 51.0986 | 82.04921 | 94.03226 | 39.80236 | 0.153467 | 1.929796 | 0.739231 |
| 1 | 12670400 | 12673399 | 57.31599 | 91.57908 | 104.893 | 44.07634 | 0.146138 | 2.091038 | 0.739247 |
| 8 | 47685200 | 47688399 | 24.63976 | 40.73818 | 62.92239 | 27.19627 | -0.24473 | 1.322287 | 0.739254 |
| 16 | 54269600 | 54272399 | 23.772 | 41.33111 | 43.46036 | 19.08981 | 0.314763 | 0.811348 | 0.739261 |
| 4 | 99208200 | 99209599 | 12.2872 | 22.11182 | 33.17137 | 14.48047 | -0.17702 | 0.383706 | 0.739274 |
| 4 | 28104000 | 28106199 | 21.7635 | 37.9654 | 41.23944 | 18.15228 | 0.268493 | 0.752273 | 0.739279 |
| 16 | 75963200 | 75966799 | 34.69323 | 56.85445 | 66.33912 | 28.55125 | 0.15444 | 1.457685 | 0.73928 |
| 5 | 110215400 | 110220799 | 82.02748 | 128.9122 | 144.848 | 60.08443 | 0.152114 | 2.549868 | 0.739283 |
| 10 | 67859200 | 67860599 | 12.48098 | 22.25478 | 33.70671 | 14.82906 | -0.19841 | 0.444176 | 0.739305 |
| 1 | 7957200 | 7960199 | 32.17706 | 53.72351 | 50.30947 | 21.88819 | 0.475508 | 1.222842 | 0.739307 |
| X | 3050000 | 3055999 | 152.5938 | 230.8927 | 202.6915 | 83.84229 | 0.465149 | 3.295286 | 0.739328 |
| 10 | 5887400 | 5889199 | 30.0086 | 50.08621 | 57.32974 | 24.93723 | 0.186949 | 1.229242 | 0.739369 |
| 13 | 44751600 | 44753399 | 16.59886 | 28.95952 | 42.55352 | 18.73216 | -0.16525 | 0.728938 | 0.739374 |
| 7 | 5577000 | 5599799 | 71.98832 | 115.2398 | 81.9338 | 34.52103 | 0.835789 | 2.044431 | 0.739392 |
| 17 | 7707600 | 7709999 | 28.1914 | 47.0574 | 55.11498 | 23.97521 | 0.153838 | 1.218198 | 0.739415 |
| 18 | 78816000 | 78820599 | 37.22899 | 61.26007 | 67.38017 | 29.04954 | 0.241754 | 1.415702 | 0.739437 |
| 18 | 74709000 | 74711999 | 19.24577 | 33.29456 | 49.01432 | 21.50371 | -0.16974 | 0.853423 | 0.739455 |
| 2 | 58504200 | 58506399 | 19.62801 | 34.2725 | 39.18595 | 17.25864 | 0.195007 | 0.725521 | 0.739472 |

| | | | | | | | | | |
|---|---|---|---|---|---|---|---|---|---|
| 4 | 16324600 | 16326599 | 19.62801 | 34.2725 | 39.18595 | 17.25864 | 0.195007 | 0.725521 | 0.739472 |
| 9 | 118643800 | 118646199 | 23.81707 | 39.64384 | 59.12612 | 25.67438 | -0.19354 | 1.240869 | 0.739476 |
| 7 | 72933200 | 72935599 | 24.06226 | 40.90291 | 47.96081 | 21.01964 | 0.15281 | 1.028718 | 0.739523 |
| 17 | 53674800 | 53675999 | 36.2575 | 58.53376 | 84.3987 | 35.99602 | -0.16765 | 1.740685 | 0.739523 |
| 3 | 127784800 | 127786199 | 10.57998 | 19.66168 | 30.465 | 12.75225 | -0.18184 | 0.180773 | 0.739526 |
| 6 | 6209600 | 6211999 | 23.41363 | 39.28954 | 57.07417 | 24.91045 | -0.15585 | 1.150179 | 0.739537 |
| 5 | 6234000 | 6234599 | 12.64861 | 23.56627 | 22.48986 | 9.374281 | 0.517536 | 0.163081 | 0.739545 |
| 4 | 121288200 | 121312999 | 456.536 | 629.5611 | 628.9863 | 261.3805 | 0.149271 | 4.795146 | 0.739592 |
| 17 | 55404400 | 55406599 | 20.25448 | 35.56199 | 38.38793 | 16.98342 | 0.277631 | 0.643008 | 0.739631 |
| 11 | 103056800 | 103057199 | 8.308258 | 16.08878 | 21.0226 | 7.988612 | 0.186433 | -0.1927 | 0.739634 |
| 7 | 3763400 | 3764799 | 9.440517 | 17.6348 | 27.89286 | 11.57541 | -0.20131 | 0.131693 | 0.739646 |
| 8 | 84585800 | 84587999 | 22.90789 | 39.02894 | 45.33569 | 19.87276 | 0.166838 | 1.001101 | 0.739656 |
| 6 | 57060400 | 57060999 | 11.93059 | 22.11065 | 26.63182 | 11.18959 | 0.175223 | 0.198222 | 0.73966 |
| 15 | 54498000 | 54498799 | 11.93059 | 22.11065 | 26.63182 | 11.18959 | 0.175223 | 0.198222 | 0.73966 |
| 15 | 67903800 | 67904999 | 12.90582 | 22.67615 | 41.96735 | 18.57853 | -0.49585 | 0.599917 | 0.739689 |
| 17 | 22927400 | 22956399 | 523.9941 | 716.844 | 684.4961 | 284.8073 | 0.202928 | 4.920487 | 0.739693 |
| 2 | 88019400 | 88024199 | 56.00102 | 89.52962 | 102.9787 | 43.30731 | 0.140977 | 2.071618 | 0.739725 |
| 4 | 109035000 | 109039999 | 49.95681 | 80.5372 | 88.15542 | 37.53057 | 0.226651 | 1.803223 | 0.739732 |
| 2 | 164779200 | 164781599 | 16.52781 | 28.46654 | 44.85174 | 19.67484 | -0.26841 | 0.875963 | 0.739735 |
| 6 | 103793400 | 103793799 | 5.945274 | 11.60037 | 21.69854 | 7.737331 | -0.2409 | -0.4384 | 0.739738 |
| 1 | 87457800 | 87460599 | 26.76347 | 45.16976 | 52.31684 | 22.88192 | 0.170011 | 1.105043 | 0.739762 |
| 5 | 21306600 | 21308799 | 34.61819 | 56.51898 | 65.91472 | 28.32792 | 0.151919 | 1.509967 | 0.739785 |
| 2 | 178805000 | 178810399 | 60.445 | 96.26194 | 111.0349 | 46.44346 | 0.130898 | 2.198282 | 0.739791 |
| 4 | 68679600 | 68681399 | 20.56612 | 35.54673 | 41.5851 | 18.24384 | 0.160222 | 0.856939 | 0.739806 |
| 2 | 172001800 | 172003999 | 20.07675 | 34.43196 | 50.07577 | 21.96224 | -0.15474 | 0.936198 | 0.739807 |
| 6 | 81830600 | 81834399 | 50.34713 | 80.77888 | 93.1633 | 39.38731 | 0.143048 | 1.953955 | 0.739813 |
| 4 | 70035400 | 70038399 | 42.94614 | 69.33589 | 80.32247 | 34.26009 | 0.147469 | 1.74851 | 0.739831 |
| 1 | 148426200 | 148430199 | 49.56108 | 79.4942 | 92.05466 | 38.90111 | 0.136677 | 1.959759 | 0.739832 |
| 10 | 13348600 | 13349999 | 13.76235 | 24.19384 | 36.81147 | 16.30094 | -0.21472 | 0.586412 | 0.739864 |
| 12 | 72259200 | 72261799 | 15.90762 | 27.75258 | 42.03406 | 18.50309 | -0.20881 | 0.723689 | 0.739867 |
| 1 | 121196600 | 121199599 | 68.44665 | 108.2708 | 124.3504 | 51.67873 | 0.127873 | 2.399378 | 0.739876 |
| 4 | 73178200 | 73184999 | 46.9352 | 75.79032 | 73.59374 | 31.39763 | 0.401821 | 1.745245 | 0.739877 |
| X | 32381800 | 32384599 | 49.14568 | 80.37561 | 55.85281 | 24.01874 | 0.898628 | 1.512086 | 0.739892 |
| 6 | 70062600 | 70063799 | 12.24515 | 23.08713 | 26.89801 | 10.98848 | 0.25466 | 0.095128 | 0.739896 |
| 18 | 4492600 | 4496799 | 53.60249 | 85.46326 | 118.676 | 49.71877 | -0.13457 | 2.156928 | 0.739917 |
| 7 | 103481800 | 103483199 | 12.18816 | 22.93061 | 26.83289 | 11.00164 | 0.244102 | 0.107162 | 0.739962 |
| 17 | 92174000 | 92175399 | 20.03995 | 35.1133 | 33.397 | 14.7498 | 0.459517 | 0.667106 | 0.739985 |
| 12 | 9968800 | 9969799 | 8.50851 | 16.10335 | 25.98454 | 10.53139 | -0.2014 | 0.033567 | 0.73999 |
| 10 | 74079000 | 74079999 | 15.61663 | 27.37278 | 40.9344 | 18.08421 | -0.18999 | 0.652719 | 0.739995 |
| 13 | 57212800 | 57213999 | 11.51049 | 21.41037 | 25.58888 | 10.69592 | 0.19161 | 0.169985 | 0.740008 |
| 12 | 99363200 | 99364799 | 17.58934 | 30.53153 | 44.75645 | 19.63886 | -0.16255 | 0.809708 | 0.740037 |
| 11 | 27562600 | 27566599 | 58.48333 | 93.20492 | 107.7494 | 45.11412 | 0.128794 | 2.172827 | 0.740051 |
| 5 | 51296800 | 51299999 | 27.41897 | 46.6674 | 51.22566 | 22.45279 | 0.247912 | 1.033754 | 0.740058 |
| 3 | 151459800 | 151460199 | 5.807177 | 11.29939 | 20.7084 | 7.63268 | -0.25591 | -0.33052 | 0.740094 |
| 14 | 22527800 | 22530799 | 37.6836 | 61.28207 | 71.26199 | 30.5821 | 0.152067 | 1.574152 | 0.740107 |
| 12 | 51338800 | 51340199 | 16.46347 | 28.63824 | 42.65336 | 18.74114 | -0.18499 | 0.771129 | 0.740118 |
| 10 | 9728800 | 9733199 | 66.38955 | 103.4163 | 178.494 | 74.08537 | -0.47628 | 2.706781 | 0.740122 |

| | | | | | | | | | |
|---|---|---|---|---|---|---|---|---|---|
| 18 | 51794600 | 51796999 | 28.30759 | 47.5478 | 54.52927 | 23.81421 | 0.184004 | 1.154107 | 0.740131 |
| 7 | 86632600 | 86634399 | 20.63345 | 35.57282 | 40.64582 | 17.83039 | 0.193467 | 0.88003 | 0.740142 |
| 13 | 33328600 | 33330999 | 22.90212 | 39.31198 | 45.22621 | 19.83672 | 0.182146 | 0.930899 | 0.740157 |
| 19 | 32330200 | 32331199 | 6.866467 | 13.25909 | 23.14824 | 8.913793 | -0.24514 | -0.15431 | 0.740157 |
| 17 | 93257800 | 93262199 | 66.2926 | 105.4697 | 118.2206 | 49.38114 | 0.170622 | 2.23781 | 0.74016 |
| 19 | 8144200 | 8145599 | 18.45097 | 32.38161 | 36.35659 | 16.07892 | 0.221056 | 0.634585 | 0.740174 |
| 8 | 124994000 | 124995399 | 13.81901 | 24.86455 | 28.56429 | 12.4712 | 0.205647 | 0.385805 | 0.740185 |
| 6 | 41340400 | 41341199 | 16.62103 | 29.03296 | 42.52556 | 18.73395 | -0.16047 | 0.712569 | 0.740195 |
| 5 | 63133800 | 63138799 | 101.8297 | 155.7925 | 209.3903 | 86.88024 | -0.13591 | 3.057798 | 0.740199 |
| 3 | 45168400 | 45174799 | 105.3608 | 162.5221 | 182.0335 | 75.61339 | 0.135917 | 2.896997 | 0.740203 |
| 10 | 49925000 | 49925799 | 9.780782 | 18.21681 | 28.38058 | 11.83657 | -0.18531 | 0.155419 | 0.740207 |
| X | 93867600 | 93869999 | 29.71974 | 50.36327 | 44.73306 | 19.58909 | 0.551841 | 1.074062 | 0.740212 |
| 16 | 77019600 | 77021199 | 18.39563 | 32.56023 | 35.13755 | 15.53851 | 0.281749 | 0.534107 | 0.740223 |
| 17 | 82237800 | 82239799 | 24.52629 | 42.23381 | 46.82101 | 20.53856 | 0.236335 | 0.913214 | 0.740232 |
| 1 | 65607600 | 65611199 | 44.69425 | 71.40232 | 118.1485 | 49.63668 | -0.38385 | 2.08691 | 0.740243 |
| 9 | 67321400 | 67322599 | 7.468702 | 14.36128 | 24.1461 | 9.420216 | -0.20956 | -0.10338 | 0.740243 |
| 1 | 111932200 | 111933799 | 21.34438 | 36.88827 | 43.10422 | 18.91023 | 0.161797 | 0.862429 | 0.740253 |
| 3 | 31816600 | 31820799 | 50.01098 | 79.74807 | 112.4819 | 47.13282 | -0.1568 | 2.147477 | 0.740257 |
| 4 | 83794400 | 83796599 | 18.82384 | 32.50575 | 47.93788 | 21.03029 | -0.17289 | 0.864419 | 0.74026 |
| X | 77898600 | 77908199 | 59.46728 | 95.63242 | 78.35706 | 33.24149 | 0.639315 | 1.890421 | 0.740279 |
| 15 | 12653800 | 12656599 | 29.15801 | 48.40642 | 56.2352 | 24.3748 | 0.165327 | 1.281319 | 0.740292 |
| X | 93730800 | 93733399 | 20.63911 | 35.95549 | 40.42997 | 17.78092 | 0.218988 | 0.769505 | 0.740294 |
| 12 | 55963200 | 55964199 | 10.25437 | 19.03327 | 29.37522 | 12.31656 | -0.17785 | 0.182349 | 0.740299 |
| 15 | 79510800 | 79512599 | 17.11601 | 29.98146 | 34.19159 | 15.09706 | 0.198939 | 0.656611 | 0.740301 |
| 4 | 5870000 | 5872799 | 29.66725 | 49.764 | 48.05896 | 20.9705 | 0.431158 | 1.166009 | 0.74031 |
| 7 | 38198800 | 38200599 | 28.75368 | 48.16256 | 55.67254 | 24.27075 | 0.172682 | 1.185015 | 0.740311 |
| 12 | 26153000 | 26160199 | 83.54482 | 130.134 | 177.184 | 73.5421 | -0.13462 | 2.714365 | 0.740363 |
| 1 | 13146000 | 13147999 | 16.15297 | 28.20962 | 41.52033 | 18.28877 | -0.16738 | 0.71203 | 0.740391 |
| 13 | 52540200 | 52548799 | 134.0079 | 200.3731 | 311.4345 | 128.4185 | -0.37356 | 3.526388 | 0.740421 |
| 3 | 18001200 | 18002999 | 19.99065 | 34.61773 | 40.00213 | 17.55039 | 0.17843 | 0.834089 | 0.740425 |
| X | 119589800 | 119591199 | 16.59164 | 29.56963 | 32.91773 | 14.48809 | 0.242816 | 0.473799 | 0.740463 |
| X | 13937000 | 13937599 | 13.39258 | 24.96254 | 28.28688 | 11.7825 | 0.272319 | 0.181829 | 0.740463 |
| 11 | 103637000 | 103639599 | 20.7008 | 35.65843 | 41.11436 | 18.03545 | 0.180093 | 0.886833 | 0.740477 |
| 1 | 117903400 | 117913999 | 117.5146 | 181.9472 | 179.8078 | 74.68743 | 0.320495 | 2.834627 | 0.740491 |
| 14 | 58805600 | 58807799 | 24.83415 | 42.84841 | 45.69319 | 20.04503 | 0.292606 | 0.893642 | 0.740526 |
| 14 | 21004200 | 21004799 | 11.36767 | 21.19284 | 25.21883 | 10.50074 | 0.202856 | 0.15658 | 0.740527 |
| 8 | 111186800 | 111189599 | 18.11562 | 31.50011 | 46.30382 | 20.3402 | -0.1663 | 0.789941 | 0.740529 |
| 14 | 62001800 | 62003199 | 16.76728 | 29.16002 | 43.16064 | 18.96129 | -0.17612 | 0.776158 | 0.74053 |
| 11 | 94724400 | 94726999 | 25.68958 | 43.48435 | 50.67417 | 22.18313 | 0.16132 | 1.073474 | 0.740546 |
| 18 | 19372400 | 19382999 | 105.1123 | 163.4475 | 170.7291 | 70.88111 | 0.245422 | 2.754043 | 0.740548 |
| 4 | 112283200 | 112284799 | 29.05367 | 48.5589 | 55.60782 | 24.20916 | 0.186272 | 1.211209 | 0.740549 |
| X | 96799800 | 96805799 | 70.44991 | 112.1506 | 118.7574 | 49.64627 | 0.253504 | 2.224603 | 0.740549 |
| 10 | 121444400 | 121446599 | 18.09416 | 31.30254 | 45.82251 | 20.10339 | -0.16153 | 0.844267 | 0.74061 |
| 4 | 144020800 | 144025399 | 54.34542 | 86.94074 | 100.4503 | 42.28388 | 0.135571 | 2.048695 | 0.740615 |
| 18 | 85020000 | 85020399 | 6.647035 | 12.94895 | 16.88139 | 6.149462 | 0.246363 | -0.37344 | 0.740622 |
| 12 | 104246200 | 104250199 | 79.57229 | 124.7983 | 142.2551 | 58.99949 | 0.129158 | 2.589109 | 0.740629 |
| 5 | 7482400 | 7486799 | 54.35717 | 87.04995 | 99.78305 | 42.0663 | 0.148651 | 2.016972 | 0.740653 |

| | | | | | | | | | |
|---|---|---|---|---|---|---|---|---|---|
| 17 | 91450200 | 91451199 | 12.78704 | 23.33766 | 27.56829 | 11.82444 | 0.182073 | 0.293487 | 0.740657 |
| X | 35151800 | 35154199 | 26.39098 | 44.73546 | 50.65405 | 22.18393 | 0.202855 | 1.066942 | 0.740675 |
| 1 | 47041400 | 47042599 | 12.49594 | 22.64823 | 34.75685 | 15.038 | -0.20166 | 0.344578 | 0.740676 |
| X | 149535400 | 149536599 | 15.401 | 27.11302 | 42.92716 | 19.00371 | -0.27105 | 0.594103 | 0.740682 |
| 1 | 11815800 | 11818799 | 43.94678 | 71.01214 | 81.20135 | 34.67321 | 0.167553 | 1.723226 | 0.740693 |
| 13 | 45441800 | 45445199 | 37.19052 | 60.13886 | 85.29495 | 36.45553 | -0.14123 | 1.693947 | 0.740701 |
| 16 | 46718000 | 46718999 | 12.61932 | 22.90159 | 35.35513 | 15.27203 | -0.20843 | 0.341786 | 0.74076 |
| 15 | 32791800 | 32804399 | 180.8396 | 269.7965 | 289.4736 | 119.3114 | 0.157868 | 3.574292 | 0.740767 |
| 9 | 76396000 | 76400199 | 51.733 | 82.87845 | 96.13816 | 40.55273 | 0.132169 | 2.001358 | 0.740793 |
| 6 | 74168400 | 74171199 | 38.40078 | 62.38532 | 72.86997 | 31.25441 | 0.144453 | 1.594986 | 0.740805 |
| 5 | 126011000 | 126022799 | 111.0297 | 169.525 | 225.9785 | 93.71923 | -0.12635 | 3.101813 | 0.740811 |
| 14 | 60232400 | 60234999 | 25.71378 | 42.90173 | 62.13146 | 27.01945 | -0.15143 | 1.222849 | 0.740814 |
| 16 | 64239800 | 64240199 | 4.934968 | 9.632869 | 18.13592 | 6.423077 | -0.24041 | -0.48953 | 0.74082 |
| 17 | 87165000 | 87168799 | 46.51142 | 74.80839 | 86.94319 | 36.89182 | 0.13627 | 1.869524 | 0.740821 |
| 14 | 105519200 | 105519799 | 6.702751 | 13.05023 | 23.80291 | 8.728497 | -0.24202 | -0.32421 | 0.740829 |
| 18 | 50797200 | 50799399 | 24.52136 | 40.17612 | 75.59942 | 32.53368 | -0.53307 | 1.445687 | 0.740831 |
| 12 | 105151600 | 105156199 | 44.80737 | 72.11962 | 83.47698 | 35.45808 | 0.143423 | 1.842206 | 0.740872 |
| 5 | 27188400 | 27190599 | 15.33299 | 26.76305 | 40.53847 | 17.85019 | -0.20866 | 0.713413 | 0.740887 |
| 1 | 21658800 | 21659399 | 11.09126 | 20.10068 | 31.19686 | 13.49912 | -0.21763 | 0.328135 | 0.7409 |
| 10 | 92584400 | 92590599 | 90.24627 | 141.6752 | 131.9256 | 54.71639 | 0.421597 | 2.574063 | 0.740909 |
| 3 | 47489600 | 47490399 | 11.03536 | 20.1992 | 31.33865 | 13.39152 | -0.20484 | 0.274359 | 0.740927 |
| 4 | 31126400 | 31129399 | 39.10552 | 63.30917 | 73.46599 | 31.42918 | 0.149249 | 1.667277 | 0.740953 |
| 11 | 120962000 | 120963199 | 11.48115 | 20.7028 | 31.72499 | 13.81435 | -0.20505 | 0.363291 | 0.740956 |
| 19 | 28486200 | 28488799 | 25.74549 | 42.79094 | 62.01928 | 26.90349 | -0.15263 | 1.265157 | 0.740989 |
| 7 | 60148800 | 60149399 | 8.988453 | 17.37386 | 22.07429 | 8.460915 | 0.214917 | -0.15198 | 0.740994 |
| 10 | 116467600 | 116468199 | 7.754464 | 14.7004 | 24.96989 | 10.085 | -0.27032 | 0.011883 | 0.741014 |
| 3 | 137877800 | 137882199 | 51.44168 | 82.06233 | 114.754 | 48.11156 | -0.14381 | 2.136013 | 0.741023 |
| 9 | 7297000 | 7302799 | 57.60774 | 91.69916 | 128.0163 | 53.51065 | -0.1448 | 2.215981 | 0.741038 |
| 19 | 27469400 | 27477799 | 115.9963 | 177.461 | 199.1417 | 82.62732 | 0.124096 | 3.062075 | 0.741039 |
| 15 | 88164600 | 88169799 | 61.11564 | 97.41594 | 111.2835 | 46.60437 | 0.1462 | 2.170976 | 0.741072 |
| 6 | 59763800 | 59766199 | 26.13832 | 43.8855 | 51.2193 | 22.36269 | 0.15885 | 1.148307 | 0.741097 |
| 5 | 89149800 | 89151199 | 22.94049 | 39.67467 | 38.34522 | 16.82278 | 0.434742 | 0.857649 | 0.741098 |
| 16 | 21146600 | 21148599 | 17.77985 | 31.07399 | 35.77642 | 15.7667 | 0.185333 | 0.699515 | 0.741102 |
| 19 | 58300600 | 58305799 | 58.05542 | 92.3902 | 129.5586 | 54.13785 | -0.15161 | 2.223492 | 0.741129 |
| 3 | 131665200 | 131666399 | 14.94565 | 27.08702 | 30.85046 | 13.34859 | 0.2259 | 0.354 | 0.741151 |
| 9 | 8624400 | 8625399 | 13.76854 | 25.17071 | 29.53356 | 12.64112 | 0.193541 | 0.295405 | 0.741157 |
| 14 | 42358400 | 42369199 | 140.8419 | 213.6433 | 205.1611 | 84.92985 | 0.338782 | 3.240892 | 0.741162 |
| 6 | 131895000 | 131897199 | 25.36842 | 43.8849 | 46.26755 | 20.29861 | 0.309946 | 0.877801 | 0.74117 |
| 13 | 87858200 | 87858599 | 6.381627 | 12.38775 | 21.68337 | 8.146413 | -0.21568 | -0.25159 | 0.741172 |
| 6 | 125819200 | 125821399 | 20.38755 | 35.54879 | 40.82518 | 17.96475 | 0.188557 | 0.752676 | 0.741189 |
| 7 | 55136600 | 55138599 | 20.38755 | 35.54879 | 40.82518 | 17.96475 | 0.188557 | 0.752676 | 0.741189 |
| 12 | 79908200 | 79912199 | 36.59622 | 58.94363 | 102.5462 | 43.60116 | -0.44203 | 1.816167 | 0.741189 |
| 6 | 117714000 | 117718199 | 44.22806 | 70.95341 | 99.94471 | 42.24728 | -0.14508 | 1.950155 | 0.74119 |
| 1 | 51956800 | 51958999 | 20.23712 | 34.86557 | 51.72384 | 22.68959 | -0.18209 | 0.897616 | 0.741208 |
| 11 | 96306400 | 96309799 | 24.21872 | 40.5855 | 58.76394 | 25.62605 | -0.15116 | 1.168349 | 0.741232 |
| 1 | 179217000 | 179221799 | 61.61078 | 97.44437 | 134.5539 | 55.91534 | -0.13767 | 2.399755 | 0.741237 |
| 5 | 143954600 | 143957199 | 20.6677 | 35.69177 | 41.56852 | 18.23609 | 0.166339 | 0.863995 | 0.741245 |

| | | | | | | | | | |
|---|---|---|---|---|---|---|---|---|---|
| 15 | 29648200 | 29648799 | 9.213424 | 17.66181 | 22.39695 | 8.815213 | 0.181394 | -0.06427 | 0.741261 |
| 1 | 48753800 | 48754999 | 15.60425 | 28.03595 | 26.29975 | 11.49658 | 0.494852 | 0.400911 | 0.741283 |
| 15 | 87629000 | 87631199 | 21.95337 | 37.77358 | 44.21682 | 19.39573 | 0.158059 | 0.907535 | 0.741293 |
| 9 | 22175800 | 22182199 | 110.2884 | 169.0659 | 190.3103 | 78.99647 | 0.121876 | 3.019311 | 0.741295 |
| 16 | 31190200 | 31193599 | 41.90524 | 67.46988 | 95.76913 | 40.69805 | -0.14977 | 1.832511 | 0.74131 |
| 9 | 36423200 | 36429999 | 86.53097 | 135.423 | 152.9076 | 63.43143 | 0.140439 | 2.631312 | 0.741312 |
| 19 | 8383000 | 8392799 | 121.3467 | 184.1111 | 243.4421 | 100.7835 | -0.12205 | 3.233867 | 0.741351 |
| 13 | 34840400 | 34842199 | 19.85068 | 34.3008 | 38.56241 | 16.91761 | 0.217316 | 0.85151 | 0.741376 |
| 6 | 105040200 | 105042199 | 24.33462 | 40.5497 | 59.68321 | 25.9342 | -0.17454 | 1.232351 | 0.741382 |
| X | 47911400 | 47913799 | 14.81988 | 26.24261 | 40.24675 | 17.79634 | -0.22213 | 0.520853 | 0.741413 |
| Y | 8441000 | 8442199 | 12.858 | 23.30028 | 22.2231 | 9.617266 | 0.480266 | 0.319567 | 0.74143 |
| 10 | 63372000 | 63376399 | 39.00721 | 62.86102 | 89.20825 | 37.95993 | -0.14778 | 1.798234 | 0.741436 |
| 8 | 82994600 | 82995199 | 5.951378 | 11.57081 | 20.65206 | 7.660757 | -0.22585 | -0.30339 | 0.741439 |
| 9 | 93922800 | 93923399 | 5.951378 | 11.57081 | 20.65206 | 7.660757 | -0.22585 | -0.30339 | 0.741439 |
| 13 | 50710200 | 50715599 | 92.18554 | 143.2536 | 162.8898 | 67.63304 | 0.12238 | 2.757455 | 0.741443 |
| 11 | 96302800 | 96303999 | 12.73586 | 23.24841 | 27.529 | 11.80494 | 0.178709 | 0.291095 | 0.741461 |
| 2 | 129614800 | 129618399 | 32.44467 | 52.68913 | 76.45876 | 32.7845 | -0.1683 | 1.593484 | 0.741464 |
| 10 | 19885000 | 19895199 | 265.9733 | 379.3506 | 475.46 | 196.5154 | -0.12875 | 4.286662 | 0.741472 |
| 8 | 11974000 | 11975399 | 14.51332 | 25.46847 | 38.30752 | 16.94031 | -0.19845 | 0.620148 | 0.741478 |
| 11 | 100051200 | 100052199 | 9.768237 | 18.23233 | 28.40849 | 11.80544 | -0.18136 | 0.141443 | 0.741497 |
| 16 | 9152400 | 9153399 | 9.427062 | 17.64793 | 27.92213 | 11.54506 | -0.19754 | 0.117484 | 0.741525 |
| 5 | 61436400 | 61439399 | 32.79591 | 54.1342 | 62.96231 | 27.18269 | 0.163089 | 1.364737 | 0.741534 |
| 7 | 80167600 | 80169599 | 24.10789 | 42.03374 | 35.43618 | 15.5929 | 0.632673 | 0.766017 | 0.741536 |
| 7 | 140083400 | 140084599 | 10.87418 | 20.05561 | 30.85385 | 13.04909 | -0.18247 | 0.225638 | 0.741554 |
| 10 | 101183000 | 101184399 | 14.98931 | 27.2267 | 25.82931 | 11.14493 | 0.490874 | 0.333052 | 0.741589 |
| Y | 88971400 | 88973999 | 16.60093 | 29.12289 | 27.38335 | 12.10628 | 0.475868 | 0.615584 | 0.741591 |
| 10 | 106027600 | 106030199 | 29.22108 | 48.97985 | 55.21705 | 24.08285 | 0.208583 | 1.175821 | 0.741592 |
| 7 | 3566600 | 3569799 | 27.10704 | 45.28829 | 52.86575 | 23.00991 | 0.158696 | 1.20468 | 0.741592 |
| 1 | 168456800 | 168457599 | 11.15512 | 19.76394 | 38.36102 | 16.96066 | -0.55929 | 0.489256 | 0.7416 |
| 17 | 69517000 | 69519799 | 21.8922 | 37.73879 | 44.18312 | 19.38206 | 0.158318 | 0.88958 | 0.74163 |
| 2 | 59397200 | 59400599 | 47.28608 | 76.09401 | 88.12087 | 37.4198 | 0.14234 | 1.853981 | 0.741649 |
| 1 | 36785600 | 36788599 | 20.7039 | 35.20296 | 51.67353 | 22.64726 | -0.17001 | 1.01751 | 0.741661 |
| 3 | 91478400 | 91483599 | 78.66416 | 124.0301 | 139.2062 | 57.76863 | 0.157349 | 2.48445 | 0.741682 |
| 12 | 62106800 | 62109199 | 33.12351 | 54.91457 | 53.61815 | 23.21853 | 0.415193 | 1.307295 | 0.741711 |
| 5 | 114083000 | 114085599 | 18.17985 | 31.1175 | 47.45006 | 20.80736 | -0.22305 | 0.944156 | 0.741721 |
| 5 | 101757200 | 101759999 | 26.68425 | 44.33981 | 64.40616 | 27.93394 | -0.15591 | 1.271594 | 0.741735 |
| 8 | 107926600 | 107928799 | 22.57126 | 38.82452 | 44.68182 | 19.59952 | 0.182196 | 0.910388 | 0.741738 |
| 14 | 59607800 | 59611399 | 62.75094 | 99.19359 | 136.6125 | 56.75517 | -0.13455 | 2.413544 | 0.741746 |
| 10 | 27329600 | 27331199 | 17.35508 | 31.05665 | 33.74493 | 14.80118 | 0.280653 | 0.451792 | 0.741753 |
| 17 | 23391200 | 23400999 | 114.4372 | 177.615 | 174.8751 | 72.6227 | 0.328569 | 2.792059 | 0.741768 |
| 9 | 103488600 | 103488999 | 8.090816 | 15.71733 | 20.04707 | 7.486228 | 0.245367 | -0.25442 | 0.741776 |
| 6 | 70733600 | 70738399 | 54.76962 | 87.10244 | 121.283 | 50.63258 | -0.14243 | 2.240252 | 0.741785 |
| 3 | 90670600 | 90673199 | 35.06107 | 58.17617 | 53.09816 | 23.00684 | 0.51233 | 1.304401 | 0.741786 |
| 3 | 10351800 | 10353799 | 15.64299 | 27.44126 | 40.31301 | 17.82191 | -0.16468 | 0.635643 | 0.741807 |
| 6 | 125365800 | 125367599 | 15.36207 | 27.15644 | 31.00995 | 13.71667 | 0.200175 | 0.523046 | 0.74181 |
| 2 | 157586400 | 157588799 | 23.93299 | 40.18431 | 58.17968 | 25.40223 | -0.15107 | 1.1475 | 0.741812 |
| 7 | 106137600 | 106138599 | 17.2524 | 30.53426 | 34.83029 | 15.40288 | 0.202309 | 0.529819 | 0.741816 |

| | | | | | | | | | |
|---|---|---|---|---|---|---|---|---|---|
| 10 | 114165800 | 114166799 | 10.47774 | 19.38719 | 29.88001 | 12.5816 | -0.1801 | 0.204386 | 0.741819 |
| 5 | 9342200 | 9343599 | 16.04488 | 28.73898 | 32.18244 | 14.1047 | 0.237995 | 0.432546 | 0.741844 |
| 7 | 106159800 | 106164799 | 114.1593 | 174.7426 | 196.6346 | 81.59699 | 0.120704 | 3.050476 | 0.741852 |
| 6 | 136489400 | 136492599 | 45.98463 | 74.43074 | 70.36958 | 30.09997 | 0.44388 | 1.682266 | 0.74186 |
| 2 | 181556400 | 181570399 | 291.7224 | 412.7148 | 512.9499 | 212.2984 | -0.12841 | 4.406214 | 0.741868 |
| Y | 38995400 | 38995999 | 8.882078 | 17.0077 | 17.27094 | 6.819413 | 0.494715 | -0.07524 | 0.741881 |
| 13 | 38282200 | 38284799 | 17.42022 | 30.60986 | 35.41748 | 15.67971 | 0.177941 | 0.609385 | 0.741921 |
| 13 | 77138200 | 77146199 | 165.1528 | 243.9273 | 326.1133 | 134.3001 | -0.17272 | 3.74632 | 0.741945 |
| 15 | 33269800 | 33271399 | 27.34249 | 45.37896 | 67.04193 | 29.05297 | -0.18034 | 1.292961 | 0.741952 |
| 15 | 18429400 | 18429999 | 10.12232 | 19.40835 | 24.04853 | 9.459338 | 0.215008 | -0.05379 | 0.741953 |
| 11 | 76444200 | 76444599 | 6.235579 | 12.12964 | 21.26719 | 7.856004 | -0.1954 | -0.31395 | 0.741957 |
| 16 | 16464800 | 16465199 | 6.235579 | 12.12964 | 21.26719 | 7.856004 | -0.1954 | -0.31395 | 0.741957 |
| 17 | 48487600 | 48488799 | 15.51692 | 26.79121 | 50.55328 | 22.17756 | -0.5266 | 0.859524 | 0.741986 |
| 1 | 13714400 | 13716999 | 23.64219 | 40.90493 | 45.17709 | 19.8203 | 0.243099 | 0.865067 | 0.741996 |
| 13 | 6316800 | 6319999 | 46.83181 | 75.31881 | 87.62178 | 37.17626 | 0.134746 | 1.872095 | 0.742015 |
| 6 | 11078400 | 11080199 | 13.58581 | 24.08916 | 35.61476 | 15.74285 | -0.16869 | 0.489466 | 0.742031 |
| 8 | 14321400 | 14325399 | 30.9285 | 50.49213 | 72.85145 | 31.30843 | -0.15437 | 1.50707 | 0.742036 |
| 18 | 7316800 | 7321399 | 106.3757 | 162.5062 | 216.4887 | 89.79469 | -0.12484 | 3.089294 | 0.742066 |
| 6 | 90144600 | 90145399 | 11.29673 | 20.84607 | 32.49933 | 13.73503 | -0.20061 | 0.231639 | 0.742105 |
| X | 126137200 | 126139399 | 24.67134 | 42.50794 | 46.82983 | 20.54278 | 0.24561 | 0.910595 | 0.742113 |
| 15 | 92709000 | 92712399 | 36.54343 | 59.08483 | 83.91059 | 35.85786 | -0.14332 | 1.696068 | 0.742118 |
| 13 | 80048200 | 80049399 | 16.17135 | 28.28966 | 41.46355 | 18.2877 | -0.16127 | 0.686076 | 0.742128 |
| 8 | 78643800 | 78647199 | 39.05063 | 63.714 | 72.26866 | 31.05125 | 0.191217 | 1.532049 | 0.74219 |
| X | 122476400 | 122477999 | 35.09894 | 58.05533 | 53.60775 | 23.17703 | 0.495309 | 1.341378 | 0.742191 |
| 16 | 68064000 | 68065999 | 16.78497 | 29.89985 | 33.8231 | 14.89259 | 0.219209 | 0.477557 | 0.742194 |
| 2 | 178901200 | 178902999 | 12.49051 | 22.34605 | 33.70411 | 14.78636 | -0.18961 | 0.423806 | 0.742228 |
| 5 | 94320600 | 94322199 | 14.12155 | 26.03618 | 24.84414 | 10.51214 | 0.501677 | 0.240135 | 0.742237 |
| 13 | 48457400 | 48461399 | 44.95972 | 72.46589 | 84.10869 | 35.79798 | 0.141919 | 1.799241 | 0.742238 |
| 1 | 150232600 | 150234799 | 23.6117 | 41.67843 | 36.55668 | 16.17492 | 0.57817 | 0.599984 | 0.742243 |
| X | 36454200 | 36458799 | 48.58766 | 78.48339 | 83.55387 | 35.66145 | 0.269933 | 1.747285 | 0.742243 |
| 12 | 62391600 | 62392799 | 13.54556 | 24.94599 | 29.09147 | 12.32825 | 0.212499 | 0.251053 | 0.742248 |
| 2 | 99133400 | 99138799 | 51.92857 | 83.23423 | 96.36498 | 40.67996 | 0.135815 | 1.986323 | 0.742256 |
| 14 | 74859200 | 74860599 | 19.40268 | 34.41198 | 35.65573 | 15.75753 | 0.34149 | 0.523374 | 0.742274 |
| 11 | 19358800 | 19359799 | 10.63362 | 18.98566 | 36.83092 | 16.18562 | -0.55228 | 0.429419 | 0.742286 |
| 16 | 22083600 | 22087999 | 27.04375 | 44.34584 | 77.36024 | 33.3032 | -0.42379 | 1.440957 | 0.742287 |
| 7 | 40668800 | 40671399 | 22.23763 | 38.80699 | 41.19625 | 18.13753 | 0.301526 | 0.749155 | 0.742294 |
| 10 | 69979800 | 69984599 | 59.26696 | 94.37221 | 108.749 | 45.47826 | 0.132082 | 2.20093 | 0.742295 |
| 2 | 11103600 | 11113799 | 140.1617 | 210.3739 | 277.1433 | 114.3719 | -0.1302 | 3.453634 | 0.742303 |
| 6 | 65929000 | 65931399 | 27.57276 | 46.0214 | 53.97461 | 23.47806 | 0.15193 | 1.216847 | 0.74232 |
| 9 | 119696600 | 119699599 | 78.94788 | 122.091 | 175.5923 | 72.93687 | -0.22155 | 2.839773 | 0.742337 |
| 12 | 83411600 | 83413199 | 22.36623 | 37.88781 | 54.93854 | 24.05545 | -0.15282 | 1.06014 | 0.742341 |
| 13 | 67180600 | 67196199 | 212.9999 | 316.3233 | 311.2448 | 128.2145 | 0.276644 | 3.661214 | 0.742345 |
| 2 | 61675600 | 61679999 | 28.11737 | 47.51773 | 45.85019 | 20.06165 | 0.432598 | 1.09125 | 0.742349 |
| 10 | 127789400 | 127790799 | 12.3504 | 21.85232 | 40.83113 | 18.05762 | -0.50604 | 0.51273 | 0.742416 |
| 4 | 62041400 | 62063199 | 550.6495 | 744.9061 | 742.9717 | 309.6747 | 0.123758 | 5.095676 | 0.742429 |
| 9 | 38344800 | 38345799 | 14.26646 | 25.43984 | 28.14741 | 12.38272 | 0.252053 | 0.443508 | 0.742432 |
| 17 | 5571600 | 5574799 | 64.93667 | 102.9461 | 118.0102 | 49.11026 | 0.133278 | 2.340483 | 0.742433 |

| | | | | | | | | | |
|---|---|---|---|---|---|---|---|---|---|
| 4 | 59984400 | 59985399 | 10.48977 | 19.36435 | 29.8585 | 12.61241 | -0.18401 | 0.217751 | 0.742435 |
| 2 | 27084200 | 27086399 | 18.94973 | 33.0791 | 38.7926 | 17.08246 | 0.158716 | 0.72744 | 0.742438 |
| 1 | 96261400 | 96263399 | 19.08026 | 32.47766 | 57.52785 | 25.21528 | -0.43975 | 1.010572 | 0.742444 |
| 11 | 13634800 | 13635599 | 11.6981 | 21.56044 | 25.078 | 10.61617 | 0.217409 | 0.222434 | 0.742448 |
| 12 | 17783800 | 17785999 | 16.82421 | 29.27273 | 43.16628 | 18.96935 | -0.17062 | 0.771183 | 0.742453 |
| 7 | 26780000 | 26782599 | 34.7464 | 56.78412 | 66.64345 | 28.65137 | 0.143804 | 1.495939 | 0.742463 |
| 13 | 44065800 | 44070199 | 57.04361 | 90.59974 | 126.3053 | 52.6593 | -0.14604 | 2.281213 | 0.742487 |
| 17 | 47553200 | 47560399 | 64.56265 | 101.9301 | 140.2513 | 58.23007 | -0.13467 | 2.444709 | 0.742511 |
| 1 | 67752800 | 67753399 | 15.68882 | 27.76838 | 32.38433 | 14.32106 | 0.17053 | 0.513973 | 0.742531 |
| 7 | 34150600 | 34152199 | 19.56331 | 34.11244 | 39.81685 | 17.52132 | 0.165261 | 0.746854 | 0.742556 |
| X | 74025400 | 74027199 | 22.32969 | 38.95119 | 41.87817 | 18.43286 | 0.283502 | 0.761401 | 0.742561 |
| 13 | 5030000 | 5034199 | 91.64766 | 140.6279 | 195.9987 | 81.36856 | -0.18494 | 2.994838 | 0.742567 |
| 9 | 19544400 | 19545399 | 10.47069 | 19.58002 | 31.19077 | 12.92068 | -0.2094 | 0.143111 | 0.742576 |
| 9 | 23355600 | 23357399 | 15.1475 | 26.49162 | 39.94653 | 17.61471 | -0.20207 | 0.680976 | 0.742576 |
| 15 | 50436000 | 50438399 | 24.77554 | 42.34838 | 48.47795 | 21.25573 | 0.188502 | 0.983044 | 0.742583 |
| 1 | 112801200 | 112803799 | 34.45806 | 56.58445 | 65.36911 | 28.15808 | 0.169919 | 1.430448 | 0.742631 |
| 2 | 150291800 | 150293599 | 16.54058 | 28.84134 | 43.07108 | 18.95317 | -0.18867 | 0.735964 | 0.742639 |
| 6 | 6138600 | 6140199 | 18.47177 | 32.25875 | 37.79267 | 16.64668 | 0.160254 | 0.717714 | 0.742702 |
| 6 | 117568800 | 117569599 | 15.4672 | 27.85107 | 25.84322 | 11.27308 | 0.512569 | 0.386086 | 0.742731 |
| 3 | 156034800 | 156036799 | 20.33146 | 35.91606 | 36.68042 | 16.22716 | 0.359913 | 0.570497 | 0.742742 |
| 1 | 152412200 | 152413999 | 10.7122 | 19.23885 | 31.30482 | 13.68905 | -0.29509 | 0.38055 | 0.742757 |
| 13 | 96279400 | 96282999 | 35.20388 | 57.5412 | 67.34676 | 28.95547 | 0.147901 | 1.494359 | 0.742762 |
| 10 | 129880400 | 129881399 | 28.16608 | 46.27118 | 67.75526 | 29.19099 | -0.17076 | 1.416186 | 0.742769 |
| 11 | 22648600 | 22651999 | 40.50339 | 65.28142 | 91.90448 | 39.11463 | -0.13603 | 1.798048 | 0.742785 |
| 15 | 38150600 | 38152399 | 27.42515 | 47.01546 | 41.39596 | 18.15485 | 0.566296 | 0.958381 | 0.742795 |
| 5 | 86213000 | 86213799 | 7.321372 | 13.89162 | 24.48078 | 9.869027 | -0.32045 | 0.000421 | 0.742801 |
| 15 | 29737200 | 29737599 | 6.999613 | 13.47229 | 23.08385 | 8.980863 | -0.23283 | -0.12136 | 0.742811 |
| 6 | 48324000 | 48324799 | 13.90348 | 24.71258 | 36.61143 | 16.15018 | -0.16967 | 0.477918 | 0.742821 |
| 15 | 91891800 | 91892599 | 12.79477 | 22.87685 | 34.1698 | 14.99813 | -0.17621 | 0.429956 | 0.742829 |
| X | 13254000 | 13254999 | 12.79477 | 22.87685 | 34.1698 | 14.99813 | -0.17621 | 0.429956 | 0.742829 |
| 12 | 48474400 | 48476399 | 20.25166 | 34.83363 | 51.16472 | 22.4432 | -0.16824 | 0.912014 | 0.742836 |
| 5 | 136196200 | 136201799 | 42.49492 | 68.49179 | 78.25217 | 33.30451 | 0.164616 | 1.793615 | 0.742844 |
| 1 | 84678200 | 84679199 | 9.364703 | 17.34067 | 27.8665 | 11.72184 | -0.23872 | 0.185989 | 0.742846 |
| 5 | 94578000 | 94579399 | 16.66562 | 29.72052 | 27.79634 | 12.2274 | 0.493116 | 0.458118 | 0.742851 |
| 7 | 26704800 | 26712799 | 215.7514 | 317.2793 | 294.2083 | 121.1603 | 0.348941 | 3.821928 | 0.742853 |
| 15 | 4721800 | 4726599 | 59.65781 | 95.61574 | 101.6655 | 42.91251 | 0.25827 | 1.999242 | 0.742871 |
| 10 | 113415000 | 113416199 | 14.02763 | 25.368 | 29.90564 | 12.97503 | 0.174088 | 0.358789 | 0.742873 |
| 11 | 31159600 | 31161999 | 41.81176 | 67.51701 | 78.21625 | 33.36973 | 0.147722 | 1.737587 | 0.742881 |
| 3 | 68705000 | 68709199 | 82.3296 | 128.0592 | 172.8827 | 71.77201 | -0.12394 | 2.738706 | 0.742899 |
| 3 | 88827200 | 88829199 | 24.35597 | 40.7642 | 59.33942 | 25.85647 | -0.15886 | 1.181855 | 0.742909 |
| 12 | 5099000 | 5114599 | 656.4574 | 865.1163 | 1026.054 | 429.208 | -0.16385 | 5.513575 | 0.742936 |
| 10 | 6889000 | 6889799 | 13.95958 | 25.06376 | 29.03187 | 12.69824 | 0.191953 | 0.400599 | 0.742976 |
| 5 | 19848000 | 19849199 | 14.19889 | 25.13113 | 37.10447 | 16.40843 | -0.16791 | 0.514162 | 0.743024 |
| X | 62032000 | 62034599 | 31.84043 | 54.29745 | 40.84744 | 17.90668 | 0.791473 | 1.024348 | 0.74303 |
| 11 | 10988200 | 10993799 | 83.48765 | 130.9008 | 148.7527 | 61.69616 | 0.133254 | 2.59666 | 0.743041 |
| 5 | 145309400 | 145327399 | 202.0978 | 300.0558 | 310.7655 | 128.0137 | 0.202395 | 3.663388 | 0.743053 |
| 5 | 109743400 | 109753199 | 139.2301 | 210.3985 | 233.9403 | 96.72855 | 0.122854 | 3.317826 | 0.743053 |

| | | | | | | | | | |
|---|---|---|---|---|---|---|---|---|---|
| 13 | 34502800 | 34504799 | 18.83148 | 32.83539 | 38.37134 | 16.88491 | 0.163707 | 0.743087 | 0.743071 |
| 1 | 90234800 | 90236199 | 15.21753 | 27.24102 | 31.85035 | 13.96607 | 0.175826 | 0.432142 | 0.74308 |
| 3 | 71459000 | 71461599 | 22.51768 | 39.01833 | 43.54218 | 19.1039 | 0.228574 | 0.843119 | 0.743086 |
| 1 | 119427200 | 119429399 | 19.02741 | 32.84838 | 47.93811 | 21.03021 | -0.15787 | 0.868697 | 0.743087 |
| 1 | 5496600 | 5501799 | 98.14772 | 151.9612 | 170.8678 | 70.97399 | 0.134433 | 2.825723 | 0.743089 |
| 8 | 63999600 | 63999999 | 7.242404 | 14.07723 | 18.42377 | 6.848392 | 0.214176 | -0.28884 | 0.74313 |
| 18 | 4698200 | 4704999 | 27.82007 | 46.90512 | 44.80858 | 19.59195 | 0.4468 | 1.107461 | 0.743134 |
| 7 | 97655600 | 97656999 | 18.7049 | 32.60119 | 37.42576 | 16.46454 | 0.189291 | 0.747743 | 0.743138 |
| 4 | 52063600 | 52063999 | 8.24384 | 15.47522 | 25.924 | 10.66927 | -0.27345 | 0.085633 | 0.743138 |
| 8 | 4898800 | 4900799 | 34.77194 | 56.96797 | 66.7871 | 28.74076 | 0.147503 | 1.462915 | 0.743141 |
| 5 | 37370600 | 37375199 | 66.94308 | 105.9436 | 121.0542 | 50.32118 | 0.135832 | 2.386118 | 0.743151 |
| 18 | 41959000 | 41961799 | 24.94038 | 41.70864 | 60.38647 | 26.29931 | -0.151 | 1.195457 | 0.743165 |
| 16 | 9110400 | 9111599 | 16.16674 | 28.29776 | 41.43809 | 18.2853 | -0.16 | 0.67663 | 0.743171 |
| 8 | 31506600 | 31515199 | 100.9708 | 155.0268 | 206.7043 | 85.82003 | -0.12045 | 2.987265 | 0.743212 |
| 6 | 63899600 | 63902599 | 50.68413 | 81.19548 | 93.61743 | 39.48749 | 0.140827 | 1.999393 | 0.743222 |
| 15 | 46914400 | 46917799 | 36.41692 | 59.43273 | 69.57792 | 29.89803 | 0.146006 | 1.521204 | 0.743236 |
| 17 | 31037200 | 31038599 | 13.5934 | 24.07822 | 35.64211 | 15.75929 | -0.17102 | 0.500289 | 0.743245 |
| 19 | 39873600 | 39874199 | 13.5934 | 24.07822 | 35.64211 | 15.75929 | -0.17102 | 0.500289 | 0.743245 |
| 1 | 11293200 | 11294399 | 12.57736 | 22.54253 | 33.6322 | 14.73212 | -0.17267 | 0.410936 | 0.743267 |
| 9 | 37364800 | 37366599 | 17.68541 | 30.71477 | 45.25736 | 19.86536 | -0.16998 | 0.801629 | 0.743309 |
| 13 | 53818600 | 53822599 | 43.15312 | 69.38978 | 97.44415 | 41.33278 | -0.13661 | 1.877254 | 0.743314 |
| 4 | 100531400 | 100534399 | 22.59917 | 38.00612 | 55.42132 | 24.2136 | -0.16127 | 1.125011 | 0.743318 |
| 4 | 69680200 | 69681599 | 14.98857 | 26.8486 | 31.31045 | 13.72181 | 0.179985 | 0.424554 | 0.743338 |
| 8 | 112135000 | 112136399 | 21.72577 | 37.30348 | 43.70881 | 19.17161 | 0.156025 | 0.923787 | 0.743348 |
| 19 | 33873800 | 33877599 | 37.07195 | 60.79714 | 59.2153 | 25.49318 | 0.414762 | 1.449344 | 0.743375 |
| 9 | 74749600 | 74750399 | 9.932018 | 18.68537 | 29.68542 | 12.17181 | -0.19373 | 0.095557 | 0.74339 |
| 3 | 15437400 | 15438399 | 19.43331 | 34.07022 | 31.52292 | 13.92897 | 0.49922 | 0.652399 | 0.743413 |
| 3 | 123492600 | 123495599 | 29.85823 | 49.15148 | 71.19131 | 30.6979 | -0.15401 | 1.394832 | 0.743416 |
| 4 | 147544800 | 147597199 | 2487.592 | 2962.331 | 3120.43 | 1320.697 | -0.14045 | 7.142372 | 0.743421 |
| 5 | 104967200 | 104977999 | 342.0022 | 481.3716 | 500.691 | 207.4004 | 0.120952 | 4.483452 | 0.743447 |
| 6 | 141544600 | 141547999 | 46.39425 | 74.39889 | 103.8337 | 43.86458 | -0.13246 | 1.967296 | 0.743465 |
| 13 | 65924800 | 65925999 | 25.69893 | 43.3556 | 50.12897 | 21.92232 | 0.172569 | 1.104472 | 0.743476 |
| 2 | 28873000 | 28874799 | 17.42853 | 30.15139 | 44.78464 | 19.64806 | -0.18243 | 0.839269 | 0.743518 |
| 7 | 95111200 | 95112799 | 17.42853 | 30.15139 | 44.78464 | 19.64806 | -0.18243 | 0.839269 | 0.743518 |
| 6 | 37758600 | 37763399 | 115.6113 | 178.1643 | 166.9767 | 69.35944 | 0.391747 | 2.916386 | 0.743523 |
| 11 | 17184000 | 17185999 | 20.19768 | 35.2695 | 40.23834 | 17.72287 | 0.198191 | 0.730022 | 0.743526 |
| 17 | 50068000 | 50070599 | 28.73525 | 47.34391 | 68.1615 | 29.40222 | -0.1448 | 1.380062 | 0.74355 |
| 9 | 113831400 | 113832199 | 14.53962 | 25.60655 | 37.7036 | 16.68996 | -0.16675 | 0.572172 | 0.743552 |
| 3 | 32758200 | 32759999 | 23.43157 | 39.77473 | 45.88894 | 20.10561 | 0.175981 | 1.037177 | 0.743552 |
| 14 | 41842800 | 41848999 | 71.92659 | 114.1619 | 109.3647 | 45.56611 | 0.393504 | 2.311187 | 0.743557 |
| X | 89171000 | 89172199 | 20.76002 | 35.93738 | 40.61611 | 17.81957 | 0.210448 | 0.846027 | 0.7436 |
| 2 | 129934000 | 129934799 | 11.3737 | 21.43227 | 25.82998 | 10.56202 | 0.205437 | 0.090153 | 0.743617 |
| 1 | 113539800 | 113547199 | 79.46864 | 126.0264 | 130.1394 | 54.17606 | 0.284225 | 2.345883 | 0.74362 |
| 17 | 94264000 | 94265999 | 20.20873 | 34.92881 | 53.42434 | 23.43784 | -0.22509 | 0.870032 | 0.743627 |
| 6 | 68999400 | 69001199 | 32.78025 | 53.96287 | 63.42126 | 27.3477 | 0.14667 | 1.397298 | 0.743651 |
| 8 | 43249000 | 43249599 | 10.412 | 19.58468 | 23.68254 | 9.713567 | 0.196817 | 0.085199 | 0.743653 |
| 13 | 46761600 | 46764999 | 30.05575 | 49.38399 | 71.02254 | 30.60085 | -0.14486 | 1.418897 | 0.743658 |

| | | | | | | | | | |
|---|---|---|---|---|---|---|---|---|---|
| 6 | 108597800 | 108600399 | 21.86489 | 37.03547 | 53.84003 | 23.57386 | -0.15649 | 1.057659 | 0.74366 |
| 12 | 62298600 | 62301599 | 32.97507 | 54.67264 | 61.95815 | 26.83145 | 0.200928 | 1.314071 | 0.743661 |
| 7 | 10937400 | 10945799 | 271.1826 | 387.9936 | 396.3845 | 163.7486 | 0.170592 | 4.23969 | 0.743668 |
| 19 | 19709200 | 19710199 | 15.39836 | 27.76456 | 23.46218 | 10.21718 | 0.648282 | 0.372407 | 0.743688 |
| 12 | 5134600 | 5136399 | 16.12695 | 28.14831 | 41.52704 | 18.28525 | -0.17086 | 0.71925 | 0.743761 |
| 5 | 126150600 | 126159799 | 120.3068 | 182.723 | 242.4782 | 100.4157 | -0.12602 | 3.212144 | 0.743764 |
| 2 | 21550400 | 21550799 | 8.010848 | 15.55194 | 20.5609 | 7.718257 | 0.186419 | -0.23827 | 0.743793 |
| X | 135827800 | 135832199 | 66.90687 | 106.2678 | 120.2396 | 50.13164 | 0.154685 | 2.290071 | 0.743804 |
| 3 | 10261800 | 10262999 | 11.72522 | 21.8508 | 25.5263 | 10.63586 | 0.227677 | 0.158177 | 0.743815 |
| 9 | 114913400 | 114916599 | 60.43665 | 95.66917 | 132.3462 | 55.02174 | -0.13933 | 2.378082 | 0.74382 |
| 15 | 14501400 | 14502799 | 20.31591 | 35.64318 | 38.98509 | 17.23672 | 0.258813 | 0.656792 | 0.74387 |
| 15 | 90570000 | 90572999 | 24.18412 | 40.06819 | 72.58669 | 31.41849 | -0.47394 | 1.304548 | 0.743881 |
| 17 | 76527400 | 76528399 | 11.585 | 21.22671 | 32.34177 | 13.80118 | -0.17786 | 0.272591 | 0.743884 |
| 3 | 94664800 | 94666599 | 13.22906 | 23.44493 | 35.10994 | 15.52239 | -0.18748 | 0.490012 | 0.743919 |
| 2 | 29185200 | 29188399 | 45.65936 | 73.58782 | 85.31143 | 36.30623 | 0.143474 | 1.803745 | 0.743922 |
| 6 | 11428600 | 11429199 | 14.5861 | 25.93935 | 29.54662 | 13.02802 | 0.208 | 0.465032 | 0.743938 |
| 18 | 3336600 | 3339599 | 18.66035 | 31.42187 | 59.55107 | 26.0279 | -0.5379 | 1.105204 | 0.743966 |
| 17 | 48323600 | 48326799 | 28.46163 | 47.2879 | 55.02814 | 23.8647 | 0.163087 | 1.271746 | 0.743994 |
| 17 | 58541800 | 58543999 | 23.91235 | 41.13818 | 46.2756 | 20.29873 | 0.215065 | 0.91676 | 0.744003 |
| 1 | 151938600 | 151941199 | 23.17434 | 39.18295 | 57.16837 | 25.01754 | -0.16186 | 1.080934 | 0.74401 |
| 9 | 37861000 | 37868399 | 155.4596 | 233.2746 | 258.4693 | 106.658 | 0.119018 | 3.460991 | 0.744028 |
| 8 | 16735400 | 16736399 | 11.03414 | 20.59179 | 24.61805 | 10.2331 | 0.197647 | 0.142821 | 0.744047 |
| 5 | 126886400 | 126887199 | 12.35955 | 23.08433 | 27.26865 | 11.31902 | 0.216765 | 0.157816 | 0.744073 |
| 2 | 89903400 | 89906399 | 33.72471 | 55.51232 | 64.718 | 27.90574 | 0.158193 | 1.400838 | 0.744081 |
| 1 | 147902400 | 147903599 | 10.53528 | 19.58226 | 30.46166 | 12.74762 | -0.18737 | 0.178261 | 0.744097 |
| 8 | 114296600 | 114309799 | 186.1442 | 270.7128 | 423.4638 | 174.5447 | -0.42046 | 3.993691 | 0.744111 |
| 13 | 117031000 | 117031799 | 8.947443 | 16.95198 | 27.08878 | 10.95443 | -0.18489 | 0.032461 | 0.744141 |
| 3 | 149109600 | 149110999 | 16.73913 | 29.40966 | 34.11166 | 15.10012 | 0.174693 | 0.607584 | 0.744243 |
| 13 | 58603000 | 58603999 | 8.52002 | 16.09077 | 25.95482 | 10.56693 | -0.20654 | 0.048671 | 0.744252 |
| 15 | 27290800 | 27294399 | 66.86371 | 105.2694 | 144.9276 | 60.13142 | -0.13859 | 2.507171 | 0.744262 |
| 8 | 48646000 | 48655799 | 163.1879 | 241.9812 | 317.5032 | 130.7724 | -0.14036 | 3.679384 | 0.744269 |
| X | 66637200 | 66638999 | 20.17724 | 35.4225 | 38.52531 | 17.04263 | 0.267075 | 0.647741 | 0.744278 |
| 6 | 138634400 | 138635799 | 17.63197 | 30.33268 | 45.86753 | 20.11971 | -0.20979 | 0.892585 | 0.744305 |
| 5 | 10405000 | 10409999 | 73.06725 | 115.7216 | 112.4614 | 46.76681 | 0.369938 | 2.372412 | 0.744307 |
| 14 | 62137000 | 62139799 | 24.69414 | 41.31605 | 59.90053 | 26.09534 | -0.15305 | 1.187354 | 0.74431 |
| 14 | 76509800 | 76512999 | 36.39513 | 59.03061 | 84.48984 | 36.1947 | -0.15023 | 1.631134 | 0.744311 |
| 17 | 58162800 | 58163199 | 10.68738 | 18.60158 | 47.35729 | 20.8156 | -0.95404 | 0.742961 | 0.744324 |
| 11 | 14509400 | 14510399 | 13.0646 | 23.73228 | 28.00404 | 12.08224 | 0.178032 | 0.322567 | 0.744372 |
| 17 | 70101800 | 70104399 | 20.07086 | 35.19484 | 39.06984 | 17.26656 | 0.237612 | 0.665034 | 0.744421 |
| 4 | 87488600 | 87490999 | 17.60697 | 30.89525 | 36.01631 | 15.92619 | 0.167265 | 0.633934 | 0.744422 |
| 10 | 56985800 | 56986799 | 8.070877 | 15.57988 | 26.64834 | 10.27673 | -0.21906 | -0.12936 | 0.744436 |
| 14 | 18625800 | 18627799 | 15.25843 | 27.17072 | 31.90202 | 14.051 | 0.165499 | 0.465866 | 0.744447 |
| 4 | 112072200 | 112072799 | 10.13491 | 18.93491 | 29.50755 | 12.24224 | -0.17959 | 0.142254 | 0.744449 |
| 6 | 15667400 | 15667999 | 9.336066 | 18.06108 | 22.70727 | 8.669119 | 0.234598 | -0.16281 | 0.744453 |
| 13 | 62914200 | 62938399 | 377.4355 | 530.764 | 525.9315 | 217.887 | 0.189584 | 4.499156 | 0.74447 |
| 6 | 90228800 | 90229199 | 6.372944 | 12.43231 | 16.94924 | 6.062541 | 0.206505 | -0.43992 | 0.744486 |
| 11 | 20690600 | 20691999 | 11.22639 | 19.6473 | 44.67989 | 19.71323 | -0.79159 | 0.657319 | 0.74449 |

| | | | | | | | | | |
|---|---|---|---|---|---|---|---|---|---|
| 9 | 17989800 | 17992199 | 22.92454 | 39.82098 | 44.08572 | 19.3533 | 0.240731 | 0.822751 | 0.744493 |
| 7 | 24373400 | 24374799 | 16.56632 | 29.11125 | 33.6194 | 14.88446 | 0.180919 | 0.602764 | 0.744513 |
| Y | 87319600 | 87321799 | 20.04345 | 34.64914 | 31.11859 | 13.65213 | 0.539897 | 0.834859 | 0.744516 |
| 1 | 26485200 | 26486199 | 18.08793 | 31.86055 | 29.41003 | 13.01831 | 0.50354 | 0.576371 | 0.744545 |
| 4 | 52556400 | 52557999 | 14.53188 | 25.61622 | 37.67996 | 16.67646 | -0.16482 | 0.56191 | 0.744546 |
| 13 | 45522600 | 45523799 | 13.39605 | 24.5662 | 29.0464 | 12.38041 | 0.186787 | 0.273559 | 0.744548 |
| 18 | 78695400 | 78699599 | 44.62333 | 72.11463 | 81.92729 | 34.98899 | 0.177134 | 1.721299 | 0.744584 |
| 11 | 25152000 | 25152999 | 11.39609 | 21.53139 | 25.24364 | 10.26837 | 0.250579 | 0.066915 | 0.744588 |
| 7 | 84623400 | 84626799 | 38.47732 | 62.74065 | 71.96315 | 30.91317 | 0.174579 | 1.540729 | 0.744596 |
| 15 | 35333400 | 35335399 | 31.87465 | 53.32338 | 49.73225 | 21.66392 | 0.481302 | 1.201182 | 0.744641 |
| 14 | 99796800 | 99798799 | 21.58612 | 37.57917 | 42.44051 | 18.657 | 0.212414 | 0.783335 | 0.744652 |
| 5 | 118647400 | 118650199 | 17.36583 | 30.12878 | 44.75113 | 19.63507 | -0.18168 | 0.811998 | 0.744671 |
| 4 | 94704600 | 94712199 | 94.15223 | 145.2846 | 194.5587 | 80.81601 | -0.12119 | 2.887684 | 0.744673 |
| 2 | 150237600 | 150283799 | 1132.317 | 1439.339 | 1624.406 | 681.985 | -0.14437 | 6.101815 | 0.744693 |
| 16 | 18573200 | 18580799 | 64.14026 | 101.8398 | 117.6786 | 49.0476 | 0.123917 | 2.295413 | 0.744729 |
| 5 | 124078400 | 124081399 | 30.9127 | 50.46916 | 72.90308 | 31.33126 | -0.156 | 1.505765 | 0.744737 |
| 1 | 13576200 | 13577399 | 12.99764 | 23.31111 | 35.10316 | 15.36794 | -0.18568 | 0.410939 | 0.744764 |
| 8 | 119383000 | 119385799 | 19.71847 | 34.4448 | 39.72915 | 17.50247 | 0.182564 | 0.722156 | 0.744765 |
| 6 | 123096800 | 123100199 | 33.27556 | 55.1246 | 62.14741 | 26.89783 | 0.208219 | 1.321495 | 0.744767 |
| 12 | 103756400 | 103763999 | 81.47581 | 128.0483 | 123.5853 | 51.26461 | 0.371198 | 2.545714 | 0.744794 |
| 11 | 40424400 | 40426399 | 12.46877 | 22.55669 | 34.70666 | 15.05153 | -0.20766 | 0.354326 | 0.744794 |
| 3 | 144438600 | 144441199 | 31.82148 | 52.55648 | 61.3495 | 26.49708 | 0.157874 | 1.351748 | 0.744806 |
| 6 | 48741200 | 48743999 | 42.01418 | 67.49956 | 95.44109 | 40.43207 | -0.148 | 1.902493 | 0.744807 |
| 6 | 51464800 | 51466999 | 33.07426 | 54.34635 | 63.97499 | 27.56492 | 0.143282 | 1.420238 | 0.744841 |
| 5 | 18550200 | 18555799 | 99.65927 | 153.0259 | 203.9078 | 84.65996 | -0.11945 | 2.98452 | 0.744851 |
| 7 | 64885600 | 64886999 | 18.72343 | 32.71365 | 38.2163 | 16.83826 | 0.164311 | 0.711739 | 0.744863 |
| 1 | 94192800 | 94193799 | 11.79498 | 21.21388 | 32.19192 | 14.05996 | -0.19394 | 0.381431 | 0.744866 |
| 18 | 6427600 | 6430399 | 29.39781 | 48.53227 | 70.65942 | 30.50883 | -0.15999 | 1.359552 | 0.744878 |
| 2 | 82172600 | 82174399 | 49.11917 | 78.76109 | 90.97675 | 38.42756 | 0.139789 | 1.969252 | 0.744879 |
| 8 | 91424800 | 91425599 | 11.30152 | 21.11364 | 25.70459 | 10.66515 | 0.173989 | 0.1426 | 0.744884 |
| 10 | 129136000 | 129141399 | 69.06339 | 108.7616 | 148.3044 | 61.53462 | -0.12448 | 2.502263 | 0.74492 |
| 9 | 113406200 | 113407999 | 14.02672 | 24.67488 | 37.22332 | 16.48114 | -0.20224 | 0.579822 | 0.744965 |
| 17 | 21619600 | 21620999 | 10.42171 | 19.11879 | 29.76264 | 12.67932 | -0.20687 | 0.249747 | 0.744996 |
| 15 | 34218200 | 34220799 | 24.72125 | 41.1717 | 60.37628 | 26.22641 | -0.16936 | 1.236874 | 0.744997 |
| 6 | 77420200 | 77421599 | 16.71854 | 29.44318 | 34.48679 | 15.26608 | 0.161599 | 0.579348 | 0.745007 |
| 8 | 95179000 | 95180999 | 24.91868 | 42.0231 | 49.07337 | 21.45818 | 0.158204 | 1.101925 | 0.745037 |
| 17 | 24047200 | 24049399 | 13.64499 | 23.93473 | 43.52393 | 19.24033 | -0.47089 | 0.631371 | 0.745048 |
| 1 | 178049600 | 178058399 | 292.5951 | 413.714 | 514.0246 | 212.7632 | -0.12877 | 4.41461 | 0.745058 |
| 13 | 54103200 | 54107599 | 48.46951 | 77.22418 | 126.1347 | 52.79143 | -0.36945 | 2.177862 | 0.745073 |
| X | 30771400 | 30801999 | 1162.603 | 1496.566 | 1243.089 | 521.2525 | 0.316079 | 5.898283 | 0.745085 |
| 4 | 105685600 | 105687599 | 21.9178 | 37.84665 | 44.17267 | 19.37847 | 0.163264 | 0.875056 | 0.745115 |
| 14 | 84013800 | 84016999 | 40.10901 | 64.68503 | 91.73906 | 39.0748 | -0.14566 | 1.778197 | 0.745127 |
| 4 | 23438000 | 23441399 | 63.63146 | 100.5231 | 137.9273 | 57.28341 | -0.12988 | 2.429851 | 0.745153 |
| 11 | 110382600 | 110384599 | 21.84064 | 37.92517 | 36.20901 | 15.89205 | 0.45329 | 0.810273 | 0.745166 |
| 8 | 72800000 | 72803799 | 39.00388 | 63.5331 | 72.77502 | 31.25007 | 0.175398 | 1.554641 | 0.745175 |
| 17 | 23065200 | 23066399 | 16.49362 | 29.21896 | 33.77246 | 14.93225 | 0.183791 | 0.511554 | 0.745197 |
| 7 | 10010000 | 10019599 | 113.811 | 176.0261 | 181.9109 | 75.56144 | 0.254884 | 2.853672 | 0.745204 |

| | | | | | | | | | |
|---|---|---|---|---|---|---|---|---|---|
| 14 | 107493200 | 107494599 | 20.06846 | 34.89227 | 40.95393 | 17.98897 | 0.156917 | 0.791826 | 0.745246 |
| 17 | 17933800 | 17935399 | 20.06846 | 34.89227 | 40.95393 | 17.98897 | 0.156917 | 0.791826 | 0.745246 |
| 10 | 83240000 | 83294399 | 1580.74 | 1949.581 | 2105.767 | 887.0185 | -0.12513 | 6.587258 | 0.745248 |
| 4 | 43801800 | 43805999 | 40.85166 | 65.91553 | 94.20794 | 40.15215 | -0.15593 | 1.767839 | 0.74525 |
| 19 | 39192400 | 39192999 | 16.03315 | 28.24032 | 32.55361 | 14.40991 | 0.184641 | 0.569951 | 0.74525 |
| 5 | 36932800 | 36939399 | 106.1033 | 162.8485 | 181.8624 | 75.5024 | 0.134657 | 2.993523 | 0.745273 |
| 12 | 115301400 | 115305599 | 60.96123 | 97.33639 | 108.6083 | 45.59305 | 0.182756 | 2.118916 | 0.745324 |
| 2 | 40088600 | 40092799 | 51.62729 | 82.58487 | 119.0483 | 50.10619 | -0.18324 | 2.056455 | 0.745326 |
| 7 | 29223200 | 29226799 | 30.42158 | 49.62995 | 72.80724 | 31.28148 | -0.179 | 1.514877 | 0.745339 |
| 13 | 48766600 | 48771599 | 50.78944 | 81.68228 | 92.21038 | 39.12205 | 0.177982 | 1.879836 | 0.745344 |
| X | 60100600 | 60103599 | 35.39899 | 58.2173 | 65.75553 | 28.34262 | 0.203329 | 1.419447 | 0.745403 |
| 1 | 46738800 | 46742199 | 19.32761 | 33.8952 | 37.93634 | 16.76716 | 0.225649 | 0.654344 | 0.745441 |
| 5 | 145510600 | 145592399 | 2376.822 | 2844.878 | 3050.678 | 1290.213 | -0.15871 | 7.062158 | 0.745442 |
| 16 | 52783800 | 52784399 | 7.566841 | 14.57642 | 24.2063 | 9.394634 | -0.18547 | -0.12193 | 0.745445 |
| X | 54745000 | 54751599 | 70.13049 | 112.4654 | 98.30376 | 41.52985 | 0.541487 | 1.998306 | 0.745461 |
| 1 | 138715600 | 138717599 | 18.7963 | 32.31496 | 47.42456 | 20.80229 | -0.16701 | 0.906357 | 0.745476 |
| 13 | 5039400 | 5040799 | 27.9259 | 46.9952 | 45.38111 | 19.83098 | 0.431286 | 1.125923 | 0.745502 |
| 11 | 93886600 | 93888399 | 12.2996 | 21.77904 | 34.22042 | 15.13172 | -0.2568 | 0.491673 | 0.745504 |
| 10 | 108051800 | 108053999 | 28.25851 | 46.82939 | 68.34451 | 29.587 | -0.16299 | 1.311778 | 0.74554 |
| 8 | 54694800 | 54702199 | 93.96331 | 146.164 | 164.6906 | 68.37044 | 0.136649 | 2.741252 | 0.745566 |
| 6 | 149191000 | 149196199 | 65.70345 | 104.7757 | 113.897 | 47.72849 | 0.21837 | 2.163255 | 0.745575 |
| 10 | 21682600 | 21685999 | 21.80332 | 37.10577 | 54.42809 | 23.85625 | -0.1689 | 1.015537 | 0.745582 |
| 19 | 22116200 | 22116999 | 7.615966 | 14.4108 | 24.95873 | 10.12054 | -0.30318 | 0.023413 | 0.74559 |
| 3 | 27083400 | 27088399 | 50.65084 | 80.95862 | 112.5187 | 47.30476 | -0.13189 | 2.075414 | 0.745613 |
| 15 | 54778000 | 54779399 | 15.65821 | 27.53397 | 40.74609 | 18.04233 | -0.175 | 0.600074 | 0.745626 |
| 10 | 113407400 | 113412999 | 76.97268 | 120.4982 | 163.3834 | 67.77519 | -0.123 | 2.622254 | 0.745639 |
| 7 | 57563200 | 57563799 | 6.51709 | 12.64897 | 22.25645 | 8.368019 | -0.22462 | -0.24708 | 0.745642 |
| 7 | 34278000 | 34292799 | 198.8305 | 290.5847 | 373.0568 | 153.6912 | -0.12871 | 3.919563 | 0.745666 |
| 16 | 94676600 | 94679599 | 23.2127 | 39.12879 | 56.57295 | 24.73418 | -0.14892 | 1.107381 | 0.74567 |
| 16 | 29332800 | 29336399 | 42.62751 | 68.40562 | 97.05978 | 41.04597 | -0.15503 | 1.938801 | 0.745702 |
| 14 | 7235600 | 7236599 | 9.041897 | 17.22994 | 28.39248 | 11.34255 | -0.21309 | -0.00042 | 0.745721 |
| 8 | 6358000 | 6358599 | 7.918076 | 15.23964 | 25.30896 | 9.847368 | -0.18868 | -0.10511 | 0.74581 |
| 8 | 21397600 | 21408599 | 124.5665 | 191.2503 | 200.2498 | 83.13999 | 0.227915 | 2.994636 | 0.745819 |
| 14 | 92269200 | 92273199 | 50.9831 | 81.95525 | 80.68558 | 34.20915 | 0.37418 | 1.889327 | 0.745831 |
| 2 | 134926800 | 134927199 | 4.220858 | 8.25691 | 16.25573 | 5.380341 | -0.21538 | -0.7459 | 0.745835 |
| 12 | 37113600 | 37116599 | 21.97508 | 38.36968 | 41.80055 | 18.40988 | 0.264551 | 0.743828 | 0.745843 |
| 12 | 70714600 | 70722199 | 77.62995 | 121.856 | 139.1598 | 57.71599 | 0.127421 | 2.570239 | 0.745869 |
| 3 | 155894800 | 155896599 | 18.14483 | 32.15155 | 35.18144 | 15.5545 | 0.2625 | 0.521852 | 0.745911 |
| 2 | 33887000 | 33887799 | 14.44383 | 25.31471 | 38.4508 | 16.98563 | -0.21226 | 0.643032 | 0.74593 |
| 9 | 36498600 | 36499399 | 19.25766 | 33.62734 | 38.83875 | 17.10621 | 0.1808 | 0.726838 | 0.745931 |
| 6 | 120897400 | 120902399 | 47.17803 | 75.62779 | 105.9061 | 44.71488 | -0.13798 | 1.979653 | 0.745997 |
| 3 | 139539800 | 139541799 | 25.73828 | 43.66434 | 50.6604 | 22.19215 | 0.167787 | 1.055326 | 0.746009 |
| 14 | 51712200 | 51714199 | 15.71489 | 27.72012 | 42.3281 | 18.73114 | -0.21826 | 0.569661 | 0.746037 |
| 3 | 102448800 | 102449999 | 12.86439 | 23.79711 | 27.50043 | 11.5837 | 0.231468 | 0.215141 | 0.746057 |
| 4 | 71536000 | 71542799 | 127.1373 | 192.4996 | 254.8098 | 105.4225 | -0.12611 | 3.27726 | 0.746074 |
| 4 | 89639800 | 89641999 | 22.93348 | 39.52989 | 45.20221 | 19.82905 | 0.192052 | 0.894526 | 0.746075 |
| X | 105881000 | 105883199 | 35.13868 | 56.97656 | 81.60728 | 34.95252 | -0.15157 | 1.634589 | 0.746157 |

| | | | | | | | | | |
|---|---|---|---|---|---|---|---|---|---|
| 2 | 177202600 | 177212799 | 144.306 | 218.7068 | 236.6624 | 97.94427 | 0.165597 | 3.2617 | 0.746167 |
| 4 | 113809600 | 113811599 | 24.55664 | 42.66645 | 44.57904 | 19.57287 | 0.323915 | 0.826972 | 0.746169 |
| 17 | 70753800 | 70755999 | 24.98235 | 42.59572 | 49.02392 | 21.49081 | 0.180167 | 1.00495 | 0.746169 |
| 4 | 145371200 | 145376799 | 116.4057 | 178.3746 | 199.1226 | 82.65001 | 0.133462 | 3.029893 | 0.746179 |
| 12 | 57283800 | 57286599 | 33.01401 | 54.52533 | 63.13073 | 27.26615 | 0.169613 | 1.356975 | 0.746202 |
| 8 | 84467200 | 84469599 | 18.58792 | 31.99692 | 46.89208 | 20.56951 | -0.16467 | 0.892445 | 0.746208 |
| 3 | 13324200 | 13325599 | 19.38778 | 33.96993 | 32.01424 | 14.13887 | 0.47275 | 0.661945 | 0.746229 |
| 6 | 7635000 | 7638399 | 36.35958 | 59.62073 | 68.29093 | 29.39869 | 0.18141 | 1.455863 | 0.74624 |
| 9 | 109486800 | 109487399 | 9.777283 | 18.75785 | 23.49617 | 9.226445 | 0.201675 | -0.06462 | 0.746278 |
| 6 | 56010200 | 56012599 | 18.08165 | 31.22787 | 45.83026 | 20.10575 | -0.16573 | 0.859062 | 0.746282 |
| 16 | 70170000 | 70170799 | 14.29659 | 24.94536 | 45.74659 | 20.13904 | -0.48329 | 0.721421 | 0.746315 |
| 5 | 30915400 | 30918799 | 30.13832 | 49.15336 | 72.03303 | 30.94552 | -0.17776 | 1.51939 | 0.746321 |
| 9 | 115368600 | 115373399 | 74.15157 | 115.6939 | 160.0184 | 66.40218 | -0.15537 | 2.679718 | 0.74634 |
| 7 | 27644400 | 27671199 | 343.4641 | 486.6127 | 491.1539 | 203.2295 | 0.173833 | 4.387949 | 0.746368 |
| 6 | 11693600 | 11693999 | 10.64826 | 19.97063 | 24.22057 | 9.983534 | 0.187135 | 0.108627 | 0.746375 |
| 4 | 21030200 | 21032599 | 11.07994 | 19.71756 | 39.84507 | 17.55885 | -0.6141 | 0.456206 | 0.746396 |
| 10 | 114371200 | 114372999 | 14.96774 | 25.98774 | 47.92216 | 21.03141 | -0.49198 | 0.799248 | 0.746403 |
| 6 | 124033600 | 124035399 | 20.297 | 34.6354 | 51.69922 | 22.66508 | -0.19339 | 0.981412 | 0.746404 |
| 11 | 68634000 | 68635399 | 14.58805 | 26.24042 | 30.90118 | 13.4949 | 0.169615 | 0.396616 | 0.746404 |
| 8 | 119411000 | 119412799 | 26.71553 | 45.27642 | 51.77237 | 22.67231 | 0.188622 | 1.068267 | 0.746426 |
| 1 | 33139000 | 33146599 | 75.27891 | 118.2403 | 162.7097 | 67.48873 | -0.14057 | 2.56095 | 0.746434 |
| 7 | 95614000 | 95615199 | 10.50296 | 19.21827 | 29.80431 | 12.74166 | -0.20475 | 0.267432 | 0.74644 |
| 6 | 145599400 | 145603199 | 33.87091 | 55.14329 | 78.87003 | 33.86086 | -0.14467 | 1.555967 | 0.746446 |
| 3 | 67325000 | 67333999 | 293.8676 | 414.217 | 521.1317 | 215.8163 | -0.15136 | 4.460404 | 0.746466 |
| 1 | 51720000 | 51722799 | 30.53023 | 50.12628 | 72.24113 | 31.1161 | -0.14832 | 1.42943 | 0.746493 |
| 8 | 62350400 | 62351399 | 8.831694 | 16.70226 | 26.50542 | 10.75986 | -0.17955 | 0.043802 | 0.746511 |
| 15 | 34856400 | 34857399 | 12.89069 | 23.42907 | 27.51581 | 11.86355 | 0.185443 | 0.316948 | 0.746527 |
| 13 | 21603800 | 21604199 | 4.223585 | 8.254374 | 16.5789 | 5.735722 | -0.30083 | -0.58801 | 0.746552 |
| 18 | 8305000 | 8305799 | 12.60048 | 22.35381 | 34.15627 | 15.09145 | -0.21552 | 0.475943 | 0.74658 |
| 15 | 47989600 | 47991199 | 17.78411 | 31.34199 | 35.83674 | 15.86141 | 0.196428 | 0.578783 | 0.746582 |
| X | 144693200 | 144694999 | 26.59629 | 45.18957 | 51.20758 | 22.44176 | 0.201802 | 1.046222 | 0.746588 |
| X | 89318400 | 89320599 | 23.26015 | 40.2514 | 45.20031 | 19.8306 | 0.219343 | 0.863213 | 0.746608 |
| 17 | 46088400 | 46091799 | 29.93397 | 49.00206 | 70.68054 | 30.40581 | -0.15118 | 1.464593 | 0.746613 |
| 6 | 147134000 | 147135399 | 13.60055 | 24.06793 | 35.67122 | 15.77516 | -0.17332 | 0.51102 | 0.746619 |
| 1 | 57170200 | 57172599 | 17.63365 | 31.16976 | 35.26969 | 15.60112 | 0.213057 | 0.544835 | 0.746644 |
| X | 148680600 | 148682999 | 51.66487 | 82.68038 | 94.46483 | 39.78339 | 0.152259 | 2.035462 | 0.746651 |
| 9 | 119986800 | 119989999 | 28.83228 | 48.11289 | 56.16554 | 24.42763 | 0.158662 | 1.226724 | 0.746663 |
| 6 | 122766200 | 122770999 | 37.85059 | 60.99809 | 87.86865 | 37.39077 | -0.16927 | 1.794794 | 0.746668 |
| 2 | 115549600 | 115552599 | 45.68904 | 73.76557 | 84.04795 | 35.84995 | 0.171348 | 1.752932 | 0.74668 |
| 5 | 37470800 | 37476399 | 66.74902 | 105.1502 | 144.4146 | 59.92315 | -0.13448 | 2.493716 | 0.746682 |
| 4 | 32566000 | 32568199 | 18.54785 | 31.95922 | 46.88352 | 20.56637 | -0.16586 | 0.882533 | 0.746683 |
| 1 | 67664400 | 67666999 | 33.17019 | 54.83424 | 52.47869 | 22.68009 | 0.443729 | 1.340214 | 0.746697 |
| 13 | 51761200 | 51762399 | 13.96011 | 24.73589 | 36.65234 | 16.2049 | -0.17224 | 0.501893 | 0.746697 |
| 14 | 11241000 | 11242399 | 15.39605 | 27.05934 | 39.76697 | 17.6087 | -0.16511 | 0.6046 | 0.746726 |
| 16 | 91802800 | 91806199 | 40.08087 | 64.87619 | 75.96094 | 32.4921 | 0.136106 | 1.673671 | 0.74674 |
| 10 | 47556000 | 47565199 | 208.1914 | 307.105 | 286.4175 | 117.951 | 0.344808 | 3.770755 | 0.746755 |
| 2 | 27768000 | 27769999 | 11.62791 | 20.84751 | 32.19976 | 14.10085 | -0.22198 | 0.399304 | 0.746767 |

| | | | | | | | | | |
|---|---|---|---|---|---|---|---|---|---|
| 3 | 97235200 | 97235999 | 9.910231 | 18.56849 | 29.01964 | 11.9813 | -0.17826 | 0.119269 | 0.746776 |
| 8 | 64451600 | 64452599 | 9.910231 | 18.56849 | 29.01964 | 11.9813 | -0.17826 | 0.119269 | 0.746776 |
| 5 | 142988800 | 142990799 | 15.67505 | 27.4026 | 40.48223 | 17.84452 | -0.17258 | 0.692944 | 0.746798 |
| 8 | 42725000 | 42725599 | 13.90422 | 24.866 | 26.71134 | 11.72222 | 0.296858 | 0.419443 | 0.746868 |
| 12 | 62191600 | 62192399 | 17.80363 | 30.25432 | 55.82768 | 24.4668 | -0.49882 | 1.019125 | 0.746915 |
| 2 | 69312800 | 69314199 | 14.93043 | 26.88653 | 31.39397 | 13.6931 | 0.182979 | 0.393494 | 0.746926 |
| 9 | 22477200 | 22479199 | 35.29473 | 57.69384 | 67.78453 | 29.14445 | 0.142499 | 1.495573 | 0.746934 |
| 3 | 55088600 | 55092199 | 38.01504 | 60.90902 | 109.3206 | 46.12175 | -0.49608 | 1.986843 | 0.746937 |
| 8 | 69796800 | 69799599 | 40.91032 | 66.07918 | 76.72712 | 32.74579 | 0.144889 | 1.729238 | 0.746959 |
| 3 | 101112600 | 101114599 | 23.47201 | 40.07835 | 46.88973 | 20.55762 | 0.15702 | 0.986084 | 0.746987 |
| 2 | 150829800 | 150830599 | 9.717836 | 18.29515 | 29.14356 | 11.93556 | -0.19638 | 0.086604 | 0.747027 |
| 2 | 98460000 | 98465199 | 60.08957 | 95.74517 | 110.0544 | 46.06615 | 0.136734 | 2.183061 | 0.747037 |
| 18 | 29477600 | 29478199 | 9.23056 | 17.63545 | 21.84182 | 8.677554 | 0.202839 | -0.03468 | 0.747048 |
| 1 | 5110800 | 5112799 | 23.04042 | 39.64589 | 45.19966 | 19.82689 | 0.195806 | 0.909441 | 0.74706 |
| X | 108096400 | 108096999 | 10.39766 | 19.76252 | 24.39601 | 9.799178 | 0.193573 | 0.019415 | 0.747078 |
| 14 | 121098800 | 121104399 | 71.9371 | 112.9696 | 153.496 | 63.66548 | -0.12248 | 2.56074 | 0.747079 |
| 18 | 90321600 | 90325599 | 29.16217 | 49.05517 | 54.10446 | 23.63998 | 0.24005 | 1.139453 | 0.747134 |
| 14 | 68648600 | 68650199 | 23.05315 | 39.48247 | 46.33303 | 20.31851 | 0.15324 | 0.955117 | 0.747143 |
| 5 | 126664800 | 126667799 | 35.0364 | 56.89322 | 81.03031 | 34.74256 | -0.14157 | 1.602382 | 0.747144 |
| Y | 90110000 | 90110599 | 8.076097 | 14.82184 | 28.45506 | 12.11548 | -0.50609 | 0.226419 | 0.747144 |
| 12 | 15185400 | 15186199 | 13.47097 | 24.64963 | 28.96534 | 12.38264 | 0.192408 | 0.284852 | 0.747149 |
| 7 | 52778200 | 52778799 | 9.807347 | 18.18411 | 28.34629 | 11.90244 | -0.19346 | 0.182915 | 0.747155 |
| 14 | 31551800 | 31553999 | 14.51737 | 25.07257 | 51.09416 | 22.41504 | -0.63724 | 0.849309 | 0.747157 |
| 16 | 7815400 | 7817599 | 22.58031 | 38.93898 | 44.66487 | 19.5936 | 0.187714 | 0.887985 | 0.74716 |
| 8 | 74741000 | 74742799 | 19.10435 | 33.60448 | 31.77485 | 14.06898 | 0.468246 | 0.601523 | 0.747169 |
| 6 | 90623000 | 90626199 | 38.8773 | 62.95285 | 73.04775 | 31.25507 | 0.149598 | 1.661101 | 0.747202 |
| 3 | 7006000 | 7007799 | 15.58731 | 28.27256 | 31.34365 | 13.54842 | 0.265494 | 0.354833 | 0.747221 |
| X | 45154200 | 45154799 | 7.029315 | 13.65413 | 24.67839 | 9.221816 | -0.25391 | -0.25031 | 0.747232 |
| 13 | 67859000 | 67870799 | 75.25726 | 119.3276 | 112.0531 | 46.639 | 0.420865 | 2.343875 | 0.747235 |
| 14 | 53014800 | 53016599 | 17.13959 | 30.06906 | 35.0831 | 15.51088 | 0.166107 | 0.633497 | 0.747244 |
| 9 | 112453800 | 112454399 | 7.135366 | 13.79188 | 23.19575 | 8.891174 | -0.18615 | -0.16938 | 0.747246 |
| 14 | 120023600 | 120024199 | 7.135366 | 13.79188 | 23.19575 | 8.891174 | -0.18615 | -0.16938 | 0.747246 |
| 4 | 126478400 | 126482999 | 52.87659 | 84.54853 | 97.81331 | 41.14465 | 0.132924 | 2.061732 | 0.747251 |
| 7 | 48530200 | 48532999 | 16.24353 | 27.59419 | 55.7551 | 24.43399 | -0.62914 | 1.016532 | 0.747253 |
| 7 | 30919600 | 30924599 | 90.94775 | 142.1331 | 157.481 | 65.33819 | 0.166072 | 2.660073 | 0.74727 |
| 7 | 116248000 | 116252799 | 34.04667 | 55.45678 | 79.51266 | 34.1429 | -0.14769 | 1.549992 | 0.747282 |
| 1 | 81169400 | 81171999 | 21.07605 | 36.61375 | 42.52532 | 18.66947 | 0.171889 | 0.810565 | 0.747314 |
| 2 | 61705000 | 61706599 | 21.28406 | 36.84568 | 43.11715 | 18.91687 | 0.160274 | 0.848843 | 0.747323 |
| 11 | 106636000 | 106638799 | 28.16231 | 47.43303 | 54.029 | 23.61604 | 0.193839 | 1.127491 | 0.747339 |
| 6 | 141529600 | 141531799 | 19.55499 | 33.00572 | 60.24192 | 26.3486 | -0.48379 | 1.086882 | 0.747344 |
| 5 | 24109400 | 24116399 | 78.77725 | 122.2955 | 193.7056 | 80.43999 | -0.3564 | 2.771778 | 0.747348 |
| 17 | 94620200 | 94621799 | 23.33855 | 40.01993 | 46.33695 | 20.32222 | 0.172891 | 0.947197 | 0.747349 |
| 7 | 111592600 | 111597599 | 60.25134 | 95.5784 | 133.7424 | 55.69229 | -0.15308 | 2.324449 | 0.74736 |
| 9 | 92308600 | 92312599 | 46.26856 | 74.72079 | 84.03092 | 35.85481 | 0.190613 | 1.747662 | 0.747361 |
| 7 | 57179200 | 57182599 | 29.20764 | 47.94376 | 69.78964 | 30.05726 | -0.16281 | 1.426828 | 0.747382 |
| 15 | 70167800 | 70171399 | 40.71745 | 65.96695 | 76.87672 | 32.90603 | 0.144066 | 1.656549 | 0.747395 |
| 5 | 117127400 | 117130399 | 35.58327 | 57.6834 | 81.80778 | 35.03211 | -0.13771 | 1.63913 | 0.747404 |

| | | | | | | | | | |
|---|---|---|---|---|---|---|---|---|---|
| 8 | 110486000 | 110492999 | 85.2854 | 133.3298 | 151.1891 | 62.7255 | 0.133064 | 2.647698 | 0.747438 |
| 10 | 99726600 | 99727199 | 8.282281 | 15.98767 | 20.39601 | 7.86626 | 0.20044 | -0.14535 | 0.747442 |
| 17 | 42626200 | 42627999 | 16.62388 | 28.68344 | 44.81318 | 19.659 | -0.25586 | 0.858379 | 0.747454 |
| 3 | 70479000 | 70484399 | 66.2298 | 104.4405 | 143.1172 | 59.3928 | -0.13009 | 2.47255 | 0.747473 |
| 19 | 39110000 | 39112599 | 19.86833 | 33.73733 | 59.2008 | 25.94061 | -0.42682 | 1.036076 | 0.747473 |
| 15 | 9771000 | 9776999 | 72.8435 | 115.1854 | 131.2962 | 54.55566 | 0.138338 | 2.411362 | 0.74748 |
| 1 | 89128000 | 89129199 | 33.49387 | 54.71244 | 61.29525 | 26.34773 | 0.210317 | 1.497929 | 0.747521 |
| 1 | 16795200 | 16798999 | 39.28505 | 63.41605 | 90.10527 | 38.42568 | -0.14674 | 1.74874 | 0.747527 |
| 2 | 104692600 | 104693399 | 12.44818 | 23.09035 | 27.66944 | 11.61154 | 0.18382 | 0.200203 | 0.747527 |
| 5 | 59109000 | 59109399 | 6.549573 | 12.73661 | 22.39916 | 8.294191 | -0.20254 | -0.29559 | 0.747532 |
| 10 | 57746000 | 57746599 | 6.549573 | 12.73661 | 22.39916 | 8.294191 | -0.20254 | -0.29559 | 0.747532 |
| 12 | 96616600 | 96618399 | 20.8453 | 36.24415 | 41.95475 | 18.42898 | 0.176782 | 0.795659 | 0.747537 |
| 4 | 16408600 | 16409799 | 11.07512 | 20.458 | 31.42777 | 13.26372 | -0.17812 | 0.220762 | 0.747542 |
| 13 | 47055800 | 47057199 | 18.16589 | 31.89776 | 36.99643 | 16.36904 | 0.174589 | 0.628664 | 0.747591 |
| 16 | 13079400 | 13081599 | 16.99428 | 29.69023 | 43.5714 | 19.19683 | -0.16317 | 0.714403 | 0.747607 |
| 6 | 82453400 | 82455199 | 21.56961 | 36.37748 | 53.81342 | 23.5313 | -0.18177 | 1.094035 | 0.747627 |
| 9 | 26384000 | 26386599 | 22.82549 | 39.14059 | 45.78562 | 20.08044 | 0.158043 | 0.94159 | 0.747646 |
| 10 | 30507400 | 30509199 | 14.68629 | 25.92934 | 38.14826 | 16.87839 | -0.16427 | 0.546124 | 0.747653 |
| 7 | 78631600 | 78632999 | 15.91558 | 28.13286 | 32.96313 | 14.58084 | 0.163197 | 0.531803 | 0.747671 |
| 9 | 19196400 | 19202399 | 92.79108 | 145.4849 | 133.9128 | 55.54134 | 0.436891 | 2.601051 | 0.747675 |
| 9 | 79935600 | 79945599 | 131.1874 | 199.2313 | 222.7693 | 92.25136 | 0.120778 | 3.21559 | 0.747678 |
| 1 | 26525400 | 26526999 | 21.22327 | 37.2291 | 33.1976 | 14.67586 | 0.552072 | 0.655419 | 0.747681 |
| 2 | 22010400 | 22012599 | 13.4943 | 24.5804 | 29.05078 | 12.49079 | 0.179104 | 0.313849 | 0.747683 |
| 5 | 7602200 | 7611999 | 247.7341 | 358.5141 | 384.2097 | 158.4559 | 0.117962 | 4.071218 | 0.747707 |
| 2 | 23674600 | 23678199 | 47.77067 | 76.75985 | 89.40303 | 37.88271 | 0.131444 | 1.901282 | 0.747714 |
| 15 | 39454800 | 39457399 | 24.00941 | 41.29125 | 46.82985 | 20.54164 | 0.20325 | 0.922026 | 0.747722 |
| 3 | 31692000 | 31694999 | 67.74396 | 107.2409 | 123.6448 | 51.40532 | 0.123173 | 2.380392 | 0.747731 |
| 14 | 15068600 | 15069599 | 9.215503 | 17.21987 | 27.42358 | 11.37463 | -0.20992 | 0.129571 | 0.747739 |
| 19 | 6873200 | 6878999 | 85.98455 | 133.5309 | 179.4673 | 74.52338 | -0.11931 | 2.767836 | 0.747764 |
| 12 | 114900600 | 114912799 | 225.5542 | 330.6534 | 351.665 | 144.8505 | 0.146929 | 3.873923 | 0.747788 |
| X | 109043200 | 109046199 | 47.42628 | 76.45313 | 86.64748 | 36.88578 | 0.176496 | 1.802172 | 0.747809 |
| 17 | 82885400 | 82889399 | 31.46237 | 53.02563 | 55.78915 | 24.3902 | 0.308092 | 1.136379 | 0.747832 |
| 9 | 36144800 | 36148199 | 43.56329 | 70.26139 | 82.00979 | 34.93433 | 0.134946 | 1.778606 | 0.747859 |
| 17 | 91987200 | 91992399 | 21.00247 | 36.57856 | 34.71325 | 15.26413 | 0.462459 | 0.764048 | 0.747869 |
| 9 | 87469000 | 87477399 | 86.58311 | 135.0596 | 154.3768 | 64.06259 | 0.119103 | 2.688501 | 0.747878 |
| 5 | 146299000 | 146300399 | 13.56052 | 24.12595 | 35.60047 | 15.69162 | -0.16315 | 0.467542 | 0.747894 |
| 14 | 19536400 | 19537599 | 18.17223 | 32.08826 | 29.83326 | 13.19913 | 0.49465 | 0.551704 | 0.747902 |
| 5 | 57804200 | 57810599 | 92.51048 | 143.6757 | 163.5355 | 67.90699 | 0.120272 | 2.767 | 0.747907 |
| 11 | 15545600 | 15547199 | 14.19694 | 25.39236 | 29.43679 | 12.91705 | 0.187379 | 0.425256 | 0.747929 |
| 4 | 86774000 | 86775999 | 17.24447 | 29.81415 | 44.78565 | 19.64813 | -0.19885 | 0.843314 | 0.74793 |
| 1 | 171477800 | 171480799 | 26.68923 | 45.11237 | 51.25909 | 22.42953 | 0.197497 | 1.087379 | 0.747937 |
| 5 | 134819600 | 134821799 | 31.45016 | 51.84671 | 61.11416 | 26.36867 | 0.143246 | 1.374883 | 0.747952 |
| 9 | 78003600 | 78005799 | 16.65751 | 29.23764 | 44.48552 | 19.67602 | -0.21517 | 0.638731 | 0.747963 |
| 2 | 10329400 | 10331599 | 17.34854 | 30.61452 | 35.36462 | 15.64834 | 0.182544 | 0.562016 | 0.747987 |
| 16 | 22164800 | 22167999 | 30.38292 | 49.82816 | 71.52731 | 30.79393 | -0.14324 | 1.442799 | 0.748058 |
| 5 | 85430000 | 85430799 | 6.684311 | 12.95592 | 22.18493 | 8.394227 | -0.19385 | -0.22133 | 0.748059 |
| 2 | 17523400 | 17526599 | 29.4307 | 49.09029 | 57.34223 | 24.93261 | 0.157731 | 1.232898 | 0.748061 |

| | | | | | | | | | |
|---|---|---|---|---|---|---|---|---|---|
| 17 | 38041800 | 38042399 | 11.57092 | 20.98905 | 31.69057 | 13.69716 | -0.17707 | 0.328002 | 0.748069 |
| 19 | 45660000 | 45660799 | 11.57092 | 20.98905 | 31.69057 | 13.69716 | -0.17707 | 0.328002 | 0.748069 |
| 13 | 21043400 | 21045399 | 15.13043 | 26.67231 | 39.67761 | 17.56045 | -0.18114 | 0.567101 | 0.748081 |
| 12 | 118754200 | 118762199 | 69.61505 | 110.4678 | 109.1638 | 45.47203 | 0.348434 | 2.314136 | 0.748096 |
| 3 | 45591800 | 45594199 | 20.06205 | 34.9023 | 40.92823 | 17.98455 | 0.1583 | 0.78307 | 0.748106 |
| 3 | 49978000 | 49979399 | 16.52221 | 29.56808 | 32.65209 | 14.32108 | 0.257271 | 0.441809 | 0.748127 |
| 4 | 135802800 | 135805599 | 16.64961 | 28.99212 | 42.62859 | 18.74248 | -0.16634 | 0.75791 | 0.748144 |
| 9 | 12552800 | 12554399 | 16.64961 | 28.99212 | 42.62859 | 18.74248 | -0.16634 | 0.75791 | 0.748144 |
| 1 | 59598400 | 59602999 | 47.87675 | 76.66369 | 106.4611 | 44.87428 | -0.12778 | 2.015481 | 0.74816 |
| 7 | 12785400 | 12789799 | 34.11586 | 55.2006 | 80.04888 | 34.23137 | -0.17225 | 1.673025 | 0.748172 |
| 2 | 87551400 | 87552799 | 16.37642 | 28.98514 | 33.84688 | 14.96786 | 0.168678 | 0.521464 | 0.748173 |
| 14 | 103474800 | 103478599 | 24.42386 | 42.45932 | 44.59644 | 19.58734 | 0.316539 | 0.820873 | 0.748187 |
| 5 | 52418200 | 52422599 | 47.048 | 75.40225 | 104.9245 | 44.28519 | -0.12924 | 1.986873 | 0.748189 |
| Y | 10385600 | 10386999 | 13.90805 | 24.86478 | 24.25697 | 10.64804 | 0.43454 | 0.413882 | 0.748196 |
| 12 | 113434200 | 113436599 | 25.28876 | 42.67303 | 49.61125 | 21.69736 | 0.164529 | 1.096383 | 0.748202 |
| 3 | 20957000 | 20957799 | 12.60127 | 22.62225 | 33.65118 | 14.72018 | -0.16699 | 0.401655 | 0.748218 |
| 11 | 101289000 | 101290999 | 13.82759 | 24.26548 | 43.51656 | 19.24409 | -0.45089 | 0.626408 | 0.748222 |
| 12 | 95900000 | 95902599 | 24.4989 | 42.30617 | 46.26507 | 20.29644 | 0.256763 | 0.887188 | 0.748223 |
| 9 | 20286000 | 20293399 | 80.7868 | 127.4754 | 140.2115 | 58.19763 | 0.187324 | 2.46956 | 0.748224 |
| 18 | 57171200 | 57171999 | 20.82359 | 35.8363 | 40.19322 | 17.63088 | 0.219686 | 0.89696 | 0.748256 |
| 8 | 41228000 | 41229599 | 33.31398 | 53.94 | 78.81553 | 33.72094 | -0.18221 | 1.657635 | 0.748256 |
| 4 | 105955600 | 105957199 | 24.20463 | 40.4219 | 59.67788 | 25.96789 | -0.17896 | 1.209485 | 0.748262 |
| 8 | 127354600 | 127360399 | 132.5325 | 199.6484 | 263.1295 | 108.6945 | -0.12618 | 3.380841 | 0.748262 |
| 14 | 119146800 | 119148399 | 11.92623 | 21.36973 | 32.65634 | 14.30792 | -0.20729 | 0.405513 | 0.748264 |
| 13 | 56415000 | 56416599 | 15.22455 | 26.94034 | 31.01101 | 13.71439 | 0.189249 | 0.511624 | 0.748308 |
| 12 | 64478600 | 64479199 | 11.81513 | 21.28742 | 32.21259 | 14.04118 | -0.18795 | 0.371943 | 0.748333 |
| X | 38883200 | 38885999 | 38.56112 | 62.98267 | 70.8256 | 30.44309 | 0.20477 | 1.51802 | 0.748348 |
| 17 | 8752800 | 8753799 | 12.30214 | 22.73054 | 27.0354 | 11.4059 | 0.188737 | 0.217113 | 0.74835 |
| 7 | 108272200 | 108279599 | 89.57826 | 140.6193 | 151.7257 | 62.92807 | 0.209274 | 2.578615 | 0.748351 |
| 3 | 76055200 | 76055599 | 5.947129 | 11.57088 | 20.70403 | 7.635489 | -0.22234 | -0.32609 | 0.748366 |
| 9 | 93505000 | 93505599 | 7.608525 | 14.71275 | 24.96206 | 9.549066 | -0.19608 | -0.16552 | 0.74837 |
| 4 | 147766800 | 147768599 | 21.13045 | 37.33229 | 37.21592 | 16.46364 | 0.394796 | 0.575983 | 0.748378 |
| 13 | 62769400 | 62772399 | 31.17996 | 51.72131 | 60.24231 | 26.09709 | 0.161505 | 1.3001 | 0.74838 |
| 10 | 47644800 | 47646599 | 26.43607 | 44.42808 | 52.30659 | 22.84798 | 0.146283 | 1.14033 | 0.748382 |
| 6 | 146979000 | 146983799 | 89.14046 | 137.3192 | 187.602 | 77.92463 | -0.15183 | 2.916103 | 0.748411 |
| 2 | 73585000 | 73587199 | 27.42005 | 46.00885 | 53.97242 | 23.55547 | 0.15146 | 1.160743 | 0.748422 |
| 17 | 92374400 | 92374799 | 4.640778 | 9.071412 | 17.16173 | 5.908798 | -0.21017 | -0.59691 | 0.748447 |
| 17 | 55419800 | 55425599 | 70.23957 | 110.1256 | 177.1201 | 73.46197 | -0.36733 | 2.593999 | 0.748482 |
| X | 8848600 | 8849199 | 6.892285 | 13.36453 | 24.46125 | 9.237744 | -0.28568 | -0.21424 | 0.748488 |
| 11 | 60140800 | 60142199 | 12.05263 | 21.79212 | 32.66077 | 14.17382 | -0.17032 | 0.352136 | 0.748502 |
| 5 | 56243000 | 56245599 | 24.70568 | 42.31574 | 40.99989 | 17.97986 | 0.428547 | 0.953901 | 0.748526 |
| 10 | 111465600 | 111467999 | 19.05436 | 33.19742 | 38.43228 | 16.90322 | 0.177144 | 0.756634 | 0.748533 |
| 4 | 82570800 | 82571999 | 15.52063 | 28.21835 | 26.28117 | 11.32655 | 0.518415 | 0.332931 | 0.748543 |
| 3 | 124093200 | 124094199 | 10.3077 | 18.79316 | 29.76447 | 12.78293 | -0.23948 | 0.284652 | 0.748567 |
| 11 | 30545400 | 30549199 | 25.9334 | 44.25123 | 49.57408 | 21.73333 | 0.21907 | 1.001765 | 0.748575 |
| 13 | 77724200 | 77728399 | 54.39122 | 86.69227 | 120.4529 | 50.4402 | -0.1359 | 2.170747 | 0.748578 |
| 10 | 69919200 | 69925399 | 50.80486 | 81.24637 | 113.6201 | 47.80244 | -0.14001 | 2.06011 | 0.748597 |

| | | | | | | | | | |
|---|---|---|---|---|---|---|---|---|---|
| X | 158197000 | 158198799 | 23.6898 | 41.09164 | 44.14334 | 19.36826 | 0.283872 | 0.846339 | 0.748606 |
| 6 | 74577400 | 74580199 | 25.13758 | 43.20188 | 46.80086 | 20.52849 | 0.26884 | 0.930325 | 0.748609 |
| 18 | 67101800 | 67102399 | 9.68569 | 18.33585 | 21.30275 | 8.629365 | 0.269899 | 0.031123 | 0.748618 |
| 18 | 30958400 | 30960399 | 17.02869 | 30.6027 | 33.2766 | 14.54342 | 0.283107 | 0.42273 | 0.748622 |
| 15 | 97332000 | 97334199 | 22.35154 | 37.75002 | 54.93495 | 24.03211 | -0.15815 | 1.085191 | 0.74864 |
| 19 | 21650600 | 21654399 | 47.88448 | 76.70089 | 106.6292 | 44.96725 | -0.1288 | 2.005639 | 0.74864 |
| 2 | 91918600 | 91920399 | 18.10087 | 31.69368 | 37.13089 | 16.38925 | 0.160201 | 0.673008 | 0.748656 |
| 17 | 35857200 | 35859199 | 19.84396 | 33.86812 | 50.06077 | 21.94709 | -0.17898 | 0.981273 | 0.748665 |
| 10 | 28960200 | 28961599 | 14.4292 | 25.33572 | 37.84961 | 16.74611 | -0.1885 | 0.611684 | 0.748685 |
| 12 | 66123800 | 66124599 | 9.895096 | 18.75251 | 22.80953 | 9.218654 | 0.207106 | 0.030515 | 0.748784 |
| 5 | 133083000 | 133086399 | 34.94469 | 57.44178 | 66.4229 | 28.62429 | 0.169178 | 1.422669 | 0.748786 |
| 4 | 90699800 | 90701999 | 20.15472 | 35.17442 | 40.24539 | 17.71963 | 0.193955 | 0.737186 | 0.748787 |
| 1 | 88936600 | 88937999 | 11.57682 | 21.67074 | 25.60539 | 10.5885 | 0.220115 | 0.128999 | 0.748798 |
| 14 | 80035400 | 80039199 | 47.56572 | 76.18614 | 105.8227 | 44.62344 | -0.12765 | 2.006901 | 0.748806 |
| 9 | 9723800 | 9725799 | 36.09814 | 59.60532 | 65.64341 | 28.34687 | 0.241619 | 1.370846 | 0.74882 |
| 11 | 7354000 | 7355399 | 13.80591 | 24.42426 | 37.07588 | 16.39731 | -0.20817 | 0.514581 | 0.748827 |
| X | 49192600 | 49193599 | 18.27101 | 32.03997 | 37.13917 | 16.41385 | 0.175619 | 0.652324 | 0.748845 |
| 1 | 152287600 | 152290199 | 28.3649 | 47.33681 | 55.66975 | 24.21325 | 0.147953 | 1.22159 | 0.748864 |
| 5 | 138452600 | 138457599 | 89.90916 | 141.5562 | 146.835 | 60.92305 | 0.269326 | 2.520633 | 0.74887 |
| 8 | 117283800 | 117286199 | 20.92203 | 36.26942 | 42.57815 | 18.68119 | 0.156085 | 0.833692 | 0.748878 |
| 6 | 141245000 | 141247399 | 24.20008 | 41.39387 | 47.93509 | 21.0189 | 0.172029 | 0.972759 | 0.748887 |
| 13 | 15985200 | 15986799 | 21.4423 | 37.02575 | 43.09069 | 18.90381 | 0.167289 | 0.869467 | 0.748895 |
| 4 | 16359200 | 16364999 | 71.42976 | 112.3838 | 152.732 | 63.36408 | -0.1207 | 2.523485 | 0.74892 |
| 8 | 97089400 | 97089799 | 7.08189 | 13.78517 | 18.4588 | 6.78168 | 0.196612 | -0.33267 | 0.748936 |
| 4 | 113437200 | 113449999 | 23.445 | 40.28994 | 35.41861 | 15.5358 | 0.569051 | 0.911364 | 0.748943 |
| 8 | 52550400 | 52554399 | 52.9298 | 85.03467 | 82.72582 | 35.04437 | 0.390287 | 1.910614 | 0.748973 |
| 7 | 110661800 | 110664399 | 32.78786 | 54.14326 | 62.59777 | 27.033 | 0.171654 | 1.356783 | 0.748975 |
| 7 | 48011800 | 48012199 | 8.556391 | 16.62807 | 21.22498 | 7.901853 | 0.247906 | -0.25702 | 0.748978 |
| 5 | 149211200 | 149212799 | 23.6228 | 40.14424 | 46.91809 | 20.56217 | 0.157494 | 1.029604 | 0.749038 |
| 6 | 92710000 | 92712199 | 42.75825 | 69.28257 | 78.89267 | 33.77227 | 0.177581 | 1.659656 | 0.749042 |
| 11 | 120205600 | 120215199 | 110.9774 | 168.7394 | 226.4765 | 93.81996 | -0.14106 | 3.186743 | 0.749047 |
| 2 | 39194000 | 39194999 | 9.311746 | 17.4004 | 27.38953 | 11.35975 | -0.19349 | 0.12806 | 0.749048 |
| 11 | 90153200 | 90154799 | 18.1123 | 31.27751 | 45.83596 | 20.10819 | -0.16361 | 0.860898 | 0.749049 |
| 7 | 29423800 | 29428199 | 58.92323 | 93.34104 | 130.0234 | 54.07585 | -0.14848 | 2.358927 | 0.749062 |
| 3 | 54840200 | 54840999 | 8.611681 | 16.36429 | 26.61167 | 10.69421 | -0.20213 | 0.006717 | 0.749136 |
| 11 | 103469200 | 103472999 | 86.81968 | 134.279 | 181.4511 | 75.37046 | -0.13163 | 2.841473 | 0.749148 |
| 2 | 144591600 | 144594799 | 27.71769 | 45.8345 | 65.9876 | 28.5243 | -0.14364 | 1.332093 | 0.74927 |
| 13 | 82138600 | 82139199 | 10.38133 | 19.07945 | 29.85651 | 12.68781 | -0.21122 | 0.243104 | 0.74929 |
| Y | 20410800 | 20411199 | 6.172471 | 11.78906 | 23.5445 | 9.360304 | -0.48126 | -0.05943 | 0.749294 |
| X | 21110600 | 21113199 | 35.48935 | 58.87946 | 51.42992 | 22.28208 | 0.57556 | 1.307189 | 0.749295 |
| 2 | 158003200 | 158004999 | 17.59926 | 30.35956 | 45.32394 | 19.88293 | -0.1906 | 0.866498 | 0.749314 |
| 9 | 109062800 | 109065399 | 17.59926 | 30.35956 | 45.32394 | 19.88293 | -0.1906 | 0.866498 | 0.749314 |
| 5 | 91302400 | 91303199 | 8.412437 | 15.81962 | 25.94336 | 10.64535 | -0.23931 | 0.077271 | 0.749352 |
| 12 | 106221200 | 106223199 | 18.17812 | 31.42011 | 46.34141 | 20.3305 | -0.17271 | 0.851269 | 0.749352 |
| 9 | 120333400 | 120337999 | 56.66844 | 90.06962 | 124.0137 | 51.74167 | -0.12707 | 2.261165 | 0.749357 |
| X | 130204600 | 130205199 | 11.71295 | 21.26538 | 32.80143 | 14.1617 | -0.20632 | 0.329846 | 0.749371 |
| 6 | 104792600 | 104793199 | 5.950715 | 11.57698 | 20.68355 | 7.632574 | -0.21992 | -0.32006 | 0.749374 |

| | | | | | | | | | |
|---|---|---|---|---|---|---|---|---|---|
| 6 | 6437800 | 6443799 | 51.8985 | 83.65166 | 89.24755 | 37.98626 | 0.263232 | 1.811434 | 0.749408 |
| 19 | 61236800 | 61240599 | 68.11574 | 107.6855 | 123.1468 | 51.16367 | 0.133418 | 2.412344 | 0.749434 |
| 6 | 129949600 | 129957199 | 68.32822 | 108.7028 | 119.9592 | 50.10384 | 0.193012 | 2.242131 | 0.749452 |
| 3 | 140291400 | 140293199 | 20.62391 | 36.15398 | 39.53092 | 17.46598 | 0.259286 | 0.671878 | 0.749464 |
| 7 | 33077800 | 33080399 | 35.64346 | 59.56764 | 59.89712 | 26.07512 | 0.373272 | 1.235922 | 0.749467 |
| 1 | 78183400 | 78186399 | 27.37539 | 45.50069 | 66.14256 | 28.69246 | -0.15697 | 1.272607 | 0.749471 |
| 17 | 22672400 | 22676599 | 41.30788 | 68.65978 | 61.32971 | 26.60516 | 0.543638 | 1.311689 | 0.749479 |
| 6 | 38185600 | 38187599 | 28.63183 | 48.13595 | 46.96752 | 20.51812 | 0.416336 | 1.138756 | 0.749507 |
| 17 | 29346600 | 29348399 | 15.2536 | 26.81436 | 40.20202 | 17.8025 | -0.194 | 0.598265 | 0.749507 |
| 8 | 5267600 | 5270399 | 40.85769 | 66.03768 | 77.06125 | 32.91637 | 0.138795 | 1.709149 | 0.749519 |
| 18 | 6367200 | 6372199 | 67.4493 | 106.8937 | 123.5576 | 51.4016 | 0.120839 | 2.353702 | 0.749523 |
| 13 | 20858600 | 20862199 | 44.90959 | 71.57944 | 123.0961 | 51.54666 | -0.44301 | 2.162124 | 0.74957 |
| 4 | 22718400 | 22721999 | 35.63928 | 58.08582 | 68.11664 | 29.25584 | 0.142241 | 1.538644 | 0.749576 |
| 18 | 61673600 | 61675799 | 22.3875 | 37.464 | 65.21219 | 28.41279 | -0.41589 | 1.184939 | 0.749601 |
| 10 | 41375000 | 41377199 | 17.44126 | 30.31411 | 44.209 | 19.4146 | -0.15493 | 0.789987 | 0.749609 |
| 2 | 52225200 | 52226399 | 15.5481 | 27.25954 | 41.32907 | 18.26246 | -0.21015 | 0.643158 | 0.749612 |
| 8 | 21428600 | 21431999 | 30.58274 | 51.11809 | 57.99225 | 25.24835 | 0.199665 | 1.214876 | 0.749614 |
| 19 | 13975800 | 13977799 | 25.00012 | 41.04589 | 75.11159 | 32.35223 | -0.49171 | 1.422541 | 0.749618 |
| X | 104597400 | 104601599 | 24.45156 | 41.24788 | 61.6095 | 26.94231 | -0.1957 | 1.111302 | 0.749618 |
| 8 | 19082800 | 19084199 | 16.99639 | 30.0057 | 34.85384 | 15.42103 | 0.174806 | 0.55414 | 0.74963 |
| 18 | 13306000 | 13306799 | 10.30653 | 19.24553 | 30.04103 | 12.47455 | -0.18298 | 0.148427 | 0.749632 |
| X | 40088400 | 40089399 | 7.775788 | 15.01607 | 25.45262 | 9.797842 | -0.20328 | -0.14075 | 0.749636 |
| 9 | 87502600 | 87503399 | 8.167929 | 15.53841 | 25.53697 | 10.23811 | -0.21405 | -0.00763 | 0.749651 |
| 8 | 9134800 | 9138599 | 65.52295 | 104.1403 | 118.9002 | 49.60684 | 0.142614 | 2.2664 | 0.749683 |
| 6 | 137134200 | 137135999 | 12.96163 | 23.69953 | 27.94236 | 11.95754 | 0.186552 | 0.282713 | 0.749697 |
| 3 | 83592000 | 83592599 | 8.75013 | 16.8175 | 21.38793 | 8.355273 | 0.187511 | -0.09588 | 0.749702 |
| 2 | 31004000 | 31004799 | 10.2822 | 19.54911 | 24.37226 | 9.783233 | 0.179835 | 0.014039 | 0.749718 |
| 19 | 40441000 | 40443599 | 25.85751 | 42.956 | 62.65201 | 27.16874 | -0.1618 | 1.27013 | 0.749731 |
| 18 | 48197400 | 48202999 | 61.26462 | 97.43245 | 112.691 | 47.04614 | 0.12485 | 2.243579 | 0.749735 |
| 4 | 35109200 | 35110599 | 11.34325 | 20.75702 | 31.7639 | 13.57835 | -0.18612 | 0.274769 | 0.749745 |
| 10 | 27765200 | 27765999 | 9.806789 | 18.45649 | 29.1312 | 11.9372 | -0.18348 | 0.091819 | 0.749761 |
| 19 | 12135400 | 12137799 | 18.21015 | 31.99295 | 36.98818 | 16.37293 | 0.179214 | 0.620842 | 0.749761 |
| 7 | 29423600 | 29428199 | 59.92393 | 94.86059 | 131.7087 | 54.75757 | -0.14457 | 2.376459 | 0.749773 |
| 3 | 153296400 | 153298599 | 21.71181 | 36.89866 | 53.29651 | 23.35765 | -0.14696 | 1.028278 | 0.749786 |
| 8 | 55826600 | 55827199 | 6.102097 | 11.86185 | 21.19337 | 7.872181 | -0.22866 | -0.29277 | 0.7498 |
| 1 | 144162800 | 144163799 | 11.01361 | 20.77494 | 25.25039 | 10.30651 | 0.194898 | 0.075903 | 0.749805 |
| 6 | 148255400 | 148260799 | 69.47219 | 108.7413 | 171.6816 | 71.21848 | -0.34247 | 2.62224 | 0.749818 |
| 5 | 75435200 | 75437799 | 16.44558 | 28.74886 | 41.97881 | 18.50379 | -0.15597 | 0.698734 | 0.749836 |
| 13 | 62399000 | 62439599 | 824.3782 | 1081.984 | 1034.809 | 433.0678 | 0.140681 | 5.580959 | 0.749853 |
| 3 | 93599200 | 93599999 | 10.59684 | 20.04642 | 24.80542 | 10.06342 | 0.177142 | 0.052316 | 0.749896 |
| 2 | 27384800 | 27393399 | 154.4898 | 229.5695 | 304.6719 | 125.5146 | -0.15398 | 3.641004 | 0.749898 |
| X | 96535200 | 96536399 | 13.79479 | 25.73203 | 20.58198 | 8.56197 | 0.773299 | 0.16278 | 0.749898 |
| 18 | 58581200 | 58584399 | 43.35523 | 70.31076 | 78.70357 | 33.71284 | 0.203379 | 1.642429 | 0.7499 |
| 10 | 111239400 | 111241999 | 23.44802 | 39.6587 | 58.8954 | 25.77587 | -0.18735 | 1.079459 | 0.749906 |
| 6 | 27077400 | 27079999 | 23.56685 | 40.61337 | 45.77561 | 20.08046 | 0.212929 | 0.903843 | 0.749924 |
| 5 | 107715800 | 107717199 | 12.0374 | 21.96391 | 34.4227 | 14.76908 | -0.22349 | 0.306331 | 0.749969 |
| 5 | 109875600 | 109894599 | 481.2118 | 659.9074 | 659.3936 | 274.258 | 0.140969 | 4.881891 | 0.749992 |

| | | | | | | | | | |
|---|---|---|---|---|---|---|---|---|---|
| 14 | 87621200 | 87624999 | 44.27602 | 71.38819 | 82.96572 | 35.32702 | 0.140605 | 1.7874 | 0.749993 |
| 5 | 17267800 | 17270799 | 28.56064 | 46.70426 | 80.07859 | 34.43526 | -0.40096 | 1.483878 | 0.750001 |
| 2 | 11370800 | 11371999 | 8.949306 | 16.84785 | 26.94787 | 11.03682 | -0.20205 | 0.074811 | 0.750005 |
| 2 | 51778600 | 51781999 | 52.75939 | 84.36766 | 97.62772 | 41.07109 | 0.132707 | 2.059897 | 0.750041 |
| 5 | 142187600 | 142189999 | 18.73517 | 32.03019 | 48.4861 | 21.25965 | -0.213 | 0.955671 | 0.750041 |
| 7 | 108140000 | 108142599 | 36.26802 | 58.68231 | 83.22845 | 35.5908 | -0.14032 | 1.679733 | 0.750043 |
| 2 | 27382800 | 27394799 | 181.8247 | 267.0719 | 349.0479 | 143.7548 | -0.14753 | 3.838374 | 0.750069 |
| 8 | 75981600 | 75984199 | 23.13606 | 39.0643 | 57.18099 | 25.01259 | -0.16664 | 1.091706 | 0.750094 |
| X | 24045200 | 24047199 | 33.10209 | 55.21223 | 59.71204 | 25.96244 | 0.268564 | 1.248597 | 0.750177 |
| 9 | 4071600 | 4080199 | 126.8345 | 193.7191 | 187.2928 | 77.68107 | 0.336981 | 3.097417 | 0.750179 |
| 15 | 33521600 | 33523999 | 20.12712 | 35.36144 | 38.37735 | 16.98811 | 0.269912 | 0.632428 | 0.750192 |
| 1 | 141129800 | 141132799 | 27.88135 | 47.85352 | 49.01936 | 21.50024 | 0.349031 | 0.958463 | 0.750216 |
| 4 | 104688800 | 104691799 | 22.40589 | 38.49825 | 45.26042 | 19.85266 | 0.151384 | 0.922427 | 0.750245 |
| 10 | 14442200 | 14442999 | 10.60633 | 19.49136 | 29.7893 | 12.66067 | -0.17789 | 0.242587 | 0.750252 |
| 7 | 6644200 | 6645399 | 16.97471 | 29.88741 | 34.91387 | 15.45585 | 0.165188 | 0.582891 | 0.750254 |
| 13 | 77223000 | 77225799 | 44.94042 | 72.98051 | 78.42496 | 33.62713 | 0.26396 | 1.619885 | 0.750282 |
| 5 | 137824400 | 137826199 | 14.57236 | 25.86271 | 27.4756 | 12.13547 | 0.306053 | 0.470425 | 0.750289 |
| 12 | 26096600 | 26099799 | 43.1764 | 69.21301 | 98.74531 | 41.69149 | -0.16472 | 1.974342 | 0.750296 |
| 7 | 17985600 | 18067599 | 1812.242 | 2227.425 | 2012.415 | 848.0581 | 0.127438 | 6.64235 | 0.7503 |
| 14 | 87048000 | 87048399 | 6.239527 | 12.15234 | 21.38275 | 7.812381 | -0.18567 | -0.35102 | 0.750312 |
| 2 | 15404600 | 15405999 | 11.41118 | 21.09826 | 33.09665 | 13.95051 | -0.2069 | 0.221821 | 0.750325 |
| 14 | 102199200 | 102202799 | 37.78532 | 61.54976 | 71.82144 | 30.84088 | 0.148804 | 1.553018 | 0.750337 |
| 15 | 8754800 | 8757599 | 41.76685 | 67.17348 | 94.48776 | 40.08985 | -0.13865 | 1.866603 | 0.750343 |
| 1 | 11778000 | 11780599 | 28.08676 | 46.77612 | 55.06415 | 23.91871 | 0.146639 | 1.243032 | 0.750348 |
| 6 | 146776200 | 146780599 | 44.87953 | 71.99816 | 100.736 | 42.58142 | -0.1354 | 1.951406 | 0.750351 |
| 18 | 47418400 | 47419599 | 10.91324 | 20.54142 | 25.24535 | 10.34268 | 0.174877 | 0.089571 | 0.750354 |
| 17 | 23301400 | 23302799 | 12.90863 | 23.92995 | 22.91766 | 9.62537 | 0.504041 | 0.195066 | 0.750358 |
| 12 | 115516800 | 115536399 | 450.9622 | 621.7626 | 627.5733 | 260.8009 | 0.134274 | 4.79793 | 0.75038 |
| 1 | 156773600 | 156774199 | 6.246413 | 12.1758 | 22.13543 | 8.000361 | -0.21807 | -0.38591 | 0.750385 |
| 14 | 120075600 | 120077999 | 20.05289 | 34.20371 | 50.0739 | 21.95176 | -0.16547 | 0.984773 | 0.750421 |
| 5 | 127609400 | 127614799 | 37.81447 | 61.05826 | 86.87243 | 37.05964 | -0.14822 | 1.737215 | 0.750437 |
| 17 | 88804800 | 88811999 | 143.9898 | 216.8186 | 239.9703 | 99.11687 | 0.125067 | 3.38991 | 0.750441 |
| 14 | 119374400 | 119377199 | 21.13447 | 35.19576 | 65.54328 | 28.46966 | -0.51287 | 1.232812 | 0.750448 |
| 11 | 72497800 | 72499399 | 14.91798 | 25.54561 | 53.69961 | 23.54869 | -0.68422 | 0.933063 | 0.750463 |
| 17 | 41345200 | 41348599 | 40.09138 | 64.69574 | 92.5929 | 39.46927 | -0.15771 | 1.764282 | 0.750464 |
| 14 | 62077800 | 62079199 | 22.55648 | 39.23602 | 36.69818 | 16.12427 | 0.483194 | 0.792592 | 0.750476 |
| X | 71788400 | 71792999 | 78.45668 | 122.464 | 166.1992 | 68.96392 | -0.12756 | 2.676119 | 0.750479 |
| 6 | 19317600 | 19323199 | 13.8107 | 24.29229 | 45.57652 | 20.18007 | -0.51547 | 0.594579 | 0.750483 |
| X | 148983400 | 148995599 | 34.23991 | 57.33856 | 48.63103 | 21.19783 | 0.61788 | 1.197726 | 0.750506 |
| 18 | 5981400 | 5983599 | 22.0234 | 36.77813 | 56.38762 | 24.53572 | -0.23324 | 1.197559 | 0.750536 |
| 5 | 146243000 | 146246199 | 35.5819 | 58.0393 | 68.29124 | 29.33949 | 0.138195 | 1.525894 | 0.75054 |
| 1 | 128953800 | 128957199 | 38.43542 | 62.99117 | 70.15202 | 30.19309 | 0.22157 | 1.469634 | 0.750565 |
| 16 | 54668800 | 54672399 | 31.65172 | 51.99953 | 77.25201 | 33.28306 | -0.19173 | 1.426937 | 0.750567 |
| 6 | 117719200 | 117720399 | 20.68346 | 35.92338 | 41.99965 | 18.436 | 0.162342 | 0.810447 | 0.750579 |
| 6 | 34838000 | 34840599 | 15.05998 | 26.46204 | 38.77062 | 17.16389 | -0.16069 | 0.604947 | 0.750588 |
| X | 19083800 | 19084999 | 13.8923 | 26.35254 | 27.73413 | 11.18668 | 0.416542 | 0.071798 | 0.750599 |
| 2 | 6269800 | 6271399 | 14.50614 | 25.5499 | 37.67739 | 16.67805 | -0.16895 | 0.570062 | 0.750625 |

| | | | | | | | | | |
|---|---|---|---|---|---|---|---|---|---|
| 5 | 127753400 | 127758199 | 58.42533 | 92.82937 | 127.2714 | 53.08219 | -0.12146 | 2.272778 | 0.75063 |
| 8 | 52296200 | 52297999 | 16.19396 | 29.09099 | 32.80258 | 14.34107 | 0.230681 | 0.418626 | 0.750634 |
| 4 | 66569200 | 66569799 | 11.23208 | 21.03336 | 25.6961 | 10.61923 | 0.173386 | 0.125859 | 0.750638 |
| 1 | 147214800 | 147215399 | 13.22078 | 23.9146 | 27.99419 | 12.14208 | 0.184588 | 0.347824 | 0.750639 |
| 5 | 126832600 | 126833999 | 16.95653 | 29.67993 | 32.35994 | 14.27929 | 0.263594 | 0.66527 | 0.75064 |
| 12 | 47450400 | 47451599 | 10.43529 | 19.88759 | 23.81563 | 9.511663 | 0.243306 | -0.00481 | 0.750645 |
| 4 | 40423000 | 40428399 | 65.24004 | 102.9725 | 140.7759 | 58.44014 | -0.12569 | 2.451741 | 0.750654 |
| 17 | 12394600 | 12397599 | 33.59372 | 54.72959 | 78.15325 | 33.56153 | -0.14164 | 1.546611 | 0.750693 |
| X | 107089800 | 107094799 | 60.97779 | 97.681 | 103.7609 | 43.7627 | 0.258808 | 2.018952 | 0.750729 |
| 3 | 15849600 | 15850599 | 20.52053 | 35.62012 | 41.56261 | 18.23763 | 0.165279 | 0.819755 | 0.75073 |
| 2 | 97510400 | 97511999 | 17.70271 | 31.22642 | 35.87686 | 15.87633 | 0.190097 | 0.569867 | 0.750747 |
| 4 | 25487400 | 25487999 | 9.968264 | 19.0507 | 23.93963 | 9.504832 | 0.182365 | -0.02532 | 0.750814 |
| 7 | 77921600 | 77921999 | 9.205695 | 17.13124 | 27.38974 | 11.43966 | -0.22368 | 0.155053 | 0.750816 |
| 11 | 65048800 | 65050399 | 15.86118 | 27.94476 | 32.06832 | 14.19434 | 0.191246 | 0.565038 | 0.750826 |
| X | 26091000 | 26100599 | 41.11785 | 67.09765 | 61.29887 | 26.33947 | 0.502804 | 1.530239 | 0.750836 |
| 16 | 63822600 | 63823599 | 14.22842 | 25.00416 | 37.2831 | 16.50717 | -0.18585 | 0.594829 | 0.750842 |
| 2 | 95181400 | 95182999 | 15.27181 | 27.14787 | 31.92479 | 14.08141 | 0.161868 | 0.476911 | 0.750875 |
| 2 | 135437800 | 135438399 | 5.072722 | 9.893257 | 18.62276 | 6.683988 | -0.25726 | -0.43731 | 0.750906 |
| 4 | 77007600 | 77010599 | 32.72891 | 54.05827 | 63.05612 | 27.23524 | 0.158966 | 1.35612 | 0.750919 |
| 7 | 96643000 | 96647399 | 47.00812 | 76.28254 | 79.72058 | 34.16425 | 0.30311 | 1.639795 | 0.75093 |
| 1 | 106099000 | 106101199 | 18.16549 | 30.28319 | 66.97256 | 29.10974 | -0.75995 | 1.213136 | 0.750952 |
| 3 | 51269000 | 51271399 | 15.45464 | 27.00807 | 40.46837 | 17.83326 | -0.19302 | 0.695612 | 0.750969 |
| 19 | 7953000 | 7954799 | 24.23488 | 41.48377 | 47.93398 | 21.01965 | 0.175357 | 0.966828 | 0.750979 |
| 7 | 6129400 | 6138399 | 115.2425 | 174.4922 | 237.1896 | 98.13292 | -0.16436 | 3.273702 | 0.751033 |
| 4 | 17332400 | 17336599 | 66.1477 | 105.139 | 119.8312 | 49.99801 | 0.145216 | 2.267017 | 0.751042 |
| 12 | 57307800 | 57310799 | 25.04117 | 43.52158 | 37.05587 | 16.27287 | 0.618038 | 0.816339 | 0.751047 |
| 6 | 39228400 | 39229599 | 14.10126 | 25.26462 | 29.50054 | 12.92628 | 0.178606 | 0.415235 | 0.75105 |
| 8 | 9988200 | 9989799 | 15.41576 | 27.01337 | 39.91773 | 17.63101 | -0.17286 | 0.653687 | 0.751059 |
| 5 | 127013000 | 127016399 | 46.03803 | 74.06191 | 86.25962 | 36.61245 | 0.133561 | 1.86265 | 0.751059 |
| 4 | 67759000 | 67760199 | 14.00066 | 24.46591 | 44.6821 | 19.6893 | -0.477 | 0.697867 | 0.751066 |
| 2 | 19688000 | 19689799 | 15.14252 | 26.92653 | 31.47959 | 13.88135 | 0.170618 | 0.473939 | 0.751099 |
| 12 | 18212200 | 18639199 | 14984.52 | 15347.73 | 13685.26 | 5936.318 | -0.12123 | 9.53472 | 0.751118 |
| 3 | 157006800 | 157008399 | 18.47945 | 32.87601 | 35.26608 | 15.54478 | 0.294354 | 0.498944 | 0.751135 |
| 17 | 61667800 | 61668599 | 11.44672 | 21.47748 | 25.66515 | 10.57118 | 0.208585 | 0.113623 | 0.751136 |
| 15 | 92663600 | 92666599 | 34.74479 | 56.724 | 66.56792 | 28.60834 | 0.142887 | 1.510204 | 0.751145 |
| 16 | 55774400 | 55784599 | 164.8096 | 243.8335 | 321.2809 | 132.3124 | -0.14946 | 3.718374 | 0.751189 |
| X | 90240800 | 90242599 | 20.35584 | 35.77641 | 38.45241 | 17.02662 | 0.284029 | 0.630832 | 0.751196 |
| 10 | 82479800 | 82480599 | 7.357918 | 14.04554 | 24.56246 | 9.776263 | -0.29219 | -0.036 | 0.751208 |
| 12 | 64505800 | 64511199 | 49.49378 | 79.89418 | 86.60379 | 36.93165 | 0.242823 | 1.765232 | 0.751212 |
| 5 | 43807000 | 43810599 | 31.55373 | 52.45882 | 60.27585 | 26.15207 | 0.181323 | 1.276486 | 0.751217 |
| 15 | 43742600 | 43749399 | 95.33821 | 146.5403 | 196.7143 | 81.6847 | -0.12898 | 2.962235 | 0.751226 |
| 14 | 20906200 | 20907399 | 13.72563 | 24.69926 | 28.95406 | 12.6402 | 0.176498 | 0.383475 | 0.751237 |
| 17 | 84214600 | 84217799 | 26.40214 | 44.37564 | 52.31473 | 22.85225 | 0.144344 | 1.138733 | 0.751242 |
| 4 | 150895400 | 150896399 | 19.1283 | 32.89942 | 47.94689 | 21.03167 | -0.15688 | 0.904811 | 0.751245 |
| X | 77004600 | 77013199 | 118.5813 | 182.4465 | 193.4497 | 80.33998 | 0.212111 | 2.953059 | 0.751248 |
| 13 | 13582000 | 13584999 | 27.14637 | 45.77164 | 52.9013 | 23.13074 | 0.172952 | 1.11338 | 0.751289 |
| 12 | 94628400 | 94628999 | 7.944744 | 15.08522 | 25.00645 | 10.06567 | -0.2306 | 0.005573 | 0.751291 |

| | | | | | | | | | |
|---|---|---|---|---|---|---|---|---|---|
| 3 | 9742000 | 9746599 | 88.22377 | 136.6186 | 183.2702 | 76.12555 | -0.11977 | 2.818448 | 0.751299 |
| 4 | 86610200 | 86613599 | 26.08399 | 43.2762 | 62.5454 | 27.09829 | -0.14876 | 1.287211 | 0.75131 |
| 14 | 106672800 | 106675399 | 37.48223 | 61.19205 | 70.78204 | 30.41921 | 0.163656 | 1.520774 | 0.751312 |
| 1 | 146749800 | 146751399 | 15.60339 | 28.0067 | 32.3683 | 14.16116 | 0.19473 | 0.41995 | 0.751326 |
| X | 81280400 | 81284799 | 27.10344 | 46.16463 | 41.62695 | 18.24665 | 0.530859 | 1.017426 | 0.751343 |
| 17 | 52849200 | 52852199 | 32.16509 | 53.21913 | 61.80917 | 26.72861 | 0.165427 | 1.335739 | 0.751352 |
| 1 | 121562000 | 121564399 | 29.18041 | 48.71994 | 56.82979 | 24.72489 | 0.159685 | 1.220095 | 0.751356 |
| 4 | 142803200 | 142805199 | 19.04621 | 33.44166 | 38.00277 | 16.81349 | 0.203952 | 0.636762 | 0.751384 |
| 1 | 20067200 | 20071199 | 45.21519 | 73.02888 | 83.42233 | 35.60092 | 0.168302 | 1.741081 | 0.751387 |
| 15 | 90071600 | 90077799 | 74.27656 | 116.9329 | 134.3622 | 55.74732 | 0.121784 | 2.509688 | 0.751395 |
| 3 | 111493400 | 111494399 | 13.0239 | 22.9632 | 41.38111 | 18.3137 | -0.45599 | 0.557995 | 0.751397 |
| 6 | 8809800 | 8811799 | 21.66212 | 37.87814 | 41.22268 | 18.17461 | 0.266115 | 0.720217 | 0.751407 |
| Y | 44500000 | 44503199 | 17.36702 | 30.36625 | 28.45276 | 12.54396 | 0.480985 | 0.675135 | 0.751425 |
| 7 | 11242400 | 11245199 | 35.36368 | 58.24246 | 56.46383 | 24.35235 | 0.423825 | 1.390845 | 0.751425 |
| 3 | 10162200 | 10163199 | 9.512834 | 17.65066 | 27.86429 | 11.68868 | -0.21037 | 0.175887 | 0.751432 |
| 8 | 111329000 | 111330199 | 9.512834 | 17.65066 | 27.86429 | 11.68868 | -0.21037 | 0.175887 | 0.751432 |
| 12 | 34080000 | 34080599 | 8.015247 | 15.54385 | 20.52126 | 7.745535 | 0.181115 | -0.21966 | 0.751468 |
| 15 | 98291600 | 98294599 | 28.24858 | 46.68617 | 67.09599 | 28.99218 | -0.14117 | 1.341718 | 0.751475 |
| X | 32527000 | 32529399 | 47.17961 | 76.84784 | 76.05653 | 32.65876 | 0.385777 | 1.576608 | 0.751486 |
| 12 | 56867200 | 56867999 | 8.976418 | 16.82386 | 26.8978 | 11.09964 | -0.21004 | 0.103947 | 0.751518 |
| 3 | 83214600 | 83216599 | 21.20344 | 36.68029 | 43.10443 | 18.91089 | 0.153965 | 0.853498 | 0.751538 |
| 6 | 65208000 | 65210599 | 36.58204 | 60.30749 | 57.08975 | 24.6324 | 0.458658 | 1.383587 | 0.751547 |
| 3 | 41737000 | 41752199 | 185.1836 | 277.2631 | 283.2288 | 116.8184 | 0.233331 | 3.509545 | 0.751548 |
| 8 | 127354400 | 127360399 | 134.079 | 201.8527 | 265.5176 | 109.6627 | -0.12416 | 3.393293 | 0.751557 |
| 9 | 35998400 | 36007799 | 143.8355 | 218.1869 | 207.8103 | 86.02684 | 0.350638 | 3.241248 | 0.751562 |
| 11 | 7054400 | 7056399 | 18.90576 | 32.50463 | 47.41909 | 20.79992 | -0.15849 | 0.905155 | 0.751565 |
| 1 | 111070200 | 111072999 | 41.27421 | 66.63854 | 77.14348 | 32.90539 | 0.14854 | 1.743109 | 0.751569 |
| 9 | 95148800 | 95153199 | 36.60944 | 59.00883 | 99.01501 | 42.14409 | -0.38865 | 1.792823 | 0.751575 |
| 7 | 87723400 | 87724399 | 10.57765 | 19.52754 | 29.82465 | 12.59727 | -0.17056 | 0.216264 | 0.751578 |
| 17 | 22229400 | 22231199 | 15.86329 | 28.93833 | 31.49117 | 13.51507 | 0.298907 | 0.326008 | 0.751578 |
| 3 | 142587000 | 142588199 | 12.68218 | 22.80507 | 34.10461 | 14.8978 | -0.1735 | 0.390831 | 0.751604 |
| 6 | 149258800 | 149261799 | 42.09357 | 68.17819 | 77.9027 | 33.33861 | 0.172047 | 1.663082 | 0.751631 |
| 8 | 86709000 | 86709799 | 17.93835 | 31.09577 | 45.28984 | 19.87092 | -0.15363 | 0.825033 | 0.75164 |
| 7 | 107491000 | 107492399 | 23.01274 | 39.54265 | 45.79768 | 20.08835 | 0.172973 | 0.926867 | 0.751644 |
| 4 | 103170800 | 103171599 | 8.038859 | 15.29814 | 25.01081 | 10.01987 | -0.20536 | -0.01157 | 0.751651 |
| 4 | 20127000 | 20129599 | 24.64241 | 41.63816 | 63.91084 | 27.96262 | -0.23483 | 1.100184 | 0.75166 |
| 10 | 67000600 | 67001599 | 16.12192 | 28.54153 | 33.42363 | 14.77998 | 0.164859 | 0.517773 | 0.751683 |
| 15 | 19495400 | 19500799 | 120.7859 | 184.83 | 178.9972 | 74.27288 | 0.336713 | 3.056553 | 0.751688 |
| 17 | 88877800 | 88880199 | 7.345227 | 13.90738 | 29.27395 | 11.85432 | -0.58086 | 0.027814 | 0.751708 |
| 17 | 50524400 | 50524999 | 13.37071 | 24.36428 | 28.5875 | 12.28592 | 0.190037 | 0.310562 | 0.751719 |
| 2 | 14078200 | 14079599 | 11.60003 | 20.95645 | 31.68655 | 13.76526 | -0.18379 | 0.352228 | 0.751731 |
| 1 | 85592600 | 85596599 | 45.07859 | 72.40577 | 101.276 | 42.88917 | -0.13289 | 1.914881 | 0.75175 |
| 8 | 30103800 | 30106399 | 18.88199 | 32.74504 | 47.93932 | 21.03364 | -0.16108 | 0.828113 | 0.751794 |
| 7 | 111002400 | 111004199 | 17.46532 | 30.8477 | 35.29986 | 15.61694 | 0.196486 | 0.552385 | 0.751798 |
| 8 | 22784800 | 22786199 | 9.49976 | 17.19555 | 34.38852 | 14.90274 | -0.58211 | 0.332639 | 0.7518 |
| 15 | 19792400 | 19792999 | 13.59856 | 24.47925 | 28.51973 | 12.44508 | 0.185861 | 0.380338 | 0.751801 |
| 13 | 24391800 | 24394799 | 21.72745 | 36.3224 | 64.04715 | 27.88724 | -0.43444 | 1.191953 | 0.751816 |

| | | | | | | | | | |
|---|---|---|---|---|---|---|---|---|---|
| 9 | 114931400 | 114932399 | 11.40286 | 20.61003 | 31.66278 | 13.74677 | -0.2062 | 0.34669 | 0.75183 |
| 17 | 33256200 | 33260799 | 53.62787 | 86.03631 | 97.08906 | 41.04244 | 0.17424 | 1.959033 | 0.75185 |
| 12 | 56199400 | 56200199 | 11.55186 | 21.52458 | 26.14711 | 10.89724 | 0.171771 | 0.161229 | 0.751864 |
| 5 | 89743600 | 89745199 | 15.67214 | 27.6665 | 31.97756 | 14.14856 | 0.181974 | 0.541431 | 0.751902 |
| 1 | 49588800 | 49591799 | 30.30439 | 50.76943 | 56.91665 | 24.81603 | 0.216641 | 1.187419 | 0.751904 |
| X | 92328200 | 92335799 | 74.825 | 117.7101 | 165.5058 | 68.66239 | -0.16988 | 2.531206 | 0.751904 |
| 6 | 143197600 | 143199799 | 19.30117 | 33.30858 | 49.01574 | 21.50272 | -0.17001 | 0.873344 | 0.751906 |
| 8 | 52658800 | 52660999 | 22.27866 | 38.93992 | 34.81973 | 15.3462 | 0.548173 | 0.719522 | 0.751906 |
| 3 | 97404800 | 97408999 | 49.00285 | 78.98542 | 88.39556 | 37.62428 | 0.19444 | 1.807416 | 0.751908 |
| 13 | 78366800 | 78374199 | 89.83955 | 140.5219 | 156.6107 | 64.97137 | 0.158585 | 2.643258 | 0.75191 |
| 4 | 73125600 | 73131199 | 51.18626 | 81.93875 | 117.4104 | 49.46757 | -0.17347 | 2.031779 | 0.751923 |
| 3 | 53270400 | 53280799 | 166.9922 | 245.4455 | 380.2948 | 156.6174 | -0.39202 | 3.827706 | 0.751925 |
| 10 | 96285600 | 96288399 | 37.30329 | 60.84487 | 70.41198 | 30.2502 | 0.162035 | 1.529735 | 0.75194 |
| 10 | 117929400 | 117934999 | 49.97995 | 79.8477 | 110.7954 | 46.54839 | -0.13032 | 2.088673 | 0.75195 |
| 1 | 151173000 | 151176199 | 26.81373 | 44.82681 | 51.82306 | 22.56514 | 0.172519 | 1.193671 | 0.751997 |
| 12 | 17931200 | 17933199 | 39.70181 | 63.72361 | 94.05854 | 39.78906 | -0.21151 | 1.927373 | 0.752009 |
| 3 | 65102600 | 65103999 | 12.26437 | 22.42063 | 26.11046 | 11.17611 | 0.20442 | 0.276321 | 0.752037 |
| 17 | 5903600 | 5906799 | 33.22461 | 54.13072 | 77.04221 | 33.08497 | -0.13683 | 1.543515 | 0.752063 |
| 14 | 101676400 | 101677199 | 13.0866 | 23.86246 | 28.52198 | 12.24756 | 0.164005 | 0.302488 | 0.752076 |
| 17 | 40774800 | 40775599 | 13.0866 | 23.86246 | 28.52198 | 12.24756 | 0.164005 | 0.302488 | 0.752076 |
| 10 | 121565400 | 121571599 | 62.42256 | 98.97184 | 135.6497 | 56.45753 | -0.12397 | 2.339523 | 0.752166 |
| 12 | 14539000 | 14544999 | 61.57792 | 98.58858 | 105.0822 | 44.28322 | 0.25297 | 2.036013 | 0.752168 |
| 18 | 70817800 | 70821599 | 38.01563 | 62.24752 | 70.16853 | 30.18878 | 0.203609 | 1.478149 | 0.752218 |
| 4 | 143418400 | 143434799 | 225.7298 | 333.4224 | 331.22 | 136.4032 | 0.255627 | 3.75445 | 0.752229 |
| 7 | 94836600 | 94838399 | 20.44503 | 35.73395 | 40.28801 | 17.75581 | 0.21549 | 0.721507 | 0.752233 |
| 6 | 139749600 | 139750999 | 15.11827 | 26.81109 | 41.8149 | 18.48076 | -0.24545 | 0.509493 | 0.752234 |
| X | 8833200 | 8839199 | 34.60804 | 57.63611 | 50.8693 | 22.0964 | 0.560897 | 1.260895 | 0.752251 |
| 12 | 74171400 | 74175399 | 57.32712 | 91.22025 | 125.9549 | 52.62328 | -0.12949 | 2.227335 | 0.752255 |
| 1 | 57619800 | 57630999 | 146.3354 | 219.5055 | 286.1676 | 118.0766 | -0.11592 | 3.467579 | 0.752286 |
| 13 | 61846000 | 61849199 | 63.34129 | 101.3734 | 106.2863 | 44.76631 | 0.276088 | 2.05087 | 0.752326 |
| 1 | 141585800 | 141591199 | 47.84367 | 77.51049 | 68.60196 | 29.36364 | 0.540032 | 1.665039 | 0.752343 |
| 10 | 68127000 | 68131399 | 52.14479 | 83.72792 | 95.34467 | 40.35703 | 0.162654 | 1.932795 | 0.752352 |
| 14 | 117528200 | 117531999 | 53.18928 | 84.71203 | 117.1192 | 48.99257 | -0.12992 | 2.192383 | 0.752356 |
| 16 | 55003400 | 55004999 | 16.05598 | 28.15233 | 41.39356 | 18.29225 | -0.16582 | 0.649191 | 0.752358 |
| 2 | 23170000 | 23171599 | 32.45937 | 53.22599 | 62.05246 | 26.71321 | 0.156078 | 1.443674 | 0.752363 |
| X | 117320800 | 117324999 | 47.30072 | 76.12554 | 86.98166 | 36.94165 | 0.161902 | 1.853258 | 0.752378 |
| 10 | 49695400 | 49696199 | 10.30724 | 19.51058 | 24.30152 | 9.846153 | 0.169525 | 0.044726 | 0.752382 |
| 16 | 19505400 | 19510999 | 63.56753 | 101.3293 | 113.5065 | 47.53864 | 0.174512 | 2.172025 | 0.7524 |
| 8 | 103091600 | 103092799 | 12.37809 | 21.88379 | 39.82422 | 17.61513 | -0.46843 | 0.52133 | 0.752416 |
| 5 | 134842600 | 134843999 | 14.48699 | 25.9727 | 30.48712 | 13.3512 | 0.171666 | 0.416706 | 0.75243 |
| 7 | 55237200 | 55238599 | 11.70108 | 20.91876 | 38.32206 | 16.82245 | -0.46933 | 0.430467 | 0.752433 |
| 17 | 20963000 | 20980199 | 373.9191 | 518.3547 | 627.142 | 260.3957 | -0.11915 | 4.712521 | 0.752441 |
| 1 | 136734000 | 136741599 | 94.3693 | 145.4426 | 194.9254 | 80.96881 | -0.12368 | 2.908954 | 0.752468 |
| 13 | 115033000 | 115033799 | 7.139772 | 13.78527 | 23.14951 | 8.919039 | -0.19093 | -0.15178 | 0.75249 |
| 16 | 82357600 | 82358199 | 7.139772 | 13.78527 | 23.14951 | 8.919039 | -0.19093 | -0.15178 | 0.75249 |
| 1 | 180886600 | 180888599 | 17.13333 | 30.29628 | 34.89723 | 15.43525 | 0.187896 | 0.539624 | 0.7525 |
| 13 | 44283200 | 44285799 | 16.69692 | 29.0175 | 43.21341 | 18.97619 | -0.18485 | 0.78807 | 0.752504 |

| | | | | | | | | | |
|---|---|---|---|---|---|---|---|---|---|
| 9 | 85949400 | 85953399 | 51.77636 | 82.81432 | 95.01039 | 39.98274 | 0.145444 | 2.051281 | 0.752508 |
| 11 | 5497400 | 5501599 | 67.1225 | 106.2883 | 122.7673 | 51.04805 | 0.120923 | 2.373366 | 0.752538 |
| 7 | 14540400 | 14572599 | 885.5078 | 1150.232 | 1102.412 | 461.8562 | 0.122705 | 5.748888 | 0.752549 |
| 14 | 92200400 | 92201399 | 10.97875 | 20.44857 | 24.20608 | 10.09508 | 0.208432 | 0.155725 | 0.752556 |
| 1 | 11735000 | 11735799 | 11.00949 | 20.98469 | 21.10542 | 8.426865 | 0.494061 | -0.00456 | 0.75256 |
| 3 | 138693000 | 138694999 | 23.0928 | 39.04378 | 56.56502 | 24.75325 | -0.15164 | 1.082191 | 0.752601 |
| 2 | 53662400 | 53666399 | 60.39985 | 96.28442 | 110.6447 | 46.34312 | 0.13783 | 2.168361 | 0.752609 |
| 11 | 14818200 | 14818999 | 12.10722 | 22.42493 | 27.1424 | 11.41314 | 0.167414 | 0.20422 | 0.752613 |
| 19 | 61300000 | 61302399 | 17.87132 | 30.98616 | 45.27785 | 19.86579 | -0.15835 | 0.821291 | 0.752615 |
| 7 | 84518600 | 84544399 | 393.7006 | 544.2849 | 656.4408 | 272.6797 | -0.11859 | 4.754825 | 0.752647 |
| 17 | 55064400 | 55069999 | 75.18652 | 118.446 | 135.7332 | 56.32116 | 0.126423 | 2.496274 | 0.752661 |
| 13 | 91680400 | 91683199 | 35.68104 | 58.68192 | 57.44258 | 24.75962 | 0.409183 | 1.411889 | 0.752675 |
| 8 | 117509800 | 117519199 | 89.16248 | 138.2621 | 186.3222 | 77.38688 | -0.12482 | 2.795001 | 0.752684 |
| 6 | 122191400 | 122194399 | 41.09938 | 66.51 | 77.78409 | 33.2665 | 0.137471 | 1.681378 | 0.752699 |
| 10 | 7055800 | 7062599 | 68.19314 | 108.1761 | 123.863 | 51.57806 | 0.135791 | 2.325624 | 0.752715 |
| 11 | 90791600 | 90793999 | 27.94108 | 45.57654 | 79.62495 | 34.20886 | -0.43048 | 1.518639 | 0.75273 |
| 1 | 190871400 | 190876399 | 56.73383 | 90.2645 | 124.2901 | 51.91842 | -0.1257 | 2.229089 | 0.752777 |
| 2 | 23896200 | 23898199 | 20.60053 | 35.77834 | 42.03031 | 18.44913 | 0.155549 | 0.811698 | 0.752789 |
| 10 | 92517000 | 92519399 | 24.31328 | 42.21783 | 45.16266 | 19.82151 | 0.290008 | 0.836033 | 0.752828 |
| 8 | 80957000 | 80959999 | 37.53344 | 61.8089 | 67.50025 | 29.11084 | 0.252455 | 1.406742 | 0.75284 |
| 5 | 144415800 | 144422199 | 59.78214 | 94.77005 | 129.9572 | 54.0795 | -0.12492 | 2.340981 | 0.75285 |
| 9 | 94395400 | 94396999 | 19.45335 | 33.85908 | 39.92452 | 17.54863 | 0.150508 | 0.772987 | 0.75287 |
| 3 | 12897800 | 12915599 | 278.2039 | 399.9664 | 422.178 | 174.2721 | 0.130463 | 4.170614 | 0.752899 |
| 12 | 115254400 | 115282199 | 384.2163 | 540.8241 | 525.088 | 217.504 | 0.220388 | 4.485371 | 0.752904 |
| 17 | 67978200 | 67980999 | 21.03527 | 36.05703 | 52.8033 | 23.15751 | -0.1649 | 0.947164 | 0.752944 |
| 10 | 121082000 | 121085999 | 48.99486 | 78.30696 | 108.9023 | 45.78038 | -0.13289 | 2.07416 | 0.752946 |
| 14 | 88933200 | 88933999 | 12.32126 | 22.05387 | 33.21031 | 14.56381 | -0.18689 | 0.418487 | 0.752947 |
| 1 | 177984800 | 177999799 | 253.8647 | 368.6518 | 387.7929 | 159.8664 | 0.149643 | 4.020899 | 0.752965 |
| Y | 89512200 | 89514399 | 12.66655 | 23.0125 | 21.713 | 9.366317 | 0.498648 | 0.301671 | 0.752987 |
| 8 | 21759800 | 21762799 | 31.38074 | 52.29468 | 50.8511 | 22.09738 | 0.421416 | 1.241129 | 0.753015 |
| 9 | 47984400 | 47985199 | 10.30837 | 19.69325 | 23.89197 | 9.493496 | 0.231204 | -0.02182 | 0.753035 |
| 10 | 23173600 | 23174799 | 11.2277 | 20.64191 | 31.33471 | 13.3099 | -0.16777 | 0.248055 | 0.753035 |
| 9 | 6746800 | 6749399 | 19.33969 | 33.87837 | 30.10119 | 13.29156 | 0.557353 | 0.661758 | 0.75305 |
| 5 | 128142200 | 128144399 | 19.05559 | 33.03622 | 48.47941 | 21.27042 | -0.16455 | 0.832374 | 0.753084 |
| 3 | 102196400 | 102198599 | 24.16988 | 40.95875 | 47.98092 | 21.01141 | 0.153878 | 1.054925 | 0.753085 |
| 6 | 121842200 | 121843199 | 12.18483 | 22.51685 | 27.07059 | 11.41862 | 0.173631 | 0.216131 | 0.753104 |
| 2 | 165208800 | 165210799 | 11.1655 | 20.11039 | 31.25278 | 13.62838 | -0.22649 | 0.36651 | 0.75312 |
| 3 | 152920000 | 152922799 | 37.30197 | 60.34343 | 86.22666 | 36.86758 | -0.15142 | 1.68388 | 0.753133 |
| 16 | 40579600 | 40582799 | 36.1449 | 58.21015 | 98.39068 | 41.82747 | -0.40065 | 1.818889 | 0.753135 |
| 2 | 167364000 | 167366399 | 23.17423 | 39.09855 | 56.60019 | 24.75274 | -0.1507 | 1.098818 | 0.753161 |
| X | 87327600 | 87328999 | 19.16727 | 33.67345 | 37.93672 | 16.79238 | 0.216302 | 0.627627 | 0.753181 |
| 13 | 62726000 | 62754799 | 396.7167 | 554.8376 | 555.0273 | 230.1366 | 0.167973 | 4.580664 | 0.753183 |
| 18 | 18080600 | 18081399 | 9.119043 | 17.04797 | 27.37988 | 11.34709 | -0.22165 | 0.121914 | 0.753202 |
| 10 | 77058800 | 77059599 | 6.102259 | 11.8354 | 21.10222 | 7.96049 | -0.24712 | -0.24166 | 0.753213 |
| 15 | 89649000 | 89650799 | 39.91725 | 64.16216 | 92.61432 | 39.26249 | -0.1768 | 1.883889 | 0.753222 |
| 15 | 19919800 | 19920599 | 23.26762 | 39.49452 | 44.81781 | 19.63602 | 0.19964 | 1.032948 | 0.753234 |
| 7 | 33281400 | 33290399 | 125.8586 | 193.5865 | 197.3136 | 81.94139 | 0.268712 | 2.960892 | 0.753268 |

| | | | | | | | | | |
|---|---|---|---|---|---|---|---|---|---|
| 9 | 103277600 | 103280199 | 32.86356 | 53.57589 | 76.56549 | 32.88772 | -0.1421 | 1.532714 | 0.753281 |
| 8 | 3055800 | 3056399 | 10.04987 | 19.31423 | 17.17029 | 6.708971 | 0.699571 | -0.0944 | 0.753289 |
| 8 | 16968600 | 16970999 | 22.58702 | 39.4161 | 35.50478 | 15.63159 | 0.537638 | 0.746314 | 0.753328 |
| 8 | 15563600 | 15566399 | 27.6249 | 45.89215 | 67.17915 | 29.13194 | -0.16703 | 1.281692 | 0.753337 |
| 4 | 35626000 | 35627399 | 12.0224 | 22.57636 | 21.55258 | 8.864792 | 0.532428 | 0.106036 | 0.753353 |
| 16 | 92027200 | 92029399 | 28.84088 | 47.20039 | 69.26153 | 29.79244 | -0.17606 | 1.466105 | 0.75336 |
| 5 | 113049200 | 113055599 | 164.2378 | 244.9337 | 265.2921 | 109.3364 | 0.143743 | 3.577938 | 0.753371 |
| 16 | 61413400 | 61419999 | 85.52359 | 133.9631 | 151.8766 | 62.99811 | 0.135545 | 2.615471 | 0.753391 |
| 10 | 106505200 | 106505999 | 7.309845 | 14.06115 | 23.61001 | 9.201064 | -0.20614 | -0.11034 | 0.753404 |
| 8 | 122663800 | 122665799 | 17.7869 | 31.06727 | 35.81786 | 15.77829 | 0.183325 | 0.708784 | 0.753406 |
| 4 | 74606200 | 74611399 | 40.48545 | 65.79146 | 75.12042 | 32.22666 | 0.177553 | 1.593446 | 0.753425 |
| 2 | 36675800 | 36677399 | 17.6031 | 31.4798 | 34.29354 | 15.05014 | 0.276466 | 0.459339 | 0.75343 |
| 9 | 93639800 | 93641999 | 33.61751 | 55.05182 | 64.91485 | 27.92974 | 0.138944 | 1.464566 | 0.753445 |
| 13 | 49951000 | 49956399 | 89.95866 | 139.587 | 156.6717 | 65.0656 | 0.139463 | 2.781512 | 0.753447 |
| 12 | 13066800 | 13068599 | 17.11287 | 30.23794 | 34.78546 | 15.38807 | 0.189109 | 0.544464 | 0.753452 |
| 1 | 133342000 | 133344199 | 14.71333 | 25.80015 | 38.36922 | 16.95687 | -0.18208 | 0.634766 | 0.75347 |
| 5 | 119707400 | 119715599 | 88.65418 | 136.1425 | 216.4478 | 89.86772 | -0.37384 | 2.979659 | 0.753475 |
| 14 | 117175600 | 117176799 | 9.399644 | 17.67479 | 28.00076 | 11.49097 | -0.1908 | 0.088575 | 0.753488 |
| 13 | 86921800 | 86922999 | 24.63719 | 42.1415 | 39.56308 | 17.34788 | 0.473703 | 0.965628 | 0.753499 |
| 8 | 45514000 | 45516999 | 34.73347 | 56.24236 | 80.40966 | 34.40479 | -0.15083 | 1.658334 | 0.753515 |
| 14 | 76288400 | 76288999 | 7.463301 | 14.49341 | 25.98822 | 9.731406 | -0.24606 | -0.23689 | 0.753516 |
| 1 | 85003200 | 85019599 | 453.7236 | 623.1612 | 633.8865 | 263.5855 | 0.117454 | 4.856777 | 0.753546 |
| 6 | 60020800 | 60024999 | 55.51235 | 89.13469 | 84.5694 | 35.79585 | 0.425432 | 1.933018 | 0.75355 |
| 1 | 59383400 | 59386799 | 74.48298 | 117.3538 | 135.372 | 56.17228 | 0.117081 | 2.494504 | 0.753574 |
| 5 | 135132600 | 135133999 | 11.13275 | 20.51311 | 31.39474 | 13.29493 | -0.17612 | 0.236453 | 0.753581 |
| 10 | 6688200 | 6689599 | 24.12874 | 40.91167 | 47.98348 | 21.01606 | 0.15227 | 1.052203 | 0.753585 |
| 4 | 134062400 | 134066999 | 50.23895 | 80.63403 | 93.72711 | 39.64715 | 0.132356 | 1.941319 | 0.753608 |
| 3 | 5056600 | 5058399 | 29.21222 | 48.5789 | 57.33249 | 24.87959 | 0.14285 | 1.263931 | 0.753608 |
| 3 | 13295400 | 13296999 | 20.88697 | 35.72746 | 51.69465 | 22.66745 | -0.14801 | 0.96478 | 0.753654 |
| 16 | 58705200 | 58707599 | 23.41642 | 39.65958 | 58.26105 | 25.50885 | -0.17156 | 1.069827 | 0.753655 |
| 4 | 50558400 | 50558999 | 8.561041 | 16.53747 | 21.47448 | 8.254857 | 0.179206 | -0.15142 | 0.753673 |
| X | 64191800 | 64197799 | 75.1578 | 119.0791 | 129.3769 | 53.81172 | 0.209739 | 2.370277 | 0.753706 |
| 3 | 69519200 | 69521999 | 36.55545 | 60.48496 | 54.90026 | 23.7414 | 0.519956 | 1.33945 | 0.753729 |
| 3 | 79168400 | 79173199 | 38.98684 | 63.02009 | 89.79623 | 38.36095 | -0.14855 | 1.709659 | 0.753729 |
| 7 | 118008000 | 118009599 | 15.94012 | 27.87755 | 40.93431 | 18.05012 | -0.16399 | 0.686871 | 0.753739 |
| 9 | 33987200 | 33988399 | 12.95431 | 23.3556 | 35.22462 | 15.34119 | -0.18261 | 0.38192 | 0.753751 |
| 7 | 52193400 | 52193799 | 9.711341 | 18.00524 | 28.38685 | 11.92025 | -0.20942 | 0.184409 | 0.753753 |
| 5 | 27500600 | 27501399 | 17.55328 | 30.70025 | 34.85623 | 15.37059 | 0.205521 | 0.686975 | 0.753789 |
| 7 | 86548200 | 86550599 | 35.77488 | 58.07528 | 85.00774 | 36.44001 | -0.1814 | 1.609482 | 0.75383 |
| 10 | 7108600 | 7111799 | 19.98817 | 34.12371 | 50.61624 | 22.1911 | -0.18421 | 0.975352 | 0.753842 |
| 13 | 12941000 | 12943599 | 59.87928 | 95.58763 | 94.44818 | 39.63627 | 0.357476 | 2.118283 | 0.75391 |
| 11 | 107180000 | 107183199 | 48.40541 | 77.67808 | 90.21219 | 38.15052 | 0.133337 | 1.942616 | 0.753933 |
| 4 | 32967800 | 32970999 | 34.1964 | 55.9496 | 65.61751 | 28.22213 | 0.145965 | 1.47662 | 0.753997 |
| 5 | 138800600 | 138807399 | 64.77694 | 102.2015 | 140.0256 | 58.11715 | -0.12926 | 2.460919 | 0.754011 |
| 3 | 92042400 | 92045599 | 31.226 | 51.02667 | 73.04971 | 31.4046 | -0.14217 | 1.493561 | 0.754023 |
| 10 | 122412600 | 122415199 | 26.75575 | 45.04991 | 52.86342 | 23.10437 | 0.151121 | 1.125605 | 0.754047 |
| 13 | 62163400 | 62164999 | 17.55276 | 30.57847 | 44.66606 | 19.64382 | -0.15701 | 0.757158 | 0.754056 |

| | | | | | | | | | |
|---|---|---|---|---|---|---|---|---|---|
| 9 | 67076800 | 67079399 | 27.5935 | 46.18088 | 54.52177 | 23.75661 | 0.142333 | 1.189056 | 0.754069 |
| 3 | 8765400 | 8768199 | 42.36547 | 67.96388 | 97.07492 | 41.03287 | -0.16509 | 1.949725 | 0.754077 |
| 1 | 164805200 | 164815599 | 265.9915 | 382.4813 | 407.9713 | 168.4027 | 0.115636 | 4.166677 | 0.754079 |
| 19 | 19239000 | 19240799 | 21.64774 | 37.63443 | 43.03365 | 18.90133 | 0.194454 | 0.804563 | 0.754097 |
| 3 | 64486800 | 64487999 | 14.89422 | 26.58824 | 31.39046 | 13.79672 | 0.159619 | 0.446399 | 0.754129 |
| 17 | 42951800 | 42953999 | 23.55737 | 39.52906 | 57.09388 | 24.91838 | -0.14763 | 1.150675 | 0.754166 |
| 16 | 49795000 | 49795799 | 11.09178 | 20.18601 | 30.72066 | 13.22359 | -0.18452 | 0.30352 | 0.754185 |
| 1 | 26464000 | 26465799 | 19.17038 | 33.45538 | 39.3343 | 17.31811 | 0.155056 | 0.733912 | 0.75419 |
| 10 | 116298800 | 116300199 | 19.13078 | 33.3675 | 39.34077 | 17.31484 | 0.150934 | 0.741038 | 0.754224 |
| 6 | 66972200 | 66972599 | 5.655152 | 11.02193 | 19.7009 | 7.131975 | -0.19568 | -0.39692 | 0.754268 |
| 15 | 97301200 | 97303599 | 19.21673 | 33.11088 | 47.94024 | 21.02991 | -0.14704 | 0.887475 | 0.754269 |
| 4 | 154521400 | 154523799 | 26.10489 | 43.30766 | 62.51383 | 27.08322 | -0.14698 | 1.288683 | 0.754282 |
| 13 | 50888800 | 50889999 | 20.06016 | 35.07316 | 32.6243 | 14.38236 | 0.491658 | 0.70109 | 0.754292 |
| 2 | 83377000 | 83380199 | 43.139 | 69.36156 | 96.84245 | 41.07282 | -0.12842 | 1.879242 | 0.754297 |
| 2 | 4526400 | 4527599 | 18.52109 | 32.32213 | 37.85617 | 16.66728 | 0.160635 | 0.728846 | 0.754307 |
| 16 | 19610400 | 19610999 | 10.99539 | 20.6001 | 25.23427 | 10.41982 | 0.170798 | 0.121142 | 0.754313 |
| 13 | 71282400 | 71283999 | 14.4229 | 25.59924 | 38.14138 | 16.84535 | -0.17905 | 0.493401 | 0.754358 |
| 3 | 68242600 | 68244399 | 14.25409 | 25.67213 | 29.78315 | 12.98826 | 0.192129 | 0.382025 | 0.754358 |
| 13 | 10810600 | 10811399 | 10.03497 | 18.79548 | 29.56046 | 12.21199 | -0.18819 | 0.122939 | 0.754396 |
| 10 | 87971200 | 87971799 | 7.125785 | 13.73857 | 23.11188 | 8.947818 | -0.20033 | -0.13726 | 0.754404 |
| 4 | 10450000 | 10453999 | 57.30553 | 90.9744 | 125.2787 | 52.20756 | -0.12896 | 2.296072 | 0.754443 |
| 13 | 31968600 | 31971199 | 19.92574 | 33.83003 | 50.59725 | 22.16976 | -0.1971 | 1.026911 | 0.754446 |
| 3 | 91070400 | 91072199 | 19.72063 | 35.29052 | 34.11616 | 14.96374 | 0.448398 | 0.467951 | 0.754502 |
| 13 | 38468400 | 38472799 | 33.21161 | 53.97495 | 77.42707 | 33.21801 | -0.15075 | 1.579362 | 0.754533 |
| 6 | 104069800 | 104072399 | 27.39691 | 46.68174 | 43.61222 | 19.11744 | 0.479832 | 1.014548 | 0.754534 |
| 9 | 18048200 | 18050399 | 24.13465 | 41.22218 | 47.9373 | 21.01741 | 0.165639 | 0.984526 | 0.754537 |
| 13 | 34355600 | 34359199 | 25.88616 | 42.65263 | 73.45784 | 31.68776 | -0.40249 | 1.376762 | 0.754598 |
| 13 | 64693400 | 64699599 | 70.25512 | 111.3068 | 127.6054 | 53.0738 | 0.132295 | 2.364425 | 0.754599 |
| 18 | 38787600 | 38791799 | 47.70883 | 76.62201 | 88.75347 | 37.5797 | 0.138463 | 1.915623 | 0.754648 |
| 13 | 52409200 | 52414599 | 13.46353 | 23.61479 | 45.09635 | 19.93427 | -0.54131 | 0.632441 | 0.754653 |
| 6 | 68728200 | 68731199 | 41.42354 | 66.72875 | 93.99236 | 39.97288 | -0.13777 | 1.815292 | 0.754653 |
| 5 | 25430400 | 25435799 | 42.85663 | 69.44621 | 79.07918 | 33.85363 | 0.177641 | 1.65759 | 0.754657 |
| 13 | 62716400 | 62719199 | 32.00603 | 52.89981 | 61.93472 | 26.7633 | 0.153667 | 1.344689 | 0.754722 |
| 1 | 10688200 | 10689199 | 13.02459 | 23.96067 | 28.55878 | 12.12029 | 0.179666 | 0.251418 | 0.754734 |
| 11 | 109179200 | 109181399 | 18.06163 | 31.16879 | 45.83149 | 20.10581 | -0.16875 | 0.865478 | 0.754737 |
| 13 | 80637800 | 80639999 | 37.69775 | 60.81513 | 86.33775 | 36.78887 | -0.14654 | 1.76406 | 0.754746 |
| 10 | 45220200 | 45221199 | 12.92164 | 23.25976 | 34.66585 | 15.12781 | -0.16747 | 0.389127 | 0.754793 |
| 6 | 11600600 | 11602199 | 18.11927 | 31.67172 | 37.24596 | 16.41631 | 0.15485 | 0.701239 | 0.754814 |
| 1 | 29776200 | 29776599 | 8.076339 | 15.43433 | 25.08902 | 9.962061 | -0.18545 | -0.03791 | 0.754816 |
| 11 | 105560000 | 105560399 | 5.941632 | 11.5938 | 21.1157 | 7.523492 | -0.20224 | -0.44507 | 0.754827 |
| 8 | 101640800 | 101643999 | 36.74523 | 59.6125 | 85.74255 | 36.73773 | -0.15699 | 1.628161 | 0.754828 |
| 13 | 107747200 | 107750199 | 27.79338 | 46.39341 | 54.03288 | 23.50462 | 0.161884 | 1.214731 | 0.754857 |
| 13 | 82842200 | 82842999 | 17.01408 | 29.02169 | 54.75024 | 24.00189 | -0.5296 | 0.981702 | 0.754861 |
| 6 | 117696000 | 117700399 | 88.23581 | 136.1513 | 184.4588 | 76.62069 | -0.13795 | 2.884743 | 0.754927 |
| 9 | 63592800 | 63595399 | 20.10819 | 33.88652 | 60.16 | 26.30032 | -0.44395 | 1.10515 | 0.754933 |
| 2 | 99319200 | 99322199 | 26.71692 | 45.44491 | 50.683 | 22.21384 | 0.224836 | 1.033371 | 0.754951 |
| 18 | 56084200 | 56086799 | 21.84 | 38.20746 | 41.19481 | 18.16967 | 0.27951 | 0.713599 | 0.754951 |

| | | | | | | | | | |
|---|---|---|---|---|---|---|---|---|---|
| 7 | 124499200 | 124503599 | 60.36422 | 96.31377 | 94.91157 | 39.80265 | 0.360545 | 2.133996 | 0.754957 |
| 3 | 61478000 | 61478999 | 9.95132 | 18.83468 | 31.08562 | 12.59793 | -0.23405 | 0.059022 | 0.754985 |
| 17 | 88585000 | 88588399 | 39.20065 | 63.5638 | 74.50638 | 31.91151 | 0.13687 | 1.634804 | 0.755031 |
| 6 | 56671200 | 56671799 | 9.119553 | 17.57544 | 22.53059 | 8.732931 | 0.186781 | -0.11298 | 0.755049 |
| 2 | 80913000 | 80913799 | 12.23624 | 22.199 | 33.22366 | 14.35728 | -0.16432 | 0.335797 | 0.755063 |
| 2 | 28482600 | 28484999 | 19.27307 | 32.95452 | 48.48663 | 21.26013 | -0.17201 | 0.958424 | 0.755068 |
| 1 | 44372000 | 44372799 | 10.22806 | 19.06852 | 29.44869 | 12.26169 | -0.17106 | 0.154762 | 0.755071 |
| 14 | 39535000 | 39540799 | 60.52761 | 96.1952 | 134.473 | 56.10075 | -0.1491 | 2.267747 | 0.75508 |
| 15 | 74884400 | 74885599 | 12.36234 | 22.23493 | 33.7266 | 14.72989 | -0.19316 | 0.39056 | 0.7551 |
| 17 | 12237600 | 12241199 | 42.03876 | 67.73114 | 96.15997 | 40.90464 | -0.14881 | 1.814409 | 0.755119 |
| 1 | 54335400 | 54338399 | 36.66152 | 60.6777 | 54.85353 | 23.72585 | 0.525826 | 1.337896 | 0.755122 |
| 3 | 87572400 | 87577799 | 66.61421 | 105.8302 | 121.0255 | 50.47229 | 0.139701 | 2.281074 | 0.755148 |
| 18 | 34674000 | 34676199 | 16.07468 | 28.10975 | 41.43748 | 18.27026 | -0.16968 | 0.68947 | 0.755153 |
| 10 | 110835200 | 110838799 | 57.38221 | 91.4712 | 106.0864 | 44.43206 | 0.124521 | 2.16466 | 0.75516 |
| 11 | 92070600 | 92073399 | 49.38219 | 79.33169 | 91.67281 | 38.83199 | 0.142422 | 1.913208 | 0.755177 |
| 14 | 43937000 | 43939799 | 20.81764 | 36.28862 | 41.32303 | 18.18065 | 0.200638 | 0.758468 | 0.755178 |
| 6 | 61431000 | 61433599 | 28.66218 | 48.73897 | 44.15924 | 19.35407 | 0.523542 | 1.039237 | 0.755183 |
| 17 | 88595400 | 88597799 | 33.8519 | 55.87385 | 64.33558 | 27.7744 | 0.177495 | 1.367343 | 0.755189 |
| 3 | 47741000 | 47741999 | 8.699475 | 16.88964 | 17.73519 | 6.654907 | 0.515972 | -0.24145 | 0.755205 |
| 1 | 130362400 | 130365199 | 26.06771 | 44.98125 | 47.35504 | 20.77413 | 0.311225 | 0.902508 | 0.755217 |
| 3 | 23774600 | 23775399 | 8.553733 | 16.25478 | 26.06719 | 10.47458 | -0.18131 | 0.008172 | 0.755282 |
| 5 | 141549200 | 141551399 | 21.89964 | 37.88417 | 44.15309 | 19.37095 | 0.165833 | 0.858504 | 0.755289 |
| 16 | 13340400 | 13341799 | 12.53668 | 22.59487 | 33.63035 | 14.65315 | -0.1642 | 0.375929 | 0.755306 |
| 19 | 11708400 | 11712399 | 42.11126 | 67.69605 | 94.8709 | 40.22541 | -0.13414 | 1.882134 | 0.755325 |
| 1 | 155998600 | 156003199 | 59.22332 | 94.37142 | 108.8989 | 45.58685 | 0.131199 | 2.177074 | 0.755351 |
| X | 157682200 | 157684799 | 32.12124 | 53.8854 | 57.42792 | 25.06013 | 0.289524 | 1.183848 | 0.755355 |
| 4 | 87532400 | 87533199 | 11.4101 | 20.30796 | 37.79412 | 16.65307 | -0.49559 | 0.455835 | 0.755403 |
| X | 8943800 | 8947799 | 48.30963 | 77.87024 | 87.25608 | 37.14062 | 0.192723 | 1.807128 | 0.755405 |
| 16 | 22832800 | 22836399 | 49.2378 | 78.68266 | 109.031 | 45.82413 | -0.12793 | 2.080851 | 0.755414 |
| 4 | 39717400 | 39720399 | 24.34261 | 41.61627 | 47.38259 | 20.77577 | 0.19617 | 0.974696 | 0.755421 |
| 7 | 121365200 | 121367399 | 20.23045 | 34.07141 | 60.21238 | 26.31827 | -0.4375 | 1.109326 | 0.755435 |
| 3 | 69527400 | 69530199 | 63.30756 | 100.3443 | 111.4098 | 46.35466 | 0.179063 | 2.343762 | 0.755476 |
| 3 | 91041200 | 91052799 | 147.4431 | 223.9034 | 234.1267 | 96.95266 | 0.217323 | 3.223129 | 0.755478 |
| X | 7810800 | 7819999 | 96.6098 | 151.347 | 155.109 | 64.3381 | 0.281173 | 2.623056 | 0.755487 |
| 5 | 109973200 | 109987799 | 194.7812 | 290.988 | 292.1162 | 120.4322 | 0.255554 | 3.552258 | 0.755504 |
| 10 | 76644200 | 76646199 | 10.32348 | 18.94235 | 29.30643 | 12.4816 | -0.19742 | 0.248788 | 0.755526 |
| 18 | 87454400 | 87454999 | 10.32348 | 18.94235 | 29.30643 | 12.4816 | -0.19742 | 0.248788 | 0.755526 |
| 8 | 21664200 | 21666799 | 32.23802 | 53.74778 | 51.40857 | 22.34605 | 0.44515 | 1.239823 | 0.755569 |
| X | 146551400 | 146554599 | 42.9618 | 69.82729 | 77.01392 | 33.04302 | 0.227671 | 1.598631 | 0.7556 |
| 18 | 23211400 | 23212999 | 15.50212 | 27.12023 | 39.94235 | 17.61748 | -0.16819 | 0.678958 | 0.755627 |
| 1 | 115487800 | 115490599 | 21.1523 | 37.13302 | 39.50416 | 17.47514 | 0.29872 | 0.654507 | 0.755639 |
| 4 | 119966200 | 119970799 | 65.53576 | 103.1884 | 142.6185 | 59.17411 | -0.14411 | 2.503878 | 0.755658 |
| 11 | 65547400 | 65551999 | 85.92074 | 134.1201 | 153.8658 | 63.84593 | 0.114606 | 2.676126 | 0.755694 |
| 12 | 23431600 | 23434999 | 60.25599 | 96.13977 | 94.03394 | 39.43383 | 0.371309 | 2.134792 | 0.755694 |
| 9 | 88992600 | 88993999 | 16.78208 | 29.4833 | 34.15298 | 15.11762 | 0.176611 | 0.609662 | 0.755728 |
| 9 | 84857000 | 84858799 | 17.80981 | 31.30035 | 36.43706 | 16.13367 | 0.169265 | 0.611129 | 0.755733 |
| 8 | 34201200 | 34206199 | 53.5128 | 85.14065 | 118.2957 | 49.41516 | -0.13863 | 2.225469 | 0.75574 |

| | | | | | | | | | |
|---|---|---|---|---|---|---|---|---|---|
| 14 | 12074800 | 12076399 | 19.32248 | 34.08178 | 31.02924 | 13.73129 | 0.523901 | 0.570446 | 0.755744 |
| 2 | 17904000 | 17905999 | 22.39842 | 38.16175 | 43.80091 | 19.20003 | 0.184065 | 0.997016 | 0.755752 |
| 6 | 130906000 | 130915799 | 144.8902 | 220.3927 | 229.3392 | 95.01742 | 0.22628 | 3.188323 | 0.755811 |
| 4 | 17743600 | 17744999 | 16.57401 | 28.98649 | 42.46655 | 18.72595 | -0.16082 | 0.691979 | 0.755814 |
| 1 | 126518800 | 126523599 | 31.67738 | 52.18133 | 61.22322 | 26.40717 | 0.149446 | 1.382873 | 0.755823 |
| 4 | 14689800 | 14693599 | 37.5098 | 61.38683 | 70.07839 | 30.14364 | 0.185027 | 1.483579 | 0.755828 |
| 10 | 127459600 | 127463199 | 43.53357 | 70.24615 | 82.25904 | 35.06154 | 0.131023 | 1.764627 | 0.755846 |
| 12 | 22955000 | 22956799 | 42.63407 | 68.6582 | 76.87658 | 32.67972 | 0.192158 | 1.818961 | 0.755847 |
| 4 | 17301800 | 17303399 | 16.074 | 28.87874 | 32.83804 | 14.35521 | 0.218777 | 0.417769 | 0.755849 |
| 3 | 153278600 | 153281399 | 22.52716 | 38.097 | 54.93832 | 24.04323 | -0.145 | 1.074905 | 0.755855 |
| 2 | 89546800 | 89549399 | 44.34639 | 71.53403 | 83.53782 | 35.5914 | 0.134415 | 1.775844 | 0.755874 |
| 3 | 8083600 | 8088399 | 80.38222 | 125.903 | 144.5674 | 59.96657 | 0.117147 | 2.612233 | 0.755876 |
| 5 | 145774600 | 145784199 | 177.2014 | 264.6677 | 286.1811 | 117.9803 | 0.148167 | 3.552868 | 0.755889 |
| 4 | 63000600 | 63003199 | 28.94195 | 47.5111 | 68.46275 | 29.4867 | -0.14813 | 1.426306 | 0.755892 |
| 14 | 83628200 | 83628599 | 6.088567 | 11.89164 | 16.48774 | 5.780454 | 0.209067 | -0.51377 | 0.755907 |
| 7 | 103727600 | 103728599 | 18.92545 | 32.97684 | 38.86259 | 17.09379 | 0.151434 | 0.752652 | 0.755915 |
| 18 | 13101400 | 13102599 | 18.9068 | 32.70181 | 47.40347 | 20.79687 | -0.14756 | 0.851624 | 0.755922 |
| 5 | 28229600 | 28231799 | 20.725 | 36.41797 | 38.90533 | 17.22427 | 0.292632 | 0.635032 | 0.75593 |
| 5 | 109349400 | 109360999 | 160.7065 | 242.4416 | 255.7368 | 105.6844 | 0.196715 | 3.358298 | 0.755943 |
| 3 | 3396200 | 3414999 | 308.949 | 438.6885 | 460.9614 | 190.6654 | 0.119036 | 4.352699 | 0.755948 |
| 8 | 21577000 | 21591999 | 196.7427 | 292.2565 | 308.8078 | 127.214 | 0.17422 | 3.653583 | 0.755949 |
| 5 | 133605600 | 133610199 | 36.86023 | 59.68799 | 84.33731 | 36.08613 | -0.13388 | 1.66374 | 0.755957 |
| 19 | 48731600 | 48734799 | 57.20856 | 91.15503 | 105.0961 | 43.99097 | 0.13238 | 2.177964 | 0.755989 |
| 5 | 138543400 | 138548199 | 36.43875 | 60.21479 | 65.70104 | 28.38708 | 0.255077 | 1.363 | 0.755994 |
| 14 | 33407400 | 33408399 | 11.98652 | 22.02134 | 25.6723 | 10.91524 | 0.209979 | 0.246299 | 0.756004 |
| 13 | 86709400 | 86712799 | 47.41458 | 76.18659 | 88.80161 | 37.62709 | 0.13034 | 1.900576 | 0.75606 |
| 17 | 71139400 | 71140199 | 11.82716 | 21.26832 | 32.22322 | 14.07555 | -0.19168 | 0.383724 | 0.756071 |
| 4 | 29185800 | 29187599 | 20.42812 | 34.98967 | 50.60994 | 22.19422 | -0.14717 | 0.950329 | 0.756089 |
| 11 | 44273000 | 44277399 | 42.33902 | 68.30495 | 79.70352 | 33.96353 | 0.135782 | 1.766632 | 0.756113 |
| 7 | 133605800 | 133610399 | 60.57905 | 95.99653 | 131.3746 | 54.64834 | -0.12256 | 2.35267 | 0.756116 |
| 18 | 76483000 | 76490199 | 72.44275 | 113.9181 | 154.5591 | 64.11774 | -0.11888 | 2.534929 | 0.756133 |
| 2 | 131004600 | 131007599 | 33.06304 | 54.44537 | 64.10048 | 27.64424 | 0.1443 | 1.393755 | 0.756156 |
| 19 | 27119600 | 27121399 | 14.71642 | 26.00449 | 38.15186 | 16.87714 | -0.15973 | 0.53779 | 0.756171 |
| 8 | 89118800 | 89122399 | 31.98266 | 51.92116 | 75.66415 | 32.43564 | -0.17482 | 1.59976 | 0.75618 |
| X | 73058600 | 73061599 | 17.90183 | 31.23746 | 47.32186 | 20.83316 | -0.209 | 0.742705 | 0.756192 |
| 18 | 6383600 | 6390399 | 76.83578 | 121.0035 | 138.9795 | 57.66563 | 0.122789 | 2.506324 | 0.756232 |
| 10 | 42347200 | 42349399 | 29.05455 | 48.20396 | 56.73135 | 24.57905 | 0.14663 | 1.28975 | 0.756237 |
| 4 | 121314000 | 121333599 | 312.6907 | 446.7094 | 456.0471 | 188.4401 | 0.16945 | 4.265717 | 0.756238 |
| 12 | 48386800 | 48388799 | 21.01703 | 35.44184 | 61.96093 | 27.09315 | -0.42196 | 1.102091 | 0.756277 |
| X | 149019800 | 149022599 | 44.44659 | 72.18811 | 78.35849 | 33.60191 | 0.249644 | 1.616349 | 0.756278 |
| 4 | 144295400 | 144306799 | 120.9761 | 186.2874 | 193.9376 | 80.55189 | 0.239426 | 2.9362 | 0.756309 |
| 8 | 14080000 | 14082999 | 27.90092 | 45.98178 | 66.40969 | 28.64986 | -0.14917 | 1.372851 | 0.756324 |
| 1 | 16035200 | 16042999 | 123.7959 | 188.7989 | 211.8381 | 87.83138 | 0.120705 | 3.127719 | 0.756359 |
| X | 153429400 | 153432399 | 31.3821 | 52.83943 | 56.3158 | 24.61325 | 0.289506 | 1.145779 | 0.756387 |
| 3 | 106522200 | 106522999 | 10.7161 | 19.55875 | 30.71112 | 13.17179 | -0.22619 | 0.281997 | 0.756444 |
| 18 | 37574800 | 37578799 | 96.42684 | 149.8411 | 146.9568 | 61.01774 | 0.335488 | 2.757555 | 0.756465 |
| 17 | 50662000 | 50663199 | 18.70416 | 32.78868 | 38.11165 | 16.83893 | 0.171584 | 0.66042 | 0.756514 |

| | | | | | | | | | |
|---|---|---|---|---|---|---|---|---|---|
| 2 | 11207000 | 11207999 | 15.06513 | 26.75195 | 31.41961 | 13.87105 | 0.162736 | 0.480508 | 0.756538 |
| 5 | 74573400 | 74574999 | 15.06513 | 26.75195 | 31.41961 | 13.87105 | 0.162736 | 0.480508 | 0.756538 |
| 14 | 6627800 | 6632399 | 50.87562 | 81.82849 | 78.74805 | 33.4149 | 0.407919 | 1.869521 | 0.756556 |
| Y | 43723400 | 43724599 | 10.35239 | 19.38159 | 20.27931 | 8.383915 | 0.395136 | 0.105047 | 0.756563 |
| 5 | 52545200 | 52548999 | 48.70876 | 78.10906 | 90.01368 | 38.02525 | 0.143258 | 1.964674 | 0.756564 |
| 6 | 68911800 | 68912599 | 10.48877 | 19.64302 | 23.23919 | 9.602549 | 0.219984 | 0.114118 | 0.756591 |
| 5 | 53059000 | 53061999 | 27.78564 | 45.32413 | 79.5642 | 34.18307 | -0.43733 | 1.518262 | 0.756597 |
| 5 | 134235800 | 134238399 | 26.74465 | 44.32605 | 63.68368 | 27.5716 | -0.14036 | 1.302075 | 0.756601 |
| 6 | 124569400 | 124570599 | 11.83212 | 21.44733 | 32.66035 | 14.12909 | -0.19041 | 0.334464 | 0.756605 |
| 1 | 177179000 | 177180999 | 16.83479 | 29.25596 | 43.203 | 18.97115 | -0.1728 | 0.788717 | 0.756624 |
| 19 | 3208000 | 3211399 | 58.62503 | 92.87018 | 129.0288 | 53.66268 | -0.14468 | 2.358638 | 0.756638 |
| 10 | 30392400 | 30396599 | 32.34971 | 52.93377 | 76.32982 | 32.83076 | -0.15126 | 1.478564 | 0.756644 |
| 7 | 29335200 | 29344799 | 27.67291 | 47.13495 | 43.64078 | 19.1294 | 0.492769 | 1.020226 | 0.756669 |
| 17 | 13731000 | 13737199 | 63.93853 | 101.6826 | 117.2499 | 48.94785 | 0.129189 | 2.248116 | 0.75667 |
| 9 | 15107600 | 15109799 | 36.56887 | 59.18042 | 83.42443 | 35.67964 | -0.1313 | 1.675828 | 0.756671 |
| 11 | 119496800 | 119500399 | 26.97757 | 44.56273 | 64.60851 | 27.90746 | -0.15377 | 1.343454 | 0.756674 |
| 15 | 52529200 | 52530999 | 16.34475 | 28.75073 | 33.53161 | 14.84678 | 0.167079 | 0.587789 | 0.756675 |
| 2 | 139914000 | 139915799 | 14.35528 | 25.33493 | 37.14803 | 16.43715 | -0.15961 | 0.546596 | 0.756692 |
| 3 | 39458000 | 39459199 | 17.65021 | 30.95777 | 35.44285 | 15.6669 | 0.193089 | 0.63836 | 0.756712 |
| 14 | 95515000 | 95517199 | 28.71642 | 48.31775 | 46.95017 | 20.51563 | 0.422276 | 1.130862 | 0.75672 |
| 2 | 71597200 | 71600199 | 22.81157 | 38.56446 | 56.06623 | 24.53424 | -0.15676 | 1.078982 | 0.756732 |
| 1 | 113804200 | 113806399 | 14.57732 | 26.47576 | 23.44355 | 10.11675 | 0.589601 | 0.324725 | 0.756734 |
| 9 | 115569400 | 115575199 | 76.34134 | 119.9243 | 137.6159 | 57.08129 | 0.121318 | 2.554788 | 0.756749 |
| 15 | 36394000 | 36394999 | 13.05508 | 24.12387 | 28.6344 | 12.07903 | 0.191926 | 0.227558 | 0.756755 |
| 1 | 174810400 | 174811399 | 12.68913 | 23.73759 | 27.79603 | 11.50692 | 0.232026 | 0.149441 | 0.756776 |
| 6 | 7021800 | 7025999 | 74.34685 | 118.3086 | 102.8594 | 42.97721 | 0.536914 | 2.234939 | 0.756786 |
| 7 | 139606800 | 139609799 | 38.53702 | 62.51891 | 73.32731 | 31.41807 | 0.136642 | 1.622845 | 0.756843 |
| 17 | 43775600 | 43779199 | 23.66237 | 39.80931 | 57.70875 | 25.21589 | -0.15287 | 1.125605 | 0.756872 |
| 14 | 25333000 | 25338199 | 47.79848 | 76.90741 | 89.33995 | 37.92949 | 0.137609 | 1.85995 | 0.756891 |
| 19 | 9267200 | 9268799 | 12.43118 | 22.32776 | 33.1501 | 14.49504 | -0.16384 | 0.396267 | 0.75691 |
| 10 | 91196800 | 91198399 | 12.4188 | 22.34608 | 33.14109 | 14.46893 | -0.16079 | 0.384575 | 0.756913 |
| 2 | 153640600 | 153641199 | 10.70025 | 20.06483 | 24.25237 | 9.999428 | 0.191861 | 0.111206 | 0.756915 |
| X | 153403600 | 153407599 | 44.51136 | 72.6641 | 74.69729 | 32.10127 | 0.333413 | 1.539139 | 0.756971 |
| 4 | 22070600 | 22074199 | 29.01736 | 48.94387 | 54.6115 | 23.88111 | 0.223585 | 1.118982 | 0.756985 |
| 16 | 70171400 | 70173799 | 26.91476 | 46.03901 | 49.57926 | 21.73991 | 0.276367 | 0.983512 | 0.757027 |
| 6 | 76472400 | 76476399 | 86.50321 | 133.2764 | 184.8787 | 76.79503 | -0.17365 | 2.912321 | 0.757036 |
| 14 | 94594800 | 94596199 | 15.81827 | 28.49288 | 32.30444 | 14.08555 | 0.225339 | 0.398412 | 0.757037 |
| 13 | 61934000 | 61944199 | 110.0786 | 169.9836 | 184.7438 | 76.73879 | 0.180504 | 2.878759 | 0.757063 |
| 1 | 128856200 | 128856799 | 11.58695 | 21.33142 | 23.25866 | 9.861182 | 0.308795 | 0.22479 | 0.757067 |
| 3 | 21206800 | 21220599 | 256.0728 | 369.6294 | 396.2416 | 163.4737 | 0.113871 | 4.11004 | 0.757073 |
| 12 | 102994200 | 102994799 | 8.050497 | 15.2865 | 24.98246 | 10.05535 | -0.21067 | 0.004031 | 0.757074 |
| 17 | 32941000 | 32942799 | 17.90218 | 30.95474 | 45.29961 | 19.87367 | -0.16127 | 0.846569 | 0.75708 |
| 10 | 119663600 | 119667599 | 42.16866 | 67.95956 | 96.47324 | 41.05452 | -0.14816 | 1.80527 | 0.757095 |
| 4 | 10693800 | 10695199 | 18.63928 | 32.10668 | 46.89457 | 20.57102 | -0.15956 | 0.887974 | 0.757097 |
| 8 | 85965800 | 85966199 | 8.189009 | 15.84244 | 20.4642 | 7.811844 | 0.197095 | -0.17907 | 0.75711 |
| X | 5992800 | 5995999 | 88.062 | 137.1975 | 155.0903 | 64.36995 | 0.133726 | 2.712303 | 0.757136 |
| 4 | 114121000 | 114124399 | 76.30565 | 120.1005 | 138.6024 | 57.50494 | 0.115279 | 2.516608 | 0.757153 |

| | | | | | | | | | |
|---|---|---|---|---|---|---|---|---|---|
| 3 | 66793200 | 66794999 | 24.03133 | 40.40849 | 58.24588 | 25.44648 | -0.14472 | 1.132893 | 0.757165 |
| 5 | 83876400 | 83876999 | 8.48844 | 16.37961 | 20.93897 | 8.086922 | 0.195906 | -0.13542 | 0.757209 |
| 12 | 115148200 | 115151999 | 51.22376 | 82.69106 | 87.43564 | 37.2889 | 0.278697 | 1.767779 | 0.757211 |
| 7 | 39435600 | 39438999 | 24.50892 | 41.21353 | 59.95893 | 26.19525 | -0.15807 | 1.13521 | 0.757219 |
| 11 | 104704000 | 104707399 | 89.11543 | 137.8393 | 184.1802 | 76.50403 | -0.11536 | 2.841153 | 0.75722 |
| 1 | 19370200 | 19371199 | 14.31144 | 24.98737 | 45.21484 | 19.91302 | -0.46392 | 0.71235 | 0.757231 |
| X | 18250600 | 18255599 | 61.9573 | 99.00128 | 108.8278 | 45.73393 | 0.205455 | 2.097206 | 0.757259 |
| 4 | 9996000 | 9996399 | 5.93948 | 11.59782 | 22.02079 | 7.750603 | -0.24608 | -0.48614 | 0.757265 |
| 1 | 35242800 | 35246599 | 51.38684 | 82.29265 | 95.90064 | 40.4292 | 0.12487 | 2.012358 | 0.757278 |
| 5 | 146296200 | 146296799 | 9.977868 | 18.86504 | 23.25266 | 9.444525 | 0.181466 | 0.04689 | 0.757293 |
| 8 | 74559200 | 74560199 | 14.49431 | 26.41541 | 24.89745 | 10.69819 | 0.504071 | 0.308491 | 0.757313 |
| 10 | 96357200 | 96358999 | 25.11386 | 42.49467 | 50.12216 | 21.93911 | 0.144015 | 1.075293 | 0.757323 |
| 17 | 50066600 | 50071199 | 42.4346 | 68.28746 | 95.47042 | 40.54122 | -0.12882 | 1.849938 | 0.757336 |
| 7 | 18593600 | 18598399 | 58.47168 | 93.16786 | 108.2844 | 45.32567 | 0.120779 | 2.179112 | 0.757345 |
| 9 | 26522200 | 26524999 | 24.43936 | 41.03645 | 59.30353 | 25.89369 | -0.14842 | 1.149946 | 0.757355 |
| X | 88575000 | 88577999 | 25.45711 | 44.40437 | 43.36831 | 19.08817 | 0.421438 | 0.778033 | 0.757364 |
| 1 | 32857400 | 32857799 | 5.497787 | 10.72003 | 19.72674 | 7.099484 | -0.22851 | -0.41648 | 0.757405 |
| 1 | 114865000 | 114869399 | 90.59636 | 141.6132 | 139.375 | 57.82485 | 0.337016 | 2.652151 | 0.757424 |
| 10 | 6587800 | 6588599 | 12.13728 | 22.77851 | 22.49716 | 9.262645 | 0.482373 | 0.112379 | 0.757441 |
| 4 | 74060000 | 74061199 | 31.65133 | 52.54085 | 50.30281 | 21.80206 | 0.443596 | 1.286496 | 0.757449 |
| 1 | 102682600 | 102683799 | 13.00988 | 24.21433 | 23.47067 | 9.800248 | 0.493241 | 0.175288 | 0.757452 |
| 14 | 41242200 | 41253399 | 205.8921 | 304.7489 | 321.7147 | 132.4917 | 0.170736 | 3.717002 | 0.757465 |
| 6 | 18824000 | 18825599 | 21.39245 | 36.76596 | 42.09037 | 18.46231 | 0.189528 | 0.909267 | 0.757482 |
| 13 | 34576800 | 34580599 | 21.72495 | 36.97563 | 53.30342 | 23.36346 | -0.14382 | 1.015246 | 0.757495 |
| 3 | 98158800 | 98164199 | 60.21935 | 95.66562 | 131.8384 | 54.97881 | -0.12912 | 2.279297 | 0.757544 |
| 6 | 146094400 | 146096199 | 28.88262 | 48.71929 | 54.59747 | 23.87538 | 0.217358 | 1.11847 | 0.757553 |
| 14 | 84628000 | 84629799 | 22.10394 | 38.12262 | 44.71233 | 19.6145 | 0.155918 | 0.887386 | 0.757556 |
| 13 | 10943400 | 10947599 | 57.08571 | 91.03663 | 105.5403 | 44.22891 | 0.125746 | 2.151325 | 0.757566 |
| 13 | 120120200 | 120123799 | 39.91123 | 64.60513 | 75.9595 | 32.49279 | 0.130179 | 1.672143 | 0.757583 |
| 11 | 63566400 | 63567799 | 18.33611 | 32.0433 | 37.78497 | 16.65061 | 0.151023 | 0.707791 | 0.757685 |
| X | 120677800 | 120678999 | 14.60532 | 26.76934 | 30.12016 | 12.84752 | 0.257413 | 0.290654 | 0.75769 |
| 18 | 59346200 | 59350199 | 40.48559 | 66.64099 | 68.87337 | 29.69748 | 0.331571 | 1.421691 | 0.757692 |
| 8 | 52811400 | 52813799 | 40.28816 | 65.14852 | 76.27677 | 32.60151 | 0.134842 | 1.695645 | 0.757731 |
| 5 | 145963600 | 145976199 | 159.6479 | 241.2878 | 250.6285 | 103.6334 | 0.221187 | 3.32189 | 0.757744 |
| 15 | 97381800 | 97386999 | 60.40614 | 96.06447 | 135.4358 | 56.53938 | -0.16042 | 2.2496 | 0.757775 |
| 12 | 101878800 | 101880199 | 9.319364 | 17.17831 | 28.80468 | 12.19179 | -0.30665 | 0.210355 | 0.757795 |
| 17 | 20076800 | 20085799 | 125.7304 | 194.0117 | 158.0665 | 65.65785 | 0.595045 | 2.897129 | 0.757833 |
| 9 | 14050800 | 14053199 | 17.36731 | 30.20784 | 44.17407 | 19.40824 | -0.15884 | 0.777421 | 0.757849 |
| 2 | 28296000 | 28297199 | 10.82746 | 19.71465 | 30.27987 | 13.02586 | -0.19684 | 0.299317 | 0.757854 |
| 18 | 79968600 | 79969999 | 14.04617 | 24.73183 | 36.72295 | 16.25661 | -0.17891 | 0.569722 | 0.757856 |
| 6 | 34381800 | 34386199 | 43.11868 | 69.45799 | 99.12732 | 42.1554 | -0.1567 | 1.821464 | 0.757858 |
| X | 33861400 | 33863199 | 19.36179 | 34.22405 | 36.84042 | 16.29595 | 0.284887 | 0.560787 | 0.757869 |
| 17 | 73180600 | 73184599 | 44.49201 | 71.77869 | 83.67087 | 35.65434 | 0.137258 | 1.771878 | 0.757891 |
| 10 | 52827000 | 52828599 | 23.06606 | 39.74531 | 35.45151 | 15.55151 | 0.549022 | 0.887247 | 0.7579 |
| 10 | 42761200 | 42763399 | 16.58149 | 29.21316 | 33.86775 | 14.99089 | 0.17637 | 0.570978 | 0.757909 |
| 7 | 57930400 | 57931599 | 15.23939 | 26.88183 | 39.72096 | 17.57736 | -0.17093 | 0.563161 | 0.757921 |
| 6 | 70591000 | 70595399 | 55.32022 | 88.0821 | 120.8323 | 50.52694 | -0.11912 | 2.203811 | 0.757934 |

| | | | | | | | | | |
|---|---|---|---|---|---|---|---|---|---|
| 2 | 89700200 | 89707399 | 74.61598 | 118.1235 | 132.2434 | 54.98112 | 0.165597 | 2.387386 | 0.757937 |
| 4 | 153426000 | 153432399 | 68.67647 | 108.64 | 125.3834 | 52.10943 | 0.120925 | 2.399286 | 0.757937 |
| 6 | 60504600 | 60506799 | 22.69259 | 39.6851 | 41.12335 | 18.13278 | 0.336423 | 0.721737 | 0.757943 |
| 6 | 7928800 | 7931199 | 22.66434 | 39.27923 | 37.83366 | 16.59941 | 0.440287 | 0.83789 | 0.757949 |
| X | 163871600 | 163872599 | 12.48541 | 22.49725 | 33.70206 | 14.68864 | -0.17345 | 0.379669 | 0.757985 |
| 5 | 118999000 | 119003799 | 60.90622 | 96.62015 | 131.8829 | 54.91962 | -0.11728 | 2.321333 | 0.757994 |
| 5 | 104704400 | 104706599 | 17.3686 | 31.01851 | 34.26335 | 15.05402 | 0.255359 | 0.465242 | 0.75801 |
| 5 | 12850600 | 12852199 | 18.62654 | 32.33879 | 47.93854 | 21.03796 | -0.1788 | 0.813301 | 0.75801 |
| 17 | 81076600 | 81078399 | 21.29421 | 36.66878 | 43.18697 | 18.94445 | 0.149526 | 0.896362 | 0.758017 |
| 19 | 7823600 | 7826199 | 53.25912 | 85.49136 | 96.86332 | 40.98102 | 0.16939 | 1.943402 | 0.758063 |
| 7 | 25672400 | 25690799 | 236.358 | 345.2975 | 366.2109 | 150.875 | 0.146231 | 3.928911 | 0.758091 |
| 12 | 72684200 | 72692799 | 244.7651 | 352.335 | 440.8987 | 181.964 | -0.11315 | 4.150568 | 0.7581 |
| 7 | 6150800 | 6152999 | 14.23412 | 24.99915 | 37.32005 | 16.51514 | -0.18753 | 0.604818 | 0.758107 |
| 3 | 47367800 | 47370199 | 21.19226 | 37.36697 | 37.72799 | 16.69682 | 0.375354 | 0.597249 | 0.75813 |
| 13 | 21291200 | 21296199 | 40.70143 | 65.26847 | 111.1245 | 46.94317 | -0.41874 | 1.962485 | 0.758141 |
| 4 | 78914000 | 78914399 | 5.376529 | 10.4802 | 19.14417 | 6.919092 | -0.22465 | -0.4095 | 0.758162 |
| 7 | 37359800 | 37364799 | 33.73517 | 54.68563 | 78.75903 | 33.72572 | -0.1599 | 1.633027 | 0.758179 |
| 18 | 53514800 | 53517599 | 23.01379 | 38.80823 | 56.03888 | 24.5035 | -0.14711 | 1.102131 | 0.758181 |
| 6 | 145654800 | 145656399 | 15.68988 | 27.65025 | 31.58645 | 13.98032 | 0.197917 | 0.560116 | 0.758185 |
| 16 | 93463400 | 93468399 | 62.16054 | 99.15156 | 111.9599 | 46.93354 | 0.164002 | 2.150334 | 0.758245 |
| 14 | 22366800 | 22367399 | 8.833866 | 16.59925 | 26.41453 | 10.85292 | -0.19812 | 0.085742 | 0.758249 |
| 1 | 15035000 | 15039399 | 65.8927 | 104.047 | 141.4 | 58.7119 | -0.11662 | 2.442299 | 0.758256 |
| 9 | 37388600 | 37389999 | 20.43399 | 35.32049 | 41.60225 | 18.25139 | 0.150506 | 0.856323 | 0.758265 |
| 15 | 25479400 | 25480999 | 17.90609 | 31.14644 | 45.23059 | 19.87303 | -0.1488 | 0.781862 | 0.758311 |
| 8 | 39427400 | 39428199 | 12.69761 | 23.48462 | 28.11777 | 11.84647 | 0.180766 | 0.218148 | 0.758394 |
| 9 | 120570600 | 120575799 | 46.97079 | 75.40652 | 86.88417 | 36.76583 | 0.145439 | 1.927793 | 0.758396 |
| 1 | 66036400 | 66038999 | 34.46983 | 56.72541 | 65.82966 | 28.38223 | 0.164683 | 1.407713 | 0.758396 |
| 5 | 78875400 | 78875999 | 9.779146 | 17.80434 | 33.91142 | 14.58966 | -0.50544 | 0.299927 | 0.758397 |
| 1 | 25987600 | 25987999 | 6.238088 | 12.15486 | 21.46417 | 7.797237 | -0.18344 | -0.37155 | 0.758401 |
| 13 | 13209000 | 13213199 | 164.737 | 245.9744 | 272.1904 | 112.2057 | 0.114419 | 3.55551 | 0.758405 |
| 12 | 119860000 | 119860799 | 11.79725 | 21.91879 | 26.67244 | 11.16824 | 0.164171 | 0.183621 | 0.758416 |
| 6 | 137026400 | 137028399 | 36.12637 | 58.69097 | 67.88326 | 29.11581 | 0.158678 | 1.588161 | 0.758423 |
| X | 114293200 | 114294599 | 36.85949 | 59.55909 | 86.71068 | 37.02495 | -0.18003 | 1.719858 | 0.75848 |
| 16 | 94219800 | 94222799 | 17.61867 | 31.41426 | 29.08393 | 12.79582 | 0.507397 | 0.467391 | 0.758511 |
| 15 | 29886400 | 29888199 | 21.04923 | 35.83197 | 51.6776 | 22.65118 | -0.14439 | 1.008918 | 0.758513 |
| 5 | 109834800 | 109836399 | 13.98819 | 25.01856 | 38.20574 | 16.76517 | -0.20818 | 0.440651 | 0.758515 |
| 17 | 9851200 | 9852199 | 9.616339 | 18.03372 | 27.95799 | 11.52576 | -0.1648 | 0.109331 | 0.758533 |
| 1 | 188991600 | 188993999 | 24.08211 | 40.26869 | 58.16039 | 25.32798 | -0.1475 | 1.191099 | 0.758544 |
| 14 | 121100200 | 121104399 | 58.05535 | 92.21696 | 126.5336 | 52.76038 | -0.123 | 2.279401 | 0.758578 |
| Y | 10365600 | 10369599 | 38.00488 | 61.53031 | 56.46383 | 24.15643 | 0.48707 | 1.650609 | 0.758578 |
| 4 | 10749000 | 10749999 | 13.8205 | 24.86187 | 29.04623 | 12.68392 | 0.181286 | 0.388068 | 0.758579 |
| 11 | 16397800 | 16398399 | 9.133859 | 17.22652 | 26.98241 | 11.01706 | -0.1683 | 0.066322 | 0.7586 |
| 4 | 75296000 | 75296999 | 13.39397 | 23.73357 | 35.10965 | 15.52273 | -0.16998 | 0.493 | 0.758614 |
| 4 | 39823000 | 39825199 | 15.77066 | 28.36663 | 27.21603 | 11.88591 | 0.463424 | 0.39691 | 0.758617 |
| 7 | 17394200 | 17472999 | 1703.777 | 2105.135 | 1908.967 | 803.946 | 0.129802 | 6.558806 | 0.758625 |
| 17 | 40436800 | 40438199 | 19.78343 | 34.5943 | 33.01789 | 14.55764 | 0.45455 | 0.696314 | 0.758636 |
| 1 | 29819200 | 29821599 | 24.87788 | 42.85777 | 47.36535 | 20.77761 | 0.24093 | 0.911669 | 0.758657 |

| | | | | | | | | | |
|---|---|---|---|---|---|---|---|---|---|
| 1 | 101139800 | 101141399 | 10.71149 | 20.41912 | 17.07878 | 6.817417 | 0.758198 | -0.02053 | 0.758665 |
| 3 | 116754200 | 116755999 | 14.75592 | 26.02531 | 39.13673 | 17.31932 | -0.19648 | 0.557148 | 0.758711 |
| 14 | 4395000 | 4398799 | 50.68078 | 81.30159 | 94.37637 | 39.89076 | 0.133408 | 1.957097 | 0.758713 |
| 14 | 62635800 | 62637599 | 17.16193 | 30.20604 | 35.06031 | 15.52181 | 0.174452 | 0.591306 | 0.75872 |
| 6 | 137056400 | 137058999 | 14.9834 | 26.20049 | 56.07722 | 24.72436 | -0.70532 | 0.707148 | 0.758722 |
| 6 | 38279400 | 38283199 | 42.63047 | 68.67268 | 97.11794 | 41.30186 | -0.14354 | 1.819634 | 0.758727 |
| 15 | 59032800 | 59034999 | 34.11755 | 55.89401 | 66.12921 | 28.45725 | 0.134351 | 1.461467 | 0.758737 |
| 13 | 106100200 | 106102399 | 28.04702 | 47.31088 | 54.01467 | 23.62068 | 0.190612 | 1.114834 | 0.758783 |
| 17 | 70363600 | 70363999 | 12.14117 | 21.7807 | 32.70006 | 14.31274 | -0.18069 | 0.401634 | 0.758807 |
| 12 | 103737000 | 103743599 | 88.24937 | 136.5945 | 182.5563 | 75.8292 | -0.11491 | 2.827139 | 0.758816 |
| 13 | 41730000 | 41730999 | 14.78264 | 27.21799 | 25.28294 | 10.71763 | 0.538283 | 0.253829 | 0.758823 |
| 8 | 89327600 | 89329399 | 21.33921 | 37.04276 | 43.05418 | 18.89263 | 0.170712 | 0.822206 | 0.758834 |
| 14 | 43062000 | 43067799 | 60.90911 | 96.96684 | 112.3206 | 46.95833 | 0.124275 | 2.209887 | 0.758854 |
| 8 | 124152000 | 124162999 | 674.1417 | 893.2522 | 845.9321 | 353.6294 | 0.168554 | 5.426191 | 0.758855 |
| 7 | 86152600 | 86154999 | 39.66216 | 64.5197 | 63.37007 | 27.19836 | 0.39541 | 1.570537 | 0.758889 |
| 7 | 84624000 | 84626799 | 33.45824 | 55.09833 | 64.63223 | 27.87403 | 0.149613 | 1.395785 | 0.758906 |
| 2 | 89288000 | 89288999 | 11.54754 | 21.29763 | 33.02689 | 13.96803 | -0.19395 | 0.236059 | 0.758909 |
| 6 | 7699600 | 7700599 | 12.50535 | 22.4489 | 33.70738 | 14.74541 | -0.18027 | 0.403077 | 0.758913 |
| 4 | 49619800 | 49621999 | 35.73953 | 57.97389 | 81.98402 | 35.12419 | -0.13272 | 1.626504 | 0.758918 |
| 6 | 148667800 | 148669399 | 22.44194 | 38.31154 | 44.81349 | 19.64702 | 0.157284 | 0.981064 | 0.758924 |
| 14 | 94168200 | 94168799 | 8.563141 | 16.23693 | 26.02945 | 10.50817 | -0.18664 | 0.023611 | 0.75893 |
| 5 | 105526200 | 105528599 | 19.2538 | 33.05981 | 47.9489 | 21.03145 | -0.1505 | 0.919543 | 0.758931 |
| 7 | 85308400 | 85326999 | 185.3891 | 277.4945 | 247.1638 | 101.9378 | 0.430546 | 3.512702 | 0.758945 |
| 12 | 52222600 | 52223399 | 22.92821 | 39.00159 | 45.35654 | 19.87936 | 0.164851 | 1.015952 | 0.758955 |
| 10 | 65907600 | 65909599 | 17.51778 | 30.62276 | 34.86263 | 15.36688 | 0.2016 | 0.694344 | 0.758957 |
| 3 | 93737600 | 93764199 | 485.0714 | 665.2124 | 667.0039 | 277.4225 | 0.135993 | 4.881246 | 0.758983 |
| X | 117042800 | 117045999 | 23.32053 | 39.79909 | 63.46983 | 27.82559 | -0.28852 | 1.007943 | 0.758992 |
| 13 | 105058400 | 105059399 | 7.86987 | 15.09919 | 24.64516 | 9.676276 | -0.17651 | -0.08009 | 0.758997 |
| 7 | 29829600 | 29867999 | 933.1272 | 1202.517 | 1348.766 | 565.5041 | -0.11556 | 5.878062 | 0.759014 |
| 11 | 63636400 | 63638999 | 22.4728 | 39.00262 | 44.09856 | 19.34895 | 0.210286 | 0.831046 | 0.759024 |
| 2 | 151312400 | 151314999 | 16.88885 | 29.40589 | 43.11233 | 18.95402 | -0.16236 | 0.75913 | 0.759027 |
| 6 | 138843000 | 138845599 | 28.81065 | 48.4178 | 55.16514 | 24.09647 | 0.193339 | 1.149234 | 0.75903 |
| 8 | 69797000 | 69827199 | 537.1418 | 733.1471 | 704.0498 | 293.0482 | 0.191399 | 4.956561 | 0.75904 |
| 6 | 124218000 | 124221799 | 42.43439 | 69.5191 | 71.85284 | 30.92025 | 0.328787 | 1.484242 | 0.759052 |
| 13 | 50962800 | 50963399 | 7.88655 | 15.07928 | 24.55495 | 9.736455 | -0.18636 | -0.04739 | 0.759064 |
| 7 | 62454800 | 62457399 | 17.45864 | 31.26319 | 27.91457 | 12.23686 | 0.562668 | 0.435986 | 0.759072 |
| 12 | 51590200 | 51602799 | 221.0745 | 320.8198 | 405.9602 | 167.3675 | -0.11772 | 4.028409 | 0.759077 |
| 11 | 37888000 | 37889199 | 10.6344 | 19.45624 | 29.77595 | 12.73179 | -0.18605 | 0.268374 | 0.759082 |
| 18 | 51308000 | 51309599 | 15.47236 | 27.58181 | 32.41688 | 14.26461 | 0.165035 | 0.462197 | 0.759083 |
| 16 | 51012800 | 51014399 | 14.12312 | 25.80258 | 29.9595 | 12.8336 | 0.207568 | 0.300591 | 0.759086 |
| 8 | 72713600 | 72715999 | 17.7178 | 30.87915 | 45.18605 | 19.87784 | -0.15954 | 0.752641 | 0.759115 |
| 6 | 13382800 | 13387399 | 51.82638 | 83.51193 | 77.50004 | 32.97357 | 0.463532 | 1.814854 | 0.759133 |
| 17 | 23370400 | 23372599 | 28.03793 | 47.51135 | 52.38047 | 22.93769 | 0.241013 | 1.07241 | 0.759136 |
| 18 | 42760400 | 42763199 | 38.95846 | 63.12461 | 74.13388 | 31.73474 | 0.133157 | 1.650629 | 0.759192 |
| 16 | 37297200 | 37298399 | 10.96053 | 19.45339 | 38.31067 | 16.92047 | -0.5791 | 0.470982 | 0.759214 |
| 9 | 40140800 | 40144799 | 31.53116 | 51.78817 | 75.31515 | 32.44506 | -0.16117 | 1.42972 | 0.759228 |
| 3 | 57986800 | 57990599 | 50.97174 | 82.13964 | 90.02516 | 38.30585 | 0.224114 | 1.817667 | 0.759239 |

| | | | | | | | | | |
|---|---|---|---|---|---|---|---|---|---|
| 6 | 142016000 | 142020199 | 50.53326 | 80.81335 | 111.9534 | 47.10232 | -0.12638 | 2.059864 | 0.759257 |
| 3 | 151719400 | 151722799 | 67.96234 | 107.5825 | 124.6827 | 51.83581 | 0.115672 | 2.382278 | 0.759295 |
| 15 | 35717400 | 35720599 | 64.17209 | 102.2688 | 113.6126 | 47.56754 | 0.186128 | 2.183227 | 0.759305 |
| 8 | 57503800 | 57504599 | 10.8301 | 20.47438 | 25.35149 | 10.29909 | 0.174666 | 0.061427 | 0.759335 |
| 18 | 24025600 | 24028799 | 22.57922 | 38.53917 | 45.3233 | 19.87025 | 0.149468 | 0.983228 | 0.759336 |
| 9 | 17551600 | 17552999 | 15.53825 | 28.49192 | 31.08555 | 13.25127 | 0.301859 | 0.29417 | 0.759341 |
| 1 | 26667000 | 26668799 | 21.982 | 38.02837 | 44.13296 | 19.36215 | 0.171898 | 0.857274 | 0.759347 |
| X | 162256200 | 162257199 | 11.4203 | 21.6111 | 25.92792 | 10.5117 | 0.222651 | 0.063003 | 0.75937 |
| 8 | 4010400 | 4012199 | 26.4693 | 44.37079 | 52.30373 | 22.81331 | 0.144566 | 1.167105 | 0.759374 |
| 2 | 64618400 | 64619999 | 19.396 | 33.80329 | 39.84484 | 17.52795 | 0.151109 | 0.753239 | 0.7594 |
| 3 | 130710400 | 130714399 | 55.18289 | 88.13479 | 102.7085 | 43.13343 | 0.120603 | 2.100696 | 0.75941 |
| 8 | 88352200 | 88353799 | 15.1377 | 26.66384 | 39.17189 | 17.3394 | -0.16359 | 0.575313 | 0.759422 |
| 17 | 17593200 | 17595199 | 12.34277 | 21.85962 | 40.82091 | 18.04966 | -0.50469 | 0.502015 | 0.75945 |
| 13 | 50860800 | 50874599 | 339.0133 | 472.8934 | 579.8084 | 240.5099 | -0.12931 | 4.617546 | 0.759452 |
| 5 | 30283800 | 30285999 | 30.08831 | 49.6413 | 57.7481 | 24.92476 | 0.162672 | 1.358321 | 0.759463 |
| 1 | 173620600 | 173624599 | 60.60955 | 96.4582 | 110.9643 | 46.37019 | 0.133722 | 2.222361 | 0.759487 |
| 10 | 120952600 | 120953599 | 10.80923 | 19.64402 | 30.25913 | 13.04798 | -0.20332 | 0.309365 | 0.759538 |
| 5 | 69496200 | 69499399 | 27.16351 | 44.19896 | 81.6037 | 35.02745 | -0.5125 | 1.552796 | 0.75954 |
| 4 | 144287400 | 144290799 | 49.64761 | 79.86619 | 90.67906 | 38.48653 | 0.170146 | 1.867608 | 0.759546 |
| 10 | 36884000 | 36884599 | 7.742816 | 14.95494 | 24.89565 | 9.575823 | -0.1766 | -0.14661 | 0.759547 |
| 16 | 30860400 | 30860999 | 7.742816 | 14.95494 | 24.89565 | 9.575823 | -0.1766 | -0.14661 | 0.759547 |
| 14 | 44590200 | 44592399 | 21.74192 | 37.54922 | 44.17283 | 19.37863 | 0.151962 | 0.872611 | 0.759567 |
| 18 | 37554200 | 37558199 | 83.13461 | 129.9528 | 148.6763 | 61.68369 | 0.120138 | 2.65033 | 0.759581 |
| X | 18404600 | 18406199 | 19.58529 | 34.08458 | 39.48062 | 17.35219 | 0.176236 | 0.776937 | 0.759607 |
| 18 | 90387800 | 90392799 | 73.68765 | 115.6687 | 156.8655 | 65.05942 | -0.12023 | 2.572312 | 0.759612 |
| 4 | 70512800 | 70513599 | 17.28918 | 29.91593 | 44.27517 | 19.42466 | -0.17715 | 0.836906 | 0.75963 |
| 6 | 97059000 | 97060199 | 12.16132 | 22.32924 | 35.68245 | 15.18356 | -0.24361 | 0.270085 | 0.759637 |
| 19 | 44456000 | 44460399 | 49.58148 | 79.57918 | 92.94959 | 39.31855 | 0.125401 | 1.940061 | 0.759641 |
| 1 | 161368000 | 161370199 | 23.05778 | 39.6211 | 46.31112 | 20.31358 | 0.159713 | 0.925546 | 0.759676 |
| 5 | 115887800 | 115890599 | 28.74123 | 47.23395 | 68.01631 | 29.30816 | -0.14652 | 1.410915 | 0.759676 |
| 13 | 87990200 | 87991999 | 20.71122 | 36.03444 | 41.9465 | 18.43288 | 0.168834 | 0.786259 | 0.759679 |
| 1 | 40993400 | 40994799 | 12.40534 | 22.36611 | 33.1484 | 14.43672 | -0.15756 | 0.372775 | 0.759688 |
| 13 | 66308600 | 66309199 | 15.10625 | 26.9968 | 31.90646 | 14.01035 | 0.159087 | 0.442646 | 0.759691 |
| 5 | 135290000 | 135291999 | 16.01874 | 28.39153 | 33.30627 | 14.72469 | 0.162812 | 0.502668 | 0.759695 |
| 4 | 121263600 | 121264199 | 10.88591 | 20.58058 | 25.31872 | 10.28505 | 0.183692 | 0.059884 | 0.759699 |
| 8 | 98629400 | 98630199 | 10.52698 | 19.75026 | 23.75552 | 9.785898 | 0.199913 | 0.104933 | 0.759701 |
| 1 | 67144400 | 67148599 | 63.11386 | 99.8677 | 135.9224 | 56.49723 | -0.11633 | 2.390759 | 0.759721 |
| 14 | 42908000 | 42911399 | 45.86786 | 73.93675 | 85.90181 | 36.56571 | 0.140652 | 1.79995 | 0.759725 |
| X | 18587200 | 18589199 | 12.30179 | 22.22283 | 33.20948 | 14.428 | -0.16698 | 0.362302 | 0.759733 |
| 12 | 91843200 | 91844199 | 13.93735 | 25.49718 | 23.43793 | 10.02281 | 0.544381 | 0.278758 | 0.759736 |
| 5 | 34918200 | 34920999 | 20.82525 | 35.19928 | 52.17414 | 22.82967 | -0.18436 | 1.072896 | 0.759747 |
| 4 | 70582200 | 70583599 | 13.56154 | 24.69819 | 29.49036 | 12.68281 | 0.163985 | 0.314753 | 0.759747 |
| 4 | 115480000 | 115480999 | 12.62677 | 23.00888 | 27.09979 | 11.64641 | 0.184524 | 0.300336 | 0.759765 |
| 1 | 16174200 | 16176999 | 20.33856 | 33.55545 | 75.78115 | 32.70796 | -0.79147 | 1.345978 | 0.759774 |
| 4 | 49466000 | 49469199 | 85.34841 | 131.9621 | 179.3549 | 74.49987 | -0.14032 | 2.845315 | 0.759777 |
| 6 | 16389400 | 16392399 | 36.22038 | 58.57762 | 82.72041 | 35.36102 | -0.13473 | 1.68763 | 0.75979 |
| 3 | 53018000 | 53020799 | 22.7158 | 39.61057 | 36.02387 | 15.85252 | 0.523741 | 0.75787 | 0.759791 |

| | | | | | | | | | |
|---|---|---|---|---|---|---|---|---|---|
| 1 | 191108800 | 191110199 | 13.95669 | 25.68128 | 29.56277 | 12.54248 | 0.229753 | 0.258841 | 0.759801 |
| 6 | 50487600 | 50489799 | 17.14387 | 29.87676 | 44.64697 | 19.63981 | -0.1898 | 0.74807 | 0.75983 |
| 13 | 63001800 | 63004999 | 34.5644 | 55.9486 | 80.35517 | 34.37242 | -0.15787 | 1.664847 | 0.759848 |
| 7 | 105911800 | 105914599 | 26.2336 | 44.50258 | 43.20344 | 18.92531 | 0.423987 | 1.04865 | 0.759863 |
| 1 | 72678600 | 72681399 | 33.6083 | 54.55291 | 77.62613 | 33.2734 | -0.14071 | 1.60683 | 0.759863 |
| 16 | 59051200 | 59052599 | 19.95441 | 34.76546 | 40.30813 | 17.72823 | 0.174676 | 0.759675 | 0.759869 |
| 1 | 61382800 | 61385599 | 33.20103 | 55.28803 | 61.00879 | 26.49929 | 0.239593 | 1.263481 | 0.759875 |
| 18 | 21400400 | 21403799 | 26.12892 | 44.36473 | 42.69759 | 18.7088 | 0.436544 | 1.040996 | 0.759887 |
| 9 | 23592600 | 23596799 | 43.82674 | 70.46947 | 98.75105 | 41.88412 | -0.13365 | 1.880519 | 0.759895 |
| 18 | 15061200 | 15061999 | 14.65197 | 25.35259 | 50.04027 | 21.95395 | -0.59076 | 0.831606 | 0.759927 |
| 19 | 5455800 | 5459199 | 17.76555 | 30.64141 | 45.32452 | 19.88308 | -0.17739 | 0.868933 | 0.759928 |
| 1 | 101976400 | 101977599 | 26.61925 | 44.69458 | 52.84903 | 23.07476 | 0.140044 | 1.150615 | 0.75995 |
| 11 | 8098000 | 8100399 | 21.06982 | 35.89198 | 51.6762 | 22.65184 | -0.14178 | 1.003186 | 0.759972 |
| 5 | 106755600 | 106757199 | 18.43321 | 32.06716 | 35.99644 | 15.81699 | 0.221193 | 0.769512 | 0.760012 |
| 13 | 94196000 | 94197599 | 11.89867 | 21.4819 | 32.15163 | 13.97874 | -0.17004 | 0.358616 | 0.760031 |
| 10 | 77943600 | 77945399 | 23.92645 | 39.87652 | 58.66294 | 25.49367 | -0.17389 | 1.225872 | 0.760033 |
| 7 | 30583600 | 30584799 | 14.57867 | 26.08628 | 30.91939 | 13.56273 | 0.15596 | 0.428956 | 0.760076 |
| 16 | 89562600 | 89571599 | 108.6648 | 165.7544 | 220.8644 | 91.57788 | -0.12687 | 3.119763 | 0.760095 |
| 6 | 52323800 | 52330599 | 89.22865 | 139.1689 | 158.7197 | 65.86565 | 0.122172 | 2.692613 | 0.760103 |
| 14 | 115450400 | 115452399 | 11.60389 | 20.47101 | 40.38317 | 17.87004 | -0.58543 | 0.54031 | 0.760145 |
| 11 | 27703000 | 27704199 | 10.12995 | 18.68198 | 28.81135 | 12.18616 | -0.18571 | 0.216724 | 0.760159 |
| 7 | 13218800 | 13247399 | 438.9982 | 607.2757 | 611.8846 | 254.147 | 0.141613 | 4.747232 | 0.760207 |
| 15 | 87115600 | 87118999 | 31.96496 | 51.97333 | 74.76564 | 32.08368 | -0.15419 | 1.569499 | 0.760226 |
| 6 | 117313800 | 117314199 | 6.848872 | 13.29858 | 22.85052 | 8.564959 | -0.18577 | -0.24823 | 0.760249 |
| 13 | 10720200 | 10723399 | 32.07402 | 52.54653 | 75.66631 | 32.56143 | -0.14837 | 1.463292 | 0.760285 |
| 7 | 61494000 | 61497799 | 61.58852 | 98.228 | 97.20632 | 40.74244 | 0.353878 | 2.149303 | 0.760297 |
| 11 | 11718800 | 11720399 | 13.17332 | 23.27706 | 35.30204 | 15.61656 | -0.20696 | 0.525643 | 0.760317 |
| 12 | 53168800 | 53169999 | 14.87626 | 27.19958 | 25.7117 | 11.00334 | 0.50333 | 0.293316 | 0.760317 |
| 13 | 94047400 | 94049399 | 14.37516 | 25.41513 | 37.12703 | 16.42202 | -0.15324 | 0.527746 | 0.760335 |
| X | 122965000 | 122966999 | 26.00332 | 44.92065 | 39.24289 | 17.21601 | 0.579625 | 0.883227 | 0.760337 |
| 15 | 33106200 | 33109799 | 55.84796 | 89.17479 | 103.9633 | 43.64501 | 0.119617 | 2.109414 | 0.760348 |
| 12 | 9116200 | 9119399 | 29.3185 | 48.37248 | 70.19 | 30.29516 | -0.15533 | 1.364121 | 0.760354 |
| 6 | 104937600 | 104938199 | 9.589838 | 17.94747 | 27.91621 | 11.54825 | -0.17365 | 0.121096 | 0.760361 |
| X | 101289800 | 101296799 | 75.34506 | 119.2638 | 130.4906 | 54.24917 | 0.198546 | 2.392716 | 0.760396 |
| 19 | 35739000 | 35739799 | 14.95545 | 26.64682 | 31.45548 | 13.84748 | 0.158341 | 0.459822 | 0.760399 |
| 7 | 80821200 | 80824799 | 47.20212 | 75.69161 | 105.3544 | 44.50449 | -0.12867 | 1.969681 | 0.760433 |
| X | 164168400 | 164170599 | 23.47111 | 40.45183 | 46.30011 | 20.31059 | 0.190785 | 0.901908 | 0.760447 |
| 12 | 72683400 | 72692799 | 250.8317 | 360.3923 | 449.7157 | 185.6574 | -0.1118 | 4.178312 | 0.760474 |
| 10 | 93362800 | 93364599 | 22.59343 | 38.17073 | 54.90488 | 24.02128 | -0.14134 | 1.085248 | 0.760515 |
| 4 | 127729000 | 127730999 | 19.58998 | 33.97883 | 38.93192 | 17.08169 | 0.191034 | 0.815226 | 0.76052 |
| 5 | 7836200 | 7837799 | 15.6537 | 28.15 | 32.29731 | 14.10767 | 0.206598 | 0.406935 | 0.7606 |
| 6 | 143152200 | 143164999 | 176.8416 | 261.7378 | 337.9067 | 139.16 | -0.11951 | 3.713544 | 0.760605 |
| 3 | 100770600 | 100772999 | 27.544 | 46.5817 | 45.30745 | 19.82843 | 0.421104 | 1.081594 | 0.760607 |
| 2 | 111212400 | 111213799 | 14.28784 | 26.39647 | 29.64064 | 12.50696 | 0.27102 | 0.239454 | 0.760614 |
| 9 | 27877200 | 27877999 | 11.15898 | 21.36473 | 25.15728 | 9.937145 | 0.281998 | -0.02868 | 0.760622 |
| 19 | 37610600 | 37618399 | 115.208 | 177.9573 | 161.5404 | 67.10045 | 0.440346 | 2.873811 | 0.760642 |
| 2 | 50962200 | 50965799 | 23.0236 | 39.53229 | 46.3113 | 20.31314 | 0.156242 | 0.931669 | 0.760669 |

| | | | | | | | | | |
|---|---|---|---|---|---|---|---|---|---|
| 17 | 88622400 | 88628399 | 42.40143 | 68.09412 | 96.49292 | 40.85303 | -0.1519 | 1.913521 | 0.760676 |
| 13 | 91960400 | 91963799 | 50.15676 | 80.47598 | 93.85222 | 39.68047 | 0.127076 | 1.952533 | 0.760688 |
| 5 | 38568800 | 38573399 | 73.24502 | 115.6265 | 133.6162 | 55.47263 | 0.11669 | 2.451889 | 0.760688 |
| 1 | 177533800 | 177534599 | 7.899983 | 15.15523 | 24.64015 | 9.677473 | -0.17107 | -0.07732 | 0.760707 |
| 7 | 24389400 | 24390199 | 7.899983 | 15.15523 | 24.64015 | 9.677473 | -0.17107 | -0.07732 | 0.760707 |
| Y | 41485800 | 41488199 | 17.55812 | 30.65589 | 29.49697 | 12.9907 | 0.44283 | 0.700228 | 0.760711 |
| 1 | 173088200 | 173091999 | 47.73759 | 76.50188 | 106.1653 | 44.80394 | -0.12545 | 1.990947 | 0.760721 |
| 10 | 19613200 | 19615599 | 17.49928 | 30.62226 | 46.18122 | 20.37382 | -0.20241 | 0.694652 | 0.760726 |
| 4 | 132241000 | 132243399 | 16.10996 | 28.39909 | 32.91288 | 14.56653 | 0.177222 | 0.560368 | 0.760727 |
| 18 | 83426800 | 83428999 | 20.01643 | 34.82532 | 40.90903 | 17.97688 | 0.155768 | 0.781154 | 0.760727 |
| 6 | 18736200 | 18737199 | 15.7264 | 27.85573 | 32.88419 | 14.53994 | 0.153501 | 0.507542 | 0.760739 |
| 4 | 89096800 | 89101999 | 59.07237 | 93.93965 | 129.6783 | 54.13345 | -0.13011 | 2.248275 | 0.76074 |
| 7 | 131872600 | 131875199 | 26.61018 | 45.33412 | 50.68098 | 22.21575 | 0.221745 | 1.019766 | 0.760765 |
| 7 | 12902400 | 12910999 | 206.6063 | 302.6806 | 324.0093 | 133.4645 | 0.136688 | 3.880941 | 0.760767 |
| 2 | 37197200 | 37199999 | 28.01713 | 47.37979 | 53.46913 | 23.40001 | 0.207413 | 1.092405 | 0.760828 |
| 16 | 3516800 | 3519799 | 22.50329 | 37.91473 | 55.48376 | 24.25444 | -0.16634 | 1.106457 | 0.760829 |
| 2 | 169753800 | 169755799 | 21.24602 | 35.10619 | 69.57131 | 30.05767 | -0.60311 | 1.328387 | 0.760855 |
| 15 | 37127800 | 37136599 | 87.69353 | 136.748 | 155.8311 | 64.6686 | 0.123132 | 2.693963 | 0.760867 |
| 17 | 8125000 | 8130599 | 100.2478 | 153.0278 | 211.2676 | 87.61362 | -0.17713 | 3.103863 | 0.760871 |
| 13 | 64931200 | 64932799 | 18.81587 | 33.28268 | 36.22552 | 16.02168 | 0.26921 | 0.546363 | 0.760885 |
| Y | 40413000 | 40413799 | 9.05136 | 16.33665 | 30.44091 | 13.23973 | -0.48405 | 0.336113 | 0.760888 |
| 1 | 190881000 | 190897799 | 292.1388 | 409.3585 | 609.4329 | 252.6203 | -0.40268 | 4.548257 | 0.760911 |
| 17 | 71447000 | 71449999 | 21.51058 | 36.57024 | 52.76421 | 23.12506 | -0.14533 | 1.022789 | 0.760925 |
| 3 | 5145600 | 5147999 | 51.84115 | 83.16867 | 95.94752 | 40.55902 | 0.14247 | 1.959498 | 0.760936 |
| 3 | 100226400 | 100228399 | 18.42504 | 32.32833 | 37.49443 | 16.57876 | 0.174584 | 0.641906 | 0.760943 |
| 13 | 63309000 | 63318199 | 102.7647 | 156.6731 | 214.3132 | 88.85085 | -0.16526 | 3.128623 | 0.760948 |
| 1 | 104182200 | 104184599 | 23.74511 | 41.34969 | 44.017 | 19.35355 | 0.297491 | 0.79329 | 0.760963 |
| 13 | 66982800 | 66984399 | 40.34604 | 65.23872 | 75.52405 | 32.27763 | 0.151048 | 1.698136 | 0.760997 |
| 2 | 98137600 | 98138199 | 10.67167 | 19.93644 | 23.25839 | 9.650615 | 0.235335 | 0.131952 | 0.761027 |
| 10 | 60388800 | 60390199 | 16.37132 | 28.84362 | 33.98866 | 15.04434 | 0.153166 | 0.571572 | 0.761036 |
| 10 | 129223800 | 129226999 | 35.11163 | 57.44096 | 67.86573 | 29.18797 | 0.135264 | 1.48336 | 0.761063 |
| 11 | 31159600 | 31165799 | 111.6427 | 171.0477 | 193.9024 | 80.48032 | 0.111109 | 3.030553 | 0.761099 |
| 18 | 62866800 | 62868399 | 15.38634 | 27.38979 | 32.3573 | 14.25534 | 0.156346 | 0.468855 | 0.761101 |
| 10 | 67143200 | 67145799 | 17.4376 | 30.22284 | 44.25427 | 19.41647 | -0.16134 | 0.822507 | 0.761105 |
| 10 | 74517800 | 74519399 | 15.44798 | 27.44798 | 32.41603 | 14.30382 | 0.155335 | 0.482085 | 0.761121 |
| 10 | 124145800 | 124149199 | 73.62647 | 115.9234 | 133.4724 | 55.37903 | 0.118998 | 2.506902 | 0.761173 |
| 4 | 16162000 | 16164399 | 21.8278 | 36.17348 | 68.18931 | 29.52033 | -0.53091 | 1.293513 | 0.761177 |
| Y | 4680200 | 4682799 | 23.17757 | 39.26579 | 37.84438 | 16.57105 | 0.434328 | 1.040854 | 0.761178 |
| 17 | 42225200 | 42225799 | 13.83154 | 24.84193 | 28.60221 | 12.50689 | 0.200966 | 0.397493 | 0.761191 |
| 5 | 109028200 | 109029799 | 32.18966 | 52.93785 | 62.09051 | 26.76259 | 0.149072 | 1.408543 | 0.761205 |
| 7 | 126884200 | 126887799 | 50.75265 | 81.37854 | 95.11653 | 40.17481 | 0.122734 | 1.97304 | 0.761248 |
| 18 | 26785400 | 26785799 | 4.641533 | 9.070484 | 17.12201 | 5.935548 | -0.21578 | -0.57275 | 0.761256 |
| 1 | 72739600 | 72740199 | 7.501733 | 14.55914 | 18.88826 | 7.103741 | 0.210636 | -0.24483 | 0.761276 |
| 15 | 96391400 | 96394399 | 34.77984 | 56.4877 | 80.18445 | 34.38487 | -0.13645 | 1.599454 | 0.761284 |
| 16 | 90178200 | 90180199 | 20.544 | 34.58817 | 60.78966 | 26.56796 | -0.42963 | 1.11344 | 0.761294 |
| 2 | 56233000 | 56233599 | 7.133674 | 13.88492 | 18.48701 | 6.796202 | 0.20485 | -0.32661 | 0.761312 |
| X | 81328400 | 81328999 | 9.994081 | 19.26913 | 23.68287 | 9.16823 | 0.248801 | -0.10377 | 0.76132 |

| | | | | | | | | | |
|---|---|---|---|---|---|---|---|---|---|
| 4 | 140669800 | 140674199 | 64.32828 | 101.3816 | 140.051 | 58.11588 | -0.14235 | 2.483466 | 0.761337 |
| 14 | 80156200 | 80157399 | 10.6004 | 19.39758 | 29.75136 | 12.71813 | -0.18908 | 0.265966 | 0.761339 |
| 5 | 95149200 | 95149599 | 12.96949 | 24.41987 | 22.3236 | 9.14056 | 0.599039 | 0.097495 | 0.761347 |
| 3 | 11063800 | 11065999 | 15.16749 | 26.73832 | 39.17657 | 17.33862 | -0.15925 | 0.567175 | 0.761347 |
| 9 | 56919400 | 56920799 | 15.16749 | 26.73832 | 39.17657 | 17.33862 | -0.15925 | 0.567175 | 0.761347 |
| 17 | 39863400 | 39870599 | 70.13581 | 111.5915 | 120.7024 | 50.42239 | 0.222087 | 2.241699 | 0.761351 |
| 7 | 43129800 | 43134999 | 39.84593 | 64.05123 | 91.82297 | 38.93028 | -0.16686 | 1.879688 | 0.761351 |
| 3 | 17040400 | 17049799 | 157.0527 | 235.347 | 261.7533 | 107.9788 | 0.111825 | 3.487203 | 0.761357 |
| 1 | 166321000 | 166323599 | 19.70192 | 33.73023 | 49.01141 | 21.49241 | -0.15386 | 0.948085 | 0.76136 |
| 16 | 17207400 | 17208599 | 15.43669 | 27.66494 | 32.3794 | 14.18439 | 0.175353 | 0.428314 | 0.761361 |
| 19 | 44704800 | 44706599 | 14.5956 | 25.47301 | 39.61409 | 17.44188 | -0.24629 | 0.711512 | 0.761379 |
| 3 | 11747200 | 11747799 | 7.561087 | 14.68307 | 19.47658 | 7.294031 | 0.185141 | -0.25459 | 0.761387 |
| 13 | 35609400 | 35610599 | 13.11883 | 24.02653 | 28.45975 | 12.15187 | 0.182152 | 0.275739 | 0.761403 |
| 13 | 51195000 | 51203799 | 145.7989 | 218.5597 | 283.9731 | 117.1529 | -0.11188 | 3.479534 | 0.761414 |
| 3 | 92391600 | 92391999 | 5.952765 | 11.56619 | 20.62137 | 7.688517 | -0.2315 | -0.28698 | 0.761417 |
| 11 | 50997400 | 50998599 | 13.84272 | 24.40148 | 36.66631 | 16.22783 | -0.19547 | 0.554461 | 0.761418 |
| 6 | 102419200 | 102420199 | 9.867705 | 18.49186 | 29.01909 | 11.97772 | -0.18401 | 0.116696 | 0.761432 |
| 11 | 110071000 | 110073799 | 21.36185 | 36.64099 | 42.69157 | 18.72388 | 0.163885 | 0.929737 | 0.761433 |
| 17 | 18602800 | 18609799 | 275.1317 | 392.9049 | 400.0655 | 165.3188 | 0.172691 | 4.267603 | 0.761441 |
| 13 | 80982600 | 80984199 | 24.34828 | 41.74597 | 47.92566 | 21.01877 | 0.184974 | 0.951916 | 0.761451 |
| 2 | 129363000 | 129363599 | 11.96051 | 22.06274 | 26.62993 | 11.26017 | 0.165719 | 0.224978 | 0.761457 |
| 1 | 8207200 | 8211799 | 62.64956 | 99.86042 | 113.501 | 47.53266 | 0.153473 | 2.173716 | 0.761467 |
| 3 | 10667600 | 10670399 | 35.17151 | 57.45361 | 67.35401 | 28.95222 | 0.144929 | 1.502154 | 0.761509 |
| 15 | 21465400 | 21466999 | 15.05592 | 27.32748 | 31.45218 | 13.58366 | 0.212739 | 0.348887 | 0.761532 |
| 5 | 25138800 | 25140999 | 23.91954 | 40.30771 | 58.22664 | 25.45478 | -0.14767 | 1.115082 | 0.761546 |
| 1 | 83047600 | 83048199 | 8.371485 | 16.12489 | 27.15124 | 10.5405 | -0.2058 | -0.10389 | 0.761551 |
| 17 | 94793800 | 94794199 | 9.53274 | 18.15598 | 22.83376 | 9.131765 | 0.172366 | -0.00813 | 0.761552 |
| 11 | 104763200 | 104763999 | 7.428302 | 14.3383 | 23.70042 | 9.143365 | -0.17044 | -0.14341 | 0.761557 |
| 5 | 21439000 | 21442199 | 31.57307 | 52.11619 | 61.79322 | 26.67812 | 0.135381 | 1.359378 | 0.761593 |
| X | 153040000 | 153043999 | 49.18869 | 79.58485 | 83.75193 | 35.81354 | 0.28915 | 1.704166 | 0.761605 |
| 17 | 72761600 | 72765799 | 63.44497 | 100.7413 | 117.0443 | 48.78585 | 0.116151 | 2.29268 | 0.761639 |
| 5 | 101200000 | 101209599 | 87.38959 | 135.4574 | 181.0177 | 75.18833 | -0.11317 | 2.801408 | 0.761645 |
| 18 | 5667400 | 5669199 | 20.55714 | 35.55892 | 42.08574 | 18.46391 | 0.143725 | 0.850345 | 0.761682 |
| 5 | 62033000 | 62038999 | 87.17358 | 135.8424 | 155.6265 | 64.59045 | 0.114682 | 2.707337 | 0.761705 |
| 12 | 103849000 | 103852399 | 35.0227 | 57.60167 | 66.58661 | 28.70152 | 0.169913 | 1.414783 | 0.76171 |
| 17 | 26461800 | 26464799 | 23.69651 | 39.36799 | 68.55888 | 29.73125 | -0.41679 | 1.269653 | 0.761735 |
| 2 | 87059200 | 87070399 | 125.1322 | 191.2538 | 210.7605 | 87.42871 | 0.149156 | 3.0847 | 0.761736 |
| 1 | 175172600 | 175174799 | 18.25006 | 31.51351 | 45.83066 | 20.10582 | -0.1527 | 0.861512 | 0.761801 |
| 13 | 44093600 | 44095799 | 25.06216 | 42.34827 | 50.13205 | 21.93419 | 0.138666 | 1.086122 | 0.761811 |
| 9 | 11844200 | 11847799 | 44.14443 | 70.87359 | 98.67586 | 41.75961 | -0.12696 | 1.927672 | 0.761813 |
| 6 | 136664000 | 136673999 | 110.9222 | 168.1103 | 264.5675 | 109.4928 | -0.37439 | 3.254671 | 0.761847 |
| 12 | 117243400 | 117243999 | 9.699039 | 18.17859 | 28.50163 | 11.76106 | -0.18171 | 0.117114 | 0.761889 |
| 10 | 23281800 | 23285199 | 46.2736 | 74.72858 | 84.45673 | 36.03627 | 0.183442 | 1.745484 | 0.761891 |
| 17 | 92281000 | 92282799 | 16.42023 | 29.3302 | 33.21254 | 14.59008 | 0.219768 | 0.454464 | 0.76191 |
| 6 | 7100000 | 7108599 | 145.6207 | 219.1645 | 244.1986 | 100.8498 | 0.114761 | 3.400459 | 0.761915 |
| 12 | 104596000 | 104597599 | 17.57712 | 31.00672 | 35.91043 | 15.891 | 0.178697 | 0.569101 | 0.761919 |
| 5 | 84011000 | 84011599 | 8.889026 | 17.02327 | 21.34032 | 8.422985 | 0.194163 | -0.06155 | 0.761929 |

| | | | | | | | | | |
|---|---|---|---|---|---|---|---|---|---|
| 10 | 11725000 | 11727799 | 22.86238 | 38.76307 | 56.05618 | 24.55151 | -0.14891 | 1.055449 | 0.76195 |
| 5 | 71049600 | 71050199 | 11.65039 | 22.09013 | 20.90031 | 8.438042 | 0.567101 | 0.034841 | 0.761969 |
| 18 | 74516400 | 74520999 | 49.31802 | 79.62542 | 86.66376 | 36.96619 | 0.237283 | 1.759444 | 0.761972 |
| 6 | 116533000 | 116534399 | 13.67747 | 24.75556 | 29.41214 | 12.7475 | 0.163849 | 0.350034 | 0.761976 |
| 6 | 86645600 | 86646799 | 10.08136 | 18.27154 | 29.87769 | 12.92732 | -0.29185 | 0.320015 | 0.762016 |
| X | 62012400 | 62015199 | 22.96325 | 40.42359 | 30.78322 | 13.6278 | 0.779352 | 0.614186 | 0.762018 |
| 13 | 34893400 | 34895999 | 19.00826 | 33.3146 | 38.60712 | 17.05474 | 0.175854 | 0.665829 | 0.762021 |
| X | 80476400 | 80478799 | 26.17944 | 44.82981 | 49.56718 | 21.73651 | 0.238816 | 0.974185 | 0.762037 |
| 8 | 60682800 | 60685599 | 24.90622 | 42.72468 | 47.91679 | 21.01575 | 0.218596 | 0.9482 | 0.762045 |
| 2 | 111730200 | 111730999 | 19.61883 | 33.50386 | 49.01317 | 21.48884 | -0.16404 | 0.972304 | 0.762048 |
| 13 | 38581200 | 38583399 | 21.92718 | 36.9651 | 53.80473 | 23.52452 | -0.15848 | 1.099943 | 0.762051 |
| 18 | 56487000 | 56490799 | 43.48786 | 70.07843 | 81.91638 | 34.85474 | 0.131357 | 1.802601 | 0.76206 |
| 1 | 188815600 | 188818399 | 40.28747 | 65.47476 | 75.42342 | 32.35848 | 0.164956 | 1.59395 | 0.762074 |
| 6 | 130348400 | 130350199 | 16.26721 | 29.39966 | 32.31229 | 14.0302 | 0.273752 | 0.382032 | 0.762078 |
| 10 | 119195800 | 119199399 | 41.27584 | 66.73354 | 77.41276 | 33.0793 | 0.1478 | 1.699989 | 0.7621 |
| 1 | 127028600 | 127031399 | 27.16551 | 46.17317 | 51.78208 | 22.69419 | 0.216741 | 1.04434 | 0.762127 |
| 13 | 13438000 | 13439399 | 23.3189 | 39.6141 | 45.89227 | 20.11178 | 0.170142 | 1.03103 | 0.762147 |
| 17 | 44632200 | 44634799 | 22.72149 | 39.2822 | 44.64145 | 19.58485 | 0.201796 | 0.865782 | 0.762148 |
| 1 | 108488600 | 108488999 | 7.42248 | 14.45377 | 19.62663 | 7.187216 | 0.181252 | -0.33416 | 0.76216 |
| 4 | 151983200 | 151984999 | 18.75637 | 32.28162 | 46.89249 | 20.56959 | -0.15199 | 0.894849 | 0.762161 |
| 4 | 57913200 | 57915599 | 22.17204 | 38.13481 | 44.75149 | 19.63003 | 0.154381 | 0.913404 | 0.762162 |
| 13 | 91481600 | 91482799 | 19.92695 | 34.68991 | 40.92034 | 17.98846 | 0.149917 | 0.773596 | 0.762171 |
| 13 | 43911600 | 43914599 | 24.03306 | 40.72918 | 47.98252 | 21.01247 | 0.145774 | 1.054392 | 0.762178 |
| 17 | 83722400 | 83729999 | 104.8637 | 163.1249 | 171.5005 | 71.19722 | 0.236521 | 2.748451 | 0.762212 |
| 9 | 61123000 | 61124399 | 13.06396 | 23.1586 | 34.61103 | 15.30099 | -0.18442 | 0.484889 | 0.762238 |
| 7 | 10395000 | 10457399 | 943.0241 | 1229.164 | 1121.845 | 469.8356 | 0.198198 | 5.694144 | 0.762239 |
| 7 | 13171800 | 13194199 | 287.432 | 412.76 | 431.7192 | 178.2427 | 0.141991 | 4.188408 | 0.762259 |
| X | 156212600 | 156214999 | 17.0584 | 29.86279 | 44.57586 | 19.67228 | -0.18756 | 0.684795 | 0.762265 |
| 19 | 31831400 | 31834599 | 58.08331 | 92.30443 | 126.1034 | 52.60559 | -0.11607 | 2.266243 | 0.762298 |
| 7 | 91719000 | 91721799 | 9.44729 | 17.25107 | 33.42378 | 14.3268 | -0.5265 | 0.278366 | 0.762302 |
| 1 | 106260200 | 106261199 | 9.917464 | 18.12728 | 33.95402 | 14.53635 | -0.47693 | 0.278088 | 0.762305 |
| 18 | 3479200 | 3481599 | 18.23461 | 31.76138 | 46.8057 | 20.58281 | -0.16976 | 0.765948 | 0.762318 |
| 11 | 116682400 | 116686799 | 40.72619 | 65.77603 | 76.73066 | 32.74362 | 0.138053 | 1.731018 | 0.762326 |
| 12 | 60645200 | 60645999 | 8.857872 | 16.78544 | 26.54983 | 10.73222 | -0.16928 | 0.031261 | 0.762326 |
| 17 | 66243200 | 66244799 | 16.20865 | 28.34084 | 41.94591 | 18.49269 | -0.17549 | 0.692066 | 0.762349 |
| 4 | 101661000 | 101661599 | 11.34311 | 20.31296 | 31.77486 | 13.9286 | -0.24102 | 0.404632 | 0.762363 |
| 1 | 7127600 | 7137599 | 197.4728 | 292.2327 | 317.4447 | 130.7329 | 0.129277 | 3.718284 | 0.762384 |
| 4 | 62816200 | 62817599 | 22.23159 | 38.63291 | 37.18797 | 16.32939 | 0.44165 | 0.803535 | 0.762392 |
| 1 | 161373000 | 161374599 | 15.43715 | 26.9623 | 50.55566 | 22.27008 | -0.51523 | 0.720115 | 0.762419 |
| X | 37574200 | 37577799 | 35.24616 | 58.02007 | 65.7834 | 28.36583 | 0.198368 | 1.407899 | 0.762432 |
| 3 | 82234800 | 82235799 | 13.42062 | 24.04361 | 35.65325 | 15.62305 | -0.16432 | 0.423946 | 0.762436 |
| 1 | 165581600 | 165582599 | 10.61453 | 19.76535 | 30.5165 | 12.7321 | -0.17288 | 0.169526 | 0.762446 |
| 3 | 141708000 | 141710199 | 29.15903 | 47.28203 | 85.67139 | 36.69395 | -0.48979 | 1.625825 | 0.762458 |
| 12 | 80349400 | 80353199 | 44.0833 | 70.92726 | 100.1038 | 42.49718 | -0.14281 | 1.859942 | 0.762469 |
| 19 | 29534400 | 29534999 | 11.14963 | 20.32677 | 30.77124 | 13.21646 | -0.17465 | 0.295793 | 0.762479 |
| X | 110186200 | 110187399 | 18.86844 | 33.40072 | 36.3244 | 16.06301 | 0.270987 | 0.541501 | 0.762484 |
| 13 | 35454200 | 35454799 | 7.941315 | 15.33223 | 26.03037 | 10.03092 | -0.20708 | -0.13362 | 0.762489 |

| | | | | | | | | | |
|---|---|---|---|---|---|---|---|---|---|
| 6 | 85858000 | 85863199 | 88.93226 | 138.8162 | 158.0728 | 65.59184 | 0.125289 | 2.679351 | 0.762503 |
| 14 | 106145400 | 106148599 | 31.0488 | 51.69362 | 59.70179 | 25.9283 | 0.174065 | 1.258395 | 0.762515 |
| 11 | 96139200 | 96139999 | 11.20868 | 20.34078 | 31.30008 | 13.52105 | -0.20329 | 0.325305 | 0.762522 |
| 12 | 59710800 | 59712999 | 23.27567 | 39.99501 | 46.29342 | 20.30581 | 0.173679 | 0.924935 | 0.762523 |
| 3 | 14375400 | 14375799 | 14.51957 | 25.90449 | 29.61845 | 13.02421 | 0.205457 | 0.446379 | 0.762523 |
| 13 | 15921600 | 15924199 | 24.97337 | 42.47725 | 49.56929 | 21.72564 | 0.159733 | 1.027072 | 0.762538 |
| 16 | 19830000 | 19833199 | 38.0154 | 61.62372 | 71.45136 | 30.59637 | 0.15212 | 1.636159 | 0.762541 |
| 15 | 31634000 | 31634999 | 13.75963 | 24.79947 | 29.46964 | 12.83996 | 0.158864 | 0.376257 | 0.762554 |
| X | 147733600 | 147735999 | 40.87584 | 66.58651 | 74.43158 | 31.96213 | 0.210683 | 1.562092 | 0.762554 |
| 9 | 37378600 | 37381399 | 21.37899 | 36.18041 | 52.74906 | 23.08999 | -0.16066 | 1.062326 | 0.762592 |
| 10 | 76532200 | 76534399 | 19.68363 | 33.48618 | 58.68351 | 25.72068 | -0.42462 | 1.017615 | 0.762618 |
| 1 | 15090400 | 15090799 | 7.239661 | 14.08065 | 18.98289 | 7.020891 | 0.178206 | -0.30511 | 0.762623 |
| X | 109171200 | 109173599 | 20.95161 | 36.60489 | 34.10535 | 15.02648 | 0.489321 | 0.720127 | 0.762629 |
| 3 | 73096600 | 73097599 | 10.34969 | 19.8294 | 20.12712 | 7.933991 | 0.497594 | -0.05523 | 0.762639 |
| 11 | 27124200 | 27127399 | 53.08191 | 85.14544 | 97.71824 | 41.29738 | 0.149669 | 1.965448 | 0.762642 |
| 9 | 32681800 | 32684199 | 27.56512 | 46.03139 | 54.52696 | 23.72572 | 0.137576 | 1.211842 | 0.762649 |
| 9 | 111529600 | 111532799 | 36.48646 | 59.31698 | 69.66765 | 29.89453 | 0.137478 | 1.57762 | 0.762668 |
| 10 | 118138200 | 118139199 | 8.325106 | 15.93157 | 25.68702 | 10.16009 | -0.16913 | -0.04884 | 0.762701 |
| 15 | 9964200 | 9964799 | 12.88513 | 22.96325 | 34.10216 | 15.00974 | -0.1709 | 0.448847 | 0.762708 |
| 14 | 4065000 | 4067799 | 33.79615 | 55.31292 | 65.44714 | 28.15223 | 0.133619 | 1.47281 | 0.762719 |
| 13 | 9205200 | 9207799 | 15.43982 | 27.0913 | 39.77176 | 17.58616 | -0.16376 | 0.628843 | 0.762735 |
| 8 | 73301200 | 73302999 | 13.61445 | 23.85031 | 37.4927 | 16.55606 | -0.2616 | 0.642682 | 0.762748 |
| 4 | 128246800 | 128248799 | 17.9542 | 31.65949 | 28.93636 | 12.80585 | 0.518393 | 0.563298 | 0.762784 |
| 14 | 117528000 | 117530999 | 46.92485 | 74.99768 | 105.4561 | 44.33096 | -0.14874 | 2.072344 | 0.762792 |
| 11 | 12100800 | 12101999 | 12.1415 | 21.90091 | 32.70216 | 14.23385 | -0.16751 | 0.368736 | 0.762811 |
| 11 | 23173600 | 23180199 | 79.59054 | 124.1506 | 166.609 | 69.13872 | -0.11216 | 2.687783 | 0.762838 |
| 5 | 21926200 | 21928199 | 16.81588 | 29.19052 | 43.2271 | 18.96843 | -0.177 | 0.804133 | 0.762889 |
| 13 | 9856200 | 9858999 | 32.57673 | 53.50163 | 63.33868 | 27.28488 | 0.13484 | 1.424692 | 0.762901 |
| 9 | 101456800 | 101459399 | 13.16863 | 24.72532 | 22.84044 | 9.396756 | 0.578711 | 0.118354 | 0.762961 |
| 4 | 78664200 | 78665599 | 14.896 | 26.44491 | 40.21981 | 17.75992 | -0.2085 | 0.497894 | 0.762978 |
| 7 | 94550800 | 94551799 | 14.77856 | 26.52166 | 31.4167 | 13.74683 | 0.159282 | 0.413629 | 0.762978 |
| 1 | 22199600 | 22200999 | 10.49126 | 19.50401 | 30.45875 | 12.74311 | -0.19285 | 0.175748 | 0.762983 |
| 3 | 37964200 | 37965599 | 14.35729 | 25.98943 | 30.50873 | 13.22059 | 0.181557 | 0.359294 | 0.763 |
| 5 | 68376000 | 68379599 | 52.95709 | 84.17047 | 119.695 | 49.93966 | -0.17355 | 2.258249 | 0.763003 |
| 9 | 116229200 | 116232999 | 35.54776 | 57.51925 | 81.59835 | 34.89477 | -0.14063 | 1.674917 | 0.763003 |
| 5 | 82801200 | 82804199 | 39.59556 | 64.00747 | 92.58194 | 39.55373 | -0.17004 | 1.712838 | 0.763017 |
| 17 | 27558400 | 27582399 | 310.0586 | 435.1825 | 542.9014 | 224.9844 | -0.14535 | 4.523746 | 0.763028 |
| 9 | 102294000 | 102295599 | 18.34909 | 32.27643 | 30.79445 | 13.63485 | 0.455268 | 0.594107 | 0.763065 |
| 5 | 135009600 | 135011199 | 12.43552 | 22.22049 | 33.16243 | 14.56372 | -0.17561 | 0.428466 | 0.76309 |
| 7 | 3631800 | 3633399 | 12.43552 | 22.22049 | 33.16243 | 14.56372 | -0.17561 | 0.428466 | 0.76309 |
| 3 | 159089600 | 159091399 | 22.02502 | 37.96071 | 44.70285 | 19.60994 | 0.149851 | 0.891881 | 0.763091 |
| 8 | 75112400 | 75112799 | 5.069247 | 9.898278 | 18.17195 | 6.398944 | -0.19701 | -0.50738 | 0.763141 |
| 3 | 117029800 | 117035799 | 62.05073 | 99.13727 | 109.0294 | 45.80995 | 0.204528 | 2.098718 | 0.763165 |
| 15 | 95650400 | 95653399 | 31.4331 | 52.06553 | 61.28414 | 26.5218 | 0.146263 | 1.319534 | 0.763176 |
| 14 | 121857000 | 121859199 | 13.99102 | 24.47837 | 44.12658 | 19.4642 | -0.45832 | 0.677222 | 0.76323 |
| 6 | 29090000 | 29097999 | 126.8871 | 191.7942 | 251.1503 | 103.8527 | -0.11264 | 3.31169 | 0.76324 |
| 12 | 21333800 | 21336999 | 28.61675 | 46.50546 | 82.8491 | 35.54305 | -0.46248 | 1.575776 | 0.763281 |

| | | | | | | | | | |
|---|---|---|---|---|---|---|---|---|---|
| 9 | 114030200 | 114038199 | 166.716 | 249.0035 | 276.43 | 113.96 | 0.11017 | 3.550351 | 0.763317 |
| 9 | 79939600 | 79943199 | 64.84136 | 102.933 | 119.4741 | 49.78629 | 0.117215 | 2.302482 | 0.763324 |
| 1 | 66578600 | 66579799 | 12.11797 | 21.94716 | 32.71623 | 14.16804 | -0.16044 | 0.344734 | 0.763332 |
| 1 | 8310000 | 8311399 | 21.38373 | 37.2273 | 42.48317 | 18.6759 | 0.197555 | 0.783948 | 0.763363 |
| 10 | 10496000 | 10497599 | 19.43866 | 33.91512 | 39.38352 | 17.33707 | 0.172934 | 0.740469 | 0.763411 |
| 2 | 144454600 | 144455599 | 17.02183 | 29.83082 | 34.71514 | 15.33446 | 0.169952 | 0.650247 | 0.763414 |
| 4 | 9561200 | 9562799 | 25.2279 | 42.04493 | 60.39234 | 26.24499 | -0.13956 | 1.231738 | 0.763417 |
| 3 | 106605000 | 106616599 | 168.5999 | 253.8847 | 263.5441 | 108.8547 | 0.217547 | 3.397007 | 0.763422 |
| 17 | 22020000 | 22030999 | 249.705 | 361.3238 | 389.0646 | 160.4606 | 0.110945 | 4.073315 | 0.763435 |
| 5 | 25741200 | 25744999 | 47.33494 | 76.14602 | 89.04119 | 37.79146 | 0.127753 | 1.866031 | 0.763447 |
| 8 | 16769000 | 16772399 | 30.24203 | 51.06816 | 55.20331 | 24.14832 | 0.269258 | 1.114682 | 0.76345 |
| 18 | 32735200 | 32736399 | 14.5416 | 25.51849 | 37.83096 | 16.72978 | -0.17757 | 0.620231 | 0.763461 |
| 7 | 29472600 | 29478599 | 57.4377 | 91.10639 | 126.1203 | 52.51334 | -0.13772 | 2.320626 | 0.763468 |
| 1 | 171149600 | 171149999 | 4.956146 | 9.674376 | 18.14059 | 6.422799 | -0.23502 | -0.49271 | 0.763484 |
| 7 | 138253800 | 138255799 | 17.42488 | 30.46551 | 45.12039 | 19.89117 | -0.17644 | 0.70557 | 0.763488 |
| 15 | 33837800 | 33838799 | 11.50778 | 21.58341 | 26.22469 | 10.80896 | 0.184093 | 0.118838 | 0.763508 |
| 4 | 144214200 | 144221799 | 75.66615 | 119.6232 | 134.7707 | 55.99329 | 0.154927 | 2.419134 | 0.763551 |
| 11 | 7261200 | 7265999 | 38.14582 | 61.96816 | 96.5813 | 41.42291 | -0.27079 | 1.610299 | 0.763562 |
| 7 | 12630000 | 12632199 | 17.34618 | 30.0701 | 44.75195 | 19.63494 | -0.18478 | 0.818656 | 0.763587 |
| 9 | 75374200 | 75376199 | 15.96597 | 27.85524 | 41.03578 | 18.06401 | -0.16884 | 0.723887 | 0.76359 |
| 6 | 91432000 | 91434199 | 23.61494 | 40.15778 | 47.43726 | 20.79082 | 0.142288 | 1.023827 | 0.763594 |
| 15 | 92625800 | 92629199 | 69.1522 | 107.766 | 177.3055 | 73.58735 | -0.40681 | 2.700861 | 0.763609 |
| 7 | 81790400 | 81791799 | 14.08874 | 25.28202 | 29.95798 | 13.10927 | 0.158773 | 0.40594 | 0.763626 |
| 5 | 23879800 | 23881599 | 17.34451 | 29.38908 | 56.86969 | 24.90307 | -0.56734 | 1.042989 | 0.763627 |
| 9 | 111022800 | 111023799 | 9.053544 | 17.03566 | 26.97391 | 11.05673 | -0.18789 | 0.082627 | 0.76363 |
| 2 | 11107600 | 11116799 | 118.8464 | 181.3346 | 204.4811 | 84.79016 | 0.114035 | 3.116011 | 0.763631 |
| 15 | 47096800 | 47098399 | 12.01457 | 21.72714 | 32.63798 | 14.1608 | -0.17351 | 0.349806 | 0.763634 |
| 13 | 61869800 | 61931399 | 1112.27 | 1427.428 | 1317.939 | 552.8011 | 0.159105 | 5.946766 | 0.763659 |
| 17 | 33070000 | 33072399 | 21.70637 | 37.32995 | 44.20589 | 19.39064 | 0.141225 | 0.909552 | 0.763676 |
| 13 | 100424800 | 100425399 | 8.668636 | 16.40194 | 25.97898 | 10.5352 | -0.17547 | 0.037325 | 0.763688 |
| 5 | 116741800 | 116744199 | 16.46641 | 28.83279 | 41.92148 | 18.5043 | -0.14983 | 0.672479 | 0.76372 |
| 5 | 150621400 | 150624599 | 29.21998 | 49.01463 | 56.26977 | 24.55357 | 0.182495 | 1.171255 | 0.763737 |
| 14 | 53800400 | 53802599 | 39.53059 | 63.90912 | 74.02524 | 31.63002 | 0.149924 | 1.697684 | 0.763749 |
| 18 | 67225000 | 67227199 | 24.59015 | 42.22323 | 40.97499 | 17.97256 | 0.426824 | 0.931632 | 0.763771 |
| 2 | 3225800 | 3227399 | 13.00045 | 22.9993 | 34.77969 | 15.38178 | -0.20213 | 0.509989 | 0.763788 |
| 14 | 61538400 | 61539799 | 14.07025 | 25.88672 | 29.53474 | 12.53291 | 0.242265 | 0.2598 | 0.763798 |
| 8 | 104033400 | 104034999 | 12.37193 | 21.83173 | 40.35029 | 17.85454 | -0.49182 | 0.542284 | 0.763817 |
| 7 | 39486600 | 39489399 | 28.09834 | 46.94725 | 55.63105 | 24.21441 | 0.137075 | 1.209371 | 0.763823 |
| 6 | 48872400 | 48875199 | 21.15894 | 36.60721 | 43.09165 | 18.90534 | 0.151519 | 0.851647 | 0.763832 |
| 19 | 3101200 | 3103599 | 28.59879 | 48.19226 | 46.93973 | 20.52043 | 0.418953 | 1.11827 | 0.763833 |
| 19 | 9356200 | 9363199 | 99.44583 | 154.5897 | 170.452 | 70.76925 | 0.167123 | 2.755843 | 0.763851 |
| 4 | 83345600 | 83347199 | 16.73299 | 29.19234 | 42.54099 | 18.72616 | -0.15337 | 0.729615 | 0.763881 |
| 12 | 36205200 | 36208199 | 20.80307 | 34.69535 | 64.44343 | 28.01816 | -0.50912 | 1.214946 | 0.763885 |
| 18 | 9911400 | 9912399 | 24.43093 | 41.14362 | 46.99372 | 20.5402 | 0.190023 | 1.110318 | 0.763888 |
| X | 104806400 | 104809999 | 44.20582 | 71.48528 | 80.826 | 34.54694 | 0.18511 | 1.705906 | 0.763908 |
| 4 | 84592000 | 84594399 | 20.48446 | 34.14614 | 66.12095 | 28.73803 | -0.56906 | 1.219942 | 0.763916 |
| 6 | 65331200 | 65333199 | 20.04079 | 35.33468 | 37.81684 | 16.73633 | 0.291666 | 0.591553 | 0.763922 |

| | | | | | | | | | |
|---|---|---|---|---|---|---|---|---|---|
| 9 | 38940400 | 38940799 | 10.29362 | 19.36449 | 23.7193 | 9.726548 | 0.178801 | 0.084124 | 0.763929 |
| 11 | 11955600 | 11962599 | 72.88944 | 115.6986 | 127.1305 | 52.97202 | 0.196012 | 2.312623 | 0.763946 |
| X | 145459200 | 145460999 | 18.03748 | 32.22651 | 34.83386 | 15.2993 | 0.286992 | 0.469943 | 0.763948 |
| 8 | 124527000 | 124528599 | 17.3853 | 30.54353 | 36.03331 | 15.95014 | 0.150235 | 0.614798 | 0.763963 |
| X | 56766600 | 56768399 | 16.23175 | 28.65219 | 45.57003 | 20.1629 | -0.27643 | 0.57003 | 0.763976 |
| 7 | 81801600 | 81804599 | 37.86303 | 62.62237 | 54.88834 | 23.72942 | 0.570236 | 1.349568 | 0.763984 |
| 13 | 86868800 | 86871199 | 24.17195 | 42.11134 | 43.97812 | 19.34175 | 0.324994 | 0.788355 | 0.763992 |
| 10 | 15509400 | 15511599 | 33.70824 | 54.44157 | 92.2094 | 39.349 | -0.39866 | 1.728439 | 0.764007 |
| 5 | 8430600 | 8434399 | 64.0219 | 101.7184 | 117.9779 | 49.20455 | 0.119459 | 2.27693 | 0.764023 |
| 8 | 4725200 | 4733599 | 116.3116 | 176.769 | 233.1442 | 96.5689 | -0.11642 | 3.197457 | 0.764026 |
| 4 | 20838000 | 20838799 | 13.26383 | 24.38223 | 29.09998 | 12.36277 | 0.176808 | 0.259707 | 0.764036 |
| 7 | 40655400 | 40657199 | 21.49147 | 37.38821 | 36.22398 | 15.91719 | 0.432554 | 0.784163 | 0.764041 |
| 5 | 125450400 | 125453399 | 41.02377 | 66.368 | 77.91538 | 33.31537 | 0.131546 | 1.685769 | 0.764045 |
| 8 | 27146000 | 27148799 | 21.96236 | 37.58669 | 43.71457 | 19.16909 | 0.165928 | 0.95268 | 0.764093 |
| 14 | 3464400 | 3465999 | 19.14992 | 32.69252 | 48.49186 | 21.25973 | -0.18393 | 0.973141 | 0.764095 |
| 16 | 86958200 | 86959399 | 11.81544 | 21.46136 | 32.6818 | 14.10191 | -0.18804 | 0.322227 | 0.764114 |
| 8 | 87073600 | 87075599 | 24.79515 | 43.03562 | 45.72895 | 20.0645 | 0.299688 | 0.846581 | 0.764121 |
| 4 | 81258200 | 81260799 | 20.98108 | 36.61584 | 41.27198 | 18.17139 | 0.215449 | 0.743035 | 0.764145 |
| 1 | 144984200 | 144986399 | 31.74116 | 51.86816 | 73.62758 | 31.65295 | -0.12998 | 1.495927 | 0.764163 |
| 11 | 5796800 | 5798999 | 24.52961 | 41.51144 | 49.05426 | 21.47252 | 0.141282 | 1.069506 | 0.764171 |
| 6 | 66316200 | 66317399 | 11.12104 | 20.15585 | 30.72665 | 13.29466 | -0.19168 | 0.328154 | 0.764193 |
| 3 | 20233800 | 20234599 | 13.01779 | 23.52214 | 27.01141 | 11.73155 | 0.210816 | 0.349705 | 0.764203 |
| 8 | 8302400 | 8304199 | 20.32025 | 35.38326 | 41.441 | 18.22015 | 0.160256 | 0.773655 | 0.764213 |
| 1 | 141679800 | 141682399 | 19.90102 | 34.58404 | 40.97812 | 17.99401 | 0.143259 | 0.79798 | 0.764219 |
| 12 | 99602200 | 99604199 | 18.88487 | 33.02777 | 38.6669 | 17.05037 | 0.161169 | 0.697707 | 0.764275 |
| 3 | 22963000 | 22966999 | 103.0187 | 158.4646 | 179.3887 | 74.49799 | 0.117489 | 2.948397 | 0.764279 |
| 1 | 108739400 | 108739999 | 14.08414 | 25.15453 | 29.0453 | 12.76067 | 0.19203 | 0.433557 | 0.764282 |
| 1 | 191670600 | 191673799 | 56.03643 | 88.78726 | 126.0676 | 52.43643 | -0.17574 | 2.351852 | 0.764295 |
| 15 | 17639000 | 17640199 | 14.84545 | 27.07531 | 30.99006 | 13.30385 | 0.226185 | 0.318624 | 0.764312 |
| 15 | 10729000 | 10732399 | 28.54455 | 47.70677 | 56.23368 | 24.48124 | 0.144619 | 1.207305 | 0.764314 |
| 11 | 34570600 | 34573199 | 26.60643 | 45.99084 | 46.2257 | 20.27977 | 0.378255 | 0.886999 | 0.764317 |
| 11 | 26369400 | 26370399 | 11.34904 | 21.68005 | 25.64118 | 10.18985 | 0.2677 | -0.0059 | 0.764317 |
| 11 | 57550400 | 57553599 | 29.9332 | 49.35279 | 56.61114 | 24.42721 | 0.182535 | 1.36542 | 0.764325 |
| 3 | 113102400 | 113105999 | 42.64614 | 68.77711 | 80.30833 | 34.2063 | 0.134248 | 1.776797 | 0.764334 |
| 5 | 108213400 | 108214199 | 8.503103 | 16.02672 | 25.92011 | 10.59471 | -0.21541 | 0.060982 | 0.764345 |
| 8 | 21082000 | 21092599 | 282.5156 | 401.7398 | 498.5485 | 206.162 | -0.11879 | 4.330646 | 0.7644 |
| 8 | 109907800 | 109909199 | 14.57544 | 26.48138 | 30.97568 | 13.36144 | 0.190704 | 0.339126 | 0.764403 |
| 12 | 18041600 | 18044199 | 44.74917 | 71.58016 | 102.6459 | 43.2007 | -0.17569 | 2.042186 | 0.764416 |
| 10 | 49964600 | 49966399 | 24.42756 | 40.80837 | 58.69978 | 25.54789 | -0.14161 | 1.203023 | 0.764451 |
| 6 | 129147200 | 129149599 | 21.3009 | 36.94899 | 43.05573 | 18.89109 | 0.166802 | 0.828856 | 0.764463 |
| 4 | 7622600 | 7625599 | 27.13209 | 45.00989 | 64.78912 | 28.06815 | -0.14296 | 1.29488 | 0.764472 |
| 7 | 29429400 | 29444399 | 222.2294 | 322.4432 | 406.0968 | 167.4267 | -0.11116 | 4.03122 | 0.764473 |
| 5 | 130043400 | 130045599 | 16.50698 | 28.76501 | 42.10557 | 18.52113 | -0.15975 | 0.746463 | 0.764481 |
| 7 | 42934600 | 42936799 | 16.8339 | 29.36423 | 43.57122 | 19.17795 | -0.17945 | 0.732048 | 0.764482 |
| 15 | 35112800 | 35113599 | 8.230745 | 15.91867 | 28.12277 | 10.7496 | -0.25335 | -0.15423 | 0.764504 |
| 10 | 98909200 | 98912799 | 33.40308 | 54.13056 | 78.70543 | 33.69508 | -0.17403 | 1.63811 | 0.764553 |
| 10 | 52097400 | 52102199 | 52.22287 | 83.72242 | 97.31748 | 41.09435 | 0.130398 | 1.980357 | 0.764558 |

| | | | | | | | | | |
|---|---|---|---|---|---|---|---|---|---|
| 10 | 119724800 | 119729599 | 71.91857 | 113.5997 | 131.6652 | 54.679 | 0.113086 | 2.436735 | 0.764559 |
| 6 | 110675200 | 110676799 | 16.96508 | 31.01799 | 32.07735 | 13.72795 | 0.374697 | 0.323338 | 0.76456 |
| 14 | 44304400 | 44305399 | 7.896931 | 15.06701 | 24.52197 | 9.770374 | -0.19173 | -0.03135 | 0.764571 |
| 1 | 134880600 | 134881199 | 7.267637 | 14.13452 | 18.99452 | 7.027338 | 0.182822 | -0.3021 | 0.764591 |
| 13 | 78256600 | 78256999 | 7.267637 | 14.13452 | 18.99452 | 7.027338 | 0.182822 | -0.3021 | 0.764591 |
| X | 41235600 | 41242599 | 66.78396 | 107.1328 | 78.98902 | 33.38623 | 0.78702 | 1.977787 | 0.764598 |
| 13 | 119000800 | 119004199 | 28.38963 | 47.20145 | 70.29126 | 30.49885 | -0.19169 | 1.275268 | 0.764604 |
| 4 | 115315400 | 115321399 | 54.98505 | 88.16428 | 99.53494 | 42.04058 | 0.172647 | 1.979172 | 0.764627 |
| 13 | 46469400 | 46471999 | 25.73031 | 42.95485 | 61.55988 | 26.78094 | -0.13635 | 1.216611 | 0.764645 |
| 13 | 51182000 | 51190799 | 121.3745 | 183.9595 | 241.4275 | 99.91755 | -0.11251 | 3.255599 | 0.764649 |
| 6 | 139079400 | 139080599 | 10.00614 | 18.70356 | 29.50367 | 12.2304 | -0.19655 | 0.134596 | 0.764655 |
| 4 | 148436400 | 148439799 | 49.35719 | 78.77452 | 109.7436 | 46.04168 | -0.13762 | 2.118622 | 0.764663 |
| 8 | 74397200 | 74397799 | 10.63633 | 19.98813 | 24.7366 | 10.16181 | 0.161989 | 0.096781 | 0.764677 |
| 3 | 29628800 | 29629799 | 9.357015 | 17.44472 | 27.3925 | 11.40542 | -0.19428 | 0.144678 | 0.764678 |
| 11 | 11560800 | 11562599 | 18.78258 | 32.77018 | 38.82337 | 17.0904 | 0.144036 | 0.733921 | 0.764679 |
| 7 | 141972200 | 141973799 | 13.7351 | 24.30458 | 35.63983 | 15.76147 | -0.15811 | 0.511812 | 0.764683 |
| 10 | 86338200 | 86339999 | 21.3171 | 36.68543 | 43.1459 | 18.92608 | 0.151237 | 0.899602 | 0.764685 |
| 18 | 87942000 | 87943399 | 18.63506 | 32.01932 | 46.90734 | 20.57506 | -0.16461 | 0.91028 | 0.764685 |
| 9 | 45945800 | 45948999 | 19.66933 | 34.01939 | 39.03518 | 17.12551 | 0.18821 | 0.842441 | 0.764707 |
| 3 | 74205800 | 74212599 | 106.3867 | 164.3241 | 159.6193 | 66.30249 | 0.342657 | 2.869421 | 0.76471 |
| 4 | 37667200 | 37667999 | 7.831869 | 15.14408 | 18.88777 | 7.22663 | 0.244033 | -0.18159 | 0.764729 |
| 13 | 52432400 | 52435799 | 36.38816 | 58.98525 | 83.18655 | 35.62122 | -0.12973 | 1.641933 | 0.764735 |
| 3 | 124759000 | 124759799 | 10.45357 | 19.54226 | 22.73419 | 9.422259 | 0.240534 | 0.12321 | 0.764738 |
| 3 | 34423000 | 34424799 | 15.38624 | 27.0435 | 40.70757 | 18.02599 | -0.19985 | 0.601015 | 0.764759 |
| 3 | 64087000 | 64087399 | 10.84518 | 20.24548 | 24.23278 | 10.06762 | 0.197051 | 0.140715 | 0.764767 |
| 12 | 90733400 | 90738799 | 56.34966 | 90.25399 | 88.28137 | 37.22572 | 0.377303 | 2.010023 | 0.764768 |
| 17 | 72307800 | 72310999 | 30.31716 | 50.23157 | 59.47351 | 25.74334 | 0.13789 | 1.312651 | 0.764786 |
| 5 | 127537000 | 127540199 | 36.74372 | 59.7354 | 70.43599 | 30.22426 | 0.131774 | 1.576945 | 0.764793 |
| 5 | 23555000 | 23557799 | 32.59687 | 53.00454 | 75.50636 | 32.40238 | -0.14006 | 1.56965 | 0.764798 |
| X | 160266400 | 160268999 | 20.8379 | 35.8572 | 53.89851 | 23.64267 | -0.2013 | 0.915177 | 0.764809 |
| 12 | 54173400 | 54175599 | 20.76552 | 35.93776 | 42.60654 | 18.69269 | 0.141456 | 0.848047 | 0.764814 |
| 18 | 89853400 | 89855399 | 24.05658 | 40.0454 | 59.80412 | 25.96996 | -0.19556 | 1.238642 | 0.764832 |
| 16 | 34232600 | 34235799 | 30.27723 | 49.63201 | 70.76053 | 30.4579 | -0.13363 | 1.449304 | 0.764835 |
| 15 | 52338800 | 52342399 | 58.825 | 93.68517 | 109.1529 | 45.65841 | 0.1165 | 2.195557 | 0.764837 |
| X | 94805600 | 94811399 | 67.26444 | 106.5945 | 153.5377 | 63.87002 | -0.19629 | 2.361754 | 0.764837 |
| 2 | 36828400 | 36831399 | 31.97502 | 53.40845 | 50.82866 | 22.11989 | 0.452332 | 1.219965 | 0.764868 |
| 6 | 105653600 | 105656199 | 25.40768 | 42.24599 | 60.90122 | 26.42562 | -0.14485 | 1.258556 | 0.76487 |
| X | 71851600 | 71853399 | 16.23115 | 29.26589 | 32.86562 | 14.3123 | 0.240285 | 0.397592 | 0.76488 |
| 10 | 119407000 | 119410199 | 38.68108 | 62.48633 | 87.5892 | 37.38761 | -0.12599 | 1.727036 | 0.764884 |
| 11 | 93685000 | 93686399 | 20.77553 | 35.67749 | 39.23442 | 17.20918 | 0.247364 | 0.914007 | 0.764896 |
| 9 | 38744400 | 38745199 | 9.339879 | 17.60283 | 27.51721 | 11.24866 | -0.16663 | 0.075333 | 0.764906 |
| 5 | 104718800 | 104758599 | 1062.165 | 1366.143 | 1278.536 | 536.159 | 0.142789 | 5.910672 | 0.76492 |
| 2 | 146731000 | 146733199 | 25.45761 | 43.31357 | 40.56265 | 17.77855 | 0.476029 | 1.015648 | 0.76496 |
| 3 | 149790600 | 149793399 | 36.37113 | 58.46112 | 101.9416 | 43.21551 | -0.44895 | 1.88338 | 0.764962 |
| 15 | 39566400 | 39570199 | 46.68232 | 75.22565 | 87.20028 | 37.10265 | 0.143389 | 1.81092 | 0.764972 |
| 10 | 121334000 | 121334799 | 6.104005 | 11.85763 | 21.15584 | 7.900768 | -0.23371 | -0.2735 | 0.76499 |
| 10 | 18215400 | 18217399 | 19.45992 | 33.75533 | 38.95621 | 17.09238 | 0.180711 | 0.814592 | 0.765002 |

| | | | | | | | | | |
|---|---|---|---|---|---|---|---|---|---|
| X | 15352000 | 15354199 | 22.36919 | 39.29058 | 32.06962 | 14.19289 | 0.679126 | 0.640922 | 0.765008 |
| 9 | 11853200 | 11857399 | 55.53632 | 89.078 | 86.44776 | 36.53153 | 0.391082 | 1.966759 | 0.765013 |
| 16 | 16769400 | 16770799 | 16.24841 | 28.48886 | 41.85349 | 18.49502 | -0.16484 | 0.6485 | 0.765017 |
| 14 | 39544600 | 39546799 | 22.99213 | 38.95705 | 57.09566 | 25.00173 | -0.16811 | 1.065181 | 0.765072 |
| 13 | 65846600 | 65853399 | 118.1906 | 181.8176 | 171.5215 | 71.2317 | 0.380365 | 2.952243 | 0.765124 |
| 8 | 14641800 | 14644999 | 39.22953 | 63.88073 | 73.8486 | 31.70746 | 0.161876 | 1.561095 | 0.765127 |
| 19 | 26659000 | 26660799 | 15.77972 | 27.92083 | 32.96558 | 14.57899 | 0.152823 | 0.520415 | 0.765151 |
| 13 | 75975400 | 75980199 | 41.24566 | 66.43012 | 93.318 | 39.67563 | -0.13419 | 1.820122 | 0.76516 |
| 17 | 76380000 | 76381199 | 14.51735 | 26.93529 | 29.76195 | 12.48537 | 0.30023 | 0.218702 | 0.765175 |
| 4 | 113934200 | 113938399 | 66.83439 | 106.391 | 118.7275 | 49.63117 | 0.17785 | 2.221123 | 0.765204 |
| 15 | 93536600 | 93541199 | 50.49949 | 81.04455 | 93.98878 | 39.75213 | 0.135479 | 1.943625 | 0.765234 |
| 7 | 106417200 | 106419199 | 14.84218 | 26.24685 | 38.62843 | 17.08533 | -0.1639 | 0.53011 | 0.765235 |
| 16 | 94635600 | 94637399 | 13.0727 | 23.1519 | 34.64374 | 15.31838 | -0.1867 | 0.495714 | 0.765237 |
| 14 | 3629800 | 3630799 | 13.16064 | 23.72991 | 26.9893 | 11.75306 | 0.222094 | 0.362541 | 0.765239 |
| 13 | 68816200 | 68818199 | 17.68297 | 31.07865 | 36.46964 | 16.14863 | 0.157819 | 0.610388 | 0.765246 |
| 11 | 27302200 | 27304599 | 26.10256 | 43.15736 | 62.99278 | 27.22698 | -0.16335 | 1.327386 | 0.765279 |
| 1 | 10374000 | 10374999 | 9.123181 | 17.13177 | 26.91451 | 11.0705 | -0.1812 | 0.093119 | 0.765282 |
| 12 | 109727800 | 109730999 | 26.81463 | 44.34254 | 64.06828 | 27.69522 | -0.14868 | 1.330034 | 0.765288 |
| X | 79665000 | 79668399 | 32.01944 | 52.75415 | 62.17424 | 26.81939 | 0.143162 | 1.385679 | 0.765291 |
| 10 | 72767600 | 72769799 | 18.83782 | 33.08661 | 37.3572 | 16.53245 | 0.213018 | 0.626508 | 0.765333 |
| X | 127859600 | 127861199 | 18.50892 | 32.53063 | 37.43814 | 16.57762 | 0.18572 | 0.616446 | 0.765339 |
| 11 | 22376600 | 22379599 | 22.4574 | 39.22337 | 42.30486 | 18.63543 | 0.278951 | 0.742543 | 0.765378 |
| 14 | 98045200 | 98047599 | 18.57034 | 32.32284 | 48.97886 | 21.52869 | -0.20977 | 0.783528 | 0.765392 |
| 4 | 86561400 | 86563399 | 21.77554 | 37.24877 | 54.98186 | 24.10892 | -0.1767 | 0.972489 | 0.765421 |
| 1 | 88527200 | 88530199 | 18.69274 | 31.87101 | 55.93987 | 24.52253 | -0.4263 | 0.990397 | 0.765429 |
| 8 | 85911600 | 85915599 | 45.13467 | 73.17496 | 80.51369 | 34.48071 | 0.227753 | 1.654257 | 0.765439 |
| 8 | 51825200 | 51826799 | 12.9226 | 23.43623 | 35.86518 | 15.50537 | -0.19675 | 0.348268 | 0.765442 |
| 5 | 73770800 | 73771999 | 10.42469 | 19.21199 | 29.29348 | 12.40198 | -0.17054 | 0.223603 | 0.765448 |
| 9 | 113203400 | 113206399 | 21.67837 | 36.70415 | 53.30277 | 23.33566 | -0.15503 | 1.059431 | 0.76546 |
| 6 | 70802600 | 70804999 | 22.64026 | 38.64906 | 45.29663 | 19.8588 | 0.154349 | 0.980332 | 0.765498 |
| 9 | 101010400 | 101014199 | 58.44376 | 93.32039 | 107.4513 | 45.10986 | 0.137533 | 2.112275 | 0.765514 |
| X | 8282600 | 8288599 | 64.96007 | 103.7089 | 112.2933 | 47.13197 | 0.225844 | 2.127723 | 0.765515 |
| 12 | 119011200 | 119011999 | 9.111305 | 17.40257 | 28.52892 | 11.34121 | -0.1999 | -0.01781 | 0.765525 |
| 8 | 109852400 | 109854599 | 20.27258 | 35.30268 | 41.42322 | 18.21312 | 0.157564 | 0.771726 | 0.765544 |
| 2 | 81243400 | 81244599 | 24.66768 | 40.88451 | 60.68491 | 26.27364 | -0.18712 | 1.294556 | 0.765556 |
| 3 | 69500800 | 69502599 | 11.7347 | 21.03561 | 32.62672 | 14.28975 | -0.22836 | 0.400123 | 0.7656 |
| 9 | 48833400 | 48835199 | 15.10983 | 26.59307 | 39.16725 | 17.34017 | -0.16774 | 0.583391 | 0.765633 |
| 1 | 42373000 | 42374199 | 12.73857 | 22.34593 | 42.02095 | 18.57673 | -0.51876 | 0.624747 | 0.765647 |
| 10 | 6957200 | 6958999 | 20.69674 | 35.47119 | 51.16269 | 22.43771 | -0.14308 | 0.945345 | 0.765648 |
| 5 | 127144800 | 127148199 | 26.50535 | 44.30876 | 65.1142 | 28.35144 | -0.17256 | 1.206301 | 0.765684 |
| 3 | 64225400 | 64227399 | 18.48157 | 32.21558 | 48.42214 | 21.30364 | -0.198 | 0.760056 | 0.765724 |
| 12 | 99366600 | 99370399 | 51.46837 | 82.14828 | 113.0491 | 47.43197 | -0.1199 | 2.119828 | 0.765727 |
| 4 | 143181400 | 143187999 | 77.23763 | 119.2668 | 200.1061 | 83.11997 | -0.44531 | 2.868233 | 0.76573 |
| 5 | 13905400 | 13909599 | 83.46534 | 131.1501 | 129.4166 | 53.68215 | 0.339044 | 2.551572 | 0.765737 |
| 3 | 85870400 | 85872999 | 52.88587 | 84.64486 | 99.00045 | 41.70087 | 0.118701 | 2.03285 | 0.765739 |
| 2 | 127408200 | 127411799 | 25.58492 | 43.47808 | 50.65231 | 22.1987 | 0.162005 | 1.040281 | 0.765774 |
| 2 | 171964400 | 171965599 | 19.28915 | 33.33994 | 48.47257 | 21.26545 | -0.15202 | 0.861857 | 0.765804 |

| | | | | | | | | | |
|---|---|---|---|---|---|---|---|---|---|
| 4 | 31527400 | 31528399 | 14.98714 | 26.01968 | 47.92193 | 21.03042 | -0.49016 | 0.801136 | 0.765804 |
| 9 | 36214200 | 36215599 | 20.54998 | 36.06695 | 39.51048 | 17.47438 | 0.256594 | 0.653856 | 0.765816 |
| 17 | 38935000 | 38948399 | 53.18535 | 85.78703 | 74.95001 | 31.9302 | 0.552047 | 1.787687 | 0.765862 |
| 2 | 5695000 | 5696399 | 14.74852 | 26.24963 | 30.95243 | 13.63849 | 0.158969 | 0.463492 | 0.765883 |
| 14 | 120194600 | 120195799 | 13.88181 | 25.22322 | 29.33365 | 12.65185 | 0.198802 | 0.327476 | 0.76589 |
| 2 | 40158200 | 40160599 | 38.82068 | 62.86192 | 73.3518 | 31.38544 | 0.141599 | 1.663174 | 0.765909 |
| 16 | 3807600 | 3810199 | 17.69139 | 30.70952 | 44.74225 | 19.63298 | -0.15383 | 0.808462 | 0.765933 |
| 7 | 23662800 | 23668999 | 91.38517 | 141.853 | 162.4383 | 67.45671 | 0.110985 | 2.775485 | 0.765933 |
| 16 | 24222000 | 24226199 | 37.31817 | 60.57314 | 88.6973 | 38.01816 | -0.18211 | 1.618355 | 0.765972 |
| 14 | 99249600 | 99250199 | 7.238743 | 14.00602 | 23.85401 | 9.099885 | -0.19838 | -0.18574 | 0.766029 |
| 1 | 102222800 | 102226199 | 38.92643 | 63.94985 | 69.81153 | 30.07721 | 0.25167 | 1.441216 | 0.766031 |
| 6 | 99623200 | 99624199 | 9.109904 | 17.14518 | 26.94232 | 11.04 | -0.17723 | 0.078495 | 0.766045 |
| 8 | 104399200 | 104400799 | 10.55592 | 19.70176 | 30.57567 | 12.70688 | -0.17592 | 0.153072 | 0.766048 |
| 7 | 134660800 | 134663399 | 37.55485 | 60.55892 | 86.0685 | 36.6541 | -0.14881 | 1.776572 | 0.766048 |
| 4 | 53799400 | 53808199 | 270.2513 | 388.5579 | 415.2042 | 171.3917 | 0.112843 | 4.168766 | 0.766065 |
| 7 | 87450400 | 87451999 | 19.60458 | 34.67808 | 36.84527 | 16.29576 | 0.304065 | 0.55539 | 0.766078 |
| 15 | 103833800 | 103838599 | 85.312 | 133.5642 | 152.5259 | 63.27073 | 0.12453 | 2.62353 | 0.766083 |
| 2 | 54246400 | 54246999 | 7.419347 | 14.27206 | 23.60732 | 9.199514 | -0.18522 | -0.11191 | 0.766103 |
| 5 | 96593200 | 96593799 | 7.410808 | 14.28128 | 23.65434 | 9.170378 | -0.18024 | -0.12896 | 0.766104 |
| 5 | 130564600 | 130568199 | 29.1946 | 48.04941 | 68.79343 | 29.66141 | -0.13744 | 1.393851 | 0.766106 |
| 15 | 14437200 | 14437999 | 8.360865 | 15.99818 | 25.67857 | 10.16007 | -0.16287 | -0.0461 | 0.766112 |
| 16 | 44881600 | 44884799 | 33.8892 | 54.60012 | 96.14696 | 40.89827 | -0.45881 | 1.801886 | 0.76614 |
| 11 | 12612600 | 12615399 | 48.96607 | 78.05149 | 111.1577 | 46.55289 | -0.17127 | 2.160612 | 0.766146 |
| 11 | 16305800 | 16310199 | 44.43981 | 71.43304 | 99.55975 | 42.20812 | -0.12641 | 1.890486 | 0.766147 |
| 5 | 131109400 | 131110999 | 16.20682 | 28.53541 | 33.53282 | 14.84448 | 0.156794 | 0.576886 | 0.766148 |
| 9 | 12097400 | 12104999 | 151.6405 | 227.9676 | 254.82 | 105.2045 | 0.108714 | 3.424056 | 0.766176 |
| 4 | 66554600 | 66557599 | 33.89685 | 54.62559 | 95.53707 | 40.653 | -0.44853 | 1.794757 | 0.766183 |
| 13 | 53319400 | 53321799 | 20.23079 | 34.74969 | 50.08008 | 21.9665 | -0.14132 | 0.922801 | 0.766231 |
| 6 | 78606400 | 78606999 | 6.537215 | 12.70517 | 21.77231 | 8.100116 | -0.17201 | -0.28239 | 0.766238 |
| 6 | 63087800 | 63091799 | 60.06967 | 95.24674 | 129.9092 | 54.07141 | -0.11686 | 2.3342 | 0.766248 |
| 9 | 117157200 | 117158799 | 15.58031 | 27.10614 | 47.90059 | 21.04871 | -0.4306 | 0.772664 | 0.766267 |
| 18 | 31623600 | 31624599 | 12.39347 | 22.78418 | 27.56391 | 11.70887 | 0.157459 | 0.248379 | 0.766279 |
| 13 | 78287000 | 78290199 | 33.52522 | 54.52736 | 77.59121 | 33.30001 | -0.1383 | 1.567818 | 0.766315 |
| 3 | 10719800 | 10720799 | 14.22817 | 25.09186 | 36.69662 | 16.23983 | -0.15621 | 0.55415 | 0.766325 |
| 9 | 113188200 | 113190799 | 28.2943 | 46.6078 | 66.85494 | 28.83611 | -0.13956 | 1.38196 | 0.766343 |
| 4 | 4126800 | 4127799 | 12.05179 | 22.52356 | 27.24617 | 11.29776 | 0.183601 | 0.149125 | 0.766346 |
| 1 | 62716000 | 62718999 | 39.34665 | 63.38711 | 89.07474 | 37.88499 | -0.1342 | 1.809402 | 0.766362 |
| 13 | 86513200 | 86515599 | 22.59196 | 38.69389 | 45.25595 | 19.84623 | 0.157969 | 0.949657 | 0.766362 |
| 12 | 113529000 | 113542799 | 169.5871 | 254.5895 | 272.4681 | 112.4484 | 0.169615 | 3.455426 | 0.766371 |
| X | 113619200 | 113619999 | 8.821188 | 17.03381 | 22.03621 | 8.48519 | 0.182839 | -0.13858 | 0.766396 |
| 12 | 49245200 | 49250399 | 44.31997 | 71.80829 | 79.11739 | 33.8668 | 0.224873 | 1.65995 | 0.766402 |
| 13 | 80213800 | 80215199 | 14.90031 | 26.15469 | 38.32958 | 16.95396 | -0.16098 | 0.619999 | 0.766404 |
| 10 | 66514600 | 66525999 | 159.8357 | 240.1161 | 264.6211 | 109.2301 | 0.128241 | 3.43968 | 0.766409 |
| 3 | 148136200 | 148137799 | 20.23496 | 33.63316 | 65.79884 | 28.54371 | -0.58376 | 1.254838 | 0.766413 |
| 2 | 20949800 | 20950799 | 10.78254 | 20.17088 | 24.69322 | 10.22249 | 0.168618 | 0.126616 | 0.76642 |
| 2 | 22785400 | 22786199 | 8.491196 | 15.96723 | 26.40966 | 10.83726 | -0.25231 | 0.075814 | 0.766432 |
| 6 | 38037400 | 38043399 | 80.52465 | 125.6485 | 168.6562 | 69.98592 | -0.11212 | 2.681945 | 0.766433 |

| | | | | | | | | | |
|---|---|---|---|---|---|---|---|---|---|
| 4 | 33110200 | 33113799 | 24.36653 | 40.59771 | 59.84355 | 26.00179 | -0.17683 | 1.230029 | 0.766435 |
| 4 | 27789800 | 27791999 | 17.04191 | 29.80808 | 44.03304 | 19.4185 | -0.17278 | 0.694399 | 0.766447 |
| 14 | 107721800 | 107726199 | 87.4727 | 136.2351 | 156.538 | 64.974 | 0.109816 | 2.715962 | 0.766456 |
| 18 | 45993800 | 45997999 | 53.46473 | 85.52448 | 99.71392 | 41.96882 | 0.122402 | 2.051626 | 0.766469 |
| 16 | 23708400 | 23709799 | 20.0598 | 34.27155 | 49.54296 | 21.72187 | -0.14692 | 0.969726 | 0.766483 |
| 9 | 9138800 | 9139599 | 11.18844 | 21.09373 | 25.78796 | 10.5359 | 0.185423 | 0.082296 | 0.766491 |
| 1 | 16622800 | 16624399 | 22.2312 | 38.20432 | 45.26038 | 19.85274 | 0.140412 | 0.920049 | 0.766511 |
| 6 | 60552000 | 60552999 | 16.13609 | 28.23182 | 40.92787 | 18.05351 | -0.14558 | 0.682162 | 0.766526 |
| 16 | 70689200 | 70689999 | 12.52198 | 23.38356 | 22.52148 | 9.351314 | 0.508882 | 0.148086 | 0.766534 |
| 1 | 54091000 | 54092599 | 18.26273 | 32.00877 | 37.57005 | 16.59697 | 0.157469 | 0.658535 | 0.766542 |
| 11 | 25500200 | 25501599 | 15.16253 | 27.00865 | 31.82055 | 14.01131 | 0.160688 | 0.46142 | 0.766549 |
| 7 | 23743200 | 23743999 | 24.207 | 41.01238 | 48.51438 | 21.24357 | 0.139873 | 1.058123 | 0.766551 |
| 1 | 28437000 | 28438799 | 53.04159 | 85.148 | 83.91539 | 35.50669 | 0.370341 | 1.936644 | 0.766578 |
| 7 | 17533600 | 17534999 | 35.87897 | 58.36843 | 67.25627 | 28.86685 | 0.165692 | 1.566758 | 0.766594 |
| 13 | 34951000 | 34952199 | 14.64115 | 25.99444 | 29.17432 | 12.88173 | 0.228019 | 0.47609 | 0.766595 |
| 11 | 101876800 | 101877199 | 4.222035 | 8.255951 | 16.07332 | 5.481351 | -0.23772 | -0.641 | 0.766669 |
| 1 | 3090000 | 3091599 | 24.05379 | 40.91606 | 48.49065 | 21.25301 | 0.137608 | 1.022933 | 0.766693 |
| 13 | 86317600 | 86321799 | 54.29684 | 87.01549 | 99.81453 | 42.12532 | 0.148857 | 1.996245 | 0.7667 |
| 15 | 95551400 | 95553199 | 16.15942 | 27.64558 | 53.19224 | 23.32444 | -0.55734 | 0.946577 | 0.766704 |
| 4 | 20206800 | 20207799 | 14.43757 | 26.07656 | 30.94274 | 13.4461 | 0.163347 | 0.372134 | 0.766777 |
| 15 | 84372200 | 84375199 | 38.18846 | 61.52229 | 87.73703 | 37.31694 | -0.15537 | 1.805977 | 0.766779 |
| 11 | 121746600 | 121757599 | 196.2192 | 293.3104 | 291.7551 | 120.2972 | 0.269578 | 3.54214 | 0.76679 |
| 13 | 79869000 | 79871199 | 30.02916 | 49.30737 | 70.45602 | 30.34819 | -0.13596 | 1.425936 | 0.766805 |
| 4 | 75786200 | 75787999 | 15.41475 | 27.34687 | 32.39714 | 14.31402 | 0.149546 | 0.490807 | 0.766806 |
| 18 | 31634800 | 31635799 | 15.09401 | 26.52022 | 38.79581 | 17.17415 | -0.1584 | 0.607026 | 0.766811 |
| 7 | 42517800 | 42521199 | 52.43168 | 84.3301 | 93.93177 | 39.85697 | 0.197421 | 1.880083 | 0.766826 |
| 15 | 51586600 | 51588399 | 15.95547 | 28.06971 | 41.77074 | 18.49433 | -0.18281 | 0.597621 | 0.766832 |
| 1 | 195044800 | 195054199 | 110.0223 | 169.5245 | 188.625 | 78.35185 | 0.144222 | 2.919309 | 0.766832 |
| 11 | 38951000 | 38951799 | 10.48057 | 19.82885 | 24.82802 | 10.07001 | 0.160679 | 0.051211 | 0.766839 |
| 2 | 176788000 | 176794799 | 43.16761 | 71.05827 | 54.61283 | 23.54885 | 0.757604 | 1.416143 | 0.766842 |
| 16 | 77326800 | 77327999 | 15.60768 | 29.03122 | 30.42184 | 12.71623 | 0.380363 | 0.216729 | 0.766872 |
| 18 | 69497400 | 69499799 | 18.54544 | 31.57438 | 55.91267 | 24.50777 | -0.43934 | 1.003086 | 0.766881 |
| X | 146575800 | 146582199 | 78.35755 | 124.3032 | 111.1246 | 46.2748 | 0.49237 | 2.331812 | 0.766905 |
| 6 | 68632400 | 68643199 | 175.6282 | 262.3286 | 286.1855 | 117.983 | 0.135397 | 3.551117 | 0.766926 |
| 19 | 9136400 | 9139999 | 45.4484 | 72.99911 | 101.8 | 43.11047 | -0.12858 | 1.916621 | 0.766978 |
| 12 | 11669600 | 11670399 | 10.41745 | 19.71313 | 24.33955 | 9.867976 | 0.181656 | 0.050029 | 0.766984 |
| 9 | 102833000 | 102833599 | 9.663276 | 17.96633 | 27.8665 | 11.65683 | -0.18216 | 0.165686 | 0.766997 |
| 5 | 3482200 | 3483799 | 10.01003 | 18.34261 | 28.81912 | 12.29645 | -0.22136 | 0.252525 | 0.767036 |
| 8 | 40182600 | 40183399 | 13.36787 | 23.34095 | 44.70543 | 19.68922 | -0.54544 | 0.705724 | 0.767053 |
| 17 | 24034000 | 24044199 | 308.1433 | 435.9738 | 448.3464 | 185.5585 | 0.144883 | 4.404966 | 0.76706 |
| 19 | 39234000 | 39238999 | 82.22623 | 128.8027 | 148.4578 | 61.57976 | 0.111791 | 2.612646 | 0.767072 |
| 12 | 99500000 | 99505199 | 67.32317 | 106.3042 | 145.0237 | 60.21605 | -0.12219 | 2.443531 | 0.767085 |
| 11 | 62547600 | 62548399 | 6.688473 | 13.00628 | 22.42062 | 8.304423 | -0.17471 | -0.29419 | 0.767104 |
| 12 | 8255000 | 8256199 | 10.92882 | 20.11168 | 30.31189 | 12.85867 | -0.15599 | 0.238786 | 0.767116 |
| 6 | 132491200 | 132511799 | 354.3534 | 498.8712 | 516.9837 | 214.1412 | 0.126603 | 4.480857 | 0.767125 |
| 7 | 16879800 | 16880399 | 17.76098 | 30.98763 | 35.3724 | 15.5712 | 0.19746 | 0.723307 | 0.767202 |
| 19 | 3208400 | 3211399 | 55.97689 | 88.81816 | 123.8634 | 51.59044 | -0.14788 | 2.3097 | 0.767203 |

| | | | | | | | | | |
|---|---|---|---|---|---|---|---|---|---|
| 16 | 90111200 | 90114999 | 31.01136 | 51.13197 | 60.08878 | 25.92811 | 0.147662 | 1.368769 | 0.767212 |
| 6 | 59883200 | 59894799 | 178.1609 | 265.5839 | 291.3586 | 120.0694 | 0.124753 | 3.588431 | 0.767257 |
| 5 | 109806000 | 109811799 | 16.88776 | 29.99877 | 28.66178 | 12.64986 | 0.459074 | 0.483986 | 0.76727 |
| X | 120463000 | 120472799 | 46.52757 | 75.09262 | 68.51678 | 29.21183 | 0.490578 | 1.759178 | 0.767298 |
| 4 | 60026200 | 60027799 | 33.0367 | 54.07703 | 62.44949 | 26.86406 | 0.168758 | 1.468882 | 0.767308 |
| 1 | 104328400 | 104336199 | 144.3728 | 217.8616 | 244.1135 | 100.8899 | 0.110001 | 3.34845 | 0.767313 |
| 6 | 136546200 | 136548799 | 21.81354 | 37.87596 | 43.56335 | 19.11917 | 0.185787 | 0.821093 | 0.767328 |
| 1 | 72160600 | 72174199 | 233.4655 | 337.3211 | 422.5995 | 174.3257 | -0.10953 | 4.094635 | 0.767332 |
| 1 | 91240000 | 91241799 | 15.02921 | 26.86341 | 31.4894 | 13.82535 | 0.17116 | 0.441856 | 0.767359 |
| 6 | 131817200 | 131838199 | 328.6611 | 467.802 | 474.7931 | 196.3073 | 0.172558 | 4.320113 | 0.767371 |
| 11 | 3215800 | 3216599 | 7.166189 | 13.69365 | 23.51902 | 9.340582 | -0.26414 | -0.05101 | 0.767384 |
| 4 | 58537000 | 58539999 | 21.53824 | 36.2389 | 63.04079 | 27.54647 | -0.41495 | 1.125197 | 0.767397 |
| 1 | 36088200 | 36092599 | 39.03116 | 63.29249 | 74.50487 | 31.91218 | 0.130815 | 1.633239 | 0.767405 |
| 3 | 44803800 | 44804799 | 18.02728 | 31.52798 | 37.26788 | 16.43349 | 0.14754 | 0.693199 | 0.767415 |
| 10 | 60258200 | 60263999 | 86.45698 | 133.835 | 179.5474 | 74.57934 | -0.12024 | 2.825147 | 0.767425 |
| 10 | 57115000 | 57116399 | 22.6103 | 39.33022 | 36.22538 | 15.91668 | 0.50532 | 0.7927 | 0.767431 |
| 2 | 88432000 | 88434999 | 53.08083 | 84.99964 | 99.23803 | 41.83179 | 0.122097 | 2.018163 | 0.767437 |
| 13 | 68030800 | 68037799 | 139.6223 | 210.3982 | 231.3099 | 95.5588 | 0.135672 | 3.374284 | 0.767449 |
| 5 | 150820600 | 150823399 | 22.17678 | 37.7258 | 54.41644 | 23.85035 | -0.14482 | 1.021732 | 0.767469 |
| 5 | 45556800 | 45557799 | 10.08999 | 18.85843 | 28.96601 | 12.00951 | -0.15819 | 0.137107 | 0.767508 |
| 11 | 8371200 | 8372399 | 12.13397 | 22.58032 | 27.12944 | 11.33117 | 0.18483 | 0.175467 | 0.767536 |
| 14 | 121423600 | 121425399 | 18.52699 | 32.22587 | 36.91885 | 16.22091 | 0.191802 | 0.771872 | 0.767612 |
| 3 | 127187400 | 127189199 | 16.54518 | 29.14221 | 34.48722 | 15.26582 | 0.1469 | 0.576481 | 0.767614 |
| 8 | 18019800 | 18020799 | 16.21036 | 28.66823 | 33.88282 | 14.98616 | 0.150939 | 0.529137 | 0.76762 |
| 1 | 138518400 | 138522199 | 33.06236 | 53.9622 | 76.60715 | 32.91946 | -0.13136 | 1.517824 | 0.767621 |
| 18 | 58813800 | 58817599 | 43.06628 | 69.37101 | 98.68956 | 41.96697 | -0.15221 | 1.821361 | 0.767622 |
| 9 | 80744200 | 80745799 | 11.80424 | 21.61215 | 32.87195 | 14.04179 | -0.17633 | 0.280778 | 0.767634 |
| 12 | 72576000 | 72577799 | 26.08143 | 44.60337 | 42.54999 | 18.65731 | 0.450582 | 0.97561 | 0.767643 |
| 6 | 76517000 | 76519399 | 24.22891 | 41.92212 | 46.27121 | 20.30035 | 0.244065 | 0.869466 | 0.767663 |
| 11 | 33111600 | 33114599 | 17.69558 | 30.33169 | 53.79899 | 23.59413 | -0.44012 | 0.935198 | 0.767681 |
| 2 | 44330200 | 44332799 | 39.8874 | 64.8638 | 74.99247 | 32.18265 | 0.160355 | 1.582203 | 0.767697 |
| 1 | 110475400 | 110475799 | 10.66142 | 19.95138 | 24.22572 | 10.02292 | 0.18121 | 0.122962 | 0.767702 |
| X | 110200000 | 110203799 | 40.35473 | 66.35739 | 69.93842 | 30.14362 | 0.30278 | 1.434826 | 0.767714 |
| 2 | 99944600 | 99947399 | 29.53032 | 48.79758 | 70.77684 | 30.57968 | -0.15445 | 1.34839 | 0.767715 |
| 17 | 12292000 | 12295999 | 61.83947 | 98.41953 | 114.4333 | 47.82245 | 0.118418 | 2.221769 | 0.767724 |
| X | 32523200 | 32553799 | 579.4825 | 790.0344 | 711.6891 | 296.2804 | 0.281959 | 4.975964 | 0.767727 |
| 17 | 27539000 | 27550399 | 131.4138 | 198.8537 | 219.6295 | 90.8499 | 0.134149 | 3.288029 | 0.76773 |
| 9 | 44648600 | 44652799 | 36.91031 | 59.76221 | 84.13618 | 35.99702 | -0.12879 | 1.667327 | 0.767741 |
| 1 | 65445400 | 65445799 | 6.089865 | 11.84774 | 20.69967 | 7.6383 | -0.18892 | -0.32165 | 0.767743 |
| 19 | 24088200 | 24091999 | 44.58893 | 71.53484 | 99.24417 | 41.95339 | -0.12313 | 1.950114 | 0.767748 |
| 19 | 59098400 | 59099599 | 9.424792 | 17.90845 | 28.90798 | 11.61832 | -0.19169 | 0.02139 | 0.767751 |
| 2 | 86799200 | 86800799 | 17.8632 | 31.09289 | 45.20961 | 19.87227 | -0.15057 | 0.771129 | 0.767769 |
| 4 | 47453600 | 47455199 | 24.41675 | 41.35171 | 49.0484 | 21.47488 | 0.135987 | 1.063508 | 0.76777 |
| 7 | 43295000 | 43297199 | 24.41675 | 41.35171 | 49.0484 | 21.47488 | 0.135987 | 1.063508 | 0.76777 |
| 5 | 8425800 | 8429599 | 45.23753 | 72.50617 | 100.7745 | 42.5385 | -0.12738 | 1.980854 | 0.767771 |
| 11 | 98528600 | 98531199 | 21.35657 | 36.87135 | 43.64412 | 19.14649 | 0.142956 | 0.872403 | 0.767776 |
| 13 | 37438000 | 37440599 | 25.06563 | 42.58684 | 50.10678 | 21.95925 | 0.147749 | 1.039336 | 0.76778 |

| | | | | | | | | | |
|---|---|---|---|---|---|---|---|---|---|
| 9 | 14947400 | 14949799 | 20.68531 | 35.10233 | 51.63855 | 22.62264 | -0.17323 | 1.038341 | 0.767793 |
| 16 | 32124800 | 32127399 | 21.98347 | 37.29475 | 53.86149 | 23.59448 | -0.14699 | 1.043792 | 0.767804 |
| 11 | 40139200 | 40140399 | 14.81026 | 25.82919 | 46.28188 | 20.36853 | -0.45001 | 0.733139 | 0.767808 |
| 2 | 126369000 | 126371999 | 27.65816 | 46.32278 | 54.56897 | 23.78809 | 0.145422 | 1.179872 | 0.767826 |
| 7 | 30923800 | 30970399 | 904.0081 | 1177.818 | 1123.472 | 470.5374 | 0.133954 | 5.7002 | 0.76783 |
| Y | 2019000 | 2021399 | 21.33469 | 37.22792 | 24.26951 | 10.68457 | 1.001425 | 0.718902 | 0.767854 |
| 12 | 114602000 | 114606399 | 60.5115 | 96.82534 | 91.74213 | 38.62969 | 0.421559 | 2.046259 | 0.767866 |
| 8 | 16245200 | 16248399 | 34.06809 | 55.8204 | 66.21274 | 28.49479 | 0.130691 | 1.458781 | 0.767871 |
| 15 | 44929400 | 44932199 | 24.49215 | 40.89649 | 59.91153 | 26.06732 | -0.16799 | 1.206577 | 0.767877 |
| 3 | 51841000 | 51843799 | 37.26792 | 60.42197 | 70.05013 | 29.99998 | 0.152543 | 1.63228 | 0.76788 |
| 15 | 99033400 | 99034999 | 21.09611 | 35.85162 | 51.66212 | 22.64134 | -0.14349 | 1.025372 | 0.767881 |
| 10 | 94798000 | 94805399 | 63.23521 | 99.52239 | 159.0086 | 65.97077 | -0.35348 | 2.512885 | 0.7679 |
| 5 | 6736400 | 6738999 | 87.77973 | 136.2547 | 151.2774 | 62.82198 | 0.155537 | 2.773199 | 0.767911 |
| 13 | 111772200 | 111776799 | 60.36166 | 96.20537 | 110.7663 | 46.38194 | 0.134747 | 2.171823 | 0.767931 |
| 9 | 89292200 | 89294599 | 36.38007 | 58.99951 | 83.30553 | 35.68438 | -0.13082 | 1.632498 | 0.767969 |
| 8 | 59828200 | 59831999 | 55.61947 | 88.88223 | 103.383 | 43.45144 | 0.12424 | 2.085628 | 0.767982 |
| 3 | 87498000 | 87498399 | 5.794817 | 11.30783 | 20.46555 | 7.286125 | -0.1916 | -0.45013 | 0.767994 |
| 2 | 83723000 | 83723599 | 9.174896 | 17.55962 | 22.35461 | 8.839378 | 0.169369 | -0.0507 | 0.767996 |
| 1 | 79597800 | 79600199 | 20.95111 | 36.22916 | 42.56381 | 18.67348 | 0.154136 | 0.853936 | 0.768022 |
| 7 | 68719400 | 68721399 | 22.21006 | 38.79469 | 42.33451 | 18.64949 | 0.262226 | 0.741186 | 0.768026 |
| 6 | 145876200 | 145880399 | 83.78845 | 130.6363 | 148.261 | 61.52864 | 0.128825 | 2.694548 | 0.768033 |
| 4 | 45149400 | 45150599 | 9.642791 | 17.88933 | 27.83992 | 11.68064 | -0.19022 | 0.176891 | 0.768038 |
| 10 | 36143400 | 36144999 | 18.69545 | 32.81257 | 38.03779 | 16.82326 | 0.175359 | 0.640992 | 0.768063 |
| 17 | 66044600 | 66049599 | 50.48435 | 80.53365 | 111.7362 | 46.84511 | -0.13251 | 2.13533 | 0.768066 |
| 13 | 13600000 | 13601999 | 12.26226 | 21.48972 | 43.07248 | 19.02614 | -0.61044 | 0.637806 | 0.768073 |
| 1 | 194337400 | 194339999 | 53.41835 | 85.3086 | 98.19353 | 41.23188 | 0.138284 | 2.101169 | 0.768086 |
| X | 134473800 | 134475999 | 15.60427 | 27.41868 | 40.30638 | 17.84391 | -0.16546 | 0.611506 | 0.768088 |
| 1 | 183221600 | 183222199 | 10.27208 | 19.25036 | 22.71148 | 9.373651 | 0.225108 | 0.105302 | 0.768118 |
| 12 | 14198200 | 14203399 | 37.86029 | 60.65458 | 109.1327 | 46.03498 | -0.49981 | 1.990228 | 0.768119 |
| 2 | 33664400 | 33670399 | 178.1001 | 261.8461 | 341.8256 | 140.7721 | -0.14458 | 3.823506 | 0.76812 |
| 13 | 20864800 | 20866599 | 15.64352 | 27.55803 | 40.72335 | 18.02573 | -0.17196 | 0.579814 | 0.768127 |
| 3 | 134520800 | 134521599 | 15.85697 | 27.17977 | 52.12067 | 22.85806 | -0.55193 | 0.928336 | 0.76816 |
| 4 | 39003600 | 39005599 | 23.22521 | 39.29696 | 56.59244 | 24.77086 | -0.14303 | 1.075643 | 0.768191 |
| 10 | 115095400 | 115097599 | 20.32618 | 35.05177 | 51.74861 | 22.7011 | -0.17492 | 0.888288 | 0.768205 |
| 2 | 20266000 | 20268199 | 20.35276 | 35.44329 | 41.38865 | 18.19824 | 0.164403 | 0.770456 | 0.768212 |
| 7 | 3284400 | 3286799 | 23.1457 | 39.4245 | 46.36053 | 20.3216 | 0.149056 | 1.003663 | 0.768221 |
| 1 | 178003000 | 178003599 | 13.47993 | 24.24444 | 26.72797 | 11.67352 | 0.264501 | 0.382167 | 0.768229 |
| 3 | 5747400 | 5748399 | 11.36001 | 21.013 | 32.55074 | 13.7121 | -0.18832 | 0.216847 | 0.768253 |
| 7 | 4858400 | 4863399 | 36.17761 | 58.4504 | 83.80906 | 35.78055 | -0.15836 | 1.715289 | 0.768255 |
| X | 91045400 | 91047599 | 24.05802 | 41.43301 | 46.84319 | 20.54837 | 0.2083 | 0.912634 | 0.768258 |
| 16 | 30279400 | 30284999 | 68.10664 | 108.1385 | 123.627 | 51.52731 | 0.139332 | 2.299677 | 0.768269 |
| 7 | 5854600 | 5863999 | 204.9638 | 301.5827 | 329.2869 | 135.6016 | 0.114125 | 3.814131 | 0.768269 |
| 12 | 97804800 | 97807399 | 18.92109 | 33.30515 | 37.32868 | 16.52577 | 0.224388 | 0.600702 | 0.768284 |
| 18 | 68153400 | 68157399 | 42.94904 | 69.47338 | 80.14325 | 34.26459 | 0.156586 | 1.692066 | 0.768296 |
| 9 | 78737200 | 78738599 | 13.60581 | 24.62452 | 29.49179 | 12.78289 | 0.152481 | 0.351477 | 0.768297 |
| 9 | 37857800 | 37867999 | 193.5154 | 286.0868 | 314.7266 | 129.6122 | 0.109662 | 3.734209 | 0.76831 |
| 10 | 25737200 | 25737799 | 6.233751 | 12.16248 | 22.43207 | 8.007507 | -0.22321 | -0.43181 | 0.768312 |

| | | | | | | | | | |
|---|---|---|---|---|---|---|---|---|---|
| 7 | 141745200 | 141746399 | 13.05027 | 23.75017 | 28.49121 | 12.26401 | 0.156464 | 0.312566 | 0.768313 |
| 8 | 72633400 | 72636199 | 21.84206 | 37.06146 | 53.30672 | 23.35272 | -0.14108 | 1.04171 | 0.76832 |
| 10 | 11314200 | 11316799 | 20.66274 | 35.41656 | 51.16831 | 22.44036 | -0.14546 | 0.943579 | 0.768322 |
| 3 | 116993800 | 117004599 | 169.9557 | 251.8279 | 323.6886 | 133.3099 | -0.11167 | 3.694345 | 0.76833 |
| 5 | 49982000 | 49983999 | 17.06353 | 30.4986 | 33.68461 | 14.78956 | 0.256074 | 0.455483 | 0.768336 |
| 6 | 119030600 | 119032199 | 16.59153 | 28.96746 | 42.01064 | 18.50151 | -0.14631 | 0.717927 | 0.768345 |
| 10 | 55591000 | 55592199 | 14.9229 | 26.54735 | 31.43883 | 13.85837 | 0.152382 | 0.468693 | 0.768348 |
| 14 | 108595800 | 108601399 | 52.37885 | 84.59171 | 87.2354 | 37.22392 | 0.315417 | 1.75734 | 0.768351 |
| 12 | 111468600 | 111471399 | 37.89104 | 61.08628 | 86.5962 | 36.86734 | -0.14553 | 1.784648 | 0.768363 |
| 14 | 53671000 | 53671599 | 10.22234 | 19.4347 | 24.40244 | 9.795997 | 0.169943 | 0.015593 | 0.768385 |
| 2 | 70189600 | 70191399 | 18.18172 | 31.86603 | 37.62842 | 16.62243 | 0.148917 | 0.659854 | 0.768385 |
| 3 | 119148400 | 119148999 | 10.45701 | 19.87225 | 24.40823 | 9.807303 | 0.200573 | 0.022102 | 0.768397 |
| 4 | 147147000 | 147157399 | 252.3601 | 368.1681 | 330.8075 | 136.3087 | 0.383403 | 3.950303 | 0.768399 |
| 8 | 42817000 | 42817799 | 7.573042 | 14.79648 | 19.06326 | 6.640372 | 0.320314 | -0.50978 | 0.768401 |
| 14 | 110473400 | 110473999 | 6.966658 | 13.51494 | 22.81543 | 8.597748 | -0.16838 | -0.2311 | 0.768437 |
| 7 | 9656200 | 9658399 | 25.86431 | 44.06697 | 41.64571 | 18.25531 | 0.463393 | 1.009031 | 0.768448 |
| 7 | 107086200 | 107088199 | 17.38179 | 30.04976 | 44.77812 | 19.64479 | -0.18736 | 0.843934 | 0.768456 |
| 17 | 5425800 | 5429199 | 22.6591 | 38.20088 | 55.49414 | 24.26353 | -0.15577 | 1.101673 | 0.768482 |
| 6 | 116475600 | 116484199 | 170.7737 | 254.8019 | 282.4128 | 116.4032 | 0.111145 | 3.570748 | 0.768487 |
| 1 | 94272000 | 94275399 | 32.33261 | 53.69813 | 61.50413 | 26.66594 | 0.185758 | 1.290514 | 0.768538 |
| 16 | 11306600 | 11310799 | 34.53151 | 56.08983 | 79.25643 | 33.98935 | -0.12978 | 1.594292 | 0.76854 |
| 6 | 140115200 | 140117399 | 20.15434 | 35.34509 | 38.94205 | 17.21167 | 0.24822 | 0.660341 | 0.768555 |
| 2 | 144961800 | 144966799 | 37.08877 | 60.21442 | 70.84067 | 30.36938 | 0.133268 | 1.604374 | 0.768587 |
| 12 | 17467400 | 17470199 | 16.60437 | 28.95346 | 42.05496 | 18.5064 | -0.14867 | 0.736165 | 0.768591 |
| 6 | 95928800 | 95931199 | 16.60853 | 28.67386 | 51.11063 | 22.42202 | -0.44498 | 0.864615 | 0.768594 |
| 7 | 117657600 | 117659999 | 29.1789 | 48.06665 | 68.87224 | 29.70666 | -0.13806 | 1.382575 | 0.768597 |
| 6 | 102317600 | 102318799 | 17.11255 | 29.82478 | 43.10005 | 18.96038 | -0.14141 | 0.74771 | 0.768598 |
| 6 | 88554800 | 88556199 | 12.73656 | 22.74504 | 33.62281 | 14.77321 | -0.1625 | 0.434588 | 0.768605 |
| 7 | 40984800 | 40989199 | 105.7577 | 162.3731 | 183.8766 | 76.34214 | 0.114937 | 2.986626 | 0.768607 |
| 8 | 34919400 | 34921199 | 17.74186 | 31.10991 | 36.53617 | 16.14641 | 0.156609 | 0.646367 | 0.768615 |
| 13 | 85172400 | 85173599 | 11.16162 | 20.3059 | 30.77181 | 13.25149 | -0.17857 | 0.308263 | 0.768625 |
| 13 | 61234200 | 61235599 | 17.18706 | 30.70272 | 34.2699 | 15.05342 | 0.240609 | 0.462169 | 0.768634 |
| 14 | 20270000 | 20271799 | 16.14648 | 28.21816 | 40.98079 | 18.05963 | -0.14808 | 0.700889 | 0.768648 |
| 5 | 6758200 | 6764799 | 59.43242 | 94.93645 | 108.0248 | 45.37592 | 0.155213 | 2.101507 | 0.76865 |
| 5 | 149227800 | 149228599 | 10.52795 | 19.18211 | 29.81897 | 12.81688 | -0.21319 | 0.292724 | 0.768694 |
| 10 | 30356400 | 30359399 | 71.65194 | 113.0993 | 131.2243 | 54.47659 | 0.110759 | 2.453625 | 0.768743 |
| 16 | 18932400 | 18933599 | 16.74961 | 28.79176 | 51.63716 | 22.65 | -0.45521 | 0.903879 | 0.768744 |
| 11 | 114080400 | 114083399 | 41.47319 | 66.63763 | 93.80062 | 39.74335 | -0.14135 | 1.896563 | 0.768755 |
| 8 | 65684200 | 65685999 | 17.01899 | 30.35692 | 34.26933 | 15.07222 | 0.223136 | 0.470276 | 0.768766 |
| Y | 27902000 | 27902399 | 7.195946 | 13.9727 | 16.35655 | 6.130137 | 0.362637 | -0.26925 | 0.768768 |
| 7 | 31969400 | 31970599 | 20.69142 | 36.07071 | 33.72001 | 14.83614 | 0.484218 | 0.748151 | 0.768772 |
| 2 | 160470200 | 160471599 | 14.90653 | 26.43819 | 30.96734 | 13.68537 | 0.165634 | 0.486277 | 0.768777 |
| 6 | 37744800 | 37752799 | 81.72255 | 128.8023 | 143.3809 | 59.49807 | 0.168735 | 2.493749 | 0.768805 |
| 4 | 139862400 | 139864799 | 31.86553 | 52.00186 | 74.22367 | 31.89368 | -0.13926 | 1.514272 | 0.768816 |
| 10 | 67492800 | 67494999 | 14.9504 | 26.29575 | 38.67044 | 17.1224 | -0.16588 | 0.588645 | 0.768818 |
| 17 | 88610800 | 88615599 | 53.00129 | 84.26295 | 118.3002 | 49.37075 | -0.15474 | 2.247991 | 0.768852 |
| 11 | 14403400 | 14407399 | 37.16337 | 61.50523 | 65.77367 | 28.44817 | 0.284167 | 1.348581 | 0.768858 |

| | | | | | | | | | |
|---|---|---|---|---|---|---|---|---|---|
| 11 | 16457600 | 16458399 | 6.105587 | 11.84105 | 21.10332 | 7.963417 | -0.24645 | -0.23874 | 0.768865 |
| 10 | 126173000 | 126176399 | 34.4116 | 57.02938 | 63.86836 | 27.65031 | 0.21788 | 1.323314 | 0.768866 |
| 7 | 38269600 | 38272599 | 47.94195 | 77.22993 | 88.74086 | 37.74005 | 0.155428 | 1.822722 | 0.768873 |
| 17 | 78918800 | 78920799 | 16.64523 | 29.47579 | 34.39441 | 15.20845 | 0.170134 | 0.520204 | 0.768874 |
| 17 | 58943400 | 58947199 | 26.34605 | 44.25687 | 52.32812 | 22.85417 | 0.140059 | 1.142299 | 0.768879 |
| 12 | 93802000 | 93802999 | 16.9494 | 30.00441 | 34.86712 | 15.4185 | 0.175819 | 0.52607 | 0.768909 |
| X | 73737000 | 73739999 | 30.53578 | 50.63248 | 59.45902 | 25.75065 | 0.149781 | 1.305501 | 0.768914 |
| 2 | 92470800 | 92472799 | 21.19865 | 35.85609 | 52.75264 | 23.08782 | -0.17376 | 1.065605 | 0.768949 |
| 7 | 4201000 | 4204399 | 29.26507 | 47.89631 | 69.49573 | 29.89356 | -0.15989 | 1.464614 | 0.768975 |
| 3 | 81681000 | 81684199 | 24.50702 | 42.0834 | 47.91979 | 21.01877 | 0.197047 | 0.938611 | 0.768992 |
| 5 | 93456800 | 93458199 | 13.20671 | 23.55953 | 34.589 | 15.21139 | -0.15365 | 0.445952 | 0.768995 |
| 18 | 70304800 | 70305999 | 10.96821 | 20.04961 | 30.2721 | 12.95863 | -0.16779 | 0.277466 | 0.769001 |
| 6 | 114296800 | 114301599 | 45.52925 | 73.38566 | 85.83277 | 36.53293 | 0.130918 | 1.801319 | 0.769025 |
| X | 109210600 | 109212799 | 33.79904 | 56.56589 | 48.63221 | 21.19021 | 0.598459 | 1.205419 | 0.769045 |
| 10 | 121404000 | 121407999 | 43.62964 | 70.04142 | 97.38185 | 41.2069 | -0.1251 | 1.928596 | 0.769061 |
| 5 | 23135000 | 23138199 | 23.52608 | 39.73332 | 57.16003 | 25.00542 | -0.14164 | 1.092291 | 0.769077 |
| 15 | 77352200 | 77353999 | 20.07819 | 35.03689 | 40.81574 | 17.96947 | 0.168169 | 0.740332 | 0.769079 |
| 5 | 51395200 | 51398799 | 65.14848 | 103.2489 | 119.4635 | 49.69915 | 0.119434 | 2.350803 | 0.769083 |
| 8 | 115520000 | 115521799 | 15.68233 | 27.49447 | 39.84777 | 17.60769 | -0.14513 | 0.645459 | 0.769094 |
| 14 | 34270400 | 34271799 | 13.27193 | 23.34721 | 35.82492 | 15.86197 | -0.22647 | 0.573849 | 0.769106 |
| 17 | 33951600 | 33953399 | 16.42579 | 28.5943 | 42.10327 | 18.50823 | -0.1685 | 0.758539 | 0.769162 |
| 4 | 17774400 | 17777599 | 26.69718 | 44.91792 | 51.82034 | 22.64339 | 0.175394 | 1.127178 | 0.769174 |
| 4 | 49464800 | 49469199 | 94.81073 | 145.813 | 194.3722 | 80.71852 | -0.11825 | 2.950817 | 0.769199 |
| 3 | 44417600 | 44417999 | 11.12896 | 20.6323 | 24.20396 | 10.16461 | 0.213772 | 0.184225 | 0.769204 |
| 8 | 109600000 | 109601599 | 26.08895 | 43.68032 | 51.24117 | 22.3328 | 0.151539 | 1.176787 | 0.769227 |
| 10 | 47496000 | 47497999 | 27.72886 | 46.99587 | 45.27446 | 19.82699 | 0.434984 | 1.064095 | 0.769227 |
| 17 | 4066600 | 4068999 | 21.23498 | 36.89193 | 42.46902 | 18.64542 | 0.184538 | 0.808059 | 0.769245 |
| 10 | 105135600 | 105137799 | 16.54456 | 29.30297 | 34.32128 | 15.17555 | 0.164715 | 0.51585 | 0.769252 |
| 10 | 103402600 | 103404199 | 16.06368 | 28.04172 | 41.49361 | 18.2721 | -0.17524 | 0.715316 | 0.769252 |
| 14 | 47844200 | 47846999 | 42.00328 | 67.43544 | 95.14405 | 40.26445 | -0.14607 | 1.923594 | 0.769268 |
| X | 57447200 | 57449599 | 19.33675 | 33.55687 | 50.10597 | 21.98474 | -0.18914 | 0.828789 | 0.769282 |
| 8 | 43038600 | 43040199 | 16.95163 | 29.63711 | 43.01095 | 18.9607 | -0.14717 | 0.70125 | 0.769294 |
| 3 | 98080600 | 98082599 | 17.67711 | 31.54367 | 34.83704 | 15.31664 | 0.255086 | 0.47497 | 0.769327 |
| 8 | 91888400 | 91894199 | 62.35006 | 99.3015 | 114.503 | 47.89787 | 0.131468 | 2.200134 | 0.769329 |
| 7 | 41111200 | 41147599 | 606.198 | 819.4488 | 779.0288 | 324.7401 | 0.191905 | 5.109281 | 0.769332 |
| 12 | 115470800 | 115498799 | 416.8365 | 582.0957 | 570.0538 | 236.4228 | 0.196423 | 4.604145 | 0.769336 |
| 1 | 89000600 | 89003599 | 23.56847 | 39.45528 | 57.07183 | 24.87202 | -0.1497 | 1.175616 | 0.769347 |
| 17 | 6251400 | 6253599 | 12.95729 | 22.9715 | 34.18643 | 15.11302 | -0.17811 | 0.484139 | 0.769371 |
| 5 | 11527400 | 11529999 | 22.43141 | 38.97157 | 31.192 | 13.69464 | 0.706678 | 0.798798 | 0.769371 |
| X | 153373400 | 153374599 | 16.738 | 30.08554 | 33.40373 | 14.59684 | 0.253562 | 0.422448 | 0.769372 |
| 11 | 110277000 | 110277399 | 5.225949 | 10.19187 | 18.62242 | 6.685758 | -0.21577 | -0.43579 | 0.769377 |
| 5 | 148671400 | 148673599 | 25.90274 | 42.96496 | 61.99299 | 26.85441 | -0.14642 | 1.288546 | 0.76938 |
| 3 | 145328200 | 145329799 | 13.35228 | 23.54358 | 35.85504 | 15.86788 | -0.21408 | 0.551533 | 0.769383 |
| 14 | 27409200 | 27412799 | 67.93708 | 107.2615 | 145.544 | 60.42864 | -0.11455 | 2.44835 | 0.769386 |
| 5 | 89469000 | 89471799 | 25.18422 | 42.99375 | 49.56865 | 21.7318 | 0.177934 | 0.995521 | 0.76942 |
| 6 | 144264200 | 144265799 | 13.38007 | 24.23333 | 28.9597 | 12.54101 | 0.156353 | 0.343592 | 0.769429 |
| 4 | 142350200 | 142352799 | 25.57819 | 42.33167 | 62.48897 | 27.02742 | -0.1795 | 1.313096 | 0.769432 |

| | | | | | | | | | |
|---|---|---|---|---|---|---|---|---|---|
| 5 | 3988400 | 3990399 | 16.62877 | 29.31091 | 34.52716 | 15.28126 | 0.154039 | 0.570467 | 0.769462 |
| 1 | 64413200 | 64414599 | 15.66383 | 27.41605 | 39.91492 | 17.6124 | -0.15164 | 0.672206 | 0.769473 |
| 3 | 133002800 | 133004199 | 18.82977 | 32.12113 | 57.0647 | 25.01666 | -0.44363 | 0.986616 | 0.769495 |
| 1 | 120272200 | 120277599 | 41.03198 | 66.12454 | 94.37255 | 40.15794 | -0.15599 | 1.800842 | 0.769545 |
| 3 | 85775600 | 85778199 | 30.9477 | 52.04765 | 56.8951 | 24.8577 | 0.252982 | 1.153378 | 0.769549 |
| 5 | 92395600 | 92405799 | 97.6812 | 148.81 | 240.2323 | 99.57602 | -0.40494 | 3.144729 | 0.76955 |
| 3 | 42803000 | 42803399 | 8.435427 | 16.3139 | 21.52855 | 8.230532 | 0.163773 | -0.17037 | 0.76957 |
| 11 | 42249400 | 42249799 | 8.435427 | 16.3139 | 21.52855 | 8.230532 | 0.163773 | -0.17037 | 0.76957 |
| 4 | 113715800 | 113722199 | 79.49516 | 125.8023 | 136.1198 | 56.58243 | 0.214301 | 2.400709 | 0.769591 |
| 17 | 95171800 | 95194799 | 298.4939 | 429.9152 | 428.9169 | 177.0058 | 0.21436 | 4.146678 | 0.769595 |
| 19 | 32806600 | 32810999 | 36.23883 | 59.33946 | 69.29069 | 29.81139 | 0.152909 | 1.473933 | 0.769604 |
| 8 | 43689600 | 43694199 | 30.64815 | 50.66564 | 59.60496 | 25.76012 | 0.146763 | 1.336289 | 0.76961 |
| 13 | 114518800 | 114522399 | 37.82801 | 61.89692 | 70.71894 | 30.41725 | 0.183662 | 1.486871 | 0.769637 |
| 15 | 55079800 | 55081799 | 25.5584 | 42.94063 | 50.68913 | 22.13951 | 0.142454 | 1.137393 | 0.769645 |
| 3 | 151388200 | 151388999 | 13.6853 | 24.8622 | 29.55221 | 12.74865 | 0.167857 | 0.331839 | 0.76966 |
| 11 | 22712200 | 22713199 | 11.73776 | 21.28528 | 31.68569 | 13.70021 | -0.15708 | 0.331255 | 0.769662 |
| 16 | 4345000 | 4346599 | 21.60649 | 36.60688 | 52.73256 | 23.0907 | -0.14323 | 1.055195 | 0.769676 |
| 3 | 3928600 | 3930999 | 35.57223 | 57.60635 | 81.05555 | 34.68372 | -0.12772 | 1.659013 | 0.769682 |
| 4 | 80175000 | 80176599 | 17.61697 | 31.20024 | 35.21321 | 15.57014 | 0.21778 | 0.523637 | 0.76971 |
| 12 | 49572400 | 49573999 | 14.99719 | 27.4717 | 25.79732 | 11.0123 | 0.515609 | 0.285443 | 0.76971 |
| 5 | 104917800 | 104919199 | 28.9429 | 48.18274 | 57.2893 | 24.87902 | 0.13228 | 1.252098 | 0.76973 |
| 9 | 71578000 | 71581399 | 26.33406 | 44.36837 | 52.33117 | 22.8762 | 0.143735 | 1.116652 | 0.769758 |
| 2 | 135585000 | 135587599 | 23.21549 | 39.44583 | 46.39004 | 20.33049 | 0.148449 | 1.027524 | 0.769773 |
| 1 | 177582400 | 177589799 | 102.7288 | 158.8369 | 154.7909 | 64.29654 | 0.339108 | 2.849127 | 0.769775 |
| 17 | 16114200 | 16115599 | 11.61858 | 21.33219 | 32.38884 | 13.78253 | -0.16983 | 0.262156 | 0.769781 |
| 17 | 31020800 | 31022999 | 24.63386 | 40.87641 | 60.20901 | 26.08873 | -0.17598 | 1.279665 | 0.76979 |
| 6 | 125698600 | 125704199 | 48.02259 | 76.91595 | 106.114 | 44.74454 | -0.11793 | 2.007692 | 0.769804 |
| 4 | 95140000 | 95150999 | 184.5462 | 270.9224 | 349.6964 | 144.0268 | -0.13031 | 3.847527 | 0.769823 |
| X | 88044800 | 88046999 | 25.14325 | 43.41603 | 47.36741 | 20.77999 | 0.260188 | 0.893399 | 0.769829 |
| 10 | 96347600 | 96348199 | 7.40264 | 14.28973 | 23.70997 | 9.144165 | -0.17578 | -0.14625 | 0.769843 |
| 18 | 90207400 | 90210599 | 29.17461 | 49.48868 | 53.50063 | 23.43557 | 0.269325 | 1.070518 | 0.769845 |
| 4 | 4932400 | 4933799 | 17.67011 | 30.94719 | 36.68171 | 16.1944 | 0.143532 | 0.666936 | 0.769849 |
| 18 | 8771200 | 8776999 | 97.67053 | 151.5671 | 141.6929 | 58.84411 | 0.403042 | 2.781467 | 0.769898 |
| 15 | 38431800 | 38435799 | 55.53116 | 88.60697 | 103.3294 | 43.33595 | 0.118133 | 2.130447 | 0.769899 |
| 1 | 3796800 | 3798799 | 27.77245 | 46.70579 | 54.53148 | 23.82475 | 0.158287 | 1.141072 | 0.769917 |
| 1 | 60386800 | 60392199 | 76.9851 | 120.7058 | 163.5743 | 67.8435 | -0.12045 | 2.595793 | 0.769926 |
| 6 | 144959800 | 144962999 | 31.24735 | 51.09556 | 72.46499 | 31.16078 | -0.12797 | 1.484543 | 0.769939 |
| 14 | 50179200 | 50181999 | 43.20043 | 69.69319 | 82.13389 | 34.99823 | 0.121474 | 1.770334 | 0.769951 |
| 9 | 85566200 | 85568999 | 31.94446 | 53.20047 | 51.94456 | 22.56354 | 0.415565 | 1.251307 | 0.769959 |
| 10 | 96945200 | 96948799 | 41.37407 | 66.47145 | 93.993 | 39.81885 | -0.14806 | 1.901355 | 0.769959 |
| 2 | 180941400 | 180944999 | 33.94086 | 55.04786 | 78.10581 | 33.45891 | -0.13768 | 1.62419 | 0.769962 |
| 5 | 18327800 | 18329799 | 25.08255 | 42.13923 | 49.09261 | 21.4419 | 0.161416 | 1.135083 | 0.769985 |
| 6 | 37775000 | 37783199 | 96.18165 | 149.442 | 168.8586 | 70.11283 | 0.131305 | 2.762871 | 0.770004 |
| 5 | 149708000 | 149719199 | 125.4586 | 191.2784 | 214.8222 | 89.06261 | 0.119015 | 3.134398 | 0.770006 |
| 13 | 11141800 | 11145399 | 46.13978 | 74.2551 | 87.28484 | 37.06894 | 0.120986 | 1.849644 | 0.77008 |
| 1 | 94154400 | 94156799 | 37.27831 | 60.5139 | 71.36714 | 30.592 | 0.129578 | 1.608292 | 0.770085 |
| 2 | 117337000 | 117339999 | 57.99006 | 92.10237 | 125.7206 | 52.41519 | -0.11565 | 2.284166 | 0.770087 |

| | | | | | | | | | |
|---|---|---|---|---|---|---|---|---|---|
| 18 | 47547600 | 47549199 | 13.93223 | 24.55544 | 36.25536 | 16.04645 | -0.1701 | 0.558772 | 0.770112 |
| 1 | 180701400 | 180702199 | 9.615929 | 18.27415 | 22.76578 | 9.14707 | 0.179699 | 0.00596 | 0.770128 |
| 4 | 69096800 | 69098199 | 15.99043 | 28.43183 | 33.27566 | 14.67442 | 0.168686 | 0.480877 | 0.770139 |
| 9 | 19263200 | 19267399 | 74.25741 | 117.2883 | 134.9928 | 56.05952 | 0.123103 | 2.440405 | 0.770146 |
| 14 | 80726400 | 80727799 | 9.968562 | 18.79394 | 29.74543 | 12.15269 | -0.18404 | 0.083472 | 0.77015 |
| 1 | 67624200 | 67627199 | 34.05663 | 55.38785 | 78.11755 | 33.52506 | -0.12555 | 1.571389 | 0.770166 |
| 1 | 68025400 | 68028199 | 28.43486 | 47.97697 | 54.56822 | 23.86455 | 0.196088 | 1.115306 | 0.770171 |
| 13 | 77570800 | 77571799 | 13.53708 | 24.41568 | 28.93972 | 12.59789 | 0.16324 | 0.368479 | 0.770177 |
| 6 | 29108600 | 29114399 | 59.10366 | 94.37329 | 108.2838 | 45.4588 | 0.142561 | 2.113451 | 0.770188 |
| 7 | 59905200 | 59906399 | 12.05226 | 22.77941 | 26.98362 | 10.96772 | 0.237147 | 0.076578 | 0.770204 |
| 9 | 79404800 | 79405599 | 12.05226 | 22.77941 | 26.98362 | 10.96772 | 0.237147 | 0.076578 | 0.770204 |
| 4 | 20596400 | 20597799 | 17.64139 | 30.9886 | 36.47772 | 16.14473 | 0.153307 | 0.618265 | 0.770247 |
| 7 | 12365600 | 12371999 | 80.2295 | 126.7077 | 121.8313 | 50.59114 | 0.382071 | 2.443813 | 0.770254 |
| 5 | 32575000 | 32577399 | 16.82172 | 29.70182 | 34.85469 | 15.42094 | 0.160227 | 0.551228 | 0.770275 |
| 3 | 51841600 | 51846599 | 48.18552 | 77.06123 | 106.4863 | 44.80412 | -0.12268 | 2.053871 | 0.770288 |
| 17 | 5572000 | 5575199 | 62.95609 | 99.9155 | 115.7743 | 48.23258 | 0.119323 | 2.306482 | 0.770306 |
| 15 | 18212600 | 18213199 | 8.627281 | 16.76491 | 21.84658 | 8.136888 | 0.217102 | -0.25542 | 0.77032 |
| 2 | 99052000 | 99053399 | 18.44919 | 32.2923 | 38.16269 | 16.83971 | 0.147721 | 0.682289 | 0.770321 |
| 8 | 58089400 | 58091599 | 16.22667 | 28.64314 | 33.95461 | 15.02355 | 0.145651 | 0.550144 | 0.770334 |
| 5 | 23056200 | 23056999 | 9.863703 | 18.80998 | 23.91238 | 9.53772 | 0.160229 | -0.01057 | 0.770338 |
| 18 | 66612400 | 66615599 | 25.4139 | 43.34994 | 49.57323 | 21.73236 | 0.189427 | 1.002435 | 0.770354 |
| 2 | 150168000 | 150195999 | 686.8923 | 917.6435 | 877.9672 | 366.6231 | 0.165618 | 5.296499 | 0.770359 |
| 7 | 31661200 | 31663599 | 49.33281 | 79.07548 | 91.87061 | 38.78409 | 0.130851 | 1.98164 | 0.770384 |
| 8 | 8840000 | 8841799 | 20.57798 | 35.53073 | 42.11918 | 18.47756 | 0.140944 | 0.866934 | 0.770393 |
| 18 | 70438000 | 70438399 | 7.396618 | 14.38403 | 19.53653 | 7.233302 | 0.166176 | -0.29744 | 0.770394 |
| 1 | 34832200 | 34834199 | 14.54738 | 25.51361 | 37.86607 | 16.73698 | -0.17915 | 0.630035 | 0.770409 |
| 10 | 99593600 | 99596399 | 30.4284 | 49.72939 | 71.17193 | 30.59869 | -0.1416 | 1.489394 | 0.77043 |
| 14 | 18556800 | 18557799 | 19.21172 | 33.66478 | 32.01286 | 14.13945 | 0.459839 | 0.659201 | 0.770433 |
| 10 | 43458400 | 43459399 | 11.12478 | 20.63733 | 32.63331 | 13.69304 | -0.21393 | 0.19741 | 0.770454 |
| 9 | 85507800 | 85509199 | 20.83673 | 34.66732 | 65.45011 | 28.41163 | -0.53257 | 1.2429 | 0.770498 |
| 8 | 76001400 | 76002399 | 13.76797 | 24.3605 | 35.67102 | 15.77555 | -0.15604 | 0.513979 | 0.770511 |
| 13 | 83341000 | 83343999 | 46.85183 | 75.61634 | 74.20615 | 31.63767 | 0.385706 | 1.75908 | 0.770524 |
| 4 | 34136200 | 34137599 | 15.6849 | 28.3135 | 32.36694 | 14.07489 | 0.216039 | 0.385749 | 0.770559 |
| 8 | 75298600 | 75299799 | 15.30878 | 26.9984 | 30.10729 | 13.32368 | 0.232859 | 0.54606 | 0.770574 |
| 8 | 97355800 | 97358599 | 85.12005 | 132.7907 | 152.9282 | 63.46078 | 0.108392 | 2.686161 | 0.770577 |
| 6 | 67108600 | 67109799 | 13.85312 | 25.05996 | 29.91147 | 12.97266 | 0.156611 | 0.355568 | 0.770579 |
| 16 | 82968600 | 82968999 | 9.75982 | 18.78334 | 19.18393 | 7.465384 | 0.506259 | -0.10474 | 0.770584 |
| 6 | 101714400 | 101715199 | 14.16829 | 25.17265 | 37.07146 | 16.35893 | -0.16161 | 0.481696 | 0.770593 |
| 3 | 63283400 | 63285799 | 28.75156 | 47.75417 | 56.71224 | 24.58958 | 0.133764 | 1.276681 | 0.770593 |
| 10 | 120697200 | 120698399 | 13.33974 | 23.78927 | 35.06604 | 15.42532 | -0.15974 | 0.448968 | 0.770616 |
| 3 | 112497800 | 112500399 | 36.29495 | 59.22829 | 70.05974 | 30.10407 | 0.131039 | 1.523279 | 0.770617 |
| X | 62060000 | 62060999 | 14.17242 | 26.39669 | 22.48523 | 9.377567 | 0.680111 | 0.18016 | 0.770621 |
| 10 | 126461600 | 126462799 | 13.27999 | 23.56533 | 34.65465 | 15.30835 | -0.16027 | 0.481381 | 0.770631 |
| 13 | 116100600 | 116104199 | 44.78727 | 71.76124 | 100.2505 | 42.29648 | -0.13528 | 1.991049 | 0.770659 |
| X | 72550800 | 72551599 | 10.50196 | 20.29345 | 24.4704 | 9.392042 | 0.287171 | -0.12747 | 0.77067 |
| 9 | 116567800 | 116569999 | 13.95183 | 24.72589 | 37.03061 | 16.37153 | -0.18785 | 0.49608 | 0.770684 |
| 4 | 73136200 | 73136999 | 9.760668 | 18.37708 | 28.58457 | 11.70524 | -0.16187 | 0.086542 | 0.7707 |

| | | | | | | | | | |
|---|---|---|---|---|---|---|---|---|---|
| 15 | 96058000 | 96059999 | 21.02489 | 35.96135 | 51.70153 | 22.67036 | -0.13886 | 0.96535 | 0.770721 |
| 7 | 17303800 | 17307199 | 78.86279 | 124.2092 | 114.1296 | 47.35556 | 0.444406 | 2.500839 | 0.770727 |
| 1 | 176444600 | 176446399 | 16.78037 | 29.34275 | 42.48247 | 18.73046 | -0.14375 | 0.696666 | 0.77074 |
| 3 | 36897800 | 36899199 | 16.78037 | 29.34275 | 42.48247 | 18.73046 | -0.14375 | 0.696666 | 0.77074 |
| 14 | 89093400 | 89095399 | 24.12355 | 40.80818 | 48.00117 | 21.00904 | 0.147983 | 1.071987 | 0.770766 |
| X | 8137600 | 8140599 | 31.55171 | 52.71699 | 57.30907 | 24.94473 | 0.261143 | 1.225892 | 0.770783 |
| 18 | 4502400 | 4507199 | 51.54935 | 82.2074 | 113.2028 | 47.43986 | -0.12213 | 2.148118 | 0.770805 |
| X | 121162000 | 121163599 | 25.84709 | 44.12169 | 50.11022 | 21.96908 | 0.199604 | 1.001623 | 0.770817 |
| 5 | 64217400 | 64218799 | 15.05398 | 27.09932 | 31.8664 | 13.90539 | 0.172362 | 0.396627 | 0.770833 |
| 1 | 186609200 | 186609799 | 11.6268 | 22.08949 | 21.1288 | 8.494333 | 0.557754 | 0.025251 | 0.770836 |
| 18 | 77948000 | 77949999 | 21.59487 | 37.20165 | 44.20506 | 19.39129 | 0.136843 | 0.894671 | 0.770852 |
| 1 | 168546600 | 168548999 | 20.24043 | 34.46316 | 50.07179 | 21.94774 | -0.15491 | 1.002034 | 0.770861 |
| 11 | 19742600 | 19745999 | 46.76174 | 74.88556 | 103.516 | 43.63936 | -0.12098 | 2.01094 | 0.770877 |
| 10 | 62713600 | 62716799 | 39.02534 | 62.91713 | 88.07259 | 37.49756 | -0.12732 | 1.785876 | 0.770878 |
| 3 | 31815800 | 31820599 | 54.44018 | 86.5694 | 119.8061 | 50.01009 | -0.13377 | 2.242763 | 0.770881 |
| 11 | 16967200 | 16971199 | 64.27198 | 101.9114 | 118.0005 | 49.11542 | 0.119105 | 2.334394 | 0.770882 |
| 6 | 64280000 | 64280599 | 7.608292 | 14.52536 | 24.01346 | 9.555142 | -0.2122 | -0.03922 | 0.770905 |
| 18 | 66486400 | 66487399 | 12.11311 | 22.94111 | 27.06562 | 10.95375 | 0.248493 | 0.064214 | 0.770905 |
| 3 | 142369200 | 142374799 | 75.08019 | 117.3258 | 158.5472 | 65.7811 | -0.12013 | 2.652483 | 0.770923 |
| 5 | 148553600 | 148556399 | 32.52619 | 53.0408 | 75.39522 | 32.38837 | -0.13409 | 1.527093 | 0.770931 |
| 2 | 93660200 | 93664199 | 38.77243 | 63.19207 | 62.37002 | 26.78765 | 0.390266 | 1.540048 | 0.770966 |
| 12 | 19373400 | 19374399 | 21.23312 | 36.89319 | 34.28423 | 15.05316 | 0.49228 | 0.796787 | 0.770969 |
| 4 | 43798800 | 43799799 | 8.692337 | 16.4799 | 26.01986 | 10.50699 | -0.16537 | 0.024733 | 0.770971 |
| 15 | 9392400 | 9392999 | 8.692337 | 16.4799 | 26.01986 | 10.50699 | -0.16537 | 0.024733 | 0.770971 |
| 14 | 106164000 | 106167599 | 36.05029 | 59.19474 | 68.31471 | 29.426 | 0.171377 | 1.438781 | 0.770975 |
| 9 | 68842200 | 68844199 | 22.44302 | 37.60827 | 55.39934 | 24.15817 | -0.17584 | 1.160076 | 0.77098 |
| 11 | 3534400 | 3536199 | 19.86227 | 34.11041 | 49.00834 | 21.49629 | -0.13698 | 0.920867 | 0.77099 |
| 18 | 62182400 | 62183399 | 8.39898 | 15.93353 | 25.50648 | 10.28628 | -0.18323 | 0.017066 | 0.771007 |
| 13 | 79706400 | 79707999 | 19.74256 | 34.80492 | 31.5802 | 13.97654 | 0.528598 | 0.580391 | 0.77101 |
| 3 | 28121600 | 28124399 | 22.56867 | 37.14455 | 72.14421 | 31.10627 | -0.57585 | 1.383903 | 0.771023 |
| 4 | 122579200 | 122581399 | 50.45858 | 80.77265 | 93.33005 | 39.32116 | 0.136482 | 2.024248 | 0.771034 |
| 9 | 101142200 | 101145999 | 31.88492 | 52.48304 | 62.2912 | 26.85913 | 0.132442 | 1.394222 | 0.771037 |
| 3 | 102173400 | 102177199 | 32.90557 | 53.5706 | 75.71638 | 32.50669 | -0.12757 | 1.553026 | 0.771061 |
| 7 | 111235400 | 111237199 | 14.85595 | 26.22494 | 38.14756 | 16.87859 | -0.14805 | 0.549037 | 0.7711 |
| 7 | 136818200 | 136818999 | 6.396135 | 12.37371 | 21.57945 | 8.238309 | -0.23212 | -0.19646 | 0.771121 |
| X | 77389800 | 77390799 | 9.310719 | 17.71569 | 30.63161 | 12.27582 | -0.28576 | 0.020024 | 0.771123 |
| 14 | 118654600 | 118662199 | 78.59061 | 122.4629 | 165.395 | 68.64512 | -0.12251 | 2.703867 | 0.771202 |
| 1 | 171184000 | 171185799 | 15.29584 | 27.10696 | 31.98438 | 14.14063 | 0.154637 | 0.498718 | 0.77121 |
| 11 | 16045800 | 16046599 | 11.71273 | 21.62805 | 25.23001 | 10.65323 | 0.216409 | 0.215582 | 0.771212 |
| 10 | 94374000 | 94379199 | 55.426 | 88.2568 | 120.6435 | 50.45297 | -0.11392 | 2.200678 | 0.771217 |
| 3 | 158153200 | 158155199 | 19.69806 | 34.19938 | 40.4897 | 17.76911 | 0.144275 | 0.809047 | 0.771219 |
| 5 | 26089600 | 26097999 | 90.42881 | 139.8317 | 185.8391 | 77.19321 | -0.10792 | 2.846192 | 0.771221 |
| 3 | 59307400 | 59309599 | 27.91636 | 45.55996 | 79.21937 | 34.04108 | -0.42315 | 1.509749 | 0.771253 |
| X | 157524400 | 157526399 | 15.57798 | 27.32701 | 39.86002 | 17.62153 | -0.15425 | 0.637054 | 0.771276 |
| 15 | 55125200 | 55127399 | 18.23607 | 31.90846 | 37.65319 | 16.61013 | 0.149819 | 0.685101 | 0.771276 |
| 3 | 93202400 | 93204999 | 21.27903 | 36.27222 | 52.79476 | 23.14337 | -0.15741 | 0.996563 | 0.771306 |
| 8 | 4849000 | 4853999 | 84.11303 | 131.4054 | 151.5243 | 62.86909 | 0.108154 | 2.661034 | 0.771311 |

| | | | | | | | | | |
|---|---|---|---|---|---|---|---|---|---|
| 6 | 131398400 | 131400199 | 14.72627 | 26.64724 | 30.8092 | 13.35714 | 0.202416 | 0.360491 | 0.771315 |
| 17 | 61326600 | 61329999 | 86.45776 | 133.8092 | 179.036 | 74.36702 | -0.11663 | 2.828342 | 0.77132 |
| 13 | 77817800 | 77823199 | 49.71441 | 79.68102 | 92.97929 | 39.24755 | 0.124402 | 1.983009 | 0.771367 |
| 11 | 47297600 | 47298399 | 11.57462 | 21.84949 | 21.4748 | 8.751568 | 0.501109 | 0.066655 | 0.771377 |
| 1 | 48505000 | 48509999 | 106.8267 | 164.7131 | 160.0679 | 66.48944 | 0.340081 | 2.902375 | 0.771389 |
| 9 | 93909400 | 93910799 | 15.79025 | 27.76456 | 40.72706 | 18.03411 | -0.16244 | 0.60082 | 0.771394 |
| 3 | 70936200 | 70941999 | 60.39378 | 96.4752 | 109.0778 | 45.82031 | 0.164463 | 2.103458 | 0.771398 |
| 8 | 126845000 | 126850199 | 55.81962 | 88.84434 | 121.268 | 50.68314 | -0.11247 | 2.216354 | 0.771424 |
| 12 | 17877000 | 18008399 | 3269.823 | 3803.06 | 3833.149 | 1628.247 | -0.11153 | 7.513158 | 0.771438 |
| 10 | 47440400 | 47441399 | 17.00979 | 30.21233 | 28.60012 | 12.62378 | 0.472189 | 0.484797 | 0.771454 |
| 5 | 130619400 | 130622799 | 43.86348 | 70.74033 | 83.33891 | 35.49711 | 0.121445 | 1.781191 | 0.771474 |
| 2 | 23463200 | 23464599 | 12.72316 | 23.00723 | 34.20291 | 14.84085 | -0.15857 | 0.359904 | 0.771477 |
| X | 86509800 | 86510599 | 7.171236 | 13.90523 | 23.926 | 9.036229 | -0.19803 | -0.21471 | 0.771494 |
| 11 | 118198400 | 118201399 | 48.96925 | 78.6398 | 92.30351 | 39.07793 | 0.119293 | 1.922775 | 0.771544 |
| 5 | 151374400 | 151376199 | 22.4687 | 39.01149 | 37.77078 | 16.57636 | 0.433277 | 0.818359 | 0.771562 |
| 1 | 119216600 | 119220399 | 28.25389 | 46.61128 | 66.43853 | 28.67474 | -0.12963 | 1.362177 | 0.771568 |
| 5 | 18877400 | 18878199 | 11.96163 | 22.26141 | 27.20571 | 11.36145 | 0.160945 | 0.175962 | 0.771595 |
| 13 | 67425600 | 67430999 | 42.36369 | 68.08676 | 96.43401 | 40.87505 | -0.14981 | 1.888214 | 0.771641 |
| 16 | 61107200 | 61115799 | 75.91154 | 120.5862 | 127.3786 | 53.11532 | 0.253877 | 2.295443 | 0.771668 |
| 6 | 111792400 | 111796599 | 63.52416 | 100.6781 | 137.3434 | 57.13953 | -0.1178 | 2.351648 | 0.771679 |
| 6 | 124575600 | 124577799 | 24.03374 | 40.94727 | 48.48601 | 21.2537 | 0.139167 | 1.008116 | 0.771692 |
| 5 | 18971200 | 18972799 | 13.35107 | 24.65308 | 29.13578 | 12.30257 | 0.197056 | 0.234468 | 0.771696 |
| 17 | 8278200 | 8278999 | 9.235902 | 17.18466 | 27.4099 | 11.45123 | -0.22039 | 0.157572 | 0.771697 |
| 8 | 45275600 | 45277599 | 21.11253 | 36.97404 | 40.61949 | 17.93199 | 0.252501 | 0.69146 | 0.771741 |
| 5 | 10212000 | 10215999 | 59.84191 | 95.36004 | 109.9412 | 46.02512 | 0.132529 | 2.176446 | 0.771779 |
| 15 | 82407600 | 82409799 | 25.70054 | 42.7262 | 61.51966 | 26.69106 | -0.14318 | 1.260823 | 0.771798 |
| 16 | 69696000 | 69698399 | 8.133604 | 15.20541 | 30.61806 | 12.68951 | -0.54541 | 0.12256 | 0.771798 |
| 19 | 21942200 | 21944199 | 17.7686 | 31.08472 | 36.69768 | 16.18607 | 0.149234 | 0.684596 | 0.771802 |
| 5 | 130802200 | 130804399 | 19.3585 | 33.71932 | 39.85138 | 17.52476 | 0.147195 | 0.760249 | 0.771804 |
| 5 | 12212800 | 12214999 | 40.42104 | 65.6648 | 64.5642 | 27.69121 | 0.392278 | 1.594195 | 0.771821 |
| 2 | 89728200 | 89736599 | 109.2148 | 168.181 | 163.8058 | 68.03691 | 0.335446 | 2.928102 | 0.771836 |
| X | 18402400 | 18406199 | 34.07968 | 56.16591 | 64.5817 | 27.86152 | 0.17884 | 1.389886 | 0.771837 |
| 3 | 3841400 | 3842199 | 9.182816 | 17.27671 | 26.96016 | 11.05348 | -0.16746 | 0.083703 | 0.771855 |
| 5 | 127899800 | 127901799 | 22.87961 | 38.73404 | 55.48988 | 24.29242 | -0.13545 | 1.068236 | 0.771856 |
| 16 | 42775800 | 42776599 | 8.205789 | 15.67975 | 25.08404 | 9.96272 | -0.16315 | -0.03674 | 0.771867 |
| 4 | 89297400 | 89297799 | 10.57853 | 19.66409 | 29.92748 | 12.52282 | -0.15574 | 0.178233 | 0.771869 |
| 3 | 63554800 | 63557199 | 24.17824 | 41.28745 | 48.47974 | 21.25486 | 0.151715 | 0.987863 | 0.771872 |
| 4 | 59450800 | 59452599 | 13.52168 | 24.18284 | 35.60131 | 15.62304 | -0.15551 | 0.433932 | 0.771875 |
| 7 | 16981000 | 16986399 | 78.25282 | 122.9934 | 141.3373 | 58.629 | 0.119883 | 2.547116 | 0.771905 |
| 8 | 64361800 | 64363599 | 37.21143 | 60.07412 | 84.677 | 36.11498 | -0.13501 | 1.741667 | 0.771921 |
| 10 | 119332000 | 119337399 | 86.14272 | 134.2217 | 154.3288 | 64.05287 | 0.109329 | 2.707728 | 0.771972 |
| 9 | 16537800 | 16539599 | 29.86031 | 50.01552 | 48.61207 | 21.19253 | 0.422013 | 1.182717 | 0.771977 |
| 17 | 23068800 | 23074399 | 91.95175 | 144.6571 | 150.5415 | 62.45447 | 0.263762 | 2.539746 | 0.772018 |
| 7 | 12524000 | 12525399 | 13.60581 | 24.24327 | 36.55496 | 16.09175 | -0.19318 | 0.457162 | 0.772039 |
| 13 | 9913400 | 9914599 | 17.49095 | 30.47041 | 44.1625 | 19.42221 | -0.14566 | 0.758535 | 0.772066 |
| 4 | 50379800 | 50381199 | 21.0973 | 35.32957 | 62.32545 | 27.167 | -0.43505 | 1.171135 | 0.772083 |
| 7 | 84828800 | 84829199 | 7.511145 | 14.64317 | 15.71448 | 5.669045 | 0.536339 | -0.40071 | 0.772165 |

| | | | | | | | | | |
|---|---|---|---|---|---|---|---|---|---|
| 13 | 61813600 | 61814199 | 17.93239 | 30.9241 | 45.3251 | 19.88324 | -0.16425 | 0.871365 | 0.772187 |
| 12 | 57285000 | 57286999 | 29.5387 | 49.26793 | 57.94715 | 25.19483 | 0.147801 | 1.233226 | 0.772198 |
| 11 | 39179400 | 39179799 | 7.117598 | 13.81865 | 23.4781 | 8.809056 | -0.17174 | -0.24235 | 0.772199 |
| 7 | 103777200 | 103779799 | 51.06264 | 81.34704 | 113.4554 | 47.47771 | -0.14209 | 2.180882 | 0.772201 |
| 7 | 20950200 | 20951999 | 31.55133 | 51.91315 | 60.35332 | 26.01913 | 0.162082 | 1.400405 | 0.772202 |
| X | 138701200 | 138703599 | 18.6567 | 32.5145 | 48.97112 | 21.54251 | -0.20096 | 0.764501 | 0.772208 |
| 4 | 114118200 | 114124199 | 102.8325 | 158.8092 | 180.3422 | 74.91042 | 0.117363 | 2.874205 | 0.772231 |
| 1 | 54363800 | 54365999 | 55.72211 | 88.84556 | 102.6954 | 43.02564 | 0.12973 | 2.154554 | 0.772232 |
| 13 | 5907200 | 5908199 | 8.22974 | 15.48667 | 25.42065 | 10.41876 | -0.2395 | 0.068328 | 0.772237 |
| 3 | 106078000 | 106108799 | 840.5299 | 1096.171 | 1272.63 | 532.955 | -0.14811 | 5.684483 | 0.772241 |
| 18 | 19108200 | 19110799 | 41.51315 | 67.44815 | 66.27437 | 28.42754 | 0.393432 | 1.596266 | 0.772263 |
| 7 | 77037800 | 77038599 | 17.42707 | 30.1527 | 44.26651 | 19.42082 | -0.1656 | 0.837529 | 0.772277 |
| 7 | 106533600 | 106537199 | 31.74706 | 53.17786 | 58.64405 | 25.56661 | 0.240341 | 1.196532 | 0.772317 |
| X | 23559000 | 23560199 | 14.4625 | 26.23106 | 30.06845 | 12.99802 | 0.218134 | 0.349566 | 0.772328 |
| 6 | 76204000 | 76205799 | 17.87334 | 31.35081 | 37.10498 | 16.40236 | 0.145635 | 0.644744 | 0.772352 |
| 8 | 67002200 | 67005199 | 25.99788 | 43.78692 | 51.78102 | 22.63335 | 0.139948 | 1.117796 | 0.772366 |
| 13 | 5088200 | 5088999 | 9.656823 | 18.10666 | 27.96468 | 11.53173 | -0.1595 | 0.11191 | 0.772377 |
| 16 | 25946800 | 25947199 | 6.533907 | 12.66732 | 21.63597 | 8.17771 | -0.18872 | -0.22869 | 0.772384 |
| 17 | 33548000 | 33554999 | 111.5822 | 173.305 | 176.6101 | 73.33555 | 0.279715 | 2.778937 | 0.77239 |
| 12 | 13598400 | 13601199 | 19.773 | 34.33886 | 52.3334 | 22.97071 | -0.21854 | 0.822561 | 0.772415 |
| 4 | 113659400 | 113663399 | 51.36635 | 82.6472 | 92.95179 | 39.46229 | 0.184171 | 1.868058 | 0.772416 |
| 10 | 34132600 | 34136199 | 34.07392 | 55.41292 | 78.05865 | 33.49911 | -0.12386 | 1.572649 | 0.772425 |
| 11 | 115345000 | 115346799 | 18.62549 | 31.95187 | 46.91435 | 20.57616 | -0.16827 | 0.924339 | 0.772436 |
| 2 | 120052000 | 120059599 | 84.56172 | 131.1448 | 175.2284 | 72.77777 | -0.11235 | 2.789926 | 0.772459 |
| 14 | 113545400 | 113548399 | 29.20808 | 48.21569 | 68.9291 | 29.76031 | -0.13374 | 1.358224 | 0.772497 |
| 2 | 134590600 | 134593399 | 29.70575 | 49.89665 | 48.03423 | 20.97777 | 0.435609 | 1.149961 | 0.772509 |
| X | 66719800 | 66722999 | 23.97752 | 41.91389 | 42.80152 | 18.86578 | 0.357635 | 0.740167 | 0.772512 |
| 2 | 112964600 | 112965199 | 5.960121 | 11.55962 | 21.10707 | 7.962636 | -0.28026 | -0.24008 | 0.772516 |
| 10 | 11430800 | 11433799 | 38.49257 | 62.43317 | 73.58578 | 31.52374 | 0.129358 | 1.629691 | 0.772517 |
| 5 | 149648600 | 149651199 | 19.65224 | 33.87309 | 49.01293 | 21.50069 | -0.14612 | 0.886371 | 0.772521 |
| 10 | 130338800 | 130339799 | 10.66972 | 20.14165 | 24.72578 | 10.07607 | 0.182607 | 0.065777 | 0.772521 |
| 10 | 123790600 | 123791799 | 12.21239 | 21.99006 | 32.61496 | 14.22717 | -0.16018 | 0.376839 | 0.772562 |
| 12 | 107663800 | 107665199 | 16.82525 | 29.55733 | 34.19408 | 15.13502 | 0.178551 | 0.611738 | 0.772563 |
| 14 | 81324400 | 81326799 | 30.68919 | 50.7303 | 59.95575 | 25.9106 | 0.140259 | 1.341342 | 0.772609 |
| 13 | 88197600 | 88198799 | 13.86605 | 25.69936 | 24.41947 | 10.25909 | 0.515214 | 0.209044 | 0.772618 |
| 16 | 53645200 | 53645599 | 6.092599 | 11.84497 | 20.65423 | 7.663648 | -0.19333 | -0.30199 | 0.772624 |
| 19 | 31352000 | 31352799 | 13.56161 | 24.91195 | 29.5908 | 12.58343 | 0.182561 | 0.266494 | 0.77263 |
| 7 | 129576600 | 129577799 | 8.352826 | 15.67678 | 26.39698 | 10.86747 | -0.28168 | 0.08676 | 0.772633 |
| 13 | 119054000 | 119056799 | 27.39126 | 47.02474 | 49.5787 | 21.74603 | 0.307666 | 0.954822 | 0.772638 |
| 1 | 63159400 | 63160599 | 10.7504 | 19.97186 | 30.45803 | 12.75499 | -0.15951 | 0.18429 | 0.772644 |
| 6 | 36937200 | 36938199 | 9.930825 | 18.68753 | 29.1293 | 11.93902 | -0.16597 | 0.092893 | 0.772653 |
| 7 | 58591000 | 58592599 | 22.89363 | 39.66747 | 45.19636 | 19.82963 | 0.198813 | 0.849889 | 0.772655 |
| 1 | 119285800 | 119288399 | 37.08036 | 60.57457 | 70.92465 | 30.48758 | 0.146783 | 1.509627 | 0.772669 |
| X | 31719200 | 31719799 | 12.85997 | 24.28744 | 20.58447 | 8.38028 | 0.715422 | 0.076599 | 0.77267 |
| 15 | 46792800 | 46796199 | 28.9092 | 48.0368 | 72.05854 | 31.25285 | -0.20225 | 1.285125 | 0.772681 |
| 9 | 25767800 | 25772199 | 94.73568 | 146.352 | 164.6008 | 68.37165 | 0.131624 | 2.86315 | 0.772683 |
| 5 | 125429600 | 125431799 | 21.37136 | 36.25576 | 52.21345 | 22.87243 | -0.14286 | 1.0409 | 0.772703 |

| | | | | | | | | | |
|---|---|---|---|---|---|---|---|---|---|
| 1 | 135289400 | 135290799 | 17.92828 | 31.39459 | 37.12952 | 16.39001 | 0.146623 | 0.670283 | 0.772706 |
| X | 116948000 | 116951999 | 46.55553 | 75.64194 | 65.59499 | 28.13647 | 0.573303 | 1.606761 | 0.772725 |
| 8 | 43174600 | 43175199 | 13.11726 | 23.18494 | 34.74512 | 15.36928 | -0.18981 | 0.51925 | 0.772738 |
| 18 | 37606800 | 37613399 | 156.2119 | 233.8975 | 258.7699 | 106.7313 | 0.118388 | 3.502282 | 0.772752 |
| 18 | 48812000 | 48812399 | 7.153232 | 13.76509 | 23.0799 | 8.984712 | -0.20258 | -0.11732 | 0.77276 |
| 7 | 94645000 | 94645799 | 8.413698 | 16.20947 | 20.88451 | 8.102966 | 0.177981 | -0.12385 | 0.772765 |
| 16 | 79417000 | 79417999 | 8.634193 | 16.44486 | 26.11373 | 10.44317 | -0.16166 | -0.00901 | 0.772792 |
| 13 | 64421800 | 64425599 | 22.40755 | 36.85459 | 74.62412 | 32.16637 | -0.63628 | 1.389984 | 0.772798 |
| 17 | 3676000 | 3677799 | 24.96718 | 42.28297 | 50.11602 | 21.94216 | 0.137067 | 1.067652 | 0.772817 |
| 5 | 109488600 | 109491999 | 52.7693 | 85.15667 | 74.26585 | 31.65825 | 0.555431 | 1.774064 | 0.772818 |
| 8 | 26538600 | 26541199 | 20.51621 | 35.4551 | 41.63709 | 18.26656 | 0.154787 | 0.860007 | 0.772823 |
| 6 | 10260600 | 10263999 | 42.50048 | 68.76659 | 80.00464 | 34.21223 | 0.144737 | 1.686514 | 0.772888 |
| 17 | 22113200 | 22121599 | 103.4774 | 159.3553 | 220.7891 | 91.7057 | -0.17247 | 2.931229 | 0.772893 |
| 5 | 68716000 | 68717199 | 15.13968 | 27.3765 | 26.31457 | 11.41831 | 0.466832 | 0.360531 | 0.772919 |
| 1 | 26104200 | 26104799 | 13.91186 | 24.85874 | 28.57219 | 12.54775 | 0.199022 | 0.428272 | 0.772933 |
| 3 | 110336200 | 110337199 | 15.15978 | 27.34135 | 31.84552 | 13.86349 | 0.18816 | 0.38573 | 0.772959 |
| 7 | 116727200 | 116728999 | 15.36857 | 27.59737 | 32.34082 | 14.14389 | 0.175088 | 0.414534 | 0.772978 |
| 16 | 87826400 | 87827199 | 9.474507 | 17.69946 | 27.38484 | 11.36332 | -0.16936 | 0.131649 | 0.773014 |
| 5 | 93982200 | 93983599 | 21.99915 | 38.16296 | 36.35113 | 15.94942 | 0.456594 | 0.82448 | 0.773019 |
| 19 | 41423800 | 41425599 | 17.15869 | 30.04984 | 35.21736 | 15.54719 | 0.159804 | 0.662503 | 0.773035 |
| 8 | 83761600 | 83763399 | 19.32316 | 33.63384 | 39.94528 | 17.5582 | 0.140216 | 0.77233 | 0.773055 |
| 6 | 9692800 | 9694599 | 14.09712 | 25.43153 | 30.44656 | 13.2501 | 0.149238 | 0.375343 | 0.773064 |
| 9 | 121147400 | 121151799 | 42.9463 | 68.77718 | 99.22288 | 41.83115 | -0.18245 | 2.002582 | 0.773067 |
| 6 | 71039000 | 71040599 | 13.89803 | 24.87241 | 29.0077 | 12.72243 | 0.179331 | 0.419097 | 0.773079 |
| 7 | 21850800 | 21851999 | 29.54488 | 49.65895 | 40.81775 | 17.83023 | 0.662991 | 1.140087 | 0.773082 |
| 9 | 91275600 | 91277799 | 19.50091 | 33.47963 | 48.47563 | 21.26225 | -0.1482 | 0.920597 | 0.773105 |
| 3 | 76669600 | 76671199 | 20.45811 | 35.41839 | 42.10055 | 18.4709 | 0.137842 | 0.843204 | 0.773125 |
| 2 | 35393000 | 35398999 | 42.55059 | 68.28318 | 97.49273 | 41.22996 | -0.16402 | 1.936481 | 0.773134 |
| 14 | 81443600 | 81444399 | 12.07622 | 22.7249 | 21.57872 | 8.844164 | 0.544281 | 0.094069 | 0.773137 |
| 6 | 100425400 | 100427199 | 28.9552 | 47.98817 | 56.65806 | 24.52932 | 0.141966 | 1.302821 | 0.773155 |
| 11 | 35494800 | 35497199 | 25.72767 | 43.14239 | 50.70233 | 22.11962 | 0.148863 | 1.158014 | 0.77316 |
| 2 | 171342000 | 171345999 | 45.09011 | 72.33916 | 99.94215 | 42.24867 | -0.11712 | 1.951091 | 0.773165 |
| 6 | 31369400 | 31371199 | 12.71977 | 22.87387 | 33.61624 | 14.68382 | -0.14832 | 0.390866 | 0.773167 |
| 11 | 20236800 | 20245399 | 161.1846 | 241.7895 | 268.2898 | 110.7012 | 0.116542 | 3.468805 | 0.773173 |
| 9 | 97277000 | 97282999 | 48.09875 | 77.76796 | 84.79694 | 36.23665 | 0.237033 | 1.717382 | 0.773185 |
| 13 | 7808400 | 7812399 | 25.78634 | 42.29677 | 74.71348 | 32.1679 | -0.44149 | 1.43427 | 0.773191 |
| 5 | 147281400 | 147283199 | 15.42516 | 27.33123 | 32.43131 | 14.33874 | 0.146367 | 0.501643 | 0.773201 |
| 3 | 76225600 | 76226199 | 11.94183 | 22.57344 | 26.99411 | 10.96916 | 0.224022 | 0.075484 | 0.773203 |
| 14 | 11187800 | 11188999 | 9.742714 | 17.78367 | 28.88336 | 12.38655 | -0.27374 | 0.273772 | 0.773218 |
| 14 | 99643000 | 99647199 | 28.36111 | 47.55007 | 55.688 | 24.29202 | 0.15382 | 1.172745 | 0.773218 |
| 10 | 44794800 | 44797999 | 22.76436 | 38.11398 | 55.94862 | 24.38448 | -0.17082 | 1.170657 | 0.773219 |
| 5 | 7409400 | 7416999 | 71.68103 | 114.033 | 122.9343 | 51.34417 | 0.226632 | 2.247091 | 0.773239 |
| 15 | 25461800 | 25465399 | 35.62841 | 57.56432 | 97.87693 | 41.78769 | -0.40371 | 1.721991 | 0.773247 |
| 18 | 56368200 | 56370799 | 29.1533 | 48.76162 | 56.8542 | 24.76338 | 0.160236 | 1.200788 | 0.773271 |
| 12 | 91847000 | 91848199 | 9.775193 | 18.45839 | 22.21527 | 9.047433 | 0.212135 | 0.050135 | 0.773278 |
| 2 | 5383200 | 5384399 | 270.7959 | 383.5501 | 488.7517 | 202.2449 | -0.16274 | 4.38814 | 0.773288 |
| 9 | 118783200 | 118785599 | 21.82085 | 37.08569 | 53.31608 | 23.36632 | -0.14026 | 1.027132 | 0.773311 |

| | | | | | | | | | |
|---|---|---|---|---|---|---|---|---|---|
| 13 | 17977800 | 17980399 | 22.40782 | 36.85792 | 73.43206 | 31.65359 | -0.61291 | 1.388882 | 0.773357 |
| 7 | 134562200 | 134564199 | 17.03754 | 29.94095 | 35.5246 | 15.72869 | 0.142022 | 0.607162 | 0.773364 |
| 13 | 37138400 | 37140599 | 15.90298 | 28.3237 | 33.34199 | 14.68294 | 0.161857 | 0.47124 | 0.773366 |
| 3 | 45997400 | 45999199 | 15.8798 | 28.18992 | 33.32778 | 14.71727 | 0.152926 | 0.491011 | 0.7734 |
| 3 | 4490000 | 4491599 | 16.96847 | 29.77181 | 35.08168 | 15.51156 | 0.151886 | 0.630717 | 0.773411 |
| X | 11011000 | 11011999 | 10.08591 | 18.67463 | 29.41506 | 12.37459 | -0.20989 | 0.198907 | 0.773427 |
| 9 | 11274600 | 11278199 | 54.4932 | 86.47027 | 122.2609 | 50.92667 | -0.1676 | 2.305708 | 0.773431 |
| 7 | 111713200 | 111715999 | 27.92326 | 46.71048 | 54.60657 | 23.7864 | 0.156411 | 1.192358 | 0.773494 |
| 2 | 106970200 | 106971399 | 11.79641 | 20.74959 | 40.44928 | 17.90981 | -0.56987 | 0.576128 | 0.773519 |
| 3 | 89455600 | 89456999 | 16.02708 | 27.95804 | 41.07185 | 18.07833 | -0.16471 | 0.727804 | 0.773531 |
| 7 | 53329400 | 53335799 | 60.56894 | 96.66064 | 96.21476 | 40.36112 | 0.346415 | 2.128601 | 0.773552 |
| 12 | 24740600 | 24744599 | 47.37711 | 75.81443 | 104.6969 | 44.08918 | -0.1208 | 2.034125 | 0.773577 |
| 14 | 8248400 | 8250599 | 19.886 | 34.17703 | 49.00784 | 21.49657 | -0.13392 | 0.914713 | 0.773584 |
| 2 | 26100400 | 26102799 | 30.18577 | 49.78849 | 57.64573 | 24.8777 | 0.169393 | 1.363981 | 0.773597 |
| 4 | 69151400 | 69153399 | 19.95258 | 35.03411 | 38.94708 | 17.23171 | 0.235405 | 0.641464 | 0.773601 |
| 13 | 4344400 | 4345799 | 13.06497 | 23.59771 | 27.5404 | 11.96744 | 0.187214 | 0.354326 | 0.773605 |
| 5 | 103189400 | 103190599 | 10.53375 | 19.58425 | 29.92157 | 12.51701 | -0.16115 | 0.175719 | 0.773613 |
| 6 | 133427800 | 133429399 | 25.03441 | 43.10815 | 47.92038 | 21.0208 | 0.232509 | 0.918773 | 0.773613 |
| 13 | 67680200 | 67692799 | 143.8287 | 218.9865 | 228.4098 | 94.65933 | 0.224028 | 3.169326 | 0.773651 |
| 16 | 45168800 | 45170199 | 18.69059 | 32.75041 | 38.0358 | 16.79914 | 0.172599 | 0.66465 | 0.773656 |
| 7 | 41070400 | 41073399 | 29.69688 | 49.70319 | 57.42181 | 25.0209 | 0.173456 | 1.196858 | 0.773668 |
| 18 | 18147200 | 18147599 | 5.659057 | 11.02979 | 19.71253 | 7.134041 | -0.19571 | -0.40001 | 0.773671 |
| 1 | 50760400 | 50760799 | 11.97458 | 22.23522 | 27.18595 | 11.39233 | 0.156556 | 0.189669 | 0.77369 |
| 12 | 29659800 | 29660799 | 12.3435 | 22.845 | 27.68009 | 11.65142 | 0.164949 | 0.21269 | 0.773704 |
| 19 | 41036400 | 41038199 | 16.79701 | 29.47618 | 34.58139 | 15.29264 | 0.158176 | 0.625963 | 0.773722 |
| 1 | 139514800 | 139515799 | 15.62466 | 26.98805 | 49.51107 | 21.72061 | -0.48616 | 0.853023 | 0.773729 |
| 10 | 66256200 | 66257999 | 30.10938 | 49.56872 | 71.04244 | 30.6347 | -0.13874 | 1.396588 | 0.773737 |
| 7 | 5517600 | 5545199 | 103.4159 | 160.8881 | 148.0231 | 61.44986 | 0.428883 | 2.74246 | 0.773752 |
| 4 | 38092200 | 38092999 | 10.75866 | 20.38681 | 25.38832 | 10.26331 | 0.172439 | 0.043658 | 0.773774 |
| 2 | 6513400 | 6516599 | 42.53206 | 68.59851 | 80.76118 | 34.40266 | 0.122554 | 1.775445 | 0.773775 |
| 19 | 31641000 | 31642999 | 16.93558 | 28.95311 | 53.7485 | 23.56702 | -0.50613 | 0.956308 | 0.773819 |
| 1 | 167830600 | 167835399 | 68.18074 | 107.5672 | 145.7022 | 60.47187 | -0.11287 | 2.464511 | 0.773827 |
| 10 | 79527200 | 79531399 | 35.27274 | 56.99535 | 81.57893 | 34.83388 | -0.15556 | 1.711431 | 0.773839 |
| 1 | 21036800 | 21038199 | 12.98828 | 23.22039 | 34.56113 | 15.17126 | -0.17162 | 0.429343 | 0.773846 |
| 4 | 127714800 | 127715799 | 10.6039 | 19.78804 | 30.57239 | 12.70907 | -0.16968 | 0.155616 | 0.773872 |
| 3 | 46875400 | 46876399 | 13.69033 | 23.85197 | 45.78172 | 20.13564 | -0.54889 | 0.740069 | 0.773874 |
| 1 | 113831400 | 113831999 | 10.46683 | 20.08437 | 20.20397 | 7.929575 | 0.516744 | -0.06612 | 0.773876 |
| 12 | 73048600 | 73051199 | 23.46041 | 39.96068 | 47.43057 | 20.79064 | 0.135775 | 1.008479 | 0.773884 |
| 7 | 36231000 | 36234599 | 24.744 | 41.25768 | 59.25429 | 25.75799 | -0.13937 | 1.224535 | 0.773898 |
| 1 | 69259400 | 69266599 | 115.5942 | 176.638 | 200.6254 | 83.22028 | 0.105359 | 3.084992 | 0.773907 |
| 2 | 99040800 | 99041399 | 7.852673 | 15.11981 | 24.7708 | 9.626901 | -0.16819 | -0.11368 | 0.773914 |
| 10 | 87814400 | 87816799 | 32.98129 | 54.18402 | 64.42772 | 27.75791 | 0.128764 | 1.423802 | 0.773919 |
| 8 | 5352400 | 5353999 | 20.20785 | 33.68603 | 64.24928 | 27.92505 | -0.54708 | 1.220941 | 0.773933 |
| 16 | 20841600 | 20844599 | 42.99011 | 69.36183 | 81.67083 | 34.80605 | 0.122929 | 1.765648 | 0.773938 |
| X | 41499600 | 41501999 | 34.65818 | 56.59113 | 67.00364 | 28.79717 | 0.130313 | 1.509733 | 0.77394 |
| 1 | 59414000 | 59416799 | 19.79235 | 33.58217 | 50.06959 | 21.93432 | -0.19258 | 1.032104 | 0.773972 |
| 8 | 85824400 | 85842999 | 212.2959 | 309.5299 | 391.1212 | 161.1711 | -0.10913 | 3.95669 | 0.773979 |

| | | | | | | | | | |
|---|---|---|---|---|---|---|---|---|---|
| 4 | 127705800 | 127710799 | 54.2115 | 87.1356 | 82.06537 | 34.78877 | 0.437812 | 1.896793 | 0.773986 |
| 4 | 93364000 | 93365199 | 18.32318 | 32.46999 | 30.10751 | 13.31101 | 0.500319 | 0.51438 | 0.773987 |
| 13 | 3052200 | 3061399 | 128.7218 | 196.907 | 185.5365 | 76.98026 | 0.375849 | 3.067017 | 0.773999 |
| 10 | 69043200 | 69044799 | 16.82349 | 29.27388 | 42.65003 | 18.74346 | -0.15325 | 0.769369 | 0.774006 |
| 4 | 70530200 | 70533799 | 32.34332 | 53.00688 | 76.47492 | 32.91443 | -0.15095 | 1.458576 | 0.774017 |
| 14 | 63877200 | 63879799 | 27.33174 | 45.92813 | 54.00011 | 23.58691 | 0.148131 | 1.144147 | 0.774043 |
| 17 | 79607600 | 79611999 | 36.67996 | 59.30808 | 83.45433 | 35.66483 | -0.12999 | 1.694193 | 0.774045 |
| 4 | 4937600 | 4940999 | 43.33831 | 70.08616 | 81.21797 | 34.71687 | 0.149726 | 1.699514 | 0.774048 |
| 1 | 26129400 | 26130999 | 14.82745 | 27.4244 | 30.22369 | 12.73336 | 0.299509 | 0.239048 | 0.774069 |
| 11 | 19691400 | 19698199 | 50.549 | 81.09702 | 94.89579 | 40.11544 | 0.122025 | 1.954957 | 0.774076 |
| 9 | 79256600 | 79257999 | 12.93243 | 23.41118 | 35.29062 | 15.29194 | -0.1772 | 0.358048 | 0.774077 |
| Y | 39731000 | 39731399 | 4.430641 | 8.639331 | 18.07922 | 6.505496 | -0.41204 | -0.44242 | 0.77408 |
| 6 | 81851800 | 81854199 | 25.65541 | 42.76907 | 61.61715 | 26.78186 | -0.14395 | 1.229581 | 0.7741 |
| 9 | 59198400 | 59199399 | 7.785905 | 14.79093 | 24.49522 | 9.849593 | -0.22787 | -0.00101 | 0.774109 |
| X | 120682000 | 120682999 | 18.0722 | 31.80255 | 36.95746 | 16.36228 | 0.172471 | 0.600082 | 0.774117 |
| 3 | 110677000 | 110678999 | 17.57128 | 31.09213 | 35.2071 | 15.57016 | 0.212523 | 0.532115 | 0.774118 |
| 10 | 120826400 | 120829199 | 19.31446 | 33.39229 | 48.47752 | 21.26779 | -0.14998 | 0.856943 | 0.774144 |
| 4 | 68893000 | 68898799 | 49.28872 | 79.71201 | 84.19338 | 35.99041 | 0.283252 | 1.710147 | 0.774173 |
| 7 | 107738600 | 107742199 | 42.07825 | 67.7319 | 94.40532 | 40.10429 | -0.12396 | 1.840172 | 0.774191 |
| X | 39387600 | 39393399 | 60.58569 | 96.90319 | 90.75095 | 38.18952 | 0.437738 | 2.061615 | 0.774196 |
| 3 | 90986000 | 90989599 | 41.59931 | 67.70309 | 75.55152 | 32.43224 | 0.212122 | 1.572747 | 0.774203 |
| 13 | 23553000 | 23553599 | 7.858602 | 14.96151 | 25.0103 | 10.01059 | -0.23637 | -0.0181 | 0.774221 |
| Y | 2523800 | 2524199 | 5.278818 | 10.23466 | 20.5777 | 7.775609 | -0.42092 | -0.24881 | 0.774225 |
| 13 | 100370400 | 100371399 | 8.21622 | 15.664 | 25.04434 | 9.994937 | -0.16834 | -0.02079 | 0.774226 |
| 18 | 8123800 | 8124799 | 8.671644 | 16.51365 | 26.10999 | 10.44508 | -0.15565 | -0.00631 | 0.774261 |
| 13 | 75441800 | 75442999 | 19.22435 | 32.72498 | 57.00002 | 24.98405 | -0.4156 | 1.009546 | 0.774263 |
| 10 | 78772400 | 78777799 | 91.13717 | 140.773 | 186.9479 | 77.65423 | -0.10803 | 2.866589 | 0.77427 |
| 18 | 81365000 | 81369399 | 37.80307 | 61.10615 | 86.05217 | 36.76114 | -0.13164 | 1.70531 | 0.774273 |
| 16 | 64223000 | 64226399 | 29.92292 | 48.60416 | 83.44566 | 35.78598 | -0.40996 | 1.589754 | 0.774278 |
| 15 | 25604200 | 25607399 | 21.00157 | 36.1188 | 53.39184 | 23.42002 | -0.17757 | 0.917285 | 0.774287 |
| 16 | 89865400 | 89868999 | 27.30359 | 44.50472 | 79.45251 | 34.12584 | -0.46232 | 1.527279 | 0.7743 |
| 9 | 19833800 | 19837799 | 61.07288 | 97.4905 | 110.6239 | 46.4229 | 0.15815 | 2.126372 | 0.774304 |
| 13 | 69306400 | 69309399 | 33.87014 | 55.94885 | 54.6708 | 23.61509 | 0.413738 | 1.352807 | 0.774315 |
| 7 | 131510000 | 131511599 | 14.27578 | 25.96052 | 30.40682 | 13.10116 | 0.189648 | 0.328779 | 0.77432 |
| 13 | 70613200 | 70615799 | 19.72243 | 34.2892 | 53.46804 | 23.48517 | -0.25135 | 0.806979 | 0.774364 |
| 9 | 15537200 | 15539599 | 16.13007 | 28.06895 | 41.62486 | 18.2936 | -0.1787 | 0.762351 | 0.77437 |
| 7 | 143056200 | 143057799 | 15.24884 | 26.61588 | 40.09248 | 17.65368 | -0.2005 | 0.714795 | 0.774381 |
| 15 | 47206400 | 47208399 | 15.56289 | 27.69082 | 32.85564 | 14.48073 | 0.149704 | 0.474056 | 0.774392 |
| 16 | 53012800 | 53013399 | 8.858896 | 16.57332 | 26.39374 | 10.92534 | -0.20773 | 0.114613 | 0.774409 |
| 3 | 128760000 | 128760599 | 7.157558 | 13.83384 | 23.18693 | 8.890604 | -0.18132 | -0.16651 | 0.774427 |
| 10 | 95269000 | 95269799 | 7.157558 | 13.83384 | 23.18693 | 8.890604 | -0.18132 | -0.16651 | 0.774427 |
| 1 | 109197000 | 109198599 | 16.88426 | 29.56537 | 43.51087 | 19.20636 | -0.16718 | 0.67823 | 0.774448 |
| 10 | 17900000 | 17904199 | 47.75936 | 76.57857 | 106.408 | 44.94367 | -0.12636 | 1.974525 | 0.774467 |
| 17 | 49552200 | 49553999 | 15.35347 | 27.28882 | 32.33565 | 14.26449 | 0.150662 | 0.47767 | 0.774474 |
| 1 | 32810000 | 32812599 | 28.815 | 47.82755 | 56.65697 | 24.55428 | 0.137345 | 1.285735 | 0.774477 |
| 11 | 66962400 | 66963799 | 14.68534 | 25.9307 | 37.63429 | 16.6506 | -0.14464 | 0.544053 | 0.774478 |
| 11 | 73951000 | 73951999 | 14.09636 | 25.63615 | 30.50265 | 13.14123 | 0.167439 | 0.329327 | 0.774478 |

| | | | | | | | | | |
|---|---|---|---|---|---|---|---|---|---|
| 1 | 24640000 | 24640799 | 7.732281 | 14.86529 | 24.14528 | 9.425033 | -0.16142 | -0.10093 | 0.774481 |
| 5 | 60036400 | 60039199 | 24.41283 | 41.65358 | 48.47924 | 21.25322 | 0.164182 | 0.994805 | 0.774507 |
| 16 | 46687600 | 46688399 | 8.969764 | 16.92529 | 26.45596 | 10.79096 | -0.16402 | 0.059915 | 0.774519 |
| 5 | 10265600 | 10266999 | 14.11183 | 26.19895 | 29.80681 | 12.49376 | 0.25933 | 0.213213 | 0.774552 |
| 14 | 35003800 | 35007799 | 42.70886 | 69.30544 | 78.68171 | 33.71887 | 0.18388 | 1.629111 | 0.774554 |
| 6 | 21198600 | 21201599 | 47.8062 | 76.67254 | 107.6259 | 45.47506 | -0.14056 | 1.968124 | 0.774588 |
| 5 | 22991200 | 22992599 | 11.89495 | 21.39477 | 32.12652 | 14.03079 | -0.17901 | 0.380006 | 0.774618 |
| 2 | 23483400 | 23487599 | 51.3265 | 82.0806 | 113.6199 | 47.80237 | -0.12529 | 2.061188 | 0.774625 |
| 2 | 156051600 | 156053599 | 40.00666 | 64.69485 | 75.52097 | 32.27946 | 0.139179 | 1.695123 | 0.77466 |
| 9 | 20264600 | 20266799 | 60.54663 | 96.27058 | 111.7385 | 46.63804 | 0.119519 | 2.250264 | 0.774684 |
| 6 | 136239400 | 136246999 | 80.28428 | 126.6433 | 141.7988 | 58.84808 | 0.161312 | 2.476464 | 0.774704 |
| 7 | 80818600 | 80820199 | 10.67425 | 19.21264 | 30.75871 | 13.42424 | -0.27004 | 0.364242 | 0.774705 |
| 11 | 29727400 | 29729799 | 17.32474 | 30.48758 | 35.92962 | 15.90719 | 0.152275 | 0.592652 | 0.774705 |
| X | 69718400 | 69721199 | 25.18481 | 42.98391 | 49.56206 | 21.72847 | 0.177839 | 1.000141 | 0.77477 |
| 8 | 20137600 | 20147199 | 163.476 | 244.4621 | 272.6568 | 112.4311 | 0.105041 | 3.527012 | 0.774803 |
| 6 | 60177000 | 60184399 | 97.38184 | 151.9392 | 143.5001 | 59.54761 | 0.394344 | 2.687994 | 0.774807 |
| 2 | 168985200 | 168986799 | 17.40008 | 30.51458 | 36.15993 | 15.98189 | 0.14393 | 0.644291 | 0.774854 |
| 3 | 80036600 | 80041799 | 112.335 | 172.0018 | 195.8209 | 81.26499 | 0.104233 | 3.04355 | 0.774858 |
| 1 | 152988000 | 152988599 | 8.787437 | 16.91466 | 21.95897 | 8.541334 | 0.164012 | -0.10702 | 0.77486 |
| 6 | 51021000 | 51022799 | 15.89372 | 27.82978 | 40.86171 | 18.03639 | -0.164 | 0.665939 | 0.774867 |
| 10 | 40855000 | 40857799 | 27.80616 | 45.72197 | 66.14555 | 28.50858 | -0.15283 | 1.401989 | 0.774899 |
| 10 | 88128400 | 88130599 | 27.18294 | 45.51696 | 53.99464 | 23.5345 | 0.135424 | 1.180429 | 0.774902 |
| 6 | 140152400 | 140155199 | 24.21644 | 40.07133 | 70.13479 | 30.33087 | -0.42448 | 1.31861 | 0.77493 |
| 3 | 143910200 | 143911799 | 13.74887 | 23.7906 | 48.96864 | 21.48382 | -0.65078 | 0.827218 | 0.774964 |
| 8 | 105091600 | 105093599 | 13.74887 | 23.7906 | 48.96864 | 21.48382 | -0.65078 | 0.827218 | 0.774964 |
| 7 | 94175600 | 94179599 | 54.71577 | 87.1995 | 119.832 | 50.1722 | -0.12023 | 2.174555 | 0.774978 |
| 1 | 117881400 | 117882799 | 30.34564 | 50.53782 | 59.00018 | 25.62836 | 0.1586 | 1.254725 | 0.774985 |
| 17 | 3053800 | 3089199 | 1221.303 | 1531.197 | 1712.687 | 720.0792 | -0.15155 | 6.324033 | 0.775023 |
| 12 | 116816600 | 116820399 | 40.96414 | 66.17401 | 77.0863 | 32.90415 | 0.140384 | 1.723794 | 0.775025 |
| 7 | 131596800 | 131597799 | 13.57955 | 23.92414 | 35.75978 | 15.82839 | -0.18801 | 0.559143 | 0.775062 |
| 7 | 108165800 | 108169599 | 64.96263 | 103.6279 | 98.65282 | 41.35897 | 0.409963 | 2.148154 | 0.775078 |
| 13 | 61813200 | 61850199 | 647.8742 | 869.8199 | 837.3839 | 349.4253 | 0.163849 | 5.217399 | 0.775083 |
| 15 | 37806000 | 37809399 | 25.63561 | 43.57067 | 50.66422 | 22.20418 | 0.16469 | 1.037447 | 0.775096 |
| 11 | 34376000 | 34377599 | 14.63283 | 25.88602 | 38.08591 | 16.84416 | -0.16333 | 0.522906 | 0.775105 |
| 7 | 61584600 | 61586599 | 26.52896 | 44.52666 | 52.87575 | 23.08109 | 0.133856 | 1.15256 | 0.775109 |
| 4 | 141784800 | 141785599 | 9.725025 | 18.43526 | 22.81363 | 9.214965 | 0.183104 | 0.026734 | 0.775156 |
| 6 | 117443800 | 117446199 | 28.95066 | 48.27528 | 57.30407 | 24.91136 | 0.134629 | 1.234776 | 0.775156 |
| 13 | 66918200 | 66930399 | 338.7781 | 476.7182 | 501.0525 | 207.5585 | 0.105549 | 4.486713 | 0.77516 |
| 1 | 60737800 | 60739399 | 13.67009 | 24.51869 | 36.66341 | 16.05016 | -0.17558 | 0.418621 | 0.775176 |
| Y | 90839000 | 90840799 | 22.70469 | 37.36505 | 62.91979 | 27.12796 | -0.37061 | 1.375858 | 0.775198 |
| 3 | 25205400 | 25212399 | 165.2639 | 246.7308 | 274.2287 | 113.0433 | 0.10793 | 3.558341 | 0.775208 |
| 5 | 53159800 | 53168199 | 71.82274 | 113.2271 | 155.9344 | 64.70942 | -0.13764 | 2.484012 | 0.77521 |
| 8 | 34382000 | 34386599 | 39.63025 | 63.86108 | 89.44064 | 38.05538 | -0.12893 | 1.800567 | 0.775314 |
| 7 | 42872200 | 42898399 | 460.9149 | 635.071 | 640.3365 | 266.1325 | 0.134832 | 4.808322 | 0.775355 |
| 1 | 39872600 | 39874799 | 17.60442 | 30.02559 | 54.26633 | 23.78955 | -0.46815 | 0.982345 | 0.775357 |
| 12 | 63960200 | 63962399 | 17.29136 | 30.72221 | 34.70943 | 15.31626 | 0.218672 | 0.496118 | 0.775363 |
| 11 | 91033800 | 91035599 | 17.98402 | 31.25886 | 45.25793 | 19.87579 | -0.1445 | 0.79442 | 0.775384 |

| | | | | | | | | | |
|---|---|---|---|---|---|---|---|---|---|
| 17 | 72605200 | 72606999 | 14.5309 | 25.61746 | 37.17224 | 16.45137 | -0.14519 | 0.55986 | 0.775387 |
| 19 | 25238200 | 25239799 | 14.5309 | 25.61746 | 37.17224 | 16.45137 | -0.14519 | 0.55986 | 0.775387 |
| 18 | 3110000 | 3122799 | 900.1185 | 1156.077 | 1328.778 | 557.3047 | -0.15571 | 5.932706 | 0.775388 |
| 13 | 103567200 | 103570399 | 22.63352 | 38.7564 | 45.79523 | 20.08235 | 0.143242 | 0.953049 | 0.775428 |
| 9 | 117450000 | 117451599 | 28.46917 | 48.06814 | 54.56148 | 23.86661 | 0.199044 | 1.109997 | 0.775432 |
| 1 | 181090800 | 181091999 | 15.84308 | 27.95809 | 32.16142 | 14.23095 | 0.188892 | 0.549976 | 0.775434 |
| 11 | 73761200 | 73766799 | 80.14863 | 126.1079 | 143.399 | 59.49247 | 0.13621 | 2.524817 | 0.775445 |
| 6 | 8601400 | 8608799 | 118.3524 | 180.9384 | 205.0582 | 85.06834 | 0.109068 | 3.077003 | 0.775458 |
| 4 | 19105800 | 19107199 | 11.2537 | 20.60235 | 30.76208 | 13.14211 | -0.14996 | 0.271421 | 0.775458 |
| 8 | 51088800 | 51090199 | 11.2537 | 20.60235 | 30.76208 | 13.14211 | -0.14996 | 0.271421 | 0.775458 |
| 13 | 72560000 | 72560999 | 11.2537 | 20.60235 | 30.76208 | 13.14211 | -0.14996 | 0.271421 | 0.775458 |
| 17 | 76297200 | 76297799 | 11.2537 | 20.60235 | 30.76208 | 13.14211 | -0.14996 | 0.271421 | 0.775458 |
| 16 | 19820800 | 19822999 | 14.07506 | 25.6779 | 24.36595 | 10.45627 | 0.495439 | 0.29711 | 0.775476 |
| 5 | 10809000 | 10810399 | 13.63624 | 24.21952 | 36.03024 | 15.9038 | -0.17646 | 0.477105 | 0.775522 |
| 14 | 87800800 | 87802399 | 12.66959 | 22.62062 | 33.7179 | 14.81766 | -0.17432 | 0.438236 | 0.775525 |
| 7 | 85767800 | 85770599 | 20.86285 | 36.56038 | 32.94442 | 14.55168 | 0.537251 | 0.67061 | 0.775525 |
| 7 | 18239400 | 18239999 | 8.741998 | 16.82912 | 21.94678 | 8.533781 | 0.15771 | -0.10983 | 0.775535 |
| X | 44318200 | 44318599 | 9.134534 | 17.541 | 22.45761 | 8.795155 | 0.174817 | -0.07924 | 0.775535 |
| 19 | 21361200 | 21361799 | 6.532375 | 12.71321 | 21.91346 | 8.062704 | -0.1658 | -0.32163 | 0.775542 |
| 8 | 126056400 | 126061199 | 56.60736 | 89.91237 | 123.1884 | 51.36294 | -0.12091 | 2.27939 | 0.775552 |
| 7 | 20931600 | 20942599 | 280.25 | 400.0705 | 421.5971 | 174.2268 | 0.122635 | 4.27344 | 0.775582 |
| X | 11943000 | 11948199 | 35.23796 | 57.19063 | 84.45351 | 36.19635 | -0.19424 | 1.620027 | 0.775584 |
| 13 | 9723800 | 9726599 | 25.19254 | 43.53641 | 47.36953 | 20.78143 | 0.264359 | 0.887037 | 0.775595 |
| 14 | 78982400 | 78983799 | 12.19706 | 22.0062 | 32.62121 | 14.19452 | -0.15711 | 0.364988 | 0.775606 |
| 4 | 58146400 | 58147399 | 14.80292 | 26.72019 | 31.34227 | 13.63007 | 0.178988 | 0.377064 | 0.775606 |
| 6 | 119849800 | 119853799 | 49.37718 | 79.29636 | 92.7162 | 39.25375 | 0.124852 | 1.921555 | 0.775607 |
| 6 | 90205000 | 90210799 | 112.5456 | 171.1628 | 226.2726 | 93.74191 | -0.11894 | 3.183074 | 0.775613 |
| 4 | 26320600 | 26321199 | 8.552765 | 16.41153 | 26.93911 | 10.57298 | -0.18396 | -0.06634 | 0.77565 |
| 3 | 30506400 | 30508799 | 14.855 | 26.03241 | 38.37831 | 16.952 | -0.16959 | 0.645186 | 0.775652 |
| 12 | 115952200 | 115959399 | 135.694 | 205.0048 | 229.1121 | 94.72491 | 0.115182 | 3.322385 | 0.775686 |
| 2 | 18560600 | 18561599 | 19.57553 | 33.85564 | 39.08451 | 17.14713 | 0.179532 | 0.843669 | 0.775697 |
| 9 | 12495400 | 12496399 | 9.716462 | 18.29688 | 28.58775 | 11.70325 | -0.16816 | 0.083932 | 0.775708 |
| 13 | 92313400 | 92314799 | 13.31925 | 23.63406 | 35.53247 | 15.69665 | -0.19239 | 0.479882 | 0.775712 |
| 3 | 29353200 | 29354999 | 18.10217 | 32.27925 | 35.2916 | 15.52578 | 0.268889 | 0.481851 | 0.775739 |
| 3 | 23544200 | 23545999 | 17.16898 | 30.50804 | 34.74625 | 15.33112 | 0.207248 | 0.495309 | 0.77574 |
| 10 | 49211800 | 49213799 | 16.80381 | 30.39707 | 32.85866 | 14.25122 | 0.298654 | 0.382043 | 0.775745 |
| 10 | 59406200 | 59410799 | 43.55569 | 70.42933 | 81.6904 | 34.91345 | 0.148224 | 1.7044 | 0.775747 |
| 17 | 49593800 | 49596799 | 36.51473 | 59.07768 | 82.97603 | 35.48117 | -0.12641 | 1.67919 | 0.775751 |
| 14 | 51017800 | 51019999 | 23.23529 | 39.91461 | 46.84814 | 20.54895 | 0.153667 | 0.929583 | 0.775754 |
| 7 | 12088800 | 12097999 | 108.7416 | 168.9356 | 149.7104 | 62.16339 | 0.48132 | 2.768794 | 0.775786 |
| 13 | 28337400 | 28338599 | 13.58427 | 24.09139 | 35.11471 | 15.51911 | -0.14807 | 0.487331 | 0.775807 |
| 3 | 144639400 | 144644399 | 49.82815 | 79.56558 | 109.7013 | 46.05626 | -0.1219 | 2.103644 | 0.775815 |
| 12 | 48578800 | 48580399 | 32.25177 | 52.93608 | 62.70858 | 27.00658 | 0.1336 | 1.430891 | 0.775829 |
| 13 | 14111600 | 14121199 | 141.932 | 215.3129 | 235.9071 | 97.65985 | 0.148789 | 3.241912 | 0.775847 |
| X | 155405600 | 155407599 | 23.19612 | 40.46312 | 35.94723 | 15.82247 | 0.557405 | 0.75477 | 0.775852 |
| 5 | 80121400 | 80122999 | 14.0583 | 25.52997 | 29.84506 | 12.88121 | 0.190732 | 0.333084 | 0.775854 |
| 6 | 51784000 | 51785799 | 16.09001 | 28.22698 | 41.38485 | 18.29661 | -0.16168 | 0.641515 | 0.775863 |

| | | | | | | | | | |
|---|---|---|---|---|---|---|---|---|---|
| 7 | 42383200 | 42384599 | 23.02841 | 39.49677 | 46.34252 | 20.32502 | 0.153872 | 0.944267 | 0.775878 |
| 8 | 70494400 | 70497199 | 14.15986 | 24.64664 | 46.31515 | 20.35813 | -0.51866 | 0.756697 | 0.775881 |
| 12 | 113829800 | 113839999 | 143.5704 | 217.666 | 237.5151 | 98.30761 | 0.153952 | 3.255314 | 0.77589 |
| 10 | 58365600 | 58372599 | 93.57111 | 146.2124 | 161.491 | 67.0029 | 0.170454 | 2.664004 | 0.775913 |
| 14 | 94834600 | 94837599 | 28.89887 | 48.06502 | 57.30667 | 24.87096 | 0.128227 | 1.26152 | 0.77593 |
| 1 | 87066200 | 87072399 | 60.3223 | 95.32295 | 132.5951 | 55.07694 | -0.14925 | 2.419268 | 0.775947 |
| 4 | 113761400 | 113769599 | 100.1368 | 156.6025 | 162.9664 | 67.60925 | 0.257222 | 2.654253 | 0.775963 |
| 4 | 34349600 | 34355999 | 93.59151 | 144.124 | 191.4357 | 79.51312 | -0.11168 | 2.924946 | 0.775976 |
| 6 | 51491200 | 51495199 | 47.12626 | 75.80664 | 89.12249 | 37.8233 | 0.119911 | 1.866544 | 0.77598 |
| 14 | 80097800 | 80101199 | 57.47108 | 91.59027 | 106.9247 | 44.76785 | 0.114665 | 2.171766 | 0.775982 |
| 1 | 49335600 | 49336599 | 10.87245 | 20.42189 | 25.20088 | 10.3611 | 0.164833 | 0.101563 | 0.776 |
| 14 | 111115800 | 111117799 | 37.64424 | 61.29664 | 72.30381 | 31.04371 | 0.132863 | 1.558791 | 0.776019 |
| 6 | 41856200 | 41859999 | 43.6526 | 70.00763 | 97.95455 | 41.3857 | -0.13595 | 1.960569 | 0.776025 |
| 5 | 44113600 | 44119399 | 55.78536 | 88.80356 | 120.9619 | 50.56605 | -0.10926 | 2.210761 | 0.776031 |
| 6 | 73052600 | 73061999 | 105.9694 | 163.5939 | 183.7577 | 76.32929 | 0.132697 | 2.882078 | 0.776061 |
| 6 | 132352400 | 132353999 | 21.61961 | 37.72992 | 42.36339 | 18.65184 | 0.221102 | 0.7495 | 0.776071 |
| 14 | 105638600 | 105640999 | 27.03325 | 44.42852 | 75.75297 | 32.64235 | -0.38959 | 1.412633 | 0.776074 |
| 4 | 147921000 | 147931399 | 292.5267 | 414.4476 | 507.8339 | 210.1311 | -0.10583 | 4.385316 | 0.776082 |
| 17 | 83658000 | 83658999 | 7.870185 | 15.01828 | 24.51574 | 9.764723 | -0.19588 | -0.03406 | 0.776102 |
| 15 | 87751000 | 87752199 | 14.24312 | 25.34815 | 29.07644 | 12.81272 | 0.198534 | 0.456795 | 0.776105 |
| 12 | 32109600 | 32111599 | 12.25477 | 21.94657 | 32.65167 | 14.31238 | -0.16929 | 0.411745 | 0.776135 |
| 18 | 59580800 | 59581799 | 13.87392 | 25.41008 | 30.05144 | 12.83105 | 0.184572 | 0.28606 | 0.77614 |
| 4 | 129997400 | 129998399 | 16.68998 | 29.34307 | 33.70016 | 14.92293 | 0.189157 | 0.598996 | 0.776174 |
| 2 | 88940800 | 88942599 | 20.93139 | 36.43034 | 42.47323 | 18.66848 | 0.166682 | 0.783765 | 0.776176 |
| 1 | 13806800 | 13807799 | 6.987975 | 13.55537 | 22.79713 | 8.593589 | -0.16301 | -0.22817 | 0.776188 |
| 7 | 36903600 | 36905399 | 17.47478 | 30.83186 | 35.84604 | 15.86192 | 0.173135 | 0.564843 | 0.776201 |
| 14 | 68658800 | 68661199 | 23.57276 | 39.57965 | 56.53827 | 24.68563 | -0.13172 | 1.144101 | 0.776217 |
| 15 | 33136800 | 33141599 | 94.57886 | 146.6355 | 168.4335 | 69.95885 | 0.105203 | 2.798232 | 0.776223 |
| 4 | 143979200 | 143994399 | 203.1781 | 301.7646 | 314.446 | 129.5344 | 0.194067 | 3.657337 | 0.776233 |
| 6 | 27524000 | 27527199 | 25.97978 | 43.57908 | 64.58848 | 28.18334 | -0.18487 | 1.167557 | 0.776313 |
| 8 | 22700800 | 22703599 | 20.38014 | 35.40179 | 42.01316 | 18.44364 | 0.140908 | 0.807275 | 0.776346 |
| 10 | 22617600 | 22617999 | 5.647684 | 11.00162 | 20.1952 | 7.361122 | -0.2414 | -0.36661 | 0.776392 |
| 4 | 6150600 | 6159799 | 120.9331 | 185.0773 | 206.7115 | 85.77495 | 0.131278 | 3.056899 | 0.776393 |
| X | 6392400 | 6393399 | 14.60378 | 26.36892 | 30.53762 | 13.27499 | 0.198198 | 0.376808 | 0.776411 |
| 3 | 26385200 | 26386399 | 13.55977 | 24.02372 | 35.10936 | 15.52307 | -0.1526 | 0.495986 | 0.77642 |
| 16 | 36805200 | 36806999 | 15.67336 | 27.66544 | 32.46684 | 14.36538 | 0.160044 | 0.5435 | 0.776442 |
| 18 | 50234600 | 50236199 | 18.55676 | 31.95058 | 46.36382 | 20.33793 | -0.15043 | 0.888949 | 0.776442 |
| 3 | 91073800 | 91079999 | 65.96228 | 105.5017 | 108.6387 | 45.71633 | 0.301043 | 2.071173 | 0.776456 |
| 1 | 171494000 | 171497599 | 42.15681 | 68.10283 | 80.15808 | 34.21674 | 0.125549 | 1.727104 | 0.776508 |
| 7 | 39004200 | 39004599 | 7.435978 | 14.25398 | 23.53818 | 9.265377 | -0.19628 | -0.07849 | 0.776531 |
| 4 | 50128800 | 50132599 | 48.70119 | 77.98096 | 107.2776 | 45.21592 | -0.11432 | 2.017938 | 0.776542 |
| 4 | 72010000 | 72013199 | 25.85701 | 44.49817 | 48.46575 | 21.25964 | 0.261588 | 0.925814 | 0.776556 |
| 9 | 30473800 | 30475599 | 15.517 | 27.31816 | 39.70969 | 17.57923 | -0.14857 | 0.585181 | 0.776566 |
| 19 | 16203000 | 16204599 | 13.79739 | 24.56999 | 36.15177 | 15.92225 | -0.15813 | 0.466111 | 0.776566 |
| 9 | 17525600 | 17527199 | 15.73865 | 27.65372 | 40.28709 | 17.83479 | -0.15257 | 0.612245 | 0.776572 |
| 17 | 52605000 | 52607199 | 19.90937 | 34.04839 | 49.01219 | 21.49087 | -0.14058 | 0.959881 | 0.776575 |
| 6 | 130402800 | 130407599 | 54.03806 | 86.97604 | 93.54919 | 39.73586 | 0.249048 | 1.859499 | 0.77658 |

| | | | | | | | | | |
|---|---|---|---|---|---|---|---|---|---|
| 2 | 162683600 | 162685399 | 21.28524 | 36.84299 | 43.64431 | 19.14808 | 0.142663 | 0.850583 | 0.776584 |
| 10 | 115718600 | 115720799 | 15.31808 | 27.06896 | 40.18214 | 17.77509 | -0.17672 | 0.543026 | 0.776601 |
| 17 | 91869400 | 91873999 | 65.68943 | 103.3113 | 142.7553 | 59.22199 | -0.14506 | 2.526221 | 0.776622 |
| X | 71785600 | 71792999 | 108.1259 | 165.1975 | 217.8129 | 90.34619 | -0.10983 | 3.088699 | 0.77664 |
| 2 | 130331200 | 130332599 | 16.70361 | 29.46755 | 34.92317 | 15.45394 | 0.145631 | 0.560846 | 0.776731 |
| 1 | 140835200 | 140838399 | 34.8014 | 57.20384 | 66.95755 | 28.85418 | 0.151653 | 1.423492 | 0.776748 |
| 9 | 71048000 | 71048799 | 10.39433 | 19.5475 | 23.78338 | 9.758273 | 0.188161 | 0.089327 | 0.776771 |
| 7 | 125117800 | 125120799 | 21.86073 | 37.65043 | 44.71227 | 19.61364 | 0.137547 | 0.897562 | 0.776785 |
| 8 | 11653200 | 11663599 | 157.7196 | 233.1304 | 355.5831 | 146.4376 | -0.36169 | 3.734137 | 0.776787 |
| 13 | 62165800 | 62232199 | 1514.679 | 1894.25 | 1720.752 | 723.6668 | 0.144949 | 6.365409 | 0.776789 |
| 2 | 70522600 | 70529599 | 62.04353 | 98.61068 | 114.6125 | 47.8182 | 0.116939 | 2.262015 | 0.77679 |
| 5 | 82045200 | 82045599 | 6.686154 | 13.01078 | 22.50724 | 8.29001 | -0.17239 | -0.31389 | 0.7768 |
| 6 | 137989000 | 137993199 | 27.98584 | 46.408 | 66.77367 | 28.92008 | -0.14252 | 1.300065 | 0.776808 |
| Y | 41160000 | 41168799 | 68.96626 | 108.7954 | 99.01553 | 41.09304 | 0.459897 | 2.455735 | 0.776825 |
| 12 | 70229200 | 70232599 | 69.75596 | 110.165 | 125.7944 | 52.23641 | 0.134399 | 2.440086 | 0.77684 |
| 4 | 10613200 | 10614999 | 15.22438 | 26.76644 | 38.74175 | 17.15541 | -0.14308 | 0.597757 | 0.776861 |
| 6 | 75054800 | 75055999 | 11.14879 | 20.67848 | 32.07043 | 13.45984 | -0.18582 | 0.201391 | 0.776864 |
| 6 | 121178000 | 121181199 | 34.11159 | 55.27072 | 78.5267 | 33.61498 | -0.14091 | 1.64447 | 0.776886 |
| 15 | 37904200 | 37904799 | 10.22605 | 19.44537 | 24.36076 | 9.775417 | 0.173096 | 0.01135 | 0.776895 |
| 7 | 125131000 | 125131599 | 5.070319 | 9.896729 | 18.13296 | 6.42583 | -0.20226 | -0.48478 | 0.776917 |
| 13 | 62560800 | 62568199 | 159.8542 | 239.3244 | 267.0314 | 110.1364 | 0.106 | 3.50528 | 0.776924 |
| 13 | 57326200 | 57327399 | 16.45637 | 28.73511 | 41.50644 | 18.28091 | -0.14046 | 0.715371 | 0.776925 |
| 10 | 41182400 | 41185599 | 36.27301 | 58.88613 | 68.41718 | 29.329 | 0.151192 | 1.602925 | 0.776942 |
| 17 | 45616800 | 45622199 | 61.24137 | 97.11036 | 131.5457 | 54.7558 | -0.1069 | 2.334062 | 0.776943 |
| 4 | 59027000 | 59036999 | 121.5216 | 184.3914 | 241.0747 | 99.80578 | -0.10568 | 3.232008 | 0.77696 |
| 1 | 128677400 | 128678799 | 8.479516 | 15.73593 | 31.02304 | 13.01086 | -0.52845 | 0.174026 | 0.776966 |
| 19 | 15341800 | 15342599 | 10.54746 | 19.90952 | 24.81646 | 10.11446 | 0.161254 | 0.068852 | 0.776969 |
| 1 | 64235200 | 64236999 | 11.73113 | 21.19072 | 31.65382 | 13.75314 | -0.1666 | 0.353115 | 0.776986 |
| 11 | 73840200 | 73841199 | 25.93878 | 43.45688 | 51.23301 | 22.33833 | 0.144388 | 1.169715 | 0.776994 |
| 8 | 85865200 | 85870199 | 77.65235 | 120.9849 | 163.3769 | 67.80884 | -0.12245 | 2.706496 | 0.776998 |
| 1 | 46328000 | 46328399 | 13.833 | 24.83925 | 29.08089 | 12.7185 | 0.176798 | 0.39974 | 0.777009 |
| 3 | 13341800 | 13344799 | 26.93022 | 45.84708 | 51.77326 | 22.69326 | 0.207185 | 1.030283 | 0.777029 |
| 7 | 116935800 | 116939599 | 44.1852 | 71.37016 | 67.63917 | 28.8679 | 0.437221 | 1.728055 | 0.777037 |
| 5 | 131167600 | 131169399 | 16.50049 | 29.06613 | 34.44921 | 15.24871 | 0.144715 | 0.57437 | 0.777041 |
| 4 | 106039400 | 106043599 | 32.39486 | 53.39107 | 63.5181 | 27.40307 | 0.129788 | 1.380978 | 0.777053 |
| 11 | 90528600 | 90529799 | 10.69793 | 18.88833 | 39.90963 | 17.65758 | -0.68331 | 0.525041 | 0.77706 |
| 3 | 49598000 | 49598999 | 12.00898 | 21.36592 | 32.78433 | 14.45014 | -0.21885 | 0.452992 | 0.777064 |
| 1 | 185388600 | 185389799 | 11.02791 | 20.3798 | 30.88551 | 13.02703 | -0.15814 | 0.215801 | 0.777076 |
| 16 | 19790400 | 19791599 | 13.76783 | 25.45464 | 29.71751 | 12.52674 | 0.216481 | 0.233067 | 0.777091 |
| 8 | 31537000 | 31538199 | 14.64837 | 25.77569 | 37.67444 | 16.67995 | -0.15678 | 0.581007 | 0.777101 |
| 6 | 44890200 | 44891199 | 16.17985 | 28.27477 | 40.99918 | 18.06682 | -0.14579 | 0.702874 | 0.777114 |
| 15 | 68963600 | 68967599 | 33.19756 | 55.00826 | 63.22396 | 27.3682 | 0.180527 | 1.320637 | 0.777127 |
| 2 | 9805000 | 9806799 | 11.78852 | 21.32991 | 31.70843 | 13.74817 | -0.15754 | 0.345738 | 0.777128 |
| 8 | 19582400 | 19593199 | 177.0293 | 261.3752 | 334.5528 | 137.7751 | -0.11043 | 3.753951 | 0.777136 |
| 9 | 85208200 | 85210799 | 23.5468 | 38.63749 | 74.49376 | 32.07829 | -0.56716 | 1.423232 | 0.777146 |
| 3 | 114380600 | 114384799 | 44.06798 | 70.92154 | 101.4809 | 43.09846 | -0.16209 | 1.854898 | 0.777161 |
| 8 | 4754600 | 4758599 | 47.27519 | 75.98808 | 89.46168 | 37.9252 | 0.11652 | 1.890498 | 0.777164 |

| | | | | | | | | | |
|---|---|---|---|---|---|---|---|---|---|
| 13 | 103457600 | 103465599 | 90.77088 | 139.449 | 218.4155 | 90.68969 | -0.35187 | 2.972397 | 0.777177 |
| 14 | 55534600 | 55535399 | 9.733999 | 17.96354 | 28.38509 | 11.99515 | -0.21914 | 0.21131 | 0.777198 |
| 13 | 113834400 | 113837399 | 25.78839 | 43.64847 | 51.22982 | 22.42588 | 0.151007 | 1.073867 | 0.777199 |
| 17 | 38074000 | 38075599 | 20.42547 | 35.33331 | 42.10131 | 18.47078 | 0.134098 | 0.849725 | 0.777222 |
| 13 | 19320000 | 19323799 | 36.36378 | 58.87414 | 82.43269 | 35.2668 | -0.12098 | 1.665215 | 0.77726 |
| 19 | 11262200 | 11263599 | 14.30141 | 26.12133 | 25.3559 | 10.86521 | 0.464306 | 0.295017 | 0.777289 |
| 9 | 94196200 | 94197999 | 16.17412 | 28.10203 | 51.16161 | 22.463 | -0.4738 | 0.796714 | 0.7773 |
| 7 | 67377600 | 67378799 | 13.48317 | 23.9108 | 35.17855 | 15.54804 | -0.16137 | 0.488718 | 0.777305 |
| 1 | 121923000 | 121924999 | 17.91528 | 31.40845 | 37.05114 | 16.37155 | 0.150232 | 0.65108 | 0.777323 |
| 1 | 159854600 | 159856999 | 17.91528 | 31.40845 | 37.05114 | 16.37155 | 0.150232 | 0.65108 | 0.777323 |
| 18 | 90603600 | 90605599 | 19.03395 | 33.2242 | 39.26185 | 17.28847 | 0.147629 | 0.72802 | 0.777356 |
| 13 | 75524800 | 75526199 | 19.14025 | 33.52238 | 39.1408 | 17.28024 | 0.16505 | 0.678037 | 0.777356 |
| 13 | 3220400 | 3235199 | 319.3029 | 450.8973 | 472.3395 | 195.5506 | 0.115455 | 4.433894 | 0.777382 |
| 9 | 37693400 | 37695999 | 19.01361 | 32.69154 | 47.41371 | 20.79759 | -0.15015 | 0.903954 | 0.777384 |
| 19 | 8415400 | 8421599 | 139.767 | 210.0574 | 272.1774 | 112.362 | -0.10462 | 3.428154 | 0.777394 |
| 8 | 21305200 | 21310799 | 105.3992 | 161.8863 | 217.558 | 90.33094 | -0.13141 | 2.982579 | 0.777425 |
| 10 | 118963800 | 118967399 | 39.15163 | 63.19541 | 88.08445 | 37.55986 | -0.11922 | 1.750911 | 0.777436 |
| 18 | 19967600 | 19969799 | 26.96212 | 44.03505 | 77.8765 | 33.47303 | -0.44672 | 1.497348 | 0.777467 |
| 18 | 81914400 | 81917199 | 31.54971 | 51.53171 | 73.0555 | 31.40177 | -0.12859 | 1.500849 | 0.77747 |
| 16 | 31032400 | 31049999 | 233.5097 | 337.8892 | 423.3282 | 174.5918 | -0.10747 | 4.07237 | 0.777488 |
| X | 158324200 | 158326199 | 15.55355 | 27.1444 | 51.67215 | 22.75073 | -0.53718 | 0.731724 | 0.777489 |
| 10 | 51764200 | 51765999 | 21.00648 | 36.07793 | 42.15981 | 18.49239 | 0.159915 | 0.91545 | 0.777516 |
| 14 | 56307000 | 56311399 | 52.75585 | 84.73366 | 96.45578 | 40.84535 | 0.163646 | 1.924329 | 0.777527 |
| 12 | 26091000 | 26092199 | 11.46374 | 21.64278 | 21.52663 | 8.770472 | 0.484525 | 0.065562 | 0.777538 |
| 15 | 8128200 | 8128799 | 5.074039 | 9.891354 | 18.59962 | 6.715311 | -0.26332 | -0.41592 | 0.777542 |
| 18 | 4452200 | 4463599 | 30.88416 | 52.43254 | 44.08641 | 19.31665 | 0.630638 | 1.06026 | 0.777545 |
| 8 | 110786600 | 110789599 | 45.23064 | 72.51379 | 100.3293 | 42.36716 | -0.12041 | 1.972611 | 0.777555 |
| 10 | 54571800 | 54575199 | 33.12917 | 55.02265 | 62.17514 | 26.9574 | 0.205189 | 1.292454 | 0.777578 |
| 16 | 51797800 | 51800199 | 15.88578 | 27.56139 | 48.99522 | 21.49722 | -0.43968 | 0.812275 | 0.777588 |
| 7 | 112882400 | 112883399 | 12.00427 | 22.38969 | 27.19123 | 11.31343 | 0.173918 | 0.16059 | 0.77759 |
| 3 | 136986600 | 136987399 | 11.47535 | 21.29965 | 25.55289 | 10.71385 | 0.182787 | 0.181187 | 0.77759 |
| 7 | 55269600 | 55271999 | 27.0908 | 45.5234 | 53.96233 | 23.57043 | 0.136462 | 1.144742 | 0.777609 |
| 8 | 108635000 | 108636799 | 16.04163 | 28.71458 | 33.32512 | 14.61347 | 0.186371 | 0.440655 | 0.77762 |
| 5 | 107731400 | 107735399 | 38.873 | 62.88833 | 89.22335 | 38.15041 | -0.14105 | 1.689244 | 0.777636 |
| 11 | 49247800 | 49249999 | 39.1077 | 63.61172 | 61.12453 | 26.23367 | 0.42682 | 1.568556 | 0.777665 |
| 6 | 129722200 | 129725399 | 28.35925 | 46.98537 | 67.28767 | 29.12479 | -0.13584 | 1.312646 | 0.777696 |
| 16 | 15422200 | 15424599 | 15.4537 | 27.00842 | 39.9573 | 17.60915 | -0.17473 | 0.693721 | 0.777697 |
| 4 | 141332800 | 141334999 | 16.46891 | 28.63288 | 42.16031 | 18.51847 | -0.1686 | 0.77811 | 0.777701 |
| 17 | 40979000 | 40980599 | 21.00717 | 36.57037 | 33.77683 | 14.84819 | 0.501396 | 0.769285 | 0.777704 |
| 5 | 65021600 | 65023399 | 15.15659 | 26.55531 | 38.86126 | 17.16225 | -0.15905 | 0.650594 | 0.777717 |
| 18 | 90364000 | 90364599 | 10.82848 | 20.11267 | 30.98897 | 12.98108 | -0.17481 | 0.185322 | 0.77776 |
| 7 | 23005200 | 23023399 | 622.5507 | 831.2188 | 821.6243 | 343.1218 | 0.117795 | 5.304118 | 0.777808 |
| 6 | 68930400 | 68930999 | 14.11146 | 25.40649 | 30.4509 | 13.28519 | 0.145327 | 0.387155 | 0.777815 |
| 3 | 27311400 | 27315799 | 66.11357 | 104.2415 | 141.1905 | 58.59207 | -0.11346 | 2.475915 | 0.777868 |
| 12 | 107026800 | 107029399 | 34.18938 | 55.71043 | 79.03968 | 33.94451 | -0.13216 | 1.544192 | 0.777892 |
| 15 | 35943200 | 35944399 | 16.94818 | 29.52574 | 42.59756 | 18.73431 | -0.13908 | 0.752239 | 0.777898 |
| 7 | 19728400 | 19729199 | 24.82851 | 41.92548 | 49.59242 | 21.69369 | 0.139846 | 1.092229 | 0.777899 |

| | | | | | | | | | |
|---|---|---|---|---|---|---|---|---|---|
| 1 | 4629200 | 4629999 | 8.37783 | 16.11065 | 26.45857 | 10.32011 | -0.17633 | -0.08993 | 0.77792 |
| 10 | 77183600 | 77187199 | 35.97488 | 58.90694 | 69.14756 | 29.74974 | 0.14538 | 1.474484 | 0.777938 |
| 2 | 25706600 | 25710199 | 45.22672 | 72.55932 | 100.1208 | 42.32509 | -0.1153 | 1.950353 | 0.777959 |
| 14 | 54478200 | 54481199 | 19.55383 | 33.99083 | 40.45567 | 17.76793 | 0.136868 | 0.791091 | 0.77796 |
| 16 | 58623000 | 58625199 | 24.73433 | 42.68998 | 47.36723 | 20.77961 | 0.235783 | 0.894868 | 0.777978 |
| 10 | 20288000 | 20289199 | 13.32502 | 23.80452 | 35.06485 | 15.40213 | -0.15734 | 0.437718 | 0.777981 |
| 6 | 107257400 | 107257799 | 6.996238 | 13.54025 | 22.68453 | 8.643464 | -0.17208 | -0.19159 | 0.777987 |
| 15 | 11051400 | 11052399 | 20.91902 | 36.66789 | 31.04672 | 13.71632 | 0.626741 | 0.664875 | 0.778002 |
| 7 | 12418600 | 12419599 | 15.57825 | 27.52227 | 32.51303 | 14.38323 | 0.151161 | 0.534389 | 0.778008 |
| 13 | 83283000 | 83286799 | 52.51852 | 83.64941 | 115.1349 | 48.16703 | -0.12338 | 2.188868 | 0.778044 |
| 3 | 133010200 | 133016999 | 75.84691 | 119.7079 | 137.3464 | 57.01501 | 0.12674 | 2.458928 | 0.778048 |
| 15 | 12952200 | 12954399 | 25.30983 | 42.73781 | 66.40798 | 29.0494 | -0.25262 | 1.107136 | 0.778048 |
| 8 | 85891600 | 85894999 | 32.30298 | 53.63608 | 53.04158 | 22.99309 | 0.39702 | 1.287242 | 0.778055 |
| 6 | 11969400 | 11976399 | 92.94422 | 145.2385 | 160.9375 | 66.77297 | 0.165817 | 2.66335 | 0.778076 |
| 19 | 40562000 | 40562799 | 8.430804 | 16.19012 | 19.86153 | 7.777758 | 0.23601 | -0.09591 | 0.778125 |
| 4 | 24850600 | 24851399 | 9.162036 | 17.09276 | 26.86838 | 11.1744 | -0.19465 | 0.135998 | 0.778164 |
| 5 | 27472000 | 27473199 | 16.64838 | 28.90518 | 42.71311 | 18.74509 | -0.17389 | 0.79983 | 0.778186 |
| 9 | 13036200 | 13038599 | 25.69926 | 43.59741 | 51.21018 | 22.43287 | 0.150034 | 1.055193 | 0.778194 |
| X | 118875400 | 118885599 | 107.1476 | 166.3277 | 151.2467 | 62.80872 | 0.443306 | 2.782106 | 0.778203 |
| 2 | 97563600 | 97564799 | 21.1864 | 36.86914 | 35.25581 | 15.49484 | 0.451468 | 0.777316 | 0.778204 |
| 10 | 60464200 | 60470399 | 120.8608 | 182.6083 | 243.6131 | 100.7226 | -0.13997 | 3.318369 | 0.778223 |
| 17 | 64054200 | 64056999 | 25.41115 | 43.2231 | 41.13876 | 18.03068 | 0.452785 | 1.020067 | 0.778231 |
| 5 | 124181600 | 124183799 | 19.252 | 32.84514 | 48.49204 | 21.25868 | -0.17748 | 0.977829 | 0.778233 |
| 9 | 113859800 | 113864199 | 52.92402 | 84.26567 | 116.84 | 48.85661 | -0.13455 | 2.199318 | 0.778259 |
| 11 | 103963600 | 103964399 | 9.486557 | 17.68407 | 27.37049 | 11.39862 | -0.174 | 0.145705 | 0.778263 |
| 18 | 3250000 | 3253399 | 45.84179 | 73.71298 | 86.73137 | 36.78864 | 0.1181 | 1.87606 | 0.778266 |
| 7 | 116389600 | 116392799 | 33.68969 | 54.50309 | 91.0253 | 38.92103 | -0.37575 | 1.681689 | 0.778271 |
| 16 | 69759000 | 69763999 | 85.11834 | 131.799 | 176.5697 | 73.34237 | -0.11793 | 2.818435 | 0.778305 |
| 10 | 126800000 | 126801799 | 18.72811 | 32.1565 | 46.90015 | 20.57141 | -0.15855 | 0.915243 | 0.778322 |
| 5 | 133038000 | 133042199 | 42.55021 | 68.44272 | 95.50167 | 40.52808 | -0.12684 | 1.863123 | 0.778344 |
| 16 | 71485000 | 71489599 | 40.97336 | 66.15425 | 77.02094 | 32.85405 | 0.140377 | 1.742468 | 0.778357 |
| 1 | 47998800 | 47999799 | 12.42003 | 22.47932 | 33.19742 | 14.38835 | -0.14789 | 0.351176 | 0.778361 |
| 12 | 88476000 | 88481199 | 74.51534 | 117.8536 | 117.3245 | 48.75532 | 0.333676 | 2.405868 | 0.778364 |
| 13 | 65384400 | 65462399 | 2155.267 | 2595.149 | 2703.054 | 1142.223 | -0.10794 | 6.967851 | 0.778381 |
| 13 | 80848200 | 80852799 | 51.36906 | 81.91712 | 112.5839 | 47.17833 | -0.11938 | 2.148575 | 0.778382 |
| 4 | 146799600 | 146805599 | 170.4683 | 258.2524 | 244.4085 | 101.137 | 0.358293 | 3.279149 | 0.778409 |
| 12 | 45364000 | 45365199 | 15.88399 | 27.85782 | 40.35576 | 17.83745 | -0.14444 | 0.642512 | 0.778413 |
| 9 | 47311800 | 47313599 | 18.12993 | 31.17418 | 45.86968 | 20.12036 | -0.17067 | 0.899772 | 0.778414 |
| 8 | 116512000 | 116513799 | 15.32788 | 27.47891 | 32.31214 | 14.15102 | 0.168764 | 0.423777 | 0.778448 |
| X | 9495000 | 9495599 | 8.863708 | 17.1531 | 22.12986 | 8.435273 | 0.200783 | -0.17104 | 0.778469 |
| 7 | 30024400 | 30027599 | 21.68216 | 36.77055 | 52.76129 | 23.11004 | -0.13764 | 1.045191 | 0.778496 |
| 1 | 130490400 | 130491199 | 7.272235 | 14.04022 | 23.14959 | 8.921449 | -0.16538 | -0.15052 | 0.778509 |
| 2 | 102325200 | 102325799 | 7.272235 | 14.04022 | 23.14959 | 8.921449 | -0.16538 | -0.15052 | 0.778509 |
| 3 | 23026800 | 23027399 | 7.272235 | 14.04022 | 23.14959 | 8.921449 | -0.16538 | -0.15052 | 0.778509 |
| 1 | 15064600 | 15065399 | 15.39157 | 26.94355 | 39.44795 | 17.40855 | -0.15958 | 0.668873 | 0.778518 |
| 15 | 78311200 | 78313399 | 17.43725 | 30.1431 | 44.27765 | 19.42517 | -0.16667 | 0.845894 | 0.778524 |
| 5 | 130791400 | 130793999 | 25.20136 | 42.57602 | 50.6751 | 22.17061 | 0.130854 | 1.088149 | 0.778536 |

| | | | | | | | | | |
|---|---|---|---|---|---|---|---|---|---|
| 5 | 67828800 | 67831599 | 23.774 | 40.93357 | 46.82175 | 20.53881 | 0.191284 | 0.909834 | 0.778553 |
| 6 | 149141800 | 149150199 | 122.5815 | 187.1105 | 211.2426 | 87.6022 | 0.112807 | 3.109807 | 0.778558 |
| 9 | 19863600 | 19875399 | 431.988 | 594.0639 | 601.7904 | 250.1889 | 0.125283 | 4.836359 | 0.778583 |
| 4 | 90150000 | 90151599 | 23.69352 | 40.47969 | 47.94126 | 21.01961 | 0.139489 | 0.981101 | 0.778629 |
| 12 | 83667600 | 83667999 | 6.383763 | 12.417 | 21.26231 | 7.858758 | -0.16229 | -0.30953 | 0.778654 |
| 13 | 77594600 | 77594999 | 6.383763 | 12.417 | 21.26231 | 7.858758 | -0.16229 | -0.30953 | 0.778654 |
| 13 | 57252800 | 57254399 | 15.01106 | 26.42443 | 38.18565 | 16.90493 | -0.14017 | 0.580687 | 0.778654 |
| 14 | 42846400 | 42847599 | 9.425636 | 17.64933 | 27.39301 | 11.32186 | -0.16952 | 0.114868 | 0.778732 |
| 9 | 118743600 | 118744999 | 14.67072 | 25.7594 | 37.76908 | 16.71027 | -0.16182 | 0.611088 | 0.778733 |
| 9 | 100112600 | 100114999 | 18.76543 | 32.95611 | 37.94338 | 16.79042 | 0.185075 | 0.62983 | 0.778737 |
| 1 | 29592800 | 29594199 | 15.54487 | 27.41175 | 40.22242 | 17.80043 | -0.16116 | 0.568872 | 0.778778 |
| 9 | 98665000 | 98666799 | 14.81907 | 25.98616 | 38.31839 | 16.93473 | -0.16999 | 0.633445 | 0.77878 |
| 1 | 109689800 | 109690399 | 7.876701 | 15.19199 | 24.82971 | 9.602604 | -0.15793 | -0.12799 | 0.778786 |
| 1 | 26606000 | 26607199 | 15.0068 | 26.55608 | 39.13493 | 17.307 | -0.16543 | 0.524546 | 0.778786 |
| 14 | 10107200 | 10109599 | 21.09171 | 36.20782 | 52.80066 | 23.15913 | -0.15846 | 0.935043 | 0.778804 |
| 6 | 147795000 | 147797599 | 72.12615 | 112.7373 | 154.7637 | 64.20971 | -0.14253 | 2.646227 | 0.778808 |
| 8 | 21436000 | 21440799 | 13.84256 | 26.20619 | 21.79554 | 8.828249 | 0.747758 | 0.06886 | 0.778821 |
| 4 | 118326000 | 118326799 | 12.67047 | 22.50967 | 33.74754 | 14.89316 | -0.18682 | 0.469433 | 0.778821 |
| 10 | 118208600 | 118209599 | 15.00338 | 26.33622 | 38.27816 | 16.93155 | -0.14924 | 0.618669 | 0.778829 |
| 1 | 64500600 | 64501199 | 7.267384 | 14.04624 | 23.19971 | 8.89509 | -0.16092 | -0.16809 | 0.778875 |
| 15 | 94276800 | 94277999 | 12.48543 | 21.77796 | 45.27286 | 19.92635 | -0.66282 | 0.719884 | 0.778875 |
| 17 | 17163200 | 17168199 | 28.49245 | 48.41368 | 44.70434 | 19.59179 | 0.496121 | 1.045862 | 0.778887 |
| 19 | 42739600 | 42742599 | 19.91359 | 34.34047 | 39.14302 | 17.17128 | 0.196988 | 0.871553 | 0.778894 |
| 7 | 47942400 | 47944599 | 18.7393 | 33.21914 | 35.61321 | 15.74405 | 0.292038 | 0.52079 | 0.778908 |
| 10 | 105409600 | 105411199 | 12.63645 | 22.39066 | 33.73324 | 14.91427 | -0.19598 | 0.486883 | 0.778914 |
| 6 | 70507600 | 70510999 | 38.74605 | 62.99698 | 74.18294 | 31.83325 | 0.133782 | 1.58211 | 0.778931 |
| 17 | 89505200 | 89510599 | 55.03033 | 87.84392 | 102.88 | 43.17257 | 0.112599 | 2.117132 | 0.778963 |
| 11 | 66436400 | 66437599 | 20.09891 | 34.84485 | 41.56491 | 18.23667 | 0.133167 | 0.828028 | 0.778973 |
| 12 | 59646600 | 59648799 | 23.69452 | 40.60728 | 47.40281 | 20.78872 | 0.161022 | 0.955167 | 0.778994 |
| 2 | 129700200 | 129702399 | 20.22249 | 34.65496 | 49.54287 | 21.72716 | -0.13027 | 0.94295 | 0.77901 |
| 8 | 54700000 | 54700599 | 10.25283 | 19.03533 | 28.84989 | 12.09205 | -0.15143 | 0.179818 | 0.779016 |
| X | 88531600 | 88532399 | 29.21931 | 48.57416 | 57.20195 | 24.81711 | 0.146051 | 1.270937 | 0.779043 |
| 6 | 26071200 | 26072199 | 10.42046 | 19.93169 | 19.66625 | 7.789562 | 0.532086 | -0.0391 | 0.779057 |
| 12 | 110030800 | 110037599 | 73.42585 | 116.2231 | 132.5253 | 55.0944 | 0.138997 | 2.389478 | 0.779061 |
| 10 | 127623200 | 127635399 | 203.1899 | 297.5934 | 376.4789 | 155.0802 | -0.10456 | 3.886529 | 0.779064 |
| 10 | 20051000 | 20052799 | 15.56171 | 27.36965 | 39.74656 | 17.59905 | -0.14768 | 0.597395 | 0.779078 |
| 18 | 21315200 | 21316599 | 17.0869 | 30.00764 | 35.60504 | 15.7554 | 0.142064 | 0.619184 | 0.779111 |
| 7 | 117876000 | 117878999 | 21.50895 | 36.42752 | 52.76206 | 23.10091 | -0.15122 | 1.055711 | 0.779115 |
| 4 | 143960200 | 143976199 | 260.4225 | 377.401 | 399.3824 | 164.6831 | 0.138121 | 4.052409 | 0.779136 |
| 2 | 78093400 | 78098599 | 38.46506 | 62.48352 | 73.08805 | 31.34565 | 0.142244 | 1.595226 | 0.77916 |
| 18 | 79686400 | 79688799 | 24.53756 | 42.20517 | 47.91872 | 21.01935 | 0.201723 | 0.924564 | 0.779165 |
| 13 | 13200600 | 13229399 | 528.6404 | 719.5107 | 720.9962 | 300.2329 | 0.125906 | 5.000058 | 0.779168 |
| 16 | 9929600 | 9932399 | 43.68573 | 71.42656 | 59.31133 | 25.50349 | 0.642156 | 1.508276 | 0.779168 |
| 12 | 113944800 | 113946199 | 17.90689 | 31.46458 | 37.01759 | 16.3879 | 0.15419 | 0.617289 | 0.77917 |
| 5 | 23327400 | 23328799 | 15.27843 | 26.77578 | 39.4313 | 17.41784 | -0.16787 | 0.651026 | 0.779194 |
| 7 | 51928000 | 51928599 | 9.488618 | 17.80058 | 27.44127 | 11.30587 | -0.15581 | 0.105639 | 0.779204 |
| 15 | 77211000 | 77212199 | 9.488618 | 17.80058 | 27.44127 | 11.30587 | -0.15581 | 0.105639 | 0.779204 |

| | | | | | | | | | |
|---|---|---|---|---|---|---|---|---|---|
| 1 | 57602600 | 57608799 | 63.11153 | 100.7104 | 112.7687 | 47.30007 | 0.176742 | 2.138043 | 0.779225 |
| 18 | 16073800 | 16077999 | 116.9226 | 178.2022 | 199.6057 | 82.74739 | 0.122453 | 3.137988 | 0.779228 |
| 6 | 92341600 | 92343199 | 10.7645 | 19.6917 | 29.74701 | 12.72169 | -0.16771 | 0.269316 | 0.77923 |
| 4 | 83319600 | 83323199 | 48.84184 | 78.67975 | 89.95849 | 38.25803 | 0.1626 | 1.825539 | 0.779241 |
| 15 | 11615600 | 11616999 | 11.26189 | 20.24591 | 31.65726 | 13.83174 | -0.23751 | 0.376845 | 0.779255 |
| 1 | 20463600 | 20465399 | 24.29565 | 41.21161 | 49.03781 | 21.48052 | 0.131529 | 1.050277 | 0.77927 |
| 16 | 67217200 | 67217599 | 7.242666 | 14.07691 | 18.96084 | 7.051357 | 0.172337 | -0.28562 | 0.77933 |
| 13 | 33906000 | 33908999 | 26.97832 | 45.89885 | 51.78759 | 22.69835 | 0.208213 | 1.034789 | 0.779379 |
| 6 | 123705200 | 123705999 | 15.4517 | 27.05902 | 39.96838 | 17.64384 | -0.17217 | 0.665305 | 0.779392 |
| 4 | 60433000 | 60443399 | 184.4989 | 274.2721 | 301.4556 | 124.1946 | 0.118373 | 3.638543 | 0.779396 |
| 19 | 25788000 | 25790199 | 19.72338 | 33.87802 | 48.47539 | 21.26264 | -0.131 | 0.918615 | 0.7794 |
| 13 | 96068600 | 96072399 | 50.75553 | 81.58457 | 93.92673 | 39.81971 | 0.148755 | 1.896276 | 0.779417 |
| 4 | 74632200 | 74636399 | 88.54463 | 138.2066 | 158.4812 | 65.76156 | 0.115187 | 2.678967 | 0.779422 |
| 14 | 104642800 | 104644599 | 16.69552 | 29.48073 | 34.89116 | 15.4369 | 0.148084 | 0.550454 | 0.779436 |
| 11 | 46495600 | 46495999 | 7.22205 | 14.02045 | 17.8828 | 6.713504 | 0.237813 | -0.25679 | 0.779447 |
| 6 | 123181600 | 123182999 | 13.72588 | 25.04896 | 29.39931 | 12.61116 | 0.190722 | 0.301731 | 0.779455 |
| 10 | 79254600 | 79262199 | 109.9387 | 168.1035 | 221.3205 | 91.81832 | -0.10681 | 3.072129 | 0.779466 |
| 5 | 66057000 | 66066399 | 143.6688 | 216.0957 | 281.6344 | 116.2912 | -0.11198 | 3.414238 | 0.779482 |
| 1 | 7418800 | 7430399 | 311.974 | 439.8423 | 536.9173 | 222.3449 | -0.10749 | 4.457662 | 0.779511 |
| 3 | 87376000 | 87377199 | 31.32916 | 51.49472 | 60.36329 | 26.01214 | 0.149588 | 1.411171 | 0.779528 |
| 17 | 38722000 | 38722399 | 9.607863 | 18.40196 | 23.47781 | 9.26368 | 0.169566 | -0.04785 | 0.77953 |
| 15 | 50506800 | 50507399 | 7.616228 | 14.69499 | 24.24364 | 9.366096 | -0.16932 | -0.13348 | 0.779541 |
| 6 | 65075600 | 65078599 | 43.24084 | 70.35206 | 65.57344 | 28.14562 | 0.470894 | 1.576103 | 0.779544 |
| 10 | 109063000 | 109063599 | 5.946172 | 11.57733 | 20.17658 | 7.397923 | -0.17643 | -0.34639 | 0.779583 |
| 18 | 8852600 | 8852999 | 5.946172 | 11.57733 | 20.17658 | 7.397923 | -0.17643 | -0.34639 | 0.779583 |
| 10 | 37076600 | 37078399 | 13.71776 | 24.44121 | 35.59362 | 15.66939 | -0.14304 | 0.45949 | 0.779611 |
| 10 | 41731000 | 41733599 | 29.64409 | 49.81095 | 56.32264 | 24.60096 | 0.20426 | 1.154282 | 0.779634 |
| 14 | 36878600 | 36884799 | 66.14416 | 105.1854 | 119.7732 | 49.9986 | 0.147223 | 2.250809 | 0.779642 |
| X | 112874600 | 112884599 | 229.0026 | 334.3526 | 361.124 | 148.7885 | 0.119122 | 3.937816 | 0.779734 |
| 10 | 71200000 | 71201999 | 16.62316 | 28.93278 | 42.11971 | 18.51341 | -0.15212 | 0.76305 | 0.77974 |
| 18 | 84335400 | 84337199 | 16.62316 | 28.93278 | 42.11971 | 18.51341 | -0.15212 | 0.76305 | 0.77974 |
| 1 | 10065600 | 10069599 | 31.08332 | 50.66078 | 72.76642 | 31.25299 | -0.14959 | 1.530466 | 0.779756 |
| 5 | 86353400 | 86355399 | 25.97454 | 43.97057 | 51.74798 | 22.65381 | 0.147221 | 1.07656 | 0.779769 |
| 9 | 40342400 | 40344799 | 12.75755 | 24.02134 | 22.94815 | 9.395948 | 0.536264 | 0.098015 | 0.779818 |
| 10 | 122812200 | 122817999 | 54.47027 | 86.85767 | 118.7751 | 49.76889 | -0.11226 | 2.15463 | 0.779819 |
| 5 | 6791200 | 6792599 | 15.55587 | 27.81953 | 32.45895 | 14.24465 | 0.178253 | 0.444218 | 0.779821 |
| 12 | 66341000 | 66342399 | 20.35981 | 35.60174 | 34.51032 | 15.2155 | 0.432343 | 0.702726 | 0.779824 |
| 1 | 149221600 | 149223599 | 23.39654 | 40.08165 | 47.39696 | 20.78553 | 0.142308 | 0.955305 | 0.77983 |
| 13 | 92266600 | 92268799 | 19.58095 | 33.47382 | 48.48667 | 21.25978 | -0.14962 | 0.961309 | 0.779834 |
| 13 | 22902800 | 22904999 | 22.19211 | 37.80837 | 54.4259 | 23.85737 | -0.14159 | 1.008617 | 0.779854 |
| 8 | 20115200 | 20120399 | 159.599 | 239.1993 | 267.3251 | 110.2806 | 0.105065 | 3.485313 | 0.77987 |
| 10 | 105675200 | 105678799 | 25.14386 | 42.00945 | 60.51087 | 26.33788 | -0.14371 | 1.20317 | 0.779899 |
| 7 | 39660200 | 39662399 | 16.23682 | 29.22386 | 32.74714 | 14.2926 | 0.241139 | 0.405574 | 0.779911 |
| 5 | 142840200 | 142841199 | 15.29231 | 26.73683 | 40.052 | 17.65673 | -0.19231 | 0.690955 | 0.779933 |
| 2 | 8132600 | 8133399 | 14.7898 | 26.08875 | 38.11988 | 16.86885 | -0.15501 | 0.555163 | 0.779954 |
| 9 | 9383000 | 9386199 | 66.02417 | 104.7011 | 122.0158 | 50.79224 | 0.109962 | 2.334661 | 0.779977 |
| 14 | 10654400 | 10657399 | 24.35747 | 41.13762 | 48.53578 | 21.23265 | 0.143453 | 1.085367 | 0.779988 |

| | | | | | | | | | |
|---|---|---|---|---|---|---|---|---|---|
| 6 | 7173800 | 7178399 | 57.47989 | 92.00788 | 90.86971 | 38.28205 | 0.36241 | 2.03269 | 0.779999 |
| 8 | 59414000 | 59414599 | 10.80873 | 20.13765 | 24.20947 | 10.09161 | 0.186799 | 0.152174 | 0.780042 |
| 10 | 112760800 | 112763799 | 28.15345 | 47.44962 | 54.59289 | 23.86732 | 0.179437 | 1.122107 | 0.780042 |
| 3 | 49356000 | 49357399 | 12.10601 | 23.09519 | 26.19276 | 10.43852 | 0.324083 | 0.010871 | 0.780044 |
| 2 | 113954800 | 113955799 | 12.25791 | 22.58953 | 27.60573 | 11.6881 | 0.146456 | 0.234324 | 0.780048 |
| 4 | 113982400 | 114006799 | 570.5885 | 770.0011 | 775.1372 | 323.2021 | 0.106881 | 5.134655 | 0.780053 |
| 6 | 86467000 | 86467999 | 11.15404 | 20.46439 | 30.82 | 13.12792 | -0.159 | 0.260056 | 0.780054 |
| 12 | 74171600 | 74176799 | 64.93943 | 102.7953 | 139.4878 | 57.99033 | -0.11159 | 2.384085 | 0.780067 |
| 7 | 42823800 | 42856199 | 1175.445 | 1501.091 | 1389.355 | 583.0348 | 0.148426 | 6.027036 | 0.780067 |
| 12 | 82246000 | 82247799 | 14.13397 | 25.60347 | 30.3821 | 13.15395 | 0.166502 | 0.349936 | 0.780079 |
| 17 | 18741600 | 18742199 | 23.00285 | 39.23261 | 46.38267 | 20.33345 | 0.141787 | 0.994444 | 0.780085 |
| 5 | 42652000 | 42652399 | 6.386281 | 12.41344 | 21.20726 | 7.881627 | -0.1662 | -0.28999 | 0.780129 |
| 10 | 6831000 | 6834399 | 30.97846 | 51.70609 | 59.20679 | 25.75406 | 0.186293 | 1.23034 | 0.780131 |
| 16 | 75622800 | 75623799 | 7.71258 | 14.80297 | 24.10038 | 9.452176 | -0.1714 | -0.08698 | 0.780134 |
| 1 | 108552200 | 108552999 | 18.01301 | 31.41431 | 36.42768 | 16.03156 | 0.17492 | 0.733568 | 0.780139 |
| 15 | 8956200 | 8959399 | 64.58093 | 101.9723 | 138.1941 | 57.3798 | -0.11262 | 2.441651 | 0.780174 |
| 4 | 146468000 | 146495399 | 1892.063 | 2310.894 | 2538.465 | 1070.606 | -0.16376 | 6.742895 | 0.780177 |
| 4 | 84850200 | 84852999 | 36.99729 | 60.06748 | 88.10926 | 37.77314 | -0.18419 | 1.61633 | 0.780179 |
| 12 | 81924000 | 81926199 | 19.31466 | 33.65254 | 52.30634 | 23.00598 | -0.24632 | 0.775038 | 0.780215 |
| 14 | 87952200 | 87952999 | 7.597338 | 14.47406 | 23.98487 | 9.586187 | -0.2214 | -0.02601 | 0.780219 |
| 2 | 88636400 | 88637799 | 22.51124 | 38.51714 | 45.81123 | 20.08812 | 0.133709 | 0.960198 | 0.780232 |
| 13 | 61330000 | 61335199 | 67.82993 | 108.2775 | 114.2306 | 47.93688 | 0.2631 | 2.13353 | 0.78025 |
| 4 | 153325400 | 153328399 | 46.50287 | 74.65162 | 86.0672 | 36.41737 | 0.144483 | 1.928861 | 0.780262 |
| 5 | 138347600 | 138352399 | 67.22912 | 106.7801 | 122.3815 | 51.02492 | 0.13618 | 2.290488 | 0.780275 |
| 16 | 92874600 | 92876799 | 15.42446 | 26.88775 | 47.34198 | 20.82894 | -0.42506 | 0.743425 | 0.780327 |
| 7 | 132052600 | 132059399 | 65.82841 | 104.0161 | 141.1084 | 58.61095 | -0.11329 | 2.425414 | 0.780331 |
| 12 | 99557200 | 99559999 | 24.5729 | 41.40436 | 59.93897 | 26.20252 | -0.1508 | 1.119539 | 0.780352 |
| 1 | 85786400 | 85794999 | 146.2886 | 218.9093 | 284.5138 | 117.336 | -0.11461 | 3.511782 | 0.780357 |
| 4 | 61673000 | 61676399 | 31.47632 | 52.34754 | 60.77675 | 26.37546 | 0.166433 | 1.274211 | 0.780358 |
| 6 | 44145400 | 44150999 | 44.29117 | 72.17217 | 76.96443 | 33.05347 | 0.27874 | 1.564773 | 0.780373 |
| 1 | 101596200 | 101599999 | 42.96518 | 69.37149 | 81.49943 | 34.76524 | 0.127383 | 1.747295 | 0.780381 |
| 10 | 81880800 | 81908799 | 764.8223 | 1010.641 | 977.1828 | 408.6345 | 0.134677 | 5.471669 | 0.780383 |
| 7 | 6588200 | 6589799 | 14.1716 | 24.97382 | 36.66564 | 16.22849 | -0.1623 | 0.560229 | 0.780388 |
| 3 | 31495400 | 31505599 | 141.2196 | 212.3353 | 274.8584 | 113.4824 | -0.10263 | 3.419815 | 0.780402 |
| 12 | 115720000 | 115723399 | 36.9758 | 60.69392 | 69.44147 | 29.90707 | 0.183661 | 1.447304 | 0.780407 |
| 6 | 123134600 | 123141999 | 168.4965 | 251.2495 | 279.3478 | 115.1262 | 0.105831 | 3.581783 | 0.780439 |
| 4 | 145002400 | 145003599 | 11.66572 | 21.58946 | 26.05862 | 10.96967 | 0.16999 | 0.200743 | 0.780443 |
| 1 | 100331000 | 100331599 | 11.66614 | 21.7254 | 26.71665 | 11.14513 | 0.153382 | 0.16885 | 0.780443 |
| 17 | 48845200 | 48846599 | 14.17356 | 24.85139 | 37.84236 | 16.72274 | -0.21615 | 0.630161 | 0.780452 |
| 10 | 67700000 | 67701199 | 14.94447 | 26.85504 | 31.9074 | 13.94591 | 0.156045 | 0.40743 | 0.780453 |
| 19 | 48210600 | 48212399 | 18.69085 | 32.62991 | 38.84124 | 17.10478 | 0.137368 | 0.726081 | 0.780479 |
| 1 | 117782200 | 117792999 | 137.0953 | 209.269 | 220.9227 | 91.61698 | 0.209583 | 3.115268 | 0.780488 |
| 18 | 48615400 | 48620199 | 38.50241 | 63.6239 | 55.49323 | 23.97636 | 0.577199 | 1.361256 | 0.780498 |
| 9 | 109504000 | 109510199 | 68.65348 | 109.0237 | 124.3753 | 51.8473 | 0.142606 | 2.296307 | 0.780516 |
| 9 | 3876600 | 3878199 | 14.96859 | 26.8013 | 31.93128 | 13.99896 | 0.149149 | 0.430453 | 0.78053 |
| 7 | 12434000 | 12436199 | 32.72001 | 54.10652 | 62.92103 | 27.19875 | 0.163523 | 1.345384 | 0.780539 |
| X | 5203000 | 5210999 | 110.0262 | 169.5807 | 163.0428 | 67.72531 | 0.355098 | 2.912395 | 0.780554 |

| | | | | | | | | | |
|---|---|---|---|---|---|---|---|---|---|
| 9 | 98785000 | 98786799 | 19.41238 | 33.46854 | 47.93982 | 21.03011 | -0.13137 | 0.88358 | 0.780557 |
| 12 | 70687400 | 70688999 | 13.17891 | 23.59266 | 34.58102 | 15.16292 | -0.14843 | 0.423307 | 0.780607 |
| 2 | 45217400 | 45220199 | 46.08918 | 74.73808 | 81.4362 | 34.88115 | 0.242047 | 1.651646 | 0.780611 |
| 11 | 74054400 | 74055199 | 15.3163 | 27.07219 | 31.15814 | 13.78256 | 0.18903 | 0.52657 | 0.780646 |
| 16 | 84912800 | 84913199 | 5.088096 | 9.931602 | 18.13721 | 6.425401 | -0.19794 | -0.48795 | 0.780646 |
| 17 | 12184200 | 12186999 | 27.87249 | 45.96226 | 65.91883 | 28.44526 | -0.13878 | 1.364467 | 0.78065 |
| 17 | 94534600 | 94542599 | 120.7008 | 183.9694 | 209.1173 | 86.69139 | 0.101314 | 3.138723 | 0.78066 |
| 7 | 80916200 | 80917399 | 10.64485 | 19.35572 | 29.76848 | 12.82775 | -0.20044 | 0.303734 | 0.780675 |
| 14 | 108421000 | 108423599 | 22.832 | 39.41887 | 45.7456 | 20.06841 | 0.171343 | 0.881137 | 0.780685 |
| 8 | 123898800 | 123908799 | 211.522 | 307.5576 | 388.6561 | 160.2009 | -0.11305 | 3.998172 | 0.780692 |
| 12 | 115052800 | 115096399 | 779.3895 | 1029.196 | 985.7577 | 412.2546 | 0.147346 | 5.482757 | 0.780697 |
| 12 | 59501000 | 59501599 | 10.13378 | 18.93717 | 28.97066 | 12.01479 | -0.15261 | 0.13966 | 0.780699 |
| 4 | 73615800 | 73616599 | 14.94497 | 27.08813 | 26.29012 | 11.37416 | 0.455634 | 0.34523 | 0.7807 |
| 4 | 123126200 | 123128799 | 15.00985 | 26.25577 | 38.95495 | 17.18016 | -0.17873 | 0.67646 | 0.780799 |
| 11 | 13611600 | 13622399 | 93.91511 | 146.6481 | 142.6001 | 59.16927 | 0.353232 | 2.672325 | 0.780804 |
| 3 | 27393200 | 27396799 | 27.04318 | 45.28485 | 53.98019 | 23.52883 | 0.128471 | 1.179943 | 0.780811 |
| 7 | 114512200 | 114512999 | 11.36229 | 21.82001 | 25.92138 | 10.15025 | 0.28155 | -0.05014 | 0.780816 |
| 13 | 43534400 | 43536799 | 15.31112 | 26.77718 | 39.43983 | 17.39095 | -0.16832 | 0.681767 | 0.780816 |
| 8 | 111237400 | 111241999 | 64.43983 | 102.1353 | 118.3126 | 49.22492 | 0.117883 | 2.346677 | 0.780834 |
| 6 | 130077200 | 130092199 | 163.5718 | 246.5496 | 261.7653 | 108.1478 | 0.186263 | 3.376396 | 0.780854 |
| 2 | 115242600 | 115243999 | 16.84441 | 29.49966 | 42.96912 | 18.96943 | -0.15242 | 0.674283 | 0.780857 |
| 16 | 60938200 | 60939399 | 16.48015 | 28.8097 | 41.4749 | 18.28189 | -0.13552 | 0.69883 | 0.780864 |
| X | 128332600 | 128334199 | 25.12941 | 43.48757 | 38.28782 | 16.79792 | 0.569031 | 0.864135 | 0.780866 |
| 7 | 84145000 | 84148199 | 39.13428 | 63.53178 | 75.05779 | 32.17592 | 0.127154 | 1.613071 | 0.780874 |
| 1 | 41084000 | 41085999 | 16.4947 | 29.21736 | 34.28445 | 15.15899 | 0.16204 | 0.513671 | 0.780935 |
| 6 | 23165000 | 23166799 | 13.03736 | 23.26855 | 34.08781 | 14.98493 | -0.15013 | 0.440694 | 0.780935 |
| 13 | 119129400 | 119131799 | 25.53148 | 44.03256 | 47.91237 | 21.01828 | 0.2636 | 0.905051 | 0.780939 |
| 6 | 109896000 | 109898199 | 25.85088 | 43.67424 | 51.77124 | 22.65032 | 0.136682 | 1.091144 | 0.780945 |
| 4 | 156228000 | 156238399 | 168.4669 | 250.0045 | 319.6731 | 131.6766 | -0.10211 | 3.666893 | 0.78097 |
| 3 | 138571000 | 138575399 | 47.11397 | 75.44174 | 104.0413 | 43.85385 | -0.11779 | 2.015237 | 0.780971 |
| X | 13500000 | 13500399 | 5.335925 | 10.43532 | 15.11888 | 4.942654 | 0.237763 | -0.77016 | 0.78098 |
| 3 | 138356200 | 138367199 | 172.785 | 256.2261 | 328.8499 | 135.4466 | -0.10845 | 3.679975 | 0.780989 |
| 2 | 14095600 | 14104399 | 119.7353 | 184.0974 | 198.8756 | 82.5857 | 0.184478 | 2.96605 | 0.780999 |
| 2 | 87322400 | 87325599 | 32.53833 | 53.23818 | 75.68698 | 32.55333 | -0.13091 | 1.480426 | 0.781001 |
| 6 | 81855600 | 81858199 | 25.66347 | 43.20893 | 51.23477 | 22.39227 | 0.136075 | 1.118937 | 0.781018 |
| 10 | 44384200 | 44385399 | 8.762118 | 16.57574 | 26.49733 | 10.74963 | -0.18963 | 0.038514 | 0.781029 |
| 2 | 80318400 | 80319399 | 20.3551 | 34.72006 | 50.5889 | 22.17757 | -0.15849 | 0.989731 | 0.781044 |
| 13 | 66157000 | 66158199 | 26.34391 | 45.20589 | 41.39561 | 18.15612 | 0.510052 | 0.944289 | 0.781044 |
| X | 146053600 | 146055199 | 17.24648 | 30.54969 | 35.3173 | 15.6156 | 0.183541 | 0.521257 | 0.781059 |
| X | 131981800 | 131983199 | 17.98834 | 32.07967 | 34.88507 | 15.34556 | 0.276933 | 0.482495 | 0.781062 |
| 19 | 7820000 | 7821799 | 37.31179 | 60.52327 | 71.17979 | 30.49498 | 0.132609 | 1.623874 | 0.781074 |
| 6 | 38067400 | 38071599 | 40.95369 | 66.27969 | 77.55928 | 33.17236 | 0.136707 | 1.676599 | 0.781103 |
| 6 | 59870400 | 59875199 | 53.93771 | 86.61035 | 97.80469 | 41.40089 | 0.174771 | 1.934257 | 0.781111 |
| 2 | 127066400 | 127067599 | 14.93081 | 26.69052 | 31.91006 | 14.00872 | 0.142761 | 0.43956 | 0.781117 |
| 19 | 30343600 | 30344999 | 12.83649 | 23.12008 | 34.11078 | 14.87464 | -0.15243 | 0.382256 | 0.781124 |
| 4 | 22698000 | 22706399 | 56.68838 | 90.83915 | 88.69484 | 37.42677 | 0.380705 | 1.998211 | 0.781128 |
| 4 | 71926800 | 71930199 | 47.98298 | 77.22796 | 90.08214 | 38.26159 | 0.132233 | 1.851214 | 0.781146 |

| | | | | | | | | | |
|---|---|---|---|---|---|---|---|---|---|
| 7 | 136715000 | 136718399 | 48.10556 | 77.09978 | 106.775 | 45.06852 | -0.1223 | 1.988975 | 0.781159 |
| 10 | 10052000 | 10053399 | 11.52467 | 21.17294 | 31.84094 | 13.53791 | -0.15544 | 0.254852 | 0.781161 |
| 11 | 59553000 | 59553999 | 6.241185 | 12.10413 | 21.098 | 7.961094 | -0.21551 | -0.24033 | 0.781164 |
| 19 | 8110200 | 8122599 | 199.6788 | 294.2763 | 324.4025 | 133.5924 | 0.102468 | 3.786404 | 0.781193 |
| 18 | 19544400 | 19546599 | 45.90851 | 73.57094 | 101.3943 | 42.79068 | -0.11546 | 1.986754 | 0.781199 |
| 10 | 116391600 | 116395399 | 33.68576 | 55.15957 | 81.36617 | 35.00687 | -0.1834 | 1.475338 | 0.781201 |
| 9 | 67382600 | 67383999 | 13.04987 | 23.61251 | 27.99792 | 12.13914 | 0.166488 | 0.344599 | 0.781209 |
| 4 | 9668400 | 9670999 | 21.26442 | 35.52005 | 63.40946 | 27.59571 | -0.45211 | 1.198958 | 0.781245 |
| 16 | 33893000 | 33894599 | 16.48503 | 28.80153 | 41.49964 | 18.28444 | -0.13679 | 0.708119 | 0.781253 |
| 6 | 47450400 | 47457399 | 96.9066 | 151.4376 | 163.5487 | 67.85605 | 0.202909 | 2.664709 | 0.781257 |
| 11 | 31893400 | 31895399 | 20.74858 | 35.38896 | 50.60547 | 22.18472 | -0.13166 | 0.989685 | 0.781272 |
| 7 | 33829000 | 33833999 | 33.27907 | 53.97836 | 76.88954 | 32.93961 | -0.14322 | 1.620495 | 0.781284 |
| 17 | 24021000 | 24023399 | 11.8012 | 20.70013 | 42.54465 | 18.80713 | -0.64641 | 0.619846 | 0.781289 |
| 10 | 35597400 | 35604199 | 102.7423 | 159.3644 | 154.2213 | 64.05146 | 0.352717 | 2.791835 | 0.78129 |
| 17 | 16888400 | 16891799 | 23.51437 | 39.65986 | 56.60283 | 24.75152 | -0.13029 | 1.103104 | 0.78131 |
| 16 | 38664600 | 38666999 | 32.73712 | 53.30377 | 75.13548 | 32.25893 | -0.12352 | 1.550253 | 0.781332 |
| 12 | 10885400 | 10887399 | 13.91262 | 25.90665 | 23.94043 | 9.988485 | 0.56266 | 0.181756 | 0.781346 |
| 8 | 43116600 | 43118399 | 19.01085 | 31.92753 | 60.04415 | 26.22026 | -0.52692 | 1.131377 | 0.78135 |
| 11 | 24512400 | 24512999 | 6.984576 | 13.56163 | 22.86749 | 8.573626 | -0.15952 | -0.24687 | 0.781381 |
| 6 | 64147800 | 64152199 | 85.36713 | 132.4039 | 176.0702 | 73.12657 | -0.1054 | 2.788597 | 0.781382 |
| 9 | 83884600 | 83886799 | 20.23756 | 35.3773 | 40.73406 | 17.95495 | 0.185081 | 0.71545 | 0.781383 |
| 4 | 4907000 | 4916199 | 179.3568 | 267.5423 | 292.7342 | 120.6517 | 0.129415 | 3.576637 | 0.781384 |
| 6 | 131942400 | 131944399 | 39.01563 | 63.51416 | 61.64042 | 26.46301 | 0.413374 | 1.558196 | 0.781387 |
| 4 | 61575600 | 61575799 | 5.20322 | 10.17701 | 14.95516 | 5.039854 | 0.177944 | -0.67066 | 0.781388 |
| 17 | 64454400 | 64461399 | 75.35838 | 118.3151 | 160.4996 | 66.56836 | -0.12043 | 2.56778 | 0.781412 |
| 1 | 166177400 | 166178999 | 14.18071 | 24.96728 | 36.69339 | 16.24371 | -0.16425 | 0.57048 | 0.781419 |
| 8 | 5501000 | 5503999 | 36.75217 | 59.6258 | 69.56504 | 29.80698 | 0.144342 | 1.616991 | 0.781428 |
| 4 | 10926400 | 10929199 | 26.67348 | 44.3526 | 63.29445 | 27.46489 | -0.13036 | 1.262737 | 0.781435 |
| 7 | 37515600 | 37518599 | 27.21781 | 45.0111 | 64.2522 | 27.7755 | -0.13135 | 1.328459 | 0.781459 |
| 7 | 89137800 | 89144399 | 82.19907 | 128.7337 | 149.2379 | 61.90466 | 0.103234 | 2.615884 | 0.781466 |
| 14 | 3219200 | 3221399 | 24.85129 | 42.32472 | 49.56087 | 21.72423 | 0.155213 | 1.018011 | 0.781482 |
| 6 | 60150000 | 60150399 | 6.996691 | 13.63278 | 18.54653 | 6.738747 | 0.189485 | -0.36874 | 0.781483 |
| X | 153590000 | 153592199 | 25.03407 | 43.23681 | 47.36381 | 20.77855 | 0.254357 | 0.889739 | 0.781514 |
| 19 | 55963400 | 55965599 | 28.28071 | 46.63105 | 66.36206 | 28.63538 | -0.12764 | 1.36932 | 0.781527 |
| 3 | 44572600 | 44573999 | 17.81906 | 31.16031 | 36.13282 | 15.93145 | 0.174879 | 0.688785 | 0.781537 |
| 4 | 60191600 | 60193199 | 80.97854 | 126.3747 | 141.101 | 58.55069 | 0.153427 | 2.676111 | 0.78154 |
| 10 | 39809400 | 39809999 | 7.804181 | 15.07649 | 18.37155 | 7.059775 | 0.271165 | -0.16975 | 0.781551 |
| 4 | 46249800 | 46251199 | 12.23323 | 22.37332 | 33.90124 | 14.50284 | -0.17267 | 0.290809 | 0.781555 |
| 1 | 111186400 | 111189199 | 35.77316 | 58.16939 | 68.5731 | 29.42708 | 0.132343 | 1.571321 | 0.781558 |
| 4 | 100871200 | 100876799 | 73.23811 | 115.1206 | 154.8336 | 64.22813 | -0.10677 | 2.54325 | 0.781567 |
| 10 | 122340400 | 122352999 | 235.8867 | 342.1839 | 370.4786 | 152.7454 | 0.106598 | 4.03478 | 0.781567 |
| 9 | 79911000 | 79913999 | 35.75442 | 57.94664 | 81.16296 | 34.7497 | -0.12009 | 1.644316 | 0.781616 |
| 1 | 15775200 | 15777999 | 28.27402 | 46.58101 | 66.3307 | 28.61171 | -0.12899 | 1.379302 | 0.781664 |
| 13 | 56552400 | 56555199 | 20.22851 | 34.55436 | 49.54333 | 21.72175 | -0.13515 | 0.972029 | 0.781667 |
| 11 | 73444000 | 73446599 | 29.50112 | 50.35548 | 52.38562 | 22.96651 | 0.325699 | 1.017818 | 0.781695 |
| 16 | 33765400 | 33767799 | 17.16167 | 29.59436 | 44.32177 | 19.44308 | -0.19514 | 0.867631 | 0.781706 |
| 16 | 3482400 | 3491199 | 94.40473 | 147.3543 | 163.7045 | 67.92889 | 0.160826 | 2.683462 | 0.781708 |

| | | | | | | | | | |
|---|---|---|---|---|---|---|---|---|---|
| 7 | 104684400 | 104689799 | 11.39893 | 21.52842 | 20.97419 | 8.538672 | 0.51498 | 0.060228 | 0.781734 |
| 12 | 114976400 | 114984599 | 85.29113 | 134.8305 | 140.7506 | 58.47576 | 0.264744 | 2.431639 | 0.781735 |
| 9 | 76335200 | 76340199 | 103.3178 | 159.0153 | 182.0546 | 75.61105 | 0.101854 | 2.937906 | 0.781739 |
| 8 | 91494800 | 91511199 | 281.1291 | 397.0281 | 566.7406 | 234.6294 | -0.33068 | 4.432403 | 0.781769 |
| 4 | 143840200 | 143844399 | 55.5858 | 89.67527 | 73.43875 | 31.29373 | 0.645498 | 1.786657 | 0.78178 |
| 12 | 76781600 | 76782599 | 16.99318 | 29.86484 | 35.48783 | 15.71323 | 0.139792 | 0.60508 | 0.781835 |
| 17 | 21563200 | 21574999 | 109.1173 | 167.2392 | 225.268 | 93.50475 | -0.13707 | 3.026567 | 0.781858 |
| 2 | 136017200 | 136019799 | 21.21392 | 36.21937 | 51.69139 | 22.66263 | -0.12868 | 0.982911 | 0.781867 |
| 5 | 115125400 | 115127199 | 21.21392 | 36.21937 | 51.69139 | 22.66263 | -0.12868 | 0.982911 | 0.781867 |
| 14 | 114114400 | 114115599 | 13.49571 | 24.577 | 29.55736 | 12.71246 | 0.153808 | 0.316213 | 0.781869 |
| 10 | 67246000 | 67247199 | 15.98663 | 28.56357 | 33.39458 | 14.66664 | 0.174447 | 0.453376 | 0.781872 |
| 9 | 20086200 | 20097799 | 23.06277 | 40.07916 | 37.64218 | 16.52932 | 0.477005 | 0.807472 | 0.781893 |
| 18 | 6991600 | 6994399 | 20.95806 | 35.85439 | 51.71493 | 22.67661 | -0.14354 | 0.961855 | 0.781906 |
| 3 | 124296800 | 124299999 | 38.26646 | 62.35774 | 73.14215 | 31.41245 | 0.14173 | 1.548897 | 0.781924 |
| 1 | 69780600 | 69781399 | 14.47729 | 25.82237 | 30.53061 | 13.42826 | 0.157028 | 0.448518 | 0.781935 |
| 13 | 96052600 | 96057999 | 55.09438 | 87.76092 | 119.5731 | 50.0307 | -0.10863 | 2.188356 | 0.781955 |
| 15 | 43950000 | 43951199 | 19.55337 | 33.43007 | 48.48588 | 21.25961 | -0.15149 | 0.959565 | 0.781961 |
| 19 | 4126200 | 4127999 | 13.07937 | 23.22396 | 34.13001 | 15.06858 | -0.15887 | 0.474025 | 0.781963 |
| 3 | 159188000 | 159192199 | 61.77774 | 97.97185 | 134.6933 | 56.07227 | -0.1281 | 2.330059 | 0.782013 |
| 12 | 26096800 | 26099799 | 42.35176 | 67.94647 | 95.95788 | 40.56486 | -0.14868 | 1.946955 | 0.782041 |
| 3 | 144533000 | 144535799 | 35.43546 | 56.83076 | 106.455 | 44.97875 | -0.55589 | 1.953585 | 0.782044 |
| 4 | 24484800 | 24485599 | 11.73665 | 21.41353 | 31.74355 | 13.62088 | -0.14395 | 0.296485 | 0.782049 |
| 7 | 17610400 | 17613399 | 39.80405 | 66.48826 | 46.44641 | 20.21335 | 0.896962 | 1.240033 | 0.782054 |
| 14 | 11730600 | 11731599 | 8.830467 | 16.52283 | 26.37388 | 10.91411 | -0.21091 | 0.112054 | 0.782099 |
| 12 | 57988200 | 57988799 | 14.40462 | 25.28102 | 37.35842 | 16.52242 | -0.17287 | 0.617524 | 0.782103 |
| 17 | 93298800 | 93299399 | 14.40462 | 25.28102 | 37.35842 | 16.52242 | -0.17287 | 0.617524 | 0.782103 |
| 6 | 143376800 | 143378199 | 10.30591 | 18.7944 | 29.26931 | 12.56652 | -0.21502 | 0.282272 | 0.782105 |
| 10 | 6276800 | 6277599 | 13.90138 | 24.99788 | 29.9348 | 13.07613 | 0.145347 | 0.391159 | 0.782105 |
| 6 | 4856800 | 4857399 | 6.538597 | 12.66193 | 21.60196 | 8.208569 | -0.19431 | -0.21036 | 0.782124 |
| 11 | 67776800 | 67778599 | 19.59228 | 34.22285 | 40.27703 | 17.74335 | 0.153628 | 0.723349 | 0.782129 |
| 5 | 25219600 | 25221799 | 19.5646 | 33.94883 | 39.09962 | 17.15548 | 0.184044 | 0.816679 | 0.782145 |
| 1 | 24915400 | 24916399 | 13.98607 | 25.02191 | 29.10722 | 12.76954 | 0.182945 | 0.423611 | 0.782182 |
| 9 | 58227800 | 58229799 | 23.60941 | 40.30106 | 47.94901 | 21.02154 | 0.132774 | 0.989849 | 0.782189 |
| 6 | 13699600 | 13700799 | 14.34384 | 26.05802 | 23.84404 | 10.28646 | 0.542601 | 0.321439 | 0.782198 |
| 4 | 40047000 | 40056599 | 98.08762 | 150.8845 | 199.4962 | 82.84879 | -0.10626 | 2.9478 | 0.782198 |
| 10 | 58487000 | 58489799 | 18.94332 | 31.99877 | 57.98715 | 25.36853 | -0.47331 | 1.077144 | 0.782282 |
| 12 | 74199800 | 74200399 | 7.454764 | 14.38841 | 23.6915 | 9.142808 | -0.16501 | -0.14057 | 0.782304 |
| 14 | 50491800 | 50497399 | 70.86299 | 112.6143 | 108.4671 | 45.24865 | 0.387445 | 2.273254 | 0.782317 |
| 6 | 56158400 | 56161599 | 33.86067 | 54.69682 | 91.75938 | 39.16534 | -0.38462 | 1.723335 | 0.782329 |
| 3 | 108844200 | 108846399 | 58.49445 | 93.18944 | 109.3369 | 45.75624 | 0.106939 | 2.1847 | 0.78235 |
| 14 | 18183800 | 18188199 | 47.02604 | 75.75607 | 87.60087 | 37.25637 | 0.146321 | 1.81881 | 0.782353 |
| 14 | 53301600 | 53309399 | 46.07716 | 74.4186 | 73.58495 | 31.40141 | 0.375997 | 1.737904 | 0.782368 |
| 6 | 37824400 | 37831999 | 122.6787 | 188.5545 | 173.5956 | 72.08427 | 0.414509 | 2.970674 | 0.782369 |
| 16 | 46137600 | 46138999 | 17.89051 | 30.96654 | 44.76891 | 19.64147 | -0.14349 | 0.836454 | 0.782376 |
| 5 | 132926200 | 132927799 | 15.37051 | 26.87782 | 39.48529 | 17.40923 | -0.16446 | 0.685761 | 0.782385 |
| 10 | 50144400 | 50146599 | 6.705426 | 12.89398 | 27.5604 | 10.75137 | -0.55365 | -0.10584 | 0.782423 |
| 1 | 97190600 | 97191199 | 6.951525 | 13.47087 | 22.76341 | 8.622029 | -0.17708 | -0.21557 | 0.782424 |

| | | | | | | | | | |
|---|---|---|---|---|---|---|---|---|---|
| 8 | 43552400 | 43553599 | 17.80115 | 31.49848 | 30.21281 | 13.3615 | 0.450912 | 0.526859 | 0.782425 |
| 3 | 63470800 | 63471799 | 16.92445 | 29.55845 | 42.51424 | 18.72717 | -0.13438 | 0.715914 | 0.78243 |
| 9 | 16771000 | 16774199 | 31.72907 | 51.43941 | 86.88367 | 37.20785 | -0.38909 | 1.63396 | 0.782443 |
| 13 | 14949600 | 14959399 | 116.3865 | 177.9589 | 202.3993 | 83.96813 | 0.10411 | 3.072573 | 0.782447 |
| 12 | 21001800 | 21022399 | 69.59482 | 110.3836 | 109.4961 | 45.58891 | 0.342297 | 2.327665 | 0.782448 |
| 6 | 61385400 | 61386399 | 9.963462 | 18.62858 | 28.44374 | 11.78588 | -0.14881 | 0.133469 | 0.78245 |
| 9 | 8384000 | 8386599 | 23.31699 | 40.23517 | 46.28134 | 20.30311 | 0.183901 | 0.888324 | 0.782454 |
| 17 | 75858400 | 75859399 | 12.61046 | 22.34663 | 33.69584 | 14.89747 | -0.19726 | 0.484698 | 0.782457 |
| 16 | 21945000 | 21948599 | 28.5804 | 47.33561 | 67.78545 | 29.33334 | -0.13577 | 1.318762 | 0.782472 |
| 15 | 72296800 | 72297399 | 10.95805 | 20.3272 | 23.72594 | 9.956027 | 0.221834 | 0.178203 | 0.782479 |
| 14 | 77359000 | 77359799 | 13.795 | 24.42936 | 35.67273 | 15.77359 | -0.15152 | 0.505444 | 0.78248 |
| 1 | 179831600 | 179832799 | 18.32102 | 32.06916 | 38.18918 | 16.85193 | 0.136806 | 0.681586 | 0.782498 |
| 1 | 170422200 | 170429599 | 116.6188 | 177.2568 | 231.9619 | 96.08257 | -0.10498 | 3.195193 | 0.782511 |
| 7 | 47861000 | 47861999 | 13.52815 | 24.31623 | 36.17441 | 15.80747 | -0.16635 | 0.40395 | 0.78252 |
| 7 | 103470800 | 103471599 | 7.472235 | 14.46792 | 24.442 | 9.293732 | -0.18136 | -0.1878 | 0.782555 |
| 17 | 85589600 | 85595199 | 59.56082 | 94.53501 | 128.1365 | 53.38731 | -0.10646 | 2.303961 | 0.782572 |
| 4 | 49011400 | 49012999 | 20.85699 | 36.27755 | 42.52341 | 18.683 | 0.158943 | 0.793806 | 0.782598 |
| 17 | 22677000 | 22679999 | 52.88288 | 85.11971 | 80.16757 | 34.05362 | 0.440316 | 1.852589 | 0.782611 |
| 15 | 14488200 | 14490199 | 22.19159 | 38.2707 | 45.22792 | 19.84063 | 0.144922 | 0.887872 | 0.782614 |
| 7 | 33784200 | 33793399 | 147.2432 | 220.5502 | 284.6709 | 117.4228 | -0.10338 | 3.494346 | 0.782643 |
| 8 | 120099200 | 120100599 | 22.43158 | 38.60079 | 45.78306 | 20.08284 | 0.139154 | 0.909945 | 0.782647 |
| 1 | 21086200 | 21089199 | 25.93162 | 43.62505 | 65.67652 | 28.69701 | -0.20728 | 1.142757 | 0.782661 |
| 4 | 58542800 | 58542999 | 4.099148 | 8.017905 | 15.56861 | 5.226563 | -0.21374 | -0.70143 | 0.782661 |
| 17 | 51409000 | 51412599 | 40.96015 | 66.62915 | 76.22863 | 32.71687 | 0.175806 | 1.581899 | 0.782687 |
| 12 | 52144400 | 52144799 | 5.514282 | 10.74824 | 19.14079 | 6.921877 | -0.18898 | -0.40491 | 0.782689 |
| 2 | 155959200 | 155961599 | 42.1953 | 67.78842 | 94.75917 | 40.14136 | -0.13157 | 1.903954 | 0.78271 |
| 13 | 8962800 | 8964399 | 19.47596 | 33.97771 | 40.35301 | 17.76303 | 0.140491 | 0.740939 | 0.78271 |
| 9 | 117165800 | 117168199 | 25.21228 | 42.79293 | 50.6566 | 22.19378 | 0.138921 | 1.048692 | 0.782719 |
| 18 | 38693000 | 38696199 | 30.47103 | 50.08335 | 71.84683 | 30.96046 | -0.14109 | 1.41445 | 0.782738 |
| 9 | 3885000 | 3888599 | 63.20251 | 100.3193 | 116.802 | 48.66692 | 0.112589 | 2.304235 | 0.782741 |
| 19 | 55875200 | 55878799 | 35.34945 | 57.46598 | 81.31608 | 34.89233 | -0.13076 | 1.581226 | 0.782773 |
| 8 | 60233800 | 60234999 | 24.90358 | 41.54272 | 59.27238 | 25.77328 | -0.12992 | 1.220178 | 0.782776 |
| 1 | 90471400 | 90472199 | 8.495679 | 16.11749 | 25.48698 | 10.27772 | -0.16606 | 0.015524 | 0.782815 |
| 6 | 124650400 | 124651799 | 15.11073 | 26.44583 | 39.38495 | 17.37735 | -0.1843 | 0.667608 | 0.782831 |
| 16 | 62346200 | 62347199 | 14.53115 | 25.88036 | 29.65937 | 13.06124 | 0.200735 | 0.457558 | 0.782851 |
| 5 | 85214600 | 85216799 | 29.61441 | 49.35281 | 58.45257 | 25.40113 | 0.137842 | 1.243947 | 0.782892 |
| 10 | 8549200 | 8549599 | 4.909793 | 9.59408 | 17.68453 | 6.141065 | -0.18403 | -0.55921 | 0.782913 |
| 2 | 163627000 | 163627999 | 10.24337 | 18.93216 | 28.79721 | 12.1443 | -0.16305 | 0.204381 | 0.782914 |
| 16 | 34842600 | 34843399 | 8.673668 | 16.30674 | 25.90213 | 10.63298 | -0.19448 | 0.07942 | 0.782918 |
| 15 | 60898000 | 60899199 | 29.35651 | 47.44865 | 89.14001 | 38.07591 | -0.54645 | 1.697963 | 0.78293 |
| 14 | 92647000 | 92650399 | 49.22822 | 78.85741 | 91.30601 | 38.5083 | 0.134643 | 1.999453 | 0.782937 |
| 15 | 44028200 | 44030799 | 26.28569 | 43.89803 | 63.23545 | 27.51621 | -0.14373 | 1.217087 | 0.78294 |
| 11 | 6676000 | 6679399 | 42.8338 | 69.13075 | 81.72253 | 34.84181 | 0.117721 | 1.756705 | 0.782962 |
| 15 | 17190000 | 17191999 | 20.36578 | 35.76916 | 39.47478 | 17.46912 | 0.245945 | 0.641274 | 0.783057 |
| 4 | 143911400 | 143913399 | 31.74753 | 53.3968 | 57.52001 | 25.13129 | 0.274065 | 1.155383 | 0.783061 |
| 5 | 83862600 | 83864799 | 21.80454 | 38.02084 | 42.92842 | 18.89068 | 0.213005 | 0.762995 | 0.783067 |
| 2 | 162216200 | 162216999 | 6.973827 | 13.45071 | 22.57549 | 8.730606 | -0.19553 | -0.14432 | 0.783091 |

| | | | | | | | | | |
|---|---|---|---|---|---|---|---|---|---|
| 7 | 140233800 | 140234399 | 6.973827 | 13.45071 | 22.57549 | 8.730606 | -0.19553 | -0.14432 | 0.783091 |
| 5 | 34103000 | 34119399 | 249.3881 | 359.436 | 448.6547 | 185.1343 | -0.10776 | 4.132805 | 0.783104 |
| 1 | 167225200 | 167227399 | 20.92762 | 36.11334 | 43.15757 | 18.93277 | 0.129146 | 0.875243 | 0.78311 |
| 3 | 31809200 | 31810799 | 15.63684 | 28.52559 | 32.08517 | 13.76978 | 0.251634 | 0.326542 | 0.783145 |
| 16 | 7330400 | 7338399 | 95.07931 | 149.0461 | 158.4848 | 65.73393 | 0.228787 | 2.609528 | 0.78315 |
| 8 | 44338600 | 44339399 | 10.80879 | 20.02879 | 30.42457 | 12.78563 | -0.15731 | 0.200381 | 0.783172 |
| 17 | 76209000 | 76210599 | 14.99795 | 27.00326 | 31.83582 | 13.88895 | 0.168765 | 0.394327 | 0.783173 |
| 1 | 177832200 | 177835599 | 22.99399 | 40.22034 | 42.2294 | 18.62367 | 0.317614 | 0.722911 | 0.783175 |
| 8 | 27084600 | 27086999 | 16.53927 | 28.77284 | 42.6144 | 18.72514 | -0.17708 | 0.767951 | 0.783186 |
| 11 | 72938400 | 72939399 | 7.918716 | 15.04141 | 24.47839 | 9.84535 | -0.20341 | 0.000127 | 0.783192 |
| 13 | 45657400 | 45659799 | 18.73848 | 32.53186 | 46.86033 | 20.56428 | -0.13743 | 0.815408 | 0.783198 |
| 3 | 78511200 | 78511799 | 11.31629 | 20.91857 | 24.70976 | 10.41824 | 0.199819 | 0.203706 | 0.783201 |
| 9 | 105219400 | 105223599 | 39.7606 | 64.207 | 90.38854 | 38.56453 | -0.13282 | 1.738887 | 0.783217 |
| 15 | 32996000 | 32998199 | 26.48264 | 44.37995 | 66.15923 | 28.85126 | -0.19314 | 1.183844 | 0.78322 |
| 13 | 65199400 | 65202799 | 46.75003 | 74.95585 | 103.046 | 43.51985 | -0.11106 | 1.973687 | 0.783251 |
| 6 | 34017200 | 34019399 | 18.87762 | 32.0226 | 57.52595 | 25.19899 | -0.46059 | 1.038239 | 0.78326 |
| 1 | 111629600 | 111633599 | 60.65712 | 96.41976 | 112.4826 | 46.93571 | 0.111796 | 2.257766 | 0.783262 |
| 13 | 105409000 | 105410999 | 31.7395 | 51.45225 | 87.40551 | 37.42915 | -0.39745 | 1.636248 | 0.783264 |
| Y | 43992400 | 43993199 | 8.359619 | 15.33919 | 27.94087 | 11.8997 | -0.43116 | 0.228327 | 0.783267 |
| 18 | 68296800 | 68300199 | 23.89035 | 40.61223 | 48.49281 | 21.25287 | 0.126695 | 1.028044 | 0.783269 |
| 2 | 174477600 | 174479199 | 14.01114 | 24.76381 | 36.10936 | 15.97288 | -0.15082 | 0.526107 | 0.78331 |
| 1 | 161015400 | 161018199 | 28.12012 | 47.51226 | 54.04783 | 23.64698 | 0.195873 | 1.099806 | 0.783315 |
| 1 | 24714400 | 24715599 | 11.39793 | 20.94819 | 31.32784 | 13.31275 | -0.14677 | 0.251454 | 0.78333 |
| 4 | 5308600 | 5311799 | 30.51235 | 51.30259 | 56.9172 | 24.86546 | 0.231637 | 1.152736 | 0.783348 |
| 7 | 20926800 | 20942799 | 338.6884 | 475.8515 | 497.6713 | 206.2078 | 0.110435 | 4.50974 | 0.783361 |
| 10 | 122176200 | 122179399 | 31.70921 | 51.85937 | 73.19767 | 31.47774 | -0.12096 | 1.482947 | 0.783361 |
| 9 | 4810800 | 4814799 | 48.71041 | 78.69461 | 73.49435 | 31.37462 | 0.458673 | 1.73386 | 0.783365 |
| X | 138861200 | 138861799 | 9.524057 | 18.34192 | 23.0751 | 8.962322 | 0.211232 | -0.09907 | 0.783372 |
| 17 | 40247400 | 40248799 | 18.35721 | 32.16793 | 38.1453 | 16.84825 | 0.142871 | 0.664542 | 0.783377 |
| 14 | 53920600 | 53934199 | 356.5142 | 494.7379 | 602.4059 | 250.0847 | -0.12686 | 4.696204 | 0.783379 |
| 18 | 20690800 | 20691599 | 7.963629 | 15.05594 | 24.98348 | 10.14867 | -0.24351 | 0.036198 | 0.783425 |
| 1 | 152192200 | 152196199 | 38.07419 | 62.78659 | 68.19954 | 29.42973 | 0.261004 | 1.392278 | 0.783428 |
| 11 | 30654000 | 30655599 | 17.02262 | 29.82993 | 35.20897 | 15.55159 | 0.149615 | 0.652192 | 0.78344 |
| 12 | 113956000 | 113958799 | 35.23487 | 57.90656 | 67.67259 | 29.16023 | 0.153813 | 1.425942 | 0.783458 |
| 1 | 88045000 | 88050399 | 40.22014 | 65.6516 | 63.03271 | 27.08732 | 0.431578 | 1.524025 | 0.783459 |
| 4 | 146500800 | 146556799 | 3626.411 | 4190.516 | 4278.21 | 1819.199 | -0.13959 | 7.615318 | 0.783463 |
| 15 | 25847800 | 25851199 | 38.99512 | 62.83773 | 88.0223 | 37.45199 | -0.12915 | 1.799386 | 0.783526 |
| 14 | 103605800 | 103609199 | 38.67659 | 62.44489 | 86.80448 | 37.0265 | -0.1149 | 1.742456 | 0.783526 |
| 6 | 133373000 | 133389599 | 333.7739 | 473.5933 | 430.1467 | 177.9415 | 0.328173 | 4.364507 | 0.783538 |
| 13 | 78408000 | 78409999 | 17.91295 | 31.55338 | 29.88672 | 13.22923 | 0.466545 | 0.575538 | 0.783541 |
| 10 | 10022400 | 10023799 | 13.2575 | 23.72565 | 34.63136 | 15.18913 | -0.14254 | 0.427825 | 0.783595 |
| 6 | 82370200 | 82374399 | 51.7702 | 83.36577 | 92.47225 | 39.30569 | 0.205535 | 1.841686 | 0.783654 |
| X | 21029800 | 21030999 | 17.15839 | 31.00287 | 33.49119 | 14.54643 | 0.298646 | 0.395977 | 0.783656 |
| 12 | 22730400 | 22841399 | 3329.163 | 3880.248 | 3371.931 | 1431.848 | 0.105512 | 7.480028 | 0.783668 |
| 5 | 66059400 | 66071199 | 170.2377 | 252.5899 | 322.9235 | 133.0132 | -0.10209 | 3.670609 | 0.78369 |
| 9 | 12916000 | 12917199 | 12.92021 | 23.26264 | 34.15192 | 14.89917 | -0.14556 | 0.38686 | 0.7837 |
| 5 | 74945800 | 74949599 | 47.08807 | 75.23292 | 105.2819 | 44.23638 | -0.14238 | 2.082887 | 0.783719 |

| | | | | | | | | | |
|---|---|---|---|---|---|---|---|---|---|
| 7 | 4265600 | 4267999 | 23.88019 | 40.62707 | 48.4904 | 21.25316 | 0.127453 | 1.020688 | 0.783765 |
| 10 | 69131800 | 69133399 | 14.18986 | 24.96104 | 36.72222 | 16.25938 | -0.16621 | 0.58065 | 0.783767 |
| 1 | 64941400 | 64942199 | 7.142334 | 13.86615 | 23.44954 | 8.801025 | -0.16512 | -0.2394 | 0.783785 |
| 15 | 50751600 | 50753999 | 17.90547 | 30.55719 | 54.28894 | 23.8006 | -0.44344 | 0.977632 | 0.783788 |
| 5 | 142288800 | 142292999 | 59.9867 | 95.44002 | 111.82 | 46.70951 | 0.106919 | 2.229298 | 0.783805 |
| 15 | 8264800 | 8270199 | 59.9867 | 95.44002 | 111.82 | 46.70951 | 0.106919 | 2.229298 | 0.783805 |
| 18 | 30868600 | 30870199 | 12.0895 | 21.86008 | 32.17677 | 13.96274 | -0.14423 | 0.35216 | 0.783862 |
| 1 | 94037600 | 94039799 | 27.89258 | 46.09951 | 65.39243 | 28.25679 | -0.12235 | 1.338526 | 0.783917 |
| 9 | 6870800 | 6872999 | 22.4411 | 38.73105 | 45.205 | 19.83105 | 0.162998 | 0.880918 | 0.783923 |
| 4 | 24768400 | 24771399 | 14.25236 | 26.2067 | 25.30714 | 10.74735 | 0.480713 | 0.256631 | 0.783945 |
| 6 | 39297800 | 39303599 | 65.17088 | 102.9812 | 139.5601 | 57.96902 | -0.11176 | 2.423749 | 0.783946 |
| 14 | 115886600 | 115888599 | 24.01204 | 39.88295 | 69.17558 | 29.9939 | -0.41103 | 1.273389 | 0.783953 |
| 6 | 121655600 | 121664199 | 101.7782 | 157.588 | 177.4356 | 73.70207 | 0.132511 | 2.827115 | 0.783959 |
| 14 | 89606600 | 89611399 | 68.38848 | 107.6881 | 145.4531 | 60.3508 | -0.11091 | 2.50307 | 0.78399 |
| 4 | 113660400 | 113662599 | 37.58447 | 61.33507 | 72.05646 | 30.96251 | 0.140915 | 1.525998 | 0.784013 |
| 8 | 5094200 | 5095799 | 17.81618 | 31.29003 | 36.99388 | 16.3707 | 0.147037 | 0.623061 | 0.784023 |
| 3 | 101164000 | 101165199 | 18.40331 | 32.21397 | 38.13469 | 16.82822 | 0.145234 | 0.680278 | 0.784036 |
| 2 | 58118600 | 58120399 | 26.22125 | 43.52836 | 62.04241 | 26.89078 | -0.12875 | 1.280251 | 0.784037 |
| 10 | 119085400 | 119093799 | 80.26501 | 125.4841 | 169.4659 | 70.30803 | -0.11876 | 2.647661 | 0.784045 |
| 1 | 87387000 | 87388399 | 21.58437 | 36.97298 | 43.75137 | 19.18662 | 0.141358 | 0.944958 | 0.78406 |
| Y | 6177800 | 6179399 | 16.63488 | 29.12547 | 28.94869 | 12.7772 | 0.396299 | 0.650538 | 0.784063 |
| 15 | 97332600 | 97335599 | 28.01151 | 46.78607 | 55.67321 | 24.2275 | 0.130998 | 1.211182 | 0.784065 |
| 5 | 111163200 | 111164799 | 20.66519 | 34.54224 | 63.27446 | 27.5486 | -0.48907 | 1.19212 | 0.78407 |
| 7 | 17392000 | 17406399 | 411.3011 | 568.3379 | 584.383 | 242.7665 | 0.110764 | 4.762474 | 0.784086 |
| 1 | 96837200 | 96839199 | 18.44991 | 32.45781 | 37.3844 | 16.55227 | 0.184967 | 0.604342 | 0.78411 |
| 7 | 36859200 | 36866399 | 94.74073 | 145.8946 | 192.6716 | 80.02653 | -0.10335 | 2.925364 | 0.784113 |
| 17 | 22484800 | 22488399 | 37.84322 | 62.35315 | 57.20116 | 24.67487 | 0.503517 | 1.392541 | 0.78412 |
| 1 | 8266400 | 8271399 | 62.03399 | 99.38496 | 88.46124 | 37.31669 | 0.513642 | 2.011431 | 0.784131 |
| 2 | 83963000 | 83964199 | 21.41115 | 36.68654 | 43.2366 | 18.9613 | 0.147252 | 0.940979 | 0.784138 |
| 4 | 56570000 | 56570599 | 9.016949 | 16.97357 | 26.44113 | 10.83103 | -0.16392 | 0.077352 | 0.784194 |
| 16 | 53734600 | 53737199 | 34.56838 | 57.07281 | 55.2295 | 23.84793 | 0.427684 | 1.361492 | 0.784195 |
| X | 60369800 | 60372399 | 33.63394 | 57.23032 | 40.32093 | 17.67275 | 0.885338 | 1.050958 | 0.784196 |
| 17 | 32882000 | 32887199 | 50.05617 | 80.26583 | 94.50328 | 39.91886 | 0.112303 | 1.969794 | 0.784208 |
| 8 | 107491600 | 107494999 | 21.3806 | 36.72271 | 43.70843 | 19.17169 | 0.133563 | 0.919044 | 0.784219 |
| 18 | 38964200 | 38967199 | 31.56979 | 51.50993 | 72.91067 | 31.32742 | -0.12741 | 1.516342 | 0.78422 |
| 7 | 65571600 | 65574599 | 33.27229 | 54.74799 | 65.1757 | 28.09883 | 0.12792 | 1.405051 | 0.784221 |
| 17 | 11515000 | 11520799 | 87.02595 | 135.7338 | 155.9037 | 64.69703 | 0.111968 | 2.692702 | 0.784237 |
| X | 54692200 | 54700799 | 84.7668 | 134.4452 | 108.0745 | 44.99617 | 0.645201 | 2.342675 | 0.784247 |
| 3 | 85297400 | 85307399 | 122.1089 | 186.8063 | 208.7263 | 86.60343 | 0.130273 | 3.065007 | 0.784278 |
| 4 | 45077800 | 45082199 | 43.93541 | 70.57441 | 97.25534 | 41.1907 | -0.1113 | 1.910056 | 0.784308 |
| 6 | 39508200 | 39513199 | 36.41873 | 59.2332 | 70.43069 | 30.22685 | 0.120181 | 1.570025 | 0.784361 |
| 14 | 8079600 | 8080799 | 18.71082 | 32.87316 | 31.82929 | 14.08955 | 0.433773 | 0.613795 | 0.784363 |
| 3 | 14809800 | 14811199 | 23.26828 | 39.70366 | 47.42037 | 20.78914 | 0.127165 | 0.991224 | 0.784364 |
| 8 | 40258800 | 40261199 | 19.10042 | 33.61185 | 37.85462 | 16.7595 | 0.217322 | 0.605568 | 0.784368 |
| 19 | 56684600 | 56687599 | 24.98752 | 41.49057 | 60.1998 | 26.0966 | -0.15421 | 1.274288 | 0.784429 |
| 18 | 72039800 | 72040799 | 14.6868 | 26.22497 | 31.02838 | 13.63451 | 0.156991 | 0.44478 | 0.784432 |
| 4 | 139373000 | 139374599 | 15.92443 | 28.11881 | 33.49731 | 14.82025 | 0.138832 | 0.544387 | 0.784474 |

| | | | | | | | | | |
|---|---|---|---|---|---|---|---|---|---|
| 5 | 77150400 | 77152599 | 28.0787 | 46.29136 | 65.84018 | 28.40847 | -0.1269 | 1.369119 | 0.784492 |
| 5 | 109572600 | 109575599 | 30.64992 | 51.72211 | 56.32635 | 24.63456 | 0.258585 | 1.123578 | 0.784508 |
| 14 | 26286600 | 26287599 | 9.486132 | 17.2099 | 33.89082 | 14.64543 | -0.55735 | 0.318109 | 0.784535 |
| 7 | 108768000 | 108770599 | 25.64667 | 42.54046 | 61.31368 | 26.56025 | -0.14477 | 1.289262 | 0.78456 |
| 8 | 93279400 | 93281199 | 15.3464 | 26.82224 | 39.49245 | 17.40503 | -0.16773 | 0.693123 | 0.784572 |
| 5 | 127080800 | 127082599 | 23.68973 | 40.25452 | 47.96148 | 21.0193 | 0.129708 | 1.029886 | 0.78458 |
| 2 | 75473800 | 75476199 | 17.69048 | 30.47708 | 53.296 | 23.37922 | -0.41835 | 0.889332 | 0.784586 |
| 7 | 58784800 | 58788199 | 59.82158 | 95.07394 | 109.5777 | 45.71508 | 0.128983 | 2.261658 | 0.784599 |
| 13 | 4357200 | 4359199 | 28.04151 | 46.47341 | 66.09617 | 28.61521 | -0.12585 | 1.307673 | 0.784602 |
| X | 66090200 | 66093199 | 28.37616 | 47.90868 | 54.53553 | 23.8549 | 0.194982 | 1.111113 | 0.784613 |
| 2 | 55518000 | 55521799 | 68.9646 | 109.054 | 127.4238 | 52.94711 | 0.102701 | 2.408282 | 0.784654 |
| 7 | 27068600 | 27073599 | 60.46884 | 95.80744 | 130.1387 | 54.12896 | -0.11196 | 2.35699 | 0.78467 |
| 4 | 112742200 | 112745799 | 45.70603 | 74.36333 | 78.68016 | 33.7714 | 0.28852 | 1.591138 | 0.784711 |
| 5 | 145038000 | 145039199 | 11.35165 | 20.64883 | 31.17023 | 13.42533 | -0.17366 | 0.307713 | 0.784746 |
| 16 | 75933400 | 75936799 | 52.84091 | 84.0534 | 116.7325 | 48.74749 | -0.13831 | 2.231155 | 0.784749 |
| 9 | 117336600 | 117340399 | 38.88969 | 62.54621 | 104.1307 | 44.18127 | -0.38133 | 1.866459 | 0.784756 |
| 6 | 18144600 | 18145399 | 10.95605 | 19.86746 | 30.23551 | 13.07339 | -0.1886 | 0.322575 | 0.784764 |
| 1 | 47719800 | 47720599 | 14.11391 | 25.07587 | 36.63067 | 16.1645 | -0.14974 | 0.483146 | 0.784777 |
| 5 | 54809000 | 54810199 | 10.44024 | 20.1425 | 20.24567 | 7.822931 | 0.538618 | -0.1167 | 0.784778 |
| 18 | 28565000 | 28567799 | 32.15014 | 53.42929 | 61.5184 | 26.68392 | 0.178264 | 1.282479 | 0.784778 |
| 1 | 157565200 | 157567799 | 23.45117 | 39.85408 | 47.44205 | 20.79186 | 0.131098 | 1.02893 | 0.784812 |
| 3 | 90973600 | 90983399 | 150.9602 | 226.3311 | 293.1864 | 120.9575 | -0.10734 | 3.478321 | 0.784838 |
| 17 | 10409600 | 10417399 | 84.57899 | 132.7773 | 150.6272 | 62.47284 | 0.137103 | 2.574239 | 0.784844 |
| 14 | 62158800 | 62159599 | 11.43737 | 20.51907 | 31.67613 | 13.86403 | -0.22081 | 0.391664 | 0.784846 |
| 16 | 49025200 | 49026999 | 16.47061 | 28.82574 | 41.42394 | 18.27697 | -0.133 | 0.680047 | 0.784849 |
| 16 | 58128600 | 58130199 | 16.67426 | 29.36447 | 34.95199 | 15.47209 | 0.138429 | 0.579272 | 0.784852 |
| 3 | 87163800 | 87164799 | 8.50734 | 16.10466 | 25.45678 | 10.31272 | -0.17125 | 0.030835 | 0.784855 |
| 11 | 105002800 | 105003799 | 8.50734 | 16.10466 | 25.45678 | 10.31272 | -0.17125 | 0.030835 | 0.784855 |
| X | 164632800 | 164634399 | 35.90325 | 58.80905 | 68.42431 | 29.44268 | 0.158344 | 1.471584 | 0.784863 |
| 13 | 93655000 | 93662199 | 62.66743 | 99.14864 | 135.2656 | 56.22059 | -0.11993 | 2.391495 | 0.78487 |
| X | 59022200 | 59025999 | 50.62806 | 82.03869 | 66.78727 | 28.59133 | 0.660689 | 1.669001 | 0.784887 |
| 3 | 51641200 | 51643399 | 20.59853 | 35.50302 | 42.15411 | 18.49189 | 0.138139 | 0.883314 | 0.784895 |
| 16 | 44118400 | 44120399 | 24.33516 | 40.66734 | 58.78915 | 25.59208 | -0.14887 | 1.196855 | 0.784932 |
| 11 | 102511800 | 102515799 | 28.27153 | 46.43877 | 66.76065 | 28.761 | -0.1444 | 1.415538 | 0.784934 |
| 1 | 53730800 | 53731999 | 20.04 | 35.01177 | 40.82007 | 17.98479 | 0.167241 | 0.725299 | 0.784939 |
| 10 | 32599400 | 32599799 | 6.247514 | 12.17295 | 21.43334 | 7.788387 | -0.17914 | -0.36847 | 0.784955 |
| 8 | 118340800 | 118350799 | 113.0696 | 172.1052 | 225.8755 | 93.60105 | -0.1075 | 3.165021 | 0.784979 |
| 6 | 78896800 | 78897999 | 8.528554 | 16.17738 | 25.4942 | 10.28396 | -0.16132 | 0.018191 | 0.785024 |
| 17 | 88199000 | 88206399 | 116.7379 | 178.0897 | 200.7925 | 83.25763 | 0.114071 | 3.117984 | 0.785045 |
| 16 | 30288200 | 30291599 | 41.72339 | 67.48965 | 79.69321 | 34.07175 | 0.122998 | 1.691581 | 0.785048 |
| 12 | 118112400 | 118114399 | 18.35739 | 31.94893 | 46.82536 | 20.58081 | -0.16188 | 0.78059 | 0.785051 |
| 9 | 88874200 | 88875999 | 28.75682 | 47.83054 | 57.28316 | 24.8614 | 0.121802 | 1.261062 | 0.785065 |
| Y | 6818200 | 6822199 | 20.18065 | 34.70492 | 34.21882 | 15.00986 | 0.404388 | 0.887953 | 0.785074 |
| 3 | 67682200 | 67685799 | 59.69521 | 95.04906 | 111.5637 | 46.65188 | 0.105497 | 2.204931 | 0.785094 |
| 7 | 108451200 | 108451599 | 6.814023 | 13.22191 | 22.24536 | 8.37383 | -0.16204 | -0.23852 | 0.785121 |
| 10 | 24139000 | 24139599 | 6.814023 | 13.22191 | 22.24536 | 8.37383 | -0.16204 | -0.23852 | 0.785121 |
| X | 77553000 | 77560199 | 111.3857 | 171.8117 | 187.8177 | 78.01608 | 0.170928 | 2.902119 | 0.785163 |

| | | | | | | | | | |
|---|---|---|---|---|---|---|---|---|---|
| 6 | 139173200 | 139175599 | 16.46387 | 29.10962 | 34.31284 | 15.17707 | 0.154519 | 0.532798 | 0.785176 |
| 14 | 35794800 | 35808999 | 172.8383 | 258.4564 | 283.919 | 117.0747 | 0.126915 | 3.530076 | 0.785178 |
| X | 55429800 | 55431399 | 18.17956 | 31.66509 | 46.27662 | 20.34999 | -0.15765 | 0.767325 | 0.785199 |
| 13 | 75095200 | 75097399 | 20.94434 | 36.50487 | 41.84386 | 18.40833 | 0.191128 | 0.760972 | 0.785205 |
| 1 | 103198400 | 103198999 | 11.478 | 20.95204 | 31.27645 | 13.41211 | -0.15312 | 0.292245 | 0.785218 |
| 3 | 12644200 | 12655599 | 195.878 | 289.4463 | 319.3544 | 131.5173 | 0.104827 | 3.742459 | 0.785219 |
| X | 95008800 | 95013999 | 33.15221 | 54.03652 | 76.80469 | 32.98826 | -0.13439 | 1.539451 | 0.785245 |
| 16 | 11704800 | 11705799 | 8.772858 | 16.79333 | 27.9687 | 11.0533 | -0.21389 | -0.0375 | 0.785262 |
| 4 | 62835200 | 62837199 | 29.38377 | 49.18744 | 57.39733 | 25.01289 | 0.159072 | 1.194336 | 0.785274 |
| 5 | 76912400 | 76914199 | 23.1131 | 39.27003 | 46.40259 | 20.33592 | 0.141683 | 1.028669 | 0.785275 |
| 10 | 128921600 | 128923599 | 19.46716 | 34.09317 | 39.04312 | 17.23636 | 0.192729 | 0.678137 | 0.785281 |
| 6 | 139508000 | 139510799 | 44.82506 | 72.15059 | 85.25755 | 36.21622 | 0.113715 | 1.842042 | 0.785283 |
| 6 | 145882200 | 145883999 | 25.80746 | 42.97685 | 61.42862 | 26.68202 | -0.13243 | 1.244958 | 0.785308 |
| 7 | 143601600 | 143604399 | 27.12518 | 44.80859 | 64.0866 | 27.68288 | -0.13419 | 1.342598 | 0.785309 |
| 9 | 30133000 | 30135999 | 25.29645 | 43.25446 | 49.57193 | 21.73603 | 0.186837 | 0.980912 | 0.78533 |
| 6 | 23930400 | 23931599 | 18.77084 | 33.1507 | 31.14129 | 13.77739 | 0.479773 | 0.554864 | 0.78533 |
| 18 | 7057200 | 7064199 | 150.6891 | 226.5029 | 254.8397 | 105.2084 | 0.099117 | 3.42671 | 0.785343 |
| X | 121253000 | 121258199 | 61.65275 | 98.71478 | 87.69588 | 36.96001 | 0.515412 | 2.030576 | 0.78538 |
| 2 | 89505800 | 89507199 | 23.09648 | 39.37616 | 46.3466 | 20.31697 | 0.147877 | 0.99447 | 0.785387 |
| 8 | 19563800 | 19567599 | 51.46991 | 82.61035 | 96.2603 | 40.71916 | 0.128842 | 1.944595 | 0.78539 |
| 12 | 28379400 | 28380199 | 13.17066 | 24.10525 | 29.09719 | 12.43541 | 0.15455 | 0.284447 | 0.785398 |
| 1 | 140108200 | 140108799 | 17.46731 | 30.56404 | 36.22424 | 15.98009 | 0.143664 | 0.679279 | 0.785415 |
| 9 | 79000000 | 79004599 | 82.86952 | 128.6471 | 171.7471 | 71.32142 | -0.11003 | 2.77155 | 0.785417 |
| 10 | 127682200 | 127683799 | 21.14886 | 36.11565 | 51.70442 | 22.66871 | -0.13318 | 0.979447 | 0.78545 |
| 12 | 108809800 | 108810399 | 10.53921 | 19.72971 | 23.76872 | 9.827922 | 0.193382 | 0.119302 | 0.785457 |
| 14 | 57580200 | 57585599 | 37.84426 | 61.14325 | 85.13007 | 36.34462 | -0.11603 | 1.72 | 0.78546 |
| 2 | 11937200 | 11939799 | 35.42794 | 58.43841 | 57.03125 | 24.61313 | 0.415166 | 1.373599 | 0.785479 |
| 6 | 84380000 | 84383799 | 37.86933 | 61.19914 | 85.11722 | 36.35092 | -0.11407 | 1.71342 | 0.785509 |
| 18 | 83542000 | 83544199 | 14.73727 | 25.85773 | 37.86046 | 16.74056 | -0.15965 | 0.625073 | 0.785513 |
| 2 | 120131000 | 120132399 | 18.01261 | 31.18916 | 45.30999 | 19.87908 | -0.1502 | 0.837219 | 0.785532 |
| 10 | 42155800 | 42163999 | 130.3599 | 196.9486 | 255.3493 | 105.5729 | -0.0989 | 3.322512 | 0.785565 |
| 3 | 88648000 | 88658999 | 141.8664 | 212.5825 | 276.3495 | 113.9993 | -0.11312 | 3.485795 | 0.785568 |
| 12 | 108689400 | 108693999 | 56.73555 | 90.18329 | 122.8022 | 51.24021 | -0.11102 | 2.257195 | 0.785602 |
| 2 | 57813400 | 57815999 | 25.97858 | 44.16966 | 51.21764 | 22.44738 | 0.16878 | 1.036157 | 0.785622 |
| 12 | 61571000 | 61573399 | 21.34871 | 36.56098 | 42.71017 | 18.72967 | 0.159771 | 0.9436 | 0.785645 |
| 5 | 77712800 | 77715799 | 25.7434 | 43.29613 | 51.76134 | 22.6149 | 0.124265 | 1.130456 | 0.785648 |
| 14 | 62515800 | 62518799 | 23.53896 | 39.94434 | 47.44625 | 20.78592 | 0.134033 | 1.040681 | 0.785686 |
| 14 | 121038200 | 121039799 | 15.62032 | 28.21169 | 27.22316 | 11.83034 | 0.459792 | 0.372432 | 0.7857 |
| 12 | 75424400 | 75427599 | 47.32829 | 75.39052 | 124.9507 | 52.28111 | -0.39078 | 2.182951 | 0.785715 |
| 1 | 15382000 | 15383799 | 14.84932 | 26.12309 | 37.70097 | 16.69248 | -0.13862 | 0.585954 | 0.785718 |
| 5 | 64699200 | 64702599 | 54.60149 | 87.25357 | 102.5752 | 43.11058 | 0.108868 | 2.084555 | 0.785722 |
| 5 | 130596200 | 130598399 | 19.85878 | 34.21985 | 49.011 | 21.49967 | -0.13146 | 0.890605 | 0.78573 |
| 2 | 12506400 | 12508999 | 21.95004 | 38.07896 | 44.10824 | 19.35294 | 0.175392 | 0.832028 | 0.785768 |
| 10 | 72091400 | 72092799 | 9.281382 | 17.10685 | 32.50905 | 13.76128 | -0.48564 | 0.223263 | 0.785795 |
| 18 | 23314000 | 23315199 | 21.53999 | 36.53594 | 52.2146 | 22.87182 | -0.13183 | 1.043113 | 0.785802 |
| 10 | 69476200 | 69477799 | 18.36632 | 31.68749 | 45.82634 | 20.10341 | -0.14497 | 0.868527 | 0.785807 |
| 5 | 117538600 | 117539999 | 12.07622 | 21.88064 | 32.18412 | 13.93036 | -0.14087 | 0.340072 | 0.785807 |

| | | | | | | | | | |
|---|---|---|---|---|---|---|---|---|---|
| 17 | 22984200 | 23001599 | 259.0003 | 378.4215 | 377.331 | 155.4546 | 0.235361 | 3.931006 | 0.785812 |
| 16 | 49850800 | 49851199 | 6.81464 | 13.28491 | 18.58437 | 6.707043 | 0.15768 | -0.39732 | 0.785814 |
| 1 | 19450000 | 19452599 | 22.89907 | 39.45462 | 46.30059 | 20.31063 | 0.154771 | 0.90094 | 0.785828 |
| 12 | 98363400 | 98363999 | 7.843833 | 15.13034 | 24.84829 | 9.606618 | -0.16467 | -0.13082 | 0.78583 |
| 17 | 19503000 | 19505599 | 20.27233 | 35.83627 | 37.7947 | 16.71781 | 0.314204 | 0.5656 | 0.785835 |
| 17 | 95095000 | 95099999 | 55.23914 | 88.82302 | 96.41546 | 40.8958 | 0.234131 | 1.892534 | 0.785836 |
| 6 | 74978000 | 74980799 | 42.50333 | 68.69568 | 80.92243 | 34.56446 | 0.125162 | 1.714479 | 0.78585 |
| 3 | 40273000 | 40275799 | 54.60125 | 87.32202 | 102.4741 | 43.11622 | 0.112655 | 2.060507 | 0.785852 |
| 18 | 15418600 | 15421799 | 15.2899 | 27.74735 | 26.7505 | 11.55559 | 0.466637 | 0.341889 | 0.785867 |
| 17 | 16658200 | 16658799 | 6.829247 | 13.26174 | 22.29237 | 8.349292 | -0.15366 | -0.25436 | 0.785881 |
| 3 | 131580400 | 131582199 | 18.02492 | 30.96163 | 53.81806 | 23.6061 | -0.41058 | 0.918064 | 0.785894 |
| 3 | 25056400 | 25059599 | 30.33626 | 49.94922 | 71.28671 | 30.74186 | -0.13266 | 1.39324 | 0.785915 |
| 12 | 62371800 | 62373599 | 21.78613 | 37.82158 | 44.13384 | 19.36736 | 0.165099 | 0.825997 | 0.785943 |
| 15 | 35946000 | 35951999 | 75.7914 | 119.5122 | 138.5657 | 57.50355 | 0.110635 | 2.479842 | 0.785947 |
| 3 | 90616200 | 90618999 | 22.58044 | 37.90638 | 54.88472 | 23.96082 | -0.15112 | 1.140671 | 0.78595 |
| 6 | 63187400 | 63188799 | 15.2213 | 26.92585 | 39.6786 | 17.54871 | -0.16553 | 0.531763 | 0.785971 |
| 13 | 29530800 | 29533599 | 39.52047 | 63.78109 | 88.54143 | 37.74733 | -0.11363 | 1.757087 | 0.785971 |
| 18 | 18387800 | 18390399 | 20.00019 | 35.16061 | 38.90489 | 17.22788 | 0.242225 | 0.623759 | 0.785981 |
| 12 | 32596600 | 32603799 | 73.9461 | 117.087 | 132.885 | 55.25358 | 0.146159 | 2.382135 | 0.785983 |
| 7 | 39431000 | 39434799 | 40.1368 | 64.74105 | 89.84497 | 38.27659 | -0.11407 | 1.773227 | 0.786017 |
| 13 | 58563200 | 58565599 | 28.0726 | 46.49779 | 66.03064 | 28.57533 | -0.12371 | 1.3151 | 0.786023 |
| 4 | 60666000 | 60669599 | 46.54036 | 74.7584 | 87.44784 | 37.03662 | 0.12469 | 1.910669 | 0.78604 |
| 7 | 66802400 | 66805199 | 27.08349 | 44.7057 | 64.13716 | 27.69018 | -0.13875 | 1.349929 | 0.7861 |
| 18 | 66787000 | 66790399 | 33.22198 | 54.57938 | 65.15715 | 28.07216 | 0.122978 | 1.423101 | 0.786109 |
| 17 | 90001200 | 90002199 | 15.55388 | 27.55435 | 32.88469 | 14.53971 | 0.137935 | 0.504564 | 0.786112 |
| 3 | 78254000 | 78255999 | 33.91328 | 55.0853 | 77.26503 | 33.13519 | -0.11913 | 1.593175 | 0.786133 |
| 7 | 30802000 | 30803199 | 8.847575 | 16.44875 | 26.84784 | 11.23186 | -0.25558 | 0.154245 | 0.786133 |
| 4 | 144017400 | 144017999 | 11.09794 | 20.28416 | 30.24463 | 12.94894 | -0.15008 | 0.278403 | 0.786142 |
| 6 | 138714600 | 138720999 | 42.82808 | 69.60392 | 77.85786 | 33.40298 | 0.207222 | 1.598367 | 0.786149 |
| 16 | 33638600 | 33639799 | 12.53497 | 22.59708 | 33.1189 | 14.42607 | -0.14178 | 0.373651 | 0.786155 |
| 10 | 72536000 | 72547999 | 119.8242 | 183.2182 | 207.2881 | 85.99664 | 0.111709 | 3.074494 | 0.786196 |
| 18 | 38506800 | 38508999 | 20.2464 | 34.72157 | 49.54147 | 21.72784 | -0.12728 | 0.936903 | 0.786211 |
| 11 | 38087000 | 38088999 | 16.67809 | 29.92764 | 33.80894 | 14.7948 | 0.227058 | 0.430221 | 0.786211 |
| 3 | 122639800 | 122640399 | 11.62809 | 21.53362 | 33.70267 | 14.17516 | -0.20129 | 0.216896 | 0.786237 |
| 3 | 158317800 | 158318599 | 10.35509 | 19.17582 | 29.31715 | 12.33097 | -0.16783 | 0.194426 | 0.786264 |
| 4 | 112100400 | 112107999 | 121.6093 | 185.595 | 210.6382 | 87.34815 | 0.105021 | 3.112752 | 0.786305 |
| 14 | 114268600 | 114269399 | 9.902254 | 18.95946 | 23.99041 | 9.475044 | 0.179252 | -0.04418 | 0.786322 |
| 3 | 21116800 | 21119599 | 32.59228 | 53.53917 | 78.91509 | 33.99808 | -0.18043 | 1.433972 | 0.786355 |
| X | 43564200 | 43567999 | 23.74495 | 40.57853 | 66.37721 | 29.10339 | -0.32474 | 1.001374 | 0.786357 |
| 13 | 70543000 | 70544799 | 26.58528 | 44.9601 | 41.7318 | 18.26354 | 0.488408 | 1.075391 | 0.786397 |
| 7 | 112185000 | 112187599 | 31.39029 | 51.46489 | 72.73983 | 31.31199 | -0.1211 | 1.450312 | 0.786398 |
| 1 | 96032400 | 96032999 | 8.988815 | 17.37277 | 22.63913 | 8.681595 | 0.177952 | -0.14894 | 0.786399 |
| 15 | 5912800 | 5917599 | 126.8483 | 192.4327 | 216.6144 | 89.67341 | 0.109841 | 3.236048 | 0.786402 |
| 1 | 29081400 | 29082199 | 10.9588 | 20.20934 | 30.34927 | 12.83771 | -0.14773 | 0.228087 | 0.786407 |
| 5 | 70566000 | 70567599 | 22.08812 | 38.62948 | 34.8923 | 15.38578 | 0.533781 | 0.71085 | 0.786421 |
| 17 | 17762600 | 17763599 | 7.160356 | 13.75442 | 23.05611 | 9.021208 | -0.20901 | -0.10044 | 0.786445 |
| 2 | 87511800 | 87514399 | 27.10206 | 44.7331 | 64.10088 | 27.67313 | -0.13705 | 1.351351 | 0.786456 |

| | | | | | | | | | |
|---|---|---|---|---|---|---|---|---|---|
| 14 | 33428800 | 33434599 | 56.62479 | 90.04871 | 122.1442 | 50.98913 | -0.10477 | 2.244453 | 0.786462 |
| 2 | 98243600 | 98246399 | 19.64613 | 34.87403 | 36.7524 | 16.2303 | 0.317921 | 0.522041 | 0.786471 |
| 7 | 137172000 | 137175999 | 36.11194 | 58.97642 | 70.14779 | 30.15063 | 0.123894 | 1.511298 | 0.786472 |
| 8 | 43755800 | 43763199 | 90.23782 | 139.5887 | 185.2421 | 76.94508 | -0.10538 | 2.838733 | 0.786476 |
| 13 | 43079200 | 43082399 | 56.12474 | 89.24453 | 121.2917 | 50.62819 | -0.10774 | 2.246614 | 0.78648 |
| 14 | 83098800 | 83099199 | 6.530104 | 12.71698 | 22.00047 | 8.048769 | -0.16362 | -0.34172 | 0.786501 |
| 10 | 70619200 | 70620999 | 22.29589 | 38.07767 | 44.77211 | 19.62952 | 0.149767 | 0.974134 | 0.786517 |
| 12 | 65428200 | 65433399 | 48.18334 | 77.58008 | 89.46655 | 38.02118 | 0.149308 | 1.837461 | 0.78652 |
| 4 | 95936200 | 95940199 | 88.5115 | 136.7233 | 182.1042 | 75.64221 | -0.11221 | 2.864589 | 0.786533 |
| 5 | 66044200 | 66046799 | 21.25607 | 36.06279 | 51.66537 | 22.63284 | -0.13529 | 1.040493 | 0.786547 |
| 12 | 66309600 | 66312199 | 19.77591 | 34.32903 | 51.2224 | 22.47693 | -0.18815 | 0.827574 | 0.786554 |
| 2 | 98131200 | 98132799 | 15.07093 | 27.29112 | 31.95177 | 13.83965 | 0.185535 | 0.363315 | 0.786557 |
| Y | 9657400 | 9657999 | 5.564303 | 10.76098 | 21.06615 | 8.054589 | -0.39925 | -0.20368 | 0.786558 |
| X | 98749600 | 98750599 | 15.92041 | 28.14196 | 32.16223 | 14.22651 | 0.199293 | 0.533308 | 0.786591 |
| 14 | 94294600 | 94295199 | 8.3079 | 15.95409 | 25.83401 | 10.11661 | -0.16189 | -0.08171 | 0.786637 |
| 14 | 122801400 | 122802799 | 13.31016 | 23.81995 | 35.0668 | 15.38009 | -0.15505 | 0.426368 | 0.786642 |
| 5 | 103519400 | 103519999 | 8.322845 | 16.06326 | 20.94299 | 8.083588 | 0.168486 | -0.13951 | 0.786644 |
| 4 | 10482800 | 10493199 | 118.4656 | 180.4675 | 203.5058 | 84.3521 | 0.112202 | 3.147973 | 0.786673 |
| 7 | 3418800 | 3421399 | 17.7517 | 30.83079 | 44.21853 | 19.40973 | -0.13108 | 0.801848 | 0.786685 |
| 3 | 47915400 | 47915999 | 12.27317 | 22.56604 | 27.60052 | 11.72228 | 0.142055 | 0.247418 | 0.786686 |
| 9 | 87901200 | 87904199 | 36.57299 | 59.01731 | 83.76198 | 35.70434 | -0.14564 | 1.753362 | 0.786687 |
| 14 | 96441400 | 96441999 | 8.036602 | 15.38921 | 24.59019 | 9.708585 | -0.15349 | -0.05969 | 0.786703 |
| 15 | 46036800 | 46037599 | 8.036602 | 15.38921 | 24.59019 | 9.708585 | -0.15349 | -0.05969 | 0.786703 |
| 17 | 68139600 | 68144999 | 137.5149 | 206.3972 | 269.3841 | 111.1692 | -0.11679 | 3.453868 | 0.786726 |
| 5 | 80432800 | 80434999 | 17.83376 | 30.71441 | 54.40041 | 23.86346 | -0.43683 | 0.893318 | 0.786729 |
| 14 | 16235400 | 16238999 | 37.65668 | 61.52915 | 61.25068 | 26.33118 | 0.380518 | 1.501707 | 0.786736 |
| 3 | 153171800 | 153173799 | 22.19136 | 38.97736 | 40.498 | 17.92266 | 0.332636 | 0.653568 | 0.786758 |
| 1 | 113202400 | 113203799 | 12.86138 | 23.83001 | 28.677 | 12.05278 | 0.17595 | 0.210713 | 0.786796 |
| 7 | 106518800 | 106524799 | 59.90559 | 94.76183 | 150.5169 | 62.55377 | -0.33924 | 2.399033 | 0.786797 |
| 2 | 136409600 | 136410599 | 14.4053 | 25.78305 | 30.92248 | 13.56098 | 0.139248 | 0.425854 | 0.786799 |
| 2 | 148915800 | 148916399 | 14.4053 | 25.78305 | 30.92248 | 13.56098 | 0.139248 | 0.425854 | 0.786799 |
| 17 | 94615200 | 94621799 | 77.57219 | 122.9103 | 134.1073 | 55.78057 | 0.203673 | 2.371 | 0.78682 |
| 19 | 42135600 | 42138199 | 19.44285 | 33.16225 | 48.49343 | 21.25885 | -0.16373 | 0.98184 | 0.78682 |
| 5 | 9929800 | 9930199 | 7.555881 | 14.69182 | 20.09928 | 7.449532 | 0.15444 | -0.28979 | 0.786834 |
| 16 | 90473800 | 90476399 | 26.77535 | 45.14334 | 53.40974 | 23.35263 | 0.139387 | 1.114953 | 0.786836 |
| 2 | 171121200 | 171122599 | 13.08551 | 23.42921 | 34.12757 | 14.96206 | -0.13914 | 0.422504 | 0.786866 |
| 18 | 47780200 | 47781999 | 12.9817 | 23.67966 | 28.45227 | 12.21215 | 0.156669 | 0.297714 | 0.786884 |
| 16 | 20390400 | 20393199 | 28.60588 | 48.05489 | 47.50384 | 20.74751 | 0.397532 | 1.145377 | 0.786889 |
| 14 | 56034000 | 56035199 | 12.65156 | 23.19433 | 27.99448 | 11.93668 | 0.156729 | 0.265651 | 0.786914 |
| 14 | 88586600 | 88588599 | 16.53735 | 29.02499 | 34.16522 | 15.11056 | 0.153526 | 0.622528 | 0.786918 |
| 11 | 4074000 | 4077999 | 58.08012 | 92.57372 | 108.5768 | 45.46828 | 0.108192 | 2.169882 | 0.786956 |
| 4 | 121184600 | 121193599 | 124.7004 | 190.9519 | 209.2723 | 86.84906 | 0.159223 | 3.047048 | 0.786964 |
| 5 | 127250200 | 127252399 | 23.24021 | 39.75365 | 47.40601 | 20.78697 | 0.129911 | 0.968406 | 0.78697 |
| 18 | 21081600 | 21087599 | 75.18457 | 118.9546 | 116.6578 | 48.48705 | 0.355677 | 2.39633 | 0.786971 |
| 5 | 128573600 | 128579599 | 67.0323 | 105.6978 | 142.8594 | 59.28521 | -0.11034 | 2.474335 | 0.78703 |
| 16 | 80797400 | 80798799 | 18.32964 | 31.69792 | 45.8336 | 20.10803 | -0.14387 | 0.850438 | 0.787033 |
| 15 | 54504200 | 54506799 | 17.28745 | 29.47057 | 54.25547 | 23.78385 | -0.49477 | 0.985319 | 0.787037 |

| | | | | | | | | | |
|---|---|---|---|---|---|---|---|---|---|
| 15 | 24181400 | 24182799 | 26.92613 | 45.22156 | 53.95655 | 23.56217 | 0.127041 | 1.149387 | 0.787041 |
| 1 | 14379800 | 14380399 | 6.689861 | 12.90916 | 22.0563 | 8.518161 | -0.21884 | -0.15265 | 0.787042 |
| 6 | 58332800 | 58333799 | 14.02321 | 25.79887 | 24.91003 | 10.57117 | 0.481863 | 0.252265 | 0.787057 |
| 1 | 171754800 | 171756199 | 23.29115 | 40.49343 | 37.63706 | 16.53153 | 0.492068 | 0.803246 | 0.787069 |
| X | 38402000 | 38402799 | 11.62796 | 21.84615 | 26.34229 | 10.82604 | 0.19908 | 0.113704 | 0.787103 |
| 15 | 27266200 | 27266999 | 6.855252 | 13.17602 | 22.54285 | 8.805637 | -0.23602 | -0.11138 | 0.78711 |
| 11 | 91161800 | 91165399 | 28.4545 | 47.49374 | 56.73285 | 24.67806 | 0.125544 | 1.222584 | 0.787122 |
| X | 67500600 | 67502799 | 34.35973 | 56.84013 | 64.85063 | 28.04039 | 0.190987 | 1.346092 | 0.787123 |
| 6 | 49486800 | 49487199 | 7.852614 | 15.23041 | 20.52616 | 7.742754 | 0.152388 | -0.22392 | 0.787134 |
| 5 | 34930200 | 34931399 | 12.33594 | 21.86538 | 33.28673 | 14.71589 | -0.21064 | 0.480956 | 0.787135 |
| 12 | 47375800 | 47378999 | 29.09032 | 47.75663 | 67.94046 | 29.26226 | -0.12968 | 1.425489 | 0.787155 |
| 8 | 15762600 | 15764199 | 21.00953 | 35.88401 | 51.15763 | 22.4293 | -0.12707 | 0.977276 | 0.787156 |
| 5 | 8306200 | 8307599 | 13.58341 | 23.99745 | 35.20355 | 15.57354 | -0.15953 | 0.527989 | 0.787172 |
| 13 | 83566200 | 83568399 | 25.42812 | 42.69601 | 50.68532 | 22.1305 | 0.134379 | 1.143638 | 0.787174 |
| 5 | 59181400 | 59182599 | 17.12747 | 29.92421 | 43.03386 | 18.96051 | -0.13408 | 0.713207 | 0.787192 |
| 9 | 78477400 | 78478199 | 12.74444 | 23.67397 | 28.2542 | 11.83371 | 0.191877 | 0.197663 | 0.787192 |
| 5 | 21382400 | 21384799 | 23.91153 | 41.18864 | 47.3751 | 20.78182 | 0.183533 | 0.908897 | 0.787192 |
| 11 | 16462800 | 16467199 | 62.32699 | 97.95934 | 158.7964 | 65.87107 | -0.37586 | 2.540339 | 0.787207 |
| 8 | 26409600 | 26413199 | 32.225 | 53.29266 | 63.06164 | 27.26119 | 0.138449 | 1.340317 | 0.787211 |
| 16 | 80562600 | 80563999 | 12.8156 | 22.83924 | 33.70255 | 14.83391 | -0.16146 | 0.450296 | 0.787233 |
| 7 | 6480000 | 6484799 | 61.41362 | 98.03467 | 97.17921 | 40.78104 | 0.352796 | 2.121859 | 0.787249 |
| 12 | 49238200 | 49240599 | 30.06033 | 48.85378 | 83.10749 | 35.65517 | -0.39604 | 1.577982 | 0.787271 |
| 6 | 13037200 | 13038399 | 10.21248 | 18.96394 | 28.83612 | 12.08323 | -0.15603 | 0.177314 | 0.787274 |
| 15 | 97318800 | 97320999 | 22.14792 | 38.06917 | 45.24324 | 19.84535 | 0.135857 | 0.916469 | 0.787284 |
| 4 | 113346600 | 113361999 | 193.5421 | 288.9456 | 298.679 | 123.122 | 0.212497 | 3.564363 | 0.787307 |
| 6 | 132240800 | 132254199 | 260.5034 | 372.8429 | 459.9246 | 189.9871 | -0.10077 | 4.235275 | 0.787318 |
| 1 | 91397800 | 91400799 | 25.17542 | 42.10906 | 59.93146 | 26.10403 | -0.12646 | 1.191907 | 0.787321 |
| 1 | 98128200 | 98129399 | 17.45171 | 30.29874 | 43.72075 | 19.18684 | -0.13982 | 0.805519 | 0.787364 |
| 8 | 5043800 | 5044199 | 10.13661 | 19.19286 | 24.30783 | 9.842995 | 0.146327 | 0.040962 | 0.787367 |
| 6 | 82242200 | 82243399 | 14.19857 | 24.95517 | 36.75312 | 16.27443 | -0.16816 | 0.59074 | 0.787372 |
| 1 | 114873800 | 114875199 | 12.46342 | 23.05783 | 28.17379 | 11.86562 | 0.152209 | 0.216177 | 0.787375 |
| 4 | 74589600 | 74590399 | 13.45967 | 24.27474 | 29.02989 | 12.63801 | 0.150599 | 0.369912 | 0.787381 |
| 13 | 75318400 | 75320799 | 30.76674 | 49.80494 | 87.02094 | 37.22569 | -0.43983 | 1.661873 | 0.787399 |
| 2 | 88039600 | 88045599 | 56.87822 | 91.06908 | 104.0082 | 43.83421 | 0.153588 | 2.027837 | 0.787436 |
| 12 | 107414000 | 107417399 | 25.72159 | 42.96954 | 61.01658 | 26.55599 | -0.12312 | 1.208803 | 0.787442 |
| 11 | 51009000 | 51010399 | 17.0314 | 30.11804 | 35.34177 | 15.63167 | 0.161154 | 0.537397 | 0.787445 |
| 3 | 16237000 | 16237999 | 11.30679 | 19.9745 | 38.88194 | 17.20096 | -0.56526 | 0.522052 | 0.787481 |
| 3 | 74461600 | 74463199 | 17.63802 | 30.83327 | 45.1294 | 19.89247 | -0.15944 | 0.71024 | 0.787494 |
| 5 | 119923200 | 119928599 | 60.43793 | 96.11945 | 112.4746 | 46.95693 | 0.108104 | 2.242126 | 0.787496 |
| 18 | 7688800 | 7693599 | 67.80347 | 107.4614 | 125.5864 | 52.24874 | 0.105088 | 2.352877 | 0.787511 |
| 6 | 17575000 | 17576399 | 21.65377 | 36.72627 | 52.76941 | 23.11434 | -0.13962 | 1.043518 | 0.787531 |
| 9 | 122584000 | 122584799 | 12.73504 | 23.70933 | 28.28818 | 11.80631 | 0.196199 | 0.183971 | 0.787538 |
| 2 | 87445200 | 87445999 | 16.78719 | 29.47223 | 34.69139 | 15.34713 | 0.153634 | 0.621567 | 0.787552 |
| 3 | 55884200 | 55884999 | 25.98785 | 42.96639 | 62.33647 | 26.94273 | -0.15461 | 1.328532 | 0.787556 |
| 14 | 17352600 | 17353199 | 8.898246 | 17.01192 | 21.33982 | 8.464089 | 0.186727 | -0.04532 | 0.78756 |
| 10 | 60377600 | 60378999 | 15.78994 | 28.07683 | 33.39174 | 14.72495 | 0.145889 | 0.481459 | 0.787577 |
| 1 | 182001400 | 182004999 | 87.33934 | 135.9522 | 157.1598 | 65.23746 | 0.100491 | 2.725576 | 0.787602 |

| | | | | | | | | | |
|---|---|---|---|---|---|---|---|---|---|
| 15 | 33172000 | 33177599 | 76.7741 | 121.0693 | 139.7265 | 57.98576 | 0.117333 | 2.47936 | 0.787605 |
| 4 | 128027800 | 128030399 | 23.10494 | 38.83856 | 55.42484 | 24.21171 | -0.13022 | 1.131329 | 0.787632 |
| 3 | 134784200 | 134785999 | 17.85436 | 31.22861 | 37.26638 | 16.43418 | 0.133884 | 0.690504 | 0.787635 |
| 10 | 47751200 | 47751799 | 12.57506 | 22.66534 | 33.14287 | 14.4397 | -0.13859 | 0.37596 | 0.78766 |
| 16 | 47825200 | 47825999 | 9.95233 | 18.64863 | 28.48122 | 11.75766 | -0.1449 | 0.11922 | 0.787691 |
| 8 | 69819400 | 69831399 | 229.1734 | 337.9627 | 342.2316 | 140.9356 | 0.225791 | 3.781741 | 0.787696 |
| 12 | 115322000 | 115323399 | 18.32481 | 32.2396 | 37.41485 | 16.56556 | 0.174186 | 0.603595 | 0.787699 |
| 18 | 47224800 | 47226999 | 25.69998 | 43.10329 | 62.22603 | 27.14992 | -0.147 | 1.16746 | 0.787716 |
| X | 49928600 | 49931199 | 19.86598 | 34.44417 | 51.74787 | 22.70452 | -0.19826 | 0.841582 | 0.787726 |
| 6 | 19188800 | 19191199 | 32.44163 | 53.32074 | 63.89459 | 27.53331 | 0.117843 | 1.415858 | 0.787728 |
| 2 | 116362400 | 116363799 | 39.75349 | 64.37744 | 76.59134 | 32.77344 | 0.113747 | 1.663628 | 0.787744 |
| 1 | 139655600 | 139658399 | 40.11171 | 65.47163 | 61.61369 | 26.47708 | 0.460251 | 1.519509 | 0.787754 |
| 19 | 44150200 | 44150599 | 5.211187 | 10.16285 | 18.62022 | 6.686973 | -0.21935 | -0.43266 | 0.787755 |
| X | 157793200 | 157793999 | 10.12307 | 18.95864 | 29.01375 | 11.98811 | -0.14886 | 0.125461 | 0.787804 |
| 4 | 4907400 | 4915999 | 174.9014 | 261.2917 | 287.376 | 118.4781 | 0.123945 | 3.547739 | 0.787809 |
| 10 | 86744000 | 86747799 | 41.63122 | 66.81741 | 94.56153 | 39.99909 | -0.15103 | 1.931273 | 0.787809 |
| 1 | 53271600 | 53283599 | 96.86369 | 150.8918 | 147.123 | 61.06424 | 0.346687 | 2.716221 | 0.787815 |
| 14 | 76141800 | 76142799 | 8.496191 | 16.22444 | 25.62087 | 10.18949 | -0.14649 | -0.02884 | 0.787817 |
| 9 | 18418800 | 18419399 | 8.236644 | 15.63329 | 24.9881 | 10.06669 | -0.17971 | 0.010508 | 0.787836 |
| 16 | 89339600 | 89340399 | 8.236644 | 15.63329 | 24.9881 | 10.06669 | -0.17971 | 0.010508 | 0.787836 |
| 6 | 149243800 | 149246399 | 18.89722 | 32.85758 | 47.91517 | 21.04738 | -0.15493 | 0.795574 | 0.787869 |
| 11 | 62585600 | 62586399 | 10.38856 | 19.78532 | 24.98501 | 9.992804 | 0.166377 | 0.006621 | 0.787871 |
| 12 | 98591400 | 98594999 | 31.59321 | 52.37425 | 61.89664 | 26.80225 | 0.140542 | 1.310813 | 0.787873 |
| 8 | 23972800 | 23978399 | 46.35562 | 74.60973 | 87.13131 | 37.00898 | 0.130526 | 1.844991 | 0.787889 |
| 6 | 44397000 | 44397799 | 12.5474 | 22.1591 | 39.83919 | 17.62557 | -0.45161 | 0.534808 | 0.787915 |
| 3 | 17104800 | 17105399 | 8.55412 | 16.41987 | 20.83692 | 8.169804 | 0.185502 | -0.08886 | 0.787932 |
| 14 | 99591800 | 99598599 | 62.99228 | 99.87063 | 135.7552 | 56.49886 | -0.11214 | 2.340113 | 0.787933 |
| 13 | 61167600 | 61177999 | 68.37425 | 108.6989 | 106.1354 | 44.29215 | 0.368263 | 2.258228 | 0.787935 |
| 12 | 61206000 | 61207799 | 17.07563 | 30.22242 | 35.34654 | 15.63107 | 0.166448 | 0.528963 | 0.787938 |
| 13 | 13196000 | 13196799 | 21.21806 | 36.42279 | 43.20478 | 18.95016 | 0.138304 | 0.922927 | 0.787942 |
| 6 | 131039400 | 131048799 | 160.4598 | 240.8073 | 266.7503 | 110.0789 | 0.11953 | 3.460916 | 0.787974 |
| Y | 4549600 | 4554999 | 39.87823 | 64.33275 | 64.20381 | 27.35752 | 0.360761 | 1.7537 | 0.787976 |
| 13 | 113406400 | 113410799 | 38.99838 | 62.83469 | 87.95468 | 37.41607 | -0.12834 | 1.803567 | 0.787977 |
| 17 | 21785800 | 21788399 | 11.45859 | 20.17139 | 42.97233 | 19.02392 | -0.69719 | 0.566172 | 0.787994 |
| 2 | 65566800 | 65567799 | 12.37714 | 22.65736 | 35.58039 | 15.20211 | -0.22239 | 0.292564 | 0.788049 |
| 18 | 19944600 | 19946999 | 28.79165 | 48.21302 | 56.77078 | 24.74528 | 0.146112 | 1.189423 | 0.788106 |
| 5 | 23438400 | 23441599 | 21.67666 | 36.70567 | 52.76323 | 23.1003 | -0.14033 | 1.057903 | 0.78813 |
| 3 | 38003400 | 38006599 | 45.6855 | 73.68829 | 86.15851 | 36.70421 | 0.132525 | 1.781599 | 0.78814 |
| 16 | 64737200 | 64740399 | 40.2626 | 64.54647 | 107.9045 | 45.56327 | -0.39286 | 1.971343 | 0.788151 |
| 4 | 35985200 | 35986799 | 15.2707 | 26.67117 | 47.31382 | 20.84183 | -0.43552 | 0.715405 | 0.788167 |
| 4 | 150104200 | 150105799 | 9.916093 | 18.34367 | 28.29277 | 11.91705 | -0.18174 | 0.194953 | 0.788171 |
| 14 | 51670400 | 51677599 | 120.6698 | 183.5345 | 205.7612 | 85.24652 | 0.118746 | 3.181622 | 0.788188 |
| 6 | 145465400 | 145467399 | 20.15027 | 35.18031 | 34.1118 | 15.02274 | 0.431833 | 0.723938 | 0.788192 |
| 10 | 117912200 | 117923999 | 142.9306 | 214.4593 | 276.8146 | 114.2261 | -0.10118 | 3.460775 | 0.788235 |
| 11 | 31327400 | 31329199 | 21.46203 | 36.97499 | 44.21587 | 19.39607 | 0.127756 | 0.89407 | 0.788237 |
| X | 97266000 | 97266799 | 7.813421 | 15.06534 | 25.34979 | 9.812353 | -0.19996 | -0.12059 | 0.788237 |
| 4 | 42818400 | 42822199 | 50.65359 | 81.00298 | 110.912 | 46.66189 | -0.10961 | 2.060062 | 0.788251 |

| | | | | | | | | | |
|---|---|---|---|---|---|---|---|---|---|
| 13 | 113640000 | 113646599 | 44.45764 | 71.12989 | 118.3784 | 49.838 | -0.38983 | 2.039623 | 0.788258 |
| 2 | 86907200 | 86908199 | 13.37045 | 24.40972 | 29.50585 | 12.65077 | 0.149159 | 0.298949 | 0.788261 |
| 17 | 69667200 | 69669599 | 25.0745 | 42.3305 | 50.67089 | 22.16381 | 0.122648 | 1.094633 | 0.788271 |
| 1 | 105185400 | 105187599 | 19.3026 | 33.86343 | 39.03924 | 17.25983 | 0.183251 | 0.651929 | 0.788285 |
| 14 | 69047400 | 69050599 | 49.12094 | 78.75866 | 92.78809 | 39.18866 | 0.111238 | 1.971854 | 0.788323 |
| 14 | 124781000 | 124782399 | 16.12955 | 28.79393 | 27.32718 | 12.01067 | 0.472892 | 0.446731 | 0.788377 |
| 11 | 88922800 | 88923599 | 9.343078 | 17.73145 | 21.78355 | 8.777668 | 0.196369 | 0.010975 | 0.788414 |
| 5 | 7021000 | 7022599 | 20.35426 | 35.61237 | 34.4619 | 15.20149 | 0.434752 | 0.693361 | 0.788428 |
| 17 | 91854800 | 91856399 | 16.28723 | 28.44533 | 40.98921 | 18.05532 | -0.13697 | 0.710837 | 0.788435 |
| 19 | 39797000 | 39802399 | 73.14641 | 116.0334 | 112.9904 | 47.05053 | 0.369158 | 2.327931 | 0.788445 |
| 4 | 26168200 | 26171999 | 47.53777 | 76.40273 | 89.70546 | 38.02319 | 0.120248 | 1.892901 | 0.788454 |
| Y | 4553200 | 4554999 | 18.3978 | 31.95982 | 32.11442 | 14.09854 | 0.37997 | 0.781267 | 0.788459 |
| 17 | 64011000 | 64015199 | 32.99821 | 53.83488 | 76.14286 | 32.71493 | -0.12647 | 1.52174 | 0.788466 |
| 6 | 149335000 | 149337399 | 39.03208 | 63.29141 | 75.09069 | 32.16213 | 0.119461 | 1.634314 | 0.788468 |
| 11 | 73761000 | 73772999 | 146.2544 | 221.5301 | 242.2067 | 100.2203 | 0.149984 | 3.273223 | 0.788476 |
| 14 | 49770400 | 49772399 | 27.25553 | 45.96647 | 53.98031 | 23.60421 | 0.150087 | 1.114313 | 0.788478 |
| 13 | 11319200 | 11322399 | 41.43464 | 67.41334 | 76.09741 | 32.66274 | 0.19526 | 1.577191 | 0.788482 |
| 16 | 93503800 | 93506199 | 28.38329 | 46.76587 | 66.45448 | 28.66636 | -0.12596 | 1.376918 | 0.788483 |
| 17 | 22186000 | 22188399 | 38.85128 | 63.10481 | 74.4042 | 31.90784 | 0.130744 | 1.602853 | 0.788522 |
| 5 | 133588000 | 133589599 | 21.64223 | 36.73668 | 52.7721 | 23.12124 | -0.13925 | 1.036266 | 0.788523 |
| 10 | 94371000 | 94372799 | 21.55452 | 37.01087 | 44.22954 | 19.40008 | 0.12768 | 0.923078 | 0.788564 |
| 2 | 22367400 | 22369599 | 21.12756 | 36.98186 | 40.58835 | 17.91067 | 0.253813 | 0.697579 | 0.788576 |
| X | 76769600 | 76772399 | 33.32333 | 54.48852 | 78.98754 | 33.96517 | -0.15953 | 1.495715 | 0.7886 |
| 14 | 59263200 | 59267999 | 56.43331 | 90.44762 | 102.4155 | 43.22899 | 0.167669 | 1.996561 | 0.788607 |
| 16 | 58666200 | 58667799 | 16.45062 | 29.21266 | 34.35426 | 15.16643 | 0.160876 | 0.495622 | 0.788614 |
| 8 | 45249000 | 45250999 | 19.66099 | 33.86254 | 48.47383 | 21.26371 | -0.13088 | 0.892786 | 0.788627 |
| 13 | 7672000 | 7686399 | 231.1951 | 334.4626 | 417.6608 | 172.2597 | -0.10306 | 4.075416 | 0.788634 |
| 2 | 156639600 | 156640199 | 10.62591 | 19.8451 | 24.19952 | 10.04613 | 0.171064 | 0.134588 | 0.788636 |
| X | 10583200 | 10585199 | 19.65744 | 34.33151 | 39.86333 | 17.55943 | 0.173218 | 0.72869 | 0.788643 |
| 4 | 89097000 | 89102399 | 60.56206 | 96.24386 | 131.5333 | 54.87087 | -0.11661 | 2.269561 | 0.788643 |
| 10 | 58648000 | 58650799 | 24.03517 | 39.39348 | 75.64957 | 32.56014 | -0.56214 | 1.439099 | 0.788653 |
| 13 | 44563000 | 44565199 | 18.01882 | 31.12982 | 45.29429 | 19.87083 | -0.15332 | 0.853664 | 0.788653 |
| 4 | 5575400 | 5578799 | 45.46781 | 73.09037 | 102.1881 | 43.32663 | -0.13079 | 1.89142 | 0.788656 |
| 6 | 86293800 | 86301999 | 55.00371 | 86.85085 | 148.5919 | 61.69737 | -0.44834 | 2.437023 | 0.788674 |
| 2 | 87030200 | 87031999 | 25.84322 | 42.62844 | 62.78038 | 27.09431 | -0.17662 | 1.355601 | 0.788675 |
| X | 60104600 | 60108799 | 44.84759 | 72.67205 | 81.47403 | 34.87873 | 0.200079 | 1.665037 | 0.788678 |
| 2 | 173477400 | 173479799 | 23.86692 | 40.20562 | 57.14566 | 24.97969 | -0.12442 | 1.115768 | 0.788698 |
| 11 | 73312800 | 73313999 | 16.41658 | 28.91738 | 34.52544 | 15.28256 | 0.134276 | 0.575723 | 0.788714 |
| 5 | 61450000 | 61451599 | 29.17728 | 47.26089 | 87.10106 | 37.2768 | -0.51569 | 1.649045 | 0.788719 |
| 10 | 6514600 | 6520199 | 64.81519 | 103.0513 | 119.6983 | 49.95722 | 0.118332 | 2.257194 | 0.788763 |
| 17 | 74834800 | 74836399 | 17.38415 | 29.32903 | 59.53663 | 26.03672 | -0.63651 | 1.082775 | 0.788788 |
| 10 | 45049000 | 45053599 | 45.16423 | 72.69122 | 103.9546 | 44.1538 | -0.16118 | 1.853841 | 0.788827 |
| 7 | 61754600 | 61761599 | 86.29589 | 135.2868 | 133.5982 | 55.41144 | 0.335435 | 2.598676 | 0.788843 |
| 2 | 128897800 | 128900799 | 55.09213 | 87.97149 | 103.4747 | 43.44237 | 0.106913 | 2.108191 | 0.788844 |
| 18 | 87967200 | 87969199 | 30.74317 | 51.40052 | 50.26745 | 21.889 | 0.413127 | 1.204444 | 0.788893 |
| 13 | 25597600 | 25599399 | 24.04131 | 41.64155 | 38.80466 | 17.02513 | 0.487635 | 0.851014 | 0.788908 |
| 4 | 146498800 | 146556799 | 3640.737 | 4205.993 | 4281.94 | 1820.86 | -0.1359 | 7.619341 | 0.788925 |

| | | | | | | | | | |
|---|---|---|---|---|---|---|---|---|---|
| 3 | 83298400 | 83303199 | 61.63986 | 98.20441 | 114.1647 | 47.77927 | 0.120254 | 2.187734 | 0.788933 |
| 3 | 55412400 | 55415999 | 27.09596 | 45.6194 | 53.46685 | 23.36759 | 0.152765 | 1.125379 | 0.788938 |
| 17 | 13333400 | 13335999 | 22.38145 | 38.18055 | 45.32428 | 19.8698 | 0.135854 | 0.98677 | 0.788958 |
| 9 | 19922400 | 19926999 | 62.41993 | 99.45414 | 114.5747 | 47.95546 | 0.133426 | 2.18779 | 0.788969 |
| 8 | 83012400 | 83013799 | 11.01314 | 20.40036 | 30.92997 | 13.00416 | -0.15547 | 0.202389 | 0.788978 |
| 13 | 6844200 | 6845399 | 13.30592 | 24.09775 | 29.04203 | 12.5775 | 0.144366 | 0.345037 | 0.788979 |
| 3 | 153544000 | 153545799 | 16.57749 | 29.11528 | 33.99344 | 15.04325 | 0.165001 | 0.609239 | 0.789001 |
| 17 | 21422000 | 21424999 | 49.16099 | 78.81003 | 92.54756 | 39.07739 | 0.115648 | 1.977551 | 0.789023 |
| 12 | 24316200 | 24379199 | 1529.592 | 1902.156 | 1769.5 | 744.6423 | 0.102427 | 6.455215 | 0.789037 |
| 8 | 33712400 | 33714399 | 31.65163 | 51.4897 | 73.56261 | 31.5732 | -0.14376 | 1.561579 | 0.789043 |
| 7 | 12407000 | 12408199 | 36.66176 | 59.46179 | 68.76619 | 29.45836 | 0.156655 | 1.623996 | 0.789068 |
| 6 | 96912400 | 96914199 | 17.61829 | 31.02234 | 36.38181 | 16.1055 | 0.159546 | 0.588147 | 0.789068 |
| 14 | 86127800 | 86129599 | 19.40214 | 33.20462 | 47.95901 | 21.03028 | -0.14526 | 0.949661 | 0.789085 |
| 14 | 89845200 | 89847199 | 25.64789 | 43.69177 | 50.66912 | 22.21043 | 0.169037 | 1.01705 | 0.789086 |
| 9 | 20142200 | 20144799 | 19.24989 | 33.83351 | 31.61709 | 14.00025 | 0.484542 | 0.608864 | 0.789099 |
| 18 | 54419000 | 54427999 | 140.7327 | 211.1255 | 274.2834 | 113.1765 | -0.11074 | 3.463284 | 0.789107 |
| 9 | 33877200 | 33879999 | 24.83556 | 42.98616 | 46.81164 | 20.53765 | 0.263457 | 0.869056 | 0.789113 |
| 5 | 143183800 | 143185399 | 30.4628 | 49.72984 | 70.98614 | 30.50635 | -0.13894 | 1.506307 | 0.789116 |
| 11 | 86818600 | 86819199 | 9.587824 | 18.46252 | 18.17499 | 7.061549 | 0.561106 | -0.1147 | 0.789122 |
| 4 | 67276000 | 67286599 | 142.8806 | 214.9597 | 278.5619 | 115.0297 | -0.10347 | 3.409246 | 0.789127 |
| 1 | 157449400 | 157450399 | 8.72278 | 16.43056 | 25.94495 | 10.61602 | -0.18186 | 0.069906 | 0.789136 |
| X | 31670000 | 31673799 | 45.32414 | 73.98617 | 76.07799 | 32.69385 | 0.332935 | 1.542103 | 0.789138 |
| 3 | 99294600 | 99296799 | 20.73791 | 35.5126 | 50.61477 | 22.19595 | -0.12606 | 0.953239 | 0.78919 |
| 14 | 7133600 | 7134799 | 10.30645 | 19.08616 | 28.84683 | 12.13278 | -0.15085 | 0.195897 | 0.789207 |
| 1 | 170426800 | 170428199 | 16.44885 | 28.65237 | 41.58269 | 18.28657 | -0.14757 | 0.749719 | 0.789237 |
| X | 21663600 | 21667399 | 55.49542 | 88.65132 | 103.5441 | 43.49643 | 0.11768 | 2.097342 | 0.789243 |
| 1 | 52348400 | 52359199 | 223.1118 | 322.367 | 407.9364 | 168.2767 | -0.12397 | 4.093115 | 0.78925 |
| 3 | 88099600 | 88101599 | 16.06342 | 28.56316 | 33.19832 | 14.63964 | 0.178553 | 0.47948 | 0.789256 |
| 9 | 84395400 | 84397999 | 28.05816 | 47.45412 | 45.33347 | 19.84018 | 0.447066 | 1.085467 | 0.789262 |
| 4 | 108809400 | 108811399 | 24.44358 | 41.55223 | 49.56273 | 21.72176 | 0.128237 | 1.032233 | 0.789265 |
| 14 | 71097000 | 71102599 | 63.72175 | 100.8193 | 135.7822 | 56.43604 | -0.10131 | 2.392865 | 0.789284 |
| 14 | 86064200 | 86066599 | 21.19623 | 36.24866 | 51.70178 | 22.67018 | -0.12746 | 0.967666 | 0.789329 |
| 4 | 35068800 | 35073199 | 59.04201 | 94.40107 | 93.38976 | 39.28047 | 0.358203 | 2.068543 | 0.789337 |
| X | 20553000 | 20556999 | 37.50957 | 61.88206 | 67.24539 | 29.02326 | 0.260716 | 1.388861 | 0.789337 |
| 2 | 26656600 | 26660199 | 40.18297 | 66.04199 | 71.08863 | 30.63289 | 0.272166 | 1.440999 | 0.789343 |
| 1 | 8374600 | 8375999 | 22.46308 | 38.3973 | 45.8423 | 20.10016 | 0.128146 | 0.970794 | 0.789359 |
| X | 155721800 | 155722799 | 11.3588 | 21.21227 | 26.24703 | 10.89751 | 0.150092 | 0.14774 | 0.789385 |
| 13 | 85845000 | 85845799 | 15.47028 | 27.06819 | 39.41119 | 17.38519 | -0.15177 | 0.67504 | 0.78942 |
| 17 | 86459800 | 86461599 | 16.79555 | 29.3184 | 42.02745 | 18.50682 | -0.12955 | 0.722545 | 0.789452 |
| 8 | 79622400 | 79623599 | 18.41713 | 31.71597 | 45.84744 | 20.11153 | -0.14482 | 0.886656 | 0.789462 |
| 13 | 67878800 | 67884399 | 80.86266 | 127.6081 | 124.0005 | 51.47157 | 0.365966 | 2.464085 | 0.789479 |
| 11 | 63658000 | 63661599 | 41.01961 | 66.3735 | 78.56228 | 33.59654 | 0.120006 | 1.682249 | 0.789497 |
| 18 | 40786400 | 40787999 | 10.8814 | 19.64021 | 35.84097 | 15.58784 | -0.45457 | 0.361519 | 0.789499 |
| 3 | 9650800 | 9652799 | 20.55556 | 35.69972 | 42.54306 | 18.67403 | 0.134916 | 0.811587 | 0.789499 |
| 1 | 28214200 | 28216199 | 33.36934 | 54.64152 | 64.7549 | 27.86008 | 0.131734 | 1.466455 | 0.789518 |
| 12 | 109037200 | 109039999 | 27.28929 | 44.89492 | 65.06191 | 28.04757 | -0.15504 | 1.392729 | 0.789567 |
| 4 | 15639800 | 15640799 | 8.934395 | 16.86112 | 26.44792 | 10.78421 | -0.16884 | 0.05729 | 0.789569 |

| | | | | | | | | | |
|---|---|---|---|---|---|---|---|---|---|
| 7 | 81887000 | 81890799 | 37.32481 | 59.89876 | 105.3963 | 44.57429 | -0.46479 | 1.933792 | 0.789574 |
| 1 | 38035600 | 38037799 | 18.44892 | 31.55557 | 46.92913 | 20.57774 | -0.18734 | 0.949658 | 0.789595 |
| 18 | 37397800 | 37400599 | 39.96391 | 64.39447 | 89.44461 | 38.0534 | -0.11713 | 1.803387 | 0.789607 |
| 14 | 7205800 | 7207199 | 16.96547 | 29.50196 | 42.6578 | 18.73929 | -0.14248 | 0.778827 | 0.789624 |
| 13 | 22602400 | 22603799 | 23.37973 | 39.74064 | 47.43441 | 20.78885 | 0.127217 | 1.025548 | 0.789646 |
| 10 | 39852000 | 39855799 | 58.20699 | 92.86185 | 108.9096 | 45.66611 | 0.109675 | 2.14004 | 0.789673 |
| 8 | 27012400 | 27014799 | 34.14735 | 55.77415 | 66.43336 | 28.55527 | 0.121995 | 1.503502 | 0.789683 |
| 5 | 102364800 | 102372999 | 83.24331 | 131.1816 | 145.7508 | 60.48045 | 0.17134 | 2.497987 | 0.789699 |
| 13 | 78459200 | 78461199 | 20.24464 | 35.18894 | 42.00516 | 18.44755 | 0.132624 | 0.797969 | 0.789719 |
| 17 | 35916200 | 35919599 | 32.53738 | 52.70097 | 88.09191 | 37.69951 | -0.37537 | 1.654285 | 0.789735 |
| 11 | 51510000 | 51510999 | 8.333188 | 15.74798 | 25.42509 | 10.33727 | -0.20636 | 0.039593 | 0.789771 |
| 11 | 39280200 | 39281799 | 13.58444 | 24.21136 | 35.11514 | 15.45461 | -0.13682 | 0.456488 | 0.789808 |
| 13 | 50266400 | 50274599 | 101.0992 | 155.3464 | 204.0634 | 84.73273 | -0.09808 | 2.971476 | 0.789831 |
| 12 | 75270800 | 75274199 | 33.96221 | 56.26133 | 64.34829 | 27.85012 | 0.187576 | 1.327796 | 0.789867 |
| 4 | 70990200 | 70995399 | 71.43397 | 111.8335 | 174.7848 | 72.50437 | -0.32767 | 2.620344 | 0.789873 |
| 19 | 6902000 | 6905799 | 24.90935 | 40.62822 | 78.51276 | 33.73011 | -0.57557 | 1.512672 | 0.789885 |
| 10 | 99051800 | 99053399 | 15.7399 | 27.26263 | 48.99045 | 21.49403 | -0.45567 | 0.826786 | 0.789886 |
| 16 | 51379400 | 51381199 | 18.83562 | 33.23088 | 31.24063 | 13.82418 | 0.478203 | 0.567383 | 0.789888 |
| X | 3361800 | 3369199 | 54.58176 | 87.62559 | 84.73422 | 35.85636 | 0.397766 | 1.937819 | 0.789891 |
| 3 | 93826800 | 93830399 | 33.99731 | 56.07565 | 54.22189 | 23.40017 | 0.42839 | 1.366928 | 0.789894 |
| 14 | 90923400 | 90927999 | 100.3482 | 154.4601 | 174.3033 | 72.39274 | 0.122781 | 2.933247 | 0.789907 |
| 3 | 151840600 | 151841999 | 12.33985 | 22.1347 | 33.07317 | 14.47739 | -0.17434 | 0.401035 | 0.78991 |
| 3 | 119628800 | 119630999 | 27.76278 | 46.19604 | 55.04152 | 23.89524 | 0.129272 | 1.251585 | 0.789956 |
| 5 | 127923600 | 127925599 | 24.19843 | 41.37558 | 48.48 | 21.25719 | 0.155166 | 0.97904 | 0.789967 |
| 12 | 72231400 | 72231999 | 6.828887 | 13.20549 | 22.10405 | 8.454751 | -0.17636 | -0.18377 | 0.789989 |
| 5 | 122228200 | 122231599 | 32.28506 | 52.44976 | 74.72596 | 32.05044 | -0.14133 | 1.584952 | 0.790008 |
| 19 | 40264200 | 40267199 | 35.67188 | 57.42297 | 97.99368 | 41.632 | -0.4152 | 1.832243 | 0.790061 |
| 13 | 93678200 | 93680999 | 28.9401 | 48.33268 | 56.86552 | 24.74495 | 0.147257 | 1.21502 | 0.790063 |
| 2 | 38433800 | 38436399 | 19.71063 | 33.7213 | 48.48405 | 21.25994 | -0.13883 | 0.954188 | 0.790064 |
| 1 | 58369400 | 58369999 | 10.85761 | 20.22542 | 24.24872 | 10.11077 | 0.190567 | 0.154703 | 0.790075 |
| 5 | 66657000 | 66659599 | 21.42612 | 37.04586 | 44.17331 | 19.37952 | 0.132894 | 0.861274 | 0.790079 |
| 8 | 36506000 | 36508599 | 25.55436 | 43.38994 | 51.20267 | 22.4356 | 0.143442 | 1.047424 | 0.790084 |
| 8 | 117568400 | 117570999 | 26.05975 | 43.44701 | 61.55593 | 26.75676 | -0.11985 | 1.231793 | 0.790107 |
| 1 | 171734400 | 171742199 | 99.99282 | 155.158 | 174.4415 | 72.44527 | 0.137036 | 2.787155 | 0.790121 |
| 11 | 59333400 | 59336399 | 20.44777 | 34.48988 | 61.27471 | 26.79497 | -0.44494 | 1.100166 | 0.790143 |
| 18 | 31556200 | 31557199 | 14.99226 | 26.74895 | 31.9669 | 14.05641 | 0.141861 | 0.453064 | 0.790155 |
| 7 | 12567000 | 12597799 | 621.525 | 835.6638 | 823.7849 | 343.6763 | 0.131797 | 5.193833 | 0.790196 |
| 7 | 27749000 | 27749799 | 9.932082 | 18.33089 | 28.28573 | 11.95156 | -0.18559 | 0.208342 | 0.790197 |
| 7 | 12309400 | 12311599 | 32.26456 | 53.71891 | 61.46779 | 26.69563 | 0.187422 | 1.263789 | 0.790232 |
| 1 | 85449000 | 85456799 | 139.451 | 210.9041 | 238.2003 | 98.51529 | 0.101299 | 3.30195 | 0.790243 |
| 17 | 36330800 | 36333999 | 34.61175 | 56.5962 | 67.78065 | 29.14631 | 0.115268 | 1.48875 | 0.790274 |
| 7 | 140048200 | 140048799 | 13.58665 | 24.495 | 29.47404 | 12.83709 | 0.141256 | 0.373072 | 0.790276 |
| 12 | 87722800 | 87729599 | 107.7162 | 165.4273 | 189.5394 | 78.69633 | 0.098387 | 2.98195 | 0.790281 |
| 6 | 117714000 | 117722599 | 85.22883 | 132.6757 | 177.0315 | 73.48555 | -0.10619 | 2.725983 | 0.790298 |
| 9 | 36582200 | 36582599 | 6.099507 | 11.85003 | 20.61661 | 7.693458 | -0.19708 | -0.27967 | 0.790303 |
| 3 | 4325200 | 4329199 | 34.8578 | 56.81052 | 80.93722 | 34.76187 | -0.13797 | 1.544126 | 0.790317 |
| 9 | 84113200 | 84114999 | 14.70208 | 26.93581 | 31.09954 | 13.27255 | 0.219466 | 0.292518 | 0.790343 |

| | | | | | | | | | |
|---|---|---|---|---|---|---|---|---|---|
| 13 | 3368400 | 3368999 | 10.45183 | 19.6917 | 24.75495 | 10.12542 | 0.144418 | 0.07844 | 0.790347 |
| 8 | 85843800 | 85847199 | 24.18186 | 40.55942 | 57.68922 | 25.17135 | -0.12557 | 1.157261 | 0.790356 |
| 7 | 88024800 | 88026799 | 17.08316 | 30.12828 | 35.3265 | 15.63333 | 0.16066 | 0.565201 | 0.790356 |
| 17 | 4493000 | 4494199 | 12.36356 | 21.90222 | 39.27905 | 17.36699 | -0.44632 | 0.497835 | 0.790379 |
| 14 | 101643600 | 101647999 | 39.95542 | 65.23714 | 74.13864 | 31.86306 | 0.189193 | 1.525991 | 0.790408 |
| 10 | 69905200 | 69908199 | 31.76968 | 52.35476 | 62.84134 | 27.1098 | 0.116959 | 1.380912 | 0.79041 |
| 8 | 79785000 | 79788999 | 27.12058 | 44.76041 | 64.06468 | 27.65611 | -0.13535 | 1.352771 | 0.790413 |
| 2 | 86272200 | 86273599 | 13.53317 | 24.16594 | 35.08561 | 15.41685 | -0.13683 | 0.443041 | 0.790418 |
| 17 | 85521200 | 85522799 | 25.00595 | 43.37085 | 38.0877 | 16.71126 | 0.573116 | 0.840517 | 0.790425 |
| X | 96638800 | 96640199 | 19.81024 | 34.82752 | 38.98207 | 17.26197 | 0.225843 | 0.624365 | 0.790436 |
| 1 | 120699000 | 120701199 | 21.12716 | 36.05293 | 51.70573 | 22.66798 | -0.13589 | 0.985292 | 0.790459 |
| 15 | 35507600 | 35508599 | 11.18257 | 20.56793 | 30.795 | 13.07287 | -0.14749 | 0.243166 | 0.790463 |
| 4 | 115026800 | 115029599 | 35.72532 | 57.94429 | 80.80984 | 34.61826 | -0.11285 | 1.627873 | 0.790488 |
| 1 | 65760200 | 65761599 | 12.52112 | 22.19669 | 39.78127 | 17.5866 | -0.44504 | 0.49213 | 0.790504 |
| 13 | 78000800 | 78002799 | 12.91058 | 23.9274 | 23.41009 | 9.835638 | 0.473034 | 0.197582 | 0.790553 |
| 1 | 105404000 | 105405599 | 22.25685 | 37.79851 | 53.83102 | 23.5887 | -0.12673 | 1.038207 | 0.790554 |
| 5 | 109166400 | 109175199 | 98.9194 | 152.6561 | 205.636 | 85.41726 | -0.12957 | 2.888311 | 0.790558 |
| 16 | 63548200 | 63548799 | 6.982374 | 13.44292 | 22.5489 | 8.765614 | -0.20158 | -0.12712 | 0.790579 |
| 17 | 49203000 | 49205199 | 14.13555 | 25.0028 | 36.56166 | 16.17048 | -0.15458 | 0.518387 | 0.790587 |
| 1 | 182854000 | 182855199 | 10.8118 | 19.85901 | 29.80463 | 12.67581 | -0.15238 | 0.248417 | 0.79059 |
| 8 | 42636800 | 42637799 | 11.90707 | 21.82907 | 26.09413 | 11.12663 | 0.170783 | 0.256817 | 0.790599 |
| 10 | 72960000 | 72960999 | 9.298572 | 17.53201 | 26.9768 | 11.02025 | -0.14336 | 0.07003 | 0.790606 |
| 8 | 41191000 | 41193799 | 62.9773 | 100.128 | 115.7147 | 48.29408 | 0.125643 | 2.256884 | 0.790622 |
| 14 | 4229200 | 4233599 | 34.65674 | 55.89801 | 95.18936 | 40.55003 | -0.40875 | 1.768932 | 0.79064 |
| 15 | 19285200 | 19294999 | 160.2309 | 240.231 | 268.9962 | 110.9776 | 0.102745 | 3.478341 | 0.790651 |
| 5 | 34487600 | 34488399 | 12.18932 | 21.93959 | 32.72939 | 14.28435 | -0.16856 | 0.382868 | 0.790653 |
| 7 | 125604400 | 125605599 | 14.37817 | 25.30893 | 36.72311 | 16.25764 | -0.14595 | 0.575446 | 0.790698 |
| 3 | 129260800 | 129264199 | 47.56279 | 76.14271 | 104.247 | 43.92515 | -0.10772 | 2.022216 | 0.7907 |
| X | 89371800 | 89374399 | 20.60511 | 35.8837 | 33.2425 | 14.61669 | 0.49705 | 0.760354 | 0.790746 |
| 7 | 43761600 | 43767799 | 59.76308 | 94.71995 | 128.5782 | 53.49442 | -0.1106 | 2.346546 | 0.790766 |
| X | 126430000 | 126432399 | 22.27644 | 38.82628 | 43.49787 | 19.13586 | 0.224216 | 0.773938 | 0.790775 |
| 14 | 101714400 | 101716999 | 18.56313 | 31.55483 | 55.88625 | 24.49308 | -0.43988 | 1.017885 | 0.790791 |
| 5 | 34098600 | 34107399 | 145.2805 | 218.5501 | 285.5442 | 117.91 | -0.1154 | 3.412156 | 0.7908 |
| 1 | 57493200 | 57494799 | 14.5518 | 26.12615 | 31.38376 | 13.72735 | 0.139486 | 0.408214 | 0.790815 |
| 10 | 40216400 | 40217399 | 9.321911 | 17.49699 | 26.9216 | 11.08195 | -0.15201 | 0.099356 | 0.79082 |
| 4 | 83324000 | 83329199 | 90.23979 | 139.6227 | 184.2978 | 76.55274 | -0.09741 | 2.835165 | 0.790827 |
| Y | 86259400 | 86264399 | 29.59301 | 48.75254 | 48.19405 | 20.78821 | 0.395972 | 1.364953 | 0.790841 |
| 18 | 45073600 | 45075599 | 19.42998 | 33.24888 | 47.96101 | 21.03099 | -0.14339 | 0.951414 | 0.790852 |
| X | 121985800 | 121992999 | 119.5152 | 184.1655 | 162.4977 | 67.49895 | 0.478606 | 2.921098 | 0.790864 |
| 18 | 57264800 | 57266999 | 13.32089 | 23.60397 | 34.71312 | 15.34743 | -0.16127 | 0.494442 | 0.790876 |
| 4 | 122412600 | 122414599 | 19.18715 | 33.419 | 39.93771 | 17.56211 | 0.131394 | 0.762791 | 0.790893 |
| 12 | 15013400 | 15013999 | 6.810479 | 13.2258 | 22.31028 | 8.353462 | -0.1587 | -0.25732 | 0.790913 |
| 13 | 105052400 | 105054399 | 19.45917 | 33.61004 | 48.4768 | 21.26685 | -0.14092 | 0.865842 | 0.79093 |
| 1 | 103336000 | 103337799 | 17.54112 | 30.10124 | 52.71839 | 23.12235 | -0.42162 | 0.923458 | 0.790939 |
| 4 | 34485200 | 34487799 | 19.89144 | 33.86153 | 61.45675 | 26.93758 | -0.47484 | 1.016339 | 0.790946 |
| 12 | 88080600 | 88105199 | 562.3095 | 761.2598 | 763.996 | 318.4006 | 0.11594 | 5.0842 | 0.790956 |
| 9 | 74641600 | 74642799 | 11.35849 | 21.0156 | 32.00226 | 13.47633 | -0.16339 | 0.21435 | 0.791008 |

| | | | | | | | | | |
|---|---|---|---|---|---|---|---|---|---|
| 14 | 38046400 | 38049199 | 19.78552 | 33.28369 | 59.58147 | 26.03341 | -0.45596 | 1.118435 | 0.791011 |
| 10 | 117581600 | 117586399 | 49.13376 | 78.60262 | 107.7706 | 45.36483 | -0.11094 | 2.045072 | 0.791032 |
| 5 | 45399200 | 45401199 | 25.44709 | 42.20224 | 60.79599 | 26.3329 | -0.1441 | 1.289125 | 0.79104 |
| 4 | 143576600 | 143599199 | 527.9117 | 717.7084 | 727.2316 | 302.8823 | 0.108225 | 5.017501 | 0.791044 |
| 14 | 117844600 | 117848799 | 37.04156 | 60.55837 | 71.06593 | 30.55699 | 0.144307 | 1.497901 | 0.791089 |
| X | 120282800 | 120285199 | 30.66827 | 51.32017 | 58.62355 | 25.54177 | 0.189698 | 1.202287 | 0.791091 |
| 12 | 79921000 | 79923799 | 22.20424 | 38.05365 | 56.72534 | 24.87719 | -0.19053 | 0.957711 | 0.791102 |
| 15 | 33122800 | 33124199 | 17.54055 | 30.72142 | 36.71272 | 16.20861 | 0.131848 | 0.666227 | 0.791103 |
| 4 | 11330600 | 11332399 | 13.99463 | 25.12131 | 29.87068 | 13.06749 | 0.153887 | 0.401384 | 0.791142 |
| 1 | 151609000 | 151611799 | 24.40476 | 41.49044 | 49.56407 | 21.72252 | 0.126055 | 1.030541 | 0.791168 |
| 1 | 182939000 | 182940999 | 24.65549 | 41.63993 | 49.59133 | 21.69426 | 0.130056 | 1.090073 | 0.791177 |
| 12 | 73096600 | 73097799 | 13.9162 | 24.97922 | 29.96059 | 13.1074 | 0.141569 | 0.402805 | 0.791183 |
| 2 | 23515200 | 23515999 | 11.97374 | 22.43348 | 22.01795 | 9.091437 | 0.488107 | 0.120491 | 0.791227 |
| 4 | 79852600 | 79853399 | 14.44676 | 25.72756 | 30.51763 | 13.4403 | 0.150999 | 0.457456 | 0.791257 |
| 3 | 67006200 | 67008799 | 72.54578 | 113.0909 | 157.2566 | 65.2633 | -0.16403 | 2.693487 | 0.791261 |
| 10 | 40203800 | 40206599 | 26.07853 | 44.30937 | 43.70044 | 19.15173 | 0.401379 | 1.035132 | 0.791274 |
| 1 | 78991200 | 78992199 | 21.98617 | 37.57871 | 44.30635 | 19.42656 | 0.146188 | 0.968409 | 0.791281 |
| 15 | 8849800 | 8857799 | 71.1429 | 112.0665 | 151.852 | 63.0088 | -0.11513 | 2.498128 | 0.791282 |
| 14 | 70965000 | 70965799 | 16.96758 | 28.54034 | 59.37789 | 25.94364 | -0.67182 | 1.110544 | 0.791287 |
| 11 | 90201600 | 90203999 | 18.2487 | 31.51514 | 45.3086 | 19.87688 | -0.13612 | 0.859792 | 0.791299 |
| 16 | 55773800 | 55782599 | 153.6841 | 228.5789 | 298.4057 | 122.9445 | -0.12905 | 3.62492 | 0.791306 |
| 17 | 31242000 | 31242999 | 12.15291 | 22.55313 | 27.70297 | 11.61825 | 0.149075 | 0.195703 | 0.791333 |
| 16 | 47650000 | 47650799 | 17.01148 | 29.84315 | 35.6197 | 15.74761 | 0.13352 | 0.634737 | 0.79137 |
| 3 | 151489200 | 151490799 | 21.10934 | 36.54467 | 43.64442 | 19.14823 | 0.131019 | 0.848105 | 0.791419 |
| 16 | 28280200 | 28281399 | 22.50095 | 37.76512 | 54.82976 | 23.9337 | -0.15494 | 1.144936 | 0.791425 |
| X | 153580000 | 153585799 | 64.58553 | 103.2911 | 110.3812 | 46.44418 | 0.247505 | 2.073744 | 0.791435 |
| 11 | 113050000 | 113052399 | 17.0995 | 29.65437 | 43.22017 | 18.9631 | -0.15445 | 0.813975 | 0.79144 |
| 14 | 10420800 | 10422799 | 15.32199 | 26.93895 | 38.69301 | 17.13371 | -0.13215 | 0.596414 | 0.791556 |
| 5 | 18914000 | 18916199 | 27.23964 | 45.7359 | 54.52322 | 23.80563 | 0.128271 | 1.153423 | 0.791557 |
| 11 | 118041400 | 118043399 | 26.05051 | 43.5711 | 51.25407 | 22.32393 | 0.147574 | 1.186753 | 0.791562 |
| 3 | 149413800 | 149416399 | 55.30753 | 88.13464 | 100.6988 | 42.15612 | 0.145555 | 2.170508 | 0.791572 |
| 10 | 92636800 | 92638399 | 12.90797 | 23.28716 | 34.17141 | 14.8704 | -0.14256 | 0.375046 | 0.791599 |
| 11 | 117188000 | 117192599 | 36.18575 | 58.47401 | 82.62421 | 35.28285 | -0.13697 | 1.709956 | 0.791619 |
| 7 | 93360600 | 93360999 | 12.72553 | 23.18057 | 27.18621 | 11.68877 | 0.190451 | 0.305097 | 0.79162 |
| 7 | 6133400 | 6141199 | 109.1923 | 165.9523 | 222.4886 | 92.15582 | -0.13997 | 3.195125 | 0.791626 |
| 12 | 72170400 | 72174399 | 57.07136 | 90.84196 | 123.4389 | 51.59466 | -0.10593 | 2.215142 | 0.791631 |
| 4 | 127482400 | 127490199 | 71.26784 | 112.0115 | 149.8599 | 62.16404 | -0.09929 | 2.543508 | 0.791653 |
| 3 | 56927800 | 56930999 | 57.33758 | 91.48382 | 107.652 | 45.14499 | 0.105019 | 2.135136 | 0.791667 |
| 2 | 20239200 | 20240799 | 19.4618 | 33.3799 | 47.9545 | 21.03202 | -0.137 | 0.93157 | 0.791675 |
| 14 | 9366400 | 9369399 | 39.40966 | 64.13913 | 74.98053 | 32.18707 | 0.145244 | 1.569963 | 0.791733 |
| 3 | 11417200 | 11417599 | 7.429965 | 14.43853 | 19.51358 | 7.26675 | 0.166121 | -0.27504 | 0.791762 |
| 3 | 11443600 | 11444199 | 7.653927 | 14.86256 | 20.00249 | 7.494231 | 0.16305 | -0.25301 | 0.791772 |
| 6 | 54804000 | 54804999 | 10.34489 | 19.45316 | 24.25872 | 9.954469 | 0.152504 | 0.089387 | 0.79178 |
| 14 | 62222200 | 62223799 | 18.45009 | 32.1596 | 38.37475 | 16.88285 | 0.133611 | 0.744925 | 0.791785 |
| 7 | 6702200 | 6703999 | 13.87523 | 24.63467 | 35.57118 | 15.70617 | -0.13378 | 0.482377 | 0.791795 |
| 5 | 131091400 | 131092799 | 17.24409 | 29.96368 | 43.1714 | 18.95586 | -0.13756 | 0.790656 | 0.791841 |
| 15 | 60844800 | 60846199 | 23.75783 | 39.24438 | 70.55504 | 30.47612 | -0.46325 | 1.337784 | 0.791844 |

| | | | | | | | | | |
|---|---|---|---|---|---|---|---|---|---|
| 10 | 35225600 | 35225999 | 4.933746 | 9.634306 | 17.61174 | 6.194907 | -0.18941 | -0.51578 | 0.791856 |
| 5 | 90143200 | 90145199 | 21.48908 | 37.09319 | 44.16245 | 19.37377 | 0.134521 | 0.874727 | 0.791868 |
| 19 | 3669600 | 3670799 | 19.45449 | 33.50801 | 47.94263 | 21.03071 | -0.13001 | 0.893514 | 0.791868 |
| 2 | 33185000 | 33187599 | 29.62062 | 49.22711 | 58.9655 | 25.57714 | 0.121481 | 1.274854 | 0.791886 |
| 1 | 99561200 | 99563799 | 50.93182 | 81.8476 | 79.63265 | 33.74766 | 0.390683 | 1.900166 | 0.791901 |
| 14 | 90925600 | 90928599 | 76.59331 | 119.4427 | 161.2326 | 66.90982 | -0.1209 | 2.689167 | 0.791914 |
| 17 | 53312800 | 53314199 | 28.30212 | 47.41175 | 56.18941 | 24.49789 | 0.136814 | 1.183417 | 0.791956 |
| 9 | 40102800 | 40104599 | 17.03581 | 29.84234 | 34.60507 | 15.28022 | 0.174861 | 0.654497 | 0.791957 |
| 16 | 22596000 | 22604199 | 106.5959 | 163.1323 | 213.3639 | 88.53496 | -0.09633 | 3.053286 | 0.791973 |
| 6 | 140692000 | 140694999 | 40.70189 | 65.92389 | 78.0651 | 33.40802 | 0.120674 | 1.662113 | 0.792017 |
| 5 | 26161400 | 26162799 | 30.8328 | 50.18069 | 72.36845 | 31.06587 | -0.15679 | 1.552425 | 0.792029 |
| 15 | 73249400 | 73254199 | 62.99244 | 100.3237 | 116.0995 | 48.56509 | 0.126255 | 2.201143 | 0.792039 |
| 8 | 72718000 | 72721999 | 46.34374 | 74.67327 | 87.73519 | 37.32534 | 0.12381 | 1.813237 | 0.792066 |
| 13 | 92188800 | 92190199 | 14.81518 | 26.06537 | 37.67402 | 16.68025 | -0.14079 | 0.583857 | 0.792075 |
| 10 | 11587600 | 11589199 | 16.64767 | 29.39895 | 34.85554 | 15.42085 | 0.145547 | 0.548312 | 0.792155 |
| 9 | 33128200 | 33132599 | 53.50609 | 85.57983 | 100.8024 | 42.41938 | 0.107474 | 2.053983 | 0.792163 |
| 16 | 67918000 | 67918799 | 12.49015 | 23.00003 | 28.14151 | 11.92728 | 0.143773 | 0.242662 | 0.792175 |
| 1 | 85518400 | 85536999 | 950.1611 | 1221.897 | 1360.085 | 570.3653 | -0.10759 | 5.91212 | 0.792188 |
| 19 | 55001600 | 55005199 | 40.49327 | 65.12713 | 91.0017 | 38.61244 | -0.12898 | 1.863369 | 0.792287 |
| 13 | 22833400 | 22843799 | 118.5475 | 182.339 | 198.6476 | 82.49525 | 0.172704 | 2.957938 | 0.79229 |
| 2 | 177435400 | 177436599 | 20.56907 | 35.5428 | 42.62177 | 18.69849 | 0.124591 | 0.860391 | 0.792294 |
| 19 | 36033800 | 36035199 | 14.38412 | 24.89634 | 48.44776 | 21.25546 | -0.57001 | 0.828835 | 0.792359 |
| X | 120976400 | 120978599 | 28.68442 | 48.48052 | 46.38091 | 20.29443 | 0.444928 | 1.093864 | 0.79238 |
| 1 | 19353400 | 19357799 | 52.09875 | 83.58348 | 81.35192 | 34.39013 | 0.387259 | 1.95183 | 0.792408 |
| 7 | 128002600 | 128004399 | 22.07966 | 37.13825 | 53.80795 | 23.50983 | -0.152 | 1.119486 | 0.792409 |
| 14 | 72858000 | 72858599 | 8.62032 | 16.59811 | 27.80975 | 10.8075 | -0.20058 | -0.09846 | 0.792411 |
| 4 | 3416600 | 3421599 | 43.18429 | 69.65692 | 82.73515 | 35.24775 | 0.109975 | 1.774474 | 0.792413 |
| 6 | 137016000 | 137017999 | 20.01741 | 34.82412 | 41.43586 | 18.20744 | 0.137318 | 0.782962 | 0.792424 |
| 8 | 34752000 | 34755199 | 30.23962 | 49.48441 | 70.16256 | 30.17887 | -0.12685 | 1.469506 | 0.792432 |
| 2 | 99136200 | 99138599 | 33.06666 | 53.76157 | 75.62034 | 32.44978 | -0.12197 | 1.571998 | 0.792433 |
| 3 | 152539000 | 152540399 | 13.9597 | 25.23778 | 30.48569 | 13.23152 | 0.138969 | 0.362533 | 0.792454 |
| 10 | 120394000 | 120398199 | 45.66005 | 73.40638 | 86.66692 | 36.75088 | 0.112818 | 1.881582 | 0.792474 |
| 10 | 22492400 | 22496399 | 49.4084 | 79.51443 | 91.79042 | 38.98169 | 0.147004 | 1.856008 | 0.792486 |
| X | 162746000 | 162751199 | 47.24158 | 76.34484 | 85.76377 | 36.62678 | 0.193312 | 1.729804 | 0.792488 |
| 8 | 37580800 | 37592799 | 186.443 | 277.7373 | 301.3668 | 124.1842 | 0.139654 | 3.602724 | 0.792495 |
| 4 | 89602600 | 89602999 | 7.067499 | 13.73038 | 17.87514 | 6.671402 | 0.216076 | -0.28012 | 0.792513 |
| 9 | 41819200 | 41820999 | 31.87143 | 51.71123 | 75.33999 | 32.28317 | -0.17518 | 1.612719 | 0.792515 |
| 12 | 85636400 | 85637599 | 9.24536 | 17.28198 | 27.36029 | 11.34135 | -0.20149 | 0.122956 | 0.79252 |
| 17 | 93435400 | 93438799 | 33.56901 | 55.65995 | 63.81845 | 27.63795 | 0.184086 | 1.315272 | 0.79254 |
| 16 | 89559400 | 89561399 | 17.69977 | 30.70178 | 44.22762 | 19.40525 | -0.13774 | 0.815256 | 0.792543 |
| 6 | 149234200 | 149239599 | 57.69931 | 92.43811 | 103.9969 | 43.86866 | 0.176233 | 2.010625 | 0.792585 |
| 10 | 53591800 | 53596799 | 39.98994 | 64.5974 | 90.29169 | 38.53863 | -0.12201 | 1.731767 | 0.792596 |
| 5 | 20575000 | 20577399 | 20.94703 | 36.08279 | 43.18635 | 18.94437 | 0.126479 | 0.891541 | 0.7926 |
| 15 | 91273200 | 91274999 | 23.68983 | 39.66006 | 57.11098 | 24.88972 | -0.14329 | 1.174541 | 0.792652 |
| 12 | 22591400 | 22591999 | 12.61486 | 23.23201 | 28.02284 | 11.87536 | 0.163907 | 0.239692 | 0.792655 |
| 8 | 23348200 | 23350599 | 17.58734 | 29.99949 | 54.27797 | 23.79484 | -0.46972 | 0.980621 | 0.792657 |
| 4 | 144266400 | 144275799 | 158.3038 | 235.3377 | 304.0339 | 125.2572 | -0.11461 | 3.63455 | 0.792661 |

| | | | | | | | | | |
|---|---|---|---|---|---|---|---|---|---|
| X | 122063200 | 122063999 | 23.6897 | 40.42985 | 47.95364 | 21.02322 | 0.137234 | 0.993303 | 0.792666 |
| 5 | 119217800 | 119218999 | 16.2647 | 28.5852 | 33.65925 | 14.90386 | 0.153078 | 0.599195 | 0.792668 |
| 2 | 104941200 | 104943399 | 16.72516 | 29.19956 | 41.99656 | 18.49482 | -0.13442 | 0.718609 | 0.792685 |
| 6 | 132499600 | 132506799 | 119.6298 | 182.553 | 208.4591 | 86.44221 | 0.096061 | 3.114576 | 0.792698 |
| 10 | 40739200 | 40741799 | 22.46021 | 38.46867 | 57.81261 | 25.35276 | -0.20232 | 0.96683 | 0.79271 |
| 12 | 51590200 | 51593399 | 34.8566 | 56.44735 | 80.01261 | 34.23742 | -0.13836 | 1.655221 | 0.792743 |
| 9 | 68092800 | 68093999 | 18.18646 | 31.708 | 37.42156 | 16.46609 | 0.149568 | 0.739893 | 0.792752 |
| 2 | 112002800 | 112006199 | 51.96953 | 83.22116 | 97.57374 | 41.13105 | 0.116006 | 2.014119 | 0.792769 |
| 10 | 93033000 | 93034999 | 25.05184 | 42.95646 | 41.04062 | 17.99933 | 0.449073 | 0.947743 | 0.792787 |
| 11 | 78789600 | 78792799 | 29.90809 | 49.59756 | 59.54103 | 25.78827 | 0.118038 | 1.299924 | 0.7928 |
| 1 | 122284200 | 122285199 | 22.58506 | 38.42975 | 45.35574 | 19.8795 | 0.143699 | 1.011455 | 0.792807 |
| 16 | 94608600 | 94619399 | 162.5708 | 243.1592 | 272.9916 | 112.5738 | 0.09584 | 3.523179 | 0.792823 |
| 6 | 14729600 | 14730199 | 11.87779 | 22.35736 | 21.93698 | 8.987006 | 0.497216 | 0.088915 | 0.792826 |
| 15 | 97869400 | 97870399 | 8.830585 | 16.70382 | 25.97341 | 10.53902 | -0.14973 | 0.041079 | 0.792832 |
| 17 | 82738400 | 82739399 | 14.49982 | 25.90783 | 30.85838 | 13.55242 | 0.1476 | 0.435803 | 0.792835 |
| 8 | 104374400 | 104375599 | 13.50462 | 24.45008 | 29.41761 | 12.74507 | 0.146099 | 0.346804 | 0.792845 |
| 18 | 79408800 | 79411199 | 22.28325 | 38.25208 | 45.80577 | 20.09102 | 0.124732 | 0.931409 | 0.792847 |
| 1 | 105542400 | 105543999 | 20.08501 | 35.02361 | 41.37199 | 18.20614 | 0.148089 | 0.751234 | 0.792862 |
| 2 | 129492000 | 129493599 | 17.69311 | 30.56684 | 44.81977 | 19.66265 | -0.16415 | 0.85561 | 0.792866 |
| 4 | 38868600 | 38872199 | 47.40728 | 76.21501 | 89.73676 | 38.05239 | 0.116764 | 1.886958 | 0.792866 |
| 6 | 60868400 | 60870799 | 49.05973 | 78.58991 | 91.30424 | 38.50934 | 0.129829 | 1.998205 | 0.792879 |
| 10 | 129346200 | 129348399 | 21.02356 | 35.96341 | 51.16034 | 22.43327 | -0.12362 | 0.963728 | 0.792882 |
| 6 | 101307400 | 101309199 | 16.20796 | 28.53413 | 34.02683 | 15.06351 | 0.135657 | 0.578917 | 0.792894 |
| 10 | 35226800 | 35227799 | 14.35442 | 25.18112 | 37.33638 | 16.50614 | -0.17783 | 0.623243 | 0.792901 |
| 11 | 90847800 | 90851199 | 47.43901 | 76.26624 | 90.21158 | 38.25389 | 0.110105 | 1.884872 | 0.79291 |
| 13 | 71341800 | 71343799 | 16.51905 | 28.58346 | 50.60575 | 22.20205 | -0.43473 | 0.840727 | 0.792929 |
| 13 | 81639200 | 81640599 | 14.58807 | 25.76112 | 37.17715 | 16.44807 | -0.13632 | 0.543346 | 0.792939 |
| X | 157460200 | 157461599 | 14.17977 | 25.92146 | 23.96931 | 10.25999 | 0.535089 | 0.286906 | 0.792957 |
| 5 | 151430400 | 151431599 | 19.99513 | 34.71928 | 41.55862 | 18.23957 | 0.128636 | 0.812189 | 0.792988 |
| 7 | 138089200 | 138091399 | 28.55886 | 48.06347 | 55.69486 | 24.33843 | 0.16914 | 1.137748 | 0.793003 |
| 13 | 75035000 | 75035399 | 7.92051 | 15.3971 | 20.65847 | 7.671721 | 0.180108 | -0.27476 | 0.79305 |
| 5 | 6430400 | 6432199 | 18.67243 | 32.20437 | 46.34753 | 20.33184 | -0.13811 | 0.873279 | 0.79307 |
| 5 | 110378600 | 110379799 | 10.1978 | 18.97913 | 28.86628 | 12.05459 | -0.15265 | 0.163559 | 0.793073 |
| 6 | 108857800 | 108860999 | 31.02032 | 50.45146 | 84.60064 | 36.30779 | -0.37458 | 1.568962 | 0.793075 |
| 2 | 144465400 | 144470399 | 59.4055 | 94.57936 | 110.7401 | 46.30181 | 0.10888 | 2.205668 | 0.793079 |
| 17 | 89271400 | 89274199 | 36.0569 | 58.71758 | 69.98758 | 30.05028 | 0.117927 | 1.550685 | 0.793106 |
| 12 | 73616000 | 73617399 | 12.79255 | 23.17591 | 34.7292 | 15.03408 | -0.16804 | 0.348714 | 0.793113 |
| 14 | 124992800 | 124995999 | 54.54018 | 87.44934 | 100.506 | 42.44954 | 0.146915 | 1.980092 | 0.793129 |
| 1 | 164485600 | 164488599 | 46.5614 | 74.78123 | 87.30317 | 36.96715 | 0.12727 | 1.915571 | 0.793162 |
| 5 | 63813400 | 63815199 | 6.913206 | 13.13126 | 27.67659 | 11.13205 | -0.57317 | 0.000789 | 0.793178 |
| 9 | 6800400 | 6815599 | 393.997 | 545.978 | 561.5995 | 233.196 | 0.114378 | 4.718429 | 0.793198 |
| 4 | 143311000 | 143312199 | 40.9153 | 65.77016 | 92.77314 | 39.33307 | -0.14353 | 1.884045 | 0.793214 |
| 5 | 115002400 | 115009399 | 95.79124 | 147.3026 | 194.5935 | 80.80902 | -0.10537 | 2.954373 | 0.793214 |
| 2 | 5152400 | 5154599 | 19.55775 | 33.68739 | 47.94047 | 21.0298 | -0.12233 | 0.892304 | 0.793241 |
| 5 | 18944000 | 18946199 | 21.38595 | 37.32894 | 42.34684 | 18.64668 | 0.206289 | 0.744903 | 0.79327 |
| 2 | 134645600 | 134646799 | 11.55171 | 21.1332 | 31.24977 | 13.3631 | -0.13714 | 0.278121 | 0.793273 |
| 15 | 100437400 | 100447999 | 198.9745 | 291.5328 | 367.2648 | 151.2811 | -0.09796 | 3.879686 | 0.793278 |

| | | | | | | | | | |
|---|---|---|---|---|---|---|---|---|---|
| 13 | 69282200 | 69283999 | 16.88153 | 29.50372 | 42.47837 | 18.72011 | -0.13583 | 0.704686 | 0.793294 |
| 14 | 83965400 | 83966399 | 15.45011 | 27.63348 | 32.89734 | 14.43568 | 0.149294 | 0.441917 | 0.793309 |
| 18 | 11998000 | 12000599 | 22.37442 | 37.96877 | 53.84804 | 23.59062 | -0.12081 | 1.044329 | 0.793334 |
| 8 | 118716000 | 118717799 | 19.75046 | 34.25672 | 41.04749 | 18.00991 | 0.126837 | 0.821257 | 0.793364 |
| 3 | 53089400 | 53093999 | 51.52321 | 82.41724 | 113.6999 | 47.85416 | -0.11998 | 2.053822 | 0.793399 |
| 10 | 130138200 | 130138999 | 8.025896 | 15.31117 | 24.51059 | 9.768394 | -0.16862 | -0.03019 | 0.793399 |
| 12 | 109780000 | 109781399 | 21.80699 | 37.42602 | 44.77331 | 19.63781 | 0.126113 | 0.930663 | 0.793413 |
| 14 | 41896800 | 41897999 | 13.1919 | 24.54042 | 23.96802 | 10.01669 | 0.481362 | 0.181372 | 0.793467 |
| 18 | 17451800 | 17455599 | 61.00607 | 97.27072 | 111.9986 | 46.92347 | 0.135277 | 2.160047 | 0.793485 |
| 11 | 61233200 | 61233999 | 6.097958 | 11.83969 | 20.5903 | 7.722136 | -0.20355 | -0.26358 | 0.793497 |
| 3 | 32562400 | 32565799 | 22.73669 | 38.00591 | 56.51054 | 24.60416 | -0.1893 | 1.187018 | 0.79352 |
| 4 | 151664000 | 151667799 | 33.94415 | 55.18477 | 77.30312 | 33.17112 | -0.11609 | 1.57736 | 0.793538 |
| 13 | 12272200 | 12275999 | 52.79359 | 84.06666 | 115.366 | 48.24715 | -0.1195 | 2.196189 | 0.793543 |
| 3 | 28911000 | 28911999 | 20.09257 | 35.10445 | 34.02063 | 14.98926 | 0.43259 | 0.71276 | 0.793556 |
| 1 | 21651800 | 21652799 | 20.91095 | 36.39772 | 31.80655 | 13.98081 | 0.580854 | 0.767017 | 0.793585 |
| 10 | 122339800 | 122350199 | 222.1496 | 323.9164 | 354.3199 | 146.0053 | 0.098968 | 3.956029 | 0.793613 |
| 18 | 61409200 | 61413799 | 52.46965 | 83.67127 | 113.838 | 47.7031 | -0.10486 | 2.149542 | 0.793619 |
| 16 | 32963800 | 32967599 | 36.71216 | 59.34953 | 82.67215 | 35.32241 | -0.1157 | 1.700016 | 0.793625 |
| 17 | 88680000 | 88686999 | 85.71649 | 133.2961 | 178.3341 | 74.03773 | -0.1112 | 2.743143 | 0.793625 |
| 12 | 15714600 | 15716599 | 17.34055 | 30.7914 | 35.35363 | 15.6083 | 0.19525 | 0.504062 | 0.793633 |
| 19 | 29058800 | 29062799 | 69.90799 | 110.2202 | 147.9169 | 61.38318 | -0.10022 | 2.480351 | 0.793635 |
| 14 | 106358800 | 106363999 | 58.07819 | 92.75438 | 107.6375 | 45.20023 | 0.126572 | 2.105649 | 0.793668 |
| 18 | 90447800 | 90451599 | 77.75308 | 122.6959 | 120.6784 | 50.09155 | 0.34849 | 2.462757 | 0.793678 |
| 18 | 37052800 | 37056199 | 116.0344 | 177.9495 | 162.3038 | 67.37506 | 0.425598 | 3.0109 | 0.793698 |
| 7 | 52698400 | 52698799 | 6.381376 | 12.38798 | 21.1279 | 7.93404 | -0.17783 | -0.25475 | 0.793727 |
| 16 | 39609200 | 39611399 | 22.20854 | 38.39986 | 45.21543 | 19.83673 | 0.150924 | 0.865046 | 0.793732 |
| 1 | 116153200 | 116154599 | 20.22671 | 35.2857 | 41.40078 | 18.22367 | 0.157971 | 0.748061 | 0.793736 |
| 10 | 112960600 | 112962999 | 17.39528 | 30.94376 | 35.29668 | 15.55973 | 0.206 | 0.491746 | 0.79374 |
| 14 | 106168800 | 106172799 | 48.84499 | 78.57669 | 91.78244 | 38.956 | 0.129366 | 1.868897 | 0.793743 |
| 13 | 62276400 | 62355399 | 1886.98 | 2317.617 | 2073.353 | 873.8164 | 0.14061 | 6.6541 | 0.793751 |
| 15 | 78387000 | 78390199 | 46.49496 | 74.71469 | 88.40745 | 37.46427 | 0.108813 | 1.89782 | 0.793765 |
| 15 | 60538200 | 60539599 | 11.33437 | 21.2644 | 26.23219 | 10.8067 | 0.162879 | 0.1152 | 0.793768 |
| 18 | 10042800 | 10045999 | 47.18819 | 75.76886 | 89.25866 | 37.78073 | 0.113823 | 1.924262 | 0.79377 |
| 4 | 12100600 | 12128799 | 673.423 | 890.8923 | 1013.064 | 423.5789 | -0.0976 | 5.452236 | 0.793784 |
| 3 | 22034800 | 22036999 | 18.21123 | 32.07865 | 36.81094 | 16.29422 | 0.190912 | 0.587242 | 0.793787 |
| X | 115892600 | 115896999 | 45.7725 | 74.3794 | 79.9213 | 34.28493 | 0.264938 | 1.609497 | 0.793843 |
| 4 | 35512200 | 35513199 | 15.16717 | 27.1495 | 32.32352 | 14.17416 | 0.149698 | 0.432097 | 0.793846 |
| 2 | 169806200 | 169807799 | 15.84014 | 28.16156 | 33.42724 | 14.74261 | 0.148628 | 0.48367 | 0.793854 |
| 1 | 9251000 | 9252999 | 20.68593 | 36.0802 | 41.85081 | 18.41965 | 0.174225 | 0.750644 | 0.793875 |
| 2 | 157795000 | 157797399 | 20.05838 | 34.27321 | 49.01402 | 21.49016 | -0.13139 | 0.968113 | 0.793916 |
| 9 | 62223000 | 62224399 | 19.2138 | 33.22524 | 47.40218 | 20.79616 | -0.1248 | 0.85471 | 0.793947 |
| 7 | 7763800 | 7766599 | 35.45325 | 58.45268 | 54.65602 | 23.58336 | 0.476542 | 1.380867 | 0.793965 |
| 6 | 103849200 | 103851399 | 26.40508 | 44.19698 | 63.91028 | 27.84946 | -0.14933 | 1.193121 | 0.794013 |
| 12 | 6025400 | 6026399 | 8.232009 | 15.75572 | 25.12768 | 9.934684 | -0.15239 | -0.05015 | 0.794019 |
| 15 | 45719800 | 45720399 | 8.232009 | 15.75572 | 25.12768 | 9.934684 | -0.15239 | -0.05015 | 0.794019 |
| 11 | 80848200 | 80849999 | 14.3373 | 25.26207 | 36.6653 | 16.22882 | -0.14589 | 0.563108 | 0.794024 |
| 9 | 104725200 | 104725599 | 7.70152 | 14.90805 | 19.42383 | 7.394492 | 0.188501 | -0.19777 | 0.794026 |

| | | | | | | | | | |
|---|---|---|---|---|---|---|---|---|---|
| 15 | 10780400 | 10781799 | 18.55607 | 32.0347 | 45.82562 | 20.10348 | -0.12907 | 0.864571 | 0.794051 |
| 4 | 22718200 | 22724399 | 65.49803 | 103.9287 | 121.9239 | 50.78424 | 0.10121 | 2.316501 | 0.794053 |
| 5 | 24419000 | 24427399 | 117.4479 | 176.8543 | 280.9556 | 116.0458 | -0.39595 | 3.386973 | 0.794066 |
| 19 | 53694800 | 53696999 | 19.31301 | 33.39405 | 47.93764 | 21.03102 | -0.13376 | 0.855213 | 0.794071 |
| 3 | 35214800 | 35217199 | 17.4579 | 30.86014 | 35.79207 | 15.83208 | 0.177616 | 0.543985 | 0.794076 |
| 16 | 54537600 | 54538599 | 13.34836 | 24.131 | 28.93019 | 12.5574 | 0.149514 | 0.353325 | 0.79408 |
| 18 | 75720000 | 75721999 | 11.82581 | 21.27045 | 31.73127 | 13.85815 | -0.16926 | 0.381463 | 0.794089 |
| 10 | 50685000 | 50687599 | 22.83464 | 39.21773 | 63.02824 | 27.64583 | -0.29823 | 0.94915 | 0.794129 |
| 7 | 144754200 | 144757199 | 34.26343 | 56.05976 | 67.24571 | 28.92251 | 0.113558 | 1.478872 | 0.794131 |
| 9 | 7426000 | 7427199 | 13.49515 | 24.22199 | 35.61789 | 15.5832 | -0.15092 | 0.41102 | 0.794154 |
| 2 | 58357400 | 58358799 | 15.31716 | 27.07054 | 39.6538 | 17.54096 | -0.15755 | 0.540944 | 0.794154 |
| 12 | 24551400 | 24552999 | 12.24386 | 22.34983 | 33.32522 | 14.29084 | -0.1518 | 0.30103 | 0.794158 |
| 1 | 28805800 | 28806799 | 11.26699 | 20.71378 | 31.83267 | 13.52174 | -0.18581 | 0.246581 | 0.79417 |
| 11 | 101685600 | 101687399 | 18.501 | 32.36001 | 38.74136 | 17.08481 | 0.129186 | 0.695621 | 0.794177 |
| 9 | 114871800 | 114877999 | 39.98726 | 63.87044 | 120.0929 | 50.44 | -0.56817 | 2.086757 | 0.794178 |
| 12 | 115480800 | 115481399 | 13.46943 | 24.48756 | 29.42029 | 12.68039 | 0.152901 | 0.322384 | 0.794189 |
| 14 | 18134000 | 18135799 | 15.46058 | 26.28465 | 55.20904 | 24.19656 | -0.68463 | 1.005462 | 0.794206 |
| 17 | 39291200 | 39295599 | 74.95456 | 118.3233 | 137.2259 | 56.97079 | 0.111483 | 2.454661 | 0.794218 |
| 4 | 112982000 | 112988199 | 75.1455 | 119.5173 | 126.0295 | 52.62035 | 0.258141 | 2.259218 | 0.794223 |
| 8 | 38046800 | 38047599 | 10.94526 | 20.34123 | 24.6746 | 10.32797 | 0.16911 | 0.169581 | 0.794236 |
| 17 | 55244200 | 55245999 | 15.48609 | 27.35254 | 40.77458 | 18.03892 | -0.18281 | 0.55369 | 0.79424 |
| 13 | 45495000 | 45497199 | 28.98863 | 48.19934 | 57.79579 | 25.07786 | 0.120031 | 1.267357 | 0.79426 |
| 8 | 71490200 | 71491199 | 11.78914 | 21.13535 | 31.70828 | 13.8863 | -0.18025 | 0.400197 | 0.79427 |
| 12 | 70405400 | 70407999 | 15.4366 | 26.79507 | 47.92716 | 21.03028 | -0.44827 | 0.804939 | 0.794275 |
| 13 | 68683000 | 68696599 | 173.3129 | 259.7896 | 245.9962 | 101.4845 | 0.344181 | 3.483677 | 0.794292 |
| 18 | 31669000 | 31671999 | 25.87183 | 43.1215 | 61.00309 | 26.51172 | -0.11769 | 1.233225 | 0.794295 |
| 19 | 11542000 | 11542599 | 11.34772 | 21.44037 | 20.10278 | 8.17692 | 0.571443 | 0.056351 | 0.794309 |
| 10 | 5336600 | 5340199 | 37.07293 | 59.98636 | 83.42912 | 35.67735 | -0.11212 | 1.680396 | 0.794332 |
| 1 | 189630600 | 189631599 | 6.251892 | 12.1102 | 21.0775 | 7.99855 | -0.2206 | -0.21898 | 0.794333 |
| 7 | 135090600 | 135094799 | 32.72171 | 53.37116 | 74.75557 | 32.11611 | -0.11265 | 1.525094 | 0.794389 |
| 7 | 29506000 | 29511999 | 69.62058 | 110.638 | 124.8566 | 52.0849 | 0.159204 | 2.276443 | 0.794412 |
| X | 157902600 | 157906399 | 79.00789 | 124.6305 | 141.6723 | 58.79562 | 0.139524 | 2.475971 | 0.794418 |
| 5 | 91524800 | 91525799 | 14.34734 | 26.019 | 31.02603 | 13.41401 | 0.161146 | 0.349517 | 0.79445 |
| 4 | 105170200 | 105172199 | 15.88909 | 28.03296 | 33.49527 | 14.82182 | 0.13402 | 0.552669 | 0.794502 |
| 10 | 71333800 | 71336999 | 35.17787 | 57.10185 | 79.86312 | 34.23279 | -0.11592 | 1.608956 | 0.794509 |
| 7 | 41156600 | 41157999 | 30.29417 | 49.9386 | 59.35225 | 25.60794 | 0.131377 | 1.371105 | 0.794547 |
| 2 | 111180800 | 111183599 | 23.4342 | 40.57282 | 46.28365 | 20.30619 | 0.196872 | 0.860818 | 0.794567 |
| 4 | 111686000 | 111688599 | 38.15754 | 62.71996 | 59.30859 | 25.55792 | 0.458609 | 1.425796 | 0.794571 |
| 2 | 32844600 | 32845599 | 17.61737 | 30.77195 | 36.31323 | 15.99676 | 0.149861 | 0.708001 | 0.79459 |
| 19 | 8229200 | 8232799 | 57.97505 | 92.03795 | 124.7629 | 51.99298 | -0.10626 | 2.296573 | 0.794627 |
| 18 | 9748800 | 9749399 | 7.004218 | 13.52568 | 22.60863 | 8.704211 | -0.18288 | -0.15606 | 0.794629 |
| 2 | 150611800 | 150614999 | 19.86265 | 33.48449 | 51.1129 | 22.34756 | -0.22674 | 1.090379 | 0.794662 |
| 15 | 82489200 | 82491799 | 35.2552 | 57.60801 | 68.98526 | 29.65673 | 0.114694 | 1.500503 | 0.794664 |
| 6 | 122242400 | 122242999 | 11.90569 | 21.58329 | 31.68081 | 13.70327 | -0.13725 | 0.334504 | 0.794688 |
| 13 | 61486800 | 61487399 | 13.48296 | 24.88086 | 29.70472 | 12.55361 | 0.181835 | 0.241881 | 0.794688 |
| X | 120793400 | 120793999 | 14.58658 | 26.44004 | 31.04744 | 13.43142 | 0.182731 | 0.355111 | 0.794692 |
| 14 | 27525800 | 27526199 | 8.45972 | 16.15729 | 25.62521 | 10.18789 | -0.15251 | -0.03157 | 0.794713 |

| | | | | | | | | | |
|---|---|---|---|---|---|---|---|---|---|
| 12 | 20464800 | 20468999 | 145.9567 | 219.2058 | 242.2244 | 99.97784 | 0.124036 | 3.441736 | 0.794717 |
| 6 | 16168200 | 16169999 | 15.84722 | 27.79719 | 39.81917 | 17.60233 | -0.12839 | 0.638503 | 0.794728 |
| 4 | 31368600 | 31368999 | 5.515847 | 10.74655 | 19.10814 | 6.950113 | -0.19414 | -0.38401 | 0.794747 |
| 10 | 110590000 | 110591799 | 25.94017 | 43.45514 | 51.76757 | 22.57007 | 0.129401 | 1.171145 | 0.794757 |
| X | 16336600 | 16338599 | 21.71904 | 37.63787 | 44.17362 | 19.381 | 0.156411 | 0.844216 | 0.79476 |
| Y | 88031600 | 88034199 | 14.88778 | 26.29268 | 26.31512 | 11.64224 | 0.388763 | 0.519416 | 0.794776 |
| 8 | 20729800 | 20730199 | 19.67665 | 33.94602 | 48.47436 | 21.26502 | -0.12685 | 0.878332 | 0.794803 |
| 1 | 43665000 | 43666599 | 22.6174 | 38.87531 | 46.3228 | 20.31888 | 0.132165 | 0.921989 | 0.794803 |
| 6 | 140715200 | 140718799 | 34.07866 | 55.1758 | 78.54382 | 33.60365 | -0.14473 | 1.65828 | 0.79483 |
| 13 | 4511000 | 4516799 | 18.51218 | 31.67546 | 54.88745 | 24.06807 | -0.40704 | 0.956158 | 0.794853 |
| 3 | 108805800 | 108807999 | 16.13272 | 28.62685 | 33.75332 | 14.91048 | 0.156318 | 0.494765 | 0.794868 |
| 10 | 83259200 | 83281199 | 638.264 | 851.223 | 850.6433 | 355.2961 | 0.100379 | 5.322343 | 0.794877 |
| 12 | 20974000 | 20982999 | 98.16717 | 150.8195 | 199.5057 | 82.83863 | -0.10831 | 2.971717 | 0.794896 |
| Y | 2521400 | 2522799 | 11.74445 | 21.49978 | 21.73846 | 9.287647 | 0.408581 | 0.251112 | 0.794904 |
| 1 | 182713200 | 182720399 | 139.7581 | 209.8317 | 271.2175 | 111.9355 | -0.10237 | 3.448197 | 0.794905 |
| 1 | 106954600 | 106955599 | 13.76142 | 24.79688 | 29.96373 | 13.05907 | 0.134569 | 0.378533 | 0.794909 |
| 2 | 50396400 | 50399599 | 25.1882 | 43.37149 | 48.46744 | 21.26075 | 0.224838 | 0.918036 | 0.794916 |
| 12 | 67266200 | 67269799 | 30.18084 | 48.70409 | 92.35124 | 39.36737 | -0.56248 | 1.74653 | 0.794959 |
| 5 | 91834200 | 91837799 | 34.4035 | 55.90991 | 78.01133 | 33.46775 | -0.11091 | 1.584464 | 0.794974 |
| 15 | 35717800 | 35720599 | 61.18646 | 97.63594 | 111.1881 | 46.6354 | 0.152407 | 2.139133 | 0.794991 |
| 7 | 85103000 | 85112799 | 312.4843 | 438.8032 | 536.6409 | 222.3729 | -0.11596 | 4.516942 | 0.795008 |
| 5 | 89153000 | 89155999 | 54.92331 | 87.27118 | 120.7778 | 50.37732 | -0.13482 | 2.268295 | 0.795027 |
| 8 | 118570400 | 118572799 | 19.81303 | 34.17493 | 49.01497 | 21.50206 | -0.13322 | 0.880646 | 0.795028 |
| 16 | 86577600 | 86580399 | 28.18575 | 47.70841 | 53.51479 | 23.42647 | 0.216093 | 1.083094 | 0.795039 |
| 1 | 102672400 | 102673199 | 10.07179 | 18.75118 | 28.35623 | 11.83458 | -0.14356 | 0.159985 | 0.795051 |
| 12 | 99339400 | 99343599 | 37.68207 | 61.12779 | 72.11545 | 30.89729 | 0.128142 | 1.621109 | 0.795052 |
| 18 | 3469600 | 3473999 | 58.65876 | 93.46536 | 110.1587 | 46.10981 | 0.100644 | 2.180189 | 0.795109 |
| 10 | 42923400 | 42930799 | 61.70582 | 98.27001 | 114.3094 | 47.81324 | 0.118761 | 2.199423 | 0.795128 |
| 6 | 105448800 | 105449399 | 8.841432 | 16.68874 | 25.94475 | 10.57493 | -0.1551 | 0.056123 | 0.795158 |
| 4 | 136947800 | 136952399 | 51.42452 | 82.4109 | 97.42704 | 41.11661 | 0.105395 | 1.991596 | 0.795175 |
| 7 | 60503800 | 60506199 | 27.34199 | 46.30076 | 53.45659 | 23.40414 | 0.174687 | 1.078283 | 0.795207 |
| 11 | 23870200 | 23871399 | 16.01378 | 28.10107 | 40.30755 | 17.82451 | -0.13031 | 0.633473 | 0.795218 |
| X | 26836800 | 26837399 | 10.36917 | 19.83392 | 25.10815 | 9.944551 | 0.175263 | -0.02512 | 0.795231 |
| 15 | 95573000 | 95574399 | 18.07317 | 31.33157 | 44.75944 | 19.63825 | -0.12586 | 0.823905 | 0.795252 |
| 12 | 91042600 | 91046999 | 55.35061 | 87.87573 | 121.3279 | 50.56392 | -0.13262 | 2.292242 | 0.795262 |
| 15 | 14318400 | 14319199 | 10.6628 | 19.94936 | 24.72876 | 10.23504 | 0.151122 | 0.125571 | 0.795327 |
| 7 | 105987800 | 105988399 | 8.612323 | 16.61133 | 21.99714 | 8.506378 | 0.143363 | -0.12818 | 0.795334 |
| 8 | 91318000 | 91318399 | 8.612323 | 16.61133 | 21.99714 | 8.506378 | 0.143363 | -0.12818 | 0.795334 |
| 2 | 12612600 | 12614599 | 22.52748 | 37.72283 | 64.70509 | 28.20344 | -0.3948 | 1.177451 | 0.795374 |
| 8 | 20140600 | 20144599 | 90.43809 | 140.3226 | 161.8638 | 67.22247 | 0.09998 | 2.782842 | 0.795379 |
| 3 | 92576000 | 92576599 | 7.582481 | 14.55708 | 23.56396 | 9.234844 | -0.16182 | -0.0911 | 0.79539 |
| X | 74344400 | 74347599 | 51.68341 | 82.94806 | 94.27167 | 39.87432 | 0.164719 | 1.949225 | 0.795405 |
| 1 | 117078200 | 117081799 | 23.30294 | 39.8339 | 47.40038 | 20.78337 | 0.132761 | 0.973142 | 0.795417 |
| 4 | 117552600 | 117555799 | 24.46349 | 41.38816 | 49.58418 | 21.70267 | 0.121611 | 1.073806 | 0.79543 |
| 1 | 19431200 | 19432399 | 15.78064 | 27.91915 | 33.46194 | 14.79885 | 0.131205 | 0.522516 | 0.795437 |
| X | 153078600 | 153086399 | 97.47987 | 152.6641 | 161.657 | 67.05623 | 0.233731 | 2.628258 | 0.795452 |
| 6 | 116498200 | 116500999 | 29.38175 | 48.41481 | 68.24916 | 29.44151 | -0.11441 | 1.38027 | 0.795472 |

| | | | | | | | | | |
|---|---|---|---|---|---|---|---|---|---|
| 11 | 30732800 | 30736399 | 32.07414 | 52.47637 | 73.77257 | 31.72942 | -0.11479 | 1.479661 | 0.795482 |
| 14 | 6896600 | 6900999 | 52.18381 | 83.19824 | 113.3246 | 47.47438 | -0.10682 | 2.156477 | 0.795523 |
| 2 | 107877200 | 107880199 | 50.25473 | 80.67356 | 95.0212 | 40.20533 | 0.113616 | 1.936511 | 0.795534 |
| X | 17847400 | 17850799 | 27.51629 | 46.88202 | 51.75175 | 22.68529 | 0.240155 | 1.028536 | 0.795544 |
| 10 | 55502200 | 55513399 | 315.119 | 442.0106 | 541.0916 | 224.2561 | -0.11898 | 4.533526 | 0.795556 |
| 11 | 81689400 | 81691999 | 17.76502 | 30.9263 | 44.16278 | 19.41382 | -0.1245 | 0.768757 | 0.795572 |
| 12 | 79524000 | 79525399 | 11.88193 | 21.50007 | 31.65047 | 13.72212 | -0.14367 | 0.344297 | 0.795573 |
| 17 | 35654800 | 35655199 | 3.981403 | 7.785933 | 15.53224 | 5.28765 | -0.27028 | -0.65494 | 0.795586 |
| 17 | 77577600 | 77578999 | 12.95926 | 24.06566 | 28.69584 | 12.02368 | 0.192165 | 0.198181 | 0.795588 |
| 19 | 37001800 | 37004399 | 29.52514 | 48.16673 | 70.13631 | 30.13375 | -0.16819 | 1.514421 | 0.795594 |
| 13 | 4863800 | 4866399 | 23.34157 | 40.2259 | 46.84425 | 20.54925 | 0.165942 | 0.903 | 0.795601 |
| 9 | 12772800 | 12775399 | 33.32556 | 55.04404 | 54.67228 | 23.61418 | 0.390215 | 1.351618 | 0.795602 |
| 1 | 29610000 | 29610599 | 7.747291 | 14.94642 | 24.25068 | 9.371424 | -0.14611 | -0.13223 | 0.795617 |
| 13 | 113623800 | 113625599 | 22.81732 | 39.28725 | 46.29445 | 20.30751 | 0.148601 | 0.905386 | 0.79562 |
| 7 | 139011000 | 139013199 | 23.43581 | 40.83963 | 44.55417 | 19.59834 | 0.262326 | 0.783019 | 0.79563 |
| 17 | 69064800 | 69066199 | 13.00195 | 23.2091 | 34.05834 | 14.96995 | -0.15252 | 0.438455 | 0.795632 |
| 10 | 82425400 | 82426599 | 15.33705 | 27.18012 | 32.34375 | 14.2995 | 0.142236 | 0.497274 | 0.795639 |
| 17 | 68511000 | 68512799 | 14.61205 | 25.806 | 37.58341 | 16.62748 | -0.14966 | 0.539769 | 0.795639 |
| 10 | 24389600 | 24393399 | 61.29176 | 97.67654 | 113.723 | 47.61242 | 0.118495 | 2.178357 | 0.795647 |
| 5 | 72791600 | 72795599 | 34.72092 | 56.81778 | 67.95369 | 29.2288 | 0.117962 | 1.477266 | 0.795721 |
| 8 | 128606600 | 128607799 | 12.4412 | 22.72039 | 27.57593 | 11.81865 | 0.1439 | 0.286838 | 0.795723 |
| 12 | 114774600 | 114815399 | 578.2488 | 783.9673 | 765.4463 | 318.9345 | 0.157693 | 5.062358 | 0.795731 |
| 2 | 173014600 | 173019199 | 38.0196 | 61.36008 | 85.35129 | 36.38822 | -0.11646 | 1.752028 | 0.795739 |
| 18 | 74167600 | 74169999 | 38.56195 | 62.10743 | 87.45378 | 37.18208 | -0.13754 | 1.816219 | 0.795759 |
| 8 | 86076200 | 86077199 | 8.475012 | 15.97639 | 25.90618 | 10.58607 | -0.21906 | 0.058367 | 0.795766 |
| 6 | 11813800 | 11815599 | 21.04558 | 36.16816 | 52.27005 | 22.92745 | -0.14524 | 0.923694 | 0.795831 |
| 9 | 103340200 | 103341999 | 15.19835 | 26.81817 | 38.66238 | 17.1078 | -0.13541 | 0.556915 | 0.795835 |
| 1 | 133442000 | 133446399 | 40.58219 | 65.63955 | 78.26338 | 33.45889 | 0.108976 | 1.690355 | 0.795851 |
| 5 | 77367200 | 77369399 | 32.40071 | 53.30432 | 63.99762 | 27.58885 | 0.115602 | 1.403361 | 0.795863 |
| 9 | 122265600 | 122272999 | 61.94931 | 97.61504 | 154.8322 | 64.24802 | -0.34167 | 2.488856 | 0.795878 |
| 17 | 43824800 | 43826799 | 27.57229 | 46.63461 | 54.00175 | 23.63419 | 0.170397 | 1.090704 | 0.795899 |
| 13 | 103101200 | 103103199 | 24.83676 | 42.67861 | 40.91038 | 17.94464 | 0.444656 | 0.924898 | 0.795911 |
| 9 | 35940800 | 35950799 | 121.7596 | 187.0576 | 202.6071 | 84.1259 | 0.179784 | 2.983013 | 0.795936 |
| 3 | 90948800 | 90951999 | 56.441 | 89.8049 | 121.406 | 50.71862 | -0.09912 | 2.226561 | 0.795989 |
| 10 | 126177400 | 126180799 | 51.38147 | 82.29943 | 97.28674 | 41.0252 | 0.104637 | 2.006831 | 0.796029 |
| 18 | 46143000 | 46143599 | 6.39118 | 12.40655 | 21.12535 | 7.935479 | -0.17546 | -0.25181 | 0.796087 |
| 3 | 103111000 | 103112799 | 17.07665 | 29.76676 | 42.56455 | 18.7266 | -0.12627 | 0.743923 | 0.796105 |
| X | 42115400 | 42116399 | 18.26874 | 32.32925 | 27.93785 | 12.35514 | 0.600933 | 0.526599 | 0.796121 |
| 6 | 7099200 | 7108599 | 152.4921 | 228.8446 | 257.8122 | 106.3897 | 0.095174 | 3.458432 | 0.796144 |
| 4 | 12342200 | 12343199 | 20.29331 | 35.10684 | 42.11674 | 18.47758 | 0.124383 | 0.849104 | 0.796149 |
| 5 | 87029600 | 87031599 | 34.905 | 56.6211 | 78.92508 | 33.81406 | -0.11194 | 1.622748 | 0.796154 |
| 9 | 23751600 | 23752199 | 11.23359 | 21.03129 | 26.22568 | 10.84235 | 0.143529 | 0.128473 | 0.796178 |
| 6 | 68728400 | 68732199 | 46.6537 | 74.84722 | 103.2197 | 43.63381 | -0.11455 | 1.952862 | 0.796201 |
| X | 114368400 | 114369999 | 21.06672 | 36.47003 | 43.15493 | 18.93346 | 0.144394 | 0.849454 | 0.796218 |
| 2 | 39188600 | 39190799 | 22.88149 | 39.0646 | 46.88523 | 20.55542 | 0.120235 | 0.983318 | 0.796223 |
| 12 | 14771600 | 14772599 | 11.27025 | 20.72154 | 30.82147 | 13.09068 | -0.13832 | 0.24805 | 0.796256 |
| 9 | 92522600 | 92523599 | 17.25588 | 30.30343 | 35.97078 | 15.91682 | 0.141247 | 0.618502 | 0.796257 |

| | | | | | | | | | |
|---|---|---|---|---|---|---|---|---|---|
| 18 | 85754400 | 85755199 | 10.14926 | 19.17547 | 24.28365 | 9.878111 | 0.140757 | 0.056058 | 0.796286 |
| 5 | 128992200 | 128994999 | 21.11803 | 35.98303 | 51.70184 | 22.66353 | -0.13891 | 0.998603 | 0.796289 |
| 13 | 77056800 | 77057999 | 16.96243 | 30.05962 | 26.94192 | 11.91137 | 0.549143 | 0.499389 | 0.796293 |
| 11 | 114306200 | 114308199 | 23.12305 | 38.98093 | 55.43416 | 24.23691 | -0.12503 | 1.107138 | 0.796304 |
| X | 103666200 | 103667799 | 13.1826 | 23.26092 | 41.36975 | 18.30585 | -0.43668 | 0.55053 | 0.796318 |
| 8 | 29083200 | 29086599 | 27.4272 | 45.34578 | 64.18284 | 27.74063 | -0.11914 | 1.333208 | 0.796326 |
| 8 | 19637200 | 19642199 | 84.30638 | 132.2726 | 151.6594 | 62.89744 | 0.121134 | 2.585972 | 0.796331 |
| X | 102163200 | 102164199 | 9.510023 | 18.38594 | 19.26701 | 7.377534 | 0.491536 | -0.1592 | 0.796348 |
| 13 | 96843000 | 96846599 | 32.90085 | 53.10785 | 91.85124 | 39.16789 | -0.42968 | 1.743012 | 0.796385 |
| 11 | 108650800 | 108655599 | 41.82675 | 68.35323 | 75.05536 | 32.2675 | 0.239562 | 1.518595 | 0.796399 |
| 13 | 45637000 | 45639399 | 31.08635 | 51.3257 | 61.75392 | 26.66578 | 0.11434 | 1.355256 | 0.7964 |
| 8 | 48177400 | 48179599 | 19.07308 | 33.07512 | 48.48835 | 21.27451 | -0.16319 | 0.82739 | 0.796417 |
| 1 | 86245000 | 86251399 | 70.77402 | 111.7743 | 131.0566 | 54.4219 | 0.096243 | 2.439597 | 0.796441 |
| 5 | 92230800 | 92231799 | 9.78642 | 18.11085 | 27.81004 | 11.70751 | -0.17464 | 0.191457 | 0.796445 |
| 4 | 145986600 | 146025399 | 1490.121 | 1851.562 | 1984.641 | 835.2556 | -0.10318 | 6.468497 | 0.796515 |
| 5 | 73369400 | 73371999 | 21.12583 | 35.73228 | 51.65766 | 22.60847 | -0.14844 | 1.066995 | 0.796527 |
| Y | 1897400 | 1900599 | 31.09717 | 50.98827 | 50.2573 | 21.63475 | 0.397494 | 1.429203 | 0.79653 |
| 10 | 42829200 | 42831999 | 33.75666 | 54.84723 | 76.67753 | 32.89046 | -0.11397 | 1.58684 | 0.796533 |
| 16 | 69758000 | 69763999 | 93.56102 | 144.1597 | 189.8948 | 78.87931 | -0.09904 | 2.913878 | 0.796549 |
| 10 | 48264000 | 48264999 | 19.58677 | 33.95142 | 40.55331 | 17.79274 | 0.13121 | 0.825497 | 0.796553 |
| 3 | 95466400 | 95467599 | 9.306938 | 17.00103 | 32.9235 | 14.10587 | -0.52519 | 0.275038 | 0.796561 |
| 14 | 107926600 | 107929799 | 42.13988 | 67.74209 | 93.50185 | 39.64496 | -0.11221 | 1.882407 | 0.796562 |
| 1 | 28346800 | 28348599 | 16.25402 | 29.19405 | 33.26539 | 14.54544 | 0.215374 | 0.419439 | 0.796566 |
| 19 | 12887800 | 12893999 | 83.44811 | 130.7243 | 151.2237 | 62.72666 | 0.106585 | 2.611436 | 0.79657 |
| 3 | 4974200 | 4981199 | 116.4927 | 179.2799 | 171.4174 | 71.1925 | 0.36143 | 2.942779 | 0.796576 |
| 17 | 86302600 | 86308399 | 35.27724 | 58.80168 | 51.46968 | 22.36946 | 0.572716 | 1.252074 | 0.796579 |
| 1 | 171371600 | 171373199 | 25.54717 | 43.56963 | 50.12707 | 21.97484 | 0.180663 | 1.004582 | 0.796597 |
| 14 | 118443600 | 118445399 | 17.39979 | 30.37543 | 43.5603 | 19.18271 | -0.13025 | 0.725647 | 0.796607 |
| X | 151686800 | 151691199 | 30.73427 | 50.83856 | 75.63947 | 32.70349 | -0.19101 | 1.344012 | 0.796631 |
| 5 | 148353200 | 148358399 | 53.10795 | 84.78451 | 115.651 | 48.53934 | -0.10684 | 2.115258 | 0.796644 |
| 16 | 90872400 | 90877799 | 55.07789 | 87.73809 | 119.0496 | 49.81437 | -0.10262 | 2.186722 | 0.796647 |
| 12 | 3108200 | 3108999 | 29.85671 | 49.29536 | 58.20626 | 25.13383 | 0.141478 | 1.35146 | 0.796655 |
| 13 | 24430400 | 24432799 | 17.08914 | 29.75185 | 42.60558 | 18.73014 | -0.12851 | 0.761823 | 0.796661 |
| 10 | 88132000 | 88135999 | 42.81892 | 68.96657 | 95.74153 | 40.70822 | -0.11708 | 1.824348 | 0.796664 |
| 17 | 47662400 | 47668199 | 64.39358 | 101.923 | 136.9115 | 56.91692 | -0.0971 | 2.384441 | 0.796673 |
| 4 | 40647200 | 40651599 | 42.39079 | 68.20745 | 94.35034 | 40.05749 | -0.11378 | 1.852927 | 0.796676 |
| X | 101121000 | 101124399 | 21.40324 | 36.56258 | 53.81713 | 23.59564 | -0.17281 | 0.98456 | 0.796679 |
| 4 | 36356600 | 36357399 | 8.883321 | 16.54817 | 26.39529 | 11.00531 | -0.21825 | 0.142839 | 0.796684 |
| 15 | 38849800 | 38856999 | 92.09682 | 141.3749 | 217.059 | 90.11675 | -0.32398 | 2.988913 | 0.796698 |
| 5 | 146084800 | 146090399 | 80.3308 | 125.9073 | 146.7787 | 60.8796 | 0.096072 | 2.600217 | 0.796708 |
| 7 | 6252600 | 6256999 | 52.80716 | 84.05416 | 115.5842 | 48.31127 | -0.12306 | 2.209488 | 0.796711 |
| 11 | 48842200 | 48842799 | 5.948403 | 11.58124 | 20.16841 | 7.39718 | -0.17522 | -0.34335 | 0.796731 |
| X | 57676000 | 57679399 | 41.86626 | 68.35996 | 75.49635 | 32.44708 | 0.230428 | 1.531369 | 0.796763 |
| 1 | 118024200 | 118026599 | 29.82907 | 49.13287 | 70.3626 | 30.34824 | -0.13728 | 1.390269 | 0.796766 |
| 8 | 129849800 | 129851599 | 30.59611 | 50.468 | 60.54825 | 26.13091 | 0.118175 | 1.364454 | 0.796768 |
| 13 | 14394200 | 14397199 | 41.97543 | 67.8194 | 80.35127 | 34.30546 | 0.11636 | 1.725551 | 0.796786 |
| 10 | 10415800 | 10419199 | 35.26655 | 58.3639 | 65.23109 | 28.21278 | 0.220587 | 1.34038 | 0.796799 |

| | | | | | | | | | |
|---|---|---|---|---|---|---|---|---|---|
| 7 | 46155600 | 46157399 | 16.23475 | 28.4051 | 40.89195 | 18.03803 | -0.13563 | 0.680858 | 0.796817 |
| 5 | 93853600 | 93862399 | 147.8563 | 225.1204 | 195.3521 | 80.95948 | 0.489219 | 3.166672 | 0.796822 |
| 4 | 73508400 | 73512199 | 28.49938 | 48.20429 | 54.61036 | 23.9008 | 0.201848 | 1.093245 | 0.796839 |
| 1 | 101138200 | 101138999 | 10.92871 | 20.51936 | 25.75243 | 10.59503 | 0.139998 | 0.106797 | 0.796845 |
| 8 | 34606600 | 34608799 | 20.33786 | 34.82516 | 49.5464 | 21.72732 | -0.12355 | 0.94954 | 0.796852 |
| 13 | 120759600 | 120763399 | 33.87395 | 55.13909 | 77.05442 | 33.07938 | -0.11142 | 1.557579 | 0.796853 |
| 10 | 50336000 | 50338199 | 16.49319 | 29.62067 | 28.18415 | 12.32468 | 0.474137 | 0.414009 | 0.796857 |
| 3 | 144195800 | 144196999 | 10.32196 | 18.94429 | 28.80633 | 12.26444 | -0.1722 | 0.246355 | 0.796872 |
| 6 | 105552800 | 105553599 | 13.15467 | 23.92585 | 29.06413 | 12.51998 | 0.137989 | 0.319664 | 0.796877 |
| 3 | 31621200 | 31626799 | 84.7648 | 133.3589 | 129.5901 | 53.7629 | 0.362562 | 2.52775 | 0.796887 |
| 1 | 93846600 | 93849599 | 25.39608 | 42.57515 | 50.16915 | 21.88339 | 0.145044 | 1.159088 | 0.796901 |
| 1 | 84429400 | 84429999 | 9.701706 | 17.91176 | 27.86419 | 11.76804 | -0.19624 | 0.20636 | 0.796901 |
| 6 | 4967800 | 4969599 | 16.84001 | 29.6756 | 35.42673 | 15.68021 | 0.134532 | 0.573958 | 0.79692 |
| 4 | 121521000 | 121527599 | 95.6275 | 149.7719 | 160.886 | 66.73598 | 0.213113 | 2.624318 | 0.796925 |
| 15 | 100119400 | 100123999 | 69.64241 | 110.0686 | 129.1138 | 53.63526 | 0.096462 | 2.421425 | 0.796927 |
| 2 | 71622200 | 71624399 | 21.05107 | 36.35217 | 43.65601 | 19.15184 | 0.122274 | 0.869348 | 0.796932 |
| 8 | 111991600 | 111994799 | 41.1784 | 66.41564 | 92.25799 | 39.30039 | -0.1156 | 1.779477 | 0.796934 |
| 10 | 76293800 | 76297999 | 67.4367 | 106.7141 | 125.0927 | 51.99729 | 0.098873 | 2.389486 | 0.796973 |
| 2 | 49137600 | 49138599 | 9.819662 | 18.16902 | 27.8334 | 11.72028 | -0.17134 | 0.193939 | 0.796984 |
| 18 | 75774000 | 75779199 | 45.20299 | 72.05885 | 122.392 | 51.26342 | -0.4249 | 2.156042 | 0.797002 |
| 9 | 31982200 | 31983399 | 11.42356 | 20.77722 | 30.71224 | 13.22999 | -0.1434 | 0.310095 | 0.797049 |
| X | 8446200 | 8499999 | 813.1169 | 1074.202 | 990.2406 | 414.1071 | 0.203166 | 5.476639 | 0.797053 |
| 15 | 97454800 | 97464999 | 164.0224 | 243.6945 | 311.2925 | 128.2399 | -0.09911 | 3.645216 | 0.797054 |
| 1 | 161568600 | 161570599 | 12.70734 | 22.54461 | 34.08578 | 15.05962 | -0.20049 | 0.474557 | 0.797064 |
| 8 | 4725200 | 4730199 | 68.45811 | 107.6405 | 145.1966 | 60.23292 | -0.11062 | 2.532618 | 0.797115 |
| 3 | 99814400 | 99816999 | 40.14665 | 64.83027 | 90.23534 | 38.49906 | -0.11646 | 1.74088 | 0.79712 |
| 7 | 8344200 | 8346199 | 53.47959 | 85.37124 | 98.27351 | 41.2414 | 0.137536 | 2.114814 | 0.797138 |
| 17 | 53423600 | 53424799 | 28.12978 | 46.2278 | 66.15186 | 28.50444 | -0.13747 | 1.409745 | 0.797141 |
| 3 | 46746400 | 46747599 | 10.77247 | 19.92156 | 30.88002 | 13.01243 | -0.18981 | 0.207388 | 0.797143 |
| 4 | 9015600 | 9016999 | 11.15114 | 20.05433 | 30.75357 | 13.43264 | -0.20901 | 0.373746 | 0.797156 |
| 18 | 15355000 | 15359799 | 56.00528 | 89.35397 | 105.3876 | 44.19252 | 0.101642 | 2.134421 | 0.797156 |
| 4 | 148061600 | 148064399 | 25.37843 | 42.34346 | 59.87285 | 26.03772 | -0.11699 | 1.219667 | 0.79716 |
| Y | 9640400 | 9641399 | 10.46118 | 19.50242 | 20.25623 | 8.434713 | 0.397185 | 0.131913 | 0.797164 |
| 9 | 20625400 | 20626999 | 18.66154 | 32.174 | 46.37107 | 20.34194 | -0.14011 | 0.87996 | 0.797164 |
| 14 | 42225400 | 42230799 | 75.85381 | 119.1475 | 138.647 | 57.50827 | 0.101098 | 2.556488 | 0.797165 |
| 1 | 16288000 | 16290799 | 20.95284 | 36.22753 | 43.09158 | 18.90494 | 0.13632 | 0.855666 | 0.797172 |
| 8 | 107171200 | 107174399 | 36.15094 | 59.07677 | 69.85303 | 30.03073 | 0.132979 | 1.500081 | 0.79719 |
| 7 | 35565800 | 35571399 | 69.92085 | 110.1583 | 147.2662 | 61.1072 | -0.09551 | 2.495028 | 0.7972 |
| 8 | 120644400 | 120645799 | 11.89181 | 21.60272 | 31.68896 | 13.67141 | -0.13395 | 0.32226 | 0.797206 |
| 8 | 60734600 | 60735399 | 8.898913 | 16.60972 | 26.4144 | 10.97681 | -0.20997 | 0.131347 | 0.79721 |
| Y | 38486200 | 38489399 | 16.82246 | 29.41358 | 29.99276 | 13.22272 | 0.359643 | 0.676058 | 0.797228 |
| 8 | 3985800 | 3996799 | 136.3059 | 206.5451 | 233.8252 | 96.76462 | 0.100283 | 3.262473 | 0.797229 |
| 4 | 13666800 | 13669599 | 39.67763 | 64.07049 | 88.80866 | 37.88854 | -0.11053 | 1.741139 | 0.797248 |
| 3 | 114217200 | 114218799 | 18.03469 | 31.68541 | 37.5042 | 16.60168 | 0.145429 | 0.620021 | 0.797251 |
| 12 | 44110400 | 44111799 | 13.71797 | 24.32596 | 35.0809 | 15.50554 | -0.13294 | 0.488138 | 0.797255 |
| 13 | 3345000 | 3351399 | 77.2851 | 122.0372 | 121.8247 | 50.58411 | 0.327851 | 2.445868 | 0.797258 |
| 11 | 10364400 | 10365999 | 18.51312 | 31.99258 | 45.81348 | 20.09876 | -0.13034 | 0.854461 | 0.797266 |

| | | | | | | | | | |
|---|---|---|---|---|---|---|---|---|---|
| 7 | 124429200 | 124433799 | 50.98341 | 81.72357 | 96.01231 | 40.53433 | 0.114787 | 1.981538 | 0.797269 |
| 5 | 88481600 | 88484399 | 24.14924 | 40.875 | 59.47047 | 26.03341 | -0.15782 | 1.077168 | 0.797282 |
| 17 | 60235600 | 60236599 | 11.5389 | 21.54181 | 26.69664 | 11.0901 | 0.146793 | 0.149843 | 0.797283 |
| X | 26121200 | 26123999 | 42.59711 | 69.48391 | 63.15412 | 27.13272 | 0.509775 | 1.539727 | 0.797297 |
| 13 | 26026400 | 26027999 | 14.86088 | 27.09045 | 31.59021 | 13.56963 | 0.199163 | 0.324847 | 0.797344 |
| 3 | 94043400 | 94046799 | 58.09216 | 92.79516 | 107.7211 | 45.24809 | 0.126401 | 2.099812 | 0.797353 |
| 8 | 22629400 | 22631399 | 23.74097 | 39.78739 | 56.53076 | 24.65316 | -0.12396 | 1.164621 | 0.797355 |
| 5 | 145861600 | 145876799 | 424.8996 | 588.4167 | 605.6277 | 251.5043 | 0.112532 | 4.730181 | 0.797375 |
| 14 | 57963200 | 57965399 | 25.24894 | 42.61313 | 51.21074 | 22.39806 | 0.116964 | 1.098281 | 0.797452 |
| 3 | 21571400 | 21575199 | 51.07645 | 82.17905 | 80.84076 | 34.31957 | 0.376949 | 1.862806 | 0.797467 |
| X | 102955600 | 102956199 | 9.560616 | 17.9656 | 28.01379 | 11.50918 | -0.16861 | 0.098962 | 0.797469 |
| 1 | 95089400 | 95090199 | 10.29607 | 19.52774 | 24.87393 | 10.03631 | 0.142587 | 0.03223 | 0.797473 |
| 14 | 3336200 | 3336799 | 10.29607 | 19.52774 | 24.87393 | 10.03631 | 0.142587 | 0.03223 | 0.797473 |
| 16 | 54844000 | 54846799 | 52.1515 | 83.7674 | 83.57494 | 35.39365 | 0.353619 | 1.91667 | 0.797478 |
| 2 | 45990000 | 45990399 | 6.392971 | 12.44308 | 21.30702 | 7.833625 | -0.15488 | -0.32635 | 0.797479 |
| 9 | 17558200 | 17559199 | 12.35605 | 21.84682 | 39.80652 | 17.60698 | -0.47026 | 0.519172 | 0.797527 |
| 18 | 89328800 | 89334599 | 101.8104 | 156.783 | 180.454 | 74.95204 | 0.09482 | 2.926617 | 0.79754 |
| 5 | 5832000 | 5836599 | 92.8674 | 143.6461 | 163.848 | 68.05849 | 0.112722 | 2.839268 | 0.797553 |
| 7 | 31878600 | 31880999 | 32.81635 | 55.42177 | 45.77633 | 20.02526 | 0.656107 | 1.113822 | 0.79757 |
| 12 | 105269400 | 105272199 | 32.85373 | 54.32335 | 51.92181 | 22.44291 | 0.445644 | 1.33662 | 0.797577 |
| 2 | 174649000 | 174651999 | 28.40411 | 46.6582 | 66.84561 | 28.79804 | -0.13945 | 1.414574 | 0.79758 |
| 13 | 61370000 | 61371799 | 33.86122 | 55.88921 | 65.28977 | 28.18632 | 0.156771 | 1.370804 | 0.797583 |
| 18 | 30901200 | 30901799 | 6.235351 | 12.12991 | 20.69531 | 7.641114 | -0.15566 | -0.31722 | 0.797591 |
| 4 | 14345000 | 14346999 | 20.78556 | 35.90255 | 43.15675 | 18.93296 | 0.12106 | 0.866404 | 0.797597 |
| 10 | 54692200 | 54693199 | 10.2951 | 19.36273 | 24.23019 | 9.939985 | 0.147643 | 0.086787 | 0.797601 |
| 6 | 143754000 | 143758599 | 37.52541 | 62.04344 | 67.52866 | 29.18743 | 0.258719 | 1.362023 | 0.797626 |
| 9 | 70370400 | 70370799 | 7.658382 | 14.85815 | 19.969 | 7.522016 | 0.157908 | -0.23426 | 0.797643 |
| 13 | 60633600 | 60636199 | 23.66537 | 40.18465 | 47.96528 | 21.0174 | 0.127009 | 1.035501 | 0.79766 |
| 13 | 66115400 | 66124199 | 252.6168 | 366.4158 | 333.4296 | 137.4718 | 0.357057 | 4.037307 | 0.797661 |
| 14 | 102554200 | 102555799 | 15.26583 | 27.57222 | 32.44207 | 14.09783 | 0.175398 | 0.380795 | 0.797673 |
| 11 | 62844200 | 62850399 | 72.89711 | 115.1078 | 134.5018 | 55.84779 | 0.100996 | 2.445361 | 0.797695 |
| 11 | 35263000 | 35264599 | 20.4634 | 35.7841 | 41.26134 | 18.19253 | 0.183058 | 0.712725 | 0.797725 |
| 3 | 138468800 | 138469999 | 16.85716 | 29.52801 | 34.75927 | 15.34771 | 0.153497 | 0.657123 | 0.797731 |
| 2 | 149126200 | 149130399 | 73.55287 | 116.063 | 135.0842 | 56.06999 | 0.105898 | 2.463948 | 0.797758 |
| 1 | 188606600 | 188607199 | 6.532547 | 12.63688 | 21.57198 | 8.237681 | -0.20225 | -0.19516 | 0.797762 |
| 4 | 115310400 | 115311999 | 15.64154 | 27.44293 | 39.28035 | 17.36764 | -0.1273 | 0.63171 | 0.797772 |
| 13 | 62508600 | 62548199 | 739.5139 | 981.5012 | 947.2071 | 395.8849 | 0.143734 | 5.401153 | 0.79778 |
| 10 | 83530200 | 83532999 | 27.84959 | 46.49402 | 55.66331 | 24.21609 | 0.122262 | 1.21558 | 0.797788 |
| 11 | 7987000 | 7988999 | 27.93803 | 46.2336 | 65.36601 | 28.27027 | -0.11742 | 1.325216 | 0.797789 |
| 3 | 85894400 | 85896399 | 17.10113 | 29.97749 | 43.97211 | 19.42769 | -0.16263 | 0.660419 | 0.797791 |
| 13 | 92755800 | 92766999 | 138.6582 | 209.5394 | 237.8387 | 98.34121 | 0.093161 | 3.317891 | 0.797798 |
| 4 | 87475000 | 87478399 | 33.10972 | 53.99861 | 75.80637 | 32.56636 | -0.116 | 1.529504 | 0.797805 |
| 6 | 143368400 | 143370599 | 18.76427 | 31.83316 | 56.39064 | 24.70234 | -0.44033 | 1.037962 | 0.797811 |
| 12 | 47977600 | 47983599 | 88.32437 | 137.4854 | 159.6496 | 66.27098 | 0.093997 | 2.726284 | 0.797812 |
| 6 | 78070000 | 78073599 | 59.45575 | 94.52915 | 109.9799 | 45.90068 | 0.115962 | 2.249616 | 0.797812 |
| 8 | 94192000 | 94198199 | 169.7153 | 252.5521 | 281.0595 | 115.8038 | 0.10178 | 3.617192 | 0.797813 |
| 5 | 57804800 | 57805599 | 14.6723 | 25.84171 | 37.16967 | 16.45342 | -0.13316 | 0.570895 | 0.797829 |

| | | | | | | | | | |
|---|---|---|---|---|---|---|---|---|---|
| 1 | 134832000 | 134834799 | 27.70911 | 46.56757 | 55.10541 | 24.07054 | 0.138898 | 1.1461 | 0.797843 |
| 11 | 25705600 | 25707599 | 16.72631 | 29.30044 | 42.39714 | 18.72104 | -0.14305 | 0.666283 | 0.797844 |
| 14 | 95982000 | 95983399 | 14.89958 | 27.22138 | 31.58289 | 13.52886 | 0.208798 | 0.310893 | 0.797873 |
| 2 | 107748800 | 107752399 | 47.61197 | 76.20586 | 104.493 | 44.01606 | -0.11023 | 2.029524 | 0.797875 |
| X | 13793400 | 13796399 | 27.91352 | 47.22233 | 53.96373 | 23.61916 | 0.189509 | 1.092416 | 0.797893 |
| Y | 4688000 | 4691999 | 31.41461 | 51.45865 | 50.78437 | 21.85264 | 0.39513 | 1.442034 | 0.797895 |
| 12 | 68766800 | 68767399 | 6.232671 | 12.13313 | 20.74993 | 7.618363 | -0.1517 | -0.33716 | 0.797896 |
| 2 | 43648000 | 43648799 | 9.223097 | 17.64691 | 22.90524 | 9.064413 | 0.140744 | -0.04509 | 0.7979 |
| 4 | 7349400 | 7351199 | 17.1147 | 30.35148 | 35.32835 | 15.61399 | 0.174498 | 0.509702 | 0.797907 |
| 2 | 173061400 | 173064399 | 23.34546 | 39.12043 | 56.56919 | 24.66828 | -0.14931 | 1.162406 | 0.797918 |
| 3 | 155457600 | 155463199 | 58.27243 | 92.93696 | 109.1406 | 45.743 | 0.107289 | 2.148885 | 0.797922 |
| 4 | 47293000 | 47300999 | 95.69929 | 147.017 | 194.8238 | 80.89378 | -0.11096 | 2.972556 | 0.797927 |
| 6 | 29090400 | 29099799 | 151.0818 | 225.7693 | 289.6352 | 119.416 | -0.09766 | 3.538867 | 0.797955 |
| 10 | 114146200 | 114146999 | 8.878464 | 16.89642 | 26.64724 | 10.6748 | -0.15354 | 0.003025 | 0.797967 |
| 8 | 105124000 | 105126199 | 26.24241 | 43.90098 | 52.32152 | 22.79178 | 0.128781 | 1.186655 | 0.797994 |
| X | 48749600 | 48750199 | 7.465533 | 14.28456 | 23.52526 | 9.308899 | -0.19911 | -0.05937 | 0.798004 |
| 17 | 69962600 | 69964799 | 26.45525 | 44.99675 | 44.20965 | 19.37653 | 0.407096 | 1.0282 | 0.798005 |
| 12 | 110270400 | 110274799 | 60.08257 | 95.26898 | 128.5149 | 53.49207 | -0.10094 | 2.333634 | 0.798016 |
| 1 | 8248000 | 8252599 | 107.2118 | 164.2501 | 185.8242 | 77.12648 | 0.113936 | 3.029351 | 0.79802 |
| 1 | 150020800 | 150022399 | 14.20464 | 25.12337 | 36.11148 | 15.97159 | -0.12976 | 0.520613 | 0.798026 |
| 12 | 80629600 | 80630799 | 12.61532 | 23.02573 | 28.06935 | 12.03838 | 0.136972 | 0.29255 | 0.798032 |
| 6 | 130679600 | 130683199 | 28.99376 | 48.29876 | 74.07386 | 32.18123 | -0.234 | 1.257267 | 0.798056 |
| 4 | 98459800 | 98460799 | 9.728288 | 18.28207 | 27.98708 | 11.49673 | -0.14277 | 0.095908 | 0.798078 |
| 7 | 63277200 | 63278399 | 9.728288 | 18.28207 | 27.98708 | 11.49673 | -0.14277 | 0.095908 | 0.798078 |
| 1 | 177698600 | 177700599 | 22.67526 | 38.40763 | 54.40699 | 23.82202 | -0.11927 | 1.061334 | 0.798104 |
| 9 | 119352600 | 119354199 | 22.89552 | 38.47564 | 54.90179 | 23.98116 | -0.13014 | 1.13135 | 0.798138 |
| 12 | 113787400 | 113798199 | 157.3847 | 237.4544 | 258.3252 | 106.7575 | 0.152355 | 3.35572 | 0.798166 |
| 3 | 113557000 | 113558399 | 12.49043 | 22.80622 | 27.61449 | 11.83782 | 0.147221 | 0.289234 | 0.798169 |
| 15 | 68233800 | 68235599 | 14.84501 | 25.97188 | 38.45166 | 16.96066 | -0.17566 | 0.671844 | 0.798176 |
| 1 | 121791400 | 121792199 | 10.36232 | 18.99074 | 36.16419 | 15.42757 | -0.4973 | 0.269853 | 0.798189 |
| 4 | 3058400 | 3092399 | 2401.198 | 2853.943 | 2953.35 | 1250.335 | -0.11767 | 7.172419 | 0.798194 |
| 6 | 137131800 | 137132599 | 9.322851 | 17.38781 | 26.86448 | 11.17818 | -0.17045 | 0.139567 | 0.798212 |
| 17 | 41393400 | 41396399 | 30.95785 | 52.37988 | 45.2102 | 19.7887 | 0.592826 | 1.089074 | 0.798228 |
| 2 | 70262200 | 70262599 | 6.681397 | 13.02007 | 18.00229 | 6.530264 | 0.168429 | -0.38125 | 0.798235 |
| 14 | 23720400 | 23721799 | 23.77703 | 40.60917 | 48.482 | 21.25613 | 0.127939 | 0.986176 | 0.798258 |
| 6 | 145348600 | 145355199 | 59.1413 | 94.52925 | 108.1923 | 45.48568 | 0.147776 | 2.085168 | 0.798273 |
| 19 | 32424200 | 32433199 | 104.7649 | 161.3792 | 185.4461 | 77.02853 | 0.097449 | 2.92274 | 0.798279 |
| 14 | 82412800 | 82416999 | 41.65678 | 67.22564 | 94.52038 | 40.29384 | -0.1321 | 1.764198 | 0.798293 |
| 5 | 21439200 | 21442199 | 31.07818 | 51.33751 | 61.77748 | 26.68504 | 0.114171 | 1.349461 | 0.798326 |
| 5 | 95765800 | 95766199 | 7.226783 | 14.09672 | 15.22182 | 5.451007 | 0.537259 | -0.43172 | 0.798333 |
| 17 | 12958400 | 12964999 | 118.771 | 178.9725 | 280.2468 | 115.7748 | -0.37425 | 3.373206 | 0.798342 |
| 10 | 26231400 | 26232799 | 15.25315 | 27.03058 | 32.43165 | 14.33845 | 0.130545 | 0.498658 | 0.798361 |
| 13 | 15275200 | 15275999 | 15.25315 | 27.03058 | 32.43165 | 14.33845 | 0.130545 | 0.498658 | 0.798361 |
| 18 | 73262600 | 73266199 | 33.84439 | 55.23819 | 78.01658 | 33.52503 | -0.124 | 1.51949 | 0.798368 |
| 4 | 115502600 | 115504999 | 20.90639 | 35.59911 | 50.60483 | 22.18148 | -0.1235 | 1.005211 | 0.798399 |
| 16 | 72588600 | 72589199 | 6.676396 | 12.96659 | 21.72454 | 8.128149 | -0.14754 | -0.26187 | 0.798409 |
| 1 | 71189400 | 71191599 | 21.59834 | 37.13754 | 55.04527 | 24.14514 | -0.18144 | 0.925079 | 0.798426 |

| | | | | | | | | | |
|---|---|---|---|---|---|---|---|---|---|
| 1 | 183293000 | 183299999 | 113.707 | 173.5428 | 195.7205 | 81.16288 | 0.114177 | 3.108555 | 0.798428 |
| 3 | 159187200 | 159191999 | 63.42861 | 100.5034 | 137.09 | 57.0221 | -0.11796 | 2.356702 | 0.798435 |
| 12 | 56690600 | 56693599 | 37.51659 | 61.06718 | 72.51928 | 31.1322 | 0.122776 | 1.561149 | 0.798445 |
| 2 | 100076800 | 100077999 | 12.22055 | 22.47345 | 27.56942 | 11.70613 | 0.137905 | 0.244977 | 0.798483 |
| 1 | 111721000 | 111721799 | 10.536 | 19.45918 | 29.28736 | 12.36294 | -0.14902 | 0.211311 | 0.798484 |
| 3 | 99768000 | 99770999 | 28.03114 | 46.41627 | 65.50788 | 28.34369 | -0.11485 | 1.316832 | 0.798487 |
| 12 | 74874600 | 74874999 | 10.57934 | 19.8406 | 24.28755 | 10.0118 | 0.174081 | 0.110149 | 0.798494 |
| 16 | 66485400 | 66485999 | 10.57934 | 19.8406 | 24.28755 | 10.0118 | 0.174081 | 0.110149 | 0.798494 |
| 5 | 132929000 | 132930799 | 16.11161 | 28.77757 | 33.87279 | 14.88072 | 0.164414 | 0.456403 | 0.798511 |
| 9 | 34550800 | 34552799 | 27.12069 | 46.09909 | 44.72649 | 19.60205 | 0.425041 | 1.037015 | 0.798521 |
| 11 | 43351800 | 43354599 | 28.26683 | 46.65485 | 65.8654 | 28.43458 | -0.11562 | 1.356562 | 0.798532 |
| X | 77450200 | 77452199 | 30.02047 | 50.43612 | 57.37683 | 25.06031 | 0.195603 | 1.160518 | 0.798559 |
| 1 | 44658600 | 44661199 | 19.18162 | 33.43067 | 50.6245 | 22.27041 | -0.20876 | 0.768881 | 0.798582 |
| 1 | 18211400 | 18211799 | 7.402323 | 14.37578 | 19.48031 | 7.29125 | 0.15531 | -0.25891 | 0.798602 |
| 8 | 54600200 | 54601799 | 27.82765 | 46.1875 | 53.89501 | 23.35685 | 0.159077 | 1.281142 | 0.798609 |
| 8 | 67729400 | 67731999 | 30.46139 | 49.62926 | 71.42571 | 30.67313 | -0.15276 | 1.534256 | 0.798611 |
| 12 | 40221600 | 40222399 | 10.5469 | 19.5672 | 29.35203 | 12.31458 | -0.13782 | 0.186856 | 0.798615 |
| 6 | 59856000 | 59869599 | 367.0342 | 508.8222 | 611.1897 | 253.7711 | -0.10872 | 4.711617 | 0.798638 |
| 16 | 79795200 | 79798399 | 43.71048 | 70.20603 | 96.82756 | 41.00329 | -0.11268 | 1.911432 | 0.798642 |
| 13 | 84626800 | 84627799 | 10.3037 | 18.87236 | 28.78466 | 12.28917 | -0.17957 | 0.256839 | 0.798645 |
| 10 | 56527200 | 56527999 | 17.46755 | 30.2754 | 43.74697 | 19.19391 | -0.14221 | 0.822619 | 0.798678 |
| 15 | 37045600 | 37047399 | 18.32492 | 31.92564 | 37.92927 | 16.68168 | 0.139898 | 0.751402 | 0.798731 |
| 15 | 30683200 | 30684399 | 7.576445 | 14.43646 | 23.97206 | 9.577974 | -0.22425 | -0.02871 | 0.798744 |
| 5 | 62295600 | 62298199 | 24.37768 | 40.47519 | 69.0984 | 29.95273 | -0.38822 | 1.280218 | 0.798763 |
| 4 | 145474400 | 145476399 | 22.48967 | 38.83079 | 45.77123 | 20.0797 | 0.149068 | 0.881141 | 0.798788 |
| 7 | 85686200 | 85688999 | 37.88652 | 61.99861 | 60.90931 | 26.1989 | 0.40098 | 1.478989 | 0.798818 |
| 6 | 70156000 | 70157199 | 14.99344 | 26.57991 | 31.544 | 13.94501 | 0.146608 | 0.492795 | 0.798832 |
| 2 | 84402400 | 84415399 | 141.6699 | 214.211 | 241.0097 | 99.67002 | 0.106542 | 3.306753 | 0.798836 |
| 2 | 142147400 | 142153799 | 132.4415 | 199.8268 | 257.8032 | 106.5426 | -0.09342 | 3.349966 | 0.798845 |
| X | 132488600 | 132489799 | 9.372106 | 17.69932 | 27.56237 | 11.22233 | -0.15608 | 0.063084 | 0.798855 |
| 8 | 20492400 | 20493799 | 17.42366 | 30.46797 | 44.04631 | 19.42014 | -0.1417 | 0.701778 | 0.798856 |
| 5 | 47509800 | 47510199 | 6.997984 | 13.62932 | 18.52122 | 6.768302 | 0.183754 | -0.34822 | 0.798856 |
| 9 | 86117800 | 86118999 | 18.05746 | 30.98789 | 53.79662 | 23.5953 | -0.40898 | 0.927753 | 0.798883 |
| 3 | 31508000 | 31509599 | 24.01055 | 40.35632 | 57.14631 | 24.96286 | -0.11919 | 1.135151 | 0.798905 |
| 1 | 58253000 | 58256999 | 33.88664 | 54.99645 | 77.22701 | 33.09866 | -0.12187 | 1.608787 | 0.798906 |
| 16 | 54687600 | 54690399 | 22.82705 | 37.68914 | 69.76608 | 30.12555 | -0.50521 | 1.341275 | 0.798924 |
| 11 | 42799400 | 42799799 | 9.24336 | 17.61583 | 22.34978 | 8.926404 | 0.161674 | -0.01594 | 0.798926 |
| 13 | 69503800 | 69506799 | 35.51888 | 57.83233 | 68.26594 | 29.30939 | 0.131736 | 1.549838 | 0.798928 |
| 3 | 110681600 | 110682999 | 25.90361 | 43.07872 | 60.92726 | 26.44036 | -0.11737 | 1.258227 | 0.798943 |
| 11 | 66363600 | 66365599 | 12.62285 | 23.60955 | 23.46518 | 9.71688 | 0.466701 | 0.140207 | 0.798974 |
| 17 | 34994600 | 35015799 | 339.917 | 476.3687 | 572.8309 | 237.4434 | -0.09453 | 4.548048 | 0.798979 |
| X | 78997000 | 79001999 | 48.68923 | 78.81649 | 85.01992 | 36.36928 | 0.254224 | 1.693784 | 0.798985 |
| 10 | 89546800 | 89547799 | 14.41809 | 26.36815 | 31.08917 | 13.30057 | 0.186942 | 0.300907 | 0.79902 |
| 10 | 115990800 | 115992199 | 21.77642 | 37.85029 | 44.09823 | 19.3664 | 0.167491 | 0.80879 | 0.799025 |
| 1 | 170058400 | 170059199 | 6.562574 | 12.66332 | 22.07386 | 8.525378 | -0.24678 | -0.15108 | 0.799036 |
| 13 | 51744200 | 51745799 | 13.56234 | 24.41965 | 36.21297 | 15.79763 | -0.1601 | 0.394561 | 0.799041 |
| 6 | 22752000 | 22755199 | 44.43773 | 71.8341 | 83.43844 | 35.64679 | 0.14569 | 1.716146 | 0.799061 |

| | | | | | | | | | |
|---|---|---|---|---|---|---|---|---|---|
| 7 | 143217800 | 143218599 | 11.38988 | 21.60029 | 26.53731 | 10.70955 | 0.193897 | 0.04637 | 0.79907 |
| 8 | 56049200 | 56050599 | 18.67233 | 32.64272 | 38.67365 | 17.04746 | 0.144039 | 0.702362 | 0.799081 |
| 15 | 11915000 | 11919199 | 65.22289 | 103.4507 | 121.4278 | 50.55736 | 0.099873 | 2.326456 | 0.799085 |
| 1 | 128747600 | 128749799 | 17.39582 | 30.52286 | 36.62368 | 16.19392 | 0.125979 | 0.636504 | 0.799104 |
| 3 | 152514000 | 152517399 | 33.28288 | 54.32988 | 63.55008 | 27.31003 | 0.147488 | 1.507341 | 0.799105 |
| 3 | 8416000 | 8417999 | 25.71093 | 43.57607 | 51.75875 | 22.66669 | 0.133966 | 1.063917 | 0.799156 |
| 6 | 140403800 | 140405999 | 17.13961 | 30.36604 | 35.25081 | 15.58562 | 0.177566 | 0.516897 | 0.79917 |
| 16 | 78547400 | 78548399 | 10.81113 | 20.24479 | 31.80814 | 13.14543 | -0.18695 | 0.138753 | 0.799177 |
| 12 | 109402800 | 109403999 | 12.3648 | 22.42624 | 25.61877 | 11.07459 | 0.222718 | 0.320025 | 0.799191 |
| 2 | 142257800 | 142260399 | 21.58799 | 37.81085 | 41.71366 | 18.41673 | 0.246551 | 0.695316 | 0.799202 |
| 2 | 8040800 | 8042799 | 19.03615 | 33.36043 | 32.01148 | 14.14002 | 0.446843 | 0.656455 | 0.799208 |
| 7 | 84520800 | 84526399 | 132.1865 | 200.0308 | 225.9344 | 93.45905 | 0.101905 | 3.28805 | 0.799211 |
| 10 | 31603000 | 31604399 | 19.76084 | 34.24301 | 41.06661 | 18.01778 | 0.125337 | 0.829751 | 0.799227 |
| 1 | 95791200 | 95792599 | 22.826 | 39.62005 | 38.28819 | 16.8003 | 0.435946 | 0.825167 | 0.799239 |
| 16 | 76819200 | 76824399 | 62.94295 | 100.2096 | 116.6576 | 48.77482 | 0.117159 | 2.211844 | 0.799249 |
| 2 | 45791000 | 45792399 | 18.29933 | 32.10651 | 38.04379 | 16.82112 | 0.143843 | 0.643147 | 0.799268 |
| 4 | 9803600 | 9805599 | 13.23233 | 23.52912 | 34.11043 | 15.04182 | -0.13813 | 0.466043 | 0.799279 |
| 3 | 43428600 | 43429199 | 9.411827 | 18.12685 | 23.64342 | 9.181503 | 0.159195 | -0.10175 | 0.799282 |
| 14 | 67793600 | 67795199 | 13.68286 | 24.65975 | 29.55586 | 12.87841 | 0.146688 | 0.37769 | 0.79931 |
| 1 | 166238800 | 166241799 | 36.69377 | 59.46735 | 82.66797 | 35.39332 | -0.10932 | 1.646248 | 0.799317 |
| 9 | 39795200 | 39797999 | 41.29933 | 66.88006 | 79.06619 | 33.83216 | 0.122859 | 1.669 | 0.799317 |
| 7 | 7297000 | 7300399 | 58.45762 | 92.67099 | 126.3118 | 52.5629 | -0.11617 | 2.338862 | 0.799318 |
| X | 8672800 | 8678199 | 54.55277 | 87.85128 | 94.23622 | 40.05861 | 0.253925 | 1.848357 | 0.799347 |
| 13 | 6316000 | 6319599 | 45.70556 | 73.34883 | 100.4599 | 42.48718 | -0.1041 | 1.941885 | 0.799374 |
| 6 | 105055000 | 105055799 | 9.038958 | 16.93944 | 26.41494 | 10.90216 | -0.17399 | 0.106466 | 0.799399 |
| 17 | 81597800 | 81600999 | 48.70813 | 77.83594 | 106.5658 | 44.78778 | -0.11058 | 2.079043 | 0.799426 |
| 6 | 95891000 | 95892999 | 22.56367 | 39.10809 | 38.28903 | 16.79924 | 0.416759 | 0.835909 | 0.799433 |
| 1 | 8106600 | 8107599 | 12.18793 | 23.07404 | 27.63278 | 11.19229 | 0.226176 | 0.069602 | 0.799442 |
| 7 | 39088400 | 39091399 | 22.40561 | 38.01767 | 53.83783 | 23.58539 | -0.11867 | 1.046006 | 0.799452 |
| X | 118495000 | 118503599 | 83.6124 | 132.1707 | 118.6504 | 49.29293 | 0.482067 | 2.426675 | 0.799481 |
| 13 | 101133400 | 101133999 | 4.806808 | 9.371583 | 18.07409 | 6.514916 | -0.29736 | -0.42856 | 0.799487 |
| 5 | 116982800 | 116988799 | 66.12204 | 104.0907 | 141.4654 | 58.69415 | -0.1201 | 2.506755 | 0.799499 |
| 8 | 43301800 | 43302399 | 8.734581 | 16.70521 | 20.84018 | 8.257122 | 0.196051 | -0.05209 | 0.799511 |
| 3 | 84516800 | 84520599 | 42.72218 | 68.95616 | 82.17617 | 35.03885 | 0.106259 | 1.755323 | 0.799513 |
| 10 | 21154000 | 21154999 | 11.04094 | 20.36183 | 30.31796 | 12.82359 | -0.13566 | 0.2266 | 0.799519 |
| 5 | 145751200 | 145780199 | 570.2595 | 772.3009 | 768.7432 | 320.3613 | 0.128299 | 5.079014 | 0.799521 |
| 15 | 49241200 | 49243199 | 14.57025 | 25.56916 | 37.36016 | 16.5218 | -0.15666 | 0.620314 | 0.79953 |
| 7 | 25265200 | 25269999 | 79.78611 | 124.2254 | 165.4882 | 68.69208 | -0.10364 | 2.718709 | 0.799537 |
| 8 | 78269000 | 78270999 | 20.72358 | 35.8644 | 43.11741 | 18.91681 | 0.12134 | 0.847932 | 0.799552 |
| 2 | 161560200 | 161563399 | 27.91366 | 46.37617 | 66.11415 | 28.67195 | -0.12896 | 1.279913 | 0.799575 |
| 10 | 8373400 | 8375599 | 24.91175 | 41.9407 | 50.14229 | 21.91439 | 0.124295 | 1.117076 | 0.799634 |
| 1 | 180057200 | 180058799 | 19.60129 | 33.61722 | 47.953 | 21.03127 | -0.12682 | 0.932153 | 0.799639 |
| 7 | 15663200 | 15665199 | 14.66401 | 25.85061 | 37.14435 | 16.43922 | -0.1312 | 0.560626 | 0.799642 |
| 10 | 69151800 | 69156399 | 41.11169 | 66.20952 | 91.27387 | 38.80243 | -0.1072 | 1.822178 | 0.799643 |
| 1 | 52360000 | 52360799 | 12.54897 | 22.95821 | 28.02854 | 11.98523 | 0.137504 | 0.277494 | 0.799651 |
| 3 | 49578600 | 49579799 | 12.54897 | 22.95821 | 28.02854 | 11.98523 | 0.137504 | 0.277494 | 0.799651 |
| 2 | 45896800 | 45898999 | 25.21485 | 42.93099 | 50.65759 | 22.20442 | 0.144034 | 1.021105 | 0.799657 |

| | | | | | | | | | |
|---|---|---|---|---|---|---|---|---|---|
| 1 | 14434600 | 14437399 | 23.45755 | 40.11005 | 47.94407 | 21.02226 | 0.126464 | 0.974083 | 0.79966 |
| 2 | 173736000 | 173736399 | 7.265213 | 14.13877 | 19.5842 | 7.208484 | 0.146117 | -0.31859 | 0.799662 |
| 7 | 100216400 | 100216799 | 7.265213 | 14.13877 | 19.5842 | 7.208484 | 0.146117 | -0.31859 | 0.799662 |
| 9 | 56506200 | 56508399 | 20.41046 | 34.91335 | 49.54248 | 21.7238 | -0.11995 | 0.960769 | 0.799671 |
| 8 | 127537200 | 127541199 | 40.4348 | 65.14824 | 90.20207 | 38.3714 | -0.11267 | 1.806369 | 0.799697 |
| 4 | 40423400 | 40428399 | 64.06064 | 101.2356 | 136.2304 | 56.58843 | -0.10148 | 2.420968 | 0.799708 |
| 17 | 89823200 | 89826599 | 27.04439 | 45.21196 | 65.6848 | 28.60073 | -0.15607 | 1.208862 | 0.799711 |
| 3 | 120749400 | 120750199 | 9.588449 | 17.94901 | 27.38751 | 11.32512 | -0.14564 | 0.118483 | 0.799732 |
| 10 | 129954400 | 129957399 | 33.11419 | 54.53869 | 65.20291 | 28.12166 | 0.12234 | 1.393415 | 0.799734 |
| 15 | 38759200 | 38761199 | 18.75776 | 32.568 | 47.39487 | 20.79993 | -0.15228 | 0.812164 | 0.799775 |
| 6 | 70254400 | 70259399 | 51.51522 | 82.89677 | 93.6068 | 39.74735 | 0.178585 | 1.862228 | 0.799808 |
| 18 | 5449800 | 5452199 | 22.51868 | 38.408 | 45.83897 | 20.09515 | 0.128091 | 0.988926 | 0.799822 |
| X | 30954400 | 30957399 | 25.23394 | 43.78868 | 38.17476 | 16.74971 | 0.583907 | 0.839535 | 0.799827 |
| X | 148624000 | 148628999 | 29.46399 | 49.68281 | 47.46362 | 20.75359 | 0.446834 | 1.116623 | 0.799828 |
| 6 | 20132600 | 20135399 | 24.4075 | 40.38041 | 69.4799 | 30.04403 | -0.3999 | 1.322079 | 0.799868 |
| 5 | 3747600 | 3748399 | 10.28347 | 19.54715 | 24.92132 | 10.00837 | 0.147152 | 0.016817 | 0.799893 |
| 13 | 14398000 | 14400799 | 37.55218 | 61.02405 | 72.86241 | 31.25773 | 0.113222 | 1.586951 | 0.799897 |
| 8 | 94199600 | 94200399 | 14.27129 | 25.96663 | 24.87911 | 10.71211 | 0.478569 | 0.315248 | 0.799904 |
| 12 | 109197600 | 109198999 | 24.58922 | 41.17163 | 58.18342 | 25.35939 | -0.1162 | 1.179183 | 0.799912 |
| 9 | 36479000 | 36483199 | 34.57599 | 55.63983 | 98.44913 | 41.80478 | -0.46785 | 1.843174 | 0.799914 |
| 19 | 21264400 | 21271599 | 72.90947 | 114.9751 | 157.9056 | 65.52999 | -0.13332 | 2.480335 | 0.799917 |
| 1 | 139576600 | 139578199 | 28.95186 | 48.27286 | 57.86146 | 25.15222 | 0.120632 | 1.236156 | 0.799929 |
| 9 | 20200000 | 20200999 | 16.8174 | 29.28355 | 42.11488 | 18.51633 | -0.13454 | 0.758676 | 0.799936 |
| 10 | 105474800 | 105476199 | 15.15189 | 27.17086 | 32.3171 | 14.14992 | 0.152575 | 0.420655 | 0.799961 |
| 13 | 11007600 | 11011799 | 74.82209 | 117.6715 | 137.4693 | 57.02889 | 0.096788 | 2.53107 | 0.799968 |
| 17 | 69643000 | 69644599 | 14.59446 | 26.06299 | 31.43758 | 13.81302 | 0.129201 | 0.44249 | 0.799996 |
| 4 | 106359800 | 106364599 | 39.86984 | 64.40034 | 75.56546 | 32.25115 | 0.129836 | 1.724877 | 0.800048 |
| 17 | 19061400 | 19062999 | 27.3171 | 44.56321 | 79.38283 | 34.10609 | -0.45829 | 1.517129 | 0.800054 |
| 13 | 98213000 | 98213999 | 7.943782 | 15.08672 | 24.49177 | 9.853909 | -0.20002 | 0.002804 | 0.80008 |
| 12 | 15212400 | 15213599 | 12.54856 | 22.70912 | 33.17178 | 14.37946 | -0.13243 | 0.352068 | 0.800145 |
| 8 | 4848600 | 4853999 | 87.16646 | 135.8552 | 157.8464 | 65.51005 | 0.094601 | 2.704548 | 0.800153 |
| 10 | 32726800 | 32728599 | 21.17831 | 36.27842 | 51.71256 | 22.67795 | -0.12624 | 0.95224 | 0.800172 |
| 18 | 76109600 | 76111199 | 13.27209 | 23.99182 | 29.01638 | 12.59566 | 0.137081 | 0.354765 | 0.800179 |
| 4 | 15235400 | 15239999 | 56.94799 | 90.25339 | 125.2253 | 52.09666 | -0.14219 | 2.344595 | 0.800187 |
| 7 | 92860200 | 92862599 | 30.96691 | 51.60664 | 51.36765 | 22.32228 | 0.387873 | 1.241107 | 0.800258 |
| 17 | 66017400 | 66020999 | 40.13414 | 64.65809 | 89.3778 | 38.01588 | -0.11047 | 1.808963 | 0.800277 |
| 15 | 103808800 | 103813599 | 82.78712 | 129.1687 | 149.0395 | 61.84626 | 0.105809 | 2.681711 | 0.800288 |
| 11 | 32875200 | 32877999 | 24.57131 | 41.05062 | 58.17407 | 25.3199 | -0.12017 | 1.202099 | 0.800326 |
| Y | 17375200 | 17379999 | 40.33371 | 66.08791 | 33.94209 | 14.60718 | 1.334333 | 1.44421 | 0.800337 |
| 5 | 79968000 | 79973399 | 114.1453 | 174.303 | 198.418 | 82.29145 | 0.101342 | 3.098668 | 0.80037 |
| 10 | 88083600 | 88085199 | 15.41648 | 27.34427 | 32.89304 | 14.53567 | 0.12742 | 0.492943 | 0.800379 |
| 4 | 8479000 | 8480799 | 21.37138 | 36.989 | 44.15403 | 19.37166 | 0.131565 | 0.851089 | 0.800397 |
| 5 | 105703800 | 105706799 | 29.56 | 49.13378 | 59.01515 | 25.60135 | 0.117524 | 1.271878 | 0.800436 |
| 2 | 85301800 | 85304799 | 76.21293 | 120.3059 | 119.4024 | 49.57033 | 0.335844 | 2.453072 | 0.800449 |
| 3 | 153258600 | 153259799 | 9.528903 | 17.48603 | 27.84498 | 11.85765 | -0.23858 | 0.237542 | 0.800461 |
| 13 | 95550400 | 95551999 | 13.08026 | 23.14605 | 34.18469 | 15.11791 | -0.16833 | 0.504343 | 0.800463 |
| 16 | 22055400 | 22060799 | 43.73535 | 70.30347 | 96.50748 | 40.91647 | -0.10446 | 1.886329 | 0.800475 |

| | | | | | | | | | |
|---|---|---|---|---|---|---|---|---|---|
| 16 | 31214600 | 31218199 | 50.08114 | 80.72814 | 77.75625 | 33.09829 | 0.410583 | 1.802747 | 0.800482 |
| X | 122320200 | 122320599 | 4.577358 | 8.949665 | 17.18721 | 5.879822 | -0.22139 | -0.61492 | 0.800483 |
| 9 | 31542000 | 31546199 | 65.38049 | 103.6193 | 121.1257 | 50.39194 | 0.104729 | 2.351363 | 0.800485 |
| 6 | 57196600 | 57198799 | 16.45018 | 29.478 | 33.77803 | 14.79847 | 0.205433 | 0.436278 | 0.800513 |
| 7 | 68215400 | 68216999 | 15.83691 | 27.96939 | 33.41604 | 14.78375 | 0.134629 | 0.540102 | 0.800516 |
| 5 | 136610200 | 136613199 | 21.66882 | 36.86693 | 52.23064 | 22.8926 | -0.11893 | 1.016095 | 0.800526 |
| 14 | 104992600 | 104994399 | 18.99871 | 33.00202 | 39.04176 | 17.13916 | 0.145631 | 0.798611 | 0.800537 |
| 3 | 110846800 | 110848199 | 18.22884 | 31.94421 | 38.1719 | 16.86054 | 0.131888 | 0.663829 | 0.800537 |
| 12 | 17926800 | 17929399 | 44.32042 | 71.57719 | 84.02967 | 35.8559 | 0.128787 | 1.741218 | 0.800558 |
| 5 | 21445200 | 21451599 | 88.69551 | 137.9276 | 160.1276 | 66.47915 | 0.093219 | 2.743362 | 0.800583 |
| 12 | 116765600 | 116769599 | 42.43911 | 68.38398 | 94.82501 | 40.34346 | -0.11469 | 1.809138 | 0.800587 |
| 2 | 60314600 | 60316399 | 18.05005 | 31.00157 | 53.80661 | 23.60101 | -0.40842 | 0.91985 | 0.800597 |
| 15 | 27281200 | 27286399 | 56.70243 | 90.34988 | 123.8472 | 51.84086 | -0.11689 | 2.180619 | 0.800609 |
| 10 | 16644600 | 16645599 | 8.852699 | 16.67316 | 25.9263 | 10.61068 | -0.16028 | 0.070989 | 0.800618 |
| X | 107493600 | 107496399 | 30.86658 | 51.4068 | 60.15004 | 26.12828 | 0.155356 | 1.257122 | 0.800625 |
| 14 | 4534800 | 4535799 | 10.13775 | 18.78461 | 28.33776 | 11.90929 | -0.14732 | 0.189899 | 0.800642 |
| 12 | 21476200 | 21478999 | 18.06898 | 31.24192 | 44.77853 | 19.64505 | -0.13137 | 0.847303 | 0.800658 |
| 5 | 52672600 | 52681799 | 105.893 | 161.7992 | 212.1514 | 87.99951 | -0.10174 | 3.084695 | 0.800678 |
| Y | 39280600 | 39281199 | 4.963851 | 9.668725 | 18.56985 | 6.776004 | -0.3079 | -0.38178 | 0.800709 |
| Y | 43590000 | 43590399 | 4.963851 | 9.668725 | 18.56985 | 6.776004 | -0.3079 | -0.38178 | 0.800709 |
| 4 | 73637200 | 73645999 | 90.6086 | 141.6735 | 159.9319 | 66.35169 | 0.139694 | 2.649592 | 0.800718 |
| 8 | 65379800 | 65381799 | 16.22843 | 28.41028 | 40.86474 | 18.03474 | -0.13444 | 0.671386 | 0.800735 |
| 13 | 72224200 | 72225999 | 15.63602 | 27.44961 | 39.24831 | 17.36221 | -0.1258 | 0.621882 | 0.800745 |
| 12 | 105893200 | 105895199 | 28.57716 | 47.56232 | 57.29367 | 24.87701 | 0.113509 | 1.252895 | 0.800762 |
| 5 | 150159200 | 150160999 | 18.48435 | 32.3566 | 38.07959 | 16.80425 | 0.153562 | 0.678969 | 0.800762 |
| 9 | 41074400 | 41076199 | 25.11673 | 42.26985 | 50.67254 | 22.14359 | 0.120414 | 1.12228 | 0.800778 |
| 11 | 98783400 | 98785399 | 24.8518 | 41.60733 | 58.8036 | 25.62816 | -0.11637 | 1.180017 | 0.800786 |
| 10 | 83445800 | 83448799 | 33.66202 | 54.92711 | 64.59157 | 27.7536 | 0.139452 | 1.515621 | 0.800816 |
| 16 | 89564200 | 89571799 | 99.90489 | 153.2357 | 201.7116 | 83.73559 | -0.10307 | 3.005141 | 0.800817 |
| 14 | 112578000 | 112579599 | 15.4822 | 27.82576 | 25.89228 | 11.31534 | 0.506826 | 0.397798 | 0.800818 |
| 5 | 138725200 | 138726199 | 12.8956 | 23.31201 | 34.19586 | 14.84339 | -0.13973 | 0.36312 | 0.800825 |
| 2 | 171418000 | 171419799 | 13.75309 | 24.38435 | 35.1117 | 15.52124 | -0.13081 | 0.49033 | 0.800833 |
| 6 | 147032000 | 147035399 | 61.88926 | 98.45151 | 115.7077 | 48.32296 | 0.102198 | 2.238039 | 0.800878 |
| 18 | 30395800 | 30397999 | 21.39551 | 36.74835 | 53.36969 | 23.40938 | -0.15247 | 0.931768 | 0.80088 |
| 5 | 144609600 | 144609999 | 7.118441 | 13.7464 | 22.60499 | 8.702392 | -0.16011 | -0.15764 | 0.800902 |
| 6 | 130218400 | 130224399 | 57.3054 | 92.17777 | 97.71474 | 41.46794 | 0.268894 | 1.887816 | 0.80091 |
| 2 | 148597600 | 148599799 | 24.78119 | 41.69597 | 49.60988 | 21.67777 | 0.131234 | 1.121941 | 0.800925 |
| 2 | 27436000 | 27440599 | 53.21456 | 85.14232 | 100.7782 | 42.4295 | 0.100977 | 2.043896 | 0.800937 |
| 4 | 83320800 | 83323199 | 41.92802 | 68.03061 | 79.03417 | 33.86708 | 0.150584 | 1.630753 | 0.800941 |
| 3 | 135602000 | 135604399 | 17.97738 | 30.8184 | 56.57349 | 24.81116 | -0.48963 | 0.936763 | 0.800952 |
| 4 | 150227600 | 150229399 | 24.07918 | 40.34031 | 57.08042 | 24.88747 | -0.11802 | 1.170172 | 0.800965 |
| 12 | 49265200 | 49268399 | 33.85667 | 55.5995 | 66.39502 | 28.60057 | 0.122294 | 1.429909 | 0.800976 |
| 18 | 3973000 | 3975199 | 29.6938 | 48.81644 | 68.76316 | 29.63435 | -0.11465 | 1.409023 | 0.801006 |
| 11 | 19761800 | 19764799 | 40.227 | 65.06266 | 77.67728 | 33.20742 | 0.107034 | 1.689653 | 0.801017 |
| 9 | 100434600 | 100435399 | 18.1402 | 31.64664 | 37.85953 | 16.66518 | 0.130073 | 0.730724 | 0.801026 |
| 16 | 70922000 | 70923399 | 15.08457 | 27.26601 | 31.84133 | 13.82334 | 0.186846 | 0.371528 | 0.801028 |
| X | 23825600 | 23828399 | 30.13361 | 51.12033 | 55.1529 | 24.16004 | 0.27234 | 1.077936 | 0.801029 |

| | | | | | | | | | |
|---|---|---|---|---|---|---|---|---|---|
| 17 | 39050400 | 39057599 | 75.84036 | 120.6188 | 126.8041 | 52.94156 | 0.262487 | 2.261173 | 0.80103 |
| 8 | 42300600 | 42301199 | 11.26176 | 20.99265 | 26.18458 | 10.90527 | 0.13475 | 0.156651 | 0.801048 |
| 9 | 77059600 | 77060399 | 17.32099 | 30.24736 | 34.91128 | 15.37696 | 0.181779 | 0.70825 | 0.801053 |
| 11 | 70981200 | 70983799 | 49.67897 | 79.95715 | 79.69889 | 33.85111 | 0.35854 | 1.848915 | 0.801068 |
| 6 | 14653400 | 14655599 | 23.94463 | 39.60202 | 69.35222 | 29.98223 | -0.42525 | 1.325227 | 0.801101 |
| 14 | 49933800 | 49935399 | 26.35549 | 45.15125 | 50.6737 | 22.22254 | 0.217436 | 0.972066 | 0.801107 |
| 3 | 111412800 | 111413799 | 14.84161 | 27.38504 | 25.32403 | 10.70387 | 0.547813 | 0.243172 | 0.801114 |
| 15 | 100118000 | 100123999 | 78.04097 | 121.938 | 161.717 | 67.09708 | -0.09324 | 2.656548 | 0.80113 |
| 13 | 47195800 | 47198399 | 16.47809 | 27.77459 | 59.99266 | 26.22876 | -0.72572 | 1.088603 | 0.801132 |
| 12 | 39288800 | 39292199 | 24.31057 | 42.23169 | 46.25356 | 20.3059 | 0.256227 | 0.831029 | 0.801152 |
| 3 | 3559600 | 3560999 | 12.10464 | 21.97352 | 32.217 | 13.91175 | -0.13398 | 0.330225 | 0.801159 |
| 1 | 145681800 | 145684599 | 29.68444 | 48.88475 | 68.87654 | 29.70474 | -0.11404 | 1.388021 | 0.801168 |
| 13 | 21775200 | 21776599 | 14.84827 | 26.03777 | 37.84171 | 16.72428 | -0.14908 | 0.63353 | 0.801168 |
| X | 126335200 | 126336199 | 16.85362 | 29.95249 | 34.88382 | 15.38974 | 0.175551 | 0.495086 | 0.80117 |
| 16 | 9366200 | 9368199 | 16.7457 | 29.39893 | 35.14731 | 15.54861 | 0.131235 | 0.621484 | 0.801194 |
| 12 | 102809400 | 102817599 | 111.7101 | 173.4086 | 180.7574 | 75.06382 | 0.246506 | 2.788086 | 0.801198 |
| 13 | 108932600 | 108933199 | 5.951165 | 11.57107 | 20.1054 | 7.45446 | -0.18672 | -0.30663 | 0.801214 |
| 3 | 7735200 | 7735599 | 6.979784 | 13.57052 | 23.03774 | 8.544384 | -0.15496 | -0.28515 | 0.801232 |
| 15 | 96176600 | 96179599 | 35.236 | 57.03619 | 79.77261 | 34.12344 | -0.11967 | 1.666505 | 0.801232 |
| 3 | 89778000 | 89782999 | 49.7582 | 79.68123 | 93.56405 | 39.4417 | 0.113896 | 2.011168 | 0.801241 |
| 5 | 99190000 | 99190799 | 8.431039 | 15.9919 | 25.51392 | 10.29261 | -0.17856 | 0.019734 | 0.801248 |
| 1 | 7194600 | 7197599 | 36.09044 | 60.04407 | 52.01889 | 22.58073 | 0.587586 | 1.27674 | 0.801248 |
| 7 | 6600400 | 6606799 | 77.25362 | 121.5696 | 141.8359 | 58.84506 | 0.099353 | 2.520845 | 0.801249 |
| 16 | 43859200 | 43862799 | 35.55762 | 57.62671 | 80.0555 | 34.27511 | -0.10831 | 1.642897 | 0.801271 |
| 10 | 31289400 | 31290199 | 11.61707 | 22.11575 | 21.64716 | 8.665524 | 0.530028 | 0.012501 | 0.801279 |
| 9 | 3501400 | 3502199 | 8.068395 | 15.44834 | 24.58965 | 9.711541 | -0.14813 | -0.05695 | 0.80129 |
| 19 | 49920600 | 49921199 | 8.068395 | 15.44834 | 24.58965 | 9.711541 | -0.14813 | -0.05695 | 0.80129 |
| 13 | 40519200 | 40522799 | 26.12488 | 43.90916 | 52.83745 | 23.08051 | 0.114882 | 1.139191 | 0.80129 |
| 16 | 5304600 | 5308599 | 33.54626 | 55.45149 | 64.91207 | 28.05196 | 0.153908 | 1.350408 | 0.801291 |
| 7 | 107535800 | 107544599 | 99.83873 | 154.7331 | 175.6952 | 72.97878 | 0.121418 | 2.807018 | 0.801294 |
| 13 | 45021800 | 45025399 | 150.5055 | 224.3197 | 292.3053 | 120.4566 | -0.1236 | 3.588347 | 0.801299 |
| 8 | 127245800 | 127249199 | 30.08907 | 49.86249 | 60.08664 | 26.01192 | 0.112509 | 1.309151 | 0.801346 |
| 18 | 54764400 | 54765399 | 8.734324 | 16.69838 | 26.77055 | 10.61838 | -0.16431 | -0.02983 | 0.801393 |
| 15 | 99414800 | 99415399 | 10.12839 | 18.68382 | 28.30511 | 11.96792 | -0.15976 | 0.214247 | 0.801407 |
| 3 | 159389800 | 159391399 | 15.13306 | 26.94472 | 32.44175 | 14.28996 | 0.1294 | 0.467213 | 0.801459 |
| 3 | 120960200 | 120961799 | 15.20364 | 26.68346 | 38.30639 | 16.94173 | -0.13142 | 0.623523 | 0.801464 |
| 5 | 85013800 | 85013999 | 5.946157 | 11.62118 | 16.00246 | 5.532683 | 0.238096 | -0.56287 | 0.801504 |
| 17 | 70101200 | 70102399 | 10.90052 | 20.32846 | 31.10988 | 12.94663 | -0.15744 | 0.162785 | 0.801516 |
| 4 | 3077800 | 3092399 | 1269.34 | 1599.056 | 1500.852 | 630.6916 | 0.10845 | 6.2472 | 0.801544 |
| 5 | 82216800 | 82217599 | 18.64244 | 32.00857 | 46.39524 | 20.3499 | -0.1495 | 0.916559 | 0.801579 |
| 17 | 22443800 | 22445399 | 12.19536 | 21.56595 | 39.80703 | 17.60667 | -0.48877 | 0.516226 | 0.8016 |
| 16 | 34704600 | 34709199 | 36.67365 | 59.88737 | 70.98076 | 30.50746 | 0.128813 | 1.512678 | 0.801616 |
| X | 46152000 | 46154999 | 49.30242 | 79.27619 | 91.85514 | 38.96095 | 0.140198 | 1.885787 | 0.801629 |
| 9 | 25767800 | 25775399 | 148.0094 | 222.0566 | 246.1037 | 101.5506 | 0.118422 | 3.462111 | 0.801652 |
| 14 | 119925200 | 119925799 | 6.237979 | 12.12675 | 20.65036 | 7.666617 | -0.16013 | -0.29761 | 0.801662 |
| 17 | 86323600 | 86328199 | 50.73951 | 81.35118 | 116.1782 | 49.06497 | -0.16596 | 1.983284 | 0.801667 |
| 8 | 45718400 | 45721199 | 31.37784 | 51.31364 | 71.99077 | 30.9579 | -0.11232 | 1.482428 | 0.801672 |

| | | | | | | | | | |
|---|---|---|---|---|---|---|---|---|---|
| 5 | 127567000 | 127568799 | 13.70142 | 24.25006 | 35.10704 | 15.5252 | -0.13966 | 0.507537 | 0.801714 |
| 17 | 29744000 | 29747999 | 50.45114 | 80.7833 | 95.38412 | 40.2032 | 0.105775 | 2.016027 | 0.801728 |
| 2 | 107349800 | 107350399 | 7.110345 | 13.82961 | 18.99829 | 7.024655 | 0.152053 | -0.30652 | 0.801737 |
| 12 | 62622600 | 62622999 | 7.110345 | 13.82961 | 18.99829 | 7.024655 | 0.152053 | -0.30652 | 0.801737 |
| 18 | 14228200 | 14232199 | 37.23099 | 60.40943 | 85.27836 | 36.5428 | -0.12986 | 1.625819 | 0.801739 |
| X | 106287000 | 106290399 | 42.82687 | 69.1889 | 81.60273 | 34.83575 | 0.122744 | 1.730557 | 0.801747 |
| 3 | 56926200 | 56930999 | 63.03652 | 99.99321 | 135.3196 | 56.34716 | -0.10501 | 2.325443 | 0.801751 |
| 19 | 13849200 | 13851799 | 17.20124 | 29.77997 | 52.78482 | 23.15844 | -0.43625 | 0.842683 | 0.801754 |
| 4 | 77931000 | 77933199 | 17.4525 | 30.70709 | 36.40917 | 16.12006 | 0.143387 | 0.595421 | 0.801758 |
| 3 | 43202000 | 43203199 | 11.94137 | 21.81784 | 32.29615 | 13.83354 | -0.14016 | 0.291999 | 0.801763 |
| 6 | 89672000 | 89677199 | 117.457 | 178.6205 | 232.4564 | 96.30188 | -0.09641 | 3.186403 | 0.801765 |
| 7 | 110998200 | 110999199 | 15.74076 | 28.18688 | 33.36603 | 14.62666 | 0.158354 | 0.43671 | 0.801794 |
| 15 | 21889200 | 21891999 | 39.83498 | 64.36212 | 75.26406 | 32.13416 | 0.135253 | 1.719354 | 0.801837 |
| 17 | 54242400 | 54244799 | 45.92104 | 74.05248 | 72.50553 | 30.87947 | 0.38765 | 1.781851 | 0.801843 |
| 19 | 21645400 | 21647599 | 21.4148 | 37.25575 | 43.53697 | 19.13089 | 0.163249 | 0.793175 | 0.801863 |
| 8 | 90645800 | 90646999 | 14.96123 | 27.06198 | 26.31858 | 11.41575 | 0.450364 | 0.357312 | 0.801882 |
| 1 | 18023000 | 18030199 | 133.9739 | 202.4884 | 227.9605 | 94.26088 | 0.105142 | 3.314007 | 0.801911 |
| 11 | 6098600 | 6100599 | 25.66895 | 43.16433 | 51.73966 | 22.60438 | 0.12054 | 1.133157 | 0.801951 |
| X | 126565600 | 126568399 | 31.14264 | 52.27623 | 58.58996 | 25.57725 | 0.217037 | 1.173221 | 0.801959 |
| 11 | 50136600 | 50138599 | 15.13731 | 26.5703 | 38.25304 | 16.91989 | -0.13565 | 0.619402 | 0.802002 |
| 14 | 66005800 | 66008399 | 26.82197 | 45.33832 | 43.74461 | 19.1416 | 0.432623 | 1.080846 | 0.802003 |
| 5 | 89829200 | 89832399 | 18.98971 | 32.71135 | 46.87701 | 20.56334 | -0.13238 | 0.886156 | 0.802056 |
| 7 | 26756600 | 26757799 | 11.85148 | 21.53329 | 31.66652 | 13.65869 | -0.13746 | 0.319899 | 0.802075 |
| 7 | 18203400 | 18204199 | 26.87612 | 44.92123 | 53.36024 | 23.23125 | 0.133656 | 1.201761 | 0.802079 |
| 18 | 8514600 | 8517599 | 33.55572 | 54.73776 | 76.6146 | 32.91616 | -0.11151 | 1.526677 | 0.802081 |
| 7 | 97584600 | 97587399 | 70.95671 | 111.6182 | 148.8204 | 61.73973 | -0.09337 | 2.526706 | 0.802096 |
| 9 | 24436800 | 24437399 | 7.278655 | 14.11511 | 23.38557 | 8.827447 | -0.1438 | -0.21944 | 0.802136 |
| 14 | 11887400 | 11888399 | 12.7843 | 23.79652 | 28.79764 | 12.02279 | 0.175251 | 0.185009 | 0.802172 |
| 6 | 133718800 | 133722799 | 55.76568 | 88.78598 | 119.9107 | 50.13737 | -0.09673 | 2.205273 | 0.802178 |
| 13 | 79093600 | 79095999 | 26.61675 | 44.11963 | 62.49591 | 27.0597 | -0.11993 | 1.300452 | 0.802183 |
| Y | 43572600 | 43576999 | 27.10587 | 43.8894 | 75.65891 | 32.37105 | -0.41998 | 1.642324 | 0.80219 |
| 17 | 3050400 | 3091199 | 1297.012 | 1619.675 | 1774.801 | 746.5189 | -0.1277 | 6.38795 | 0.802202 |
| 8 | 90011400 | 90013999 | 19.20469 | 33.67571 | 39.02272 | 17.24562 | 0.175802 | 0.657532 | 0.802209 |
| 2 | 121686800 | 121688399 | 17.27569 | 30.98666 | 28.55744 | 12.50127 | 0.518811 | 0.427348 | 0.802229 |
| 11 | 110053000 | 110054599 | 15.38373 | 27.5686 | 32.85976 | 14.39545 | 0.149079 | 0.42827 | 0.802241 |
| 2 | 149741400 | 149743599 | 21.74924 | 37.27161 | 44.77598 | 19.63664 | 0.119721 | 0.942792 | 0.802242 |
| 4 | 78691000 | 78693199 | 28.34249 | 47.19896 | 56.66765 | 24.61424 | 0.118338 | 1.247376 | 0.802244 |
| Y | 84725000 | 84727399 | 8.581788 | 16.66267 | 17.25539 | 6.470092 | 0.53722 | -0.24597 | 0.802251 |
| 7 | 29588600 | 29595799 | 269.4866 | 385.8538 | 412.0407 | 170.1964 | 0.107824 | 4.229273 | 0.802265 |
| 14 | 54768200 | 54771999 | 46.90031 | 74.54482 | 130.2055 | 54.32563 | -0.46969 | 2.253671 | 0.802274 |
| 15 | 94990000 | 94992599 | 24.40709 | 41.22974 | 59.40093 | 25.98736 | -0.14376 | 1.096722 | 0.802289 |
| 4 | 84856600 | 84857799 | 15.08574 | 26.86478 | 32.40334 | 14.2711 | 0.12689 | 0.464991 | 0.802294 |
| 11 | 95441800 | 95446399 | 42.18248 | 68.25419 | 80.53528 | 34.43993 | 0.124499 | 1.685702 | 0.802332 |
| 6 | 95685800 | 95686599 | 10.06241 | 18.65191 | 28.30005 | 11.88746 | -0.15539 | 0.184915 | 0.802339 |
| 14 | 6957600 | 6958599 | 10.06241 | 18.65191 | 28.30005 | 11.88746 | -0.15539 | 0.184915 | 0.802339 |
| 5 | 90970000 | 90971399 | 16.16553 | 28.29994 | 40.40383 | 17.8312 | -0.12356 | 0.672787 | 0.802361 |
| 5 | 132807400 | 132809399 | 22.39588 | 38.6771 | 45.75917 | 20.07454 | 0.143744 | 0.877476 | 0.80238 |

| | | | | | | | | | |
|---|---|---|---|---|---|---|---|---|---|
| 2 | 86554800 | 86558599 | 42.06327 | 68.20062 | 79.59311 | 34.08838 | 0.143055 | 1.64515 | 0.80241 |
| 14 | 3288400 | 3291799 | 42.63978 | 68.78325 | 81.61306 | 34.77272 | 0.111562 | 1.77022 | 0.802418 |
| 16 | 14401800 | 14405199 | 32.08187 | 52.60862 | 74.5362 | 32.08782 | -0.12447 | 1.451479 | 0.802449 |
| 8 | 74104600 | 74109999 | 61.25422 | 97.61087 | 113.8363 | 47.65591 | 0.115995 | 2.17979 | 0.802461 |
| 2 | 16029600 | 16030199 | 8.090538 | 14.84739 | 31.9787 | 13.61695 | -0.67078 | 0.23774 | 0.802469 |
| 12 | 115139800 | 115151599 | 144.4216 | 220.2349 | 227.949 | 94.50924 | 0.236961 | 3.138018 | 0.802477 |
| 9 | 114500600 | 114501599 | 12.98028 | 23.81771 | 29.11679 | 12.40052 | 0.139931 | 0.268288 | 0.802482 |
| 8 | 44162800 | 44163999 | 10.07818 | 18.63754 | 28.2882 | 11.92032 | -0.15915 | 0.198421 | 0.802483 |
| 3 | 74631800 | 74636799 | 60.07029 | 96.19079 | 106.7644 | 45.00351 | 0.194914 | 2.029396 | 0.802505 |
| 9 | 108977800 | 108979199 | 19.00199 | 33.14142 | 39.87931 | 17.55138 | 0.12166 | 0.742218 | 0.802524 |
| 3 | 77060800 | 77061399 | 10.81019 | 20.13579 | 24.70355 | 10.30154 | 0.157228 | 0.154739 | 0.802567 |
| 10 | 22460800 | 22466399 | 55.08341 | 88.26892 | 101.3718 | 42.77731 | 0.146987 | 1.997408 | 0.802575 |
| 11 | 54773600 | 54775999 | 17.45528 | 30.57581 | 36.64764 | 16.18154 | 0.127457 | 0.662179 | 0.802595 |
| 13 | 33573200 | 33575599 | 31.2606 | 52.17825 | 49.71956 | 21.62758 | 0.450637 | 1.226055 | 0.802598 |
| 14 | 14752600 | 14753399 | 8.455199 | 16.28478 | 21.43365 | 8.322839 | 0.146605 | -0.11807 | 0.802604 |
| X | 6694800 | 6699199 | 44.81441 | 72.51514 | 83.06198 | 35.52127 | 0.167272 | 1.693404 | 0.802617 |
| 18 | 56411800 | 56412399 | 10.77545 | 20.03661 | 24.18652 | 10.11532 | 0.177027 | 0.163512 | 0.802618 |
| 13 | 70633800 | 70636799 | 42.82526 | 69.08643 | 82.20346 | 35.02687 | 0.107617 | 1.769487 | 0.802625 |
| 10 | 38100200 | 38103799 | 40.19701 | 65.7124 | 62.59514 | 26.914 | 0.444434 | 1.502156 | 0.802628 |
| 4 | 142334800 | 142338799 | 48.46201 | 77.81318 | 90.90351 | 38.47617 | 0.125785 | 1.923563 | 0.802633 |
| 13 | 37856400 | 37856799 | 7.538314 | 14.61062 | 19.43132 | 7.35369 | 0.167044 | -0.22022 | 0.802643 |
| 8 | 129769800 | 129771399 | 18.52861 | 32.48167 | 38.62988 | 17.06823 | 0.138658 | 0.659614 | 0.802666 |
| 14 | 53805200 | 53815999 | 125.3066 | 192.1038 | 179.7479 | 74.61291 | 0.388408 | 3.02247 | 0.802715 |
| 14 | 96861400 | 96862999 | 18.15283 | 31.75967 | 38.22323 | 16.86019 | 0.121614 | 0.688317 | 0.802742 |
| X | 66512400 | 66515199 | 27.00017 | 46.4259 | 39.89013 | 17.49747 | 0.60228 | 0.931092 | 0.802753 |
| 14 | 8019800 | 8022399 | 50.90908 | 81.13819 | 111.7849 | 46.80726 | -0.12374 | 2.166434 | 0.802758 |
| 3 | 9703400 | 9706399 | 23.01051 | 38.81191 | 54.94523 | 24.02703 | -0.1186 | 1.099147 | 0.802769 |
| 2 | 80094400 | 80094999 | 6.088154 | 11.83677 | 20.66041 | 7.663617 | -0.19484 | -0.30498 | 0.802828 |
| 2 | 120665000 | 120667799 | 38.66381 | 62.63241 | 74.55038 | 31.90749 | 0.113525 | 1.655668 | 0.802836 |
| 7 | 13630600 | 13633199 | 55.49421 | 88.29208 | 119.4777 | 49.90808 | -0.10062 | 2.228208 | 0.802838 |
| 6 | 146600000 | 146604999 | 53.40897 | 85.39108 | 99.69024 | 41.92811 | 0.119643 | 2.064061 | 0.802841 |
| 4 | 7233200 | 7235999 | 27.78349 | 46.2479 | 55.58026 | 24.13528 | 0.116914 | 1.248219 | 0.802846 |
| 10 | 58760600 | 58761999 | 16.12911 | 28.06959 | 41.11582 | 18.07082 | -0.16096 | 0.760533 | 0.802865 |
| 2 | 60008000 | 60008999 | 9.938397 | 18.86466 | 23.87132 | 9.615263 | 0.154792 | 0.023346 | 0.802888 |
| 7 | 81905600 | 81910199 | 39.9899 | 64.33487 | 89.87598 | 38.14968 | -0.12819 | 1.853414 | 0.802898 |
| 2 | 64961400 | 64962599 | 11.89861 | 22.39007 | 27.41585 | 11.23707 | 0.17955 | 0.105076 | 0.802901 |
| 2 | 101231200 | 101232399 | 13.03579 | 23.7683 | 28.99592 | 12.45205 | 0.134492 | 0.302477 | 0.802916 |
| 1 | 185328400 | 185330999 | 22.08855 | 37.56176 | 53.89251 | 23.62002 | -0.13732 | 1.022441 | 0.802926 |
| 2 | 92394200 | 92397599 | 46.22252 | 74.54046 | 87.30316 | 37.18281 | 0.129773 | 1.788968 | 0.802944 |
| 2 | 150485800 | 150487599 | 20.0303 | 34.58368 | 41.09058 | 18.0263 | 0.137606 | 0.862574 | 0.802956 |
| 18 | 15508800 | 15509799 | 20.15514 | 35.05361 | 42.01306 | 18.45772 | 0.126958 | 0.790507 | 0.802967 |
| 8 | 44243000 | 44245399 | 27.03056 | 45.13677 | 53.98243 | 23.48814 | 0.123765 | 1.209292 | 0.80297 |
| 13 | 11315000 | 11317199 | 19.60078 | 34.66738 | 37.82816 | 16.73085 | 0.265608 | 0.556025 | 0.802999 |
| 3 | 11969200 | 11970199 | 18.85509 | 32.7931 | 39.00435 | 17.13606 | 0.138061 | 0.780531 | 0.803003 |
| 11 | 52407400 | 52409599 | 12.59342 | 22.15245 | 40.94024 | 18.12704 | -0.49329 | 0.582253 | 0.80303 |
| 4 | 117933000 | 117934399 | 11.13531 | 19.79352 | 37.33648 | 16.46918 | -0.51573 | 0.463105 | 0.803045 |
| 10 | 71897200 | 71898599 | 21.54762 | 37.31247 | 44.18324 | 19.38477 | 0.143403 | 0.850138 | 0.803067 |

| | | | | | | | | | |
|---|---|---|---|---|---|---|---|---|---|
| 17 | 5604400 | 5607199 | 23.02168 | 39.09507 | 46.39269 | 20.32886 | 0.135453 | 1.030918 | 0.803069 |
| 10 | 85167200 | 85171199 | 44.78072 | 71.90859 | 98.40631 | 41.65737 | -0.10184 | 1.922288 | 0.803071 |
| 3 | 22210200 | 22210999 | 14.24015 | 25.18388 | 36.14132 | 15.98507 | -0.12747 | 0.522774 | 0.803093 |
| 2 | 178491600 | 178492599 | 12.73488 | 22.74683 | 33.12496 | 14.55189 | -0.14078 | 0.43239 | 0.803101 |
| 2 | 118101600 | 118102399 | 7.601026 | 14.61682 | 23.60144 | 9.205755 | -0.15158 | -0.10509 | 0.803111 |
| 9 | 53331600 | 53332799 | 18.1473 | 31.76821 | 38.18778 | 16.85272 | 0.123305 | 0.67887 | 0.803118 |
| 7 | 98680000 | 98683599 | 38.81355 | 62.62236 | 86.69506 | 36.94659 | -0.11016 | 1.762181 | 0.80313 |
| 3 | 106157600 | 106175999 | 386.6005 | 538.4357 | 668.2074 | 277.2456 | -0.14904 | 4.643831 | 0.803144 |
| 9 | 9467200 | 9468599 | 21.41269 | 36.68474 | 43.751 | 19.1867 | 0.13015 | 0.942619 | 0.803148 |
| 13 | 58730200 | 58734799 | 50.34814 | 80.51937 | 109.4589 | 46.05448 | -0.09913 | 2.057915 | 0.803152 |
| 1 | 29180400 | 29180799 | 5.364078 | 10.46079 | 18.61947 | 6.688599 | -0.17901 | -0.43114 | 0.803162 |
| 10 | 120099400 | 120102999 | 57.48232 | 91.57738 | 107.8849 | 45.14886 | 0.101068 | 2.182885 | 0.803163 |
| 13 | 79881200 | 79883799 | 21.95207 | 37.63637 | 45.29764 | 19.86614 | 0.11719 | 0.940742 | 0.80317 |
| 19 | 21445600 | 21447199 | 16.89864 | 29.91375 | 35.34756 | 15.63104 | 0.151741 | 0.526004 | 0.803182 |
| 10 | 62696200 | 62699799 | 21.2785 | 36.69742 | 44.20456 | 19.39173 | 0.117584 | 0.883522 | 0.803182 |
| 12 | 79380000 | 79382199 | 25.57337 | 44.23822 | 39.19318 | 17.19491 | 0.559769 | 0.86822 | 0.803191 |
| 1 | 105067600 | 105068199 | 10.71329 | 20.04053 | 24.75968 | 10.25069 | 0.155723 | 0.128132 | 0.803209 |
| 7 | 88742600 | 88743399 | 9.144752 | 17.21196 | 26.42444 | 10.82653 | -0.14346 | 0.078431 | 0.803219 |
| 18 | 47489000 | 47489399 | 5.940225 | 11.57662 | 20.30296 | 7.357563 | -0.17096 | -0.39054 | 0.803222 |
| 1 | 31810400 | 31811799 | 14.74233 | 26.40769 | 31.89885 | 13.9796 | 0.129638 | 0.425081 | 0.803265 |
| 19 | 9959000 | 9960799 | 13.96848 | 24.71394 | 36.05207 | 15.94425 | -0.15096 | 0.513333 | 0.80328 |
| 14 | 76396200 | 76398799 | 24.84366 | 41.54744 | 58.80012 | 25.60865 | -0.11834 | 1.191519 | 0.803299 |
| 18 | 45038400 | 45042599 | 60.38186 | 96.2907 | 95.1031 | 39.85359 | 0.356484 | 2.151187 | 0.803324 |
| 12 | 87623200 | 87625399 | 28.86073 | 48.42267 | 47.51921 | 20.7381 | 0.408022 | 1.158197 | 0.803338 |
| 1 | 117999600 | 118002599 | 58.12704 | 92.75677 | 108.6858 | 45.58782 | 0.111398 | 2.13353 | 0.803343 |
| 14 | 8314600 | 8319399 | 96.89133 | 149.6266 | 172.5636 | 71.67937 | 0.095014 | 2.871544 | 0.803346 |
| 2 | 81952800 | 81953399 | 9.042733 | 17.05778 | 26.47306 | 10.80489 | -0.15416 | 0.065163 | 0.803352 |
| 3 | 97039800 | 97044199 | 56.62881 | 90.19214 | 122.2287 | 51.13165 | -0.10113 | 2.195869 | 0.803355 |
| 3 | 49962200 | 49962599 | 6.234146 | 12.16181 | 16.97083 | 6.068523 | 0.174133 | -0.44146 | 0.803356 |
| X | 93584200 | 93586599 | 24.12908 | 41.1249 | 47.9628 | 21.02498 | 0.16125 | 1.008798 | 0.803359 |
| 1 | 171010200 | 171011399 | 12.28958 | 22.16914 | 33.07808 | 14.39656 | -0.1669 | 0.36584 | 0.80336 |
| 5 | 41796200 | 41801599 | 67.96734 | 108.4113 | 117.7296 | 49.35053 | 0.2202 | 2.159972 | 0.803367 |
| 16 | 29574800 | 29587799 | 121.9769 | 184.8859 | 240.3439 | 99.4714 | -0.0987 | 3.253371 | 0.803389 |
| 7 | 128203800 | 128206399 | 45.95514 | 73.58972 | 101.3179 | 42.70908 | -0.11529 | 2.011368 | 0.80339 |
| 2 | 80050400 | 80053599 | 39.56447 | 64.45276 | 75.1359 | 32.26503 | 0.150249 | 1.556629 | 0.803414 |
| 19 | 14936800 | 14938399 | 17.35715 | 30.4388 | 36.63127 | 16.19 | 0.121693 | 0.644204 | 0.803418 |
| 5 | 89044400 | 89044999 | 11.20727 | 21.07171 | 26.34057 | 10.81148 | 0.148714 | 0.103778 | 0.803418 |
| 3 | 16111600 | 16117799 | 72.13619 | 114.6099 | 127.4132 | 53.13942 | 0.180501 | 2.285862 | 0.803451 |
| 19 | 35576600 | 35577399 | 20.47083 | 34.8156 | 50.06612 | 21.94323 | -0.14029 | 1.015106 | 0.803506 |
| 13 | 99427200 | 99432399 | 60.61516 | 96.47605 | 113.7988 | 47.56075 | 0.097771 | 2.217901 | 0.803538 |
| 13 | 95278000 | 95279599 | 28.87021 | 48.71263 | 55.66888 | 24.34607 | 0.189291 | 1.118897 | 0.80356 |
| 1 | 52351400 | 52360799 | 214.5736 | 310.8597 | 394.197 | 162.5517 | -0.12319 | 4.052783 | 0.803562 |
| 17 | 22545600 | 22548199 | 30.84547 | 51.60531 | 59.19848 | 25.78867 | 0.183639 | 1.20573 | 0.803566 |
| 3 | 14517600 | 14521199 | 57.21179 | 91.89676 | 100.3615 | 42.50363 | 0.223478 | 1.931146 | 0.803569 |
| 4 | 22594400 | 22596599 | 25.78174 | 43.44721 | 52.29477 | 22.86179 | 0.114651 | 1.11461 | 0.803592 |
| 11 | 22529000 | 22530799 | 13.11789 | 23.81345 | 29.03572 | 12.53764 | 0.130345 | 0.329606 | 0.803615 |
| 12 | 114330000 | 114343999 | 204.7622 | 302.3909 | 330.13 | 135.9541 | 0.119243 | 3.754229 | 0.803617 |

| | | | | | | | | | |
|---|---|---|---|---|---|---|---|---|---|
| 15 | 85666600 | 85669399 | 18.36546 | 31.77206 | 45.28625 | 19.86813 | -0.12329 | 0.843773 | 0.803621 |
| 11 | 58540200 | 58542799 | 35.93428 | 58.09995 | 81.23331 | 34.71378 | -0.1208 | 1.695128 | 0.803665 |
| 3 | 34577400 | 34583799 | 62.19448 | 99.04996 | 115.1608 | 48.17052 | 0.119484 | 2.199452 | 0.803675 |
| 17 | 69847800 | 69850799 | 25.08801 | 41.79705 | 59.24154 | 25.73915 | -0.12035 | 1.234823 | 0.803685 |
| 6 | 143166000 | 143167799 | 16.98141 | 29.2693 | 43.8148 | 19.2204 | -0.19465 | 0.869859 | 0.8037 |
| 6 | 63899800 | 63902799 | 51.02525 | 81.68286 | 96.35894 | 40.59945 | 0.106656 | 2.02313 | 0.803736 |
| 3 | 104354400 | 104357999 | 31.30779 | 51.33987 | 59.74187 | 25.71899 | 0.158812 | 1.438755 | 0.80374 |
| 16 | 29494600 | 29497599 | 32.19469 | 52.93469 | 63.43713 | 27.34052 | 0.1179 | 1.408704 | 0.803782 |
| 4 | 10136600 | 10138599 | 52.78203 | 83.32829 | 148.4398 | 61.62879 | -0.50667 | 2.44156 | 0.803795 |
| 5 | 37366800 | 37375399 | 91.76596 | 141.3712 | 187.215 | 77.76443 | -0.10688 | 2.915655 | 0.803813 |
| 4 | 124762400 | 124767599 | 53.08192 | 84.99766 | 100.4743 | 42.35008 | 0.104158 | 2.019847 | 0.803817 |
| 2 | 132554800 | 132556399 | 13.44663 | 24.29431 | 29.5057 | 12.81666 | 0.130441 | 0.360278 | 0.803839 |
| 18 | 13342600 | 13344599 | 22.74644 | 38.9609 | 46.87041 | 20.55432 | 0.117584 | 0.953842 | 0.803844 |
| 15 | 3204800 | 3206399 | 13.60749 | 24.62172 | 29.99261 | 13.00383 | 0.127867 | 0.353788 | 0.803875 |
| 4 | 111787800 | 111788199 | 10.91469 | 21.01381 | 25.47728 | 9.903788 | 0.261904 | -0.08059 | 0.803882 |
| X | 39455000 | 39461799 | 168.1853 | 250.7854 | 277.4719 | 114.3531 | 0.112882 | 3.58248 | 0.803913 |
| 12 | 28847200 | 28850199 | 36.1542 | 58.76411 | 70.19282 | 30.11695 | 0.112955 | 1.580414 | 0.803917 |
| 16 | 84153800 | 84154199 | 6.239408 | 12.15255 | 20.79485 | 7.59397 | -0.14507 | -0.35436 | 0.803928 |
| 2 | 19917600 | 19921599 | 35.34535 | 57.37047 | 79.86474 | 34.23207 | -0.10926 | 1.610543 | 0.803958 |
| 3 | 91620400 | 91623999 | 57.62671 | 91.61477 | 123.4257 | 51.50878 | -0.09539 | 2.254499 | 0.803987 |
| 16 | 62198800 | 62202599 | 70.42125 | 111.1019 | 129.5958 | 53.78707 | 0.102507 | 2.463065 | 0.803991 |
| 7 | 26807400 | 26809599 | 28.40035 | 46.91968 | 65.96736 | 28.49723 | -0.10962 | 1.345401 | 0.803993 |
| 7 | 4571000 | 4574999 | 23.0652 | 39.10125 | 55.49294 | 24.30375 | -0.12188 | 1.057783 | 0.804001 |
| 16 | 30715400 | 30739599 | 379.3377 | 526.6007 | 626.9796 | 260.2717 | -0.09398 | 4.691401 | 0.80402 |
| 3 | 131995600 | 131997199 | 17.20887 | 30.42359 | 35.85833 | 15.861 | 0.154669 | 0.542424 | 0.804027 |
| 9 | 4957600 | 4959799 | 26.70823 | 45.24105 | 52.84204 | 23.13717 | 0.158144 | 1.075779 | 0.804036 |
| 16 | 21990000 | 21994599 | 48.66968 | 77.92838 | 106.1388 | 44.73407 | -0.09997 | 2.017896 | 0.804039 |
| 8 | 29682200 | 29684999 | 35.93866 | 58.23976 | 81.10147 | 34.72097 | -0.11185 | 1.647079 | 0.804039 |
| 7 | 119861000 | 119863199 | 31.05052 | 50.98801 | 71.65914 | 30.86734 | -0.11205 | 1.429404 | 0.804055 |
| 19 | 54315800 | 54318399 | 19.22165 | 33.76154 | 39.01951 | 17.26845 | 0.179682 | 0.633674 | 0.804094 |
| 5 | 99411800 | 99413199 | 12.97942 | 23.33057 | 34.07245 | 14.8886 | -0.13984 | 0.394851 | 0.804146 |
| 3 | 52353400 | 52355399 | 15.82246 | 27.87265 | 32.99837 | 14.60642 | 0.146316 | 0.567006 | 0.80415 |
| 19 | 54899400 | 54900599 | 15.82246 | 27.87265 | 32.99837 | 14.60642 | 0.146316 | 0.567006 | 0.80415 |
| 14 | 8420600 | 8424399 | 72.99349 | 115.0099 | 134.8721 | 55.96503 | 0.093372 | 2.492853 | 0.804161 |
| 6 | 131945400 | 131949799 | 51.03022 | 82.33566 | 90.40109 | 38.52764 | 0.223606 | 1.783046 | 0.804173 |
| 15 | 59659800 | 59660999 | 17.08273 | 30.01615 | 36.0654 | 15.96624 | 0.123982 | 0.611242 | 0.804174 |
| 2 | 44824200 | 44826999 | 57.3048 | 91.01521 | 123.0766 | 51.31352 | -0.10211 | 2.282261 | 0.804196 |
| 7 | 103172400 | 103173999 | 18.27301 | 31.83143 | 45.73226 | 20.11203 | -0.13318 | 0.764224 | 0.804214 |
| 13 | 113252000 | 113254199 | 28.41326 | 48.2558 | 54.02868 | 23.67561 | 0.219015 | 1.058837 | 0.804215 |
| 15 | 49274400 | 49275599 | 11.43139 | 21.05378 | 31.37291 | 13.29393 | -0.13856 | 0.240844 | 0.804251 |
| 2 | 100963400 | 100963999 | 12.2333 | 22.07287 | 32.14198 | 13.98497 | -0.13134 | 0.365016 | 0.804257 |
| 12 | 87844600 | 87862599 | 259.1831 | 375.7205 | 401.3726 | 165.497 | 0.124954 | 4.047496 | 0.804292 |
| 1 | 117609800 | 117613799 | 81.43809 | 127.588 | 148.5802 | 61.62956 | 0.097127 | 2.610017 | 0.804301 |
| 7 | 138535200 | 138539399 | 40.56831 | 65.66125 | 78.42974 | 33.54664 | 0.107289 | 1.676663 | 0.804311 |
| X | 147505200 | 147506199 | 10.64328 | 19.90159 | 30.62937 | 12.68938 | -0.16008 | 0.144109 | 0.804324 |
| 8 | 85809800 | 85811199 | 10.40824 | 19.23072 | 28.79228 | 12.14751 | -0.1408 | 0.207839 | 0.80436 |
| 4 | 59566000 | 59567399 | 19.38415 | 33.8598 | 40.33814 | 17.77043 | 0.136251 | 0.723958 | 0.804394 |

| | | | | | | | | | |
|---|---|---|---|---|---|---|---|---|---|
| 12 | 104984400 | 104992199 | 94.84422 | 144.3711 | 240.3063 | 99.58254 | -0.44998 | 3.161363 | 0.804431 |
| 19 | 25445800 | 25450799 | 59.2726 | 93.6763 | 148.5891 | 61.72452 | -0.33829 | 2.419714 | 0.804435 |
| 8 | 92156400 | 92164399 | 132.567 | 199.402 | 259.7107 | 107.2393 | -0.11101 | 3.410267 | 0.804443 |
| 4 | 69872400 | 69873599 | 16.83028 | 29.8568 | 27.8784 | 12.31797 | 0.491144 | 0.492005 | 0.804445 |
| 1 | 146775000 | 146777799 | 34.42065 | 56.51785 | 67.50748 | 29.07808 | 0.1219 | 1.434914 | 0.804447 |
| 5 | 125699000 | 125702199 | 43.29073 | 69.47271 | 95.93862 | 40.57432 | -0.11594 | 1.938424 | 0.804467 |
| 15 | 70529600 | 70532399 | 39.85224 | 65.41548 | 59.9274 | 25.80956 | 0.503455 | 1.447466 | 0.804484 |
| 13 | 98521000 | 98524199 | 25.79221 | 43.89466 | 51.22181 | 22.45093 | 0.159876 | 1.026571 | 0.804508 |
| 8 | 69471200 | 69474199 | 37.90726 | 61.89816 | 61.32687 | 26.35766 | 0.386643 | 1.510227 | 0.804509 |
| 10 | 39447400 | 39449399 | 17.89273 | 31.05615 | 44.22656 | 19.40557 | -0.121 | 0.811093 | 0.804514 |
| X | 93584400 | 93587399 | 25.96536 | 43.87815 | 51.18406 | 22.39505 | 0.15997 | 1.093585 | 0.80452 |
| 4 | 129307000 | 129312599 | 43.7458 | 70.38811 | 96.95048 | 41.16177 | -0.1076 | 1.856572 | 0.804564 |
| 19 | 39632800 | 39633399 | 9.451914 | 18.03885 | 29.09516 | 11.58568 | -0.1787 | -0.00722 | 0.804592 |
| 3 | 150925000 | 150926999 | 16.51288 | 29.19014 | 34.8605 | 15.41996 | 0.135694 | 0.537041 | 0.804594 |
| 1 | 45728600 | 45729599 | 8.933619 | 17.10467 | 27.47155 | 10.85026 | -0.16189 | -0.0435 | 0.804614 |
| 11 | 3761200 | 3765599 | 56.0882 | 89.10261 | 121.4038 | 50.62729 | -0.11269 | 2.273282 | 0.804628 |
| 3 | 95455200 | 95457599 | 21.32713 | 37.06314 | 43.55232 | 19.12463 | 0.155096 | 0.806777 | 0.804632 |
| 8 | 76437200 | 76439999 | 35.55455 | 57.48028 | 80.78357 | 34.51716 | -0.12843 | 1.695421 | 0.804632 |
| 18 | 4743000 | 4744999 | 12.0413 | 21.61661 | 32.10717 | 14.04497 | -0.165 | 0.392559 | 0.804633 |
| X | 81083000 | 81085399 | 20.05096 | 35.2182 | 39.56941 | 17.51182 | 0.220363 | 0.641151 | 0.804633 |
| 8 | 62994600 | 62998799 | 63.29343 | 100.6079 | 118.5121 | 49.45042 | 0.09778 | 2.262031 | 0.804642 |
| 14 | 91987600 | 91987999 | 7.552669 | 14.59348 | 23.76687 | 9.124692 | -0.14291 | -0.15965 | 0.804648 |
| 16 | 49519200 | 49519999 | 8.075543 | 15.43572 | 24.55306 | 9.744609 | -0.15371 | -0.04075 | 0.804657 |
| 4 | 74430200 | 74432799 | 38.59529 | 62.40621 | 73.22247 | 31.29295 | 0.131636 | 1.692317 | 0.804659 |
| 4 | 153005200 | 153007799 | 29.49273 | 48.70615 | 69.05494 | 29.82333 | -0.12178 | 1.353296 | 0.804673 |
| 4 | 134412800 | 134416599 | 49.54684 | 79.21403 | 107.6432 | 45.27121 | -0.09904 | 2.066074 | 0.804678 |
| 12 | 5663600 | 5664599 | 21.62252 | 37.03006 | 44.27962 | 19.41793 | 0.126305 | 0.948355 | 0.804687 |
| 14 | 118730400 | 118732599 | 24.28576 | 40.72012 | 57.61351 | 25.13303 | -0.1179 | 1.16357 | 0.804693 |
| 9 | 62904800 | 62906599 | 21.91334 | 37.57385 | 45.2889 | 19.86247 | 0.115067 | 0.938973 | 0.804703 |
| 2 | 51778800 | 51781999 | 51.61823 | 82.58316 | 97.59839 | 41.08717 | 0.103124 | 2.043793 | 0.804716 |
| 13 | 36905800 | 36909599 | 33.54434 | 54.04088 | 98.80139 | 42.0233 | -0.513 | 1.802691 | 0.804718 |
| 11 | 36134800 | 36136199 | 14.37413 | 25.68603 | 30.9041 | 13.57126 | 0.1333 | 0.434965 | 0.80472 |
| 14 | 121014800 | 121017199 | 23.88493 | 40.097 | 56.5588 | 24.69196 | -0.11358 | 1.146686 | 0.804726 |
| 1 | 81598800 | 81600399 | 24.04589 | 41.21281 | 40.50488 | 17.76372 | 0.408193 | 0.944668 | 0.804737 |
| 17 | 33444000 | 33462799 | 275.1535 | 398.315 | 411.6717 | 169.775 | 0.170651 | 4.070789 | 0.804741 |
| 15 | 33890800 | 33894399 | 34.38297 | 55.72783 | 78.24229 | 33.50094 | -0.12345 | 1.636981 | 0.80476 |
| 11 | 80904200 | 80908999 | 43.76611 | 70.41017 | 96.79103 | 41.08509 | -0.10504 | 1.861665 | 0.804819 |
| 6 | 123299800 | 123302599 | 28.9504 | 48.63857 | 56.30729 | 24.59321 | 0.170379 | 1.152644 | 0.804834 |
| 9 | 115807200 | 115811399 | 47.78614 | 76.79669 | 91.28594 | 38.6893 | 0.1024 | 1.897237 | 0.804844 |
| 8 | 25613400 | 25621799 | 189.2223 | 277.3136 | 352.8626 | 145.3463 | -0.11203 | 3.874776 | 0.804853 |
| 9 | 17999600 | 18006399 | 83.39536 | 130.5047 | 152.2188 | 63.14625 | 0.093543 | 2.630934 | 0.8049 |
| X | 22271800 | 22274399 | 32.76182 | 54.54203 | 61.94876 | 26.90302 | 0.198143 | 1.268874 | 0.804901 |
| 5 | 138573200 | 138578199 | 42.4968 | 68.39321 | 94.00628 | 39.92427 | -0.10417 | 1.848315 | 0.804905 |
| 3 | 23137000 | 23138399 | 14.30225 | 25.29384 | 36.56101 | 16.17068 | -0.13801 | 0.521336 | 0.804906 |
| 5 | 93055200 | 93059199 | 42.96784 | 69.21487 | 95.89935 | 40.78254 | -0.11406 | 1.820518 | 0.804911 |
| 2 | 111709000 | 111722999 | 191.2086 | 283.8851 | 311.6778 | 128.3894 | 0.118271 | 3.661352 | 0.804916 |
| 17 | 13951600 | 13953999 | 17.37819 | 30.40739 | 36.29028 | 16.00891 | 0.133773 | 0.680583 | 0.804917 |

| | | | | | | | | | |
|---|---|---|---|---|---|---|---|---|---|
| 10 | 120585400 | 120588199 | 23.04251 | 38.77571 | 54.92044 | 23.99927 | -0.1194 | 1.119836 | 0.804926 |
| 14 | 71205800 | 71206999 | 11.07473 | 21.11095 | 26.14769 | 10.43812 | 0.196018 | 0.007687 | 0.804932 |
| 4 | 150375000 | 150377999 | 46.32652 | 74.50049 | 88.7501 | 37.65083 | 0.100454 | 1.873868 | 0.804932 |
| 9 | 54828400 | 54830999 | 38.26147 | 62.01137 | 74.18299 | 31.76184 | 0.106927 | 1.643212 | 0.804944 |
| 16 | 92411800 | 92416399 | 37.77679 | 61.77 | 72.23773 | 31.0626 | 0.149456 | 1.499242 | 0.804945 |
| 13 | 5199000 | 5200799 | 18.64582 | 32.72494 | 38.53326 | 17.04216 | 0.152876 | 0.640905 | 0.804955 |
| X | 20767800 | 20769399 | 20.09027 | 35.51133 | 30.59304 | 13.53252 | 0.604611 | 0.554488 | 0.804975 |
| 5 | 109991800 | 109993199 | 17.64947 | 30.65943 | 44.19064 | 19.39995 | -0.13829 | 0.796168 | 0.804982 |
| 13 | 50154200 | 50156599 | 17.64947 | 30.65943 | 44.19064 | 19.39995 | -0.13829 | 0.796168 | 0.804982 |
| 7 | 12452600 | 12454799 | 24.25997 | 41.27926 | 49.56263 | 21.72337 | 0.118956 | 1.022642 | 0.804983 |
| 11 | 102601400 | 102603999 | 34.92478 | 56.63781 | 78.80994 | 33.75797 | -0.10976 | 1.62872 | 0.804995 |
| 12 | 103017200 | 103018999 | 16.82281 | 29.70025 | 35.36737 | 15.64814 | 0.139115 | 0.553296 | 0.805028 |
| 14 | 29599600 | 29602199 | 30.24637 | 49.9288 | 60.01087 | 25.90953 | 0.115819 | 1.353736 | 0.805029 |
| 13 | 38489400 | 38492799 | 29.54696 | 48.39766 | 69.10494 | 29.73559 | -0.13628 | 1.453519 | 0.805042 |
| 15 | 96016000 | 96020199 | 45.73052 | 73.68528 | 86.75307 | 36.90758 | 0.120818 | 1.809923 | 0.805049 |
| 6 | 60887200 | 60888999 | 17.45741 | 30.76522 | 36.48301 | 16.14756 | 0.14427 | 0.580661 | 0.805066 |
| 12 | 115659600 | 115661399 | 31.9479 | 52.70778 | 63.46701 | 27.39276 | 0.113014 | 1.368298 | 0.805068 |
| 12 | 7199000 | 7202599 | 68.80785 | 108.817 | 128.0588 | 53.21367 | 0.09252 | 2.40602 | 0.80507 |
| 4 | 145296000 | 145302599 | 168.6348 | 251.603 | 282.2468 | 116.3341 | 0.093735 | 3.570071 | 0.805076 |
| 19 | 3609400 | 3611999 | 32.24203 | 52.82421 | 74.49624 | 32.05854 | -0.11833 | 1.462633 | 0.805079 |
| 14 | 122865200 | 122866199 | 18.6735 | 32.27771 | 46.35753 | 20.3376 | -0.1343 | 0.857096 | 0.805092 |
| 7 | 39145800 | 39149599 | 49.35401 | 79.63487 | 89.47108 | 38.13359 | 0.19055 | 1.778476 | 0.805098 |
| 2 | 80065800 | 80067999 | 17.87109 | 31.20186 | 45.14233 | 19.88086 | -0.14299 | 0.728073 | 0.805118 |
| 5 | 30489800 | 30495399 | 51.66209 | 82.68593 | 97.66449 | 41.13797 | 0.104566 | 2.032532 | 0.805142 |
| 7 | 75056200 | 75060199 | 34.85882 | 57.40723 | 67.23435 | 28.99677 | 0.151859 | 1.398849 | 0.805145 |
| 4 | 27140600 | 27141399 | 11.65551 | 21.9446 | 22.03854 | 9.024687 | 0.464717 | 0.086762 | 0.805147 |
| 12 | 28104000 | 28104999 | 13.25589 | 23.87656 | 28.5265 | 12.4388 | 0.150409 | 0.373989 | 0.805154 |
| 1 | 8708200 | 8712799 | 47.8657 | 77.08661 | 90.19492 | 38.34368 | 0.128897 | 1.832679 | 0.80516 |
| 8 | 113852600 | 113853199 | 9.275178 | 17.36339 | 27.46359 | 11.35568 | -0.19615 | 0.120468 | 0.805161 |
| 17 | 80305800 | 80310599 | 55.71479 | 88.8556 | 104.8195 | 43.93041 | 0.100776 | 2.147086 | 0.805175 |
| 15 | 56818600 | 56818999 | 7.768131 | 15.03648 | 25.04722 | 9.536779 | -0.16339 | -0.17922 | 0.805187 |
| 17 | 85840400 | 85842799 | 18.89313 | 32.51912 | 46.36462 | 20.33808 | -0.12519 | 0.893762 | 0.805199 |
| 3 | 34571400 | 34571799 | 6.376305 | 12.42755 | 16.93926 | 6.131642 | 0.191687 | -0.3972 | 0.805205 |
| 14 | 45012400 | 45014399 | 19.51626 | 34.32215 | 38.93621 | 17.24051 | 0.206619 | 0.617362 | 0.805223 |
| 8 | 19841800 | 19849799 | 146.0242 | 222.1337 | 235.0764 | 97.39649 | 0.20209 | 3.187989 | 0.805259 |
| 10 | 8364600 | 8365399 | 7.877469 | 15.09012 | 24.05363 | 9.486569 | -0.14853 | -0.06651 | 0.80531 |
| 3 | 93107800 | 93111199 | 55.7744 | 88.73162 | 119.8179 | 50.04531 | -0.09768 | 2.230025 | 0.80532 |
| 16 | 11947600 | 11950399 | 29.71553 | 49.66528 | 58.53522 | 25.48368 | 0.144777 | 1.212622 | 0.805332 |
| 9 | 87373600 | 87376799 | 36.90888 | 59.86024 | 70.71269 | 30.29133 | 0.125988 | 1.624267 | 0.805366 |
| 18 | 64557600 | 64561799 | 37.37852 | 60.37937 | 83.68101 | 35.71727 | -0.1097 | 1.724698 | 0.805374 |
| 9 | 13974600 | 13978799 | 85.72678 | 132.8965 | 175.3639 | 72.83859 | -0.0948 | 2.79751 | 0.805386 |
| 1 | 157651800 | 157653599 | 12.11603 | 22.12473 | 32.81448 | 14.06527 | -0.14362 | 0.297717 | 0.805398 |
| 13 | 78098600 | 78102799 | 58.29877 | 92.68787 | 124.9218 | 52.13575 | -0.0959 | 2.254139 | 0.805406 |
| 2 | 55924000 | 55926199 | 26.44921 | 44.62454 | 52.89614 | 23.13295 | 0.136609 | 1.104825 | 0.80542 |
| 10 | 65501600 | 65501999 | 10.08871 | 18.86041 | 28.43015 | 11.78274 | -0.13079 | 0.134509 | 0.805431 |
| 15 | 98202000 | 98206599 | 39.60261 | 64.69517 | 74.18613 | 31.8896 | 0.176792 | 1.517469 | 0.805487 |
| 18 | 56352400 | 56357399 | 51.21594 | 81.59381 | 112.3217 | 47.00533 | -0.12321 | 2.179601 | 0.805491 |

| | | | | | | | | | |
|---|---|---|---|---|---|---|---|---|---|
| 13 | 22681600 | 22683999 | 10.59729 | 20.25896 | 20.67898 | 8.203431 | 0.481375 | -0.0295 | 0.805514 |
| 1 | 120218400 | 120222199 | 53.71619 | 85.96486 | 101.6601 | 42.81487 | 0.102637 | 2.037785 | 0.805527 |
| 2 | 21626200 | 21632799 | 62.1884 | 99.56771 | 90.52677 | 38.15035 | 0.481929 | 2.0302 | 0.805528 |
| X | 108824800 | 108825999 | 15.91721 | 28.14982 | 42.32519 | 18.72015 | -0.19461 | 0.543436 | 0.805542 |
| 17 | 51058000 | 51059999 | 16.33354 | 28.65708 | 33.68114 | 14.89199 | 0.155666 | 0.625538 | 0.805567 |
| 10 | 111747000 | 111751399 | 57.98102 | 92.44364 | 109.275 | 45.77976 | 0.097381 | 2.158954 | 0.805573 |
| 9 | 117240000 | 117241399 | 13.27924 | 24.14356 | 29.5296 | 12.72633 | 0.127655 | 0.322928 | 0.805593 |
| 7 | 23986000 | 23988199 | 25.2165 | 42.12822 | 59.32236 | 25.81957 | -0.11105 | 1.204989 | 0.8056 |
| 12 | 112738000 | 112742399 | 66.36337 | 104.2366 | 143.4025 | 59.48049 | -0.14016 | 2.552129 | 0.805601 |
| 12 | 99072800 | 99075399 | 33.92754 | 55.44576 | 66.55657 | 28.61399 | 0.111361 | 1.493974 | 0.805602 |
| 5 | 55362600 | 55366599 | 48.32319 | 77.62787 | 92.28083 | 39.08767 | 0.101568 | 1.911859 | 0.805605 |
| 12 | 113864400 | 113872199 | 94.58588 | 148.169 | 161.1017 | 66.82417 | 0.195896 | 2.62162 | 0.805629 |
| 5 | 141810800 | 141813199 | 19.80981 | 34.44247 | 41.51002 | 18.2332 | 0.11896 | 0.792328 | 0.805654 |
| 3 | 39340800 | 39342399 | 22.72746 | 38.51917 | 54.38974 | 23.81881 | -0.11457 | 1.057496 | 0.805656 |
| 6 | 132659600 | 132659999 | 9.510521 | 18.03634 | 22.78911 | 9.196748 | 0.154183 | 0.020284 | 0.805669 |
| 7 | 29622800 | 29638199 | 228.8258 | 330.7863 | 411.1334 | 169.5844 | -0.09735 | 4.086518 | 0.805675 |
| 17 | 63285800 | 63289199 | 22.31372 | 37.96115 | 53.88043 | 23.61555 | -0.12165 | 1.020711 | 0.805713 |
| 8 | 68234800 | 68236599 | 10.897 | 19.70457 | 35.87786 | 15.56803 | -0.44911 | 0.35186 | 0.805718 |
| 9 | 86775800 | 86778799 | 27.94138 | 46.53265 | 67.31458 | 29.23951 | -0.14981 | 1.254718 | 0.805749 |
| 6 | 3786600 | 3795599 | 167.2168 | 250.3552 | 238.097 | 98.20307 | 0.336355 | 3.504028 | 0.805749 |
| 5 | 6305000 | 6305799 | 13.00233 | 23.81035 | 29.01499 | 12.39094 | 0.14139 | 0.277208 | 0.805753 |
| 6 | 85822400 | 85823399 | 13.00233 | 23.81035 | 29.01499 | 12.39094 | 0.14139 | 0.277208 | 0.805753 |
| 12 | 10294200 | 10294599 | 7.869062 | 15.10017 | 24.09471 | 9.455588 | -0.14337 | -0.083 | 0.805771 |
| 11 | 62133600 | 62136999 | 34.24697 | 55.67437 | 77.42092 | 33.22108 | -0.10563 | 1.578084 | 0.805773 |
| 15 | 33007600 | 33008399 | 19.13778 | 33.21993 | 39.55335 | 17.35598 | 0.136252 | 0.809651 | 0.805774 |
| 3 | 56040000 | 56040799 | 8.310712 | 15.46606 | 30.53296 | 12.75201 | -0.52588 | 0.154194 | 0.80578 |
| 18 | 6291000 | 6292999 | 17.03824 | 29.93984 | 36.03056 | 15.95165 | 0.121654 | 0.609162 | 0.805796 |
| 4 | 144660200 | 144661599 | 11.87733 | 21.4094 | 31.62177 | 13.77566 | -0.15295 | 0.365928 | 0.805809 |
| 3 | 56767400 | 56768399 | 10.93209 | 20.35572 | 25.16519 | 10.50026 | 0.145402 | 0.158322 | 0.80583 |
| 6 | 38050600 | 38057999 | 62.52341 | 98.46961 | 161.2634 | 66.91224 | -0.38834 | 2.498191 | 0.805845 |
| 19 | 35316200 | 35317199 | 17.66412 | 29.92469 | 56.3355 | 24.66766 | -0.52781 | 1.045854 | 0.805855 |
| 14 | 62263200 | 62263999 | 9.679181 | 18.22727 | 28.04378 | 11.47993 | -0.14546 | 0.085402 | 0.805879 |
| 8 | 4116200 | 4116999 | 28.11998 | 46.65776 | 55.55138 | 24.06937 | 0.130027 | 1.286042 | 0.80589 |
| 16 | 92449200 | 92451199 | 15.51655 | 27.03247 | 40.11411 | 17.64242 | -0.17935 | 0.741434 | 0.805902 |
| 7 | 29506400 | 29511999 | 66.79979 | 106.301 | 121.2146 | 50.64779 | 0.146255 | 2.231657 | 0.805908 |
| 10 | 35550200 | 35553199 | 34.15135 | 55.16654 | 92.66463 | 39.55189 | -0.38648 | 1.722689 | 0.805922 |
| 18 | 77842600 | 77845799 | 43.04142 | 69.10938 | 95.98524 | 40.62689 | -0.12331 | 1.919662 | 0.805927 |
| 3 | 52709800 | 52711199 | 14.47319 | 25.69242 | 30.11473 | 13.29862 | 0.165538 | 0.47737 | 0.805935 |
| 8 | 72703200 | 72712799 | 101.1425 | 156.3993 | 179.4526 | 74.54039 | 0.103864 | 2.850371 | 0.805936 |
| 1 | 103908200 | 103908999 | 10.08069 | 19.0443 | 29.83788 | 12.13964 | -0.16448 | 0.069703 | 0.805946 |
| 13 | 88921400 | 88925599 | 70.21998 | 110.9107 | 130.0162 | 53.99273 | 0.096666 | 2.435871 | 0.80595 |
| 1 | 102728400 | 102734599 | 111.7924 | 171.2971 | 197.0534 | 81.79038 | 0.090109 | 3.028961 | 0.805969 |
| 8 | 11132400 | 11133999 | 16.91871 | 29.69739 | 35.64701 | 15.7673 | 0.125468 | 0.626258 | 0.805996 |
| 10 | 114886000 | 114887999 | 23.48378 | 40.00573 | 47.9503 | 21.01869 | 0.121642 | 1.005483 | 0.806009 |
| 5 | 8962600 | 8963999 | 24.26493 | 40.52535 | 57.58873 | 25.0599 | -0.12411 | 1.204644 | 0.806029 |
| 4 | 117949400 | 117952599 | 26.98437 | 45.02413 | 53.92044 | 23.4495 | 0.121927 | 1.219807 | 0.806044 |
| 11 | 56762600 | 56765199 | 18.33468 | 32.43959 | 36.79071 | 16.27084 | 0.210295 | 0.541781 | 0.806048 |

| | | | | | | | | | |
|---|---|---|---|---|---|---|---|---|---|
| 6 | 69469000 | 69471599 | 44.38763 | 71.46536 | 85.09712 | 36.16191 | 0.103175 | 1.834067 | 0.806052 |
| 17 | 77104600 | 77105399 | 9.897328 | 18.74847 | 23.83447 | 9.641883 | 0.142558 | 0.036 | 0.806057 |
| 17 | 48062000 | 48062999 | 12.46617 | 22.27636 | 32.70312 | 14.3614 | -0.15183 | 0.42851 | 0.806058 |
| 1 | 83151200 | 83152799 | 13.04191 | 24.15034 | 29.20993 | 12.28657 | 0.167881 | 0.2167 | 0.806082 |
| X | 14251800 | 14252199 | 7.488117 | 14.58728 | 19.66766 | 7.165496 | 0.198795 | -0.34835 | 0.806115 |
| 16 | 4046800 | 4049999 | 28.23035 | 46.48024 | 65.81524 | 28.38192 | -0.12126 | 1.385166 | 0.806183 |
| 13 | 55960600 | 55961799 | 8.568815 | 15.97496 | 25.90655 | 10.78704 | -0.24044 | 0.133153 | 0.80619 |
| 4 | 95049400 | 95049999 | 7.566276 | 14.5024 | 23.52724 | 9.263542 | -0.17151 | -0.0773 | 0.806198 |
| 5 | 122415800 | 122417199 | 7.566276 | 14.5024 | 23.52724 | 9.263542 | -0.17151 | -0.0773 | 0.806198 |
| 3 | 9633400 | 9636199 | 19.2707 | 32.64222 | 57.46422 | 25.16092 | -0.43146 | 1.054866 | 0.806206 |
| 3 | 4090400 | 4093799 | 45.27263 | 73.59055 | 66.25355 | 28.42808 | 0.51991 | 1.594128 | 0.806212 |
| 9 | 124078800 | 124079999 | 17.13908 | 30.06145 | 35.72733 | 15.79274 | 0.139838 | 0.640886 | 0.80624 |
| 3 | 153699200 | 153700599 | 12.9522 | 23.19801 | 33.65457 | 14.75068 | -0.13297 | 0.419436 | 0.806246 |
| 3 | 90978200 | 90983199 | 110.0692 | 167.9988 | 218.6011 | 90.6522 | -0.09194 | 3.108197 | 0.806266 |
| 9 | 9605200 | 9607199 | 29.43588 | 48.3925 | 68.01207 | 29.31068 | -0.11134 | 1.409726 | 0.806275 |
| 11 | 30853400 | 30857799 | 49.02377 | 79.0338 | 90.29299 | 38.44109 | 0.16505 | 1.803036 | 0.806285 |
| X | 75087000 | 75087999 | 15.15707 | 27.32358 | 32.45418 | 14.13679 | 0.159739 | 0.391754 | 0.806297 |
| Y | 75154400 | 75155199 | 4.989981 | 9.707796 | 19.06619 | 7.036326 | -0.35537 | -0.34102 | 0.806302 |
| 1 | 178366200 | 178368199 | 15.17304 | 26.72984 | 38.16337 | 16.89251 | -0.12242 | 0.573351 | 0.806311 |
| 6 | 33717000 | 33723599 | 126.1325 | 189.4936 | 290.1511 | 119.7698 | -0.34583 | 3.434339 | 0.806312 |
| 3 | 70478600 | 70484399 | 68.72399 | 108.2112 | 144.8075 | 60.08256 | -0.09757 | 2.504271 | 0.806322 |
| 10 | 8276800 | 8282199 | 46.48541 | 74.62488 | 102.9189 | 43.54893 | -0.11354 | 1.932771 | 0.806327 |
| 6 | 135190800 | 135191799 | 12.56552 | 22.93894 | 28.03296 | 12.02008 | 0.133502 | 0.290157 | 0.806334 |
| 2 | 77788200 | 77789599 | 16.49168 | 28.6901 | 41.64415 | 18.29883 | -0.1479 | 0.769423 | 0.806345 |
| 12 | 67659400 | 67660399 | 15.61375 | 27.77101 | 33.39578 | 14.7234 | 0.130205 | 0.47843 | 0.806353 |
| 15 | 34747800 | 34749599 | 16.20348 | 28.43088 | 41.28093 | 18.25328 | -0.14806 | 0.634756 | 0.806371 |
| 11 | 114119600 | 114121799 | 32.57966 | 52.98563 | 74.40746 | 31.93279 | -0.1193 | 1.566153 | 0.806396 |
| 6 | 144949800 | 144951399 | 9.121806 | 17.13316 | 26.39083 | 10.85083 | -0.15243 | 0.090471 | 0.806403 |
| 6 | 117057800 | 117060799 | 13.01811 | 22.84803 | 41.97063 | 18.56254 | -0.48532 | 0.616376 | 0.806409 |
| 4 | 148737200 | 148739799 | 36.76413 | 59.28609 | 84.02629 | 35.78612 | -0.14472 | 1.77409 | 0.806438 |
| 15 | 93852200 | 93856799 | 59.96638 | 95.97736 | 91.27888 | 38.4484 | 0.41657 | 2.037273 | 0.806439 |
| 2 | 4613200 | 4614399 | 12.39528 | 22.78176 | 28.07637 | 11.93065 | 0.130514 | 0.250822 | 0.806444 |
| 1 | 45007000 | 45011399 | 51.20903 | 82.36135 | 94.46872 | 40.08322 | 0.155138 | 1.878441 | 0.806445 |
| 10 | 37049200 | 37050799 | 14.1752 | 25.72066 | 30.94467 | 13.36984 | 0.148709 | 0.342464 | 0.806459 |
| 15 | 8945600 | 8951599 | 157.2448 | 235.6884 | 266.3083 | 109.8627 | 0.089337 | 3.483032 | 0.806473 |
| 5 | 133073800 | 133076999 | 21.16926 | 36.29348 | 51.7181 | 22.68192 | -0.12562 | 0.944458 | 0.806474 |
| 2 | 23673200 | 23678799 | 60.48389 | 96.3792 | 113.3024 | 47.42961 | 0.104417 | 2.180943 | 0.806511 |
| 8 | 109459200 | 109461199 | 12.02672 | 21.50604 | 32.29006 | 14.1717 | -0.18329 | 0.421527 | 0.806548 |
| 14 | 95591600 | 95592399 | 10.18621 | 18.17088 | 36.36223 | 15.99148 | -0.5974 | 0.43153 | 0.806566 |
| 5 | 69308800 | 69309399 | 7.103059 | 13.76277 | 22.75737 | 8.624855 | -0.14678 | -0.21134 | 0.806567 |
| 2 | 77828000 | 77828599 | 9.957987 | 18.74842 | 23.22173 | 9.509323 | 0.164415 | 0.074091 | 0.80659 |
| 16 | 11198200 | 11199599 | 14.4609 | 25.70875 | 30.55245 | 13.47504 | 0.146923 | 0.468533 | 0.806597 |
| 2 | 100973600 | 100974999 | 18.73268 | 32.7033 | 39.28377 | 17.29978 | 0.124177 | 0.724693 | 0.806601 |
| 5 | 45086400 | 45088199 | 24.50905 | 40.88666 | 58.60217 | 25.48267 | -0.13635 | 1.220431 | 0.806603 |
| 5 | 24457400 | 24464399 | 68.39402 | 108.0553 | 145.2628 | 60.33238 | -0.10037 | 2.432457 | 0.80661 |
| 5 | 110228200 | 110234599 | 112.4078 | 172.3791 | 197.8561 | 82.13284 | 0.09421 | 3.013505 | 0.806621 |
| 17 | 40069200 | 40072399 | 36.86851 | 60.72422 | 58.12151 | 25.06787 | 0.44233 | 1.397771 | 0.806653 |

| | | | | | | | | | |
|---|---|---|---|---|---|---|---|---|---|
| 5 | 18949000 | 18952799 | 37.83031 | 61.80285 | 72.35627 | 31.10355 | 0.146932 | 1.509992 | 0.806666 |
| 17 | 57248200 | 57249399 | 20.66597 | 36.01609 | 42.45603 | 18.67648 | 0.151052 | 0.764697 | 0.806672 |
| 12 | 14594800 | 14595799 | 20.14285 | 34.91278 | 42.09936 | 18.47102 | 0.117572 | 0.831654 | 0.806697 |
| 5 | 109055800 | 109059399 | 50.97916 | 81.74766 | 96.85707 | 40.91449 | 0.103268 | 1.972498 | 0.806718 |
| 8 | 47524600 | 47526399 | 16.75183 | 29.54645 | 35.46565 | 15.69465 | 0.127278 | 0.56503 | 0.806731 |
| 14 | 52953800 | 52958399 | 26.27779 | 44.98171 | 41.94852 | 18.39498 | 0.483292 | 0.965705 | 0.806779 |
| 17 | 46672200 | 46684199 | 190.1346 | 281.9023 | 312.7884 | 128.8288 | 0.101208 | 3.685416 | 0.806781 |
| 16 | 57662000 | 57662599 | 8.694706 | 16.63083 | 20.80556 | 8.240729 | 0.192197 | -0.05482 | 0.806791 |
| 1 | 38183600 | 38184799 | 12.68384 | 23.07944 | 34.30915 | 14.77142 | -0.15134 | 0.323367 | 0.806829 |
| 14 | 114904200 | 114905599 | 16.96669 | 30.05284 | 28.82052 | 12.74314 | 0.451776 | 0.507876 | 0.806871 |
| 5 | 101599200 | 101600399 | 12.31613 | 23.1049 | 27.87599 | 11.48504 | 0.194593 | 0.127969 | 0.806871 |
| 1 | 18027000 | 18030199 | 87.33742 | 135.4785 | 150.6202 | 62.55526 | 0.152844 | 2.784913 | 0.806875 |
| 10 | 51480800 | 51483399 | 18.24274 | 31.90605 | 47.82227 | 21.0882 | -0.19371 | 0.707845 | 0.806876 |
| 5 | 14358000 | 14359199 | 13.22692 | 23.44668 | 34.11461 | 15.08163 | -0.14593 | 0.485751 | 0.806897 |
| 16 | 97546600 | 97551399 | 27.05478 | 44.68695 | 63.64006 | 27.48773 | -0.12811 | 1.341419 | 0.806902 |
| 7 | 58562000 | 58563199 | 16.81221 | 29.57553 | 35.45351 | 15.69747 | 0.127622 | 0.592196 | 0.806911 |
| 7 | 124370400 | 124372799 | 16.81221 | 29.57553 | 35.45351 | 15.69747 | 0.127622 | 0.592196 | 0.806911 |
| 8 | 112418800 | 112420199 | 20.96619 | 36.05276 | 43.21663 | 18.95664 | 0.12379 | 0.907636 | 0.806926 |
| 18 | 74441800 | 74449999 | 119.234 | 182.2342 | 206.7234 | 85.75352 | 0.107364 | 3.081706 | 0.806954 |
| 6 | 135981400 | 135983399 | 14.71442 | 26.45329 | 31.87829 | 13.92802 | 0.135989 | 0.401991 | 0.80697 |
| 7 | 18644400 | 18646199 | 15.70204 | 27.64033 | 32.59816 | 14.43137 | 0.151513 | 0.574432 | 0.806978 |
| 11 | 20062200 | 20063399 | 9.201823 | 17.05319 | 26.87505 | 11.29206 | -0.20951 | 0.177472 | 0.806997 |
| 14 | 22013400 | 22014799 | 15.90962 | 28.14232 | 33.92257 | 15.0031 | 0.122838 | 0.525403 | 0.807007 |
| 19 | 33733200 | 33734799 | 19.08474 | 33.64629 | 37.82005 | 16.73802 | 0.221092 | 0.585264 | 0.807008 |
| 14 | 83402400 | 83402799 | 9.046484 | 17.51868 | 18.26057 | 6.938412 | 0.509846 | -0.19291 | 0.807018 |
| 4 | 78351800 | 78355399 | 33.03473 | 54.32511 | 65.2249 | 28.11304 | 0.115312 | 1.410251 | 0.807044 |
| 13 | 73324000 | 73328599 | 34.55848 | 56.01991 | 78.23902 | 33.5029 | -0.11567 | 1.635044 | 0.807054 |
| X | 44341600 | 44342799 | 19.24871 | 33.98208 | 30.75042 | 13.60547 | 0.533435 | 0.562902 | 0.807055 |
| 4 | 145986600 | 146015599 | 1333.066 | 1670.849 | 1794.955 | 754.6911 | -0.09373 | 6.329078 | 0.807071 |
| 11 | 12106800 | 12109399 | 28.05052 | 47.15722 | 55.67366 | 24.32123 | 0.142257 | 1.144999 | 0.807083 |
| 14 | 118734400 | 118737599 | 36.88507 | 59.92977 | 71.60186 | 30.7131 | 0.112072 | 1.588075 | 0.807098 |
| 6 | 70163200 | 70164199 | 10.75932 | 19.45801 | 35.89331 | 15.57234 | -0.46752 | 0.350969 | 0.807104 |
| 1 | 96021200 | 96021599 | 10.93027 | 20.47729 | 25.29004 | 10.44354 | 0.159303 | 0.122664 | 0.807127 |
| 6 | 8240200 | 8250999 | 217.7764 | 318.3125 | 351.2627 | 144.7114 | 0.089684 | 3.91878 | 0.807137 |
| 2 | 107061800 | 107062599 | 8.311399 | 15.84594 | 25.03236 | 9.989522 | -0.15152 | -0.02234 | 0.807147 |
| 5 | 68544600 | 68546199 | 24.56809 | 41.56217 | 50.12371 | 21.93834 | 0.112028 | 1.074199 | 0.807153 |
| 5 | 6639800 | 6643799 | 50.40808 | 80.73645 | 95.71579 | 40.35979 | 0.100395 | 2.006867 | 0.807161 |
| 2 | 176927200 | 176935199 | 161.9148 | 240.6982 | 306.8387 | 126.4123 | -0.09542 | 3.635086 | 0.807161 |
| 18 | 83485200 | 83488199 | 32.2688 | 52.50342 | 73.77391 | 31.66603 | -0.11968 | 1.559337 | 0.807162 |
| 11 | 7608000 | 7610199 | 19.3875 | 33.97014 | 39.58974 | 17.48484 | 0.167623 | 0.671843 | 0.807214 |
| 1 | 191311600 | 191312999 | 19.28916 | 33.69858 | 40.29604 | 17.75344 | 0.130822 | 0.720004 | 0.807245 |
| 4 | 81382800 | 81385799 | 24.38042 | 40.95622 | 57.68191 | 25.19277 | -0.1114 | 1.142401 | 0.807268 |
| 10 | 113618600 | 113619399 | 12.79825 | 23.16841 | 27.61265 | 11.96488 | 0.159902 | 0.340497 | 0.807269 |
| 2 | 150330600 | 150334199 | 43.58336 | 70.37485 | 82.839 | 35.34006 | 0.124645 | 1.743037 | 0.80727 |
| 7 | 9483000 | 9485399 | 47.12822 | 75.94199 | 74.10152 | 31.52796 | 0.391337 | 1.807335 | 0.807276 |
| 7 | 33794200 | 33807399 | 231.2408 | 333.918 | 414.83 | 171.1336 | -0.0982 | 4.102471 | 0.807293 |
| 14 | 52636200 | 52637599 | 16.11336 | 28.39242 | 34.06778 | 15.07901 | 0.127367 | 0.570051 | 0.807307 |

| | | | | | | | | | |
|---|---|---|---|---|---|---|---|---|---|
| 19 | 45161000 | 45163399 | 32.81054 | 53.28806 | 75.19477 | 32.24465 | -0.12788 | 1.593357 | 0.807332 |
| 17 | 61701200 | 61703199 | 27.32862 | 45.68247 | 55.02454 | 23.95745 | 0.113597 | 1.202496 | 0.807356 |
| 1 | 75303200 | 75305199 | 21.14737 | 36.11742 | 51.16325 | 22.43166 | -0.11795 | 0.977841 | 0.807359 |
| 4 | 130948000 | 130953399 | 46.97295 | 75.51134 | 89.97813 | 38.15077 | 0.099389 | 1.8869 | 0.807435 |
| 1 | 124123600 | 124124199 | 12.16327 | 22.73734 | 21.55475 | 8.934161 | 0.53361 | 0.135724 | 0.807435 |
| 10 | 29316600 | 29317199 | 12.16327 | 22.73734 | 21.55475 | 8.934161 | 0.53361 | 0.135724 | 0.807435 |
| 6 | 68697800 | 68700799 | 67.25957 | 106.4153 | 124.9796 | 51.94571 | 0.095935 | 2.393232 | 0.807468 |
| 1 | 103669600 | 103675399 | 94.0517 | 146.3824 | 166.7729 | 69.22915 | 0.121321 | 2.733645 | 0.807469 |
| 11 | 14790800 | 14797399 | 94.28515 | 146.1282 | 169.5605 | 70.43062 | 0.090203 | 2.804766 | 0.8075 |
| 13 | 35187200 | 35189799 | 47.71115 | 76.68896 | 91.03309 | 38.59147 | 0.104691 | 1.893299 | 0.80751 |
| 14 | 5087400 | 5092599 | 82.74863 | 129.2918 | 150.8772 | 62.59971 | 0.091108 | 2.657144 | 0.807513 |
| 7 | 19662600 | 19668399 | 42.86052 | 68.82266 | 96.06714 | 40.66492 | -0.13044 | 1.917321 | 0.807538 |
| 10 | 108139400 | 108141399 | 24.33354 | 40.78725 | 57.70062 | 25.16598 | -0.1178 | 1.1644 | 0.807549 |
| 15 | 23489800 | 23491599 | 22.73636 | 38.85077 | 46.88371 | 20.5562 | 0.112594 | 0.975155 | 0.807551 |
| 19 | 39236600 | 39238999 | 62.65889 | 99.42164 | 116.5971 | 48.5645 | 0.101676 | 2.313276 | 0.807594 |
| 3 | 38846200 | 38848599 | 18.75405 | 32.67901 | 39.37487 | 17.31939 | 0.119575 | 0.751883 | 0.807626 |
| 4 | 59446400 | 59447199 | 13.74661 | 25.0273 | 29.90564 | 12.8665 | 0.162261 | 0.316572 | 0.807631 |
| 4 | 5286800 | 5290399 | 40.33528 | 65.56078 | 65.65019 | 28.16816 | 0.366734 | 1.585146 | 0.807636 |
| 1 | 120205200 | 120206999 | 17.13699 | 29.90529 | 42.55076 | 18.73366 | -0.1189 | 0.729698 | 0.807637 |
| 9 | 83305000 | 83307599 | 22.05939 | 37.36403 | 52.75233 | 23.09745 | -0.1144 | 1.058454 | 0.807701 |
| 16 | 48995600 | 49001799 | 58.78411 | 93.7415 | 110.4064 | 46.26566 | 0.102906 | 2.15384 | 0.807714 |
| 4 | 19569400 | 19570399 | 10.14254 | 18.66739 | 28.30638 | 12.00558 | -0.16426 | 0.227457 | 0.807734 |
| 2 | 171856200 | 171860399 | 45.47812 | 73.20758 | 87.27003 | 37.07537 | 0.101196 | 1.841159 | 0.807739 |
| 3 | 89735400 | 89736999 | 16.17721 | 28.41698 | 40.77206 | 18.04555 | -0.13076 | 0.61853 | 0.807751 |
| 5 | 130753000 | 130754999 | 21.2271 | 36.44261 | 52.26199 | 22.92299 | -0.13443 | 0.933972 | 0.807752 |
| 18 | 70440000 | 70444999 | 58.46979 | 93.06961 | 126.7392 | 52.97544 | -0.10901 | 2.217703 | 0.807763 |
| 13 | 7596200 | 7600199 | 39.2372 | 64.06637 | 74.13886 | 31.8635 | 0.163142 | 1.523109 | 0.807783 |
| 3 | 33569600 | 33571399 | 25.8344 | 43.46401 | 52.28106 | 22.84434 | 0.115537 | 1.129928 | 0.807817 |
| 6 | 102426200 | 102426999 | 8.076182 | 15.33631 | 24.47455 | 9.849505 | -0.17608 | 0.003937 | 0.807818 |
| 10 | 9351600 | 9352399 | 18.03844 | 31.50813 | 37.84589 | 16.67216 | 0.124519 | 0.713686 | 0.807831 |
| 18 | 6370800 | 6372199 | 31.51551 | 51.74315 | 61.41441 | 26.45257 | 0.131044 | 1.426431 | 0.807833 |
| 3 | 104027600 | 104030599 | 19.87787 | 33.13335 | 63.73401 | 27.69973 | -0.55932 | 1.218878 | 0.807833 |
| 11 | 31064400 | 31064999 | 9.338799 | 17.60494 | 26.98312 | 11.02592 | -0.13787 | 0.072649 | 0.807867 |
| 10 | 113360000 | 113360199 | 4.220886 | 8.256867 | 15.64659 | 5.176023 | -0.16006 | -0.74992 | 0.807889 |
| 13 | 119087200 | 119088799 | 11.461 | 21.8754 | 21.59928 | 8.598973 | 0.523943 | -0.00866 | 0.807892 |
| 5 | 120041200 | 120045799 | 89.16805 | 137.7328 | 181.8762 | 75.54752 | -0.09984 | 2.865246 | 0.80791 |
| 6 | 132013600 | 132023199 | 123.4554 | 189.4337 | 206.8702 | 85.88176 | 0.166637 | 3.006988 | 0.807916 |
| 2 | 71742200 | 71745799 | 25.43446 | 42.40542 | 59.8064 | 25.99663 | -0.11322 | 1.229064 | 0.807926 |
| 11 | 95832400 | 95834399 | 15.60015 | 27.38831 | 39.2244 | 17.35253 | -0.1282 | 0.619818 | 0.807979 |
| 13 | 62136200 | 62139399 | 34.0506 | 57.10182 | 60.45617 | 26.3757 | 0.298972 | 1.194685 | 0.807994 |
| 15 | 91028400 | 91030399 | 13.97543 | 24.58404 | 44.50987 | 19.70746 | -0.46424 | 0.593391 | 0.807998 |
| 5 | 107724800 | 107727399 | 40.86004 | 66.36759 | 66.15075 | 28.3684 | 0.372491 | 1.600819 | 0.808023 |
| 13 | 33838800 | 33843199 | 33.86181 | 56.26969 | 63.78912 | 27.66744 | 0.200692 | 1.292879 | 0.808055 |
| 12 | 59545200 | 59545599 | 7.897747 | 15.28335 | 19.96174 | 7.60981 | 0.183425 | -0.18762 | 0.808064 |
| 6 | 25539400 | 25543399 | 37.53945 | 60.65541 | 83.80564 | 35.7827 | -0.10484 | 1.717312 | 0.808064 |
| 15 | 20274000 | 20278799 | 69.18623 | 109.8379 | 126.279 | 52.62561 | 0.131022 | 2.304092 | 0.808065 |
| 8 | 18514800 | 18516199 | 15.14553 | 26.92064 | 32.46221 | 14.31969 | 0.12581 | 0.47825 | 0.808068 |

| | | | | | | | | | |
|---|---|---|---|---|---|---|---|---|---|
| 5 | 35321800 | 35328399 | 63.54403 | 99.64417 | 163.0017 | 67.60511 | -0.3915 | 2.586096 | 0.80807 |
| 17 | 61089400 | 61091599 | 31.59761 | 51.11872 | 88.06986 | 37.65744 | -0.42042 | 1.677385 | 0.808078 |
| 19 | 12922000 | 12926199 | 44.45671 | 71.88522 | 83.48916 | 35.68154 | 0.146313 | 1.708344 | 0.808079 |
| 7 | 26806600 | 26810199 | 38.77707 | 62.87259 | 75.4483 | 32.31329 | 0.102981 | 1.637127 | 0.808101 |
| 12 | 102770800 | 102772799 | 18.93465 | 33.27867 | 32.31752 | 14.31013 | 0.429558 | 0.610724 | 0.80813 |
| 11 | 27519200 | 27521399 | 19.14087 | 33.52089 | 39.66714 | 17.51148 | 0.145795 | 0.679964 | 0.808139 |
| 7 | 40638600 | 40648999 | 138.4945 | 211.2193 | 193.8005 | 80.35305 | 0.410766 | 3.130142 | 0.808141 |
| 15 | 81342000 | 81348599 | 79.85064 | 124.6888 | 164.8726 | 68.41071 | -0.08964 | 2.668369 | 0.808143 |
| 1 | 106756000 | 106756599 | 6.663825 | 12.93347 | 21.68237 | 8.153062 | -0.15542 | -0.24595 | 0.808149 |
| 10 | 122670600 | 122671799 | 16.33707 | 29.10596 | 27.66296 | 12.17882 | 0.468962 | 0.459527 | 0.808151 |
| 5 | 148112600 | 148121199 | 98.59261 | 151.6326 | 198.9854 | 82.63449 | -0.09564 | 2.951443 | 0.808155 |
| 3 | 153949600 | 153953399 | 39.82387 | 64.61339 | 76.77457 | 32.89765 | 0.11812 | 1.627439 | 0.808164 |
| 7 | 31461000 | 31463199 | 52.07827 | 83.15886 | 95.42133 | 40.06123 | 0.142594 | 2.102121 | 0.808172 |
| 2 | 83709800 | 83710799 | 15.75205 | 27.81862 | 33.49763 | 14.81995 | 0.123481 | 0.541469 | 0.80819 |
| 5 | 4182800 | 4185399 | 39.51756 | 63.84333 | 88.32919 | 37.70783 | -0.10703 | 1.726601 | 0.808201 |
| 9 | 25993600 | 25994199 | 10.29765 | 19.48879 | 24.36378 | 9.875276 | 0.164323 | 0.048922 | 0.80822 |
| 17 | 93582200 | 93582799 | 10.29765 | 19.48879 | 24.36378 | 9.875276 | 0.164323 | 0.048922 | 0.80822 |
| 7 | 5966800 | 5970999 | 64.98132 | 103.9043 | 93.20256 | 39.20547 | 0.499358 | 2.074831 | 0.808237 |
| 7 | 26657600 | 26660999 | 60.88143 | 96.35989 | 130.8052 | 54.37725 | -0.11226 | 2.380664 | 0.808245 |
| 18 | 8180600 | 8183799 | 55.43074 | 88.39491 | 103.8165 | 43.50484 | 0.107014 | 2.150005 | 0.80825 |
| 16 | 91224800 | 91228399 | 33.83578 | 55.1916 | 77.37694 | 33.24302 | -0.11397 | 1.527285 | 0.808266 |
| 1 | 162797400 | 162805599 | 150.2422 | 226.6215 | 253.8828 | 104.9133 | 0.109738 | 3.359384 | 0.808267 |
| 4 | 109963600 | 109964599 | 11.91997 | 21.69536 | 31.72226 | 13.65307 | -0.12701 | 0.312271 | 0.808287 |
| X | 37985000 | 37986599 | 13.61042 | 24.61219 | 39.06482 | 16.95 | -0.25234 | 0.378854 | 0.808319 |
| 13 | 76142600 | 76143399 | 10.85894 | 20.22329 | 24.73968 | 10.31945 | 0.1612 | 0.157266 | 0.808327 |
| 3 | 108456800 | 108459799 | 23.11954 | 39.08142 | 55.53259 | 24.30003 | -0.12388 | 1.080162 | 0.808333 |
| 7 | 53758600 | 53759799 | 12.66367 | 22.98901 | 34.30903 | 14.81456 | -0.15938 | 0.339474 | 0.808341 |
| 3 | 46415600 | 46419999 | 124.9063 | 189.7426 | 216.481 | 89.65442 | 0.092024 | 3.208271 | 0.808346 |
| 2 | 21798200 | 21799799 | 21.84145 | 37.18721 | 42.78497 | 18.75365 | 0.18042 | 0.998565 | 0.808355 |
| 10 | 7425400 | 7434799 | 139.4687 | 210.1481 | 235.9869 | 97.48865 | 0.104983 | 3.376175 | 0.808362 |
| Y | 86754600 | 86755199 | 8.473684 | 15.62075 | 26.93437 | 11.3989 | -0.3459 | 0.200509 | 0.808376 |
| X | 13239000 | 13242999 | 25.89473 | 43.58309 | 63.98689 | 27.96265 | -0.17115 | 1.136751 | 0.808419 |
| 15 | 84200000 | 84203399 | 30.60183 | 50.61415 | 61.18767 | 26.45318 | 0.107706 | 1.33376 | 0.808425 |
| X | 60133800 | 60135599 | 19.02049 | 33.43198 | 38.53484 | 17.0642 | 0.18365 | 0.622375 | 0.808437 |
| 2 | 167398200 | 167399799 | 16.78775 | 29.47093 | 35.18488 | 15.56442 | 0.133277 | 0.623548 | 0.808498 |
| 5 | 19056600 | 19058999 | 45.89641 | 73.74073 | 86.76681 | 36.76026 | 0.116623 | 1.902668 | 0.808502 |
| 8 | 14240400 | 14246799 | 63.35988 | 100.1508 | 135.1911 | 56.16314 | -0.10571 | 2.414313 | 0.808511 |
| 8 | 57477000 | 57480399 | 47.22673 | 75.75999 | 103.5638 | 43.77351 | -0.10202 | 1.957882 | 0.808513 |
| 5 | 138876800 | 138878599 | 20.62359 | 35.12551 | 50.06856 | 21.94682 | -0.12732 | 1.002525 | 0.808526 |
| 5 | 61187800 | 61188999 | 13.1764 | 23.27158 | 41.35956 | 18.29794 | -0.43517 | 0.540092 | 0.808531 |
| 6 | 68759400 | 68760199 | 18.99357 | 33.01006 | 39.51746 | 17.3536 | 0.128625 | 0.791721 | 0.808555 |
| 4 | 61233200 | 61234799 | 26.55789 | 44.88272 | 41.73514 | 18.26117 | 0.485766 | 1.080806 | 0.808591 |
| 10 | 58066000 | 58069799 | 121.0424 | 182.7534 | 242.5322 | 100.2533 | -0.13329 | 3.332886 | 0.808608 |
| 2 | 93671600 | 93672999 | 11.51723 | 20.65676 | 31.29914 | 13.70215 | -0.19388 | 0.396248 | 0.808614 |
| 9 | 82732400 | 82735399 | 28.52827 | 47.26407 | 66.71548 | 28.87656 | -0.11505 | 1.31323 | 0.808649 |
| 7 | 118705000 | 118707199 | 18.41767 | 31.80168 | 45.30905 | 19.87697 | -0.12317 | 0.862241 | 0.808688 |
| 14 | 80236000 | 80239599 | 26.06017 | 43.90645 | 52.85851 | 23.10665 | 0.1143 | 1.117113 | 0.808714 |

| | | | | | | | | | |
|---|---|---|---|---|---|---|---|---|---|
| 18 | 40842000 | 40848399 | 74.57115 | 117.8134 | 136.785 | 56.811 | 0.110824 | 2.434836 | 0.808716 |
| 5 | 73015800 | 73018999 | 36.84905 | 59.70428 | 82.67552 | 35.39013 | -0.10407 | 1.651269 | 0.808727 |
| 17 | 55942200 | 55943799 | 19.56177 | 33.98148 | 41.00075 | 17.99962 | 0.117159 | 0.801564 | 0.808728 |
| 11 | 80384600 | 80385999 | 12.70895 | 22.379 | 40.37969 | 17.87512 | -0.45847 | 0.568704 | 0.808743 |
| 13 | 11103200 | 11105599 | 44.51714 | 71.73921 | 85.62585 | 36.43466 | 0.101368 | 1.805411 | 0.808748 |
| 3 | 10067600 | 10070199 | 20.28525 | 35.47607 | 41.26005 | 18.19348 | 0.170673 | 0.71004 | 0.808797 |
| 2 | 169589000 | 169590399 | 19.40279 | 33.48602 | 47.40607 | 20.79661 | -0.11424 | 0.873696 | 0.808808 |
| X | 133573200 | 133574799 | 18.79089 | 33.01352 | 38.54257 | 17.06111 | 0.165237 | 0.627411 | 0.808815 |
| 4 | 86387400 | 86387799 | 7.608284 | 14.60496 | 23.5672 | 9.239127 | -0.15744 | -0.08833 | 0.808818 |
| 10 | 18624400 | 18626999 | 15.52353 | 26.92344 | 52.79893 | 23.1659 | -0.58106 | 0.815864 | 0.808822 |
| 15 | 15082200 | 15084999 | 40.4312 | 65.54293 | 78.03815 | 33.4181 | 0.114021 | 1.645168 | 0.808848 |
| 11 | 10031000 | 10035599 | 43.12305 | 69.67788 | 82.2674 | 35.12632 | 0.121393 | 1.723437 | 0.808863 |
| 7 | 11378000 | 11379199 | 22.68772 | 38.59281 | 45.3828 | 19.89088 | 0.148958 | 1.016555 | 0.808867 |
| 4 | 27414200 | 27417799 | 85.20297 | 132.7405 | 153.9118 | 63.88103 | 0.0971 | 2.71015 | 0.808891 |
| 1 | 139771000 | 139772399 | 19.80969 | 34.88376 | 32.17621 | 14.24354 | 0.504475 | 0.594737 | 0.808928 |
| 13 | 67554600 | 67555999 | 14.71673 | 26.15561 | 38.12391 | 16.81871 | -0.14649 | 0.486301 | 0.80893 |
| 5 | 128203400 | 128205399 | 21.13628 | 36.04365 | 51.16056 | 22.42774 | -0.12116 | 0.991233 | 0.808938 |
| Y | 6827800 | 6828599 | 7.828434 | 14.61627 | 25.45602 | 10.57292 | -0.33962 | 0.115247 | 0.808939 |
| 10 | 6604400 | 6605199 | 10.64005 | 19.82904 | 24.21285 | 10.08815 | 0.164991 | 0.14862 | 0.808962 |
| 15 | 69383200 | 69387799 | 35.85414 | 58.50455 | 85.50603 | 36.74021 | -0.17353 | 1.531637 | 0.808979 |
| 7 | 43797200 | 43799799 | 17.17805 | 30.53078 | 34.64823 | 15.28507 | 0.21237 | 0.491706 | 0.808985 |
| 14 | 106780800 | 106782799 | 29.32981 | 49.45927 | 47.46623 | 20.75507 | 0.440291 | 1.116111 | 0.809 |
| 10 | 12238200 | 12241399 | 41.79544 | 67.26383 | 92.20051 | 39.15684 | -0.10024 | 1.845962 | 0.809015 |
| 7 | 41323400 | 41325799 | 26.97085 | 46.21882 | 51.2453 | 22.47373 | 0.234935 | 0.971788 | 0.809023 |
| 19 | 56014800 | 56019199 | 12.39409 | 21.81143 | 40.39642 | 17.88467 | -0.49627 | 0.573181 | 0.809034 |
| 14 | 51670800 | 51674199 | 93.43892 | 144.3223 | 162.7451 | 67.60089 | 0.127641 | 2.866282 | 0.809045 |
| 1 | 107454400 | 107454799 | 9.478052 | 17.94254 | 22.25194 | 9.014109 | 0.175978 | 0.030176 | 0.809057 |
| 3 | 149410600 | 149416399 | 70.90991 | 111.3524 | 149.471 | 61.99574 | -0.10501 | 2.559599 | 0.809075 |
| 10 | 38454800 | 38455799 | 10.16193 | 19.15813 | 24.2678 | 9.915098 | 0.134979 | 0.070973 | 0.809094 |
| 5 | 45985600 | 45989599 | 40.48141 | 66.42461 | 72.86019 | 31.37473 | 0.244128 | 1.461775 | 0.809113 |
| 5 | 77148000 | 77153599 | 60.43293 | 95.9094 | 128.661 | 53.60037 | -0.09169 | 2.308522 | 0.809134 |
| 18 | 11321200 | 11321999 | 9.086146 | 17.44712 | 22.98997 | 9.005064 | 0.133065 | -0.07911 | 0.809146 |
| 4 | 79638000 | 79640599 | 42.04427 | 67.38289 | 109.5417 | 46.23389 | -0.35315 | 1.982996 | 0.809155 |
| 14 | 94882600 | 94884999 | 33.63152 | 55.57088 | 65.00246 | 28.0835 | 0.154929 | 1.353794 | 0.809167 |
| 1 | 8710800 | 8712199 | 27.28382 | 45.56119 | 55.04026 | 23.94892 | 0.109362 | 1.212289 | 0.809191 |
| 7 | 108185800 | 108192199 | 68.15539 | 108.3592 | 107.6015 | 44.90742 | 0.344111 | 2.257274 | 0.809202 |
| 15 | 90231000 | 90233599 | 30.93147 | 51.02656 | 61.7006 | 26.62942 | 0.106941 | 1.364933 | 0.809213 |
| 7 | 26487400 | 26489999 | 31.1428 | 51.01545 | 71.86703 | 30.92508 | -0.11679 | 1.461761 | 0.80923 |
| 6 | 131112600 | 131138399 | 494.5371 | 671.6789 | 780.1035 | 324.9518 | -0.09003 | 5.032331 | 0.809246 |
| 12 | 6874200 | 6876999 | 26.53067 | 45.37317 | 51.24255 | 22.46889 | 0.207995 | 0.986658 | 0.80927 |
| 17 | 81233600 | 81235399 | 22.60219 | 38.36778 | 54.37711 | 23.82475 | -0.11978 | 1.044198 | 0.809276 |
| 16 | 79531400 | 79531999 | 6.839051 | 13.19428 | 22.0531 | 8.522057 | -0.18813 | -0.14857 | 0.80928 |
| 15 | 56736000 | 56739999 | 54.24233 | 86.3313 | 117.4945 | 49.10403 | -0.10832 | 2.213351 | 0.809345 |
| 4 | 68132400 | 68133399 | 14.12281 | 25.80308 | 25.36 | 10.86277 | 0.446825 | 0.291688 | 0.809351 |
| 1 | 155986600 | 155987799 | 14.18889 | 25.28847 | 29.94133 | 13.17764 | 0.154135 | 0.447777 | 0.809352 |
| 1 | 124901600 | 124904599 | 29.57655 | 49.43508 | 58.51076 | 25.4737 | 0.138708 | 1.212145 | 0.809359 |
| 7 | 106624800 | 106625599 | 9.234533 | 17.489 | 27.07604 | 10.95844 | -0.14073 | 0.037346 | 0.80937 |

| | | | | | | | | | |
|---|---|---|---|---|---|---|---|---|---|
| 1 | 97097600 | 97103399 | 91.87717 | 142.5673 | 165.3145 | 68.65469 | 0.092537 | 2.78155 | 0.80938 |
| 7 | 38098400 | 38100399 | 49.3131 | 79.04967 | 93.84167 | 39.62061 | 0.099927 | 1.979546 | 0.809403 |
| 19 | 61317200 | 61318799 | 14.67798 | 26.53822 | 24.33617 | 10.56156 | 0.534002 | 0.352237 | 0.809417 |
| 3 | 10911400 | 10912799 | 17.71006 | 30.89021 | 44.10182 | 19.41057 | -0.12401 | 0.73993 | 0.809433 |
| 1 | 100946400 | 100947599 | 13.25148 | 24.40935 | 29.65952 | 12.56551 | 0.153919 | 0.249128 | 0.809437 |
| 14 | 92306800 | 92311399 | 39.61996 | 64.42094 | 76.0039 | 32.61523 | 0.131105 | 1.584962 | 0.809457 |
| 16 | 70578400 | 70582599 | 55.07066 | 87.59477 | 119.0915 | 49.72832 | -0.10781 | 2.234717 | 0.809484 |
| 6 | 50097400 | 50105599 | 74.21318 | 116.7149 | 156.4328 | 64.89594 | -0.10118 | 2.533601 | 0.809503 |
| 14 | 72036400 | 72041999 | 99.95054 | 154.0743 | 178.1136 | 73.98541 | 0.089629 | 2.906638 | 0.809516 |
| 8 | 126085800 | 126090199 | 48.21507 | 77.37059 | 107.2026 | 45.33391 | -0.12092 | 1.950178 | 0.809516 |
| 1 | 78701600 | 78702599 | 6.966271 | 13.41344 | 22.54195 | 8.760056 | -0.20421 | -0.12993 | 0.809518 |
| 15 | 82182800 | 82183599 | 12.27313 | 22.0197 | 32.17127 | 14.08007 | -0.14162 | 0.400252 | 0.809524 |
| 13 | 61336600 | 61338199 | 22.26133 | 38.94803 | 42.81839 | 18.8872 | 0.251595 | 0.717356 | 0.809551 |
| 1 | 34909200 | 34911599 | 19.54071 | 34.00641 | 40.92459 | 17.98663 | 0.121087 | 0.775353 | 0.809564 |
| 10 | 70932400 | 70934799 | 16.56657 | 28.99437 | 41.39063 | 18.26263 | -0.12357 | 0.67874 | 0.809567 |
| 10 | 73514000 | 73514799 | 12.30318 | 22.52008 | 27.60606 | 11.79635 | 0.132604 | 0.273206 | 0.809572 |
| 6 | 68496400 | 68497999 | 18.11855 | 31.90662 | 37.48071 | 16.5916 | 0.157395 | 0.594089 | 0.809575 |
| X | 152709600 | 152721399 | 97.78036 | 150.9038 | 200.1501 | 83.13851 | -0.10719 | 2.887557 | 0.809583 |
| 18 | 68106200 | 68108799 | 26.92838 | 44.52068 | 63.02899 | 27.24175 | -0.11944 | 1.330873 | 0.809583 |
| 18 | 74426000 | 74426999 | 10.3782 | 19.26543 | 28.8304 | 12.08616 | -0.13357 | 0.180823 | 0.809585 |
| 5 | 106721800 | 106723399 | 16.8938 | 29.49072 | 41.9979 | 18.49413 | -0.12022 | 0.721258 | 0.809611 |
| 12 | 100975200 | 100977599 | 17.02185 | 29.63873 | 42.57809 | 18.71965 | -0.13314 | 0.757966 | 0.809614 |
| 17 | 78066000 | 78067599 | 26.01612 | 43.36024 | 61.03262 | 26.52387 | -0.11046 | 1.233692 | 0.80963 |
| 4 | 147174600 | 147177599 | 48.68959 | 78.81388 | 85.63253 | 36.63024 | 0.243785 | 1.694827 | 0.809634 |
| 4 | 16268200 | 16270599 | 22.56702 | 39.20195 | 45.17447 | 19.83189 | 0.183146 | 0.821249 | 0.809638 |
| 11 | 95388600 | 95389199 | 8.368218 | 15.98566 | 25.07836 | 9.966347 | -0.13587 | -0.03284 | 0.809654 |
| 5 | 100885400 | 100890999 | 72.9082 | 115.0461 | 135.1909 | 56.11436 | 0.09207 | 2.461673 | 0.809674 |
| 17 | 41963400 | 41966799 | 44.87836 | 72.52001 | 83.71436 | 35.74789 | 0.154043 | 1.729244 | 0.809674 |
| 10 | 14955800 | 14961199 | 77.01844 | 121.9449 | 117.6152 | 48.90302 | 0.380524 | 2.382673 | 0.809675 |
| 2 | 149971800 | 149978999 | 88.93041 | 138.0875 | 183.1625 | 76.0593 | -0.10048 | 2.771309 | 0.809677 |
| 19 | 43415200 | 43416399 | 15.31024 | 26.77764 | 38.93408 | 17.16902 | -0.14974 | 0.679861 | 0.809678 |
| 3 | 105153000 | 105154399 | 13.72636 | 24.22614 | 35.20238 | 15.57609 | -0.14646 | 0.539257 | 0.809689 |
| 19 | 29058400 | 29062599 | 71.56734 | 112.6963 | 150.2674 | 62.34852 | -0.09231 | 2.506225 | 0.809697 |
| 19 | 36607000 | 36611199 | 28.97816 | 46.97168 | 85.30011 | 36.52515 | -0.49357 | 1.630071 | 0.809725 |
| 16 | 93608200 | 93610599 | 45.7642 | 73.35913 | 100.1577 | 42.28589 | -0.10148 | 1.978406 | 0.809725 |
| 3 | 57854000 | 57855599 | 17.86417 | 31.36991 | 37.54155 | 16.61109 | 0.129645 | 0.627115 | 0.809726 |
| 7 | 114009800 | 114010999 | 9.99652 | 18.72834 | 28.48563 | 11.76276 | -0.13916 | 0.121793 | 0.809731 |
| 16 | 61708200 | 61709399 | 12.05322 | 22.52099 | 27.79008 | 11.52784 | 0.154633 | 0.151717 | 0.809737 |
| 16 | 64624400 | 64627399 | 46.54941 | 74.59214 | 101.5836 | 42.86546 | -0.09844 | 1.989803 | 0.80975 |
| 10 | 50172800 | 50175399 | 31.08329 | 51.18594 | 61.65362 | 26.58914 | 0.111506 | 1.386712 | 0.809752 |
| 13 | 50322800 | 50328399 | 58.95341 | 93.02258 | 149.7806 | 62.16901 | -0.36174 | 2.457206 | 0.809758 |
| 16 | 70155200 | 70156999 | 22.48749 | 38.97429 | 38.34419 | 16.82342 | 0.409829 | 0.837149 | 0.809768 |
| 2 | 77542600 | 77545599 | 19.79547 | 34.00695 | 48.47451 | 21.26235 | -0.12557 | 0.915631 | 0.809782 |
| 6 | 102484600 | 102485799 | 11.9298 | 21.84462 | 32.32725 | 13.80688 | -0.13692 | 0.279309 | 0.809792 |
| X | 153686000 | 153690999 | 50.54104 | 81.52688 | 90.80024 | 38.68581 | 0.202656 | 1.789373 | 0.809796 |
| 2 | 73064200 | 73064999 | 13.37719 | 24.6317 | 29.69388 | 12.58646 | 0.164961 | 0.254034 | 0.8098 |
| 5 | 101761000 | 101768999 | 91.91377 | 142.0684 | 187.3611 | 77.82557 | -0.09727 | 2.853918 | 0.809829 |

| | | | | | | | | | |
|---|---|---|---|---|---|---|---|---|---|
| 11 | 3996200 | 4001199 | 77.95603 | 121.9058 | 137.671 | 57.11546 | 0.139292 | 2.638256 | 0.809838 |
| 15 | 89287800 | 89296799 | 127.1651 | 193.2223 | 221.1942 | 91.61324 | 0.087468 | 3.204319 | 0.809844 |
| 9 | 30559400 | 30563199 | 39.64646 | 64.46832 | 64.50673 | 27.68217 | 0.368267 | 1.576451 | 0.809917 |
| 8 | 77887200 | 77888799 | 13.71625 | 24.44365 | 35.08471 | 15.44261 | -0.12203 | 0.457314 | 0.809932 |
| 10 | 126099200 | 126103599 | 37.44647 | 61.15269 | 71.77243 | 30.84834 | 0.142974 | 1.513187 | 0.809933 |
| 15 | 47319000 | 47320999 | 25.20373 | 43.12472 | 50.11978 | 21.97744 | 0.166896 | 0.97606 | 0.809942 |
| 4 | 112746800 | 112751999 | 60.27909 | 96.64929 | 105.0195 | 44.35458 | 0.227704 | 1.992161 | 0.809947 |
| X | 61312200 | 61314199 | 20.13137 | 35.24882 | 40.74459 | 17.98423 | 0.179611 | 0.691971 | 0.80995 |
| 11 | 67535400 | 67538799 | 24.82503 | 41.82064 | 50.1307 | 21.91344 | 0.120598 | 1.112922 | 0.809989 |
| 9 | 12134200 | 12139799 | 273.6848 | 388.3693 | 480.4262 | 198.7378 | -0.11723 | 4.360668 | 0.810009 |
| 19 | 60632000 | 60632799 | 11.31244 | 20.33889 | 30.75211 | 13.43508 | -0.18894 | 0.376907 | 0.81002 |
| 6 | 70029400 | 70031599 | 19.71056 | 32.94913 | 62.63486 | 27.27148 | -0.54227 | 1.188127 | 0.810024 |
| 8 | 19697000 | 19712999 | 198.3546 | 294.5408 | 317.2906 | 130.7035 | 0.145875 | 3.660376 | 0.810044 |
| 12 | 115517000 | 115547999 | 621.7246 | 834.3493 | 836.7548 | 349.1853 | 0.10425 | 5.223722 | 0.810054 |
| 17 | 13472600 | 13474399 | 25.72602 | 43.72907 | 43.17875 | 18.92375 | 0.399771 | 1.02911 | 0.810075 |
| 5 | 128066600 | 128069599 | 30.1856 | 50.07574 | 60.1818 | 26.07227 | 0.116448 | 1.296112 | 0.81008 |
| 10 | 30736000 | 30737999 | 16.70487 | 30.14505 | 33.918 | 14.75527 | 0.23844 | 0.397605 | 0.810081 |
| 2 | 159299400 | 159305799 | 62.91268 | 100.2391 | 116.06 | 48.57701 | 0.126192 | 2.188574 | 0.810092 |
| 5 | 95493000 | 95499999 | 84.6462 | 133.8841 | 116.848 | 48.55871 | 0.523332 | 2.410854 | 0.810107 |
| 10 | 48881200 | 48889199 | 134.773 | 205.2963 | 198.0991 | 82.10946 | 0.336755 | 3.153712 | 0.810117 |
| 8 | 13860200 | 13861799 | 20.6825 | 35.49426 | 51.14304 | 22.43151 | -0.14117 | 0.936194 | 0.810128 |
| 6 | 21949200 | 21950399 | 11.241 | 20.49934 | 30.21301 | 12.97195 | -0.13615 | 0.291957 | 0.810128 |
| 2 | 172058800 | 172060199 | 12.54016 | 22.89292 | 27.653 | 11.85699 | 0.15058 | 0.291628 | 0.810137 |
| 1 | 104377400 | 104378399 | 13.27637 | 24.34421 | 29.16077 | 12.43091 | 0.168396 | 0.276515 | 0.810145 |
| 3 | 33980600 | 33981799 | 12.67881 | 22.80885 | 33.07988 | 14.44493 | -0.12909 | 0.386312 | 0.810153 |
| 2 | 139281200 | 139282399 | 15.2133 | 26.66895 | 38.37431 | 16.95488 | -0.13469 | 0.643055 | 0.810176 |
| 12 | 50941600 | 50943199 | 18.43722 | 32.17473 | 38.82087 | 17.09167 | 0.117798 | 0.728646 | 0.810192 |
| 15 | 43862600 | 43864399 | 22.68934 | 39.6604 | 43.40585 | 19.13147 | 0.258138 | 0.735863 | 0.810228 |
| 11 | 107849200 | 107853399 | 46.22308 | 74.30344 | 88.31065 | 37.44198 | 0.103072 | 1.887569 | 0.810229 |
| X | 84091800 | 84092399 | 7.975013 | 15.4133 | 26.0945 | 10.00691 | -0.19614 | -0.14826 | 0.810245 |
| 12 | 83553400 | 83554799 | 14.2882 | 25.41376 | 30.07605 | 13.25963 | 0.153313 | 0.463342 | 0.810247 |
| 4 | 6609000 | 6609999 | 15.68409 | 27.6557 | 33.00014 | 14.60437 | 0.13559 | 0.555948 | 0.810257 |
| 16 | 30657600 | 30661199 | 57.20369 | 91.40896 | 107.2861 | 45.08846 | 0.111134 | 2.08905 | 0.810269 |
| 10 | 20789000 | 20791799 | 20.19392 | 34.69855 | 49.00898 | 21.49692 | -0.11227 | 0.917684 | 0.810282 |
| 16 | 38578000 | 38579599 | 13.08391 | 23.57928 | 28.04912 | 12.22254 | 0.157264 | 0.368532 | 0.810311 |
| 6 | 94359800 | 94361399 | 23.91853 | 40.1254 | 56.53386 | 24.67021 | -0.11188 | 1.154967 | 0.810331 |
| 1 | 50593400 | 50594999 | 18.56552 | 32.39193 | 38.89689 | 17.12291 | 0.124774 | 0.734529 | 0.810346 |
| X | 40865400 | 40867599 | 24.29375 | 42.06812 | 39.27328 | 17.2306 | 0.484897 | 0.854003 | 0.810388 |
| 9 | 50194000 | 50196999 | 23.08881 | 38.49448 | 66.78996 | 29.03216 | -0.41117 | 1.229342 | 0.810389 |
| 5 | 127702200 | 127705999 | 40.45927 | 65.20343 | 89.58795 | 38.12351 | -0.10117 | 1.799249 | 0.810409 |
| 13 | 112113000 | 112114799 | 14.09881 | 25.42857 | 30.95064 | 13.47344 | 0.125223 | 0.377625 | 0.810415 |
| 18 | 78899200 | 78905199 | 124.0602 | 185.8799 | 298.795 | 123.2558 | -0.41937 | 3.486163 | 0.810422 |
| 10 | 95996800 | 96000399 | 36.33926 | 59.01 | 82.47284 | 35.36157 | -0.11438 | 1.605186 | 0.810453 |
| 7 | 94177200 | 94179599 | 44.46517 | 71.33694 | 97.76621 | 41.32918 | -0.10548 | 1.950107 | 0.810461 |
| 18 | 60661200 | 60661999 | 13.57012 | 24.11194 | 34.60268 | 15.26841 | -0.12419 | 0.474251 | 0.810484 |
| 12 | 28210200 | 28212199 | 14.30166 | 24.94073 | 45.23567 | 19.90737 | -0.46747 | 0.728672 | 0.810495 |
| 6 | 68105800 | 68108599 | 58.72784 | 93.46757 | 110.1287 | 46.02378 | 0.099305 | 2.218112 | 0.81053 |

| | | | | | | | | | |
|---|---|---|---|---|---|---|---|---|---|
| 2 | 109006200 | 109008599 | 37.60259 | 61.75198 | 70.80465 | 30.50027 | 0.180807 | 1.443171 | 0.810533 |
| 11 | 74117200 | 74119399 | 17.45219 | 30.56647 | 44.53417 | 19.66166 | -0.1529 | 0.677416 | 0.810535 |
| X | 56649600 | 56651199 | 17.5858 | 31.23123 | 26.49171 | 11.69447 | 0.630016 | 0.488909 | 0.810551 |
| 2 | 74124400 | 74125399 | 11.40979 | 20.92805 | 30.77618 | 13.11399 | -0.12544 | 0.261958 | 0.810561 |
| 10 | 55501600 | 55513199 | 318.5704 | 446.5773 | 542.7681 | 224.9724 | -0.10951 | 4.542808 | 0.810568 |
| 18 | 19543600 | 19551399 | 100.999 | 156.0525 | 180.0579 | 74.79211 | 0.094928 | 2.864704 | 0.810634 |
| 10 | 47187800 | 47188399 | 8.116463 | 15.68511 | 20.95209 | 8.038616 | 0.141537 | -0.16398 | 0.810652 |
| X | 77704600 | 77706799 | 26.51312 | 45.25535 | 51.76974 | 22.69651 | 0.189201 | 1.006299 | 0.81067 |
| 17 | 12283000 | 12284199 | 12.41143 | 22.76012 | 28.06927 | 11.96424 | 0.126377 | 0.263754 | 0.810699 |
| 8 | 45610400 | 45618599 | 103.5861 | 159.1138 | 208.2843 | 86.48135 | -0.09343 | 2.979235 | 0.810704 |
| 1 | 84361200 | 84361999 | 10.33539 | 19.18149 | 28.87807 | 12.10999 | -0.14207 | 0.184826 | 0.810763 |
| 7 | 52060600 | 52062599 | 21.97182 | 37.902 | 45.22061 | 19.83759 | 0.13128 | 0.883622 | 0.810773 |
| 14 | 92604400 | 92605999 | 19.30846 | 33.29203 | 47.41291 | 20.79902 | -0.12299 | 0.883093 | 0.810793 |
| 12 | 7195600 | 7202799 | 138.3475 | 207.9192 | 267.4496 | 110.4096 | -0.09414 | 3.427918 | 0.810795 |
| 9 | 98744600 | 98748199 | 55.49384 | 88.0711 | 121.2308 | 50.50507 | -0.12878 | 2.302547 | 0.810807 |
| 10 | 129189600 | 129191599 | 39.54903 | 63.98504 | 76.4984 | 32.71058 | 0.105401 | 1.68151 | 0.810848 |
| 8 | 21367600 | 21374599 | 128.9911 | 194.8963 | 251.632 | 104.0387 | -0.09277 | 3.319703 | 0.810872 |
| X | 128303200 | 128304199 | 12.0624 | 21.54046 | 39.36567 | 17.29696 | -0.46674 | 0.443283 | 0.810893 |
| 2 | 39406800 | 39409199 | 26.5463 | 44.56228 | 53.36075 | 23.29495 | 0.121984 | 1.154743 | 0.810895 |
| 10 | 22470200 | 22474399 | 41.31914 | 66.68112 | 92.61698 | 39.4827 | -0.1145 | 1.762638 | 0.810898 |
| 6 | 145397400 | 145398399 | 13.39834 | 24.21135 | 29.46699 | 12.797 | 0.127518 | 0.357953 | 0.81091 |
| 13 | 70079400 | 70082399 | 27.7198 | 45.63931 | 65.25197 | 28.13894 | -0.13518 | 1.382725 | 0.810914 |
| 3 | 24072400 | 24073999 | 14.3955 | 26.16858 | 30.90002 | 13.3198 | 0.177507 | 0.332036 | 0.81092 |
| X | 89356000 | 89358999 | 44.59505 | 72.57322 | 64.90622 | 27.86173 | 0.530651 | 1.577848 | 0.810949 |
| X | 12107200 | 12113399 | 34.53339 | 57.31035 | 54.20249 | 23.48809 | 0.461157 | 1.301307 | 0.810953 |
| 13 | 69276800 | 69279399 | 22.17175 | 38.04011 | 45.79225 | 20.08422 | 0.116966 | 0.933946 | 0.81096 |
| 14 | 91411600 | 91414799 | 33.45359 | 54.64918 | 65.30218 | 28.07058 | 0.117529 | 1.49828 | 0.810967 |
| 8 | 128286800 | 128288399 | 15.36565 | 27.15324 | 32.46916 | 14.36314 | 0.133765 | 0.529309 | 0.810982 |
| 6 | 41212200 | 41212799 | 6.674197 | 12.92377 | 21.59761 | 8.209151 | -0.16547 | -0.20905 | 0.810984 |
| 7 | 135481600 | 135484999 | 36.74371 | 59.48937 | 82.3007 | 35.21038 | -0.10368 | 1.667251 | 0.810988 |
| 4 | 12145400 | 12152999 | 120.1656 | 183.712 | 209.443 | 86.88755 | 0.100507 | 3.077692 | 0.810998 |
| 6 | 89980600 | 89988199 | 124.4741 | 190.2528 | 186.3287 | 77.29424 | 0.319223 | 3.08225 | 0.811008 |
| 1 | 84888600 | 84890999 | 36.71519 | 59.1892 | 84.17704 | 35.83629 | -0.15019 | 1.781248 | 0.811029 |
| 10 | 13479400 | 13482399 | 39.38677 | 64.09694 | 62.72774 | 26.92653 | 0.401032 | 1.564989 | 0.811037 |
| 10 | 7188600 | 7195799 | 106.9815 | 164.9829 | 163.0701 | 67.73642 | 0.315867 | 2.899213 | 0.811057 |
| 12 | 50770800 | 50772999 | 15.72541 | 27.54195 | 39.82161 | 17.581 | -0.14189 | 0.659347 | 0.811064 |
| 4 | 40294000 | 40297399 | 50.58067 | 81.11134 | 96.4183 | 40.73118 | 0.098598 | 1.969791 | 0.811083 |
| 13 | 73399400 | 73400399 | 12.25309 | 21.9482 | 32.15773 | 14.09335 | -0.14711 | 0.409521 | 0.811086 |
| 9 | 4390200 | 4393999 | 57.53696 | 91.94327 | 106.685 | 44.83704 | 0.127675 | 2.09088 | 0.81109 |
| 12 | 104873400 | 104874799 | 13.33462 | 23.90755 | 34.57728 | 15.14195 | -0.12799 | 0.414957 | 0.811106 |
| 12 | 115720800 | 115723199 | 30.97333 | 51.58198 | 60.77355 | 26.39828 | 0.145387 | 1.257442 | 0.811176 |
| 6 | 143471000 | 143472599 | 15.33979 | 27.3056 | 32.82272 | 14.46098 | 0.1313 | 0.468801 | 0.811195 |
| 18 | 9202400 | 9202999 | 8.3308 | 16.04947 | 20.93104 | 8.121451 | 0.161093 | -0.12228 | 0.811199 |
| 4 | 112109000 | 112111399 | 31.89336 | 53.0505 | 51.90815 | 22.52948 | 0.412525 | 1.266281 | 0.811208 |
| 1 | 118274800 | 118277199 | 22.48249 | 38.15402 | 53.86985 | 23.60052 | -0.11444 | 1.04319 | 0.811239 |
| 3 | 37424200 | 37427199 | 24.98716 | 42.45713 | 50.65686 | 22.20047 | 0.12763 | 1.037519 | 0.81124 |
| 17 | 83222200 | 83224399 | 19.69556 | 34.48232 | 40.13497 | 17.71367 | 0.169473 | 0.686722 | 0.811294 |

| | | | | | | | | | |
|---|---|---|---|---|---|---|---|---|---|
| 17 | 11907200 | 11908399 | 17.04913 | 29.92284 | 36.11519 | 15.97283 | 0.117532 | 0.628964 | 0.811295 |
| 8 | 16265800 | 16266799 | 16.64808 | 28.99428 | 41.60066 | 18.29245 | -0.13112 | 0.754253 | 0.811301 |
| 16 | 71713200 | 71715599 | 22.70394 | 39.19516 | 46.29328 | 20.30863 | 0.146061 | 0.88224 | 0.811306 |
| 6 | 143883200 | 143884399 | 13.88767 | 25.01546 | 30.41884 | 13.26333 | 0.125022 | 0.381679 | 0.811308 |
| 19 | 57635800 | 57637999 | 25.56883 | 42.06885 | 72.75004 | 31.36322 | -0.4092 | 1.393063 | 0.811324 |
| 11 | 90249200 | 90249999 | 7.899253 | 15.15638 | 24.09404 | 9.458419 | -0.13816 | -0.08023 | 0.811335 |
| 11 | 24121800 | 24123199 | 12.4186 | 22.48206 | 32.6841 | 14.16166 | -0.12506 | 0.348863 | 0.811355 |
| X | 76085800 | 76086999 | 17.28107 | 30.17901 | 44.13076 | 19.43831 | -0.15796 | 0.724868 | 0.811392 |
| 11 | 113026800 | 113028999 | 17.95483 | 31.36642 | 37.7888 | 16.64846 | 0.120125 | 0.70973 | 0.811425 |
| 10 | 126250600 | 126252599 | 26.60026 | 45.19696 | 52.33895 | 22.93761 | 0.17053 | 1.044813 | 0.811428 |
| 1 | 162818000 | 162819399 | 17.3138 | 30.36465 | 36.59842 | 16.17632 | 0.11942 | 0.642159 | 0.811433 |
| 4 | 91963600 | 91964599 | 11.07223 | 20.46314 | 30.36098 | 12.80457 | -0.12746 | 0.215785 | 0.811433 |
| 16 | 23134800 | 23135399 | 8.82157 | 17.03281 | 22.59192 | 8.703325 | 0.146396 | -0.13557 | 0.811433 |
| 6 | 89924200 | 89925999 | 35.72007 | 57.74571 | 80.78517 | 34.51618 | -0.12189 | 1.696923 | 0.811439 |
| 10 | 90688400 | 90690599 | 16.43329 | 28.76435 | 40.8852 | 18.04135 | -0.11731 | 0.676118 | 0.81144 |
| 15 | 97289400 | 97290399 | 14.05899 | 25.18225 | 30.43368 | 13.33812 | 0.128887 | 0.417418 | 0.811461 |
| 3 | 71132600 | 71133199 | 10.27527 | 19.24745 | 23.69793 | 9.788427 | 0.162984 | 0.110549 | 0.811481 |
| 14 | 8731600 | 8733999 | 20.92435 | 36.61789 | 41.78183 | 18.43381 | 0.198196 | 0.705328 | 0.81151 |
| 17 | 53586400 | 53587799 | 19.85898 | 34.50484 | 41.55096 | 18.2435 | 0.120107 | 0.802921 | 0.811524 |
| 8 | 80111600 | 80113599 | 20.57885 | 34.94917 | 50.07328 | 21.94213 | -0.13532 | 1.026957 | 0.811531 |
| 5 | 102345200 | 102350799 | 95.39111 | 148.4673 | 147.8833 | 61.3879 | 0.314931 | 2.730297 | 0.811537 |
| 4 | 36327000 | 36327599 | 6.846607 | 13.3038 | 22.33973 | 8.324747 | -0.1451 | -0.27045 | 0.811555 |
| 6 | 29029200 | 29031399 | 24.73169 | 41.33782 | 58.18936 | 25.334 | -0.11054 | 1.197662 | 0.811611 |
| 2 | 140189000 | 140189799 | 10.2305 | 19.28811 | 24.21145 | 9.891459 | 0.147711 | 0.069441 | 0.811628 |
| 3 | 44145600 | 44154599 | 164.1747 | 245.5747 | 276.3896 | 113.9765 | 0.092346 | 3.52297 | 0.811639 |
| 3 | 44972600 | 44973999 | 17.47199 | 30.51672 | 43.56415 | 19.19044 | -0.12356 | 0.720411 | 0.811649 |
| 6 | 123855000 | 123861799 | 105.9891 | 164.1821 | 158.2453 | 65.73078 | 0.357048 | 2.817783 | 0.811659 |
| 3 | 34965400 | 34969999 | 48.16633 | 77.12397 | 104.8052 | 44.17335 | -0.09666 | 2.016095 | 0.811693 |
| 6 | 143447200 | 143453799 | 74.95835 | 117.5173 | 155.696 | 64.57653 | -0.08799 | 2.594455 | 0.811707 |
| 18 | 53638400 | 53638799 | 12.87524 | 24.0571 | 20.13108 | 8.35063 | 0.711726 | 0.142304 | 0.811709 |
| 10 | 20908200 | 20912399 | 25.51184 | 43.09954 | 51.73582 | 22.63452 | 0.118721 | 1.092863 | 0.811723 |
| 3 | 17007600 | 17009599 | 15.0122 | 26.54662 | 38.10743 | 16.85501 | -0.1281 | 0.530962 | 0.811736 |
| 3 | 122533600 | 122535799 | 20.2844 | 35.38759 | 34.61648 | 15.23772 | 0.419052 | 0.735606 | 0.811773 |
| 6 | 54168000 | 54170399 | 18.97627 | 33.03723 | 39.94266 | 17.55945 | 0.114611 | 0.767163 | 0.811774 |
| 7 | 79123600 | 79126199 | 18.97627 | 33.03723 | 39.94266 | 17.55945 | 0.114611 | 0.767163 | 0.811774 |
| 12 | 26305400 | 26306599 | 16.20616 | 29.04089 | 33.9323 | 14.86598 | 0.177774 | 0.438275 | 0.811788 |
| 14 | 108087000 | 108087999 | 8.773945 | 16.56639 | 25.92237 | 10.55902 | -0.16404 | 0.050867 | 0.811799 |
| 10 | 33620800 | 33630999 | 180.7086 | 268.0842 | 300.1794 | 123.6429 | 0.088848 | 3.673086 | 0.811824 |
| 10 | 50997200 | 51000199 | 29.75136 | 49.22175 | 59.47237 | 25.71675 | 0.108513 | 1.326367 | 0.811831 |
| 6 | 9943200 | 9946599 | 43.29874 | 69.88367 | 83.11809 | 35.43846 | 0.108985 | 1.756454 | 0.811853 |
| X | 106286800 | 106290399 | 44.31755 | 71.50495 | 84.4357 | 35.98526 | 0.118874 | 1.771409 | 0.811854 |
| 6 | 71014800 | 71020199 | 56.28638 | 89.66392 | 121.0571 | 50.65492 | -0.09542 | 2.18898 | 0.811873 |
| 5 | 26378800 | 26379799 | 18.11484 | 31.6771 | 38.23091 | 16.85633 | 0.117561 | 0.695701 | 0.811876 |
| 13 | 45538800 | 45540999 | 22.02415 | 37.25236 | 52.76204 | 23.09194 | -0.11908 | 1.069396 | 0.811893 |
| 3 | 108598400 | 108600399 | 16.72839 | 29.42711 | 35.52436 | 15.72895 | 0.117626 | 0.593542 | 0.81192 |
| 18 | 24191200 | 24193999 | 23.7771 | 40.92837 | 40.45551 | 17.74606 | 0.401118 | 0.905704 | 0.811928 |
| 14 | 53024000 | 53025199 | 10.53093 | 19.36477 | 29.23507 | 12.41468 | -0.15969 | 0.235263 | 0.811933 |

| | | | | | | | | | |
|---|---|---|---|---|---|---|---|---|---|
| X | 117109600 | 117110799 | 15.85539 | 27.92778 | 40.21438 | 17.80088 | -0.13476 | 0.582759 | 0.81196 |
| X | 102030400 | 102031799 | 16.22528 | 29.36196 | 33.53871 | 14.5391 | 0.220286 | 0.379924 | 0.811983 |
| X | 125218200 | 125264599 | 645.5006 | 868.7649 | 826.3127 | 344.68 | 0.184853 | 5.179293 | 0.811984 |
| 9 | 95002400 | 95004599 | 15.85044 | 27.85364 | 32.64774 | 14.45443 | 0.159556 | 0.595391 | 0.811994 |
| 1 | 121977400 | 121978199 | 9.109878 | 17.2588 | 26.53452 | 10.73152 | -0.12964 | 0.033494 | 0.812018 |
| 19 | 5232200 | 5235599 | 56.20655 | 89.74235 | 106.4515 | 44.68606 | 0.094601 | 2.11248 | 0.81206 |
| 14 | 91796000 | 91797799 | 13.92462 | 25.54935 | 30.63971 | 13.04989 | 0.167233 | 0.278117 | 0.812071 |
| 13 | 8802600 | 8803599 | 15.00237 | 26.43616 | 37.65251 | 16.6655 | -0.11878 | 0.568436 | 0.812139 |
| 7 | 134671200 | 134672799 | 18.10872 | 31.68441 | 38.19561 | 16.84887 | 0.119191 | 0.686313 | 0.812147 |
| 1 | 6096600 | 6099399 | 36.9694 | 61.06937 | 55.32695 | 23.89877 | 0.522585 | 1.362854 | 0.812155 |
| 1 | 145059600 | 145061399 | 21.41035 | 36.57514 | 51.69031 | 22.66317 | -0.11445 | 0.97935 | 0.812167 |
| 5 | 151424400 | 151425799 | 32.13253 | 52.79634 | 63.82184 | 27.49836 | 0.105099 | 1.419194 | 0.8122 |
| 9 | 12445400 | 12452599 | 98.39782 | 151.3996 | 197.583 | 82.05708 | -0.08717 | 2.943899 | 0.812204 |
| X | 118962200 | 118969999 | 89.80199 | 141.1031 | 154.6118 | 64.13294 | 0.188123 | 2.56287 | 0.812218 |
| 13 | 33125800 | 33128399 | 27.74894 | 45.74296 | 64.67641 | 27.90511 | -0.11844 | 1.368639 | 0.812221 |
| 10 | 82037000 | 82041399 | 46.2546 | 74.68253 | 86.29314 | 36.81007 | 0.151239 | 1.751715 | 0.812232 |
| 10 | 73971000 | 73972599 | 18.62423 | 32.7716 | 31.82662 | 14.0909 | 0.430094 | 0.595086 | 0.812248 |
| 18 | 41746400 | 41747199 | 9.004968 | 17.13505 | 26.64668 | 10.67736 | -0.13395 | 0.004167 | 0.812254 |
| 15 | 90591000 | 90592199 | 12.35882 | 23.2835 | 23.04051 | 9.424889 | 0.487141 | 0.092172 | 0.812269 |
| 1 | 121434000 | 121435599 | 24.25615 | 41.04826 | 49.58441 | 21.70451 | 0.109731 | 1.069973 | 0.812274 |
| 11 | 52567000 | 52579799 | 189.2596 | 279.6178 | 311.5541 | 128.3048 | 0.090436 | 3.743554 | 0.812293 |
| 1 | 4067000 | 4068999 | 20.4537 | 35.69854 | 35.07387 | 15.4433 | 0.412806 | 0.730997 | 0.812302 |
| 14 | 6358400 | 6359799 | 23.55266 | 39.15773 | 66.80663 | 28.98606 | -0.38711 | 1.260484 | 0.812304 |
| 12 | 6708400 | 6710199 | 22.21518 | 37.98734 | 45.82696 | 20.09402 | 0.113207 | 0.964996 | 0.812318 |
| 12 | 104173400 | 104176399 | 7.316596 | 13.98392 | 27.87043 | 11.06379 | -0.47718 | -0.04047 | 0.812352 |
| 8 | 21324800 | 21325799 | 19.40537 | 33.93973 | 40.21631 | 17.73432 | 0.143856 | 0.702131 | 0.812363 |
| 12 | 74588600 | 74590399 | 19.40537 | 33.93973 | 40.21631 | 17.73432 | 0.143856 | 0.702131 | 0.812363 |
| 14 | 39713800 | 39714199 | 9.344409 | 17.73007 | 22.2853 | 8.984031 | 0.163 | 0.013736 | 0.812375 |
| 8 | 69787000 | 69789399 | 23.93845 | 41.65395 | 37.10884 | 16.30816 | 0.553164 | 0.795487 | 0.812405 |
| 5 | 11830000 | 11831199 | 9.318011 | 17.30679 | 26.83819 | 11.24266 | -0.18347 | 0.164808 | 0.812429 |
| 10 | 105108000 | 105111799 | 29.51491 | 49.43024 | 57.9927 | 25.27985 | 0.15128 | 1.189705 | 0.812434 |
| 1 | 125726800 | 125729399 | 38.10245 | 61.78321 | 74.22662 | 31.79175 | 0.101418 | 1.633446 | 0.812444 |
| 5 | 89682000 | 89683199 | 11.616 | 21.61222 | 34.48351 | 14.40748 | -0.22225 | 0.18828 | 0.812452 |
| 11 | 8299800 | 8302199 | 30.95308 | 51.25333 | 61.79236 | 26.73559 | 0.111737 | 1.322561 | 0.812482 |
| 15 | 66086800 | 66092399 | 76.50439 | 120.1234 | 140.7053 | 58.3591 | 0.091189 | 2.564537 | 0.812498 |
| 3 | 62546000 | 62546399 | 7.432372 | 14.4336 | 19.49374 | 7.298493 | 0.160068 | -0.25595 | 0.812535 |
| 1 | 104364600 | 104365599 | 12.3593 | 21.98956 | 39.30363 | 17.32337 | -0.43808 | 0.467264 | 0.812539 |
| 16 | 80079800 | 80081199 | 14.73533 | 26.11337 | 37.59944 | 16.62314 | -0.13115 | 0.505906 | 0.812545 |
| 8 | 127245600 | 127248599 | 28.10309 | 46.84061 | 56.7163 | 24.64876 | 0.106075 | 1.236093 | 0.812568 |
| 8 | 125901800 | 125903599 | 16.87979 | 29.41307 | 42.04911 | 18.49566 | -0.12602 | 0.746524 | 0.812592 |
| 4 | 151427200 | 151431199 | 33.31294 | 54.45668 | 65.29505 | 28.07452 | 0.113314 | 1.488765 | 0.812598 |
| 13 | 58826800 | 58828799 | 20.69396 | 36.06602 | 35.2028 | 15.48607 | 0.422165 | 0.753676 | 0.812599 |
| 7 | 143231400 | 143231999 | 8.055409 | 15.61904 | 21.08053 | 7.963059 | 0.148723 | -0.2136 | 0.812626 |
| 4 | 123692000 | 123693799 | 16.77282 | 29.5026 | 35.5593 | 15.7447 | 0.119905 | 0.595637 | 0.812635 |
| 13 | 62153800 | 62156799 | 28.46132 | 47.13968 | 66.80466 | 28.90946 | -0.12083 | 1.314824 | 0.812655 |
| 6 | 35372000 | 35379399 | 77.41263 | 121.3099 | 161.3587 | 66.92833 | -0.09422 | 2.604138 | 0.812667 |
| 8 | 78584200 | 78586799 | 24.68326 | 42.00642 | 50.09851 | 21.95854 | 0.128683 | 1.024729 | 0.812704 |

| | | | | | | | | | |
|---|---|---|---|---|---|---|---|---|---|
| 5 | 57303800 | 57306599 | 30.07443 | 50.73415 | 56.89858 | 24.88232 | 0.216269 | 1.123935 | 0.812711 |
| 4 | 72310200 | 72310599 | 4.954705 | 9.675404 | 17.61316 | 6.193538 | -0.18386 | -0.51899 | 0.812739 |
| 3 | 68464600 | 68465399 | 11.76137 | 21.37355 | 31.20855 | 13.45802 | -0.12678 | 0.319004 | 0.812764 |
| 5 | 87457200 | 87459599 | 33.67712 | 55.82905 | 54.72403 | 23.69644 | 0.409598 | 1.311117 | 0.812775 |
| 10 | 15775400 | 15775799 | 6.797503 | 13.24005 | 18.50569 | 6.754659 | 0.144018 | -0.35885 | 0.81278 |
| 8 | 69049800 | 69050799 | 19.35173 | 33.58004 | 40.08236 | 17.58668 | 0.132617 | 0.815822 | 0.812808 |
| 13 | 15608400 | 15608599 | 6.508778 | 12.69607 | 18.05947 | 6.467817 | 0.143532 | -0.43143 | 0.812809 |
| 9 | 55948800 | 55949599 | 8.359681 | 15.79908 | 24.92259 | 10.13157 | -0.17265 | 0.039514 | 0.812816 |
| 1 | 97174000 | 97175199 | 14.79202 | 26.13635 | 37.68766 | 16.67201 | -0.13478 | 0.540008 | 0.812818 |
| 4 | 143670400 | 143698599 | 573.9542 | 778.0985 | 770.5945 | 321.0825 | 0.137099 | 5.06278 | 0.812842 |
| 12 | 65620800 | 65621799 | 10.12483 | 18.59637 | 28.28448 | 12.03003 | -0.17165 | 0.238098 | 0.812866 |
| 1 | 69660200 | 69665399 | 27.91303 | 46.90363 | 42.32401 | 18.48671 | 0.528708 | 1.13592 | 0.812879 |
| 9 | 102998400 | 103001399 | 29.33099 | 47.81754 | 69.62862 | 29.90822 | -0.16893 | 1.521773 | 0.812918 |
| 1 | 109469600 | 109471599 | 27.0319 | 45.39671 | 54.5333 | 23.81321 | 0.11727 | 1.149747 | 0.812919 |
| 9 | 104243800 | 104248999 | 48.82631 | 78.21283 | 106.4913 | 44.912 | -0.09876 | 2.004715 | 0.81295 |
| 5 | 139065400 | 139066799 | 16.31797 | 28.41052 | 41.11135 | 18.07375 | -0.14336 | 0.756157 | 0.812986 |
| 11 | 84952200 | 84960599 | 45.55558 | 73.60876 | 73.09066 | 31.20829 | 0.370669 | 1.723629 | 0.813031 |
| 1 | 109269400 | 109277999 | 73.74018 | 116.8234 | 133.488 | 55.51932 | 0.137001 | 2.369211 | 0.813046 |
| 3 | 108453800 | 108455199 | 18.91266 | 32.49652 | 46.37819 | 20.34296 | -0.1271 | 0.909793 | 0.813067 |
| 18 | 74968000 | 74977999 | 121.6529 | 184.2609 | 238.8031 | 98.81104 | -0.0952 | 3.26825 | 0.813092 |
| 5 | 8088400 | 8089399 | 10.07665 | 18.8792 | 28.47036 | 11.75568 | -0.12714 | 0.120271 | 0.813103 |
| 17 | 29744400 | 29751599 | 79.45007 | 123.7305 | 164.1913 | 68.1514 | -0.09779 | 2.714516 | 0.81312 |
| 5 | 58834800 | 58835599 | 8.686707 | 16.22722 | 25.87055 | 10.73557 | -0.21231 | 0.120085 | 0.813133 |
| 2 | 77859000 | 77860399 | 17.8939 | 31.7418 | 36.2897 | 16.03526 | 0.200388 | 0.513741 | 0.813142 |
| 13 | 33225200 | 33227799 | 26.27288 | 43.94424 | 52.85016 | 23.01952 | 0.115762 | 1.189622 | 0.813147 |
| 10 | 42307600 | 42309999 | 26.69819 | 45.16169 | 53.43519 | 23.38709 | 0.139406 | 1.085391 | 0.813148 |
| 7 | 14137200 | 14138999 | 64.01889 | 101.5469 | 119.7595 | 49.86562 | 0.093142 | 2.324906 | 0.81316 |
| 17 | 91670600 | 91672199 | 18.59562 | 32.36936 | 38.92254 | 17.10927 | 0.122388 | 0.765296 | 0.813173 |
| X | 116527000 | 116530799 | 38.09824 | 62.95442 | 68.68379 | 29.67495 | 0.255243 | 1.371581 | 0.813181 |
| 12 | 72322000 | 72322799 | 10.79796 | 20.32225 | 25.80347 | 10.57326 | 0.128076 | 0.091163 | 0.813186 |
| 5 | 6639600 | 6643799 | 50.90523 | 81.50478 | 96.86861 | 40.82462 | 0.096209 | 2.018495 | 0.813189 |
| 9 | 68035800 | 68038399 | 31.29619 | 52.07731 | 61.3609 | 26.63933 | 0.145278 | 1.267437 | 0.813196 |
| 9 | 32032400 | 32035599 | 36.26903 | 59.68178 | 58.69791 | 25.3066 | 0.402662 | 1.406253 | 0.813206 |
| 8 | 102687600 | 102693799 | 96.35918 | 148.9148 | 172.606 | 71.69671 | 0.088601 | 2.857689 | 0.813297 |
| 1 | 30122400 | 30130999 | 211.8379 | 307.7034 | 386.4611 | 159.313 | -0.10563 | 4.013277 | 0.813319 |
| 16 | 33466200 | 33473999 | 78.78487 | 124.5961 | 139.5693 | 57.99696 | 0.163602 | 2.41559 | 0.813324 |
| 15 | 15108000 | 15108599 | 10.67276 | 20.31282 | 20.60158 | 8.251442 | 0.478149 | 0.000645 | 0.813333 |
| 12 | 114943000 | 114953799 | 144.3792 | 219.6614 | 235.0686 | 97.39725 | 0.186163 | 3.184359 | 0.813379 |
| 17 | 85217200 | 85218799 | 13.76992 | 24.95333 | 30.49466 | 13.19651 | 0.124971 | 0.347249 | 0.813385 |
| 6 | 85858800 | 85862999 | 82.04815 | 128.6873 | 150.0476 | 62.23107 | 0.096573 | 2.590711 | 0.813402 |
| 5 | 121523000 | 121524999 | 14.18864 | 24.96174 | 36.22379 | 16.03833 | -0.14647 | 0.578629 | 0.813405 |
| 5 | 57529000 | 57529999 | 10.78482 | 20.34377 | 25.83674 | 10.54539 | 0.132266 | 0.076419 | 0.813422 |
| 17 | 10409600 | 10412199 | 29.84609 | 49.7924 | 59.11672 | 25.70718 | 0.134264 | 1.232117 | 0.813425 |
| 16 | 47273400 | 47274599 | 12.06174 | 22.26482 | 34.16995 | 14.43483 | -0.17788 | 0.238121 | 0.813448 |
| 13 | 44869800 | 44872199 | 18.4058 | 31.72819 | 45.31927 | 19.88044 | -0.12733 | 0.876829 | 0.813463 |
| 12 | 88136200 | 88138799 | 24.78925 | 42.7692 | 48.47163 | 21.26387 | 0.205238 | 0.900097 | 0.813483 |
| 10 | 110523800 | 110524399 | 8.810387 | 16.87232 | 21.39844 | 8.446428 | 0.177943 | -0.05999 | 0.813504 |

| | | | | | | | | | |
|---|---|---|---|---|---|---|---|---|---|
| 5 | 110273000 | 110274799 | 18.16718 | 31.89505 | 38.03398 | 16.82588 | 0.134736 | 0.632624 | 0.813526 |
| 1 | 144128200 | 144131599 | 55.83701 | 88.99255 | 104.5631 | 43.78347 | 0.105513 | 2.167357 | 0.813531 |
| 15 | 86428800 | 86435599 | 39.69606 | 64.62636 | 64.53403 | 27.70598 | 0.372406 | 1.560403 | 0.813542 |
| 1 | 9046600 | 9047399 | 13.06378 | 23.24042 | 33.6174 | 14.81849 | -0.13455 | 0.460848 | 0.813587 |
| 2 | 107876400 | 107880199 | 56.54894 | 90.41427 | 106.1854 | 44.66195 | 0.111127 | 2.070791 | 0.813594 |
| X | 22053000 | 22053999 | 15.16611 | 27.7492 | 31.68495 | 13.54677 | 0.233903 | 0.307928 | 0.813607 |
| 13 | 104032200 | 104035599 | 38.67852 | 63.32132 | 72.64185 | 31.25046 | 0.178261 | 1.483165 | 0.813618 |
| 4 | 110574400 | 110577199 | 25.25479 | 41.5023 | 73.80839 | 31.80333 | -0.45019 | 1.407297 | 0.813624 |
| 13 | 67049000 | 67053599 | 36.34505 | 59.6436 | 69.88594 | 30.0953 | 0.14923 | 1.44963 | 0.813631 |
| 1 | 85101200 | 85105799 | 78.21028 | 122.797 | 166.755 | 69.16314 | -0.12192 | 2.570313 | 0.813633 |
| 17 | 56222400 | 56223799 | 16.44818 | 29.02035 | 34.99523 | 15.48546 | 0.120888 | 0.559309 | 0.813642 |
| 5 | 130244600 | 130248599 | 46.48305 | 74.67715 | 88.94244 | 37.67752 | 0.09896 | 1.90548 | 0.813642 |
| 1 | 87036000 | 87037999 | 16.95536 | 29.68141 | 35.22262 | 15.54406 | 0.141831 | 0.667297 | 0.813661 |
| 4 | 140894600 | 140895599 | 9.892175 | 18.46561 | 27.86339 | 11.57685 | -0.13527 | 0.139892 | 0.813685 |
| 7 | 120155800 | 120158399 | 33.41131 | 54.82913 | 66.23407 | 28.52266 | 0.105329 | 1.438265 | 0.813705 |
| 18 | 22545400 | 22546199 | 12.2294 | 21.64774 | 38.80358 | 17.15942 | -0.44592 | 0.505677 | 0.813719 |
| 4 | 92932000 | 92933399 | 18.73491 | 32.3094 | 45.83306 | 20.10609 | -0.11733 | 0.875219 | 0.813731 |
| 18 | 14230800 | 14232799 | 23.92971 | 40.19027 | 56.53395 | 24.68812 | -0.10958 | 1.143139 | 0.813749 |
| 3 | 73335200 | 73335999 | 8.315742 | 15.94383 | 25.19032 | 9.909583 | -0.13273 | -0.06806 | 0.81375 |
| 3 | 83498600 | 83501999 | 26.28347 | 44.11052 | 53.38731 | 23.30303 | 0.106562 | 1.1546 | 0.813805 |
| 2 | 114302200 | 114304399 | 19.17478 | 33.32412 | 39.94187 | 17.53968 | 0.1267 | 0.791004 | 0.813806 |
| 2 | 108298200 | 108303999 | 80.12175 | 125.7358 | 147.1895 | 61.04426 | 0.091465 | 2.577954 | 0.813821 |
| 15 | 33391000 | 33399999 | 102.7035 | 159.5215 | 176.9284 | 73.46725 | 0.157716 | 2.773245 | 0.813836 |
| 12 | 82185600 | 82186399 | 12.08086 | 22.12967 | 27.07597 | 11.55808 | 0.136256 | 0.265023 | 0.813869 |
| 3 | 15994600 | 15995399 | 10.84785 | 20.24118 | 25.23139 | 10.49064 | 0.137896 | 0.145881 | 0.813895 |
| 14 | 42637800 | 42639799 | 16.66768 | 30.47264 | 33.21795 | 14.21701 | 0.2991 | 0.323857 | 0.813906 |
| 4 | 80205000 | 80206399 | 19.75449 | 33.93211 | 47.94706 | 21.03091 | -0.1129 | 0.918736 | 0.813916 |
| 8 | 26538000 | 26540999 | 19.75449 | 33.93211 | 47.94706 | 21.03091 | -0.1129 | 0.918736 | 0.813916 |
| 3 | 141908000 | 141909199 | 17.85922 | 31.3803 | 37.50293 | 16.60253 | 0.13157 | 0.617206 | 0.813925 |
| 2 | 87421000 | 87424199 | 31.05845 | 51.45597 | 61.87874 | 26.78308 | 0.115432 | 1.315603 | 0.81394 |
| 3 | 6761200 | 6763599 | 32.63266 | 53.83227 | 64.64716 | 27.90098 | 0.116823 | 1.372454 | 0.813945 |
| X | 23398600 | 23401999 | 42.75145 | 69.12814 | 80.44097 | 34.37899 | 0.143609 | 1.706076 | 0.813954 |
| 6 | 116790400 | 116792999 | 29.9854 | 49.83113 | 59.6634 | 25.87907 | 0.122021 | 1.274587 | 0.813974 |
| 1 | 97097800 | 97103399 | 90.35019 | 140.3337 | 162.9901 | 67.6796 | 0.09127 | 2.763981 | 0.813974 |
| 1 | 45647600 | 45652799 | 76.43343 | 120.3401 | 141.1818 | 58.57756 | 0.091933 | 2.510331 | 0.814094 |
| 14 | 24433800 | 24435199 | 14.89501 | 26.45138 | 31.91316 | 14.08824 | 0.124168 | 0.47964 | 0.814106 |
| 17 | 91880000 | 91882599 | 21.73956 | 37.35017 | 54.48854 | 23.90035 | -0.15887 | 0.93193 | 0.814113 |
| 3 | 100165800 | 100167999 | 22.66844 | 39.12791 | 46.30782 | 20.31492 | 0.143187 | 0.885289 | 0.814114 |
| 15 | 24648200 | 24648999 | 10.33327 | 19.47082 | 24.78256 | 10.13451 | 0.127064 | 0.077358 | 0.814116 |
| 1 | 149352600 | 149353799 | 11.74833 | 21.39461 | 31.21524 | 13.42568 | -0.12327 | 0.306615 | 0.814118 |
| 19 | 44617400 | 44618399 | 11.25216 | 20.60468 | 30.24876 | 12.91843 | -0.12529 | 0.26901 | 0.814122 |
| 10 | 73418200 | 73419799 | 14.87801 | 25.66588 | 49.49647 | 21.71337 | -0.55809 | 0.862804 | 0.814126 |
| 18 | 50915800 | 50916599 | 10.04402 | 18.48514 | 28.36153 | 12.02979 | -0.18088 | 0.228935 | 0.814138 |
| 8 | 35768000 | 35772399 | 35.6051 | 58.1423 | 70.10024 | 30.12899 | 0.104258 | 1.511407 | 0.814155 |
| 6 | 122974600 | 122979399 | 64.33113 | 102.1895 | 120.6395 | 50.30464 | 0.093693 | 2.283184 | 0.814182 |
| 7 | 42112200 | 42115799 | 52.798 | 84.48398 | 100.3994 | 42.27579 | 0.095388 | 2.041518 | 0.814183 |
| 13 | 51153800 | 51155599 | 15.14151 | 27.3733 | 26.7998 | 11.63232 | 0.440166 | 0.362833 | 0.814191 |

| | | | | | | | | | |
|---|---|---|---|---|---|---|---|---|---|
| 9 | 11273800 | 11278199 | 60.926 | 96.27196 | 131.937 | 54.80213 | -0.12788 | 2.419917 | 0.814229 |
| 7 | 137445000 | 137446199 | 13.87658 | 24.76054 | 35.58925 | 15.64806 | -0.12302 | 0.451376 | 0.814268 |
| 11 | 17651800 | 17653599 | 21.55202 | 36.2152 | 61.88524 | 27.03093 | -0.38933 | 1.135861 | 0.814275 |
| 17 | 55605600 | 55608999 | 37.43331 | 61.10801 | 72.63797 | 31.21603 | 0.124337 | 1.521643 | 0.814275 |
| 8 | 75993400 | 75996399 | 51.41753 | 82.20644 | 96.67151 | 40.658 | 0.109216 | 2.063373 | 0.814287 |
| 14 | 50770800 | 50773399 | 37.11669 | 60.29055 | 72.60795 | 31.13874 | 0.100308 | 1.595624 | 0.814287 |
| 15 | 96899600 | 96901399 | 17.03697 | 29.8421 | 35.1043 | 15.49967 | 0.154272 | 0.656434 | 0.814288 |
| 9 | 26548800 | 26549599 | 10.48178 | 19.82678 | 25.36618 | 10.29309 | 0.129186 | 0.053933 | 0.814355 |
| 6 | 57174600 | 57177199 | 35.43115 | 58.16 | 57.99397 | 24.97714 | 0.381474 | 1.433406 | 0.814359 |
| 5 | 85904200 | 85904999 | 10.29849 | 19.73529 | 25.19319 | 9.925242 | 0.170207 | -0.04406 | 0.814367 |
| 8 | 52729800 | 52733399 | 48.47403 | 77.6048 | 105.531 | 44.4689 | -0.09792 | 2.021043 | 0.81441 |
| 8 | 52730000 | 52733399 | 48.47403 | 77.6048 | 105.531 | 44.4689 | -0.09792 | 2.021043 | 0.81441 |
| 7 | 40304000 | 40305399 | 13.21041 | 23.81711 | 34.65829 | 15.0947 | -0.13143 | 0.381392 | 0.814416 |
| 12 | 44287200 | 44290199 | 23.90093 | 41.02035 | 48.47057 | 21.25947 | 0.14376 | 0.942233 | 0.814444 |
| 10 | 125471600 | 125473799 | 25.18948 | 42.35108 | 50.66352 | 22.13313 | 0.123437 | 1.132306 | 0.8145 |
| 2 | 156627600 | 156629799 | 28.23635 | 47.42247 | 56.22663 | 24.55364 | 0.136083 | 1.155283 | 0.814521 |
| 17 | 41010800 | 41012199 | 18.40893 | 32.23529 | 38.72404 | 17.09344 | 0.124321 | 0.678052 | 0.814527 |
| X | 100752000 | 100753399 | 13.13245 | 23.79075 | 29.04615 | 12.57454 | 0.126102 | 0.341799 | 0.814559 |
| X | 66715600 | 66717799 | 27.6518 | 47.69598 | 39.23893 | 17.21347 | 0.665566 | 0.905125 | 0.81456 |
| 13 | 68044000 | 68048599 | 84.61052 | 132.0206 | 154.3601 | 64.05378 | 0.086788 | 2.682594 | 0.814572 |
| X | 120287600 | 120292599 | 75.8675 | 120.2395 | 134.6167 | 56.0045 | 0.166887 | 2.362515 | 0.814597 |
| 12 | 47282400 | 47282999 | 7.547477 | 14.59969 | 23.83982 | 9.105204 | -0.13962 | -0.17739 | 0.814598 |
| 10 | 40946200 | 40948799 | 25.00561 | 40.94609 | 75.03585 | 32.28334 | -0.49533 | 1.453734 | 0.814624 |
| 6 | 24342000 | 24345199 | 28.86708 | 47.055 | 82.34241 | 35.36759 | -0.43364 | 1.529298 | 0.81464 |
| 12 | 26597600 | 26600199 | 30.56324 | 50.37482 | 60.55703 | 26.12615 | 0.114849 | 1.373068 | 0.814641 |
| 12 | 86909800 | 86911399 | 17.59249 | 30.77517 | 36.82326 | 16.24082 | 0.130158 | 0.687202 | 0.814647 |
| 17 | 92725000 | 92725799 | 9.26511 | 17.58264 | 27.13538 | 10.93529 | -0.13062 | 0.024712 | 0.814677 |
| 14 | 63449400 | 63451199 | 14.44148 | 25.55494 | 36.64849 | 16.20749 | -0.1261 | 0.519355 | 0.814699 |
| 18 | 51227400 | 51228999 | 16.11964 | 29.26782 | 27.16104 | 11.72557 | 0.521856 | 0.345958 | 0.814724 |
| 8 | 84742000 | 84744799 | 29.7619 | 49.02711 | 57.68322 | 24.88052 | 0.145506 | 1.376109 | 0.814735 |
| 14 | 42401800 | 42402999 | 17.172 | 30.03656 | 35.79883 | 15.78765 | 0.135704 | 0.683393 | 0.814744 |
| 13 | 45734200 | 45737399 | 38.58554 | 62.37295 | 73.17761 | 31.26247 | 0.131284 | 1.700163 | 0.814793 |
| 17 | 95171800 | 95175799 | 38.25505 | 61.83467 | 86.26839 | 36.85192 | -0.11826 | 1.706297 | 0.814812 |
| 1 | 186691400 | 186693999 | 25.09036 | 41.73505 | 59.21402 | 25.70062 | -0.12186 | 1.252073 | 0.814817 |
| 8 | 48467400 | 48469599 | 22.09001 | 37.91233 | 45.24425 | 19.84439 | 0.129444 | 0.928739 | 0.814823 |
| 10 | 124379600 | 124382999 | 60.89831 | 96.88082 | 114.3585 | 47.76338 | 0.095998 | 2.232839 | 0.814845 |
| X | 48582200 | 48584599 | 26.20555 | 43.97845 | 52.8151 | 23.05284 | 0.117813 | 1.155826 | 0.814845 |
| 1 | 63286000 | 63288999 | 28.75426 | 47.75071 | 57.8183 | 25.06627 | 0.105851 | 1.279362 | 0.814862 |
| 6 | 89671800 | 89676599 | 113.8878 | 173.4938 | 225.3404 | 93.4024 | -0.09152 | 3.149771 | 0.814923 |
| 17 | 91869600 | 91873999 | 65.34033 | 102.8316 | 139.9414 | 58.05975 | -0.12232 | 2.512086 | 0.814926 |
| 10 | 31281200 | 31284199 | 28.77094 | 48.00514 | 57.87094 | 25.16757 | 0.112372 | 1.22783 | 0.81493 |
| 11 | 119819400 | 119822199 | 32.23264 | 52.50426 | 73.4 | 31.5186 | -0.11121 | 1.541911 | 0.814931 |
| 11 | 104206200 | 104214999 | 70.01599 | 108.9631 | 178.6831 | 74.17425 | -0.40359 | 2.724468 | 0.814936 |
| 18 | 5785200 | 5786999 | 14.23269 | 25.48103 | 30.92557 | 13.55924 | 0.122404 | 0.422749 | 0.814946 |
| 11 | 75833400 | 75835599 | 23.98492 | 40.89638 | 49.02353 | 21.49069 | 0.121659 | 1.000515 | 0.81496 |
| 4 | 128350000 | 128351999 | 21.84864 | 37.13784 | 52.2249 | 22.88834 | -0.10844 | 1.025701 | 0.814964 |
| Y | 3982000 | 3982599 | 7.462038 | 14.44038 | 16.84336 | 6.42063 | 0.344358 | -0.20792 | 0.81501 |

| | | | | | | | | | |
|---|---|---|---|---|---|---|---|---|---|
| 5 | 109665000 | 109682999 | 349.8574 | 491.0149 | 521.7963 | 216.2401 | 0.085842 | 4.526328 | 0.815013 |
| 11 | 7394400 | 7395399 | 9.365039 | 17.8418 | 28.45122 | 11.37128 | -0.16738 | 0.000321 | 0.815013 |
| 6 | 130627800 | 130639399 | 120.7697 | 186.325 | 195.8648 | 81.35861 | 0.22757 | 2.901449 | 0.815044 |
| 14 | 39212400 | 39220199 | 110.3867 | 169.9569 | 167.8475 | 69.71399 | 0.315272 | 2.932496 | 0.815061 |
| 2 | 143880800 | 143891799 | 112.3838 | 171.5364 | 222.9585 | 92.45979 | -0.09033 | 3.108329 | 0.815105 |
| 10 | 94160600 | 94163599 | 28.29926 | 46.85654 | 66.19875 | 28.64104 | -0.11642 | 1.317568 | 0.815131 |
| 13 | 86347000 | 86350599 | 35.87279 | 59.09875 | 68.30416 | 29.46273 | 0.171191 | 1.400497 | 0.815136 |
| 2 | 171646200 | 171647999 | 16.19689 | 28.43632 | 41.25343 | 18.25035 | -0.14685 | 0.624934 | 0.815153 |
| 9 | 92005000 | 92005999 | 17.85507 | 31.22749 | 37.77318 | 16.65661 | 0.114422 | 0.692408 | 0.815157 |
| 11 | 46583800 | 46587399 | 41.77961 | 67.45635 | 80.77745 | 34.45666 | 0.099867 | 1.744781 | 0.815182 |
| 8 | 113205000 | 113210399 | 60.62747 | 96.69694 | 112.9175 | 47.3287 | 0.115474 | 2.151444 | 0.815212 |
| X | 31380800 | 31411799 | 1151.088 | 1478.724 | 1319.279 | 553.3093 | 0.210019 | 5.931444 | 0.815221 |
| 11 | 68193600 | 68194999 | 17.53601 | 30.72899 | 37.17985 | 16.42186 | 0.114005 | 0.658577 | 0.815225 |
| 2 | 23167600 | 23173199 | 55.53479 | 88.7114 | 105.3701 | 44.26151 | 0.093404 | 2.097006 | 0.815249 |
| 14 | 83152000 | 83153399 | 16.82992 | 29.76256 | 35.44324 | 15.67646 | 0.140173 | 0.537989 | 0.815254 |
| 1 | 187333000 | 187333999 | 23.12111 | 39.97011 | 39.37342 | 17.27365 | 0.407261 | 0.86357 | 0.815273 |
| 12 | 46008800 | 46009799 | 17.326 | 29.55099 | 54.23089 | 23.77403 | -0.48998 | 0.982949 | 0.81528 |
| 3 | 78394000 | 78396199 | 23.23897 | 38.85443 | 55.9671 | 24.37109 | -0.14363 | 1.186748 | 0.815287 |
| 13 | 29009200 | 29010999 | 22.4947 | 38.34015 | 45.84024 | 20.09457 | 0.125338 | 0.994719 | 0.815296 |
| 5 | 88592200 | 88592599 | 5.376392 | 10.48032 | 18.59093 | 6.715929 | -0.18198 | -0.41292 | 0.815321 |
| 1 | 141116400 | 141118999 | 49.70532 | 79.44891 | 107.5603 | 45.2214 | -0.09403 | 2.073237 | 0.815333 |
| 10 | 32212400 | 32214599 | 37.34704 | 61.46146 | 58.27403 | 25.12485 | 0.455385 | 1.406471 | 0.815333 |
| 15 | 16664400 | 16664799 | 6.231811 | 12.16593 | 17.57911 | 6.211111 | 0.139412 | -0.4822 | 0.815353 |
| 6 | 140200200 | 140202799 | 17.25241 | 30.32846 | 36.53432 | 16.1783 | 0.120332 | 0.606076 | 0.815354 |
| 10 | 127656600 | 127657399 | 17.25241 | 30.32846 | 36.53432 | 16.1783 | 0.120332 | 0.606076 | 0.815354 |
| 8 | 77912200 | 77913999 | 13.95962 | 24.39711 | 44.66614 | 19.68426 | -0.48067 | 0.693896 | 0.815374 |
| 4 | 60707200 | 60708799 | 10.71818 | 20.0932 | 25.18861 | 10.39005 | 0.138517 | 0.112366 | 0.815419 |
| 10 | 98341400 | 98342199 | 6.971069 | 13.50922 | 22.18382 | 8.401097 | -0.13543 | -0.21577 | 0.815425 |
| X | 8340400 | 8342999 | 41.95818 | 68.6431 | 60.97937 | 26.22646 | 0.545635 | 1.496065 | 0.815461 |
| X | 39453000 | 39462999 | 199.6006 | 294.2382 | 323.4421 | 133.1973 | 0.106914 | 3.783529 | 0.815462 |
| 15 | 58362800 | 58364399 | 15.44798 | 27.88608 | 32.9574 | 14.33148 | 0.168403 | 0.38628 | 0.815476 |
| 11 | 50760600 | 50774999 | 215.8152 | 317.0363 | 348.5375 | 143.5436 | 0.102032 | 3.840643 | 0.815483 |
| 10 | 21325600 | 21327999 | 17.32717 | 30.31814 | 43.51318 | 19.19519 | -0.1312 | 0.691805 | 0.815513 |
| 1 | 182822000 | 182825199 | 39.39531 | 63.61411 | 74.19328 | 31.65284 | 0.138036 | 1.7328 | 0.815517 |
| 6 | 124593000 | 124607199 | 261.4646 | 376.7723 | 410.3475 | 169.3312 | 0.088617 | 4.135178 | 0.815524 |
| 12 | 77925600 | 77927999 | 12.36939 | 22.57607 | 33.3082 | 14.28584 | -0.13686 | 0.301957 | 0.815526 |
| 6 | 143461000 | 143461199 | 4.928775 | 9.640069 | 18.00934 | 6.077401 | -0.16727 | -0.66154 | 0.815541 |
| 12 | 21790800 | 21876199 | 1959.523 | 2386.57 | 2195.307 | 926.2193 | 0.088121 | 6.787081 | 0.815542 |
| 10 | 8038400 | 8048999 | 254.0148 | 366.8576 | 401.1194 | 165.4742 | 0.086143 | 4.101972 | 0.815563 |
| 9 | 11416200 | 11417399 | 19.64678 | 33.98953 | 40.61596 | 17.81917 | 0.130112 | 0.844216 | 0.815566 |
| 4 | 145611400 | 145616199 | 201.759 | 295.9445 | 323.7951 | 133.366 | 0.106886 | 3.861918 | 0.815585 |
| 10 | 117932000 | 117936399 | 43.39508 | 69.77291 | 95.36162 | 40.44444 | -0.09774 | 1.877855 | 0.815595 |
| 11 | 34819000 | 34819599 | 10.16723 | 19.32859 | 24.93632 | 10.01178 | 0.130674 | 0.015683 | 0.815658 |
| 7 | 27982600 | 27984599 | 18.54759 | 31.88677 | 45.851 | 20.11206 | -0.13777 | 0.901557 | 0.81566 |
| 1 | 148149600 | 148155199 | 60.6751 | 96.8967 | 96.28211 | 40.42876 | 0.349988 | 2.107071 | 0.815661 |
| 13 | 62441200 | 62444399 | 75.61101 | 118.4706 | 133.6159 | 55.42196 | 0.143384 | 2.602219 | 0.815671 |
| 14 | 79252400 | 79252999 | 8.253339 | 15.90388 | 20.88903 | 8.099826 | 0.151184 | -0.12791 | 0.815703 |

| | | | | | | | | | |
|---|---|---|---|---|---|---|---|---|---|
| 13 | 52509800 | 52514799 | 13.7434 | 25.41468 | 24.84823 | 10.47129 | 0.470881 | 0.219989 | 0.815741 |
| 9 | 7890400 | 7893399 | 36.26283 | 59.28283 | 69.99543 | 30.09625 | 0.135434 | 1.493927 | 0.815787 |
| 5 | 93536000 | 93541799 | 104.0788 | 160.3851 | 185.3585 | 76.99495 | 0.089652 | 2.914594 | 0.81581 |
| 18 | 6802800 | 6803399 | 17.24161 | 30.20865 | 36.10572 | 15.94544 | 0.131438 | 0.655475 | 0.815847 |
| 8 | 116953600 | 116955399 | 21.83085 | 37.52292 | 45.28569 | 19.86395 | 0.113876 | 0.918278 | 0.815878 |
| 6 | 25174200 | 25175999 | 19.20809 | 33.37298 | 47.92628 | 21.0423 | -0.13302 | 0.807445 | 0.815919 |
| 4 | 71715000 | 71716399 | 15.71546 | 28.2441 | 33.37252 | 14.58471 | 0.163958 | 0.413726 | 0.815954 |
| 2 | 17042200 | 17045199 | 33.90871 | 55.8691 | 66.51536 | 28.69246 | 0.128542 | 1.39091 | 0.815964 |
| 7 | 72299600 | 72301199 | 19.76289 | 33.9176 | 47.95003 | 21.03112 | -0.11381 | 0.926646 | 0.816011 |
| X | 112207200 | 112209199 | 15.28715 | 26.9204 | 38.74654 | 17.15197 | -0.13403 | 0.581717 | 0.816034 |
| 1 | 189280600 | 189285799 | 53.9087 | 86.1395 | 102.4851 | 43.06779 | 0.091494 | 2.085722 | 0.816035 |
| 12 | 63955600 | 63956999 | 21.50779 | 37.35831 | 34.85803 | 15.30197 | 0.486498 | 0.804902 | 0.816086 |
| 14 | 15983000 | 15983999 | 17.67101 | 30.82462 | 36.42464 | 16.03243 | 0.147833 | 0.728307 | 0.816095 |
| 5 | 44980400 | 44983399 | 36.69364 | 59.41751 | 81.90457 | 35.04491 | -0.09829 | 1.66261 | 0.816107 |
| 18 | 82277200 | 82279399 | 19.19065 | 32.96631 | 46.89154 | 20.56798 | -0.12244 | 0.912643 | 0.816111 |
| 1 | 60152200 | 60154199 | 32.60388 | 53.78851 | 64.68552 | 27.91835 | 0.114826 | 1.371041 | 0.816127 |
| 17 | 48564000 | 48569799 | 66.33241 | 105.373 | 122.7766 | 51.19796 | 0.112631 | 2.282411 | 0.816141 |
| 1 | 40959400 | 40960199 | 6.975832 | 13.50306 | 22.13743 | 8.428286 | -0.1403 | -0.19756 | 0.816157 |
| 2 | 111575000 | 111578799 | 46.43396 | 74.80157 | 88.50942 | 37.64226 | 0.113207 | 1.819511 | 0.816163 |
| 2 | 53662200 | 53666399 | 60.90543 | 97.03406 | 114.1493 | 47.77262 | 0.103188 | 2.188214 | 0.816188 |
| 10 | 15105800 | 15106399 | 11.46934 | 21.69496 | 27.02419 | 10.96569 | 0.167412 | 0.066897 | 0.816192 |
| 7 | 107740800 | 107742799 | 34.11897 | 55.69121 | 67.00731 | 28.79508 | 0.106793 | 1.512396 | 0.81624 |
| 2 | 178342600 | 178344799 | 18.05054 | 31.51442 | 37.76755 | 16.63273 | 0.127534 | 0.717654 | 0.816248 |
| Y | 90113600 | 90115999 | 23.38402 | 38.35683 | 63.91879 | 27.52041 | -0.35752 | 1.418858 | 0.816266 |
| 5 | 124816200 | 124821799 | 63.16079 | 99.53264 | 136.4921 | 56.63837 | -0.13188 | 2.483488 | 0.816272 |
| 17 | 13060400 | 13063399 | 25.57123 | 42.92632 | 51.76412 | 22.59821 | 0.111806 | 1.14704 | 0.816275 |
| 16 | 93866400 | 93875199 | 84.59559 | 130.1744 | 206.6771 | 85.83493 | -0.36962 | 2.938008 | 0.816323 |
| 1 | 45009000 | 45011599 | 35.19945 | 57.87374 | 68.33934 | 29.453 | 0.139123 | 1.420275 | 0.816351 |
| 13 | 9933800 | 9937399 | 27.93741 | 46.69762 | 55.71355 | 24.25663 | 0.127172 | 1.20166 | 0.816354 |
| 2 | 7450400 | 7453399 | 34.38027 | 56.82232 | 66.15819 | 28.58777 | 0.161666 | 1.355035 | 0.816355 |
| 6 | 66545000 | 66547199 | 28.20954 | 46.83978 | 66.16775 | 28.68337 | -0.1159 | 1.286916 | 0.816356 |
| 8 | 37043000 | 37044399 | 12.296 | 22.09689 | 32.18499 | 14.06664 | -0.13577 | 0.390903 | 0.816385 |
| 4 | 9055200 | 9056599 | 11.58081 | 20.81238 | 31.16104 | 13.61871 | -0.17547 | 0.3809 | 0.816415 |
| 2 | 145843800 | 145844999 | 14.77933 | 26.20993 | 31.49975 | 13.92185 | 0.128691 | 0.487606 | 0.816415 |
| 5 | 142367800 | 142368999 | 10.71736 | 19.74243 | 29.26407 | 12.39716 | -0.1308 | 0.228001 | 0.816419 |
| 9 | 41234600 | 41235399 | 12.6636 | 22.82618 | 33.08891 | 14.4152 | -0.12623 | 0.374528 | 0.816424 |
| 13 | 92539600 | 92540999 | 15.26071 | 27.80321 | 32.00127 | 13.75616 | 0.216823 | 0.328494 | 0.816435 |
| X | 11628000 | 11630999 | 18.38856 | 32.06073 | 46.79522 | 20.5908 | -0.15567 | 0.755811 | 0.816437 |
| 18 | 86412800 | 86414799 | 19.89743 | 34.35633 | 41.10926 | 18.03453 | 0.127528 | 0.861961 | 0.816438 |
| 18 | 74963000 | 74966399 | 34.03073 | 55.48253 | 77.26629 | 33.18958 | -0.10482 | 1.535334 | 0.816442 |
| 7 | 107459600 | 107463199 | 47.39756 | 76.38958 | 76.73112 | 32.65488 | 0.349881 | 1.802338 | 0.816448 |
| 5 | 58468800 | 58470199 | 20.65091 | 35.97113 | 35.20832 | 15.48313 | 0.418052 | 0.760686 | 0.816454 |
| 19 | 11022400 | 11022999 | 8.166674 | 15.73928 | 20.419 | 7.914106 | 0.170111 | -0.12931 | 0.816457 |
| 10 | 66640400 | 66643799 | 33.91624 | 55.75567 | 66.95607 | 28.85503 | 0.114839 | 1.418727 | 0.816466 |
| 5 | 128519800 | 128522199 | 19.5684 | 33.24939 | 49.02328 | 21.48453 | -0.1765 | 1.016089 | 0.816497 |
| X | 32991600 | 32992399 | 11.94267 | 22.57111 | 22.1041 | 8.98538 | 0.510008 | 0.06605 | 0.816514 |
| 6 | 65611000 | 65616799 | 66.27077 | 105.2949 | 123.0548 | 51.32369 | 0.108576 | 2.278129 | 0.81652 |

| | | | | | | | | | |
|---|---|---|---|---|---|---|---|---|---|
| 10 | 61092400 | 61094799 | 29.26063 | 48.07363 | 67.40901 | 29.04292 | -0.10841 | 1.416406 | 0.816526 |
| 10 | 30667200 | 30668999 | 19.85492 | 35.03255 | 38.94798 | 17.23441 | 0.23752 | 0.586001 | 0.816531 |
| 2 | 35919400 | 35923599 | 41.2247 | 66.72133 | 79.73919 | 34.10586 | 0.106455 | 1.679084 | 0.816532 |
| 1 | 89054600 | 89055199 | 9.563019 | 18.51381 | 23.98263 | 9.124435 | 0.196758 | -0.16746 | 0.816549 |
| 15 | 62263200 | 62266399 | 27.06355 | 46.29537 | 51.80789 | 22.71721 | 0.221208 | 0.987575 | 0.816591 |
| 13 | 74765200 | 74766199 | 13.89669 | 24.43416 | 35.79438 | 15.85001 | -0.16024 | 0.583135 | 0.816619 |
| 5 | 70235400 | 70236799 | 11.32335 | 20.77714 | 30.7439 | 13.09368 | -0.13405 | 0.257106 | 0.816619 |
| 11 | 108463600 | 108464399 | 11.35697 | 21.0182 | 31.45683 | 13.242 | -0.13818 | 0.21185 | 0.81662 |
| 9 | 116028600 | 116031799 | 32.71559 | 53.33013 | 74.12898 | 31.84017 | -0.10224 | 1.532843 | 0.816624 |
| 13 | 44488600 | 44490999 | 27.50977 | 46.45828 | 54.57369 | 23.87351 | 0.149692 | 1.102948 | 0.816639 |
| 19 | 9205800 | 9206799 | 15.27505 | 27.1423 | 32.91671 | 14.52404 | 0.117283 | 0.481216 | 0.816665 |
| 13 | 27731000 | 27731399 | 7.16794 | 13.91534 | 23.42198 | 8.793365 | -0.1584 | -0.23645 | 0.816674 |
| X | 30168000 | 30169199 | 10.96333 | 20.90079 | 20.67376 | 8.251104 | 0.5189 | -0.00581 | 0.816675 |
| 7 | 129964800 | 129968399 | 41.35752 | 67.14037 | 78.6058 | 33.69663 | 0.140133 | 1.617693 | 0.816678 |
| 4 | 93581400 | 93582399 | 10.12237 | 18.19117 | 35.8208 | 15.65536 | -0.5683 | 0.381097 | 0.816693 |
| 7 | 102803600 | 102804999 | 13.8688 | 25.69224 | 30.30427 | 12.73963 | 0.204436 | 0.223202 | 0.816696 |
| 10 | 83573200 | 83581199 | 113.5672 | 173.637 | 200.0198 | 82.97975 | 0.08565 | 3.073585 | 0.816704 |
| 5 | 115095800 | 115098399 | 26.55942 | 44.0156 | 62.00886 | 26.84503 | -0.11214 | 1.300749 | 0.816725 |
| 7 | 40720200 | 40730799 | 136.7257 | 207.704 | 233.142 | 96.55542 | 0.115559 | 3.212154 | 0.816727 |
| 5 | 141181400 | 141183399 | 16.27471 | 28.4593 | 40.42124 | 17.82208 | -0.11618 | 0.690354 | 0.816728 |
| 9 | 115984000 | 115986199 | 17.53038 | 30.84514 | 36.34377 | 16.091 | 0.152324 | 0.594073 | 0.816807 |
| 2 | 29456600 | 29457599 | 13.3134 | 24.57644 | 29.23669 | 12.35001 | 0.187761 | 0.239927 | 0.816811 |
| 3 | 64366200 | 64366599 | 7.615431 | 14.59331 | 23.54078 | 9.274908 | -0.16364 | -0.07178 | 0.816812 |
| 19 | 34291600 | 34292199 | 7.615431 | 14.59331 | 23.54078 | 9.274908 | -0.16364 | -0.07178 | 0.816812 |
| 2 | 138674600 | 138675999 | 23.22104 | 39.6547 | 47.94628 | 21.02105 | 0.109664 | 0.983535 | 0.816812 |
| 5 | 8960800 | 8964199 | 33.42111 | 54.3127 | 75.51626 | 32.39703 | -0.10583 | 1.581404 | 0.816831 |
| 1 | 26009200 | 26010399 | 19.51891 | 33.544 | 47.41431 | 20.79754 | -0.1133 | 0.911113 | 0.816835 |
| 2 | 176743600 | 176744399 | 7.530201 | 14.53514 | 23.71372 | 9.148048 | -0.15227 | -0.14499 | 0.816845 |
| 2 | 142904800 | 142905399 | 6.679248 | 12.91905 | 21.56847 | 8.241151 | -0.17116 | -0.19099 | 0.816854 |
| 14 | 87842800 | 87848399 | 89.84039 | 139.3464 | 159.6028 | 66.28693 | 0.109811 | 2.789534 | 0.816876 |
| 7 | 5029200 | 5041999 | 297.9733 | 414.8097 | 641.2864 | 266.0949 | -0.4666 | 4.647019 | 0.816926 |
| X | 153351000 | 153356799 | 68.88106 | 109.8921 | 118.5698 | 49.71733 | 0.229832 | 2.15517 | 0.816947 |
| 13 | 68157400 | 68215199 | 1319.447 | 1663.62 | 1579.917 | 663.8574 | 0.092751 | 6.233439 | 0.816994 |
| 16 | 11881400 | 11883599 | 19.78137 | 34.49939 | 41.38763 | 18.21476 | 0.125927 | 0.747943 | 0.81701 |
| 9 | 67095400 | 67096799 | 19.50867 | 33.5554 | 47.41081 | 20.79656 | -0.11246 | 0.90312 | 0.817036 |
| 10 | 71143200 | 71144599 | 19.50867 | 33.5554 | 47.41081 | 20.79656 | -0.11246 | 0.90312 | 0.817036 |
| 7 | 102382800 | 102384599 | 24.62538 | 41.1101 | 58.11736 | 25.28325 | -0.11662 | 1.211598 | 0.817085 |
| 11 | 20463000 | 20466399 | 46.01356 | 73.67867 | 101.2651 | 42.68277 | -0.11294 | 2.011154 | 0.81709 |
| 5 | 109671000 | 109677999 | 208.1576 | 304.747 | 335.7471 | 138.3053 | 0.094256 | 3.892304 | 0.817127 |
| 4 | 17654800 | 17658999 | 65.52813 | 103.8173 | 122.2878 | 50.86198 | 0.093278 | 2.360585 | 0.817144 |
| 13 | 87589000 | 87591199 | 35.25889 | 58.39274 | 54.74293 | 23.68823 | 0.473616 | 1.328181 | 0.817145 |
| 12 | 96804400 | 96810999 | 78.83388 | 124.4795 | 123.3404 | 51.21278 | 0.338561 | 2.448438 | 0.817156 |
| 17 | 11516000 | 11520799 | 79.78906 | 125.0501 | 146.1173 | 60.6057 | 0.092675 | 2.602515 | 0.817164 |
| 15 | 9044400 | 9046199 | 16.12067 | 28.37953 | 34.10622 | 15.09876 | 0.124617 | 0.580266 | 0.817179 |
| 4 | 73120200 | 73120999 | 14.97545 | 27.03919 | 25.87386 | 11.24804 | 0.471585 | 0.366989 | 0.817181 |
| 10 | 93176400 | 93177599 | 16.78484 | 29.14332 | 42.18999 | 18.51601 | -0.14443 | 0.798691 | 0.817183 |
| 12 | 54917200 | 54920799 | 32.69128 | 53.31204 | 74.23268 | 31.8894 | -0.10434 | 1.526464 | 0.817185 |

| | | | | | | | | | |
|---|---|---|---|---|---|---|---|---|---|
| 1 | 144368000 | 144369599 | 17.46856 | 30.38277 | 43.15998 | 18.96224 | -0.11702 | 0.779547 | 0.817275 |
| 19 | 12027200 | 12029199 | 27.92243 | 47.44049 | 54.01414 | 23.67164 | 0.194936 | 1.053862 | 0.817276 |
| 17 | 69630000 | 69631199 | 27.54647 | 45.31197 | 65.42201 | 28.20122 | -0.14967 | 1.398495 | 0.817302 |
| 9 | 71405800 | 71408399 | 19.25802 | 33.46149 | 40.49991 | 17.7823 | 0.11273 | 0.796594 | 0.817342 |
| 1 | 23420800 | 23422799 | 16.06524 | 27.76598 | 49.52143 | 21.72556 | -0.44556 | 0.848407 | 0.817364 |
| 6 | 96907200 | 96910199 | 29.02613 | 48.51286 | 57.94975 | 25.22873 | 0.125461 | 1.209584 | 0.817418 |
| 17 | 52048800 | 52050999 | 17.73441 | 30.59854 | 44.28254 | 19.42619 | -0.1459 | 0.863685 | 0.817429 |
| 7 | 106267000 | 106268799 | 19.16388 | 33.68291 | 38.95806 | 17.25113 | 0.178519 | 0.621686 | 0.817433 |
| 10 | 125550400 | 125551199 | 20.133 | 34.33806 | 49.01574 | 21.48776 | -0.12903 | 0.98675 | 0.817439 |
| 10 | 42031600 | 42033999 | 35.57855 | 58.04379 | 70.06317 | 30.10264 | 0.101607 | 1.524234 | 0.817455 |
| 3 | 84981400 | 84981999 | 10.15536 | 19.3496 | 24.98898 | 9.985191 | 0.135175 | 6.38E-05 | 0.81747 |
| 11 | 7819600 | 7820399 | 15.10946 | 26.63891 | 38.21565 | 16.91297 | -0.12866 | 0.566558 | 0.817477 |
| 11 | 53924600 | 53932799 | 87.81533 | 134.4862 | 220.3732 | 91.45848 | -0.42041 | 3.033999 | 0.81749 |
| 6 | 109002200 | 109005799 | 14.49776 | 26.40888 | 25.86732 | 11.12168 | 0.448317 | 0.313226 | 0.817492 |
| 1 | 151723800 | 151726199 | 25.16346 | 42.40029 | 51.19778 | 22.38142 | 0.110105 | 1.113254 | 0.817515 |
| 4 | 24089600 | 24091399 | 17.3701 | 29.89374 | 51.65741 | 22.6597 | -0.4015 | 0.895609 | 0.817555 |
| 16 | 47907400 | 47910599 | 32.28985 | 53.81429 | 62.04695 | 26.964 | 0.176463 | 1.254185 | 0.817602 |
| 5 | 100269600 | 100271799 | 25.44233 | 42.76604 | 51.74996 | 22.60612 | 0.106862 | 1.13464 | 0.817606 |
| 4 | 155999400 | 156002999 | 52.03198 | 83.12137 | 97.14283 | 40.80931 | 0.116867 | 2.088307 | 0.817665 |
| X | 35912600 | 35917199 | 49.09971 | 79.00508 | 92.29274 | 39.18598 | 0.129647 | 1.863425 | 0.817666 |
| 18 | 79977000 | 79978599 | 21.76921 | 37.36049 | 45.28731 | 19.86323 | 0.107143 | 0.93054 | 0.817668 |
| 19 | 6873000 | 6878599 | 84.42863 | 131.353 | 172.6039 | 71.6539 | -0.08478 | 2.735185 | 0.817693 |
| 13 | 3910800 | 3913799 | 20.38485 | 34.57614 | 50.08048 | 21.93691 | -0.15107 | 1.035936 | 0.817699 |
| 13 | 25535000 | 25535999 | 16.71741 | 29.48553 | 35.06503 | 15.51738 | 0.140812 | 0.566464 | 0.817707 |
| 3 | 122539200 | 122542199 | 37.06985 | 61.25054 | 68.18746 | 29.459 | 0.226142 | 1.3669 | 0.817765 |
| 18 | 64872600 | 64874599 | 27.24091 | 45.73366 | 55.07601 | 24.04554 | 0.113692 | 1.154872 | 0.817765 |
| 17 | 27924600 | 27925999 | 15.70281 | 27.36893 | 40.10987 | 17.64541 | -0.16133 | 0.736982 | 0.817783 |
| 11 | 20397400 | 20402999 | 37.90194 | 61.01264 | 101.2787 | 43.02744 | -0.37546 | 1.835533 | 0.817808 |
| X | 35946600 | 35949799 | 45.96267 | 74.42497 | 68.42302 | 29.27598 | 0.484719 | 1.675551 | 0.817812 |
| 4 | 60283800 | 60290799 | 105.9365 | 162.0949 | 210.3208 | 87.26875 | -0.08501 | 3.056675 | 0.817817 |
| 3 | 117569600 | 117570799 | 13.99267 | 24.78099 | 35.5474 | 15.71798 | -0.12622 | 0.502684 | 0.817836 |
| 3 | 118741600 | 118743199 | 17.2399 | 30.06677 | 42.54506 | 18.72407 | -0.11114 | 0.737507 | 0.817859 |
| 5 | 24510400 | 24514399 | 35.06373 | 56.81423 | 78.77027 | 33.71958 | -0.10573 | 1.645476 | 0.817861 |
| 8 | 24543400 | 24544599 | 16.02683 | 28.1417 | 33.18519 | 14.68273 | 0.150925 | 0.610171 | 0.817873 |
| 2 | 87223400 | 87225399 | 25.21098 | 42.96376 | 37.12072 | 16.27209 | 0.592268 | 0.997762 | 0.817891 |
| 4 | 105860800 | 105863399 | 26.41443 | 43.63047 | 62.45332 | 26.97566 | -0.13542 | 1.338663 | 0.817893 |
| 16 | 76472600 | 76474599 | 22.75174 | 39.39257 | 45.72824 | 20.06254 | 0.171699 | 0.854442 | 0.817896 |
| 12 | 104246600 | 104249999 | 75.89889 | 119.2692 | 139.9279 | 58.04278 | 0.089795 | 2.548932 | 0.817927 |
| 6 | 3163000 | 3164199 | 20.60044 | 35.97317 | 34.64701 | 15.26029 | 0.441612 | 0.725883 | 0.817957 |
| 11 | 27153000 | 27156399 | 26.00307 | 43.249 | 61.01345 | 26.47983 | -0.11377 | 1.255692 | 0.817963 |
| 2 | 4710600 | 4712599 | 21.04567 | 36.60639 | 35.68673 | 15.67943 | 0.423543 | 0.782174 | 0.817972 |
| 9 | 63567800 | 63568999 | 13.63448 | 24.01935 | 35.33523 | 15.64068 | -0.16501 | 0.561329 | 0.817984 |
| 13 | 96351400 | 96352799 | 10.80756 | 20.03168 | 29.89556 | 12.55891 | -0.13155 | 0.197866 | 0.817997 |
| 7 | 16877600 | 16898599 | 328.8073 | 464.1555 | 494.4214 | 204.7045 | 0.090845 | 4.439442 | 0.818015 |
| 1 | 113528000 | 113528599 | 6.668367 | 12.97743 | 21.93457 | 8.072718 | -0.13852 | -0.3202 | 0.818054 |
| 14 | 92443600 | 92443999 | 6.973683 | 13.58181 | 19.06901 | 6.969699 | 0.136293 | -0.34794 | 0.818058 |
| 3 | 47551000 | 47552599 | 17.64013 | 31.36587 | 35.75638 | 15.76807 | 0.206417 | 0.495522 | 0.818064 |

| | | | | | | | | | |
|---|---|---|---|---|---|---|---|---|---|
| 2 | 62895600 | 62896199 | 6.523321 | 12.67939 | 21.21386 | 7.886222 | -0.13709 | -0.2886 | 0.818069 |
| 7 | 42916000 | 42923599 | 86.82046 | 134.7633 | 176.6925 | 73.37649 | -0.08416 | 2.776155 | 0.81807 |
| 11 | 100728200 | 100728999 | 15.60756 | 27.61713 | 33.46235 | 14.79858 | 0.115646 | 0.51956 | 0.81807 |
| 7 | 27405600 | 27414599 | 102.147 | 158.3598 | 156.2729 | 64.90852 | 0.32399 | 2.801177 | 0.818073 |
| 16 | 78641600 | 78647399 | 73.04851 | 114.6717 | 152.7352 | 63.34683 | -0.0942 | 2.568077 | 0.8181 |
| 6 | 67644000 | 67646399 | 39.66141 | 64.44724 | 76.19365 | 32.68516 | 0.127315 | 1.593745 | 0.818104 |
| 4 | 41080000 | 41080599 | 10.99535 | 20.42526 | 25.20707 | 10.55721 | 0.143922 | 0.174629 | 0.818123 |
| 7 | 12969800 | 12973399 | 47.65413 | 76.33168 | 103.7347 | 43.7464 | -0.09613 | 2.003469 | 0.818126 |
| 10 | 106174200 | 106175399 | 36.85041 | 59.96459 | 71.83538 | 30.83519 | 0.109883 | 1.563443 | 0.818226 |
| 8 | 110245800 | 110248399 | 33.5021 | 55.54509 | 54.72193 | 23.69735 | 0.402326 | 1.309212 | 0.818236 |
| 8 | 34361800 | 34365799 | 28.25915 | 46.04773 | 79.63795 | 34.20155 | -0.417 | 1.53338 | 0.818284 |
| 12 | 112428800 | 112429999 | 12.69035 | 22.9163 | 33.12692 | 14.39876 | -0.12002 | 0.364954 | 0.818312 |
| 4 | 134300600 | 134302999 | 24.00077 | 40.45218 | 47.51185 | 20.77181 | 0.149982 | 1.103364 | 0.818332 |
| 4 | 88689000 | 88690799 | 15.6351 | 26.93419 | 50.03656 | 21.94936 | -0.50503 | 0.877545 | 0.818338 |
| 4 | 111106800 | 111107599 | 6.092216 | 11.86082 | 20.17267 | 7.400804 | -0.14227 | -0.34191 | 0.81838 |
| 8 | 101319400 | 101319799 | 6.092216 | 11.86082 | 20.17267 | 7.400804 | -0.14227 | -0.34191 | 0.81838 |
| 1 | 85197200 | 85202599 | 259.9462 | 369.7939 | 463.3758 | 191.6043 | -0.13218 | 4.32384 | 0.818381 |
| 16 | 8372000 | 8378599 | 129.4404 | 196.5724 | 224.4389 | 92.94198 | 0.090615 | 3.216948 | 0.818408 |
| 1 | 24898600 | 24899399 | 10.50501 | 19.62174 | 24.23962 | 10.06067 | 0.152861 | 0.133551 | 0.818411 |
| 5 | 73355200 | 73356599 | 13.83593 | 25.01712 | 30.52405 | 13.24612 | 0.12479 | 0.361646 | 0.818412 |
| 14 | 118468400 | 118469999 | 17.26178 | 30.30698 | 36.61457 | 16.19862 | 0.116161 | 0.625954 | 0.818429 |
| 3 | 98679800 | 98681599 | 16.91664 | 30.34228 | 34.2768 | 15.00513 | 0.226603 | 0.434149 | 0.818446 |
| 5 | 117733400 | 117735199 | 16.73792 | 29.18759 | 41.52268 | 18.27266 | -0.11883 | 0.734987 | 0.818451 |
| 5 | 74884600 | 74887599 | 23.8154 | 40.19059 | 56.63255 | 24.76881 | -0.11192 | 1.098811 | 0.818461 |
| 1 | 22065400 | 22066799 | 16.31809 | 29.08424 | 26.44136 | 11.63692 | 0.533343 | 0.456626 | 0.818478 |
| 3 | 52565400 | 52566999 | 16.73109 | 29.19395 | 41.49913 | 18.26949 | -0.11763 | 0.725894 | 0.818479 |
| 10 | 99163600 | 99164599 | 8.094379 | 15.40249 | 24.50273 | 9.818068 | -0.16585 | -0.00898 | 0.818479 |
| 2 | 43596200 | 43598599 | 16.13053 | 28.52274 | 34.46253 | 15.24302 | 0.119312 | 0.532604 | 0.818515 |
| 5 | 17105600 | 17108999 | 36.57259 | 59.28343 | 81.93962 | 35.08694 | -0.10074 | 1.642564 | 0.818545 |
| 15 | 83850800 | 83851999 | 17.27894 | 30.2678 | 36.27478 | 16.01731 | 0.127825 | 0.662872 | 0.818549 |
| 4 | 147978000 | 147979599 | 14.12637 | 25.38041 | 30.96623 | 13.53658 | 0.117889 | 0.401108 | 0.818554 |
| 2 | 144068600 | 144074199 | 48.96269 | 78.23089 | 106.9695 | 44.94782 | -0.10899 | 2.082646 | 0.81858 |
| 4 | 22695400 | 22696999 | 16.31831 | 28.79589 | 42.83411 | 18.95357 | -0.18054 | 0.568714 | 0.818581 |
| 5 | 110592400 | 110593199 | 7.426938 | 14.26382 | 23.03446 | 9.0176 | -0.15693 | -0.09801 | 0.818629 |
| 8 | 113580600 | 113581199 | 7.426938 | 14.26382 | 23.03446 | 9.0176 | -0.15693 | -0.09801 | 0.818629 |
| 1 | 94620800 | 94622199 | 20.44286 | 35.84949 | 33.84293 | 14.9574 | 0.470278 | 0.65733 | 0.818645 |
| 6 | 130608600 | 130613199 | 39.46979 | 64.0932 | 97.94628 | 41.99589 | -0.24305 | 1.61817 | 0.81867 |
| 12 | 114646200 | 114646799 | 10.45756 | 19.86808 | 25.46741 | 10.23854 | 0.1381 | 0.023408 | 0.81871 |
| 11 | 16331200 | 16336399 | 46.88206 | 75.23341 | 102.4884 | 43.34202 | -0.09644 | 1.946021 | 0.818758 |
| 7 | 107838400 | 107838999 | 11.64923 | 21.25809 | 31.27182 | 13.41527 | -0.13245 | 0.295566 | 0.818759 |
| 1 | 169673600 | 169674399 | 8.691305 | 16.48158 | 25.48906 | 10.28785 | -0.13505 | 0.021981 | 0.818778 |
| 17 | 90508400 | 90509799 | 18.04927 | 31.65134 | 38.13442 | 16.8538 | 0.120031 | 0.651427 | 0.818785 |
| 5 | 54617600 | 54619199 | 17.72737 | 30.86543 | 43.62548 | 19.17946 | -0.10976 | 0.765008 | 0.818788 |
| 16 | 15832000 | 15835999 | 29.41903 | 49.85169 | 55.70088 | 24.39172 | 0.22192 | 1.084959 | 0.81879 |
| 11 | 46616200 | 46620199 | 47.71321 | 76.43729 | 103.535 | 43.67173 | -0.0911 | 2.000143 | 0.818806 |
| 15 | 4881000 | 4882399 | 16.68415 | 29.35194 | 35.48937 | 15.71317 | 0.115356 | 0.591447 | 0.818818 |
| 6 | 136647200 | 136656199 | 107.4226 | 163.5287 | 248.3979 | 102.9378 | -0.31807 | 3.162933 | 0.818829 |

| | | | | | | | | | |
|---|---|---|---|---|---|---|---|---|---|
| 7 | 12481400 | 12487999 | 101.2615 | 155.1704 | 202.5159 | 84.0586 | -0.09175 | 3.02455 | 0.818844 |
| 2 | 116313600 | 116316199 | 33.72909 | 55.33946 | 66.85415 | 28.78723 | 0.105128 | 1.441641 | 0.818924 |
| X | 37583200 | 37585799 | 21.15113 | 36.63511 | 55.70006 | 24.4378 | -0.21585 | 0.859724 | 0.81897 |
| 8 | 22196400 | 22197799 | 25.65205 | 43.19738 | 52.28296 | 22.85161 | 0.106673 | 1.120976 | 0.819018 |
| 13 | 51495000 | 51496799 | 19.61133 | 34.37441 | 40.11497 | 17.72207 | 0.165717 | 0.668971 | 0.81903 |
| 8 | 88904000 | 88908999 | 55.38391 | 88.4711 | 105.271 | 44.22038 | 0.090869 | 2.096759 | 0.81903 |
| 4 | 80501200 | 80503599 | 40.63497 | 65.34923 | 90.48955 | 38.39019 | -0.11611 | 1.865667 | 0.819038 |
| 18 | 50636000 | 50637599 | 59.67411 | 93.92814 | 154.3972 | 64.05848 | -0.39427 | 2.50861 | 0.819038 |
| 1 | 63653200 | 63653999 | 7.105263 | 13.7029 | 22.56796 | 8.730098 | -0.16914 | -0.14307 | 0.819045 |
| 4 | 133196000 | 133198799 | 24.78285 | 41.69387 | 50.13907 | 21.90823 | 0.115893 | 1.123413 | 0.819065 |
| 2 | 75949200 | 75951199 | 25.59845 | 43.17896 | 52.29613 | 22.86892 | 0.105745 | 1.105559 | 0.819073 |
| 5 | 21449800 | 21450599 | 13.23434 | 23.94746 | 35.26434 | 15.28873 | -0.1442 | 0.362164 | 0.819086 |
| Y | 43575600 | 43577199 | 12.86626 | 22.45939 | 38.1789 | 16.8121 | -0.37405 | 0.695144 | 0.819086 |
| 12 | 81390400 | 81406599 | 206.5407 | 305.8989 | 328.3127 | 135.2099 | 0.147734 | 3.707634 | 0.819087 |
| 3 | 24863200 | 24865599 | 14.80278 | 25.94692 | 47.78134 | 21.10953 | -0.48912 | 0.656657 | 0.819131 |
| 13 | 4918000 | 4927199 | 159.7628 | 238.453 | 302.9886 | 124.8922 | -0.08543 | 3.563683 | 0.819221 |
| 6 | 64238200 | 64241199 | 31.24449 | 51.962 | 60.95654 | 26.45349 | 0.151452 | 1.269922 | 0.819227 |
| 7 | 105311800 | 105316799 | 48.16133 | 77.76784 | 88.34145 | 37.688 | 0.175902 | 1.754508 | 0.819253 |
| 15 | 14737400 | 14741999 | 90.6657 | 141.7531 | 138.9747 | 57.65747 | 0.342824 | 2.647229 | 0.819268 |
| 4 | 151220000 | 151222399 | 19.41804 | 33.77372 | 40.96789 | 17.99923 | 0.109676 | 0.783498 | 0.819272 |
| 15 | 52707000 | 52708799 | 33.93226 | 54.58207 | 98.72669 | 41.89831 | -0.50025 | 1.854784 | 0.819279 |
| 13 | 52046000 | 52050799 | 40.0032 | 65.15866 | 75.90461 | 32.59352 | 0.15112 | 1.558868 | 0.81929 |
| X | 40794000 | 40796999 | 23.10176 | 40.00039 | 46.30146 | 20.31406 | 0.176027 | 0.860812 | 0.819304 |
| 4 | 143294600 | 143312199 | 357.5776 | 503.0533 | 528.6017 | 218.9787 | 0.105562 | 4.491834 | 0.819311 |
| 4 | 110001800 | 110003199 | 12.33498 | 21.86625 | 32.80668 | 14.50332 | -0.18964 | 0.478822 | 0.819342 |
| 11 | 7342200 | 7343999 | 17.31624 | 30.25309 | 43.51894 | 19.17387 | -0.13468 | 0.712503 | 0.819351 |
| 7 | 54804400 | 54805399 | 12.69557 | 23.05783 | 33.73956 | 14.56005 | -0.13079 | 0.333326 | 0.819362 |
| 4 | 134065600 | 134068799 | 38.21616 | 61.71772 | 84.87086 | 36.21398 | -0.09887 | 1.734094 | 0.819365 |
| 17 | 67736600 | 67737399 | 14.94556 | 26.6643 | 32.4276 | 14.25958 | 0.116639 | 0.453014 | 0.819366 |
| 17 | 23740200 | 23743199 | 37.15522 | 60.2352 | 83.51819 | 35.7661 | -0.10516 | 1.640171 | 0.819417 |
| 4 | 118364600 | 118365399 | 10.78497 | 19.77406 | 29.27051 | 12.47922 | -0.13522 | 0.256494 | 0.819448 |
| X | 39089200 | 39092399 | 28.50689 | 48.15366 | 55.67351 | 24.35639 | 0.172618 | 1.107663 | 0.819472 |
| 1 | 96042600 | 96045799 | 51.33498 | 81.96608 | 110.5303 | 46.39979 | -0.09004 | 2.108216 | 0.819474 |
| 13 | 79650000 | 79651599 | 15.42821 | 27.33461 | 40.1993 | 17.77344 | -0.16172 | 0.517677 | 0.819505 |
| 6 | 19714000 | 19714799 | 12.60021 | 22.51345 | 32.66681 | 14.34677 | -0.13522 | 0.42935 | 0.819516 |
| 12 | 23461400 | 23464999 | 49.36661 | 79.37052 | 78.56561 | 33.31871 | 0.366695 | 1.881317 | 0.819521 |
| 13 | 68039000 | 68048799 | 235.0179 | 340.3362 | 369.309 | 152.2972 | 0.100905 | 4.060249 | 0.819542 |
| 4 | 37996600 | 38001399 | 40.88251 | 66.40992 | 77.9877 | 33.44678 | 0.136578 | 1.604654 | 0.819563 |
| 1 | 112858200 | 112858799 | 13.18639 | 24.28697 | 21.48263 | 9.10256 | 0.60888 | 0.229612 | 0.819565 |
| 14 | 123869200 | 123871599 | 38.93976 | 63.03422 | 75.78384 | 32.41827 | 0.098123 | 1.669949 | 0.81957 |
| 14 | 34955800 | 34956399 | 9.631214 | 18.25899 | 23.27326 | 9.398914 | 0.140505 | 0.024157 | 0.819582 |
| Y | 5704200 | 5704799 | 3.867351 | 7.561008 | 15.52802 | 5.320371 | -0.32007 | -0.63491 | 0.819592 |
| 3 | 158806400 | 158806999 | 8.038658 | 15.50108 | 24.82238 | 9.605617 | -0.12945 | -0.12391 | 0.819597 |
| 3 | 45065200 | 45070999 | 67.76292 | 107.3604 | 148.0292 | 61.5691 | -0.1336 | 2.366986 | 0.819606 |
| 7 | 75449600 | 75450599 | 12.24418 | 22.61074 | 28.13757 | 11.87996 | 0.12328 | 0.223575 | 0.819627 |
| 1 | 64713000 | 64713799 | 11.35934 | 21.0818 | 26.10362 | 10.94655 | 0.137414 | 0.182712 | 0.819638 |
| 1 | 194895000 | 194895799 | 10.26541 | 19.01856 | 28.31424 | 11.90162 | -0.12873 | 0.190896 | 0.819657 |

| | | | | | | | | | |
|---|---|---|---|---|---|---|---|---|---|
| 1 | 46592000 | 46594599 | 21.41059 | 36.02656 | 62.42736 | 27.27898 | -0.40928 | 1.124827 | 0.819664 |
| 5 | 64849600 | 64851399 | 24.08554 | 40.98884 | 49.5624 | 21.72354 | 0.108841 | 1.020394 | 0.819696 |
| 6 | 69971800 | 69971999 | 6.089713 | 11.88907 | 17.02164 | 6.009383 | 0.154247 | -0.48736 | 0.819716 |
| 13 | 34724800 | 34726999 | 24.05054 | 40.98965 | 49.02191 | 21.4892 | 0.125041 | 1.00854 | 0.819723 |
| 16 | 42859200 | 42860599 | 15.47577 | 27.39573 | 40.15928 | 17.75874 | -0.15772 | 0.524812 | 0.819728 |
| X | 78572800 | 78584199 | 117.7348 | 181.9827 | 159.0739 | 66.07569 | 0.495505 | 2.863432 | 0.819748 |
| 16 | 19925400 | 19926399 | 11.29064 | 20.9533 | 26.16997 | 10.97509 | 0.125223 | 0.184211 | 0.819755 |
| 10 | 58973600 | 58975999 | 21.64076 | 36.55251 | 52.18659 | 22.83036 | -0.13053 | 1.082082 | 0.819755 |
| 6 | 81830400 | 81834399 | 51.34882 | 82.30051 | 97.76237 | 41.26645 | 0.098758 | 1.988822 | 0.819756 |
| 6 | 50421400 | 50422399 | 10.25129 | 19.03739 | 28.32855 | 11.86937 | -0.12473 | 0.177285 | 0.819783 |
| 16 | 47801200 | 47802799 | 24.81353 | 41.74119 | 50.13584 | 21.90613 | 0.117636 | 1.125019 | 0.819795 |
| 11 | 3072800 | 3077799 | 603.4308 | 805.6769 | 784.7876 | 327.7389 | 0.138917 | 5.30408 | 0.819823 |
| 12 | 36447200 | 36449799 | 30.30137 | 50.86061 | 58.54827 | 25.55795 | 0.178579 | 1.171085 | 0.819833 |
| 2 | 132458800 | 132459999 | 14.82132 | 26.28021 | 31.54501 | 13.94373 | 0.130424 | 0.489799 | 0.819884 |
| 17 | 89556000 | 89556399 | 5.647295 | 11.02097 | 19.27387 | 6.851029 | -0.14038 | -0.4619 | 0.819885 |
| 14 | 99000400 | 99001999 | 15.3434 | 26.91178 | 38.31568 | 16.93682 | -0.11956 | 0.634051 | 0.819887 |
| 4 | 95992600 | 95995399 | 30.52528 | 50.13272 | 69.92551 | 30.1224 | -0.10108 | 1.424539 | 0.819895 |
| 10 | 11497800 | 11500399 | 34.37474 | 56.08343 | 78.55177 | 33.75055 | -0.11234 | 1.526967 | 0.819896 |
| 17 | 21833000 | 21838399 | 67.19047 | 106.0021 | 141.3023 | 58.64499 | -0.08983 | 2.463722 | 0.819913 |
| 1 | 138580000 | 138580599 | 9.954322 | 18.8942 | 24.3465 | 9.80742 | 0.128064 | 0.021884 | 0.819926 |
| 4 | 144685000 | 144687599 | 16.9504 | 29.63941 | 41.95937 | 18.49948 | -0.11149 | 0.697456 | 0.819946 |
| X | 38894200 | 38895199 | 16.15794 | 28.65483 | 34.40405 | 15.20544 | 0.130082 | 0.502678 | 0.819968 |
| 4 | 70452200 | 70454399 | 19.37928 | 32.03468 | 71.71687 | 30.99335 | -0.77842 | 1.318431 | 0.819987 |
| 14 | 31645400 | 31648599 | 66.9504 | 105.5574 | 140.6762 | 58.37841 | -0.09017 | 2.477319 | 0.820001 |
| 13 | 38762800 | 38764999 | 21.12573 | 35.29019 | 62.7951 | 27.32922 | -0.44738 | 1.198619 | 0.820006 |
| 1 | 188342200 | 188343599 | 9.423459 | 17.76519 | 26.96481 | 11.01775 | -0.12422 | 0.071116 | 0.820011 |
| 7 | 8713600 | 8714399 | 16.78993 | 29.74785 | 28.34511 | 12.53226 | 0.46115 | 0.502776 | 0.820043 |
| 16 | 56139400 | 56146399 | 97.99798 | 151.5092 | 176.2045 | 73.19127 | 0.084207 | 2.851604 | 0.82005 |
| 9 | 75305600 | 75306799 | 9.450987 | 17.62464 | 26.83975 | 11.1702 | -0.15014 | 0.140597 | 0.820054 |
| 3 | 139234800 | 139237599 | 33.01231 | 53.38417 | 89.77585 | 38.37225 | -0.38633 | 1.688205 | 0.820065 |
| 4 | 34940400 | 34942199 | 14.07443 | 25.16532 | 30.46048 | 13.36979 | 0.125222 | 0.428831 | 0.820071 |
| Y | 5190800 | 5191599 | 8.2086 | 15.75069 | 18.34892 | 7.202097 | 0.306524 | -0.09749 | 0.820104 |
| 6 | 112416800 | 112417599 | 8.670599 | 16.51535 | 25.56438 | 10.22197 | -0.1246 | -0.00911 | 0.820112 |
| X | 67702800 | 67703599 | 8.76716 | 16.8146 | 27.48387 | 10.80212 | -0.17959 | -0.05669 | 0.820132 |
| 10 | 124043600 | 124043999 | 8.159575 | 15.75011 | 20.94687 | 8.080378 | 0.14079 | -0.14361 | 0.820141 |
| 11 | 81431800 | 81438599 | 67.99506 | 106.4196 | 168.3677 | 69.84443 | -0.34557 | 2.622411 | 0.820179 |
| 14 | 37862000 | 37862599 | 11.05893 | 20.48665 | 30.39723 | 12.77894 | -0.12425 | 0.202369 | 0.820197 |
| 17 | 8211000 | 8213599 | 14.8323 | 26.1434 | 37.14387 | 16.4395 | -0.11508 | 0.563511 | 0.820211 |
| 17 | 71578800 | 71584399 | 51.31456 | 81.36408 | 137.4395 | 57.20535 | -0.42494 | 2.329422 | 0.820228 |
| 15 | 97307600 | 97312799 | 54.20618 | 86.57517 | 103.0366 | 43.27147 | 0.090304 | 2.099693 | 0.82024 |
| 4 | 25815400 | 25818199 | 23.99437 | 41.00882 | 49.02098 | 21.49365 | 0.126187 | 0.979578 | 0.820247 |
| 5 | 52249400 | 52251599 | 19.473 | 33.82633 | 41.03676 | 18.01511 | 0.109398 | 0.802816 | 0.820249 |
| 4 | 135368800 | 135370599 | 17.09807 | 29.65568 | 42.70708 | 18.73805 | -0.13718 | 0.812209 | 0.820266 |
| 11 | 102980200 | 102980999 | 11.58123 | 20.64341 | 37.30157 | 16.41531 | -0.45189 | 0.447785 | 0.820272 |
| 3 | 41229600 | 41234999 | 64.64829 | 102.7739 | 120.8673 | 50.43926 | 0.100286 | 2.260599 | 0.820299 |
| 3 | 156014200 | 156018599 | 70.60405 | 111.0173 | 147.2673 | 61.09211 | -0.08649 | 2.53403 | 0.820307 |
| 7 | 26803800 | 26804799 | 9.660809 | 18.27258 | 28.15532 | 11.42657 | -0.13704 | 0.055624 | 0.820307 |

| | | | | | | | | | |
|---|---|---|---|---|---|---|---|---|---|
| 9 | 110830400 | 110832199 | 12.7889 | 22.68784 | 33.21391 | 14.67522 | -0.15394 | 0.476862 | 0.820316 |
| 5 | 10410800 | 10425599 | 189.0222 | 280.8242 | 310.604 | 127.9556 | 0.108493 | 3.650143 | 0.820317 |
| 2 | 144625600 | 144626799 | 12.03451 | 21.81411 | 31.65092 | 13.69246 | -0.12088 | 0.335404 | 0.820327 |
| 1 | 155279400 | 155282999 | 43.89456 | 70.75239 | 84.87084 | 36.12455 | 0.094583 | 1.797917 | 0.82034 |
| 12 | 113444600 | 113450199 | 64.93836 | 102.6785 | 138.4896 | 57.54269 | -0.10418 | 2.407748 | 0.820349 |
| 11 | 15794400 | 15795199 | 9.323781 | 17.89699 | 23.52226 | 9.223068 | 0.134899 | -0.07378 | 0.820423 |
| 3 | 40830200 | 40842399 | 207.7226 | 308.8248 | 269.7213 | 111.1217 | 0.449962 | 3.63876 | 0.820426 |
| 6 | 74977200 | 74981199 | 49.57124 | 79.65723 | 94.22734 | 39.92993 | 0.109194 | 1.903512 | 0.820437 |
| 1 | 111942400 | 111942999 | 9.859628 | 18.63964 | 23.79833 | 9.668185 | 0.130882 | 0.048504 | 0.820451 |
| 5 | 142325400 | 142327599 | 21.69738 | 36.57501 | 52.75684 | 23.06585 | -0.14552 | 1.096889 | 0.820455 |
| 16 | 70902200 | 70903999 | 18.38184 | 31.96577 | 45.22796 | 19.86867 | -0.11148 | 0.787638 | 0.820458 |
| 4 | 99333200 | 99334799 | 18.53492 | 32.05989 | 45.2865 | 19.86813 | -0.11038 | 0.846245 | 0.820472 |
| 2 | 177655200 | 177657999 | 53.65699 | 85.99281 | 85.80371 | 36.214 | 0.349691 | 1.988785 | 0.820492 |
| 5 | 131161600 | 131163399 | 16.40457 | 29.61587 | 33.95783 | 14.76465 | 0.211879 | 0.39354 | 0.8205 |
| 8 | 48939200 | 48941399 | 15.74503 | 27.83292 | 40.73129 | 18.01679 | -0.15595 | 0.544767 | 0.82052 |
| 3 | 37113200 | 37118399 | 56.63517 | 90.23789 | 121.8979 | 51.02111 | -0.09589 | 2.183287 | 0.820541 |
| X | 79182000 | 79184399 | 31.353 | 51.91846 | 62.32315 | 26.96629 | 0.118013 | 1.324617 | 0.820554 |
| X | 14355200 | 14356599 | 19.31559 | 33.65664 | 39.9754 | 17.58325 | 0.140396 | 0.759671 | 0.820558 |
| 11 | 94505600 | 94509599 | 38.28183 | 62.03947 | 74.70871 | 31.98507 | 0.097315 | 1.645489 | 0.820573 |
| 10 | 121976000 | 121978399 | 40.95828 | 66.06369 | 90.6025 | 38.5975 | -0.09711 | 1.775996 | 0.820578 |
| 1 | 111229200 | 111230399 | 22.03757 | 37.74346 | 45.32524 | 19.87657 | 0.120251 | 0.952055 | 0.820581 |
| 2 | 94540800 | 94541999 | 18.94537 | 32.90485 | 46.85419 | 20.56702 | -0.12077 | 0.811226 | 0.820584 |
| 4 | 117356400 | 117357199 | 9.222055 | 17.50363 | 27.14174 | 10.9343 | -0.1372 | 0.022039 | 0.820586 |
| 10 | 90100000 | 90103199 | 35.80957 | 58.94061 | 68.5513 | 29.55788 | 0.161546 | 1.407997 | 0.820601 |
| 7 | 94167600 | 94170199 | 40.97191 | 66.38506 | 79.11253 | 33.86524 | 0.112033 | 1.657841 | 0.820602 |
| 10 | 78303800 | 78307199 | 26.36576 | 43.77028 | 61.51573 | 26.66336 | -0.10852 | 1.279365 | 0.820603 |
| 1 | 57347600 | 57348199 | 13.10743 | 23.39101 | 33.64861 | 14.79325 | -0.12388 | 0.442979 | 0.82061 |
| 9 | 37502800 | 37503199 | 6.783396 | 13.19764 | 17.90779 | 6.605464 | 0.172462 | -0.32492 | 0.82063 |
| 3 | 33411600 | 33413399 | 25.81416 | 44.38426 | 41.43575 | 18.17541 | 0.482891 | 0.926165 | 0.820639 |
| 14 | 37506200 | 37507599 | 14.38065 | 25.40694 | 36.13871 | 15.98712 | -0.11531 | 0.534123 | 0.820677 |
| 9 | 31542800 | 31546199 | 59.5686 | 94.70827 | 111.3171 | 46.45863 | 0.101292 | 2.251634 | 0.82068 |
| 14 | 58259000 | 58262399 | 40.49966 | 65.63261 | 78.23268 | 33.49339 | 0.11194 | 1.650048 | 0.820692 |
| X | 92215000 | 92217799 | 27.685 | 47.15691 | 53.45029 | 23.42933 | 0.201988 | 1.030944 | 0.8207 |
| X | 155583600 | 155585599 | 24.62544 | 42.60548 | 47.92761 | 21.02701 | 0.216843 | 0.876215 | 0.820706 |
| 5 | 105791400 | 105792399 | 12.29413 | 22.46931 | 26.95833 | 11.54271 | 0.160987 | 0.277251 | 0.82071 |
| 11 | 116606600 | 116614599 | 97.49649 | 149.3207 | 227.0619 | 94.23557 | -0.31266 | 3.036249 | 0.820718 |
| 9 | 7866400 | 7869599 | 31.07033 | 50.83803 | 70.78478 | 30.44525 | -0.10084 | 1.474001 | 0.820723 |
| 1 | 74093800 | 74095799 | 16.74115 | 29.40634 | 35.6021 | 15.75678 | 0.113092 | 0.613568 | 0.820723 |
| 13 | 45801000 | 45801999 | 12.85447 | 23.39556 | 28.60955 | 12.31381 | 0.129333 | 0.313107 | 0.820744 |
| 9 | 92933800 | 92935799 | 17.34207 | 30.27433 | 43.06783 | 18.96571 | -0.11856 | 0.727035 | 0.820768 |
| 3 | 91340800 | 91342999 | 30.88822 | 51.6568 | 50.24561 | 21.88317 | 0.42097 | 1.204164 | 0.820771 |
| 7 | 26531000 | 26536999 | 99.49392 | 153.2413 | 176.7634 | 73.42078 | 0.091855 | 2.921609 | 0.820809 |
| 5 | 119926600 | 119927999 | 15.23055 | 26.74849 | 38.68032 | 17.11674 | -0.14216 | 0.610158 | 0.820827 |
| 10 | 119971800 | 119972599 | 13.19093 | 23.70039 | 34.09207 | 14.90267 | -0.11836 | 0.400289 | 0.820836 |
| 3 | 4157400 | 4157799 | 8.555061 | 16.41892 | 21.35677 | 8.377668 | 0.149431 | -0.08595 | 0.820851 |
| 3 | 4127000 | 4134399 | 133.8995 | 204.1047 | 196.6802 | 81.5372 | 0.339521 | 3.139677 | 0.820857 |
| 2 | 80633400 | 80635599 | 20.98942 | 36.44041 | 43.58802 | 19.12849 | 0.129444 | 0.81894 | 0.820858 |

| | | | | | | | | | |
|---|---|---|---|---|---|---|---|---|---|
| 4 | 18518600 | 18519199 | 10.03131 | 19.12071 | 23.94714 | 9.56099 | 0.181024 | -0.00243 | 0.820869 |
| 16 | 20592400 | 20594399 | 15.89861 | 27.82661 | 39.85303 | 17.58488 | -0.12822 | 0.671604 | 0.820872 |
| 5 | 60105000 | 60110799 | 47.29417 | 76.14915 | 90.21188 | 38.33847 | 0.110596 | 1.837558 | 0.820873 |
| 1 | 63348000 | 63349199 | 28.55199 | 47.50783 | 57.18656 | 24.82595 | 0.114608 | 1.258441 | 0.820874 |
| 13 | 94399800 | 94401799 | 16.24637 | 28.3955 | 40.42903 | 17.81796 | -0.11971 | 0.697704 | 0.820882 |
| 12 | 99149000 | 99149999 | 11.23248 | 20.79604 | 30.92699 | 13.01195 | -0.12836 | 0.208358 | 0.82091 |
| 12 | 99282200 | 99285999 | 26.00582 | 42.99111 | 73.17211 | 31.62352 | -0.38447 | 1.336968 | 0.82092 |
| X | 99908200 | 99909599 | 16.57887 | 28.73409 | 50.06121 | 21.96424 | -0.41102 | 0.827699 | 0.82093 |
| 1 | 65435000 | 65436599 | 15.69417 | 27.60421 | 39.20883 | 17.36075 | -0.11587 | 0.596129 | 0.820931 |
| 6 | 96754400 | 96757599 | 17.63985 | 31.15867 | 30.31745 | 13.41245 | 0.429469 | 0.545021 | 0.820944 |
| 18 | 3096400 | 3123399 | 1138.931 | 1439.328 | 1570.929 | 659.9236 | -0.10457 | 6.195812 | 0.820956 |
| 3 | 142370200 | 142374799 | 67.99537 | 106.9184 | 143.3967 | 59.48667 | -0.10227 | 2.531588 | 0.820962 |
| 4 | 52103800 | 52106199 | 17.39669 | 30.93971 | 35.82151 | 15.79391 | 0.184443 | 0.493899 | 0.820963 |
| 14 | 17673200 | 17674999 | 17.39669 | 30.93971 | 35.82151 | 15.79391 | 0.184443 | 0.493899 | 0.820963 |
| 1 | 73686000 | 73686399 | 5.648357 | 11.01886 | 19.21684 | 6.873035 | -0.14431 | -0.43997 | 0.820968 |
| 11 | 92787200 | 92788599 | 16.80033 | 29.31075 | 41.53287 | 18.28276 | -0.11297 | 0.729815 | 0.820991 |
| 6 | 117656000 | 117659599 | 35.01396 | 56.92372 | 78.89031 | 33.85212 | -0.10075 | 1.578457 | 0.820991 |
| 6 | 122079800 | 122098999 | 479.0594 | 652.061 | 756.201 | 314.8902 | -0.08476 | 4.997947 | 0.821025 |
| 4 | 28486000 | 28486399 | 6.227747 | 12.13899 | 16.42497 | 5.939191 | 0.203597 | -0.40519 | 0.821026 |
| 4 | 74781800 | 74782999 | 16.37345 | 28.72316 | 34.21135 | 15.12466 | 0.1366 | 0.629559 | 0.821034 |
| 12 | 74992400 | 74993399 | 10.15548 | 19.02321 | 29.00274 | 11.97942 | -0.14352 | 0.121371 | 0.821069 |
| X | 40337200 | 40338999 | 23.3378 | 39.38559 | 55.96841 | 24.47853 | -0.12389 | 1.100142 | 0.821074 |
| 6 | 6625800 | 6627999 | 32.78306 | 53.61333 | 64.20632 | 27.6107 | 0.11547 | 1.479958 | 0.821082 |
| 1 | 120041000 | 120044199 | 35.23947 | 57.56146 | 69.56717 | 29.90295 | 0.101068 | 1.505557 | 0.821087 |
| 4 | 149304200 | 149306999 | 23.85306 | 40.36486 | 48.5227 | 21.23957 | 0.116669 | 1.066368 | 0.821092 |
| 4 | 23082000 | 23084599 | 29.6309 | 49.46595 | 46.0257 | 20.0228 | 0.484841 | 1.214369 | 0.821114 |
| X | 126895800 | 126900399 | 47.45807 | 76.91031 | 84.57219 | 36.20787 | 0.228076 | 1.668179 | 0.821156 |
| 5 | 87041800 | 87043199 | 21.11057 | 36.54209 | 44.17379 | 19.3804 | 0.113548 | 0.849846 | 0.821172 |
| 15 | 97763800 | 97767599 | 46.98879 | 75.43431 | 89.86808 | 38.02765 | 0.09732 | 1.929449 | 0.821173 |
| 5 | 8829600 | 8831799 | 18.78001 | 33.00471 | 37.81731 | 16.74449 | 0.191795 | 0.6165 | 0.821192 |
| 11 | 10860000 | 10862599 | 36.75002 | 59.42596 | 82.00796 | 35.0502 | -0.10179 | 1.693479 | 0.821201 |
| 2 | 149584600 | 149586199 | 23.00304 | 37.9548 | 69.6852 | 30.07801 | -0.49354 | 1.350338 | 0.821216 |
| 7 | 12347800 | 12350199 | 34.08939 | 56.38092 | 54.66987 | 23.63492 | 0.424976 | 1.34004 | 0.821242 |
| 17 | 14719400 | 14720199 | 8.735491 | 16.70419 | 21.34351 | 8.460568 | 0.161087 | -0.04923 | 0.82127 |
| 1 | 5874400 | 5875799 | 19.60827 | 34.38324 | 33.43192 | 14.77478 | 0.427879 | 0.653291 | 0.821295 |
| 6 | 90963600 | 90970999 | 82.23761 | 128.7075 | 151.1691 | 62.71005 | 0.083662 | 2.628171 | 0.821295 |
| 5 | 70497400 | 70501199 | 44.16929 | 71.0715 | 97.91151 | 41.57148 | -0.10782 | 1.85847 | 0.821311 |
| 5 | 84972200 | 84974399 | 50.63411 | 81.08711 | 96.67787 | 40.75679 | 0.092006 | 2.012734 | 0.821363 |
| 9 | 18054000 | 18056399 | 24.11103 | 41.25621 | 40.06747 | 17.56955 | 0.425029 | 0.960292 | 0.821387 |
| 6 | 127294600 | 127296399 | 14.87568 | 26.19365 | 37.19747 | 16.46651 | -0.11487 | 0.575871 | 0.821393 |
| 2 | 30573400 | 30576399 | 17.06787 | 29.60644 | 42.6942 | 18.73246 | -0.13915 | 0.810333 | 0.821425 |
| 2 | 50886600 | 50890799 | 57.27095 | 90.84266 | 123.4809 | 51.41481 | -0.11141 | 2.319731 | 0.821431 |
| 3 | 41745200 | 41746399 | 25.03088 | 42.15144 | 50.6585 | 22.14161 | 0.11687 | 1.118143 | 0.821464 |
| 8 | 20121200 | 20121799 | 15.5803 | 27.897 | 26.33785 | 11.5466 | 0.482796 | 0.421707 | 0.821481 |
| 8 | 100873800 | 100875999 | 11.35182 | 21.43404 | 21.57721 | 8.788857 | 0.467766 | 0.064469 | 0.821483 |
| 2 | 176777200 | 176777799 | 10.22633 | 18.94967 | 28.29511 | 11.89059 | -0.13286 | 0.188405 | 0.821484 |
| 1 | 85400000 | 85437599 | 1186.096 | 1502.757 | 1629.283 | 684.2572 | -0.09127 | 6.154913 | 0.821498 |

| | | | | | | | | | |
|---|---|---|---|---|---|---|---|---|---|
| 1 | 101161200 | 101162199 | 19.97474 | 34.87773 | 34.1106 | 15.02326 | 0.419479 | 0.721287 | 0.821505 |
| 6 | 121876800 | 121885599 | 111.6001 | 171.6837 | 195.3802 | 81.14034 | 0.110014 | 2.95188 | 0.821517 |
| 13 | 68774600 | 68779799 | 48.20528 | 77.44123 | 92.08206 | 39.0055 | 0.101245 | 1.908606 | 0.821523 |
| 4 | 51295800 | 51300199 | 76.28175 | 119.713 | 139.3104 | 57.77674 | 0.100066 | 2.575623 | 0.821526 |
| 7 | 40915600 | 40918599 | 39.35024 | 65.26702 | 52.19935 | 22.61115 | 0.702187 | 1.317717 | 0.821551 |
| 11 | 19981000 | 19982399 | 12.37671 | 23.16511 | 22.91479 | 9.477904 | 0.474625 | 0.131434 | 0.821567 |
| 7 | 114285000 | 114286999 | 21.72678 | 37.92949 | 43.45308 | 19.13559 | 0.192295 | 0.748065 | 0.821574 |
| 8 | 76323600 | 76327199 | 31.12262 | 52.35561 | 58.61991 | 25.61332 | 0.218524 | 1.153104 | 0.82158 |
| 5 | 57012200 | 57012599 | 9.959329 | 19.06075 | 24.56477 | 9.713576 | 0.152186 | -0.03424 | 0.821596 |
| 4 | 113252000 | 113253599 | 31.31643 | 51.88826 | 62.37732 | 27.00063 | 0.11597 | 1.31722 | 0.8216 |
| 9 | 80452200 | 80452799 | 7.756254 | 15.06868 | 20.61474 | 7.692964 | 0.145866 | -0.25991 | 0.821619 |
| 4 | 20918600 | 20920599 | 28.55494 | 48.07347 | 46.96245 | 20.52456 | 0.414739 | 1.127849 | 0.821628 |
| 5 | 62240400 | 62241399 | 7.00317 | 13.61547 | 23.00461 | 8.534715 | -0.14818 | -0.28215 | 0.821697 |
| 9 | 124193400 | 124204599 | 326.5095 | 450.4602 | 704.0968 | 292.5721 | -0.49558 | 4.784965 | 0.821711 |
| 14 | 112371200 | 112371599 | 9.433482 | 17.89361 | 22.34946 | 9.015727 | 0.171691 | 0.019072 | 0.821724 |
| 8 | 16345600 | 16352399 | 67.79226 | 107.6128 | 125.86 | 52.44509 | 0.10618 | 2.307099 | 0.821726 |
| 9 | 115460200 | 115463399 | 31.25412 | 51.26928 | 71.52372 | 30.79552 | -0.10199 | 1.443436 | 0.821742 |
| 2 | 36301400 | 36303399 | 33.27455 | 54.84302 | 65.84846 | 28.4089 | 0.116581 | 1.385178 | 0.821751 |
| 3 | 22368400 | 22371599 | 24.54303 | 41.84792 | 50.1098 | 21.96688 | 0.123244 | 1.004743 | 0.821752 |
| 14 | 12001200 | 12005399 | 48.68278 | 78.01082 | 105.9375 | 44.70277 | -0.09436 | 1.993669 | 0.821781 |
| 5 | 92084600 | 92085599 | 8.196987 | 15.69309 | 24.58452 | 9.712103 | -0.12592 | -0.05576 | 0.8218 |
| 8 | 119660200 | 119663199 | 34.26017 | 56.13785 | 67.93514 | 29.23664 | 0.101929 | 1.460098 | 0.821806 |
| 15 | 95665200 | 95666999 | 16.34806 | 28.46098 | 41.12913 | 18.0808 | -0.14139 | 0.758085 | 0.821816 |
| 6 | 131581800 | 131583199 | 36.76238 | 60.36763 | 59.79584 | 25.75723 | 0.391262 | 1.433433 | 0.821817 |
| 12 | 96488000 | 96490199 | 18.99212 | 33.6054 | 37.88467 | 16.75439 | 0.219085 | 0.548577 | 0.821864 |
| 6 | 129698000 | 129699399 | 16.78663 | 29.75263 | 26.94179 | 11.91095 | 0.534483 | 0.4964 | 0.821865 |
| 3 | 17016200 | 17017399 | 16.04006 | 28.19775 | 40.24135 | 17.8198 | -0.12296 | 0.605821 | 0.821869 |
| 11 | 108540800 | 108543199 | 15.20008 | 27.48083 | 26.37009 | 11.44495 | 0.469168 | 0.362861 | 0.82187 |
| 11 | 10477000 | 10479999 | 20.55535 | 35.87099 | 35.15176 | 15.47625 | 0.416629 | 0.734923 | 0.821898 |
| X | 48053200 | 48066199 | 216.4424 | 317.1618 | 349.7141 | 144.0507 | 0.094301 | 3.880305 | 0.821898 |
| 7 | 12969600 | 12976999 | 81.7886 | 128.2815 | 150.0621 | 62.23702 | 0.091885 | 2.589241 | 0.821934 |
| 6 | 85819400 | 85820999 | 14.89704 | 25.99933 | 46.26007 | 20.36819 | -0.43999 | 0.720784 | 0.821938 |
| 9 | 22082400 | 22090399 | 115.0265 | 176.457 | 173.4028 | 71.98543 | 0.318444 | 3.004545 | 0.821944 |
| 1 | 172092000 | 172093799 | 29.45239 | 48.68955 | 58.84597 | 25.43233 | 0.108049 | 1.334519 | 0.821964 |
| Y | 5959200 | 5959599 | 7.152992 | 13.93241 | 17.8685 | 6.52243 | 0.265323 | -0.36089 | 0.821965 |
| 4 | 90545000 | 90547999 | 57.86352 | 92.06012 | 107.8763 | 45.06087 | 0.106684 | 2.227343 | 0.821977 |
| 13 | 113438000 | 113447799 | 200.163 | 293.6628 | 366.594 | 150.9922 | -0.08301 | 3.859442 | 0.821978 |
| 1 | 182758600 | 182759199 | 6.520528 | 12.68259 | 21.27324 | 7.864959 | -0.13353 | -0.30812 | 0.821982 |
| 17 | 88360200 | 88361399 | 8.701443 | 16.46511 | 25.46283 | 10.32444 | -0.14075 | 0.037244 | 0.822009 |
| 17 | 87896800 | 87899399 | 27.09282 | 45.40693 | 55.06127 | 24.01305 | 0.103824 | 1.17276 | 0.822016 |
| 19 | 54054600 | 54059599 | 49.68648 | 79.27496 | 108.3279 | 45.42735 | -0.11029 | 2.128411 | 0.822034 |
| 2 | 116364600 | 116370799 | 192.8593 | 284.3512 | 316.9484 | 130.5234 | 0.087111 | 3.777928 | 0.822034 |
| 12 | 8661600 | 8666599 | 79.54491 | 124.894 | 146.6574 | 60.82761 | 0.087575 | 2.568639 | 0.822036 |
| 5 | 132779200 | 132781599 | 22.69325 | 37.8894 | 64.66177 | 28.13258 | -0.38745 | 1.208837 | 0.822049 |
| 17 | 34210600 | 34212599 | 13.99504 | 25.00779 | 36.08389 | 15.84636 | -0.12748 | 0.443151 | 0.822052 |
| 1 | 27198000 | 27200399 | 37.98585 | 61.08205 | 101.4179 | 43.01954 | -0.37767 | 1.87203 | 0.822054 |
| 2 | 34719000 | 34722199 | 71.29828 | 112.4191 | 132.4288 | 54.95628 | 0.087675 | 2.476772 | 0.822057 |

| | | | | | | | | | |
|---|---|---|---|---|---|---|---|---|---|
| 7 | 66117800 | 66119199 | 15.42514 | 27.1022 | 38.67341 | 17.12006 | -0.12299 | 0.605091 | 0.822071 |
| 7 | 125539200 | 125540599 | 14.24767 | 25.46283 | 30.94922 | 13.58983 | 0.118845 | 0.434126 | 0.822077 |
| 2 | 22080600 | 22082999 | 33.05738 | 54.62345 | 54.17418 | 23.40565 | 0.392338 | 1.34419 | 0.822079 |
| 17 | 83825000 | 83826599 | 13.62859 | 24.66561 | 29.88227 | 12.95223 | 0.135691 | 0.35001 | 0.822082 |
| 3 | 66052800 | 66053799 | 14.88273 | 26.18248 | 37.22436 | 16.48138 | -0.117 | 0.586031 | 0.822151 |
| 10 | 35308000 | 35310399 | 22.56147 | 37.79273 | 65.226 | 28.43666 | -0.40364 | 1.175429 | 0.822151 |
| 1 | 140650000 | 140651599 | 24.56179 | 41.06272 | 57.63825 | 25.09749 | -0.10646 | 1.194228 | 0.822157 |
| 10 | 11228200 | 11235399 | 71.3356 | 112.4086 | 149.8881 | 62.19669 | -0.0916 | 2.49171 | 0.822161 |
| 7 | 129532600 | 129538599 | 50.42448 | 80.88092 | 96.48713 | 40.77563 | 0.093897 | 1.962065 | 0.822182 |
| 4 | 80482200 | 80484199 | 21.88113 | 37.01695 | 52.21687 | 22.85455 | -0.11322 | 1.06735 | 0.822255 |
| 5 | 145676400 | 145693799 | 469.8063 | 643.8904 | 668.1982 | 277.9457 | 0.085546 | 4.890078 | 0.822273 |
| 17 | 43275400 | 43278999 | 29.5095 | 48.1229 | 81.87627 | 35.17289 | -0.39274 | 1.524626 | 0.822308 |
| X | 87469400 | 87471799 | 37.48078 | 61.23253 | 71.15224 | 30.58611 | 0.157862 | 1.513902 | 0.822372 |
| 10 | 110962400 | 110964199 | 13.31966 | 24.03493 | 28.9011 | 12.57362 | 0.143031 | 0.362971 | 0.822385 |
| 2 | 14386800 | 14391599 | 103.1145 | 158.3484 | 181.5736 | 75.38818 | 0.097162 | 2.981398 | 0.822387 |
| 6 | 126239600 | 126240399 | 10.47509 | 19.39222 | 28.84221 | 12.13615 | -0.12826 | 0.199377 | 0.822391 |
| X | 112025800 | 112026799 | 10.47509 | 19.39222 | 28.84221 | 12.13615 | -0.12826 | 0.199377 | 0.822391 |
| 16 | 16053000 | 16055199 | 25.41069 | 42.71724 | 51.75689 | 22.60986 | 0.105009 | 1.13304 | 0.822397 |
| 2 | 71147200 | 71148999 | 14.95704 | 26.38288 | 37.60388 | 16.64062 | -0.11934 | 0.556003 | 0.822429 |
| 14 | 110154800 | 110159799 | 71.99986 | 112.4606 | 174.0924 | 72.23423 | -0.31647 | 2.661099 | 0.82244 |
| 7 | 117325400 | 117329799 | 51.23112 | 82.13991 | 97.12656 | 41.01938 | 0.105933 | 1.978634 | 0.82245 |
| 10 | 7199600 | 7200799 | 11.53776 | 21.02638 | 30.68651 | 13.18633 | -0.12294 | 0.298566 | 0.822473 |
| X | 113665600 | 113668599 | 42.63529 | 69.34034 | 64.58587 | 27.71787 | 0.471575 | 1.583147 | 0.822478 |
| 3 | 11723200 | 11724599 | 16.50684 | 28.88959 | 40.92297 | 18.05616 | -0.11228 | 0.68014 | 0.822485 |
| 14 | 35300800 | 35303399 | 20.35641 | 34.8035 | 49.01418 | 21.49128 | -0.10923 | 0.963337 | 0.822514 |
| 4 | 35865800 | 35868199 | 31.89637 | 52.0557 | 72.38503 | 31.10438 | -0.10158 | 1.513402 | 0.822524 |
| 9 | 118702400 | 118703799 | 15.3629 | 26.99266 | 38.23878 | 16.92756 | -0.11238 | 0.60651 | 0.822538 |
| 10 | 83284000 | 83294599 | 285.6958 | 409.1025 | 442.6405 | 182.8331 | 0.089032 | 4.229526 | 0.822541 |
| 5 | 143190000 | 143196599 | 82.53873 | 128.2706 | 169.6678 | 70.44668 | -0.0955 | 2.754119 | 0.822563 |
| 17 | 47702200 | 47704799 | 33.57032 | 54.39274 | 76.91326 | 32.92388 | -0.13412 | 1.641152 | 0.822564 |
| 14 | 80324800 | 80328399 | 67.48786 | 106.4056 | 141.833 | 58.85854 | -0.09042 | 2.478373 | 0.822577 |
| 18 | 76663000 | 76664799 | 17.8023 | 30.95862 | 44.15926 | 19.39945 | -0.12319 | 0.781289 | 0.82259 |
| 7 | 15901400 | 15903999 | 43.10023 | 69.27579 | 94.36004 | 40.00036 | -0.09341 | 1.888207 | 0.822591 |
| 1 | 37288400 | 37288799 | 3.936611 | 7.679681 | 12.29169 | 3.849097 | 0.151636 | -0.9612 | 0.822597 |
| 15 | 93912200 | 93913599 | 11.67779 | 21.36106 | 31.22628 | 13.35545 | -0.12097 | 0.279059 | 0.822598 |
| 9 | 85950200 | 85953399 | 47.99329 | 76.9565 | 91.24944 | 38.5433 | 0.102128 | 1.96793 | 0.822631 |
| 1 | 153242600 | 153243799 | 13.91797 | 24.97704 | 30.45388 | 13.32561 | 0.117817 | 0.405045 | 0.822636 |
| 2 | 111709400 | 111716999 | 118.0411 | 180.4505 | 207.5069 | 86.08289 | 0.087986 | 3.077877 | 0.822663 |
| 11 | 53440400 | 53448999 | 77.91497 | 121.4609 | 161.5009 | 67.02473 | -0.09955 | 2.695875 | 0.822667 |
| 15 | 9461200 | 9462799 | 15.1171 | 26.92642 | 32.2836 | 14.21579 | 0.135445 | 0.461368 | 0.822677 |
| 6 | 69854200 | 69855599 | 14.67433 | 26.47889 | 31.98793 | 13.9169 | 0.136617 | 0.382227 | 0.822684 |
| 4 | 15858000 | 15862799 | 53.10815 | 84.8901 | 101.1377 | 42.52268 | 0.090073 | 2.074113 | 0.822692 |
| 6 | 97074200 | 97075999 | 16.88775 | 29.75355 | 35.97001 | 15.92133 | 0.116385 | 0.578113 | 0.822694 |
| 4 | 114634600 | 114636199 | 9.087525 | 17.07158 | 26.37598 | 10.84172 | -0.15667 | 0.087882 | 0.822704 |
| 15 | 49282000 | 49283799 | 17.44479 | 31.07443 | 35.85869 | 15.78921 | 0.19055 | 0.485121 | 0.822711 |
| 12 | 5413800 | 5414999 | 10.8239 | 20.11848 | 24.71671 | 10.34365 | 0.151218 | 0.168569 | 0.822715 |
| 3 | 16916000 | 16917399 | 14.03235 | 24.94941 | 35.56225 | 15.68325 | -0.11402 | 0.474423 | 0.822722 |

| | | | | | | | | | |
|---|---|---|---|---|---|---|---|---|---|
| 7 | 85293400 | 85296399 | 28.88984 | 48.89842 | 55.68329 | 24.37548 | 0.194548 | 1.092586 | 0.822733 |
| 5 | 64870400 | 64872399 | 13.37952 | 23.74814 | 34.09112 | 15.0561 | -0.12557 | 0.47785 | 0.822774 |
| 6 | 69395000 | 69395799 | 10.25689 | 18.91832 | 28.27735 | 11.95837 | -0.1408 | 0.215225 | 0.822785 |
| 1 | 13174000 | 13176399 | 17.15566 | 29.79838 | 43.15374 | 18.94345 | -0.14511 | 0.793646 | 0.82282 |
| 3 | 49257000 | 49257999 | 11.17124 | 20.18597 | 36.89528 | 16.02389 | -0.45571 | 0.358239 | 0.822823 |
| 15 | 36400600 | 36402599 | 13.23783 | 23.77951 | 35.06037 | 15.32873 | -0.15423 | 0.401075 | 0.822832 |
| 3 | 57848400 | 57850599 | 19.93587 | 33.46376 | 60.69446 | 26.49561 | -0.47497 | 1.139015 | 0.822835 |
| 17 | 81551600 | 81554999 | 53.80978 | 85.9861 | 102.1959 | 42.94957 | 0.093095 | 2.084859 | 0.822864 |
| 15 | 27388800 | 27392799 | 36.51597 | 59.16235 | 81.42971 | 34.85587 | -0.09538 | 1.650714 | 0.822869 |
| 3 | 55783600 | 55786799 | 32.97117 | 54.53611 | 64.77761 | 28.00635 | 0.133047 | 1.34284 | 0.822878 |
| 5 | 43504400 | 43507399 | 14.58504 | 25.64404 | 36.78306 | 16.28786 | -0.12994 | 0.59052 | 0.82288 |
| 3 | 12048600 | 12048999 | 6.685955 | 13.01117 | 21.9075 | 8.06523 | -0.13301 | -0.31717 | 0.82288 |
| 3 | 106273000 | 106273999 | 15.48564 | 27.81995 | 26.83908 | 11.73337 | 0.454617 | 0.402298 | 0.822922 |
| 16 | 70773000 | 70775599 | 21.02299 | 36.72738 | 42.32128 | 18.64563 | 0.183903 | 0.730469 | 0.822934 |
| 15 | 36405400 | 36408799 | 28.36926 | 46.89877 | 65.46609 | 28.29337 | -0.09923 | 1.336842 | 0.822945 |
| 6 | 89589200 | 89592199 | 21.71996 | 36.66688 | 52.21423 | 22.83878 | -0.12693 | 1.084706 | 0.822946 |
| 1 | 102532200 | 102533799 | 15.36759 | 26.98462 | 38.27357 | 16.93433 | -0.11409 | 0.616449 | 0.822952 |
| 12 | 115657200 | 115663199 | 67.91119 | 108.0919 | 122.5927 | 51.23829 | 0.154365 | 2.227602 | 0.822954 |
| 11 | 14161600 | 14164399 | 25.59292 | 43.84058 | 50.67467 | 22.22279 | 0.174999 | 0.968704 | 0.82296 |
| 5 | 131493200 | 131495799 | 24.55842 | 41.22433 | 57.67506 | 25.17801 | -0.10182 | 1.151214 | 0.822967 |
| 4 | 62785400 | 62788399 | 49.16955 | 78.74125 | 92.53878 | 39.0115 | 0.112753 | 2.010249 | 0.82298 |
| 16 | 3825600 | 3827999 | 38.99839 | 62.95377 | 86.2188 | 36.76864 | -0.09375 | 1.747596 | 0.822998 |
| 18 | 5530200 | 5533199 | 20.0858 | 35.02202 | 41.90368 | 18.43913 | 0.129661 | 0.75308 | 0.823003 |
| 10 | 24601400 | 24604799 | 29.62939 | 49.20686 | 59.08739 | 25.6175 | 0.11776 | 1.280126 | 0.823012 |
| 5 | 46146400 | 46148199 | 23.67529 | 40.16904 | 57.20688 | 25.06074 | -0.1269 | 1.053474 | 0.823015 |
| 17 | 44462000 | 44464399 | 19.09443 | 33.27992 | 40.3849 | 17.76628 | 0.109382 | 0.751802 | 0.823018 |
| 18 | 54454400 | 54455799 | 15.6481 | 27.55056 | 39.16875 | 17.33956 | -0.11672 | 0.583907 | 0.823021 |
| 1 | 27767000 | 27767599 | 17.52085 | 30.619 | 36.83705 | 16.23308 | 0.122259 | 0.701913 | 0.823029 |
| 5 | 25917600 | 25920199 | 26.52258 | 44.27936 | 62.20916 | 27.06412 | -0.1078 | 1.218576 | 0.823059 |
| 12 | 44194600 | 44204599 | 138.1555 | 209.2559 | 238.204 | 98.56306 | 0.091793 | 3.271934 | 0.823074 |
| 9 | 113051000 | 113060399 | 135.1817 | 203.9236 | 261.1404 | 107.9155 | -0.08293 | 3.353936 | 0.823079 |
| 5 | 97402200 | 97402999 | 7.875957 | 15.19318 | 24.25742 | 9.37664 | -0.12371 | -0.13098 | 0.823081 |
| 3 | 54673000 | 54674599 | 18.36608 | 31.99248 | 38.45914 | 16.91311 | 0.122992 | 0.755173 | 0.823116 |
| 15 | 55270800 | 55272399 | 14.57735 | 26.26027 | 31.91824 | 13.91543 | 0.125775 | 0.389415 | 0.823144 |
| 16 | 12234800 | 12235599 | 15.25958 | 26.82898 | 38.25913 | 16.94162 | -0.12164 | 0.597836 | 0.823145 |
| 4 | 149679800 | 149681199 | 18.96314 | 32.60454 | 46.38243 | 20.34523 | -0.1222 | 0.905396 | 0.82315 |
| X | 101975200 | 101977999 | 20.15482 | 34.82535 | 50.09186 | 21.97564 | -0.13671 | 0.871027 | 0.82317 |
| 4 | 74367600 | 74371599 | 48.87978 | 78.57554 | 93.17071 | 39.50361 | 0.106442 | 1.892559 | 0.823194 |
| 2 | 22190800 | 22198999 | 108.7482 | 168.357 | 158.4204 | 65.80372 | 0.39099 | 2.828556 | 0.823221 |
| 10 | 3618600 | 3619999 | 19.51492 | 33.89686 | 41.05531 | 18.02248 | 0.111775 | 0.804709 | 0.823223 |
| 10 | 48941400 | 48942199 | 19.51492 | 33.89686 | 41.05531 | 18.02248 | 0.111775 | 0.804709 | 0.823223 |
| 7 | 61668000 | 61673599 | 72.95138 | 115.6683 | 114.593 | 47.69487 | 0.343655 | 2.341701 | 0.823261 |
| 13 | 30153800 | 30157599 | 31.70705 | 51.86216 | 72.03962 | 30.98111 | -0.0978 | 1.480584 | 0.823285 |
| 3 | 96243200 | 96246799 | 32.51963 | 53.14641 | 73.84096 | 31.74603 | -0.09902 | 1.495261 | 0.823328 |
| 6 | 99530600 | 99533599 | 25.36207 | 42.44178 | 59.36636 | 25.86578 | -0.1015 | 1.187566 | 0.823341 |
| 9 | 122704800 | 122709599 | 45.86586 | 74.02728 | 87.0165 | 37.10079 | 0.125941 | 1.763706 | 0.82337 |
| 2 | 102376800 | 102378599 | 14.28927 | 26.14228 | 31.05527 | 13.27935 | 0.176496 | 0.296093 | 0.823382 |

| | | | | | | | | | |
|---|---|---|---|---|---|---|---|---|---|
| 10 | 6132000 | 6132599 | 8.323539 | 16.06205 | 21.4753 | 8.293114 | 0.131723 | -0.13651 | 0.823389 |
| 7 | 16787600 | 16790799 | 26.08027 | 43.28074 | 60.87553 | 26.37931 | -0.10964 | 1.283214 | 0.823396 |
| 5 | 60831000 | 60833599 | 46.9673 | 75.41076 | 90.01876 | 38.09981 | 0.094698 | 1.924596 | 0.82343 |
| 5 | 68073600 | 68075999 | 25.90147 | 43.70996 | 52.85512 | 23.11655 | 0.108029 | 1.102764 | 0.82344 |
| 11 | 12518400 | 12526799 | 98.31739 | 151.9794 | 176.6272 | 73.36693 | 0.085045 | 2.854109 | 0.823442 |
| 12 | 115057200 | 115060599 | 80.75537 | 126.3395 | 148.2976 | 61.52128 | 0.084114 | 2.633541 | 0.823444 |
| 5 | 52362800 | 52364599 | 15.44605 | 27.17987 | 38.64805 | 17.11093 | -0.1173 | 0.587072 | 0.823453 |
| 1 | 46125400 | 46129799 | 50.57126 | 81.12539 | 96.65388 | 40.85302 | 0.095941 | 1.958596 | 0.823465 |
| 2 | 105652200 | 105656399 | 47.06947 | 76.0654 | 86.98293 | 37.14662 | 0.16768 | 1.730354 | 0.823477 |
| 12 | 71811000 | 71813999 | 25.4016 | 42.38254 | 59.2942 | 25.78624 | -0.10167 | 1.219807 | 0.823487 |
| 1 | 9055400 | 9056399 | 12.09376 | 22.18191 | 32.89228 | 14.01628 | -0.13748 | 0.272293 | 0.82352 |
| 17 | 87791800 | 87794799 | 32.51355 | 53.10152 | 73.79566 | 31.71894 | -0.10002 | 1.504357 | 0.823526 |
| 11 | 97773200 | 97775999 | 18.00307 | 30.8521 | 53.8108 | 23.5989 | -0.41607 | 0.938125 | 0.823527 |
| Y | 8706600 | 8711599 | 44.09806 | 70.82892 | 69.80066 | 29.55852 | 0.371275 | 1.89831 | 0.823579 |
| 2 | 33780400 | 33781399 | 11.79467 | 21.4788 | 31.66186 | 13.61795 | -0.13839 | 0.305146 | 0.823642 |
| 1 | 104414400 | 104422999 | 81.41137 | 128.5537 | 144.11 | 59.83762 | 0.160771 | 2.453348 | 0.823643 |
| 2 | 96261200 | 96261799 | 12.36383 | 21.98214 | 32.8196 | 14.47408 | -0.18005 | 0.461276 | 0.823654 |
| 12 | 114842600 | 114860599 | 250.516 | 366.0227 | 383.1033 | 157.8328 | 0.165418 | 3.929609 | 0.823657 |
| 9 | 115341800 | 115348599 | 89.83813 | 139.1641 | 182.7276 | 75.90005 | -0.08853 | 2.812652 | 0.823658 |
| 3 | 18349600 | 18350199 | 6.961744 | 13.46162 | 22.09982 | 8.455413 | -0.1493 | -0.18248 | 0.823677 |
| 5 | 64454000 | 64457399 | 21.06287 | 35.61291 | 51.14446 | 22.3814 | -0.13899 | 1.067621 | 0.823691 |
| 3 | 10875000 | 10880399 | 46.89486 | 75.33212 | 103.6637 | 43.90808 | -0.1092 | 1.914544 | 0.823706 |
| 1 | 94520600 | 94521999 | 16.79245 | 29.92449 | 29.18829 | 12.84774 | 0.432101 | 0.465478 | 0.823706 |
| 11 | 17913600 | 17915599 | 44.49348 | 71.66317 | 84.93213 | 36.11179 | 0.110657 | 1.824158 | 0.823715 |
| X | 106683600 | 106685199 | 14.27047 | 25.64266 | 38.31221 | 16.74596 | -0.17275 | 0.41621 | 0.823728 |
| 17 | 57997000 | 58002199 | 61.5087 | 98.07096 | 114.6345 | 48.02703 | 0.11357 | 2.163578 | 0.823743 |
| 1 | 7777200 | 7785399 | 198.1143 | 291.84 | 326.1851 | 134.3259 | 0.081976 | 3.794037 | 0.823744 |
| 9 | 39141800 | 39144599 | 31.72559 | 52.74943 | 51.90328 | 22.52088 | 0.404434 | 1.270501 | 0.823753 |
| 8 | 83229400 | 83231599 | 26.85138 | 45.8111 | 43.59671 | 19.11262 | 0.45352 | 1.000259 | 0.823766 |
| 10 | 90306400 | 90307799 | 16.05862 | 28.48787 | 34.3371 | 15.17173 | 0.124618 | 0.498291 | 0.823776 |
| 5 | 138492200 | 138495799 | 58.50013 | 93.13113 | 110.6339 | 46.25271 | 0.087938 | 2.208182 | 0.823789 |
| 1 | 84634200 | 84637799 | 31.00492 | 51.275 | 61.88749 | 26.75363 | 0.109942 | 1.334919 | 0.82379 |
| 17 | 46153400 | 46158199 | 56.00466 | 89.40699 | 106.4627 | 44.68154 | 0.088823 | 2.11618 | 0.823806 |
| X | 84388200 | 84395399 | 70.82874 | 112.6777 | 125.3479 | 52.35255 | 0.181415 | 2.248048 | 0.823816 |
| 11 | 43527400 | 43534199 | 65.59294 | 104.0419 | 123.3354 | 51.3537 | 0.085684 | 2.326414 | 0.823829 |
| 9 | 112428000 | 112429399 | 15.05444 | 27.63384 | 25.83479 | 10.9972 | 0.524961 | 0.275065 | 0.823845 |
| 3 | 47514200 | 47514799 | 12.92967 | 23.528 | 29.03454 | 12.49967 | 0.115829 | 0.314009 | 0.823859 |
| 14 | 80353800 | 80355399 | 12.92967 | 23.528 | 29.03454 | 12.49967 | 0.115829 | 0.314009 | 0.823859 |
| 9 | 113859000 | 113864199 | 58.36075 | 92.63387 | 124.8382 | 52.01514 | -0.0981 | 2.300384 | 0.823867 |
| 12 | 33493400 | 33496999 | 29.03909 | 48.73834 | 57.39964 | 25.05539 | 0.145745 | 1.165413 | 0.823871 |
| X | 74713400 | 74718199 | 68.29707 | 108.8082 | 120.0345 | 50.23423 | 0.195723 | 2.199929 | 0.823884 |
| 7 | 29834800 | 29866999 | 859.449 | 1115.82 | 1233.146 | 516.6528 | -0.0835 | 5.756997 | 0.823893 |
| 14 | 42972800 | 42986999 | 62.09136 | 97.83604 | 153.805 | 63.82155 | -0.32885 | 2.488905 | 0.82391 |
| 9 | 75549400 | 75554799 | 54.43448 | 87.04426 | 103.6253 | 43.59278 | 0.091775 | 2.064422 | 0.823937 |
| X | 111293800 | 111298599 | 56.31565 | 90.01456 | 106.1413 | 44.62463 | 0.104894 | 2.080828 | 0.823943 |
| 8 | 88834000 | 88835199 | 12.84907 | 23.23755 | 33.65812 | 14.60239 | -0.12117 | 0.358488 | 0.82395 |
| 11 | 20331000 | 20344999 | 176.7317 | 262.2461 | 332.5588 | 136.9832 | -0.09029 | 3.669959 | 0.823951 |

| | | | | | | | | | |
|---|---|---|---|---|---|---|---|---|---|
| 14 | 3945400 | 3948599 | 44.72213 | 72.0145 | 86.42657 | 36.73468 | 0.092133 | 1.830537 | 0.823953 |
| 18 | 44868200 | 44869199 | 11.90422 | 22.15315 | 27.71697 | 11.57623 | 0.126977 | 0.175991 | 0.823989 |
| 5 | 20387800 | 20388999 | 12.6681 | 23.50209 | 22.37625 | 9.388165 | 0.513341 | 0.186551 | 0.823997 |
| 14 | 118999200 | 119001799 | 15.38771 | 27.60899 | 26.28859 | 11.505 | 0.472227 | 0.407087 | 0.824005 |
| 7 | 40698800 | 40699599 | 9.765549 | 18.23567 | 27.35959 | 11.36074 | -0.1262 | 0.13627 | 0.824015 |
| 14 | 78812200 | 78813799 | 11.30458 | 20.52566 | 30.24959 | 13.05862 | -0.14091 | 0.319179 | 0.82402 |
| 15 | 36912800 | 36917199 | 34.22654 | 55.60561 | 76.79571 | 32.93893 | -0.09656 | 1.589165 | 0.824043 |
| 7 | 105460200 | 105464199 | 30.65854 | 50.45429 | 70.71413 | 30.4884 | -0.10687 | 1.399826 | 0.824049 |
| 13 | 40092400 | 40093599 | 11.76978 | 21.62313 | 26.61473 | 11.31596 | 0.131892 | 0.245353 | 0.82405 |
| 6 | 50448800 | 50451399 | 32.37791 | 53.50815 | 64.22588 | 27.7511 | 0.117851 | 1.349169 | 0.82405 |
| 6 | 123026000 | 123027999 | 21.5026 | 37.20905 | 44.6797 | 19.60218 | 0.122992 | 0.853448 | 0.824051 |
| 6 | 33335400 | 33335999 | 5.955281 | 11.56415 | 20.06018 | 7.515571 | -0.19851 | -0.27106 | 0.824079 |
| 9 | 101435000 | 101437199 | 20.89975 | 35.96001 | 51.17654 | 22.44861 | -0.12277 | 0.911785 | 0.824088 |
| 7 | 57544600 | 57545799 | 19.01253 | 32.98042 | 39.09946 | 17.15525 | 0.142327 | 0.815808 | 0.824091 |
| 2 | 22908400 | 22909599 | 10.17532 | 19.13989 | 24.26294 | 9.951275 | 0.129529 | 0.085713 | 0.824097 |
| 7 | 106600000 | 106601199 | 16.7389 | 29.3262 | 34.64181 | 15.2982 | 0.148318 | 0.651042 | 0.824122 |
| 3 | 53213200 | 53214599 | 18.60845 | 32.72014 | 38.51178 | 17.05315 | 0.153727 | 0.614636 | 0.824127 |
| 6 | 65238000 | 65238599 | 12.65296 | 22.31766 | 33.88508 | 14.9951 | -0.20963 | 0.537842 | 0.824134 |
| 6 | 43761600 | 43764399 | 69.21979 | 108.6169 | 146.3481 | 60.69455 | -0.11123 | 2.574753 | 0.824175 |
| 15 | 93979200 | 93979999 | 13.66257 | 25.15152 | 24.91933 | 10.56589 | 0.445738 | 0.245444 | 0.824201 |
| 8 | 21094000 | 21099199 | 109.9252 | 168.0882 | 217.5262 | 90.24484 | -0.08198 | 3.071542 | 0.824222 |
| 8 | 21081800 | 21092199 | 283.6723 | 403.497 | 490.186 | 202.6944 | -0.08769 | 4.326541 | 0.824228 |
| 9 | 35210000 | 35212199 | 19.72344 | 34.28727 | 50.1148 | 22.01081 | -0.15831 | 0.804904 | 0.824246 |
| 6 | 108010200 | 108010999 | 20.27828 | 34.77689 | 49.00816 | 21.49397 | -0.10939 | 0.937026 | 0.824247 |
| 7 | 7036400 | 7050999 | 229.7341 | 334.6423 | 369.7477 | 152.3784 | 0.083095 | 3.971858 | 0.824249 |
| X | 65121000 | 65121599 | 8.524816 | 16.48283 | 22.09173 | 8.455285 | 0.140588 | -0.16157 | 0.824249 |
| 7 | 42730800 | 42735199 | 65.5048 | 104.1162 | 122.1571 | 50.96802 | 0.103428 | 2.267336 | 0.824256 |
| 1 | 5497800 | 5501599 | 88.71284 | 138.1235 | 161.1994 | 66.91191 | 0.087025 | 2.723643 | 0.824267 |
| 5 | 137992800 | 137997399 | 70.09417 | 110.6424 | 129.3712 | 53.70776 | 0.099644 | 2.452975 | 0.824273 |
| 5 | 61660000 | 61661399 | 11.6452 | 21.40246 | 26.16477 | 11.11884 | 0.142321 | 0.24195 | 0.824311 |
| X | 120389000 | 120390199 | 12.42773 | 23.19703 | 28.35569 | 11.78284 | 0.166565 | 0.165531 | 0.824332 |
| 13 | 14076400 | 14078599 | 48.51123 | 77.56498 | 105.1727 | 44.23758 | -0.09577 | 2.060658 | 0.824343 |
| 13 | 65456800 | 65462399 | 144.6948 | 216.7777 | 276.8602 | 114.2055 | -0.08787 | 3.48998 | 0.824344 |
| 7 | 24298600 | 24300799 | 12.10514 | 21.72883 | 31.681 | 13.85983 | -0.13825 | 0.394874 | 0.824348 |
| 10 | 119268800 | 119269799 | 12.10514 | 21.72883 | 31.681 | 13.85983 | -0.13825 | 0.394874 | 0.824348 |
| 10 | 47273200 | 47277799 | 45.34188 | 73.22667 | 85.98349 | 36.68949 | 0.128493 | 1.744281 | 0.824352 |
| 3 | 106595600 | 106603999 | 106.329 | 165.5496 | 176.7239 | 73.35602 | 0.215529 | 2.734699 | 0.824368 |
| 16 | 37392000 | 37393799 | 14.1275 | 24.80315 | 43.4877 | 19.23877 | -0.41855 | 0.620082 | 0.82439 |
| 13 | 9569200 | 9579999 | 123.7727 | 190.0463 | 207.4641 | 86.13612 | 0.167886 | 2.994842 | 0.824419 |
| 10 | 48409800 | 48417999 | 114.0457 | 174.4753 | 201.4248 | 83.57443 | 0.083193 | 3.060763 | 0.824433 |
| 10 | 21305800 | 21307599 | 18.4024 | 32.09598 | 38.82719 | 17.08843 | 0.113935 | 0.735785 | 0.824441 |
| 13 | 39170200 | 39171399 | 15.63001 | 27.99771 | 33.30622 | 14.59641 | 0.151383 | 0.432184 | 0.824442 |
| 15 | 18248400 | 18248999 | 8.945045 | 16.95491 | 25.99921 | 10.50402 | -0.12456 | 0.026973 | 0.824455 |
| 3 | 144350600 | 144353399 | 33.39829 | 54.57958 | 76.1761 | 32.74934 | -0.10558 | 1.50064 | 0.824457 |
| 10 | 49999800 | 50002199 | 23.34574 | 39.68223 | 47.96093 | 21.01958 | 0.109196 | 1.025418 | 0.824459 |
| 3 | 48960800 | 48961399 | 11.1909 | 20.02611 | 38.32349 | 16.80931 | -0.53128 | 0.418922 | 0.82448 |
| 8 | 39871800 | 39873199 | 27.01535 | 45.66672 | 44.83237 | 19.61778 | 0.407811 | 1.085043 | 0.824492 |

| | | | | | | | | | |
|---|---|---|---|---|---|---|---|---|---|
| 3 | 54484200 | 54485599 | 17.5626 | 30.56572 | 43.11788 | 18.9513 | -0.10701 | 0.769469 | 0.8245 |
| 2 | 162122800 | 162134199 | 130.5871 | 199.524 | 219.9012 | 91.21338 | 0.1486 | 3.094885 | 0.824501 |
| 9 | 77722400 | 77728799 | 59.66024 | 94.75877 | 127.2467 | 53.05354 | -0.09205 | 2.282735 | 0.824507 |
| 12 | 119787600 | 119798399 | 122.2271 | 187.1359 | 210.6473 | 87.41659 | 0.120512 | 3.049957 | 0.824546 |
| 2 | 51801000 | 51801799 | 11.61078 | 21.62091 | 27.21892 | 11.3559 | 0.119471 | 0.168881 | 0.824559 |
| 8 | 43013200 | 43015199 | 18.76703 | 32.62091 | 46.30539 | 20.33622 | -0.11618 | 0.798247 | 0.824616 |
| X | 72444200 | 72445399 | 16.75654 | 29.6072 | 35.03506 | 15.4986 | 0.14896 | 0.547799 | 0.824618 |
| 14 | 38327000 | 38328799 | 24.49972 | 41.73409 | 50.09968 | 21.96075 | 0.119493 | 1.015072 | 0.824626 |
| 8 | 65962200 | 65964199 | 24.2702 | 41.57529 | 49.02028 | 21.4973 | 0.14638 | 0.95943 | 0.824631 |
| X | 131731000 | 131734599 | 48.7887 | 78.73372 | 89.76341 | 38.26508 | 0.169717 | 1.775045 | 0.824652 |
| X | 54307200 | 54307999 | 16.50642 | 30.22087 | 26.15146 | 11.17122 | 0.632064 | 0.300497 | 0.8247 |
| 2 | 87254400 | 87255399 | 14.78353 | 26.44008 | 31.797 | 13.95336 | 0.134477 | 0.432806 | 0.824723 |
| 17 | 34616400 | 34619799 | 24.50425 | 40.99761 | 57.69011 | 25.1321 | -0.1101 | 1.18462 | 0.824735 |
| 15 | 88207800 | 88210799 | 24.02844 | 40.26576 | 56.56899 | 24.66852 | -0.10778 | 1.165602 | 0.824754 |
| 19 | 33429600 | 33431599 | 24.02844 | 40.26576 | 56.56899 | 24.66852 | -0.10778 | 1.165602 | 0.824754 |
| 7 | 132749200 | 132751799 | 22.64668 | 38.27973 | 53.86749 | 23.57097 | -0.10987 | 1.077822 | 0.82479 |
| 17 | 67067200 | 67069199 | 17.18683 | 29.93415 | 42.62654 | 18.74416 | -0.12025 | 0.758709 | 0.824813 |
| 4 | 38382800 | 38387999 | 66.23501 | 105.1131 | 124.1231 | 51.70766 | 0.091967 | 2.312315 | 0.82484 |
| 3 | 39522600 | 39525199 | 33.37991 | 54.32047 | 75.1982 | 32.27938 | -0.09811 | 1.559264 | 0.824858 |
| 15 | 14764200 | 14768999 | 74.97097 | 117.887 | 138.8831 | 57.61422 | 0.084503 | 2.534227 | 0.824865 |
| 9 | 36293400 | 36294199 | 35.70517 | 57.97743 | 68.18253 | 29.23329 | 0.13411 | 1.599149 | 0.824883 |
| 11 | 119214800 | 119216199 | 14.01103 | 25.1359 | 30.53595 | 13.36518 | 0.122972 | 0.409599 | 0.824891 |
| 5 | 15885600 | 15887599 | 38.79609 | 62.78347 | 87.09891 | 37.2506 | -0.10836 | 1.680337 | 0.824902 |
| 17 | 19801400 | 19802399 | 12.40739 | 23.47846 | 28.20622 | 11.43592 | 0.220143 | 0.074528 | 0.824909 |
| X | 123086400 | 123094399 | 57.11336 | 91.70878 | 84.64123 | 35.82827 | 0.465284 | 1.935355 | 0.824923 |
| X | 47783200 | 47784199 | 10.65165 | 19.87847 | 30.02646 | 12.48072 | -0.137 | 0.15558 | 0.824934 |
| 7 | 47716600 | 47717799 | 13.08726 | 24.05925 | 29.63433 | 12.58864 | 0.131644 | 0.258752 | 0.82494 |
| 15 | 101813400 | 101815199 | 18.50477 | 32.09602 | 45.26596 | 19.86079 | -0.10734 | 0.820926 | 0.82494 |
| 15 | 32869800 | 32877199 | 185.384 | 271.5062 | 348.0922 | 143.3874 | -0.12384 | 3.8844 | 0.824942 |
| 11 | 103094800 | 103095999 | 11.05845 | 20.22049 | 29.71295 | 12.71461 | -0.12876 | 0.273601 | 0.824985 |
| 14 | 86107400 | 86107999 | 12.64482 | 22.98246 | 28.09701 | 12.11396 | 0.128044 | 0.317474 | 0.824993 |
| 12 | 107740400 | 107742599 | 26.46491 | 44.49034 | 53.93416 | 23.56113 | 0.104251 | 1.139544 | 0.825051 |
| 1 | 12510200 | 12511599 | 24.76485 | 41.53262 | 58.19964 | 25.39228 | -0.10408 | 1.162993 | 0.825066 |
| 3 | 84054200 | 84054999 | 17.37552 | 30.65563 | 28.87565 | 12.7776 | 0.475322 | 0.552483 | 0.825078 |
| 10 | 129045000 | 129053999 | 224.0991 | 326.3613 | 360.2334 | 148.4608 | 0.084242 | 3.974563 | 0.825084 |
| 1 | 65396200 | 65401599 | 89.99321 | 140.46 | 140.4175 | 58.26638 | 0.312822 | 2.678269 | 0.825087 |
| 14 | 103555400 | 103560199 | 54.05797 | 86.94634 | 97.73579 | 41.47147 | 0.184276 | 1.88506 | 0.825105 |
| 8 | 4940000 | 4943399 | 38.18796 | 62.24944 | 61.66211 | 26.48515 | 0.385261 | 1.537414 | 0.825116 |
| 10 | 122704400 | 122705999 | 11.47891 | 21.13737 | 31.38168 | 13.30101 | -0.13342 | 0.243296 | 0.825128 |
| 3 | 33830800 | 33834799 | 39.56427 | 64.07442 | 77.23586 | 33.05053 | 0.094954 | 1.661948 | 0.825149 |
| 5 | 133541800 | 133544799 | 24.39559 | 41.27697 | 50.12257 | 21.93891 | 0.102164 | 1.072021 | 0.825155 |
| 1 | 52659600 | 52664399 | 49.85883 | 80.029 | 95.11617 | 40.2386 | 0.100421 | 1.938454 | 0.825165 |
| 1 | 156243200 | 156246599 | 32.19298 | 53.14179 | 63.98758 | 27.62634 | 0.11326 | 1.364168 | 0.825166 |
| 10 | 50101400 | 50103799 | 22.93377 | 38.89965 | 54.99507 | 24.08969 | -0.11642 | 1.049563 | 0.825169 |
| X | 161100400 | 161108199 | 146.5651 | 222.8704 | 200.6949 | 83.14197 | 0.434351 | 3.193503 | 0.82517 |
| X | 75334800 | 75337599 | 26.553 | 44.87074 | 53.37991 | 23.35575 | 0.131655 | 1.096315 | 0.825184 |
| 4 | 48677800 | 48679999 | 23.41469 | 39.69845 | 47.98367 | 21.01574 | 0.108946 | 1.048919 | 0.825192 |

| | | | | | | | | | |
|---|---|---|---|---|---|---|---|---|---|
| 1 | 36729000 | 36731399 | 15.80903 | 27.63993 | 39.44484 | 17.38889 | -0.12293 | 0.689843 | 0.825198 |
| 10 | 128568400 | 128570399 | 19.7559 | 34.04956 | 47.94005 | 21.03 | -0.10674 | 0.888425 | 0.825211 |
| 9 | 123144800 | 123147199 | 31.30996 | 51.90417 | 62.39891 | 27.01956 | 0.115964 | 1.311246 | 0.825213 |
| 14 | 10437200 | 10439999 | 30.17302 | 50.18224 | 60.23873 | 26.14276 | 0.118406 | 1.267063 | 0.825222 |
| 2 | 72817600 | 72826399 | 104.6547 | 160.517 | 208.1314 | 86.40032 | -0.08135 | 3.008355 | 0.825248 |
| 3 | 106755800 | 106766999 | 147.1406 | 222.875 | 246.8205 | 102.1298 | 0.131522 | 3.27447 | 0.825272 |
| 1 | 58513200 | 58515199 | 18.72711 | 32.59423 | 39.41092 | 17.32261 | 0.114354 | 0.767774 | 0.825275 |
| 3 | 138189400 | 138193399 | 30.36672 | 50.04524 | 70.11217 | 30.24708 | -0.10544 | 1.382224 | 0.82528 |
| 7 | 34474400 | 34481799 | 73.2918 | 115.0417 | 151.9341 | 63.01373 | -0.08209 | 2.57139 | 0.825297 |
| 12 | 3275400 | 3277199 | 24.57003 | 41.05224 | 57.62575 | 25.0827 | -0.10649 | 1.200696 | 0.825308 |
| 7 | 90512600 | 90516599 | 28.22849 | 46.80243 | 65.51737 | 28.37264 | -0.10298 | 1.303724 | 0.825321 |
| 9 | 13067600 | 13069599 | 14.81602 | 26.60033 | 31.18462 | 13.64033 | 0.174037 | 0.406226 | 0.825344 |
| 7 | 3358600 | 3360999 | 27.53727 | 45.71165 | 63.82736 | 27.66386 | -0.09918 | 1.286029 | 0.825362 |
| 1 | 4566400 | 4568399 | 18.60453 | 31.43965 | 58.00353 | 25.37892 | -0.49907 | 1.071169 | 0.825366 |
| 7 | 91191800 | 91192399 | 7.731526 | 14.86636 | 23.59685 | 9.206472 | -0.12771 | -0.10387 | 0.825369 |
| 7 | 29622600 | 29631999 | 167.777 | 248.7896 | 314.4818 | 129.5282 | -0.08655 | 3.680019 | 0.825398 |
| X | 73379600 | 73382599 | 19.2528 | 33.32939 | 48.46927 | 21.26502 | -0.15179 | 0.850116 | 0.825411 |
| 3 | 52991000 | 52992999 | 17.01827 | 29.83518 | 36.16428 | 15.97937 | 0.11137 | 0.646426 | 0.825419 |
| 5 | 82419800 | 82420399 | 8.305862 | 15.76757 | 24.95239 | 10.04839 | -0.16565 | 0.006294 | 0.82547 |
| 15 | 3191200 | 3195999 | 74.37283 | 117.5966 | 118.2578 | 49.13649 | 0.318811 | 2.412413 | 0.825472 |
| 11 | 114188200 | 114190199 | 17.59851 | 30.78785 | 37.20365 | 16.40955 | 0.115764 | 0.683803 | 0.825477 |
| 1 | 165274600 | 165282199 | 78.13582 | 122.8813 | 169.4306 | 70.28789 | -0.14204 | 2.539706 | 0.825487 |
| 14 | 57582200 | 57583999 | 19.47891 | 33.81652 | 41.06142 | 18.01921 | 0.108051 | 0.811432 | 0.825489 |
| 4 | 89763800 | 89768199 | 33.75861 | 55.64901 | 66.50759 | 28.69505 | 0.123315 | 1.384827 | 0.825495 |
| 9 | 19653800 | 19655199 | 21.88378 | 37.91695 | 45.21189 | 19.83643 | 0.133454 | 0.845238 | 0.825498 |
| 5 | 64720200 | 64723399 | 28.10624 | 46.25703 | 65.1373 | 28.08472 | -0.11349 | 1.390524 | 0.82553 |
| 9 | 22663400 | 22665799 | 19.91646 | 34.56206 | 42.06862 | 18.45822 | 0.104406 | 0.818766 | 0.825541 |
| 3 | 106563800 | 106564399 | 15.33358 | 27.19575 | 32.97487 | 14.5725 | 0.11613 | 0.494332 | 0.825581 |
| 11 | 99883400 | 99885399 | 17.02794 | 29.56268 | 42.66449 | 18.72317 | -0.14013 | 0.799864 | 0.825589 |
| 12 | 81377800 | 81380999 | 28.13514 | 45.7215 | 82.20335 | 35.26529 | -0.47571 | 1.574731 | 0.82559 |
| 4 | 130167400 | 130169799 | 18.56444 | 32.02469 | 45.31085 | 19.87719 | -0.11349 | 0.871079 | 0.825632 |
| 3 | 24481800 | 24483199 | 14.52933 | 25.73588 | 36.56728 | 16.16838 | -0.1124 | 0.50727 | 0.825637 |
| 5 | 135494600 | 135500199 | 52.64099 | 83.46498 | 136.9287 | 56.99117 | -0.38289 | 2.331905 | 0.825678 |
| 5 | 139858400 | 139861999 | 30.26463 | 49.848 | 59.91394 | 25.8411 | 0.114713 | 1.380925 | 0.825686 |
| 14 | 70152600 | 70157799 | 40.22831 | 64.48821 | 108.8109 | 45.94248 | -0.40632 | 1.971915 | 0.825701 |
| 5 | 92767400 | 92771399 | 28.31577 | 47.14854 | 56.79736 | 24.66862 | 0.113399 | 1.245046 | 0.825718 |
| 10 | 8376400 | 8379199 | 24.10833 | 40.73092 | 57.78747 | 25.28259 | -0.12173 | 1.090503 | 0.825732 |
| 9 | 92629200 | 92631999 | 33.49432 | 54.92657 | 66.64403 | 28.69063 | 0.098641 | 1.448971 | 0.825745 |
| 8 | 43701600 | 43703399 | 23.57223 | 40.51114 | 38.34832 | 16.82091 | 0.462666 | 0.914009 | 0.825758 |
| 10 | 30348200 | 30349599 | 18.42036 | 32.03056 | 45.23476 | 19.87081 | -0.10875 | 0.789547 | 0.825764 |
| 12 | 115840600 | 115841799 | 12.58065 | 23.85254 | 28.30518 | 11.42906 | 0.24296 | 0.063213 | 0.825764 |
| X | 115917000 | 115920399 | 37.20024 | 61.57277 | 68.12511 | 29.46735 | 0.235224 | 1.349344 | 0.825785 |
| 14 | 52744600 | 52747799 | 36.49952 | 59.69055 | 71.1129 | 30.58061 | 0.12288 | 1.491463 | 0.825807 |
| 13 | 61941000 | 61942999 | 23.31298 | 39.94189 | 47.93123 | 21.01998 | 0.12113 | 0.953419 | 0.825808 |
| 7 | 92795000 | 92797199 | 24.59929 | 40.95836 | 58.10729 | 25.23683 | -0.12166 | 1.238713 | 0.825832 |
| 11 | 28251000 | 28252399 | 10.6945 | 19.65932 | 29.22991 | 12.41784 | -0.13822 | 0.238665 | 0.825862 |
| 12 | 19407200 | 19409199 | 47.99745 | 77.12677 | 92.34365 | 39.13058 | 0.091776 | 1.902475 | 0.825876 |

| | | | | | | | | | |
|---|---|---|---|---|---|---|---|---|---|
| 8 | 3889200 | 3890799 | 21.2075 | 36.66845 | 44.20333 | 19.3927 | 0.117302 | 0.861885 | 0.825899 |
| 12 | 68815800 | 68818199 | 21.91694 | 37.45906 | 45.32057 | 19.8712 | 0.108983 | 0.968157 | 0.825922 |
| 1 | 177954000 | 177957399 | 81.62485 | 127.5682 | 149.573 | 62.05696 | 0.084568 | 2.652501 | 0.825977 |
| X | 80283400 | 80287399 | 37.49189 | 61.56343 | 71.29618 | 30.71058 | 0.166404 | 1.44535 | 0.825984 |
| 4 | 91570600 | 91573999 | 37.47479 | 61.01841 | 73.46901 | 31.54355 | 0.103263 | 1.559408 | 0.825994 |
| X | 156599800 | 156603399 | 29.04838 | 48.28733 | 58.30388 | 25.29422 | 0.11007 | 1.271689 | 0.826026 |
| 13 | 84453800 | 84457799 | 57.18736 | 91.7027 | 103.8462 | 43.86675 | 0.168387 | 1.981366 | 0.826029 |
| 10 | 22000200 | 22001999 | 19.85724 | 34.02724 | 47.95139 | 21.02912 | -0.10963 | 0.939372 | 0.826035 |
| 6 | 13328800 | 13332199 | 70.49295 | 111.1669 | 130.7837 | 54.27471 | 0.089727 | 2.473113 | 0.826048 |
| 13 | 100976400 | 100980799 | 39.23515 | 63.5776 | 76.69271 | 32.83189 | 0.094672 | 1.649693 | 0.826066 |
| X | 64615800 | 64623999 | 144.5355 | 220.6942 | 186.2806 | 77.25636 | 0.532418 | 3.10932 | 0.826077 |
| 6 | 81670000 | 81671599 | 15.50036 | 27.626 | 33.31726 | 14.66337 | 0.127601 | 0.462918 | 0.82608 |
| 7 | 106757000 | 106758399 | 20.75626 | 35.70131 | 43.20209 | 18.95042 | 0.11023 | 0.903447 | 0.826121 |
| 3 | 54394800 | 54395799 | 16.28298 | 28.96422 | 27.36314 | 12.06273 | 0.476518 | 0.469844 | 0.826121 |
| 9 | 36035000 | 36038399 | 52.55017 | 83.82521 | 112.9828 | 47.36473 | -0.09086 | 2.14108 | 0.826122 |
| 18 | 57971400 | 57974199 | 19.60202 | 33.74268 | 47.93752 | 21.02811 | -0.1202 | 0.895557 | 0.826131 |
| 16 | 28764000 | 28766999 | 19.93522 | 34.85087 | 41.28542 | 18.1989 | 0.14425 | 0.713974 | 0.826141 |
| 10 | 11352000 | 11371799 | 52.78144 | 84.66684 | 85.00915 | 35.92847 | 0.342246 | 1.958473 | 0.826203 |
| 16 | 36263200 | 36266599 | 41.86444 | 67.86717 | 79.81378 | 34.17878 | 0.131834 | 1.650379 | 0.826224 |
| 18 | 60659000 | 60662999 | 9.445843 | 17.25258 | 32.90758 | 14.10129 | -0.50374 | 0.275974 | 0.826237 |
| 7 | 57706600 | 57708199 | 24.14198 | 40.39032 | 57.02162 | 24.84042 | -0.11467 | 1.186418 | 0.82626 |
| 16 | 55204200 | 55205799 | 14.12506 | 25.232 | 36.58082 | 16.06894 | -0.13465 | 0.446184 | 0.826264 |
| 18 | 4514200 | 4515399 | 9.173327 | 17.08149 | 26.36404 | 10.99982 | -0.17205 | 0.147413 | 0.826271 |
| 14 | 123338400 | 123339799 | 11.80459 | 21.20195 | 31.23185 | 13.65649 | -0.15237 | 0.388622 | 0.82628 |
| 16 | 51653400 | 51655199 | 18.08404 | 31.76763 | 38.05264 | 16.84244 | 0.128351 | 0.624088 | 0.826283 |
| 3 | 74494600 | 74495399 | 8.34473 | 16.02569 | 25.24694 | 9.885152 | -0.12204 | -0.08184 | 0.826286 |
| 1 | 31410200 | 31410799 | 9.498785 | 18.04973 | 23.31845 | 9.371106 | 0.127521 | 0.007601 | 0.826324 |
| 1 | 102903000 | 102904399 | 19.64698 | 34.8091 | 30.4403 | 13.45866 | 0.584237 | 0.525231 | 0.826324 |
| 4 | 7758400 | 7765799 | 136.5137 | 207.6704 | 202.1133 | 83.73976 | 0.322842 | 3.181525 | 0.826336 |
| 8 | 21572400 | 21575999 | 54.87091 | 88.11572 | 100.2068 | 42.42276 | 0.164766 | 1.933012 | 0.826392 |
| 7 | 57843200 | 57844399 | 11.6795 | 21.69739 | 27.22004 | 11.39939 | 0.120407 | 0.185151 | 0.826402 |
| 15 | 46514400 | 46515199 | 11.6795 | 21.69739 | 27.22004 | 11.39939 | 0.120407 | 0.185151 | 0.826402 |
| 1 | 19448200 | 19452999 | 43.40942 | 70.27819 | 82.51026 | 35.29886 | 0.132438 | 1.678991 | 0.826412 |
| 3 | 78961400 | 78963199 | 31.70219 | 52.90498 | 51.40136 | 22.35639 | 0.422573 | 1.227518 | 0.826415 |
| 10 | 20688600 | 20689399 | 10.72571 | 20.01749 | 30.01871 | 12.47665 | -0.12684 | 0.154065 | 0.826428 |
| 12 | 39468800 | 39470799 | 16.3407 | 28.73957 | 41.2543 | 18.26667 | -0.13115 | 0.603383 | 0.826434 |
| 17 | 84368200 | 84370199 | 20.36944 | 35.25876 | 42.6482 | 18.71107 | 0.112794 | 0.84618 | 0.826449 |
| 6 | 14879400 | 14879999 | 6.532165 | 12.71356 | 21.32215 | 7.84137 | -0.12593 | -0.32492 | 0.826467 |
| 16 | 9571000 | 9575399 | 40.24423 | 64.98992 | 77.77699 | 33.18504 | 0.101054 | 1.734167 | 0.826479 |
| 4 | 8460400 | 8469199 | 139.2553 | 211.1836 | 237.9226 | 98.48462 | 0.108226 | 3.248544 | 0.826502 |
| 17 | 79446800 | 79449999 | 34.87734 | 56.66116 | 78.08014 | 33.48907 | -0.09343 | 1.591674 | 0.826509 |
| 8 | 104810400 | 104812599 | 24.85297 | 42.38999 | 41.58481 | 18.23021 | 0.409825 | 0.99252 | 0.826523 |
| 5 | 138612000 | 138616799 | 108.4984 | 166.1657 | 216.4045 | 89.81191 | -0.08933 | 3.040392 | 0.826537 |
| 13 | 54328400 | 54330999 | 27.07257 | 44.71664 | 63.04521 | 27.23093 | -0.11362 | 1.341561 | 0.82654 |
| 14 | 32119800 | 32121399 | 7.969469 | 15.13319 | 24.50535 | 9.862555 | -0.19654 | 0.00548 | 0.826554 |
| 8 | 70217000 | 70223199 | 69.72186 | 110.481 | 129.775 | 53.98117 | 0.097471 | 2.359859 | 0.82656 |
| 1 | 114735600 | 114738199 | 13.99213 | 24.55368 | 42.99565 | 19.01336 | -0.41662 | 0.627221 | 0.826614 |

| | | | | | | | | | |
|---|---|---|---|---|---|---|---|---|---|
| X | 116672400 | 116673399 | 12.34118 | 23.14165 | 28.45497 | 11.73237 | 0.166914 | 0.13471 | 0.826626 |
| 2 | 22174600 | 22177199 | 28.90643 | 48.14483 | 58.43977 | 25.38622 | 0.102508 | 1.247075 | 0.826626 |
| 3 | 4935200 | 4937399 | 12.79677 | 23.71973 | 23.46516 | 9.856928 | 0.457491 | 0.196586 | 0.826647 |
| 1 | 74168000 | 74168999 | 7.45014 | 14.30676 | 23.04165 | 9.023354 | -0.15319 | -0.09523 | 0.826648 |
| 11 | 6729800 | 6733199 | 33.02032 | 53.76835 | 74.72183 | 32.08208 | -0.10305 | 1.548585 | 0.826653 |
| 5 | 63883000 | 63885399 | 25.55263 | 43.1402 | 52.30613 | 22.87949 | 0.104194 | 1.096969 | 0.826657 |
| 5 | 60860800 | 60861799 | 12.56033 | 22.68965 | 32.6441 | 14.18254 | -0.11269 | 0.361775 | 0.826663 |
| 2 | 6146400 | 6147999 | 18.64905 | 32.42093 | 39.00869 | 17.13338 | 0.121406 | 0.78482 | 0.826674 |
| 8 | 91524400 | 91526399 | 21.07314 | 36.32081 | 44.20488 | 19.3915 | 0.102537 | 0.887421 | 0.826677 |
| 6 | 49869400 | 49874399 | 70.57819 | 111.6436 | 131.8348 | 54.78945 | 0.087864 | 2.401803 | 0.826702 |
| 16 | 28665000 | 28668199 | 27.97654 | 46.29329 | 64.99436 | 28.10784 | -0.10746 | 1.323032 | 0.826707 |
| 16 | 19829200 | 19831399 | 25.33734 | 42.26113 | 59.35054 | 25.80527 | -0.1072 | 1.22159 | 0.826714 |
| 13 | 84382000 | 84386599 | 80.56392 | 126.3326 | 148.5982 | 61.63128 | 0.083714 | 2.593328 | 0.826724 |
| 4 | 54490600 | 54490999 | 5.658911 | 11.02991 | 19.14194 | 6.924229 | -0.15298 | -0.40342 | 0.826725 |
| 13 | 104637400 | 104637799 | 5.658911 | 11.02991 | 19.14194 | 6.924229 | -0.15298 | -0.40342 | 0.826725 |
| 10 | 65515600 | 65516199 | 8.768524 | 16.79126 | 21.88564 | 8.640056 | 0.138268 | -0.05987 | 0.826735 |
| 9 | 3743200 | 3745799 | 22.63661 | 39.15307 | 38.88425 | 17.05931 | 0.395717 | 0.856297 | 0.826738 |
| 15 | 23522200 | 23526999 | 66.61449 | 106.0638 | 104.0424 | 43.50539 | 0.363936 | 2.210588 | 0.826745 |
| 8 | 37775200 | 37780799 | 76.49569 | 120.6207 | 118.964 | 49.36894 | 0.343672 | 2.476582 | 0.826753 |
| 7 | 55982800 | 55986799 | 37.95785 | 61.59541 | 86.96622 | 37.26902 | -0.12999 | 1.623743 | 0.826764 |
| 8 | 93125000 | 93126399 | 12.38544 | 23.22557 | 28.44148 | 11.72603 | 0.172614 | 0.13318 | 0.826773 |
| 3 | 40527800 | 40527999 | 5.793765 | 11.30876 | 15.91693 | 5.641037 | 0.173244 | -0.48471 | 0.826775 |
| 11 | 113238200 | 113242599 | 64.8988 | 102.555 | 136.3568 | 56.63952 | -0.08422 | 2.423059 | 0.826829 |
| 13 | 72517800 | 72519599 | 16.71886 | 29.12033 | 41.55239 | 18.27232 | -0.12335 | 0.751049 | 0.826831 |
| 4 | 39855400 | 39857399 | 24.85242 | 41.68466 | 58.26164 | 25.42132 | -0.10042 | 1.16033 | 0.826851 |
| 1 | 33660400 | 33662399 | 18.99927 | 32.70331 | 46.35453 | 20.33368 | -0.11683 | 0.892556 | 0.826861 |
| 2 | 125801400 | 125805999 | 103.9382 | 159.1281 | 206.3419 | 85.62905 | -0.08345 | 3.044061 | 0.826899 |
| 13 | 3558200 | 3567799 | 87.44362 | 137.1692 | 156.3122 | 64.82832 | 0.129759 | 2.591527 | 0.826919 |
| 1 | 29371600 | 29372399 | 12.59795 | 22.90135 | 28.05673 | 12.0939 | 0.125118 | 0.315109 | 0.826969 |
| 9 | 26861600 | 26864399 | 53.76573 | 85.78194 | 115.383 | 48.38476 | -0.08762 | 2.134674 | 0.826989 |
| 2 | 17981400 | 17986199 | 49.86834 | 79.83279 | 108.0137 | 45.51159 | -0.09072 | 2.026434 | 0.827013 |
| 18 | 25800000 | 25802399 | 16.33561 | 28.59487 | 40.40544 | 17.83043 | -0.10872 | 0.675506 | 0.827046 |
| 1 | 167666000 | 167666199 | 4.052764 | 7.925204 | 15.54345 | 5.296301 | -0.24587 | -0.64655 | 0.827046 |
| X | 12050800 | 12052999 | 22.54951 | 39.14091 | 45.75361 | 20.07626 | 0.162593 | 0.83471 | 0.827056 |
| 10 | 42614800 | 42620999 | 100.8977 | 155.642 | 181.1227 | 75.23507 | 0.080834 | 2.894195 | 0.827057 |
| 16 | 65056200 | 65057599 | 17.52562 | 30.61336 | 43.53144 | 19.1772 | -0.11806 | 0.717144 | 0.827063 |
| 3 | 46746600 | 46750999 | 29.32203 | 49.05494 | 44.45175 | 19.36414 | 0.522834 | 1.189833 | 0.827072 |
| 6 | 141985400 | 141988399 | 27.50861 | 46.01405 | 55.66421 | 24.24617 | 0.107259 | 1.193738 | 0.827118 |
| 3 | 113140600 | 113148799 | 143.8628 | 217.8159 | 213.115 | 88.17108 | 0.309558 | 3.279229 | 0.827125 |
| 4 | 149529600 | 149530599 | 6.985115 | 13.49107 | 22.07118 | 8.490609 | -0.15133 | -0.16193 | 0.827188 |
| 10 | 76053000 | 76054999 | 15.99482 | 27.90836 | 40.03818 | 17.62597 | -0.13065 | 0.722124 | 0.827203 |
| 12 | 49620800 | 49622599 | 12.77368 | 23.99132 | 23.46679 | 9.649295 | 0.49761 | 0.115224 | 0.827207 |
| 17 | 62088000 | 62090199 | 13.73532 | 23.88182 | 46.34139 | 20.35581 | -0.56468 | 0.7724 | 0.827207 |
| 1 | 8752600 | 8755999 | 38.44959 | 62.70599 | 74.41604 | 31.96881 | 0.125698 | 1.538941 | 0.827209 |
| 1 | 79103000 | 79105399 | 20.50415 | 34.69268 | 59.2107 | 25.91683 | -0.38717 | 1.068655 | 0.827232 |
| 7 | 125113200 | 125116399 | 24.75177 | 41.73336 | 50.6699 | 22.15496 | 0.102193 | 1.104258 | 0.827299 |
| 1 | 8445600 | 8446399 | 10.96476 | 20.31977 | 30.38348 | 12.76668 | -0.13508 | 0.197386 | 0.827303 |

| | | | | | | | | | |
|---|---|---|---|---|---|---|---|---|---|
| 14 | 15113200 | 15113599 | 5.648746 | 11.03026 | 15.42107 | 5.426475 | 0.192637 | -0.51251 | 0.827341 |
| 15 | 74727000 | 74735799 | 115.8449 | 176.2071 | 227.1156 | 94.09539 | -0.08225 | 3.180752 | 0.827347 |
| 1 | 142651000 | 142651599 | 10.28226 | 19.37792 | 24.76038 | 10.12259 | 0.121625 | 0.074742 | 0.827377 |
| 1 | 181267200 | 181268999 | 15.17671 | 26.87422 | 32.3856 | 14.32032 | 0.123699 | 0.505071 | 0.827429 |
| 3 | 140430800 | 140432599 | 21.56751 | 36.08791 | 62.36755 | 27.17154 | -0.40551 | 1.180716 | 0.827443 |
| 6 | 105392200 | 105394799 | 23.07846 | 39.74708 | 47.38169 | 20.78461 | 0.132087 | 0.906727 | 0.827458 |
| 8 | 78391200 | 78391999 | 8.701977 | 16.36109 | 25.41108 | 10.43019 | -0.16188 | 0.079356 | 0.82746 |
| 11 | 79205600 | 79208199 | 17.60487 | 30.74754 | 43.55412 | 19.18551 | -0.11245 | 0.721095 | 0.827471 |
| 8 | 99570800 | 99574199 | 57.42638 | 91.51072 | 108.8038 | 45.54891 | 0.088196 | 2.176686 | 0.827478 |
| 18 | 59065400 | 59068799 | 45.67215 | 73.64153 | 86.79214 | 36.9578 | 0.120531 | 1.794191 | 0.827485 |
| 11 | 10574000 | 10576399 | 55.64328 | 88.64198 | 118.8253 | 49.72303 | -0.08508 | 2.187213 | 0.827492 |
| 5 | 148112800 | 148123199 | 115.4545 | 175.9752 | 227.8774 | 94.46886 | -0.08656 | 3.13775 | 0.827494 |
| 17 | 18770800 | 18791999 | 449.0935 | 622.8084 | 537.2613 | 223.0628 | 0.369295 | 4.708605 | 0.827495 |
| 15 | 18323600 | 18324999 | 14.65976 | 27.04481 | 31.28276 | 13.22561 | 0.226511 | 0.253759 | 0.827504 |
| 1 | 182596400 | 182599599 | 43.3166 | 69.91919 | 83.71081 | 35.69558 | 0.099649 | 1.754252 | 0.82752 |
| 3 | 122581400 | 122588399 | 45.48131 | 73.48178 | 73.45517 | 31.36008 | 0.360892 | 1.727791 | 0.827523 |
| 9 | 11570800 | 11573799 | 35.03333 | 57.00689 | 78.93796 | 33.88378 | -0.09859 | 1.566149 | 0.827545 |
| 2 | 89955600 | 89958999 | 38.38414 | 62.65458 | 74.03358 | 31.81475 | 0.132807 | 1.524989 | 0.827546 |
| 14 | 110727800 | 110730999 | 37.21641 | 60.7442 | 72.69351 | 31.23808 | 0.114464 | 1.522501 | 0.827554 |
| 16 | 55606000 | 55612199 | 50.24374 | 80.26044 | 109.2432 | 45.8892 | -0.10275 | 2.090775 | 0.827565 |
| 1 | 33083000 | 33085999 | 29.56998 | 48.04392 | 83.04954 | 35.62323 | -0.41946 | 1.581694 | 0.827573 |
| 1 | 161865400 | 161867999 | 16.12941 | 28.24056 | 39.86853 | 17.59698 | -0.10747 | 0.668846 | 0.82759 |
| 18 | 34219800 | 34228199 | 85.96143 | 133.4518 | 175.1506 | 72.7344 | -0.08544 | 2.771941 | 0.827595 |
| 7 | 144632800 | 144636199 | 27.72815 | 46.66119 | 55.66441 | 24.32446 | 0.127344 | 1.135608 | 0.827618 |
| 2 | 36466200 | 36470799 | 43.62829 | 70.73123 | 69.79466 | 29.88972 | 0.38445 | 1.644666 | 0.827638 |
| 11 | 4082000 | 4083999 | 20.63794 | 35.99033 | 35.62375 | 15.67726 | 0.402075 | 0.744567 | 0.82764 |
| 7 | 16504000 | 16507199 | 57.25165 | 90.86031 | 122.7717 | 51.14671 | -0.10209 | 2.303821 | 0.827659 |
| 3 | 111800600 | 111801599 | 12.80436 | 23.16082 | 33.64029 | 14.5914 | -0.12504 | 0.356157 | 0.827676 |
| 1 | 73921600 | 73925399 | 62.8417 | 99.87979 | 118.7018 | 49.52459 | 0.084877 | 2.265107 | 0.827693 |
| 8 | 85828800 | 85829599 | 10.22533 | 19.07219 | 28.37995 | 11.80753 | -0.11683 | 0.149615 | 0.827698 |
| 2 | 26767000 | 26777799 | 146.0318 | 220.6191 | 249.7007 | 103.2365 | 0.096925 | 3.326331 | 0.827753 |
| 4 | 89505800 | 89507599 | 23.43549 | 38.82088 | 67.72009 | 29.3084 | -0.41946 | 1.303771 | 0.82776 |
| 4 | 143485000 | 143526999 | 738.1513 | 980.2667 | 954.2561 | 398.8022 | 0.132066 | 5.391854 | 0.827763 |
| 17 | 16499400 | 16500199 | 8.933082 | 16.8623 | 25.91646 | 10.56265 | -0.13903 | 0.054591 | 0.827795 |
| 6 | 119270600 | 119272199 | 13.81182 | 24.87746 | 30.49792 | 13.29872 | 0.113472 | 0.383111 | 0.827803 |
| 9 | 48080200 | 48082199 | 18.05397 | 31.48046 | 44.63813 | 19.64285 | -0.11422 | 0.747027 | 0.827804 |
| 13 | 3507200 | 3507799 | 8.494502 | 16.11881 | 24.95751 | 10.05952 | -0.13524 | 0.012765 | 0.827816 |
| 2 | 150023200 | 150024799 | 23.66753 | 40.78949 | 39.86671 | 17.48843 | 0.417567 | 0.892321 | 0.827825 |
| 10 | 22423800 | 22431399 | 92.80177 | 144.7894 | 165.6077 | 68.72171 | 0.118378 | 2.690928 | 0.827829 |
| 8 | 126515200 | 126518399 | 20.17262 | 34.39003 | 49.01868 | 21.48826 | -0.12719 | 0.987952 | 0.827868 |
| X | 7452000 | 7458199 | 55.02392 | 88.25595 | 100.3593 | 42.4102 | 0.162884 | 1.973997 | 0.827873 |
| 1 | 117373600 | 117375399 | 26.75715 | 44.62786 | 62.69935 | 27.26077 | -0.10775 | 1.231379 | 0.827884 |
| 4 | 145025200 | 145028399 | 31.48828 | 51.70573 | 61.12618 | 26.33 | 0.136736 | 1.420169 | 0.827887 |
| 5 | 138001200 | 138006199 | 67.85327 | 107.7548 | 125.4912 | 52.31337 | 0.112883 | 2.293802 | 0.827888 |
| 14 | 58286800 | 58292799 | 60.17764 | 95.84333 | 113.9609 | 47.67232 | 0.087273 | 2.197005 | 0.827894 |
| 8 | 30496000 | 30496999 | 16.0295 | 28.3764 | 34.35639 | 15.19261 | 0.116826 | 0.517646 | 0.827897 |
| 3 | 32902200 | 32903599 | 13.36018 | 23.87357 | 34.07223 | 14.96282 | -0.11163 | 0.435564 | 0.827904 |

| | | | | | | | | | |
|---|---|---|---|---|---|---|---|---|---|
| 3 | 112567200 | 112569399 | 20.49476 | 35.5016 | 43.14204 | 18.92816 | 0.106273 | 0.840144 | 0.827955 |
| 2 | 76614000 | 76617199 | 40.58115 | 65.70663 | 79.07123 | 33.83037 | 0.09706 | 1.669704 | 0.82796 |
| Y | 257600 | 259399 | 11.73773 | 20.66579 | 33.9989 | 15.05093 | -0.32582 | 0.55155 | 0.827962 |
| 9 | 8358000 | 8363199 | 46.88715 | 75.70214 | 87.89731 | 37.49327 | 0.144232 | 1.756096 | 0.827984 |
| 1 | 16935200 | 16936399 | 13.85192 | 24.46894 | 35.13645 | 15.54423 | -0.12907 | 0.529551 | 0.828019 |
| 1 | 120227600 | 120228999 | 16.12792 | 28.36677 | 34.14548 | 15.11886 | 0.121845 | 0.590401 | 0.828036 |
| 18 | 70277600 | 70282599 | 58.70854 | 93.84595 | 108.5694 | 45.65003 | 0.132422 | 2.078765 | 0.828048 |
| 1 | 69879000 | 69879799 | 8.292587 | 15.77712 | 24.98697 | 10.0155 | -0.16093 | -0.00928 | 0.828049 |
| 2 | 47089800 | 47090599 | 9.740565 | 17.85755 | 32.94442 | 14.04729 | -0.45097 | 0.257125 | 0.82808 |
| 6 | 102708000 | 102716799 | 103.5365 | 159.1982 | 208.5063 | 86.58549 | -0.09309 | 2.960103 | 0.828106 |
| 4 | 26034200 | 26034999 | 10.70413 | 20.2783 | 25.95973 | 10.49997 | 0.132527 | 0.047946 | 0.828115 |
| 3 | 87472600 | 87473799 | 9.497188 | 17.67053 | 26.85753 | 11.22078 | -0.15159 | 0.157054 | 0.828123 |
| 13 | 64396000 | 64396999 | 9.497188 | 17.67053 | 26.85753 | 11.22078 | -0.15159 | 0.157054 | 0.828123 |
| 16 | 70953400 | 70953799 | 10.64557 | 20.17119 | 25.43728 | 10.28453 | 0.154947 | 0.046756 | 0.82814 |
| 10 | 37727400 | 37732799 | 113.4291 | 172.8305 | 222.7814 | 92.34738 | -0.08033 | 3.1447 | 0.82814 |
| 14 | 103294400 | 103299399 | 57.06523 | 90.76103 | 121.4388 | 50.70362 | -0.08489 | 2.239756 | 0.828152 |
| 2 | 40008600 | 40009599 | 16.10586 | 28.16889 | 39.90586 | 17.5974 | -0.11247 | 0.685757 | 0.828153 |
| 18 | 38637200 | 38640399 | 26.10679 | 43.94109 | 53.39083 | 23.33228 | 0.100999 | 1.127059 | 0.828164 |
| 1 | 16033400 | 16044199 | 146.0636 | 220.3886 | 250.8292 | 103.6618 | 0.087364 | 3.351135 | 0.828171 |
| 10 | 104329400 | 104329999 | 7.702977 | 14.81315 | 23.59748 | 9.203703 | -0.13274 | -0.10666 | 0.828196 |
| 3 | 98618600 | 98619599 | 13.07499 | 23.88074 | 29.5461 | 12.66111 | 0.116353 | 0.294695 | 0.828235 |
| 12 | 93896000 | 93897399 | 15.45001 | 27.29412 | 33.04611 | 14.61923 | 0.115881 | 0.53569 | 0.828244 |
| 7 | 85526600 | 85530799 | 58.18438 | 93.3345 | 86.92794 | 36.7399 | 0.450458 | 1.965358 | 0.828285 |
| 11 | 48641200 | 48645199 | 36.407 | 59.24744 | 71.67938 | 30.76906 | 0.095793 | 1.562344 | 0.828289 |
| 13 | 40982000 | 40985399 | 49.09374 | 78.74139 | 94.31563 | 39.85408 | 0.087967 | 1.961426 | 0.828295 |
| 7 | 82317000 | 82319599 | 27.93446 | 46.39918 | 56.12038 | 24.33373 | 0.107469 | 1.273262 | 0.828334 |
| 8 | 57280200 | 57281599 | 13.67544 | 23.90574 | 43.64045 | 19.23584 | -0.47635 | 0.688721 | 0.828336 |
| 7 | 67315200 | 67316799 | 15.32772 | 26.83908 | 38.39067 | 16.94586 | -0.12624 | 0.660986 | 0.828343 |
| 6 | 103831800 | 103833199 | 15.5916 | 27.68582 | 41.28468 | 18.22679 | -0.18001 | 0.503324 | 0.828346 |
| 4 | 143951400 | 143956999 | 119.8863 | 183.1017 | 210.8284 | 87.44204 | 0.085071 | 3.098029 | 0.828386 |
| 7 | 31721600 | 31723399 | 65.69651 | 103.8967 | 120.9532 | 50.26108 | 0.108121 | 2.402715 | 0.828391 |
| 19 | 9245200 | 9246399 | 10.66419 | 19.94735 | 25.23705 | 10.4495 | 0.121331 | 0.128178 | 0.828401 |
| 19 | 60429600 | 60431599 | 23.28502 | 39.72513 | 57.78073 | 25.33077 | -0.15619 | 1.005299 | 0.828404 |
| 5 | 54460000 | 54460599 | 11.68732 | 21.71702 | 27.14555 | 11.36457 | 0.125554 | 0.181138 | 0.828414 |
| 12 | 25686000 | 25687799 | 11.68732 | 21.71702 | 27.14555 | 11.36457 | 0.125554 | 0.181138 | 0.828414 |
| 10 | 73538400 | 73538999 | 7.726387 | 14.98648 | 20.53704 | 7.7447 | 0.129219 | -0.22525 | 0.828458 |
| 12 | 59370600 | 59371399 | 14.78868 | 26.19805 | 31.54303 | 13.94658 | 0.125418 | 0.49844 | 0.82846 |
| 15 | 46409200 | 46412399 | 50.01203 | 80.59765 | 78.42986 | 33.37477 | 0.39543 | 1.808774 | 0.828528 |
| 12 | 21043400 | 21124999 | 2172.6 | 2614.415 | 2671.552 | 1129.008 | -0.08126 | 6.977468 | 0.828555 |
| 8 | 24815600 | 24818599 | 22.53275 | 38.08985 | 53.30414 | 23.32496 | -0.10184 | 1.077432 | 0.828558 |
| 7 | 139363000 | 139364199 | 9.754504 | 18.14079 | 27.31164 | 11.41956 | -0.13925 | 0.161638 | 0.82856 |
| 14 | 5086800 | 5091599 | 80.21172 | 125.522 | 147.3996 | 61.1471 | 0.083801 | 2.627897 | 0.82857 |
| 1 | 37784400 | 37786799 | 25.98963 | 43.79671 | 52.87676 | 23.11608 | 0.110171 | 1.113877 | 0.828575 |
| 15 | 7292800 | 7294999 | 16.57744 | 28.98263 | 40.92197 | 18.03967 | -0.10774 | 0.695636 | 0.828577 |
| 17 | 40230200 | 40232599 | 40.37385 | 65.65971 | 65.64355 | 28.17136 | 0.369631 | 1.577825 | 0.828583 |
| 13 | 78466000 | 78467599 | 18.19799 | 32.09653 | 31.18822 | 13.8018 | 0.430434 | 0.564771 | 0.828615 |
| 5 | 15801400 | 15801799 | 12.56327 | 22.84397 | 27.21683 | 11.72822 | 0.165799 | 0.314225 | 0.828632 |

| | | | | | | | | | |
|---|---|---|---|---|---|---|---|---|---|
| 13 | 113437600 | 113447799 | 203.6004 | 298.3718 | 371.3699 | 152.9701 | -0.08027 | 3.877368 | 0.828634 |
| 12 | 3571000 | 3574199 | 27.22582 | 45.26076 | 53.95461 | 23.40846 | 0.128422 | 1.25861 | 0.828645 |
| 9 | 20302200 | 20313399 | 232.0896 | 338.2622 | 373.2801 | 153.8252 | 0.085678 | 3.96436 | 0.828654 |
| 6 | 56200800 | 56203199 | 24.02509 | 41.69155 | 38.75286 | 17.00337 | 0.491789 | 0.834116 | 0.828665 |
| 9 | 80407800 | 80408599 | 11.42062 | 20.29639 | 37.28928 | 16.45129 | -0.47831 | 0.464741 | 0.828703 |
| 8 | 8964600 | 8966199 | 22.68635 | 39.72481 | 43.33293 | 19.12766 | 0.26282 | 0.707939 | 0.828739 |
| 4 | 41481000 | 41484399 | 22.35402 | 37.68304 | 53.30261 | 23.30467 | -0.11741 | 1.099996 | 0.828748 |
| 12 | 90942400 | 90943599 | 13.10018 | 23.68817 | 29.02047 | 12.59271 | 0.118936 | 0.351547 | 0.828752 |
| 9 | 32821400 | 32822399 | 10.89932 | 20.33092 | 30.55851 | 12.7122 | -0.13115 | 0.160212 | 0.828753 |
| 16 | 35597200 | 35598799 | 15.23687 | 26.92725 | 32.51339 | 14.38253 | 0.11991 | 0.528526 | 0.82878 |
| 9 | 31726000 | 31726799 | 12.91282 | 23.13615 | 33.1239 | 14.5135 | -0.11375 | 0.414947 | 0.828783 |
| 5 | 83285600 | 83286399 | 14.15117 | 25.76347 | 24.27882 | 10.44583 | 0.502775 | 0.308142 | 0.828809 |
| 8 | 83030800 | 83031799 | 17.46633 | 30.27717 | 43.23308 | 18.96851 | -0.12509 | 0.82086 | 0.828814 |
| 3 | 157061400 | 157061999 | 8.261167 | 15.83706 | 25.17218 | 9.906276 | -0.14107 | -0.06373 | 0.828828 |
| 8 | 31224400 | 31225799 | 11.21059 | 20.66809 | 30.31147 | 12.82644 | -0.11438 | 0.230033 | 0.828862 |
| 9 | 72726600 | 72728199 | 16.70465 | 29.32639 | 35.60948 | 15.75286 | 0.108854 | 0.621401 | 0.828883 |
| 18 | 83284200 | 83284799 | 19.30861 | 33.85028 | 32.05698 | 14.16449 | 0.465839 | 0.653632 | 0.828896 |
| 10 | 88496800 | 88498599 | 15.79994 | 27.88415 | 40.17296 | 17.77566 | -0.13453 | 0.560069 | 0.828897 |
| 12 | 87612200 | 87615799 | 48.89656 | 78.84846 | 90.88715 | 38.70655 | 0.152646 | 1.796381 | 0.828899 |
| 1 | 91606800 | 91607599 | 11.7898 | 21.46631 | 31.23766 | 13.43833 | -0.11959 | 0.308991 | 0.828909 |
| 1 | 139709400 | 139712599 | 45.58954 | 73.10192 | 99.10077 | 41.85948 | -0.09073 | 1.968003 | 0.828931 |
| 12 | 61080000 | 61080799 | 10.38227 | 19.39669 | 28.94732 | 12.01017 | -0.11786 | 0.141733 | 0.828936 |
| 4 | 143349200 | 143372799 | 471.3177 | 647.3715 | 667.1444 | 277.4119 | 0.098777 | 4.856963 | 0.82894 |
| 6 | 57319000 | 57320799 | 38.65723 | 62.38325 | 85.49817 | 36.44639 | -0.0953 | 1.755568 | 0.828944 |
| 6 | 9329600 | 9330599 | 12.52456 | 23.16062 | 28.71918 | 12.10265 | 0.130574 | 0.221154 | 0.828959 |
| 12 | 69517400 | 69518599 | 15.28197 | 26.99176 | 38.59893 | 17.07643 | -0.12339 | 0.545129 | 0.82896 |
| Y | 43763200 | 43763799 | 5.264446 | 10.24069 | 19.05247 | 7.037198 | -0.27953 | -0.33518 | 0.828962 |
| 14 | 76595600 | 76597399 | 21.64746 | 36.89369 | 51.70132 | 22.66372 | -0.10288 | 0.999647 | 0.829014 |
| 3 | 108434800 | 108436799 | 19.5985 | 34.06055 | 41.51524 | 18.23062 | 0.102663 | 0.796581 | 0.829016 |
| 5 | 109030600 | 109034999 | 80.34655 | 126.2571 | 147.8373 | 61.32356 | 0.092584 | 2.552792 | 0.829035 |
| 5 | 46559600 | 46561599 | 14.70657 | 26.3854 | 31.99454 | 14.00283 | 0.125657 | 0.417288 | 0.829038 |
| 3 | 20438400 | 20440199 | 24.1963 | 40.22718 | 68.01653 | 29.51068 | -0.37421 | 1.26332 | 0.829072 |
| 12 | 48577200 | 48580399 | 39.87843 | 64.53473 | 77.78399 | 33.26667 | 0.094067 | 1.677496 | 0.829091 |
| X | 103852400 | 103856399 | 41.28795 | 68.01307 | 56.65862 | 24.43842 | 0.642192 | 1.40738 | 0.829096 |
| 11 | 33510200 | 33513199 | 37.13592 | 60.88073 | 60.85254 | 26.19415 | 0.377322 | 1.455664 | 0.829097 |
| 12 | 44055400 | 44057199 | 20.68545 | 36.22688 | 41.73545 | 18.42499 | 0.184292 | 0.691177 | 0.82913 |
| 12 | 88761200 | 88762999 | 22.81045 | 38.16921 | 54.83455 | 23.89026 | -0.13973 | 1.177495 | 0.829143 |
| 10 | 32137200 | 32137999 | 7.142141 | 13.86658 | 22.86161 | 8.576338 | -0.12805 | -0.24257 | 0.829152 |
| 9 | 25410600 | 25412399 | 13.48342 | 24.40024 | 30.03222 | 13.01898 | 0.113239 | 0.352896 | 0.829197 |
| 10 | 96554000 | 96555599 | 16.27367 | 28.37253 | 40.5346 | 17.83566 | -0.12487 | 0.734423 | 0.829249 |
| 9 | 53107800 | 53109999 | 28.49289 | 48.20803 | 55.68779 | 24.37461 | 0.173943 | 1.093645 | 0.829271 |
| 2 | 69018000 | 69018799 | 5.807935 | 11.29404 | 19.57112 | 7.248779 | -0.18142 | -0.31428 | 0.829289 |
| 6 | 142658200 | 142662999 | 55.71831 | 88.95906 | 106.2554 | 44.60107 | 0.084568 | 2.113262 | 0.829296 |
| 9 | 23654600 | 23656799 | 35.5873 | 58.4574 | 58.60271 | 25.24682 | 0.37418 | 1.425436 | 0.8293 |
| 3 | 16389800 | 16399399 | 21.59784 | 37.63819 | 36.5793 | 16.09077 | 0.428233 | 0.760044 | 0.829318 |
| 4 | 86554800 | 86557999 | 53.92982 | 85.96467 | 115.3329 | 48.30227 | -0.08534 | 2.164173 | 0.829368 |
| 5 | 88492400 | 88497599 | 66.09554 | 104.8841 | 122.7928 | 51.14976 | 0.104233 | 2.312433 | 0.829372 |

| | | | | | | | | | |
|---|---|---|---|---|---|---|---|---|---|
| 2 | 88608400 | 88610999 | 23.66901 | 41.42231 | 43.86726 | 19.35437 | 0.305322 | 0.722058 | 0.829374 |
| 16 | 25178600 | 25179599 | 13.44544 | 25.39273 | 29.31154 | 11.93365 | 0.272169 | 0.099726 | 0.829386 |
| 18 | 75898600 | 75900599 | 21.09588 | 36.19739 | 51.16049 | 22.43885 | -0.11358 | 0.937944 | 0.82939 |
| 8 | 100401800 | 100402799 | 16.40583 | 28.72866 | 40.93207 | 18.0686 | -0.12054 | 0.671955 | 0.829422 |
| 3 | 12473800 | 12474799 | 24.38696 | 41.68489 | 37.08021 | 16.25828 | 0.551024 | 0.967063 | 0.829452 |
| 17 | 18296000 | 18299599 | 59.9009 | 95.60011 | 112.1189 | 47.03661 | 0.110289 | 2.131527 | 0.829458 |
| 3 | 66187400 | 66188999 | 19.74378 | 34.7736 | 39.46358 | 17.46887 | 0.206909 | 0.6042 | 0.829469 |
| 14 | 66589800 | 66594199 | 49.84078 | 79.85315 | 94.8258 | 40.00381 | 0.098532 | 1.994847 | 0.829476 |
| 1 | 55334600 | 55337799 | 66.26415 | 104.5269 | 139.0015 | 57.68713 | -0.0865 | 2.466549 | 0.829483 |
| 12 | 81403600 | 81404799 | 37.83016 | 61.38197 | 74.15486 | 31.77639 | 0.094334 | 1.620942 | 0.829486 |
| 13 | 51735000 | 51740999 | 52.18398 | 83.40341 | 112.6276 | 47.34594 | -0.09054 | 2.080187 | 0.829519 |
| 17 | 67485400 | 67486799 | 17.57985 | 30.9258 | 36.89419 | 16.33537 | 0.134428 | 0.598186 | 0.829528 |
| 16 | 93580200 | 93581599 | 17.52022 | 30.51271 | 43.08618 | 18.94547 | -0.10841 | 0.75866 | 0.829547 |
| 18 | 76886600 | 76893799 | 66.16299 | 104.7054 | 139.74 | 58.08768 | -0.08797 | 2.389758 | 0.829554 |
| 6 | 123654000 | 123656999 | 43.71349 | 70.38176 | 84.43037 | 35.87989 | 0.092548 | 1.831674 | 0.829595 |
| 10 | 18430600 | 18432799 | 19.64162 | 34.13187 | 49.57478 | 21.76835 | -0.14938 | 0.80798 | 0.829602 |
| 5 | 126089600 | 126090999 | 14.63591 | 26.00216 | 31.51593 | 13.90797 | 0.118063 | 0.475865 | 0.829636 |
| 2 | 169841600 | 169842999 | 11.28117 | 20.70009 | 30.28251 | 12.8969 | -0.11727 | 0.258579 | 0.829637 |
| 14 | 123946000 | 123947199 | 17.97845 | 31.27323 | 44.19033 | 19.41633 | -0.10942 | 0.782076 | 0.829642 |
| 2 | 21984800 | 21985199 | 6.649529 | 12.94665 | 17.94107 | 6.578791 | 0.15043 | -0.34637 | 0.829643 |
| 3 | 138372600 | 138379199 | 71.10242 | 111.2226 | 174.3505 | 72.33036 | -0.33291 | 2.633912 | 0.829646 |
| 6 | 10637200 | 10638799 | 12.72016 | 23.01274 | 33.16547 | 14.38215 | -0.11344 | 0.355293 | 0.82965 |
| 7 | 133428200 | 133430199 | 19.9779 | 34.60076 | 42.11521 | 18.47756 | 0.103918 | 0.83761 | 0.829655 |
| 4 | 19188400 | 19189199 | 12.61449 | 23.40756 | 22.01659 | 9.234627 | 0.531421 | 0.185524 | 0.829679 |
| 17 | 53737200 | 53738399 | 13.76321 | 24.79429 | 30.46109 | 13.27972 | 0.11051 | 0.380806 | 0.829695 |
| 14 | 40219600 | 40219999 | 5.944264 | 11.6033 | 20.55424 | 7.276105 | -0.15356 | -0.46756 | 0.829722 |
| 3 | 51247200 | 51251799 | 33.8949 | 54.92512 | 76.96949 | 32.95082 | -0.121 | 1.639571 | 0.829726 |
| 5 | 149238000 | 149239199 | 18.87666 | 32.63598 | 45.81314 | 20.0989 | -0.10154 | 0.852913 | 0.829728 |
| 17 | 49856600 | 49858199 | 15.81464 | 27.63427 | 39.4749 | 17.39394 | -0.12429 | 0.699167 | 0.829766 |
| 19 | 18786000 | 18786599 | 8.094741 | 15.52385 | 24.63033 | 9.682566 | -0.13702 | -0.07058 | 0.829804 |
| 4 | 110727400 | 110729799 | 38.19148 | 61.81648 | 85.01475 | 36.36421 | -0.09556 | 1.675852 | 0.829861 |
| 4 | 40624200 | 40625199 | 8.50617 | 16.10597 | 24.93442 | 10.09644 | -0.1408 | 0.028101 | 0.829873 |
| 6 | 102381000 | 102382199 | 16.85268 | 29.5328 | 35.70427 | 15.77066 | 0.115154 | 0.651388 | 0.829874 |
| 6 | 43742600 | 43745999 | 29.1455 | 48.49278 | 58.5108 | 25.3999 | 0.111044 | 1.257456 | 0.829889 |
| 13 | 33936200 | 33938999 | 36.6082 | 59.60076 | 72.25365 | 31.02048 | 0.093318 | 1.557673 | 0.829907 |
| 9 | 12840000 | 12842799 | 37.37456 | 61.13416 | 61.23949 | 26.33673 | 0.372619 | 1.48655 | 0.82992 |
| 3 | 96396400 | 96403799 | 68.90997 | 108.697 | 143.8902 | 59.71574 | -0.07999 | 2.469934 | 0.829966 |
| 5 | 145471800 | 145473599 | 35.39301 | 57.50967 | 79.30743 | 34.02016 | -0.09423 | 1.590198 | 0.829975 |
| 1 | 94028600 | 94029599 | 22.98953 | 38.55027 | 54.84074 | 23.92484 | -0.12564 | 1.156274 | 0.829978 |
| 9 | 16782200 | 16785399 | 50.27408 | 80.47447 | 95.8342 | 40.37417 | 0.09296 | 2.025399 | 0.830006 |
| 6 | 3189600 | 3215199 | 701.2642 | 924.0802 | 1035.455 | 433.1451 | -0.08204 | 5.515594 | 0.830007 |
| 2 | 98924600 | 98926199 | 19.86533 | 34.20954 | 47.9395 | 21.02923 | -0.10029 | 0.895315 | 0.830017 |
| 7 | 105967600 | 105979399 | 279.0586 | 396.4619 | 481.0617 | 198.9612 | -0.08769 | 4.344046 | 0.83003 |
| 12 | 40903600 | 40909199 | 60.90841 | 96.96773 | 114.9522 | 48.05991 | 0.090957 | 2.209732 | 0.830042 |
| 2 | 85178000 | 85178999 | 14.58208 | 26.08127 | 31.91725 | 14.00547 | 0.109721 | 0.433377 | 0.830043 |
| 12 | 103546400 | 103548199 | 18.61955 | 32.83125 | 31.08169 | 13.7554 | 0.467597 | 0.569191 | 0.830054 |
| 5 | 43806400 | 43808399 | 20.72502 | 35.8623 | 43.64461 | 19.14803 | 0.103749 | 0.84967 | 0.830061 |

| | | | | | | | | | |
|---|---|---|---|---|---|---|---|---|---|
| X | 109136200 | 109137399 | 17.63685 | 31.74057 | 27.65783 | 12.07285 | 0.602256 | 0.409718 | 0.830112 |
| 1 | 49973400 | 49975399 | 27.30697 | 45.61877 | 55.57997 | 24.19009 | 0.097143 | 1.208803 | 0.83015 |
| 12 | 63166000 | 63167399 | 23.68525 | 38.96602 | 71.73323 | 30.92361 | -0.49908 | 1.391122 | 0.830166 |
| 1 | 29878400 | 29879599 | 13.3069 | 23.9446 | 34.58784 | 15.10029 | -0.12324 | 0.391731 | 0.830233 |
| 12 | 23516600 | 23518799 | 40.71876 | 66.11931 | 60.3939 | 25.89418 | 0.49778 | 1.604008 | 0.830238 |
| 5 | 126023600 | 126025599 | 22.58368 | 38.53304 | 46.89573 | 20.55903 | 0.100119 | 0.988028 | 0.830249 |
| 4 | 72596000 | 72598599 | 25.18623 | 41.22695 | 75.44697 | 32.45527 | -0.49353 | 1.460678 | 0.830261 |
| Y | 41496200 | 41496799 | 8.339613 | 15.97316 | 18.33197 | 7.232912 | 0.320781 | -0.07975 | 0.830262 |
| X | 146480400 | 146483399 | 32.61911 | 54.47835 | 62.11231 | 27.02851 | 0.19253 | 1.232898 | 0.830265 |
| 1 | 96111000 | 96112199 | 8.817185 | 16.85997 | 26.22663 | 10.397 | -0.12079 | -0.03424 | 0.830279 |
| 9 | 100383600 | 100384599 | 8.817185 | 16.85997 | 26.22663 | 10.397 | -0.12079 | -0.03424 | 0.830279 |
| 12 | 4778800 | 4779399 | 8.817185 | 16.85997 | 26.22663 | 10.397 | -0.12079 | -0.03424 | 0.830279 |
| 11 | 70014400 | 70016599 | 26.74468 | 44.95058 | 54.50984 | 23.811 | 0.103733 | 1.142008 | 0.830323 |
| 15 | 7451800 | 7454199 | 31.27509 | 51.30024 | 71.47457 | 30.77349 | -0.10016 | 1.444791 | 0.830324 |
| 10 | 112480200 | 112482199 | 18.56131 | 32.89258 | 37.30996 | 16.49527 | 0.210989 | 0.527742 | 0.830337 |
| 11 | 27091600 | 27092999 | 11.67168 | 21.7352 | 27.17252 | 11.33576 | 0.129289 | 0.167372 | 0.830373 |
| 17 | 92086200 | 92087799 | 17.43655 | 30.60838 | 37.12421 | 16.42161 | 0.110523 | 0.630727 | 0.830381 |
| 13 | 65498400 | 65500799 | 56.7573 | 90.42203 | 107.1729 | 44.8507 | 0.092333 | 2.18349 | 0.830386 |
| 13 | 77605000 | 77606799 | 16.38679 | 29.14802 | 34.83531 | 15.35714 | 0.139026 | 0.484619 | 0.830395 |
| 1 | 11139800 | 11141399 | 16.90029 | 30.13561 | 35.35193 | 15.55253 | 0.167767 | 0.473807 | 0.830405 |
| 13 | 112900400 | 112904399 | 35.04647 | 57.11698 | 79.6889 | 34.2255 | -0.10786 | 1.544786 | 0.83041 |
| 1 | 98816800 | 98817999 | 23.12645 | 39.25097 | 47.45551 | 20.79104 | 0.108659 | 1.039066 | 0.830422 |
| 4 | 97670800 | 97671999 | 19.37519 | 33.70149 | 40.94898 | 17.9917 | 0.107228 | 0.781585 | 0.830425 |
| 3 | 134726600 | 134726999 | 6.70002 | 13.05806 | 22.78746 | 8.252044 | -0.16249 | -0.37183 | 0.830448 |
| 10 | 63385000 | 63386799 | 15.81339 | 27.85958 | 39.67239 | 17.56025 | -0.11871 | 0.578612 | 0.83046 |
| 6 | 60897400 | 60898599 | 16.79436 | 30.14332 | 28.62119 | 12.52217 | 0.476598 | 0.42024 | 0.830476 |
| 9 | 46289800 | 46291399 | 16.39618 | 28.59368 | 41.09307 | 18.08441 | -0.13314 | 0.73502 | 0.830482 |
| X | 153376800 | 153378999 | 23.34737 | 40.49124 | 46.83578 | 20.54949 | 0.177466 | 0.847005 | 0.830487 |
| 2 | 171197200 | 171198599 | 18.67622 | 32.50705 | 46.27312 | 20.33936 | -0.12012 | 0.776898 | 0.830489 |
| 18 | 90109200 | 90111399 | 23.44945 | 38.80194 | 68.21696 | 29.50135 | -0.43079 | 1.316982 | 0.830499 |
| X | 77008400 | 77013199 | 69.7707 | 110.8624 | 126.7367 | 52.86244 | 0.140424 | 2.282151 | 0.830526 |
| 13 | 10753400 | 10756399 | 33.43199 | 55.43717 | 54.75699 | 23.71499 | 0.398591 | 1.306286 | 0.830552 |
| 10 | 14859200 | 14861399 | 21.77215 | 37.86278 | 44.62286 | 19.60327 | 0.151029 | 0.801897 | 0.830602 |
| 12 | 11548000 | 11548999 | 16.63027 | 29.01969 | 41.01702 | 18.05789 | -0.10932 | 0.725316 | 0.830603 |
| X | 75667200 | 75671599 | 69.21571 | 110.3068 | 121.6629 | 50.93814 | 0.196499 | 2.190229 | 0.830608 |
| 8 | 126813800 | 126818799 | 58.27628 | 92.78931 | 125.1434 | 52.33006 | -0.09464 | 2.207259 | 0.830611 |
| 4 | 144156800 | 144160399 | 43.09269 | 69.3855 | 94.63184 | 40.2179 | -0.09215 | 1.833207 | 0.830621 |
| 17 | 59000400 | 59003199 | 27.26156 | 46.11347 | 54.54489 | 23.8726 | 0.13986 | 1.089524 | 0.830634 |
| 8 | 25709600 | 25710999 | 12.42824 | 22.33207 | 32.15055 | 14.0529 | -0.11923 | 0.391767 | 0.830645 |
| 7 | 89256800 | 89259799 | 25.21393 | 41.63271 | 70.65807 | 30.51207 | -0.38043 | 1.347253 | 0.830662 |
| 10 | 30175800 | 30179199 | 30.72722 | 51.07411 | 61.35861 | 26.61804 | 0.117208 | 1.277094 | 0.830663 |
| 3 | 18342800 | 18346799 | 61.81095 | 98.13873 | 116.3811 | 48.50549 | 0.086525 | 2.293615 | 0.830667 |
| 17 | 53619200 | 53620799 | 22.2575 | 37.9362 | 45.86246 | 20.10433 | 0.109502 | 0.995323 | 0.830737 |
| 4 | 147281000 | 147288799 | 134.0292 | 203.5072 | 194.2273 | 80.42706 | 0.348804 | 3.218508 | 0.830744 |
| 14 | 52870600 | 52872599 | 20.18751 | 34.60697 | 48.4788 | 21.2611 | -0.10099 | 0.937898 | 0.830745 |
| 15 | 43161600 | 43161999 | 10.91902 | 20.37014 | 25.67828 | 10.67986 | 0.121126 | 0.14695 | 0.83075 |
| X | 46513200 | 46515199 | 15.26815 | 27.21926 | 32.53535 | 14.31608 | 0.141065 | 0.462775 | 0.830763 |

| | | | | | | | | | |
|---|---|---|---|---|---|---|---|---|---|
| X | 31662000 | 31663999 | 50.84419 | 83.45763 | 51.88993 | 22.34791 | 1.061056 | 1.470275 | 0.83078 |
| 19 | 8079000 | 8088199 | 125.5253 | 191.2303 | 219.4275 | 90.95275 | 0.087167 | 3.150237 | 0.830782 |
| X | 153323800 | 153324399 | 15.43953 | 27.65812 | 33.40593 | 14.63939 | 0.129828 | 0.432757 | 0.830789 |
| 6 | 7189000 | 7189599 | 7.628512 | 14.668 | 23.60121 | 9.208683 | -0.14668 | -0.1023 | 0.830817 |
| 17 | 59874000 | 59875999 | 15.07897 | 26.26498 | 46.29556 | 20.35855 | -0.42673 | 0.750556 | 0.830819 |
| 3 | 15451600 | 15454599 | 21.8246 | 37.85676 | 37.35528 | 16.38995 | 0.405825 | 0.825498 | 0.830834 |
| 11 | 116683000 | 116687399 | 39.76006 | 64.20732 | 76.49491 | 32.63755 | 0.107533 | 1.733282 | 0.83086 |
| 8 | 81823600 | 81824399 | 7.271785 | 14.04077 | 22.60054 | 8.705737 | -0.13025 | -0.15353 | 0.830877 |
| 11 | 23170200 | 23170599 | 7.271785 | 14.04077 | 22.60054 | 8.705737 | -0.13025 | -0.15353 | 0.830877 |
| 5 | 102274400 | 102274799 | 5.645224 | 11.0251 | 19.42271 | 6.816276 | -0.13449 | -0.50699 | 0.830891 |
| 7 | 40568400 | 40572399 | 35.91714 | 57.95462 | 98.93253 | 42.16684 | -0.41164 | 1.759425 | 0.830901 |
| 15 | 94750000 | 94753799 | 38.79445 | 62.73314 | 86.31424 | 36.89168 | -0.09754 | 1.696309 | 0.830917 |
| 5 | 133537000 | 133538199 | 11.14652 | 20.68502 | 30.98602 | 12.99552 | -0.13526 | 0.196346 | 0.830919 |
| 10 | 33686600 | 33688199 | 24.56379 | 40.99716 | 57.62006 | 25.063 | -0.10826 | 1.212006 | 0.830951 |
| Y | 68112400 | 68112999 | 4.986956 | 9.709154 | 18.55361 | 6.806014 | -0.30819 | -0.3644 | 0.830957 |
| Y | 69246000 | 69246599 | 4.986956 | 9.709154 | 18.55361 | 6.806014 | -0.30819 | -0.3644 | 0.830957 |
| X | 76535600 | 76537999 | 28.60577 | 48.54967 | 55.10812 | 24.14363 | 0.19934 | 1.068627 | 0.83097 |
| 17 | 83727200 | 83729399 | 26.01607 | 44.0255 | 52.83216 | 23.12598 | 0.119199 | 1.081253 | 0.830974 |
| X | 105854800 | 105855399 | 7.329479 | 14.23652 | 24.09949 | 9.008884 | -0.16094 | -0.2478 | 0.830981 |
| 11 | 5566200 | 5567599 | 15.34342 | 27.02596 | 38.16269 | 16.89271 | -0.10665 | 0.576222 | 0.830998 |
| 16 | 94872200 | 94873599 | 10.81044 | 19.86151 | 29.2971 | 12.45571 | -0.12718 | 0.245977 | 0.83102 |
| 8 | 9993400 | 9994999 | 11.89517 | 21.48565 | 31.15012 | 13.53536 | -0.12379 | 0.354027 | 0.831046 |
| 16 | 69130600 | 69132599 | 24.27835 | 41.11453 | 58.83924 | 25.76107 | -0.13396 | 1.075164 | 0.831062 |
| 15 | 61817600 | 61819199 | 17.99213 | 31.59092 | 38.03271 | 16.82675 | 0.121012 | 0.629829 | 0.831065 |
| 4 | 83274000 | 83280599 | 71.45719 | 113.3004 | 130.7958 | 54.43924 | 0.123338 | 2.342755 | 0.831103 |
| 15 | 60911600 | 60912199 | 8.60422 | 16.48278 | 21.37522 | 8.429513 | 0.146969 | -0.06669 | 0.831105 |
| 8 | 87137600 | 87140599 | 25.88084 | 43.74464 | 52.86114 | 23.13033 | 0.109073 | 1.088776 | 0.831107 |
| 5 | 145666600 | 145720399 | 1313.487 | 1651.332 | 1775.784 | 746.3846 | -0.09074 | 6.280111 | 0.831127 |
| 15 | 93057200 | 93061199 | 34.36815 | 55.86152 | 76.9581 | 33.01985 | -0.09239 | 1.579738 | 0.831128 |
| 12 | 40335600 | 40342599 | 78.01276 | 121.4736 | 188.0162 | 78.04226 | -0.31978 | 2.71973 | 0.831146 |
| X | 160560600 | 160561999 | 13.31566 | 23.74741 | 34.17404 | 15.03236 | -0.12487 | 0.450469 | 0.831159 |
| X | 126452400 | 126453399 | 16.53933 | 29.45616 | 34.92271 | 15.37944 | 0.151746 | 0.480252 | 0.831169 |
| 7 | 84466000 | 84467599 | 16.84111 | 29.67408 | 35.93751 | 15.90665 | 0.113832 | 0.575999 | 0.831171 |
| 16 | 20764400 | 20769199 | 66.24995 | 104.991 | 124.716 | 51.88254 | 0.081488 | 2.352915 | 0.831172 |
| 9 | 41194000 | 41195199 | 14.12684 | 25.79671 | 31.02497 | 13.29817 | 0.15653 | 0.305363 | 0.831226 |
| 17 | 47709400 | 47712199 | 24.56752 | 40.88904 | 58.13643 | 25.24301 | -0.12485 | 1.241957 | 0.831233 |
| 5 | 32688800 | 32700199 | 217.0754 | 317.4446 | 353.3625 | 145.572 | 0.077831 | 3.911181 | 0.831242 |
| X | 75666400 | 75669799 | 49.53212 | 80.18886 | 87.06844 | 37.2483 | 0.244931 | 1.694206 | 0.831267 |
| 18 | 3785200 | 3785799 | 8.985418 | 17.17524 | 26.7848 | 10.62969 | -0.12588 | -0.02749 | 0.831267 |
| 10 | 87882200 | 87885199 | 23.69802 | 40.35079 | 49.02468 | 21.48878 | 0.102051 | 1.01261 | 0.831284 |
| 3 | 101929200 | 101931399 | 16.08811 | 28.18624 | 39.81654 | 17.58352 | -0.10844 | 0.657258 | 0.831311 |
| 7 | 73518400 | 73519399 | 10.30507 | 19.08851 | 28.33351 | 11.91272 | -0.12454 | 0.193386 | 0.831327 |
| 18 | 7932200 | 7933399 | 13.65012 | 24.55152 | 30.02357 | 13.11243 | 0.115596 | 0.389427 | 0.831328 |
| 6 | 10707000 | 10708599 | 17.15714 | 30.01095 | 42.4769 | 18.73294 | -0.11117 | 0.694685 | 0.831346 |
| 9 | 36402000 | 36437199 | 491.4946 | 673.161 | 690.5025 | 287.254 | 0.101325 | 4.90157 | 0.831379 |
| 8 | 62660600 | 62662799 | 22.75328 | 38.30207 | 53.81487 | 23.51859 | -0.10775 | 1.11587 | 0.83139 |
| 12 | 25261800 | 25263599 | 14.69096 | 25.9226 | 36.6401 | 16.21299 | -0.10699 | 0.550344 | 0.831398 |

| | | | | | | | | | |
|---|---|---|---|---|---|---|---|---|---|
| 17 | 56147200 | 56149599 | 40.99629 | 66.47026 | 65.40684 | 28.01252 | 0.388616 | 1.635892 | 0.831421 |
| 13 | 60828400 | 60831799 | 27.43329 | 45.70218 | 55.58687 | 24.15056 | 0.099637 | 1.238011 | 0.831447 |
| 7 | 74736800 | 74738799 | 19.39915 | 33.95056 | 32.12748 | 14.17502 | 0.466944 | 0.681 | 0.831457 |
| 18 | 16073400 | 16077999 | 119.1648 | 181.3469 | 207.67 | 86.05177 | 0.088807 | 3.169721 | 0.831472 |
| 6 | 129833600 | 129834999 | 20.97312 | 34.97823 | 63.30648 | 27.52348 | -0.4718 | 1.215695 | 0.831478 |
| 4 | 139404000 | 139411599 | 103.8958 | 159.2029 | 205.9838 | 85.49773 | -0.07928 | 3.026047 | 0.831485 |
| 18 | 66396200 | 66399199 | 26.18762 | 44.04991 | 53.4183 | 23.33991 | 0.103731 | 1.13117 | 0.831494 |
| 13 | 114748600 | 114751599 | 27.11202 | 45.60039 | 54.57219 | 23.84337 | 0.12269 | 1.135138 | 0.831498 |
| 17 | 12392400 | 12393799 | 10.19914 | 18.80742 | 28.35258 | 11.994 | -0.1529 | 0.219169 | 0.831518 |
| 18 | 90570800 | 90573799 | 32.61138 | 54.2792 | 53.06804 | 23.04298 | 0.413674 | 1.260393 | 0.831521 |
| 10 | 122666000 | 122669799 | 53.32902 | 85.22245 | 101.5309 | 42.67334 | 0.089708 | 2.080231 | 0.831532 |
| 1 | 101588000 | 101589199 | 14.23912 | 25.62135 | 31.41485 | 13.71535 | 0.111932 | 0.392552 | 0.831546 |
| 6 | 145432400 | 145436199 | 25.934 | 43.46294 | 52.84507 | 23.04592 | 0.100006 | 1.167401 | 0.83158 |
| 11 | 66407600 | 66410399 | 19.3896 | 33.68602 | 40.99947 | 18.00027 | 0.104663 | 0.799026 | 0.831582 |
| 18 | 55571200 | 55571799 | 8.890818 | 17.02102 | 22.3839 | 8.843354 | 0.12421 | -0.05581 | 0.831588 |
| 1 | 103452800 | 103454199 | 14.68968 | 26.96753 | 31.70185 | 13.49246 | 0.196456 | 0.28216 | 0.831615 |
| 3 | 17061400 | 17063399 | 19.02131 | 32.96674 | 46.32928 | 20.32686 | -0.10247 | 0.831952 | 0.83163 |
| 5 | 88583200 | 88585799 | 60.49989 | 96.13708 | 114.0511 | 47.57398 | 0.087142 | 2.267895 | 0.831642 |
| 12 | 49598200 | 49602599 | 49.83951 | 80.03327 | 95.43613 | 40.40104 | 0.096439 | 1.927151 | 0.83167 |
| 6 | 41439600 | 41441799 | 15.6763 | 27.50435 | 38.79301 | 17.15239 | -0.10609 | 0.631794 | 0.831677 |
| 9 | 8221400 | 8223799 | 31.09291 | 51.17509 | 62.18616 | 26.81289 | 0.098533 | 1.393592 | 0.831681 |
| 10 | 58593000 | 58598799 | 58.52973 | 93.09308 | 124.6649 | 52.05275 | -0.08608 | 2.240596 | 0.831684 |
| 2 | 19378200 | 19379799 | 12.5039 | 22.63771 | 32.63517 | 14.1395 | -0.11314 | 0.347419 | 0.831704 |
| 12 | 26853800 | 26855399 | 11.20101 | 20.34841 | 30.15944 | 13.01106 | -0.14884 | 0.312103 | 0.831706 |
| 9 | 62990000 | 62991399 | 12.99372 | 23.591 | 29.07633 | 12.55321 | 0.115115 | 0.328701 | 0.831716 |
| 2 | 109986800 | 109989799 | 49.9906 | 79.68359 | 127.9361 | 53.5788 | -0.34403 | 2.166437 | 0.831738 |
| 5 | 73432400 | 73435799 | 27.38855 | 45.40343 | 63.18816 | 27.3613 | -0.09456 | 1.300913 | 0.831759 |
| 3 | 113647800 | 113649799 | 20.77279 | 36.08629 | 43.60122 | 19.1416 | 0.115147 | 0.810955 | 0.831778 |
| 7 | 64257000 | 64261599 | 41.99844 | 67.87505 | 81.45575 | 34.78921 | 0.098289 | 1.715491 | 0.831783 |
| 8 | 97939200 | 97942799 | 27.26675 | 45.62526 | 65.66094 | 28.60678 | -0.14246 | 1.201039 | 0.831784 |
| 15 | 95535800 | 95537799 | 15.02462 | 26.6732 | 32.36273 | 14.29048 | 0.115737 | 0.482597 | 0.831805 |
| 6 | 149254200 | 149257599 | 51.4934 | 82.29827 | 110.6526 | 46.51456 | -0.0843 | 2.078935 | 0.831838 |
| 5 | 18546800 | 18556199 | 126.3338 | 191.5041 | 245.4636 | 101.5927 | -0.07829 | 3.253806 | 0.831852 |
| 19 | 13923800 | 13930399 | 78.4608 | 123.4332 | 145.092 | 60.19364 | 0.088249 | 2.52836 | 0.831853 |
| 14 | 14754400 | 14756199 | 23.70195 | 40.24327 | 57.19963 | 25.06304 | -0.12401 | 1.047915 | 0.831855 |
| 14 | 124236600 | 124237799 | 16.95146 | 29.42949 | 42.1915 | 18.51544 | -0.13045 | 0.801219 | 0.831864 |
| 1 | 8330800 | 8332999 | 19.57916 | 34.08904 | 41.443 | 18.21976 | 0.106575 | 0.770248 | 0.831867 |
| 4 | 36205000 | 36206599 | 17.68429 | 31.07608 | 37.50165 | 16.60339 | 0.117616 | 0.614387 | 0.831877 |
| 15 | 94594200 | 94598199 | 38.11589 | 62.12824 | 74.31323 | 31.91837 | 0.113821 | 1.545755 | 0.831902 |
| 15 | 89509400 | 89509999 | 8.539675 | 16.44371 | 21.46656 | 8.34111 | 0.158021 | -0.11244 | 0.831909 |
| 8 | 124580400 | 124582399 | 17.0566 | 29.79055 | 41.97668 | 18.49088 | -0.10489 | 0.714722 | 0.831913 |
| 15 | 40136600 | 40137599 | 17.5102 | 30.63263 | 37.26336 | 16.43556 | 0.106303 | 0.685106 | 0.831988 |
| X | 131905400 | 131906799 | 19.367 | 34.76829 | 27.50815 | 12.03302 | 0.738713 | 0.4366 | 0.831992 |
| 13 | 26556200 | 26558399 | 18.4976 | 32.46718 | 31.40388 | 13.8899 | 0.435256 | 0.626723 | 0.832018 |
| 19 | 19140000 | 19140399 | 7.715306 | 14.88899 | 23.69794 | 9.150275 | -0.11776 | -0.13806 | 0.832021 |
| X | 129854000 | 129856799 | 45.28545 | 73.25849 | 68.77835 | 29.40734 | 0.453258 | 1.693375 | 0.832028 |
| 1 | 115235200 | 115236199 | 15.52955 | 27.94529 | 26.85257 | 11.72281 | 0.461781 | 0.392917 | 0.832041 |

| | | | | | | | | | |
|---|---|---|---|---|---|---|---|---|---|
| 5 | 136842000 | 136846799 | 52.64882 | 84.06064 | 112.8161 | 47.35677 | -0.08325 | 2.11136 | 0.832072 |
| 9 | 13137600 | 13141599 | 44.35186 | 71.52488 | 85.87625 | 36.57583 | 0.094064 | 1.78403 | 0.832074 |
| 9 | 115365800 | 115371999 | 86.12141 | 133.2789 | 175.4079 | 72.86004 | -0.09293 | 2.828424 | 0.832076 |
| 2 | 148792800 | 148799199 | 86.30892 | 135.5781 | 133.3871 | 55.327 | 0.343023 | 2.560021 | 0.832114 |
| 14 | 9028200 | 9028999 | 10.86026 | 20.22115 | 25.23581 | 10.53046 | 0.132105 | 0.159827 | 0.83212 |
| 10 | 50291600 | 50292599 | 13.20041 | 24.0307 | 35.95343 | 15.47024 | -0.16005 | 0.327937 | 0.832143 |
| 15 | 68175800 | 68178999 | 42.37072 | 68.49764 | 82.10118 | 35.07871 | 0.100587 | 1.708414 | 0.832158 |
| 18 | 38772200 | 38775599 | 36.07413 | 58.69858 | 71.05163 | 30.49824 | 0.094937 | 1.562928 | 0.832158 |
| 10 | 112799200 | 112802199 | 46.32912 | 73.92342 | 119.678 | 50.19197 | -0.35435 | 2.126595 | 0.832173 |
| 6 | 119492600 | 119494799 | 19.29069 | 33.89489 | 32.81691 | 14.5278 | 0.433914 | 0.618363 | 0.832176 |
| 5 | 109910200 | 109918599 | 139.6141 | 212.2363 | 235.6972 | 97.63357 | 0.131767 | 3.199973 | 0.832181 |
| 16 | 48359400 | 48363199 | 42.42342 | 68.10911 | 93.76386 | 39.67959 | -0.11076 | 1.922505 | 0.832192 |
| 10 | 121528600 | 121530999 | 20.36024 | 34.99133 | 41.66564 | 18.276 | 0.133123 | 0.906051 | 0.832245 |
| 9 | 6726000 | 6726999 | 12.70643 | 23.13851 | 28.50088 | 12.25882 | 0.119217 | 0.305981 | 0.832253 |
| 5 | 70467600 | 70471599 | 46.99228 | 75.75881 | 89.37145 | 38.04881 | 0.118884 | 1.798778 | 0.832313 |
| 5 | 80196800 | 80198799 | 23.1572 | 39.94558 | 47.38377 | 20.78651 | 0.139687 | 0.894168 | 0.832327 |
| 8 | 65609000 | 65609599 | 6.248592 | 12.17015 | 20.77622 | 7.589452 | -0.14159 | -0.3513 | 0.832331 |
| 5 | 141843600 | 141844399 | 9.644921 | 18.24548 | 23.26246 | 9.43762 | 0.1344 | 0.039392 | 0.832336 |
| 4 | 80604000 | 80604399 | 7.450392 | 14.39718 | 23.18573 | 8.897752 | -0.12558 | -0.16109 | 0.83234 |
| 1 | 29121800 | 29122199 | 5.509656 | 10.75323 | 18.70802 | 6.640668 | -0.13175 | -0.47334 | 0.832351 |
| 10 | 10381800 | 10382199 | 5.509656 | 10.75323 | 18.70802 | 6.640668 | -0.13175 | -0.47334 | 0.832351 |
| 10 | 84919200 | 84922199 | 28.02028 | 47.62303 | 53.95692 | 23.64722 | 0.202167 | 1.054484 | 0.832364 |
| 17 | 78876200 | 78879199 | 26.7086 | 44.53235 | 62.17492 | 27.02718 | -0.09877 | 1.233283 | 0.832408 |
| X | 12937200 | 12940199 | 45.26896 | 74.082 | 57.71833 | 24.82668 | 0.734772 | 1.499168 | 0.832431 |
| 14 | 118756800 | 118758799 | 13.12353 | 24.18176 | 20.94897 | 8.871166 | 0.639178 | 0.224694 | 0.832441 |
| 2 | 16843000 | 16844399 | 20.20126 | 34.98254 | 42.59241 | 18.68684 | 0.103339 | 0.838747 | 0.832466 |
| 6 | 131517200 | 131517999 | 9.689677 | 18.33061 | 28.17098 | 11.4284 | -0.13333 | 0.051434 | 0.832479 |
| 16 | 23567600 | 23568999 | 21.72641 | 37.76421 | 36.80349 | 16.16303 | 0.424086 | 0.799166 | 0.832481 |
| 10 | 53127200 | 53128999 | 23.87496 | 40.03832 | 56.01286 | 24.43746 | -0.10172 | 1.157034 | 0.832493 |
| 8 | 122850200 | 122853799 | 36.21872 | 58.62084 | 80.48511 | 34.42559 | -0.09317 | 1.670938 | 0.832505 |
| 3 | 85270400 | 85274199 | 33.90531 | 55.0529 | 76.0573 | 32.60876 | -0.09777 | 1.599433 | 0.832539 |
| 8 | 86958600 | 86958999 | 7.399747 | 14.37951 | 20.0609 | 7.471824 | 0.119988 | -0.27481 | 0.83255 |
| 13 | 95358200 | 95359999 | 14.5171 | 25.46594 | 36.84057 | 16.28664 | -0.14245 | 0.624054 | 0.832564 |
| 18 | 42619800 | 42620799 | 12.28994 | 22.54032 | 28.1056 | 11.97757 | 0.110897 | 0.262805 | 0.832614 |
| 8 | 129971600 | 129972399 | 11.0692 | 20.55812 | 25.63521 | 10.7396 | 0.128448 | 0.175666 | 0.832622 |
| 4 | 106066800 | 106067599 | 7.44333 | 14.24605 | 22.99661 | 9.092441 | -0.1696 | -0.06498 | 0.832639 |
| 2 | 87718800 | 87723799 | 55.81736 | 89.44895 | 103.9231 | 43.85681 | 0.130421 | 2.000725 | 0.832644 |
| 12 | 44620400 | 44620999 | 5.942717 | 11.58163 | 19.70391 | 7.1392 | -0.12665 | -0.39085 | 0.832672 |
| 14 | 121900400 | 121901799 | 13.70284 | 24.34518 | 34.56838 | 15.25464 | -0.10911 | 0.475065 | 0.832675 |
| 8 | 29027600 | 29029599 | 37.16609 | 60.40469 | 73.17239 | 31.39379 | 0.092653 | 1.585839 | 0.832698 |
| 4 | 81318000 | 81325999 | 102.8198 | 159.0635 | 182.006 | 75.60084 | 0.108339 | 2.844097 | 0.832708 |
| 5 | 106551200 | 106552599 | 18.48609 | 32.0257 | 45.29712 | 19.87374 | -0.11165 | 0.83439 | 0.83271 |
| 11 | 13755000 | 13758799 | 39.47927 | 64.53191 | 74.45604 | 32.01253 | 0.168493 | 1.507037 | 0.832714 |
| 8 | 83474000 | 83482799 | 181.8938 | 269.7659 | 304.4462 | 125.3968 | 0.077138 | 3.67866 | 0.832747 |
| 4 | 144308800 | 144316599 | 122.7482 | 186.3222 | 238.8571 | 98.89881 | -0.07689 | 3.22513 | 0.832748 |
| 6 | 149163800 | 149170999 | 74.53745 | 117.2633 | 156.1818 | 64.79461 | -0.09175 | 2.527305 | 0.832752 |
| 12 | 84130200 | 84144399 | 264.9533 | 379.8033 | 462.3503 | 190.9425 | -0.07943 | 4.213157 | 0.832778 |

| | | | | | | | | | |
|---|---|---|---|---|---|---|---|---|---|
| 2 | 118381800 | 118385999 | 37.99459 | 61.38877 | 84.01342 | 35.86988 | -0.09118 | 1.71829 | 0.832783 |
| 12 | 44771000 | 44772199 | 17.16299 | 29.86511 | 42.58613 | 18.71555 | -0.1226 | 0.767498 | 0.832809 |
| 3 | 72186200 | 72187199 | 14.25314 | 25.5998 | 31.42766 | 13.74208 | 0.108653 | 0.404215 | 0.832839 |
| 5 | 86490600 | 86491599 | 14.25314 | 25.5998 | 31.42766 | 13.74208 | 0.108653 | 0.404215 | 0.832839 |
| 6 | 79923400 | 79924599 | 19.77479 | 34.71999 | 40.01897 | 17.70512 | 0.183461 | 0.642624 | 0.83286 |
| 4 | 90590800 | 90591399 | 11.30605 | 20.93241 | 26.17263 | 11.01152 | 0.120368 | 0.197768 | 0.832872 |
| 19 | 9293600 | 9294599 | 17.9901 | 32.23946 | 28.37499 | 12.4307 | 0.583978 | 0.436128 | 0.832894 |
| 11 | 32120000 | 32121399 | 13.6337 | 24.76097 | 30.53738 | 13.17863 | 0.114471 | 0.33416 | 0.832925 |
| 18 | 7322400 | 7327799 | 92.85052 | 143.6571 | 166.983 | 69.36061 | 0.085798 | 2.834638 | 0.832955 |
| X | 138232800 | 138236799 | 36.33889 | 59.41031 | 71.34524 | 30.67722 | 0.111176 | 1.498466 | 0.832972 |
| 1 | 139711200 | 139712799 | 39.6294 | 63.98753 | 75.74523 | 32.31196 | 0.116567 | 1.737487 | 0.832975 |
| 7 | 16981600 | 16986799 | 78.44315 | 123.2083 | 144.7033 | 60.01992 | 0.087729 | 2.561367 | 0.832976 |
| 12 | 8134200 | 8136199 | 19.48294 | 33.68654 | 47.39942 | 20.79487 | -0.105 | 0.855946 | 0.83298 |
| 5 | 44282000 | 44284399 | 25.76372 | 43.47989 | 52.84501 | 23.11251 | 0.100736 | 1.102248 | 0.832988 |
| 3 | 67621600 | 67623599 | 18.80357 | 32.74013 | 39.94135 | 17.56008 | 0.101688 | 0.764576 | 0.833001 |
| 16 | 52374200 | 52375399 | 10.58147 | 19.41544 | 28.74574 | 12.24304 | -0.1346 | 0.248265 | 0.833009 |
| 9 | 105075400 | 105077599 | 17.87949 | 31.07254 | 43.66928 | 19.17632 | -0.10186 | 0.792035 | 0.833016 |
| 7 | 12173000 | 12173999 | 12.81787 | 23.90963 | 23.97407 | 9.973128 | 0.448865 | 0.160431 | 0.833051 |
| 11 | 51332600 | 51333799 | 17.52714 | 30.61091 | 36.87475 | 16.24297 | 0.12036 | 0.71117 | 0.833066 |
| 1 | 85205200 | 85269799 | 3817.052 | 4387.041 | 4281.882 | 1822.37 | -0.08266 | 7.700077 | 0.833066 |
| 5 | 75600400 | 75602399 | 17.51335 | 30.51959 | 43.06664 | 18.94422 | -0.10736 | 0.749712 | 0.833075 |
| 1 | 33209600 | 33212599 | 22.25568 | 38.44726 | 45.74167 | 20.06706 | 0.135722 | 0.871972 | 0.833082 |
| 3 | 41851800 | 41853399 | 12.21446 | 21.60282 | 38.80142 | 17.1614 | -0.44938 | 0.514192 | 0.833087 |
| 14 | 76473000 | 76475199 | 16.69295 | 28.37894 | 54.73288 | 23.98782 | -0.56216 | 1.009373 | 0.833094 |
| 15 | 46889400 | 46893199 | 34.66511 | 57.55574 | 65.20097 | 28.26325 | 0.201552 | 1.302839 | 0.833103 |
| 2 | 58802000 | 58803599 | 22.90198 | 38.20123 | 65.13559 | 28.32157 | -0.3861 | 1.221677 | 0.833112 |
| 3 | 128582600 | 128583599 | 9.25782 | 17.46203 | 26.43511 | 10.79134 | -0.11935 | 0.06472 | 0.833116 |
| 1 | 13748000 | 13748599 | 8.447284 | 16.29646 | 22.0027 | 8.50335 | 0.116378 | -0.13227 | 0.833117 |
| 7 | 79188000 | 79188799 | 8.447284 | 16.29646 | 22.0027 | 8.50335 | 0.116378 | -0.13227 | 0.833117 |
| 9 | 87419400 | 87419999 | 8.447284 | 16.29646 | 22.0027 | 8.50335 | 0.116378 | -0.13227 | 0.833117 |
| 16 | 21772800 | 21775999 | 29.48338 | 48.59095 | 67.80703 | 29.25299 | -0.09977 | 1.376623 | 0.83315 |
| 6 | 103730200 | 103731799 | 25.15071 | 42.90254 | 42.65411 | 18.69914 | 0.390716 | 0.995634 | 0.833181 |
| 17 | 49233400 | 49235399 | 17.67719 | 30.93706 | 37.73534 | 16.64932 | 0.102402 | 0.680289 | 0.833187 |
| 6 | 130930600 | 130934199 | 31.34303 | 51.82301 | 62.85355 | 27.16722 | 0.103007 | 1.342139 | 0.833197 |
| 14 | 25857800 | 25860799 | 20.54079 | 34.91988 | 49.54833 | 21.71555 | -0.12124 | 1.01637 | 0.833238 |
| 4 | 73962200 | 73963399 | 12.3627 | 21.90329 | 38.76507 | 17.13933 | -0.42726 | 0.495713 | 0.833249 |
| 14 | 62648800 | 62650599 | 18.0392 | 31.50676 | 38.35224 | 16.89432 | 0.105343 | 0.715564 | 0.83325 |
| 11 | 6889000 | 6891799 | 22.75971 | 38.46807 | 54.43736 | 23.81982 | -0.118 | 1.078212 | 0.833262 |
| 9 | 114968400 | 114969799 | 13.49917 | 24.21856 | 28.92657 | 12.66036 | 0.147007 | 0.399088 | 0.83328 |
| 3 | 25205600 | 25206999 | 42.96793 | 69.21513 | 82.42704 | 35.05334 | 0.103958 | 1.816034 | 0.833389 |
| X | 56453800 | 56455799 | 20.3264 | 35.45163 | 41.96198 | 18.46809 | 0.145413 | 0.75382 | 0.833394 |
| 10 | 73530000 | 73531799 | 17.41997 | 30.87494 | 36.33515 | 16.06377 | 0.158256 | 0.517741 | 0.83341 |
| 5 | 148662800 | 148664999 | 23.38796 | 39.40137 | 54.92628 | 24.01002 | -0.09646 | 1.111961 | 0.833438 |
| 6 | 120520200 | 120523399 | 32.38268 | 52.83625 | 73.01595 | 31.37266 | -0.09291 | 1.518292 | 0.833439 |
| X | 4285200 | 4286399 | 26.47344 | 44.26237 | 51.71974 | 22.52159 | 0.157401 | 1.197649 | 0.833443 |
| 7 | 128277200 | 128280399 | 20.70184 | 35.46307 | 49.54562 | 21.72771 | -0.09732 | 0.948209 | 0.833459 |
| 3 | 108641800 | 108643399 | 18.85779 | 32.6629 | 45.80172 | 20.09501 | -0.09948 | 0.836157 | 0.833466 |

| | | | | | | | | | |
|---|---|---|---|---|---|---|---|---|---|
| 5 | 145534800 | 145552199 | 584.3808 | 788.6623 | 799.7978 | 333.4815 | 0.096337 | 5.134002 | 0.833486 |
| 13 | 22414000 | 22417599 | 52.34826 | 83.46981 | 112.1789 | 47.00173 | -0.08731 | 2.152818 | 0.833488 |
| 7 | 7040800 | 7049999 | 163.8799 | 244.7634 | 278.1265 | 114.6591 | 0.076565 | 3.550352 | 0.83351 |
| 4 | 117308600 | 117310399 | 18.18299 | 31.86337 | 38.66202 | 17.0768 | 0.109865 | 0.663739 | 0.833528 |
| 8 | 119688200 | 119692999 | 42.63637 | 68.46772 | 93.66759 | 39.6536 | -0.10129 | 1.916409 | 0.833538 |
| 4 | 73289800 | 73291399 | 11.70697 | 22.08635 | 27.49186 | 11.21762 | 0.161198 | 0.086692 | 0.833542 |
| 8 | 26471800 | 26473399 | 15.492 | 27.36581 | 33.0878 | 14.63794 | 0.11786 | 0.537836 | 0.833546 |
| 19 | 9603600 | 9606399 | 55.93921 | 89.11774 | 104.5883 | 43.76843 | 0.106527 | 2.180076 | 0.833546 |
| 6 | 100373200 | 100375599 | 19.23759 | 33.27493 | 47.39748 | 20.79426 | -0.12258 | 0.849785 | 0.83355 |
| 11 | 27591800 | 27592799 | 12.93196 | 23.38317 | 34.14046 | 14.81504 | -0.13303 | 0.35935 | 0.833553 |
| X | 141954600 | 141955799 | 24.15732 | 41.29955 | 49.55982 | 21.73042 | 0.120787 | 0.980527 | 0.833556 |
| 12 | 113676000 | 113681199 | 35.3838 | 58.7707 | 53.74424 | 23.30294 | 0.509583 | 1.292016 | 0.8336 |
| 13 | 50866600 | 50874599 | 288.5527 | 407.8104 | 497.1214 | 205.7824 | -0.10194 | 4.42112 | 0.833611 |
| 18 | 14815000 | 14816999 | 10.66312 | 18.90428 | 38.3188 | 16.93983 | -0.62142 | 0.475905 | 0.833612 |
| 2 | 103338400 | 103340399 | 25.37047 | 43.1146 | 51.75081 | 22.68016 | 0.119056 | 1.039476 | 0.833626 |
| 14 | 25095800 | 25103599 | 90.12522 | 139.2942 | 181.561 | 75.41635 | -0.08044 | 2.854797 | 0.833628 |
| 11 | 31496400 | 31505999 | 67.92962 | 106.5305 | 165.4827 | 68.63349 | -0.31692 | 2.586708 | 0.833628 |
| 4 | 107388200 | 107391999 | 38.49996 | 62.49381 | 75.4751 | 32.35168 | 0.0953 | 1.615085 | 0.833635 |
| 7 | 104416000 | 104426799 | 164.9137 | 246.1859 | 315.097 | 129.8879 | -0.09568 | 3.560908 | 0.833642 |
| 10 | 78512600 | 78514799 | 22.07903 | 37.9779 | 55.63373 | 24.40354 | -0.16449 | 0.924153 | 0.833649 |
| 1 | 159541600 | 159542199 | 10.20458 | 18.86873 | 28.2622 | 11.91156 | -0.14048 | 0.19941 | 0.833717 |
| 13 | 31885400 | 31890199 | 31.41772 | 51.67665 | 73.26305 | 31.58038 | -0.12373 | 1.40824 | 0.833803 |
| 10 | 58523200 | 58524599 | 18.89171 | 32.52071 | 45.8424 | 20.10908 | -0.1088 | 0.892075 | 0.833814 |
| 2 | 42933000 | 42933599 | 10.08366 | 18.73698 | 27.81594 | 11.64538 | -0.12052 | 0.171232 | 0.833815 |
| 1 | 7286800 | 7288399 | 18.1203 | 31.98822 | 31.24285 | 13.8235 | 0.423659 | 0.55577 | 0.83384 |
| 13 | 62574000 | 62615199 | 911.5558 | 1185.793 | 1160.501 | 486.1226 | 0.094654 | 5.725694 | 0.833844 |
| 10 | 29945000 | 29945599 | 10.1379 | 19.19107 | 24.84166 | 10.06385 | 0.114448 | 0.043695 | 0.833845 |
| 3 | 11463000 | 11463599 | 9.502397 | 18.20107 | 24.01966 | 9.475825 | 0.120897 | -0.05049 | 0.833852 |
| 1 | 130327800 | 130328599 | 9.939868 | 18.67114 | 27.98416 | 11.50445 | -0.11313 | 0.102161 | 0.833862 |
| 1 | 87589400 | 87592599 | 35.27463 | 57.51379 | 69.92399 | 30.03633 | 0.09068 | 1.532392 | 0.833868 |
| 2 | 53929600 | 53930399 | 12.62507 | 22.88645 | 33.22851 | 14.37348 | -0.12156 | 0.344646 | 0.833882 |
| 2 | 26745800 | 26748199 | 24.09363 | 40.31983 | 56.51465 | 24.62357 | -0.10438 | 1.181888 | 0.833891 |
| 7 | 8839400 | 8840799 | 13.99038 | 25.6207 | 25.39277 | 10.84349 | 0.437904 | 0.278057 | 0.833977 |
| 10 | 84386800 | 84388199 | 14.59615 | 26.0605 | 31.93467 | 14.03393 | 0.10636 | 0.444684 | 0.833986 |
| 1 | 9245400 | 9246199 | 13.45594 | 23.84465 | 34.18648 | 15.11441 | -0.12479 | 0.493117 | 0.834054 |
| 12 | 80934000 | 80938599 | 36.49359 | 59.37171 | 72.05469 | 30.92701 | 0.091045 | 1.56853 | 0.834055 |
| 10 | 123359000 | 123361799 | 30.12907 | 50.76828 | 58.01015 | 25.35963 | 0.189373 | 1.135385 | 0.834072 |
| 13 | 110285200 | 110285799 | 9.235424 | 17.48474 | 27.07598 | 10.96629 | -0.14118 | 0.044199 | 0.8341 |
| 13 | 34642800 | 34645399 | 23.87548 | 40.90617 | 49.01812 | 21.49665 | 0.123221 | 0.957705 | 0.834109 |
| 11 | 48357800 | 48358199 | 5.944312 | 11.57965 | 19.65674 | 7.163266 | -0.13083 | -0.37011 | 0.83413 |
| 15 | 34550400 | 34552799 | 28.41236 | 47.06329 | 56.66606 | 24.52371 | 0.113681 | 1.30617 | 0.83416 |
| 16 | 32331800 | 32332999 | 16.09441 | 28.28657 | 34.1467 | 15.11998 | 0.117329 | 0.598372 | 0.834161 |
| 4 | 20408600 | 20411799 | 61.44598 | 97.17824 | 130.4954 | 54.22725 | -0.09754 | 2.400309 | 0.834168 |
| 10 | 41846800 | 41847999 | 13.60679 | 24.81694 | 30.56509 | 13.12094 | 0.121263 | 0.309415 | 0.834184 |
| 3 | 103746400 | 103754199 | 98.99624 | 152.9937 | 178.4576 | 74.12729 | 0.07952 | 2.859037 | 0.834197 |
| X | 64691000 | 64691999 | 12.98659 | 23.46814 | 34.22869 | 14.86427 | -0.13175 | 0.367774 | 0.83422 |
| 8 | 9208600 | 9209599 | 8.986929 | 16.81462 | 25.86273 | 10.70363 | -0.15782 | 0.113051 | 0.834247 |

| | | | | | | | | | |
|---|---|---|---|---|---|---|---|---|---|
| 10 | 81510600 | 81513399 | 32.60046 | 53.88245 | 53.58485 | 23.15519 | 0.388475 | 1.340176 | 0.834303 |
| 9 | 118001800 | 118002599 | 13.86079 | 24.96125 | 30.53478 | 13.31774 | 0.116455 | 0.385414 | 0.83433 |
| 14 | 122180600 | 122182999 | 29.7102 | 50.50169 | 55.74553 | 24.43135 | 0.239612 | 1.058966 | 0.834331 |
| 13 | 21830600 | 21832599 | 24.55321 | 41.69709 | 50.64831 | 22.19383 | 0.101892 | 1.041498 | 0.834367 |
| 9 | 16782000 | 16785399 | 51.09873 | 81.74261 | 97.51524 | 41.04547 | 0.089434 | 2.046229 | 0.834381 |
| 13 | 12545400 | 12553999 | 86.51695 | 135.4504 | 157.3907 | 65.28874 | 0.099492 | 2.625399 | 0.834396 |
| 7 | 86816400 | 86817399 | 17.57149 | 30.46082 | 43.21035 | 18.95856 | -0.11573 | 0.81962 | 0.834409 |
| 14 | 82806600 | 82807399 | 11.18237 | 20.71157 | 25.71157 | 10.81176 | 0.131359 | 0.194264 | 0.83441 |
| 4 | 73121800 | 73123799 | 27.23157 | 45.84716 | 42.8061 | 18.70826 | 0.479737 | 1.116925 | 0.834443 |
| 1 | 190873000 | 190875999 | 47.45674 | 76.24215 | 91.66986 | 38.83344 | 0.085336 | 1.907187 | 0.834464 |
| 7 | 18909600 | 18913999 | 44.12965 | 70.72245 | 97.12994 | 40.99301 | -0.11036 | 1.982726 | 0.834489 |
| 11 | 11339000 | 11352199 | 227.2814 | 329.7045 | 405.9428 | 167.3668 | -0.07879 | 4.034517 | 0.834515 |
| Y | 42192200 | 42194399 | 22.44414 | 38.71055 | 34.3936 | 15.08787 | 0.554807 | 0.87293 | 0.834518 |
| 5 | 76858400 | 76860399 | 17.42712 | 30.46136 | 43.00265 | 18.95333 | -0.10752 | 0.707302 | 0.83452 |
| 1 | 111751000 | 111754199 | 38.57016 | 62.79688 | 74.5834 | 32.02107 | 0.122889 | 1.565328 | 0.834557 |
| 2 | 150965200 | 150966799 | 36.48775 | 59.38247 | 72.10226 | 30.95127 | 0.090699 | 1.563563 | 0.834584 |
| 4 | 12720800 | 12724199 | 27.18619 | 45.71028 | 55.1014 | 24.07222 | 0.112294 | 1.139804 | 0.834608 |
| 12 | 14709200 | 14710799 | 29.97747 | 49.57506 | 60.53512 | 26.16874 | 0.093304 | 1.333722 | 0.834622 |
| 8 | 25787400 | 25787999 | 9.309105 | 17.51618 | 26.4197 | 10.82994 | -0.11865 | 0.082113 | 0.834629 |
| 14 | 41703400 | 41704999 | 14.40194 | 26.41189 | 25.41351 | 10.83138 | 0.482476 | 0.274409 | 0.834674 |
| X | 147017600 | 147019199 | 37.74795 | 61.73166 | 54.54656 | 23.45653 | 0.552972 | 1.487661 | 0.834708 |
| 7 | 41772000 | 41773599 | 25.66533 | 44.00416 | 50.68746 | 22.23001 | 0.180133 | 0.959776 | 0.834709 |
| 7 | 38527800 | 38529599 | 20.15063 | 35.06425 | 42.51841 | 18.68593 | 0.110267 | 0.78354 | 0.834709 |
| 12 | 116015200 | 116030999 | 217.8222 | 321.2084 | 346.7481 | 142.7955 | 0.133502 | 3.78014 | 0.834735 |
| 8 | 101341600 | 101345199 | 39.59277 | 64.48754 | 76.26773 | 32.74897 | 0.129334 | 1.560168 | 0.834742 |
| 13 | 120773200 | 120783199 | 133.0997 | 201.1277 | 257.7851 | 106.5867 | -0.08208 | 3.318503 | 0.834744 |
| 1 | 112706000 | 112709999 | 36.00669 | 59.36753 | 69.01339 | 29.77888 | 0.163303 | 1.390867 | 0.83476 |
| 12 | 25439000 | 25440999 | 16.15558 | 28.31751 | 39.83049 | 17.59654 | -0.10218 | 0.651691 | 0.834797 |
| X | 13169000 | 13170599 | 14.06031 | 25.03807 | 35.58551 | 15.67292 | -0.10851 | 0.465586 | 0.834803 |
| 2 | 44823000 | 44826799 | 67.75495 | 106.8533 | 141.312 | 58.64424 | -0.07881 | 2.472697 | 0.834827 |
| 10 | 8026000 | 8038199 | 18.84304 | 33.20927 | 31.78167 | 14.06654 | 0.452068 | 0.579717 | 0.83483 |
| 9 | 10172800 | 10174199 | 14.39653 | 25.50524 | 36.08681 | 15.95536 | -0.10615 | 0.504353 | 0.834843 |
| 2 | 148965600 | 148968599 | 35.7166 | 58.23737 | 82.24881 | 35.3312 | -0.12493 | 1.540799 | 0.834853 |
| 1 | 175222800 | 175226999 | 47.02133 | 76.22693 | 69.51094 | 29.76673 | 0.497945 | 1.65353 | 0.834854 |
| 2 | 146644400 | 146646199 | 15.31531 | 27.07368 | 38.60373 | 17.07565 | -0.11869 | 0.536774 | 0.834864 |
| 18 | 76056800 | 76057799 | 12.5841 | 22.79159 | 27.56908 | 11.93888 | 0.13906 | 0.334923 | 0.834875 |
| 10 | 6593000 | 6593999 | 12.52249 | 23.33001 | 28.20458 | 11.75741 | 0.177933 | 0.172344 | 0.834877 |
| 5 | 142690400 | 142692199 | 15.82096 | 27.62775 | 39.50385 | 17.39975 | -0.12576 | 0.708425 | 0.834913 |
| 5 | 30125400 | 30127199 | 26.44018 | 45.39503 | 41.94503 | 18.39788 | 0.497269 | 0.941647 | 0.834922 |
| 10 | 24944400 | 24947799 | 27.98364 | 46.48465 | 65.09107 | 28.225 | -0.10323 | 1.279346 | 0.834924 |
| 15 | 41359400 | 41361199 | 13.14085 | 24.00225 | 29.48444 | 12.63387 | 0.126488 | 0.293228 | 0.834933 |
| 11 | 36198000 | 36198799 | 10.998 | 20.59541 | 26.27669 | 10.85881 | 0.111485 | 0.126378 | 0.834958 |
| 5 | 131557800 | 131558999 | 13.04793 | 23.49524 | 33.61166 | 14.66141 | -0.10834 | 0.385463 | 0.834961 |
| 1 | 7775200 | 7786999 | 238.8537 | 346.9145 | 383.1058 | 157.9313 | 0.079784 | 4.017331 | 0.834981 |
| 13 | 87757400 | 87758599 | 16.00573 | 28.28702 | 41.23203 | 18.23915 | -0.1505 | 0.548407 | 0.834988 |
| 1 | 48329600 | 48330199 | 7.567874 | 14.67169 | 23.9892 | 9.067678 | -0.1271 | -0.21073 | 0.834989 |
| 4 | 145611600 | 145616399 | 202.2939 | 296.6143 | 327.3561 | 134.8359 | 0.093844 | 3.868098 | 0.83499 |

| | | | | | | | | | |
|---|---|---|---|---|---|---|---|---|---|
| 8 | 125870600 | 125873399 | 37.70816 | 61.20086 | 73.89789 | 31.67269 | 0.095395 | 1.61248 | 0.834993 |
| 14 | 32461600 | 32462799 | 18.48632 | 32.38797 | 39.16052 | 17.29386 | 0.114883 | 0.66913 | 0.834995 |
| 11 | 75610400 | 75612799 | 34.65794 | 56.52391 | 68.63274 | 29.48464 | 0.092829 | 1.527307 | 0.834998 |
| 14 | 118091800 | 118093999 | 34.36422 | 56.18454 | 68.17217 | 29.31335 | 0.096388 | 1.491795 | 0.835007 |
| 12 | 87232000 | 87234399 | 16.53729 | 28.71492 | 49.53289 | 21.73928 | -0.3963 | 0.806967 | 0.83503 |
| 5 | 119574200 | 119578999 | 50.73515 | 81.14534 | 96.22261 | 40.49032 | 0.097783 | 2.050769 | 0.835097 |
| 10 | 129226400 | 129226999 | 17.46264 | 30.56874 | 37.19483 | 16.4141 | 0.105862 | 0.673643 | 0.835126 |
| 4 | 114009200 | 114010199 | 14.62706 | 26.61374 | 32.07788 | 13.8118 | 0.149422 | 0.333904 | 0.835136 |
| 4 | 145560400 | 145584199 | 954.8892 | 1233.055 | 1212.287 | 508.1824 | 0.077433 | 5.845225 | 0.83515 |
| 2 | 5778600 | 5792599 | 199.8342 | 295.1253 | 329.3884 | 135.6487 | 0.087472 | 3.752348 | 0.835189 |
| 8 | 5191400 | 5193999 | 26.64636 | 44.64514 | 54.47439 | 23.75256 | 0.094951 | 1.174894 | 0.835239 |
| 2 | 65601800 | 65606599 | 65.63981 | 104.3193 | 122.9205 | 51.28088 | 0.097102 | 2.270437 | 0.835245 |
| 7 | 102203600 | 102204799 | 10.0704 | 18.75285 | 27.83337 | 11.61188 | -0.11627 | 0.157426 | 0.835258 |
| 15 | 44230600 | 44231799 | 21.28101 | 37.14169 | 36.01581 | 15.85784 | 0.431643 | 0.736845 | 0.835258 |
| 16 | 30371600 | 30380799 | 117.2658 | 179.1585 | 207.1309 | 85.91507 | 0.079565 | 3.089346 | 0.835292 |
| 9 | 102364200 | 102366399 | 32.09358 | 52.24634 | 72.67594 | 31.20085 | -0.10462 | 1.550455 | 0.835299 |
| 4 | 7302200 | 7303199 | 30.20357 | 49.7579 | 59.7896 | 25.78977 | 0.115287 | 1.380441 | 0.83532 |
| X | 84312800 | 84314399 | 18.1525 | 31.7727 | 38.29639 | 16.89857 | 0.11963 | 0.684204 | 0.835341 |
| 4 | 29888600 | 29890599 | 19.2526 | 33.95653 | 38.9459 | 17.23483 | 0.192513 | 0.585475 | 0.83537 |
| 1 | 181263200 | 181266199 | 25.50393 | 42.60489 | 59.37366 | 25.84055 | -0.09614 | 1.205942 | 0.835373 |
| 2 | 163021000 | 163021999 | 10.89939 | 20.25452 | 25.13505 | 10.52173 | 0.136181 | 0.169611 | 0.835379 |
| 7 | 12894400 | 12896199 | 16.3214 | 28.91138 | 34.80023 | 15.38694 | 0.125488 | 0.512812 | 0.835383 |
| 16 | 68532400 | 68532799 | 6.521522 | 12.64547 | 21.08934 | 7.964729 | -0.15383 | -0.23474 | 0.835387 |
| 19 | 11142600 | 11142999 | 5.941089 | 11.58367 | 19.76128 | 7.117961 | -0.12303 | -0.41195 | 0.835389 |
| 3 | 59235200 | 59238799 | 37.26569 | 60.37853 | 83.60564 | 35.78793 | -0.10406 | 1.65095 | 0.835397 |
| 14 | 83419200 | 83419599 | 12.19177 | 22.23767 | 27.02041 | 11.59828 | 0.140656 | 0.288948 | 0.835416 |
| 13 | 96606000 | 96607999 | 14.32281 | 25.19634 | 36.19213 | 16.02449 | -0.13188 | 0.579381 | 0.835437 |
| 7 | 64109400 | 64109999 | 9.199288 | 17.53062 | 22.84373 | 9.125035 | 0.122937 | -0.01583 | 0.835441 |
| 14 | 119989800 | 119991399 | 18.91178 | 32.69323 | 45.8186 | 20.10123 | -0.09918 | 0.854758 | 0.835493 |
| 6 | 127040600 | 127042199 | 12.33833 | 22.06207 | 32.56629 | 14.29358 | -0.15955 | 0.419512 | 0.835517 |
| 13 | 91206400 | 91207999 | 19.18837 | 33.54644 | 40.22382 | 17.73148 | 0.126811 | 0.706776 | 0.835533 |
| 8 | 73571600 | 73571999 | 5.236743 | 10.22158 | 18.13817 | 6.427606 | -0.1579 | -0.48637 | 0.835539 |
| 8 | 21132200 | 21134399 | 92.23874 | 142.1489 | 185.6813 | 77.12903 | -0.08674 | 2.909301 | 0.835544 |
| 14 | 22003600 | 22011399 | 64.62858 | 102.1627 | 136.7046 | 56.79373 | -0.09304 | 2.413415 | 0.835556 |
| 11 | 29203400 | 29205399 | 20.3597 | 35.12363 | 49.55632 | 21.73965 | -0.10958 | 0.883297 | 0.83556 |
| 6 | 8561200 | 8565999 | 36.55341 | 60.2053 | 69.76468 | 30.0896 | 0.167249 | 1.403679 | 0.8356 |
| 1 | 44192800 | 44194599 | 18.46821 | 31.28704 | 56.35194 | 24.6749 | -0.46425 | 1.049658 | 0.835623 |
| 5 | 120848400 | 120851799 | 27.97418 | 45.81169 | 65.98105 | 28.38883 | -0.14877 | 1.451608 | 0.835633 |
| 3 | 91730400 | 91732599 | 38.3626 | 62.92604 | 60.91268 | 26.22609 | 0.423775 | 1.453064 | 0.835639 |
| 2 | 177064800 | 177066399 | 11.60579 | 21.18773 | 30.74017 | 13.18025 | -0.11222 | 0.290789 | 0.835689 |
| 8 | 19100600 | 19101199 | 11.60579 | 21.18773 | 30.74017 | 13.18025 | -0.11222 | 0.290789 | 0.835689 |
| 10 | 111782400 | 111784599 | 20.70933 | 35.7059 | 50.6442 | 22.21649 | -0.11733 | 0.891554 | 0.8357 |
| 10 | 36653000 | 36657599 | 64.42705 | 101.7939 | 135.2239 | 56.16456 | -0.0831 | 2.426437 | 0.835717 |
| 10 | 90940600 | 90951599 | 114.026 | 174.4649 | 202.449 | 84.00161 | 0.075925 | 3.059021 | 0.835718 |
| 12 | 49564200 | 49565799 | 19.29815 | 33.08471 | 46.9162 | 20.57592 | -0.11838 | 0.933762 | 0.835739 |
| 6 | 130416400 | 130418199 | 17.51824 | 31.4338 | 35.37399 | 15.48026 | 0.23253 | 0.438794 | 0.83574 |
| 14 | 8663800 | 8664799 | 13.20985 | 23.92658 | 29.46497 | 12.7588 | 0.113302 | 0.342674 | 0.835742 |

| | | | | | | | | | |
|---|---|---|---|---|---|---|---|---|---|
| 9 | 60248400 | 60249599 | 10.01013 | 18.43078 | 27.84025 | 11.8017 | -0.15801 | 0.224027 | 0.835755 |
| 18 | 11328800 | 11330199 | 13.18563 | 24.47279 | 29.20558 | 12.24628 | 0.190016 | 0.202706 | 0.835758 |
| 1 | 17717800 | 17718399 | 7.805859 | 15.07516 | 19.88511 | 7.652236 | 0.155563 | -0.16068 | 0.83577 |
| 6 | 23064600 | 23067199 | 24.00299 | 41.13816 | 40.06456 | 17.57058 | 0.421395 | 0.945979 | 0.835789 |
| 15 | 100793000 | 100798399 | 68.65503 | 107.8927 | 143.9135 | 59.6965 | -0.09504 | 2.543511 | 0.835796 |
| 11 | 16911200 | 16916399 | 58.13275 | 92.86046 | 92.92618 | 39.03379 | 0.340225 | 2.096837 | 0.835816 |
| 3 | 7926600 | 7929799 | 34.29858 | 56.00957 | 68.18597 | 29.30639 | 0.090358 | 1.507377 | 0.835883 |
| 7 | 93170000 | 93171799 | 15.61484 | 27.22946 | 47.86954 | 21.06583 | -0.42276 | 0.74081 | 0.8359 |
| 1 | 88648800 | 88654599 | 48.75718 | 78.06742 | 105.9087 | 44.63614 | -0.09431 | 2.01701 | 0.835902 |
| 12 | 115551000 | 115553199 | 47.66979 | 76.78376 | 90.94989 | 38.67387 | 0.111512 | 1.827031 | 0.835919 |
| 13 | 85337400 | 85338199 | 8.202355 | 15.50931 | 24.41596 | 9.915381 | -0.16842 | 0.033052 | 0.835935 |
| 14 | 5822000 | 5833999 | 229.4927 | 332.3271 | 408.2575 | 168.3595 | -0.07806 | 4.060744 | 0.835949 |
| X | 113157000 | 113158999 | 29.2171 | 49.34912 | 56.81364 | 24.85453 | 0.178732 | 1.112297 | 0.835971 |
| 12 | 110057000 | 110066599 | 129.9185 | 196.039 | 251.2509 | 103.8379 | -0.08377 | 3.346561 | 0.835981 |
| 9 | 117942000 | 117942599 | 6.99065 | 13.48392 | 22.0499 | 8.525958 | -0.15761 | -0.14448 | 0.835982 |
| 9 | 10971000 | 10971399 | 6.09375 | 11.88007 | 20.3259 | 7.331378 | -0.12843 | -0.40078 | 0.835998 |
| 11 | 5497600 | 5502599 | 69.45809 | 109.7785 | 130.5142 | 54.21731 | 0.077126 | 2.421 | 0.836012 |
| 3 | 80036400 | 80041799 | 112.8662 | 172.7042 | 200.5281 | 83.20612 | 0.07514 | 3.056988 | 0.836022 |
| 18 | 70679800 | 70681599 | 12.45447 | 23.51302 | 23.0121 | 9.375849 | 0.507569 | 0.078471 | 0.836049 |
| 4 | 53946000 | 53949199 | 15.83373 | 27.17195 | 51.07927 | 22.40347 | -0.52312 | 0.91538 | 0.836066 |
| 5 | 149824600 | 149826799 | 18.82024 | 32.71654 | 46.29454 | 20.33269 | -0.11174 | 0.795087 | 0.836099 |
| 3 | 54379400 | 54379999 | 6.669232 | 12.97624 | 17.44295 | 6.431128 | 0.187485 | -0.3266 | 0.836118 |
| 16 | 94148800 | 94151599 | 16.90859 | 29.14039 | 50.58049 | 22.18826 | -0.40739 | 0.880511 | 0.836122 |
| X | 153370400 | 153372399 | 26.64395 | 45.05745 | 53.95627 | 23.61317 | 0.122171 | 1.089675 | 0.836169 |
| 18 | 38660800 | 38663399 | 26.9623 | 45.05829 | 55.02855 | 23.95523 | 0.093693 | 1.203365 | 0.836181 |
| 5 | 51162200 | 51165999 | 66.88191 | 105.7012 | 123.882 | 51.46083 | 0.097695 | 2.420066 | 0.836194 |
| 15 | 37089800 | 37092799 | 30.39277 | 49.6858 | 69.55595 | 29.90715 | -0.10953 | 1.483229 | 0.836236 |
| 16 | 54872800 | 54876999 | 74.03759 | 116.2867 | 135.9749 | 56.39945 | 0.094048 | 2.556244 | 0.83625 |
| 3 | 56121400 | 56123199 | 19.80315 | 34.90441 | 32.13222 | 14.22144 | 0.507768 | 0.584552 | 0.836257 |
| 4 | 10354200 | 10354599 | 6.813567 | 13.22241 | 21.67737 | 8.156042 | -0.12424 | -0.24167 | 0.83626 |
| 9 | 6886800 | 6887999 | 10.60662 | 20.24119 | 19.68214 | 7.838565 | 0.546062 | -0.01919 | 0.836262 |
| 11 | 74010600 | 74012999 | 41.30456 | 66.78479 | 80.50861 | 34.40539 | 0.092631 | 1.702875 | 0.836273 |
| 16 | 92745400 | 92748599 | 18.01811 | 31.44011 | 37.7737 | 16.62956 | 0.123798 | 0.724844 | 0.836274 |
| 15 | 9991600 | 9992599 | 18.95034 | 32.45186 | 46.40675 | 20.35064 | -0.1307 | 0.941277 | 0.836278 |
| 9 | 74823400 | 74825199 | 19.64208 | 34.30155 | 41.35404 | 18.21504 | 0.119019 | 0.72907 | 0.836302 |
| 10 | 41712400 | 41713399 | 15.18251 | 26.98653 | 32.84076 | 14.4866 | 0.112449 | 0.476801 | 0.836307 |
| 6 | 80288400 | 80289799 | 15.00208 | 27.45897 | 32.15539 | 13.74117 | 0.197746 | 0.301617 | 0.836325 |
| 15 | 54313600 | 54314399 | 11.81459 | 21.57347 | 25.71903 | 11.02446 | 0.169652 | 0.278999 | 0.836332 |
| Y | 42995200 | 42999199 | 35.69597 | 57.26035 | 90.66881 | 38.32084 | -0.31359 | 1.937984 | 0.836333 |
| 13 | 77745000 | 77746799 | 18.32407 | 32.46765 | 30.60378 | 13.53074 | 0.47668 | 0.5165 | 0.836345 |
| 13 | 23184200 | 23184599 | 6.663197 | 12.98443 | 18.54914 | 6.729832 | 0.12089 | -0.38095 | 0.836359 |
| 8 | 4731000 | 4733399 | 39.20943 | 63.553 | 76.80387 | 32.88599 | 0.09238 | 1.643799 | 0.836362 |
| 2 | 76080600 | 76082399 | 18.14227 | 31.16078 | 54.38737 | 23.85576 | -0.4166 | 0.918532 | 0.836365 |
| 8 | 84251400 | 84252199 | 13.60786 | 24.34343 | 35.03376 | 15.37054 | -0.12287 | 0.430311 | 0.836393 |
| 3 | 5093800 | 5094399 | 7.139899 | 13.87143 | 22.94056 | 8.559954 | -0.12544 | -0.26151 | 0.836398 |
| 11 | 110915600 | 110916799 | 11.36928 | 20.73064 | 30.18262 | 12.96096 | -0.11887 | 0.292883 | 0.836401 |
| 3 | 145567600 | 145571599 | 32.19988 | 52.10729 | 87.97445 | 37.62421 | -0.39099 | 1.670732 | 0.836453 |

| | | | | | | | | | |
|---|---|---|---|---|---|---|---|---|---|
| 5 | 77165400 | 77167599 | 18.32478 | 31.81633 | 44.72516 | 19.62832 | -0.10241 | 0.808077 | 0.836453 |
| 6 | 40883800 | 40885799 | 21.66945 | 37.07138 | 44.8189 | 19.65275 | 0.11032 | 0.95947 | 0.836454 |
| 18 | 68550400 | 68550799 | 6.805514 | 13.23114 | 17.93272 | 6.65324 | 0.167015 | -0.30223 | 0.836478 |
| 19 | 8638800 | 8640399 | 14.80405 | 26.1786 | 31.15757 | 13.78106 | 0.14109 | 0.517742 | 0.83648 |
| 1 | 122891400 | 122893199 | 17.58702 | 30.78591 | 44.00753 | 19.42025 | -0.12555 | 0.685687 | 0.836499 |
| 15 | 92976800 | 92979799 | 21.11321 | 36.06705 | 50.62804 | 22.1974 | -0.10481 | 0.974497 | 0.836502 |
| 11 | 6155200 | 6157999 | 33.95406 | 55.53606 | 67.7322 | 29.1284 | 0.089369 | 1.483229 | 0.836505 |
| 6 | 123651400 | 123656999 | 57.74755 | 91.91019 | 122.8112 | 51.32603 | -0.08188 | 2.218762 | 0.836507 |
| 18 | 19763000 | 19764999 | 21.05949 | 36.12142 | 43.73598 | 19.18191 | 0.108626 | 0.930099 | 0.836521 |
| 14 | 48188000 | 48192999 | 59.76249 | 94.68587 | 126.9696 | 52.81013 | -0.09344 | 2.355699 | 0.836532 |
| 5 | 94443000 | 94443399 | 5.238132 | 10.22062 | 18.10004 | 6.453952 | -0.16293 | -0.46418 | 0.836538 |
| 13 | 72539000 | 72543599 | 52.83677 | 84.12552 | 113.7011 | 47.54312 | -0.09772 | 2.200459 | 0.836543 |
| 12 | 106724600 | 106726599 | 22.15246 | 38.39854 | 45.75872 | 20.0766 | 0.134497 | 0.844781 | 0.836561 |
| 4 | 71827000 | 71827799 | 15.47145 | 27.40418 | 33.46517 | 14.79682 | 0.105014 | 0.508065 | 0.836561 |
| 4 | 101730600 | 101735399 | 46.87039 | 75.25317 | 102.1975 | 43.25263 | -0.0911 | 1.929926 | 0.836587 |
| 19 | 19803800 | 19804199 | 9.727327 | 18.43149 | 23.83947 | 9.63831 | 0.11853 | 0.032226 | 0.836594 |
| 15 | 93255600 | 93258599 | 24.23449 | 40.89024 | 49.59959 | 21.69166 | 0.103578 | 1.094844 | 0.83661 |
| X | 90173400 | 90177999 | 91.57899 | 143.4673 | 134.0915 | 55.62011 | 0.413942 | 2.61559 | 0.836621 |
| 5 | 21298600 | 21307599 | 123.8211 | 188.287 | 217.5286 | 90.11594 | 0.075172 | 3.187399 | 0.836627 |
| 13 | 15410800 | 15411399 | 10.96334 | 20.32139 | 25.18391 | 10.57888 | 0.134745 | 0.18577 | 0.836681 |
| 16 | 61339000 | 61339599 | 10.96334 | 20.32139 | 25.18391 | 10.57888 | 0.134745 | 0.18577 | 0.836681 |
| 7 | 88921800 | 88926199 | 90.23931 | 139.6315 | 181.5293 | 75.40273 | -0.07543 | 2.833781 | 0.836721 |
| 11 | 103309400 | 103310799 | 16.04726 | 28.03141 | 40.02549 | 17.63286 | -0.12364 | 0.707683 | 0.83674 |
| 3 | 97405800 | 97408799 | 37.73994 | 61.55429 | 73.82582 | 31.71632 | 0.110561 | 1.534931 | 0.836745 |
| 16 | 21562200 | 21566199 | 38.74861 | 62.73982 | 87.36824 | 37.38035 | -0.11311 | 1.668765 | 0.83675 |
| 10 | 66507200 | 66508199 | 19.78282 | 34.2141 | 41.618 | 18.25862 | 0.104408 | 0.84831 | 0.836758 |
| 7 | 44585400 | 44589599 | 38.9536 | 63.46379 | 75.55578 | 32.44648 | 0.120087 | 1.555006 | 0.836782 |
| 1 | 146313400 | 146316199 | 26.67714 | 45.77198 | 51.26235 | 22.48345 | 0.220696 | 0.95816 | 0.83681 |
| 15 | 15476000 | 15479799 | 31.87268 | 52.74386 | 63.65453 | 27.52969 | 0.110151 | 1.331989 | 0.83688 |
| 19 | 54281400 | 54284599 | 24.30067 | 40.89804 | 57.19115 | 24.99248 | -0.10111 | 1.124151 | 0.836888 |
| 5 | 149021200 | 149025999 | 40.94281 | 65.98669 | 89.7382 | 38.19043 | -0.0863 | 1.798189 | 0.836889 |
| 7 | 29485000 | 29493999 | 124.6924 | 188.5976 | 242.0885 | 100.1258 | -0.0831 | 3.297875 | 0.836889 |
| 15 | 68419800 | 68421799 | 18.59554 | 32.64012 | 30.55003 | 13.51261 | 0.482615 | 0.626892 | 0.836953 |
| X | 169066400 | 169073999 | 501.0205 | 665.6757 | 1041.701 | 435.3211 | -0.55196 | 5.381452 | 0.836954 |
| 15 | 27990200 | 27991599 | 15.18558 | 26.53256 | 38.51999 | 16.97218 | -0.14748 | 0.695975 | 0.836995 |
| 11 | 24209800 | 24212999 | 29.76458 | 49.0291 | 68.16449 | 29.40079 | -0.09465 | 1.386708 | 0.836996 |
| 9 | 88543000 | 88548599 | 64.15671 | 102.6848 | 111.8815 | 47.12711 | 0.220802 | 2.05109 | 0.837005 |
| 2 | 154479600 | 154481399 | 27.78484 | 45.85937 | 64.10146 | 27.67309 | -0.10127 | 1.354384 | 0.837012 |
| 16 | 19166600 | 19171999 | 56.50225 | 89.28818 | 145.8796 | 60.59562 | -0.381 | 2.420184 | 0.83702 |
| 8 | 35771200 | 35772599 | 17.87456 | 31.34811 | 38.13304 | 16.85462 | 0.106247 | 0.648663 | 0.837022 |
| 1 | 130675200 | 130677199 | 16.66268 | 29.22535 | 41.33767 | 18.27243 | -0.11026 | 0.646457 | 0.837025 |
| 1 | 69869600 | 69871799 | 22.04307 | 38.19464 | 37.80454 | 16.58653 | 0.401004 | 0.835126 | 0.837057 |
| 8 | 106125800 | 106127999 | 12.84202 | 22.6325 | 40.34645 | 17.85737 | -0.44074 | 0.559204 | 0.837095 |
| 7 | 108437200 | 108438799 | 19.7211 | 34.4405 | 41.31899 | 18.19992 | 0.125969 | 0.727779 | 0.837105 |
| 10 | 67316400 | 67323799 | 73.52438 | 116.1487 | 136.6622 | 56.75729 | 0.091502 | 2.43563 | 0.837116 |
| 19 | 3157800 | 3164999 | 82.21731 | 130.3968 | 111.3785 | 46.3701 | 0.557719 | 2.341808 | 0.83712 |
| 18 | 11839800 | 11842199 | 25.75404 | 44.20193 | 41.51609 | 18.20924 | 0.473796 | 0.943154 | 0.837133 |

| | | | | | | | | | |
|---|---|---|---|---|---|---|---|---|---|
| 7 | 18606400 | 18611399 | 43.3776 | 69.67202 | 94.92426 | 40.19787 | -0.09507 | 1.909963 | 0.837143 |
| 5 | 146147000 | 146151199 | 45.86733 | 73.30824 | 117.9191 | 49.5708 | -0.34243 | 2.075725 | 0.837171 |
| 6 | 5849400 | 5852399 | 35.30636 | 58.07267 | 58.58684 | 25.25411 | 0.365836 | 1.409398 | 0.837188 |
| 12 | 91910200 | 91913799 | 31.5395 | 51.54275 | 71.37296 | 30.68457 | -0.0942 | 1.49213 | 0.837195 |
| 6 | 111792200 | 111796399 | 63.68386 | 100.9578 | 134.3476 | 55.90792 | -0.08165 | 2.343577 | 0.837267 |
| 1 | 28286800 | 28290799 | 55.714 | 88.80788 | 105.6893 | 44.26289 | 0.087294 | 2.16398 | 0.837293 |
| 8 | 9806000 | 9807399 | 19.84698 | 34.23851 | 47.93734 | 21.02941 | -0.0985 | 0.879058 | 0.837309 |
| 14 | 84095800 | 84096999 | 20.22575 | 34.55787 | 48.48744 | 21.25922 | -0.10397 | 0.968806 | 0.837313 |
| 13 | 7943400 | 7946199 | 22.91232 | 39.8628 | 45.68087 | 20.07351 | 0.191342 | 0.80214 | 0.83732 |
| 7 | 35358600 | 35361999 | 40.9116 | 66.02497 | 79.42899 | 33.86166 | 0.092472 | 1.754953 | 0.83733 |
| 18 | 30138600 | 30141599 | 26.70994 | 44.89684 | 54.52266 | 23.81734 | 0.101659 | 1.14041 | 0.837349 |
| 1 | 39304000 | 39306999 | 32.10182 | 52.76882 | 64.45981 | 27.77835 | 0.090264 | 1.413521 | 0.837402 |
| 13 | 67902800 | 67903399 | 7.256071 | 13.9908 | 22.56335 | 8.733581 | -0.13985 | -0.13899 | 0.837409 |
| 13 | 78506600 | 78507399 | 15.23212 | 26.64078 | 38.50648 | 16.98033 | -0.14104 | 0.681254 | 0.83742 |
| 17 | 29700200 | 29702799 | 26.26814 | 43.73913 | 60.93418 | 26.46539 | -0.09554 | 1.246488 | 0.837422 |
| 3 | 91140000 | 91140999 | 11.24271 | 21.46979 | 26.82224 | 10.66668 | 0.188467 | -0.0014 | 0.837448 |
| 2 | 171361600 | 171362799 | 15.19838 | 26.96892 | 32.85732 | 14.51467 | 0.109403 | 0.487746 | 0.837452 |
| 17 | 38638800 | 38640999 | 24.80271 | 42.45661 | 50.12291 | 21.97956 | 0.144415 | 0.969672 | 0.837464 |
| 10 | 68128000 | 68131399 | 45.58496 | 73.50657 | 87.68716 | 37.3426 | 0.103222 | 1.790707 | 0.837479 |
| 4 | 98574600 | 98578399 | 36.92113 | 60.12101 | 72.9386 | 31.3166 | 0.092418 | 1.55581 | 0.837492 |
| 4 | 49681600 | 49682199 | 6.82357 | 13.21134 | 21.59362 | 8.212464 | -0.13447 | -0.20484 | 0.837497 |
| 14 | 106694600 | 106700999 | 132.3784 | 200.5212 | 230.9793 | 95.57601 | 0.074825 | 3.268663 | 0.837504 |
| 14 | 59313400 | 59315199 | 18.96064 | 32.93273 | 40.032 | 17.57281 | 0.106545 | 0.800287 | 0.837525 |
| 7 | 139600800 | 139604599 | 81.10249 | 126.7321 | 148.9266 | 61.78975 | 0.081157 | 2.654342 | 0.837536 |
| 13 | 120772400 | 120783199 | 137.9007 | 207.9373 | 265.7068 | 109.788 | -0.08035 | 3.362968 | 0.83754 |
| 2 | 97076400 | 97076999 | 13.93042 | 24.55791 | 35.26874 | 15.60906 | -0.13018 | 0.554674 | 0.837545 |
| 17 | 40364600 | 40365799 | 22.26666 | 37.58922 | 52.73822 | 23.06787 | -0.10552 | 1.088941 | 0.837557 |
| X | 22240800 | 22241999 | 15.6021 | 28.3854 | 33.15551 | 14.27731 | 0.194503 | 0.345083 | 0.837564 |
| 6 | 132028800 | 132033799 | 29.23057 | 49.54111 | 45.80705 | 20.06024 | 0.494081 | 1.072206 | 0.837592 |
| 11 | 15472000 | 15472999 | 10.00487 | 18.83201 | 28.57497 | 11.7064 | -0.12715 | 0.088695 | 0.837614 |
| 12 | 113941400 | 113943999 | 30.55127 | 50.58794 | 72.70992 | 31.45924 | -0.14116 | 1.329749 | 0.837627 |
| 12 | 60408000 | 60410599 | 48.60782 | 77.82335 | 91.35394 | 38.49822 | 0.114013 | 2.015334 | 0.837651 |
| 4 | 74566800 | 74567799 | 16.94072 | 29.53599 | 41.5409 | 18.27863 | -0.10238 | 0.73956 | 0.837659 |
| 8 | 86901000 | 86902399 | 18.92874 | 32.66384 | 45.83223 | 20.10611 | -0.10141 | 0.871285 | 0.837681 |
| 14 | 107880600 | 107883199 | 24.54304 | 41.71348 | 50.6485 | 22.19728 | 0.102554 | 1.034204 | 0.837687 |
| 15 | 22648600 | 22648999 | 10.61464 | 19.85793 | 25.2094 | 10.43515 | 0.116607 | 0.125618 | 0.837702 |
| 9 | 6440000 | 6440799 | 8.027654 | 15.4013 | 24.08904 | 9.459003 | -0.11552 | -0.07902 | 0.837739 |
| 15 | 50492600 | 50495799 | 39.82446 | 64.39377 | 77.1635 | 32.97727 | 0.101251 | 1.694481 | 0.837747 |
| X | 77415400 | 77424799 | 52.27197 | 83.93049 | 83.0875 | 35.16784 | 0.364255 | 1.929516 | 0.837753 |
| 2 | 77502600 | 77502799 | 5.346253 | 10.45276 | 15.47795 | 5.298335 | 0.147027 | -0.61102 | 0.837754 |
| 3 | 43807200 | 43816999 | 100.4582 | 155.9222 | 154.6007 | 64.20282 | 0.318433 | 2.780621 | 0.83776 |
| 16 | 33628200 | 33632999 | 57.65733 | 91.99469 | 109.7336 | 46.01843 | 0.085461 | 2.135565 | 0.837776 |
| 19 | 48502600 | 48504799 | 16.80783 | 29.42706 | 41.4011 | 18.27273 | -0.10256 | 0.676001 | 0.837776 |
| 1 | 72162600 | 72175999 | 224.5085 | 325.9595 | 400.5815 | 165.1418 | -0.0749 | 4.0217 | 0.837793 |
| 3 | 61766000 | 61766999 | 10.86749 | 19.65933 | 35.34834 | 15.33035 | -0.43069 | 0.347221 | 0.837832 |
| 14 | 89528800 | 89532199 | 42.91768 | 69.12573 | 94.16597 | 40.0384 | -0.08988 | 1.822638 | 0.837835 |
| 14 | 51716400 | 51720799 | 57.00384 | 90.83512 | 108.4917 | 45.41666 | 0.081609 | 2.175671 | 0.837844 |

| | | | | | | | | | |
|---|---|---|---|---|---|---|---|---|---|
| Y | 41922000 | 41926399 | 38.51124 | 61.5516 | 98.68094 | 41.4838 | -0.33753 | 2.058709 | 0.837852 |
| 2 | 30081200 | 30082199 | 14.48669 | 25.67462 | 30.63256 | 13.54281 | 0.138832 | 0.490402 | 0.837853 |
| 7 | 24028800 | 24033399 | 57.93702 | 91.97043 | 122.9894 | 51.24989 | -0.0868 | 2.297876 | 0.837857 |
| 12 | 99499800 | 99503199 | 58.84328 | 93.52392 | 124.8872 | 52.10476 | -0.08303 | 2.262173 | 0.837864 |
| 19 | 5490000 | 5491199 | 13.94847 | 24.63278 | 35.23672 | 15.58943 | -0.12349 | 0.535986 | 0.837867 |
| 2 | 8223800 | 8227999 | 59.63844 | 95.0832 | 112.9451 | 47.31529 | 0.090294 | 2.162579 | 0.837872 |
| 13 | 51548800 | 51551399 | 31.78429 | 52.41518 | 64.00398 | 27.61947 | 0.092636 | 1.373456 | 0.837874 |
| 11 | 28008600 | 28010199 | 18.5246 | 32.49143 | 39.11817 | 17.29152 | 0.121016 | 0.651892 | 0.83792 |
| 2 | 66258400 | 66260799 | 34.39837 | 55.86826 | 76.74518 | 32.91009 | -0.08922 | 1.595633 | 0.837921 |
| 5 | 98387600 | 98389399 | 15.62395 | 27.46424 | 38.66731 | 17.12303 | -0.10363 | 0.599993 | 0.837939 |
| 14 | 121624600 | 121627799 | 30.97097 | 50.97256 | 62.11647 | 26.78244 | 0.094447 | 1.394568 | 0.837944 |
| 5 | 106143600 | 106145199 | 13.66644 | 24.2844 | 34.53716 | 15.2388 | -0.11136 | 0.472859 | 0.837976 |
| 11 | 55009400 | 55011999 | 30.74244 | 50.39384 | 69.71325 | 30.00691 | -0.09028 | 1.449436 | 0.837994 |
| 14 | 53019000 | 53021199 | 15.66167 | 27.52834 | 38.69218 | 17.13392 | -0.10118 | 0.602078 | 0.838019 |
| 11 | 112768600 | 112770799 | 16.83595 | 29.52403 | 41.86507 | 18.50278 | -0.11387 | 0.65118 | 0.838027 |
| 3 | 70759400 | 70762999 | 38.5355 | 62.67911 | 75.12627 | 32.24264 | 0.108748 | 1.578797 | 0.83804 |
| 10 | 116593000 | 116595599 | 22.1843 | 38.02485 | 46.33193 | 20.31933 | 0.099347 | 0.943421 | 0.838057 |
| 10 | 122812400 | 122817999 | 53.96257 | 86.10827 | 115.3709 | 48.38945 | -0.08177 | 2.13103 | 0.838061 |
| 14 | 17509800 | 17512399 | 16.18478 | 28.88785 | 27.38807 | 12.03907 | 0.474332 | 0.44897 | 0.838075 |
| 10 | 3080400 | 3080999 | 8.733716 | 16.70001 | 26.21165 | 10.3917 | -0.13325 | -0.03267 | 0.838079 |
| X | 148289800 | 148290199 | 7.630638 | 14.80373 | 19.49586 | 7.345172 | 0.188265 | -0.22994 | 0.838082 |
| 1 | 150096200 | 150097199 | 9.275345 | 17.34315 | 26.34137 | 10.91354 | -0.14119 | 0.120307 | 0.83811 |
| 18 | 5769600 | 5773999 | 52.33045 | 83.69055 | 99.91436 | 42.03932 | 0.087901 | 2.055196 | 0.838125 |
| Y | 88740200 | 88741799 | 7.489728 | 14.12208 | 23.96309 | 9.788475 | -0.2829 | 0.044421 | 0.83816 |
| 2 | 26572400 | 26574599 | 17.09793 | 30.16877 | 36.45988 | 16.13335 | 0.11771 | 0.564597 | 0.838187 |
| 9 | 38790000 | 38792799 | 46.43088 | 74.68823 | 89.72694 | 38.07995 | 0.088772 | 1.864977 | 0.838211 |
| 9 | 77639400 | 77643999 | 54.79165 | 87.66289 | 104.0231 | 43.79369 | 0.097284 | 2.046564 | 0.838235 |
| 13 | 45846800 | 45850199 | 28.73232 | 47.29259 | 65.81895 | 28.37994 | -0.09658 | 1.390594 | 0.838242 |
| 1 | 192635400 | 192636199 | 12.26474 | 22.30387 | 32.22599 | 13.88464 | -0.11092 | 0.321198 | 0.83826 |
| 2 | 20258400 | 20261199 | 52.08923 | 83.39281 | 100.1189 | 42.18851 | 0.0815 | 2.024163 | 0.838287 |
| 2 | 10261800 | 10263599 | 18.10444 | 31.47679 | 44.68545 | 19.62777 | -0.11643 | 0.7863 | 0.838357 |
| 1 | 39845600 | 39847199 | 13.72714 | 24.42607 | 34.58758 | 15.24324 | -0.1038 | 0.466251 | 0.838368 |
| 1 | 104664400 | 104667799 | 42.58434 | 68.58128 | 81.85167 | 34.79692 | 0.100371 | 1.82141 | 0.838373 |
| 3 | 29099600 | 29101999 | 15.9655 | 27.9424 | 39.38544 | 17.37789 | -0.10518 | 0.673706 | 0.838406 |
| 9 | 57211000 | 57216999 | 78.61928 | 123.8585 | 124.5146 | 51.66637 | 0.315143 | 2.496064 | 0.838412 |
| 5 | 4213600 | 4214599 | 9.87163 | 18.62268 | 23.79096 | 9.708758 | 0.1244 | 0.063486 | 0.83845 |
| 1 | 117976800 | 117979799 | 53.11725 | 85.14798 | 101.0302 | 42.65579 | 0.100633 | 1.986474 | 0.838465 |
| 12 | 98139000 | 98144999 | 68.08336 | 106.785 | 165.7613 | 68.74818 | -0.31576 | 2.584487 | 0.838476 |
| 1 | 179092000 | 179098199 | 118.6604 | 181.9952 | 177.9977 | 73.89091 | 0.325071 | 3.009564 | 0.838497 |
| 1 | 33849200 | 33850799 | 13.69674 | 24.63136 | 30.06483 | 13.13226 | 0.118246 | 0.39172 | 0.838511 |
| 5 | 128491000 | 128493399 | 21.28602 | 36.55377 | 51.73688 | 22.69306 | -0.11555 | 0.929338 | 0.838511 |
| 5 | 103341400 | 103342999 | 14.28462 | 25.22068 | 36.07537 | 15.9612 | -0.12392 | 0.538183 | 0.838529 |
| X | 77929200 | 77938399 | 178.2501 | 264.8549 | 298.4409 | 122.9466 | 0.08185 | 3.646368 | 0.83854 |
| 14 | 25864400 | 25866799 | 15.75967 | 27.52451 | 39.4586 | 17.38132 | -0.12959 | 0.704475 | 0.83854 |
| X | 34871600 | 34873199 | 24.30755 | 42.28745 | 37.53695 | 16.49416 | 0.558244 | 0.799242 | 0.838559 |
| 9 | 7087800 | 7092999 | 70.72071 | 112.359 | 110.0575 | 45.90018 | 0.362849 | 2.281155 | 0.838573 |
| Y | 20735400 | 20736599 | 8.760142 | 16.17936 | 26.42886 | 11.15625 | -0.26565 | 0.191559 | 0.838606 |

| | | | | | | | | | |
|---|---|---|---|---|---|---|---|---|---|
| 5 | 86153000 | 86155399 | 29.88982 | 49.23716 | 68.25335 | 29.43957 | -0.09043 | 1.385727 | 0.83862 |
| 7 | 70386400 | 70387399 | 13.51249 | 24.34569 | 35.14889 | 15.32098 | -0.12095 | 0.387721 | 0.838645 |
| 6 | 108024400 | 108024999 | 7.256967 | 14.05919 | 22.75134 | 8.627684 | -0.11666 | -0.20711 | 0.838652 |
| 3 | 66888200 | 66890399 | 27.18253 | 44.85913 | 63.69283 | 27.4941 | -0.1239 | 1.350597 | 0.838665 |
| 13 | 85292800 | 85295199 | 24.10912 | 40.94641 | 58.95569 | 25.83475 | -0.14265 | 1.04692 | 0.838675 |
| 5 | 60829600 | 60833999 | 60.93963 | 96.88145 | 115.5288 | 48.21308 | 0.080356 | 2.254193 | 0.838723 |
| 4 | 5375000 | 5376399 | 14.7561 | 25.75878 | 45.20638 | 19.90688 | -0.42012 | 0.716367 | 0.838734 |
| 15 | 50487000 | 50491399 | 86.79419 | 134.8538 | 156.4159 | 64.94659 | 0.093579 | 2.757801 | 0.838735 |
| Y | 87297800 | 87298399 | 7.626695 | 14.72196 | 17.87342 | 6.893793 | 0.271159 | -0.16216 | 0.838737 |
| 3 | 15959400 | 15959999 | 10.60221 | 19.62487 | 28.82033 | 12.12921 | -0.1104 | 0.200369 | 0.838745 |
| 6 | 104150000 | 104150799 | 10.60221 | 19.62487 | 28.82033 | 12.12921 | -0.1104 | 0.200369 | 0.838745 |
| 10 | 43337600 | 43339999 | 19.54877 | 33.99691 | 41.47403 | 18.2202 | 0.101438 | 0.785953 | 0.838746 |
| 11 | 11277400 | 11278799 | 19.54877 | 33.99691 | 41.47403 | 18.2202 | 0.101438 | 0.785953 | 0.838746 |
| 9 | 19045000 | 19046399 | 19.37646 | 33.88372 | 40.80624 | 17.99202 | 0.12069 | 0.707326 | 0.838756 |
| 2 | 171639200 | 171643599 | 35.22756 | 57.12923 | 78.34927 | 33.56082 | -0.0889 | 1.628339 | 0.838758 |
| 9 | 119588600 | 119589999 | 16.94783 | 29.82542 | 43.92158 | 19.44534 | -0.16749 | 0.606047 | 0.838821 |
| 13 | 51025400 | 51030399 | 39.33658 | 64.66465 | 72.82303 | 31.38309 | 0.207264 | 1.437168 | 0.838828 |
| 5 | 145461000 | 145461999 | 17.05603 | 30.06967 | 36.45473 | 16.13377 | 0.112685 | 0.572783 | 0.838829 |
| 10 | 118102400 | 118102599 | 4.510141 | 8.820841 | 16.09544 | 5.454593 | -0.13806 | -0.65951 | 0.838852 |
| 1 | 5437800 | 5439599 | 23.6576 | 40.6894 | 40.50091 | 17.76553 | 0.390897 | 0.913122 | 0.838878 |
| 16 | 80692400 | 80698199 | 85.11852 | 133.263 | 129.1756 | 53.5845 | 0.360729 | 2.62015 | 0.838928 |
| 4 | 18710400 | 18710799 | 7.264988 | 13.98278 | 22.53355 | 8.767285 | -0.14568 | -0.12185 | 0.838938 |
| 13 | 92227600 | 92231999 | 37.32979 | 60.95286 | 72.84801 | 31.30889 | 0.116716 | 1.516433 | 0.838942 |
| 11 | 99822800 | 99823999 | 22.80623 | 38.23127 | 54.3111 | 23.68923 | -0.12363 | 1.157842 | 0.838944 |
| 10 | 58521400 | 58524399 | 29.44538 | 48.69824 | 59.42092 | 25.68821 | 0.094354 | 1.329944 | 0.838944 |
| 13 | 97400800 | 97406799 | 80.77351 | 127.5854 | 123.2313 | 51.17624 | 0.375728 | 2.44147 | 0.83898 |
| 5 | 39877400 | 39878599 | 16.86324 | 28.66121 | 54.72123 | 23.98222 | -0.54764 | 1.013309 | 0.838987 |
| 8 | 33656800 | 33658599 | 21.1847 | 36.2678 | 50.62411 | 22.19956 | -0.09619 | 0.956718 | 0.838996 |
| 2 | 8274200 | 8274799 | 12.45825 | 22.69777 | 28.08485 | 12.07242 | 0.113515 | 0.301773 | 0.839001 |
| 14 | 41478000 | 41480399 | 25.52746 | 43.46721 | 43.14548 | 18.91182 | 0.392584 | 1.012111 | 0.839013 |
| 16 | 12347200 | 12349599 | 35.87137 | 58.14673 | 79.59022 | 34.0809 | -0.08673 | 1.639032 | 0.839014 |
| 12 | 70464400 | 70465799 | 15.39345 | 27.1272 | 32.59772 | 14.42798 | 0.125264 | 0.560562 | 0.839028 |
| 13 | 60534600 | 60535799 | 14.3077 | 25.67372 | 37.71208 | 16.50305 | -0.1497 | 0.42181 | 0.839061 |
| 4 | 85301200 | 85302199 | 9.245612 | 17.4761 | 26.47656 | 10.75979 | -0.11481 | 0.049755 | 0.839077 |
| 12 | 59299200 | 59300199 | 12.25311 | 21.80363 | 32.18413 | 14.18375 | -0.16337 | 0.450254 | 0.839095 |
| 7 | 47232000 | 47232199 | 6.515209 | 12.68873 | 21.42374 | 7.831686 | -0.12819 | -0.34811 | 0.839102 |
| 7 | 87177400 | 87178599 | 14.04529 | 25.19736 | 30.90806 | 13.52805 | 0.108815 | 0.408139 | 0.839103 |
| 5 | 24499200 | 24500999 | 21.24251 | 36.76349 | 44.69683 | 19.60976 | 0.105222 | 0.852207 | 0.839141 |
| 4 | 23517400 | 23519599 | 36.78823 | 59.73531 | 72.51127 | 31.08895 | 0.088427 | 1.60212 | 0.839173 |
| 7 | 12845800 | 12846399 | 13.46257 | 23.83361 | 34.22164 | 15.13285 | -0.12744 | 0.503897 | 0.839181 |
| 14 | 33624600 | 33625599 | 11.86201 | 22.02872 | 27.66839 | 11.5927 | 0.117819 | 0.187187 | 0.8392 |
| 17 | 19649800 | 19651799 | 37.63841 | 61.40502 | 73.71274 | 31.67081 | 0.109563 | 1.532044 | 0.83921 |
| 10 | 78920200 | 78928999 | 97.78183 | 151.9235 | 174.3722 | 72.40252 | 0.10869 | 2.763882 | 0.839214 |
| 4 | 145080000 | 145080599 | 9.698902 | 18.19348 | 27.40118 | 11.29103 | -0.1231 | 0.105177 | 0.839254 |
| 7 | 5193400 | 5199599 | 75.38962 | 119.9418 | 104.78 | 43.76647 | 0.529687 | 2.243611 | 0.839261 |
| X | 7744200 | 7745199 | 13.12297 | 24.46515 | 28.35454 | 11.8065 | 0.240878 | 0.181312 | 0.839273 |
| 3 | 27049800 | 27051599 | 15.85668 | 27.94938 | 33.98962 | 15.04342 | 0.108119 | 0.562935 | 0.839297 |

| | | | | | | | | | |
|---|---|---|---|---|---|---|---|---|---|
| 14 | 6550200 | 6550799 | 9.286356 | 17.55043 | 26.48177 | 10.76539 | -0.10921 | 0.052394 | 0.839299 |
| 1 | 161499800 | 161500199 | 7.710217 | 14.89603 | 23.76036 | 9.127539 | -0.1139 | -0.15551 | 0.839301 |
| 5 | 78560400 | 78560999 | 7.710217 | 14.89603 | 23.76036 | 9.127539 | -0.1139 | -0.15551 | 0.839301 |
| 4 | 21373800 | 21374999 | 21.38502 | 36.1748 | 51.14748 | 22.38596 | -0.11655 | 1.064877 | 0.839318 |
| 4 | 6145000 | 6146999 | 23.36859 | 39.63937 | 55.51666 | 24.31857 | -0.10284 | 1.053322 | 0.839319 |
| 5 | 115917400 | 115928199 | 170.7996 | 254.7647 | 289.079 | 119.144 | 0.076906 | 3.576056 | 0.839333 |
| 1 | 90354800 | 90356999 | 31.72997 | 51.67139 | 72.20394 | 31.00193 | -0.11085 | 1.543484 | 0.839361 |
| 12 | 113786200 | 113791999 | 63.7657 | 101.4193 | 141.8745 | 59.23708 | -0.14902 | 2.250726 | 0.839383 |
| 4 | 28622200 | 28625599 | 34.50991 | 56.61999 | 68.49269 | 29.49266 | 0.103236 | 1.446873 | 0.839394 |
| 17 | 61332600 | 61335799 | 43.88836 | 71.02284 | 70.85519 | 30.303 | 0.36621 | 1.680963 | 0.839408 |
| 7 | 4281800 | 4286399 | 36.65325 | 59.33446 | 81.38922 | 34.81677 | -0.09164 | 1.667208 | 0.83941 |
| 8 | 121160000 | 121162799 | 21.63958 | 36.63843 | 51.65871 | 22.61589 | -0.1124 | 1.060957 | 0.839411 |
| 8 | 8710000 | 8711999 | 20.09736 | 34.10058 | 49.02984 | 21.479 | -0.1404 | 1.031764 | 0.839415 |
| 8 | 117426000 | 117428199 | 35.46612 | 58.05776 | 70.26487 | 30.2269 | 0.101254 | 1.476999 | 0.83943 |
| 6 | 82486400 | 82488399 | 21.03002 | 35.9534 | 50.0817 | 21.95927 | -0.09349 | 0.96816 | 0.839446 |
| 12 | 111890600 | 111893799 | 26.93251 | 45.01222 | 55.04375 | 23.9631 | 0.091803 | 1.201822 | 0.839476 |
| 12 | 85340000 | 85343399 | 37.731 | 61.74667 | 72.70806 | 31.27437 | 0.140452 | 1.489569 | 0.839487 |
| 6 | 101723200 | 101723599 | 7.286429 | 14.02245 | 22.54117 | 8.773118 | -0.1422 | -0.11904 | 0.839494 |
| 8 | 19570000 | 19611599 | 656.6247 | 871.7268 | 980.0016 | 409.5775 | -0.07605 | 5.395616 | 0.839512 |
| X | 52775800 | 52778599 | 41.40275 | 66.95732 | 79.71731 | 34.07627 | 0.110937 | 1.698687 | 0.839513 |
| 15 | 60316400 | 60317799 | 19.17716 | 33.28726 | 46.85933 | 20.56144 | -0.1045 | 0.8192 | 0.83955 |
| 6 | 123751200 | 123753999 | 19.11448 | 33.81575 | 38.39508 | 16.98074 | 0.208684 | 0.551451 | 0.839561 |
| 2 | 89916000 | 89918599 | 33.48885 | 54.1937 | 89.45152 | 38.25619 | -0.35846 | 1.673476 | 0.839589 |
| 8 | 26823400 | 26825999 | 16.70977 | 29.27168 | 41.40856 | 18.28462 | -0.11035 | 0.66778 | 0.839593 |
| 6 | 10225800 | 10226199 | 7.853063 | 15.22969 | 21.08493 | 7.957416 | 0.11316 | -0.22079 | 0.839598 |
| 2 | 110328000 | 110328599 | 12.05114 | 22.03083 | 27.04872 | 11.57771 | 0.128432 | 0.275377 | 0.839599 |
| X | 125508400 | 125511399 | 28.59382 | 48.28441 | 56.20403 | 24.586 | 0.162896 | 1.112428 | 0.8396 |
| 16 | 8316800 | 8327999 | 139.88 | 209.0849 | 311.679 | 128.5111 | -0.31376 | 3.533246 | 0.839606 |
| 14 | 60882000 | 60885599 | 25.68861 | 43.02526 | 59.9558 | 26.1368 | -0.09611 | 1.179979 | 0.83961 |
| 6 | 129752400 | 129755199 | 21.23061 | 37.14303 | 42.85032 | 18.90057 | 0.18235 | 0.712529 | 0.839616 |
| 15 | 74900800 | 74905999 | 61.95462 | 98.59844 | 117.2979 | 49.01661 | 0.08534 | 2.223223 | 0.839641 |
| 16 | 22808200 | 22810799 | 37.47 | 60.94775 | 73.95757 | 31.74138 | 0.09101 | 1.574174 | 0.83966 |
| X | 163533200 | 163533799 | 6.832034 | 13.25734 | 21.66987 | 8.155814 | -0.11999 | -0.23874 | 0.839689 |
| X | 117057200 | 117060399 | 28.33698 | 47.83021 | 56.22525 | 24.59209 | 0.148704 | 1.113965 | 0.839699 |
| 7 | 42215800 | 42294399 | 1757.506 | 2167.998 | 2004.221 | 844.225 | 0.09963 | 6.584832 | 0.8397 |
| 1 | 23395000 | 23395399 | 6.810223 | 13.22607 | 21.73172 | 8.133023 | -0.12039 | -0.2605 | 0.839723 |
| 15 | 3917000 | 3917999 | 16.09408 | 28.45811 | 34.07452 | 15.07142 | 0.132515 | 0.534045 | 0.839724 |
| 4 | 146806400 | 146831799 | 1362.71 | 1715.285 | 1630.56 | 685.2554 | 0.08891 | 6.260762 | 0.839727 |
| 5 | 143333000 | 143340599 | 113.4062 | 173.4213 | 201.419 | 83.56362 | 0.074007 | 3.069656 | 0.839752 |
| 11 | 47432200 | 47432999 | 8.829476 | 16.70537 | 25.44658 | 10.32047 | -0.11961 | 0.038354 | 0.839764 |
| 9 | 19511200 | 19514799 | 46.7141 | 75.23604 | 89.95593 | 38.24507 | 0.09783 | 1.829236 | 0.839765 |
| 17 | 75752000 | 75752599 | 6.829029 | 13.26209 | 21.71972 | 8.130999 | -0.11569 | -0.25754 | 0.839807 |
| 7 | 143219800 | 143223599 | 36.33925 | 58.81944 | 80.67525 | 34.50844 | -0.09164 | 1.668894 | 0.839826 |
| 12 | 55938200 | 55939799 | 11.93806 | 21.43931 | 31.19359 | 13.64103 | -0.13483 | 0.389485 | 0.839835 |
| 19 | 59589600 | 59596399 | 65.95567 | 104.9153 | 122.6091 | 51.20184 | 0.110249 | 2.243838 | 0.839838 |
| 17 | 27451200 | 27481399 | 349.8349 | 485.9787 | 587.3843 | 243.8088 | -0.11468 | 4.679263 | 0.839843 |
| 5 | 13779800 | 13780799 | 14.99599 | 26.34308 | 37.23168 | 16.47914 | -0.10906 | 0.604777 | 0.839845 |

| | | | | | | | | | |
|---|---|---|---|---|---|---|---|---|---|
| 7 | 18605600 | 18609599 | 36.83762 | 59.51273 | 82.04861 | 35.02592 | -0.10197 | 1.718173 | 0.839868 |
| X | 48505600 | 48506599 | 10.80249 | 20.59656 | 26.1473 | 10.43308 | 0.161917 | 0.006956 | 0.839894 |
| 3 | 27378800 | 27383199 | 36.52238 | 59.3183 | 81.8725 | 35.10894 | -0.09611 | 1.602088 | 0.839898 |
| X | 16484800 | 16486399 | 15.53234 | 27.82124 | 32.55094 | 14.26607 | 0.175785 | 0.436533 | 0.839914 |
| 3 | 97628200 | 97632399 | 48.01769 | 77.41346 | 90.74671 | 38.63529 | 0.12804 | 1.800814 | 0.839917 |
| 12 | 24770800 | 24780399 | 130.8415 | 197.3167 | 252.3254 | 104.2628 | -0.08127 | 3.359179 | 0.839927 |
| 14 | 99886800 | 99887399 | 8.486159 | 16.23913 | 25.12344 | 9.938305 | -0.11005 | -0.04778 | 0.83994 |
| 6 | 131091600 | 131094399 | 45.5506 | 73.34249 | 88.38005 | 37.56048 | 0.086106 | 1.835129 | 0.839952 |
| 2 | 86955600 | 86956399 | 11.78698 | 21.93767 | 27.74125 | 11.58395 | 0.112064 | 0.174977 | 0.839954 |
| 6 | 50879000 | 50880599 | 20.61019 | 35.48772 | 50.1021 | 21.97779 | -0.11114 | 0.90061 | 0.839957 |
| 14 | 35153400 | 35154999 | 17.8451 | 31.24345 | 38.21262 | 16.86562 | 0.098479 | 0.675445 | 0.839973 |
| 9 | 70459200 | 70461199 | 27.0958 | 45.94188 | 53.94354 | 23.62638 | 0.150641 | 1.070603 | 0.839991 |
| 19 | 33798600 | 33803799 | 59.11674 | 94.02447 | 125.5924 | 52.4383 | -0.08244 | 2.243411 | 0.840002 |
| 7 | 28542600 | 28544399 | 20.67119 | 35.79658 | 43.66658 | 19.15812 | 0.100709 | 0.844377 | 0.84001 |
| 1 | 64438000 | 64439999 | 22.93942 | 39.54521 | 47.38325 | 20.78589 | 0.125014 | 0.898048 | 0.840017 |
| 13 | 36742800 | 36744599 | 15.21097 | 26.95511 | 32.88569 | 14.53927 | 0.106471 | 0.498598 | 0.840045 |
| 12 | 48968400 | 48969799 | 16.7527 | 29.54454 | 35.97517 | 15.9205 | 0.10663 | 0.567083 | 0.840059 |
| 14 | 37360800 | 37361799 | 14.96814 | 26.48354 | 32.07214 | 14.184 | 0.116445 | 0.514249 | 0.84009 |
| 3 | 40271400 | 40274999 | 55.48712 | 88.52579 | 119.3881 | 50.06223 | -0.09146 | 2.139447 | 0.84011 |
| 8 | 71642400 | 71643399 | 14.92783 | 26.41463 | 32.027 | 14.16377 | 0.114698 | 0.512082 | 0.840112 |
| 8 | 45894600 | 45900399 | 96.15097 | 147.8027 | 192.265 | 79.83814 | -0.08354 | 2.960599 | 0.840127 |
| 16 | 75430400 | 75433399 | 56.80964 | 90.54975 | 108.1582 | 45.29372 | 0.081917 | 2.16647 | 0.840131 |
| 10 | 55100400 | 55101199 | 15.07523 | 26.62386 | 32.07018 | 14.18836 | 0.123007 | 0.534041 | 0.840141 |
| 1 | 27165400 | 27166199 | 11.52628 | 21.55582 | 27.31781 | 11.31466 | 0.118525 | 0.142419 | 0.840171 |
| 13 | 66906000 | 66906399 | 11.52628 | 21.55582 | 27.31781 | 11.31466 | 0.118525 | 0.142419 | 0.840171 |
| 2 | 89305600 | 89306399 | 18.17063 | 31.73228 | 38.83734 | 17.10664 | 0.097463 | 0.718124 | 0.840177 |
| 17 | 75128800 | 75133399 | 57.78536 | 92.81937 | 102.8701 | 43.56698 | 0.20234 | 1.933043 | 0.840209 |
| 11 | 33096600 | 33098599 | 15.91893 | 27.74715 | 40.02968 | 17.61074 | -0.13895 | 0.734432 | 0.840218 |
| 8 | 14316400 | 14320399 | 28.02891 | 46.35369 | 64.32639 | 27.8081 | -0.09074 | 1.330511 | 0.840223 |
| 16 | 38876600 | 38877799 | 9.600161 | 17.93609 | 26.84643 | 11.13792 | -0.12178 | 0.130093 | 0.840242 |
| 6 | 37551200 | 37552199 | 11.56954 | 21.50115 | 27.17454 | 11.37455 | 0.110016 | 0.18013 | 0.840259 |
| 3 | 102508800 | 102511199 | 23.72734 | 40.21003 | 49.04401 | 21.47715 | 0.095878 | 1.054729 | 0.840266 |
| 2 | 116680200 | 116681399 | 10.92822 | 19.98988 | 29.24036 | 12.50612 | -0.12157 | 0.270258 | 0.840285 |
| 12 | 105273200 | 105276399 | 53.23229 | 85.11659 | 102.2026 | 42.99049 | 0.079332 | 2.063855 | 0.840292 |
| 2 | 23310200 | 23311599 | 18.43057 | 32.49508 | 31.73369 | 14.0442 | 0.423144 | 0.571834 | 0.840297 |
| 18 | 39080600 | 39084599 | 37.0458 | 60.32588 | 73.04515 | 31.3627 | 0.095228 | 1.554905 | 0.840303 |
| 8 | 61074200 | 61075599 | 14.14236 | 24.99719 | 35.57319 | 15.73553 | -0.1159 | 0.524803 | 0.840311 |
| 15 | 87419000 | 87421999 | 36.14427 | 59.52802 | 58.68677 | 25.31107 | 0.399727 | 1.396015 | 0.840329 |
| 5 | 76818800 | 76823599 | 44.34368 | 71.26988 | 96.77583 | 41.02068 | -0.0891 | 1.891413 | 0.840389 |
| 1 | 90779800 | 90780999 | 12.31971 | 22.49494 | 32.76543 | 14.0456 | -0.11805 | 0.297179 | 0.840408 |
| 3 | 91307600 | 91309799 | 21.71759 | 37.98308 | 43.38614 | 19.13183 | 0.196618 | 0.720496 | 0.840423 |
| 9 | 19815000 | 19818799 | 63.57151 | 100.9153 | 120.0586 | 50.02893 | 0.081649 | 2.302653 | 0.840425 |
| 15 | 25533000 | 25535799 | 19.97308 | 34.63331 | 42.03887 | 18.44473 | 0.108051 | 0.824172 | 0.840432 |
| 6 | 140663400 | 140666199 | 31.95545 | 52.34722 | 72.96317 | 31.3969 | -0.10159 | 1.459962 | 0.840433 |
| 17 | 8131400 | 8133999 | 28.91563 | 47.89363 | 58.34193 | 25.24779 | 0.096897 | 1.309834 | 0.840434 |
| 1 | 73418400 | 73420999 | 17.11474 | 30.09464 | 36.43841 | 16.13495 | 0.112967 | 0.599811 | 0.840438 |
| 9 | 5861000 | 5865799 | 112.264 | 172.6032 | 172.0182 | 71.43244 | 0.300615 | 2.960542 | 0.840451 |

| | | | | | | | | | |
|---|---|---|---|---|---|---|---|---|---|
| X | 55198800 | 55201999 | 32.8338 | 54.62387 | 63.75639 | 27.6751 | 0.158839 | 1.276585 | 0.840454 |
| 9 | 89965200 | 89968799 | 38.4487 | 62.44524 | 75.68831 | 32.45635 | 0.090842 | 1.602972 | 0.840463 |
| 15 | 94564400 | 94566399 | 16.34841 | 28.74731 | 35.02068 | 15.50716 | 0.104298 | 0.593833 | 0.840486 |
| 8 | 128011800 | 128013199 | 14.50773 | 26.13341 | 31.98426 | 13.94518 | 0.115957 | 0.390849 | 0.840491 |
| 12 | 31033600 | 31034199 | 7.69734 | 14.91386 | 20.49914 | 7.769921 | 0.117667 | -0.20977 | 0.840512 |
| 3 | 6487600 | 6491399 | 45.98793 | 74.08343 | 88.88406 | 37.80187 | 0.093256 | 1.819559 | 0.840516 |
| 4 | 148453600 | 148456399 | 37.99625 | 61.31077 | 84.18031 | 35.88022 | -0.09805 | 1.756424 | 0.840525 |
| 17 | 21464800 | 21467599 | 29.94397 | 50.08727 | 50.2485 | 21.88708 | 0.376452 | 1.197401 | 0.840533 |
| 4 | 47428000 | 47430799 | 33.94497 | 55.48426 | 67.69478 | 29.10525 | 0.088144 | 1.492409 | 0.84055 |
| 17 | 31877200 | 31878199 | 7.43155 | 14.20038 | 22.97112 | 9.123453 | -0.17892 | -0.0515 | 0.840601 |
| 3 | 57476000 | 57478399 | 23.82336 | 40.84537 | 49.01565 | 21.49678 | 0.121375 | 0.952886 | 0.840604 |
| 11 | 30652000 | 30655399 | 25.08837 | 42.31055 | 51.74869 | 22.62822 | 0.091653 | 1.104583 | 0.84061 |
| 14 | 121088600 | 121097999 | 113.8408 | 173.4211 | 223.0497 | 92.45275 | -0.07741 | 3.148248 | 0.840733 |
| 1 | 88771400 | 88772799 | 17.83426 | 31.26087 | 38.1412 | 16.85068 | 0.101911 | 0.656299 | 0.84074 |
| 19 | 24183400 | 24187799 | 39.30588 | 63.7239 | 77.20562 | 33.06359 | 0.089074 | 1.641529 | 0.840787 |
| 10 | 37727200 | 37732799 | 113.9313 | 173.572 | 222.7855 | 92.34531 | -0.07435 | 3.14762 | 0.840793 |
| 13 | 7577400 | 7579199 | 18.61951 | 32.09549 | 45.81577 | 20.09823 | -0.12662 | 0.876105 | 0.840799 |
| 7 | 66524200 | 66527199 | 27.10484 | 44.98294 | 62.57319 | 27.11575 | -0.09371 | 1.288087 | 0.840813 |
| 18 | 73482400 | 73482799 | 8.292402 | 15.9746 | 21.43828 | 8.319611 | 0.11953 | -0.12212 | 0.840839 |
| 4 | 117666400 | 117667799 | 11.09407 | 20.60098 | 25.27854 | 10.5925 | 0.151945 | 0.179644 | 0.840848 |
| 1 | 50854400 | 50855599 | 20.23516 | 34.54584 | 48.48979 | 21.25885 | -0.1047 | 0.97642 | 0.840872 |
| 7 | 3313800 | 3315199 | 12.22347 | 21.97838 | 31.62379 | 13.81509 | -0.11799 | 0.384068 | 0.840891 |
| 1 | 58253800 | 58257199 | 31.52866 | 51.50956 | 71.37654 | 30.68275 | -0.09551 | 1.496034 | 0.840952 |
| 4 | 95142600 | 95152199 | 183.2109 | 269.3791 | 338.7042 | 139.486 | -0.09036 | 3.822897 | 0.841038 |
| 12 | 67579200 | 67580399 | 13.37472 | 23.98067 | 34.08914 | 14.92753 | -0.10306 | 0.414997 | 0.841054 |
| X | 21900400 | 21903599 | 39.89149 | 64.60779 | 77.45913 | 33.14737 | 0.102859 | 1.664866 | 0.84106 |
| 3 | 95280200 | 95285199 | 79.16229 | 124.1969 | 147.0492 | 60.98568 | 0.074802 | 2.582641 | 0.84106 |
| 1 | 45051000 | 45052399 | 14.73458 | 27.17065 | 31.84524 | 13.47164 | 0.20707 | 0.258688 | 0.841065 |
| 12 | 22469600 | 22469799 | 5.944085 | 11.58924 | 19.74551 | 7.114425 | -0.12102 | -0.40883 | 0.841073 |
| 9 | 105220200 | 105223199 | 36.19423 | 59.09112 | 71.46652 | 30.71371 | 0.099511 | 1.515249 | 0.841086 |
| 5 | 141032600 | 141039999 | 80.50987 | 126.1218 | 149.1077 | 61.84897 | 0.075249 | 2.609772 | 0.841114 |
| 7 | 42087800 | 42117999 | 556.9804 | 752.7312 | 774.5943 | 322.9038 | 0.077285 | 5.111625 | 0.841134 |
| Y | 6742200 | 6742999 | 11.67374 | 21.26729 | 23.27663 | 10.00629 | 0.289262 | 0.282745 | 0.841142 |
| 15 | 3972200 | 3973199 | 11.61744 | 21.30185 | 31.24143 | 13.31826 | -0.1224 | 0.263858 | 0.84115 |
| 17 | 37617200 | 37618799 | 15.61033 | 27.29886 | 38.913 | 17.15779 | -0.12132 | 0.683255 | 0.841193 |
| 12 | 105857600 | 105867599 | 201.861 | 296.8252 | 333.1397 | 137.1937 | 0.073534 | 3.82357 | 0.841199 |
| 9 | 102272800 | 102277199 | 45.88933 | 73.69555 | 99.53967 | 42.14302 | -0.08285 | 1.920552 | 0.841209 |
| 6 | 142177600 | 142181599 | 63.92046 | 101.5931 | 120.7686 | 50.38624 | 0.084481 | 2.2659 | 0.841217 |
| 14 | 8546000 | 8546799 | 11.80034 | 21.91314 | 27.71505 | 11.613 | 0.108048 | 0.18869 | 0.841236 |
| 9 | 18065600 | 18066199 | 9.314753 | 17.84596 | 23.49682 | 9.263143 | 0.12581 | -0.05298 | 0.841248 |
| 13 | 68217600 | 68338599 | 3127.952 | 3654.378 | 3611.814 | 1533.137 | -0.07666 | 7.442811 | 0.841256 |
| 7 | 133462600 | 133464999 | 20.75265 | 34.77044 | 61.24516 | 26.70484 | -0.43286 | 1.162595 | 0.841265 |
| 6 | 97215600 | 97217399 | 19.86453 | 34.49105 | 42.09317 | 18.47368 | 0.100816 | 0.813327 | 0.84128 |
| 19 | 52605400 | 52605999 | 7.867312 | 15.20693 | 20.45737 | 7.840839 | 0.133449 | -0.16958 | 0.841295 |
| 6 | 124524200 | 124528399 | 52.29789 | 83.54553 | 111.9379 | 47.02416 | -0.08 | 2.093911 | 0.841306 |
| 5 | 52596400 | 52601999 | 63.16205 | 100.1351 | 133.057 | 55.37335 | -0.07947 | 2.34124 | 0.841309 |
| 1 | 148272600 | 148273799 | 10.589 | 19.64672 | 28.83917 | 12.09822 | -0.10645 | 0.186831 | 0.841309 |

| | | | | | | | | | |
|---|---|---|---|---|---|---|---|---|---|
| 14 | 109947600 | 109948599 | 10.589 | 19.64672 | 28.83917 | 12.09822 | -0.10645 | 0.186831 | 0.841309 |
| 1 | 10957600 | 10959599 | 20.83555 | 35.33111 | 62.03921 | 27.17302 | -0.42796 | 1.053733 | 0.841349 |
| 13 | 11437600 | 11443999 | 76.62859 | 121.068 | 140.553 | 58.37692 | 0.110974 | 2.43686 | 0.841375 |
| 16 | 70578800 | 70582599 | 53.43075 | 85.08699 | 114.4363 | 47.86279 | -0.09037 | 2.194199 | 0.841405 |
| 10 | 83111400 | 83112999 | 19.54992 | 33.82598 | 47.40073 | 20.79589 | -0.09883 | 0.851307 | 0.841411 |
| 13 | 111205600 | 111208399 | 39.64907 | 64.39593 | 63.24114 | 27.11916 | 0.393715 | 1.597435 | 0.841416 |
| 11 | 76474600 | 76475999 | 14.42372 | 25.53137 | 36.50716 | 16.144 | -0.12207 | 0.51147 | 0.841432 |
| 5 | 143327600 | 143332199 | 48.87191 | 78.36345 | 106.2673 | 44.88511 | -0.09127 | 1.974846 | 0.841439 |
| 15 | 91001200 | 91007199 | 41.07669 | 66.48021 | 80.26913 | 34.33193 | 0.091681 | 1.679794 | 0.841451 |
| 17 | 55844600 | 55845199 | 7.467422 | 14.36369 | 23.07201 | 8.992421 | -0.14271 | -0.10926 | 0.841466 |
| 16 | 55596200 | 55598999 | 22.438 | 38.11863 | 53.33676 | 23.37444 | -0.10145 | 1.03268 | 0.841498 |
| 3 | 4685400 | 4685799 | 7.291333 | 14.09302 | 22.63759 | 8.678172 | -0.12041 | -0.16827 | 0.841503 |
| 3 | 20100000 | 20107199 | 54.44934 | 87.1824 | 86.82069 | 36.59198 | 0.350943 | 2.018169 | 0.841523 |
| 15 | 39093600 | 39096599 | 30.03331 | 49.77341 | 60.70785 | 26.28205 | 0.095114 | 1.30755 | 0.841559 |
| X | 36813800 | 36816399 | 25.44368 | 43.6843 | 41.01752 | 17.99106 | 0.474324 | 0.938556 | 0.841572 |
| 5 | 133747400 | 133750599 | 31.36595 | 51.4395 | 71.05034 | 30.58838 | -0.08784 | 1.445337 | 0.841581 |
| 18 | 68461600 | 68462999 | 41.52372 | 66.68462 | 92.06416 | 38.97803 | -0.1143 | 1.9134 | 0.841598 |
| 11 | 63839200 | 63840999 | 15.51159 | 27.20408 | 38.31731 | 16.93612 | -0.1041 | 0.636822 | 0.841614 |
| X | 125260800 | 125264599 | 54.54882 | 87.45278 | 102.3535 | 43.22202 | 0.12056 | 1.986566 | 0.841621 |
| 9 | 80314800 | 80315999 | 16.8644 | 29.42535 | 41.48357 | 18.26216 | -0.1058 | 0.726572 | 0.841632 |
| 5 | 138598200 | 138616999 | 356.2942 | 500.008 | 612.4583 | 253.815 | -0.11916 | 4.529507 | 0.841648 |
| 3 | 85930600 | 85933199 | 21.84793 | 37.05471 | 51.68709 | 22.64009 | -0.09698 | 1.044317 | 0.84168 |
| 15 | 89287600 | 89296799 | 127.1837 | 193.1934 | 223.4071 | 92.52158 | 0.072553 | 3.210491 | 0.841692 |
| 13 | 33418400 | 33424599 | 76.85918 | 121.647 | 138.7382 | 57.67473 | 0.138596 | 2.394636 | 0.841708 |
| 4 | 88493200 | 88493599 | 6.681518 | 13.01983 | 18.55405 | 6.73364 | 0.124521 | -0.37786 | 0.841727 |
| 6 | 66740400 | 66743599 | 28.05131 | 46.38734 | 64.29051 | 27.79118 | -0.08889 | 1.33195 | 0.841741 |
| 9 | 31150800 | 31152599 | 21.81627 | 37.09086 | 51.69294 | 22.6556 | -0.09552 | 1.022445 | 0.841756 |
| 10 | 125454600 | 125455999 | 13.82256 | 25.2842 | 23.97508 | 10.25416 | 0.499787 | 0.280222 | 0.841762 |
| 18 | 65912400 | 65913799 | 13.3839 | 24.22875 | 29.96336 | 12.98336 | 0.106621 | 0.348224 | 0.841799 |
| 16 | 4675800 | 4678799 | 23.10765 | 39.27783 | 47.96779 | 21.0226 | 0.094342 | 1.026021 | 0.841819 |
| 5 | 75886400 | 75889199 | 20.47371 | 35.06002 | 49.01337 | 21.49369 | -0.09835 | 0.948482 | 0.84184 |
| 12 | 22107600 | 22108999 | 27.65378 | 45.91633 | 55.52142 | 24.068 | 0.10786 | 1.27713 | 0.841864 |
| X | 14388000 | 14391799 | 78.42353 | 123.5013 | 144.2993 | 59.87278 | 0.098084 | 2.509606 | 0.841882 |
| 5 | 49512000 | 49512599 | 6.378874 | 12.42392 | 20.80903 | 7.601252 | -0.11526 | -0.3529 | 0.841884 |
| 5 | 80505800 | 80506199 | 6.378874 | 12.42392 | 20.80903 | 7.601252 | -0.11526 | -0.3529 | 0.841884 |
| 13 | 4918600 | 4927199 | 156.4785 | 233.8928 | 295.2816 | 121.75 | -0.0742 | 3.535786 | 0.841885 |
| 13 | 108133000 | 108135199 | 44.15306 | 70.97114 | 96.17069 | 40.77052 | -0.08588 | 1.890325 | 0.841895 |
| 1 | 40571600 | 40572599 | 15.70784 | 27.57442 | 38.78507 | 17.15662 | -0.1021 | 0.624029 | 0.841898 |
| 5 | 105536200 | 105543799 | 106.9901 | 164.5383 | 191.4217 | 79.49284 | 0.077889 | 2.95613 | 0.841908 |
| 19 | 8233600 | 8234999 | 25.34731 | 42.61944 | 51.77085 | 22.61736 | 0.10129 | 1.129838 | 0.841908 |
| 3 | 27307400 | 27315799 | 97.68211 | 150.4841 | 194.7813 | 80.90719 | -0.07397 | 2.919403 | 0.841911 |
| 16 | 59192800 | 59194999 | 22.89071 | 38.94526 | 46.9301 | 20.56935 | 0.113872 | 1.017827 | 0.841925 |
| 3 | 32705200 | 32708599 | 30.90359 | 51.1489 | 62.40264 | 26.99144 | 0.09459 | 1.32689 | 0.841939 |
| 14 | 82864000 | 82867399 | 40.85169 | 65.8527 | 89.49772 | 38.09769 | -0.08501 | 1.793905 | 0.841943 |
| 2 | 131239000 | 131240999 | 17.57085 | 30.82211 | 37.6445 | 16.64172 | 0.100503 | 0.643211 | 0.841953 |
| 14 | 9447400 | 9447799 | 5.049708 | 9.863951 | 17.63962 | 6.170082 | -0.15002 | -0.53081 | 0.84198 |
| 1 | 161597000 | 161598799 | 15.34818 | 27.16464 | 33.04932 | 14.61522 | 0.110031 | 0.515874 | 0.841983 |

| | | | | | | | | | |
|---|---|---|---|---|---|---|---|---|---|
| 1 | 46753200 | 46755999 | 29.33918 | 49.02712 | 47.5811 | 20.7123 | 0.424213 | 1.204767 | 0.841983 |
| 19 | 49019000 | 49021199 | 21.19441 | 36.85326 | 44.08441 | 19.36514 | 0.129578 | 0.799241 | 0.841991 |
| 8 | 53583800 | 53584799 | 13.57908 | 24.87505 | 30.61841 | 13.06958 | 0.127571 | 0.284173 | 0.84201 |
| 8 | 93347200 | 93348199 | 11.65327 | 21.55032 | 27.17692 | 11.45206 | 0.106454 | 0.209717 | 0.842021 |
| 2 | 56657400 | 56657799 | 5.939424 | 11.58574 | 19.82964 | 7.099737 | -0.12002 | -0.43341 | 0.842037 |
| 13 | 112977400 | 112979399 | 17.41096 | 30.49327 | 37.24726 | 16.44407 | 0.100419 | 0.667449 | 0.842063 |
| 7 | 58987200 | 58987399 | 4.510779 | 8.820258 | 16.06306 | 5.482902 | -0.14447 | -0.6342 | 0.842066 |
| 6 | 143282400 | 143284999 | 22.22799 | 37.97213 | 53.94165 | 23.65026 | -0.12203 | 0.981566 | 0.842073 |
| 5 | 145769800 | 145771799 | 42.48325 | 68.74406 | 82.05924 | 35.09285 | 0.107904 | 1.68833 | 0.842103 |
| 1 | 182460400 | 182466999 | 66.03391 | 104.2396 | 138.0645 | 57.31848 | -0.07989 | 2.448379 | 0.842112 |
| 8 | 59539000 | 59539999 | 15.51604 | 27.48064 | 33.50456 | 14.8145 | 0.107347 | 0.510244 | 0.842113 |
| 8 | 35570200 | 35571999 | 17.93472 | 31.10146 | 43.72649 | 19.18523 | -0.10275 | 0.819751 | 0.842128 |
| 10 | 7361200 | 7362599 | 31.77763 | 51.98462 | 71.88475 | 30.91603 | -0.09113 | 1.480613 | 0.842129 |
| 5 | 45553800 | 45555999 | 36.11439 | 58.38903 | 80.58915 | 34.43689 | -0.10224 | 1.696774 | 0.842145 |
| 3 | 9930400 | 9931799 | 10.09936 | 18.84387 | 27.87091 | 11.58882 | -0.10728 | 0.146008 | 0.842148 |
| 14 | 106251400 | 106252799 | 10.09936 | 18.84387 | 27.87091 | 11.58882 | -0.10728 | 0.146008 | 0.842148 |
| 13 | 93721200 | 93723799 | 26.69419 | 44.39485 | 61.57342 | 26.72135 | -0.08929 | 1.26017 | 0.842157 |
| X | 54474200 | 54476599 | 27.17645 | 46.33857 | 44.16857 | 19.36241 | 0.451214 | 1.009284 | 0.842192 |
| 3 | 92086800 | 92088399 | 17.62709 | 30.57894 | 43.22023 | 18.96331 | -0.1102 | 0.814843 | 0.842218 |
| 18 | 85844800 | 85847199 | 35.48915 | 57.62066 | 68.83962 | 29.51276 | 0.111252 | 1.59977 | 0.842219 |
| 4 | 105913200 | 105915599 | 19.2315 | 33.49387 | 40.91546 | 17.99125 | 0.099706 | 0.763244 | 0.842223 |
| 15 | 37591200 | 37595999 | 58.57174 | 93.08217 | 123.6782 | 51.59471 | -0.07598 | 2.265707 | 0.842231 |
| 4 | 143361600 | 143373999 | 278.9879 | 399.3102 | 436.5657 | 180.3371 | 0.073349 | 4.235818 | 0.842242 |
| 8 | 41099600 | 41103799 | 42.49609 | 68.13728 | 110.1945 | 46.54087 | -0.34494 | 1.969036 | 0.842251 |
| 17 | 30111200 | 30114399 | 31.36669 | 51.19762 | 71.28682 | 30.63387 | -0.10338 | 1.50859 | 0.842258 |
| 7 | 40330600 | 40332999 | 19.61982 | 33.91147 | 41.12565 | 18.04228 | 0.10858 | 0.852445 | 0.842264 |
| 10 | 69897800 | 69898599 | 7.760904 | 14.92114 | 23.59663 | 9.209404 | -0.12256 | -0.10107 | 0.842282 |
| 3 | 121309600 | 121311999 | 25.29171 | 42.45664 | 59.44784 | 25.95246 | -0.10309 | 1.152929 | 0.842328 |
| 15 | 93574200 | 93575799 | 15.43518 | 27.31626 | 33.46275 | 14.79831 | 0.099976 | 0.516602 | 0.84233 |
| 16 | 55309600 | 55311399 | 21.50016 | 37.4894 | 44.02401 | 19.37257 | 0.156584 | 0.763398 | 0.842352 |
| 3 | 116986400 | 116987999 | 15.20992 | 26.79546 | 37.67823 | 16.67768 | -0.10043 | 0.573426 | 0.842354 |
| 1 | 27315200 | 27316199 | 17.33664 | 30.13887 | 42.67429 | 18.7432 | -0.11234 | 0.785853 | 0.842393 |
| X | 113674000 | 113674599 | 9.584396 | 18.24338 | 23.90474 | 9.572251 | 0.112122 | 0.000168 | 0.842395 |
| 17 | 43522800 | 43524999 | 26.67773 | 44.34749 | 61.58252 | 26.71698 | -0.0911 | 1.264853 | 0.842404 |
| 1 | 35523800 | 35524999 | 19.50493 | 33.91742 | 41.52469 | 18.24135 | 0.09643 | 0.789123 | 0.842409 |
| 17 | 82107400 | 82116599 | 182.552 | 273.4832 | 247.7701 | 102.205 | 0.407142 | 3.495507 | 0.842441 |
| 10 | 129044800 | 129053999 | 225.5155 | 328.2153 | 364.3096 | 150.1507 | 0.075297 | 3.98435 | 0.842453 |
| 10 | 84412000 | 84414399 | 13.71422 | 24.10217 | 35.30704 | 15.63316 | -0.16042 | 0.58628 | 0.842472 |
| 11 | 80191400 | 80192999 | 15.16097 | 26.55169 | 37.88732 | 16.72613 | -0.12274 | 0.656323 | 0.842473 |
| 8 | 68165200 | 68166199 | 12.30486 | 22.51753 | 28.10589 | 12.01401 | 0.106331 | 0.275614 | 0.842475 |
| 2 | 98067800 | 98075799 | 83.96741 | 132.082 | 151.6829 | 62.92713 | 0.121699 | 2.535337 | 0.842497 |
| 4 | 3660600 | 3661799 | 12.23906 | 22.45114 | 28.07475 | 11.96143 | 0.106919 | 0.26038 | 0.842524 |
| 3 | 42260600 | 42261599 | 11.32936 | 21.06556 | 26.71366 | 11.17315 | 0.106293 | 0.175569 | 0.842524 |
| 9 | 120307800 | 120309199 | 14.3273 | 25.63665 | 31.50078 | 13.81769 | 0.104413 | 0.429482 | 0.84254 |
| X | 98863000 | 98869999 | 68.82323 | 109.2287 | 126.967 | 52.89637 | 0.114789 | 2.31586 | 0.842544 |
| 14 | 118406000 | 118407799 | 20.04835 | 34.37454 | 47.95032 | 21.02963 | -0.09484 | 0.935671 | 0.842546 |
| 8 | 4042800 | 4048799 | 101.9727 | 156.0825 | 202.7345 | 84.1275 | -0.08615 | 3.048072 | 0.842572 |

| | | | | | | | | | |
|---|---|---|---|---|---|---|---|---|---|
| 8 | 18117000 | 18118599 | 14.78405 | 26.95499 | 32.04789 | 13.76329 | 0.171181 | 0.320978 | 0.842595 |
| 6 | 128746800 | 128758599 | 202.0368 | 296.7819 | 370.4308 | 152.5606 | -0.08108 | 3.840612 | 0.842668 |
| 16 | 10039200 | 10042799 | 43.44785 | 70.17953 | 84.19825 | 35.93545 | 0.097816 | 1.73705 | 0.842681 |
| 8 | 124851400 | 124856999 | 69.63783 | 108.9829 | 168.9246 | 70.076 | -0.31612 | 2.624801 | 0.842682 |
| 6 | 48740200 | 48742999 | 46.56116 | 74.72255 | 88.73822 | 37.53025 | 0.101337 | 1.940166 | 0.842698 |
| 16 | 88555800 | 88556199 | 5.514151 | 10.74838 | 18.58802 | 6.718763 | -0.14633 | -0.40832 | 0.842718 |
| 11 | 30484000 | 30488199 | 43.49675 | 70.55777 | 81.89811 | 35.08765 | 0.151474 | 1.638701 | 0.842735 |
| 9 | 89932800 | 89935199 | 23.23101 | 38.53431 | 67.29625 | 29.1522 | -0.42098 | 1.285575 | 0.84274 |
| Y | 43829000 | 43829399 | 4.706308 | 9.17659 | 17.54356 | 6.315141 | -0.28381 | -0.43972 | 0.84276 |
| 6 | 57201000 | 57204199 | 24.98181 | 42.5638 | 51.1934 | 22.44053 | 0.116761 | 1.015873 | 0.842774 |
| 18 | 36974200 | 36975199 | 14.50315 | 26.03743 | 31.85675 | 13.93485 | 0.112997 | 0.408175 | 0.842792 |
| 1 | 25965600 | 25977599 | 156.3121 | 235.0523 | 266.6285 | 110.0875 | 0.087865 | 3.420749 | 0.842798 |
| 10 | 88508000 | 88509799 | 15.00137 | 26.43747 | 37.14342 | 16.43979 | -0.09908 | 0.566392 | 0.842808 |
| 16 | 20658400 | 20659599 | 14.81858 | 26.44062 | 32.4678 | 14.27596 | 0.102928 | 0.452184 | 0.842828 |
| 14 | 41996000 | 41998199 | 31.37207 | 52.47724 | 50.79725 | 22.12642 | 0.427909 | 1.203632 | 0.842858 |
| 4 | 70003800 | 70006999 | 38.0056 | 61.71662 | 74.88138 | 32.10688 | 0.089208 | 1.605644 | 0.842865 |
| 15 | 90074000 | 90077799 | 60.52064 | 96.15627 | 114.4662 | 47.7403 | 0.081998 | 2.27244 | 0.842865 |
| 1 | 88681600 | 88683399 | 25.98906 | 43.47315 | 52.82596 | 23.01019 | 0.100955 | 1.188658 | 0.842881 |
| 15 | 33005400 | 33009599 | 46.80298 | 75.47852 | 89.04103 | 37.92427 | 0.119456 | 1.791054 | 0.842881 |
| 1 | 67134600 | 67136799 | 21.76426 | 37.50632 | 45.78376 | 20.08402 | 0.098211 | 0.894067 | 0.842886 |
| 8 | 62504200 | 62504599 | 5.800533 | 11.2944 | 19.61858 | 7.190065 | -0.1699 | -0.34811 | 0.842898 |
| 13 | 72993600 | 72995599 | 21.981 | 37.47614 | 52.24981 | 22.90485 | -0.09546 | 0.998291 | 0.842924 |
| 12 | 93875400 | 93876999 | 14.39779 | 25.65891 | 36.60559 | 16.11167 | -0.11331 | 0.464917 | 0.842929 |
| 9 | 94274800 | 94275599 | 11.4263 | 21.06556 | 24.82758 | 10.50604 | 0.199635 | 0.221956 | 0.842936 |
| 1 | 190880800 | 190884199 | 47.79989 | 76.72076 | 92.44992 | 39.10801 | 0.080525 | 1.937462 | 0.842967 |
| 4 | 61979800 | 61984799 | 53.65281 | 85.99908 | 101.5965 | 42.88926 | 0.106738 | 1.988241 | 0.84298 |
| 10 | 121151200 | 121152599 | 15.62465 | 26.9431 | 49.51145 | 21.71968 | -0.48909 | 0.867678 | 0.842983 |
| 7 | 18206600 | 18206999 | 9.330629 | 18.06607 | 23.95115 | 9.107381 | 0.164779 | -0.17006 | 0.843048 |
| 14 | 111116200 | 111125999 | 131.9663 | 201.4356 | 224.6518 | 93.16303 | 0.130432 | 3.114371 | 0.843074 |
| 11 | 115987400 | 115987999 | 8.601473 | 16.6295 | 22.65786 | 8.675739 | 0.115857 | -0.16011 | 0.843079 |
| 3 | 50962400 | 50966599 | 47.47811 | 76.32986 | 91.94844 | 38.99096 | 0.083869 | 1.885783 | 0.843121 |
| 5 | 98682400 | 98683199 | 7.573561 | 14.56813 | 23.05998 | 8.98724 | -0.12228 | -0.11083 | 0.843124 |
| 12 | 42589800 | 42591999 | 25.64423 | 43.21268 | 52.82782 | 23.09437 | 0.092296 | 1.115582 | 0.843155 |
| 11 | 31383600 | 31389799 | 94.85143 | 146.2214 | 189.2237 | 78.60033 | -0.07292 | 2.90494 | 0.843164 |
| 6 | 24491800 | 24492599 | 9.682633 | 17.48314 | 34.39528 | 14.952 | -0.56131 | 0.350249 | 0.84319 |
| 19 | 29942000 | 29944999 | 38.83072 | 63.65145 | 61.52634 | 26.48283 | 0.425457 | 1.4622 | 0.843202 |
| 12 | 5232000 | 5234999 | 21.33603 | 36.37922 | 50.61865 | 22.18993 | -0.09251 | 0.993638 | 0.843217 |
| 3 | 104083200 | 104086399 | 30.92766 | 51.42046 | 61.92137 | 26.86692 | 0.113835 | 1.275732 | 0.843246 |
| 4 | 147394800 | 147395999 | 8.798691 | 17.0923 | 17.80392 | 6.649079 | 0.534415 | -0.25443 | 0.843249 |
| 3 | 81463200 | 81463999 | 11.43444 | 20.88639 | 30.23612 | 12.95523 | -0.1084 | 0.285072 | 0.843255 |
| 6 | 59860000 | 59869599 | 330.8193 | 462.2345 | 554.3865 | 229.905 | -0.0951 | 4.591695 | 0.843266 |
| 4 | 14198600 | 14198999 | 7.565269 | 14.57837 | 23.09787 | 8.955659 | -0.11673 | -0.12788 | 0.843267 |
| 3 | 62815400 | 62815799 | 6.955476 | 13.52936 | 19.00206 | 7.021975 | 0.121079 | -0.31093 | 0.843277 |
| X | 51701200 | 51703599 | 17.94133 | 31.31493 | 45.18408 | 19.89543 | -0.1389 | 0.737227 | 0.843308 |
| 17 | 40690800 | 40691599 | 17.60372 | 30.9338 | 37.56058 | 16.62916 | 0.108873 | 0.615728 | 0.84332 |
| 15 | 60426600 | 60428599 | 21.5498 | 37.28195 | 45.22418 | 19.84091 | 0.108359 | 0.857035 | 0.843337 |
| 1 | 176545800 | 176547399 | 17.02178 | 29.51283 | 51.16065 | 22.44688 | -0.40382 | 0.82649 | 0.843384 |

| | | | | | | | | | |
|---|---|---|---|---|---|---|---|---|---|
| 1 | 180699000 | 180700799 | 14.66818 | 25.95539 | 36.56314 | 16.16983 | -0.10065 | 0.518768 | 0.843408 |
| 8 | 89152400 | 89154199 | 23.72099 | 39.97616 | 55.48186 | 24.25541 | -0.0901 | 1.110809 | 0.843416 |
| 11 | 11640200 | 11644399 | 26.43884 | 44.08084 | 61.70421 | 26.82191 | -0.10256 | 1.230157 | 0.843435 |
| 14 | 58224800 | 58231399 | 64.43944 | 102.5102 | 120.9749 | 50.52404 | 0.096274 | 2.239685 | 0.843452 |
| 10 | 112445400 | 112447199 | 14.77134 | 26.3606 | 32.43117 | 14.25791 | 0.100251 | 0.449947 | 0.843503 |
| 9 | 3646800 | 3652999 | 57.71812 | 92.06742 | 126.751 | 53.13547 | -0.12147 | 2.150623 | 0.843526 |
| 8 | 99594400 | 99597399 | 37.89785 | 62.0662 | 61.32919 | 26.38789 | 0.393303 | 1.471858 | 0.84354 |
| 6 | 107855000 | 107855399 | 6.384988 | 12.45962 | 18.10889 | 6.44581 | 0.121605 | -0.45164 | 0.843569 |
| 18 | 23102000 | 23102599 | 10.18503 | 18.20112 | 35.87572 | 15.75389 | -0.57406 | 0.420251 | 0.843583 |
| 19 | 40471400 | 40474199 | 32.96268 | 53.9891 | 66.00516 | 28.40099 | 0.087028 | 1.461362 | 0.843594 |
| 11 | 20456000 | 20461999 | 68.05602 | 107.8603 | 128.1097 | 53.29787 | 0.081742 | 2.352932 | 0.843625 |
| 13 | 20835800 | 20843799 | 180.8097 | 265.7648 | 336.2396 | 138.4738 | -0.09974 | 3.827267 | 0.843629 |
| 12 | 113835200 | 113836599 | 36.59056 | 59.14467 | 81.24478 | 34.70628 | -0.09581 | 1.704462 | 0.843629 |
| 15 | 92950200 | 92954599 | 37.81414 | 61.60141 | 73.8985 | 31.73367 | 0.109001 | 1.550401 | 0.84366 |
| 18 | 37061400 | 37065799 | 111.6698 | 170.0031 | 219.5251 | 90.97408 | -0.0839 | 3.161657 | 0.843666 |
| 2 | 89485200 | 89486199 | 11.69429 | 22.11017 | 27.57774 | 11.20134 | 0.164011 | 0.07181 | 0.843704 |
| 2 | 9046600 | 9047599 | 13.53277 | 24.05441 | 34.07204 | 15.0296 | -0.1052 | 0.469889 | 0.843704 |
| 3 | 3384400 | 3387799 | 34.71357 | 57.29666 | 56.92043 | 24.57337 | 0.389888 | 1.365745 | 0.843724 |
| 14 | 7446600 | 7448399 | 40.41185 | 65.31215 | 79.22454 | 33.83711 | 0.082925 | 1.712259 | 0.843725 |
| 12 | 21519400 | 21522199 | 29.58358 | 49.01609 | 60.06077 | 25.99752 | 0.088478 | 1.309428 | 0.843744 |
| 12 | 118083200 | 118083799 | 6.532068 | 12.63724 | 21.04015 | 8.030769 | -0.16575 | -0.19823 | 0.843756 |
| 5 | 102528200 | 102529999 | 24.73348 | 41.85875 | 51.20186 | 22.41296 | 0.091659 | 1.07232 | 0.843756 |
| 4 | 90544800 | 90547599 | 55.87798 | 88.99118 | 105.0431 | 43.93894 | 0.097748 | 2.192101 | 0.843783 |
| 19 | 29736600 | 29741399 | 72.70396 | 115.471 | 130.9821 | 54.60853 | 0.150994 | 2.297862 | 0.84379 |
| 19 | 15154000 | 15155199 | 13.28088 | 24.14073 | 30.03779 | 12.94924 | 0.102713 | 0.325277 | 0.843796 |
| 2 | 82742600 | 82747399 | 71.32537 | 112.5049 | 133.8297 | 55.5404 | 0.074037 | 2.469864 | 0.843798 |
| 4 | 72409200 | 72412399 | 67.75571 | 107.1015 | 127.076 | 52.79241 | 0.080168 | 2.415498 | 0.843801 |
| 10 | 9721400 | 9723999 | 38.38501 | 61.85475 | 85.29648 | 36.29235 | -0.10651 | 1.798483 | 0.843811 |
| 7 | 10664200 | 10664999 | 16.73425 | 29.05077 | 42.24678 | 18.53904 | -0.15075 | 0.803712 | 0.84385 |
| 2 | 23656000 | 23657799 | 25.82864 | 43.03957 | 59.92194 | 26.03806 | -0.09472 | 1.233745 | 0.843854 |
| 1 | 151398200 | 151403199 | 56.35458 | 89.82725 | 120.8543 | 50.61272 | -0.08946 | 2.168152 | 0.843869 |
| 18 | 79742800 | 79744599 | 9.713716 | 18.30034 | 27.48875 | 11.24436 | -0.11072 | 0.078582 | 0.843884 |
| 18 | 74385000 | 74386999 | 25.04582 | 41.78592 | 58.19536 | 25.30722 | -0.09516 | 1.217874 | 0.843901 |
| 13 | 95513000 | 95514599 | 17.96287 | 31.35782 | 38.33166 | 16.87997 | 0.099154 | 0.720812 | 0.843921 |
| 14 | 76744400 | 76749799 | 52.26672 | 83.44619 | 111.6746 | 46.87412 | -0.07926 | 2.111396 | 0.843927 |
| 13 | 69483200 | 69484399 | 10.85935 | 19.95604 | 29.22476 | 12.42099 | -0.11691 | 0.242064 | 0.843931 |
| 6 | 23773400 | 23774599 | 16.36065 | 28.687 | 40.33989 | 17.82686 | -0.10179 | 0.648764 | 0.843938 |
| 4 | 52890600 | 52891399 | 15.6194 | 27.46937 | 32.73373 | 14.49408 | 0.136293 | 0.585922 | 0.843943 |
| X | 103899200 | 103900999 | 16.71362 | 29.45117 | 35.97095 | 15.92127 | 0.101753 | 0.575239 | 0.843949 |
| X | 148592800 | 148594799 | 22.81706 | 39.82126 | 45.08117 | 19.84892 | 0.209365 | 0.760384 | 0.843982 |
| 10 | 54762600 | 54767199 | 126.1595 | 191.3305 | 219.8808 | 91.01748 | 0.079634 | 3.242632 | 0.844005 |
| 1 | 117480400 | 117481599 | 20.60284 | 35.77351 | 43.61853 | 19.1433 | 0.102006 | 0.817029 | 0.844035 |
| 14 | 102875800 | 102877599 | 21.76095 | 37.37365 | 45.80522 | 20.09136 | 0.091461 | 0.924304 | 0.844041 |
| 12 | 94668400 | 94669599 | 12.1542 | 22.55498 | 28.16952 | 11.81431 | 0.124815 | 0.194224 | 0.844044 |
| 6 | 71647400 | 71649799 | 25.84571 | 43.33159 | 60.54568 | 26.41057 | -0.10002 | 1.170273 | 0.84406 |
| 2 | 103900000 | 103904599 | 57.3103 | 91.31999 | 122.7527 | 51.38386 | -0.08868 | 2.182118 | 0.844079 |
| 1 | 82680400 | 82681799 | 18.98319 | 33.02636 | 40.48488 | 17.79006 | 0.094679 | 0.777818 | 0.84408 |

| | | | | | | | | | |
|---|---|---|---|---|---|---|---|---|---|
| 12 | 103936400 | 103940799 | 46.94892 | 75.21952 | 102.0589 | 43.05488 | -0.09342 | 1.993954 | 0.844094 |
| 4 | 55912000 | 55913199 | 16.31316 | 28.63961 | 40.26002 | 17.81042 | -0.10142 | 0.627166 | 0.84411 |
| 7 | 107550400 | 107552199 | 35.0576 | 56.7829 | 78.13786 | 33.43957 | -0.09543 | 1.652541 | 0.844126 |
| 18 | 6857800 | 6861399 | 46.02076 | 74.25047 | 88.17927 | 37.57904 | 0.110562 | 1.774606 | 0.844153 |
| 13 | 78462600 | 78464799 | 29.23364 | 47.81205 | 81.89721 | 35.21924 | -0.3994 | 1.485063 | 0.844164 |
| 6 | 79274800 | 79275599 | 14.82267 | 26.27805 | 32.02966 | 14.15961 | 0.108273 | 0.491932 | 0.844187 |
| 10 | 106069000 | 106069999 | 11.03757 | 20.93954 | 21.61785 | 8.717724 | 0.444226 | 0.029324 | 0.844192 |
| 8 | 112966400 | 112968599 | 24.41726 | 41.46857 | 50.64385 | 22.19226 | 0.094142 | 1.040958 | 0.844195 |
| 12 | 103723200 | 103736199 | 170.5685 | 253.2412 | 317.1671 | 130.6506 | -0.0716 | 3.658948 | 0.844222 |
| 7 | 14320000 | 14321199 | 22.67372 | 38.32603 | 53.31501 | 23.32933 | -0.09327 | 1.077955 | 0.84423 |
| 3 | 120062800 | 120066599 | 45.78668 | 73.62429 | 88.75685 | 37.64773 | 0.083105 | 1.87565 | 0.844238 |
| 8 | 108663400 | 108665399 | 20.59162 | 35.86007 | 43.03683 | 18.92319 | 0.12527 | 0.776702 | 0.844242 |
| 6 | 97523200 | 97526199 | 23.92375 | 41.28399 | 40.38857 | 17.71804 | 0.41662 | 0.883801 | 0.844252 |
| 7 | 47801400 | 47802799 | 26.34368 | 43.93008 | 52.81554 | 22.96074 | 0.116373 | 1.223292 | 0.84428 |
| 3 | 153473200 | 153474199 | 12.56744 | 23.55434 | 28.99521 | 11.96508 | 0.164102 | 0.13943 | 0.844296 |
| 1 | 175707000 | 175709799 | 15.60486 | 26.96008 | 48.98641 | 21.49059 | -0.47231 | 0.849458 | 0.844299 |
| X | 155517200 | 155518199 | 11.31341 | 21.03285 | 31.57263 | 13.2087 | -0.1357 | 0.188758 | 0.844305 |
| 5 | 145721600 | 145750399 | 513.599 | 700.3991 | 721.3332 | 300.2829 | 0.089231 | 4.969268 | 0.844313 |
| 3 | 36407800 | 36409199 | 18.06105 | 31.83392 | 37.95595 | 16.79897 | 0.136445 | 0.583824 | 0.844314 |
| 5 | 90028000 | 90028599 | 15.56764 | 27.22029 | 39.00852 | 17.19782 | -0.12874 | 0.688546 | 0.844319 |
| 5 | 150334800 | 150336799 | 23.11131 | 38.82884 | 54.33873 | 23.73347 | -0.10213 | 1.135183 | 0.844325 |
| 11 | 63407600 | 63409999 | 22.17953 | 37.83093 | 52.79329 | 23.1439 | -0.09668 | 0.996373 | 0.844375 |
| 17 | 76345000 | 76357399 | 187.2669 | 276.2274 | 344.3453 | 141.8064 | -0.07363 | 3.771041 | 0.844383 |
| 5 | 9985200 | 9986199 | 18.27649 | 31.82316 | 44.67983 | 19.64363 | -0.10013 | 0.769467 | 0.844421 |
| 16 | 57866800 | 57869399 | 22.53791 | 38.87506 | 46.83892 | 20.54746 | 0.117182 | 0.889685 | 0.844422 |
| 7 | 6517400 | 6521999 | 94.96436 | 146.5225 | 169.3343 | 70.33828 | 0.091066 | 2.88654 | 0.844438 |
| 4 | 14758400 | 14759599 | 17.85577 | 31.22637 | 38.2827 | 16.88022 | 0.095117 | 0.69431 | 0.844438 |
| 10 | 39157200 | 39160599 | 33.40796 | 54.91214 | 66.91482 | 28.83549 | 0.093705 | 1.418906 | 0.844479 |
| 4 | 121071400 | 121074199 | 22.70246 | 38.37062 | 53.30531 | 23.32435 | -0.09132 | 1.0796 | 0.844488 |
| 3 | 12368000 | 12370799 | 25.54361 | 43.57249 | 51.77161 | 22.69615 | 0.134471 | 1.006463 | 0.844506 |
| 15 | 77557400 | 77559799 | 24.87437 | 42.32561 | 51.20098 | 22.44151 | 0.108141 | 1.025562 | 0.844509 |
| X | 100135200 | 100139199 | 57.40711 | 92.51097 | 75.53785 | 32.14193 | 0.647958 | 1.823788 | 0.844549 |
| 4 | 119780000 | 119788399 | 126.6304 | 191.2946 | 245.0154 | 101.2972 | -0.08146 | 3.322977 | 0.844551 |
| 17 | 80561000 | 80562999 | 17.32079 | 30.35673 | 37.14746 | 16.40977 | 0.097684 | 0.653796 | 0.844558 |
| 11 | 98503200 | 98506199 | 25.82252 | 43.27436 | 52.82388 | 23.03609 | 0.094371 | 1.168482 | 0.844575 |
| 7 | 17135000 | 17136799 | 29.91224 | 49.27074 | 68.20955 | 29.41983 | -0.08855 | 1.387125 | 0.844583 |
| 1 | 44298000 | 44299799 | 18.01954 | 31.4056 | 44.11202 | 19.40558 | -0.1007 | 0.752239 | 0.844606 |
| 7 | 56021200 | 56025199 | 45.20345 | 72.96705 | 86.93444 | 37.07163 | 0.106733 | 1.759032 | 0.844612 |
| 2 | 19841400 | 19842999 | 15.3126 | 27.07842 | 33.04779 | 14.61705 | 0.105009 | 0.52435 | 0.844637 |
| 11 | 31281400 | 31282399 | 15.89958 | 28.5494 | 26.42168 | 11.55573 | 0.513985 | 0.408623 | 0.844639 |
| X | 18175400 | 18195799 | 261.9291 | 379.7083 | 408.4057 | 168.3965 | 0.115157 | 4.048676 | 0.84469 |
| 2 | 48948800 | 48951999 | 70.48089 | 111.5288 | 132.3439 | 55.01077 | 0.081238 | 2.394404 | 0.844695 |
| X | 91615800 | 91616599 | 10.9954 | 20.54326 | 31.16465 | 12.92898 | -0.14103 | 0.153887 | 0.844733 |
| 12 | 84048600 | 84050399 | 29.2597 | 48.51532 | 59.48753 | 25.76254 | 0.087589 | 1.299766 | 0.844764 |
| 13 | 45355600 | 45357399 | 21.91562 | 37.97616 | 37.83367 | 16.59934 | 0.391707 | 0.834499 | 0.844772 |
| 1 | 34918800 | 34923799 | 61.26888 | 97.15763 | 128.7495 | 53.59389 | -0.07518 | 2.332928 | 0.844775 |
| 1 | 151391600 | 151394799 | 26.30653 | 43.77028 | 61.06504 | 26.50914 | -0.09773 | 1.251901 | 0.844795 |

| | | | | | | | | | |
|---|---|---|---|---|---|---|---|---|---|
| 16 | 19495200 | 19495799 | 7.654227 | 14.86228 | 20.56987 | 7.710425 | 0.122333 | -0.24983 | 0.844805 |
| 8 | 49438200 | 49438799 | 14.84315 | 26.24534 | 31.20608 | 13.80277 | 0.142543 | 0.519901 | 0.844812 |
| 14 | 113345800 | 113349399 | 39.3558 | 64.2542 | 75.44354 | 32.42316 | 0.14214 | 1.524501 | 0.844816 |
| 15 | 82307400 | 82320799 | 156.2127 | 234.3006 | 301.9307 | 124.5777 | -0.09931 | 3.47242 | 0.844817 |
| 4 | 112143000 | 112146999 | 66.41016 | 105.3734 | 125.2979 | 52.18832 | 0.081726 | 2.317636 | 0.844828 |
| 5 | 91562400 | 91563399 | 12.02014 | 21.96824 | 31.73654 | 13.58882 | -0.10508 | 0.288138 | 0.844838 |
| 1 | 143771800 | 143772799 | 13.11539 | 23.69797 | 34.20545 | 14.85647 | -0.11702 | 0.368658 | 0.844838 |
| 17 | 48154400 | 48155199 | 9.343471 | 17.46483 | 26.38311 | 10.93687 | -0.13368 | 0.125415 | 0.844838 |
| 1 | 179218200 | 179221799 | 56.93744 | 90.41156 | 121.0858 | 50.47223 | -0.08852 | 2.28742 | 0.844846 |
| 14 | 61480400 | 61490999 | 122.9318 | 187.5506 | 216.5005 | 89.77222 | 0.080181 | 3.120641 | 0.844849 |
| 19 | 60919200 | 60923799 | 74.27147 | 115.1095 | 186.9427 | 77.65067 | -0.39428 | 2.798527 | 0.84489 |
| 7 | 107241400 | 107244399 | 30.70424 | 50.241 | 69.68862 | 29.9742 | -0.09536 | 1.470823 | 0.844893 |
| 4 | 43296800 | 43297799 | 11.36441 | 20.63903 | 30.15528 | 13.01417 | -0.12863 | 0.315373 | 0.844898 |
| 19 | 44517000 | 44517799 | 11.36441 | 20.63903 | 30.15528 | 13.01417 | -0.12863 | 0.315373 | 0.844898 |
| 1 | 46575600 | 46577199 | 21.4467 | 37.00507 | 45.25204 | 19.85153 | 0.096045 | 0.881194 | 0.844928 |
| X | 30134200 | 30134599 | 9.095712 | 17.63317 | 18.2927 | 6.915405 | 0.523813 | -0.20824 | 0.844949 |
| 10 | 40482800 | 40485999 | 43.7987 | 70.59672 | 85.16043 | 36.24705 | 0.086453 | 1.796577 | 0.844958 |
| 1 | 8247600 | 8252399 | 108.7697 | 166.4286 | 192.0353 | 79.68133 | 0.084118 | 3.055606 | 0.844962 |
| 4 | 82403800 | 82406399 | 25.42302 | 43.51438 | 51.22528 | 22.4628 | 0.148552 | 0.975701 | 0.844963 |
| 12 | 109951400 | 109956599 | 44.74354 | 71.866 | 96.82691 | 41.00364 | -0.079 | 1.913678 | 0.844971 |
| 7 | 110333600 | 110338399 | 48.00476 | 76.94065 | 103.4474 | 43.66607 | -0.07963 | 1.985209 | 0.844983 |
| 5 | 42702200 | 42704799 | 11.48561 | 20.61568 | 38.35083 | 16.77851 | -0.48817 | 0.402147 | 0.845006 |
| 3 | 138139200 | 138145999 | 85.95896 | 133.9607 | 157.9803 | 65.56657 | 0.073032 | 2.70458 | 0.845017 |
| 2 | 95242600 | 95245399 | 37.00133 | 60.05658 | 72.93824 | 31.26251 | 0.087165 | 1.61209 | 0.845043 |
| 4 | 26751800 | 26752199 | 9.819109 | 18.89204 | 19.73947 | 7.687883 | 0.472799 | -0.09899 | 0.845051 |
| 16 | 28768600 | 28770399 | 18.63096 | 32.5812 | 39.75819 | 17.53044 | 0.101705 | 0.700118 | 0.845054 |
| 13 | 102958000 | 102964199 | 87.9516 | 136.6993 | 178.7879 | 74.23201 | -0.07912 | 2.754963 | 0.845074 |
| 10 | 59076000 | 59080399 | 45.75071 | 73.33376 | 98.94563 | 41.77091 | -0.08453 | 1.979372 | 0.845091 |
| 9 | 86081400 | 86082399 | 11.24389 | 20.77371 | 30.35449 | 12.80745 | -0.106 | 0.219242 | 0.845103 |
| 18 | 10858400 | 10861599 | 28.33704 | 46.87611 | 64.96236 | 28.08833 | -0.08876 | 1.328235 | 0.845121 |
| 2 | 10800800 | 10802199 | 12.17792 | 22.02635 | 31.61342 | 13.71331 | -0.10787 | 0.348414 | 0.845134 |
| 16 | 38820400 | 38821199 | 7.707003 | 14.90045 | 19.93945 | 7.634222 | 0.142491 | -0.17677 | 0.845142 |
| 3 | 146600800 | 146602199 | 15.97887 | 28.44839 | 34.2988 | 15.10898 | 0.127099 | 0.474173 | 0.845155 |
| 11 | 80162800 | 80164799 | 15.97887 | 28.44839 | 34.2988 | 15.10898 | 0.127099 | 0.474173 | 0.845155 |
| 1 | 30121400 | 30130799 | 215.6666 | 312.98 | 387.6394 | 159.8139 | -0.08677 | 4.027326 | 0.845232 |
| 15 | 31157400 | 31161799 | 39.79269 | 64.77954 | 76.51072 | 32.84709 | 0.130693 | 1.565801 | 0.845253 |
| 4 | 89763600 | 89764599 | 9.637601 | 18.0031 | 26.86493 | 11.14868 | -0.11756 | 0.132643 | 0.84527 |
| 5 | 6476800 | 6479599 | 54.12439 | 86.74212 | 87.84088 | 37.07395 | 0.328392 | 1.989487 | 0.845324 |
| 3 | 6672600 | 6673999 | 19.5076 | 32.74672 | 60.02675 | 26.20505 | -0.49004 | 1.139243 | 0.845337 |
| 4 | 142326800 | 142329599 | 31.43681 | 51.5979 | 62.63579 | 26.9751 | 0.098432 | 1.428891 | 0.845352 |
| 2 | 16270600 | 16276999 | 54.69371 | 87.41005 | 104.9083 | 44.09766 | 0.079204 | 2.080988 | 0.845379 |
| 1 | 8951600 | 8952399 | 12.07897 | 22.02457 | 31.73216 | 13.62644 | -0.10376 | 0.303111 | 0.845399 |
| 6 | 70210400 | 70212199 | 22.38012 | 38.28573 | 46.87714 | 20.55521 | 0.09196 | 0.962875 | 0.845416 |
| 5 | 127444800 | 127447999 | 28.73665 | 47.69634 | 58.38224 | 25.30156 | 0.090179 | 1.28531 | 0.845418 |
| 14 | 52752200 | 52756399 | 50.88275 | 81.45519 | 110.082 | 46.38184 | -0.0891 | 2.029185 | 0.845429 |
| 7 | 15646000 | 15648999 | 21.86548 | 36.58006 | 62.98429 | 27.43718 | -0.40024 | 1.183102 | 0.845448 |
| 15 | 32805200 | 32807599 | 28.8504 | 48.80389 | 56.31853 | 24.6493 | 0.17534 | 1.095076 | 0.84546 |

| | | | | | | | | | |
|---|---|---|---|---|---|---|---|---|---|
| 12 | 103434200 | 103434599 | 8.247546 | 15.72482 | 24.51987 | 9.78418 | -0.13197 | -0.0209 | 0.845497 |
| 3 | 105958800 | 105960399 | 20.37966 | 34.48451 | 58.61882 | 25.65831 | -0.3813 | 1.068261 | 0.845504 |
| X | 53222600 | 53227199 | 68.7003 | 109.5009 | 100.6766 | 42.1598 | 0.458877 | 2.179143 | 0.845518 |
| 14 | 21770200 | 21774599 | 44.11895 | 70.92118 | 95.72965 | 40.58759 | -0.08016 | 1.887309 | 0.84552 |
| 16 | 91826200 | 91830599 | 45.86317 | 73.43328 | 99.86804 | 42.08992 | -0.09783 | 2.012802 | 0.845536 |
| 4 | 154602000 | 154602799 | 11.96804 | 21.60967 | 31.21219 | 13.56831 | -0.11857 | 0.358669 | 0.845588 |
| 6 | 90261400 | 90262999 | 21.83018 | 36.99846 | 51.69195 | 22.63737 | -0.09936 | 1.049859 | 0.845604 |
| 6 | 82112800 | 82113799 | 10.9224 | 19.89806 | 29.22595 | 12.56415 | -0.13233 | 0.293125 | 0.845617 |
| X | 88740200 | 88742399 | 27.25255 | 46.35215 | 53.96757 | 23.65328 | 0.163111 | 1.043876 | 0.845658 |
| 10 | 118992800 | 118994599 | 22.65369 | 38.79553 | 47.41071 | 20.79098 | 0.095009 | 0.956658 | 0.845671 |
| 1 | 131236000 | 131236999 | 11.98704 | 22.41439 | 21.55802 | 8.931144 | 0.513419 | 0.132127 | 0.845704 |
| 6 | 11691000 | 11696199 | 14.10744 | 26.06423 | 25.34936 | 10.69566 | 0.477214 | 0.229393 | 0.845714 |
| 6 | 63182200 | 63182999 | 15.2034 | 26.80852 | 37.65498 | 16.66434 | -0.09836 | 0.563149 | 0.845759 |
| 7 | 90459600 | 90460199 | 8.453047 | 16.28792 | 26.01231 | 10.08781 | -0.12902 | -0.1114 | 0.8458 |
| 2 | 174825600 | 174828199 | 35.09478 | 57.5983 | 57.41193 | 24.72473 | 0.38193 | 1.434119 | 0.845819 |
| 4 | 147271400 | 147275199 | 109.3959 | 167.3879 | 193.4202 | 80.25612 | 0.082055 | 3.056632 | 0.84582 |
| 19 | 37806200 | 37806999 | 10.76563 | 19.99173 | 30.48175 | 12.77225 | -0.15923 | 0.190812 | 0.845825 |
| 6 | 57648800 | 57652399 | 38.86244 | 63.69917 | 61.49213 | 26.46735 | 0.427309 | 1.463546 | 0.84583 |
| 15 | 31940000 | 31942599 | 32.91229 | 54.55409 | 64.91081 | 28.10498 | 0.130765 | 1.317736 | 0.845839 |
| 1 | 53757000 | 53759199 | 16.52092 | 29.04306 | 35.52213 | 15.73002 | 0.098437 | 0.598678 | 0.845877 |
| 6 | 6159600 | 6161799 | 16.52092 | 29.04306 | 35.52213 | 15.73002 | 0.098437 | 0.598678 | 0.845877 |
| 9 | 18753400 | 18754399 | 16.52092 | 29.04306 | 35.52213 | 15.73002 | 0.098437 | 0.598678 | 0.845877 |
| 19 | 39116400 | 39119999 | 106.2447 | 162.7264 | 187.0721 | 77.63978 | 0.090562 | 3.03429 | 0.84589 |
| 15 | 20011800 | 20019999 | 107.0786 | 164.9278 | 191.2326 | 79.43103 | 0.084407 | 2.925901 | 0.845902 |
| 12 | 119736000 | 119739599 | 42.73935 | 68.68935 | 93.16078 | 39.48667 | -0.08741 | 1.889469 | 0.845911 |
| Y | 4003000 | 4005199 | 24.68822 | 41.26775 | 44.53116 | 19.38837 | 0.271716 | 1.181572 | 0.845912 |
| 3 | 129058600 | 129062199 | 45.36646 | 73.09831 | 88.11178 | 37.48534 | 0.086949 | 1.812131 | 0.845917 |
| 19 | 21581800 | 21582999 | 11.91099 | 21.82538 | 27.08024 | 11.55495 | 0.116594 | 0.261659 | 0.845942 |
| 4 | 151798800 | 151800199 | 13.20228 | 23.68303 | 33.58342 | 14.70005 | -0.09912 | 0.409638 | 0.845986 |
| 13 | 53999800 | 54000599 | 6.70611 | 13.04085 | 21.81137 | 8.077378 | -0.13076 | -0.29444 | 0.845993 |
| 5 | 131185600 | 131187799 | 15.59296 | 27.64118 | 33.89205 | 14.9832 | 0.099518 | 0.500203 | 0.846023 |
| 4 | 18479200 | 18483399 | 67.17108 | 106.1327 | 125.3849 | 52.07881 | 0.085906 | 2.425599 | 0.846026 |
| 9 | 9231000 | 9234999 | 40.62044 | 65.77752 | 79.59609 | 34.0577 | 0.089247 | 1.668532 | 0.846035 |
| 1 | 141301400 | 141301999 | 11.08933 | 21.08124 | 21.65688 | 8.695393 | 0.456951 | 0.016571 | 0.846036 |
| 18 | 56839800 | 56843199 | 22.89462 | 37.61997 | 72.82604 | 31.37872 | -0.57214 | 1.402797 | 0.846062 |
| 1 | 55587800 | 55588399 | 12.47589 | 22.3738 | 32.20277 | 14.099 | -0.12038 | 0.405658 | 0.846097 |
| 4 | 20809800 | 20810999 | 18.64484 | 32.2107 | 45.31521 | 19.87999 | -0.10477 | 0.860097 | 0.846122 |
| 15 | 27278400 | 27289199 | 111.5104 | 171.2388 | 198.4333 | 82.38803 | 0.081956 | 2.986077 | 0.846123 |
| 7 | 12899800 | 12910799 | 218.5285 | 318.4377 | 352.276 | 145.1725 | 0.08184 | 3.96484 | 0.846134 |
| 6 | 50247400 | 50252799 | 46.8258 | 75.32902 | 90.85317 | 38.56202 | 0.083255 | 1.864739 | 0.846141 |
| 1 | 46390600 | 46391399 | 7.557298 | 14.58793 | 23.14387 | 8.92681 | -0.11166 | -0.14515 | 0.846141 |
| 5 | 70233600 | 70237399 | 25.19513 | 42.82328 | 51.75067 | 22.68038 | 0.109341 | 1.037246 | 0.846145 |
| 18 | 41160200 | 41160599 | 8.352279 | 16.16956 | 22.14781 | 8.427048 | 0.117428 | -0.18362 | 0.846155 |
| 3 | 30590000 | 30590799 | 9.032572 | 17.07863 | 25.97635 | 10.54969 | -0.11899 | 0.047472 | 0.846167 |
| 4 | 92926200 | 92933599 | 66.09856 | 105.2525 | 121.6733 | 50.88457 | 0.127535 | 2.212071 | 0.846169 |
| 18 | 66520400 | 66522599 | 24.01477 | 41.27978 | 49.01883 | 21.50149 | 0.137028 | 0.928702 | 0.846231 |
| 18 | 46721400 | 46723999 | 27.54339 | 45.70195 | 63.23828 | 27.40033 | -0.08618 | 1.290764 | 0.846234 |

| | | | | | | | | | |
|---|---|---|---|---|---|---|---|---|---|
| 13 | 52202000 | 52205199 | 42.58311 | 68.34826 | 93.69501 | 39.63527 | -0.10509 | 1.930425 | 0.846236 |
| 3 | 91025000 | 91035399 | 157.3619 | 236.2757 | 268.9374 | 110.9962 | 0.081001 | 3.450445 | 0.846246 |
| 3 | 108857200 | 108859999 | 33.93227 | 55.92033 | 56.35868 | 24.3135 | 0.368356 | 1.380085 | 0.846293 |
| 10 | 111532400 | 111534999 | 25.54748 | 43.40355 | 52.30085 | 22.92071 | 0.113402 | 1.043258 | 0.846322 |
| 4 | 51067200 | 51067799 | 15.74417 | 27.70566 | 33.61496 | 14.88247 | 0.11067 | 0.580587 | 0.846325 |
| 13 | 64199000 | 64205399 | 69.72545 | 110.2961 | 148.8797 | 61.87355 | -0.10481 | 2.405046 | 0.846327 |
| 3 | 36758400 | 36761799 | 33.95913 | 56.2712 | 66.12014 | 28.62225 | 0.148769 | 1.325787 | 0.846331 |
| 6 | 135833000 | 135833799 | 9.779167 | 18.75509 | 24.64093 | 9.685403 | 0.131978 | -0.05882 | 0.846357 |
| 14 | 53922400 | 53938599 | 396.9365 | 547.4072 | 640.9354 | 266.3435 | -0.07946 | 4.792781 | 0.846358 |
| 19 | 51579800 | 51580999 | 14.99365 | 26.44812 | 37.11867 | 16.42582 | -0.09705 | 0.556074 | 0.846379 |
| 18 | 34214000 | 34216799 | 33.75289 | 54.80861 | 75.42244 | 32.33796 | -0.09202 | 1.59798 | 0.8464 |
| 3 | 95279600 | 95285399 | 85.58969 | 133.592 | 157.7751 | 65.46867 | 0.072667 | 2.677788 | 0.846405 |
| 3 | 66139400 | 66141399 | 38.64353 | 62.05358 | 103.2982 | 43.72825 | -0.38387 | 1.916982 | 0.846434 |
| 10 | 120539000 | 120541399 | 18.28666 | 31.85582 | 38.91389 | 17.11367 | 0.099853 | 0.753133 | 0.846436 |
| X | 31373800 | 31430799 | 2204.861 | 2678.331 | 2340.205 | 987.7016 | 0.158443 | 6.829504 | 0.846444 |
| 9 | 84528000 | 84530399 | 19.82183 | 33.81629 | 59.87308 | 26.24801 | -0.4388 | 0.993225 | 0.846451 |
| 9 | 8790000 | 8796799 | 69.81592 | 111.0995 | 106.2615 | 44.39926 | 0.399488 | 2.230587 | 0.846458 |
| 16 | 54874800 | 54876999 | 64.30989 | 101.7875 | 119.6207 | 49.72806 | 0.095472 | 2.38119 | 0.846464 |
| 2 | 177382000 | 177383599 | 15.46758 | 27.41109 | 39.61558 | 17.5145 | -0.13674 | 0.511988 | 0.846467 |
| 3 | 45168200 | 45169399 | 20.26933 | 34.93935 | 49.00913 | 21.49907 | -0.10134 | 0.890957 | 0.846478 |
| 6 | 26000400 | 26001799 | 15.38317 | 27.25285 | 33.38578 | 14.76117 | 0.100428 | 0.503704 | 0.846479 |
| 1 | 41277600 | 41279199 | 17.67438 | 30.72399 | 43.13265 | 18.94373 | -0.1003 | 0.785787 | 0.846491 |
| X | 152722200 | 152731599 | 96.5124 | 148.7577 | 193.3913 | 80.33157 | -0.07972 | 2.91017 | 0.846491 |
| 12 | 112847200 | 112851599 | 48.18796 | 77.09815 | 103.5768 | 43.60638 | -0.08141 | 2.040172 | 0.846506 |
| 18 | 90503200 | 90504799 | 20.65961 | 35.81897 | 43.61268 | 19.13516 | 0.103571 | 0.830967 | 0.846542 |
| 5 | 109688400 | 109691799 | 26.40453 | 44.37209 | 64.13051 | 28.0122 | -0.14874 | 1.149107 | 0.846561 |
| 4 | 120787600 | 120791199 | 47.69671 | 76.57871 | 92.34767 | 39.08303 | 0.079987 | 1.928817 | 0.846563 |
| 4 | 44617800 | 44624599 | 47.12012 | 75.09793 | 124.4472 | 52.10788 | -0.38977 | 2.16485 | 0.84658 |
| 6 | 67664800 | 67668599 | 44.58859 | 72.02584 | 72.82461 | 31.0833 | 0.344166 | 1.730334 | 0.846595 |
| 1 | 43203000 | 43204799 | 19.02183 | 32.85533 | 45.81404 | 20.0987 | -0.09225 | 0.861825 | 0.846605 |
| 7 | 66979400 | 66983399 | 57.13386 | 91.86814 | 101.3361 | 42.98189 | 0.21099 | 1.89989 | 0.846616 |
| 11 | 74130400 | 74130999 | 9.746253 | 18.58413 | 24.47063 | 9.762467 | 0.109746 | -0.00891 | 0.846656 |
| 10 | 44098000 | 44100599 | 24.49487 | 41.33367 | 57.75973 | 25.26113 | -0.09994 | 1.103427 | 0.846662 |
| 1 | 68628000 | 68629999 | 32.02606 | 51.8541 | 87.51065 | 37.44082 | -0.38966 | 1.65893 | 0.846671 |
| Y | 76191400 | 76191999 | 4.83178 | 9.420826 | 17.54188 | 6.318179 | -0.24678 | -0.4351 | 0.846698 |
| 8 | 91378400 | 91381399 | 25.49826 | 43.22194 | 52.31345 | 22.91062 | 0.106882 | 1.061587 | 0.846699 |
| 5 | 82202000 | 82202399 | 5.388582 | 10.49968 | 18.56309 | 6.742778 | -0.18484 | -0.39501 | 0.846744 |
| 18 | 15355400 | 15359799 | 53.70949 | 85.8066 | 103.1133 | 43.32132 | 0.076843 | 2.090302 | 0.846807 |
| X | 112410200 | 112411399 | 11.8696 | 21.98521 | 27.71968 | 11.6556 | 0.109076 | 0.204744 | 0.84681 |
| 1 | 143020200 | 143022399 | 22.64148 | 38.36374 | 53.32775 | 23.35194 | -0.0921 | 1.056608 | 0.846853 |
| 6 | 9683400 | 9684199 | 13.03389 | 23.27214 | 33.08655 | 14.53963 | -0.10674 | 0.436306 | 0.846869 |
| 11 | 22667400 | 22670599 | 35.89755 | 58.43585 | 71.09696 | 30.52238 | 0.087984 | 1.556309 | 0.846895 |
| 4 | 141994800 | 141997199 | 32.25158 | 52.48441 | 72.6801 | 31.19849 | -0.09853 | 1.55582 | 0.84694 |
| 3 | 49560200 | 49562799 | 53.89065 | 86.44737 | 86.73398 | 36.65826 | 0.343335 | 1.961591 | 0.846948 |
| 16 | 14993400 | 14998799 | 81.74938 | 128.3814 | 150.6077 | 62.46766 | 0.089173 | 2.56523 | 0.846963 |
| 12 | 44012600 | 44017799 | 51.3085 | 82.31632 | 98.48964 | 41.63531 | 0.090013 | 1.958139 | 0.846982 |
| 7 | 29293800 | 29297999 | 62.48664 | 99.43143 | 118.4689 | 49.49653 | 0.082938 | 2.228811 | 0.84706 |

| | | | | | | | | | |
|---|---|---|---|---|---|---|---|---|---|
| 6 | 37497600 | 37503999 | 62.70463 | 99.49978 | 132.1977 | 55.06548 | -0.07804 | 2.314962 | 0.847077 |
| 12 | 91519600 | 91521599 | 18.77583 | 32.65429 | 39.9755 | 17.5624 | 0.096515 | 0.780318 | 0.847087 |
| 14 | 87133200 | 87133999 | 7.251852 | 14.06557 | 22.82134 | 8.607514 | -0.11307 | -0.2255 | 0.847113 |
| 10 | 48299400 | 48300199 | 20.03577 | 34.63406 | 42.16427 | 18.49799 | 0.103131 | 0.856199 | 0.847116 |
| 5 | 91709200 | 91710999 | 19.83747 | 34.53309 | 41.94066 | 18.43664 | 0.107921 | 0.773384 | 0.847139 |
| 7 | 97130000 | 97130599 | 5.948479 | 11.57443 | 19.58959 | 7.219814 | -0.14119 | -0.32963 | 0.847157 |
| 14 | 108537400 | 108541599 | 81.49288 | 126.6953 | 165.9514 | 68.89957 | -0.08095 | 2.745124 | 0.847164 |
| 11 | 23454800 | 23457599 | 35.36388 | 57.9806 | 69.95079 | 30.11161 | 0.106635 | 1.455737 | 0.847166 |
| 9 | 26570200 | 26571399 | 11.70561 | 21.32576 | 30.68159 | 13.18925 | -0.10277 | 0.301869 | 0.84717 |
| 1 | 69263200 | 69266599 | 45.32277 | 72.91897 | 88.19909 | 37.44139 | 0.079323 | 1.857857 | 0.847196 |
| 2 | 59016400 | 59019199 | 18.95462 | 32.73432 | 45.83101 | 20.10606 | -0.09801 | 0.864884 | 0.847227 |
| 12 | 72700800 | 72703199 | 22.40107 | 38.25859 | 46.88927 | 20.55781 | 0.090236 | 0.978164 | 0.847231 |
| 10 | 56384600 | 56386599 | 30.71132 | 50.23439 | 69.64502 | 29.9515 | -0.09501 | 1.476117 | 0.847241 |
| 7 | 111494000 | 111495799 | 15.89767 | 27.82705 | 39.33882 | 17.3591 | -0.10954 | 0.669683 | 0.847243 |
| 16 | 53514800 | 53515199 | 6.514307 | 12.68974 | 17.97461 | 6.552105 | 0.127112 | -0.36819 | 0.847261 |
| 10 | 120842200 | 120855799 | 148.1834 | 223.2471 | 255.8531 | 105.6915 | 0.075442 | 3.381761 | 0.847273 |
| 5 | 28205600 | 28206599 | 10.14189 | 19.03752 | 24.23313 | 9.97398 | 0.119424 | 0.097698 | 0.847305 |
| 2 | 44557800 | 44559399 | 18.54822 | 32.27119 | 45.21503 | 19.86922 | -0.09734 | 0.781431 | 0.847326 |
| 14 | 4045800 | 4047999 | 13.78708 | 24.44575 | 34.64706 | 15.31524 | -0.10794 | 0.490388 | 0.847328 |
| 10 | 19137600 | 19140799 | 30.55634 | 50.44589 | 61.7559 | 26.66491 | 0.089392 | 1.354068 | 0.847383 |
| 13 | 26429000 | 26430399 | 28.72507 | 48.0956 | 42.89206 | 18.69436 | 0.545711 | 1.17633 | 0.847422 |
| Y | 39839200 | 39842599 | 17.77017 | 30.08002 | 48.01675 | 21.01997 | -0.29073 | 1.042287 | 0.847426 |
| 17 | 95097400 | 95099399 | 29.98867 | 50.26075 | 59.17841 | 25.80874 | 0.146098 | 1.184014 | 0.847439 |
| 3 | 121162200 | 121163999 | 19.77002 | 33.90531 | 47.42917 | 20.80146 | -0.09877 | 0.932859 | 0.847449 |
| 18 | 37775400 | 37798599 | 384.4295 | 535.6229 | 570.178 | 236.5578 | 0.072801 | 4.636241 | 0.847468 |
| Y | 39983600 | 39987199 | 32.01847 | 52.21959 | 55.39288 | 23.79686 | 0.287227 | 1.509017 | 0.847469 |
| 6 | 117481800 | 117483999 | 17.68966 | 30.80446 | 43.08753 | 18.94476 | -0.09479 | 0.761253 | 0.847479 |
| 6 | 115928000 | 115931199 | 36.5038 | 59.01534 | 81.04207 | 34.62794 | -0.0951 | 1.696414 | 0.847486 |
| 13 | 59427000 | 59428199 | 12.4675 | 22.36598 | 32.5344 | 14.24031 | -0.13584 | 0.399654 | 0.847496 |
| 10 | 58592400 | 58596599 | 55.46883 | 88.58255 | 106.1121 | 44.55646 | 0.080775 | 2.104247 | 0.847542 |
| 12 | 84007800 | 84010999 | 50.00896 | 80.03865 | 95.40519 | 40.1853 | 0.091394 | 2.030374 | 0.847545 |
| 14 | 45080600 | 45082199 | 13.7622 | 24.54492 | 29.56503 | 13.00445 | 0.130187 | 0.440872 | 0.847558 |
| 6 | 140956000 | 140961599 | 41.12729 | 66.54643 | 79.93998 | 34.18521 | 0.098678 | 1.682301 | 0.847582 |
| 12 | 65415600 | 65417599 | 16.3898 | 28.7087 | 40.31685 | 17.80128 | -0.10005 | 0.662646 | 0.847589 |
| 10 | 42615800 | 42623999 | 116.1253 | 177.354 | 206.2971 | 85.5623 | 0.070393 | 3.096298 | 0.847591 |
| 13 | 47324800 | 47326999 | 24.96788 | 41.95677 | 51.20318 | 22.36537 | 0.094675 | 1.135251 | 0.84763 |
| 7 | 34182000 | 34184999 | 27.10517 | 45.29099 | 55.59435 | 24.19949 | 0.086372 | 1.205246 | 0.847673 |
| 13 | 100687800 | 100689799 | 18.40691 | 31.27332 | 55.33461 | 24.25032 | -0.43859 | 1.021432 | 0.847701 |
| 6 | 58681000 | 58682599 | 14.90103 | 26.3355 | 37.53491 | 16.60361 | -0.11831 | 0.532967 | 0.847708 |
| 10 | 57506800 | 57509399 | 22.64496 | 38.28142 | 53.32473 | 23.33432 | -0.09523 | 1.076309 | 0.847726 |
| 2 | 162368200 | 162370599 | 19.84274 | 34.40943 | 42.05397 | 18.45139 | 0.098268 | 0.82354 | 0.847739 |
| X | 8159200 | 8160599 | 18.89109 | 33.23238 | 39.06428 | 17.29621 | 0.155921 | 0.613644 | 0.847739 |
| 7 | 10310200 | 10312399 | 38.16387 | 62.84369 | 58.80553 | 25.36049 | 0.474715 | 1.403687 | 0.847773 |
| 5 | 5318400 | 5319599 | 13.53383 | 23.9627 | 34.11284 | 15.08437 | -0.11529 | 0.500345 | 0.847818 |
| 4 | 141610400 | 141613199 | 22.72236 | 38.34371 | 53.29553 | 23.30891 | -0.09213 | 1.093628 | 0.847837 |
| 3 | 67320200 | 67334199 | 361.2714 | 502.243 | 592.0242 | 245.706 | -0.07745 | 4.669198 | 0.847839 |
| 2 | 59454200 | 59454999 | 8.796053 | 16.75224 | 25.55768 | 10.22196 | -0.10436 | -0.00797 | 0.847867 |

| | | | | | | | | | |
|---|---|---|---|---|---|---|---|---|---|
| 1 | 173845400 | 173846199 | 11.16955 | 20.73144 | 26.20467 | 10.98757 | 0.108388 | 0.183207 | 0.847869 |
| 15 | 97454600 | 97464399 | 161.6871 | 240.6118 | 302.4448 | 124.6154 | -0.07375 | 3.616443 | 0.847869 |
| 7 | 102715400 | 102716799 | 20.1707 | 34.86076 | 42.66419 | 18.71721 | 0.095471 | 0.858544 | 0.847885 |
| 19 | 3795600 | 3801199 | 73.43594 | 115.7692 | 137.8046 | 57.18587 | 0.072446 | 2.482429 | 0.847891 |
| 1 | 44519400 | 44521199 | 26.69347 | 43.44253 | 79.62469 | 34.18039 | -0.50173 | 1.549582 | 0.847917 |
| 1 | 150354600 | 150357199 | 26.05799 | 44.33936 | 44.19672 | 19.37151 | 0.386405 | 1.022014 | 0.847936 |
| 8 | 22745800 | 22750799 | 76.9674 | 119.9155 | 158.9885 | 65.98767 | -0.09603 | 2.705159 | 0.84796 |
| 8 | 79711800 | 79714399 | 24.16589 | 40.44907 | 56.58719 | 24.65838 | -0.10171 | 1.177698 | 0.847981 |
| 8 | 118377800 | 118380599 | 32.20995 | 52.52801 | 72.39404 | 31.09974 | -0.08953 | 1.524409 | 0.847984 |
| 12 | 29001000 | 29003599 | 47.46509 | 76.08577 | 90.50513 | 38.21075 | 0.097001 | 1.97607 | 0.847992 |
| 19 | 53713400 | 53714399 | 9.604638 | 18.28529 | 23.8561 | 9.549039 | 0.118188 | -0.00408 | 0.84801 |
| 1 | 140001600 | 140006199 | 60.66808 | 96.62427 | 115.4406 | 48.29118 | 0.080374 | 2.197784 | 0.848032 |
| 5 | 65094800 | 65097399 | 31.32365 | 51.5778 | 63.2695 | 27.28513 | 0.085243 | 1.390827 | 0.848049 |
| 3 | 31410800 | 31413199 | 24.83191 | 41.63223 | 57.71716 | 25.17686 | -0.08873 | 1.163962 | 0.848052 |
| 8 | 37258800 | 37260399 | 15.02524 | 26.80224 | 32.85594 | 14.44986 | 0.10492 | 0.453794 | 0.848087 |
| 6 | 10171600 | 10172799 | 15.30518 | 27.09687 | 32.84016 | 14.52183 | 0.115363 | 0.50798 | 0.84809 |
| 5 | 142693400 | 142698799 | 63.46157 | 100.2628 | 133.3343 | 55.37814 | -0.08472 | 2.427067 | 0.8481 |
| 15 | 22296800 | 22298799 | 18.18582 | 31.70905 | 44.63096 | 19.63918 | -0.10364 | 0.747697 | 0.848106 |
| 15 | 17700800 | 17703199 | 39.34057 | 63.94046 | 63.82791 | 27.38517 | 0.371163 | 1.583192 | 0.84811 |
| 10 | 58056600 | 58062399 | 147.9421 | 220.871 | 281.9866 | 116.2414 | -0.09192 | 3.555762 | 0.848117 |
| 5 | 74147600 | 74150399 | 21.89714 | 36.58676 | 62.91414 | 27.38478 | -0.39831 | 1.197638 | 0.848166 |
| 1 | 87385400 | 87389399 | 34.4868 | 56.14241 | 67.90145 | 29.15111 | 0.096657 | 1.552266 | 0.848176 |
| 10 | 39078200 | 39080399 | 28.90598 | 47.61308 | 65.85113 | 28.40256 | -0.08715 | 1.382519 | 0.848178 |
| 5 | 138599400 | 138601199 | 22.72695 | 39.33344 | 46.83509 | 20.54789 | 0.135132 | 0.861006 | 0.84821 |
| 13 | 31921000 | 31924199 | 24.83183 | 41.56016 | 57.70553 | 25.14434 | -0.09089 | 1.182166 | 0.84822 |
| 1 | 117937800 | 117956399 | 308.1582 | 438.878 | 474.0716 | 195.9895 | 0.083562 | 4.309941 | 0.848243 |
| 8 | 102942400 | 102944999 | 24.41155 | 41.97508 | 49.57021 | 21.74356 | 0.145081 | 0.928984 | 0.848259 |
| 13 | 21679200 | 21680799 | 14.47852 | 25.48933 | 36.16149 | 16.00853 | -0.11366 | 0.572068 | 0.848277 |
| 6 | 11748400 | 11749999 | 19.71431 | 32.90315 | 62.62426 | 27.23958 | -0.54404 | 1.204403 | 0.848281 |
| 12 | 88519000 | 88521199 | 20.32889 | 35.04732 | 49.0142 | 21.5014 | -0.0971 | 0.887925 | 0.8483 |
| 16 | 20290000 | 20291399 | 14.38615 | 25.97597 | 31.93981 | 13.88453 | 0.111836 | 0.374394 | 0.848304 |
| 3 | 95316400 | 95335999 | 359.3178 | 501.2156 | 590.0966 | 244.7776 | -0.0708 | 4.618179 | 0.848309 |
| 11 | 46710800 | 46712599 | 16.45587 | 29.00374 | 35.53103 | 15.72582 | 0.097684 | 0.571675 | 0.848316 |
| 15 | 34295400 | 34297999 | 26.19359 | 43.70176 | 60.4831 | 26.30243 | -0.08618 | 1.223155 | 0.848317 |
| 4 | 152679200 | 152681199 | 28.59623 | 47.53575 | 58.37859 | 25.32664 | 0.085596 | 1.267991 | 0.848332 |
| 12 | 113392200 | 113395199 | 32.52836 | 53.03848 | 73.02417 | 31.36854 | -0.08827 | 1.5276 | 0.848334 |
| 9 | 105480200 | 105482199 | 19.72276 | 34.29295 | 42.00145 | 18.44969 | 0.095714 | 0.790313 | 0.848334 |
| 11 | 121832000 | 121835399 | 53.85143 | 86.77375 | 96.37997 | 41.00543 | 0.204821 | 1.829194 | 0.84835 |
| 8 | 78121800 | 78127599 | 104.7457 | 160.4311 | 206.3565 | 85.64334 | -0.07133 | 3.035653 | 0.848368 |
| 8 | 78122000 | 78127599 | 104.7457 | 160.4311 | 206.3565 | 85.64334 | -0.07133 | 3.035653 | 0.848368 |
| 17 | 41275600 | 41276999 | 30.91146 | 50.13395 | 84.45519 | 36.17854 | -0.38472 | 1.623039 | 0.84837 |
| 13 | 4263800 | 4264199 | 8.416186 | 16.3423 | 17.25851 | 6.467552 | 0.50988 | -0.25028 | 0.848374 |
| 7 | 28993800 | 28995599 | 13.56801 | 23.86864 | 34.81655 | 15.41543 | -0.15378 | 0.573408 | 0.848415 |
| 9 | 25898400 | 25900599 | 32.08707 | 52.95219 | 64.68083 | 27.92027 | 0.092533 | 1.365525 | 0.848419 |
| X | 24304800 | 24308999 | 40.89478 | 67.15478 | 74.69654 | 32.17602 | 0.224493 | 1.455384 | 0.848432 |
| X | 92339400 | 92341199 | 15.67709 | 27.92276 | 33.0612 | 14.55868 | 0.153996 | 0.473278 | 0.848435 |
| 2 | 158690000 | 158692799 | 16.85427 | 29.33032 | 41.64118 | 18.30122 | -0.116 | 0.767658 | 0.848437 |

| | | | | | | | | | |
|---|---|---|---|---|---|---|---|---|---|
| 5 | 104862000 | 104871199 | 248.5897 | 359.48 | 397.8892 | 164.114 | 0.070315 | 4.082821 | 0.848444 |
| 4 | 3058400 | 3076999 | 1047.407 | 1332.442 | 1452.232 | 609.6529 | -0.09293 | 6.088432 | 0.848587 |
| 19 | 57830000 | 57835999 | 68.90345 | 109.325 | 128.7131 | 53.60857 | 0.095959 | 2.321718 | 0.848613 |
| 3 | 32836400 | 32837999 | 22.58341 | 38.65082 | 47.40855 | 20.78898 | 0.089516 | 0.960873 | 0.848617 |
| 3 | 14992000 | 14994399 | 24.20584 | 41.01046 | 57.75943 | 25.29181 | -0.11098 | 1.068792 | 0.848617 |
| 16 | 35183600 | 35185199 | 15.95611 | 28.06873 | 34.14523 | 15.1183 | 0.106761 | 0.587556 | 0.848627 |
| 12 | 114827000 | 114829199 | 43.71525 | 70.64405 | 84.59144 | 36.12477 | 0.10135 | 1.723277 | 0.848648 |
| 3 | 76304800 | 76307199 | 24.01864 | 41.10947 | 49.56581 | 21.73505 | 0.114139 | 0.96763 | 0.848659 |
| 4 | 35583800 | 35587599 | 34.92175 | 57.67319 | 57.03493 | 24.63251 | 0.396424 | 1.357987 | 0.848661 |
| 5 | 93739000 | 93749599 | 354.2107 | 494.8384 | 523.7614 | 217.2101 | 0.084923 | 4.594582 | 0.848671 |
| 3 | 127729200 | 127731599 | 34.13995 | 55.93486 | 68.42103 | 29.4444 | 0.086418 | 1.462245 | 0.848677 |
| 18 | 85865000 | 85867799 | 41.10884 | 66.62332 | 66.02519 | 28.26815 | 0.377775 | 1.644292 | 0.848677 |
| 2 | 32405800 | 32407399 | 13.76527 | 24.48997 | 34.61761 | 15.25854 | -0.10135 | 0.468468 | 0.848707 |
| 13 | 44040200 | 44042799 | 28.59271 | 47.63006 | 58.41221 | 25.37685 | 0.087731 | 1.244638 | 0.848712 |
| 7 | 110248400 | 110250599 | 25.42214 | 43.2088 | 51.77038 | 22.68901 | 0.121616 | 1.036642 | 0.848728 |
| X | 169176600 | 169177599 | 9.064172 | 16.35612 | 34.38772 | 14.9605 | -0.65745 | 0.347099 | 0.848749 |
| 5 | 104619800 | 104624999 | 46.27389 | 74.2683 | 88.60699 | 37.47994 | 0.094809 | 1.934641 | 0.848751 |
| 18 | 49508600 | 49511799 | 33.94694 | 55.89452 | 67.59798 | 29.15117 | 0.105595 | 1.400442 | 0.848757 |
| 1 | 49129600 | 49130799 | 16.61246 | 29.44572 | 35.25468 | 15.58589 | 0.133439 | 0.507951 | 0.848765 |
| 5 | 110018800 | 110051399 | 674.6685 | 900.0415 | 904.0771 | 377.599 | 0.093373 | 5.318807 | 0.848842 |
| 14 | 122731800 | 122733599 | 24.8746 | 41.5788 | 57.66232 | 25.10535 | -0.08911 | 1.196734 | 0.848846 |
| 17 | 21090800 | 21091799 | 17.85879 | 31.82871 | 29.46767 | 12.96935 | 0.50691 | 0.471214 | 0.848856 |
| 11 | 121830400 | 121835599 | 68.60189 | 109.5276 | 119.5674 | 50.18805 | 0.214141 | 2.133268 | 0.848906 |
| 17 | 74779800 | 74782999 | 28.60934 | 47.37317 | 57.74378 | 24.98415 | 0.095995 | 1.312145 | 0.848913 |
| 7 | 117327400 | 117329399 | 33.78272 | 54.90299 | 75.45925 | 32.37371 | -0.08911 | 1.582234 | 0.848983 |
| 3 | 24565400 | 24566199 | 11.17671 | 21.14987 | 27.04216 | 10.96373 | 0.131163 | 0.062043 | 0.849024 |
| 12 | 20881600 | 20973199 | 3010.74 | 3538.127 | 3173.182 | 1345.712 | 0.072113 | 7.350664 | 0.849037 |
| 10 | 42486400 | 42493599 | 44.30988 | 70.87757 | 114.8542 | 48.33889 | -0.35176 | 2.04579 | 0.849051 |
| 4 | 116004400 | 116006599 | 16.18447 | 28.39995 | 39.79173 | 17.59607 | -0.0966 | 0.634286 | 0.84907 |
| 4 | 35854400 | 35854799 | 7.742025 | 14.95647 | 23.74715 | 9.125355 | -0.10742 | -0.15266 | 0.849094 |
| 15 | 24945000 | 24945599 | 7.742025 | 14.95647 | 23.74715 | 9.125355 | -0.10742 | -0.15266 | 0.849094 |
| 16 | 48019200 | 48019999 | 7.742025 | 14.95647 | 23.74715 | 9.125355 | -0.10742 | -0.15266 | 0.849094 |
| 8 | 119243000 | 119245799 | 21.42864 | 37.04075 | 45.23783 | 19.84638 | 0.098388 | 0.864728 | 0.84912 |
| 5 | 117019400 | 117022599 | 33.24983 | 54.47535 | 66.69981 | 28.70344 | 0.085005 | 1.457993 | 0.849124 |
| 17 | 66957200 | 66959799 | 29.10436 | 48.14864 | 58.86163 | 25.45167 | 0.091661 | 1.324892 | 0.849141 |
| 14 | 61572400 | 61573799 | 12.28559 | 21.61784 | 40.45031 | 17.90897 | -0.51078 | 0.576627 | 0.849143 |
| 1 | 58676600 | 58677999 | 10.38402 | 19.15023 | 28.24901 | 11.94857 | -0.12222 | 0.216202 | 0.849179 |
| 15 | 14951200 | 14951799 | 8.010943 | 15.42394 | 24.19622 | 9.404668 | -0.10614 | -0.1126 | 0.849203 |
| 14 | 89922800 | 89926999 | 62.6195 | 99.47473 | 119.1405 | 49.68166 | 0.072974 | 2.279389 | 0.849209 |
| 8 | 48680000 | 48683599 | 59.73995 | 94.97211 | 113.1887 | 47.23546 | 0.081153 | 2.254866 | 0.849229 |
| X | 143369000 | 143370199 | 13.61671 | 24.47799 | 35.16366 | 15.36442 | -0.11588 | 0.404027 | 0.84923 |
| 9 | 16970200 | 16971199 | 11.47202 | 20.82339 | 30.24641 | 13.06218 | -0.12045 | 0.322443 | 0.849231 |
| 1 | 72940000 | 72943999 | 29.14844 | 48.76911 | 58.55844 | 25.51083 | 0.117958 | 1.198531 | 0.849244 |
| 9 | 92506800 | 92507399 | 14.73948 | 26.13805 | 31.94221 | 14.11796 | 0.104592 | 0.487549 | 0.849251 |
| 5 | 57222000 | 57224999 | 73.49552 | 116.1392 | 115.8151 | 48.10681 | 0.330217 | 2.424961 | 0.849265 |
| 15 | 39617000 | 39621199 | 33.03331 | 53.756 | 74.04168 | 31.78284 | -0.09084 | 1.557458 | 0.849275 |
| 3 | 94648200 | 94659799 | 145.3348 | 218.4142 | 276.4008 | 114.104 | -0.07064 | 3.430541 | 0.849281 |

| | | | | | | | | | |
|---|---|---|---|---|---|---|---|---|---|
| 12 | 30869200 | 30870999 | 14.9264 | 26.11369 | 45.15432 | 19.91676 | -0.39867 | 0.683832 | 0.849284 |
| 5 | 127034000 | 127048799 | 180.2552 | 268.0255 | 302.9982 | 124.8331 | 0.078101 | 3.633036 | 0.849285 |
| 12 | 60407000 | 60410799 | 56.00789 | 89.21494 | 106.0475 | 44.37068 | 0.08794 | 2.185864 | 0.849293 |
| 13 | 106866600 | 106867599 | 12.75665 | 22.94635 | 32.63067 | 14.25009 | -0.10072 | 0.388641 | 0.849322 |
| 15 | 64072000 | 64073399 | 21.49952 | 37.37333 | 37.2771 | 16.37198 | 0.39072 | 0.796508 | 0.849324 |
| 16 | 80833000 | 80834399 | 23.57135 | 40.39549 | 39.0294 | 17.1165 | 0.432752 | 0.942274 | 0.849326 |
| 17 | 88805200 | 88815399 | 165.6373 | 245.9546 | 308.9011 | 127.2474 | -0.07543 | 3.655168 | 0.849331 |
| 11 | 40700600 | 40702799 | 29.91917 | 49.43405 | 68.95125 | 29.78839 | -0.09818 | 1.351108 | 0.849382 |
| 3 | 97313800 | 97314799 | 11.33497 | 20.74869 | 30.29139 | 12.94309 | -0.11757 | 0.273836 | 0.849406 |
| 3 | 3028000 | 3100599 | 3284.745 | 3803.478 | 3780.483 | 1607.019 | -0.09795 | 7.582809 | 0.849412 |
| 8 | 84219200 | 84219599 | 7.110581 | 13.82925 | 19.55045 | 7.232262 | 0.110329 | -0.30325 | 0.849434 |
| 5 | 114799400 | 114800199 | 10.63912 | 19.98403 | 25.78356 | 10.6003 | 0.101312 | 0.10208 | 0.849451 |
| 17 | 48657600 | 48660999 | 32.61412 | 53.45511 | 65.47052 | 28.17918 | 0.084816 | 1.451312 | 0.849491 |
| 6 | 137824400 | 137834599 | 114.4206 | 175.2677 | 233.402 | 96.8703 | -0.12126 | 3.040684 | 0.849492 |
| 14 | 108537200 | 108541399 | 82.51021 | 128.1695 | 167.6168 | 69.59969 | -0.07961 | 2.760089 | 0.849501 |
| 6 | 44723600 | 44725199 | 18.61817 | 32.46103 | 39.88328 | 17.54941 | 0.091624 | 0.74406 | 0.849543 |
| 6 | 120103600 | 120107799 | 31.07265 | 51.34552 | 62.93794 | 27.19257 | 0.087647 | 1.346217 | 0.849556 |
| 10 | 105523400 | 105523999 | 12.59872 | 23.23855 | 29.17634 | 12.33795 | 0.108946 | 0.235414 | 0.849583 |
| 1 | 83929600 | 83931399 | 13.54268 | 24.55338 | 30.47027 | 13.17801 | 0.103379 | 0.341659 | 0.849593 |
| 11 | 96735000 | 96737999 | 26.72964 | 44.49455 | 61.55511 | 26.73014 | -0.08552 | 1.250746 | 0.849651 |
| 13 | 68713600 | 68715199 | 13.13628 | 23.53015 | 33.533 | 14.69611 | -0.10764 | 0.416605 | 0.849655 |
| 11 | 107190600 | 107194399 | 41.36373 | 66.79676 | 81.03293 | 34.57357 | 0.081441 | 1.737268 | 0.849672 |
| 6 | 64237600 | 64241399 | 33.73314 | 55.33396 | 79.37028 | 34.17347 | -0.14215 | 1.450807 | 0.849677 |
| 3 | 40520800 | 40522999 | 27.00148 | 44.91914 | 62.19008 | 26.99445 | -0.08673 | 1.257745 | 0.849733 |
| 2 | 89350000 | 89350799 | 21.15777 | 35.92864 | 50.59301 | 22.16919 | -0.11035 | 1.032945 | 0.849756 |
| 11 | 77038400 | 77041799 | 29.73528 | 48.68461 | 67.88365 | 29.20461 | -0.10234 | 1.459555 | 0.849763 |
| 1 | 4422200 | 4425199 | 20.7521 | 35.60624 | 43.2336 | 18.96209 | 0.104689 | 0.925503 | 0.849789 |
| 12 | 88492400 | 88499799 | 66.42061 | 105.3052 | 125.8024 | 52.35628 | 0.073881 | 2.341206 | 0.849792 |
| 1 | 117130800 | 117132599 | 28.86063 | 47.50548 | 65.90061 | 28.41558 | -0.09191 | 1.389617 | 0.849797 |
| 11 | 61520200 | 61525599 | 63.94238 | 101.8094 | 102.7225 | 42.95357 | 0.323357 | 2.20744 | 0.849803 |
| 3 | 33653600 | 33654399 | 15.82015 | 27.72309 | 38.83529 | 17.15349 | -0.09628 | 0.651946 | 0.849813 |
| 2 | 68062200 | 68062999 | 7.547559 | 14.51977 | 23.05643 | 8.982918 | -0.1267 | -0.11363 | 0.84982 |
| 6 | 130143200 | 130146999 | 27.12161 | 45.58695 | 67.05002 | 29.28936 | -0.17392 | 1.152883 | 0.849843 |
| 11 | 31483400 | 31488599 | 62.23369 | 99.22613 | 116.5412 | 48.82534 | 0.106668 | 2.165707 | 0.849859 |
| 11 | 73465800 | 73466399 | 10.73605 | 19.99834 | 29.42714 | 12.27063 | -0.10338 | 0.165425 | 0.849889 |
| 2 | 17734800 | 17737199 | 17.66776 | 29.85478 | 58.50568 | 25.59816 | -0.58579 | 1.06838 | 0.849896 |
| 14 | 107540600 | 107541399 | 16.62901 | 29.58993 | 29.22671 | 12.88052 | 0.412884 | 0.473516 | 0.849908 |
| 14 | 94441200 | 94442999 | 21.25543 | 36.90916 | 44.66895 | 19.60516 | 0.112697 | 0.820208 | 0.849917 |
| 15 | 95673600 | 95676799 | 25.04043 | 41.73211 | 58.18833 | 25.28703 | -0.09682 | 1.229039 | 0.849925 |
| 13 | 87654000 | 87658399 | 86.2854 | 134.4899 | 158.9044 | 65.94856 | 0.070478 | 2.702306 | 0.849943 |
| 2 | 177213600 | 177220799 | 138.4957 | 209.9344 | 239.9991 | 99.33021 | 0.086594 | 3.257429 | 0.849955 |
| 19 | 10543800 | 10544999 | 16.30378 | 28.52672 | 39.89976 | 17.6008 | -0.09408 | 0.681043 | 0.849955 |
| 19 | 39100000 | 39100799 | 14.34435 | 25.46114 | 30.63946 | 13.5315 | 0.127739 | 0.478703 | 0.849968 |
| 2 | 69457400 | 69460599 | 29.83994 | 49.3248 | 60.48163 | 26.13726 | 0.087264 | 1.339148 | 0.849969 |
| 5 | 55365000 | 55366199 | 30.95352 | 50.76487 | 60.87222 | 26.20691 | 0.115679 | 1.436313 | 0.850005 |
| 17 | 83910000 | 83914399 | 53.99522 | 86.10532 | 114.6744 | 48.0546 | -0.07408 | 2.151651 | 0.850009 |
| 19 | 11003400 | 11004399 | 15.0316 | 26.51264 | 37.14611 | 16.43826 | -0.0946 | 0.558203 | 0.850011 |

| | | | | | | | | | |
|---|---|---|---|---|---|---|---|---|---|
| 11 | 48215000 | 48215599 | 7.774786 | 14.8956 | 23.53687 | 9.278865 | -0.13482 | -0.06783 | 0.850011 |
| 7 | 102908000 | 102924599 | 228.5195 | 333.9248 | 370.3639 | 152.5832 | 0.082001 | 3.924482 | 0.850017 |
| 6 | 140315000 | 140317199 | 21.80618 | 38.01393 | 44.5621 | 19.60641 | 0.159035 | 0.768514 | 0.850025 |
| 9 | 8360400 | 8362199 | 23.81512 | 40.28677 | 56.09239 | 24.55045 | -0.09457 | 1.077878 | 0.850042 |
| 6 | 131065000 | 131066999 | 22.08881 | 38.3548 | 45.73939 | 20.07448 | 0.133852 | 0.825913 | 0.850052 |
| 12 | 60931600 | 60934199 | 25.61354 | 42.95904 | 60.02713 | 26.19079 | -0.10011 | 1.163618 | 0.850054 |
| 12 | 118189600 | 118192199 | 27.77856 | 46.91146 | 46.92485 | 20.52739 | 0.380785 | 1.096947 | 0.850055 |
| 15 | 6973200 | 6975199 | 21.26962 | 36.48962 | 51.16073 | 22.43867 | -0.10207 | 0.940301 | 0.850056 |
| 13 | 81343000 | 81343599 | 11.25378 | 20.48444 | 29.71502 | 12.78791 | -0.11546 | 0.302088 | 0.85006 |
| 6 | 127255200 | 127257999 | 28.51266 | 46.66209 | 77.973 | 33.53932 | -0.36315 | 1.469871 | 0.850063 |
| 2 | 86040800 | 86042199 | 29.83345 | 49.53312 | 60.6079 | 26.27209 | 0.090787 | 1.288401 | 0.850072 |
| 14 | 116125800 | 116127999 | 19.6414 | 34.53219 | 39.98008 | 17.70375 | 0.177181 | 0.622111 | 0.850087 |
| 3 | 7146200 | 7148599 | 22.51905 | 37.5843 | 64.66888 | 28.12894 | -0.39925 | 1.2117 | 0.850096 |
| 9 | 60950600 | 60951199 | 11.81946 | 21.70496 | 27.14782 | 11.54945 | 0.108368 | 0.250226 | 0.8501 |
| 6 | 116498200 | 116507399 | 130.0395 | 197.1677 | 253.4674 | 104.9128 | -0.08226 | 3.250964 | 0.850148 |
| 13 | 13893200 | 13897999 | 73.05013 | 114.8866 | 135.8257 | 56.34719 | 0.079634 | 2.529911 | 0.850158 |
| 5 | 93987400 | 93989999 | 44.98844 | 72.88387 | 70.98084 | 30.38158 | 0.402398 | 1.662667 | 0.850166 |
| 4 | 6439200 | 6441199 | 13.49716 | 23.99495 | 34.03924 | 15.01312 | -0.10732 | 0.467679 | 0.850171 |
| 14 | 98818800 | 98821999 | 26.9518 | 44.98621 | 62.87234 | 27.34891 | -0.10032 | 1.221385 | 0.850194 |
| 11 | 61184800 | 61186799 | 17.21803 | 30.22968 | 37.09153 | 16.40941 | 0.09384 | 0.625869 | 0.850195 |
| 3 | 16933800 | 16934399 | 8.3141 | 16.07854 | 22.09796 | 8.449984 | 0.105534 | -0.1686 | 0.850198 |
| 5 | 26867800 | 26869599 | 24.72725 | 42.28489 | 50.6643 | 22.21517 | 0.122979 | 0.980133 | 0.850201 |
| 13 | 79365800 | 79366599 | 10.00752 | 18.98804 | 24.89911 | 10.03801 | 0.102177 | 0.027316 | 0.850209 |
| 1 | 107061400 | 107071599 | 101.9689 | 158.2919 | 179.8661 | 74.69336 | 0.122197 | 2.781421 | 0.850218 |
| 13 | 53413200 | 53417999 | 33.84211 | 54.6048 | 93.3729 | 39.7957 | -0.41406 | 1.755585 | 0.850232 |
| 17 | 18975600 | 18987799 | 207.9098 | 308.6032 | 277.9591 | 114.4967 | 0.403403 | 3.667534 | 0.850233 |
| 13 | 120701000 | 120703799 | 26.5693 | 44.32751 | 53.951 | 23.46144 | 0.098628 | 1.217596 | 0.850239 |
| 7 | 102982400 | 102984199 | 14.19003 | 25.90105 | 31.5792 | 13.54295 | 0.136515 | 0.310139 | 0.850261 |
| 9 | 88921400 | 88927599 | 54.75401 | 87.55795 | 105.0579 | 44.1974 | 0.080519 | 2.064037 | 0.850279 |
| 11 | 40699600 | 40703599 | 41.56445 | 67.15481 | 93.46034 | 39.90185 | -0.11542 | 1.731514 | 0.850279 |
| 7 | 61135000 | 61136799 | 17.54096 | 31.01957 | 36.96096 | 16.34779 | 0.139352 | 0.545476 | 0.85029 |
| 16 | 44208800 | 44213199 | 52.5075 | 84.10083 | 100.8234 | 42.5149 | 0.084336 | 2.009602 | 0.850296 |
| 2 | 64529000 | 64530999 | 18.04984 | 31.6499 | 38.65232 | 17.08158 | 0.100585 | 0.653381 | 0.850356 |
| 16 | 61845600 | 61847399 | 15.32177 | 27.06276 | 38.10147 | 16.85627 | -0.1009 | 0.545198 | 0.850384 |
| 8 | 6978200 | 6979599 | 13.19906 | 24.0341 | 35.40978 | 15.23167 | -0.13768 | 0.325579 | 0.850396 |
| 9 | 60918400 | 60921399 | 30.3692 | 49.8451 | 68.69359 | 29.58387 | -0.08412 | 1.431285 | 0.850432 |
| 17 | 12635600 | 12637999 | 18.20892 | 31.67374 | 44.1644 | 19.40479 | -0.09042 | 0.781467 | 0.850436 |
| 1 | 29657800 | 29660999 | 36.76945 | 60.5142 | 70.86428 | 30.55384 | 0.151682 | 1.414955 | 0.850456 |
| 4 | 15189000 | 15190199 | 14.92948 | 26.81651 | 32.51683 | 14.21746 | 0.126697 | 0.413389 | 0.850468 |
| 16 | 85846400 | 85847999 | 15.29584 | 27.73577 | 32.99089 | 14.26566 | 0.164197 | 0.357661 | 0.850483 |
| 19 | 33426000 | 33431599 | 66.25492 | 105.1383 | 125.2906 | 52.19085 | 0.078742 | 2.314522 | 0.850487 |
| 13 | 55924200 | 55929599 | 51.3164 | 82.09807 | 98.63736 | 41.52298 | 0.079352 | 2.044934 | 0.850503 |
| 2 | 91819800 | 91821399 | 17.40943 | 30.4116 | 36.77979 | 16.20325 | 0.114541 | 0.70524 | 0.85051 |
| 11 | 13563800 | 13566799 | 40.64502 | 64.97022 | 113.3567 | 47.66182 | -0.4592 | 2.064226 | 0.850537 |
| 7 | 9395400 | 9396599 | 23.35459 | 40.16219 | 38.52257 | 16.89766 | 0.444137 | 0.911429 | 0.850542 |
| X | 14352200 | 14356599 | 40.42916 | 65.73045 | 77.60098 | 33.29943 | 0.130077 | 1.59122 | 0.850549 |
| 7 | 15991000 | 15993399 | 17.96493 | 31.25276 | 37.50162 | 16.47846 | 0.125384 | 0.769148 | 0.85055 |

| | | | | | | | | | |
|---|---|---|---|---|---|---|---|---|---|
| 2 | 5738400 | 5740399 | 22.64938 | 38.66408 | 47.42637 | 20.79249 | 0.088975 | 0.985527 | 0.850567 |
| 1 | 139675000 | 139677599 | 23.79884 | 41.2629 | 47.37841 | 20.7874 | 0.187836 | 0.849396 | 0.850575 |
| 2 | 54786200 | 54786999 | 14.42831 | 25.58681 | 36.09295 | 15.95222 | -0.10114 | 0.495727 | 0.850577 |
| 6 | 131965200 | 131967799 | 25.38312 | 43.78988 | 50.13661 | 21.99422 | 0.190598 | 0.906171 | 0.850587 |
| 5 | 141657800 | 141658999 | 17.7775 | 31.04558 | 43.55541 | 19.18474 | -0.09864 | 0.723751 | 0.850591 |
| 2 | 43283400 | 43285799 | 25.29739 | 42.37002 | 58.8658 | 25.66163 | -0.09182 | 1.176471 | 0.850638 |
| 1 | 153694400 | 153695399 | 9.120189 | 17.24486 | 25.95367 | 10.53975 | -0.10419 | 0.045934 | 0.850672 |
| 13 | 120323000 | 120323799 | 9.120189 | 17.24486 | 25.95367 | 10.53975 | -0.10419 | 0.045934 | 0.850672 |
| 14 | 58885800 | 58887999 | 21.07251 | 35.60535 | 50.6167 | 22.14595 | -0.1244 | 1.07318 | 0.850695 |
| 7 | 47773400 | 47775399 | 18.82309 | 32.35665 | 45.83822 | 20.10638 | -0.11645 | 0.902745 | 0.850711 |
| 3 | 44146200 | 44147999 | 48.60838 | 77.68241 | 104.9466 | 44.112 | -0.09123 | 2.077496 | 0.85072 |
| 4 | 26607800 | 26610199 | 28.6763 | 47.58122 | 66.22477 | 28.69411 | -0.09465 | 1.295801 | 0.850736 |
| 4 | 91438400 | 91439599 | 13.80864 | 24.83384 | 30.35166 | 13.25699 | 0.115939 | 0.388832 | 0.850762 |
| 4 | 37922800 | 37923399 | 11.77318 | 21.4965 | 26.14729 | 11.20892 | 0.140601 | 0.278995 | 0.850778 |
| 11 | 32227600 | 32229199 | 11.77663 | 21.23663 | 30.70361 | 13.36878 | -0.12156 | 0.362795 | 0.850799 |
| 10 | 3531800 | 3533599 | 25.99503 | 43.55389 | 53.36466 | 23.26869 | 0.088986 | 1.171972 | 0.850828 |
| 11 | 101694000 | 101695199 | 13.19603 | 23.84497 | 34.16502 | 14.8379 | -0.10656 | 0.367217 | 0.85085 |
| 3 | 93680600 | 93683199 | 34.35344 | 55.93169 | 76.58324 | 32.87969 | -0.08184 | 1.5551 | 0.850903 |
| 3 | 130039200 | 130042599 | 32.14404 | 52.71169 | 72.99843 | 31.42609 | -0.09169 | 1.448253 | 0.85091 |
| 9 | 93830000 | 93830799 | 9.424743 | 17.90847 | 23.36611 | 9.39088 | 0.113679 | 0.009149 | 0.850927 |
| 18 | 37813200 | 37826599 | 500.5507 | 679.9021 | 703.9566 | 293.2291 | 0.075788 | 5.030709 | 0.850935 |
| 18 | 9247600 | 9250399 | 49.49374 | 79.07061 | 106.1637 | 44.60218 | -0.08289 | 2.087256 | 0.850939 |
| 5 | 114680200 | 114684399 | 40.58461 | 65.57714 | 79.75899 | 34.05585 | 0.078718 | 1.719224 | 0.850943 |
| 5 | 76296800 | 76301799 | 46.35385 | 74.20367 | 100.4887 | 42.3386 | -0.09207 | 2.019955 | 0.85098 |
| 17 | 62991800 | 62992599 | 9.549028 | 17.88227 | 26.84784 | 11.09454 | -0.12183 | 0.113302 | 0.850985 |
| 13 | 11008000 | 11011599 | 71.15059 | 112.1668 | 133.4625 | 55.38402 | 0.073044 | 2.480416 | 0.850985 |
| 8 | 4123400 | 4125599 | 44.41537 | 71.3187 | 95.98254 | 40.62856 | -0.07787 | 1.921711 | 0.85101 |
| 14 | 3870800 | 3872999 | 17.24227 | 29.96543 | 42.13798 | 18.50401 | -0.10262 | 0.78707 | 0.851014 |
| 17 | 80910800 | 80913399 | 23.74161 | 40.03799 | 55.47653 | 24.25806 | -0.08768 | 1.105514 | 0.851023 |
| 9 | 92503600 | 92508399 | 48.57693 | 78.18362 | 92.9283 | 39.47027 | 0.105125 | 1.852103 | 0.851138 |
| 16 | 55461400 | 55463999 | 32.71416 | 53.57022 | 73.97539 | 31.82748 | -0.08836 | 1.469302 | 0.851191 |
| 18 | 37544600 | 37547399 | 73.5915 | 116.3697 | 108.0949 | 44.91541 | 0.433248 | 2.409017 | 0.851213 |
| 12 | 106593000 | 106594799 | 20.36486 | 34.79363 | 48.48697 | 21.25891 | -0.09422 | 0.969372 | 0.851223 |
| 11 | 9391000 | 9401599 | 178.809 | 264.6944 | 297.5041 | 122.5212 | 0.080529 | 3.709924 | 0.851276 |
| 11 | 15998400 | 16002599 | 74.10731 | 116.7919 | 139.1004 | 57.7213 | 0.071294 | 2.489184 | 0.85128 |
| 2 | 23269800 | 23272599 | 26.97656 | 44.81376 | 62.1888 | 26.96734 | -0.09025 | 1.273255 | 0.851297 |
| 11 | 67198200 | 67200799 | 28.68171 | 47.57171 | 58.26673 | 25.23925 | 0.08926 | 1.295334 | 0.851313 |
| 4 | 112765600 | 112767399 | 30.85705 | 51.1365 | 62.30615 | 26.97325 | 0.096658 | 1.31444 | 0.851331 |
| 6 | 140245600 | 140249199 | 26.2125 | 44.116 | 53.96202 | 23.58141 | 0.091363 | 1.127429 | 0.851368 |
| 8 | 62995400 | 62998799 | 58.83409 | 93.72563 | 112.5766 | 47.10861 | 0.073058 | 2.187516 | 0.851467 |
| 12 | 114335400 | 114343999 | 145.9092 | 220.2202 | 252.189 | 104.2334 | 0.078831 | 3.346069 | 0.851475 |
| 5 | 114608600 | 114610799 | 28.88804 | 47.52627 | 65.82466 | 28.37668 | -0.0899 | 1.396618 | 0.851494 |
| 18 | 77143600 | 77144999 | 17.22319 | 30.22065 | 37.13061 | 16.41838 | 0.091928 | 0.635692 | 0.851513 |
| 4 | 67607600 | 67610999 | 24.75345 | 41.73124 | 51.20637 | 22.38872 | 0.086965 | 1.105747 | 0.851516 |
| 12 | 59412800 | 59413599 | 8.323463 | 15.93375 | 24.57894 | 9.712489 | -0.10439 | -0.05457 | 0.851529 |
| 15 | 54044400 | 54045199 | 8.323463 | 15.93375 | 24.57894 | 9.712489 | -0.10439 | -0.05457 | 0.851529 |
| 13 | 119622800 | 119629199 | 360.4924 | 494.6447 | 739.3041 | 307.4667 | -0.43894 | 4.869841 | 0.851535 |

| | | | | | | | | | |
|---|---|---|---|---|---|---|---|---|---|
| 15 | 88065600 | 88067599 | 17.26907 | 30.02794 | 42.13181 | 18.50747 | -0.09931 | 0.780236 | 0.851559 |
| 3 | 11263400 | 11263799 | 9.378744 | 17.8238 | 23.34496 | 9.37933 | 0.108368 | 0.006466 | 0.851592 |
| 13 | 97305000 | 97306799 | 15.29325 | 27.49913 | 33.44243 | 14.60907 | 0.123122 | 0.408856 | 0.851596 |
| 6 | 134576000 | 134576599 | 8.105435 | 15.69745 | 21.59043 | 8.200876 | 0.113204 | -0.19672 | 0.851606 |
| 7 | 61880400 | 61884399 | 49.31351 | 79.37601 | 79.76106 | 33.88188 | 0.347064 | 1.845087 | 0.851619 |
| 3 | 12430600 | 12432399 | 14.23673 | 25.76209 | 31.56098 | 13.68262 | 0.119935 | 0.363022 | 0.851638 |
| 1 | 122907800 | 122910799 | 26.27101 | 44.37449 | 53.95507 | 23.60422 | 0.100164 | 1.097746 | 0.851641 |
| 5 | 71407800 | 71411999 | 43.47701 | 70.19117 | 84.54369 | 36.05938 | 0.091334 | 1.752325 | 0.851645 |
| 14 | 33700800 | 33703599 | 26.08672 | 43.27278 | 60.3014 | 26.12275 | -0.09631 | 1.287947 | 0.851649 |
| 18 | 7774000 | 7776999 | 34.63027 | 56.39054 | 68.55727 | 29.43543 | 0.089416 | 1.548754 | 0.851651 |
| 6 | 38342600 | 38348799 | 63.10023 | 100.0353 | 132.3224 | 55.06663 | -0.07294 | 2.34299 | 0.851672 |
| 15 | 88728600 | 88730799 | 20.3242 | 34.75613 | 48.48396 | 21.25917 | -0.09553 | 0.959971 | 0.851683 |
| 9 | 114648800 | 114650399 | 17.28651 | 30.11111 | 42.08248 | 18.50622 | -0.0934 | 0.755607 | 0.85173 |
| 13 | 68643000 | 68643599 | 11.9738 | 21.88833 | 27.1329 | 11.61448 | 0.114852 | 0.276841 | 0.851756 |
| 9 | 5612400 | 5617599 | 78.53966 | 123.3391 | 146.5242 | 60.77385 | 0.071043 | 2.564102 | 0.85176 |
| 12 | 115439400 | 115463999 | 291.6284 | 420.6516 | 439.7941 | 181.4589 | 0.148942 | 4.124776 | 0.851768 |
| 9 | 78798800 | 78800999 | 19.083 | 33.12559 | 46.3272 | 20.32863 | -0.095 | 0.818676 | 0.85177 |
| 4 | 16655200 | 16656199 | 10.03073 | 18.80063 | 27.92526 | 11.52285 | -0.10477 | 0.115075 | 0.851779 |
| 7 | 33474600 | 33572999 | 2318.353 | 2790.806 | 2549.884 | 1077.537 | 0.080695 | 6.972041 | 0.851796 |
| 12 | 115932800 | 115937199 | 110.4546 | 168.8952 | 195.8399 | 81.24758 | 0.076296 | 3.06976 | 0.851815 |
| 17 | 55423000 | 55425599 | 28.62258 | 47.49922 | 66.29993 | 28.72965 | -0.09877 | 1.292858 | 0.851851 |
| 2 | 11749000 | 11749999 | 12.06662 | 22.04923 | 31.75243 | 13.59868 | -0.10061 | 0.290539 | 0.851861 |
| 9 | 72851400 | 72854799 | 47.97675 | 76.92826 | 92.53436 | 39.08472 | 0.081418 | 1.968664 | 0.851871 |
| 2 | 39338400 | 39341399 | 29.59519 | 50.28176 | 45.19775 | 19.80784 | 0.534705 | 1.050656 | 0.851882 |
| 10 | 66601800 | 66612799 | 149.7255 | 225.4064 | 258.6077 | 106.8086 | 0.072953 | 3.396321 | 0.851904 |
| 8 | 100831400 | 100832399 | 12.57073 | 23.03373 | 28.66634 | 12.23211 | 0.11254 | 0.273438 | 0.851971 |
| 8 | 74148600 | 74149199 | 8.757091 | 16.68061 | 25.55752 | 10.21857 | -0.11032 | -0.01067 | 0.851972 |
| 13 | 62276200 | 62294999 | 355.472 | 500.2523 | 532.8175 | 220.7138 | 0.08652 | 4.486917 | 0.852015 |
| 6 | 79998400 | 80000199 | 25.09931 | 42.20261 | 51.74772 | 22.60728 | 0.087862 | 1.130432 | 0.852016 |
| 10 | 45046600 | 45048199 | 14.34418 | 25.34778 | 35.6048 | 15.75027 | -0.09723 | 0.52989 | 0.852027 |
| 1 | 177131200 | 177132199 | 12.5857 | 22.53182 | 32.15354 | 14.09794 | -0.10969 | 0.415744 | 0.852046 |
| 5 | 83018200 | 83022799 | 66.53 | 105.66 | 106.0995 | 44.23235 | 0.326514 | 2.287347 | 0.852047 |
| X | 48189600 | 48191799 | 20.92507 | 36.06304 | 43.74084 | 19.1879 | 0.107627 | 0.889966 | 0.85206 |
| 8 | 86674400 | 86675999 | 23.20686 | 38.68426 | 65.1827 | 28.33027 | -0.3691 | 1.228955 | 0.852065 |
| 12 | 91736600 | 91738599 | 23.6559 | 39.68627 | 55.49632 | 24.218 | -0.10114 | 1.150356 | 0.852077 |
| 13 | 22179400 | 22180199 | 8.560442 | 16.31306 | 25.56525 | 10.21234 | -0.14179 | -0.01722 | 0.852097 |
| 11 | 67484400 | 67485399 | 14.15835 | 25.31137 | 30.86503 | 13.54918 | 0.114279 | 0.429628 | 0.852104 |
| 17 | 15588000 | 15591399 | 23.4587 | 38.68198 | 71.57912 | 30.88716 | -0.5053 | 1.358357 | 0.852119 |
| Y | 8893400 | 8893999 | 6.264378 | 12.07791 | 21.02653 | 8.14019 | -0.24884 | -0.15351 | 0.852122 |
| 8 | 30005400 | 30006799 | 17.80911 | 31.14449 | 37.85759 | 16.69265 | 0.107547 | 0.695707 | 0.852132 |
| 9 | 87368400 | 87370199 | 27.49859 | 45.7732 | 56.08658 | 24.35499 | 0.089058 | 1.24866 | 0.852142 |
| 13 | 50710200 | 50724799 | 170.1968 | 252.9908 | 316.8971 | 130.5552 | -0.07022 | 3.637845 | 0.852146 |
| 4 | 69772200 | 69773799 | 22.60296 | 38.00129 | 53.26655 | 23.2701 | -0.10441 | 1.12595 | 0.852174 |
| 5 | 103388000 | 103388599 | 24.46183 | 41.03729 | 49.09987 | 21.42632 | 0.123166 | 1.148504 | 0.852213 |
| 15 | 56736400 | 56739799 | 51.75716 | 82.50749 | 111.1606 | 46.55962 | -0.09127 | 2.157858 | 0.852244 |
| 8 | 19294000 | 19297799 | 46.67596 | 75.23951 | 89.83355 | 38.23942 | 0.101353 | 1.802228 | 0.852245 |
| 14 | 20170000 | 20171399 | 15.31526 | 26.85035 | 37.81873 | 16.71081 | -0.10412 | 0.639782 | 0.852283 |

| | | | | | | | | | |
|---|---|---|---|---|---|---|---|---|---|
| 5 | 109289000 | 109294799 | 97.44103 | 150.971 | 176.5858 | 73.34885 | 0.078325 | 2.812867 | 0.852289 |
| 2 | 43090600 | 43090999 | 6.999912 | 13.62422 | 19.04379 | 7.0023 | 0.135354 | -0.32473 | 0.852296 |
| 9 | 92901600 | 92901999 | 6.999912 | 13.62422 | 19.04379 | 7.0023 | 0.135354 | -0.32473 | 0.852296 |
| 2 | 131562600 | 131567999 | 70.94033 | 112.2502 | 133.3627 | 55.43284 | 0.079417 | 2.395659 | 0.852307 |
| 14 | 7939600 | 7940799 | 10.82652 | 19.98665 | 29.2588 | 12.35854 | -0.11002 | 0.215741 | 0.852314 |
| 7 | 71553800 | 71554599 | 9.566256 | 18.32377 | 24.00722 | 9.470352 | 0.130921 | -0.05205 | 0.852322 |
| 6 | 42138600 | 42140599 | 61.50745 | 97.56557 | 115.3222 | 48.01919 | 0.089955 | 2.320723 | 0.852335 |
| 8 | 113876800 | 113878799 | 19.3114 | 34.05304 | 39.50182 | 17.48159 | 0.176163 | 0.589643 | 0.852343 |
| 5 | 120145400 | 120148999 | 39.77493 | 64.2371 | 87.29832 | 37.25129 | -0.08187 | 1.734999 | 0.852363 |
| 13 | 68292200 | 68333599 | 991.7786 | 1278.58 | 1259.777 | 528.1739 | 0.071917 | 5.872252 | 0.852383 |
| 12 | 99587800 | 99588399 | 8.393044 | 16.26612 | 22.1988 | 8.403752 | 0.129831 | -0.19887 | 0.852412 |
| 5 | 122547600 | 122549599 | 20.88403 | 35.54475 | 49.5476 | 21.71727 | -0.09545 | 1.007781 | 0.852499 |
| 15 | 96088800 | 96097199 | 89.65474 | 139.0423 | 180.3904 | 74.9218 | -0.06994 | 2.792697 | 0.8525 |
| 8 | 37809400 | 37812399 | 46.6905 | 74.89508 | 89.27754 | 37.73151 | 0.095128 | 1.954373 | 0.852517 |
| 7 | 80596800 | 80597799 | 10.0746 | 18.5538 | 27.75173 | 11.76046 | -0.14405 | 0.220099 | 0.852518 |
| 3 | 85210600 | 85219399 | 97.18482 | 149.7226 | 192.9491 | 80.14648 | -0.06763 | 2.918035 | 0.852551 |
| 1 | 142538600 | 142547799 | 138.755 | 209.7869 | 242.1494 | 100.1388 | 0.06956 | 3.308611 | 0.85256 |
| 6 | 130726800 | 130727999 | 12.36135 | 23.0264 | 28.80654 | 12.01134 | 0.128917 | 0.175399 | 0.85265 |
| 14 | 3701200 | 3702199 | 20.58705 | 35.44322 | 49.54684 | 21.73414 | -0.09683 | 0.903745 | 0.852656 |
| 5 | 101809400 | 101813799 | 38.15405 | 61.16061 | 106.7261 | 45.05945 | -0.45476 | 1.970788 | 0.852669 |
| 10 | 62196000 | 62203799 | 111.2762 | 169.7333 | 217.6453 | 90.24257 | -0.07154 | 3.121128 | 0.852671 |
| 16 | 27232400 | 27234799 | 22.69133 | 38.44886 | 54.46701 | 23.85094 | -0.11942 | 1.055212 | 0.852704 |
| 10 | 24905000 | 24907599 | 23.04437 | 38.93143 | 53.84822 | 23.55877 | -0.08505 | 1.085438 | 0.852709 |
| 7 | 14551800 | 14561799 | 437.7032 | 600.1238 | 619.581 | 257.707 | 0.093537 | 4.883162 | 0.852731 |
| 6 | 140672000 | 140675799 | 48.19365 | 77.35594 | 92.9459 | 39.32027 | 0.084736 | 1.935231 | 0.852766 |
| 16 | 35215400 | 35216799 | 19.91306 | 34.54722 | 42.12787 | 18.48407 | 0.101849 | 0.823767 | 0.852769 |
| 13 | 15091800 | 15092999 | 15.88271 | 28.04114 | 34.45723 | 15.24538 | 0.094126 | 0.546389 | 0.852777 |
| 8 | 26363000 | 26363999 | 16.04551 | 28.73789 | 28.1511 | 12.33867 | 0.429991 | 0.426215 | 0.852778 |
| 13 | 114866800 | 114870399 | 37.82962 | 61.20816 | 83.17611 | 35.56805 | -0.07882 | 1.681031 | 0.852788 |
| 8 | 79953600 | 79956199 | 25.13028 | 42.25057 | 58.88861 | 25.7262 | -0.09646 | 1.137495 | 0.852796 |
| 2 | 66679400 | 66682399 | 43.08834 | 69.78955 | 70.25015 | 30.06373 | 0.354519 | 1.662017 | 0.852818 |
| 13 | 51147400 | 51148599 | 18.28902 | 31.97376 | 39.26581 | 17.309 | 0.092582 | 0.70218 | 0.852821 |
| 17 | 66044800 | 66049799 | 50.95899 | 81.31931 | 109.024 | 45.73047 | -0.08259 | 2.124039 | 0.85284 |
| 7 | 6678000 | 6681999 | 41.32359 | 66.55578 | 89.91481 | 38.22884 | -0.07794 | 1.820814 | 0.852864 |
| 12 | 64345000 | 64346199 | 16.93542 | 29.83135 | 36.5123 | 16.16203 | 0.098585 | 0.582261 | 0.852874 |
| 3 | 102842400 | 102844999 | 43.51062 | 69.78522 | 95.37517 | 40.30057 | -0.10196 | 1.957812 | 0.852901 |
| 5 | 12193600 | 12195199 | 13.86665 | 24.54167 | 34.60819 | 15.30469 | -0.10195 | 0.508392 | 0.852903 |
| 6 | 68198200 | 68199999 | 20.82962 | 35.70991 | 49.54831 | 21.73026 | -0.08729 | 0.940996 | 0.852915 |
| 13 | 68547800 | 68551399 | 61.57959 | 98.2343 | 115.4363 | 48.39728 | 0.106722 | 2.148384 | 0.852936 |
| 7 | 136267400 | 136268599 | 13.08618 | 23.70552 | 29.50619 | 12.77466 | 0.098213 | 0.341778 | 0.852938 |
| 8 | 23148800 | 23150399 | 11.69411 | 21.483 | 26.69353 | 11.35025 | 0.118516 | 0.246829 | 0.85294 |
| 3 | 35911000 | 35911799 | 10.56407 | 19.68815 | 28.89887 | 12.03981 | -0.0988 | 0.159309 | 0.852956 |
| 3 | 83249600 | 83252399 | 26.87869 | 44.11131 | 74.45494 | 32.0636 | -0.37583 | 1.429042 | 0.85296 |
| 8 | 41879000 | 41880399 | 14.80398 | 26.07513 | 36.63496 | 16.21654 | -0.09946 | 0.569618 | 0.852985 |
| 7 | 107082200 | 107083599 | 21.12441 | 36.36872 | 44.75066 | 19.63026 | 0.086512 | 0.899019 | 0.852989 |
| 2 | 48468600 | 48469199 | 11.26807 | 20.46785 | 29.72477 | 12.8254 | -0.11947 | 0.314477 | 0.853011 |
| 6 | 137181600 | 137182599 | 11.66756 | 21.52671 | 27.17476 | 11.4874 | 0.101764 | 0.223039 | 0.853039 |

| | | | | | | | | | |
|---|---|---|---|---|---|---|---|---|---|
| X | 140820000 | 140820799 | 12.3628 | 23.07429 | 28.89142 | 12.00681 | 0.131845 | 0.165564 | 0.85304 |
| 3 | 98851200 | 98853399 | 16.7784 | 29.34594 | 40.91544 | 18.043 | -0.08958 | 0.690961 | 0.853056 |
| 9 | 10197200 | 10199199 | 16.7784 | 29.34594 | 40.91544 | 18.043 | -0.08958 | 0.690961 | 0.853056 |
| 6 | 8325800 | 8332599 | 76.18073 | 120.5748 | 139.021 | 57.79267 | 0.122934 | 2.395322 | 0.85307 |
| 12 | 96022800 | 96024399 | 16.51457 | 28.87245 | 40.46308 | 17.83724 | -0.09684 | 0.69703 | 0.85308 |
| 14 | 44450200 | 44451399 | 10.89879 | 20.15942 | 29.31384 | 12.34891 | -0.09736 | 0.207333 | 0.853093 |
| 4 | 112746200 | 112748599 | 34.20605 | 56.66076 | 66.63105 | 28.83652 | 0.147608 | 1.331892 | 0.853103 |
| 14 | 73853200 | 73854799 | 13.57461 | 24.51095 | 30.46297 | 13.24232 | 0.09656 | 0.365751 | 0.853112 |
| 18 | 10396800 | 10397599 | 9.404527 | 18.14306 | 24.30067 | 9.391601 | 0.127584 | -0.11587 | 0.853114 |
| 7 | 23872600 | 23881399 | 86.00959 | 132.9581 | 175.3621 | 72.84143 | -0.09726 | 2.848086 | 0.853117 |
| X | 164627600 | 164630399 | 34.34189 | 56.91308 | 66.6984 | 28.87524 | 0.152586 | 1.326396 | 0.853119 |
| 7 | 63635000 | 63637399 | 19.85577 | 34.02884 | 47.42463 | 20.79831 | -0.09365 | 0.93773 | 0.853137 |
| 9 | 83626200 | 83626999 | 11.54336 | 21.30585 | 26.70792 | 11.28407 | 0.112524 | 0.219583 | 0.853143 |
| 10 | 34795800 | 34796599 | 11.54336 | 21.30585 | 26.70792 | 11.28407 | 0.112524 | 0.219583 | 0.853143 |
| 18 | 85561800 | 85562399 | 7.58577 | 14.64191 | 23.13968 | 8.92801 | -0.10622 | -0.14232 | 0.85316 |
| 7 | 3158000 | 3158999 | 17.54708 | 30.71049 | 37.76278 | 16.66195 | 0.090846 | 0.679586 | 0.853161 |
| 17 | 64466400 | 64468399 | 18.08852 | 31.75373 | 38.60878 | 17.0787 | 0.106906 | 0.635934 | 0.853184 |
| 7 | 57812400 | 57813199 | 14.29177 | 25.40954 | 31.03581 | 13.68755 | 0.107612 | 0.467662 | 0.85319 |
| 8 | 58438400 | 58439399 | 10.27608 | 18.57633 | 34.35244 | 14.91161 | -0.47148 | 0.346435 | 0.853202 |
| X | 105240000 | 105243999 | 19.75141 | 34.51157 | 52.28103 | 23.03601 | -0.20897 | 0.73905 | 0.853208 |
| 13 | 77050800 | 77051799 | 13.03059 | 23.38644 | 33.07998 | 14.47374 | -0.09512 | 0.404267 | 0.853212 |
| 1 | 103825600 | 103826199 | 10.15857 | 19.06873 | 28.52026 | 11.73859 | -0.11133 | 0.111047 | 0.853232 |
| 7 | 65251600 | 65256399 | 50.42992 | 80.71034 | 97.05828 | 40.88013 | 0.078628 | 2.030888 | 0.853235 |
| 10 | 79248400 | 79263199 | 265.5102 | 382.2718 | 421.0025 | 173.7479 | 0.07132 | 4.14858 | 0.853239 |
| 6 | 67636000 | 67649599 | 253.5466 | 367.2189 | 404.4233 | 166.7747 | 0.079751 | 4.060517 | 0.853258 |
| Y | 998400 | 999399 | 7.948226 | 14.90644 | 24.44892 | 10.07803 | -0.24482 | 0.084954 | 0.853274 |
| X | 98175600 | 98184599 | 65.32171 | 103.6064 | 142.8575 | 59.4792 | -0.13211 | 2.335689 | 0.853288 |
| 2 | 54464600 | 54465199 | 13.53003 | 24.17054 | 34.06865 | 14.96466 | -0.09395 | 0.438648 | 0.853305 |
| X | 144069800 | 144070999 | 12.17963 | 22.42756 | 33.54098 | 14.21632 | -0.14398 | 0.25371 | 0.853312 |
| 10 | 129752800 | 129754599 | 25.16265 | 42.12468 | 58.27445 | 25.39636 | -0.08561 | 1.181126 | 0.853336 |
| 5 | 3534000 | 3536999 | 33.52699 | 55.04044 | 67.24432 | 28.96249 | 0.089351 | 1.436596 | 0.85334 |
| 6 | 117608200 | 117611399 | 44.02626 | 71.02389 | 85.92938 | 36.61448 | 0.083616 | 1.773754 | 0.853352 |
| X | 168102000 | 168104199 | 22.97741 | 39.28757 | 58.27735 | 25.55306 | -0.18382 | 0.987745 | 0.853355 |
| 11 | 62506800 | 62508799 | 19.27344 | 33.4427 | 41.06455 | 18.01761 | 0.091824 | 0.8156 | 0.853393 |
| 13 | 65145800 | 65148799 | 60.5592 | 96.24889 | 115.1852 | 48.05584 | 0.074824 | 2.263665 | 0.853402 |
| 4 | 59369800 | 59370999 | 10.40564 | 19.36165 | 28.34451 | 11.84114 | -0.09811 | 0.167064 | 0.853405 |
| 9 | 39896800 | 39897199 | 5.797279 | 11.30393 | 19.17506 | 6.901513 | -0.11332 | -0.4169 | 0.853441 |
| 8 | 48681400 | 48684799 | 52.03501 | 82.99164 | 110.922 | 46.49161 | -0.07896 | 2.144266 | 0.853442 |
| 4 | 115087600 | 115090999 | 50.82995 | 81.54077 | 97.62239 | 41.26268 | 0.08893 | 1.957999 | 0.853453 |
| 6 | 20865000 | 20865599 | 8.210029 | 15.79941 | 24.74354 | 9.631123 | -0.10538 | -0.10279 | 0.853469 |
| 13 | 7949000 | 7950599 | 28.98338 | 47.88276 | 58.22373 | 25.15202 | 0.099259 | 1.340754 | 0.85347 |
| 16 | 10008400 | 10011399 | 28.90736 | 47.55484 | 65.78452 | 28.35857 | -0.08818 | 1.398005 | 0.853491 |
| 7 | 90106200 | 90106799 | 6.240202 | 12.09042 | 20.53432 | 7.783794 | -0.1849 | -0.22789 | 0.853506 |
| 7 | 23790000 | 23812799 | 839.0442 | 1089.057 | 1195.38 | 500.8422 | -0.07401 | 5.760921 | 0.853528 |
| 14 | 89059400 | 89060199 | 11.48429 | 20.80296 | 30.25847 | 13.10094 | -0.12478 | 0.334659 | 0.853531 |
| 12 | 103436600 | 103439199 | 32.9555 | 53.82961 | 65.26209 | 28.05223 | 0.096557 | 1.498397 | 0.853552 |
| 9 | 78509600 | 78510799 | 16.04696 | 28.22624 | 34.56039 | 15.30237 | 0.097261 | 0.588229 | 0.853568 |

| | | | | | | | | | |
|---|---|---|---|---|---|---|---|---|---|
| 6 | 5423200 | 5423999 | 10.97796 | 20.03436 | 29.27682 | 12.55845 | -0.12275 | 0.285335 | 0.853606 |
| 18 | 86084400 | 86084999 | 12.6465 | 22.98002 | 28.58896 | 12.32975 | 0.102692 | 0.319822 | 0.853607 |
| 7 | 144894000 | 144894599 | 5.506636 | 10.73348 | 18.59047 | 6.721664 | -0.14824 | -0.40523 | 0.853612 |
| 3 | 53599000 | 53601599 | 27.05958 | 44.98525 | 62.1095 | 26.94684 | -0.08278 | 1.266918 | 0.853632 |
| 5 | 83818600 | 83820199 | 16.9287 | 29.84694 | 36.48619 | 16.14751 | 0.100853 | 0.57202 | 0.85365 |
| 7 | 132517200 | 132520799 | 49.47086 | 79.54912 | 94.95714 | 40.27822 | 0.09733 | 1.883706 | 0.853652 |
| 15 | 20919600 | 20921199 | 15.20407 | 26.96889 | 33.01677 | 14.58971 | 0.102437 | 0.493526 | 0.853656 |
| 3 | 28927800 | 28937599 | 290.1752 | 413.4462 | 450.896 | 186.3947 | 0.070387 | 4.301868 | 0.853667 |
| 10 | 120939400 | 120943599 | 52.31319 | 83.80559 | 100.3737 | 42.33755 | 0.086029 | 2.002907 | 0.85369 |
| 14 | 121254400 | 121255999 | 24.10647 | 40.6447 | 49.59932 | 21.68676 | 0.09492 | 1.101281 | 0.853703 |
| 4 | 130243600 | 130244799 | 9.700893 | 18.31647 | 27.54307 | 11.21935 | -0.10722 | 0.063735 | 0.853709 |
| 13 | 29904000 | 29904799 | 9.700893 | 18.31647 | 27.54307 | 11.21935 | -0.10722 | 0.063735 | 0.853709 |
| 5 | 24212200 | 24213799 | 20.22536 | 35.62903 | 40.56199 | 17.95436 | 0.202589 | 0.608068 | 0.853722 |
| 14 | 40317600 | 40318999 | 19.0834 | 32.87062 | 45.84149 | 20.10905 | -0.09318 | 0.888205 | 0.853729 |
| 1 | 3805000 | 3805799 | 13.48354 | 24.82991 | 24.92398 | 10.56325 | 0.427441 | 0.242028 | 0.853731 |
| 1 | 183321000 | 183324999 | 34.32543 | 55.81934 | 76.54171 | 32.84755 | -0.08518 | 1.571185 | 0.853743 |
| 11 | 30363200 | 30364399 | 15.90766 | 27.60352 | 47.93056 | 21.03018 | -0.40579 | 0.810616 | 0.85375 |
| X | 144715000 | 144716599 | 48.85471 | 78.55689 | 77.97288 | 33.07307 | 0.362972 | 1.876357 | 0.853789 |
| X | 9215800 | 9218399 | 29.95437 | 50.25757 | 48.05701 | 20.9728 | 0.445465 | 1.166165 | 0.853801 |
| 4 | 106253000 | 106254599 | 27.91292 | 46.48924 | 56.60337 | 24.5882 | 0.098222 | 1.245987 | 0.853805 |
| 19 | 13900800 | 13911199 | 190.008 | 278.8066 | 347.0484 | 142.9405 | -0.07865 | 3.855431 | 0.85381 |
| 4 | 121359800 | 121419399 | 1158.285 | 1479.224 | 1422.45 | 596.9212 | 0.093344 | 6.026363 | 0.853812 |
| 1 | 72798000 | 72799799 | 14.01244 | 25.13304 | 31.0283 | 13.58313 | 0.09966 | 0.411834 | 0.853814 |
| X | 130490400 | 130491799 | 13.83372 | 24.89207 | 36.23006 | 15.81752 | -0.13384 | 0.402325 | 0.853834 |
| 14 | 44864800 | 44865799 | 10.18637 | 19.12483 | 28.50917 | 11.72998 | -0.10665 | 0.106933 | 0.853853 |
| 14 | 83169600 | 83170399 | 10.18637 | 19.12483 | 28.50917 | 11.72998 | -0.10665 | 0.106933 | 0.853853 |
| 15 | 48760800 | 48761999 | 10.18637 | 19.12483 | 28.50917 | 11.72998 | -0.10665 | 0.106933 | 0.853853 |
| 13 | 59255400 | 59258599 | 26.93119 | 44.02875 | 76.36698 | 32.83623 | -0.41776 | 1.48074 | 0.853855 |
| 6 | 43528800 | 43529599 | 7.894301 | 15.29091 | 21.04822 | 7.989191 | 0.113995 | -0.19953 | 0.853859 |
| 10 | 95144800 | 95147599 | 24.73531 | 41.21039 | 57.60074 | 25.02662 | -0.10028 | 1.231521 | 0.853884 |
| 3 | 104499200 | 104505999 | 65.70697 | 103.9241 | 136.8115 | 56.85386 | -0.0689 | 2.402856 | 0.853929 |
| 17 | 12406600 | 12413199 | 52.87268 | 84.11705 | 132.3225 | 55.26981 | -0.31759 | 2.229751 | 0.853943 |
| 3 | 78032800 | 78033599 | 7.876666 | 15.0911 | 23.5178 | 9.270798 | -0.11547 | -0.06939 | 0.853985 |
| 7 | 46883800 | 46887799 | 41.45693 | 67.15924 | 67.12711 | 28.73103 | 0.364929 | 1.654193 | 0.853993 |
| 9 | 30551800 | 30554999 | 38.93749 | 62.93656 | 76.05883 | 32.48983 | 0.088502 | 1.704973 | 0.853996 |
| 8 | 44288600 | 44291199 | 26.6691 | 45.5708 | 43.62266 | 19.12639 | 0.445516 | 0.987492 | 0.854023 |
| 4 | 121471800 | 121566999 | 1521.946 | 1895.129 | 2041.831 | 859.1171 | -0.1075 | 6.434733 | 0.854039 |
| 14 | 111222000 | 111223999 | 23.42825 | 40.02136 | 49.01975 | 21.49224 | 0.091149 | 0.983189 | 0.854051 |
| X | 153329400 | 153348999 | 297.3542 | 426.883 | 453.1106 | 187.0831 | 0.120344 | 4.194348 | 0.854074 |
| 3 | 70761600 | 70762399 | 18.76166 | 32.27469 | 45.33943 | 19.88803 | -0.104 | 0.897847 | 0.854083 |
| 5 | 64807400 | 64808199 | 8.621556 | 16.59602 | 22.50666 | 8.753371 | 0.101571 | -0.1081 | 0.854085 |
| 17 | 17351400 | 17352399 | 8.621556 | 16.59602 | 22.50666 | 8.753371 | 0.101571 | -0.1081 | 0.854085 |
| 7 | 19669200 | 19675399 | 54.21845 | 86.42608 | 102.2263 | 42.81404 | 0.096166 | 2.1606 | 0.854105 |
| 17 | 55201000 | 55201999 | 10.07104 | 19.06388 | 24.87708 | 10.0789 | 0.103157 | 0.045238 | 0.854106 |
| 14 | 5401000 | 5403399 | 30.86133 | 51.31798 | 61.98118 | 26.89569 | 0.109563 | 1.272749 | 0.854109 |
| 1 | 73448800 | 73449999 | 12.50732 | 22.51696 | 32.07288 | 13.99393 | -0.10242 | 0.378615 | 0.854112 |
| 3 | 129537800 | 129539599 | 17.56788 | 30.5589 | 42.61636 | 18.7247 | -0.09057 | 0.776478 | 0.854132 |

| | | | | | | | | | |
|---|---|---|---|---|---|---|---|---|---|
| X | 5539200 | 5548799 | 41.72292 | 68.61366 | 58.39683 | 25.17017 | 0.610407 | 1.429041 | 0.854184 |
| 7 | 12776400 | 12780799 | 48.2314 | 77.0843 | 103.9135 | 43.68123 | -0.08803 | 2.074117 | 0.854184 |
| 5 | 90513000 | 90514999 | 16.64258 | 28.39217 | 53.72047 | 23.55072 | -0.53394 | 0.974589 | 0.854188 |
| 8 | 15204400 | 15205999 | 19.10847 | 33.26049 | 40.95883 | 18.00383 | 0.088111 | 0.771461 | 0.854192 |
| 6 | 123158000 | 123160999 | 31.88202 | 52.72844 | 64.20649 | 27.75719 | 0.097248 | 1.339165 | 0.854211 |
| 19 | 46885600 | 46886999 | 21.19604 | 36.51652 | 44.75947 | 19.63452 | 0.092313 | 0.89456 | 0.854217 |
| X | 44099600 | 44100799 | 13.53179 | 24.73216 | 30.63597 | 13.11679 | 0.115634 | 0.298328 | 0.854233 |
| X | 26531000 | 26541599 | 156.2038 | 238.1761 | 183.5559 | 76.10162 | 0.661965 | 3.141451 | 0.854275 |
| 8 | 4152200 | 4155999 | 27.13929 | 45.2421 | 55.55847 | 24.14823 | 0.08583 | 1.230774 | 0.854282 |
| 8 | 75520200 | 75521199 | 27.8159 | 46.13253 | 56.05884 | 24.28173 | 0.100631 | 1.291058 | 0.854287 |
| 19 | 33537400 | 33538199 | 9.327934 | 17.62631 | 26.48736 | 10.77115 | -0.10353 | 0.055032 | 0.854317 |
| 3 | 94589600 | 94590599 | 9.947179 | 18.40714 | 27.30295 | 11.49507 | -0.12503 | 0.192427 | 0.854338 |
| 5 | 9457600 | 9458999 | 13.88506 | 24.62082 | 34.57892 | 15.28441 | -0.09498 | 0.489001 | 0.854341 |
| 12 | 110964000 | 110966599 | 40.96328 | 66.05604 | 89.26051 | 38.01412 | -0.07618 | 1.782551 | 0.854349 |
| 8 | 110811400 | 110818199 | 69.30228 | 108.5856 | 167.3926 | 69.43206 | -0.30693 | 2.604353 | 0.85443 |
| 15 | 47919400 | 47919799 | 6.804359 | 13.23246 | 21.87002 | 8.096374 | -0.11453 | -0.29905 | 0.854434 |
| 13 | 51315800 | 51317799 | 25.9354 | 43.4612 | 53.38641 | 23.28062 | 0.085289 | 1.168834 | 0.854465 |
| 10 | 21541800 | 21545999 | 36.29503 | 58.91046 | 80.70806 | 34.5929 | -0.08632 | 1.612647 | 0.854471 |
| 17 | 82921800 | 82923399 | 19.1033 | 33.50907 | 40.16702 | 17.75632 | 0.127354 | 0.656454 | 0.854478 |
| 10 | 104595400 | 104599599 | 38.56779 | 63.12514 | 74.32328 | 31.97081 | 0.140696 | 1.487664 | 0.85449 |
| 4 | 113496000 | 113509599 | 205.4928 | 304.155 | 288.3559 | 118.7535 | 0.325752 | 3.715755 | 0.854505 |
| 6 | 38594600 | 38597199 | 47.95571 | 76.90539 | 92.69231 | 39.15983 | 0.078767 | 1.963941 | 0.854528 |
| 7 | 39985800 | 39988399 | 16.3514 | 28.30529 | 49.53178 | 21.73115 | -0.41781 | 0.834762 | 0.854533 |
| 13 | 35404000 | 35406399 | 38.95752 | 63.3569 | 76.53355 | 32.84499 | 0.097347 | 1.580857 | 0.854535 |
| 2 | 36406400 | 36408199 | 18.14325 | 31.80977 | 38.70492 | 17.10304 | 0.105974 | 0.657463 | 0.854542 |
| 11 | 16402400 | 16403399 | 14.39416 | 25.79809 | 31.90661 | 13.97657 | 0.096196 | 0.418851 | 0.854553 |
| 2 | 88293800 | 88296999 | 39.05547 | 63.60936 | 76.26893 | 32.74891 | 0.109517 | 1.55897 | 0.854565 |
| 6 | 112412600 | 112416199 | 23.21135 | 38.16409 | 71.64131 | 30.87636 | -0.52747 | 1.396914 | 0.854579 |
| 3 | 15050800 | 15052999 | 23.8747 | 39.66651 | 66.82036 | 28.97858 | -0.36893 | 1.269044 | 0.854621 |
| 13 | 21542800 | 21543999 | 13.30917 | 24.09218 | 30.03704 | 13.01616 | 0.095115 | 0.349668 | 0.854636 |
| X | 93814800 | 93817799 | 36.82791 | 59.79261 | 72.98459 | 31.28913 | 0.080298 | 1.605691 | 0.854648 |
| 8 | 37808600 | 37812399 | 49.82263 | 79.74465 | 95.48936 | 40.22376 | 0.084903 | 2.028182 | 0.854651 |
| 2 | 19370400 | 19372199 | 13.52388 | 24.32436 | 34.6039 | 15.1129 | -0.10159 | 0.397188 | 0.854656 |
| 11 | 27876600 | 27878399 | 19.81391 | 34.6171 | 41.86251 | 18.44418 | 0.114668 | 0.724415 | 0.854658 |
| 1 | 49308400 | 49310199 | 11.43637 | 21.68 | 22.04608 | 8.904392 | 0.463603 | 0.0381 | 0.854668 |
| 1 | 176861600 | 176862399 | 7.600328 | 14.61795 | 23.06386 | 8.991696 | -0.11775 | -0.10803 | 0.854681 |
| 1 | 177755600 | 177756599 | 26.11461 | 43.35721 | 60.28752 | 26.13261 | -0.09305 | 1.278745 | 0.854685 |
| 9 | 109367600 | 109375999 | 150.6072 | 227.5551 | 222.6057 | 92.03742 | 0.307266 | 3.323105 | 0.85471 |
| 2 | 169070200 | 169073199 | 25.93939 | 43.89966 | 53.39918 | 23.37482 | 0.09969 | 1.079466 | 0.854738 |
| 1 | 7685000 | 7686799 | 21.45599 | 37.27939 | 44.62902 | 19.59489 | 0.1283 | 0.810991 | 0.854752 |
| 8 | 108290800 | 108293399 | 23.25679 | 39.49274 | 48.50003 | 21.25202 | 0.086167 | 1.035432 | 0.854754 |
| 9 | 4391200 | 4393999 | 42.88749 | 69.08293 | 94.27495 | 40.08962 | -0.09225 | 1.822681 | 0.854759 |
| 1 | 152958400 | 152961199 | 37.82211 | 61.2718 | 74.49311 | 31.88408 | 0.083078 | 1.652598 | 0.854789 |
| 6 | 49542000 | 49545599 | 24.44129 | 41.1325 | 57.21046 | 25.00051 | -0.09339 | 1.124658 | 0.85486 |
| 10 | 126713000 | 126717399 | 50.10204 | 80.3058 | 109.2176 | 46.10498 | -0.09608 | 1.990008 | 0.854868 |
| 18 | 54419200 | 54428599 | 143.2898 | 214.8408 | 272.4385 | 112.3986 | -0.07658 | 3.474511 | 0.854869 |
| 3 | 63467800 | 63471799 | 32.32227 | 52.91192 | 72.86847 | 31.34708 | -0.08466 | 1.470102 | 0.854872 |

| | | | | | | | | | |
|---|---|---|---|---|---|---|---|---|---|
| 7 | 98048200 | 98052599 | 36.78762 | 59.55327 | 81.51906 | 34.87287 | -0.08862 | 1.666353 | 0.854888 |
| 7 | 56523000 | 56524799 | 15.95715 | 28.65329 | 34.44341 | 15.06387 | 0.138617 | 0.423637 | 0.854899 |
| X | 36909000 | 36911199 | 19.04784 | 33.21307 | 47.87177 | 21.06565 | -0.13766 | 0.759022 | 0.854918 |
| 11 | 16678600 | 16679199 | 7.21965 | 14.02254 | 19.47114 | 7.281439 | 0.121078 | -0.2662 | 0.85492 |
| 2 | 97231000 | 97232999 | 30.22983 | 50.80991 | 47.47697 | 20.73911 | 0.478416 | 1.143761 | 0.854935 |
| 18 | 72904800 | 72907999 | 75.22446 | 117.6148 | 154.4276 | 64.06833 | -0.07804 | 2.642535 | 0.854953 |
| 8 | 45217400 | 45219799 | 14.37239 | 25.41923 | 35.60708 | 15.7486 | -0.09275 | 0.521465 | 0.854958 |
| 2 | 150289000 | 150290999 | 15.23145 | 26.64197 | 53.87846 | 23.76074 | -0.62395 | 0.698944 | 0.854975 |
| 4 | 112516200 | 112519199 | 35.46568 | 58.4125 | 69.52628 | 29.98651 | 0.12869 | 1.402213 | 0.855036 |
| 10 | 97163400 | 97166999 | 64.47636 | 101.6956 | 135.4481 | 56.21164 | -0.08886 | 2.4672 | 0.855044 |
| 6 | 132235000 | 132255199 | 340.8891 | 477.8309 | 564.0507 | 233.7963 | -0.06753 | 4.545095 | 0.855048 |
| 10 | 70268600 | 70270799 | 17.75013 | 31.11267 | 38.19542 | 16.8743 | 0.093144 | 0.657619 | 0.855053 |
| 8 | 21433000 | 21438799 | 605.263 | 808.7585 | 820.24 | 342.5076 | 0.081825 | 5.29136 | 0.855064 |
| 4 | 112671000 | 112675599 | 49.83351 | 80.34364 | 93.06908 | 39.62579 | 0.145198 | 1.803377 | 0.855065 |
| 10 | 23376800 | 23377399 | 10.4652 | 19.16692 | 28.28792 | 12.07817 | -0.13243 | 0.257784 | 0.855086 |
| 1 | 65562400 | 65563399 | 19.14967 | 33.20343 | 40.59108 | 17.80947 | 0.098016 | 0.821732 | 0.855115 |
| 19 | 20248400 | 20250799 | 30.85402 | 51.65263 | 60.81542 | 26.50375 | 0.14624 | 1.202711 | 0.855118 |
| 17 | 34992200 | 34993399 | 23.32801 | 39.29029 | 54.87229 | 23.98455 | -0.09902 | 1.1163 | 0.855123 |
| 17 | 80824600 | 80826599 | 17.07492 | 29.76874 | 41.52546 | 18.27137 | -0.09062 | 0.740232 | 0.855123 |
| 13 | 17516800 | 17517799 | 12.69491 | 23.25234 | 29.14145 | 12.44134 | 0.101845 | 0.276797 | 0.855132 |
| 6 | 81252400 | 81254399 | 16.19939 | 28.76666 | 34.93634 | 15.42335 | 0.114778 | 0.496148 | 0.85515 |
| 1 | 71894000 | 71894399 | 5.799309 | 11.30306 | 19.13813 | 6.926865 | -0.1184 | -0.39883 | 0.855155 |
| 4 | 7733000 | 7734399 | 17.97681 | 31.34278 | 38.39747 | 16.89561 | 0.095864 | 0.739019 | 0.855161 |
| 18 | 53636000 | 53637399 | 13.07721 | 22.83919 | 43.6897 | 19.24491 | -0.54341 | 0.700594 | 0.855174 |
| 4 | 73883400 | 73885399 | 17.99223 | 32.01669 | 36.82516 | 16.2288 | 0.194268 | 0.494924 | 0.8552 |
| X | 115904800 | 115906799 | 17.16424 | 30.30091 | 45.56699 | 20.1612 | -0.19588 | 0.576337 | 0.855214 |
| 4 | 24886400 | 24894399 | 81.59393 | 128.0794 | 150.8334 | 62.5576 | 0.083135 | 2.576028 | 0.855223 |
| 19 | 39000000 | 39002199 | 27.31127 | 45.85314 | 55.67709 | 24.30841 | 0.101776 | 1.153644 | 0.85524 |
| 11 | 33696600 | 33697799 | 9.167355 | 17.18188 | 25.88377 | 10.68102 | -0.12445 | 0.104895 | 0.855258 |
| 16 | 88574800 | 88575399 | 9.167355 | 17.18188 | 25.88377 | 10.68102 | -0.12445 | 0.104895 | 0.855258 |
| 10 | 50221400 | 50222799 | 16.8963 | 29.73805 | 36.50822 | 16.16287 | 0.093753 | 0.59032 | 0.855276 |
| 1 | 85043200 | 85105799 | 2782.891 | 3287.007 | 3280.681 | 1390.219 | -0.07469 | 7.27085 | 0.855315 |
| 11 | 110638000 | 110641399 | 26.03859 | 43.40206 | 59.89002 | 26.02897 | -0.08187 | 1.232455 | 0.855336 |
| 3 | 124558800 | 124561199 | 45.41863 | 73.06305 | 88.42299 | 37.52888 | 0.078293 | 1.864129 | 0.855336 |
| 2 | 46580600 | 46581999 | 31.29707 | 51.47205 | 63.24695 | 27.26152 | 0.082105 | 1.404858 | 0.855345 |
| 18 | 46764200 | 46764999 | 6.673536 | 12.97044 | 21.20904 | 7.888975 | -0.105 | -0.28423 | 0.855348 |
| 5 | 64270000 | 64275199 | 58.34403 | 93.19732 | 110.6657 | 46.48483 | 0.093792 | 2.101921 | 0.855365 |
| 5 | 148098800 | 148100399 | 18.60117 | 32.17933 | 45.26692 | 19.85964 | -0.10449 | 0.843165 | 0.855367 |
| 10 | 71815600 | 71817199 | 18.58699 | 32.65432 | 32.3714 | 14.32905 | 0.399981 | 0.615051 | 0.855385 |
| 15 | 32865400 | 32878799 | 240.167 | 345.5618 | 421.8318 | 174.1071 | -0.07803 | 4.156216 | 0.855385 |
| 18 | 36619400 | 36622599 | 29.15838 | 48.29623 | 59.39442 | 25.70345 | 0.083259 | 1.312758 | 0.855386 |
| 11 | 99600800 | 99601399 | 8.784596 | 16.76841 | 25.61106 | 10.1943 | -0.09996 | -0.02379 | 0.855388 |
| 15 | 44155400 | 44156199 | 10.57344 | 20.08128 | 26.01761 | 10.46742 | 0.121668 | 0.027305 | 0.855443 |
| 5 | 32689000 | 32692599 | 99.02089 | 152.4624 | 177.8818 | 73.88198 | 0.075059 | 2.928247 | 0.855456 |
| Y | 8707000 | 8711599 | 41.20575 | 66.30046 | 68.73826 | 29.18345 | 0.30138 | 1.838082 | 0.855463 |
| 1 | 40249600 | 40253399 | 54.02689 | 85.84903 | 134.8842 | 56.26264 | -0.31762 | 2.267861 | 0.855472 |
| 3 | 103746600 | 103750599 | 57.96959 | 92.45928 | 110.9157 | 46.49069 | 0.076718 | 2.14746 | 0.855477 |

| | | | | | | | | | |
|---|---|---|---|---|---|---|---|---|---|
| 16 | 97797000 | 97834999 | 739.962 | 979.4609 | 978.4117 | 409.0623 | 0.090115 | 5.444291 | 0.855487 |
| 7 | 27808800 | 27811599 | 25.26414 | 42.00089 | 58.65018 | 25.44618 | -0.09902 | 1.258867 | 0.855491 |
| 5 | 46844800 | 46845199 | 8.011322 | 15.55105 | 21.70785 | 8.156463 | 0.107315 | -0.23195 | 0.855505 |
| 3 | 92626000 | 92626799 | 7.134547 | 13.79311 | 22.09606 | 8.461585 | -0.11505 | -0.17546 | 0.855547 |
| 2 | 30107200 | 30111399 | 42.0811 | 67.93619 | 82.68999 | 35.26783 | 0.076198 | 1.748425 | 0.855578 |
| 6 | 7499200 | 7499599 | 9.281388 | 17.71783 | 23.42554 | 9.32312 | 0.107135 | -0.02346 | 0.855587 |
| 14 | 81738800 | 81739199 | 14.20956 | 24.94842 | 35.84377 | 15.85777 | -0.13249 | 0.606716 | 0.855597 |
| 14 | 50904200 | 50910399 | 71.88538 | 113.5526 | 135.2603 | 56.1733 | 0.073721 | 2.435767 | 0.855602 |
| 14 | 88045000 | 88049399 | 54.49099 | 86.93124 | 115.8044 | 48.55543 | -0.07383 | 2.139464 | 0.85562 |
| 2 | 88436600 | 88448799 | 180.3101 | 267.4978 | 336.7521 | 138.7073 | -0.08006 | 3.674518 | 0.855685 |
| 10 | 68742200 | 68743399 | 13.80545 | 24.73352 | 30.52353 | 13.37503 | 0.099195 | 0.415719 | 0.85571 |
| 14 | 91683800 | 91684599 | 13.80545 | 24.73352 | 30.52353 | 13.37503 | 0.099195 | 0.415719 | 0.85571 |
| 1 | 173391600 | 173394599 | 49.35205 | 79.64315 | 75.98016 | 32.3896 | 0.426255 | 1.769614 | 0.855723 |
| 10 | 34435600 | 34437199 | 17.08119 | 29.76124 | 41.54878 | 18.27452 | -0.09186 | 0.749233 | 0.855742 |
| 3 | 117308400 | 117309399 | 11.5224 | 20.5808 | 37.26855 | 16.37239 | -0.45349 | 0.43206 | 0.855754 |
| 13 | 32719000 | 32720599 | 18.07918 | 31.59646 | 38.75414 | 17.07868 | 0.094342 | 0.704924 | 0.855794 |
| 5 | 141053400 | 141054999 | 18.46666 | 32.03747 | 44.73001 | 19.62578 | -0.09281 | 0.817285 | 0.855798 |
| 6 | 79132400 | 79135599 | 23.32681 | 38.96977 | 65.87925 | 28.67274 | -0.3739 | 1.205298 | 0.855802 |
| 4 | 50723600 | 50724999 | 18.11188 | 31.68294 | 38.8143 | 17.11859 | 0.096238 | 0.693531 | 0.855839 |
| X | 122470600 | 122471799 | 17.95567 | 31.83226 | 29.30201 | 12.95378 | 0.511199 | 0.507734 | 0.855843 |
| 11 | 78939400 | 78943399 | 49.43832 | 79.22008 | 95.68491 | 40.37534 | 0.074353 | 1.99142 | 0.855843 |
| 4 | 113829600 | 113832799 | 57.49701 | 91.97401 | 92.92465 | 39.11086 | 0.328636 | 2.053654 | 0.855899 |
| 16 | 81589200 | 81591599 | 21.0746 | 36.31876 | 44.73139 | 19.62239 | 0.085397 | 0.889119 | 0.855903 |
| 18 | 23074800 | 23075199 | 13.28844 | 23.84188 | 29.07482 | 12.72462 | 0.117503 | 0.399648 | 0.855908 |
| 4 | 82020800 | 82022999 | 25.99725 | 43.31526 | 59.92411 | 26.037 | -0.08559 | 1.235727 | 0.855958 |
| 3 | 91189000 | 91195599 | 138.2922 | 210.0729 | 206.4283 | 85.49067 | 0.307276 | 3.21112 | 0.85597 |
| 15 | 41431000 | 41432199 | 13.34976 | 24.02303 | 34.08994 | 14.88221 | -0.09758 | 0.391766 | 0.85598 |
| 3 | 10032000 | 10032599 | 6.983421 | 13.40145 | 21.98934 | 8.627968 | -0.18272 | -0.0991 | 0.856022 |
| 1 | 48840400 | 48843399 | 46.27168 | 74.58324 | 74.72816 | 31.80698 | 0.353702 | 1.797049 | 0.856029 |
| 11 | 28094800 | 28095799 | 25.30438 | 42.29276 | 50.69486 | 22.07036 | 0.120617 | 1.192831 | 0.856032 |
| 11 | 24730200 | 24730999 | 8.254278 | 15.90297 | 21.42233 | 8.310688 | 0.114299 | -0.12494 | 0.856056 |
| X | 121209000 | 121209399 | 7.817176 | 15.16113 | 21.08482 | 7.954837 | 0.106795 | -0.22371 | 0.856065 |
| 9 | 19505600 | 19508799 | 39.53773 | 64.32589 | 65.028 | 27.9113 | 0.35406 | 1.569842 | 0.856066 |
| 16 | 94610200 | 94611199 | 12.02489 | 21.71731 | 31.11475 | 13.52206 | -0.10702 | 0.354913 | 0.856091 |
| 18 | 40297000 | 40297999 | 14.95216 | 26.48107 | 31.25342 | 13.81922 | 0.154288 | 0.505041 | 0.856141 |
| 6 | 134665800 | 134669199 | 54.32995 | 86.45564 | 115.4929 | 48.25581 | -0.08174 | 2.220159 | 0.856149 |
| 16 | 54343400 | 54345399 | 19.93042 | 33.58155 | 58.52733 | 25.58532 | -0.41739 | 1.103851 | 0.856154 |
| 12 | 111182400 | 111189599 | 77.6016 | 121.9621 | 144.9666 | 60.13397 | 0.071132 | 2.54648 | 0.856181 |
| 3 | 21759200 | 21765999 | 93.79072 | 145.2721 | 171.2442 | 71.13022 | 0.066809 | 2.815356 | 0.856195 |
| 14 | 110524600 | 110525999 | 16.88871 | 29.75178 | 36.48158 | 16.14814 | 0.095949 | 0.580149 | 0.856236 |
| 8 | 125741200 | 125742999 | 17.78376 | 31.22554 | 38.1151 | 16.86343 | 0.101339 | 0.630431 | 0.856239 |
| 11 | 91252800 | 91253399 | 9.320128 | 17.75473 | 23.34679 | 9.332835 | 0.108766 | -0.01188 | 0.856256 |
| 3 | 17166400 | 17168599 | 23.79846 | 41.26553 | 39.27512 | 17.23211 | 0.457532 | 0.839981 | 0.856262 |
| 1 | 15572200 | 15573799 | 15.36699 | 27.80842 | 32.89939 | 14.26217 | 0.169462 | 0.368261 | 0.856274 |
| 1 | 162716400 | 162717399 | 13.91973 | 24.97486 | 30.95045 | 13.54534 | 0.09429 | 0.407284 | 0.856275 |
| 3 | 74669800 | 74673199 | 28.6647 | 48.10755 | 57.88148 | 25.26492 | 0.115045 | 1.166957 | 0.856336 |
| 10 | 17779600 | 17782599 | 31.89357 | 53.19778 | 62.66428 | 27.24647 | 0.145601 | 1.243794 | 0.85635 |

| | | | | | | | | | |
|---|---|---|---|---|---|---|---|---|---|
| 7 | 18243800 | 18249999 | 60.72231 | 97.30829 | 89.50495 | 37.7716 | 0.466774 | 2.00246 | 0.856354 |
| 5 | 128220800 | 128227199 | 82.40835 | 128.3252 | 166.7284 | 69.20548 | -0.06746 | 2.717915 | 0.856402 |
| 12 | 94043800 | 94046999 | 23.03474 | 39.85176 | 47.38016 | 20.7868 | 0.137191 | 0.865826 | 0.856409 |
| 8 | 71656600 | 71659599 | 21.29869 | 36.18294 | 50.6178 | 22.18284 | -0.1011 | 1.0256 | 0.856418 |
| 4 | 142813600 | 142815599 | 18.35606 | 32.00937 | 39.41194 | 17.34382 | 0.088686 | 0.740869 | 0.856456 |
| 4 | 50327000 | 50329599 | 21.68279 | 37.66367 | 45.18404 | 19.83534 | 0.125281 | 0.817199 | 0.856476 |
| 13 | 33586200 | 33588599 | 20.79452 | 35.88655 | 50.6624 | 22.22511 | -0.11039 | 0.882182 | 0.856479 |
| 18 | 33809200 | 33809799 | 8.318955 | 16.07005 | 22.05234 | 8.476465 | 0.100699 | -0.15094 | 0.856498 |
| 12 | 118953600 | 118955999 | 34.96489 | 57.70704 | 57.55941 | 24.8479 | 0.384051 | 1.366451 | 0.856501 |
| 8 | 71528400 | 71530199 | 16.35747 | 28.86221 | 35.43347 | 15.67917 | 0.095071 | 0.557106 | 0.856526 |
| 14 | 63321400 | 63322399 | 12.09941 | 22.10976 | 27.58358 | 11.81286 | 0.105098 | 0.280175 | 0.856544 |
| 4 | 32777200 | 32782799 | 63.24713 | 100.414 | 120.3291 | 50.14846 | 0.0714 | 2.296457 | 0.856546 |
| X | 127694800 | 127695999 | 16.75582 | 29.98153 | 35.48208 | 15.56433 | 0.158206 | 0.454222 | 0.85655 |
| 10 | 94076000 | 94077599 | 15.51447 | 27.32318 | 38.16201 | 16.89291 | -0.09099 | 0.579089 | 0.856555 |
| X | 137393400 | 137395199 | 18.85054 | 33.05109 | 39.7156 | 17.55021 | 0.123929 | 0.662737 | 0.856561 |
| 14 | 95167200 | 95168199 | 10.97112 | 20.4593 | 26.22058 | 10.91267 | 0.096797 | 0.152065 | 0.856572 |
| 5 | 34666400 | 34666999 | 10.47625 | 20.06391 | 20.72121 | 8.177507 | 0.471725 | -0.04684 | 0.856581 |
| 1 | 57134800 | 57136399 | 15.7529 | 27.69627 | 33.65656 | 14.90355 | 0.107937 | 0.590699 | 0.856584 |
| 13 | 50855000 | 50856399 | 10.92182 | 19.81825 | 29.22093 | 12.62445 | -0.14252 | 0.315556 | 0.856613 |
| 8 | 20506600 | 20517399 | 187.1671 | 277.7976 | 313.6517 | 129.1984 | 0.077496 | 3.666554 | 0.856614 |
| 5 | 51153400 | 51154199 | 14.46897 | 25.83624 | 31.97613 | 14.05086 | 0.092255 | 0.44385 | 0.856629 |
| 2 | 58378600 | 58381799 | 33.83442 | 55.09058 | 75.41529 | 32.37909 | -0.08146 | 1.551647 | 0.856644 |
| 2 | 6704400 | 6706999 | 17.02555 | 30.12937 | 36.35536 | 16.07767 | 0.121463 | 0.530998 | 0.856663 |
| 3 | 12198800 | 12199999 | 19.83184 | 34.81126 | 33.37365 | 14.76192 | 0.448148 | 0.638629 | 0.856669 |
| 5 | 128232000 | 128233599 | 11.67607 | 21.36323 | 30.70613 | 13.12853 | -0.09635 | 0.276656 | 0.856671 |
| 12 | 7060600 | 7062599 | 20.14505 | 34.6703 | 48.4738 | 21.26321 | -0.09707 | 0.904823 | 0.856678 |
| 14 | 47207400 | 47209799 | 23.53253 | 39.95155 | 49.03262 | 21.48386 | 0.087008 | 1.038064 | 0.856695 |
| 3 | 121370000 | 121373199 | 20.3845 | 34.7708 | 48.49166 | 21.2582 | -0.09564 | 0.98455 | 0.856699 |
| 5 | 111274600 | 111275799 | 11.34711 | 20.96445 | 26.07027 | 10.99978 | 0.124845 | 0.207222 | 0.856722 |
| 17 | 85415400 | 85419799 | 41.26509 | 66.49413 | 89.74908 | 38.18547 | -0.07576 | 1.804096 | 0.856735 |
| 10 | 95065600 | 95067999 | 18.03724 | 31.38112 | 43.6352 | 19.17458 | -0.0864 | 0.777089 | 0.856741 |
| 14 | 79232200 | 79234999 | 5.545837 | 10.71409 | 25.92868 | 9.96216 | -0.70915 | -0.17059 | 0.856752 |
| 12 | 114911000 | 114921999 | 39.22741 | 64.71429 | 57.18301 | 24.68003 | 0.558219 | 1.383948 | 0.85676 |
| 9 | 38104600 | 38104999 | 7.082437 | 13.7846 | 19.58889 | 7.203791 | 0.110168 | -0.32607 | 0.856765 |
| 8 | 73984400 | 73989399 | 70.09536 | 110.8915 | 149.531 | 62.14712 | -0.10323 | 2.403725 | 0.856774 |
| 15 | 13034000 | 13034999 | 10.15255 | 18.89966 | 27.86614 | 11.63199 | -0.10708 | 0.162475 | 0.856775 |
| 14 | 111282200 | 111284999 | 27.41848 | 45.72063 | 64.06759 | 27.85171 | -0.10412 | 1.23344 | 0.856779 |
| 15 | 101012600 | 101014399 | 29.95797 | 49.38695 | 60.45904 | 26.08605 | 0.088833 | 1.369976 | 0.856799 |
| 13 | 22641600 | 22642999 | 11.47778 | 21.14068 | 30.85528 | 13.07337 | -0.10855 | 0.240836 | 0.856869 |
| 2 | 140945000 | 140945799 | 14.50425 | 25.17866 | 46.87813 | 20.57098 | -0.50569 | 0.798618 | 0.856883 |
| 7 | 30692800 | 30697199 | 48.19361 | 77.13377 | 103.1307 | 43.4398 | -0.07398 | 2.02968 | 0.856896 |
| 11 | 40905200 | 40907799 | 52.16446 | 83.22627 | 110.7949 | 46.46011 | -0.07273 | 2.132954 | 0.856897 |
| 10 | 129396600 | 129397999 | 15.23668 | 27.40192 | 33.41622 | 14.59559 | 0.119223 | 0.406563 | 0.856912 |
| X | 90968200 | 90971999 | 43.40299 | 70.44997 | 82.07847 | 35.18123 | 0.147002 | 1.629866 | 0.856925 |
| 16 | 89506000 | 89515399 | 160.6785 | 241.1131 | 274.7399 | 113.3731 | 0.078689 | 3.462209 | 0.856939 |
| 18 | 7240600 | 7241399 | 12.42772 | 22.59874 | 28.06051 | 12.09095 | 0.106176 | 0.31183 | 0.856948 |
| 19 | 53577200 | 53577799 | 8.407126 | 16.21962 | 21.50057 | 8.308358 | 0.143738 | -0.13084 | 0.856953 |

| | | | | | | | | | |
|---|---|---|---|---|---|---|---|---|---|
| 12 | 17544800 | 17547799 | 20.88883 | 35.83443 | 43.73562 | 19.182 | 0.097201 | 0.927741 | 0.856954 |
| 14 | 84092000 | 84096799 | 52.51942 | 84.01834 | 113.3747 | 47.72339 | -0.08801 | 2.052295 | 0.856974 |
| 13 | 69422200 | 69425799 | 39.89701 | 64.76389 | 78.30148 | 33.56409 | 0.093744 | 1.616718 | 0.856975 |
| 11 | 69228400 | 69229399 | 12.39208 | 22.5326 | 32.20115 | 13.87613 | -0.09542 | 0.32211 | 0.856978 |
| 18 | 65045000 | 65046599 | 16.36885 | 28.72768 | 35.1514 | 15.54601 | 0.097845 | 0.62369 | 0.856986 |
| 3 | 62431600 | 62431999 | 5.798728 | 11.30167 | 19.13403 | 6.927458 | -0.11807 | -0.39574 | 0.857008 |
| 1 | 38957400 | 38959399 | 25.8824 | 43.742 | 53.4115 | 23.37029 | 0.094083 | 1.090284 | 0.857028 |
| 17 | 38556000 | 38557999 | 34.25592 | 55.93113 | 77.30383 | 33.22335 | -0.09243 | 1.516607 | 0.857029 |
| 12 | 48471800 | 48472999 | 17.64566 | 30.74889 | 43.05537 | 18.93878 | -0.09628 | 0.750379 | 0.857035 |
| 2 | 33739400 | 33741799 | 21.20418 | 36.04951 | 50.07814 | 21.94784 | -0.09076 | 1.019536 | 0.857052 |
| 16 | 82437000 | 82438399 | 18.50733 | 32.34787 | 39.81035 | 17.54559 | 0.089562 | 0.708739 | 0.857054 |
| 2 | 160463400 | 160465199 | 17.06218 | 29.78388 | 41.47812 | 18.26495 | -0.0881 | 0.722038 | 0.857072 |
| X | 72810000 | 72814199 | 27.03632 | 45.09116 | 63.7282 | 27.70719 | -0.11626 | 1.235893 | 0.857073 |
| 7 | 65461400 | 65469399 | 89.89424 | 139.124 | 179.8467 | 74.70378 | -0.06705 | 2.829653 | 0.857104 |
| 1 | 58968400 | 58975199 | 151.0834 | 225.6727 | 284.8845 | 117.4473 | -0.075 | 3.546529 | 0.857121 |
| 15 | 46596600 | 46598799 | 21.66633 | 37.36912 | 45.79039 | 20.08745 | 0.093035 | 0.887153 | 0.857126 |
| 6 | 121389200 | 121392399 | 35.69975 | 57.70141 | 79.62941 | 34.01367 | -0.10259 | 1.70152 | 0.857129 |
| 14 | 116392800 | 116393799 | 17.06579 | 29.89692 | 36.74042 | 16.22406 | 0.09172 | 0.660044 | 0.857136 |
| 10 | 121017600 | 121022399 | 45.08336 | 72.67766 | 87.40152 | 37.20772 | 0.091096 | 1.794877 | 0.857145 |
| 14 | 59108200 | 59109799 | 14.58375 | 25.94272 | 37.02204 | 16.32017 | -0.11555 | 0.475358 | 0.857145 |
| X | 20878800 | 20883599 | 70.64017 | 112.1818 | 129.6509 | 54.04799 | 0.123944 | 2.299804 | 0.857148 |
| 9 | 122711000 | 122713799 | 28.11549 | 47.15391 | 57.31654 | 25.00765 | 0.100358 | 1.17102 | 0.857161 |
| 7 | 27954200 | 27955599 | 20.60278 | 35.88674 | 35.67214 | 15.68695 | 0.39582 | 0.760582 | 0.857189 |
| 17 | 20738400 | 20741799 | 47.52002 | 76.73871 | 75.06191 | 32.02617 | 0.391353 | 1.744163 | 0.857202 |
| 16 | 92009800 | 92013199 | 21.84884 | 36.91582 | 51.69196 | 22.61617 | -0.10273 | 1.076693 | 0.85721 |
| 18 | 7861200 | 7864199 | 33.38527 | 54.94831 | 67.02513 | 28.89938 | 0.093032 | 1.402215 | 0.857212 |
| X | 153498400 | 153503199 | 62.57193 | 100.2968 | 88.13248 | 37.20619 | 0.533003 | 1.997113 | 0.857216 |
| 6 | 132287600 | 132290599 | 42.67498 | 68.77709 | 93.43425 | 39.76242 | -0.08481 | 1.804675 | 0.857238 |
| 7 | 81356800 | 81361199 | 45.3924 | 72.82941 | 97.68085 | 41.29731 | -0.07431 | 1.948126 | 0.857243 |
| 6 | 138018200 | 138022599 | 60.97615 | 96.65206 | 128.497 | 53.4659 | -0.08048 | 2.343193 | 0.857276 |
| 12 | 114293800 | 114296999 | 70.54453 | 111.2557 | 132.4509 | 54.96707 | 0.072704 | 2.472756 | 0.857277 |
| 10 | 127796200 | 127798999 | 31.0501 | 50.40918 | 83.59072 | 35.83462 | -0.3607 | 1.602061 | 0.857307 |
| 3 | 20742200 | 20746599 | 110.8353 | 169.2608 | 194.5409 | 80.68484 | 0.08756 | 3.096249 | 0.857311 |
| 3 | 33510200 | 33511199 | 17.08742 | 29.75381 | 41.57187 | 18.27761 | -0.09309 | 0.758173 | 0.85732 |
| 18 | 47051200 | 47052199 | 12.01658 | 22.0596 | 27.53913 | 11.72123 | 0.110238 | 0.252141 | 0.85732 |
| 1 | 147306400 | 147307599 | 14.22332 | 26.03385 | 25.78344 | 11.01804 | 0.438009 | 0.282347 | 0.857331 |
| 10 | 55276800 | 55279799 | 26.08441 | 43.86897 | 53.94324 | 23.56808 | 0.083776 | 1.133736 | 0.857372 |
| 4 | 117358200 | 117361199 | 33.64671 | 54.89208 | 66.88141 | 28.73558 | 0.088193 | 1.518941 | 0.857424 |
| 4 | 7831400 | 7838599 | 115.8889 | 177.6645 | 175.4157 | 72.81457 | 0.310932 | 3.018474 | 0.857425 |
| 11 | 44243600 | 44245999 | 17.97489 | 31.46711 | 38.70317 | 17.08049 | 0.090353 | 0.678073 | 0.857446 |
| 14 | 9424000 | 9424799 | 8.684335 | 16.6389 | 25.72845 | 10.14161 | -0.10433 | -0.0545 | 0.857453 |
| 13 | 92122400 | 92124999 | 29.19013 | 48.93882 | 49.13331 | 21.43243 | 0.375346 | 1.170893 | 0.857475 |
| 4 | 72407000 | 72419199 | 187.9287 | 278.1241 | 314.7455 | 129.6214 | 0.070314 | 3.715424 | 0.857479 |
| 13 | 27927400 | 27930399 | 26.46052 | 45.4746 | 51.81962 | 22.72988 | 0.196244 | 0.94475 | 0.857483 |
| 14 | 9249800 | 9251399 | 19.90638 | 34.81474 | 41.84447 | 18.45174 | 0.123532 | 0.70986 | 0.857491 |
| 11 | 74149800 | 74150199 | 5.492984 | 10.73237 | 18.87082 | 6.573572 | -0.12151 | -0.53548 | 0.857525 |
| 5 | 61489200 | 61491999 | 34.19168 | 55.19595 | 91.56147 | 39.04841 | -0.36951 | 1.742696 | 0.857541 |

| | | | | | | | | | |
|---|---|---|---|---|---|---|---|---|---|
| 4 | 122638400 | 122642999 | 76.8409 | 120.4719 | 157.7868 | 65.44352 | -0.0714 | 2.596442 | 0.857593 |
| 1 | 49185000 | 49186999 | 20.22739 | 35.05347 | 49.57237 | 21.74969 | -0.11157 | 0.843028 | 0.857626 |
| X | 109192200 | 109195199 | 41.40924 | 67.55851 | 56.2906 | 24.18556 | 0.635077 | 1.530854 | 0.857641 |
| 8 | 124150200 | 124151199 | 11.42997 | 21.01741 | 26.12811 | 11.09571 | 0.118835 | 0.236109 | 0.857697 |
| 7 | 136784800 | 136786199 | 13.14836 | 24.16094 | 30.18165 | 12.82882 | 0.110921 | 0.263632 | 0.857729 |
| 5 | 87430400 | 87432399 | 32.11287 | 52.82635 | 65.05834 | 28.04512 | 0.078959 | 1.405019 | 0.85774 |
| 1 | 112449200 | 112450599 | 15.47791 | 28.10293 | 33.62107 | 14.51422 | 0.157664 | 0.354912 | 0.857762 |
| 2 | 150309400 | 150322999 | 185.7205 | 275.5008 | 352.0789 | 145.0191 | -0.10184 | 3.677598 | 0.857764 |
| Y | 2519200 | 2524199 | 31.99743 | 52.19431 | 54.35519 | 23.35254 | 0.313906 | 1.50541 | 0.857766 |
| 15 | 4383600 | 4386999 | 30.09265 | 50.17538 | 60.77533 | 26.41901 | 0.105526 | 1.241291 | 0.857768 |
| 4 | 15881000 | 15881999 | 11.551 | 21.01141 | 30.17765 | 12.99786 | -0.10208 | 0.308654 | 0.857768 |
| 5 | 4430200 | 4434999 | 68.37043 | 107.77 | 141.5724 | 58.74846 | -0.06971 | 2.48231 | 0.85777 |
| 9 | 88921400 | 88923599 | 36.81765 | 59.75006 | 72.94146 | 31.26065 | 0.079542 | 1.614071 | 0.857788 |
| 7 | 140317800 | 140321399 | 26.91408 | 44.48542 | 61.92225 | 26.75858 | -0.09513 | 1.332712 | 0.857803 |
| 1 | 189601800 | 189603399 | 16.50487 | 28.76855 | 40.57515 | 17.85075 | -0.10631 | 0.740937 | 0.857815 |
| 8 | 22588400 | 22590399 | 8.116723 | 15.28285 | 29.13441 | 11.92915 | -0.45267 | 0.073807 | 0.857834 |
| 10 | 19672800 | 19675199 | 66.12781 | 105.1091 | 97.35745 | 40.6218 | 0.444024 | 2.262125 | 0.857838 |
| 13 | 13972400 | 13976799 | 40.23434 | 64.87183 | 88.14528 | 37.53266 | -0.08437 | 1.783148 | 0.857843 |
| 10 | 27479200 | 27480599 | 14.27043 | 25.03821 | 43.49875 | 19.23335 | -0.40541 | 0.630648 | 0.857873 |
| 3 | 14267400 | 14267999 | 6.389609 | 12.45 | 20.85905 | 7.578186 | -0.10824 | -0.37037 | 0.857875 |
| 10 | 17958600 | 17964799 | 77.4287 | 122.1894 | 143.3534 | 59.50489 | 0.094486 | 2.464775 | 0.857923 |
| 18 | 33790400 | 33790799 | 7.249074 | 14.06899 | 19.46788 | 7.321217 | 0.119097 | -0.24437 | 0.857927 |
| 2 | 27466600 | 27468799 | 19.20424 | 33.3821 | 41.04642 | 18.02703 | 0.090219 | 0.792849 | 0.857969 |
| 1 | 33344400 | 33346199 | 27.87056 | 46.01296 | 63.68561 | 27.49856 | -0.08713 | 1.349267 | 0.857975 |
| 12 | 92169600 | 92172199 | 26.35449 | 44.3212 | 62.88571 | 27.47472 | -0.12209 | 1.143723 | 0.857975 |
| 16 | 3050000 | 3202399 | 7989.62 | 8635.827 | 7910.062 | 3401.474 | -0.08125 | 8.670397 | 0.857994 |
| 5 | 97634800 | 97639199 | 67.47101 | 106.6699 | 127.4682 | 52.95997 | 0.070114 | 2.41188 | 0.857998 |
| 1 | 111388200 | 111388999 | 18.7887 | 32.74934 | 39.97126 | 17.58507 | 0.101313 | 0.751846 | 0.858001 |
| 5 | 110147000 | 110171999 | 473.1029 | 650.5302 | 673.9956 | 280.2127 | 0.092645 | 4.840158 | 0.858014 |
| 14 | 48188400 | 48193199 | 58.42015 | 92.67847 | 123.3064 | 51.34934 | -0.08036 | 2.316697 | 0.858022 |
| 13 | 43173800 | 43174999 | 13.59033 | 24.49016 | 30.46618 | 13.27699 | 0.092892 | 0.377627 | 0.858036 |
| 17 | 61327200 | 61329999 | 84.05669 | 130.3994 | 169.888 | 70.55671 | -0.07563 | 2.784198 | 0.858047 |
| 14 | 61067800 | 61069799 | 20.39085 | 35.54246 | 42.96263 | 18.90128 | 0.11492 | 0.760043 | 0.858056 |
| 15 | 68979200 | 68980599 | 9.45972 | 17.71842 | 26.3669 | 10.89226 | -0.10834 | 0.112199 | 0.85806 |
| 15 | 63245200 | 63245799 | 6.663351 | 12.9339 | 21.12236 | 7.938612 | -0.11719 | -0.24911 | 0.85807 |
| 19 | 45986400 | 45987199 | 6.663351 | 12.9339 | 21.12236 | 7.938612 | -0.11719 | -0.24911 | 0.85807 |
| 18 | 26930000 | 26930999 | 11.85905 | 22.22657 | 21.58419 | 8.906925 | 0.504224 | 0.116815 | 0.858073 |
| 9 | 20126800 | 20135799 | 133.7546 | 204.3352 | 191.4609 | 79.41401 | 0.382423 | 3.095386 | 0.858076 |
| 5 | 66427800 | 66429799 | 16.05747 | 28.2163 | 39.66125 | 17.55899 | -0.10143 | 0.608309 | 0.858081 |
| 3 | 124806200 | 124807399 | 14.84067 | 26.04375 | 36.80571 | 16.27661 | -0.10889 | 0.6198 | 0.858087 |
| 11 | 15710400 | 15711599 | 14.84067 | 26.04375 | 36.80571 | 16.27661 | -0.10889 | 0.6198 | 0.858087 |
| 7 | 6394600 | 6397799 | 33.97459 | 55.50625 | 68.18061 | 29.30905 | 0.077955 | 1.500113 | 0.858108 |
| 4 | 150167000 | 150170199 | 28.7365 | 47.34217 | 65.28508 | 28.16049 | -0.08283 | 1.379444 | 0.858114 |
| 4 | 60108800 | 60110599 | 63.49025 | 100.1616 | 133.9299 | 55.58439 | -0.09428 | 2.461849 | 0.858115 |
| X | 55200600 | 55202199 | 20.1294 | 35.26064 | 41.78271 | 18.4492 | 0.143919 | 0.686337 | 0.858145 |
| 13 | 59577200 | 59579599 | 26.65271 | 44.45174 | 54.4501 | 23.67346 | 0.089474 | 1.22356 | 0.858148 |
| 18 | 21034600 | 21037399 | 15.76168 | 27.68792 | 38.67631 | 17.11844 | -0.09237 | 0.610724 | 0.858152 |

| | | | | | | | | | |
|---|---|---|---|---|---|---|---|---|---|
| 16 | 47921200 | 47925999 | 49.86699 | 80.13727 | 95.70765 | 40.56075 | 0.095497 | 1.901995 | 0.858158 |
| X | 54686400 | 54691399 | 68.37014 | 108.6729 | 104.6061 | 43.64614 | 0.388614 | 2.266555 | 0.858158 |
| 19 | 58580200 | 58585799 | 101.9319 | 156.6036 | 201.0533 | 83.48124 | -0.0652 | 2.974061 | 0.858192 |
| 2 | 125397200 | 125397799 | 12.27159 | 22.41208 | 28.0777 | 12.03273 | 0.098473 | 0.2859 | 0.858199 |
| 14 | 52581200 | 52584199 | 31.53834 | 51.54398 | 70.79848 | 30.43827 | -0.08247 | 1.490957 | 0.858205 |
| 2 | 176439800 | 176445999 | 12.56678 | 23.55581 | 23.54827 | 9.715469 | 0.462849 | 0.127496 | 0.858213 |
| 18 | 7722400 | 7728199 | 63.41908 | 100.2847 | 132.3777 | 55.00556 | -0.07301 | 2.405465 | 0.858215 |
| X | 152304800 | 152305399 | 7.274312 | 14.12272 | 22.87803 | 8.584802 | -0.10358 | -0.24121 | 0.858228 |
| X | 156890000 | 156893799 | 25.62563 | 44.67027 | 35.72526 | 15.71794 | 0.708227 | 0.775274 | 0.858271 |
| 10 | 121919400 | 121924599 | 104.4771 | 161.703 | 185.1484 | 76.90594 | 0.107903 | 2.836851 | 0.858315 |
| 7 | 3220200 | 3221399 | 19.55613 | 33.77817 | 41.10832 | 18.0343 | 0.103255 | 0.857062 | 0.858372 |
| 12 | 72707400 | 72708599 | 8.540264 | 15.99737 | 24.90015 | 10.28512 | -0.17237 | 0.099535 | 0.858374 |
| 13 | 33504600 | 33507999 | 33.60452 | 55.00308 | 76.67151 | 32.98124 | -0.10225 | 1.476417 | 0.858379 |
| 12 | 32522000 | 32523599 | 18.51843 | 32.32697 | 39.86895 | 17.5567 | 0.08642 | 0.727169 | 0.858383 |
| 7 | 110937400 | 110940399 | 28.55037 | 47.51332 | 58.43092 | 25.36918 | 0.083721 | 1.254115 | 0.858394 |
| 4 | 8483600 | 8487799 | 37.28068 | 60.51083 | 83.28371 | 35.69711 | -0.09296 | 1.615684 | 0.858421 |
| 2 | 33605000 | 33606199 | 9.961826 | 18.3917 | 27.30827 | 11.53351 | -0.12973 | 0.20584 | 0.858426 |
| 13 | 17634400 | 17635199 | 10.81005 | 20.00269 | 29.28434 | 12.33026 | -0.10688 | 0.202371 | 0.858433 |
| 2 | 52843800 | 52846199 | 24.22656 | 40.09485 | 68.38507 | 29.57764 | -0.38723 | 1.316142 | 0.858472 |
| 12 | 112848800 | 112854199 | 67.27924 | 106.4338 | 127.4109 | 52.95307 | 0.068291 | 2.396641 | 0.858502 |
| 11 | 7031000 | 7031999 | 10.70925 | 19.64731 | 28.71453 | 12.23195 | -0.11633 | 0.249219 | 0.858515 |
| 15 | 60678200 | 60680599 | 27.3311 | 46.17382 | 46.40153 | 20.30087 | 0.374176 | 1.09225 | 0.858526 |
| 3 | 21061000 | 21062199 | 19.84859 | 34.65891 | 34.15172 | 15.04176 | 0.408759 | 0.720605 | 0.858533 |
| 10 | 89894400 | 89896199 | 20.11326 | 34.99583 | 42.56056 | 18.70331 | 0.106139 | 0.786722 | 0.858544 |
| 9 | 98613800 | 98616199 | 20.35494 | 35.00417 | 48.47387 | 21.26274 | -0.08364 | 0.910427 | 0.858551 |
| 13 | 67833200 | 67834999 | 20.35494 | 35.00417 | 48.47387 | 21.26274 | -0.08364 | 0.910427 | 0.858551 |
| 14 | 100635200 | 100638599 | 31.53339 | 52.37921 | 63.15689 | 27.38555 | 0.11186 | 1.287835 | 0.858571 |
| 9 | 38875000 | 38880199 | 81.42292 | 127.4597 | 151.5545 | 62.86855 | 0.066194 | 2.624228 | 0.858612 |
| 2 | 97505200 | 97509399 | 70.94878 | 112.2348 | 133.7694 | 55.59471 | 0.074539 | 2.401794 | 0.858639 |
| 5 | 138427600 | 138429799 | 20.62276 | 35.91475 | 43.53513 | 19.14245 | 0.110805 | 0.775324 | 0.858715 |
| 3 | 38659200 | 38664399 | 57.68381 | 92.01239 | 110.699 | 46.40609 | 0.072713 | 2.14461 | 0.858719 |
| 5 | 70159200 | 70159999 | 12.56055 | 23.06076 | 34.00044 | 14.46912 | -0.12802 | 0.271991 | 0.858721 |
| 12 | 23352200 | 23355799 | 115.9031 | 177.6396 | 170.8184 | 70.90369 | 0.348721 | 3.023428 | 0.858732 |
| 6 | 35324200 | 35325999 | 16.70543 | 29.32549 | 36.11229 | 15.97424 | 0.088666 | 0.623381 | 0.858743 |
| 16 | 86585800 | 86587599 | 18.57926 | 32.51136 | 39.70385 | 17.51584 | 0.100511 | 0.688734 | 0.858757 |
| 19 | 35932800 | 35935799 | 18.57926 | 32.51136 | 39.70385 | 17.51584 | 0.100511 | 0.688734 | 0.858757 |
| 7 | 50744800 | 50745799 | 9.358948 | 17.71532 | 22.79222 | 9.234905 | 0.123234 | 0.031796 | 0.858764 |
| X | 8256200 | 8264399 | 83.36145 | 131.8489 | 117.351 | 48.76714 | 0.495076 | 2.413762 | 0.858769 |
| 12 | 113268400 | 113270999 | 23.51643 | 39.34194 | 55.47129 | 24.16434 | -0.11295 | 1.17978 | 0.85877 |
| 3 | 42131600 | 42136199 | 90.37887 | 140.4488 | 166.1239 | 68.97569 | 0.065573 | 2.756689 | 0.858776 |
| X | 17531200 | 17532399 | 18.59691 | 32.50048 | 39.85533 | 17.56361 | 0.0948 | 0.712709 | 0.858822 |
| 14 | 121265000 | 121269199 | 55.23485 | 88.01869 | 116.8295 | 48.90888 | -0.07037 | 2.17523 | 0.858873 |
| 17 | 47208200 | 47212799 | 27.97397 | 45.94637 | 78.32524 | 33.742 | -0.38973 | 1.422737 | 0.858875 |
| 2 | 45588800 | 45589199 | 6.378068 | 12.42506 | 17.46339 | 6.360973 | 0.139772 | -0.37301 | 0.858895 |
| 3 | 84698000 | 84701799 | 79.27705 | 123.4671 | 161.5982 | 67.07461 | -0.07787 | 2.71314 | 0.858898 |
| 10 | 49164400 | 49165399 | 15.60515 | 27.48736 | 33.61496 | 14.87962 | 0.099879 | 0.569636 | 0.858926 |
| 8 | 27646000 | 27647199 | 14.85321 | 26.2293 | 36.61343 | 16.19867 | -0.08864 | 0.542825 | 0.858927 |

| | | | | | | | | | |
|---|---|---|---|---|---|---|---|---|---|
| 4 | 56433000 | 56435999 | 27.40201 | 45.38332 | 62.57802 | 27.07963 | -0.08133 | 1.311563 | 0.858939 |
| 9 | 15888600 | 15889999 | 11.07451 | 20.64532 | 26.2698 | 10.93923 | 0.106763 | 0.157137 | 0.858943 |
| 2 | 174686200 | 174688799 | 21.52189 | 36.85964 | 51.16941 | 22.4394 | -0.08824 | 0.955275 | 0.858956 |
| 7 | 23055800 | 23058399 | 87.3438 | 135.4572 | 157.1416 | 65.26594 | 0.091273 | 2.789391 | 0.858988 |
| 17 | 23412000 | 23412799 | 7.234267 | 13.95035 | 22.5599 | 8.729385 | -0.14368 | -0.14181 | 0.85899 |
| X | 105374000 | 105375399 | 14.83451 | 26.18757 | 37.23034 | 16.4727 | -0.11484 | 0.550542 | 0.858993 |
| 17 | 90413800 | 90417999 | 49.98428 | 80.48696 | 80.00738 | 34.00566 | 0.363325 | 1.834285 | 0.859018 |
| 9 | 21944600 | 21948399 | 35.77038 | 58.12336 | 70.82094 | 30.37994 | 0.083938 | 1.584341 | 0.859047 |
| 13 | 56418200 | 56422599 | 66.20803 | 104.6018 | 137.3815 | 57.05903 | -0.06679 | 2.429053 | 0.859049 |
| 4 | 98930800 | 98931999 | 11.65373 | 20.98566 | 38.40836 | 16.7528 | -0.46177 | 0.38423 | 0.85906 |
| 13 | 68587400 | 68612599 | 807.7608 | 1052.319 | 1155.601 | 483.9987 | -0.06938 | 5.70085 | 0.859067 |
| 7 | 51990800 | 51991999 | 13.06756 | 23.62137 | 33.66952 | 14.6161 | -0.0986 | 0.364034 | 0.859076 |
| 19 | 18344800 | 18347199 | 16.47409 | 29.24841 | 35.28707 | 15.58091 | 0.123801 | 0.49631 | 0.859084 |
| 14 | 124988400 | 124989599 | 25.35039 | 42.1934 | 58.7574 | 25.51321 | -0.09501 | 1.243902 | 0.859102 |
| 19 | 52826400 | 52829999 | 32.18008 | 52.12627 | 87.04849 | 37.25509 | -0.37392 | 1.650124 | 0.859128 |
| 3 | 48889200 | 48889799 | 10.96961 | 20.37395 | 29.92151 | 12.53463 | -0.10562 | 0.187779 | 0.859144 |
| 19 | 24153800 | 24156399 | 18.19705 | 31.69185 | 44.13058 | 19.40485 | -0.08835 | 0.763781 | 0.859156 |
| 6 | 9429800 | 9431599 | 21.50831 | 36.8857 | 45.31403 | 19.87377 | 0.087824 | 0.937229 | 0.85916 |
| 17 | 28542800 | 28544399 | 20.65387 | 35.25065 | 49.01763 | 21.48978 | -0.09139 | 0.979712 | 0.859173 |
| 1 | 184328400 | 184331399 | 22.11208 | 38.29196 | 46.29925 | 20.31317 | 0.113319 | 0.853074 | 0.859187 |
| 3 | 89828000 | 89829399 | 15.64008 | 27.4446 | 38.25773 | 16.91769 | -0.08931 | 0.627769 | 0.859193 |
| 14 | 122941000 | 122943199 | 21.62662 | 37.70512 | 44.56096 | 19.60728 | 0.147359 | 0.765905 | 0.859197 |
| 17 | 55065000 | 55069599 | 68.20003 | 107.8824 | 128.7751 | 53.518 | 0.072327 | 2.398486 | 0.859203 |
| 3 | 35433000 | 35434599 | 19.31405 | 33.651 | 41.45122 | 18.23139 | 0.087863 | 0.760022 | 0.859211 |
| 8 | 25667800 | 25670799 | 22.04677 | 37.38198 | 51.68242 | 22.63622 | -0.08419 | 1.048197 | 0.859247 |
| 13 | 34745600 | 34746399 | 13.82904 | 25.50204 | 25.40505 | 10.74651 | 0.440298 | 0.238202 | 0.859251 |
| 17 | 35516600 | 35521399 | 55.88054 | 89.12853 | 107.4895 | 45.05543 | 0.069107 | 2.144777 | 0.859254 |
| 3 | 79112600 | 79114399 | 14.14085 | 25.35512 | 31.48394 | 13.78637 | 0.091001 | 0.41492 | 0.859256 |
| 10 | 94303800 | 94305399 | 15.88479 | 27.9234 | 34.10482 | 15.10256 | 0.100501 | 0.593456 | 0.859279 |
| 11 | 54352800 | 54354399 | 19.96072 | 35.01237 | 33.8834 | 14.97831 | 0.434644 | 0.651079 | 0.859291 |
| 3 | 61735400 | 61737999 | 42.88458 | 69.4241 | 69.63264 | 29.78826 | 0.35898 | 1.670898 | 0.859313 |
| 9 | 82472600 | 82474799 | 31.38787 | 52.43594 | 51.34729 | 22.3478 | 0.411336 | 1.217927 | 0.859338 |
| 4 | 50345200 | 50347799 | 43.94657 | 70.60585 | 94.82287 | 40.17159 | -0.07384 | 1.903599 | 0.859347 |
| X | 139674600 | 139677999 | 29.86112 | 49.87805 | 60.14097 | 26.17264 | 0.112072 | 1.222969 | 0.859348 |
| 12 | 115361600 | 115368399 | 105.1458 | 163.1793 | 155.1491 | 64.43166 | 0.378772 | 2.785533 | 0.85935 |
| 11 | 81327800 | 81328999 | 18.30057 | 31.84114 | 38.97547 | 17.12759 | 0.096786 | 0.770907 | 0.859358 |
| 2 | 34715800 | 34722399 | 97.26628 | 149.6185 | 192.6849 | 80.01993 | -0.06834 | 2.947489 | 0.859359 |
| 2 | 58977800 | 58979799 | 26.81613 | 44.74069 | 55.00271 | 23.91926 | 0.084252 | 1.220584 | 0.859375 |
| 10 | 129980000 | 129980799 | 14.57391 | 25.7908 | 36.11319 | 15.97057 | -0.09181 | 0.518042 | 0.85938 |
| 15 | 85191200 | 85193799 | 24.92805 | 41.73244 | 57.63803 | 25.1192 | -0.08319 | 1.181854 | 0.859437 |
| 7 | 104194200 | 104194799 | 11.77929 | 21.68056 | 26.75036 | 11.34511 | 0.131502 | 0.238662 | 0.859443 |
| X | 31331400 | 31331799 | 10.49813 | 20.30488 | 25.85642 | 9.881084 | 0.214974 | -0.13895 | 0.859447 |
| 17 | 18203200 | 18204999 | 36.50992 | 59.84409 | 60.75397 | 26.14981 | 0.354862 | 1.456764 | 0.859448 |
| X | 26767600 | 26771799 | 53.59167 | 86.3892 | 76.15898 | 32.41948 | 0.53812 | 1.809381 | 0.859468 |
| 3 | 158888600 | 158890999 | 26.85732 | 45.48624 | 54.5591 | 23.88782 | 0.119821 | 1.076433 | 0.859485 |
| 1 | 146265000 | 146268199 | 36.84409 | 60.27971 | 72.65951 | 31.25055 | 0.106537 | 1.490206 | 0.859506 |
| 17 | 7187200 | 7200599 | 212.1488 | 310.4235 | 348.6743 | 143.6353 | 0.065652 | 3.901245 | 0.859514 |

| | | | | | | | | | |
|---|---|---|---|---|---|---|---|---|---|
| 9 | 13919800 | 13920999 | 13.48558 | 24.39601 | 35.22627 | 15.27698 | -0.11647 | 0.363941 | 0.85954 |
| 1 | 23956000 | 23957799 | 19.50034 | 34.11604 | 41.29068 | 18.2127 | 0.113488 | 0.700575 | 0.859543 |
| 7 | 95057600 | 95060799 | 21.40886 | 35.74192 | 63.44997 | 27.60389 | -0.44411 | 1.20433 | 0.859568 |
| 15 | 96832200 | 96833399 | 10.48564 | 19.24344 | 28.30726 | 12.05303 | -0.12472 | 0.247305 | 0.859603 |
| 7 | 133605400 | 133609799 | 61.5511 | 97.55299 | 128.661 | 53.52843 | -0.06909 | 2.348035 | 0.859655 |
| 10 | 103722800 | 103723399 | 10.75639 | 20.15272 | 25.80724 | 10.65607 | 0.107444 | 0.121661 | 0.85969 |
| 17 | 72734800 | 72735799 | 15.74324 | 27.98657 | 34.37346 | 15.16117 | 0.099367 | 0.483556 | 0.859695 |
| 2 | 171710400 | 171712999 | 17.9935 | 31.45287 | 38.18346 | 16.82997 | 0.109008 | 0.69906 | 0.859723 |
| 10 | 25426200 | 25427599 | 17.7588 | 31.09161 | 38.26604 | 16.88886 | 0.089573 | 0.676756 | 0.859744 |
| 4 | 143333200 | 143348399 | 194.9661 | 289.8256 | 319.7142 | 131.7165 | 0.113084 | 3.640891 | 0.859763 |
| 18 | 79659800 | 79660399 | 6.683493 | 12.95702 | 21.0874 | 7.97123 | -0.11963 | -0.22757 | 0.859772 |
| 14 | 5396200 | 5397999 | 22.00495 | 38.00407 | 46.3138 | 20.31789 | 0.101291 | 0.87736 | 0.859775 |
| 1 | 29734200 | 29740199 | 100.8687 | 154.7624 | 199.1504 | 82.6759 | -0.07001 | 2.999212 | 0.859778 |
| X | 153762000 | 153765799 | 42.75948 | 69.65613 | 80.09454 | 34.39411 | 0.170021 | 1.569052 | 0.859787 |
| 15 | 82400200 | 82402999 | 22.72593 | 37.39363 | 71.07153 | 30.64038 | -0.54486 | 1.386056 | 0.85979 |
| 4 | 79859000 | 79860199 | 12.01793 | 21.83432 | 31.14925 | 13.43665 | -0.09395 | 0.320825 | 0.859797 |
| X | 14323600 | 14326399 | 21.74213 | 37.53627 | 55.06396 | 24.15632 | -0.16531 | 0.892022 | 0.859797 |
| 1 | 7305800 | 7313399 | 105.406 | 162.4294 | 189.757 | 78.822 | 0.074098 | 2.915929 | 0.859803 |
| 1 | 96047600 | 96049399 | 24.49009 | 42.57006 | 37.66791 | 16.54317 | 0.562849 | 0.813658 | 0.859812 |
| 19 | 10422400 | 10424199 | 15.16333 | 26.63437 | 37.23327 | 16.47846 | -0.09332 | 0.607591 | 0.859845 |
| 5 | 75862200 | 75864999 | 34.73404 | 56.45912 | 76.89866 | 32.99547 | -0.07588 | 1.580568 | 0.859868 |
| 3 | 3005000 | 3111599 | 4596.485 | 5181.367 | 4952.345 | 2114.544 | -0.08099 | 8.005285 | 0.859887 |
| 10 | 116474800 | 116475599 | 10.50443 | 19.62247 | 28.93497 | 12.00758 | -0.10103 | 0.142769 | 0.859897 |
| 18 | 71299200 | 71300199 | 10.50443 | 19.62247 | 28.93497 | 12.00758 | -0.10103 | 0.142769 | 0.859897 |
| 8 | 65905600 | 65907399 | 27.59544 | 45.84629 | 56.11674 | 24.3359 | 0.090338 | 1.269385 | 0.859907 |
| 3 | 18037000 | 18039799 | 27.03822 | 45.29117 | 55.63815 | 24.25644 | 0.08514 | 1.177647 | 0.859909 |
| 9 | 7836400 | 7837999 | 15.66104 | 27.49231 | 32.70562 | 14.46739 | 0.138254 | 0.610424 | 0.85991 |
| 9 | 91530800 | 91531199 | 7.138958 | 13.78649 | 22.06766 | 8.494188 | -0.12085 | -0.15781 | 0.859915 |
| 14 | 5742800 | 5743199 | 7.825906 | 15.15106 | 24.53237 | 9.33283 | -0.12247 | -0.18679 | 0.859919 |
| 8 | 125531200 | 125533599 | 22.25599 | 37.60532 | 52.20879 | 22.84258 | -0.09035 | 1.080415 | 0.859948 |
| 17 | 57891200 | 57892399 | 11.66587 | 21.52947 | 31.38607 | 13.26267 | -0.10449 | 0.229611 | 0.859948 |
| 1 | 154531000 | 154533399 | 18.0311 | 31.38958 | 43.61713 | 19.17399 | -0.08535 | 0.768253 | 0.859995 |
| 15 | 68637600 | 68638999 | 18.0311 | 31.38958 | 43.61713 | 19.17399 | -0.08535 | 0.768253 | 0.859995 |
| 2 | 16899400 | 16899999 | 10.22465 | 18.9514 | 27.78042 | 11.67023 | -0.10602 | 0.185888 | 0.860007 |
| 14 | 29460200 | 29461399 | 13.50564 | 23.99723 | 34.20563 | 15.09329 | -0.11421 | 0.475698 | 0.86002 |
| 13 | 107752400 | 107753599 | 11.22305 | 20.64901 | 29.76484 | 12.63004 | -0.09242 | 0.240708 | 0.860041 |
| 3 | 38417400 | 38419199 | 16.90299 | 30.07471 | 35.94673 | 15.84333 | 0.13929 | 0.490647 | 0.860048 |
| 17 | 39432600 | 39446599 | 143.1883 | 216.6388 | 213.176 | 88.17628 | 0.300454 | 3.294162 | 0.860113 |
| 12 | 17680200 | 17681799 | 20.20536 | 34.584 | 47.95549 | 21.02897 | -0.08665 | 0.951814 | 0.860144 |
| 3 | 91353600 | 91354599 | 20.66326 | 36.12202 | 35.48689 | 15.64228 | 0.413073 | 0.709889 | 0.860151 |
| 11 | 94635000 | 94638399 | 49.31749 | 78.98877 | 95.19418 | 40.13913 | 0.076748 | 2.006678 | 0.860179 |
| X | 77547200 | 77549199 | 23.72825 | 41.00688 | 48.4769 | 21.26726 | 0.145151 | 0.882177 | 0.860183 |
| 2 | 59849200 | 59850599 | 15.7938 | 27.75949 | 38.66788 | 17.1226 | -0.08832 | 0.602822 | 0.860195 |
| 14 | 87841800 | 87844199 | 31.30669 | 51.22744 | 70.63701 | 30.38227 | -0.08676 | 1.475745 | 0.860224 |
| 5 | 58790800 | 58795399 | 109.5634 | 167.1601 | 214.127 | 88.78865 | -0.06979 | 3.116428 | 0.860251 |
| 17 | 82134000 | 82135199 | 11.38144 | 20.97633 | 30.30499 | 12.82929 | -0.09328 | 0.233462 | 0.860271 |
| 7 | 118460400 | 118461199 | 9.906275 | 18.57402 | 27.40489 | 11.30126 | -0.09431 | 0.111393 | 0.86028 |

| | | | | | | | | | |
|---|---|---|---|---|---|---|---|---|---|
| 15 | 47010200 | 47011999 | 17.95629 | 31.65511 | 37.90488 | 16.77575 | 0.130282 | 0.579586 | 0.860283 |
| 8 | 96589600 | 96591799 | 14.75521 | 25.83343 | 44.60661 | 19.68756 | -0.39665 | 0.669731 | 0.860295 |
| 14 | 87822800 | 87825399 | 20.53161 | 35.91943 | 42.28227 | 18.64971 | 0.153267 | 0.70589 | 0.860305 |
| 18 | 19337600 | 19339799 | 23.30013 | 40.21052 | 39.32268 | 17.25048 | 0.417126 | 0.877611 | 0.860306 |
| 14 | 116206600 | 116208999 | 27.50915 | 47.01254 | 43.59421 | 19.11412 | 0.491249 | 0.989662 | 0.860319 |
| 17 | 72660000 | 72665399 | 49.28624 | 79.15607 | 95.59682 | 40.47781 | 0.078377 | 1.921154 | 0.860325 |
| 6 | 46276400 | 46276799 | 5.523074 | 10.76126 | 18.56021 | 6.745602 | -0.15009 | -0.39046 | 0.860331 |
| 15 | 8955800 | 8962799 | 96.35176 | 148.9337 | 175.5015 | 72.89934 | 0.065032 | 2.854279 | 0.86034 |
| 5 | 143132600 | 143141799 | 98.23334 | 151.158 | 194.5042 | 80.77929 | -0.06676 | 2.941101 | 0.860347 |
| 3 | 112490600 | 112494799 | 70.48292 | 111.9915 | 110.3742 | 46.03653 | 0.354112 | 2.277604 | 0.860353 |
| 6 | 132938400 | 132948399 | 208.331 | 303.6988 | 373.6682 | 153.9818 | -0.07095 | 3.958644 | 0.86036 |
| 18 | 40747200 | 40749599 | 37.95252 | 62.042 | 60.02691 | 25.80956 | 0.422036 | 1.497208 | 0.860397 |
| 6 | 36556200 | 36557199 | 8.359382 | 16.00069 | 24.57884 | 9.715624 | -0.09856 | -0.05183 | 0.860415 |
| Y | 5886600 | 5887399 | 9.004702 | 17.03868 | 20.34323 | 8.248861 | 0.229344 | 0.023193 | 0.860439 |
| 14 | 46319200 | 46324999 | 39.26201 | 63.59896 | 77.75639 | 33.2787 | 0.074869 | 1.657683 | 0.860446 |
| 6 | 32756000 | 32758199 | 24.58335 | 41.36409 | 50.67365 | 22.14277 | 0.089198 | 1.121164 | 0.860447 |
| 17 | 66780000 | 66785599 | 53.3906 | 85.46322 | 102.9767 | 43.38331 | 0.076033 | 2.030869 | 0.860453 |
| 19 | 23146200 | 23151599 | 57.15137 | 91.12982 | 109.7897 | 46.00167 | 0.070146 | 2.154959 | 0.860459 |
| 17 | 48775800 | 48777199 | 13.00784 | 23.56585 | 33.67838 | 14.57926 | -0.09988 | 0.349657 | 0.860467 |
| 4 | 112870800 | 112881799 | 131.928 | 201.4908 | 225.4619 | 93.51115 | 0.12628 | 3.103813 | 0.860478 |
| 4 | 5388600 | 5390399 | 22.4694 | 38.89252 | 46.84145 | 20.55076 | 0.118822 | 0.859767 | 0.86048 |
| 1 | 153790400 | 153792599 | 25.07991 | 42.14221 | 51.7523 | 22.60477 | 0.085626 | 1.135616 | 0.860485 |
| X | 43455400 | 43457999 | 22.64666 | 38.97487 | 46.86956 | 20.5595 | 0.119471 | 0.914298 | 0.86052 |
| 3 | 23407200 | 23408399 | 13.22779 | 23.05585 | 44.74716 | 19.68586 | -0.56464 | 0.730487 | 0.86053 |
| 9 | 65181600 | 65183599 | 19.29257 | 33.22979 | 46.34405 | 20.32949 | -0.0934 | 0.887479 | 0.860531 |
| 2 | 48995800 | 48999199 | 47.44152 | 76.51943 | 90.55692 | 38.57716 | 0.115101 | 1.788002 | 0.860563 |
| 6 | 51995800 | 51997399 | 18.81682 | 32.88208 | 47.29179 | 20.83927 | -0.1343 | 0.725248 | 0.860584 |
| 5 | 104989600 | 104991999 | 25.32388 | 42.93332 | 52.31209 | 22.91116 | 0.09729 | 1.059387 | 0.860586 |
| 18 | 83791000 | 83792399 | 18.3774 | 31.85575 | 44.22875 | 19.40535 | -0.08494 | 0.825279 | 0.860595 |
| X | 115924000 | 115924999 | 13.59068 | 24.83529 | 30.65726 | 13.12888 | 0.120498 | 0.30073 | 0.860606 |
| 13 | 33239000 | 33242199 | 41.0487 | 66.18929 | 79.85585 | 34.00591 | 0.086902 | 1.779729 | 0.86066 |
| 14 | 79190800 | 79192199 | 15.09264 | 26.524 | 37.28397 | 16.50815 | -0.10107 | 0.601024 | 0.86066 |
| 2 | 7926600 | 7927399 | 11.93074 | 21.10601 | 38.33802 | 16.95564 | -0.46518 | 0.510511 | 0.860669 |
| 11 | 4825800 | 4827599 | 11.93074 | 21.10601 | 38.33802 | 16.95564 | -0.46518 | 0.510511 | 0.860669 |
| 11 | 19450800 | 19453399 | 27.33337 | 46.52377 | 54.56109 | 23.91483 | 0.152811 | 1.036956 | 0.86068 |
| 11 | 76580600 | 76582599 | 16.95755 | 28.97105 | 52.66549 | 23.0909 | -0.47605 | 0.962481 | 0.860693 |
| 2 | 14631600 | 14632199 | 9.169884 | 17.29934 | 25.93468 | 10.58295 | -0.10445 | 0.06356 | 0.860711 |
| 9 | 19129600 | 19134599 | 66.9272 | 106.79 | 99.48727 | 41.72415 | 0.441529 | 2.141827 | 0.860718 |
| 18 | 90294200 | 90300199 | 53.13403 | 85.51153 | 98.11743 | 41.66959 | 0.155755 | 1.864138 | 0.860724 |
| 15 | 15778200 | 15780199 | 22.10474 | 38.25727 | 38.37252 | 16.83513 | 0.381655 | 0.847145 | 0.860761 |
| 7 | 86604800 | 86609599 | 57.37839 | 91.42232 | 121.9398 | 51.03884 | -0.07761 | 2.182807 | 0.860763 |
| 3 | 37444600 | 37447199 | 20.73171 | 35.62758 | 43.72044 | 19.17728 | 0.089827 | 0.911319 | 0.86077 |
| 8 | 78825000 | 78826599 | 15.78248 | 27.91579 | 34.46361 | 15.24257 | 0.088534 | 0.526712 | 0.86077 |
| 1 | 102890600 | 102893599 | 29.23732 | 48.32014 | 66.60963 | 28.78198 | -0.08122 | 1.344341 | 0.860775 |
| 6 | 44760400 | 44763799 | 36.62087 | 59.02499 | 95.82917 | 40.78606 | -0.34121 | 1.793289 | 0.860776 |
| 13 | 15031000 | 15035999 | 66.33415 | 105.1177 | 126.1596 | 52.47976 | 0.066566 | 2.356095 | 0.860778 |
| 14 | 18249800 | 18251599 | 5.651931 | 11.01177 | 25.14474 | 9.144843 | -0.55209 | -0.36336 | 0.86079 |

| | | | | | | | | | |
|---|---|---|---|---|---|---|---|---|---|
| 6 | 79919600 | 79920999 | 14.61309 | 26.03569 | 37.05488 | 16.31308 | -0.11033 | 0.466512 | 0.860809 |
| 8 | 27814400 | 27816599 | 19.72805 | 34.60987 | 33.83822 | 14.9604 | 0.419881 | 0.646259 | 0.86083 |
| 5 | 113580600 | 113581999 | 14.42334 | 25.75892 | 31.93799 | 14.0322 | 0.089715 | 0.441607 | 0.860838 |
| 8 | 72071600 | 72077599 | 111.5056 | 169.5232 | 219.1918 | 90.79911 | -0.08738 | 3.188583 | 0.860853 |
| 7 | 122695400 | 122697599 | 24.2974 | 40.79403 | 56.52165 | 24.67749 | -0.08779 | 1.148907 | 0.860861 |
| 8 | 30650800 | 30654599 | 44.18241 | 71.25232 | 86.47675 | 36.83206 | 0.078549 | 1.783579 | 0.86087 |
| 7 | 78813600 | 78818999 | 52.25311 | 83.41894 | 111.0556 | 46.60993 | -0.07183 | 2.113037 | 0.860873 |
| 2 | 105398000 | 105402399 | 50.37731 | 80.30943 | 128.3314 | 53.75309 | -0.33705 | 2.161024 | 0.860916 |
| 15 | 4843800 | 4846799 | 39.4132 | 63.89918 | 77.9743 | 33.39327 | 0.078771 | 1.640518 | 0.860941 |
| 14 | 13726200 | 13728399 | 37.20529 | 60.04555 | 82.20861 | 35.04815 | -0.09361 | 1.749585 | 0.860944 |
| 19 | 7730000 | 7732599 | 27.13562 | 44.65404 | 77.23526 | 33.29915 | -0.40952 | 1.396045 | 0.860945 |
| 9 | 109245600 | 109252999 | 75.92669 | 119.182 | 155.7483 | 64.59594 | -0.06683 | 2.572764 | 0.860948 |
| 4 | 153189800 | 153191199 | 20.90737 | 35.80575 | 49.54412 | 21.72659 | -0.08347 | 0.952305 | 0.860997 |
| 14 | 50907200 | 50913599 | 84.93542 | 132.0621 | 171.0065 | 70.99714 | -0.06429 | 2.745331 | 0.861037 |
| 9 | 11810800 | 11819599 | 327.2907 | 458.2611 | 544.3398 | 225.6652 | -0.07814 | 4.560577 | 0.861058 |
| 11 | 34484800 | 34490999 | 84.2727 | 132.2383 | 133.7276 | 55.46055 | 0.302239 | 2.580862 | 0.861064 |
| 10 | 121161400 | 121163399 | 16.87329 | 29.77964 | 36.43081 | 16.1197 | 0.100278 | 0.559562 | 0.861065 |
| 12 | 85295000 | 85296799 | 19.88573 | 33.99571 | 47.43882 | 20.80038 | -0.09598 | 0.960954 | 0.86107 |
| 5 | 150394400 | 150397799 | 26.93013 | 45.10826 | 55.60489 | 24.24132 | 0.080217 | 1.178725 | 0.861074 |
| 13 | 88921600 | 88925599 | 68.22481 | 107.8572 | 128.8291 | 53.5243 | 0.070703 | 2.412743 | 0.861074 |
| 8 | 48783600 | 48787599 | 40.20254 | 65.22405 | 78.91408 | 33.813 | 0.091974 | 1.628059 | 0.861076 |
| X | 144991800 | 144992599 | 9.560981 | 18.20378 | 28.36607 | 11.35216 | -0.13518 | 0.011138 | 0.861076 |
| 18 | 17788000 | 17789999 | 26.77947 | 45.1341 | 55.065 | 24.07375 | 0.095174 | 1.119406 | 0.861106 |
| 1 | 3601600 | 3604399 | 38.72516 | 62.51416 | 84.75004 | 36.14336 | -0.07903 | 1.747239 | 0.861112 |
| 4 | 19071600 | 19073399 | 19.85694 | 33.94993 | 47.43549 | 20.79908 | -0.09783 | 0.95921 | 0.861116 |
| 1 | 192450600 | 192453399 | 21.73256 | 37.09307 | 45.33178 | 19.87393 | 0.094203 | 0.979346 | 0.861137 |
| 5 | 51395600 | 51398999 | 63.51054 | 100.6978 | 120.5872 | 50.18899 | 0.070529 | 2.336782 | 0.861157 |
| X | 109961600 | 109962999 | 19.35639 | 33.94775 | 40.65887 | 17.97139 | 0.128476 | 0.659849 | 0.861166 |
| 16 | 48150000 | 48156999 | 54.00396 | 86.20509 | 115.6734 | 48.54086 | -0.08338 | 2.118353 | 0.861173 |
| 12 | 90048200 | 90049799 | 19.77878 | 33.25435 | 59.04228 | 25.79358 | -0.4442 | 1.122186 | 0.861182 |
| 3 | 47093000 | 47093799 | 14.67013 | 25.84383 | 36.16274 | 16.00696 | -0.09345 | 0.566819 | 0.861183 |
| 2 | 77302800 | 77305999 | 31.23452 | 51.55612 | 63.49544 | 27.41256 | 0.080725 | 1.359516 | 0.861197 |
| 14 | 41145800 | 41148599 | 27.406 | 44.81253 | 76.47807 | 32.88601 | -0.39431 | 1.479287 | 0.86121 |
| 8 | 63953200 | 63954399 | 16.05591 | 28.33936 | 34.96117 | 15.46918 | 0.088322 | 0.551367 | 0.861262 |
| 2 | 92783600 | 92784399 | 6.394953 | 12.40122 | 20.56463 | 7.763123 | -0.14415 | -0.23628 | 0.861285 |
| 18 | 23288200 | 23288799 | 7.862941 | 15.2139 | 21.01081 | 8.014897 | 0.102226 | -0.18436 | 0.861331 |
| 19 | 7873400 | 7876599 | 28.80552 | 48.05003 | 58.96871 | 25.64067 | 0.086704 | 1.232124 | 0.861368 |
| 2 | 66972200 | 66973199 | 10.55206 | 19.7082 | 28.94011 | 12.01307 | -0.0952 | 0.145318 | 0.861371 |
| 5 | 109003400 | 109004199 | 10.55206 | 19.7082 | 28.94011 | 12.01307 | -0.0952 | 0.145318 | 0.861371 |
| 11 | 50780800 | 50810399 | 508.6255 | 694.0246 | 719.6249 | 299.5445 | 0.08032 | 4.960096 | 0.861374 |
| 13 | 96921600 | 96922599 | 12.93142 | 23.52553 | 29.54003 | 12.72112 | 0.090621 | 0.316365 | 0.861427 |
| 5 | 102143400 | 102146599 | 22.83637 | 38.538 | 53.3164 | 23.31839 | -0.08547 | 1.092513 | 0.861429 |
| 19 | 55876400 | 55878999 | 30.67219 | 50.50344 | 69.54788 | 29.99228 | -0.08123 | 1.393108 | 0.861431 |
| 8 | 15980000 | 15982799 | 32.54928 | 53.46797 | 65.6898 | 28.30025 | 0.081587 | 1.422726 | 0.861433 |
| 10 | 40832400 | 40834999 | 17.75138 | 31.1012 | 38.12135 | 16.83533 | 0.095222 | 0.661853 | 0.861435 |
| Y | 29218000 | 29219199 | 9.398947 | 17.14884 | 27.40546 | 11.75905 | -0.25126 | 0.267839 | 0.861436 |
| 6 | 44672600 | 44673599 | 12.55436 | 23.58543 | 23.55064 | 9.679571 | 0.468822 | 0.1131 | 0.861451 |

| | | | | | | | | | |
|---|---|---|---|---|---|---|---|---|---|
| 7 | 108255000 | 108257199 | 21.52596 | 37.38912 | 45.21048 | 19.84624 | 0.114046 | 0.819703 | 0.86151 |
| 11 | 30903800 | 30916799 | 163.5883 | 246.1872 | 274.5548 | 113.384 | 0.113362 | 3.407822 | 0.861516 |
| 12 | 113484800 | 113499799 | 209.5623 | 308.3642 | 345.5321 | 142.2914 | 0.076816 | 3.81338 | 0.861517 |
| 11 | 28232600 | 28234399 | 31.64071 | 52.51363 | 63.61386 | 27.56781 | 0.105129 | 1.299668 | 0.86153 |
| 8 | 76665600 | 76666399 | 12.86 | 23.8325 | 23.5155 | 9.880578 | 0.461114 | 0.19908 | 0.861535 |
| 2 | 171384600 | 171385599 | 16.81594 | 29.28589 | 41.0919 | 18.06845 | -0.09905 | 0.75502 | 0.861567 |
| X | 51350600 | 51352999 | 14.50161 | 25.74678 | 36.64676 | 16.18109 | -0.11297 | 0.491378 | 0.861575 |
| 13 | 78524600 | 78527199 | 19.35729 | 33.85494 | 33.10686 | 14.59899 | 0.419736 | 0.692232 | 0.861624 |
| 4 | 33237800 | 33240799 | 26.3206 | 43.81786 | 60.49566 | 26.27167 | -0.08264 | 1.245807 | 0.861625 |
| 9 | 114614800 | 114616999 | 19.66677 | 33.85548 | 46.8776 | 20.56319 | -0.08316 | 0.895796 | 0.86163 |
| 1 | 85445400 | 85484199 | 805.8276 | 1057.079 | 1060.326 | 443.773 | 0.071193 | 5.589008 | 0.861646 |
| Y | 36421000 | 36421399 | 4.548228 | 8.879638 | 16.52651 | 5.834132 | -0.21955 | -0.51451 | 0.861647 |
| 14 | 31581600 | 31590399 | 232.5653 | 337.686 | 376.679 | 155.2865 | 0.064908 | 4.020696 | 0.861681 |
| 12 | 58653400 | 58654799 | 23.91618 | 40.12905 | 55.44636 | 24.19853 | -0.08381 | 1.152077 | 0.861696 |
| 5 | 60713800 | 60717199 | 37.21739 | 61.21354 | 59.36198 | 25.58781 | 0.422653 | 1.414111 | 0.861705 |
| 15 | 15747600 | 15749599 | 17.58079 | 30.65456 | 42.54778 | 18.72268 | -0.08344 | 0.742754 | 0.861707 |
| 8 | 18317400 | 18318799 | 19.12778 | 33.23379 | 41.03409 | 18.01636 | 0.084116 | 0.797739 | 0.861723 |
| 1 | 183380800 | 183382999 | 25.11345 | 42.08096 | 58.19282 | 25.37549 | -0.08502 | 1.173744 | 0.861737 |
| 13 | 85118600 | 85122199 | 42.4384 | 68.67402 | 82.78981 | 35.40623 | 0.093711 | 1.687004 | 0.86174 |
| 12 | 87872200 | 87872599 | 9.221623 | 17.83718 | 17.76187 | 6.785527 | 0.56812 | -0.17381 | 0.861742 |
| 4 | 37179800 | 37181599 | 19.06965 | 32.71282 | 45.86896 | 20.11761 | -0.10211 | 0.924561 | 0.861749 |
| 4 | 28412200 | 28415199 | 93.29375 | 144.0295 | 167.1213 | 69.41883 | 0.085943 | 2.875034 | 0.861761 |
| 1 | 21271600 | 21272399 | 11.59495 | 20.96204 | 30.20279 | 13.10913 | -0.1137 | 0.345322 | 0.861789 |
| 3 | 33641600 | 33645399 | 28.4619 | 47.28754 | 65.73371 | 28.50718 | -0.09273 | 1.278896 | 0.86179 |
| 13 | 119258200 | 119261999 | 49.58805 | 78.92099 | 127.7682 | 53.3952 | -0.35837 | 2.213263 | 0.861809 |
| 2 | 97583800 | 97584799 | 10.49531 | 20.02277 | 26.17188 | 10.42976 | 0.121279 | -0.00235 | 0.861809 |
| 11 | 20232000 | 20249999 | 240.5376 | 348.1284 | 424.4416 | 175.0464 | -0.06792 | 4.070356 | 0.861813 |
| 13 | 80069000 | 80071599 | 24.68918 | 41.25681 | 57.11958 | 24.86446 | -0.08669 | 1.198228 | 0.861823 |
| 13 | 29244800 | 29248799 | 35.44411 | 57.62197 | 79.13544 | 33.95709 | -0.08772 | 1.578287 | 0.86183 |
| 5 | 94539000 | 94540599 | 22.73391 | 39.31802 | 38.42862 | 16.85937 | 0.418756 | 0.85826 | 0.86187 |
| 7 | 144113800 | 144114799 | 9.159781 | 17.31708 | 25.96231 | 10.54666 | -0.09887 | 0.048576 | 0.861879 |
| 10 | 77287800 | 77289399 | 20.0101 | 34.54511 | 47.93675 | 21.02963 | -0.08547 | 0.873295 | 0.861879 |
| 5 | 75685800 | 75686799 | 12.5409 | 23.1362 | 29.1582 | 12.32716 | 0.103649 | 0.232951 | 0.86188 |
| 2 | 30651600 | 30653399 | 16.67143 | 29.40025 | 36.03948 | 15.94906 | 0.097132 | 0.568443 | 0.861885 |
| 7 | 42772200 | 42822799 | 1890.241 | 2319.782 | 2138.901 | 901.5307 | 0.095894 | 6.666722 | 0.861887 |
| 3 | 17567800 | 17568199 | 8.769361 | 16.79006 | 22.40699 | 8.85012 | 0.10375 | -0.057 | 0.861889 |
| Y | 41163800 | 41168399 | 44.66597 | 71.62229 | 75.40487 | 31.85137 | 0.273331 | 1.95386 | 0.861904 |
| 11 | 108593000 | 108593599 | 8.202698 | 15.81404 | 24.81508 | 9.608647 | -0.1012 | -0.11984 | 0.861921 |
| 4 | 156366800 | 156373399 | 205.6403 | 296.0418 | 459.1401 | 189.49 | -0.42252 | 4.14705 | 0.861932 |
| 5 | 144603200 | 144606599 | 31.048 | 50.68894 | 70.09235 | 30.12306 | -0.09319 | 1.504202 | 0.861976 |
| 10 | 110950600 | 110953799 | 23.79265 | 39.98809 | 55.51889 | 24.2516 | -0.09086 | 1.132244 | 0.861987 |
| 5 | 68570000 | 68572199 | 11.27909 | 21.38851 | 21.65147 | 8.739242 | 0.471498 | 0.035783 | 0.862014 |
| 16 | 97798200 | 97802199 | 48.34207 | 77.66066 | 94.10099 | 39.86047 | 0.074107 | 1.911727 | 0.862015 |
| 13 | 55581000 | 55589799 | 125.8431 | 190.4637 | 241.7949 | 100.0251 | -0.06635 | 3.284489 | 0.862029 |
| 1 | 98385200 | 98387599 | 28.34985 | 47.5697 | 47.52776 | 20.74299 | 0.382301 | 1.1578 | 0.862075 |
| X | 16456400 | 16457599 | 17.73804 | 31.36333 | 29.41329 | 13.00986 | 0.483013 | 0.534641 | 0.862089 |
| 11 | 91950000 | 91952599 | 35.00606 | 57.5352 | 58.50901 | 25.2125 | 0.35396 | 1.418296 | 0.862103 |

| | | | | | | | | | |
|---|---|---|---|---|---|---|---|---|---|
| 6 | 8194400 | 8198599 | 91.39301 | 141.9368 | 167.9401 | 69.73621 | 0.064397 | 2.767961 | 0.862115 |
| 11 | 11968000 | 11968799 | 7.898523 | 15.15753 | 23.55443 | 9.242123 | -0.1049 | -0.08313 | 0.862126 |
| 6 | 137286800 | 137292399 | 59.49976 | 94.65881 | 125.9801 | 52.61952 | -0.0768 | 2.233801 | 0.862128 |
| 5 | 88192600 | 88194599 | 35.89132 | 58.21742 | 79.0494 | 33.86576 | -0.07428 | 1.626206 | 0.862131 |
| 11 | 100702000 | 100703399 | 17.39707 | 30.52032 | 37.64308 | 16.6425 | 0.086415 | 0.640437 | 0.862141 |
| 5 | 47298400 | 47301999 | 38.50077 | 62.28954 | 84.43373 | 36.1039 | -0.07531 | 1.684629 | 0.862173 |
| 5 | 68571600 | 68572199 | 11.24403 | 20.77341 | 25.76326 | 10.87066 | 0.129408 | 0.21018 | 0.862175 |
| 4 | 78587000 | 78589599 | 15.76326 | 28.37869 | 33.86304 | 14.77498 | 0.151248 | 0.402902 | 0.862181 |
| 2 | 48126800 | 48127399 | 8.034251 | 15.50909 | 20.96752 | 8.08619 | 0.11795 | -0.14487 | 0.862194 |
| 12 | 8144800 | 8145399 | 8.034251 | 15.50909 | 20.96752 | 8.08619 | 0.11795 | -0.14487 | 0.862194 |
| 13 | 56431200 | 56432399 | 13.36811 | 23.67567 | 33.62012 | 14.8657 | -0.11155 | 0.495253 | 0.862201 |
| 18 | 57211800 | 57214999 | 40.35344 | 65.39145 | 79.36004 | 33.97443 | 0.085964 | 1.651834 | 0.862233 |
| 11 | 48389200 | 48389999 | 7.478437 | 14.45039 | 23.17785 | 8.897543 | -0.1199 | -0.15823 | 0.862269 |
| 6 | 13978400 | 13982799 | 99.15324 | 152.7579 | 179.3714 | 74.50684 | 0.066489 | 2.916752 | 0.862282 |
| 4 | 142974600 | 142981799 | 70.70581 | 111.3699 | 145.7162 | 60.46226 | -0.06475 | 2.4992 | 0.862282 |
| 10 | 73525200 | 73526999 | 24.05153 | 40.70786 | 50.12028 | 21.94004 | 0.082276 | 1.067658 | 0.862318 |
| 14 | 47844200 | 47850399 | 71.82753 | 113.1416 | 135.0167 | 56.02302 | 0.067924 | 2.497297 | 0.862327 |
| 5 | 114979200 | 114980199 | 13.46855 | 24.09803 | 34.1483 | 14.98006 | -0.1001 | 0.43097 | 0.862351 |
| 10 | 54762400 | 54767199 | 126.6769 | 192.0497 | 222.2377 | 91.98371 | 0.06927 | 3.24986 | 0.862383 |
| 10 | 31704200 | 31704999 | 11.27947 | 20.36773 | 29.74052 | 12.92749 | -0.13417 | 0.348616 | 0.862391 |
| 18 | 86873600 | 86874599 | 13.74974 | 24.81387 | 30.9623 | 13.4684 | 0.090236 | 0.371239 | 0.862403 |
| 4 | 107985400 | 107988199 | 27.32707 | 45.49577 | 62.80091 | 27.27367 | -0.0824 | 1.250588 | 0.86241 |
| 5 | 119111400 | 119113199 | 9.69685 | 18.03211 | 37.25007 | 15.57715 | -0.59314 | 0.183964 | 0.862425 |
| 5 | 72232800 | 72234599 | 20.63131 | 34.46275 | 62.24611 | 27.08955 | -0.46888 | 1.196071 | 0.862439 |
| 1 | 61794000 | 61794799 | 12.0597 | 21.90632 | 31.17427 | 13.45044 | -0.09048 | 0.323186 | 0.862442 |
| 10 | 83511800 | 83519399 | 73.86353 | 115.7201 | 151.4954 | 62.8388 | -0.07155 | 2.606181 | 0.862461 |
| 8 | 116007600 | 116008799 | 11.19687 | 20.86607 | 26.75584 | 11.14828 | 0.094873 | 0.160729 | 0.862498 |
| 16 | 19413000 | 19422599 | 299.4164 | 422.2152 | 506.4002 | 209.701 | -0.08181 | 4.45465 | 0.86251 |
| 3 | 31242200 | 31246799 | 41.65528 | 67.23601 | 81.64937 | 34.8155 | 0.079176 | 1.750861 | 0.862521 |
| 6 | 9278000 | 9281399 | 43.43027 | 70.5029 | 68.83933 | 29.50921 | 0.401374 | 1.617185 | 0.86253 |
| 4 | 144460400 | 144462599 | 27.43287 | 47.07611 | 41.3118 | 18.11941 | 0.571182 | 0.947624 | 0.862549 |
| 6 | 123105200 | 123105799 | 9.533795 | 18.00982 | 27.00195 | 10.9875 | -0.10166 | 0.057302 | 0.862553 |
| X | 61673400 | 61674799 | 14.13427 | 26.12109 | 24.47314 | 10.32203 | 0.531575 | 0.228391 | 0.862635 |
| 1 | 165884800 | 165886799 | 17.88484 | 31.06587 | 43.165 | 18.94876 | -0.08552 | 0.798923 | 0.862637 |
| 9 | 7298600 | 7301999 | 50.9317 | 81.57575 | 98.85781 | 41.68516 | 0.068785 | 2.008583 | 0.862639 |
| 2 | 70428400 | 70430199 | 13.46175 | 23.93583 | 34.00649 | 14.99665 | -0.10944 | 0.465468 | 0.862653 |
| 5 | 33489200 | 33491199 | 44.73967 | 71.7413 | 96.5882 | 40.80003 | -0.08075 | 1.964592 | 0.862664 |
| 1 | 109965200 | 109972599 | 123.8009 | 188.7089 | 219.3032 | 90.91565 | 0.069508 | 3.139149 | 0.862678 |
| 19 | 57978000 | 57978799 | 7.890875 | 15.16958 | 23.59161 | 9.209955 | -0.09925 | -0.09985 | 0.862702 |
| 8 | 75993000 | 75995599 | 50.29149 | 80.45256 | 96.63952 | 40.67779 | 0.079549 | 2.044766 | 0.862719 |
| 5 | 126160600 | 126168799 | 101.9618 | 157.0717 | 184.8121 | 76.76588 | 0.063465 | 2.920448 | 0.862748 |
| 10 | 23790800 | 23793799 | 33.65264 | 55.56915 | 67.09485 | 28.97431 | 0.109292 | 1.365612 | 0.862766 |
| 17 | 87849200 | 87856199 | 78.74447 | 124.178 | 125.1454 | 51.93478 | 0.312675 | 2.478306 | 0.862767 |
| 14 | 113845000 | 113845999 | 17.27917 | 30.18533 | 36.8102 | 16.21707 | 0.102679 | 0.704553 | 0.86277 |
| 2 | 132215200 | 132216199 | 14.10433 | 24.83829 | 35.30302 | 15.62752 | -0.11583 | 0.567834 | 0.86277 |
| 8 | 21105800 | 21105999 | 5.936654 | 11.58919 | 20.00301 | 7.073132 | -0.11609 | -0.4775 | 0.862772 |
| 7 | 17051400 | 17053799 | 33.74553 | 55.19466 | 76.27793 | 32.80345 | -0.0905 | 1.489665 | 0.862776 |

| | | | | | | | | | |
|---|---|---|---|---|---|---|---|---|---|
| 7 | 34022400 | 34027799 | 50.31546 | 80.46851 | 107.0746 | 45.05238 | -0.0683 | 2.056219 | 0.86279 |
| 6 | 146607000 | 146614199 | 77.33946 | 121.566 | 145.0918 | 60.18693 | 0.065352 | 2.544616 | 0.862833 |
| 8 | 22746400 | 22750799 | 73.91378 | 115.4542 | 152.8522 | 63.42101 | -0.09106 | 2.657877 | 0.862837 |
| 6 | 123131600 | 123141599 | 191.9892 | 283.4221 | 321.4311 | 132.3712 | 0.06383 | 3.758186 | 0.862838 |
| 13 | 24117200 | 24119399 | 32.90888 | 53.08202 | 92.1671 | 39.26556 | -0.43654 | 1.764481 | 0.86291 |
| 8 | 112478800 | 112479399 | 8.828254 | 17.01637 | 23.09977 | 8.950026 | 0.105452 | -0.11532 | 0.862931 |
| 14 | 97964200 | 97967599 | 38.93266 | 63.92699 | 73.99059 | 31.87072 | 0.167192 | 1.449746 | 0.862939 |
| 7 | 136560400 | 136562199 | 21.7256 | 37.19483 | 45.82479 | 20.09502 | 0.083233 | 0.952073 | 0.862945 |
| 9 | 51789000 | 51790799 | 21.7256 | 37.19483 | 45.82479 | 20.09502 | 0.083233 | 0.952073 | 0.862945 |
| 5 | 30023800 | 30024399 | 12.11733 | 22.2308 | 28.11071 | 11.97458 | 0.091233 | 0.259429 | 0.862958 |
| 7 | 84603400 | 84604799 | 21.68493 | 37.0445 | 51.1627 | 22.43186 | -0.08141 | 0.97887 | 0.862973 |
| 7 | 77571800 | 77572999 | 15.83218 | 27.82523 | 38.69368 | 17.13313 | -0.08582 | 0.604906 | 0.862992 |
| 17 | 31730000 | 31731799 | 11.23262 | 20.32745 | 29.68441 | 12.86792 | -0.13192 | 0.334231 | 0.863088 |
| 1 | 120090600 | 120092799 | 17.70224 | 30.90136 | 43.02087 | 18.94436 | -0.08772 | 0.727106 | 0.863105 |
| 9 | 88871000 | 88876199 | 60.69971 | 96.4088 | 127.2472 | 53.05322 | -0.06719 | 2.284726 | 0.863117 |
| 14 | 7932400 | 7935799 | 35.67284 | 58.73929 | 70.06947 | 30.21765 | 0.125391 | 1.406659 | 0.863153 |
| X | 13351800 | 13355599 | 66.37509 | 105.2024 | 123.8191 | 51.51584 | 0.094994 | 2.352777 | 0.863158 |
| 6 | 143394200 | 143401399 | 86.21226 | 133.6226 | 173.1726 | 71.93053 | -0.06905 | 2.800305 | 0.863176 |
| 18 | 4414000 | 4415599 | 18.46284 | 32.27206 | 39.78776 | 17.53653 | 0.086957 | 0.706752 | 0.863195 |
| 11 | 48205400 | 48208399 | 23.1174 | 39.81517 | 47.92146 | 21.0214 | 0.118185 | 0.905396 | 0.863206 |
| 13 | 14952600 | 14958999 | 86.16045 | 134.077 | 158.3205 | 65.72222 | 0.069533 | 2.729448 | 0.863215 |
| 1 | 90841000 | 90848799 | 74.40846 | 116.8626 | 152.4816 | 63.2456 | -0.064 | 2.560241 | 0.863217 |
| 1 | 35505400 | 35514799 | 122.9326 | 186.6384 | 236.9641 | 98.12068 | -0.06275 | 3.220812 | 0.863234 |
| 8 | 78771000 | 78773599 | 36.58082 | 59.40653 | 72.61361 | 31.13592 | 0.078614 | 1.598044 | 0.863275 |
| 1 | 26542800 | 26544399 | 20.11674 | 34.96223 | 43.07608 | 18.91638 | 0.087167 | 0.800908 | 0.863339 |
| 1 | 58907400 | 58919599 | 194.201 | 286.1705 | 355.0773 | 146.2244 | -0.06824 | 3.788907 | 0.863352 |
| 3 | 139767000 | 139767999 | 15.07638 | 26.6228 | 32.55597 | 14.40362 | 0.101289 | 0.536117 | 0.863357 |
| X | 20878400 | 20883399 | 73.29292 | 116.2212 | 134.4713 | 55.97393 | 0.120227 | 2.345825 | 0.863404 |
| 5 | 122650800 | 122652799 | 14.69747 | 25.91325 | 36.16441 | 16.00507 | -0.08916 | 0.558647 | 0.863404 |
| 2 | 156169800 | 156171599 | 16.71129 | 29.3183 | 36.154 | 15.98465 | 0.086687 | 0.6332 | 0.863408 |
| X | 80578400 | 80579999 | 19.84018 | 35.29436 | 39.01482 | 17.19818 | 0.251261 | 0.515531 | 0.863412 |
| 15 | 103561600 | 103564599 | 26.76213 | 44.30228 | 61.37676 | 26.551 | -0.08814 | 1.314725 | 0.863426 |
| 3 | 93795800 | 93798399 | 15.49977 | 27.51832 | 33.9502 | 14.99073 | 0.092016 | 0.490728 | 0.86343 |
| 4 | 127400600 | 127403199 | 23.70263 | 39.83537 | 54.93005 | 23.99413 | -0.08095 | 1.133476 | 0.863452 |
| 5 | 98191000 | 98191399 | 6.237849 | 12.15529 | 20.27465 | 7.358059 | -0.10045 | -0.37832 | 0.863454 |
| 2 | 111554600 | 111555399 | 12.54295 | 22.88692 | 28.64201 | 12.2885 | 0.099188 | 0.296396 | 0.863475 |
| 7 | 93292200 | 93292599 | 7.113301 | 13.82514 | 19.52111 | 7.261175 | 0.104885 | -0.28378 | 0.863475 |
| 11 | 24228000 | 24230199 | 15.34581 | 27.06726 | 33.00064 | 14.60372 | 0.10483 | 0.550159 | 0.863477 |
| 9 | 79019800 | 79023399 | 43.67916 | 70.4298 | 85.73887 | 36.51047 | 0.073914 | 1.786803 | 0.863501 |
| 1 | 170923400 | 170927999 | 32.74779 | 53.84874 | 66.2906 | 28.57137 | 0.079288 | 1.411904 | 0.863505 |
| 2 | 68217600 | 68220799 | 19.70643 | 33.8073 | 46.89811 | 20.56909 | -0.08674 | 0.92766 | 0.863511 |
| 4 | 38346200 | 38346799 | 10.46157 | 19.41799 | 28.34068 | 11.8827 | -0.09751 | 0.1833 | 0.863524 |
| 15 | 23776000 | 23778799 | 35.55748 | 58.52031 | 58.55093 | 25.24495 | 0.378134 | 1.40419 | 0.863534 |
| 10 | 56499000 | 56500599 | 16.64125 | 29.48677 | 35.90358 | 15.87384 | 0.109307 | 0.51583 | 0.863539 |
| 16 | 89408800 | 89409999 | 12.86632 | 23.32394 | 28.97404 | 12.53303 | 0.101762 | 0.333868 | 0.863584 |
| X | 164374400 | 164376399 | 15.82338 | 28.04749 | 40.73407 | 18.00824 | -0.14343 | 0.517198 | 0.863594 |
| 7 | 104743400 | 104745199 | 18.28287 | 32.27355 | 38.47276 | 17.02257 | 0.137581 | 0.570311 | 0.863602 |

| | | | | | | | | | |
|---|---|---|---|---|---|---|---|---|---|
| 8 | 121732200 | 121735999 | 41.6361 | 67.13749 | 90.54216 | 38.55735 | -0.07342 | 1.785975 | 0.863603 |
| 3 | 8510200 | 8511799 | 12.4375 | 22.89575 | 34.07962 | 14.45033 | -0.13812 | 0.25415 | 0.863604 |
| X | 13610600 | 13614999 | 38.21378 | 62.59163 | 74.0881 | 31.87935 | 0.133546 | 1.479891 | 0.863617 |
| 4 | 35707000 | 35710799 | 41.27755 | 66.4792 | 89.4732 | 38.03906 | -0.07258 | 1.820712 | 0.863617 |
| 6 | 91049400 | 91051399 | 27.79074 | 46.32333 | 57.24534 | 24.87981 | 0.076756 | 1.235019 | 0.86362 |
| 4 | 78507600 | 78508599 | 13.4098 | 23.9216 | 33.61154 | 14.78165 | -0.09049 | 0.44688 | 0.863629 |
| 16 | 37984400 | 37985999 | 13.4098 | 23.9216 | 33.61154 | 14.78165 | -0.09049 | 0.44688 | 0.863629 |
| 13 | 35739000 | 35742199 | 36.93238 | 60.34138 | 73.26684 | 31.49602 | 0.094616 | 1.508721 | 0.863638 |
| 5 | 73051400 | 73054999 | 25.22175 | 42.28992 | 58.24618 | 25.40919 | -0.07927 | 1.166046 | 0.86364 |
| 3 | 64140600 | 64145399 | 59.30629 | 94.32809 | 113.3985 | 47.34849 | 0.069362 | 2.238624 | 0.863668 |
| 2 | 170158600 | 170159999 | 14.11609 | 24.86368 | 35.62926 | 15.77134 | -0.12786 | 0.562191 | 0.863669 |
| 13 | 111746200 | 111747799 | 19.00347 | 33.13839 | 40.91854 | 18.00684 | 0.084532 | 0.745917 | 0.863671 |
| 5 | 146605800 | 146612199 | 65.37531 | 103.2729 | 135.7268 | 56.3681 | -0.06789 | 2.431031 | 0.863676 |
| 2 | 78342800 | 78343999 | 27.58224 | 45.52154 | 63.02346 | 27.20629 | -0.08754 | 1.352917 | 0.863696 |
| 2 | 131804000 | 131807399 | 25.9954 | 43.77557 | 61.76144 | 26.99442 | -0.11389 | 1.12976 | 0.863712 |
| 15 | 65544800 | 65546599 | 16.39292 | 28.77607 | 40.30203 | 17.82714 | -0.09598 | 0.631292 | 0.863727 |
| 6 | 66742600 | 66743599 | 17.19225 | 30.01616 | 36.41421 | 16.0368 | 0.1102 | 0.713192 | 0.86373 |
| 17 | 13632600 | 13635399 | 60.3678 | 94.9367 | 155.5365 | 64.52383 | -0.39045 | 2.52728 | 0.863736 |
| 9 | 90231000 | 90232199 | 21.93609 | 37.54086 | 46.35812 | 20.32827 | 0.079861 | 0.957784 | 0.863743 |
| 5 | 122437800 | 122439599 | 14.39158 | 25.29424 | 35.79714 | 15.85104 | -0.11083 | 0.591641 | 0.863746 |
| X | 158038200 | 158040399 | 28.82613 | 48.27287 | 57.77943 | 25.18562 | 0.122656 | 1.19304 | 0.863787 |
| 13 | 110549200 | 110551599 | 17.40194 | 30.51079 | 37.68033 | 16.65062 | 0.084575 | 0.650157 | 0.863796 |
| 7 | 57715800 | 57716599 | 10.87839 | 20.37398 | 26.30123 | 10.86663 | 0.095039 | 0.12533 | 0.863801 |
| 15 | 71476600 | 71476999 | 8.652459 | 16.54506 | 21.40631 | 8.485981 | 0.143602 | -0.04768 | 0.86381 |
| 17 | 12987600 | 12996999 | 102.6624 | 157.9899 | 185.4822 | 77.03346 | 0.065526 | 2.939549 | 0.86382 |
| 4 | 144129800 | 144137799 | 97.87093 | 150.6912 | 194.0537 | 80.59882 | -0.06721 | 2.928904 | 0.863848 |
| 18 | 55504400 | 55504999 | 7.906053 | 15.14569 | 23.52524 | 9.276756 | -0.1109 | -0.06664 | 0.863867 |
| 14 | 16270200 | 16271399 | 13.54568 | 24.2838 | 29.64584 | 12.98336 | 0.115474 | 0.40956 | 0.863868 |
| 3 | 158455800 | 158456799 | 20.06157 | 34.73382 | 42.66147 | 18.71702 | 0.090888 | 0.843076 | 0.863883 |
| 2 | 88799200 | 88801799 | 38.0273 | 61.74687 | 75.40941 | 32.33136 | 0.079697 | 1.607977 | 0.863901 |
| 7 | 47204200 | 47206199 | 45.7876 | 73.86067 | 74.87206 | 31.90038 | 0.338206 | 1.773797 | 0.863909 |
| Y | 29263000 | 29264799 | 11.51136 | 20.97776 | 23.27751 | 10.0029 | 0.269869 | 0.279439 | 0.863916 |
| 14 | 108883800 | 108888799 | 63.09471 | 100.2723 | 120.4311 | 50.24127 | 0.069547 | 2.267468 | 0.863917 |
| 1 | 24717600 | 24720199 | 17.86544 | 30.9919 | 43.18667 | 18.94858 | -0.08995 | 0.814235 | 0.863925 |
| 5 | 89140400 | 89148999 | 101.2095 | 156.2305 | 183.6694 | 76.29266 | 0.066909 | 2.882472 | 0.863934 |
| 9 | 117068200 | 117071199 | 40.82991 | 66.01593 | 80.24185 | 34.29046 | 0.080659 | 1.699644 | 0.863942 |
| Y | 5645200 | 5649399 | 36.85098 | 60.67764 | 35.9643 | 15.50953 | 1.131108 | 1.378121 | 0.863959 |
| 19 | 54320400 | 54321199 | 21.81687 | 37.13393 | 51.66533 | 22.64557 | -0.09268 | 1.015635 | 0.863979 |
| 3 | 41946000 | 41946799 | 12.12516 | 21.80517 | 31.2036 | 13.62965 | -0.10978 | 0.383233 | 0.863994 |
| 16 | 65997400 | 65998799 | 16.11262 | 28.26273 | 39.26112 | 17.35566 | -0.08436 | 0.637988 | 0.864005 |
| 11 | 108870600 | 108872799 | 19.16366 | 32.6133 | 55.94385 | 24.52061 | -0.39376 | 1.008702 | 0.864015 |
| 3 | 89504800 | 89507599 | 41.82112 | 67.18879 | 91.25712 | 38.6591 | -0.0901 | 1.900534 | 0.864033 |
| 12 | 101411000 | 101413399 | 31.94729 | 53.19548 | 53.04712 | 23.03982 | 0.385224 | 1.254038 | 0.864081 |
| 2 | 151055200 | 151068399 | 146.0723 | 221.191 | 250.2333 | 103.5322 | 0.100423 | 3.279672 | 0.864093 |
| 10 | 94992000 | 94992599 | 8.171374 | 15.62182 | 24.00373 | 9.521477 | -0.10388 | -0.04515 | 0.864095 |
| 2 | 55665600 | 55668399 | 18.19564 | 31.94903 | 30.01426 | 13.27943 | 0.477192 | 0.617254 | 0.864102 |
| 13 | 87144200 | 87144799 | 9.445376 | 17.87346 | 22.85436 | 9.266056 | 0.131709 | 0.037094 | 0.864119 |

| | | | | | | | | | |
|---|---|---|---|---|---|---|---|---|---|
| 2 | 13797600 | 13798999 | 14.33218 | 24.9615 | 45.26852 | 19.90555 | -0.46741 | 0.748646 | 0.86414 |
| 9 | 81708400 | 81709399 | 10.18653 | 18.83229 | 27.8504 | 11.74061 | -0.12177 | 0.203355 | 0.864155 |
| 6 | 45401600 | 45402399 | 12.89218 | 23.31917 | 28.66315 | 12.43291 | 0.114862 | 0.34979 | 0.864162 |
| 1 | 144463600 | 144466999 | 48.43425 | 77.64718 | 93.92122 | 39.6591 | 0.073056 | 1.974913 | 0.864181 |
| 16 | 12979400 | 12986599 | 84.95196 | 133.4597 | 154.6823 | 64.16077 | 0.107025 | 2.562967 | 0.864184 |
| 6 | 83980600 | 83984599 | 61.4272 | 97.23386 | 128.6566 | 53.48689 | -0.07526 | 2.379202 | 0.864185 |
| 4 | 11462600 | 11463999 | 17.76361 | 31.28734 | 38.00394 | 16.82261 | 0.109416 | 0.590394 | 0.864186 |
| 1 | 127802000 | 127803599 | 26.03864 | 44.20381 | 43.24416 | 18.94731 | 0.413077 | 1.043687 | 0.864197 |
| 6 | 59212200 | 59221399 | 143.3705 | 216.695 | 249.352 | 103.1069 | 0.073613 | 3.316688 | 0.864203 |
| 3 | 91788400 | 91789599 | 15.45879 | 27.28105 | 33.57674 | 14.85709 | 0.091756 | 0.548222 | 0.86421 |
| 3 | 16503200 | 16504799 | 26.79698 | 44.68645 | 55.00979 | 23.91481 | 0.082322 | 1.225414 | 0.864217 |
| 14 | 89607000 | 89612399 | 71.72512 | 112.7032 | 147.5355 | 61.19796 | -0.06816 | 2.547632 | 0.864266 |
| 6 | 71266200 | 71268599 | 19.39061 | 33.65015 | 46.86202 | 20.5614 | -0.08901 | 0.823596 | 0.864276 |
| 7 | 6146000 | 6148599 | 20.94556 | 35.86801 | 43.77696 | 19.19738 | 0.096869 | 0.945142 | 0.864287 |
| 9 | 92521200 | 92530199 | 131.0856 | 198.5576 | 230.5054 | 95.37911 | 0.063566 | 3.268696 | 0.864309 |
| 16 | 55579800 | 55581199 | 16.23939 | 28.49874 | 39.7258 | 17.56818 | -0.0894 | 0.630901 | 0.864323 |
| 6 | 5315200 | 5319999 | 54.41198 | 86.62896 | 115.1848 | 48.15998 | -0.07426 | 2.202956 | 0.864328 |
| 7 | 99112000 | 99114599 | 21.01383 | 35.79211 | 49.54938 | 21.71934 | -0.08541 | 1.000893 | 0.864348 |
| 4 | 100878000 | 100880999 | 24.24158 | 41.78026 | 40.91148 | 17.94675 | 0.414938 | 0.896671 | 0.864354 |
| 16 | 22023600 | 22026799 | 37.53909 | 61.53427 | 73.15877 | 31.4898 | 0.127587 | 1.465591 | 0.864415 |
| 4 | 72272800 | 72273799 | 13.48499 | 24.39728 | 30.5344 | 13.24061 | 0.08898 | 0.355205 | 0.864472 |
| 11 | 53681000 | 53681799 | 13.48499 | 24.39728 | 30.5344 | 13.24061 | 0.08898 | 0.355205 | 0.864472 |
| 14 | 29679800 | 29680999 | 13.32419 | 23.72658 | 33.69631 | 14.8412 | -0.10714 | 0.45945 | 0.864486 |
| 16 | 24775200 | 24776399 | 17.7201 | 31.18504 | 37.99771 | 16.82261 | 0.104455 | 0.598424 | 0.864488 |
| X | 72891600 | 72891999 | 12.21811 | 22.40743 | 28.17188 | 12.00665 | 0.099199 | 0.26428 | 0.864489 |
| 4 | 27681200 | 27682599 | 15.68047 | 27.49754 | 38.31894 | 16.93542 | -0.08874 | 0.639591 | 0.864489 |
| 2 | 111765800 | 111768599 | 51.97837 | 83.14337 | 100.5722 | 42.32756 | 0.06933 | 2.051302 | 0.864509 |
| 2 | 30616200 | 30618799 | 24.72175 | 41.28361 | 57.08827 | 24.84035 | -0.08493 | 1.20621 | 0.86452 |
| 6 | 43843000 | 43843599 | 11.83433 | 21.68379 | 27.15929 | 11.58949 | 0.103255 | 0.26313 | 0.86454 |
| 10 | 74669000 | 74669799 | 8.979497 | 16.91377 | 25.39015 | 10.38938 | -0.10974 | 0.069394 | 0.864559 |
| 18 | 70442800 | 70444999 | 42.12746 | 67.80566 | 91.07306 | 38.68421 | -0.07077 | 1.842771 | 0.86458 |
| 14 | 110648200 | 110651399 | 27.73112 | 46.80268 | 56.24376 | 24.5994 | 0.116951 | 1.111215 | 0.8646 |
| Y | 89212600 | 89213199 | 7.346894 | 14.1998 | 17.89065 | 6.859793 | 0.226304 | -0.18518 | 0.864619 |
| 5 | 120449600 | 120451999 | 36.49546 | 59.09805 | 80.20416 | 34.31796 | -0.07583 | 1.660141 | 0.864625 |
| 13 | 3551600 | 3552399 | 8.037937 | 15.50236 | 24.25063 | 9.379725 | -0.09525 | -0.1269 | 0.864648 |
| 5 | 139825200 | 139834199 | 103.2181 | 156.9065 | 243.6896 | 100.9439 | -0.35172 | 3.190684 | 0.864652 |
| 6 | 53141400 | 53142599 | 10.34295 | 19.65644 | 25.46063 | 10.22912 | 0.123577 | 0.01804 | 0.864656 |
| 18 | 19592400 | 19592999 | 10.34295 | 19.65644 | 25.46063 | 10.22912 | 0.123577 | 0.01804 | 0.864656 |
| 9 | 79247600 | 79250199 | 48.72672 | 77.85172 | 104.4463 | 43.88584 | -0.08162 | 2.084405 | 0.864674 |
| 3 | 64140200 | 64145399 | 62.11819 | 98.61915 | 118.4974 | 49.38383 | 0.067489 | 2.296151 | 0.864679 |
| 4 | 34503400 | 34511799 | 150.3512 | 225.6718 | 260.011 | 107.3022 | 0.063027 | 3.454598 | 0.86471 |
| 17 | 22975400 | 22977399 | 26.04679 | 44.35574 | 44.18317 | 19.36682 | 0.387535 | 1.014615 | 0.864739 |
| 10 | 48523800 | 48528199 | 41.57791 | 67.59963 | 80.00433 | 34.33101 | 0.126499 | 1.59128 | 0.864757 |
| 2 | 71607200 | 71609199 | 19.52344 | 33.74337 | 41.16244 | 18.05839 | 0.10022 | 0.853671 | 0.864758 |
| 17 | 75196200 | 75199199 | 31.42327 | 52.08539 | 63.61228 | 27.54254 | 0.093228 | 1.312863 | 0.864761 |
| 5 | 120955200 | 120966599 | 94.33088 | 144.224 | 225.4432 | 93.52657 | -0.35435 | 3.069801 | 0.864774 |
| 6 | 10261800 | 10264599 | 35.96555 | 58.52174 | 80.43156 | 34.52445 | -0.08789 | 1.568152 | 0.864783 |

| | | | | | | | | | |
|---|---|---|---|---|---|---|---|---|---|
| 8 | 65126600 | 65128399 | 15.99835 | 28.13345 | 39.18466 | 17.35076 | -0.08788 | 0.599715 | 0.86479 |
| 2 | 3241600 | 3245399 | 38.65036 | 62.73168 | 76.70562 | 32.87672 | 0.077347 | 1.617614 | 0.864792 |
| 13 | 94490000 | 94493599 | 41.1503 | 66.55368 | 92.18801 | 39.40982 | -0.10697 | 1.699594 | 0.8648 |
| X | 64199200 | 64204399 | 47.77255 | 77.46047 | 86.92399 | 37.22911 | 0.199573 | 1.661361 | 0.864812 |
| 18 | 33690200 | 33692399 | 17.66703 | 30.82921 | 37.88368 | 16.67855 | 0.091629 | 0.724772 | 0.864818 |
| 1 | 85494800 | 85523399 | 729.8771 | 961.0734 | 1064.922 | 445.5083 | -0.06696 | 5.52751 | 0.864829 |
| 3 | 144045000 | 144046199 | 12.36113 | 22.30604 | 31.60612 | 13.74988 | -0.09208 | 0.363609 | 0.864835 |
| 11 | 103243000 | 103244399 | 21.64509 | 37.11362 | 45.82921 | 20.09943 | 0.080354 | 0.939457 | 0.864848 |
| 10 | 75230200 | 75231999 | 13.72453 | 24.31768 | 34.11417 | 15.08281 | -0.09378 | 0.494721 | 0.864857 |
| 5 | 7076200 | 7077599 | 25.66844 | 43.65412 | 43.72616 | 19.16447 | 0.379375 | 1.026173 | 0.864866 |
| 15 | 57103200 | 57104199 | 8.82571 | 16.84428 | 25.60763 | 10.19627 | -0.09348 | -0.02107 | 0.864885 |
| 12 | 113535800 | 113538599 | 37.24158 | 60.62926 | 74.1653 | 31.84069 | 0.080294 | 1.560076 | 0.864902 |
| 2 | 96802800 | 96804199 | 17.91396 | 30.50162 | 54.27034 | 23.78805 | -0.44608 | 0.99846 | 0.864904 |
| 3 | 39472600 | 39476999 | 44.31557 | 71.58523 | 85.96867 | 36.69376 | 0.096466 | 1.735454 | 0.864921 |
| X | 72549000 | 72553199 | 25.36851 | 42.8798 | 60.61245 | 26.52226 | -0.11639 | 1.095393 | 0.864921 |
| 2 | 173558400 | 173564999 | 58.61179 | 92.35538 | 154.5645 | 64.1369 | -0.41898 | 2.484836 | 0.864952 |
| 2 | 48377400 | 48381199 | 39.7705 | 64.3511 | 78.77842 | 33.6885 | 0.07151 | 1.681622 | 0.864994 |
| 1 | 95195600 | 95197199 | 12.97936 | 23.14207 | 33.03511 | 14.53432 | -0.11388 | 0.443083 | 0.86502 |
| 1 | 183485600 | 183487599 | 14.56958 | 26.09978 | 32.4096 | 14.20273 | 0.090032 | 0.42419 | 0.865033 |
| 3 | 9237200 | 9237599 | 9.149523 | 17.33508 | 25.99769 | 10.51291 | -0.09361 | 0.033411 | 0.865034 |
| 6 | 60168800 | 60174599 | 48.88051 | 78.48798 | 107.0945 | 45.33172 | -0.09774 | 1.931688 | 0.86504 |
| 1 | 146950000 | 146950399 | 6.086635 | 11.8691 | 16.96745 | 6.100449 | 0.132633 | -0.42452 | 0.865045 |
| 1 | 78240400 | 78241599 | 11.08667 | 20.1912 | 29.22279 | 12.56755 | -0.11153 | 0.296426 | 0.865047 |
| 3 | 55517200 | 55520199 | 27.65815 | 45.47326 | 63.84264 | 27.51467 | -0.10971 | 1.402194 | 0.865052 |
| 18 | 89984400 | 89985599 | 11.92034 | 22.12715 | 28.21884 | 11.83051 | 0.095458 | 0.192222 | 0.865066 |
| 1 | 14516000 | 14518199 | 30.99492 | 51.38526 | 62.93167 | 27.25147 | 0.089292 | 1.310218 | 0.865068 |
| X | 7938400 | 7941399 | 29.53484 | 49.80979 | 57.88003 | 25.30935 | 0.165329 | 1.129217 | 0.865077 |
| 5 | 32678600 | 32685199 | 73.30402 | 116.1661 | 135.0729 | 56.18913 | 0.112238 | 2.364176 | 0.865079 |
| 16 | 50923400 | 50924399 | 11.0808 | 20.30739 | 29.23689 | 12.47247 | -0.09614 | 0.260814 | 0.865112 |
| X | 118591200 | 118593399 | 34.66583 | 57.40333 | 53.57219 | 23.17962 | 0.480157 | 1.32829 | 0.865112 |
| 5 | 129719000 | 129721799 | 42.57637 | 68.45673 | 92.09585 | 39.05953 | -0.07497 | 1.875506 | 0.865154 |
| 14 | 7713600 | 7716999 | 36.22629 | 59.65105 | 70.65641 | 30.47086 | 0.135567 | 1.409107 | 0.865172 |
| 10 | 116722000 | 116724799 | 34.17572 | 55.53419 | 75.80105 | 32.51712 | -0.0794 | 1.583206 | 0.86518 |
| 9 | 7757200 | 7757999 | 16.24963 | 28.58505 | 40.24481 | 17.81898 | -0.10308 | 0.600703 | 0.865227 |
| 16 | 10820000 | 10823399 | 36.20974 | 58.5519 | 79.73774 | 34.07961 | -0.08273 | 1.690341 | 0.865281 |
| 2 | 61173600 | 61175599 | 15.72115 | 27.86339 | 34.36194 | 15.19148 | 0.091077 | 0.503131 | 0.865306 |
| 5 | 130787600 | 130790199 | 27.31884 | 45.5082 | 62.82089 | 27.29224 | -0.08248 | 1.244333 | 0.86531 |
| 19 | 38512400 | 38514399 | 32.03874 | 51.88652 | 86.25407 | 36.90942 | -0.36761 | 1.654577 | 0.86534 |
| 1 | 16905000 | 16905999 | 16.40889 | 29.32049 | 27.87286 | 12.24068 | 0.471467 | 0.445226 | 0.865356 |
| 2 | 24631200 | 24633199 | 9.972122 | 18.61302 | 27.37075 | 11.37405 | -0.09811 | 0.142394 | 0.865396 |
| 2 | 149542400 | 149545399 | 35.86747 | 58.93517 | 70.74614 | 30.48221 | 0.115107 | 1.430941 | 0.865406 |
| 11 | 109815400 | 109816199 | 10.43177 | 19.56442 | 25.22468 | 10.39759 | 0.099171 | 0.107676 | 0.865432 |
| 9 | 110732200 | 110738799 | 67.54804 | 106.7777 | 140.1832 | 58.2389 | -0.06558 | 2.417899 | 0.86544 |
| 4 | 35074000 | 35075599 | 10.88719 | 20.79713 | 21.68545 | 8.618398 | 0.448294 | -0.01875 | 0.865444 |
| 3 | 78934800 | 78940999 | 69.81049 | 110.201 | 144.9057 | 60.15859 | -0.06947 | 2.452116 | 0.865456 |
| 11 | 109073600 | 109074999 | 20.20473 | 34.50147 | 47.96398 | 21.0287 | -0.09085 | 0.97308 | 0.865458 |
| 13 | 47310200 | 47312999 | 18.66282 | 32.54201 | 39.81767 | 17.52159 | 0.097447 | 0.740832 | 0.865467 |

| | | | | | | | | | |
|---|---|---|---|---|---|---|---|---|---|
| 3 | 133038000 | 133040199 | 18.31704 | 31.8244 | 44.1798 | 19.39722 | -0.08432 | 0.79766 | 0.865499 |
| 1 | 90743600 | 90744199 | 6.518831 | 12.68452 | 17.445 | 6.42816 | 0.1555 | -0.33104 | 0.865506 |
| 16 | 68982400 | 68983399 | 10.29164 | 18.98373 | 27.79947 | 11.75518 | -0.11112 | 0.215209 | 0.865509 |
| 6 | 13532800 | 13533999 | 14.05891 | 25.04085 | 35.0771 | 15.44626 | -0.0875 | 0.463415 | 0.865511 |
| 4 | 112834800 | 112837199 | 68.46879 | 107.9464 | 127.0113 | 52.70737 | 0.089249 | 2.475553 | 0.865512 |
| 4 | 118822600 | 118826399 | 33.39364 | 54.39904 | 74.08315 | 31.81272 | -0.07346 | 1.544523 | 0.865531 |
| 5 | 59430200 | 59430799 | 11.61624 | 21.03939 | 30.22288 | 13.08754 | -0.10713 | 0.335558 | 0.865555 |
| 5 | 78332400 | 78339399 | 140.6329 | 213.4528 | 209.6934 | 86.82146 | 0.306695 | 3.229032 | 0.865568 |
| 4 | 9919000 | 9920399 | 16.87771 | 29.62266 | 36.61937 | 16.19612 | 0.083123 | 0.62815 | 0.865608 |
| 7 | 18140600 | 18143399 | 61.07299 | 97.10117 | 116.8619 | 48.77331 | 0.067218 | 2.254023 | 0.865611 |
| 11 | 98194200 | 98195799 | 19.15709 | 33.08698 | 45.80852 | 20.09625 | -0.08203 | 0.862439 | 0.865638 |
| 7 | 6950800 | 6952799 | 22.89249 | 38.71239 | 53.30152 | 23.32645 | -0.07842 | 1.076354 | 0.865656 |
| 4 | 115216600 | 115218399 | 19.31352 | 33.27966 | 46.36805 | 20.34027 | -0.09165 | 0.887831 | 0.865657 |
| X | 169286400 | 169291599 | 134.3977 | 198.0471 | 367.7282 | 151.4342 | -0.64941 | 3.782263 | 0.865664 |
| 1 | 182085600 | 182087399 | 16.44959 | 27.81833 | 56.74553 | 24.83399 | -0.64301 | 1.059621 | 0.865675 |
| 14 | 35610800 | 35616199 | 56.17595 | 89.57115 | 108.1335 | 45.30537 | 0.067141 | 2.154919 | 0.86568 |
| 2 | 101031400 | 101033599 | 26.42524 | 44.27061 | 61.6303 | 26.87121 | -0.09467 | 1.182447 | 0.865709 |
| 18 | 8251800 | 8252199 | 11.68169 | 21.50341 | 27.17809 | 11.5247 | 0.096845 | 0.236222 | 0.865716 |
| 17 | 58475800 | 58476399 | 9.982993 | 18.59355 | 27.35763 | 11.4098 | -0.10306 | 0.156369 | 0.86572 |
| 6 | 57422600 | 57424199 | 21.24949 | 36.60954 | 45.25997 | 19.85409 | 0.079889 | 0.89323 | 0.865751 |
| 17 | 74846800 | 74852199 | 50.49034 | 80.96037 | 110.0716 | 46.49359 | -0.09495 | 1.976767 | 0.865751 |
| 18 | 54807400 | 54808399 | 12.95089 | 23.25093 | 33.02158 | 14.44423 | -0.10084 | 0.399709 | 0.865751 |
| 10 | 32848000 | 32849399 | 17.89944 | 31.04758 | 43.19822 | 18.95358 | -0.08774 | 0.816112 | 0.865752 |
| 10 | 78650200 | 78650599 | 7.275864 | 14.12003 | 19.48444 | 7.329775 | 0.123034 | -0.24143 | 0.865784 |
| 6 | 132424400 | 132426599 | 47.67097 | 76.87313 | 76.70479 | 32.66718 | 0.360404 | 1.788239 | 0.865816 |
| 2 | 36510000 | 36520999 | 150.2277 | 228.4192 | 202.1509 | 83.74291 | 0.459276 | 3.196026 | 0.865825 |
| 8 | 36854200 | 36854799 | 6.666961 | 12.97933 | 18.49702 | 6.787975 | 0.10976 | -0.34001 | 0.86585 |
| 10 | 75336200 | 75337799 | 15.68653 | 27.65359 | 33.99 | 15.04314 | 0.092898 | 0.56005 | 0.865851 |
| 13 | 99181200 | 99184199 | 23.52448 | 39.9633 | 55.53192 | 24.33625 | -0.09146 | 1.041042 | 0.865911 |
| 17 | 12500400 | 12509199 | 167.7316 | 248.918 | 310.2565 | 127.7981 | -0.06526 | 3.666019 | 0.865926 |
| 14 | 29952200 | 29953799 | 19.59094 | 34.33041 | 41.24366 | 18.217 | 0.124142 | 0.676202 | 0.865937 |
| 18 | 89330800 | 89333999 | 67.32978 | 106.3325 | 126.4263 | 52.4986 | 0.076147 | 2.437001 | 0.865972 |
| 1 | 25267400 | 25268599 | 11.02546 | 20.25489 | 29.21961 | 12.42415 | -0.09577 | 0.245459 | 0.865995 |
| 6 | 135571400 | 135573599 | 17.46651 | 30.53064 | 42.48875 | 18.72705 | -0.08693 | 0.707399 | 0.866008 |
| 8 | 34933200 | 34935799 | 25.37209 | 42.59692 | 52.30949 | 22.83796 | 0.085624 | 1.144817 | 0.866029 |
| 9 | 5612200 | 5617599 | 79.55221 | 124.8355 | 148.8671 | 61.73961 | 0.06473 | 2.579957 | 0.866039 |
| 10 | 70672000 | 70675799 | 40.48335 | 65.79479 | 66.84103 | 28.67697 | 0.345847 | 1.587724 | 0.866048 |
| 1 | 14030600 | 14034999 | 63.22121 | 100.186 | 131.5861 | 54.73779 | -0.06331 | 2.354298 | 0.866048 |
| 4 | 133560400 | 133563399 | 23.36167 | 39.58001 | 48.51495 | 21.24378 | 0.088731 | 1.054313 | 0.866122 |
| 9 | 78610800 | 78615799 | 47.8729 | 77.16559 | 91.82537 | 39.0851 | 0.106102 | 1.806645 | 0.866129 |
| X | 48053800 | 48061999 | 174.7918 | 259.8062 | 296.0285 | 121.9571 | 0.066261 | 3.639662 | 0.866148 |
| 16 | 19629000 | 19630599 | 15.14333 | 27.3701 | 27.28893 | 11.84815 | 0.413766 | 0.365133 | 0.866149 |
| 1 | 119265200 | 119266199 | 10.9441 | 20.23947 | 29.3289 | 12.35838 | -0.09254 | 0.209813 | 0.866153 |
| 14 | 54317800 | 54319199 | 17.40416 | 30.26165 | 42.11821 | 18.50103 | -0.08775 | 0.780887 | 0.866164 |
| 17 | 73361200 | 73363999 | 29.65053 | 48.92875 | 67.30211 | 29.05083 | -0.07831 | 1.361437 | 0.866175 |
| 15 | 81173000 | 81174799 | 19.13412 | 33.22505 | 41.05871 | 18.02044 | 0.082809 | 0.806381 | 0.866218 |
| 13 | 11092800 | 11095999 | 30.17738 | 49.88846 | 61.62797 | 26.63482 | 0.076603 | 1.339617 | 0.866221 |

| | | | | | | | | | |
|---|---|---|---|---|---|---|---|---|---|
| 14 | 6395400 | 6402199 | 126.3189 | 191.1117 | 242.1846 | 100.1745 | -0.06423 | 3.292539 | 0.866221 |
| 12 | 103798400 | 103804399 | 64.91866 | 102.6444 | 134.666 | 55.95319 | -0.06433 | 2.409906 | 0.866241 |
| 1 | 174841200 | 174844799 | 69.68722 | 110.4782 | 99.12179 | 41.25013 | 0.48628 | 2.340643 | 0.866262 |
| 14 | 96174400 | 96175999 | 14.58378 | 26.07877 | 32.42069 | 14.22893 | 0.086934 | 0.435584 | 0.86627 |
| 1 | 79041000 | 79041599 | 8.424468 | 16.09185 | 20.36989 | 8.096512 | 0.17104 | -0.04661 | 0.866276 |
| 1 | 41409200 | 41410399 | 13.4912 | 24.46441 | 30.44006 | 13.16201 | 0.099699 | 0.339311 | 0.866278 |
| 3 | 103105200 | 103109999 | 46.31033 | 74.36383 | 99.90542 | 42.29112 | -0.07533 | 1.923178 | 0.866305 |
| 17 | 77320200 | 77321599 | 22.73154 | 38.62618 | 53.30648 | 23.36176 | -0.08147 | 1.034352 | 0.866308 |
| 15 | 63644000 | 63647199 | 25.98623 | 43.20876 | 59.90646 | 25.99432 | -0.0888 | 1.257678 | 0.866311 |
| X | 98227400 | 98230399 | 19.64369 | 34.09741 | 47.93748 | 21.03466 | -0.10254 | 0.824868 | 0.866322 |
| 13 | 119395000 | 119395799 | 11.8115 | 21.42158 | 30.73499 | 13.28714 | -0.10379 | 0.331293 | 0.866332 |
| 5 | 115005400 | 115009399 | 64.3229 | 101.7125 | 134.0458 | 55.69828 | -0.0707 | 2.407943 | 0.866363 |
| 1 | 127109200 | 127110999 | 29.07308 | 48.43555 | 59.58088 | 25.88635 | 0.083337 | 1.245523 | 0.866363 |
| Y | 40347600 | 40347999 | 6.214588 | 12.10331 | 16.38741 | 5.990219 | 0.187794 | -0.36982 | 0.866378 |
| 17 | 17829800 | 17831999 | 9.76691 | 17.90257 | 32.95157 | 14.05384 | -0.44778 | 0.259546 | 0.86639 |
| 15 | 28290000 | 28291599 | 16.07519 | 28.19881 | 39.23718 | 17.34597 | -0.08679 | 0.635939 | 0.866405 |
| 11 | 32126400 | 32130999 | 33.26796 | 54.22715 | 74.02159 | 31.79331 | -0.07618 | 1.53661 | 0.866408 |
| 5 | 85708000 | 85708399 | 10.21932 | 18.85725 | 27.74646 | 11.7269 | -0.11777 | 0.210285 | 0.866409 |
| 7 | 68274800 | 68275199 | 8.306236 | 15.9563 | 24.69174 | 9.659883 | -0.09553 | -0.08755 | 0.866421 |
| 4 | 33120200 | 33120999 | 10.58704 | 19.88949 | 25.76493 | 10.58968 | 0.095698 | 0.099488 | 0.866427 |
| 3 | 14657000 | 14660399 | 43.3168 | 70.35233 | 82.38628 | 35.32881 | 0.140399 | 1.616576 | 0.866427 |
| 19 | 13017000 | 13029599 | 389.8678 | 539.9711 | 569.9126 | 236.6678 | 0.076459 | 4.726258 | 0.866429 |
| 13 | 20832200 | 20843399 | 216.467 | 314.2501 | 385.6626 | 158.9932 | -0.07307 | 4.022118 | 0.866439 |
| 17 | 71589200 | 71590999 | 20.98406 | 35.74499 | 49.55085 | 21.72016 | -0.08736 | 0.999182 | 0.866447 |
| 9 | 6837000 | 6838799 | 15.30695 | 27.0879 | 38.0634 | 16.83304 | -0.09711 | 0.524088 | 0.866477 |
| 7 | 25195800 | 25197599 | 11.40714 | 20.31112 | 36.78086 | 16.20093 | -0.45597 | 0.451493 | 0.866478 |
| 3 | 34655400 | 34655999 | 10.87109 | 20.20376 | 29.37167 | 12.29369 | -0.09042 | 0.180222 | 0.86649 |
| 8 | 45289800 | 45292999 | 28.8586 | 48.12036 | 59.05215 | 25.67082 | 0.086725 | 1.234355 | 0.866493 |
| 5 | 130437400 | 130446599 | 115.2665 | 175.6219 | 223.3539 | 92.58343 | -0.061 | 3.14588 | 0.866496 |
| 6 | 142734200 | 142737199 | 23.72933 | 39.87616 | 54.91648 | 23.98745 | -0.07911 | 1.135074 | 0.866508 |
| 7 | 24030800 | 24033399 | 47.84089 | 76.66601 | 92.06934 | 38.85406 | 0.082781 | 1.986839 | 0.866521 |
| 9 | 95173400 | 95175799 | 13.25179 | 23.06611 | 45.29475 | 19.90964 | -0.58159 | 0.752205 | 0.866525 |
| 2 | 24013000 | 24015999 | 21.3947 | 37.3081 | 36.63892 | 16.12349 | 0.413427 | 0.751672 | 0.866528 |
| 1 | 184126000 | 184128999 | 28.87766 | 47.72365 | 65.47315 | 28.29005 | -0.07434 | 1.342439 | 0.866528 |
| 5 | 145457000 | 145460199 | 43.58619 | 70.21924 | 95.00397 | 40.40868 | -0.07961 | 1.816768 | 0.866531 |
| 6 | 38230800 | 38231999 | 10.86342 | 19.71457 | 29.64365 | 12.80523 | -0.17081 | 0.313189 | 0.866533 |
| 4 | 21186400 | 21188799 | 23.39897 | 39.03142 | 66.29225 | 28.82505 | -0.38052 | 1.22408 | 0.866554 |
| 2 | 173888400 | 173891799 | 37.96692 | 61.37707 | 83.07642 | 35.49445 | -0.07443 | 1.70167 | 0.866555 |
| 4 | 115136400 | 115140999 | 40.08884 | 64.80648 | 88.3547 | 37.74894 | -0.0849 | 1.707903 | 0.866592 |
| 8 | 55104000 | 55106599 | 40.01106 | 65.2665 | 63.80754 | 27.41359 | 0.404782 | 1.530161 | 0.866607 |
| 1 | 195001000 | 195005999 | 77.69803 | 121.8701 | 145.4822 | 60.33731 | 0.062719 | 2.585961 | 0.866608 |
| 17 | 36489800 | 36490599 | 9.764206 | 18.23734 | 26.84177 | 11.1414 | -0.0982 | 0.13368 | 0.866611 |
| 3 | 36302200 | 36303999 | 17.7378 | 31.11402 | 38.04848 | 16.81987 | 0.098505 | 0.642522 | 0.86664 |
| 5 | 33808400 | 33810399 | 12.92273 | 22.92756 | 32.69016 | 14.44258 | -0.11602 | 0.475531 | 0.866667 |
| 4 | 113723200 | 113729599 | 39.46697 | 65.10972 | 57.26171 | 24.71403 | 0.564974 | 1.38338 | 0.8667 |
| 4 | 70611400 | 70613799 | 24.08287 | 41.6229 | 48.48219 | 21.26963 | 0.166272 | 0.87911 | 0.866711 |
| 13 | 86042800 | 86044999 | 19.22076 | 33.10526 | 45.83543 | 20.10636 | -0.08282 | 0.888807 | 0.86673 |

| | | | | | | | | | |
|---|---|---|---|---|---|---|---|---|---|
| 4 | 15803000 | 15803799 | 9.377239 | 17.41483 | 26.39719 | 11.05961 | -0.15053 | 0.167281 | 0.866736 |
| 9 | 4327000 | 4328399 | 12.48317 | 22.97007 | 34.07095 | 14.45501 | -0.1333 | 0.260535 | 0.866736 |
| 11 | 22335600 | 22336999 | 19.19773 | 33.25259 | 40.63199 | 17.82686 | 0.098453 | 0.832077 | 0.866737 |
| 7 | 108464000 | 108466599 | 34.26344 | 55.93527 | 68.8081 | 29.57053 | 0.07505 | 1.511097 | 0.866773 |
| 7 | 12681600 | 12683999 | 22.22833 | 38.34886 | 36.92605 | 16.19895 | 0.439389 | 0.873887 | 0.866783 |
| 3 | 135203600 | 135204999 | 21.23438 | 36.01359 | 50.08151 | 21.93672 | -0.0925 | 1.041428 | 0.866804 |
| 7 | 144541400 | 144542199 | 10.79558 | 20.01687 | 25.18734 | 10.57569 | 0.11333 | 0.182276 | 0.866826 |
| X | 160962200 | 160963999 | 37.4147 | 60.57616 | 72.35616 | 30.95601 | 0.107808 | 1.664721 | 0.866845 |
| 18 | 73334600 | 73337199 | 41.17312 | 66.57974 | 81.02853 | 34.6326 | 0.079133 | 1.70024 | 0.866862 |
| 9 | 7477200 | 7479399 | 18.81393 | 32.89578 | 47.27826 | 20.84335 | -0.13331 | 0.715997 | 0.86688 |
| 12 | 105994400 | 105996399 | 20.34093 | 35.09444 | 43.14441 | 18.92689 | 0.088318 | 0.872481 | 0.86689 |
| 8 | 5825000 | 5826999 | 50.05287 | 80.17066 | 97.02015 | 40.91223 | 0.070834 | 2.005095 | 0.866901 |
| 5 | 89130600 | 89132199 | 18.89455 | 32.86387 | 45.75325 | 20.10193 | -0.08823 | 0.78837 | 0.866905 |
| 13 | 32485600 | 32490599 | 49.26436 | 78.5612 | 123.4123 | 51.71632 | -0.31191 | 2.148938 | 0.866926 |
| 14 | 111112000 | 111113399 | 13.05554 | 23.64805 | 33.69442 | 14.58947 | -0.09568 | 0.351999 | 0.866938 |
| 1 | 118174400 | 118177999 | 53.91047 | 86.13679 | 104.3414 | 43.84387 | 0.065492 | 2.088138 | 0.866956 |
| 9 | 70205600 | 70210199 | 49.2938 | 78.97053 | 105.5311 | 44.51465 | -0.07165 | 2.001164 | 0.867035 |
| 14 | 31644800 | 31648599 | 70.62777 | 111.0817 | 145.199 | 60.23601 | -0.065 | 2.529384 | 0.867094 |
| X | 79605800 | 79608999 | 40.05332 | 65.3153 | 77.43759 | 33.26572 | 0.127014 | 1.546973 | 0.867096 |
| 4 | 54632600 | 54633199 | 11.05918 | 20.4862 | 25.74994 | 10.82587 | 0.113863 | 0.193269 | 0.867108 |
| 12 | 92333800 | 92336399 | 26.16196 | 44.80139 | 43.07601 | 18.89002 | 0.439613 | 0.965369 | 0.867109 |
| 1 | 34464000 | 34472999 | 210.0458 | 306.5063 | 375.2473 | 154.6164 | -0.06236 | 3.943809 | 0.867133 |
| 3 | 85205400 | 85209399 | 26.38542 | 43.80689 | 60.37391 | 26.17009 | -0.08032 | 1.278168 | 0.867163 |
| 4 | 121323000 | 121334599 | 179.9439 | 268.5417 | 301.8314 | 124.4122 | 0.091267 | 3.569049 | 0.86718 |
| 2 | 5794600 | 5800199 | 69.29749 | 109.5955 | 131.6242 | 54.6963 | 0.063265 | 2.405978 | 0.86721 |
| 16 | 81213000 | 81214199 | 9.381326 | 17.69286 | 26.41972 | 10.7877 | -0.10018 | 0.065809 | 0.867217 |
| 5 | 128131200 | 128131799 | 7.422776 | 14.34625 | 22.63657 | 8.680118 | -0.09553 | -0.16699 | 0.867269 |
| 5 | 52369800 | 52372199 | 18.39155 | 31.94952 | 44.19873 | 19.40388 | -0.07923 | 0.801451 | 0.867271 |
| X | 121116200 | 121118199 | 23.91358 | 40.98649 | 49.55584 | 21.73311 | 0.110556 | 0.958055 | 0.867283 |
| 3 | 82015800 | 82016599 | 7.434257 | 14.32972 | 22.56306 | 8.741605 | -0.10643 | -0.1321 | 0.867288 |
| 9 | 100524000 | 100526799 | 20.70326 | 34.35361 | 68.20762 | 29.55425 | -0.60539 | 1.274045 | 0.867316 |
| 8 | 101628200 | 101629399 | 13.16191 | 23.74508 | 33.6014 | 14.62252 | -0.09064 | 0.374518 | 0.86735 |
| 13 | 78976800 | 78980199 | 30.28028 | 51.1725 | 58.63 | 25.65335 | 0.185627 | 1.111383 | 0.867351 |
| 6 | 110346000 | 110347199 | 10.33491 | 19.31586 | 28.40153 | 11.77584 | -0.09592 | 0.136587 | 0.867363 |
| 1 | 108936600 | 108937999 | 25.79427 | 43.22655 | 53.37368 | 23.27569 | 0.077864 | 1.168344 | 0.867374 |
| 2 | 31403400 | 31406399 | 42.1806 | 67.81607 | 91.53854 | 38.81958 | -0.07984 | 1.878863 | 0.867387 |
| 5 | 70446200 | 70446799 | 7.593362 | 14.74808 | 24.16093 | 9.036763 | -0.11542 | -0.24508 | 0.867388 |
| 12 | 113624400 | 113626999 | 27.5434 | 46.05984 | 56.75872 | 24.71872 | 0.080704 | 1.19811 | 0.86739 |
| 3 | 3389400 | 3392399 | 44.74533 | 71.6668 | 97.16181 | 40.96965 | -0.09276 | 2.001038 | 0.867433 |
| 5 | 20216400 | 20218399 | 22.42958 | 38.33018 | 47.41771 | 20.79049 | 0.076956 | 0.973896 | 0.867434 |
| 7 | 110603400 | 110610199 | 64.74139 | 102.3933 | 134.2643 | 55.79434 | -0.06323 | 2.402848 | 0.867445 |
| 15 | 33867000 | 33869999 | 23.77796 | 39.33698 | 68.22097 | 29.49866 | -0.4112 | 1.320747 | 0.867453 |
| 10 | 11509200 | 11509599 | 7.244496 | 14.0747 | 22.99033 | 8.578235 | -0.10838 | -0.26312 | 0.867455 |
| 17 | 55443600 | 55445199 | 17.52175 | 31.14553 | 36.84846 | 16.25399 | 0.152886 | 0.49901 | 0.867463 |
| 1 | 81556600 | 81557199 | 7.913469 | 15.13405 | 23.50363 | 9.313755 | -0.11724 | -0.05036 | 0.867465 |
| 4 | 153223800 | 153233999 | 150.9808 | 225.518 | 283.5232 | 116.8859 | -0.06909 | 3.546863 | 0.867495 |
| 17 | 34970800 | 34979599 | 270.2178 | 385.8758 | 417.2258 | 172.4108 | 0.086046 | 4.267945 | 0.867559 |

| | | | | | | | | | |
|---|---|---|---|---|---|---|---|---|---|
| 4 | 115228600 | 115231199 | 42.20889 | 68.09316 | 83.1833 | 35.4451 | 0.069747 | 1.769077 | 0.867572 |
| 5 | 47571400 | 47572799 | 15.74597 | 27.82742 | 34.46068 | 15.24387 | 0.083585 | 0.535125 | 0.867587 |
| 5 | 4763000 | 4766399 | 34.6808 | 57.13622 | 58.00759 | 25.02157 | 0.357673 | 1.388272 | 0.867613 |
| X | 141959800 | 141960599 | 14.66237 | 26.53899 | 25.90155 | 11.22555 | 0.446133 | 0.350887 | 0.867657 |
| 1 | 169771400 | 169774199 | 28.97801 | 48.12074 | 66.89696 | 29.00155 | -0.09289 | 1.287375 | 0.867664 |
| 14 | 41237200 | 41238999 | 32.3805 | 53.50375 | 65.38236 | 28.24763 | 0.092035 | 1.351741 | 0.867668 |
| 19 | 51726000 | 51728799 | 31.7903 | 52.32258 | 72.66382 | 31.33047 | -0.09366 | 1.402198 | 0.867669 |
| 4 | 144144800 | 144148999 | 49.02513 | 78.46411 | 106.0393 | 44.66449 | -0.08947 | 2.028908 | 0.86767 |
| 1 | 85420000 | 85431799 | 359.897 | 505.5399 | 542.2544 | 224.6881 | 0.07367 | 4.514134 | 0.86769 |
| 2 | 177527600 | 177530999 | 60.62613 | 96.69949 | 98.15243 | 41.14277 | 0.317445 | 2.14672 | 0.867702 |
| 1 | 50025800 | 50027799 | 18.30376 | 31.92817 | 44.64024 | 19.64848 | -0.09387 | 0.744517 | 0.867703 |
| 11 | 49276200 | 49278199 | 22.29888 | 38.56829 | 46.84342 | 20.55115 | 0.106514 | 0.865623 | 0.867715 |
| 6 | 104354200 | 104355999 | 20.26297 | 35.41399 | 34.98217 | 15.41676 | 0.405166 | 0.709942 | 0.867725 |
| 14 | 39319800 | 39320199 | 9.245654 | 17.61232 | 23.38739 | 9.349664 | 0.095095 | -0.01032 | 0.867747 |
| 8 | 119326000 | 119329599 | 36.63029 | 59.50578 | 73.1308 | 31.36503 | 0.071246 | 1.593163 | 0.867747 |
| 8 | 39655200 | 39657999 | 20.34153 | 35.16015 | 49.02031 | 21.50578 | -0.09189 | 0.86513 | 0.86775 |
| 1 | 46112800 | 46117399 | 40.22396 | 65.18601 | 91.03898 | 38.97618 | -0.11609 | 1.658729 | 0.867767 |
| 13 | 64693400 | 64694199 | 15.38533 | 26.95425 | 37.90957 | 16.741 | -0.10186 | 0.653514 | 0.867769 |
| 4 | 142003000 | 142006199 | 28.15371 | 46.45453 | 64.33388 | 27.76854 | -0.08811 | 1.355755 | 0.867779 |
| 2 | 16026000 | 16028399 | 30.9751 | 50.43998 | 70.20339 | 30.1425 | -0.1051 | 1.543259 | 0.867782 |
| 6 | 77878200 | 77881199 | 27.3675 | 46.24641 | 55.6865 | 24.36485 | 0.114128 | 1.098444 | 0.867787 |
| 6 | 94269800 | 94272399 | 18.17862 | 31.87203 | 39.14926 | 17.2995 | 0.092235 | 0.656051 | 0.867788 |
| 10 | 129067600 | 129071199 | 38.36693 | 62.51198 | 75.66996 | 32.49628 | 0.096202 | 1.551252 | 0.867803 |
| 16 | 17805800 | 17820199 | 300.49 | 426.6263 | 465.935 | 192.7265 | 0.063177 | 4.354213 | 0.867805 |
| 6 | 74113200 | 74114199 | 9.642224 | 18.39146 | 24.45757 | 9.750864 | 0.095973 | -0.01433 | 0.867861 |
| 17 | 41295800 | 41297599 | 25.36615 | 42.35937 | 58.22693 | 25.33567 | -0.07636 | 1.209096 | 0.867961 |
| 11 | 73995600 | 73999199 | 50.67517 | 81.1343 | 108.3989 | 45.68218 | -0.07237 | 2.0235 | 0.867972 |
| 18 | 54525600 | 54528199 | 20.96824 | 36.55099 | 44.03948 | 19.37574 | 0.119603 | 0.764529 | 0.867989 |
| 19 | 7365200 | 7374999 | 123.9677 | 188.1514 | 238.6354 | 98.80354 | -0.06161 | 3.227262 | 0.867996 |
| 8 | 46610600 | 46612999 | 27.87538 | 45.60418 | 76.64188 | 32.96281 | -0.37167 | 1.472545 | 0.868002 |
| X | 81807400 | 81811999 | 29.21676 | 48.88397 | 72.15489 | 31.43431 | -0.17892 | 1.214518 | 0.868017 |
| 9 | 120537800 | 120542999 | 62.08013 | 98.71164 | 118.7173 | 49.55227 | 0.068329 | 2.250714 | 0.86803 |
| 13 | 37838400 | 37838999 | 6.690348 | 13.00262 | 21.19855 | 7.887473 | -0.10071 | -0.28125 | 0.86803 |
| 6 | 101865800 | 101866799 | 10.22397 | 19.07402 | 27.85164 | 11.58323 | -0.08928 | 0.147038 | 0.868035 |
| 1 | 17132800 | 17133399 | 7.541709 | 14.6066 | 23.33413 | 8.860673 | -0.10035 | -0.19848 | 0.868045 |
| 5 | 127461800 | 127464199 | 27.74074 | 46.63525 | 56.77359 | 24.80155 | 0.098118 | 1.14532 | 0.868057 |
| 5 | 110202200 | 110206399 | 59.98005 | 95.5596 | 114.9976 | 48.12754 | 0.070452 | 2.186254 | 0.868061 |
| 7 | 22460400 | 22461399 | 10.44885 | 20.12404 | 26.10242 | 10.13652 | 0.166656 | -0.08696 | 0.868075 |
| 11 | 58852200 | 58854799 | 38.07338 | 61.40947 | 83.71228 | 35.66144 | -0.08836 | 1.770658 | 0.86808 |
| 8 | 48570000 | 48574999 | 66.81717 | 105.8034 | 139.8337 | 58.14409 | -0.07323 | 2.376414 | 0.868081 |
| 8 | 81690600 | 81693599 | 29.08863 | 48.51382 | 59.58453 | 25.90559 | 0.085551 | 1.234443 | 0.868126 |
| 5 | 58752600 | 58753399 | 9.751484 | 18.25314 | 26.85824 | 11.10735 | -0.09363 | 0.119483 | 0.868129 |
| 7 | 89598400 | 89601599 | 34.47684 | 56.20894 | 76.85455 | 33.01253 | -0.07845 | 1.535312 | 0.868195 |
| 2 | 110911600 | 110911999 | 5.220636 | 10.19784 | 17.64701 | 6.172636 | -0.10456 | -0.53241 | 0.86822 |
| 12 | 17780400 | 17782799 | 22.01634 | 37.19294 | 51.69304 | 22.61556 | -0.09201 | 1.078858 | 0.868224 |
| 3 | 98806600 | 98808599 | 19.88244 | 34.92048 | 41.10948 | 18.19263 | 0.153181 | 0.640888 | 0.868232 |
| 17 | 54309600 | 54311199 | 18.32755 | 31.92221 | 39.44771 | 17.34677 | 0.083289 | 0.756901 | 0.868232 |

| | | | | | | | | | |
|---|---|---|---|---|---|---|---|---|---|
| 4 | 61493200 | 61497799 | 56.71331 | 90.69934 | 108.3444 | 45.58587 | 0.087053 | 2.064029 | 0.868233 |
| 17 | 66057400 | 66064399 | 102.9529 | 158.0789 | 202.2363 | 83.96566 | -0.06079 | 2.985869 | 0.868238 |
| 6 | 130149400 | 130152599 | 40.25066 | 65.37209 | 79.00174 | 33.8773 | 0.095084 | 1.607783 | 0.868243 |
| 1 | 105461000 | 105468599 | 128.3364 | 195.8335 | 224.0258 | 92.89665 | 0.093441 | 3.119804 | 0.868264 |
| 11 | 32717000 | 32726999 | 121.709 | 183.5404 | 278.3374 | 115.0104 | -0.32709 | 3.358443 | 0.868264 |
| 6 | 60865200 | 60870799 | 81.4095 | 127.2611 | 151.5287 | 62.86689 | 0.062628 | 2.647858 | 0.868272 |
| 12 | 4591000 | 4592599 | 16.88527 | 29.49961 | 40.93272 | 18.03421 | -0.08276 | 0.708298 | 0.868281 |
| 7 | 44314200 | 44314999 | 11.27981 | 20.70279 | 29.77013 | 12.67436 | -0.09223 | 0.256148 | 0.868308 |
| 13 | 43428600 | 43431599 | 29.03293 | 47.72422 | 65.83478 | 28.37088 | -0.08464 | 1.406798 | 0.868313 |
| 11 | 4643200 | 4645999 | 39.4399 | 63.74602 | 77.86338 | 33.25892 | 0.07307 | 1.708055 | 0.868323 |
| 11 | 74191400 | 74193599 | 29.84171 | 49.11224 | 59.76858 | 25.76971 | 0.096412 | 1.390954 | 0.868329 |
| 11 | 109194800 | 109195199 | 5.221885 | 10.19644 | 17.60644 | 6.198558 | -0.10974 | -0.50937 | 0.86833 |
| 1 | 126827000 | 126828399 | 13.67749 | 24.88231 | 31.09092 | 13.38933 | 0.09772 | 0.326575 | 0.868348 |
| 15 | 95721200 | 95723999 | 23.2638 | 39.39157 | 54.44744 | 23.83736 | -0.08405 | 1.065114 | 0.868352 |
| 13 | 66689600 | 66700999 | 296.7041 | 424.3866 | 395.8034 | 163.5159 | 0.30159 | 4.245856 | 0.868356 |
| 6 | 140069200 | 140070999 | 27.47522 | 46.01935 | 56.70896 | 24.72198 | 0.080715 | 1.182722 | 0.868358 |
| 9 | 113497200 | 113499599 | 27.47522 | 46.01935 | 56.70896 | 24.72198 | 0.080715 | 1.182722 | 0.868358 |
| X | 77674400 | 77675199 | 15.77836 | 27.95947 | 34.04165 | 15.05045 | 0.109713 | 0.50895 | 0.868406 |
| 6 | 10531400 | 10532399 | 10.90107 | 20.55505 | 31.52809 | 12.87779 | -0.1391 | 0.093746 | 0.868432 |
| 9 | 14441400 | 14445399 | 41.7123 | 67.22686 | 81.5686 | 34.71365 | 0.07794 | 1.795674 | 0.868433 |
| 11 | 26848000 | 26851199 | 26.50945 | 45.27012 | 53.44781 | 23.43307 | 0.143892 | 1.003567 | 0.868478 |
| 17 | 37048600 | 37049399 | 8.972206 | 16.8276 | 25.35862 | 10.45222 | -0.12379 | 0.096054 | 0.868481 |
| 11 | 116964400 | 116966799 | 22.36084 | 37.99088 | 52.2258 | 22.88789 | -0.07591 | 1.032399 | 0.868509 |
| 8 | 97823400 | 97823799 | 5.20585 | 10.17224 | 17.68771 | 6.147745 | -0.10258 | -0.55263 | 0.86854 |
| 1 | 114110200 | 114111799 | 35.64637 | 58.3822 | 54.5738 | 23.48183 | 0.473192 | 1.460352 | 0.868583 |
| 15 | 27354800 | 27355999 | 12.94533 | 23.36801 | 29.04111 | 12.62875 | 0.096418 | 0.360324 | 0.868597 |
| 1 | 185809000 | 185816199 | 79.5738 | 125.0882 | 168.0121 | 69.69545 | -0.10477 | 2.54835 | 0.868601 |
| 10 | 70036400 | 70038999 | 19.89758 | 34.59199 | 49.01193 | 21.52721 | -0.11359 | 0.803867 | 0.868617 |
| 5 | 138602200 | 138611199 | 187.8734 | 280.5456 | 307.7522 | 126.8677 | 0.127106 | 3.560175 | 0.868633 |
| 3 | 93623600 | 93625599 | 34.5512 | 56.87737 | 57.37917 | 24.74216 | 0.366383 | 1.397872 | 0.868646 |
| 1 | 146037600 | 146039399 | 15.95369 | 27.95605 | 38.81182 | 17.14258 | -0.08344 | 0.652661 | 0.868666 |
| 11 | 27073600 | 27074199 | 15.95369 | 27.95605 | 38.81182 | 17.14258 | -0.08344 | 0.652661 | 0.868666 |
| 5 | 93529400 | 93532399 | 45.57849 | 73.5103 | 74.29689 | 31.64806 | 0.342159 | 1.777672 | 0.868692 |
| 13 | 31680800 | 31683399 | 23.98811 | 40.89209 | 50.11049 | 21.96677 | 0.089922 | 1.003692 | 0.868718 |
| 18 | 84773600 | 84775199 | 14.10806 | 25.25529 | 31.46525 | 13.79706 | 0.084782 | 0.42412 | 0.868754 |
| 10 | 124199200 | 124200199 | 8.613892 | 16.27565 | 24.87887 | 10.11887 | -0.1286 | 0.04172 | 0.868764 |
| 6 | 129004200 | 129004999 | 7.572151 | 14.66456 | 23.31577 | 8.856524 | -0.09366 | -0.19558 | 0.868783 |
| 12 | 24076000 | 24187399 | 3122.186 | 3655.639 | 3286.598 | 1394.612 | 0.06318 | 7.408371 | 0.868795 |
| 7 | 53492800 | 53494599 | 18.21369 | 31.80256 | 39.37427 | 17.34145 | 0.080937 | 0.721969 | 0.868798 |
| X | 70710200 | 70711799 | 18.21369 | 31.80256 | 39.37427 | 17.34145 | 0.080937 | 0.721969 | 0.868798 |
| 10 | 14518400 | 14524399 | 125.1901 | 189.5224 | 240.5056 | 99.49958 | -0.06547 | 3.280244 | 0.868812 |
| 17 | 92541000 | 92541999 | 13.29564 | 24.11528 | 34.74651 | 15.01979 | -0.11038 | 0.346311 | 0.868813 |
| 5 | 67648200 | 67650599 | 21.85473 | 37.6592 | 46.30734 | 20.31365 | 0.087626 | 0.894549 | 0.868836 |
| 12 | 91448000 | 91452199 | 46.37305 | 74.48538 | 99.7931 | 42.26154 | -0.07087 | 1.915753 | 0.868839 |
| 2 | 86609800 | 86611799 | 15.77682 | 28.33735 | 34.44953 | 15.06302 | 0.122665 | 0.420499 | 0.868873 |
| 10 | 5207200 | 5208399 | 18.21678 | 31.97514 | 39.10668 | 17.29724 | 0.098435 | 0.638637 | 0.868909 |
| 3 | 73781800 | 73783799 | 21.35897 | 37.3389 | 43.95399 | 19.3748 | 0.15336 | 0.726295 | 0.868915 |

| | | | | | | | | | |
|---|---|---|---|---|---|---|---|---|---|
| 16 | 48360200 | 48363199 | 40.28297 | 64.80296 | 88.2523 | 37.45764 | -0.09161 | 1.854161 | 0.868947 |
| 15 | 99732000 | 99733399 | 8.700751 | 16.36258 | 24.90804 | 10.2197 | -0.13259 | 0.076695 | 0.868974 |
| 4 | 122651400 | 122667599 | 207.2493 | 304.2975 | 378.8022 | 156.0129 | -0.07788 | 3.84895 | 0.868979 |
| 1 | 61608000 | 61612599 | 34.14083 | 56.12421 | 68.64747 | 29.5838 | 0.088429 | 1.421429 | 0.869 |
| 9 | 116908400 | 116909999 | 17.85726 | 31.09978 | 37.98662 | 16.70316 | 0.100075 | 0.754482 | 0.869007 |
| 3 | 158883600 | 158885999 | 23.57385 | 40.7532 | 48.48069 | 21.26913 | 0.136127 | 0.876659 | 0.869024 |
| 6 | 132508400 | 132509399 | 17.64368 | 30.98535 | 38.03017 | 16.8285 | 0.093286 | 0.624229 | 0.869038 |
| 10 | 66903800 | 66908199 | 49.6304 | 79.45913 | 105.8182 | 44.59251 | -0.06776 | 2.02154 | 0.869066 |
| 5 | 95617600 | 95621199 | 39.84702 | 65.43095 | 60.52736 | 26.072 | 0.489667 | 1.443237 | 0.86907 |
| 12 | 16620600 | 16621799 | 15.47234 | 27.40249 | 33.96496 | 15.01816 | 0.083566 | 0.510182 | 0.869074 |
| X | 37070000 | 37073999 | 33.02288 | 54.52238 | 66.39269 | 28.6686 | 0.097074 | 1.365555 | 0.869076 |
| 11 | 113961000 | 113965799 | 34.56929 | 56.18316 | 76.2633 | 32.71947 | -0.07117 | 1.58275 | 0.86909 |
| 5 | 115243200 | 115246999 | 37.01405 | 59.84113 | 81.04153 | 34.62837 | -0.07506 | 1.697587 | 0.86909 |
| 8 | 93998600 | 94000599 | 16.34032 | 28.75557 | 35.49071 | 15.71283 | 0.085969 | 0.585749 | 0.869096 |
| 4 | 156561400 | 156592799 | 584.2761 | 786.4878 | 904.0442 | 377.0954 | -0.08812 | 5.175262 | 0.869108 |
| 14 | 35543400 | 35544199 | 12.43906 | 22.31611 | 31.67728 | 13.86455 | -0.10025 | 0.401147 | 0.86912 |
| 12 | 95973000 | 95975199 | 30.11863 | 48.74833 | 86.36835 | 36.94235 | -0.46006 | 1.662725 | 0.869127 |
| 1 | 190530200 | 190531999 | 16.12983 | 28.14516 | 39.51325 | 17.39535 | -0.09959 | 0.720939 | 0.869137 |
| 6 | 70658000 | 70658999 | 14.31642 | 25.65742 | 31.98107 | 14.0098 | 0.085121 | 0.420269 | 0.869174 |
| 6 | 129984800 | 129987999 | 33.06957 | 54.7475 | 66.06772 | 28.58165 | 0.110303 | 1.332667 | 0.869178 |
| 8 | 10240000 | 10240999 | 7.791146 | 14.97757 | 23.59681 | 9.212498 | -0.1173 | -0.09828 | 0.869207 |
| 6 | 11421200 | 11421799 | 10.30535 | 18.96581 | 27.80613 | 11.79446 | -0.11599 | 0.228413 | 0.869221 |
| 14 | 87321400 | 87326399 | 42.945 | 69.25676 | 84.07852 | 35.81061 | 0.078128 | 1.779046 | 0.869301 |
| 13 | 74564000 | 74567999 | 66.72981 | 105.4109 | 137.9034 | 57.27158 | -0.06131 | 2.433305 | 0.869313 |
| 7 | 58628000 | 58636999 | 107.817 | 166.4194 | 192.1672 | 79.82248 | 0.092706 | 2.887055 | 0.869314 |
| 12 | 64911600 | 64913599 | 15.13609 | 26.93879 | 33.44242 | 14.73595 | 0.085105 | 0.471545 | 0.86932 |
| 10 | 17770800 | 17774399 | 57.86851 | 92.67832 | 108.3786 | 45.69386 | 0.119982 | 2.024237 | 0.869339 |
| X | 72333400 | 72339799 | 53.11917 | 84.88959 | 114.2016 | 48.00078 | -0.08524 | 2.086378 | 0.86934 |
| 4 | 4969800 | 4970599 | 9.480865 | 17.94001 | 23.26736 | 9.434173 | 0.110597 | 0.035638 | 0.869359 |
| 5 | 11982800 | 11983199 | 6.224148 | 12.11772 | 20.18516 | 7.40519 | -0.11343 | -0.34351 | 0.869372 |
| 13 | 19249200 | 19251999 | 41.60666 | 66.97871 | 89.94358 | 38.21379 | -0.07017 | 1.835668 | 0.869378 |
| 15 | 103826800 | 103836999 | 132.7871 | 201.6659 | 233.4638 | 96.68083 | 0.070722 | 3.216591 | 0.869412 |
| X | 5217000 | 5233399 | 242.3955 | 356.6158 | 300.6336 | 123.8029 | 0.486924 | 3.82004 | 0.869417 |
| 5 | 122824600 | 122826199 | 17.20266 | 30.11589 | 37.25307 | 16.44083 | 0.082271 | 0.672232 | 0.869465 |
| 15 | 10755800 | 10756799 | 13.54648 | 24.28201 | 34.08525 | 14.92927 | -0.08519 | 0.418119 | 0.869488 |
| 2 | 36485000 | 36491599 | 160.7116 | 239.9538 | 275.0727 | 113.393 | 0.063449 | 3.555924 | 0.869498 |
| 1 | 95755000 | 95756599 | 14.46585 | 25.70244 | 36.00759 | 15.88888 | -0.08991 | 0.481228 | 0.869519 |
| 14 | 120222800 | 120226599 | 25.0054 | 41.99166 | 57.72788 | 25.20752 | -0.07666 | 1.147581 | 0.869546 |
| 17 | 31271000 | 31274199 | 31.49707 | 52.44598 | 63.09015 | 27.40184 | 0.115467 | 1.262567 | 0.869567 |
| 6 | 7612600 | 7614999 | 49.77793 | 79.61631 | 95.26484 | 40.08867 | 0.084845 | 2.049056 | 0.869569 |
| 5 | 104944800 | 105008999 | 1572.106 | 1946.726 | 2026.429 | 853.1899 | -0.06597 | 6.523495 | 0.869578 |
| 12 | 19750600 | 19944199 | 7556.319 | 8213.285 | 6848.547 | 2942.24 | 0.062464 | 8.581371 | 0.869607 |
| Y | 21025800 | 21028199 | 11.22761 | 19.8951 | 31.4306 | 13.89611 | -0.264 | 0.471056 | 0.869639 |
| 18 | 45744600 | 45747399 | 24.04352 | 40.39102 | 55.47576 | 24.22936 | -0.07527 | 1.139132 | 0.869652 |
| 10 | 21524000 | 21531999 | 93.4183 | 144.8777 | 171.5734 | 71.25995 | 0.061524 | 2.793182 | 0.86966 |
| 10 | 14931400 | 14934399 | 15.11889 | 26.45736 | 46.20955 | 20.3867 | -0.41306 | 0.682935 | 0.86967 |
| 1 | 75979200 | 75984599 | 76.79449 | 120.638 | 144.1878 | 59.80733 | 0.062657 | 2.555058 | 0.869678 |

| | | | | | | | | | |
|---|---|---|---|---|---|---|---|---|---|
| 8 | 15673400 | 15675199 | 38.20305 | 61.92681 | 75.69193 | 32.41181 | 0.076235 | 1.642737 | 0.869678 |
| 18 | 24761400 | 24763599 | 16.61222 | 29.30483 | 35.8791 | 15.87711 | 0.098737 | 0.560701 | 0.869685 |
| 12 | 80002000 | 80003599 | 13.30473 | 23.82543 | 33.52914 | 14.69778 | -0.08979 | 0.419717 | 0.869688 |
| 2 | 89011200 | 89015199 | 43.72637 | 70.41822 | 94.95013 | 40.36347 | -0.07541 | 1.828373 | 0.869691 |
| 18 | 72926600 | 72929199 | 27.06823 | 45.86302 | 55.11368 | 24.13376 | 0.117178 | 1.074683 | 0.869698 |
| 7 | 136499200 | 136503199 | 42.80882 | 68.92408 | 92.49463 | 39.30601 | -0.06895 | 1.834947 | 0.869723 |
| 18 | 64943800 | 64945399 | 13.88574 | 24.74424 | 34.57478 | 15.21892 | -0.0835 | 0.458191 | 0.86974 |
| 14 | 115616600 | 115618399 | 15.83912 | 28.45074 | 34.39747 | 15.0396 | 0.130478 | 0.419071 | 0.869745 |
| 2 | 46728000 | 46731799 | 85.17296 | 132.5536 | 156.6221 | 65.01618 | 0.068723 | 2.727106 | 0.869747 |
| 10 | 117159800 | 117161599 | 20.6465 | 35.6877 | 44.20354 | 19.39261 | 0.078206 | 0.86096 | 0.869811 |
| 16 | 93268400 | 93273999 | 62.46955 | 99.53269 | 118.4126 | 49.56148 | 0.087077 | 2.187394 | 0.869814 |
| 2 | 39308000 | 39311399 | 46.96734 | 75.36327 | 100.9242 | 42.67451 | -0.07198 | 1.947143 | 0.869836 |
| 16 | 35412200 | 35414599 | 23.76848 | 40.00818 | 48.5559 | 21.21982 | 0.102715 | 1.113136 | 0.869836 |
| 3 | 49680800 | 49682599 | 14.74624 | 25.84573 | 45.10255 | 19.92469 | -0.4119 | 0.652499 | 0.869845 |
| 3 | 39243400 | 39244599 | 10.11324 | 18.36326 | 33.88629 | 14.62786 | -0.46313 | 0.318872 | 0.869855 |
| 6 | 57293600 | 57295799 | 28.60884 | 48.05044 | 48.57515 | 21.21297 | 0.365331 | 1.148287 | 0.869886 |
| 4 | 12712800 | 12713599 | 10.99541 | 20.28641 | 25.22286 | 10.6624 | 0.123619 | 0.212635 | 0.86992 |
| 9 | 88968600 | 88969799 | 15.58972 | 27.50731 | 34.0313 | 15.05874 | 0.084137 | 0.551057 | 0.869927 |
| 1 | 7341800 | 7342599 | 20.49615 | 35.22756 | 43.24468 | 18.96867 | 0.089473 | 0.910255 | 0.869938 |
| 2 | 118475800 | 118477199 | 18.14319 | 31.77457 | 39.19104 | 17.3017 | 0.086305 | 0.673214 | 0.869945 |
| 16 | 8639400 | 8641399 | 18.14319 | 31.77457 | 39.19104 | 17.3017 | 0.086305 | 0.673214 | 0.869945 |
| 16 | 90228400 | 90231199 | 35.88835 | 58.12342 | 78.78191 | 33.71303 | -0.07378 | 1.656664 | 0.869952 |
| 11 | 78816000 | 78817399 | 13.84427 | 24.67823 | 30.13868 | 13.26269 | 0.110114 | 0.447519 | 0.869958 |
| 7 | 29448800 | 29452199 | 41.53018 | 67.07782 | 91.58424 | 39.08462 | -0.08842 | 1.739397 | 0.869982 |
| 4 | 43488200 | 43490999 | 22.29176 | 36.96066 | 66.58342 | 28.83497 | -0.46561 | 1.287782 | 0.870014 |
| 8 | 4023000 | 4023799 | 11.86636 | 21.7923 | 31.29499 | 13.31276 | -0.08977 | 0.259195 | 0.87003 |
| 1 | 49872200 | 49873799 | 32.95669 | 53.9352 | 66.54074 | 28.6219 | 0.073395 | 1.471876 | 0.870032 |
| 6 | 43642400 | 43643399 | 11.32178 | 20.33151 | 37.82921 | 16.54351 | -0.48801 | 0.396764 | 0.870037 |
| 8 | 69723400 | 69772999 | 867.7895 | 1131.125 | 1264.046 | 529.3894 | -0.09383 | 5.69334 | 0.870052 |
| 7 | 7273800 | 7276799 | 52.00637 | 83.03908 | 110.0105 | 46.18238 | -0.0645 | 2.108727 | 0.870056 |
| 13 | 91212200 | 91215599 | 28.1907 | 47.42968 | 57.30583 | 25.04015 | 0.10908 | 1.142243 | 0.870056 |
| 7 | 6463200 | 6464399 | 23.27219 | 38.9807 | 54.33463 | 23.68736 | -0.09636 | 1.167508 | 0.87009 |
| 8 | 27367800 | 27369999 | 13.39463 | 23.65593 | 33.73309 | 14.92404 | -0.11905 | 0.527201 | 0.870097 |
| 1 | 119905400 | 119908799 | 31.15238 | 51.10841 | 69.98891 | 30.13619 | -0.07527 | 1.438448 | 0.870112 |
| 1 | 173811400 | 173811999 | 6.975401 | 13.50361 | 21.58963 | 8.21578 | -0.10365 | -0.20064 | 0.870119 |
| 14 | 11395600 | 11397399 | 16.66609 | 29.6661 | 35.92785 | 15.8292 | 0.120693 | 0.485398 | 0.87013 |
| 16 | 16452400 | 16459199 | 107.3341 | 164.9995 | 193.8491 | 80.49633 | 0.063306 | 2.96369 | 0.870147 |
| 12 | 91412400 | 91412799 | 4.911472 | 9.590919 | 17.06815 | 6.002536 | -0.14982 | -0.51617 | 0.870149 |
| 10 | 63199000 | 63201599 | 24.35269 | 41.14239 | 50.67037 | 22.16856 | 0.081725 | 1.084368 | 0.870154 |
| 3 | 46414600 | 46420199 | 127.1522 | 192.8777 | 224.6027 | 92.97808 | 0.060866 | 3.238521 | 0.870167 |
| 5 | 102361200 | 102362799 | 23.18773 | 39.38378 | 48.49666 | 21.2519 | 0.08225 | 1.032064 | 0.870192 |
| 11 | 6648800 | 6652399 | 34.02949 | 55.32857 | 75.20948 | 32.27381 | -0.07276 | 1.573162 | 0.870198 |
| 12 | 109756600 | 109758799 | 21.79234 | 37.21333 | 45.8586 | 20.10568 | 0.082401 | 0.976826 | 0.870225 |
| 12 | 103521200 | 103522399 | 17.92444 | 31.39889 | 38.64262 | 17.063 | 0.089405 | 0.666528 | 0.870236 |
| 19 | 58581400 | 58585799 | 93.60512 | 144.6002 | 185.6951 | 77.13377 | -0.05957 | 2.866083 | 0.870247 |
| 13 | 71100000 | 71104399 | 49.8858 | 79.99748 | 97.03072 | 40.99024 | 0.06957 | 1.968279 | 0.870258 |
| 8 | 16507800 | 16512399 | 40.86426 | 66.27987 | 80.17899 | 34.3482 | 0.091836 | 1.635278 | 0.870286 |

| | | | | | | | | | |
|---|---|---|---|---|---|---|---|---|---|
| 3 | 5279800 | 5281599 | 16.23345 | 28.5054 | 39.69439 | 17.56318 | -0.08795 | 0.621053 | 0.870291 |
| 6 | 45612200 | 45612599 | 7.441908 | 14.31867 | 22.53727 | 8.776861 | -0.1128 | -0.11501 | 0.870297 |
| 15 | 18240800 | 18241599 | 12.81431 | 22.93494 | 32.54135 | 14.2712 | -0.10196 | 0.417398 | 0.87032 |
| 9 | 93205400 | 93210199 | 36.4655 | 59.589 | 58.95002 | 25.34311 | 0.389602 | 1.495564 | 0.870339 |
| 9 | 18983200 | 18986199 | 25.0087 | 42.3298 | 59.49223 | 26.04295 | -0.10805 | 1.080974 | 0.870339 |
| 7 | 118148400 | 118154799 | 48.87944 | 77.99172 | 126.1645 | 52.91139 | -0.3533 | 2.129835 | 0.870343 |
| 18 | 30147200 | 30149799 | 50.3066 | 80.35023 | 106.8939 | 44.89395 | -0.07002 | 2.095486 | 0.870353 |
| 6 | 129771200 | 129772399 | 14.5991 | 26.05627 | 37.07034 | 16.29585 | -0.10836 | 0.455355 | 0.870373 |
| 15 | 98683400 | 98683999 | 9.968821 | 18.87618 | 24.84847 | 10.06077 | 0.091049 | 0.039931 | 0.870375 |
| 1 | 112749800 | 112751599 | 16.02139 | 28.45821 | 34.94088 | 15.42186 | 0.099328 | 0.493136 | 0.870383 |
| 18 | 37079000 | 37091599 | 238.173 | 344.9435 | 383.4238 | 158.1156 | 0.066366 | 4.05976 | 0.870398 |
| X | 107383200 | 107384799 | 19.46501 | 34.53781 | 30.06506 | 13.28186 | 0.592005 | 0.512939 | 0.870429 |
| 18 | 4634400 | 4636599 | 16.22619 | 28.26115 | 40.03863 | 17.60624 | -0.11304 | 0.746703 | 0.870472 |
| 3 | 152347600 | 152350199 | 12.16407 | 21.66051 | 38.25879 | 16.85084 | -0.4204 | 0.457669 | 0.870495 |
| 10 | 94101600 | 94102399 | 7.778822 | 15.00811 | 23.68115 | 9.149831 | -0.10559 | -0.13239 | 0.870513 |
| 4 | 107441600 | 107442199 | 12.86056 | 23.21918 | 28.64532 | 12.45383 | 0.107374 | 0.359467 | 0.87052 |
| 3 | 19654800 | 19655399 | 12.59793 | 22.08882 | 41.54913 | 18.36084 | -0.51904 | 0.631829 | 0.870534 |
| 6 | 90337000 | 90340599 | 26.21748 | 43.42834 | 60.32376 | 26.10698 | -0.09187 | 1.303512 | 0.870537 |
| 9 | 44201400 | 44203399 | 14.63702 | 25.88111 | 36.04972 | 15.94513 | -0.08491 | 0.525133 | 0.870538 |
| 13 | 4119400 | 4120999 | 22.19403 | 38.63124 | 37.68452 | 16.56457 | 0.422818 | 0.782285 | 0.870571 |
| 12 | 84036200 | 84037999 | 16.56437 | 29.11417 | 36.06114 | 15.96844 | 0.080304 | 0.602771 | 0.870577 |
| 14 | 24466000 | 24467999 | 26.81351 | 44.9384 | 55.57824 | 24.23837 | 0.075497 | 1.173213 | 0.87058 |
| 5 | 59349600 | 59351199 | 16.82499 | 29.69712 | 36.4006 | 16.10604 | 0.097475 | 0.557427 | 0.870585 |
| 4 | 23113400 | 23114999 | 16.13994 | 28.3455 | 34.72395 | 15.36593 | 0.096335 | 0.612423 | 0.870585 |
| 2 | 49125600 | 49127199 | 19.27916 | 33.83185 | 40.66848 | 17.98442 | 0.123283 | 0.651468 | 0.870622 |
| 10 | 42754000 | 42756399 | 20.75368 | 35.60463 | 43.74808 | 19.18755 | 0.087588 | 0.927157 | 0.87063 |
| 5 | 51171800 | 51173399 | 14.90135 | 27.15869 | 32.56737 | 13.99293 | 0.158407 | 0.324263 | 0.870632 |
| 12 | 52782000 | 52782999 | 9.463752 | 17.84241 | 26.43902 | 10.80163 | -0.08941 | 0.071049 | 0.870634 |
| 5 | 97460400 | 97462399 | 11.62899 | 22.08373 | 22.16005 | 8.917371 | 0.487865 | 0.030788 | 0.870673 |
| X | 77878800 | 77885399 | 70.95894 | 112.8625 | 129.3828 | 54.02295 | 0.137823 | 2.254836 | 0.870682 |
| 8 | 26751200 | 26753999 | 24.55612 | 41.2283 | 56.57492 | 24.70071 | -0.07398 | 1.148314 | 0.870698 |
| 11 | 32867600 | 32869399 | 13.73793 | 24.4087 | 34.10208 | 15.04826 | -0.08574 | 0.47512 | 0.870698 |
| 9 | 56909000 | 56910399 | 12.8316 | 23.44223 | 29.61204 | 12.68529 | 0.087057 | 0.292832 | 0.870711 |
| 18 | 56596000 | 56596399 | 5.945077 | 11.60098 | 19.85691 | 7.070158 | -0.11185 | -0.44893 | 0.870723 |
| 6 | 149080000 | 149081799 | 20.13885 | 35.32229 | 41.67548 | 18.42172 | 0.149953 | 0.66381 | 0.870809 |
| 18 | 68518000 | 68521599 | 33.13058 | 53.89644 | 73.7228 | 31.64214 | -0.08115 | 1.565057 | 0.870818 |
| 11 | 119256200 | 119258399 | 18.06181 | 31.83153 | 38.48217 | 17.03227 | 0.116494 | 0.585863 | 0.870831 |
| 13 | 50722400 | 50723799 | 18.66764 | 32.20821 | 44.77421 | 19.64187 | -0.08817 | 0.868152 | 0.870863 |
| 11 | 23080600 | 23082199 | 15.41663 | 27.71181 | 33.93496 | 14.82836 | 0.11285 | 0.411967 | 0.870864 |
| 3 | 106644000 | 106657199 | 192.0279 | 286.4497 | 313.3466 | 129.1477 | 0.129792 | 3.579984 | 0.870864 |
| 10 | 111607400 | 111609999 | 18.56606 | 32.40067 | 46.21317 | 20.34671 | -0.12272 | 0.737157 | 0.870865 |
| 16 | 5273400 | 5275599 | 19.1318 | 33.11516 | 40.61411 | 17.81864 | 0.092856 | 0.836784 | 0.870874 |
| 5 | 87018200 | 87019999 | 18.18146 | 31.49977 | 43.70796 | 19.17655 | -0.08424 | 0.827635 | 0.870891 |
| 14 | 105443200 | 105445599 | 32.74447 | 53.93627 | 66.39231 | 28.6361 | 0.080397 | 1.390898 | 0.8709 |
| 7 | 4568600 | 4569599 | 11.84793 | 21.53713 | 30.64564 | 13.21045 | -0.0896 | 0.315203 | 0.870931 |
| X | 102086200 | 102095599 | 116.4337 | 179.0719 | 203.9111 | 84.68006 | 0.108854 | 2.960476 | 0.870935 |
| X | 108816000 | 108817399 | 18.92794 | 33.51898 | 30.65444 | 13.55465 | 0.519866 | 0.5276 | 0.870976 |

| | | | | | | | | | |
|---|---|---|---|---|---|---|---|---|---|
| 7 | 7212200 | 7214199 | 15.72011 | 27.4761 | 38.89737 | 17.14313 | -0.11154 | 0.69132 | 0.870987 |
| 14 | 35613800 | 35616199 | 30.82769 | 50.55309 | 61.35842 | 26.41503 | 0.09818 | 1.438479 | 0.870987 |
| 4 | 8884600 | 8889399 | 109.5534 | 167.6861 | 196.5541 | 81.56292 | 0.061838 | 3.049614 | 0.870989 |
| 6 | 40581800 | 40582599 | 11.92985 | 21.54957 | 30.68717 | 13.33322 | -0.09784 | 0.354052 | 0.870998 |
| 8 | 48595400 | 48600399 | 50.78417 | 81.13678 | 96.59885 | 40.58701 | 0.090342 | 2.084455 | 0.871009 |
| 12 | 99366400 | 99370199 | 51.77887 | 82.66909 | 109.5989 | 46.00426 | -0.06567 | 2.109801 | 0.871011 |
| 13 | 87202400 | 87203999 | 16.2376 | 28.78656 | 35.35653 | 15.63034 | 0.097056 | 0.505549 | 0.871033 |
| 3 | 10416000 | 10416799 | 8.600455 | 16.28502 | 24.90103 | 10.08223 | -0.1234 | 0.026564 | 0.871034 |
| 4 | 59299800 | 59306399 | 64.22352 | 101.3082 | 157.4947 | 65.36204 | -0.31154 | 2.469189 | 0.871052 |
| 5 | 105565000 | 105568799 | 45.63055 | 73.3907 | 89.42175 | 37.94285 | 0.068208 | 1.868318 | 0.871053 |
| 3 | 132829400 | 132831999 | 41.28121 | 67.0257 | 64.85629 | 27.80533 | 0.414655 | 1.610366 | 0.871076 |
| 8 | 77405200 | 77405599 | 7.533884 | 14.61596 | 20.53596 | 7.737199 | 0.0943 | -0.23245 | 0.871088 |
| 6 | 122034600 | 122035799 | 15.00324 | 26.56243 | 37.05178 | 16.38419 | -0.08624 | 0.516123 | 0.871119 |
| 11 | 15283800 | 15284799 | 14.67162 | 27.21686 | 24.43351 | 10.25244 | 0.598102 | 0.212018 | 0.871122 |
| 10 | 24896200 | 24898599 | 21.50262 | 36.77357 | 50.62593 | 22.19849 | -0.07657 | 0.967311 | 0.871125 |
| 2 | 87344200 | 87348999 | 64.38004 | 101.962 | 135.2931 | 56.26102 | -0.07875 | 2.367914 | 0.871139 |
| 3 | 87120400 | 87125199 | 13.15449 | 23.21884 | 40.31705 | 17.83888 | -0.40214 | 0.544265 | 0.871143 |
| 5 | 20183400 | 20184799 | 21.40531 | 36.15625 | 50.61866 | 22.14469 | -0.10239 | 1.07751 | 0.871157 |
| 2 | 71384800 | 71390199 | 58.4976 | 92.62481 | 144.2741 | 59.98887 | -0.30995 | 2.372526 | 0.871172 |
| X | 79664800 | 79668399 | 33.54523 | 54.99478 | 67.70229 | 29.14279 | 0.077616 | 1.454085 | 0.871222 |
| 2 | 48131000 | 48132799 | 17.43814 | 30.31867 | 42.13327 | 18.50686 | -0.08552 | 0.782796 | 0.871231 |
| 7 | 41308400 | 41320399 | 264.3337 | 379.7093 | 420.6386 | 173.6593 | 0.059579 | 4.181705 | 0.871249 |
| 10 | 36475400 | 36477999 | 20.80048 | 36.2172 | 44.07534 | 19.37761 | 0.10511 | 0.779731 | 0.871253 |
| 15 | 11436400 | 11437799 | 13.77948 | 24.77075 | 30.97251 | 13.53306 | 0.083135 | 0.394806 | 0.871263 |
| 2 | 39250000 | 39252799 | 42.85254 | 69.29483 | 68.02679 | 29.07514 | 0.388403 | 1.696026 | 0.871284 |
| 4 | 24288200 | 24291199 | 48.37104 | 77.48505 | 103.0202 | 43.4494 | -0.06445 | 2.0033 | 0.871299 |
| 15 | 87329400 | 87341599 | 143.7379 | 217.1028 | 250.7812 | 103.6757 | 0.06725 | 3.330567 | 0.87133 |
| 14 | 113255800 | 113259999 | 66.1312 | 104.5082 | 124.234 | 51.60538 | 0.07717 | 2.41974 | 0.871345 |
| 4 | 128529800 | 128533599 | 22.62263 | 38.30342 | 46.44409 | 20.33305 | 0.104389 | 1.056538 | 0.871365 |
| 15 | 9048000 | 9048399 | 6.965266 | 13.59749 | 15.28995 | 5.404434 | 0.496789 | -0.47894 | 0.871369 |
| 13 | 80636000 | 80640599 | 52.59931 | 83.86201 | 111.3592 | 46.65204 | -0.07016 | 2.15467 | 0.871372 |
| 4 | 113363800 | 113366799 | 34.80051 | 57.65282 | 67.96523 | 29.41652 | 0.144041 | 1.331785 | 0.871373 |
| X | 36384800 | 36386399 | 15.95394 | 28.6413 | 34.51859 | 15.09952 | 0.134909 | 0.427345 | 0.87138 |
| 12 | 3407000 | 3408999 | 14.8236 | 26.27656 | 32.51857 | 14.37609 | 0.086366 | 0.494063 | 0.871388 |
| 6 | 136542600 | 136545399 | 34.58105 | 57.33451 | 67.43668 | 29.20368 | 0.147376 | 1.321174 | 0.871394 |
| 16 | 53637200 | 53637999 | 6.245581 | 12.14178 | 20.09941 | 7.460816 | -0.11886 | -0.29785 | 0.871403 |
| 15 | 43154800 | 43157399 | 33.21213 | 54.14845 | 73.60275 | 31.61603 | -0.0699 | 1.531604 | 0.871403 |
| 14 | 109950200 | 109950999 | 7.875213 | 15.19436 | 23.69171 | 9.15344 | -0.08909 | -0.13397 | 0.871405 |
| 7 | 122231600 | 122233199 | 9.738603 | 18.26919 | 26.8809 | 11.07537 | -0.08933 | 0.105123 | 0.871406 |
| 10 | 114148000 | 114150599 | 23.3079 | 39.52209 | 54.43514 | 23.8422 | -0.07886 | 1.054107 | 0.871412 |
| 10 | 7201600 | 7203999 | 20.96592 | 35.94151 | 49.55049 | 21.73111 | -0.0781 | 0.941576 | 0.871417 |
| 13 | 69242400 | 69244199 | 15.87078 | 27.89134 | 38.7194 | 17.14361 | -0.0833 | 0.606989 | 0.871424 |
| 12 | 45900000 | 45901399 | 13.06326 | 24.0881 | 23.89395 | 10.10832 | 0.446172 | 0.227791 | 0.871458 |
| 7 | 128077400 | 128091599 | 165.5375 | 247.1548 | 311.1273 | 128.2553 | -0.07153 | 3.558577 | 0.871464 |
| 14 | 116504600 | 116505799 | 20.15929 | 35.1678 | 35.15507 | 15.47438 | 0.387958 | 0.736787 | 0.871472 |
| 3 | 155407400 | 155408399 | 11.34688 | 21.5084 | 27.67924 | 11.18176 | 0.126395 | 0.053415 | 0.871477 |
| 16 | 21292000 | 21293399 | 16.393 | 28.61757 | 39.95157 | 17.59343 | -0.09159 | 0.715005 | 0.871486 |

| | | | | | | | | | |
|---|---|---|---|---|---|---|---|---|---|
| X | 147640400 | 147641399 | 11.38286 | 21.20606 | 27.20634 | 11.34197 | 0.093066 | 0.162806 | 0.87149 |
| X | 71260800 | 71262399 | 16.03155 | 28.42778 | 34.94682 | 15.44666 | 0.096152 | 0.504006 | 0.871534 |
| 8 | 52756600 | 52758799 | 31.6756 | 51.42496 | 72.09685 | 30.90742 | -0.11902 | 1.59731 | 0.871551 |
| 17 | 17108400 | 17111799 | 44.87374 | 72.31972 | 88.09779 | 37.49073 | 0.072135 | 1.804832 | 0.871579 |
| 13 | 4173200 | 4176599 | 50.27197 | 80.71183 | 97.55341 | 41.28498 | 0.076647 | 1.933465 | 0.871591 |
| X | 93833200 | 93834599 | 12.64319 | 23.06036 | 33.32116 | 14.30337 | -0.10753 | 0.310053 | 0.871597 |
| 3 | 77913800 | 77915199 | 18.92921 | 32.85824 | 40.54935 | 17.79314 | 0.08472 | 0.808793 | 0.871607 |
| 16 | 47644800 | 47645799 | 23.49997 | 39.59671 | 54.40972 | 23.78543 | -0.07579 | 1.109371 | 0.871612 |
| 2 | 179883000 | 179884599 | 18.94679 | 33.08375 | 40.84703 | 17.99048 | 0.084783 | 0.725758 | 0.871623 |
| 7 | 104997000 | 104999599 | 28.80558 | 48.92315 | 45.76723 | 20.05553 | 0.477328 | 1.051725 | 0.871631 |
| 12 | 106490800 | 106491199 | 5.647667 | 11.03167 | 16.49759 | 5.773471 | 0.103066 | -0.52833 | 0.871632 |
| 4 | 58708800 | 58711399 | 26.46064 | 44.78548 | 54.51385 | 23.86334 | 0.098711 | 1.081953 | 0.871667 |
| 9 | 11104200 | 11104799 | 11.88859 | 21.60722 | 30.67257 | 13.225 | -0.08628 | 0.317567 | 0.871668 |
| 2 | 115549000 | 115552199 | 43.7442 | 70.66354 | 85.60657 | 36.54006 | 0.08394 | 1.735313 | 0.871671 |
| 16 | 11328800 | 11331399 | 19.66305 | 33.96586 | 46.86598 | 20.56025 | -0.07712 | 0.864968 | 0.871687 |
| 1 | 28210000 | 28215799 | 93.29894 | 144.7582 | 170.8825 | 70.96821 | 0.066664 | 2.786646 | 0.871693 |
| 6 | 122556600 | 122558599 | 18.32945 | 32.04715 | 39.19405 | 17.2786 | 0.098361 | 0.698891 | 0.871716 |
| 4 | 65491400 | 65492199 | 13.26319 | 23.75811 | 29.06401 | 12.73725 | 0.111575 | 0.408919 | 0.871721 |
| 1 | 189978800 | 189984199 | 63.92491 | 101.347 | 121.7985 | 50.6892 | 0.065268 | 2.338534 | 0.871732 |
| 17 | 59840200 | 59842799 | 23.91339 | 41.30339 | 40.36699 | 17.70906 | 0.41831 | 0.875597 | 0.87174 |
| 15 | 47885400 | 47887599 | 26.67401 | 44.75119 | 55.08923 | 24.04089 | 0.082094 | 1.158837 | 0.871743 |
| 6 | 118541600 | 118542999 | 10.52927 | 19.95629 | 25.94741 | 10.4847 | 0.110961 | 0.039961 | 0.871767 |
| 5 | 109749000 | 109755599 | 77.99896 | 121.8787 | 158.222 | 65.64666 | -0.0624 | 2.654482 | 0.871792 |
| 11 | 71406800 | 71409999 | 19.75132 | 33.8887 | 47.41066 | 20.79411 | -0.09902 | 0.924571 | 0.871813 |
| 4 | 101715000 | 101723999 | 83.82097 | 129.7461 | 199.0402 | 82.67617 | -0.31359 | 2.825967 | 0.871824 |
| 2 | 73878400 | 73880199 | 24.81224 | 41.58134 | 57.17372 | 24.93303 | -0.07688 | 1.167581 | 0.871853 |
| 1 | 85129800 | 85204399 | 2363.375 | 2827.23 | 2832.898 | 1198.202 | -0.06165 | 7.070639 | 0.871916 |
| 11 | 51049800 | 51052199 | 27.42745 | 46.10727 | 45.40324 | 19.83425 | 0.403111 | 1.133213 | 0.87193 |
| 11 | 50721200 | 50725799 | 70.96604 | 111.4743 | 145.7182 | 60.44157 | -0.06645 | 2.554215 | 0.871939 |
| 13 | 113986200 | 113989199 | 29.51073 | 48.68097 | 66.6491 | 28.76468 | -0.07177 | 1.365486 | 0.87198 |
| 4 | 69831600 | 69831999 | 5.219526 | 10.19908 | 17.69848 | 6.14962 | -0.10006 | -0.55589 | 0.871982 |
| 5 | 4843200 | 4844599 | 15.42816 | 27.32677 | 33.92724 | 15.0012 | 0.081167 | 0.508003 | 0.872002 |
| 14 | 3706800 | 3707999 | 15.42816 | 27.32677 | 33.92724 | 15.0012 | 0.081167 | 0.508003 | 0.872002 |
| 14 | 66784600 | 66785999 | 14.03146 | 25.21267 | 31.3955 | 13.72296 | 0.088512 | 0.398784 | 0.872023 |
| 11 | 80194000 | 80198399 | 41.54442 | 67.05254 | 82.04245 | 34.98006 | 0.068234 | 1.753814 | 0.872041 |
| 5 | 146084800 | 146122999 | 611.8278 | 810.795 | 1073.859 | 448.9036 | -0.31509 | 5.423986 | 0.872083 |
| 4 | 13929600 | 13931999 | 30.99582 | 51.55892 | 52.45084 | 22.76544 | 0.356516 | 1.262555 | 0.872091 |
| 19 | 8411600 | 8421999 | 175.4743 | 260.4028 | 323.5686 | 133.2812 | -0.06083 | 3.668198 | 0.8721 |
| 1 | 31583800 | 31586599 | 15.27847 | 28.17565 | 25.12178 | 10.62632 | 0.59893 | 0.247053 | 0.872124 |
| 4 | 140187200 | 140189799 | 41.3392 | 66.37206 | 90.81047 | 38.43295 | -0.10174 | 1.919778 | 0.872146 |
| 15 | 44253400 | 44253799 | 5.223104 | 10.19507 | 17.57589 | 6.22717 | -0.11552 | -0.48674 | 0.87216 |
| 17 | 55256400 | 55256999 | 11.08514 | 20.6212 | 26.26558 | 10.97696 | 0.1012 | 0.170997 | 0.872209 |
| 4 | 113825200 | 113829799 | 52.65027 | 84.69151 | 98.97267 | 42.00299 | 0.128512 | 1.87901 | 0.872218 |
| 7 | 54433200 | 54434599 | 18.55644 | 32.25636 | 39.55916 | 17.37395 | 0.094026 | 0.787875 | 0.872222 |
| 8 | 50834200 | 50835199 | 11.6453 | 21.40251 | 30.76045 | 13.07191 | -0.09 | 0.250928 | 0.872227 |
| 5 | 120040200 | 120045599 | 92.02722 | 141.9534 | 182.8994 | 75.97307 | -0.06595 | 2.891799 | 0.872228 |
| 16 | 30843800 | 30846399 | 12.2065 | 21.61172 | 38.27341 | 16.92427 | -0.42852 | 0.501391 | 0.872245 |

| | | | | | | | | | |
|---|---|---|---|---|---|---|---|---|---|
| 8 | 48822800 | 48826199 | 26.64149 | 44.55015 | 54.55131 | 23.7569 | 0.089794 | 1.192202 | 0.872273 |
| X | 45215800 | 45217399 | 21.04076 | 36.5588 | 44.67602 | 19.61582 | 0.098932 | 0.812217 | 0.872302 |
| 15 | 5766800 | 5767999 | 17.39079 | 30.18666 | 42.16646 | 18.50231 | -0.09321 | 0.805022 | 0.872311 |
| 2 | 48673600 | 48677599 | 42.38945 | 68.17006 | 91.64387 | 38.8793 | -0.07357 | 1.868195 | 0.872332 |
| 13 | 22756000 | 22758399 | 28.05798 | 46.81508 | 57.87136 | 25.16759 | 0.076231 | 1.224671 | 0.87235 |
| 4 | 150852600 | 150860999 | 89.41849 | 137.0106 | 217.1108 | 90.11503 | -0.37154 | 3.024135 | 0.872361 |
| 12 | 115930200 | 115936999 | 133.7415 | 202.3839 | 235.6155 | 97.46369 | 0.058279 | 3.28955 | 0.872363 |
| 16 | 27489000 | 27491399 | 11.88505 | 21.17705 | 37.27633 | 16.40938 | -0.4147 | 0.451669 | 0.87237 |
| 11 | 5551000 | 5553399 | 17.96383 | 31.35685 | 38.84241 | 17.10397 | 0.080074 | 0.722681 | 0.872374 |
| 2 | 78390600 | 78390999 | 7.114473 | 13.82337 | 22.37212 | 8.341349 | -0.0938 | -0.2677 | 0.872427 |
| 1 | 96366000 | 96366799 | 12.26341 | 22.30682 | 31.71166 | 13.65893 | -0.08734 | 0.318845 | 0.872434 |
| 13 | 92970000 | 92970999 | 16.77185 | 29.35657 | 35.85723 | 15.82249 | 0.100594 | 0.671835 | 0.872445 |
| 14 | 94302800 | 94305799 | 33.90778 | 56.52678 | 52.56058 | 22.84556 | 0.485786 | 1.246779 | 0.87245 |
| 3 | 93864200 | 93865599 | 11.4897 | 21.08099 | 30.21473 | 12.86946 | -0.08838 | 0.258056 | 0.872456 |
| 10 | 54337000 | 54337999 | 9.563949 | 18.28234 | 24.5244 | 9.730707 | 0.089991 | -0.02871 | 0.872463 |
| 10 | 23522000 | 23522599 | 6.694992 | 12.99355 | 21.11834 | 7.942143 | -0.11035 | -0.24323 | 0.872478 |
| 3 | 41036200 | 41038399 | 17.95022 | 30.43412 | 55.84141 | 24.45735 | -0.49091 | 1.038172 | 0.872483 |
| 6 | 70041000 | 70042199 | 19.5194 | 33.63748 | 46.34853 | 20.33177 | -0.07579 | 0.885407 | 0.872485 |
| 16 | 61441800 | 61443199 | 23.20811 | 39.97266 | 39.50124 | 17.32777 | 0.401728 | 0.89813 | 0.87249 |
| 8 | 18644200 | 18646199 | 20.91075 | 35.73172 | 43.28404 | 18.97794 | 0.107186 | 0.963145 | 0.872498 |
| 18 | 24528000 | 24530599 | 31.19396 | 51.79988 | 63.131 | 27.36867 | 0.096416 | 1.290064 | 0.8725 |
| 9 | 44648200 | 44650999 | 27.97784 | 46.67562 | 57.80539 | 25.13695 | 0.073634 | 1.227253 | 0.872528 |
| X | 42516000 | 42519599 | 34.70219 | 57.09449 | 69.26122 | 29.85879 | 0.100774 | 1.414149 | 0.872551 |
| 6 | 43759200 | 43764399 | 86.22899 | 133.6458 | 172.6168 | 71.69992 | -0.06419 | 2.801254 | 0.872564 |
| 3 | 62414000 | 62414799 | 9.003728 | 16.99418 | 25.4202 | 10.3623 | -0.09991 | 0.057145 | 0.872591 |
| 17 | 23593000 | 23601399 | 85.86826 | 134.0887 | 176.8671 | 73.38738 | -0.08609 | 2.673727 | 0.8726 |
| 15 | 75204000 | 75206199 | 22.1769 | 37.63121 | 51.69019 | 22.64484 | -0.07485 | 1.041501 | 0.872603 |
| 12 | 12516800 | 12519199 | 19.84742 | 34.75883 | 33.09041 | 14.60831 | 0.458383 | 0.675333 | 0.87261 |
| 7 | 53181000 | 53181599 | 8.993238 | 17.01058 | 25.44148 | 10.32435 | -0.09408 | 0.042107 | 0.872617 |
| 3 | 115275400 | 115279199 | 43.08457 | 69.45856 | 93.98187 | 40.01132 | -0.07855 | 1.794978 | 0.872633 |
| 2 | 92416000 | 92418199 | 19.69453 | 34.09531 | 42.08507 | 18.46409 | 0.083559 | 0.837964 | 0.872636 |
| 9 | 99095800 | 99096799 | 11.03147 | 20.53547 | 29.9469 | 12.5009 | -0.09187 | 0.172545 | 0.87265 |
| 3 | 20747400 | 20750799 | 32.31437 | 52.86619 | 72.24334 | 31.07101 | -0.0741 | 1.478188 | 0.872676 |
| 7 | 127036600 | 127038599 | 30.16252 | 49.75022 | 61.56706 | 26.57001 | 0.073562 | 1.365723 | 0.872682 |
| 5 | 4748600 | 4749399 | 10.3037 | 19.09086 | 27.82433 | 11.69457 | -0.09796 | 0.190872 | 0.872684 |
| 9 | 20307200 | 20318999 | 221.0732 | 323.688 | 363.2636 | 149.6394 | 0.067754 | 3.893127 | 0.872684 |
| 1 | 10932800 | 10934199 | 15.98434 | 28.02442 | 38.80404 | 17.14672 | -0.07961 | 0.645033 | 0.872697 |
| 6 | 62685400 | 62687199 | 13.57488 | 23.60732 | 45.82393 | 20.1309 | -0.5651 | 0.768029 | 0.872707 |
| 6 | 38593800 | 38596999 | 48.67342 | 77.92333 | 103.4904 | 43.60843 | -0.0639 | 2.021699 | 0.872718 |
| 2 | 56758000 | 56758999 | 21.46576 | 36.96752 | 45.79714 | 20.08949 | 0.076835 | 0.899142 | 0.87272 |
| 9 | 18447400 | 18448599 | 18.98401 | 33.02506 | 41.00157 | 18.01617 | 0.076382 | 0.779627 | 0.872734 |
| 2 | 16576000 | 16576999 | 10.02083 | 18.81266 | 24.26769 | 9.986069 | 0.100764 | 0.096634 | 0.872758 |
| 13 | 33745200 | 33753399 | 154.3822 | 230.7933 | 288.2842 | 118.8682 | -0.05867 | 3.532294 | 0.872765 |
| 12 | 52637600 | 52639199 | 18.53393 | 32.3186 | 39.83884 | 17.53136 | 0.086831 | 0.740161 | 0.872776 |
| 5 | 149190400 | 149192399 | 16.10166 | 28.27727 | 39.19621 | 17.34446 | -0.08129 | 0.618333 | 0.872791 |
| 1 | 26548800 | 26549799 | 13.98967 | 25.17846 | 31.49216 | 13.74545 | 0.083787 | 0.390817 | 0.872805 |
| 7 | 26336800 | 26337599 | 13.98967 | 25.17846 | 31.49216 | 13.74545 | 0.083787 | 0.390817 | 0.872805 |

| | | | | | | | | | |
|---|---|---|---|---|---|---|---|---|---|
| 10 | 93465200 | 93466999 | 13.98967 | 25.17846 | 31.49216 | 13.74545 | 0.083787 | 0.390817 | 0.872805 |
| 3 | 76757400 | 76760399 | 57.0527 | 90.86821 | 109.7714 | 45.92106 | 0.064491 | 2.192441 | 0.872813 |
| X | 10916600 | 10918999 | 37.0715 | 62.06619 | 48.08267 | 20.95578 | 0.748251 | 1.212031 | 0.872822 |
| 15 | 34973800 | 34974999 | 16.60258 | 29.31511 | 35.85162 | 15.862 | 0.100836 | 0.550313 | 0.872843 |
| 11 | 7876800 | 7878799 | 52.01382 | 83.07905 | 99.45527 | 41.77128 | 0.082028 | 2.096109 | 0.872843 |
| 10 | 83989200 | 83991799 | 26.71669 | 44.30115 | 60.91966 | 26.38374 | -0.07728 | 1.293999 | 0.872844 |
| 8 | 86315400 | 86317199 | 18.0221 | 31.53431 | 38.66035 | 17.05454 | 0.094888 | 0.684214 | 0.87287 |
| 16 | 62484400 | 62485799 | 38.67039 | 62.50295 | 75.82578 | 32.38902 | 0.082948 | 1.706649 | 0.872919 |
| 17 | 63044400 | 63047399 | 38.70663 | 62.78985 | 76.89856 | 32.94658 | 0.074319 | 1.626148 | 0.872928 |
| 18 | 37720600 | 37722999 | 38.30153 | 62.37113 | 75.982 | 32.62375 | 0.086508 | 1.560857 | 0.873012 |
| 12 | 60692800 | 60694599 | 18.76092 | 32.56487 | 45.24727 | 19.85413 | -0.08578 | 0.81365 | 0.873018 |
| 5 | 43167400 | 43167999 | 9.591555 | 18.06965 | 22.84393 | 9.344854 | 0.13687 | 0.068167 | 0.873025 |
| 7 | 45989000 | 45991399 | 22.70151 | 38.46205 | 52.76843 | 23.1064 | -0.07321 | 1.058405 | 0.873025 |
| 5 | 39674200 | 39675199 | 12.47907 | 23.28296 | 29.42805 | 12.23724 | 0.117542 | 0.169165 | 0.873029 |
| 9 | 14168200 | 14169199 | 13.97703 | 25.20321 | 36.2079 | 15.7665 | -0.11302 | 0.387711 | 0.873036 |
| 8 | 92869600 | 92874599 | 43.37171 | 70.18224 | 84.33661 | 36.06696 | 0.098276 | 1.688905 | 0.873037 |
| 8 | 45401600 | 45402399 | 11.94333 | 21.53086 | 30.70047 | 13.37237 | -0.102 | 0.365984 | 0.873065 |
| 4 | 25710400 | 25713199 | 35.90237 | 59.25793 | 57.66478 | 24.89445 | 0.419555 | 1.367501 | 0.873094 |
| 10 | 127650800 | 127655599 | 59.16106 | 94.3434 | 113.5569 | 47.58644 | 0.071612 | 2.155247 | 0.873095 |
| 3 | 118892000 | 118893999 | 17.86541 | 31.36718 | 38.58076 | 17.06866 | 0.090294 | 0.631077 | 0.873105 |
| 1 | 131573000 | 131574999 | 41.28047 | 66.89403 | 60.36101 | 25.84096 | 0.51257 | 1.644551 | 0.873108 |
| 1 | 27764400 | 27781999 | 408.0612 | 561.9632 | 649.6002 | 270.0043 | -0.06302 | 4.814139 | 0.873109 |
| 17 | 63846200 | 63847599 | 8.236707 | 15.63317 | 30.54177 | 12.30444 | -0.46782 | 0.022703 | 0.873121 |
| 4 | 55651000 | 55654199 | 18.00719 | 31.06936 | 43.74702 | 19.19127 | -0.10653 | 0.863195 | 0.873121 |
| 12 | 30715600 | 30716399 | 15.14854 | 26.91476 | 33.45647 | 14.76347 | 0.081819 | 0.482554 | 0.87313 |
| 7 | 18649200 | 18650599 | 15.21212 | 26.95393 | 33.38635 | 14.76097 | 0.084642 | 0.500722 | 0.873135 |
| 4 | 79287200 | 79292399 | 99.55361 | 154.1645 | 154.6588 | 64.24116 | 0.299006 | 2.820176 | 0.873187 |
| 10 | 59233000 | 59233999 | 17.71479 | 30.77631 | 42.64424 | 18.72237 | -0.08151 | 0.794598 | 0.873197 |
| 8 | 121961200 | 121961999 | 10.6157 | 19.47955 | 28.27483 | 12.04139 | -0.10591 | 0.24826 | 0.873216 |
| 13 | 104797000 | 104797999 | 16.44396 | 28.87096 | 35.67825 | 15.7843 | 0.083712 | 0.619935 | 0.873233 |
| 13 | 111726400 | 111730199 | 31.34543 | 51.54085 | 63.73698 | 27.47032 | 0.072828 | 1.408859 | 0.873292 |
| 10 | 39175600 | 39176799 | 14.30272 | 26.51659 | 32.03947 | 13.45493 | 0.171068 | 0.226774 | 0.873294 |
| 6 | 110409200 | 110411799 | 20.44698 | 35.83348 | 42.23336 | 18.65546 | 0.151493 | 0.678833 | 0.873308 |
| 4 | 8442000 | 8445599 | 62.05158 | 98.54941 | 118.8833 | 49.56276 | 0.06231 | 2.286837 | 0.873322 |
| X | 71709600 | 71712799 | 25.10409 | 42.7886 | 51.72636 | 22.67486 | 0.109588 | 1.015149 | 0.873338 |
| 3 | 38658400 | 38660999 | 39.14408 | 63.59551 | 77.56098 | 33.26744 | 0.082376 | 1.599363 | 0.873338 |
| 12 | 115577400 | 115605199 | 69.90776 | 110.9132 | 112.408 | 46.81533 | 0.311815 | 2.321301 | 0.873381 |
| 1 | 50216600 | 50217399 | 10.1899 | 19.12006 | 24.77084 | 10.20214 | 0.093553 | 0.102922 | 0.873402 |
| 7 | 17666600 | 17754799 | 1404.881 | 1769.739 | 1665.654 | 699.9478 | 0.104405 | 6.248708 | 0.873434 |
| 18 | 22391800 | 22393399 | 17.49429 | 30.23625 | 51.6949 | 22.67909 | -0.38495 | 0.857153 | 0.873435 |
| 3 | 74672200 | 74673199 | 14.87826 | 26.32069 | 36.59082 | 16.18287 | -0.08173 | 0.523889 | 0.873481 |
| 14 | 119386000 | 119388999 | 33.47407 | 55.4487 | 66.63157 | 28.8369 | 0.116468 | 1.32881 | 0.873486 |
| 4 | 29758800 | 29760199 | 14.66722 | 25.95676 | 36.0529 | 15.9438 | -0.08032 | 0.516664 | 0.873496 |
| 1 | 68649800 | 68653399 | 77.52797 | 122.1014 | 121.5532 | 50.4355 | 0.328903 | 2.501667 | 0.873547 |
| 15 | 102294800 | 102297799 | 16.49791 | 28.86558 | 40.33497 | 17.79236 | -0.09292 | 0.680347 | 0.873559 |
| 6 | 76705000 | 76706199 | 13.53778 | 23.70833 | 42.57287 | 18.79422 | -0.45257 | 0.657812 | 0.873568 |
| X | 74707200 | 74720799 | 153.6569 | 233.1321 | 252.8781 | 104.6898 | 0.163439 | 3.245487 | 0.873599 |

| | | | | | | | | | |
|---|---|---|---|---|---|---|---|---|---|
| 15 | 86599400 | 86603799 | 57.57779 | 91.39636 | 121.1031 | 50.46178 | -0.07362 | 2.297931 | 0.873639 |
| 6 | 37774800 | 37778199 | 39.35131 | 63.97245 | 77.60288 | 33.29917 | 0.090914 | 1.588828 | 0.873642 |
| 17 | 44826000 | 44829799 | 34.38356 | 55.88908 | 75.7365 | 32.49675 | -0.06857 | 1.57877 | 0.873651 |
| 17 | 79429000 | 79430799 | 26.88404 | 45.02173 | 55.5411 | 24.21034 | 0.079187 | 1.182929 | 0.873656 |
| 5 | 55311600 | 55312399 | 13.20824 | 23.8215 | 29.60704 | 12.88912 | 0.095482 | 0.370438 | 0.87366 |
| 13 | 24409000 | 24411199 | 37.16747 | 60.40189 | 74.33492 | 31.8906 | 0.069783 | 1.588056 | 0.873662 |
| 14 | 110297600 | 110299599 | 40.85067 | 66.20971 | 61.60259 | 26.37605 | 0.468664 | 1.638736 | 0.873668 |
| 7 | 62056200 | 62056999 | 15.42645 | 27.20141 | 33.57528 | 14.8589 | 0.087113 | 0.556467 | 0.873704 |
| 6 | 140119200 | 140124599 | 38.37508 | 62.19313 | 84.6523 | 36.24332 | -0.07894 | 1.650927 | 0.873705 |
| 14 | 8866600 | 8867199 | 9.295174 | 17.41927 | 25.85887 | 10.673 | -0.10383 | 0.105951 | 0.873738 |
| 1 | 16486800 | 16498399 | 115.1856 | 176.7468 | 235.5906 | 97.80516 | -0.12068 | 3.006203 | 0.87374 |
| 11 | 51018200 | 51018999 | 12.90553 | 23.9882 | 22.09183 | 9.244861 | 0.564334 | 0.180386 | 0.873747 |
| 5 | 16879200 | 16879799 | 10.14074 | 18.919 | 23.73279 | 9.870008 | 0.12842 | 0.135354 | 0.873791 |
| 14 | 64487000 | 64487799 | 13.22615 | 23.64663 | 33.10086 | 14.52973 | -0.08327 | 0.430317 | 0.873794 |
| 5 | 63381000 | 63386599 | 66.38156 | 105.0463 | 125.8472 | 52.3114 | 0.067529 | 2.387462 | 0.873799 |
| 9 | 62517000 | 62521199 | 66.81805 | 105.7534 | 127.3968 | 52.95967 | 0.059751 | 2.385824 | 0.873833 |
| 10 | 7236200 | 7240799 | 51.78734 | 82.79814 | 109.8844 | 46.21541 | -0.06498 | 2.067462 | 0.873851 |
| 2 | 156861800 | 156865999 | 30.30775 | 49.00544 | 87.14478 | 37.24258 | -0.46676 | 1.686979 | 0.873852 |
| 9 | 9298400 | 9300599 | 27.47854 | 46.37798 | 46.93753 | 20.52934 | 0.363946 | 1.101256 | 0.873866 |
| 15 | 65050200 | 65052799 | 31.75061 | 51.72307 | 71.00954 | 30.49298 | -0.08503 | 1.538732 | 0.873868 |
| 3 | 158817800 | 158819399 | 20.63491 | 35.67324 | 50.10124 | 21.9801 | -0.10229 | 0.868282 | 0.873893 |
| 4 | 137531800 | 137533199 | 18.93421 | 32.93799 | 40.54388 | 17.81491 | 0.088854 | 0.780996 | 0.873897 |
| 14 | 7348200 | 7348799 | 9.035148 | 17.2226 | 22.84872 | 9.121675 | 0.097933 | -0.01969 | 0.873897 |
| 3 | 23317800 | 23318599 | 11.6044 | 21.19034 | 30.23187 | 12.95838 | -0.08781 | 0.288401 | 0.873924 |
| 6 | 51202800 | 51209399 | 64.88593 | 103.2785 | 122.4278 | 51.17059 | 0.09071 | 2.223767 | 0.873927 |
| 3 | 37298200 | 37306799 | 128.1931 | 193.8185 | 244.727 | 101.2048 | -0.05983 | 3.306415 | 0.873937 |
| 4 | 64211400 | 64212399 | 11.42036 | 20.78135 | 29.71151 | 12.79125 | -0.09498 | 0.305379 | 0.873942 |
| 13 | 33931400 | 33933599 | 28.58934 | 47.54535 | 58.95194 | 25.5832 | 0.071867 | 1.263173 | 0.873943 |
| 6 | 128587800 | 128595399 | 104.5341 | 161.34 | 188.6451 | 78.35951 | 0.074613 | 2.887316 | 0.87395 |
| 3 | 11079000 | 11081199 | 18.51462 | 32.10551 | 44.77399 | 19.64479 | -0.09106 | 0.826028 | 0.873976 |
| 4 | 18216800 | 18218599 | 17.51243 | 30.85986 | 29.82931 | 13.2028 | 0.437505 | 0.567676 | 0.873994 |
| 15 | 38586600 | 38594799 | 81.85165 | 128.0618 | 152.7347 | 63.36165 | 0.061196 | 2.632893 | 0.874007 |
| 5 | 24759400 | 24762199 | 26.78949 | 44.32211 | 61.32227 | 26.51692 | -0.08626 | 1.322095 | 0.874021 |
| 9 | 123018600 | 123020399 | 19.12562 | 33.12272 | 45.79188 | 20.09062 | -0.07919 | 0.837409 | 0.874027 |
| 4 | 36611400 | 36612599 | 35.6211 | 58.34034 | 58.46139 | 25.15448 | 0.373119 | 1.462038 | 0.874044 |
| 12 | 88482800 | 88486599 | 51.81428 | 82.66868 | 109.554 | 45.94058 | -0.06616 | 2.131489 | 0.874076 |
| 15 | 50457600 | 50460599 | 50.43091 | 80.62135 | 107.1036 | 45.03945 | -0.06659 | 2.067529 | 0.874098 |
| 13 | 72253000 | 72255399 | 19.46269 | 33.84346 | 42.03054 | 18.46339 | 0.075843 | 0.789015 | 0.874129 |
| 7 | 53425000 | 53426199 | 10.49111 | 19.76152 | 29.72029 | 12.16073 | -0.11256 | 0.094567 | 0.874152 |
| 7 | 40940600 | 40948599 | 100.9678 | 156.8138 | 180.5434 | 74.96919 | 0.103801 | 2.772907 | 0.874166 |
| 9 | 46271000 | 46272999 | 22.11978 | 37.81338 | 46.89219 | 20.56063 | 0.073563 | 0.969407 | 0.874195 |
| 2 | 169247200 | 169249799 | 24.32926 | 40.86334 | 56.05606 | 24.4802 | -0.07356 | 1.14159 | 0.874211 |
| 12 | 88048800 | 88069799 | 636.4605 | 844.5341 | 942.1133 | 393.767 | -0.06561 | 5.403249 | 0.874212 |
| 2 | 33804400 | 33809199 | 28.41143 | 47.00788 | 65.11289 | 28.15702 | -0.08809 | 1.324342 | 0.874216 |
| 4 | 84209800 | 84212199 | 33.37829 | 53.82052 | 93.52715 | 39.82718 | -0.43835 | 1.774805 | 0.874227 |
| 16 | 14869600 | 14871999 | 21.91999 | 37.45533 | 46.36685 | 20.32956 | 0.075967 | 0.971385 | 0.874242 |
| 10 | 39853400 | 39855799 | 47.34063 | 76.06812 | 92.78851 | 39.3167 | 0.064881 | 1.903012 | 0.874255 |

| | | | | | | | | | |
|---|---|---|---|---|---|---|---|---|---|
| 8 | 127722400 | 127724399 | 19.47434 | 33.59308 | 46.33838 | 20.32793 | -0.07713 | 0.87543 | 0.874258 |
| 2 | 12645800 | 12647199 | 21.90457 | 37.14747 | 51.14297 | 22.40116 | -0.07811 | 1.046121 | 0.874273 |
| 11 | 12947400 | 12949599 | 18.81318 | 32.97083 | 29.15889 | 12.88036 | 0.563992 | 0.649643 | 0.874296 |
| 15 | 14763200 | 14767399 | 66.5161 | 105.0539 | 137.3924 | 57.05403 | -0.06107 | 2.436686 | 0.874319 |
| 1 | 94396000 | 94397999 | 20.66398 | 35.65367 | 44.22504 | 19.40098 | 0.075633 | 0.877445 | 0.87434 |
| 6 | 122976800 | 122978399 | 35.729 | 57.80572 | 78.56118 | 33.59304 | -0.079 | 1.676978 | 0.874341 |
| 16 | 44956800 | 44960599 | 41.77243 | 67.54209 | 82.48572 | 35.25085 | 0.07394 | 1.703627 | 0.874343 |
| 7 | 16773000 | 16775399 | 17.90654 | 30.95006 | 43.24101 | 18.97031 | -0.09471 | 0.847978 | 0.874343 |
| 2 | 165938200 | 165939999 | 14.9685 | 26.28269 | 36.73724 | 16.25397 | -0.09319 | 0.610655 | 0.874375 |
| 12 | 55531600 | 55533799 | 14.9685 | 26.28269 | 36.73724 | 16.25397 | -0.09319 | 0.610655 | 0.874375 |
| 7 | 5183800 | 5184399 | 10.23721 | 19.20568 | 24.79812 | 10.21622 | 0.098245 | 0.105504 | 0.87438 |
| Y | 42994800 | 42998799 | 36.95886 | 59.27376 | 89.16269 | 37.6723 | -0.2401 | 1.945357 | 0.87439 |
| 7 | 6412400 | 6413599 | 8.188313 | 15.70627 | 24.08337 | 9.462421 | -0.08787 | -0.07505 | 0.874407 |
| 9 | 6428000 | 6428599 | 8.188313 | 15.70627 | 24.08337 | 9.462421 | -0.08787 | -0.07505 | 0.874407 |
| 12 | 67793800 | 67794399 | 8.188313 | 15.70627 | 24.08337 | 9.462421 | -0.08787 | -0.07505 | 0.874407 |
| 11 | 53643200 | 53644399 | 14.53563 | 25.61131 | 35.69599 | 15.79981 | -0.0876 | 0.564016 | 0.874417 |
| 3 | 91928600 | 91932399 | 41.99537 | 67.69862 | 91.0043 | 38.74073 | -0.0692 | 1.795136 | 0.874431 |
| 10 | 125628000 | 125630999 | 40.62149 | 65.70239 | 80.55803 | 34.43961 | 0.068745 | 1.692325 | 0.874448 |
| 10 | 118627200 | 118628999 | 15.13612 | 26.80164 | 32.84074 | 14.52165 | 0.09968 | 0.50501 | 0.874557 |
| 4 | 45322200 | 45324599 | 21.49345 | 36.78842 | 50.62889 | 22.20127 | -0.0759 | 0.959619 | 0.874567 |
| 11 | 99564600 | 99565199 | 7.145206 | 13.86005 | 22.22207 | 8.377727 | -0.09468 | -0.22704 | 0.874585 |
| 4 | 66600000 | 66601399 | 21.05424 | 35.74731 | 49.55201 | 21.71202 | -0.08784 | 1.030397 | 0.87459 |
| 1 | 64633600 | 64634599 | 11.87455 | 21.62696 | 30.67667 | 13.19216 | -0.08276 | 0.305169 | 0.874593 |
| 7 | 66003800 | 66014599 | 133.8067 | 201.8448 | 254.0967 | 105.004 | -0.05831 | 3.354114 | 0.874598 |
| 13 | 119320000 | 119323399 | 43.73349 | 70.2219 | 94.08918 | 39.82549 | -0.07154 | 1.920438 | 0.874601 |
| 7 | 108799800 | 108803399 | 26.77331 | 44.90224 | 55.59907 | 24.25809 | 0.073757 | 1.165014 | 0.874622 |
| 5 | 77567000 | 77572799 | 84.87116 | 132.9071 | 157.5036 | 65.3339 | 0.071427 | 2.620244 | 0.874636 |
| 7 | 26811200 | 26815199 | 33.49969 | 54.70438 | 74.48857 | 32.01512 | -0.07083 | 1.509168 | 0.874638 |
| 3 | 80243400 | 80244199 | 8.094114 | 15.52527 | 24.08898 | 9.465175 | -0.10435 | -0.07348 | 0.874669 |
| 17 | 93711400 | 93711999 | 10.33846 | 19.31175 | 24.74088 | 10.26771 | 0.100747 | 0.132558 | 0.874693 |
| 1 | 108063000 | 108067799 | 59.76749 | 95.23866 | 114.6738 | 48.00441 | 0.069948 | 2.179424 | 0.874693 |
| 7 | 38811800 | 38876999 | 1133.776 | 1448.319 | 1414.982 | 593.7717 | 0.070875 | 6.021766 | 0.874723 |
| 11 | 65077800 | 65078599 | 11.27855 | 20.70423 | 25.66449 | 10.92372 | 0.120668 | 0.243298 | 0.874725 |
| 7 | 22125200 | 22126399 | 23.79472 | 40.75277 | 40.57368 | 17.79286 | 0.389537 | 0.951289 | 0.874736 |
| 11 | 109764200 | 109766799 | 29.09624 | 48.8498 | 49.12033 | 21.44572 | 0.37301 | 1.15441 | 0.874784 |
| 12 | 8115800 | 8117999 | 25.35905 | 42.88334 | 52.85731 | 23.13203 | 0.080595 | 1.082274 | 0.874792 |
| 3 | 14389800 | 14392999 | 32.60215 | 53.62715 | 74.80975 | 32.24768 | -0.10039 | 1.415696 | 0.874802 |
| 9 | 99250200 | 99252399 | 17.7376 | 30.8508 | 42.61777 | 18.72405 | -0.07696 | 0.779045 | 0.874808 |
| X | 133567200 | 133571999 | 32.05761 | 52.76266 | 73.09325 | 31.51566 | -0.09004 | 1.405201 | 0.874814 |
| 9 | 95515200 | 95517199 | 18.59497 | 32.26965 | 39.55418 | 17.35499 | 0.094416 | 0.808789 | 0.87485 |
| 16 | 90209000 | 90213999 | 41.28984 | 66.46449 | 89.19558 | 37.89252 | -0.06935 | 1.837125 | 0.874865 |
| 18 | 9417800 | 9420399 | 46.8615 | 75.26834 | 91.80041 | 38.87545 | 0.064385 | 1.914705 | 0.874866 |
| 3 | 70728000 | 70728799 | 13.45847 | 23.96313 | 33.661 | 14.82698 | -0.09157 | 0.460273 | 0.874879 |
| 3 | 59574600 | 59575599 | 13.29566 | 24.31252 | 30.62171 | 13.10263 | 0.092204 | 0.292588 | 0.874926 |
| Y | 82132200 | 82132599 | 5.93898 | 11.57758 | 15.9242 | 5.748706 | 0.182257 | -0.4178 | 0.874939 |
| 10 | 39774200 | 39778999 | 41.02288 | 66.2135 | 80.85864 | 34.47715 | 0.071093 | 1.750657 | 0.87495 |
| 14 | 56346000 | 56348799 | 26.99374 | 45.20678 | 62.88593 | 27.41246 | -0.09365 | 1.18724 | 0.874964 |

| | | | | | | | | | |
|---|---|---|---|---|---|---|---|---|---|
| 6 | 3671200 | 3674599 | 35.41512 | 57.4683 | 77.81211 | 33.34566 | -0.06965 | 1.616996 | 0.874993 |
| 7 | 39193000 | 39194599 | 17.1621 | 30.04671 | 37.22258 | 16.42822 | 0.080087 | 0.670217 | 0.875012 |
| 13 | 47156400 | 47158999 | 32.69369 | 53.46898 | 65.77035 | 28.28362 | 0.077027 | 1.481817 | 0.875033 |
| 2 | 150063400 | 150066399 | 43.72211 | 70.62971 | 85.67679 | 36.57128 | 0.082108 | 1.734161 | 0.875035 |
| 3 | 131661000 | 131663399 | 23.96788 | 40.49957 | 50.11799 | 21.92809 | 0.074946 | 1.082937 | 0.875052 |
| 8 | 20685200 | 20685599 | 7.449427 | 14.30779 | 22.51876 | 8.814332 | -0.1195 | -0.09815 | 0.875059 |
| 5 | 96631000 | 96634199 | 26.45215 | 44.80762 | 54.51887 | 23.87144 | 0.099325 | 1.074849 | 0.875079 |
| 17 | 72085000 | 72090399 | 29.73885 | 49.97253 | 49.11343 | 21.45238 | 0.405871 | 1.14773 | 0.875107 |
| 3 | 57975600 | 57979999 | 36.4596 | 59.51635 | 73.06506 | 31.39916 | 0.07789 | 1.520465 | 0.875129 |
| X | 51859000 | 51859999 | 9.860931 | 18.76303 | 28.94173 | 11.59801 | -0.12249 | 0.018909 | 0.875168 |
| 6 | 24014800 | 24018999 | 43.78996 | 70.63613 | 96.71737 | 41.20631 | -0.0947 | 1.781389 | 0.875187 |
| 3 | 76599400 | 76601599 | 18.1165 | 31.67523 | 39.2644 | 17.30981 | 0.079149 | 0.699497 | 0.875212 |
| 6 | 18968200 | 18969199 | 10.98686 | 20.61547 | 26.83677 | 11.05301 | 0.086461 | 0.114662 | 0.875218 |
| 10 | 42159200 | 42163399 | 101.5426 | 156.1861 | 183.999 | 76.41206 | 0.059988 | 2.950012 | 0.875317 |
| 4 | 15911000 | 15919599 | 358.4004 | 503.726 | 457.5553 | 189.5754 | 0.314258 | 4.504567 | 0.875333 |
| 12 | 21328200 | 21330599 | 16.92948 | 29.69837 | 36.56273 | 16.1641 | 0.088873 | 0.634566 | 0.875392 |
| 7 | 6181600 | 6185599 | 29.89226 | 48.96103 | 67.35829 | 28.98345 | -0.08279 | 1.45474 | 0.875435 |
| 8 | 46523400 | 46525799 | 21.28322 | 36.55803 | 50.64207 | 22.21302 | -0.08455 | 0.926016 | 0.875435 |
| 5 | 151393400 | 151394799 | 35.16659 | 57.16261 | 70.09371 | 30.0757 | 0.075309 | 1.576884 | 0.875436 |
| 2 | 52017000 | 52024199 | 25.50654 | 43.48549 | 43.69484 | 19.15454 | 0.375354 | 1.003369 | 0.875441 |
| 13 | 4584200 | 4584799 | 9.048301 | 17.34305 | 23.48976 | 9.248746 | 0.086465 | -0.06246 | 0.875463 |
| 8 | 57635000 | 57636799 | 17.5386 | 30.10423 | 51.64662 | 22.65271 | -0.39188 | 0.920143 | 0.8755 |
| 12 | 119091800 | 119093199 | 19.18395 | 33.4257 | 41.46745 | 18.23901 | 0.077691 | 0.759358 | 0.875503 |
| 3 | 37473200 | 37475199 | 18.44169 | 31.17311 | 57.46013 | 25.14328 | -0.49775 | 1.067484 | 0.875511 |
| 3 | 62988400 | 62990399 | 22.80847 | 39.30144 | 39.47533 | 17.31664 | 0.378405 | 0.891493 | 0.875514 |
| 13 | 94566800 | 94570999 | 43.66849 | 70.65562 | 71.75984 | 30.68625 | 0.340305 | 1.685934 | 0.875557 |
| X | 36926600 | 36928599 | 19.39983 | 33.85692 | 40.96366 | 18.03586 | 0.114078 | 0.740202 | 0.875567 |
| 7 | 132753200 | 132754799 | 17.4826 | 30.54945 | 37.81336 | 16.66207 | 0.081345 | 0.703705 | 0.875575 |
| 6 | 120110000 | 120112199 | 31.32309 | 51.12875 | 70.37217 | 30.24131 | -0.0866 | 1.511012 | 0.875576 |
| 9 | 100289400 | 100291399 | 24.31085 | 40.81063 | 56.06506 | 24.47588 | -0.07565 | 1.146728 | 0.875623 |
| 10 | 85084800 | 85085599 | 13.97996 | 25.69164 | 31.7804 | 13.50653 | 0.125057 | 0.27254 | 0.875628 |
| 4 | 60665600 | 60669599 | 48.36439 | 77.55302 | 94.25687 | 39.81476 | 0.066584 | 1.969391 | 0.875635 |
| 6 | 34201200 | 34204399 | 32.90491 | 53.6192 | 72.87389 | 31.29692 | -0.07023 | 1.53857 | 0.875646 |
| 15 | 6729200 | 6730999 | 16.00201 | 28.58748 | 25.99111 | 11.41629 | 0.535228 | 0.439309 | 0.875682 |
| X | 125051000 | 125052999 | 20.37469 | 35.4419 | 43.59438 | 19.15464 | 0.089737 | 0.791638 | 0.8757 |
| 15 | 96200600 | 96201999 | 14.93246 | 26.5433 | 32.58547 | 14.37343 | 0.100369 | 0.478831 | 0.875723 |
| 17 | 6389000 | 6392799 | 21.66442 | 36.8724 | 50.61805 | 22.18643 | -0.07354 | 1.011378 | 0.875733 |
| 10 | 58877000 | 58878199 | 15.70536 | 27.96143 | 34.46564 | 15.18248 | 0.095668 | 0.476156 | 0.875734 |
| 9 | 31990800 | 31992599 | 15.76213 | 27.80411 | 34.52884 | 15.27972 | 0.078558 | 0.556017 | 0.875741 |
| 9 | 117839600 | 117841599 | 16.7205 | 29.08351 | 41.17353 | 18.09021 | -0.1119 | 0.774014 | 0.875756 |
| 9 | 9231600 | 9234999 | 36.0218 | 58.78887 | 72.27425 | 31.05685 | 0.07566 | 1.52331 | 0.875821 |
| 7 | 43546200 | 43547799 | 11.36976 | 20.49 | 30.12856 | 13.12875 | -0.14684 | 0.359101 | 0.875834 |
| 12 | 15864400 | 15867799 | 26.85478 | 43.66182 | 82.13035 | 35.24327 | -0.54018 | 1.563032 | 0.87584 |
| 15 | 79501600 | 79505199 | 31.41709 | 51.19098 | 70.43939 | 30.25056 | -0.08808 | 1.534284 | 0.875844 |
| 15 | 96753200 | 96755199 | 20.80356 | 35.64328 | 49.01322 | 21.49451 | -0.07452 | 0.945387 | 0.875863 |
| 12 | 23682200 | 23685799 | 126.9178 | 193.502 | 178.0776 | 73.82862 | 0.406079 | 3.133361 | 0.875907 |
| 14 | 86324400 | 86325799 | 16.20969 | 28.22284 | 48.42857 | 21.29102 | -0.38832 | 0.765179 | 0.875916 |

| | | | | | | | | | |
|---|---|---|---|---|---|---|---|---|---|
| 1 | 43355400 | 43356999 | 21.80377 | 37.71001 | 35.82216 | 15.71588 | 0.45948 | 0.847156 | 0.875919 |
| 10 | 97360000 | 97361999 | 19.38502 | 33.97457 | 33.9374 | 14.99175 | 0.389118 | 0.6601 | 0.875939 |
| 6 | 13701800 | 13708399 | 92.20248 | 143.5522 | 144.0094 | 59.77682 | 0.305037 | 2.722013 | 0.875939 |
| 1 | 138736200 | 138738199 | 18.63818 | 32.74223 | 32.42052 | 14.35004 | 0.40173 | 0.617126 | 0.875968 |
| 6 | 97653200 | 97656599 | 32.31543 | 53.31753 | 75.24435 | 32.47681 | -0.11537 | 1.375864 | 0.875984 |
| 4 | 156403600 | 156410999 | 408.5655 | 553.817 | 835.279 | 348.041 | -0.46977 | 5.061789 | 0.875999 |
| 3 | 12675400 | 12675799 | 4.543628 | 8.882177 | 16.03115 | 5.508908 | -0.14119 | -0.61269 | 0.876006 |
| 14 | 88044200 | 88049399 | 57.61397 | 91.74446 | 121.0895 | 50.6423 | -0.06335 | 2.202396 | 0.876012 |
| 2 | 94763000 | 94764999 | 19.02376 | 33.11757 | 40.92094 | 17.98872 | 0.083183 | 0.767612 | 0.876014 |
| 9 | 116457400 | 116460199 | 19.07719 | 33.46203 | 33.42806 | 14.77682 | 0.389 | 0.644993 | 0.876021 |
| 5 | 8223400 | 8226199 | 24.26132 | 41.9284 | 40.34498 | 17.69972 | 0.440845 | 0.872251 | 0.87605 |
| X | 107724200 | 107724999 | 12.52583 | 23.46988 | 29.56458 | 12.20573 | 0.130756 | 0.143579 | 0.876101 |
| 14 | 87914200 | 87914999 | 11.10821 | 20.39942 | 29.26573 | 12.44999 | -0.08807 | 0.250327 | 0.876101 |
| 8 | 40613200 | 40613999 | 10.26779 | 19.15263 | 27.86542 | 11.59218 | -0.08424 | 0.149577 | 0.876132 |
| 8 | 110479200 | 110483999 | 40.88409 | 66.00808 | 80.84503 | 34.48388 | 0.067338 | 1.742769 | 0.876138 |
| 4 | 95397800 | 95399799 | 23.58612 | 39.07219 | 67.1992 | 29.08388 | -0.39908 | 1.302898 | 0.876141 |
| 13 | 110472800 | 110477399 | 65.41962 | 103.3238 | 135.2353 | 56.15874 | -0.06216 | 2.435965 | 0.876143 |
| 1 | 108994000 | 108994999 | 16.32684 | 29.06928 | 28.80294 | 12.68749 | 0.408913 | 0.467492 | 0.876167 |
| 8 | 118500400 | 118502999 | 21.33694 | 36.2188 | 50.08495 | 21.94397 | -0.08446 | 1.03303 | 0.876174 |
| 4 | 41353400 | 41355599 | 25.09349 | 42.67737 | 52.29164 | 22.91863 | 0.08965 | 1.032632 | 0.876178 |
| 13 | 5293400 | 5295799 | 35.60935 | 58.00278 | 71.61646 | 30.75238 | 0.067448 | 1.547953 | 0.876217 |
| 17 | 57999800 | 58003399 | 46.70132 | 75.04317 | 100.734 | 42.68529 | -0.07292 | 1.903777 | 0.876223 |
| 6 | 58796400 | 58796999 | 12.4388 | 23.08901 | 22.01955 | 9.231798 | 0.512009 | 0.182019 | 0.876254 |
| 1 | 44530200 | 44530999 | 8.16823 | 15.7369 | 21.45854 | 8.325178 | 0.09735 | -0.12337 | 0.876256 |
| 7 | 56673600 | 56674599 | 11.592 | 21.21361 | 30.23845 | 12.9251 | -0.08389 | 0.275719 | 0.876277 |
| 15 | 35437200 | 35437599 | 11.592 | 21.21361 | 30.23845 | 12.9251 | -0.08389 | 0.275719 | 0.876277 |
| 15 | 92664200 | 92668999 | 17.04388 | 28.84627 | 56.30885 | 24.64883 | -0.57992 | 1.051425 | 0.876287 |
| 7 | 88019400 | 88020999 | 14.12657 | 24.92811 | 35.10021 | 15.53143 | -0.10159 | 0.541579 | 0.876339 |
| 2 | 73997200 | 73999599 | 18.63168 | 32.58 | 40.2826 | 17.76054 | 0.082823 | 0.702016 | 0.876348 |
| 7 | 36966600 | 36971399 | 52.93983 | 84.81253 | 102.5694 | 43.26607 | 0.072145 | 2.005062 | 0.876378 |
| 7 | 106620800 | 106621599 | 10.82723 | 20.11428 | 25.70935 | 10.76661 | 0.093679 | 0.17365 | 0.876393 |
| 19 | 35051800 | 35056599 | 53.26824 | 85.0645 | 103.0846 | 43.28251 | 0.064059 | 2.10392 | 0.876403 |
| 13 | 113409200 | 113411799 | 27.26938 | 44.9559 | 62.37468 | 26.91289 | -0.09119 | 1.365207 | 0.876432 |
| 17 | 40329800 | 40331199 | 27.4912 | 46.23486 | 45.92908 | 20.06664 | 0.390587 | 1.131087 | 0.876439 |
| X | 16305200 | 16307199 | 38.77612 | 62.82099 | 76.0541 | 32.55362 | 0.089287 | 1.656758 | 0.876455 |
| 14 | 108227200 | 108228199 | 16.70151 | 30.15672 | 28.16925 | 12.24539 | 0.506212 | 0.384325 | 0.876464 |
| 4 | 26004400 | 26004999 | 7.730771 | 14.86742 | 23.05522 | 8.990777 | -0.09364 | -0.10681 | 0.876499 |
| 5 | 89143800 | 89148999 | 63.18841 | 100.4687 | 121.0372 | 50.51804 | 0.065804 | 2.255125 | 0.876506 |
| 9 | 6264600 | 6266799 | 27.22663 | 45.20539 | 62.86389 | 27.25019 | -0.09338 | 1.278649 | 0.876526 |
| X | 68327200 | 68328399 | 16.02698 | 28.71531 | 34.98097 | 15.32744 | 0.117744 | 0.439603 | 0.876533 |
| 9 | 116770800 | 116771599 | 8.486425 | 16.0402 | 24.38758 | 9.912013 | -0.11975 | 0.037902 | 0.876546 |
| 2 | 60844000 | 60845599 | 14.52018 | 26.36657 | 32.53814 | 14.045 | 0.113115 | 0.343779 | 0.876547 |
| 13 | 28013600 | 28016199 | 36.14827 | 58.25977 | 94.98698 | 40.42449 | -0.34736 | 1.794492 | 0.87658 |
| 2 | 74497400 | 74499399 | 19.26249 | 32.5094 | 57.45286 | 25.12811 | -0.4373 | 1.085338 | 0.876585 |
| 16 | 3345200 | 3345999 | 14.82004 | 26.43764 | 32.96853 | 14.49871 | 0.080611 | 0.45437 | 0.876587 |
| Y | 43874000 | 43878999 | 43.11576 | 69.21572 | 78.7896 | 33.3391 | 0.162864 | 1.918557 | 0.876596 |
| 9 | 55920600 | 55924399 | 37.2031 | 61.30555 | 72.07454 | 31.09232 | 0.146751 | 1.402821 | 0.876597 |

| | | | | | | | | | |
|---|---|---|---|---|---|---|---|---|---|
| 4 | 96961800 | 96963799 | 16.508 | 29.05919 | 40.71575 | 18.02506 | -0.09591 | 0.594124 | 0.876647 |
| 18 | 37093400 | 37098999 | 109.4671 | 167.3285 | 194.8926 | 80.84828 | 0.069472 | 3.076745 | 0.876664 |
| 13 | 81750800 | 81751599 | 10.70288 | 19.89114 | 25.24916 | 10.56819 | 0.104314 | 0.1701 | 0.876681 |
| 10 | 126235600 | 126238999 | 31.36584 | 52.33706 | 62.65448 | 27.24863 | 0.122342 | 1.237824 | 0.876684 |
| X | 77489800 | 77492399 | 20.61391 | 36.11648 | 42.82389 | 18.91208 | 0.14297 | 0.68685 | 0.876707 |
| 1 | 96817600 | 96819399 | 16.22746 | 28.51214 | 39.66263 | 17.5581 | -0.08649 | 0.611129 | 0.876734 |
| 4 | 20081400 | 20084999 | 20.82612 | 35.34845 | 61.49549 | 26.94258 | -0.41455 | 1.044964 | 0.87675 |
| 4 | 79545800 | 79546799 | 17.95512 | 31.18373 | 43.18931 | 18.95782 | -0.08082 | 0.802708 | 0.876776 |
| 5 | 87531800 | 87534199 | 30.64639 | 50.95232 | 62.42712 | 27.08618 | 0.088996 | 1.276171 | 0.876787 |
| 6 | 128450200 | 128453999 | 40.1074 | 64.68663 | 86.86 | 36.99974 | -0.06682 | 1.774608 | 0.876816 |
| 10 | 70633400 | 70637199 | 31.16165 | 51.00108 | 69.82993 | 30.03748 | -0.07646 | 1.46802 | 0.87683 |
| X | 102491000 | 102494599 | 36.6428 | 59.47152 | 81.37891 | 34.87896 | -0.08459 | 1.619133 | 0.876884 |
| 10 | 96713600 | 96715999 | 9.686637 | 17.84209 | 33.0152 | 13.98801 | -0.44889 | 0.228645 | 0.876885 |
| 17 | 79427800 | 79430999 | 33.54804 | 55.09142 | 67.81339 | 29.21127 | 0.078731 | 1.433712 | 0.876904 |
| 7 | 56601200 | 56601999 | 9.814805 | 18.28968 | 26.85279 | 11.18952 | -0.09896 | 0.150251 | 0.876909 |
| 7 | 85738200 | 85750999 | 182.5167 | 271.6024 | 308.5251 | 127.1203 | 0.072201 | 3.620436 | 0.876932 |
| 10 | 22065200 | 22069599 | 43.37737 | 69.70907 | 93.33013 | 39.55377 | -0.06896 | 1.892712 | 0.876932 |
| 10 | 8039000 | 8052399 | 271.402 | 389.2726 | 431.4347 | 178.1587 | 0.056721 | 4.204462 | 0.876935 |
| 2 | 60123000 | 60128999 | 73.59791 | 116.044 | 139.3515 | 57.82907 | 0.059964 | 2.479435 | 0.876951 |
| 17 | 84647200 | 84648399 | 12.19161 | 22.01186 | 31.11029 | 13.52516 | -0.08782 | 0.358118 | 0.876969 |
| 17 | 36386600 | 36387599 | 28.32748 | 47.35066 | 58.39324 | 25.42366 | 0.079684 | 1.208601 | 0.876983 |
| 17 | 34230200 | 34234199 | 72.34079 | 113.3794 | 148.2775 | 61.50153 | -0.06962 | 2.597331 | 0.877017 |
| 5 | 69201400 | 69209599 | 85.18319 | 131.716 | 200.2849 | 83.19367 | -0.30202 | 2.846157 | 0.87703 |
| 4 | 153205000 | 153208599 | 30.24314 | 49.86905 | 61.67182 | 26.61203 | 0.074335 | 1.367638 | 0.877035 |
| X | 87286800 | 87288599 | 14.46347 | 26.96091 | 24.4383 | 10.17795 | 0.592421 | 0.181152 | 0.877051 |
| 10 | 47455800 | 47457599 | 25.90964 | 43.49369 | 53.93947 | 23.54764 | 0.071475 | 1.150299 | 0.877065 |
| 3 | 5192400 | 5193599 | 22.10793 | 38.4132 | 38.24954 | 16.7942 | 0.393032 | 0.806466 | 0.877098 |
| 10 | 86831400 | 86834599 | 22.11429 | 37.61935 | 51.70471 | 22.66182 | -0.07541 | 1.017869 | 0.877099 |
| 6 | 56137200 | 56137999 | 10.36035 | 19.42973 | 25.27853 | 10.42047 | 0.086472 | 0.109197 | 0.877112 |
| 10 | 34773000 | 34776199 | 69.62283 | 109.5065 | 143.0106 | 59.32852 | -0.06364 | 2.527203 | 0.87715 |
| 12 | 80460000 | 80465599 | 68.96296 | 109.2757 | 131.1002 | 54.5314 | 0.067004 | 2.360866 | 0.877159 |
| 7 | 22462600 | 22559599 | 3343.442 | 3901.796 | 3483.347 | 1478.863 | 0.068456 | 7.460457 | 0.877167 |
| 9 | 78632200 | 78633199 | 12.7152 | 22.88044 | 32.09824 | 14.01311 | -0.081 | 0.384083 | 0.877175 |
| 4 | 4169000 | 4173599 | 40.26267 | 65.26539 | 79.81703 | 34.17819 | 0.075374 | 1.646818 | 0.877179 |
| 4 | 39381200 | 39396599 | 262.8639 | 377.422 | 418.2516 | 172.6865 | 0.058401 | 4.188307 | 0.877187 |
| 18 | 6412600 | 6426199 | 239.8803 | 347.5173 | 388.9404 | 160.3844 | 0.056898 | 4.055307 | 0.877195 |
| 1 | 168394200 | 168395999 | 20.06991 | 35.05401 | 31.77289 | 13.9962 | 0.528877 | 0.71586 | 0.877231 |
| 5 | 116978800 | 116981799 | 42.79113 | 68.71035 | 92.47664 | 39.1443 | -0.07792 | 1.918747 | 0.877278 |
| 9 | 17600000 | 17602799 | 33.12707 | 54.20619 | 67.11163 | 28.86594 | 0.068167 | 1.474786 | 0.877354 |
| 5 | 70438400 | 70440599 | 26.1782 | 44.04175 | 54.46129 | 23.79688 | 0.075754 | 1.133213 | 0.877359 |
| 4 | 61806800 | 61816999 | 177.9631 | 263.2774 | 326.0251 | 134.2661 | -0.06006 | 3.720765 | 0.87737 |
| 15 | 9469200 | 9470199 | 11.6815 | 21.36421 | 30.79041 | 13.17207 | -0.10034 | 0.28294 | 0.877396 |
| 16 | 75937600 | 75937999 | 6.092011 | 11.86113 | 19.61557 | 7.192967 | -0.10145 | -0.34522 | 0.877409 |
| 13 | 39770000 | 39772799 | 29.80552 | 49.40872 | 60.76379 | 26.31099 | 0.083204 | 1.302723 | 0.877417 |
| X | 157370200 | 157371799 | 13.92756 | 25.54176 | 31.7169 | 13.51804 | 0.116505 | 0.28296 | 0.877444 |
| 9 | 92529200 | 92530399 | 14.99267 | 26.44947 | 36.61016 | 16.20042 | -0.0771 | 0.554016 | 0.877463 |
| 15 | 81098200 | 81101599 | 26.16279 | 43.47474 | 59.89031 | 25.97593 | -0.07965 | 1.265795 | 0.877503 |

| | | | | | | | | | |
|---|---|---|---|---|---|---|---|---|---|
| 10 | 129879000 | 129885799 | 84.43409 | 132.0657 | 157.3023 | 65.25834 | 0.062792 | 2.641109 | 0.877546 |
| 10 | 40789200 | 40791599 | 21.42991 | 37.0382 | 45.77312 | 20.08114 | 0.08134 | 0.866453 | 0.877576 |
| 10 | 46852800 | 46853599 | 12.66836 | 23.96288 | 22.60222 | 9.171456 | 0.566111 | 0.068744 | 0.87758 |
| 4 | 112760400 | 112766399 | 66.37309 | 105.735 | 123.0251 | 51.48046 | 0.118875 | 2.200133 | 0.877589 |
| 2 | 125475400 | 125476399 | 22.53942 | 39.05504 | 38.88324 | 17.0598 | 0.392713 | 0.840766 | 0.877649 |
| 17 | 65474800 | 65475599 | 14.06274 | 25.17824 | 31.42683 | 13.77833 | 0.082195 | 0.42186 | 0.877665 |
| 9 | 14588800 | 14592199 | 29.35629 | 48.61264 | 66.76807 | 28.88979 | -0.07577 | 1.321243 | 0.877666 |
| 6 | 89835800 | 89845999 | 161.6651 | 241.086 | 299.933 | 123.6121 | -0.0561 | 3.579335 | 0.877673 |
| 4 | 10733800 | 10735999 | 32.4511 | 52.88681 | 72.14426 | 30.98508 | -0.07537 | 1.536311 | 0.877703 |
| 3 | 70733400 | 70735399 | 28.73563 | 47.65011 | 58.76865 | 25.4526 | 0.079295 | 1.299593 | 0.877717 |
| 2 | 67154200 | 67156599 | 23.27413 | 40.20745 | 48.47793 | 21.26748 | 0.116656 | 0.881836 | 0.877729 |
| 1 | 118175600 | 118177799 | 43.86983 | 70.56615 | 85.89451 | 36.45168 | 0.069859 | 1.862074 | 0.877745 |
| 2 | 28189000 | 28190999 | 16.76653 | 28.72667 | 51.62309 | 22.63907 | -0.45887 | 0.933711 | 0.877753 |
| 14 | 47244400 | 47248399 | 39.61592 | 63.9574 | 85.90597 | 36.64023 | -0.06559 | 1.744658 | 0.877758 |
| 18 | 7314600 | 7329999 | 266.8239 | 381.8311 | 456.6592 | 188.6427 | -0.05636 | 4.237916 | 0.877761 |
| X | 153754000 | 153758799 | 38.73252 | 63.11198 | 90.60912 | 38.91284 | -0.14931 | 1.561339 | 0.877764 |
| 3 | 130857400 | 130859599 | 23.37471 | 39.62698 | 54.41211 | 23.83058 | -0.0744 | 1.057446 | 0.877769 |
| 10 | 68835000 | 68838399 | 47.56395 | 76.37773 | 92.91565 | 39.33295 | 0.067611 | 1.921618 | 0.877772 |
| 2 | 93843600 | 93844799 | 19.17488 | 33.72189 | 32.67228 | 14.46691 | 0.433035 | 0.604201 | 0.877789 |
| 13 | 34105000 | 34106799 | 21.76819 | 37.14573 | 45.85998 | 20.10514 | 0.079569 | 0.982676 | 0.877796 |
| 10 | 81743000 | 81744599 | 46.2753 | 74.64814 | 76.16253 | 32.45061 | 0.328883 | 1.774678 | 0.877798 |
| 12 | 21553400 | 21558599 | 51.77846 | 82.59126 | 109.5698 | 45.93164 | -0.06811 | 2.138657 | 0.877805 |
| X | 106242800 | 106243599 | 16.6709 | 29.50776 | 35.54542 | 15.71884 | 0.124406 | 0.530147 | 0.877845 |
| 5 | 7227600 | 7230799 | 35.59911 | 57.70754 | 78.17873 | 33.47741 | -0.0718 | 1.638625 | 0.877855 |
| 1 | 101301000 | 101302399 | 23.85189 | 40.14131 | 54.94839 | 24.01214 | -0.07037 | 1.121966 | 0.87791 |
| 9 | 58415600 | 58416999 | 14.43422 | 25.57268 | 35.61171 | 15.74521 | -0.08323 | 0.504414 | 0.877911 |
| 5 | 139402400 | 139404399 | 23.76139 | 39.82952 | 54.89024 | 23.94073 | -0.08004 | 1.161888 | 0.877945 |
| 4 | 30928800 | 30930399 | 10.96182 | 19.5316 | 36.33042 | 15.99326 | -0.49362 | 0.444612 | 0.877955 |
| 16 | 16487600 | 16489399 | 16.71535 | 29.20847 | 40.41362 | 17.80771 | -0.07864 | 0.703744 | 0.878009 |
| 7 | 9453800 | 9454199 | 6.53091 | 12.71566 | 17.98282 | 6.594273 | 0.122274 | -0.34477 | 0.878017 |
| 10 | 68543600 | 68546799 | 26.76924 | 45.36143 | 45.83221 | 20.07017 | 0.366506 | 1.062943 | 0.878027 |
| 8 | 56259000 | 56259799 | 9.780541 | 18.34485 | 26.89402 | 11.08386 | -0.08424 | 0.107704 | 0.878034 |
| X | 164773600 | 164775599 | 15.74314 | 27.83952 | 39.15073 | 17.31642 | -0.09854 | 0.538505 | 0.87806 |
| 16 | 43799600 | 43800599 | 10.90531 | 19.91367 | 28.72005 | 12.31043 | -0.10344 | 0.278161 | 0.878072 |
| 18 | 14705400 | 14706399 | 8.64997 | 16.54918 | 25.11682 | 9.941588 | -0.08335 | -0.04386 | 0.878073 |
| 15 | 35390200 | 35392999 | 53.11301 | 85.00669 | 103.2266 | 43.47975 | 0.064612 | 2.035705 | 0.878099 |
| 5 | 145193800 | 145201399 | 103.3677 | 158.504 | 202.2406 | 83.95207 | -0.05842 | 3.011711 | 0.878103 |
| 11 | 18286400 | 18287599 | 16.13638 | 27.85524 | 49.50971 | 21.71962 | -0.4408 | 0.862207 | 0.878109 |
| 14 | 113493800 | 113495399 | 23.52524 | 40.10853 | 49.55602 | 21.72394 | 0.078281 | 1.003316 | 0.87811 |
| 15 | 97116800 | 97118199 | 17.23206 | 30.07712 | 41.47947 | 18.26427 | -0.07409 | 0.724684 | 0.878112 |
| 2 | 95680000 | 95680999 | 13.08987 | 23.57214 | 33.12628 | 14.44864 | -0.0826 | 0.385496 | 0.878149 |
| 3 | 39874200 | 39877199 | 34.08095 | 55.45631 | 75.1437 | 32.25493 | -0.06735 | 1.563357 | 0.878155 |
| X | 77683800 | 77685199 | 13.80259 | 25.43317 | 31.33293 | 13.26831 | 0.134894 | 0.257633 | 0.878188 |
| 16 | 94516000 | 94518599 | 38.67069 | 62.45307 | 83.89757 | 35.79965 | -0.06519 | 1.733044 | 0.878188 |
| 2 | 21651200 | 21653199 | 24.89004 | 41.55953 | 57.09011 | 24.83923 | -0.07539 | 1.208222 | 0.87819 |
| 1 | 59427000 | 59429399 | 20.82825 | 35.7119 | 49.0119 | 21.49521 | -0.07153 | 0.939351 | 0.878222 |
| 14 | 112834800 | 112836799 | 16.43459 | 28.88672 | 40.17952 | 17.79247 | -0.0863 | 0.60805 | 0.87823 |

| | | | | | | | | | |
|---|---|---|---|---|---|---|---|---|---|
| 17 | 85801000 | 85804199 | 41.9132 | 67.72716 | 82.98181 | 35.43415 | 0.068201 | 1.721708 | 0.878238 |
| Y | 82851200 | 82851599 | 4.264583 | 8.334921 | 15.50488 | 5.357143 | -0.19031 | -0.59869 | 0.878251 |
| 13 | 72505800 | 72507599 | 16.5799 | 28.81654 | 40.6255 | 17.8407 | -0.10623 | 0.780205 | 0.878256 |
| 18 | 45036400 | 45037199 | 7.722738 | 14.87868 | 23.09269 | 8.959059 | -0.08799 | -0.12381 | 0.878261 |
| 6 | 59601200 | 59605399 | 11.76322 | 22.23316 | 22.62287 | 9.195203 | 0.45524 | 0.065023 | 0.87827 |
| 7 | 134657600 | 134660399 | 34.46615 | 57.04573 | 56.45047 | 24.41703 | 0.395746 | 1.332037 | 0.878293 |
| 1 | 3413600 | 3415599 | 29.55678 | 48.75025 | 66.56675 | 28.72747 | -0.068 | 1.368312 | 0.878298 |
| 9 | 110812600 | 110819199 | 62.35809 | 98.80174 | 129.4338 | 53.83602 | -0.05981 | 2.356972 | 0.878302 |
| 18 | 84665400 | 84666799 | 17.95084 | 31.07627 | 43.23712 | 18.96952 | -0.0882 | 0.834935 | 0.878315 |
| 8 | 19672200 | 19692399 | 403.6032 | 559.177 | 598.2209 | 248.4105 | 0.057425 | 4.721818 | 0.878329 |
| 2 | 32394400 | 32397399 | 24.24125 | 41.46945 | 50.65966 | 22.21381 | 0.095208 | 0.974941 | 0.878333 |
| 16 | 19450400 | 19452999 | 34.03139 | 55.42411 | 67.80685 | 29.11491 | 0.080579 | 1.546142 | 0.878335 |
| 4 | 47900600 | 47901599 | 12.7936 | 23.327 | 29.5784 | 12.70199 | 0.079264 | 0.303004 | 0.878347 |
| 13 | 6038000 | 6039199 | 12.01856 | 21.97072 | 31.21352 | 13.36079 | -0.08078 | 0.285743 | 0.878354 |
| 4 | 142662200 | 142663199 | 10.9993 | 20.59306 | 26.80507 | 11.08152 | 0.082304 | 0.128993 | 0.878365 |
| 4 | 17655400 | 17659399 | 61.91485 | 98.1296 | 128.5413 | 53.47865 | -0.05926 | 2.348799 | 0.878365 |
| 11 | 46376400 | 46377399 | 13.38326 | 24.12427 | 30.09213 | 13.10904 | 0.089658 | 0.375907 | 0.878396 |
| 9 | 116349000 | 116350799 | 17.55129 | 30.8646 | 38.0197 | 16.8349 | 0.088379 | 0.605649 | 0.878408 |
| 8 | 6547000 | 6551399 | 97.18802 | 149.7062 | 175.4129 | 72.86096 | 0.069391 | 2.918235 | 0.87842 |
| 15 | 98382200 | 98388399 | 69.33858 | 109.5775 | 132.0725 | 54.8616 | 0.057264 | 2.424272 | 0.878442 |
| 3 | 124835800 | 124836799 | 13.95049 | 24.82113 | 30.55794 | 13.4684 | 0.09686 | 0.459434 | 0.87845 |
| 4 | 23207200 | 23214199 | 78.62006 | 123.7163 | 147.4437 | 61.17137 | 0.068671 | 2.523566 | 0.878455 |
| 9 | 82658400 | 82665999 | 84.30509 | 131.6569 | 171.5742 | 71.19068 | -0.0686 | 2.670005 | 0.878462 |
| 15 | 59170400 | 59172399 | 22.62991 | 38.46959 | 52.79047 | 23.1364 | -0.07331 | 1.027723 | 0.878483 |
| 8 | 5009200 | 5016599 | 238.4724 | 342.7433 | 417.5741 | 172.3797 | -0.07681 | 4.172233 | 0.878523 |
| 14 | 95714400 | 95715399 | 10.85335 | 19.59764 | 35.31152 | 15.34969 | -0.43595 | 0.356908 | 0.878549 |
| 6 | 147959800 | 147964799 | 78.16761 | 122.8274 | 147.1767 | 61.04917 | 0.059301 | 2.551869 | 0.87857 |
| 5 | 127263800 | 127265199 | 17.29829 | 30.17405 | 41.97763 | 18.47644 | -0.08684 | 0.732417 | 0.878619 |
| 3 | 139116400 | 139118599 | 16.80755 | 29.7224 | 36.35095 | 16.07801 | 0.101662 | 0.536478 | 0.878621 |
| 12 | 112034000 | 112040199 | 70.24943 | 110.5511 | 143.9081 | 59.70502 | -0.05839 | 2.516837 | 0.878627 |
| 3 | 93829200 | 93830399 | 41.84045 | 67.20186 | 91.03155 | 38.54835 | -0.08672 | 1.909337 | 0.87863 |
| 10 | 90676400 | 90678399 | 24.81126 | 42.81289 | 40.8909 | 17.93838 | 0.451126 | 0.889499 | 0.878638 |
| 4 | 135424800 | 135425999 | 15.44382 | 27.1822 | 37.62081 | 16.65537 | -0.07835 | 0.583025 | 0.878642 |
| 10 | 14519200 | 14524399 | 121.1143 | 183.6835 | 232.8821 | 96.39995 | -0.06195 | 3.242394 | 0.878646 |
| 2 | 104387000 | 104387799 | 13.53272 | 23.96358 | 33.62005 | 14.86611 | -0.09427 | 0.498229 | 0.878671 |
| 18 | 82715200 | 82717399 | 14.72096 | 26.84115 | 23.33912 | 10.02267 | 0.620072 | 0.303219 | 0.87868 |
| 4 | 79599200 | 79599799 | 10.64268 | 19.90307 | 25.3235 | 10.48892 | 0.113344 | 0.132254 | 0.878705 |
| 18 | 45489600 | 45491799 | 22.77464 | 39.42545 | 37.92223 | 16.63769 | 0.442039 | 0.850026 | 0.878705 |
| 8 | 124100200 | 124101199 | 8.213124 | 15.50027 | 23.91381 | 9.750457 | -0.14443 | 0.045407 | 0.878746 |
| X | 121233000 | 121235999 | 38.44475 | 64.0622 | 50.95114 | 22.1399 | 0.710578 | 1.265964 | 0.878764 |
| 18 | 31859800 | 31861999 | 16.25361 | 28.60204 | 35.56044 | 15.74381 | 0.075563 | 0.587096 | 0.878767 |
| 8 | 79333600 | 79338599 | 38.5389 | 62.63824 | 76.60294 | 32.86655 | 0.078957 | 1.587336 | 0.878771 |
| 6 | 134404400 | 134407399 | 24.42085 | 41.24402 | 51.19969 | 22.39791 | 0.070358 | 1.089151 | 0.878791 |
| 8 | 98531200 | 98532399 | 13.07298 | 23.59003 | 29.48724 | 12.82847 | 0.087589 | 0.363496 | 0.878831 |
| 6 | 72268400 | 72270999 | 25.04233 | 42.25013 | 52.26082 | 22.85499 | 0.075536 | 1.1035 | 0.878878 |
| 15 | 52510800 | 52512599 | 13.15743 | 23.30428 | 41.33296 | 18.27578 | -0.43069 | 0.508257 | 0.87888 |
| 3 | 55483200 | 55486599 | 36.20383 | 59.0787 | 72.59097 | 31.19155 | 0.07626 | 1.522704 | 0.878882 |

| | | | | | | | | | |
|---|---|---|---|---|---|---|---|---|---|
| 1 | 152154800 | 152155999 | 11.64232 | 21.76951 | 27.79666 | 11.51597 | 0.1069 | 0.141979 | 0.878887 |
| 5 | 40962200 | 40963199 | 12.04881 | 22.16465 | 32.37723 | 13.74161 | -0.11224 | 0.250572 | 0.878908 |
| Y | 90111600 | 90116399 | 33.53952 | 53.98015 | 81.84489 | 34.76173 | -0.24523 | 1.826145 | 0.878917 |
| 4 | 143961200 | 143977199 | 254.3825 | 368.9244 | 407.8244 | 168.1474 | 0.076263 | 4.04115 | 0.878922 |
| 9 | 26535400 | 26539599 | 36.80624 | 59.89603 | 82.34714 | 35.34806 | -0.08826 | 1.569745 | 0.878922 |
| X | 65493800 | 65496199 | 25.35431 | 43.31165 | 52.30805 | 22.93397 | 0.111395 | 0.99534 | 0.878993 |
| X | 103849000 | 103851199 | 19.1551 | 33.59523 | 40.71778 | 17.99766 | 0.111523 | 0.660456 | 0.878997 |
| 12 | 11578600 | 11580399 | 25.1799 | 42.03621 | 57.68223 | 25.09409 | -0.07386 | 1.210567 | 0.878999 |
| 12 | 102391400 | 102397599 | 69.83861 | 108.7271 | 176.2258 | 73.15013 | -0.38634 | 2.717845 | 0.878999 |
| 8 | 84521000 | 84523599 | 24.62337 | 41.69895 | 51.73156 | 22.6492 | 0.071426 | 1.067854 | 0.879009 |
| 2 | 134346800 | 134348799 | 12.67932 | 22.35164 | 39.83417 | 17.62971 | -0.44016 | 0.55427 | 0.87901 |
| 6 | 5296400 | 5298599 | 17.96006 | 31.1739 | 43.20764 | 18.95883 | -0.08195 | 0.811313 | 0.879028 |
| 11 | 16547400 | 16551999 | 54.1284 | 86.73473 | 104.1741 | 43.95687 | 0.082417 | 2.000973 | 0.879038 |
| 2 | 23737400 | 23738399 | 9.626112 | 18.35733 | 24.4837 | 9.765103 | 0.091913 | -0.01006 | 0.879061 |
| 12 | 46326800 | 46327799 | 15.05116 | 26.47792 | 36.72345 | 16.25971 | -0.08143 | 0.586859 | 0.879087 |
| X | 38592200 | 38598199 | 69.17695 | 109.9956 | 128.2119 | 53.51509 | 0.113422 | 2.262517 | 0.879104 |
| 1 | 118062800 | 118087199 | 645.5 | 856.7284 | 951.0783 | 397.5026 | -0.05826 | 5.399659 | 0.879112 |
| 13 | 38745800 | 38746999 | 14.77298 | 26.35795 | 32.93388 | 14.4815 | 0.077841 | 0.452134 | 0.879114 |
| 7 | 134661400 | 134663999 | 38.52044 | 62.12117 | 83.92658 | 35.74577 | -0.07573 | 1.774744 | 0.879124 |
| 6 | 132291600 | 132294199 | 30.90361 | 52.17582 | 47.43418 | 20.74862 | 0.518143 | 1.110223 | 0.879131 |
| 9 | 4291200 | 4294599 | 28.63788 | 48.04522 | 69.89552 | 30.50204 | -0.15812 | 1.181509 | 0.879144 |
| 8 | 66161400 | 66165199 | 52.29707 | 83.54676 | 101.1273 | 42.48454 | 0.066373 | 2.090481 | 0.879157 |
| 4 | 153357000 | 153361599 | 47.8666 | 76.67852 | 101.6984 | 42.89322 | -0.06089 | 2.001487 | 0.879183 |
| 7 | 34136000 | 34140799 | 29.47985 | 48.49021 | 66.56088 | 28.69164 | -0.07718 | 1.399512 | 0.879196 |
| 15 | 61159400 | 61161199 | 19.57003 | 34.43426 | 32.31963 | 14.30931 | 0.479156 | 0.603345 | 0.8792 |
| 3 | 57583800 | 57590199 | 65.98646 | 104.6537 | 126.3004 | 52.58205 | 0.059747 | 2.331433 | 0.879218 |
| 14 | 51116800 | 51119199 | 35.87324 | 58.73595 | 58.37791 | 25.11543 | 0.384775 | 1.467785 | 0.879222 |
| 3 | 91164400 | 91165999 | 21.18859 | 36.70323 | 45.21937 | 19.83956 | 0.086373 | 0.843704 | 0.879232 |
| 6 | 48212000 | 48212799 | 22.44032 | 38.11579 | 46.92918 | 20.56525 | 0.08253 | 1.028621 | 0.879233 |
| Y | 38887400 | 38888199 | 8.368316 | 15.96056 | 19.35102 | 7.716054 | 0.228415 | -0.04168 | 0.879242 |
| 6 | 22750200 | 22753599 | 45.3701 | 73.25135 | 88.28228 | 37.65653 | 0.090523 | 1.753801 | 0.879242 |
| 5 | 146801000 | 146802799 | 18.62063 | 32.34446 | 44.70475 | 19.62488 | -0.07808 | 0.802626 | 0.879248 |
| X | 58178800 | 58180799 | 20.79604 | 35.81118 | 43.76645 | 19.19865 | 0.096517 | 0.897461 | 0.87925 |
| 6 | 84309800 | 84311199 | 16.79396 | 29.45675 | 40.79608 | 18.03362 | -0.08003 | 0.645219 | 0.87926 |
| 6 | 122766400 | 122771399 | 40.10814 | 64.60845 | 87.00327 | 36.99947 | -0.07302 | 1.811242 | 0.879262 |
| 5 | 7089600 | 7090399 | 9.150454 | 17.50462 | 23.46956 | 9.292757 | 0.093972 | -0.04067 | 0.879287 |
| 1 | 46524000 | 46524999 | 15.95638 | 28.48045 | 28.28579 | 12.43359 | 0.407543 | 0.44804 | 0.87929 |
| 18 | 8698000 | 8699799 | 27.38808 | 45.6652 | 62.76915 | 27.28547 | -0.07632 | 1.236294 | 0.879293 |
| 11 | 19433400 | 19436599 | 39.15445 | 63.77081 | 77.27964 | 33.18294 | 0.094248 | 1.561323 | 0.879293 |
| 15 | 95281200 | 95282399 | 14.74143 | 26.1361 | 32.43303 | 14.33739 | 0.082412 | 0.489683 | 0.879295 |
| 1 | 182046400 | 182048799 | 39.26745 | 63.59074 | 78.29049 | 33.50067 | 0.064471 | 1.663379 | 0.8793 |
| 3 | 130389200 | 130389799 | 6.536805 | 12.70586 | 20.64264 | 7.672581 | -0.09429 | -0.28885 | 0.879329 |
| 17 | 39512600 | 39513799 | 24.4097 | 42.10063 | 40.89094 | 17.93814 | 0.426842 | 0.890971 | 0.879329 |
| X | 47551600 | 47553599 | 16.29174 | 28.65651 | 40.2641 | 17.82785 | -0.10017 | 0.602798 | 0.879338 |
| 7 | 78980000 | 78981199 | 11.84901 | 21.66297 | 27.17578 | 11.63098 | 0.098006 | 0.275904 | 0.879362 |
| 9 | 119045400 | 119047999 | 45.26265 | 73.13309 | 74.15167 | 31.66 | 0.340547 | 1.725012 | 0.879375 |
| 2 | 8411000 | 8412999 | 23.25311 | 39.48378 | 54.39886 | 23.83631 | -0.07914 | 1.044134 | 0.879391 |

| | | | | | | | | | |
|---|---|---|---|---|---|---|---|---|---|
| 8 | 87916000 | 87917999 | 19.25514 | 33.27572 | 45.81912 | 20.10125 | -0.07391 | 0.859678 | 0.879405 |
| 15 | 61153400 | 61156599 | 18.64754 | 33.72824 | 26.73967 | 11.59907 | 0.743084 | 0.383673 | 0.879407 |
| 3 | 46934400 | 46935199 | 21.723 | 37.10245 | 45.84362 | 20.09944 | 0.078559 | 0.973357 | 0.879407 |
| 18 | 7545600 | 7547199 | 14.29355 | 25.40744 | 31.52053 | 13.90376 | 0.08506 | 0.469819 | 0.879409 |
| 10 | 59214000 | 59217999 | 44.66046 | 71.66672 | 96.10996 | 40.64323 | -0.07388 | 1.94003 | 0.879412 |
| 5 | 127646600 | 127648399 | 15.77873 | 27.67049 | 38.26707 | 16.91283 | -0.07791 | 0.638269 | 0.879424 |
| 3 | 32366200 | 32368999 | 14.00813 | 24.52447 | 43.08016 | 19.01322 | -0.42107 | 0.665908 | 0.879438 |
| 7 | 51117800 | 51118999 | 12.77833 | 22.79076 | 32.2005 | 14.16339 | -0.09845 | 0.441515 | 0.87947 |
| 9 | 18102600 | 18103599 | 16.68615 | 29.67936 | 26.04071 | 11.48041 | 0.58272 | 0.471642 | 0.87949 |
| 2 | 122974600 | 122975399 | 8.504978 | 16.21164 | 24.49514 | 9.779428 | -0.08798 | -0.01859 | 0.879497 |
| 1 | 53733200 | 53740199 | 132.9621 | 201.0563 | 253.3536 | 104.7768 | -0.05676 | 3.305109 | 0.879508 |
| 1 | 125053000 | 125055199 | 19.86633 | 34.10499 | 46.8926 | 20.56781 | -0.07383 | 0.922136 | 0.879529 |
| 1 | 122699200 | 122703199 | 36.23017 | 58.16111 | 102.922 | 43.54954 | -0.47233 | 1.9236 | 0.87954 |
| 12 | 81751000 | 81759999 | 93.87392 | 145.6239 | 172.5572 | 71.66564 | 0.060987 | 2.788272 | 0.879542 |
| 16 | 46960800 | 46961999 | 15.92737 | 28.11414 | 34.99681 | 15.48476 | 0.075493 | 0.550595 | 0.879554 |
| 3 | 37316400 | 37318599 | 19.15186 | 33.33096 | 41.49826 | 18.2392 | 0.07237 | 0.775201 | 0.879564 |
| 3 | 14305200 | 14308599 | 66.08304 | 103.9006 | 159.3967 | 66.12319 | -0.29607 | 2.535488 | 0.879575 |
| 12 | 71122000 | 71123599 | 12.36029 | 22.0953 | 31.80574 | 13.96299 | -0.12305 | 0.425507 | 0.879585 |
| 6 | 22874800 | 22876599 | 12.77228 | 22.70584 | 32.21947 | 14.21073 | -0.10741 | 0.461573 | 0.8796 |
| X | 12213400 | 12219399 | 48.15796 | 77.49408 | 93.1798 | 39.56583 | 0.088179 | 1.861383 | 0.879613 |
| 12 | 114069200 | 114097199 | 398.7304 | 558.1327 | 583.158 | 241.7684 | 0.106452 | 4.568654 | 0.879622 |
| 4 | 28412000 | 28415199 | 93.81184 | 144.7597 | 169.4339 | 70.37964 | 0.072893 | 2.884338 | 0.879631 |
| 2 | 102379600 | 102382199 | 13.88107 | 24.13626 | 45.82244 | 20.12839 | -0.53332 | 0.771247 | 0.879631 |
| 14 | 109850800 | 109851599 | 12.34784 | 22.45231 | 28.1435 | 12.1275 | 0.092779 | 0.313283 | 0.879631 |
| 6 | 145055400 | 145057599 | 19.48053 | 33.8138 | 42.09681 | 18.47168 | 0.072114 | 0.814978 | 0.879639 |
| 1 | 29734800 | 29739799 | 93.56087 | 144.1657 | 185.2296 | 76.94144 | -0.06309 | 2.911915 | 0.879641 |
| 2 | 46494200 | 46495399 | 17.95967 | 31.25735 | 38.46792 | 16.90773 | 0.089068 | 0.763936 | 0.879642 |
| 1 | 105747200 | 105748199 | 10.05594 | 18.77022 | 27.33304 | 11.35762 | -0.08433 | 0.140884 | 0.879642 |
| 6 | 32520200 | 32522799 | 20.48118 | 35.24994 | 48.47351 | 21.26311 | -0.07338 | 0.903007 | 0.879655 |
| 8 | 3386000 | 3391599 | 52.11807 | 83.56274 | 81.61977 | 34.47089 | 0.381127 | 1.97137 | 0.879661 |
| 5 | 38010400 | 38011999 | 12.56926 | 22.93465 | 29.04395 | 12.46113 | 0.081817 | 0.294924 | 0.879666 |
| 16 | 89758600 | 89760199 | 10.62675 | 19.58444 | 28.29392 | 11.97875 | -0.09299 | 0.22455 | 0.879666 |
| 18 | 70058400 | 70059199 | 6.533052 | 12.66828 | 20.55463 | 7.761487 | -0.11375 | -0.23495 | 0.879679 |
| 1 | 166862200 | 166862799 | 7.607466 | 14.71525 | 23.23461 | 8.875423 | -0.09102 | -0.17469 | 0.87968 |
| 12 | 33883800 | 33885399 | 20.14429 | 34.77467 | 47.93617 | 21.02933 | -0.07598 | 0.873905 | 0.879688 |
| X | 129692000 | 129694599 | 22.03279 | 37.92147 | 53.41309 | 23.4299 | -0.10789 | 0.915651 | 0.879699 |
| 4 | 79815200 | 79816399 | 13.60484 | 24.47124 | 34.55663 | 15.0916 | -0.09111 | 0.395759 | 0.8797 |
| 5 | 126661400 | 126680999 | 251.0131 | 361.5542 | 435.0759 | 179.5458 | -0.05584 | 4.141632 | 0.879714 |
| 15 | 96050400 | 96051199 | 9.307893 | 17.518 | 25.89842 | 10.61205 | -0.08944 | 0.079445 | 0.879715 |
| 5 | 150273400 | 150274599 | 13.95412 | 25.07241 | 31.47049 | 13.75633 | 0.077209 | 0.400209 | 0.879726 |
| 16 | 48126600 | 48136399 | 91.83127 | 142.6205 | 169.0513 | 70.19743 | 0.061834 | 2.766835 | 0.879735 |
| 9 | 42869400 | 42871999 | 30.6109 | 50.5205 | 62.82135 | 27.11922 | 0.066899 | 1.359463 | 0.879735 |
| 15 | 35384000 | 35395199 | 159.201 | 239.6352 | 271.8344 | 112.2669 | 0.0891 | 3.403134 | 0.879742 |
| 11 | 113753200 | 113754799 | 12.21697 | 21.8963 | 31.14069 | 13.64091 | -0.10396 | 0.402798 | 0.879814 |
| 14 | 42448400 | 42451399 | 16.51776 | 29.83399 | 28.17539 | 12.24341 | 0.490794 | 0.381125 | 0.879817 |
| 6 | 6247600 | 6248399 | 13.50429 | 24.20836 | 30.0833 | 13.17552 | 0.089827 | 0.40952 | 0.879869 |
| 9 | 23625200 | 23626599 | 18.99057 | 32.89507 | 45.26635 | 19.86016 | -0.07242 | 0.835034 | 0.879873 |

| | | | | | | | | | |
|---|---|---|---|---|---|---|---|---|---|
| 6 | 124524400 | 124528399 | 51.29613 | 82.022 | 108.5286 | 45.65243 | -0.06048 | 2.063177 | 0.879881 |
| 14 | 49078800 | 49081199 | 18.78893 | 31.79807 | 56.35856 | 24.6702 | -0.44121 | 1.059542 | 0.879889 |
| 14 | 28342200 | 28343399 | 22.33631 | 37.48471 | 52.70175 | 23.00323 | -0.1087 | 1.143765 | 0.87989 |
| 4 | 62172800 | 62173399 | 7.570045 | 14.66809 | 20.51319 | 7.773104 | 0.093599 | -0.21108 | 0.879899 |
| 15 | 26221200 | 26221999 | 7.570045 | 14.66809 | 20.51319 | 7.773104 | 0.093599 | -0.21108 | 0.879899 |
| 13 | 13863400 | 13865199 | 16.22433 | 29.28535 | 35.04511 | 15.24073 | 0.150315 | 0.394911 | 0.879911 |
| 7 | 51159000 | 51161399 | 29.13488 | 47.9571 | 66.02471 | 28.46914 | -0.08104 | 1.389056 | 0.879919 |
| 11 | 110250200 | 110251399 | 17.24724 | 30.33967 | 37.53229 | 16.618 | 0.082497 | 0.600052 | 0.879987 |
| 17 | 37475400 | 37476799 | 12.02145 | 21.96618 | 27.66249 | 11.84742 | 0.091835 | 0.281639 | 0.879989 |
| 1 | 177954400 | 177957199 | 76.07622 | 119.3341 | 142.6429 | 59.15934 | 0.0609 | 2.58384 | 0.88001 |
| 15 | 97413400 | 97417199 | 32.71225 | 53.33419 | 72.39704 | 31.09828 | -0.06791 | 1.529408 | 0.880014 |
| 9 | 62236200 | 62239199 | 21.51316 | 37.34851 | 34.41515 | 15.10269 | 0.504454 | 0.811734 | 0.880036 |
| 5 | 24883000 | 24884799 | 22.26904 | 38.27955 | 47.39364 | 20.78863 | 0.07712 | 0.920563 | 0.88004 |
| 18 | 11130600 | 11132399 | 17.56361 | 30.94367 | 30.36719 | 13.44142 | 0.415698 | 0.571832 | 0.88004 |
| 11 | 24736200 | 24737599 | 12.0556 | 22.77173 | 23.09842 | 9.400743 | 0.457943 | 0.072561 | 0.880087 |
| 7 | 139276800 | 139279799 | 24.6869 | 41.39868 | 56.59451 | 24.69076 | -0.06854 | 1.160594 | 0.880092 |
| 15 | 37263200 | 37265599 | 21.66782 | 36.96319 | 50.62191 | 22.19224 | -0.06961 | 0.990676 | 0.880108 |
| 19 | 58149800 | 58155399 | 54.27612 | 86.40487 | 114.2439 | 47.76063 | -0.06631 | 2.205989 | 0.880109 |
| 4 | 93351800 | 93353199 | 13.20075 | 23.68536 | 33.0766 | 14.47559 | -0.07696 | 0.407401 | 0.880114 |
| 14 | 14798800 | 14799599 | 11.96881 | 21.49565 | 30.74629 | 13.44484 | -0.11016 | 0.389519 | 0.880125 |
| 1 | 17609200 | 17610999 | 15.72302 | 28.44494 | 27.64983 | 11.99053 | 0.451114 | 0.363654 | 0.880127 |
| 4 | 55248800 | 55253399 | 40.88174 | 66.06224 | 81.13089 | 34.64501 | 0.064885 | 1.717467 | 0.880142 |
| 12 | 91448800 | 91451599 | 37.66813 | 60.98012 | 82.04329 | 35.09774 | -0.06371 | 1.669425 | 0.880157 |
| 15 | 86881600 | 86884399 | 29.4689 | 49.2688 | 60.17318 | 26.20205 | 0.093575 | 1.210977 | 0.880177 |
| 4 | 131192400 | 131194199 | 27.50107 | 46.2325 | 56.71918 | 24.77781 | 0.087072 | 1.145916 | 0.880185 |
| 1 | 130963400 | 130966799 | 37.69909 | 61.21005 | 75.16782 | 32.22627 | 0.0716 | 1.60504 | 0.880212 |
| 16 | 21500600 | 21503199 | 25.32784 | 42.9897 | 52.82498 | 23.1439 | 0.085301 | 1.052667 | 0.880271 |
| 3 | 72933400 | 72933799 | 6.405056 | 12.46687 | 20.71778 | 7.615658 | -0.11122 | -0.32662 | 0.880275 |
| 12 | 5792000 | 5798199 | 65.11632 | 102.9659 | 135.1393 | 56.15234 | -0.0648 | 2.406216 | 0.88028 |
| 15 | 96392000 | 96394799 | 35.52345 | 57.93193 | 71.55053 | 30.73743 | 0.068257 | 1.532479 | 0.880286 |
| 13 | 53972800 | 53976399 | 33.15882 | 54.54319 | 67.39215 | 29.05032 | 0.074233 | 1.410986 | 0.880354 |
| 1 | 8603000 | 8608599 | 47.09869 | 75.99631 | 90.54475 | 38.59152 | 0.106032 | 1.773445 | 0.880384 |
| 16 | 69003800 | 69006599 | 32.69642 | 53.05371 | 87.2787 | 37.40095 | -0.35001 | 1.61842 | 0.880387 |
| 7 | 12498600 | 12499199 | 13.05818 | 23.47738 | 28.70531 | 12.54096 | 0.115747 | 0.38631 | 0.880397 |
| 7 | 5162000 | 5164199 | 20.82554 | 35.97332 | 50.1146 | 21.98542 | -0.09099 | 0.87409 | 0.880404 |
| 6 | 111732600 | 111733999 | 23.75353 | 39.77168 | 54.8894 | 23.92341 | -0.0821 | 1.17353 | 0.880428 |
| 1 | 181319400 | 181323399 | 44.5962 | 71.61561 | 95.37212 | 40.37578 | -0.06254 | 1.919313 | 0.880472 |
| 14 | 123959200 | 123962999 | 67.75915 | 107.0142 | 127.848 | 53.08988 | 0.069291 | 2.438118 | 0.880507 |
| 13 | 56531600 | 56533199 | 16.74027 | 29.28322 | 40.37743 | 17.80779 | -0.07368 | 0.687056 | 0.880509 |
| 14 | 24658600 | 24660999 | 20.0592 | 34.94441 | 43.03303 | 18.92546 | 0.088259 | 0.768926 | 0.880542 |
| 14 | 18371400 | 18372199 | 18.61887 | 31.49349 | 56.36181 | 24.66764 | -0.45518 | 1.062827 | 0.880554 |
| 3 | 134395000 | 134396399 | 14.65477 | 26.77411 | 32.6665 | 13.99303 | 0.136513 | 0.309939 | 0.880584 |
| 4 | 117298400 | 117300799 | 18.01894 | 31.29756 | 43.15421 | 18.94355 | -0.07453 | 0.799565 | 0.880596 |
| 2 | 40882000 | 40884599 | 26.41984 | 44.2554 | 61.2086 | 26.685 | -0.08538 | 1.180257 | 0.8806 |
| 12 | 46060400 | 46061399 | 11.18054 | 20.71441 | 29.84702 | 12.54558 | -0.08212 | 0.200851 | 0.880606 |
| 5 | 149630800 | 149632799 | 41.82829 | 67.42378 | 90.33002 | 38.44964 | -0.06449 | 1.796957 | 0.880609 |
| 17 | 66315000 | 66315999 | 11.71027 | 21.21084 | 30.15374 | 13.05673 | -0.09237 | 0.334115 | 0.880625 |

| | | | | | | | | | |
|---|---|---|---|---|---|---|---|---|---|
| 14 | 37538400 | 37538999 | 7.833879 | 15.14179 | 20.97419 | 8.040218 | 0.090964 | -0.1694 | 0.880628 |
| 8 | 67637800 | 67639599 | 12.44073 | 23.08665 | 22.49842 | 9.435878 | 0.480636 | 0.184547 | 0.880678 |
| 3 | 67604800 | 67605599 | 12.51586 | 23.50426 | 29.64356 | 12.19158 | 0.133418 | 0.129271 | 0.880683 |
| 3 | 52026200 | 52028199 | 20.51575 | 35.31446 | 43.74267 | 19.18799 | 0.077004 | 0.8977 | 0.880696 |
| 6 | 123004200 | 123006799 | 41.91643 | 67.50078 | 90.23395 | 38.35643 | -0.06302 | 1.825054 | 0.880712 |
| 10 | 78693200 | 78695599 | 41.97774 | 67.55823 | 90.32315 | 38.36037 | -0.06429 | 1.845595 | 0.880737 |
| 12 | 100896600 | 100898399 | 14.89845 | 26.3371 | 32.51374 | 14.38182 | 0.088224 | 0.522651 | 0.880741 |
| 1 | 102554800 | 102556399 | 16.69858 | 29.47573 | 36.43178 | 16.11965 | 0.085585 | 0.556656 | 0.880743 |
| 3 | 102157000 | 102160199 | 33.63792 | 55.02162 | 68.25851 | 29.35526 | 0.064892 | 1.482308 | 0.880746 |
| 2 | 178240200 | 178242999 | 34.7286 | 56.41402 | 76.21861 | 32.6883 | -0.0651 | 1.592876 | 0.880747 |
| 10 | 18544600 | 18552599 | 146.0223 | 218.9123 | 273.9612 | 113.0243 | -0.0577 | 3.477482 | 0.880782 |
| 12 | 60821600 | 60822199 | 9.296207 | 17.53557 | 25.91701 | 10.57839 | -0.08455 | 0.06465 | 0.880785 |
| 3 | 69429400 | 69430199 | 10.25628 | 19.17255 | 27.89251 | 11.56105 | -0.07996 | 0.135505 | 0.880804 |
| 1 | 24842000 | 24848399 | 126.3115 | 191.9401 | 224.7304 | 93.08 | 0.055098 | 3.202984 | 0.880804 |
| 8 | 69848000 | 69850199 | 16.6641 | 29.15667 | 35.79408 | 15.78944 | 0.093159 | 0.675282 | 0.880805 |
| X | 140504400 | 140509199 | 48.99393 | 78.48394 | 105.1009 | 44.32786 | -0.07476 | 2.006436 | 0.880806 |
| 2 | 75329200 | 75330599 | 26.57294 | 45.25549 | 44.21986 | 19.38303 | 0.415346 | 1.018538 | 0.880859 |
| X | 153308000 | 153328599 | 226.8293 | 335.2282 | 356.1749 | 146.681 | 0.159406 | 3.746689 | 0.880866 |
| 12 | 54307000 | 54309799 | 31.1582 | 51.87416 | 51.41322 | 22.32816 | 0.39399 | 1.250034 | 0.880881 |
| 8 | 19070200 | 19071799 | 13.47689 | 24.24905 | 34.06236 | 14.87162 | -0.08316 | 0.392634 | 0.880889 |
| 3 | 20685000 | 20693999 | 218.4187 | 315.2651 | 451.3447 | 186.208 | -0.30337 | 4.108895 | 0.880893 |
| 1 | 103022600 | 103023999 | 13.26816 | 24.16266 | 30.56141 | 13.14426 | 0.081343 | 0.315266 | 0.880896 |
| 14 | 55462800 | 55469199 | 78.75589 | 123.3074 | 147.5256 | 61.19611 | 0.057505 | 2.618615 | 0.880915 |
| 17 | 17926400 | 17928399 | 21.71785 | 37.70417 | 37.77208 | 16.57593 | 0.384175 | 0.814961 | 0.880923 |
| 4 | 61657600 | 61658999 | 43.37077 | 69.75578 | 84.57148 | 35.88453 | 0.075389 | 1.86415 | 0.880937 |
| 7 | 50021400 | 50022599 | 12.896 | 23.50529 | 29.63697 | 12.73286 | 0.087155 | 0.307773 | 0.880939 |
| 15 | 17933800 | 17934399 | 8.332311 | 16.0469 | 21.9854 | 8.538926 | 0.089056 | -0.11634 | 0.880952 |
| 8 | 122568000 | 122568399 | 6.976037 | 13.57746 | 19.59372 | 7.205767 | 0.088947 | -0.32447 | 0.880991 |
| 17 | 36368600 | 36388199 | 339.9302 | 470.3441 | 674.1323 | 279.9911 | -0.36631 | 4.740675 | 0.881007 |
| 6 | 79224800 | 79226399 | 10.50047 | 19.99048 | 20.60633 | 8.248498 | 0.455601 | -0.0032 | 0.881037 |
| 8 | 55161600 | 55165799 | 63.76791 | 100.8734 | 131.9738 | 54.84793 | -0.05975 | 2.397377 | 0.881052 |
| 10 | 103188000 | 103189199 | 13.25364 | 24.18777 | 30.57864 | 13.11663 | 0.08447 | 0.302786 | 0.881055 |
| 3 | 53339600 | 53342599 | 21.60172 | 36.7731 | 50.62586 | 22.19021 | -0.07765 | 1.007969 | 0.881091 |
| 12 | 11037600 | 11039399 | 23.17477 | 39.52815 | 49.02051 | 21.48994 | 0.073037 | 0.997174 | 0.881139 |
| 7 | 88481400 | 88482999 | 22.0652 | 37.58363 | 46.38492 | 20.33227 | 0.079622 | 1.000405 | 0.881198 |
| 7 | 66979200 | 66983399 | 57.13911 | 91.81699 | 104.8119 | 44.41437 | 0.160465 | 1.920967 | 0.88121 |
| 11 | 84538000 | 84539599 | 14.70937 | 26.37558 | 37.70971 | 16.51198 | -0.1114 | 0.430333 | 0.881224 |
| 17 | 39997000 | 39998799 | 25.00764 | 42.86937 | 42.58035 | 18.67422 | 0.393123 | 0.950902 | 0.88125 |
| 16 | 46462800 | 46463799 | 8.792864 | 16.64228 | 24.91025 | 10.09502 | -0.0937 | 0.032977 | 0.881252 |
| 7 | 65252000 | 65256399 | 48.63516 | 77.92518 | 94.73915 | 39.97009 | 0.064761 | 1.993573 | 0.881294 |
| 3 | 123753400 | 123754799 | 20.24717 | 35.18459 | 43.59747 | 19.14386 | 0.078934 | 0.803341 | 0.881296 |
| 4 | 145358000 | 145380599 | 618.7426 | 828.024 | 856.3227 | 357.4995 | 0.055873 | 5.267889 | 0.881303 |
| 5 | 26249200 | 26252599 | 44.67858 | 71.85375 | 87.84019 | 37.26738 | 0.063301 | 1.869509 | 0.881334 |
| 17 | 41923400 | 41925999 | 32.67211 | 54.05675 | 55.21372 | 23.8761 | 0.350178 | 1.330776 | 0.88134 |
| 2 | 157495600 | 157499399 | 41.90889 | 67.61063 | 83.04365 | 35.38657 | 0.061975 | 1.768448 | 0.881345 |
| 11 | 73473400 | 73475799 | 10.00164 | 19.20486 | 19.73942 | 7.732052 | 0.488753 | -0.07821 | 0.881345 |
| 3 | 16105800 | 16106999 | 31.09892 | 50.79964 | 69.42308 | 29.84134 | -0.07565 | 1.497895 | 0.881352 |

| | | | | | | | | | |
|---|---|---|---|---|---|---|---|---|---|
| 15 | 9736400 | 9738399 | 24.30461 | 41.52235 | 39.11694 | 17.15059 | 0.468494 | 0.97469 | 0.88136 |
| 15 | 37554800 | 37558599 | 40.24056 | 65.23208 | 79.87927 | 34.20582 | 0.073562 | 1.645607 | 0.881362 |
| 10 | 65326400 | 65327399 | 11.08246 | 20.44501 | 29.28829 | 12.38262 | -0.07962 | 0.224113 | 0.881387 |
| 4 | 10732200 | 10735399 | 40.32111 | 64.98655 | 87.04845 | 37.04709 | -0.0644 | 1.796176 | 0.881389 |
| 12 | 27809000 | 27810599 | 20.7455 | 36.1465 | 44.04654 | 19.37338 | 0.103287 | 0.76892 | 0.881398 |
| 1 | 173443200 | 173444199 | 15.31258 | 27.23986 | 33.94284 | 14.96249 | 0.07934 | 0.476742 | 0.881406 |
| 19 | 57609000 | 57610599 | 13.63182 | 24.80944 | 30.98947 | 13.33876 | 0.098298 | 0.320373 | 0.881433 |
| 11 | 90397400 | 90402199 | 61.1189 | 97.41049 | 98.34521 | 41.17971 | 0.323989 | 2.171811 | 0.881441 |
| 5 | 14727600 | 14732599 | 61.00948 | 96.84292 | 126.9303 | 52.88946 | -0.05797 | 2.302015 | 0.881444 |
| 17 | 64064600 | 64070999 | 66.74616 | 105.3105 | 137.4516 | 57.04939 | -0.05933 | 2.461492 | 0.881468 |
| 4 | 116374800 | 116375799 | 8.396253 | 16.06944 | 24.57916 | 9.718927 | -0.09261 | -0.04908 | 0.881482 |
| 11 | 52047400 | 52051999 | 56.68889 | 90.36777 | 119.6869 | 50.12969 | -0.06671 | 2.165645 | 0.881498 |
| 7 | 3896800 | 3898799 | 18.4938 | 32.10898 | 44.19748 | 19.39644 | -0.07218 | 0.808826 | 0.881505 |
| 19 | 37783800 | 37784999 | 9.59878 | 17.93762 | 26.33285 | 10.92061 | -0.09349 | 0.127498 | 0.881523 |
| 15 | 91168800 | 91173999 | 83.15986 | 129.7848 | 167.8066 | 69.63203 | -0.05796 | 2.680119 | 0.881558 |
| 2 | 90670800 | 90672799 | 18.14297 | 31.64345 | 39.38709 | 17.33483 | 0.073043 | 0.736291 | 0.881568 |
| 7 | 47533000 | 47534199 | 22.36018 | 38.23741 | 47.42307 | 20.79397 | 0.073534 | 0.969167 | 0.881573 |
| 9 | 20270600 | 20278999 | 148.4063 | 225.2479 | 210.3489 | 87.09249 | 0.379717 | 3.231183 | 0.881575 |
| 2 | 74015400 | 74017199 | 20.86316 | 35.76807 | 49.01126 | 21.49475 | -0.06924 | 0.94112 | 0.881577 |
| 1 | 107057800 | 107059199 | 13.01978 | 23.54477 | 33.13234 | 14.37543 | -0.07978 | 0.359403 | 0.881579 |
| 16 | 39382400 | 39382999 | 10.17635 | 19.02238 | 27.92001 | 11.57323 | -0.0924 | 0.137021 | 0.881602 |
| 2 | 45534000 | 45534599 | 7.111763 | 13.82746 | 22.45666 | 8.327187 | -0.09159 | -0.287 | 0.881626 |
| 14 | 70195800 | 70196399 | 8.358803 | 16.15389 | 25.57392 | 9.810561 | -0.10128 | -0.14747 | 0.881628 |
| 6 | 89832200 | 89834999 | 33.71772 | 55.26819 | 68.27691 | 29.38803 | 0.072493 | 1.457034 | 0.881714 |
| 5 | 115920400 | 115928799 | 139.9675 | 211.4674 | 246.2492 | 101.8111 | 0.055972 | 3.323572 | 0.88173 |
| 4 | 16544000 | 16545999 | 25.95464 | 43.43701 | 53.92061 | 23.49431 | 0.070212 | 1.18341 | 0.881753 |
| 5 | 47444400 | 47445199 | 11.03997 | 19.48168 | 39.48152 | 17.47006 | -0.62325 | 0.53735 | 0.881774 |
| 12 | 13238200 | 13239999 | 15.80906 | 27.73835 | 38.25929 | 16.91696 | -0.07407 | 0.630552 | 0.881788 |
| 8 | 76206200 | 76206799 | 6.531956 | 12.71391 | 20.74019 | 7.623685 | -0.08559 | -0.32822 | 0.881789 |
| 12 | 3631400 | 3633399 | 18.62571 | 32.45126 | 40.42886 | 17.78158 | 0.071583 | 0.754889 | 0.881804 |
| 1 | 10435200 | 10435999 | 13.0518 | 23.87888 | 30.15745 | 12.89502 | 0.089215 | 0.288317 | 0.881807 |
| 1 | 112259200 | 112262999 | 32.91756 | 54.21189 | 66.97208 | 28.88403 | 0.075121 | 1.393963 | 0.881812 |
| 11 | 17876000 | 17878399 | 20.78457 | 36.05964 | 51.2649 | 22.49305 | -0.11893 | 0.835757 | 0.881824 |
| 12 | 47979600 | 47983599 | 73.47694 | 115.6358 | 139.0029 | 57.6703 | 0.056418 | 2.516537 | 0.881834 |
| 4 | 11035000 | 11036599 | 20.59414 | 35.50904 | 44.21522 | 19.39627 | 0.069849 | 0.881971 | 0.881836 |
| X | 156971600 | 156974999 | 34.5991 | 56.57731 | 78.10654 | 33.58693 | -0.08923 | 1.496493 | 0.881859 |
| 13 | 85263400 | 85266399 | 16.33548 | 28.9715 | 26.80572 | 11.8466 | 0.503789 | 0.486029 | 0.88189 |
| 16 | 81500200 | 81502599 | 25.13684 | 42.08156 | 51.74915 | 22.55263 | 0.08376 | 1.175612 | 0.881895 |
| 2 | 96599800 | 96601199 | 15.07773 | 26.62197 | 37.03895 | 16.3874 | -0.08417 | 0.544308 | 0.881897 |
| 18 | 24169200 | 24170799 | 20.50955 | 34.62713 | 58.63684 | 25.64888 | -0.376 | 1.086666 | 0.8819 |
| 3 | 124617800 | 124628599 | 133.2406 | 202.1533 | 235.4022 | 97.45448 | 0.061106 | 3.236465 | 0.881942 |
| 1 | 176429400 | 176430199 | 6.539155 | 12.70195 | 20.60782 | 7.701043 | -0.09938 | -0.26962 | 0.881942 |
| 1 | 23824000 | 23825999 | 33.86757 | 55.05437 | 74.57781 | 32.00043 | -0.06797 | 1.575355 | 0.881972 |
| 1 | 27233000 | 27234599 | 20.98103 | 36.45208 | 35.34656 | 15.51876 | 0.431219 | 0.799076 | 0.881977 |
| 3 | 32235600 | 32237399 | 19.35756 | 33.3389 | 45.82931 | 20.10364 | -0.07257 | 0.889408 | 0.882 |
| 7 | 55595000 | 55606399 | 203.9023 | 298.6148 | 364.8686 | 150.2984 | -0.05455 | 3.887807 | 0.882016 |
| 9 | 57692600 | 57694799 | 22.23794 | 37.72682 | 51.67878 | 22.63839 | -0.07084 | 1.044854 | 0.88203 |

| | | | | | | | | | |
|---|---|---|---|---|---|---|---|---|---|
| 5 | 145168200 | 145169199 | 9.611729 | 17.92336 | 26.32766 | 10.95816 | -0.09849 | 0.141586 | 0.882059 |
| 14 | 15661800 | 15665399 | 21.40443 | 36.80514 | 45.79761 | 20.08875 | 0.069996 | 0.911594 | 0.882096 |
| 10 | 97163600 | 97167599 | 68.31483 | 107.4908 | 140.7022 | 58.37382 | -0.06652 | 2.518581 | 0.882116 |
| 12 | 58648400 | 58650399 | 22.2599 | 38.29729 | 47.39059 | 20.78785 | 0.078125 | 0.912586 | 0.882122 |
| 7 | 32040200 | 32056199 | 86.73567 | 136.6236 | 129.0236 | 53.53623 | 0.405062 | 2.506936 | 0.882151 |
| 8 | 105157800 | 105159799 | 18.57283 | 32.28748 | 39.98024 | 17.55978 | 0.080018 | 0.784606 | 0.882159 |
| 9 | 123880000 | 123883199 | 37.85895 | 61.95324 | 61.51403 | 26.45897 | 0.385695 | 1.478845 | 0.882168 |
| 14 | 93047200 | 93047799 | 6.95955 | 13.52411 | 21.78973 | 8.115682 | -0.08597 | -0.27526 | 0.88218 |
| 7 | 81279000 | 81286199 | 82.04149 | 127.8721 | 164.8183 | 68.40322 | -0.05492 | 2.701663 | 0.882284 |
| 10 | 34623600 | 34627199 | 28.59104 | 48.20549 | 48.00571 | 20.98991 | 0.387075 | 1.114344 | 0.882293 |
| 5 | 4085200 | 4085999 | 8.494025 | 16.22762 | 24.52997 | 9.746339 | -0.08259 | -0.03451 | 0.882318 |
| 16 | 89224400 | 89224999 | 7.539283 | 14.60947 | 20.50352 | 7.766976 | 0.088608 | -0.21399 | 0.882322 |
| 6 | 60665800 | 60673799 | 192.0567 | 279.7092 | 410.4998 | 169.1761 | -0.32649 | 3.972478 | 0.882327 |
| 1 | 158707000 | 158707999 | 15.5892 | 27.0935 | 46.85788 | 20.5771 | -0.39975 | 0.786592 | 0.882345 |
| 14 | 9248200 | 9248999 | 14.81169 | 26.06817 | 36.16134 | 16.00941 | -0.08156 | 0.577787 | 0.882349 |
| 12 | 5851600 | 5852399 | 9.200204 | 17.39084 | 25.97127 | 10.55372 | -0.09347 | 0.051217 | 0.882352 |
| 13 | 87475800 | 87480999 | 96.46481 | 148.7617 | 175.2225 | 72.78435 | 0.063091 | 2.896566 | 0.882355 |
| 18 | 29480000 | 29481399 | 13.28252 | 24.13789 | 30.54928 | 13.17372 | 0.078025 | 0.327623 | 0.882374 |
| 17 | 20627600 | 20633399 | 51.34889 | 82.79218 | 78.19358 | 33.29507 | 0.43925 | 1.797028 | 0.882382 |
| 11 | 13394800 | 13395999 | 15.47435 | 27.25815 | 37.62427 | 16.65418 | -0.07397 | 0.574944 | 0.882412 |
| 10 | 46251200 | 46251999 | 12.20163 | 22.11562 | 31.13876 | 13.47198 | -0.07782 | 0.336324 | 0.882432 |
| 9 | 115493400 | 115495399 | 15.81624 | 27.87741 | 34.45152 | 15.24792 | 0.084944 | 0.562863 | 0.882481 |
| 4 | 115228000 | 115231199 | 46.00254 | 73.99611 | 90.622 | 38.45748 | 0.061016 | 1.86702 | 0.882488 |
| 5 | 120034800 | 120036599 | 17.51061 | 30.77159 | 38.02024 | 16.83331 | 0.083735 | 0.613565 | 0.88249 |
| 2 | 5815800 | 5817799 | 14.84623 | 26.12661 | 36.19318 | 16.0238 | -0.07959 | 0.579888 | 0.88252 |
| 2 | 65198600 | 65200999 | 22.09309 | 37.64316 | 51.71494 | 22.66921 | -0.07445 | 1.003024 | 0.882523 |
| 1 | 35534800 | 35536199 | 11.82114 | 21.70258 | 27.64725 | 11.76594 | 0.081689 | 0.252662 | 0.882541 |
| 1 | 51958000 | 51959999 | 19.34899 | 33.58468 | 46.32202 | 20.32535 | -0.07514 | 0.819947 | 0.882571 |
| 3 | 107037800 | 107038599 | 7.271335 | 14.04133 | 22.059 | 8.493121 | -0.09476 | -0.15654 | 0.882628 |
| 16 | 38697000 | 38699399 | 18.15814 | 31.91306 | 39.01226 | 17.27416 | 0.099009 | 0.616799 | 0.882637 |
| 1 | 138519800 | 138520999 | 17.82014 | 31.14602 | 38.80251 | 17.10106 | 0.07201 | 0.703544 | 0.882641 |
| 8 | 47911400 | 47915399 | 53.96103 | 86.46082 | 88.29902 | 37.25454 | 0.315842 | 1.996397 | 0.882641 |
| 11 | 114368000 | 114372199 | 33.1123 | 54.16543 | 67.11509 | 28.86421 | 0.066693 | 1.478747 | 0.882652 |
| 9 | 20098800 | 20106599 | 145.2903 | 219.5081 | 215.6624 | 89.16491 | 0.300968 | 3.323164 | 0.882692 |
| 4 | 5573600 | 5578999 | 63.66377 | 101.4148 | 120.3129 | 50.34267 | 0.090833 | 2.197752 | 0.882699 |
| 12 | 16918600 | 16922399 | 26.3353 | 43.86782 | 59.92844 | 26.03489 | -0.06745 | 1.239683 | 0.882713 |
| 10 | 79043400 | 79048599 | 48.32356 | 77.96364 | 76.77276 | 32.71678 | 0.380169 | 1.772665 | 0.882739 |
| 17 | 76830000 | 76830399 | 5.655177 | 11.03309 | 18.66081 | 6.666893 | -0.10024 | -0.44974 | 0.882743 |
| 16 | 46146200 | 46148399 | 23.37858 | 40.72526 | 47.31959 | 20.81009 | 0.171672 | 0.790209 | 0.882794 |
| 10 | 8419200 | 8429199 | 225.4359 | 325.8627 | 397.4461 | 163.9402 | -0.07025 | 4.086734 | 0.882806 |
| 5 | 91971200 | 91973399 | 24.11669 | 41.22435 | 50.65203 | 22.20907 | 0.086764 | 0.982012 | 0.882815 |
| 18 | 17325600 | 17326599 | 11.00053 | 20.29286 | 29.32922 | 12.40076 | -0.09214 | 0.225601 | 0.882829 |
| 2 | 21806200 | 21807999 | 15.04239 | 26.78065 | 33.37344 | 14.70156 | 0.079732 | 0.467107 | 0.882847 |
| 11 | 19742200 | 19745799 | 51.86397 | 82.83049 | 109.374 | 45.9299 | -0.0594 | 2.101869 | 0.882904 |
| 9 | 109413200 | 109421199 | 96.67489 | 150.2447 | 176.1739 | 73.14764 | 0.078171 | 2.759772 | 0.882921 |
| 8 | 40595200 | 40599599 | 42.56587 | 68.60726 | 92.0124 | 39.16175 | -0.06617 | 1.799326 | 0.882933 |
| 7 | 103141200 | 103142199 | 15.97822 | 28.17523 | 35.06414 | 15.51771 | 0.075375 | 0.563093 | 0.88296 |

| | | | | | | | | | |
|---|---|---|---|---|---|---|---|---|---|
| 8 | 107203000 | 107206199 | 30.25888 | 49.91518 | 68.5938 | 29.604 | -0.07719 | 1.367094 | 0.882966 |
| 5 | 129366000 | 129368199 | 21.21329 | 36.12259 | 49.54832 | 21.71843 | -0.07219 | 1.004864 | 0.882973 |
| 10 | 129720800 | 129722399 | 29.45648 | 48.68453 | 60.51494 | 26.14927 | 0.067631 | 1.338395 | 0.882997 |
| 1 | 64353600 | 64356399 | 24.03813 | 40.19653 | 55.46073 | 24.15285 | -0.08173 | 1.187441 | 0.883044 |
| 15 | 47218800 | 47221599 | 42.36527 | 68.58005 | 69.28711 | 29.63938 | 0.348518 | 1.673123 | 0.883079 |
| 10 | 108225800 | 108227999 | 16.72406 | 29.11644 | 40.55121 | 17.82708 | -0.08853 | 0.756316 | 0.883115 |
| 5 | 81775400 | 81779399 | 65.47083 | 103.0578 | 136.4996 | 56.63328 | -0.08306 | 2.507166 | 0.883131 |
| 5 | 81775600 | 81779399 | 65.47083 | 103.0578 | 136.4996 | 56.63328 | -0.08306 | 2.507166 | 0.883131 |
| 1 | 83494600 | 83495799 | 21.01097 | 36.56378 | 32.38278 | 14.23214 | 0.561682 | 0.77268 | 0.883197 |
| 14 | 83381000 | 83383999 | 23.408 | 39.61767 | 49.05102 | 21.47181 | 0.074278 | 1.064652 | 0.883199 |
| 19 | 57490000 | 57491599 | 15.76688 | 27.68255 | 38.19724 | 16.89891 | -0.07472 | 0.618652 | 0.883235 |
| X | 53180000 | 53182999 | 52.50153 | 84.72899 | 73.20536 | 31.20829 | 0.569021 | 1.774612 | 0.883236 |
| 1 | 179717000 | 179723999 | 78.90285 | 122.1554 | 190.4836 | 79.12192 | -0.33692 | 2.820056 | 0.883249 |
| 12 | 57385000 | 57386199 | 11.25817 | 20.94321 | 30.48406 | 12.73982 | -0.09037 | 0.181153 | 0.883263 |
| 8 | 112421600 | 112425999 | 42.22579 | 67.9784 | 90.85387 | 38.60302 | -0.06328 | 1.834807 | 0.883266 |
| X | 71150200 | 71151799 | 20.96321 | 36.56897 | 35.69307 | 15.71085 | 0.422377 | 0.744006 | 0.883269 |
| 10 | 48078200 | 48081999 | 34.49348 | 56.78633 | 69.34513 | 29.90282 | 0.091581 | 1.40538 | 0.883273 |
| X | 110140400 | 110142799 | 15.7587 | 28.03527 | 40.24102 | 17.73786 | -0.12366 | 0.491125 | 0.883288 |
| 13 | 87708600 | 87710199 | 17.60596 | 31.04027 | 30.85274 | 13.6544 | 0.397824 | 0.565722 | 0.883295 |
| 15 | 32850000 | 32850399 | 6.085172 | 11.87128 | 17.51923 | 6.263991 | 0.094065 | -0.44266 | 0.883301 |
| 6 | 143454600 | 143459999 | 163.3701 | 243.3253 | 277.9792 | 114.5449 | 0.065187 | 3.601351 | 0.88332 |
| 7 | 29633400 | 29637199 | 44.06235 | 70.85243 | 86.28191 | 36.59856 | 0.06863 | 1.873399 | 0.883324 |
| 10 | 38202200 | 38204999 | 35.88211 | 58.43812 | 79.69931 | 34.22127 | -0.07585 | 1.556782 | 0.883326 |
| 5 | 10613800 | 10615999 | 19.35813 | 33.85385 | 33.59843 | 14.81481 | 0.398517 | 0.694134 | 0.883368 |
| 6 | 132412200 | 132413799 | 21.31064 | 36.51632 | 50.09044 | 21.96713 | -0.07097 | 0.947905 | 0.883391 |
| 10 | 48633200 | 48634999 | 17.37572 | 30.99791 | 36.92089 | 16.23634 | 0.146266 | 0.476159 | 0.883417 |
| 2 | 107190400 | 107192399 | 18.82369 | 32.84925 | 40.89129 | 18.00324 | 0.072979 | 0.734218 | 0.883424 |
| X | 60846800 | 60848999 | 17.75192 | 31.12162 | 43.98795 | 19.43644 | -0.1092 | 0.665155 | 0.88348 |
| 4 | 130747600 | 130751599 | 40.81094 | 65.81452 | 87.98467 | 37.47382 | -0.06061 | 1.780282 | 0.883494 |
| 18 | 30319400 | 30320199 | 17.34721 | 30.82181 | 30.19254 | 13.32295 | 0.42358 | 0.48993 | 0.883505 |
| 17 | 49948000 | 49950999 | 28.89152 | 47.91903 | 59.50987 | 25.77749 | 0.06927 | 1.294474 | 0.883505 |
| 13 | 88119800 | 88121399 | 18.04249 | 31.37386 | 43.12875 | 18.94581 | -0.07001 | 0.784063 | 0.883526 |
| 19 | 28412200 | 28414399 | 19.44233 | 33.62846 | 46.32526 | 20.32384 | -0.07445 | 0.850635 | 0.883532 |
| 14 | 3608000 | 3609199 | 18.71864 | 32.86188 | 32.87477 | 14.54299 | 0.386911 | 0.627668 | 0.883533 |
| 15 | 30894200 | 30894799 | 7.139705 | 13.87187 | 22.35007 | 8.335676 | -0.08739 | -0.26472 | 0.883539 |
| 16 | 22109800 | 22113999 | 45.65783 | 73.52397 | 89.88857 | 38.20823 | 0.065491 | 1.831051 | 0.883569 |
| 9 | 60644200 | 60646599 | 40.19019 | 64.63213 | 87.73963 | 37.22202 | -0.08758 | 1.866142 | 0.883573 |
| 17 | 60873200 | 60873399 | 5.500204 | 10.74413 | 15.97128 | 5.584819 | 0.113475 | -0.53359 | 0.883592 |
| 1 | 114484800 | 114486399 | 30.86754 | 51.25923 | 45.56972 | 19.75585 | 0.550196 | 1.274492 | 0.883592 |
| 5 | 8805600 | 8810199 | 57.61951 | 91.7882 | 111.2184 | 46.5385 | 0.060408 | 2.186013 | 0.883625 |
| 7 | 23895600 | 23905799 | 168.5368 | 250.2146 | 309.7616 | 127.5995 | -0.05492 | 3.659278 | 0.883625 |
| 18 | 53412600 | 53415599 | 21.62785 | 37.09722 | 45.7928 | 20.08403 | 0.080776 | 0.932954 | 0.883639 |
| 6 | 137598200 | 137598599 | 7.26418 | 13.98359 | 22.00494 | 8.557456 | -0.11089 | -0.1248 | 0.883643 |
| 9 | 61794400 | 61797399 | 23.18444 | 38.89402 | 53.84688 | 23.49771 | -0.08671 | 1.148057 | 0.88365 |
| 4 | 99396400 | 99397199 | 13.54143 | 24.15381 | 33.5766 | 14.76762 | -0.0753 | 0.447711 | 0.883663 |
| 7 | 38077600 | 38083599 | 62.66715 | 99.74878 | 120.0519 | 50.17885 | 0.068873 | 2.219828 | 0.883667 |
| 3 | 5652200 | 5653999 | 17.22385 | 30.21947 | 37.64166 | 16.64327 | 0.072232 | 0.63766 | 0.88368 |

| | | | | | | | | | |
|---|---|---|---|---|---|---|---|---|---|
| 5 | 6724800 | 6726599 | 17.22385 | 30.21947 | 37.64166 | 16.64327 | 0.072232 | 0.63766 | 0.88368 |
| 2 | 35276000 | 35279599 | 39.28626 | 63.68939 | 78.4435 | 33.59266 | 0.06533 | 1.642457 | 0.8837 |
| 10 | 129114400 | 129115599 | 22.50385 | 38.15036 | 52.72885 | 23.09332 | -0.08367 | 1.055158 | 0.883704 |
| X | 113684000 | 113685199 | 19.07289 | 33.15038 | 45.76773 | 20.09927 | -0.0763 | 0.799647 | 0.883718 |
| 17 | 42112200 | 42113199 | 11.16107 | 20.67548 | 26.07402 | 10.96174 | 0.108393 | 0.190306 | 0.883745 |
| 17 | 68529400 | 68533599 | 51.41724 | 82.61433 | 83.17453 | 35.24043 | 0.341123 | 1.904526 | 0.883762 |
| 5 | 75995800 | 75996999 | 13.30595 | 23.94577 | 30.02895 | 13.10827 | 0.079947 | 0.383098 | 0.88377 |
| 11 | 26414000 | 26417999 | 47.71381 | 76.76372 | 93.35934 | 39.63035 | 0.071387 | 1.865366 | 0.883787 |
| 6 | 95932600 | 95937199 | 36.62203 | 59.5174 | 80.98503 | 34.7444 | -0.07478 | 1.588014 | 0.88379 |
| 10 | 122920400 | 122923199 | 27.16044 | 45.13812 | 61.61874 | 26.72787 | -0.06657 | 1.2692 | 0.883811 |
| 6 | 70263600 | 70264599 | 15.41475 | 27.17467 | 37.68037 | 16.67638 | -0.07991 | 0.568159 | 0.883828 |
| 8 | 11930400 | 11937599 | 89.0132 | 139.3044 | 139.6475 | 57.93083 | 0.311853 | 2.631647 | 0.883829 |
| 18 | 49797600 | 49802999 | 54.23265 | 86.7027 | 105.5036 | 44.36927 | 0.059891 | 2.070712 | 0.883871 |
| 1 | 21654600 | 21656599 | 31.87982 | 52.83404 | 52.98268 | 22.93776 | 0.3768 | 1.30857 | 0.883878 |
| 1 | 75918200 | 75921799 | 32.63396 | 53.84309 | 66.04125 | 28.50459 | 0.08643 | 1.36835 | 0.88389 |
| 4 | 82953400 | 82957399 | 34.93762 | 56.91295 | 77.51468 | 33.28602 | -0.07371 | 1.547676 | 0.883921 |
| 3 | 151629400 | 151633399 | 58.70512 | 92.96508 | 143.3676 | 59.61575 | -0.29543 | 2.370414 | 0.883922 |
| 14 | 34378800 | 34385199 | 56.88187 | 90.47176 | 118.9121 | 49.65046 | -0.05916 | 2.237977 | 0.883954 |
| 18 | 38706200 | 38709799 | 23.10186 | 38.89463 | 53.88593 | 23.55071 | -0.08771 | 1.112559 | 0.883958 |
| 9 | 120184800 | 120187399 | 29.11553 | 48.20409 | 59.98674 | 25.95191 | 0.06616 | 1.317111 | 0.883981 |
| 17 | 23248600 | 23249399 | 11.05577 | 19.91395 | 35.30109 | 15.39257 | -0.41527 | 0.374302 | 0.883994 |
| 1 | 46658200 | 46660199 | 24.47362 | 41.69442 | 51.21026 | 22.44777 | 0.086498 | 1.011945 | 0.884023 |
| 1 | 118979800 | 118981799 | 18.93568 | 32.28282 | 45.92406 | 20.13175 | -0.12418 | 0.980378 | 0.88404 |
| 3 | 142397600 | 142402599 | 77.32624 | 121.9765 | 124.7428 | 51.7696 | 0.291853 | 2.470653 | 0.884085 |
| 4 | 83592200 | 83595199 | 34.21411 | 56.13057 | 69.26112 | 29.82262 | 0.074593 | 1.444975 | 0.88409 |
| 7 | 119031000 | 119034199 | 29.68697 | 48.88143 | 66.54729 | 28.69833 | -0.06471 | 1.388652 | 0.884111 |
| 9 | 73244400 | 73246199 | 27.51506 | 46.05371 | 64.03976 | 27.90543 | -0.09309 | 1.196206 | 0.884115 |
| 15 | 38863400 | 38866599 | 34.20728 | 56.31499 | 68.90222 | 29.71179 | 0.088766 | 1.402373 | 0.884137 |
| 1 | 179643400 | 179649599 | 84.13381 | 131.4019 | 157.2387 | 65.24188 | 0.054461 | 2.666662 | 0.884143 |
| 2 | 122245800 | 122246599 | 14.77635 | 26.10145 | 32.11561 | 14.20787 | 0.092697 | 0.530337 | 0.884145 |
| 1 | 72785200 | 72787799 | 22.5689 | 38.81538 | 47.92855 | 21.0236 | 0.081088 | 0.917246 | 0.884147 |
| 7 | 30905400 | 30911199 | 55.41529 | 88.22756 | 116.0267 | 48.51295 | -0.05839 | 2.204011 | 0.884162 |
| 9 | 39856400 | 39858399 | 19.35588 | 33.25293 | 45.84614 | 20.10949 | -0.07755 | 0.911687 | 0.884165 |
| X | 122168000 | 122169799 | 32.34835 | 53.55863 | 48.19513 | 20.85114 | 0.532537 | 1.317415 | 0.884172 |
| 10 | 95209000 | 95217999 | 118.0941 | 179.9136 | 227.8944 | 94.46293 | -0.05532 | 3.148454 | 0.884176 |
| 10 | 4097600 | 4099999 | 17.18387 | 29.85268 | 41.61457 | 18.26983 | -0.0902 | 0.790179 | 0.884183 |
| X | 110121400 | 110123399 | 22.51427 | 38.75465 | 55.11709 | 24.17745 | -0.1218 | 0.91807 | 0.8842 |
| 15 | 36995000 | 37000799 | 52.10237 | 83.70069 | 84.38799 | 35.74561 | 0.338777 | 1.911629 | 0.884211 |
| 7 | 112717400 | 112718999 | 18.02994 | 31.52314 | 39.31113 | 17.33006 | 0.070619 | 0.700796 | 0.884216 |
| 14 | 4815600 | 4816599 | 15.81924 | 28.49131 | 28.22395 | 12.30816 | 0.418957 | 0.392014 | 0.884217 |
| 16 | 33129400 | 33134599 | 51.80744 | 82.67718 | 109.1224 | 45.77485 | -0.05996 | 2.124078 | 0.884237 |
| 4 | 27274400 | 27275199 | 8.960891 | 17.06144 | 25.55112 | 10.22534 | -0.07851 | -0.00412 | 0.884266 |
| 3 | 3304000 | 3309599 | 72.6023 | 115.8819 | 96.03944 | 40.29617 | 0.610509 | 2.137647 | 0.884277 |
| 7 | 12470200 | 12471199 | 14.23632 | 25.47669 | 31.92446 | 14.00221 | 0.07616 | 0.42718 | 0.884328 |
| 5 | 60226600 | 60227399 | 10.84147 | 20.4507 | 30.96998 | 12.64048 | -0.12002 | 0.088436 | 0.88433 |
| 3 | 78358200 | 78359399 | 8.784433 | 16.99987 | 17.16603 | 6.54287 | 0.550477 | -0.191 | 0.884335 |
| 18 | 6419000 | 6424799 | 152.5465 | 228.5646 | 264.1563 | 108.9673 | 0.05632 | 3.48914 | 0.884372 |

| | | | | | | | | | |
|---|---|---|---|---|---|---|---|---|---|
| 17 | 31988400 | 31991599 | 38.26492 | 62.00695 | 76.51473 | 32.75629 | 0.062054 | 1.647479 | 0.884407 |
| 15 | 29128000 | 29129599 | 27.82512 | 46.71587 | 46.47041 | 20.28882 | 0.388582 | 1.14866 | 0.884429 |
| 1 | 7405200 | 7409799 | 82.61456 | 129.818 | 131.5202 | 54.555 | 0.301155 | 2.552093 | 0.884483 |
| 1 | 138379600 | 138380199 | 8.074494 | 15.57107 | 24.23811 | 9.377984 | -0.08833 | -0.12407 | 0.88449 |
| 2 | 39593400 | 39594999 | 24.63777 | 41.56172 | 51.73214 | 22.62281 | 0.066486 | 1.101412 | 0.884511 |
| 6 | 29576400 | 29578199 | 15.86645 | 28.0619 | 34.89883 | 15.43532 | 0.077827 | 0.527433 | 0.884526 |
| 5 | 109898600 | 109901399 | 14.92918 | 25.88329 | 48.45827 | 21.26116 | -0.51429 | 0.812533 | 0.884566 |
| 3 | 7154400 | 7155199 | 9.285227 | 17.55214 | 25.9477 | 10.5434 | -0.07925 | 0.049679 | 0.884568 |
| 13 | 25438600 | 25439399 | 8.434834 | 16.08288 | 24.48023 | 9.767124 | -0.09822 | -0.024 | 0.884574 |
| 4 | 33132400 | 33133599 | 10.47541 | 19.40589 | 28.24603 | 11.87439 | -0.09691 | 0.190399 | 0.884583 |
| 14 | 54133400 | 54134799 | 15.18904 | 26.83623 | 37.09719 | 16.411 | -0.07437 | 0.540265 | 0.884618 |
| 4 | 109412000 | 109414799 | 40.21095 | 65.67562 | 64.24783 | 27.61548 | 0.40522 | 1.516042 | 0.884648 |
| 6 | 76075000 | 76078599 | 36.23829 | 59.63084 | 59.29465 | 25.56381 | 0.386864 | 1.406109 | 0.884677 |
| 7 | 121361600 | 121362999 | 11.34417 | 20.72545 | 29.78744 | 12.75993 | -0.09755 | 0.284144 | 0.884681 |
| 2 | 72914800 | 72916799 | 17.11613 | 30.23314 | 28.46684 | 12.59378 | 0.47661 | 0.538428 | 0.88471 |
| 5 | 109177600 | 109191999 | 175.3465 | 262.7913 | 294.3476 | 121.4274 | 0.101797 | 3.490218 | 0.884713 |
| 13 | 98481800 | 98483399 | 11.70394 | 21.32785 | 30.1735 | 12.96693 | -0.07837 | 0.299499 | 0.884722 |
| 16 | 23260000 | 23262599 | 41.59407 | 66.95079 | 80.30609 | 34.11352 | 0.092239 | 1.835804 | 0.884741 |
| 4 | 67276200 | 67278199 | 22.87257 | 39.59912 | 47.93372 | 21.03011 | 0.11166 | 0.861696 | 0.88475 |
| 13 | 4260600 | 4262199 | 15.02677 | 27.13422 | 33.54912 | 14.58308 | 0.104118 | 0.383614 | 0.884759 |
| 1 | 107166000 | 107167599 | 15.84611 | 28.14358 | 40.18291 | 17.73729 | -0.11754 | 0.500657 | 0.884769 |
| 17 | 8498200 | 8500199 | 14.14784 | 25.20059 | 35.01831 | 15.41968 | -0.07607 | 0.462015 | 0.88479 |
| 3 | 65907600 | 65911399 | 44.67678 | 71.72611 | 95.64441 | 40.47484 | -0.0649 | 1.925961 | 0.884794 |
| 2 | 108299600 | 108303999 | 70.26724 | 111.0218 | 133.9824 | 55.64904 | 0.055192 | 2.429407 | 0.884795 |
| 9 | 50312200 | 50314399 | 22.83134 | 38.62171 | 52.77985 | 23.10046 | -0.06768 | 1.071537 | 0.8848 |
| 5 | 99268200 | 99268799 | 10.78065 | 20.03147 | 25.67679 | 10.75026 | 0.089678 | 0.171141 | 0.884801 |
| 17 | 93819800 | 93822199 | 20.79371 | 36.16804 | 36.25966 | 15.93113 | 0.383429 | 0.782677 | 0.884818 |
| 2 | 24071000 | 24074799 | 23.81184 | 39.60583 | 68.12866 | 29.56852 | -0.39898 | 1.254928 | 0.884819 |
| 10 | 89005200 | 89007799 | 26.65812 | 44.73476 | 55.5725 | 24.25529 | 0.069087 | 1.15944 | 0.884852 |
| 17 | 3098800 | 3099199 | 7.27321 | 14.12469 | 20.03776 | 7.501397 | 0.089742 | -0.25709 | 0.884862 |
| X | 79103200 | 79105399 | 21.59271 | 37.15583 | 51.72937 | 22.69117 | -0.09116 | 0.915036 | 0.88487 |
| 2 | 22396400 | 22397399 | 17.58447 | 30.64861 | 37.50986 | 16.50162 | 0.097419 | 0.742342 | 0.884875 |
| 12 | 21457600 | 21459599 | 19.65965 | 34.26118 | 42.48572 | 18.68915 | 0.078283 | 0.759953 | 0.884878 |
| 7 | 116397800 | 116408599 | 172.0761 | 255.4039 | 315.8225 | 130.0928 | -0.05361 | 3.664002 | 0.884889 |
| 4 | 83543400 | 83544599 | 16.4974 | 28.91091 | 39.83363 | 17.59498 | -0.07249 | 0.657192 | 0.884896 |
| 14 | 3521400 | 3522599 | 13.00342 | 23.30464 | 32.5569 | 14.26189 | -0.0785 | 0.411303 | 0.884905 |
| 10 | 116954000 | 116954599 | 7.261536 | 14.0535 | 22.13155 | 8.433313 | -0.0842 | -0.19207 | 0.884911 |
| 5 | 149692800 | 149694799 | 21.31723 | 36.50629 | 45.32514 | 19.87638 | 0.072289 | 0.948713 | 0.884911 |
| 13 | 82355800 | 82360199 | 93.24833 | 144.1305 | 184.2972 | 76.55292 | -0.05274 | 2.855302 | 0.884946 |
| 3 | 155274000 | 155274999 | 11.25075 | 21.00771 | 27.26196 | 11.32143 | 0.081197 | 0.147818 | 0.88496 |
| 15 | 45661600 | 45662599 | 13.74812 | 24.6737 | 30.95356 | 13.54359 | 0.076947 | 0.40415 | 0.884974 |
| 17 | 14860000 | 14861399 | 13.74812 | 24.6737 | 30.95356 | 13.54359 | 0.076947 | 0.40415 | 0.884974 |
| 2 | 19215800 | 19223399 | 79.7681 | 125.7801 | 148.2374 | 61.51707 | 0.087038 | 2.484378 | 0.88502 |
| 8 | 115054400 | 115057199 | 48.83456 | 78.30895 | 95.72373 | 40.43594 | 0.058347 | 1.969228 | 0.88503 |
| 9 | 39656000 | 39656399 | 7.445947 | 14.40612 | 22.67821 | 8.65361 | -0.08522 | -0.18193 | 0.885045 |
| 1 | 190246600 | 190247799 | 12.69405 | 23.30267 | 29.60036 | 12.59898 | 0.085299 | 0.262479 | 0.885047 |
| 4 | 16623000 | 16624199 | 10.79232 | 20.30272 | 26.38473 | 10.81949 | 0.094372 | 0.097966 | 0.88506 |

| | | | | | | | | | |
|---|---|---|---|---|---|---|---|---|---|
| 5 | 120052400 | 120053799 | 16.80435 | 29.30435 | 40.53279 | 17.8374 | -0.0783 | 0.735188 | 0.885084 |
| 16 | 9936200 | 9937199 | 10.31218 | 19.34258 | 25.25427 | 10.40757 | 0.081543 | 0.106617 | 0.8851 |
| 17 | 30977200 | 30981999 | 38.17308 | 62.19596 | 63.35317 | 27.20684 | 0.344606 | 1.544606 | 0.885102 |
| 2 | 150269200 | 150283599 | 326.5946 | 463.569 | 501.3626 | 207.3896 | 0.076882 | 4.360676 | 0.885104 |
| 5 | 59716600 | 59717199 | 9.994858 | 18.84424 | 24.79266 | 10.12854 | 0.080692 | 0.069954 | 0.885127 |
| 9 | 109337200 | 109346799 | 172.526 | 256.6277 | 319.0268 | 131.4417 | -0.05839 | 3.626498 | 0.885141 |
| 13 | 22855200 | 22857199 | 27.36219 | 45.78993 | 56.7646 | 24.73257 | 0.072075 | 1.189573 | 0.885154 |
| 4 | 24649200 | 24650599 | 14.4356 | 25.19112 | 44.69343 | 19.67699 | -0.43576 | 0.718372 | 0.885175 |
| 3 | 116782400 | 116783799 | 12.75528 | 22.94884 | 32.12968 | 14.02875 | -0.07816 | 0.38638 | 0.885191 |
| 5 | 24713400 | 24714199 | 13.96752 | 25.05118 | 31.48705 | 13.78456 | 0.073766 | 0.411794 | 0.885205 |
| 11 | 28980200 | 28981399 | 22.4568 | 38.93532 | 36.42007 | 15.9794 | 0.482609 | 0.833236 | 0.885227 |
| 16 | 23159800 | 23162199 | 29.30953 | 48.60762 | 60.57182 | 26.23573 | 0.064381 | 1.299325 | 0.885227 |
| 4 | 142441400 | 142441599 | 6.677164 | 13.02836 | 19.29266 | 6.880048 | 0.091907 | -0.43835 | 0.885239 |
| X | 36880600 | 36886199 | 49.70489 | 79.79116 | 109.0349 | 46.13499 | -0.10031 | 1.943814 | 0.885265 |
| 1 | 193497800 | 193498799 | 16.67766 | 29.36028 | 36.47806 | 16.14874 | 0.076644 | 0.585362 | 0.885267 |
| 5 | 80148600 | 80149799 | 18.83871 | 32.58209 | 44.75479 | 19.6348 | -0.07024 | 0.8476 | 0.885308 |
| 7 | 118161000 | 118164199 | 26.05325 | 43.31407 | 59.2753 | 25.71793 | -0.07002 | 1.260807 | 0.88538 |
| 11 | 19816800 | 19820399 | 44.56238 | 71.72656 | 88.0202 | 37.38884 | 0.059296 | 1.844245 | 0.88543 |
| 1 | 85796000 | 85812799 | 405.6402 | 559.5588 | 643.9325 | 267.5776 | -0.05414 | 4.788705 | 0.885437 |
| 12 | 78005000 | 78006599 | 21.89441 | 37.26468 | 45.39828 | 19.89858 | 0.098292 | 1.008705 | 0.885442 |
| 2 | 28928200 | 28930199 | 16.19691 | 28.34702 | 39.31843 | 17.34812 | -0.08224 | 0.673104 | 0.885444 |
| 6 | 84535600 | 84537799 | 19.83198 | 33.79904 | 57.06644 | 25.01554 | -0.37067 | 1.000277 | 0.885488 |
| 14 | 110064600 | 110069199 | 33.46028 | 55.50399 | 55.30681 | 23.95905 | 0.385999 | 1.303061 | 0.885495 |
| 3 | 93998000 | 93998799 | 9.585474 | 18.24135 | 24.43823 | 9.790514 | 0.079704 | 0.002961 | 0.885522 |
| 1 | 13664400 | 13665799 | 12.84583 | 23.24273 | 32.62258 | 14.14472 | -0.07542 | 0.353875 | 0.885529 |
| 18 | 73076600 | 73082799 | 95.5344 | 147.7008 | 175.595 | 72.93812 | 0.052516 | 2.850672 | 0.885531 |
| 12 | 61679800 | 61680399 | 8.035035 | 15.39135 | 23.51308 | 9.274432 | -0.08774 | -0.06545 | 0.885532 |
| 17 | 68003600 | 68004999 | 13.16143 | 23.34509 | 33.05572 | 14.60723 | -0.10657 | 0.477096 | 0.88555 |
| 5 | 142694800 | 142698999 | 55.27009 | 88.06376 | 106.2468 | 44.47086 | 0.066968 | 2.17766 | 0.885562 |
| 6 | 89180000 | 89182599 | 19.07673 | 33.14192 | 41.08802 | 18.03896 | 0.078365 | 0.800888 | 0.885587 |
| 3 | 152189600 | 152194599 | 40.59095 | 65.46887 | 87.60292 | 37.31793 | -0.06172 | 1.774505 | 0.885593 |
| 4 | 25030600 | 25032799 | 23.7307 | 40.55296 | 50.10677 | 21.96949 | 0.078632 | 0.981669 | 0.885632 |
| 5 | 146163600 | 146165199 | 14.3671 | 25.32103 | 35.20103 | 15.57991 | -0.08376 | 0.55909 | 0.885644 |
| 13 | 84301600 | 84303799 | 20.6557 | 35.42603 | 48.47711 | 21.26115 | -0.06725 | 0.935352 | 0.885651 |
| 10 | 37659000 | 37661799 | 34.7961 | 57.22502 | 58.55966 | 25.24084 | 0.345261 | 1.409585 | 0.885684 |
| 7 | 55314800 | 55318599 | 43.91025 | 70.5626 | 93.85313 | 39.77311 | -0.05954 | 1.895796 | 0.885704 |
| 8 | 96679600 | 96683999 | 37.26421 | 60.47716 | 82.09371 | 35.18408 | -0.0732 | 1.617069 | 0.885717 |
| 5 | 109196600 | 109200399 | 72.50207 | 114.4221 | 137.9095 | 57.24706 | 0.055723 | 2.459385 | 0.88572 |
| 14 | 103538400 | 103540199 | 18.93397 | 33.27902 | 40.00318 | 17.71402 | 0.123107 | 0.620779 | 0.885737 |
| 12 | 116138800 | 116141599 | 27.25385 | 45.60676 | 56.72615 | 24.71518 | 0.06732 | 1.190645 | 0.885738 |
| 3 | 69934600 | 69935399 | 11.3626 | 20.56016 | 29.64488 | 12.85275 | -0.11393 | 0.335127 | 0.885756 |
| 2 | 66369800 | 66371599 | 20.47372 | 35.54779 | 44.16097 | 19.38085 | 0.07517 | 0.818166 | 0.885772 |
| 12 | 87565400 | 87567199 | 16.05327 | 28.24337 | 34.40573 | 15.23324 | 0.104433 | 0.582693 | 0.885779 |
| 18 | 74549400 | 74556199 | 99.65758 | 153.4958 | 181.6197 | 75.43809 | 0.055198 | 2.921781 | 0.885783 |
| Y | 88322800 | 88323599 | 8.557486 | 15.93012 | 24.9214 | 10.40302 | -0.19188 | 0.139474 | 0.885784 |
| 1 | 37015400 | 37017599 | 18.5482 | 32.43324 | 40.32346 | 17.77819 | 0.074969 | 0.703317 | 0.885799 |
| 4 | 17513000 | 17513399 | 7.993664 | 15.44712 | 21.51246 | 8.255318 | 0.081807 | -0.16223 | 0.885802 |

| | | | | | | | | | |
|---|---|---|---|---|---|---|---|---|---|
| 6 | 49459800 | 49468799 | 155.7558 | 233.4106 | 270.5627 | 111.6135 | 0.052227 | 3.486652 | 0.885815 |
| 4 | 90004600 | 90007399 | 24.03203 | 39.82347 | 67.92163 | 29.40296 | -0.3871 | 1.299553 | 0.885817 |
| 1 | 16254200 | 16257199 | 17.18128 | 30.02805 | 41.41365 | 18.25114 | -0.07407 | 0.704273 | 0.885849 |
| 13 | 12468200 | 12470199 | 15.80749 | 27.64136 | 38.43609 | 16.94633 | -0.08562 | 0.686042 | 0.88585 |
| X | 145745400 | 145746999 | 17.01509 | 30.04622 | 37.01022 | 16.3743 | 0.090872 | 0.557645 | 0.885942 |
| 5 | 33484000 | 33485999 | 21.34671 | 36.57436 | 50.08731 | 21.96557 | -0.06859 | 0.949668 | 0.885962 |
| 3 | 16061800 | 16068999 | 131.7656 | 199.439 | 233.1201 | 96.43837 | 0.052776 | 3.28427 | 0.885964 |
| 3 | 73129600 | 73130799 | 12.89503 | 23.61663 | 29.48805 | 12.59094 | 0.106402 | 0.274791 | 0.885975 |
| 19 | 4326000 | 4327199 | 14.4574 | 25.85561 | 32.45937 | 14.24459 | 0.07305 | 0.434744 | 0.886001 |
| 15 | 34877800 | 34880399 | 21.43484 | 36.76281 | 45.82159 | 20.09651 | 0.06696 | 0.935334 | 0.886023 |
| 4 | 71816000 | 71816599 | 7.246136 | 14.07264 | 20.03085 | 7.496531 | 0.084952 | -0.26005 | 0.886024 |
| 3 | 20740600 | 20746599 | 119.1726 | 181.2318 | 211.3067 | 87.54035 | 0.062045 | 3.184681 | 0.886025 |
| 2 | 25607400 | 25609399 | 24.264 | 41.27154 | 51.18588 | 22.43421 | 0.072265 | 1.027344 | 0.886028 |
| 6 | 76470400 | 76476399 | 104.0668 | 158.9734 | 203.9776 | 84.60468 | -0.07075 | 3.088706 | 0.886061 |
| 3 | 155475600 | 155478199 | 22.82113 | 37.75737 | 67.53302 | 29.20297 | -0.4555 | 1.316955 | 0.886068 |
| X | 45253600 | 45255399 | 16.5968 | 29.54115 | 35.98238 | 15.85396 | 0.112528 | 0.4868 | 0.886071 |
| 15 | 35161800 | 35162399 | 13.80708 | 24.73119 | 31.01482 | 13.59274 | 0.075953 | 0.417943 | 0.886125 |
| 18 | 38677800 | 38684599 | 90.91075 | 141.0348 | 167.8888 | 69.72622 | 0.054432 | 2.785545 | 0.886129 |
| 12 | 103845200 | 103847799 | 30.81319 | 50.91252 | 63.39897 | 27.39018 | 0.064988 | 1.348073 | 0.886154 |
| 2 | 69627200 | 69630799 | 28.43066 | 47.33438 | 58.95532 | 25.60402 | 0.065474 | 1.25029 | 0.886175 |
| 9 | 6159800 | 6161399 | 16.6967 | 29.22545 | 40.32818 | 17.79556 | -0.07484 | 0.675607 | 0.88619 |
| 9 | 118297200 | 118299399 | 16.6967 | 29.22545 | 40.32818 | 17.79556 | -0.07484 | 0.675607 | 0.88619 |
| 17 | 88637600 | 88639399 | 28.84685 | 47.57226 | 64.86689 | 27.99213 | -0.06607 | 1.365797 | 0.886207 |
| 7 | 25257600 | 25260399 | 41.00935 | 66.21297 | 88.96546 | 37.94956 | -0.06592 | 1.747341 | 0.886244 |
| 17 | 12177000 | 12179599 | 33.64819 | 54.94546 | 74.544 | 32.03865 | -0.06561 | 1.509549 | 0.886284 |
| 13 | 45023000 | 45025399 | 146.6298 | 219.1145 | 276.5681 | 114.0282 | -0.07424 | 3.538753 | 0.886288 |
| 12 | 116654600 | 116656799 | 23.41961 | 39.60567 | 49.05521 | 21.46844 | 0.073688 | 1.071756 | 0.88632 |
| 10 | 87931000 | 87931999 | 14.46825 | 25.16662 | 45.80304 | 20.12458 | -0.47282 | 0.766972 | 0.886324 |
| 3 | 112171400 | 112172199 | 10.95057 | 20.33537 | 26.17898 | 10.96919 | 0.082676 | 0.177194 | 0.886325 |
| 4 | 119293000 | 119294599 | 18.8989 | 32.85425 | 45.23207 | 19.86642 | -0.07224 | 0.795305 | 0.886338 |
| 5 | 109069000 | 109079599 | 128.5499 | 197.2628 | 179.4854 | 74.51365 | 0.429749 | 3.004244 | 0.886355 |
| 2 | 177229000 | 177233199 | 53.57934 | 85.76696 | 104.2176 | 43.90726 | 0.063922 | 2.031175 | 0.886355 |
| 6 | 130576200 | 130581399 | 101.5649 | 156.78 | 185.0726 | 76.87552 | 0.061011 | 2.883498 | 0.886375 |
| 8 | 42116000 | 42116599 | 7.439922 | 14.4183 | 19.99662 | 7.571601 | 0.106969 | -0.21552 | 0.886386 |
| 3 | 15693200 | 15694399 | 25.86371 | 43.21008 | 53.33945 | 23.21576 | 0.078372 | 1.202545 | 0.886416 |
| 8 | 72801400 | 72803599 | 25.89209 | 43.85453 | 53.96589 | 23.62859 | 0.083039 | 1.072225 | 0.886421 |
| 17 | 92366000 | 92366599 | 8.978605 | 17.15642 | 22.85106 | 9.075884 | 0.098428 | -0.03835 | 0.886432 |
| 18 | 47321200 | 47323999 | 20.64955 | 34.47035 | 62.19616 | 27.05648 | -0.46736 | 1.204045 | 0.886446 |
| 8 | 18883600 | 18886999 | 67.00224 | 106.0957 | 128.3805 | 53.38153 | 0.053877 | 2.37518 | 0.886452 |
| 9 | 18451200 | 18453799 | 39.47368 | 63.80394 | 85.59958 | 36.56585 | -0.06192 | 1.71028 | 0.886474 |
| 4 | 132296000 | 132299199 | 23.86788 | 39.55579 | 67.91934 | 29.4042 | -0.39674 | 1.297648 | 0.886521 |
| 16 | 68712600 | 68713199 | 18.04102 | 31.3757 | 38.55176 | 16.9367 | 0.091387 | 0.776595 | 0.886531 |
| 8 | 73760200 | 73761199 | 10.57466 | 19.67052 | 28.33866 | 11.84442 | -0.07567 | 0.170597 | 0.886562 |
| 15 | 5909000 | 5917599 | 188.4765 | 278.07 | 316.7495 | 130.4423 | 0.056687 | 3.766691 | 0.88657 |
| X | 153761800 | 153763399 | 20.16098 | 35.05973 | 49.55192 | 21.76821 | -0.10994 | 0.803328 | 0.886627 |
| 2 | 4810200 | 4814799 | 56.8741 | 90.7205 | 110.3876 | 46.27531 | 0.056403 | 2.143988 | 0.886632 |
| 10 | 126091600 | 126093599 | 19.18144 | 33.42686 | 41.39838 | 18.21052 | 0.079989 | 0.754204 | 0.886653 |

| | | | | | | | | | |
|---|---|---|---|---|---|---|---|---|---|
| 12 | 16824000 | 16826199 | 17.0613 | 29.78488 | 40.95938 | 18.03748 | -0.06995 | 0.720169 | 0.886661 |
| 9 | 113049000 | 113059799 | 147.6562 | 221.6976 | 276.9478 | 114.3015 | -0.05316 | 3.448618 | 0.886673 |
| 5 | 67371800 | 67374599 | 31.68013 | 51.6735 | 70.72248 | 30.3837 | -0.07934 | 1.518471 | 0.886701 |
| 4 | 78359400 | 78360999 | 13.87971 | 25.13129 | 31.57286 | 13.67738 | 0.084505 | 0.356567 | 0.886704 |
| 3 | 120516600 | 120516999 | 7.39419 | 14.38756 | 20.74753 | 7.646161 | 0.086418 | -0.31059 | 0.886766 |
| 3 | 140361200 | 140362199 | 10.08789 | 19.02968 | 28.07525 | 11.45836 | -0.08216 | 0.076496 | 0.88678 |
| X | 74346400 | 74347599 | 30.84366 | 50.44616 | 69.31602 | 29.809 | -0.08214 | 1.481449 | 0.886784 |
| 14 | 26446800 | 26447799 | 15.69804 | 27.76555 | 34.53616 | 15.27475 | 0.077879 | 0.528088 | 0.886788 |
| 14 | 123959400 | 123962999 | 67.25783 | 106.2565 | 127.3257 | 52.88156 | 0.065323 | 2.430053 | 0.88681 |
| 2 | 111797400 | 111797999 | 12.80058 | 22.86645 | 32.21125 | 14.1493 | -0.09279 | 0.432479 | 0.886822 |
| 2 | 163495400 | 163498399 | 37.46719 | 60.70431 | 81.68974 | 34.96752 | -0.06294 | 1.65172 | 0.88683 |
| 2 | 173656800 | 173659599 | 29.62358 | 49.46167 | 60.74331 | 26.42816 | 0.085656 | 1.225676 | 0.886845 |
| 3 | 152654200 | 152656999 | 24.6636 | 41.22124 | 56.5908 | 24.63699 | -0.07457 | 1.195278 | 0.886856 |
| 15 | 100439600 | 100449399 | 195.8243 | 287.4559 | 351.93 | 144.9474 | -0.05419 | 3.84963 | 0.88686 |
| 12 | 82204600 | 82206399 | 17.55288 | 30.70081 | 38.30679 | 16.89277 | 0.069867 | 0.690921 | 0.886866 |
| 8 | 105761200 | 105762599 | 21.88194 | 37.92181 | 36.91725 | 16.19731 | 0.425045 | 0.834097 | 0.88688 |
| 1 | 48851800 | 48854999 | 31.3646 | 51.50833 | 69.97411 | 30.14264 | -0.06315 | 1.427912 | 0.886886 |
| 7 | 43482000 | 43483199 | 11.25213 | 20.48634 | 29.21962 | 12.57095 | -0.0909 | 0.299724 | 0.886889 |
| 6 | 95697800 | 95700599 | 24.28589 | 41.37073 | 51.18865 | 22.43813 | 0.075971 | 1.014261 | 0.8869 |
| 1 | 89518600 | 89520599 | 34.47497 | 56.33993 | 69.99316 | 30.09136 | 0.061932 | 1.498295 | 0.886901 |
| 9 | 19643400 | 19644799 | 13.35294 | 24.64964 | 24.95958 | 10.54318 | 0.418447 | 0.227876 | 0.886906 |
| 8 | 97982200 | 97985399 | 40.9275 | 66.11086 | 81.4771 | 34.77567 | 0.059196 | 1.72966 | 0.886911 |
| 1 | 27320400 | 27322599 | 29.90407 | 49.28694 | 67.10365 | 28.9502 | -0.06422 | 1.378942 | 0.886919 |
| 15 | 74295400 | 74296799 | 15.36171 | 26.99477 | 37.22723 | 16.48194 | -0.07374 | 0.60251 | 0.886937 |
| 2 | 7632600 | 7639399 | 85.31506 | 133.6403 | 135.2877 | 56.1169 | 0.298774 | 2.612414 | 0.886944 |
| 19 | 56106400 | 56111199 | 70.84277 | 111.7181 | 134.7405 | 55.91675 | 0.053901 | 2.474269 | 0.886961 |
| 17 | 15084200 | 15085599 | 27.94863 | 47.15851 | 45.38193 | 19.84731 | 0.436484 | 1.105952 | 0.886973 |
| 4 | 96427200 | 96427999 | 10.92437 | 20.11873 | 28.76816 | 12.19122 | -0.07937 | 0.231411 | 0.886974 |
| 8 | 21502000 | 21502599 | 10.24462 | 19.19279 | 27.92613 | 11.5321 | -0.07592 | 0.121274 | 0.887003 |
| 15 | 33785600 | 33787999 | 26.40435 | 44.79977 | 45.28578 | 19.83849 | 0.365878 | 1.049805 | 0.887043 |
| 6 | 22521000 | 22522199 | 13.41832 | 24.18741 | 30.47128 | 13.27426 | 0.075134 | 0.374444 | 0.887119 |
| 5 | 74946600 | 74949399 | 42.63631 | 68.43058 | 92.17475 | 38.98951 | -0.07985 | 1.932739 | 0.887126 |
| 11 | 31127200 | 31128399 | 11.40682 | 20.93311 | 29.74391 | 12.66548 | -0.0754 | 0.257101 | 0.887131 |
| 16 | 90486400 | 90487399 | 11.40682 | 20.93311 | 29.74391 | 12.66548 | -0.0754 | 0.257101 | 0.887131 |
| 17 | 90182200 | 90182799 | 11.40682 | 20.93311 | 29.74391 | 12.66548 | -0.0754 | 0.257101 | 0.887131 |
| X | 65218000 | 65218999 | 14.31398 | 26.16205 | 32.23369 | 13.80067 | 0.123427 | 0.308641 | 0.887143 |
| 3 | 145678600 | 145681999 | 55.79124 | 89.08842 | 108.0944 | 45.38233 | 0.062155 | 2.108047 | 0.887153 |
| 8 | 107287200 | 107290199 | 21.48341 | 36.5203 | 50.08395 | 21.9502 | -0.07235 | 1.020628 | 0.887156 |
| 4 | 112921800 | 112925199 | 46.36285 | 74.88096 | 89.39602 | 38.1492 | 0.104773 | 1.748164 | 0.887192 |
| 10 | 41562400 | 41564199 | 18.8436 | 32.98061 | 40.77335 | 17.99075 | 0.082988 | 0.689855 | 0.887194 |
| 18 | 16274800 | 16278199 | 46.01597 | 73.82562 | 97.94845 | 41.40679 | -0.0588 | 1.950239 | 0.887195 |
| 8 | 12730200 | 12731599 | 17.57917 | 30.5444 | 42.14089 | 18.50279 | -0.07524 | 0.792163 | 0.887214 |
| 13 | 18219400 | 18220799 | 14.72618 | 26.27871 | 32.89924 | 14.46429 | 0.075083 | 0.449898 | 0.887214 |
| 17 | 49269800 | 49271199 | 12.54432 | 22.58487 | 31.60399 | 13.78836 | -0.07678 | 0.378646 | 0.88723 |
| 11 | 67192000 | 67192999 | 9.756962 | 18.3831 | 26.9528 | 11.02382 | -0.07568 | 0.078517 | 0.887235 |
| 11 | 10336000 | 10336799 | 8.485359 | 16.24031 | 24.57283 | 9.715837 | -0.07751 | -0.05064 | 0.887239 |
| 8 | 123290200 | 123292799 | 20.37492 | 34.70651 | 47.97322 | 21.02858 | -0.08315 | 0.996259 | 0.88725 |

| | | | | | | | | | |
|---|---|---|---|---|---|---|---|---|---|
| 13 | 87475600 | 87480999 | 97.65748 | 150.4865 | 177.5516 | 73.75212 | 0.059833 | 2.911212 | 0.8873 |
| 7 | 47759200 | 47760599 | 16.07832 | 28.45343 | 35.41363 | 15.66116 | 0.077073 | 0.524026 | 0.887311 |
| 2 | 55854600 | 55855999 | 17.37181 | 30.34539 | 37.25384 | 16.41192 | 0.092948 | 0.705144 | 0.887403 |
| 1 | 30714600 | 30714999 | 5.189555 | 10.14932 | 15.51284 | 5.270622 | 0.112109 | -0.6378 | 0.887403 |
| 11 | 101894800 | 101895799 | 12.75916 | 23.08537 | 29.02865 | 12.58681 | 0.082212 | 0.345101 | 0.887444 |
| 13 | 72830800 | 72833399 | 21.05873 | 36.48054 | 45.22624 | 19.84261 | 0.077456 | 0.843079 | 0.887445 |
| 8 | 21095000 | 21099399 | 105.5729 | 162.1376 | 191.8938 | 79.67401 | 0.051658 | 2.98188 | 0.887452 |
| 3 | 5787600 | 5789399 | 19.36434 | 33.71262 | 42.01574 | 18.47073 | 0.070999 | 0.772679 | 0.88746 |
| 2 | 8407000 | 8412399 | 91.18853 | 141.761 | 168.9265 | 70.13533 | 0.055288 | 2.750855 | 0.887461 |
| 6 | 121544800 | 121545399 | 14.42468 | 25.63248 | 31.96405 | 14.10313 | 0.077315 | 0.47279 | 0.887481 |
| 10 | 65007800 | 65008599 | 10.05854 | 17.91154 | 36.42021 | 16.0408 | -0.62148 | 0.444146 | 0.887502 |
| 14 | 7244600 | 7247599 | 43.69986 | 70.40082 | 86.54708 | 36.81049 | 0.058311 | 1.813908 | 0.887513 |
| 8 | 41645200 | 41648199 | 25.86877 | 43.51253 | 60.00335 | 26.21691 | -0.08105 | 1.138473 | 0.887516 |
| X | 34844200 | 34846399 | 31.24202 | 51.88094 | 63.49468 | 27.52679 | 0.090492 | 1.293739 | 0.887527 |
| 2 | 21589800 | 21592199 | 40.42432 | 65.64281 | 63.69001 | 27.30789 | 0.410932 | 1.60504 | 0.887537 |
| 8 | 44237800 | 44239199 | 23.94316 | 40.64536 | 50.63924 | 22.18726 | 0.065442 | 1.043246 | 0.887547 |
| 8 | 68069800 | 68070799 | 12.50461 | 22.78236 | 32.19909 | 13.84086 | -0.07733 | 0.310631 | 0.887584 |
| 2 | 165458000 | 165460399 | 27.97296 | 46.42407 | 57.73173 | 25.01789 | 0.067444 | 1.284854 | 0.88762 |
| 10 | 121875400 | 121877599 | 14.48407 | 25.44527 | 45.58557 | 20.17764 | -0.44953 | 0.6159 | 0.887621 |
| 6 | 142948800 | 142955399 | 64.23346 | 101.059 | 156.2769 | 64.83455 | -0.30681 | 2.520156 | 0.887629 |
| 9 | 85992600 | 85993999 | 20.7217 | 35.67739 | 49.01041 | 21.49887 | -0.07172 | 0.902657 | 0.887645 |
| 15 | 58624400 | 58626999 | 25.42824 | 42.88843 | 53.38516 | 23.34458 | 0.066383 | 1.106401 | 0.887652 |
| 8 | 15207400 | 15210599 | 31.25946 | 51.39066 | 63.85307 | 27.51835 | 0.065863 | 1.408897 | 0.88766 |
| 14 | 80073200 | 80074199 | 14.0974 | 24.9655 | 34.64437 | 15.3188 | -0.07838 | 0.504983 | 0.887664 |
| 4 | 143699400 | 143700399 | 21.70546 | 37.21815 | 46.33724 | 20.32228 | 0.068457 | 0.938145 | 0.887689 |
| 3 | 134857600 | 134859999 | 19.93593 | 35.03106 | 41.64049 | 18.43502 | 0.139306 | 0.63508 | 0.887733 |
| 3 | 103130800 | 103138199 | 107.9126 | 165.616 | 195.7114 | 81.25133 | 0.053099 | 2.996888 | 0.88775 |
| 4 | 16557000 | 16558799 | 20.10254 | 35.09343 | 35.07168 | 15.44449 | 0.388346 | 0.725708 | 0.887751 |
| X | 148634400 | 148637999 | 42.24909 | 68.37672 | 95.37224 | 40.79127 | -0.11604 | 1.691341 | 0.887797 |
| 5 | 14326200 | 14327599 | 22.94741 | 38.9759 | 48.49955 | 21.25322 | 0.067274 | 1.032666 | 0.887807 |
| 14 | 44456200 | 44459799 | 50.99522 | 81.43698 | 107.4226 | 45.10569 | -0.05803 | 2.101903 | 0.887841 |
| 15 | 76448000 | 76449799 | 15.66051 | 27.0477 | 48.98026 | 21.48769 | -0.4674 | 0.854975 | 0.887856 |
| 4 | 90901200 | 90904399 | 26.88964 | 45.41964 | 55.66006 | 24.35015 | 0.088861 | 1.100607 | 0.887877 |
| 5 | 116991400 | 116991999 | 6.81816 | 13.21731 | 21.08138 | 7.971153 | -0.09151 | -0.22624 | 0.887911 |
| 17 | 55605400 | 55607399 | 28.11276 | 46.92194 | 58.39905 | 25.40235 | 0.066473 | 1.222749 | 0.887914 |
| 10 | 126067000 | 126072599 | 49.26121 | 79.01335 | 105.7619 | 44.69372 | -0.07197 | 1.963471 | 0.88793 |
| 14 | 37992400 | 37992999 | 7.709758 | 14.89666 | 23.19142 | 8.904733 | -0.07844 | -0.15854 | 0.88796 |
| 15 | 26484000 | 26484599 | 7.709758 | 14.89666 | 23.19142 | 8.904733 | -0.07844 | -0.15854 | 0.88796 |
| 4 | 34905200 | 34905799 | 7.9092 | 15.25818 | 23.6835 | 9.15329 | -0.08274 | -0.13114 | 0.887965 |
| 5 | 16504800 | 16505799 | 7.9092 | 15.25818 | 23.6835 | 9.15329 | -0.08274 | -0.13114 | 0.887965 |
| 3 | 45121800 | 45123999 | 19.73828 | 34.58755 | 41.7117 | 18.42327 | 0.11849 | 0.6755 | 0.887969 |
| 4 | 79704600 | 79705399 | 9.728477 | 18.62181 | 25.13641 | 9.935198 | 0.086183 | -0.03806 | 0.887986 |
| 15 | 91005400 | 91006999 | 13.88183 | 24.52527 | 34.18385 | 15.12237 | -0.08612 | 0.527535 | 0.887995 |
| 3 | 21858200 | 21858999 | 8.768356 | 16.66852 | 24.96897 | 10.0258 | -0.08326 | 0.002204 | 0.888007 |
| 1 | 191189400 | 191190399 | 8.804884 | 16.88249 | 22.97476 | 9.036731 | 0.081528 | -0.06774 | 0.888026 |
| 14 | 48622200 | 48623399 | 26.30191 | 44.9952 | 43.60742 | 19.12148 | 0.427965 | 0.975252 | 0.888032 |
| 15 | 37866600 | 37886199 | 308.8833 | 437.7817 | 518.8347 | 214.6701 | -0.05721 | 4.3811 | 0.888046 |

| | | | | | | | | | |
|---|---|---|---|---|---|---|---|---|---|
| 1 | 40289600 | 40291399 | 18.739 | 33.00448 | 39.44575 | 17.46092 | 0.132441 | 0.59582 | 0.888056 |
| 11 | 61280400 | 61288999 | 133.0769 | 200.5745 | 251.9961 | 104.1102 | -0.0565 | 3.37049 | 0.88807 |
| 3 | 99796800 | 99805799 | 186.2883 | 275.9709 | 316.2703 | 130.2519 | 0.053484 | 3.699728 | 0.888081 |
| 1 | 115137400 | 115141399 | 41.71768 | 67.76699 | 81.38469 | 34.90381 | 0.104367 | 1.610562 | 0.888146 |
| 1 | 75978800 | 75984799 | 80.00449 | 125.3381 | 150.4085 | 62.38813 | 0.05382 | 2.608373 | 0.88815 |
| 10 | 69922400 | 69925199 | 29.39204 | 48.84782 | 60.70526 | 26.332 | 0.068485 | 1.273975 | 0.888177 |
| 13 | 3166000 | 3171199 | 63.4296 | 100.8154 | 121.9285 | 50.87142 | 0.059675 | 2.265002 | 0.888179 |
| 17 | 71204600 | 71208199 | 36.00309 | 58.52486 | 72.48542 | 31.10242 | 0.060949 | 1.579586 | 0.888179 |
| 7 | 16702200 | 16705599 | 29.59462 | 48.61985 | 66.28083 | 28.55623 | -0.06789 | 1.417507 | 0.88819 |
| 19 | 60917200 | 60917599 | 9.096127 | 17.63212 | 18.82198 | 7.118978 | 0.48219 | -0.20512 | 0.888226 |
| X | 91122600 | 91124599 | 17.34596 | 30.47792 | 42.8323 | 18.96156 | -0.10109 | 0.624262 | 0.888233 |
| X | 77865000 | 77867199 | 23.59493 | 40.81136 | 49.02954 | 21.51027 | 0.12225 | 0.875031 | 0.888234 |
| 17 | 68871000 | 68872399 | 11.98119 | 22.08923 | 28.07666 | 11.88044 | 0.090391 | 0.228448 | 0.888242 |
| 15 | 20154600 | 20158199 | 50.36378 | 80.90874 | 82.60722 | 34.99195 | 0.32068 | 1.909272 | 0.888256 |
| 17 | 54179400 | 54198599 | 381.2355 | 531.7391 | 573.6926 | 237.9765 | 0.054992 | 4.620216 | 0.888257 |
| 17 | 4939800 | 4942799 | 27.37599 | 45.24083 | 61.99223 | 26.78553 | -0.07258 | 1.335444 | 0.888258 |
| 10 | 18549400 | 18553599 | 82.22579 | 128.035 | 164.9094 | 68.45093 | -0.05497 | 2.717987 | 0.888268 |
| 1 | 121665000 | 121666399 | 18.59259 | 32.19033 | 44.24846 | 19.41329 | -0.07085 | 0.838087 | 0.88827 |
| 5 | 51878400 | 51879399 | 10.38742 | 19.77791 | 26.1036 | 10.44604 | 0.101791 | 0.007953 | 0.888297 |
| 10 | 123974800 | 123979399 | 49.32844 | 79.21655 | 96.65665 | 40.92084 | 0.06345 | 1.925009 | 0.888297 |
| 5 | 83540200 | 83542799 | 34.98654 | 56.42214 | 93.32045 | 39.7468 | -0.36699 | 1.773126 | 0.888331 |
| 6 | 131314000 | 131314599 | 9.216 | 17.65783 | 24.03627 | 9.474162 | 0.077833 | -0.05562 | 0.88834 |
| 1 | 177743800 | 177756799 | 176.8905 | 262.8122 | 302.3821 | 124.5691 | 0.051654 | 3.647193 | 0.888343 |
| 9 | 19836000 | 19837999 | 40.81777 | 66.15897 | 80.94451 | 34.65858 | 0.074644 | 1.651349 | 0.888346 |
| 4 | 17741800 | 17744999 | 32.80066 | 53.64815 | 72.88026 | 31.34204 | -0.06587 | 1.483191 | 0.888348 |
| X | 146626200 | 146627999 | 43.90265 | 70.96793 | 66.81505 | 28.54427 | 0.448058 | 1.70865 | 0.888368 |
| 17 | 80290000 | 80293399 | 41.74973 | 67.11787 | 90.07009 | 38.19234 | -0.07163 | 1.877904 | 0.888369 |
| 8 | 125085000 | 125086999 | 20.41423 | 35.01657 | 47.94959 | 21.03002 | -0.0681 | 0.93431 | 0.888417 |
| 3 | 6200400 | 6204199 | 71.38566 | 111.7375 | 146.9734 | 60.96876 | -0.07936 | 2.619136 | 0.888461 |
| 5 | 151274200 | 151277999 | 51.83546 | 82.72728 | 109.0168 | 45.73474 | -0.05759 | 2.12258 | 0.888463 |
| 11 | 113346800 | 113353199 | 72.57131 | 114.2091 | 148.0571 | 61.42666 | -0.05241 | 2.518465 | 0.888468 |
| 1 | 145811200 | 145814199 | 41.01052 | 66.28675 | 81.44546 | 34.79044 | 0.064648 | 1.713247 | 0.888481 |
| 8 | 58549600 | 58549999 | 5.225703 | 10.19214 | 17.54177 | 6.291717 | -0.12877 | -0.44271 | 0.888511 |
| 9 | 38437200 | 38437399 | 5.366269 | 10.48909 | 15.98953 | 5.511359 | 0.094993 | -0.58933 | 0.888514 |
| 12 | 54767600 | 54769199 | 20.43545 | 35.45944 | 44.16588 | 19.37836 | 0.071285 | 0.824835 | 0.888518 |
| 1 | 107196000 | 107198199 | 22.3958 | 38.72262 | 47.39002 | 20.79074 | 0.095537 | 0.871041 | 0.888522 |
| 1 | 58403000 | 58409599 | 83.94659 | 130.9741 | 156.8253 | 65.07726 | 0.052396 | 2.683776 | 0.888542 |
| 13 | 62644200 | 62706999 | 1509.307 | 1883.081 | 1806.244 | 759.8227 | 0.06302 | 6.403142 | 0.888546 |
| 9 | 20792400 | 20796799 | 80.11229 | 125.4828 | 150.6438 | 62.48697 | 0.053025 | 2.613264 | 0.888554 |
| 6 | 65368800 | 65369999 | 10.85245 | 20.08258 | 28.77504 | 12.11465 | -0.07573 | 0.202351 | 0.888572 |
| 19 | 47049200 | 47052799 | 27.99865 | 46.27197 | 63.21377 | 27.3142 | -0.06826 | 1.336774 | 0.888574 |
| 7 | 12776600 | 12780999 | 48.70823 | 77.83578 | 103.4063 | 43.45966 | -0.06723 | 2.078433 | 0.888584 |
| 5 | 103912800 | 103915999 | 35.72913 | 58.4166 | 72.08339 | 30.99523 | 0.072102 | 1.494486 | 0.888593 |
| 4 | 49423200 | 49424199 | 25.95439 | 43.94732 | 41.7444 | 18.27589 | 0.455301 | 1.062863 | 0.888612 |
| 13 | 34771200 | 34776799 | 88.98585 | 138.534 | 165.427 | 68.66781 | 0.053867 | 2.724993 | 0.888633 |
| 7 | 91967800 | 91968999 | 10.08617 | 18.86437 | 27.37102 | 11.33492 | -0.0751 | 0.129306 | 0.888658 |
| 10 | 55637800 | 55639999 | 21.60062 | 37.35746 | 52.88859 | 23.20323 | -0.11378 | 0.871413 | 0.888664 |

| | | | | | | | | | |
|---|---|---|---|---|---|---|---|---|---|
| 8 | 47637800 | 47640599 | 30.24556 | 50.07812 | 62.36028 | 26.97994 | 0.065174 | 1.320796 | 0.888671 |
| 17 | 14296000 | 14298999 | 39.53356 | 63.66725 | 85.83984 | 36.4917 | -0.07516 | 1.819542 | 0.88869 |
| 5 | 109201200 | 109212199 | 113.0185 | 174.1798 | 200.6239 | 83.33591 | 0.094626 | 2.91821 | 0.888691 |
| 13 | 69317600 | 69319599 | 15.8546 | 28.12152 | 34.95109 | 15.44506 | 0.08064 | 0.50101 | 0.888694 |
| 15 | 92369000 | 92372999 | 61.69689 | 97.80603 | 127.7334 | 53.15443 | -0.05463 | 2.341199 | 0.888696 |
| 9 | 114780800 | 114782399 | 18.11785 | 31.5609 | 43.57774 | 19.16442 | -0.07624 | 0.758085 | 0.888718 |
| 12 | 94502000 | 94503399 | 14.17797 | 25.458 | 32.00472 | 13.99756 | 0.0745 | 0.407884 | 0.888725 |
| 6 | 114814800 | 114815999 | 13.99354 | 24.80467 | 34.6801 | 15.33175 | -0.08849 | 0.495573 | 0.888726 |
| 4 | 96346000 | 96349599 | 31.98745 | 53.08587 | 77.30184 | 33.49825 | -0.15975 | 1.309902 | 0.88873 |
| 12 | 85453000 | 85455799 | 54.42766 | 86.72973 | 104.3011 | 43.66222 | 0.071753 | 2.172997 | 0.888731 |
| 1 | 19884600 | 19885599 | 12.15501 | 22.34049 | 28.65592 | 12.17667 | 0.073374 | 0.251348 | 0.888761 |
| 18 | 37092600 | 37096799 | 89.3956 | 137.9111 | 178.0958 | 73.97771 | -0.06908 | 2.888462 | 0.888783 |
| 6 | 42080000 | 42081799 | 37.91036 | 61.34741 | 74.6444 | 31.923 | 0.080507 | 1.676914 | 0.888788 |
| 15 | 93059000 | 93061199 | 18.3611 | 31.77665 | 43.7201 | 19.18124 | -0.07236 | 0.838553 | 0.888789 |
| 17 | 55899400 | 55902199 | 28.53664 | 47.61913 | 59.03639 | 25.67609 | 0.072034 | 1.225552 | 0.888805 |
| 2 | 18504400 | 18510399 | 61.61566 | 98.11771 | 118.4356 | 49.53305 | 0.065314 | 2.2039 | 0.888807 |
| 3 | 37267000 | 37272599 | 78.34162 | 122.3645 | 144.5454 | 59.97468 | 0.07317 | 2.660309 | 0.888808 |
| 13 | 67742600 | 67744799 | 29.08437 | 48.00916 | 65.37861 | 28.22644 | -0.06378 | 1.357326 | 0.888818 |
| 1 | 19529200 | 19530599 | 11.30323 | 20.79756 | 29.72237 | 12.61116 | -0.08027 | 0.239227 | 0.888821 |
| 11 | 56123400 | 56125399 | 19.4083 | 33.78684 | 42.03158 | 18.4769 | 0.073651 | 0.774605 | 0.888853 |
| X | 36377800 | 36381999 | 27.34402 | 45.86395 | 64.49161 | 28.1397 | -0.10912 | 1.176646 | 0.888857 |
| 3 | 102504000 | 102507999 | 36.00895 | 58.36229 | 78.48604 | 33.60492 | -0.06148 | 1.640727 | 0.888866 |
| 17 | 65465600 | 65470599 | 49.93218 | 80.04571 | 97.92621 | 41.34799 | 0.056661 | 1.979418 | 0.888872 |
| 4 | 42825000 | 42826999 | 22.52065 | 38.29664 | 47.45119 | 20.79692 | 0.073711 | 1.020582 | 0.888878 |
| 12 | 17852600 | 17855999 | 28.87573 | 47.59502 | 64.7949 | 27.95489 | -0.06406 | 1.372919 | 0.888904 |
| 17 | 39319000 | 39323199 | 58.76287 | 93.64639 | 113.6976 | 47.60176 | 0.058142 | 2.177248 | 0.888911 |
| 10 | 7045200 | 7056399 | 199.1219 | 292.0851 | 356.7349 | 146.9331 | -0.0517 | 3.861513 | 0.888951 |
| 12 | 5661800 | 5668999 | 91.57381 | 142.1529 | 169.6206 | 70.43883 | 0.051729 | 2.775955 | 0.888953 |
| 5 | 103329400 | 103335599 | 61.53203 | 98.09649 | 117.7161 | 49.30996 | 0.07555 | 2.168305 | 0.888975 |
| 9 | 22050000 | 22053199 | 30.23279 | 49.58708 | 67.42983 | 29.03116 | -0.06524 | 1.439347 | 0.888985 |
| 5 | 127155200 | 127159999 | 49.09337 | 78.85702 | 107.2026 | 45.40133 | -0.09187 | 1.919545 | 0.888991 |
| 14 | 61597400 | 61598399 | 7.994902 | 15.2612 | 23.44199 | 9.330665 | -0.10802 | -0.03855 | 0.888993 |
| 9 | 9500800 | 9502199 | 15.07754 | 26.62174 | 33.04496 | 14.6203 | 0.079751 | 0.538191 | 0.888996 |
| 6 | 12592200 | 12592999 | 9.105755 | 17.14941 | 25.36743 | 10.38232 | -0.08907 | 0.070477 | 0.889 |
| 18 | 21209400 | 21210199 | 9.105755 | 17.14941 | 25.36743 | 10.38232 | -0.08907 | 0.070477 | 0.889 |
| 3 | 65422800 | 65423399 | 7.4418 | 14.41446 | 22.73269 | 8.628473 | -0.08062 | -0.19999 | 0.889045 |
| 11 | 79704400 | 79706799 | 21.59196 | 37.07553 | 51.19446 | 22.45507 | -0.08007 | 0.930633 | 0.889064 |
| 2 | 23798200 | 23798799 | 10.67387 | 19.93322 | 28.92735 | 12.0103 | -0.07867 | 0.146352 | 0.88908 |
| 17 | 29173600 | 29177799 | 34.33888 | 55.89851 | 75.41845 | 32.37761 | -0.0608 | 1.55657 | 0.889096 |
| 9 | 105487200 | 105492799 | 70.96976 | 111.4693 | 144.9464 | 60.1207 | -0.05896 | 2.556068 | 0.889115 |
| 13 | 8050200 | 8052199 | 25.52824 | 42.93029 | 58.86387 | 25.71733 | -0.07285 | 1.13812 | 0.889128 |
| 9 | 85152000 | 85152799 | 12.96933 | 23.63444 | 30.08428 | 12.92811 | 0.073082 | 0.308682 | 0.889132 |
| 5 | 76094200 | 76096199 | 18.75154 | 32.57255 | 40.52209 | 17.78898 | 0.073213 | 0.797642 | 0.889185 |
| 5 | 117524400 | 117535799 | 131.7284 | 198.9208 | 249.4929 | 103.136 | -0.05168 | 3.332129 | 0.889197 |
| 1 | 104645600 | 104646999 | 6.537756 | 12.66297 | 26.88728 | 10.20691 | -0.50678 | -0.2007 | 0.889203 |
| 12 | 119573600 | 119575599 | 27.85455 | 45.93205 | 63.10553 | 27.2311 | -0.07698 | 1.362441 | 0.889205 |
| 15 | 29130600 | 29134199 | 52.81629 | 84.33088 | 110.9409 | 46.57186 | -0.05443 | 2.110147 | 0.889206 |

| | | | | | | | | | |
|---|---|---|---|---|---|---|---|---|---|
| 11 | 45967000 | 45969599 | 17.6514 | 30.45131 | 51.68672 | 22.67422 | -0.37511 | 0.874259 | 0.889206 |
| 12 | 56096400 | 56098999 | 25.66009 | 43.66947 | 44.23325 | 19.38778 | 0.363436 | 1.020371 | 0.889213 |
| Y | 43172800 | 43174399 | 25.24727 | 41.98196 | 47.61493 | 20.66149 | 0.20012 | 1.243683 | 0.889219 |
| 7 | 64850200 | 64857999 | 83.65313 | 130.6751 | 169.3157 | 70.25158 | -0.05997 | 2.664782 | 0.889229 |
| 2 | 111552400 | 111553799 | 22.30805 | 37.79412 | 51.70134 | 22.6389 | -0.06906 | 1.054791 | 0.889234 |
| 6 | 18997200 | 18998199 | 14.88793 | 26.17426 | 36.25682 | 16.05512 | -0.07995 | 0.592093 | 0.889263 |
| 19 | 6697200 | 6698399 | 15.29153 | 27.11426 | 33.93251 | 14.99893 | 0.070416 | 0.496406 | 0.889263 |
| 11 | 5789000 | 5794999 | 76.03936 | 119.4217 | 143.798 | 59.64371 | 0.051685 | 2.560357 | 0.889288 |
| 11 | 5789200 | 5794999 | 76.03936 | 119.4217 | 143.798 | 59.64371 | 0.051685 | 2.560357 | 0.889288 |
| 19 | 29736200 | 29744799 | 127.2407 | 194.5842 | 222.8735 | 92.46533 | 0.094117 | 3.07599 | 0.889293 |
| 2 | 120654200 | 120656799 | 22.00566 | 37.6694 | 51.72034 | 22.68009 | -0.07253 | 0.962138 | 0.889323 |
| 11 | 94956600 | 94958799 | 18.85513 | 32.66667 | 44.74547 | 19.63174 | -0.0657 | 0.832656 | 0.889323 |
| 4 | 72769400 | 72771199 | 21.96054 | 36.45998 | 65.48522 | 28.38545 | -0.46113 | 1.27059 | 0.889384 |
| 11 | 51813200 | 51816799 | 39.96906 | 64.63503 | 79.78352 | 34.10205 | 0.058804 | 1.695487 | 0.889393 |
| 10 | 126506400 | 126508399 | 23.37036 | 39.86071 | 49.56218 | 21.72736 | 0.06917 | 0.99644 | 0.889461 |
| 3 | 33756000 | 33756999 | 13.98523 | 24.82836 | 34.65982 | 15.30461 | -0.08506 | 0.484691 | 0.889491 |
| 6 | 104423400 | 104425399 | 15.79303 | 27.76053 | 38.15393 | 16.8958 | -0.06903 | 0.600817 | 0.889497 |
| 3 | 77504600 | 77508199 | 44.68097 | 72.16556 | 73.18844 | 31.23328 | 0.339593 | 1.73595 | 0.889499 |
| 9 | 72776200 | 72777399 | 9.648033 | 17.98813 | 26.35181 | 10.97123 | -0.09476 | 0.14412 | 0.889518 |
| 9 | 25669000 | 25673599 | 59.09535 | 94.0083 | 123.2615 | 51.47857 | -0.05541 | 2.236576 | 0.889523 |
| 9 | 79020200 | 79023199 | 36.63214 | 59.40009 | 79.95667 | 34.24638 | -0.06223 | 1.632546 | 0.889534 |
| 6 | 131992800 | 132000199 | 126.4544 | 192.6957 | 224.9334 | 93.2422 | 0.062754 | 3.146375 | 0.889551 |
| 6 | 102066600 | 102068199 | 15.72023 | 27.73251 | 38.56841 | 17.06702 | -0.08413 | 0.55999 | 0.889574 |
| 3 | 6545400 | 6546199 | 10.18499 | 19.12671 | 27.96219 | 11.50037 | -0.07825 | 0.10429 | 0.889575 |
| 3 | 60262800 | 60263599 | 8.806227 | 16.73794 | 24.97687 | 10.03222 | -0.07793 | 0.004888 | 0.889627 |
| 9 | 43837000 | 43841999 | 62.22477 | 98.53317 | 128.7298 | 53.52744 | -0.05654 | 2.370777 | 0.889656 |
| 1 | 88846200 | 88847599 | 20.91233 | 36.73041 | 33.64417 | 14.8893 | 0.513468 | 0.634311 | 0.88969 |
| Y | 75648400 | 75649199 | 7.611104 | 14.41411 | 22.95803 | 9.292209 | -0.18058 | 0.012473 | 0.889705 |
| 4 | 75225400 | 75230599 | 91.79249 | 142.3808 | 169.8084 | 70.52509 | 0.05157 | 2.789979 | 0.889721 |
| 12 | 65552800 | 65558599 | 92.37034 | 143.7618 | 143.878 | 59.72547 | 0.308052 | 2.728048 | 0.889734 |
| 18 | 47045600 | 47048599 | 19.86974 | 34.20375 | 47.39792 | 20.79141 | -0.08447 | 0.895068 | 0.889788 |
| 1 | 12109200 | 12112799 | 60.12662 | 95.43996 | 114.5906 | 47.74694 | 0.068333 | 2.298669 | 0.889791 |
| 5 | 130623600 | 130624399 | 11.15043 | 20.95843 | 27.41376 | 11.25779 | 0.082944 | 0.106487 | 0.889802 |
| 14 | 95802200 | 95806999 | 99.28961 | 153.0833 | 181.7575 | 75.49893 | 0.051342 | 2.903744 | 0.889869 |
| 7 | 48804200 | 48806599 | 24.31828 | 41.05699 | 56.0936 | 24.53631 | -0.06747 | 1.096788 | 0.88987 |
| 17 | 47621200 | 47622399 | 14.1754 | 25.1627 | 31.11754 | 13.74196 | 0.088611 | 0.477837 | 0.889872 |
| 18 | 61212000 | 61215199 | 51.18029 | 81.76221 | 98.35658 | 41.32022 | 0.075259 | 2.086487 | 0.889902 |
| 6 | 14569800 | 14571999 | 24.12991 | 40.70212 | 50.65711 | 22.15222 | 0.066635 | 1.098947 | 0.88991 |
| 13 | 12383800 | 12385199 | 15.61608 | 27.37401 | 37.79381 | 16.69787 | -0.07548 | 0.643262 | 0.889921 |
| 13 | 3339000 | 3342199 | 75.24716 | 117.6735 | 152.8741 | 63.42255 | -0.06253 | 2.63947 | 0.889924 |
| 3 | 9451000 | 9463999 | 202.0977 | 296.8218 | 363.3418 | 149.6426 | -0.05325 | 3.842822 | 0.889925 |
| X | 42767800 | 42769599 | 28.63322 | 47.14943 | 64.57287 | 27.84764 | -0.07314 | 1.386225 | 0.889933 |
| 17 | 49572800 | 49575599 | 25.32215 | 42.59985 | 58.32456 | 25.48463 | -0.07072 | 1.132945 | 0.889936 |
| 9 | 62485400 | 62492599 | 100.0809 | 155.0126 | 182.4963 | 75.80402 | 0.068601 | 2.819993 | 0.889938 |
| 1 | 96652800 | 96653999 | 14.19527 | 25.28399 | 31.40757 | 13.83026 | 0.084705 | 0.454308 | 0.889944 |
| 4 | 21710400 | 21712599 | 18.68987 | 32.3678 | 39.64832 | 17.3952 | 0.09489 | 0.829552 | 0.88995 |
| 7 | 27715200 | 27718799 | 35.5089 | 57.69485 | 77.67021 | 33.31652 | -0.05967 | 1.589624 | 0.889963 |

| | | | | | | | | | |
|---|---|---|---|---|---|---|---|---|---|
| 11 | 14391600 | 14395199 | 68.27591 | 107.4817 | 140.0531 | 58.10822 | -0.05945 | 2.508369 | 0.889964 |
| 15 | 65542000 | 65545599 | 31.36913 | 52.00833 | 64.06956 | 27.7452 | 0.080865 | 1.312137 | 0.889968 |
| 14 | 8419200 | 8424999 | 82.77856 | 129.3064 | 155.1522 | 64.37508 | 0.050728 | 2.662334 | 0.88999 |
| 13 | 49485000 | 49486599 | 12.52984 | 22.60374 | 31.60131 | 13.75292 | -0.07317 | 0.366805 | 0.889991 |
| 13 | 8714600 | 8717399 | 62.71143 | 99.31068 | 118.5292 | 49.28765 | 0.073816 | 2.367826 | 0.890004 |
| 16 | 55669000 | 55669999 | 9.930749 | 18.42454 | 26.7985 | 11.24288 | -0.09328 | 0.176344 | 0.890014 |
| 7 | 81810800 | 81814999 | 70.21051 | 110.5686 | 143.3871 | 59.49438 | -0.05215 | 2.502259 | 0.890065 |
| 16 | 20909400 | 20912999 | 34.38486 | 56.33207 | 69.76911 | 30.02363 | 0.068439 | 1.462747 | 0.890124 |
| 4 | 129293200 | 129296199 | 22.14732 | 36.77675 | 65.40876 | 28.35586 | -0.44693 | 1.270851 | 0.89013 |
| 5 | 138492000 | 138495599 | 58.0255 | 92.38299 | 112.3967 | 46.99496 | 0.053675 | 2.20573 | 0.890131 |
| 14 | 122785600 | 122789799 | 49.01099 | 78.62881 | 104.8335 | 44.31578 | -0.06592 | 1.957589 | 0.890133 |
| 5 | 146574000 | 146576199 | 31.72296 | 51.89475 | 70.26366 | 30.21875 | -0.06083 | 1.478362 | 0.890145 |
| 9 | 69870000 | 69872599 | 18.37606 | 31.85752 | 43.7072 | 19.17659 | -0.06776 | 0.82352 | 0.890161 |
| 11 | 85181400 | 85185999 | 53.54145 | 85.53705 | 104.3498 | 43.84004 | 0.05512 | 2.090728 | 0.890165 |
| 10 | 29119000 | 29120999 | 19.69082 | 33.30197 | 57.53088 | 25.17822 | -0.40459 | 1.067406 | 0.890205 |
| 3 | 90727800 | 90736399 | 129.8394 | 198.1611 | 194.5805 | 80.68982 | 0.313798 | 3.114161 | 0.890209 |
| 17 | 21254400 | 21257999 | 121.7581 | 184.7033 | 232.9971 | 96.4543 | -0.0544 | 3.237772 | 0.890271 |
| 13 | 21446400 | 21448399 | 16.69038 | 29.23122 | 40.29907 | 17.79138 | -0.07355 | 0.666089 | 0.890333 |
| 9 | 69989600 | 69993599 | 36.95972 | 59.94897 | 74.10801 | 31.74825 | 0.060767 | 1.624908 | 0.890346 |
| 4 | 112370400 | 112372199 | 43.3169 | 69.66897 | 84.14177 | 35.70209 | 0.080967 | 1.866244 | 0.890364 |
| 17 | 63971800 | 63972199 | 7.130509 | 13.89169 | 20.27628 | 7.379933 | 0.086017 | -0.35715 | 0.890365 |
| 4 | 92806800 | 92810399 | 32.04735 | 53.42167 | 64.27907 | 27.93783 | 0.115106 | 1.253988 | 0.890385 |
| 3 | 120982600 | 120984799 | 27.70593 | 46.65671 | 46.94855 | 20.52016 | 0.372056 | 1.119773 | 0.890387 |
| 14 | 44626400 | 44627599 | 13.83289 | 24.57873 | 34.03272 | 15.01696 | -0.07296 | 0.473728 | 0.890419 |
| 3 | 19392600 | 19393599 | 16.48702 | 28.96419 | 36.03418 | 15.94998 | 0.073867 | 0.608598 | 0.890443 |
| 5 | 109024600 | 109038799 | 201.5976 | 297.5374 | 337.7845 | 139.1055 | 0.062042 | 3.763824 | 0.890449 |
| 1 | 35087200 | 35090399 | 50.57083 | 80.91925 | 107.036 | 45.06956 | -0.05889 | 2.042107 | 0.890475 |
| 3 | 25373000 | 25375999 | 26.52823 | 45.17859 | 44.73459 | 19.60863 | 0.39623 | 1.018408 | 0.890494 |
| 8 | 54754000 | 54754799 | 9.073406 | 16.73923 | 31.98064 | 13.52032 | -0.49229 | 0.212435 | 0.890499 |
| 5 | 73346400 | 73347399 | 11.64686 | 21.27278 | 30.15371 | 12.92115 | -0.07862 | 0.284541 | 0.890504 |
| 6 | 38918600 | 38922799 | 42.11572 | 67.87765 | 90.49323 | 38.5122 | -0.05767 | 1.800664 | 0.890531 |
| 1 | 184159200 | 184160399 | 10.80087 | 20.14759 | 26.22594 | 10.90959 | 0.075031 | 0.148499 | 0.890545 |
| 10 | 40318600 | 40319199 | 10.80087 | 20.14759 | 26.22594 | 10.90959 | 0.075031 | 0.148499 | 0.890545 |
| 5 | 149365200 | 149366199 | 8.321295 | 15.83635 | 23.93624 | 9.58844 | -0.09289 | -0.01217 | 0.890564 |
| 11 | 45659000 | 45660799 | 23.65252 | 40.54556 | 40.57208 | 17.79344 | 0.382472 | 0.942931 | 0.890586 |
| 13 | 73172200 | 73173599 | 13.9205 | 25.01439 | 35.13475 | 15.35681 | -0.08398 | 0.408037 | 0.890613 |
| 13 | 6752800 | 6753399 | 9.525978 | 18.01869 | 23.80821 | 9.661053 | 0.083133 | 0.041009 | 0.890627 |
| 7 | 49722600 | 49725799 | 24.31661 | 41.30296 | 57.24858 | 25.08745 | -0.08789 | 1.046173 | 0.890669 |
| 14 | 58823800 | 58824999 | 14.47055 | 25.83359 | 32.47445 | 14.27238 | 0.069724 | 0.446043 | 0.890675 |
| X | 147437400 | 147439999 | 28.49558 | 47.64973 | 58.96797 | 25.68005 | 0.074637 | 1.205572 | 0.890684 |
| 2 | 151134000 | 151134799 | 19.99048 | 34.35406 | 42.24245 | 18.52898 | 0.087066 | 0.907737 | 0.890701 |
| 2 | 159437600 | 159438799 | 12.0458 | 21.80034 | 30.64272 | 13.28234 | -0.07709 | 0.342901 | 0.890716 |
| 5 | 146274000 | 146275399 | 12.0458 | 21.80034 | 30.64272 | 13.28234 | -0.07709 | 0.342901 | 0.890716 |
| 13 | 33717800 | 33719999 | 26.37723 | 45.33118 | 41.90773 | 18.38218 | 0.496751 | 0.931977 | 0.890728 |
| 9 | 62312600 | 62316599 | 55.27745 | 88.41952 | 107.201 | 45.11669 | 0.065973 | 2.057117 | 0.89075 |
| X | 43962000 | 43963799 | 20.00111 | 35.05532 | 42.30836 | 18.69004 | 0.117577 | 0.678023 | 0.890763 |
| 5 | 61369200 | 61371199 | 19.66078 | 34.37335 | 34.55306 | 15.23208 | 0.380113 | 0.701312 | 0.890791 |

| | | | | | | | | | |
|---|---|---|---|---|---|---|---|---|---|
| 15 | 43327200 | 43331199 | 101.796 | 155.8869 | 199.0129 | 82.59234 | -0.0607 | 3.03769 | 0.890794 |
| 13 | 62556400 | 62569599 | 215.1991 | 315.2339 | 357.6461 | 147.321 | 0.052705 | 3.885014 | 0.890804 |
| 13 | 54116400 | 54127599 | 105.7721 | 162.3223 | 206.3693 | 85.67543 | -0.05235 | 2.997973 | 0.890812 |
| 1 | 32626800 | 32628199 | 25.03909 | 42.10508 | 52.30222 | 22.85015 | 0.069161 | 1.128694 | 0.890812 |
| 11 | 8097200 | 8100399 | 27.4688 | 45.82077 | 57.23268 | 24.88598 | 0.061373 | 1.226232 | 0.890889 |
| 15 | 87011600 | 87013999 | 18.02407 | 31.39928 | 43.07062 | 18.94206 | -0.06669 | 0.757525 | 0.890915 |
| 8 | 36085600 | 36088599 | 51.99828 | 83.00262 | 109.2069 | 45.82661 | -0.05502 | 2.11707 | 0.890925 |
| 8 | 126813600 | 126818599 | 58.26631 | 92.80972 | 122.2639 | 51.15364 | -0.06019 | 2.194055 | 0.890938 |
| 15 | 7947800 | 7949199 | 14.51627 | 25.91104 | 32.51057 | 14.2902 | 0.072374 | 0.448283 | 0.890939 |
| 15 | 95906000 | 95915399 | 41.51785 | 66.59356 | 106.9862 | 45.2114 | -0.33473 | 1.955301 | 0.890965 |
| 3 | 129454800 | 129462599 | 116.2587 | 177.2256 | 224.1738 | 92.93647 | -0.05258 | 3.135088 | 0.890996 |
| 2 | 22199800 | 22202599 | 39.09218 | 64.16443 | 61.10837 | 26.31758 | 0.447531 | 1.445575 | 0.890997 |
| 4 | 10426200 | 10429799 | 36.94495 | 59.96717 | 74.25623 | 31.82824 | 0.05926 | 1.610572 | 0.891018 |
| 11 | 84952200 | 84955399 | 94.60226 | 145.5959 | 186.511 | 77.46148 | -0.06011 | 2.93678 | 0.89103 |
| 14 | 50932200 | 50933199 | 11.27855 | 20.96974 | 27.20877 | 11.38147 | 0.073128 | 0.175598 | 0.891057 |
| 9 | 94511400 | 94513799 | 22.74103 | 38.97019 | 48.47561 | 21.25907 | 0.0696 | 0.950988 | 0.891059 |
| 19 | 26203800 | 26205999 | 14.54792 | 26.54564 | 26.38709 | 11.3209 | 0.429232 | 0.305502 | 0.891064 |
| 4 | 15222000 | 15222599 | 7.890409 | 15.29947 | 21.64704 | 8.176159 | 0.081234 | -0.21473 | 0.891069 |
| 2 | 152022400 | 152024599 | 16.20904 | 28.26844 | 39.47561 | 17.37302 | -0.09216 | 0.726815 | 0.891092 |
| X | 78673400 | 78677199 | 54.00811 | 86.6543 | 102.9022 | 43.50592 | 0.10101 | 1.963347 | 0.891096 |
| 1 | 116136600 | 116138399 | 17.43332 | 30.6164 | 38.11232 | 16.86508 | 0.07313 | 0.62483 | 0.891135 |
| 18 | 7838200 | 7839999 | 24.13132 | 40.90608 | 50.66303 | 22.18803 | 0.073784 | 1.053747 | 0.891142 |
| 17 | 33358200 | 33361799 | 37.26889 | 60.5267 | 74.99097 | 32.15638 | 0.059195 | 1.600379 | 0.891165 |
| 4 | 93844200 | 93845999 | 24.52695 | 41.1867 | 56.04278 | 24.47114 | -0.06186 | 1.145272 | 0.891185 |
| 6 | 27013600 | 27014799 | 15.06904 | 26.57935 | 36.66829 | 16.22673 | -0.07231 | 0.558274 | 0.891188 |
| 15 | 3974200 | 3974999 | 13.76323 | 24.65662 | 30.97487 | 13.57338 | 0.07351 | 0.415679 | 0.891195 |
| 10 | 39096600 | 39099599 | 38.37064 | 62.58939 | 63.98461 | 27.48946 | 0.34057 | 1.529693 | 0.891229 |
| 5 | 143782400 | 143789599 | 68.42082 | 107.2971 | 164.3156 | 68.14971 | -0.29642 | 2.587438 | 0.891262 |
| 13 | 40961000 | 40964199 | 43.99794 | 70.83333 | 87.03698 | 36.98275 | 0.057794 | 1.834976 | 0.89127 |
| 2 | 145657200 | 145659799 | 31.46141 | 52.00807 | 64.6392 | 27.93508 | 0.067777 | 1.34501 | 0.89127 |
| X | 156931800 | 156934599 | 26.4062 | 44.32471 | 61.10683 | 26.67564 | -0.0807 | 1.16113 | 0.891274 |
| 2 | 111950800 | 111954199 | 32.35198 | 53.78432 | 65.01009 | 28.20555 | 0.108413 | 1.281721 | 0.891295 |
| 1 | 185631800 | 185635399 | 35.4608 | 57.65265 | 77.894 | 33.42513 | -0.06412 | 1.577317 | 0.89132 |
| 14 | 50314400 | 50315599 | 20.21369 | 34.91853 | 47.93673 | 21.03002 | -0.0698 | 0.869338 | 0.891337 |
| 6 | 75702600 | 75703399 | 13.61727 | 25.00938 | 25.38582 | 10.79809 | 0.407618 | 0.258459 | 0.891349 |
| 7 | 66208200 | 66211399 | 27.43805 | 46.23921 | 56.76754 | 24.81729 | 0.086145 | 1.123866 | 0.891356 |
| 19 | 19787200 | 19787999 | 8.067691 | 15.58806 | 21.52843 | 8.26668 | 0.093528 | -0.15651 | 0.891359 |
| 5 | 143767800 | 143773199 | 60.61664 | 96.25202 | 125.8013 | 52.43684 | -0.05342 | 2.29303 | 0.891379 |
| 1 | 113486200 | 113486999 | 8.01857 | 15.41361 | 23.58614 | 9.210331 | -0.0767 | -0.09863 | 0.89138 |
| 4 | 71206000 | 71215999 | 158.5203 | 238.4367 | 273.5754 | 112.9639 | 0.071732 | 3.415867 | 0.89138 |
| 12 | 57624200 | 57624799 | 11.13812 | 20.98311 | 27.46602 | 11.23575 | 0.086317 | 0.091871 | 0.891401 |
| 13 | 114799200 | 114800999 | 15.82105 | 28.12052 | 34.82098 | 15.36057 | 0.087387 | 0.484312 | 0.891482 |
| X | 49600000 | 49601599 | 22.11712 | 38.5234 | 37.19385 | 16.35584 | 0.437785 | 0.772882 | 0.891493 |
| 1 | 18283200 | 18284799 | 18.32202 | 31.92963 | 39.93112 | 17.56525 | 0.066173 | 0.74978 | 0.891497 |
| 8 | 43626000 | 43638599 | 122.0951 | 186.3416 | 218.4437 | 90.58553 | 0.058388 | 3.11298 | 0.89151 |
| 17 | 76201200 | 76204399 | 25.5302 | 43.29638 | 53.41186 | 23.395 | 0.079512 | 1.059143 | 0.891512 |
| 1 | 123195800 | 123196799 | 12.25523 | 22.30109 | 31.62199 | 13.61332 | -0.0836 | 0.312647 | 0.891519 |

| | | | | | | | | | |
|---|---|---|---|---|---|---|---|---|---|
| 2 | 24054400 | 24063999 | 171.4375 | 254.5768 | 314.1126 | 129.3947 | -0.04983 | 3.655737 | 0.89154 |
| 13 | 78109800 | 78112599 | 23.92555 | 40.80761 | 50.64703 | 22.20294 | 0.071817 | 1.000671 | 0.891546 |
| 10 | 3207000 | 3209599 | 30.02729 | 49.78069 | 61.86347 | 26.78877 | 0.068225 | 1.304194 | 0.891565 |
| 1 | 184516200 | 184520399 | 41.77819 | 67.28763 | 89.54834 | 38.07276 | -0.05633 | 1.820767 | 0.891568 |
| 16 | 16508200 | 16513799 | 51.23388 | 82.24041 | 99.69122 | 42.17741 | 0.072111 | 1.940868 | 0.891612 |
| 2 | 45216800 | 45219399 | 40.80837 | 66.54578 | 79.53909 | 34.16833 | 0.115298 | 1.548486 | 0.891613 |
| 19 | 25433400 | 25436199 | 29.27887 | 48.48973 | 60.56895 | 26.20947 | 0.060791 | 1.314368 | 0.891624 |
| 5 | 116205600 | 116206399 | 9.849955 | 18.65336 | 24.86904 | 10.06652 | 0.073341 | 0.038819 | 0.891719 |
| 18 | 42521000 | 42522199 | 11.39345 | 20.95584 | 29.75923 | 12.63314 | -0.07143 | 0.244115 | 0.891732 |
| 4 | 52599600 | 52601799 | 23.59959 | 39.54305 | 54.34467 | 23.697 | -0.07604 | 1.165016 | 0.891745 |
| 6 | 82491800 | 82493199 | 12.17265 | 22.15499 | 31.15402 | 13.4087 | -0.0712 | 0.311745 | 0.891749 |
| 18 | 4052400 | 4053399 | 12.17265 | 22.15499 | 31.15402 | 13.4087 | -0.0712 | 0.311745 | 0.891749 |
| 11 | 53786800 | 53787399 | 5.510829 | 10.75197 | 18.09334 | 6.459152 | -0.09132 | -0.45478 | 0.891761 |
| 2 | 75767800 | 75768799 | 9.145159 | 17.10968 | 25.35737 | 10.4955 | -0.10464 | 0.114048 | 0.891773 |
| 14 | 50144800 | 50146199 | 14.48007 | 25.6834 | 32.0336 | 14.15585 | 0.075619 | 0.485935 | 0.891803 |
| 13 | 59541000 | 59542599 | 10.72926 | 19.72951 | 28.23838 | 11.9914 | -0.08297 | 0.236208 | 0.891805 |
| 14 | 13726400 | 13729999 | 49.62829 | 79.46176 | 97.0319 | 40.89462 | 0.05727 | 2.016938 | 0.891827 |
| 12 | 63227200 | 63234999 | 117.2931 | 179.1139 | 211.0423 | 87.52771 | 0.051694 | 3.099438 | 0.891831 |
| 18 | 15106000 | 15110399 | 35.91451 | 58.08762 | 78.44607 | 33.53606 | -0.0703 | 1.68424 | 0.891839 |
| 10 | 118222400 | 118223199 | 9.536337 | 18.15064 | 24.42818 | 9.783295 | 0.073345 | 0.000265 | 0.891865 |
| 6 | 20083600 | 20084199 | 13.51744 | 24.18896 | 30.11586 | 13.20953 | 0.085682 | 0.420991 | 0.89187 |
| 16 | 8360000 | 8368599 | 54.84669 | 87.08834 | 135.8452 | 56.62228 | -0.30823 | 2.289876 | 0.891885 |
| 6 | 81823000 | 81826999 | 53.30435 | 85.17115 | 103.9515 | 43.68209 | 0.05471 | 2.087144 | 0.89191 |
| 7 | 43892800 | 43893799 | 11.66401 | 21.37845 | 27.16283 | 11.58603 | 0.083155 | 0.259763 | 0.891953 |
| 18 | 76073000 | 76075799 | 26.16473 | 43.59295 | 59.36968 | 25.7956 | -0.06302 | 1.236332 | 0.891978 |
| 5 | 126808200 | 126809199 | 8.744812 | 16.58977 | 24.92461 | 10.05141 | -0.09314 | 0.015018 | 0.891981 |
| 12 | 101677400 | 101680999 | 32.35482 | 53.1377 | 72.44168 | 31.20638 | -0.0682 | 1.431096 | 0.891983 |
| 1 | 79889200 | 79889799 | 11.17458 | 20.91027 | 27.33126 | 11.30908 | 0.075324 | 0.135213 | 0.891991 |
| 5 | 109159200 | 109162599 | 46.74193 | 75.74857 | 70.80026 | 30.31136 | 0.461957 | 1.658626 | 0.892002 |
| 9 | 19862000 | 19882799 | 602.3565 | 805.5395 | 833.2597 | 347.905 | 0.054548 | 5.278342 | 0.892011 |
| 5 | 58790400 | 58795399 | 112.2607 | 171.1016 | 216.4918 | 89.74812 | -0.05319 | 3.137859 | 0.892013 |
| 10 | 83742600 | 83744399 | 13.64813 | 24.415 | 30.55422 | 13.40543 | 0.077923 | 0.42405 | 0.892039 |
| 5 | 29266600 | 29267799 | 18.10011 | 31.47888 | 38.97991 | 17.12494 | 0.080084 | 0.77523 | 0.892052 |
| 8 | 124969400 | 124972399 | 46.68309 | 74.85803 | 99.02559 | 41.82996 | -0.05541 | 1.967321 | 0.89208 |
| X | 18675800 | 18677799 | 20.13711 | 35.20727 | 42.93141 | 18.92604 | 0.10284 | 0.718324 | 0.89208 |
| 8 | 35573800 | 35575599 | 20.22649 | 34.95359 | 43.66753 | 19.15732 | 0.065732 | 0.858928 | 0.892105 |
| 9 | 82348200 | 82348799 | 9.984542 | 18.75634 | 27.45456 | 11.28541 | -0.07888 | 0.102106 | 0.892118 |
| 17 | 39058400 | 39058999 | 13.9808 | 25.03021 | 31.50648 | 13.81386 | 0.070224 | 0.423274 | 0.892122 |
| 7 | 49298400 | 49300799 | 26.08837 | 43.68112 | 54.4849 | 23.74715 | 0.063258 | 1.178765 | 0.892122 |
| 6 | 67711200 | 67713599 | 41.49054 | 67.24667 | 68.87966 | 29.4918 | 0.330367 | 1.645779 | 0.892141 |
| 5 | 149652600 | 149664999 | 151.549 | 226.9339 | 282.1354 | 116.3703 | -0.04973 | 3.500413 | 0.892141 |
| 3 | 142355800 | 142357199 | 15.12198 | 26.58207 | 36.66089 | 16.2345 | -0.07395 | 0.593508 | 0.892175 |
| X | 102863200 | 102867799 | 34.50108 | 56.25747 | 76.97266 | 33.06519 | -0.07917 | 1.537519 | 0.892177 |
| 17 | 14572400 | 14574199 | 19.08406 | 33.31363 | 41.4278 | 18.2429 | 0.074536 | 0.733531 | 0.89218 |
| 7 | 81054600 | 81054999 | 5.942584 | 11.5818 | 19.13432 | 6.929503 | -0.08397 | -0.39425 | 0.892183 |
| 15 | 86781400 | 86784799 | 27.02958 | 44.66287 | 61.43875 | 26.54412 | -0.0782 | 1.334837 | 0.892197 |
| 3 | 68549200 | 68551599 | 19.16596 | 33.07688 | 45.28914 | 19.86793 | -0.06629 | 0.868964 | 0.892204 |

| | | | | | | | | | |
|---|---|---|---|---|---|---|---|---|---|
| 4 | 114144400 | 114146799 | 22.14551 | 37.08284 | 64.20933 | 27.98717 | -0.4084 | 1.17381 | 0.892211 |
| 2 | 83265800 | 83267599 | 10.80824 | 20.00446 | 28.75762 | 12.10427 | -0.08034 | 0.199867 | 0.892229 |
| 1 | 79921600 | 79922599 | 12.29903 | 22.95374 | 29.43747 | 12.23512 | 0.097212 | 0.165596 | 0.892255 |
| 10 | 47778200 | 47779399 | 16.51307 | 29.04511 | 40.20391 | 17.80128 | -0.07885 | 0.602198 | 0.892257 |
| 17 | 33424200 | 33424999 | 9.472696 | 17.99282 | 23.90377 | 9.614608 | 0.086995 | 0.014603 | 0.892264 |
| 7 | 52450000 | 52450999 | 15.45977 | 27.27961 | 34.06995 | 15.07567 | 0.070666 | 0.550286 | 0.892275 |
| 8 | 128587800 | 128593799 | 26.21011 | 44.41488 | 44.77644 | 19.60796 | 0.369663 | 1.058792 | 0.892298 |
| 5 | 16684000 | 16684999 | 13.85132 | 24.98242 | 31.53104 | 13.72595 | 0.073319 | 0.378153 | 0.89231 |
| 10 | 63229600 | 63231999 | 28.35385 | 47.1751 | 58.86571 | 25.55378 | 0.062833 | 1.259058 | 0.892335 |
| 1 | 44305800 | 44306799 | 15.14885 | 26.65065 | 36.6626 | 16.23266 | -0.06982 | 0.585513 | 0.892361 |
| 13 | 26549800 | 26552599 | 21.6641 | 37.52961 | 35.92688 | 15.7626 | 0.449043 | 0.834685 | 0.892415 |
| 10 | 62206600 | 62209199 | 29.49409 | 48.63825 | 66.08301 | 28.51658 | -0.06095 | 1.368534 | 0.892417 |
| 2 | 174893800 | 174896199 | 112.2806 | 170.5832 | 218.534 | 90.50784 | -0.07487 | 3.203748 | 0.892422 |
| 4 | 8296600 | 8298199 | 18.91103 | 32.58717 | 44.80275 | 19.65364 | -0.07255 | 0.882397 | 0.89244 |
| X | 83202800 | 83205199 | 18.78528 | 31.99029 | 55.86739 | 24.48839 | -0.41918 | 1.004735 | 0.892476 |
| 8 | 115052800 | 115053399 | 10.08186 | 18.9659 | 24.72702 | 10.13997 | 0.089057 | 0.083174 | 0.892487 |
| 11 | 24831800 | 24832799 | 10.61023 | 20.23438 | 21.19403 | 8.452775 | 0.437662 | -0.01073 | 0.892508 |
| 11 | 121743400 | 121752999 | 143.1889 | 218.8112 | 236.6034 | 98.14477 | 0.176265 | 3.098413 | 0.892524 |
| 17 | 38535400 | 38537199 | 40.13788 | 64.78953 | 86.43855 | 36.85964 | -0.05614 | 1.751175 | 0.892536 |
| 8 | 118468000 | 118471599 | 28.7893 | 47.2244 | 64.97411 | 27.9751 | -0.08198 | 1.433619 | 0.892538 |
| 6 | 70476800 | 70477599 | 16.62871 | 29.02189 | 39.99571 | 17.61013 | -0.07294 | 0.721586 | 0.892551 |
| 17 | 73117600 | 73119399 | 15.70095 | 27.55374 | 38.17807 | 16.88367 | -0.08079 | 0.624385 | 0.892555 |
| 16 | 66082600 | 66083799 | 13.199 | 24.09239 | 30.55584 | 13.1039 | 0.079976 | 0.300394 | 0.892563 |
| 15 | 28504800 | 28506399 | 12.48383 | 22.88259 | 29.03632 | 12.38403 | 0.084603 | 0.267124 | 0.892576 |
| 7 | 127738400 | 127739399 | 15.61442 | 27.47575 | 34.14472 | 15.11718 | 0.076269 | 0.581858 | 0.892594 |
| 8 | 34516600 | 34518799 | 38.16181 | 61.61535 | 82.59507 | 35.23214 | -0.06253 | 1.738638 | 0.892608 |
| 17 | 48503400 | 48515799 | 235.935 | 341.4098 | 411.3105 | 169.6347 | -0.05095 | 4.071805 | 0.89261 |
| 7 | 23849800 | 23851399 | 15.59763 | 27.39026 | 37.68527 | 16.67491 | -0.07058 | 0.613864 | 0.892614 |
| 6 | 79184600 | 79185199 | 9.517251 | 18.09069 | 27.17876 | 10.91497 | -0.08733 | 0.018438 | 0.892621 |
| 3 | 69287800 | 69288799 | 10.71716 | 19.87053 | 25.26531 | 10.60912 | 0.09862 | 0.183771 | 0.892622 |
| 11 | 114102400 | 114106399 | 39.66435 | 63.86366 | 85.92259 | 36.51511 | -0.07252 | 1.825899 | 0.892625 |
| 7 | 29561200 | 29565199 | 34.47335 | 56.06311 | 68.99909 | 29.61129 | 0.070493 | 1.566908 | 0.892627 |
| 7 | 96715000 | 96716999 | 20.32373 | 35.61806 | 34.43948 | 15.21293 | 0.435962 | 0.668198 | 0.892643 |
| 5 | 55808400 | 55810799 | 30.49885 | 50.77794 | 51.89349 | 22.53716 | 0.349987 | 1.250596 | 0.892644 |
| 16 | 12181400 | 12181999 | 6.707787 | 13.03602 | 21.18871 | 7.886214 | -0.09634 | -0.27826 | 0.892668 |
| 2 | 37184400 | 37187999 | 47.94247 | 76.96215 | 102.6432 | 43.43073 | -0.06529 | 1.937894 | 0.892691 |
| 18 | 41553000 | 41556999 | 32.18267 | 54.17548 | 48.55525 | 21.22003 | 0.538555 | 1.14184 | 0.89271 |
| 1 | 110594200 | 110601199 | 84.27735 | 133.2384 | 121.1043 | 50.31559 | 0.464296 | 2.423198 | 0.892736 |
| 3 | 6122800 | 6129799 | 132.3233 | 201.9584 | 195.5457 | 81.09066 | 0.334061 | 3.114013 | 0.892763 |
| 14 | 119119200 | 119121199 | 15.16817 | 26.62899 | 36.7704 | 16.26643 | -0.07561 | 0.615522 | 0.892768 |
| 5 | 142289800 | 142294599 | 60.54537 | 96.12183 | 125.4107 | 52.26475 | -0.05115 | 2.297041 | 0.892777 |
| 6 | 143455000 | 143459999 | 159.4236 | 237.7836 | 272.7215 | 112.4011 | 0.061369 | 3.575155 | 0.892778 |
| 1 | 106849800 | 106852999 | 58.65557 | 93.23765 | 112.6775 | 47.01681 | 0.060953 | 2.255657 | 0.892788 |
| 15 | 35548000 | 35549999 | 25.36848 | 43.25932 | 52.84623 | 23.16664 | 0.094568 | 1.011967 | 0.892794 |
| 12 | 44015000 | 44018199 | 36.53581 | 59.35942 | 80.08577 | 34.35098 | -0.0629 | 1.595175 | 0.892803 |
| 7 | 58530000 | 58533999 | 44.29679 | 71.23324 | 94.66223 | 40.15642 | -0.05703 | 1.876 | 0.892827 |
| 5 | 24280200 | 24280999 | 9.787131 | 18.49792 | 24.36331 | 9.902986 | 0.086 | 0.052758 | 0.89283 |

| | | | | | | | | | |
|---|---|---|---|---|---|---|---|---|---|
| 8 | 114139400 | 114140399 | 12.83185 | 23.26519 | 32.6416 | 14.11664 | -0.07251 | 0.341771 | 0.89286 |
| 1 | 64097800 | 64099199 | 15.1482 | 26.91512 | 37.54726 | 16.56813 | -0.08384 | 0.488793 | 0.892945 |
| 7 | 8343000 | 8346199 | 60.61693 | 96.30893 | 116.5319 | 48.60165 | 0.058525 | 2.274322 | 0.892954 |
| 6 | 103383200 | 103384199 | 13.37034 | 23.86006 | 33.08092 | 14.54374 | -0.07111 | 0.442455 | 0.892972 |
| 1 | 13959200 | 13962399 | 25.96109 | 43.72177 | 54.47947 | 23.81231 | 0.06481 | 1.122657 | 0.892985 |
| 13 | 56811600 | 56813199 | 12.102 | 21.73245 | 30.70605 | 13.42849 | -0.09276 | 0.390382 | 0.892991 |
| 2 | 30338200 | 30340999 | 38.13999 | 61.63869 | 75.25449 | 32.1403 | 0.073593 | 1.707763 | 0.893002 |
| 1 | 11436200 | 11444199 | 96.71992 | 149.4963 | 191.6804 | 79.6197 | -0.05668 | 2.857859 | 0.893017 |
| 6 | 74365800 | 74366599 | 14.80616 | 26.17622 | 36.04914 | 15.94536 | -0.06869 | 0.528075 | 0.893018 |
| 16 | 16793000 | 16795999 | 51.00607 | 81.51552 | 107.2501 | 45.08119 | -0.05314 | 2.079131 | 0.893037 |
| 1 | 14030000 | 14034399 | 67.32557 | 106.504 | 129.0856 | 53.64832 | 0.050398 | 2.398198 | 0.893069 |
| 3 | 77193000 | 77194999 | 25.35833 | 42.4474 | 57.70309 | 25.14555 | -0.06043 | 1.183309 | 0.893083 |
| 8 | 49525400 | 49526799 | 22.20391 | 38.41019 | 35.96668 | 15.77939 | 0.480413 | 0.850524 | 0.893102 |
| 10 | 92740000 | 92741599 | 15.83159 | 27.82637 | 38.18098 | 16.90723 | -0.06659 | 0.602902 | 0.893107 |
| 15 | 103810400 | 103813599 | 71.19055 | 111.9524 | 133.5435 | 55.39978 | 0.066675 | 2.529319 | 0.893125 |
| 12 | 111486000 | 111491799 | 61.16066 | 97.1782 | 127.2929 | 53.09265 | -0.05573 | 2.273579 | 0.893147 |
| 11 | 64392600 | 64395599 | 23.02857 | 40.02137 | 47.91918 | 21.04266 | 0.128215 | 0.821797 | 0.89319 |
| 3 | 21244000 | 21247799 | 41.27108 | 66.70809 | 82.30363 | 35.15716 | 0.058658 | 1.712596 | 0.893203 |
| 6 | 127445000 | 127445999 | 13.19006 | 23.7017 | 29.61458 | 12.9441 | 0.084059 | 0.391686 | 0.89322 |
| 1 | 106848000 | 106852999 | 76.16563 | 119.5076 | 143.5469 | 59.53256 | 0.054193 | 2.578161 | 0.893225 |
| 4 | 16602000 | 16603399 | 21.26764 | 36.49516 | 50.08126 | 21.96565 | -0.07112 | 0.936999 | 0.893246 |
| 7 | 41006200 | 41008399 | 26.23186 | 44.04191 | 60.54009 | 26.43196 | -0.07651 | 1.157602 | 0.893271 |
| 5 | 99603200 | 99604399 | 14.02573 | 25.1066 | 31.54457 | 13.83245 | 0.072818 | 0.425534 | 0.893284 |
| 15 | 33168800 | 33170999 | 28.99626 | 48.6869 | 59.13202 | 25.81844 | 0.101423 | 1.161489 | 0.893296 |
| 8 | 126113800 | 126116199 | 22.31384 | 37.71727 | 51.70342 | 22.62405 | -0.07217 | 1.074495 | 0.893313 |
| 18 | 90390200 | 90392799 | 55.81442 | 88.71942 | 116.879 | 48.76885 | -0.06334 | 2.256163 | 0.893319 |
| 8 | 83371200 | 83371399 | 5.085304 | 9.934122 | 17.07937 | 5.967513 | -0.09383 | -0.54091 | 0.893331 |
| 9 | 26034800 | 26036199 | 13.73002 | 25.02627 | 31.58819 | 13.57058 | 0.085258 | 0.31508 | 0.893347 |
| 1 | 37349200 | 37350399 | 11.13911 | 20.4987 | 29.29502 | 12.42726 | -0.07951 | 0.239735 | 0.893359 |
| 4 | 156005000 | 156016999 | 148.6308 | 222.8031 | 277.1821 | 114.3512 | -0.04927 | 3.479834 | 0.893368 |
| 19 | 31779000 | 31779799 | 13.24657 | 24.02679 | 30.51237 | 13.18915 | 0.070858 | 0.33751 | 0.893371 |
| 6 | 77725200 | 77726799 | 17.57435 | 30.66518 | 42.00324 | 18.49135 | -0.06434 | 0.731829 | 0.893391 |
| 3 | 62071600 | 62074399 | 43.2935 | 69.69852 | 85.24666 | 36.2215 | 0.06444 | 1.832616 | 0.893413 |
| 6 | 89886200 | 89889399 | 39.30396 | 63.89881 | 78.31508 | 33.60542 | 0.076142 | 1.587431 | 0.893418 |
| 10 | 52378600 | 52381399 | 34.78856 | 56.67267 | 70.36207 | 30.21507 | 0.059625 | 1.543748 | 0.893468 |
| 7 | 41311800 | 41319999 | 197.2471 | 290.0218 | 330.6459 | 136.1673 | 0.050844 | 3.823805 | 0.893472 |
| 4 | 15745200 | 15747799 | 23.3064 | 39.52437 | 53.88677 | 23.60289 | -0.0642 | 1.052569 | 0.893473 |
| 18 | 45203200 | 45204799 | 19.37909 | 33.82621 | 41.87525 | 18.43911 | 0.080825 | 0.733498 | 0.893475 |
| 7 | 3459400 | 3465799 | 60.06643 | 95.38082 | 114.6028 | 47.7703 | 0.06781 | 2.286661 | 0.893503 |
| 8 | 46774400 | 46776599 | 20.9629 | 35.94538 | 44.79438 | 19.64565 | 0.067226 | 0.934587 | 0.893507 |
| 14 | 18183600 | 18187399 | 41.49206 | 67.11614 | 82.27294 | 35.17494 | 0.069135 | 1.69068 | 0.893522 |
| 17 | 42212000 | 42216799 | 65.5307 | 103.7621 | 135.0723 | 56.1631 | -0.05141 | 2.376005 | 0.893529 |
| 1 | 178958200 | 178962399 | 51.27877 | 82.10236 | 100.5427 | 42.37265 | 0.053107 | 2.020785 | 0.89353 |
| 11 | 50153800 | 50154399 | 6.37806 | 12.39109 | 20.06501 | 7.488138 | -0.09528 | -0.28012 | 0.893563 |
| 7 | 20133200 | 20133999 | 14.57046 | 26.49341 | 24.9929 | 10.77003 | 0.500335 | 0.32327 | 0.893565 |
| 17 | 29655800 | 29659199 | 22.38835 | 38.00436 | 46.92783 | 20.56086 | 0.078281 | 1.032549 | 0.89357 |
| 2 | 19958000 | 19961199 | 28.6194 | 47.58775 | 59.4866 | 25.81239 | 0.060158 | 1.26602 | 0.893585 |

| | | | | | | | | | |
|---|---|---|---|---|---|---|---|---|---|
| 11 | 52966400 | 52966999 | 10.58784 | 20.05598 | 26.52044 | 10.72904 | 0.085965 | 0.049572 | 0.893593 |
| 3 | 40539600 | 40541799 | 19.27681 | 33.67682 | 41.28089 | 18.18728 | 0.095023 | 0.718513 | 0.893597 |
| 4 | 68683800 | 68684799 | 8.342228 | 15.90938 | 23.97101 | 9.56004 | -0.0826 | -0.02527 | 0.893598 |
| 17 | 72920600 | 72925199 | 62.50663 | 99.17879 | 120.3739 | 50.137 | 0.052207 | 2.314328 | 0.893599 |
| 6 | 111187400 | 111189399 | 18.07879 | 31.78025 | 46.04116 | 20.38829 | -0.14489 | 0.62421 | 0.89362 |
| 6 | 138019000 | 138022799 | 58.98899 | 93.65989 | 122.8663 | 51.20932 | -0.05887 | 2.291256 | 0.893625 |
| 6 | 5226200 | 5227999 | 17.74554 | 31.10679 | 38.60654 | 17.05693 | 0.077283 | 0.654173 | 0.893634 |
| 6 | 36931600 | 36936399 | 45.5202 | 73.33027 | 89.93986 | 38.25114 | 0.061555 | 1.819467 | 0.893637 |
| 4 | 117964200 | 117966599 | 26.32601 | 43.69907 | 59.88141 | 25.95284 | -0.07215 | 1.278491 | 0.893668 |
| 7 | 128662600 | 128668599 | 71.09947 | 112.2782 | 135.8095 | 56.39305 | 0.05129 | 2.442555 | 0.893675 |
| 12 | 44873800 | 44879199 | 61.19452 | 97.71228 | 116.3128 | 48.82975 | 0.089625 | 2.118648 | 0.893675 |
| 2 | 78447000 | 78447799 | 7.564471 | 14.57932 | 22.55414 | 8.740556 | -0.08181 | -0.13085 | 0.893685 |
| 6 | 9392000 | 9392999 | 12.98317 | 23.60975 | 30.07823 | 12.95969 | 0.069451 | 0.321088 | 0.893688 |
| 17 | 40487600 | 40488599 | 12.98317 | 23.60975 | 30.07823 | 12.95969 | 0.069451 | 0.321088 | 0.893688 |
| 14 | 76991400 | 76991799 | 6.514457 | 12.68957 | 18.52248 | 6.755021 | 0.083454 | -0.36483 | 0.893691 |
| 19 | 12076200 | 12076799 | 6.514457 | 12.68957 | 18.52248 | 6.755021 | 0.083454 | -0.36483 | 0.893691 |
| 14 | 109737200 | 109739199 | 23.16368 | 39.19802 | 53.32745 | 23.34268 | -0.06121 | 1.070316 | 0.893697 |
| 6 | 90057000 | 90062599 | 69.66788 | 110.2372 | 133.2638 | 55.39184 | 0.054424 | 2.395149 | 0.893714 |
| 11 | 10577000 | 10581399 | 75.72837 | 119.9912 | 117.5924 | 48.91188 | 0.358387 | 2.364744 | 0.893744 |
| 4 | 132545800 | 132549799 | 57.91418 | 92.23699 | 112.3725 | 47.00684 | 0.052238 | 2.195522 | 0.893771 |
| 6 | 34399400 | 34401599 | 17.15399 | 30.18114 | 37.47617 | 16.59255 | 0.07711 | 0.595861 | 0.893772 |
| 8 | 94176400 | 94177599 | 14.1616 | 25.33929 | 35.6571 | 15.64128 | -0.09008 | 0.43863 | 0.893784 |
| 14 | 36982600 | 36983199 | 15.95048 | 28.07761 | 35.09615 | 15.53725 | 0.068098 | 0.581463 | 0.893794 |
| 17 | 75909600 | 75910599 | 14.32069 | 26.08802 | 26.36317 | 11.33396 | 0.403588 | 0.312293 | 0.893795 |
| 7 | 62306600 | 62313999 | 288.9747 | 407.9852 | 488.1804 | 202.1157 | -0.07665 | 4.435779 | 0.893805 |
| 10 | 4960000 | 4960799 | 9.15426 | 17.19929 | 25.3799 | 10.42987 | -0.09005 | 0.08776 | 0.893813 |
| 19 | 58395400 | 58400199 | 15.2968 | 27.9937 | 26.75717 | 11.43693 | 0.488786 | 0.294547 | 0.893814 |
| X | 160068200 | 160071199 | 26.98515 | 45.22425 | 62.33024 | 27.18231 | -0.08034 | 1.179243 | 0.893859 |
| 12 | 80471400 | 80473799 | 19.27004 | 33.94479 | 32.68157 | 14.46628 | 0.443062 | 0.588178 | 0.89386 |
| X | 29895400 | 29896599 | 47.76278 | 77.03453 | 67.21681 | 28.63283 | 0.553831 | 1.7813 | 0.893861 |
| 9 | 57228000 | 57228799 | 12.51522 | 22.62285 | 31.60306 | 13.7191 | -0.06973 | 0.354852 | 0.893893 |
| 2 | 170629600 | 170631399 | 29.08269 | 48.09808 | 65.45707 | 28.29756 | -0.06264 | 1.335473 | 0.893914 |
| 5 | 77365200 | 77369199 | 39.59471 | 63.98265 | 85.68597 | 36.59027 | -0.0598 | 1.717208 | 0.893978 |
| 11 | 95303400 | 95305799 | 9.403512 | 17.29785 | 31.93919 | 13.55634 | -0.44741 | 0.232478 | 0.893982 |
| 5 | 14601800 | 14602999 | 21.74859 | 37.16831 | 46.36652 | 20.32968 | 0.064949 | 0.969079 | 0.893983 |
| 13 | 22398400 | 22401399 | 27.83214 | 46.18152 | 62.74308 | 27.18594 | -0.05988 | 1.290329 | 0.894031 |
| 5 | 40696800 | 40697399 | 8.477067 | 16.25249 | 24.62421 | 9.687988 | -0.07279 | -0.06699 | 0.894038 |
| 1 | 68360000 | 68363199 | 33.66325 | 55.86288 | 67.28159 | 29.15455 | 0.113401 | 1.309591 | 0.894054 |
| 11 | 32624000 | 32624599 | 5.940954 | 11.58383 | 19.17958 | 6.905097 | -0.07958 | -0.41538 | 0.894082 |
| 6 | 100351400 | 100354999 | 32.37262 | 52.9014 | 71.42409 | 30.70595 | -0.05784 | 1.49541 | 0.894111 |
| 17 | 47365600 | 47371799 | 57.64623 | 91.64544 | 120.4284 | 50.2577 | -0.05952 | 2.251879 | 0.894121 |
| 9 | 15483400 | 15485199 | 19.92494 | 34.33005 | 42.68341 | 18.72388 | 0.071804 | 0.883464 | 0.894125 |
| 16 | 6809200 | 6810399 | 20.69694 | 35.56714 | 44.22881 | 19.40025 | 0.070722 | 0.911198 | 0.89413 |
| 6 | 78966600 | 78967399 | 12.49516 | 22.79826 | 29.11116 | 12.49077 | 0.070102 | 0.296387 | 0.894144 |
| 8 | 117798000 | 117799399 | 17.25286 | 29.87518 | 41.70958 | 18.29948 | -0.0931 | 0.826319 | 0.894149 |
| X | 49039200 | 49040199 | 9.012684 | 17.11919 | 25.51154 | 10.26111 | -0.07754 | 0.014161 | 0.894158 |
| 8 | 60403400 | 60412199 | 150.0699 | 226.4766 | 224.972 | 92.98122 | 0.283729 | 3.346626 | 0.894192 |

| | | | | | | | | | |
|---|---|---|---|---|---|---|---|---|---|
| 12 | 57625600 | 57627999 | 23.29207 | 39.66141 | 49.55831 | 21.72276 | 0.061787 | 1.012535 | 0.894196 |
| 8 | 9819000 | 9822999 | 37.3408 | 59.87453 | 105.4935 | 44.55894 | -0.46812 | 1.959985 | 0.894201 |
| 8 | 96876800 | 96878199 | 13.85721 | 24.66004 | 34.05188 | 15.00585 | -0.06769 | 0.464907 | 0.894209 |
| 6 | 137788600 | 137790399 | 19.20055 | 33.13309 | 45.29645 | 19.87107 | -0.06407 | 0.870793 | 0.894223 |
| 13 | 35536400 | 35537599 | 11.27367 | 20.97636 | 30.47189 | 12.7303 | -0.08751 | 0.177125 | 0.89424 |
| 13 | 52472200 | 52473399 | 10.47782 | 19.15295 | 27.81786 | 11.90822 | -0.112 | 0.268162 | 0.894242 |
| 9 | 119052800 | 119056199 | 42.60826 | 69.05022 | 83.74483 | 35.85346 | 0.087026 | 1.658491 | 0.894261 |
| 5 | 110056600 | 110059399 | 47.39108 | 76.34922 | 92.6867 | 39.42317 | 0.076446 | 1.822119 | 0.894279 |
| 16 | 79294000 | 79295199 | 8.794924 | 16.75384 | 25.01418 | 9.999855 | -0.07279 | -0.01077 | 0.894297 |
| 19 | 29782200 | 29784199 | 15.03068 | 25.81393 | 51.60932 | 22.63717 | -0.61151 | 0.903397 | 0.894311 |
| 12 | 59874000 | 59876199 | 20.25841 | 35.33234 | 43.4932 | 19.14176 | 0.088862 | 0.752226 | 0.894314 |
| 5 | 118696000 | 118699999 | 27.39278 | 45.2242 | 61.94104 | 26.74529 | -0.07201 | 1.347087 | 0.89432 |
| 8 | 45834000 | 45835399 | 13.93697 | 24.83442 | 31.00905 | 13.6499 | 0.077798 | 0.450479 | 0.894326 |
| 7 | 13876200 | 13876999 | 10.83333 | 20.26466 | 29.52015 | 12.22199 | -0.0809 | 0.138426 | 0.894328 |
| 18 | 62144400 | 62145199 | 8.21655 | 15.78638 | 24.12809 | 9.435313 | -0.07665 | -0.0889 | 0.894333 |
| 4 | 26107800 | 26109999 | 27.43578 | 45.87372 | 57.27538 | 24.94162 | 0.061881 | 1.200587 | 0.894337 |
| 4 | 115150200 | 115152399 | 19.42676 | 33.33943 | 45.86832 | 20.11801 | -0.07469 | 0.923179 | 0.894381 |
| 8 | 81610800 | 81613199 | 22.35951 | 37.99304 | 51.6877 | 22.65227 | -0.0609 | 1.030842 | 0.894394 |
| 5 | 47572400 | 47574799 | 24.33003 | 41.60767 | 51.19883 | 22.44963 | 0.084749 | 0.980048 | 0.89443 |
| 1 | 85494800 | 85516999 | 527.1056 | 715.5281 | 751.9898 | 313.2671 | 0.053345 | 5.040934 | 0.894436 |
| X | 60110200 | 60115199 | 67.13582 | 106.7427 | 126.977 | 52.99596 | 0.084023 | 2.261985 | 0.894498 |
| 8 | 67628200 | 67632999 | 88.62213 | 137.4573 | 176.2171 | 73.1872 | -0.05259 | 2.788671 | 0.894523 |
| 6 | 85296000 | 85297999 | 19.19101 | 33.14626 | 45.2883 | 19.86804 | -0.06299 | 0.862555 | 0.894557 |
| 11 | 10621800 | 10624799 | 32.41209 | 53.28767 | 73.00759 | 31.46414 | -0.07472 | 1.419914 | 0.894562 |
| 8 | 19723000 | 19725799 | 55.20423 | 88.34814 | 88.24771 | 37.16725 | 0.345832 | 2.035631 | 0.894573 |
| 14 | 95495200 | 95495999 | 8.862408 | 16.9346 | 22.43621 | 8.910882 | 0.10732 | -0.03535 | 0.894595 |
| 4 | 6834400 | 6835399 | 9.819969 | 18.89047 | 20.26214 | 7.895319 | 0.434561 | -0.09604 | 0.894617 |
| 13 | 42693800 | 42695599 | 22.37758 | 38.79916 | 47.39027 | 20.79261 | 0.099157 | 0.845962 | 0.894645 |
| 17 | 63917600 | 63919599 | 23.22597 | 38.50918 | 67.30471 | 29.1471 | -0.42216 | 1.290152 | 0.894696 |
| 10 | 22205000 | 22205399 | 6.380758 | 12.42125 | 20.1723 | 7.407049 | -0.07775 | -0.336 | 0.894729 |
| 7 | 42138200 | 42155599 | 245.4333 | 356.4322 | 398.8535 | 164.425 | 0.060572 | 4.019088 | 0.894748 |
| 14 | 71564200 | 71568599 | 34.17956 | 56.10206 | 69.34842 | 29.86655 | 0.072321 | 1.438177 | 0.894775 |
| Y | 84429400 | 84429799 | 5.963363 | 11.55854 | 19.51879 | 7.385414 | -0.17367 | -0.23967 | 0.894782 |
| 5 | 103516200 | 103517199 | 9.953993 | 18.50771 | 26.8225 | 11.21922 | -0.08483 | 0.165134 | 0.894808 |
| 10 | 6674800 | 6678799 | 47.43007 | 76.16935 | 93.43374 | 39.55747 | 0.055824 | 1.919417 | 0.894827 |
| 12 | 92680200 | 92680599 | 5.942126 | 11.5929 | 16.99389 | 6.071454 | 0.105338 | -0.44769 | 0.894828 |
| 6 | 134845600 | 134847599 | 21.7547 | 36.75991 | 50.61688 | 22.14638 | -0.07849 | 1.076436 | 0.894844 |
| 8 | 21142200 | 21142799 | 23.79709 | 39.25354 | 69.21742 | 29.87242 | -0.43566 | 1.355444 | 0.894869 |
| 15 | 39604800 | 39607399 | 22.08629 | 37.75031 | 51.72978 | 22.68147 | -0.0701 | 0.974559 | 0.894924 |
| 13 | 90574600 | 90578199 | 50.37994 | 80.77623 | 107.8085 | 45.53155 | -0.06845 | 1.9796 | 0.894933 |
| 13 | 47102600 | 47104999 | 22.78026 | 38.78623 | 48.49185 | 21.25514 | 0.06097 | 1.010033 | 0.894934 |
| 8 | 110754800 | 110758599 | 53.47375 | 85.11417 | 112.4789 | 47.01257 | -0.06579 | 2.210149 | 0.894947 |
| 8 | 98004800 | 98006599 | 17.16892 | 30.04046 | 41.3614 | 18.24556 | -0.07171 | 0.685561 | 0.89498 |
| 3 | 87703200 | 87704199 | 11.5271 | 20.93316 | 29.65262 | 12.79826 | -0.08425 | 0.316297 | 0.894982 |
| 15 | 51601400 | 51603399 | 16.05696 | 28.3379 | 35.46843 | 15.69399 | 0.067492 | 0.553432 | 0.895001 |
| 5 | 77997000 | 78000799 | 42.74195 | 68.92533 | 84.153 | 35.83967 | 0.06984 | 1.779631 | 0.895013 |
| 6 | 14790800 | 14791399 | 10.00864 | 18.82746 | 24.78085 | 10.16269 | 0.075723 | 0.084701 | 0.895032 |

| | | | | | | | | | |
|---|---|---|---|---|---|---|---|---|---|
| 15 | 75232600 | 75233199 | 10.00864 | 18.82746 | 24.78085 | 10.16269 | 0.075723 | 0.084701 | 0.895032 |
| 3 | 94497000 | 94507599 | 150.3223 | 224.869 | 279.6046 | 115.3038 | -0.05124 | 3.518958 | 0.895034 |
| 13 | 51734800 | 51738599 | 36.03465 | 58.375 | 78.31603 | 33.51987 | -0.05869 | 1.651294 | 0.895073 |
| 1 | 40592000 | 40595599 | 33.04654 | 54.94397 | 54.14944 | 23.49497 | 0.402103 | 1.27618 | 0.895117 |
| 19 | 3089600 | 3090599 | 12.00373 | 21.67538 | 30.75354 | 13.36795 | -0.09277 | 0.358696 | 0.895143 |
| 12 | 26744400 | 26746599 | 19.83629 | 34.6769 | 42.32872 | 18.65834 | 0.101116 | 0.713777 | 0.895175 |
| 6 | 147789800 | 147799799 | 146.1779 | 219.9067 | 256.5079 | 105.9209 | 0.048173 | 3.409493 | 0.895203 |
| 17 | 81368600 | 81370599 | 21.78439 | 37.33635 | 51.18542 | 22.44773 | -0.07012 | 0.948663 | 0.895222 |
| 13 | 61748000 | 61812399 | 1147.613 | 1468.189 | 1427.992 | 599.1439 | 0.079497 | 5.997808 | 0.895234 |
| 15 | 74789800 | 74791199 | 13.82069 | 24.71178 | 31.04008 | 13.62412 | 0.072217 | 0.429356 | 0.895237 |
| 18 | 60614000 | 60616399 | 15.59138 | 27.39579 | 37.64842 | 16.66709 | -0.06891 | 0.603914 | 0.895245 |
| 4 | 55303800 | 55311999 | 93.88771 | 146.2317 | 148.5398 | 61.65381 | 0.287481 | 2.71757 | 0.895286 |
| 17 | 62714600 | 62717599 | 26.51679 | 45.54918 | 53.50556 | 23.46905 | 0.152448 | 0.951503 | 0.895287 |
| 1 | 151393000 | 151394799 | 17.8588 | 31.0982 | 42.56423 | 18.71438 | -0.06359 | 0.762019 | 0.895298 |
| 5 | 18602800 | 18603399 | 5.503826 | 10.73812 | 18.09448 | 6.461505 | -0.09297 | -0.45163 | 0.895309 |
| 8 | 110699600 | 110700999 | 13.79558 | 24.51636 | 33.99947 | 15.00027 | -0.07516 | 0.47152 | 0.895365 |
| 9 | 73082000 | 73084799 | 22.59869 | 38.25967 | 52.21213 | 22.8577 | -0.06553 | 1.065157 | 0.895374 |
| 13 | 99272400 | 99274399 | 35.71182 | 57.41527 | 98.92423 | 41.95107 | -0.43122 | 1.873418 | 0.895375 |
| 8 | 127907200 | 127909399 | 19.3552 | 33.72312 | 41.92997 | 18.44211 | 0.074312 | 0.758644 | 0.895392 |
| 5 | 92941800 | 92944599 | 30.83417 | 50.79174 | 69.01046 | 29.76587 | -0.06163 | 1.388028 | 0.8954 |
| 11 | 51770400 | 51774199 | 35.20457 | 57.11442 | 76.71475 | 32.87021 | -0.0584 | 1.619119 | 0.895416 |
| 4 | 101727600 | 101735199 | 80.33391 | 125.4689 | 161.2307 | 66.89815 | -0.04823 | 2.664597 | 0.895436 |
| 10 | 15352400 | 15352799 | 6.513262 | 12.69096 | 20.90252 | 7.593814 | -0.08468 | -0.37198 | 0.895443 |
| 17 | 58241600 | 58242199 | 8.140722 | 15.65753 | 21.42532 | 8.352544 | 0.085591 | -0.10916 | 0.895445 |
| 9 | 11570600 | 11573399 | 34.97745 | 57.10246 | 71.11131 | 30.56064 | 0.057427 | 1.51476 | 0.895446 |
| 5 | 67523600 | 67526399 | 31.96544 | 52.45044 | 65.45635 | 28.18639 | 0.058471 | 1.436172 | 0.895471 |
| 1 | 97054200 | 97055999 | 23.26442 | 39.71047 | 49.55915 | 21.72737 | 0.064036 | 0.990072 | 0.895486 |
| X | 97539800 | 97541199 | 12.87677 | 23.83428 | 30.3424 | 12.77038 | 0.094116 | 0.222218 | 0.895495 |
| 15 | 96379200 | 96382399 | 31.52806 | 51.60509 | 69.7809 | 30.01731 | -0.0584 | 1.468822 | 0.895514 |
| 17 | 40103200 | 40104199 | 17.65605 | 30.9675 | 38.62321 | 17.07248 | 0.070287 | 0.645807 | 0.895521 |
| 7 | 15761400 | 15771199 | 69.25296 | 108.0765 | 171.1912 | 71.03679 | -0.3505 | 2.674779 | 0.895563 |
| 1 | 79232200 | 79232999 | 13.92145 | 24.57027 | 34.25707 | 15.15784 | -0.08702 | 0.540168 | 0.895585 |
| 10 | 46795000 | 46795799 | 8.331066 | 15.92384 | 23.99859 | 9.525048 | -0.0769 | -0.04125 | 0.895608 |
| 10 | 8025600 | 8030399 | 116.9544 | 178.068 | 224.2442 | 92.93286 | -0.04762 | 3.160802 | 0.895609 |
| 5 | 142267600 | 142280199 | 156.6763 | 234.8022 | 272.9366 | 112.594 | 0.048271 | 3.486163 | 0.895628 |
| 9 | 26861000 | 26864399 | 56.07107 | 89.36573 | 117.1434 | 49.05133 | -0.05211 | 2.171198 | 0.895658 |
| 3 | 31404800 | 31405999 | 18.76499 | 32.6274 | 40.56843 | 17.82013 | 0.074314 | 0.7872 | 0.895704 |
| 6 | 88827800 | 88829799 | 25.71756 | 43.23684 | 53.94621 | 23.56687 | 0.062784 | 1.134716 | 0.895706 |
| 18 | 30943000 | 30944399 | 15.78595 | 27.77031 | 34.63654 | 15.33573 | 0.070897 | 0.586728 | 0.895731 |
| 4 | 126441200 | 126443399 | 17.08167 | 29.87233 | 40.89964 | 18.03398 | -0.06352 | 0.694351 | 0.89574 |
| 7 | 33701800 | 33740399 | 799.0163 | 1043.55 | 1133.188 | 474.4878 | -0.04957 | 5.66001 | 0.895759 |
| 10 | 116531200 | 116531999 | 9.996651 | 19.00533 | 25.49514 | 10.23497 | 0.074841 | 0.014639 | 0.895773 |
| 1 | 79913800 | 79915599 | 21.33226 | 35.89884 | 60.2912 | 26.34645 | -0.36438 | 1.119339 | 0.895789 |
| 5 | 29624200 | 29625399 | 12.66268 | 22.95642 | 29.09594 | 12.58465 | 0.073338 | 0.334413 | 0.895825 |
| 7 | 103921400 | 103924799 | 23.6729 | 40.16501 | 55.47132 | 24.30044 | -0.08262 | 1.05501 | 0.895836 |
| 11 | 13875800 | 13879799 | 64.64944 | 102.3943 | 124.0787 | 51.599 | 0.052096 | 2.366484 | 0.895869 |
| 3 | 8856000 | 8857599 | 15.35982 | 27.15889 | 33.92126 | 15.00372 | 0.071522 | 0.525006 | 0.895916 |

| | | | | | | | | | |
|---|---|---|---|---|---|---|---|---|---|
| 6 | 53088800 | 53089399 | 8.352663 | 16.16863 | 22.7194 | 8.648729 | 0.080159 | -0.18054 | 0.895916 |
| 16 | 30657000 | 30661199 | 59.20158 | 94.40044 | 114.6136 | 48.02392 | 0.059022 | 2.158582 | 0.895953 |
| 13 | 3605000 | 3605599 | 8.37581 | 16.11504 | 24.73679 | 9.634325 | -0.07745 | -0.09876 | 0.895962 |
| 4 | 17450600 | 17451799 | 26.65271 | 44.98189 | 43.31814 | 18.94602 | 0.435414 | 1.095605 | 0.895992 |
| 7 | 42516200 | 42516999 | 22.48646 | 38.35627 | 47.95877 | 21.02459 | 0.061269 | 0.992386 | 0.896005 |
| 5 | 128497000 | 128501999 | 60.07652 | 95.63262 | 116.394 | 48.65518 | 0.052878 | 2.214811 | 0.896011 |
| 5 | 149011800 | 149013999 | 29.74347 | 48.86286 | 66.32311 | 28.57406 | -0.06166 | 1.417914 | 0.896017 |
| 10 | 26715800 | 26716799 | 9.166118 | 17.18352 | 25.37926 | 10.46874 | -0.0956 | 0.102266 | 0.896028 |
| 3 | 51519200 | 51526399 | 82.97153 | 128.9316 | 165.8453 | 68.8605 | -0.05536 | 2.753614 | 0.89605 |
| 1 | 3299000 | 3301599 | 24.36153 | 41.98396 | 50.67998 | 22.23188 | 0.114266 | 0.911352 | 0.896063 |
| 4 | 21482000 | 21483999 | 34.88426 | 57.35835 | 58.47572 | 25.20249 | 0.350599 | 1.413725 | 0.896078 |
| 17 | 64005600 | 64007799 | 24.60902 | 41.5056 | 56.67258 | 24.78192 | -0.0667 | 1.106272 | 0.896096 |
| 8 | 116922600 | 116923599 | 9.929777 | 18.54126 | 26.83711 | 11.14488 | -0.0748 | 0.137263 | 0.896121 |
| 2 | 30074400 | 30076599 | 43.17137 | 69.25992 | 93.05879 | 39.33803 | -0.07696 | 1.947988 | 0.896136 |
| 9 | 4248600 | 4251599 | 85.05443 | 131.8885 | 169.373 | 70.34749 | -0.05539 | 2.792016 | 0.896159 |
| 19 | 13011000 | 13036599 | 591.5448 | 792.5827 | 823.8095 | 343.8644 | 0.050354 | 5.248236 | 0.896168 |
| 3 | 61408400 | 61408999 | 7.544588 | 14.60311 | 20.4798 | 7.799387 | 0.082454 | -0.19581 | 0.896177 |
| 15 | 19121800 | 19122399 | 7.544588 | 14.60311 | 20.4798 | 7.799387 | 0.082454 | -0.19581 | 0.896177 |
| 6 | 13507200 | 13507799 | 6.832787 | 13.20117 | 21.02221 | 8.074757 | -0.11051 | -0.1721 | 0.89619 |
| 12 | 50069200 | 50071199 | 14.9793 | 26.71897 | 33.33181 | 14.65976 | 0.079686 | 0.453744 | 0.89619 |
| 8 | 87279200 | 87281799 | 24.22865 | 40.5068 | 55.44655 | 24.14354 | -0.07028 | 1.191021 | 0.896205 |
| 15 | 37866800 | 37871999 | 105.0426 | 161.435 | 206.0687 | 85.56751 | -0.05657 | 2.970304 | 0.89621 |
| 3 | 115214000 | 115215399 | 19.82313 | 34.41584 | 43.12537 | 18.92554 | 0.062799 | 0.814991 | 0.896237 |
| 17 | 21472400 | 21473799 | 20.82963 | 36.26935 | 36.7201 | 16.14428 | 0.369467 | 0.768759 | 0.896246 |
| 10 | 61353000 | 61355599 | 33.18981 | 54.24055 | 67.47275 | 29.00796 | 0.060115 | 1.494393 | 0.896256 |
| 2 | 63127600 | 63136399 | 86.65591 | 135.088 | 162.0343 | 67.246 | 0.048938 | 2.700971 | 0.896273 |
| 14 | 15042200 | 15044399 | 25.75328 | 43.69371 | 44.25641 | 19.38737 | 0.363011 | 1.046778 | 0.896282 |
| 14 | 13989200 | 13990799 | 21.20886 | 36.34467 | 49.54696 | 21.72892 | -0.06198 | 0.947452 | 0.896283 |
| 9 | 63615600 | 63617599 | 23.98972 | 40.23998 | 54.95561 | 23.97966 | -0.06712 | 1.153064 | 0.89629 |
| Y | 89523400 | 89523999 | 7.14435 | 13.63436 | 21.97478 | 8.751071 | -0.17692 | -0.04888 | 0.896318 |
| 8 | 18315000 | 18315399 | 5.241369 | 10.21831 | 17.52147 | 6.319887 | -0.13131 | -0.42436 | 0.896319 |
| 1 | 170876800 | 170879799 | 30.1395 | 49.71193 | 62.16088 | 26.82621 | 0.058622 | 1.36558 | 0.896328 |
| 7 | 66264800 | 66266999 | 21.3233 | 36.39554 | 49.55052 | 21.72351 | -0.06098 | 0.982905 | 0.896418 |
| 15 | 9771800 | 9780799 | 133.2248 | 201.5275 | 235.7885 | 97.52387 | 0.050651 | 3.296085 | 0.896468 |
| 18 | 14821000 | 14821599 | 7.231519 | 14.01229 | 22.18967 | 8.407945 | -0.08488 | -0.21313 | 0.896504 |
| 1 | 108962000 | 108963399 | 22.79092 | 39.1921 | 34.97012 | 15.33939 | 0.547867 | 0.901936 | 0.896529 |
| 17 | 6172000 | 6173199 | 16.4588 | 28.84582 | 35.80544 | 15.81686 | 0.077445 | 0.649238 | 0.896537 |
| 19 | 39704800 | 39707399 | 31.23291 | 52.44515 | 49.66374 | 21.68508 | 0.459369 | 1.162352 | 0.896545 |
| 18 | 73956600 | 73957999 | 12.77573 | 23.06875 | 29.04485 | 12.62567 | 0.07807 | 0.357121 | 0.896564 |
| 3 | 22996200 | 22997799 | 14.28904 | 26.14275 | 26.33221 | 11.25922 | 0.413295 | 0.287161 | 0.896576 |
| 19 | 52973000 | 52975599 | 40.35891 | 64.97578 | 87.03222 | 36.98156 | -0.0663 | 1.831412 | 0.896587 |
| 3 | 147786600 | 147787399 | 7.556823 | 14.58849 | 22.59091 | 8.709409 | -0.07628 | -0.14816 | 0.896619 |
| 7 | 95004000 | 95004599 | 7.556823 | 14.58849 | 22.59091 | 8.709409 | -0.07628 | -0.14816 | 0.896619 |
| 10 | 64780000 | 64780599 | 7.556823 | 14.58849 | 22.59091 | 8.709409 | -0.07628 | -0.14816 | 0.896619 |
| Y | 32537000 | 32537799 | 8.710992 | 16.21132 | 24.91968 | 10.40726 | -0.16723 | 0.143016 | 0.896627 |
| 18 | 28913400 | 28913999 | 10.32326 | 19.48608 | 25.87114 | 10.54458 | 0.070497 | 0.067897 | 0.896654 |
| 2 | 87546600 | 87547999 | 20.33157 | 35.65598 | 34.26411 | 15.14414 | 0.444627 | 0.655197 | 0.896659 |

| | | | | | | | | | |
|---|---|---|---|---|---|---|---|---|---|
| 11 | 27124000 | 27126999 | 49.79364 | 79.9776 | 97.60586 | 41.33369 | 0.063434 | 1.920083 | 0.896684 |
| 1 | 96493600 | 96493799 | 5.20492 | 10.17391 | 15.47065 | 5.329784 | 0.100883 | -0.59148 | 0.896704 |
| 13 | 12462600 | 12463799 | 10.81719 | 19.80251 | 28.65207 | 12.24127 | -0.1052 | 0.260624 | 0.89672 |
| 1 | 23924200 | 23924999 | 8.789286 | 16.76184 | 22.38672 | 8.928339 | 0.090087 | -0.02482 | 0.896724 |
| 5 | 133495000 | 133498399 | 62.71487 | 99.34639 | 119.3727 | 49.64604 | 0.064468 | 2.360888 | 0.896732 |
| 13 | 75825400 | 75826799 | 26.69222 | 45.17382 | 40.76307 | 17.84353 | 0.529112 | 1.070214 | 0.896744 |
| 2 | 148881600 | 148883599 | 20.37326 | 35.54702 | 51.18617 | 22.53324 | -0.1363 | 0.76015 | 0.896746 |
| 3 | 36832800 | 36833999 | 10.31789 | 19.20902 | 27.78899 | 11.59818 | -0.08029 | 0.159465 | 0.896748 |
| 2 | 26765200 | 26772799 | 102.3738 | 157.9723 | 187.4026 | 77.8435 | 0.053498 | 2.889477 | 0.896761 |
| 11 | 108741000 | 108742399 | 12.71716 | 23.01825 | 32.13895 | 13.92872 | -0.06739 | 0.350673 | 0.896773 |
| 9 | 110128000 | 110131599 | 51.59523 | 82.46358 | 108.4628 | 45.59604 | -0.05258 | 2.076565 | 0.896774 |
| 11 | 5073600 | 5076799 | 24.5699 | 41.20469 | 56.01268 | 24.43775 | -0.06039 | 1.160238 | 0.896793 |
| 7 | 18511800 | 18513399 | 21.82228 | 37.39741 | 51.17974 | 22.44505 | -0.06761 | 0.950427 | 0.896801 |
| 1 | 76246000 | 76246999 | 10.07211 | 19.06171 | 25.40919 | 10.29929 | 0.072042 | 0.047967 | 0.896808 |
| 1 | 55335000 | 55337799 | 64.74714 | 102.2705 | 133.1951 | 55.31391 | -0.05492 | 2.432913 | 0.896813 |
| 2 | 166173600 | 166176199 | 22.46502 | 38.2799 | 47.41939 | 20.78639 | 0.074289 | 0.997766 | 0.896841 |
| 7 | 9960800 | 9964399 | 53.60214 | 86.55698 | 73.69104 | 31.4396 | 0.591197 | 1.758612 | 0.896851 |
| 8 | 45941800 | 45945799 | 68.05307 | 107.4401 | 139.2096 | 57.79742 | -0.04808 | 2.447849 | 0.896856 |
| 7 | 18573800 | 18575799 | 12.89102 | 22.87963 | 32.1674 | 14.20726 | -0.09566 | 0.471189 | 0.896874 |
| 5 | 10554800 | 10558399 | 63.97448 | 101.817 | 102.8748 | 42.99182 | 0.320681 | 2.221534 | 0.896893 |
| 5 | 133959200 | 133961399 | 23.39853 | 40.10086 | 39.5335 | 17.33763 | 0.403784 | 0.941571 | 0.896894 |
| 5 | 38568800 | 38577399 | 101.0338 | 155.4481 | 197.514 | 82.0279 | -0.04869 | 2.944987 | 0.896902 |
| 1 | 40497000 | 40498599 | 26.79894 | 44.85911 | 56.13125 | 24.46066 | 0.058736 | 1.18622 | 0.896907 |
| 13 | 14009600 | 14013999 | 66.15691 | 104.6285 | 135.902 | 56.47392 | -0.04964 | 2.404092 | 0.896924 |
| 3 | 9794000 | 9804199 | 169.6703 | 252.4846 | 291.609 | 120.1504 | 0.048264 | 3.617646 | 0.896929 |
| 14 | 85899400 | 85899799 | 7.645866 | 14.78274 | 20.42579 | 7.81922 | 0.096077 | -0.17946 | 0.896929 |
| 4 | 61657600 | 61666599 | 128.0306 | 194.9687 | 194.2492 | 80.50666 | 0.290296 | 3.15756 | 0.896933 |
| 7 | 4039600 | 4039999 | 7.09486 | 13.77145 | 19.47131 | 7.318145 | 0.089017 | -0.24866 | 0.896936 |
| 3 | 146583600 | 146596599 | 173.8991 | 259.1482 | 298.858 | 123.1551 | 0.052319 | 3.59366 | 0.896951 |
| 4 | 60573600 | 60578599 | 114.5615 | 174.6357 | 204.8444 | 84.92273 | 0.05618 | 3.134561 | 0.896966 |
| 19 | 33800200 | 33803999 | 47.86605 | 76.67933 | 101.0848 | 42.63583 | -0.05211 | 2.000637 | 0.89699 |
| 3 | 89059800 | 89060599 | 8.644127 | 16.33385 | 24.38343 | 9.915992 | -0.09421 | 0.041642 | 0.89706 |
| 18 | 56774000 | 56776199 | 21.89564 | 38.08978 | 46.24635 | 20.3206 | 0.108262 | 0.799222 | 0.897068 |
| 11 | 68247600 | 68249599 | 13.9603 | 25.67484 | 25.86991 | 10.98334 | 0.420125 | 0.254716 | 0.89707 |
| 7 | 102931800 | 102932799 | 29.18273 | 47.68348 | 66.30851 | 28.50558 | -0.1003 | 1.488666 | 0.897089 |
| 3 | 145261600 | 145262999 | 14.33903 | 25.61401 | 35.5551 | 15.61849 | -0.07215 | 0.447116 | 0.897122 |
| 14 | 70100200 | 70103799 | 29.83942 | 49.24679 | 61.55945 | 26.5743 | 0.059399 | 1.357709 | 0.897131 |
| 9 | 24681000 | 24682999 | 16.88389 | 29.40948 | 40.52692 | 17.82203 | -0.07322 | 0.749984 | 0.897134 |
| 5 | 128004600 | 128015799 | 143.7207 | 216.7893 | 253.278 | 104.6641 | 0.049255 | 3.35705 | 0.897172 |
| 8 | 109070400 | 109072799 | 27.13876 | 45.09593 | 56.07243 | 24.3198 | 0.067786 | 1.266535 | 0.897176 |
| 1 | 163557000 | 163560199 | 53.43902 | 85.44306 | 104.5823 | 43.98851 | 0.051623 | 2.066745 | 0.897188 |
| 18 | 25886000 | 25889199 | 28.87403 | 47.49234 | 64.70196 | 27.88999 | -0.06631 | 1.398339 | 0.89721 |
| 3 | 6112200 | 6114199 | 23.75766 | 40.96227 | 40.93294 | 17.95639 | 0.385893 | 0.891229 | 0.897225 |
| 4 | 53291400 | 53295399 | 74.65724 | 118.3082 | 117.7011 | 48.96165 | 0.336853 | 2.359997 | 0.897273 |
| X | 159845200 | 159846399 | 13.1234 | 23.67569 | 33.16016 | 14.43043 | -0.07558 | 0.375986 | 0.897283 |
| 2 | 150914600 | 150916399 | 39.48069 | 63.69449 | 84.91979 | 36.18693 | -0.05604 | 1.76499 | 0.897286 |
| 4 | 35376200 | 35378199 | 30.71897 | 49.85771 | 82.94737 | 35.55153 | -0.36579 | 1.606189 | 0.897296 |

| | | | | | | | | | |
|---|---|---|---|---|---|---|---|---|---|
| 3 | 6200200 | 6204199 | 71.71756 | 112.2367 | 146.9729 | 60.96969 | -0.07313 | 2.622476 | 0.897324 |
| 9 | 115585800 | 115588799 | 31.40011 | 51.841 | 64.67044 | 27.92414 | 0.062289 | 1.358146 | 0.897325 |
| 2 | 5686600 | 5689399 | 31.07025 | 51.02809 | 63.26341 | 27.25259 | 0.068439 | 1.41972 | 0.897334 |
| 15 | 33267400 | 33271199 | 46.10777 | 74.12329 | 98.6774 | 41.84154 | -0.06014 | 1.889185 | 0.897352 |
| 8 | 48746000 | 48748999 | 30.5929 | 50.54574 | 69.27138 | 29.92609 | -0.07293 | 1.350399 | 0.897373 |
| X | 69315400 | 69317999 | 28.44773 | 48.50433 | 56.86493 | 24.9281 | 0.153677 | 1.028932 | 0.89743 |
| 18 | 64583000 | 64585599 | 29.25246 | 48.44957 | 60.59807 | 26.22341 | 0.0589 | 1.312914 | 0.897445 |
| 6 | 7061600 | 7062799 | 13.50213 | 24.21163 | 33.54814 | 14.68935 | -0.06632 | 0.413619 | 0.897468 |
| 1 | 45618000 | 45618999 | 10.34719 | 19.30151 | 27.83184 | 11.57742 | -0.07164 | 0.148067 | 0.897489 |
| 15 | 65186400 | 65187199 | 10.34719 | 19.30151 | 27.83184 | 11.57742 | -0.07164 | 0.148067 | 0.897489 |
| 3 | 106517600 | 106519199 | 34.88437 | 56.70351 | 69.94848 | 30.01332 | 0.066711 | 1.577416 | 0.897492 |
| 4 | 71431400 | 71433799 | 20.72162 | 36.01558 | 44.64897 | 19.60781 | 0.078217 | 0.803982 | 0.897511 |
| 19 | 13499000 | 13505199 | 41.56287 | 67.43469 | 68.32868 | 29.27856 | 0.347368 | 1.625945 | 0.897513 |
| 6 | 51160800 | 51163399 | 22.16551 | 38.04779 | 47.39135 | 20.78638 | 0.067998 | 0.93104 | 0.897516 |
| 18 | 22099200 | 22100999 | 21.34857 | 37.20371 | 37.15753 | 16.3451 | 0.389094 | 0.76055 | 0.897518 |
| 3 | 75636600 | 75637199 | 13.55691 | 24.26257 | 33.58409 | 14.72926 | -0.06635 | 0.427343 | 0.897534 |
| 8 | 69022600 | 69023399 | 11.37994 | 20.9789 | 29.7803 | 12.6028 | -0.06767 | 0.230994 | 0.897539 |
| 17 | 49589400 | 49589999 | 11.37994 | 20.9789 | 29.7803 | 12.6028 | -0.06767 | 0.230994 | 0.897539 |
| 18 | 53356200 | 53358399 | 25.81598 | 42.93766 | 58.79509 | 25.517 | -0.07088 | 1.253012 | 0.897571 |
| 2 | 39192000 | 39193199 | 14.69325 | 26.04708 | 32.56124 | 14.39478 | 0.071995 | 0.493259 | 0.897574 |
| 3 | 100038800 | 100040199 | 14.69325 | 26.04708 | 32.56124 | 14.39478 | 0.071995 | 0.493259 | 0.897574 |
| 7 | 68484800 | 68487199 | 16.65351 | 29.24889 | 40.23286 | 17.80445 | -0.07023 | 0.622893 | 0.897592 |
| 7 | 7270000 | 7276999 | 82.98898 | 129.5301 | 166.4944 | 69.08687 | -0.04937 | 2.678214 | 0.897598 |
| 10 | 14127400 | 14129399 | 18.63751 | 32.74243 | 40.05621 | 17.7304 | 0.097908 | 0.627157 | 0.897599 |
| 7 | 50919000 | 50921399 | 27.40108 | 45.21598 | 61.91471 | 26.72517 | -0.07172 | 1.35287 | 0.897603 |
| 6 | 32350600 | 32351399 | 8.955413 | 17.0701 | 22.89529 | 9.140849 | 0.082656 | -0.01814 | 0.897611 |
| 13 | 42283000 | 42284799 | 27.76967 | 45.95561 | 62.52483 | 27.04174 | -0.06214 | 1.322728 | 0.897635 |
| 13 | 67774800 | 67781399 | 157.9995 | 238.0738 | 232.4177 | 96.01457 | 0.306852 | 3.380098 | 0.897646 |
| 8 | 98309000 | 98312399 | 23.63428 | 40.80539 | 49.58049 | 21.75078 | 0.10538 | 0.890142 | 0.897655 |
| 14 | 47027000 | 47033399 | 62.97915 | 99.88692 | 130.1829 | 54.19995 | -0.051 | 2.328215 | 0.897656 |
| 10 | 99503800 | 99504199 | 8.30208 | 15.96184 | 21.94844 | 8.565783 | 0.077135 | -0.1022 | 0.897658 |
| 10 | 68202200 | 68204599 | 28.39793 | 46.99762 | 63.74842 | 27.57187 | -0.0578 | 1.324063 | 0.897667 |
| 4 | 14740800 | 14741199 | 8.034668 | 15.50832 | 21.49588 | 8.293876 | 0.081537 | -0.14187 | 0.89767 |
| 16 | 51651800 | 51655199 | 23.84216 | 40.61503 | 56.14152 | 24.60919 | -0.08333 | 1.01768 | 0.897721 |
| 10 | 119000400 | 119003199 | 28.97472 | 47.89327 | 65.01863 | 28.09739 | -0.05923 | 1.338714 | 0.897743 |
| 9 | 61770600 | 61772199 | 14.34574 | 25.4591 | 35.02594 | 15.47103 | -0.06475 | 0.487062 | 0.897756 |
| 17 | 84930800 | 84932799 | 22.70021 | 38.90425 | 48.47495 | 21.25896 | 0.067178 | 0.949224 | 0.897778 |
| 1 | 110484400 | 110489199 | 45.81678 | 74.04702 | 88.88583 | 37.96324 | 0.097919 | 1.725132 | 0.89778 |
| 14 | 91219200 | 91223599 | 49.57049 | 79.88961 | 95.04137 | 40.44594 | 0.106204 | 1.813009 | 0.89779 |
| 9 | 15069800 | 15080799 | 235.8113 | 342.336 | 386.466 | 159.3242 | 0.047381 | 4.023627 | 0.897798 |
| 13 | 78439000 | 78441599 | 21.5525 | 37.27705 | 46.30253 | 20.31385 | 0.07422 | 0.860499 | 0.897802 |
| X | 113594400 | 113595399 | 12.91327 | 24.06383 | 30.51817 | 12.71669 | 0.110442 | 0.183956 | 0.897802 |
| 8 | 38875400 | 38875999 | 12.11142 | 21.63797 | 30.76232 | 13.51147 | -0.10593 | 0.422624 | 0.897816 |
| X | 64091200 | 64095399 | 45.60525 | 73.79369 | 87.82692 | 37.55858 | 0.112124 | 1.698873 | 0.897819 |
| 10 | 121362200 | 121365599 | 36.69495 | 59.41567 | 79.495 | 34.01224 | -0.05543 | 1.663017 | 0.897822 |
| 12 | 49610600 | 49612999 | 15.99083 | 28.14886 | 38.64949 | 17.11031 | -0.06678 | 0.587596 | 0.897842 |
| 7 | 58166000 | 58168999 | 32.60243 | 53.4091 | 66.61446 | 28.66544 | 0.058353 | 1.457745 | 0.897856 |

| | | | | | | | | | |
|---|---|---|---|---|---|---|---|---|---|
| 10 | 117893600 | 117895599 | 24.77119 | 41.48704 | 51.22143 | 22.32862 | 0.078004 | 1.168521 | 0.897867 |
| 7 | 62375400 | 62377599 | 63.89756 | 101.3016 | 122.8886 | 51.14184 | 0.051773 | 2.341085 | 0.897889 |
| 3 | 50005200 | 50008799 | 45.85785 | 73.8096 | 90.81823 | 38.57577 | 0.055386 | 1.847653 | 0.897899 |
| 3 | 67030600 | 67031599 | 12.15794 | 22.17509 | 31.16897 | 13.37968 | -0.06816 | 0.299273 | 0.897914 |
| 3 | 10224200 | 10228599 | 70.21635 | 110.8002 | 134.0882 | 55.65711 | 0.049725 | 2.460606 | 0.897924 |
| 4 | 24965400 | 24966999 | 13.32548 | 23.62986 | 33.05321 | 14.60936 | -0.08926 | 0.480098 | 0.897939 |
| 16 | 27552200 | 27555399 | 29.54785 | 48.94571 | 66.75975 | 28.89265 | -0.06572 | 1.318645 | 0.897989 |
| 3 | 148287400 | 148287999 | 6.826305 | 13.26641 | 21.20422 | 7.89173 | -0.07311 | -0.27986 | 0.898022 |
| 7 | 29806800 | 29816399 | 155.6161 | 232.4525 | 287.9616 | 118.7168 | -0.04777 | 3.54684 | 0.898045 |
| 1 | 28494400 | 28500399 | 94.44416 | 146.841 | 149.1553 | 61.92552 | 0.285558 | 2.747697 | 0.898053 |
| 5 | 98153600 | 98156599 | 61.79096 | 97.29286 | 152.8793 | 63.43149 | -0.32896 | 2.503107 | 0.898054 |
| 12 | 33070000 | 33072799 | 25.68021 | 42.80562 | 58.23302 | 25.30916 | -0.06141 | 1.229146 | 0.898068 |
| 13 | 54295600 | 54298399 | 21.15924 | 36.74869 | 36.84359 | 16.17243 | 0.383214 | 0.806964 | 0.898071 |
| 16 | 16231400 | 16235199 | 41.8396 | 67.9637 | 82.11666 | 35.21722 | 0.095611 | 1.610774 | 0.898079 |
| 7 | 47786800 | 47788399 | 17.89492 | 30.9618 | 42.71434 | 18.73984 | -0.07626 | 0.837884 | 0.898099 |
| 8 | 56044400 | 56050399 | 58.8849 | 93.98853 | 96.02235 | 40.28977 | 0.309175 | 2.121284 | 0.898115 |
| 2 | 170035800 | 170038199 | 16.91728 | 29.46564 | 40.54657 | 17.82991 | -0.07114 | 0.751919 | 0.898131 |
| 5 | 14727400 | 14732599 | 62.00355 | 98.37514 | 128.118 | 53.35955 | -0.04942 | 2.316683 | 0.898152 |
| X | 36454000 | 36455999 | 25.50091 | 43.47172 | 52.81968 | 23.15442 | 0.102336 | 1.017113 | 0.898212 |
| 8 | 104446000 | 104449599 | 31.68619 | 52.48573 | 64.85499 | 28.06751 | 0.076302 | 1.320877 | 0.898247 |
| 15 | 16585800 | 16588399 | 21.26921 | 36.92845 | 52.36659 | 22.98183 | -0.11503 | 0.832922 | 0.898258 |
| 12 | 87814600 | 87815999 | 18.00005 | 31.66821 | 31.28367 | 13.85096 | 0.40539 | 0.590435 | 0.898265 |
| 2 | 178959000 | 178967999 | 103.5406 | 159.1888 | 189.0909 | 78.52189 | 0.047598 | 2.959107 | 0.898292 |
| 4 | 112340000 | 112354599 | 340.3112 | 483.5506 | 434.108 | 179.5376 | 0.347008 | 4.34461 | 0.898303 |
| 16 | 27736200 | 27739599 | 43.997 | 70.88581 | 87.36469 | 37.16302 | 0.054802 | 1.811319 | 0.898309 |
| 15 | 14677000 | 14678399 | 14.31731 | 25.90296 | 32.46009 | 14.07576 | 0.08679 | 0.36241 | 0.898321 |
| 8 | 5163400 | 5164399 | 11.38108 | 21.06157 | 27.1273 | 11.42037 | 0.076845 | 0.201286 | 0.898334 |
| 14 | 60097400 | 60098999 | 12.37429 | 22.29056 | 31.11224 | 13.56482 | -0.07247 | 0.373199 | 0.89834 |
| 6 | 62458600 | 62459599 | 12.24793 | 22.81268 | 29.37272 | 12.24963 | 0.087523 | 0.17694 | 0.898367 |
| X | 153035000 | 153038599 | 37.08526 | 61.18974 | 72.6941 | 31.37661 | 0.13245 | 1.388068 | 0.898378 |
| 5 | 88406400 | 88407599 | 18.05207 | 31.48403 | 39.43269 | 17.3546 | 0.064214 | 0.737572 | 0.898383 |
| 1 | 162655600 | 162658999 | 28.44714 | 47.63015 | 59.01212 | 25.7202 | 0.072911 | 1.191017 | 0.898398 |
| 4 | 63366000 | 63369199 | 34.65324 | 56.46999 | 70.30922 | 30.19584 | 0.055885 | 1.53961 | 0.898416 |
| 6 | 28928000 | 28930599 | 20.9477 | 35.53999 | 49.03103 | 21.4789 | -0.08097 | 1.037262 | 0.898429 |
| 6 | 69637400 | 69638799 | 17.40846 | 30.36254 | 41.50465 | 18.26686 | -0.06147 | 0.736428 | 0.898438 |
| 13 | 95695800 | 95696599 | 8.605936 | 16.48056 | 22.4116 | 8.846694 | 0.07757 | -0.06093 | 0.898443 |
| X | 137292000 | 137299799 | 131.0936 | 200.2864 | 190.5808 | 79.05048 | 0.360311 | 3.093806 | 0.898473 |
| 11 | 29910800 | 29913999 | 42.25424 | 68.2694 | 84.18314 | 35.94113 | 0.058774 | 1.726708 | 0.898484 |
| 13 | 77809200 | 77810199 | 13.67755 | 24.27317 | 33.55588 | 14.82176 | -0.07165 | 0.479526 | 0.898491 |
| 10 | 11775600 | 11781999 | 80.33422 | 126.3093 | 128.68 | 53.38096 | 0.293774 | 2.537247 | 0.898501 |
| 3 | 10939200 | 10948799 | 344.7815 | 482.3768 | 521.3801 | 216.173 | 0.056849 | 4.572485 | 0.898502 |
| 13 | 33287000 | 33288199 | 12.00896 | 21.98586 | 28.14782 | 12.02462 | 0.070642 | 0.27132 | 0.89853 |
| 16 | 86336400 | 86339399 | 60.43695 | 95.95209 | 115.9065 | 48.3051 | 0.059899 | 2.295342 | 0.89853 |
| 2 | 173345600 | 173348999 | 34.59368 | 56.25476 | 75.67465 | 32.47529 | -0.05755 | 1.574312 | 0.898532 |
| 10 | 126718600 | 126726399 | 82.9937 | 129.5502 | 166.4954 | 69.08657 | -0.04904 | 2.676619 | 0.898536 |
| Y | 41922000 | 41927399 | 46.64336 | 74.14179 | 107.9041 | 45.02408 | -0.20675 | 2.243548 | 0.898544 |
| 2 | 123586400 | 123586999 | 9.56707 | 17.67046 | 32.00168 | 13.50787 | -0.41379 | 0.211929 | 0.898556 |

| | | | | | | | | | |
|---|---|---|---|---|---|---|---|---|---|
| 9 | 18119800 | 18121999 | 23.24527 | 39.61752 | 49.56002 | 21.72509 | 0.060304 | 1.003376 | 0.898609 |
| 17 | 51018800 | 51025799 | 62.89708 | 100.1926 | 120.5399 | 50.437 | 0.070627 | 2.196149 | 0.898611 |
| 2 | 130585000 | 130590399 | 56.20947 | 89.73754 | 109.5817 | 45.99314 | 0.052613 | 2.116437 | 0.89862 |
| 8 | 105696400 | 105698199 | 15.28699 | 26.98492 | 37.06572 | 16.40004 | -0.06583 | 0.549359 | 0.898645 |
| 11 | 35512600 | 35515799 | 30.25219 | 49.3987 | 67.68862 | 29.09196 | -0.07985 | 1.498899 | 0.898651 |
| 5 | 109957000 | 109959399 | 35.79928 | 58.53891 | 72.56224 | 31.20262 | 0.065739 | 1.494382 | 0.89868 |
| X | 148175600 | 148178399 | 49.11005 | 78.96503 | 75.3946 | 31.97803 | 0.418818 | 1.878252 | 0.898704 |
| 4 | 73247600 | 73248999 | 16.61687 | 29.0239 | 40.38015 | 17.78813 | -0.08677 | 0.707064 | 0.898704 |
| 12 | 59438000 | 59438999 | 10.54018 | 19.92537 | 26.4619 | 10.75124 | 0.074229 | 0.062014 | 0.898713 |
| 9 | 23717000 | 23719199 | 18.7577 | 32.67542 | 45.16719 | 19.86339 | -0.0779 | 0.763109 | 0.898723 |
| 9 | 56827800 | 56829799 | 14.71913 | 25.72763 | 45.17083 | 19.90872 | -0.42068 | 0.695891 | 0.898726 |
| 16 | 89506000 | 89519199 | 185.9426 | 275.942 | 316.5621 | 130.3953 | 0.05433 | 3.668717 | 0.898733 |
| 11 | 107934800 | 107936799 | 17.44162 | 30.81286 | 37.97662 | 16.8004 | 0.090159 | 0.555823 | 0.898741 |
| 1 | 8318800 | 8319999 | 25.70497 | 42.9146 | 58.23099 | 25.33351 | -0.05772 | 1.213115 | 0.898744 |
| 15 | 38112800 | 38114399 | 18.49486 | 32.2246 | 40.44795 | 17.79043 | 0.060881 | 0.754224 | 0.898758 |
| 6 | 130408600 | 130410999 | 18.54178 | 32.98996 | 39.04131 | 17.2075 | 0.153408 | 0.505287 | 0.898774 |
| 1 | 115126200 | 115128199 | 19.01997 | 33.28347 | 41.30501 | 18.22264 | 0.077514 | 0.694475 | 0.898792 |
| 15 | 4191000 | 4191599 | 9.003015 | 17.139 | 25.55215 | 10.22909 | -0.07225 | -0.00142 | 0.8988 |
| 18 | 3358400 | 3361599 | 34.69467 | 56.85261 | 70.44659 | 30.31809 | 0.067888 | 1.46072 | 0.898807 |
| 13 | 27317400 | 27318799 | 21.88226 | 37.56103 | 46.88026 | 20.56219 | 0.065254 | 0.932966 | 0.898808 |
| 2 | 53213400 | 53217199 | 53.62448 | 85.43659 | 112.1263 | 46.92788 | -0.05434 | 2.179047 | 0.898837 |
| X | 126366200 | 126367999 | 23.14725 | 40.12907 | 48.485 | 21.27286 | 0.114773 | 0.852878 | 0.898856 |
| 3 | 119037400 | 119038599 | 12.87875 | 23.34607 | 32.66156 | 14.12843 | -0.06852 | 0.344118 | 0.89886 |
| 9 | 47080800 | 47082399 | 12.87875 | 23.34607 | 32.66156 | 14.12843 | -0.06852 | 0.344118 | 0.89886 |
| 18 | 62096800 | 62099599 | 20.25328 | 34.98319 | 47.93708 | 21.03011 | -0.06715 | 0.871175 | 0.898876 |
| 16 | 43859400 | 43862799 | 34.39167 | 55.87766 | 75.05632 | 32.19448 | -0.05646 | 1.587458 | 0.898888 |
| 9 | 103646000 | 103646799 | 9.905129 | 18.7343 | 27.59655 | 11.20029 | -0.0729 | 0.05511 | 0.898894 |
| 15 | 26870000 | 26871799 | 19.20531 | 33.3792 | 45.7636 | 20.09695 | -0.06632 | 0.800293 | 0.898905 |
| 6 | 102388600 | 102390999 | 29.11935 | 48.69659 | 60.16761 | 26.20377 | 0.076893 | 1.206929 | 0.898925 |
| 13 | 102733800 | 102736999 | 22.93554 | 39.22676 | 49.01792 | 21.49352 | 0.062637 | 0.970381 | 0.898929 |
| 8 | 89515000 | 89517599 | 21.3926 | 36.52427 | 45.33104 | 19.8741 | 0.07207 | 0.974758 | 0.898972 |
| X | 85417200 | 85420199 | 30.82628 | 50.66867 | 69.15312 | 29.80003 | -0.06924 | 1.418154 | 0.898977 |
| 4 | 21312000 | 21312799 | 10.86811 | 20.39525 | 26.85584 | 11.05849 | 0.070435 | 0.113603 | 0.898994 |
| 1 | 5881000 | 5882399 | 17.4031 | 30.77337 | 30.70267 | 13.57997 | 0.393942 | 0.531497 | 0.899011 |
| 2 | 15305800 | 15306999 | 12.62548 | 23.39279 | 29.76233 | 12.50903 | 0.095951 | 0.209756 | 0.899054 |
| 7 | 80420000 | 80421399 | 12.55865 | 23.10969 | 29.65735 | 12.58194 | 0.074072 | 0.248542 | 0.89909 |
| 10 | 82233200 | 82243999 | 96.30938 | 150.676 | 172.8047 | 71.68793 | 0.117584 | 2.646481 | 0.899092 |
| 14 | 61975000 | 61976999 | 24.32784 | 40.79304 | 55.50423 | 24.21377 | -0.06177 | 1.15865 | 0.899106 |
| 5 | 18747400 | 18748399 | 12.95855 | 23.35349 | 32.54009 | 14.1794 | -0.06973 | 0.376309 | 0.899114 |
| 1 | 44925200 | 44925999 | 11.65904 | 21.38492 | 30.20836 | 12.8723 | -0.06797 | 0.261433 | 0.899118 |
| 16 | 68887600 | 68889399 | 15.72219 | 27.86189 | 34.87321 | 15.41789 | 0.069722 | 0.505256 | 0.899121 |
| 2 | 128346600 | 128349799 | 29.44499 | 48.6989 | 66.26704 | 28.64838 | -0.06255 | 1.333754 | 0.899129 |
| X | 103391200 | 103392599 | 15.51115 | 27.492 | 34.47069 | 15.23947 | 0.067485 | 0.503717 | 0.899139 |
| 6 | 7160600 | 7163999 | 35.599 | 58.44252 | 59.70456 | 25.71517 | 0.346686 | 1.433364 | 0.899156 |
| 10 | 120978600 | 120979999 | 11.04255 | 20.51573 | 29.36514 | 12.29642 | -0.06857 | 0.183741 | 0.899161 |
| 4 | 17339400 | 17340199 | 14.39489 | 25.50842 | 35.07707 | 15.50817 | -0.06507 | 0.500107 | 0.89918 |
| 10 | 119694400 | 119698599 | 41.95555 | 67.6512 | 90.01865 | 38.33339 | -0.05413 | 1.786736 | 0.899185 |

| | | | | | | | | | |
|---|---|---|---|---|---|---|---|---|---|
| 2 | 48693200 | 48695399 | 17.707 | 31.21565 | 38.48609 | 17.03297 | 0.088438 | 0.580108 | 0.89919 |
| 14 | 121026600 | 121037599 | 126.1827 | 190.1386 | 281.7484 | 116.3948 | -0.29473 | 3.374958 | 0.8992 |
| X | 44576000 | 44577799 | 24.32032 | 42.1875 | 38.80482 | 17.02597 | 0.506833 | 0.840373 | 0.899202 |
| 3 | 144140000 | 144141599 | 13.73127 | 24.30925 | 33.65829 | 14.88374 | -0.07539 | 0.503376 | 0.899245 |
| 1 | 11443200 | 11444199 | 17.88061 | 31.33538 | 39.20446 | 17.31728 | 0.065879 | 0.662222 | 0.899255 |
| 3 | 20104400 | 20106599 | 26.03951 | 44.06058 | 54.52982 | 23.86834 | 0.074767 | 1.081376 | 0.899276 |
| 6 | 131801000 | 131802399 | 30.16971 | 50.12004 | 62.34908 | 27.03751 | 0.067035 | 1.284091 | 0.899289 |
| 13 | 21238600 | 21239799 | 15.12832 | 26.77646 | 37.13101 | 16.42013 | -0.07773 | 0.522638 | 0.899294 |
| 15 | 59509800 | 59512199 | 20.82224 | 35.62028 | 48.4828 | 21.25926 | -0.06011 | 0.959236 | 0.899303 |
| X | 16014800 | 16015399 | 7.595475 | 14.74309 | 23.45721 | 8.816447 | -0.07997 | -0.22946 | 0.899306 |
| 5 | 133976800 | 133978399 | 13.80378 | 24.61129 | 34.0099 | 14.96356 | -0.0673 | 0.451554 | 0.899311 |
| 1 | 54409800 | 54410799 | 14.05905 | 25.4422 | 32.07942 | 13.90626 | 0.078676 | 0.362105 | 0.899323 |
| 3 | 90751400 | 90752599 | 12.89519 | 23.42431 | 33.15279 | 14.30383 | -0.08305 | 0.330438 | 0.899335 |
| 11 | 105272800 | 105276199 | 38.16972 | 61.6916 | 82.29537 | 35.15095 | -0.05381 | 1.708961 | 0.899342 |
| 10 | 22373400 | 22376599 | 51.3193 | 82.21882 | 100.7717 | 42.50969 | 0.052945 | 2.002074 | 0.899357 |
| 12 | 24771400 | 24780999 | 132.0096 | 199.1066 | 249.2704 | 103.0051 | -0.05052 | 3.356268 | 0.89939 |
| 15 | 70605600 | 70606799 | 15.69747 | 27.64452 | 34.5246 | 15.28287 | 0.069466 | 0.572357 | 0.899427 |
| 1 | 6496200 | 6500799 | 63.60941 | 100.5937 | 130.9812 | 54.42745 | -0.05321 | 2.403941 | 0.899435 |
| 3 | 48764800 | 48765399 | 7.824091 | 15.05992 | 20.39987 | 7.935531 | 0.103049 | -0.12312 | 0.899552 |
| 1 | 17992400 | 17993399 | 10.9584 | 20.40329 | 29.42654 | 12.27977 | -0.07533 | 0.171499 | 0.899568 |
| 6 | 90017600 | 90034399 | 208.4196 | 307.8627 | 346.3812 | 142.647 | 0.07612 | 3.751 | 0.899574 |
| 1 | 10435400 | 10438399 | 6.74196 | 12.84179 | 27.75873 | 11.09936 | -0.60263 | -0.02409 | 0.89961 |
| 11 | 74471200 | 74474199 | 19.12412 | 33.12437 | 45.26006 | 19.85737 | -0.06229 | 0.835661 | 0.899643 |
| 11 | 62516800 | 62518399 | 19.26972 | 33.73703 | 33.58397 | 14.82246 | 0.394201 | 0.676578 | 0.899644 |
| 4 | 156500200 | 156504799 | 81.02825 | 127.3655 | 152.2967 | 63.17573 | 0.062676 | 2.548302 | 0.899647 |
| 3 | 112798600 | 112800999 | 18.21287 | 31.98515 | 39.59956 | 17.52305 | 0.080865 | 0.630781 | 0.899681 |
| 9 | 89809000 | 89811399 | 28.89294 | 48.39121 | 59.64266 | 25.99998 | 0.080386 | 1.188909 | 0.899685 |
| 7 | 84709400 | 84710599 | 27.641 | 45.8635 | 57.1632 | 24.76792 | 0.064187 | 1.285745 | 0.899688 |
| 14 | 72038200 | 72041399 | 80.66912 | 126.0141 | 151.231 | 62.74794 | 0.050485 | 2.660871 | 0.899688 |
| 7 | 99407800 | 99414599 | 94.27414 | 146.2753 | 174.347 | 72.40671 | 0.05279 | 2.786635 | 0.899709 |
| 3 | 144814000 | 144821399 | 69.15318 | 108.9527 | 130.5749 | 54.1815 | 0.062011 | 2.489939 | 0.899757 |
| 10 | 42717600 | 42721999 | 56.02406 | 89.19905 | 108.1404 | 45.21714 | 0.058814 | 2.201507 | 0.899768 |
| 4 | 101681000 | 101683599 | 29.04009 | 48.60478 | 49.14883 | 21.41627 | 0.365109 | 1.18857 | 0.89977 |
| 16 | 73131200 | 73131999 | 13.46287 | 24.94239 | 31.45463 | 13.2217 | 0.108776 | 0.222757 | 0.89977 |
| 8 | 61188600 | 61192399 | 42.78767 | 68.96689 | 91.85142 | 39.0948 | -0.05605 | 1.799866 | 0.899772 |
| 9 | 113976200 | 113976599 | 6.083981 | 11.85362 | 16.94956 | 6.163784 | 0.117241 | -0.38546 | 0.899774 |
| 2 | 83831800 | 83836999 | 51.48183 | 82.50563 | 101.023 | 42.63648 | 0.054916 | 1.992388 | 0.899776 |
| 3 | 21219800 | 21220599 | 25.88512 | 43.10344 | 52.77988 | 22.92445 | 0.090095 | 1.237787 | 0.899803 |
| 1 | 58545400 | 58553599 | 178.0245 | 265.4947 | 304.2319 | 125.385 | 0.062492 | 3.581093 | 0.899808 |
| 4 | 61561400 | 61561999 | 20.75893 | 36.07252 | 35.30745 | 15.5033 | 0.417735 | 0.794636 | 0.899808 |
| 1 | 88983600 | 88985799 | 27.06146 | 44.90561 | 60.93727 | 26.40467 | -0.05811 | 1.287209 | 0.89985 |
| 10 | 110728200 | 110732599 | 40.52822 | 65.72411 | 80.71857 | 34.5752 | 0.06984 | 1.637951 | 0.899853 |
| 10 | 48937600 | 48940399 | 32.90327 | 54.07083 | 67.45101 | 29.06387 | 0.059876 | 1.420315 | 0.89986 |
| 14 | 41218600 | 41241399 | 328.4794 | 465.47 | 562.1355 | 232.593 | -0.08464 | 4.385484 | 0.899867 |
| 12 | 49600600 | 49602599 | 31.87206 | 52.1373 | 70.31268 | 30.23954 | -0.05516 | 1.478751 | 0.899869 |
| 16 | 51219800 | 51221399 | 18.81566 | 33.04809 | 32.91509 | 14.56674 | 0.393353 | 0.621927 | 0.899891 |
| 13 | 62259400 | 62264999 | 58.26914 | 92.94127 | 123.4892 | 51.76416 | -0.0704 | 2.150767 | 0.899892 |

| | | | | | | | | | |
|---|---|---|---|---|---|---|---|---|---|
| 14 | 83733800 | 83736199 | 20.78414 | 36.09408 | 44.70644 | 19.62274 | 0.079572 | 0.819546 | 0.899894 |
| 7 | 72925400 | 72928799 | 31.38843 | 51.70898 | 64.58813 | 27.85938 | 0.059427 | 1.383913 | 0.899896 |
| 3 | 108691000 | 108694199 | 54.02477 | 86.27308 | 105.5204 | 44.30555 | 0.050732 | 2.104161 | 0.899901 |
| 1 | 102151200 | 102151999 | 10.01479 | 18.97651 | 28.23451 | 11.41481 | -0.08256 | 0.043808 | 0.89991 |
| 1 | 23180800 | 23184799 | 49.9065 | 80.02529 | 98.17429 | 41.4691 | 0.053085 | 1.971018 | 0.899919 |
| 8 | 9170600 | 9172199 | 19.01979 | 33.38333 | 33.33005 | 14.74169 | 0.389734 | 0.633166 | 0.899922 |
| 8 | 28674400 | 28677799 | 35.61043 | 57.94475 | 72.10439 | 30.95026 | 0.055189 | 1.56281 | 0.899925 |
| 2 | 40274400 | 40275799 | 25.72802 | 43.85172 | 43.70884 | 19.16027 | 0.386931 | 1.007355 | 0.899931 |
| 15 | 96407400 | 96412999 | 58.03508 | 92.53199 | 112.9327 | 47.31368 | 0.051371 | 2.159761 | 0.899972 |
| 13 | 44099000 | 44103199 | 44.05099 | 70.86807 | 93.91946 | 39.86586 | -0.05236 | 1.860545 | 0.89999 |
| 19 | 18171800 | 18174799 | 28.47915 | 47.9388 | 48.55807 | 21.22095 | 0.362602 | 1.12905 | 0.899997 |
| 19 | 42415600 | 42423799 | 45.17634 | 72.02349 | 121.4675 | 50.88315 | -0.41454 | 2.15182 | 0.900018 |
| X | 62973200 | 62978399 | 60.48874 | 96.85788 | 113.1328 | 47.6857 | 0.121389 | 2.035274 | 0.90004 |
| 1 | 54940600 | 54943799 | 31.82822 | 52.03687 | 70.36811 | 30.2573 | -0.05965 | 1.485312 | 0.900056 |
| 18 | 71277000 | 71279199 | 20.37721 | 35.40624 | 44.12398 | 19.37356 | 0.070654 | 0.805772 | 0.900067 |
| 9 | 13664200 | 13671599 | 142.3458 | 214.0564 | 266.379 | 109.9853 | -0.04565 | 3.417231 | 0.900078 |
| 10 | 82045800 | 82078999 | 200.3977 | 299.2383 | 255.3945 | 105.2832 | 0.488946 | 3.560751 | 0.900083 |
| 12 | 26549000 | 26550399 | 13.38101 | 24.12883 | 33.61566 | 14.63746 | -0.0687 | 0.380021 | 0.900114 |
| Y | 82270200 | 82270799 | 5.815092 | 11.29401 | 19.02452 | 7.116923 | -0.15435 | -0.28174 | 0.900166 |
| 6 | 89804600 | 89815199 | 197.1532 | 289.9877 | 354.6902 | 146.0659 | -0.05005 | 3.819655 | 0.900207 |
| 8 | 53718000 | 53720799 | 71.86488 | 112.6164 | 146.4736 | 60.75429 | -0.06187 | 2.599726 | 0.900223 |
| 8 | 8341200 | 8342199 | 11.90079 | 22.38553 | 23.09198 | 9.471009 | 0.424678 | 0.098381 | 0.900229 |
| 13 | 67523200 | 67525199 | 17.15032 | 30.05864 | 37.65744 | 16.63537 | 0.063909 | 0.652991 | 0.900252 |
| 2 | 120052000 | 120055999 | 58.24199 | 92.63109 | 112.5432 | 46.98843 | 0.054065 | 2.240131 | 0.90029 |
| 7 | 7531200 | 7531799 | 18.43755 | 31.87333 | 43.75437 | 19.19562 | -0.06935 | 0.850616 | 0.900292 |
| 11 | 90475800 | 90477999 | 30.95955 | 50.98817 | 63.86892 | 27.54589 | 0.055223 | 1.387059 | 0.90031 |
| X | 156308800 | 156311399 | 23.64065 | 40.56334 | 50.10719 | 21.97686 | 0.079898 | 0.94602 | 0.900315 |
| 2 | 148819600 | 148823199 | 44.14103 | 71.47586 | 72.00595 | 30.80919 | 0.35302 | 1.670976 | 0.900373 |
| 9 | 109337400 | 109346999 | 173.5388 | 258.0746 | 319.0313 | 131.4404 | -0.05061 | 3.630692 | 0.90038 |
| 16 | 72801400 | 72801799 | 7.276262 | 14.11933 | 20.01518 | 7.533107 | 0.083758 | -0.23828 | 0.900384 |
| 10 | 70218200 | 70221599 | 33.01191 | 53.82006 | 72.50164 | 31.14264 | -0.05701 | 1.530157 | 0.900392 |
| 8 | 104948200 | 104952799 | 63.15953 | 100.2314 | 122.0891 | 50.85985 | 0.047261 | 2.310283 | 0.900426 |
| 8 | 39495800 | 39497399 | 13.7003 | 25.0811 | 31.66422 | 13.52773 | 0.090223 | 0.289929 | 0.900431 |
| 9 | 108042400 | 108044199 | 16.91567 | 29.57044 | 40.40733 | 17.811 | -0.06069 | 0.699108 | 0.900431 |
| 3 | 131099400 | 131104599 | 96.85706 | 149.7562 | 178.7805 | 74.26126 | 0.046599 | 2.850479 | 0.900433 |
| 9 | 110823800 | 110824999 | 12.42906 | 22.22763 | 31.20675 | 13.69513 | -0.08713 | 0.419637 | 0.900438 |
| 2 | 146767800 | 146769599 | 24.88452 | 42.94127 | 40.84891 | 17.91998 | 0.456854 | 0.888281 | 0.900442 |
| 9 | 57303000 | 57310399 | 82.57367 | 129.2701 | 167.8725 | 69.63717 | -0.06078 | 2.625484 | 0.900463 |
| 16 | 96216200 | 96218399 | 16.43108 | 28.76703 | 39.3357 | 17.36093 | -0.06165 | 0.670351 | 0.900471 |
| 12 | 64436600 | 64438199 | 17.6897 | 30.91919 | 38.82644 | 17.11214 | 0.060668 | 0.702853 | 0.900479 |
| 8 | 43814400 | 43817199 | 25.57123 | 43.01363 | 58.95006 | 25.75699 | -0.07223 | 1.133816 | 0.900482 |
| 18 | 30357600 | 30357999 | 5.945343 | 11.58691 | 19.13064 | 6.930252 | -0.08288 | -0.39116 | 0.900504 |
| 1 | 75750400 | 75753799 | 25.79342 | 43.14679 | 53.91358 | 23.48425 | 0.060773 | 1.187898 | 0.90052 |
| 13 | 57109000 | 57116399 | 119.1909 | 181.6497 | 229.176 | 95.00437 | -0.04913 | 3.142058 | 0.900546 |
| 9 | 120340800 | 120346799 | 55.99419 | 89.31829 | 109.136 | 45.75159 | 0.05041 | 2.141481 | 0.900604 |
| 5 | 93353400 | 93354799 | 14.0151 | 24.4471 | 44.21046 | 19.45924 | -0.46305 | 0.723608 | 0.900609 |
| 15 | 3147200 | 3161599 | 252.1009 | 365.8575 | 350.1253 | 144.3476 | 0.285129 | 4.029689 | 0.90061 |

| | | | | | | | | | |
|---|---|---|---|---|---|---|---|---|---|
| 7 | 29795800 | 29798999 | 39.92086 | 64.43855 | 79.35641 | 33.83917 | 0.059098 | 1.747361 | 0.900613 |
| 5 | 119291200 | 119292399 | 16.14485 | 28.33655 | 38.73888 | 17.13363 | -0.06134 | 0.628283 | 0.90063 |
| 11 | 14214600 | 14217399 | 50.4075 | 80.97531 | 82.46704 | 34.93027 | 0.32424 | 1.911348 | 0.90063 |
| 3 | 114969800 | 114970799 | 15.85915 | 27.94727 | 34.99111 | 15.48738 | 0.066168 | 0.567025 | 0.900637 |
| 17 | 34988600 | 34993199 | 55.77145 | 88.41163 | 138.3217 | 57.55996 | -0.31482 | 2.340784 | 0.900642 |
| 4 | 114573000 | 114575599 | 31.03041 | 50.89188 | 68.70378 | 29.5788 | -0.05487 | 1.442409 | 0.900649 |
| 13 | 71179600 | 71180399 | 8.783461 | 16.77002 | 25.05966 | 9.970064 | -0.06801 | -0.02662 | 0.900668 |
| 12 | 54674800 | 54678399 | 46.48084 | 74.62999 | 91.74175 | 38.82929 | 0.052416 | 1.924379 | 0.900673 |
| 14 | 12339800 | 12340999 | 19.40211 | 33.94546 | 34.08659 | 15.03556 | 0.381699 | 0.688652 | 0.900689 |
| 13 | 79098600 | 79103199 | 46.82172 | 75.21542 | 92.64041 | 39.23964 | 0.050498 | 1.909662 | 0.900702 |
| 16 | 55547800 | 55548599 | 11.65816 | 21.74221 | 28.33353 | 11.79048 | 0.072992 | 0.162661 | 0.900716 |
| 7 | 42341800 | 42342799 | 15.96402 | 27.94402 | 38.37105 | 16.93247 | -0.06759 | 0.669865 | 0.900725 |
| 12 | 71363800 | 71366399 | 63.08269 | 98.90143 | 162.0457 | 67.20995 | -0.39401 | 2.590027 | 0.90073 |
| 5 | 122718400 | 122719599 | 24.99703 | 41.91775 | 52.26995 | 22.80371 | 0.063724 | 1.158292 | 0.90077 |
| 12 | 108538800 | 108542999 | 41.5448 | 67.30391 | 67.55833 | 28.91633 | 0.358887 | 1.65541 | 0.900777 |
| 12 | 81408600 | 81410199 | 15.51144 | 27.98214 | 34.5001 | 15.01473 | 0.106761 | 0.391683 | 0.900797 |
| 4 | 9759000 | 9759999 | 9.107543 | 17.26199 | 25.45608 | 10.28576 | -0.06867 | 0.028007 | 0.900809 |
| X | 11152400 | 11155999 | 28.0402 | 47.24733 | 57.87074 | 25.2985 | 0.089511 | 1.129568 | 0.900818 |
| 2 | 71776400 | 71780399 | 101.6379 | 155.6649 | 198.1382 | 82.2318 | -0.05626 | 3.034636 | 0.900867 |
| 14 | 45108200 | 45110399 | 19.37468 | 33.68749 | 46.29069 | 20.33371 | -0.06953 | 0.795937 | 0.900869 |
| 11 | 22019600 | 22020999 | 13.48655 | 24.39432 | 31.03977 | 13.46376 | 0.06496 | 0.357512 | 0.900872 |
| 19 | 16244200 | 16244999 | 8.676408 | 16.39236 | 24.40465 | 9.927811 | -0.0905 | 0.044276 | 0.900879 |
| 7 | 28747400 | 28748799 | 18.20755 | 31.67579 | 39.52761 | 17.36902 | 0.069162 | 0.773982 | 0.900889 |
| 8 | 110949400 | 110949999 | 10.80271 | 19.65084 | 28.30721 | 12.19178 | -0.10595 | 0.299948 | 0.900899 |
| 3 | 130064600 | 130068799 | 47.34421 | 76.37759 | 92.16366 | 39.27165 | 0.087413 | 1.781219 | 0.900912 |
| 8 | 46421400 | 46426199 | 46.99243 | 75.54162 | 93.00603 | 39.43383 | 0.052281 | 1.888588 | 0.900918 |
| 3 | 70625600 | 70626599 | 11.63215 | 20.62196 | 37.83767 | 16.72718 | -0.47824 | 0.485249 | 0.900965 |
| 18 | 27374000 | 27375199 | 17.94816 | 31.57147 | 39.06584 | 17.29714 | 0.082115 | 0.607396 | 0.900971 |
| 15 | 22055200 | 22059399 | 59.51294 | 95.14337 | 95.42214 | 40.12893 | 0.338298 | 2.072688 | 0.900976 |
| 8 | 62919000 | 62919599 | 7.551712 | 14.59461 | 22.63514 | 8.681909 | -0.07154 | -0.16571 | 0.900981 |
| 2 | 92469200 | 92471999 | 29.49569 | 48.47708 | 65.91954 | 28.40547 | -0.06409 | 1.409404 | 0.901009 |
| 19 | 48955000 | 48956999 | 15.05315 | 26.28996 | 45.18873 | 19.90513 | -0.39024 | 0.710504 | 0.901013 |
| 6 | 19209200 | 19210599 | 12.45739 | 22.56824 | 31.58364 | 13.67276 | -0.06988 | 0.340444 | 0.901014 |
| 2 | 163872800 | 163875999 | 29.56678 | 48.6636 | 65.86075 | 28.39735 | -0.05641 | 1.394006 | 0.901024 |
| 4 | 80499600 | 80500399 | 13.193 | 23.47758 | 32.59442 | 14.36382 | -0.07541 | 0.457338 | 0.901044 |
| 18 | 80612200 | 80614199 | 20.4702 | 35.26627 | 44.23703 | 19.40535 | 0.059083 | 0.889498 | 0.90106 |
| 12 | 49123000 | 49124799 | 18.78153 | 32.7725 | 40.9473 | 18.0268 | 0.06777 | 0.737464 | 0.901073 |
| 3 | 5307800 | 5309399 | 13.99163 | 24.88517 | 31.08479 | 13.70485 | 0.075792 | 0.463812 | 0.901076 |
| 5 | 31487200 | 31499399 | 209.5094 | 306.1514 | 370.9665 | 152.8313 | -0.04554 | 3.92253 | 0.901077 |
| 6 | 107325400 | 107326599 | 15.62256 | 27.46555 | 34.18437 | 15.13744 | 0.073597 | 0.591962 | 0.90108 |
| 4 | 74429000 | 74432999 | 42.08695 | 67.72232 | 89.89119 | 38.16737 | -0.05422 | 1.849826 | 0.901081 |
| 10 | 4545400 | 4549799 | 37.59542 | 60.83539 | 81.59708 | 34.89543 | -0.05991 | 1.676206 | 0.901083 |
| 17 | 67830400 | 67834799 | 51.09475 | 81.70034 | 107.3845 | 45.17148 | -0.05087 | 2.06192 | 0.901107 |
| 6 | 35695000 | 35701999 | 159.0723 | 237.5356 | 293.7638 | 121.101 | -0.04578 | 3.554367 | 0.901108 |
| 1 | 110288400 | 110289199 | 11.45233 | 21.28076 | 27.72713 | 11.60776 | 0.066311 | 0.181661 | 0.901112 |
| 16 | 48126800 | 48129999 | 25.9853 | 43.32818 | 58.82762 | 25.57281 | -0.05861 | 1.226656 | 0.901153 |
| 17 | 63736800 | 63737799 | 7.878746 | 15.01088 | 22.93594 | 9.166066 | -0.10517 | -0.02662 | 0.901161 |

| | | | | | | | | | |
|---|---|---|---|---|---|---|---|---|---|
| 14 | 122815200 | 122815599 | 9.774062 | 18.59435 | 24.94488 | 10.00018 | 0.076317 | 0.005338 | 0.901186 |
| 7 | 79995000 | 79996799 | 15.43062 | 26.74325 | 47.40552 | 20.79905 | -0.43578 | 0.818394 | 0.901206 |
| 2 | 9031600 | 9033399 | 37.38132 | 61.03135 | 62.2424 | 26.74987 | 0.344949 | 1.513636 | 0.901211 |
| 3 | 87008400 | 87010999 | 23.52431 | 39.49929 | 53.84757 | 23.5112 | -0.06453 | 1.140384 | 0.90124 |
| 7 | 47003200 | 47013399 | 166.668 | 248.4945 | 308.0021 | 126.9325 | -0.05101 | 3.585022 | 0.901243 |
| 9 | 15549600 | 15552199 | 22.79547 | 39.36033 | 48.47645 | 21.2665 | 0.085964 | 0.884549 | 0.901252 |
| 11 | 58745600 | 58756199 | 152.3337 | 228.8972 | 266.869 | 110.1646 | 0.047346 | 3.434242 | 0.90126 |
| 7 | 48438400 | 48439399 | 9.674601 | 18.0734 | 26.37603 | 10.94457 | -0.08529 | 0.132599 | 0.901263 |
| 12 | 9693200 | 9693799 | 7.529131 | 14.53601 | 22.59062 | 8.706556 | -0.08132 | -0.151 | 0.901324 |
| Y | 6724800 | 6727599 | 26.42362 | 45.09129 | 27.05256 | 11.85995 | 1.116088 | 0.972659 | 0.901326 |
| 10 | 117089800 | 117096399 | 78.86962 | 124.0309 | 149.1354 | 61.86819 | 0.055199 | 2.537213 | 0.901338 |
| 7 | 56569400 | 56574599 | 88.10872 | 134.9727 | 216.5656 | 89.88398 | -0.38977 | 3.028495 | 0.901359 |
| 14 | 13059200 | 13060399 | 13.71966 | 24.58254 | 30.93486 | 13.554 | 0.071087 | 0.413413 | 0.901359 |
| 3 | 70745000 | 70747999 | 36.4608 | 59.29439 | 73.85126 | 31.69345 | 0.053293 | 1.574309 | 0.901361 |
| 3 | 70745200 | 70748199 | 36.4608 | 59.29439 | 73.85126 | 31.69345 | 0.053293 | 1.574309 | 0.901361 |
| 13 | 13540800 | 13543199 | 18.39522 | 32.00142 | 39.95114 | 17.55483 | 0.068351 | 0.773269 | 0.901372 |
| 3 | 131411200 | 131414199 | 25.60018 | 43.29934 | 53.94366 | 23.60881 | 0.065267 | 1.083707 | 0.901374 |
| X | 86974800 | 86976199 | 21.16863 | 35.44575 | 63.86122 | 27.8348 | -0.46523 | 1.176653 | 0.901406 |
| 2 | 109857000 | 109859999 | 27.27978 | 46.74317 | 54.65618 | 23.96938 | 0.158464 | 0.976761 | 0.901409 |
| 10 | 8420600 | 8435399 | 311.3541 | 438.5555 | 514.4376 | 213.0689 | -0.05139 | 4.471277 | 0.901436 |
| 4 | 25281400 | 25281999 | 7.884156 | 15.08219 | 22.97123 | 9.095498 | -0.08849 | -0.05595 | 0.901452 |
| 3 | 95861800 | 95864199 | 26.01068 | 43.36698 | 58.80434 | 25.56137 | -0.05673 | 1.228177 | 0.901454 |
| 12 | 44275400 | 44278599 | 33.32239 | 53.99276 | 87.2518 | 37.35034 | -0.32625 | 1.646956 | 0.901467 |
| 12 | 41418800 | 41422399 | 34.32121 | 56.19036 | 70.11625 | 30.16451 | 0.057335 | 1.4729 | 0.901493 |
| 17 | 83111000 | 83111599 | 9.653453 | 18.11297 | 26.39286 | 10.87072 | -0.0746 | 0.104008 | 0.901506 |
| X | 80578800 | 80582399 | 34.04194 | 56.63984 | 67.1948 | 29.16978 | 0.135466 | 1.283894 | 0.901519 |
| 13 | 9243800 | 9246599 | 20.60438 | 35.7703 | 44.68841 | 19.61082 | 0.067022 | 0.820575 | 0.901523 |
| 17 | 23967200 | 23969199 | 14.28186 | 25.31177 | 31.54312 | 13.94535 | 0.076228 | 0.489482 | 0.901565 |
| 2 | 149239000 | 149240999 | 28.61837 | 47.79557 | 59.55524 | 25.91545 | 0.064785 | 1.218775 | 0.901571 |
| 9 | 12739600 | 12740799 | 21.57497 | 36.49526 | 50.09041 | 21.92315 | -0.07379 | 1.065668 | 0.901573 |
| Y | 6543000 | 6543999 | 13.40063 | 23.74924 | 27.78628 | 12.28451 | 0.166718 | 0.474833 | 0.901603 |
| 18 | 24977600 | 24977999 | 6.665067 | 12.9819 | 19.07742 | 6.964728 | 0.07243 | -0.35696 | 0.901603 |
| 3 | 117172000 | 117173599 | 14.73715 | 26.29544 | 36.60356 | 16.09438 | -0.07705 | 0.459881 | 0.901613 |
| 17 | 50719600 | 50726399 | 83.87283 | 130.5286 | 167.4129 | 69.49573 | -0.0495 | 2.728008 | 0.901619 |
| 3 | 56050200 | 56052599 | 27.8336 | 47.06261 | 57.34911 | 25.09692 | 0.097057 | 1.098186 | 0.90163 |
| 11 | 90026400 | 90026799 | 7.219947 | 14.0223 | 20.0242 | 7.491769 | 0.080298 | -0.26301 | 0.901684 |
| 8 | 81134800 | 81135399 | 9.225781 | 17.4994 | 23.33712 | 9.414421 | 0.077424 | 0.018125 | 0.901688 |
| 6 | 146624200 | 146627199 | 24.55161 | 41.39688 | 51.75022 | 22.62749 | 0.060181 | 1.103474 | 0.90169 |
| 5 | 102535600 | 102537999 | 21.8544 | 36.16639 | 66.93111 | 28.94736 | -0.5045 | 1.310265 | 0.901694 |
| 2 | 150964000 | 150967399 | 44.57482 | 71.83762 | 88.54878 | 37.68241 | 0.055153 | 1.805185 | 0.901711 |
| 9 | 78836800 | 78844999 | 88.60258 | 138.3554 | 165.0601 | 68.48846 | 0.058596 | 2.673264 | 0.901717 |
| 10 | 4409200 | 4414199 | 55.95586 | 89.06724 | 116.3436 | 48.6296 | -0.04904 | 2.212905 | 0.901758 |
| 17 | 69451200 | 69453799 | 17.70096 | 30.79171 | 42.08436 | 18.49145 | -0.06164 | 0.775323 | 0.901789 |
| 12 | 66373600 | 66376199 | 31.31978 | 51.38174 | 69.41331 | 29.88803 | -0.05573 | 1.438993 | 0.901817 |
| 7 | 132956200 | 132957999 | 18.94764 | 33.0824 | 41.37466 | 18.22189 | 0.066268 | 0.727628 | 0.901818 |
| 4 | 96833000 | 96842799 | 200.3316 | 294.4491 | 336.8683 | 138.733 | 0.045301 | 3.829933 | 0.901842 |
| 12 | 12009600 | 12010199 | 7.277857 | 14.11652 | 22.22871 | 8.382569 | -0.06959 | -0.22571 | 0.901845 |

| | | | | | | | | | |
|---|---|---|---|---|---|---|---|---|---|
| 13 | 33123200 | 33124999 | 17.9648 | 31.35587 | 39.35579 | 17.32911 | 0.061147 | 0.724547 | 0.901846 |
| 13 | 22381600 | 22385799 | 58.51457 | 93.39751 | 113.3733 | 47.57007 | 0.060848 | 2.126269 | 0.901855 |
| 18 | 47183200 | 47185199 | 15.21683 | 26.8411 | 37.03424 | 16.38855 | -0.07281 | 0.555483 | 0.901864 |
| 4 | 117605800 | 117608599 | 39.20204 | 63.38378 | 78.24674 | 33.43778 | 0.058412 | 1.699623 | 0.901869 |
| 17 | 13370600 | 13374399 | 49.30642 | 78.86415 | 95.69098 | 40.26944 | 0.064793 | 2.048159 | 0.901894 |
| X | 117905600 | 117909199 | 43.58684 | 70.4848 | 85.46704 | 36.53094 | 0.084455 | 1.707154 | 0.901902 |
| 2 | 32361800 | 32374599 | 163.1633 | 243.2063 | 300.2946 | 123.7501 | -0.04583 | 3.587942 | 0.901904 |
| 4 | 63721000 | 63726399 | 53.28753 | 85.6128 | 102.3878 | 43.37609 | 0.093076 | 1.918104 | 0.901916 |
| 3 | 133014400 | 133016599 | 26.47511 | 43.9872 | 59.92104 | 25.98659 | -0.06356 | 1.268192 | 0.901926 |
| 4 | 71928200 | 71929999 | 22.62844 | 38.4716 | 52.24712 | 22.89848 | -0.05833 | 1.026161 | 0.901937 |
| 18 | 37533400 | 37536599 | 69.03775 | 108.9022 | 131.5672 | 54.60182 | 0.051779 | 2.467123 | 0.901943 |
| 9 | 9726600 | 9731399 | 64.71775 | 103.0927 | 123.147 | 51.5278 | 0.080899 | 2.19845 | 0.901961 |
| 12 | 32607600 | 32609399 | 25.65486 | 42.59031 | 58.56913 | 25.38627 | -0.07708 | 1.277997 | 0.901978 |
| 1 | 98516000 | 98522799 | 58.22835 | 93.05177 | 95.34916 | 40.07511 | 0.306755 | 2.083994 | 0.902006 |
| 16 | 96219600 | 96221999 | 30.05327 | 49.70218 | 62.14764 | 26.86646 | 0.059201 | 1.336056 | 0.902007 |
| 2 | 156865400 | 156870399 | 64.95603 | 102.3913 | 134.059 | 55.63085 | -0.06485 | 2.478726 | 0.902022 |
| 16 | 56142200 | 56146199 | 78.05135 | 122.338 | 147.6725 | 61.25015 | 0.045919 | 2.598839 | 0.902084 |
| 3 | 84614000 | 84616799 | 28.07689 | 47.10052 | 47.52693 | 20.73952 | 0.368013 | 1.157654 | 0.902084 |
| 10 | 6580400 | 6580999 | 10.95488 | 20.21754 | 28.79782 | 12.1688 | -0.07074 | 0.220658 | 0.902105 |
| 13 | 96572800 | 96575199 | 17.89904 | 31.29667 | 42.97186 | 18.9451 | -0.06768 | 0.704049 | 0.902127 |
| 12 | 25302600 | 25303999 | 20.81248 | 35.46655 | 48.4923 | 21.25682 | -0.06746 | 0.993742 | 0.902137 |
| 11 | 89409000 | 89410799 | 16.75241 | 29.38796 | 40.25216 | 17.79527 | -0.06416 | 0.641208 | 0.902144 |
| 13 | 69071000 | 69079999 | 79.27954 | 124.2302 | 160.1844 | 66.44153 | -0.04946 | 2.606616 | 0.902155 |
| 2 | 57172000 | 57174999 | 34.37009 | 56.09275 | 75.60922 | 32.48982 | -0.05684 | 1.521201 | 0.902168 |
| 5 | 137993800 | 137996999 | 59.00525 | 93.64265 | 122.5432 | 51.05113 | -0.05595 | 2.307341 | 0.902169 |
| 7 | 48120600 | 48122199 | 13.87577 | 24.8947 | 31.41332 | 13.74851 | 0.068225 | 0.407245 | 0.902222 |
| 10 | 66824000 | 66828799 | 49.60831 | 79.00959 | 124.2726 | 51.98566 | -0.31568 | 2.190281 | 0.902231 |
| 9 | 71103000 | 71104599 | 11.94924 | 21.07749 | 38.90219 | 17.21514 | -0.48969 | 0.544189 | 0.902239 |
| 1 | 31259800 | 31262799 | 28.03821 | 46.75437 | 58.44112 | 25.40587 | 0.060272 | 1.230875 | 0.902248 |
| 7 | 108093600 | 108097399 | 50.76071 | 81.25581 | 99.7595 | 42.02961 | 0.049098 | 2.026005 | 0.902259 |
| 4 | 92958400 | 92960199 | 20.09415 | 35.00573 | 43.53155 | 19.14485 | 0.074111 | 0.767547 | 0.90227 |
| 3 | 54777600 | 54783999 | 84.25515 | 131.5552 | 158.2955 | 65.68222 | 0.046174 | 2.670883 | 0.902302 |
| 9 | 34856800 | 34858999 | 18.99468 | 32.79082 | 44.76517 | 19.63824 | -0.06196 | 0.864807 | 0.902348 |
| 5 | 59070200 | 59071799 | 20.72194 | 36.07639 | 44.63283 | 19.62132 | 0.0815 | 0.782819 | 0.902376 |
| 6 | 31883000 | 31886599 | 43.70061 | 70.34688 | 86.81902 | 36.88423 | 0.051297 | 1.838461 | 0.902385 |
| 11 | 14390000 | 14395599 | 78.69513 | 122.9244 | 157.8145 | 65.47988 | -0.04675 | 2.661357 | 0.902393 |
| 6 | 48781200 | 48782399 | 14.99964 | 26.29717 | 49.40217 | 21.82925 | -0.51778 | 0.657643 | 0.902402 |
| 15 | 72350400 | 72352399 | 16.42558 | 28.77401 | 39.30416 | 17.35548 | -0.06017 | 0.6608 | 0.902406 |
| 16 | 34398400 | 34403999 | 59.77706 | 95.13888 | 115.9701 | 48.46594 | 0.050408 | 2.220587 | 0.902458 |
| 2 | 20494000 | 20501199 | 74.21133 | 117.0157 | 141.2922 | 58.63472 | 0.052086 | 2.478844 | 0.902462 |
| 19 | 33619200 | 33621199 | 26.23509 | 44.38549 | 44.80705 | 19.61176 | 0.367673 | 1.073057 | 0.902481 |
| 13 | 60054600 | 60058199 | 43.55273 | 70.46876 | 71.38988 | 30.52818 | 0.343906 | 1.683534 | 0.90251 |
| 5 | 75845200 | 75846999 | 16.78873 | 29.65374 | 36.98325 | 16.36157 | 0.073106 | 0.552586 | 0.902519 |
| 6 | 81997400 | 81999999 | 21.89837 | 37.25729 | 50.61078 | 22.1826 | -0.05842 | 1.017032 | 0.902527 |
| 12 | 64480200 | 64482399 | 15.61617 | 27.60294 | 38.05646 | 16.83389 | -0.07048 | 0.538406 | 0.902542 |
| 12 | 115322600 | 115329599 | 45.95147 | 74.40485 | 72.27619 | 30.92402 | 0.405386 | 1.675035 | 0.902552 |
| 15 | 90958800 | 90960999 | 17.63027 | 30.86989 | 38.70039 | 17.08211 | 0.062913 | 0.67263 | 0.902556 |

| | | | | | | | | | |
|---|---|---|---|---|---|---|---|---|---|
| 6 | 61477200 | 61479399 | 28.54258 | 47.41881 | 64.48179 | 27.96312 | -0.06103 | 1.280664 | 0.902578 |
| 4 | 116597400 | 116600199 | 28.73585 | 47.47512 | 64.27161 | 27.76515 | -0.05521 | 1.34526 | 0.902583 |
| 2 | 66062600 | 66063599 | 10.22086 | 19.23425 | 28.01414 | 11.48101 | -0.06866 | 0.092317 | 0.902597 |
| 2 | 176939000 | 176941399 | 29.19658 | 49.1281 | 48.56084 | 21.21976 | 0.397677 | 1.133184 | 0.902597 |
| 2 | 143648000 | 143648999 | 10.02065 | 18.96723 | 25.39855 | 10.29163 | 0.065738 | 0.045316 | 0.902605 |
| 17 | 63931000 | 63931999 | 13.11954 | 23.66431 | 30.02045 | 13.06716 | 0.065997 | 0.368096 | 0.90261 |
| 10 | 102159400 | 102161399 | 26.54111 | 44.98065 | 45.32949 | 19.8508 | 0.370321 | 1.062013 | 0.902611 |
| 16 | 50807800 | 50808799 | 18.70451 | 32.60158 | 40.96066 | 18.02001 | 0.059572 | 0.75164 | 0.902629 |
| X | 49148200 | 49150399 | 22.20467 | 38.46984 | 47.39032 | 20.79215 | 0.086684 | 0.851892 | 0.902633 |
| 12 | 107291200 | 107292799 | 15.65264 | 27.54315 | 37.64726 | 16.66787 | -0.06047 | 0.587978 | 0.902665 |
| 14 | 31568800 | 31569399 | 7.219771 | 13.95895 | 22.08997 | 8.456199 | -0.09809 | -0.17991 | 0.902691 |
| 14 | 61719200 | 61721799 | 43.58836 | 70.63348 | 70.97427 | 30.38443 | 0.357702 | 1.653752 | 0.902725 |
| 11 | 88172600 | 88174999 | 23.47281 | 39.27511 | 66.51867 | 28.97965 | -0.37663 | 1.189297 | 0.902734 |
| 2 | 58972400 | 58973399 | 10.97972 | 20.44718 | 29.39986 | 12.26437 | -0.07096 | 0.167463 | 0.90274 |
| 9 | 22930800 | 22933199 | 30.96483 | 51.23011 | 64.04784 | 27.69553 | 0.059412 | 1.333681 | 0.902741 |
| 14 | 111104000 | 111106999 | 52.44078 | 84.06746 | 84.66849 | 35.75026 | 0.336731 | 1.977234 | 0.902788 |
| 6 | 122969800 | 122973399 | 43.75934 | 71.43306 | 64.11699 | 27.55389 | 0.528512 | 1.529864 | 0.902796 |
| 13 | 113903600 | 113906399 | 27.50343 | 46.02006 | 57.34487 | 24.98326 | 0.064643 | 1.191472 | 0.902799 |
| 5 | 145712000 | 145719399 | 128.2642 | 194.7835 | 246.0787 | 101.9028 | -0.05528 | 3.218286 | 0.902818 |
| 10 | 82292200 | 82296799 | 57.67606 | 92.02483 | 112.2481 | 47.07307 | 0.053304 | 2.136294 | 0.902821 |
| 9 | 41002800 | 41003999 | 12.54389 | 22.71687 | 31.63526 | 13.70118 | -0.063 | 0.345126 | 0.90284 |
| X | 14437400 | 14438599 | 8.925856 | 16.95373 | 25.52706 | 10.26804 | -0.09197 | 0.01571 | 0.902848 |
| 16 | 75968200 | 75968999 | 16.54144 | 29.54163 | 29.10696 | 12.78838 | 0.419293 | 0.448995 | 0.902853 |
| 8 | 14853200 | 14857399 | 66.67131 | 105.6011 | 128.4921 | 53.43548 | 0.046213 | 2.368411 | 0.902873 |
| 19 | 21778400 | 21782199 | 33.74996 | 55.24346 | 69.01148 | 29.68664 | 0.055572 | 1.471968 | 0.902875 |
| 13 | 87429400 | 87434799 | 53.95465 | 86.56243 | 104.2872 | 44.08676 | 0.080075 | 1.9629 | 0.902906 |
| 15 | 54820400 | 54822599 | 22.67954 | 38.93497 | 48.47153 | 21.26037 | 0.068762 | 0.933566 | 0.902909 |
| 1 | 170011400 | 170014399 | 20.41689 | 35.12036 | 47.93574 | 21.02687 | -0.06285 | 0.90575 | 0.902909 |
| 2 | 89962400 | 89962999 | 12.67016 | 23.10488 | 29.61256 | 12.71464 | 0.064128 | 0.30208 | 0.902926 |
| 14 | 50266600 | 50269799 | 46.28584 | 74.63055 | 76.5867 | 32.61185 | 0.319789 | 1.788422 | 0.902936 |
| 12 | 44698000 | 44700199 | 13.65975 | 24.07213 | 41.36443 | 18.30816 | -0.38805 | 0.567413 | 0.902946 |
| 13 | 84569000 | 84572799 | 51.93578 | 83.30526 | 84.26407 | 35.61028 | 0.331471 | 1.958677 | 0.902963 |
| 1 | 26644000 | 26645199 | 12.04164 | 21.28235 | 38.79959 | 17.16298 | -0.47123 | 0.519704 | 0.902963 |
| 3 | 66175000 | 66178199 | 43.71641 | 70.72776 | 71.6557 | 30.63999 | 0.343778 | 1.688301 | 0.902969 |
| 11 | 32826800 | 32833999 | 66.59769 | 105.5123 | 137.8514 | 57.33533 | -0.05601 | 2.362782 | 0.902984 |
| 2 | 135505800 | 135506999 | 16.51607 | 29.18541 | 36.40762 | 16.10544 | 0.072898 | 0.543292 | 0.902991 |
| 15 | 100266000 | 100267599 | 22.65195 | 38.65925 | 48.48659 | 21.25689 | 0.056949 | 0.988728 | 0.902996 |
| 3 | 20334800 | 20334999 | 6.226437 | 12.14056 | 18.03695 | 6.493079 | 0.075234 | -0.41612 | 0.903059 |
| 10 | 34861800 | 34862599 | 9.385424 | 17.97567 | 24.58795 | 9.702097 | 0.069428 | -0.0488 | 0.903065 |
| 14 | 115621400 | 115625399 | 44.32898 | 71.06232 | 112.8921 | 47.66615 | -0.31958 | 1.979052 | 0.903076 |
| 2 | 142627800 | 142631199 | 38.43181 | 62.13401 | 76.68787 | 32.76861 | 0.058558 | 1.697006 | 0.903084 |
| 17 | 31992600 | 31993599 | 13.95239 | 24.81934 | 31.03993 | 13.68324 | 0.074105 | 0.461598 | 0.903103 |
| 2 | 18333400 | 18335799 | 48.14575 | 77.60568 | 78.49481 | 33.41116 | 0.340139 | 1.801635 | 0.903105 |
| 1 | 185512400 | 185513999 | 12.52652 | 23.15777 | 29.73361 | 12.5349 | 0.079844 | 0.222165 | 0.903114 |
| 16 | 93771600 | 93773399 | 24.22391 | 40.96823 | 51.17249 | 22.39544 | 0.061647 | 1.078957 | 0.903138 |
| 3 | 35930800 | 35932599 | 16.89169 | 29.79839 | 41.76439 | 18.48138 | -0.09516 | 0.575422 | 0.903142 |
| 1 | 133718000 | 133719199 | 13.68589 | 24.36675 | 33.55541 | 14.78114 | -0.06338 | 0.459698 | 0.903174 |

| | | | | | | | | | |
|---|---|---|---|---|---|---|---|---|---|
| 2 | 130318800 | 130323999 | 51.40023 | 82.32535 | 109.1875 | 46.04015 | -0.06119 | 2.014255 | 0.903213 |
| 18 | 82295800 | 82297999 | 24.53915 | 41.36096 | 51.71617 | 22.61013 | 0.059978 | 1.109453 | 0.903214 |
| 1 | 39083800 | 39089999 | 69.35859 | 109.0017 | 168.7825 | 70.01266 | -0.30998 | 2.547309 | 0.903215 |
| 13 | 65425600 | 65430999 | 96.58695 | 149.7181 | 178.3333 | 74.07298 | 0.052642 | 2.803997 | 0.903215 |
| 15 | 66889400 | 66892999 | 23.57381 | 38.9728 | 69.49431 | 30.03519 | -0.45136 | 1.326048 | 0.903232 |
| 3 | 5464000 | 5469199 | 59.92699 | 95.23906 | 115.7364 | 48.28326 | 0.05266 | 2.262807 | 0.903243 |
| 6 | 41162800 | 41163799 | 12.55555 | 22.81742 | 29.01371 | 12.51111 | 0.071234 | 0.317443 | 0.903248 |
| 5 | 14596400 | 14598599 | 18.0497 | 31.86599 | 38.9847 | 17.24864 | 0.100351 | 0.567332 | 0.903249 |
| 1 | 8730600 | 8733199 | 45.22742 | 72.55828 | 87.77372 | 37.10379 | 0.074199 | 1.94784 | 0.903249 |
| 17 | 75728800 | 75730199 | 14.96762 | 26.28316 | 36.23801 | 16.03414 | -0.0735 | 0.608668 | 0.903252 |
| 14 | 113184800 | 113185599 | 14.22298 | 25.36775 | 31.92182 | 14.04063 | 0.067316 | 0.447541 | 0.903262 |
| 19 | 38333800 | 38337999 | 46.45654 | 74.71134 | 92.00784 | 39.0342 | 0.052641 | 1.873737 | 0.903267 |
| 8 | 49719800 | 49721199 | 11.79261 | 21.7434 | 28.16383 | 11.91594 | 0.06379 | 0.228961 | 0.903297 |
| 7 | 52089200 | 52090999 | 30.06246 | 49.78901 | 62.18606 | 26.90981 | 0.060978 | 1.319786 | 0.903307 |
| 13 | 65873800 | 65875199 | 50.41584 | 80.55594 | 96.57411 | 40.58206 | 0.080546 | 2.083286 | 0.903312 |
| 12 | 17922200 | 17925199 | 62.23291 | 98.60202 | 118.6296 | 49.34182 | 0.062856 | 2.356 | 0.903328 |
| 6 | 49324400 | 49327599 | 32.44653 | 53.24313 | 66.67841 | 28.71338 | 0.053457 | 1.437079 | 0.903352 |
| 1 | 89431000 | 89434399 | 39.27237 | 63.42952 | 78.22901 | 33.38406 | 0.05797 | 1.728641 | 0.903359 |
| 7 | 35357400 | 35361799 | 44.22697 | 70.99181 | 94.02482 | 39.77911 | -0.05539 | 1.930983 | 0.90336 |
| 8 | 104261200 | 104262399 | 11.02369 | 20.25671 | 28.70907 | 12.20291 | -0.06998 | 0.243017 | 0.90339 |
| 6 | 20728400 | 20729799 | 11.95821 | 21.78046 | 30.62175 | 13.16768 | -0.0702 | 0.303712 | 0.903403 |
| 6 | 20818800 | 20820599 | 15.98409 | 28.16265 | 38.63054 | 17.09881 | -0.06489 | 0.577408 | 0.903421 |
| 16 | 15660800 | 15663999 | 38.84599 | 62.81623 | 77.72099 | 33.21865 | 0.055376 | 1.694404 | 0.903421 |
| 18 | 29845600 | 29846599 | 12.45479 | 22.78424 | 32.30547 | 13.81546 | -0.07686 | 0.28936 | 0.903436 |
| 9 | 59280200 | 59283799 | 61.9906 | 98.30664 | 119.204 | 49.62275 | 0.052794 | 2.32917 | 0.903441 |
| 18 | 33527800 | 33528799 | 19.8197 | 33.21012 | 60.0499 | 26.18507 | -0.47043 | 1.158695 | 0.903463 |
| 17 | 43218600 | 43219199 | 7.247174 | 14.07138 | 22.31009 | 8.365211 | -0.0721 | -0.24734 | 0.903476 |
| 8 | 81362200 | 81365199 | 32.17843 | 53.79117 | 51.92074 | 22.60654 | 0.432016 | 1.213474 | 0.903498 |
| 4 | 25007000 | 25009799 | 30.48343 | 50.126 | 67.75023 | 29.20063 | -0.0551 | 1.407183 | 0.903501 |
| 8 | 21349800 | 21350199 | 6.683102 | 13.01673 | 19.07949 | 6.967575 | 0.076206 | -0.3539 | 0.903515 |
| 12 | 42387800 | 42389599 | 14.5274 | 26.16259 | 32.92322 | 14.35885 | 0.075494 | 0.391649 | 0.903517 |
| 11 | 89444400 | 89446399 | 13.61459 | 24.48481 | 34.1319 | 14.90815 | -0.07234 | 0.399519 | 0.903522 |
| 3 | 104424400 | 104424999 | 6.367087 | 12.4017 | 20.23758 | 7.386776 | -0.07741 | -0.35931 | 0.903539 |
| 11 | 24546400 | 24547399 | 14.78347 | 26.20472 | 32.96973 | 14.57562 | 0.062549 | 0.494004 | 0.903575 |
| 1 | 13880200 | 13890999 | 177.3411 | 263.1113 | 303.8833 | 125.1696 | 0.044285 | 3.671638 | 0.903593 |
| 8 | 28675200 | 28677599 | 31.29204 | 51.41441 | 64.34786 | 27.72479 | 0.055107 | 1.417042 | 0.903693 |
| 7 | 47898600 | 47903199 | 50.17829 | 80.30673 | 105.4586 | 44.41754 | -0.04816 | 2.034285 | 0.903706 |
| 2 | 38356400 | 38357599 | 12.66265 | 22.82707 | 28.63049 | 12.47111 | 0.081821 | 0.365871 | 0.903727 |
| 5 | 75673400 | 75674799 | 16.19367 | 28.43785 | 35.60509 | 15.75491 | 0.064814 | 0.612997 | 0.903738 |
| 16 | 88592400 | 88594599 | 48.07907 | 76.83784 | 101.9005 | 42.83269 | -0.06452 | 2.074876 | 0.903739 |
| 15 | 56782000 | 56784399 | 19.17959 | 33.43445 | 46.23904 | 20.34398 | -0.07844 | 0.757974 | 0.903744 |
| 4 | 3352200 | 3353399 | 17.91519 | 30.94121 | 42.74617 | 18.75278 | -0.07879 | 0.854557 | 0.903763 |
| 7 | 52174600 | 52175999 | 17.13701 | 30.0664 | 37.69877 | 16.6682 | 0.062867 | 0.638987 | 0.903807 |
| X | 36491600 | 36494599 | 19.3106 | 33.6742 | 46.7823 | 20.58737 | -0.08481 | 0.75667 | 0.90383 |
| 9 | 33303000 | 33303399 | 7.012867 | 13.58979 | 21.62185 | 8.189511 | -0.08954 | -0.21306 | 0.903849 |
| 5 | 76820400 | 76823199 | 32.58877 | 53.36762 | 66.61918 | 28.66313 | 0.056921 | 1.461757 | 0.903869 |
| 13 | 86560200 | 86563799 | 65.53442 | 104.1241 | 106.182 | 44.28599 | 0.304904 | 2.273802 | 0.903893 |

| | | | | | | | | | |
|---|---|---|---|---|---|---|---|---|---|
| 8 | 6547200 | 6551399 | 96.18416 | 148.2092 | 175.4101 | 72.86258 | 0.055295 | 2.911394 | 0.90391 |
| 3 | 14513800 | 14521999 | 84.29121 | 133.1343 | 153.2397 | 63.63649 | 0.123004 | 2.45288 | 0.903917 |
| 7 | 108402000 | 108404599 | 29.35463 | 48.45544 | 65.49844 | 28.27823 | -0.05314 | 1.356758 | 0.903923 |
| 5 | 93498000 | 93504599 | 93.71982 | 145.464 | 173.8818 | 72.20987 | 0.048996 | 2.779686 | 0.903958 |
| 3 | 68443600 | 68444799 | 15.37895 | 27.40102 | 34.38056 | 15.13535 | 0.070554 | 0.466442 | 0.903963 |
| 18 | 13545600 | 13547199 | 18.66828 | 32.67698 | 40.77198 | 17.99165 | 0.06974 | 0.687144 | 0.903999 |
| 3 | 51834000 | 51838799 | 44.99734 | 72.27377 | 88.56518 | 37.5051 | 0.057581 | 1.910138 | 0.904009 |
| 15 | 14141600 | 14147999 | 97.33744 | 150.901 | 151.9569 | 63.11598 | 0.294801 | 2.798359 | 0.904026 |
| 6 | 140643000 | 140647199 | 47.83601 | 76.58345 | 100.8511 | 42.49695 | -0.05165 | 2.019337 | 0.904028 |
| 13 | 3581600 | 3589399 | 72.60033 | 114.3967 | 138.5925 | 57.50921 | 0.046515 | 2.491197 | 0.904045 |
| 12 | 25332400 | 25336999 | 31.5452 | 52.28323 | 72.87626 | 31.55184 | -0.09693 | 1.321834 | 0.904077 |
| 18 | 82256400 | 82258199 | 19.37338 | 33.99447 | 33.83587 | 14.96189 | 0.394186 | 0.640705 | 0.904083 |
| 13 | 87850800 | 87852799 | 27.12308 | 45.18452 | 56.66181 | 24.61728 | 0.05571 | 1.238313 | 0.90409 |
| 4 | 86611200 | 86613599 | 21.69267 | 36.83704 | 50.08262 | 21.94555 | -0.06001 | 1.031872 | 0.904155 |
| 16 | 7857200 | 7862999 | 85.00086 | 131.9718 | 168.6744 | 70.04414 | -0.04711 | 2.770249 | 0.904184 |
| 16 | 72392600 | 72393799 | 10.33345 | 19.31758 | 27.86453 | 11.54868 | -0.06795 | 0.13399 | 0.904196 |
| 12 | 118304200 | 118306399 | 19.35907 | 33.85312 | 41.80055 | 18.43034 | 0.084628 | 0.705831 | 0.904224 |
| 12 | 40228600 | 40236399 | 108.7075 | 167.4231 | 169.4245 | 70.37196 | 0.280479 | 2.925423 | 0.904234 |
| 2 | 58821800 | 58823799 | 22.99665 | 39.58936 | 49.01595 | 21.50122 | 0.077548 | 0.912988 | 0.904235 |
| 19 | 3453600 | 3455399 | 21.23347 | 36.31384 | 45.31917 | 19.8716 | 0.064511 | 0.958904 | 0.904251 |
| 2 | 29444000 | 29446199 | 18.44746 | 31.97449 | 43.72796 | 19.18557 | -0.06315 | 0.827261 | 0.904263 |
| 14 | 59475200 | 59481799 | 85.96378 | 135.075 | 159.1144 | 66.00079 | 0.083851 | 2.558876 | 0.904275 |
| 10 | 63553400 | 63554599 | 11.36627 | 21.00231 | 29.80733 | 12.57454 | -0.06414 | 0.217735 | 0.904283 |
| 10 | 103778200 | 103778799 | 5.227199 | 10.19045 | 17.53683 | 6.327314 | -0.13615 | -0.42126 | 0.904287 |
| 5 | 60466400 | 60467599 | 12.8154 | 23.69236 | 23.98018 | 10.1091 | 0.421083 | 0.212543 | 0.9043 |
| 9 | 120736000 | 120738199 | 21.35775 | 36.19967 | 49.56221 | 21.70504 | -0.07018 | 1.045973 | 0.904311 |
| 3 | 114038200 | 114038999 | 9.743806 | 17.94531 | 32.48901 | 13.76724 | -0.41739 | 0.231085 | 0.904336 |
| 7 | 66513200 | 66515199 | 20.14306 | 34.77655 | 43.70249 | 19.1721 | 0.057092 | 0.868415 | 0.904357 |
| 6 | 122927200 | 122929999 | 34.19057 | 55.75364 | 68.8685 | 29.58436 | 0.067931 | 1.525131 | 0.904361 |
| 1 | 12292200 | 12293399 | 15.9371 | 28.45538 | 35.43819 | 15.57325 | 0.083062 | 0.459902 | 0.904395 |
| 9 | 19949800 | 19950599 | 10.78747 | 20.33942 | 26.929 | 11.00059 | 0.071548 | 0.081793 | 0.904411 |
| 7 | 13183400 | 13187599 | 52.79909 | 84.71701 | 102.7455 | 43.44165 | 0.070665 | 1.957928 | 0.904417 |
| 5 | 130509600 | 130510599 | 12.39896 | 22.3756 | 31.13772 | 13.54512 | -0.06599 | 0.363637 | 0.904419 |
| 2 | 140382000 | 140383599 | 19.26844 | 32.64627 | 56.38365 | 24.6896 | -0.40396 | 1.0518 | 0.904474 |
| 15 | 46078800 | 46080799 | 33.30764 | 54.52802 | 68.29412 | 29.37995 | 0.051986 | 1.469602 | 0.904489 |
| 2 | 50567600 | 50571599 | 35.21827 | 57.38602 | 77.2741 | 33.18606 | -0.05699 | 1.546933 | 0.904494 |
| 9 | 105030600 | 105031199 | 11.20748 | 20.67286 | 29.26478 | 12.37496 | -0.06289 | 0.225089 | 0.904505 |
| 3 | 14523800 | 14529799 | 64.192 | 102.409 | 100.2232 | 42.02321 | 0.370338 | 2.145105 | 0.904513 |
| 4 | 23539200 | 23543599 | 59.1915 | 94.23464 | 115.2338 | 48.17782 | 0.046269 | 2.209246 | 0.904543 |
| 10 | 86973200 | 86975199 | 28.63828 | 47.40603 | 59.37649 | 25.68494 | 0.056854 | 1.317522 | 0.904563 |
| 15 | 25381000 | 25383399 | 25.65963 | 43.18226 | 58.90086 | 25.73909 | -0.06532 | 1.131824 | 0.904567 |
| 3 | 16306600 | 16308599 | 27.28841 | 45.4695 | 61.71684 | 26.81705 | -0.05822 | 1.240076 | 0.90463 |
| 5 | 147895600 | 147898199 | 19.45747 | 33.85214 | 42.53482 | 18.69108 | 0.059127 | 0.782053 | 0.904644 |
| X | 152749600 | 152751999 | 23.13443 | 39.34822 | 53.84758 | 23.60071 | -0.06912 | 1.02911 | 0.904667 |
| 10 | 69218000 | 69218599 | 8.943632 | 16.85293 | 24.85672 | 10.16424 | -0.08388 | 0.064102 | 0.904675 |
| 19 | 25939800 | 25940599 | 9.392138 | 17.80614 | 23.84947 | 9.631173 | 0.069876 | 0.024665 | 0.904679 |
| 10 | 22462600 | 22465999 | 31.39516 | 51.63907 | 70.33652 | 30.31877 | -0.06612 | 1.406232 | 0.904697 |

| | | | | | | | | | |
|---|---|---|---|---|---|---|---|---|---|
| 17 | 32287600 | 32291199 | 44.73657 | 71.82812 | 87.45266 | 37.01297 | 0.06627 | 1.9223 | 0.90471 |
| 14 | 13264000 | 13265599 | 14.49299 | 25.79611 | 35.52099 | 15.65088 | -0.06362 | 0.47027 | 0.90472 |
| 16 | 44958800 | 44960599 | 26.86876 | 44.66415 | 60.43848 | 26.22028 | -0.05386 | 1.265591 | 0.904733 |
| 7 | 15005400 | 15033999 | 1110.035 | 1426.222 | 1179.962 | 494.8716 | 0.319098 | 5.927556 | 0.904762 |
| 10 | 67711600 | 67712399 | 9.854629 | 18.50601 | 24.27152 | 9.982735 | 0.077522 | 0.092981 | 0.904775 |
| 13 | 3573400 | 3574999 | 14.06747 | 25.05903 | 31.45279 | 13.84892 | 0.069976 | 0.453482 | 0.90479 |
| 11 | 7875600 | 7878199 | 53.53591 | 85.32147 | 111.8978 | 46.85197 | -0.05285 | 2.170421 | 0.904793 |
| 4 | 35437400 | 35438999 | 18.21781 | 31.81232 | 39.80095 | 17.53026 | 0.065708 | 0.71861 | 0.904814 |
| 1 | 171715200 | 171716399 | 18.71144 | 32.34641 | 44.73105 | 19.6241 | -0.08021 | 0.848552 | 0.904815 |
| X | 48445200 | 48446199 | 8.949422 | 17.10697 | 26.19494 | 10.39467 | -0.09885 | -0.02603 | 0.904829 |
| 6 | 130681400 | 130682999 | 16.88298 | 29.76176 | 41.71423 | 18.46182 | -0.09578 | 0.580087 | 0.90484 |
| 10 | 5241000 | 5243999 | 25.24482 | 42.61943 | 53.38831 | 23.35267 | 0.057289 | 1.097299 | 0.904852 |
| 7 | 115870800 | 115874399 | 34.99301 | 56.77004 | 76.25108 | 32.67105 | -0.05843 | 1.617552 | 0.904853 |
| 13 | 67999200 | 68001599 | 26.67447 | 44.59338 | 55.63926 | 24.2266 | 0.062808 | 1.196569 | 0.90486 |
| 10 | 82468400 | 82470599 | 21.86365 | 37.42003 | 46.87639 | 20.55567 | 0.059194 | 0.955907 | 0.904876 |
| 3 | 7904600 | 7905599 | 23.9898 | 40.46798 | 50.65188 | 22.1503 | 0.058513 | 1.098431 | 0.904881 |
| 5 | 87937000 | 87938999 | 19.17044 | 33.66307 | 41.13467 | 18.20064 | 0.099607 | 0.639533 | 0.904895 |
| 13 | 11517400 | 11518399 | 24.0103 | 40.28451 | 49.61417 | 21.65235 | 0.081591 | 1.146879 | 0.904905 |
| 11 | 57300800 | 57302799 | 13.83416 | 24.69261 | 31.06794 | 13.65649 | 0.068391 | 0.440667 | 0.904918 |
| 5 | 109507400 | 109508999 | 15.49501 | 27.22795 | 37.19855 | 16.4687 | -0.06034 | 0.603252 | 0.904926 |
| 5 | 60623000 | 60625399 | 25.69665 | 43.60211 | 53.97171 | 23.64408 | 0.074659 | 1.055685 | 0.904931 |
| 7 | 60000400 | 60002599 | 47.27422 | 75.85105 | 92.84883 | 39.2563 | 0.057361 | 1.94976 | 0.904939 |
| 11 | 88637800 | 88639599 | 15.93597 | 27.97662 | 38.19873 | 16.89814 | -0.05959 | 0.62145 | 0.904943 |
| 4 | 112734600 | 112741199 | 72.27226 | 114.8914 | 109.505 | 45.69658 | 0.403028 | 2.266066 | 0.904969 |
| 6 | 140288600 | 140292199 | 33.62613 | 55.03907 | 68.91755 | 29.6459 | 0.052187 | 1.47291 | 0.904984 |
| 7 | 18106200 | 18183399 | 1607.419 | 1990.4 | 1917.435 | 807.3027 | 0.045752 | 6.523893 | 0.904996 |
| 14 | 78127000 | 78129199 | 27.01065 | 45.17277 | 56.65881 | 24.67653 | 0.055348 | 1.197738 | 0.905009 |
| 5 | 37836800 | 37839399 | 24.38954 | 41.28494 | 51.74193 | 22.65074 | 0.056724 | 1.069477 | 0.905018 |
| X | 63416400 | 63419399 | 30.46391 | 50.60812 | 70.36038 | 30.51131 | -0.09293 | 1.292406 | 0.905044 |
| 13 | 39697000 | 39698199 | 11.10441 | 20.57768 | 29.34648 | 12.33167 | -0.06679 | 0.199836 | 0.90506 |
| 9 | 101005400 | 101013799 | 98.33377 | 152.762 | 180.0593 | 74.76529 | 0.070231 | 2.767581 | 0.905076 |
| 6 | 23919000 | 23919599 | 10.68839 | 19.77451 | 25.2467 | 10.63244 | 0.089579 | 0.194814 | 0.905107 |
| 16 | 31409600 | 31418199 | 57.9901 | 91.3515 | 151.7017 | 62.94676 | -0.40807 | 2.489926 | 0.905107 |
| 18 | 73078800 | 73082799 | 71.40669 | 112.3464 | 144.8145 | 60.07928 | -0.04448 | 2.522296 | 0.905175 |
| 6 | 82360000 | 82362799 | 34.12359 | 55.88356 | 76.12648 | 32.7543 | -0.06877 | 1.471109 | 0.905177 |
| 13 | 66394200 | 66396399 | 45.58584 | 73.50077 | 67.7872 | 28.86432 | 0.473511 | 1.783907 | 0.905187 |
| 12 | 17705800 | 17709999 | 54.07596 | 86.35689 | 105.9409 | 44.48363 | 0.046447 | 2.104358 | 0.905188 |
| 6 | 77592800 | 77594599 | 21.32235 | 35.58446 | 63.26588 | 27.51744 | -0.44611 | 1.211522 | 0.90521 |
| 10 | 82039200 | 82041199 | 28.15273 | 47.10632 | 58.51102 | 25.49126 | 0.069227 | 1.19438 | 0.905213 |
| 6 | 67503000 | 67503599 | 9.91133 | 18.92841 | 25.62494 | 10.18929 | 0.07409 | -0.01525 | 0.90522 |
| 14 | 90434800 | 90435599 | 12.22623 | 22.32783 | 28.54636 | 12.23455 | 0.06911 | 0.285893 | 0.905261 |
| 10 | 47399600 | 47400999 | 18.66216 | 32.53001 | 40.94176 | 18.01247 | 0.057042 | 0.749696 | 0.905272 |
| 9 | 42232000 | 42233199 | 11.11669 | 20.51298 | 29.22898 | 12.3536 | -0.07202 | 0.220163 | 0.905294 |
| 2 | 143598200 | 143600599 | 22.81886 | 38.21467 | 63.64608 | 27.74364 | -0.35247 | 1.175574 | 0.905296 |
| 11 | 88851600 | 88853599 | 15.11578 | 26.68983 | 36.55434 | 16.17291 | -0.06151 | 0.544377 | 0.905303 |
| 15 | 80065200 | 80066599 | 15.11578 | 26.68983 | 36.55434 | 16.17291 | -0.06151 | 0.544377 | 0.905303 |
| 12 | 106616200 | 106617599 | 14.77425 | 26.18934 | 36.08872 | 15.95438 | -0.06785 | 0.501715 | 0.90532 |

| | | | | | | | | | |
|---|---|---|---|---|---|---|---|---|---|
| 4 | 29270000 | 29272399 | 32.52271 | 53.1998 | 71.56354 | 30.77711 | -0.05155 | 1.481375 | 0.905337 |
| 16 | 38774800 | 38779399 | 47.57541 | 76.46642 | 93.95947 | 39.82874 | 0.054807 | 1.893533 | 0.905359 |
| 16 | 21431200 | 21432199 | 16.9117 | 29.38695 | 40.6285 | 17.83963 | -0.07819 | 0.785305 | 0.905415 |
| 10 | 37387800 | 37389399 | 17.78176 | 31.49116 | 38.45375 | 17.00311 | 0.104826 | 0.53215 | 0.905437 |
| 4 | 106104000 | 106105799 | 15.60541 | 27.48742 | 37.59689 | 16.64226 | -0.06099 | 0.575702 | 0.905456 |
| 3 | 97897600 | 97899999 | 16.81117 | 29.3766 | 40.3746 | 17.79087 | -0.06911 | 0.70245 | 0.905459 |
| 18 | 74427800 | 74432999 | 71.83464 | 113.4438 | 137.6198 | 57.14591 | 0.047144 | 2.443304 | 0.905494 |
| 15 | 48899600 | 48900799 | 9.524416 | 17.63598 | 32.04053 | 13.47807 | -0.41498 | 0.196035 | 0.905523 |
| 9 | 15683000 | 15685399 | 19.56547 | 33.57556 | 45.8617 | 20.11502 | -0.06439 | 0.923765 | 0.905533 |
| 11 | 63378200 | 63381399 | 38.93308 | 63.76896 | 76.81686 | 33.05332 | 0.108264 | 1.4813 | 0.90561 |
| 2 | 36276600 | 36279399 | 16.7796 | 29.94761 | 29.65065 | 13.03435 | 0.411889 | 0.456498 | 0.905654 |
| 14 | 92471000 | 92472799 | 13.13302 | 22.79901 | 46.88226 | 20.57322 | -0.64804 | 0.79393 | 0.905655 |
| 14 | 105738400 | 105740399 | 14.476 | 25.49072 | 35.1658 | 15.56703 | -0.07334 | 0.568003 | 0.905672 |
| 5 | 85659600 | 85660599 | 16.5706 | 29.10517 | 36.60879 | 16.20157 | 0.058237 | 0.61474 | 0.905672 |
| 6 | 6850400 | 6852199 | 16.5706 | 29.10517 | 36.60879 | 16.20157 | 0.058237 | 0.61474 | 0.905672 |
| 4 | 143845600 | 143875799 | 855.4309 | 1111.145 | 1195.446 | 500.8335 | -0.04406 | 5.749073 | 0.905693 |
| 3 | 52344200 | 52345199 | 11.32579 | 20.61312 | 29.28489 | 12.60463 | -0.0854 | 0.304473 | 0.905705 |
| 9 | 27167200 | 27168599 | 14.22376 | 25.21202 | 34.5767 | 15.28612 | -0.06106 | 0.494991 | 0.905742 |
| 7 | 50818200 | 50819399 | 12.38 | 22.75392 | 32.38167 | 13.75993 | -0.07568 | 0.261297 | 0.905752 |
| 2 | 125497000 | 125497799 | 8.834898 | 16.8274 | 25.01868 | 10.00491 | -0.06695 | -0.00807 | 0.905776 |
| 12 | 81488000 | 81489799 | 20.72474 | 36.42683 | 33.21865 | 14.70982 | 0.519933 | 0.623016 | 0.905783 |
| 10 | 48626800 | 48632399 | 79.83463 | 125.7777 | 128.5979 | 53.36095 | 0.290849 | 2.497395 | 0.905825 |
| 13 | 91643200 | 91644199 | 22.90535 | 39.4305 | 39.96323 | 17.53015 | 0.365128 | 0.900054 | 0.905845 |
| 2 | 68548800 | 68551999 | 36.72472 | 60.24274 | 73.68561 | 31.72624 | 0.087149 | 1.45538 | 0.905861 |
| 1 | 13505000 | 13506799 | 16.72823 | 29.19637 | 39.95442 | 17.59225 | -0.06295 | 0.720301 | 0.905875 |
| 9 | 26317600 | 26319799 | 23.60128 | 39.79801 | 49.06398 | 21.45352 | 0.080412 | 1.100983 | 0.905892 |
| 10 | 60085800 | 60089999 | 32.83795 | 53.10873 | 88.65581 | 37.89924 | -0.37551 | 1.683439 | 0.905905 |
| 2 | 149629000 | 149630799 | 17.28925 | 30.17984 | 41.42937 | 18.24332 | -0.06758 | 0.721432 | 0.905906 |
| 1 | 130307600 | 130308399 | 13.46943 | 24.23708 | 30.35884 | 13.254 | 0.081115 | 0.382507 | 0.905906 |
| 15 | 83943800 | 83948199 | 39.74915 | 64.26926 | 85.93609 | 36.72438 | -0.05661 | 1.700786 | 0.905916 |
| 3 | 34711000 | 34712399 | 13.89178 | 24.8789 | 31.43017 | 13.77664 | 0.065088 | 0.41875 | 0.905917 |
| 19 | 59416400 | 59418399 | 22.65276 | 38.53953 | 52.24583 | 22.89918 | -0.05559 | 1.020496 | 0.905963 |
| 16 | 91803200 | 91807199 | 43.23867 | 69.58311 | 92.00268 | 39.07006 | -0.04844 | 1.84902 | 0.905984 |
| 16 | 5341800 | 5343599 | 23.26002 | 39.48853 | 49.56432 | 21.71676 | 0.054782 | 1.038535 | 0.905987 |
| 7 | 80723200 | 80724999 | 15.43553 | 27.01713 | 37.36948 | 16.48957 | -0.07809 | 0.665357 | 0.906024 |
| 8 | 73674000 | 73675399 | 16.40827 | 28.45416 | 48.46125 | 21.26262 | -0.3782 | 0.819918 | 0.906039 |
| 3 | 40754200 | 40755199 | 7.866517 | 14.95668 | 22.91146 | 9.196333 | -0.11455 | -0.01357 | 0.906042 |
| 12 | 58909200 | 58911199 | 34.78087 | 57.08635 | 57.93921 | 24.9524 | 0.355969 | 1.43317 | 0.906049 |
| 11 | 47500800 | 47501799 | 11.065 | 20.32889 | 28.73459 | 12.21681 | -0.06627 | 0.245453 | 0.906099 |
| 14 | 10796000 | 10796799 | 11.065 | 20.32889 | 28.73459 | 12.21681 | -0.06627 | 0.245453 | 0.906099 |
| 16 | 11501400 | 11503799 | 16.38191 | 28.71599 | 39.24838 | 17.34028 | -0.06111 | 0.649141 | 0.906106 |
| 13 | 33057400 | 33059799 | 23.82113 | 40.85259 | 41.57586 | 18.23431 | 0.358268 | 0.940754 | 0.90611 |
| 1 | 114340600 | 114341999 | 17.6927 | 31.24897 | 38.45564 | 17.01312 | 0.092113 | 0.559443 | 0.906202 |
| 4 | 113882600 | 113902399 | 371.8266 | 519.8571 | 564.8014 | 234.2012 | 0.048399 | 4.5846 | 0.906208 |
| 3 | 111505000 | 111507199 | 18.39032 | 32.54653 | 31.20067 | 13.79791 | 0.451886 | 0.531468 | 0.906214 |
| 1 | 6498000 | 6500799 | 50.37284 | 80.38978 | 105.9578 | 44.44889 | -0.05792 | 2.121756 | 0.906214 |
| X | 9211200 | 9212399 | 13.90991 | 25.58738 | 25.45793 | 10.80544 | 0.438784 | 0.253753 | 0.906217 |

| | | | | | | | | | |
|---|---|---|---|---|---|---|---|---|---|
| X | 122019200 | 122022599 | 34.92768 | 57.71354 | 70.08298 | 30.27883 | 0.101165 | 1.364701 | 0.90623 |
| 5 | 79724400 | 79729199 | 96.96489 | 149.5505 | 189.6422 | 78.774 | -0.04314 | 2.896878 | 0.906231 |
| 19 | 15811600 | 15819199 | 66.19181 | 104.4196 | 135.6614 | 56.30191 | -0.05287 | 2.463542 | 0.906231 |
| 11 | 45550200 | 45552999 | 56.33352 | 89.63087 | 117.0781 | 48.9083 | -0.04965 | 2.227592 | 0.906232 |
| 4 | 60573600 | 60581199 | 145.2661 | 217.6622 | 270.602 | 111.6269 | -0.04886 | 3.487643 | 0.90624 |
| 7 | 57718000 | 57721399 | 54.91754 | 87.46248 | 114.2345 | 47.78432 | -0.04803 | 2.193032 | 0.906249 |
| 1 | 92406800 | 92407999 | 8.36622 | 15.98905 | 24.00646 | 9.531274 | -0.07168 | -0.03852 | 0.906252 |
| 14 | 23583600 | 23586799 | 54.84349 | 87.50238 | 106.8655 | 44.81367 | 0.051488 | 2.135046 | 0.906262 |
| 8 | 94189200 | 94198599 | 203.8603 | 299.1952 | 342.1384 | 140.9163 | 0.043953 | 3.853158 | 0.906268 |
| 6 | 57271200 | 57283799 | 382.0497 | 534.6713 | 494.7302 | 205.1143 | 0.28138 | 4.569472 | 0.906276 |
| 6 | 37731400 | 37737999 | 84.52715 | 132.6962 | 157.9891 | 65.52623 | 0.067466 | 2.576292 | 0.90628 |
| 16 | 61346000 | 61346399 | 6.235896 | 12.15873 | 18.04396 | 6.49756 | 0.076991 | -0.413 | 0.906309 |
| 8 | 75533600 | 75535599 | 20.31262 | 34.69141 | 47.43472 | 20.79693 | -0.06703 | 0.970302 | 0.906334 |
| 10 | 49830400 | 49831799 | 16.13067 | 28.36187 | 35.62801 | 15.77637 | 0.060339 | 0.596441 | 0.906358 |
| 1 | 5318200 | 5324599 | 76.69669 | 120.3541 | 154.9867 | 64.27754 | -0.04601 | 2.578316 | 0.906404 |
| 1 | 101230000 | 101230399 | 6.368331 | 12.40048 | 17.96653 | 6.582454 | 0.088267 | -0.35229 | 0.906409 |
| 11 | 58747200 | 58755999 | 138.6313 | 209.5472 | 246.0421 | 101.7406 | 0.044611 | 3.313669 | 0.906409 |
| 12 | 87932000 | 87938999 | 73.64862 | 116.7239 | 138.3988 | 57.57851 | 0.084208 | 2.358844 | 0.906413 |
| 16 | 21516600 | 21520999 | 46.61315 | 74.9989 | 92.29903 | 39.18477 | 0.054467 | 1.859086 | 0.906438 |
| 13 | 62900600 | 62903799 | 30.50595 | 50.22333 | 67.77767 | 29.22731 | -0.05219 | 1.393176 | 0.906444 |
| 13 | 67335000 | 67338599 | 40.50416 | 65.33412 | 86.70761 | 36.94035 | -0.04973 | 1.773428 | 0.906479 |
| 3 | 29609600 | 29611599 | 33.00668 | 54.22217 | 67.85313 | 29.23289 | 0.055195 | 1.427028 | 0.906482 |
| 18 | 23111600 | 23112199 | 7.436066 | 14.42607 | 20.5513 | 7.743285 | 0.075171 | -0.23086 | 0.906512 |
| 13 | 51068200 | 51069599 | 13.15824 | 24.35509 | 24.95456 | 10.50288 | 0.40517 | 0.21108 | 0.906514 |
| 16 | 47574200 | 47576599 | 22.89789 | 39.65764 | 39.39753 | 17.2852 | 0.395712 | 0.847489 | 0.906543 |
| 17 | 74353800 | 74355999 | 19.2257 | 32.50555 | 56.95415 | 24.92354 | -0.42488 | 1.068103 | 0.906556 |
| 17 | 49045600 | 49047399 | 17.51753 | 30.51474 | 37.98374 | 16.70417 | 0.072918 | 0.749284 | 0.90657 |
| 15 | 89463400 | 89464199 | 8.978209 | 16.9153 | 24.87854 | 10.176 | -0.07994 | 0.066714 | 0.90658 |
| X | 39728000 | 39728999 | 12.55043 | 23.08911 | 29.71934 | 12.61229 | 0.069801 | 0.252473 | 0.906583 |
| 7 | 127064600 | 127068399 | 35.24948 | 57.48581 | 71.75523 | 30.8255 | 0.052984 | 1.529451 | 0.906608 |
| X | 135376800 | 135378799 | 20.11411 | 34.99145 | 43.61478 | 19.16457 | 0.070764 | 0.790337 | 0.906637 |
| 1 | 109127600 | 109128199 | 9.42085 | 17.76858 | 25.89874 | 10.57358 | -0.06512 | 0.06574 | 0.906674 |
| 12 | 56465200 | 56465999 | 11.89751 | 21.83843 | 28.08573 | 11.9558 | 0.067523 | 0.25362 | 0.906681 |
| 7 | 110627800 | 110628599 | 8.930455 | 16.86466 | 24.8689 | 10.1264 | -0.07846 | 0.049184 | 0.90671 |
| 6 | 57063200 | 57072399 | 236.8233 | 345.0431 | 329.6863 | 135.8655 | 0.292899 | 3.968496 | 0.906724 |
| 14 | 121131800 | 121133199 | 13.793 | 24.6231 | 31.02563 | 13.63605 | 0.066343 | 0.438427 | 0.906756 |
| 11 | 21463600 | 21464399 | 14.16787 | 25.30817 | 32.01989 | 14.0656 | 0.061031 | 0.439936 | 0.906769 |
| 6 | 21198000 | 21201599 | 53.0868 | 84.91442 | 112.4491 | 47.32461 | -0.06113 | 2.057963 | 0.906792 |
| 5 | 56135400 | 56135999 | 13.26186 | 23.59258 | 32.67009 | 14.40104 | -0.07181 | 0.461771 | 0.906807 |
| 8 | 47157600 | 47159199 | 14.24724 | 26.02105 | 25.46334 | 10.91195 | 0.452891 | 0.296484 | 0.906815 |
| 16 | 75430200 | 75433199 | 56.00047 | 89.26518 | 109.1944 | 45.73133 | 0.047695 | 2.164155 | 0.906845 |
| 9 | 19377800 | 19380799 | 23.43744 | 40.81792 | 48.44102 | 21.30003 | 0.14127 | 0.795775 | 0.906852 |
| 6 | 20629000 | 20629399 | 9.0959 | 17.2778 | 25.4957 | 10.25406 | -0.06377 | 0.012613 | 0.906879 |
| 18 | 19563400 | 19565799 | 30.85768 | 51.44291 | 52.44005 | 22.79342 | 0.3536 | 1.238611 | 0.906883 |
| X | 101910600 | 101912999 | 16.74583 | 29.35473 | 40.36378 | 17.83349 | -0.06952 | 0.65632 | 0.906891 |
| 14 | 121074800 | 121076599 | 14.93515 | 26.22776 | 36.20471 | 16.02023 | -0.07528 | 0.606598 | 0.9069 |
| 17 | 82837200 | 82837599 | 7.129648 | 13.80047 | 21.58564 | 8.2191 | -0.07302 | -0.19645 | 0.906902 |

| | | | | | | | | | |
|---|---|---|---|---|---|---|---|---|---|
| 2 | 76608200 | 76609999 | 16.58033 | 29.09124 | 39.72869 | 17.56648 | -0.05993 | 0.636466 | 0.906907 |
| 4 | 36269600 | 36271199 | 20.38303 | 34.85798 | 47.43523 | 20.79938 | -0.05979 | 0.960221 | 0.906917 |
| X | 72866400 | 72868799 | 20.4057 | 35.2866 | 49.00857 | 21.50091 | -0.08607 | 0.865441 | 0.906927 |
| 6 | 17394200 | 17395599 | 17.01823 | 29.97125 | 37.48341 | 16.59255 | 0.067376 | 0.584973 | 0.906929 |
| 9 | 36513200 | 36516599 | 36.33223 | 59.02119 | 73.60163 | 31.56681 | 0.050245 | 1.59355 | 0.906942 |
| 1 | 35238600 | 35241999 | 26.84625 | 44.94759 | 56.20069 | 24.49412 | 0.059712 | 1.182026 | 0.906944 |
| 19 | 57544800 | 57545999 | 13.92903 | 25.53797 | 32.25973 | 13.75415 | 0.09158 | 0.285378 | 0.906971 |
| 18 | 25535200 | 25536799 | 18.09349 | 31.86122 | 45.52518 | 20.15282 | -0.12393 | 0.607683 | 0.906971 |
| 17 | 20296400 | 20299999 | 40.73179 | 66.33695 | 80.38512 | 34.51586 | 0.094287 | 1.568019 | 0.906975 |
| 19 | 39115600 | 39119999 | 110.5216 | 168.8413 | 199.283 | 82.65834 | 0.049664 | 3.088576 | 0.906975 |
| 8 | 82766800 | 82767599 | 11.36706 | 21.41057 | 22.09169 | 9.039978 | 0.426912 | 0.081993 | 0.906996 |
| 1 | 107808800 | 107809799 | 15.80286 | 27.24651 | 48.98521 | 21.48873 | -0.45762 | 0.870225 | 0.906999 |
| 5 | 119208000 | 119210799 | 18.08873 | 31.40322 | 43.13771 | 18.93041 | -0.06933 | 0.805052 | 0.907011 |
| X | 149181000 | 149181799 | 9.108289 | 17.51351 | 23.59631 | 9.20644 | 0.107384 | -0.08573 | 0.907022 |
| 9 | 42491400 | 42494399 | 30.37907 | 50.00455 | 62.22026 | 26.82858 | 0.06447 | 1.389204 | 0.907027 |
| 3 | 3516800 | 3518999 | 24.60506 | 42.19723 | 42.5633 | 18.66738 | 0.371028 | 0.94444 | 0.907038 |
| 5 | 33397800 | 33399999 | 19.77189 | 33.80761 | 46.41079 | 20.34994 | -0.07234 | 0.958889 | 0.907039 |
| 7 | 26739200 | 26739799 | 7.546513 | 14.60083 | 22.68867 | 8.657216 | -0.06724 | -0.18351 | 0.90704 |
| 12 | 113682000 | 113683799 | 22.95094 | 39.51486 | 49.01717 | 21.50182 | 0.074806 | 0.911184 | 0.907073 |
| 6 | 139651200 | 139651799 | 6.668117 | 12.92942 | 20.54405 | 7.75963 | -0.08458 | -0.23361 | 0.907074 |
| 14 | 67676400 | 67679199 | 18.17585 | 31.60373 | 39.53343 | 17.36574 | 0.065572 | 0.780856 | 0.907075 |
| 16 | 88603400 | 88604199 | 11.90714 | 22.14843 | 28.79213 | 12.03462 | 0.071175 | 0.181088 | 0.907084 |
| 12 | 103927800 | 103930199 | 21.85594 | 37.3352 | 50.6236 | 22.19543 | -0.05495 | 0.979561 | 0.907128 |
| 3 | 97286000 | 97291399 | 54.89762 | 87.81116 | 107.3014 | 45.15839 | 0.054659 | 2.056291 | 0.907133 |
| 4 | 17843200 | 17845399 | 21.33132 | 37.14286 | 45.69361 | 20.08955 | 0.089563 | 0.784098 | 0.907167 |
| 14 | 25095600 | 25100799 | 73.26618 | 114.9865 | 148.0513 | 61.40229 | -0.04562 | 2.573042 | 0.907188 |
| 2 | 168939400 | 168941199 | 21.69563 | 36.75613 | 50.0911 | 21.93373 | -0.06354 | 1.05191 | 0.90723 |
| 2 | 72580000 | 72581199 | 13.35417 | 24.40911 | 31.16642 | 13.34282 | 0.072191 | 0.297387 | 0.907258 |
| 19 | 15918400 | 15923199 | 43.4038 | 69.97171 | 86.65493 | 36.89098 | 0.0489 | 1.793473 | 0.907259 |
| 9 | 117661800 | 117663399 | 13.9981 | 24.77575 | 34.07552 | 15.06896 | -0.06608 | 0.507077 | 0.907291 |
| X | 75382600 | 75383999 | 21.96161 | 37.68942 | 46.88945 | 20.56587 | 0.069897 | 0.936521 | 0.907297 |
| X | 111363400 | 111365799 | 24.3215 | 42.02313 | 50.68613 | 22.23633 | 0.116239 | 0.890001 | 0.907324 |
| 4 | 143799600 | 143804399 | 58.80175 | 93.97043 | 113.2534 | 47.60193 | 0.073091 | 2.088988 | 0.907324 |
| 10 | 22443000 | 22443799 | 10.47221 | 19.84309 | 26.51965 | 10.7217 | 0.071183 | 0.04425 | 0.907324 |
| 1 | 37136600 | 37138399 | 15.47516 | 27.25684 | 34.14465 | 15.11425 | 0.065355 | 0.570914 | 0.907324 |
| 7 | 40423000 | 40423799 | 10.53861 | 19.73074 | 28.4446 | 11.75912 | -0.06409 | 0.128558 | 0.907362 |
| 1 | 11436400 | 11440599 | 61.03722 | 96.94325 | 125.9717 | 52.52058 | -0.04474 | 2.285397 | 0.907389 |
| 14 | 13422000 | 13422999 | 12.84791 | 23.85456 | 24.49181 | 10.26471 | 0.406646 | 0.190554 | 0.907407 |
| 6 | 138769200 | 138770199 | 13.14269 | 23.43192 | 32.53534 | 14.31514 | -0.07416 | 0.443961 | 0.907452 |
| 2 | 57850200 | 57857199 | 90.62501 | 140.5205 | 179.1252 | 74.39845 | -0.04477 | 2.795597 | 0.907463 |
| 15 | 48261200 | 48262599 | 13.38469 | 24.12119 | 30.584 | 13.32722 | 0.065918 | 0.378184 | 0.907478 |
| 16 | 12148800 | 12149399 | 13.23716 | 23.62991 | 32.62305 | 14.33859 | -0.06465 | 0.439429 | 0.907479 |
| 17 | 41708800 | 41709999 | 19.12895 | 33.25845 | 41.46092 | 18.21156 | 0.070215 | 0.785767 | 0.907482 |
| 5 | 72784000 | 72784999 | 9.775437 | 18.22343 | 26.32379 | 10.96184 | -0.07501 | 0.145148 | 0.907511 |
| 13 | 95642200 | 95643999 | 16.0558 | 28.21778 | 38.61879 | 17.09973 | -0.06308 | 0.604308 | 0.907536 |
| X | 94805400 | 94810399 | 64.07711 | 102.059 | 122.7019 | 51.33238 | 0.071404 | 2.202103 | 0.907554 |
| 4 | 73637600 | 73638599 | 16.74721 | 29.3964 | 40.22203 | 17.79094 | -0.06269 | 0.63142 | 0.907564 |

| | | | | | | | | | |
|---|---|---|---|---|---|---|---|---|---|
| 4 | 138563000 | 138573999 | 149.659 | 224.552 | 278.3361 | 114.8536 | -0.0428 | 3.462424 | 0.907575 |
| 12 | 109298400 | 109300199 | 24.30405 | 40.89372 | 51.19456 | 22.37031 | 0.058083 | 1.121623 | 0.907618 |
| 16 | 12893600 | 12894999 | 14.51148 | 25.64256 | 35.10358 | 15.52865 | -0.06027 | 0.530738 | 0.907645 |
| 8 | 12793000 | 12797399 | 61.19717 | 97.12838 | 126.0646 | 52.52198 | -0.04404 | 2.306091 | 0.90767 |
| 5 | 102160600 | 102164399 | 54.37638 | 86.35739 | 133.6713 | 55.72637 | -0.29686 | 2.284156 | 0.907696 |
| 13 | 49951400 | 49956999 | 88.86164 | 137.7026 | 164.2228 | 68.21353 | 0.05056 | 2.804618 | 0.907701 |
| 1 | 26081600 | 26082199 | 7.474751 | 14.46087 | 22.67091 | 8.653569 | -0.07941 | -0.17904 | 0.90771 |
| 14 | 60967200 | 60970599 | 28.75234 | 47.52241 | 64.26334 | 27.76967 | -0.05353 | 1.340861 | 0.907716 |
| 11 | 92317400 | 92318199 | 12.70456 | 23.22925 | 32.79847 | 14.03584 | -0.07173 | 0.293629 | 0.907719 |
| 6 | 90963600 | 90968199 | 49.3073 | 78.90627 | 103.5181 | 43.59518 | -0.04688 | 2.034384 | 0.907744 |
| 8 | 25777600 | 25781199 | 45.1501 | 73.08554 | 72.65516 | 31.07936 | 0.371718 | 1.680014 | 0.907745 |
| 11 | 20465000 | 20468599 | 63.40256 | 100.1707 | 130.5239 | 54.21117 | -0.05534 | 2.425405 | 0.90776 |
| 15 | 41203200 | 41205199 | 22.14732 | 38.52569 | 38.20892 | 16.78844 | 0.398921 | 0.790965 | 0.907785 |
| 1 | 102935600 | 102936999 | 25.85165 | 43.70311 | 54.47324 | 23.83702 | 0.064584 | 1.090679 | 0.907787 |
| 18 | 83066000 | 83069599 | 36.3234 | 59.03521 | 73.69773 | 31.61923 | 0.049347 | 1.583784 | 0.907802 |
| 13 | 53874400 | 53876599 | 10.27975 | 18.51235 | 34.32054 | 14.96887 | -0.47962 | 0.367993 | 0.907813 |
| 17 | 80094000 | 80094999 | 10.04948 | 18.92176 | 27.47504 | 11.24996 | -0.06325 | 0.085964 | 0.907816 |
| 13 | 85017800 | 85019399 | 16.13516 | 28.35375 | 38.67081 | 17.12097 | -0.05799 | 0.608472 | 0.907819 |
| 14 | 119717000 | 119719999 | 38.33809 | 61.95935 | 76.34663 | 32.60709 | 0.060345 | 1.709161 | 0.907821 |
| 4 | 156576800 | 156589399 | 253.3894 | 366.8681 | 411.9835 | 169.8999 | 0.051187 | 4.065589 | 0.907834 |
| 1 | 86702800 | 86704399 | 16.56612 | 29.34916 | 36.48108 | 16.12965 | 0.079705 | 0.519334 | 0.907836 |
| 17 | 67964800 | 67968199 | 31.07844 | 51.26048 | 69.85106 | 30.14496 | -0.06514 | 1.371669 | 0.90785 |
| 4 | 51298800 | 51300599 | 53.15002 | 84.7257 | 102.1622 | 42.78909 | 0.068522 | 2.159721 | 0.907861 |
| 2 | 148575800 | 148577999 | 30.59468 | 50.46983 | 63.39436 | 27.36017 | 0.052169 | 1.36507 | 0.907876 |
| 10 | 4159000 | 4164599 | 56.3926 | 89.92757 | 117.7519 | 49.34361 | -0.0497 | 2.15403 | 0.907881 |
| 8 | 97982000 | 97983599 | 29.13345 | 48.18151 | 60.50323 | 26.15579 | 0.053043 | 1.33023 | 0.907885 |
| 11 | 11097400 | 11099599 | 35.75684 | 58.5173 | 58.46953 | 25.15007 | 0.376727 | 1.47175 | 0.907898 |
| 19 | 56106200 | 56110999 | 70.01059 | 110.4626 | 134.2157 | 55.70672 | 0.043821 | 2.462674 | 0.907906 |
| 12 | 86504400 | 86505399 | 12.32907 | 22.34384 | 31.10616 | 13.45996 | -0.06183 | 0.337224 | 0.907934 |
| 16 | 42606000 | 42607999 | 16.75063 | 29.27055 | 39.91988 | 17.59036 | -0.05787 | 0.70384 | 0.907937 |
| 13 | 11099800 | 11105599 | 83.36881 | 130.6774 | 156.9937 | 65.11609 | 0.052773 | 2.602979 | 0.907937 |
| 12 | 114116600 | 114141599 | 410.429 | 570.949 | 611.4645 | 253.749 | 0.061803 | 4.661714 | 0.90794 |
| 7 | 134260400 | 134262799 | 31.82225 | 52.5324 | 65.73922 | 28.38356 | 0.05779 | 1.362886 | 0.907942 |
| 7 | 61485000 | 61487399 | 29.17032 | 49.16632 | 48.0034 | 20.98685 | 0.41546 | 1.11801 | 0.907966 |
| 13 | 44306400 | 44308399 | 22.7207 | 38.6619 | 48.49203 | 21.25421 | 0.056173 | 1.014038 | 0.907968 |
| 12 | 49823000 | 49824199 | 15.87796 | 27.92989 | 35.05834 | 15.52274 | 0.061562 | 0.587402 | 0.907995 |
| X | 127564200 | 127565599 | 15.35241 | 27.58892 | 34.51216 | 15.08401 | 0.08217 | 0.415644 | 0.908005 |
| 5 | 8585800 | 8586199 | 6.663039 | 12.93417 | 20.57105 | 7.727779 | -0.07862 | -0.25227 | 0.90805 |
| 1 | 160646400 | 160648799 | 17.2346 | 30.45843 | 26.10205 | 11.54637 | 0.612025 | 0.528799 | 0.908086 |
| X | 117798000 | 117807599 | 150.0482 | 228.083 | 203.4722 | 84.28325 | 0.447439 | 3.202291 | 0.908087 |
| 8 | 3223800 | 3226799 | 36.63467 | 59.44594 | 79.47146 | 34.05619 | -0.05141 | 1.618427 | 0.90809 |
| 7 | 40856800 | 40860199 | 35.6208 | 58.75679 | 71.46252 | 30.84133 | 0.098468 | 1.385172 | 0.908093 |
| 3 | 115162200 | 115163999 | 12.01937 | 22.14466 | 28.69816 | 12.15457 | 0.062118 | 0.237314 | 0.908102 |
| 17 | 65858200 | 65859599 | 14.98581 | 26.45892 | 33.12839 | 14.65731 | 0.067446 | 0.539555 | 0.908141 |
| 5 | 115642200 | 115642999 | 10.34891 | 19.44706 | 25.81325 | 10.60497 | 0.061566 | 0.097355 | 0.908165 |
| 17 | 74111600 | 74112599 | 10.18197 | 18.79652 | 27.21442 | 11.49521 | -0.09415 | 0.205339 | 0.90818 |
| 15 | 43326200 | 43330799 | 106.2112 | 162.2899 | 205.249 | 85.13698 | -0.04966 | 3.08395 | 0.908181 |

| | | | | | | | | | |
|---|---|---|---|---|---|---|---|---|---|
| 13 | 86319400 | 86320199 | 22.90967 | 38.90165 | 48.50512 | 21.25398 | 0.064464 | 1.037177 | 0.90821 |
| 13 | 13512600 | 13515599 | 66.39575 | 104.8394 | 126.2338 | 52.4141 | 0.05778 | 2.441046 | 0.908211 |
| 12 | 17551600 | 17553599 | 19.09557 | 33.26069 | 46.27589 | 20.34971 | -0.08695 | 0.773331 | 0.908264 |
| X | 56961800 | 56967799 | 48.11434 | 77.31994 | 103.2885 | 43.77286 | -0.0658 | 1.905563 | 0.908286 |
| X | 145135600 | 145136799 | 13.11592 | 24.31616 | 30.93511 | 12.99127 | 0.097438 | 0.217278 | 0.908312 |
| 10 | 25545400 | 25545999 | 9.453956 | 18.03407 | 24.48976 | 9.761741 | 0.066827 | -0.01393 | 0.908312 |
| 19 | 47793200 | 47793999 | 12.1589 | 21.867 | 30.75439 | 13.43284 | -0.08474 | 0.383266 | 0.908358 |
| 3 | 51834400 | 51838799 | 43.36533 | 69.7614 | 85.88929 | 36.45458 | 0.053603 | 1.857992 | 0.90837 |
| 11 | 105956600 | 105958199 | 10.79548 | 19.88906 | 28.29028 | 11.9824 | -0.07111 | 0.227977 | 0.908396 |
| 1 | 14790400 | 14792199 | 23.61278 | 40.88121 | 49.59981 | 21.76109 | 0.108275 | 0.865372 | 0.908396 |
| 4 | 28658800 | 28664799 | 69.73482 | 110.7193 | 112.3906 | 46.84215 | 0.31053 | 2.29998 | 0.908403 |
| 3 | 11201400 | 11215999 | 261.016 | 371.0209 | 520.7411 | 215.3534 | -0.29688 | 4.336043 | 0.908409 |
| 15 | 49331800 | 49333799 | 33.28549 | 54.56146 | 68.41416 | 29.44751 | 0.051057 | 1.453538 | 0.908412 |
| 10 | 81119200 | 81121599 | 17.93495 | 30.74193 | 43.35614 | 19.01431 | -0.11054 | 0.923402 | 0.908419 |
| 6 | 8193600 | 8198399 | 94.28076 | 146.1009 | 175.0815 | 72.72393 | 0.043597 | 2.807589 | 0.908429 |
| X | 52074000 | 52077599 | 37.80286 | 61.96486 | 75.14821 | 32.34572 | 0.099063 | 1.471771 | 0.908507 |
| 12 | 33172000 | 33174199 | 19.54128 | 33.89011 | 42.59217 | 18.68758 | 0.058193 | 0.822164 | 0.90852 |
| 1 | 172712200 | 172712799 | 8.192849 | 15.59727 | 23.43897 | 9.382406 | -0.0834 | -0.01612 | 0.908534 |
| 11 | 29551800 | 29552599 | 8.192849 | 15.59727 | 23.43897 | 9.382406 | -0.0834 | -0.01612 | 0.908534 |
| 16 | 88479000 | 88479599 | 8.192849 | 15.59727 | 23.43897 | 9.382406 | -0.0834 | -0.01612 | 0.908534 |
| 10 | 83678800 | 83681399 | 35.47943 | 57.94192 | 72.36998 | 31.10547 | 0.053522 | 1.511811 | 0.908559 |
| 5 | 64996400 | 65004799 | 95.6149 | 148.0579 | 177.1942 | 73.60178 | 0.044659 | 2.82164 | 0.908561 |
| 18 | 79610200 | 79612799 | 31.51216 | 52.11173 | 71.6275 | 30.96332 | -0.07704 | 1.345847 | 0.908561 |
| 2 | 170387800 | 170388999 | 10.53079 | 19.4644 | 27.74093 | 11.698 | -0.06969 | 0.20384 | 0.908563 |
| 1 | 85034800 | 85035799 | 53.27292 | 84.88388 | 102.0867 | 42.73121 | 0.071579 | 2.173391 | 0.908573 |
| 7 | 102920200 | 102922999 | 51.68583 | 82.77361 | 109.3973 | 46.12085 | -0.05633 | 2.017404 | 0.908578 |
| 6 | 118882600 | 118883799 | 12.13448 | 22.2072 | 28.61464 | 12.23038 | 0.060749 | 0.274671 | 0.908588 |
| 10 | 70175000 | 70176999 | 26.32612 | 43.95486 | 59.40859 | 25.84703 | -0.05212 | 1.214465 | 0.908637 |
| 3 | 140218000 | 140218999 | 14.49208 | 25.19132 | 45.81929 | 20.12331 | -0.47194 | 0.77767 | 0.908654 |
| 2 | 139881200 | 139882199 | 13.89496 | 25.36737 | 32.14903 | 13.78398 | 0.081266 | 0.308275 | 0.908676 |
| 14 | 26575000 | 26575399 | 6.390857 | 12.44743 | 20.21252 | 7.382062 | -0.07016 | -0.35319 | 0.908681 |
| 18 | 69786200 | 69786999 | 6.390857 | 12.44743 | 20.21252 | 7.382062 | -0.07016 | -0.35319 | 0.908681 |
| 1 | 194336000 | 194339599 | 62.1137 | 98.30729 | 127.908 | 53.17207 | -0.05119 | 2.383242 | 0.908694 |
| X | 149143000 | 149146799 | 39.46084 | 64.32273 | 78.69939 | 33.8027 | 0.081268 | 1.550538 | 0.9087 |
| 9 | 31532800 | 31533999 | 13.52286 | 23.97189 | 33.09131 | 14.62917 | -0.07043 | 0.485287 | 0.90871 |
| 9 | 118906600 | 118908999 | 20.58385 | 35.23417 | 43.77757 | 19.19701 | 0.0711 | 0.946476 | 0.908757 |
| X | 45300800 | 45301399 | 11.70233 | 21.54336 | 30.86117 | 13.08487 | -0.08234 | 0.246706 | 0.908761 |
| 4 | 94593200 | 94594399 | 16.26677 | 28.46798 | 38.85527 | 17.14311 | -0.05898 | 0.675205 | 0.908781 |
| 6 | 14250400 | 14251599 | 30.09628 | 49.32601 | 66.74425 | 28.72683 | -0.05853 | 1.448561 | 0.908789 |
| 6 | 145121600 | 145122999 | 13.53162 | 23.96445 | 33.13033 | 14.64922 | -0.07307 | 0.496111 | 0.908833 |
| 4 | 115508400 | 115513599 | 49.47402 | 79.2237 | 103.9471 | 43.81658 | -0.04602 | 2.015167 | 0.908842 |
| 6 | 131039400 | 131041799 | 68.66485 | 108.0576 | 140.062 | 58.10935 | -0.05221 | 2.516183 | 0.908905 |
| 11 | 26585000 | 26585599 | 9.307039 | 17.66734 | 26.03442 | 10.48518 | -0.06333 | 0.021864 | 0.908918 |
| 14 | 6015800 | 6017999 | 24.62931 | 41.57518 | 52.27365 | 22.86425 | 0.052 | 1.095944 | 0.908927 |
| 9 | 6804800 | 6815999 | 335.3076 | 469.8657 | 509.2135 | 211.0765 | 0.055301 | 4.548661 | 0.908931 |
| 1 | 118737600 | 118743199 | 46.07478 | 74.22249 | 91.02249 | 38.7107 | 0.061683 | 1.819446 | 0.908932 |
| 16 | 34936200 | 34941799 | 49.16221 | 78.0652 | 131.7511 | 54.91787 | -0.4216 | 2.283249 | 0.908974 |

| | | | | | | | | | |
|---|---|---|---|---|---|---|---|---|---|
| 8 | 81581800 | 81583199 | 11.47898 | 21.09307 | 27.15569 | 11.54285 | 0.067795 | 0.24344 | 0.908987 |
| 5 | 135430800 | 135432199 | 15.43716 | 27.3133 | 34.4647 | 15.24212 | 0.057312 | 0.520808 | 0.908988 |
| 4 | 121212400 | 121214199 | 52.70414 | 84.25255 | 103.2086 | 43.3994 | 0.050191 | 2.072536 | 0.908989 |
| 19 | 53166000 | 53167999 | 10.69315 | 19.47605 | 34.40565 | 14.79503 | -0.39732 | 0.304859 | 0.908997 |
| 5 | 79268000 | 79270599 | 31.13155 | 52.181 | 50.78858 | 22.15107 | 0.419911 | 1.180419 | 0.909001 |
| 17 | 74193000 | 74197199 | 62.93974 | 99.46424 | 129.6427 | 53.85198 | -0.05548 | 2.419567 | 0.909015 |
| 9 | 120750400 | 120751999 | 14.44834 | 25.51185 | 35.06744 | 15.51499 | -0.06667 | 0.536954 | 0.909015 |
| 5 | 68153800 | 68154999 | 15.31824 | 27.43207 | 34.47377 | 15.11141 | 0.072667 | 0.436348 | 0.909025 |
| 11 | 16180400 | 16182799 | 30.76502 | 51.24106 | 50.79158 | 22.06403 | 0.393962 | 1.248105 | 0.909043 |
| 4 | 142037600 | 142042599 | 58.66884 | 93.30656 | 121.4924 | 50.72225 | -0.04577 | 2.242443 | 0.909048 |
| 1 | 12896400 | 12899999 | 35.9219 | 58.65643 | 73.11456 | 31.42392 | 0.056264 | 1.514115 | 0.909066 |
| 7 | 29567800 | 29600199 | 802.0692 | 1049.441 | 1071.062 | 448.3903 | 0.042492 | 5.63064 | 0.909066 |
| 1 | 101219000 | 101222399 | 37.88635 | 61.57436 | 76.48771 | 32.81582 | 0.056401 | 1.587035 | 0.909083 |
| 1 | 187230600 | 187232799 | 16.38339 | 28.82821 | 39.62005 | 17.5414 | -0.06855 | 0.593895 | 0.909122 |
| 12 | 54404400 | 54411799 | 100.2751 | 154.1033 | 194.8156 | 80.89446 | -0.04265 | 2.966955 | 0.909128 |
| X | 119199600 | 119203199 | 29.10275 | 48.39011 | 66.20879 | 28.72872 | -0.06978 | 1.2751 | 0.909163 |
| 5 | 53269200 | 53271599 | 23.81012 | 40.29465 | 50.64433 | 22.16867 | 0.052709 | 1.06978 | 0.909165 |
| 1 | 57054400 | 57055599 | 13.66942 | 25.13848 | 31.77618 | 13.49193 | 0.094791 | 0.264263 | 0.909192 |
| 7 | 54623800 | 54625399 | 24.901 | 42.15415 | 52.83876 | 23.13147 | 0.056521 | 1.07196 | 0.909213 |
| 13 | 43869200 | 43872399 | 27.15362 | 45.02077 | 61.01348 | 26.42239 | -0.05637 | 1.295336 | 0.909223 |
| 19 | 8229200 | 8235399 | 92.11116 | 142.324 | 180.6777 | 75.04958 | -0.04271 | 2.860742 | 0.909241 |
| 2 | 149888000 | 149888999 | 11.69103 | 21.48802 | 30.24987 | 12.85375 | -0.06003 | 0.250924 | 0.90926 |
| 11 | 13161200 | 13162799 | 30.69353 | 51.26631 | 51.88601 | 22.57958 | 0.363874 | 1.217688 | 0.909265 |
| 1 | 186793200 | 186795399 | 24.71851 | 41.35566 | 56.01266 | 24.40117 | -0.05509 | 1.185518 | 0.909274 |
| 1 | 53649800 | 53655799 | 90.09781 | 139.889 | 178.5403 | 74.14076 | -0.04505 | 2.773052 | 0.909371 |
| 8 | 40704400 | 40706199 | 19.76665 | 34.23549 | 43.14213 | 18.92812 | 0.053993 | 0.836757 | 0.909384 |
| 14 | 88313600 | 88317199 | 26.63916 | 45.41845 | 44.13145 | 19.346 | 0.423454 | 1.00535 | 0.909493 |
| X | 130510000 | 130510799 | 11.94819 | 22.31255 | 28.92692 | 12.01052 | 0.08322 | 0.159932 | 0.909493 |
| 14 | 67179000 | 67180399 | 21.38732 | 36.50247 | 45.37462 | 19.89265 | 0.06992 | 0.981117 | 0.909499 |
| 3 | 139525000 | 139528199 | 28.06116 | 46.838 | 63.97573 | 27.82891 | -0.06727 | 1.228017 | 0.909503 |
| 18 | 56958600 | 56963999 | 83.67492 | 130.5727 | 157.4531 | 65.33662 | 0.042414 | 2.681724 | 0.909523 |
| 6 | 129021000 | 129021399 | 5.360184 | 10.46612 | 17.60307 | 6.201072 | -0.07286 | -0.50456 | 0.909535 |
| 13 | 66296000 | 66296599 | 20.3489 | 34.81199 | 42.81946 | 18.776 | 0.085589 | 0.951556 | 0.909555 |
| 6 | 131210200 | 131212799 | 34.04035 | 55.6808 | 75.21637 | 32.34775 | -0.05765 | 1.488081 | 0.90959 |
| 6 | 99011800 | 99013399 | 14.589 | 25.82613 | 35.48148 | 15.69016 | -0.06459 | 0.510216 | 0.909598 |
| 6 | 147036400 | 147036999 | 14.589 | 25.82613 | 35.48148 | 15.69016 | -0.06459 | 0.510216 | 0.909598 |
| 12 | 30237200 | 30237999 | 10.05045 | 18.37691 | 32.89687 | 14.07748 | -0.41161 | 0.274226 | 0.909604 |
| 13 | 13517000 | 13519599 | 15.89654 | 28.16316 | 35.38626 | 15.64557 | 0.064048 | 0.510359 | 0.909611 |
| 8 | 126923200 | 126924599 | 18.95023 | 32.94588 | 41.52721 | 18.24003 | 0.054507 | 0.788283 | 0.909616 |
| 7 | 41284200 | 41285599 | 18.40036 | 32.21762 | 40.19965 | 17.74356 | 0.069607 | 0.678479 | 0.90963 |
| 7 | 134987000 | 134990399 | 29.96844 | 50.04538 | 61.90588 | 26.93665 | 0.075254 | 1.226088 | 0.909642 |
| 10 | 55123400 | 55124199 | 10.0324 | 18.94865 | 25.36253 | 10.32392 | 0.060666 | 0.060376 | 0.909672 |
| 16 | 35016800 | 35017799 | 11.62726 | 21.14768 | 29.73899 | 12.81078 | -0.07166 | 0.311037 | 0.909684 |
| 7 | 28430400 | 28433399 | 35.26227 | 57.22593 | 76.59095 | 32.82474 | -0.05282 | 1.61447 | 0.909687 |
| 4 | 35833000 | 35834399 | 13.10826 | 23.82849 | 30.55524 | 13.17126 | 0.059575 | 0.324349 | 0.909715 |
| 13 | 46710000 | 46714199 | 46.00887 | 73.79104 | 97.18586 | 41.06479 | -0.04875 | 1.959602 | 0.909716 |
| 8 | 74420800 | 74423199 | 19.19676 | 33.58695 | 41.83892 | 18.45546 | 0.072108 | 0.699085 | 0.909726 |

| | | | | | | | | | |
|---|---|---|---|---|---|---|---|---|---|
| 17 | 48362400 | 48365599 | 17.61411 | 30.68221 | 42.00958 | 18.47433 | -0.06414 | 0.753729 | 0.909745 |
| 3 | 26932200 | 26934599 | 23.62522 | 39.93327 | 50.11581 | 21.92925 | 0.054771 | 1.078604 | 0.90976 |
| 7 | 65831400 | 65833799 | 23.05318 | 39.01381 | 52.76758 | 23.09812 | -0.05275 | 1.069921 | 0.909765 |
| 1 | 64981400 | 64985599 | 34.07013 | 55.89205 | 69.79591 | 30.05236 | 0.05738 | 1.445786 | 0.909769 |
| 5 | 99325600 | 99331799 | 72.27787 | 114.2572 | 138.3313 | 57.47023 | 0.051144 | 2.419922 | 0.909787 |
| 10 | 10999400 | 11000199 | 7.898878 | 15.28083 | 21.56168 | 8.231603 | 0.070647 | -0.17841 | 0.909787 |
| 12 | 7195600 | 7198199 | 65.04663 | 102.6468 | 133.4906 | 55.41024 | -0.05391 | 2.454737 | 0.909832 |
| 7 | 6954600 | 6956399 | 20.39675 | 34.93571 | 47.42899 | 20.79925 | -0.05605 | 0.946518 | 0.909849 |
| 11 | 28923200 | 28926799 | 41.13206 | 67.39147 | 61.56709 | 26.495 | 0.506434 | 1.475425 | 0.909901 |
| 12 | 80350600 | 80352999 | 38.08372 | 61.87165 | 76.97177 | 33.01426 | 0.053755 | 1.595876 | 0.909917 |
| 10 | 120999000 | 121001999 | 41.5273 | 67.26008 | 68.13229 | 29.15705 | 0.345459 | 1.659916 | 0.909932 |
| 11 | 110430800 | 110431999 | 13.84035 | 24.82583 | 31.52674 | 13.80065 | 0.059162 | 0.41094 | 0.909955 |
| 8 | 39417400 | 39417799 | 6.21531 | 12.12472 | 18.06881 | 6.463546 | 0.078322 | -0.44078 | 0.909972 |
| 16 | 19407800 | 19425399 | 365.5457 | 507.9241 | 586.2707 | 243.3376 | -0.04789 | 4.677219 | 0.909979 |
| 1 | 33574200 | 33576999 | 25.03968 | 42.27719 | 57.85254 | 25.30611 | -0.06984 | 1.098527 | 0.90998 |
| 13 | 4272000 | 4275199 | 33.98905 | 55.79815 | 69.68833 | 30.01494 | 0.057614 | 1.438581 | 0.909992 |
| 11 | 116626800 | 116627599 | 12.63832 | 23.16403 | 32.83942 | 14.00747 | -0.07431 | 0.278499 | 0.910029 |
| 16 | 8369400 | 8387599 | 226.3014 | 328.9509 | 395.4805 | 163.0189 | -0.0416 | 4.004113 | 0.910047 |
| 3 | 135247800 | 135249399 | 13.68539 | 24.74637 | 31.45171 | 13.64767 | 0.065972 | 0.359263 | 0.910054 |
| 17 | 94200400 | 94203199 | 25.77904 | 43.7804 | 53.9973 | 23.6616 | 0.079844 | 1.047307 | 0.910069 |
| 18 | 84711800 | 84712599 | 6.678322 | 12.91989 | 20.51421 | 7.83087 | -0.09787 | -0.19712 | 0.910101 |
| 4 | 143404200 | 143409999 | 164.7083 | 244.5751 | 301.3557 | 124.1392 | -0.04788 | 3.655341 | 0.910108 |
| 7 | 68320600 | 68321999 | 17.32378 | 30.66137 | 30.75802 | 13.60194 | 0.386733 | 0.522257 | 0.910125 |
| 8 | 61893400 | 61897599 | 58.59925 | 93.23891 | 121.3395 | 50.69081 | -0.04428 | 2.228914 | 0.91013 |
| 17 | 71975600 | 71976399 | 13.45183 | 24.28651 | 31.00589 | 13.48024 | 0.057951 | 0.36716 | 0.910141 |
| 3 | 78947800 | 78949799 | 23.29271 | 40.48938 | 38.6833 | 16.98929 | 0.452509 | 0.806831 | 0.910151 |
| 4 | 56870400 | 56873999 | 40.13169 | 64.9891 | 80.71481 | 34.53756 | 0.05186 | 1.667229 | 0.910219 |
| 15 | 65049600 | 65052799 | 37.13802 | 60.11329 | 74.34411 | 31.8007 | 0.057357 | 1.667213 | 0.91029 |
| 18 | 68519600 | 68521799 | 28.4765 | 46.79885 | 63.88473 | 27.52786 | -0.06958 | 1.409759 | 0.91029 |
| X | 147436600 | 147439999 | 33.83454 | 55.69498 | 69.25334 | 29.86176 | 0.065504 | 1.405158 | 0.910291 |
| 8 | 60030400 | 60031399 | 11.81068 | 21.29448 | 39.52032 | 17.21131 | -0.48039 | 0.380156 | 0.91031 |
| 7 | 136258800 | 136260199 | 22.24618 | 37.95128 | 47.44104 | 20.79783 | 0.061731 | 0.991396 | 0.910357 |
| 11 | 35493400 | 35496999 | 32.59928 | 53.30723 | 66.50645 | 28.59874 | 0.056453 | 1.481643 | 0.910361 |
| 11 | 64598400 | 64603799 | 44.06408 | 71.02403 | 94.76927 | 40.33657 | -0.0588 | 1.802234 | 0.910364 |
| 3 | 102551000 | 102553399 | 19.48971 | 33.34789 | 45.86769 | 20.11291 | -0.07509 | 0.947945 | 0.910373 |
| 5 | 79462200 | 79463599 | 21.76527 | 37.24994 | 46.88553 | 20.5596 | 0.052414 | 0.957087 | 0.910407 |
| 9 | 5795800 | 5796999 | 14.50681 | 26.39163 | 33.11778 | 14.26179 | 0.091284 | 0.333864 | 0.910417 |
| 5 | 131032000 | 131039399 | 63.78707 | 100.5645 | 159.49 | 66.1854 | -0.34093 | 2.477883 | 0.910458 |
| 13 | 67844200 | 67848799 | 49.23469 | 78.89549 | 104.032 | 43.89893 | -0.052 | 1.992002 | 0.910466 |
| 13 | 79252400 | 79253599 | 11.35014 | 20.74987 | 29.17174 | 12.48601 | -0.06563 | 0.275481 | 0.910492 |
| 3 | 111345000 | 111346399 | 13.03168 | 23.94936 | 30.68461 | 13.04052 | 0.074533 | 0.261195 | 0.91052 |
| 8 | 117592400 | 117594399 | 16.73951 | 29.28409 | 39.86075 | 17.58104 | -0.05513 | 0.685148 | 0.910521 |
| X | 153035400 | 153059199 | 302.1361 | 434.1284 | 469.3749 | 193.7718 | 0.095021 | 4.18251 | 0.910534 |
| 9 | 16559200 | 16560799 | 15.88676 | 28.17877 | 35.36592 | 15.62828 | 0.066423 | 0.49956 | 0.91054 |
| 13 | 85774800 | 85776199 | 15.88676 | 28.17877 | 35.36592 | 15.62828 | 0.066423 | 0.49956 | 0.91054 |
| 11 | 10189200 | 10190399 | 12.37148 | 22.41672 | 31.1339 | 13.47495 | -0.05853 | 0.339568 | 0.910549 |
| 6 | 69990800 | 69994199 | 30.51342 | 50.4595 | 68.57482 | 29.64345 | -0.06071 | 1.341309 | 0.910554 |

| | | | | | | | | | |
|---|---|---|---|---|---|---|---|---|---|
| 3 | 19969600 | 19974399 | 63.43255 | 100.6177 | 122.7513 | 51.11184 | 0.044351 | 2.32345 | 0.91057 |
| 1 | 87246800 | 87250199 | 41.95182 | 67.96745 | 83.55073 | 35.76301 | 0.067245 | 1.662822 | 0.910578 |
| 17 | 5408200 | 5410599 | 22.55667 | 38.23419 | 47.47895 | 20.79311 | 0.070243 | 1.04971 | 0.910609 |
| 3 | 46890000 | 46890799 | 18.56026 | 32.02953 | 43.75854 | 19.19643 | -0.06303 | 0.865946 | 0.910623 |
| 16 | 76918000 | 76920399 | 31.59849 | 52.33586 | 53.02259 | 22.94537 | 0.361966 | 1.312786 | 0.910643 |
| 18 | 74463400 | 74483599 | 270.2498 | 387.5229 | 433.8183 | 179.1501 | 0.041917 | 4.207764 | 0.910661 |
| 7 | 40115600 | 40119799 | 44.21112 | 71.09799 | 93.78075 | 39.78452 | -0.04626 | 1.872832 | 0.910671 |
| 8 | 127715800 | 127720599 | 49.88537 | 79.90131 | 98.2439 | 41.42743 | 0.047911 | 2.005547 | 0.91069 |
| 18 | 45278000 | 45280399 | 20.69429 | 34.87906 | 58.60871 | 25.62322 | -0.36496 | 1.10274 | 0.910726 |
| 9 | 19511600 | 19514799 | 43.91734 | 70.85005 | 87.58485 | 37.32115 | 0.052669 | 1.774366 | 0.910749 |
| 5 | 58563400 | 58563799 | 6.234894 | 12.13045 | 19.58074 | 7.222716 | -0.07498 | -0.32377 | 0.910769 |
| X | 113926000 | 113928399 | 35.56536 | 58.36411 | 72.07527 | 31.0381 | 0.073647 | 1.449765 | 0.91077 |
| 7 | 98701600 | 98703999 | 25.76363 | 43.36691 | 54.48386 | 23.81051 | 0.05293 | 1.125773 | 0.91078 |
| 13 | 29144800 | 29146399 | 17.92424 | 31.28723 | 39.33212 | 17.31944 | 0.058823 | 0.722582 | 0.910797 |
| 3 | 7740000 | 7740599 | 9.148473 | 17.33691 | 25.4647 | 10.29258 | -0.06314 | 0.030668 | 0.910802 |
| 15 | 82180000 | 82183799 | 30.54979 | 50.1006 | 67.47552 | 29.04925 | -0.05147 | 1.441502 | 0.910803 |
| X | 52712800 | 52717599 | 34.59255 | 56.87806 | 70.292 | 30.29505 | 0.073709 | 1.423139 | 0.910809 |
| 7 | 48034400 | 48040199 | 71.91641 | 113.884 | 137.185 | 57.04424 | 0.06042 | 2.378549 | 0.91082 |
| 16 | 78642400 | 78647199 | 69.48104 | 109.4849 | 141.4433 | 58.69241 | -0.04602 | 2.488859 | 0.910853 |
| 12 | 115043600 | 115046599 | 38.57713 | 63.23087 | 61.99176 | 26.68229 | 0.405058 | 1.463955 | 0.91086 |
| 5 | 17253400 | 17254399 | 15.92051 | 28.12502 | 35.47254 | 15.69269 | 0.057084 | 0.542216 | 0.910876 |
| 7 | 7063400 | 7101199 | 873.2088 | 1130.978 | 1213.225 | 508.4264 | -0.04399 | 5.795818 | 0.910893 |
| 5 | 93828400 | 93829399 | 26.32284 | 43.88293 | 54.97939 | 23.89718 | 0.057045 | 1.227299 | 0.910919 |
| 2 | 132737800 | 132738999 | 8.936289 | 17.10042 | 23.46485 | 9.28234 | 0.061761 | -0.04734 | 0.910945 |
| 6 | 132283600 | 132290399 | 83.41627 | 130.8207 | 156.9995 | 65.11495 | 0.054894 | 2.594513 | 0.910946 |
| 12 | 7641000 | 7642399 | 20.16025 | 35.16649 | 35.65729 | 15.69461 | 0.367513 | 0.738642 | 0.910964 |
| 4 | 94663800 | 94666599 | 24.99057 | 41.92649 | 56.62368 | 24.71039 | -0.0511 | 1.156497 | 0.910981 |
| 14 | 93157400 | 93160199 | 23.52865 | 39.39508 | 68.17185 | 29.7125 | -0.40753 | 1.185156 | 0.911006 |
| 13 | 80849200 | 80851999 | 41.98964 | 67.49373 | 89.68738 | 38.02204 | -0.05769 | 1.884512 | 0.911017 |
| 19 | 26325000 | 26328399 | 23.68508 | 40.36874 | 55.64334 | 24.39185 | -0.07924 | 1.009255 | 0.911029 |
| 16 | 48725200 | 48725799 | 8.824692 | 16.84614 | 25.06028 | 9.973572 | -0.06173 | -0.0239 | 0.911049 |
| 10 | 66468000 | 66469199 | 16.45343 | 28.85479 | 39.26414 | 17.35407 | -0.05468 | 0.643529 | 0.911063 |
| 16 | 19201000 | 19203199 | 30.82105 | 50.73891 | 63.79829 | 27.51057 | 0.049543 | 1.392279 | 0.911079 |
| 1 | 79466000 | 79466399 | 6.683084 | 12.95757 | 20.55157 | 7.764926 | -0.08199 | -0.2307 | 0.911079 |
| 8 | 5009600 | 5018799 | 259.4801 | 370.8219 | 441.3963 | 182.3762 | -0.05146 | 4.256824 | 0.911081 |
| 1 | 24929200 | 24933199 | 39.12882 | 63.39475 | 84.93936 | 36.35779 | -0.05672 | 1.657309 | 0.911085 |
| Y | 1717800 | 1718399 | 4.980887 | 9.711696 | 17.00667 | 6.124202 | -0.15929 | -0.43702 | 0.911095 |
| 6 | 53733800 | 53735799 | 19.54358 | 34.22408 | 42.32588 | 18.68425 | 0.082449 | 0.68756 | 0.911148 |
| 9 | 53864200 | 53866799 | 22.4588 | 37.64345 | 64.25347 | 28.02333 | -0.38784 | 1.16472 | 0.911155 |
| 13 | 17112000 | 17114799 | 23.81123 | 40.4989 | 50.65172 | 22.19998 | 0.060089 | 1.023563 | 0.911187 |
| 6 | 93615400 | 93617799 | 24.66952 | 41.51128 | 56.07287 | 24.50244 | -0.05128 | 1.127043 | 0.911203 |
| X | 67205000 | 67206599 | 18.41515 | 32.26839 | 45.06075 | 19.90215 | -0.09188 | 0.677987 | 0.911233 |
| 6 | 60554000 | 60555799 | 34.51005 | 56.06985 | 69.3908 | 29.76427 | 0.061439 | 1.58403 | 0.911255 |
| 8 | 53228000 | 53229199 | 17.30776 | 30.37875 | 38.17496 | 16.88474 | 0.059654 | 0.633949 | 0.911275 |
| 16 | 63828000 | 63828799 | 10.94834 | 21.10009 | 28.0141 | 10.85811 | 0.13623 | -0.08123 | 0.911278 |
| X | 30807800 | 30855599 | 2674.552 | 3188.35 | 2888.488 | 1222.339 | 0.078604 | 7.1261 | 0.911284 |
| 7 | 57537400 | 57538199 | 12.72653 | 22.86417 | 31.61638 | 13.82139 | -0.0616 | 0.393529 | 0.911289 |

| | | | | | | | | | |
|---|---|---|---|---|---|---|---|---|---|
| 16 | 26665200 | 26667199 | 16.12516 | 28.53383 | 39.60349 | 17.51437 | -0.0795 | 0.532371 | 0.911319 |
| 4 | 40043600 | 40056999 | 124.7013 | 189.1964 | 236.795 | 98.03094 | -0.04289 | 3.234302 | 0.91133 |
| 12 | 43263000 | 43263599 | 6.531119 | 12.71531 | 18.51287 | 6.7919 | 0.079951 | -0.34145 | 0.911348 |
| 5 | 127660600 | 127665399 | 43.98597 | 70.67585 | 93.58814 | 39.65395 | -0.0534 | 1.897767 | 0.911356 |
| 2 | 177208400 | 177220799 | 214.8358 | 315.8369 | 358.802 | 147.7639 | 0.055755 | 3.829258 | 0.911374 |
| 12 | 72378600 | 72380599 | 22.29548 | 38.0979 | 51.72325 | 22.67813 | -0.05677 | 0.978602 | 0.911377 |
| 10 | 76863200 | 76865599 | 20.30655 | 34.96384 | 47.40251 | 20.79357 | -0.05285 | 0.897233 | 0.911379 |
| 3 | 106515200 | 106519599 | 56.33197 | 89.96695 | 109.9715 | 46.18136 | 0.051784 | 2.106882 | 0.911387 |
| 11 | 121197400 | 121201999 | 57.43519 | 91.29222 | 118.7967 | 49.56713 | -0.04568 | 2.258804 | 0.911427 |
| 4 | 34425800 | 34430599 | 43.82457 | 70.61971 | 87.42903 | 37.19933 | 0.048621 | 1.805742 | 0.911437 |
| 2 | 13861000 | 13861599 | 6.673304 | 12.97074 | 20.64317 | 7.674864 | -0.06558 | -0.28746 | 0.911463 |
| 11 | 104702200 | 104707399 | 104.6149 | 160.5446 | 202.6754 | 84.14176 | -0.0422 | 2.997827 | 0.91148 |
| 12 | 5535000 | 5539399 | 41.06714 | 66.13462 | 87.76306 | 37.30733 | -0.05244 | 1.821434 | 0.911483 |
| 16 | 15123600 | 15134199 | 131.2693 | 198.7202 | 248.1872 | 102.6767 | -0.04252 | 3.281523 | 0.911505 |
| 6 | 58162600 | 58164999 | 25.73102 | 43.41963 | 59.45606 | 26.00302 | -0.07076 | 1.110651 | 0.911521 |
| 10 | 71373200 | 71374799 | 17.40204 | 30.51018 | 41.82129 | 18.48008 | -0.0653 | 0.656097 | 0.911531 |
| 1 | 33790200 | 33795199 | 39.68494 | 64.105 | 79.18487 | 33.79817 | 0.055933 | 1.723904 | 0.911583 |
| 7 | 44168200 | 44171799 | 31.33923 | 51.35868 | 64.14452 | 27.60689 | 0.056653 | 1.449764 | 0.911616 |
| 3 | 71060000 | 71060799 | 11.3524 | 21.02601 | 29.84015 | 12.54823 | -0.06082 | 0.204335 | 0.911618 |
| 16 | 40805400 | 40806799 | 11.3524 | 21.02601 | 29.84015 | 12.54823 | -0.06082 | 0.204335 | 0.911618 |
| 17 | 17001800 | 17003799 | 23.56671 | 39.18469 | 65.64487 | 28.4838 | -0.36084 | 1.259271 | 0.91163 |
| X | 15662000 | 15662999 | 18.99591 | 33.18011 | 41.43718 | 18.25407 | 0.068596 | 0.725661 | 0.911631 |
| 14 | 79477400 | 79478999 | 20.24445 | 34.83369 | 43.71863 | 19.17721 | 0.057872 | 0.897937 | 0.911635 |
| 1 | 126406400 | 126408999 | 16.9593 | 29.67937 | 37.22844 | 16.42506 | 0.062153 | 0.674979 | 0.911641 |
| 13 | 119324000 | 119326799 | 35.61422 | 57.5482 | 92.43462 | 39.47017 | -0.32153 | 1.716112 | 0.911648 |
| 13 | 83567000 | 83569599 | 28.73225 | 47.24325 | 64.06868 | 27.61308 | -0.05989 | 1.402266 | 0.911653 |
| 1 | 116766200 | 116767399 | 21.76115 | 37.79616 | 38.28142 | 16.80415 | 0.368515 | 0.810029 | 0.911659 |
| 17 | 49980600 | 49982999 | 27.71121 | 45.78491 | 61.99603 | 26.78317 | -0.05549 | 1.339171 | 0.91169 |
| 6 | 96738800 | 96739599 | 13.0885 | 23.57519 | 32.62065 | 14.22393 | -0.06005 | 0.383226 | 0.911693 |
| 2 | 28009600 | 28014799 | 72.63473 | 113.978 | 146.7534 | 60.86282 | -0.04579 | 2.575894 | 0.911699 |
| 8 | 51607200 | 51608199 | 12.57567 | 23.27288 | 22.95508 | 9.663539 | 0.459391 | 0.201598 | 0.9117 |
| 4 | 8485000 | 8486999 | 25.77748 | 43.0917 | 58.3233 | 25.39451 | -0.05418 | 1.197636 | 0.911738 |
| 9 | 96807800 | 96810799 | 27.02808 | 44.66396 | 60.88362 | 26.30573 | -0.06509 | 1.333546 | 0.911747 |
| 18 | 52538800 | 52539599 | 8.784927 | 16.87805 | 22.97751 | 8.985905 | 0.08766 | -0.09146 | 0.911777 |
| 9 | 83858800 | 83861599 | 35.45562 | 57.65836 | 72.00812 | 30.9022 | 0.04938 | 1.571095 | 0.91178 |
| 15 | 96662400 | 96664999 | 30.01614 | 49.32166 | 61.37178 | 26.44334 | 0.063547 | 1.411812 | 0.911807 |
| 4 | 21649600 | 21651799 | 37.84197 | 61.24121 | 76.02085 | 32.51341 | 0.051771 | 1.674255 | 0.911813 |
| 18 | 36605400 | 36607599 | 16.52917 | 29.03418 | 36.57603 | 16.18791 | 0.055955 | 0.612667 | 0.911852 |
| 9 | 39650800 | 39651599 | 11.48236 | 21.23567 | 27.69496 | 11.6693 | 0.058336 | 0.208748 | 0.911863 |
| 19 | 57783400 | 57784599 | 11.48236 | 21.23567 | 27.69496 | 11.6693 | 0.058336 | 0.208748 | 0.911863 |
| 14 | 51363200 | 51365799 | 28.04177 | 46.47303 | 62.66699 | 27.13028 | -0.04919 | 1.305753 | 0.911863 |
| 7 | 12476400 | 12485199 | 118.0567 | 179.5628 | 224.9879 | 93.21254 | -0.04155 | 3.181433 | 0.911898 |
| 8 | 67832200 | 67833399 | 15.23134 | 26.93332 | 33.96006 | 15.02071 | 0.058001 | 0.52422 | 0.91191 |
| 11 | 25559600 | 25564399 | 45.11531 | 72.13373 | 115.511 | 48.58463 | -0.33539 | 2.060266 | 0.911912 |
| 8 | 64131200 | 64132599 | 17.96817 | 31.30056 | 39.03326 | 17.166 | 0.070419 | 0.755013 | 0.911914 |
| 19 | 30154400 | 30156999 | 27.18755 | 45.25178 | 61.05783 | 26.51233 | -0.04961 | 1.252271 | 0.911915 |
| 13 | 84330400 | 84331999 | 12.75876 | 22.88464 | 31.99121 | 14.00455 | -0.07897 | 0.401405 | 0.911918 |

| | | | | | | | | | |
|---|---|---|---|---|---|---|---|---|---|
| 2 | 166008200 | 166009599 | 13.57225 | 24.09703 | 29.58842 | 13.06441 | 0.098762 | 0.46672 | 0.911924 |
| 1 | 128245600 | 128248199 | 22.84399 | 38.52987 | 52.23724 | 22.84264 | -0.05638 | 1.09649 | 0.911935 |
| 11 | 51424000 | 51426599 | 16.19594 | 28.34746 | 38.8052 | 17.12276 | -0.06334 | 0.671184 | 0.911946 |
| 15 | 9275000 | 9276999 | 22.4553 | 38.18873 | 47.96888 | 21.02395 | 0.054007 | 1.018756 | 0.911966 |
| 10 | 41648200 | 41668799 | 517.3238 | 701.6569 | 788.0079 | 328.3144 | -0.0436 | 5.054511 | 0.911982 |
| 1 | 33058800 | 33060799 | 25.36877 | 42.51889 | 53.336 | 23.26113 | 0.055196 | 1.166873 | 0.912024 |
| 7 | 102861200 | 102862999 | 14.06461 | 25.17624 | 31.92806 | 14.00059 | 0.059183 | 0.424077 | 0.912034 |
| 6 | 129073000 | 129073399 | 7.255114 | 14.15661 | 16.34316 | 5.824158 | 0.448231 | -0.44336 | 0.912085 |
| 3 | 61168000 | 61168599 | 6.237526 | 12.12729 | 19.55665 | 7.253718 | -0.08079 | -0.30408 | 0.912086 |
| 18 | 60074600 | 60075399 | 10.76003 | 20.05171 | 28.85889 | 12.02483 | -0.07116 | 0.158822 | 0.91209 |
| 1 | 162251200 | 162253999 | 25.34024 | 42.47461 | 53.34671 | 23.26701 | 0.053384 | 1.165301 | 0.912095 |
| 9 | 84733600 | 84734599 | 18.27096 | 31.83758 | 39.995 | 17.59228 | 0.059854 | 0.752996 | 0.912119 |
| 1 | 67704000 | 67704399 | 5.794195 | 11.30876 | 17.00792 | 6.03544 | 0.077256 | -0.47446 | 0.912132 |
| 1 | 182326400 | 182329799 | 26.36039 | 43.77583 | 59.29393 | 25.70621 | -0.05536 | 1.274008 | 0.912168 |
| 13 | 15061200 | 15061799 | 8.259962 | 15.84008 | 24.08416 | 9.468936 | -0.0761 | -0.06951 | 0.912178 |
| 1 | 53064400 | 53065799 | 15.21124 | 27.4653 | 33.94061 | 14.75461 | 0.104347 | 0.381576 | 0.912185 |
| 3 | 91865800 | 91866999 | 12.91297 | 23.55186 | 33.24582 | 14.27145 | -0.07452 | 0.308046 | 0.912198 |
| 1 | 18957400 | 18964399 | 80.02513 | 125.7424 | 164.0369 | 68.04256 | -0.06323 | 2.55619 | 0.912229 |
| 7 | 86748800 | 86750799 | 20.39549 | 35.62942 | 44.00974 | 19.38894 | 0.084231 | 0.732684 | 0.912232 |
| 1 | 112413800 | 112415199 | 20.24752 | 35.04546 | 44.18391 | 19.3848 | 0.053081 | 0.845836 | 0.912297 |
| X | 81514200 | 81518799 | 35.89203 | 58.8495 | 81.7182 | 35.17787 | -0.09568 | 1.465829 | 0.912336 |
| 12 | 17076800 | 17079399 | 28.84961 | 47.56899 | 59.23154 | 25.55852 | 0.064543 | 1.364567 | 0.912338 |
| 9 | 32864000 | 32864999 | 13.368 | 24.62046 | 24.97479 | 10.58284 | 0.412676 | 0.241061 | 0.912344 |
| 4 | 10486800 | 10492799 | 93.16664 | 143.7312 | 170.3169 | 70.74648 | 0.054874 | 2.888263 | 0.912347 |
| 12 | 65080600 | 65083599 | 33.18601 | 54.39281 | 68.2779 | 29.38758 | 0.049446 | 1.455836 | 0.91236 |
| 3 | 117449800 | 117450199 | 4.929636 | 9.639092 | 14.94101 | 5.1025 | 0.084681 | -0.62995 | 0.912372 |
| X | 46560800 | 46562999 | 22.77 | 39.72074 | 37.0502 | 16.3082 | 0.487455 | 0.754505 | 0.912384 |
| 2 | 65409200 | 65413999 | 87.5851 | 136.0174 | 163.3064 | 67.81253 | 0.043283 | 2.765859 | 0.912408 |
| 13 | 99107600 | 99111999 | 46.26113 | 74.36019 | 91.83492 | 38.93284 | 0.047719 | 1.891222 | 0.912418 |
| 5 | 32827000 | 32828999 | 17.72256 | 31.1779 | 43.85687 | 19.41805 | -0.10212 | 0.610526 | 0.912429 |
| 2 | 150099000 | 150100399 | 25.54587 | 42.95489 | 53.94497 | 23.56745 | 0.053413 | 1.132612 | 0.912436 |
| 5 | 127660400 | 127665199 | 44.30877 | 71.18127 | 94.20562 | 39.90482 | -0.05295 | 1.904179 | 0.912445 |
| 8 | 67457600 | 67459799 | 25.76373 | 44.16988 | 42.58006 | 18.67424 | 0.436157 | 0.95426 | 0.912451 |
| 15 | 26280400 | 26281199 | 16.35784 | 28.64098 | 35.78411 | 15.79444 | 0.067939 | 0.66236 | 0.912453 |
| 5 | 85833600 | 85834199 | 8.508889 | 16.3403 | 24.67658 | 9.662937 | -0.06155 | -0.08077 | 0.912479 |
| 10 | 124519800 | 124523399 | 56.06427 | 89.01812 | 137.6476 | 57.37124 | -0.2957 | 2.293711 | 0.912499 |
| 4 | 10428000 | 10429799 | 25.94609 | 43.37061 | 54.46518 | 23.71363 | 0.053577 | 1.196285 | 0.912499 |
| 8 | 65466000 | 65472399 | 77.42192 | 121.1162 | 155.5447 | 64.5282 | -0.04557 | 2.632868 | 0.912527 |
| 14 | 8252400 | 8253399 | 14.07485 | 24.88354 | 34.18474 | 15.12066 | -0.06488 | 0.522057 | 0.912531 |
| 13 | 51229800 | 51231599 | 29.691 | 48.7687 | 65.84406 | 28.36572 | -0.05413 | 1.418086 | 0.912547 |
| 1 | 13879200 | 13890999 | 183.7333 | 271.929 | 314.289 | 129.4335 | 0.039997 | 3.714701 | 0.912563 |
| 5 | 62032800 | 62033999 | 18.05371 | 31.35841 | 42.64708 | 18.7211 | -0.0547 | 0.799674 | 0.912571 |
| 9 | 63722800 | 63724199 | 14.76101 | 26.37731 | 33.43174 | 14.68128 | 0.058608 | 0.443078 | 0.912579 |
| 7 | 74501000 | 74502199 | 18.26255 | 32.17031 | 32.27616 | 14.28681 | 0.383909 | 0.581243 | 0.912591 |
| 9 | 19519000 | 19520799 | 24.43868 | 41.88316 | 42.58083 | 18.67405 | 0.359557 | 0.949893 | 0.912597 |
| 15 | 8640000 | 8642199 | 20.04036 | 34.9363 | 43.5001 | 19.13938 | 0.072341 | 0.756662 | 0.9126 |
| X | 7240200 | 7243199 | 39.5282 | 64.38248 | 78.10488 | 33.53785 | 0.092801 | 1.565414 | 0.912604 |

| | | | | | | | | | |
|---|---|---|---|---|---|---|---|---|---|
| 16 | 18857400 | 18861999 | 55.92436 | 89.43123 | 109.2033 | 45.94261 | 0.055294 | 2.06598 | 0.912618 |
| 19 | 24947000 | 24947999 | 11.65039 | 21.39588 | 27.65158 | 11.76271 | 0.06148 | 0.249274 | 0.912626 |
| 3 | 59670400 | 59671999 | 28.01271 | 46.52068 | 62.75812 | 27.20843 | -0.0496 | 1.281574 | 0.912628 |
| 18 | 42511800 | 42513199 | 15.7285 | 27.53986 | 37.81016 | 16.68911 | -0.06749 | 0.661184 | 0.912663 |
| 11 | 87784200 | 87785999 | 17.26871 | 30.14156 | 40.97931 | 18.04422 | -0.05352 | 0.724783 | 0.912668 |
| 13 | 78936000 | 78937599 | 13.04412 | 23.9296 | 34.51753 | 14.70501 | -0.09809 | 0.27821 | 0.912675 |
| 13 | 91860200 | 91862999 | 36.23599 | 59.04258 | 79.75366 | 34.25056 | -0.06153 | 1.551311 | 0.912695 |
| 6 | 129142400 | 129143799 | 15.5176 | 28.33848 | 34.90305 | 14.95901 | 0.122672 | 0.325038 | 0.912702 |
| 3 | 87506000 | 87516799 | 315.0141 | 444.3738 | 518.0931 | 214.53 | -0.04043 | 4.449811 | 0.912706 |
| 6 | 131497400 | 131498999 | 20.78749 | 36.17623 | 36.72588 | 16.1413 | 0.365446 | 0.775686 | 0.912719 |
| 7 | 92995000 | 93006599 | 268.3633 | 387.2015 | 370.0189 | 152.664 | 0.278967 | 4.11669 | 0.912726 |
| 15 | 98524400 | 98526999 | 20.62995 | 35.36037 | 44.28868 | 19.42323 | 0.059884 | 0.936068 | 0.912732 |
| 17 | 34971000 | 34981599 | 281.4573 | 400.4795 | 441.9328 | 182.7312 | 0.051416 | 4.321309 | 0.912753 |
| 2 | 62543400 | 62545599 | 23.1739 | 39.87291 | 40.52288 | 17.77538 | 0.361112 | 0.907742 | 0.912761 |
| 14 | 56808800 | 56809599 | 6.665052 | 12.89508 | 20.49944 | 7.822778 | -0.09958 | -0.19999 | 0.912785 |
| 7 | 23926800 | 23934799 | 91.08529 | 140.8362 | 167.8008 | 69.70052 | 0.049479 | 2.846088 | 0.912797 |
| 2 | 149018200 | 149022599 | 85.61587 | 133.9956 | 160.7544 | 66.68631 | 0.053095 | 2.632027 | 0.912812 |
| 18 | 54835800 | 54841399 | 59.01145 | 94.21128 | 126.1061 | 52.92906 | -0.07963 | 2.125001 | 0.912825 |
| 14 | 120279600 | 120282599 | 34.34506 | 56.0024 | 74.98109 | 32.20953 | -0.04805 | 1.532791 | 0.912838 |
| 14 | 21408400 | 21412599 | 58.57114 | 93.08294 | 113.2892 | 47.26162 | 0.050467 | 2.263257 | 0.912845 |
| 14 | 105454800 | 105456599 | 16.29741 | 28.53621 | 38.84773 | 17.1472 | -0.05523 | 0.667717 | 0.912849 |
| 7 | 10212800 | 10215799 | 30.93605 | 51.28685 | 64.14454 | 27.77638 | 0.059037 | 1.309932 | 0.912854 |
| 10 | 28217600 | 28221599 | 45.17113 | 72.62775 | 95.86968 | 40.65928 | -0.04769 | 1.880513 | 0.912866 |
| 17 | 19218200 | 19219399 | 15.69268 | 27.77929 | 35.00659 | 15.48023 | 0.059595 | 0.519549 | 0.912867 |
| 18 | 33461400 | 33463999 | 24.37461 | 40.54076 | 67.44082 | 29.26943 | -0.35075 | 1.259231 | 0.912873 |
| Y | 43876800 | 43878799 | 30.33492 | 49.25538 | 68.78968 | 29.49276 | -0.1133 | 1.587529 | 0.912879 |
| 6 | 140250000 | 140252199 | 25.78666 | 43.15492 | 58.3253 | 25.41317 | -0.05214 | 1.186182 | 0.912909 |
| 13 | 71295600 | 71298399 | 17.18044 | 30.02883 | 40.89042 | 18.02168 | -0.05576 | 0.702384 | 0.912922 |
| Y | 4951600 | 4952999 | 15.99821 | 27.55572 | 40.83736 | 17.9139 | -0.18042 | 0.865975 | 0.912937 |
| 8 | 61134600 | 61140199 | 74.31986 | 116.6469 | 149.8585 | 62.15227 | -0.04237 | 2.573811 | 0.912956 |
| 18 | 41690000 | 41691599 | 15.71911 | 27.73377 | 38.04585 | 16.8354 | -0.0644 | 0.557932 | 0.912971 |
| 5 | 15635000 | 15651999 | 206.5283 | 304.9535 | 346.4998 | 142.6953 | 0.061423 | 3.756637 | 0.912977 |
| 8 | 85836200 | 85840399 | 85.74133 | 132.811 | 169.8803 | 70.56365 | -0.05085 | 2.810647 | 0.912977 |
| 4 | 100865600 | 100876599 | 160.2382 | 239.7564 | 279.7846 | 115.3831 | 0.040552 | 3.516325 | 0.913006 |
| 17 | 59109800 | 59111199 | 15.8136 | 27.73207 | 37.78792 | 16.70128 | -0.05655 | 0.638355 | 0.913007 |
| 6 | 3241200 | 3245799 | 41.48215 | 67.32589 | 82.55214 | 35.38183 | 0.073285 | 1.62667 | 0.913033 |
| 8 | 25992200 | 25995599 | 23.7775 | 40.24198 | 54.45146 | 23.83563 | -0.05346 | 1.071668 | 0.913056 |
| 13 | 45733000 | 45737199 | 39.13927 | 63.08931 | 84.01708 | 35.76249 | -0.05582 | 1.788707 | 0.913056 |
| 13 | 35146600 | 35148999 | 20.14008 | 34.4876 | 46.89755 | 20.56579 | -0.05856 | 0.945035 | 0.913061 |
| 5 | 24986600 | 24991199 | 58.81122 | 93.70416 | 114.7748 | 48.03909 | 0.045112 | 2.18253 | 0.913077 |
| 7 | 12866800 | 12867399 | 8.037225 | 15.50366 | 23.68549 | 9.156608 | -0.06065 | -0.12988 | 0.913079 |
| 15 | 23978000 | 23978599 | 8.037225 | 15.50366 | 23.68549 | 9.156608 | -0.06065 | -0.12988 | 0.913079 |
| 8 | 55159400 | 55165399 | 72.61212 | 114.1056 | 146.6266 | 60.82154 | -0.04134 | 2.54759 | 0.913089 |
| 1 | 18192000 | 18192399 | 7.082711 | 13.78431 | 20.16958 | 7.420903 | 0.067663 | -0.32278 | 0.913118 |
| 4 | 108136600 | 108141399 | 72.03181 | 113.8496 | 137.9424 | 57.30392 | 0.049855 | 2.422583 | 0.913135 |
| 13 | 29092800 | 29093999 | 18.18228 | 31.596 | 39.56199 | 17.37175 | 0.064118 | 0.789587 | 0.913142 |
| 7 | 87858200 | 87860199 | 23.57554 | 40.84686 | 39.79965 | 17.46183 | 0.423555 | 0.845812 | 0.913149 |

| | | | | | | | | | |
|---|---|---|---|---|---|---|---|---|---|
| 10 | 39879000 | 39881199 | 16.20679 | 28.42707 | 35.69107 | 15.77709 | 0.060869 | 0.632679 | 0.913177 |
| 16 | 57604200 | 57605199 | 8.476565 | 16.37317 | 22.60565 | 8.692461 | 0.090326 | -0.15108 | 0.913202 |
| 1 | 102236600 | 102238799 | 29.81658 | 49.36585 | 66.85505 | 28.92378 | -0.05556 | 1.324008 | 0.913219 |
| 17 | 72920400 | 72925199 | 63.33663 | 100.4385 | 122.705 | 51.07877 | 0.041959 | 2.332103 | 0.913242 |
| 19 | 6190600 | 6193799 | 23.20238 | 39.23445 | 49.05223 | 21.4665 | 0.060333 | 1.074004 | 0.913244 |
| 8 | 11633400 | 11637599 | 55.74874 | 88.764 | 115.504 | 48.29859 | -0.04306 | 2.202321 | 0.913244 |
| 6 | 116704200 | 116704799 | 12.43889 | 22.4437 | 31.16968 | 13.56202 | -0.0632 | 0.365953 | 0.91326 |
| 9 | 88948400 | 88951599 | 56.57592 | 89.9099 | 117.3299 | 48.94614 | -0.05002 | 2.262448 | 0.913279 |
| 2 | 68104000 | 68105799 | 16.30745 | 28.52124 | 38.91232 | 17.15917 | -0.05831 | 0.686642 | 0.913288 |
| 13 | 108138000 | 108141199 | 33.21137 | 53.61219 | 90.22494 | 38.47727 | -0.39027 | 1.740649 | 0.913312 |
| 7 | 93348800 | 93352199 | 37.58484 | 61.24864 | 76.08689 | 32.67745 | 0.059105 | 1.547974 | 0.913312 |
| 18 | 36304000 | 36306199 | 16.894 | 29.6003 | 40.29201 | 17.7946 | -0.05523 | 0.661275 | 0.913331 |
| 6 | 121954600 | 121957799 | 45.15497 | 72.64773 | 96.24724 | 40.85687 | -0.05183 | 1.860567 | 0.913341 |
| 7 | 20033600 | 20033799 | 10.71881 | 20.52874 | 19.73596 | 7.788373 | 0.574721 | -0.04589 | 0.913374 |
| 3 | 112497200 | 112499599 | 36.31911 | 59.18055 | 73.9885 | 31.77518 | 0.049861 | 1.546551 | 0.913379 |
| 2 | 112877800 | 112878599 | 9.188941 | 17.54282 | 23.93812 | 9.525434 | 0.061619 | -0.02607 | 0.913407 |
| 5 | 145190200 | 145191999 | 26.62169 | 44.41462 | 60.00674 | 26.09476 | -0.05157 | 1.22318 | 0.913425 |
| 11 | 87507600 | 87509599 | 31.96162 | 51.55121 | 91.7063 | 39.06644 | -0.47148 | 1.765756 | 0.913448 |
| 10 | 55632600 | 55633799 | 18.39895 | 32.07655 | 40.4605 | 17.8026 | 0.053974 | 0.746514 | 0.913452 |
| 1 | 35349000 | 35349399 | 6.375053 | 12.42933 | 18.59793 | 6.702182 | 0.063653 | -0.40806 | 0.913453 |
| 8 | 18181600 | 18184399 | 27.89148 | 46.97574 | 47.49188 | 20.75851 | 0.365426 | 1.122458 | 0.913455 |
| 2 | 143730000 | 143732999 | 27.90463 | 46.24281 | 62.83893 | 27.20352 | -0.06037 | 1.302983 | 0.913491 |
| 10 | 3108800 | 3110399 | 15.76608 | 27.68329 | 37.68677 | 16.67416 | -0.05535 | 0.616668 | 0.913497 |
| 3 | 92224000 | 92225199 | 11.32461 | 20.90734 | 27.1836 | 11.48112 | 0.060297 | 0.216151 | 0.913516 |
| X | 103967400 | 103969399 | 22.73862 | 39.40134 | 48.4849 | 21.2725 | 0.088297 | 0.854444 | 0.913531 |
| 10 | 77948000 | 77952199 | 48.14958 | 77.10008 | 101.1151 | 42.62048 | -0.04542 | 2.013658 | 0.913537 |
| 2 | 88441200 | 88448399 | 143.7102 | 216.545 | 254.3774 | 105.0867 | 0.040057 | 3.377495 | 0.913551 |
| 19 | 48732600 | 48734799 | 49.49585 | 79.19391 | 97.05731 | 40.86399 | 0.050917 | 2.03887 | 0.913554 |
| 9 | 93258600 | 93261399 | 27.0881 | 44.96494 | 61.11337 | 26.48713 | -0.06043 | 1.280281 | 0.913595 |
| 2 | 150377600 | 150378999 | 14.04821 | 25.33896 | 35.64249 | 15.51463 | -0.08252 | 0.383996 | 0.913599 |
| 8 | 6687600 | 6687999 | 6.657047 | 12.93963 | 20.64809 | 7.674361 | -0.06939 | -0.29044 | 0.913608 |
| 2 | 164514800 | 164515199 | 8.15233 | 15.76119 | 24.41795 | 9.349581 | -0.06796 | -0.16056 | 0.913621 |
| 5 | 34108800 | 34118599 | 131.9585 | 200.4525 | 235.792 | 97.65152 | 0.04798 | 3.211807 | 0.913625 |
| 1 | 37812800 | 37814199 | 20.55407 | 35.37238 | 47.93969 | 21.02889 | -0.05243 | 0.90493 | 0.913632 |
| 15 | 73248600 | 73252399 | 50.984 | 81.8133 | 100.4129 | 42.46221 | 0.053699 | 1.951689 | 0.913648 |
| 13 | 9079800 | 9082599 | 29.38856 | 48.69397 | 61.21424 | 26.49718 | 0.051606 | 1.308702 | 0.913654 |
| 18 | 4502200 | 4505399 | 38.21615 | 61.67096 | 82.21292 | 35.04558 | -0.05539 | 1.755105 | 0.913659 |
| 2 | 9679200 | 9680399 | 17.97225 | 31.28934 | 42.6061 | 18.73038 | -0.05609 | 0.767805 | 0.913689 |
| 13 | 22352200 | 22353399 | 38.86495 | 62.78178 | 83.29644 | 35.55408 | -0.04687 | 1.726995 | 0.913708 |
| 12 | 91439800 | 91442399 | 61.41226 | 96.37895 | 159.0939 | 65.98174 | -0.40362 | 2.570267 | 0.913711 |
| 3 | 133485200 | 133488399 | 34.53504 | 55.67836 | 92.86059 | 39.53659 | -0.37941 | 1.781812 | 0.913724 |
| 8 | 24902000 | 24905599 | 29.66641 | 49.0525 | 66.19164 | 28.6107 | -0.0505 | 1.338501 | 0.913727 |
| 3 | 87507000 | 87516999 | 303.7557 | 429.6845 | 502.3558 | 207.9181 | -0.04039 | 4.408488 | 0.913732 |
| 13 | 11626000 | 11630599 | 86.10751 | 133.8948 | 170.4927 | 70.78302 | -0.04 | 2.745071 | 0.913733 |
| 8 | 81883000 | 81883999 | 12.12143 | 22.28814 | 28.54819 | 12.12397 | 0.075518 | 0.244989 | 0.913734 |
| 13 | 114315000 | 114316599 | 14.47686 | 26.50334 | 33.31946 | 14.23445 | 0.096396 | 0.300269 | 0.913752 |
| 10 | 48901200 | 48902799 | 20.7802 | 35.68548 | 48.47567 | 21.26262 | -0.05632 | 0.926715 | 0.913772 |

| | | | | | | | | | |
|---|---|---|---|---|---|---|---|---|---|
| 11 | 73507800 | 73510599 | 22.61985 | 39.64012 | 34.69993 | 15.32733 | 0.578712 | 0.681252 | 0.913788 |
| 5 | 150382400 | 150384599 | 18.93016 | 32.85782 | 44.71455 | 19.62006 | -0.05588 | 0.814397 | 0.913794 |
| 17 | 16950200 | 16952599 | 18.93016 | 32.85782 | 44.71455 | 19.62006 | -0.05588 | 0.814397 | 0.913794 |
| 1 | 13410600 | 13412999 | 17.3406 | 30.33432 | 38.27822 | 16.88242 | 0.053625 | 0.686222 | 0.913794 |
| 3 | 71672600 | 71673799 | 12.78077 | 23.34786 | 30.10128 | 12.8959 | 0.057561 | 0.292824 | 0.913795 |
| 8 | 105294600 | 105296399 | 12.19846 | 21.62071 | 37.7448 | 16.68693 | -0.40736 | 0.488471 | 0.9138 |
| 16 | 62024600 | 62025799 | 23.0405 | 39.65562 | 39.52737 | 17.33886 | 0.389134 | 0.903783 | 0.913807 |
| 7 | 77125200 | 77125999 | 14.45774 | 25.59833 | 35.00738 | 15.47995 | -0.05783 | 0.50731 | 0.913833 |
| 8 | 109741400 | 109742399 | 9.293843 | 17.42079 | 25.3516 | 10.45954 | -0.07482 | 0.103324 | 0.913842 |
| 16 | 23761800 | 23762399 | 8.762186 | 16.67528 | 22.35919 | 8.956844 | 0.078669 | -0.01173 | 0.913859 |
| 10 | 33156800 | 33158399 | 17.78464 | 31.22221 | 39.15264 | 17.32018 | 0.062599 | 0.634389 | 0.913864 |
| 4 | 120247800 | 120249199 | 14.09864 | 25.64797 | 25.8496 | 11.13243 | 0.405718 | 0.316674 | 0.913886 |
| 1 | 181487600 | 181489399 | 20.28031 | 35.09006 | 47.937 | 21.03117 | -0.06224 | 0.856432 | 0.913922 |
| 7 | 67003200 | 67006799 | 41.40347 | 67.05189 | 82.89771 | 35.4729 | 0.058465 | 1.66865 | 0.91393 |
| 3 | 38715800 | 38718199 | 19.40066 | 33.79716 | 42.47722 | 18.68101 | 0.058849 | 0.762413 | 0.91396 |
| 15 | 9557200 | 9560599 | 34.97787 | 56.86869 | 76.01805 | 32.62039 | -0.0487 | 1.575298 | 0.913964 |
| 18 | 65915000 | 65916399 | 13.00981 | 23.20235 | 32.08245 | 14.11211 | -0.06781 | 0.440945 | 0.913988 |
| 15 | 74687400 | 74690999 | 23.24205 | 39.18969 | 53.35507 | 23.32936 | -0.06253 | 1.099394 | 0.914002 |
| 7 | 79922400 | 79926199 | 42.22617 | 68.18454 | 84.67459 | 36.12394 | 0.047637 | 1.743023 | 0.914028 |
| 9 | 52973000 | 52974799 | 15.58344 | 27.65684 | 34.89354 | 15.41198 | 0.059425 | 0.493618 | 0.914028 |
| 15 | 34165200 | 34168199 | 42.94696 | 69.2593 | 85.96494 | 36.61365 | 0.046266 | 1.7831 | 0.91407 |
| 6 | 76041800 | 76042999 | 14.09153 | 25.27835 | 35.03283 | 15.33441 | -0.06645 | 0.416755 | 0.914093 |
| 14 | 18478200 | 18479799 | 15.67571 | 26.93162 | 50.0295 | 21.94447 | -0.50567 | 0.903516 | 0.914099 |
| 17 | 41963200 | 41965799 | 35.94504 | 58.53426 | 78.92666 | 33.88817 | -0.05954 | 1.559474 | 0.91411 |
| 11 | 35049600 | 35050399 | 8.925832 | 16.97827 | 22.86443 | 9.168882 | 0.071111 | -0.00511 | 0.914129 |
| 7 | 48347200 | 48348599 | 26.23121 | 43.83953 | 55.04393 | 23.96296 | 0.053806 | 1.19865 | 0.914155 |
| 13 | 23000200 | 23003599 | 50.45947 | 80.88582 | 99.77281 | 42.12359 | 0.044686 | 1.983108 | 0.914201 |
| 1 | 140086600 | 140087599 | 9.737245 | 18.27089 | 26.35565 | 10.85468 | -0.06042 | 0.102487 | 0.914212 |
| 6 | 100763000 | 100770799 | 83.05345 | 129.2774 | 165.0428 | 68.50996 | -0.04262 | 2.724381 | 0.914221 |
| 7 | 6637400 | 6643399 | 87.18934 | 135.7138 | 163.6686 | 67.93966 | 0.039438 | 2.727012 | 0.914229 |
| X | 79021400 | 79022199 | 15.14639 | 27.33389 | 34.11024 | 14.83808 | 0.090286 | 0.390536 | 0.91423 |
| 10 | 53362600 | 53376399 | 235.9803 | 342.5385 | 388.3892 | 160.1192 | 0.040901 | 4.025325 | 0.914235 |
| 14 | 87652800 | 87655199 | 17.36178 | 29.78342 | 51.64778 | 22.65216 | -0.40747 | 0.923892 | 0.914235 |
| 7 | 6434400 | 6437599 | 32.17008 | 52.44464 | 70.72874 | 30.38047 | -0.0587 | 1.527288 | 0.914279 |
| 6 | 130331200 | 130335399 | 31.4753 | 52.20669 | 72.40804 | 31.36543 | -0.08971 | 1.311614 | 0.914318 |
| 6 | 95049600 | 95050199 | 7.541223 | 14.60717 | 22.7521 | 8.635474 | -0.06343 | -0.20158 | 0.91432 |
| 16 | 44208600 | 44213199 | 53.01831 | 84.85284 | 104.2333 | 43.90211 | 0.047966 | 2.035263 | 0.914322 |
| 16 | 84191600 | 84192399 | 9.306447 | 17.40644 | 25.35423 | 10.4993 | -0.08036 | 0.117653 | 0.91433 |
| 14 | 51753400 | 51754199 | 14.57353 | 26.48623 | 25.94789 | 11.18829 | 0.445566 | 0.327966 | 0.914376 |
| 6 | 143157400 | 143165199 | 131.2297 | 198.4115 | 247.1525 | 102.2082 | -0.04028 | 3.306544 | 0.914377 |
| 8 | 116757400 | 116759599 | 18.79009 | 32.45049 | 44.23337 | 19.40518 | -0.0597 | 0.858807 | 0.914442 |
| X | 31611400 | 31614399 | 41.06781 | 67.93801 | 55.04528 | 23.80094 | 0.68336 | 1.355944 | 0.914469 |
| 5 | 80178400 | 80178999 | 8.196196 | 15.82735 | 24.31503 | 9.361532 | -0.06311 | -0.14013 | 0.914504 |
| 5 | 77752600 | 77754799 | 29.59602 | 48.99982 | 61.72972 | 26.70606 | 0.048486 | 1.319316 | 0.914507 |
| 8 | 16476600 | 16480199 | 30.29656 | 49.93579 | 67.31927 | 29.04371 | -0.05005 | 1.377393 | 0.914513 |
| 11 | 45491800 | 45493199 | 18.81859 | 32.85844 | 41.39124 | 18.22967 | 0.055977 | 0.726947 | 0.914519 |
| 6 | 50401400 | 50403599 | 16.78777 | 29.48385 | 37.02603 | 16.38492 | 0.060359 | 0.61613 | 0.91452 |

| | | | | | | | | | |
|---|---|---|---|---|---|---|---|---|---|
| 19 | 44351800 | 44352399 | 9.496168 | 17.92681 | 23.7727 | 9.687083 | 0.072506 | 0.053423 | 0.914537 |
| 5 | 49348800 | 49350199 | 32.04837 | 52.45653 | 70.35449 | 30.26401 | -0.04663 | 1.471207 | 0.91454 |
| 17 | 7196600 | 7197799 | 12.17428 | 22.03015 | 30.60689 | 13.26885 | -0.06061 | 0.343795 | 0.914542 |
| 1 | 85208400 | 85238399 | 2334.316 | 2798.011 | 2770.571 | 1171.508 | -0.04163 | 7.039412 | 0.914558 |
| 13 | 57579200 | 57580399 | 20.2591 | 34.91656 | 47.39794 | 20.79219 | -0.05441 | 0.887325 | 0.914559 |
| 10 | 10393000 | 10396199 | 42.04823 | 67.81309 | 84.24724 | 35.8842 | 0.045006 | 1.778155 | 0.914611 |
| 12 | 108713400 | 108719199 | 46.42683 | 74.46997 | 98.31693 | 41.55 | -0.05205 | 1.955538 | 0.914626 |
| 7 | 113495000 | 113496799 | 20.71282 | 35.69814 | 48.47353 | 21.26402 | -0.05476 | 0.892884 | 0.914635 |
| 12 | 93886400 | 93887799 | 20.71282 | 35.69814 | 48.47353 | 21.26402 | -0.05476 | 0.892884 | 0.914635 |
| 5 | 96420000 | 96423999 | 76.52509 | 119.5803 | 153.5329 | 63.70087 | -0.04643 | 2.65256 | 0.914643 |
| 3 | 22134800 | 22136199 | 11.70846 | 21.21257 | 29.65986 | 12.83909 | -0.06826 | 0.331793 | 0.914651 |
| 1 | 67145200 | 67148599 | 61.43562 | 97.42698 | 126.2049 | 52.53733 | -0.04236 | 2.331196 | 0.914659 |
| 17 | 69876800 | 69878799 | 20.95436 | 35.95671 | 45.30436 | 19.87047 | 0.051544 | 0.928383 | 0.914665 |
| 8 | 99174400 | 99176599 | 20.82771 | 36.12393 | 45.21536 | 19.83857 | 0.063984 | 0.830258 | 0.914704 |
| 7 | 36626400 | 36628399 | 17.51048 | 30.52228 | 41.49564 | 18.25606 | -0.05378 | 0.744189 | 0.914732 |
| 1 | 25158400 | 25159399 | 10.23869 | 18.63184 | 33.37066 | 14.36585 | -0.4179 | 0.305069 | 0.914743 |
| 5 | 75474000 | 75475399 | 11.4154 | 21.08918 | 29.82056 | 12.5833 | -0.05899 | 0.220209 | 0.914743 |
| 3 | 61418400 | 61420199 | 28.48523 | 47.17813 | 59.36723 | 25.69051 | 0.050213 | 1.311137 | 0.914749 |
| 16 | 6068200 | 6071399 | 34.44689 | 56.3141 | 70.66512 | 30.38419 | 0.047866 | 1.492925 | 0.914749 |
| 6 | 57205400 | 57208199 | 31.67738 | 52.60423 | 54.10496 | 23.45617 | 0.340557 | 1.285157 | 0.914755 |
| 8 | 49157400 | 49162399 | 58.01388 | 92.39882 | 113.4225 | 47.44826 | 0.041403 | 2.193875 | 0.914762 |
| Y | 12752600 | 12752999 | 6.090012 | 11.8265 | 19.50757 | 7.304894 | -0.12663 | -0.27582 | 0.914785 |
| 1 | 31720800 | 31721399 | 12.5688 | 22.83217 | 29.15908 | 12.58022 | 0.064961 | 0.323619 | 0.914824 |
| 13 | 19653600 | 19655199 | 13.40268 | 24.55938 | 31.10627 | 13.27233 | 0.086682 | 0.279333 | 0.914833 |
| 4 | 112392800 | 112395999 | 58.29071 | 92.75996 | 120.5829 | 50.3837 | -0.04248 | 2.224637 | 0.91484 |
| 2 | 117628400 | 117631599 | 22.62467 | 38.58445 | 52.26477 | 22.91193 | -0.05393 | 0.998173 | 0.914846 |
| 1 | 32811200 | 32813599 | 25.56656 | 42.59385 | 57.62408 | 25.03622 | -0.05337 | 1.235126 | 0.914859 |
| 2 | 116411000 | 116412599 | 13.10575 | 23.72561 | 30.14728 | 13.06278 | 0.068229 | 0.350219 | 0.914863 |
| 4 | 84849400 | 84852999 | 41.11993 | 66.51324 | 89.90525 | 38.43765 | -0.07153 | 1.698693 | 0.914884 |
| X | 36166000 | 36167399 | 12.66403 | 23.28945 | 29.75678 | 12.63457 | 0.080095 | 0.257362 | 0.914904 |
| 10 | 112517400 | 112521599 | 43.73428 | 70.35278 | 87.00359 | 36.92589 | 0.047151 | 1.860697 | 0.914918 |
| 5 | 23462200 | 23463999 | 15.1882 | 26.8379 | 36.58653 | 16.18471 | -0.0543 | 0.538184 | 0.914924 |
| 15 | 11966800 | 11968399 | 20.06559 | 34.6501 | 43.67933 | 19.16207 | 0.052603 | 0.864748 | 0.914929 |
| 5 | 79969000 | 79973199 | 105.2108 | 161.2149 | 191.355 | 79.42547 | 0.044993 | 3.025889 | 0.914934 |
| 17 | 13280000 | 13282399 | 17.57228 | 30.55319 | 41.60525 | 18.27542 | -0.05636 | 0.781644 | 0.914956 |
| 10 | 34133200 | 34136199 | 31.33397 | 51.56026 | 64.93442 | 27.9952 | 0.046996 | 1.400185 | 0.914956 |
| 14 | 122549200 | 122550599 | 16.2365 | 28.8681 | 28.73098 | 12.6707 | 0.40124 | 0.474108 | 0.914963 |
| 1 | 188929400 | 188933799 | 35.75446 | 58.30855 | 72.972 | 31.34806 | 0.04922 | 1.53165 | 0.914976 |
| 7 | 36038800 | 36041599 | 24.23636 | 41.19122 | 51.73171 | 22.67196 | 0.054017 | 1.034547 | 0.914991 |
| 17 | 22659400 | 22662999 | 59.49228 | 95.17439 | 114.0913 | 48.02587 | 0.08249 | 2.060392 | 0.914994 |
| 9 | 22173800 | 22202399 | 505.2595 | 689.1741 | 730.5682 | 304.1172 | 0.047891 | 4.965869 | 0.914997 |
| 5 | 138450200 | 138451599 | 16.29214 | 29.38831 | 36.13785 | 15.72895 | 0.110576 | 0.401782 | 0.914998 |
| 16 | 36050600 | 36051799 | 17.01763 | 29.64125 | 40.5086 | 17.81352 | -0.06135 | 0.750648 | 0.915035 |
| 16 | 61187400 | 61190799 | 33.49212 | 54.93084 | 68.98213 | 29.699 | 0.049202 | 1.448395 | 0.915035 |
| 13 | 90752400 | 90754999 | 38.52237 | 63.11781 | 62.60456 | 26.94184 | 0.388101 | 1.467825 | 0.915039 |
| 7 | 46838400 | 46840199 | 20.71202 | 35.99344 | 35.26941 | 15.48725 | 0.416089 | 0.792736 | 0.915049 |
| 1 | 13860600 | 13862799 | 30.60525 | 50.80282 | 63.57532 | 27.554 | 0.058358 | 1.294196 | 0.915056 |

| | | | | | | | | | |
|---|---|---|---|---|---|---|---|---|---|
| 12 | 92375200 | 92376199 | 10.08054 | 18.74072 | 26.79386 | 11.20954 | -0.06576 | 0.166147 | 0.91513 |
| 12 | 16485400 | 16487599 | 20.15383 | 34.65041 | 46.87863 | 20.56286 | -0.05022 | 0.90921 | 0.91517 |
| 10 | 7287400 | 7287999 | 10.1928 | 19.28352 | 25.93223 | 10.51889 | 0.058315 | 0.051799 | 0.91519 |
| 14 | 33670000 | 33672599 | 22.34659 | 37.82956 | 51.16004 | 22.39651 | -0.05258 | 1.062078 | 0.915197 |
| X | 164611800 | 164616799 | 61.56247 | 98.50028 | 116.5184 | 49.05746 | 0.101871 | 2.059875 | 0.915201 |
| 2 | 161187600 | 161189599 | 25.00291 | 41.9955 | 56.62157 | 24.72741 | -0.0487 | 1.144673 | 0.915206 |
| 8 | 115919400 | 115923399 | 39.92246 | 64.53314 | 80.39256 | 34.34416 | 0.044888 | 1.706516 | 0.915224 |
| 9 | 88531200 | 88534799 | 100.4466 | 154.3138 | 194.8218 | 80.89321 | -0.0411 | 2.974666 | 0.915234 |
| 6 | 131113200 | 131121199 | 282.5662 | 400.679 | 473.2171 | 195.7797 | -0.05149 | 4.371514 | 0.915285 |
| 1 | 189281800 | 189285199 | 41.61472 | 67.11615 | 83.37252 | 35.51312 | 0.045239 | 1.777358 | 0.915334 |
| 13 | 14748400 | 14748999 | 9.462574 | 17.8444 | 25.91182 | 10.58202 | -0.05988 | 0.068361 | 0.915377 |
| 14 | 102651400 | 102653199 | 28.37731 | 47.23171 | 59.41358 | 25.79824 | 0.051279 | 1.255706 | 0.915383 |
| 4 | 96557800 | 96558399 | 9.31887 | 17.8358 | 26.85049 | 10.61603 | -0.07027 | -0.03576 | 0.915426 |
| 13 | 87211400 | 87214399 | 39.69632 | 64.82398 | 64.78402 | 27.84421 | 0.3743 | 1.517309 | 0.915455 |
| 4 | 60579400 | 60580799 | 26.0258 | 43.42034 | 54.41407 | 23.66267 | 0.056684 | 1.218625 | 0.915469 |
| 10 | 82065600 | 82075199 | 180.3448 | 269.3226 | 307.308 | 126.6864 | 0.070416 | 3.55785 | 0.915473 |
| 7 | 116397600 | 116404399 | 144.3007 | 217.2004 | 255.1386 | 105.3708 | 0.03873 | 3.398599 | 0.915491 |
| 15 | 87251600 | 87261799 | 114.2828 | 174.1816 | 218.3281 | 90.50918 | -0.03966 | 3.138109 | 0.915494 |
| X | 78663000 | 78665599 | 23.18752 | 39.8408 | 55.61664 | 24.39522 | -0.09549 | 0.940213 | 0.915502 |
| 8 | 24113800 | 24116199 | 27.51706 | 45.54053 | 61.50919 | 26.60367 | -0.05168 | 1.318366 | 0.915531 |
| 7 | 86627600 | 86629799 | 24.75732 | 42.24406 | 43.15735 | 18.92021 | 0.351638 | 0.990941 | 0.915538 |
| 10 | 66357400 | 66358799 | 19.3163 | 33.76368 | 33.60487 | 14.81134 | 0.394389 | 0.701477 | 0.915541 |
| X | 69580200 | 69583999 | 28.6948 | 47.74304 | 65.5092 | 28.43801 | -0.07374 | 1.267703 | 0.915558 |
| 13 | 106803800 | 106809399 | 31.88057 | 53.18625 | 52.5109 | 22.83453 | 0.399498 | 1.232284 | 0.915572 |
| 10 | 49835800 | 49836599 | 16.79001 | 29.46577 | 37.18802 | 16.44623 | 0.053468 | 0.631453 | 0.915606 |
| 9 | 116705200 | 116712399 | 67.08798 | 106.393 | 129.5946 | 53.94959 | 0.046285 | 2.334377 | 0.915609 |
| 5 | 116449600 | 116452399 | 19.83523 | 34.25715 | 43.19508 | 18.94971 | 0.052389 | 0.863637 | 0.915619 |
| 11 | 15757400 | 15758399 | 13.46658 | 24.26441 | 31.00647 | 13.51435 | 0.054317 | 0.379027 | 0.915652 |
| 13 | 42044200 | 42047799 | 32.36588 | 53.20311 | 71.89511 | 30.98258 | -0.05502 | 1.420229 | 0.91566 |
| 4 | 109552200 | 109557199 | 109.0105 | 167.5043 | 199.2295 | 82.7257 | 0.045091 | 2.974054 | 0.915668 |
| 9 | 81341800 | 81342599 | 10.67992 | 19.92443 | 26.25514 | 10.91945 | 0.05785 | 0.14747 | 0.915683 |
| 10 | 59605000 | 59606399 | 10.91041 | 20.01139 | 28.23819 | 12.0331 | -0.06616 | 0.252593 | 0.915713 |
| 13 | 95675600 | 95676999 | 11.89763 | 21.71999 | 30.61235 | 13.12584 | -0.07123 | 0.288793 | 0.915731 |
| 5 | 79177200 | 79180999 | 21.56095 | 37.43273 | 37.38137 | 16.40473 | 0.388902 | 0.815683 | 0.915733 |
| 17 | 54049200 | 54054599 | 73.01588 | 114.5198 | 147.3867 | 61.12752 | -0.0457 | 2.58566 | 0.915739 |
| 1 | 55562200 | 55563399 | 13.27456 | 23.69286 | 32.65846 | 14.3561 | -0.06243 | 0.441668 | 0.915752 |
| 8 | 52314600 | 52316999 | 31.64202 | 52.01382 | 65.51519 | 28.23334 | 0.046185 | 1.413384 | 0.915757 |
| 5 | 48801200 | 48802199 | 14.114 | 25.62849 | 25.39768 | 10.96312 | 0.427848 | 0.326649 | 0.915757 |
| 11 | 81022200 | 81024399 | 19.66184 | 33.77115 | 45.8397 | 20.10653 | -0.05522 | 0.914655 | 0.915766 |
| 7 | 104686200 | 104689599 | 44.71467 | 71.81056 | 88.26243 | 37.36963 | 0.053053 | 1.914471 | 0.915781 |
| 12 | 8337200 | 8338399 | 16.84786 | 29.43757 | 36.86887 | 16.24489 | 0.064476 | 0.700507 | 0.915791 |
| 4 | 49428200 | 49436599 | 102.3444 | 159.3155 | 151.4428 | 62.86392 | 0.382472 | 2.731522 | 0.915822 |
| 10 | 54879600 | 54880799 | 15.73657 | 27.66103 | 37.72718 | 16.70685 | -0.05784 | 0.602238 | 0.915835 |
| 5 | 57808400 | 57811999 | 43.03194 | 69.48309 | 86.09186 | 36.72683 | 0.050826 | 1.745695 | 0.915838 |
| 2 | 103688400 | 103691799 | 40.89 | 66.05195 | 82.20179 | 35.08612 | 0.045191 | 1.728366 | 0.915839 |
| 16 | 18232400 | 18235999 | 21.06452 | 36.11659 | 49.01446 | 21.49632 | -0.0555 | 0.938743 | 0.91584 |
| 19 | 34002800 | 34003199 | 6.228174 | 12.13849 | 18.01214 | 6.522466 | 0.069463 | -0.39498 | 0.915849 |

| | | | | | | | | | |
|---|---|---|---|---|---|---|---|---|---|
| 16 | 87406800 | 87409199 | 19.87323 | 34.31916 | 43.20806 | 18.95534 | 0.054565 | 0.865473 | 0.915857 |
| 16 | 12162200 | 12165199 | 44.57941 | 71.90567 | 88.87773 | 37.86359 | 0.052573 | 1.780956 | 0.915873 |
| 7 | 97348200 | 97348799 | 11.87081 | 21.7843 | 28.18058 | 12.00001 | 0.05917 | 0.257524 | 0.915874 |
| 18 | 37427800 | 37431799 | 89.75261 | 139.9512 | 135.5874 | 56.26811 | 0.356979 | 2.695011 | 0.915905 |
| 17 | 85848200 | 85850199 | 29.00534 | 47.99179 | 60.46424 | 26.14726 | 0.048368 | 1.325346 | 0.91593 |
| 1 | 94176000 | 94176799 | 9.138062 | 17.35522 | 25.50063 | 10.25938 | -0.05783 | 0.015292 | 0.915959 |
| 2 | 36688800 | 36696199 | 122.7793 | 188.3549 | 182.7244 | 75.85539 | 0.337055 | 3.007837 | 0.915964 |
| 1 | 5264800 | 5267999 | 27.31047 | 45.98518 | 57.29862 | 25.04302 | 0.064805 | 1.131686 | 0.915967 |
| 2 | 3357000 | 3358799 | 13.95572 | 25.06988 | 31.9747 | 13.97927 | 0.054062 | 0.402458 | 0.915973 |
| 13 | 93260600 | 93262999 | 17.77861 | 31.19744 | 39.08793 | 17.28492 | 0.063659 | 0.638744 | 0.915978 |
| 1 | 16216400 | 16217599 | 15.49463 | 27.36083 | 34.56815 | 15.29392 | 0.054526 | 0.544063 | 0.916 |
| 5 | 95963400 | 95966399 | 19.36432 | 33.84557 | 33.64899 | 14.83003 | 0.396043 | 0.703465 | 0.916019 |
| 5 | 88557200 | 88557799 | 11.94069 | 21.65833 | 30.20544 | 13.05636 | -0.06304 | 0.329864 | 0.916033 |
| 3 | 41786800 | 41790999 | 67.25182 | 106.6708 | 129.8251 | 54.05468 | 0.047717 | 2.329649 | 0.916058 |
| 9 | 8466800 | 8468599 | 34.88345 | 56.48951 | 75.77463 | 32.42342 | -0.05894 | 1.654858 | 0.916065 |
| 4 | 75410200 | 75410799 | 9.268156 | 17.45017 | 25.36312 | 10.38569 | -0.06445 | 0.074167 | 0.916098 |
| 2 | 108880000 | 108881799 | 20.12375 | 34.94839 | 47.93434 | 21.03997 | -0.06712 | 0.816962 | 0.916098 |
| 17 | 95144600 | 95152399 | 88.10041 | 138.5761 | 162.0733 | 67.23722 | 0.095332 | 2.541707 | 0.916109 |
| 7 | 130575800 | 130577799 | 15.06868 | 26.34868 | 44.6365 | 19.67957 | -0.36926 | 0.692016 | 0.916116 |
| 3 | 8084600 | 8089199 | 74.77209 | 117.3457 | 150.3891 | 62.3716 | -0.03897 | 2.57536 | 0.916121 |
| 5 | 110346400 | 110348599 | 28.04579 | 46.54792 | 62.70422 | 27.17391 | -0.04757 | 1.289136 | 0.916146 |
| 5 | 43364400 | 43366199 | 11.17964 | 20.17477 | 35.29962 | 15.35596 | -0.3945 | 0.363311 | 0.916153 |
| 3 | 27080200 | 27082599 | 20.27344 | 34.92865 | 47.40951 | 20.79704 | -0.0542 | 0.89398 | 0.916156 |
| 2 | 35365600 | 35366799 | 14.82926 | 26.14712 | 35.63198 | 15.76926 | -0.05498 | 0.557362 | 0.916159 |
| 15 | 74177200 | 74181199 | 41.08238 | 66.24286 | 82.13508 | 34.97599 | 0.047538 | 1.781629 | 0.916179 |
| 1 | 164700600 | 164701999 | 14.08002 | 25.15936 | 31.94464 | 14.02873 | 0.056037 | 0.435444 | 0.916181 |
| 5 | 70533400 | 70534399 | 14.08002 | 25.15936 | 31.94464 | 14.02873 | 0.056037 | 0.435444 | 0.916181 |
| 7 | 39286800 | 39287999 | 11.51821 | 21.18245 | 29.73467 | 12.62502 | -0.05513 | 0.245078 | 0.916182 |
| 13 | 102746200 | 102748799 | 30.16698 | 50.19086 | 50.80581 | 22.05467 | 0.363703 | 1.256396 | 0.91619 |
| 12 | 119529600 | 119532599 | 76.69393 | 119.8789 | 153.6672 | 63.7547 | -0.04377 | 2.647848 | 0.9162 |
| 6 | 126134800 | 126137399 | 18.84189 | 32.81656 | 41.489 | 18.24389 | 0.050466 | 0.763087 | 0.916204 |
| 7 | 40940800 | 40943199 | 34.76442 | 57.2984 | 70.73846 | 30.51853 | 0.0765 | 1.391824 | 0.91622 |
| 4 | 89524400 | 89525599 | 12.80806 | 23.03649 | 29.09137 | 12.70553 | 0.069442 | 0.380825 | 0.916226 |
| 2 | 161510000 | 161510599 | 7.598114 | 14.73702 | 21.09546 | 7.95695 | 0.066773 | -0.22345 | 0.916228 |
| 9 | 5287200 | 5289799 | 27.25806 | 45.42955 | 57.24755 | 24.87873 | 0.048659 | 1.233857 | 0.916236 |
| 12 | 111354000 | 111356199 | 22.58539 | 38.18461 | 51.68132 | 22.61588 | -0.05373 | 1.076185 | 0.916256 |
| 13 | 28397600 | 28399999 | 25.34702 | 42.31389 | 57.13499 | 24.85554 | -0.05067 | 1.21118 | 0.916272 |
| 8 | 47658200 | 47660199 | 18.76656 | 32.46829 | 44.21262 | 19.39714 | -0.05773 | 0.842162 | 0.916273 |
| 7 | 53406800 | 53408599 | 17.25778 | 30.16035 | 40.92822 | 18.0365 | -0.05071 | 0.706355 | 0.91631 |
| 9 | 19731800 | 19733199 | 21.5106 | 37.44773 | 46.24307 | 20.32863 | 0.084109 | 0.788494 | 0.916315 |
| X | 60723600 | 60727399 | 33.68038 | 56.00106 | 67.92591 | 29.47364 | 0.103452 | 1.288118 | 0.916322 |
| 3 | 75646800 | 75649599 | 25.14906 | 43.10771 | 52.89467 | 23.19754 | 0.089276 | 0.964022 | 0.916345 |
| 16 | 90527400 | 90530199 | 41.39447 | 66.92736 | 83.15347 | 35.53375 | 0.049067 | 1.703811 | 0.916352 |
| 17 | 20525600 | 20526599 | 12.01342 | 22.15659 | 31.36956 | 13.26733 | -0.06344 | 0.236486 | 0.916372 |
| 3 | 144585400 | 144586799 | 23.52866 | 40.46497 | 41.03719 | 18.00073 | 0.364049 | 0.914204 | 0.916377 |
| 4 | 8238400 | 8240399 | 16.65612 | 28.82622 | 49.53623 | 21.73295 | -0.39185 | 0.837868 | 0.916389 |
| 4 | 96968400 | 96968599 | 6.502345 | 12.66024 | 18.4879 | 6.778607 | 0.075476 | -0.34753 | 0.9164 |

| | | | | | | | | | |
|---|---|---|---|---|---|---|---|---|---|
| 3 | 153036200 | 153036599 | 6.844119 | 13.30956 | 21.24768 | 7.867038 | -0.06418 | -0.29629 | 0.916413 |
| 19 | 9484600 | 9488999 | 51.57251 | 82.54081 | 101.8019 | 42.87943 | 0.042223 | 2.032709 | 0.916413 |
| 2 | 38786200 | 38788599 | 29.89744 | 49.38882 | 66.91443 | 28.90467 | -0.05646 | 1.348024 | 0.916431 |
| 8 | 19364000 | 19365799 | 32.68389 | 53.42344 | 71.52216 | 30.75092 | -0.04545 | 1.492271 | 0.916473 |
| X | 19103000 | 19104599 | 15.81529 | 28.0936 | 35.0391 | 15.46453 | 0.077019 | 0.493722 | 0.916475 |
| 5 | 26088400 | 26094999 | 79.70673 | 124.893 | 151.445 | 62.81691 | 0.038994 | 2.60582 | 0.916481 |
| 1 | 48391600 | 48397399 | 65.22538 | 103.8503 | 105.8194 | 44.24741 | 0.309127 | 2.209448 | 0.916481 |
| 1 | 48550800 | 48551599 | 10.29511 | 19.22952 | 25.21447 | 10.46542 | 0.06711 | 0.132602 | 0.91649 |
| 8 | 102947600 | 102948799 | 14.65587 | 26.29535 | 36.6316 | 16.03183 | -0.07369 | 0.423601 | 0.916503 |
| 12 | 105285000 | 105288799 | 39.10299 | 63.20834 | 83.87888 | 35.83342 | -0.0461 | 1.706358 | 0.916524 |
| 6 | 140815000 | 140820799 | 55.59609 | 88.66959 | 115.5206 | 48.41781 | -0.04221 | 2.148726 | 0.916533 |
| 10 | 51616400 | 51623399 | 87.25945 | 135.6674 | 163.4545 | 67.86236 | 0.039556 | 2.747159 | 0.916551 |
| X | 87874800 | 87885799 | 124.3861 | 191.2289 | 174.2068 | 72.34067 | 0.43004 | 2.968118 | 0.916578 |
| 19 | 53390800 | 53393999 | 32.04219 | 51.88246 | 85.60194 | 36.62535 | -0.35704 | 1.658158 | 0.916588 |
| 4 | 45080800 | 45082199 | 19.04893 | 33.04943 | 41.6282 | 18.26486 | 0.055315 | 0.817977 | 0.916596 |
| 3 | 21508000 | 21512399 | 32.45974 | 53.66471 | 74.20089 | 32.06996 | -0.08564 | 1.352062 | 0.916637 |
| 15 | 33160400 | 33166599 | 70.0607 | 111.0758 | 147.0587 | 61.20241 | -0.0742 | 2.34968 | 0.916646 |
| 6 | 130149200 | 130152199 | 37.26622 | 60.82183 | 75.58705 | 32.48078 | 0.060072 | 1.525742 | 0.916653 |
| 10 | 94004600 | 94006399 | 19.72628 | 34.16921 | 43.12991 | 18.92282 | 0.051606 | 0.834895 | 0.916671 |
| 2 | 121333600 | 121337199 | 27.03852 | 45.12932 | 56.73346 | 24.67798 | 0.052032 | 1.216262 | 0.916675 |
| Y | 87905600 | 87905999 | 6.392199 | 12.35703 | 20.00826 | 7.6647 | -0.13063 | -0.19058 | 0.916694 |
| 18 | 66863200 | 66865399 | 21.82717 | 37.53053 | 51.19437 | 22.45597 | -0.06209 | 0.920732 | 0.916697 |
| 1 | 17418000 | 17419599 | 13.03544 | 23.62699 | 33.09411 | 14.31789 | -0.07076 | 0.343546 | 0.916701 |
| 17 | 63461600 | 63463999 | 19.97693 | 34.75799 | 47.90643 | 21.05232 | -0.07388 | 0.790276 | 0.916715 |
| 17 | 86288000 | 86291399 | 34.13658 | 55.94124 | 70.19944 | 30.21245 | 0.049768 | 1.460482 | 0.916722 |
| 13 | 74651000 | 74652999 | 19.67397 | 33.84678 | 45.83097 | 20.10369 | -0.05124 | 0.900476 | 0.916724 |
| 1 | 37203600 | 37204999 | 15.56767 | 27.41752 | 34.56234 | 15.2994 | 0.056148 | 0.571599 | 0.916751 |
| 3 | 49365400 | 49365999 | 9.661265 | 18.22958 | 24.28287 | 9.905188 | 0.064775 | 0.059841 | 0.916758 |
| 13 | 30080600 | 30083599 | 50.7006 | 81.21427 | 106.6862 | 44.99346 | -0.04718 | 2.00853 | 0.91676 |
| 8 | 39867800 | 39868799 | 10.36169 | 19.42772 | 25.79559 | 10.6378 | 0.056814 | 0.111828 | 0.916769 |
| 5 | 132951400 | 132953399 | 14.29764 | 25.53181 | 32.47777 | 14.27059 | 0.052924 | 0.442968 | 0.916789 |
| 10 | 51327000 | 51328199 | 12.24369 | 22.35056 | 28.68261 | 12.29924 | 0.063736 | 0.292146 | 0.916841 |
| Y | 38806600 | 38806999 | 4.423141 | 8.642471 | 15.48554 | 5.388008 | -0.14661 | -0.57304 | 0.916884 |
| 13 | 43513400 | 43515599 | 16.74839 | 29.39454 | 37.15723 | 16.43347 | 0.051121 | 0.629395 | 0.916886 |
| 9 | 81312200 | 81312999 | 12.57572 | 22.81743 | 31.66792 | 13.68308 | -0.05588 | 0.335311 | 0.916898 |
| 3 | 27003400 | 27007199 | 33.66283 | 55.53784 | 77.20187 | 33.32101 | -0.09366 | 1.381495 | 0.91694 |
| 5 | 144174400 | 144175799 | 15.02285 | 26.52799 | 36.1068 | 15.97463 | -0.05237 | 0.54367 | 0.91695 |
| 6 | 85794600 | 85795999 | 14.97606 | 26.4726 | 36.04855 | 15.94558 | -0.05257 | 0.531014 | 0.916982 |
| 11 | 73608000 | 73608999 | 11.33549 | 21.53985 | 21.68731 | 8.715591 | 0.484868 | 0.023078 | 0.916996 |
| X | 91111200 | 91116999 | 79.55712 | 125.9277 | 120.0081 | 49.89057 | 0.397479 | 2.392282 | 0.917052 |
| 2 | 46619600 | 46620999 | 41.19462 | 66.22491 | 88.26843 | 37.42792 | -0.06179 | 1.879129 | 0.91708 |
| 1 | 88498400 | 88501599 | 41.92878 | 67.63741 | 84.16373 | 35.86023 | 0.043129 | 1.771546 | 0.917093 |
| 19 | 4329000 | 4330799 | 19.66386 | 33.85889 | 45.82337 | 20.1009 | -0.05024 | 0.892419 | 0.917143 |
| 7 | 115064200 | 115066599 | 31.34232 | 51.28619 | 69.40482 | 29.85237 | -0.05974 | 1.476033 | 0.917171 |
| 13 | 87662400 | 87662999 | 10.54883 | 19.56473 | 25.26811 | 10.60565 | 0.076658 | 0.180273 | 0.917173 |
| 8 | 25898000 | 25899199 | 12.25598 | 22.53474 | 31.86013 | 13.53114 | -0.06561 | 0.257897 | 0.917173 |
| 19 | 43963400 | 43965399 | 17.50094 | 30.64459 | 38.72489 | 17.09347 | 0.051526 | 0.671924 | 0.917223 |

| | | | | | | | | | |
|---|---|---|---|---|---|---|---|---|---|
| X | 11657000 | 11657999 | 10.46021 | 19.60597 | 25.83767 | 10.6611 | 0.067279 | 0.116981 | 0.917224 |
| 5 | 23229400 | 23230199 | 9.551947 | 17.98957 | 23.80075 | 9.741269 | 0.070807 | 0.070913 | 0.917225 |
| 5 | 132343200 | 132345199 | 17.19366 | 30.27214 | 38.00231 | 16.82314 | 0.061834 | 0.589863 | 0.917245 |
| 13 | 98223800 | 98227799 | 39.64109 | 64.20534 | 80.05398 | 34.25901 | 0.046458 | 1.662757 | 0.917271 |
| 3 | 94628000 | 94631999 | 26.0618 | 43.71205 | 55.04818 | 24.01881 | 0.049418 | 1.160367 | 0.917306 |
| 2 | 63725800 | 63726599 | 9.024577 | 16.96172 | 24.90241 | 10.22743 | -0.08182 | 0.084035 | 0.917328 |
| 10 | 112497000 | 112498999 | 23.36082 | 39.95668 | 50.09606 | 21.96638 | 0.057925 | 0.974078 | 0.917341 |
| 16 | 54784800 | 54786599 | 20.52592 | 35.74886 | 36.24601 | 15.93819 | 0.367435 | 0.763682 | 0.917347 |
| 5 | 93510400 | 93516799 | 74.78785 | 119.0241 | 105.2665 | 43.98998 | 0.512452 | 2.231638 | 0.917359 |
| 1 | 49920400 | 49922199 | 23.61056 | 40.64815 | 39.52739 | 17.33901 | 0.424787 | 0.904751 | 0.917359 |
| 1 | 71877000 | 71879199 | 21.89902 | 37.5947 | 47.39741 | 20.78923 | 0.050672 | 0.929869 | 0.917392 |
| 13 | 97693600 | 97695199 | 14.97424 | 26.78849 | 33.96532 | 14.90145 | 0.059073 | 0.439303 | 0.917399 |
| 6 | 117928200 | 117931599 | 34.88448 | 56.60385 | 75.49636 | 32.35185 | -0.04806 | 1.61502 | 0.917403 |
| 12 | 29414000 | 29416199 | 16.68754 | 29.34779 | 37.01673 | 16.39055 | 0.054429 | 0.597492 | 0.917413 |
| X | 76210000 | 76210799 | 16.22608 | 28.67622 | 36.04759 | 15.94582 | 0.062255 | 0.543174 | 0.91742 |
| 16 | 87098800 | 87099799 | 8.480311 | 16.13452 | 23.9316 | 9.592169 | -0.06659 | -0.00834 | 0.917434 |
| 13 | 50374200 | 50376399 | 25.05484 | 42.00162 | 56.58163 | 24.67991 | -0.04739 | 1.166303 | 0.91744 |
| 1 | 117769000 | 117774999 | 113.0471 | 173.4239 | 205.7525 | 85.41511 | 0.046921 | 3.0076 | 0.917455 |
| 13 | 113416200 | 113416799 | 7.116899 | 13.8197 | 21.72115 | 8.13838 | -0.05828 | -0.25187 | 0.917461 |
| 1 | 31471200 | 31472799 | 17.14788 | 30.03619 | 40.85537 | 18.03886 | -0.05407 | 0.670001 | 0.917481 |
| 5 | 145781200 | 145831399 | 885.3621 | 1155.417 | 1165.505 | 488.0593 | 0.055629 | 5.673725 | 0.917483 |
| 8 | 87383000 | 87386599 | 44.81673 | 72.26778 | 89.19405 | 37.98433 | 0.054233 | 1.790969 | 0.917525 |
| 2 | 149165600 | 149177599 | 78.46955 | 124.1939 | 120.6547 | 50.15686 | 0.369642 | 2.393009 | 0.917534 |
| 1 | 16688200 | 16689999 | 13.69325 | 24.00787 | 42.01661 | 18.56648 | -0.41568 | 0.639326 | 0.91754 |
| X | 122665000 | 122665599 | 12.10211 | 22.6665 | 22.11809 | 9.137891 | 0.496142 | 0.125634 | 0.917546 |
| 15 | 9777800 | 9780999 | 56.33168 | 89.64271 | 109.2031 | 45.62901 | 0.051077 | 2.218197 | 0.917559 |
| 6 | 102660000 | 102662399 | 23.21026 | 38.8004 | 64.67204 | 28.15893 | -0.35351 | 1.198643 | 0.917572 |
| 10 | 96141000 | 96151399 | 102.1752 | 157.7842 | 188.2742 | 78.20521 | 0.04592 | 2.875735 | 0.91758 |
| 6 | 132939800 | 132947399 | 194.8724 | 285.553 | 346.8584 | 142.8744 | -0.04527 | 3.877112 | 0.917587 |
| 12 | 97529800 | 97531599 | 29.6041 | 48.91892 | 61.66982 | 26.64471 | 0.047303 | 1.341406 | 0.917589 |
| 6 | 66141000 | 66143599 | 37.65599 | 61.45093 | 63.45126 | 27.26473 | 0.326628 | 1.520554 | 0.917609 |
| X | 96093200 | 96094799 | 17.82327 | 31.6537 | 29.31888 | 12.94225 | 0.503788 | 0.496076 | 0.917651 |
| 12 | 113986600 | 114007999 | 277.2229 | 399.7786 | 443.424 | 182.9623 | 0.063356 | 4.126834 | 0.917652 |
| 5 | 128413400 | 128419399 | 58.07053 | 92.36803 | 119.7621 | 50.00963 | -0.03944 | 2.237888 | 0.917715 |
| 7 | 113810600 | 113811599 | 9.46389 | 18.0122 | 24.45411 | 9.794317 | 0.061184 | 0.001819 | 0.917737 |
| 16 | 89124600 | 89127399 | 23.87019 | 39.72522 | 65.7378 | 28.54195 | -0.3431 | 1.249273 | 0.917744 |
| 10 | 82106200 | 82107399 | 14.93795 | 26.86931 | 33.9761 | 14.83798 | 0.067393 | 0.404799 | 0.917768 |
| 9 | 66781200 | 66784199 | 32.83519 | 53.94579 | 67.85174 | 29.23352 | 0.047925 | 1.425252 | 0.917772 |
| 6 | 62422200 | 62422799 | 6.960767 | 13.46248 | 21.02055 | 8.03483 | -0.0761 | -0.18859 | 0.917803 |
| 12 | 14840400 | 14841799 | 14.14422 | 24.77794 | 42.53059 | 18.7804 | -0.38793 | 0.655119 | 0.917806 |
| 6 | 48668800 | 48672799 | 34.38648 | 55.88496 | 74.52764 | 31.97435 | -0.04572 | 1.581457 | 0.91785 |
| 3 | 111361800 | 111362999 | 12.4552 | 22.70192 | 31.65752 | 13.60089 | -0.05767 | 0.30588 | 0.917865 |
| 19 | 34107600 | 34111199 | 50.62796 | 80.94225 | 99.43836 | 41.82034 | 0.046224 | 2.064298 | 0.917872 |
| 6 | 76453800 | 76454599 | 9.404143 | 17.79036 | 23.83676 | 9.67019 | 0.06365 | 0.039901 | 0.917876 |
| 7 | 12523200 | 12532599 | 138.4243 | 209.119 | 246.2526 | 101.8099 | 0.039751 | 3.324222 | 0.917891 |
| 14 | 91439600 | 91441199 | 12.50988 | 22.73847 | 28.97826 | 12.49305 | 0.068105 | 0.31508 | 0.917892 |
| 16 | 18179800 | 18184999 | 40.68977 | 65.63248 | 81.21233 | 34.59848 | 0.051028 | 1.769175 | 0.917926 |

| | | | | | | | | | |
|---|---|---|---|---|---|---|---|---|---|
| 6 | 101421200 | 101423799 | 26.27203 | 44.12915 | 55.58736 | 24.27646 | 0.049044 | 1.146981 | 0.917937 |
| 14 | 78016600 | 78018599 | 16.8086 | 29.46781 | 37.14322 | 16.41202 | 0.055129 | 0.645532 | 0.91795 |
| 18 | 52464400 | 52465599 | 17.58433 | 30.53776 | 41.65023 | 18.28149 | -0.05877 | 0.799041 | 0.917969 |
| 4 | 113888400 | 113892399 | 58.29625 | 93.12235 | 113.4035 | 47.63582 | 0.057467 | 2.103537 | 0.917993 |
| 3 | 93108200 | 93111199 | 54.43906 | 86.71588 | 112.868 | 47.22433 | -0.04276 | 2.186789 | 0.918004 |
| 4 | 150374200 | 150378199 | 48.98079 | 78.55796 | 97.1908 | 41.06721 | 0.041321 | 1.963793 | 0.918017 |
| 15 | 92513800 | 92514799 | 13.8573 | 24.46432 | 33.69467 | 14.90811 | -0.06943 | 0.533821 | 0.918024 |
| 11 | 20397400 | 20398599 | 14.69057 | 26.0518 | 35.52328 | 15.70049 | -0.05257 | 0.495203 | 0.918025 |
| 17 | 75216800 | 75224199 | 133.083 | 202.3843 | 203.4868 | 84.30045 | 0.275519 | 3.187262 | 0.918045 |
| 8 | 48725800 | 48733599 | 93.19413 | 143.2051 | 212.8987 | 88.40188 | -0.27643 | 2.96907 | 0.918071 |
| 3 | 127212600 | 127214999 | 20.1818 | 34.99097 | 44.19303 | 19.3898 | 0.05113 | 0.832075 | 0.918081 |
| X | 169281600 | 169282599 | 10.32231 | 19.67875 | 18.1612 | 7.248308 | 0.617858 | -0.02107 | 0.918156 |
| 12 | 69603800 | 69605399 | 15.84988 | 28.08311 | 35.35372 | 15.63089 | 0.061265 | 0.508177 | 0.918167 |
| 7 | 61395600 | 61397599 | 26.66047 | 44.52418 | 59.98698 | 26.10276 | -0.04755 | 1.213472 | 0.918168 |
| 15 | 99846200 | 99851199 | 46.15805 | 74.12385 | 91.73143 | 38.83459 | 0.04306 | 1.918946 | 0.918185 |
| 12 | 107829000 | 107831599 | 19.97287 | 34.3747 | 43.22771 | 18.96201 | 0.055183 | 0.895057 | 0.918198 |
| 18 | 56438600 | 56441199 | 29.58874 | 49.18311 | 61.83285 | 26.82427 | 0.051876 | 1.273959 | 0.918204 |
| 14 | 63882200 | 63892199 | 142.9046 | 214.7071 | 250.7518 | 103.5085 | 0.044762 | 3.433317 | 0.918208 |
| 17 | 68312600 | 68318999 | 50.26269 | 80.44074 | 105.228 | 44.31959 | -0.04264 | 2.0334 | 0.918211 |
| 1 | 84976400 | 84983999 | 185.8892 | 277.1048 | 268.4817 | 110.6418 | 0.303883 | 3.588527 | 0.918211 |
| 11 | 34482200 | 34483799 | 13.83078 | 24.4862 | 33.5813 | 14.84961 | -0.06183 | 0.501998 | 0.918218 |
| 14 | 110044000 | 110050799 | 135.0881 | 205.0367 | 204.3854 | 84.62374 | 0.285662 | 3.226626 | 0.918223 |
| 14 | 41403400 | 41406999 | 46.43463 | 74.8677 | 77.23808 | 32.88766 | 0.312098 | 1.789707 | 0.918231 |
| 4 | 117786200 | 117788199 | 16.54064 | 28.84046 | 39.47856 | 17.37193 | -0.06351 | 0.732068 | 0.918235 |
| 15 | 4155800 | 4156999 | 19.23586 | 32.70439 | 55.34712 | 24.25716 | -0.37443 | 1.019616 | 0.918237 |
| 8 | 3988000 | 3996799 | 115.2275 | 176.1734 | 209.6209 | 86.96261 | 0.038953 | 3.07457 | 0.918244 |
| 17 | 14064000 | 14065399 | 17.86324 | 30.11407 | 57.36657 | 25.08183 | -0.54514 | 1.090205 | 0.918272 |
| 14 | 5775200 | 5776599 | 18.07691 | 31.5866 | 42.99563 | 18.94502 | -0.0552 | 0.716006 | 0.918321 |
| 1 | 182637600 | 182645399 | 79.20609 | 122.6901 | 187.8647 | 78.03387 | -0.31008 | 2.810913 | 0.918333 |
| 16 | 56806000 | 56808999 | 29.67824 | 48.69099 | 65.81646 | 28.33986 | -0.0565 | 1.431798 | 0.918345 |
| 6 | 67685000 | 67688799 | 71.4177 | 112.4029 | 144.2955 | 59.86665 | -0.0382 | 2.515591 | 0.918349 |
| 8 | 110104200 | 110106399 | 20.2446 | 34.05501 | 58.53924 | 25.5778 | -0.39769 | 1.118619 | 0.918383 |
| 5 | 51160000 | 51160399 | 12.25039 | 22.78747 | 22.96117 | 9.596971 | 0.436295 | 0.169864 | 0.918387 |
| 3 | 9450000 | 9460399 | 184.7975 | 274.1059 | 317.175 | 130.6412 | 0.041253 | 3.677701 | 0.918392 |
| 19 | 26061600 | 26062199 | 10.26657 | 19.15474 | 27.34438 | 11.37101 | -0.05654 | 0.146999 | 0.918395 |
| 17 | 78950200 | 78951799 | 14.42008 | 25.53401 | 34.97368 | 15.46477 | -0.06008 | 0.505135 | 0.918445 |
| 7 | 104098400 | 104100799 | 26.10914 | 43.65717 | 58.89777 | 25.64873 | -0.04953 | 1.19662 | 0.918462 |
| 6 | 90087400 | 90087999 | 11.02147 | 20.38877 | 28.76938 | 12.11758 | -0.05419 | 0.205822 | 0.918486 |
| 12 | 113555000 | 113555399 | 8.206547 | 15.80621 | 22.00097 | 8.54262 | 0.067041 | -0.11758 | 0.918504 |
| 1 | 12031000 | 12032199 | 14.68307 | 26.06553 | 33.0251 | 14.58359 | 0.053802 | 0.48451 | 0.918505 |
| 13 | 113892600 | 113893999 | 13.13749 | 23.78292 | 30.54475 | 13.23511 | 0.052573 | 0.34875 | 0.918513 |
| 2 | 47040800 | 47041799 | 11.85975 | 22.00791 | 28.78845 | 12.07437 | 0.058849 | 0.196251 | 0.918547 |
| 1 | 184894600 | 184896599 | 16.09758 | 28.17676 | 38.34624 | 16.92099 | -0.0548 | 0.67057 | 0.918562 |
| 9 | 15717600 | 15718599 | 16.09758 | 28.17676 | 38.34624 | 16.92099 | -0.0548 | 0.67057 | 0.918562 |
| 5 | 121807200 | 121809399 | 20.13504 | 34.6741 | 43.72949 | 19.18246 | 0.051216 | 0.891078 | 0.918569 |
| 17 | 93953400 | 93954799 | 12.07859 | 22.22136 | 31.33512 | 13.29776 | -0.06104 | 0.252047 | 0.918573 |
| 1 | 164906400 | 164915599 | 104.5722 | 160.6505 | 191.9587 | 79.70433 | 0.038226 | 2.974745 | 0.918596 |

| | | | | | | | | | |
|---|---|---|---|---|---|---|---|---|---|
| 4 | 21142600 | 21143999 | 15.59545 | 27.52011 | 34.38023 | 15.21285 | 0.069933 | 0.547493 | 0.918599 |
| 14 | 45996600 | 45997599 | 12.649 | 22.84269 | 29.0898 | 12.64327 | 0.06196 | 0.356236 | 0.918632 |
| 12 | 41004400 | 41006199 | 22.3192 | 38.14865 | 47.94303 | 21.02114 | 0.054238 | 0.97029 | 0.918632 |
| 8 | 34839800 | 34841799 | 18.10345 | 31.57402 | 39.8795 | 17.55138 | 0.051979 | 0.73619 | 0.91867 |
| 5 | 150763200 | 150764999 | 13.40406 | 24.20454 | 30.97396 | 13.46337 | 0.054686 | 0.36484 | 0.91867 |
| 18 | 85153200 | 85154599 | 15.54283 | 27.86635 | 35.02043 | 15.33614 | 0.07364 | 0.432529 | 0.918671 |
| X | 161409400 | 161409799 | 7.629328 | 14.80778 | 23.43 | 8.809047 | -0.07214 | -0.22651 | 0.918716 |
| 5 | 85657000 | 85660799 | 40.63771 | 66.0087 | 81.38783 | 34.90361 | 0.066372 | 1.608173 | 0.918765 |
| 18 | 74283800 | 74284799 | 16.38091 | 29.16085 | 28.85827 | 12.7136 | 0.410625 | 0.469713 | 0.918768 |
| X | 103181600 | 103182199 | 8.104585 | 15.66766 | 24.3594 | 9.33057 | -0.0727 | -0.1561 | 0.918789 |
| 10 | 126665000 | 126670199 | 48.70464 | 78.11286 | 96.32153 | 40.69816 | 0.046006 | 1.9622 | 0.9188 |
| 3 | 68705600 | 68709399 | 81.7639 | 127.4427 | 162.5571 | 67.46448 | -0.03982 | 2.701904 | 0.918804 |
| 4 | 72592200 | 72594599 | 20.32441 | 35.37066 | 49.00691 | 21.53901 | -0.08138 | 0.790253 | 0.918819 |
| 6 | 63219000 | 63219599 | 9.790891 | 18.32813 | 26.3583 | 10.89937 | -0.06047 | 0.119434 | 0.91882 |
| 18 | 71343000 | 71343999 | 9.790891 | 18.32813 | 26.3583 | 10.89937 | -0.06047 | 0.119434 | 0.91882 |
| 1 | 163632400 | 163632999 | 11.57786 | 21.24027 | 29.73873 | 12.66874 | -0.05469 | 0.260481 | 0.918831 |
| 1 | 165271600 | 165273799 | 19.70017 | 34.62367 | 33.31215 | 14.75039 | 0.443005 | 0.618125 | 0.918837 |
| 4 | 138600400 | 138603399 | 26.17408 | 43.52024 | 58.77482 | 25.50293 | -0.05101 | 1.258417 | 0.918844 |
| 2 | 43406400 | 43409599 | 39.84878 | 64.33561 | 85.0962 | 36.2965 | -0.04338 | 1.745324 | 0.918876 |
| 3 | 15956400 | 15959999 | 37.53866 | 61.04065 | 82.16376 | 35.25939 | -0.05857 | 1.583706 | 0.918883 |
| 11 | 118198800 | 118201599 | 46.51174 | 74.74645 | 92.6973 | 39.28582 | 0.041304 | 1.897408 | 0.918892 |
| 9 | 79652400 | 79652999 | 7.376089 | 14.24376 | 22.04493 | 8.487296 | -0.07394 | -0.15812 | 0.918929 |
| 8 | 40860000 | 40860799 | 7.828915 | 15.14754 | 21.55482 | 8.223763 | 0.05876 | -0.18417 | 0.918937 |
| 7 | 6600400 | 6645399 | 971.7995 | 1249.558 | 1325.568 | 555.8975 | -0.03841 | 5.914498 | 0.918955 |
| 8 | 86124200 | 86126399 | 16.61109 | 29.17848 | 39.68834 | 17.56557 | -0.05416 | 0.618832 | 0.918958 |
| 4 | 38401000 | 38402799 | 28.20739 | 47.43589 | 48.02746 | 20.98297 | 0.363134 | 1.134276 | 0.918979 |
| 6 | 23714800 | 23720999 | 89.07022 | 138.491 | 167.0244 | 69.34388 | 0.038167 | 2.747367 | 0.918985 |
| 12 | 60925000 | 60926599 | 15.71025 | 27.744 | 38.02639 | 16.82365 | -0.06263 | 0.547526 | 0.918994 |
| 16 | 3136400 | 3202399 | 3802.091 | 4384.242 | 3934.1 | 1673.448 | 0.043465 | 7.648536 | 0.919011 |
| 1 | 191723200 | 191724799 | 11.82173 | 21.56572 | 27.64493 | 11.86429 | 0.064339 | 0.288543 | 0.919048 |
| 12 | 58240400 | 58241199 | 11.44423 | 20.86993 | 29.23723 | 12.55302 | -0.06316 | 0.2929 | 0.919055 |
| 6 | 121809000 | 121813799 | 53.63811 | 86.14304 | 104.1205 | 44.08509 | 0.077134 | 1.931135 | 0.919062 |
| 5 | 129115400 | 129116999 | 16.42116 | 28.7794 | 36.26681 | 16.02046 | 0.055673 | 0.64915 | 0.919083 |
| 15 | 20537000 | 20538199 | 12.56569 | 22.13309 | 39.38322 | 17.433 | -0.43817 | 0.56387 | 0.919094 |
| 1 | 118068600 | 118089799 | 580.6471 | 777.4349 | 859.3408 | 358.7328 | -0.03941 | 5.259271 | 0.919111 |
| 17 | 45457800 | 45459999 | 20.06184 | 34.47327 | 46.8707 | 20.55903 | -0.0576 | 0.911809 | 0.919111 |
| 6 | 89428200 | 89428599 | 6.239171 | 12.15297 | 19.64909 | 7.168737 | -0.06256 | -0.36105 | 0.919111 |
| 15 | 97530400 | 97534599 | 63.1635 | 100.0912 | 129.2727 | 53.77361 | -0.03917 | 2.354059 | 0.919137 |
| 13 | 3079800 | 3080999 | 13.61774 | 24.73031 | 34.80935 | 15.02351 | -0.07513 | 0.344445 | 0.919146 |
| 17 | 44988800 | 44992799 | 32.80608 | 53.83454 | 67.82366 | 29.20687 | 0.044868 | 1.438945 | 0.919147 |
| 11 | 114011600 | 114012599 | 15.84945 | 27.79323 | 37.81622 | 16.71292 | -0.0544 | 0.6404 | 0.919171 |
| 3 | 67264200 | 67268199 | 28.00951 | 47.24025 | 48.00811 | 20.99302 | 0.357984 | 1.11067 | 0.919184 |
| 8 | 71553400 | 71554599 | 20.16596 | 34.39485 | 46.92335 | 20.57051 | -0.06406 | 0.981573 | 0.919214 |
| 5 | 90316200 | 90317999 | 25.10378 | 42.28845 | 53.37328 | 23.33057 | 0.046388 | 1.115947 | 0.919243 |
| 14 | 5023200 | 5024599 | 19.18119 | 33.63593 | 30.67276 | 13.55594 | 0.520173 | 0.647052 | 0.919246 |
| 10 | 12680200 | 12681999 | 15.876 | 27.93231 | 38.07927 | 16.85961 | -0.05697 | 0.589403 | 0.919274 |
| 19 | 48240000 | 48241999 | 26.71412 | 44.30417 | 59.81164 | 25.9068 | -0.05075 | 1.291346 | 0.919305 |

| | | | | | | | | | |
|---|---|---|---|---|---|---|---|---|---|
| 5 | 125013200 | 125016799 | 30.67129 | 50.58203 | 63.85178 | 27.55431 | 0.044899 | 1.370576 | 0.919319 |
| 4 | 72806200 | 72808199 | 28.23869 | 47.62655 | 48.00348 | 20.99096 | 0.369773 | 1.110072 | 0.919346 |
| 11 | 57236000 | 57236999 | 13.25212 | 23.73623 | 32.6195 | 14.29648 | -0.05522 | 0.418952 | 0.919386 |
| 13 | 36817400 | 36818999 | 19.12955 | 33.23139 | 42.07229 | 18.47029 | 0.048072 | 0.801312 | 0.919396 |
| 5 | 78611800 | 78613599 | 18.03623 | 32.17893 | 30.09552 | 13.23389 | 0.493444 | 0.46896 | 0.919397 |
| 14 | 15572200 | 15575999 | 50.452 | 80.68407 | 105.3189 | 44.31149 | -0.04069 | 2.056549 | 0.919401 |
| 14 | 15572400 | 15575999 | 50.452 | 80.68407 | 105.3189 | 44.31149 | -0.04069 | 2.056549 | 0.919401 |
| 5 | 46450200 | 46452999 | 18.43499 | 31.2448 | 55.30876 | 24.22142 | -0.43935 | 1.043251 | 0.919421 |
| 1 | 103625400 | 103626599 | 14.64092 | 25.99487 | 32.98694 | 14.56477 | 0.051619 | 0.48231 | 0.919453 |
| 13 | 8714800 | 8721399 | 94.34837 | 145.4707 | 173.1411 | 71.91983 | 0.047874 | 2.899972 | 0.919484 |
| 3 | 36689800 | 36691999 | 18.2809 | 32.00497 | 40.30372 | 17.78786 | 0.056609 | 0.683116 | 0.919486 |
| 3 | 106318400 | 106319399 | 12.40783 | 22.61963 | 31.63878 | 13.58979 | -0.06193 | 0.303497 | 0.919504 |
| 5 | 8807200 | 8810199 | 44.28945 | 71.19446 | 87.79018 | 37.22079 | 0.049996 | 1.882419 | 0.919508 |
| 7 | 102491600 | 102493799 | 29.58963 | 48.94823 | 61.51794 | 26.59905 | 0.051919 | 1.332016 | 0.919567 |
| 4 | 97719000 | 97722599 | 34.09517 | 56.12004 | 69.98098 | 30.17461 | 0.061333 | 1.40584 | 0.919573 |
| 10 | 118348200 | 118349999 | 12.19996 | 22.11786 | 30.63498 | 13.25008 | -0.05397 | 0.333998 | 0.919592 |
| 11 | 94635400 | 94638399 | 43.61377 | 70.06479 | 92.25927 | 39.08038 | -0.0456 | 1.90511 | 0.919611 |
| 3 | 57180000 | 57180999 | 9.610972 | 18.19315 | 26.51357 | 10.74173 | -0.05509 | 0.044797 | 0.919627 |
| 2 | 170258000 | 170260799 | 19.62152 | 33.65184 | 45.84229 | 20.10554 | -0.06078 | 0.926898 | 0.919628 |
| 14 | 94684000 | 94686199 | 23.62885 | 40.4463 | 40.09052 | 17.58019 | 0.3958 | 0.955068 | 0.919671 |
| 8 | 33980600 | 33983199 | 22.74221 | 38.84216 | 49.01971 | 21.4919 | 0.048106 | 0.981591 | 0.919675 |
| 5 | 127522400 | 127528799 | 74.70987 | 117.2416 | 150.2719 | 62.32255 | -0.03919 | 2.575443 | 0.919676 |
| 9 | 84647800 | 84649599 | 27.18443 | 45.34744 | 61.10152 | 26.56861 | -0.04765 | 1.228637 | 0.919692 |
| 12 | 58338000 | 58339399 | 13.87719 | 25.02899 | 31.94585 | 13.90658 | 0.056955 | 0.376686 | 0.91974 |
| Y | 6166000 | 6166799 | 7.132202 | 13.64139 | 21.47107 | 8.50569 | -0.13615 | -0.06786 | 0.919741 |
| 8 | 32477600 | 32479999 | 22.37057 | 38.37018 | 52.27939 | 22.92891 | -0.06105 | 0.946925 | 0.919801 |
| 15 | 81980200 | 81982399 | 18.18335 | 31.71281 | 43.032 | 18.93861 | -0.05099 | 0.742151 | 0.919806 |
| 8 | 114398400 | 114399799 | 11.71678 | 21.31161 | 29.67324 | 12.78162 | -0.05759 | 0.309583 | 0.919876 |
| 5 | 71476200 | 71476999 | 16.66778 | 29.52617 | 29.35778 | 12.98045 | 0.399982 | 0.506219 | 0.919879 |
| 1 | 88096200 | 88107399 | 69.84739 | 110.9603 | 111.854 | 46.64471 | 0.321333 | 2.283171 | 0.919889 |
| 3 | 45847200 | 45847799 | 7.972853 | 15.36547 | 21.44742 | 8.31311 | 0.064791 | -0.13025 | 0.919902 |
| 2 | 33673200 | 33674599 | 13.36584 | 23.67722 | 32.64369 | 14.43327 | -0.06898 | 0.49101 | 0.919949 |
| 5 | 150207600 | 150208999 | 15.47131 | 27.40422 | 37.52108 | 16.59012 | -0.05932 | 0.514259 | 0.919956 |
| 9 | 4975400 | 4979599 | 60.93446 | 96.53822 | 125.1938 | 52.07032 | -0.04529 | 2.357832 | 0.919958 |
| 13 | 28381000 | 28387799 | 87.04323 | 136.6499 | 136.7775 | 56.72886 | 0.317548 | 2.573308 | 0.919986 |
| 17 | 55858000 | 55860199 | 17.51538 | 30.94051 | 38.45713 | 17.01326 | 0.077886 | 0.556526 | 0.919986 |
| 5 | 131502800 | 131513199 | 136.7344 | 206.186 | 255.7524 | 105.676 | -0.03744 | 3.361638 | 0.920012 |
| 14 | 73597600 | 73599599 | 18.84425 | 32.68223 | 44.21087 | 19.39782 | -0.04743 | 0.822415 | 0.920018 |
| 4 | 91490000 | 91493199 | 42.0722 | 67.83401 | 89.5751 | 38.14068 | -0.04328 | 1.788074 | 0.920068 |
| 1 | 3219800 | 3223399 | 52.92647 | 84.92127 | 86.56448 | 36.6001 | 0.32092 | 1.953162 | 0.920068 |
| 14 | 53720000 | 53720999 | 11.16875 | 20.58406 | 26.69075 | 11.29945 | 0.061806 | 0.223492 | 0.920098 |
| 10 | 129175200 | 129178199 | 45.80152 | 73.73784 | 98.03151 | 41.65514 | -0.0556 | 1.843718 | 0.920105 |
| 2 | 33278400 | 33283199 | 61.35992 | 97.6174 | 119.7196 | 50.00406 | 0.040958 | 2.235267 | 0.920107 |
| 3 | 8908200 | 8908999 | 10.1449 | 19.03434 | 25.25927 | 10.40451 | 0.058773 | 0.102977 | 0.920123 |
| 6 | 139508200 | 139512199 | 52.92017 | 84.59441 | 104.4048 | 43.89983 | 0.039336 | 2.07259 | 0.920152 |
| 1 | 35177000 | 35178799 | 16.89062 | 29.74824 | 37.51212 | 16.60507 | 0.055632 | 0.584217 | 0.920162 |
| 13 | 65403000 | 65420599 | 675.0184 | 893.616 | 923.007 | 385.8919 | 0.042081 | 5.44094 | 0.92017 |

| | | | | | | | | | |
|---|---|---|---|---|---|---|---|---|---|
| 10 | 61811400 | 61812399 | 12.56948 | 22.55241 | 31.1492 | 13.63224 | -0.06099 | 0.399765 | 0.920183 |
| Y | 4458000 | 4460799 | 33.14851 | 53.7589 | 62.51422 | 26.77838 | 0.148604 | 1.612261 | 0.920202 |
| 16 | 92982200 | 92982999 | 8.195461 | 15.69541 | 23.50836 | 9.278068 | -0.0602 | -0.0615 | 0.920221 |
| 5 | 27225600 | 27227399 | 30.42517 | 50.26481 | 63.43853 | 27.40479 | 0.045628 | 1.347314 | 0.920222 |
| 9 | 39048200 | 39048799 | 10.25014 | 19.1832 | 25.30394 | 10.46765 | 0.062883 | 0.12248 | 0.920257 |
| 18 | 81477600 | 81479199 | 19.07241 | 33.35149 | 41.88016 | 18.46528 | 0.060645 | 0.707741 | 0.920306 |
| 19 | 25076200 | 25077399 | 11.91565 | 21.70316 | 30.19613 | 12.98449 | -0.05485 | 0.305181 | 0.920311 |
| 19 | 23807000 | 23810399 | 26.45191 | 44.06787 | 59.42328 | 25.81757 | -0.04876 | 1.237264 | 0.920331 |
| 12 | 106743200 | 106745399 | 20.51073 | 35.02074 | 47.43852 | 20.79817 | -0.05359 | 0.974335 | 0.920332 |
| 5 | 143568800 | 143569999 | 14.86585 | 26.20925 | 35.66457 | 15.78397 | -0.05286 | 0.559485 | 0.920335 |
| 5 | 61388800 | 61390999 | 13.34967 | 23.39965 | 41.509 | 18.33831 | -0.43511 | 0.639517 | 0.920341 |
| 5 | 106810800 | 106813199 | 13.46407 | 23.83095 | 43.44825 | 19.21373 | -0.47078 | 0.520463 | 0.920348 |
| 2 | 125797000 | 125800599 | 92.14497 | 142.8112 | 171.9959 | 71.44222 | 0.03656 | 2.804073 | 0.92045 |
| 15 | 25633600 | 25635799 | 19.5858 | 33.67646 | 45.81823 | 20.09777 | -0.05835 | 0.9031 | 0.920457 |
| 16 | 95583000 | 95584799 | 18.35868 | 31.88086 | 43.157 | 18.94223 | -0.04811 | 0.80463 | 0.920457 |
| 6 | 60233800 | 60236799 | 25.70212 | 43.3463 | 58.90128 | 25.75597 | -0.05974 | 1.11446 | 0.920458 |
| 15 | 8623600 | 8626599 | 24.95227 | 41.89353 | 52.81882 | 23.06083 | 0.047798 | 1.144775 | 0.920473 |
| 1 | 46952600 | 46954399 | 17.97643 | 31.46503 | 39.74428 | 17.5377 | 0.052118 | 0.681912 | 0.920475 |
| 8 | 49238000 | 49239399 | 17.97643 | 31.46503 | 39.74428 | 17.5377 | 0.052118 | 0.681912 | 0.920475 |
| 18 | 57212600 | 57215999 | 40.52573 | 65.59909 | 81.58477 | 34.89879 | 0.049266 | 1.674932 | 0.920496 |
| 4 | 156234000 | 156236599 | 32.65836 | 53.52414 | 71.79046 | 30.89849 | -0.04598 | 1.456475 | 0.920501 |
| 10 | 53472000 | 53473999 | 15.79544 | 27.13804 | 50.56562 | 22.17964 | -0.51018 | 0.9022 | 0.920513 |
| 14 | 76087800 | 76089599 | 17.0973 | 29.84821 | 40.46312 | 17.81912 | -0.04933 | 0.720267 | 0.920521 |
| 12 | 109810600 | 109815599 | 47.26959 | 75.39584 | 120.6591 | 50.57711 | -0.33831 | 2.138712 | 0.920548 |
| 5 | 129719200 | 129722399 | 44.66457 | 71.77745 | 88.76908 | 37.62062 | 0.045287 | 1.892952 | 0.920559 |
| 1 | 113991800 | 113992399 | 11.19819 | 20.43967 | 28.68077 | 12.30023 | -0.06468 | 0.28242 | 0.920575 |
| 4 | 150741000 | 150744199 | 33.20365 | 54.02003 | 72.26685 | 31.01823 | -0.04911 | 1.560697 | 0.9206 |
| 6 | 146364600 | 146368199 | 29.50687 | 49.04144 | 61.74118 | 26.7824 | 0.049891 | 1.276481 | 0.920619 |
| 15 | 25233000 | 25235399 | 20.4888 | 35.51964 | 44.68325 | 19.60482 | 0.05662 | 0.831981 | 0.920619 |
| 1 | 12662000 | 12673399 | 91.50343 | 142.9004 | 145.2002 | 60.24735 | 0.290034 | 2.669926 | 0.920634 |
| 10 | 82124400 | 82125799 | 14.07482 | 25.4706 | 32.52089 | 14.09774 | 0.060387 | 0.360632 | 0.920647 |
| 14 | 116143200 | 116147399 | 12.24825 | 21.52439 | 39.93077 | 17.67947 | -0.49925 | 0.588545 | 0.920676 |
| 17 | 84050400 | 84054399 | 69.16775 | 109.2228 | 133.4038 | 55.39282 | 0.037247 | 2.442419 | 0.920682 |
| 10 | 76227000 | 76228799 | 17.26185 | 29.86746 | 41.22599 | 18.08688 | -0.07692 | 0.833 | 0.920705 |
| 6 | 130901800 | 130903199 | 27.22856 | 45.75522 | 57.31875 | 25.03655 | 0.056947 | 1.147241 | 0.920705 |
| 10 | 46876000 | 46876799 | 11.90845 | 21.71618 | 27.72308 | 11.90313 | 0.070108 | 0.293318 | 0.92071 |
| 17 | 89728400 | 89729999 | 16.61283 | 29.34752 | 36.98421 | 16.3615 | 0.058238 | 0.549665 | 0.92076 |
| 17 | 66438200 | 66440999 | 31.86468 | 52.14738 | 69.7829 | 30.01634 | -0.04357 | 1.472271 | 0.920765 |
| 5 | 101892800 | 101894199 | 19.68886 | 33.92818 | 45.82251 | 20.10099 | -0.04702 | 0.886128 | 0.920771 |
| 7 | 104579600 | 104579999 | 6.650066 | 12.94616 | 19.03831 | 6.98784 | 0.064048 | -0.33974 | 0.92079 |
| 1 | 94061400 | 94061999 | 10.59934 | 19.62961 | 27.79386 | 11.68906 | -0.05727 | 0.195353 | 0.920791 |
| 8 | 89532400 | 89533999 | 17.51252 | 30.62977 | 38.79191 | 17.10648 | 0.048401 | 0.690826 | 0.920827 |
| 19 | 12197200 | 12198599 | 16.2395 | 28.62612 | 39.17496 | 17.33815 | -0.06118 | 0.576317 | 0.920848 |
| 2 | 124946200 | 124950199 | 45.29767 | 73.23079 | 74.23859 | 31.72008 | 0.341715 | 1.709497 | 0.920848 |
| 1 | 189841000 | 189844199 | 42.9721 | 69.14501 | 91.01152 | 38.64175 | -0.04215 | 1.853681 | 0.920869 |
| 7 | 140318800 | 140322399 | 29.80849 | 49.2195 | 62.17817 | 26.85035 | 0.044243 | 1.351787 | 0.920879 |
| 14 | 106587200 | 106591399 | 74.86017 | 117.523 | 142.64 | 59.16033 | 0.0398 | 2.566854 | 0.920891 |

| | | | | | | | | | |
|---|---|---|---|---|---|---|---|---|---|
| X | 88177000 | 88177599 | 10.18968 | 19.21563 | 28.07064 | 11.46359 | -0.0684 | 0.081744 | 0.920917 |
| 10 | 123664400 | 123664999 | 8.570531 | 16.40367 | 24.56642 | 9.712716 | -0.06319 | -0.05221 | 0.920936 |
| 10 | 59665600 | 59669399 | 34.13273 | 55.6224 | 69.73642 | 29.94996 | 0.045895 | 1.537277 | 0.920975 |
| 4 | 144308000 | 144323399 | 219.5567 | 321.2725 | 367.2897 | 151.3036 | 0.040225 | 3.902446 | 0.920979 |
| 2 | 99378600 | 99383999 | 84.78286 | 131.8742 | 158.6242 | 65.8527 | 0.04241 | 2.738237 | 0.921092 |
| 2 | 164191000 | 164193399 | 18.6435 | 32.56047 | 44.57614 | 19.63531 | -0.06364 | 0.726353 | 0.921103 |
| 7 | 60833000 | 60833199 | 5.358723 | 10.46812 | 15.97369 | 5.582309 | 0.076778 | -0.53846 | 0.921107 |
| 18 | 6764800 | 6765599 | 10.05454 | 18.77191 | 26.813 | 11.1372 | -0.05618 | 0.138298 | 0.921112 |
| 14 | 50255800 | 50256799 | 20.74021 | 35.50838 | 47.95517 | 21.02925 | -0.04864 | 0.952821 | 0.921116 |
| 18 | 62408600 | 62413599 | 59.5525 | 94.73508 | 116.4171 | 48.61983 | 0.037977 | 2.235345 | 0.921155 |
| 5 | 30717200 | 30719399 | 17.14392 | 30.04683 | 40.82737 | 18.03541 | -0.0526 | 0.660413 | 0.921172 |
| 16 | 19327400 | 19329399 | 21.33301 | 36.77843 | 46.30842 | 20.31443 | 0.053705 | 0.887293 | 0.921206 |
| 10 | 107829600 | 107829999 | 7.088997 | 13.77761 | 20.07032 | 7.466717 | 0.059549 | -0.28354 | 0.92121 |
| 8 | 89659800 | 89661799 | 21.80511 | 37.19615 | 46.91004 | 20.56493 | 0.048921 | 0.987598 | 0.921225 |
| 6 | 109402800 | 109405199 | 37.35012 | 61.08642 | 62.97358 | 27.08125 | 0.331173 | 1.488766 | 0.921227 |
| 1 | 66018400 | 66020399 | 21.74672 | 36.30076 | 62.31626 | 27.10821 | -0.39582 | 1.207092 | 0.921229 |
| 11 | 61280000 | 61289199 | 138.1997 | 207.8868 | 257.6264 | 106.3803 | -0.03924 | 3.40997 | 0.921229 |
| 7 | 123564000 | 123568799 | 59.76516 | 95.09215 | 116.653 | 48.73162 | 0.040767 | 2.227697 | 0.92125 |
| 5 | 77901200 | 77910599 | 126.055 | 191.773 | 227.2343 | 94.1498 | 0.039236 | 3.179666 | 0.921262 |
| 5 | 109554400 | 109563399 | 101.5125 | 157.8159 | 185.1567 | 76.87344 | 0.077729 | 2.757871 | 0.921265 |
| 11 | 44274800 | 44277399 | 29.30505 | 48.38791 | 61.02597 | 26.35267 | 0.046643 | 1.349463 | 0.921307 |
| 13 | 18981000 | 18982199 | 9.597399 | 17.93915 | 25.82408 | 10.70546 | -0.06491 | 0.124899 | 0.921312 |
| 4 | 56535000 | 56537999 | 20.43344 | 35.72594 | 35.44767 | 15.62709 | 0.398756 | 0.705218 | 0.921336 |
| 1 | 127620800 | 127621799 | 10.52811 | 19.59159 | 27.79982 | 11.61152 | -0.05327 | 0.165527 | 0.921336 |
| 5 | 9145800 | 9147599 | 17.02822 | 30.3345 | 37.44737 | 16.48777 | 0.093677 | 0.483312 | 0.921345 |
| 5 | 66331400 | 66334799 | 44.66177 | 71.82502 | 89.11593 | 37.80734 | 0.041914 | 1.870297 | 0.921345 |
| 15 | 79163200 | 79165799 | 23.26782 | 38.59313 | 66.05874 | 28.61497 | -0.39201 | 1.287328 | 0.921354 |
| 13 | 90459400 | 90460799 | 15.81575 | 28.01914 | 35.45171 | 15.67461 | 0.054141 | 0.511744 | 0.921358 |
| 3 | 75089800 | 75092399 | 19.08403 | 33.31203 | 45.67694 | 20.11343 | -0.06589 | 0.739322 | 0.921359 |
| 2 | 176780400 | 176787199 | 95.8214 | 148.3263 | 178.3173 | 74.06843 | 0.037786 | 2.828852 | 0.921393 |
| 5 | 119921600 | 119925799 | 46.14366 | 74.09332 | 91.73771 | 38.83133 | 0.042195 | 1.92176 | 0.921395 |
| 5 | 138588800 | 138589399 | 7.701315 | 14.81497 | 22.51205 | 8.77182 | -0.06372 | -0.11255 | 0.921406 |
| 14 | 86275400 | 86275799 | 7.701315 | 14.81497 | 22.51205 | 8.77182 | -0.06372 | -0.11255 | 0.921406 |
| 8 | 74259600 | 74262399 | 30.06214 | 50.11478 | 62.52131 | 27.17466 | 0.063028 | 1.243902 | 0.921432 |
| 3 | 101729400 | 101730999 | 13.35987 | 23.76821 | 32.59212 | 14.36602 | -0.05786 | 0.460377 | 0.921451 |
| 8 | 110480400 | 110483599 | 30.15729 | 49.62553 | 66.59778 | 28.71269 | -0.04443 | 1.396909 | 0.921454 |
| Y | 20800800 | 20801399 | 5.258612 | 10.24318 | 17.50451 | 6.384763 | -0.14135 | -0.38555 | 0.921496 |
| 16 | 82933600 | 82936599 | 25.12591 | 41.8071 | 74.0627 | 32.15224 | -0.44135 | 1.259354 | 0.921503 |
| 7 | 137242800 | 137243599 | 10.19677 | 19.11074 | 27.38219 | 11.29987 | -0.05323 | 0.116078 | 0.92151 |
| 9 | 53926600 | 53928599 | 35.21633 | 57.31775 | 71.84372 | 30.84117 | 0.045047 | 1.560537 | 0.921525 |
| 17 | 4653000 | 4654199 | 15.89343 | 28.02756 | 35.49923 | 15.71005 | 0.050016 | 0.560895 | 0.921527 |
| 7 | 34300200 | 34301799 | 16.12729 | 28.24327 | 38.33883 | 16.92506 | -0.0511 | 0.663059 | 0.921556 |
| 10 | 20558800 | 20560399 | 26.54475 | 44.56603 | 56.09674 | 24.49157 | 0.050164 | 1.155288 | 0.921574 |
| 6 | 123905000 | 123905999 | 14.01037 | 25.13686 | 34.56208 | 15.12627 | -0.05479 | 0.415943 | 0.92159 |
| 16 | 92084800 | 92087999 | 19.40981 | 33.66419 | 42.60548 | 18.69346 | 0.048186 | 0.821532 | 0.921598 |
| 3 | 57256400 | 57258399 | 17.34478 | 30.82626 | 37.92365 | 16.73041 | 0.096654 | 0.500287 | 0.921619 |
| 6 | 123788800 | 123797999 | 123.9036 | 189.0347 | 223.505 | 92.67678 | 0.045652 | 3.121623 | 0.921635 |

| | | | | | | | | | |
|---|---|---|---|---|---|---|---|---|---|
| 6 | 54056400 | 54058199 | 17.09152 | 29.85712 | 40.43644 | 17.8145 | -0.04788 | 0.711061 | 0.921639 |
| 12 | 21820200 | 21836399 | 390.9883 | 541.7816 | 587.1934 | 243.8255 | 0.0389 | 4.719115 | 0.921654 |
| 17 | 70088600 | 70091799 | 46.03318 | 74.15558 | 72.32439 | 30.7587 | 0.391468 | 1.81244 | 0.921704 |
| 3 | 87484600 | 87491199 | 83.08036 | 129.8782 | 157.4075 | 65.30562 | 0.037153 | 2.649132 | 0.921711 |
| 6 | 70510000 | 70510799 | 16.38307 | 28.7144 | 36.23325 | 16.00646 | 0.053692 | 0.647115 | 0.921735 |
| 18 | 34029800 | 34031799 | 26.4064 | 43.78149 | 59.21107 | 25.64169 | -0.0533 | 1.293694 | 0.92184 |
| 1 | 181662000 | 181664599 | 32.85131 | 53.26091 | 85.98089 | 36.82232 | -0.32395 | 1.634385 | 0.921846 |
| 12 | 118380200 | 118382199 | 19.13588 | 33.22267 | 42.09412 | 18.47298 | 0.046881 | 0.809935 | 0.92185 |
| 2 | 80691400 | 80692799 | 19.05736 | 33.03341 | 41.6461 | 18.27219 | 0.053744 | 0.826491 | 0.921914 |
| 10 | 62198000 | 62205599 | 122.3225 | 186.0701 | 221.003 | 91.5644 | 0.035646 | 3.180342 | 0.921936 |
| Y | 1898200 | 1900799 | 26.61034 | 43.91837 | 51.22504 | 22.11339 | 0.15904 | 1.337463 | 0.92196 |
| 16 | 93029600 | 93030399 | 15.50163 | 27.34781 | 34.60252 | 15.31197 | 0.051923 | 0.554479 | 0.921965 |
| 3 | 58566000 | 58568399 | 20.23897 | 34.93931 | 44.2042 | 19.39216 | 0.047296 | 0.868909 | 0.921997 |
| X | 169254000 | 169261999 | 260.8857 | 363.9504 | 630.9024 | 261.6999 | -0.6286 | 4.613354 | 0.922029 |
| 7 | 113989600 | 113992199 | 35.08999 | 57.37245 | 77.57691 | 33.35757 | -0.05949 | 1.499114 | 0.922039 |
| 16 | 18794000 | 18795799 | 59.56287 | 94.57406 | 115.3874 | 48.09369 | 0.045665 | 2.290748 | 0.922057 |
| 1 | 72336600 | 72337199 | 6.690156 | 13.00301 | 20.63878 | 7.675567 | -0.06168 | -0.28447 | 0.922074 |
| 13 | 33787400 | 33788799 | 25.6243 | 43.13471 | 54.47005 | 23.80487 | 0.045595 | 1.125267 | 0.922076 |
| 11 | 48118400 | 48119399 | 15.94062 | 28.70648 | 27.83315 | 12.13955 | 0.44982 | 0.394353 | 0.922082 |
| 19 | 32991000 | 32992599 | 15.99669 | 28.13699 | 38.1566 | 16.89477 | -0.04937 | 0.595689 | 0.922093 |
| 12 | 86285800 | 86287199 | 15.54954 | 27.56016 | 34.87409 | 15.4178 | 0.054127 | 0.50227 | 0.922103 |
| 1 | 27768800 | 27777399 | 316.9246 | 446.0333 | 487.5864 | 201.9752 | 0.049104 | 4.483168 | 0.922124 |
| 3 | 53639800 | 53641599 | 15.45787 | 27.28246 | 37.065 | 16.40021 | -0.05013 | 0.552268 | 0.922172 |
| 6 | 143409600 | 143410999 | 12.0181 | 21.63288 | 30.11493 | 13.14218 | -0.06962 | 0.371815 | 0.922186 |
| 16 | 30668000 | 30668599 | 7.535839 | 14.61364 | 22.82605 | 8.616837 | -0.06011 | -0.21991 | 0.922186 |
| 11 | 22225600 | 22227999 | 30.33283 | 50.10667 | 63.33142 | 27.35641 | 0.043555 | 1.349741 | 0.922192 |
| 6 | 115485200 | 115486399 | 15.13522 | 26.78792 | 34.00088 | 15.03601 | 0.049126 | 0.514955 | 0.922204 |
| 10 | 48879800 | 48880399 | 16.61261 | 28.94412 | 39.5526 | 17.39627 | -0.06102 | 0.74498 | 0.922209 |
| 13 | 55584600 | 55589199 | 56.59407 | 90.09442 | 116.7715 | 48.81694 | -0.03762 | 2.209296 | 0.922249 |
| 6 | 123341400 | 123345199 | 43.94384 | 71.16323 | 72.05706 | 30.83319 | 0.345804 | 1.668245 | 0.922261 |
| 13 | 34058000 | 34065399 | 148.0939 | 223.8605 | 222.9434 | 92.19039 | 0.282045 | 3.313336 | 0.922287 |
| 3 | 26976400 | 26978599 | 39.80956 | 64.1941 | 84.80087 | 36.11378 | -0.04357 | 1.77956 | 0.922296 |
| 9 | 121355600 | 121361399 | 72.06164 | 113.285 | 145.2074 | 60.23593 | -0.0373 | 2.538555 | 0.922329 |
| 11 | 16874200 | 16875399 | 14.74735 | 26.03138 | 35.53621 | 15.72341 | -0.05704 | 0.542716 | 0.922342 |
| 19 | 37000600 | 37003999 | 38.06367 | 61.38244 | 81.90392 | 34.88274 | -0.05786 | 1.775125 | 0.922348 |
| Y | 35139200 | 35139799 | 7.795805 | 14.9355 | 19.38919 | 7.64428 | 0.145609 | -0.08427 | 0.922357 |
| 10 | 38832800 | 38833599 | 9.571742 | 18.09853 | 24.34739 | 9.887613 | 0.056562 | 0.046374 | 0.922375 |
| 10 | 33536600 | 33539799 | 42.75088 | 68.90928 | 85.84557 | 36.53968 | 0.040153 | 1.79636 | 0.922427 |
| 1 | 85118400 | 85122999 | 202.6644 | 295.5667 | 358.2749 | 147.6318 | -0.04882 | 3.951243 | 0.922433 |
| 10 | 104646200 | 104647199 | 13.94411 | 25.70415 | 25.86889 | 10.94857 | 0.425007 | 0.24164 | 0.922441 |
| X | 118449400 | 118451199 | 19.56415 | 34.33691 | 42.28356 | 18.70092 | 0.08869 | 0.654511 | 0.922449 |
| 12 | 54587600 | 54589799 | 22.17812 | 38.03244 | 47.93416 | 21.02282 | 0.050844 | 0.940542 | 0.92245 |
| 15 | 44863600 | 44864199 | 8.485259 | 16.3859 | 23.16792 | 8.919006 | 0.055747 | -0.14078 | 0.922452 |
| 12 | 115385200 | 115392599 | 46.68905 | 75.39376 | 75.76675 | 32.32523 | 0.352419 | 1.746537 | 0.922467 |
| 15 | 84529200 | 84532799 | 38.95146 | 63.13646 | 78.97628 | 33.8168 | 0.042818 | 1.645111 | 0.922486 |
| 13 | 99735400 | 99737999 | 23.43935 | 39.51207 | 53.34614 | 23.32361 | -0.0505 | 1.103156 | 0.922505 |
| 7 | 96876200 | 96877199 | 22.6978 | 37.42499 | 68.99655 | 29.77196 | -0.49983 | 1.358308 | 0.922515 |

| | | | | | | | | | |
|---|---|---|---|---|---|---|---|---|---|
| 2 | 24051400 | 24059799 | 157.9958 | 236.1772 | 290.3563 | 119.7208 | -0.03585 | 3.533978 | 0.922542 |
| 3 | 78449000 | 78450599 | 20.84852 | 36.23902 | 36.33405 | 15.95707 | 0.383281 | 0.793327 | 0.922552 |
| 12 | 78054400 | 78056599 | 19.1258 | 32.80565 | 56.60582 | 24.82647 | -0.40052 | 0.937564 | 0.922565 |
| 5 | 128452400 | 128458599 | 93.7807 | 145.3 | 174.9407 | 72.6655 | 0.036635 | 2.810295 | 0.922579 |
| Y | 40643600 | 40643999 | 5.109521 | 9.962068 | 17.00503 | 6.127142 | -0.12343 | -0.4324 | 0.92259 |
| Y | 78948600 | 78948999 | 5.109521 | 9.962068 | 17.00503 | 6.127142 | -0.12343 | -0.4324 | 0.92259 |
| 13 | 19146400 | 19149199 | 25.34639 | 42.93991 | 53.91768 | 23.60909 | 0.054103 | 1.070109 | 0.922605 |
| 6 | 128748800 | 128758599 | 189.3478 | 279.8913 | 341.62 | 140.6873 | -0.0403 | 3.73701 | 0.922631 |
| 15 | 46989000 | 46991599 | 24.51742 | 41.75487 | 52.30552 | 22.92725 | 0.058101 | 1.016811 | 0.922652 |
| 7 | 131638400 | 131639999 | 11.50362 | 21.20404 | 29.75806 | 12.59571 | -0.0516 | 0.231966 | 0.922661 |
| 3 | 57143800 | 57150999 | 116.4251 | 177.759 | 211.5801 | 87.74738 | 0.036897 | 3.103795 | 0.922699 |
| 13 | 38375400 | 38381999 | 160.9373 | 237.155 | 359.2782 | 147.9544 | -0.35608 | 3.785153 | 0.922709 |
| 11 | 85626800 | 85628399 | 15.01649 | 26.69515 | 36.53554 | 16.11454 | -0.05589 | 0.48368 | 0.922713 |
| 5 | 81477400 | 81478199 | 11.41856 | 20.91322 | 29.22817 | 12.47944 | -0.05441 | 0.267541 | 0.922719 |
| 2 | 125477200 | 125478399 | 17.79162 | 30.96565 | 38.97134 | 17.12927 | 0.056905 | 0.763173 | 0.922723 |
| 2 | 63443600 | 63444199 | 10.44237 | 19.70424 | 26.39665 | 10.76647 | 0.056553 | 0.071684 | 0.922745 |
| 1 | 101566800 | 101568799 | 21.24644 | 36.75614 | 46.30744 | 20.31614 | 0.053932 | 0.857414 | 0.922763 |
| 1 | 181607000 | 181616199 | 102.4029 | 157.8688 | 188.9506 | 78.48688 | 0.039656 | 2.906156 | 0.922774 |
| 12 | 8805200 | 8811999 | 70.30683 | 110.6873 | 142.4854 | 59.11793 | -0.04185 | 2.507015 | 0.922783 |
| 11 | 41600200 | 41601399 | 10.18378 | 18.20268 | 35.37974 | 15.53335 | -0.55375 | 0.418044 | 0.922816 |
| 8 | 61137000 | 61140399 | 60.10485 | 95.40924 | 116.3177 | 48.46854 | 0.046396 | 2.299849 | 0.922822 |
| 1 | 101195400 | 101198199 | 28.85026 | 48.3471 | 49.67249 | 21.66143 | 0.342167 | 1.174815 | 0.922824 |
| 15 | 16454600 | 16457799 | 54.30877 | 86.9323 | 88.3977 | 37.24089 | 0.320418 | 2.027575 | 0.922829 |
| 7 | 124635400 | 124639599 | 40.80649 | 65.8977 | 81.7588 | 34.88382 | 0.049092 | 1.734813 | 0.922858 |
| 18 | 61537200 | 61539199 | 32.14588 | 52.77599 | 66.62864 | 28.69798 | 0.042151 | 1.430813 | 0.922873 |
| 8 | 122040200 | 122041599 | 13.91331 | 24.58152 | 33.73386 | 14.92322 | -0.06369 | 0.527601 | 0.922882 |
| 9 | 109552600 | 109556799 | 59.94221 | 95.7656 | 116.0369 | 48.75213 | 0.064938 | 2.101347 | 0.922928 |
| 2 | 162981400 | 162983999 | 33.34343 | 54.53943 | 68.73853 | 29.5616 | 0.042149 | 1.482712 | 0.922951 |
| 9 | 9598400 | 9601999 | 60.34978 | 95.99266 | 118.0195 | 49.28142 | 0.0371 | 2.239626 | 0.922961 |
| 8 | 21362800 | 21392599 | 436.1547 | 600.932 | 685.1563 | 284.7619 | -0.0425 | 4.808074 | 0.922964 |
| 7 | 25564200 | 25566399 | 25.25457 | 42.34157 | 53.37885 | 23.28469 | 0.047934 | 1.160579 | 0.923027 |
| 16 | 97770000 | 97770999 | 14.2531 | 25.45637 | 32.4419 | 14.25288 | 0.050313 | 0.440725 | 0.923035 |
| 16 | 35491600 | 35494799 | 28.14496 | 46.62861 | 58.80979 | 25.45458 | 0.046962 | 1.306028 | 0.923053 |
| 2 | 154986600 | 154995599 | 118.3691 | 181.4492 | 213.962 | 88.81382 | 0.054886 | 3.025081 | 0.923054 |
| 1 | 70524800 | 70525799 | 10.25505 | 19.17468 | 27.36553 | 11.33819 | -0.05197 | 0.132905 | 0.923057 |
| 19 | 18501600 | 18504599 | 27.28872 | 45.65029 | 61.77693 | 26.91026 | -0.054 | 1.198417 | 0.923077 |
| 11 | 15177600 | 15180599 | 27.86245 | 46.77019 | 46.99995 | 20.5177 | 0.373986 | 1.151698 | 0.923086 |
| 1 | 85400000 | 85405399 | 196.9516 | 288.2725 | 349.3154 | 143.8967 | -0.04334 | 3.895348 | 0.923124 |
| 14 | 81808400 | 81808999 | 10.95317 | 20.41695 | 26.80915 | 11.16583 | 0.061582 | 0.158683 | 0.923149 |
| 1 | 164484400 | 164487799 | 49.87321 | 79.76764 | 97.93036 | 41.20959 | 0.047833 | 2.051891 | 0.923169 |
| 10 | 43731800 | 43734799 | 30.98348 | 50.95553 | 64.33628 | 27.73087 | 0.043018 | 1.405131 | 0.923169 |
| 6 | 131945800 | 131951399 | 53.56561 | 85.99017 | 117.1559 | 49.57241 | -0.09603 | 1.948799 | 0.923179 |
| 4 | 75250400 | 75251599 | 11.91528 | 21.96103 | 28.65335 | 12.12948 | 0.052681 | 0.232402 | 0.923216 |
| 19 | 35067400 | 35069999 | 27.84564 | 45.98619 | 62.03768 | 26.79278 | -0.05022 | 1.343995 | 0.923235 |
| 2 | 23812200 | 23817199 | 114.1228 | 173.5113 | 217.7717 | 90.21226 | -0.04443 | 3.188179 | 0.923245 |
| 5 | 50117800 | 50119599 | 15.19968 | 26.76173 | 33.57316 | 14.86238 | 0.062876 | 0.569907 | 0.923311 |
| X | 111305400 | 111308999 | 31.9423 | 53.26583 | 65.38913 | 28.42678 | 0.086279 | 1.251508 | 0.923363 |

| | | | | | | | | | |
|---|---|---|---|---|---|---|---|---|---|
| 13 | 40960800 | 40963199 | 37.61798 | 60.81644 | 80.71521 | 34.48815 | -0.04603 | 1.69788 | 0.923378 |
| 1 | 110509000 | 110509999 | 9.228219 | 17.63902 | 26.22496 | 10.40806 | -0.05787 | -0.02799 | 0.923393 |
| 8 | 51849200 | 51851399 | 30.21006 | 49.90152 | 62.91073 | 27.17382 | 0.047172 | 1.347129 | 0.923408 |
| 9 | 106695000 | 106697799 | 14.50121 | 26.40236 | 26.85426 | 11.55302 | 0.39367 | 0.317952 | 0.923451 |
| 15 | 31795000 | 31797199 | 22.6593 | 38.94798 | 53.42294 | 23.43327 | -0.07019 | 0.929574 | 0.923472 |
| 9 | 38467600 | 38475199 | 104.5338 | 162.2198 | 156.8797 | 65.15094 | 0.354209 | 2.784803 | 0.923477 |
| 19 | 20040600 | 20041199 | 11.22141 | 20.52216 | 28.70388 | 12.27994 | -0.05767 | 0.27214 | 0.923487 |
| 5 | 109187400 | 109190799 | 53.85203 | 86.3907 | 105.2611 | 44.49293 | 0.063696 | 1.966394 | 0.923497 |
| 13 | 95006800 | 95007799 | 8.791642 | 16.64361 | 24.39076 | 9.879921 | -0.06276 | 0.030245 | 0.923534 |
| Y | 41651400 | 41652399 | 9.169639 | 16.89901 | 25.38744 | 10.74829 | -0.14907 | 0.202888 | 0.923624 |
| X | 80194400 | 80194999 | 11.06711 | 20.98257 | 27.6718 | 11.17371 | 0.092455 | 0.052744 | 0.923689 |
| 9 | 86636800 | 86640199 | 56.46835 | 89.90857 | 110.2023 | 46.0808 | 0.043073 | 2.203358 | 0.923696 |
| 2 | 90432400 | 90435599 | 28.95218 | 48.1662 | 65.23601 | 28.31736 | -0.05517 | 1.264924 | 0.92373 |
| 19 | 37453000 | 37453999 | 10.87567 | 20.15602 | 26.11656 | 10.97278 | 0.070317 | 0.185821 | 0.923748 |
| 9 | 80932200 | 80932799 | 11.55658 | 21.712 | 22.10533 | 9.084784 | 0.44119 | 0.100305 | 0.923762 |
| 5 | 72293600 | 72294399 | 9.037602 | 17.24119 | 25.59851 | 10.20201 | -0.06051 | -0.01448 | 0.923772 |
| 18 | 46843800 | 46844399 | 7.417595 | 14.35371 | 22.1266 | 8.436393 | -0.05438 | -0.18788 | 0.923774 |
| 3 | 34237000 | 34238999 | 15.77626 | 27.80626 | 34.91976 | 15.45524 | 0.061801 | 0.56279 | 0.923814 |
| 11 | 26619200 | 26620199 | 12.08216 | 21.29327 | 38.87998 | 17.20832 | -0.47491 | 0.553191 | 0.92383 |
| 3 | 120651400 | 120652599 | 10.58611 | 19.6515 | 27.80213 | 11.655 | -0.0528 | 0.181774 | 0.923842 |
| 9 | 10375400 | 10375999 | 10.58611 | 19.6515 | 27.80213 | 11.655 | -0.0528 | 0.181774 | 0.923842 |
| 4 | 60288800 | 60290599 | 38.50525 | 62.28218 | 77.77257 | 33.24983 | 0.042534 | 1.686846 | 0.92385 |
| 14 | 46016800 | 46019599 | 30.98783 | 51.28605 | 64.61018 | 27.94543 | 0.04845 | 1.330533 | 0.923865 |
| 19 | 21048200 | 21050199 | 28.50045 | 47.77746 | 49.13246 | 21.43062 | 0.340826 | 1.169261 | 0.92389 |
| 19 | 8058600 | 8064799 | 30.15886 | 50.30727 | 51.88643 | 22.56109 | 0.336727 | 1.229274 | 0.92392 |
| 14 | 65420400 | 65425399 | 60.67784 | 96.34695 | 124.4804 | 51.88505 | -0.03684 | 2.292079 | 0.92393 |
| 13 | 51018800 | 51026999 | 129.3048 | 196.8069 | 232.104 | 96.18061 | 0.046558 | 3.171939 | 0.923971 |
| 14 | 26707400 | 26710999 | 26.53315 | 44.46583 | 59.9937 | 26.163 | -0.04966 | 1.177083 | 0.923976 |
| 3 | 62071200 | 62074399 | 45.44899 | 72.96092 | 95.99209 | 40.6178 | -0.04568 | 1.928383 | 0.92398 |
| 1 | 154841200 | 154842799 | 29.02927 | 47.8285 | 64.23696 | 27.70937 | -0.04482 | 1.377765 | 0.924031 |
| 8 | 33686000 | 33688399 | 18.81247 | 32.72747 | 41.51942 | 18.244 | 0.045332 | 0.778867 | 0.924082 |
| 2 | 83516400 | 83520999 | 90.16613 | 140.54 | 141.7013 | 58.80884 | 0.299017 | 2.701308 | 0.924082 |
| 3 | 42945800 | 42946399 | 9.000862 | 17.3394 | 24.22044 | 9.399263 | 0.062331 | -0.10531 | 0.924084 |
| 3 | 101247600 | 101251999 | 37.16407 | 60.22249 | 79.78708 | 34.15696 | -0.04026 | 1.647841 | 0.924089 |
| 4 | 74409600 | 74410599 | 13.69502 | 25.09108 | 25.45029 | 10.8614 | 0.405416 | 0.273754 | 0.924109 |
| 17 | 56203400 | 56209799 | 82.13256 | 129.7035 | 153.1137 | 63.5789 | 0.086373 | 2.453683 | 0.92411 |
| 11 | 6909800 | 6912799 | 48.68458 | 78.34638 | 80.69228 | 34.26649 | 0.31117 | 1.851734 | 0.924116 |
| 12 | 15921800 | 15923399 | 18.11945 | 31.65874 | 43.00859 | 18.9499 | -0.05232 | 0.717987 | 0.924151 |
| 10 | 89799800 | 89800999 | 15.34074 | 27.03061 | 36.63617 | 16.21587 | -0.04755 | 0.570106 | 0.924169 |
| 2 | 88797600 | 88801799 | 45.45072 | 73.13568 | 96.50535 | 40.97628 | -0.04572 | 1.858283 | 0.9242 |
| 9 | 116707800 | 116712599 | 54.59715 | 87.26166 | 114.2992 | 48.04969 | -0.04662 | 2.081954 | 0.924235 |
| 18 | 74427800 | 74439999 | 158.5833 | 237.0851 | 291.3795 | 120.1456 | -0.03522 | 3.53197 | 0.924242 |
| 1 | 117554200 | 117571799 | 229.1339 | 335.1485 | 325.6444 | 134.1514 | 0.274168 | 3.908369 | 0.924273 |
| 13 | 9066200 | 9067999 | 12.63323 | 22.47214 | 39.26761 | 17.31061 | -0.40595 | 0.468902 | 0.924278 |
| 6 | 19496400 | 19498999 | 20.48117 | 35.73098 | 44.57112 | 19.61966 | 0.069872 | 0.751474 | 0.924346 |
| 17 | 64976800 | 64979599 | 19.90015 | 34.46019 | 43.65593 | 19.15343 | 0.046285 | 0.83913 | 0.924363 |
| 1 | 101655200 | 101656799 | 13.24038 | 23.89359 | 33.0326 | 14.36977 | -0.05702 | 0.370797 | 0.924366 |

| | | | | | | | | | |
|---|---|---|---|---|---|---|---|---|---|
| 10 | 47681000 | 47681999 | 26.48178 | 44.68985 | 44.34474 | 19.39455 | 0.392357 | 1.096451 | 0.924386 |
| 4 | 86648200 | 86652599 | 44.54749 | 71.69165 | 89.22841 | 37.89367 | 0.038686 | 1.849104 | 0.924399 |
| 1 | 111465200 | 111468399 | 38.62506 | 62.70665 | 84.07663 | 36.04279 | -0.05499 | 1.61506 | 0.924417 |
| 4 | 25137200 | 25140599 | 33.15108 | 54.37817 | 73.09892 | 31.47304 | -0.04872 | 1.447942 | 0.924471 |
| 15 | 42805200 | 42806199 | 13.76504 | 24.65462 | 31.47144 | 13.79346 | 0.050378 | 0.417902 | 0.924478 |
| 17 | 32502600 | 32503999 | 10.75394 | 19.61118 | 27.76255 | 11.91981 | -0.07822 | 0.28269 | 0.924495 |
| 18 | 66515400 | 66516399 | 8.080506 | 15.55696 | 23.62806 | 9.18538 | -0.0594 | -0.10993 | 0.924515 |
| 12 | 108739400 | 108742599 | 31.78092 | 52.89508 | 65.48105 | 28.43221 | 0.074198 | 1.268924 | 0.924526 |
| 9 | 62241200 | 62241999 | 10.62376 | 19.8475 | 28.37892 | 11.77471 | -0.05677 | 0.141213 | 0.924534 |
| 17 | 54695200 | 54696399 | 15.17442 | 26.86458 | 36.53634 | 16.15637 | -0.04989 | 0.51698 | 0.924538 |
| 1 | 146153800 | 146156999 | 47.10335 | 75.72095 | 93.86294 | 39.79792 | 0.042498 | 1.887688 | 0.924545 |
| 6 | 3137600 | 3141999 | 212.2224 | 308.8263 | 346.707 | 142.8978 | 0.058719 | 3.985626 | 0.924566 |
| 2 | 76614200 | 76617199 | 38.94706 | 63.14395 | 79.03141 | 33.84619 | 0.042292 | 1.640409 | 0.924606 |
| 4 | 47109400 | 47112599 | 37.84342 | 61.14421 | 80.93887 | 34.55696 | -0.04325 | 1.717318 | 0.92465 |
| 10 | 52489800 | 52491799 | 26.41622 | 44.87232 | 55.60324 | 24.36693 | 0.073272 | 1.053654 | 0.924675 |
| 1 | 31982400 | 31982799 | 8.144124 | 15.65347 | 23.58022 | 9.210526 | -0.05484 | -0.09741 | 0.924676 |
| 8 | 69922400 | 69923599 | 13.67062 | 24.6846 | 34.12526 | 14.83411 | -0.05613 | 0.374509 | 0.924685 |
| X | 118544600 | 118546199 | 17.28865 | 30.56617 | 42.84009 | 18.94902 | -0.09391 | 0.556345 | 0.924702 |
| 5 | 111857000 | 111860799 | 34.32169 | 56.50078 | 70.51524 | 30.40677 | 0.060216 | 1.406068 | 0.924718 |
| 1 | 17459400 | 17460799 | 14.20648 | 24.7419 | 45.27959 | 19.91008 | -0.48033 | 0.749916 | 0.924727 |
| 15 | 14531400 | 14532199 | 16.12113 | 28.64109 | 26.95337 | 11.89441 | 0.481089 | 0.475674 | 0.924737 |
| 15 | 20015600 | 20019399 | 54.31605 | 86.96243 | 106.7922 | 45.00413 | 0.049019 | 2.028083 | 0.924739 |
| 1 | 163504000 | 163506199 | 14.3585 | 25.43992 | 32.11747 | 14.20013 | 0.057561 | 0.495979 | 0.924766 |
| 15 | 66910600 | 66912599 | 14.3585 | 25.43992 | 32.11747 | 14.20013 | 0.057561 | 0.495979 | 0.924766 |
| 7 | 113355800 | 113357999 | 20.87094 | 35.85151 | 45.27766 | 19.86007 | 0.048356 | 0.916912 | 0.924784 |
| 10 | 55046200 | 55047999 | 18.85768 | 32.94696 | 45.11123 | 19.87572 | -0.06375 | 0.723669 | 0.924802 |
| 17 | 84347000 | 84349799 | 32.47994 | 53.19881 | 71.1692 | 30.62309 | -0.0426 | 1.462918 | 0.924838 |
| X | 14386800 | 14391199 | 78.9482 | 124.1898 | 150.1728 | 62.30082 | 0.047376 | 2.532739 | 0.924841 |
| 1 | 35944200 | 35945999 | 15.34395 | 27.17351 | 34.50384 | 15.25661 | 0.048939 | 0.511501 | 0.92485 |
| 10 | 105760600 | 105761599 | 12.1097 | 21.63943 | 30.28745 | 13.30054 | -0.0833 | 0.420419 | 0.924885 |
| 12 | 113351400 | 113352799 | 25.6048 | 42.60636 | 57.57277 | 24.99458 | -0.05164 | 1.249156 | 0.924886 |
| 4 | 60108800 | 60116599 | 121.5014 | 184.6235 | 230.0856 | 95.29694 | -0.03492 | 3.203173 | 0.924901 |
| 5 | 37713000 | 37715999 | 28.15685 | 46.75833 | 62.79224 | 27.22253 | -0.04307 | 1.282025 | 0.924917 |
| 3 | 106621600 | 106622399 | 19.09608 | 33.05326 | 44.73055 | 19.62464 | -0.04868 | 0.840471 | 0.924926 |
| 11 | 21842200 | 21844999 | 24.81618 | 42.05354 | 52.85785 | 23.14701 | 0.052564 | 1.0615 | 0.92494 |
| 10 | 31648200 | 31649199 | 15.59053 | 27.39644 | 37.13911 | 16.4427 | -0.04931 | 0.601914 | 0.92496 |
| 7 | 136631200 | 136633399 | 26.27992 | 43.93564 | 55.56803 | 24.19623 | 0.043348 | 1.196676 | 0.924963 |
| 11 | 57652400 | 57655199 | 23.48466 | 39.70087 | 53.33526 | 23.33891 | -0.04316 | 1.080097 | 0.924998 |
| 12 | 82556600 | 82559199 | 30.48999 | 50.11834 | 67.13963 | 28.93295 | -0.04249 | 1.411501 | 0.925021 |
| 1 | 53730600 | 53738199 | 151.5488 | 227.6213 | 267.5793 | 110.4452 | 0.034913 | 3.442362 | 0.925035 |
| 1 | 28288000 | 28290599 | 47.18491 | 75.61906 | 93.18138 | 39.32674 | 0.045788 | 1.984512 | 0.925051 |
| 5 | 131338600 | 131342999 | 30.32781 | 50.09794 | 62.98193 | 27.20529 | 0.051174 | 1.346131 | 0.925057 |
| 18 | 26351000 | 26351799 | 8.918483 | 16.98712 | 25.00355 | 9.998214 | -0.05291 | -0.00962 | 0.925068 |
| 18 | 90312600 | 90315399 | 22.30232 | 37.5356 | 63.15374 | 27.5983 | -0.36703 | 1.124114 | 0.925093 |
| 3 | 94148200 | 94150399 | 35.8848 | 58.17567 | 77.19544 | 33.05844 | -0.04186 | 1.63596 | 0.925108 |
| 7 | 39057000 | 39063799 | 12.00328 | 22.39113 | 22.0723 | 9.181326 | 0.473523 | 0.148805 | 0.925145 |
| 1 | 170225200 | 170226199 | 11.75113 | 22.00394 | 28.97445 | 11.97533 | 0.065738 | 0.138143 | 0.925209 |

| | | | | | | | | | |
|---|---|---|---|---|---|---|---|---|---|
| 9 | 8463800 | 8468599 | 59.86051 | 95.21288 | 117.1672 | 48.92443 | 0.03577 | 2.239589 | 0.92522 |
| 3 | 26750800 | 26752399 | 14.93112 | 26.41112 | 33.50131 | 14.81683 | 0.049664 | 0.518388 | 0.92524 |
| 16 | 91172600 | 91174999 | 24.06796 | 40.43737 | 54.37833 | 23.75099 | -0.04485 | 1.13781 | 0.925254 |
| 17 | 46659600 | 46671399 | 255.4793 | 367.5089 | 414.583 | 171.1214 | 0.035429 | 4.160443 | 0.925256 |
| 3 | 3353200 | 3375399 | 150.7896 | 227.5075 | 226.6401 | 93.6635 | 0.279328 | 3.351495 | 0.925259 |
| 5 | 52826600 | 52827199 | 6.520549 | 12.6462 | 20.01475 | 7.551589 | -0.07742 | -0.241 | 0.925301 |
| 7 | 12643600 | 12644599 | 10.42179 | 19.49432 | 25.80779 | 10.68548 | 0.056651 | 0.128696 | 0.925304 |
| 3 | 144120600 | 144123999 | 36.48673 | 59.06656 | 78.39453 | 33.53639 | -0.04413 | 1.665029 | 0.925305 |
| 7 | 29829600 | 29833999 | 50.65354 | 80.95491 | 105.3959 | 44.30397 | -0.03791 | 2.076822 | 0.925312 |
| 10 | 61862000 | 61867199 | 41.22167 | 66.45284 | 87.63057 | 37.30558 | -0.04154 | 1.789603 | 0.925316 |
| 2 | 89545200 | 89549399 | 52.6146 | 84.31082 | 103.8864 | 43.83688 | 0.045616 | 1.997976 | 0.925321 |
| 17 | 79223000 | 79224399 | 13.38202 | 23.84461 | 32.60633 | 14.35348 | -0.05252 | 0.451476 | 0.925322 |
| 11 | 15231000 | 15238999 | 103.3197 | 158.4805 | 199.2233 | 82.70315 | -0.03661 | 3.006587 | 0.925322 |
| 7 | 11335400 | 11347399 | 222.1325 | 323.889 | 370.4565 | 152.6548 | 0.034615 | 3.956905 | 0.92533 |
| 5 | 104324800 | 104341999 | 309.6679 | 438.719 | 487.7677 | 201.8292 | 0.034345 | 4.386382 | 0.925332 |
| 2 | 80647400 | 80647999 | 8.333067 | 16.04561 | 22.52407 | 8.752432 | 0.053566 | -0.11338 | 0.925408 |
| 14 | 79025600 | 79026199 | 8.333067 | 16.04561 | 22.52407 | 8.752432 | 0.053566 | -0.11338 | 0.925408 |
| 4 | 142958000 | 142961599 | 42.52099 | 68.46857 | 85.01745 | 36.1346 | 0.043058 | 1.825928 | 0.925415 |
| 8 | 57267000 | 57267999 | 8.456132 | 16.16136 | 23.98545 | 9.522342 | -0.05549 | -0.04008 | 0.925424 |
| 7 | 13331000 | 13331799 | 17.09235 | 29.66096 | 40.67894 | 17.84847 | -0.06682 | 0.805132 | 0.925455 |
| 5 | 139666000 | 139668399 | 26.49732 | 43.89337 | 59.63138 | 25.80793 | -0.05983 | 1.30686 | 0.92546 |
| 1 | 9310600 | 9311799 | 24.85378 | 40.68739 | 73.79523 | 31.74624 | -0.48057 | 1.456298 | 0.925481 |
| 2 | 83055600 | 83056799 | 11.22826 | 20.80431 | 29.32761 | 12.32606 | -0.05048 | 0.200831 | 0.925482 |
| 13 | 112488800 | 112489999 | 14.35223 | 25.85489 | 32.93127 | 14.35697 | 0.058498 | 0.388476 | 0.925488 |
| 10 | 95209200 | 95211399 | 22.99148 | 38.83338 | 52.24423 | 22.85547 | -0.04523 | 1.084664 | 0.9255 |
| 19 | 34082400 | 34083199 | 8.011988 | 15.32532 | 22.95207 | 9.090313 | -0.065 | -0.05477 | 0.925504 |
| 10 | 42334600 | 42337199 | 37.98556 | 62.10827 | 63.62328 | 27.35785 | 0.33998 | 1.494134 | 0.925516 |
| 14 | 86656600 | 86657199 | 11.99621 | 22.00599 | 28.65291 | 12.20751 | 0.049175 | 0.260905 | 0.925528 |
| 9 | 36724200 | 36726399 | 15.49901 | 27.35256 | 37.09255 | 16.4127 | -0.04749 | 0.554403 | 0.925562 |
| 6 | 35690800 | 35703599 | 209.1729 | 306.6793 | 370.8756 | 152.7621 | -0.03812 | 3.871115 | 0.92557 |
| 14 | 73896800 | 73897399 | 9.295084 | 17.53727 | 25.39504 | 10.36057 | -0.05463 | 0.061956 | 0.925581 |
| 1 | 33272400 | 33272799 | 10.74809 | 20.17257 | 26.87435 | 11.06373 | 0.054102 | 0.112544 | 0.925595 |
| 3 | 15120000 | 15126599 | 69.58336 | 110.2434 | 143.9712 | 59.88091 | -0.05538 | 2.368026 | 0.925613 |
| X | 73582600 | 73582999 | 6.669928 | 12.9753 | 19.01967 | 7.022475 | 0.061418 | -0.31677 | 0.925613 |
| 16 | 68247600 | 68250599 | 30.65866 | 50.45405 | 63.76465 | 27.49203 | 0.042007 | 1.396089 | 0.92564 |
| 5 | 93439400 | 93446199 | 80.05875 | 125.394 | 152.4762 | 63.24729 | 0.034532 | 2.613372 | 0.925658 |
| 5 | 57418600 | 57419399 | 8.649192 | 16.55046 | 24.56672 | 9.719187 | -0.05083 | -0.04672 | 0.925662 |
| 7 | 110334200 | 110338599 | 48.6072 | 77.95958 | 96.6672 | 40.84661 | 0.038099 | 1.962085 | 0.925687 |
| 16 | 81509400 | 81512399 | 27.582 | 46.52459 | 57.84029 | 25.29332 | 0.068176 | 1.119538 | 0.925699 |
| 12 | 37001600 | 37002599 | 13.14335 | 24.16983 | 24.46044 | 10.38642 | 0.413786 | 0.245825 | 0.925701 |
| 15 | 17681800 | 17683599 | 14.03903 | 26.06034 | 25.43979 | 10.66522 | 0.478662 | 0.204056 | 0.925704 |
| X | 92328000 | 92331199 | 31.7023 | 52.34614 | 70.94322 | 30.63519 | -0.05692 | 1.36384 | 0.925801 |
| 3 | 91241600 | 91242999 | 16.7097 | 29.55113 | 28.81785 | 12.74597 | 0.426771 | 0.521937 | 0.925836 |
| 2 | 165598800 | 165599999 | 12.86191 | 23.08796 | 29.17308 | 12.76233 | 0.067131 | 0.394794 | 0.925885 |
| 17 | 30386000 | 30390799 | 49.74316 | 80.02692 | 98.08072 | 41.63327 | 0.060248 | 1.871097 | 0.925887 |
| 10 | 46627600 | 46631199 | 36.56832 | 60.31205 | 58.69715 | 25.33077 | 0.419129 | 1.381199 | 0.925911 |
| 17 | 81106200 | 81106799 | 11.45122 | 20.99888 | 27.12972 | 11.56408 | 0.059756 | 0.253971 | 0.925918 |

| | | | | | | | | | |
|---|---|---|---|---|---|---|---|---|---|
| 6 | 137824600 | 137828199 | 49.81077 | 79.99316 | 106.3603 | 45.03073 | -0.06022 | 1.929957 | 0.925925 |
| 5 | 65042600 | 65047799 | 75.45029 | 118.257 | 151.016 | 62.63715 | -0.03527 | 2.59874 | 0.925942 |
| 4 | 143004000 | 143007199 | 25.10756 | 41.97939 | 56.64349 | 24.66591 | -0.04975 | 1.191783 | 0.925949 |
| 4 | 145179200 | 145200599 | 539.7416 | 729.5478 | 773.5962 | 322.48 | 0.034249 | 5.108846 | 0.925976 |
| 12 | 19568800 | 19573999 | 114.2357 | 175.1443 | 173.4141 | 71.98451 | 0.306969 | 3.016203 | 0.925988 |
| 13 | 98464800 | 98467199 | 23.42292 | 39.21533 | 53.2762 | 23.21927 | -0.05943 | 1.17173 | 0.926002 |
| 14 | 10754000 | 10755199 | 14.42933 | 26.33905 | 25.94673 | 11.12679 | 0.442705 | 0.302194 | 0.926037 |
| 1 | 117991400 | 118012199 | 302.9685 | 432.1389 | 480.0593 | 198.4145 | 0.045336 | 4.288388 | 0.926042 |
| 13 | 83377600 | 83379799 | 18.78957 | 33.01262 | 46.59934 | 20.62825 | -0.10757 | 0.637465 | 0.926055 |
| X | 108647200 | 108648199 | 21.15455 | 36.19009 | 49.01193 | 21.49134 | -0.05275 | 0.964231 | 0.926072 |
| 5 | 84852400 | 84855999 | 39.17242 | 63.3791 | 83.906 | 35.87946 | -0.04126 | 1.6851 | 0.926076 |
| 13 | 90944600 | 90945999 | 14.39769 | 25.79339 | 32.92373 | 14.42732 | 0.05052 | 0.4233 | 0.92609 |
| 7 | 60425200 | 60425999 | 11.99173 | 21.70263 | 30.24994 | 13.112 | -0.06466 | 0.344365 | 0.926119 |
| 8 | 13893400 | 13908599 | 220.1696 | 321.2956 | 386.2711 | 159.1577 | -0.03609 | 3.944474 | 0.926121 |
| 2 | 41101400 | 41112599 | 299.1614 | 418.5528 | 599.5907 | 248.6002 | -0.34941 | 4.571698 | 0.926123 |
| 3 | 69423200 | 69425399 | 19.91306 | 34.25342 | 46.33975 | 20.32678 | -0.05009 | 0.901548 | 0.926142 |
| 16 | 84385200 | 84386399 | 11.74013 | 22.0781 | 29.11375 | 11.94558 | 0.072422 | 0.113521 | 0.926154 |
| 12 | 90531000 | 90531999 | 16.83397 | 29.36331 | 39.94075 | 17.5796 | -0.05443 | 0.728145 | 0.926165 |
| 2 | 91117400 | 91119599 | 20.10966 | 34.60379 | 43.73069 | 19.18254 | 0.048008 | 0.89734 | 0.926166 |
| 3 | 4826400 | 4834999 | 140.7758 | 211.7352 | 261.6396 | 108.0339 | -0.03525 | 3.412684 | 0.92617 |
| 5 | 149652000 | 149659399 | 97.69409 | 150.4581 | 189.3734 | 78.65774 | -0.03396 | 2.924227 | 0.926178 |
| 3 | 10661000 | 10664799 | 29.17655 | 49.1119 | 60.3108 | 26.35702 | 0.085626 | 1.141119 | 0.926205 |
| 13 | 116583800 | 116586199 | 31.35346 | 51.39685 | 64.70173 | 27.85011 | 0.045446 | 1.446923 | 0.926223 |
| 1 | 31590600 | 31591599 | 15.25782 | 26.90496 | 34.06846 | 15.07706 | 0.050439 | 0.555628 | 0.926253 |
| 18 | 82304600 | 82306799 | 20.84069 | 35.59764 | 47.95979 | 21.02735 | -0.04574 | 0.972933 | 0.926254 |
| X | 96959000 | 96960999 | 17.91792 | 31.43363 | 39.68537 | 17.54521 | 0.052857 | 0.646868 | 0.92627 |
| 1 | 125424600 | 125426599 | 20.18846 | 34.97537 | 47.3975 | 20.79528 | -0.05038 | 0.842594 | 0.926271 |
| X | 17174800 | 17175999 | 16.44256 | 29.10379 | 36.53492 | 16.1564 | 0.065103 | 0.529224 | 0.926281 |
| 17 | 83624200 | 83628399 | 35.49752 | 57.15086 | 95.67083 | 40.65399 | -0.38723 | 1.825986 | 0.926286 |
| 14 | 106701800 | 106702599 | 12.99292 | 23.43878 | 30.06227 | 13.08337 | 0.050491 | 0.367217 | 0.926307 |
| 7 | 99155800 | 99156599 | 11.58918 | 21.47523 | 28.20571 | 11.85289 | 0.050799 | 0.198788 | 0.92633 |
| X | 36887800 | 36892199 | 44.87198 | 72.40445 | 89.3058 | 38.06418 | 0.056248 | 1.773019 | 0.926348 |
| 11 | 13871600 | 13874399 | 33.36604 | 55.13954 | 55.87706 | 24.1432 | 0.36134 | 1.344518 | 0.926387 |
| 10 | 124145400 | 124149199 | 74.22942 | 116.5499 | 148.9694 | 61.78664 | -0.03458 | 2.564757 | 0.926433 |
| 18 | 51204400 | 51207599 | 40.67256 | 65.54833 | 86.38595 | 36.76177 | -0.04112 | 1.799144 | 0.926445 |
| 4 | 136964400 | 136969399 | 64.75554 | 102.6498 | 125.8576 | 52.37129 | 0.036242 | 2.344012 | 0.92646 |
| 1 | 98773600 | 98776199 | 24.06234 | 41.35732 | 41.55399 | 18.22708 | 0.377251 | 0.923029 | 0.926474 |
| 16 | 53498200 | 53499199 | 25.81272 | 43.12403 | 54.44236 | 23.69549 | 0.046018 | 1.202267 | 0.926477 |
| 18 | 7534000 | 7536599 | 35.89775 | 58.98061 | 60.19436 | 25.93499 | 0.348627 | 1.426739 | 0.926489 |
| 5 | 128466000 | 128475799 | 122.3679 | 186.5353 | 221.4136 | 91.7918 | 0.039062 | 3.136675 | 0.926505 |
| 4 | 60656000 | 60656399 | 6.51577 | 12.64991 | 20.03218 | 7.517586 | -0.07104 | -0.25973 | 0.926511 |
| 14 | 8870600 | 8872599 | 16.85288 | 29.65748 | 37.50677 | 16.60542 | 0.050955 | 0.592263 | 0.926512 |
| 3 | 139001600 | 139002199 | 6.98922 | 13.48576 | 21.01532 | 8.117966 | -0.08714 | -0.15047 | 0.926515 |
| 17 | 54254000 | 54257199 | 27.71635 | 46.25221 | 58.42642 | 25.41145 | 0.04509 | 1.222061 | 0.92654 |
| 15 | 24296400 | 24297199 | 8.600435 | 16.48747 | 24.62136 | 9.689475 | -0.05265 | -0.06579 | 0.926561 |
| 2 | 55150600 | 55150999 | 6.641353 | 12.90961 | 20.65357 | 7.674065 | -0.07313 | -0.29343 | 0.926564 |
| 16 | 46574200 | 46577999 | 28.22253 | 47.15741 | 64.01642 | 27.86528 | -0.05844 | 1.217242 | 0.926565 |

| | | | | | | | | | |
|---|---|---|---|---|---|---|---|---|---|
| 12 | 78012000 | 78014399 | 22.08383 | 37.86902 | 47.93783 | 21.02435 | 0.044591 | 0.941731 | 0.926571 |
| X | 120564000 | 120565599 | 18.772 | 32.88453 | 45.60559 | 20.13805 | -0.08197 | 0.685388 | 0.926589 |
| 9 | 93526200 | 93529799 | 105.4978 | 161.7767 | 193.4547 | 80.30538 | 0.035103 | 3.011232 | 0.926605 |
| X | 55834400 | 55837199 | 22.19902 | 37.94923 | 51.6823 | 22.66095 | -0.06098 | 0.978545 | 0.926609 |
| 16 | 36280400 | 36283999 | 30.82355 | 51.27385 | 52.99542 | 23.00226 | 0.333682 | 1.261956 | 0.926612 |
| 1 | 88982400 | 88990399 | 75.66188 | 119.1242 | 145.2109 | 60.2492 | 0.036747 | 2.51117 | 0.926634 |
| 5 | 53119800 | 53122199 | 20.39078 | 35.05302 | 47.40006 | 20.79149 | -0.04966 | 0.910275 | 0.92665 |
| 12 | 20974000 | 20989799 | 181.1148 | 267.7783 | 326.0509 | 134.2759 | -0.03655 | 3.730933 | 0.926663 |
| 5 | 4507200 | 4509399 | 17.85149 | 31.10727 | 42.01963 | 18.48307 | -0.04457 | 0.751251 | 0.926681 |
| 12 | 103866000 | 103874199 | 105.8896 | 162.2409 | 193.5396 | 80.33062 | 0.037672 | 3.027448 | 0.926689 |
| 11 | 107291800 | 107293999 | 17.31347 | 30.25459 | 38.32004 | 16.88565 | 0.048275 | 0.702919 | 0.926695 |
| 9 | 31875000 | 31879199 | 49.84728 | 80.01498 | 98.97194 | 41.87337 | 0.043064 | 1.940128 | 0.926697 |
| 7 | 73921000 | 73922199 | 10.73358 | 20.0029 | 28.36105 | 11.81691 | -0.04919 | 0.160297 | 0.926701 |
| 3 | 8943000 | 8945199 | 23.47753 | 40.50016 | 56.84868 | 24.93866 | -0.10228 | 0.906584 | 0.926714 |
| 15 | 19264400 | 19265999 | 17.62236 | 30.71098 | 41.54655 | 18.2761 | -0.04657 | 0.750021 | 0.926715 |
| 5 | 134216200 | 134218999 | 22.10332 | 37.83881 | 47.9443 | 21.02434 | 0.042907 | 0.957276 | 0.92674 |
| 6 | 71180400 | 71181399 | 12.04409 | 21.8024 | 30.14613 | 13.06323 | -0.05321 | 0.340587 | 0.926749 |
| 17 | 83258800 | 83259799 | 12.04409 | 21.8024 | 30.14613 | 13.06323 | -0.05321 | 0.340587 | 0.926749 |
| 8 | 44749600 | 44752999 | 31.73754 | 52.87319 | 65.37983 | 28.40539 | 0.075855 | 1.260544 | 0.926753 |
| 5 | 49347800 | 49354999 | 94.0852 | 145.8893 | 175.794 | 73.01459 | 0.036353 | 2.796559 | 0.926768 |
| 16 | 13727400 | 13741399 | 193.1058 | 285.2243 | 347.8606 | 143.2561 | -0.04024 | 3.750346 | 0.926787 |
| 14 | 114762400 | 114763199 | 13.5013 | 24.79972 | 25.43669 | 10.81776 | 0.39297 | 0.257504 | 0.926792 |
| 14 | 81756600 | 81758599 | 24.85067 | 41.77745 | 56.0639 | 24.49166 | -0.0419 | 1.135954 | 0.926817 |
| 5 | 17556200 | 17562799 | 65.84497 | 104.2362 | 127.788 | 53.12823 | 0.03477 | 2.380181 | 0.926823 |
| 12 | 87334200 | 87335799 | 18.65899 | 32.40823 | 41.07295 | 18.02956 | 0.046606 | 0.800677 | 0.926828 |
| 16 | 80559600 | 80559999 | 6.521956 | 12.7306 | 19.38123 | 6.864211 | 0.06105 | -0.46517 | 0.92683 |
| X | 73519600 | 73521399 | 12.39329 | 22.5786 | 31.78383 | 13.66307 | -0.0711 | 0.313567 | 0.926846 |
| 18 | 10972200 | 10977199 | 40.60011 | 65.45866 | 86.20949 | 36.70628 | -0.03948 | 1.785302 | 0.926869 |
| 6 | 142148400 | 142149199 | 12.59881 | 23.05012 | 32.24351 | 13.787 | -0.05771 | 0.286417 | 0.926885 |
| 1 | 131406200 | 131408599 | 26.76868 | 45.49936 | 56.23802 | 24.64709 | 0.076937 | 1.047871 | 0.926898 |
| 6 | 82766800 | 82771599 | 44.3876 | 71.60075 | 88.95015 | 37.89749 | 0.045378 | 1.778399 | 0.926941 |
| 17 | 52632800 | 52636799 | 54.89565 | 87.95628 | 89.91845 | 37.93645 | 0.314313 | 1.998027 | 0.926944 |
| 8 | 118109200 | 118111999 | 33.64466 | 54.95002 | 69.28148 | 29.77888 | 0.04009 | 1.504255 | 0.926953 |
| 1 | 129701400 | 129701799 | 6.94453 | 13.47715 | 21.11521 | 7.942873 | -0.05933 | -0.24348 | 0.92698 |
| 10 | 39477200 | 39485599 | 120.6137 | 184.5358 | 218.3079 | 90.58094 | 0.048051 | 3.063607 | 0.927 |
| 16 | 77824800 | 77827199 | 33.95636 | 55.46953 | 69.94963 | 30.06813 | 0.039995 | 1.502305 | 0.927006 |
| 8 | 118711400 | 118714199 | 26.68666 | 44.79588 | 60.59556 | 26.4526 | -0.05344 | 1.160674 | 0.927013 |
| 8 | 124145800 | 124164999 | 743.2074 | 974.5371 | 992.6316 | 415.4601 | 0.048493 | 5.594827 | 0.927023 |
| 3 | 122717600 | 122718399 | 8.779465 | 16.65663 | 24.40918 | 9.842148 | -0.05698 | 0.014928 | 0.927032 |
| 15 | 69321800 | 69322799 | 8.779465 | 16.65663 | 24.40918 | 9.842148 | -0.05698 | 0.014928 | 0.927032 |
| 1 | 59012200 | 59013399 | 13.11427 | 23.45139 | 29.58202 | 12.98289 | 0.066164 | 0.419477 | 0.927065 |
| 7 | 104905400 | 104906799 | 12.89705 | 23.04485 | 31.6474 | 13.89854 | -0.05624 | 0.42887 | 0.927082 |
| 19 | 9431800 | 9436599 | 61.3289 | 97.4467 | 126.0122 | 52.55914 | -0.03717 | 2.273204 | 0.927089 |
| 14 | 9352200 | 9354399 | 33.37847 | 54.4217 | 72.44876 | 31.12077 | -0.03987 | 1.530976 | 0.927092 |
| 12 | 58540000 | 58541999 | 32.35792 | 53.64981 | 55.21649 | 23.91205 | 0.339458 | 1.306112 | 0.927104 |
| 14 | 4265800 | 4268799 | 40.01442 | 64.67721 | 85.52941 | 36.53531 | -0.04117 | 1.712619 | 0.927125 |
| 17 | 94336000 | 94339199 | 32.62613 | 53.74405 | 72.80337 | 31.40197 | -0.05736 | 1.39438 | 0.927131 |

| | | | | | | | | | |
|---|---|---|---|---|---|---|---|---|---|
| 8 | 78493800 | 78497799 | 39.27516 | 63.52314 | 84.05453 | 35.93372 | -0.04103 | 1.690989 | 0.927134 |
| 18 | 20037400 | 20039199 | 19.5258 | 34.04779 | 43.00865 | 18.92613 | 0.051827 | 0.752145 | 0.927137 |
| 19 | 39428600 | 39431199 | 26.49772 | 45.04031 | 62.98546 | 27.60432 | -0.10053 | 1.055403 | 0.927151 |
| 1 | 44681400 | 44684399 | 29.43225 | 48.87717 | 61.79164 | 26.78906 | 0.043765 | 1.28426 | 0.927155 |
| 3 | 97039800 | 97053199 | 145.5942 | 219.3722 | 272.3361 | 112.5041 | -0.03966 | 3.380174 | 0.927177 |
| 1 | 49360200 | 49368599 | 269.1105 | 383.2431 | 452.3551 | 187.0134 | -0.04439 | 4.308167 | 0.927207 |
| 18 | 67511200 | 67515199 | 32.50965 | 53.05775 | 70.81494 | 30.42995 | -0.04258 | 1.512747 | 0.927237 |
| 18 | 21064000 | 21067399 | 31.23225 | 51.2555 | 69.02068 | 29.72323 | -0.05096 | 1.432341 | 0.92725 |
| X | 160689600 | 160692399 | 23.96323 | 40.78675 | 51.15865 | 22.42346 | 0.056318 | 1.023431 | 0.927279 |
| 10 | 68271800 | 68273399 | 15.39189 | 27.06934 | 36.72361 | 16.26074 | -0.04987 | 0.592547 | 0.927319 |
| 9 | 120701200 | 120704799 | 39.27445 | 63.4734 | 83.82041 | 35.79964 | -0.03938 | 1.712328 | 0.927319 |
| 18 | 84010600 | 84011999 | 14.46238 | 25.707 | 34.98524 | 15.43232 | -0.04793 | 0.476924 | 0.927355 |
| 13 | 34346600 | 34347599 | 9.646792 | 17.98991 | 25.85173 | 10.75888 | -0.06667 | 0.141545 | 0.92737 |
| 15 | 6830000 | 6842799 | 239.008 | 349.5604 | 331.1312 | 136.4125 | 0.310667 | 3.911239 | 0.927398 |
| 16 | 10073800 | 10075199 | 15.1429 | 26.77607 | 34.0363 | 15.05453 | 0.046494 | 0.525593 | 0.927404 |
| 15 | 32817200 | 32819999 | 37.29539 | 60.54796 | 76.05489 | 32.60395 | 0.038949 | 1.608637 | 0.927426 |
| 8 | 102723200 | 102723799 | 8.027241 | 15.52185 | 23.81854 | 9.1132 | -0.05296 | -0.16489 | 0.927428 |
| 1 | 109255000 | 109257599 | 33.86436 | 55.51936 | 69.80633 | 30.04881 | 0.047161 | 1.452148 | 0.927492 |
| 10 | 96544400 | 96545799 | 17.41936 | 30.34637 | 41.03705 | 18.04792 | -0.04608 | 0.752606 | 0.927513 |
| 7 | 77862800 | 77865999 | 29.69804 | 49.14218 | 66.41789 | 28.72359 | -0.05277 | 1.327296 | 0.927517 |
| 6 | 45155000 | 45155999 | 10.5758 | 19.76115 | 28.37095 | 11.76812 | -0.06245 | 0.138662 | 0.927521 |
| 1 | 187170200 | 187173399 | 22.66986 | 38.51229 | 51.69462 | 22.65474 | -0.04164 | 1.033614 | 0.927528 |
| 6 | 7611400 | 7614799 | 53.6082 | 85.41565 | 110.9852 | 46.45407 | -0.03987 | 2.176648 | 0.927547 |
| 14 | 33478600 | 33482199 | 32.47237 | 53.14958 | 67.06442 | 28.84854 | 0.041138 | 1.469225 | 0.927551 |
| 5 | 33951400 | 33952199 | 13.11101 | 24.21535 | 24.92068 | 10.51947 | 0.395743 | 0.221959 | 0.927571 |
| 2 | 5110600 | 5113199 | 45.97757 | 74.17474 | 91.39011 | 38.94331 | 0.057448 | 1.777282 | 0.927581 |
| 3 | 54434600 | 54437199 | 25.5804 | 43.09872 | 54.48941 | 23.81968 | 0.043901 | 1.116793 | 0.927583 |
| 11 | 109196200 | 109198399 | 21.68417 | 37.17193 | 50.0896 | 21.96751 | -0.04525 | 0.946568 | 0.927614 |
| 6 | 46219200 | 46221199 | 13.92087 | 25.3384 | 25.36755 | 10.91738 | 0.41592 | 0.311029 | 0.927623 |
| 4 | 83184200 | 83185399 | 13.01408 | 23.40923 | 32.06042 | 14.00115 | -0.04692 | 0.388116 | 0.927643 |
| 3 | 96768000 | 96769199 | 15.40507 | 27.03888 | 36.83485 | 16.29188 | -0.05605 | 0.622443 | 0.927648 |
| 7 | 112213000 | 112215999 | 26.57409 | 44.39341 | 56.14705 | 24.43654 | 0.043334 | 1.205495 | 0.927653 |
| 4 | 142959600 | 142964399 | 49.608 | 79.35154 | 103.4538 | 43.54016 | -0.0386 | 2.048042 | 0.927665 |
| 7 | 99025400 | 99026399 | 11.72265 | 22.05694 | 23.09689 | 9.468298 | 0.403723 | 0.094704 | 0.92769 |
| 15 | 11222000 | 11227199 | 18.54839 | 31.69345 | 53.23568 | 23.34116 | -0.36252 | 0.968395 | 0.927703 |
| X | 77319400 | 77321999 | 26.65277 | 45.03197 | 61.8565 | 27.06324 | -0.07525 | 1.10375 | 0.927707 |
| X | 141924000 | 141926799 | 30.66841 | 51.64191 | 48.01747 | 20.98691 | 0.485796 | 1.1357 | 0.927712 |
| 7 | 26211400 | 26212599 | 22.20388 | 37.90926 | 47.95903 | 21.02626 | 0.044652 | 0.983713 | 0.92772 |
| 5 | 3815400 | 3816999 | 24.98286 | 42.07248 | 52.76483 | 23.06256 | 0.055657 | 1.121524 | 0.927728 |
| 13 | 98161000 | 98163599 | 22.81087 | 38.74145 | 49.02871 | 21.48467 | 0.043013 | 1.033679 | 0.927845 |
| 4 | 144433600 | 144436599 | 27.98891 | 47.14035 | 58.54486 | 25.58981 | 0.069471 | 1.130332 | 0.927851 |
| 9 | 95631400 | 95632599 | 9.635105 | 18.00669 | 25.84868 | 10.71862 | -0.06102 | 0.127452 | 0.927861 |
| 17 | 20937600 | 20939599 | 44.77104 | 72.05161 | 89.67325 | 38.08273 | 0.038755 | 1.850591 | 0.927878 |
| 12 | 66247000 | 66250799 | 44.12041 | 70.66536 | 111.944 | 47.20222 | -0.31702 | 2.004878 | 0.927906 |
| 3 | 151884600 | 151888399 | 26.89868 | 44.97107 | 60.6597 | 26.41368 | -0.04932 | 1.201407 | 0.927912 |
| 9 | 72169200 | 72177799 | 92.83637 | 144.7993 | 147.5262 | 61.2204 | 0.284744 | 2.692624 | 0.927932 |
| 7 | 26651400 | 26669399 | 230.2969 | 334.7477 | 400.7097 | 165.1749 | -0.03539 | 4.00465 | 0.927963 |

| | | | | | | | | | |
|---|---|---|---|---|---|---|---|---|---|
| 10 | 71009600 | 71016599 | 64.46226 | 102.2439 | 132.0747 | 54.99006 | -0.03813 | 2.328489 | 0.927964 |
| 17 | 80401400 | 80403999 | 20.80903 | 35.73981 | 45.30287 | 19.87104 | 0.043128 | 0.919895 | 0.92799 |
| 17 | 27047000 | 27050199 | 37.14905 | 60.10192 | 74.8172 | 31.99159 | 0.047307 | 1.677435 | 0.927999 |
| 8 | 51583200 | 51586399 | 43.54963 | 70.41022 | 71.83878 | 30.69775 | 0.332578 | 1.701527 | 0.928078 |
| 7 | 98012600 | 98014599 | 27.62671 | 46.2301 | 58.39441 | 25.44182 | 0.045165 | 1.194005 | 0.92809 |
| 19 | 7791800 | 7794599 | 41.72738 | 67.82841 | 68.0346 | 29.19239 | 0.364428 | 1.589044 | 0.928091 |
| 12 | 119483800 | 119485999 | 16.22587 | 28.51381 | 38.62017 | 17.09888 | -0.04814 | 0.607132 | 0.928103 |
| 6 | 124260400 | 124261199 | 13.56724 | 24.24337 | 33.09519 | 14.53386 | -0.04769 | 0.436494 | 0.928114 |
| 1 | 21695400 | 21697599 | 29.05369 | 48.27293 | 65.08491 | 28.22669 | -0.04873 | 1.2817 | 0.928126 |
| 12 | 98059800 | 98062199 | 30.10167 | 50.03676 | 63.00261 | 27.33213 | 0.049669 | 1.27634 | 0.92813 |
| 9 | 11662000 | 11662599 | 10.05673 | 18.91037 | 25.32073 | 10.39112 | 0.05051 | 0.089963 | 0.928166 |
| X | 35043400 | 35047199 | 33.30071 | 54.8746 | 74.99389 | 32.35161 | -0.06981 | 1.396297 | 0.928185 |
| 18 | 74160800 | 74164399 | 36.65244 | 59.47488 | 78.82086 | 33.77744 | -0.03887 | 1.618569 | 0.928196 |
| 10 | 119173600 | 119176199 | 23.11099 | 39.27312 | 52.7998 | 23.13981 | -0.04393 | 1.032733 | 0.928206 |
| 14 | 13553800 | 13554799 | 13.1222 | 23.67912 | 32.65245 | 14.2053 | -0.05294 | 0.373701 | 0.928207 |
| 2 | 111757800 | 111761799 | 54.14723 | 86.6848 | 106.7017 | 44.96029 | 0.045492 | 2.030421 | 0.928238 |
| 1 | 180443200 | 180452199 | 96.60982 | 148.6225 | 187.579 | 77.90049 | -0.03916 | 2.94448 | 0.928249 |
| 11 | 6676200 | 6680799 | 46.19043 | 74.13172 | 96.88148 | 40.97778 | -0.03646 | 1.939449 | 0.928287 |
| 1 | 171801400 | 171803199 | 17.80795 | 31.16137 | 42.387 | 18.6994 | -0.05433 | 0.688776 | 0.92829 |
| Y | 9634600 | 9635399 | 9.020249 | 16.95235 | 21.34453 | 8.767295 | 0.137478 | 0.070787 | 0.928328 |
| 11 | 78223400 | 78224799 | 12.36218 | 22.11978 | 30.64193 | 13.44124 | -0.06765 | 0.412914 | 0.928345 |
| 11 | 97956000 | 97960799 | 77.38829 | 120.8372 | 154.2882 | 64.01925 | -0.03928 | 2.666848 | 0.928369 |
| 10 | 37351400 | 37352799 | 15.71864 | 27.43544 | 46.25567 | 20.36727 | -0.36261 | 0.724858 | 0.92837 |
| 5 | 133278600 | 133280999 | 28.09669 | 47.19683 | 59.03031 | 25.78105 | 0.05923 | 1.155332 | 0.928378 |
| 1 | 189452600 | 189454799 | 21.10157 | 36.17232 | 45.82959 | 20.09918 | 0.043324 | 0.93846 | 0.928413 |
| 17 | 22128000 | 22129599 | 18.21513 | 31.79964 | 40.40522 | 17.7936 | 0.043634 | 0.72571 | 0.928439 |
| 2 | 92453200 | 92455599 | 28.48464 | 47.43513 | 60.04243 | 26.07992 | 0.042266 | 1.24979 | 0.928452 |
| 9 | 70093400 | 70096999 | 32.78246 | 53.51894 | 71.58829 | 30.76561 | -0.04549 | 1.508206 | 0.92846 |
| 12 | 88048400 | 88067999 | 636.0296 | 844.3969 | 924.5005 | 386.381 | -0.03788 | 5.394961 | 0.92846 |
| 15 | 4338000 | 4341599 | 34.29995 | 55.50794 | 89.00167 | 38.06464 | -0.31668 | 1.674281 | 0.928461 |
| 1 | 181320400 | 181322999 | 36.05651 | 58.34689 | 77.46055 | 33.12713 | -0.04502 | 1.673355 | 0.928477 |
| 7 | 3973800 | 3978399 | 52.96912 | 84.75191 | 104.8195 | 44.13266 | 0.037782 | 2.04501 | 0.928479 |
| 4 | 79222000 | 79223199 | 17.21269 | 30.34146 | 29.40915 | 13.01599 | 0.433779 | 0.561986 | 0.928483 |
| 3 | 137374000 | 137375399 | 28.26161 | 46.73762 | 58.64794 | 25.35298 | 0.054177 | 1.329611 | 0.928524 |
| 13 | 68057600 | 68087599 | 824.1495 | 1074.453 | 1098.156 | 459.9044 | 0.035246 | 5.689056 | 0.928529 |
| 17 | 66091800 | 66093599 | 15.77623 | 27.78417 | 37.59604 | 16.64238 | -0.04561 | 0.57857 | 0.928538 |
| 3 | 66601000 | 66601799 | 11.74384 | 21.40189 | 29.69802 | 12.76159 | -0.0504 | 0.299509 | 0.928583 |
| 7 | 17840200 | 17842399 | 57.33125 | 91.27204 | 112.0713 | 46.85489 | 0.040354 | 2.208813 | 0.928606 |
| 1 | 186627400 | 186631799 | 37.81292 | 61.28598 | 81.1519 | 34.74628 | -0.03923 | 1.645609 | 0.928636 |
| 8 | 31359000 | 31359599 | 12.70532 | 22.89463 | 29.15436 | 12.70579 | 0.059659 | 0.370458 | 0.928639 |
| 16 | 5679600 | 5682799 | 32.37531 | 52.71764 | 70.54472 | 30.28854 | -0.04864 | 1.545393 | 0.928654 |
| 15 | 91779400 | 91780999 | 14.11229 | 25.4049 | 35.08281 | 15.30835 | -0.05824 | 0.395804 | 0.928688 |
| 10 | 38203600 | 38209799 | 70.27267 | 111.499 | 134.8224 | 56.15386 | 0.057548 | 2.325737 | 0.92869 |
| 1 | 133270200 | 133272399 | 23.64829 | 39.99514 | 50.64412 | 22.16427 | 0.04196 | 1.074697 | 0.928709 |
| 17 | 46652600 | 46658799 | 136.488 | 205.8732 | 242.9781 | 100.4068 | 0.03459 | 3.35558 | 0.92873 |
| 10 | 82028600 | 82030599 | 26.99641 | 45.0965 | 60.56836 | 26.35991 | -0.04307 | 1.212478 | 0.928756 |
| 16 | 69492600 | 69494999 | 25.56979 | 43.11455 | 54.49601 | 23.82822 | 0.044289 | 1.109893 | 0.928782 |

| | | | | | | | | | |
|---|---|---|---|---|---|---|---|---|---|
| 7 | 139600200 | 139604399 | 82.66777 | 128.9184 | 156.2615 | 64.84666 | 0.034268 | 2.691595 | 0.9288 |
| 18 | 89152200 | 89155599 | 37.34562 | 60.42317 | 75.33435 | 32.21399 | 0.045149 | 1.677799 | 0.928817 |
| 2 | 87526200 | 87528599 | 19.31559 | 33.83031 | 42.30894 | 18.67932 | 0.066346 | 0.682816 | 0.928825 |
| 4 | 12213000 | 12213399 | 4.799177 | 9.378809 | 16.00928 | 5.545886 | -0.07165 | -0.57848 | 0.92885 |
| Y | 43609000 | 43609599 | 6.939736 | 13.42954 | 18.43812 | 7.028783 | 0.1117 | -0.20679 | 0.928873 |
| 3 | 85167600 | 85168399 | 10.34136 | 19.45852 | 28.00544 | 11.47994 | -0.052 | 0.09339 | 0.928877 |
| X | 64991200 | 64993999 | 26.22965 | 44.38396 | 55.58216 | 24.32923 | 0.057889 | 1.08559 | 0.928911 |
| 8 | 4474600 | 4475799 | 16.8385 | 29.7354 | 37.529 | 16.60371 | 0.055953 | 0.5568 | 0.928929 |
| 11 | 75104400 | 75107399 | 15.56889 | 27.18818 | 37.45604 | 16.49801 | -0.07245 | 0.701564 | 0.928959 |
| 5 | 93535800 | 93562999 | 392.8994 | 548.5717 | 595.3697 | 246.929 | 0.047944 | 4.605568 | 0.928977 |
| 6 | 100863800 | 100865199 | 16.93831 | 29.53924 | 40.00505 | 17.60563 | -0.04803 | 0.734009 | 0.928985 |
| 18 | 73618400 | 73620799 | 33.87875 | 55.73025 | 55.66266 | 23.99468 | 0.380368 | 1.403726 | 0.928987 |
| 15 | 95586600 | 95588599 | 23.94976 | 40.43153 | 51.1906 | 22.39111 | 0.042005 | 1.091323 | 0.928991 |
| 14 | 62619600 | 62622599 | 25.48081 | 42.63261 | 57.18152 | 24.91095 | -0.04113 | 1.187352 | 0.929002 |
| 9 | 5917600 | 5919199 | 14.68221 | 26.23511 | 33.49828 | 14.71116 | 0.048099 | 0.444488 | 0.929006 |
| 15 | 47246000 | 47250399 | 49.17258 | 78.74163 | 96.94789 | 40.86672 | 0.045583 | 2.010392 | 0.929019 |
| 5 | 113825000 | 113826199 | 14.36656 | 25.56199 | 32.55412 | 14.34794 | 0.048371 | 0.467526 | 0.929028 |
| 11 | 95884800 | 95885799 | 15.66471 | 27.52339 | 37.20012 | 16.46799 | -0.0449 | 0.606074 | 0.92905 |
| 17 | 76915400 | 76915999 | 9.368861 | 17.83684 | 24.43462 | 9.780125 | 0.04869 | -0.00358 | 0.929124 |
| 2 | 142629000 | 142630999 | 32.18795 | 52.50756 | 70.15024 | 30.13906 | -0.04448 | 1.518511 | 0.929151 |
| 3 | 25545800 | 25547999 | 18.68348 | 32.64487 | 41.38295 | 18.23386 | 0.047015 | 0.717066 | 0.929162 |
| 14 | 37288600 | 37291599 | 41.08768 | 66.70408 | 67.7419 | 29.04008 | 0.344917 | 1.613002 | 0.929173 |
| 15 | 80152000 | 80154799 | 41.4766 | 66.85117 | 87.86453 | 37.39607 | -0.03706 | 1.797544 | 0.929191 |
| 8 | 59828600 | 59835199 | 81.85377 | 127.7337 | 162.2547 | 67.32894 | -0.03253 | 2.680723 | 0.929192 |
| 7 | 97584000 | 97587799 | 79.3268 | 124.1422 | 158.152 | 65.60734 | -0.03354 | 2.629471 | 0.929205 |
| 1 | 99344800 | 99346599 | 21.74971 | 37.53403 | 47.38876 | 20.78898 | 0.050128 | 0.882886 | 0.929213 |
| 4 | 9723200 | 9725199 | 20.86103 | 35.49358 | 47.97599 | 21.02784 | -0.05126 | 1.008769 | 0.929225 |
| 8 | 93243600 | 93246399 | 37.52738 | 60.77521 | 76.06237 | 32.54819 | 0.040947 | 1.659742 | 0.929273 |
| 4 | 69228000 | 69228599 | 11.44966 | 21.22012 | 27.77913 | 11.67106 | 0.056289 | 0.19927 | 0.929274 |
| 14 | 98069200 | 98069999 | 11.44966 | 21.22012 | 27.77913 | 11.67106 | 0.056289 | 0.19927 | 0.929274 |
| 13 | 60387800 | 60388999 | 19.53588 | 33.61536 | 45.31617 | 19.87799 | -0.04476 | 0.898197 | 0.929275 |
| X | 91959800 | 91961199 | 17.69659 | 31.10486 | 39.13354 | 17.32925 | 0.057935 | 0.615921 | 0.929341 |
| 16 | 7974600 | 7976799 | 21.36146 | 37.17697 | 37.74499 | 16.58972 | 0.365383 | 0.780142 | 0.929349 |
| Y | 41429800 | 41430199 | 4.701326 | 9.178642 | 15.98738 | 5.642685 | -0.12519 | -0.51649 | 0.929368 |
| 14 | 39599200 | 39600399 | 24.06077 | 40.54559 | 54.39602 | 23.78014 | -0.04132 | 1.112102 | 0.929412 |
| 11 | 8217800 | 8225399 | 84.16472 | 131.1682 | 166.5768 | 69.13404 | -0.03365 | 2.704328 | 0.929473 |
| 10 | 22520600 | 22522399 | 22.46483 | 38.08907 | 51.16604 | 22.40973 | -0.04288 | 1.048263 | 0.929474 |
| 5 | 27379000 | 27380399 | 14.27856 | 25.5609 | 35.01909 | 15.35517 | -0.05163 | 0.431328 | 0.929483 |
| 8 | 53909000 | 53910199 | 10.90395 | 20.54728 | 27.48716 | 11.23947 | 0.055908 | 0.089714 | 0.929486 |
| 13 | 73178600 | 73181599 | 22.37417 | 38.29349 | 48.47832 | 21.25812 | 0.044048 | 0.96005 | 0.929496 |
| 5 | 105391800 | 105392599 | 10.69457 | 20.07545 | 26.86218 | 11.05561 | 0.047992 | 0.109957 | 0.929511 |
| 17 | 48337800 | 48338199 | 7.988881 | 15.4535 | 22.10979 | 8.444534 | 0.049532 | -0.17694 | 0.929559 |
| 4 | 8555800 | 8558799 | 35.47024 | 58.39811 | 72.4808 | 31.25583 | 0.068228 | 1.408449 | 0.929562 |
| 13 | 58638800 | 58642399 | 39.50222 | 63.81297 | 84.09183 | 35.89486 | -0.03707 | 1.727194 | 0.929586 |
| 5 | 12716800 | 12720999 | 42.13427 | 67.79535 | 89.0652 | 37.81415 | -0.03944 | 1.85211 | 0.929591 |
| 1 | 163599200 | 163599799 | 8.388885 | 16.08581 | 24.07832 | 9.469191 | -0.05432 | -0.06831 | 0.929622 |
| 3 | 84181200 | 84182599 | 16.9167 | 29.56889 | 37.39129 | 16.47999 | 0.050749 | 0.697067 | 0.929627 |

| | | | | | | | | | |
|---|---|---|---|---|---|---|---|---|---|
| 6 | 136062800 | 136065199 | 31.23032 | 51.40379 | 68.88385 | 29.70136 | -0.04217 | 1.400063 | 0.929644 |
| 12 | 109782600 | 109785599 | 20.84442 | 35.68843 | 47.95146 | 21.02771 | -0.04131 | 0.951639 | 0.929676 |
| 16 | 20257000 | 20259199 | 28.47029 | 47.36266 | 60.03554 | 26.05926 | 0.040182 | 1.260733 | 0.929683 |
| X | 142854000 | 142855599 | 16.00144 | 28.51683 | 39.70349 | 17.47519 | -0.07855 | 0.479573 | 0.929693 |
| 7 | 105644600 | 105645399 | 6.40238 | 12.47506 | 20.25341 | 7.356843 | -0.0624 | -0.37067 | 0.929699 |
| 1 | 134198200 | 134199199 | 9.505133 | 17.92169 | 25.92525 | 10.59047 | -0.05455 | 0.070981 | 0.929708 |
| 16 | 38762800 | 38764199 | 15.87995 | 28.04987 | 35.57138 | 15.73681 | 0.049369 | 0.545731 | 0.92971 |
| 8 | 67628000 | 67632999 | 89.79464 | 139.2013 | 176.2096 | 73.18995 | -0.03496 | 2.798325 | 0.92971 |
| 8 | 4429400 | 4430599 | 32.89593 | 53.46241 | 71.50532 | 30.67765 | -0.04991 | 1.576099 | 0.929742 |
| 17 | 15853600 | 15856999 | 47.21304 | 76.29102 | 75.14612 | 32.08817 | 0.382357 | 1.725891 | 0.929807 |
| 4 | 62835000 | 62843399 | 96.03171 | 149.0404 | 178.9791 | 74.32758 | 0.042266 | 2.782595 | 0.929812 |
| 5 | 56087000 | 56087999 | 14.45634 | 26.27 | 26.82419 | 11.56792 | 0.385766 | 0.32793 | 0.929849 |
| 18 | 33767200 | 33768399 | 13.80286 | 24.89357 | 32.0091 | 13.93507 | 0.046438 | 0.378121 | 0.929859 |
| X | 31616800 | 31620599 | 77.23073 | 122.4909 | 113.0002 | 47.04635 | 0.446836 | 2.338879 | 0.929863 |
| 4 | 146799600 | 146833399 | 1572.194 | 1955.56 | 1896.047 | 797.8764 | 0.043042 | 6.452182 | 0.929883 |
| 4 | 78635000 | 78635799 | 9.159622 | 17.48362 | 23.97286 | 9.542978 | 0.054884 | -0.02179 | 0.929902 |
| X | 40832800 | 40833599 | 9.159622 | 17.48362 | 23.97286 | 9.542978 | 0.054884 | -0.02179 | 0.929902 |
| 5 | 109207600 | 109208999 | 22.41601 | 38.23933 | 48.48663 | 21.25614 | 0.041096 | 0.990535 | 0.929938 |
| 14 | 120764000 | 120765799 | 16.39934 | 28.80724 | 36.60735 | 16.2023 | 0.043508 | 0.611929 | 0.929952 |
| 15 | 60909000 | 60909999 | 16.39934 | 28.80724 | 36.60735 | 16.2023 | 0.043508 | 0.611929 | 0.929952 |
| X | 105260800 | 105266199 | 52.07486 | 83.54477 | 102.7252 | 43.42502 | 0.050694 | 1.962198 | 0.929969 |
| 2 | 77818000 | 77818599 | 8.482159 | 16.24503 | 22.43356 | 8.853 | 0.056149 | -0.06212 | 0.929975 |
| 16 | 43891800 | 43899799 | 103.6574 | 159.5369 | 191.489 | 79.52926 | 0.033746 | 2.939938 | 0.929992 |
| 4 | 74463200 | 74465799 | 27.56607 | 46.37386 | 63.51083 | 27.75057 | -0.07118 | 1.146359 | 0.930027 |
| 5 | 145256800 | 145268999 | 199.4557 | 295.448 | 288.161 | 118.6743 | 0.285893 | 3.702265 | 0.930044 |
| 10 | 46870400 | 46872799 | 19.53754 | 34.01036 | 43.0192 | 18.91054 | 0.049538 | 0.773782 | 0.930052 |
| 1 | 99596800 | 99597799 | 8.707196 | 16.58993 | 24.49831 | 9.789621 | -0.056 | -0.01202 | 0.930063 |
| 11 | 6701400 | 6702799 | 12.41404 | 22.24462 | 30.69352 | 13.44751 | -0.0606 | 0.405957 | 0.930064 |
| 1 | 95095600 | 95097799 | 21.76512 | 37.367 | 47.4004 | 20.79064 | 0.041888 | 0.929284 | 0.930064 |
| 14 | 97956600 | 97959799 | 40.85221 | 67.55389 | 55.62927 | 24.0467 | 0.65997 | 1.359762 | 0.930075 |
| 1 | 26301000 | 26302599 | 12.65075 | 23.14087 | 32.25487 | 13.79532 | -0.05274 | 0.288824 | 0.93011 |
| 1 | 176946200 | 176947199 | 11.62854 | 21.29078 | 29.65146 | 12.66584 | -0.05022 | 0.269429 | 0.930137 |
| 3 | 130502600 | 130502999 | 6.229876 | 12.13646 | 17.99597 | 6.554299 | 0.063169 | -0.37419 | 0.93015 |
| 7 | 112435400 | 112438999 | 33.29317 | 54.48326 | 68.87303 | 29.6246 | 0.038311 | 1.474769 | 0.930156 |
| 6 | 21841400 | 21846199 | 62.03404 | 97.83526 | 150.3302 | 62.38711 | -0.29492 | 2.469441 | 0.93019 |
| 9 | 14278800 | 14280799 | 16.26199 | 28.73986 | 36.46877 | 16.13211 | 0.048571 | 0.541733 | 0.930191 |
| 8 | 15468200 | 15468999 | 9.284098 | 17.55385 | 25.41874 | 10.32368 | -0.04897 | 0.046961 | 0.930195 |
| 11 | 13896000 | 13898999 | 37.46314 | 60.36909 | 96.36192 | 41.0004 | -0.31715 | 1.802433 | 0.930219 |
| 7 | 3775400 | 3775999 | 12.13447 | 22.57974 | 22.52844 | 9.410529 | 0.451284 | 0.166298 | 0.930239 |
| 4 | 23455200 | 23456799 | 15.79171 | 27.89913 | 35.50607 | 15.70734 | 0.044239 | 0.541438 | 0.930255 |
| 17 | 12157600 | 12159399 | 17.72415 | 31.01483 | 39.20497 | 17.29566 | 0.050959 | 0.682719 | 0.930264 |
| 8 | 126422400 | 126426199 | 40.40903 | 65.50876 | 81.73072 | 35.00005 | 0.046739 | 1.645674 | 0.930273 |
| 7 | 59028800 | 59031999 | 47.9265 | 76.99193 | 95.57749 | 40.48505 | 0.039188 | 1.910614 | 0.930275 |
| 1 | 107227000 | 107229199 | 26.43005 | 44.33358 | 56.14764 | 24.49988 | 0.041293 | 1.162313 | 0.930285 |
| 10 | 53365800 | 53376399 | 200.7429 | 294.9233 | 339.9683 | 140.0135 | 0.033805 | 3.836516 | 0.930289 |
| 8 | 67272200 | 67273599 | 18.22031 | 31.46112 | 42.73361 | 18.74714 | -0.05452 | 0.857623 | 0.930305 |
| 14 | 92833400 | 92833999 | 11.10249 | 21.05476 | 22.16884 | 8.947045 | 0.415213 | 0.034746 | 0.930329 |

| | | | | | | | | | |
|---|---|---|---|---|---|---|---|---|---|
| 4 | 98752800 | 98757199 | 58.469 | 92.98947 | 120.337 | 50.24044 | -0.0369 | 2.241271 | 0.93033 |
| 13 | 92930600 | 92931599 | 9.508379 | 17.81728 | 25.78541 | 10.64423 | -0.06808 | 0.10549 | 0.930386 |
| 10 | 3096000 | 3096599 | 11.18705 | 20.70433 | 27.21814 | 11.45724 | 0.048041 | 0.201788 | 0.930403 |
| 6 | 115576800 | 115579199 | 25.93899 | 43.45681 | 58.34913 | 25.4409 | -0.04273 | 1.175107 | 0.93042 |
| 9 | 26066800 | 26079599 | 359.0115 | 500.667 | 547.1046 | 226.9527 | 0.036376 | 4.621572 | 0.930424 |
| 15 | 27290200 | 27298399 | 109.6406 | 168.0062 | 201.1573 | 83.49445 | 0.032479 | 3.028202 | 0.930472 |
| 5 | 52723400 | 52735999 | 161.6999 | 241.9687 | 283.7057 | 117.0025 | 0.033866 | 3.514804 | 0.930493 |
| 8 | 91033000 | 91033799 | 21.01962 | 36.54847 | 37.30465 | 16.38673 | 0.357646 | 0.790753 | 0.930495 |
| 18 | 50935400 | 50936599 | 12.99463 | 23.00829 | 39.25798 | 17.35946 | -0.37519 | 0.507561 | 0.930497 |
| 3 | 59682000 | 59682999 | 11.33131 | 20.89844 | 29.24073 | 12.36706 | -0.04645 | 0.226064 | 0.930521 |
| 10 | 71659600 | 71661199 | 37.85139 | 61.22671 | 76.49281 | 32.70354 | 0.041873 | 1.684438 | 0.930521 |
| 17 | 94971200 | 94980399 | 176.9142 | 262.2004 | 304.8653 | 125.5606 | 0.033227 | 3.689934 | 0.930615 |
| 8 | 55572600 | 55574999 | 13.90667 | 24.70705 | 33.60469 | 14.8295 | -0.04723 | 0.475989 | 0.930627 |
| 1 | 85526000 | 85536999 | 720.8858 | 949.5009 | 1028.887 | 430.4197 | -0.03437 | 5.522766 | 0.930635 |
| 10 | 39390800 | 39392399 | 19.69957 | 33.91252 | 42.74416 | 18.75004 | 0.052045 | 0.892172 | 0.930656 |
| 9 | 53634000 | 53636399 | 24.47948 | 41.36033 | 55.5514 | 24.30471 | -0.04288 | 1.090441 | 0.93066 |
| 19 | 6413600 | 6414799 | 10.23209 | 18.84734 | 26.75637 | 11.33579 | -0.06882 | 0.218593 | 0.930662 |
| 3 | 7797800 | 7798399 | 15.17044 | 26.54401 | 36.47699 | 16.09108 | -0.06857 | 0.669601 | 0.930692 |
| 14 | 6996000 | 6998199 | 33.14436 | 54.07927 | 68.01262 | 29.22293 | 0.042705 | 1.516223 | 0.930698 |
| X | 71786800 | 71787599 | 18.09807 | 31.53976 | 39.9936 | 17.59287 | 0.046412 | 0.750388 | 0.930703 |
| 1 | 107250800 | 107251399 | 8.597842 | 16.49082 | 22.97016 | 9.026571 | 0.049141 | -0.07447 | 0.930729 |
| 3 | 142453600 | 142456799 | 32.21598 | 52.5658 | 70.09913 | 30.11972 | -0.04158 | 1.515976 | 0.930743 |
| 12 | 59615800 | 59617799 | 17.11965 | 30.22849 | 37.92242 | 16.77815 | 0.064266 | 0.556905 | 0.930756 |
| 15 | 5398400 | 5399599 | 9.113585 | 17.39844 | 23.96317 | 9.535999 | 0.048643 | -0.02451 | 0.930767 |
| 11 | 75801000 | 75802999 | 22.3269 | 37.94262 | 47.97481 | 21.02532 | 0.044412 | 1.025562 | 0.930789 |
| 4 | 112051000 | 112066599 | 206.3696 | 305.4061 | 292.986 | 120.6602 | 0.308497 | 3.718439 | 0.930802 |
| 4 | 127338000 | 127340599 | 23.80748 | 40.20124 | 53.88705 | 23.57231 | -0.04002 | 1.091315 | 0.93081 |
| 14 | 40980000 | 40982999 | 8.935462 | 16.64489 | 30.54673 | 12.73675 | -0.41932 | 0.154485 | 0.930842 |
| 16 | 90462400 | 90466399 | 46.6925 | 75.05912 | 93.44854 | 39.62117 | 0.036219 | 1.888927 | 0.930892 |
| 7 | 48280600 | 48282599 | 21.75951 | 37.68444 | 47.38943 | 20.79153 | 0.056889 | 0.851597 | 0.930908 |
| Y | 88287200 | 88288399 | 9.50897 | 17.73491 | 22.34201 | 9.296426 | 0.121799 | 0.127401 | 0.930911 |
| 7 | 10624000 | 10626999 | 38.38106 | 63.33657 | 58.29207 | 25.16183 | 0.499784 | 1.378726 | 0.930941 |
| 5 | 110178000 | 110236199 | 985.2862 | 1271.864 | 1362.436 | 571.143 | -0.04681 | 5.851527 | 0.930949 |
| 9 | 70987200 | 70990399 | 41.96937 | 67.68776 | 84.55632 | 36.01712 | 0.037143 | 1.779046 | 0.930975 |
| 4 | 122456400 | 122456999 | 13.73794 | 24.30091 | 33.20999 | 14.68794 | -0.05705 | 0.511966 | 0.930986 |
| 1 | 59382400 | 59391199 | 151.3256 | 226.6792 | 278.3891 | 114.833 | -0.03161 | 3.493771 | 0.930999 |
| 12 | 25642000 | 25642599 | 9.774438 | 18.35472 | 24.33447 | 10.00927 | 0.062345 | 0.094503 | 0.93102 |
| 10 | 68999000 | 69001399 | 20.12868 | 34.93898 | 47.39629 | 20.79709 | -0.05134 | 0.823756 | 0.931022 |
| 7 | 49325400 | 49327199 | 16.67231 | 29.24721 | 37.1341 | 16.41669 | 0.044709 | 0.635081 | 0.931045 |
| 4 | 71021400 | 71022999 | 17.69044 | 30.91816 | 39.33978 | 17.33729 | 0.041744 | 0.704743 | 0.931047 |
| 1 | 116067800 | 116068599 | 10.9917 | 20.43037 | 28.82086 | 12.06117 | -0.04722 | 0.178691 | 0.931052 |
| 16 | 7554000 | 7561399 | 138.5429 | 210.1262 | 211.415 | 87.51506 | 0.2714 | 3.243514 | 0.931063 |
| 6 | 126477200 | 126479999 | 25.1674 | 41.93279 | 56.57619 | 24.5824 | -0.04952 | 1.230169 | 0.931071 |
| 9 | 39243800 | 39245799 | 18.1501 | 31.86361 | 31.40125 | 13.89109 | 0.408398 | 0.621136 | 0.931071 |
| 2 | 150059600 | 150061799 | 21.12839 | 36.05151 | 48.48835 | 21.25731 | -0.04356 | 0.983251 | 0.931076 |
| 16 | 76660800 | 76661399 | 8.959795 | 17.06317 | 25.00813 | 10.00331 | -0.04695 | -0.00692 | 0.931085 |
| 8 | 34488800 | 34490999 | 19.10256 | 33.26885 | 45.16971 | 19.86176 | -0.05213 | 0.768297 | 0.931093 |

| | | | | | | | | | |
|---|---|---|---|---|---|---|---|---|---|
| 5 | 7736800 | 7738199 | 15.16075 | 27.33973 | 27.80088 | 12.10459 | 0.382918 | 0.379326 | 0.931094 |
| 4 | 79687400 | 79688599 | 11.47295 | 20.96314 | 29.2583 | 12.53147 | -0.05544 | 0.282684 | 0.931098 |
| 17 | 83491600 | 83495799 | 41.08651 | 66.33915 | 87.45973 | 37.30843 | -0.03858 | 1.744447 | 0.931107 |
| X | 72453200 | 72456399 | 25.16359 | 42.52081 | 57.70768 | 25.24895 | -0.05776 | 1.097118 | 0.931132 |
| 8 | 102690400 | 102694199 | 74.39242 | 116.7276 | 141.874 | 58.83864 | 0.037199 | 2.577027 | 0.931139 |
| 3 | 76041000 | 76041599 | 8.52087 | 16.31732 | 22.45099 | 8.862769 | 0.061232 | -0.05937 | 0.931156 |
| 16 | 93348800 | 93349999 | 12.83555 | 23.12112 | 29.59252 | 12.90246 | 0.051851 | 0.373609 | 0.931159 |
| 1 | 165354000 | 165355399 | 11.37055 | 20.72908 | 26.28067 | 11.28803 | 0.079625 | 0.28642 | 0.931203 |
| 19 | 21650400 | 21652999 | 44.69834 | 71.87886 | 89.56382 | 37.99355 | 0.03566 | 1.873342 | 0.931204 |
| 13 | 23822600 | 23826199 | 49.5873 | 79.51486 | 98.6976 | 41.69126 | 0.036246 | 1.970342 | 0.931207 |
| 3 | 112056600 | 112059799 | 49.83497 | 79.83493 | 99.09367 | 41.79719 | 0.03464 | 2.001999 | 0.931222 |
| 1 | 95625600 | 95627999 | 16.3051 | 28.79253 | 36.40376 | 16.10598 | 0.053149 | 0.548694 | 0.931224 |
| 3 | 22389400 | 22392599 | 48.38458 | 77.64188 | 96.45675 | 40.78913 | 0.036244 | 1.947818 | 0.931232 |
| 13 | 58732400 | 58734799 | 34.03036 | 55.37406 | 73.51983 | 31.5579 | -0.03806 | 1.56116 | 0.931235 |
| 7 | 122049000 | 122051199 | 18.74467 | 32.39692 | 43.74442 | 19.19113 | -0.04571 | 0.853696 | 0.931292 |
| 6 | 130931600 | 130934199 | 23.01109 | 38.98282 | 52.23912 | 22.87395 | -0.03939 | 1.0596 | 0.931298 |
| 13 | 93667800 | 93675599 | 114.5883 | 174.7183 | 217.9092 | 90.34397 | -0.032 | 3.129891 | 0.931318 |
| 12 | 44000200 | 44010599 | 197.8563 | 290.4087 | 334.0984 | 137.6041 | 0.035383 | 3.851028 | 0.931322 |
| 12 | 51627400 | 51628999 | 14.26458 | 25.32842 | 34.45888 | 15.21379 | -0.04841 | 0.480519 | 0.931323 |
| 1 | 5277000 | 5278199 | 13.58215 | 24.37331 | 31.11231 | 13.61423 | 0.052296 | 0.406992 | 0.931335 |
| 1 | 16228000 | 16230799 | 28.4806 | 47.25955 | 59.97329 | 25.9866 | 0.038276 | 1.289885 | 0.931369 |
| 1 | 30337400 | 30338799 | 20.05071 | 34.47581 | 46.36169 | 20.33615 | -0.04144 | 0.908738 | 0.931371 |
| 3 | 97008800 | 97018999 | 103.8035 | 160.4973 | 191.3581 | 79.48572 | 0.048443 | 2.854712 | 0.931394 |
| 9 | 95451400 | 95452199 | 7.270886 | 14.04188 | 21.52523 | 8.283696 | -0.05892 | -0.15956 | 0.931401 |
| 4 | 64031200 | 64032999 | 15.5127 | 27.77451 | 26.46683 | 11.60366 | 0.469615 | 0.423121 | 0.931412 |
| 14 | 17659000 | 17660799 | 15.25729 | 27.02503 | 34.43245 | 15.22448 | 0.044004 | 0.507144 | 0.931418 |
| 1 | 125235000 | 125247599 | 141.4869 | 213.389 | 252.2985 | 104.2546 | 0.031872 | 3.359089 | 0.931445 |
| 2 | 80835600 | 80838399 | 31.57119 | 51.19259 | 84.46109 | 36.17515 | -0.35504 | 1.629315 | 0.931448 |
| 10 | 95212400 | 95217799 | 84.13503 | 131.282 | 166.9094 | 69.26109 | -0.03391 | 2.683528 | 0.931465 |
| 13 | 45278600 | 45280799 | 25.18 | 42.79429 | 58.40658 | 25.59726 | -0.06559 | 1.045848 | 0.931483 |
| 9 | 67555000 | 67558799 | 45.04334 | 72.46229 | 90.37626 | 38.36016 | 0.035001 | 1.862063 | 0.9315 |
| 3 | 111997200 | 111999999 | 39.50133 | 64.15911 | 66.06474 | 28.331 | 0.325614 | 1.597724 | 0.931532 |
| 9 | 113348200 | 113350199 | 20.19912 | 34.69059 | 43.78313 | 19.20441 | 0.049422 | 0.916908 | 0.931546 |
| 4 | 60349600 | 60353199 | 20.86841 | 36.23617 | 49.58457 | 21.76226 | -0.06376 | 0.825693 | 0.931552 |
| 3 | 37998400 | 38007399 | 94.10379 | 146.7828 | 175.2176 | 72.71143 | 0.056498 | 2.696748 | 0.931564 |
| 3 | 10463400 | 10464399 | 14.20522 | 25.24383 | 32.0933 | 14.15953 | 0.049866 | 0.473367 | 0.931575 |
| X | 39000600 | 39001399 | 14.00613 | 25.3449 | 32.57254 | 14.12109 | 0.051114 | 0.362078 | 0.931621 |
| 15 | 14954200 | 14955399 | 8.933947 | 17.28201 | 24.49673 | 9.355197 | 0.062778 | -0.16038 | 0.931636 |
| 8 | 66036400 | 66037599 | 19.035 | 32.74917 | 44.29104 | 19.42824 | -0.04942 | 0.895677 | 0.931652 |
| 3 | 15564400 | 15569799 | 69.62672 | 110.5372 | 133.9419 | 55.81905 | 0.055296 | 2.306946 | 0.931656 |
| 11 | 15788600 | 15790799 | 26.21296 | 43.93274 | 55.61146 | 24.25315 | 0.042024 | 1.168931 | 0.931688 |
| 12 | 98901600 | 98903199 | 20.62108 | 35.37056 | 44.79364 | 19.64583 | 0.044138 | 0.929878 | 0.931705 |
| 9 | 3807600 | 3810399 | 32.35951 | 52.98115 | 66.97314 | 28.81289 | 0.038773 | 1.466179 | 0.93171 |
| 6 | 22853600 | 22854399 | 10.0849 | 18.86634 | 26.84809 | 11.11402 | -0.04681 | 0.1267 | 0.931751 |
| 8 | 120062400 | 120072199 | 125.7562 | 190.1638 | 236.3999 | 97.76608 | -0.03717 | 3.302035 | 0.931766 |
| 3 | 159316200 | 159320199 | 34.08681 | 56.55501 | 69.58923 | 30.15054 | 0.082595 | 1.315634 | 0.931794 |
| 5 | 146744000 | 146747399 | 23.8835 | 40.42019 | 51.19622 | 22.41048 | 0.041522 | 1.06852 | 0.931813 |

| | | | | | | | | | |
|---|---|---|---|---|---|---|---|---|---|
| 2 | 6174600 | 6176999 | 27.17533 | 45.19076 | 60.4887 | 26.24889 | -0.03818 | 1.261825 | 0.931816 |
| 5 | 119251200 | 119253399 | 16.78327 | 29.39102 | 37.18907 | 16.417 | 0.049609 | 0.662745 | 0.931877 |
| 2 | 160468600 | 160483399 | 197.8709 | 290.62 | 335.0214 | 137.9786 | 0.033352 | 3.840915 | 0.931881 |
| 3 | 129551000 | 129555599 | 44.05754 | 71.40755 | 87.44239 | 37.43984 | 0.07333 | 1.662221 | 0.931888 |
| 2 | 95838600 | 95840999 | 25.17394 | 42.27737 | 56.62381 | 24.7264 | -0.03913 | 1.146763 | 0.931906 |
| 8 | 37854600 | 37857599 | 59.31821 | 94.27251 | 115.9239 | 48.36195 | 0.035657 | 2.262651 | 0.931912 |
| 5 | 127714000 | 127722599 | 131.3552 | 198.6383 | 245.9616 | 101.7217 | -0.03144 | 3.302906 | 0.931917 |
| 18 | 82044000 | 82047999 | 40.64394 | 65.54323 | 86.1261 | 36.68081 | -0.03587 | 1.780165 | 0.931919 |
| 18 | 90428400 | 90430799 | 25.49424 | 42.48754 | 57.13932 | 24.83084 | -0.04487 | 1.229203 | 0.931921 |
| 7 | 99404400 | 99415599 | 127.9967 | 194.8131 | 230.9585 | 95.70554 | 0.038995 | 3.171599 | 0.931921 |
| 4 | 100461000 | 100464599 | 32.14955 | 53.02429 | 71.60557 | 30.90143 | -0.05217 | 1.377856 | 0.931933 |
| 3 | 22768600 | 22771399 | 19.55738 | 33.00351 | 57.4911 | 25.14401 | -0.41664 | 1.086376 | 0.931934 |
| 10 | 22423800 | 22425399 | 32.04194 | 52.53525 | 66.43205 | 28.59716 | 0.0391 | 1.448085 | 0.931955 |
| 3 | 4828000 | 4838399 | 167.5817 | 249.1057 | 303.8401 | 125.1761 | -0.03185 | 3.636882 | 0.931967 |
| 6 | 142965000 | 142967399 | 18.04526 | 31.49585 | 42.4949 | 18.71115 | -0.04267 | 0.730529 | 0.931995 |
| 19 | 39509400 | 39512999 | 38.55364 | 62.44356 | 82.68637 | 35.38534 | -0.04016 | 1.66326 | 0.932005 |
| 4 | 88278600 | 88279799 | 11.76158 | 20.83454 | 37.3117 | 16.49733 | -0.44404 | 0.492579 | 0.932029 |
| 1 | 127795400 | 127796999 | 15.26655 | 26.89582 | 34.10564 | 15.09626 | 0.047904 | 0.565929 | 0.932034 |
| 1 | 120842000 | 120843399 | 16.57949 | 29.29185 | 40.17932 | 17.77459 | -0.06332 | 0.555178 | 0.932042 |
| 2 | 19838200 | 19839399 | 18.17311 | 31.72849 | 40.38471 | 17.78535 | 0.041107 | 0.723741 | 0.932064 |
| 9 | 43304800 | 43306799 | 24.60678 | 41.41915 | 55.55317 | 24.27773 | -0.04111 | 1.122353 | 0.932086 |
| 5 | 73623200 | 73626599 | 41.43362 | 66.7173 | 87.61926 | 37.24014 | -0.03764 | 1.826396 | 0.932089 |
| 15 | 73244400 | 73245599 | 6.169139 | 11.76608 | 26.13192 | 10.42002 | -0.63669 | -0.03986 | 0.932093 |
| 7 | 26357800 | 26363999 | 104.1674 | 162.0219 | 153.0101 | 63.52223 | 0.391004 | 2.746026 | 0.932103 |
| 9 | 10977400 | 10978599 | 11.70731 | 21.87273 | 28.92985 | 12.00244 | 0.055272 | 0.153798 | 0.932108 |
| X | 161522200 | 161524399 | 17.44119 | 30.60897 | 41.851 | 18.50861 | -0.06148 | 0.644131 | 0.932109 |
| 1 | 108065200 | 108067999 | 40.23129 | 65.00503 | 85.83442 | 36.64908 | -0.03964 | 1.720833 | 0.93211 |
| 1 | 150162000 | 150164199 | 30.45871 | 50.31347 | 67.76401 | 29.27064 | -0.04782 | 1.355535 | 0.932132 |
| 12 | 65079400 | 65083399 | 41.94147 | 67.61686 | 84.5043 | 35.97719 | 0.035879 | 1.789539 | 0.93215 |
| 9 | 115434800 | 115436799 | 17.61792 | 30.60106 | 41.58582 | 18.25508 | -0.05374 | 0.794007 | 0.932151 |
| 13 | 68548000 | 68570999 | 360.241 | 509.9488 | 449.8483 | 186.1642 | 0.366848 | 4.4001 | 0.932152 |
| 19 | 34687200 | 34688599 | 12.08334 | 21.8696 | 30.17974 | 13.08069 | -0.05049 | 0.342923 | 0.932172 |
| 10 | 43510200 | 43514199 | 44.57282 | 71.77122 | 94.35102 | 40.10057 | -0.03914 | 1.834366 | 0.932174 |
| 5 | 93750400 | 93761399 | 413.3456 | 569.0032 | 612.2528 | 254.4979 | 0.039735 | 4.820367 | 0.932196 |
| 7 | 90222600 | 90223399 | 9.189463 | 17.41782 | 25.43289 | 10.27569 | -0.05481 | 0.026464 | 0.932207 |
| 8 | 52754800 | 52758799 | 45.22709 | 72.60382 | 89.91109 | 38.044 | 0.041528 | 1.928759 | 0.932221 |
| 15 | 92882400 | 92886399 | 34.25394 | 56.0065 | 70.71321 | 30.40643 | 0.039135 | 1.489899 | 0.932234 |
| 17 | 14294800 | 14299999 | 56.29392 | 89.65073 | 110.3965 | 46.17617 | 0.036741 | 2.195957 | 0.93224 |
| 6 | 111984400 | 111987199 | 22.36719 | 38.83413 | 38.75082 | 17.00617 | 0.389831 | 0.818135 | 0.932297 |
| 13 | 24572000 | 24575399 | 27.02583 | 45.14191 | 60.54276 | 26.34739 | -0.04099 | 1.214015 | 0.932303 |
| 18 | 64181200 | 64183599 | 14.93069 | 26.03032 | 44.71305 | 19.67386 | -0.38944 | 0.735368 | 0.932331 |
| 14 | 52972400 | 52978599 | 132.1535 | 199.8999 | 237.0336 | 98.03777 | 0.031327 | 3.297214 | 0.932332 |
| 2 | 99379800 | 99383199 | 67.98324 | 107.0254 | 137.6334 | 57.10459 | -0.04041 | 2.507614 | 0.932332 |
| 1 | 147603000 | 147605399 | 25.52309 | 43.22697 | 54.46991 | 23.84884 | 0.049016 | 1.073825 | 0.932334 |
| 13 | 100551600 | 100556999 | 85.89056 | 133.6151 | 161.9183 | 67.21888 | 0.03183 | 2.735967 | 0.932386 |
| 16 | 46887600 | 46888599 | 10.86424 | 20.21475 | 26.75355 | 11.1767 | 0.046494 | 0.167494 | 0.932389 |
| 14 | 108884800 | 108888799 | 57.32735 | 91.3897 | 112.8392 | 47.26455 | 0.034433 | 2.163177 | 0.932397 |

| | | | | | | | | | |
|---|---|---|---|---|---|---|---|---|---|
| 3 | 91403400 | 91407199 | 56.01854 | 89.42925 | 116.6043 | 48.93797 | -0.04187 | 2.120906 | 0.932403 |
| 10 | 29170200 | 29173199 | 19.00253 | 33.32867 | 41.73001 | 18.44555 | 0.06466 | 0.658231 | 0.932448 |
| 17 | 64753000 | 64753599 | 10.31631 | 19.21045 | 27.26524 | 11.3752 | -0.05244 | 0.156906 | 0.932478 |
| 4 | 92812400 | 92813799 | 24.26546 | 41.7371 | 42.00105 | 18.42359 | 0.375118 | 0.916125 | 0.932503 |
| 9 | 25328600 | 25331999 | 36.42022 | 59.23147 | 74.54361 | 31.99118 | 0.038367 | 1.572905 | 0.932512 |
| 6 | 84741000 | 84742599 | 32.80705 | 53.38818 | 71.38966 | 30.64452 | -0.04813 | 1.558565 | 0.93252 |
| 12 | 119528600 | 119539599 | 213.5182 | 312.4857 | 359.5649 | 148.1199 | 0.031105 | 3.898958 | 0.932528 |
| 8 | 118906200 | 118908599 | 24.13293 | 40.61472 | 54.43213 | 23.78661 | -0.03997 | 1.121507 | 0.932554 |
| 3 | 56821600 | 56822199 | 11.77041 | 22.14764 | 23.01673 | 9.434832 | 0.414322 | 0.093172 | 0.932557 |
| 11 | 20978000 | 20982199 | 51.60054 | 82.45722 | 107.0346 | 44.98419 | -0.03388 | 2.081814 | 0.932579 |
| 2 | 17896200 | 17900599 | 66.67395 | 105.9094 | 109.6311 | 45.71362 | 0.283008 | 2.282203 | 0.932605 |
| 5 | 40527200 | 40528199 | 12.56557 | 22.70003 | 29.00734 | 12.60188 | 0.057228 | 0.351592 | 0.932621 |
| 9 | 71877800 | 71879599 | 11.31779 | 19.99677 | 37.90479 | 16.76821 | -0.52689 | 0.520032 | 0.932632 |
| 1 | 108255000 | 108256599 | 31.69282 | 52.47532 | 50.29137 | 21.75865 | 0.441902 | 1.316541 | 0.932638 |
| 5 | 87107200 | 87107599 | 7.688388 | 14.92631 | 21.72036 | 8.151801 | 0.049231 | -0.24054 | 0.932652 |
| 16 | 25008600 | 25010399 | 17.39576 | 30.38136 | 40.9367 | 18.03219 | -0.04062 | 0.716286 | 0.932662 |
| 13 | 69372000 | 69372799 | 8.034252 | 15.39242 | 22.98433 | 9.061533 | -0.05436 | -0.06833 | 0.932672 |
| 10 | 52333800 | 52334399 | 6.090219 | 11.86378 | 19.0953 | 6.959371 | -0.05471 | -0.36893 | 0.932691 |
| 5 | 39721200 | 39721799 | 7.872918 | 15.19801 | 21.4998 | 8.290561 | 0.053099 | -0.14597 | 0.93273 |
| 5 | 121586600 | 121587399 | 7.872918 | 15.19801 | 21.4998 | 8.290561 | 0.053099 | -0.14597 | 0.93273 |
| 8 | 95449400 | 95451799 | 16.98781 | 29.89311 | 40.69353 | 18.01651 | -0.05472 | 0.600574 | 0.932733 |
| 3 | 9720800 | 9724399 | 37.93885 | 61.36285 | 76.73126 | 32.80339 | 0.040436 | 1.684599 | 0.932738 |
| 13 | 52359800 | 52361599 | 27.72238 | 45.89158 | 61.47282 | 26.59244 | -0.03972 | 1.317221 | 0.932768 |
| 15 | 53712800 | 53713599 | 10.25189 | 18.92345 | 26.77693 | 11.31218 | -0.06106 | 0.207786 | 0.932775 |
| 12 | 19387400 | 19388399 | 22.9285 | 39.40172 | 40.01246 | 17.55082 | 0.361835 | 0.916061 | 0.932793 |
| 6 | 59534400 | 59537199 | 29.11717 | 48.57242 | 61.27417 | 26.644 | 0.047063 | 1.233781 | 0.932808 |
| 8 | 65014400 | 65016599 | 17.02186 | 30.137 | 30.32256 | 13.40848 | 0.382675 | 0.516394 | 0.932813 |
| X | 104014600 | 104016599 | 14.63838 | 26.21646 | 36.14666 | 15.84375 | -0.06013 | 0.436516 | 0.932814 |
| 9 | 13812600 | 13813799 | 11.02501 | 20.54699 | 27.28185 | 11.36633 | 0.044934 | 0.159741 | 0.932825 |
| 11 | 54695400 | 54696199 | 13.4069 | 23.92649 | 32.62072 | 14.34082 | -0.04687 | 0.442502 | 0.932845 |
| 3 | 94652600 | 94659799 | 118.7244 | 180.8081 | 225.1679 | 93.32265 | -0.03124 | 3.156607 | 0.932851 |
| 2 | 4092400 | 4094399 | 13.70304 | 24.34469 | 30.69214 | 13.54449 | 0.061708 | 0.468842 | 0.932891 |
| 12 | 28846400 | 28848799 | 30.50756 | 50.29414 | 63.81855 | 27.53584 | 0.037237 | 1.374455 | 0.932927 |
| 7 | 3914400 | 3918799 | 53.73108 | 85.73356 | 111.0958 | 46.59227 | -0.03376 | 2.130691 | 0.932927 |
| 14 | 84058600 | 84061199 | 29.04108 | 48.30598 | 61.1302 | 26.53205 | 0.042571 | 1.266114 | 0.932927 |
| 4 | 15467400 | 15469199 | 24.63742 | 41.46643 | 55.53844 | 24.27054 | -0.03907 | 1.123963 | 0.93293 |
| 17 | 95143600 | 95145399 | 17.51328 | 30.82127 | 42.27422 | 18.71592 | -0.0655 | 0.605086 | 0.932969 |
| 6 | 3280000 | 3280799 | 10.96285 | 20.39156 | 26.80144 | 11.20291 | 0.05609 | 0.172533 | 0.932985 |
| 12 | 114730400 | 114768799 | 1085.834 | 1394.419 | 1389.99 | 582.985 | 0.049528 | 5.93613 | 0.932994 |
| 1 | 117871600 | 117875399 | 60.50103 | 96.33356 | 125.4882 | 52.47572 | -0.04457 | 2.209849 | 0.932997 |
| 13 | 28315400 | 28317199 | 19.34749 | 33.58757 | 45.25075 | 19.85734 | -0.04135 | 0.816405 | 0.933 |
| 17 | 81415400 | 81417799 | 16.96185 | 29.76833 | 40.24473 | 17.79854 | -0.04538 | 0.636267 | 0.933002 |
| 12 | 54613800 | 54621599 | 71.09254 | 112.2019 | 144.1333 | 59.83176 | -0.03608 | 2.457238 | 0.933045 |
| 16 | 31374400 | 31378599 | 47.63759 | 76.49619 | 100.1735 | 42.40552 | -0.03846 | 1.924951 | 0.933052 |
| 8 | 4736400 | 4738399 | 28.45688 | 47.29505 | 63.44119 | 27.51933 | -0.04138 | 1.273518 | 0.933139 |
| 9 | 49250800 | 49253399 | 27.89842 | 45.98417 | 61.89262 | 26.70274 | -0.04763 | 1.368445 | 0.933141 |
| 5 | 85128400 | 85128999 | 10.41576 | 19.34653 | 27.29419 | 11.433 | -0.04796 | 0.175729 | 0.933144 |

| | | | | | | | | | |
|---|---|---|---|---|---|---|---|---|---|
| X | 107333000 | 107336999 | 49.72685 | 80.38081 | 74.05594 | 31.63674 | 0.479178 | 1.722631 | 0.933153 |
| 17 | 21198600 | 21201999 | 31.10032 | 51.70725 | 64.82387 | 28.12618 | 0.055923 | 1.278271 | 0.933155 |
| 1 | 166938200 | 166938599 | 8.231448 | 15.9169 | 22.65868 | 8.668963 | 0.055002 | -0.16396 | 0.933159 |
| 3 | 7281800 | 7283199 | 26.35066 | 43.78605 | 58.7599 | 25.48529 | -0.04195 | 1.266532 | 0.933165 |
| 2 | 139381800 | 139383199 | 12.43276 | 22.73168 | 29.59635 | 12.66594 | 0.044442 | 0.28376 | 0.933177 |
| X | 72683000 | 72685999 | 27.15566 | 45.6443 | 57.18284 | 24.97911 | 0.057023 | 1.149068 | 0.933201 |
| 13 | 7277600 | 7278399 | 13.62984 | 24.68503 | 31.63639 | 13.70062 | 0.056371 | 0.354781 | 0.933287 |
| 12 | 75820800 | 75827999 | 91.6596 | 142.4063 | 171.9675 | 71.40442 | 0.035431 | 2.760919 | 0.933293 |
| 12 | 35871400 | 35872799 | 16.25964 | 28.59294 | 38.61899 | 17.10029 | -0.04382 | 0.599196 | 0.933335 |
| 9 | 5928000 | 5930999 | 36.68539 | 59.53772 | 78.75503 | 33.75315 | -0.03592 | 1.616243 | 0.933357 |
| 1 | 88703800 | 88705999 | 24.16577 | 40.50937 | 54.40932 | 23.73172 | -0.04313 | 1.160352 | 0.933389 |
| 1 | 133232400 | 133234999 | 19.81015 | 34.07906 | 45.82448 | 20.10081 | -0.04125 | 0.901072 | 0.933413 |
| 13 | 50361000 | 50362399 | 11.82122 | 21.18213 | 36.75904 | 16.1051 | -0.38975 | 0.413242 | 0.933416 |
| 5 | 113349000 | 113353199 | 44.27222 | 70.94473 | 93.54306 | 39.47395 | -0.0518 | 1.984189 | 0.933427 |
| 16 | 94889000 | 94892399 | 26.88243 | 44.17874 | 72.76584 | 31.3546 | -0.33979 | 1.411599 | 0.933429 |
| 4 | 111288600 | 111290999 | 17.17064 | 29.94147 | 40.45752 | 17.80366 | -0.04493 | 0.735255 | 0.933451 |
| 13 | 46220400 | 46221799 | 10.72198 | 20.0244 | 28.39581 | 11.78829 | -0.04523 | 0.146312 | 0.933464 |
| 13 | 34103400 | 34106799 | 29.41681 | 48.66311 | 61.714 | 26.68412 | 0.038837 | 1.327812 | 0.933508 |
| X | 49435200 | 49437999 | 21.55928 | 37.55012 | 46.78406 | 20.57253 | 0.071404 | 0.783408 | 0.933512 |
| X | 73883400 | 73885999 | 22.8012 | 38.89347 | 49.01875 | 21.48929 | 0.049935 | 0.997223 | 0.933535 |
| 1 | 183467000 | 183469799 | 32.84017 | 54.73421 | 53.61104 | 23.29871 | 0.411024 | 1.248009 | 0.933538 |
| 13 | 43886400 | 43886999 | 7.266024 | 14.04791 | 21.5482 | 8.250079 | -0.05288 | -0.17722 | 0.93355 |
| 2 | 15988600 | 15990599 | 26.66339 | 46.05796 | 40.31877 | 17.68796 | 0.576759 | 0.889447 | 0.933555 |
| 3 | 74806600 | 74809199 | 24.59732 | 41.74697 | 52.8259 | 23.14383 | 0.043064 | 1.04937 | 0.93356 |
| 6 | 81979800 | 81982799 | 29.97875 | 49.42556 | 62.69185 | 27.05044 | 0.037829 | 1.370736 | 0.933563 |
| 9 | 11076600 | 11082399 | 8.077154 | 15.43302 | 31.46044 | 12.49686 | -0.5108 | -0.0227 | 0.933566 |
| 16 | 8615800 | 8619199 | 45.21681 | 72.72538 | 95.2358 | 40.40992 | -0.0356 | 1.870844 | 0.933568 |
| 5 | 36929400 | 36939199 | 128.4157 | 194.1142 | 240.5521 | 99.47176 | -0.03309 | 3.310249 | 0.933685 |
| 2 | 144068000 | 144070599 | 31.59951 | 51.82235 | 65.46303 | 28.18291 | 0.040665 | 1.440814 | 0.933712 |
| 4 | 128157600 | 128162199 | 46.35752 | 74.20057 | 97.2854 | 40.98194 | -0.04565 | 2.022908 | 0.933763 |
| 11 | 50702200 | 50710999 | 172.9994 | 257.9805 | 301.0565 | 124.0744 | 0.036124 | 3.582287 | 0.933767 |
| 2 | 169213000 | 169221199 | 99.1038 | 152.8215 | 184.0679 | 76.4586 | 0.030842 | 2.900039 | 0.933788 |
| 6 | 9891600 | 9894599 | 38.92927 | 63.17426 | 79.251 | 33.96368 | 0.040235 | 1.621434 | 0.933797 |
| 13 | 103693200 | 103693599 | 5.942238 | 11.59269 | 17.53229 | 6.266862 | 0.059975 | -0.4442 | 0.933834 |
| 1 | 32211400 | 32212199 | 9.899687 | 18.58325 | 24.80585 | 10.20936 | 0.051666 | 0.098213 | 0.933887 |
| 10 | 4445400 | 4447399 | 17.45039 | 30.41716 | 41.03071 | 18.05138 | -0.0424 | 0.745567 | 0.933892 |
| 10 | 100646000 | 100646999 | 12.05381 | 22.51993 | 23.07536 | 9.573639 | 0.421096 | 0.142473 | 0.933906 |
| 8 | 125709800 | 125712399 | 22.21766 | 37.76286 | 50.61287 | 22.18174 | -0.03933 | 1.027179 | 0.93392 |
| 3 | 31586000 | 31588399 | 22.36237 | 38.4321 | 52.32887 | 22.95327 | -0.05972 | 0.926428 | 0.933944 |
| 13 | 35922200 | 35925999 | 31.90374 | 52.04676 | 69.43728 | 29.83334 | -0.04237 | 1.517899 | 0.933947 |
| 5 | 57719000 | 57719599 | 9.353859 | 16.91037 | 33.38325 | 14.48966 | -0.5649 | 0.336895 | 0.933953 |
| 1 | 76206800 | 76209399 | 34.20084 | 56.07956 | 70.72125 | 30.44461 | 0.043023 | 1.454966 | 0.933978 |
| 8 | 120500200 | 120501199 | 13.83579 | 24.69032 | 31.55585 | 13.87342 | 0.045702 | 0.442858 | 0.933997 |
| 10 | 16520200 | 16521199 | 17.23305 | 29.97475 | 40.58107 | 17.82979 | -0.04795 | 0.772906 | 0.933997 |
| 2 | 176967200 | 176970999 | 31.01853 | 51.32538 | 69.22715 | 29.93775 | -0.04979 | 1.336488 | 0.934001 |
| 1 | 103643800 | 103644199 | 5.650481 | 11.02802 | 18.12671 | 6.434972 | -0.05053 | -0.47213 | 0.934002 |
| 5 | 17956400 | 17957399 | 10.23668 | 18.93905 | 26.76808 | 11.27296 | -0.0562 | 0.194388 | 0.934016 |

| | | | | | | | | | |
|---|---|---|---|---|---|---|---|---|---|
| 1 | 113460200 | 113461199 | 12.21564 | 22.23461 | 30.66683 | 13.19799 | -0.04323 | 0.311758 | 0.934035 |
| 1 | 190741000 | 190741799 | 12.21564 | 22.23461 | 30.66683 | 13.19799 | -0.04323 | 0.311758 | 0.934035 |
| 10 | 109396200 | 109396999 | 11.17584 | 20.90776 | 27.86809 | 11.53575 | 0.04679 | 0.137819 | 0.93404 |
| 7 | 44160400 | 44164599 | 42.94525 | 68.95347 | 90.69332 | 38.38577 | -0.0449 | 1.920185 | 0.934074 |
| 6 | 57910600 | 57912599 | 25.45427 | 43.31683 | 43.1311 | 18.90461 | 0.387906 | 1.016112 | 0.934084 |
| 7 | 103855000 | 103856599 | 17.97276 | 31.34737 | 39.90022 | 17.56095 | 0.040931 | 0.735517 | 0.93409 |
| 11 | 41449200 | 41451199 | 19.2967 | 33.87953 | 33.87128 | 14.98488 | 0.387888 | 0.63223 | 0.934096 |
| 13 | 107568800 | 107569599 | 14.29203 | 25.30259 | 32.07224 | 14.18233 | 0.051247 | 0.502395 | 0.934111 |
| 1 | 137248200 | 137248799 | 7.139511 | 13.8723 | 21.76797 | 8.114734 | -0.0489 | -0.26793 | 0.934113 |
| 9 | 88945800 | 88951599 | 74.04537 | 116.2473 | 141.6117 | 58.73405 | 0.034536 | 2.565729 | 0.934127 |
| 4 | 111874200 | 111874999 | 9.860182 | 18.81654 | 27.93891 | 11.12712 | -0.06058 | -0.00667 | 0.934144 |
| 14 | 52843400 | 52853599 | 188.2996 | 279.7625 | 276.1922 | 113.7799 | 0.272391 | 3.64735 | 0.934146 |
| 5 | 112809200 | 112811399 | 24.93218 | 41.83948 | 56.10517 | 24.48816 | -0.0409 | 1.151887 | 0.934175 |
| 11 | 48212400 | 48213799 | 19.70065 | 34.02677 | 43.2076 | 18.95524 | 0.042329 | 0.863024 | 0.934189 |
| 10 | 7037000 | 7054399 | 325.7966 | 459.0861 | 531.031 | 219.9267 | -0.0306 | 4.46627 | 0.934213 |
| 16 | 6069400 | 6072399 | 34.13994 | 55.85575 | 70.65517 | 30.38854 | 0.037107 | 1.481632 | 0.934248 |
| 8 | 70095200 | 70109199 | 49.70152 | 79.20118 | 126.0008 | 52.74835 | -0.33126 | 2.171799 | 0.934252 |
| 6 | 140875400 | 140876999 | 22.45421 | 38.19022 | 51.16535 | 22.42507 | -0.03871 | 1.02079 | 0.934268 |
| 2 | 85766600 | 85769799 | 27.92799 | 47.11261 | 48.00355 | 20.99235 | 0.354193 | 1.107418 | 0.934268 |
| 10 | 4390400 | 4391799 | 14.59708 | 25.81893 | 35.00391 | 15.48155 | -0.04597 | 0.518782 | 0.934282 |
| 10 | 87694800 | 87697999 | 24.19177 | 40.62064 | 54.39921 | 23.75229 | -0.03893 | 1.143462 | 0.934309 |
| 1 | 111113800 | 111114599 | 15.35721 | 27.14523 | 34.57047 | 15.2919 | 0.043663 | 0.53278 | 0.934312 |
| 6 | 89738000 | 89762799 | 603.326 | 808.2413 | 851.6185 | 355.4803 | 0.030459 | 5.251011 | 0.934312 |
| 4 | 17875600 | 17876999 | 16.50418 | 28.59843 | 48.46635 | 21.26436 | -0.37145 | 0.825306 | 0.934324 |
| 4 | 144081000 | 144085399 | 53.61495 | 85.70137 | 111.4602 | 46.8636 | -0.0362 | 2.076891 | 0.934333 |
| 16 | 44151000 | 44153399 | 26.57431 | 44.05502 | 59.2128 | 25.64049 | -0.0444 | 1.295605 | 0.934334 |
| 2 | 40445000 | 40447399 | 23.52786 | 39.19402 | 64.6348 | 28.08069 | -0.33812 | 1.236334 | 0.934348 |
| 1 | 93697200 | 93699199 | 16.28376 | 28.69802 | 36.54607 | 16.17534 | 0.041947 | 0.572882 | 0.934349 |
| 16 | 64624800 | 64627799 | 48.81263 | 78.18218 | 97.01028 | 40.9065 | 0.03479 | 2.00549 | 0.934355 |
| 2 | 68395400 | 68396599 | 11.01514 | 20.1599 | 28.17821 | 12.04287 | -0.05579 | 0.263977 | 0.934358 |
| 4 | 91867600 | 91869199 | 14.99419 | 26.57843 | 35.99877 | 15.91471 | -0.04331 | 0.50113 | 0.934408 |
| 9 | 116894600 | 116899999 | 49.72563 | 79.63375 | 103.6554 | 43.69958 | -0.03436 | 2.014089 | 0.934414 |
| 6 | 73368200 | 73370999 | 22.4467 | 38.1998 | 48.49364 | 21.2551 | 0.038912 | 1.012939 | 0.934423 |
| 3 | 46095600 | 46096399 | 9.647839 | 18.12349 | 23.87378 | 9.813817 | 0.072497 | 0.090787 | 0.934439 |
| 17 | 82290000 | 82290799 | 11.17566 | 20.57599 | 28.71247 | 12.17207 | -0.04499 | 0.233349 | 0.934454 |
| 13 | 50445600 | 50447599 | 20.78304 | 35.66845 | 45.30403 | 19.87058 | 0.040015 | 0.92602 | 0.934468 |
| 6 | 101001800 | 101002999 | 11.21289 | 20.64416 | 29.16001 | 12.3621 | -0.06287 | 0.231878 | 0.93447 |
| 3 | 33196200 | 33198199 | 18.47476 | 31.86775 | 43.22148 | 18.96057 | -0.0528 | 0.866968 | 0.934488 |
| 7 | 50768200 | 50768599 | 7.634774 | 14.78996 | 21.06616 | 7.991591 | 0.066496 | -0.20216 | 0.934517 |
| 7 | 61966400 | 61967999 | 21.70748 | 37.61611 | 37.91994 | 16.6372 | 0.374865 | 0.835109 | 0.934532 |
| 10 | 78692400 | 78705199 | 288.197 | 408.4421 | 477.1847 | 197.4389 | -0.03664 | 4.38008 | 0.934543 |
| 11 | 75719600 | 75720799 | 17.44517 | 30.42704 | 41.00586 | 18.04752 | -0.04099 | 0.736529 | 0.934564 |
| 9 | 107022200 | 107024599 | 36.88252 | 59.75781 | 74.84181 | 32.03655 | 0.040552 | 1.649195 | 0.934567 |
| 1 | 183025600 | 183026599 | 15.06195 | 27.08339 | 34.47406 | 15.05965 | 0.057552 | 0.407915 | 0.934579 |
| 1 | 51720600 | 51723399 | 31.9609 | 52.52358 | 70.20468 | 30.25073 | -0.03945 | 1.419898 | 0.934626 |
| 1 | 43264200 | 43265999 | 12.83553 | 22.71107 | 38.76094 | 17.14201 | -0.37591 | 0.513188 | 0.934636 |
| 13 | 56161200 | 56163399 | 19.4372 | 34.06131 | 34.48171 | 15.23043 | 0.369985 | 0.664075 | 0.934644 |

| | | | | | | | | | |
|---|---|---|---|---|---|---|---|---|---|
| 7 | 11335400 | 11340799 | 161.0042 | 239.9172 | 280.0099 | 115.3874 | 0.034995 | 3.59224 | 0.934646 |
| 3 | 104354000 | 104357999 | 33.02566 | 53.76298 | 67.15983 | 28.83256 | 0.050145 | 1.548921 | 0.934656 |
| 14 | 123758400 | 123759799 | 17.98251 | 31.60874 | 31.00952 | 13.73206 | 0.415064 | 0.601924 | 0.934689 |
| 3 | 33945800 | 33947999 | 16.89034 | 29.60523 | 37.71278 | 16.66088 | 0.040065 | 0.651558 | 0.934722 |
| 1 | 51953800 | 51955999 | 16.69245 | 29.13839 | 39.42152 | 17.35928 | -0.04653 | 0.716466 | 0.934729 |
| 13 | 50636400 | 50637799 | 16.69245 | 29.13839 | 39.42152 | 17.35928 | -0.04653 | 0.716466 | 0.934729 |
| 12 | 41713600 | 41714799 | 10.9772 | 20.17163 | 28.29019 | 12.02459 | -0.05447 | 0.244487 | 0.934739 |
| 5 | 71545800 | 71548599 | 43.12523 | 69.67499 | 70.86551 | 30.25438 | 0.335973 | 1.721864 | 0.934749 |
| 8 | 110487000 | 110487999 | 9.04901 | 17.01386 | 24.80042 | 10.17924 | -0.07167 | 0.077324 | 0.934767 |
| 18 | 82760800 | 82764399 | 41.26906 | 66.48934 | 87.21702 | 37.09912 | -0.03495 | 1.808588 | 0.934779 |
| 15 | 82406000 | 82414399 | 104.4533 | 160.3369 | 192.4889 | 79.91555 | 0.03054 | 2.99096 | 0.934784 |
| 16 | 32740800 | 32744399 | 53.0106 | 84.6664 | 109.7757 | 46.10218 | -0.03303 | 2.101461 | 0.934785 |
| 4 | 117358000 | 117359799 | 23.3742 | 39.34004 | 52.768 | 23.05979 | -0.04109 | 1.118173 | 0.934786 |
| 18 | 22368400 | 22370199 | 20.94699 | 35.99713 | 48.47479 | 21.26269 | -0.04359 | 0.921166 | 0.934788 |
| 2 | 111411800 | 111412599 | 23.08914 | 39.11316 | 49.57548 | 21.70731 | 0.040611 | 1.056314 | 0.934798 |
| 12 | 106411600 | 106420399 | 100.0724 | 152.9506 | 226.7911 | 94.07837 | -0.27866 | 3.078761 | 0.9348 |
| 3 | 44618600 | 44620599 | 6.344815 | 12.01524 | 27.09461 | 10.96789 | -0.67794 | 0.013722 | 0.934821 |
| 6 | 90531000 | 90531999 | 11.46477 | 21.10939 | 27.65539 | 11.7231 | 0.045528 | 0.232815 | 0.934826 |
| 13 | 72052800 | 72054199 | 11.46477 | 21.10939 | 27.65539 | 11.7231 | 0.045528 | 0.232815 | 0.934826 |
| 11 | 15280000 | 15280399 | 8.327317 | 16.05551 | 24.24933 | 9.38755 | -0.04638 | -0.12157 | 0.93483 |
| 15 | 47490600 | 47491199 | 6.527923 | 12.72063 | 19.1215 | 6.938874 | 0.048187 | -0.37904 | 0.934834 |
| 2 | 56500800 | 56502399 | 17.16381 | 30.04515 | 38.24198 | 16.87597 | 0.041189 | 0.674063 | 0.934842 |
| 6 | 117105600 | 117107999 | 18.25627 | 31.60185 | 42.71376 | 18.73985 | -0.04666 | 0.836298 | 0.93485 |
| X | 62553000 | 62553599 | 6.863035 | 13.35748 | 19.59841 | 7.20769 | 0.066074 | -0.32286 | 0.934861 |
| 1 | 191016800 | 191019999 | 26.36014 | 43.98633 | 58.81466 | 25.57933 | -0.03659 | 1.22099 | 0.934872 |
| 2 | 44295200 | 44299199 | 42.53898 | 68.66456 | 90.86516 | 38.74671 | -0.04443 | 1.759114 | 0.934916 |
| X | 164482000 | 164484799 | 30.97237 | 51.33616 | 69.71251 | 30.18304 | -0.05931 | 1.319462 | 0.934951 |
| 7 | 89141200 | 89145999 | 71.78617 | 113.1956 | 138.7132 | 57.56471 | 0.030855 | 2.476663 | 0.934953 |
| 5 | 110052600 | 110057999 | 186.9984 | 274.9397 | 332.9642 | 137.1224 | -0.0363 | 3.823677 | 0.934959 |
| 13 | 66785400 | 66792599 | 235.4005 | 343.0784 | 329.1602 | 135.6442 | 0.287409 | 3.963796 | 0.934964 |
| 17 | 35500600 | 35501799 | 13.47324 | 23.81593 | 32.85177 | 14.53155 | -0.0706 | 0.518922 | 0.934969 |
| 11 | 50447200 | 50453599 | 61.72092 | 97.94935 | 125.9532 | 52.47005 | -0.03079 | 2.309994 | 0.935 |
| 15 | 49647600 | 49652999 | 153.6476 | 231.5574 | 231.6087 | 95.68494 | 0.272091 | 3.375026 | 0.935053 |
| 1 | 6677200 | 6678199 | 11.62093 | 21.43372 | 28.16942 | 11.91311 | 0.043365 | 0.225527 | 0.935064 |
| 7 | 74542600 | 74543199 | 11.62093 | 21.43372 | 28.16942 | 11.91311 | 0.043365 | 0.225527 | 0.935064 |
| 18 | 87954600 | 87959199 | 22.21198 | 38.58552 | 38.8217 | 17.04307 | 0.378184 | 0.812039 | 0.935065 |
| 2 | 176450600 | 176453199 | 30.10446 | 49.83889 | 66.93994 | 28.95906 | -0.0437 | 1.324884 | 0.935067 |
| 11 | 3541200 | 3543199 | 17.23954 | 29.96806 | 40.60639 | 17.8342 | -0.04923 | 0.78168 | 0.935099 |
| 16 | 16624400 | 16625799 | 17.23954 | 29.96806 | 40.60639 | 17.8342 | -0.04923 | 0.78168 | 0.935099 |
| 14 | 13627200 | 13627999 | 13.00748 | 23.4184 | 30.07637 | 13.12212 | 0.04629 | 0.379057 | 0.935141 |
| 16 | 87025200 | 87030399 | 78.50941 | 123.2261 | 150.4335 | 62.39102 | 0.031188 | 2.574544 | 0.935175 |
| 4 | 73312200 | 73313799 | 30.83188 | 50.79553 | 67.85619 | 29.26985 | -0.03716 | 1.385518 | 0.935178 |
| 10 | 82608400 | 82609799 | 16.39406 | 28.59962 | 39.0843 | 17.20402 | -0.06074 | 0.727597 | 0.935187 |
| Y | 38150600 | 38151199 | 6.531959 | 12.61278 | 19.99164 | 7.699344 | -0.10772 | -0.1716 | 0.935197 |
| 13 | 93242600 | 93252599 | 112.8498 | 172.6892 | 216.1077 | 89.67185 | -0.03278 | 3.057304 | 0.935209 |
| 8 | 113224400 | 113229199 | 37.5616 | 61.21056 | 76.73771 | 32.95692 | 0.045937 | 1.54783 | 0.935236 |
| 7 | 10205000 | 10206199 | 17.82589 | 31.44971 | 27.14924 | 12.0137 | 0.600759 | 0.554306 | 0.935263 |

| | | | | | | | | | |
|---|---|---|---|---|---|---|---|---|---|
| 6 | 131684800 | 131706799 | 344.2892 | 486.0002 | 535.1893 | 221.5843 | 0.042872 | 4.442031 | 0.935271 |
| X | 40274400 | 40276999 | 19.87783 | 34.79413 | 43.44595 | 19.17157 | 0.068785 | 0.700151 | 0.935281 |
| 16 | 51032400 | 51032999 | 6.97625 | 13.57707 | 20.1611 | 7.41799 | 0.047364 | -0.32117 | 0.935286 |
| 2 | 162292400 | 162294799 | 24.66803 | 41.51365 | 55.52377 | 24.26337 | -0.03703 | 1.125572 | 0.935317 |
| 6 | 41295400 | 41297399 | 20.32815 | 35.60215 | 34.99223 | 15.44842 | 0.412486 | 0.679688 | 0.935317 |
| 8 | 119437000 | 119438999 | 20.32304 | 35.48853 | 36.08126 | 15.89194 | 0.363736 | 0.724902 | 0.935321 |
| 6 | 130745200 | 130753799 | 89.6258 | 140.8573 | 166.6182 | 69.11517 | 0.078077 | 2.55984 | 0.935329 |
| 13 | 7675400 | 7681999 | 155.3719 | 232.2106 | 272.0025 | 112.1489 | 0.03363 | 3.536012 | 0.93533 |
| 4 | 101802600 | 101805799 | 30.81533 | 51.00618 | 64.60755 | 27.94642 | 0.040646 | 1.328651 | 0.935332 |
| 4 | 21583600 | 21590399 | 97.97727 | 152.0579 | 155.6183 | 64.62631 | 0.27271 | 2.780947 | 0.935333 |
| 6 | 148479200 | 148479599 | 8.644147 | 16.72627 | 23.89058 | 9.111337 | 0.05393 | -0.16891 | 0.93534 |
| 4 | 5541400 | 5542199 | 9.589789 | 18.23322 | 26.62831 | 10.68814 | -0.04643 | 0.014105 | 0.935379 |
| 1 | 139553600 | 139554199 | 10.01006 | 18.82572 | 25.29993 | 10.37968 | 0.045391 | 0.087363 | 0.935384 |
| 17 | 56083800 | 56084399 | 10.01006 | 18.82572 | 25.29993 | 10.37968 | 0.045391 | 0.087363 | 0.935384 |
| 8 | 67056000 | 67060999 | 44.31176 | 71.34896 | 89.0583 | 37.84963 | 0.03543 | 1.831312 | 0.935392 |
| 14 | 39939800 | 39941599 | 20.26185 | 35.15962 | 44.69957 | 19.61255 | 0.041822 | 0.82409 | 0.935395 |
| 9 | 61259600 | 61265799 | 60.67075 | 96.27248 | 123.9364 | 51.62439 | -0.03256 | 2.310943 | 0.935402 |
| 17 | 45695400 | 45697199 | 14.65459 | 25.86094 | 32.68546 | 14.46281 | 0.053693 | 0.542107 | 0.935423 |
| 13 | 29008000 | 29011599 | 32.73508 | 53.66288 | 67.88356 | 29.21997 | 0.038382 | 1.449789 | 0.935436 |
| 10 | 125423600 | 125425399 | 16.56581 | 29.11213 | 37.07795 | 16.41653 | 0.040255 | 0.606776 | 0.935452 |
| 19 | 36129200 | 36132799 | 26.47734 | 44.35686 | 59.46602 | 25.92719 | -0.0405 | 1.179107 | 0.935452 |
| 12 | 37846400 | 37847399 | 11.14125 | 20.49057 | 26.66675 | 11.32139 | 0.053523 | 0.23419 | 0.935502 |
| 2 | 148251000 | 148253999 | 29.70918 | 48.79883 | 65.23989 | 28.10551 | -0.03996 | 1.418229 | 0.93551 |
| 1 | 113293200 | 113294199 | 12.65824 | 23.127 | 30.13148 | 12.90677 | 0.042733 | 0.291892 | 0.935525 |
| 1 | 117851000 | 117853999 | 35.39259 | 58.22004 | 59.64891 | 25.71296 | 0.343721 | 1.410457 | 0.935567 |
| 17 | 70630400 | 70631599 | 15.7021 | 27.58739 | 37.23043 | 16.48056 | -0.04267 | 0.608152 | 0.935574 |
| X | 20482600 | 20484799 | 21.09541 | 36.79864 | 45.68261 | 20.10758 | 0.076944 | 0.76117 | 0.935601 |
| 4 | 135481200 | 135482599 | 20.58496 | 35.30805 | 47.42322 | 20.79909 | -0.04032 | 0.935015 | 0.935613 |
| 6 | 139714000 | 139716599 | 17.82638 | 30.72474 | 51.13976 | 22.43365 | -0.34722 | 0.883148 | 0.935622 |
| 1 | 179156800 | 179157399 | 7.445573 | 14.40688 | 22.12395 | 8.438058 | -0.049 | -0.185 | 0.935647 |
| 10 | 99248000 | 99248399 | 7.445573 | 14.40688 | 22.12395 | 8.438058 | -0.049 | -0.185 | 0.935647 |
| 3 | 18022800 | 18023999 | 11.26805 | 20.8339 | 29.24959 | 12.32872 | -0.04796 | 0.210293 | 0.935652 |
| 18 | 29347000 | 29348399 | 13.40266 | 24.54147 | 31.743 | 13.55743 | 0.056109 | 0.289528 | 0.935693 |
| 12 | 112313200 | 112314399 | 20.66962 | 35.31519 | 47.43406 | 20.79731 | -0.04128 | 0.969019 | 0.935693 |
| 1 | 125132600 | 125135399 | 42.67169 | 69.0729 | 85.7822 | 36.69434 | 0.051356 | 1.683592 | 0.935738 |
| 13 | 79711200 | 79712999 | 24.38658 | 41.6681 | 41.15091 | 18.04256 | 0.400738 | 0.97699 | 0.93575 |
| 7 | 6512600 | 6535199 | 325.8115 | 460.2076 | 511.1978 | 211.629 | 0.031288 | 4.430885 | 0.935768 |
| 16 | 13834200 | 13834799 | 7.225282 | 13.95568 | 21.50593 | 8.27107 | -0.06631 | -0.16525 | 0.935782 |
| 12 | 5839400 | 5841799 | 18.03954 | 31.50595 | 42.47328 | 18.70913 | -0.0414 | 0.721368 | 0.935788 |
| 7 | 11571800 | 11573399 | 23.93758 | 41.11983 | 40.56887 | 17.79466 | 0.403345 | 0.927039 | 0.935815 |
| 17 | 94998000 | 95000999 | 35.0073 | 56.93279 | 75.37459 | 32.34764 | -0.03454 | 1.571766 | 0.935823 |
| 1 | 53003000 | 53004199 | 8.549917 | 16.26197 | 23.96183 | 9.610512 | -0.05744 | -0.00296 | 0.935842 |
| 16 | 18348800 | 18351199 | 23.18814 | 39.97385 | 40.41098 | 17.72732 | 0.369141 | 0.888576 | 0.935846 |
| 11 | 21633200 | 21634199 | 11.79742 | 21.45065 | 29.73581 | 12.81427 | -0.05145 | 0.314319 | 0.935849 |
| 4 | 56434600 | 56436199 | 19.70885 | 33.89905 | 42.76144 | 18.75712 | 0.050652 | 0.900223 | 0.93586 |
| 7 | 35906200 | 35908199 | 15.11472 | 26.69109 | 36.04484 | 15.94711 | -0.04122 | 0.542304 | 0.93589 |
| 6 | 56788200 | 56790799 | 19.9773 | 34.7577 | 44.11517 | 19.38594 | 0.044634 | 0.786398 | 0.935891 |

| | | | | | | | | | |
|---|---|---|---|---|---|---|---|---|---|
| 4 | 61489200 | 61492399 | 94.17728 | 146.5978 | 143.6271 | 59.61988 | 0.338916 | 2.728549 | 0.935895 |
| 18 | 14337000 | 14338599 | 18.90022 | 33.03483 | 41.90423 | 18.46854 | 0.046129 | 0.714353 | 0.935905 |
| 19 | 30495200 | 30496599 | 9.272968 | 17.57071 | 25.45008 | 10.28931 | -0.04364 | 0.031785 | 0.935973 |
| 9 | 116484800 | 116486799 | 16.49937 | 28.955 | 39.16039 | 17.31814 | -0.04609 | 0.630385 | 0.936 |
| 1 | 27263600 | 27264399 | 18.0581 | 31.90758 | 31.64396 | 13.99841 | 0.402121 | 0.545253 | 0.936021 |
| 13 | 107494800 | 107497399 | 21.9533 | 37.77351 | 47.93197 | 21.02539 | 0.042049 | 0.911276 | 0.936027 |
| 9 | 5075800 | 5079399 | 47.55159 | 76.32544 | 99.57962 | 42.12669 | -0.03382 | 1.939204 | 0.936031 |
| 1 | 130252000 | 130252799 | 7.908562 | 15.25957 | 23.1295 | 8.934757 | -0.04799 | -0.13413 | 0.936036 |
| 16 | 12362000 | 12363599 | 18.90541 | 33.28171 | 32.36935 | 14.32987 | 0.428358 | 0.594044 | 0.936072 |
| 17 | 42514000 | 42515599 | 18.55323 | 32.26219 | 41.05935 | 18.03683 | 0.04081 | 0.784534 | 0.936085 |
| 9 | 3609000 | 3609599 | 8.17134 | 15.73214 | 23.62922 | 9.185334 | -0.04392 | -0.11151 | 0.936087 |
| 1 | 49356400 | 49368799 | 319.0933 | 448.5717 | 518.8175 | 214.9509 | -0.03391 | 4.500744 | 0.936112 |
| 1 | 75640600 | 75643199 | 25.34245 | 42.67268 | 57.20499 | 25.00232 | -0.04032 | 1.124782 | 0.936117 |
| 11 | 65786600 | 65788399 | 18.50921 | 32.09051 | 43.17647 | 18.94377 | -0.03975 | 0.822392 | 0.936128 |
| 1 | 85020400 | 85027399 | 144.0556 | 217.7714 | 217.2884 | 89.85394 | 0.279416 | 3.311297 | 0.936139 |
| 11 | 109698400 | 109702199 | 44.0113 | 70.92257 | 88.63162 | 37.71252 | 0.035164 | 1.806005 | 0.936146 |
| 12 | 38093600 | 38094399 | 15.46564 | 27.15795 | 34.27556 | 15.16594 | 0.053548 | 0.609132 | 0.936154 |
| 7 | 104597600 | 104600999 | 50.99256 | 81.79361 | 101.4104 | 42.8574 | 0.038422 | 1.9647 | 0.936174 |
| 2 | 95095000 | 95097199 | 26.65716 | 43.9972 | 72.00827 | 31.08608 | -0.32831 | 1.356406 | 0.936192 |
| X | 107610400 | 107610999 | 9.054121 | 17.42794 | 26.52087 | 10.31585 | -0.06336 | -0.09065 | 0.936196 |
| 11 | 110377600 | 110379399 | 18.89626 | 33.04672 | 41.87872 | 18.46618 | 0.047498 | 0.705054 | 0.936196 |
| 7 | 73310400 | 73314199 | 31.50283 | 51.63633 | 68.75801 | 29.59478 | -0.03546 | 1.450364 | 0.936197 |
| X | 99338200 | 99340199 | 26.72834 | 45.24833 | 45.86849 | 20.08016 | 0.361816 | 1.072871 | 0.936204 |
| 1 | 48017600 | 48019399 | 18.09391 | 31.73671 | 40.21177 | 17.77526 | 0.047716 | 0.651598 | 0.936225 |
| 15 | 37887400 | 37894399 | 113.6685 | 174.4939 | 177.4605 | 73.67687 | 0.269736 | 2.991325 | 0.936277 |
| 2 | 124313800 | 124314999 | 13.95518 | 24.74748 | 33.66208 | 14.87798 | -0.04877 | 0.489117 | 0.936306 |
| 5 | 142530600 | 142533999 | 22.75138 | 38.47413 | 48.5282 | 21.23755 | 0.047539 | 1.068626 | 0.936327 |
| 1 | 71288600 | 71289999 | 8.931325 | 16.56568 | 30.48031 | 12.79237 | -0.43036 | 0.179509 | 0.936341 |
| 5 | 96208200 | 96214399 | 83.36491 | 130.5977 | 158.6854 | 65.82154 | 0.035818 | 2.612784 | 0.936344 |
| 11 | 107590800 | 107592799 | 19.18073 | 33.28125 | 44.72976 | 19.62576 | -0.03798 | 0.820673 | 0.936349 |
| 4 | 7082600 | 7091199 | 124.3476 | 189.9956 | 191.5379 | 79.44902 | 0.277144 | 3.088454 | 0.936362 |
| X | 43618400 | 43619799 | 12.57421 | 23.35525 | 33.7981 | 14.15772 | -0.08455 | 0.209885 | 0.936414 |
| 4 | 34534800 | 34536599 | 19.73238 | 34.28054 | 43.61221 | 19.14701 | 0.040998 | 0.804378 | 0.936417 |
| 8 | 74193800 | 74194799 | 10.70598 | 19.88665 | 26.24635 | 10.99328 | 0.047665 | 0.175179 | 0.936486 |
| 7 | 107879800 | 107880599 | 9.694536 | 18.3245 | 24.84578 | 10.09724 | 0.044186 | 0.050194 | 0.936498 |
| 19 | 13726400 | 13727199 | 9.694536 | 18.3245 | 24.84578 | 10.09724 | 0.044186 | 0.050194 | 0.936498 |
| 1 | 77645600 | 77648799 | 30.34912 | 49.86996 | 63.07231 | 27.17632 | 0.040129 | 1.413539 | 0.936502 |
| 10 | 65431400 | 65433599 | 32.12564 | 53.60617 | 53.04825 | 23.0712 | 0.396195 | 1.23237 | 0.936555 |
| 15 | 81851200 | 81861999 | 128.6389 | 194.8762 | 241.3244 | 99.85979 | -0.02938 | 3.267141 | 0.936569 |
| 3 | 3638400 | 3639199 | 18.75698 | 32.6477 | 41.54579 | 18.26148 | 0.041099 | 0.773231 | 0.936574 |
| 6 | 89965400 | 89972799 | 129.0203 | 195.6583 | 232.5093 | 96.24331 | 0.031309 | 3.246072 | 0.936583 |
| 9 | 32184400 | 32186199 | 18.06704 | 31.73369 | 31.93312 | 14.13235 | 0.378489 | 0.614558 | 0.93662 |
| X | 76120600 | 76122999 | 33.97351 | 56.07503 | 55.14885 | 23.80852 | 0.404422 | 1.364132 | 0.936632 |
| 13 | 54998400 | 54999999 | 16.68517 | 29.23545 | 37.21071 | 16.43426 | 0.04123 | 0.65447 | 0.936635 |
| 14 | 119408000 | 119410199 | 16.68517 | 29.23545 | 37.21071 | 16.43426 | 0.04123 | 0.65447 | 0.936635 |
| 6 | 23177400 | 23179199 | 15.03241 | 26.39573 | 35.81113 | 15.84465 | -0.05019 | 0.610817 | 0.936639 |
| 10 | 24713600 | 24715199 | 24.33783 | 41.60834 | 42.65026 | 18.7008 | 0.347296 | 0.972458 | 0.936647 |

| | | | | | | | | | |
|---|---|---|---|---|---|---|---|---|---|
| 19 | 59415000 | 59418799 | 34.773 | 56.81368 | 71.83408 | 30.88017 | 0.036359 | 1.502572 | 0.936647 |
| 10 | 5912800 | 5914599 | 26.90357 | 44.76623 | 59.86602 | 25.98966 | -0.03675 | 1.25485 | 0.93665 |
| 9 | 89447400 | 89448399 | 13.20673 | 23.93019 | 33.08595 | 14.31979 | -0.05245 | 0.346787 | 0.93666 |
| 5 | 92531800 | 92535599 | 27.18726 | 44.9893 | 60.37037 | 26.10894 | -0.04231 | 1.317908 | 0.93666 |
| 2 | 170042400 | 170045799 | 23.6553 | 39.7477 | 53.32554 | 23.29181 | -0.04153 | 1.132672 | 0.936677 |
| 12 | 7442200 | 7443799 | 15.84865 | 27.94513 | 35.59937 | 15.75476 | 0.041862 | 0.564378 | 0.936746 |
| 8 | 24165600 | 24167599 | 19.95031 | 34.76889 | 34.70594 | 15.26721 | 0.389896 | 0.748436 | 0.93676 |
| 5 | 79723800 | 79729199 | 100.0021 | 153.9968 | 193.1696 | 80.23336 | -0.02918 | 2.927318 | 0.936761 |
| 1 | 73647400 | 73649599 | 41.0611 | 66.58499 | 68.19997 | 29.21189 | 0.331121 | 1.635468 | 0.936767 |
| 18 | 11884200 | 11888199 | 79.26303 | 124.9035 | 127.9886 | 53.10957 | 0.287886 | 2.493732 | 0.936792 |
| 2 | 168131000 | 168133799 | 38.00162 | 61.38064 | 80.88706 | 34.52092 | -0.03729 | 1.726501 | 0.936793 |
| 10 | 7259400 | 7260999 | 15.41272 | 27.11178 | 36.60067 | 16.20561 | -0.04294 | 0.587015 | 0.936801 |
| 12 | 96567200 | 96568599 | 16.48504 | 29.1235 | 37.01287 | 16.37398 | 0.046208 | 0.548889 | 0.936829 |
| 15 | 3948000 | 3950799 | 59.02858 | 94.20576 | 94.79109 | 39.76591 | 0.330911 | 2.126897 | 0.936846 |
| 4 | 69428000 | 69431199 | 25.4433 | 43.44769 | 44.17928 | 19.36942 | 0.358543 | 0.988187 | 0.93687 |
| 2 | 152305600 | 152307799 | 22.23573 | 37.0388 | 63.42182 | 27.55243 | -0.39217 | 1.231852 | 0.936884 |
| 14 | 116470200 | 116470399 | 4.771515 | 9.332806 | 14.9816 | 5.040918 | 0.054516 | -0.6828 | 0.936895 |
| 14 | 93087400 | 93088399 | 18.01903 | 31.29709 | 39.58885 | 17.37823 | 0.049429 | 0.795726 | 0.936924 |
| 8 | 73884200 | 73885199 | 17.09928 | 29.98196 | 38.21712 | 16.88869 | 0.039176 | 0.64863 | 0.936934 |
| 10 | 81413200 | 81415199 | 19.29212 | 33.20048 | 44.82441 | 19.66236 | -0.04677 | 0.897121 | 0.93694 |
| 9 | 62303000 | 62303399 | 6.803331 | 13.23359 | 19.55896 | 7.227022 | 0.048139 | -0.31209 | 0.936991 |
| 18 | 3540400 | 3544999 | 79.99694 | 124.8424 | 151.3438 | 62.80105 | 0.034854 | 2.676809 | 0.936993 |
| 9 | 120147200 | 120148599 | 13.79479 | 24.62113 | 31.51553 | 13.85382 | 0.043552 | 0.44062 | 0.93701 |
| 1 | 7650600 | 7652399 | 20.66379 | 35.6537 | 47.93548 | 21.02871 | -0.04021 | 0.881216 | 0.937015 |
| 10 | 83831000 | 83832999 | 15.82706 | 27.71308 | 37.75517 | 16.66516 | -0.05651 | 0.659875 | 0.937033 |
| 6 | 67552200 | 67553599 | 16.67758 | 29.14858 | 39.36187 | 17.34733 | -0.04375 | 0.697982 | 0.937034 |
| 2 | 43333000 | 43334399 | 17.38117 | 30.98439 | 30.18514 | 13.28263 | 0.434237 | 0.470095 | 0.937035 |
| 9 | 20863800 | 20865999 | 19.07075 | 32.89011 | 44.27541 | 19.42281 | -0.04199 | 0.874963 | 0.937048 |
| 17 | 30614600 | 30618199 | 36.1652 | 58.6441 | 77.42691 | 33.16334 | -0.03434 | 1.630713 | 0.937055 |
| 8 | 95250400 | 95251999 | 12.55757 | 22.81564 | 29.51777 | 12.73227 | 0.04612 | 0.31979 | 0.937073 |
| 2 | 27709200 | 27712599 | 32.6636 | 53.45158 | 71.08165 | 30.57431 | -0.03464 | 1.475293 | 0.937074 |
| 17 | 53295400 | 53296199 | 8.222967 | 15.77359 | 23.5449 | 9.249415 | -0.04882 | -0.07518 | 0.937092 |
| 6 | 38923800 | 38926599 | 24.48623 | 41.0824 | 54.96714 | 23.98823 | -0.03763 | 1.152613 | 0.937138 |
| 10 | 112518600 | 112521999 | 39.80839 | 64.2415 | 80.27552 | 34.22045 | 0.037731 | 1.755645 | 0.937148 |
| 13 | 49909000 | 49910399 | 11.87488 | 21.50763 | 29.6564 | 12.8424 | -0.04861 | 0.335033 | 0.93715 |
| 10 | 47096000 | 47098399 | 30.57768 | 50.85929 | 50.26004 | 21.81334 | 0.398284 | 1.261949 | 0.937163 |
| 1 | 124969200 | 124970799 | 17.8198 | 31.28416 | 39.60629 | 17.52097 | 0.048746 | 0.63297 | 0.937165 |
| 4 | 57932200 | 57934399 | 25.10289 | 42.28977 | 56.68298 | 24.7778 | -0.04011 | 1.117954 | 0.937171 |
| 17 | 77231600 | 77232799 | 18.13819 | 31.95312 | 32.32399 | 14.30781 | 0.372142 | 0.580485 | 0.937172 |
| X | 19884600 | 19885399 | 10.02031 | 19.16555 | 26.24453 | 10.39632 | 0.062984 | -0.02293 | 0.937177 |
| 14 | 66997000 | 66999999 | 31.78286 | 52.41758 | 70.43585 | 30.39616 | -0.04506 | 1.375961 | 0.937178 |
| 4 | 105367400 | 105368999 | 28.4556 | 46.89313 | 62.93299 | 27.14929 | -0.04353 | 1.374969 | 0.937188 |
| 2 | 159541600 | 159542799 | 17.68506 | 31.50367 | 39.09642 | 17.21426 | 0.086198 | 0.492347 | 0.937194 |
| 4 | 146431800 | 146447999 | 932.9722 | 1212.235 | 1227.902 | 514.4158 | 0.043344 | 5.744973 | 0.937221 |
| 6 | 104680800 | 104682999 | 41.3968 | 66.71373 | 87.49868 | 37.23395 | -0.03426 | 1.799435 | 0.937242 |
| 1 | 101598400 | 101599999 | 28.87326 | 47.66041 | 60.34941 | 26.05503 | 0.040906 | 1.352688 | 0.937247 |
| 1 | 55006800 | 55009399 | 29.45066 | 48.69185 | 65.09095 | 28.13338 | -0.03702 | 1.33703 | 0.937247 |

| | | | | | | | | | |
|---|---|---|---|---|---|---|---|---|---|
| 1 | 110355200 | 110358799 | 45.449 | 73.06483 | 91.06476 | 38.61405 | 0.034796 | 1.885712 | 0.937273 |
| 9 | 63771800 | 63779199 | 96.71518 | 149.3852 | 180.3244 | 74.90298 | 0.029506 | 2.868291 | 0.93731 |
| 7 | 90067600 | 90069399 | 26.69687 | 44.55363 | 56.61381 | 24.62404 | 0.036712 | 1.219435 | 0.937319 |
| 2 | 89741400 | 89751999 | 132.6352 | 201.1521 | 238.821 | 98.85769 | 0.032678 | 3.244626 | 0.93732 |
| 6 | 51269600 | 51271399 | 16.23175 | 28.63536 | 36.49018 | 16.14733 | 0.041487 | 0.560454 | 0.937336 |
| 14 | 51987400 | 51992399 | 70.20935 | 110.5364 | 141.1984 | 58.58415 | -0.03071 | 2.507303 | 0.93734 |
| 6 | 144916200 | 144920999 | 46.47982 | 74.63129 | 97.36997 | 41.21388 | -0.03324 | 1.924749 | 0.93735 |
| 11 | 23173400 | 23177799 | 67.28182 | 106.2577 | 136.1262 | 56.52676 | -0.03136 | 2.439794 | 0.937353 |
| 15 | 63053800 | 63055599 | 17.25317 | 30.32584 | 41.19535 | 18.24247 | -0.05239 | 0.614108 | 0.937355 |
| 14 | 44549000 | 44550399 | 13.46855 | 24.02015 | 30.57512 | 13.44924 | 0.050954 | 0.441348 | 0.937363 |
| 15 | 33814600 | 33818399 | 54.37029 | 86.85047 | 107.662 | 45.22368 | 0.031861 | 2.096297 | 0.937374 |
| 12 | 87946200 | 87946999 | 11.64427 | 20.4715 | 39.466 | 17.47623 | -0.55413 | 0.579372 | 0.937388 |
| X | 148979400 | 148982399 | 21.76385 | 37.88972 | 37.74124 | 16.59149 | 0.392899 | 0.778393 | 0.937397 |
| 2 | 47210800 | 47211999 | 19.49658 | 34.12876 | 33.62961 | 14.84061 | 0.408904 | 0.681311 | 0.93741 |
| 1 | 117366200 | 117367999 | 18.62214 | 32.59573 | 41.28395 | 18.21734 | 0.048186 | 0.687049 | 0.937416 |
| 5 | 149789200 | 149791599 | 17.85787 | 31.22299 | 39.82697 | 17.5578 | 0.038133 | 0.700007 | 0.937422 |
| 9 | 101173000 | 101176599 | 35.49245 | 57.57366 | 72.16658 | 30.91835 | 0.040915 | 1.619469 | 0.937436 |
| 7 | 135030000 | 135032999 | 30.88669 | 50.94059 | 64.61164 | 27.88299 | 0.038011 | 1.368456 | 0.937458 |
| X | 43096800 | 43097599 | 13.68247 | 24.77592 | 31.66243 | 13.71497 | 0.06035 | 0.357122 | 0.937474 |
| 18 | 20937400 | 20938999 | 15.13912 | 26.93286 | 34.45398 | 15.18706 | 0.041642 | 0.47587 | 0.937475 |
| 14 | 53623000 | 53623799 | 24.72622 | 41.27832 | 55.44829 | 24.12206 | -0.04316 | 1.208438 | 0.937541 |
| 3 | 69527600 | 69534399 | 88.68221 | 137.4217 | 165.9245 | 68.92037 | 0.032735 | 2.804528 | 0.937546 |
| 12 | 3464600 | 3466399 | 20.69216 | 35.599 | 45.30118 | 19.87131 | 0.038045 | 0.905148 | 0.937552 |
| 7 | 63440600 | 63441599 | 11.04108 | 20.51833 | 28.83349 | 12.06953 | -0.0418 | 0.181203 | 0.937575 |
| 1 | 147241200 | 147246399 | 89.48849 | 139.4608 | 168.9072 | 70.10159 | 0.034631 | 2.708642 | 0.937584 |
| 6 | 37604000 | 37606999 | 21.09054 | 36.42617 | 46.3148 | 20.31835 | 0.040274 | 0.871529 | 0.937588 |
| 12 | 114142400 | 114145399 | 30.66623 | 51.18995 | 52.44226 | 22.813 | 0.346397 | 1.223938 | 0.937591 |
| 18 | 66610800 | 66614399 | 25.41676 | 42.71003 | 57.24287 | 25.0033 | -0.04018 | 1.140883 | 0.937608 |
| 6 | 103204600 | 103208199 | 28.34453 | 47.5787 | 59.66184 | 26.04497 | 0.05549 | 1.162883 | 0.937613 |
| 9 | 68059200 | 68060199 | 11.57216 | 21.25113 | 27.71946 | 11.7924 | 0.048411 | 0.25075 | 0.937675 |
| 2 | 23300400 | 23300799 | 7.54747 | 14.70597 | 16.83139 | 6.115452 | 0.435653 | -0.36989 | 0.937686 |
| 11 | 13237400 | 13238199 | 9.749202 | 17.72342 | 33.35509 | 14.37706 | -0.49023 | 0.305256 | 0.937703 |
| X | 130681000 | 130683199 | 25.96412 | 45.05567 | 39.19791 | 17.19864 | 0.586914 | 0.844598 | 0.937743 |
| 9 | 86569000 | 86569999 | 9.10654 | 17.41096 | 25.65806 | 10.17548 | -0.04397 | -0.03201 | 0.937745 |
| 2 | 4916000 | 4919399 | 39.95388 | 64.49216 | 84.65461 | 36.09866 | -0.03275 | 1.74993 | 0.937752 |
| 3 | 138100000 | 138100799 | 11.11492 | 20.55402 | 28.80543 | 12.13889 | -0.04465 | 0.210782 | 0.937755 |
| 13 | 22782400 | 22783799 | 17.99171 | 31.44404 | 42.4365 | 18.7017 | -0.04298 | 0.710156 | 0.937776 |
| 2 | 102217600 | 102221199 | 37.38332 | 61.56825 | 75.66805 | 32.63485 | 0.082565 | 1.412125 | 0.937788 |
| 15 | 50486800 | 50491199 | 84.46885 | 131.3267 | 158.731 | 65.90145 | 0.034943 | 2.745779 | 0.937792 |
| 12 | 28477600 | 28481999 | 29.52421 | 48.10827 | 79.36079 | 34.0823 | -0.34905 | 1.535294 | 0.937793 |
| 10 | 116508800 | 116511399 | 27.70993 | 46.76268 | 47.46134 | 20.75765 | 0.359797 | 1.102153 | 0.937794 |
| 13 | 11392400 | 11396599 | 53.87111 | 86.32257 | 106.3811 | 44.88821 | 0.045375 | 2.002185 | 0.937818 |
| 14 | 9566600 | 9567799 | 11.36021 | 21.01264 | 27.7243 | 11.67949 | 0.041993 | 0.207761 | 0.937838 |
| 17 | 89918400 | 89919399 | 16.13454 | 28.35488 | 38.1565 | 16.89439 | -0.0387 | 0.606471 | 0.937838 |
| 19 | 5056200 | 5058999 | 25.16123 | 42.40442 | 53.90227 | 23.56503 | 0.036201 | 1.113778 | 0.937864 |
| 13 | 24531800 | 24534799 | 31.26189 | 52.17291 | 64.93126 | 28.24198 | 0.066487 | 1.238079 | 0.937886 |
| 12 | 109348800 | 109349399 | 6.691615 | 13.00019 | 18.5016 | 6.904962 | 0.090235 | -0.27808 | 0.937891 |

| | | | | | | | | | |
|---|---|---|---|---|---|---|---|---|---|
| X | 92675400 | 92676199 | 11.12406 | 20.7503 | 29.46238 | 12.25606 | -0.04927 | 0.161169 | 0.937907 |
| 3 | 123384600 | 123385399 | 9.748352 | 18.57974 | 25.56977 | 10.21071 | 0.045188 | -0.00329 | 0.937916 |
| 12 | 64863400 | 64865199 | 14.13271 | 25.10214 | 33.99246 | 15.0039 | -0.04139 | 0.47756 | 0.937925 |
| 1 | 11931200 | 11932399 | 17.87227 | 31.20002 | 41.9674 | 18.48116 | -0.03825 | 0.726018 | 0.937947 |
| 2 | 89674600 | 89675199 | 8.948698 | 17.08073 | 25.05309 | 9.973346 | -0.04205 | -0.02274 | 0.937963 |
| 10 | 4577000 | 4577599 | 8.948698 | 17.08073 | 25.05309 | 9.973346 | -0.04205 | -0.02274 | 0.937963 |
| 14 | 106354400 | 106356199 | 27.33128 | 44.92859 | 73.5438 | 31.69347 | -0.33077 | 1.408516 | 0.937969 |
| 12 | 114866400 | 114910199 | 730.754 | 968.9229 | 1000.591 | 418.2474 | 0.044629 | 5.417 | 0.937975 |
| 1 | 46095200 | 46096999 | 15.33678 | 27.46531 | 34.91873 | 15.30645 | 0.05578 | 0.434884 | 0.937975 |
| 5 | 71712800 | 71714399 | 23.89356 | 39.63965 | 66.17056 | 28.66773 | -0.35599 | 1.286006 | 0.937978 |
| 1 | 137083400 | 137087799 | 54.53027 | 86.90341 | 112.4317 | 47.07346 | -0.03333 | 2.17007 | 0.938004 |
| 15 | 91357400 | 91358399 | 11.39887 | 21.36184 | 28.43636 | 11.73662 | 0.051721 | 0.128288 | 0.938031 |
| X | 69709000 | 69710599 | 15.15414 | 27.28598 | 34.60222 | 15.0948 | 0.064589 | 0.40457 | 0.938057 |
| 2 | 169169600 | 169172399 | 25.27584 | 42.25552 | 56.64554 | 24.66488 | -0.04037 | 1.193811 | 0.938062 |
| 19 | 6686400 | 6689399 | 20.79656 | 35.75802 | 47.93711 | 21.02719 | -0.03711 | 0.912328 | 0.938082 |
| 6 | 82163400 | 82170599 | 75.08493 | 118.3944 | 144.6815 | 60.04131 | 0.03482 | 2.480295 | 0.93809 |
| 5 | 28743400 | 28745599 | 30.82919 | 50.66209 | 64.268 | 27.69227 | 0.035798 | 1.414445 | 0.938118 |
| 8 | 19677000 | 19693599 | 337.8206 | 474.0596 | 545.5359 | 226.0827 | -0.0292 | 4.527795 | 0.938138 |
| 16 | 66037400 | 66039399 | 19.03321 | 33.36463 | 33.93487 | 14.99316 | 0.363179 | 0.654597 | 0.938147 |
| 3 | 144032400 | 144037399 | 42.69213 | 68.82707 | 90.41685 | 38.49443 | -0.03599 | 1.791401 | 0.938177 |
| 16 | 19277400 | 19278999 | 23.29932 | 40.47301 | 39.24213 | 17.22727 | 0.431246 | 0.817254 | 0.938195 |
| 18 | 18898000 | 18899999 | 48.38942 | 77.56616 | 96.44079 | 40.7157 | 0.033252 | 1.981248 | 0.938201 |
| 6 | 37575000 | 37577199 | 26.47218 | 44.59635 | 56.22971 | 24.57963 | 0.04773 | 1.12065 | 0.938226 |
| 8 | 60388000 | 60389199 | 16.34762 | 28.72241 | 38.69818 | 17.13073 | -0.04039 | 0.61337 | 0.938247 |
| 13 | 60824000 | 60826399 | 20.69141 | 35.37778 | 47.43263 | 20.79789 | -0.03851 | 0.963107 | 0.938258 |
| 2 | 95037000 | 95038599 | 23.64557 | 39.90748 | 53.33711 | 23.32814 | -0.03583 | 1.094651 | 0.938266 |
| 16 | 13254800 | 13256399 | 11.35848 | 20.56234 | 28.67035 | 12.42294 | -0.06523 | 0.330496 | 0.938269 |
| 1 | 185755600 | 185757799 | 39.1267 | 63.22739 | 83.09642 | 35.4852 | -0.03266 | 1.71544 | 0.938275 |
| 5 | 93439600 | 93441399 | 41.82366 | 67.43392 | 84.35782 | 35.91963 | 0.034665 | 1.787228 | 0.938297 |
| 6 | 124498600 | 124500599 | 22.5622 | 37.98335 | 51.15914 | 22.35849 | -0.04697 | 1.111614 | 0.938301 |
| 3 | 73982800 | 73983799 | 11.5262 | 21.17063 | 27.68765 | 11.77593 | 0.044737 | 0.248316 | 0.938304 |
| 7 | 42041400 | 42049999 | 362.3573 | 503.9435 | 547.947 | 227.3994 | 0.03956 | 4.663022 | 0.938305 |
| 4 | 60434400 | 60436399 | 80.84793 | 126.23 | 153.5465 | 63.71198 | 0.030492 | 2.670029 | 0.938323 |
| Y | 39984800 | 39986799 | 24.79767 | 41.17082 | 49.66304 | 21.52729 | 0.11132 | 1.261765 | 0.938347 |
| 5 | 130656200 | 130658799 | 26.09331 | 44.23961 | 55.63806 | 24.36814 | 0.051786 | 1.066376 | 0.938349 |
| 4 | 153124200 | 153127999 | 39.98225 | 64.37584 | 84.81959 | 36.04663 | -0.04241 | 1.825688 | 0.938374 |
| 8 | 61887200 | 61888799 | 21.28875 | 36.54961 | 49.01219 | 21.49762 | -0.03796 | 0.928939 | 0.938376 |
| 10 | 105565800 | 105568399 | 22.23497 | 38.34662 | 48.47485 | 21.26502 | 0.048129 | 0.891759 | 0.938378 |
| 9 | 124067800 | 124070199 | 16.89876 | 29.59371 | 39.80187 | 17.57063 | -0.03794 | 0.66891 | 0.938435 |
| 1 | 5637200 | 5638399 | 15.6796 | 27.81274 | 35.46357 | 15.67076 | 0.04392 | 0.500146 | 0.938444 |
| 7 | 120050600 | 120051799 | 12.94471 | 23.60576 | 32.76857 | 14.06956 | -0.05029 | 0.311911 | 0.93845 |
| 1 | 180422000 | 180424399 | 18.01708 | 31.29946 | 42.11502 | 18.48928 | -0.03938 | 0.795994 | 0.938488 |
| 3 | 6253400 | 6254799 | 16.3495 | 28.96068 | 27.338 | 12.08755 | 0.474297 | 0.49899 | 0.938498 |
| 18 | 11291000 | 11296599 | 139.9206 | 210.4117 | 259.1598 | 107.0048 | -0.03078 | 3.416759 | 0.938498 |
| 10 | 8539000 | 8541399 | 22.23689 | 38.06942 | 51.19278 | 22.44895 | -0.04266 | 0.959902 | 0.938512 |
| 7 | 103452800 | 103454399 | 17.34952 | 30.46615 | 38.68104 | 17.11527 | 0.044886 | 0.62815 | 0.93855 |
| 1 | 35243400 | 35246599 | 45.50555 | 73.11965 | 91.25445 | 38.66682 | 0.031972 | 1.899333 | 0.938561 |

| | | | | | | | | | |
|---|---|---|---|---|---|---|---|---|---|
| 4 | 120949800 | 120950999 | 11.68375 | 21.23646 | 27.21319 | 11.73236 | 0.060708 | 0.307834 | 0.938572 |
| 6 | 115336400 | 115339199 | 20.00776 | 34.98771 | 35.51969 | 15.66112 | 0.365917 | 0.701147 | 0.938579 |
| 1 | 99716800 | 99719599 | 21.59058 | 36.96531 | 49.55456 | 21.73067 | -0.03816 | 0.955179 | 0.938589 |
| 4 | 6255800 | 6256399 | 6.983959 | 13.56279 | 21.15141 | 7.91907 | -0.04569 | -0.25641 | 0.93861 |
| 13 | 72169000 | 72170399 | 15.88995 | 28.57127 | 36.04468 | 15.74621 | 0.070295 | 0.416514 | 0.938617 |
| 15 | 46939800 | 46940399 | 7.599186 | 14.73447 | 22.79101 | 8.609109 | -0.04631 | -0.21406 | 0.93862 |
| 1 | 79665000 | 79666999 | 17.74494 | 31.61208 | 39.06434 | 17.19934 | 0.092256 | 0.490941 | 0.938665 |
| 12 | 81469600 | 81479999 | 392.0799 | 542.3608 | 588.86 | 244.5845 | 0.033847 | 4.745408 | 0.938668 |
| X | 79600000 | 79602599 | 28.07702 | 47.82487 | 45.23446 | 19.82803 | 0.462133 | 1.023766 | 0.938681 |
| 16 | 59190400 | 59193399 | 47.16371 | 75.59996 | 98.35288 | 41.52176 | -0.03205 | 1.979369 | 0.938693 |
| 2 | 172376200 | 172380399 | 47.33765 | 75.98298 | 94.1629 | 39.83593 | 0.04023 | 1.933289 | 0.938704 |
| 7 | 47897800 | 47901599 | 45.40216 | 72.91223 | 95.01927 | 40.22839 | -0.03132 | 1.919069 | 0.938708 |
| 3 | 17731800 | 17732999 | 16.23977 | 28.6231 | 36.51594 | 16.16171 | 0.039366 | 0.570764 | 0.938755 |
| 13 | 44395000 | 44397599 | 19.76809 | 34.23361 | 43.66672 | 19.1582 | 0.036501 | 0.838505 | 0.938757 |
| 19 | 16450600 | 16453399 | 31.35441 | 51.79499 | 65.68226 | 28.37217 | 0.038787 | 1.354401 | 0.938767 |
| 8 | 21167200 | 21167799 | 11.93626 | 21.82985 | 30.22814 | 12.93178 | -0.04319 | 0.282413 | 0.938792 |
| Y | 83779200 | 83779599 | 6.374193 | 12.39476 | 17.96554 | 6.65595 | 0.073176 | -0.31242 | 0.938795 |
| 14 | 42227000 | 42229599 | 51.31824 | 81.82574 | 106.4214 | 44.58813 | -0.0401 | 2.150307 | 0.938806 |
| 16 | 79111800 | 79112399 | 9.406274 | 17.66832 | 25.3284 | 10.41316 | -0.04944 | 0.089898 | 0.938819 |
| 1 | 85838600 | 85840399 | 16.9094 | 29.71409 | 40.16209 | 17.78111 | -0.04515 | 0.614437 | 0.938873 |
| 6 | 62901400 | 62903199 | 15.21441 | 26.95156 | 34.3965 | 15.2083 | 0.041574 | 0.504965 | 0.938892 |
| 12 | 62444400 | 62445999 | 17.25949 | 30.1575 | 38.39841 | 16.91908 | 0.040872 | 0.706196 | 0.938908 |
| 16 | 76504200 | 76506199 | 21.50274 | 37.47786 | 46.76801 | 20.57452 | 0.069182 | 0.772586 | 0.938925 |
| 16 | 28462600 | 28463399 | 10.20573 | 18.97095 | 26.76469 | 11.19951 | -0.04705 | 0.16716 | 0.93895 |
| 6 | 56983400 | 56984599 | 11.20311 | 20.85378 | 29.39856 | 12.27322 | -0.04339 | 0.17353 | 0.938968 |
| 3 | 66950200 | 66953199 | 34.75664 | 57.81833 | 54.20416 | 23.52745 | 0.47406 | 1.275698 | 0.938976 |
| X | 25459600 | 25460599 | 12.33365 | 22.8769 | 32.62726 | 13.69292 | -0.06567 | 0.212361 | 0.938979 |
| 5 | 133320200 | 133327799 | 126.5478 | 191.5445 | 237.2338 | 98.14062 | -0.03064 | 3.282565 | 0.938983 |
| 18 | 11938200 | 11940199 | 19.28868 | 33.32067 | 44.7478 | 19.63102 | -0.03821 | 0.85961 | 0.938984 |
| 18 | 28761800 | 28763599 | 15.18462 | 26.38955 | 47.38541 | 20.80836 | -0.45345 | 0.788347 | 0.939 |
| 19 | 7562600 | 7568399 | 60.21492 | 95.72319 | 123.1973 | 51.41006 | -0.02964 | 2.258107 | 0.939001 |
| 19 | 18432400 | 18436199 | 47.56754 | 76.35864 | 99.57188 | 42.1298 | -0.03291 | 1.936422 | 0.939002 |
| 14 | 53620400 | 53623799 | 45.76154 | 73.67529 | 91.93015 | 39.06398 | 0.035756 | 1.840269 | 0.939017 |
| 1 | 194648800 | 194653399 | 73.65842 | 115.8471 | 141.8486 | 58.84602 | 0.029138 | 2.529758 | 0.939039 |
| 17 | 22411800 | 22435799 | 320.7666 | 457.6542 | 503.0145 | 207.8924 | 0.061116 | 4.286488 | 0.939068 |
| 17 | 94709200 | 94710599 | 17.32326 | 30.3541 | 38.6895 | 17.08765 | 0.039121 | 0.659625 | 0.939079 |
| 11 | 113809400 | 113813199 | 38.95702 | 62.87401 | 82.68103 | 35.25054 | -0.0356 | 1.751227 | 0.939088 |
| 11 | 70022000 | 70023599 | 16.09096 | 28.29136 | 36.09472 | 15.98311 | 0.037988 | 0.596362 | 0.939112 |
| X | 147554800 | 147555399 | 13.84362 | 25.24987 | 25.44797 | 10.92374 | 0.408929 | 0.300015 | 0.939113 |
| 12 | 17992600 | 17995399 | 40.05869 | 64.74019 | 85.29355 | 36.42834 | -0.03606 | 1.713873 | 0.939124 |
| 15 | 35356400 | 35358799 | 20.82199 | 35.98097 | 48.47764 | 21.26752 | -0.04282 | 0.86893 | 0.939166 |
| 4 | 99879600 | 99880399 | 11.61873 | 20.58273 | 37.33641 | 16.50806 | -0.46231 | 0.491778 | 0.939178 |
| 15 | 17691200 | 17694999 | 46.9114 | 75.29824 | 98.09057 | 41.49707 | -0.03162 | 1.938079 | 0.93918 |
| 1 | 177632800 | 177633799 | 10.7825 | 20.09076 | 28.37302 | 11.82518 | -0.04368 | 0.162829 | 0.939189 |
| 19 | 60333400 | 60335199 | 14.31936 | 25.65171 | 32.99048 | 14.45721 | 0.03982 | 0.424717 | 0.939193 |
| 10 | 43501600 | 43504399 | 22.59969 | 38.51103 | 49.02225 | 21.48908 | 0.035256 | 1.001903 | 0.939204 |
| 1 | 123230200 | 123230599 | 7.095341 | 13.77095 | 20.00835 | 7.523656 | 0.049292 | -0.2455 | 0.939273 |

| | | | | | | | | | |
|---|---|---|---|---|---|---|---|---|---|
| 6 | 108836600 | 108840399 | 38.2279 | 61.99845 | 82.18303 | 35.20448 | -0.03994 | 1.632367 | 0.939306 |
| 9 | 87375000 | 87376599 | 26.81002 | 44.59932 | 56.6208 | 24.57653 | 0.037976 | 1.253119 | 0.939308 |
| 15 | 14618600 | 14619999 | 14.76816 | 26.54234 | 36.63465 | 16.00997 | -0.05904 | 0.412905 | 0.93933 |
| 17 | 44551000 | 44552199 | 13.98494 | 25.26528 | 32.44965 | 14.09656 | 0.049805 | 0.367957 | 0.939333 |
| 16 | 34399800 | 34403999 | 52.67418 | 84.18581 | 104.4539 | 43.90911 | 0.031417 | 2.079165 | 0.939334 |
| 6 | 131373600 | 131375799 | 20.13873 | 34.57288 | 46.35786 | 20.33328 | -0.03781 | 0.921651 | 0.939371 |
| 16 | 23781200 | 23783799 | 19.81427 | 33.67662 | 55.94666 | 24.51927 | -0.34801 | 1.02336 | 0.939396 |
| 2 | 170835000 | 170836999 | 25.87327 | 43.50197 | 58.30873 | 25.47327 | -0.0402 | 1.140862 | 0.9394 |
| 4 | 101849600 | 101850999 | 24.69162 | 42.19472 | 43.16985 | 18.92796 | 0.349923 | 0.978685 | 0.939402 |
| 5 | 87027600 | 87030999 | 45.76869 | 73.58052 | 96.14653 | 40.77027 | -0.03325 | 1.886782 | 0.939404 |
| 4 | 98753000 | 98757199 | 57.80784 | 91.99308 | 118.7246 | 49.6084 | -0.03206 | 2.221068 | 0.939415 |
| 3 | 80357600 | 80360199 | 19.726 | 34.51469 | 43.43113 | 19.15862 | 0.057587 | 0.703524 | 0.939435 |
| 11 | 107197600 | 107199199 | 19.12959 | 33.36945 | 42.46003 | 18.68912 | 0.041361 | 0.743097 | 0.939439 |
| 1 | 18163000 | 18170599 | 167.1899 | 250.3919 | 249.514 | 102.9182 | 0.269443 | 3.498713 | 0.939449 |
| 17 | 12467400 | 12478399 | 170.2335 | 250.3612 | 367.5434 | 151.3603 | -0.31342 | 3.818469 | 0.939492 |
| 13 | 55581000 | 55583799 | 59.66313 | 94.75562 | 116.6618 | 48.63636 | 0.032921 | 2.283272 | 0.939505 |
| 10 | 92875000 | 92876999 | 17.06006 | 29.90583 | 40.26381 | 17.78946 | -0.03949 | 0.654396 | 0.939531 |
| 1 | 108566200 | 108567399 | 12.30004 | 22.52491 | 31.21094 | 13.32822 | -0.04264 | 0.277313 | 0.939534 |
| 5 | 133901800 | 133903399 | 17.91805 | 31.25799 | 42.00592 | 18.48996 | -0.03694 | 0.737098 | 0.939541 |
| 6 | 142139600 | 142140799 | 16.74434 | 29.27816 | 39.37769 | 17.36077 | -0.03787 | 0.692629 | 0.939544 |
| 13 | 57432400 | 57434799 | 21.6526 | 37.23649 | 47.39958 | 20.79152 | 0.037405 | 0.91461 | 0.939563 |
| 5 | 27734800 | 27735999 | 14.21354 | 25.38028 | 32.55234 | 14.30399 | 0.041204 | 0.444375 | 0.939581 |
| 1 | 141007000 | 141008999 | 15.03873 | 27.12941 | 27.36867 | 11.91106 | 0.394829 | 0.376168 | 0.939629 |
| 5 | 148434000 | 148439599 | 58.4579 | 93.00001 | 120.0165 | 50.12765 | -0.03243 | 2.231335 | 0.939675 |
| 5 | 26765400 | 26770399 | 49.70333 | 79.68363 | 103.8024 | 43.83313 | -0.03368 | 1.980428 | 0.939677 |
| 8 | 69995800 | 69997799 | 37.44603 | 60.68045 | 76.39698 | 32.70608 | 0.03316 | 1.645608 | 0.939716 |
| 2 | 79676000 | 79678399 | 27.20336 | 45.66758 | 57.79425 | 25.23123 | 0.04238 | 1.159681 | 0.939739 |
| 1 | 58928800 | 58932399 | 33.6271 | 54.81494 | 69.16161 | 29.70635 | 0.037065 | 1.531157 | 0.939742 |
| 4 | 99585600 | 99587999 | 18.3726 | 32.13308 | 43.47631 | 19.17172 | -0.0466 | 0.702682 | 0.93975 |
| 11 | 11863400 | 11865399 | 18.06067 | 31.62448 | 40.28108 | 17.78028 | 0.040179 | 0.67838 | 0.939788 |
| 2 | 51872400 | 51875799 | 30.94553 | 50.87762 | 67.84175 | 29.23807 | -0.03574 | 1.409794 | 0.939802 |
| 14 | 50545400 | 50546399 | 16.45517 | 28.85193 | 36.75893 | 16.24399 | 0.039951 | 0.643459 | 0.939817 |
| 9 | 9526200 | 9526799 | 9.227031 | 17.49791 | 23.85446 | 9.627609 | 0.045258 | 0.020879 | 0.93984 |
| 15 | 48914000 | 48914999 | 12.80785 | 23.491 | 30.69312 | 13.07967 | 0.04381 | 0.272188 | 0.939847 |
| 6 | 24289000 | 24290399 | 13.05993 | 23.60284 | 30.47939 | 13.23456 | 0.042613 | 0.356127 | 0.939855 |
| 6 | 45714200 | 45715199 | 9.268241 | 17.74393 | 26.29228 | 10.38687 | -0.04666 | -0.04137 | 0.939875 |
| X | 109777800 | 109780199 | 42.95858 | 69.58443 | 68.70133 | 29.40249 | 0.382537 | 1.662981 | 0.939891 |
| 17 | 58540400 | 58540799 | 8.902634 | 17.15414 | 24.24302 | 9.401783 | 0.045999 | -0.11095 | 0.939897 |
| 8 | 98703200 | 98704599 | 14.82111 | 26.28052 | 35.48999 | 15.68717 | -0.03882 | 0.496007 | 0.939927 |
| 2 | 23353600 | 23354399 | 11.43509 | 20.90624 | 29.10954 | 12.4586 | -0.05198 | 0.274016 | 0.939935 |
| 7 | 6263200 | 6270999 | 77.19446 | 121.052 | 147.9387 | 61.3576 | 0.028621 | 2.589799 | 0.939945 |
| 9 | 81271600 | 81273199 | 20.60771 | 35.38603 | 47.40631 | 20.79351 | -0.03654 | 0.922388 | 0.939952 |
| 5 | 106303600 | 106305999 | 28.37827 | 46.8861 | 62.58825 | 27.03865 | -0.03498 | 1.342074 | 0.939955 |
| 7 | 83553800 | 83558599 | 56.18136 | 89.51908 | 110.781 | 46.37111 | 0.030415 | 2.177208 | 0.939963 |
| 4 | 97179600 | 97180399 | 14.24411 | 25.61368 | 32.95171 | 14.39421 | 0.042388 | 0.399327 | 0.939995 |
| 3 | 16472800 | 16474399 | 25.17742 | 42.37365 | 56.63855 | 24.751 | -0.03614 | 1.12803 | 0.940001 |
| 6 | 140310000 | 140313399 | 28.45326 | 47.14872 | 59.97047 | 25.9605 | 0.0348 | 1.305021 | 0.940016 |

| | | | | | | | | | |
|---|---|---|---|---|---|---|---|---|---|
| 11 | 93690800 | 93693799 | 31.03018 | 51.29651 | 65.09362 | 28.13196 | 0.037912 | 1.345006 | 0.940024 |
| 7 | 85092000 | 85114599 | 516.2538 | 699.7106 | 746.9836 | 311.2558 | 0.028482 | 5.065441 | 0.940048 |
| X | 41261800 | 41263399 | 16.78745 | 29.80179 | 37.52722 | 16.57143 | 0.062592 | 0.507194 | 0.940069 |
| 18 | 35206800 | 35207599 | 9.674586 | 18.46833 | 27.32068 | 10.87346 | -0.05386 | -0.00908 | 0.940071 |
| 14 | 92188000 | 92192999 | 68.23085 | 108.2142 | 132.5698 | 55.19265 | 0.038161 | 2.33519 | 0.940073 |
| 4 | 137783400 | 137786199 | 24.52198 | 41.04296 | 54.94121 | 23.94034 | -0.03825 | 1.179083 | 0.940084 |
| 8 | 31111000 | 31112799 | 27.49631 | 46.22717 | 47.50588 | 20.75334 | 0.341786 | 1.135483 | 0.940113 |
| 5 | 145482000 | 145495399 | 341.156 | 482.6184 | 453.1614 | 187.5554 | 0.275877 | 4.408808 | 0.940114 |
| 16 | 81893000 | 81893999 | 13.35547 | 24.40548 | 31.69349 | 13.57264 | 0.04762 | 0.299779 | 0.940127 |
| 6 | 146268600 | 146268999 | 7.752433 | 14.93673 | 22.55468 | 8.748919 | -0.04806 | -0.12397 | 0.940141 |
| 9 | 11126400 | 11127999 | 16.28921 | 28.44282 | 38.4801 | 16.94785 | -0.04631 | 0.710673 | 0.940142 |
| 6 | 142617200 | 142621399 | 36.81317 | 59.65388 | 78.61387 | 33.65471 | -0.03259 | 1.646079 | 0.940145 |
| 2 | 38451000 | 38451999 | 12.23242 | 21.96893 | 30.18126 | 13.19784 | -0.05248 | 0.388996 | 0.940149 |
| 13 | 36716800 | 36718199 | 15.43015 | 27.32379 | 34.93922 | 15.44939 | 0.038663 | 0.512234 | 0.940173 |
| 1 | 53726200 | 53727599 | 15.10635 | 26.70132 | 36.01732 | 15.93195 | -0.03907 | 0.531797 | 0.940178 |
| 3 | 151458000 | 151458999 | 17.52818 | 30.74203 | 39.1686 | 17.31299 | 0.039689 | 0.647044 | 0.940181 |
| 3 | 148332200 | 148334399 | 22.72013 | 38.65088 | 51.69175 | 22.65618 | -0.03607 | 1.022325 | 0.940217 |
| 3 | 78330800 | 78335199 | 52.80896 | 84.53041 | 110.1702 | 46.41249 | -0.03705 | 2.033708 | 0.940231 |
| X | 130426000 | 130430799 | 33.04009 | 54.70947 | 75.16851 | 32.52318 | -0.0763 | 1.34061 | 0.940233 |
| 17 | 65466600 | 65470399 | 40.29082 | 65.02953 | 85.25362 | 36.3494 | -0.03112 | 1.753839 | 0.940255 |
| 13 | 91704600 | 91707799 | 29.6279 | 49.11708 | 65.75237 | 28.47219 | -0.03877 | 1.307568 | 0.940257 |
| 13 | 93250000 | 93252599 | 37.34418 | 60.59133 | 76.42253 | 32.74727 | 0.032267 | 1.621772 | 0.940262 |
| 9 | 71681600 | 71688999 | 80.7344 | 126.2007 | 160.267 | 66.49253 | -0.03028 | 2.648849 | 0.940291 |
| 12 | 114619800 | 114622399 | 39.48988 | 64.88141 | 61.128 | 26.33706 | 0.463636 | 1.435449 | 0.940294 |
| 12 | 6617200 | 6623399 | 91.79651 | 143.1761 | 146.2271 | 60.68136 | 0.28123 | 2.691449 | 0.940339 |
| 4 | 4340000 | 4344799 | 75.77735 | 117.7967 | 179.8597 | 74.6752 | -0.302 | 2.746166 | 0.940366 |
| X | 45007200 | 45008999 | 17.23503 | 30.36649 | 38.53157 | 17.05502 | 0.046865 | 0.583826 | 0.940368 |
| 13 | 52207000 | 52209599 | 25.80688 | 43.20941 | 54.99903 | 23.97109 | 0.034167 | 1.179216 | 0.940403 |
| 1 | 8094600 | 8102199 | 126.2786 | 192.498 | 192.2489 | 79.70107 | 0.288034 | 3.136836 | 0.940409 |
| 6 | 7537200 | 7537999 | 8.633014 | 16.5771 | 24.67745 | 9.665908 | -0.04159 | -0.07955 | 0.940422 |
| 7 | 72297800 | 72301199 | 30.69099 | 50.63043 | 64.41363 | 27.80047 | 0.033856 | 1.36753 | 0.940423 |
| 3 | 47560600 | 47562799 | 23.22567 | 39.64759 | 53.39912 | 23.41102 | -0.04558 | 0.991992 | 0.940431 |
| 8 | 83472800 | 83482199 | 177.7982 | 263.5972 | 319.7991 | 131.7156 | -0.02763 | 3.684831 | 0.940438 |
| 7 | 86180400 | 86183799 | 29.46149 | 49.17729 | 50.79346 | 22.09529 | 0.334641 | 1.218504 | 0.940443 |
| 3 | 152321200 | 152322799 | 17.22416 | 30.08757 | 40.41765 | 17.80575 | -0.03622 | 0.711747 | 0.94046 |
| 3 | 36413200 | 36414199 | 10.76898 | 20.12279 | 26.82778 | 11.12738 | 0.044507 | 0.14109 | 0.94047 |
| 2 | 97819000 | 97820999 | 28.71879 | 48.20256 | 49.1249 | 21.44391 | 0.353679 | 1.155356 | 0.940488 |
| 3 | 9793800 | 9808599 | 209.4685 | 306.6388 | 367.6199 | 151.4356 | -0.02772 | 3.894523 | 0.940495 |
| 8 | 39443800 | 39444799 | 16.17882 | 28.4137 | 38.21945 | 16.9127 | -0.038 | 0.618498 | 0.940498 |
| 5 | 15868000 | 15869799 | 27.84532 | 46.16357 | 61.57326 | 26.6632 | -0.03344 | 1.299745 | 0.940507 |
| 15 | 38899400 | 38900399 | 11.03698 | 20.52563 | 27.26329 | 11.39956 | 0.040302 | 0.173576 | 0.94052 |
| 3 | 52506200 | 52511199 | 32.86343 | 53.10771 | 88.25766 | 37.69065 | -0.37035 | 1.707153 | 0.940554 |
| 7 | 36270400 | 36272799 | 21.42771 | 36.77274 | 46.87189 | 20.55815 | 0.034881 | 0.93078 | 0.940586 |
| 13 | 90426000 | 90426399 | 6.514908 | 12.68907 | 20.23235 | 7.389126 | -0.04496 | -0.35479 | 0.9406 |
| 11 | 42790200 | 42791399 | 16.57678 | 29.09627 | 37.15577 | 16.43421 | 0.036521 | 0.626605 | 0.940634 |
| 6 | 14183400 | 14184399 | 15.15156 | 26.90888 | 34.46167 | 15.21239 | 0.038641 | 0.486849 | 0.94064 |
| 7 | 8764400 | 8764799 | 16.0588 | 28.67824 | 26.51812 | 11.65144 | 0.510748 | 0.443751 | 0.94064 |

| | | | | | | | | | |
|---|---|---|---|---|---|---|---|---|---|
| 8 | 33976800 | 33978999 | 49.80914 | 79.66736 | 103.2683 | 43.45753 | -0.0304 | 2.051161 | 0.940641 |
| 7 | 87541400 | 87547599 | 79.03088 | 124.6972 | 127.2589 | 52.81981 | 0.29516 | 2.466942 | 0.940655 |
| 6 | 71667800 | 71668999 | 13.62491 | 24.77911 | 32.11343 | 13.83495 | 0.044852 | 0.328909 | 0.940668 |
| 4 | 12934000 | 12941999 | 104.1324 | 160.3426 | 193.0176 | 80.16925 | 0.030055 | 2.931228 | 0.940669 |
| 12 | 87895600 | 87900799 | 68.80335 | 108.6475 | 133.5273 | 55.44417 | 0.028303 | 2.441677 | 0.940715 |
| 13 | 43670200 | 43671399 | 12.8837 | 23.18229 | 31.59561 | 13.79436 | -0.03957 | 0.384985 | 0.940724 |
| X | 35913600 | 35917199 | 41.06182 | 66.51689 | 83.08158 | 35.55879 | 0.044157 | 1.665418 | 0.940726 |
| 7 | 35885800 | 35889399 | 31.72891 | 51.787 | 68.91252 | 29.61318 | -0.03816 | 1.509923 | 0.940802 |
| 15 | 91170200 | 91174999 | 71.30389 | 112.5837 | 144.2467 | 59.8918 | -0.03182 | 2.447999 | 0.940861 |
| 19 | 43308200 | 43309799 | 13.66498 | 24.06225 | 40.85925 | 18.08744 | -0.37141 | 0.575612 | 0.940921 |
| 1 | 114043000 | 114044199 | 18.13198 | 31.62284 | 42.5449 | 18.7241 | -0.03858 | 0.743534 | 0.940929 |
| 8 | 37849400 | 37857599 | 131.496 | 198.9956 | 236.6356 | 97.88716 | 0.02777 | 3.287626 | 0.94095 |
| 13 | 92222200 | 92222999 | 14.13761 | 25.78503 | 33.22898 | 14.26453 | 0.056239 | 0.318734 | 0.940951 |
| 15 | 47374400 | 47375999 | 22.17555 | 38.28151 | 38.37447 | 16.83467 | 0.381683 | 0.868981 | 0.940998 |
| X | 74501000 | 74502199 | 17.34809 | 30.47022 | 41.26099 | 18.26015 | -0.04772 | 0.628197 | 0.941003 |
| X | 152755800 | 152759599 | 28.20462 | 46.95126 | 62.80308 | 27.27259 | -0.03713 | 1.25569 | 0.941005 |
| 7 | 33276600 | 33277599 | 31.10892 | 50.94884 | 67.88081 | 29.20708 | -0.03666 | 1.461403 | 0.941006 |
| 2 | 43406600 | 43409599 | 39.02735 | 63.06258 | 82.82888 | 35.36787 | -0.03186 | 1.718595 | 0.941007 |
| 9 | 67590800 | 67595199 | 79.82933 | 124.6259 | 151.4773 | 62.8541 | 0.031488 | 2.670553 | 0.941066 |
| 9 | 111579200 | 111582399 | 44.88953 | 72.11695 | 93.95287 | 39.79991 | -0.03018 | 1.905577 | 0.941071 |
| 15 | 34295000 | 34298199 | 29.5009 | 48.76773 | 65.01634 | 28.09828 | -0.03312 | 1.3399 | 0.941091 |
| 3 | 5619200 | 5624599 | 70.72903 | 112.32 | 136.1208 | 56.74369 | 0.055513 | 2.300932 | 0.941118 |
| 16 | 21991200 | 21994599 | 41.57572 | 67.06468 | 84.16615 | 35.85906 | 0.03076 | 1.771825 | 0.941118 |
| 8 | 62001200 | 62007199 | 89.23019 | 137.1836 | 206.9207 | 85.92559 | -0.29672 | 2.957211 | 0.941127 |
| 10 | 95192600 | 95194599 | 17.25949 | 30.31182 | 38.62847 | 17.0938 | 0.039434 | 0.624012 | 0.94113 |
| 14 | 72846200 | 72858599 | 134.6239 | 204.6529 | 258.1642 | 106.9405 | -0.05193 | 3.200014 | 0.94113 |
| 2 | 150545000 | 150546999 | 10.90117 | 20.77144 | 22.19927 | 8.869736 | 0.406324 | -9.68E-05 | 0.941131 |
| 18 | 66675600 | 66676199 | 9.117868 | 17.24802 | 24.90014 | 10.10256 | -0.04329 | 0.04049 | 0.941149 |
| 1 | 55576200 | 55579999 | 51.82705 | 82.90102 | 107.4558 | 45.22443 | -0.03024 | 2.053102 | 0.941153 |
| X | 115322000 | 115323399 | 16.34293 | 29.04193 | 29.3242 | 12.9381 | 0.38025 | 0.48068 | 0.941156 |
| 8 | 66162600 | 66164999 | 43.46898 | 69.80866 | 91.35791 | 38.67908 | -0.03731 | 1.915538 | 0.941175 |
| 12 | 104121800 | 104122399 | 7.545069 | 14.60254 | 21.01959 | 8.008805 | 0.04444 | -0.19273 | 0.941189 |
| 4 | 112106000 | 112107999 | 43.90857 | 70.56476 | 92.0436 | 39.01015 | -0.03134 | 1.893065 | 0.94119 |
| 7 | 142328200 | 142328599 | 7.904128 | 15.26927 | 23.17778 | 8.907536 | -0.04311 | -0.15156 | 0.941199 |
| 8 | 96959600 | 96960399 | 7.904128 | 15.26927 | 23.17778 | 8.907536 | -0.04311 | -0.15156 | 0.941199 |
| 18 | 29295000 | 29297799 | 37.66843 | 61.59648 | 63.58323 | 27.34172 | 0.329096 | 1.49204 | 0.941204 |
| 10 | 6464600 | 6465199 | 12.16402 | 21.85312 | 30.0986 | 13.15811 | -0.05608 | 0.384428 | 0.941204 |
| 4 | 5263000 | 5264199 | 17.17959 | 30.0296 | 40.36957 | 17.79326 | -0.0373 | 0.700492 | 0.941214 |
| 6 | 101119200 | 101121599 | 31.64258 | 51.94615 | 66.0177 | 28.43401 | 0.032564 | 1.429762 | 0.941235 |
| 9 | 34437200 | 34440799 | 34.42502 | 56.27812 | 71.29591 | 30.65538 | 0.034141 | 1.492781 | 0.941244 |
| 18 | 17562600 | 17563599 | 15.19077 | 27.50065 | 27.81311 | 12.05113 | 0.395242 | 0.35776 | 0.941245 |
| 19 | 12712000 | 12713199 | 13.40744 | 24.07541 | 30.9598 | 13.54008 | 0.041855 | 0.397872 | 0.94125 |
| 15 | 85481800 | 85483799 | 21.8691 | 37.17049 | 46.91717 | 20.56205 | 0.046829 | 1.018543 | 0.941255 |
| 18 | 88605400 | 88606799 | 15.11881 | 26.68597 | 34.07041 | 15.07497 | 0.039205 | 0.544471 | 0.941259 |
| 10 | 77537400 | 77545399 | 113.248 | 173.1446 | 207.7488 | 86.18565 | 0.026874 | 3.074251 | 0.941262 |
| 14 | 118059200 | 118060999 | 16.0169 | 28.26038 | 36.07314 | 15.96233 | 0.039327 | 0.558989 | 0.941264 |
| 1 | 14067600 | 14070599 | 24.28564 | 41.1103 | 55.01267 | 24.08278 | -0.0375 | 1.072628 | 0.941266 |

| | | | | | | | | | |
|---|---|---|---|---|---|---|---|---|---|
| 15 | 97028600 | 97030799 | 18.95775 | 32.93649 | 42.07096 | 18.47095 | 0.035318 | 0.798774 | 0.941267 |
| 8 | 113660400 | 113662199 | 17.82 | 31.46101 | 31.78784 | 14.06437 | 0.375061 | 0.554153 | 0.941274 |
| 14 | 107444400 | 107445399 | 8.749752 | 15.83331 | 33.89536 | 14.69443 | -0.68008 | 0.326303 | 0.941281 |
| 8 | 45895600 | 45903599 | 114.7842 | 174.9076 | 217.3636 | 90.10351 | -0.02754 | 3.14274 | 0.941294 |
| 7 | 119820400 | 119823599 | 22.07128 | 37.66756 | 47.95365 | 21.02322 | 0.035418 | 0.984366 | 0.941301 |
| 1 | 194649000 | 194653399 | 72.64689 | 114.3415 | 140.1632 | 58.1525 | 0.028339 | 2.513971 | 0.941302 |
| 14 | 17709200 | 17710999 | 11.70341 | 21.46773 | 29.71156 | 12.65945 | -0.0384 | 0.261432 | 0.941316 |
| 5 | 34095800 | 34101799 | 82.35569 | 129.1831 | 157.2758 | 65.22819 | 0.034427 | 2.589687 | 0.941336 |
| X | 141715200 | 141716999 | 20.41794 | 35.76531 | 44.48919 | 19.64389 | 0.074218 | 0.693246 | 0.941354 |
| X | 148712000 | 148715999 | 42.93569 | 69.78072 | 68.08463 | 29.21061 | 0.40401 | 1.595432 | 0.941365 |
| 7 | 21136000 | 21138799 | 55.42007 | 88.76115 | 90.41725 | 38.12378 | 0.318823 | 2.013695 | 0.941383 |
| 2 | 178792400 | 178794199 | 20.36593 | 35.11165 | 44.7675 | 19.63848 | 0.035895 | 0.884282 | 0.941385 |
| 14 | 15821200 | 15823399 | 30.9159 | 51.44748 | 52.41903 | 22.7582 | 0.354364 | 1.258894 | 0.941406 |
| 11 | 120209000 | 120214999 | 91.94125 | 142.0703 | 171.2774 | 71.14486 | 0.031803 | 2.858573 | 0.941412 |
| 18 | 57660600 | 57661999 | 18.7978 | 32.74853 | 44.09871 | 19.39471 | -0.04022 | 0.761383 | 0.94142 |
| Y | 83351200 | 83351599 | 5.241226 | 10.2184 | 17.00337 | 6.130085 | -0.08763 | -0.42779 | 0.94143 |
| X | 43357200 | 43358999 | 35.81909 | 58.96379 | 57.93456 | 24.98151 | 0.404386 | 1.403316 | 0.941438 |
| 15 | 42224000 | 42224599 | 12.18442 | 22.28894 | 29.15028 | 12.46666 | 0.038909 | 0.279485 | 0.941485 |
| 7 | 35330400 | 35332199 | 15.31873 | 26.84694 | 36.36147 | 16.06209 | -0.0478 | 0.643618 | 0.941523 |
| 6 | 8927400 | 8928399 | 18.27493 | 31.05482 | 54.24025 | 23.77107 | -0.41982 | 1.019456 | 0.941551 |
| 7 | 75446200 | 75448399 | 28.11433 | 46.53851 | 62.19655 | 26.90476 | -0.03651 | 1.316945 | 0.94156 |
| 4 | 92699000 | 92700999 | 20.34201 | 35.0256 | 46.87366 | 20.5616 | -0.03414 | 0.897353 | 0.941563 |
| X | 23401000 | 23402199 | 21.84031 | 37.21295 | 50.07102 | 21.94862 | -0.04429 | 1.005174 | 0.941566 |
| 8 | 81373800 | 81375999 | 22.14443 | 38.03444 | 51.19405 | 22.45508 | -0.04324 | 0.931626 | 0.941574 |
| X | 147680000 | 147682199 | 27.59883 | 47.11756 | 44.65395 | 19.57716 | 0.459676 | 0.999261 | 0.941576 |
| 3 | 126735000 | 126742599 | 95.48992 | 146.0344 | 224.1688 | 93.00351 | -0.32791 | 3.064363 | 0.941584 |
| 1 | 7305800 | 7308599 | 36.19834 | 58.81348 | 77.67087 | 33.31615 | -0.03206 | 1.592421 | 0.941587 |
| 10 | 126034400 | 126035399 | 12.55169 | 23.08556 | 30.24363 | 12.83924 | 0.04412 | 0.254921 | 0.941603 |
| 7 | 10018800 | 10019399 | 22.57573 | 38.582 | 49.01642 | 21.49155 | 0.038805 | 0.978064 | 0.941618 |
| 8 | 16843800 | 16847199 | 38.38592 | 62.65129 | 64.58678 | 27.75414 | 0.329111 | 1.521904 | 0.941645 |
| 8 | 15439000 | 15439799 | 9.916133 | 18.56026 | 26.33446 | 10.89181 | -0.04154 | 0.120482 | 0.94166 |
| 9 | 6920400 | 6921799 | 14.46537 | 25.59051 | 34.53648 | 15.27427 | -0.03925 | 0.515893 | 0.941661 |
| 8 | 92718800 | 92719399 | 9.949112 | 18.75668 | 25.30053 | 10.33725 | 0.044595 | 0.070022 | 0.941683 |
| 9 | 5363400 | 5365399 | 20.37121 | 35.25346 | 47.39771 | 20.79468 | -0.03931 | 0.853464 | 0.941685 |
| 18 | 47904000 | 47908199 | 55.18847 | 88.3069 | 90.66328 | 38.17482 | 0.305981 | 2.04153 | 0.941707 |
| 1 | 145811800 | 145814599 | 38.72208 | 62.64448 | 82.45472 | 35.25638 | -0.03303 | 1.686572 | 0.941757 |
| 5 | 117888600 | 117890199 | 15.88766 | 27.92119 | 35.59474 | 15.76021 | 0.038912 | 0.599474 | 0.941778 |
| 13 | 30452000 | 30454799 | 36.52004 | 59.26117 | 78.11208 | 33.47418 | -0.031 | 1.618361 | 0.941789 |
| 2 | 129009200 | 129011199 | 18.87968 | 32.89774 | 42.00498 | 18.4762 | 0.036345 | 0.757924 | 0.941796 |
| 3 | 142761000 | 142761999 | 9.419218 | 17.91791 | 26.08034 | 10.45708 | -0.04029 | 0.007493 | 0.941822 |
| 9 | 28246400 | 28248799 | 19.31455 | 33.39191 | 44.74718 | 19.63123 | -0.03486 | 0.853152 | 0.941843 |
| 12 | 115108600 | 115118599 | 48.33363 | 77.93756 | 78.60509 | 33.47419 | 0.344879 | 1.791521 | 0.941844 |
| 2 | 52471400 | 52473199 | 22.34209 | 37.32559 | 63.0089 | 27.42389 | -0.37179 | 1.199072 | 0.94185 |
| 14 | 98478000 | 98479599 | 17.66628 | 30.95287 | 41.81698 | 18.4678 | -0.04452 | 0.667146 | 0.941865 |
| 9 | 4245200 | 4251599 | 114.1464 | 174.2148 | 208.6004 | 86.50418 | 0.027889 | 3.110302 | 0.941875 |
| 2 | 150062600 | 150102399 | 724.3301 | 960.5382 | 995.7647 | 416.2228 | 0.039266 | 5.41462 | 0.941885 |
| 14 | 89688600 | 89688999 | 5.50141 | 10.74213 | 16.49991 | 5.81016 | 0.05766 | -0.50693 | 0.941909 |

| | | | | | | | | | |
|---|---|---|---|---|---|---|---|---|---|
| 10 | 40546200 | 40547999 | 15.63696 | 27.5689 | 37.08852 | 16.41415 | -0.03661 | 0.565517 | 0.941913 |
| 5 | 104682400 | 104683399 | 13.63827 | 24.7514 | 32.08827 | 13.86044 | 0.04196 | 0.341168 | 0.941973 |
| 3 | 67488600 | 67491399 | 20.03357 | 34.94697 | 35.69279 | 15.71064 | 0.357135 | 0.737968 | 0.94198 |
| 3 | 6663600 | 6664799 | 20.5096 | 35.20751 | 44.80174 | 19.65065 | 0.037465 | 0.92319 | 0.941984 |
| 8 | 63107400 | 63108799 | 17.70277 | 30.38486 | 51.11383 | 22.41895 | -0.36364 | 0.920851 | 0.941993 |
| 3 | 141084400 | 141084999 | 11.93172 | 21.05092 | 38.37977 | 16.98326 | -0.47205 | 0.539996 | 0.942008 |
| 16 | 7841200 | 7844599 | 31.98374 | 52.83703 | 66.95491 | 28.92283 | 0.039812 | 1.355734 | 0.94202 |
| 7 | 28155000 | 28156599 | 13.84913 | 24.67141 | 31.58303 | 13.90564 | 0.041956 | 0.454055 | 0.942037 |
| X | 23182800 | 23184999 | 19.25646 | 33.80166 | 42.29679 | 18.709 | 0.0657 | 0.651004 | 0.942049 |
| 4 | 147809400 | 147832999 | 943.2028 | 1216.498 | 1284.273 | 538.421 | -0.02702 | 5.864838 | 0.942058 |
| 16 | 84739200 | 84742199 | 31.94199 | 52.62982 | 66.87878 | 28.84949 | 0.03479 | 1.38587 | 0.942077 |
| 5 | 89140800 | 89142799 | 24.92216 | 41.92865 | 53.3722 | 23.32101 | 0.034052 | 1.125918 | 0.942099 |
| 16 | 14454800 | 14461799 | 60.66293 | 96.77187 | 118.843 | 49.82544 | 0.043208 | 2.147126 | 0.942114 |
| 4 | 87225000 | 87225599 | 9.578734 | 17.9598 | 23.82213 | 9.823881 | 0.058561 | 0.100124 | 0.942123 |
| 4 | 121111400 | 121113799 | 24.52379 | 41.51817 | 52.8013 | 23.11551 | 0.035769 | 1.072033 | 0.942123 |
| 17 | 32893000 | 32895799 | 27.86087 | 46.14243 | 61.52854 | 26.62515 | -0.03316 | 1.311715 | 0.94215 |
| 14 | 65567600 | 65570199 | 31.08323 | 51.18611 | 65.0845 | 28.06882 | 0.033577 | 1.388585 | 0.942157 |
| 5 | 150745800 | 150754199 | 100.9668 | 156.0238 | 187.8215 | 78.01691 | 0.033963 | 2.862136 | 0.942161 |
| 17 | 88793800 | 88794599 | 11.31739 | 20.67453 | 26.68083 | 11.43056 | 0.056336 | 0.273792 | 0.942175 |
| 1 | 166183600 | 166188999 | 48.03101 | 76.74708 | 100.1846 | 42.10056 | -0.04206 | 2.078997 | 0.942199 |
| 9 | 91919800 | 91921199 | 20.21298 | 35.09497 | 44.70961 | 19.62159 | 0.039063 | 0.818666 | 0.942199 |
| 13 | 23060000 | 23063799 | 40.71837 | 65.89952 | 82.71842 | 35.37921 | 0.035752 | 1.679741 | 0.9422 |
| 3 | 5024200 | 5024999 | 11.67327 | 21.91817 | 23.06211 | 9.487561 | 0.392645 | 0.106662 | 0.942201 |
| 2 | 124204400 | 124207399 | 27.18192 | 45.72765 | 61.79906 | 27.00263 | -0.05208 | 1.142934 | 0.942286 |
| 17 | 43294000 | 43296999 | 22.76278 | 38.93277 | 49.55781 | 21.73032 | 0.036075 | 0.969686 | 0.942294 |
| 3 | 158332200 | 158336199 | 82.2352 | 128.0581 | 155.2951 | 64.4596 | 0.03194 | 2.716159 | 0.942311 |
| 3 | 62584200 | 62585599 | 16.61529 | 29.1705 | 39.20056 | 17.34197 | -0.03676 | 0.626734 | 0.942317 |
| 6 | 35120000 | 35124599 | 51.75156 | 82.91023 | 108.1797 | 45.63376 | -0.03721 | 2.003982 | 0.942344 |
| 12 | 94336600 | 94338399 | 16.13729 | 28.5077 | 38.57634 | 17.06581 | -0.04396 | 0.549271 | 0.942359 |
| 4 | 51350800 | 51351399 | 9.531181 | 18.01276 | 25.9168 | 10.53618 | -0.04142 | 0.051882 | 0.942361 |
| 1 | 37544000 | 37549599 | 53.89233 | 86.16463 | 111.9556 | 47.08731 | -0.03447 | 2.069544 | 0.942382 |
| 2 | 126814000 | 126815999 | 19.85098 | 34.23219 | 45.81566 | 20.09843 | -0.03378 | 0.880408 | 0.942405 |
| 9 | 114784600 | 114785399 | 12.47205 | 22.38121 | 30.74271 | 13.45251 | -0.05272 | 0.398944 | 0.942409 |
| 5 | 50082200 | 50085399 | 33.68467 | 55.16229 | 70.01523 | 30.1243 | 0.032998 | 1.468656 | 0.942411 |
| 19 | 10155600 | 10157599 | 22.02251 | 37.63634 | 47.95511 | 21.0262 | 0.03458 | 0.97381 | 0.94242 |
| 6 | 121960800 | 121961199 | 8.33025 | 15.92492 | 23.46786 | 9.309971 | -0.04408 | -0.0441 | 0.942429 |
| 16 | 63042800 | 63043599 | 8.33025 | 15.92492 | 23.46786 | 9.309971 | -0.04408 | -0.0441 | 0.942429 |
| 14 | 123267000 | 123268799 | 18.03286 | 31.68913 | 40.09595 | 17.75222 | 0.049578 | 0.620055 | 0.942433 |
| 4 | 113813400 | 113816399 | 32.73128 | 54.44792 | 67.78688 | 29.42262 | 0.066004 | 1.280923 | 0.942444 |
| 3 | 107901200 | 107902199 | 14.58878 | 25.91919 | 35.0732 | 15.47741 | -0.03991 | 0.483547 | 0.942448 |
| 17 | 27752200 | 27752799 | 10.82387 | 19.8847 | 25.71602 | 10.93432 | 0.061031 | 0.235557 | 0.942451 |
| 14 | 35691400 | 35693799 | 18.61513 | 31.96469 | 53.85294 | 23.62116 | -0.3659 | 0.922202 | 0.942507 |
| 7 | 6476600 | 6481999 | 67.5928 | 107.0448 | 131.7951 | 54.80498 | 0.029026 | 2.372303 | 0.942561 |
| 3 | 14915800 | 14916199 | 6.092019 | 11.88392 | 18.07994 | 6.469579 | 0.050023 | -0.43603 | 0.942577 |
| 3 | 32705000 | 32707999 | 27.13033 | 45.34521 | 60.61128 | 26.38777 | -0.03621 | 1.206519 | 0.94261 |
| 4 | 156228000 | 156233199 | 111.1292 | 169.3981 | 211.0467 | 87.49336 | -0.03075 | 3.134328 | 0.942612 |
| 14 | 120885800 | 120891199 | 46.81392 | 74.93715 | 120.2304 | 50.65343 | -0.33632 | 2.029385 | 0.942614 |

| | | | | | | | | | |
|---|---|---|---|---|---|---|---|---|---|
| 12 | 82555200 | 82558599 | 36.29061 | 58.91541 | 77.75396 | 33.33177 | -0.03223 | 1.607192 | 0.942627 |
| 15 | 4832600 | 4835599 | 31.53891 | 52.26178 | 66.0013 | 28.57116 | 0.044987 | 1.317108 | 0.942629 |
| 17 | 61684400 | 61685399 | 17.1261 | 30.21912 | 28.99653 | 12.83079 | 0.448931 | 0.550941 | 0.942648 |
| 8 | 102861600 | 102862599 | 9.392441 | 17.68192 | 25.33941 | 10.37822 | -0.04463 | 0.075247 | 0.94266 |
| 7 | 119329200 | 119336199 | 80.45437 | 124.7755 | 188.7858 | 78.40856 | -0.2915 | 2.788815 | 0.942704 |
| 12 | 70795200 | 70795599 | 6.981581 | 13.5672 | 21.19941 | 7.894487 | -0.04143 | -0.2755 | 0.942705 |
| X | 94840200 | 94841799 | 16.09531 | 28.82054 | 36.52294 | 16.01112 | 0.060434 | 0.44483 | 0.942734 |
| 14 | 120122200 | 120123399 | 17.21827 | 30.09539 | 40.39004 | 17.80137 | -0.03484 | 0.70248 | 0.942736 |
| 12 | 117065600 | 117067599 | 15.55969 | 27.42523 | 35.02749 | 15.50284 | 0.037763 | 0.563382 | 0.942738 |
| 11 | 87697400 | 87700199 | 23.99019 | 40.3814 | 53.87551 | 23.54459 | -0.03344 | 1.119525 | 0.942749 |
| 10 | 57551800 | 57554799 | 20.95781 | 35.85449 | 47.9511 | 21.02622 | -0.03485 | 0.958183 | 0.942757 |
| 6 | 65163400 | 65165599 | 45.86239 | 73.54255 | 95.72542 | 40.43637 | -0.03211 | 1.965046 | 0.942762 |
| 10 | 39473000 | 39488399 | 172.2209 | 256.5781 | 313.4013 | 129.1447 | -0.03092 | 3.599104 | 0.94278 |
| 19 | 57456400 | 57459399 | 29.4845 | 48.88026 | 62.2649 | 26.96238 | 0.032722 | 1.305202 | 0.94278 |
| 2 | 152261600 | 152263399 | 27.66055 | 45.84155 | 58.22789 | 25.20859 | 0.036829 | 1.300462 | 0.942787 |
| 9 | 12607600 | 12608399 | 12.11111 | 21.96082 | 30.20194 | 13.05977 | -0.04332 | 0.333117 | 0.942811 |
| 5 | 112945000 | 112946999 | 17.50569 | 30.52675 | 38.92599 | 17.13006 | 0.038371 | 0.734951 | 0.942838 |
| 10 | 10460200 | 10461199 | 8.433537 | 16.14165 | 24.04736 | 9.507192 | -0.05433 | -0.0492 | 0.94285 |
| 13 | 109989000 | 109989999 | 36.35906 | 58.83769 | 73.50119 | 31.43433 | 0.042594 | 1.671289 | 0.942862 |
| 17 | 14674400 | 14677999 | 32.06478 | 52.32355 | 69.48178 | 29.85538 | -0.03539 | 1.514441 | 0.942906 |
| 3 | 8875600 | 8882599 | 73.63836 | 116.9071 | 140.5841 | 58.58769 | 0.066304 | 2.313099 | 0.942931 |
| 2 | 173985400 | 173988999 | 42.2937 | 68.52118 | 85.42757 | 36.56631 | 0.04694 | 1.66444 | 0.942932 |
| 1 | 16940400 | 16954999 | 179.8345 | 267.6937 | 328.6571 | 135.4195 | -0.03943 | 3.615671 | 0.942945 |
| 5 | 142030600 | 142048799 | 295.1937 | 421.3915 | 473.1735 | 195.5418 | 0.03103 | 4.275396 | 0.942947 |
| X | 127778200 | 127778999 | 14.66022 | 26.12211 | 35.62187 | 15.68096 | -0.04848 | 0.468075 | 0.942982 |
| X | 132376800 | 132378399 | 16.39361 | 29.33058 | 37.08204 | 16.26749 | 0.063258 | 0.454651 | 0.942986 |
| 5 | 61932800 | 61935999 | 55.58503 | 88.49191 | 114.0465 | 47.6806 | -0.02939 | 2.20601 | 0.942986 |
| 8 | 101029400 | 101029799 | 6.978738 | 13.57249 | 20.11538 | 7.442746 | 0.042808 | -0.3014 | 0.943003 |
| 4 | 48033800 | 48034599 | 8.813407 | 16.86683 | 23.47982 | 9.285788 | 0.041728 | -0.04852 | 0.943088 |
| 10 | 5422600 | 5424199 | 13.97707 | 24.79556 | 33.51379 | 14.80778 | -0.03965 | 0.483341 | 0.943094 |
| 15 | 24024800 | 24025799 | 11.45397 | 21.27829 | 28.25552 | 11.83314 | 0.038691 | 0.184197 | 0.943097 |
| 3 | 25365800 | 25366999 | 12.46347 | 22.6909 | 29.58634 | 12.72934 | 0.037471 | 0.308871 | 0.943105 |
| 6 | 10578000 | 10578599 | 12.46347 | 22.6909 | 29.58634 | 12.72934 | 0.037471 | 0.308871 | 0.943105 |
| 18 | 49902000 | 49903199 | 16.17407 | 28.61077 | 36.50301 | 16.14429 | 0.041367 | 0.532555 | 0.943136 |
| 11 | 98509800 | 98517399 | 113.1138 | 171.8593 | 251.5359 | 104.1778 | -0.26683 | 3.20438 | 0.943139 |
| 13 | 45761400 | 45763199 | 17.26797 | 30.14284 | 40.46454 | 17.81851 | -0.03528 | 0.722916 | 0.943155 |
| 5 | 90248400 | 90251799 | 28.92935 | 48.10919 | 61.22548 | 26.56923 | 0.034376 | 1.266312 | 0.943164 |
| 6 | 67511600 | 67513199 | 21.82341 | 37.00491 | 49.56344 | 21.70843 | -0.0386 | 1.043704 | 0.943166 |
| 8 | 108465400 | 108471599 | 71.62151 | 112.9665 | 138.838 | 57.61854 | 0.026939 | 2.47129 | 0.943181 |
| 9 | 31547800 | 31548799 | 13.8873 | 24.90067 | 32.06021 | 14.03863 | 0.039161 | 0.415446 | 0.943224 |
| 14 | 114046000 | 114049199 | 28.93307 | 48.01836 | 61.17872 | 26.51197 | 0.032543 | 1.289304 | 0.943266 |
| 6 | 62971600 | 62973999 | 18.22299 | 31.12557 | 52.69283 | 23.10244 | -0.37384 | 0.969805 | 0.943266 |
| 7 | 142889400 | 142892999 | 34.74313 | 56.78852 | 71.93185 | 30.9269 | 0.034173 | 1.496044 | 0.943271 |
| 18 | 4487800 | 4496999 | 91.36986 | 142.2216 | 172.2797 | 71.51409 | 0.033046 | 2.730808 | 0.943275 |
| 13 | 69896400 | 69897999 | 14.42839 | 25.52742 | 34.50082 | 15.25822 | -0.04134 | 0.513728 | 0.943298 |
| 18 | 55688400 | 55689799 | 17.8195 | 31.34998 | 42.79409 | 18.94731 | -0.05885 | 0.610691 | 0.943327 |
| 10 | 119394400 | 119397199 | 22.70414 | 38.68545 | 51.70158 | 22.66341 | -0.03473 | 1.007499 | 0.943339 |

| | | | | | | | | | |
|---|---|---|---|---|---|---|---|---|---|
| 7 | 118061000 | 118063199 | 15.25294 | 26.8138 | 36.12705 | 15.99792 | -0.04043 | 0.592245 | 0.943357 |
| 8 | 14860800 | 14862599 | 18.4215 | 32.0787 | 40.95375 | 18.00652 | 0.036342 | 0.761116 | 0.943365 |
| 2 | 140581400 | 140583199 | 24.68976 | 41.47929 | 52.79041 | 23.05696 | 0.034329 | 1.138627 | 0.943369 |
| 1 | 34815400 | 34817399 | 30.42653 | 50.26301 | 64.00892 | 27.64966 | 0.032687 | 1.348599 | 0.943372 |
| 3 | 50525800 | 50527199 | 18.19795 | 31.83318 | 42.93875 | 18.93746 | -0.0422 | 0.698087 | 0.943377 |
| X | 104072800 | 104075199 | 24.36999 | 41.69075 | 56.77802 | 24.8967 | -0.06105 | 0.982678 | 0.943381 |
| X | 17116000 | 17120399 | 32.77119 | 54.25297 | 68.13837 | 29.47723 | 0.052912 | 1.336453 | 0.943395 |
| 1 | 142624600 | 142625199 | 9.122466 | 17.56892 | 24.80131 | 9.632131 | 0.045927 | -0.10111 | 0.943408 |
| 6 | 112422800 | 112425599 | 22.7231 | 38.57288 | 49.03265 | 21.48311 | 0.036527 | 1.035915 | 0.943461 |
| 9 | 86777400 | 86778999 | 19.00689 | 32.98923 | 44.20159 | 19.39433 | -0.03348 | 0.816389 | 0.943464 |
| 8 | 70042600 | 70044399 | 24.75293 | 41.55789 | 55.46921 | 24.21765 | -0.03411 | 1.149085 | 0.943468 |
| 9 | 117966400 | 117968599 | 15.26143 | 27.01949 | 36.47858 | 16.13072 | -0.03947 | 0.515599 | 0.943492 |
| 4 | 153206600 | 153222399 | 199.006 | 289.2327 | 419.4234 | 172.8908 | -0.31211 | 4.004 | 0.943519 |
| 11 | 90199400 | 90204599 | 39.46996 | 63.7563 | 83.57235 | 35.66982 | -0.02956 | 1.728291 | 0.943522 |
| X | 75571600 | 75572999 | 14.20219 | 25.60906 | 35.15079 | 15.3063 | -0.04746 | 0.388653 | 0.943543 |
| 15 | 26904800 | 26906199 | 10.51059 | 19.11725 | 33.85205 | 14.58203 | -0.40241 | 0.309261 | 0.94358 |
| 13 | 103413400 | 103416399 | 35.29779 | 57.68795 | 73.04909 | 31.40583 | 0.034498 | 1.500901 | 0.943648 |
| 16 | 18667200 | 18671199 | 33.63907 | 55.55897 | 57.49219 | 24.83127 | 0.331235 | 1.354514 | 0.943657 |
| 4 | 62636200 | 62637999 | 20.68762 | 35.48576 | 47.41713 | 20.7965 | -0.03297 | 0.933825 | 0.943693 |
| 8 | 99618600 | 99619199 | 6.374943 | 12.39404 | 19.5467 | 7.252003 | -0.04973 | -0.30269 | 0.943718 |
| 3 | 106623400 | 106626799 | 38.95783 | 63.26997 | 84.19855 | 36.10477 | -0.04372 | 1.608607 | 0.943723 |
| X | 56098000 | 56099399 | 13.18519 | 23.97564 | 33.27161 | 14.33672 | -0.05306 | 0.332336 | 0.943744 |
| 2 | 126782600 | 126785399 | 26.89124 | 45.16895 | 57.30496 | 25.02648 | 0.03873 | 1.149792 | 0.943777 |
| 6 | 143740200 | 143741799 | 12.83449 | 23.49276 | 30.59848 | 13.07492 | 0.045239 | 0.281081 | 0.943786 |
| 1 | 177961800 | 177962799 | 13.1297 | 23.79503 | 32.60776 | 14.10956 | -0.03928 | 0.345913 | 0.943802 |
| 15 | 33831600 | 33833399 | 58.52969 | 92.96818 | 119.5729 | 49.8568 | -0.02983 | 2.27902 | 0.943859 |
| 13 | 12791600 | 12792199 | 16.45836 | 28.99801 | 37.05627 | 16.40203 | 0.036823 | 0.577798 | 0.943869 |
| 5 | 76977400 | 76978999 | 16.07883 | 28.30797 | 38.08294 | 16.859 | -0.0375 | 0.584219 | 0.943885 |
| 13 | 68459400 | 68529399 | 1631.043 | 2015.7 | 1963.839 | 827.0334 | 0.02661 | 6.555048 | 0.943894 |
| 13 | 108783400 | 108784199 | 9.428294 | 17.75896 | 23.834 | 9.752774 | 0.050751 | 0.06983 | 0.943899 |
| 14 | 70118000 | 70122399 | 52.88769 | 84.53486 | 105.1279 | 44.19824 | 0.028232 | 2.074966 | 0.943904 |
| 1 | 85938600 | 85948199 | 111.9573 | 170.1209 | 249.0344 | 103.1451 | -0.26694 | 3.201677 | 0.943959 |
| 4 | 143607400 | 143609399 | 47.38417 | 76.05158 | 98.88331 | 41.82783 | -0.02901 | 1.940818 | 0.943982 |
| 1 | 85073800 | 85100399 | 1267.027 | 1598.681 | 1589.914 | 668.0079 | 0.027267 | 6.22152 | 0.943994 |
| 15 | 50284200 | 50285799 | 21.7286 | 37.75861 | 38.32997 | 16.83078 | 0.365438 | 0.804537 | 0.944054 |
| 18 | 23171600 | 23175199 | 36.29041 | 58.44464 | 95.46688 | 40.58466 | -0.35142 | 1.817459 | 0.94408 |
| 2 | 149018200 | 149025399 | 105.8925 | 163.3744 | 195.8 | 81.33169 | 0.038597 | 2.896044 | 0.944082 |
| 1 | 35548000 | 35550399 | 29.34446 | 48.74238 | 61.88222 | 26.83251 | 0.037663 | 1.279816 | 0.944091 |
| 4 | 145181200 | 145200599 | 511.1157 | 693.602 | 742.7648 | 309.4393 | 0.025796 | 5.045768 | 0.944126 |
| 9 | 94810400 | 94811599 | 19.03392 | 33.7824 | 31.02208 | 13.7014 | 0.51541 | 0.509011 | 0.94413 |
| 6 | 99718600 | 99720799 | 16.99378 | 29.6574 | 39.91409 | 17.57358 | -0.03909 | 0.721661 | 0.944148 |
| 13 | 65375400 | 65376199 | 14.86324 | 26.35107 | 35.51937 | 15.70221 | -0.03622 | 0.498198 | 0.94419 |
| 6 | 91071400 | 91073999 | 20.04488 | 34.61464 | 46.33776 | 20.32832 | -0.03368 | 0.869981 | 0.944197 |
| 17 | 22257600 | 22263999 | 59.6426 | 94.77695 | 121.8356 | 50.82227 | -0.02848 | 2.26737 | 0.944226 |
| 16 | 73364000 | 73365399 | 20.56446 | 35.50317 | 45.28191 | 19.86498 | 0.035829 | 0.873882 | 0.944245 |
| 6 | 81006200 | 81014999 | 158.9482 | 239.0373 | 238.9792 | 98.67377 | 0.270052 | 3.41766 | 0.944264 |
| 6 | 140067000 | 140071199 | 46.34195 | 74.53696 | 97.49498 | 41.37008 | -0.0339 | 1.871935 | 0.944274 |

| | | | | | | | | | |
|---|---|---|---|---|---|---|---|---|---|
| 1 | 18023000 | 18023799 | 19.09935 | 33.04986 | 42.13101 | 18.48399 | 0.037178 | 0.840155 | 0.944309 |
| X | 67673000 | 67674599 | 16.6607 | 29.23091 | 39.25742 | 17.35743 | -0.03579 | 0.638605 | 0.944314 |
| 2 | 77651000 | 77652199 | 14.64281 | 26.32575 | 36.12128 | 15.78123 | -0.0502 | 0.409796 | 0.944321 |
| 14 | 96353400 | 96354599 | 15.76528 | 27.68403 | 37.17895 | 16.45056 | -0.03581 | 0.614682 | 0.944324 |
| 8 | 90178600 | 90182799 | 33.95258 | 55.68909 | 74.43715 | 32.0483 | -0.0406 | 1.451004 | 0.944329 |
| 13 | 105643800 | 105644999 | 10.81487 | 19.73228 | 34.95921 | 15.00018 | -0.3996 | 0.295646 | 0.944363 |
| 6 | 11831800 | 11833599 | 37.92882 | 61.14808 | 81.06996 | 34.51525 | -0.04906 | 1.783448 | 0.944372 |
| 13 | 86348400 | 86349399 | 15.96283 | 28.05815 | 37.64329 | 16.66968 | -0.03436 | 0.601641 | 0.944386 |
| 14 | 51455000 | 51456599 | 21.63388 | 37.42776 | 37.92227 | 16.6374 | 0.367042 | 0.848204 | 0.944415 |
| 1 | 46418600 | 46422799 | 45.64744 | 73.68392 | 76.19213 | 32.49087 | 0.310702 | 1.755054 | 0.944416 |
| 12 | 63228400 | 63230399 | 37.99394 | 61.43369 | 80.64343 | 34.46431 | -0.02994 | 1.695383 | 0.944434 |
| 13 | 75556200 | 75559199 | 38.32537 | 62.08945 | 81.6922 | 34.96637 | -0.03061 | 1.655996 | 0.944448 |
| 3 | 39526200 | 39531399 | 69.28967 | 109.984 | 134.3591 | 55.98368 | 0.043356 | 2.31143 | 0.94447 |
| X | 107029400 | 107032199 | 38.13202 | 61.90343 | 82.28508 | 35.2736 | -0.04254 | 1.617635 | 0.944489 |
| 5 | 96420400 | 96424199 | 75.16142 | 117.6052 | 149.5563 | 62.0425 | -0.03112 | 2.627878 | 0.944495 |
| 19 | 59597800 | 59599399 | 13.07699 | 23.58619 | 30.48147 | 13.2688 | 0.039198 | 0.368067 | 0.944499 |
| 3 | 139679600 | 139681799 | 42.52982 | 69.0453 | 69.75796 | 29.90433 | 0.352334 | 1.616014 | 0.944535 |
| 7 | 71202400 | 71202799 | 13.02886 | 24.17634 | 24.99411 | 10.48358 | 0.395939 | 0.196586 | 0.944554 |
| 17 | 31646600 | 31650399 | 37.64005 | 61.08115 | 77.17892 | 33.07529 | 0.029875 | 1.618783 | 0.944556 |
| 12 | 13836400 | 13842399 | 78.11196 | 122.2294 | 154.8967 | 64.25755 | -0.02605 | 2.629049 | 0.944583 |
| 7 | 5258600 | 5264599 | 36.94348 | 61.11982 | 57.16975 | 24.72083 | 0.476717 | 1.343955 | 0.944601 |
| 14 | 115430200 | 115431199 | 7.999326 | 15.43958 | 22.02549 | 8.499559 | 0.039739 | -0.1417 | 0.944611 |
| 2 | 9851800 | 9854199 | 21.50246 | 36.77386 | 46.90038 | 20.56493 | 0.033381 | 0.963642 | 0.944634 |
| 2 | 151212200 | 151218999 | 161.6318 | 240.6741 | 293.0744 | 120.7619 | -0.0272 | 3.605973 | 0.944635 |
| 7 | 137544400 | 137545399 | 12.52885 | 22.74188 | 31.13353 | 13.44516 | -0.03582 | 0.330622 | 0.94465 |
| 2 | 132853600 | 132855799 | 20.16371 | 35.03277 | 44.68554 | 19.61925 | 0.03733 | 0.808169 | 0.944652 |
| 14 | 58517000 | 58519599 | 22.43023 | 38.60339 | 49.01635 | 21.50121 | 0.041166 | 0.912009 | 0.944657 |
| 8 | 51677200 | 51679999 | 26.74102 | 45.7283 | 44.11517 | 19.34351 | 0.434434 | 0.980316 | 0.944666 |
| 2 | 142418400 | 142419799 | 10.86822 | 20.20835 | 28.31186 | 11.84139 | -0.03655 | 0.175139 | 0.944696 |
| 4 | 79274600 | 79275599 | 10.86822 | 20.20835 | 28.31186 | 11.84139 | -0.03655 | 0.175139 | 0.944696 |
| 14 | 82191600 | 82192199 | 11.0634 | 20.33105 | 28.23042 | 11.99836 | -0.04035 | 0.24301 | 0.944707 |
| 15 | 96989000 | 96989799 | 8.286667 | 15.87041 | 23.48708 | 9.269118 | -0.04346 | -0.06306 | 0.944713 |
| X | 69701400 | 69706399 | 59.78857 | 95.95512 | 87.42109 | 36.97606 | 0.483064 | 1.955589 | 0.94472 |
| 12 | 49568400 | 49568999 | 6.689579 | 13.00417 | 19.01729 | 7.062514 | 0.057664 | -0.29416 | 0.94476 |
| 14 | 118880600 | 118881799 | 16.44782 | 28.86431 | 38.71623 | 17.12161 | -0.03404 | 0.631799 | 0.944783 |
| 10 | 53076400 | 53078799 | 16.91742 | 29.87222 | 37.99928 | 16.81206 | 0.044501 | 0.5575 | 0.944791 |
| 10 | 78696400 | 78704399 | 223.0286 | 323.394 | 367.7941 | 151.6464 | 0.035045 | 4.04012 | 0.944823 |
| 5 | 148972400 | 148974799 | 30.65457 | 50.53108 | 64.42366 | 27.79567 | 0.030345 | 1.376139 | 0.944827 |
| 4 | 74476000 | 74477199 | 12.41628 | 23.15888 | 30.54458 | 12.70796 | 0.05575 | 0.171769 | 0.944846 |
| X | 38256000 | 38261399 | 51.74798 | 83.1801 | 102.5035 | 43.45598 | 0.05083 | 1.902398 | 0.944906 |
| 2 | 138127600 | 138128799 | 15.10628 | 26.83277 | 34.42964 | 15.19619 | 0.035969 | 0.484645 | 0.944914 |
| 12 | 59449200 | 59450599 | 14.45663 | 25.59941 | 34.50348 | 15.25677 | -0.03687 | 0.505196 | 0.944956 |
| 15 | 84255600 | 84257799 | 18.90565 | 32.50445 | 43.80376 | 19.21409 | -0.04444 | 0.901429 | 0.944964 |
| 17 | 52253400 | 52260599 | 87.51799 | 136.0947 | 165.4789 | 68.70095 | 0.026546 | 2.743007 | 0.944984 |
| 7 | 59448600 | 59449199 | 13.57654 | 24.10256 | 32.65476 | 14.41936 | -0.0422 | 0.476626 | 0.944985 |
| 3 | 28063600 | 28064599 | 11.4222 | 21.07435 | 27.74506 | 11.72821 | 0.041767 | 0.223563 | 0.945004 |
| 2 | 99294800 | 99297399 | 16.08408 | 28.4433 | 38.53727 | 17.04498 | -0.04525 | 0.536606 | 0.945005 |

| | | | | | | | | | |
|---|---|---|---|---|---|---|---|---|---|
| 5 | 68294400 | 68295799 | 14.0597 | 24.90694 | 33.62102 | 14.86533 | -0.03857 | 0.498565 | 0.945009 |
| 18 | 7388000 | 7392999 | 64.60954 | 102.8527 | 105.9472 | 44.29071 | 0.293237 | 2.214672 | 0.945016 |
| 9 | 105851800 | 105852199 | 7.134881 | 13.8823 | 21.91521 | 8.07977 | -0.043 | -0.30686 | 0.945018 |
| 4 | 143846600 | 143862599 | 566.3403 | 759.7643 | 834.471 | 348.2499 | -0.02738 | 5.228034 | 0.945031 |
| 12 | 75478000 | 75479799 | 24.18308 | 41.06468 | 54.99767 | 24.09901 | -0.0385 | 1.046697 | 0.945043 |
| 4 | 115033800 | 115034599 | 10.07132 | 18.64382 | 26.2749 | 11.05662 | -0.05124 | 0.188407 | 0.945063 |
| 5 | 104457200 | 104458399 | 12.87017 | 23.20383 | 31.5917 | 13.759 | -0.03581 | 0.373186 | 0.945076 |
| 3 | 155961200 | 155963999 | 19.9379 | 34.53147 | 46.31785 | 20.32142 | -0.03576 | 0.841307 | 0.945084 |
| 2 | 30593000 | 30595599 | 23.16063 | 39.20234 | 52.2454 | 22.87072 | -0.03155 | 1.067271 | 0.945093 |
| 10 | 66793200 | 66798199 | 34.00832 | 55.58548 | 70.23665 | 30.19766 | 0.037618 | 1.49222 | 0.945097 |
| 10 | 81669800 | 81671199 | 15.72596 | 28.22107 | 28.33818 | 12.40007 | 0.396358 | 0.414447 | 0.945123 |
| 19 | 30075400 | 30084999 | 131.249 | 198.2919 | 244.4038 | 101.0476 | -0.02594 | 3.321803 | 0.945125 |
| 12 | 36321200 | 36322199 | 10.96353 | 20.20067 | 28.18027 | 11.93419 | -0.04364 | 0.225013 | 0.945138 |
| 16 | 18888000 | 18888599 | 10.77227 | 20.115 | 28.40472 | 11.79528 | -0.03938 | 0.148861 | 0.945141 |
| 10 | 7760200 | 7768999 | 95.52778 | 147.6612 | 185.5059 | 77.05494 | -0.02723 | 2.854584 | 0.945182 |
| 1 | 8062800 | 8067599 | 46.80611 | 75.606 | 72.65986 | 31.01198 | 0.417279 | 1.738472 | 0.945191 |
| 16 | 38861600 | 38864199 | 19.6624 | 33.8606 | 45.29937 | 19.87111 | -0.03359 | 0.890729 | 0.945195 |
| 19 | 59758800 | 59759799 | 12.4916 | 22.66901 | 31.2163 | 13.4848 | -0.04419 | 0.334428 | 0.94522 |
| 5 | 138207200 | 138217199 | 253.7271 | 365.2571 | 430.0409 | 177.4812 | -0.02516 | 4.149994 | 0.945231 |
| 8 | 96011600 | 96014199 | 27.3031 | 45.45004 | 60.55388 | 26.29569 | -0.03143 | 1.249864 | 0.945265 |
| 5 | 87622400 | 87624399 | 15.9956 | 28.16655 | 38.03218 | 16.83591 | -0.0428 | 0.580007 | 0.945272 |
| 15 | 49193400 | 49196799 | 26.25772 | 44.26833 | 59.55722 | 26.04012 | -0.04549 | 1.117994 | 0.945275 |
| 17 | 61704200 | 61705199 | 11.72211 | 21.19929 | 29.18883 | 12.66353 | -0.04822 | 0.3416 | 0.945279 |
| 9 | 79723000 | 79727999 | 68.39334 | 106.8273 | 168.6417 | 69.97263 | -0.34466 | 2.658264 | 0.945314 |
| 12 | 77425000 | 77431999 | 99.42284 | 154.6767 | 153.3576 | 63.66435 | 0.321126 | 2.740725 | 0.945318 |
| 15 | 86751200 | 86756799 | 56.34293 | 90.00539 | 111.5078 | 46.84196 | 0.032757 | 2.098053 | 0.945342 |
| 4 | 72320400 | 72321999 | 25.59633 | 42.78626 | 57.09464 | 24.85858 | -0.03366 | 1.200047 | 0.945343 |
| 8 | 79691200 | 79691799 | 12.10298 | 21.89033 | 28.08686 | 12.18388 | 0.052862 | 0.33966 | 0.945343 |
| 3 | 98343000 | 98344399 | 17.21274 | 30.10271 | 40.36149 | 17.79727 | -0.0335 | 0.693145 | 0.945376 |
| 6 | 70643200 | 70643799 | 9.013411 | 17.24548 | 23.98894 | 9.493638 | 0.041161 | -0.04604 | 0.945381 |
| 2 | 36469600 | 36493199 | 365.7475 | 511.8687 | 564.767 | 234.1506 | 0.027601 | 4.569397 | 0.945402 |
| 3 | 112055600 | 112059599 | 53.64068 | 85.71515 | 106.6146 | 44.80604 | 0.027563 | 2.085898 | 0.945435 |
| 15 | 88106000 | 88110399 | 42.2208 | 68.13262 | 89.37753 | 38.09912 | -0.03235 | 1.762897 | 0.945444 |
| 4 | 113933800 | 113938599 | 70.1924 | 111.3786 | 135.968 | 56.63456 | 0.043884 | 2.323182 | 0.94545 |
| 4 | 53933200 | 53942599 | 89.84534 | 139.1974 | 174.9416 | 72.66589 | -0.02525 | 2.809533 | 0.945469 |
| 6 | 129768000 | 129770399 | 17.78693 | 31.52672 | 31.10576 | 13.75174 | 0.41125 | 0.509673 | 0.945479 |
| 15 | 45557000 | 45559599 | 39.36698 | 63.68129 | 80.0921 | 34.24358 | 0.032255 | 1.685057 | 0.945523 |
| 14 | 124968800 | 124969399 | 11.61383 | 21.17275 | 29.23403 | 12.55644 | -0.04268 | 0.296211 | 0.945529 |
| 9 | 35785600 | 35786399 | 9.733086 | 18.42208 | 26.50386 | 10.74054 | -0.03714 | 0.045906 | 0.94556 |
| 12 | 113799200 | 113802799 | 44.91319 | 72.47094 | 95.96396 | 40.90204 | -0.04601 | 1.77328 | 0.945561 |
| 3 | 96219600 | 96222999 | 36.6281 | 59.45415 | 78.3459 | 33.58172 | -0.03025 | 1.61146 | 0.945566 |
| 4 | 21205400 | 21211199 | 58.0732 | 92.52462 | 119.291 | 49.92661 | -0.02878 | 2.185117 | 0.945591 |
| 11 | 37643200 | 37644599 | 23.19176 | 38.16206 | 70.091 | 30.21831 | -0.49564 | 1.384806 | 0.945597 |
| 2 | 97408000 | 97408999 | 12.76891 | 23.07575 | 31.53565 | 13.69405 | -0.03869 | 0.356627 | 0.945601 |
| 2 | 89560000 | 89566799 | 55.93319 | 89.49169 | 117.3679 | 49.41376 | -0.04678 | 2.052338 | 0.94562 |
| 8 | 115693000 | 115697399 | 48.97001 | 78.46527 | 101.7152 | 42.91624 | -0.02755 | 1.994531 | 0.945644 |
| 18 | 7686200 | 7696599 | 114.6538 | 175.4937 | 210.6512 | 87.41279 | 0.027592 | 3.052302 | 0.945652 |

| | | | | | | | | | |
|---|---|---|---|---|---|---|---|---|---|
| 16 | 9305200 | 9306599 | 12.4687 | 22.68403 | 31.11911 | 13.4009 | -0.03632 | 0.315948 | 0.94566 |
| 1 | 8205200 | 8205599 | 6.077124 | 11.84403 | 19.1385 | 6.930874 | -0.05328 | -0.39585 | 0.945671 |
| X | 26324200 | 26324799 | 8.70092 | 16.79195 | 25.45846 | 9.828496 | -0.04793 | -0.12181 | 0.945674 |
| 17 | 3673400 | 3674799 | 13.91827 | 25.02122 | 34.10018 | 14.89917 | -0.0403 | 0.403537 | 0.945683 |
| 2 | 145267800 | 145268999 | 15.89412 | 27.91568 | 35.63796 | 15.7713 | 0.036918 | 0.609452 | 0.945684 |
| 17 | 12285000 | 12296599 | 179.7745 | 267.1069 | 312.5336 | 128.7516 | 0.027463 | 3.647426 | 0.945702 |
| 19 | 35053600 | 35056599 | 41.4488 | 66.74261 | 83.33974 | 35.42192 | 0.035048 | 1.824943 | 0.945722 |
| 8 | 14856400 | 14858599 | 33.88319 | 55.30234 | 73.17059 | 31.4428 | -0.02998 | 1.51798 | 0.945734 |
| 7 | 3159800 | 3161999 | 30.89285 | 50.48836 | 63.31183 | 27.21941 | 0.048635 | 1.485901 | 0.945738 |
| 15 | 15908200 | 15910199 | 18.25225 | 32.0224 | 31.08546 | 13.74394 | 0.430239 | 0.633105 | 0.945763 |
| 1 | 48776800 | 48778399 | 28.91691 | 47.90517 | 60.96716 | 26.38745 | 0.03403 | 1.312764 | 0.945781 |
| 1 | 90113000 | 90119399 | 66.53866 | 104.111 | 164.6607 | 68.30864 | -0.34539 | 2.626249 | 0.9458 |
| 2 | 55124600 | 55125599 | 16.14178 | 28.34214 | 36.20842 | 16.01968 | 0.036015 | 0.618384 | 0.945804 |
| 1 | 111071200 | 111073599 | 38.48779 | 62.20501 | 78.20086 | 33.40159 | 0.03158 | 1.708088 | 0.945808 |
| 17 | 41497600 | 41500599 | 21.82474 | 38.04418 | 37.60908 | 16.54664 | 0.403836 | 0.759309 | 0.945829 |
| 5 | 109902400 | 109936399 | 688.5802 | 916.4849 | 956.3153 | 399.5427 | 0.035136 | 5.355714 | 0.945832 |
| 4 | 101727600 | 101729599 | 25.01696 | 41.75504 | 52.79249 | 22.96369 | 0.04395 | 1.208821 | 0.945846 |
| 5 | 93742200 | 93766399 | 814.8444 | 1059.657 | 1131.723 | 474.0825 | -0.03181 | 5.72966 | 0.945855 |
| 2 | 177502800 | 177505999 | 33.30158 | 54.53716 | 72.43625 | 31.16649 | -0.03233 | 1.467294 | 0.945866 |
| 18 | 52158800 | 52159399 | 7.304193 | 14.17108 | 20.56054 | 7.744605 | 0.049738 | -0.2322 | 0.945878 |
| 5 | 79491200 | 79494599 | 32.94361 | 54.35937 | 68.81653 | 29.7037 | 0.041034 | 1.372144 | 0.945879 |
| 14 | 121248200 | 121249399 | 23.64027 | 39.83301 | 50.65412 | 22.14363 | 0.035635 | 1.106394 | 0.945885 |
| 2 | 177538000 | 177540399 | 16.86212 | 29.52327 | 37.75675 | 16.66482 | 0.034408 | 0.668712 | 0.945906 |
| 14 | 90709800 | 90711199 | 16.86212 | 29.52327 | 37.75675 | 16.66482 | 0.034408 | 0.668712 | 0.945906 |
| 18 | 67773800 | 67776199 | 17.49797 | 29.98247 | 53.27001 | 23.36148 | -0.4427 | 0.935491 | 0.945908 |
| 11 | 20711600 | 20712999 | 19.97036 | 34.88342 | 35.57053 | 15.67002 | 0.359532 | 0.717714 | 0.945923 |
| 18 | 78456600 | 78457199 | 7.606206 | 14.60833 | 22.00008 | 8.612285 | -0.05641 | -0.09708 | 0.945948 |
| 6 | 89966200 | 89977199 | 163.0221 | 243.5865 | 297.7175 | 122.7458 | -0.02797 | 3.543857 | 0.945969 |
| 5 | 109856000 | 109897799 | 928.6482 | 1203.172 | 1280.357 | 536.542 | -0.03195 | 5.791398 | 0.945982 |
| 10 | 94660400 | 94661599 | 12.18798 | 22.01564 | 30.11856 | 13.08633 | -0.0405 | 0.353527 | 0.945985 |
| 10 | 130305400 | 130306599 | 12.18798 | 22.01564 | 30.11856 | 13.08633 | -0.0405 | 0.353527 | 0.945985 |
| 13 | 3528000 | 3528799 | 12.18798 | 22.01564 | 30.11856 | 13.08633 | -0.0405 | 0.353527 | 0.945985 |
| 2 | 42845600 | 42847399 | 28.11577 | 47.15289 | 48.59448 | 21.20173 | 0.337694 | 1.162115 | 0.945986 |
| 7 | 105983000 | 105983599 | 16.42189 | 28.7785 | 36.76616 | 16.24 | 0.035977 | 0.651096 | 0.945987 |
| 12 | 36462800 | 36467999 | 50.55208 | 81.23047 | 107.2582 | 45.44957 | -0.04923 | 1.91206 | 0.94599 |
| 9 | 119575000 | 119576799 | 17.75175 | 31.10836 | 39.74733 | 17.5565 | 0.035718 | 0.66345 | 0.946016 |
| 11 | 31430800 | 31432599 | 12.1496 | 21.95016 | 30.08276 | 13.06789 | -0.04297 | 0.351204 | 0.946028 |
| 1 | 101657600 | 101658399 | 9.706901 | 18.30893 | 24.82373 | 10.13372 | 0.038613 | 0.065158 | 0.946039 |
| 2 | 58880600 | 58882799 | 23.32769 | 39.59849 | 52.78937 | 23.12895 | -0.03185 | 1.043925 | 0.946041 |
| 1 | 140886600 | 140902599 | 387.3952 | 535.92 | 607.501 | 252.3241 | -0.02823 | 4.744236 | 0.946049 |
| 6 | 130739000 | 130743999 | 69.43901 | 110.3919 | 133.9987 | 55.92012 | 0.054722 | 2.266676 | 0.946098 |
| 2 | 8422000 | 8422599 | 11.29159 | 21.36079 | 22.16251 | 8.991122 | 0.429845 | 0.053711 | 0.946098 |
| 10 | 85896600 | 85898799 | 20.98991 | 36.48593 | 46.27858 | 20.32487 | 0.045417 | 0.807029 | 0.9461 |
| X | 57064000 | 57065999 | 16.27354 | 28.75405 | 36.58619 | 16.18475 | 0.044787 | 0.547405 | 0.946118 |
| 7 | 13820600 | 13822799 | 31.8045 | 51.77598 | 69.08401 | 29.65883 | -0.04462 | 1.549218 | 0.946158 |
| 10 | 94202000 | 94204199 | 33.91456 | 55.00403 | 72.94158 | 31.24616 | -0.04038 | 1.621534 | 0.946159 |
| 16 | 62198000 | 62202599 | 72.48019 | 114.0434 | 139.9107 | 58.04538 | 0.026829 | 2.519894 | 0.946189 |

| | | | | | | | | | |
|---|---|---|---|---|---|---|---|---|---|
| 14 | 117057400 | 117060399 | 24.27916 | 41.76045 | 52.35315 | 22.96449 | 0.059281 | 0.928796 | 0.946197 |
| 5 | 41547200 | 41555399 | 34.15674 | 56.21338 | 58.52258 | 25.23242 | 0.320922 | 1.399584 | 0.946234 |
| X | 153056000 | 153058399 | 28.14078 | 47.40212 | 59.61146 | 26.05702 | 0.051536 | 1.132891 | 0.94624 |
| 4 | 39344400 | 39344799 | 5.346349 | 10.45253 | 16.61373 | 5.692991 | 0.044257 | -0.60346 | 0.946244 |
| 8 | 48244200 | 48252199 | 149.677 | 224.3629 | 274.467 | 113.2302 | -0.02507 | 3.480439 | 0.946295 |
| 1 | 113946600 | 113949399 | 28.48417 | 47.69308 | 60.24381 | 26.25735 | 0.045029 | 1.189663 | 0.946303 |
| 8 | 106567600 | 106568599 | 12.73535 | 22.98512 | 29.6563 | 12.89896 | 0.042722 | 0.363167 | 0.946321 |
| 4 | 86562600 | 86566999 | 48.1662 | 77.33407 | 100.7115 | 42.62405 | -0.03072 | 1.930795 | 0.94636 |
| 4 | 98522000 | 98524399 | 23.12383 | 39.25473 | 50.0962 | 21.94937 | 0.030913 | 1.039545 | 0.946383 |
| 10 | 72536000 | 72539799 | 38.79399 | 62.73367 | 79.07132 | 33.79685 | 0.028701 | 1.692945 | 0.946399 |
| 1 | 178579400 | 178581199 | 19.11763 | 33.24788 | 42.55342 | 18.69466 | 0.032566 | 0.785697 | 0.946401 |
| 5 | 84620400 | 84621399 | 13.0159 | 23.73682 | 32.73501 | 14.05423 | -0.04101 | 0.310447 | 0.946428 |
| 12 | 119715800 | 119716599 | 17.39254 | 30.38611 | 38.88088 | 17.1303 | 0.033708 | 0.708768 | 0.946436 |
| 14 | 44232600 | 44234199 | 19.20909 | 33.37124 | 44.70979 | 19.6288 | -0.03322 | 0.805379 | 0.946441 |
| 11 | 106619400 | 106622599 | 33.42661 | 54.74014 | 69.58241 | 29.93819 | 0.030832 | 1.465714 | 0.946446 |
| 4 | 27473000 | 27475599 | 40.2329 | 64.80343 | 80.54201 | 34.24712 | 0.042261 | 1.811262 | 0.946457 |
| 1 | 39119200 | 39121799 | 33.65294 | 54.99944 | 72.83758 | 31.31483 | -0.03 | 1.496968 | 0.946497 |
| X | 104531400 | 104532799 | 16.82643 | 29.55193 | 39.71268 | 17.57392 | -0.03672 | 0.623711 | 0.946498 |
| 16 | 8920800 | 8925199 | 61.90263 | 98.14826 | 125.6994 | 52.3166 | -0.02623 | 2.336858 | 0.946547 |
| 7 | 106316200 | 106316999 | 10.33731 | 19.46467 | 26.37234 | 10.79755 | 0.036034 | 0.085386 | 0.94655 |
| 6 | 4092000 | 4092799 | 10.76491 | 19.94514 | 26.28332 | 11.04707 | 0.046466 | 0.1913 | 0.946567 |
| 3 | 62015600 | 62015999 | 9.106376 | 17.26358 | 24.9248 | 10.0664 | -0.03769 | 0.025259 | 0.946582 |
| 6 | 123792600 | 123797999 | 95.11395 | 147.3693 | 178.6537 | 74.20779 | 0.026772 | 2.811179 | 0.946583 |
| 6 | 68464200 | 68465399 | 10.47255 | 19.82236 | 28.75648 | 11.65255 | -0.04885 | 0.059392 | 0.946594 |
| 12 | 86974000 | 86975599 | 12.36559 | 22.08287 | 38.30467 | 16.83013 | -0.39185 | 0.442805 | 0.946605 |
| 2 | 72632200 | 72633599 | 25.04501 | 42.62677 | 42.21324 | 18.50254 | 0.395972 | 1.015877 | 0.946638 |
| 10 | 121972000 | 121972799 | 12.78649 | 23.53507 | 32.95082 | 13.97424 | -0.05043 | 0.256067 | 0.946646 |
| 12 | 45244600 | 45246799 | 32.69625 | 54.70208 | 51.92425 | 22.61989 | 0.456048 | 1.20663 | 0.946675 |
| 8 | 18158000 | 18158599 | 7.132109 | 13.88823 | 20.80003 | 7.617658 | 0.041176 | -0.33344 | 0.946681 |
| 4 | 67199400 | 67199799 | 7.216903 | 14.02491 | 20.6368 | 7.683179 | 0.043794 | -0.27893 | 0.946693 |
| 1 | 190252000 | 190254599 | 30.15734 | 49.62539 | 63.16643 | 27.23326 | 0.031703 | 1.395041 | 0.946713 |
| 6 | 20269000 | 20271999 | 24.49982 | 41.65762 | 52.8516 | 23.1636 | 0.039454 | 1.03143 | 0.946714 |
| 16 | 15316400 | 15316999 | 12.15127 | 22.05444 | 28.61502 | 12.35897 | 0.041046 | 0.322317 | 0.946715 |
| 18 | 47868000 | 47870399 | 17.48995 | 30.65873 | 39.17715 | 17.30903 | 0.035451 | 0.654687 | 0.946756 |
| 3 | 62192000 | 62193799 | 24.91372 | 42.71199 | 43.07503 | 18.8913 | 0.371186 | 0.949011 | 0.946777 |
| 6 | 130693400 | 130695599 | 22.11238 | 38.10344 | 48.47316 | 21.26374 | 0.038825 | 0.899269 | 0.946786 |
| 3 | 44067200 | 44068399 | 16.20486 | 28.53809 | 36.51183 | 16.16255 | 0.034757 | 0.578883 | 0.946794 |
| 18 | 34171200 | 34172399 | 14.6876 | 26.22243 | 35.54534 | 15.62135 | -0.03849 | 0.453267 | 0.946795 |
| 5 | 57595000 | 57596399 | 19.62915 | 32.94363 | 59.01641 | 25.76047 | -0.45711 | 1.138801 | 0.9468 |
| 4 | 78429600 | 78432199 | 59.66898 | 95.1595 | 95.77594 | 40.13883 | 0.329419 | 2.150081 | 0.946803 |
| 16 | 44298600 | 44300599 | 15.12705 | 26.67592 | 34.10698 | 15.09393 | 0.036637 | 0.554861 | 0.946823 |
| 5 | 79149000 | 79150999 | 25.26399 | 42.49309 | 56.67969 | 24.76434 | -0.03322 | 1.132138 | 0.946869 |
| 9 | 67347400 | 67349599 | 18.4623 | 31.40761 | 54.77019 | 24.00544 | -0.41727 | 1.011915 | 0.946881 |
| 7 | 91596400 | 91598799 | 14.39158 | 26.4394 | 26.40727 | 11.228 | 0.431365 | 0.26639 | 0.946907 |
| 7 | 16810200 | 16811399 | 13.96185 | 24.8104 | 33.49611 | 14.77892 | -0.03666 | 0.47239 | 0.946916 |
| 12 | 11151600 | 11154399 | 23.02246 | 39.40048 | 50.10819 | 21.97271 | 0.037434 | 0.964767 | 0.946918 |
| 5 | 125748400 | 125750199 | 19.35551 | 33.44598 | 42.69185 | 18.72924 | 0.034978 | 0.856393 | 0.946958 |

| | | | | | | | | | |
|---|---|---|---|---|---|---|---|---|---|
| 4 | 33086400 | 33088999 | 29.36618 | 49.09551 | 61.96721 | 26.98271 | 0.046195 | 1.210556 | 0.946978 |
| 5 | 57784000 | 57784799 | 10.83055 | 20.11009 | 26.72134 | 11.19827 | 0.037242 | 0.178721 | 0.946987 |
| 11 | 23768800 | 23770199 | 11.84979 | 21.82257 | 30.27116 | 12.82851 | -0.03613 | 0.241277 | 0.946994 |
| 9 | 112196800 | 112197599 | 10.50787 | 19.48737 | 25.72541 | 10.79864 | 0.04496 | 0.1803 | 0.947005 |
| 4 | 102118200 | 102119399 | 13.51952 | 24.33266 | 33.05787 | 14.42857 | -0.03485 | 0.390406 | 0.947021 |
| 17 | 77062600 | 77066799 | 53.24312 | 85.21845 | 105.9989 | 44.6501 | 0.03008 | 2.034784 | 0.947032 |
| X | 114036600 | 114037799 | 20.51198 | 35.7083 | 45.1925 | 19.86647 | 0.048941 | 0.785248 | 0.947049 |
| 6 | 3189600 | 3191999 | 44.31406 | 71.19603 | 88.6041 | 37.53685 | 0.035849 | 1.903173 | 0.947055 |
| 10 | 22070400 | 22071599 | 10.23394 | 19.06063 | 26.79593 | 11.17586 | -0.03806 | 0.15585 | 0.947092 |
| 8 | 34683400 | 34687999 | 43.63083 | 70.19348 | 91.73978 | 38.94174 | -0.0323 | 1.856787 | 0.947098 |
| 6 | 48098800 | 48101999 | 32.13576 | 52.86841 | 67.34618 | 29.03238 | 0.030421 | 1.405788 | 0.947101 |
| 1 | 71629200 | 71630199 | 13.45357 | 24.28391 | 31.51223 | 13.70442 | 0.034266 | 0.369449 | 0.947121 |
| 4 | 121208400 | 121239199 | 515.3641 | 701.8431 | 751.7937 | 313.0296 | 0.030549 | 4.99072 | 0.947127 |
| 10 | 117831600 | 117833599 | 26.55344 | 44.58944 | 56.73437 | 24.77304 | 0.03456 | 1.150884 | 0.947134 |
| 1 | 102741000 | 102743399 | 20.25134 | 34.98875 | 46.86869 | 20.56156 | -0.03443 | 0.867735 | 0.947135 |
| 14 | 63882600 | 63892199 | 141.1893 | 212.2312 | 250.7397 | 103.5167 | 0.02872 | 3.423769 | 0.947144 |
| 5 | 146278400 | 146282599 | 43.60434 | 70.07871 | 91.2811 | 38.6893 | -0.02923 | 1.89158 | 0.947155 |
| 4 | 147809400 | 147847199 | 1376.183 | 1724.924 | 1770.442 | 744.3827 | -0.02792 | 6.328964 | 0.947161 |
| 8 | 72011000 | 72012999 | 17.67345 | 30.72417 | 39.05464 | 17.15332 | 0.042508 | 0.780136 | 0.947177 |
| 16 | 9634200 | 9636199 | 15.31688 | 26.94538 | 36.16111 | 16.01073 | -0.03425 | 0.586341 | 0.947196 |
| 4 | 45846400 | 45849399 | 31.86429 | 52.20786 | 66.39789 | 28.5741 | 0.030446 | 1.455766 | 0.947199 |
| 4 | 12396200 | 12396599 | 6.243527 | 12.18343 | 18.68634 | 6.6527 | 0.044854 | -0.44968 | 0.9472 |
| 8 | 7829400 | 7829999 | 9.819763 | 18.62933 | 25.47642 | 10.27362 | 0.042171 | 0.030499 | 0.947201 |
| 1 | 195034400 | 195035599 | 32.90408 | 53.49039 | 70.9333 | 30.43708 | -0.03725 | 1.57132 | 0.947216 |
| 10 | 100887600 | 100888799 | 9.655209 | 18.28618 | 24.92181 | 10.086 | 0.042617 | 0.03925 | 0.947271 |
| 1 | 50851400 | 50851799 | 7.837049 | 15.04804 | 22.47064 | 8.802615 | -0.04613 | -0.09451 | 0.947277 |
| 14 | 76553800 | 76555199 | 23.31309 | 40.24336 | 40.95405 | 17.96629 | 0.360085 | 0.879421 | 0.947288 |
| 5 | 100461800 | 100463399 | 18.81974 | 32.60392 | 41.63069 | 18.26513 | 0.035234 | 0.828733 | 0.947296 |
| 1 | 132610600 | 132612399 | 16.18619 | 28.57806 | 36.55381 | 16.17287 | 0.036717 | 0.553567 | 0.947317 |
| 3 | 112439400 | 112441599 | 35.69877 | 58.46408 | 60.631 | 26.08736 | 0.323731 | 1.465495 | 0.947338 |
| 3 | 43978400 | 43979799 | 19.17659 | 33.28838 | 42.62934 | 18.7061 | 0.031579 | 0.813594 | 0.947366 |
| 7 | 106673200 | 106673799 | 8.155985 | 15.7556 | 22.61556 | 8.692004 | 0.036361 | -0.15204 | 0.947379 |
| 19 | 53801000 | 53802799 | 16.57677 | 28.81816 | 39.12153 | 17.18288 | -0.05188 | 0.774806 | 0.947397 |
| 7 | 12120600 | 12121999 | 30.19287 | 49.64032 | 65.91534 | 28.40787 | -0.02965 | 1.408211 | 0.947397 |
| 1 | 131575800 | 131578799 | 32.56039 | 53.6979 | 50.33607 | 21.72188 | 0.472996 | 1.362953 | 0.947407 |
| 7 | 127224400 | 127227799 | 26.79968 | 44.61088 | 59.41158 | 25.79922 | -0.03083 | 1.247475 | 0.947417 |
| 3 | 82877200 | 82881599 | 41.10866 | 66.30718 | 83.3672 | 35.51584 | 0.02805 | 1.773053 | 0.947428 |
| 11 | 17910800 | 17920599 | 112.2275 | 172.1298 | 216.0368 | 89.68186 | -0.03442 | 3.012988 | 0.947436 |
| 8 | 112866600 | 112870999 | 59.35849 | 94.26944 | 120.9557 | 50.42523 | -0.0266 | 2.284831 | 0.94746 |
| 18 | 40941800 | 40942399 | 9.378648 | 17.69552 | 25.35776 | 10.34463 | -0.03995 | 0.060424 | 0.947468 |
| 2 | 171341800 | 171344399 | 35.4935 | 57.66361 | 72.96891 | 31.29742 | 0.029294 | 1.589294 | 0.947501 |
| 16 | 15561200 | 15564599 | 87.05768 | 134.8599 | 162.7688 | 67.6097 | 0.032929 | 2.809644 | 0.947512 |
| 13 | 118297200 | 118298399 | 14.1027 | 25.13478 | 33.97119 | 14.95127 | -0.03582 | 0.455439 | 0.947515 |
| 2 | 53965400 | 53965799 | 7.111989 | 13.75323 | 21.01708 | 8.038199 | -0.04603 | -0.18443 | 0.947528 |
| 12 | 104547000 | 104550999 | 33.76 | 54.84235 | 72.55025 | 31.11576 | -0.03469 | 1.588864 | 0.947531 |
| 9 | 17999800 | 18001199 | 31.76282 | 51.83654 | 68.82109 | 29.57268 | -0.03496 | 1.512527 | 0.947571 |
| 15 | 13252600 | 13252999 | 6.953577 | 13.53184 | 21.34583 | 7.859045 | -0.04051 | -0.31762 | 0.947576 |

| | | | | | | | | | |
|---|---|---|---|---|---|---|---|---|---|
| 4 | 113972200 | 114001199 | 775.0058 | 1016.784 | 1053.588 | 440.9463 | 0.024521 | 5.586618 | 0.947588 |
| 4 | 153166000 | 153171599 | 73.76408 | 116.188 | 142.737 | 59.22618 | 0.026036 | 2.499829 | 0.947593 |
| 16 | 36373400 | 36374199 | 15.01545 | 26.81472 | 34.39203 | 15.11118 | 0.040883 | 0.44914 | 0.947595 |
| 3 | 47752000 | 47754399 | 26.31962 | 44.15272 | 58.90229 | 25.70348 | -0.03346 | 1.163948 | 0.947598 |
| 17 | 24061000 | 24063399 | 20.74164 | 34.71615 | 61.26091 | 26.69432 | -0.4355 | 1.172649 | 0.947611 |
| 6 | 142081600 | 142084599 | 26.54655 | 44.50549 | 59.54739 | 25.9747 | -0.03773 | 1.169856 | 0.947611 |
| X | 157910600 | 157913199 | 23.61025 | 40.42636 | 51.18032 | 22.44374 | 0.044203 | 0.967781 | 0.947616 |
| 4 | 80501400 | 80505199 | 54.70351 | 87.2047 | 108.1579 | 45.30216 | 0.027926 | 2.160281 | 0.947624 |
| 1 | 35877600 | 35879199 | 17.02146 | 29.83163 | 38.19759 | 16.87343 | 0.032605 | 0.654221 | 0.947628 |
| 9 | 38985200 | 38988799 | 68.00214 | 107.3161 | 131.9225 | 54.7607 | 0.027255 | 2.45556 | 0.947643 |
| 2 | 33800200 | 33802399 | 25.0165 | 41.8918 | 53.33522 | 23.24795 | 0.033858 | 1.174391 | 0.947671 |
| 12 | 51595000 | 51601399 | 162.6485 | 242.31 | 294.6341 | 121.4108 | -0.02442 | 3.59717 | 0.9477 |
| 16 | 19745200 | 19748199 | 42.23201 | 68.4832 | 70.06896 | 30.01225 | 0.332651 | 1.638977 | 0.947719 |
| 7 | 127036800 | 127039799 | 37.49661 | 60.71547 | 79.68238 | 34.09315 | -0.02764 | 1.66368 | 0.947769 |
| 16 | 68689600 | 68690399 | 18.25144 | 32.02345 | 30.60929 | 13.53429 | 0.452459 | 0.631136 | 0.94777 |
| 6 | 55708600 | 55709999 | 16.12905 | 28.24079 | 35.88508 | 15.8392 | 0.043854 | 0.66296 | 0.947775 |
| X | 21365000 | 21366599 | 19.61175 | 34.23348 | 43.52677 | 19.16673 | 0.042631 | 0.739795 | 0.947801 |
| 1 | 111021200 | 111026399 | 98.99196 | 152.5078 | 190.6713 | 79.20045 | -0.02394 | 2.91809 | 0.947866 |
| 6 | 62878600 | 62878999 | 6.380098 | 12.42219 | 18.51579 | 6.789011 | 0.047086 | -0.34593 | 0.947885 |
| 13 | 93342400 | 93344199 | 18.10447 | 31.57313 | 40.39804 | 17.77865 | 0.03336 | 0.738042 | 0.947897 |
| 6 | 76195400 | 76197399 | 30.89331 | 50.93259 | 67.80161 | 29.25502 | -0.03169 | 1.378373 | 0.947907 |
| 6 | 59762800 | 59764999 | 20.74724 | 35.49524 | 47.43456 | 20.79977 | -0.03362 | 0.958914 | 0.947931 |
| 2 | 23812200 | 23820399 | 154.9886 | 231.7177 | 273.0112 | 112.5724 | 0.025682 | 3.530092 | 0.947947 |
| 10 | 89964400 | 89972199 | 114.695 | 173.5681 | 258.1328 | 106.7793 | -0.29467 | 3.286206 | 0.94796 |
| 5 | 51163000 | 51165599 | 60.92528 | 96.59205 | 118.7159 | 49.40666 | 0.033011 | 2.336992 | 0.947963 |
| 7 | 118926800 | 118927399 | 7.895061 | 15.28921 | 22.15946 | 8.419139 | 0.038749 | -0.19334 | 0.947977 |
| 8 | 97391200 | 97394399 | 49.51713 | 79.3184 | 102.689 | 43.30756 | -0.0262 | 2.005586 | 0.947982 |
| 2 | 86380800 | 86382199 | 15.65531 | 27.68595 | 35.50985 | 15.70585 | 0.033667 | 0.530116 | 0.948006 |
| X | 109940400 | 109941799 | 16.95107 | 29.79484 | 38.09971 | 16.87163 | 0.03454 | 0.608518 | 0.948029 |
| 14 | 34497600 | 34500599 | 32.20815 | 52.68155 | 69.7412 | 29.99261 | -0.02857 | 1.481 | 0.948054 |
| 4 | 64270800 | 64273799 | 22.63423 | 39.01336 | 52.89889 | 23.20537 | -0.0528 | 0.903879 | 0.948062 |
| Y | 90745200 | 90759399 | 6948.871 | 7460.657 | 9303.998 | 4005.373 | -0.53627 | 8.777125 | 0.9481 |
| 13 | 30575600 | 30578199 | 31.83086 | 52.51323 | 66.85965 | 28.85671 | 0.032757 | 1.371177 | 0.948108 |
| 13 | 61610600 | 61612199 | 34.42503 | 56.04901 | 74.0358 | 31.78615 | -0.03007 | 1.555674 | 0.948121 |
| 19 | 54081600 | 54084399 | 22.14518 | 37.50757 | 50.09917 | 21.93545 | -0.03473 | 1.05517 | 0.94813 |
| 1 | 182532000 | 182536799 | 61.86674 | 98.3677 | 126.5545 | 52.82103 | -0.02885 | 2.254799 | 0.948153 |
| 14 | 52883200 | 52889399 | 81.19753 | 127.4718 | 162.9662 | 67.59067 | -0.03509 | 2.574232 | 0.948159 |
| 8 | 26762200 | 26765199 | 28.50103 | 47.15753 | 62.69929 | 27.11386 | -0.0291 | 1.324911 | 0.948164 |
| 1 | 93816800 | 93820599 | 33.0719 | 54.09866 | 71.87035 | 30.90917 | -0.03365 | 1.480279 | 0.948184 |
| 2 | 80891600 | 80892599 | 11.71007 | 21.69067 | 28.71161 | 12.07198 | 0.038933 | 0.202294 | 0.948191 |
| 3 | 159408400 | 159410399 | 23.12294 | 40.05285 | 39.25009 | 17.22059 | 0.415172 | 0.843781 | 0.948259 |
| 14 | 95568400 | 95571399 | 42.6921 | 69.07765 | 71.60033 | 30.61863 | 0.311039 | 1.68375 | 0.948312 |
| 14 | 63445800 | 63448399 | 25.16658 | 42.05022 | 56.05005 | 24.3973 | -0.03211 | 1.19994 | 0.948325 |
| 1 | 96408000 | 96409399 | 13.3565 | 24.26194 | 31.52894 | 13.60345 | 0.039384 | 0.330855 | 0.948332 |
| 14 | 62882400 | 62883399 | 11.26478 | 20.73243 | 28.75935 | 12.19806 | -0.03671 | 0.238245 | 0.948358 |
| Y | 39628400 | 39628999 | 6.081755 | 11.8296 | 16.93119 | 6.257762 | 0.093952 | -0.33075 | 0.948365 |
| 15 | 65538000 | 65540799 | 41.67556 | 67.43093 | 66.88032 | 28.59961 | 0.374437 | 1.679986 | 0.948405 |

| | | | | | | | | | |
|---|---|---|---|---|---|---|---|---|---|
| 10 | 3820600 | 3824199 | 39.84072 | 64.24275 | 84.0719 | 35.80219 | -0.03008 | 1.780817 | 0.948443 |
| 7 | 89513200 | 89516999 | 34.61514 | 56.40751 | 71.56002 | 30.73294 | 0.028881 | 1.539323 | 0.948456 |
| X | 35151000 | 35153599 | 25.82409 | 43.44161 | 58.24161 | 25.44791 | -0.04044 | 1.138381 | 0.948493 |
| 5 | 45314000 | 45314399 | 6.080692 | 11.85691 | 18.04068 | 6.490632 | 0.04182 | -0.42075 | 0.948496 |
| 15 | 65030000 | 65030399 | 6.080692 | 11.85691 | 18.04068 | 6.490632 | 0.04182 | -0.42075 | 0.948496 |
| 19 | 51458000 | 51458599 | 13.01665 | 23.5503 | 32.10305 | 13.92057 | -0.03357 | 0.354791 | 0.948537 |
| 1 | 136720200 | 136721999 | 17.31053 | 30.196 | 40.51105 | 17.83115 | -0.03443 | 0.734 | 0.948561 |
| 11 | 51003200 | 51011399 | 38.9528 | 63.85742 | 61.98936 | 26.68317 | 0.419411 | 1.463415 | 0.948576 |
| 11 | 114837400 | 114840799 | 45.47262 | 73.21505 | 91.71996 | 38.97845 | 0.03012 | 1.835523 | 0.948577 |
| 11 | 89196200 | 89198999 | 32.73284 | 53.35576 | 70.58248 | 30.31634 | -0.03101 | 1.532345 | 0.94858 |
| 8 | 19139600 | 19142399 | 27.86445 | 45.6858 | 74.46844 | 32.05656 | -0.32619 | 1.443672 | 0.948583 |
| 7 | 129536400 | 129540399 | 32.76808 | 53.81884 | 68.5236 | 29.51911 | 0.030147 | 1.427867 | 0.94859 |
| 1 | 23822600 | 23825999 | 45.84539 | 73.5609 | 95.51062 | 40.38369 | -0.02746 | 1.945431 | 0.948606 |
| 2 | 50887200 | 50890599 | 55.41433 | 88.11589 | 113.6701 | 47.44739 | -0.03257 | 2.243274 | 0.948616 |
| 5 | 116335600 | 116338199 | 32.58492 | 52.98008 | 70.31659 | 30.17422 | -0.03831 | 1.56822 | 0.948622 |
| 18 | 21595400 | 21598199 | 26.65714 | 44.62543 | 59.45662 | 25.91113 | -0.03157 | 1.187714 | 0.948672 |
| 6 | 22817600 | 22818199 | 8.771037 | 16.78766 | 23.47041 | 9.27903 | 0.035748 | -0.05126 | 0.948681 |
| 8 | 113135000 | 113138999 | 40.122 | 64.76115 | 84.65649 | 36.09775 | -0.02685 | 1.751387 | 0.94869 |
| 5 | 78242200 | 78245399 | 39.53865 | 64.31514 | 80.68798 | 34.63047 | 0.043084 | 1.582522 | 0.948712 |
| 1 | 189840800 | 189844199 | 44.28796 | 71.18994 | 92.6317 | 39.27177 | -0.02741 | 1.888012 | 0.948716 |
| 7 | 72846600 | 72850399 | 34.00452 | 55.35717 | 73.04389 | 31.35881 | -0.0287 | 1.553775 | 0.948743 |
| 9 | 48659400 | 48661199 | 17.05307 | 29.79455 | 39.90104 | 17.58033 | -0.03176 | 0.707199 | 0.94876 |
| 7 | 113570000 | 113573399 | 42.57394 | 68.54579 | 85.99183 | 36.54252 | 0.02808 | 1.831087 | 0.948789 |
| 16 | 82070600 | 82072599 | 18.22718 | 31.9478 | 40.74971 | 18.00226 | 0.038128 | 0.66393 | 0.948799 |
| 12 | 116725800 | 116727999 | 23.00302 | 39.23203 | 50.09206 | 21.95956 | 0.031107 | 0.998081 | 0.948807 |
| 6 | 131330200 | 131331399 | 10.65058 | 19.81707 | 27.77658 | 11.60421 | -0.03604 | 0.166543 | 0.948813 |
| 9 | 16945000 | 16946399 | 12.94246 | 24.35069 | 24.00956 | 9.8434 | 0.489512 | 0.107025 | 0.948819 |
| 4 | 45524200 | 45526799 | 20.10033 | 34.61363 | 44.23739 | 19.40524 | 0.032059 | 0.890959 | 0.948832 |
| 7 | 6698600 | 6699799 | 10.92913 | 20.26989 | 28.30863 | 11.88197 | -0.03554 | 0.191307 | 0.948845 |
| 17 | 72761600 | 72770399 | 112.8701 | 172.7143 | 207.7553 | 86.2054 | 0.024211 | 3.056785 | 0.948854 |
| 1 | 137663800 | 137665999 | 19.59856 | 34.293 | 43.44215 | 19.16409 | 0.047999 | 0.702828 | 0.948859 |
| 18 | 68667800 | 68668999 | 14.24276 | 25.49678 | 34.58046 | 15.16285 | -0.0369 | 0.432789 | 0.948866 |
| 11 | 50900400 | 50903199 | 20.99498 | 36.43254 | 46.29811 | 20.31395 | 0.042338 | 0.827783 | 0.94888 |
| 1 | 149957200 | 149959599 | 38.54497 | 62.86107 | 65.1251 | 27.97739 | 0.321175 | 1.533656 | 0.948887 |
| 2 | 100186200 | 100187999 | 19.82937 | 33.89734 | 57.66589 | 25.28449 | -0.38086 | 0.973368 | 0.948914 |
| 8 | 3843200 | 3844399 | 31.71483 | 51.71906 | 64.91296 | 27.88565 | 0.045128 | 1.519782 | 0.948917 |
| 6 | 67528800 | 67529999 | 12.19787 | 22.2655 | 29.15067 | 12.50093 | 0.03483 | 0.292138 | 0.94893 |
| 9 | 7705600 | 7706999 | 12.19787 | 22.2655 | 29.15067 | 12.50093 | 0.03483 | 0.292138 | 0.94893 |
| 11 | 71398800 | 71399199 | 7.866639 | 15.10314 | 22.48305 | 8.81035 | -0.04182 | -0.09173 | 0.948939 |
| 13 | 104032000 | 104045199 | 171.7703 | 256.7364 | 316.2104 | 130.3779 | -0.03865 | 3.540213 | 0.948943 |
| 10 | 126753200 | 126757199 | 45.33696 | 72.85106 | 91.25251 | 38.66777 | 0.02676 | 1.898003 | 0.948946 |
| 11 | 34501400 | 34514199 | 121.2033 | 186.9168 | 217.2361 | 90.23604 | 0.082414 | 2.91235 | 0.948984 |
| 4 | 6733400 | 6734799 | 16.76642 | 29.36529 | 39.30722 | 17.35394 | -0.03105 | 0.66627 | 0.948999 |
| X | 37388400 | 37390799 | 21.92021 | 37.48589 | 50.6004 | 22.18721 | -0.04808 | 0.974179 | 0.949002 |
| 19 | 23989000 | 23990599 | 16.59984 | 29.37705 | 37.4627 | 16.56725 | 0.042172 | 0.530668 | 0.94901 |
| 2 | 7933600 | 7935999 | 18.60922 | 32.46159 | 43.5694 | 19.17737 | -0.03519 | 0.744112 | 0.949024 |
| 18 | 46160200 | 46162999 | 25.15759 | 41.65942 | 68.58909 | 29.67753 | -0.33645 | 1.311246 | 0.94907 |

| | | | | | | | | | |
|---|---|---|---|---|---|---|---|---|---|
| 13 | 72696800 | 72698199 | 13.98099 | 24.89844 | 32.02879 | 14.10566 | 0.034632 | 0.457054 | 0.949112 |
| 8 | 31905600 | 31906399 | 11.44484 | 20.89966 | 27.12167 | 11.62476 | 0.047794 | 0.27712 | 0.949135 |
| 1 | 66506600 | 66508199 | 16.01889 | 27.93146 | 37.97041 | 16.70857 | -0.05358 | 0.725758 | 0.949151 |
| 6 | 81023200 | 81023599 | 5.793253 | 11.31393 | 18.78861 | 6.596424 | -0.04982 | -0.50254 | 0.949151 |
| 1 | 155280200 | 155283199 | 41.44684 | 66.90328 | 84.27077 | 35.93584 | 0.026664 | 1.752217 | 0.949151 |
| 8 | 34382600 | 34387799 | 43.33294 | 69.70832 | 90.90144 | 38.58114 | -0.02921 | 1.859287 | 0.949159 |
| 5 | 133496000 | 133498399 | 55.80096 | 88.69803 | 114.3905 | 47.7305 | -0.03269 | 2.255462 | 0.949174 |
| X | 147453400 | 147455799 | 17.83049 | 31.305 | 42.31152 | 18.71876 | -0.04506 | 0.638489 | 0.949208 |
| 11 | 107158400 | 107160599 | 27.48338 | 45.89242 | 61.16626 | 26.6137 | -0.03204 | 1.218369 | 0.949221 |
| 5 | 100159200 | 100163799 | 35.76008 | 58.04581 | 76.58619 | 32.8277 | -0.03206 | 1.6074 | 0.949232 |
| 10 | 34362400 | 34370599 | 88.05907 | 138.6455 | 165.3354 | 68.59917 | 0.068407 | 2.52281 | 0.949239 |
| 12 | 93881400 | 93887799 | 66.88219 | 106.2527 | 130.553 | 54.44397 | 0.03623 | 2.285032 | 0.949273 |
| 1 | 32003600 | 32005199 | 19.85496 | 34.22592 | 43.73942 | 19.1874 | 0.03254 | 0.881754 | 0.949282 |
| 6 | 9319800 | 9320399 | 9.099412 | 17.37726 | 23.9197 | 9.51189 | 0.049488 | -0.03151 | 0.949288 |
| 15 | 8306800 | 8307599 | 10.20762 | 19.25745 | 27.51773 | 11.22845 | -0.03621 | 0.074892 | 0.949291 |
| 13 | 81364000 | 81366799 | 19.48891 | 33.95014 | 45.70618 | 20.10078 | -0.0399 | 0.768347 | 0.949314 |
| 10 | 13329000 | 13330999 | 20.7811 | 35.78047 | 45.80403 | 20.09237 | 0.029636 | 0.897582 | 0.94933 |
| 10 | 97151200 | 97152799 | 10.21357 | 18.66945 | 32.89184 | 14.08072 | -0.38909 | 0.277567 | 0.949332 |
| 3 | 9073800 | 9076599 | 39.15607 | 63.48393 | 80.15012 | 34.32574 | 0.029823 | 1.641832 | 0.949332 |
| 7 | 60187600 | 60191399 | 46.11974 | 74.09345 | 92.66483 | 39.25438 | 0.028649 | 1.908383 | 0.949336 |
| 14 | 8456000 | 8458399 | 17.14408 | 29.96147 | 38.31853 | 16.88636 | 0.034338 | 0.700245 | 0.949356 |
| 14 | 114973400 | 114974999 | 21.1176 | 35.45073 | 60.66856 | 26.48588 | -0.39135 | 1.144938 | 0.949357 |
| 5 | 127724600 | 127729199 | 44.29689 | 71.22817 | 89.12436 | 37.80318 | 0.029502 | 1.873433 | 0.949359 |
| 19 | 49671400 | 49681799 | 89.97422 | 140.6336 | 144.1695 | 59.81438 | 0.278222 | 2.653661 | 0.949375 |
| 15 | 33105800 | 33109599 | 55.54967 | 88.63333 | 110.2618 | 46.24105 | 0.024966 | 2.133343 | 0.949375 |
| 6 | 17439000 | 17441399 | 24.5285 | 40.77714 | 66.88613 | 29.01908 | -0.33057 | 1.264515 | 0.949388 |
| 15 | 52406800 | 52414399 | 97.6134 | 150.56 | 188.3886 | 78.25324 | -0.02382 | 2.895442 | 0.949392 |
| 3 | 10181400 | 10182599 | 14.48664 | 25.67487 | 34.50617 | 15.25531 | -0.03222 | 0.496597 | 0.94942 |
| 1 | 106045400 | 106048599 | 34.19277 | 55.5182 | 73.29027 | 31.42181 | -0.03237 | 1.59826 | 0.94942 |
| 3 | 92510600 | 92512999 | 18.48898 | 32.38269 | 41.27326 | 18.22215 | 0.039153 | 0.676859 | 0.949436 |
| 9 | 109288800 | 109292199 | 51.38975 | 82.14828 | 106.1649 | 44.64015 | -0.027 | 2.070243 | 0.949477 |
| 18 | 37125200 | 37127999 | 85.41462 | 132.3597 | 159.2655 | 66.1544 | 0.037704 | 2.802925 | 0.949481 |
| 5 | 101306600 | 101307199 | 9.505206 | 18.12493 | 25.03581 | 9.987438 | 0.041598 | -0.0084 | 0.949486 |
| 4 | 75521200 | 75523599 | 25.91658 | 43.57815 | 55.59325 | 24.28753 | 0.030857 | 1.136027 | 0.949488 |
| 2 | 112679800 | 112682399 | 23.87991 | 41.12285 | 51.77967 | 22.71376 | 0.053407 | 0.917356 | 0.949498 |
| 13 | 11140600 | 11144599 | 47.85646 | 76.80404 | 99.68016 | 42.16026 | -0.02652 | 1.943551 | 0.949502 |
| 15 | 45669200 | 45670399 | 14.4204 | 25.91791 | 33.46463 | 14.62434 | 0.03676 | 0.404732 | 0.949512 |
| 9 | 86102600 | 86104999 | 27.06899 | 44.96879 | 59.83958 | 25.94923 | -0.0298 | 1.27382 | 0.949516 |
| 6 | 113140000 | 113141199 | 11.21531 | 20.66084 | 27.22135 | 11.53093 | 0.03842 | 0.228414 | 0.949538 |
| 2 | 84298800 | 84300599 | 17.53411 | 30.60226 | 40.94478 | 18.02808 | -0.03062 | 0.726141 | 0.949578 |
| 1 | 73921400 | 73925399 | 63.35832 | 100.6123 | 124.5386 | 51.91402 | 0.025019 | 2.292422 | 0.94958 |
| 15 | 42611400 | 42613799 | 22.14518 | 38.19988 | 48.47654 | 21.2659 | 0.042569 | 0.888116 | 0.949607 |
| 16 | 20074400 | 20076799 | 20.98553 | 35.9155 | 45.85391 | 20.10723 | 0.032024 | 0.953427 | 0.949628 |
| 17 | 78382400 | 78383399 | 16.33392 | 27.97039 | 51.07387 | 22.39724 | -0.4818 | 0.938131 | 0.949662 |
| 9 | 8203800 | 8204399 | 8.467836 | 16.26609 | 24.12309 | 9.438497 | -0.0347 | -0.08647 | 0.949675 |
| 16 | 95643200 | 95645399 | 30.01994 | 49.50555 | 63.21456 | 27.2788 | 0.028369 | 1.369106 | 0.949677 |
| 5 | 104997800 | 105010199 | 202.3138 | 299.1139 | 346.2838 | 142.6083 | 0.036177 | 3.735793 | 0.949695 |

| | | | | | | | | | |
|---|---|---|---|---|---|---|---|---|---|
| 11 | 18247200 | 18247599 | 7.579648 | 14.65209 | 21.00666 | 8.048325 | 0.04303 | -0.17197 | 0.949702 |
| 4 | 112838200 | 112839599 | 28.09625 | 46.62856 | 62.07981 | 26.90206 | -0.03065 | 1.290752 | 0.949737 |
| 6 | 101575400 | 101578199 | 22.20581 | 38.46267 | 52.37516 | 22.97905 | -0.05746 | 0.859202 | 0.949751 |
| 4 | 104919800 | 104921399 | 20.84062 | 35.17914 | 58.09139 | 25.40877 | -0.33982 | 1.089473 | 0.949755 |
| 4 | 7095800 | 7101999 | 65.80557 | 104.116 | 132.9221 | 55.24719 | -0.02429 | 2.393396 | 0.949781 |
| 3 | 111187800 | 111189999 | 21.49949 | 37.23874 | 47.39235 | 20.79288 | 0.039792 | 0.85035 | 0.949783 |
| 3 | 86242400 | 86243799 | 15.46934 | 26.28264 | 54.13619 | 23.72514 | -0.65659 | 1.011399 | 0.949784 |
| 13 | 79899800 | 79900399 | 8.916306 | 16.87759 | 24.3674 | 9.872961 | -0.0419 | 0.031357 | 0.949791 |
| 4 | 112139800 | 112146799 | 90.61946 | 141.3132 | 171.5082 | 71.17431 | 0.032334 | 2.698584 | 0.949799 |
| 10 | 70944200 | 70952799 | 114.6693 | 173.8586 | 257.5777 | 106.6132 | -0.28684 | 3.247127 | 0.94981 |
| 6 | 27204800 | 27205199 | 6.828593 | 13.26278 | 20.60071 | 7.704861 | -0.03854 | -0.26392 | 0.949813 |
| 1 | 141341400 | 141341999 | 7.70337 | 14.9055 | 21.56296 | 8.224629 | 0.035825 | -0.18547 | 0.949814 |
| 1 | 40815600 | 40819799 | 51.12945 | 81.80234 | 105.6768 | 44.48891 | -0.02508 | 2.044821 | 0.94982 |
| X | 159043400 | 159045799 | 27.79478 | 46.65244 | 62.85972 | 27.4397 | -0.04777 | 1.16874 | 0.949824 |
| X | 71906800 | 71910199 | 29.41154 | 48.80043 | 62.18129 | 26.94185 | 0.032444 | 1.297114 | 0.949834 |
| 16 | 47604000 | 47604799 | 8.992222 | 17.16115 | 25.05416 | 9.977034 | -0.03556 | -0.02002 | 0.949834 |
| 4 | 45386400 | 45389599 | 31.16866 | 51.35533 | 65.56594 | 28.2831 | 0.02808 | 1.384127 | 0.949834 |
| 4 | 12110600 | 12119599 | 292.12 | 414.7005 | 464.8484 | 192.2804 | 0.025508 | 4.355493 | 0.949836 |
| 8 | 61458600 | 61459799 | 33.76448 | 55.75419 | 56.83798 | 24.54509 | 0.352783 | 1.358929 | 0.949845 |
| 13 | 100978000 | 100980799 | 29.64522 | 48.93561 | 65.00949 | 28.06733 | -0.02819 | 1.356334 | 0.949873 |
| 15 | 40170800 | 40171199 | 7.2819 | 14.10946 | 21.61744 | 8.192355 | -0.037 | -0.21042 | 0.9499 |
| 18 | 20435200 | 20435799 | 5.94245 | 11.58196 | 18.57589 | 6.724047 | -0.04088 | -0.39765 | 0.949908 |
| 13 | 85616400 | 85620399 | 57.11526 | 90.92383 | 116.7718 | 48.81684 | -0.02443 | 2.210319 | 0.949934 |
| 14 | 56431600 | 56433199 | 19.632 | 33.03876 | 57.93475 | 25.31699 | -0.4262 | 1.113506 | 0.949935 |
| 3 | 119499000 | 119499599 | 7.123833 | 13.80923 | 20.00266 | 7.564917 | 0.046307 | -0.224 | 0.949953 |
| 19 | 4075000 | 4078599 | 50.81411 | 81.18695 | 100.8395 | 42.37051 | 0.029397 | 2.085127 | 0.950016 |
| X | 54643400 | 54643799 | 7.134478 | 13.88318 | 20.73311 | 7.636592 | 0.037807 | -0.31343 | 0.950033 |
| 3 | 16397200 | 16400799 | 39.00429 | 63.08003 | 83.13814 | 35.53967 | -0.03547 | 1.689407 | 0.950048 |
| 16 | 68471800 | 68472799 | 13.34192 | 24.21235 | 33.16848 | 14.32794 | -0.03716 | 0.341674 | 0.950048 |
| 4 | 147178600 | 147220399 | 1193.997 | 1518.655 | 1590.806 | 667.8387 | -0.03596 | 6.092363 | 0.950056 |
| 14 | 50083000 | 50083999 | 22.08167 | 37.64165 | 47.96838 | 21.02771 | 0.033896 | 0.998223 | 0.950069 |
| 3 | 119431400 | 119432799 | 13.30927 | 23.94129 | 31.0186 | 13.54566 | 0.03273 | 0.387624 | 0.950102 |
| 7 | 106936000 | 106940199 | 46.67016 | 75.09935 | 94.03806 | 39.92979 | 0.029732 | 1.857043 | 0.950133 |
| 13 | 69431000 | 69433399 | 29.45268 | 49.48422 | 49.10557 | 21.44797 | 0.392088 | 1.151014 | 0.95014 |
| 6 | 59945000 | 59945999 | 12.35915 | 22.59594 | 29.65403 | 12.69144 | 0.033217 | 0.285228 | 0.950144 |
| 6 | 87251400 | 87253599 | 23.11965 | 39.26072 | 52.25187 | 22.89617 | -0.02945 | 1.038475 | 0.950147 |
| 13 | 33878600 | 33880599 | 26.80467 | 44.75746 | 57.21574 | 24.89424 | 0.028006 | 1.213377 | 0.950154 |
| 8 | 8130600 | 8132799 | 33.7199 | 55.496 | 70.32511 | 30.32152 | 0.038166 | 1.408621 | 0.950169 |
| 18 | 18894400 | 18899999 | 23.14026 | 39.77955 | 41.03746 | 18.0006 | 0.339323 | 0.915613 | 0.95017 |
| 2 | 177492200 | 177493999 | 24.50562 | 41.91819 | 43.15337 | 18.92224 | 0.341212 | 0.968731 | 0.950171 |
| 5 | 89852200 | 89855399 | 32.83912 | 53.71478 | 71.03717 | 30.5502 | -0.02712 | 1.482297 | 0.950183 |
| 17 | 52701800 | 52704799 | 21.82686 | 37.93623 | 47.92028 | 21.0443 | 0.051295 | 0.814554 | 0.950186 |
| 15 | 42738000 | 42741399 | 30.82167 | 50.61261 | 67.10948 | 28.90765 | -0.0283 | 1.424722 | 0.950214 |
| X | 88122600 | 88125399 | 21.76121 | 38.22443 | 33.68635 | 14.90897 | 0.569065 | 0.639805 | 0.950234 |
| 7 | 6502400 | 6505999 | 42.27643 | 68.29789 | 85.91586 | 36.67579 | 0.029866 | 1.730906 | 0.950236 |
| 10 | 111699200 | 111700799 | 14.58784 | 25.54871 | 43.0288 | 18.9964 | -0.3608 | 0.661215 | 0.950336 |
| 1 | 51705000 | 51706799 | 16.62049 | 29.33005 | 37.52543 | 16.60444 | 0.035946 | 0.562145 | 0.950338 |

| | | | | | | | | | |
|---|---|---|---|---|---|---|---|---|---|
| 1 | 52976600 | 52979399 | 47.76188 | 76.89933 | 79.93667 | 33.96983 | 0.298765 | 1.835723 | 0.950341 |
| 3 | 38934000 | 38935399 | 15.29664 | 27.10548 | 36.48302 | 16.12987 | -0.03455 | 0.507042 | 0.950364 |
| 13 | 93242600 | 93245799 | 41.71144 | 67.27397 | 87.90324 | 37.44434 | -0.02756 | 1.776914 | 0.950373 |
| 18 | 37592400 | 37633599 | 811.845 | 1061.586 | 1096.776 | 459.1841 | 0.023994 | 5.640864 | 0.950374 |
| 8 | 39831600 | 39834399 | 14.1424 | 25.9948 | 26.37402 | 11.20613 | 0.409418 | 0.260559 | 0.950384 |
| 10 | 118881200 | 118883599 | 25.31189 | 42.61782 | 54.45507 | 23.79984 | 0.028681 | 1.122641 | 0.950412 |
| 1 | 102806800 | 102808399 | 16.08485 | 28.18912 | 37.77018 | 16.68412 | -0.03248 | 0.649488 | 0.950435 |
| 9 | 57576800 | 57578999 | 19.20195 | 33.13102 | 42.21406 | 18.51879 | 0.037149 | 0.868737 | 0.950436 |
| 18 | 30202600 | 30204999 | 22.42406 | 38.33851 | 49.01803 | 21.49296 | 0.029628 | 0.9711 | 0.950444 |
| 15 | 82383400 | 82384999 | 50.55369 | 80.89277 | 100.7912 | 42.44078 | 0.027242 | 2.039912 | 0.950456 |
| 7 | 103121600 | 103124399 | 31.68114 | 51.84177 | 68.64076 | 29.52397 | -0.02838 | 1.472146 | 0.950477 |
| 4 | 23856800 | 23858199 | 14.81716 | 27.26808 | 25.92555 | 10.99743 | 0.504579 | 0.26022 | 0.95049 |
| 3 | 54211200 | 54213199 | 24.44283 | 41.88865 | 52.8862 | 23.19344 | 0.048209 | 0.962421 | 0.950501 |
| 7 | 65498800 | 65503599 | 35.47069 | 57.55336 | 75.78196 | 32.47354 | -0.02963 | 1.615198 | 0.950528 |
| 12 | 94008000 | 94009399 | 12.08499 | 22.20505 | 29.22252 | 12.42256 | 0.036354 | 0.25527 | 0.950537 |
| 8 | 3575400 | 3576999 | 12.90217 | 23.41633 | 30.52419 | 13.18426 | 0.034049 | 0.331 | 0.95055 |
| 1 | 187470600 | 187471999 | 11.33226 | 20.76795 | 28.66858 | 12.23026 | -0.03606 | 0.26029 | 0.950552 |
| 10 | 100656800 | 100659199 | 32.52232 | 53.26496 | 70.49282 | 30.33126 | -0.02708 | 1.464402 | 0.95056 |
| 13 | 76472600 | 76475999 | 34.12851 | 57.05265 | 51.97858 | 22.63213 | 0.515048 | 1.21866 | 0.950566 |
| 18 | 86210200 | 86219199 | 217.8353 | 316.6672 | 362.5197 | 149.4311 | 0.029118 | 4.001456 | 0.950604 |
| 4 | 4878600 | 4883999 | 47.78626 | 76.92916 | 95.90106 | 40.74681 | 0.036915 | 1.84504 | 0.950605 |
| 10 | 41922600 | 41923599 | 12.42522 | 22.46957 | 30.66652 | 13.30558 | -0.03559 | 0.351684 | 0.950609 |
| 9 | 71701200 | 71703799 | 24.1306 | 40.90827 | 52.26513 | 22.89025 | 0.029202 | 1.057304 | 0.950642 |
| 15 | 99003200 | 99006399 | 36.27984 | 58.75801 | 73.86699 | 31.61002 | 0.034557 | 1.652652 | 0.950653 |
| 13 | 15248000 | 15248799 | 9.243097 | 17.37089 | 24.82131 | 10.19565 | -0.04376 | 0.083585 | 0.95066 |
| 11 | 19861200 | 19865799 | 47.91617 | 76.85512 | 96.05353 | 40.59041 | 0.026804 | 1.958823 | 0.950666 |
| 7 | 133420800 | 133423799 | 27.94259 | 46.74211 | 59.54631 | 25.93789 | 0.032901 | 1.199347 | 0.950704 |
| 10 | 17829000 | 17836599 | 82.08509 | 129.2898 | 156.8169 | 65.06804 | 0.044395 | 2.510391 | 0.950731 |
| 2 | 97090600 | 97091199 | 10.23162 | 19.43854 | 26.60509 | 10.69626 | 0.04386 | 0.022453 | 0.950732 |
| 3 | 112473400 | 112473999 | 8.731526 | 16.70836 | 24.55942 | 9.715708 | -0.03718 | -0.04828 | 0.950735 |
| 5 | 44115000 | 44119599 | 51.62636 | 82.57053 | 103.0367 | 43.35754 | 0.024312 | 2.054723 | 0.950756 |
| 3 | 23366800 | 23370999 | 34.63531 | 56.80305 | 72.08653 | 31.03491 | 0.03403 | 1.453176 | 0.950772 |
| 1 | 44348200 | 44350599 | 25.85333 | 42.87452 | 71.50495 | 30.97062 | -0.35482 | 1.299884 | 0.950781 |
| 4 | 85325200 | 85327599 | 20.10164 | 34.99118 | 44.6283 | 19.61747 | 0.037654 | 0.78006 | 0.95079 |
| X | 88041200 | 88042799 | 26.73767 | 45.22764 | 45.8807 | 20.08059 | 0.360747 | 1.079964 | 0.950801 |
| 4 | 156506800 | 156509399 | 42.47378 | 68.50932 | 89.55064 | 38.15025 | -0.02793 | 1.778412 | 0.950807 |
| 3 | 11988200 | 11989999 | 29.09893 | 48.90523 | 49.10338 | 21.44905 | 0.375233 | 1.146826 | 0.95081 |
| 15 | 10533200 | 10534999 | 16.29912 | 28.53362 | 36.37853 | 16.05465 | 0.03905 | 0.667612 | 0.95083 |
| 3 | 10938600 | 10947799 | 338.1014 | 473.5068 | 522.965 | 216.7962 | 0.027165 | 4.557324 | 0.950836 |
| X | 142911600 | 142912199 | 11.25278 | 21.23033 | 28.64481 | 11.68706 | 0.046234 | 0.087732 | 0.950839 |
| 15 | 50418400 | 50420399 | 16.86343 | 29.99029 | 37.96485 | 16.73941 | 0.056186 | 0.493438 | 0.950859 |
| 2 | 73107400 | 73110199 | 19.20259 | 32.08985 | 62.75806 | 27.31968 | -0.58326 | 1.185743 | 0.950873 |
| 7 | 140630200 | 140631799 | 13.9124 | 24.58277 | 33.25012 | 14.70889 | -0.0428 | 0.525516 | 0.950875 |
| 15 | 98360600 | 98361599 | 9.419639 | 17.77016 | 25.37379 | 10.3545 | -0.03502 | 0.063048 | 0.950876 |
| 1 | 16807000 | 16808599 | 25.69805 | 43.00934 | 57.1561 | 24.90473 | -0.02781 | 1.185975 | 0.950886 |
| 17 | 44470800 | 44472599 | 23.71914 | 40.08247 | 53.31708 | 23.32835 | -0.02888 | 1.085551 | 0.95089 |
| 6 | 128998400 | 128999999 | 17.95269 | 31.55448 | 42.78432 | 18.94565 | -0.04971 | 0.621433 | 0.95091 |

| | | | | | | | | | |
|---|---|---|---|---|---|---|---|---|---|
| 10 | 51310600 | 51317999 | 80.9641 | 126.8107 | 161.1778 | 66.85677 | -0.02928 | 2.613938 | 0.950935 |
| 7 | 102748600 | 102751199 | 58.76456 | 93.31386 | 115.0562 | 47.95945 | 0.030598 | 2.288027 | 0.950969 |
| 2 | 51403200 | 51403999 | 12.57449 | 23.02327 | 30.19779 | 12.8988 | 0.036108 | 0.280677 | 0.950974 |
| 1 | 146744800 | 146745799 | 12.17525 | 22.30493 | 30.70994 | 13.10744 | -0.03284 | 0.273957 | 0.950979 |
| 4 | 108137400 | 108141399 | 68.55028 | 108.5806 | 133.823 | 55.65331 | 0.027749 | 2.368839 | 0.950994 |
| 9 | 81105600 | 81107399 | 24.6862 | 41.39957 | 52.79273 | 23.03286 | 0.031535 | 1.157168 | 0.951007 |
| 4 | 143740800 | 143762599 | 667.1121 | 884.978 | 928.0702 | 387.9099 | 0.023154 | 5.407745 | 0.951007 |
| 4 | 121345000 | 121353799 | 144.899 | 219.9452 | 213.0302 | 88.20515 | 0.327225 | 3.22897 | 0.951016 |
| 4 | 51017800 | 51019399 | 22.19491 | 37.17368 | 61.33234 | 26.73686 | -0.33885 | 1.171369 | 0.951024 |
| 11 | 81522000 | 81523999 | 16.5387 | 28.96906 | 38.86683 | 17.16081 | -0.03421 | 0.664941 | 0.951053 |
| 18 | 63591800 | 63593199 | 14.55436 | 26.00949 | 33.53304 | 14.72507 | 0.034374 | 0.443652 | 0.95107 |
| 16 | 70950400 | 70952399 | 26.195 | 43.88575 | 56.08097 | 24.45197 | 0.028496 | 1.176149 | 0.951092 |
| 1 | 153273800 | 153275199 | 21.96966 | 37.4924 | 47.95781 | 21.02496 | 0.028641 | 0.985532 | 0.951092 |
| 3 | 32650400 | 32652399 | 19.45088 | 33.37258 | 44.80226 | 19.65004 | -0.03967 | 0.921231 | 0.951105 |
| 3 | 12422200 | 12422799 | 11.32627 | 20.90509 | 27.69183 | 11.69982 | 0.033197 | 0.218633 | 0.951142 |
| 14 | 88633600 | 88634999 | 11.32627 | 20.90509 | 27.69183 | 11.69982 | 0.033197 | 0.218633 | 0.951142 |
| 14 | 124307800 | 124308799 | 12.86536 | 23.2115 | 30.10346 | 13.0993 | 0.034779 | 0.366336 | 0.951163 |
| 4 | 145112800 | 145114599 | 32.21698 | 53.41809 | 55.19601 | 23.90378 | 0.333781 | 1.305669 | 0.951166 |
| 11 | 10387000 | 10395599 | 264.3455 | 376.901 | 441.6332 | 182.5446 | -0.03215 | 4.29051 | 0.951167 |
| 12 | 82339600 | 82341799 | 16.0641 | 28.20922 | 36.14062 | 15.99135 | 0.031827 | 0.614248 | 0.951184 |
| 17 | 65223800 | 65225599 | 16.0641 | 28.20922 | 36.14062 | 15.99135 | 0.031827 | 0.614248 | 0.951184 |
| 13 | 64069600 | 64072999 | 23.45696 | 39.96699 | 53.37111 | 23.39519 | -0.03365 | 1.010982 | 0.951231 |
| 3 | 97149200 | 97152399 | 23.62954 | 40.02289 | 51.17889 | 22.40841 | 0.027909 | 1.061988 | 0.951255 |
| 12 | 103225400 | 103226799 | 19.6754 | 34.07155 | 43.68819 | 19.16759 | 0.029037 | 0.83974 | 0.951264 |
| 12 | 64518600 | 64519999 | 18.93299 | 32.85457 | 42.09786 | 18.47103 | 0.030614 | 0.814108 | 0.951266 |
| 1 | 155388400 | 155389399 | 11.12861 | 20.63268 | 28.77065 | 12.0816 | -0.03423 | 0.193328 | 0.951275 |
| 3 | 6888200 | 6889199 | 12.36182 | 23.02557 | 24.03535 | 10.02318 | 0.3885 | 0.166199 | 0.951283 |
| 18 | 59603400 | 59603999 | 10.46125 | 19.60337 | 26.35318 | 10.87809 | 0.038281 | 0.119606 | 0.951292 |
| 14 | 88897200 | 88899199 | 19.57403 | 34.25735 | 35.04062 | 15.46061 | 0.355159 | 0.685755 | 0.951306 |
| 16 | 30906400 | 30911199 | 57.92347 | 91.66296 | 141.4737 | 58.80736 | -0.29739 | 2.386589 | 0.95133 |
| 12 | 80505200 | 80507799 | 27.84714 | 46.09542 | 58.74232 | 25.41023 | 0.032001 | 1.315607 | 0.951358 |
| 6 | 63087800 | 63094799 | 85.17786 | 132.7558 | 167.3721 | 69.46346 | -0.02311 | 2.704322 | 0.951363 |
| 17 | 94629200 | 94631399 | 18.84772 | 32.97023 | 44.5304 | 19.63965 | -0.04412 | 0.703214 | 0.95137 |
| 3 | 18906800 | 18908199 | 18.66429 | 32.29813 | 43.19114 | 18.94911 | -0.03153 | 0.839913 | 0.951395 |
| X | 40319200 | 40321799 | 31.31873 | 52.58034 | 49.66063 | 21.6814 | 0.463206 | 1.165523 | 0.951395 |
| 14 | 91908200 | 91909999 | 18.74521 | 32.8997 | 41.73453 | 18.45815 | 0.045947 | 0.647061 | 0.951405 |
| 19 | 20497000 | 20498799 | 14.98752 | 26.75907 | 34.48019 | 15.15267 | 0.034263 | 0.45279 | 0.951406 |
| 12 | 67498000 | 67502199 | 36.60631 | 60.15743 | 60.95596 | 26.26547 | 0.35903 | 1.424491 | 0.951455 |
| 1 | 152666000 | 152668799 | 23.74779 | 40.1011 | 51.19474 | 22.3947 | 0.030074 | 1.087532 | 0.951477 |
| X | 89655400 | 89655799 | 8.137184 | 15.78402 | 24.60068 | 9.27512 | -0.05441 | -0.20738 | 0.951477 |
| 14 | 46053000 | 46054799 | 20.50181 | 35.33937 | 45.28323 | 19.8646 | 0.02863 | 0.886586 | 0.951483 |
| 6 | 40848800 | 40850799 | 23.40781 | 39.68043 | 50.62447 | 22.17128 | 0.031352 | 1.055551 | 0.951506 |
| 16 | 3350000 | 3355999 | 81.29764 | 128.1823 | 129.1701 | 53.6038 | 0.312597 | 2.483032 | 0.951507 |
| 8 | 6064000 | 6066399 | 20.20618 | 35.372 | 34.57621 | 15.25873 | 0.42043 | 0.686547 | 0.951524 |
| 1 | 166867600 | 166871999 | 47.5351 | 76.3583 | 95.67976 | 40.52463 | 0.025659 | 1.913778 | 0.951531 |
| 2 | 11410000 | 11412599 | 18.20422 | 32.00703 | 40.60594 | 17.98071 | 0.045902 | 0.614908 | 0.951534 |
| 12 | 59293000 | 59294399 | 17.98866 | 31.44815 | 40.32741 | 17.77674 | 0.030469 | 0.702594 | 0.951553 |

| | | | | | | | | | |
|---|---|---|---|---|---|---|---|---|---|
| 18 | 61742000 | 61750599 | 75.10075 | 117.9657 | 144.5762 | 59.96769 | 0.02639 | 2.555258 | 0.951558 |
| 15 | 17430200 | 17430999 | 10.76907 | 19.93752 | 27.78964 | 11.69241 | -0.03517 | 0.198834 | 0.951566 |
| 3 | 69265200 | 69268399 | 33.0945 | 54.12947 | 68.87944 | 29.62164 | 0.028249 | 1.480968 | 0.951568 |
| 8 | 28522000 | 28523199 | 21.62126 | 37.03195 | 47.41374 | 20.79252 | 0.028051 | 0.950405 | 0.95158 |
| 5 | 77903400 | 77913399 | 136.4202 | 206.2496 | 253.9955 | 105.0365 | -0.02386 | 3.30953 | 0.951586 |
| 17 | 61780400 | 61790599 | 183.9744 | 272.0664 | 328.1603 | 135.1449 | -0.02264 | 3.728112 | 0.951595 |
| 10 | 37431600 | 37432399 | 12.42595 | 22.91551 | 30.25033 | 12.79541 | 0.036708 | 0.236921 | 0.951608 |
| 6 | 49465200 | 49470199 | 71.85693 | 113.1032 | 143.6504 | 59.59966 | -0.02271 | 2.513479 | 0.951644 |
| 1 | 125682800 | 125683999 | 15.19145 | 26.86209 | 34.4259 | 15.22683 | 0.034542 | 0.524155 | 0.951649 |
| 6 | 136351200 | 136358999 | 74.04474 | 116.7904 | 143.5034 | 59.55484 | 0.027288 | 2.47276 | 0.951652 |
| 1 | 89984000 | 89987799 | 29.37351 | 48.36428 | 64.20635 | 27.68856 | -0.02846 | 1.387055 | 0.951652 |
| 10 | 38631000 | 38632199 | 16.18026 | 28.09545 | 47.39649 | 20.80186 | -0.36428 | 0.801523 | 0.951669 |
| 4 | 37007200 | 37007999 | 13.67753 | 24.67055 | 32.04333 | 13.94783 | 0.032213 | 0.377245 | 0.951674 |
| 14 | 100292200 | 100294199 | 35.70547 | 57.4451 | 96.83697 | 41.10744 | -0.39851 | 1.850775 | 0.951675 |
| 15 | 74249000 | 74255999 | 71.51415 | 112.6211 | 138.6168 | 57.51503 | 0.023099 | 2.502438 | 0.951726 |
| 1 | 26186200 | 26188799 | 27.27325 | 46.77342 | 43.04283 | 18.87771 | 0.502989 | 0.957904 | 0.951739 |
| 9 | 4054600 | 4057399 | 32.98932 | 53.95271 | 68.74011 | 29.56078 | 0.0264 | 1.48323 | 0.951742 |
| 6 | 134667200 | 134669199 | 47.05927 | 75.3859 | 97.68191 | 41.19972 | -0.0273 | 1.999372 | 0.951751 |
| 8 | 69760200 | 69767399 | 117.7052 | 180.442 | 227.4003 | 94.3927 | -0.04088 | 3.025009 | 0.951753 |
| 16 | 42695200 | 42697199 | 13.20374 | 23.01514 | 45.25089 | 19.90812 | -0.58342 | 0.728441 | 0.951765 |
| 11 | 97784800 | 97785599 | 16.59289 | 29.07308 | 37.26991 | 16.46008 | 0.031058 | 0.655786 | 0.951785 |
| 4 | 140455400 | 140457199 | 15.16366 | 26.54921 | 35.94949 | 15.867 | -0.04738 | 0.658164 | 0.951785 |
| 16 | 23157400 | 23162399 | 49.16242 | 78.86867 | 102.3588 | 43.26709 | -0.02719 | 1.95811 | 0.951802 |
| 18 | 24389200 | 24392599 | 32.2824 | 52.5263 | 69.69761 | 29.91652 | -0.0372 | 1.557714 | 0.951839 |
| 10 | 38405800 | 38406399 | 10.2967 | 19.22808 | 25.72582 | 10.68181 | 0.037745 | 0.135192 | 0.951843 |
| 18 | 53268800 | 53269999 | 11.46658 | 21.30671 | 28.19972 | 11.80601 | 0.043368 | 0.180185 | 0.95185 |
| 14 | 90581400 | 90583599 | 38.67348 | 62.61376 | 79.16182 | 33.86749 | 0.025955 | 1.66875 | 0.951855 |
| 14 | 57094200 | 57095199 | 15.88597 | 28.03707 | 35.97913 | 15.91985 | 0.03164 | 0.55259 | 0.951863 |
| 8 | 78446200 | 78446999 | 7.551234 | 14.59517 | 22.08041 | 8.465106 | -0.03525 | -0.16875 | 0.951905 |
| 17 | 69261000 | 69263399 | 19.85945 | 34.64367 | 44.02212 | 19.37706 | 0.043183 | 0.746874 | 0.95191 |
| 17 | 54182600 | 54184399 | 22.47107 | 38.45646 | 49.01924 | 21.49518 | 0.034127 | 0.960497 | 0.951927 |
| 1 | 30330400 | 30330999 | 10.22125 | 19.07769 | 26.80797 | 11.14053 | -0.03329 | 0.141875 | 0.951985 |
| 17 | 35618200 | 35619799 | 12.51693 | 22.50705 | 30.59632 | 13.36573 | -0.03649 | 0.383576 | 0.951987 |
| 13 | 52640800 | 52645799 | 54.89375 | 87.63485 | 109.1747 | 45.82075 | 0.023786 | 2.115672 | 0.952011 |
| 2 | 151072000 | 151079999 | 109.8858 | 168.6666 | 203.2987 | 84.40279 | 0.024722 | 2.995374 | 0.952018 |
| 4 | 19834400 | 19837399 | 34.91448 | 57.06039 | 72.5178 | 31.17719 | 0.029225 | 1.498917 | 0.952018 |
| X | 21719000 | 21719799 | 18.00218 | 31.38075 | 42.05961 | 18.5047 | -0.03314 | 0.750039 | 0.952019 |
| 6 | 9580600 | 9583799 | 21.54703 | 37.46226 | 47.36293 | 20.80387 | 0.050017 | 0.806417 | 0.952022 |
| 13 | 98749200 | 98751999 | 87.96093 | 136.0386 | 171.0988 | 71.07037 | -0.02825 | 2.840954 | 0.952046 |
| 16 | 28263400 | 28266999 | 20.95072 | 34.97343 | 61.70739 | 26.84402 | -0.43523 | 1.203599 | 0.952054 |
| 1 | 93883800 | 93886599 | 34.40557 | 55.81471 | 73.61314 | 31.53982 | -0.03217 | 1.617813 | 0.952063 |
| 15 | 3922600 | 3924799 | 27.38042 | 45.85908 | 58.44393 | 25.47781 | 0.03229 | 1.182305 | 0.952069 |
| X | 7778200 | 7781599 | 27.22917 | 46.11333 | 57.79983 | 25.30628 | 0.056678 | 1.086274 | 0.95207 |
| 18 | 59067000 | 59068199 | 28.81646 | 47.76249 | 63.7182 | 27.5876 | -0.03369 | 1.309867 | 0.952076 |
| 3 | 85200200 | 85200799 | 6.380538 | 12.42156 | 19.6096 | 7.196977 | -0.03653 | -0.3393 | 0.95208 |
| 3 | 115154400 | 115154799 | 6.380538 | 12.42156 | 19.6096 | 7.196977 | -0.03653 | -0.3393 | 0.95208 |
| 12 | 63934600 | 63935599 | 13.20102 | 24.29026 | 31.84258 | 13.51013 | 0.043665 | 0.260431 | 0.952091 |

| | | | | | | | | | |
|---|---|---|---|---|---|---|---|---|---|
| 17 | 20610200 | 20611199 | 17.7174 | 31.01395 | 39.78679 | 17.55757 | 0.029926 | 0.680506 | 0.952093 |
| 12 | 75922600 | 75924199 | 17.52769 | 30.6105 | 40.91927 | 18.02404 | -0.02926 | 0.71697 | 0.952104 |
| 13 | 113422200 | 113422799 | 10.98418 | 20.33586 | 26.80624 | 11.27857 | 0.045094 | 0.199784 | 0.952132 |
| 3 | 50489200 | 50491599 | 24.54114 | 42.42911 | 40.89303 | 17.94037 | 0.4387 | 0.871845 | 0.952156 |
| 13 | 33741800 | 33753599 | 193.157 | 284.5914 | 341.9244 | 140.8077 | -0.02199 | 3.790689 | 0.952192 |
| 8 | 14916400 | 14919599 | 42.39907 | 68.51796 | 86.18356 | 36.80528 | 0.030535 | 1.721209 | 0.9522 |
| 7 | 13546200 | 13551599 | 66.82813 | 106.0893 | 109.8643 | 45.78279 | 0.281539 | 2.300017 | 0.952209 |
| 5 | 127238200 | 127240799 | 32.86502 | 53.91674 | 68.5873 | 29.53235 | 0.030744 | 1.440568 | 0.952209 |
| 8 | 5500200 | 5503399 | 34.82904 | 56.5009 | 74.46034 | 31.90248 | -0.03113 | 1.617497 | 0.952226 |
| 9 | 11069800 | 11071999 | 32.72609 | 54.05235 | 55.26545 | 23.86991 | 0.348481 | 1.348861 | 0.952226 |
| 6 | 133737000 | 133740199 | 65.85856 | 103.953 | 132.7013 | 55.08926 | -0.02688 | 2.449518 | 0.952257 |
| 13 | 67260200 | 67265999 | 49.24515 | 78.98055 | 98.77317 | 41.73471 | 0.025741 | 1.96407 | 0.952265 |
| 9 | 71029600 | 71031399 | 19.62161 | 33.81074 | 43.21169 | 18.95568 | 0.03227 | 0.882139 | 0.952266 |
| 19 | 43449800 | 43451399 | 14.78971 | 26.33093 | 33.95498 | 14.95768 | 0.030749 | 0.467632 | 0.952274 |
| 6 | 50353000 | 50354599 | 15.9477 | 28.08198 | 37.5952 | 16.6425 | -0.03034 | 0.581434 | 0.95228 |
| 8 | 14917400 | 14919799 | 33.61385 | 55.05727 | 70.15841 | 30.18845 | 0.027397 | 1.46464 | 0.952293 |
| 19 | 42148000 | 42151399 | 32.18278 | 52.60516 | 66.86051 | 28.74681 | 0.029376 | 1.488233 | 0.952313 |
| 2 | 48198400 | 48200199 | 30.55574 | 50.57638 | 64.49457 | 27.8975 | 0.031027 | 1.329224 | 0.952327 |
| 9 | 61372200 | 61374799 | 26.48149 | 44.73016 | 60.12567 | 26.30375 | -0.0441 | 1.104075 | 0.952386 |
| 2 | 70204200 | 70206799 | 21.26362 | 36.58796 | 46.86163 | 20.5559 | 0.028684 | 0.90635 | 0.952405 |
| 14 | 14782600 | 14787399 | 45.23824 | 72.81411 | 94.90655 | 40.31375 | -0.02752 | 1.847907 | 0.952415 |
| 15 | 32813400 | 32819999 | 99.79256 | 153.819 | 192.202 | 79.83754 | -0.02257 | 2.909298 | 0.952417 |
| 8 | 73230600 | 73233399 | 29.66661 | 49.0522 | 62.74798 | 27.12197 | 0.026377 | 1.336597 | 0.952432 |
| 18 | 7539200 | 7547599 | 76.85766 | 121.3289 | 148.3205 | 61.57715 | 0.035612 | 2.458202 | 0.952448 |
| 19 | 12129600 | 12130599 | 29.0575 | 47.85 | 60.7991 | 26.22061 | 0.034719 | 1.382953 | 0.952454 |
| 4 | 53938000 | 53942599 | 52.44808 | 83.74206 | 107.8346 | 45.26718 | -0.02364 | 2.108726 | 0.952458 |
| 4 | 23539800 | 23543799 | 54.08577 | 86.29268 | 110.9982 | 46.54611 | -0.02325 | 2.133899 | 0.952482 |
| 18 | 26628400 | 26630799 | 17.4078 | 30.25732 | 40.60787 | 17.83365 | -0.03549 | 0.784234 | 0.952487 |
| 3 | 74691600 | 74694399 | 22.91884 | 38.83092 | 49.57439 | 21.70789 | 0.030234 | 1.054117 | 0.952495 |
| 6 | 86121800 | 86122799 | 20.39739 | 34.89114 | 44.29257 | 19.42178 | 0.040043 | 0.951757 | 0.95251 |
| 6 | 63150400 | 63155999 | 91.59804 | 143.2111 | 144.5637 | 59.97622 | 0.300764 | 2.649563 | 0.952512 |
| 4 | 156493800 | 156509399 | 223.7816 | 327.8199 | 395.5246 | 162.9235 | -0.03595 | 3.886584 | 0.952521 |
| 1 | 140837400 | 140838199 | 15.62458 | 27.46312 | 35.1667 | 15.57274 | 0.032655 | 0.59598 | 0.952523 |
| 13 | 74654000 | 74656399 | 26.12978 | 44.0175 | 56.14582 | 24.54262 | 0.031177 | 1.122353 | 0.952523 |
| 3 | 67006200 | 67011999 | 119.4555 | 181.0627 | 224.3313 | 92.84399 | -0.02943 | 3.252536 | 0.952525 |
| 13 | 24412200 | 24417199 | 64.39434 | 101.8005 | 129.962 | 53.9947 | -0.02517 | 2.411762 | 0.952583 |
| 9 | 66808800 | 66809599 | 12.69393 | 23.72752 | 24.5367 | 10.17145 | 0.408751 | 0.147963 | 0.952595 |
| 8 | 44115600 | 44116399 | 13.65151 | 24.41071 | 31.53383 | 13.84016 | 0.031996 | 0.42847 | 0.952633 |
| 12 | 84074200 | 84075999 | 13.65151 | 24.41071 | 31.53383 | 13.84016 | 0.031996 | 0.42847 | 0.952633 |
| 18 | 58351600 | 58352799 | 10.79291 | 20.15767 | 28.37135 | 11.77734 | -0.0347 | 0.144797 | 0.952647 |
| 13 | 82322200 | 82327999 | 107.9834 | 165.5387 | 199.7607 | 82.91966 | 0.021669 | 3.018334 | 0.952661 |
| 11 | 89127200 | 89128599 | 15.30097 | 26.96612 | 36.10102 | 15.97834 | -0.02979 | 0.565946 | 0.952683 |
| 17 | 22653400 | 22665999 | 146.0951 | 221.7986 | 260.0111 | 107.6631 | 0.052225 | 3.229924 | 0.952692 |
| 14 | 118217200 | 118220599 | 39.74936 | 64.32358 | 81.26543 | 34.7547 | 0.026274 | 1.68004 | 0.952697 |
| 5 | 150040200 | 150041199 | 15.16109 | 26.63665 | 35.73613 | 15.81897 | -0.03423 | 0.601603 | 0.952727 |
| 18 | 18956200 | 18958399 | 49.44302 | 79.45203 | 76.96991 | 32.61739 | 0.39682 | 1.898371 | 0.952727 |
| 13 | 88677800 | 88679599 | 20.56848 | 35.49212 | 47.40201 | 20.79473 | -0.03056 | 0.880727 | 0.952749 |

| | | | | | | | | | |
|---|---|---|---|---|---|---|---|---|---|
| 1 | 178288800 | 178290399 | 19.5338 | 33.79681 | 45.24685 | 19.8509 | -0.0334 | 0.845933 | 0.952753 |
| 10 | 51619800 | 51620999 | 33.05889 | 53.91005 | 68.35712 | 29.36505 | 0.03041 | 1.526 | 0.952779 |
| 4 | 4785400 | 4788399 | 42.91188 | 69.19354 | 90.32775 | 38.46524 | -0.02662 | 1.787709 | 0.952783 |
| 7 | 42756800 | 42856399 | 3517.148 | 4084.111 | 3884.944 | 1650.674 | -0.03041 | 7.538777 | 0.952791 |
| 5 | 135388000 | 135388599 | 10.8891 | 20.03719 | 26.25718 | 11.13967 | 0.044653 | 0.229805 | 0.952806 |
| 9 | 47370600 | 47371999 | 13.35809 | 23.76999 | 32.10363 | 14.14823 | -0.03589 | 0.458211 | 0.952837 |
| 11 | 58598800 | 58604599 | 58.00894 | 92.40557 | 114.8829 | 48.0697 | 0.023321 | 2.189039 | 0.95284 |
| 4 | 21124600 | 21132599 | 89.95162 | 140.9587 | 140.678 | 58.3504 | 0.319741 | 2.609038 | 0.952886 |
| 5 | 115512000 | 115515599 | 22.67808 | 38.49922 | 51.15719 | 22.41444 | -0.02713 | 1.039346 | 0.952914 |
| 16 | 59548000 | 59550399 | 21.21348 | 36.66483 | 46.84719 | 20.55236 | 0.0334 | 0.865604 | 0.952918 |
| 1 | 25429000 | 25431399 | 31.57193 | 52.20618 | 66.47748 | 28.73538 | 0.032898 | 1.342037 | 0.952921 |
| 14 | 78182600 | 78185199 | 29.33252 | 48.56817 | 64.48704 | 27.90078 | -0.02708 | 1.320495 | 0.952922 |
| 2 | 19081200 | 19082399 | 14.69694 | 25.81581 | 34.83041 | 15.41533 | -0.04221 | 0.601215 | 0.952949 |
| 3 | 129186000 | 129190399 | 42.57596 | 68.59308 | 89.31493 | 37.99002 | -0.02439 | 1.812182 | 0.952962 |
| 3 | 91662000 | 91664999 | 38.2252 | 62.26654 | 64.40731 | 27.65721 | 0.322277 | 1.549437 | 0.952982 |
| 10 | 70682000 | 70684799 | 35.78322 | 58.40364 | 74.18487 | 31.87868 | 0.028683 | 1.521201 | 0.952998 |
| 13 | 54140000 | 54145199 | 46.48758 | 74.67143 | 96.90457 | 41.04002 | -0.02494 | 1.913331 | 0.953005 |
| 13 | 12272600 | 12277599 | 74.02776 | 116.2986 | 142.8917 | 59.27019 | 0.022955 | 2.552385 | 0.953022 |
| 1 | 121196800 | 121200599 | 72.68328 | 114.4816 | 141.0581 | 58.52928 | 0.021738 | 2.500474 | 0.953034 |
| 5 | 115442600 | 115444799 | 20.35381 | 35.13571 | 46.86574 | 20.55973 | -0.02863 | 0.87475 | 0.953044 |
| 15 | 100735200 | 100737199 | 35.69201 | 58.39581 | 56.1408 | 24.14607 | 0.432011 | 1.475903 | 0.953057 |
| 10 | 114690400 | 114691799 | 16.09559 | 28.42332 | 36.49495 | 16.14624 | 0.03121 | 0.549286 | 0.953064 |
| 16 | 3497000 | 3499399 | 23.22835 | 39.64347 | 50.64633 | 22.20376 | 0.030346 | 0.991544 | 0.953092 |
| 3 | 153743600 | 153748399 | 43.50048 | 70.01868 | 91.18175 | 38.73285 | -0.02621 | 1.841521 | 0.953104 |
| 10 | 4005200 | 4006599 | 12.15119 | 22.18427 | 29.12027 | 12.4851 | 0.031171 | 0.289745 | 0.953127 |
| 14 | 6931800 | 6933199 | 11.49712 | 21.33279 | 29.90293 | 12.54355 | -0.04013 | 0.198301 | 0.953142 |
| 3 | 78245400 | 78249399 | 38.59304 | 61.94599 | 102.9001 | 43.5314 | -0.38157 | 1.928055 | 0.95315 |
| 3 | 154176200 | 154179799 | 41.59916 | 67.03364 | 84.31443 | 35.87412 | 0.025858 | 1.802414 | 0.953166 |
| 2 | 17340000 | 17341399 | 16.12126 | 28.2512 | 37.79859 | 16.69581 | -0.03033 | 0.651521 | 0.953178 |
| 9 | 13483600 | 13484399 | 12.24378 | 22.56996 | 31.3684 | 13.27604 | -0.03746 | 0.242377 | 0.953199 |
| 8 | 8597800 | 8601799 | 40.51509 | 65.50293 | 82.55914 | 35.27796 | 0.028344 | 1.701933 | 0.953203 |
| 10 | 42712400 | 42718799 | 68.31792 | 107.8314 | 132.9245 | 55.18104 | 0.023434 | 2.452063 | 0.953203 |
| 7 | 55319400 | 55320399 | 10.53388 | 19.93911 | 28.77397 | 11.65882 | -0.04153 | 0.057843 | 0.953207 |
| 2 | 132935000 | 132936199 | 12.97546 | 23.84441 | 31.21309 | 13.26635 | 0.043524 | 0.261196 | 0.953209 |
| X | 60654000 | 60658399 | 46.42085 | 74.88009 | 99.34782 | 42.32712 | -0.04932 | 1.788884 | 0.953246 |
| 5 | 132090400 | 132091399 | 10.71749 | 20.24606 | 27.5559 | 11.2127 | 0.037112 | 0.07117 | 0.953261 |
| 2 | 70463800 | 70469999 | 62.69567 | 98.94554 | 149.6787 | 62.13244 | -0.27164 | 2.455108 | 0.95327 |
| 14 | 112626800 | 112627999 | 20.53108 | 35.36763 | 45.26027 | 19.85415 | 0.030196 | 0.889862 | 0.953284 |
| 9 | 22057600 | 22064999 | 76.2638 | 119.992 | 147.4364 | 61.16728 | 0.024545 | 2.524142 | 0.95333 |
| 6 | 140928600 | 140933199 | 45.80237 | 73.9093 | 76.73482 | 32.70819 | 0.304328 | 1.765011 | 0.953337 |
| 14 | 98991200 | 98996999 | 124.7587 | 190.3992 | 193.2545 | 80.13918 | 0.266004 | 3.112411 | 0.953369 |
| 5 | 103474000 | 103476799 | 28.73177 | 47.70206 | 63.50145 | 27.52562 | -0.03056 | 1.285165 | 0.95337 |
| 13 | 53184600 | 53186799 | 26.31362 | 43.89433 | 58.29147 | 25.34655 | -0.02677 | 1.222913 | 0.953371 |
| 1 | 82127000 | 82128999 | 14.58586 | 25.93997 | 33.432 | 14.74071 | 0.030604 | 0.471271 | 0.953374 |
| 7 | 35680400 | 35683399 | 27.99435 | 46.27601 | 61.53026 | 26.5912 | -0.02927 | 1.332897 | 0.953378 |
| 2 | 170960800 | 170964399 | 48.38027 | 77.52421 | 96.8941 | 40.88521 | 0.025119 | 1.992121 | 0.95338 |
| 15 | 14730400 | 14735999 | 82.15467 | 128.9327 | 130.8196 | 54.25551 | 0.2976 | 2.576828 | 0.953396 |

| | | | | | | | | | |
|---|---|---|---|---|---|---|---|---|---|
| 15 | 89646800 | 89650599 | 57.83816 | 91.95752 | 118.045 | 49.2668 | -0.02568 | 2.252318 | 0.953405 |
| 13 | 94830600 | 94834199 | 35.36638 | 57.71051 | 76.31132 | 32.78982 | -0.02937 | 1.525983 | 0.953438 |
| 18 | 71120400 | 71121799 | 17.91422 | 31.44356 | 40.17955 | 17.77153 | 0.035504 | 0.639067 | 0.953443 |
| 9 | 115726800 | 115729599 | 20.62551 | 35.90741 | 45.70325 | 20.09151 | 0.04061 | 0.782603 | 0.953445 |
| 1 | 24190800 | 24193199 | 29.03483 | 48.10711 | 63.86844 | 27.64578 | -0.02684 | 1.312352 | 0.953454 |
| 1 | 108117000 | 108118199 | 22.54957 | 38.35879 | 49.02386 | 21.48677 | 0.029239 | 1.019611 | 0.953495 |
| Y | 75680000 | 75680599 | 6.672144 | 12.86332 | 19.97491 | 7.734406 | -0.08592 | -0.1529 | 0.953496 |
| 19 | 13473600 | 13474599 | 18.18753 | 31.49353 | 42.17363 | 18.50307 | -0.03329 | 0.830833 | 0.953514 |
| 16 | 3408000 | 3410799 | 27.29796 | 45.74311 | 58.38879 | 25.46155 | 0.030041 | 1.178372 | 0.953578 |
| 1 | 77901600 | 77902399 | 11.95298 | 21.91421 | 30.17578 | 12.86628 | -0.03208 | 0.265754 | 0.953599 |
| 9 | 94432600 | 94433999 | 12.40118 | 22.70176 | 31.24259 | 13.34844 | -0.03317 | 0.28214 | 0.953646 |
| 6 | 100677800 | 100680599 | 23.08278 | 39.59684 | 40.03875 | 17.56054 | 0.367555 | 0.932457 | 0.953655 |
| 15 | 6548200 | 6549999 | 14.28757 | 25.30688 | 34.00284 | 15.03459 | -0.03224 | 0.500104 | 0.953676 |
| 3 | 58571600 | 58576399 | 63.08617 | 100.0093 | 127.7329 | 53.15474 | -0.02251 | 2.34311 | 0.953683 |
| 2 | 67959400 | 67969399 | 286.961 | 401.9826 | 590.889 | 244.945 | -0.38477 | 4.552437 | 0.953687 |
| 7 | 4415200 | 4416999 | 26.56809 | 44.20201 | 58.73954 | 25.49823 | -0.02768 | 1.254424 | 0.953691 |
| 2 | 10173000 | 10175599 | 21.61775 | 36.83899 | 49.02126 | 21.4887 | -0.02851 | 0.99748 | 0.953724 |
| 9 | 109004000 | 109007599 | 41.68526 | 67.91488 | 84.39745 | 36.24355 | 0.058363 | 1.565832 | 0.953727 |
| 13 | 44043000 | 44045799 | 16.52163 | 29.34551 | 29.80468 | 13.15537 | 0.371461 | 0.485847 | 0.953739 |
| 13 | 110473400 | 110477399 | 63.39518 | 100.3561 | 128.1035 | 53.25884 | -0.02345 | 2.379 | 0.953747 |
| 5 | 109177400 | 109183799 | 71.6145 | 113.625 | 138.8218 | 57.81812 | 0.042591 | 2.326685 | 0.953778 |
| 13 | 6997000 | 6998799 | 25.52707 | 43.40135 | 55.04458 | 24.1246 | 0.040027 | 1.040969 | 0.953786 |
| 1 | 53391400 | 53391999 | 7.858087 | 15.1134 | 22.50735 | 8.775248 | -0.03563 | -0.10852 | 0.953797 |
| 4 | 145390000 | 145435399 | 1099.818 | 1410.773 | 1496.989 | 627.9315 | -0.04228 | 5.954845 | 0.953822 |
| 5 | 147775000 | 147776999 | 17.23934 | 30.06727 | 38.46948 | 16.93093 | 0.033659 | 0.73175 | 0.953834 |
| 2 | 124695200 | 124697799 | 19.16183 | 33.47317 | 45.10608 | 19.87152 | -0.04087 | 0.727012 | 0.95386 |
| 8 | 93863800 | 93865399 | 17.10748 | 30.14353 | 38.55679 | 17.06599 | 0.035422 | 0.583068 | 0.95386 |
| 5 | 119419600 | 119421399 | 18.4686 | 31.82222 | 53.87718 | 23.63427 | -0.37191 | 0.891369 | 0.953866 |
| 15 | 100609600 | 100610599 | 13.409 | 24.06916 | 31.11443 | 13.61203 | 0.034395 | 0.403856 | 0.953927 |
| 7 | 39903400 | 39906399 | 19.26587 | 33.58182 | 43.00184 | 18.91862 | 0.032125 | 0.754632 | 0.953959 |
| 1 | 78642200 | 78644199 | 18.25312 | 31.8814 | 40.91758 | 18.02472 | 0.029311 | 0.720373 | 0.953997 |
| 18 | 82338200 | 82341399 | 29.95938 | 49.69761 | 63.48688 | 27.50302 | 0.02866 | 1.300084 | 0.954022 |
| 5 | 10419200 | 10420999 | 30.28862 | 49.82491 | 66.07469 | 28.48312 | -0.02752 | 1.400286 | 0.954024 |
| 2 | 55517800 | 55521399 | 65.3282 | 103.4528 | 128.0175 | 53.23268 | 0.021701 | 2.371667 | 0.954031 |
| 16 | 58138800 | 58140599 | 25.02773 | 42.56846 | 43.17768 | 18.92422 | 0.361243 | 1.020152 | 0.954036 |
| 5 | 117168200 | 117174199 | 65.3411 | 103.4803 | 131.9757 | 54.88046 | -0.02171 | 2.370891 | 0.954047 |
| 2 | 150339400 | 150345599 | 62.36007 | 99.14351 | 127.3202 | 53.13601 | -0.02635 | 2.25787 | 0.954059 |
| 3 | 114401800 | 114404199 | 22.90096 | 39.29014 | 50.10792 | 21.97699 | 0.033972 | 0.942659 | 0.95407 |
| 6 | 122891200 | 122894799 | 31.07566 | 50.94476 | 67.43606 | 29.02787 | -0.02677 | 1.448108 | 0.954089 |
| 11 | 8187800 | 8191599 | 35.70727 | 58.17537 | 76.7474 | 32.95833 | -0.02762 | 1.549469 | 0.95409 |
| 8 | 60101800 | 60103999 | 27.61418 | 45.04828 | 76.68391 | 32.94664 | -0.39295 | 1.514652 | 0.954101 |
| 17 | 19015800 | 19021399 | 65.34105 | 103.7238 | 127.922 | 53.30536 | 0.029791 | 2.307085 | 0.954143 |
| 1 | 141495800 | 141498199 | 31.95621 | 53.03039 | 67.15907 | 29.10157 | 0.041155 | 1.303964 | 0.954159 |
| 9 | 52924200 | 52925399 | 19.4908 | 33.57477 | 44.77713 | 19.6422 | -0.02902 | 0.886624 | 0.954171 |
| 10 | 28771400 | 28771799 | 6.526604 | 12.72283 | 20.35471 | 7.350238 | -0.03483 | -0.39317 | 0.954171 |
| 10 | 17810200 | 17818999 | 109.3195 | 168.041 | 202.5572 | 84.11169 | 0.026225 | 2.96674 | 0.954172 |
| 5 | 116035800 | 116039599 | 30.54616 | 50.1636 | 66.41182 | 28.60796 | -0.02601 | 1.424055 | 0.954193 |

| | | | | | | | | | |
|---|---|---|---|---|---|---|---|---|---|
| 5 | 140994400 | 140995599 | 12.14601 | 22.40337 | 31.32687 | 13.24747 | -0.04589 | 0.233507 | 0.954194 |
| 2 | 59862800 | 59864199 | 14.11061 | 25.40787 | 34.55769 | 15.07476 | -0.03614 | 0.393539 | 0.954222 |
| 6 | 116502000 | 116506999 | 87.51209 | 136.4512 | 172.4498 | 71.56633 | -0.02614 | 2.699514 | 0.954251 |
| 13 | 75236400 | 75239399 | 28.29189 | 46.80388 | 62.17509 | 26.88415 | -0.02789 | 1.324744 | 0.954325 |
| 4 | 7230400 | 7235999 | 46.50288 | 74.70349 | 96.8972 | 41.0429 | -0.02403 | 1.910495 | 0.954364 |
| 8 | 65767800 | 65771199 | 36.37154 | 59.07736 | 77.61977 | 33.28695 | -0.02507 | 1.598887 | 0.95437 |
| 1 | 125243800 | 125247799 | 59.93401 | 95.14642 | 117.6737 | 49.0379 | 0.025845 | 2.297282 | 0.954377 |
| X | 15459400 | 15462999 | 21.58263 | 37.86542 | 34.29871 | 15.16466 | 0.529706 | 0.65855 | 0.954401 |
| 14 | 8041000 | 8041999 | 22.47581 | 38.68259 | 38.52028 | 16.89708 | 0.390477 | 0.899452 | 0.954423 |
| 5 | 124009400 | 124009999 | 9.472751 | 17.82723 | 25.37835 | 10.39997 | -0.03534 | 0.080459 | 0.954431 |
| 13 | 24151400 | 24151999 | 9.472751 | 17.82723 | 25.37835 | 10.39997 | -0.03534 | 0.080459 | 0.954431 |
| 14 | 80810400 | 80810999 | 9.472751 | 17.82723 | 25.37835 | 10.39997 | -0.03534 | 0.080459 | 0.954431 |
| 10 | 70434000 | 70436199 | 21.46686 | 35.69534 | 63.83507 | 27.69964 | -0.4547 | 1.249882 | 0.954437 |
| 2 | 157466200 | 157467999 | 18.99117 | 32.70964 | 43.75183 | 19.19236 | -0.03327 | 0.884103 | 0.95446 |
| 12 | 76537000 | 76538999 | 22.47423 | 38.07875 | 50.63176 | 22.1711 | -0.0282 | 1.053922 | 0.954462 |
| X | 77065800 | 77070999 | 39.50577 | 64.58749 | 87.91536 | 37.80202 | -0.06919 | 1.516411 | 0.95448 |
| 15 | 40544200 | 40545999 | 20.35127 | 35.30261 | 45.24702 | 19.85255 | 0.030129 | 0.829607 | 0.954483 |
| 3 | 115699600 | 115701399 | 19.96514 | 34.58381 | 44.23034 | 19.40563 | 0.032949 | 0.841138 | 0.954526 |
| 17 | 74432000 | 74432799 | 13.55741 | 24.65503 | 32.15839 | 13.85523 | 0.035759 | 0.330367 | 0.954561 |
| 7 | 125206800 | 125210799 | 32.60392 | 53.40759 | 70.68784 | 30.41734 | -0.02716 | 1.460775 | 0.954571 |
| 5 | 86320000 | 86320999 | 11.55532 | 20.90877 | 28.70871 | 12.44693 | -0.04359 | 0.336063 | 0.954571 |
| 6 | 6735000 | 6736999 | 21.60709 | 36.8504 | 49.02024 | 21.48964 | -0.02787 | 0.989975 | 0.954579 |
| 8 | 8628400 | 8631199 | 21.60709 | 36.8504 | 49.02024 | 21.48964 | -0.02787 | 0.989975 | 0.954579 |
| 17 | 22266400 | 22275599 | 129.5973 | 196.2046 | 234.4843 | 97.01012 | 0.021092 | 3.279015 | 0.954585 |
| 11 | 42897600 | 42897999 | 5.50158 | 10.74184 | 17.59971 | 6.203589 | -0.03613 | -0.49976 | 0.954606 |
| 17 | 64066800 | 64070999 | 55.76872 | 88.78178 | 110.1069 | 46.03168 | 0.025928 | 2.206641 | 0.954628 |
| X | 78245000 | 78248599 | 37.2521 | 61.05748 | 82.45778 | 35.49082 | -0.05584 | 1.474243 | 0.954648 |
| 19 | 43337800 | 43339799 | 38.02735 | 61.48342 | 77.54516 | 33.13722 | 0.027532 | 1.697084 | 0.95465 |
| X | 77897800 | 77901999 | 91.45883 | 141.9904 | 172.856 | 71.7812 | 0.022981 | 2.775394 | 0.954671 |
| 3 | 59699400 | 59700599 | 16.6258 | 29.29155 | 37.45915 | 16.58052 | 0.035501 | 0.577127 | 0.954675 |
| 6 | 100369400 | 100371999 | 30.54735 | 50.10378 | 66.34219 | 28.56307 | -0.02692 | 1.43908 | 0.954701 |
| 4 | 129655400 | 129656599 | 10.36569 | 19.42167 | 27.37586 | 11.30282 | -0.03031 | 0.119704 | 0.954712 |
| 10 | 50239000 | 50239599 | 12.55472 | 22.57133 | 30.6349 | 13.38441 | -0.03423 | 0.385868 | 0.954714 |
| 6 | 36917600 | 36920399 | 27.69772 | 46.72437 | 48.00821 | 20.99442 | 0.342217 | 1.108017 | 0.954738 |
| 15 | 17661200 | 17662999 | 16.76584 | 29.51648 | 39.62332 | 17.54103 | -0.03444 | 0.591579 | 0.954751 |
| 1 | 97204600 | 97207999 | 55.30517 | 88.2194 | 113.4107 | 47.54378 | -0.02274 | 2.14355 | 0.954762 |
| 8 | 41195800 | 41199799 | 78.93563 | 124.0554 | 152.1638 | 63.11925 | 0.025798 | 2.548953 | 0.954774 |
| 7 | 37466800 | 37472399 | 38.50951 | 61.62342 | 109.4161 | 46.07565 | -0.48267 | 2.02616 | 0.954823 |
| 5 | 103491800 | 103500799 | 134.0782 | 202.5415 | 248.7185 | 102.8275 | -0.02076 | 3.324973 | 0.95484 |
| 18 | 85232200 | 85233199 | 16.4358 | 28.88478 | 37.10765 | 16.43022 | 0.027898 | 0.606034 | 0.954877 |
| 19 | 37418400 | 37425999 | 66.19501 | 103.6009 | 164.2882 | 68.15223 | -0.34891 | 2.621008 | 0.954885 |
| 12 | 114128000 | 114129799 | 28.84606 | 48.3779 | 61.28689 | 26.7395 | 0.040883 | 1.177661 | 0.954888 |
| X | 164156600 | 164160999 | 49.07407 | 78.874 | 103.1152 | 43.70884 | -0.03446 | 1.901689 | 0.95489 |
| 15 | 83638600 | 83639599 | 13.60477 | 24.6262 | 33.61439 | 14.56715 | -0.03513 | 0.357908 | 0.95492 |
| 10 | 102557800 | 102559799 | 23.09495 | 39.48249 | 52.80692 | 23.15414 | -0.03509 | 0.981023 | 0.954921 |
| 5 | 146374200 | 146377599 | 41.81925 | 67.38047 | 87.73404 | 37.32282 | -0.02415 | 1.808736 | 0.954926 |
| 8 | 52032200 | 52034199 | 20.56387 | 35.70014 | 47.93726 | 21.03659 | -0.0366 | 0.825777 | 0.954966 |

| | | | | | | | | | |
|---|---|---|---|---|---|---|---|---|---|
| 14 | 60804400 | 60807799 | 29.03181 | 48.50356 | 49.69678 | 21.63047 | 0.346215 | 1.207879 | 0.954969 |
| 18 | 37078800 | 37082799 | 86.55444 | 134.2226 | 168.7251 | 70.07774 | -0.02451 | 2.792426 | 0.954977 |
| 7 | 25633600 | 25637399 | 47.57626 | 76.41034 | 99.18733 | 41.9986 | -0.02552 | 1.92124 | 0.954979 |
| 2 | 83017800 | 83018199 | 9.297528 | 17.93247 | 19.75716 | 7.640739 | 0.406108 | -0.12822 | 0.954989 |
| 7 | 67393400 | 67396399 | 31.34993 | 51.1735 | 67.87164 | 29.16694 | -0.03334 | 1.505488 | 0.955 |
| 2 | 103649200 | 103651799 | 24.67469 | 41.41532 | 54.96093 | 23.99163 | -0.02586 | 1.149583 | 0.955006 |
| 10 | 66601400 | 66612399 | 149.0703 | 224.3353 | 266.4457 | 110.0347 | 0.022557 | 3.40397 | 0.95502 |
| 14 | 72579000 | 72579999 | 8.222428 | 15.65466 | 22.95125 | 9.185839 | -0.0475 | -0.01622 | 0.955052 |
| 3 | 37268000 | 37272999 | 75.17706 | 117.6092 | 143.2741 | 59.43747 | 0.030574 | 2.630428 | 0.955055 |
| 6 | 80834000 | 80834399 | 6.95881 | 13.52505 | 20.07687 | 7.467124 | 0.033336 | -0.28492 | 0.955062 |
| 17 | 95153400 | 95154799 | 18.45778 | 32.04538 | 41.0776 | 18.02687 | 0.030171 | 0.804882 | 0.955063 |
| 6 | 114442800 | 114444399 | 20.10711 | 34.83213 | 44.71433 | 19.61802 | 0.027421 | 0.83855 | 0.95507 |
| 1 | 19692400 | 19695399 | 40.38616 | 65.63804 | 58.9734 | 25.30185 | 0.522636 | 1.581885 | 0.955104 |
| 11 | 11549000 | 11549599 | 10.0336 | 18.94676 | 25.89619 | 10.54603 | 0.030066 | 0.06308 | 0.95511 |
| 2 | 132239600 | 132243199 | 56.99982 | 91.06214 | 113.0767 | 47.50649 | 0.029568 | 2.096642 | 0.955122 |
| 12 | 42713000 | 42714799 | 16.57229 | 28.65811 | 48.46246 | 21.26134 | -0.36882 | 0.845848 | 0.955129 |
| 6 | 129891400 | 129914599 | 646.1075 | 859.5181 | 932.0272 | 389.4223 | -0.02106 | 5.36275 | 0.955129 |
| 13 | 60297800 | 60300599 | 29.10794 | 48.47779 | 64.70554 | 28.1071 | -0.03411 | 1.253092 | 0.955138 |
| 2 | 107482200 | 107482799 | 13.28462 | 23.97482 | 32.54025 | 14.15455 | -0.03042 | 0.370827 | 0.9552 |
| 18 | 30741600 | 30743199 | 13.28462 | 23.97482 | 32.54025 | 14.15455 | -0.03042 | 0.370827 | 0.9552 |
| 15 | 94263200 | 94264399 | 10.83749 | 19.97618 | 27.70217 | 11.72554 | -0.03443 | 0.221572 | 0.955217 |
| X | 60301800 | 60302799 | 12.83027 | 23.98559 | 23.58734 | 9.775688 | 0.481435 | 0.147885 | 0.955223 |
| 15 | 49190200 | 49191999 | 16.93446 | 29.83271 | 40.13605 | 17.76571 | -0.03725 | 0.586305 | 0.955239 |
| 1 | 52661400 | 52663599 | 32.09207 | 52.71392 | 67.32492 | 29.00399 | 0.025751 | 1.423816 | 0.955249 |
| 19 | 44160200 | 44161599 | 19.54589 | 33.99565 | 43.61793 | 19.16337 | 0.029102 | 0.789493 | 0.955256 |
| 2 | 148798000 | 148798599 | 13.73813 | 24.72595 | 32.0682 | 13.99673 | 0.031905 | 0.391344 | 0.955263 |
| 10 | 14752400 | 14753799 | 15.76147 | 27.95368 | 35.91438 | 15.87202 | 0.032779 | 0.500889 | 0.955267 |
| 4 | 75226400 | 75230599 | 82.34511 | 128.5878 | 157.4582 | 65.33402 | 0.02108 | 2.668833 | 0.955269 |
| 2 | 73577800 | 73582999 | 88.05079 | 136.5997 | 171.8795 | 71.38347 | -0.02544 | 2.787084 | 0.955282 |
| 8 | 90551400 | 90552999 | 19.91213 | 34.34281 | 45.79738 | 20.0905 | -0.02863 | 0.87737 | 0.955285 |
| 3 | 86023400 | 86024599 | 16.63983 | 29.28328 | 37.59181 | 16.64298 | 0.02964 | 0.592861 | 0.955289 |
| 17 | 65555200 | 65556599 | 12.50171 | 22.52261 | 30.57921 | 13.33055 | -0.03276 | 0.371799 | 0.955306 |
| 13 | 40770800 | 40773199 | 22.45204 | 38.10308 | 50.62909 | 22.18087 | -0.02714 | 1.039484 | 0.955307 |
| 6 | 128665400 | 128672999 | 107.3856 | 164.9215 | 199.3079 | 82.7523 | 0.02155 | 2.982765 | 0.95532 |
| 4 | 143039400 | 143041199 | 12.80651 | 22.85908 | 31.06732 | 13.65585 | -0.04243 | 0.431258 | 0.955329 |
| 13 | 39218400 | 39224199 | 77.43311 | 121.3763 | 149.0883 | 61.83703 | 0.020867 | 2.598194 | 0.955345 |
| 4 | 123204800 | 123205999 | 13.88025 | 24.62753 | 33.09886 | 14.62257 | -0.03123 | 0.482589 | 0.955358 |
| 7 | 56103000 | 56103599 | 7.277458 | 14.11722 | 21.66234 | 8.164999 | -0.03185 | -0.22885 | 0.955358 |
| 4 | 83226400 | 83227799 | 12.67106 | 22.71465 | 29.16582 | 12.7737 | 0.042888 | 0.400973 | 0.955421 |
| 13 | 58683200 | 58686999 | 24.65964 | 40.89343 | 67.46477 | 29.21997 | -0.33919 | 1.29237 | 0.955442 |
| 3 | 97889800 | 97891199 | 12.07141 | 21.76926 | 29.68862 | 12.92605 | -0.03777 | 0.362298 | 0.955446 |
| 18 | 23927400 | 23928999 | 12.07141 | 21.76926 | 29.68862 | 12.92605 | -0.03777 | 0.362298 | 0.955446 |
| 3 | 71311800 | 71312199 | 7.136832 | 13.87814 | 20.67671 | 7.658539 | 0.033865 | -0.29374 | 0.955463 |
| 2 | 59666600 | 59668399 | 19.63254 | 33.79999 | 43.22667 | 18.96174 | 0.031072 | 0.890246 | 0.955463 |
| 8 | 109805400 | 109809399 | 31.8757 | 52.64794 | 67.08915 | 28.97678 | 0.031704 | 1.355827 | 0.955466 |
| 1 | 102385400 | 102386999 | 14.7298 | 26.27406 | 33.91691 | 14.91703 | 0.030732 | 0.45427 | 0.955468 |
| 15 | 54421400 | 54423199 | 18.72673 | 32.96328 | 41.65794 | 18.44235 | 0.05207 | 0.607372 | 0.955478 |

| | | | | | | | | | |
|---|---|---|---|---|---|---|---|---|---|
| 16 | 48138200 | 48141399 | 28.25755 | 46.90164 | 60.0256 | 26.01395 | 0.026079 | 1.285013 | 0.955489 |
| 12 | 81462000 | 81479999 | 473.4199 | 644.89 | 716.4131 | 298.2859 | -0.02161 | 4.98539 | 0.955528 |
| 6 | 101764000 | 101766199 | 20.59451 | 35.79014 | 45.73966 | 20.0852 | 0.034516 | 0.806917 | 0.955553 |
| 18 | 23332000 | 23332799 | 9.144692 | 17.51789 | 24.59774 | 9.700738 | 0.033036 | -0.05117 | 0.955616 |
| 16 | 47955600 | 47958599 | 29.78168 | 49.78992 | 63.0833 | 27.46861 | 0.040748 | 1.214047 | 0.955636 |
| 6 | 31884200 | 31886599 | 33.01544 | 53.86404 | 68.48968 | 29.42694 | 0.026834 | 1.518296 | 0.955644 |
| 3 | 16119200 | 16122399 | 35.1946 | 57.36091 | 75.57122 | 32.45752 | -0.02531 | 1.542235 | 0.955652 |
| 1 | 21128000 | 21130799 | 34.47363 | 56.72325 | 59.11445 | 25.4854 | 0.31931 | 1.403042 | 0.955668 |
| 3 | 138240800 | 138242399 | 13.13917 | 23.78031 | 31.05084 | 13.4584 | 0.028545 | 0.351063 | 0.955701 |
| 18 | 18691800 | 18692799 | 13.13917 | 23.78031 | 31.05084 | 13.4584 | 0.028545 | 0.351063 | 0.955701 |
| 13 | 98224600 | 98227999 | 35.84276 | 58.30096 | 76.65414 | 32.89872 | -0.02443 | 1.572239 | 0.955724 |
| 10 | 106226400 | 106227999 | 31.59959 | 51.61101 | 68.29464 | 29.35494 | -0.02938 | 1.500642 | 0.955754 |
| 3 | 100628400 | 100633999 | 68.70922 | 108.6764 | 138.8331 | 57.69498 | -0.02559 | 2.403047 | 0.955763 |
| 1 | 14175200 | 14177799 | 22.25416 | 38.32092 | 37.49171 | 16.44615 | 0.415982 | 0.891866 | 0.955777 |
| 12 | 7639400 | 7640199 | 8.569267 | 16.37 | 24.01029 | 9.541539 | -0.03906 | -0.0319 | 0.955789 |
| 3 | 46191800 | 46192999 | 23.49515 | 39.78301 | 52.79102 | 23.11211 | -0.02534 | 1.066041 | 0.955804 |
| 2 | 39415200 | 39424399 | 119.3899 | 181.91 | 218.6973 | 90.65403 | 0.02012 | 3.146011 | 0.955813 |
| 12 | 84006600 | 84011599 | 59.10428 | 93.9375 | 116.5149 | 48.6111 | 0.023295 | 2.262666 | 0.955829 |
| 8 | 53192600 | 53192999 | 6.662773 | 12.9344 | 20.02887 | 7.520608 | -0.03968 | -0.25543 | 0.955831 |
| 14 | 98544600 | 98545799 | 15.12351 | 26.80603 | 35.96808 | 15.90134 | -0.02994 | 0.501932 | 0.955851 |
| 5 | 65207600 | 65208799 | 11.70978 | 21.63418 | 28.74673 | 12.1293 | 0.030217 | 0.219669 | 0.955886 |
| 1 | 18109400 | 18111799 | 25.6189 | 42.95203 | 55.00889 | 23.99562 | 0.025269 | 1.16396 | 0.955889 |
| 15 | 12003800 | 12005599 | 15.25893 | 26.90381 | 34.56446 | 15.29692 | 0.029549 | 0.557681 | 0.955897 |
| 11 | 103104000 | 103105199 | 13.8953 | 25.00574 | 32.44587 | 14.16376 | 0.030755 | 0.390747 | 0.955906 |
| 7 | 43525000 | 43529399 | 32.34341 | 52.88531 | 69.79307 | 30.01131 | -0.02444 | 1.485329 | 0.955917 |
| 15 | 93861200 | 93862799 | 18.95727 | 32.73792 | 43.71287 | 19.17681 | -0.03001 | 0.859537 | 0.955921 |
| 6 | 130153800 | 130156599 | 22.93919 | 39.77 | 50.16614 | 22.01049 | 0.052748 | 0.850663 | 0.955935 |
| 9 | 48068800 | 48070599 | 20.33913 | 34.82355 | 46.37998 | 20.33859 | -0.02865 | 0.947493 | 0.955941 |
| 15 | 36796400 | 36798999 | 25.50901 | 42.59863 | 54.41719 | 23.67823 | 0.029086 | 1.20621 | 0.955956 |
| 5 | 21165200 | 21174199 | 87.5548 | 136.086 | 166.0908 | 68.95997 | 0.020595 | 2.751703 | 0.955993 |
| 2 | 127153200 | 127154599 | 9.553982 | 17.69198 | 31.50804 | 13.2528 | -0.3861 | 0.196013 | 0.956024 |
| 13 | 19299000 | 19300199 | 11.27069 | 20.98043 | 29.37709 | 12.2634 | -0.03384 | 0.172022 | 0.956035 |
| 3 | 7332800 | 7333999 | 17.64581 | 30.74818 | 39.44231 | 17.34915 | 0.029685 | 0.746489 | 0.956059 |
| 3 | 144468000 | 144469199 | 17.08339 | 29.86966 | 38.36313 | 16.91284 | 0.02839 | 0.694208 | 0.95606 |
| 13 | 63585200 | 63585799 | 7.740444 | 14.95952 | 21.53634 | 8.260031 | 0.035621 | -0.16478 | 0.956073 |
| 4 | 16521800 | 16522199 | 7.163521 | 13.92951 | 20.66912 | 7.658071 | 0.039688 | -0.29073 | 0.956079 |
| 17 | 86325200 | 86327999 | 41.92316 | 68.04304 | 85.23986 | 36.53434 | 0.042421 | 1.627289 | 0.956095 |
| 9 | 41960600 | 41962599 | 15.29299 | 26.97679 | 36.07236 | 15.96273 | -0.02759 | 0.555626 | 0.956104 |
| 7 | 121388000 | 121396999 | 161.1697 | 240.2827 | 291.5623 | 120.156 | -0.02045 | 3.584041 | 0.956186 |
| 4 | 144970800 | 144973999 | 34.27958 | 55.8184 | 70.89209 | 30.43768 | 0.026501 | 1.548718 | 0.956217 |
| 14 | 8134400 | 8141599 | 109.3607 | 166.9365 | 207.1555 | 85.90805 | -0.02343 | 3.105687 | 0.956221 |
| 11 | 103467200 | 103473199 | 105.5503 | 161.6421 | 200.8481 | 83.3547 | -0.02164 | 3.038946 | 0.956223 |
| 11 | 67850200 | 67851599 | 18.48407 | 32.2406 | 41.43802 | 18.23827 | 0.026954 | 0.739894 | 0.956232 |
| 13 | 52647400 | 52650599 | 19.36023 | 33.60631 | 43.10441 | 18.91427 | 0.029147 | 0.812874 | 0.956234 |
| 1 | 97250000 | 97250399 | 7.108285 | 13.83273 | 20.74626 | 7.639092 | 0.03168 | -0.31646 | 0.956317 |
| 17 | 61780800 | 61790599 | 182.2125 | 269.7043 | 325.1057 | 133.8882 | -0.02051 | 3.712961 | 0.95633 |
| 1 | 51438200 | 51442399 | 36.60499 | 59.67737 | 78.98879 | 33.92903 | -0.03174 | 1.541251 | 0.956351 |

| | | | | | | | | | |
|---|---|---|---|---|---|---|---|---|---|
| 2 | 55382200 | 55382599 | 5.508143 | 10.75484 | 17.59983 | 6.201754 | -0.03472 | -0.50296 | 0.956355 |
| 11 | 40304200 | 40304599 | 7.40696 | 14.36908 | 21.10035 | 7.951507 | 0.031097 | -0.23065 | 0.956366 |
| 1 | 187779000 | 187781599 | 13.5171 | 24.77296 | 25.45452 | 10.85852 | 0.387301 | 0.27039 | 0.95637 |
| 6 | 49591600 | 49592199 | 9.405354 | 17.78876 | 24.35239 | 9.883946 | 0.03223 | 0.042625 | 0.956382 |
| 10 | 24900800 | 24904199 | 42.45559 | 68.2594 | 88.79836 | 37.6585 | -0.02668 | 1.876693 | 0.956386 |
| 4 | 146667200 | 146667999 | 12.1983 | 22.1201 | 30.13449 | 13.02974 | -0.0299 | 0.33167 | 0.956403 |
| 19 | 19938000 | 19938999 | 11.79724 | 21.59286 | 29.64556 | 12.66875 | -0.03015 | 0.272783 | 0.956405 |
| 2 | 49021000 | 49021399 | 6.378605 | 12.4243 | 19.65057 | 7.171259 | -0.03191 | -0.35959 | 0.956408 |
| 19 | 29083400 | 29087599 | 40.9952 | 66.1128 | 86.12381 | 36.68206 | -0.02329 | 1.779893 | 0.956476 |
| 12 | 63904600 | 63906199 | 13.3231 | 23.92273 | 31.03876 | 13.57521 | 0.029209 | 0.399294 | 0.956484 |
| 10 | 37583600 | 37585999 | 25.5831 | 43.48279 | 42.74275 | 18.73255 | 0.406368 | 1.031535 | 0.956551 |
| 8 | 22329000 | 22330399 | 15.77026 | 27.9378 | 35.93432 | 15.88807 | 0.030463 | 0.511681 | 0.956564 |
| 9 | 4209200 | 4211199 | 30.81614 | 50.74609 | 64.96459 | 28.01706 | 0.023842 | 1.389148 | 0.956584 |
| 5 | 80372200 | 80372999 | 15.49813 | 27.35421 | 36.58229 | 16.18655 | -0.02744 | 0.552339 | 0.956589 |
| 10 | 11496800 | 11500599 | 45.09369 | 72.71954 | 91.3174 | 38.89214 | 0.029428 | 1.789166 | 0.956594 |
| 5 | 37918200 | 37921799 | 30.03161 | 49.77554 | 63.57888 | 27.52689 | 0.028694 | 1.308152 | 0.956635 |
| 12 | 84133400 | 84146199 | 264.6624 | 379.8198 | 444.2483 | 183.4338 | -0.02013 | 4.196242 | 0.956643 |
| 18 | 10374600 | 10377399 | 25.39487 | 42.65188 | 56.69459 | 24.75859 | -0.02832 | 1.144578 | 0.956655 |
| 7 | 102647200 | 102649799 | 22.40006 | 38.37482 | 51.19744 | 22.45222 | -0.03116 | 0.954495 | 0.956676 |
| 4 | 91077800 | 91078399 | 6.39459 | 12.43979 | 19.58065 | 7.229066 | -0.03957 | -0.31633 | 0.956686 |
| 3 | 84056400 | 84060399 | 36.7652 | 59.57178 | 78.15249 | 33.45539 | -0.02622 | 1.646071 | 0.956713 |
| 4 | 96833600 | 96843399 | 198.4444 | 291.7363 | 339.7667 | 139.9248 | 0.01989 | 3.826575 | 0.956715 |
| 7 | 25266200 | 25270199 | 77.15928 | 120.6561 | 152.3956 | 63.22444 | -0.02202 | 2.640442 | 0.95673 |
| 4 | 46246600 | 46247799 | 16.3574 | 28.69676 | 36.78188 | 16.26192 | 0.031395 | 0.635041 | 0.956758 |
| 10 | 65410800 | 65411599 | 10.93833 | 20.34882 | 27.22109 | 11.37561 | 0.030451 | 0.168541 | 0.956763 |
| 15 | 50493800 | 50495799 | 29.43506 | 48.4772 | 64.39634 | 27.7733 | -0.0293 | 1.381887 | 0.956782 |
| 3 | 67659400 | 67661599 | 21.67127 | 37.51795 | 38.42672 | 16.85906 | 0.35173 | 0.84342 | 0.956787 |
| 10 | 7770600 | 7773599 | 26.43743 | 44.63882 | 45.88463 | 20.07128 | 0.341613 | 1.092332 | 0.956788 |
| 9 | 6793400 | 6794399 | 15.71375 | 27.72481 | 37.11946 | 16.4254 | -0.02919 | 0.559439 | 0.956804 |
| 3 | 57285200 | 57286399 | 14.5631 | 26.51029 | 26.90024 | 11.5754 | 0.396973 | 0.320328 | 0.956811 |
| 12 | 109281400 | 109287799 | 69.15735 | 109.2381 | 135.0505 | 56.08484 | 0.020104 | 2.435726 | 0.956827 |
| 6 | 85797600 | 85799999 | 22.65452 | 38.65442 | 49.55467 | 21.72473 | 0.025382 | 0.991925 | 0.956832 |
| 2 | 123909000 | 123909399 | 7.517021 | 14.54625 | 22.12753 | 8.436377 | -0.03596 | -0.18947 | 0.956863 |
| 4 | 98113600 | 98115199 | 13.01581 | 23.42115 | 31.92847 | 13.9391 | -0.04034 | 0.382097 | 0.956864 |
| 7 | 54197200 | 54198599 | 14.85046 | 26.23367 | 35.103 | 15.52933 | -0.02768 | 0.536596 | 0.956898 |
| 12 | 14546000 | 14546799 | 14.85046 | 26.23367 | 35.103 | 15.52933 | -0.02768 | 0.536596 | 0.956898 |
| 7 | 117584000 | 117586799 | 22.03714 | 37.5264 | 47.95976 | 21.02216 | 0.029331 | 1.003439 | 0.956918 |
| Y | 987400 | 990799 | 18.16008 | 31.23712 | 38.81185 | 17.02468 | 0.072415 | 0.88572 | 0.956922 |
| 8 | 52577800 | 52578399 | 9.052262 | 17.50803 | 25.12708 | 9.603231 | 0.043728 | -0.15603 | 0.956927 |
| 12 | 108431400 | 108432599 | 13.52531 | 24.17744 | 32.56366 | 14.29594 | -0.02812 | 0.432042 | 0.956935 |
| 8 | 33543600 | 33547799 | 54.94501 | 87.78632 | 109.5185 | 46.01673 | 0.023 | 2.09236 | 0.956939 |
| 7 | 98012800 | 98018399 | 48.35832 | 77.62812 | 97.38858 | 41.20608 | 0.022757 | 1.936307 | 0.95698 |
| 5 | 147083400 | 147086999 | 33.06764 | 54.8876 | 69.12128 | 29.95667 | 0.049239 | 1.304884 | 0.956986 |
| 13 | 105830200 | 105832599 | 20.03555 | 35.02601 | 44.53117 | 19.62953 | 0.042768 | 0.720014 | 0.956998 |
| 12 | 105856600 | 105868199 | 209.0242 | 305.8781 | 364.5697 | 150.1826 | -0.01978 | 3.900378 | 0.957 |
| 14 | 118116800 | 118117399 | 9.719369 | 18.44455 | 25.46339 | 10.26167 | 0.029038 | 0.025154 | 0.957022 |
| 12 | 68453400 | 68453799 | 5.502505 | 10.74031 | 16.49504 | 5.845398 | 0.049785 | -0.48433 | 0.957024 |

| | | | | | | | | | |
|---|---|---|---|---|---|---|---|---|---|
| 8 | 45941400 | 45945399 | 69.19288 | 109.228 | 134.965 | 56.03212 | 0.020192 | 2.450368 | 0.957029 |
| 16 | 31031400 | 31041399 | 144.263 | 216.9224 | 265.1768 | 109.4866 | -0.02005 | 3.418671 | 0.957036 |
| 7 | 71473800 | 71475999 | 17.41857 | 30.34753 | 40.525 | 17.82366 | -0.02796 | 0.750771 | 0.957037 |
| 11 | 61334200 | 61336199 | 21.65005 | 36.98936 | 47.42793 | 20.79458 | 0.025461 | 0.973477 | 0.95704 |
| 14 | 91482200 | 91483599 | 15.34476 | 27.17172 | 35.00766 | 15.47976 | 0.027955 | 0.513615 | 0.957062 |
| 14 | 41224800 | 41225999 | 31.4027 | 51.51145 | 68.08555 | 29.3148 | -0.02432 | 1.441993 | 0.957064 |
| 2 | 13841200 | 13842799 | 16.02837 | 28.14166 | 37.63173 | 16.64873 | -0.02977 | 0.618136 | 0.957069 |
| 7 | 85026400 | 85028999 | 65.26313 | 103.1615 | 127.1917 | 52.84051 | 0.024882 | 2.415496 | 0.95711 |
| 10 | 126987800 | 126989599 | 22.43 | 38.1273 | 50.62856 | 22.18878 | -0.02605 | 1.024885 | 0.957112 |
| 5 | 89949000 | 89951999 | 23.14551 | 39.59403 | 50.64254 | 22.20626 | 0.029214 | 0.971636 | 0.957131 |
| 11 | 19933400 | 19934599 | 10.5405 | 19.57551 | 27.26694 | 11.42374 | -0.02999 | 0.176736 | 0.95716 |
| 11 | 15116000 | 15117999 | 32.35062 | 52.93242 | 69.83329 | 30.03591 | -0.02331 | 1.476105 | 0.957195 |
| 5 | 61557800 | 61558999 | 11.11262 | 20.65225 | 28.8016 | 12.0554 | -0.03109 | 0.1797 | 0.957205 |
| 2 | 150826000 | 150826999 | 9.407781 | 17.78557 | 25.40049 | 10.31889 | -0.02966 | 0.048065 | 0.957252 |
| 8 | 72016400 | 72019399 | 32.66156 | 53.39648 | 70.41785 | 30.27809 | -0.0236 | 1.488602 | 0.957265 |
| 9 | 80383800 | 80385799 | 18.86752 | 32.77073 | 43.6667 | 19.16824 | -0.02541 | 0.80538 | 0.957284 |
| 17 | 87566400 | 87569599 | 59.04716 | 93.92806 | 120.2069 | 50.20025 | -0.02055 | 2.236501 | 0.957292 |
| 10 | 38651200 | 38652799 | 18.08727 | 32.01705 | 31.25181 | 13.81993 | 0.42615 | 0.52772 | 0.957307 |
| 17 | 93761200 | 93762199 | 11.88416 | 22.00032 | 29.23959 | 12.30371 | 0.032317 | 0.208279 | 0.957329 |
| 16 | 33846000 | 33847999 | 24.14574 | 40.77409 | 52.26254 | 22.86197 | 0.024362 | 1.091119 | 0.957343 |
| X | 8421000 | 8429599 | 103.2185 | 160.8611 | 149.5037 | 62.04836 | 0.416246 | 2.713164 | 0.957349 |
| 6 | 78329600 | 78330599 | 20.89745 | 35.39292 | 47.49079 | 20.79316 | -0.04117 | 1.049286 | 0.957349 |
| 3 | 12604600 | 12607799 | 45.25353 | 72.78278 | 94.46202 | 40.08028 | -0.02274 | 1.872044 | 0.957357 |
| 7 | 64303200 | 64305599 | 21.63 | 36.91556 | 49.01892 | 21.49029 | -0.02511 | 0.984156 | 0.957367 |
| X | 18200800 | 18202199 | 12.73448 | 23.39041 | 32.36439 | 13.7646 | -0.03622 | 0.26808 | 0.957426 |
| 1 | 164744000 | 164744599 | 8.21212 | 15.6702 | 22.95408 | 9.143421 | -0.04032 | -0.03207 | 0.957433 |
| 10 | 121738200 | 121738999 | 8.445413 | 16.07415 | 23.39365 | 9.369116 | -0.0386 | -0.01382 | 0.957434 |
| 17 | 58946000 | 58946999 | 8.445413 | 16.07415 | 23.39365 | 9.369116 | -0.0386 | -0.01382 | 0.957434 |
| 13 | 57375600 | 57378399 | 22.95688 | 39.06708 | 50.09321 | 21.95616 | 0.024511 | 1.017011 | 0.957453 |
| 12 | 38097000 | 38097999 | 10.0033 | 19.20153 | 20.77416 | 8.145301 | 0.413947 | -0.07238 | 0.957511 |
| 13 | 86838000 | 86838799 | 9.037735 | 17.21983 | 23.90843 | 9.553764 | 0.031486 | -0.01408 | 0.95752 |
| 1 | 117731200 | 117737799 | 33.72683 | 55.71924 | 57.43496 | 24.81126 | 0.336879 | 1.352993 | 0.95754 |
| 10 | 82642400 | 82643399 | 15.30589 | 26.95958 | 34.63672 | 15.33197 | 0.029061 | 0.57008 | 0.957541 |
| 1 | 90952600 | 90954999 | 26.95889 | 45.16308 | 57.81506 | 25.20719 | 0.025899 | 1.179419 | 0.95755 |
| 10 | 8275400 | 8277999 | 17.99725 | 31.65793 | 42.79173 | 18.94683 | -0.04481 | 0.613529 | 0.957591 |
| 10 | 72526600 | 72527599 | 10.20841 | 19.09501 | 26.82601 | 11.10722 | -0.02876 | 0.127744 | 0.957602 |
| 6 | 123619200 | 123620599 | 20.62889 | 35.11616 | 59.87715 | 26.24517 | -0.38519 | 1.016068 | 0.957613 |
| 1 | 142050800 | 142055799 | 81.14016 | 126.8783 | 155.6761 | 64.58555 | 0.019717 | 2.643941 | 0.957622 |
| 14 | 122762400 | 122764399 | 37.27891 | 60.00016 | 96.53739 | 41.00365 | -0.33075 | 1.838885 | 0.957635 |
| 8 | 102920000 | 102920399 | 8.72195 | 16.71885 | 24.61809 | 9.690796 | -0.03308 | -0.0646 | 0.957661 |
| 2 | 36730400 | 36732799 | 34.08913 | 55.84614 | 74.07593 | 31.87662 | -0.03018 | 1.468489 | 0.957666 |
| 2 | 71467800 | 71472999 | 62.0532 | 98.4927 | 122.2519 | 50.93647 | 0.020383 | 2.304947 | 0.957683 |
| 7 | 73377600 | 73381999 | 32.30246 | 53.00071 | 67.78074 | 29.18667 | 0.023236 | 1.439108 | 0.957709 |
| 14 | 95132200 | 95132999 | 12.0963 | 22.17584 | 29.21396 | 12.45704 | 0.031742 | 0.268177 | 0.957717 |
| 5 | 71731800 | 71733999 | 31.05538 | 51.23348 | 65.54885 | 28.29087 | 0.025783 | 1.369425 | 0.957723 |
| 6 | 149271000 | 149272999 | 19.02448 | 33.09681 | 44.14455 | 19.3977 | -0.02663 | 0.783604 | 0.95773 |
| 15 | 82729400 | 82732399 | 15.7231 | 27.42826 | 45.73707 | 20.13202 | -0.34684 | 0.732116 | 0.957734 |

| | | | | | | | | | |
|---|---|---|---|---|---|---|---|---|---|
| X | 74593200 | 74595399 | 20.57939 | 35.91411 | 45.6611 | 20.1036 | 0.042601 | 0.750045 | 0.957749 |
| 5 | 99954000 | 99962999 | 118.9701 | 181.5382 | 218.4509 | 90.5846 | 0.020484 | 3.115963 | 0.957767 |
| 6 | 149159400 | 149162999 | 36.37427 | 59.59312 | 75.38581 | 32.44107 | 0.037994 | 1.470271 | 0.957777 |
| 8 | 5689400 | 5694799 | 67.60282 | 106.6824 | 135.6517 | 56.30776 | -0.02147 | 2.457899 | 0.957799 |
| 12 | 17860400 | 17862999 | 24.9657 | 42.05316 | 53.90637 | 23.56313 | 0.024085 | 1.116918 | 0.957804 |
| X | 130861400 | 130863599 | 32.07893 | 52.81377 | 70.12742 | 30.24076 | -0.02884 | 1.400362 | 0.957822 |
| 3 | 22128000 | 22131799 | 24.07013 | 39.78712 | 67.75889 | 29.28198 | -0.38521 | 1.329213 | 0.957832 |
| 3 | 62905000 | 62906799 | 19.76198 | 34.53833 | 43.98486 | 19.38425 | 0.040291 | 0.720735 | 0.957839 |
| 3 | 26451200 | 26451799 | 9.648961 | 18.12138 | 24.36673 | 10.0204 | 0.042537 | 0.093437 | 0.957843 |
| 4 | 142045400 | 142048599 | 21.46357 | 36.62499 | 46.90931 | 20.56512 | 0.026758 | 0.983027 | 0.957844 |
| Y | 82696600 | 82697199 | 5.943429 | 11.56888 | 16.94805 | 6.22767 | 0.068677 | -0.35214 | 0.957846 |
| 5 | 52633400 | 52636999 | 45.96679 | 73.88112 | 92.85238 | 39.3596 | 0.022369 | 1.891311 | 0.957848 |
| 16 | 38096200 | 38098599 | 23.69684 | 40.89838 | 40.89241 | 17.93915 | 0.385314 | 0.881247 | 0.957868 |
| 5 | 26418200 | 26420599 | 19.21523 | 33.22293 | 44.24173 | 19.40953 | -0.02572 | 0.852649 | 0.957878 |
| 15 | 8268400 | 8269399 | 29.15396 | 48.09385 | 61.53155 | 26.55636 | 0.025756 | 1.360436 | 0.957931 |
| 11 | 54778000 | 54779999 | 18.10067 | 31.5337 | 42.05211 | 18.49449 | -0.02606 | 0.757733 | 0.957995 |
| 13 | 97363800 | 97365599 | 23.88538 | 40.17686 | 53.30737 | 23.29136 | -0.02548 | 1.125988 | 0.95805 |
| 14 | 50044200 | 50052799 | 132.0884 | 199.8993 | 239.158 | 98.93172 | 0.019061 | 3.287621 | 0.958054 |
| 1 | 6587600 | 6588599 | 24.1108 | 40.29551 | 53.80029 | 23.42147 | -0.03437 | 1.194345 | 0.958065 |
| 5 | 109081400 | 109088199 | 85.4264 | 133.7822 | 163.4322 | 67.79282 | 0.027685 | 2.621231 | 0.958094 |
| 12 | 87595000 | 87609799 | 199.5403 | 292.5889 | 349.4669 | 143.9429 | -0.02005 | 3.867219 | 0.958101 |
| 4 | 60283800 | 60287999 | 64.18735 | 101.7145 | 126.0851 | 52.44813 | 0.019991 | 2.35464 | 0.958122 |
| 12 | 26155800 | 26160199 | 69.64346 | 110.02 | 136.0652 | 56.50989 | 0.019741 | 2.433124 | 0.958158 |
| 3 | 148370800 | 148371799 | 15.42121 | 27.20771 | 35.03047 | 15.50113 | 0.026786 | 0.552277 | 0.958175 |
| 10 | 60259000 | 60263599 | 81.24797 | 126.7162 | 159.5889 | 66.22663 | -0.02081 | 2.690797 | 0.958181 |
| 18 | 70625200 | 70628599 | 51.51488 | 82.43512 | 106.2816 | 44.75596 | -0.02189 | 2.040381 | 0.958186 |
| 2 | 18460200 | 18467999 | 133.5567 | 202.2683 | 249.1678 | 103.0961 | -0.02217 | 3.274837 | 0.958197 |
| 11 | 40903600 | 40907799 | 65.04507 | 102.9314 | 131.0234 | 54.46303 | -0.01983 | 2.389109 | 0.958214 |
| 7 | 35731000 | 35733799 | 29.34725 | 48.46653 | 62.10609 | 26.82252 | 0.023776 | 1.349078 | 0.95822 |
| 18 | 63324000 | 63334799 | 45.82537 | 73.94451 | 76.67449 | 32.68138 | 0.30611 | 1.766141 | 0.958226 |
| 9 | 112376400 | 112378399 | 19.74148 | 34.45552 | 44.05308 | 19.39693 | 0.034593 | 0.742332 | 0.958231 |
| 5 | 147057600 | 147059999 | 26.71881 | 44.60862 | 57.15462 | 24.86572 | 0.02481 | 1.215971 | 0.958261 |
| X | 100892800 | 100894599 | 27.38987 | 47.39783 | 37.23086 | 16.33418 | 0.733086 | 0.875508 | 0.958295 |
| 5 | 12328600 | 12329599 | 12.74472 | 23.62211 | 23.5688 | 9.901112 | 0.445479 | 0.198085 | 0.958316 |
| 5 | 138450000 | 138457999 | 112.981 | 174.3986 | 207.9649 | 86.38453 | 0.046386 | 2.890295 | 0.958317 |
| 1 | 73931800 | 73934999 | 42.54864 | 68.57764 | 86.51993 | 36.8228 | 0.021792 | 1.7979 | 0.958326 |
| 5 | 116433000 | 116435999 | 33.48356 | 54.86532 | 70.10367 | 30.16988 | 0.023734 | 1.460232 | 0.958356 |
| 14 | 43143000 | 43143599 | 10.9769 | 20.63348 | 27.97183 | 11.48582 | 0.031726 | 0.105475 | 0.958356 |
| 4 | 145109000 | 145111399 | 31.53078 | 52.09381 | 66.50132 | 28.72881 | 0.029173 | 1.350834 | 0.958358 |
| 4 | 118180600 | 118182199 | 16.26555 | 28.584 | 38.13034 | 16.88225 | -0.02621 | 0.607213 | 0.958364 |
| 15 | 63666000 | 63672599 | 63.58381 | 101.2991 | 124.8854 | 52.26429 | 0.035603 | 2.192086 | 0.958394 |
| 18 | 74308000 | 74309999 | 19.63963 | 34.00014 | 45.25992 | 19.857 | -0.02491 | 0.843115 | 0.958411 |
| 17 | 30155000 | 30157399 | 23.25914 | 39.24231 | 50.12404 | 21.92063 | 0.02931 | 1.092058 | 0.958411 |
| 6 | 39290600 | 39291999 | 14.85893 | 26.35976 | 34.01314 | 15.02852 | 0.027013 | 0.491797 | 0.958424 |
| 17 | 22265000 | 22275599 | 139.6736 | 210.598 | 251.1806 | 103.784 | 0.018968 | 3.364151 | 0.958436 |
| 10 | 82869200 | 82871599 | 21.95917 | 37.31991 | 49.55559 | 21.7182 | -0.02603 | 1.024084 | 0.958436 |
| 10 | 53714200 | 53715799 | 27.15168 | 45.76342 | 47.48426 | 20.75982 | 0.328053 | 1.112332 | 0.95845 |

| | | | | | | | | | |
|---|---|---|---|---|---|---|---|---|---|
| 16 | 97255600 | 97260399 | 56.43231 | 89.97414 | 112.1953 | 47.00303 | 0.020401 | 2.159418 | 0.95846 |
| 13 | 100389400 | 100391999 | 21.11493 | 36.38465 | 48.474 | 21.26411 | -0.02753 | 0.89469 | 0.958464 |
| 8 | 55822200 | 55824799 | 17.28222 | 29.56875 | 51.61346 | 22.63229 | -0.41733 | 0.950242 | 0.958465 |
| X | 9193400 | 9196799 | 50.29817 | 80.58108 | 104.3371 | 44.01224 | -0.02612 | 2.004512 | 0.958467 |
| 12 | 82228200 | 82229399 | 22.18676 | 37.81808 | 48.49137 | 21.25685 | 0.02495 | 0.998651 | 0.958479 |
| 4 | 50932200 | 50932999 | 8.780057 | 16.77486 | 23.44082 | 9.312678 | 0.030003 | -0.03512 | 0.958522 |
| 5 | 129109000 | 129113999 | 45.83147 | 73.57593 | 95.21227 | 40.28866 | -0.0218 | 1.929357 | 0.958554 |
| 2 | 145008200 | 145010799 | 16.78905 | 28.67074 | 52.13775 | 22.85871 | -0.47667 | 0.964466 | 0.958558 |
| 8 | 80529600 | 80531399 | 20.08034 | 34.74412 | 46.31706 | 20.32046 | -0.02722 | 0.850316 | 0.958559 |
| 6 | 43127800 | 43128999 | 13.91025 | 24.98689 | 32.4506 | 14.18796 | 0.027955 | 0.40242 | 0.958563 |
| 5 | 148348600 | 148349799 | 19.13883 | 33.14628 | 44.24842 | 19.41345 | -0.02867 | 0.838985 | 0.958604 |
| 4 | 125578200 | 125579999 | 15.88019 | 27.83701 | 37.23352 | 16.45034 | -0.03004 | 0.642749 | 0.958607 |
| Y | 4709400 | 4709999 | 7.269445 | 13.92687 | 20.96431 | 8.265644 | -0.06585 | -0.08316 | 0.958615 |
| 7 | 11394600 | 11395399 | 16.59555 | 29.20802 | 37.56431 | 16.6305 | 0.026987 | 0.590761 | 0.958624 |
| X | 122060400 | 122063999 | 51.25252 | 82.16558 | 102.6615 | 43.35002 | 0.026381 | 1.983712 | 0.958629 |
| 13 | 68018000 | 68022799 | 55.80513 | 88.98137 | 110.8353 | 46.43845 | 0.022094 | 2.15394 | 0.95863 |
| 1 | 149499800 | 149500799 | 14.6521 | 26.55598 | 27.33237 | 11.82581 | 0.37139 | 0.345735 | 0.958654 |
| 1 | 12762200 | 12763799 | 14.23992 | 25.25134 | 33.93365 | 14.99856 | -0.03188 | 0.487093 | 0.958684 |
| 2 | 138623200 | 138624799 | 16.56018 | 29.00075 | 37.27721 | 16.4561 | 0.027168 | 0.663347 | 0.958713 |
| X | 8312000 | 8334599 | 80.3015 | 127.3954 | 114.0114 | 47.48009 | 0.490927 | 2.332978 | 0.958726 |
| 8 | 105726800 | 105728999 | 13.18114 | 23.1842 | 39.86937 | 17.65321 | -0.39013 | 0.583315 | 0.958728 |
| 1 | 90908800 | 90909599 | 12.24465 | 22.44449 | 29.61983 | 12.63265 | 0.028541 | 0.26765 | 0.958738 |
| 5 | 38802400 | 38805199 | 29.22501 | 49.30843 | 61.9522 | 27.09352 | 0.052941 | 1.125724 | 0.958749 |
| 17 | 36516000 | 36517999 | 15.11886 | 26.13991 | 47.93419 | 21.02946 | -0.48525 | 0.838607 | 0.958754 |
| 6 | 76512200 | 76514799 | 27.23289 | 45.73742 | 58.342 | 25.47748 | 0.03101 | 1.156019 | 0.958766 |
| 4 | 143138400 | 143141599 | 34.24906 | 55.84582 | 71.07617 | 30.53209 | 0.024861 | 1.528497 | 0.95877 |
| 12 | 65219000 | 65220999 | 15.2187 | 26.40688 | 47.9313 | 21.03048 | -0.46961 | 0.80536 | 0.958772 |
| 2 | 24091800 | 24092399 | 6.823528 | 13.27081 | 20.68635 | 7.653214 | -0.02938 | -0.30255 | 0.958795 |
| 4 | 13730200 | 13732599 | 24.02391 | 40.96261 | 52.30404 | 22.92877 | 0.030858 | 1.004278 | 0.958805 |
| 7 | 90413600 | 90414999 | 22.54638 | 38.85932 | 39.01972 | 17.1169 | 0.37897 | 0.888591 | 0.958813 |
| 1 | 84843800 | 84845599 | 18.13636 | 31.54317 | 42.0578 | 18.4776 | -0.02619 | 0.779196 | 0.958813 |
| 17 | 9997800 | 9999999 | 18.13636 | 31.54317 | 42.0578 | 18.4776 | -0.02619 | 0.779196 | 0.958813 |
| 6 | 4819200 | 4821599 | 19.50953 | 34.04374 | 43.47675 | 19.14071 | 0.036025 | 0.739757 | 0.958875 |
| 12 | 38372000 | 38372999 | 11.49875 | 21.21124 | 28.19905 | 11.92344 | 0.027202 | 0.224552 | 0.958888 |
| 3 | 14510000 | 14512999 | 31.78808 | 52.18719 | 68.96806 | 29.70527 | -0.02369 | 1.430795 | 0.958907 |
| 2 | 27288800 | 27290799 | 38.69243 | 62.51995 | 81.58745 | 34.83674 | -0.0226 | 1.718032 | 0.95892 |
| 17 | 13365600 | 13374199 | 88.36726 | 136.9414 | 171.4959 | 71.23456 | -0.01985 | 2.804407 | 0.958921 |
| 6 | 132817600 | 132819999 | 31.55045 | 51.27194 | 83.17481 | 35.68221 | -0.32777 | 1.583948 | 0.958952 |
| 10 | 81128000 | 81129199 | 12.12673 | 21.96873 | 28.59311 | 12.37846 | 0.0343 | 0.332188 | 0.958967 |
| 17 | 63867200 | 63869199 | 18.41606 | 31.19043 | 55.86377 | 24.45918 | -0.45632 | 1.048651 | 0.959003 |
| 5 | 148179800 | 148183399 | 45.48432 | 73.05124 | 91.72332 | 38.83869 | 0.022414 | 1.913688 | 0.959007 |
| 2 | 53487800 | 53488999 | 10.36302 | 19.4257 | 26.31762 | 10.85735 | 0.027463 | 0.114456 | 0.959022 |
| 8 | 106139400 | 106140599 | 10.36302 | 19.4257 | 26.31762 | 10.85735 | 0.027463 | 0.114456 | 0.959022 |
| 15 | 5816400 | 5818599 | 26.9429 | 45.19155 | 57.83251 | 25.23427 | 0.026331 | 1.166195 | 0.959054 |
| 5 | 59341400 | 59344999 | 28.70769 | 48.00737 | 48.62464 | 21.17643 | 0.362778 | 1.196105 | 0.959059 |
| 8 | 12001400 | 12002599 | 12.13148 | 22.08559 | 30.21954 | 13.00242 | -0.03134 | 0.310854 | 0.959096 |
| 6 | 43720400 | 43721799 | 20.00402 | 33.45197 | 60.54047 | 26.36567 | -0.47164 | 1.181891 | 0.959105 |

| | | | | | | | | | |
|---|---|---|---|---|---|---|---|---|---|
| 3 | 10227800 | 10228599 | 31.59371 | 51.4839 | 68.13655 | 29.26266 | -0.03191 | 1.532661 | 0.95912 |
| 3 | 153457600 | 153459999 | 22.14294 | 37.77769 | 48.48798 | 21.25692 | 0.023671 | 0.989416 | 0.959139 |
| 10 | 113408000 | 113412999 | 75.88797 | 119.1004 | 150.537 | 62.43318 | -0.01897 | 2.574964 | 0.959142 |
| 8 | 57575400 | 57576999 | 11.45648 | 20.73472 | 36.91785 | 16.00147 | -0.41635 | 0.350323 | 0.959142 |
| 3 | 18343800 | 18346399 | 50.13479 | 80.08836 | 103.1967 | 43.35081 | -0.02377 | 2.089504 | 0.959236 |
| 11 | 46770400 | 46772999 | 24.88573 | 42.45246 | 53.98812 | 23.66795 | 0.03676 | 1.005441 | 0.959254 |
| 5 | 8432000 | 8433799 | 40.95493 | 65.97187 | 85.8192 | 36.49541 | -0.02323 | 1.813275 | 0.95926 |
| 15 | 52407200 | 52410199 | 55.55929 | 88.61951 | 110.6014 | 46.36226 | 0.019806 | 2.143998 | 0.959326 |
| 14 | 87845200 | 87847999 | 46.10499 | 73.88631 | 95.67534 | 40.37759 | -0.02566 | 1.985939 | 0.959337 |
| 8 | 129921000 | 129931399 | 221.7166 | 323.1982 | 383.5866 | 158.0697 | -0.01906 | 3.96072 | 0.95936 |
| 5 | 128083400 | 128087199 | 37.90407 | 61.5511 | 78.10765 | 33.48995 | 0.024561 | 1.604929 | 0.959361 |
| X | 159063400 | 159063999 | 7.891557 | 15.29689 | 23.38056 | 8.843881 | -0.03143 | -0.20536 | 0.959372 |
| 5 | 142815000 | 142820399 | 94.38435 | 145.4266 | 181.5941 | 75.43123 | -0.02203 | 2.912806 | 0.959407 |
| 15 | 33174200 | 33177599 | 54.38781 | 86.8272 | 108.4496 | 45.51644 | 0.020052 | 2.114742 | 0.959409 |
| 5 | 9958000 | 9958999 | 10.14943 | 19.02957 | 26.83649 | 11.06723 | -0.02985 | 0.110876 | 0.959427 |
| 5 | 136742800 | 136746599 | 30.2506 | 49.5634 | 65.62583 | 28.24172 | -0.02749 | 1.451936 | 0.959428 |
| 14 | 10884400 | 10889799 | 56.5635 | 90.19379 | 115.7537 | 48.50157 | -0.02089 | 2.156107 | 0.959436 |
| 1 | 128426400 | 128434999 | 74.76554 | 117.79 | 145.2085 | 60.25329 | 0.02126 | 2.496946 | 0.959443 |
| X | 168754800 | 168772799 | 2530.268 | 2993.406 | 2905.455 | 1230.918 | -0.03209 | 7.245458 | 0.959448 |
| 3 | 154856000 | 154857999 | 32.53327 | 53.31733 | 68.18816 | 29.34765 | 0.022551 | 1.455589 | 0.95945 |
| Y | 41989200 | 41989599 | 5.250516 | 10.2464 | 15.42561 | 5.485327 | 0.07266 | -0.48488 | 0.959451 |
| 5 | 116749400 | 116751799 | 18.64814 | 32.34756 | 43.21039 | 18.95895 | -0.0291 | 0.821373 | 0.959465 |
| 8 | 61124000 | 61125399 | 12.93489 | 23.37973 | 30.51803 | 13.2493 | 0.027399 | 0.35524 | 0.959465 |
| 3 | 156014600 | 156018599 | 69.05855 | 108.8356 | 138.1052 | 57.30838 | -0.02 | 2.486184 | 0.959469 |
| 10 | 36856400 | 36861399 | 17.14834 | 30.76168 | 28.96008 | 12.67634 | 0.488178 | 0.42503 | 0.959508 |
| 9 | 114034200 | 114038799 | 110.7576 | 169.4043 | 204.4889 | 84.84108 | 0.018646 | 3.065172 | 0.959545 |
| 11 | 51849600 | 51851999 | 22.33855 | 38.14542 | 50.62848 | 22.1969 | -0.02432 | 0.984718 | 0.959547 |
| 1 | 89697200 | 89697799 | 6.237299 | 12.12755 | 19.02479 | 7.05312 | -0.04059 | -0.30733 | 0.95957 |
| 4 | 5923600 | 5924599 | 12.63069 | 23.10448 | 31.77381 | 13.58932 | -0.03309 | 0.290303 | 0.959573 |
| 18 | 19124600 | 19127599 | 36.88472 | 60.04875 | 79.17454 | 33.99136 | -0.02758 | 1.564222 | 0.959592 |
| X | 9360400 | 9363799 | 25.61566 | 43.51189 | 58.36856 | 25.57868 | -0.04079 | 1.053663 | 0.959599 |
| 17 | 15026600 | 15029399 | 31.74264 | 52.24304 | 66.85241 | 28.82451 | 0.024094 | 1.398171 | 0.959609 |
| 9 | 120577400 | 120578999 | 16.92679 | 29.68247 | 38.21563 | 16.88946 | 0.024804 | 0.645867 | 0.959645 |
| 2 | 39124400 | 39125799 | 10.82112 | 19.99193 | 27.70642 | 11.6907 | -0.03031 | 0.208283 | 0.959647 |
| 8 | 120050400 | 120058199 | 103.7516 | 159.3812 | 198.1217 | 82.26269 | -0.01882 | 2.976229 | 0.959661 |
| 13 | 14970800 | 14972999 | 20.81305 | 36.18243 | 46.27729 | 20.3257 | 0.033463 | 0.804482 | 0.959662 |
| 1 | 121438200 | 121438999 | 10.38684 | 19.47431 | 26.25886 | 10.82938 | 0.03415 | 0.110359 | 0.959679 |
| 18 | 17094800 | 17096399 | 23.92754 | 40.54514 | 53.85945 | 23.58517 | -0.02677 | 1.064173 | 0.959709 |
| 6 | 26751600 | 26754599 | 52.86116 | 84.3585 | 108.4031 | 45.47289 | -0.02143 | 2.126313 | 0.959725 |
| 2 | 92600200 | 92601999 | 15.23585 | 27.0544 | 34.90308 | 15.40925 | 0.027838 | 0.487601 | 0.959744 |
| 4 | 73849200 | 73850999 | 24.29604 | 41.21075 | 52.8132 | 23.13403 | 0.024814 | 1.053848 | 0.959754 |
| 16 | 46986200 | 46986799 | 7.849402 | 15.12384 | 22.53958 | 8.742598 | -0.02981 | -0.12554 | 0.959757 |
| 6 | 137509600 | 137510999 | 17.344 | 30.45856 | 39.10014 | 17.30172 | 0.028833 | 0.624662 | 0.959776 |
| 1 | 178028800 | 178035799 | 86.9632 | 136.1247 | 166.0697 | 68.89024 | 0.0291 | 2.631066 | 0.95978 |
| 3 | 94355000 | 94355599 | 7.40067 | 14.29171 | 21.51154 | 8.280635 | -0.03344 | -0.15829 | 0.959782 |
| 11 | 10397600 | 10399599 | 39.19118 | 63.63764 | 64.25081 | 27.54747 | 0.353581 | 1.602163 | 0.959803 |
| 12 | 88191200 | 88197599 | 83.36498 | 130.4462 | 159.9572 | 66.35701 | 0.021339 | 2.632814 | 0.959828 |

| | | | | | | | | | |
|---|---|---|---|---|---|---|---|---|---|
| 12 | 63660000 | 63662599 | 29.97158 | 50.03663 | 63.63053 | 27.6822 | 0.035475 | 1.230258 | 0.959851 |
| 14 | 106687800 | 106702599 | 319.3007 | 450.1764 | 503.285 | 208.417 | 0.019314 | 4.457613 | 0.959861 |
| 3 | 4154200 | 4155199 | 12.43738 | 21.92625 | 39.40443 | 17.43934 | -0.45177 | 0.55493 | 0.95988 |
| 9 | 95971200 | 95986799 | 242.5936 | 351.6173 | 415.7181 | 171.4156 | -0.02152 | 4.048465 | 0.9599 |
| 10 | 65225200 | 65225799 | 8.50816 | 16.34168 | 24.11963 | 9.440245 | -0.02803 | -0.08368 | 0.959903 |
| 19 | 40116800 | 40119399 | 25.96155 | 43.05885 | 71.64322 | 31.03292 | -0.35147 | 1.297396 | 0.959915 |
| 10 | 93616600 | 93617799 | 13.91699 | 24.68879 | 33.13698 | 14.64138 | -0.02937 | 0.484785 | 0.959926 |
| 7 | 11721400 | 11730399 | 185.1108 | 276.0203 | 269.6673 | 111.1363 | 0.292236 | 3.583592 | 0.959934 |
| 6 | 43890200 | 43891999 | 16.3926 | 28.70573 | 38.28856 | 16.9018 | -0.02599 | 0.664478 | 0.959944 |
| 7 | 100725800 | 100726399 | 8.319846 | 16.06848 | 23.17438 | 8.916257 | 0.028106 | -0.1449 | 0.959945 |
| 8 | 77991000 | 77998599 | 86.92263 | 135.0318 | 169.3122 | 70.30306 | -0.01889 | 2.760843 | 0.959951 |
| 2 | 113645600 | 113650999 | 63.79239 | 100.2316 | 155.9698 | 64.69592 | -0.31729 | 2.546704 | 0.959968 |
| 15 | 61973400 | 61975399 | 28.35415 | 46.91998 | 62.10253 | 26.85796 | -0.02257 | 1.323172 | 0.959988 |
| 14 | 37746400 | 37750399 | 58.52957 | 93.30538 | 116.2093 | 48.67555 | 0.021909 | 2.16609 | 0.959996 |
| 15 | 47410600 | 47411999 | 17.35614 | 30.31685 | 40.39873 | 17.79695 | -0.02469 | 0.712433 | 0.960012 |
| 18 | 45418800 | 45426999 | 79.9542 | 125.6345 | 154.2719 | 63.99225 | 0.024006 | 2.55357 | 0.960023 |
| 15 | 28425200 | 28426799 | 18.99508 | 33.50978 | 42.25512 | 18.69844 | 0.056659 | 0.589471 | 0.960025 |
| 4 | 50197000 | 50197799 | 9.190212 | 17.54129 | 24.48478 | 9.747604 | 0.028514 | -0.02324 | 0.96005 |
| 15 | 94506000 | 94507399 | 17.20492 | 30.11299 | 38.78121 | 17.11196 | 0.024365 | 0.677995 | 0.960052 |
| 4 | 61474200 | 61484399 | 159.1296 | 237.8501 | 281.5073 | 116.07 | 0.018832 | 3.535192 | 0.960074 |
| 8 | 11986600 | 11987999 | 13.4289 | 24.32104 | 33.09633 | 14.33292 | -0.0301 | 0.352365 | 0.96009 |
| 16 | 34550000 | 34550999 | 13.4289 | 24.32104 | 33.09633 | 14.33292 | -0.0301 | 0.352365 | 0.96009 |
| 15 | 93770400 | 93772399 | 19.331 | 32.95367 | 54.32766 | 23.81546 | -0.33627 | 0.993009 | 0.960129 |
| 12 | 107452200 | 107455599 | 36.17179 | 58.8634 | 77.35144 | 33.20358 | -0.02318 | 1.566704 | 0.960137 |
| 8 | 107530400 | 107532799 | 24.66653 | 41.51603 | 54.97633 | 24.025 | -0.02269 | 1.124099 | 0.960137 |
| 10 | 37399400 | 37402399 | 24.66653 | 41.51603 | 54.97633 | 24.025 | -0.02269 | 1.124099 | 0.960137 |
| 13 | 81780800 | 81782599 | 13.81482 | 24.59882 | 33.01991 | 14.544 | -0.02665 | 0.458351 | 0.960183 |
| 3 | 69857400 | 69858999 | 26.65493 | 44.44874 | 58.85076 | 25.58432 | -0.02245 | 1.228284 | 0.960183 |
| X | 75936000 | 75944199 | 178.2712 | 263.9569 | 318.1119 | 131.0085 | -0.01976 | 3.707911 | 0.960204 |
| 14 | 31637400 | 31643799 | 109.3604 | 168.3994 | 170.7728 | 70.93033 | 0.277241 | 2.928525 | 0.960251 |
| 4 | 24890200 | 24890799 | 17.0342 | 29.74783 | 38.00918 | 16.74192 | 0.035969 | 0.714161 | 0.960266 |
| 11 | 52626200 | 52628199 | 14.36801 | 25.55923 | 33.04725 | 14.56785 | 0.026446 | 0.469688 | 0.960269 |
| 13 | 54869800 | 54872999 | 42.77256 | 68.73211 | 89.2642 | 37.82661 | -0.02524 | 1.893938 | 0.960274 |
| 1 | 15045200 | 15048199 | 60.03104 | 95.33443 | 121.7036 | 50.73537 | -0.01934 | 2.286747 | 0.960281 |
| 4 | 10448600 | 10453999 | 71.54679 | 112.6888 | 139.1317 | 57.72976 | 0.018787 | 2.500144 | 0.960283 |
| 3 | 152275800 | 152277599 | 16.69131 | 29.33595 | 39.18331 | 17.35032 | -0.02795 | 0.611039 | 0.960322 |
| 13 | 35813000 | 35814999 | 21.44864 | 36.47757 | 48.4965 | 21.25524 | -0.02764 | 1.014234 | 0.960349 |
| 6 | 41878000 | 41880199 | 22.17408 | 38.55249 | 38.1591 | 16.76115 | 0.401622 | 0.79653 | 0.960353 |
| 5 | 18819000 | 18821799 | 26.32698 | 44.76521 | 45.8055 | 20.07549 | 0.348653 | 1.032345 | 0.960366 |
| 12 | 36422400 | 36424399 | 21.4715 | 36.45849 | 48.50365 | 21.25314 | -0.02884 | 1.028913 | 0.960367 |
| 6 | 140267000 | 140279199 | 143.5809 | 216.5113 | 258.2109 | 106.6921 | 0.019216 | 3.363471 | 0.960378 |
| 14 | 24942600 | 24943199 | 8.551529 | 16.54932 | 23.93529 | 9.122546 | 0.036346 | -0.17469 | 0.960407 |
| 5 | 151276400 | 151277799 | 35.77294 | 57.8665 | 76.02558 | 32.50454 | -0.03046 | 1.680926 | 0.960428 |
| 11 | 114615800 | 114618599 | 19.76305 | 34.03548 | 45.28386 | 19.86434 | -0.02575 | 0.88952 | 0.960469 |
| 6 | 11975600 | 11976399 | 13.0785 | 23.4416 | 31.63329 | 13.85609 | -0.02848 | 0.411383 | 0.96047 |
| 15 | 4844000 | 4847799 | 42.3975 | 68.45314 | 86.48026 | 36.88868 | 0.02272 | 1.747495 | 0.960474 |
| 13 | 114419800 | 114422399 | 20.84593 | 36.24311 | 46.26553 | 20.32177 | 0.03615 | 0.80132 | 0.960476 |

| | | | | | | | | | |
|---|---|---|---|---|---|---|---|---|---|
| 2 | 157040000 | 157041399 | 16.97247 | 29.74327 | 38.27794 | 16.90772 | 0.025488 | 0.657572 | 0.960478 |
| 9 | 110791600 | 110795399 | 31.82729 | 52.13928 | 68.76557 | 29.59111 | -0.02199 | 1.457878 | 0.960497 |
| 1 | 58678800 | 58685999 | 177.3723 | 261.7839 | 315.3685 | 129.8742 | -0.02354 | 3.759477 | 0.960527 |
| 6 | 19539800 | 19541199 | 13.61411 | 24.45911 | 33.00036 | 14.42625 | -0.0266 | 0.400661 | 0.960528 |
| 5 | 50737200 | 50741999 | 72.31874 | 114.4047 | 118.9922 | 49.45387 | 0.270803 | 2.398739 | 0.960542 |
| 2 | 139275000 | 139278199 | 41.03555 | 66.63505 | 69.02654 | 29.5959 | 0.316679 | 1.607899 | 0.960545 |
| 10 | 71006800 | 71007799 | 13.17111 | 23.73087 | 32.05441 | 13.97196 | -0.0253 | 0.379522 | 0.960555 |
| 1 | 105688600 | 105692399 | 90.728 | 140.4503 | 175.6414 | 72.95687 | -0.01898 | 2.825835 | 0.960562 |
| 12 | 49186800 | 49189399 | 32.37452 | 53.61805 | 55.74768 | 24.12367 | 0.324728 | 1.319373 | 0.96057 |
| 5 | 83121000 | 83121799 | 11.16367 | 20.47052 | 28.1694 | 12.00809 | -0.03075 | 0.254497 | 0.960579 |
| 1 | 26239600 | 26242999 | 42.52587 | 68.50737 | 86.40113 | 36.74675 | 0.02143 | 1.812401 | 0.960589 |
| 9 | 86001800 | 86002999 | 20.06699 | 33.89757 | 56.94323 | 24.91199 | -0.36433 | 1.080589 | 0.960612 |
| 9 | 23771200 | 23771999 | 8.304573 | 15.95852 | 23.57411 | 9.213673 | -0.0276 | -0.0934 | 0.960621 |
| 4 | 72790400 | 72791399 | 13.64293 | 24.5633 | 32.00779 | 13.96418 | 0.025368 | 0.386744 | 0.960626 |
| 6 | 16437400 | 16438799 | 13.64293 | 24.5633 | 32.00779 | 13.96418 | 0.025368 | 0.386744 | 0.960626 |
| 16 | 56253000 | 56254799 | 14.7959 | 26.18911 | 35.00631 | 15.48042 | -0.02517 | 0.513244 | 0.960648 |
| 3 | 65962400 | 65965199 | 22.12534 | 37.88145 | 48.47678 | 21.25806 | 0.028555 | 0.954214 | 0.960665 |
| 4 | 113062600 | 113064199 | 41.78443 | 67.3848 | 87.61022 | 37.31459 | -0.02053 | 1.781395 | 0.960666 |
| 18 | 5951400 | 5954399 | 31.71741 | 52.42797 | 69.79247 | 30.16086 | -0.03101 | 1.35111 | 0.960666 |
| 2 | 56652800 | 56657799 | 56.32658 | 89.98246 | 112.139 | 47.10945 | 0.024336 | 2.099127 | 0.960673 |
| 4 | 129275000 | 129276599 | 17.60665 | 30.74414 | 40.95674 | 18.03898 | -0.02424 | 0.720916 | 0.960704 |
| 6 | 65874000 | 65874999 | 8.285328 | 15.95156 | 22.54828 | 8.765202 | 0.043949 | -0.10899 | 0.960725 |
| 18 | 24389400 | 24392599 | 32.25797 | 52.55876 | 66.46351 | 28.54264 | 0.03347 | 1.535866 | 0.960752 |
| 8 | 119639800 | 119648399 | 120.0977 | 182.8158 | 225.4948 | 93.4453 | -0.01796 | 3.165971 | 0.960763 |
| 14 | 30002600 | 30003399 | 11.06952 | 20.65673 | 28.91851 | 12.02175 | -0.02838 | 0.15605 | 0.960767 |
| 1 | 140849000 | 140853599 | 43.26818 | 69.70322 | 90.52262 | 38.49965 | -0.02074 | 1.816653 | 0.960788 |
| 18 | 37898800 | 37902399 | 204.7968 | 298.8312 | 343.6163 | 141.5839 | 0.027984 | 3.943885 | 0.960805 |
| 8 | 53230600 | 53231399 | 9.936309 | 18.67757 | 26.38671 | 10.83483 | -0.02692 | 0.094206 | 0.960839 |
| 11 | 10561400 | 10564999 | 27.02308 | 45.80086 | 57.96643 | 25.38537 | 0.042636 | 1.073526 | 0.960847 |
| 7 | 63148400 | 63149199 | 9.993822 | 18.71504 | 25.2554 | 10.43666 | 0.031031 | 0.113735 | 0.960849 |
| 9 | 83872400 | 83874799 | 32.34407 | 53.46783 | 55.73702 | 24.08804 | 0.320763 | 1.340066 | 0.960849 |
| 14 | 53347400 | 53352999 | 100.6099 | 155.6194 | 158.9058 | 66.00564 | 0.272224 | 2.842631 | 0.960884 |
| 6 | 12775400 | 12777599 | 26.00943 | 43.8815 | 56.13189 | 24.54786 | 0.027187 | 1.109664 | 0.960889 |
| 17 | 91394800 | 91397199 | 58.54511 | 92.21329 | 150.6511 | 62.50972 | -0.38466 | 2.494836 | 0.96089 |
| 15 | 45339600 | 45345399 | 87.57923 | 137.7685 | 133.6036 | 55.42702 | 0.365354 | 2.534271 | 0.960916 |
| X | 36440000 | 36443599 | 25.45708 | 43.1384 | 54.98904 | 24.08007 | 0.032506 | 1.071012 | 0.960926 |
| 3 | 101242800 | 101244599 | 17.89133 | 31.31296 | 40.30926 | 17.78551 | 0.02497 | 0.685137 | 0.960983 |
| 11 | 27531800 | 27533799 | 18.01353 | 31.18466 | 57.82696 | 25.37056 | -0.5013 | 0.85213 | 0.961056 |
| 14 | 26701600 | 26703999 | 39.46178 | 63.25293 | 105.0023 | 44.32744 | -0.38298 | 1.974634 | 0.961062 |
| 13 | 37797400 | 37797799 | 7.995117 | 15.4452 | 23.18729 | 8.910766 | -0.02781 | -0.15314 | 0.961072 |
| 15 | 81856600 | 81860599 | 76.45378 | 119.8948 | 147.5352 | 61.19002 | 0.018768 | 2.589877 | 0.961148 |
| 3 | 37589600 | 37595199 | 65.61078 | 104.6188 | 137.4722 | 57.60065 | -0.05493 | 2.173325 | 0.96115 |
| 1 | 145737200 | 145739599 | 23.46361 | 40.26587 | 53.99255 | 23.68126 | -0.03805 | 0.947378 | 0.961151 |
| 13 | 54609200 | 54613199 | 45.67562 | 73.49627 | 92.57613 | 39.30693 | 0.021164 | 1.856109 | 0.961162 |
| 1 | 47696600 | 47699999 | 36.71219 | 59.66357 | 75.78467 | 32.51319 | 0.02426 | 1.584435 | 0.961166 |
| 7 | 18419400 | 18422399 | 27.06167 | 45.10729 | 57.82139 | 25.1299 | 0.024046 | 1.231733 | 0.961176 |
| 11 | 113797800 | 113805199 | 49.52134 | 78.61207 | 130.5397 | 54.39674 | -0.39864 | 2.292998 | 0.961184 |

| | | | | | | | | | |
|---|---|---|---|---|---|---|---|---|---|
| 3 | 99796600 | 99808999 | 212.341 | 311.1343 | 371.2698 | 152.9303 | -0.0197 | 3.880656 | 0.961197 |
| 14 | 96431800 | 96432799 | 16.02516 | 27.38253 | 51.58971 | 22.61952 | -0.52727 | 0.956536 | 0.961234 |
| 18 | 6594600 | 6596199 | 20.00119 | 34.44134 | 44.25862 | 19.41453 | 0.024256 | 0.892168 | 0.961237 |
| 4 | 88294400 | 88294999 | 10.66428 | 19.65991 | 27.32575 | 11.5638 | -0.03687 | 0.222067 | 0.961255 |
| 10 | 59045000 | 59045799 | 10.35293 | 19.44096 | 27.41269 | 11.27548 | -0.02651 | 0.105298 | 0.961294 |
| 10 | 72544600 | 72545199 | 12.83732 | 23.1189 | 30.08012 | 13.11888 | 0.027973 | 0.375882 | 0.961298 |
| X | 6461400 | 6462599 | 12.78598 | 23.23608 | 31.69576 | 13.66772 | -0.02911 | 0.328679 | 0.961327 |
| 9 | 59279400 | 59283799 | 66.50932 | 105.1383 | 130.1743 | 54.08139 | 0.018893 | 2.414368 | 0.961331 |
| 10 | 9559600 | 9560799 | 12.04238 | 21.80446 | 29.65293 | 12.84572 | -0.02912 | 0.33827 | 0.961348 |
| 4 | 48418600 | 48420599 | 21.24308 | 35.54262 | 60.69155 | 26.44021 | -0.3882 | 1.178918 | 0.96137 |
| 2 | 91949800 | 91952799 | 34.79696 | 56.66293 | 74.41619 | 31.95118 | -0.02168 | 1.551685 | 0.961385 |
| 8 | 85857200 | 85863999 | 94.83859 | 146.2337 | 182.3018 | 75.725 | -0.01885 | 2.89998 | 0.961389 |
| 15 | 35991000 | 35992399 | 14.34602 | 25.08081 | 43.63976 | 19.23737 | -0.40736 | 0.692105 | 0.961395 |
| 6 | 46851400 | 46853199 | 17.80111 | 31.42342 | 40.09605 | 17.74077 | 0.03991 | 0.569918 | 0.961401 |
| 5 | 80754000 | 80756599 | 27.73296 | 46.15615 | 61.20775 | 26.57606 | -0.02476 | 1.252611 | 0.961407 |
| X | 99301800 | 99302999 | 20.91619 | 36.08926 | 46.33807 | 20.32795 | 0.026046 | 0.882179 | 0.961455 |
| 10 | 111888600 | 111889599 | 11.43415 | 21.14502 | 28.19076 | 11.87935 | 0.026496 | 0.208766 | 0.961482 |
| 11 | 10573000 | 10576199 | 63.92503 | 101.4203 | 129.3994 | 53.89137 | -0.01988 | 2.320975 | 0.961593 |
| 7 | 63998600 | 64000799 | 18.22656 | 31.64688 | 40.62101 | 17.82754 | 0.028403 | 0.803712 | 0.961598 |
| 14 | 19174800 | 19176599 | 27.08399 | 45.66947 | 44.88547 | 19.62627 | 0.406134 | 1.107629 | 0.961622 |
| 2 | 32387200 | 32389599 | 34.98895 | 57.31958 | 58.40061 | 25.13104 | 0.349377 | 1.457776 | 0.961654 |
| 16 | 18792600 | 18795799 | 64.73394 | 102.3632 | 130.1965 | 54.099 | -0.01953 | 2.406318 | 0.961664 |
| 15 | 49085200 | 49085799 | 8.631436 | 16.5797 | 23.5619 | 9.219974 | 0.026282 | -0.08539 | 0.961702 |
| 4 | 11436400 | 11442599 | 69.34419 | 109.7292 | 139.8598 | 58.13466 | -0.02179 | 2.395082 | 0.961709 |
| 17 | 12810600 | 12812199 | 18.09041 | 31.74713 | 40.70961 | 18.00326 | 0.030484 | 0.643822 | 0.961738 |
| 14 | 121265600 | 121269199 | 53.07368 | 84.73461 | 108.7633 | 45.6522 | -0.01914 | 2.111417 | 0.961784 |
| 4 | 108762000 | 108763599 | 17.48552 | 30.54678 | 39.34546 | 17.33435 | 0.024105 | 0.709358 | 0.961827 |
| 8 | 4147000 | 4151399 | 48.08531 | 77.55728 | 78.85097 | 33.59015 | 0.333749 | 1.782493 | 0.961853 |
| 7 | 29429000 | 29435999 | 140.381 | 211.1816 | 257.9278 | 106.5066 | -0.01818 | 3.408954 | 0.961864 |
| 18 | 59506600 | 59506999 | 8.990257 | 17.16514 | 23.97801 | 9.539385 | 0.028963 | -0.02568 | 0.9619 |
| 16 | 88592600 | 88595799 | 61.8877 | 98.08316 | 121.4669 | 50.53366 | 0.021571 | 2.348237 | 0.961921 |
| 10 | 45681200 | 45683999 | 21.4572 | 36.5462 | 48.49271 | 21.25611 | -0.02449 | 1.001099 | 0.961935 |
| 8 | 13676000 | 13677799 | 28.9013 | 47.68473 | 63.09969 | 27.23501 | -0.02261 | 1.360486 | 0.961949 |
| 12 | 59409800 | 59411799 | 26.97755 | 44.87782 | 59.39734 | 25.78163 | -0.02196 | 1.255661 | 0.961974 |
| 1 | 8137800 | 8143599 | 131.692 | 199.399 | 238.9487 | 98.86074 | 0.017325 | 3.277676 | 0.961991 |
| 14 | 101827000 | 101828999 | 16.54325 | 29.14515 | 37.51469 | 16.60521 | 0.026269 | 0.578481 | 0.962019 |
| 16 | 41395400 | 41396199 | 17.93637 | 31.52013 | 29.12581 | 12.89546 | 0.501011 | 0.601839 | 0.962038 |
| 2 | 148302000 | 148302999 | 35.94081 | 58.23531 | 76.33784 | 32.67829 | -0.02499 | 1.647914 | 0.962053 |
| 5 | 92621400 | 92628399 | 91.20735 | 141.1725 | 176.3423 | 73.24807 | -0.01749 | 2.827942 | 0.96209 |
| 2 | 41690200 | 41690799 | 5.800699 | 11.30161 | 18.01817 | 6.553025 | -0.03828 | -0.38474 | 0.96209 |
| 11 | 63306600 | 63307799 | 9.722828 | 18.28893 | 25.85173 | 10.60273 | -0.02653 | 0.085277 | 0.962105 |
| 8 | 63069800 | 63070199 | 6.234666 | 12.13072 | 19.0387 | 7.019434 | -0.03406 | -0.32705 | 0.962116 |
| 7 | 59970400 | 59972399 | 29.86644 | 50.00223 | 51.31249 | 22.36313 | 0.343888 | 1.189829 | 0.962123 |
| 5 | 125395400 | 125396999 | 16.97791 | 29.59036 | 39.54973 | 17.39863 | -0.0291 | 0.743148 | 0.962124 |
| 7 | 57704600 | 57708399 | 40.70402 | 65.81789 | 83.26659 | 35.58689 | 0.023264 | 1.702201 | 0.96213 |
| 10 | 70076200 | 70077799 | 16.76947 | 29.36038 | 37.81233 | 16.68902 | 0.024456 | 0.670022 | 0.962143 |
| 2 | 181387400 | 181388799 | 11.17426 | 20.72414 | 27.74083 | 11.64393 | 0.025032 | 0.190784 | 0.962159 |

| | | | | | | | | | |
|---|---|---|---|---|---|---|---|---|---|
| 4 | 88205600 | 88206799 | 15.78422 | 27.26944 | 47.93718 | 21.03028 | -0.42473 | 0.850304 | 0.96216 |
| 19 | 31698600 | 31701599 | 48.1813 | 76.72538 | 122.6412 | 51.29262 | -0.3391 | 2.194325 | 0.962178 |
| 1 | 16822600 | 16824799 | 20.49957 | 35.67758 | 47.90101 | 21.05374 | -0.03614 | 0.789204 | 0.962213 |
| 13 | 33516400 | 33517399 | 12.20983 | 22.42578 | 29.71067 | 12.63673 | 0.026103 | 0.258664 | 0.962228 |
| 19 | 20732400 | 20733399 | 12.20983 | 22.42578 | 29.71067 | 12.63673 | 0.026103 | 0.258664 | 0.962228 |
| 4 | 54446600 | 54449199 | 41.25282 | 66.16668 | 105.1969 | 44.45347 | -0.31971 | 1.957039 | 0.962262 |
| 15 | 64400600 | 64401399 | 22.05234 | 37.29331 | 49.57131 | 21.69428 | -0.02774 | 1.069031 | 0.962318 |
| 16 | 15560200 | 15564799 | 90.98377 | 140.5025 | 170.3159 | 70.7457 | 0.023232 | 2.869412 | 0.962342 |
| 9 | 31580400 | 31580999 | 7.440673 | 14.41676 | 21.06545 | 7.984037 | 0.030816 | -0.2093 | 0.96235 |
| 12 | 87664800 | 87666599 | 13.41325 | 24.07037 | 32.47053 | 14.20849 | -0.02739 | 0.404599 | 0.962367 |
| 4 | 36029000 | 36029999 | 10.38676 | 19.25553 | 26.7601 | 11.2387 | -0.02919 | 0.184343 | 0.962419 |
| 13 | 106426600 | 106428199 | 16.68848 | 28.86034 | 48.4653 | 21.26261 | -0.35888 | 0.84462 | 0.962426 |
| 10 | 40482600 | 40485399 | 39.89015 | 64.47091 | 81.64807 | 34.87355 | 0.020918 | 1.710711 | 0.962432 |
| X | 169298600 | 169298799 | 4.098781 | 8.018038 | 13.93347 | 4.635028 | -0.04307 | -0.73997 | 0.962432 |
| 1 | 28776000 | 28776799 | 8.569594 | 16.40462 | 24.01371 | 9.489646 | -0.02983 | -0.05507 | 0.96244 |
| 12 | 32080200 | 32083199 | 16.93167 | 29.67358 | 38.25078 | 16.8966 | 0.023081 | 0.655539 | 0.962523 |
| 7 | 3350400 | 3355599 | 30.42782 | 49.29563 | 83.63165 | 35.79688 | -0.39635 | 1.643144 | 0.962525 |
| 6 | 75666200 | 75667599 | 15.23587 | 26.92864 | 35.98582 | 15.91821 | -0.02574 | 0.532579 | 0.962534 |
| 8 | 34774400 | 34776999 | 26.45224 | 44.13589 | 56.65749 | 24.63969 | 0.022004 | 1.219222 | 0.962536 |
| Y | 6821600 | 6822199 | 7.563012 | 14.39698 | 21.43356 | 8.581205 | -0.07035 | -0.02961 | 0.962577 |
| 2 | 6517800 | 6519599 | 14.67814 | 25.98783 | 34.90429 | 15.43445 | -0.03213 | 0.506727 | 0.962585 |
| 18 | 75681800 | 75683999 | 22.93161 | 38.99713 | 50.09449 | 21.9555 | 0.021729 | 1.022689 | 0.962586 |
| 8 | 59810000 | 59810399 | 7.562027 | 14.68148 | 21.71561 | 8.147828 | 0.02654 | -0.24189 | 0.962592 |
| 16 | 34232200 | 34235599 | 32.06671 | 52.49003 | 69.16102 | 29.75133 | -0.021 | 1.470161 | 0.962664 |
| 18 | 35213000 | 35214799 | 20.13689 | 34.49043 | 45.85199 | 20.10768 | -0.02596 | 0.941762 | 0.962688 |
| X | 167405200 | 167408399 | 34.56411 | 56.99978 | 72.34594 | 31.21935 | 0.036999 | 1.387632 | 0.962698 |
| 15 | 17570600 | 17581599 | 119.9898 | 184.0131 | 184.0784 | 76.41394 | 0.292425 | 3.013217 | 0.962709 |
| 13 | 57154400 | 57155999 | 14.6138 | 25.51789 | 43.64727 | 19.22388 | -0.38306 | 0.710066 | 0.962713 |
| 11 | 101513600 | 101515999 | 17.16214 | 29.94791 | 39.91544 | 17.57296 | -0.02515 | 0.724303 | 0.962729 |
| 9 | 60204200 | 60205799 | 14.46083 | 25.50175 | 32.68496 | 14.46455 | 0.033196 | 0.547475 | 0.962777 |
| 17 | 57877600 | 57881799 | 36.88014 | 60.26397 | 62.83347 | 27.01221 | 0.313988 | 1.50067 | 0.96278 |
| 5 | 5864800 | 5867199 | 22.26025 | 38.15608 | 49.01529 | 21.4962 | 0.023403 | 0.947365 | 0.962787 |
| 11 | 45548800 | 45552999 | 63.26587 | 100.206 | 127.5724 | 53.05266 | -0.01899 | 2.366562 | 0.962802 |
| 10 | 114170200 | 114172199 | 18.68616 | 32.63122 | 41.87111 | 18.44244 | 0.029323 | 0.722906 | 0.962824 |
| 5 | 141164800 | 141165999 | 23.13318 | 38.93175 | 51.71832 | 22.60061 | -0.02726 | 1.115126 | 0.962826 |
| 15 | 87033000 | 87038599 | 99.39487 | 154.459 | 155.688 | 64.64269 | 0.296096 | 2.759193 | 0.962834 |
| 17 | 66776800 | 66782999 | 68.49541 | 108.1605 | 133.9211 | 55.60747 | 0.017584 | 2.441933 | 0.96287 |
| 17 | 17563800 | 17564599 | 13.58017 | 24.50364 | 31.98742 | 13.91732 | 0.025203 | 0.372616 | 0.962908 |
| 5 | 61979400 | 61979799 | 5.935956 | 11.58011 | 18.6585 | 6.671903 | -0.03246 | -0.44351 | 0.962941 |
| 1 | 100700000 | 100716799 | 236.5995 | 342.5994 | 403.5012 | 166.3993 | -0.01734 | 4.061957 | 0.962968 |
| 8 | 67610800 | 67618999 | 75.609 | 119.575 | 153.9249 | 63.96271 | -0.03672 | 2.416528 | 0.962969 |
| 4 | 144503000 | 144503799 | 13.62539 | 24.32531 | 31.5181 | 13.85181 | 0.026298 | 0.437546 | 0.962973 |
| 7 | 10228200 | 10230199 | 18.86897 | 33.00699 | 30.59058 | 13.49151 | 0.496668 | 0.680705 | 0.96302 |
| 14 | 14859000 | 14859799 | 9.677077 | 18.21439 | 24.79012 | 10.15644 | 0.028824 | 0.077341 | 0.963026 |
| 11 | 73537600 | 73538599 | 12.32756 | 22.48301 | 30.64613 | 13.1563 | -0.02415 | 0.300209 | 0.963034 |
| 14 | 122065800 | 122079799 | 207.087 | 303.266 | 361.0608 | 148.7302 | -0.01719 | 3.888146 | 0.963054 |
| 10 | 123508000 | 123509399 | 14.45212 | 26.27793 | 34.23133 | 14.7513 | 0.037149 | 0.34002 | 0.963078 |

| | | | | | | | | | |
|---|---|---|---|---|---|---|---|---|---|
| 5 | 97144800 | 97146199 | 12.22658 | 21.88873 | 29.72928 | 13.03509 | -0.03874 | 0.407627 | 0.963079 |
| 1 | 111732600 | 111735599 | 39.30776 | 63.85084 | 66.10575 | 28.3506 | 0.317905 | 1.59488 | 0.963099 |
| 12 | 118180200 | 118181799 | 12.87254 | 23.3186 | 30.48772 | 13.19955 | 0.027503 | 0.340837 | 0.963145 |
| 6 | 101366000 | 101367799 | 17.99095 | 31.44507 | 41.90581 | 18.46899 | -0.02484 | 0.708268 | 0.963155 |
| 14 | 79788200 | 79793199 | 48.29254 | 77.61484 | 100.8206 | 42.73377 | -0.02532 | 1.899228 | 0.963174 |
| 18 | 84846600 | 84849599 | 31.41181 | 51.65501 | 68.23241 | 29.40903 | -0.02199 | 1.411779 | 0.963181 |
| 5 | 149668400 | 149669399 | 12.66937 | 22.94671 | 31.09576 | 13.46561 | -0.02382 | 0.343704 | 0.963182 |
| 14 | 44777800 | 44778599 | 11.20462 | 20.54184 | 28.20018 | 12.0242 | -0.02745 | 0.256919 | 0.963182 |
| 5 | 82213600 | 82217599 | 48.27423 | 77.63149 | 97.42118 | 41.32845 | 0.025476 | 1.882026 | 0.963197 |
| 16 | 19511800 | 19513599 | 16.69827 | 29.33458 | 39.11967 | 17.31536 | -0.02587 | 0.615503 | 0.963206 |
| 9 | 90532800 | 90534599 | 15.90548 | 27.6065 | 46.87693 | 20.56829 | -0.3736 | 0.807073 | 0.963214 |
| 16 | 79462800 | 79468799 | 65.81769 | 104.4716 | 103.1406 | 42.97525 | 0.350197 | 2.301471 | 0.963219 |
| 4 | 133371200 | 133374799 | 37.67043 | 60.9276 | 79.46116 | 33.97 | -0.02014 | 1.686938 | 0.963234 |
| 5 | 147800400 | 147801599 | 13.90523 | 24.70963 | 33.11289 | 14.60991 | -0.02573 | 0.473836 | 0.963242 |
| 9 | 116273200 | 116277599 | 33.45242 | 54.84147 | 70.19361 | 30.21493 | 0.021531 | 1.453509 | 0.963247 |
| 11 | 50867200 | 50869999 | 32.00348 | 53.21323 | 54.6848 | 23.72833 | 0.341901 | 1.272004 | 0.963256 |
| 1 | 140891200 | 140900999 | 326.8725 | 458.2026 | 524.0452 | 217.2128 | -0.02186 | 4.543063 | 0.963298 |
| 1 | 109275200 | 109277599 | 21.54587 | 36.946 | 49.01164 | 21.49527 | -0.02263 | 0.942701 | 0.963308 |
| 6 | 75436800 | 75439799 | 24.70632 | 41.54675 | 54.95446 | 24.00909 | -0.02107 | 1.132522 | 0.963345 |
| 11 | 103191200 | 103192399 | 15.17615 | 27.03543 | 34.9896 | 15.4055 | 0.026124 | 0.469194 | 0.963349 |
| 5 | 133933000 | 133937599 | 46.5252 | 75.00522 | 94.18703 | 40.09857 | 0.02891 | 1.800301 | 0.96341 |
| 3 | 18501000 | 18504199 | 34.82242 | 57.26884 | 72.88482 | 31.4155 | 0.031516 | 1.420598 | 0.963418 |
| 7 | 81309000 | 81312199 | 26.47047 | 44.11522 | 56.63466 | 24.61179 | 0.021951 | 1.231903 | 0.963441 |
| 10 | 36011600 | 36014399 | 28.1797 | 47.27672 | 49.12157 | 21.43645 | 0.325977 | 1.160055 | 0.963443 |
| 3 | 78474200 | 78474999 | 11.83933 | 21.67672 | 28.65445 | 12.23996 | 0.024798 | 0.270375 | 0.963448 |
| X | 9198800 | 9199599 | 9.771138 | 18.5317 | 26.55909 | 10.7168 | -0.02604 | 0.033336 | 0.963451 |
| 12 | 25460800 | 25463399 | 35.64363 | 58.55066 | 60.82729 | 26.20534 | 0.322893 | 1.427579 | 0.963467 |
| 7 | 29799800 | 29818799 | 295.9816 | 421.439 | 476.4893 | 196.9966 | 0.017416 | 4.31253 | 0.963519 |
| 12 | 72758800 | 72762999 | 33.45576 | 54.91462 | 70.2718 | 30.26437 | 0.02255 | 1.4386 | 0.963522 |
| 9 | 56540800 | 56542399 | 15.28169 | 26.72624 | 35.96429 | 15.85868 | -0.0385 | 0.675835 | 0.963528 |
| 6 | 39716200 | 39718199 | 22.31118 | 37.45796 | 61.42454 | 26.81751 | -0.33015 | 1.145242 | 0.963556 |
| 4 | 25134200 | 25134799 | 9.193329 | 16.99299 | 32.54597 | 13.72351 | -0.49336 | 0.202562 | 0.963586 |
| 3 | 139143800 | 139144599 | 12.38484 | 22.54656 | 29.65203 | 12.75844 | 0.02529 | 0.310328 | 0.963624 |
| 11 | 8189400 | 8190799 | 21.40306 | 36.90547 | 47.39948 | 20.79317 | 0.025414 | 0.891049 | 0.963627 |
| 14 | 8200800 | 8202799 | 16.47946 | 29.13784 | 37.50984 | 16.59108 | 0.028197 | 0.540466 | 0.963648 |
| 13 | 13330000 | 13330599 | 10.41411 | 19.50918 | 27.38708 | 11.31062 | -0.02461 | 0.122276 | 0.963655 |
| 6 | 123144600 | 123145599 | 16.2476 | 28.59468 | 38.1848 | 16.90532 | -0.02683 | 0.59263 | 0.963657 |
| 3 | 63564000 | 63566399 | 32.23229 | 54.0083 | 51.34363 | 22.38852 | 0.45383 | 1.189163 | 0.963739 |
| 8 | 4496600 | 4497999 | 16.96212 | 29.61849 | 39.46268 | 17.38086 | -0.02436 | 0.71584 | 0.963741 |
| 8 | 57481800 | 57483599 | 26.60477 | 44.61253 | 57.26111 | 24.98076 | 0.022075 | 1.16726 | 0.963806 |
| 13 | 104036400 | 104039799 | 63.26961 | 100.5643 | 124.9456 | 52.13189 | 0.020932 | 2.266559 | 0.963817 |
| X | 60873600 | 60875799 | 21.74714 | 37.52983 | 50.10715 | 21.98152 | -0.03012 | 0.88812 | 0.963825 |
| 11 | 55256800 | 55258799 | 12.86845 | 23.08003 | 31.10717 | 13.61822 | -0.02627 | 0.403754 | 0.963834 |
| 7 | 117883400 | 117887999 | 51.88243 | 82.91107 | 106.5378 | 44.778 | -0.01918 | 2.082705 | 0.963858 |
| 7 | 117883600 | 117887999 | 51.88243 | 82.91107 | 106.5378 | 44.778 | -0.01918 | 2.082705 | 0.963858 |
| 13 | 106668800 | 106670599 | 8.667399 | 15.93521 | 30.93483 | 13.13963 | -0.51958 | 0.228522 | 0.963886 |
| 2 | 174883200 | 174898399 | 323.6665 | 455.8263 | 509.5755 | 211.0606 | 0.017779 | 4.474583 | 0.963899 |

| | | | | | | | | | |
|---|---|---|---|---|---|---|---|---|---|
| 6 | 129068000 | 129069399 | 16.90888 | 29.57973 | 39.34812 | 17.35461 | -0.02209 | 0.68594 | 0.963899 |
| 1 | 161115800 | 161119999 | 48.81237 | 78.28676 | 98.41495 | 41.5836 | 0.01829 | 1.964259 | 0.963911 |
| 15 | 50458000 | 50461399 | 54.71435 | 87.28047 | 111.8445 | 46.88978 | -0.01807 | 2.139838 | 0.963917 |
| 18 | 83185800 | 83190799 | 68.13489 | 107.5486 | 133.046 | 55.23303 | 0.018396 | 2.450377 | 0.963918 |
| 2 | 161282400 | 161285599 | 38.32583 | 62.22881 | 79.10968 | 33.91671 | 0.021849 | 1.610428 | 0.96393 |
| 11 | 86868200 | 86870399 | 16.11975 | 28.75521 | 37.0123 | 16.27753 | 0.034949 | 0.469521 | 0.963947 |
| 5 | 138584800 | 138587999 | 65.50358 | 104.1676 | 108.3216 | 45.21854 | 0.278001 | 2.248079 | 0.963978 |
| 4 | 32777800 | 32790399 | 148.9442 | 223.8277 | 266.5373 | 110.0307 | 0.016982 | 3.431194 | 0.963979 |
| 4 | 56873000 | 56873999 | 13.65933 | 24.53635 | 33.02914 | 14.44082 | -0.02337 | 0.402946 | 0.963981 |
| X | 70705400 | 70708999 | 59.46682 | 94.66858 | 117.9506 | 49.31053 | 0.019259 | 2.213008 | 0.963997 |
| 5 | 70371200 | 70372799 | 22.40423 | 38.85438 | 39.39711 | 17.28572 | 0.366709 | 0.833433 | 0.964011 |
| 16 | 30720200 | 30741399 | 361.3852 | 504.5092 | 572.6859 | 237.5255 | -0.01697 | 4.605743 | 0.964012 |
| 4 | 128303400 | 128304199 | 8.320179 | 15.9381 | 22.43777 | 8.849416 | 0.029326 | -0.06607 | 0.964018 |
| 16 | 62363600 | 62366199 | 19.69809 | 33.99992 | 43.67962 | 19.16193 | 0.025198 | 0.866267 | 0.964041 |
| 11 | 13306200 | 13306999 | 15.21959 | 26.7764 | 35.72997 | 15.81958 | -0.02587 | 0.585667 | 0.964049 |
| 10 | 27770400 | 27771199 | 13.14524 | 24.16719 | 24.95088 | 10.59831 | 0.384791 | 0.248272 | 0.96406 |
| 14 | 51987200 | 51992399 | 71.54298 | 112.5495 | 142.3644 | 59.06203 | -0.01741 | 2.523629 | 0.964068 |
| 4 | 143739000 | 143789599 | 1268.872 | 1599.753 | 1639.352 | 688.8354 | -0.01708 | 6.234375 | 0.964126 |
| 6 | 81457200 | 81458599 | 14.44644 | 25.72819 | 34.47119 | 15.18088 | -0.02391 | 0.463713 | 0.964134 |
| 15 | 18558000 | 18559399 | 14.00024 | 24.99962 | 33.52269 | 14.73027 | -0.02276 | 0.443445 | 0.964153 |
| 18 | 63639600 | 63640999 | 14.49913 | 26.20517 | 34.04184 | 14.7805 | 0.03397 | 0.372527 | 0.964153 |
| 2 | 165272200 | 165273599 | 16.44369 | 28.75088 | 36.91964 | 16.27631 | 0.028732 | 0.689107 | 0.964168 |
| 8 | 57818400 | 57819399 | 16.2083 | 28.53425 | 38.0563 | 16.8474 | -0.02515 | 0.584973 | 0.9642 |
| Y | 28970200 | 28970799 | 5.813509 | 11.3001 | 17.98213 | 6.68303 | -0.06472 | -0.31041 | 0.964212 |
| 8 | 92951600 | 92953199 | 20.35028 | 35.14296 | 45.27442 | 19.86186 | 0.02145 | 0.869605 | 0.964236 |
| 11 | 6404200 | 6405999 | 16.75619 | 29.53688 | 38.03817 | 16.83503 | 0.025896 | 0.575274 | 0.964251 |
| 4 | 34876600 | 34880199 | 53.63853 | 85.71943 | 107.3279 | 45.11157 | 0.01817 | 2.083281 | 0.964252 |
| 14 | 63144000 | 63146399 | 21.46687 | 36.83163 | 47.40761 | 20.79274 | 0.020824 | 0.93421 | 0.96426 |
| 15 | 83415600 | 83417399 | 16.20093 | 28.34433 | 37.82215 | 16.6831 | -0.02659 | 0.676819 | 0.964297 |
| 2 | 173339200 | 173344799 | 71.50351 | 112.7132 | 142.7309 | 59.22951 | -0.0168 | 2.483969 | 0.964298 |
| 4 | 39382600 | 39387599 | 113.5553 | 172.8453 | 213.4737 | 88.46197 | -0.0199 | 3.164442 | 0.964307 |
| 12 | 62934800 | 62935199 | 7.540738 | 14.60775 | 22.17771 | 8.41339 | -0.02606 | -0.20468 | 0.964341 |
| 6 | 57571600 | 57572199 | 6.656777 | 12.93986 | 20.08538 | 7.46169 | -0.02914 | -0.29367 | 0.964355 |
| 10 | 43688000 | 43689399 | 16.41665 | 28.91709 | 38.58015 | 17.07741 | -0.02529 | 0.581795 | 0.964358 |
| 12 | 44001200 | 44010399 | 186.5315 | 274.629 | 329.0261 | 135.4951 | -0.01891 | 3.801134 | 0.964383 |
| X | 35135200 | 35136199 | 14.47257 | 26.17991 | 35.77843 | 15.51763 | -0.0374 | 0.375022 | 0.964421 |
| 6 | 12869000 | 12869599 | 11.53012 | 20.84443 | 26.80685 | 11.63544 | 0.049296 | 0.336585 | 0.964423 |
| 7 | 41042400 | 41043199 | 10.6941 | 19.90385 | 26.75854 | 11.17338 | 0.024543 | 0.163957 | 0.964436 |
| 15 | 20348400 | 20349799 | 15.74817 | 28.41896 | 28.30859 | 12.31484 | 0.413422 | 0.381647 | 0.964445 |
| 1 | 170298000 | 170300399 | 17.67934 | 30.81582 | 40.97803 | 18.02805 | -0.02199 | 0.74496 | 0.964463 |
| 12 | 111406400 | 111408199 | 18.86365 | 32.46767 | 43.26615 | 18.97891 | -0.02804 | 0.889905 | 0.964493 |
| 11 | 10982400 | 10986399 | 29.63326 | 49.20277 | 65.21411 | 28.26958 | -0.02426 | 1.2896 | 0.964502 |
| 12 | 105593000 | 105597999 | 76.67422 | 120.0071 | 151.2416 | 62.73977 | -0.01781 | 2.62461 | 0.964526 |
| 18 | 74435800 | 74438199 | 56.72333 | 90.32217 | 112.6931 | 47.12781 | 0.017602 | 2.200296 | 0.964551 |
| 12 | 114665200 | 114667999 | 37.35138 | 61.02289 | 80.99633 | 34.81814 | -0.03362 | 1.515631 | 0.964576 |
| 1 | 27089200 | 27090999 | 17.47924 | 30.95462 | 39.54958 | 17.48774 | 0.039812 | 0.532609 | 0.964578 |
| 4 | 112874200 | 112879799 | 77.24674 | 122.0583 | 149.736 | 62.1945 | 0.031652 | 2.437238 | 0.964583 |

| | | | | | | | | | |
|---|---|---|---|---|---|---|---|---|---|
| 14 | 110849200 | 110849999 | 10.83064 | 20.26904 | 28.42438 | 11.75906 | -0.02541 | 0.133214 | 0.964589 |
| 6 | 109465400 | 109466599 | 13.92818 | 25.85819 | 25.48099 | 10.68045 | 0.465484 | 0.203057 | 0.964594 |
| 2 | 43298400 | 43298999 | 6.954958 | 13.46771 | 20.50719 | 7.793918 | -0.03234 | -0.20971 | 0.964603 |
| 3 | 47236400 | 47238599 | 13.81606 | 24.28383 | 40.87203 | 18.0989 | -0.35973 | 0.596601 | 0.964696 |
| 4 | 25511800 | 25513399 | 20.98842 | 35.74077 | 47.44754 | 20.79765 | -0.0252 | 1.000312 | 0.964701 |
| 6 | 70043000 | 70046399 | 54.03089 | 86.26615 | 107.9202 | 45.30078 | 0.017918 | 2.110703 | 0.96471 |
| 3 | 79691000 | 79693799 | 22.26893 | 37.83409 | 50.09072 | 21.95216 | -0.0219 | 1.02842 | 0.96472 |
| 7 | 45416400 | 45417599 | 16.24643 | 28.49049 | 36.72542 | 16.23138 | 0.023085 | 0.63865 | 0.964736 |
| 17 | 24071800 | 24073599 | 16.24643 | 28.49049 | 36.72542 | 16.23138 | 0.023085 | 0.63865 | 0.964736 |
| 9 | 5440600 | 5447799 | 104.3941 | 161.4228 | 164.7071 | 68.41547 | 0.272971 | 2.849889 | 0.964762 |
| 11 | 3590000 | 3600199 | 108.1186 | 166.4828 | 208.4653 | 86.58554 | -0.02674 | 2.934719 | 0.964767 |
| 16 | 86711600 | 86714199 | 23.30235 | 40.12191 | 51.22459 | 22.47016 | 0.033403 | 0.914689 | 0.96479 |
| 6 | 142943600 | 142945999 | 16.00612 | 28.27052 | 36.43567 | 16.11942 | 0.025778 | 0.545 | 0.964831 |
| 9 | 18810800 | 18811799 | 17.65073 | 30.79943 | 41.00072 | 18.05036 | -0.02323 | 0.73203 | 0.964917 |
| X | 18167200 | 18168599 | 21.16572 | 36.56723 | 46.86846 | 20.56144 | 0.029023 | 0.873572 | 0.964924 |
| 4 | 61606400 | 61607799 | 27.47265 | 45.8146 | 58.89938 | 25.60625 | 0.019886 | 1.230382 | 0.964931 |
| 1 | 35988200 | 35993399 | 44.5014 | 71.54814 | 90.17572 | 38.24225 | 0.018857 | 1.878737 | 0.96495 |
| 4 | 112937200 | 112939399 | 25.10433 | 43.01962 | 54.59279 | 23.94175 | 0.040815 | 0.967164 | 0.964979 |
| 17 | 4170000 | 4170199 | 5.499399 | 10.74546 | 17.68979 | 6.154054 | -0.02622 | -0.54607 | 0.964979 |
| 19 | 9314200 | 9315399 | 14.59778 | 25.53691 | 43.07068 | 18.99654 | -0.36282 | 0.68021 | 0.964999 |
| X | 168193400 | 168195199 | 14.29516 | 25.55731 | 33.10109 | 14.53011 | 0.028332 | 0.439774 | 0.965026 |
| 17 | 85417200 | 85419199 | 25.06376 | 41.90187 | 55.46436 | 24.15089 | -0.02204 | 1.194676 | 0.965055 |
| 11 | 104175600 | 104177999 | 27.55206 | 45.68822 | 60.39523 | 26.15735 | -0.02048 | 1.296208 | 0.965069 |
| 6 | 51325800 | 51326199 | 5.799028 | 11.30334 | 18.03185 | 6.520212 | -0.03179 | -0.40564 | 0.965074 |
| 2 | 32365400 | 32373999 | 119.4478 | 181.8795 | 224.1037 | 92.87693 | -0.01609 | 3.159395 | 0.965078 |
| 9 | 35207200 | 35208599 | 16.48581 | 29.12142 | 37.5309 | 16.60354 | 0.026073 | 0.550964 | 0.965089 |
| 3 | 17503400 | 17504799 | 15.70019 | 28.06448 | 38.10677 | 16.72981 | -0.0402 | 0.454315 | 0.965105 |
| X | 153484600 | 153486599 | 21.12582 | 36.78661 | 49.54881 | 21.7851 | -0.04063 | 0.788609 | 0.965109 |
| 10 | 129727800 | 129731399 | 38.15324 | 62.13426 | 78.94383 | 33.89636 | 0.026063 | 1.561136 | 0.965116 |
| 15 | 99711600 | 99714599 | 23.91289 | 40.30976 | 51.73719 | 22.62006 | 0.022282 | 1.103477 | 0.965129 |
| 10 | 93049200 | 93054399 | 44.00961 | 70.82046 | 91.72297 | 38.94864 | -0.01878 | 1.850637 | 0.965149 |
| 5 | 149699600 | 149707199 | 91.60591 | 142.0248 | 173.493 | 72.06042 | 0.016492 | 2.797266 | 0.965178 |
| 10 | 67971600 | 67974599 | 21.73511 | 36.87126 | 49.0359 | 21.48021 | -0.02847 | 1.037138 | 0.965188 |
| 6 | 89810800 | 89817399 | 168.8035 | 250.9382 | 302.8078 | 124.7516 | -0.01629 | 3.636419 | 0.965189 |
| 2 | 149951600 | 149977399 | 450.5634 | 620.9035 | 699.2817 | 290.6238 | -0.0245 | 4.806161 | 0.965203 |
| 5 | 132154400 | 132155999 | 10.25624 | 19.0308 | 25.30985 | 10.61653 | 0.0354 | 0.175767 | 0.965208 |
| 8 | 21222000 | 21223799 | 12.02764 | 21.8224 | 29.64119 | 12.8078 | -0.02502 | 0.326084 | 0.96521 |
| X | 96326600 | 96328799 | 17.50089 | 30.87461 | 39.59726 | 17.52222 | 0.032302 | 0.574526 | 0.965243 |
| 4 | 145450400 | 145451799 | 18.28666 | 32.12868 | 32.91114 | 14.56872 | 0.352966 | 0.613491 | 0.965254 |
| 12 | 48468400 | 48472799 | 37.95385 | 61.92696 | 78.62287 | 33.78191 | 0.029029 | 1.533079 | 0.965257 |
| 1 | 42830000 | 42831599 | 16.55962 | 29.12116 | 37.55994 | 16.63127 | 0.022386 | 0.598758 | 0.965299 |
| 4 | 90089000 | 90091999 | 24.49961 | 41.57395 | 53.35567 | 23.37471 | 0.022775 | 1.052054 | 0.965302 |
| 10 | 100938200 | 100938799 | 8.513637 | 16.19905 | 23.43137 | 9.390301 | -0.03012 | -0.00844 | 0.965318 |
| 17 | 93258800 | 93262999 | 60.03923 | 95.71157 | 119.1897 | 49.92361 | 0.022095 | 2.168707 | 0.965324 |
| 6 | 47831400 | 47832599 | 10.15557 | 19.07652 | 26.92009 | 11.06681 | -0.0265 | 0.103115 | 0.965328 |
| 1 | 46684600 | 46685199 | 7.411883 | 14.36196 | 21.06154 | 7.980024 | 0.025693 | -0.21221 | 0.965365 |
| 1 | 15142800 | 15145999 | 49.89788 | 79.80791 | 99.87338 | 42.02782 | 0.020243 | 2.049664 | 0.9654 |

| | | | | | | | | | |
|---|---|---|---|---|---|---|---|---|---|
| 6 | 135690800 | 135691199 | 6.519558 | 12.68369 | 19.02302 | 7.019644 | 0.02936 | -0.3212 | 0.965415 |
| 11 | 61195200 | 61196799 | 23.18393 | 39.37604 | 50.62367 | 22.18377 | 0.020355 | 1.037184 | 0.965422 |
| 14 | 22385600 | 22387599 | 16.76759 | 29.24963 | 38.99244 | 17.16299 | -0.0253 | 0.727668 | 0.96543 |
| 4 | 34503400 | 34507999 | 114.1051 | 173.9004 | 209.4655 | 86.83368 | 0.017678 | 3.138993 | 0.965445 |
| 15 | 31223400 | 31225399 | 16.38321 | 28.18382 | 50.58299 | 22.18812 | -0.45607 | 0.89214 | 0.965448 |
| 15 | 95709600 | 95711399 | 25.50671 | 42.7622 | 54.97896 | 23.98195 | 0.019722 | 1.165043 | 0.965453 |
| 13 | 19769600 | 19770999 | 12.37734 | 22.76061 | 31.30989 | 13.29529 | -0.02623 | 0.256411 | 0.965461 |
| 13 | 77946800 | 77947799 | 31.26413 | 51.07179 | 64.89691 | 27.89623 | 0.02914 | 1.495448 | 0.965482 |
| 6 | 136485400 | 136488199 | 33.44272 | 54.71984 | 70.07554 | 30.14021 | 0.019472 | 1.477533 | 0.965487 |
| 3 | 85141800 | 85144799 | 23.97076 | 40.32662 | 53.33915 | 23.30632 | -0.02105 | 1.123282 | 0.965501 |
| 3 | 131111000 | 131112599 | 17.78676 | 30.38554 | 52.66623 | 23.09109 | -0.40755 | 0.968107 | 0.965515 |
| 14 | 50318400 | 50323599 | 45.32883 | 73.34616 | 91.64732 | 39.19224 | 0.041997 | 1.695616 | 0.96555 |
| 14 | 124651400 | 124653799 | 39.52826 | 63.6746 | 82.97523 | 35.28669 | -0.02567 | 1.81062 | 0.965571 |
| 13 | 17763800 | 17765399 | 14.63608 | 26.15137 | 33.97699 | 14.92203 | 0.023009 | 0.44444 | 0.965581 |
| 5 | 52620800 | 52629199 | 84.99594 | 132.519 | 166.3435 | 69.03291 | -0.0164 | 2.698028 | 0.965584 |
| 10 | 69082200 | 69083799 | 14.82638 | 26.27209 | 34.00407 | 15.03389 | 0.021759 | 0.500445 | 0.965612 |
| 14 | 11114400 | 11115799 | 22.18465 | 38.28256 | 38.99107 | 17.10497 | 0.358859 | 0.875572 | 0.965628 |
| 9 | 25260200 | 25261799 | 14.42859 | 25.75196 | 33.44755 | 14.70329 | 0.022479 | 0.448186 | 0.965644 |
| 5 | 16888800 | 16890199 | 13.40545 | 23.92893 | 32.12975 | 14.12247 | -0.02476 | 0.44031 | 0.965674 |
| 1 | 35506800 | 35517599 | 140.7829 | 212.2839 | 259.4313 | 107.1951 | -0.01606 | 3.363695 | 0.965689 |
| 10 | 68060600 | 68061599 | 9.453254 | 17.72675 | 23.85936 | 9.837151 | 0.038031 | 0.099065 | 0.965694 |
| 14 | 90270400 | 90271599 | 10.94146 | 20.34537 | 28.27535 | 11.8253 | -0.02521 | 0.173635 | 0.965695 |
| 17 | 27505000 | 27529999 | 466.9922 | 636.4591 | 687.4869 | 286.2196 | 0.019599 | 4.976871 | 0.965729 |
| 1 | 3601000 | 3603799 | 38.70181 | 62.55937 | 79.29688 | 33.87564 | 0.019959 | 1.706173 | 0.965744 |
| 14 | 74361200 | 74362399 | 13.84291 | 24.24125 | 42.04959 | 18.56228 | -0.40294 | 0.659322 | 0.965746 |
| 12 | 58531800 | 58532599 | 9.52687 | 17.9009 | 24.27918 | 9.976075 | 0.030498 | 0.085665 | 0.96575 |
| 1 | 34449200 | 34451799 | 19.12671 | 33.23539 | 44.16029 | 19.39022 | -0.02136 | 0.7998 | 0.965761 |
| 18 | 62886600 | 62888199 | 17.11028 | 29.95936 | 39.82569 | 17.57878 | -0.0211 | 0.673655 | 0.965786 |
| 18 | 89741000 | 89742199 | 10.2053 | 19.26152 | 26.42742 | 10.77357 | 0.023259 | 0.069504 | 0.965794 |
| 10 | 62115200 | 62117399 | 22.47885 | 38.15855 | 49.03144 | 21.48263 | 0.021114 | 1.036517 | 0.965802 |
| 16 | 7394200 | 7395199 | 12.86799 | 22.74497 | 38.80022 | 17.16293 | -0.3757 | 0.52597 | 0.965807 |
| X | 114490000 | 114490999 | 16.71939 | 30.19636 | 27.33133 | 11.87752 | 0.551986 | 0.383513 | 0.96582 |
| 7 | 92877000 | 92877599 | 6.66743 | 12.9787 | 19.60233 | 7.200513 | 0.025241 | -0.3334 | 0.965824 |
| 11 | 42880800 | 42881199 | 6.66743 | 12.9787 | 19.60233 | 7.200513 | 0.025241 | -0.3334 | 0.965824 |
| 8 | 61196000 | 61198199 | 14.69485 | 26.20607 | 34.013 | 14.962 | 0.022996 | 0.457908 | 0.965876 |
| 5 | 101199400 | 101203599 | 42.81396 | 68.91377 | 89.27619 | 37.92348 | -0.01864 | 1.842467 | 0.965888 |
| 5 | 145832800 | 145882199 | 1097.348 | 1402.652 | 1459.672 | 612.4882 | -0.0193 | 6.011694 | 0.965888 |
| 11 | 104479400 | 104480399 | 11.82783 | 21.69309 | 29.68382 | 12.64991 | -0.02238 | 0.262382 | 0.965947 |
| 9 | 44281000 | 44283999 | 24.54572 | 41.08313 | 54.3874 | 23.69918 | -0.02225 | 1.17922 | 0.965955 |
| 6 | 70356000 | 70357799 | 20.30371 | 34.9677 | 46.33908 | 20.32724 | -0.01999 | 0.893855 | 0.965975 |
| 6 | 38908000 | 38911799 | 46.02011 | 73.96926 | 93.29153 | 39.54764 | 0.017363 | 1.891443 | 0.965984 |
| 6 | 41217600 | 41220599 | 29.32911 | 48.07205 | 78.18378 | 33.65143 | -0.32323 | 1.453142 | 0.965992 |
| 10 | 35525600 | 35526399 | 8.608513 | 16.47724 | 24.01803 | 9.494452 | -0.02395 | -0.05233 | 0.965995 |
| 11 | 19303400 | 19307799 | 33.21662 | 54.58673 | 69.86444 | 30.10402 | 0.022945 | 1.421902 | 0.966005 |
| X | 4365400 | 4368799 | 32.56487 | 53.91699 | 68.76064 | 29.74856 | 0.030888 | 1.334427 | 0.966005 |
| 5 | 9597600 | 9598399 | 14.06541 | 25.6903 | 26.30351 | 11.27125 | 0.387651 | 0.294069 | 0.966046 |
| 4 | 4168600 | 4171999 | 33.82429 | 55.41667 | 73.06578 | 31.44299 | -0.02152 | 1.464343 | 0.96605 |

| | | | | | | | | | |
|---|---|---|---|---|---|---|---|---|---|
| 14 | 58649600 | 58651399 | 16.41676 | 29.08567 | 37.44435 | 16.55542 | 0.029157 | 0.517024 | 0.966056 |
| 6 | 136892000 | 136893199 | 13.37629 | 24.10458 | 31.44499 | 13.70345 | 0.024611 | 0.376689 | 0.966066 |
| 5 | 21544600 | 21547199 | 24.46777 | 42.03031 | 53.46437 | 23.45099 | 0.037971 | 0.942411 | 0.966069 |
| 8 | 117403800 | 117408999 | 52.92223 | 84.53978 | 108.4026 | 45.53661 | -0.0168 | 2.094421 | 0.966075 |
| 8 | 41617200 | 41617599 | 8.194699 | 15.69657 | 22.98007 | 9.065224 | -0.02684 | -0.06438 | 0.966086 |
| 14 | 87177200 | 87178599 | 10.81682 | 19.27932 | 35.85844 | 15.7813 | -0.4931 | 0.441602 | 0.966086 |
| 17 | 93804600 | 93808599 | 12.47475 | 23.29443 | 22.62383 | 9.394452 | 0.497146 | 0.149602 | 0.96609 |
| 10 | 126288400 | 126297399 | 84.92187 | 132.6655 | 162.9159 | 67.59576 | 0.017409 | 2.662932 | 0.966091 |
| 4 | 126964000 | 126969999 | 72.83923 | 114.9057 | 145.715 | 60.47424 | -0.01802 | 2.470676 | 0.966098 |
| 17 | 68268200 | 68269399 | 11.07671 | 20.45522 | 27.24975 | 11.50481 | 0.026101 | 0.21419 | 0.966108 |
| 18 | 45616200 | 45616999 | 9.992372 | 18.73799 | 26.38419 | 10.87785 | -0.02661 | 0.111289 | 0.966108 |
| 7 | 47003600 | 47051199 | 759.5601 | 1001.898 | 1046.524 | 437.7266 | 0.020686 | 5.500357 | 0.966114 |
| 9 | 88191800 | 88192399 | 8.016191 | 15.41687 | 21.97165 | 8.567361 | 0.027276 | -0.10744 | 0.966117 |
| 15 | 28476600 | 28478599 | 19.46592 | 33.7094 | 44.7303 | 19.62481 | -0.02025 | 0.838887 | 0.966125 |
| 4 | 38905200 | 38908599 | 59.43335 | 94.74798 | 95.09786 | 39.83386 | 0.332822 | 2.160853 | 0.966136 |
| 7 | 84856800 | 84858799 | 29.47854 | 50.02606 | 46.84795 | 20.52413 | 0.475789 | 1.061991 | 0.966148 |
| 15 | 83093600 | 83095799 | 16.66983 | 29.15358 | 38.81737 | 17.11657 | -0.02351 | 0.686736 | 0.966184 |
| 19 | 15929800 | 15933799 | 39.8066 | 64.2953 | 83.57612 | 35.66802 | -0.01763 | 1.731242 | 0.966198 |
| 12 | 114851400 | 114855399 | 58.22246 | 93.03747 | 120.4605 | 50.62558 | -0.03024 | 2.095445 | 0.966268 |
| 5 | 39912800 | 39913199 | 5.798493 | 11.30204 | 18.04182 | 6.525773 | -0.03252 | -0.40255 | 0.966281 |
| 11 | 28482800 | 28484399 | 32.26602 | 53.3175 | 68.23782 | 29.48241 | 0.025529 | 1.353356 | 0.966306 |
| 15 | 87162600 | 87164199 | 15.07425 | 26.62516 | 35.50531 | 15.70776 | -0.02306 | 0.538086 | 0.966326 |
| 5 | 31487200 | 31494199 | 158.3714 | 236.5135 | 286.4414 | 118.0847 | -0.01547 | 3.551733 | 0.96633 |
| 12 | 85546800 | 85549999 | 47.82253 | 76.74314 | 96.62751 | 40.86391 | 0.016988 | 1.944671 | 0.966357 |
| 3 | 43718200 | 43719399 | 16.28207 | 28.45083 | 37.94776 | 16.72096 | -0.0258 | 0.69954 | 0.966365 |
| 6 | 23861600 | 23863199 | 15.2986 | 27.5239 | 28.2588 | 12.33891 | 0.366421 | 0.39424 | 0.966372 |
| 5 | 126334200 | 126336199 | 22.06108 | 37.78366 | 50.10095 | 21.97087 | -0.0224 | 0.954309 | 0.966383 |
| 4 | 149643400 | 149646799 | 33.03852 | 53.88581 | 70.7236 | 30.38349 | -0.01907 | 1.524199 | 0.966386 |
| 9 | 9852000 | 9853799 | 18.58743 | 32.52973 | 41.8667 | 18.47197 | 0.025266 | 0.692282 | 0.96641 |
| 11 | 13092600 | 13092999 | 7.840755 | 15.13388 | 22.57987 | 8.712543 | -0.02443 | -0.1428 | 0.966416 |
| 18 | 31166800 | 31167799 | 17.90545 | 31.28379 | 40.3959 | 17.79836 | 0.0206 | 0.713169 | 0.966456 |
| 10 | 34649000 | 34653799 | 50.30614 | 80.0521 | 125.3101 | 52.35731 | -0.3101 | 2.219713 | 0.966483 |
| 1 | 115952400 | 115952999 | 11.88816 | 21.49316 | 29.18604 | 12.66704 | -0.02866 | 0.344821 | 0.9665 |
| 11 | 31452000 | 31456199 | 29.01144 | 47.84957 | 63.17615 | 27.26394 | -0.01963 | 1.363825 | 0.96651 |
| 16 | 82520800 | 82525599 | 53.62975 | 86.03349 | 88.85998 | 37.5599 | 0.301625 | 1.958897 | 0.966512 |
| 9 | 22623000 | 22627399 | 34.57996 | 56.34312 | 73.98984 | 31.77502 | -0.02098 | 1.541314 | 0.96655 |
| 13 | 53145600 | 53148599 | 33.03896 | 53.94042 | 70.81095 | 30.4321 | -0.01838 | 1.510087 | 0.966554 |
| 1 | 140726800 | 140728999 | 30.82489 | 51.27175 | 53.54379 | 23.23893 | 0.318806 | 1.263311 | 0.966556 |
| 11 | 107177600 | 107183199 | 59.13782 | 94.06095 | 117.2802 | 48.96982 | 0.0168 | 2.238861 | 0.96656 |
| 1 | 101129000 | 101130399 | 11.90214 | 22.38278 | 23.60713 | 9.686165 | 0.392401 | 0.10104 | 0.966571 |
| 17 | 65656600 | 65670199 | 195.1924 | 287.9169 | 345.0406 | 142.0928 | -0.01678 | 3.772302 | 0.966573 |
| 3 | 38178800 | 38181199 | 38.40235 | 62.06213 | 80.86399 | 34.53564 | -0.02004 | 1.713064 | 0.96661 |
| 4 | 5095000 | 5097999 | 48.02141 | 77.14934 | 97.15076 | 41.15528 | 0.01875 | 1.910757 | 0.966613 |
| 13 | 7096000 | 7097799 | 16.73449 | 29.41001 | 39.2078 | 17.36025 | -0.02517 | 0.613122 | 0.96662 |
| 6 | 11450000 | 11452199 | 56.60937 | 89.94641 | 115.0292 | 47.97748 | -0.02115 | 2.267981 | 0.966633 |
| 12 | 87868000 | 87868599 | 16.63415 | 29.41658 | 28.93805 | 12.79921 | 0.414379 | 0.523311 | 0.966633 |
| 4 | 141127600 | 141130199 | 23.7234 | 39.88481 | 52.76551 | 23.05118 | -0.02129 | 1.12922 | 0.966715 |

| | | | | | | | | | |
|---|---|---|---|---|---|---|---|---|---|
| 8 | 49331000 | 49332399 | 12.76254 | 23.21418 | 30.56715 | 13.16635 | 0.02222 | 0.317791 | 0.966741 |
| 3 | 16222800 | 16224599 | 34.91041 | 57.80938 | 56.97732 | 24.65326 | 0.401604 | 1.329632 | 0.966749 |
| 17 | 53636200 | 53638399 | 21.32929 | 35.90226 | 59.18842 | 25.86639 | -0.33766 | 1.116389 | 0.966758 |
| 1 | 132739000 | 132740999 | 15.7995 | 27.64922 | 36.90606 | 16.2826 | -0.02686 | 0.67088 | 0.966761 |
| 5 | 123613000 | 123613999 | 9.417217 | 17.77331 | 24.33973 | 9.923403 | 0.026094 | 0.057655 | 0.966764 |
| 7 | 48790800 | 48792799 | 22.05984 | 37.69177 | 48.48591 | 21.25855 | 0.020847 | 0.97716 | 0.966797 |
| 8 | 39646200 | 39647399 | 18.43708 | 32.15552 | 41.48889 | 18.25955 | 0.021514 | 0.743129 | 0.966798 |
| 3 | 145705400 | 145708199 | 30.70735 | 50.28233 | 66.27716 | 28.51495 | -0.02134 | 1.459683 | 0.96682 |
| 4 | 143400200 | 143409999 | 244.1133 | 352.4273 | 404.4726 | 166.8657 | 0.015743 | 4.106927 | 0.966834 |
| 4 | 115456600 | 115464199 | 93.78469 | 143.9375 | 215.1125 | 89.30622 | -0.28533 | 2.992305 | 0.966836 |
| 13 | 112001400 | 112004599 | 22.20231 | 37.81414 | 50.09765 | 21.95956 | -0.02237 | 1.004576 | 0.96687 |
| 11 | 14678000 | 14680199 | 19.94472 | 34.65841 | 44.67583 | 19.61716 | 0.022184 | 0.803733 | 0.966912 |
| 14 | 122338400 | 122340599 | 21.40884 | 36.79902 | 47.39693 | 20.79004 | 0.020293 | 0.91661 | 0.96693 |
| 5 | 130674800 | 130676199 | 15.95253 | 28.22579 | 36.47494 | 16.13109 | 0.023084 | 0.52744 | 0.966952 |
| 17 | 36357000 | 36359999 | 24.77883 | 41.61749 | 54.99463 | 24.01631 | -0.0198 | 1.141769 | 0.966957 |
| 4 | 19277800 | 19278799 | 14.54503 | 25.85102 | 34.53997 | 15.23735 | -0.02145 | 0.479188 | 0.966961 |
| 10 | 34772200 | 34776399 | 77.31843 | 121.0037 | 152.2846 | 63.17309 | -0.01591 | 2.626517 | 0.966964 |
| 12 | 119763800 | 119764399 | 9.569609 | 17.9699 | 25.32414 | 10.41652 | -0.02548 | 0.093554 | 0.966969 |
| 13 | 67446200 | 67447199 | 13.48635 | 23.91865 | 32.07608 | 14.17571 | -0.02858 | 0.478824 | 0.966978 |
| 12 | 26579200 | 26581999 | 34.51619 | 56.0636 | 73.38924 | 31.47263 | -0.01976 | 1.591757 | 0.966978 |
| 12 | 80539200 | 80540999 | 16.80068 | 29.12861 | 49.00636 | 21.50166 | -0.36087 | 0.821704 | 0.966994 |
| 5 | 77420600 | 77421399 | 8.876754 | 16.91402 | 24.45393 | 9.770632 | -0.02643 | -0.01513 | 0.96701 |
| 17 | 63277600 | 63282199 | 48.01019 | 77.17213 | 100.0096 | 42.39886 | -0.02166 | 1.89487 | 0.96702 |
| 14 | 52971800 | 52986799 | 213.0782 | 311.8869 | 362.8652 | 149.4781 | 0.015363 | 3.89664 | 0.96702 |
| 8 | 97875800 | 97877199 | 13.94025 | 25.27639 | 33.10345 | 14.31527 | 0.026236 | 0.349974 | 0.967053 |
| 7 | 107945400 | 107946199 | 11.01258 | 20.61255 | 27.8406 | 11.5147 | 0.028328 | 0.130129 | 0.967068 |
| 18 | 34217800 | 34228599 | 115.349 | 176.0431 | 217.2168 | 90.07642 | -0.01546 | 3.111109 | 0.967079 |
| 16 | 89761200 | 89765599 | 58.68768 | 93.28488 | 118.9498 | 49.63203 | -0.01642 | 2.258499 | 0.967106 |
| 15 | 68456200 | 68456799 | 6.223621 | 12.11813 | 19.05573 | 6.984138 | -0.0296 | -0.35014 | 0.967121 |
| Y | 6191600 | 6192199 | 8.17419 | 15.40861 | 22.40334 | 9.155413 | -0.06251 | 0.04704 | 0.967122 |
| 14 | 11419200 | 11422399 | 33.50525 | 54.20257 | 86.87352 | 37.14472 | -0.31654 | 1.679635 | 0.96713 |
| 10 | 93836800 | 93837799 | 10.35365 | 19.14117 | 37.02627 | 15.60888 | -0.50742 | 0.222473 | 0.967149 |
| 6 | 81509600 | 81510399 | 9.070931 | 17.31188 | 25.04543 | 9.972928 | -0.02289 | -0.02158 | 0.967154 |
| 14 | 42215800 | 42217999 | 38.90769 | 63.28884 | 65.47292 | 28.10033 | 0.320678 | 1.571929 | 0.967166 |
| 14 | 83675800 | 83676799 | 9.13725 | 17.222 | 23.37511 | 9.554418 | 0.036108 | 0.062364 | 0.967179 |
| 17 | 65854800 | 65856799 | 12.53259 | 22.02067 | 40.40098 | 17.88533 | -0.48352 | 0.592039 | 0.967207 |
| 3 | 14706400 | 14707199 | 7.253512 | 13.99303 | 20.9743 | 8.106588 | -0.03289 | -0.14795 | 0.967209 |
| 19 | 57074400 | 57075599 | 20.78122 | 36.1845 | 37.19797 | 16.35423 | 0.347473 | 0.768653 | 0.967215 |
| 2 | 41542600 | 41543799 | 11.48724 | 21.22837 | 29.2587 | 12.34108 | -0.02207 | 0.216228 | 0.967218 |
| 2 | 123752800 | 123754199 | 16.04503 | 28.22876 | 37.56289 | 16.63109 | -0.0222 | 0.590231 | 0.967218 |
| 18 | 48201000 | 48202999 | 27.53888 | 45.46698 | 58.09786 | 25.08646 | 0.027863 | 1.344342 | 0.967241 |
| 7 | 27715400 | 27718799 | 35.15586 | 57.23161 | 74.94707 | 32.1759 | -0.01781 | 1.55737 | 0.967245 |
| 6 | 65162200 | 65165599 | 52.44433 | 83.74699 | 107.3061 | 45.05362 | -0.01627 | 2.104566 | 0.967245 |
| 18 | 81750800 | 81758199 | 83.5654 | 130.146 | 163.0948 | 67.6958 | -0.01524 | 2.715887 | 0.967261 |
| 8 | 129919600 | 129939799 | 385.8646 | 536.9322 | 607.2747 | 251.9989 | -0.01646 | 4.656208 | 0.967263 |
| 7 | 29694200 | 29696399 | 19.20183 | 33.387 | 43.1065 | 18.93503 | 0.020057 | 0.791316 | 0.967298 |
| 1 | 28893600 | 28895399 | 23.9082 | 40.33175 | 53.30871 | 23.31239 | -0.01981 | 1.101688 | 0.967354 |

| | | | | | | | | | |
|---|---|---|---|---|---|---|---|---|---|
| 1 | 132591600 | 132593199 | 15.92715 | 27.98669 | 36.13162 | 15.99604 | 0.020835 | 0.603558 | 0.967355 |
| 14 | 48984200 | 48985599 | 13.16778 | 23.736 | 31.05009 | 13.52626 | 0.021266 | 0.37497 | 0.967368 |
| 9 | 72304400 | 72306799 | 27.85586 | 46.1492 | 59.30621 | 25.66927 | 0.020028 | 1.30646 | 0.967389 |
| 1 | 85595200 | 85597599 | 28.08215 | 46.56448 | 61.50676 | 26.63767 | -0.01938 | 1.300034 | 0.967396 |
| 7 | 144565800 | 144567599 | 16.85726 | 29.43031 | 37.90453 | 16.69278 | 0.024215 | 0.713504 | 0.967409 |
| 15 | 49167600 | 49168799 | 23.42555 | 38.87354 | 65.53134 | 28.396 | -0.37001 | 1.281854 | 0.967409 |
| 4 | 89654000 | 89658199 | 62.86065 | 99.98202 | 100.6704 | 42.03437 | 0.324714 | 2.238548 | 0.967415 |
| 2 | 61167800 | 61168199 | 5.940094 | 11.5967 | 18.16506 | 6.4156 | 0.024965 | -0.485 | 0.967439 |
| 15 | 29927800 | 29928599 | 10.0399 | 18.78956 | 26.31 | 10.88399 | -0.02301 | 0.121529 | 0.967472 |
| 14 | 34372200 | 34375199 | 26.29695 | 43.72152 | 56.06369 | 24.32539 | 0.023538 | 1.256797 | 0.967506 |
| 3 | 13195200 | 13199799 | 48.65323 | 78.26756 | 80.1689 | 34.02636 | 0.318482 | 1.864844 | 0.967517 |
| 1 | 145325400 | 145326999 | 20.64956 | 36.0703 | 36.14697 | 15.9241 | 0.384665 | 0.724293 | 0.967521 |
| 7 | 34233800 | 34235999 | 25.61836 | 42.80416 | 54.9982 | 23.93913 | 0.02067 | 1.200109 | 0.967529 |
| 12 | 98557600 | 98565199 | 84.45238 | 132.7526 | 162.4325 | 67.38061 | 0.029554 | 2.550419 | 0.967531 |
| X | 128212400 | 128216199 | 33.8014 | 55.84658 | 74.65612 | 32.25226 | -0.03712 | 1.366257 | 0.967537 |
| 16 | 21437200 | 21438399 | 12.41865 | 22.75052 | 30.13468 | 12.86215 | 0.022452 | 0.273431 | 0.967576 |
| 19 | 57522600 | 57524199 | 18.03218 | 31.38796 | 40.51692 | 17.80989 | 0.020565 | 0.766197 | 0.967585 |
| 18 | 46825400 | 46825999 | 14.54458 | 25.71075 | 33.17014 | 14.67225 | 0.025228 | 0.524542 | 0.967597 |
| 9 | 11810400 | 11824999 | 471.5772 | 641.8164 | 710.2527 | 295.7611 | -0.01732 | 4.999017 | 0.967609 |
| 16 | 87645000 | 87646199 | 14.82824 | 26.26912 | 35.00927 | 15.47905 | -0.02038 | 0.504689 | 0.967611 |
| 9 | 9625600 | 9627199 | 10.71051 | 19.74975 | 27.23217 | 11.52053 | -0.02553 | 0.218125 | 0.967612 |
| 11 | 61496600 | 61502199 | 60.02497 | 95.60509 | 122.4285 | 51.21812 | -0.01948 | 2.197049 | 0.967614 |
| 12 | 47147200 | 47147799 | 10.8901 | 20.26239 | 27.20019 | 11.36374 | 0.025614 | 0.166022 | 0.967643 |
| 1 | 142051600 | 142055599 | 74.90489 | 117.6625 | 145.3445 | 60.28663 | 0.015101 | 2.556358 | 0.967646 |
| 13 | 55739600 | 55741599 | 5.560664 | 10.73507 | 24.21892 | 9.336359 | -0.61352 | -0.16491 | 0.967647 |
| 13 | 8070200 | 8077199 | 66.03744 | 104.8846 | 109.214 | 45.53371 | 0.274315 | 2.28562 | 0.967656 |
| 15 | 28982200 | 28983199 | 12.67775 | 22.41676 | 38.27031 | 16.92732 | -0.37669 | 0.51878 | 0.967708 |
| 15 | 59174200 | 59176799 | 26.1873 | 43.71132 | 57.70692 | 25.10242 | -0.01827 | 1.216208 | 0.967725 |
| 2 | 170883600 | 170886199 | 23.99917 | 40.37026 | 53.32889 | 23.30111 | -0.01919 | 1.124888 | 0.967751 |
| 6 | 136495200 | 136496399 | 9.537547 | 17.54207 | 31.92586 | 13.55313 | -0.42705 | 0.233443 | 0.967777 |
| 10 | 51618200 | 51618999 | 22.22113 | 37.75673 | 48.50161 | 21.25591 | 0.021665 | 1.028267 | 0.967784 |
| 5 | 5990000 | 5991399 | 15.90848 | 27.96243 | 37.22435 | 16.48407 | -0.023 | 0.603062 | 0.967785 |
| 13 | 82322000 | 82327999 | 108.8378 | 166.7592 | 202.1182 | 83.89138 | 0.014755 | 3.029295 | 0.967792 |
| 8 | 79117400 | 79118999 | 12.72406 | 23.00569 | 30.13092 | 13.07625 | 0.023239 | 0.353488 | 0.967822 |
| 2 | 40720800 | 40722799 | 18.92319 | 32.96852 | 42.54434 | 18.71164 | 0.021116 | 0.761696 | 0.967847 |
| 9 | 99576200 | 99581199 | 42.63488 | 68.46978 | 108.7486 | 46.04259 | -0.31617 | 1.918674 | 0.967855 |
| 4 | 143805200 | 143822199 | 545.1346 | 734.7854 | 787.1288 | 328.2614 | 0.015504 | 5.152688 | 0.967857 |
| 14 | 18570600 | 18571999 | 10.70714 | 19.75352 | 26.24165 | 11.09386 | 0.028648 | 0.213172 | 0.967871 |
| 13 | 116433000 | 116434399 | 11.52743 | 21.03792 | 27.68968 | 11.87708 | 0.027077 | 0.284294 | 0.967873 |
| 17 | 75679800 | 75682799 | 19.21471 | 33.39476 | 43.07678 | 18.91676 | 0.021173 | 0.794964 | 0.967881 |
| X | 168755800 | 168772799 | 2524.271 | 2987.294 | 2887.241 | 1223.14 | -0.02548 | 7.240284 | 0.967919 |
| 5 | 123448800 | 123450599 | 14.923 | 26.3163 | 35.06409 | 15.51747 | -0.02269 | 0.553951 | 0.967944 |
| 5 | 145559800 | 145592399 | 1127.647 | 1439.929 | 1497.726 | 628.5173 | -0.02005 | 6.028116 | 0.96796 |
| X | 49363200 | 49365599 | 23.5353 | 40.47105 | 51.76443 | 22.70608 | 0.030446 | 0.928537 | 0.967964 |
| 11 | 49543600 | 49547599 | 39.30952 | 63.38956 | 79.97913 | 34.06161 | 0.022618 | 1.77594 | 0.967973 |
| 5 | 100263200 | 100265599 | 24.0323 | 40.20163 | 53.31051 | 23.22185 | -0.02467 | 1.181783 | 0.967982 |
| 2 | 89338400 | 89342999 | 28.23727 | 47.56885 | 48.55262 | 21.22361 | 0.351712 | 1.122799 | 0.968 |

| | | | | | | | | | |
|---|---|---|---|---|---|---|---|---|---|
| 4 | 38581400 | 38583999 | 29.48914 | 48.87183 | 64.54022 | 27.9413 | -0.01921 | 1.310476 | 0.968004 |
| 17 | 40729400 | 40732199 | 35.06794 | 57.00608 | 74.57796 | 32.00058 | -0.01781 | 1.579354 | 0.968057 |
| 11 | 28320400 | 28323199 | 33.44092 | 55.03265 | 70.4728 | 30.38422 | 0.022996 | 1.40686 | 0.968132 |
| 1 | 15624000 | 15627199 | 26.81452 | 45.42226 | 46.90365 | 20.53718 | 0.335215 | 1.067665 | 0.968145 |
| 8 | 24199600 | 24202199 | 16.16174 | 27.98816 | 49.0004 | 21.49828 | -0.41842 | 0.828669 | 0.968164 |
| 5 | 149680600 | 149684199 | 52.04571 | 82.73088 | 129.4036 | 53.99617 | -0.31063 | 2.254108 | 0.968164 |
| 7 | 113569200 | 113572399 | 39.48975 | 63.77872 | 82.85223 | 35.35564 | -0.01683 | 1.732839 | 0.968175 |
| 13 | 67632000 | 67650199 | 227.2228 | 332.2035 | 384.9097 | 158.5706 | 0.019716 | 3.917059 | 0.968204 |
| 10 | 66101200 | 66102999 | 18.03862 | 31.1263 | 52.21959 | 22.90814 | -0.35821 | 0.876296 | 0.968214 |
| 7 | 4914400 | 4915799 | 13.74385 | 24.56154 | 32.95686 | 14.47182 | -0.02321 | 0.433676 | 0.968228 |
| 6 | 92595600 | 92596999 | 18.19977 | 31.68769 | 42.0446 | 18.48435 | -0.01896 | 0.765374 | 0.968231 |
| 10 | 15476400 | 15478799 | 27.52421 | 46.29938 | 48.03151 | 20.98644 | 0.328247 | 1.131757 | 0.96824 |
| 10 | 55595000 | 55597999 | 30.02181 | 50.19507 | 51.86964 | 22.58685 | 0.333926 | 1.204724 | 0.968278 |
| 18 | 76904000 | 76905199 | 12.22223 | 22.39811 | 29.69489 | 12.66872 | 0.021954 | 0.271543 | 0.968292 |
| 1 | 190300600 | 190302599 | 22.77354 | 38.9143 | 50.09793 | 21.96548 | 0.019609 | 0.978941 | 0.968308 |
| 6 | 136390000 | 136391399 | 12.27703 | 22.17768 | 30.04256 | 13.05242 | -0.0265 | 0.352091 | 0.96831 |
| 1 | 111465000 | 111467999 | 38.43051 | 62.47874 | 79.50498 | 34.116 | 0.022072 | 1.585227 | 0.968336 |
| 19 | 24119200 | 24126599 | 60.21894 | 95.00102 | 147.9958 | 61.42216 | -0.31438 | 2.461425 | 0.968347 |
| X | 55925200 | 55926399 | 11.38021 | 21.41397 | 29.13216 | 11.94112 | 0.028851 | 0.106174 | 0.968347 |
| 1 | 59491800 | 59492199 | 7.29731 | 14.18834 | 21.2465 | 7.877921 | 0.025173 | -0.28609 | 0.968352 |
| 6 | 46455800 | 46457399 | 14.87582 | 26.32572 | 35.0736 | 15.51075 | -0.02042 | 0.517583 | 0.968356 |
| 5 | 27436000 | 27437999 | 18.73874 | 32.5317 | 43.17208 | 18.9455 | -0.01956 | 0.811571 | 0.968362 |
| 11 | 34582600 | 34586399 | 42.69472 | 68.77545 | 87.00181 | 36.99916 | 0.016975 | 1.81476 | 0.968374 |
| 18 | 66361600 | 66363999 | 23.39511 | 39.86042 | 52.79489 | 23.14255 | -0.02181 | 1.012271 | 0.968404 |
| 11 | 21736400 | 21737599 | 22.28021 | 37.90903 | 50.09072 | 21.95476 | -0.01872 | 1.015406 | 0.968411 |
| 3 | 51004800 | 51014799 | 192.3361 | 283.4063 | 331.8253 | 136.649 | 0.015344 | 3.788229 | 0.968415 |
| 1 | 20170800 | 20174399 | 28.72794 | 48.37174 | 49.09277 | 21.45654 | 0.359837 | 1.129026 | 0.968465 |
| 17 | 28714400 | 28716799 | 22.70679 | 38.67706 | 51.18407 | 22.43595 | -0.02076 | 1.007077 | 0.968468 |
| 1 | 46789600 | 46792199 | 7.30826 | 13.94158 | 27.25661 | 10.86368 | -0.45491 | -0.03002 | 0.968516 |
| 18 | 89850800 | 89855399 | 38.71177 | 62.55581 | 81.52411 | 34.813 | -0.02059 | 1.712505 | 0.968522 |
| 6 | 88672800 | 88674199 | 12.31315 | 22.50491 | 30.6611 | 13.1275 | -0.02099 | 0.287618 | 0.968524 |
| 7 | 3821600 | 3828799 | 85.22039 | 132.5882 | 165.9648 | 68.89753 | -0.01473 | 2.733847 | 0.96853 |
| 2 | 146837200 | 146838199 | 10.9704 | 20.31365 | 27.19639 | 11.44019 | 0.022417 | 0.195825 | 0.968534 |
| 13 | 3100800 | 3102199 | 24.563 | 41.1333 | 54.38075 | 23.70401 | -0.02031 | 1.174217 | 0.968555 |
| 13 | 86931600 | 86931999 | 6.839551 | 13.31989 | 20.21036 | 7.391854 | 0.024684 | -0.34272 | 0.968561 |
| 10 | 83758600 | 83760399 | 14.62536 | 25.89425 | 34.50302 | 15.25703 | -0.02049 | 0.508167 | 0.968579 |
| 1 | 5321200 | 5324599 | 53.40992 | 85.37364 | 107.0864 | 45.02465 | 0.015943 | 2.075027 | 0.968582 |
| 10 | 110463200 | 110463999 | 13.87236 | 24.4518 | 32.81282 | 14.52257 | -0.0329 | 0.55051 | 0.968613 |
| 11 | 63262400 | 63265599 | 23.49433 | 40.18703 | 53.39562 | 23.41328 | -0.02556 | 0.976702 | 0.968621 |
| 10 | 100339400 | 100342199 | 22.40583 | 39.01344 | 49.56277 | 21.79052 | 0.043357 | 0.79466 | 0.968625 |
| 6 | 125719200 | 125720599 | 15.44386 | 27.08719 | 34.81467 | 15.38887 | 0.027374 | 0.628748 | 0.968694 |
| 9 | 81919400 | 81920799 | 20.00779 | 33.86237 | 56.45652 | 24.71341 | -0.35344 | 1.061611 | 0.968702 |
| 3 | 130434600 | 130435399 | 9.12644 | 17.37571 | 25.00207 | 10.00676 | -0.02131 | -0.00307 | 0.968764 |
| 7 | 103578200 | 103578999 | 9.12644 | 17.37571 | 25.00207 | 10.00676 | -0.02131 | -0.00307 | 0.968764 |
| 7 | 57717800 | 57721199 | 56.22781 | 89.50914 | 114.266 | 47.7679 | -0.0158 | 2.208655 | 0.968776 |
| 3 | 94182200 | 94185199 | 24.42243 | 40.22233 | 70.45799 | 30.38778 | -0.42713 | 1.375387 | 0.968842 |
| 6 | 10397200 | 10405799 | 203.7004 | 297.8713 | 345.7701 | 142.4446 | 0.017317 | 3.910598 | 0.968845 |

| | | | | | | | | | |
|---|---|---|---|---|---|---|---|---|---|
| 1 | 24114600 | 24115599 | 26.37097 | 44.54108 | 43.83832 | 19.17806 | 0.40416 | 1.089075 | 0.968866 |
| 8 | 53415200 | 53417599 | 78.26746 | 122.0671 | 149.2528 | 61.93734 | 0.021823 | 2.687802 | 0.968867 |
| 9 | 87727600 | 87728799 | 10.92055 | 20.53097 | 27.96954 | 11.48165 | 0.024872 | 0.102876 | 0.968875 |
| 16 | 86128600 | 86130599 | 18.63512 | 32.60374 | 43.48009 | 19.17911 | -0.02576 | 0.700015 | 0.968879 |
| 8 | 26386800 | 26390399 | 38.52668 | 62.2467 | 80.94009 | 34.55643 | -0.01757 | 1.719972 | 0.968883 |
| 1 | 124899400 | 124904599 | 60.69872 | 96.09029 | 146.3508 | 60.84605 | -0.27786 | 2.380556 | 0.968885 |
| 13 | 82287600 | 82288199 | 11.08381 | 20.68766 | 28.89198 | 12.00655 | -0.02494 | 0.151992 | 0.968935 |
| 6 | 39895400 | 39897799 | 17.91736 | 31.25944 | 41.48486 | 18.26158 | -0.01892 | 0.735241 | 0.96895 |
| 8 | 3865400 | 3867199 | 25.63742 | 42.78591 | 54.98155 | 23.91422 | 0.020534 | 1.212953 | 0.968975 |
| 17 | 22937600 | 22941199 | 36.11179 | 59.91139 | 75.02258 | 32.50337 | 0.057386 | 1.327466 | 0.96899 |
| 8 | 69723400 | 69735799 | 203.5753 | 300.2194 | 350.9392 | 144.5215 | 0.018824 | 3.776981 | 0.968992 |
| 9 | 12533600 | 12534999 | 19.96581 | 34.62157 | 44.71458 | 19.61872 | 0.019011 | 0.829461 | 0.968993 |
| 5 | 122850000 | 122855199 | 50.14727 | 80.24918 | 102.9988 | 43.37549 | -0.01535 | 2.036117 | 0.96901 |
| 15 | 48851000 | 48851599 | 8.342476 | 16.02952 | 23.5748 | 9.21697 | -0.02145 | -0.09061 | 0.969011 |
| 19 | 21883400 | 21883799 | 8.342476 | 16.02952 | 23.5748 | 9.21697 | -0.02145 | -0.09061 | 0.969011 |
| X | 84293400 | 84294599 | 8.342476 | 16.02952 | 23.5748 | 9.21697 | -0.02145 | -0.09061 | 0.969011 |
| X | 75935000 | 75944799 | 193.6236 | 285.2955 | 333.9533 | 137.5253 | 0.01567 | 3.790303 | 0.969012 |
| 2 | 68011200 | 68012399 | 14.38277 | 25.53133 | 34.05693 | 15.03978 | -0.02009 | 0.484977 | 0.969028 |
| 7 | 39275600 | 39277799 | 19.01886 | 33.28551 | 44.51186 | 19.63941 | -0.02986 | 0.696531 | 0.96904 |
| 12 | 103879800 | 103887599 | 139.543 | 210.0616 | 250.8128 | 103.5872 | 0.015342 | 3.39525 | 0.969044 |
| 5 | 63675000 | 63679199 | 44.46888 | 71.58178 | 92.7119 | 39.38632 | -0.01828 | 1.841371 | 0.969067 |
| 8 | 65689600 | 65691599 | 26.14669 | 43.77328 | 57.76416 | 25.17442 | -0.01776 | 1.183865 | 0.969088 |
| 13 | 49898000 | 49903199 | 102.2637 | 157.1002 | 194.6742 | 80.83158 | -0.01425 | 2.975531 | 0.969093 |
| 17 | 82349000 | 82349999 | 11.18862 | 20.70191 | 27.7292 | 11.67644 | 0.020812 | 0.204291 | 0.969113 |
| 3 | 97664200 | 97666399 | 17.42846 | 30.32381 | 38.97443 | 17.12716 | 0.026606 | 0.764948 | 0.969137 |
| 17 | 66941000 | 66943999 | 29.47462 | 48.89806 | 64.5891 | 27.98233 | -0.01942 | 1.298461 | 0.969146 |
| 3 | 7919200 | 7921999 | 34.04989 | 56.10756 | 58.50812 | 25.23952 | 0.319297 | 1.385056 | 0.969155 |
| 14 | 58860200 | 58861599 | 13.53936 | 24.47384 | 32.08529 | 13.92863 | 0.0214 | 0.364448 | 0.969158 |
| 17 | 13309000 | 13313799 | 62.21338 | 98.47411 | 125.2185 | 52.05644 | -0.01803 | 2.379766 | 0.969175 |
| 4 | 6105400 | 6106599 | 14.77739 | 26.21262 | 34.95259 | 15.44484 | -0.02039 | 0.49168 | 0.969217 |
| 1 | 145978800 | 145990199 | 167.7894 | 250.4274 | 303.2572 | 125.0024 | -0.01609 | 3.565115 | 0.969218 |
| 17 | 51939000 | 51941199 | 22.44702 | 38.10853 | 49.03112 | 21.48318 | 0.019214 | 1.034836 | 0.96922 |
| Y | 2761800 | 2762599 | 8.744081 | 16.43633 | 21.8666 | 8.97896 | 0.059017 | 0.068704 | 0.969233 |
| 9 | 38875800 | 38880199 | 72.37805 | 113.9452 | 141.0667 | 58.52914 | 0.014344 | 2.51014 | 0.969235 |
| 19 | 27724800 | 27726399 | 20.04013 | 34.79573 | 46.30087 | 20.32026 | -0.02366 | 0.816446 | 0.969258 |
| 2 | 82745400 | 82747399 | 41.61389 | 67.06308 | 84.90119 | 36.12823 | 0.01666 | 1.800298 | 0.969265 |
| 2 | 36914400 | 36916399 | 32.39863 | 53.12895 | 69.89912 | 30.09177 | -0.01785 | 1.449371 | 0.969272 |
| 15 | 93243600 | 93245599 | 18.8752 | 32.90771 | 42.5085 | 18.70403 | 0.019714 | 0.750786 | 0.969279 |
| 17 | 81551400 | 81554599 | 50.52197 | 80.83453 | 101.5012 | 42.73413 | 0.015942 | 2.04235 | 0.96928 |
| 12 | 74017000 | 74018999 | 32.3839 | 53.16244 | 68.25602 | 29.3979 | 0.017904 | 1.434862 | 0.96929 |
| 7 | 100346800 | 100348799 | 14.92432 | 26.03759 | 44.17661 | 19.44646 | -0.37159 | 0.72443 | 0.969305 |
| 7 | 92209400 | 92211199 | 16.05482 | 28.04525 | 37.42565 | 16.48775 | -0.02666 | 0.700301 | 0.969313 |
| 9 | 115226200 | 115228399 | 29.25381 | 48.20912 | 63.59881 | 27.43667 | -0.01888 | 1.376855 | 0.96932 |
| 10 | 31859400 | 31859999 | 7.713305 | 14.89174 | 21.50371 | 8.287249 | 0.02445 | -0.15007 | 0.969339 |
| 13 | 80660400 | 80660799 | 7.713305 | 14.89174 | 21.50371 | 8.287249 | 0.02445 | -0.15007 | 0.969339 |
| 16 | 39715800 | 39716199 | 5.797323 | 11.30511 | 18.0539 | 6.489759 | -0.02579 | -0.42691 | 0.96938 |
| 8 | 37947400 | 37948799 | 25.33115 | 42.33167 | 54.42303 | 23.69138 | 0.019864 | 1.197748 | 0.969381 |

| | | | | | | | | | |
|---|---|---|---|---|---|---|---|---|---|
| 8 | 14233600 | 14235999 | 23.58953 | 39.80887 | 51.19306 | 22.3897 | 0.019577 | 1.092373 | 0.969397 |
| 5 | 123340200 | 123344799 | 49.9772 | 80.03482 | 102.8027 | 43.33861 | -0.01529 | 2.013597 | 0.969425 |
| 14 | 42274200 | 42275399 | 50.34054 | 80.43403 | 100.6506 | 42.2939 | 0.018759 | 2.083446 | 0.969431 |
| 9 | 19800000 | 19800799 | 12.30648 | 22.69829 | 30.2706 | 12.80162 | 0.022379 | 0.23595 | 0.969454 |
| 9 | 43307600 | 43312799 | 41.40621 | 66.79638 | 86.61767 | 36.90719 | -0.01623 | 1.770045 | 0.96946 |
| X | 78672600 | 78676999 | 55.51321 | 88.86062 | 115.2014 | 48.53327 | -0.02938 | 2.040651 | 0.969478 |
| 9 | 61127200 | 61132799 | 44.65115 | 71.79058 | 92.74793 | 39.33442 | -0.01674 | 1.878743 | 0.969478 |
| 2 | 128374400 | 128375799 | 15.62133 | 27.75534 | 35.94916 | 15.86315 | 0.022711 | 0.489194 | 0.96949 |
| 12 | 23946000 | 23946999 | 11.11426 | 21.0311 | 22.1673 | 8.987844 | 0.407947 | 0.049956 | 0.969495 |
| 7 | 64916400 | 64918399 | 23.26351 | 39.8556 | 51.20258 | 22.45454 | 0.02315 | 0.958422 | 0.969495 |
| Y | 10667400 | 10667999 | 7.821848 | 14.92354 | 20.40364 | 8.129732 | 0.057539 | -0.04611 | 0.969503 |
| 5 | 127919600 | 127922799 | 30.18912 | 49.53628 | 65.25186 | 28.09846 | -0.01924 | 1.431774 | 0.969511 |
| 4 | 86555000 | 86558599 | 58.36514 | 92.86712 | 118.4013 | 49.46384 | -0.01471 | 2.226902 | 0.969538 |
| X | 149114800 | 149119599 | 41.49112 | 67.29663 | 88.55723 | 37.9374 | -0.02931 | 1.642218 | 0.969539 |
| 17 | 30920600 | 30921199 | 10.57212 | 19.67475 | 27.29967 | 11.40137 | -0.02085 | 0.165495 | 0.969554 |
| 8 | 109858200 | 109858999 | 9.580121 | 17.95826 | 24.31575 | 10.03136 | 0.028549 | 0.102755 | 0.969566 |
| 10 | 90936000 | 90939399 | 26.28554 | 44.01212 | 56.67697 | 24.70287 | 0.017386 | 1.181506 | 0.969571 |
| 18 | 18616800 | 18617199 | 6.653717 | 12.94273 | 20.12706 | 7.436165 | -0.02461 | -0.31324 | 0.969599 |
| 2 | 106956000 | 106956799 | 8.755221 | 16.81017 | 24.67795 | 9.668737 | -0.02221 | -0.07835 | 0.969617 |
| 9 | 112318200 | 112319199 | 15.30016 | 26.86381 | 35.74493 | 15.81433 | -0.02242 | 0.612289 | 0.969633 |
| 14 | 21408800 | 21412799 | 55.95978 | 89.06127 | 111.0487 | 46.40764 | 0.017762 | 2.216328 | 0.969642 |
| 5 | 115076400 | 115077799 | 16.16867 | 28.6253 | 36.99643 | 16.35976 | 0.023256 | 0.524025 | 0.969659 |
| Y | 689600 | 690399 | 11.13497 | 20.23223 | 26.21036 | 11.30474 | 0.044454 | 0.300959 | 0.969677 |
| 1 | 133370400 | 133372599 | 26.60865 | 44.60635 | 58.9398 | 25.70849 | -0.01969 | 1.171567 | 0.969681 |
| 1 | 125251400 | 125253599 | 23.27829 | 39.86372 | 39.58579 | 17.35948 | 0.39324 | 0.948773 | 0.969695 |
| 1 | 33588000 | 33590999 | 30.32079 | 50.12848 | 64.52179 | 27.88661 | 0.017464 | 1.34066 | 0.969712 |
| 4 | 20131800 | 20133799 | 23.23866 | 40.08832 | 51.24853 | 22.48197 | 0.032042 | 0.896725 | 0.969721 |
| 1 | 46331600 | 46331999 | 4.930602 | 9.637977 | 16.02256 | 5.516009 | -0.02586 | -0.59771 | 0.969724 |
| 15 | 34443000 | 34443999 | 11.32963 | 20.90066 | 28.72059 | 12.14287 | -0.02018 | 0.223589 | 0.969724 |
| X | 153492200 | 153497399 | 73.29614 | 115.7041 | 146.9907 | 61.02439 | -0.01982 | 2.456956 | 0.969731 |
| 10 | 81878000 | 81961599 | 2028.693 | 2467.096 | 2439.915 | 1029.63 | -0.01862 | 6.81124 | 0.969735 |
| 11 | 51012800 | 51017399 | 51.51509 | 82.76883 | 103.7028 | 43.93601 | 0.026135 | 1.915342 | 0.969769 |
| 16 | 15769200 | 15770399 | 10.05801 | 18.90836 | 25.84156 | 10.60906 | 0.020685 | 0.092621 | 0.969788 |
| 1 | 133116600 | 133118599 | 23.33571 | 39.5844 | 52.24498 | 22.88558 | -0.01748 | 1.049619 | 0.969793 |
| 19 | 61163800 | 61165999 | 41.69212 | 67.54332 | 70.51852 | 30.18359 | 0.302274 | 1.6564 | 0.969801 |
| 18 | 59398400 | 59401399 | 35.05474 | 56.86829 | 74.2945 | 31.83194 | -0.01852 | 1.619206 | 0.969809 |
| 1 | 67727200 | 67728199 | 7.981391 | 14.96903 | 28.53342 | 11.76117 | -0.46012 | 0.096703 | 0.969813 |
| 18 | 68614200 | 68615799 | 22.29564 | 37.83058 | 50.07947 | 21.93883 | -0.0216 | 1.043572 | 0.969841 |
| 18 | 33245800 | 33245999 | 5.349581 | 10.44554 | 17.05384 | 6.006428 | -0.02925 | -0.50494 | 0.969847 |
| X | 44560200 | 44562399 | 15.08758 | 26.41216 | 44.08246 | 19.45425 | -0.34783 | 0.67325 | 0.969852 |
| 2 | 163490400 | 163494399 | 58.69758 | 93.3159 | 116.425 | 48.5865 | 0.015204 | 2.253433 | 0.969854 |
| 5 | 64379200 | 64382999 | 29.02579 | 47.52704 | 76.80146 | 33.04238 | -0.31451 | 1.464694 | 0.969871 |
| 11 | 19434400 | 19437199 | 31.49015 | 51.94997 | 68.59199 | 29.60411 | -0.01959 | 1.371313 | 0.969878 |
| 14 | 31580800 | 31590799 | 235.9155 | 341.726 | 393.9629 | 162.4584 | 0.013965 | 4.05712 | 0.969879 |
| 7 | 62438200 | 62440599 | 20.6793 | 35.97335 | 46.26765 | 20.32978 | 0.025544 | 0.795119 | 0.969896 |
| 6 | 43303400 | 43304599 | 13.78117 | 24.18009 | 41.43091 | 18.31924 | -0.38547 | 0.624226 | 0.969903 |
| 2 | 171578600 | 171580999 | 28.11459 | 46.59778 | 59.96633 | 25.96266 | 0.01801 | 1.301214 | 0.969912 |

| | | | | | | | | | |
|---|---|---|---|---|---|---|---|---|---|
| 11 | 20338400 | 20341799 | 70.58008 | 111.4927 | 138.2694 | 57.42351 | 0.015681 | 2.436977 | 0.969913 |
| 15 | 32809000 | 32811599 | 27.13426 | 45.69877 | 60.70168 | 26.53218 | -0.02717 | 1.131578 | 0.969926 |
| 13 | 55815200 | 55816799 | 16.06175 | 28.33361 | 36.60997 | 16.20055 | 0.021768 | 0.563178 | 0.969937 |
| 10 | 24872000 | 24874999 | 27.18969 | 45.44113 | 59.99857 | 26.11983 | -0.01852 | 1.208201 | 0.969944 |
| 10 | 90815000 | 90818599 | 34.82662 | 56.86758 | 72.84898 | 31.3097 | 0.016953 | 1.512701 | 0.970017 |
| 3 | 58130000 | 58130999 | 10.18222 | 19.13048 | 26.88121 | 11.04701 | -0.02049 | 0.098995 | 0.97002 |
| 4 | 62132800 | 62136399 | 39.94951 | 64.66985 | 84.30064 | 36.06197 | -0.01832 | 1.675422 | 0.970057 |
| 2 | 8444200 | 8444999 | 14.76797 | 26.76923 | 26.45689 | 11.44529 | 0.429951 | 0.345781 | 0.970088 |
| 6 | 136275400 | 136276799 | 18.80344 | 32.88592 | 42.3804 | 18.68813 | 0.023288 | 0.702961 | 0.970114 |
| 6 | 82013800 | 82015999 | 17.70954 | 30.89082 | 39.94124 | 17.57991 | 0.018475 | 0.734168 | 0.970117 |
| 13 | 73481600 | 73483599 | 17.70954 | 30.89082 | 39.94124 | 17.57991 | 0.018475 | 0.734168 | 0.970117 |
| 10 | 97695200 | 97697399 | 17.01868 | 30.03127 | 40.15092 | 17.76635 | -0.02725 | 0.570023 | 0.970124 |
| 14 | 50318000 | 50322199 | 40.43088 | 65.82747 | 87.39336 | 37.52103 | -0.03748 | 1.572417 | 0.970132 |
| 12 | 8804400 | 8808399 | 44.47357 | 71.48882 | 90.188 | 38.23616 | 0.017096 | 1.884531 | 0.970139 |
| 3 | 26504200 | 26504799 | 8.168033 | 15.62577 | 21.91893 | 8.677987 | 0.029411 | -0.05657 | 0.970151 |
| 7 | 12268200 | 12270999 | 40.60523 | 66.10234 | 66.81973 | 28.68663 | 0.354772 | 1.565664 | 0.970158 |
| 9 | 48406200 | 48410199 | 29.85638 | 49.29583 | 64.96494 | 28.05268 | -0.0166 | 1.355247 | 0.970162 |
| 18 | 69617800 | 69630199 | 126.7393 | 192.8082 | 232.2054 | 96.2086 | 0.015762 | 3.180418 | 0.97018 |
| 7 | 106245200 | 106247599 | 24.02487 | 40.64737 | 52.2894 | 22.88695 | 0.019156 | 1.072088 | 0.970194 |
| 2 | 154000200 | 154005599 | 20.69735 | 36.05976 | 37.22231 | 16.37101 | 0.341718 | 0.761019 | 0.970242 |
| 13 | 96364600 | 96366999 | 20.48701 | 35.04579 | 46.3818 | 20.33794 | -0.01978 | 0.955797 | 0.970251 |
| 1 | 178440800 | 178443599 | 18.68347 | 32.02897 | 52.72976 | 23.12544 | -0.33297 | 0.938179 | 0.970265 |
| 3 | 77719200 | 77720999 | 16.17987 | 27.68815 | 51.06999 | 22.39434 | -0.4964 | 0.943564 | 0.970285 |
| 17 | 34527000 | 34529799 | 27.05908 | 44.78274 | 59.2102 | 25.60938 | -0.02096 | 1.316307 | 0.970286 |
| 1 | 179222600 | 179225599 | 23.6888 | 40.14248 | 51.70651 | 22.64272 | 0.017513 | 1.059685 | 0.9703 |
| 3 | 101814000 | 101817599 | 34.29078 | 55.90427 | 71.55441 | 30.73561 | 0.01656 | 1.532222 | 0.970325 |
| 4 | 129758600 | 129761399 | 24.78699 | 41.68877 | 54.99657 | 24.0286 | -0.01732 | 1.12982 | 0.970339 |
| X | 22585400 | 22588599 | 21.54425 | 37.40921 | 50.1244 | 21.99901 | -0.03344 | 0.832129 | 0.970345 |
| 9 | 46506800 | 46507799 | 10.99014 | 20.43255 | 28.29037 | 11.83485 | -0.02001 | 0.176149 | 0.970366 |
| 10 | 55905800 | 55907799 | 28.2531 | 46.20397 | 76.4203 | 32.86281 | -0.34905 | 1.481285 | 0.970387 |
| 2 | 110850400 | 110852599 | 31.48344 | 51.96348 | 68.62775 | 29.62693 | -0.01978 | 1.365574 | 0.970387 |
| 11 | 107161400 | 107163199 | 14.95794 | 26.38163 | 34.10713 | 15.09355 | 0.020776 | 0.551966 | 0.9704 |
| 8 | 33973800 | 33979599 | 75.85729 | 119.1427 | 147.2952 | 61.09471 | 0.01375 | 2.559299 | 0.970414 |
| Y | 89315000 | 89315599 | 6.935045 | 13.39063 | 18.9361 | 7.294047 | 0.054901 | -0.17098 | 0.970418 |
| 5 | 126670800 | 126679599 | 145.2768 | 218.2543 | 265.4007 | 109.5532 | -0.01357 | 3.436266 | 0.970452 |
| 4 | 132306000 | 132308799 | 33.04713 | 53.74277 | 70.46084 | 30.23842 | -0.0205 | 1.565627 | 0.970453 |
| 11 | 63581200 | 63583199 | 17.26005 | 30.31043 | 39.14849 | 17.32278 | 0.020156 | 0.625997 | 0.970454 |
| 2 | 132914800 | 132916799 | 22.97845 | 39.14853 | 51.71481 | 22.669 | -0.01806 | 1.008605 | 0.970466 |
| 2 | 116880400 | 116886399 | 58.71856 | 93.74567 | 120.6725 | 50.64997 | -0.02329 | 2.121253 | 0.970472 |
| 4 | 8442800 | 8444999 | 50.65503 | 80.90876 | 103.7212 | 43.563 | -0.0166 | 2.094767 | 0.970473 |
| Y | 40447000 | 40447399 | 4.977178 | 9.713201 | 15.96745 | 5.675809 | -0.05215 | -0.48765 | 0.970479 |
| 18 | 65102600 | 65105599 | 22.85917 | 38.76856 | 51.15875 | 22.40845 | -0.01719 | 1.048804 | 0.970488 |
| 2 | 165337200 | 165338599 | 13.89047 | 24.73592 | 32.11038 | 14.14223 | 0.021701 | 0.458453 | 0.970495 |
| 2 | 176927200 | 176930999 | 92.82967 | 143.2424 | 178.2448 | 74.03953 | -0.01537 | 2.885164 | 0.970525 |
| 18 | 86208800 | 86219599 | 227.88 | 330.1004 | 379.9567 | 156.6819 | 0.016315 | 4.055446 | 0.970572 |
| 16 | 76438400 | 76441599 | 35.17737 | 57.10224 | 72.77617 | 31.1955 | 0.017911 | 1.605481 | 0.97059 |
| 2 | 172760600 | 172762799 | 21.31605 | 36.40479 | 46.90755 | 20.56579 | 0.018348 | 0.97488 | 0.970656 |

| | | | | | | | | | |
|---|---|---|---|---|---|---|---|---|---|
| 14 | 21536600 | 21539599 | 20.96722 | 35.93971 | 46.36088 | 20.33207 | 0.017492 | 0.939524 | 0.970677 |
| 4 | 122143200 | 122143599 | 11.34152 | 21.52271 | 19.72767 | 7.950359 | 0.616101 | 0.027435 | 0.970724 |
| 15 | 31702600 | 31703599 | 18.61297 | 32.25452 | 41.61377 | 18.25786 | 0.020366 | 0.824371 | 0.970733 |
| 4 | 22493800 | 22495799 | 16.05098 | 27.72641 | 48.46392 | 21.26127 | -0.41676 | 0.851423 | 0.970733 |
| 8 | 52795600 | 52797399 | 16.05098 | 27.72641 | 48.46392 | 21.26127 | -0.41676 | 0.851423 | 0.970733 |
| 18 | 21175600 | 21176799 | 15.76074 | 28.38438 | 28.32603 | 12.35353 | 0.40849 | 0.393448 | 0.970789 |
| 1 | 97438600 | 97444599 | 23.986 | 39.48732 | 69.61841 | 30.02046 | -0.43654 | 1.381331 | 0.970797 |
| 12 | 94002200 | 94003999 | 15.33976 | 27.46065 | 27.40915 | 12.01835 | 0.402935 | 0.42445 | 0.970832 |
| 2 | 4424400 | 4429199 | 29.7542 | 49.37975 | 65.26496 | 28.2822 | -0.02029 | 1.29464 | 0.970842 |
| 18 | 68558800 | 68561999 | 31.3622 | 51.33177 | 67.45565 | 29.01669 | -0.01729 | 1.467652 | 0.970875 |
| 7 | 26477400 | 26480399 | 34.51492 | 56.50613 | 72.44096 | 31.1643 | 0.018447 | 1.475407 | 0.970884 |
| 16 | 75932800 | 75936799 | 60.717 | 96.25918 | 119.5681 | 49.75993 | 0.017687 | 2.337116 | 0.970905 |
| 1 | 129630400 | 129631399 | 9.008347 | 17.31941 | 24.72027 | 9.648064 | 0.023754 | -0.08546 | 0.970924 |
| 11 | 57870800 | 57873799 | 15.02481 | 26.19033 | 46.27673 | 20.35989 | -0.43025 | 0.737685 | 0.970933 |
| 9 | 73312400 | 73315199 | 30.3694 | 49.37866 | 80.30806 | 34.46197 | -0.33082 | 1.569472 | 0.970968 |
| 14 | 80860000 | 80861599 | 13.23814 | 24.03841 | 32.6078 | 14.07519 | -0.02252 | 0.334628 | 0.970998 |
| 16 | 41775600 | 41777999 | 51.83264 | 82.73143 | 105.9264 | 44.44485 | -0.01592 | 2.118645 | 0.971006 |
| 12 | 7362800 | 7366199 | 43.81675 | 70.41038 | 88.69478 | 37.58557 | 0.018683 | 1.893076 | 0.971007 |
| 4 | 109960600 | 109961599 | 17.10243 | 29.73577 | 39.61914 | 17.40283 | -0.02495 | 0.777427 | 0.971015 |
| 15 | 94659600 | 94661599 | 44.45056 | 71.77432 | 75.00513 | 31.99757 | 0.296012 | 1.74283 | 0.971025 |
| 2 | 3424800 | 3425799 | 8.002219 | 15.33534 | 22.4462 | 8.842256 | -0.0249 | -0.07396 | 0.971076 |
| 3 | 49114000 | 49120599 | 170.893 | 255.3229 | 255.0398 | 105.1483 | 0.262836 | 3.544192 | 0.971081 |
| 6 | 68078400 | 68079799 | 16.092 | 28.28997 | 36.59879 | 16.20669 | 0.017927 | 0.598371 | 0.971082 |
| 1 | 72935200 | 72937799 | 18.67731 | 32.50532 | 43.08294 | 18.93579 | -0.01751 | 0.774836 | 0.9711 |
| 2 | 129226400 | 129228199 | 18.67731 | 32.50532 | 43.08294 | 18.93579 | -0.01751 | 0.774836 | 0.9711 |
| 10 | 125998600 | 126000399 | 13.20082 | 23.70454 | 31.93382 | 13.96599 | -0.02501 | 0.396939 | 0.971112 |
| 10 | 89518800 | 89519799 | 8.969645 | 17.04763 | 24.44156 | 9.818328 | -0.0207 | 0.00593 | 0.971112 |
| 6 | 95647400 | 95649599 | 26.30298 | 44.18379 | 58.35995 | 25.48829 | -0.01918 | 1.149115 | 0.971115 |
| 7 | 75542400 | 75543799 | 14.66434 | 25.96089 | 34.53845 | 15.27298 | -0.01825 | 0.510341 | 0.971162 |
| 10 | 98667000 | 98671399 | 44.8361 | 72.31716 | 94.15159 | 40.10854 | -0.0223 | 1.782928 | 0.971178 |
| X | 15644400 | 15645599 | 18.21832 | 32.1094 | 41.18468 | 18.22817 | 0.031645 | 0.592195 | 0.9712 |
| 5 | 136843000 | 136846799 | 48.01095 | 77.03959 | 97.15585 | 41.08254 | 0.01456 | 1.94786 | 0.971206 |
| 2 | 99374600 | 99377599 | 32.16899 | 53.28751 | 55.76568 | 24.13454 | 0.315368 | 1.316026 | 0.971233 |
| 12 | 15531000 | 15531799 | 16.55733 | 28.9086 | 38.43122 | 16.92544 | -0.02126 | 0.71204 | 0.971244 |
| 6 | 10869600 | 10871399 | 20.43931 | 35.45232 | 45.77208 | 20.08287 | 0.01953 | 0.830125 | 0.971249 |
| 5 | 13914000 | 13915199 | 14.57261 | 25.56684 | 42.9637 | 18.9958 | -0.35767 | 0.632187 | 0.971267 |
| 11 | 84443800 | 84445399 | 26.06683 | 43.52045 | 56.02439 | 24.37413 | 0.018045 | 1.213087 | 0.971296 |
| 5 | 98960000 | 98961599 | 19.92026 | 33.54991 | 57.44884 | 25.11056 | -0.39203 | 1.106169 | 0.971298 |
| 17 | 64568800 | 64576199 | 89.74311 | 139.2569 | 173.9263 | 72.23105 | -0.0145 | 2.781278 | 0.971303 |
| 7 | 51825800 | 51826599 | 14.6317 | 25.88696 | 33.54133 | 14.8339 | 0.019345 | 0.514644 | 0.971332 |
| 4 | 73779000 | 73780399 | 25.37461 | 42.5087 | 56.04662 | 24.43605 | -0.01646 | 1.17403 | 0.971337 |
| 2 | 3843200 | 3845599 | 12.57422 | 22.18907 | 38.86063 | 17.19533 | -0.41426 | 0.543263 | 0.971348 |
| 13 | 54025400 | 54029799 | 56.5924 | 90.09691 | 114.8861 | 48.03253 | -0.01404 | 2.207064 | 0.971375 |
| 1 | 107044600 | 107045399 | 12.38669 | 22.73761 | 30.22418 | 12.86514 | 0.020519 | 0.264484 | 0.971383 |
| 14 | 122486200 | 122487199 | 10.36747 | 19.59995 | 27.01961 | 10.97657 | 0.020833 | 0.060986 | 0.971392 |
| 2 | 75537800 | 75540599 | 22.43554 | 38.44569 | 49.55687 | 21.73322 | 0.018414 | 0.951347 | 0.971411 |
| 12 | 105592200 | 105597799 | 79.22212 | 123.6643 | 155.1981 | 64.39876 | -0.01476 | 2.673451 | 0.971436 |

| | | | | | | | | | |
|---|---|---|---|---|---|---|---|---|---|
| 12 | 11577800 | 11580999 | 33.14851 | 54.17889 | 71.04681 | 30.54572 | -0.01573 | 1.493571 | 0.971446 |
| 17 | 44933000 | 44933999 | 12.43452 | 22.72935 | 30.11336 | 12.89113 | 0.019154 | 0.286157 | 0.971453 |
| 10 | 127673000 | 127677799 | 65.68609 | 104.0516 | 129.5267 | 53.8686 | 0.013485 | 2.365384 | 0.971455 |
| 4 | 111102000 | 111102599 | 8.872329 | 16.75907 | 23.92267 | 9.735968 | -0.03045 | 0.047933 | 0.97149 |
| 14 | 69396200 | 69401199 | 81.97773 | 128.4772 | 158.3071 | 65.66186 | 0.016123 | 2.604057 | 0.971529 |
| 13 | 91197600 | 91205199 | 85.93012 | 134.7633 | 170.9773 | 70.91217 | -0.02523 | 2.59793 | 0.971543 |
| 5 | 115513600 | 115516999 | 23.78059 | 40.14752 | 51.7264 | 22.62574 | 0.016891 | 1.09067 | 0.97155 |
| 3 | 57044200 | 57045799 | 14.27331 | 25.43203 | 33.96714 | 14.95287 | -0.01899 | 0.458492 | 0.971557 |
| 12 | 39442800 | 39443799 | 14.27331 | 25.43203 | 33.96714 | 14.95287 | -0.01899 | 0.458492 | 0.971557 |
| 7 | 28928400 | 28933599 | 40.92233 | 65.92403 | 85.36869 | 36.30756 | -0.01658 | 1.810121 | 0.971559 |
| 2 | 14420800 | 14422999 | 27.3461 | 45.76012 | 58.90628 | 25.66469 | 0.017943 | 1.194456 | 0.971578 |
| 11 | 13859400 | 13861199 | 20.57387 | 35.66959 | 47.39466 | 20.7945 | -0.02193 | 0.836031 | 0.971603 |
| 3 | 99030400 | 99031399 | 10.66654 | 19.68223 | 26.20602 | 11.07603 | 0.025512 | 0.21071 | 0.97162 |
| 6 | 75113200 | 75116199 | 33.31577 | 54.75506 | 70.29983 | 30.29288 | 0.01853 | 1.423107 | 0.971623 |
| 2 | 21067600 | 21070199 | 25.55487 | 42.94486 | 56.70562 | 24.76912 | -0.01875 | 1.139924 | 0.971638 |
| 17 | 16147000 | 16150799 | 47.66486 | 76.36225 | 98.12112 | 41.39011 | -0.01511 | 1.996798 | 0.971638 |
| 4 | 74578600 | 74579599 | 15.20005 | 26.81509 | 35.63336 | 15.7682 | -0.01846 | 0.554928 | 0.971645 |
| 4 | 122452000 | 122455599 | 44.15835 | 71.29537 | 90.17802 | 38.46546 | 0.021003 | 1.754108 | 0.971707 |
| 8 | 19431200 | 19450999 | 329.6254 | 465.1842 | 532.9909 | 220.6832 | -0.01469 | 4.444657 | 0.971711 |
| 5 | 59129400 | 59130199 | 9.000086 | 17.14494 | 23.95228 | 9.575019 | 0.022808 | -0.00982 | 0.971715 |
| 2 | 56655600 | 56656599 | 21.53347 | 36.72377 | 48.49097 | 21.25757 | -0.01712 | 0.991266 | 0.971733 |
| 12 | 64771600 | 64773799 | 24.8692 | 41.66031 | 54.94471 | 23.955 | -0.01688 | 1.171599 | 0.971757 |
| 13 | 117593800 | 117597199 | 43.08239 | 69.34774 | 89.58347 | 38.05553 | -0.01453 | 1.840903 | 0.97177 |
| 7 | 118153800 | 118156999 | 20.66268 | 35.6976 | 47.39018 | 20.79031 | -0.02176 | 0.866397 | 0.971772 |
| 5 | 72998200 | 72998799 | 11.24347 | 20.98011 | 28.37555 | 11.79721 | 0.020991 | 0.157042 | 0.971778 |
| 4 | 121138600 | 121142399 | 39.94923 | 64.74096 | 84.53716 | 36.19264 | -0.01929 | 1.653452 | 0.971789 |
| 18 | 8240400 | 8242399 | 21.912 | 37.57583 | 48.47533 | 21.26015 | 0.017389 | 0.941027 | 0.971796 |
| 5 | 59242200 | 59245199 | 28.13287 | 47.11386 | 62.40627 | 27.20314 | -0.02322 | 1.190471 | 0.971809 |
| 3 | 16794000 | 16795999 | 25.21329 | 43.11893 | 55.10414 | 24.16207 | 0.030342 | 0.987637 | 0.971842 |
| 7 | 85645800 | 85647199 | 34.75972 | 56.4695 | 73.79839 | 31.65236 | -0.01709 | 1.590956 | 0.97185 |
| 7 | 107018400 | 107020999 | 32.53192 | 53.18613 | 69.74671 | 29.98985 | -0.01545 | 1.488359 | 0.97186 |
| 3 | 51005400 | 51014999 | 189.3324 | 279.2846 | 327.7322 | 134.965 | 0.013589 | 3.769889 | 0.971861 |
| 1 | 188898200 | 188900599 | 19.21353 | 33.3618 | 44.19614 | 19.39703 | -0.01708 | 0.812218 | 0.971903 |
| 3 | 85956400 | 85959199 | 21.77072 | 37.95264 | 37.57844 | 16.53389 | 0.401509 | 0.757365 | 0.971914 |
| 15 | 36711200 | 36713599 | 19.98428 | 34.58964 | 44.73032 | 19.62452 | 0.016661 | 0.846332 | 0.971917 |
| 2 | 84041000 | 84041799 | 17.70507 | 30.89724 | 40.9461 | 18.02741 | -0.01689 | 0.728783 | 0.971938 |
| 17 | 36411000 | 36418199 | 122.8543 | 186.0005 | 228.184 | 94.40432 | -0.0166 | 3.276591 | 0.971943 |
| 2 | 163798800 | 163802599 | 48.99982 | 78.30182 | 100.5503 | 42.25939 | -0.0183 | 2.077385 | 0.971946 |
| 1 | 29624800 | 29626599 | 21.53926 | 37.30127 | 37.44831 | 16.42996 | 0.380605 | 0.839305 | 0.971951 |
| 5 | 63381400 | 63386599 | 62.92204 | 99.74643 | 124.1101 | 51.64574 | 0.015097 | 2.340921 | 0.971951 |
| 8 | 43776200 | 43787399 | 126.2268 | 191.6857 | 230.9723 | 95.64654 | 0.012916 | 3.216274 | 0.971954 |
| 5 | 88648800 | 88650399 | 17.68005 | 30.24325 | 52.71599 | 23.11537 | -0.41557 | 0.951714 | 0.971955 |
| 12 | 37412400 | 37413399 | 16.21502 | 28.42051 | 36.73273 | 16.22742 | 0.019234 | 0.64631 | 0.971965 |
| Y | 38001800 | 38002599 | 7.408059 | 14.16716 | 20.94639 | 8.30094 | -0.04723 | -0.06562 | 0.971976 |
| 9 | 90809200 | 90809599 | 9.13975 | 17.21865 | 24.36929 | 9.968681 | -0.02487 | 0.06773 | 0.972006 |
| 1 | 111021400 | 111026399 | 97.9427 | 151.0749 | 184.2961 | 76.55328 | 0.012811 | 2.894575 | 0.972015 |
| 14 | 13424400 | 13425799 | 13.02691 | 23.71212 | 32.17873 | 13.84804 | -0.02006 | 0.320515 | 0.972015 |

| | | | | | | | | | |
|---|---|---|---|---|---|---|---|---|---|
| 1 | 52632400 | 52634399 | 24.28524 | 41.55211 | 43.13809 | 18.91594 | 0.32913 | 0.964664 | 0.972078 |
| 3 | 22299200 | 22322599 | 396.3635 | 550.3577 | 608.2834 | 252.5018 | 0.013644 | 4.688904 | 0.972101 |
| 16 | 13733400 | 13742399 | 126.235 | 192.07 | 231.5271 | 95.93193 | 0.014628 | 3.177168 | 0.972111 |
| 3 | 76756800 | 76760399 | 58.72927 | 93.36431 | 118.8301 | 49.58913 | -0.01353 | 2.25498 | 0.972135 |
| 6 | 62367000 | 62368599 | 19.94841 | 34.82244 | 35.22363 | 15.51086 | 0.371185 | 0.728283 | 0.972139 |
| 14 | 76098400 | 76101999 | 28.0801 | 47.09699 | 48.59421 | 21.20272 | 0.335972 | 1.160537 | 0.972139 |
| 7 | 12526600 | 12533999 | 115.4592 | 176.3382 | 213.4536 | 88.53064 | 0.012964 | 3.09654 | 0.972146 |
| 16 | 32025400 | 32030199 | 68.98371 | 108.8971 | 135.2143 | 56.13546 | 0.013148 | 2.449819 | 0.972241 |
| Y | 40105800 | 40106599 | 9.739321 | 18.06371 | 23.86786 | 10.01765 | 0.044039 | 0.16884 | 0.97225 |
| 9 | 7758800 | 7759799 | 23.9109 | 40.99806 | 41.61465 | 18.25102 | 0.362061 | 0.944294 | 0.972257 |
| 1 | 85375000 | 85386599 | 418.6722 | 582.3728 | 536.5504 | 222.6632 | 0.278769 | 4.661906 | 0.972275 |
| X | 81497400 | 81499599 | 16.57839 | 29.29687 | 39.12521 | 17.30745 | -0.02474 | 0.551023 | 0.972281 |
| 18 | 5769200 | 5773999 | 52.68358 | 84.16798 | 105.5873 | 44.36102 | 0.014923 | 2.088908 | 0.972297 |
| 13 | 34323600 | 34327599 | 46.48587 | 74.22867 | 115.8004 | 48.61689 | -0.2998 | 2.103094 | 0.972301 |
| 17 | 12768000 | 12769599 | 15.73229 | 27.61769 | 35.67928 | 15.78294 | 0.019863 | 0.616557 | 0.972305 |
| 17 | 57359000 | 57360599 | 15.04018 | 26.49767 | 34.22054 | 15.14705 | 0.021874 | 0.56652 | 0.972321 |
| 9 | 45739800 | 45741399 | 20.17585 | 35.00426 | 45.24722 | 19.85274 | 0.017971 | 0.827099 | 0.972325 |
| X | 118160200 | 118161799 | 13.10359 | 23.99257 | 32.83533 | 14.02497 | -0.02491 | 0.290051 | 0.972334 |
| 18 | 16271800 | 16277999 | 68.67945 | 108.4448 | 137.1345 | 56.94006 | -0.01287 | 2.444171 | 0.972336 |
| 2 | 8583000 | 8583399 | 7.116667 | 13.82005 | 21.15225 | 7.921513 | -0.01958 | -0.25505 | 0.972336 |
| 5 | 88130400 | 88132199 | 20.13475 | 34.92764 | 46.31538 | 20.32131 | -0.0188 | 0.82875 | 0.972346 |
| 14 | 3946600 | 3948599 | 35.18739 | 57.13621 | 74.54891 | 31.9627 | -0.01546 | 1.600349 | 0.972353 |
| 9 | 115258600 | 115261999 | 30.80371 | 50.6951 | 66.68141 | 28.75015 | -0.0155 | 1.397126 | 0.972387 |
| 1 | 98799600 | 98800599 | 14.6168 | 25.90398 | 34.47042 | 15.23969 | -0.01808 | 0.497399 | 0.972436 |
| 4 | 29574400 | 29574999 | 11.87528 | 21.77611 | 29.706 | 12.66254 | -0.01811 | 0.264806 | 0.972441 |
| 8 | 26872000 | 26873399 | 14.5095 | 25.64508 | 34.11048 | 15.08863 | -0.01877 | 0.526566 | 0.972442 |
| X | 21060800 | 21066399 | 47.5 | 76.71083 | 101.1755 | 43.17113 | -0.03884 | 1.758765 | 0.972447 |
| 12 | 79898800 | 79900599 | 17.86988 | 31.20383 | 40.40263 | 17.79494 | 0.016572 | 0.720407 | 0.972453 |
| 3 | 153160400 | 153169799 | 115.9392 | 176.8089 | 213.602 | 88.56183 | 0.014119 | 3.12571 | 0.972473 |
| 17 | 70964400 | 70964999 | 7.444828 | 14.4084 | 21.0396 | 8.01678 | 0.024543 | -0.19114 | 0.972481 |
| 10 | 95219000 | 95220599 | 13.7276 | 24.57584 | 32.94758 | 14.44623 | -0.02052 | 0.4223 | 0.972483 |
| 8 | 89975200 | 89980599 | 38.04278 | 61.74584 | 78.71538 | 33.73824 | 0.017279 | 1.614765 | 0.972494 |
| 12 | 102820600 | 102822199 | 17.63472 | 31.16795 | 31.74314 | 14.04153 | 0.364205 | 0.540765 | 0.972495 |
| 13 | 66289800 | 66296599 | 231.6734 | 337.7151 | 309.3367 | 127.4724 | 0.354537 | 3.959765 | 0.972565 |
| 12 | 40010400 | 40011799 | 15.96146 | 28.20805 | 37.53376 | 16.60329 | -0.01957 | 0.542187 | 0.97257 |
| 11 | 105708000 | 105710199 | 19.01719 | 33.29025 | 42.88534 | 18.92543 | 0.023833 | 0.690789 | 0.972579 |
| 14 | 104019000 | 104020399 | 16.52461 | 29.03926 | 37.56078 | 16.63007 | 0.018008 | 0.606695 | 0.972606 |
| 6 | 138323600 | 138325599 | 17.85997 | 31.21963 | 41.39359 | 18.24497 | -0.01747 | 0.705688 | 0.97261 |
| 9 | 115553400 | 115554999 | 19.77316 | 34.72603 | 34.28739 | 15.17265 | 0.405739 | 0.630685 | 0.972649 |
| 2 | 21774600 | 21774999 | 6.986108 | 13.55883 | 20.03223 | 7.532923 | 0.0261 | -0.24391 | 0.972655 |
| 3 | 66218800 | 66222599 | 48.72834 | 78.35462 | 78.29881 | 33.21189 | 0.353242 | 1.873978 | 0.972659 |
| 17 | 52027000 | 52028999 | 17.90833 | 31.69804 | 40.59212 | 17.95059 | 0.036155 | 0.541353 | 0.97266 |
| 13 | 84381200 | 84387199 | 84.8303 | 132.433 | 162.9785 | 67.62644 | 0.013577 | 2.675525 | 0.972662 |
| 11 | 59326800 | 59329799 | 35.23389 | 57.90896 | 60.1584 | 25.92311 | 0.323291 | 1.420994 | 0.972676 |
| 9 | 52659400 | 52661799 | 21.36631 | 35.81493 | 60.21876 | 26.26485 | -0.36605 | 1.158203 | 0.972693 |
| 5 | 145374000 | 145414199 | 565.3487 | 765.4411 | 845.1855 | 352.2321 | -0.02171 | 5.083546 | 0.972699 |
| X | 55167600 | 55170199 | 28.13234 | 47.30463 | 62.91459 | 27.4862 | -0.02903 | 1.152798 | 0.972701 |

| | | | | | | | | | |
|---|---|---|---|---|---|---|---|---|---|
| 14 | 37747200 | 37750199 | 51.92035 | 83.1218 | 104.5403 | 44.05113 | 0.014553 | 2.025229 | 0.972719 |
| 18 | 7548800 | 7549599 | 9.32368 | 17.82337 | 25.09036 | 9.954745 | 0.021311 | -0.02821 | 0.97272 |
| 3 | 63527400 | 63528999 | 18.07462 | 31.59222 | 40.88969 | 18.0216 | 0.017237 | 0.708463 | 0.972741 |
| 6 | 26082600 | 26082999 | 5.79544 | 11.30681 | 17.53517 | 6.264207 | 0.024748 | -0.44889 | 0.972746 |
| 8 | 117400400 | 117401799 | 11.98341 | 21.75635 | 28.61847 | 12.3558 | 0.021683 | 0.319054 | 0.972768 |
| 10 | 45132200 | 45134199 | 19.02163 | 33.27876 | 42.90906 | 18.92665 | 0.022564 | 0.700198 | 0.972771 |
| 5 | 56736800 | 56738599 | 15.01681 | 26.89685 | 35.02818 | 15.35224 | 0.021669 | 0.434614 | 0.972784 |
| 17 | 22984400 | 22985399 | 18.60757 | 32.46702 | 41.98843 | 18.48446 | 0.018212 | 0.738546 | 0.972798 |
| 7 | 96989000 | 96990599 | 16.98494 | 29.66322 | 39.37146 | 17.34248 | -0.01906 | 0.710604 | 0.972802 |
| 1 | 85923000 | 85925599 | 31.11334 | 51.15218 | 67.26821 | 28.99057 | -0.01579 | 1.410347 | 0.97281 |
| 7 | 141707200 | 141708999 | 16.72754 | 29.42842 | 39.09325 | 17.30879 | -0.01981 | 0.597569 | 0.972831 |
| 3 | 17839400 | 17843199 | 49.61538 | 79.30593 | 101.7052 | 42.76037 | -0.01601 | 2.069813 | 0.972834 |
| 5 | 92889000 | 92890999 | 17.17258 | 30.03656 | 39.82107 | 17.56099 | -0.01734 | 0.689243 | 0.972849 |
| 6 | 128896800 | 128903999 | 146.5155 | 221.2964 | 223.9563 | 92.58573 | 0.257915 | 3.329197 | 0.972853 |
| 6 | 26279800 | 26280799 | 19.42911 | 33.43841 | 44.30138 | 19.43297 | -0.01965 | 0.896034 | 0.97288 |
| 18 | 45304200 | 45305599 | 17.98934 | 31.32949 | 41.50626 | 18.25067 | -0.01678 | 0.759026 | 0.972898 |
| 2 | 67360000 | 67361199 | 14.02284 | 24.84686 | 33.09106 | 14.63039 | -0.01918 | 0.494258 | 0.972903 |
| 10 | 119650000 | 119651199 | 15.77045 | 27.6787 | 36.71338 | 16.23742 | -0.01795 | 0.622576 | 0.972908 |
| 10 | 19558200 | 19560799 | 26.81543 | 44.66303 | 58.84666 | 25.56328 | -0.01541 | 1.241357 | 0.97292 |
| 19 | 7991800 | 8004599 | 214.6508 | 317.3608 | 298.0197 | 122.7319 | 0.338022 | 3.73502 | 0.972959 |
| 9 | 76083600 | 76085199 | 13.6291 | 24.32169 | 32.50764 | 14.29176 | -0.01866 | 0.441931 | 0.972966 |
| 18 | 89153600 | 89155599 | 28.74377 | 47.23108 | 62.3376 | 26.85952 | -0.0212 | 1.409667 | 0.972998 |
| 7 | 30494800 | 30496999 | 17.76775 | 30.89733 | 41.02339 | 18.0214 | -0.02018 | 0.777325 | 0.973025 |
| 17 | 89785400 | 89788799 | 48.24843 | 77.43766 | 97.66242 | 41.31054 | 0.01488 | 1.942757 | 0.973047 |
| 5 | 5125200 | 5129599 | 32.75673 | 54.00636 | 69.36812 | 29.93132 | 0.019693 | 1.381589 | 0.973072 |
| 8 | 13551000 | 13551599 | 8.798527 | 16.89425 | 24.10986 | 9.444669 | 0.018797 | -0.07848 | 0.973074 |
| 11 | 14003600 | 14004199 | 8.798527 | 16.89425 | 24.10986 | 9.444669 | 0.018797 | -0.07848 | 0.973074 |
| 17 | 73600400 | 73602999 | 18.35334 | 32.01389 | 41.45418 | 18.24586 | 0.016301 | 0.739221 | 0.973122 |
| 10 | 118856800 | 118858399 | 20.79652 | 35.48031 | 46.914 | 20.56688 | -0.01908 | 0.981436 | 0.973131 |
| 13 | 65564400 | 65565599 | 14.46047 | 26.02833 | 34.01828 | 14.84622 | 0.020674 | 0.397611 | 0.973139 |
| 8 | 20072600 | 20073999 | 20.92455 | 35.89176 | 47.40811 | 20.79253 | -0.01644 | 0.933217 | 0.97314 |
| 5 | 149513400 | 149520199 | 72.12749 | 112.7046 | 170.2196 | 70.62444 | -0.28048 | 2.653136 | 0.973148 |
| 8 | 33836000 | 33837999 | 15.9673 | 28.05113 | 37.17271 | 16.45393 | -0.01661 | 0.609631 | 0.973203 |
| 16 | 93974000 | 93976399 | 11.54528 | 20.55918 | 36.25213 | 15.96667 | -0.41776 | 0.450146 | 0.973213 |
| 6 | 18859600 | 18862199 | 44.43032 | 71.95544 | 72.75084 | 31.13146 | 0.348025 | 1.668089 | 0.973227 |
| 3 | 81686000 | 81688599 | 32.95113 | 53.93867 | 70.87867 | 30.49045 | -0.01711 | 1.4744 | 0.973227 |
| 4 | 60024600 | 60027799 | 40.98218 | 66.13751 | 84.00296 | 35.81066 | 0.014294 | 1.758793 | 0.973245 |
| 17 | 94855200 | 94857199 | 21.22525 | 36.90251 | 37.72723 | 16.57136 | 0.355177 | 0.791942 | 0.973251 |
| 13 | 4850200 | 4850999 | 36.3553 | 58.80306 | 74.5969 | 31.89152 | 0.019789 | 1.681336 | 0.973257 |
| 1 | 91639800 | 91641199 | 17.0697 | 29.77467 | 38.46796 | 16.93143 | 0.019676 | 0.729119 | 0.973288 |
| 19 | 49833600 | 49834999 | 13.03174 | 24.45315 | 24.55311 | 10.11196 | 0.458448 | 0.126737 | 0.973295 |
| 1 | 101263600 | 101265199 | 17.10911 | 29.96187 | 38.79741 | 17.1271 | 0.016614 | 0.669801 | 0.973307 |
| 1 | 101299400 | 101302599 | 36.3177 | 59.10982 | 75.64682 | 32.47367 | 0.015058 | 1.563698 | 0.973317 |
| 3 | 159471600 | 159473599 | 19.70058 | 34.62166 | 33.8151 | 14.9721 | 0.421402 | 0.620123 | 0.973319 |
| 4 | 17469800 | 17470999 | 14.94741 | 26.51268 | 34.4429 | 15.22037 | 0.017012 | 0.492544 | 0.973367 |
| 18 | 9144200 | 9145799 | 14.94741 | 26.51268 | 34.4429 | 15.22037 | 0.017012 | 0.492544 | 0.973367 |
| 1 | 120503800 | 120508199 | 61.14988 | 97.19292 | 123.671 | 51.59893 | -0.01346 | 2.26328 | 0.973368 |

| | | | | | | | | | |
|---|---|---|---|---|---|---|---|---|---|
| 17 | 35892400 | 35894199 | 19.72943 | 34.16591 | 44.18418 | 19.38521 | 0.016675 | 0.838391 | 0.973371 |
| 17 | 47353400 | 47356199 | 25.58322 | 42.02764 | 71.47909 | 30.7952 | -0.38613 | 1.411472 | 0.973377 |
| 17 | 46136200 | 46143599 | 75.29628 | 118.0992 | 148.5654 | 61.61626 | -0.01282 | 2.586396 | 0.973397 |
| 5 | 148515400 | 148516599 | 13.54916 | 24.40311 | 31.94783 | 13.93186 | 0.018893 | 0.38214 | 0.973402 |
| 12 | 48141600 | 48143999 | 22.25155 | 37.9379 | 50.08106 | 21.95415 | -0.01679 | 0.99936 | 0.973418 |
| 16 | 27515200 | 27516999 | 19.33577 | 33.74428 | 43.47563 | 19.14155 | 0.023375 | 0.737121 | 0.97342 |
| 3 | 146755600 | 146757799 | 26.44399 | 45.18709 | 44.67553 | 19.58806 | 0.399045 | 0.986672 | 0.973423 |
| 4 | 105770800 | 105774599 | 26.9862 | 45.11459 | 59.45931 | 25.88997 | -0.01593 | 1.203231 | 0.973434 |
| 1 | 191232000 | 191234599 | 39.42542 | 63.55577 | 82.45967 | 35.10329 | -0.01816 | 1.785295 | 0.973453 |
| 15 | 14245000 | 14245799 | 9.215766 | 17.51113 | 24.40552 | 9.808923 | 0.0188 | 0.0082 | 0.973471 |
| 17 | 89083600 | 89086199 | 24.2431 | 40.96808 | 52.82514 | 23.11314 | 0.01577 | 1.084487 | 0.973492 |
| X | 114052400 | 114053799 | 20.32909 | 35.12905 | 45.29527 | 19.87139 | 0.02054 | 0.866247 | 0.973509 |
| 5 | 15686200 | 15689399 | 57.33815 | 91.54285 | 114.6152 | 48.10835 | 0.016659 | 2.117309 | 0.973514 |
| 1 | 90496200 | 90499599 | 38.4857 | 62.37695 | 81.2162 | 34.77404 | -0.01495 | 1.647138 | 0.973514 |
| 16 | 67730800 | 67731599 | 11.46925 | 20.2445 | 37.90481 | 16.77151 | -0.50985 | 0.531333 | 0.973523 |
| 2 | 118627800 | 118629999 | 22.75509 | 38.8232 | 50.09747 | 21.96256 | 0.01591 | 0.992356 | 0.97354 |
| 2 | 154748400 | 154750799 | 22.75509 | 38.8232 | 50.09747 | 21.96256 | 0.01591 | 0.992356 | 0.97354 |
| 12 | 47376600 | 47379399 | 27.68136 | 45.88095 | 60.43311 | 26.16522 | -0.0154 | 1.301203 | 0.973555 |
| 2 | 91990200 | 91992199 | 16.73337 | 29.41897 | 38.0738 | 16.85968 | 0.017338 | 0.603602 | 0.973603 |
| 10 | 3441400 | 3441999 | 6.96447 | 13.51776 | 20.02215 | 7.526743 | 0.022492 | -0.24685 | 0.973615 |
| 18 | 42760000 | 42762999 | 37.8472 | 61.28799 | 79.68024 | 34.09421 | -0.01398 | 1.663324 | 0.973632 |
| 5 | 89754400 | 89754999 | 10.2526 | 19.26794 | 26.3128 | 10.80844 | 0.020634 | 0.094818 | 0.973633 |
| 7 | 30512000 | 30512999 | 10.2526 | 19.26794 | 26.3128 | 10.80844 | 0.020634 | 0.094818 | 0.973633 |
| 9 | 67917400 | 67918599 | 13.15567 | 23.51176 | 30.59064 | 13.43213 | 0.02132 | 0.426143 | 0.973653 |
| 17 | 18401800 | 18403399 | 18.16054 | 32.04051 | 41.1639 | 18.21523 | 0.029795 | 0.579828 | 0.97369 |
| 3 | 7461800 | 7465399 | 39.3373 | 63.73179 | 83.01061 | 35.5319 | -0.01602 | 1.658079 | 0.973743 |
| 17 | 19634200 | 19637999 | 39.76853 | 65.78254 | 56.67473 | 24.50393 | 0.595023 | 1.355026 | 0.973744 |
| 7 | 140236600 | 140236999 | 8.333823 | 16.04432 | 23.07004 | 8.968987 | 0.018444 | -0.11042 | 0.97377 |
| 16 | 53927200 | 53930199 | 33.0755 | 54.3446 | 71.57239 | 30.8375 | -0.01831 | 1.425179 | 0.973775 |
| 8 | 45171400 | 45173599 | 16.79856 | 29.47597 | 39.1386 | 17.30647 | -0.01971 | 0.633896 | 0.9738 |
| 7 | 3404400 | 3405199 | 9.913743 | 18.71857 | 26.44887 | 10.77337 | -0.0177 | 0.064688 | 0.973802 |
| 13 | 114171200 | 114175999 | 49.70565 | 79.74031 | 100.4374 | 42.45504 | 0.015907 | 1.956422 | 0.973817 |
| 5 | 139580600 | 139582599 | 14.43422 | 25.52265 | 33.04713 | 14.61704 | 0.019933 | 0.518072 | 0.973845 |
| 18 | 67361200 | 67364999 | 28.10834 | 46.54372 | 59.95427 | 25.94062 | 0.016523 | 1.31171 | 0.973878 |
| 14 | 121459800 | 121461599 | 15.16645 | 26.73987 | 34.63761 | 15.32939 | 0.017853 | 0.559038 | 0.973879 |
| 7 | 56675600 | 56677399 | 29.67382 | 48.69422 | 64.12892 | 27.61573 | -0.0189 | 1.428147 | 0.97388 |
| 16 | 79028800 | 79031399 | 16.8534 | 29.81503 | 38.46986 | 17.01388 | 0.025018 | 0.536437 | 0.973902 |
| X | 107028600 | 107031799 | 40.6005 | 65.71707 | 85.74976 | 36.67916 | -0.01983 | 1.682296 | 0.973906 |
| 4 | 51040000 | 51042199 | 25.98853 | 43.47394 | 56.08757 | 24.43149 | 0.01478 | 1.190631 | 0.973917 |
| 14 | 107062400 | 107066599 | 57.88933 | 92.16887 | 117.42 | 49.09725 | -0.01261 | 2.20606 | 0.973935 |
| 11 | 97904000 | 97906399 | 18.87006 | 32.64531 | 43.20389 | 18.95455 | -0.01663 | 0.844242 | 0.973942 |
| 9 | 3936400 | 3952599 | 434.2149 | 594.978 | 661.399 | 275.1271 | -0.01398 | 4.892692 | 0.973948 |
| 3 | 39845600 | 39847599 | 24.6839 | 41.84032 | 53.90176 | 23.60602 | 0.017265 | 1.062996 | 0.973968 |
| 8 | 86711000 | 86713599 | 29.44158 | 49.17143 | 65.20593 | 28.37471 | -0.02476 | 1.227701 | 0.973972 |
| 10 | 90023800 | 90024999 | 12.64027 | 22.98915 | 31.11313 | 13.40344 | -0.01721 | 0.319231 | 0.973976 |
| 13 | 114125200 | 114126599 | 16.18824 | 28.55788 | 36.97705 | 16.36286 | 0.018479 | 0.560382 | 0.973977 |
| 14 | 62836200 | 62838199 | 18.34311 | 32.03122 | 42.46126 | 18.70203 | -0.0173 | 0.724705 | 0.973983 |

| | | | | | | | | | |
|---|---|---|---|---|---|---|---|---|---|
| 16 | 9848000 | 9849399 | 18.34311 | 32.03122 | 42.46126 | 18.70203 | -0.0173 | 0.724705 | 0.973983 |
| 17 | 82932600 | 82933599 | 10.2743 | 19.14138 | 25.81068 | 10.75962 | 0.02252 | 0.153249 | 0.974 |
| 16 | 68416200 | 68423999 | 127.4585 | 193.3441 | 232.9381 | 96.42894 | 0.011827 | 3.239029 | 0.974007 |
| 8 | 104370200 | 104371799 | 17.26903 | 30.28984 | 39.21483 | 17.33455 | 0.016801 | 0.645601 | 0.974017 |
| 4 | 48801200 | 48801999 | 11.33619 | 20.66022 | 27.18957 | 11.68282 | 0.025538 | 0.28879 | 0.974017 |
| 12 | 112220200 | 112225199 | 45.35265 | 72.78924 | 93.68148 | 39.626 | -0.01435 | 1.936129 | 0.974019 |
| 19 | 28660400 | 28661799 | 14.45717 | 25.71385 | 33.47058 | 14.75639 | 0.016538 | 0.470453 | 0.974029 |
| 3 | 57143800 | 57148599 | 93.09973 | 143.8716 | 178.8575 | 74.29347 | -0.01237 | 2.85885 | 0.974055 |
| 3 | 159345800 | 159348799 | 38.93167 | 62.8212 | 81.51457 | 34.74474 | -0.01667 | 1.755626 | 0.974059 |
| 4 | 83207800 | 83208599 | 15.13054 | 26.5748 | 34.27216 | 15.16699 | 0.0225 | 0.603514 | 0.974084 |
| 16 | 76438200 | 76440999 | 31.38207 | 51.30858 | 67.33469 | 28.9532 | -0.01645 | 1.483515 | 0.974086 |
| 10 | 66819000 | 66820399 | 14.11316 | 25.2483 | 32.97163 | 14.46537 | 0.016699 | 0.430761 | 0.974091 |
| 12 | 91645000 | 91647399 | 17.89682 | 31.21131 | 41.42386 | 18.23021 | -0.01929 | 0.737168 | 0.97411 |
| 3 | 35690000 | 35691799 | 30.29936 | 50.00146 | 64.48623 | 27.83606 | 0.014399 | 1.361019 | 0.974122 |
| 15 | 19997400 | 20005999 | 222.0665 | 322.7105 | 373.4383 | 153.9372 | 0.01312 | 4.006892 | 0.974129 |
| 5 | 33484800 | 33491799 | 89.51906 | 138.6687 | 172.7027 | 71.736 | -0.01235 | 2.812071 | 0.97416 |
| 5 | 43475800 | 43476999 | 19.29258 | 33.41912 | 44.20266 | 19.3935 | -0.01554 | 0.834651 | 0.974185 |
| 2 | 178841600 | 178843199 | 13.94928 | 24.72692 | 32.0289 | 14.15959 | 0.020797 | 0.485628 | 0.974188 |
| 2 | 158486400 | 158488999 | 21.94183 | 37.53259 | 48.48115 | 21.2584 | 0.015048 | 0.964283 | 0.974197 |
| 18 | 60137600 | 60138999 | 15.2925 | 26.87065 | 34.71576 | 15.36881 | 0.019894 | 0.598346 | 0.974198 |
| 5 | 48883400 | 48884399 | 13.55131 | 24.44589 | 32.07701 | 13.96035 | 0.017803 | 0.376367 | 0.97421 |
| 2 | 136811200 | 136814599 | 24.47809 | 41.62374 | 55.0316 | 24.11919 | -0.01975 | 1.03488 | 0.974247 |
| 2 | 21175800 | 21177199 | 12.70115 | 23.04764 | 31.12829 | 13.448 | -0.01676 | 0.333881 | 0.974251 |
| 5 | 17813000 | 17814999 | 23.7607 | 40.00055 | 52.75695 | 23.05671 | -0.01679 | 1.11878 | 0.974255 |
| 12 | 83122400 | 83122999 | 7.268533 | 14.13293 | 21.2607 | 7.880769 | 0.018628 | -0.28909 | 0.974256 |
| 7 | 106068400 | 106071599 | 29.07858 | 48.20897 | 62.1325 | 26.90569 | 0.01595 | 1.305334 | 0.974264 |
| 9 | 93525000 | 93530799 | 122.584 | 186.2483 | 224.6663 | 93.04934 | 0.01194 | 3.204469 | 0.974269 |
| 3 | 112647000 | 112647799 | 12.1184 | 22.40516 | 30.8629 | 13.0095 | -0.02073 | 0.221723 | 0.974271 |
| 18 | 45340200 | 45341599 | 25.03407 | 42.14462 | 55.52082 | 24.26469 | -0.01523 | 1.124576 | 0.974287 |
| X | 48800800 | 48802799 | 19.29054 | 33.77054 | 45.05638 | 19.8844 | -0.02648 | 0.695752 | 0.97429 |
| 3 | 9526200 | 9527999 | 17.61099 | 30.60515 | 40.62875 | 17.84086 | -0.01978 | 0.788687 | 0.974301 |
| 15 | 55843000 | 55843999 | 10.1749 | 19.00298 | 25.75669 | 10.69238 | 0.019537 | 0.134156 | 0.974306 |
| 10 | 18544600 | 18548399 | 56.39775 | 89.77373 | 112.2912 | 46.93803 | 0.013463 | 2.210647 | 0.974311 |
| 17 | 55028400 | 55029599 | 16.83275 | 29.53439 | 38.23357 | 16.90553 | 0.017026 | 0.63745 | 0.974324 |
| 1 | 109972000 | 109972799 | 11.1771 | 20.90526 | 28.40911 | 11.76433 | 0.018577 | 0.140423 | 0.974339 |
| 14 | 111377200 | 111379599 | 21.70328 | 37.69694 | 48.48057 | 21.28166 | 0.025435 | 0.824307 | 0.974347 |
| 2 | 18461000 | 18474199 | 175.038 | 260.419 | 313.8817 | 129.3247 | -0.01352 | 3.623662 | 0.974349 |
| 13 | 67501800 | 67503999 | 16.34522 | 28.56787 | 36.86412 | 16.24663 | 0.021568 | 0.69248 | 0.974364 |
| 17 | 10337800 | 10338999 | 13.97388 | 25.04113 | 33.50775 | 14.67978 | -0.01684 | 0.420711 | 0.974376 |
| 8 | 115399200 | 115400999 | 17.46927 | 30.46276 | 39.41154 | 17.34356 | 0.017485 | 0.73485 | 0.9744 |
| 13 | 98736200 | 98738399 | 18.70828 | 32.51096 | 42.03869 | 18.4596 | 0.017674 | 0.792435 | 0.97442 |
| 11 | 11236600 | 11238799 | 54.96705 | 86.934 | 140.1029 | 58.22044 | -0.36032 | 2.39728 | 0.974421 |
| 9 | 6763200 | 6764199 | 20.66801 | 35.5171 | 46.87901 | 20.56271 | -0.01485 | 0.916362 | 0.974437 |
| 11 | 100555000 | 100556199 | 16.2252 | 28.04843 | 47.93643 | 21.03034 | -0.38417 | 0.845679 | 0.974439 |
| 1 | 64680800 | 64682399 | 15.86173 | 27.7706 | 36.93017 | 16.29943 | -0.0214 | 0.665399 | 0.974462 |
| 10 | 104075000 | 104075799 | 9.901793 | 18.74038 | 25.95349 | 10.51871 | 0.017394 | 0.046937 | 0.974476 |
| 3 | 129934000 | 129938599 | 48.09936 | 77.15459 | 99.19612 | 41.92308 | -0.01383 | 1.957998 | 0.974483 |

| | | | | | | | | | |
|---|---|---|---|---|---|---|---|---|---|
| 14 | 9302000 | 9310599 | 157.5541 | 237.0242 | 238.6427 | 98.54411 | 0.26034 | 3.410747 | 0.974501 |
| 3 | 5991600 | 5993399 | 32.23581 | 53.27326 | 54.04109 | 23.35038 | 0.35998 | 1.343037 | 0.974563 |
| 7 | 34413000 | 34415399 | 21.66289 | 36.7178 | 48.5155 | 21.24682 | -0.01907 | 1.045636 | 0.974607 |
| 7 | 106802600 | 106804599 | 16.45919 | 28.9961 | 37.56971 | 16.62967 | 0.016894 | 0.579836 | 0.974609 |
| 9 | 122609200 | 122609599 | 5.358907 | 10.46787 | 17.07269 | 5.97235 | -0.01988 | -0.53118 | 0.974637 |
| X | 115395200 | 115396599 | 16.66976 | 29.40393 | 39.11302 | 17.30846 | -0.02016 | 0.570445 | 0.974645 |
| 2 | 6203000 | 6205799 | 47.55396 | 76.49119 | 77.24369 | 32.77984 | 0.33874 | 1.863804 | 0.97466 |
| 8 | 120576000 | 120577799 | 21.38704 | 36.31391 | 47.98689 | 21.02651 | -0.01906 | 1.030186 | 0.974662 |
| 3 | 146409800 | 146411199 | 16.53081 | 29.03254 | 37.59856 | 16.63828 | 0.016261 | 0.616647 | 0.974674 |
| 12 | 68980400 | 68983799 | 68.388 | 106.951 | 166.1906 | 68.94687 | -0.3205 | 2.637473 | 0.974677 |
| 19 | 33610600 | 33611599 | 12.63704 | 22.85643 | 30.06123 | 13.04023 | 0.017418 | 0.348827 | 0.974685 |
| 15 | 28259200 | 28259599 | 8.328619 | 15.92708 | 22.42613 | 8.888246 | 0.022541 | -0.04979 | 0.974686 |
| 13 | 74716200 | 74741799 | 344.9167 | 486.9056 | 559.821 | 231.781 | -0.01928 | 4.441634 | 0.974697 |
| 7 | 110910200 | 110911999 | 24.67616 | 41.71993 | 54.99557 | 24.06629 | -0.01591 | 1.085708 | 0.974699 |
| 13 | 82758400 | 82758799 | 6.841549 | 13.31537 | 20.15922 | 7.415234 | 0.020424 | -0.3226 | 0.974764 |
| 10 | 29259800 | 29260999 | 14.96967 | 26.28122 | 33.83441 | 14.96795 | 0.025046 | 0.608652 | 0.974778 |
| 13 | 37653400 | 37656199 | 22.00574 | 37.54569 | 48.48864 | 21.25731 | 0.014834 | 0.988857 | 0.974778 |
| 3 | 35235800 | 35239199 | 31.56456 | 52.12892 | 68.7927 | 29.71056 | -0.01864 | 1.35739 | 0.974811 |
| 10 | 60911200 | 60913199 | 30.29137 | 50.01281 | 64.514 | 27.85748 | 0.014155 | 1.355249 | 0.97482 |
| 14 | 8691000 | 8691999 | 8.765887 | 16.67122 | 23.90664 | 9.590233 | -0.01947 | -0.00336 | 0.974851 |
| 11 | 61883200 | 61886399 | 31.63647 | 52.09814 | 68.52903 | 29.55457 | -0.0152 | 1.393048 | 0.974854 |
| 11 | 98985600 | 98986399 | 10.50015 | 19.62807 | 27.32547 | 11.32638 | -0.01715 | 0.134986 | 0.974891 |
| X | 9213400 | 9214199 | 12.30559 | 23.1233 | 24.04861 | 9.880467 | 0.410696 | 0.109493 | 0.974893 |
| 13 | 13893400 | 13898599 | 72.75945 | 114.2979 | 143.921 | 59.69659 | -0.01246 | 2.553799 | 0.974894 |
| 10 | 98767800 | 98770599 | 22.77481 | 38.79423 | 50.09436 | 21.95835 | 0.014595 | 1.007334 | 0.974901 |
| 13 | 39041400 | 39046999 | 67.11519 | 106.4047 | 135.1884 | 56.26164 | -0.01485 | 2.344194 | 0.974912 |
| 16 | 14642200 | 14643999 | 15.19331 | 26.82809 | 35.60301 | 15.75185 | -0.01604 | 0.544515 | 0.974913 |
| 3 | 83299200 | 83302799 | 52.27154 | 83.70411 | 105.334 | 44.40177 | 0.014123 | 2.018764 | 0.974917 |
| X | 138961800 | 138964399 | 50.50925 | 81.62884 | 73.30452 | 31.30748 | 0.515728 | 1.731047 | 0.974921 |
| 16 | 75523200 | 75525399 | 23.81351 | 40.10916 | 51.72068 | 22.60719 | 0.015579 | 1.11149 | 0.974939 |
| 2 | 39415000 | 39417399 | 30.52685 | 50.43075 | 65.03098 | 28.09192 | 0.014739 | 1.351069 | 0.974941 |
| 19 | 27473200 | 27477399 | 61.38824 | 97.5647 | 121.9416 | 50.87367 | 0.012235 | 2.266261 | 0.974943 |
| 11 | 100343000 | 100344599 | 17.82253 | 31.14321 | 40.35492 | 17.78187 | 0.015508 | 0.709206 | 0.974955 |
| 6 | 44832400 | 44833599 | 13.77037 | 24.78393 | 32.48088 | 14.17689 | 0.016638 | 0.389879 | 0.974957 |
| 9 | 26014200 | 26016199 | 28.06096 | 47.27933 | 48.55152 | 21.22415 | 0.342969 | 1.120674 | 0.974965 |
| 6 | 66227400 | 66230999 | 37.00667 | 60.11198 | 78.3851 | 33.61639 | -0.01415 | 1.599048 | 0.974972 |
| 17 | 47568400 | 47574999 | 65.43079 | 102.6664 | 160.6521 | 66.62638 | -0.32685 | 2.574493 | 0.974983 |
| 15 | 9641400 | 9644799 | 44.878 | 71.9995 | 92.68796 | 39.18263 | -0.01536 | 1.947515 | 0.975009 |
| 3 | 4461200 | 4465199 | 39.79967 | 64.58322 | 82.29478 | 35.2667 | 0.017663 | 1.626429 | 0.97503 |
| 10 | 7330200 | 7334999 | 22.58134 | 37.74964 | 62.90587 | 27.3902 | -0.3531 | 1.195884 | 0.975064 |
| 11 | 90129600 | 90130999 | 12.51209 | 22.76987 | 30.12228 | 12.96672 | 0.016309 | 0.313605 | 0.975073 |
| 13 | 6939400 | 6940399 | 12.51209 | 22.76987 | 30.12228 | 12.96672 | 0.016309 | 0.313605 | 0.975073 |
| 6 | 142660000 | 142662599 | 43.40725 | 69.90569 | 88.66206 | 37.69149 | 0.01279 | 1.825397 | 0.975077 |
| 16 | 86132600 | 86135599 | 24.03349 | 41.24011 | 52.90434 | 23.2038 | 0.02552 | 0.948179 | 0.975108 |
| 13 | 42138800 | 42142399 | 36.62494 | 59.51329 | 76.14503 | 32.66436 | 0.013642 | 1.588934 | 0.975153 |
| 4 | 29025000 | 29026199 | 18.16007 | 31.91 | 32.46517 | 14.37273 | 0.362806 | 0.610755 | 0.975153 |
| 1 | 97206200 | 97207999 | 43.58116 | 70.06368 | 90.3054 | 38.29336 | -0.0135 | 1.880821 | 0.975155 |

| | | | | | | | | | |
|---|---|---|---|---|---|---|---|---|---|
| 17 | 82421000 | 82423199 | 23.52969 | 40.09703 | 51.71422 | 22.66917 | 0.016457 | 1.009659 | 0.975161 |
| 1 | 182014400 | 182017399 | 34.95937 | 57.71709 | 58.7089 | 25.34995 | 0.355884 | 1.363244 | 0.97517 |
| 1 | 142145200 | 142146199 | 16.37853 | 28.71886 | 37.20023 | 16.43964 | 0.01604 | 0.641329 | 0.975185 |
| 2 | 17616000 | 17617999 | 18.80336 | 32.62322 | 43.15268 | 18.93374 | -0.01514 | 0.817009 | 0.97519 |
| 1 | 63345400 | 63348999 | 42.63554 | 68.79989 | 89.15072 | 38.00101 | -0.01468 | 1.766007 | 0.975226 |
| 1 | 147714800 | 147716599 | 16.10489 | 28.52122 | 36.8593 | 16.29812 | 0.023223 | 0.51612 | 0.975241 |
| X | 79859200 | 79859999 | 9.494766 | 17.94453 | 25.4263 | 10.33606 | -0.01876 | 0.053345 | 0.97525 |
| 10 | 13912200 | 13913999 | 15.74291 | 27.8319 | 36.98905 | 16.36115 | -0.01771 | 0.535014 | 0.975257 |
| 7 | 40004800 | 40005999 | 12.57694 | 22.79822 | 30.02356 | 12.98782 | 0.017991 | 0.334373 | 0.975257 |
| 11 | 15310000 | 15311199 | 10.54692 | 19.7168 | 26.8151 | 11.11358 | 0.01667 | 0.134943 | 0.975288 |
| 14 | 5825400 | 5835199 | 205.8464 | 301.4081 | 357.3696 | 147.211 | -0.01142 | 3.890181 | 0.975325 |
| 15 | 28286400 | 28286999 | 7.682815 | 14.83388 | 21.49286 | 8.280461 | 0.019694 | -0.15291 | 0.975362 |
| 17 | 23811400 | 23811999 | 7.682815 | 14.83388 | 21.49286 | 8.280461 | 0.019694 | -0.15291 | 0.975362 |
| 3 | 111814600 | 111815799 | 18.11843 | 31.66211 | 41.95426 | 18.4875 | -0.01648 | 0.714221 | 0.975372 |
| 3 | 42131200 | 42144599 | 253.7317 | 365.528 | 426.5609 | 176.0282 | -0.01135 | 4.139313 | 0.97539 |
| 13 | 42012200 | 42015199 | 31.54749 | 51.99741 | 66.9602 | 28.88875 | 0.015809 | 1.379423 | 0.975443 |
| 13 | 19181200 | 19185399 | 121.321 | 184.0924 | 221.6383 | 91.7599 | 0.013206 | 3.230645 | 0.975458 |
| 1 | 137083200 | 137087799 | 55.34663 | 88.17846 | 112.4531 | 47.06324 | -0.01309 | 2.180296 | 0.975478 |
| 1 | 97415200 | 97417999 | 29.64589 | 49.28424 | 63.50261 | 27.55102 | 0.016529 | 1.274835 | 0.975501 |
| 10 | 15655800 | 15657799 | 19.30682 | 33.85409 | 34.4733 | 15.23479 | 0.36158 | 0.653717 | 0.975511 |
| 14 | 90749600 | 90750999 | 10.34302 | 19.65666 | 20.12404 | 8.084991 | 0.460873 | 0.005802 | 0.97558 |
| 10 | 47479000 | 47479799 | 10.15441 | 19.1684 | 26.42019 | 10.76713 | 0.016933 | 0.066874 | 0.975594 |
| 9 | 38617200 | 38618799 | 23.21195 | 39.59136 | 52.25646 | 22.90853 | -0.01658 | 1.000941 | 0.975606 |
| 11 | 75609600 | 75612599 | 37.0025 | 59.94378 | 77.91517 | 33.34866 | -0.01313 | 1.652441 | 0.975625 |
| 14 | 124650600 | 124653799 | 41.97446 | 67.5156 | 85.23714 | 36.17131 | 0.017286 | 1.861241 | 0.975628 |
| 7 | 123470400 | 123472799 | 13.82045 | 24.27685 | 40.89918 | 18.10167 | -0.36107 | 0.606599 | 0.97563 |
| 4 | 35384000 | 35387399 | 29.82953 | 48.83274 | 79.3172 | 34.12168 | -0.32206 | 1.47011 | 0.975641 |
| 6 | 46416600 | 46418199 | 29.36908 | 48.58799 | 62.70173 | 27.11267 | 0.013842 | 1.329853 | 0.97565 |
| 1 | 20556400 | 20557999 | 19.19922 | 32.12885 | 59.99079 | 26.13795 | -0.51656 | 1.170271 | 0.975669 |
| 11 | 20261600 | 20262999 | 9.785202 | 18.21138 | 25.33513 | 10.57984 | -0.02421 | 0.153945 | 0.975679 |
| 1 | 191240200 | 191242399 | 23.18114 | 39.26935 | 51.70215 | 22.63856 | -0.014 | 1.060263 | 0.975681 |
| 7 | 102807600 | 102808199 | 12.44995 | 22.70884 | 30.10118 | 12.9205 | 0.015984 | 0.298757 | 0.975702 |
| 16 | 86336600 | 86339999 | 60.95767 | 96.72633 | 120.607 | 50.2368 | 0.013445 | 2.312799 | 0.975727 |
| 3 | 23732800 | 23733199 | 7.570887 | 14.66668 | 21.60962 | 8.196515 | 0.01748 | -0.20487 | 0.975747 |
| 4 | 50367000 | 50368399 | 14.68626 | 26.2256 | 35.02977 | 15.39201 | -0.01716 | 0.451075 | 0.975756 |
| 3 | 6630000 | 6630599 | 10.56023 | 19.69455 | 27.3167 | 11.3674 | -0.01618 | 0.151595 | 0.975769 |
| 9 | 12127800 | 12132599 | 83.64287 | 130.5686 | 163.4948 | 67.84125 | -0.01151 | 2.675827 | 0.975775 |
| 16 | 82668000 | 82669399 | 14.62635 | 26.16922 | 35.00872 | 15.35768 | -0.01787 | 0.437529 | 0.975818 |
| X | 94896400 | 94897799 | 25.3864 | 42.58479 | 54.95519 | 23.98021 | 0.014385 | 1.159478 | 0.975869 |
| 4 | 118052600 | 118055199 | 29.40263 | 48.52324 | 62.57386 | 27.01118 | 0.014617 | 1.358983 | 0.975893 |
| 14 | 118560000 | 118561999 | 15.33086 | 27.04743 | 35.10015 | 15.53207 | 0.015562 | 0.553638 | 0.975939 |
| 5 | 74362200 | 74364599 | 22.64404 | 38.4502 | 50.62524 | 22.1829 | -0.01394 | 1.036114 | 0.975942 |
| 10 | 8432200 | 8435799 | 44.84919 | 72.07252 | 91.16769 | 38.63589 | 0.012761 | 1.895055 | 0.975958 |
| 7 | 26527000 | 26548599 | 261.1663 | 375.1972 | 436.6893 | 180.2829 | -0.01152 | 4.180003 | 0.975963 |
| 5 | 42596200 | 42597999 | 15.53598 | 27.45007 | 36.46804 | 16.13258 | -0.01755 | 0.538374 | 0.975986 |
| 15 | 91781800 | 91785399 | 40.9225 | 66.06238 | 85.52847 | 36.47607 | -0.01275 | 1.748702 | 0.975991 |
| 2 | 64703800 | 64705199 | 11.14099 | 20.61761 | 27.70544 | 11.66349 | 0.016305 | 0.201808 | 0.976037 |

| | | | | | | | | | |
|---|---|---|---|---|---|---|---|---|---|
| 15 | 45505200 | 45505799 | 7.417157 | 14.35434 | 21.57502 | 8.222136 | -0.01746 | -0.19095 | 0.976042 |
| 4 | 33035200 | 33036799 | 14.37809 | 25.98964 | 34.0674 | 14.7893 | 0.021254 | 0.371643 | 0.976044 |
| 16 | 61410200 | 61411399 | 16.302 | 29.32647 | 29.15897 | 12.73293 | 0.412348 | 0.403738 | 0.976058 |
| 10 | 45114400 | 45116599 | 19.64813 | 34.13599 | 44.12654 | 19.37346 | 0.018001 | 0.802373 | 0.976089 |
| 5 | 24503600 | 24506799 | 25.19138 | 42.16367 | 55.49903 | 24.18352 | -0.01404 | 1.18213 | 0.976105 |
| 4 | 28399000 | 28399399 | 8.746269 | 16.82296 | 24.75129 | 9.647935 | -0.01852 | -0.09508 | 0.976113 |
| 14 | 29478200 | 29500199 | 251.575 | 363.4711 | 418.6196 | 172.685 | 0.011722 | 4.096448 | 0.976129 |
| 15 | 93474200 | 93480199 | 64.16192 | 101.7531 | 126.9885 | 52.86491 | 0.01133 | 2.331878 | 0.976129 |
| X | 153384800 | 153388999 | 50.64957 | 81.47545 | 102.4317 | 43.47267 | 0.023365 | 1.878509 | 0.97613 |
| 16 | 19647600 | 19649999 | 25.59415 | 43.07823 | 55.58473 | 24.29103 | 0.014544 | 1.126604 | 0.976131 |
| 18 | 15838600 | 15839199 | 13.53028 | 24.31248 | 32.55482 | 14.22952 | -0.0154 | 0.399837 | 0.976142 |
| 13 | 72366600 | 72367799 | 13.86929 | 24.77371 | 33.05633 | 14.52143 | -0.01533 | 0.440414 | 0.976185 |
| 8 | 23951800 | 23957199 | 58.85139 | 93.87815 | 117.5486 | 49.27993 | 0.015194 | 2.14634 | 0.976191 |
| Y | 8685000 | 8685599 | 5.239344 | 10.21973 | 16.48466 | 5.905102 | -0.03487 | -0.45277 | 0.976206 |
| 10 | 115402800 | 115404999 | 18.81107 | 32.38944 | 55.52961 | 24.35861 | -0.39017 | 0.901262 | 0.976219 |
| 3 | 12898000 | 12900399 | 50.42964 | 80.71077 | 103.3701 | 43.53926 | -0.0121 | 2.032124 | 0.97625 |
| 16 | 81347600 | 81348199 | 8.33458 | 16.04304 | 23.62305 | 9.188492 | -0.0163 | -0.10747 | 0.97625 |
| 10 | 66690000 | 66696399 | 61.945 | 98.40253 | 122.9531 | 51.27114 | 0.011969 | 2.279463 | 0.976259 |
| 17 | 42202400 | 42203999 | 18.84343 | 32.81374 | 42.53701 | 18.70276 | 0.014463 | 0.76674 | 0.976277 |
| 14 | 108928200 | 108930399 | 19.28221 | 33.43202 | 44.19332 | 19.38994 | -0.01442 | 0.826184 | 0.976284 |
| 8 | 45257200 | 45258799 | 14.25852 | 25.44991 | 33.96062 | 14.92809 | -0.01628 | 0.447305 | 0.976292 |
| 15 | 47676000 | 47677399 | 11.37694 | 20.98405 | 28.74169 | 12.15486 | -0.01565 | 0.226051 | 0.976294 |
| 8 | 78058000 | 78058999 | 10.1206 | 18.69306 | 25.83833 | 10.90616 | -0.02634 | 0.201829 | 0.976295 |
| 4 | 74963200 | 74963799 | 12.32874 | 22.23313 | 29.11762 | 12.67759 | 0.020156 | 0.361774 | 0.976296 |
| 13 | 110883600 | 110886199 | 22.65543 | 38.97081 | 40.03523 | 17.56133 | 0.345546 | 0.906893 | 0.976303 |
| 13 | 114142800 | 114144599 | 12.92502 | 23.27271 | 30.49267 | 13.30178 | 0.017522 | 0.376724 | 0.976327 |
| 8 | 69663600 | 69692999 | 595.0003 | 800.4451 | 856.6573 | 357.361 | 0.014054 | 5.184822 | 0.976369 |
| 15 | 44290600 | 44291999 | 11.95633 | 20.95035 | 41.55683 | 18.35425 | -0.59543 | 0.636138 | 0.976396 |
| 17 | 20372800 | 20387399 | 181.0069 | 272.4782 | 233.7229 | 96.52882 | 0.492153 | 3.403831 | 0.976403 |
| 4 | 122717400 | 122721599 | 54.08944 | 86.28772 | 108.3132 | 45.41233 | 0.011748 | 2.137375 | 0.976406 |
| 13 | 17706000 | 17706799 | 10.95224 | 20.22295 | 27.78297 | 11.73184 | -0.01813 | 0.215694 | 0.976407 |
| 10 | 19410800 | 19411399 | 8.858172 | 16.94491 | 24.56183 | 9.726205 | -0.01777 | -0.04006 | 0.976408 |
| 14 | 6481200 | 6482199 | 13.36385 | 23.86855 | 31.11571 | 13.67005 | 0.018069 | 0.433676 | 0.97642 |
| 8 | 13858800 | 13865599 | 74.23269 | 116.8077 | 147.1546 | 61.05111 | -0.01133 | 2.52229 | 0.976426 |
| 3 | 105426400 | 105428799 | 17.27367 | 30.10581 | 39.92955 | 17.5657 | -0.01833 | 0.741166 | 0.976433 |
| 10 | 55065400 | 55067599 | 16.56807 | 29.35146 | 29.72744 | 13.14372 | 0.373367 | 0.503976 | 0.976443 |
| 4 | 121567800 | 121598199 | 379.7736 | 532.9167 | 610.6244 | 253.058 | -0.02301 | 4.531548 | 0.976484 |
| X | 30807800 | 30876799 | 3743.82 | 4331.841 | 3963.469 | 1685.006 | 0.020424 | 7.595021 | 0.976491 |
| 3 | 120973600 | 120974799 | 11.20281 | 20.68002 | 27.72282 | 11.71076 | 0.016385 | 0.217654 | 0.976512 |
| 6 | 56942800 | 56943399 | 11.20186 | 20.85623 | 28.86198 | 12.04453 | -0.01634 | 0.170975 | 0.976516 |
| 3 | 145675400 | 145680799 | 65.54295 | 103.872 | 129.4369 | 53.84723 | 0.012535 | 2.352067 | 0.976545 |
| 7 | 3843600 | 3846599 | 23.81684 | 40.10533 | 52.79382 | 23.07462 | -0.01411 | 1.11445 | 0.976564 |
| 7 | 33824600 | 33827799 | 40.10224 | 64.73783 | 82.35249 | 35.12132 | 0.012578 | 1.746252 | 0.976565 |
| 1 | 106915800 | 106916399 | 7.65378 | 14.77732 | 22.02976 | 8.48865 | -0.0217 | -0.15274 | 0.976567 |
| 10 | 130391000 | 130398199 | 109.6017 | 169.8472 | 161.0759 | 66.90641 | 0.380835 | 2.812095 | 0.976578 |
| 12 | 8403800 | 8407399 | 21.38384 | 36.71907 | 48.47306 | 21.26142 | -0.01543 | 0.926084 | 0.976583 |
| 4 | 19488000 | 19488799 | 8.913703 | 16.88004 | 23.33934 | 9.447679 | 0.021322 | 0.025887 | 0.976608 |

| | | | | | | | | | |
|---|---|---|---|---|---|---|---|---|---|
| 7 | 103830600 | 103831799 | 8.913703 | 16.88004 | 23.33934 | 9.447679 | 0.021322 | 0.025887 | 0.976608 |
| 6 | 7230200 | 7232999 | 39.35098 | 63.63133 | 82.53876 | 35.27673 | -0.0127 | 1.698153 | 0.976617 |
| 11 | 27087400 | 27088199 | 9.18114 | 17.71179 | 25.5255 | 9.865469 | 0.022319 | -0.11681 | 0.976632 |
| 3 | 139726800 | 139727599 | 28.09254 | 46.39023 | 59.70291 | 25.78303 | 0.017629 | 1.34642 | 0.976652 |
| 6 | 8497200 | 8501599 | 14.53286 | 26.35556 | 26.73929 | 11.56059 | 0.392468 | 0.338716 | 0.976657 |
| 5 | 138465600 | 138467399 | 17.85048 | 31.53379 | 40.64101 | 17.97925 | 0.025998 | 0.563739 | 0.976687 |
| 8 | 118366200 | 118370199 | 42.56238 | 68.51343 | 88.38441 | 37.54832 | -0.01248 | 1.837874 | 0.976696 |
| 4 | 155460400 | 155463399 | 25.41267 | 42.6332 | 56.12775 | 24.49344 | -0.01446 | 1.156537 | 0.976714 |
| 11 | 105308600 | 105310399 | 14.1781 | 24.90779 | 42.44486 | 18.78749 | -0.3775 | 0.61033 | 0.976724 |
| 5 | 69092800 | 69095799 | 55.33802 | 88.4651 | 86.50765 | 36.3778 | 0.374664 | 2.066162 | 0.976728 |
| 2 | 150153000 | 150200999 | 1059.685 | 1360.392 | 1417.885 | 594.7036 | -0.01525 | 5.941294 | 0.976756 |
| 4 | 33557000 | 33557999 | 11.36218 | 20.73766 | 28.18292 | 12.08768 | -0.01884 | 0.283357 | 0.976766 |
| 1 | 100589600 | 100590399 | 9.936268 | 18.76871 | 25.87573 | 10.53088 | 0.018104 | 0.057814 | 0.976767 |
| 15 | 11972600 | 11976199 | 33.53424 | 54.84753 | 71.97555 | 30.95288 | -0.01612 | 1.48277 | 0.976774 |
| 17 | 68132200 | 68138599 | 118.7871 | 180.9014 | 222.0961 | 92.0492 | -0.01071 | 3.155832 | 0.976803 |
| 7 | 75977400 | 75979799 | 28.01097 | 47.79525 | 45.20334 | 19.81716 | 0.462553 | 1.007146 | 0.976806 |
| 15 | 33810000 | 33819999 | 106.2299 | 163.6875 | 199.2748 | 82.77561 | 0.014595 | 2.91811 | 0.976818 |
| 10 | 103829200 | 103829799 | 9.551874 | 17.52907 | 31.8994 | 13.58328 | -0.43007 | 0.246467 | 0.976823 |
| 17 | 22232000 | 22256799 | 307.0051 | 436.8806 | 495.0319 | 204.6828 | 0.013409 | 4.321773 | 0.976823 |
| 3 | 25007800 | 25008999 | 16.79113 | 29.46319 | 38.20619 | 16.8943 | 0.014531 | 0.635395 | 0.976827 |
| 2 | 7752000 | 7753599 | 25.40451 | 42.93807 | 56.65256 | 24.78908 | -0.01722 | 1.093505 | 0.976828 |
| 11 | 109682400 | 109685999 | 41.86925 | 67.35024 | 85.08111 | 36.10811 | 0.016456 | 1.856855 | 0.976848 |
| 18 | 52337400 | 52341799 | 95.54291 | 147.1856 | 182.4766 | 75.79595 | -0.01188 | 2.917603 | 0.976848 |
| 6 | 122902600 | 122904399 | 19.19996 | 33.30511 | 43.11781 | 18.91833 | 0.015587 | 0.818912 | 0.976856 |
| 6 | 37716400 | 37719999 | 52.63257 | 84.4294 | 88.01868 | 37.20134 | 0.288133 | 1.958828 | 0.97687 |
| 5 | 19026400 | 19029399 | 36.38257 | 59.05558 | 76.87732 | 32.95222 | -0.01265 | 1.612272 | 0.976897 |
| 8 | 122946400 | 122948399 | 19.07631 | 32.97943 | 42.64905 | 18.71074 | 0.016243 | 0.848402 | 0.976927 |
| 4 | 144518200 | 144520199 | 20.73544 | 35.51738 | 45.86621 | 20.11399 | 0.015883 | 0.946283 | 0.976954 |
| 8 | 16605200 | 16605799 | 9.927169 | 18.54489 | 25.81634 | 10.71266 | -0.01796 | 0.132081 | 0.976957 |
| 6 | 70064800 | 70067399 | 26.8534 | 44.17937 | 71.75738 | 30.93439 | -0.31908 | 1.396606 | 0.976961 |
| 13 | 57589000 | 57593399 | 35.01148 | 57.04421 | 73.1928 | 31.43233 | 0.012426 | 1.544842 | 0.976981 |
| 16 | 15320200 | 15322199 | 32.54665 | 53.70011 | 54.5872 | 23.56102 | 0.356746 | 1.362546 | 0.976993 |
| 9 | 88531200 | 88540799 | 155.7054 | 233.5251 | 278.6919 | 114.9874 | 0.01131 | 3.47214 | 0.977 |
| 14 | 114180600 | 114181399 | 9.490597 | 18.16682 | 25.70198 | 10.15819 | 0.019289 | -0.03755 | 0.977011 |
| 14 | 59185600 | 59185999 | 10.23124 | 19.06427 | 25.78219 | 10.74477 | 0.01846 | 0.150719 | 0.977059 |
| 6 | 42037800 | 42038999 | 17.30007 | 30.64207 | 39.55124 | 17.48793 | 0.025256 | 0.529642 | 0.977118 |
| 5 | 108291800 | 108294399 | 24.67181 | 41.34211 | 54.41481 | 23.72849 | -0.014 | 1.166542 | 0.977136 |
| 4 | 142258400 | 142265199 | 64.95963 | 102.4759 | 155.3132 | 64.45782 | -0.27487 | 2.466629 | 0.977142 |
| 2 | 126055200 | 126055799 | 7.384481 | 14.30835 | 21.61671 | 8.191736 | -0.01755 | -0.21201 | 0.977142 |
| 1 | 143288000 | 143289399 | 10.57838 | 19.8994 | 27.98671 | 11.47724 | -0.0197 | 0.095535 | 0.977145 |
| 8 | 57271800 | 57274599 | 23.28345 | 39.98384 | 41.5545 | 18.22682 | 0.328368 | 0.925806 | 0.977183 |
| 9 | 80062600 | 80065999 | 29.69693 | 48.931 | 64.27483 | 27.7263 | -0.01285 | 1.379374 | 0.977209 |
| 1 | 112184400 | 112185199 | 17.02281 | 30.1351 | 30.81036 | 13.62451 | 0.359603 | 0.518503 | 0.977286 |
| 9 | 26068000 | 26075799 | 288.8533 | 409.2759 | 470.4518 | 194.6606 | -0.01355 | 4.383399 | 0.97729 |
| X | 111638800 | 111642399 | 26.70957 | 45.03711 | 59.50883 | 26.02019 | -0.01949 | 1.120226 | 0.9773 |
| X | 77967200 | 77967999 | 11.62662 | 21.85187 | 30.81148 | 12.65469 | -0.02468 | 0.120413 | 0.977312 |
| 14 | 50042800 | 50053599 | 143.0706 | 215.4161 | 261.8885 | 108.1675 | -0.01041 | 3.392789 | 0.977318 |

| | | | | | | | | | |
|---|---|---|---|---|---|---|---|---|---|
| 3 | 16731600 | 16731999 | 6.667664 | 12.97838 | 20.17072 | 7.412862 | -0.01639 | -0.33009 | 0.977322 |
| 5 | 141721400 | 141725199 | 34.13076 | 55.67706 | 72.75634 | 31.25878 | -0.01266 | 1.523945 | 0.977335 |
| 7 | 61157000 | 61157799 | 15.09527 | 27.23299 | 26.94021 | 11.72373 | 0.42312 | 0.376199 | 0.977338 |
| 4 | 61323000 | 61324399 | 32.39716 | 52.86681 | 67.78574 | 29.12679 | 0.015063 | 1.51433 | 0.977339 |
| 8 | 101343200 | 101344999 | 21.43206 | 36.76673 | 48.47531 | 21.26079 | -0.0138 | 0.935738 | 0.977375 |
| 1 | 26240200 | 26242799 | 38.09946 | 61.59627 | 78.43468 | 33.51472 | 0.013606 | 1.697093 | 0.977384 |
| 15 | 88469400 | 88474799 | 81.66004 | 127.3729 | 156.9115 | 65.1148 | 0.011133 | 2.68731 | 0.977394 |
| 3 | 66556800 | 66558799 | 16.5307 | 28.92991 | 38.29762 | 16.89709 | -0.01518 | 0.674695 | 0.977403 |
| 17 | 21285800 | 21294599 | 111.8555 | 170.8101 | 210.1559 | 87.16003 | -0.01066 | 3.097985 | 0.977416 |
| 4 | 45734200 | 45737399 | 26.30338 | 43.90687 | 57.75067 | 25.12209 | -0.01297 | 1.215149 | 0.977424 |
| 13 | 4652200 | 4653799 | 11.94301 | 21.68683 | 28.58 | 12.33646 | 0.019105 | 0.316698 | 0.977443 |
| 18 | 24662400 | 24668799 | 75.65025 | 118.4214 | 148.6641 | 61.66962 | -0.01205 | 2.620621 | 0.97746 |
| 5 | 150658000 | 150658599 | 7.534765 | 14.53116 | 20.94907 | 8.10395 | 0.021422 | -0.14262 | 0.977465 |
| 14 | 116562400 | 116563199 | 12.85323 | 23.23088 | 30.58899 | 13.28101 | 0.015036 | 0.356679 | 0.97747 |
| X | 43960400 | 43961199 | 12.85323 | 23.23088 | 30.58899 | 13.28101 | 0.015036 | 0.356679 | 0.97747 |
| 3 | 5463800 | 5468999 | 59.44424 | 94.46064 | 118.0403 | 49.23855 | 0.012277 | 2.266349 | 0.977483 |
| 5 | 149899400 | 149901999 | 35.96845 | 58.51749 | 75.01137 | 32.19609 | 0.012287 | 1.567982 | 0.977516 |
| X | 43189800 | 43190399 | 10.30301 | 19.47313 | 27.57652 | 11.20884 | -0.01809 | 0.065243 | 0.97753 |
| 2 | 73577600 | 73582999 | 88.88596 | 137.8442 | 171.8758 | 71.3859 | -0.01276 | 2.79379 | 0.977537 |
| 4 | 124237400 | 124238599 | 13.71664 | 24.78093 | 32.59116 | 14.15778 | 0.016214 | 0.369955 | 0.977557 |
| 3 | 85281600 | 85284999 | 30.81066 | 50.91718 | 65.71297 | 28.3933 | 0.013546 | 1.347399 | 0.977571 |
| 7 | 24834200 | 24835399 | 17.10719 | 29.83299 | 39.49971 | 17.38296 | -0.01554 | 0.734783 | 0.977582 |
| 5 | 66448000 | 66449999 | 18.40971 | 32.08979 | 42.4844 | 18.69109 | -0.01579 | 0.748719 | 0.977598 |
| 3 | 132722200 | 132726199 | 43.41556 | 70.20197 | 91.59275 | 39.14503 | -0.02115 | 1.711453 | 0.977614 |
| 1 | 50035400 | 50038399 | 21.15616 | 36.61309 | 48.48144 | 21.27016 | -0.01747 | 0.857246 | 0.977621 |
| 18 | 69173800 | 69175799 | 19.79465 | 33.70029 | 55.42817 | 24.29535 | -0.33325 | 1.00702 | 0.97763 |
| 9 | 109398600 | 109400199 | 16.9516 | 29.79343 | 38.61706 | 17.09959 | 0.015095 | 0.61052 | 0.977633 |
| 9 | 70076200 | 70077199 | 15.16486 | 26.88326 | 34.97778 | 15.46308 | 0.014449 | 0.499864 | 0.97764 |
| 13 | 98747000 | 98751799 | 101.0261 | 154.985 | 191.5632 | 79.5248 | -0.01208 | 3.001672 | 0.977648 |
| 1 | 29714600 | 29715399 | 8.492075 | 16.37252 | 23.6737 | 9.165756 | 0.015941 | -0.12048 | 0.977653 |
| 7 | 29524800 | 29529799 | 69.9162 | 109.962 | 136.1043 | 56.46302 | 0.013604 | 2.52698 | 0.977656 |
| 6 | 64813200 | 64817999 | 40.06653 | 64.89807 | 82.75739 | 35.41741 | 0.014138 | 1.662171 | 0.977667 |
| 11 | 74171000 | 74172399 | 20.86993 | 36.23212 | 46.84899 | 20.55925 | 0.017652 | 0.82539 | 0.977685 |
| 3 | 119364000 | 119364999 | 15.66859 | 27.94039 | 27.44298 | 12.07272 | 0.422816 | 0.450984 | 0.977697 |
| 18 | 37552600 | 37556999 | 81.02777 | 126.5808 | 158.4911 | 65.75992 | -0.01057 | 2.661484 | 0.977774 |
| 17 | 77861000 | 77862599 | 14.33316 | 25.62532 | 33.50599 | 14.70781 | 0.014444 | 0.438315 | 0.977788 |
| 7 | 87064400 | 87065199 | 11.45601 | 20.22297 | 37.88078 | 16.76057 | -0.51053 | 0.529195 | 0.977794 |
| X | 35374800 | 35376999 | 14.01175 | 25.95289 | 25.00872 | 10.51573 | 0.494475 | 0.216541 | 0.977796 |
| 8 | 41191000 | 41199799 | 157.8838 | 236.6261 | 286.7291 | 118.2851 | -0.01157 | 3.486152 | 0.977816 |
| 8 | 31511000 | 31514199 | 68.77809 | 108.454 | 134.6542 | 55.88038 | 0.012032 | 2.47532 | 0.977839 |
| 16 | 64675400 | 64676399 | 8.873093 | 17.12056 | 25.45004 | 9.831286 | -0.0205 | -0.11772 | 0.977841 |
| 17 | 94971400 | 94981999 | 182.436 | 269.5003 | 322.0217 | 132.6153 | -0.01053 | 3.745552 | 0.977841 |
| 13 | 3788400 | 3789199 | 10.07027 | 18.88918 | 25.82591 | 10.64279 | 0.015785 | 0.107134 | 0.977847 |
| 5 | 128183600 | 128189199 | 63.79822 | 101.2559 | 126.4908 | 52.69918 | 0.011029 | 2.308197 | 0.977851 |
| 17 | 5666200 | 5668799 | 30.28485 | 49.95044 | 65.63346 | 28.32455 | -0.01275 | 1.367889 | 0.977856 |
| 5 | 142254600 | 142257599 | 32.31952 | 52.44445 | 83.86955 | 35.94122 | -0.30923 | 1.613396 | 0.977892 |
| 4 | 112606600 | 112607399 | 15.44382 | 28.12291 | 27.73623 | 11.93015 | 0.437913 | 0.325282 | 0.977895 |

| | | | | | | | | | |
|---|---|---|---|---|---|---|---|---|---|
| 18 | 37847200 | 37862999 | 662.8755 | 877.3576 | 940.7503 | 393.3202 | -0.01217 | 5.444294 | 0.977897 |
| 8 | 61693600 | 61698399 | 66.20344 | 104.7356 | 130.5149 | 54.24408 | 0.010488 | 2.396465 | 0.977898 |
| 2 | 52644600 | 52645799 | 10.70774 | 20.05088 | 27.3632 | 11.30873 | 0.014964 | 0.126945 | 0.9779 |
| 13 | 24722000 | 24722999 | 10.70774 | 20.05088 | 27.3632 | 11.30873 | 0.014964 | 0.126945 | 0.9779 |
| 19 | 23274400 | 23275999 | 14.80205 | 26.18129 | 34.03692 | 15.05389 | 0.014366 | 0.519698 | 0.977903 |
| 13 | 68504400 | 68513399 | 379.2957 | 525.2487 | 578.9194 | 240.4139 | 0.013753 | 4.728292 | 0.977904 |
| 16 | 13227400 | 13230999 | 18.64393 | 32.80277 | 32.67563 | 14.4671 | 0.393411 | 0.595677 | 0.977928 |
| 2 | 114848200 | 114849599 | 15.03272 | 26.79293 | 34.90746 | 15.36305 | 0.016746 | 0.462515 | 0.977937 |
| 1 | 22287400 | 22288799 | 14.75409 | 26.12369 | 33.96415 | 15.01857 | 0.014754 | 0.50685 | 0.977947 |
| 5 | 64147600 | 64148799 | 14.75409 | 26.12369 | 33.96415 | 15.01857 | 0.014754 | 0.50685 | 0.977947 |
| 8 | 114936800 | 114938799 | 22.26419 | 38.03078 | 50.09556 | 21.96383 | -0.01342 | 0.979646 | 0.977952 |
| 8 | 65076800 | 65079599 | 22.83955 | 38.8217 | 50.10105 | 21.95751 | 0.01488 | 1.024957 | 0.978009 |
| 2 | 47999200 | 47999799 | 9.58871 | 18.23526 | 26.07282 | 10.46022 | -0.01544 | 0.011323 | 0.978019 |
| 8 | 19825000 | 19837599 | 344.1045 | 479.5398 | 647.1339 | 268.4767 | -0.26901 | 4.632345 | 0.978033 |
| 8 | 125705800 | 125708199 | 32.9815 | 53.44257 | 85.47551 | 36.59104 | -0.31132 | 1.644129 | 0.978043 |
| 17 | 16889200 | 16891799 | 22.62403 | 38.47761 | 50.62755 | 22.18929 | -0.01274 | 1.021466 | 0.97805 |
| 19 | 21213200 | 21215999 | 33.1959 | 54.23077 | 70.90278 | 30.47856 | -0.01189 | 1.501413 | 0.978061 |
| 4 | 135119400 | 135121399 | 16.28585 | 28.55349 | 37.76421 | 16.68747 | -0.01376 | 0.644606 | 0.978062 |
| 3 | 82938200 | 82939399 | 11.2163 | 20.52807 | 27.22624 | 11.63602 | 0.019678 | 0.26492 | 0.978072 |
| Y | 20887800 | 20890599 | 13.99254 | 24.528 | 33.02558 | 14.59117 | -0.03932 | 0.625264 | 0.978122 |
| 6 | 130290600 | 130293199 | 32.34165 | 53.64042 | 68.88335 | 29.83737 | 0.021088 | 1.313226 | 0.978129 |
| 4 | 94702800 | 94708599 | 79.86464 | 124.669 | 153.6603 | 63.76054 | 0.011262 | 2.673104 | 0.978142 |
| 3 | 146730200 | 146731399 | 13.50312 | 24.36327 | 32.0492 | 13.94515 | 0.014302 | 0.374052 | 0.978152 |
| 17 | 22009600 | 22030399 | 358.0548 | 501.7027 | 560.6824 | 232.4139 | 0.010874 | 4.551345 | 0.978162 |
| 4 | 12933600 | 12936399 | 62.09221 | 98.42696 | 124.7261 | 51.89998 | -0.01125 | 2.343683 | 0.978178 |
| 1 | 100704600 | 100712399 | 154.9354 | 231.4083 | 275.0639 | 113.3969 | 0.011657 | 3.548135 | 0.97821 |
| 1 | 79413000 | 79414599 | 14.98194 | 26.46436 | 35.06861 | 15.5143 | -0.01379 | 0.537375 | 0.978211 |
| 18 | 54943200 | 54946199 | 19.32849 | 33.62349 | 43.60531 | 19.15996 | 0.013679 | 0.785016 | 0.978231 |
| 14 | 120336200 | 120336999 | 11.09509 | 20.84804 | 28.51843 | 11.71749 | 0.017955 | 0.109191 | 0.978237 |
| 4 | 80177600 | 80184999 | 88.30125 | 137.5708 | 172.0592 | 71.41282 | -0.01201 | 2.712625 | 0.978239 |
| 17 | 22687800 | 22690999 | 67.43239 | 106.4217 | 134.3723 | 55.77885 | -0.01125 | 2.455699 | 0.978311 |
| 9 | 86326400 | 86329199 | 23.77327 | 40.47441 | 52.27504 | 22.91329 | 0.014149 | 1.018061 | 0.978314 |
| 15 | 57641000 | 57647199 | 57.8482 | 92.49957 | 115.8445 | 48.73052 | 0.018689 | 2.071636 | 0.978322 |
| 11 | 95633200 | 95635999 | 26.46652 | 44.27885 | 57.2235 | 24.92819 | 0.012298 | 1.191501 | 0.978327 |
| 3 | 58571400 | 58575599 | 61.44158 | 97.51897 | 123.6907 | 51.53438 | -0.01082 | 2.306196 | 0.978331 |
| 7 | 34777800 | 34778799 | 8.53939 | 16.28201 | 23.45949 | 9.360142 | -0.01875 | -0.0215 | 0.97834 |
| 8 | 93018600 | 93019999 | 14.9045 | 26.4408 | 34.40984 | 15.20371 | 0.014552 | 0.49035 | 0.978344 |
| 18 | 65974800 | 65978999 | 71.02901 | 111.998 | 139.2407 | 57.78337 | 0.010031 | 2.477715 | 0.978393 |
| 7 | 92847600 | 92850599 | 25.33279 | 43.18579 | 44.74392 | 19.6143 | 0.331277 | 1.004272 | 0.978417 |
| 4 | 69700400 | 69705799 | 47.96176 | 76.94728 | 97.25626 | 41.11413 | 0.011039 | 1.952354 | 0.978446 |
| 15 | 51751400 | 51752199 | 9.297521 | 17.77781 | 25.09306 | 9.948672 | 0.017591 | -0.03524 | 0.978452 |
| 5 | 42744400 | 42745199 | 13.36884 | 23.99068 | 32.06881 | 14.0327 | -0.014 | 0.406058 | 0.97846 |
| 11 | 114248400 | 114250199 | 18.72113 | 32.74118 | 42.40911 | 18.70041 | 0.01606 | 0.704264 | 0.978465 |
| 2 | 89298600 | 89301399 | 34.98459 | 57.39868 | 73.62968 | 31.7046 | 0.018726 | 1.447319 | 0.978473 |
| 6 | 142099600 | 142102199 | 36.59319 | 60.08259 | 61.50469 | 26.49198 | 0.343838 | 1.435327 | 0.978479 |
| 15 | 82190600 | 82192199 | 13.75541 | 24.66565 | 32.97172 | 14.43704 | -0.01493 | 0.413093 | 0.978484 |
| 9 | 100999600 | 101001199 | 15.56625 | 27.12611 | 45.23501 | 19.89787 | -0.34701 | 0.745728 | 0.978504 |

| | | | | | | | | | |
|---|---|---|---|---|---|---|---|---|---|
| 16 | 87587000 | 87589599 | 20.18991 | 34.99601 | 46.30776 | 20.31748 | -0.01608 | 0.834121 | 0.978508 |
| 1 | 185944800 | 185948199 | 51.78028 | 82.8639 | 106.0479 | 44.66066 | -0.01118 | 2.038142 | 0.978538 |
| 4 | 35612800 | 35615199 | 25.02228 | 42.67059 | 44.22983 | 19.38945 | 0.330728 | 0.999854 | 0.978562 |
| 2 | 76280000 | 76283599 | 31.07514 | 51.34195 | 67.5739 | 29.19254 | -0.01475 | 1.350688 | 0.978565 |
| 5 | 26376200 | 26377999 | 18.72682 | 32.31974 | 42.7358 | 18.74782 | -0.01603 | 0.864959 | 0.978572 |
| 5 | 137266200 | 137273399 | 71.84029 | 113.0552 | 140.2981 | 58.20733 | 0.010491 | 2.516548 | 0.978593 |
| 7 | 49402000 | 49402799 | 6.99066 | 13.55048 | 20.54545 | 7.771813 | -0.01908 | -0.2222 | 0.978595 |
| X | 80237400 | 80240399 | 41.04883 | 67.3115 | 60.94964 | 26.23858 | 0.519724 | 1.46581 | 0.978607 |
| 13 | 33280600 | 33282599 | 16.60461 | 29.19092 | 38.61686 | 17.10036 | -0.01417 | 0.604864 | 0.97861 |
| 11 | 113047000 | 113049199 | 22.68626 | 38.48623 | 50.62346 | 22.17589 | -0.01255 | 1.045052 | 0.978614 |
| 8 | 80480800 | 80486599 | 69.30479 | 110.2025 | 136.7205 | 57.06838 | 0.023583 | 2.261114 | 0.978616 |
| 8 | 23833600 | 23834999 | 13.08193 | 24.32768 | 25.02967 | 10.46632 | 0.406181 | 0.185453 | 0.978623 |
| X | 91109000 | 91111199 | 21.06635 | 36.74463 | 47.31091 | 20.82311 | 0.024345 | 0.761523 | 0.978628 |
| 4 | 19750200 | 19751199 | 10.22909 | 19.21985 | 26.36191 | 10.83234 | 0.014492 | 0.098939 | 0.978637 |
| 5 | 141006000 | 141008999 | 44.56604 | 71.7843 | 92.66469 | 39.40295 | -0.01231 | 1.823283 | 0.978638 |
| 4 | 100740400 | 100741799 | 20.59118 | 35.51316 | 46.85889 | 20.55615 | -0.01341 | 0.882328 | 0.978647 |
| 6 | 127207800 | 127212999 | 50.87296 | 81.39947 | 104.1988 | 43.87243 | -0.01178 | 2.039848 | 0.978648 |
| 10 | 82174000 | 82174999 | 12.67685 | 23.29482 | 31.81938 | 13.5247 | -0.0172 | 0.263207 | 0.978661 |
| 8 | 97143000 | 97143799 | 11.15835 | 20.59651 | 28.20945 | 11.91745 | -0.01464 | 0.217669 | 0.978674 |
| 8 | 76630200 | 76630999 | 15.60749 | 27.3813 | 36.25168 | 16.02759 | -0.01526 | 0.627639 | 0.978677 |
| 17 | 78766800 | 78769199 | 17.97797 | 31.46294 | 40.79507 | 17.99907 | 0.014527 | 0.685744 | 0.978692 |
| 2 | 7591000 | 7592799 | 32.0353 | 52.41601 | 67.4595 | 29.01482 | 0.012369 | 1.474545 | 0.978696 |
| 17 | 75238600 | 75240399 | 16.39584 | 28.81183 | 38.104 | 16.87003 | -0.01386 | 0.607954 | 0.978696 |
| 14 | 69047600 | 69051399 | 50.48138 | 80.79177 | 101.857 | 42.90059 | 0.010562 | 2.033351 | 0.978723 |
| 9 | 13853200 | 13855599 | 30.15548 | 50.91813 | 47.99406 | 20.9942 | 0.466316 | 1.110258 | 0.978738 |
| 9 | 73842600 | 73843799 | 15.38069 | 26.79099 | 45.24053 | 19.89482 | -0.36515 | 0.750155 | 0.978788 |
| 9 | 19838600 | 19842999 | 58.52105 | 93.6189 | 95.6333 | 40.25505 | 0.312902 | 2.053699 | 0.978804 |
| 18 | 18562800 | 18567199 | 45.66246 | 73.68217 | 93.34236 | 39.78619 | 0.017811 | 1.774192 | 0.978863 |
| 9 | 54821400 | 54824999 | 34.8095 | 56.55622 | 72.39181 | 31.05145 | 0.012896 | 1.585393 | 0.978874 |
| 1 | 15044600 | 15048399 | 64.50606 | 102.1823 | 129.2849 | 53.76806 | -0.00994 | 2.364043 | 0.978891 |
| 1 | 53652200 | 53655599 | 75.90853 | 119.2941 | 150.0212 | 62.23022 | -0.01018 | 2.548736 | 0.978901 |
| 2 | 179209600 | 179210599 | 13.70871 | 24.45589 | 32.57993 | 14.32743 | -0.01404 | 0.446404 | 0.97891 |
| 16 | 81888200 | 81889399 | 23.27481 | 38.54893 | 66.00534 | 28.56331 | -0.39266 | 1.303904 | 0.978911 |
| 14 | 25369400 | 25371199 | 20.15027 | 34.89867 | 45.25315 | 19.85428 | 0.012912 | 0.84216 | 0.978928 |
| 4 | 79388200 | 79392599 | 43.37715 | 70.20147 | 72.021 | 30.80351 | 0.326082 | 1.678483 | 0.978967 |
| 8 | 14904200 | 14906799 | 23.79147 | 40.44102 | 52.26241 | 22.90482 | 0.012981 | 1.032787 | 0.978978 |
| 5 | 109581000 | 109581599 | 9.431352 | 18.08395 | 26.37133 | 10.37227 | -0.01783 | -0.05373 | 0.978997 |
| 9 | 14888000 | 14888599 | 9.841392 | 18.66554 | 26.55359 | 10.71385 | -0.01571 | 0.031788 | 0.979002 |
| 2 | 98213400 | 98220399 | 68.23081 | 108.136 | 137.1817 | 57.07141 | -0.01316 | 2.356917 | 0.979003 |
| 6 | 60489600 | 60492799 | 34.57269 | 56.74091 | 74.64575 | 32.14643 | -0.01738 | 1.444469 | 0.979013 |
| 8 | 77278400 | 77280799 | 18.00044 | 31.09552 | 51.69461 | 22.67862 | -0.34482 | 0.864518 | 0.979039 |
| 14 | 115517400 | 115519599 | 26.49961 | 44.22942 | 57.18039 | 24.87224 | 0.011856 | 1.217293 | 0.979056 |
| 8 | 9885000 | 9885399 | 6.086161 | 11.85147 | 18.52161 | 6.783263 | -0.01922 | -0.3549 | 0.979075 |
| 15 | 63312200 | 63314199 | 19.70618 | 34.46498 | 35.55455 | 15.67828 | 0.342914 | 0.697762 | 0.979082 |
| 17 | 68661000 | 68662999 | 18.99113 | 33.20354 | 42.97164 | 18.94371 | 0.017366 | 0.71283 | 0.979121 |
| 1 | 48673000 | 48674199 | 11.94323 | 22.33153 | 31.20105 | 12.92702 | -0.02171 | 0.156252 | 0.97913 |
| 14 | 86635400 | 86636399 | 29.18229 | 47.96463 | 63.03253 | 27.16206 | -0.01488 | 1.406584 | 0.979135 |

| | | | | | | | | | |
|---|---|---|---|---|---|---|---|---|---|
| 9 | 42201600 | 42202999 | 13.79207 | 25.11647 | 33.20208 | 14.28021 | 0.01793 | 0.324554 | 0.979147 |
| 12 | 68603800 | 68604999 | 9.38082 | 18.19809 | 19.91972 | 7.506918 | 0.450818 | -0.21169 | 0.979153 |
| 7 | 66142400 | 66143799 | 15.74542 | 27.60811 | 35.7639 | 15.80459 | 0.016026 | 0.636164 | 0.979154 |
| 1 | 182053200 | 182071599 | 214.4876 | 311.4488 | 438.7396 | 180.8703 | -0.27161 | 4.019598 | 0.979166 |
| 17 | 54729000 | 54731599 | 33.51493 | 54.74272 | 71.531 | 30.74672 | -0.01124 | 1.504656 | 0.979172 |
| 6 | 118831000 | 118832999 | 19.1965 | 33.13861 | 43.73613 | 19.1868 | -0.01332 | 0.865652 | 0.979195 |
| 5 | 13810400 | 13811599 | 11.78065 | 21.76061 | 29.2292 | 12.33634 | 0.014147 | 0.220664 | 0.979236 |
| 14 | 60190400 | 60191799 | 7.600685 | 14.61743 | 29.44395 | 11.48176 | -0.46849 | -0.09254 | 0.97924 |
| 19 | 15316200 | 15318799 | 19.61975 | 33.91186 | 44.73912 | 19.62756 | -0.01249 | 0.856232 | 0.979261 |
| 1 | 57627400 | 57629399 | 39.1623 | 63.33554 | 80.82777 | 34.55201 | 0.010989 | 1.692365 | 0.979262 |
| 8 | 111631600 | 111634599 | 20.99579 | 36.00655 | 47.41453 | 20.795 | -0.01202 | 0.936762 | 0.979266 |
| 7 | 81810800 | 81821799 | 190.602 | 281.3233 | 335.6696 | 138.2346 | -0.00963 | 3.761228 | 0.979306 |
| 11 | 5405800 | 5410199 | 64.14796 | 101.6342 | 126.847 | 52.75916 | 0.009955 | 2.358571 | 0.979341 |
| 6 | 25931000 | 25932199 | 15.35991 | 26.99793 | 35.73055 | 15.82243 | -0.01461 | 0.596497 | 0.979342 |
| 5 | 131188800 | 131194999 | 59.6351 | 94.82751 | 120.4626 | 50.28271 | -0.01046 | 2.247915 | 0.979389 |
| 13 | 34390200 | 34390599 | 5.347747 | 10.44947 | 17.10817 | 5.946912 | -0.01652 | -0.5514 | 0.979394 |
| 18 | 75168800 | 75171599 | 26.63657 | 44.32804 | 58.2834 | 25.30499 | -0.01241 | 1.249018 | 0.979409 |
| 15 | 34452000 | 34454199 | 20.82785 | 35.61334 | 46.90504 | 20.56666 | -0.0129 | 0.962057 | 0.979441 |
| 1 | 4303400 | 4304799 | 14.64593 | 25.98758 | 34.48348 | 15.23353 | -0.01303 | 0.488736 | 0.979462 |
| 8 | 4699800 | 4702799 | 41.58881 | 67.10016 | 85.41589 | 36.40156 | 0.010656 | 1.766348 | 0.979509 |
| 13 | 75611000 | 75612399 | 17.23491 | 30.62576 | 30.58905 | 13.49663 | 0.395627 | 0.487666 | 0.979542 |
| 17 | 53610600 | 53616399 | 59.08658 | 94.22591 | 120.3177 | 50.42054 | -0.01346 | 2.157315 | 0.979557 |
| 10 | 81474000 | 81497599 | 412.9955 | 570.0486 | 636.3835 | 264.4154 | -0.00946 | 4.779625 | 0.979584 |
| 16 | 38541800 | 38543399 | 19.10006 | 33.42153 | 34.12608 | 15.0547 | 0.357727 | 0.685247 | 0.979596 |
| 13 | 38522000 | 38524599 | 37.32592 | 59.93048 | 100.6204 | 42.58681 | -0.39637 | 1.917261 | 0.979627 |
| 3 | 158331400 | 158336199 | 85.29985 | 132.4883 | 162.7351 | 67.57377 | 0.010623 | 2.762675 | 0.979633 |
| X | 68223200 | 68224399 | 14.36628 | 25.96222 | 34.12487 | 14.81858 | 0.017284 | 0.375413 | 0.979649 |
| 1 | 132775400 | 132778399 | 54.25524 | 86.72566 | 109.1581 | 45.89633 | 0.010986 | 2.076531 | 0.979668 |
| 15 | 21876200 | 21877199 | 17.46043 | 30.54656 | 40.37553 | 17.80872 | -0.01288 | 0.690456 | 0.979677 |
| 1 | 185558600 | 185562799 | 41.88137 | 67.49437 | 85.78311 | 36.5037 | 0.010992 | 1.799667 | 0.979706 |
| 16 | 21284200 | 21286799 | 19.08142 | 33.06667 | 43.66656 | 19.15852 | -0.01316 | 0.82854 | 0.979724 |
| 16 | 9912400 | 9914799 | 31.61331 | 52.22767 | 68.73672 | 29.69353 | -0.01469 | 1.354447 | 0.979727 |
| Y | 40583000 | 40583799 | 17.65189 | 29.53098 | 60.40458 | 26.31346 | -0.64754 | 1.169046 | 0.979784 |
| 11 | 117398600 | 117401399 | 31.50591 | 51.53119 | 67.46322 | 29.0126 | -0.0126 | 1.477311 | 0.979807 |
| 10 | 120905000 | 120907199 | 19.02756 | 32.92842 | 42.6204 | 18.69875 | 0.015224 | 0.838187 | 0.979844 |
| X | 127929800 | 127931799 | 19.21111 | 33.63845 | 43.45254 | 19.17995 | 0.020053 | 0.69131 | 0.979848 |
| 15 | 88920400 | 88921999 | 16.58655 | 29.33163 | 29.35499 | 12.98369 | 0.389588 | 0.523246 | 0.979868 |
| 14 | 45707000 | 45707599 | 13.65489 | 24.40643 | 32.52596 | 14.28069 | -0.0131 | 0.432882 | 0.979872 |
| 18 | 15732000 | 15733799 | 15.11003 | 27.06687 | 26.87775 | 11.77892 | 0.41073 | 0.416857 | 0.979879 |
| 14 | 115173600 | 115176799 | 37.47699 | 60.93412 | 78.07257 | 33.50246 | 0.012325 | 1.58198 | 0.979888 |
| 6 | 9596000 | 9597999 | 20.25291 | 34.92344 | 45.26778 | 19.85805 | 0.012137 | 0.880523 | 0.9799 |
| 1 | 100501000 | 100506799 | 87.78492 | 136.9299 | 168.6833 | 69.99966 | 0.011162 | 2.691895 | 0.979905 |
| 17 | 12281800 | 12291199 | 130.7078 | 198.2896 | 239.2264 | 99.03454 | 0.009933 | 3.238847 | 0.979915 |
| 15 | 19996600 | 20005999 | 227.237 | 329.686 | 381.7606 | 157.3944 | 0.009895 | 4.032123 | 0.979935 |
| 13 | 4529200 | 4530999 | 28.14175 | 46.8713 | 60.60013 | 26.3247 | 0.011755 | 1.247214 | 0.979974 |
| X | 24085200 | 24088799 | 39.6418 | 64.43536 | 82.16912 | 35.25656 | 0.018864 | 1.595784 | 0.979986 |
| 8 | 18473000 | 18474399 | 13.29623 | 23.959 | 31.51403 | 13.73448 | 0.012887 | 0.378121 | 0.980021 |

| | | | | | | | | | |
|---|---|---|---|---|---|---|---|---|---|
| 10 | 43575000 | 43575999 | 13.29623 | 23.959 | 31.51403 | 13.73448 | 0.012887 | 0.378121 | 0.980021 |
| 12 | 75744200 | 75746799 | 21.70028 | 37.46729 | 48.47554 | 21.26606 | 0.015196 | 0.881163 | 0.980031 |
| 12 | 98323600 | 98324799 | 16.23624 | 28.50203 | 37.66227 | 16.65996 | -0.01259 | 0.622998 | 0.980032 |
| 11 | 53117000 | 53121399 | 46.50354 | 74.60363 | 94.31235 | 39.86625 | 0.010684 | 1.95158 | 0.980032 |
| 13 | 57895600 | 57897199 | 23.58166 | 40.66043 | 41.45783 | 18.18666 | 0.356955 | 0.890506 | 0.980032 |
| 4 | 147351600 | 147361599 | 485.4218 | 659.1576 | 724.4207 | 301.7671 | -0.01067 | 5.035031 | 0.980035 |
| 11 | 10385000 | 10395799 | 291.4986 | 412.6973 | 473.4977 | 195.9466 | -0.012 | 4.394938 | 0.980063 |
| 19 | 45241600 | 45244399 | 28.02891 | 46.41943 | 60.9725 | 26.38402 | -0.01148 | 1.312297 | 0.980075 |
| 10 | 93192800 | 93194399 | 15.90533 | 28.01314 | 37.05488 | 16.40264 | -0.01302 | 0.577293 | 0.980078 |
| 4 | 131864000 | 131865399 | 12.5941 | 22.78279 | 30.02644 | 13.02226 | 0.014545 | 0.346496 | 0.980091 |
| 1 | 117753200 | 117780599 | 378.5982 | 530.3299 | 590.9471 | 244.9705 | 0.014652 | 4.557156 | 0.980116 |
| 10 | 62008600 | 62015199 | 58.20989 | 92.67965 | 116.2687 | 48.61613 | 0.009554 | 2.205894 | 0.980186 |
| 7 | 43696000 | 43699399 | 29.44001 | 48.5775 | 62.75333 | 27.08644 | 0.011964 | 1.35808 | 0.98022 |
| 13 | 87781800 | 87782199 | 9.274537 | 17.73145 | 25.09244 | 9.952302 | 0.014003 | -0.03094 | 0.980226 |
| 13 | 99907800 | 99913199 | 58.3343 | 92.96126 | 116.6556 | 48.83863 | 0.010516 | 2.176827 | 0.98024 |
| 7 | 56206800 | 56209199 | 27.30969 | 45.52201 | 58.88287 | 25.59169 | 0.011087 | 1.234715 | 0.980249 |
| 8 | 101194000 | 101196199 | 27.69137 | 46.48006 | 48.05256 | 20.97637 | 0.33318 | 1.152513 | 0.980255 |
| 11 | 29989800 | 29990399 | 8.806406 | 16.87973 | 24.05915 | 9.47241 | 0.013844 | -0.06196 | 0.980257 |
| 15 | 21477800 | 21478399 | 7.092337 | 13.77412 | 20.5919 | 7.705706 | 0.014762 | -0.26119 | 0.98027 |
| 15 | 19917000 | 19920799 | 41.87178 | 67.6649 | 86.18476 | 36.80512 | 0.012448 | 1.720036 | 0.980286 |
| 2 | 18104600 | 18105999 | 12.72104 | 23.4314 | 31.33656 | 13.27668 | 0.016549 | 0.250191 | 0.980304 |
| 3 | 91033400 | 91035399 | 75.57296 | 118.1572 | 148.2493 | 61.50523 | -0.01259 | 2.642784 | 0.980332 |
| 8 | 23016800 | 23018999 | 27.16127 | 45.0717 | 58.23292 | 25.23304 | 0.01248 | 1.284116 | 0.980335 |
| 18 | 62946400 | 62948399 | 21.37668 | 36.62642 | 47.40689 | 20.79012 | 0.01244 | 0.944544 | 0.98034 |
| 3 | 140981800 | 140982999 | 10.70899 | 20.04856 | 27.89817 | 11.53437 | -0.01344 | 0.129559 | 0.980378 |
| 15 | 8067800 | 8068399 | 8.142422 | 15.6555 | 22.48834 | 8.775489 | 0.014647 | -0.10328 | 0.980382 |
| 3 | 54367400 | 54368999 | 18.50495 | 32.09549 | 41.6282 | 18.26468 | 0.013092 | 0.817104 | 0.980417 |
| 14 | 26061600 | 26062799 | 18.50495 | 32.09549 | 41.6282 | 18.26468 | 0.013092 | 0.817104 | 0.980417 |
| 5 | 146079200 | 146083399 | 56.55962 | 90.14581 | 114.7313 | 48.04235 | -0.00959 | 2.170626 | 0.980418 |
| 18 | 51933200 | 51935199 | 20.51712 | 35.95091 | 35.341 | 15.60893 | 0.41207 | 0.671469 | 0.980428 |
| 7 | 85065600 | 85066199 | 10.7218 | 19.86385 | 26.75242 | 11.24532 | 0.014954 | 0.191324 | 0.980431 |
| 10 | 66528000 | 66535799 | 92.0108 | 142.4036 | 174.5672 | 72.51078 | 0.009503 | 2.828945 | 0.980432 |
| 4 | 58522400 | 58525399 | 25.31128 | 42.51836 | 54.992 | 24.01751 | 0.011087 | 1.142897 | 0.980439 |
| 2 | 45061000 | 45062799 | 13.38171 | 23.89933 | 31.93834 | 14.03201 | -0.01797 | 0.432181 | 0.980482 |
| 11 | 23263000 | 23264799 | 11.67156 | 20.8132 | 36.73115 | 16.15853 | -0.41795 | 0.44197 | 0.980502 |
| 6 | 21800000 | 21803399 | 30.10744 | 50.02528 | 65.8841 | 28.57389 | -0.01506 | 1.283061 | 0.980528 |
| 6 | 62339200 | 62339599 | 5.794206 | 11.30833 | 18.12597 | 6.436581 | -0.01553 | -0.47056 | 0.980533 |
| 10 | 22066200 | 22071799 | 53.06562 | 84.65805 | 106.3531 | 44.59313 | 0.010655 | 2.133844 | 0.980548 |
| 15 | 27642400 | 27643399 | 11.63835 | 21.41135 | 28.6726 | 12.16792 | 0.012908 | 0.24114 | 0.980551 |
| 6 | 144161800 | 144162199 | 8.803984 | 16.74111 | 23.92232 | 9.599635 | -0.01458 | -0.00067 | 0.980593 |
| 12 | 115760800 | 115770399 | 108.8222 | 168.2248 | 204.5201 | 84.95296 | 0.020129 | 2.862315 | 0.9806 |
| X | 65029600 | 65031599 | 21.64739 | 37.42887 | 49.56336 | 21.74422 | -0.0177 | 0.871182 | 0.98063 |
| 12 | 36623200 | 36623999 | 9.050946 | 17.33936 | 25.16873 | 9.924192 | -0.01478 | -0.05381 | 0.980636 |
| 4 | 107560200 | 107561999 | 23.75052 | 40.18281 | 52.81586 | 23.11731 | -0.01174 | 1.072555 | 0.980637 |
| 8 | 52742200 | 52744199 | 18.06818 | 31.89165 | 32.19012 | 14.24303 | 0.376282 | 0.557763 | 0.980688 |
| 12 | 31418600 | 31421599 | 24.07651 | 40.59637 | 53.366 | 23.33408 | -0.01211 | 1.103153 | 0.980703 |
| 10 | 46321800 | 46322399 | 8.081334 | 15.25577 | 28.64839 | 11.67406 | -0.42558 | 0.053642 | 0.980748 |

| | | | | | | | | | |
|---|---|---|---|---|---|---|---|---|---|
| 8 | 84493000 | 84494999 | 21.9261 | 37.44787 | 48.48454 | 21.25725 | 0.011355 | 0.977826 | 0.980752 |
| 4 | 105272000 | 105272999 | 18.60561 | 32.62329 | 33.52061 | 14.81329 | 0.348632 | 0.648451 | 0.980786 |
| 16 | 19021400 | 19029399 | 99.3047 | 153.5994 | 190.9978 | 79.33587 | -0.01174 | 2.8453 | 0.980798 |
| 5 | 85058800 | 85059399 | 7.766463 | 15.04106 | 22.15877 | 8.416555 | 0.01605 | -0.19465 | 0.980826 |
| 11 | 31384600 | 31394599 | 129.2379 | 195.6469 | 238.6968 | 98.75073 | -0.00877 | 3.280787 | 0.980841 |
| 6 | 44769800 | 44770199 | 8.222281 | 15.77493 | 23.01294 | 9.036072 | -0.01526 | -0.07808 | 0.980843 |
| 15 | 91216200 | 91217399 | 15.46368 | 27.27382 | 36.07186 | 15.96299 | -0.01193 | 0.558522 | 0.980874 |
| 15 | 58165800 | 58168199 | 18.49823 | 32.21966 | 42.53441 | 18.70418 | -0.01168 | 0.761553 | 0.98089 |
| 15 | 103684200 | 103685599 | 17.08692 | 30.00707 | 39.66954 | 17.55367 | -0.0133 | 0.625179 | 0.980903 |
| 12 | 79149800 | 79152799 | 20.92977 | 35.98942 | 47.39981 | 20.79155 | -0.01164 | 0.911247 | 0.980908 |
| Y | 4437400 | 4438399 | 12.55492 | 22.21208 | 29.9011 | 13.22405 | -0.03537 | 0.49359 | 0.98093 |
| 5 | 125022200 | 125024999 | 26.17982 | 43.96854 | 57.86772 | 25.27028 | -0.01409 | 1.148023 | 0.980945 |
| 12 | 34986600 | 34989599 | 61.21851 | 97.19119 | 121.6059 | 50.67946 | 0.009191 | 2.296969 | 0.980954 |
| 15 | 12236200 | 12236799 | 9.389562 | 17.68477 | 24.31038 | 9.948877 | 0.016 | 0.069898 | 0.980979 |
| 10 | 127621800 | 127622399 | 10.21711 | 19.44667 | 27.24312 | 10.91031 | 0.015869 | 0.013957 | 0.981015 |
| 18 | 6345600 | 6356399 | 364.1882 | 505.6156 | 568.794 | 236.1144 | -0.01203 | 4.689275 | 0.981021 |
| 4 | 84635200 | 84635599 | 5.357847 | 10.46932 | 17.10949 | 5.945563 | -0.01428 | -0.55466 | 0.98104 |
| 13 | 28715600 | 28716599 | 11.6144 | 21.19263 | 28.10392 | 12.05762 | 0.015796 | 0.285219 | 0.981053 |
| 1 | 147242000 | 147245999 | 79.22363 | 124.2925 | 156.0482 | 64.71789 | -0.00969 | 2.584644 | 0.981062 |
| 14 | 48194200 | 48197599 | 29.70185 | 49.20667 | 64.69943 | 28.00327 | -0.01303 | 1.312959 | 0.981084 |
| 4 | 22023800 | 22024599 | 11.34654 | 20.75344 | 28.17037 | 12.04978 | -0.01461 | 0.270677 | 0.981093 |
| 18 | 6674000 | 6678199 | 52.10141 | 83.84693 | 83.50697 | 35.46298 | 0.359558 | 1.858626 | 0.981094 |
| 1 | 105688400 | 105692399 | 92.07198 | 142.4341 | 176.807 | 73.44132 | -0.00905 | 2.83897 | 0.981141 |
| 12 | 66306600 | 66307199 | 9.443453 | 17.73937 | 24.84294 | 10.21239 | -0.01557 | 0.089838 | 0.981152 |
| 7 | 85409000 | 85410399 | 28.2668 | 46.72735 | 61.39996 | 26.53526 | -0.01218 | 1.336086 | 0.981155 |
| 2 | 43347600 | 43349199 | 15.9875 | 28.156 | 36.60456 | 16.20351 | 0.011945 | 0.579478 | 0.981165 |
| 17 | 30893400 | 30901199 | 67.32734 | 105.7618 | 163.632 | 67.87219 | -0.3093 | 2.55291 | 0.981167 |
| 3 | 122240400 | 122243599 | 36.70691 | 59.44699 | 75.94627 | 32.49872 | 0.011358 | 1.65788 | 0.981173 |
| 8 | 20011400 | 20017199 | 68.21524 | 107.0433 | 161.7474 | 67.08128 | -0.27642 | 2.575083 | 0.981174 |
| 9 | 35728400 | 35732599 | 67.19423 | 106.3758 | 132.753 | 55.19337 | 0.009161 | 2.381612 | 0.981179 |
| 15 | 70321400 | 70321999 | 8.852878 | 16.79448 | 23.92639 | 9.649222 | -0.01626 | 0.017507 | 0.981214 |
| 15 | 19625200 | 19626999 | 19.91094 | 34.71857 | 34.779 | 15.30441 | 0.384992 | 0.742648 | 0.98122 |
| 15 | 95300200 | 95301399 | 11.0618 | 20.33322 | 27.73022 | 11.78174 | -0.01418 | 0.240565 | 0.98124 |
| 1 | 116634600 | 116635799 | 15.19269 | 26.86111 | 34.93282 | 15.45141 | 0.013429 | 0.526247 | 0.981241 |
| 9 | 12093400 | 12104999 | 176.8769 | 262.9067 | 311.6639 | 128.3986 | 0.009158 | 3.639764 | 0.98125 |
| 3 | 158111000 | 158112199 | 16.56406 | 29.35651 | 39.08188 | 17.27827 | -0.01867 | 0.519192 | 0.981257 |
| 1 | 15142600 | 15145799 | 49.40811 | 79.03557 | 101.0886 | 42.54757 | -0.01096 | 2.045129 | 0.981257 |
| 14 | 36899600 | 36900399 | 13.20895 | 23.67282 | 31.59439 | 13.84033 | -0.0128 | 0.412235 | 0.98126 |
| 18 | 74125200 | 74129599 | 42.88184 | 69.18082 | 89.39468 | 38.09326 | -0.01109 | 1.771843 | 0.98128 |
| 8 | 71120600 | 71122199 | 13.89952 | 24.87281 | 32.57524 | 14.28821 | 0.013031 | 0.42912 | 0.981294 |
| 16 | 17400800 | 17402599 | 13.89952 | 24.87281 | 32.57524 | 14.28821 | 0.013031 | 0.42912 | 0.981294 |
| X | 77793200 | 77795599 | 22.53641 | 38.98473 | 51.77475 | 22.71472 | -0.02178 | 0.873824 | 0.981294 |
| 4 | 53339400 | 53343399 | 44.64219 | 71.79923 | 91.07457 | 38.64273 | 0.010223 | 1.868021 | 0.981305 |
| X | 103765000 | 103765599 | 11.5421 | 21.14725 | 28.19228 | 12.03114 | 0.013756 | 0.263655 | 0.981328 |
| 8 | 105825800 | 105828599 | 28.87183 | 47.73329 | 62.6455 | 27.0758 | -0.01047 | 1.335982 | 0.981402 |
| 7 | 26667200 | 26668199 | 20.023 | 34.41053 | 45.31851 | 19.87826 | -0.01162 | 0.91158 | 0.981428 |
| 5 | 52196200 | 52198399 | 17.20377 | 30.00476 | 38.94788 | 17.14131 | 0.012847 | 0.73165 | 0.981447 |

| | | | | | | | | | |
|---|---|---|---|---|---|---|---|---|---|
| 13 | 29121200 | 29124199 | 25.93182 | 43.38624 | 56.12146 | 24.4488 | 0.010963 | 1.187524 | 0.981458 |
| 3 | 120547600 | 120548599 | 14.56876 | 26.69542 | 26.78532 | 11.4291 | 0.420866 | 0.281131 | 0.981462 |
| 1 | 15595600 | 15600799 | 57.46062 | 91.63576 | 115.1125 | 48.24126 | 0.010155 | 2.153931 | 0.981473 |
| 10 | 99115000 | 99116799 | 17.84235 | 31.11857 | 40.43621 | 17.79637 | 0.011265 | 0.736696 | 0.981474 |
| 5 | 14930800 | 14934399 | 35.6574 | 57.99441 | 75.48641 | 32.39652 | -0.01001 | 1.572061 | 0.981488 |
| 6 | 38994400 | 38996999 | 18.94551 | 32.84159 | 42.63236 | 18.70495 | 0.011613 | 0.824397 | 0.981497 |
| 18 | 85844200 | 85847399 | 38.48568 | 62.16223 | 79.2023 | 33.80172 | 0.01122 | 1.726742 | 0.981535 |
| 9 | 113454400 | 113456999 | 26.37168 | 43.89088 | 57.66166 | 25.03649 | -0.01117 | 1.248236 | 0.981542 |
| 7 | 18185000 | 18187799 | 30.01643 | 49.51126 | 64.94305 | 28.02745 | -0.01016 | 1.367186 | 0.981553 |
| 13 | 10541600 | 10542199 | 9.22828 | 17.49642 | 24.37776 | 9.843396 | 0.013416 | 0.02363 | 0.981554 |
| X | 125859000 | 125861999 | 26.14783 | 44.7678 | 57.39519 | 25.16894 | 0.025983 | 0.984501 | 0.981561 |
| 3 | 11032600 | 11033999 | 10.127 | 18.95073 | 25.83937 | 10.69106 | 0.015309 | 0.124045 | 0.981565 |
| 16 | 20596600 | 20597799 | 13.07257 | 23.61046 | 31.11563 | 13.52172 | 0.013063 | 0.364511 | 0.981604 |
| X | 105588400 | 105589999 | 16.95765 | 30.18121 | 39.14143 | 17.24692 | 0.022372 | 0.494826 | 0.981604 |
| 9 | 13003800 | 13004999 | 19.97345 | 34.48071 | 44.7484 | 19.63089 | 0.010779 | 0.869402 | 0.981652 |
| 18 | 14855000 | 14856399 | 12.77105 | 23.36369 | 31.7016 | 13.5565 | -0.01408 | 0.28736 | 0.981662 |
| 2 | 53402200 | 53403399 | 13.45704 | 24.27871 | 32.53367 | 14.15699 | -0.01239 | 0.374022 | 0.981676 |
| 14 | 46844400 | 46846399 | 21.56516 | 36.99746 | 47.93722 | 21.0249 | 0.01123 | 0.93467 | 0.981702 |
| 3 | 85869600 | 85872399 | 51.78623 | 82.85432 | 105.8706 | 44.57142 | -0.0093 | 2.045201 | 0.981723 |
| 6 | 26750000 | 26754599 | 62.93466 | 99.8165 | 124.7655 | 51.94455 | 0.009245 | 2.32884 | 0.981748 |
| 12 | 69967200 | 69968399 | 9.493866 | 17.79178 | 24.87441 | 10.26651 | -0.01754 | 0.106899 | 0.981759 |
| 3 | 131099400 | 131107599 | 119.0674 | 181.3907 | 222.3942 | 92.1822 | -0.00835 | 3.149312 | 0.981779 |
| 16 | 90348400 | 90352599 | 60.20109 | 95.69327 | 119.9174 | 50.03723 | 0.008673 | 2.259707 | 0.981782 |
| 4 | 146198200 | 146201399 | 35.08011 | 56.335 | 99.99477 | 42.3339 | -0.47609 | 1.908918 | 0.981784 |
| 14 | 55930000 | 55933199 | 31.82843 | 52.08462 | 68.11613 | 29.2987 | -0.01059 | 1.471349 | 0.981794 |
| 4 | 135818400 | 135820999 | 28.70127 | 47.62605 | 61.61861 | 26.69965 | 0.010474 | 1.292686 | 0.981812 |
| X | 95576400 | 95577999 | 16.58001 | 29.62383 | 39.7606 | 17.46252 | -0.02375 | 0.468722 | 0.981818 |
| 14 | 100892400 | 100892799 | 5.939017 | 11.58624 | 18.08123 | 6.464282 | 0.014146 | -0.44386 | 0.981828 |
| 3 | 12532800 | 12533999 | 14.97159 | 26.86989 | 26.89156 | 11.76611 | 0.401126 | 0.404479 | 0.981834 |
| 9 | 119513400 | 119516599 | 29.70242 | 49.0715 | 64.38526 | 27.81406 | -0.01019 | 1.347737 | 0.98185 |
| 17 | 61836400 | 61837599 | 17.63415 | 30.66875 | 40.52641 | 17.80444 | -0.01327 | 0.774867 | 0.981851 |
| 11 | 22984600 | 22990599 | 67.79782 | 107.2602 | 135.4763 | 56.30673 | -0.00894 | 2.397273 | 0.981861 |
| 5 | 18491400 | 18493999 | 26.28821 | 44.00835 | 57.78628 | 25.18339 | -0.01062 | 1.18435 | 0.981878 |
| 4 | 38870000 | 38872599 | 36.38819 | 58.93267 | 76.46439 | 32.72119 | -0.01089 | 1.657056 | 0.981891 |
| 3 | 129157800 | 129159199 | 17.24625 | 30.05811 | 39.00296 | 17.15812 | 0.013344 | 0.742644 | 0.981902 |
| 5 | 151194600 | 151195399 | 9.912638 | 18.72058 | 25.9133 | 10.55033 | 0.012395 | 0.061985 | 0.981903 |
| 16 | 47921000 | 47924799 | 43.48501 | 70.19243 | 89.28687 | 38.07428 | 0.012443 | 1.756004 | 0.981912 |
| 2 | 18970000 | 18972199 | 16.26949 | 28.57804 | 37.14502 | 16.43973 | 0.01114 | 0.613183 | 0.981918 |
| 3 | 91619800 | 91624199 | 66.03484 | 104.5672 | 130.5753 | 54.29504 | 0.008586 | 2.374289 | 0.981922 |
| 2 | 71770600 | 71780399 | 145.5789 | 218.5793 | 264.7519 | 109.2681 | -0.00865 | 3.447463 | 0.981929 |
| X | 59477800 | 59479399 | 17.5486 | 30.98138 | 40.13537 | 17.75774 | 0.018311 | 0.56838 | 0.981953 |
| 1 | 10285200 | 10287799 | 28.32795 | 47.48978 | 49.66703 | 21.66423 | 0.316572 | 1.168632 | 0.981993 |
| 16 | 55965200 | 55966999 | 16.31428 | 28.63772 | 37.21402 | 16.46104 | 0.011561 | 0.625144 | 0.982014 |
| 7 | 120749000 | 120750399 | 18.26869 | 31.8427 | 42.03116 | 18.49122 | -0.01127 | 0.751375 | 0.982022 |
| 3 | 52329000 | 52332599 | 30.80753 | 51.29899 | 53.55462 | 23.26062 | 0.319326 | 1.25088 | 0.982043 |
| 3 | 104488600 | 104489799 | 14.63061 | 26.00995 | 33.96906 | 14.98217 | 0.011355 | 0.47564 | 0.982053 |
| 9 | 46594200 | 46598199 | 33.85235 | 55.22591 | 72.02081 | 30.94338 | -0.00966 | 1.523335 | 0.982055 |

| | | | | | | | | | |
|---|---|---|---|---|---|---|---|---|---|
| 1 | 18008400 | 18015799 | 98.93991 | 152.5386 | 186.4733 | 77.45784 | 0.009235 | 2.903114 | 0.982082 |
| 2 | 68736800 | 68737599 | 9.200593 | 17.68162 | 25.91056 | 10.12478 | -0.01629 | -0.07996 | 0.982085 |
| 2 | 80432400 | 80436599 | 40.85391 | 65.99177 | 84.12711 | 35.90657 | 0.010483 | 1.729097 | 0.982114 |
| 8 | 89728200 | 89730199 | 17.69076 | 30.80291 | 40.00303 | 17.58732 | 0.011911 | 0.7592 | 0.982116 |
| 5 | 136930400 | 136934799 | 41.64413 | 67.11945 | 86.66806 | 36.8856 | -0.01165 | 1.794792 | 0.982129 |
| 16 | 9704200 | 9705399 | 9.485289 | 17.96534 | 25.45356 | 10.30003 | -0.01292 | 0.038217 | 0.982163 |
| 7 | 121026800 | 121027999 | 16.3239 | 28.61846 | 37.79273 | 16.69921 | -0.01152 | 0.646647 | 0.982204 |
| 8 | 48066000 | 48074199 | 89.74278 | 139.4495 | 171.6008 | 71.25057 | 0.008447 | 2.758631 | 0.982216 |
| 2 | 170634800 | 170643999 | 112.3791 | 171.954 | 211.5967 | 87.7983 | -0.00862 | 3.057947 | 0.982225 |
| 11 | 90750600 | 90751199 | 7.096057 | 13.70963 | 20.46526 | 7.861308 | -0.01867 | -0.17257 | 0.982287 |
| 6 | 144292800 | 144293799 | 12.1875 | 22.28354 | 30.17177 | 12.91198 | -0.01154 | 0.284287 | 0.982288 |
| Y | 21022400 | 21026999 | 20.49685 | 34.61177 | 43.96542 | 19.23234 | 0.037403 | 1.067064 | 0.982299 |
| 12 | 42921600 | 42922799 | 15.17964 | 26.85452 | 35.54376 | 15.71971 | -0.01123 | 0.523431 | 0.982309 |
| 3 | 145893400 | 145896799 | 31.82599 | 52.14085 | 68.19254 | 29.34541 | -0.00986 | 1.456679 | 0.982328 |
| 5 | 17559800 | 17563199 | 41.80944 | 67.29044 | 86.67924 | 36.81523 | -0.01062 | 1.842641 | 0.98233 |
| 18 | 32602600 | 32606199 | 46.33882 | 74.42737 | 95.51826 | 40.44868 | -0.00901 | 1.915114 | 0.982383 |
| 6 | 79097000 | 79098599 | 18.94163 | 32.76219 | 43.22873 | 18.96533 | -0.01229 | 0.847944 | 0.982389 |
| 14 | 53296400 | 53297999 | 21.72754 | 36.87565 | 48.50913 | 21.25247 | -0.01258 | 1.036181 | 0.982389 |
| 5 | 151436000 | 151442799 | 125.0374 | 190.8743 | 193.9814 | 80.44546 | 0.264486 | 3.108306 | 0.982398 |
| 6 | 141717400 | 141718999 | 17.6562 | 30.99309 | 40.22007 | 17.79062 | 0.013416 | 0.637611 | 0.9824 |
| 5 | 65106200 | 65106999 | 11.15664 | 20.59859 | 27.6967 | 11.69678 | 0.012196 | 0.215187 | 0.982404 |
| 8 | 47932800 | 47933999 | 15.75189 | 27.60339 | 35.8057 | 15.81528 | 0.014134 | 0.645857 | 0.982409 |
| 18 | 45225200 | 45225799 | 7.212436 | 13.96323 | 21.02146 | 7.997562 | -0.0182 | -0.204 | 0.982414 |
| 10 | 94202600 | 94208399 | 65.77887 | 103.2145 | 159.0819 | 65.97535 | -0.305 | 2.574722 | 0.982419 |
| Y | 2072800 | 2073999 | 11.5212 | 20.71434 | 26.83954 | 11.72569 | 0.032342 | 0.367735 | 0.982423 |
| 10 | 126331000 | 126333999 | 22.95529 | 38.9654 | 51.16851 | 22.41863 | -0.01019 | 1.040424 | 0.982438 |
| 7 | 43549600 | 43551799 | 18.80791 | 32.73402 | 42.54396 | 18.6994 | 0.010627 | 0.773676 | 0.98244 |
| 15 | 53824200 | 53824799 | 8.29491 | 15.97124 | 23.0687 | 8.965513 | 0.012146 | -0.11323 | 0.982456 |
| 12 | 67996800 | 67997999 | 15.49613 | 27.12602 | 35.95308 | 15.86597 | -0.01664 | 0.663613 | 0.982459 |
| 4 | 130793800 | 130797199 | 19.43468 | 33.63667 | 43.67782 | 19.16273 | 0.010688 | 0.841945 | 0.982471 |
| 13 | 53385600 | 53389399 | 24.51486 | 40.88057 | 53.85272 | 23.4117 | -0.01509 | 1.216831 | 0.982496 |
| 10 | 121910600 | 121913799 | 31.47888 | 51.80108 | 67.8732 | 29.26276 | -0.00999 | 1.401222 | 0.982521 |
| 13 | 19438000 | 19438599 | 6.536395 | 12.70655 | 19.55077 | 7.259884 | -0.01504 | -0.29531 | 0.982546 |
| 17 | 65660600 | 65670399 | 170.078 | 253.3537 | 304.6269 | 125.5274 | -0.00838 | 3.60032 | 0.982607 |
| 1 | 33266600 | 33268199 | 17.05321 | 30.07497 | 39.04429 | 17.27866 | 0.014529 | 0.572739 | 0.982659 |
| 12 | 98283600 | 98285799 | 17.05321 | 30.07497 | 39.04429 | 17.27866 | 0.014529 | 0.572739 | 0.982659 |
| 10 | 121072200 | 121080799 | 67.92725 | 107.4028 | 133.9578 | 55.65931 | 0.008546 | 2.410202 | 0.982675 |
| 1 | 114203000 | 114205599 | 21.37667 | 36.98927 | 47.93576 | 21.03066 | 0.013487 | 0.859833 | 0.982693 |
| 4 | 78591000 | 78591399 | 7.412324 | 14.36132 | 21.61458 | 8.193531 | -0.01219 | -0.2091 | 0.982705 |
| 13 | 111309400 | 111309999 | 7.412324 | 14.36132 | 21.61458 | 8.193531 | -0.01219 | -0.2091 | 0.982705 |
| 1 | 30612000 | 30616999 | 59.01368 | 93.30996 | 143.1668 | 59.48588 | -0.28986 | 2.406123 | 0.982706 |
| 17 | 31340400 | 31345199 | 43.59298 | 70.04796 | 90.01735 | 38.1438 | -0.0101 | 1.896546 | 0.982726 |
| 15 | 16815400 | 16816399 | 10.97542 | 20.45324 | 27.79154 | 11.57996 | 0.011535 | 0.159775 | 0.982728 |
| 13 | 37220800 | 37222999 | 34.08082 | 55.35312 | 70.85148 | 30.38298 | 0.012508 | 1.592385 | 0.982748 |
| 10 | 100487400 | 100491199 | 33.55097 | 55.23654 | 71.20062 | 30.70337 | 0.013743 | 1.401532 | 0.982757 |
| 13 | 51545800 | 51551799 | 61.02064 | 96.91892 | 122.7893 | 51.19481 | -0.0082 | 2.283994 | 0.982759 |
| 2 | 98462400 | 98466199 | 26.60772 | 44.67918 | 57.72881 | 25.20677 | 0.012581 | 1.15611 | 0.982777 |

| | | | | | | | | | |
|---|---|---|---|---|---|---|---|---|---|
| 5 | 97635800 | 97639199 | 59.52789 | 94.53386 | 118.3261 | 49.32657 | 0.00904 | 2.286007 | 0.982793 |
| 1 | 191652800 | 191654399 | 26.74077 | 44.56332 | 57.66521 | 25.05879 | 0.010614 | 1.2364 | 0.982794 |
| 8 | 72637400 | 72660199 | 339.0182 | 478.5503 | 547.1366 | 226.5316 | -0.01128 | 4.442168 | 0.982803 |
| 17 | 21098800 | 21101799 | 42.12869 | 68.95003 | 63.34529 | 27.24804 | 0.497689 | 1.491976 | 0.982811 |
| 12 | 72630400 | 72633599 | 48.27874 | 77.32354 | 98.92913 | 41.71274 | -0.00941 | 2.005958 | 0.982812 |
| 15 | 64370400 | 64371599 | 14.07823 | 25.46459 | 33.57207 | 14.56212 | 0.013824 | 0.365243 | 0.982822 |
| 15 | 35315200 | 35317599 | 17.48451 | 30.35231 | 51.73026 | 22.70022 | -0.37916 | 0.816849 | 0.982823 |
| X | 54347400 | 54349799 | 28.87927 | 48.45028 | 62.41806 | 27.23895 | 0.016725 | 1.175485 | 0.982836 |
| 15 | 71038400 | 71038999 | 6.863332 | 13.35647 | 20.71781 | 7.626707 | -0.01494 | -0.31627 | 0.982881 |
| 11 | 109062200 | 109064199 | 18.77471 | 32.65586 | 43.0786 | 18.92697 | -0.01088 | 0.782376 | 0.9829 |
| 7 | 19553600 | 19554999 | 13.94458 | 24.82721 | 32.9809 | 14.52813 | -0.01186 | 0.459175 | 0.982913 |
| 4 | 70258200 | 70258999 | 10.49873 | 19.62994 | 26.79751 | 11.1032 | 0.011436 | 0.132386 | 0.982962 |
| 3 | 87794600 | 87796999 | 28.12879 | 46.8012 | 60.59278 | 26.30372 | 0.009745 | 1.258176 | 0.982978 |
| 3 | 48275400 | 48275999 | 10.27827 | 18.51397 | 33.82054 | 14.74651 | -0.45813 | 0.365719 | 0.982985 |
| 9 | 82491000 | 82493199 | 19.11207 | 32.95302 | 42.69587 | 18.72975 | 0.012773 | 0.873165 | 0.983019 |
| 3 | 32294600 | 32295399 | 15.38354 | 27.08834 | 35.20228 | 15.58324 | 0.012607 | 0.576335 | 0.983032 |
| 8 | 118494200 | 118494999 | 11.98077 | 21.87983 | 29.12483 | 12.48227 | 0.011465 | 0.286418 | 0.983035 |
| 5 | 107832800 | 107836199 | 35.98019 | 58.39538 | 75.83262 | 32.50153 | -0.00933 | 1.611594 | 0.983039 |
| 9 | 24184800 | 24188199 | 48.43658 | 77.93147 | 81.54723 | 34.61937 | 0.28801 | 1.859191 | 0.983058 |
| 10 | 83239000 | 83256999 | 540.9638 | 728.9481 | 784.3966 | 327.1368 | 0.008586 | 5.157241 | 0.983063 |
| 2 | 17939200 | 17941799 | 32.38081 | 52.88983 | 69.04475 | 29.67794 | -0.00986 | 1.501148 | 0.983063 |
| 9 | 58209000 | 58209799 | 9.858985 | 18.50129 | 25.81072 | 10.62834 | -0.01243 | 0.10091 | 0.983072 |
| 15 | 6700800 | 6702399 | 19.51463 | 33.89705 | 44.70425 | 19.62455 | -0.01063 | 0.808555 | 0.98308 |
| 6 | 12748000 | 12749599 | 11.59453 | 21.11057 | 28.5759 | 12.29339 | -0.01629 | 0.297746 | 0.983088 |
| 5 | 148975600 | 148980399 | 51.05903 | 81.74192 | 103.2059 | 43.48808 | 0.008924 | 2.023574 | 0.983121 |
| 6 | 65177000 | 65179199 | 27.13035 | 45.70858 | 58.9988 | 25.79067 | 0.014088 | 1.127149 | 0.983144 |
| 5 | 133487600 | 133490399 | 33.86955 | 55.84149 | 58.53004 | 25.25492 | 0.312227 | 1.377551 | 0.983158 |
| 15 | 35135200 | 35137199 | 23.55343 | 39.78675 | 52.23545 | 22.85236 | -0.01006 | 1.087402 | 0.983165 |
| 15 | 9227200 | 9227799 | 9.105747 | 17.41236 | 25.09951 | 9.949192 | -0.01164 | -0.03485 | 0.983195 |
| 14 | 56149000 | 56149599 | 7.91412 | 15.24765 | 22.0198 | 8.542412 | 0.015739 | -0.12288 | 0.983204 |
| X | 94487000 | 94489399 | 22.40195 | 38.3687 | 49.5422 | 21.7259 | 0.016076 | 0.960758 | 0.983246 |
| 4 | 156389400 | 156391799 | 27.90603 | 45.52717 | 76.91106 | 33.04497 | -0.38199 | 1.512813 | 0.983258 |
| 11 | 70639600 | 70640199 | 7.378893 | 14.31411 | 21.6703 | 8.167673 | -0.01319 | -0.23044 | 0.983268 |
| 4 | 73970800 | 73973199 | 14.47804 | 25.26596 | 43.67372 | 19.22842 | -0.39818 | 0.718568 | 0.983316 |
| 6 | 57443800 | 57446399 | 24.74149 | 41.65838 | 53.92186 | 23.56697 | 0.010046 | 1.118434 | 0.983329 |
| 13 | 16781400 | 16783399 | 20.37176 | 33.92838 | 62.58842 | 27.18757 | -0.49907 | 1.229911 | 0.983332 |
| 9 | 11296200 | 11297599 | 18.53269 | 32.3002 | 42.00236 | 18.4776 | 0.010117 | 0.75271 | 0.983344 |
| 16 | 25545600 | 25546999 | 12.51521 | 22.50878 | 30.10739 | 13.14977 | -0.01307 | 0.381316 | 0.983349 |
| 13 | 119358000 | 119358599 | 11.48434 | 21.08703 | 28.6683 | 12.19829 | -0.01149 | 0.250759 | 0.983401 |
| 9 | 30673400 | 30675999 | 23.37178 | 39.76563 | 52.24554 | 22.89919 | -0.0105 | 1.023805 | 0.983436 |
| 8 | 51368200 | 51370399 | 26.95043 | 44.67191 | 57.70202 | 24.98372 | 0.012683 | 1.294597 | 0.983465 |
| 2 | 3903200 | 3904199 | 13.3564 | 24.0118 | 32.05411 | 14.00487 | -0.0106 | 0.394432 | 0.983466 |
| 13 | 111807600 | 111813999 | 65.61545 | 104.1143 | 130.1994 | 54.23087 | 0.009151 | 2.317608 | 0.983481 |
| 10 | 96968400 | 96969799 | 14.43552 | 25.89236 | 33.98046 | 14.87525 | 0.011714 | 0.418544 | 0.983495 |
| 1 | 107021200 | 107021799 | 7.540253 | 14.60832 | 21.61127 | 8.194536 | 0.011745 | -0.20778 | 0.983498 |
| 2 | 88042200 | 88044199 | 26.09561 | 43.5986 | 57.22695 | 24.90817 | -0.01005 | 1.203786 | 0.983553 |
| 7 | 61274600 | 61277799 | 33.39781 | 55.1443 | 71.0893 | 30.69772 | 0.015043 | 1.369353 | 0.983553 |

| | | | | | | | | | |
|---|---|---|---|---|---|---|---|---|---|
| 10 | 27657000 | 27658599 | 15.89673 | 28.0236 | 37.03078 | 16.38897 | -0.01106 | 0.567045 | 0.983587 |
| 16 | 77913800 | 77917599 | 47.12358 | 75.55629 | 95.55854 | 40.35885 | 0.009092 | 1.969352 | 0.983592 |
| 5 | 109457600 | 109462399 | 83.75794 | 131.0206 | 162.0307 | 67.21932 | 0.008767 | 2.639404 | 0.983593 |
| 1 | 136332000 | 136334199 | 23.47974 | 39.92419 | 51.71173 | 22.66407 | 0.009812 | 1.028446 | 0.983626 |
| 6 | 26932200 | 26933399 | 15.45362 | 27.49595 | 36.57655 | 16.1209 | -0.01421 | 0.48317 | 0.983646 |
| 17 | 39966400 | 39968999 | 36.70233 | 59.90746 | 78.31391 | 33.65377 | -0.01261 | 1.526174 | 0.983664 |
| 16 | 69965200 | 69967799 | 23.65191 | 40.7 | 42.03687 | 18.43955 | 0.337898 | 0.910152 | 0.983666 |
| 13 | 82268400 | 82269799 | 16.80066 | 29.31026 | 38.01961 | 16.73584 | 0.014069 | 0.725908 | 0.983695 |
| 10 | 45471600 | 45474199 | 28.28971 | 47.59605 | 61.33442 | 26.80043 | 0.016348 | 1.144615 | 0.983717 |
| 11 | 25920200 | 25920999 | 8.91606 | 16.9901 | 23.9268 | 9.558605 | 0.011678 | -0.01523 | 0.983735 |
| 15 | 87120400 | 87121999 | 15.55185 | 27.33441 | 36.09456 | 15.98318 | -0.01164 | 0.595823 | 0.983748 |
| 7 | 93594800 | 93595599 | 9.531452 | 18.34421 | 19.78321 | 7.698134 | 0.428879 | -0.10425 | 0.983764 |
| 16 | 32740600 | 32744399 | 54.64799 | 87.22517 | 109.8238 | 46.0802 | 0.008201 | 2.122853 | 0.983771 |
| 13 | 75754400 | 75756199 | 22.7035 | 38.68058 | 50.09993 | 21.96115 | 0.010205 | 1.003913 | 0.98378 |
| 5 | 12719000 | 12720999 | 22.24152 | 37.7205 | 49.55917 | 21.70771 | -0.01084 | 1.045914 | 0.983783 |
| 13 | 19181600 | 19185399 | 117.4283 | 178.464 | 215.7472 | 89.36267 | 0.00912 | 3.197418 | 0.9838 |
| 5 | 149016000 | 149017599 | 15.79 | 27.76472 | 36.63079 | 16.2202 | -0.01008 | 0.59479 | 0.983803 |
| 4 | 60628000 | 60629799 | 18.14945 | 31.53075 | 41.58378 | 18.25654 | -0.01059 | 0.79485 | 0.983822 |
| 11 | 37374400 | 37376999 | 27.21309 | 45.21029 | 59.31422 | 25.72224 | -0.00942 | 1.27293 | 0.983831 |
| 3 | 22620600 | 22622799 | 33.37489 | 54.81657 | 70.74461 | 30.47616 | 0.010723 | 1.432573 | 0.983838 |
| 2 | 160478000 | 160483599 | 104.4656 | 159.8857 | 196.838 | 81.67918 | -0.00899 | 3.04951 | 0.983841 |
| 14 | 24574800 | 24576999 | 14.08561 | 25.15339 | 33.45572 | 14.70019 | -0.0112 | 0.442038 | 0.983853 |
| 6 | 10396200 | 10404399 | 194.5308 | 285.297 | 333.9037 | 137.5309 | 0.009507 | 3.863636 | 0.983856 |
| 2 | 169865600 | 169867599 | 20.80431 | 35.47347 | 45.88283 | 20.1139 | 0.012499 | 0.985787 | 0.983862 |
| 7 | 33718200 | 33733599 | 425.6628 | 584.5047 | 648.6815 | 269.7462 | -0.00848 | 4.85927 | 0.983874 |
| 17 | 30021800 | 30026599 | 58.07338 | 92.46661 | 117.4158 | 49.09897 | -0.00783 | 2.204888 | 0.983901 |
| 7 | 143514000 | 143516999 | 31.74244 | 52.05857 | 68.08814 | 29.31349 | -0.00928 | 1.445502 | 0.983902 |
| X | 149951400 | 149953399 | 17.40038 | 30.50986 | 40.31991 | 17.81792 | -0.01258 | 0.655647 | 0.983908 |
| 4 | 73722400 | 73722999 | 9.962597 | 18.80793 | 26.45591 | 10.77973 | -0.01147 | 0.06732 | 0.983943 |
| 3 | 116960800 | 116965599 | 40.08012 | 65.09547 | 83.26228 | 35.70443 | 0.013367 | 1.609029 | 0.983954 |
| 15 | 7945400 | 7948999 | 24.45985 | 41.19346 | 53.35327 | 23.32006 | 0.009206 | 1.115938 | 0.983996 |
| 17 | 26195600 | 26197599 | 14.57002 | 25.7992 | 33.62078 | 14.86658 | 0.011733 | 0.507495 | 0.984019 |
| X | 101929800 | 101930799 | 12.62705 | 23.71244 | 24.10486 | 9.91412 | 0.442549 | 0.118338 | 0.984038 |
| 2 | 55510800 | 55511399 | 7.713761 | 14.89112 | 22.041 | 8.49833 | -0.01165 | -0.14706 | 0.98404 |
| 5 | 81525200 | 81527599 | 27.90429 | 46.21882 | 59.80416 | 25.88069 | 0.010174 | 1.310687 | 0.98407 |
| 13 | 4791600 | 4792199 | 7.575514 | 14.65896 | 21.57057 | 8.225289 | 0.012124 | -0.18677 | 0.984081 |
| 11 | 77028000 | 77028999 | 14.70849 | 25.89756 | 34.25857 | 15.16573 | -0.01294 | 0.571016 | 0.984099 |
| 15 | 19649800 | 19650399 | 14.48464 | 25.67744 | 33.50649 | 14.81264 | 0.010284 | 0.492338 | 0.984122 |
| X | 163673000 | 163678199 | 50.82033 | 81.80112 | 82.60819 | 35.09114 | 0.339926 | 1.851019 | 0.984141 |
| 3 | 154683600 | 154685199 | 18.81755 | 32.51087 | 42.17933 | 18.5043 | 0.0115 | 0.853754 | 0.984177 |
| 15 | 22320200 | 22322999 | 30.67932 | 50.36581 | 65.89206 | 28.38014 | -0.00912 | 1.427356 | 0.984192 |
| 9 | 8795600 | 8796799 | 26.03279 | 43.5849 | 57.17029 | 24.91643 | -0.00905 | 1.183518 | 0.9842 |
| 1 | 103081800 | 103085599 | 41.86458 | 67.89483 | 70.69552 | 30.28307 | 0.307553 | 1.635705 | 0.984221 |
| 11 | 3227200 | 3236399 | 99.26079 | 153.1181 | 187.5037 | 77.88561 | 0.007375 | 2.894405 | 0.984239 |
| 12 | 63674600 | 63675799 | 11.37544 | 20.98662 | 28.22806 | 11.93351 | 0.010772 | 0.223576 | 0.984245 |
| 5 | 106307200 | 106308999 | 20.00394 | 34.43749 | 45.29997 | 19.87118 | -0.00942 | 0.895545 | 0.984253 |
| 12 | 60487800 | 60490399 | 26.9006 | 44.15093 | 72.62962 | 31.27863 | -0.33873 | 1.42811 | 0.984261 |

| | | | | | | | | | |
|---|---|---|---|---|---|---|---|---|---|
| 4 | 4463800 | 4465799 | 19.56759 | 33.97487 | 44.16168 | 19.38129 | 0.010057 | 0.812246 | 0.98427 |
| 1 | 63044600 | 63049199 | 34.16818 | 55.80767 | 72.79274 | 31.28864 | -0.00873 | 1.507256 | 0.984271 |
| 3 | 15605200 | 15605799 | 9.999087 | 19.00145 | 26.61355 | 10.69407 | 0.011787 | 0.020192 | 0.984294 |
| 6 | 89738600 | 89743799 | 137.7368 | 207.7967 | 249.9948 | 103.3126 | 0.007189 | 3.352578 | 0.984296 |
| 8 | 110702200 | 110703599 | 15.72677 | 28.55246 | 37.51007 | 16.19475 | 0.023283 | 0.364689 | 0.9843 |
| 16 | 47001200 | 47008799 | 78.92599 | 124.3282 | 154.2638 | 64.00966 | 0.011791 | 2.505691 | 0.984332 |
| 12 | 48478400 | 48478999 | 9.967457 | 18.62138 | 25.34169 | 10.51453 | 0.014895 | 0.132037 | 0.984345 |
| 13 | 27942200 | 27942599 | 6.084425 | 11.8532 | 18.54104 | 6.752137 | -0.01331 | -0.37536 | 0.984354 |
| 4 | 122406800 | 122408999 | 17.90595 | 31.30264 | 41.34181 | 18.22345 | -0.01199 | 0.702402 | 0.984354 |
| 1 | 176727000 | 176727599 | 8.064875 | 15.45449 | 21.96824 | 8.65546 | 0.017432 | -0.07134 | 0.98437 |
| 6 | 137809000 | 137812599 | 30.34772 | 50.00178 | 65.48501 | 28.24797 | -0.00857 | 1.382098 | 0.984385 |
| X | 109076000 | 109076799 | 14.61166 | 26.05632 | 34.0776 | 14.9911 | 0.01215 | 0.459313 | 0.984411 |
| 18 | 28729200 | 28729799 | 8.338909 | 15.91372 | 22.42133 | 8.929323 | 0.015481 | -0.03373 | 0.984417 |
| 17 | 50555000 | 50561199 | 131.8286 | 199.0346 | 242.1164 | 100.0808 | -0.00782 | 3.335144 | 0.98442 |
| 4 | 24288800 | 24292599 | 54.43943 | 86.90983 | 109.4922 | 45.95418 | 0.007652 | 2.115732 | 0.984424 |
| 13 | 11626400 | 11631799 | 92.50739 | 143.2751 | 176.017 | 73.11273 | 0.007174 | 2.817242 | 0.984426 |
| 15 | 100271000 | 100273999 | 31.90213 | 52.27774 | 67.48557 | 29.04403 | 0.00909 | 1.455486 | 0.984488 |
| 2 | 14770800 | 14772199 | 14.28643 | 25.30786 | 33.50469 | 14.81398 | -0.01092 | 0.497983 | 0.98449 |
| 1 | 105465200 | 105468399 | 71.27378 | 112.6575 | 142.2522 | 59.09588 | -0.00956 | 2.424105 | 0.984515 |
| 1 | 70736200 | 70738399 | 24.21622 | 40.75667 | 52.77812 | 23.06412 | 0.009508 | 1.121914 | 0.984529 |
| X | 109617200 | 109619799 | 21.66352 | 37.7308 | 50.10474 | 22.03242 | -0.02022 | 0.789349 | 0.98454 |
| 6 | 113590200 | 113592199 | 28.7165 | 47.50956 | 61.51555 | 26.60022 | 0.009052 | 1.327076 | 0.984551 |
| 16 | 48549400 | 48552199 | 24.98728 | 42.01968 | 54.4382 | 23.78362 | 0.008745 | 1.132063 | 0.984565 |
| 6 | 36237600 | 36239999 | 26.82429 | 45.17257 | 58.3978 | 25.52446 | 0.011846 | 1.129893 | 0.984599 |
| 15 | 14888200 | 14888999 | 11.07861 | 20.5491 | 28.22487 | 11.84526 | -0.01209 | 0.18831 | 0.984623 |
| 10 | 3799400 | 3801199 | 17.4064 | 30.6229 | 29.52789 | 13.07365 | 0.440259 | 0.585276 | 0.984696 |
| 17 | 33296400 | 33299199 | 19.64798 | 33.98787 | 44.73868 | 19.62784 | -0.009 | 0.84975 | 0.984728 |
| 10 | 89708400 | 89710399 | 15.46615 | 26.63365 | 48.98788 | 21.48911 | -0.49083 | 0.878052 | 0.984732 |
| 3 | 28928000 | 28943399 | 363.0988 | 507.1612 | 566.3796 | 234.8912 | 0.007304 | 4.597989 | 0.98474 |
| 16 | 15209200 | 15212999 | 61.47312 | 97.60594 | 123.5601 | 51.49946 | -0.00751 | 2.293388 | 0.984744 |
| 1 | 99339200 | 99341199 | 19.33085 | 33.92502 | 43.89199 | 19.41111 | 0.01766 | 0.655573 | 0.984747 |
| 6 | 122084000 | 122098599 | 413.2875 | 570.5233 | 635.9357 | 264.224 | -0.00706 | 4.777691 | 0.984802 |
| 4 | 118534400 | 118537199 | 31.65914 | 51.38645 | 83.27389 | 35.69291 | -0.32795 | 1.605968 | 0.984803 |
| 7 | 12317600 | 12324399 | 93.96757 | 147.1284 | 142.7517 | 59.21563 | 0.359451 | 2.626406 | 0.984829 |
| 4 | 61685000 | 61689399 | 144.1179 | 215.9041 | 257.6481 | 106.2796 | 0.00994 | 3.489733 | 0.984847 |
| Y | 1962000 | 1962599 | 6.228324 | 12.09633 | 17.96229 | 6.720699 | 0.025182 | -0.28106 | 0.98488 |
| 9 | 20492000 | 20498599 | 70.46802 | 111.2872 | 138.7862 | 57.63066 | 0.007342 | 2.441578 | 0.984887 |
| 15 | 39106400 | 39108799 | 19.89139 | 34.42729 | 44.74639 | 19.63154 | 0.009442 | 0.847565 | 0.984908 |
| 7 | 143549200 | 143551999 | 18.95651 | 33.06838 | 42.99182 | 18.9235 | 0.010447 | 0.742322 | 0.984922 |
| 16 | 83301600 | 83303599 | 16.31038 | 28.81592 | 37.53186 | 16.60346 | 0.01097 | 0.548042 | 0.984939 |
| 14 | 46100200 | 46103799 | 71.98 | 112.9998 | 140.0854 | 58.1007 | 0.00929 | 2.566221 | 0.984962 |
| 13 | 79949800 | 79950799 | 19.15652 | 33.194 | 43.16571 | 18.93873 | 0.009007 | 0.830519 | 0.985012 |
| 4 | 110467600 | 110470399 | 22.86457 | 39.74937 | 39.25764 | 17.24268 | 0.404899 | 0.804786 | 0.985056 |
| 10 | 120020600 | 120022999 | 22.86457 | 39.74937 | 39.25764 | 17.24268 | 0.404899 | 0.804786 | 0.985056 |
| 10 | 18190200 | 18191999 | 24.90015 | 41.894 | 54.95057 | 24.01109 | -0.00896 | 1.129428 | 0.985075 |
| 10 | 118115000 | 118117199 | 17.29776 | 30.40822 | 40.15759 | 17.78277 | -0.01172 | 0.612188 | 0.985086 |
| 5 | 151317000 | 151317999 | 10.56183 | 19.91847 | 27.48921 | 11.22515 | 0.012286 | 0.078152 | 0.985093 |

| | | | | | | | | | |
|---|---|---|---|---|---|---|---|---|---|
| 8 | 102736800 | 102737399 | 9.064915 | 17.1914 | 24.38322 | 9.839897 | -0.0114 | 0.019844 | 0.985097 |
| 18 | 9940400 | 9940999 | 8.85188 | 16.79629 | 23.41905 | 9.440212 | 0.015248 | 0.014748 | 0.985097 |
| X | 17645400 | 17649999 | 62.8141 | 99.75484 | 126.3805 | 52.68541 | -0.00835 | 2.291785 | 0.985126 |
| 16 | 22342000 | 22343999 | 21.93615 | 37.34395 | 49.02322 | 21.48782 | -0.00922 | 1.007752 | 0.985131 |
| 13 | 75217000 | 75217599 | 10.78031 | 20.09591 | 27.32917 | 11.38129 | 0.011383 | 0.157694 | 0.985137 |
| 1 | 11050200 | 11052399 | 30.58219 | 50.72483 | 50.83978 | 22.02243 | 0.377603 | 1.291536 | 0.985142 |
| 12 | 109194800 | 109199599 | 55.6022 | 88.65935 | 112.7084 | 47.22438 | -0.00722 | 2.155423 | 0.985142 |
| 7 | 116702200 | 116704399 | 17.80416 | 31.05148 | 40.93534 | 18.01586 | -0.00952 | 0.736595 | 0.98515 |
| 13 | 10797400 | 10798999 | 19.09804 | 33.1538 | 43.12676 | 18.92501 | 0.009004 | 0.811599 | 0.985233 |
| 5 | 109454400 | 109456399 | 25.77393 | 43.58955 | 57.293 | 25.07404 | -0.01178 | 1.087901 | 0.985234 |
| 12 | 9521800 | 9523799 | 21.47705 | 36.52648 | 47.97286 | 21.02576 | -0.01006 | 1.014363 | 0.985261 |
| 4 | 10093600 | 10095599 | 29.29466 | 49.22751 | 50.19564 | 21.92527 | 0.352956 | 1.147551 | 0.985262 |
| 2 | 46727600 | 46731999 | 88.4525 | 137.2052 | 168.7413 | 70.08148 | 0.007331 | 2.786473 | 0.985262 |
| 6 | 60523400 | 60531599 | 104.8628 | 161.8178 | 197.8725 | 82.19247 | 0.009836 | 2.89128 | 0.985307 |
| 13 | 48084200 | 48085399 | 20.90152 | 35.7028 | 46.92064 | 20.5718 | -0.00992 | 0.973185 | 0.985317 |
| 14 | 108027600 | 108030199 | 34.63412 | 56.56113 | 72.90011 | 31.33326 | 0.008296 | 1.509708 | 0.985332 |
| 8 | 103591400 | 103593199 | 21.59896 | 35.59005 | 68.86812 | 29.70838 | -0.56981 | 1.362654 | 0.98534 |
| 13 | 23449400 | 23453199 | 39.42925 | 63.71039 | 81.39544 | 34.75466 | 0.007907 | 1.717485 | 0.985345 |
| 11 | 35605200 | 35608799 | 36.08264 | 58.59291 | 76.1063 | 32.63166 | -0.00897 | 1.59984 | 0.985349 |
| 13 | 9487600 | 9489999 | 23.49775 | 39.89046 | 51.70626 | 22.65497 | 0.008539 | 1.043047 | 0.985411 |
| 3 | 21526800 | 21528199 | 11.43045 | 21.52132 | 22.63617 | 9.269083 | 0.3988 | 0.087278 | 0.985411 |
| 4 | 46206200 | 46209799 | 42.75 | 68.85471 | 88.57914 | 37.66218 | -0.00748 | 1.820913 | 0.985447 |
| 14 | 45742000 | 45743599 | 13.75214 | 24.45829 | 32.46865 | 14.31544 | -0.01183 | 0.462837 | 0.985459 |
| X | 55615800 | 55616999 | 12.15849 | 22.52302 | 30.93964 | 13.0074 | -0.01359 | 0.214804 | 0.985508 |
| 4 | 50406000 | 50410799 | 100.081 | 153.8953 | 189.905 | 78.86174 | -0.00715 | 2.954521 | 0.985529 |
| 15 | 46730800 | 46732999 | 35.19186 | 57.23525 | 73.57981 | 31.57786 | 0.007834 | 1.571299 | 0.98553 |
| 18 | 85901600 | 85902599 | 16.75556 | 29.38287 | 38.72734 | 17.11593 | -0.00889 | 0.645059 | 0.98553 |
| 1 | 18007800 | 18013399 | 77.66334 | 121.7863 | 151.0674 | 62.65526 | 0.007165 | 2.590245 | 0.985541 |
| 13 | 3407800 | 3411799 | 54.93014 | 86.51803 | 149.2285 | 61.91933 | -0.46284 | 2.491298 | 0.985543 |
| 13 | 88082000 | 88083999 | 19.58984 | 34.33287 | 44.46364 | 19.64136 | 0.016207 | 0.678259 | 0.985579 |
| 6 | 101274000 | 101275999 | 21.12523 | 36.14386 | 46.895 | 20.56326 | 0.008756 | 0.957254 | 0.98558 |
| 4 | 11022400 | 11027399 | 47.36943 | 76.01807 | 96.3654 | 40.75472 | 0.007287 | 1.943842 | 0.985596 |
| 3 | 101360200 | 101362799 | 25.06164 | 41.83185 | 54.91266 | 23.88672 | -0.01002 | 1.211059 | 0.985597 |
| 8 | 23201800 | 23203199 | 11.83854 | 22.05694 | 29.95724 | 12.48691 | 0.011741 | 0.173918 | 0.985598 |
| 18 | 10156400 | 10159199 | 45.82833 | 73.85482 | 95.30946 | 40.55723 | -0.01098 | 1.812262 | 0.98563 |
| 16 | 22800400 | 22805999 | 24.45713 | 41.64587 | 40.58387 | 17.78904 | 0.419003 | 1.003826 | 0.985636 |
| 4 | 125759800 | 125763399 | 36.04676 | 58.51048 | 75.07981 | 32.18174 | 0.00806 | 1.610633 | 0.98565 |
| 4 | 33187600 | 33189599 | 15.63163 | 27.45465 | 36.20367 | 16.02165 | -0.00951 | 0.609967 | 0.985651 |
| Y | 75641600 | 75642399 | 6.232482 | 12.09432 | 18.47656 | 6.95292 | -0.0231 | -0.25896 | 0.985656 |
| 4 | 142181800 | 142186599 | 43.88715 | 70.46363 | 89.22909 | 37.76426 | 0.00967 | 1.921244 | 0.985657 |
| 1 | 23736000 | 23737399 | 17.46282 | 30.74604 | 40.6603 | 17.9998 | -0.01288 | 0.59692 | 0.985692 |
| 2 | 146830200 | 146831199 | 10.80971 | 20.51682 | 21.20079 | 8.541107 | 0.44456 | 0.024317 | 0.985701 |
| 15 | 90912800 | 90915599 | 29.75693 | 49.29059 | 64.62755 | 27.96929 | -0.00897 | 1.315869 | 0.985719 |
| 15 | 82388200 | 82394199 | 83.34771 | 130.0967 | 160.8467 | 66.74302 | 0.006717 | 2.677409 | 0.985737 |
| 1 | 66086400 | 66087999 | 18.75986 | 32.6727 | 42.50842 | 18.6918 | 0.009171 | 0.762877 | 0.985747 |
| 13 | 111873800 | 111875399 | 21.42237 | 37.33021 | 37.61338 | 16.54504 | 0.376361 | 0.76111 | 0.985769 |
| X | 57265200 | 57266199 | 9.820673 | 18.62691 | 26.01478 | 10.49565 | 0.011388 | 0.033253 | 0.98587 |

| | | | | | | | | | |
|---|---|---|---|---|---|---|---|---|---|
| 12 | 50953000 | 50955599 | 30.61041 | 50.83026 | 65.70811 | 28.48567 | 0.011765 | 1.294327 | 0.98588 |
| 5 | 105536200 | 105539399 | 51.04791 | 81.60409 | 104.0544 | 43.75445 | -0.00758 | 2.07101 | 0.985885 |
| 15 | 42737800 | 42741399 | 31.9724 | 52.38287 | 67.66601 | 29.11932 | 0.007973 | 1.455875 | 0.985886 |
| 17 | 32918800 | 32927799 | 90.90926 | 141.2658 | 173.994 | 72.24393 | 0.007166 | 2.759217 | 0.98589 |
| 2 | 14387800 | 14391799 | 93.18537 | 143.7122 | 175.8743 | 73.05506 | 0.008036 | 2.895026 | 0.985922 |
| 1 | 32953800 | 32955399 | 23.07418 | 39.36326 | 51.70582 | 22.66746 | -0.00959 | 0.998791 | 0.985946 |
| 11 | 11339000 | 11345599 | 164.3442 | 244.4471 | 293.2496 | 120.8208 | -0.00708 | 3.625138 | 0.985948 |
| 4 | 79725000 | 79726199 | 17.03986 | 29.81 | 38.8128 | 17.11906 | 0.008687 | 0.684731 | 0.985955 |
| 2 | 24125800 | 24128199 | 26.28096 | 44.22489 | 57.26718 | 25.02468 | 0.009423 | 1.132647 | 0.985965 |
| 6 | 132032200 | 132033999 | 47.28263 | 75.76611 | 95.76743 | 40.41041 | 0.008969 | 1.991077 | 0.985968 |
| 1 | 19796600 | 19796999 | 10.24092 | 19.19923 | 26.3404 | 10.86443 | 0.009813 | 0.113401 | 0.985969 |
| 17 | 87470800 | 87472799 | 19.74821 | 34.04835 | 44.2255 | 19.40074 | 0.009112 | 0.877997 | 0.985976 |
| 7 | 83553600 | 83557999 | 53.73568 | 85.72819 | 109.0319 | 45.71722 | -0.00707 | 2.134955 | 0.985997 |
| 6 | 67683600 | 67689599 | 86.56778 | 134.7424 | 166.2458 | 69.00975 | 0.006541 | 2.728131 | 0.986039 |
| 10 | 8278800 | 8282199 | 36.09047 | 58.63127 | 76.15175 | 32.66168 | -0.0083 | 1.591374 | 0.986041 |
| 12 | 36392200 | 36393599 | 24.86936 | 40.65597 | 74.27048 | 31.93421 | -0.49232 | 1.476046 | 0.986043 |
| 5 | 142062400 | 142063999 | 12.55315 | 22.57335 | 30.14805 | 13.16933 | -0.01092 | 0.383609 | 0.986058 |
| 4 | 34339400 | 34348799 | 221.6521 | 321.7248 | 373.2426 | 153.8755 | 0.007801 | 4.024784 | 0.986086 |
| 5 | 25070800 | 25072399 | 16.13896 | 28.61851 | 27.47228 | 12.14277 | 0.45105 | 0.488744 | 0.986124 |
| 1 | 88777400 | 88780599 | 12.19138 | 21.56903 | 37.76425 | 16.70158 | -0.4126 | 0.507826 | 0.986152 |
| 16 | 64842800 | 64847999 | 54.71387 | 87.49845 | 111.7173 | 47.00253 | -0.00891 | 2.063367 | 0.986173 |
| 9 | 15064600 | 15080799 | 305.4977 | 433.3613 | 491.8288 | 203.4667 | 0.00651 | 4.367449 | 0.986178 |
| 3 | 8815400 | 8816599 | 11.67341 | 21.24666 | 28.64766 | 12.32974 | -0.01082 | 0.302494 | 0.986183 |
| 13 | 78385600 | 78386799 | 21.67084 | 36.71058 | 47.48131 | 20.79035 | 0.01162 | 1.04976 | 0.986199 |
| 8 | 82618600 | 82620799 | 27.11138 | 45.37632 | 59.51012 | 25.93101 | -0.00889 | 1.190723 | 0.986228 |
| 17 | 82988600 | 82989399 | 15.47936 | 27.14366 | 35.83016 | 15.83489 | -0.01088 | 0.634719 | 0.986239 |
| 8 | 47189400 | 47190799 | 19.77699 | 34.35004 | 44.69032 | 19.61746 | 0.008813 | 0.809836 | 0.98627 |
| 13 | 43978800 | 43980599 | 19.77699 | 34.35004 | 44.69032 | 19.61746 | 0.008813 | 0.809836 | 0.98627 |
| X | 164656200 | 164658599 | 23.0349 | 39.9956 | 38.36622 | 16.83796 | 0.44677 | 0.824149 | 0.986282 |
| 11 | 99891600 | 99894199 | 26.02707 | 43.6677 | 57.29012 | 25.00204 | -0.0095 | 1.157891 | 0.986296 |
| 8 | 77194600 | 77195199 | 11.63177 | 21.17969 | 28.11777 | 12.09537 | 0.011809 | 0.297756 | 0.986315 |
| 12 | 67678000 | 67680199 | 19.35261 | 33.57814 | 44.20186 | 19.39411 | -0.00815 | 0.821419 | 0.986322 |
| 3 | 23433800 | 23434799 | 13.74754 | 24.52639 | 32.12557 | 14.12703 | 0.010392 | 0.446452 | 0.986327 |
| 9 | 3939200 | 3950199 | 369.0194 | 512.0331 | 574.1761 | 238.3695 | -0.00824 | 4.697857 | 0.986363 |
| X | 157079200 | 157082599 | 28.50645 | 47.94235 | 61.88339 | 27.03717 | 0.013956 | 1.149781 | 0.98639 |
| 6 | 130163000 | 130165599 | 20.2333 | 35.49545 | 47.18493 | 20.86078 | -0.02127 | 0.67149 | 0.98641 |
| 19 | 49856600 | 49857399 | 9.96576 | 18.74112 | 25.80699 | 10.58894 | 0.010141 | 0.087424 | 0.986412 |
| 13 | 33796800 | 33799999 | 36.53727 | 60.11401 | 60.90628 | 26.25701 | 0.359857 | 1.411592 | 0.986433 |
| 5 | 143774200 | 143775799 | 20.26846 | 35.31224 | 34.82656 | 15.31724 | 0.407315 | 0.756863 | 0.986472 |
| 15 | 83903800 | 83904999 | 13.67903 | 24.66744 | 32.55298 | 14.17386 | 0.009099 | 0.379525 | 0.986478 |
| 19 | 4074600 | 4077999 | 48.7151 | 77.91081 | 98.26779 | 41.34816 | 0.008871 | 2.052208 | 0.986478 |
| 13 | 14009400 | 14013799 | 67.957 | 107.408 | 134.1176 | 55.7147 | 0.006453 | 2.419728 | 0.986505 |
| 3 | 17048400 | 17049799 | 51.21631 | 81.71496 | 102.7277 | 43.07829 | 0.009827 | 2.130233 | 0.986516 |
| 3 | 133502800 | 133503799 | 17.24016 | 29.63154 | 50.04766 | 21.9527 | -0.36895 | 0.906019 | 0.986525 |
| 15 | 91918800 | 91920199 | 13.44217 | 24.30104 | 32.55067 | 14.12856 | -0.0095 | 0.362092 | 0.986544 |
| 13 | 112832200 | 112832999 | 10.43649 | 19.90413 | 21.71798 | 8.660751 | 0.379182 | -0.01178 | 0.986548 |
| 1 | 5658600 | 5659599 | 13.8504 | 24.39607 | 32.33876 | 14.31476 | -0.0157 | 0.556653 | 0.986549 |

| | | | | | | | | | |
|---|---|---|---|---|---|---|---|---|---|
| 2 | 23480200 | 23487599 | 102.3401 | 157.4915 | 192.6645 | 80.01622 | 0.006198 | 2.941194 | 0.986565 |
| 9 | 50311200 | 50314399 | 28.67707 | 47.42377 | 62.11473 | 26.85128 | -0.00757 | 1.331378 | 0.986569 |
| 11 | 58518400 | 58519799 | 23.97822 | 39.61577 | 68.6321 | 29.64931 | -0.40979 | 1.338531 | 0.986578 |
| 2 | 177163000 | 177165199 | 25.83698 | 44.32444 | 43.59025 | 19.11829 | 0.407545 | 0.948558 | 0.986602 |
| 11 | 63368200 | 63373999 | 72.23867 | 113.5677 | 142.4997 | 59.11299 | -0.00654 | 2.537753 | 0.986624 |
| 9 | 81793200 | 81795199 | 54.80108 | 87.21497 | 110.7363 | 46.27546 | -0.00839 | 2.215789 | 0.986627 |
| 6 | 30395800 | 30397199 | 14.81825 | 26.16089 | 34.10872 | 15.09129 | 0.009231 | 0.540783 | 0.986646 |
| 14 | 11520200 | 11523199 | 29.72754 | 49.45711 | 64.01884 | 27.78939 | 0.010032 | 1.268338 | 0.986663 |
| 1 | 88713600 | 88715599 | 24.37353 | 41.6946 | 43.16809 | 18.92878 | 0.333071 | 0.968161 | 0.98667 |
| 10 | 112743600 | 112745399 | 14.91688 | 26.42699 | 34.94132 | 15.44891 | -0.00911 | 0.503307 | 0.986682 |
| 7 | 8959400 | 8960799 | 10.18084 | 19.13236 | 26.34756 | 10.82342 | 0.008876 | 0.096344 | 0.986715 |
| 17 | 40701400 | 40705399 | 36.46307 | 59.7051 | 78.20781 | 33.64749 | -0.0125 | 1.480246 | 0.986729 |
| 2 | 169809200 | 169814199 | 76.15005 | 119.656 | 148.7334 | 61.69485 | 0.006434 | 2.550452 | 0.986766 |
| 4 | 50130800 | 50132599 | 34.52229 | 56.14844 | 72.94937 | 31.30774 | -0.00733 | 1.568625 | 0.986768 |
| 4 | 12121600 | 12128799 | 171.8948 | 254.9199 | 302.0815 | 124.4219 | 0.00668 | 3.678564 | 0.986789 |
| 2 | 111347000 | 111350999 | 61.05689 | 97.20056 | 123.4119 | 51.59417 | -0.00793 | 2.214568 | 0.986799 |
| 7 | 84548400 | 84556399 | 122.3645 | 189.0175 | 164.9517 | 68.51743 | 0.497173 | 2.876965 | 0.986804 |
| Y | 75919400 | 75920599 | 9.390012 | 17.4346 | 23.35519 | 9.788126 | 0.025654 | 0.156797 | 0.986835 |
| 5 | 125797800 | 125798199 | 10.08846 | 19.25983 | 20.69578 | 8.235501 | 0.40378 | -0.02518 | 0.986842 |
| 13 | 23528000 | 23530199 | 36.61834 | 59.28796 | 76.76507 | 32.84274 | -0.00833 | 1.661014 | 0.986845 |
| 3 | 13110800 | 13111999 | 15.41374 | 27.2166 | 35.50412 | 15.70818 | 0.008389 | 0.543925 | 0.986877 |
| 18 | 56960600 | 56963799 | 64.24465 | 101.7169 | 128.3383 | 53.36074 | -0.0065 | 2.376375 | 0.986893 |
| 7 | 126024800 | 126027799 | 22.38505 | 38.1768 | 50.09826 | 21.96221 | -0.00836 | 0.993596 | 0.986902 |
| X | 155894000 | 155896599 | 23.86909 | 40.89864 | 53.92366 | 23.64811 | -0.014 | 0.966337 | 0.986915 |
| 1 | 148093800 | 148094399 | 9.284489 | 17.71176 | 25.03997 | 9.979917 | 0.009239 | -0.01499 | 0.986937 |
| 14 | 85484000 | 85484799 | 9.284489 | 17.71176 | 25.03997 | 9.979917 | 0.009239 | -0.01499 | 0.986937 |
| 8 | 116056200 | 116056599 | 8.017394 | 15.53988 | 23.39542 | 8.851972 | -0.01044 | -0.20404 | 0.986943 |
| 14 | 52981600 | 52983799 | 19.58659 | 33.95194 | 44.18425 | 19.38583 | 0.007954 | 0.829315 | 0.986973 |
| 11 | 78081600 | 78085999 | 36.03722 | 58.46977 | 75.03401 | 32.15188 | 0.007356 | 1.618982 | 0.986982 |
| 9 | 44442400 | 44443599 | 13.56453 | 24.41507 | 32.57713 | 14.21772 | -0.0088 | 0.390441 | 0.986986 |
| 13 | 16229600 | 16231399 | 15.88815 | 28.03435 | 37.00701 | 16.37543 | -0.00909 | 0.556714 | 0.987014 |
| X | 114568800 | 114571999 | 32.8108 | 54.95203 | 51.3442 | 22.38219 | 0.478773 | 1.196868 | 0.987032 |
| 10 | 75858400 | 75863399 | 67.56559 | 107.1924 | 111.759 | 46.54295 | 0.270936 | 2.320202 | 0.987048 |
| 9 | 119693400 | 119699599 | 111.4744 | 169.7825 | 205.671 | 85.24433 | 0.008738 | 3.151596 | 0.98707 |
| 11 | 32882200 | 32883799 | 17.51484 | 30.62692 | 40.34382 | 17.78746 | -0.0081 | 0.696531 | 0.987073 |
| 17 | 94914400 | 94917599 | 41.73368 | 67.34942 | 86.81662 | 37.00952 | -0.00705 | 1.760029 | 0.987076 |
| 11 | 28668800 | 28669399 | 7.243711 | 14.07565 | 21.209 | 7.902356 | 0.009142 | -0.27274 | 0.987083 |
| 6 | 130736000 | 130744199 | 100.6851 | 156.4189 | 195.4249 | 81.14517 | -0.01367 | 2.770173 | 0.987097 |
| 13 | 46986600 | 46988599 | 28.24188 | 46.56921 | 60.23678 | 25.99283 | 0.009834 | 1.364001 | 0.987108 |
| 13 | 45231200 | 45234999 | 44.97344 | 72.29675 | 92.75898 | 39.32963 | -0.00709 | 1.884329 | 0.987124 |
| 1 | 132186400 | 132189399 | 23.81459 | 40.28761 | 52.25742 | 22.87226 | 0.00734 | 1.07434 | 0.987139 |
| 2 | 96791000 | 96792199 | 10.33307 | 18.84413 | 33.36176 | 14.32338 | -0.39905 | 0.29112 | 0.987145 |
| 6 | 3131600 | 3143599 | 451.1888 | 615.4822 | 670.4962 | 279.1065 | 0.008683 | 4.962799 | 0.987148 |
| 2 | 65885400 | 65886599 | 12.66809 | 22.82092 | 30.08387 | 13.11565 | 0.009511 | 0.372704 | 0.987161 |
| 6 | 143745600 | 143753199 | 106.6689 | 164.0072 | 202.3229 | 84.01722 | -0.00686 | 2.96096 | 0.987166 |
| 18 | 83186200 | 83190799 | 65.97161 | 104.2706 | 130.1793 | 54.07882 | 0.006699 | 2.417297 | 0.987178 |
| 16 | 15124600 | 15134199 | 126.9813 | 192.6673 | 235.0633 | 97.31583 | -0.00603 | 3.233705 | 0.987185 |

| | | | | | | | | | |
|---|---|---|---|---|---|---|---|---|---|
| 9 | 115246400 | 115257799 | 127.3082 | 193.1069 | 233.6098 | 96.7056 | 0.00585 | 3.2399 | 0.987196 |
| 14 | 123657000 | 123657599 | 9.001062 | 17.14296 | 24.4797 | 9.79049 | -0.00921 | -0.00702 | 0.987247 |
| 1 | 10891800 | 10897399 | 12.65077 | 23.35908 | 22.57234 | 9.531638 | 0.485879 | 0.214962 | 0.987287 |
| 5 | 37841400 | 37858199 | 198.3596 | 292.1168 | 344.0507 | 141.6827 | 0.00606 | 3.799412 | 0.987342 |
| 3 | 158344800 | 158345599 | 12.51366 | 22.76724 | 30.63089 | 13.18964 | -0.00818 | 0.315963 | 0.987342 |
| 11 | 101530400 | 101532399 | 9.546964 | 18.34368 | 19.66998 | 7.690532 | 0.429938 | -0.09178 | 0.987354 |
| 3 | 17031000 | 17032999 | 23.74716 | 39.77955 | 52.22447 | 22.7686 | -0.01022 | 1.16639 | 0.987365 |
| 10 | 83659800 | 83661799 | 19.16628 | 33.18044 | 43.17881 | 18.94394 | 0.007728 | 0.838954 | 0.987375 |
| 8 | 29250600 | 29251999 | 13.18834 | 23.82075 | 31.45944 | 13.67027 | 0.009588 | 0.361607 | 0.987382 |
| 5 | 145190200 | 145198799 | 125.0189 | 189.6944 | 231.4317 | 95.81302 | -0.00598 | 3.232407 | 0.987403 |
| 3 | 43493000 | 43496399 | 64.7733 | 102.271 | 127.4149 | 52.9028 | 0.00856 | 2.441835 | 0.987418 |
| 12 | 59126800 | 59129399 | 26.39208 | 44.01846 | 57.67501 | 25.07608 | -0.00735 | 1.226102 | 0.987442 |
| 4 | 88681600 | 88685799 | 32.17281 | 52.6728 | 68.72747 | 29.56699 | -0.00689 | 1.466691 | 0.987449 |
| 13 | 35177600 | 35178399 | 9.905068 | 18.00563 | 32.83967 | 14.15602 | -0.44526 | 0.306183 | 0.987497 |
| 3 | 40370000 | 40372199 | 18.92449 | 33.20058 | 43.90053 | 19.40931 | -0.01367 | 0.657694 | 0.987506 |
| 12 | 111980400 | 111994199 | 215.375 | 311.4379 | 437.0959 | 180.2861 | -0.27214 | 4.081573 | 0.987518 |
| 9 | 49012000 | 49014199 | 20.08983 | 34.52247 | 44.81915 | 19.65917 | 0.009033 | 0.913491 | 0.987519 |
| 4 | 12132400 | 12141999 | 201.8944 | 293.979 | 414.3594 | 170.7702 | -0.26852 | 3.976084 | 0.987527 |
| 5 | 24431200 | 24432399 | 10.98824 | 20.29418 | 27.70156 | 11.69374 | -0.00898 | 0.211737 | 0.987542 |
| 1 | 112893400 | 112894399 | 12.38049 | 22.40321 | 29.64601 | 12.85236 | 0.009413 | 0.344733 | 0.987555 |
| 11 | 107395400 | 107396199 | 13.42785 | 23.89145 | 31.70178 | 13.97244 | -0.01038 | 0.460479 | 0.987579 |
| 9 | 10218600 | 10220199 | 13.61593 | 24.60775 | 32.54368 | 14.13086 | 0.008473 | 0.365305 | 0.987599 |
| 18 | 35276800 | 35281999 | 50.95687 | 81.96645 | 103.7976 | 44.04977 | 0.012837 | 1.877919 | 0.987617 |
| 9 | 87679600 | 87680599 | 10.92779 | 20.27263 | 27.79293 | 11.66142 | -0.00859 | 0.188784 | 0.987627 |
| 16 | 7967600 | 7968399 | 10.92779 | 20.27263 | 27.79293 | 11.66142 | -0.00859 | 0.188784 | 0.987627 |
| 3 | 45393800 | 45394599 | 10.13065 | 18.79552 | 25.83458 | 10.8384 | -0.0119 | 0.177291 | 0.987671 |
| 9 | 19268600 | 19271199 | 38.18465 | 62.65397 | 62.65869 | 26.98145 | 0.377048 | 1.449526 | 0.987693 |
| 8 | 63606800 | 63607999 | 12.82438 | 23.45693 | 31.71578 | 13.56597 | -0.00916 | 0.289767 | 0.987742 |
| 15 | 93470000 | 93472599 | 28.37182 | 47.04527 | 61.01181 | 26.42349 | 0.006993 | 1.2996 | 0.987769 |
| X | 7425200 | 7428399 | 27.56114 | 46.49179 | 60.02005 | 26.24693 | 0.014011 | 1.122942 | 0.987802 |
| X | 121661400 | 121668199 | 82.11708 | 129.5745 | 127.8639 | 53.07252 | 0.343685 | 2.46878 | 0.987819 |
| 1 | 16484000 | 16485999 | 28.04169 | 46.65096 | 60.50904 | 26.2653 | 0.007145 | 1.260715 | 0.98782 |
| 6 | 75841600 | 75844599 | 25.16543 | 42.39639 | 55.5456 | 24.28084 | -0.00729 | 1.118225 | 0.987823 |
| 1 | 57416400 | 57417399 | 8.792668 | 16.75704 | 23.94453 | 9.56319 | -0.00863 | -0.01638 | 0.987839 |
| 7 | 87551800 | 87552199 | 8.792668 | 16.75704 | 23.94453 | 9.56319 | -0.00863 | -0.01638 | 0.987839 |
| 2 | 36811200 | 36815599 | 65.17705 | 103.881 | 104.2662 | 43.6605 | 0.332458 | 2.177332 | 0.987859 |
| 1 | 3823200 | 3827599 | 57.03726 | 91.20373 | 94.95388 | 39.94333 | 0.28482 | 2.065685 | 0.987878 |
| 4 | 10018800 | 10020599 | 17.77321 | 31.25751 | 40.64877 | 17.99872 | 0.010576 | 0.610589 | 0.987883 |
| 3 | 130829400 | 130830999 | 18.36613 | 32.14759 | 41.84864 | 18.4665 | 0.008855 | 0.687562 | 0.987884 |
| X | 121698600 | 121699799 | 10.47792 | 20.06034 | 21.77645 | 8.602272 | 0.398966 | -0.0411 | 0.987893 |
| 4 | 77417000 | 77418399 | 11.4707 | 21.25314 | 28.75779 | 12.08686 | 0.008023 | 0.200307 | 0.987935 |
| 2 | 18538200 | 18542399 | 26.45816 | 44.28133 | 57.34069 | 24.9851 | 0.009315 | 1.184208 | 0.987935 |
| 3 | 137289000 | 137292599 | 37.28012 | 60.57099 | 78.6209 | 33.7236 | -0.00723 | 1.593656 | 0.98797 |
| 18 | 8347400 | 8349199 | 18.47947 | 32.24745 | 42.46974 | 18.69779 | -0.00804 | 0.734524 | 0.987974 |
| 4 | 38321600 | 38322399 | 8.771008 | 16.27773 | 29.96403 | 12.56581 | -0.43016 | 0.173478 | 0.987982 |
| 15 | 85474400 | 85478599 | 38.48 | 62.08856 | 80.08606 | 34.13314 | -0.00818 | 1.757009 | 0.987987 |
| 1 | 130847800 | 130848399 | 10.29892 | 19.26244 | 26.34542 | 10.91004 | 0.009856 | 0.130258 | 0.987991 |

| | | | | | | | | | |
|---|---|---|---|---|---|---|---|---|---|
| 15 | 23973000 | 23973599 | 15.43284 | 27.09573 | 35.71146 | 15.79901 | -0.00879 | 0.613024 | 0.987997 |
| 3 | 99813000 | 99816799 | 49.66145 | 79.70217 | 102.0738 | 43.17376 | -0.00734 | 1.946029 | 0.987998 |
| 5 | 127365600 | 127368199 | 31.31983 | 51.44165 | 66.58013 | 28.68223 | 0.006597 | 1.423323 | 0.988021 |
| 16 | 9009400 | 9011599 | 21.5495 | 36.91217 | 47.94159 | 21.02414 | 0.007428 | 0.948521 | 0.988052 |
| 10 | 3269800 | 3273599 | 38.71445 | 62.73947 | 80.43656 | 34.4369 | 0.007122 | 1.65125 | 0.988069 |
| 16 | 59840800 | 59842999 | 21.3468 | 36.87404 | 47.93435 | 21.02894 | 0.008525 | 0.874561 | 0.988069 |
| 5 | 68672400 | 68673399 | 16.92478 | 29.55783 | 38.93249 | 17.149 | -0.0079 | 0.711991 | 0.988071 |
| 7 | 144370600 | 144374399 | 42.06921 | 67.83807 | 86.56689 | 36.86603 | 0.006215 | 1.783191 | 0.988077 |
| 2 | 134404000 | 134406199 | 46.36039 | 74.80672 | 77.26313 | 32.93166 | 0.311773 | 1.768294 | 0.98808 |
| 10 | 13768400 | 13771199 | 36.86035 | 59.19044 | 100.3492 | 42.4802 | -0.41013 | 1.9136 | 0.988081 |
| 12 | 107778600 | 107781399 | 21.5979 | 36.95602 | 48.48064 | 21.25872 | -0.00713 | 0.959643 | 0.988087 |
| 12 | 86390400 | 86393199 | 30.47654 | 50.0264 | 65.36947 | 28.1535 | -0.00748 | 1.427115 | 0.988095 |
| 11 | 31545000 | 31551999 | 87.65099 | 135.061 | 200.9964 | 83.49072 | -0.27484 | 2.911388 | 0.988129 |
| 18 | 37534000 | 37536599 | 64.89378 | 102.5615 | 129.2147 | 53.67667 | -0.00645 | 2.418629 | 0.988149 |
| X | 72464800 | 72466599 | 24.83265 | 41.86721 | 54.92076 | 24.01317 | -0.00892 | 1.113174 | 0.988159 |
| 6 | 149251800 | 149256799 | 66.71182 | 105.757 | 132.3879 | 55.09182 | 0.006413 | 2.346776 | 0.988174 |
| 8 | 97774200 | 97776799 | 24.18148 | 41.43026 | 54.54163 | 23.91895 | -0.01195 | 0.966173 | 0.988223 |
| 19 | 14960200 | 14962199 | 18.47613 | 32.52218 | 42.23053 | 18.69584 | 0.013184 | 0.607894 | 0.988239 |
| 1 | 134615800 | 134616599 | 11.58774 | 21.22168 | 28.72194 | 12.26439 | -0.00841 | 0.268487 | 0.988278 |
| 16 | 12998800 | 13000799 | 17.13234 | 29.91495 | 38.95859 | 17.15856 | 0.008426 | 0.716622 | 0.988281 |
| 2 | 174966000 | 174967199 | 8.107826 | 15.59076 | 22.47896 | 8.768956 | 0.009456 | -0.10607 | 0.98836 |
| 5 | 104848000 | 104872799 | 503.4861 | 686.2898 | 748.2025 | 311.4906 | 0.006453 | 4.976413 | 0.988372 |
| 18 | 60566800 | 60567599 | 9.652093 | 18.12548 | 25.27428 | 10.39534 | -0.01059 | 0.092033 | 0.988388 |
| 3 | 16310800 | 16315999 | 81.90897 | 128.9146 | 132.7451 | 55.07452 | 0.279493 | 2.520148 | 0.988405 |
| 19 | 47158600 | 47160599 | 17.65886 | 30.77765 | 40.5401 | 17.83448 | -0.00814 | 0.748283 | 0.988417 |
| 9 | 78182800 | 78188399 | 51.57696 | 82.43936 | 104.1212 | 43.77466 | 0.005938 | 2.073392 | 0.988431 |
| 6 | 131643000 | 131648799 | 73.8776 | 116.7768 | 145.6401 | 60.50428 | 0.008297 | 2.425258 | 0.988444 |
| 1 | 4246200 | 4247199 | 18.92502 | 32.79761 | 42.68563 | 18.72818 | 0.007984 | 0.829374 | 0.988448 |
| 7 | 84455000 | 84459599 | 60.72403 | 96.3739 | 121.7975 | 50.74232 | -0.00571 | 2.306291 | 0.988487 |
| 6 | 85633200 | 85634599 | 10.01115 | 18.98227 | 26.54515 | 10.71671 | 0.00815 | 0.035575 | 0.98849 |
| 14 | 36836000 | 36838399 | 25.89448 | 43.16229 | 56.55041 | 24.57769 | -0.00725 | 1.230034 | 0.988491 |
| 4 | 12430800 | 12431999 | 13.82755 | 24.58455 | 32.54639 | 14.35358 | -0.00796 | 0.467262 | 0.988515 |
| 14 | 46099800 | 46103599 | 73.67657 | 115.4619 | 143.1515 | 59.37598 | 0.007204 | 2.599548 | 0.988517 |
| 6 | 117762000 | 117763399 | 17.56065 | 30.68785 | 40.39311 | 17.7998 | -0.00692 | 0.70783 | 0.988523 |
| 3 | 13176800 | 13177999 | 18.17945 | 31.59957 | 41.57757 | 18.25988 | -0.00714 | 0.788053 | 0.988586 |
| 12 | 50836400 | 50839599 | 32.37395 | 53.18496 | 69.48745 | 29.93753 | -0.00681 | 1.426343 | 0.9886 |
| 4 | 5369400 | 5371199 | 15.83433 | 28.263 | 36.98325 | 16.25654 | 0.011647 | 0.461986 | 0.988602 |
| 9 | 115468800 | 115471599 | 24.77504 | 42.35494 | 42.55091 | 18.65721 | 0.37604 | 0.971244 | 0.988615 |
| 2 | 144591000 | 144593599 | 25.26902 | 42.40206 | 55.00985 | 24.00913 | 0.006677 | 1.153152 | 0.98862 |
| X | 112972000 | 112972799 | 13.63191 | 24.67696 | 32.65779 | 14.15143 | 0.009715 | 0.359408 | 0.988668 |
| 3 | 106569600 | 106570799 | 13.2731 | 24.1541 | 32.12845 | 13.8308 | 0.008211 | 0.322344 | 0.988676 |
| 10 | 26494400 | 26494999 | 11.5562 | 21.12658 | 28.20063 | 12.07037 | 0.008938 | 0.276425 | 0.988678 |
| 13 | 16672800 | 16675199 | 29.92099 | 49.88601 | 51.314 | 22.30529 | 0.340661 | 1.234807 | 0.98868 |
| 16 | 97332800 | 97333799 | 10.53602 | 19.73507 | 27.36953 | 11.30578 | -0.00758 | 0.123326 | 0.988681 |
| 4 | 155994200 | 156002799 | 85.34957 | 132.8009 | 163.9055 | 68.04172 | 0.005669 | 2.73154 | 0.988698 |
| 8 | 64059000 | 64060399 | 18.71556 | 32.44352 | 42.67174 | 18.72224 | -0.00711 | 0.825006 | 0.98871 |
| 1 | 45225800 | 45227199 | 17.93187 | 31.39533 | 41.34957 | 18.25014 | -0.00785 | 0.681521 | 0.988716 |

| | | | | | | | | | |
|---|---|---|---|---|---|---|---|---|---|
| 13 | 29713800 | 29716799 | 32.46145 | 53.29108 | 69.59308 | 29.97409 | -0.00657 | 1.433614 | 0.988728 |
| 9 | 43069600 | 43069799 | 5.941558 | 11.61118 | 18.97356 | 6.574695 | -0.01058 | -0.54761 | 0.988732 |
| 3 | 10426800 | 10427799 | 14.25501 | 25.81569 | 26.33465 | 11.4055 | 0.383163 | 0.344399 | 0.988736 |
| 6 | 36017800 | 36019599 | 16.62851 | 29.52158 | 38.52235 | 17.00991 | 0.011312 | 0.504871 | 0.988751 |
| 15 | 5118800 | 5119999 | 10.36074 | 19.17085 | 26.23574 | 11.04699 | -0.0099 | 0.192876 | 0.988759 |
| 4 | 152409800 | 152412799 | 22.1349 | 37.44763 | 48.52617 | 21.23942 | 0.008783 | 1.063215 | 0.988777 |
| 2 | 95627000 | 95631199 | 48.30607 | 77.51959 | 99.2388 | 41.96862 | -0.00688 | 1.948459 | 0.988777 |
| 6 | 9394200 | 9396399 | 21.54525 | 36.61073 | 47.46274 | 20.80073 | 0.008509 | 1.023544 | 0.988785 |
| 1 | 107613600 | 107616599 | 30.6255 | 50.66331 | 66.34228 | 28.68558 | -0.00726 | 1.335122 | 0.988793 |
| 14 | 77386800 | 77387999 | 11.57634 | 21.43535 | 29.3154 | 12.33181 | -0.00791 | 0.207796 | 0.988793 |
| 9 | 64489200 | 64490599 | 12.69622 | 22.90779 | 30.60046 | 13.3145 | -0.00757 | 0.365427 | 0.988794 |
| 12 | 97527400 | 97528799 | 20.09097 | 34.52017 | 45.3351 | 19.88541 | -0.00756 | 0.91516 | 0.9888 |
| 13 | 86994200 | 86996199 | 25.30325 | 42.89071 | 55.59596 | 24.34808 | 0.008367 | 1.065885 | 0.988812 |
| 15 | 8645200 | 8647999 | 23.79617 | 40.3166 | 52.8046 | 23.12222 | -0.00655 | 1.061615 | 0.988843 |
| X | 133389600 | 133391599 | 21.83479 | 37.3743 | 49.0148 | 21.49355 | -0.00658 | 0.959387 | 0.988848 |
| 3 | 38500600 | 38502999 | 19.62847 | 34.01653 | 44.72237 | 19.62178 | -0.00675 | 0.832938 | 0.988852 |
| 5 | 109554400 | 109559999 | 68.66187 | 109.0018 | 138.1351 | 57.5646 | -0.00932 | 2.308112 | 0.988886 |
| 17 | 19092600 | 19094999 | 30.8397 | 52.45812 | 45.71088 | 20.03463 | 0.579665 | 1.045952 | 0.988896 |
| 7 | 25635800 | 25637399 | 29.88607 | 49.11342 | 63.55501 | 27.38533 | 0.007236 | 1.412171 | 0.988896 |
| 12 | 58188600 | 58193599 | 54.16102 | 86.69079 | 90.12467 | 37.96534 | 0.288461 | 2.029823 | 0.988917 |
| 2 | 114384000 | 114386599 | 28.95206 | 47.99252 | 62.26729 | 26.96146 | 0.006272 | 1.304021 | 0.988922 |
| 7 | 122627200 | 122629199 | 24.66353 | 41.52078 | 53.88656 | 23.55045 | 0.006287 | 1.12115 | 0.988945 |
| 18 | 52337200 | 52341599 | 94.68616 | 145.9688 | 180.2895 | 74.88923 | -0.00568 | 2.903856 | 0.988958 |
| 10 | 7419400 | 7434799 | 200.4357 | 293.8143 | 344.6618 | 141.9663 | 0.005529 | 3.871661 | 0.988977 |
| 17 | 33530400 | 33575599 | 735.3024 | 975.2885 | 1032.045 | 431.377 | 0.009922 | 5.412 | 0.988979 |
| 5 | 103492400 | 103501599 | 138.1081 | 208.3948 | 251.1316 | 103.7884 | 0.005014 | 3.348717 | 0.988983 |
| 10 | 119932400 | 119933199 | 11.11107 | 20.51825 | 27.67062 | 11.68283 | 0.008057 | 0.212719 | 0.989001 |
| 15 | 5667200 | 5669199 | 20.33792 | 35.56203 | 46.15709 | 20.34983 | 0.013013 | 0.71978 | 0.989017 |
| 16 | 13368200 | 13373599 | 77.72767 | 122.8177 | 125.2427 | 52.02082 | 0.297942 | 2.432301 | 0.989059 |
| 13 | 90153000 | 90153799 | 9.902321 | 18.57956 | 25.83058 | 10.63948 | -0.00762 | 0.103492 | 0.989062 |
| 9 | 35299800 | 35302599 | 24.69582 | 41.56592 | 54.41819 | 23.78119 | -0.00626 | 1.124235 | 0.989077 |
| 7 | 53516800 | 53519399 | 27.7745 | 47.27582 | 45.78739 | 20.06924 | 0.427938 | 1.029926 | 0.989084 |
| 3 | 22101400 | 22103199 | 22.27115 | 38.13803 | 49.55601 | 21.73165 | 0.006747 | 0.956766 | 0.989103 |
| 17 | 37792200 | 37795599 | 47.12713 | 75.9439 | 79.49605 | 33.82554 | 0.290282 | 1.807658 | 0.989112 |
| 3 | 45539600 | 45540599 | 20.76431 | 35.36416 | 46.43214 | 20.3528 | -0.00916 | 0.999461 | 0.989112 |
| 10 | 68826400 | 68828399 | 17.55378 | 30.47871 | 39.63479 | 17.39471 | 0.009695 | 0.798661 | 0.989145 |
| 7 | 42924800 | 42927399 | 28.1238 | 46.65345 | 60.53672 | 26.22529 | 0.006204 | 1.291957 | 0.98915 |
| 6 | 131628800 | 131651799 | 282.0822 | 406.306 | 467.2159 | 192.8092 | 0.009661 | 4.145033 | 0.989167 |
| 10 | 64770000 | 64771799 | 19.49844 | 33.9305 | 44.65682 | 19.62612 | -0.00747 | 0.782414 | 0.989197 |
| 1 | 73037200 | 73037599 | 6.543742 | 12.7507 | 19.68667 | 7.14955 | 0.009317 | -0.37257 | 0.989237 |
| 15 | 58730800 | 58732199 | 15.13739 | 26.66368 | 35.13189 | 15.55105 | -0.00717 | 0.569247 | 0.98924 |
| 17 | 73941800 | 73945599 | 29.2937 | 48.40127 | 63.32663 | 27.35869 | -0.00615 | 1.342978 | 0.989257 |
| 2 | 129082000 | 129083199 | 18.3933 | 31.94665 | 41.61852 | 18.26901 | 0.007105 | 0.801176 | 0.989327 |
| 11 | 71403000 | 71405199 | 17.47977 | 30.5525 | 39.832 | 17.55544 | 0.006723 | 0.70197 | 0.989327 |
| 5 | 12788600 | 12789399 | 11.9364 | 21.66602 | 28.7362 | 12.4102 | 0.009912 | 0.322858 | 0.989328 |
| 6 | 106819000 | 106820599 | 16.63501 | 29.29466 | 38.60249 | 17.08812 | -0.00756 | 0.586769 | 0.989365 |
| 16 | 68419200 | 68423999 | 99.90647 | 153.6459 | 188.0136 | 78.07763 | 0.005075 | 2.952514 | 0.989385 |

| | | | | | | | | | |
|---|---|---|---|---|---|---|---|---|---|
| 3 | 154176000 | 154179799 | 42.4299 | 68.29366 | 86.97272 | 36.94349 | 0.005921 | 1.840844 | 0.989389 |
| 11 | 31602200 | 31603999 | 16.11154 | 28.39544 | 37.05803 | 16.40182 | 0.006751 | 0.572046 | 0.9894 |
| 3 | 114965400 | 114970799 | 55.60205 | 88.66183 | 111.6679 | 46.79043 | 0.006198 | 2.151782 | 0.9894 |
| 8 | 22192000 | 22193999 | 24.51787 | 41.53173 | 53.89602 | 23.59882 | 0.006744 | 1.067956 | 0.989425 |
| 19 | 56833200 | 56838799 | 74.86196 | 117.6533 | 147.4209 | 61.15183 | -0.00491 | 2.546324 | 0.989454 |
| 17 | 47547600 | 47550999 | 27.03309 | 44.87672 | 58.75006 | 25.46406 | -0.00641 | 1.280395 | 0.989471 |
| 9 | 5062800 | 5063999 | 27.8878 | 46.10589 | 59.77448 | 25.83473 | 0.007157 | 1.331454 | 0.989474 |
| 2 | 153117600 | 153120199 | 21.58756 | 36.96909 | 48.47866 | 21.25928 | -0.00639 | 0.951915 | 0.989493 |
| 17 | 76345000 | 76352999 | 160.9541 | 240.2061 | 286.6373 | 118.1498 | 0.004987 | 3.564324 | 0.989496 |
| 9 | 76874200 | 76875599 | 14.90494 | 26.24217 | 34.59867 | 15.31643 | -0.00846 | 0.570351 | 0.989497 |
| 4 | 67279600 | 67284399 | 96.18082 | 148.2214 | 181.6483 | 75.45377 | 0.005179 | 2.910491 | 0.989515 |
| 8 | 19228600 | 19231199 | 40.15844 | 64.86861 | 82.98072 | 35.41745 | 0.005581 | 1.729925 | 0.989518 |
| 3 | 52500200 | 52501399 | 12.47267 | 23.30032 | 31.75906 | 13.17845 | 0.011686 | 0.165527 | 0.989542 |
| 17 | 89800000 | 89801799 | 19.95653 | 34.39396 | 44.75872 | 19.63442 | 0.006323 | 0.88394 | 0.989545 |
| 18 | 88423200 | 88424399 | 17.96066 | 31.36054 | 40.88526 | 18.00686 | 0.006602 | 0.720991 | 0.989549 |
| 3 | 63574200 | 63575999 | 14.36128 | 25.43579 | 33.58233 | 14.84885 | -0.00694 | 0.502343 | 0.989557 |
| 18 | 30592000 | 30592799 | 10.58094 | 19.52333 | 26.27999 | 11.10804 | 0.010076 | 0.212193 | 0.989616 |
| 6 | 71522800 | 71523799 | 9.816321 | 18.70142 | 26.70097 | 10.67192 | -0.00903 | 0.000766 | 0.989641 |
| 1 | 132990400 | 132992999 | 29.51514 | 49.20094 | 51.3255 | 22.30788 | 0.320415 | 1.234111 | 0.989698 |
| 16 | 61360200 | 61361999 | 15.50533 | 27.50524 | 35.97896 | 15.89771 | 0.007275 | 0.499281 | 0.989702 |
| 1 | 139969200 | 139970999 | 41.96791 | 67.91332 | 71.07185 | 30.3951 | 0.297361 | 1.67882 | 0.989715 |
| 9 | 62529600 | 62531399 | 17.13444 | 30.07288 | 39.20492 | 17.33948 | 0.006859 | 0.635109 | 0.989725 |
| 9 | 71678200 | 71680199 | 18.03909 | 31.27504 | 41.15211 | 18.05558 | -0.00765 | 0.818217 | 0.98973 |
| 8 | 89492800 | 89497999 | 71.62394 | 112.9653 | 140.918 | 58.48141 | 0.005463 | 2.474575 | 0.989745 |
| 18 | 16987000 | 16987999 | 18.66046 | 32.13842 | 41.72898 | 18.30497 | 0.009466 | 0.880613 | 0.989764 |
| 18 | 70464000 | 70465799 | 21.77964 | 37.65522 | 49.56666 | 21.74563 | -0.00918 | 0.871796 | 0.989773 |
| 7 | 60667000 | 60668199 | 13.86783 | 24.77635 | 32.558 | 14.29997 | 0.006819 | 0.438215 | 0.989797 |
| 1 | 195050800 | 195054199 | 34.91596 | 56.99454 | 74.18369 | 31.87948 | -0.00634 | 1.516645 | 0.98982 |
| X | 153655600 | 153656199 | 10.11813 | 19.17492 | 27.09521 | 10.95107 | -0.00785 | 0.04365 | 0.98985 |
| 10 | 98704600 | 98706399 | 16.66015 | 29.73133 | 30.2005 | 13.27722 | 0.375092 | 0.457854 | 0.989863 |
| 14 | 108487600 | 108487999 | 6.084077 | 11.87291 | 18.701 | 6.651941 | 0.007351 | -0.45757 | 0.98987 |
| 10 | 67178600 | 67179799 | 12.09636 | 21.99374 | 29.56533 | 12.74113 | -0.00791 | 0.312327 | 0.989889 |
| 3 | 49385000 | 49390199 | 121.2573 | 184.2033 | 224.7578 | 93.0828 | -0.00475 | 3.207603 | 0.989896 |
| 1 | 177818800 | 177819999 | 9.240626 | 17.48197 | 24.35739 | 9.880281 | 0.007706 | 0.038873 | 0.989916 |
| 18 | 37124600 | 37127999 | 89.4862 | 138.2698 | 169.6311 | 70.46105 | 0.006597 | 2.858897 | 0.989918 |
| 6 | 13117200 | 13119599 | 31.93322 | 52.42763 | 68.43746 | 29.47724 | -0.00595 | 1.429978 | 0.98992 |
| 4 | 143175000 | 143177799 | 26.34799 | 44.72305 | 46.35802 | 20.31082 | 0.329852 | 1.048474 | 0.989925 |
| 7 | 64901800 | 64905799 | 60.17753 | 95.67764 | 120.2436 | 50.18476 | 0.00484 | 2.252809 | 0.989927 |
| 2 | 39473600 | 39476199 | 30.357 | 50.0629 | 64.94568 | 28.02618 | 0.005612 | 1.370855 | 0.989934 |
| 6 | 124417400 | 124421999 | 30.72054 | 50.72881 | 66.38994 | 28.67042 | -0.00666 | 1.354586 | 0.989948 |
| 19 | 47742600 | 47745999 | 39.29629 | 63.49459 | 81.26427 | 34.69797 | 0.005333 | 1.718318 | 0.989959 |
| 5 | 20918400 | 20919599 | 13.57609 | 24.22701 | 32.12766 | 14.12475 | -0.00711 | 0.443383 | 0.989982 |
| 2 | 181603400 | 181606399 | 45.91574 | 73.7062 | 93.5874 | 39.59702 | 0.005443 | 1.9327 | 0.989992 |
| 5 | 30281400 | 30285999 | 41.94064 | 67.61819 | 86.9534 | 37.02176 | -0.00517 | 1.789209 | 0.989995 |
| 2 | 46619800 | 46623199 | 56.99438 | 90.67439 | 113.9635 | 47.60141 | 0.005729 | 2.230474 | 0.990008 |
| X | 131646000 | 131647199 | 14.33587 | 25.66059 | 34.1242 | 14.96421 | -0.00856 | 0.435094 | 0.990026 |
| 7 | 121387200 | 121397199 | 169.4405 | 251.8895 | 299.5845 | 123.4239 | 0.004621 | 3.635817 | 0.990029 |

| | | | | | | | | | |
|---|---|---|---|---|---|---|---|---|---|
| 1 | 114050800 | 114052199 | 21.88774 | 37.39915 | 49.01619 | 21.49112 | -0.006 | 0.976468 | 0.99006 |
| 18 | 84188800 | 84190199 | 16.25321 | 28.38914 | 36.96164 | 16.28238 | 0.008838 | 0.703062 | 0.99008 |
| 12 | 43082400 | 43084799 | 25.459 | 41.85023 | 71.32499 | 30.73709 | -0.38866 | 1.405483 | 0.990091 |
| 1 | 155602400 | 155603199 | 9.731934 | 18.42396 | 25.96042 | 10.51541 | -0.00668 | 0.043175 | 0.990097 |
| 10 | 38794600 | 38795199 | 12.66129 | 23.12129 | 31.16972 | 13.35964 | -0.00682 | 0.296671 | 0.990108 |
| 8 | 84068000 | 84069199 | 9.681182 | 18.10476 | 24.75608 | 10.25408 | 0.009486 | 0.118184 | 0.990117 |
| 10 | 43657200 | 43663799 | 67.12509 | 106.4453 | 133.3512 | 55.51005 | 0.00574 | 2.337237 | 0.990126 |
| 13 | 113600000 | 113602199 | 17.11624 | 29.89128 | 39.3494 | 17.33231 | -0.0073 | 0.711289 | 0.99016 |
| 18 | 11155800 | 11157599 | 14.80467 | 26.30687 | 34.46971 | 15.20909 | 0.00625 | 0.480816 | 0.990164 |
| 8 | 75720200 | 75721399 | 13.35051 | 23.61583 | 31.2516 | 13.82176 | -0.01093 | 0.503992 | 0.990174 |
| 1 | 10415200 | 10415999 | 16.91884 | 29.56602 | 38.90135 | 17.14231 | -0.00629 | 0.702738 | 0.990178 |
| 5 | 99336200 | 99337799 | 19.89747 | 34.3568 | 44.73109 | 19.62342 | 0.006147 | 0.865736 | 0.990209 |
| 4 | 109549600 | 109559199 | 140.6511 | 212.6038 | 256.5082 | 106.0689 | 0.005452 | 3.315118 | 0.990244 |
| 1 | 136746400 | 136748799 | 24.07161 | 40.68477 | 52.82391 | 23.11372 | 0.00583 | 1.082322 | 0.990272 |
| 5 | 131102600 | 131104999 | 21.86809 | 37.3129 | 48.48929 | 21.25768 | 0.005913 | 0.988297 | 0.990285 |
| 4 | 109427800 | 109429199 | 13.06149 | 23.73629 | 31.56602 | 13.61212 | 0.006838 | 0.326678 | 0.990295 |
| 2 | 109641200 | 109641999 | 8.974345 | 16.9199 | 23.37753 | 9.550686 | 0.011144 | 0.05866 | 0.990295 |
| 9 | 35130600 | 35130999 | 5.068912 | 9.898762 | 15.9996 | 5.622827 | -0.01187 | -0.52176 | 0.99031 |
| 9 | 109195800 | 109196799 | 16.34811 | 27.99184 | 51.0688 | 22.39483 | -0.48053 | 0.939891 | 0.990323 |
| 10 | 86420400 | 86422399 | 21.61719 | 36.51311 | 59.82338 | 26.17136 | -0.3286 | 1.087479 | 0.990335 |
| 11 | 43549800 | 43550999 | 15.47042 | 27.26384 | 35.59797 | 15.7558 | 0.006125 | 0.566782 | 0.990349 |
| 12 | 115609800 | 115656399 | 700.2668 | 931.99 | 990.9632 | 414.0214 | 0.007922 | 5.357072 | 0.990364 |
| 19 | 44312400 | 44322799 | 110.2103 | 169.1836 | 206.7087 | 85.81983 | 0.005278 | 2.993282 | 0.990385 |
| 1 | 68444800 | 68445599 | 10.83566 | 19.97784 | 27.19791 | 11.50815 | -0.00758 | 0.2191 | 0.990389 |
| 4 | 74903400 | 74906799 | 31.40334 | 52.12573 | 68.36083 | 29.62735 | -0.00911 | 1.299395 | 0.99039 |
| 5 | 137841000 | 137848799 | 91.86569 | 142.171 | 175.969 | 73.09306 | -0.00444 | 2.830631 | 0.990402 |
| 9 | 102450200 | 102454999 | 44.81948 | 72.10611 | 91.81986 | 38.97599 | 0.005166 | 1.859358 | 0.990405 |
| 7 | 58531600 | 58533999 | 36.54791 | 59.21506 | 76.57196 | 32.7769 | -0.0055 | 1.649287 | 0.99042 |
| 10 | 48409800 | 48414999 | 81.7974 | 127.7137 | 159.1653 | 66.04336 | -0.00446 | 2.670338 | 0.990442 |
| 8 | 21447200 | 21448399 | 20.62925 | 35.57954 | 46.32869 | 20.32358 | 0.005674 | 0.882446 | 0.990444 |
| 4 | 37756000 | 37758199 | 27.84499 | 46.23865 | 60.48722 | 26.22252 | -0.00532 | 1.280388 | 0.990447 |
| 17 | 81033200 | 81033999 | 11.32795 | 20.90287 | 28.20424 | 11.92044 | 0.006369 | 0.221112 | 0.990456 |
| 13 | 34066400 | 34067599 | 14.35825 | 25.58474 | 33.9 | 14.92264 | -0.00766 | 0.457091 | 0.990471 |
| 12 | 6748600 | 6749999 | 20.34296 | 35.15761 | 45.79679 | 20.09149 | 0.005772 | 0.863064 | 0.990484 |
| 11 | 114834400 | 114836599 | 24.68305 | 41.48988 | 53.87373 | 23.53388 | 0.005494 | 1.13478 | 0.990508 |
| 3 | 47636200 | 47636799 | 9.394575 | 17.8029 | 24.90084 | 10.06513 | 0.00655 | 0.030159 | 0.990512 |
| 10 | 16813600 | 16819999 | 24.02653 | 41.09549 | 38.15422 | 16.73006 | 0.489762 | 0.962669 | 0.990513 |
| 2 | 36276200 | 36277399 | 18.85557 | 32.79204 | 43.10383 | 18.93643 | -0.00567 | 0.786199 | 0.990547 |
| 7 | 108982600 | 108984399 | 25.11113 | 42.38081 | 55.02476 | 24.06624 | 0.005791 | 1.101274 | 0.990549 |
| 1 | 126348200 | 126354399 | 72.95326 | 115.3329 | 120.1662 | 49.92523 | 0.267544 | 2.414436 | 0.990563 |
| 2 | 76844800 | 76846599 | 15.62757 | 27.58428 | 36.04309 | 15.94778 | 0.00589 | 0.551045 | 0.99057 |
| 11 | 15402400 | 15403999 | 15.62757 | 27.58428 | 36.04309 | 15.94778 | 0.00589 | 0.551045 | 0.99057 |
| 2 | 65408600 | 65413999 | 88.80822 | 137.7026 | 169.6266 | 70.45331 | 0.004564 | 2.793744 | 0.990599 |
| 9 | 25361800 | 25363599 | 19.09295 | 32.89052 | 43.21168 | 18.95551 | -0.00751 | 0.881199 | 0.990614 |
| 14 | 114329800 | 114334999 | 79.90917 | 124.8909 | 155.8069 | 64.64314 | -0.00481 | 2.65053 | 0.990616 |
| 11 | 20853000 | 20854799 | 21.10278 | 36.40531 | 47.39338 | 20.79079 | 0.006025 | 0.883156 | 0.99065 |
| 19 | 41398000 | 41413399 | 131.7916 | 198.6612 | 290.5625 | 120.0475 | -0.27548 | 3.370118 | 0.990656 |

| | | | | | | | | | |
|---|---|---|---|---|---|---|---|---|---|
| 14 | 75300400 | 75302599 | 19.84138 | 34.1377 | 44.7885 | 19.64647 | -0.00582 | 0.899511 | 0.99069 |
| 3 | 62224600 | 62224999 | 6.803588 | 13.2333 | 20.1223 | 7.438717 | 0.006789 | -0.30882 | 0.990712 |
| 7 | 108158200 | 108161599 | 56.00778 | 89.46092 | 113.8287 | 47.81004 | -0.00579 | 2.103491 | 0.990714 |
| 10 | 57216400 | 57218599 | 22.82191 | 38.95632 | 50.6333 | 22.19835 | 0.005685 | 0.989863 | 0.990722 |
| 13 | 9916000 | 9918799 | 32.53102 | 52.99441 | 68.88661 | 29.58135 | -0.00642 | 1.538601 | 0.990723 |
| 3 | 46603600 | 46607799 | 82.20339 | 128.965 | 133.4784 | 55.35759 | 0.268578 | 2.582157 | 0.990727 |
| 8 | 28551800 | 28553199 | 19.54925 | 34.52155 | 32.64984 | 14.44512 | 0.470065 | 0.564457 | 0.990735 |
| 7 | 3667800 | 3669199 | 13.32475 | 23.92056 | 31.53534 | 13.79488 | 0.006107 | 0.401538 | 0.990739 |
| 5 | 100741000 | 100744599 | 39.95577 | 64.59637 | 82.75074 | 35.35822 | 0.004921 | 1.703983 | 0.990756 |
| 8 | 18082600 | 18084999 | 45.29046 | 72.76942 | 93.17771 | 39.47812 | -0.00506 | 1.903704 | 0.990763 |
| 4 | 46363200 | 46364799 | 12.57487 | 22.92821 | 30.58687 | 13.13504 | 0.006273 | 0.302058 | 0.990814 |
| 18 | 11961600 | 11967199 | 49.64388 | 79.60258 | 100.9801 | 42.65296 | 0.004851 | 1.973105 | 0.990816 |
| 5 | 8594600 | 8595799 | 10.06397 | 18.28897 | 32.83542 | 14.15953 | -0.42302 | 0.30946 | 0.990844 |
| 5 | 28718200 | 28718799 | 12.50672 | 22.14119 | 37.74194 | 16.68956 | -0.37382 | 0.503044 | 0.990845 |
| 7 | 81750600 | 81752599 | 24.08298 | 40.67173 | 52.82058 | 23.10671 | 0.005426 | 1.089348 | 0.990847 |
| 6 | 51468400 | 51470399 | 16.39508 | 28.81276 | 37.58898 | 16.64311 | 0.005735 | 0.605954 | 0.990849 |
| 4 | 147351800 | 147361799 | 488.5034 | 663.1163 | 725.5956 | 302.2725 | -0.00486 | 5.040437 | 0.990863 |
| 3 | 18439200 | 18440599 | 18.17071 | 32.13512 | 31.54866 | 13.95377 | 0.417056 | 0.535506 | 0.990899 |
| 8 | 53414400 | 53417599 | 80.30669 | 125.0291 | 154.3507 | 64.06957 | 0.006015 | 2.719451 | 0.990906 |
| 9 | 86242400 | 86244999 | 17.25608 | 30.20053 | 39.74713 | 17.53718 | -0.00697 | 0.678513 | 0.990917 |
| 8 | 43609200 | 43617799 | 86.22702 | 134.0389 | 166.4695 | 69.11593 | -0.00439 | 2.748787 | 0.990918 |
| 6 | 140293200 | 140295199 | 17.55557 | 30.69628 | 40.3645 | 17.79566 | -0.00554 | 0.698525 | 0.990925 |
| 3 | 92364000 | 92364799 | 7.635346 | 14.788 | 22.1577 | 8.41383 | -0.00747 | -0.19596 | 0.990929 |
| 1 | 131305400 | 131305999 | 9.747434 | 18.2582 | 25.31554 | 10.45709 | -0.00699 | 0.111632 | 0.990953 |
| 17 | 50897000 | 50897599 | 7.708839 | 14.89792 | 22.07551 | 8.468181 | -0.0063 | -0.16461 | 0.99102 |
| 10 | 7354600 | 7356999 | 28.83411 | 47.98826 | 62.30663 | 27.05298 | 0.005688 | 1.259175 | 0.991028 |
| X | 70705400 | 70711799 | 86.23038 | 134.3658 | 166.1588 | 68.96225 | 0.004503 | 2.709556 | 0.991038 |
| 1 | 67481000 | 67481599 | 9.10521 | 17.26518 | 24.39908 | 9.849485 | -0.00638 | 0.022509 | 0.991054 |
| 3 | 111913400 | 111914399 | 15.88705 | 28.02966 | 36.61046 | 16.20028 | 0.006339 | 0.56028 | 0.991058 |
| 5 | 4182000 | 4185599 | 46.58738 | 74.93716 | 95.39653 | 40.48391 | 0.005294 | 1.870774 | 0.991074 |
| 9 | 83852400 | 83854599 | 19.76548 | 34.36856 | 45.19773 | 19.85418 | -0.00651 | 0.794284 | 0.991108 |
| 3 | 81523600 | 81524199 | 6.088572 | 11.86623 | 18.57256 | 6.726741 | -0.00664 | -0.39307 | 0.991123 |
| 5 | 109772800 | 109773399 | 14.20822 | 25.23866 | 33.07087 | 14.59595 | 0.006124 | 0.477671 | 0.991135 |
| 16 | 96058000 | 96058999 | 9.772862 | 18.34996 | 25.24619 | 10.38601 | 0.008233 | 0.093099 | 0.991144 |
| 19 | 59707600 | 59710799 | 28.93486 | 47.52025 | 79.09362 | 34.07168 | -0.3554 | 1.426373 | 0.991148 |
| 18 | 40598400 | 40599399 | 11.63956 | 21.49788 | 29.30761 | 12.3711 | -0.00676 | 0.223647 | 0.991151 |
| 3 | 112632800 | 112633599 | 11.60859 | 21.44971 | 29.23605 | 12.33382 | -0.00639 | 0.217211 | 0.991153 |
| 15 | 33848000 | 33850199 | 27.46021 | 46.40963 | 47.99008 | 20.99829 | 0.333086 | 1.09027 | 0.991158 |
| 16 | 8841600 | 8842799 | 10.41278 | 19.35097 | 26.28254 | 11.00152 | 0.007317 | 0.170651 | 0.991167 |
| 14 | 8950000 | 8954799 | 55.44482 | 88.5498 | 112.583 | 47.27781 | -0.0049 | 2.105739 | 0.991175 |
| 5 | 105244800 | 105246199 | 14.86037 | 26.35685 | 34.51261 | 15.25188 | 0.005741 | 0.493933 | 0.991179 |
| 19 | 46217000 | 46218199 | 14.86037 | 26.35685 | 34.51261 | 15.25188 | 0.005741 | 0.493933 | 0.991179 |
| 17 | 83567400 | 83569399 | 17.73825 | 30.97529 | 40.42167 | 17.8038 | 0.005384 | 0.719725 | 0.99119 |
| 10 | 83611400 | 83617199 | 69.46827 | 109.6009 | 137.6795 | 57.14281 | -0.00425 | 2.463413 | 0.99122 |
| 4 | 61684800 | 61688799 | 141.067 | 211.5408 | 255.4434 | 105.3911 | -0.00582 | 3.470714 | 0.991234 |
| 3 | 77106800 | 77111799 | 45.15399 | 72.37528 | 113.0048 | 47.70414 | -0.29489 | 1.984375 | 0.991265 |
| 9 | 23631400 | 23633399 | 23.48105 | 39.92172 | 52.25451 | 22.90174 | -0.00532 | 1.030008 | 0.991299 |

| | | | | | | | | | |
|---|---|---|---|---|---|---|---|---|---|
| 3 | 153649600 | 153650599 | 10.71639 | 19.64129 | 26.26803 | 11.20461 | 0.009762 | 0.250002 | 0.991319 |
| 2 | 108444600 | 108444999 | 7.430801 | 14.43677 | 21.77224 | 8.123221 | 0.006467 | -0.2622 | 0.991323 |
| 7 | 9002800 | 9003199 | 9.83686 | 18.85925 | 19.75398 | 7.772208 | 0.455767 | -0.06565 | 0.991328 |
| 14 | 11784200 | 11785399 | 17.48002 | 30.37263 | 39.56192 | 17.37063 | 0.007312 | 0.786206 | 0.99136 |
| 16 | 53927400 | 53930599 | 35.22236 | 57.5848 | 75.02746 | 32.26049 | -0.00622 | 1.495246 | 0.991399 |
| 10 | 93200600 | 93202199 | 17.30309 | 30.26371 | 39.80025 | 17.55126 | -0.00581 | 0.689941 | 0.991444 |
| 3 | 91132200 | 91133799 | 21.87977 | 37.41636 | 49.01521 | 21.49211 | -0.00514 | 0.968827 | 0.991456 |
| 3 | 94494600 | 94503999 | 133.8197 | 201.9021 | 244.8441 | 101.1864 | -0.00419 | 3.349782 | 0.991465 |
| 10 | 68892000 | 68892399 | 6.675476 | 12.96782 | 19.52171 | 7.290297 | 0.008377 | -0.27464 | 0.991485 |
| 9 | 12912400 | 12912799 | 6.976462 | 13.57667 | 20.73845 | 7.634092 | 0.006228 | -0.31788 | 0.991546 |
| 14 | 27374800 | 27375399 | 6.385413 | 12.41466 | 19.02227 | 7.056243 | -0.00764 | -0.30294 | 0.991549 |
| 19 | 9352200 | 9353199 | 18.24933 | 32.02645 | 32.97162 | 14.5814 | 0.345767 | 0.631188 | 0.991576 |
| 19 | 33514800 | 33515799 | 13.7787 | 24.83249 | 33.11355 | 14.4286 | -0.00636 | 0.386422 | 0.991637 |
| 11 | 81053400 | 81057799 | 45.22878 | 72.90961 | 93.78362 | 39.92289 | -0.00583 | 1.8007 | 0.991685 |
| 7 | 67054600 | 67055599 | 16.77013 | 29.35931 | 38.3185 | 16.91135 | 0.005293 | 0.671946 | 0.9917 |
| 16 | 84235800 | 84237199 | 19.72044 | 33.99179 | 44.26706 | 19.41882 | 0.005448 | 0.88285 | 0.991732 |
| 2 | 152485400 | 152489799 | 44.04345 | 71.18566 | 73.60697 | 31.44025 | 0.312811 | 1.712417 | 0.991749 |
| 11 | 114837600 | 114844799 | 82.2527 | 129.1942 | 160.5937 | 66.61108 | 0.005955 | 2.565329 | 0.991784 |
| X | 62933600 | 62936599 | 23.20611 | 40.14703 | 52.90738 | 23.2117 | -0.01066 | 0.876995 | 0.991839 |
| 1 | 35833400 | 35834999 | 15.16257 | 27.33655 | 28.29289 | 12.32245 | 0.357299 | 0.381605 | 0.991856 |
| 3 | 29135800 | 29137799 | 16.64855 | 28.36336 | 53.18291 | 23.31248 | -0.52125 | 0.986348 | 0.991859 |
| 1 | 28340600 | 28344399 | 22.63066 | 39.16357 | 40.41231 | 17.72999 | 0.340844 | 0.853158 | 0.991861 |
| 5 | 9619600 | 9621199 | 24.30793 | 40.68729 | 52.80564 | 23.0107 | 0.006227 | 1.174787 | 0.991873 |
| 10 | 35655000 | 35656799 | 31.08555 | 51.74295 | 53.51624 | 23.23857 | 0.332818 | 1.257361 | 0.991889 |
| 19 | 55421200 | 55423799 | 22.8617 | 39.39946 | 51.19378 | 22.45728 | 0.00856 | 0.90787 | 0.991898 |
| 11 | 53134000 | 53136199 | 33.04266 | 53.88018 | 69.53753 | 29.87144 | 0.004893 | 1.527005 | 0.9919 |
| 14 | 108450000 | 108450599 | 9.60884 | 18.19714 | 25.43809 | 10.2964 | 0.005816 | 0.03933 | 0.991911 |
| 13 | 25499000 | 25500799 | 20.69457 | 35.59429 | 46.35064 | 20.3315 | 0.00485 | 0.908505 | 0.991917 |
| 2 | 38074000 | 38075599 | 15.57127 | 27.41439 | 36.06586 | 15.96605 | -0.00538 | 0.577697 | 0.991927 |
| 2 | 76139800 | 76141999 | 32.02725 | 52.96769 | 68.70521 | 29.70167 | 0.006363 | 1.344989 | 0.991943 |
| 3 | 27455200 | 27456799 | 11.31415 | 20.78632 | 28.16488 | 11.97483 | -0.00587 | 0.24493 | 0.991959 |
| 2 | 151018000 | 151021599 | 46.73787 | 75.26254 | 96.68335 | 41.09593 | -0.0058 | 1.836668 | 0.992002 |
| 1 | 177796600 | 177798799 | 52.81537 | 84.23449 | 106.9056 | 44.80689 | -0.00446 | 2.144209 | 0.992038 |
| 12 | 113497800 | 113499999 | 40.48624 | 65.43123 | 83.82074 | 35.79936 | 0.004344 | 1.716164 | 0.992056 |
| 16 | 3490400 | 3491999 | 18.65249 | 32.54578 | 42.47096 | 18.69702 | 0.005142 | 0.73716 | 0.992058 |
| 8 | 65688000 | 65690399 | 28.85857 | 48.04792 | 62.86416 | 27.30201 | -0.00533 | 1.255818 | 0.992085 |
| 4 | 14170200 | 14170999 | 13.0771 | 23.44408 | 31.14361 | 13.63915 | -0.00574 | 0.409153 | 0.992106 |
| 13 | 35769800 | 35772799 | 29.99753 | 49.7233 | 64.57287 | 27.95889 | 0.004934 | 1.3102 | 0.992108 |
| 6 | 85689400 | 85692799 | 28.61219 | 47.51359 | 61.72924 | 26.76134 | 0.00452 | 1.282182 | 0.992132 |
| 17 | 86137800 | 86138999 | 17.21418 | 30.91775 | 29.64273 | 12.96171 | 0.463321 | 0.42173 | 0.992178 |
| 18 | 14260000 | 14261799 | 22.59259 | 38.52081 | 50.09563 | 21.96061 | 0.004585 | 0.997593 | 0.992201 |
| 14 | 52945800 | 52950199 | 15.324 | 26.65105 | 46.33006 | 20.35418 | -0.40686 | 0.776645 | 0.992209 |
| 7 | 6307600 | 6310599 | 35.12976 | 57.58418 | 74.50288 | 32.06819 | 0.005929 | 1.461883 | 0.992223 |
| 6 | 114871800 | 114873999 | 23.94906 | 40.63541 | 52.81059 | 23.13517 | 0.00468 | 1.049429 | 0.992241 |
| 6 | 118728200 | 118729599 | 14.01076 | 24.86162 | 32.56891 | 14.38722 | 0.005542 | 0.481257 | 0.992296 |
| 1 | 194933800 | 194939599 | 74.99514 | 117.967 | 147.0344 | 60.99855 | 0.00369 | 2.528659 | 0.992302 |
| X | 141464200 | 141466199 | 17.1305 | 30.09798 | 39.23529 | 17.36853 | 0.007105 | 0.62084 | 0.992308 |

| | | | | | | | | | |
|---|---|---|---|---|---|---|---|---|---|
| 16 | 54289800 | 54294199 | 34.94849 | 57.69175 | 75.66823 | 32.66993 | -0.00988 | 1.376692 | 0.992309 |
| 9 | 92621400 | 92625599 | 97.12264 | 148.4169 | 222.8041 | 92.42086 | -0.29665 | 3.081121 | 0.992324 |
| 5 | 136236600 | 136237599 | 11.54912 | 21.29479 | 28.73648 | 12.15795 | 0.005257 | 0.229486 | 0.992363 |
| 9 | 78841400 | 78844399 | 33.01247 | 53.9809 | 69.84093 | 30.03221 | 0.004114 | 1.486912 | 0.992419 |
| 5 | 127436400 | 127439799 | 41.23799 | 66.39822 | 84.66062 | 35.97945 | 0.004805 | 1.826299 | 0.992431 |
| 3 | 37298200 | 37302799 | 96.52548 | 148.711 | 182.3895 | 75.76059 | 0.003756 | 2.916395 | 0.992432 |
| 13 | 67014800 | 67017199 | 27.16678 | 45.28563 | 58.85724 | 25.58121 | 0.00424 | 1.234248 | 0.992436 |
| 16 | 93208400 | 93210599 | 27.16678 | 45.28563 | 58.85724 | 25.58121 | 0.00424 | 1.234248 | 0.992436 |
| 1 | 152850800 | 152855399 | 43.67788 | 70.37552 | 90.37643 | 38.4472 | -0.0043 | 1.810252 | 0.992437 |
| 13 | 65278800 | 65284599 | 64.50896 | 102.5791 | 128.9027 | 53.80649 | 0.005185 | 2.25474 | 0.992443 |
| 15 | 86781200 | 86784599 | 27.65 | 45.68958 | 59.29018 | 25.61646 | 0.005703 | 1.334268 | 0.99245 |
| 15 | 26552200 | 26554199 | 21.0943 | 35.38089 | 59.59083 | 26.00113 | -0.36841 | 1.152271 | 0.992453 |
| 14 | 112529600 | 112532599 | 25.27402 | 42.81067 | 55.60303 | 24.34592 | 0.005446 | 1.07137 | 0.992472 |
| 10 | 68340800 | 68342599 | 17.41403 | 30.48705 | 39.80634 | 17.56822 | 0.004642 | 0.677083 | 0.992484 |
| 13 | 119221200 | 119222799 | 16.88066 | 29.61868 | 38.6688 | 17.09808 | 0.004691 | 0.636036 | 0.992486 |
| 7 | 106255800 | 106263999 | 113.2099 | 173.0505 | 212.0646 | 87.97188 | -0.00378 | 3.080008 | 0.992529 |
| 10 | 72853200 | 72854399 | 10.43631 | 19.43594 | 26.7175 | 11.14469 | -0.00652 | 0.155366 | 0.992536 |
| 4 | 130913200 | 130918799 | 50.84471 | 81.14763 | 103.094 | 43.25119 | -0.00485 | 2.119465 | 0.99254 |
| 6 | 147066400 | 147068399 | 18.4331 | 32.16615 | 41.98711 | 18.48517 | 0.004892 | 0.73591 | 0.992546 |
| 1 | 148208000 | 148208999 | 13.92107 | 24.82407 | 32.61373 | 14.34749 | 0.005654 | 0.451702 | 0.992564 |
| 14 | 11152800 | 11154199 | 22.29923 | 37.88618 | 49.55626 | 21.71796 | -0.00446 | 1.02855 | 0.992596 |
| 1 | 156438600 | 156439799 | 14.94196 | 26.39925 | 34.53302 | 15.27672 | 0.004821 | 0.533148 | 0.992612 |
| 6 | 102151600 | 102152399 | 15.12914 | 26.67343 | 35.09848 | 15.53344 | -0.0048 | 0.558966 | 0.992637 |
| 12 | 72734400 | 72735799 | 14.89382 | 26.34134 | 34.46338 | 15.24277 | 0.005041 | 0.520422 | 0.992643 |
| 3 | 19720600 | 19722199 | 24.60074 | 40.49422 | 69.80476 | 30.0992 | -0.40431 | 1.382997 | 0.992648 |
| 8 | 64165800 | 64167799 | 20.06666 | 34.72036 | 45.27325 | 19.86251 | 0.004712 | 0.851778 | 0.99266 |
| 7 | 24398600 | 24400199 | 13.4688 | 24.12809 | 32.01314 | 14.02888 | -0.0049 | 0.416162 | 0.992662 |
| 17 | 7601800 | 7603799 | 15.14481 | 26.77314 | 35.03651 | 15.49765 | 0.004599 | 0.529775 | 0.992669 |
| 3 | 71655600 | 71657199 | 17.2265 | 30.21476 | 39.71161 | 17.55391 | -0.00491 | 0.645513 | 0.992682 |
| 14 | 85764600 | 85765999 | 16.46672 | 28.9796 | 38.10645 | 16.87071 | -0.00486 | 0.592032 | 0.992687 |
| 13 | 74983600 | 74984599 | 14.46319 | 26.08525 | 25.92324 | 11.28518 | 0.416018 | 0.369302 | 0.992694 |
| 15 | 14169000 | 14170799 | 9.021491 | 16.49173 | 31.9416 | 13.67259 | -0.52409 | 0.268367 | 0.992694 |
| 13 | 3582000 | 3582999 | 10.84749 | 20.08881 | 27.2272 | 11.45102 | 0.005158 | 0.194831 | 0.99276 |
| 12 | 93525800 | 93526399 | 9.73352 | 18.16064 | 24.79725 | 10.31138 | 0.007256 | 0.13488 | 0.992768 |
| 12 | 5952800 | 5954399 | 21.27958 | 36.88327 | 48.48176 | 21.2713 | -0.00639 | 0.846047 | 0.992771 |
| 10 | 42328200 | 42331799 | 31.66163 | 51.92126 | 67.6538 | 29.12536 | -0.00397 | 1.444295 | 0.992811 |
| 19 | 30073600 | 30084399 | 144.4764 | 217.0972 | 262.1793 | 108.2295 | -0.00336 | 3.43175 | 0.99283 |
| 5 | 145458200 | 145476199 | 279.9014 | 401.8824 | 465.259 | 192.0949 | -0.00465 | 4.190882 | 0.992831 |
| 12 | 39588800 | 39589399 | 12.27543 | 22.13191 | 29.24651 | 12.73737 | 0.007335 | 0.365516 | 0.992857 |
| 3 | 10003600 | 10005599 | 13.20142 | 23.83462 | 31.58235 | 13.7307 | 0.004734 | 0.367687 | 0.992924 |
| X | 6494800 | 6496999 | 20.46784 | 35.60295 | 46.82041 | 20.57174 | -0.00631 | 0.797535 | 0.992925 |
| 4 | 73931000 | 73933199 | 19.03389 | 32.21915 | 56.43864 | 24.70677 | -0.4245 | 1.055628 | 0.992941 |
| 13 | 78679200 | 78679799 | 9.550688 | 18.12945 | 25.47033 | 10.25848 | 0.004666 | 0.021355 | 0.99297 |
| 2 | 8124600 | 8125999 | 19.40918 | 33.93342 | 35.13076 | 15.48672 | 0.337879 | 0.701823 | 0.992979 |
| 12 | 98898600 | 98900799 | 17.83009 | 31.13385 | 40.90297 | 18.01513 | -0.00439 | 0.720308 | 0.993003 |
| Y | 43288200 | 43288799 | 6.806607 | 13.14206 | 19.46348 | 7.498717 | -0.01127 | -0.16924 | 0.993006 |
| 10 | 85958200 | 85959399 | 16.76056 | 29.25821 | 38.45881 | 16.93569 | -0.00499 | 0.716662 | 0.993007 |

| | | | | | | | | | |
|---|---|---|---|---|---|---|---|---|---|
| 11 | 63544600 | 63545799 | 14.66282 | 26.10234 | 34.49686 | 15.1977 | -0.00452 | 0.468978 | 0.993048 |
| 6 | 111546000 | 111546799 | 10.89332 | 20.34321 | 27.85648 | 11.56553 | 0.004837 | 0.147306 | 0.993085 |
| 4 | 91354600 | 91355799 | 16.58361 | 29.08646 | 38.21267 | 16.89099 | -0.00423 | 0.640332 | 0.993105 |
| 8 | 25640400 | 25644199 | 51.48188 | 82.21732 | 104.4401 | 43.85577 | -0.00367 | 2.100507 | 0.993141 |
| 7 | 144637400 | 144639799 | 22.48525 | 38.48987 | 50.09504 | 21.96735 | 0.004341 | 0.962479 | 0.993142 |
| 5 | 21910400 | 21911799 | 16.55252 | 29.13157 | 38.05381 | 16.8473 | 0.00455 | 0.590683 | 0.993169 |
| 1 | 7809400 | 7813599 | 68.96894 | 109.2987 | 113.59 | 47.2535 | 0.274017 | 2.35271 | 0.993195 |
| 17 | 82272400 | 82278199 | 55.55982 | 88.9383 | 112.5498 | 47.41849 | 0.00544 | 2.035227 | 0.993203 |
| 11 | 39169000 | 39169799 | 9.081732 | 17.29713 | 24.45623 | 9.780473 | 0.004637 | -0.00857 | 0.993233 |
| 13 | 63362600 | 63363399 | 9.081732 | 17.29713 | 24.45623 | 9.780473 | 0.004637 | -0.00857 | 0.993233 |
| 4 | 86066200 | 86068599 | 23.53758 | 39.29836 | 64.10308 | 27.88916 | -0.32244 | 1.212613 | 0.993237 |
| 1 | 163955800 | 163957799 | 19.13924 | 33.10697 | 43.17985 | 18.94393 | 0.004243 | 0.845485 | 0.993237 |
| 7 | 48928800 | 48930799 | 14.40935 | 25.45423 | 33.53802 | 14.83702 | -0.00569 | 0.528578 | 0.993268 |
| 16 | 88965200 | 88966599 | 22.10389 | 38.56999 | 37.59724 | 16.55201 | 0.424208 | 0.7478 | 0.993301 |
| 8 | 52591200 | 52595599 | 71.9831 | 114.217 | 116.1573 | 48.38156 | 0.30696 | 2.320089 | 0.993305 |
| 10 | 88799200 | 88801399 | 26.75611 | 45.1866 | 45.90446 | 20.08127 | 0.358639 | 1.094035 | 0.993316 |
| 13 | 9885800 | 9888799 | 40.4835 | 65.43684 | 83.8812 | 35.83227 | 0.003681 | 1.711697 | 0.993324 |
| 16 | 54671600 | 54672599 | 12.56081 | 22.84894 | 30.657 | 13.20383 | -0.00436 | 0.318333 | 0.993364 |
| 2 | 119498000 | 119499999 | 17.55046 | 30.70479 | 40.33559 | 17.79146 | -0.00413 | 0.689151 | 0.993367 |
| 6 | 146097600 | 146099999 | 23.30848 | 39.76259 | 51.72566 | 22.67614 | 0.004291 | 0.998184 | 0.993383 |
| 19 | 45980800 | 45984599 | 40.62847 | 65.6889 | 84.66982 | 36.18175 | -0.00382 | 1.704078 | 0.99339 |
| 3 | 90708400 | 90708999 | 9.162408 | 17.18846 | 23.89786 | 9.846126 | -0.0075 | 0.094364 | 0.993393 |
| 7 | 34257800 | 34258799 | 13.82558 | 24.70486 | 32.52288 | 14.28259 | 0.004267 | 0.435969 | 0.993393 |
| 1 | 185475200 | 185477199 | 17.57377 | 30.81532 | 40.23804 | 17.78245 | 0.004458 | 0.653 | 0.993398 |
| 2 | 177368800 | 177369799 | 15.20266 | 27.26689 | 27.42299 | 12.0052 | 0.393661 | 0.412124 | 0.993437 |
| 11 | 8887600 | 8889399 | 21.98212 | 37.47449 | 49.02056 | 21.48907 | -0.00378 | 0.996249 | 0.993465 |
| 1 | 32627000 | 32629999 | 34.27075 | 55.92705 | 72.27558 | 31.0566 | 0.003706 | 1.518043 | 0.99347 |
| 17 | 70149800 | 70150199 | 5.644985 | 11.0256 | 17.63548 | 6.179749 | 0.005765 | -0.51784 | 0.993474 |
| 12 | 115925600 | 115927799 | 24.29208 | 41.52185 | 53.97393 | 23.6655 | 0.005739 | 0.986372 | 0.993484 |
| 18 | 38668200 | 38670199 | 15.04535 | 26.65299 | 34.90661 | 15.43375 | 0.004444 | 0.504105 | 0.993499 |
| 1 | 120497000 | 120498399 | 24.71807 | 41.63128 | 54.41307 | 23.78379 | -0.00382 | 1.118994 | 0.993506 |
| 10 | 59572000 | 59573599 | 13.02425 | 23.3951 | 31.08134 | 13.58944 | -0.00437 | 0.395289 | 0.993516 |
| 2 | 48947800 | 48951999 | 74.20032 | 116.9688 | 146.764 | 60.90346 | -0.00351 | 2.486339 | 0.993539 |
| 2 | 163798600 | 163802599 | 49.81508 | 79.57864 | 100.568 | 42.2473 | 0.004226 | 2.088271 | 0.993543 |
| 2 | 125798000 | 125803799 | 193.3694 | 284.2999 | 335.0027 | 137.9621 | 0.003138 | 3.824701 | 0.993571 |
| 5 | 8256200 | 8256999 | 13.36547 | 23.86643 | 31.60237 | 13.88632 | -0.00453 | 0.435874 | 0.993575 |
| 4 | 71536000 | 71539999 | 109.3716 | 167.4606 | 205.362 | 85.22405 | -0.00294 | 3.04374 | 0.993583 |
| 13 | 9884400 | 9884799 | 13.5608 | 24.12544 | 31.65843 | 13.95458 | 0.0054 | 0.4613 | 0.993584 |
| X | 118452600 | 118453799 | 14.96686 | 26.99874 | 35.69651 | 15.53613 | 0.006817 | 0.394186 | 0.993598 |
| 6 | 99783200 | 99784999 | 17.01708 | 29.8369 | 39.19746 | 17.32147 | -0.0043 | 0.648455 | 0.993601 |
| 8 | 3222200 | 3227799 | 50.58671 | 81.04566 | 102.8097 | 43.36984 | 0.003408 | 2.000313 | 0.993612 |
| 13 | 113648200 | 113651799 | 33.4384 | 54.79076 | 71.36442 | 30.71233 | -0.00391 | 1.459944 | 0.993632 |
| 19 | 13199000 | 13199399 | 8.458461 | 16.27993 | 23.61993 | 9.189805 | 0.004266 | -0.10623 | 0.993643 |
| 10 | 41185000 | 41185599 | 12.87088 | 22.97625 | 30.19275 | 13.27032 | 0.006212 | 0.431347 | 0.993654 |
| 18 | 73857200 | 73862199 | 48.49602 | 77.71639 | 99.06971 | 41.80871 | -0.00316 | 1.98932 | 0.993675 |
| 9 | 124030200 | 124045599 | 336.8632 | 477.9605 | 547.4267 | 226.4587 | -0.00635 | 4.367843 | 0.99368 |
| 3 | 130710600 | 130715799 | 60.18469 | 95.62026 | 120.238 | 50.14636 | 0.003039 | 2.273767 | 0.993694 |

| | | | | | | | | | |
|---|---|---|---|---|---|---|---|---|---|
| 12 | 28539400 | 28539999 | 8.367415 | 16.13374 | 23.67303 | 9.162948 | -0.00443 | -0.12172 | 0.993695 |
| 11 | 37371000 | 37376799 | 60.06143 | 95.43088 | 120.0078 | 50.05374 | 0.003044 | 2.272229 | 0.993697 |
| 10 | 122182000 | 122188399 | 58.11069 | 92.89666 | 117.365 | 49.35311 | 0.005683 | 2.078789 | 0.993782 |
| 8 | 68015000 | 68016599 | 11.35284 | 20.56069 | 35.35757 | 15.31257 | -0.36543 | 0.344823 | 0.993791 |
| 6 | 11448400 | 11452199 | 68.2502 | 107.7002 | 134.6232 | 55.87981 | 0.003135 | 2.458882 | 0.99382 |
| Y | 41515200 | 41515799 | 8.015604 | 15.24604 | 21.42709 | 8.593491 | 0.010106 | -0.0181 | 0.993822 |
| 6 | 100365400 | 100368399 | 34.27922 | 55.86582 | 72.13675 | 30.98178 | 0.00354 | 1.537151 | 0.993827 |
| 2 | 169118200 | 169119999 | 16.7641 | 29.52027 | 38.57743 | 17.07723 | 0.004307 | 0.587524 | 0.993852 |
| 10 | 41648800 | 41663999 | 446.6726 | 613.3142 | 677.9732 | 281.9297 | -0.00289 | 4.861275 | 0.993854 |
| 5 | 138207000 | 138216599 | 253.4396 | 365.0904 | 421.8677 | 174.0925 | 0.002811 | 4.13988 | 0.993864 |
| 2 | 29670600 | 29673199 | 27.8708 | 46.27803 | 60.45945 | 26.20913 | -0.00343 | 1.281868 | 0.993909 |
| 4 | 3163000 | 3165599 | 65.824 | 102.7618 | 167.9616 | 69.69407 | -0.39529 | 2.667323 | 0.993909 |
| 10 | 51319200 | 51321399 | 26.85882 | 45.2198 | 46.45036 | 20.301 | 0.342509 | 1.120491 | 0.993914 |
| 2 | 111381400 | 111384599 | 44.0661 | 70.89905 | 90.81205 | 38.55212 | -0.0031 | 1.856954 | 0.993921 |
| 7 | 78120200 | 78122399 | 18.02687 | 31.39532 | 41.00575 | 18.03076 | 0.00372 | 0.75913 | 0.993928 |
| 2 | 5553400 | 5557999 | 37.95078 | 61.48992 | 79.39049 | 33.98417 | -0.00328 | 1.650907 | 0.993954 |
| 1 | 167357200 | 167362199 | 56.02084 | 89.43939 | 113.5585 | 47.66456 | -0.00348 | 2.115752 | 0.993959 |
| 8 | 40525600 | 40526999 | 14.16852 | 25.17217 | 33.03344 | 14.57746 | 0.004013 | 0.475465 | 0.993972 |
| 13 | 67375200 | 67380999 | 66.0985 | 104.6581 | 131.1884 | 54.54739 | 0.002967 | 2.375347 | 0.993974 |
| 11 | 105527200 | 105531199 | 50.42982 | 81.00389 | 102.9848 | 43.61521 | 0.004651 | 1.922891 | 0.993983 |
| 6 | 12106200 | 12107999 | 16.96226 | 29.67784 | 38.76201 | 17.09841 | 0.004074 | 0.680723 | 0.993997 |
| 2 | 92825800 | 92827399 | 16.52981 | 28.93052 | 37.78967 | 16.67404 | 0.004034 | 0.672776 | 0.994031 |
| 7 | 33874800 | 33881599 | 74.5474 | 117.0335 | 145.7349 | 60.44404 | 0.00287 | 2.56702 | 0.994062 |
| 5 | 70497800 | 70501799 | 48.61666 | 78.07233 | 99.80978 | 42.2548 | -0.00374 | 1.927194 | 0.994136 |
| 8 | 19719400 | 19722199 | 49.02785 | 78.88806 | 82.48879 | 35.02248 | 0.289147 | 1.857276 | 0.994147 |
| 5 | 110052600 | 110055399 | 137.8228 | 206.79 | 249.5987 | 102.995 | -0.00449 | 3.459334 | 0.994149 |
| 1 | 8137600 | 8143599 | 131.7056 | 199.3774 | 241.3224 | 99.83612 | 0.002658 | 3.282077 | 0.994173 |
| 8 | 65413200 | 65415199 | 23.65284 | 39.98841 | 52.25398 | 22.86632 | -0.00331 | 1.079241 | 0.994201 |
| 15 | 93986000 | 93987999 | 24.72697 | 40.85521 | 67.767 | 29.27676 | -0.34733 | 1.336678 | 0.99422 |
| 4 | 105642800 | 105645599 | 24.71608 | 42.17297 | 43.66642 | 19.14336 | 0.332323 | 0.990813 | 0.994239 |
| 1 | 182373600 | 182374799 | 25.19965 | 42.01523 | 54.92631 | 23.87582 | -0.00409 | 1.222745 | 0.994267 |
| 15 | 93128600 | 93137999 | 136.7776 | 206.5399 | 250.3861 | 103.5044 | -0.00268 | 3.333199 | 0.994275 |
| 6 | 130885600 | 130889999 | 49.33895 | 79.57025 | 101.4816 | 43.22433 | 0.00718 | 1.799128 | 0.994282 |
| 5 | 26345400 | 26346399 | 13.85566 | 24.97257 | 33.0588 | 14.40375 | 0.003987 | 0.384987 | 0.994288 |
| 14 | 96533400 | 96535599 | 20.02475 | 34.82913 | 45.73788 | 20.09499 | -0.00435 | 0.793727 | 0.994289 |
| X | 75196200 | 75201399 | 72.06485 | 114.4245 | 111.7561 | 46.58008 | 0.366192 | 2.302161 | 0.994327 |
| 11 | 34676400 | 34677599 | 10.46847 | 19.40481 | 26.32032 | 11.05571 | 0.00585 | 0.186812 | 0.994334 |
| 7 | 4844400 | 4851799 | 66.3066 | 104.6995 | 131.5124 | 54.60609 | -0.00317 | 2.44025 | 0.994357 |
| 1 | 13502600 | 13506799 | 29.77951 | 49.18761 | 63.93682 | 27.61597 | 0.003189 | 1.34841 | 0.994361 |
| 4 | 94998400 | 95002399 | 42.08553 | 68.20918 | 88.22253 | 37.77298 | -0.00546 | 1.659914 | 0.99437 |
| 5 | 129054800 | 129058399 | 33.55175 | 54.74485 | 70.76854 | 30.40724 | 0.003186 | 1.520275 | 0.994375 |
| 3 | 67682000 | 67685399 | 55.44677 | 88.42288 | 112.0734 | 46.96667 | -0.00273 | 2.151156 | 0.994406 |
| 14 | 113255400 | 113258199 | 56.56074 | 89.88943 | 113.587 | 47.38665 | -0.00358 | 2.260313 | 0.994411 |
| X | 164760000 | 164761199 | 11.71117 | 21.6312 | 29.26869 | 12.35384 | 0.00383 | 0.222156 | 0.994495 |
| X | 100102400 | 100105999 | 33.84701 | 55.37258 | 71.71458 | 30.84339 | 0.003129 | 1.483221 | 0.994496 |
| 9 | 74639600 | 74640799 | 13.02551 | 23.71531 | 31.65727 | 13.61951 | 0.003869 | 0.318156 | 0.994498 |
| 18 | 18565000 | 18568399 | 39.02262 | 63.37824 | 81.60126 | 34.99228 | 0.003937 | 1.607942 | 0.994502 |

| | | | | | | | | | |
|---|---|---|---|---|---|---|---|---|---|
| 4 | 143799400 | 143830799 | 771.6297 | 1012.751 | 1069.727 | 447.681 | -0.00258 | 5.580384 | 0.994509 |
| 14 | 47466000 | 47468199 | 22.44645 | 38.30478 | 50.09414 | 21.96148 | -0.00325 | 0.989512 | 0.994569 |
| 8 | 70006400 | 70008199 | 25.49577 | 42.69164 | 55.52612 | 24.20195 | 0.003122 | 1.178105 | 0.99457 |
| 3 | 11439200 | 11440199 | 8.901991 | 16.89077 | 23.86446 | 9.620489 | -0.00454 | 0.013296 | 0.994589 |
| 10 | 71510000 | 71511599 | 15.81515 | 27.85661 | 36.60503 | 16.20324 | -0.00335 | 0.576611 | 0.99459 |
| 12 | 118130200 | 118131399 | 15.09587 | 26.84898 | 35.44878 | 15.62896 | -0.00401 | 0.477973 | 0.994594 |
| 4 | 72363600 | 72366199 | 24.56407 | 41.67959 | 54.49109 | 23.87147 | -0.00393 | 1.052326 | 0.99461 |
| 10 | 4409000 | 4410399 | 16.24721 | 28.48967 | 37.22913 | 16.45294 | 0.003469 | 0.640608 | 0.994613 |
| 8 | 67888000 | 67889199 | 12.29785 | 22.26154 | 29.57316 | 12.81511 | 0.004025 | 0.340062 | 0.994621 |
| 12 | 3250400 | 3257399 | 71.00594 | 111.7377 | 139.4009 | 57.83478 | 0.002801 | 2.51647 | 0.994622 |
| 18 | 4640600 | 4642599 | 14.56129 | 25.85102 | 33.91308 | 14.97489 | 0.003645 | 0.482197 | 0.994623 |
| 15 | 14902000 | 14903399 | 16.83821 | 30.01372 | 27.97336 | 12.31031 | 0.497959 | 0.464812 | 0.994624 |
| 12 | 54824200 | 54825799 | 27.61642 | 45.06785 | 76.1076 | 32.70325 | -0.3811 | 1.509653 | 0.994647 |
| 16 | 59141200 | 59141599 | 8.315446 | 16.07617 | 23.8212 | 9.12209 | -0.00428 | -0.15945 | 0.994662 |
| 6 | 23889400 | 23889999 | 10.52917 | 19.95663 | 20.63469 | 8.337497 | 0.439595 | 0.027786 | 0.994662 |
| 13 | 56665400 | 56667799 | 22.21216 | 37.89238 | 49.55864 | 21.72614 | -0.00348 | 0.989721 | 0.994664 |
| 15 | 37859400 | 37861799 | 51.33834 | 82.17964 | 104.5928 | 44.06529 | -0.00277 | 2.029562 | 0.994665 |
| 6 | 101802000 | 101802399 | 7.999923 | 15.43879 | 22.57443 | 8.715578 | 0.003714 | -0.13871 | 0.994679 |
| 4 | 58404000 | 58407199 | 34.16717 | 55.80939 | 72.2116 | 31.03953 | 0.002913 | 1.506091 | 0.994686 |
| 8 | 22092200 | 22093799 | 18.8442 | 32.81234 | 43.06448 | 18.93368 | -0.00333 | 0.768564 | 0.994691 |
| 7 | 62993600 | 62995999 | 17.14598 | 29.96026 | 39.34289 | 17.33564 | -0.00364 | 0.704021 | 0.994691 |
| 6 | 33128200 | 33129199 | 14.19355 | 25.42369 | 33.53259 | 14.69606 | 0.003586 | 0.426091 | 0.994692 |
| 16 | 13890200 | 13890999 | 8.168868 | 15.62478 | 22.42991 | 8.884513 | -0.00439 | -0.05371 | 0.994734 |
| 8 | 38754000 | 38757599 | 64.68276 | 102.6431 | 128.9458 | 53.71329 | 0.002714 | 2.314838 | 0.994747 |
| 13 | 80366000 | 80367999 | 14.90649 | 26.57082 | 34.93985 | 15.3759 | 0.003618 | 0.461689 | 0.994758 |
| 7 | 39629400 | 39630599 | 11.69893 | 21.33413 | 28.67027 | 12.30979 | -0.00373 | 0.292375 | 0.994774 |
| 2 | 176783800 | 176786599 | 54.15446 | 86.33734 | 109.3983 | 45.82751 | -0.00274 | 2.158736 | 0.994803 |
| 5 | 73621800 | 73626399 | 50.81343 | 81.14507 | 103.0255 | 43.25821 | -0.00303 | 2.100984 | 0.994806 |
| 9 | 113587800 | 113589199 | 23.04525 | 38.93006 | 50.633 | 22.15161 | 0.003465 | 1.083424 | 0.994815 |
| 5 | 17811800 | 17814999 | 29.71497 | 49.04784 | 63.76406 | 27.5285 | 0.002975 | 1.35804 | 0.994819 |
| 7 | 62298600 | 62315999 | 438.2446 | 600.5715 | 662.4677 | 275.5665 | -0.00264 | 4.89098 | 0.994828 |
| 6 | 49289000 | 49289799 | 8.616558 | 16.46708 | 23.44591 | 9.309203 | 0.003877 | -0.03902 | 0.994847 |
| X | 31343000 | 31345199 | 34.00842 | 57.18157 | 48.56325 | 21.21595 | 0.615841 | 1.155867 | 0.994865 |
| 3 | 145795400 | 145802199 | 71.42945 | 111.7271 | 168.3115 | 69.8255 | -0.27565 | 2.634474 | 0.994873 |
| 8 | 126295800 | 126298199 | 15.55294 | 27.00434 | 46.34582 | 20.33987 | -0.38891 | 0.798361 | 0.994876 |
| 7 | 7073200 | 7081799 | 257.3433 | 368.2226 | 424.3987 | 175.3192 | -0.0032 | 4.238345 | 0.994908 |
| 4 | 3152400 | 3154199 | 23.03511 | 39.30533 | 51.16998 | 22.43295 | 0.003292 | 0.995473 | 0.994932 |
| 6 | 58206200 | 58207399 | 19.8095 | 34.44275 | 45.20475 | 19.85641 | -0.0036 | 0.796191 | 0.994962 |
| 10 | 57726400 | 57729599 | 44.88154 | 72.18097 | 92.04375 | 39.05143 | 0.002554 | 1.871486 | 0.994979 |
| 6 | 70095600 | 70096199 | 10.10652 | 18.99195 | 26.38158 | 10.83809 | -0.00327 | 0.097871 | 0.995029 |
| 3 | 138767200 | 138768599 | 19.14322 | 33.3458 | 43.56659 | 19.15906 | 0.003194 | 0.764748 | 0.995073 |
| 19 | 30373800 | 30375199 | 19.65993 | 34.09906 | 44.72229 | 19.62224 | -0.00299 | 0.826325 | 0.995087 |
| 13 | 36823200 | 36824599 | 12.45803 | 22.28838 | 38.28961 | 16.79779 | -0.37667 | 0.430069 | 0.9951 |
| 1 | 181563800 | 181565799 | 23.64275 | 40.00333 | 52.25623 | 22.87285 | -0.0028 | 1.072162 | 0.995103 |
| 8 | 13894800 | 13907399 | 203.5093 | 299.0412 | 352.3853 | 145.1257 | 0.002318 | 3.835126 | 0.995119 |
| 7 | 70366600 | 70368599 | 14.88486 | 26.25446 | 34.52845 | 15.2798 | -0.00391 | 0.549768 | 0.995131 |
| 13 | 53145400 | 53148599 | 33.86349 | 55.20874 | 71.35132 | 30.64859 | 0.002691 | 1.532387 | 0.995138 |

| | | | | | | | | | |
|---|---|---|---|---|---|---|---|---|---|
| 13 | 68055600 | 68081999 | 762.5838 | 999.3386 | 1053.835 | 441.1062 | -0.00243 | 5.604961 | 0.995138 |
| 17 | 48842800 | 48846799 | 28.59336 | 47.17145 | 61.59808 | 26.58564 | -0.00368 | 1.356419 | 0.99514 |
| 6 | 89838000 | 89844399 | 132.9734 | 200.8115 | 243.3988 | 100.6186 | -0.00232 | 3.330489 | 0.995153 |
| 17 | 16777600 | 16778199 | 7.101147 | 13.76476 | 20.525 | 7.763759 | 0.004095 | -0.22379 | 0.995158 |
| 8 | 41685000 | 41685799 | 7.773813 | 15.02146 | 22.08007 | 8.475292 | 0.005025 | -0.15889 | 0.995192 |
| 9 | 88982200 | 88987199 | 54.69932 | 87.29871 | 110.345 | 46.29258 | 0.002424 | 2.124266 | 0.995195 |
| 5 | 88583800 | 88586599 | 61.52606 | 97.62755 | 122.6705 | 51.09597 | 0.002337 | 2.312587 | 0.995197 |
| 4 | 23014800 | 23016199 | 14.54679 | 26.54749 | 35.44422 | 15.20389 | 0.00598 | 0.320917 | 0.995218 |
| 15 | 33233200 | 33234599 | 9.828111 | 18.2684 | 32.67518 | 13.67335 | -0.38722 | 0.178353 | 0.99524 |
| 2 | 5517400 | 5520799 | 29.64256 | 48.93906 | 63.87163 | 27.57912 | -0.00266 | 1.353777 | 0.99525 |
| 1 | 163372600 | 163373399 | 10.49852 | 19.79834 | 27.50009 | 11.23029 | 0.003286 | 0.079689 | 0.995254 |
| 19 | 38640400 | 38644399 | 36.14867 | 58.83597 | 76.23269 | 32.72542 | -0.00271 | 1.563188 | 0.995279 |
| 8 | 7154600 | 7156199 | 16.18673 | 28.80714 | 38.06852 | 16.77514 | -0.00474 | 0.487155 | 0.995283 |
| 19 | 9730000 | 9732199 | 42.727 | 68.70081 | 87.91326 | 37.28711 | -0.00286 | 1.873317 | 0.995296 |
| 3 | 47242800 | 47245399 | 26.72222 | 44.84113 | 58.31695 | 25.45271 | 0.003188 | 1.163152 | 0.995297 |
| 19 | 55822000 | 55824799 | 24.80775 | 41.84974 | 54.45258 | 23.8127 | 0.002693 | 1.104073 | 0.995357 |
| Y | 87888400 | 87888799 | 6.944676 | 13.38694 | 19.44694 | 7.532873 | 0.008833 | -0.15056 | 0.995362 |
| 2 | 150251200 | 150257399 | 125.9396 | 191.7358 | 234.071 | 97.00341 | -0.00292 | 3.166491 | 0.995371 |
| 2 | 101769400 | 101771999 | 23.45664 | 40.65155 | 52.95804 | 23.23517 | 0.006368 | 0.86157 | 0.995376 |
| 13 | 45277400 | 45281199 | 36.33292 | 59.5454 | 77.59097 | 33.39478 | -0.00443 | 1.469337 | 0.995402 |
| 16 | 87125600 | 87126799 | 11.03411 | 20.24571 | 27.21255 | 11.59026 | 0.00404 | 0.248687 | 0.995425 |
| 15 | 46907800 | 46908399 | 7.698738 | 14.91193 | 22.17173 | 8.416116 | 0.003064 | -0.20046 | 0.995428 |
| 12 | 49317200 | 49317799 | 13.52416 | 24.06409 | 31.61637 | 13.93417 | 0.003695 | 0.459084 | 0.995428 |
| 9 | 26258200 | 26259399 | 14.6021 | 26.28031 | 25.90154 | 11.30552 | 0.425418 | 0.381997 | 0.995429 |
| 9 | 92207800 | 92209199 | 11.52526 | 20.52253 | 36.73511 | 16.18005 | -0.43948 | 0.450097 | 0.99543 |
| 5 | 83218200 | 83218999 | 12.99636 | 23.43398 | 31.05488 | 13.52329 | 0.003004 | 0.371783 | 0.995448 |
| 9 | 86950600 | 86951999 | 37.04882 | 59.97581 | 77.05264 | 32.96207 | 0.002731 | 1.667685 | 0.995466 |
| 7 | 23707200 | 23710799 | 67.84642 | 107.7657 | 112.4758 | 46.89713 | 0.271058 | 2.285586 | 0.995467 |
| 3 | 36152400 | 36157999 | 57.45748 | 91.80488 | 116.6539 | 49.0181 | -0.00338 | 2.095796 | 0.995477 |
| 17 | 22184600 | 22227199 | 667.7697 | 892.9844 | 962.1015 | 401.7015 | -0.00424 | 5.281136 | 0.995483 |
| 16 | 16419600 | 16421399 | 18.93845 | 33.09698 | 43.46791 | 19.15542 | -0.00394 | 0.716729 | 0.995485 |
| 4 | 20174400 | 20175399 | 15.89509 | 27.75641 | 36.4725 | 16.06428 | -0.00448 | 0.703127 | 0.995494 |
| 6 | 122191000 | 122194199 | 40.63419 | 65.67789 | 84.54826 | 36.11543 | -0.0025 | 1.713049 | 0.995534 |
| 9 | 103268800 | 103275599 | 63.491 | 100.5925 | 126.6092 | 52.66004 | -0.00219 | 2.358193 | 0.995539 |
| 9 | 26610800 | 26611999 | 12.86682 | 23.08201 | 30.61851 | 13.40188 | -0.00323 | 0.401517 | 0.995558 |
| 9 | 100339400 | 100341199 | 12.86682 | 23.08201 | 30.61851 | 13.40188 | -0.00323 | 0.401517 | 0.995558 |
| 19 | 17918800 | 17919799 | 15.45632 | 27.16801 | 35.6616 | 15.7922 | -0.00295 | 0.595156 | 0.99556 |
| 17 | 69627600 | 69631199 | 46.42525 | 74.45089 | 94.94691 | 40.11223 | -0.00252 | 1.96457 | 0.995565 |
| 19 | 50304400 | 50308599 | 41.27716 | 66.69677 | 85.83239 | 36.64953 | -0.00256 | 1.723176 | 0.995592 |
| 5 | 31591400 | 31592799 | 16.45151 | 28.85809 | 37.72944 | 16.67901 | 0.002782 | 0.637646 | 0.995595 |
| X | 6316000 | 6316999 | 11.79783 | 21.88056 | 29.88156 | 12.54253 | -0.00351 | 0.202798 | 0.995597 |
| 12 | 36192400 | 36196999 | 37.26579 | 60.66105 | 78.29659 | 33.6128 | 0.002895 | 1.564245 | 0.995598 |
| 12 | 82114800 | 82116599 | 20.65006 | 35.43363 | 46.35885 | 20.33292 | -0.00259 | 0.928745 | 0.995623 |
| 4 | 8391800 | 8396199 | 45.7131 | 72.79026 | 121.4633 | 50.79547 | -0.40111 | 2.192812 | 0.995631 |
| 16 | 17766800 | 17771999 | 42.88465 | 69.02041 | 88.39434 | 37.54325 | -0.00234 | 1.843618 | 0.995644 |
| 2 | 148857000 | 148860999 | 45.49018 | 73.31397 | 73.28691 | 31.18826 | 0.356831 | 1.799757 | 0.995645 |
| 12 | 19380600 | 19383199 | 20.75276 | 36.13145 | 37.29063 | 16.39397 | 0.341911 | 0.77185 | 0.995659 |

| | | | | | | | | | |
|---|---|---|---|---|---|---|---|---|---|
| 10 | 119954800 | 119958799 | 41.29768 | 66.54591 | 85.35527 | 36.31546 | -0.00236 | 1.803731 | 0.995671 |
| 18 | 6338400 | 6359799 | 478.3164 | 650.9921 | 712.8753 | 296.8529 | -0.00215 | 4.999306 | 0.995683 |
| 3 | 45062200 | 45062799 | 14.66338 | 25.96226 | 34.03931 | 15.0519 | 0.002808 | 0.508229 | 0.995696 |
| 18 | 74126000 | 74129399 | 36.62337 | 59.57702 | 76.88596 | 32.99948 | 0.002472 | 1.571636 | 0.995714 |
| 1 | 170909000 | 170909799 | 21.85212 | 36.16844 | 65.82073 | 28.47029 | -0.48032 | 1.307648 | 0.995727 |
| 10 | 83682400 | 83683399 | 11.41955 | 21.16231 | 28.74176 | 12.07706 | 0.002806 | 0.197815 | 0.99575 |
| X | 39672400 | 39673999 | 17.21035 | 31.04547 | 28.25166 | 12.29607 | 0.542799 | 0.395345 | 0.995752 |
| 3 | 49384600 | 49390199 | 123.6422 | 187.6372 | 228.3197 | 94.52942 | -0.00198 | 3.228888 | 0.99576 |
| 7 | 9337000 | 9339799 | 69.90831 | 110.6665 | 110.5486 | 45.96042 | 0.329768 | 2.380399 | 0.995772 |
| 8 | 112865200 | 112870999 | 67.99189 | 107.2928 | 134.6035 | 55.87169 | -0.00213 | 2.457344 | 0.995795 |
| 9 | 18008800 | 18011599 | 49.98354 | 80.3709 | 83.22347 | 35.29945 | 0.30201 | 1.882152 | 0.995798 |
| 3 | 47004800 | 47007199 | 22.05271 | 37.0479 | 60.78296 | 26.54759 | -0.33079 | 1.139233 | 0.995806 |
| 19 | 9728200 | 9731999 | 55.88225 | 89.02981 | 112.349 | 47.01821 | 0.002059 | 2.182211 | 0.995856 |
| 1 | 73325800 | 73332399 | 131.9037 | 199.5644 | 241.5617 | 99.91732 | 0.001887 | 3.293566 | 0.995863 |
| 3 | 34720400 | 34721599 | 10.36997 | 19.27499 | 26.25403 | 10.98681 | 0.003326 | 0.16814 | 0.995868 |
| 7 | 13931400 | 13931799 | 10.81932 | 19.99368 | 27.19708 | 11.47181 | -0.0032 | 0.205792 | 0.99587 |
| 4 | 135336800 | 135341799 | 70.38084 | 110.8642 | 138.8773 | 57.62501 | -0.00198 | 2.497045 | 0.995891 |
| 16 | 68326200 | 68327199 | 16.5782 | 29.09426 | 38.17725 | 16.88381 | -0.00255 | 0.630555 | 0.995933 |
| 15 | 59103200 | 59107599 | 58.22259 | 92.66043 | 117.1228 | 48.94447 | -0.00194 | 2.219867 | 0.995941 |
| 8 | 19533000 | 19537799 | 61.1907 | 97.82177 | 97.7172 | 41.09182 | 0.344017 | 2.078538 | 0.995989 |
| 1 | 140583000 | 140584799 | 20.68582 | 35.76156 | 46.85962 | 20.55796 | -0.00264 | 0.863142 | 0.99599 |
| 1 | 102919000 | 102923999 | 51.1242 | 82.15817 | 104.5889 | 44.325 | 0.003585 | 1.907627 | 0.996009 |
| 10 | 107681200 | 107682999 | 14.66375 | 25.4132 | 46.36961 | 20.34452 | -0.47718 | 0.814613 | 0.996018 |
| 6 | 146000800 | 146004599 | 34.46006 | 56.44385 | 73.17628 | 31.48713 | 0.002578 | 1.468601 | 0.996032 |
| 4 | 106568200 | 106568999 | 10.00989 | 18.70016 | 25.76127 | 10.68909 | -0.00321 | 0.13057 | 0.996042 |
| 4 | 71533200 | 71534999 | 13.95815 | 24.61545 | 41.33808 | 18.29391 | -0.35477 | 0.560762 | 0.996044 |
| 8 | 38754200 | 38757799 | 64.01253 | 101.6132 | 127.7602 | 53.23706 | 0.001959 | 2.304312 | 0.99616 |
| 19 | 30376400 | 30379399 | 22.35644 | 38.44112 | 50.11469 | 21.98238 | 0.002928 | 0.919755 | 0.99616 |
| 19 | 49327400 | 49329999 | 27.77267 | 46.70585 | 47.50906 | 20.75662 | 0.356584 | 1.135603 | 0.996163 |
| 13 | 58203600 | 58205199 | 13.98554 | 25.02273 | 33.00755 | 14.47979 | 0.002422 | 0.429919 | 0.996169 |
| 15 | 99848800 | 99849599 | 13.98554 | 25.02273 | 33.00755 | 14.47979 | 0.002422 | 0.429919 | 0.996169 |
| 2 | 77792000 | 77793399 | 13.92679 | 24.96614 | 32.96758 | 14.43855 | 0.002416 | 0.416216 | 0.996191 |
| 3 | 10465200 | 10468599 | 32.95334 | 54.14987 | 70.30293 | 30.29196 | 0.002363 | 1.423689 | 0.9962 |
| 2 | 148755000 | 148765399 | 151.9907 | 228.9772 | 232.3829 | 95.99543 | 0.250663 | 3.381984 | 0.996206 |
| 10 | 15934400 | 15935199 | 10.93989 | 20.34709 | 27.74617 | 11.59951 | 0.00247 | 0.171089 | 0.996225 |
| 16 | 56778000 | 56780199 | 26.39358 | 43.99381 | 57.24561 | 24.87969 | 0.002488 | 1.228728 | 0.996293 |
| 1 | 191235400 | 191238599 | 28.37483 | 47.23743 | 61.67426 | 26.78341 | -0.00229 | 1.254929 | 0.996332 |
| 1 | 97453800 | 97454799 | 13.4777 | 24.4109 | 32.59367 | 14.11417 | -0.00268 | 0.35239 | 0.996335 |
| 10 | 6896600 | 6898399 | 18.84973 | 33.05138 | 43.40397 | 19.18093 | -0.00369 | 0.666737 | 0.996375 |
| 14 | 83195000 | 83196199 | 14.69519 | 26.48488 | 26.45113 | 11.52522 | 0.408202 | 0.377079 | 0.996413 |
| 1 | 164802400 | 164814599 | 277.1853 | 396.0635 | 455.3661 | 188.1528 | -0.00165 | 4.260139 | 0.99642 |
| 18 | 49887000 | 49889399 | 22.57719 | 37.49824 | 64.47687 | 27.95581 | -0.39842 | 1.266559 | 0.996421 |
| 5 | 141069600 | 141071199 | 20.02607 | 34.65384 | 45.2665 | 19.85961 | 0.002163 | 0.849925 | 0.996465 |
| 4 | 18946400 | 18947399 | 13.70351 | 24.617 | 32.55143 | 14.23133 | 0.002394 | 0.402984 | 0.996479 |
| 2 | 158583200 | 158584599 | 16.13571 | 28.23209 | 36.90949 | 16.28144 | 0.002989 | 0.676302 | 0.996481 |
| 13 | 6785600 | 6786199 | 6.371795 | 12.39711 | 19.02636 | 7.016815 | -0.00288 | -0.32563 | 0.996504 |
| 18 | 36678400 | 36679999 | 18.43823 | 32.06346 | 42.045 | 18.47026 | -0.0023 | 0.78241 | 0.996512 |

| | | | | | | | | | |
|---|---|---|---|---|---|---|---|---|---|
| 4 | 146271600 | 146279599 | 86.93384 | 135.9213 | 141.0668 | 58.5253 | 0.260785 | 2.645714 | 0.996526 |
| 7 | 107819600 | 107821799 | 44.52412 | 71.48435 | 91.25785 | 38.62127 | -0.00206 | 1.923266 | 0.996549 |
| 1 | 35163600 | 35166999 | 39.47394 | 63.92786 | 82.4285 | 35.27244 | -0.002 | 1.665443 | 0.996581 |
| 1 | 39119600 | 39127399 | 99.49655 | 153.2426 | 188.1854 | 78.16489 | 0.001595 | 2.923078 | 0.996588 |
| 10 | 126629200 | 126632399 | 23.39282 | 39.72497 | 51.72323 | 22.66459 | 0.002029 | 1.038003 | 0.996594 |
| 9 | 7057200 | 7057599 | 7.397623 | 14.38259 | 21.86545 | 8.111034 | 0.002173 | -0.28445 | 0.996596 |
| 3 | 31729600 | 31730199 | 10.52529 | 19.59527 | 26.76353 | 11.17006 | 0.002411 | 0.160416 | 0.996623 |
| 3 | 46870400 | 46871999 | 16.83321 | 29.53291 | 38.74497 | 17.13053 | -0.00214 | 0.639417 | 0.996628 |
| 10 | 124380200 | 124391199 | 139.4508 | 211.1164 | 255.6504 | 105.7659 | 0.002063 | 3.283221 | 0.996657 |
| 5 | 28224200 | 28225199 | 12.45171 | 22.57402 | 30.06115 | 13.00206 | 0.002256 | 0.333473 | 0.99666 |
| 8 | 108900200 | 108902599 | 16.04384 | 28.22991 | 37.05001 | 16.40364 | -0.00233 | 0.588211 | 0.996671 |
| 1 | 10893400 | 10895399 | 31.46328 | 51.82583 | 67.37203 | 29.05866 | 0.001909 | 1.38873 | 0.996672 |
| 16 | 25909200 | 25911399 | 21.20622 | 36.91092 | 48.45055 | 21.29654 | -0.00367 | 0.792516 | 0.996706 |
| 10 | 61349600 | 61351999 | 19.94258 | 34.31338 | 44.76976 | 19.63828 | 0.002101 | 0.898306 | 0.996707 |
| 10 | 21522800 | 21529999 | 83.64189 | 130.5152 | 161.9075 | 67.18536 | 0.001519 | 2.682405 | 0.996722 |
| 6 | 56372000 | 56373199 | 9.249096 | 17.60706 | 24.99055 | 10.00476 | -0.00223 | -0.00193 | 0.996724 |
| 10 | 37899600 | 37901599 | 19.99821 | 35.07307 | 46.08617 | 20.37013 | -0.00463 | 0.672426 | 0.996767 |
| 2 | 16031600 | 16033799 | 39.50103 | 63.62995 | 81.30043 | 34.57382 | 0.002649 | 1.809702 | 0.996772 |
| 10 | 66299400 | 66299799 | 8.791194 | 17.11188 | 26.17672 | 9.619553 | 0.00528 | -0.28984 | 0.996781 |
| 19 | 9694800 | 9696999 | 28.78997 | 47.70476 | 62.24261 | 26.94334 | -0.00185 | 1.308108 | 0.996786 |
| 3 | 12650000 | 12655799 | 121.002 | 184.0186 | 223.7696 | 92.70419 | 0.001459 | 3.185093 | 0.996793 |
| 17 | 22538600 | 22544599 | 58.27267 | 93.42981 | 90.97489 | 38.42137 | 0.385603 | 1.984764 | 0.996818 |
| 5 | 78287200 | 78287599 | 8.200615 | 15.8182 | 23.12152 | 8.939602 | 0.002547 | -0.12879 | 0.996871 |
| 19 | 61288400 | 61289199 | 8.200615 | 15.8182 | 23.12152 | 8.939602 | 0.002547 | -0.12879 | 0.996871 |
| 5 | 126004400 | 126007999 | 30.94967 | 50.87313 | 66.08495 | 28.47826 | 0.001773 | 1.411638 | 0.996894 |
| 14 | 44456400 | 44459799 | 50.1513 | 80.19508 | 101.8686 | 42.8537 | -0.00162 | 2.058913 | 0.99691 |
| 13 | 76083800 | 76086799 | 28.65594 | 47.97164 | 62.41543 | 27.20062 | 0.002539 | 1.196598 | 0.996916 |
| 1 | 129724000 | 129725799 | 22.6774 | 38.74357 | 50.63179 | 22.19925 | -0.00194 | 0.981722 | 0.996928 |
| 2 | 144607200 | 144608999 | 22.44618 | 38.42694 | 50.09814 | 21.96889 | 0.001893 | 0.960726 | 0.996945 |
| 5 | 118693200 | 118695199 | 19.62477 | 33.80776 | 44.24716 | 19.40976 | -0.00198 | 0.885532 | 0.996946 |
| X | 97626200 | 97627799 | 18.15157 | 31.86204 | 41.83069 | 18.50275 | -0.00315 | 0.64549 | 0.996971 |
| 3 | 143735000 | 143736399 | 17.6513 | 30.847 | 40.35345 | 17.78268 | 0.001822 | 0.706535 | 0.996974 |
| 1 | 53321800 | 53324599 | 42.93155 | 69.35603 | 89.22986 | 38.09032 | -0.00246 | 1.736247 | 0.996975 |
| 11 | 79119800 | 79129999 | 116.8296 | 177.1901 | 256.0351 | 105.9848 | -0.25044 | 3.24196 | 0.996992 |
| 1 | 91403800 | 91404599 | 8.666422 | 16.52218 | 23.43949 | 9.354606 | 0.002887 | -0.02035 | 0.997001 |
| 8 | 90326800 | 90332799 | 45.54605 | 73.42826 | 93.93985 | 39.99445 | 0.00214 | 1.798218 | 0.997009 |
| 15 | 30251600 | 30252799 | 14.39737 | 26.16146 | 26.92407 | 11.61176 | 0.374638 | 0.329384 | 0.997029 |
| 5 | 127468200 | 127471199 | 34.78288 | 56.79917 | 73.5383 | 31.60657 | 0.001698 | 1.511251 | 0.997037 |
| 16 | 47363200 | 47364399 | 15.40648 | 27.19793 | 35.63503 | 15.76678 | 0.002265 | 0.54957 | 0.997045 |
| 14 | 106587000 | 106591399 | 75.39082 | 118.2458 | 147.2832 | 61.08455 | 0.001437 | 2.585485 | 0.997073 |
| 12 | 17873200 | 17875199 | 29.05214 | 48.58218 | 50.76006 | 22.10608 | 0.318075 | 1.199317 | 0.997085 |
| 7 | 20025600 | 20025999 | 10.44276 | 19.89334 | 19.22955 | 7.686965 | 0.549958 | -0.01004 | 0.997122 |
| 16 | 20923400 | 20924399 | 21.42218 | 36.68317 | 47.95312 | 21.02869 | -0.00174 | 0.954405 | 0.997153 |
| 11 | 107993800 | 107995599 | 19.54152 | 33.32346 | 54.85488 | 24.04721 | -0.33399 | 0.992805 | 0.997178 |
| 3 | 9721200 | 9724399 | 35.97547 | 58.31323 | 75.05675 | 32.13873 | 0.001736 | 1.636586 | 0.997178 |
| 8 | 7640800 | 7641599 | 9.035884 | 17.2457 | 24.51857 | 9.762291 | 0.002909 | -0.02013 | 0.997194 |
| 3 | 22389000 | 22390799 | 37.85445 | 61.22201 | 78.81685 | 33.69325 | -0.00154 | 1.688593 | 0.997194 |

| | | | | | | | | | |
|---|---|---|---|---|---|---|---|---|---|
| 1 | 46418200 | 46420799 | 23.69195 | 40.42316 | 52.81915 | 23.15581 | -0.00205 | 1.00014 | 0.997247 |
| 8 | 67502600 | 67503199 | 10.58623 | 19.82584 | 27.38115 | 11.31374 | -0.00177 | 0.125897 | 0.99725 |
| 7 | 45980800 | 45982599 | 20.13351 | 34.93001 | 45.77735 | 20.08532 | -0.00187 | 0.826986 | 0.997279 |
| 1 | 63877600 | 63883599 | 48.13362 | 77.54352 | 99.45048 | 42.29851 | -0.0027 | 1.824922 | 0.997282 |
| 4 | 88728800 | 88730399 | 13.7836 | 24.63381 | 32.48775 | 14.26523 | 0.00173 | 0.433721 | 0.997286 |
| X | 132245600 | 132249399 | 42.0296 | 68.50027 | 65.21492 | 28.00946 | 0.44223 | 1.552927 | 0.997304 |
| 8 | 84604600 | 84605599 | 15.19921 | 26.81674 | 35.13476 | 15.5472 | 0.001943 | 0.552868 | 0.997309 |
| 3 | 130607800 | 130609999 | 16.61408 | 29.17291 | 38.16912 | 16.88789 | 0.001668 | 0.622771 | 0.997318 |
| X | 70860000 | 70860799 | 9.857768 | 18.77397 | 26.67282 | 10.66837 | -0.00213 | 0.007736 | 0.997339 |
| 10 | 67095600 | 67099399 | 40.52276 | 65.40476 | 83.82188 | 35.73958 | 0.001537 | 1.751276 | 0.997374 |
| 11 | 61482200 | 61485999 | 41.78612 | 67.23183 | 86.03057 | 36.52312 | -0.00162 | 1.851699 | 0.997396 |
| 19 | 31831600 | 31836799 | 85.97649 | 134.0626 | 166.2447 | 68.99077 | 0.001266 | 2.696583 | 0.997405 |
| 9 | 70793800 | 70799599 | 44.18736 | 71.07868 | 90.90509 | 38.57945 | -0.00133 | 1.863145 | 0.997471 |
| 1 | 51423400 | 51424799 | 27.80413 | 46.02355 | 59.9017 | 25.91148 | 0.001596 | 1.315323 | 0.997531 |
| 2 | 4914400 | 4919199 | 46.4944 | 74.61427 | 95.15047 | 40.24127 | -0.00128 | 1.941141 | 0.997535 |
| 6 | 143905600 | 143907999 | 27.11109 | 45.12605 | 58.88336 | 25.5687 | -0.00148 | 1.248539 | 0.997539 |
| 6 | 69424200 | 69424799 | 11.1613 | 20.74424 | 28.28413 | 11.83763 | 0.00155 | 0.179672 | 0.997562 |
| 5 | 104507000 | 104517599 | 87.40094 | 134.9056 | 199.1034 | 82.70366 | -0.26097 | 2.879577 | 0.997569 |
| 4 | 145154600 | 145178399 | 791.311 | 1043.977 | 1108.279 | 463.547 | -0.0024 | 5.498168 | 0.997574 |
| 12 | 97350200 | 97350999 | 12.82697 | 23.27304 | 31.09646 | 13.43635 | -0.00157 | 0.334784 | 0.997604 |
| 16 | 56775800 | 56776799 | 12.82697 | 23.27304 | 31.09646 | 13.43635 | -0.00157 | 0.334784 | 0.997604 |
| 10 | 96368400 | 96370399 | 16.02555 | 28.24904 | 37.00149 | 16.3762 | 0.001497 | 0.567808 | 0.997614 |
| 4 | 135226800 | 135229799 | 32.12127 | 52.62141 | 68.28483 | 29.38436 | 0.00134 | 1.45747 | 0.997629 |
| 8 | 4038600 | 4049599 | 165.908 | 247.1991 | 295.347 | 121.7062 | 0.001079 | 3.595409 | 0.99763 |
| 7 | 57605200 | 57606799 | 13.3463 | 24.43046 | 32.80173 | 14.0157 | 0.001959 | 0.291927 | 0.997641 |
| 4 | 50405200 | 50410799 | 103.4742 | 158.8337 | 194.6101 | 80.79617 | 0.001161 | 2.990324 | 0.997649 |
| 1 | 122012600 | 122015199 | 31.19772 | 50.6683 | 81.37274 | 34.89367 | -0.3141 | 1.593186 | 0.997677 |
| 3 | 127425200 | 127428599 | 32.21847 | 52.04177 | 88.64871 | 37.83544 | -0.40666 | 1.718864 | 0.997686 |
| 15 | 65395800 | 65396399 | 6.087017 | 11.86854 | 18.60543 | 6.699066 | -0.00138 | -0.4142 | 0.997714 |
| 18 | 64261400 | 64263799 | 28.51039 | 47.40488 | 61.85779 | 26.84066 | -0.00169 | 1.265455 | 0.997744 |
| 15 | 24137200 | 24139199 | 38.25101 | 62.30142 | 64.93387 | 27.88215 | 0.311258 | 1.551848 | 0.997744 |
| 10 | 20167800 | 20170799 | 21.53216 | 36.72589 | 47.9649 | 21.02715 | -0.00132 | 0.989671 | 0.997752 |
| 4 | 37611000 | 37612799 | 22.09142 | 37.61924 | 49.02124 | 21.48746 | 0.001597 | 1.008854 | 0.997764 |
| 15 | 4233200 | 4234799 | 24.05435 | 41.19689 | 42.64723 | 18.70215 | 0.33352 | 0.955926 | 0.997766 |
| 14 | 80673200 | 80678399 | 65.36505 | 103.9114 | 109.1742 | 45.5587 | 0.262681 | 2.258411 | 0.997778 |
| 17 | 15753000 | 15754399 | 16.33095 | 29.26525 | 29.70694 | 13.01454 | 0.379497 | 0.429086 | 0.997779 |
| 10 | 78771800 | 78776999 | 94.45505 | 145.7858 | 179.6523 | 74.62406 | -0.00108 | 2.881076 | 0.997784 |
| 8 | 48250400 | 48252799 | 39.76012 | 64.16346 | 82.23313 | 35.05485 | 0.001264 | 1.755225 | 0.997795 |
| 9 | 11016600 | 11017999 | 19.2077 | 33.37434 | 43.65161 | 19.16622 | 0.001366 | 0.801807 | 0.997798 |
| 2 | 36352600 | 36355599 | 38.94959 | 63.20927 | 81.62907 | 34.98366 | -0.00148 | 1.622226 | 0.997802 |
| 9 | 50305600 | 50309799 | 40.60769 | 65.49501 | 83.89128 | 35.7362 | 0.001179 | 1.774 | 0.997836 |
| 6 | 68388200 | 68390799 | 32.2896 | 53.26961 | 69.31067 | 29.91447 | 0.001493 | 1.373209 | 0.997844 |
| 1 | 81656600 | 81657399 | 9.146374 | 17.34058 | 24.41511 | 9.859148 | -0.00126 | 0.025173 | 0.997849 |
| 9 | 21816200 | 21818999 | 26.61659 | 44.34889 | 57.75876 | 25.09677 | 0.00126 | 1.235122 | 0.997856 |
| 10 | 38689200 | 38704599 | 179.126 | 266.4883 | 318.4606 | 131.2106 | -0.00125 | 3.624582 | 0.997868 |
| 16 | 15263600 | 15268199 | 44.09856 | 71.09742 | 91.01911 | 38.7529 | 0.00126 | 1.794423 | 0.997891 |
| 7 | 96965600 | 96967199 | 17.18415 | 30.02487 | 39.36631 | 17.3452 | -0.00135 | 0.706003 | 0.9979 |

| | | | | | | | | | |
|---|---|---|---|---|---|---|---|---|---|
| 10 | 117842800 | 117846999 | 47.97133 | 76.98708 | 98.02034 | 41.45716 | 0.001055 | 1.942928 | 0.997944 |
| 12 | 57600000 | 57601199 | 12.22575 | 22.21715 | 29.66172 | 12.7912 | 0.001649 | 0.319421 | 0.997957 |
| 17 | 66776200 | 66779199 | 33.55149 | 54.58381 | 70.506 | 30.26201 | 0.001319 | 1.561858 | 0.99801 |
| 7 | 85995600 | 85999599 | 46.11112 | 74.19561 | 94.76284 | 40.23427 | 0.001168 | 1.859875 | 0.998058 |
| X | 80921800 | 80925399 | 46.78257 | 75.3211 | 96.44307 | 40.98321 | -0.0014 | 1.842927 | 0.998061 |
| 16 | 19017400 | 19027799 | 115.0683 | 176.3851 | 215.877 | 89.60844 | 0.001211 | 3.023898 | 0.9981 |
| 4 | 5927400 | 5928199 | 14.14385 | 25.08997 | 33.01704 | 14.58901 | -0.0013 | 0.484206 | 0.998101 |
| 17 | 53524200 | 53525399 | 14.09239 | 24.76565 | 41.35667 | 18.3111 | -0.34859 | 0.600094 | 0.998106 |
| X | 153384600 | 153388999 | 50.97662 | 81.95896 | 104.7917 | 44.44078 | -0.00185 | 1.895711 | 0.998146 |
| 13 | 58372200 | 58375599 | 39.1736 | 63.37686 | 81.47153 | 34.83676 | 0.000999 | 1.685493 | 0.998159 |
| 1 | 72202800 | 72203999 | 12.69695 | 22.78831 | 30.13482 | 13.18479 | 0.001658 | 0.396141 | 0.998163 |
| 13 | 20881600 | 20882199 | 9.678575 | 18.21295 | 25.31002 | 10.37355 | -0.00153 | 0.080009 | 0.998184 |
| 17 | 77552000 | 77552799 | 9.678575 | 18.21295 | 25.31002 | 10.37355 | -0.00153 | 0.080009 | 0.998184 |
| 8 | 55828600 | 55830199 | 14.49592 | 25.31138 | 43.66689 | 19.2322 | -0.39525 | 0.711354 | 0.998196 |
| 12 | 17287800 | 17289199 | 13.69451 | 24.35566 | 32.0995 | 14.15275 | -0.00119 | 0.464283 | 0.998198 |
| 1 | 141217800 | 141219399 | 18.62934 | 32.44669 | 42.52146 | 18.69801 | -0.0012 | 0.762216 | 0.998216 |
| 14 | 13900000 | 13900799 | 10.0955 | 19.01423 | 26.40698 | 10.80703 | 0.00141 | 0.083228 | 0.998218 |
| 3 | 93195200 | 93196599 | 16.15339 | 27.76601 | 49.5106 | 21.71719 | -0.4467 | 0.90101 | 0.998255 |
| 13 | 32586200 | 32587599 | 18.07518 | 31.59083 | 41.41275 | 18.25097 | -0.00109 | 0.710351 | 0.998306 |
| 11 | 73932400 | 73936399 | 56.54505 | 90.28418 | 114.2876 | 47.97651 | 0.000978 | 2.114251 | 0.998316 |
| 5 | 138697800 | 138699199 | 18.43093 | 32.07017 | 42.02281 | 18.46754 | -0.00117 | 0.773618 | 0.998323 |
| 16 | 86590000 | 86591399 | 14.82902 | 26.87037 | 35.78038 | 15.48551 | 0.001638 | 0.365679 | 0.998326 |
| 8 | 129755400 | 129757399 | 22.48272 | 38.36276 | 50.09129 | 21.96005 | -0.00099 | 0.991237 | 0.998331 |
| 17 | 7724400 | 7726599 | 18.64851 | 32.22415 | 42.17875 | 18.50414 | -0.00117 | 0.851296 | 0.998366 |
| 2 | 4214200 | 4216999 | 21.70869 | 37.11662 | 48.48139 | 21.25771 | -0.00114 | 0.96615 | 0.998377 |
| 18 | 12839600 | 12847999 | 61.17179 | 97.60531 | 100.9304 | 42.3286 | 0.291217 | 2.135043 | 0.998383 |
| 11 | 103563200 | 103564799 | 15.64985 | 27.54775 | 36.13017 | 15.9951 | -0.00094 | 0.581906 | 0.998412 |
| 4 | 143701400 | 143712799 | 280.5491 | 400.171 | 458.521 | 189.5117 | 0.000736 | 4.285714 | 0.998418 |
| 1 | 80329600 | 80329999 | 7.740839 | 14.95875 | 22.07758 | 8.471657 | -0.0007 | -0.16175 | 0.998471 |
| 7 | 39456800 | 39458799 | 16.56724 | 29.11009 | 38.10539 | 16.8692 | 0.000886 | 0.610774 | 0.998488 |
| 4 | 10770000 | 10772599 | 23.11656 | 39.26513 | 51.16882 | 22.42327 | 0.000845 | 1.035369 | 0.998492 |
| 13 | 72545200 | 72546399 | 10.70882 | 19.75163 | 26.73467 | 11.30618 | 0.001432 | 0.21565 | 0.9985 |
| 9 | 58401200 | 58402399 | 14.35514 | 25.33483 | 33.20986 | 14.69458 | 0.001428 | 0.540491 | 0.998503 |
| 1 | 22911200 | 22914199 | 26.74909 | 44.64727 | 58.23498 | 25.33148 | -0.00086 | 1.220287 | 0.998503 |
| X | 6311000 | 6313799 | 18.86752 | 33.41089 | 30.75945 | 13.60117 | 0.51043 | 0.528984 | 0.998511 |
| 7 | 137431200 | 137432599 | 12.70043 | 23.23911 | 31.2122 | 13.34362 | 0.001067 | 0.286429 | 0.998526 |
| 4 | 16234200 | 16235199 | 10.80281 | 20.00978 | 27.20126 | 11.43723 | 0.000979 | 0.192343 | 0.998546 |
| 5 | 86007600 | 86008799 | 15.57327 | 27.75412 | 36.55663 | 16.08995 | 0.001265 | 0.472936 | 0.998547 |
| 6 | 142094600 | 142096799 | 33.54505 | 54.59183 | 70.55137 | 30.28522 | 0.000945 | 1.55688 | 0.998548 |
| 14 | 114329600 | 114334999 | 80.24188 | 125.3886 | 155.8055 | 64.64374 | 0.000731 | 2.653802 | 0.998565 |
| 18 | 72904200 | 72908199 | 80.91319 | 126.1311 | 156.5933 | 64.9883 | -0.00073 | 2.698401 | 0.99859 |
| 18 | 32767600 | 32769399 | 18.42439 | 32.50674 | 32.69198 | 14.4663 | 0.381294 | 0.565627 | 0.998592 |
| X | 92064000 | 92065799 | 19.04068 | 33.266 | 43.53765 | 19.18217 | 0.001298 | 0.725947 | 0.998595 |
| 3 | 124596000 | 124596799 | 9.303438 | 17.87513 | 25.86963 | 10.11549 | 0.001129 | -0.07437 | 0.998609 |
| 14 | 95802600 | 95806999 | 94.2494 | 145.6328 | 179.5633 | 74.58667 | -0.00063 | 2.860998 | 0.998624 |
| 4 | 61475400 | 61485399 | 158.6719 | 237.1168 | 284.4032 | 117.2593 | -0.00065 | 3.539024 | 0.998644 |
| 6 | 9602600 | 9604799 | 27.15012 | 44.72154 | 71.93963 | 31.02881 | -0.30451 | 1.381943 | 0.998649 |

| | | | | | | | | | |
|---|---|---|---|---|---|---|---|---|---|
| 12 | 115875800 | 115876799 | 13.83904 | 24.68574 | 32.54097 | 14.3116 | 0.000932 | 0.447235 | 0.998661 |
| 18 | 8236000 | 8236799 | 13.0109 | 23.41364 | 31.06074 | 13.5595 | -0.00079 | 0.38359 | 0.998711 |
| 3 | 90779800 | 90782599 | 35.31699 | 57.45664 | 74.31824 | 31.89839 | -0.00069 | 1.566568 | 0.998741 |
| 7 | 109803800 | 109809799 | 66.30797 | 105.2243 | 132.3722 | 55.14171 | 0.000731 | 2.315893 | 0.998761 |
| 3 | 27975000 | 27976799 | 20.91227 | 35.90553 | 46.87113 | 20.55862 | 0.00069 | 0.923684 | 0.998771 |
| 15 | 55538000 | 55541199 | 26.42545 | 44.34051 | 57.82495 | 25.23703 | -0.00084 | 1.159976 | 0.998796 |
| 9 | 82658400 | 82662999 | 61.45946 | 97.57845 | 122.928 | 51.23293 | -0.00061 | 2.294806 | 0.998803 |
| 19 | 48185600 | 48187999 | 20.1495 | 34.65584 | 45.30125 | 19.87118 | -0.00067 | 0.904182 | 0.998863 |
| 14 | 110303000 | 110305199 | 23.07166 | 39.22399 | 51.17655 | 22.42923 | -0.00075 | 1.026362 | 0.998873 |
| 3 | 144813800 | 144817199 | 47.83602 | 76.50748 | 97.14809 | 40.87899 | -0.00076 | 2.048661 | 0.99888 |
| 5 | 38582400 | 38583199 | 9.093553 | 17.28102 | 24.42385 | 9.813752 | -0.00071 | 0.007066 | 0.998885 |
| 12 | 17608200 | 17610599 | 23.89979 | 40.49416 | 52.82252 | 23.13041 | -0.00076 | 1.060483 | 0.998893 |
| 8 | 56532600 | 56533199 | 6.529595 | 12.71784 | 19.6463 | 7.173841 | 0.001109 | -0.35508 | 0.998897 |
| 16 | 70316600 | 70317199 | 10.05174 | 18.77529 | 25.78685 | 10.70267 | 0.000977 | 0.133119 | 0.998898 |
| 18 | 51204600 | 51207599 | 40.16995 | 64.7911 | 83.02339 | 35.36798 | 0.000676 | 1.773201 | 0.998902 |
| 6 | 67921600 | 67922599 | 17.47708 | 30.50498 | 39.97531 | 17.60203 | -0.00059 | 0.725661 | 0.998903 |
| 16 | 7857000 | 7862199 | 83.27482 | 129.5503 | 160.5648 | 66.65664 | -0.00055 | 2.734316 | 0.998919 |
| 12 | 20489800 | 20554999 | 2109.665 | 2563.073 | 2086.75 | 880.711 | 0.260572 | 6.826357 | 0.998948 |
| 2 | 70726600 | 70727599 | 11.50037 | 21.20884 | 28.71498 | 12.1458 | 0.000657 | 0.227026 | 0.998978 |
| 13 | 68343600 | 68458199 | 2770.822 | 3278.625 | 3114.005 | 1319.233 | -0.00051 | 7.242697 | 0.998979 |
| 10 | 89777600 | 89780599 | 25.92577 | 43.47342 | 56.68822 | 24.73064 | -0.00066 | 1.164192 | 0.998985 |
| 5 | 46537000 | 46538799 | 25.20243 | 42.24924 | 55.04166 | 24.00828 | 0.000626 | 1.161742 | 0.998989 |
| 6 | 132611200 | 132614599 | 28.91617 | 48.34424 | 63.05182 | 27.45538 | -0.00093 | 1.210388 | 0.999031 |
| 4 | 119259400 | 119265399 | 78.88586 | 123.8511 | 154.4857 | 64.0739 | 0.000482 | 2.570484 | 0.999063 |
| 9 | 81792400 | 81795799 | 62.23141 | 98.60415 | 123.882 | 51.52766 | 0.000518 | 2.355769 | 0.99907 |
| 1 | 112791200 | 112792999 | 20.3361 | 35.34096 | 46.29022 | 20.32692 | -0.00077 | 0.807419 | 0.999091 |
| 15 | 29474800 | 29477199 | 18.61871 | 32.67384 | 42.82578 | 18.9389 | -0.001 | 0.650271 | 0.999103 |
| 12 | 7950600 | 7951999 | 10.96493 | 20.38614 | 27.82435 | 11.63946 | 0.00105 | 0.177625 | 0.99911 |
| 2 | 73194200 | 73195599 | 12.32593 | 22.48549 | 30.13257 | 12.93191 | 0.000556 | 0.297831 | 0.999119 |
| 17 | 72746400 | 72749199 | 12.77644 | 22.47163 | 40.45174 | 17.91118 | -0.45548 | 0.585175 | 0.99913 |
| 7 | 49378600 | 49382799 | 32.623 | 53.50816 | 69.55894 | 29.94799 | -0.00053 | 1.444902 | 0.999132 |
| 3 | 77792600 | 77793799 | 14.20442 | 25.87323 | 26.90917 | 11.57035 | 0.362275 | 0.31374 | 0.999134 |
| 5 | 47735000 | 47738199 | 44.68544 | 72.33395 | 73.71886 | 31.53455 | 0.335936 | 1.679908 | 0.999164 |
| 3 | 5509800 | 5511799 | 20.6759 | 35.50448 | 46.35752 | 20.33351 | 0.000516 | 0.922632 | 0.999192 |
| 1 | 19038800 | 19040799 | 22.52734 | 38.40388 | 50.08706 | 21.95659 | 0.00051 | 1.000472 | 0.999193 |
| 7 | 5126800 | 5127799 | 8.467025 | 16.26729 | 23.56801 | 9.216822 | -0.00056 | -0.08939 | 0.999216 |
| 12 | 12998400 | 13001399 | 27.16206 | 45.13603 | 58.81107 | 25.50815 | 0.000467 | 1.268611 | 0.999222 |
| 7 | 44160200 | 44163199 | 37.20544 | 60.11986 | 77.26495 | 32.99294 | -0.00049 | 1.712025 | 0.999229 |
| 9 | 121415400 | 121418999 | 46.08255 | 73.97898 | 94.26938 | 39.88967 | 0.00042 | 1.931077 | 0.999249 |
| 12 | 17795800 | 17797999 | 19.06919 | 33.01086 | 43.20295 | 18.95451 | -0.00037 | 0.840188 | 0.99925 |
| 11 | 107141800 | 107143199 | 13.89191 | 24.73332 | 32.59938 | 14.36013 | -0.00033 | 0.460624 | 0.999265 |
| 15 | 5253200 | 5255599 | 23.64201 | 40.71883 | 42.01926 | 18.43232 | 0.339408 | 0.902112 | 0.999269 |
| 5 | 136801400 | 136806599 | 65.63282 | 103.8559 | 130.3549 | 54.18367 | 0.000351 | 2.389781 | 0.999272 |
| 19 | 16471600 | 16472999 | 12.89228 | 23.30134 | 30.99146 | 13.45583 | 0.000304 | 0.355214 | 0.999292 |
| 4 | 142280200 | 142286999 | 79.76652 | 124.7514 | 155.1615 | 64.37094 | 0.000314 | 2.639347 | 0.999324 |
| X | 80707200 | 80708799 | 17.69915 | 31.35161 | 41.21879 | 18.22494 | -0.00095 | 0.536986 | 0.999327 |
| 4 | 15340400 | 15344199 | 52.03921 | 83.40863 | 85.41588 | 36.05593 | 0.312461 | 1.983825 | 0.999341 |

| | | | | | | | | | |
|---|---|---|---|---|---|---|---|---|---|
| 9 | 25328200 | 25332199 | 40.54026 | 65.52644 | 84.22959 | 35.97957 | -0.00041 | 1.710671 | 0.999384 |
| 5 | 37496800 | 37500199 | 29.4475 | 48.62972 | 63.33769 | 27.3535 | 0.000306 | 1.349221 | 0.999425 |
| 3 | 138893600 | 138894799 | 13.48845 | 23.6011 | 42.0616 | 18.55467 | -0.44189 | 0.668893 | 0.999426 |
| 16 | 49866000 | 49868199 | 25.18191 | 42.17708 | 54.96264 | 23.96029 | 0.000367 | 1.174142 | 0.999429 |
| 13 | 111785400 | 111785999 | 8.15645 | 15.75489 | 23.18083 | 8.91351 | 0.000252 | -0.14902 | 0.999437 |
| 3 | 95320200 | 95335999 | 335.1675 | 471.2128 | 532.2977 | 220.5326 | 0.000246 | 4.50014 | 0.999466 |
| 15 | 39566600 | 39570399 | 44.72752 | 72.07238 | 92.26952 | 39.257 | 0.000297 | 1.811276 | 0.999467 |
| 6 | 69572400 | 69573399 | 22.52533 | 38.02299 | 49.58186 | 21.68677 | -0.00031 | 1.086335 | 0.999473 |
| 8 | 83236600 | 83239999 | 27.32945 | 44.89252 | 72.92633 | 31.41759 | -0.32022 | 1.416975 | 0.999488 |
| 6 | 99549200 | 99550799 | 15.15993 | 26.74988 | 35.10044 | 15.53173 | -0.00026 | 0.550736 | 0.999492 |
| 1 | 129912200 | 129913999 | 19.43668 | 33.74891 | 44.17002 | 19.38543 | 0.000236 | 0.811608 | 0.999492 |
| 16 | 26738200 | 26739599 | 12.40435 | 22.76967 | 30.69297 | 13.06497 | 9.27E-05 | 0.263006 | 0.999511 |
| 10 | 71895600 | 71900399 | 11.49836 | 21.59607 | 23.06697 | 9.484862 | 0.371662 | 0.103009 | 0.999537 |
| 2 | 129018200 | 129020599 | 19.84838 | 34.23631 | 44.76663 | 19.63839 | -0.00023 | 0.876997 | 0.999544 |
| 12 | 52936200 | 52938599 | 19.84838 | 34.23631 | 44.76663 | 19.63839 | -0.00023 | 0.876997 | 0.999544 |
| 16 | 49724800 | 49726199 | 12.6072 | 22.89078 | 30.56431 | 13.19452 | 0.00017 | 0.326834 | 0.999561 |
| 8 | 126302600 | 126306799 | 50.08471 | 80.09733 | 101.6246 | 42.75766 | 0.000238 | 2.054291 | 0.999565 |
| 1 | 99750800 | 99752399 | 16.45914 | 29.11492 | 28.46711 | 12.59024 | 0.423278 | 0.518264 | 0.999579 |
| 1 | 104207200 | 104209599 | 34.99187 | 57.31619 | 59.51221 | 25.60784 | 0.322079 | 1.460172 | 0.999579 |
| 10 | 68139600 | 68144999 | 45.54324 | 73.2882 | 93.70527 | 39.79024 | -0.00025 | 1.854721 | 0.999583 |
| 15 | 17816200 | 17828999 | 246.6553 | 359.0552 | 349.5818 | 144.0776 | 0.264591 | 3.982777 | 0.999617 |
| 12 | 20228800 | 20370399 | 5382.964 | 6011.763 | 5381.754 | 2301.503 | -0.00018 | 8.152023 | 0.999631 |
| 9 | 22198800 | 22202199 | 33.13991 | 54.58608 | 71.04247 | 30.64142 | 0.000206 | 1.39606 | 0.999653 |
| 3 | 58391400 | 58393599 | 22.39716 | 38.37921 | 50.10329 | 21.97282 | 0.000125 | 0.951222 | 0.999663 |
| 2 | 9239800 | 9242399 | 26.84049 | 44.70519 | 58.26941 | 25.31267 | 0.000153 | 1.241498 | 0.999756 |
| 14 | 91693200 | 91693999 | 10.25814 | 19.35404 | 26.97439 | 11.00522 | -7.85E-05 | 0.074826 | 0.999759 |
| 3 | 53027200 | 53029599 | 18.4769 | 32.12853 | 42.06047 | 18.47627 | 0.00012 | 0.784319 | 0.999761 |
| 16 | 3359600 | 3361199 | 21.99174 | 38.37964 | 37.53555 | 16.5263 | 0.419387 | 0.74388 | 0.999771 |
| 3 | 112880600 | 112882199 | 13.5527 | 24.27052 | 32.08378 | 14.06368 | 0.000296 | 0.42069 | 0.999791 |
| 3 | 153879000 | 153882399 | 36.17544 | 58.67922 | 75.67513 | 32.42057 | -9.85E-05 | 1.623955 | 0.999794 |
| 12 | 56499000 | 56499799 | 9.562474 | 18.11211 | 25.42848 | 10.28917 | -0.00015 | 0.036674 | 0.999801 |
| 10 | 5337800 | 5340399 | 29.07908 | 48.01967 | 62.56268 | 27.01816 | -5.86E-05 | 1.350928 | 0.999803 |
| 11 | 96117400 | 96119999 | 30.26553 | 49.65201 | 64.53534 | 27.78763 | -4.78E-05 | 1.435994 | 0.999807 |
| 2 | 47419400 | 47425799 | 106.5952 | 162.0327 | 242.896 | 100.6138 | -0.30074 | 3.191743 | 0.999812 |
| 5 | 117401000 | 117403199 | 26.68177 | 44.2265 | 57.62829 | 24.95175 | 0.000159 | 1.295171 | 0.999847 |
| 7 | 104456800 | 104460999 | 40.82421 | 66.24098 | 85.43837 | 36.61192 | 8.11E-05 | 1.631771 | 0.999853 |
| 10 | 121400000 | 121408399 | 93.73018 | 144.9104 | 178.6779 | 74.21866 | -6.26E-05 | 2.850524 | 0.999859 |
| 3 | 68739400 | 68741999 | 20.38567 | 35.05725 | 45.80463 | 20.0919 | -0.00015 | 0.905252 | 0.999907 |
| 18 | 50208000 | 50209399 | 17.05332 | 29.91631 | 39.18898 | 17.32549 | -0.00014 | 0.640779 | 0.999913 |
| 11 | 61481400 | 61485999 | 45.05745 | 72.29382 | 92.08462 | 38.93283 | 7.82E-05 | 1.945169 | 0.99992 |
| 5 | 29248600 | 29250199 | 28.47384 | 47.35658 | 61.72619 | 26.78843 | 1.06E-05 | 1.264799 | 0.999928 |
| 1 | 58543200 | 58552999 | 199.5422 | 294.3237 | 348.6187 | 143.5662 | 3.09E-05 | 3.773361 | 0.999939 |
| 6 | 129903200 | 129914599 | 373.6273 | 519.3769 | 580.1733 | 240.7907 | -2.91E-05 | 4.670465 | 0.99994 |
| 2 | 151017800 | 151021599 | 47.23444 | 76.03469 | 97.24305 | 41.31219 | -5.80E-05 | 1.848835 | 0.999959 |
| 14 | 3613800 | 3614999 | 19.01356 | 32.97882 | 43.1646 | 18.93881 | 1.15E-05 | 0.821396 | 0.999962 |
| 2 | 63119200 | 63119999 | 9.619207 | 18.17753 | 25.40779 | 10.33115 | 0.000264 | 0.054445 | 0.999981 |
| 1 | 3213000 | 3218599 | 93.65957 | 145.6163 | 168.7293 | 70.05246 | 0.095747 | 2.750409 | 1 |

| | | | | | | | | | |
|---|---|---|---|---|---|---|---|---|---|
| 1 | 3370800 | 3374599 | 32.29163 | 53.63909 | 62.54821 | 27.12176 | 0.160022 | 1.291091 | 1 |
| 1 | 3853200 | 3855999 | 39.89279 | 65.04651 | 68.13567 | 29.26864 | 0.305125 | 1.541236 | 1 |
| 1 | 3960600 | 3962799 | 24.92622 | 41.95373 | 59.9258 | 26.18807 | -0.13157 | 1.13063 | 1 |
| 1 | 4051000 | 4059999 | 111.6598 | 171.0124 | 201.3987 | 83.58476 | 0.055728 | 3.038515 | 1 |
| 1 | 4155200 | 4156799 | 16.40017 | 29.12072 | 30.83165 | 13.61223 | 0.311589 | 0.489327 | 1 |
| 1 | 4323800 | 4330799 | 106.1063 | 161.6907 | 209.3029 | 86.76318 | -0.08627 | 3.138657 | 1 |
| 1 | 4468400 | 4471999 | 59.35115 | 95.06662 | 95.98731 | 40.47846 | 0.33172 | 2.016331 | 1 |
| 1 | 4637000 | 4638399 | 14.39795 | 25.46084 | 33.00568 | 14.59844 | 0.018225 | 0.515913 | 1 |
| 1 | 4679400 | 4682599 | 32.28168 | 53.29736 | 55.75859 | 24.07636 | 0.31545 | 1.353229 | 1 |
| 1 | 5001800 | 5004399 | 46.73156 | 75.21885 | 82.78289 | 35.16472 | 0.216058 | 1.843315 | 1 |
| 1 | 5053000 | 5060999 | 117.4416 | 178.6505 | 236.5774 | 98.01789 | -0.12115 | 3.180503 | 1 |
| 1 | 5178600 | 5180999 | 27.98211 | 46.0273 | 65.58254 | 28.27009 | -0.13074 | 1.398348 | 1 |
| 1 | 5235600 | 5237399 | 20.81932 | 35.2072 | 57.02344 | 24.95543 | -0.31189 | 1.072379 | 1 |
| 1 | 5257800 | 5260199 | 35.60119 | 57.56614 | 87.84156 | 37.54169 | -0.24657 | 1.694443 | 1 |
| 1 | 5380400 | 5384199 | 19.29534 | 33.41605 | 49.02674 | 21.50986 | -0.16467 | 0.841957 | 1 |
| 1 | 5493400 | 5494599 | 13.54943 | 24.54433 | 27.96596 | 12.10804 | 0.224991 | 0.342192 | 1 |
| 1 | 5814000 | 5820799 | 104.4089 | 160.8615 | 184.3279 | 76.56606 | 0.10174 | 2.917944 | 1 |
| 1 | 5889000 | 5892399 | 35.68091 | 58.01729 | 70.70981 | 30.34361 | 0.084404 | 1.574264 | 1 |
| 1 | 5919400 | 5921399 | 23.8313 | 40.83513 | 45.24604 | 19.84261 | 0.235855 | 0.95225 | 1 |
| 1 | 6046600 | 6047799 | 13.02115 | 23.78668 | 29.51859 | 12.64647 | 0.112126 | 0.292297 | 1 |
| 1 | 6062600 | 6063999 | 18.05257 | 31.17699 | 46.35018 | 20.33383 | -0.18434 | 0.858947 | 1 |
| 1 | 6081600 | 6086399 | 62.3083 | 99.11436 | 121.4579 | 50.72175 | 0.041922 | 2.239774 | 1 |
| 1 | 6241200 | 6247399 | 120.84 | 184.28 | 202.719 | 84.04931 | 0.149146 | 3.127398 | 1 |
| 1 | 6504800 | 6511999 | 128.3078 | 192.4179 | 276.8486 | 114.2242 | -0.25836 | 3.468444 | 1 |
| 1 | 6637200 | 6639599 | 28.15578 | 47.19747 | 56.177 | 24.50259 | 0.13061 | 1.176342 | 1 |
| 1 | 6759400 | 6766199 | 102.3267 | 157.0143 | 196.5159 | 81.5833 | -0.02989 | 2.998331 | 1 |
| 1 | 6980800 | 6982199 | 19.57292 | 34.09818 | 37.81979 | 16.63301 | 0.23852 | 0.757457 | 1 |
| 1 | 6986000 | 6988199 | 26.76602 | 43.78886 | 73.49805 | 31.61058 | -0.36978 | 1.469157 | 1 |
| 1 | 7083600 | 7084399 | 8.943632 | 16.85293 | 24.85672 | 10.16424 | -0.08388 | 0.064102 | 1 |
| 1 | 7199400 | 7203799 | 21.94123 | 36.94957 | 58.70038 | 25.65675 | -0.28443 | 1.113307 | 1 |
| 1 | 7357200 | 7358999 | 25.50991 | 42.59768 | 58.25061 | 25.34532 | -0.0689 | 1.209571 | 1 |
| 1 | 7676000 | 7683199 | 156.8253 | 235.4807 | 245.226 | 101.2112 | 0.209267 | 3.447348 | 1 |
| 1 | 7704400 | 7704999 | 8.835557 | 17.08886 | 19.8262 | 7.577381 | 0.348181 | -0.17279 | 1 |
| 1 | 7754000 | 7754999 | 16.43238 | 29.05371 | 30.05517 | 13.29387 | 0.342265 | 0.528085 | 1 |
| 1 | 7889400 | 7891399 | 38.28795 | 62.31159 | 65.38433 | 28.06737 | 0.30075 | 1.564243 | 1 |
| 1 | 7913400 | 7914999 | 32.19878 | 52.76075 | 64.14635 | 27.60623 | 0.095312 | 1.454444 | 1 |
| 1 | 8012800 | 8015399 | 50.34461 | 80.38594 | 101.4347 | 42.58241 | 0.005655 | 2.104476 | 1 |
| 1 | 8288600 | 8292399 | 22.05252 | 36.97866 | 54.25338 | 23.66845 | -0.16992 | 1.1538 | 1 |
| 1 | 8469400 | 8470399 | 19.09353 | 33.28116 | 39.86479 | 17.5384 | 0.128076 | 0.749961 | 1 |
| 1 | 8579200 | 8580599 | 21.77318 | 37.14022 | 51.17088 | 22.43277 | -0.07832 | 0.991108 | 1 |
| 1 | 8756800 | 8759799 | 44.54518 | 71.59019 | 89.20714 | 37.8098 | 0.034565 | 1.89175 | 1 |
| 1 | 8779800 | 8784599 | 70.74754 | 112.1506 | 127.0062 | 52.85038 | 0.150708 | 2.349134 | 1 |
| 1 | 8856600 | 8862599 | 121.0683 | 184.4995 | 206.1081 | 85.44006 | 0.126165 | 3.141482 | 1 |
| 1 | 9163200 | 9164599 | 15.63123 | 27.93602 | 29.91861 | 13.13707 | 0.30061 | 0.443099 | 1 |
| 1 | 9190400 | 9191599 | 15.52219 | 27.18888 | 38.88926 | 17.17089 | -0.12614 | 0.658103 | 1 |
| 1 | 9470400 | 9472599 | 17.83535 | 31.42941 | 35.86975 | 15.87631 | 0.199113 | 0.580899 | 1 |
| 1 | 9706400 | 9708199 | 25.27422 | 42.8852 | 49.0382 | 21.48277 | 0.188635 | 1.048478 | 1 |

| | | | | | | | | | |
|---|---|---|---|---|---|---|---|---|---|
| 1 | 9978600 | 9979799 | 14.21787 | 25.01377 | 35.76139 | 15.83352 | -0.12493 | 0.578704 | 1 |
| 1 | 10452000 | 10455399 | 38.54262 | 62.27215 | 90.57657 | 38.67038 | -0.17897 | 1.724088 | 1 |
| 1 | 10620200 | 10621199 | 12.24077 | 22.61602 | 27.09954 | 11.43372 | 0.178274 | 0.218605 | 1 |
| 1 | 10726400 | 10727799 | 20.34303 | 35.32449 | 40.0153 | 17.56258 | 0.207882 | 0.811911 | 1 |
| 1 | 10804400 | 10808799 | 65.48335 | 103.1733 | 135.5957 | 56.26505 | -0.07083 | 2.489213 | 1 |
| 1 | 10874400 | 10877799 | 41.05703 | 65.84838 | 96.65699 | 40.84068 | -0.20482 | 1.95547 | 1 |
| 1 | 10993000 | 10996999 | 51.85273 | 83.1018 | 92.80003 | 39.16725 | 0.187577 | 1.988211 | 1 |
| 1 | 11057800 | 11059799 | 28.05477 | 46.5311 | 63.81048 | 27.64031 | -0.07332 | 1.297842 | 1 |
| 1 | 11099200 | 11105599 | 57.46965 | 91.64134 | 107.3909 | 44.99929 | 0.110096 | 2.153246 | 1 |
| 1 | 11163600 | 11164799 | 15.54501 | 27.91254 | 31.82131 | 13.91738 | 0.214599 | 0.415431 | 1 |
| 1 | 11555200 | 11559399 | 49.90341 | 80.22757 | 92.72063 | 39.31741 | 0.143524 | 1.891807 | 1 |
| 1 | 11689400 | 11691199 | 17.1415 | 30.05414 | 38.2445 | 16.89992 | 0.041658 | 0.650672 | 1 |
| 1 | 11696600 | 11697399 | 9.832381 | 18.40278 | 26.37762 | 10.91038 | -0.05583 | 0.121999 | 1 |
| 1 | 11743000 | 11745199 | 15.35859 | 26.81146 | 43.10185 | 18.98056 | -0.29415 | 0.71555 | 1 |
| 1 | 11790400 | 11792599 | 22.2455 | 37.5422 | 58.19361 | 25.45175 | -0.24898 | 1.095316 | 1 |
| 1 | 11852800 | 11854199 | 28.76222 | 47.30702 | 64.02753 | 27.59893 | -0.05682 | 1.399454 | 1 |
| 1 | 12016200 | 12021399 | 91.66461 | 142.4199 | 154.1682 | 64.01341 | 0.193083 | 2.75865 | 1 |
| 1 | 12190800 | 12191999 | 10.36493 | 19.6334 | 22.6685 | 9.171037 | 0.279745 | 0.03498 | 1 |
| 1 | 12239800 | 12243199 | 44.92096 | 72.45208 | 78.54129 | 33.45739 | 0.241398 | 1.779923 | 1 |
| 1 | 12412600 | 12415399 | 28.17022 | 46.91762 | 67.47125 | 29.30909 | -0.14136 | 1.252627 | 1 |
| 1 | 12539800 | 12544999 | 55.66233 | 88.87353 | 107.1749 | 44.98978 | 0.07083 | 2.111292 | 1 |
| 1 | 12635800 | 12636799 | 15.82182 | 27.62693 | 40.00924 | 17.62142 | -0.14409 | 0.710301 | 1 |
| 1 | 13611600 | 13619799 | 109.4484 | 167.8569 | 188.8325 | 78.38806 | 0.123279 | 3.008861 | 1 |
| 1 | 13620600 | 13623999 | 55.43459 | 87.77237 | 135.3433 | 56.27467 | -0.29539 | 2.370207 | 1 |
| 1 | 13683400 | 13693199 | 126.2119 | 190.7596 | 271.0524 | 112.0797 | -0.23047 | 3.314068 | 1 |
| 1 | 14149600 | 14151399 | 25.82203 | 43.71672 | 49.59331 | 21.7112 | 0.199979 | 1.070956 | 1 |
| 1 | 14182000 | 14184599 | 40.51593 | 65.21478 | 95.56011 | 40.59277 | -0.19603 | 1.840851 | 1 |
| 1 | 14576400 | 14578799 | 18.05251 | 31.1765 | 42.73288 | 18.74691 | -0.06751 | 0.855173 | 1 |
| 1 | 14589000 | 14590399 | 11.06509 | 20.32888 | 33.4804 | 14.23471 | -0.28577 | 0.254428 | 1 |
| 1 | 15148200 | 15154799 | 103.437 | 158.4245 | 194.7872 | 80.84153 | -0.00632 | 3.03459 | 1 |
| 1 | 15241800 | 15243999 | 44.81941 | 72.15765 | 82.94136 | 35.24378 | 0.153914 | 1.834186 | 1 |
| 1 | 15334400 | 15337599 | 39.35807 | 64.1522 | 71.0558 | 30.51913 | 0.224437 | 1.545745 | 1 |
| 1 | 15566400 | 15569999 | 47.66845 | 76.78723 | 82.35136 | 35.02131 | 0.254714 | 1.821925 | 1 |
| 1 | 15927600 | 15935199 | 80.52463 | 126.1684 | 146.3831 | 60.71763 | 0.102568 | 2.606481 | 1 |
| 1 | 16163000 | 16164199 | 12.43813 | 22.0716 | 34.15855 | 15.08911 | -0.23365 | 0.472935 | 1 |
| 1 | 16416600 | 16420599 | 57.46071 | 91.89343 | 97.94143 | 41.20807 | 0.251255 | 2.061815 | 1 |
| 1 | 16521400 | 16526199 | 70.34377 | 111.7124 | 118.2006 | 49.27526 | 0.25112 | 2.295876 | 1 |
| 1 | 16542800 | 16546999 | 42.13292 | 68.21655 | 78.54338 | 33.60357 | 0.160628 | 1.67442 | 1 |
| 1 | 16780000 | 16787399 | 52.23521 | 83.12398 | 121.7061 | 50.85332 | -0.21379 | 2.219317 | 1 |
| 1 | 17011600 | 17013599 | 23.9261 | 40.85372 | 43.17112 | 18.9273 | 0.303218 | 0.981221 | 1 |
| 1 | 17141800 | 17147799 | 33.73692 | 54.92821 | 88.46641 | 37.9816 | -0.31554 | 1.559934 | 1 |
| 1 | 17292000 | 17294599 | 17.00983 | 29.84579 | 42.31081 | 18.70871 | -0.11374 | 0.640759 | 1 |
| 1 | 17542600 | 17544999 | 34.86424 | 57.05825 | 59.98295 | 25.80134 | 0.3037 | 1.471635 | 1 |
| 1 | 17644800 | 17645599 | 10.89406 | 20.03122 | 27.73381 | 11.77819 | -0.03539 | 0.237163 | 1 |
| 1 | 17973600 | 17975999 | 18.99118 | 32.65743 | 45.3566 | 19.89534 | -0.08763 | 0.903839 | 1 |
| 1 | 18145200 | 18150999 | 81.54697 | 127.4522 | 142.3689 | 59.06776 | 0.154434 | 2.649681 | 1 |
| 1 | 18542200 | 18543999 | 32.5839 | 52.67862 | 86.19352 | 36.82956 | -0.34718 | 1.695784 | 1 |

| | | | | | | | | | |
|---|---|---|---|---|---|---|---|---|---|
| 1 | 18562400 | 18564799 | 30.04875 | 49.39398 | 64.14877 | 27.6443 | 0.002197 | 1.409648 | 1 |
| 1 | 18606000 | 18608799 | 41.37419 | 66.56612 | 92.2569 | 39.16392 | -0.11663 | 1.856239 | 1 |
| 1 | 18728800 | 18730199 | 29.62948 | 48.47325 | 70.54102 | 30.33876 | -0.16453 | 1.473101 | 1 |
| 1 | 18942200 | 18942999 | 20.95861 | 36.20002 | 42.63671 | 18.7048 | 0.150341 | 0.867134 | 1 |
| 1 | 18997800 | 18998399 | 10.31327 | 19.50135 | 21.73584 | 8.826358 | 0.326134 | 0.046225 | 1 |
| 1 | 19468200 | 19469799 | 16.51427 | 28.47529 | 47.41809 | 20.80132 | -0.34769 | 0.868147 | 1 |
| 1 | 19762600 | 19764199 | 24.38898 | 41.39371 | 43.30318 | 18.97183 | 0.316708 | 1.039385 | 1 |
| 1 | 19907200 | 19910199 | 50.34799 | 80.46963 | 104.5843 | 43.96518 | -0.03536 | 2.075511 | 1 |
| 1 | 20166800 | 20169799 | 22.91872 | 39.84171 | 42.98154 | 18.87776 | 0.278026 | 0.810557 | 1 |
| 1 | 20510600 | 20512399 | 12.0382 | 22.10642 | 30.76074 | 13.08667 | -0.04461 | 0.260147 | 1 |
| 1 | 20594400 | 20596399 | 17.19595 | 29.91103 | 43.18141 | 18.97261 | -0.14011 | 0.778241 | 1 |
| 1 | 20643400 | 20645599 | 31.7922 | 52.49401 | 57.81678 | 24.96665 | 0.241546 | 1.355121 | 1 |
| 1 | 20717200 | 20719199 | 22.8253 | 38.42542 | 59.23193 | 25.88643 | -0.24106 | 1.121756 | 1 |
| 1 | 21191400 | 21192599 | 15.27499 | 26.78543 | 39.4006 | 17.41289 | -0.16626 | 0.641341 | 1 |
| 1 | 21426200 | 21435199 | 128.9848 | 195.661 | 228.0175 | 94.39219 | 0.059787 | 3.238572 | 1 |
| 1 | 22131000 | 22131799 | 8.633033 | 16.44633 | 25.56424 | 10.21858 | -0.13042 | -0.01182 | 1 |
| 1 | 22227800 | 22230599 | 41.4097 | 66.74402 | 80.2608 | 34.16029 | 0.090968 | 1.793119 | 1 |
| 1 | 23340000 | 23342599 | 29.79724 | 48.69983 | 69.3227 | 29.80409 | -0.13374 | 1.486928 | 1 |
| 1 | 23346000 | 23354599 | 127.3747 | 193.7314 | 229.2547 | 94.97971 | 0.040823 | 3.185506 | 1 |
| 1 | 23462200 | 23468799 | 76.19523 | 120.0402 | 125.9916 | 52.27927 | 0.253048 | 2.495049 | 1 |
| 1 | 23636000 | 23638599 | 14.77709 | 26.52472 | 30.91473 | 13.52472 | 0.182753 | 0.411391 | 1 |
| 1 | 23651000 | 23653999 | 44.43053 | 71.67642 | 83.21517 | 35.45819 | 0.142981 | 1.771799 | 1 |
| 1 | 23702200 | 23702799 | 12.41143 | 22.76012 | 28.06927 | 11.96424 | 0.126377 | 0.263754 | 1 |
| 1 | 23819400 | 23823199 | 35.0761 | 56.79937 | 80.99294 | 34.65534 | -0.14699 | 1.659031 | 1 |
| 1 | 24107400 | 24108999 | 19.67114 | 34.01821 | 39.53994 | 17.34689 | 0.169654 | 0.844176 | 1 |
| 1 | 24457800 | 24463199 | 91.67477 | 142.4475 | 147.3351 | 61.17542 | 0.258791 | 2.756818 | 1 |
| 1 | 24535400 | 24536399 | 11.52446 | 20.64786 | 33.74602 | 14.78654 | -0.30352 | 0.407412 | 1 |
| 1 | 24759400 | 24760799 | 19.47061 | 33.19844 | 49.53808 | 21.71622 | -0.19285 | 0.986754 | 1 |
| 1 | 24834800 | 24847599 | 16.08676 | 28.29693 | 34.10557 | 15.10091 | 0.119967 | 0.588306 | 1 |
| 1 | 25262600 | 25263999 | 12.15119 | 22.18427 | 29.12027 | 12.4851 | 0.031171 | 0.289745 | 1 |
| 1 | 25629200 | 25630599 | 25.9403 | 43.54361 | 50.15821 | 21.89641 | 0.177703 | 1.148445 | 1 |
| 1 | 25953400 | 25956199 | 41.96865 | 67.92347 | 78.43132 | 33.54576 | 0.155955 | 1.682073 | 1 |
| 1 | 25957200 | 25958399 | 17.86417 | 31.37052 | 33.54413 | 14.84259 | 0.291438 | 0.621121 | 1 |
| 1 | 26001200 | 26002399 | 11.53087 | 21.03407 | 28.67377 | 12.3066 | -0.02389 | 0.28906 | 1 |
| 1 | 26049800 | 26050599 | 12.53457 | 22.73239 | 28.64751 | 12.38508 | 0.082094 | 0.331183 | 1 |
| 1 | 26452200 | 26455199 | 55.19546 | 88.11788 | 98.57962 | 41.37487 | 0.178772 | 2.112762 | 1 |
| 1 | 26513200 | 26516199 | 32.58346 | 53.1631 | 71.19154 | 30.58861 | -0.0476 | 1.5191 | 1 |
| 1 | 26628800 | 26634399 | 65.84734 | 104.9233 | 109.4171 | 45.80175 | 0.277002 | 2.18699 | 1 |
| 1 | 27070200 | 27072399 | 25.50501 | 42.22922 | 67.41514 | 29.16696 | -0.29206 | 1.313862 | 1 |
| 1 | 27147800 | 27149199 | 16.08359 | 28.62428 | 30.36601 | 13.3807 | 0.31069 | 0.47235 | 1 |
| 1 | 27174400 | 27181599 | 64.30982 | 102.1082 | 120.3249 | 50.15017 | 0.095658 | 2.29632 | 1 |
| 1 | 27215200 | 27222599 | 128.5937 | 196.0096 | 204.3184 | 84.70302 | 0.226222 | 3.140018 | 1 |
| 1 | 27345000 | 27347599 | 29.92053 | 49.88681 | 54.58451 | 23.72739 | 0.251795 | 1.236765 | 1 |
| 1 | 28018000 | 28019199 | 15.51225 | 27.20315 | 38.82435 | 17.15912 | -0.12304 | 0.638782 | 1 |
| 1 | 28326600 | 28327999 | 15.10205 | 26.70622 | 30.1606 | 13.34004 | 0.216405 | 0.517285 | 1 |
| 1 | 28946200 | 28952799 | 23.16263 | 39.40812 | 47.95698 | 21.01945 | 0.099537 | 1.015813 | 1 |
| 1 | 29407200 | 29408599 | 35.72272 | 57.81302 | 77.45798 | 33.12868 | -0.05806 | 1.670304 | 1 |

| | | | | | | | | | |
|---|---|---|---|---|---|---|---|---|---|
| 1 | 29523600 | 29524999 | 23.47757 | 39.11039 | 59.14059 | 25.69378 | -0.2134 | 1.230642 | 1 |
| 1 | 29642600 | 29645399 | 42.52935 | 68.8667 | 74.07157 | 31.6932 | 0.258804 | 1.669242 | 1 |
| 1 | 29650200 | 29652999 | 43.87952 | 70.77943 | 85.19088 | 36.29468 | 0.090906 | 1.776732 | 1 |
| 1 | 29689000 | 29700599 | 85.04726 | 133.3238 | 141.856 | 58.83671 | 0.227898 | 2.598575 | 1 |
| 1 | 29985800 | 29991399 | 104.0304 | 159.9544 | 183.4062 | 76.16209 | 0.098601 | 2.957836 | 1 |
| 1 | 30039400 | 30040599 | 9.983559 | 18.467 | 32.55954 | 13.71573 | -0.37361 | 0.206478 | 1 |
| 1 | 30086000 | 30087199 | 17.05351 | 29.69678 | 41.55519 | 18.27106 | -0.09531 | 0.756242 | 1 |
| 1 | 30204400 | 30204999 | 8.824692 | 16.84614 | 25.06028 | 9.973572 | -0.06173 | -0.0239 | 1 |
| 1 | 30312600 | 30314599 | 27.56888 | 46.11834 | 50.73418 | 22.09981 | 0.243963 | 1.190629 | 1 |
| 1 | 30349400 | 30351199 | 27.66032 | 45.78273 | 57.10477 | 24.70059 | 0.062872 | 1.314283 | 1 |
| 1 | 30371800 | 30373399 | 23.9498 | 40.07709 | 57.03841 | 24.85098 | -0.12631 | 1.182823 | 1 |
| 1 | 30463600 | 30464399 | 12.67969 | 23.28845 | 28.1288 | 11.96414 | 0.158684 | 0.259088 | 1 |
| 1 | 30499000 | 30499599 | 9.803733 | 18.7196 | 21.25169 | 8.453517 | 0.325668 | -0.0274 | 1 |
| 1 | 30610200 | 30610999 | 11.05456 | 20.4944 | 28.80755 | 12.0979 | -0.04551 | 0.194826 | 1 |
| 1 | 30705600 | 30709399 | 46.73976 | 74.92118 | 100.4311 | 42.39981 | -0.07511 | 1.980316 | 1 |
| 1 | 30786000 | 30788799 | 17.566 | 30.56148 | 45.22781 | 19.87451 | -0.17593 | 0.776725 | 1 |
| 1 | 30987600 | 30989399 | 13.82492 | 24.19919 | 41.54644 | 18.33342 | -0.38818 | 0.664918 | 1 |
| 1 | 31016200 | 31017199 | 10.83528 | 20.26151 | 25.23246 | 10.45271 | 0.143452 | 0.131779 | 1 |
| 1 | 31242400 | 31243399 | 12.80254 | 22.67349 | 35.20606 | 15.56702 | -0.23964 | 0.496235 | 1 |
| 1 | 31315400 | 31316799 | 17.55838 | 30.47487 | 45.80514 | 20.09781 | -0.19858 | 0.81137 | 1 |
| 1 | 31339800 | 31341599 | 26.44612 | 44.52744 | 53.94613 | 23.5774 | 0.105196 | 1.125929 | 1 |
| 1 | 31422200 | 31424999 | 17.37977 | 30.55479 | 32.07118 | 14.20169 | 0.318204 | 0.598785 | 1 |
| 1 | 31454600 | 31456199 | 23.79383 | 39.78258 | 59.21135 | 25.784 | -0.1907 | 1.192958 | 1 |
| 1 | 31530600 | 31531799 | 20.57798 | 35.53073 | 42.11918 | 18.47756 | 0.140944 | 0.866934 | 1 |
| 1 | 31545400 | 31546599 | 16.28397 | 28.09749 | 47.4229 | 20.80387 | -0.36659 | 0.864196 | 1 |
| 1 | 31768800 | 31776599 | 137.7843 | 207.2931 | 271.2779 | 112.0219 | -0.11762 | 3.407117 | 1 |
| 1 | 31894400 | 31895599 | 23.38042 | 39.74534 | 49.02568 | 21.4864 | 0.080079 | 1.02912 | 1 |
| 1 | 32107000 | 32108399 | 13.59789 | 24.07173 | 34.1856 | 15.1171 | -0.11177 | 0.504693 | 1 |
| 1 | 32112200 | 32114399 | 30.14695 | 49.145 | 69.67711 | 29.92843 | -0.13059 | 1.525003 | 1 |
| 1 | 32408800 | 32413199 | 42.26445 | 68.59297 | 74.03368 | 31.72994 | 0.256798 | 1.621913 | 1 |
| 1 | 32416800 | 32418799 | 25.92232 | 43.56978 | 53.93581 | 23.56047 | 0.074105 | 1.138416 | 1 |
| 1 | 32475400 | 32476399 | 20.86259 | 35.85926 | 42.6728 | 18.71785 | 0.134038 | 0.908569 | 1 |
| 1 | 33201000 | 33203399 | 28.99499 | 47.93424 | 59.89389 | 25.88555 | 0.060185 | 1.334371 | 1 |
| 1 | 33221600 | 33222399 | 14.88731 | 26.07953 | 40.43087 | 17.85314 | -0.24181 | 0.654986 | 1 |
| 1 | 33295600 | 33297599 | 30.39399 | 50.16224 | 57.79727 | 24.9503 | 0.176581 | 1.354828 | 1 |
| 1 | 33445200 | 33447999 | 49.98591 | 79.40485 | 123.0633 | 51.31704 | -0.29824 | 2.271288 | 1 |
| 1 | 33645800 | 33646799 | 19.00544 | 32.9915 | 39.57482 | 17.36583 | 0.125603 | 0.809013 | 1 |
| 1 | 33678200 | 33680799 | 33.31764 | 54.6619 | 63.80189 | 27.4724 | 0.154674 | 1.441163 | 1 |
| 1 | 34097200 | 34099799 | 12.75251 | 23.60609 | 26.06222 | 10.96782 | 0.298402 | 0.210633 | 1 |
| 1 | 34127000 | 34129799 | 15.13491 | 27.38479 | 29.281 | 12.69569 | 0.314651 | 0.362351 | 1 |
| 1 | 34456400 | 34457599 | 19.67757 | 33.68072 | 48.48524 | 21.26128 | -0.14033 | 0.952902 | 1 |
| 1 | 34965400 | 34973399 | 172.9691 | 255.5881 | 332.2642 | 136.8339 | -0.13172 | 3.7405 | 1 |
| 1 | 35308600 | 35316599 | 92.9379 | 143.5666 | 193.5261 | 80.38663 | -0.12947 | 2.866217 | 1 |
| 1 | 35353000 | 35362399 | 90.51444 | 140.2376 | 168.7948 | 70.1127 | 0.037065 | 2.809195 | 1 |
| 1 | 35645800 | 35649199 | 19.24007 | 33.1811 | 47.40852 | 20.79723 | -0.12766 | 0.87945 | 1 |
| 1 | 36055800 | 36062399 | 64.79441 | 102.1515 | 150.096 | 62.28697 | -0.23089 | 2.478005 | 1 |
| 1 | 36076800 | 36084599 | 109.7821 | 168.8565 | 181.659 | 75.44014 | 0.190838 | 2.953148 | 1 |

| | | | | | | | | | |
|---|---|---|---|---|---|---|---|---|---|
| 1 | 36297600 | 36300999 | 30.26019 | 49.55079 | 73.65385 | 31.68887 | -0.1943 | 1.463353 | 1 |
| 1 | 37089400 | 37092399 | 29.93243 | 49.56208 | 59.48575 | 25.73715 | 0.118197 | 1.317862 | 1 |
| 1 | 37313200 | 37320399 | 78.36015 | 122.3633 | 162.364 | 67.36968 | -0.09465 | 2.666803 | 1 |
| 1 | 37321600 | 37324399 | 46.15822 | 74.17357 | 83.50843 | 35.38822 | 0.180444 | 1.895707 | 1 |
| 1 | 37614000 | 37615799 | 27.32015 | 44.98689 | 70.17642 | 30.26412 | -0.26047 | 1.385647 | 1 |
| 1 | 37809000 | 37814399 | 23.97663 | 40.90746 | 42.68624 | 18.71351 | 0.321223 | 0.988911 | 1 |
| 1 | 37913600 | 37915999 | 41.39485 | 66.81087 | 78.96828 | 33.66917 | 0.118025 | 1.75285 | 1 |
| 1 | 38168000 | 38169399 | 12.77205 | 22.96561 | 30.02066 | 13.11434 | 0.019535 | 0.383052 | 1 |
| 1 | 38234200 | 38236599 | 17.72275 | 31.17947 | 35.45317 | 15.69711 | 0.203605 | 0.598425 | 1 |
| 1 | 38657800 | 38658799 | 10.99464 | 19.91039 | 32.76206 | 14.18991 | -0.30223 | 0.336545 | 1 |
| 1 | 38744800 | 38745399 | 12.06869 | 21.3613 | 35.82299 | 15.84166 | -0.35021 | 0.500769 | 1 |
| 1 | 38794400 | 38795999 | 16.72227 | 29.2277 | 45.15864 | 19.90264 | -0.23706 | 0.709425 | 1 |
| 1 | 38989800 | 38992999 | 34.58229 | 57.038 | 62.54939 | 26.99351 | 0.247389 | 1.378022 | 1 |
| 1 | 39751000 | 39753399 | 16.85583 | 29.07367 | 47.94163 | 21.03119 | -0.33346 | 0.868347 | 1 |
| 1 | 39849000 | 39851999 | 19.85839 | 33.6235 | 54.82145 | 24.0007 | -0.32129 | 1.056502 | 1 |
| 1 | 39910200 | 39911199 | 11.47586 | 21.24541 | 25.66868 | 10.80046 | 0.169484 | 0.198739 | 1 |
| 1 | 40233200 | 40234799 | 15.9398 | 27.97162 | 33.32223 | 14.73568 | 0.136467 | 0.621378 | 1 |
| 1 | 40701400 | 40703199 | 13.81528 | 24.38763 | 37.25648 | 16.48436 | -0.2181 | 0.543501 | 1 |
| 1 | 40708600 | 40710999 | 11.38437 | 20.82845 | 31.77787 | 13.58771 | -0.18195 | 0.277178 | 1 |
| 1 | 40733400 | 40740799 | 43.32007 | 69.47342 | 106.6699 | 45.06702 | -0.26978 | 1.962753 | 1 |
| 1 | 41108400 | 41114999 | 81.1718 | 126.624 | 171.7019 | 71.25177 | -0.12707 | 2.686999 | 1 |
| 1 | 41124000 | 41131399 | 86.28684 | 134.0429 | 175.441 | 72.84801 | -0.08074 | 2.761798 | 1 |
| 1 | 41887600 | 41889799 | 29.977 | 49.84107 | 52.97563 | 22.98585 | 0.293397 | 1.264556 | 1 |
| 1 | 42061200 | 42062999 | 28.24821 | 47.54203 | 52.89653 | 23.11577 | 0.227616 | 1.134595 | 1 |
| 1 | 43549800 | 43552799 | 20.91937 | 35.58053 | 52.20106 | 22.87905 | -0.16916 | 1.017341 | 1 |
| 1 | 43719400 | 43721799 | 20.11683 | 34.36486 | 56.57941 | 24.80672 | -0.33401 | 0.978899 | 1 |
| 1 | 44283800 | 44284799 | 14.0707 | 25.1693 | 29.48728 | 12.93802 | 0.172429 | 0.424412 | 1 |
| 1 | 44387600 | 44389799 | 14.25469 | 24.98151 | 39.93489 | 17.64021 | -0.28579 | 0.644089 | 1 |
| 1 | 44413800 | 44416799 | 32.28899 | 53.27127 | 58.38184 | 25.19812 | 0.248506 | 1.366273 | 1 |
| 1 | 44457000 | 44458599 | 20.90112 | 36.31 | 39.40965 | 17.30168 | 0.269286 | 0.805963 | 1 |
| 1 | 44535600 | 44538199 | 32.9294 | 54.28475 | 62.37814 | 26.91392 | 0.179881 | 1.379648 | 1 |
| 1 | 44647000 | 44649599 | 75.16526 | 117.2428 | 156.3974 | 64.90143 | -0.10353 | 2.685368 | 1 |
| 1 | 44731600 | 44732999 | 10.62564 | 19.37399 | 32.81709 | 14.09289 | -0.33597 | 0.292981 | 1 |
| 1 | 44765800 | 44768799 | 48.26106 | 77.16894 | 111.0467 | 46.71169 | -0.18133 | 2.061694 | 1 |
| 1 | 45048400 | 45049399 | 15.55678 | 27.77494 | 29.21149 | 12.83748 | 0.32542 | 0.444038 | 1 |
| 1 | 45128600 | 45139599 | 172.6146 | 258.0956 | 282.7264 | 116.5805 | 0.130836 | 3.531578 | 1 |
| 1 | 45162400 | 45163799 | 14.35058 | 25.45184 | 36.54921 | 16.15234 | -0.12683 | 0.493475 | 1 |
| 1 | 45201200 | 45202599 | 20.95131 | 36.38273 | 41.50805 | 18.21836 | 0.197587 | 0.814976 | 1 |
| 1 | 45243600 | 45245599 | 18.65425 | 31.91721 | 51.12079 | 22.41631 | -0.29388 | 0.953059 | 1 |
| 1 | 45289800 | 45292199 | 11.01811 | 20.77453 | 25.84271 | 10.55661 | 0.161164 | 0.08275 | 1 |
| 1 | 45671000 | 45676999 | 55.38239 | 88.47362 | 97.07804 | 40.78207 | 0.207703 | 2.093139 | 1 |
| 1 | 45785400 | 45788399 | 43.81379 | 70.86401 | 80.35356 | 34.35159 | 0.181242 | 1.699509 | 1 |
| 1 | 45789200 | 45791399 | 28.04393 | 46.88761 | 53.93533 | 23.4862 | 0.17983 | 1.202861 | 1 |
| 1 | 46488800 | 46489399 | 11.25048 | 20.48823 | 28.7279 | 12.3557 | -0.06612 | 0.297358 | 1 |
| 1 | 46595200 | 46596999 | 17.79409 | 30.77035 | 44.79374 | 19.65179 | -0.1536 | 0.846063 | 1 |
| 1 | 46601800 | 46602799 | 14.42433 | 25.91133 | 30.4222 | 13.30068 | 0.172842 | 0.402867 | 1 |
| 1 | 46621200 | 46622999 | 24.86638 | 42.11017 | 50.11013 | 21.93926 | 0.131391 | 1.068782 | 1 |

| | | | | | | | | | |
|---|---|---|---|---|---|---|---|---|---|
| 1 | 46744800 | 46747399 | 45.28254 | 72.97089 | 78.05873 | 33.21363 | 0.259044 | 1.803865 | 1 |
| 1 | 46943400 | 46944599 | 18.81746 | 32.86078 | 36.7844 | 16.20205 | 0.225685 | 0.719277 | 1 |
| 1 | 47036400 | 47038199 | 19.74964 | 33.94052 | 45.85272 | 20.11258 | -0.04817 | 0.912067 | 1 |
| 1 | 47231800 | 47233999 | 25.79891 | 43.68032 | 51.74672 | 22.65437 | 0.137738 | 1.074377 | 1 |
| 1 | 47264800 | 47265799 | 16.08858 | 28.29425 | 31.26416 | 13.8434 | 0.244725 | 0.586309 | 1 |
| 1 | 47289400 | 47290399 | 10.50688 | 19.04873 | 32.26933 | 13.95666 | -0.34264 | 0.324491 | 1 |
| 1 | 47404800 | 47406599 | 15.55661 | 27.38225 | 40.24875 | 17.8186 | -0.16464 | 0.58931 | 1 |
| 1 | 47758600 | 47760999 | 32.8726 | 53.19296 | 84.31077 | 36.05587 | -0.30011 | 1.672929 | 1 |
| 1 | 47776800 | 47779999 | 27.69643 | 46.36805 | 52.90589 | 23.05721 | 0.191384 | 1.183723 | 1 |
| 1 | 47790600 | 47791999 | 18.27426 | 32.02921 | 36.67132 | 16.20005 | 0.193381 | 0.660034 | 1 |
| 1 | 47993600 | 47999799 | 34.05974 | 55.76318 | 62.73843 | 26.9907 | 0.206342 | 1.466305 | 1 |
| 1 | 48007000 | 48012799 | 120.9023 | 184.6217 | 189.3667 | 78.53759 | 0.251548 | 3.100016 | 1 |
| 1 | 48270200 | 48271399 | 29.21968 | 47.60692 | 73.95442 | 31.75924 | -0.26263 | 1.534408 | 1 |
| 1 | 48371200 | 48372599 | 11.77582 | 21.23777 | 34.7287 | 15.11905 | -0.2983 | 0.369174 | 1 |
| 1 | 48418800 | 48419799 | 13.92397 | 24.48217 | 39.29747 | 17.4012 | -0.29155 | 0.589231 | 1 |
| 1 | 48451800 | 48453199 | 14.73702 | 26.60609 | 30.89781 | 13.43372 | 0.193724 | 0.376216 | 1 |
| 1 | 48518800 | 48519799 | 9.034465 | 16.84916 | 27.91308 | 11.61543 | -0.27046 | 0.140125 | 1 |
| 1 | 48533400 | 48535199 | 17.72012 | 30.51326 | 47.42718 | 20.80445 | -0.24892 | 0.888725 | 1 |
| 1 | 48584600 | 48586599 | 45.02878 | 71.89722 | 110.147 | 46.23906 | -0.27374 | 2.102464 | 1 |
| 1 | 48734000 | 48735999 | 26.1492 | 44.26447 | 51.20558 | 22.41609 | 0.171957 | 1.07722 | 1 |
| 1 | 48736800 | 48738599 | 25.50558 | 42.53655 | 59.35266 | 25.8042 | -0.09791 | 1.223585 | 1 |
| 1 | 48776800 | 48784399 | 49.12583 | 78.86863 | 89.74311 | 37.9774 | 0.163501 | 1.93098 | 1 |
| 1 | 48966400 | 48967399 | 16.9558 | 29.51515 | 40.08645 | 17.62098 | -0.05233 | 0.760934 | 1 |
| 1 | 49295600 | 49297599 | 20.25116 | 35.51109 | 34.47266 | 15.23492 | 0.430322 | 0.659889 | 1 |
| 1 | 49300000 | 49301399 | 13.81584 | 24.86999 | 36.17107 | 15.78418 | -0.13265 | 0.396296 | 1 |
| 1 | 49342200 | 49344999 | 45.39678 | 73.1375 | 75.15188 | 31.9657 | 0.316531 | 1.809668 | 1 |
| 1 | 49513600 | 49515799 | 12.75251 | 23.60609 | 26.06222 | 10.96782 | 0.298402 | 0.210633 | 1 |
| 1 | 49631600 | 49632199 | 13.28958 | 24.18693 | 23.59529 | 10.15591 | 0.453135 | 0.303221 | 1 |
| 1 | 49649200 | 49650599 | 12.57503 | 22.12001 | 37.44209 | 16.57818 | -0.36707 | 0.576149 | 1 |
| 1 | 49758400 | 49768399 | 153.2033 | 230.9778 | 244.22 | 100.9056 | 0.192539 | 3.368391 | 1 |
| 1 | 49882200 | 49884999 | 20.87001 | 36.52065 | 38.60264 | 17.03033 | 0.308056 | 0.701257 | 1 |
| 1 | 50017200 | 50018799 | 14.95276 | 26.74502 | 26.92934 | 11.81646 | 0.389596 | 0.42518 | 1 |
| 1 | 50059800 | 50062799 | 40.6959 | 65.93275 | 73.94529 | 31.65152 | 0.199297 | 1.654696 | 1 |
| 1 | 50121400 | 50124199 | 22.53554 | 38.72835 | 44.19481 | 19.38536 | 0.194231 | 0.919684 | 1 |
| 1 | 50421400 | 50426399 | 89.67869 | 138.4727 | 189.0797 | 78.53987 | -0.1485 | 2.872691 | 1 |
| 1 | 51295800 | 51296799 | 14.37649 | 25.82166 | 27.01495 | 11.81245 | 0.338231 | 0.396479 | 1 |
| 1 | 51409400 | 51411199 | 16.53834 | 28.84328 | 45.71314 | 20.11837 | -0.27414 | 0.736162 | 1 |
| 1 | 51608200 | 51610399 | 14.59577 | 25.53916 | 41.52223 | 18.31707 | -0.31006 | 0.674472 | 1 |
| 1 | 51745200 | 51747799 | 22.72194 | 38.42972 | 64.47103 | 28.21587 | -0.36267 | 1.08327 | 1 |
| 1 | 52628200 | 52630599 | 22.07036 | 38.1976 | 41.51451 | 18.21361 | 0.266393 | 0.852297 | 1 |
| 1 | 52884000 | 52884799 | 9.020005 | 16.78558 | 28.39803 | 11.8609 | -0.30503 | 0.154106 | 1 |
| 1 | 53020400 | 53022199 | 21.01045 | 36.28081 | 40.53937 | 17.78456 | 0.22589 | 0.865453 | 1 |
| 1 | 53171800 | 53174599 | 46.06545 | 74.12452 | 77.163 | 32.76328 | 0.295655 | 1.849645 | 1 |
| 1 | 53540200 | 53548799 | 103.9277 | 159.6153 | 194.2366 | 80.64705 | 0.011596 | 2.980636 | 1 |
| 1 | 53636800 | 53639599 | 19.5699 | 33.78627 | 49.01131 | 21.50105 | -0.1492 | 0.87306 | 1 |
| 1 | 53744800 | 53746399 | 13.178 | 23.01898 | 43.65021 | 19.22949 | -0.53114 | 0.69535 | 1 |
| 1 | 54063600 | 54065799 | 30.24211 | 50.00411 | 58.37237 | 25.2308 | 0.157996 | 1.333838 | 1 |

| | | | | | | | | | |
|---|---|---|---|---|---|---|---|---|---|
| 1 | 55166000 | 55167799 | 15.91856 | 27.89483 | 35.31537 | 15.59841 | 0.049042 | 0.646661 | 1 |
| 1 | 55400600 | 55403199 | 57.96413 | 91.54185 | 143.4671 | 59.57492 | -0.32178 | 2.435 | 1 |
| 1 | 55511400 | 55515599 | 41.1009 | 66.26763 | 89.94661 | 38.29862 | -0.08287 | 1.786963 | 1 |
| 1 | 55534200 | 55537999 | 55.81364 | 88.90718 | 113.5965 | 47.53041 | -0.01605 | 2.188647 | 1 |
| 1 | 56753400 | 56754399 | 15.07402 | 26.26947 | 43.16725 | 18.98883 | -0.32581 | 0.739531 | 1 |
| 1 | 56809400 | 56812399 | 13.45302 | 24.50486 | 25.43282 | 10.93539 | 0.365164 | 0.30352 | 1 |
| 1 | 57219000 | 57221399 | 32.70871 | 54.33955 | 60.92291 | 26.41958 | 0.216549 | 1.288427 | 1 |
| 1 | 57411400 | 57413399 | 19.24282 | 33.61707 | 40.80328 | 17.97673 | 0.109319 | 0.719905 | 1 |
| 1 | 58361200 | 58364199 | 31.22933 | 51.06924 | 69.03136 | 29.68495 | -0.05878 | 1.481404 | 1 |
| 1 | 58367200 | 58368799 | 35.75163 | 57.8562 | 73.85131 | 31.58466 | 0.011606 | 1.670988 | 1 |
| 1 | 58644800 | 58649799 | 58.19769 | 92.69813 | 112.7898 | 47.18816 | 0.054233 | 2.19194 | 1 |
| 1 | 58741800 | 58748199 | 75.76877 | 118.6899 | 155.6897 | 64.57939 | -0.07455 | 2.60883 | 1 |
| 1 | 58922400 | 58927399 | 62.47909 | 99.26857 | 119.7642 | 49.95024 | 0.062704 | 2.274752 | 1 |
| 1 | 58968400 | 58986199 | 22.06493 | 38.03958 | 39.54839 | 17.34895 | 0.329079 | 0.884842 | 1 |
| 1 | 59341400 | 59344599 | 10.57629 | 19.90188 | 22.17187 | 9.087643 | 0.315229 | 0.080882 | 1 |
| 1 | 60086000 | 60088599 | 35.27648 | 57.81906 | 64.53718 | 27.77752 | 0.218403 | 1.457969 | 1 |
| 1 | 60584000 | 60589199 | 100.3885 | 153.8446 | 217.4827 | 90.27189 | -0.20681 | 3.024993 | 1 |
| 1 | 61209000 | 61209999 | 17.28109 | 30.26327 | 34.35426 | 15.16577 | 0.205811 | 0.664705 | 1 |
| 1 | 61406000 | 61408399 | 15.49526 | 26.986 | 42.63812 | 18.7484 | -0.26967 | 0.748671 | 1 |
| 1 | 61552600 | 61554599 | 25.73471 | 42.7459 | 59.78614 | 25.92342 | -0.10145 | 1.270572 | 1 |
| 1 | 61644200 | 61645799 | 13.30255 | 23.53167 | 38.22614 | 16.90632 | -0.3052 | 0.517788 | 1 |
| 1 | 61765200 | 61768599 | 38.72156 | 63.76765 | 62.28291 | 26.86115 | 0.413123 | 1.406764 | 1 |
| 1 | 62333800 | 62335599 | 14.66974 | 26.2099 | 30.36027 | 13.33432 | 0.18757 | 0.436687 | 1 |
| 1 | 62580000 | 62582399 | 21.48298 | 37.40368 | 38.71912 | 17.02223 | 0.337506 | 0.775597 | 1 |
| 1 | 63177200 | 63180199 | 29.17532 | 47.69337 | 75.55211 | 32.4848 | -0.28742 | 1.48651 | 1 |
| 1 | 63238400 | 63239999 | 23.19566 | 38.76062 | 56.93702 | 24.78481 | -0.17167 | 1.196817 | 1 |
| 1 | 63312000 | 63313599 | 14.5971 | 25.62402 | 38.85637 | 17.18786 | -0.20997 | 0.618451 | 1 |
| 1 | 63403600 | 63405199 | 24.00205 | 39.94182 | 62.98131 | 27.34379 | -0.27368 | 1.24765 | 1 |
| 1 | 63438000 | 63439399 | 25.0023 | 42.56222 | 45.29247 | 19.8525 | 0.292557 | 1.016213 | 1 |
| 1 | 63932400 | 63935399 | 18.45718 | 32.28046 | 39.75681 | 17.53128 | 0.088423 | 0.697428 | 1 |
| 1 | 64217000 | 64219199 | 33.03304 | 54.59311 | 61.40057 | 26.53109 | 0.211465 | 1.348745 | 1 |
| 1 | 64224000 | 64225799 | 21.06999 | 36.2105 | 45.80491 | 20.09162 | 0.046197 | 0.914797 | 1 |
| 1 | 64298800 | 64300399 | 18.87006 | 32.64531 | 43.20389 | 18.95455 | -0.01663 | 0.844242 | 1 |
| 1 | 64421600 | 64424399 | 31.6355 | 52.09983 | 61.05971 | 26.33398 | 0.150977 | 1.388047 | 1 |
| 1 | 64621000 | 64622999 | 16.38268 | 29.38729 | 31.24915 | 13.67866 | 0.313702 | 0.426574 | 1 |
| 1 | 65005200 | 65007999 | 49.82118 | 80.07632 | 91.60228 | 38.82979 | 0.157844 | 1.897568 | 1 |
| 1 | 66062000 | 66065399 | 52.37051 | 83.90189 | 90.15215 | 38.02919 | 0.24247 | 1.999506 | 1 |
| 1 | 66619800 | 66620799 | 11.29415 | 21.12644 | 27.85104 | 11.53115 | 0.06221 | 0.13886 | 1 |
| 1 | 67315600 | 67317199 | 13.25289 | 23.73437 | 37.2188 | 16.31414 | -0.24494 | 0.427497 | 1 |
| 1 | 68307200 | 68307799 | 15.9455 | 27.87229 | 36.95203 | 16.28688 | -0.0171 | 0.690401 | 1 |
| 1 | 68508000 | 68509199 | 22.99413 | 38.33085 | 60.22441 | 26.17577 | -0.2685 | 1.222642 | 1 |
| 1 | 68636400 | 68638399 | 37.89877 | 61.19961 | 81.15369 | 34.62384 | -0.04665 | 1.735039 | 1 |
| 1 | 68663800 | 68664799 | 9.044971 | 16.83692 | 27.39328 | 11.43564 | -0.24817 | 0.151512 | 1 |
| 1 | 69953800 | 69955599 | 14.78163 | 26.00325 | 41.34052 | 18.30404 | -0.27751 | 0.593417 | 1 |
| 1 | 70008000 | 70008599 | 10.63649 | 20.18467 | 22.21746 | 8.953874 | 0.353516 | 0.026039 | 1 |
| 1 | 70276200 | 70278799 | 10.18716 | 19.1238 | 23.7624 | 9.779107 | 0.154382 | 0.097641 | 1 |
| 1 | 70673800 | 70675999 | 23.80419 | 39.91513 | 55.40784 | 24.17178 | -0.09043 | 1.159832 | 1 |

| | | | | | | | | | |
|---|---|---|---|---|---|---|---|---|---|
| 1 | 70963400 | 70964799 | 15.70732 | 27.57565 | 38.27678 | 16.93289 | -0.08308 | 0.622049 | 1 |
| 1 | 71555800 | 71556599 | 13.41415 | 23.91427 | 35.1187 | 15.45267 | -0.15447 | 0.453431 | 1 |
| 1 | 71791800 | 71793199 | 22.07402 | 38.18921 | 43.0909 | 18.905 | 0.212394 | 0.857506 | 1 |
| 1 | 71907600 | 71909999 | 39.99112 | 64.80805 | 76.57199 | 32.78224 | 0.124675 | 1.651374 | 1 |
| 1 | 72848400 | 72850999 | 18.42818 | 31.78782 | 49.5415 | 21.73307 | -0.25251 | 0.875944 | 1 |
| 1 | 73148800 | 73152799 | 49.17246 | 78.5623 | 108.627 | 45.64119 | -0.12512 | 2.085631 | 1 |
| 1 | 73362400 | 73364399 | 23.02523 | 38.38462 | 62.39763 | 27.12119 | -0.31746 | 1.224874 | 1 |
| 1 | 73746200 | 73747799 | 26.68117 | 45.13054 | 45.38433 | 19.86315 | 0.373372 | 1.081223 | 1 |
| 1 | 74696200 | 74698799 | 13.88886 | 25.64097 | 27.51417 | 11.62102 | 0.335339 | 0.237366 | 1 |
| 1 | 74865200 | 74867199 | 17.89319 | 31.31228 | 38.16653 | 16.8383 | 0.10325 | 0.6816 | 1 |
| 1 | 76243200 | 76245199 | 9.786296 | 18.95375 | 20.9854 | 7.969004 | 0.423676 | -0.18029 | 1 |
| 1 | 76287600 | 76289599 | 21.61516 | 36.44928 | 57.04579 | 24.94369 | -0.26281 | 1.099294 | 1 |
| 1 | 76564000 | 76570399 | 140.1217 | 210.8636 | 255.2221 | 105.399 | -0.00471 | 3.402428 | 1 |
| 1 | 76591400 | 76592199 | 15.27435 | 26.8001 | 36.74779 | 16.24859 | -0.06562 | 0.624046 | 1 |
| 1 | 76593200 | 76595399 | 24.04374 | 40.47747 | 52.24179 | 22.83164 | 0.014329 | 1.11832 | 1 |
| 1 | 78805400 | 78808399 | 46.32863 | 73.98441 | 113.2109 | 47.53595 | -0.27181 | 2.099664 | 1 |
| 1 | 78967000 | 78968199 | 22.0142 | 37.13499 | 51.69535 | 22.60654 | -0.0944 | 1.091233 | 1 |
| 1 | 78996400 | 79000799 | 94.10428 | 144.3888 | 211.4133 | 87.7672 | -0.2561 | 2.998089 | 1 |
| 1 | 79022400 | 79023399 | 11.29503 | 21.35315 | 23.71347 | 9.633814 | 0.330562 | 0.061868 | 1 |
| 1 | 79059800 | 79060799 | 10.5469 | 19.5672 | 29.35203 | 12.31458 | -0.13782 | 0.186856 | 1 |
| 1 | 79229200 | 79233599 | 12.82452 | 23.69212 | 29.75074 | 12.55406 | 0.111008 | 0.230571 | 1 |
| 1 | 79916600 | 79917599 | 25.26393 | 42.40762 | 49.62593 | 21.66385 | 0.15503 | 1.145369 | 1 |
| 1 | 80116600 | 80125199 | 171.1518 | 255.3651 | 267.2859 | 110.1683 | 0.193693 | 3.569821 | 1 |
| 1 | 80960800 | 80962199 | 23.99635 | 40.07083 | 61.9364 | 26.94927 | -0.245 | 1.209989 | 1 |
| 1 | 80988400 | 80991999 | 27.87702 | 47.59136 | 46.82845 | 20.53046 | 0.405755 | 1.003861 | 1 |
| 1 | 81374200 | 81377199 | 23.93961 | 40.78017 | 49.0235 | 21.48903 | 0.117449 | 1.010855 | 1 |
| 1 | 81396400 | 81398599 | 37.35274 | 60.46184 | 74.80655 | 31.99889 | 0.056797 | 1.668765 | 1 |
| 1 | 81476200 | 81476599 | 9.428294 | 17.75896 | 23.834 | 9.752774 | 0.050751 | 0.06983 | 1 |
| 1 | 81546600 | 81549799 | 37.07398 | 60.1081 | 76.67279 | 32.83646 | 0.015067 | 1.637603 | 1 |
| 1 | 81574200 | 81575399 | 14.00732 | 24.86579 | 31.59929 | 13.95403 | 0.049542 | 0.47697 | 1 |
| 1 | 82478000 | 82482399 | 33.59785 | 54.56741 | 83.12745 | 35.64662 | -0.23815 | 1.597365 | 1 |
| 1 | 82936600 | 82938599 | 22.71434 | 38.27899 | 52.75912 | 23.06496 | -0.08009 | 1.104402 | 1 |
| 1 | 83210000 | 83213999 | 63.44358 | 100.219 | 132.703 | 55.11165 | -0.07869 | 2.430255 | 1 |
| 1 | 83346800 | 83349199 | 21.94997 | 38.17328 | 39.8464 | 17.50529 | 0.325496 | 0.796984 | 1 |
| 1 | 83384600 | 83386199 | 12.64067 | 22.57381 | 30.58428 | 13.43813 | -0.03756 | 0.425983 | 1 |
| 1 | 83593000 | 83593799 | 8.778237 | 16.65797 | 23.88833 | 9.627645 | -0.02533 | 0.012168 | 1 |
| 1 | 83761200 | 83762999 | 11.01237 | 20.56964 | 32.31059 | 13.41027 | -0.19191 | 0.154992 | 1 |
| 1 | 83888400 | 83891199 | 12.06383 | 22.05488 | 30.7143 | 13.14605 | -0.05195 | 0.285749 | 1 |
| 1 | 83961000 | 83961399 | 7.714217 | 14.89049 | 22.586 | 8.712583 | -0.04739 | -0.14406 | 1 |
| 1 | 84928600 | 84948799 | 329.9285 | 467.132 | 460.4674 | 190.5525 | 0.207008 | 4.39568 | 1 |
| 1 | 84962800 | 84975599 | 363.1106 | 506.6857 | 548.0454 | 227.3219 | 0.051999 | 4.612513 | 1 |
| 1 | 84985000 | 84993599 | 313.5795 | 447.7611 | 428.5235 | 177.0817 | 0.261908 | 4.271648 | 1 |
| 1 | 84997200 | 85002399 | 81.34135 | 127.7386 | 138.2408 | 57.33813 | 0.205399 | 2.564767 | 1 |
| 1 | 85111000 | 85123599 | 61.85051 | 98.41332 | 98.90697 | 41.3201 | 0.327981 | 2.224575 | 1 |
| 1 | 85124200 | 85126199 | 104.6257 | 159.5098 | 215.8355 | 89.4819 | -0.1496 | 3.130534 | 1 |
| 1 | 85282000 | 85285599 | 46.03835 | 74.2058 | 77.84204 | 33.13159 | 0.287571 | 1.797154 | 1 |
| 1 | 85387800 | 85396599 | 255.1354 | 371.1594 | 356.4028 | 146.8987 | 0.283635 | 3.993707 | 1 |

| | | | | | | | | | |
|---|---|---|---|---|---|---|---|---|---|
| 1 | 85396800 | 85399199 | 68.69727 | 108.6893 | 111.3743 | 46.29075 | 0.292435 | 2.391847 | 1 |
| 1 | 85485200 | 85493599 | 106.6974 | 165.2533 | 168.0935 | 69.82149 | 0.279241 | 2.822477 | 1 |
| 1 | 86001800 | 86003199 | 19.89891 | 34.73979 | 38.71505 | 17.04949 | 0.231914 | 0.730239 | 1 |
| 1 | 86393200 | 86395799 | 21.25359 | 35.99092 | 53.27102 | 23.32294 | -0.1823 | 1.05784 | 1 |
| 1 | 88131000 | 88133999 | 18.24558 | 31.04307 | 52.66748 | 23.08409 | -0.37781 | 1.005676 | 1 |
| 1 | 88853200 | 88856199 | 27.73114 | 45.76293 | 70.52625 | 30.44314 | -0.24185 | 1.358398 | 1 |
| 1 | 88858800 | 88866999 | 71.52841 | 112.1009 | 162.0141 | 67.19942 | -0.21369 | 2.599411 | 1 |
| 1 | 88890400 | 88893399 | 16.47313 | 28.82143 | 43.5262 | 19.19961 | -0.20437 | 0.6877 | 1 |
| 1 | 89125600 | 89129199 | 14.03949 | 24.66848 | 39.28002 | 17.38913 | -0.28032 | 0.597934 | 1 |
| 1 | 92213800 | 92216599 | 20.09651 | 34.16059 | 54.88776 | 24.05544 | -0.30004 | 1.021326 | 1 |
| 1 | 92413800 | 92419399 | 80.82223 | 126.2168 | 153.2997 | 63.60819 | 0.032895 | 2.665912 | 1 |
| 1 | 94169400 | 94171999 | 29.42924 | 48.24439 | 74.85094 | 32.21925 | -0.25531 | 1.447657 | 1 |
| 1 | 94409000 | 94409999 | 8.883107 | 16.73943 | 24.4254 | 9.987319 | -0.06771 | 0.06556 | 1 |
| 1 | 94697000 | 94702399 | 55.09234 | 87.68099 | 120.1239 | 50.2004 | -0.11791 | 2.216398 | 1 |
| 1 | 94733400 | 94735999 | 27.9565 | 46.2772 | 69.96454 | 30.26498 | -0.21388 | 1.325269 | 1 |
| 1 | 94849200 | 94849999 | 16.41336 | 28.7892 | 34.74097 | 15.35701 | 0.117892 | 0.633575 | 1 |
| 1 | 95822000 | 95826999 | 101.6654 | 155.5455 | 221.918 | 92.07882 | -0.22193 | 3.057951 | 1 |
| 1 | 95885400 | 95887399 | 23.40218 | 40.09698 | 44.17494 | 19.37277 | 0.243997 | 0.948543 | 1 |
| 1 | 95961800 | 95966599 | 86.00272 | 133.9959 | 157.3038 | 65.28817 | 0.079324 | 2.70933 | 1 |
| 1 | 96118800 | 96119999 | 21.76508 | 37.25011 | 43.25319 | 18.96682 | 0.16839 | 0.95341 | 1 |
| 1 | 96660600 | 96662199 | 11.30163 | 20.34841 | 31.70681 | 13.83246 | -0.23085 | 0.369658 | 1 |
| 1 | 96676800 | 96678599 | 12.44295 | 22.71808 | 28.07352 | 12.03556 | 0.117801 | 0.289227 | 1 |
| 1 | 96894400 | 96898999 | 63.43646 | 100.8615 | 113.6592 | 47.43786 | 0.162001 | 2.253456 | 1 |
| 1 | 96950800 | 96953799 | 31.20323 | 51.8954 | 55.751 | 24.19507 | 0.278093 | 1.267947 | 1 |
| 1 | 97037200 | 97039999 | 7.584791 | 14.55417 | 25.20497 | 9.892279 | -0.26064 | -0.08237 | 1 |
| 1 | 97156800 | 97158799 | 15.27179 | 26.79295 | 44.01638 | 19.46088 | -0.32513 | 0.639511 | 1 |
| 1 | 97324600 | 97327199 | 29.15286 | 48.56712 | 53.50288 | 23.24521 | 0.241997 | 1.24043 | 1 |
| 1 | 97449800 | 97451999 | 24.61877 | 40.93737 | 63.0606 | 27.36403 | -0.24021 | 1.257199 | 1 |
| 1 | 97706000 | 97708999 | 20.68583 | 34.79361 | 54.81009 | 23.94765 | -0.27221 | 1.116672 | 1 |
| 1 | 97793200 | 97794399 | 14.08895 | 24.69909 | 38.42151 | 16.97638 | -0.2466 | 0.635478 | 1 |
| 1 | 97904600 | 97905199 | 9.2846 | 17.79167 | 18.27768 | 7.201562 | 0.481485 | -0.0725 | 1 |
| 1 | 98122200 | 98123399 | 14.43863 | 25.96495 | 28.30622 | 12.36434 | 0.279919 | 0.390143 | 1 |
| 1 | 98211200 | 98225999 | 323.464 | 454.2945 | 543.4268 | 225.1792 | -0.08381 | 4.514871 | 1 |
| 1 | 98312000 | 98313599 | 18.04379 | 31.49889 | 33.9941 | 14.96979 | 0.278327 | 0.717093 | 1 |
| 1 | 98367800 | 98368599 | 10.77959 | 20.17465 | 23.68665 | 9.798128 | 0.229501 | 0.121398 | 1 |
| 1 | 98379400 | 98381999 | 34.11499 | 55.98247 | 69.74272 | 30.03328 | 0.060992 | 1.443069 | 1 |
| 1 | 98527200 | 98533399 | 98.48044 | 151.755 | 168.8247 | 70.12707 | 0.144579 | 2.911067 | 1 |
| 1 | 98548000 | 98548999 | 12.17865 | 21.72383 | 33.74224 | 14.84001 | -0.23418 | 0.44028 | 1 |
| 1 | 98578200 | 98579199 | 13.92736 | 24.96514 | 29.01971 | 12.7104 | 0.185479 | 0.409888 | 1 |
| 1 | 98872600 | 98876799 | 36.11572 | 58.89183 | 71.7565 | 30.82477 | 0.087635 | 1.533978 | 1 |
| 1 | 99027400 | 99029199 | 17.23947 | 30.45269 | 31.74128 | 14.04224 | 0.330586 | 0.543259 | 1 |
| 1 | 99053200 | 99059599 | 77.91643 | 122.6047 | 137.302 | 56.96356 | 0.158313 | 2.52198 | 1 |
| 1 | 99431000 | 99431799 | 10.55376 | 19.34885 | 30.7673 | 13.11857 | -0.2384 | 0.255552 | 1 |
| 1 | 100210000 | 100212999 | 27.36001 | 45.52714 | 65.61827 | 28.4875 | -0.14456 | 1.258901 | 1 |
| 1 | 100575200 | 100576399 | 11.08598 | 20.61938 | 31.58533 | 13.20388 | -0.164 | 0.182716 | 1 |
| 1 | 100622800 | 100625799 | 28.5327 | 47.84078 | 54.57718 | 23.80841 | 0.191571 | 1.171939 | 1 |
| 1 | 100677600 | 100683399 | 60.11001 | 95.77304 | 106.8269 | 44.71218 | 0.179934 | 2.182147 | 1 |

| | | | | | | | | | |
|---|---|---|---|---|---|---|---|---|---|
| 1 | 101015800 | 101024199 | 134.6754 | 203.4108 | 250.0937 | 103.3908 | -0.02273 | 3.328881 | 1 |
| 1 | 101313400 | 101314399 | 24.03584 | 40.82755 | 41.28589 | 18.09249 | 0.365541 | 1.028729 | 1 |
| 1 | 101383000 | 101385399 | 17.80029 | 30.7616 | 47.40917 | 20.79883 | -0.2358 | 0.854716 | 1 |
| 1 | 101463600 | 101465799 | 21.07174 | 36.83363 | 38.06931 | 16.77964 | 0.340239 | 0.717399 | 1 |
| 1 | 101578200 | 101582199 | 53.6142 | 86.10768 | 90.23429 | 38.20767 | 0.282742 | 1.925778 | 1 |
| 1 | 101692000 | 101693199 | 10.34388 | 19.64593 | 21.25284 | 8.549867 | 0.380829 | 0.015578 | 1 |
| 1 | 101825200 | 101831599 | 87.41561 | 137.2224 | 140.4127 | 58.23591 | 0.28567 | 2.574429 | 1 |
| 1 | 101929400 | 101930199 | 16.05386 | 28.49636 | 28.94881 | 12.78452 | 0.370696 | 0.485653 | 1 |
| 1 | 102019800 | 102020999 | 31.80634 | 51.73409 | 74.76146 | 32.0861 | -0.16038 | 1.564187 | 1 |
| 1 | 102039000 | 102039599 | 12.13578 | 21.69306 | 33.19897 | 14.57514 | -0.21127 | 0.424536 | 1 |
| 1 | 102109400 | 102111799 | 24.43774 | 41.51563 | 46.8889 | 20.54867 | 0.206478 | 1.032859 | 1 |
| 1 | 102182800 | 102184599 | 45.03776 | 72.47128 | 80.78905 | 34.30319 | 0.197107 | 1.85116 | 1 |
| 1 | 102316000 | 102317199 | 19.81834 | 34.60707 | 37.22823 | 16.39708 | 0.282805 | 0.72592 | 1 |
| 1 | 102453400 | 102456399 | 35.41772 | 57.97459 | 61.65468 | 26.52218 | 0.287273 | 1.472421 | 1 |
| 1 | 102533000 | 102535599 | 44.47589 | 71.52864 | 83.09288 | 35.25388 | 0.136793 | 1.866607 | 1 |
| 1 | 102745400 | 102747199 | 15.95977 | 28.401 | 30.90424 | 13.61905 | 0.274202 | 0.473694 | 1 |
| 1 | 102868600 | 102870199 | 19.38636 | 33.58233 | 40.55413 | 17.79275 | 0.115251 | 0.829605 | 1 |
| 1 | 102926400 | 102930799 | 65.73152 | 104.4006 | 125.87 | 52.47898 | 0.06321 | 2.289964 | 1 |
| 1 | 102953000 | 102954999 | 15.33989 | 27.0319 | 40.2182 | 17.80098 | -0.18148 | 0.574143 | 1 |
| 1 | 103044400 | 103047399 | 14.38566 | 25.52585 | 35.05635 | 15.48655 | -0.06229 | 0.489259 | 1 |
| 1 | 103105400 | 103109799 | 67.55488 | 106.1092 | 156.0225 | 64.7151 | -0.23604 | 2.555215 | 1 |
| 1 | 103132200 | 103133399 | 28.89454 | 47.98885 | 53.44536 | 23.17218 | 0.226162 | 1.275201 | 1 |
| 1 | 103137600 | 103144799 | 96.93034 | 149.6081 | 170.6688 | 70.89251 | 0.110173 | 2.881726 | 1 |
| 1 | 103403600 | 103411399 | 161.3364 | 241.0339 | 273.534 | 112.7721 | 0.078809 | 3.544561 | 1 |
| 1 | 103467200 | 103471399 | 71.79109 | 113.6546 | 127.2689 | 52.91367 | 0.16531 | 2.383679 | 1 |
| 1 | 104083600 | 104088999 | 72.42181 | 114.4591 | 122.305 | 50.8063 | 0.230844 | 2.422654 | 1 |
| 1 | 104316000 | 104318199 | 35.29239 | 57.62084 | 69.42873 | 29.83834 | 0.104967 | 1.513344 | 1 |
| 1 | 104412600 | 104413399 | 9.703437 | 18.31325 | 28.65416 | 11.68275 | -0.16538 | 0.069132 | 1 |
| 1 | 104439400 | 104442199 | 31.05477 | 51.37141 | 62.41514 | 26.9868 | 0.100482 | 1.33322 | 1 |
| 1 | 106173400 | 106178199 | 48.24747 | 78.55727 | 66.58496 | 28.58768 | 0.608441 | 1.578626 | 1 |
| 1 | 106686000 | 106688199 | 15.33763 | 26.65481 | 45.79951 | 20.11163 | -0.39057 | 0.781565 | 1 |
| 1 | 106727200 | 106734599 | 99.64849 | 153.5279 | 181.9873 | 75.59388 | 0.052918 | 2.916693 | 1 |
| 1 | 106900600 | 106901599 | 21.4044 | 36.80511 | 42.15872 | 18.49258 | 0.18904 | 0.907863 | 1 |
| 1 | 107075400 | 107076599 | 13.73002 | 24.57318 | 29.49146 | 12.93396 | 0.138244 | 0.418209 | 1 |
| 1 | 107146600 | 107147999 | 13.75654 | 25.01681 | 27.84468 | 11.99692 | 0.262836 | 0.319523 | 1 |
| 1 | 107724200 | 107726799 | 23.72243 | 40.39026 | 43.20567 | 18.93808 | 0.285037 | 1.005546 | 1 |
| 1 | 107853200 | 107854399 | 16.87255 | 29.62966 | 33.14167 | 14.66471 | 0.22682 | 0.61428 | 1 |
| 1 | 108641400 | 108642799 | 21.48172 | 37.09444 | 41.60429 | 18.25173 | 0.220603 | 0.87104 | 1 |
| 1 | 109262400 | 109267199 | 25.9096 | 44.29442 | 48.47132 | 21.25312 | 0.253204 | 0.98343 | 1 |
| 1 | 109338000 | 109339399 | 24.68478 | 42.0593 | 45.8181 | 20.08434 | 0.259077 | 1.00755 | 1 |
| 1 | 109494600 | 109496199 | 17.30175 | 30.37799 | 33.05547 | 14.62438 | 0.266283 | 0.618553 | 1 |
| 1 | 109795800 | 109799999 | 55.47556 | 88.27601 | 121.6917 | 50.84375 | -0.1271 | 2.223371 | 1 |
| 1 | 110219600 | 110222599 | 37.62831 | 61.82191 | 66.64533 | 28.71402 | 0.269822 | 1.435989 | 1 |
| 1 | 110546200 | 110548999 | 31.41249 | 52.10737 | 61.91413 | 26.82145 | 0.132846 | 1.302899 | 1 |
| 1 | 111204000 | 111206799 | 13.0094 | 22.9855 | 36.28706 | 16.05257 | -0.26466 | 0.52688 | 1 |
| 1 | 111338400 | 111340399 | 23.55228 | 40.13799 | 45.31757 | 19.86527 | 0.207731 | 1.000987 | 1 |
| 1 | 111437200 | 111438199 | 12.84587 | 23.9348 | 26.14633 | 10.89762 | 0.324181 | 0.174756 | 1 |

| | | | | | | | | | |
|---|---|---|---|---|---|---|---|---|---|
| 1 | 111508400 | 111509399 | 31.20179 | 51.22355 | 59.70084 | 25.71354 | 0.157215 | 1.426073 | 1 |
| 1 | 111553600 | 111554799 | 21.32117 | 36.5003 | 43.28049 | 18.97921 | 0.138369 | 0.949932 | 1 |
| 1 | 112195000 | 112199599 | 90.12424 | 139.2737 | 167.63 | 69.62979 | 0.034261 | 2.855534 | 1 |
| 1 | 113483800 | 113485599 | 17.05701 | 30.06822 | 32.95686 | 14.58603 | 0.257472 | 0.568737 | 1 |
| 1 | 113522600 | 113524799 | 25.43985 | 43.46663 | 44.21464 | 19.38581 | 0.358243 | 0.985241 | 1 |
| 1 | 113677800 | 113679999 | 29.23847 | 48.63114 | 55.65145 | 24.15304 | 0.187246 | 1.261363 | 1 |
| 1 | 113743000 | 113744999 | 30.63758 | 50.4833 | 61.02063 | 26.32306 | 0.106855 | 1.378541 | 1 |
| 1 | 113825400 | 113825999 | 6.081294 | 11.85634 | 20.37225 | 7.343659 | -0.13485 | -0.40701 | 1 |
| 1 | 113841000 | 113843599 | 12.30415 | 22.90616 | 25.53694 | 10.65881 | 0.292945 | 0.171347 | 1 |
| 1 | 114427000 | 114428399 | 14.21795 | 25.10563 | 39.20616 | 17.34601 | -0.24981 | 0.54366 | 1 |
| 1 | 114503800 | 114507599 | 52.25686 | 83.80121 | 97.99881 | 41.40031 | 0.122203 | 1.972115 | 1 |
| 1 | 114658800 | 114659999 | 14.42284 | 25.45545 | 36.19715 | 16.01624 | -0.11536 | 0.546735 | 1 |
| 1 | 114698000 | 114699199 | 16.08361 | 28.30111 | 32.64422 | 14.45288 | 0.183287 | 0.582245 | 1 |
| 1 | 115009000 | 115010399 | 19.18607 | 33.42161 | 36.31026 | 15.96833 | 0.268205 | 0.753764 | 1 |
| 1 | 115267400 | 115269199 | 17.33375 | 30.6425 | 31.79214 | 14.06261 | 0.337741 | 0.537049 | 1 |
| 1 | 115434000 | 115434999 | 12.91764 | 22.86174 | 34.18468 | 15.11745 | -0.186 | 0.50138 | 1 |
| 1 | 116434200 | 116434999 | 11.05456 | 20.4944 | 28.80755 | 12.0979 | -0.04551 | 0.194826 | 1 |
| 1 | 116647600 | 116649599 | 18.44279 | 32.47165 | 33.77985 | 14.9539 | 0.331643 | 0.590199 | 1 |
| 1 | 117484800 | 117489799 | 22.88817 | 38.62933 | 55.47029 | 24.26116 | -0.13899 | 1.094792 | 1 |
| 1 | 117512600 | 117523799 | 46.4718 | 74.95253 | 78.465 | 33.42439 | 0.291593 | 1.781166 | 1 |
| 1 | 117808400 | 117809799 | 41.03059 | 66.25248 | 71.50289 | 30.50373 | 0.249651 | 1.735773 | 1 |
| 1 | 118133800 | 118136799 | 41.38075 | 67.01855 | 79.31474 | 33.94099 | 0.121424 | 1.665748 | 1 |
| 1 | 118638200 | 118644599 | 65.87335 | 104.1499 | 146.6664 | 60.93812 | -0.16641 | 2.412855 | 1 |
| 1 | 118728600 | 118731199 | 14.06461 | 25.48864 | 29.43509 | 12.73718 | 0.206179 | 0.346744 | 1 |
| 1 | 119509200 | 119511599 | 35.39278 | 57.8592 | 67.92007 | 29.20367 | 0.143802 | 1.493322 | 1 |
| 1 | 120814600 | 120816599 | 9.500669 | 17.57664 | 29.37048 | 12.3696 | -0.29625 | 0.194504 | 1 |
| 1 | 120867000 | 120869199 | 29.34524 | 48.63427 | 61.08353 | 26.44514 | 0.053013 | 1.308114 | 1 |
| 1 | 121596800 | 121598599 | 25.47048 | 42.45139 | 57.15971 | 24.84097 | -0.04663 | 1.227685 | 1 |
| 1 | 121650800 | 121652799 | 16.53798 | 29.1528 | 34.95606 | 15.47088 | 0.128444 | 0.568273 | 1 |
| 1 | 121702000 | 121706999 | 42.50346 | 68.69547 | 74.32105 | 31.74456 | 0.247663 | 1.71216 | 1 |
| 1 | 121825400 | 121825999 | 9.535138 | 18.15242 | 23.89061 | 9.563471 | 0.105997 | -0.00253 | 1 |
| 1 | 122572600 | 122573399 | 10.08841 | 18.7313 | 29.38097 | 12.31354 | -0.20062 | 0.178841 | 1 |
| 1 | 122706000 | 122707199 | 23.22248 | 39.78544 | 44.22201 | 19.39327 | 0.231351 | 0.950913 | 1 |
| 1 | 122880600 | 122881599 | 13.37722 | 24.10342 | 27.51513 | 11.99262 | 0.216115 | 0.370315 | 1 |
| 1 | 123186600 | 123187599 | 15.30234 | 27.35151 | 31.27836 | 13.73273 | 0.206276 | 0.439872 | 1 |
| 1 | 123784000 | 123785599 | 41.28452 | 66.47081 | 86.88166 | 36.92148 | -0.03096 | 1.829252 | 1 |
| 1 | 123795000 | 123796799 | 18.65147 | 32.71348 | 35.01118 | 15.47591 | 0.290126 | 0.646658 | 1 |
| 1 | 123800000 | 123801999 | 18.98921 | 32.47667 | 47.9673 | 21.03281 | -0.1774 | 0.95566 | 1 |
| 1 | 124067800 | 124069599 | 22.95321 | 38.53603 | 55.40658 | 24.19018 | -0.141 | 1.14275 | 1 |
| 1 | 124403000 | 124406199 | 76.22313 | 120.0464 | 129.581 | 53.76644 | 0.212303 | 2.502018 | 1 |
| 1 | 124813600 | 124816399 | 27.6633 | 47.10799 | 45.7378 | 20.04824 | 0.424327 | 1.02196 | 1 |
| 1 | 125022200 | 125028799 | 54.1729 | 86.4126 | 114.2402 | 47.891 | -0.06305 | 2.143101 | 1 |
| 1 | 125062200 | 125062999 | 9.548284 | 17.82736 | 27.92993 | 11.60043 | -0.18804 | 0.139335 | 1 |
| 1 | 125167200 | 125176999 | 100.7691 | 154.3813 | 212.6396 | 88.25644 | -0.16967 | 3.029879 | 1 |
| 1 | 125410200 | 125414599 | 39.31828 | 63.38059 | 88.22897 | 37.55741 | -0.11956 | 1.789608 | 1 |
| 1 | 126556000 | 126558799 | 50.36908 | 80.35529 | 110.8026 | 46.4581 | -0.1235 | 2.133734 | 1 |
| 1 | 126863200 | 126865199 | 23.3583 | 40.00566 | 49.01827 | 21.49609 | 0.091133 | 0.958447 | 1 |

| | | | | | | | | | |
|---|---|---|---|---|---|---|---|---|---|
| 1 | 127375000 | 127377599 | 24.91531 | 41.47042 | 59.22976 | 25.71881 | -0.13133 | 1.243881 | 1 |
| 1 | 127515200 | 127523999 | 78.97152 | 122.9945 | 174.6281 | 72.48262 | -0.19512 | 2.714477 | 1 |
| 1 | 127808800 | 127812799 | 10.96766 | 20.65018 | 24.2173 | 9.916366 | 0.242854 | 0.08684 | 1 |
| 1 | 127823200 | 127824999 | 14.38604 | 25.67325 | 29.95318 | 13.16838 | 0.176396 | 0.441857 | 1 |
| 1 | 127865000 | 127869399 | 46.83699 | 75.2492 | 92.0278 | 38.98711 | 0.060922 | 1.905915 | 1 |
| 1 | 128780400 | 128781399 | 14.21124 | 25.11435 | 31.7734 | 14.05523 | 0.0532 | 0.522975 | 1 |
| 1 | 128812600 | 128820199 | 85.9456 | 133.6093 | 178.5059 | 74.11264 | -0.10956 | 2.749944 | 1 |
| 1 | 129333000 | 129335199 | 18.45986 | 32.43875 | 38.98896 | 17.26178 | 0.123304 | 0.620358 | 1 |
| 1 | 129363400 | 129364799 | 13.9573 | 24.522 | 39.34812 | 17.41803 | -0.29115 | 0.601349 | 1 |
| 1 | 129965200 | 129966599 | 16.80707 | 29.60741 | 32.59995 | 14.43004 | 0.250705 | 0.575429 | 1 |
| 1 | 130038600 | 130039999 | 14.53762 | 25.60893 | 36.69456 | 16.24255 | -0.12751 | 0.568098 | 1 |
| 1 | 130179800 | 130181799 | 25.45441 | 43.08836 | 46.94061 | 20.54903 | 0.25831 | 1.070529 | 1 |
| 1 | 130417800 | 130419399 | 38.41327 | 61.93456 | 87.18178 | 37.12156 | -0.1352 | 1.783659 | 1 |
| 1 | 130680200 | 130680999 | 11.86486 | 21.64056 | 27.68404 | 11.88372 | 0.067204 | 0.290931 | 1 |
| 1 | 130994000 | 130996999 | 20.97325 | 35.75514 | 49.55101 | 21.72163 | -0.08679 | 0.991694 | 1 |
| 1 | 131319800 | 131320999 | 14.38355 | 25.81222 | 28.92947 | 12.65873 | 0.238688 | 0.405453 | 1 |
| 1 | 133356200 | 133358999 | 30.50739 | 51.51298 | 51.28946 | 22.4358 | 0.387452 | 1.112194 | 1 |
| 1 | 134057800 | 134058799 | 15.52894 | 27.58757 | 32.83272 | 14.48958 | 0.144003 | 0.482841 | 1 |
| 1 | 134182000 | 134184599 | 34.32896 | 56.20006 | 66.88586 | 28.77413 | 0.125372 | 1.468774 | 1 |
| 1 | 134185800 | 134191599 | 65.86901 | 103.629 | 153.5719 | 63.71208 | -0.24551 | 2.521439 | 1 |
| 1 | 134726800 | 134727799 | 33.072 | 54.43086 | 54.52943 | 23.51124 | 0.376327 | 1.39501 | 1 |
| 1 | 135623600 | 135627599 | 30.88583 | 50.1181 | 81.72232 | 35.02109 | -0.33714 | 1.610396 | 1 |
| 1 | 136008600 | 136012199 | 26.79964 | 45.44912 | 49.04267 | 21.48123 | 0.271979 | 1.059463 | 1 |
| 1 | 136689000 | 136690799 | 20.50498 | 35.0269 | 49.01592 | 21.49053 | -0.10029 | 0.97155 | 1 |
| 1 | 138874000 | 138879399 | 61.83173 | 97.77362 | 146.0022 | 60.66646 | -0.25071 | 2.407417 | 1 |
| 1 | 139473800 | 139475399 | 16.72691 | 28.81675 | 45.87977 | 20.12595 | -0.28327 | 0.877267 | 1 |
| 1 | 139539800 | 139542599 | 29.91809 | 49.5492 | 69.64585 | 30.13759 | -0.109 | 1.324488 | 1 |
| 1 | 139568400 | 139569999 | 25.68253 | 42.32287 | 64.41003 | 27.78637 | -0.22441 | 1.367617 | 1 |
| 1 | 139671000 | 139674599 | 24.23168 | 41.19898 | 46.37647 | 20.3256 | 0.211533 | 1.026269 | 1 |
| 1 | 139792800 | 139793599 | 11.55957 | 21.00187 | 32.72967 | 14.11814 | -0.2207 | 0.320408 | 1 |
| 1 | 139855600 | 139857599 | 19.20666 | 33.03098 | 45.32479 | 19.88144 | -0.0704 | 0.901435 | 1 |
| 1 | 139932600 | 139934399 | 17.29581 | 30.41598 | 34.56087 | 15.30328 | 0.204513 | 0.60013 | 1 |
| 1 | 139966800 | 139968399 | 26.32894 | 43.89214 | 59.23249 | 25.74546 | -0.04974 | 1.234341 | 1 |
| 1 | 139975400 | 139975999 | 11.80554 | 21.20106 | 35.73315 | 15.62636 | -0.34597 | 0.397209 | 1 |
| 1 | 140229000 | 140231599 | 74.90196 | 117.7868 | 134.7986 | 55.92033 | 0.126381 | 2.533533 | 1 |
| 1 | 140245000 | 140247799 | 17.39758 | 30.18066 | 44.75824 | 19.63812 | -0.17942 | 0.813876 | 1 |
| 1 | 140720000 | 140725999 | 87.54231 | 136.5463 | 148.286 | 61.53551 | 0.192807 | 2.689129 | 1 |
| 1 | 140946800 | 140949599 | 36.80158 | 60.41392 | 65.90752 | 28.38625 | 0.252232 | 1.443692 | 1 |
| 1 | 141158200 | 141159199 | 11.63137 | 20.84489 | 33.1888 | 14.53915 | -0.26597 | 0.403769 | 1 |
| 1 | 141189400 | 141190799 | 28.89672 | 47.9132 | 51.2909 | 22.21271 | 0.282927 | 1.292685 | 1 |
| 1 | 141257000 | 141257799 | 11.23078 | 20.74178 | 27.12514 | 11.45067 | 0.052213 | 0.211215 | 1 |
| 1 | 141963000 | 141963799 | 16.12124 | 28.54349 | 29.11355 | 12.87377 | 0.363344 | 0.509481 | 1 |
| 1 | 142429400 | 142432199 | 25.52294 | 43.45264 | 46.34215 | 20.31275 | 0.289407 | 1.018717 | 1 |
| 1 | 142593800 | 142595599 | 20.37383 | 35.41127 | 38.38094 | 16.85522 | 0.271331 | 0.794758 | 1 |
| 1 | 143038200 | 143038999 | 14.34159 | 24.94971 | 44.779 | 19.67634 | -0.45259 | 0.764608 | 1 |
| 1 | 143319600 | 143321199 | 12.75646 | 22.82169 | 33.14087 | 14.53933 | -0.13536 | 0.423282 | 1 |
| 1 | 143456800 | 143458599 | 10.68225 | 19.49373 | 30.30413 | 12.99994 | -0.21146 | 0.285853 | 1 |

| | | | | | | | | | |
|---|---|---|---|---|---|---|---|---|---|
| 1 | 143540200 | 143542399 | 20.72742 | 35.42243 | 52.77205 | 23.13818 | -0.1902 | 0.97336 | 1 |
| 1 | 143592600 | 143593599 | 8.811021 | 16.62303 | 27.53344 | 11.23433 | -0.2476 | 0.06159 | 1 |
| 1 | 143707600 | 143713599 | 77.53548 | 122.1837 | 128.5077 | 53.3251 | 0.250349 | 2.491424 | 1 |
| 1 | 143759800 | 143761399 | 19.73238 | 34.28054 | 43.61221 | 19.14701 | 0.040998 | 0.804378 | 1 |
| 1 | 143846400 | 143848399 | 13.79568 | 24.23153 | 37.87023 | 16.76002 | -0.25321 | 0.602521 | 1 |
| 1 | 143921800 | 143922799 | 9.841322 | 18.06119 | 30.84317 | 13.13259 | -0.33889 | 0.245939 | 1 |
| 1 | 144196400 | 144197399 | 20.18597 | 34.71068 | 42.24112 | 18.52873 | 0.102159 | 0.903926 | 1 |
| 1 | 144256800 | 144268999 | 141.0667 | 211.9091 | 260.3717 | 107.4745 | -0.02865 | 3.436885 | 1 |
| 1 | 144349600 | 144350799 | 20.27234 | 35.0109 | 42.13464 | 18.48448 | 0.11933 | 0.863874 | 1 |
| 1 | 144680800 | 144683799 | 21.49155 | 36.37992 | 55.4911 | 24.2921 | -0.2257 | 1.06169 | 1 |
| 1 | 144720600 | 144722399 | 27.48462 | 45.79546 | 60.56385 | 26.3157 | -0.02079 | 1.240823 | 1 |
| 1 | 145031200 | 145033599 | 29.36865 | 47.81264 | 75.73048 | 32.51279 | -0.2914 | 1.545705 | 1 |
| 1 | 145040600 | 145042199 | 16.277 | 28.45692 | 41.97723 | 18.50461 | -0.17058 | 0.696051 | 1 |
| 1 | 145385000 | 145386199 | 15.68654 | 27.77093 | 31.39962 | 13.88494 | 0.21514 | 0.509755 | 1 |
| 1 | 145700200 | 145702799 | 50.40707 | 80.42289 | 115.7673 | 48.54561 | -0.18529 | 2.132799 | 1 |
| 1 | 145841400 | 145841799 | 8.094647 | 15.70945 | 16.78115 | 6.312008 | 0.488664 | -0.24325 | 1 |
| 1 | 146149000 | 146159599 | 22.09797 | 38.23095 | 40.87697 | 17.93369 | 0.289539 | 0.850897 | 1 |
| 1 | 146162800 | 146165799 | 44.05982 | 70.80816 | 88.45769 | 37.49063 | 0.030855 | 1.891806 | 1 |
| 1 | 146341200 | 146341999 | 12.87163 | 23.56856 | 29.13592 | 12.44434 | 0.121061 | 0.280154 | 1 |
| 1 | 146373000 | 146374599 | 13.4492 | 24.03742 | 29.57881 | 12.98736 | 0.101324 | 0.425663 | 1 |
| 1 | 146536600 | 146540399 | 32.76922 | 54.0102 | 63.70695 | 27.485 | 0.142071 | 1.379429 | 1 |
| 1 | 147839200 | 147841199 | 22.31072 | 38.37611 | 45.23449 | 19.84194 | 0.147879 | 0.910876 | 1 |
| 1 | 147864400 | 147867599 | 15.14466 | 26.15807 | 44.35676 | 19.45939 | -0.37291 | 0.845011 | 1 |
| 1 | 147954800 | 147961199 | 55.99489 | 88.77397 | 131.53 | 54.73873 | -0.23627 | 2.335146 | 1 |
| 1 | 148006000 | 148008399 | 33.70178 | 55.71585 | 64.75205 | 27.98533 | 0.164373 | 1.351742 | 1 |
| 1 | 148132800 | 148134999 | 20.04551 | 34.9282 | 38.78786 | 17.06082 | 0.236738 | 0.75805 | 1 |
| 1 | 148177600 | 148178599 | 18.41752 | 32.0815 | 35.4259 | 15.57901 | 0.244931 | 0.749914 | 1 |
| 1 | 148252800 | 148255199 | 36.2223 | 59.05467 | 64.31816 | 27.62758 | 0.248919 | 1.532137 | 1 |
| 1 | 148360800 | 148362599 | 26.01224 | 43.68223 | 48.57416 | 21.21038 | 0.228354 | 1.141408 | 1 |
| 1 | 148397400 | 148399799 | 36.81873 | 60.12609 | 63.77746 | 27.41179 | 0.288647 | 1.511738 | 1 |
| 1 | 148759200 | 148761799 | 32.65957 | 53.03724 | 76.86268 | 32.95729 | -0.16639 | 1.596298 | 1 |
| 1 | 149378600 | 149380199 | 15.70497 | 27.88702 | 33.33205 | 14.71584 | 0.137443 | 0.488002 | 1 |
| 1 | 149395000 | 149395599 | 9.664094 | 18.09303 | 26.38156 | 10.90667 | -0.07983 | 0.118385 | 1 |
| 1 | 149453600 | 149460599 | 95.83329 | 148.2638 | 164.2835 | 68.23933 | 0.15472 | 2.837321 | 1 |
| 1 | 150104800 | 150109799 | 92.62125 | 143.056 | 181.1903 | 75.26264 | -0.03983 | 2.868122 | 1 |
| 1 | 150141200 | 150142199 | 16.54565 | 29.30113 | 30.86231 | 13.64648 | 0.317133 | 0.511772 | 1 |
| 1 | 150144800 | 150149199 | 31.79451 | 52.51099 | 58.53149 | 25.27966 | 0.224274 | 1.349709 | 1 |
| 1 | 150159200 | 150163599 | 32.47144 | 53.5549 | 59.08617 | 25.49862 | 0.238632 | 1.36847 | 1 |
| 1 | 150235600 | 150238599 | 31.25423 | 51.79631 | 58.46357 | 25.31028 | 0.206652 | 1.31107 | 1 |
| 1 | 150349600 | 150352599 | 14.06682 | 24.79406 | 35.72854 | 15.8131 | -0.13522 | 0.557489 | 1 |
| 1 | 151011800 | 151014199 | 23.44835 | 39.24531 | 58.12411 | 25.32699 | -0.18361 | 1.179535 | 1 |
| 1 | 151275000 | 151276199 | 17.60319 | 31.17568 | 34.79781 | 15.38648 | 0.233863 | 0.525087 | 1 |
| 1 | 151395600 | 151396999 | 14.326 | 25.73753 | 30.84448 | 13.48404 | 0.143299 | 0.400544 | 1 |
| 1 | 151835800 | 151840999 | 50.34309 | 80.34871 | 121.6337 | 51.03067 | -0.25731 | 2.122159 | 1 |
| 1 | 152011800 | 152012799 | 20.78982 | 36.33369 | 37.00033 | 16.30576 | 0.36144 | 0.716241 | 1 |
| 1 | 152116600 | 152118799 | 30.313 | 49.75204 | 60.2813 | 25.96081 | 0.101168 | 1.425315 | 1 |
| 1 | 152174400 | 152176199 | 24.1738 | 40.94881 | 51.71598 | 22.64426 | 0.04581 | 1.064605 | 1 |

| | | | | | | | | | | |
|---|---|---|---|---|---|---|---|---|---|---|
| 1 | 152538600 | 152540399 | 15.54255 | 27.86955 | 30.44551 | 13.3312 | 0.274963 | 0.423949 | | 1 |
| 1 | 153755800 | 153756999 | 13.75583 | 24.27877 | 34.7449 | 15.37359 | -0.12446 | 0.539325 | | 1 |
| 1 | 154043600 | 154044799 | 15.28756 | 27.22845 | 31.77567 | 13.99281 | 0.174191 | 0.462243 | | 1 |
| 1 | 154061800 | 154063599 | 17.78947 | 30.69124 | 47.41308 | 20.79948 | -0.23972 | 0.869393 | | 1 |
| 1 | 154778600 | 154782199 | 33.67301 | 54.91272 | 66.79872 | 28.69581 | 0.09012 | 1.52535 | | 1 |
| 1 | 155640400 | 155650399 | 183.3244 | 271.9432 | 304.9685 | 125.6145 | 0.086529 | 3.675676 | | 1 |
| 1 | 155974600 | 155981599 | 105.0533 | 160.544 | 217.4488 | 90.20047 | -0.14832 | 3.081378 | | 1 |
| 1 | 156161000 | 156163999 | 21.47246 | 37.32045 | 41.4632 | 18.20872 | 0.235819 | 0.805319 | | 1 |
| 1 | 156676600 | 156677999 | 13.86073 | 25.43688 | 30.62396 | 13.03988 | 0.161801 | 0.275688 | | 1 |
| 1 | 156881200 | 156882999 | 24.5872 | 41.0305 | 61.48006 | 26.73963 | -0.20038 | 1.216888 | | 1 |
| 1 | 157541200 | 157543599 | 17.70583 | 30.6965 | 46.33606 | 20.33003 | -0.2051 | 0.822308 | | 1 |
| 1 | 158159000 | 158160199 | 22.61649 | 38.23608 | 46.4569 | 20.32948 | 0.101379 | 1.069121 | | 1 |
| 1 | 159205400 | 159207799 | 14.65948 | 25.96794 | 36.02632 | 15.929 | -0.07813 | 0.505949 | | 1 |
| 1 | 159795800 | 159799999 | 44.98482 | 72.62816 | 77.95005 | 33.24867 | 0.257401 | 1.749009 | | 1 |
| 1 | 159801200 | 159805799 | 43.9353 | 70.48923 | 94.15126 | 39.80306 | -0.06837 | 1.946475 | | 1 |
| 1 | 160595000 | 160595599 | 8.169007 | 15.73571 | 21.9901 | 8.535589 | 0.061487 | -0.1204 | | 1 |
| 1 | 161891000 | 161891599 | 7.777069 | 15.01282 | 21.52608 | 8.3011 | 0.034363 | -0.14439 | | 1 |
| 1 | 162097400 | 162099199 | 18.43397 | 31.62779 | 47.43048 | 20.80276 | -0.19867 | 0.921856 | | 1 |
| 1 | 162355600 | 162357599 | 26.12344 | 43.41374 | 60.3005 | 26.15566 | -0.09138 | 1.26798 | | 1 |
| 1 | 162588400 | 162589999 | 16.57152 | 29.25334 | 32.46734 | 14.36535 | 0.240302 | 0.549764 | | 1 |
| 1 | 163263600 | 163264999 | 18.68364 | 31.88182 | 52.17085 | 22.87182 | -0.32528 | 0.979219 | | 1 |
| 1 | 163299400 | 163300999 | 17.82034 | 31.28332 | 36.00995 | 15.92926 | 0.185485 | 0.628964 | | 1 |
| 1 | 163537600 | 163538599 | 19.99909 | 34.51569 | 39.56788 | 17.35808 | 0.188917 | 0.863837 | | 1 |
| 1 | 163626400 | 163628799 | 24.13189 | 41.71863 | 43.00662 | 18.86759 | 0.341937 | 0.871987 | | 1 |
| 1 | 164557000 | 164557999 | 32.75342 | 53.24434 | 68.02575 | 29.18924 | 0.016355 | 1.570315 | | 1 |
| 1 | 165441600 | 165443799 | 24.70129 | 41.90642 | 51.19927 | 22.42842 | 0.093469 | 1.05083 | | 1 |
| 1 | 165913000 | 165914399 | 15.70296 | 27.55302 | 39.21029 | 17.3379 | -0.11916 | 0.628329 | | 1 |
| 1 | 166315600 | 166317799 | 36.45663 | 59.7517 | 66.01754 | 28.41435 | 0.233 | 1.459515 | | 1 |
| 1 | 166355000 | 166356799 | 18.07544 | 31.79416 | 34.94641 | 15.4733 | 0.25224 | 0.602094 | | 1 |
| 1 | 166401000 | 166413799 | 374.0027 | 523.0647 | 522.4322 | 216.6217 | 0.170194 | 4.578959 | | 1 |
| 1 | 166424400 | 166429799 | 100.1974 | 153.7519 | 195.2737 | 81.06794 | -0.05098 | 2.996491 | | 1 |
| 1 | 166739800 | 166740799 | 17.02354 | 29.35059 | 45.35251 | 19.89513 | -0.2401 | 0.870436 | | 1 |
| 1 | 168444600 | 168447399 | 50.21764 | 80.10749 | 106.8111 | 44.77973 | -0.07521 | 2.135411 | | 1 |
| 1 | 168476200 | 168478799 | 22.25345 | 38.31015 | 44.7217 | 19.61751 | 0.162129 | 0.904181 | | 1 |
| 1 | 168810800 | 168813199 | 11.55716 | 20.60703 | 32.8312 | 14.44418 | -0.27081 | 0.437113 | | 1 |
| 1 | 169001400 | 169003199 | 24.52055 | 40.63619 | 65.10935 | 28.18707 | -0.29717 | 1.29867 | | 1 |
| 1 | 169110800 | 169112999 | 19.44674 | 33.8702 | 35.88413 | 15.77932 | 0.304416 | 0.7584 | | 1 |
| 1 | 169369200 | 169371399 | 30.43229 | 50.25528 | 59.4474 | 25.67315 | 0.138824 | 1.349941 | | 1 |
| 1 | 169841400 | 169854399 | 316.272 | 442.6987 | 582.7842 | 241.6144 | -0.22684 | 4.565067 | | 1 |
| 1 | 169944600 | 169944999 | 8.551055 | 16.05747 | 23.93843 | 9.846316 | -0.10539 | 0.082352 | | 1 |
| 1 | 170411600 | 170414599 | 27.32233 | 44.98386 | 71.32263 | 30.75651 | -0.28397 | 1.388151 | | 1 |
| 1 | 170766800 | 170779599 | 397.3254 | 540.8808 | 762.1951 | 317.3798 | -0.36559 | 4.994357 | | 1 |
| 1 | 170837000 | 170840799 | 43.49978 | 70.11636 | 78.37685 | 33.36063 | 0.196192 | 1.793704 | | 1 |
| 1 | 170911000 | 170911799 | 26.82224 | 43.75929 | 74.75255 | 32.11763 | -0.39788 | 1.511836 | | 1 |
| 1 | 171450800 | 171454799 | 69.36102 | 109.2669 | 130.2123 | 54.03018 | 0.070039 | 2.493039 | | 1 |
| 1 | 171635200 | 171636799 | 14.56294 | 25.68052 | 35.69729 | 15.79779 | -0.08327 | 0.555828 | | 1 |
| 1 | 171730000 | 171732399 | 31.46079 | 51.75515 | 60.45019 | 26.05897 | 0.155238 | 1.400451 | | 1 |

| | | | | | | | | | |
|---|---|---|---|---|---|---|---|---|---|
| 1 | 172504600 | 172505399 | 10.39877 | 19.04024 | 32.92182 | 14.06174 | -0.35978 | 0.270582 | 1 |
| 1 | 173243600 | 173246199 | 20.98769 | 36.4427 | 38.87735 | 17.06278 | 0.293944 | 0.81155 | 1 |
| 1 | 173389000 | 173391199 | 38.59731 | 62.27104 | 87.37197 | 37.23319 | -0.12947 | 1.766002 | 1 |
| 1 | 173491400 | 173494199 | 30.98947 | 51.37473 | 58.53651 | 25.34771 | 0.193088 | 1.304784 | 1 |
| 1 | 173548600 | 173549799 | 14.88239 | 26.08478 | 36.91271 | 16.30693 | -0.11067 | 0.641375 | 1 |
| 1 | 173639000 | 173641399 | 22.82598 | 39.27231 | 48.47152 | 21.26202 | 0.08188 | 0.91692 | 1 |
| 1 | 173660400 | 173661599 | 16.76698 | 28.82575 | 46.92651 | 20.58382 | -0.31581 | 0.898599 | 1 |
| 1 | 174109000 | 174112799 | 31.38875 | 52.01594 | 54.14211 | 23.43839 | 0.323116 | 1.306088 | 1 |
| 1 | 174222000 | 174222799 | 12.50112 | 22.78828 | 26.66352 | 11.45366 | 0.194543 | 0.297039 | 1 |
| 1 | 174663600 | 174668799 | 23.36366 | 38.87255 | 59.59795 | 25.8717 | -0.23323 | 1.245587 | 1 |
| 1 | 174769400 | 174771199 | 21.82417 | 37.27346 | 41.31105 | 18.11213 | 0.234985 | 0.971693 | 1 |
| 1 | 175057200 | 175060399 | 48.09669 | 77.38497 | 86.39981 | 36.6797 | 0.194651 | 1.859899 | 1 |
| 1 | 175519400 | 175521599 | 18.85537 | 31.95916 | 54.82706 | 24.0109 | -0.3944 | 1.041649 | 1 |
| 1 | 176375400 | 176377799 | 9.098997 | 17.50082 | 18.21161 | 7.099495 | 0.477102 | -0.10983 | 1 |
| 1 | 176407200 | 176416199 | 176.3456 | 260.7845 | 334.1807 | 137.6243 | -0.10992 | 3.727374 | 1 |
| 1 | 177108400 | 177109399 | 11.46097 | 20.99058 | 29.65737 | 12.66294 | -0.07044 | 0.266071 | 1 |
| 1 | 177614200 | 177618599 | 23.14323 | 38.66334 | 56.44144 | 24.56521 | -0.16276 | 1.197245 | 1 |
| 1 | 177836800 | 177839999 | 40.63465 | 65.55769 | 82.69789 | 35.24115 | 0.023682 | 1.765714 | 1 |
| 1 | 177920200 | 177921199 | 12.65977 | 23.18609 | 30.07777 | 12.84249 | 0.051802 | 0.275317 | 1 |
| 1 | 177958200 | 177962599 | 19.31112 | 33.06339 | 49.00815 | 21.4911 | -0.18232 | 0.948116 | 1 |
| 1 | 178000400 | 178003599 | 36.20039 | 58.74855 | 81.04811 | 34.73503 | -0.09649 | 1.617076 | 1 |
| 1 | 178014800 | 178015599 | 9.446053 | 17.73601 | 25.86346 | 10.64027 | -0.07453 | 0.095134 | 1 |
| 1 | 178092400 | 178094999 | 28.56837 | 47.89869 | 57.92451 | 25.26799 | 0.107701 | 1.175501 | 1 |
| 1 | 178109400 | 178113599 | 37.88094 | 61.52167 | 73.40699 | 31.47753 | 0.113396 | 1.599751 | 1 |
| 1 | 178122000 | 178131199 | 108.0709 | 165.4373 | 209.4075 | 86.89907 | -0.0488 | 3.047342 | 1 |
| 1 | 179117600 | 179118999 | 16.92791 | 29.68063 | 35.23781 | 15.57172 | 0.141313 | 0.643839 | 1 |
| 1 | 179255600 | 179257799 | 27.06728 | 44.89673 | 63.73824 | 27.61049 | -0.12311 | 1.293857 | 1 |
| 1 | 179405800 | 179408199 | 17.20389 | 29.50323 | 50.05186 | 21.95165 | -0.37597 | 0.924446 | 1 |
| 1 | 179672600 | 179673799 | 8.822832 | 16.85147 | 19.30866 | 7.675958 | 0.313204 | -0.04605 | 1 |
| 1 | 179724800 | 179729199 | 60.46073 | 96.41949 | 107.5736 | 45.08138 | 0.180991 | 2.153917 | 1 |
| 1 | 180432400 | 180440999 | 86.39273 | 134.3134 | 148.9936 | 61.85901 | 0.158542 | 2.745146 | 1 |
| 1 | 181096600 | 181096999 | 8.629462 | 16.03366 | 27.94202 | 11.69934 | -0.34964 | 0.162325 | 1 |
| 1 | 181159600 | 181166799 | 66.53327 | 104.5995 | 153.9588 | 63.86666 | -0.23649 | 2.535931 | 1 |
| 1 | 181324600 | 181325799 | 16.86735 | 29.51776 | 37.28754 | 16.45081 | 0.052139 | 0.676285 | 1 |
| 1 | 181540400 | 181543999 | 58.4648 | 93.23069 | 95.59966 | 40.05996 | 0.302435 | 2.151017 | 1 |
| 1 | 182011200 | 182013399 | 18.94889 | 33.08044 | 38.27875 | 16.85709 | 0.177948 | 0.725573 | 1 |
| 1 | 182089600 | 182091799 | 15.0839 | 26.2045 | 40.75223 | 17.89124 | -0.247 | 0.781667 | 1 |
| 1 | 182482400 | 182485999 | 11.96231 | 22.26067 | 23.01991 | 9.614881 | 0.400007 | 0.165314 | 1 |
| 1 | 183344600 | 183347199 | 9.343185 | 17.36482 | 29.95428 | 12.53692 | -0.33532 | 0.168812 | 1 |
| 1 | 183886400 | 183888199 | 18.68811 | 31.94237 | 47.97205 | 21.03387 | -0.20147 | 0.960458 | 1 |
| 1 | 184027200 | 184028199 | 14.09941 | 24.69194 | 41.4925 | 18.31731 | -0.35752 | 0.656789 | 1 |
| 1 | 184229000 | 184231399 | 10.16182 | 18.88127 | 32.11243 | 13.44504 | -0.3155 | 0.182853 | 1 |
| 1 | 184685200 | 184696399 | 149.4097 | 223.4554 | 293.7039 | 121.113 | -0.13154 | 3.523645 | 1 |
| 1 | 185442000 | 185443599 | 18.97173 | 32.91918 | 45.7876 | 20.08991 | -0.08734 | 0.819815 | 1 |
| 1 | 185547200 | 185548799 | 16.02959 | 28.53878 | 29.82948 | 13.14017 | 0.332217 | 0.467964 | 1 |
| 1 | 186379600 | 186382199 | 23.87488 | 39.84803 | 64.66857 | 28.12919 | -0.31509 | 1.218017 | 1 |
| 1 | 186697400 | 186702199 | 123.3051 | 185.244 | 278.9991 | 115.1662 | -0.32202 | 3.433678 | 1 |

| | | | | | | | | | |
|---|---|---|---|---|---|---|---|---|---|
| 1 | 187509800 | 187511199 | 13.98077 | 24.79254 | 38.60522 | 17.06294 | -0.24334 | 0.493822 | 1 |
| 1 | 188972000 | 188973799 | 18.86142 | 32.9394 | 39.84253 | 17.54965 | 0.11439 | 0.723303 | 1 |
| 1 | 189183600 | 189191599 | 94.83326 | 145.832 | 199.3463 | 82.78293 | -0.15461 | 2.952334 | 1 |
| 1 | 190213000 | 190215999 | 47.40106 | 75.82469 | 101.1943 | 42.59184 | -0.07216 | 2.045053 | 1 |
| 1 | 190477200 | 190483399 | 72.71137 | 113.8751 | 159.6448 | 66.22148 | -0.17062 | 2.612394 | 1 |
| 1 | 191307000 | 191308399 | 10.8613 | 20.4344 | 21.62746 | 8.867288 | 0.38823 | 0.082972 | 1 |
| 1 | 191485800 | 191487999 | 24.34357 | 41.59814 | 44.74021 | 19.61642 | 0.278062 | 0.978912 | 1 |
| 1 | 191667200 | 191668399 | 16.23496 | 28.40427 | 37.34815 | 16.47422 | -0.00537 | 0.676911 | 1 |
| 1 | 191834000 | 191840999 | 59.46593 | 94.56115 | 117.1855 | 48.91618 | 0.02526 | 2.246265 | 1 |
| 1 | 191909200 | 191912599 | 42.84077 | 69.24417 | 74.95931 | 32.01928 | 0.24689 | 1.711699 | 1 |
| 1 | 192308800 | 192321399 | 183.0594 | 270.0512 | 338.6056 | 139.4429 | -0.08185 | 3.768186 | 1 |
| 1 | 192637200 | 192643799 | 82.30631 | 128.8442 | 151.5865 | 62.88175 | 0.081463 | 2.62307 | 1 |
| 1 | 193715000 | 193717599 | 19.53908 | 33.52127 | 51.15532 | 22.43632 | -0.22392 | 0.930657 | 1 |
| 1 | 194150000 | 194158799 | 124.3284 | 188.1329 | 255.6144 | 105.7352 | -0.16448 | 3.289232 | 1 |
| 1 | 194737600 | 194740399 | 40.47441 | 65.45564 | 76.27292 | 32.60369 | 0.141841 | 1.693804 | 1 |
| 1 | 194934400 | 194952999 | 415.0697 | 569.8458 | 688.9002 | 286.4795 | -0.13217 | 4.862261 | 1 |
| 1 | 195020600 | 195029799 | 171.8953 | 256.3925 | 290.9743 | 119.9245 | 0.076631 | 3.576497 | 1 |
| 1 | 195034400 | 195043799 | 194.9924 | 283.9254 | 395.9091 | 163.1665 | -0.25302 | 3.975545 | 1 |
| 2 | 3283600 | 3286199 | 26.18382 | 43.00236 | 63.18997 | 27.22082 | -0.17594 | 1.409863 | 1 |
| 2 | 3474600 | 3477199 | 14.30873 | 24.85324 | 44.79744 | 19.66489 | -0.45913 | 0.785172 | 1 |
| 2 | 3548000 | 3554599 | 17.09972 | 30.16508 | 33.45399 | 14.80393 | 0.241022 | 0.5626 | 1 |
| 2 | 3558000 | 3560999 | 35.18382 | 58.10179 | 63.70673 | 27.51258 | 0.247995 | 1.367385 | 1 |
| 2 | 3973400 | 3978199 | 52.7156 | 84.64917 | 93.92368 | 39.75893 | 0.200116 | 1.930209 | 1 |
| 2 | 3979600 | 3987199 | 84.10071 | 130.848 | 175.8865 | 73.01621 | -0.11744 | 2.734727 | 1 |
| 2 | 4553600 | 4556599 | 21.85914 | 36.58641 | 60.2167 | 26.23924 | -0.33533 | 1.176005 | 1 |
| 2 | 4938400 | 4951599 | 181.3614 | 268.7822 | 308.1594 | 126.9169 | 0.053432 | 3.691551 | 1 |
| 2 | 4951800 | 4958999 | 64.64316 | 101.9585 | 150.8732 | 62.61312 | -0.24057 | 2.469554 | 1 |
| 2 | 5028800 | 5031399 | 13.6917 | 25.08463 | 25.80071 | 11.01117 | 0.385039 | 0.272278 | 1 |
| 2 | 5147400 | 5149799 | 24.72456 | 41.768 | 49.06428 | 21.46548 | 0.149781 | 1.085811 | 1 |
| 2 | 5279600 | 5282399 | 35.95996 | 58.83995 | 69.11017 | 29.72523 | 0.143935 | 1.483752 | 1 |
| 2 | 5721200 | 5722999 | 15.80517 | 27.74377 | 39.24983 | 17.36141 | -0.11054 | 0.624677 | 1 |
| 2 | 6183400 | 6185399 | 14.07075 | 24.78958 | 37.73027 | 16.70058 | -0.21407 | 0.565668 | 1 |
| 2 | 6776000 | 6778399 | 27.65256 | 45.48093 | 67.82938 | 29.23744 | -0.19646 | 1.39972 | 1 |
| 2 | 6781400 | 6783799 | 26.14746 | 43.54672 | 58.82296 | 25.55159 | -0.05124 | 1.239731 | 1 |
| 2 | 6876600 | 6877799 | 13.16255 | 23.14093 | 38.35721 | 16.98521 | -0.33755 | 0.585278 | 1 |
| 2 | 7080400 | 7083399 | 38.69875 | 62.62807 | 91.77705 | 39.25232 | -0.18719 | 1.682769 | 1 |
| 2 | 7419600 | 7420599 | 12.03108 | 21.81826 | 30.63949 | 13.24704 | -0.07341 | 0.330748 | 1 |
| 2 | 7582200 | 7583199 | 13.81538 | 24.28492 | 40.35902 | 17.8722 | -0.34148 | 0.594593 | 1 |
| 2 | 7605600 | 7606399 | 16.28264 | 28.45062 | 42.00093 | 18.50641 | -0.1717 | 0.705354 | 1 |
| 2 | 8025000 | 8026199 | 13.57628 | 24.71435 | 26.35284 | 11.33959 | 0.325572 | 0.309168 | 1 |
| 2 | 8170000 | 8170799 | 8.369218 | 15.98397 | 25.62321 | 10.18773 | -0.16746 | -0.03001 | 1 |
| 2 | 8380600 | 8384199 | 74.84346 | 118.0463 | 125.8236 | 52.21734 | 0.232106 | 2.471242 | 1 |
| 2 | 8539400 | 8541199 | 28.46683 | 47.58693 | 50.73662 | 22.09105 | 0.289034 | 1.202325 | 1 |
| 2 | 9028800 | 9030599 | 18.51063 | 32.08885 | 43.69717 | 19.17215 | -0.05709 | 0.824151 | 1 |
| 2 | 9217400 | 9217999 | 16.1246 | 28.09547 | 43.1822 | 18.99335 | -0.22969 | 0.75008 | 1 |
| 2 | 9519000 | 9520999 | 14.98679 | 26.45757 | 33.61694 | 14.87381 | 0.046276 | 0.541627 | 1 |
| 2 | 10892200 | 10894799 | 32.96965 | 53.45562 | 76.72354 | 32.85904 | -0.15462 | 1.628069 | 1 |

| | | | | | | | | | |
|---|---|---|---|---|---|---|---|---|---|
| 2 | 11142400 | 11144199 | 29.95254 | 49.53283 | 57.76509 | 24.97092 | 0.159464 | 1.331675 | 1 |
| 2 | 11352400 | 11364999 | 203.1455 | 296.0971 | 390.8826 | 161.0779 | -0.17265 | 3.960725 | 1 |
| 2 | 11583600 | 11586799 | 57.14955 | 91.24125 | 103.0252 | 43.24324 | 0.16546 | 2.114387 | 1 |
| 2 | 11814800 | 11816399 | 13.84534 | 24.56516 | 37.5913 | 16.60752 | -0.21783 | 0.488744 | 1 |
| 2 | 11947800 | 11949799 | 35.75642 | 58.30576 | 67.10686 | 28.82737 | 0.169659 | 1.530366 | 1 |
| 2 | 11981600 | 11984999 | 48.06998 | 77.47235 | 80.01548 | 34.05208 | 0.30982 | 1.807549 | 1 |
| 2 | 12047800 | 12049599 | 26.50946 | 44.65091 | 45.92655 | 20.0747 | 0.340627 | 1.114767 | 1 |
| 2 | 12050400 | 12052999 | 32.05518 | 52.28937 | 70.05372 | 30.09725 | -0.0485 | 1.51943 | 1 |
| 2 | 12165600 | 12167199 | 13.63095 | 24.31989 | 33.00685 | 14.51381 | -0.04083 | 0.44412 | 1 |
| 2 | 12949400 | 12951199 | 40.34734 | 64.99408 | 86.11827 | 36.62337 | -0.04969 | 1.813018 | 1 |
| 2 | 13250600 | 13257599 | 83.8893 | 130.1024 | 172.7617 | 71.75326 | -0.10341 | 2.788675 | 1 |
| 2 | 13835000 | 13836799 | 26.97524 | 45.13412 | 54.48118 | 23.73493 | 0.110471 | 1.190535 | 1 |
| 2 | 13868400 | 13871199 | 41.13674 | 66.11026 | 90.21119 | 38.23359 | -0.09624 | 1.888439 | 1 |
| 2 | 13914000 | 13920799 | 89.36978 | 140.2558 | 142.1175 | 58.94111 | 0.299527 | 2.581946 | 1 |
| 2 | 14038200 | 14040199 | 20.76001 | 35.94739 | 44.17412 | 19.38075 | 0.090044 | 0.844886 | 1 |
| 2 | 14089800 | 14092999 | 72.80933 | 114.3911 | 146.237 | 60.65824 | -0.03414 | 2.552335 | 1 |
| 2 | 14129600 | 14131199 | 12.23617 | 21.53217 | 37.4128 | 16.56395 | -0.4045 | 0.568345 | 1 |
| 2 | 14293200 | 14294199 | 18.74448 | 32.52034 | 42.67035 | 18.7221 | -0.0034 | 0.818364 | 1 |
| 2 | 14300200 | 14301399 | 13.12169 | 23.36621 | 34.65891 | 15.26504 | -0.16967 | 0.458524 | 1 |
| 2 | 14354400 | 14357199 | 29.6056 | 49.06587 | 61.17727 | 26.48576 | 0.063475 | 1.307539 | 1 |
| 2 | 14369800 | 14370999 | 11.5469 | 21.50599 | 26.74503 | 11.15463 | 0.137675 | 0.167831 | 1 |
| 2 | 14445800 | 14447999 | 21.62665 | 36.66243 | 53.26372 | 23.32742 | -0.15542 | 1.051209 | 1 |
| 2 | 14494200 | 14495199 | 10.55537 | 19.4404 | 30.30886 | 12.84303 | -0.20355 | 0.229505 | 1 |
| 2 | 14730400 | 14732799 | 38.72913 | 62.41192 | 85.29483 | 36.29387 | -0.0935 | 1.798222 | 1 |
| 2 | 15156600 | 15157199 | 8.884772 | 17.18288 | 19.84926 | 7.588608 | 0.354261 | -0.16992 | 1 |
| 2 | 15249200 | 15252399 | 28.78128 | 48.399 | 54.55888 | 23.83622 | 0.208772 | 1.145311 | 1 |
| 2 | 15338600 | 15340399 | 16.42917 | 28.67146 | 41.51318 | 18.2774 | -0.14399 | 0.722575 | 1 |
| 2 | 15444200 | 15444999 | 10.55488 | 19.65882 | 26.16816 | 10.9136 | 0.039703 | 0.153826 | 1 |
| 2 | 15525600 | 15529599 | 82.49924 | 128.3111 | 164.6582 | 68.35857 | -0.05096 | 2.739628 | 1 |
| 2 | 16021600 | 16023199 | 20.61273 | 36.02773 | 36.99916 | 16.3066 | 0.349324 | 0.713569 | 1 |
| 2 | 16031800 | 16034999 | 49.22994 | 78.5023 | 110.7281 | 46.3977 | -0.15683 | 2.147111 | 1 |
| 2 | 16113200 | 16114199 | 14.02465 | 24.58096 | 39.01711 | 17.23634 | -0.2753 | 0.642826 | 1 |
| 2 | 16202800 | 16203399 | 9.432887 | 18.12621 | 19.71336 | 7.705395 | 0.410315 | -0.09299 | 1 |
| 2 | 16264600 | 16266799 | 42.66099 | 68.81097 | 73.19873 | 31.18386 | 0.268665 | 1.770396 | 1 |
| 2 | 16321800 | 16324799 | 23.4431 | 40.08789 | 57.83376 | 25.35876 | -0.14395 | 0.984911 | 1 |
| 2 | 16386000 | 16387799 | 45.84409 | 73.38512 | 94.87547 | 39.97165 | -0.02531 | 2.018469 | 1 |
| 2 | 16584400 | 16586399 | 33.73328 | 55.4109 | 57.244 | 24.66128 | 0.330988 | 1.422268 | 1 |
| 2 | 16982400 | 16984799 | 18.36383 | 31.87416 | 46.32893 | 20.32817 | -0.15045 | 0.815283 | 1 |
| 2 | 17099600 | 17100799 | 18.91797 | 32.8696 | 42.05548 | 18.46494 | 0.032891 | 0.796877 | 1 |
| 2 | 17101800 | 17105199 | 93.79746 | 143.9021 | 203.8679 | 84.63345 | -0.2087 | 2.999263 | 1 |
| 2 | 17134800 | 17135199 | 8.140722 | 15.65753 | 21.42532 | 8.352544 | 0.085591 | -0.10916 | 1 |
| 2 | 17157000 | 17159399 | 29.12139 | 47.87134 | 73.17476 | 31.53446 | -0.23228 | 1.413093 | 1 |
| 2 | 17491000 | 17491799 | 12.24996 | 22.43795 | 26.5773 | 11.34622 | 0.182708 | 0.265968 | 1 |
| 2 | 17617200 | 17619199 | 8.428019 | 16.32481 | 19.36345 | 7.349465 | 0.326418 | -0.20067 | 1 |
| 2 | 17658400 | 17661199 | 25.03895 | 41.27931 | 67.19524 | 28.99296 | -0.32101 | 1.362319 | 1 |
| 2 | 17715800 | 17716599 | 10.51839 | 19.19593 | 30.80915 | 13.21553 | -0.25721 | 0.284917 | 1 |
| 2 | 17958000 | 17963999 | 101.4742 | 156.7101 | 176.017 | 73.11382 | 0.133159 | 2.872257 | 1 |

| | | | | | | | | | | |
|---|---|---|---|---|---|---|---|---|---|---|
| 2 | 18020400 | 18024599 | 45.87472 | 73.98556 | 78.00442 | 33.22591 | 0.281261 | 1.779923 | | 1 |
| 2 | 18255400 | 18259199 | 71.29023 | 113.0389 | 121.2143 | 50.45271 | 0.229673 | 2.341925 | | 1 |
| 2 | 18342600 | 18344599 | 17.32366 | 29.81626 | 48.47185 | 21.26239 | -0.31373 | 0.889948 | | 1 |
| 2 | 18359400 | 18361199 | 30.47021 | 50.81629 | 57.36688 | 24.94088 | 0.206834 | 1.238512 | | 1 |
| 2 | 18369400 | 18372999 | 23.96017 | 41.21609 | 46.28138 | 20.3012 | 0.217605 | 0.918557 | | 1 |
| 2 | 18476000 | 18482399 | 93.55318 | 144.6451 | 174.2423 | 72.3762 | 0.03359 | 2.848607 | | 1 |
| 2 | 19235600 | 19237999 | 37.80596 | 61.42837 | 71.4983 | 30.66955 | 0.149793 | 1.591345 | | 1 |
| 2 | 19318200 | 19325999 | 100.9249 | 154.5667 | 224.6582 | 93.23731 | -0.24759 | 3.038343 | | 1 |
| 2 | 19360600 | 19369199 | 118.167 | 178.5245 | 262.5788 | 108.5614 | -0.28076 | 3.320577 | | 1 |
| 2 | 20177800 | 20186199 | 115.0656 | 177.0401 | 195.1532 | 81.04762 | 0.156128 | 2.950384 | | 1 |
| 2 | 20387400 | 20388599 | 60.80755 | 96.29245 | 123.8771 | 51.51059 | -0.03425 | 2.368952 | | 1 |
| 2 | 20389400 | 20394999 | 298.238 | 416.4126 | 589.464 | 244.4789 | -0.33519 | 4.601539 | | 1 |
| 2 | 20848600 | 20852599 | 39.0245 | 63.24563 | 83.47239 | 35.73815 | -0.03464 | 1.648087 | | 1 |
| 2 | 20904200 | 20905199 | 27.21679 | 44.67056 | 68.10564 | 29.33052 | -0.22932 | 1.425994 | | 1 |
| 2 | 21108200 | 21110799 | 26.28788 | 43.45293 | 66.92513 | 28.92165 | -0.2407 | 1.333889 | | 1 |
| 2 | 21172200 | 21173999 | 21.41445 | 37.06911 | 43.10439 | 18.91124 | 0.169339 | 0.849465 | | 1 |
| 2 | 21304600 | 21305799 | 18.94197 | 32.54351 | 46.38372 | 20.34532 | -0.12519 | 0.911583 | | 1 |
| 2 | 21328000 | 21330399 | 35.35303 | 57.51902 | 72.47465 | 31.10783 | 0.037117 | 1.565733 | | 1 |
| 2 | 21368400 | 21369599 | 13.48089 | 23.7174 | 37.83257 | 16.75051 | -0.28204 | 0.578834 | | 1 |
| 2 | 21497200 | 21498599 | 15.70997 | 27.44946 | 43.60066 | 19.21104 | -0.27675 | 0.706274 | | 1 |
| 2 | 21594600 | 21595599 | 12.29327 | 21.99335 | 34.68193 | 15.21524 | -0.25362 | 0.422876 | | 1 |
| 2 | 21606800 | 21609799 | 46.29009 | 74.49374 | 84.89204 | 36.05047 | 0.165552 | 1.850238 | | 1 |
| 2 | 21662000 | 21662999 | 12.19223 | 22.1284 | 32.68693 | 14.11796 | -0.14512 | 0.331111 | | 1 |
| 2 | 21837400 | 21843799 | 132.6545 | 200.9443 | 223.5094 | 92.48577 | 0.125288 | 3.268301 | | 1 |
| 2 | 21998400 | 22000599 | 32.48347 | 53.25816 | 62.57594 | 26.937 | 0.144791 | 1.446235 | | 1 |
| 2 | 22006800 | 22007399 | 9.291181 | 17.42384 | 24.35287 | 10.03964 | -0.01597 | 0.098061 | | 1 |
| 2 | 22060000 | 22061599 | 31.91487 | 52.76889 | 58.45444 | 25.26592 | 0.233395 | 1.339379 | | 1 |
| 2 | 22079000 | 22079799 | 11.62001 | 21.63111 | 23.52211 | 9.818924 | 0.328529 | 0.160791 | | 1 |
| 2 | 22168800 | 22171799 | 28.29335 | 47.00575 | 56.11701 | 24.33601 | 0.126278 | 1.272495 | | 1 |
| 2 | 22335400 | 22336399 | 17.07332 | 30.13909 | 31.29361 | 13.84604 | 0.33577 | 0.548722 | | 1 |
| 2 | 23210800 | 23212599 | 13.7615 | 24.2719 | 37.73289 | 16.69801 | -0.2438 | 0.551703 | | 1 |
| 2 | 23286000 | 23287799 | 23.58257 | 40.09272 | 49.02645 | 21.4869 | 0.092485 | 1.026824 | | 1 |
| 2 | 23510200 | 23512999 | 31.29027 | 51.5534 | 62.7718 | 27.07729 | 0.096246 | 1.382524 | | 1 |
| 2 | 23590800 | 23595199 | 58.05234 | 92.77948 | 101.7611 | 42.77589 | 0.20897 | 2.084294 | | 1 |
| 2 | 23765200 | 23766599 | 17.21512 | 29.99337 | 41.53338 | 18.26727 | -0.08021 | 0.749899 | | 1 |
| 2 | 23828000 | 23829599 | 18.83183 | 33.00955 | 34.55817 | 15.26775 | 0.321822 | 0.657148 | | 1 |
| 2 | 23921200 | 23929199 | 165.9526 | 248.2814 | 271.6196 | 112.0136 | 0.133498 | 3.519115 | | 1 |
| 2 | 24141600 | 24144399 | 39.653 | 64.39101 | 68.97826 | 29.57553 | 0.268383 | 1.60357 | | 1 |
| 2 | 25636800 | 25641199 | 46.68142 | 74.81674 | 100.6233 | 42.47152 | -0.08014 | 1.983698 | | 1 |
| 2 | 26605200 | 26607199 | 9.754614 | 18.56638 | 23.37226 | 9.360098 | 0.169464 | 0.001215 | | 1 |
| 2 | 26857000 | 26858199 | 13.65328 | 24.64272 | 31.4096 | 13.66095 | 0.059657 | 0.368893 | | 1 |
| 2 | 27945800 | 27946999 | 16.83819 | 29.54826 | 38.1248 | 16.85952 | 0.021571 | 0.632011 | | 1 |
| 2 | 28380600 | 28387199 | 59.17226 | 93.70249 | 140.4046 | 58.38293 | -0.25399 | 2.370462 | | 1 |
| 2 | 29224000 | 29225199 | 18.10189 | 31.71411 | 36.20749 | 15.9896 | 0.197491 | 0.664955 | | 1 |
| 2 | 30054600 | 30059999 | 61.78063 | 97.6426 | 145.3738 | 60.38967 | -0.24703 | 2.418406 | | 1 |
| 2 | 30116000 | 30126599 | 251.5539 | 358.4971 | 481.2742 | 198.9453 | -0.22898 | 4.297347 | | 1 |
| 2 | 33266400 | 33270399 | 43.45083 | 69.94758 | 97.6097 | 41.47095 | -0.12552 | 1.841784 | | 1 |

| | | | | | | | | | |
|---|---|---|---|---|---|---|---|---|---|
| 2 | 33433600 | 33436199 | 16.74409 | 28.66725 | 48.48418 | 21.26126 | -0.37184 | 0.935943 | 1 |
| 2 | 34930200 | 34932799 | 30.35821 | 50.06077 | 62.09444 | 26.79484 | 0.070117 | 1.370237 | 1 |
| 2 | 36188800 | 36190799 | 33.01396 | 53.57716 | 77.27844 | 33.11935 | -0.16045 | 1.611453 | 1 |
| 2 | 36243600 | 36251999 | 218.8699 | 320.4997 | 321.741 | 132.534 | 0.228673 | 3.890056 | 1 |
| 2 | 36337000 | 36339399 | 35.24611 | 57.78562 | 62.86958 | 27.063 | 0.255394 | 1.453009 | 1 |
| 2 | 36348400 | 36351199 | 35.41078 | 57.76834 | 67.71714 | 29.09434 | 0.143753 | 1.523072 | 1 |
| 2 | 36438000 | 36440199 | 23.65801 | 40.72567 | 43.13134 | 18.91986 | 0.302082 | 0.910304 | 1 |
| 2 | 36442600 | 36446999 | 28.95351 | 48.26967 | 55.08343 | 23.94276 | 0.191332 | 1.235577 | 1 |
| 2 | 36532000 | 36533799 | 19.00888 | 33.31402 | 35.06002 | 15.48715 | 0.31431 | 0.661843 | 1 |
| 2 | 36654600 | 36662799 | 114.4564 | 176.479 | 179.6577 | 74.6267 | 0.273142 | 2.907764 | 1 |
| 2 | 36783200 | 36794199 | 78.71998 | 123.9032 | 137.2871 | 56.9594 | 0.173952 | 2.516896 | 1 |
| 2 | 37135000 | 37142999 | 113.3435 | 175.1617 | 167.883 | 69.73486 | 0.362549 | 2.86437 | 1 |
| 2 | 37205400 | 37207999 | 16.77775 | 29.347 | 40.39784 | 17.81588 | -0.07123 | 0.689056 | 1 |
| 2 | 37231000 | 37231799 | 18.2299 | 31.6418 | 39.14228 | 17.17485 | 0.081455 | 0.806986 | 1 |
| 2 | 38346200 | 38348199 | 19.79878 | 34.19307 | 40.13688 | 17.60806 | 0.155237 | 0.859744 | 1 |
| 2 | 38873600 | 38875999 | 38.98041 | 63.39585 | 68.80563 | 29.52878 | 0.251493 | 1.577475 | 1 |
| 2 | 39136400 | 39137199 | 12.20489 | 21.55765 | 36.81048 | 16.28509 | -0.37746 | 0.524855 | 1 |
| 2 | 39498000 | 39498599 | 9.715894 | 18.45025 | 23.85835 | 9.601301 | 0.124652 | 0.016864 | 1 |
| 2 | 39531400 | 39538199 | 89.31739 | 139.5355 | 159.0093 | 65.97471 | 0.125178 | 2.6658 | 1 |
| 2 | 39581400 | 39583999 | 31.48414 | 51.8721 | 65.03763 | 28.05451 | 0.054086 | 1.385107 | 1 |
| 2 | 39612400 | 39620199 | 98.61475 | 152.6123 | 167.3608 | 69.51746 | 0.169959 | 2.83317 | 1 |
| 2 | 39663000 | 39665799 | 54.90787 | 87.66844 | 104.7165 | 43.95748 | 0.084597 | 2.111051 | 1 |
| 2 | 39674400 | 39676999 | 26.49873 | 45.07013 | 47.40311 | 20.77617 | 0.309177 | 1.031935 | 1 |
| 2 | 39879000 | 39881799 | 29.71246 | 49.97182 | 54.61212 | 23.86048 | 0.253331 | 1.146352 | 1 |
| 2 | 39905400 | 39911599 | 137.0311 | 207.4943 | 218.0602 | 90.22002 | 0.206636 | 3.276293 | 1 |
| 2 | 39968000 | 39968799 | 11.58625 | 21.89503 | 23.67468 | 9.62611 | 0.367641 | 0.066706 | 1 |
| 2 | 40125000 | 40129599 | 53.15706 | 85.22208 | 89.66946 | 37.86615 | 0.273901 | 1.980315 | 1 |
| 2 | 40252200 | 40253599 | 21.80774 | 37.72772 | 40.97465 | 17.97653 | 0.267231 | 0.852738 | 1 |
| 2 | 40264000 | 40264999 | 19.79084 | 33.7803 | 50.59537 | 22.18161 | -0.19804 | 0.979417 | 1 |
| 2 | 40471000 | 40472199 | 15.76345 | 28.04501 | 28.79544 | 12.69227 | 0.357075 | 0.467228 | 1 |
| 2 | 40473200 | 40477599 | 91.4861 | 142.2415 | 154.5732 | 64.17488 | 0.188276 | 2.74697 | 1 |
| 2 | 41461400 | 41462599 | 11.56982 | 21.34882 | 27.62796 | 11.67714 | 0.066008 | 0.220589 | 1 |
| 2 | 41497200 | 41499799 | 26.49836 | 44.91668 | 48.50614 | 21.24314 | 0.270852 | 1.062376 | 1 |
| 2 | 41665000 | 41666399 | 17.34676 | 29.92442 | 45.34448 | 19.89195 | -0.21187 | 0.867105 | 1 |
| 2 | 41973200 | 41974399 | 27.21139 | 44.57856 | 67.47807 | 29.03736 | -0.22005 | 1.447856 | 1 |
| 2 | 42090000 | 42093999 | 67.87339 | 107.511 | 113.8698 | 47.35589 | 0.246147 | 2.364325 | 1 |
| 2 | 42448000 | 42450599 | 24.12985 | 41.10805 | 49.02482 | 21.48977 | 0.128844 | 1.00854 | 1 |
| 2 | 43196600 | 43197999 | 22.38269 | 38.53989 | 40.05693 | 17.57138 | 0.329123 | 0.89766 | 1 |
| 2 | 43237600 | 43239599 | 23.66195 | 41.08036 | 41.37016 | 18.15521 | 0.376713 | 0.8294 | 1 |
| 2 | 44667600 | 44673599 | 13.52063 | 24.76699 | 26.44566 | 11.28904 | 0.331434 | 0.275222 | 1 |
| 2 | 44770600 | 44772599 | 25.46233 | 43.61986 | 46.84562 | 20.54372 | 0.28075 | 0.964437 | 1 |
| 2 | 44878400 | 44880199 | 24.71005 | 41.23582 | 58.19149 | 25.30945 | -0.11413 | 1.213871 | 1 |
| 2 | 45828600 | 45830599 | 24.94836 | 41.61444 | 58.67553 | 25.51259 | -0.11283 | 1.221843 | 1 |
| 2 | 46186600 | 46187199 | 33.91288 | 55.04375 | 74.80176 | 32.06125 | -0.07456 | 1.606892 | 1 |
| 2 | 46586200 | 46589399 | 46.0897 | 74.39849 | 78.80726 | 33.60645 | 0.276011 | 1.755658 | 1 |
| 2 | 47531600 | 47532999 | 29.87955 | 48.70053 | 70.00265 | 30.06634 | -0.15049 | 1.529264 | 1 |
| 2 | 48117400 | 48118999 | 10.26164 | 19.40245 | 22.19422 | 9.013752 | 0.288515 | 0.046302 | 1 |

| | | | | | | | | | |
|---|---|---|---|---|---|---|---|---|---|
| 2 | 48374600 | 48378399 | 17.97468 | 31.56553 | 33.82646 | 14.96789 | 0.288006 | 0.61837 | 1 |
| 2 | 48742000 | 48746999 | 74.16767 | 117.4697 | 128.8657 | 53.58973 | 0.195431 | 2.375907 | 1 |
| 2 | 49054200 | 49056599 | 25.83723 | 43.81822 | 52.31679 | 22.91552 | 0.126519 | 1.058805 | 1 |
| 2 | 50489200 | 50491199 | 31.0117 | 51.05526 | 58.72844 | 25.32492 | 0.177696 | 1.38986 | 1 |
| 2 | 50563400 | 50565599 | 17.77311 | 30.71009 | 48.47061 | 21.26439 | -0.27013 | 0.856327 | 1 |
| 2 | 50576200 | 50578399 | 20.12275 | 34.95098 | 39.48036 | 17.33059 | 0.211939 | 0.805711 | 1 |
| 2 | 50732600 | 50734799 | 13.62143 | 24.14316 | 36.65318 | 16.20428 | -0.20693 | 0.495919 | 1 |
| 2 | 51511200 | 51511599 | 8.496855 | 16.11617 | 26.02189 | 10.49832 | -0.19657 | 0.01828 | 1 |
| 2 | 51557200 | 51557999 | 9.311746 | 17.4004 | 27.38953 | 11.35975 | -0.19349 | 0.12806 | 1 |
| 2 | 52228200 | 52230399 | 19.96058 | 34.21417 | 46.89129 | 20.56469 | -0.06961 | 0.934899 | 1 |
| 2 | 52757000 | 52758799 | 27.30491 | 45.52924 | 53.39094 | 23.20967 | 0.15218 | 1.227241 | 1 |
| 2 | 53255200 | 53258199 | 32.35195 | 53.23511 | 63.84189 | 27.52412 | 0.117414 | 1.402905 | 1 |
| 2 | 53598000 | 53598799 | 12.96324 | 23.69762 | 24.03 | 10.28595 | 0.402712 | 0.276216 | 1 |
| 2 | 53734600 | 53735599 | 15.81742 | 27.72676 | 36.82289 | 16.2688 | -0.01962 | 0.644154 | 1 |
| 2 | 53923200 | 53925799 | 29.88502 | 50.46897 | 50.184 | 21.95314 | 0.389295 | 1.106448 | 1 |
| 2 | 54148400 | 54148999 | 9.659879 | 18.23093 | 23.76784 | 9.69065 | 0.096216 | 0.057144 | 1 |
| 2 | 54187000 | 54187599 | 11.18879 | 21.11878 | 22.7124 | 9.258135 | 0.372599 | 0.070315 | 1 |
| 2 | 54274200 | 54276199 | 29.51494 | 48.74549 | 57.65568 | 24.90118 | 0.139045 | 1.346471 | 1 |
| 2 | 54489600 | 54494799 | 59.8121 | 95.17926 | 110.0393 | 45.97709 | 0.126377 | 2.222917 | 1 |
| 2 | 54607800 | 54608399 | 12.77471 | 22.96327 | 26.72537 | 11.67786 | 0.186146 | 0.378969 | 1 |
| 2 | 54630600 | 54632199 | 18.08454 | 31.71259 | 31.14381 | 13.76402 | 0.413615 | 0.640087 | 1 |
| 2 | 54775600 | 54776399 | 16.79062 | 29.06774 | 44.27325 | 19.42411 | -0.21828 | 0.829461 | 1 |
| 2 | 54916400 | 54917199 | 11.77663 | 21.23663 | 30.70361 | 13.36878 | -0.12156 | 0.362795 | 1 |
| 2 | 55182000 | 55184399 | 27.89139 | 46.72172 | 51.81234 | 22.58894 | 0.232357 | 1.177922 | 1 |
| 2 | 55213000 | 55214599 | 8.706171 | 16.2144 | 27.89222 | 11.63402 | -0.32735 | 0.144459 | 1 |
| 2 | 55236600 | 55238399 | 34.82746 | 56.26168 | 83.58305 | 35.67805 | -0.20954 | 1.720128 | 1 |
| 2 | 55627000 | 55627799 | 9.023212 | 17.23512 | 23.4035 | 9.302763 | 0.069968 | -0.03279 | 1 |
| 2 | 55667600 | 55682199 | 433.086 | 591.5927 | 710.941 | 295.88 | -0.13091 | 4.940882 | 1 |
| 2 | 55790800 | 55791999 | 16.18473 | 28.56189 | 31.49117 | 13.93429 | 0.249571 | 0.548094 | 1 |
| 2 | 55829000 | 55831999 | 33.46353 | 54.98787 | 64.50375 | 27.79315 | 0.148366 | 1.422255 | 1 |
| 2 | 55953200 | 55955799 | 51.24301 | 82.0987 | 92.92016 | 39.19912 | 0.167701 | 1.997971 | 1 |
| 2 | 55956600 | 55959199 | 49.50229 | 79.36055 | 85.25171 | 35.99912 | 0.243967 | 1.972226 | 1 |
| 2 | 56360800 | 56361999 | 36.55003 | 59.07702 | 77.25506 | 33.0004 | -0.02503 | 1.702746 | 1 |
| 2 | 56510800 | 56513599 | 13.70968 | 24.33636 | 32.61323 | 14.40231 | -0.02669 | 0.477438 | 1 |
| 2 | 56689400 | 56691399 | 20.94943 | 35.62188 | 45.9397 | 20.13433 | 0.01629 | 1.014976 | 1 |
| 2 | 56735400 | 56737999 | 27.72723 | 46.13087 | 61.03684 | 26.49561 | -0.02146 | 1.259656 | 1 |
| 2 | 56742400 | 56745999 | 91.5543 | 141.6035 | 163.3084 | 67.83439 | 0.096745 | 2.840018 | 1 |
| 2 | 56836400 | 56837199 | 11.65294 | 20.89787 | 31.26076 | 13.68661 | -0.17567 | 0.397105 | 1 |
| 2 | 57105400 | 57106999 | 21.97453 | 36.97107 | 55.38792 | 24.20207 | -0.20004 | 1.119775 | 1 |
| 2 | 57160600 | 57161799 | 15.122 | 27.05375 | 29.78349 | 13.06639 | 0.261609 | 0.43016 | 1 |
| 2 | 57570200 | 57575199 | 99.69255 | 153.4623 | 201.9951 | 83.89474 | -0.099 | 2.935401 | 1 |
| 2 | 57651200 | 57653799 | 21.95609 | 37.62953 | 46.87855 | 20.55888 | 0.067522 | 0.94567 | 1 |
| 2 | 58227800 | 58235199 | 97.12378 | 150.3208 | 164.8578 | 68.47773 | 0.16999 | 2.830287 | 1 |
| 2 | 58562800 | 58568999 | 111.3341 | 170.064 | 199.7952 | 82.8687 | 0.05624 | 3.091411 | 1 |
| 2 | 58581400 | 58582999 | 19.32739 | 33.03675 | 49.01039 | 21.48923 | -0.18388 | 0.963532 | 1 |
| 2 | 58611800 | 58613199 | 23.42026 | 39.21818 | 52.21811 | 22.76079 | -0.03047 | 1.168877 | 1 |
| 2 | 58930800 | 58934199 | 24.64852 | 40.90522 | 62.40945 | 27.04424 | -0.2266 | 1.280396 | 1 |

| | | | | | | | | | |
|---|---|---|---|---|---|---|---|---|---|
| 2 | 59148800 | 59153199 | 60.78707 | 97.07154 | 105.4627 | 44.28111 | 0.221379 | 2.113789 | 1 |
| 2 | 59413600 | 59418399 | 39.05422 | 62.87831 | 97.79424 | 41.55742 | -0.28136 | 1.83136 | 1 |
| 2 | 59558800 | 59560599 | 15.81986 | 27.62929 | 42.5533 | 18.74438 | -0.23257 | 0.710465 | 1 |
| 2 | 59893800 | 59896599 | 33.95893 | 55.35855 | 77.44503 | 33.26485 | -0.11153 | 1.535261 | 1 |
| 2 | 59940000 | 59943199 | 35.09032 | 57.14849 | 69.26716 | 29.7419 | 0.09394 | 1.547251 | 1 |
| 2 | 60056000 | 60058399 | 16.82006 | 29.27927 | 47.89699 | 21.05447 | -0.31977 | 0.769846 | 1 |
| 2 | 60087800 | 60089799 | 27.95681 | 46.61292 | 58.3712 | 25.37351 | 0.057688 | 1.233448 | 1 |
| 2 | 60393400 | 60405599 | 264.3931 | 378.7017 | 448.6435 | 185.3047 | -0.04136 | 4.224272 | 1 |
| 2 | 60598400 | 60603199 | 62.24937 | 98.50284 | 134.6407 | 55.96536 | -0.12247 | 2.388506 | 1 |
| 2 | 60895000 | 60896599 | 15.37331 | 27.11012 | 40.74864 | 18.03319 | -0.19572 | 0.568033 | 1 |
| 2 | 61166000 | 61168199 | 18.2713 | 31.57917 | 45.30723 | 19.87676 | -0.13289 | 0.853342 | 1 |
| 2 | 61237600 | 61238599 | 11.83649 | 22.06158 | 24.17317 | 10.06943 | 0.320655 | 0.159582 | 1 |
| 2 | 61644600 | 61646799 | 24.40396 | 40.57407 | 55.94182 | 24.27244 | -0.08072 | 1.251722 | 1 |
| 2 | 61681600 | 61684199 | 26.41513 | 43.37016 | 68.01319 | 29.2951 | -0.26964 | 1.419875 | 1 |
| 2 | 62004600 | 62007199 | 31.37481 | 51.22912 | 70.71041 | 30.39012 | -0.09044 | 1.503564 | 1 |
| 2 | 62030000 | 62031599 | 24.57082 | 41.12174 | 54.37401 | 23.69211 | -0.02052 | 1.180779 | 1 |
| 2 | 62656000 | 62657599 | 16.03763 | 27.94177 | 41.12006 | 18.08545 | -0.16738 | 0.745886 | 1 |
| 2 | 63314800 | 63315999 | 12.47629 | 22.26165 | 36.20382 | 15.91847 | -0.3001 | 0.443885 | 1 |
| 2 | 63389800 | 63390999 | 12.90552 | 23.1477 | 30.65883 | 13.42136 | -0.00107 | 0.403791 | 1 |
| 2 | 63554400 | 63556399 | 22.81815 | 37.76158 | 65.8624 | 28.4853 | -0.41926 | 1.313039 | 1 |
| 2 | 63668000 | 63670799 | 33.5422 | 54.17949 | 83.82935 | 35.77814 | -0.26821 | 1.721623 | 1 |
| 2 | 63780600 | 63780999 | 8.000739 | 15.43771 | 23.1306 | 8.93472 | -0.03195 | -0.13571 | 1 |
| 2 | 64463600 | 64465199 | 17.06198 | 30.25574 | 31.75748 | 14.03666 | 0.32287 | 0.503416 | 1 |
| 2 | 66102200 | 66104599 | 31.05359 | 51.46398 | 56.24446 | 24.34963 | 0.253054 | 1.308423 | 1 |
| 2 | 66242000 | 66243199 | 18.61765 | 32.27156 | 39.19481 | 17.1967 | 0.107534 | 0.822429 | 1 |
| 2 | 66547400 | 66549199 | 13.28508 | 24.33586 | 27.03549 | 11.5406 | 0.274818 | 0.275749 | 1 |
| 2 | 66675600 | 66677799 | 37.73377 | 60.72511 | 86.46028 | 36.71722 | -0.15491 | 1.838968 | 1 |
| 2 | 66786000 | 66787599 | 16.35313 | 28.86805 | 33.4146 | 14.78445 | 0.179882 | 0.54885 | 1 |
| 2 | 67768400 | 67771199 | 25.23521 | 42.28914 | 49.64144 | 21.64793 | 0.150577 | 1.162223 | 1 |
| 2 | 68031800 | 68035599 | 18.64264 | 32.73214 | 35.93742 | 15.89934 | 0.25337 | 0.631029 | 1 |
| 2 | 68289400 | 68290199 | 14.81223 | 26.16852 | 35.06898 | 15.514 | -0.02988 | 0.534449 | 1 |
| 2 | 68486200 | 68487199 | 12.60677 | 23.22345 | 27.08396 | 11.47323 | 0.213027 | 0.238708 | 1 |
| 2 | 68503200 | 68506599 | 24.1372 | 40.20183 | 60.81488 | 26.41811 | -0.21399 | 1.236386 | 1 |
| 2 | 68527800 | 68530799 | 30.02666 | 49.2319 | 78.17796 | 33.6537 | -0.28861 | 1.452043 | 1 |
| 2 | 68566400 | 68567799 | 14.28804 | 25.92235 | 30.48399 | 13.17274 | 0.181501 | 0.344839 | 1 |
| 2 | 68600200 | 68601399 | 22.20259 | 37.21987 | 59.12404 | 25.78875 | -0.2843 | 1.160593 | 1 |
| 2 | 69856600 | 69863399 | 35.24837 | 57.69531 | 59.39657 | 25.55046 | 0.333986 | 1.47125 | 1 |
| 2 | 70092200 | 70099199 | 87.38266 | 135.8913 | 151.4389 | 62.87155 | 0.152233 | 2.741296 | 1 |
| 2 | 70100000 | 70101999 | 29.68478 | 48.8266 | 64.78069 | 27.9241 | -0.02819 | 1.400354 | 1 |
| 2 | 70714200 | 70715799 | 27.63899 | 46.2065 | 53.40887 | 23.25566 | 0.172859 | 1.20078 | 1 |
| 2 | 70893200 | 70894999 | 16.97757 | 29.4854 | 43.21067 | 18.96697 | -0.16196 | 0.798039 | 1 |
| 2 | 71637800 | 71640799 | 14.27419 | 25.56751 | 29.49843 | 12.92831 | 0.1956 | 0.41835 | 1 |
| 2 | 71735600 | 71739999 | 34.71601 | 56.25916 | 78.82771 | 33.74786 | -0.12072 | 1.641162 | 1 |
| 2 | 72492000 | 72495399 | 31.03001 | 50.95401 | 68.77923 | 29.62673 | -0.05399 | 1.427409 | 1 |
| 2 | 72637800 | 72643399 | 61.09916 | 97.10866 | 120.3306 | 50.20336 | 0.024701 | 2.265229 | 1 |
| 2 | 72668400 | 72672799 | 12.62289 | 22.5868 | 37.24002 | 16.33521 | -0.31752 | 0.428745 | 1 |
| 2 | 74274800 | 74276199 | 17.02636 | 29.77087 | 41.47168 | 18.28375 | -0.08822 | 0.699546 | 1 |

| | | | | | | | | | |
|---|---|---|---|---|---|---|---|---|---|
| 2 | 74419800 | 74422599 | 27.82963 | 46.26256 | 63.93496 | 27.73765 | -0.08424 | 1.270069 | 1 |
| 2 | 74428400 | 74429999 | 13.30578 | 24.66531 | 26.40428 | 11.09028 | 0.344039 | 0.204636 | 1 |
| 2 | 74509400 | 74514599 | 39.12374 | 62.62983 | 108.2506 | 45.6091 | -0.44335 | 2.013988 | 1 |
| 2 | 75127800 | 75129999 | 22.99881 | 39.94034 | 41.91995 | 18.39975 | 0.317341 | 0.822975 | 1 |
| 2 | 76118000 | 76119999 | 20.02927 | 34.51227 | 48.47361 | 21.2639 | -0.10339 | 0.891326 | 1 |
| 2 | 76538800 | 76557999 | 533.505 | 716.3075 | 879.4137 | 366.9608 | -0.1868 | 5.214484 | 1 |
| 2 | 76558600 | 76593799 | 453.8601 | 622.1878 | 644.0993 | 267.9111 | 0.08943 | 4.885369 | 1 |
| 2 | 77073800 | 77076199 | 13.60588 | 24.06034 | 38.71634 | 17.12425 | -0.29187 | 0.523622 | 1 |
| 2 | 77124800 | 77126399 | 16.08419 | 28.44334 | 30.53112 | 13.50389 | 0.289294 | 0.524289 | 1 |
| 2 | 77173400 | 77175199 | 14.75679 | 25.83115 | 42.01445 | 18.54043 | -0.31069 | 0.669622 | 1 |
| 2 | 77842400 | 77844199 | 12.23277 | 22.20507 | 27.72847 | 11.97425 | 0.096391 | 0.32235 | 1 |
| 2 | 77959200 | 77960799 | 25.13948 | 41.63587 | 68.05998 | 29.45175 | -0.32612 | 1.30848 | 1 |
| 2 | 78920400 | 78922199 | 16.58377 | 29.4143 | 32.80864 | 14.49853 | 0.235841 | 0.500885 | 1 |
| 2 | 79765800 | 79768399 | 29.53284 | 48.15683 | 71.21776 | 30.59318 | -0.19098 | 1.521845 | 1 |
| 2 | 80060200 | 80061799 | 18.34704 | 31.39454 | 51.64664 | 22.64707 | -0.33228 | 0.951788 | 1 |
| 2 | 80123000 | 80124599 | 18.69485 | 32.62176 | 37.85754 | 16.66672 | 0.173831 | 0.731486 | 1 |
| 2 | 80206800 | 80212599 | 45.32965 | 72.69556 | 100.2397 | 42.34975 | -0.115 | 1.961576 | 1 |
| 2 | 80439000 | 80442799 | 32.65453 | 53.89841 | 62.53748 | 26.99844 | 0.166485 | 1.362128 | 1 |
| 2 | 80562600 | 80563799 | 17.36265 | 30.43057 | 36.67142 | 16.19969 | 0.119762 | 0.653885 | 1 |
| 2 | 80571600 | 80572799 | 16.28197 | 28.88377 | 33.35679 | 14.73872 | 0.186101 | 0.499216 | 1 |
| 2 | 80641800 | 80644599 | 20.82757 | 35.34607 | 51.11746 | 22.39479 | -0.14871 | 1.037331 | 1 |
| 2 | 80661200 | 80665199 | 68.52824 | 107.7447 | 153.8453 | 63.82025 | -0.19265 | 2.536073 | 1 |
| 2 | 80803000 | 80810599 | 118.6636 | 180.126 | 233.6089 | 96.72642 | -0.09359 | 3.223096 | 1 |
| 2 | 80905000 | 80907399 | 27.86712 | 46.13398 | 64.34202 | 27.83479 | -0.09781 | 1.318263 | 1 |
| 2 | 81013200 | 81014199 | 10.75739 | 20.38892 | 24.83943 | 10.03671 | 0.204519 | 0.040916 | 1 |
| 2 | 81055200 | 81056799 | 24.88502 | 42.30636 | 47.95581 | 21.01764 | 0.201474 | 1.030873 | 1 |
| 2 | 81141600 | 81143399 | 34.5896 | 56.7114 | 61.67064 | 26.54732 | 0.25613 | 1.450352 | 1 |
| 2 | 81339000 | 81341399 | 26.39035 | 44.29385 | 49.63709 | 21.66731 | 0.217259 | 1.150073 | 1 |
| 2 | 81424000 | 81426799 | 29.70878 | 49.77573 | 52.44134 | 22.86597 | 0.305956 | 1.179867 | 1 |
| 2 | 81472600 | 81475599 | 21.56267 | 36.58705 | 53.82688 | 23.58052 | -0.17352 | 1.04218 | 1 |
| 2 | 81585800 | 81588199 | 21.48465 | 36.93071 | 50.09702 | 21.97443 | -0.05417 | 0.920583 | 1 |
| 2 | 81818400 | 81822399 | 58.40175 | 92.59894 | 123.6164 | 51.45046 | -0.08598 | 2.332233 | 1 |
| 2 | 81823400 | 81825799 | 14.13338 | 25.36791 | 37.71898 | 16.50223 | -0.16694 | 0.41868 | 1 |
| 2 | 82236200 | 82243599 | 196.5808 | 289.7582 | 319.4007 | 131.5329 | 0.102785 | 3.782898 | 1 |
| 2 | 82260800 | 82261799 | 14.78661 | 26.2006 | 30.58247 | 13.52124 | 0.170111 | 0.494193 | 1 |
| 2 | 82503600 | 82507199 | 60.44073 | 96.11617 | 102.1551 | 42.64426 | 0.246571 | 2.242707 | 1 |
| 2 | 82592400 | 82593599 | 15.25394 | 27.02677 | 40.73186 | 18.00878 | -0.19688 | 0.516816 | 1 |
| 2 | 82718000 | 82719799 | 27.52476 | 45.8172 | 55.52371 | 24.11072 | 0.104971 | 1.248801 | 1 |
| 2 | 82945800 | 82953599 | 155.4237 | 232.4328 | 272.3268 | 112.2962 | 0.034107 | 3.525074 | 1 |
| 2 | 82970800 | 82973199 | 22.44629 | 38.04542 | 58.21445 | 25.49432 | -0.23008 | 1.059992 | 1 |
| 2 | 83026200 | 83028399 | 18.11755 | 31.49563 | 47.91656 | 21.04535 | -0.21576 | 0.795335 | 1 |
| 2 | 83202400 | 83206199 | 33.23573 | 54.45518 | 73.43648 | 31.6032 | -0.05392 | 1.463962 | 1 |
| 2 | 83243800 | 83245199 | 24.28973 | 40.34831 | 58.04315 | 25.16876 | -0.14186 | 1.264328 | 1 |
| 2 | 83260000 | 83261599 | 15.63935 | 27.44544 | 37.75032 | 16.6944 | -0.07008 | 0.625796 | 1 |
| 2 | 83297600 | 83300799 | 37.0859 | 59.96169 | 89.85749 | 38.39899 | -0.22071 | 1.700883 | 1 |
| 2 | 83386000 | 83387999 | 31.66701 | 51.43528 | 72.1908 | 30.95847 | -0.11998 | 1.587606 | 1 |
| 2 | 83760000 | 83761599 | 26.58243 | 44.10669 | 59.77587 | 25.8998 | -0.05623 | 1.286318 | 1 |

| | | | | | | | | | |
|---|---|---|---|---|---|---|---|---|---|
| 2 | 83916000 | 83917599 | 10.90149 | 19.61266 | 34.28479 | 14.97032 | -0.39647 | 0.378382 | 1 |
| 2 | 84025000 | 84028599 | 28.29119 | 47.19345 | 53.96839 | 23.46686 | 0.188313 | 1.226373 | 1 |
| 2 | 84391200 | 84398599 | 74.64616 | 116.9644 | 157.1034 | 65.16391 | -0.10836 | 2.603963 | 1 |
| 2 | 84416200 | 84421599 | 77.5059 | 121.6399 | 145.3577 | 60.28514 | 0.06188 | 2.574899 | 1 |
| 2 | 85524200 | 85525399 | 24.58095 | 40.97837 | 57.59228 | 25.03264 | -0.10819 | 1.224735 | 1 |
| 2 | 85531200 | 85533399 | 22.27644 | 38.56357 | 40.41263 | 17.73031 | 0.318728 | 0.848206 | 1 |
| 2 | 85785600 | 85791399 | 71.7102 | 113.7562 | 120.1919 | 50.04978 | 0.251611 | 2.328212 | 1 |
| 2 | 86196200 | 86198799 | 26.55058 | 44.21904 | 55.04558 | 23.91109 | 0.066231 | 1.236294 | 1 |
| 2 | 86671400 | 86674399 | 40.54592 | 65.43466 | 80.14439 | 34.16655 | 0.066679 | 1.757337 | 1 |
| 2 | 86776000 | 86778599 | 31.94597 | 52.54525 | 63.31994 | 27.2943 | 0.110191 | 1.40508 | 1 |
| 2 | 87267400 | 87269199 | 15.08067 | 26.8727 | 30.91828 | 13.60981 | 0.195207 | 0.458472 | 1 |
| 2 | 87276000 | 87279599 | 43.01686 | 69.95443 | 69.47827 | 29.8165 | 0.379043 | 1.585708 | 1 |
| 2 | 87356200 | 87356799 | 8.634293 | 16.34303 | 25.4408 | 10.30886 | -0.14983 | 0.031948 | 1 |
| 2 | 87537000 | 87537799 | 11.68857 | 20.93622 | 33.74392 | 14.78899 | -0.28375 | 0.410528 | 1 |
| 2 | 87599200 | 87601999 | 56.44668 | 89.79671 | 117.936 | 49.25588 | -0.05776 | 2.232144 | 1 |
| 2 | 87620600 | 87621599 | 15.1754 | 26.86292 | 33.04949 | 14.61485 | 0.094064 | 0.51291 | 1 |
| 2 | 87701000 | 87707599 | 92.55625 | 144.0483 | 164.0753 | 68.10984 | 0.121594 | 2.730809 | 1 |
| 2 | 88071000 | 88071999 | 16.65819 | 29.43885 | 34.96571 | 15.46735 | 0.143592 | 0.543499 | 1 |
| 2 | 88303400 | 88305999 | 28.88195 | 47.79495 | 66.09125 | 28.58364 | -0.08556 | 1.327568 | 1 |
| 2 | 88305800 | 88307999 | 34.73466 | 57.1356 | 62.44256 | 26.91679 | 0.250787 | 1.410859 | 1 |
| 2 | 88500400 | 88501399 | 12.14562 | 22.35711 | 25.64688 | 10.87559 | 0.235815 | 0.236647 | 1 |
| 2 | 88541200 | 88542199 | 16.72251 | 29.20121 | 36.93556 | 16.26821 | 0.050289 | 0.709071 | 1 |
| 2 | 88752600 | 88754999 | 18.82527 | 33.02445 | 37.00967 | 16.36186 | 0.223954 | 0.647544 | 1 |
| 2 | 88929400 | 88933799 | 74.23366 | 116.9087 | 126.9453 | 52.67259 | 0.20371 | 2.500889 | 1 |
| 2 | 89364200 | 89365799 | 19.01589 | 32.97519 | 47.93997 | 21.03389 | -0.15109 | 0.828744 | 1 |
| 2 | 89390800 | 89394399 | 28.58033 | 47.05454 | 67.51356 | 29.1138 | -0.14028 | 1.388299 | 1 |
| 2 | 89521800 | 89523599 | 19.04355 | 33.05936 | 46.32323 | 20.32775 | -0.09778 | 0.816793 | 1 |
| 2 | 89617200 | 89617999 | 14.7686 | 25.69161 | 44.24451 | 19.44102 | -0.39342 | 0.764763 | 1 |
| 2 | 89627200 | 89629399 | 21.49199 | 37.38698 | 40.3249 | 17.71854 | 0.278374 | 0.789842 | 1 |
| 2 | 90079000 | 90080199 | 17.59155 | 30.63697 | 38.53607 | 16.94474 | 0.057985 | 0.754986 | 1 |
| 2 | 90102600 | 90104799 | 21.58113 | 36.48821 | 52.20451 | 22.84511 | -0.13356 | 1.07174 | 1 |
| 2 | 90219400 | 90220599 | 10.39832 | 18.84645 | 32.82017 | 14.20004 | -0.38233 | 0.328263 | 1 |
| 2 | 90224600 | 90226999 | 22.1733 | 37.34302 | 59.75675 | 26.11902 | -0.29479 | 1.11636 | 1 |
| 2 | 93772200 | 93775199 | 35.60861 | 57.48943 | 89.11597 | 38.01068 | -0.27179 | 1.739628 | 1 |
| 2 | 94684200 | 94686199 | 24.53566 | 41.72539 | 47.95532 | 21.01796 | 0.181666 | 1.026397 | 1 |
| 2 | 94720400 | 94722199 | 18.46169 | 32.27365 | 35.24301 | 15.53488 | 0.261339 | 0.699028 | 1 |
| 2 | 95128400 | 95129399 | 18.54811 | 32.41698 | 33.24877 | 14.65288 | 0.351325 | 0.699475 | 1 |
| 2 | 95131400 | 95133399 | 28.11183 | 46.30104 | 65.72983 | 28.34922 | -0.12472 | 1.381892 | 1 |
| 2 | 95163400 | 95165999 | 35.49833 | 59.40799 | 53.08519 | 23.13004 | 0.542969 | 1.214475 | 1 |
| 2 | 95386000 | 95386999 | 16.47694 | 28.95757 | 36.63222 | 16.21993 | 0.050101 | 0.606115 | 1 |
| 2 | 95478200 | 95482199 | 19.49349 | 33.17008 | 47.98731 | 21.03297 | -0.14874 | 1.004422 | 1 |
| 2 | 95764200 | 95765199 | 18.32044 | 31.60026 | 45.84991 | 20.1136 | -0.14958 | 0.873396 | 1 |
| 2 | 95900800 | 95903199 | 52.14891 | 83.24881 | 100.5127 | 42.18239 | 0.068859 | 2.112945 | 1 |
| 2 | 95925200 | 95926999 | 22.43848 | 37.86514 | 60.9691 | 26.66507 | -0.30368 | 1.099609 | 1 |
| 2 | 96037000 | 96039399 | 23.44835 | 39.75134 | 47.98634 | 21.01624 | 0.110803 | 1.050589 | 1 |
| 2 | 96864600 | 96866199 | 30.26807 | 49.54031 | 70.68275 | 30.40471 | -0.13565 | 1.468048 | 1 |
| 2 | 97015000 | 97017399 | 43.44752 | 70.06962 | 77.49954 | 33.00999 | 0.212305 | 1.776815 | 1 |

| | | | | | | | | | |
|---|---|---|---|---|---|---|---|---|---|
| 2 | 97018400 | 97021199 | 24.11729 | 40.53328 | 55.44168 | 24.2178 | -0.06925 | 1.137158 | 1 |
| 2 | 97157000 | 97158399 | 18.55889 | 31.75316 | 50.57701 | 22.17782 | -0.2858 | 0.954361 | 1 |
| 2 | 97663400 | 97664999 | 13.53317 | 24.16594 | 35.08561 | 15.41685 | -0.13683 | 0.443041 | 1 |
| 2 | 97716000 | 97717399 | 13.76251 | 25.18477 | 27.48454 | 11.74854 | 0.298861 | 0.285808 | 1 |
| 2 | 97721800 | 97726799 | 21.17629 | 36.58474 | 41.55062 | 18.22846 | 0.202593 | 0.863083 | 1 |
| 2 | 97784600 | 97786599 | 35.63125 | 59.04356 | 57.67503 | 24.96697 | 0.414508 | 1.322717 | 1 |
| 2 | 97970400 | 97971399 | 15.41285 | 27.01993 | 40.91397 | 18.07763 | -0.20797 | 0.647926 | 1 |
| 2 | 98011200 | 98015599 | 51.6593 | 82.3495 | 122.3352 | 51.23837 | -0.23205 | 2.162047 | 1 |
| 2 | 98159600 | 98169799 | 286.4259 | 409.5814 | 418.8594 | 173.0478 | 0.168366 | 4.249233 | 1 |
| 2 | 98239600 | 98240399 | 14.1332 | 25.36855 | 29.03 | 12.70035 | 0.209179 | 0.403738 | 1 |
| 2 | 98258400 | 98263399 | 31.65022 | 51.93615 | 65.98134 | 28.41314 | 0.032837 | 1.435224 | 1 |
| 2 | 98307200 | 98308199 | 13.57461 | 24.51095 | 30.46297 | 13.24232 | 0.09656 | 0.365751 | 1 |
| 2 | 98580000 | 98589399 | 55.35441 | 88.37978 | 119.1224 | 50.00341 | -0.08946 | 2.116993 | 1 |
| 2 | 98595600 | 98596799 | 15.83301 | 28.10464 | 29.85208 | 13.18331 | 0.306615 | 0.484729 | 1 |
| 2 | 98810400 | 98811799 | 18.39298 | 32.00281 | 38.92717 | 17.10667 | 0.105756 | 0.769645 | 1 |
| 2 | 98885800 | 98886599 | 14.1561 | 25.5755 | 27.38882 | 11.89749 | 0.310792 | 0.360164 | 1 |
| 2 | 98928000 | 98929399 | 17.18464 | 30.52215 | 32.05883 | 14.15071 | 0.322934 | 0.4919 | 1 |
| 2 | 98933400 | 98934399 | 16.28637 | 28.44625 | 40.47585 | 17.83014 | -0.11879 | 0.708959 | 1 |
| 2 | 98996200 | 98997999 | 31.24192 | 51.62672 | 55.08871 | 23.80176 | 0.287186 | 1.341703 | 1 |
| 2 | 99189000 | 99191999 | 30.07323 | 49.48662 | 75.81363 | 32.68581 | -0.23498 | 1.403775 | 1 |
| 2 | 99196000 | 99197999 | 22.61718 | 39.1874 | 43.02677 | 18.87772 | 0.251947 | 0.845097 | 1 |
| 2 | 99266000 | 99269399 | 55.0385 | 87.94113 | 96.54782 | 40.57034 | 0.207196 | 2.086566 | 1 |
| 2 | 99452800 | 99454199 | 13.96217 | 25.46153 | 29.94471 | 12.85689 | 0.187061 | 0.309809 | 1 |
| 2 | 99476200 | 99479399 | 31.95078 | 53.04585 | 56.83166 | 24.63431 | 0.281871 | 1.292116 | 1 |
| 2 | 99974600 | 99975999 | 22.36319 | 38.30755 | 41.16547 | 18.05262 | 0.279883 | 0.944941 | 1 |
| 2 | 100733400 | 100735599 | 21.46022 | 36.29186 | 58.15576 | 25.45114 | -0.29658 | 1.075379 | 1 |
| 2 | 100975800 | 100978799 | 33.37606 | 54.87922 | 57.83718 | 24.92742 | 0.302834 | 1.40887 | 1 |
| 2 | 101174400 | 101175599 | 14.93115 | 26.21073 | 41.39105 | 18.30923 | -0.2683 | 0.624078 | 1 |
| 2 | 101311800 | 101312399 | 11.72918 | 20.95981 | 34.77883 | 15.27269 | -0.32736 | 0.428629 | 1 |
| 2 | 103359400 | 103359999 | 9.768828 | 18.36382 | 26.92009 | 11.05275 | -0.07983 | 0.093195 | 1 |
| 2 | 103694200 | 103697199 | 29.34482 | 48.82343 | 57.3686 | 24.90372 | 0.149199 | 1.258197 | 1 |
| 2 | 103845600 | 103847199 | 12.41614 | 21.83628 | 38.9256 | 17.2357 | -0.44157 | 0.579361 | 1 |
| 2 | 103942600 | 103944199 | 26.0882 | 43.27088 | 57.54676 | 24.92769 | -0.02909 | 1.287353 | 1 |
| 2 | 104071000 | 104073599 | 23.02546 | 39.28097 | 46.35779 | 20.32296 | 0.144267 | 0.989835 | 1 |
| 2 | 104523000 | 104528599 | 53.36671 | 84.88491 | 127.6489 | 53.30196 | -0.25302 | 2.236233 | 1 |
| 2 | 104708800 | 104711399 | 19.14688 | 33.50794 | 36.62862 | 16.16052 | 0.259791 | 0.687387 | 1 |
| 2 | 105675400 | 105676599 | 47.42206 | 75.92712 | 93.58568 | 39.44048 | 0.043817 | 2.015807 | 1 |
| 2 | 105688800 | 105691399 | 53.11577 | 84.87749 | 99.65227 | 41.88082 | 0.110715 | 2.082802 | 1 |
| 2 | 105958400 | 105962399 | 59.00747 | 94.24913 | 103.6729 | 43.54233 | 0.203845 | 2.103673 | 1 |
| 2 | 106223200 | 106224999 | 15.10586 | 26.29976 | 41.19237 | 18.10919 | -0.25687 | 0.752056 | 1 |
| 2 | 106397000 | 106401399 | 55.97886 | 89.28832 | 108.5163 | 45.48794 | 0.058031 | 2.143083 | 1 |
| 2 | 107308800 | 107311599 | 31.85197 | 52.29676 | 70.02546 | 30.16184 | -0.04249 | 1.431856 | 1 |
| 2 | 107577800 | 107579399 | 14.6055 | 25.72345 | 40.80928 | 18.0646 | -0.27382 | 0.578386 | 1 |
| 2 | 107806400 | 107807799 | 23.47145 | 39.27708 | 58.65068 | 25.55361 | -0.1954 | 1.182512 | 1 |
| 2 | 108163000 | 108165399 | 20.8279 | 35.34536 | 54.8583 | 24.03341 | -0.25036 | 1.040906 | 1 |
| 2 | 108293800 | 108294799 | 17.99249 | 31.41409 | 39.95152 | 17.5949 | 0.042491 | 0.724448 | 1 |
| 2 | 108383200 | 108383999 | 10.20405 | 18.79746 | 30.38116 | 12.86984 | -0.25433 | 0.229047 | 1 |

| | | | | | | | | | |
|---|---|---|---|---|---|---|---|---|---|
| 2 | 108916400 | 108916999 | 11.02501 | 20.54699 | 27.28185 | 11.36633 | 0.044934 | 0.159741 | 1 |
| 2 | 109465400 | 109466399 | 15.42822 | 27.32661 | 30.05507 | 13.28912 | 0.255243 | 0.501817 | 1 |
| 2 | 109894600 | 109895399 | 12.22807 | 22.85166 | 26.69895 | 11.07196 | 0.233345 | 0.150122 | 1 |
| 2 | 109956400 | 109958999 | 29.19824 | 48.6926 | 56.81101 | 24.69845 | 0.159394 | 1.232811 | 1 |
| 2 | 110265600 | 110266999 | 12.00735 | 21.98839 | 27.64978 | 11.80786 | 0.096788 | 0.268907 | 1 |
| 2 | 110835200 | 110836599 | 17.63313 | 30.86649 | 36.68901 | 16.19057 | 0.139469 | 0.67444 | 1 |
| 2 | 110918000 | 110918999 | 24.02373 | 40.33978 | 48.05414 | 20.98227 | 0.129519 | 1.137392 | 1 |
| 2 | 110952600 | 110953599 | 20.33881 | 34.35814 | 53.22513 | 23.28566 | -0.24786 | 1.076474 | 1 |
| 2 | 111010000 | 111011399 | 25.95735 | 43.51892 | 50.16393 | 21.88201 | 0.176771 | 1.161797 | 1 |
| 2 | 111363400 | 111364399 | 13.29347 | 24.11909 | 29.53146 | 12.76036 | 0.123724 | 0.3352 | 1 |
| 2 | 111374800 | 111384599 | 78.83077 | 123.8617 | 142.6315 | 59.16335 | 0.116519 | 2.552978 | 1 |
| 2 | 111480000 | 111480999 | 21.88684 | 37.74827 | 38.54295 | 16.9081 | 0.355165 | 0.879009 | 1 |
| 2 | 111680600 | 111682199 | 27.09593 | 44.92159 | 63.11796 | 27.33227 | -0.10827 | 1.300025 | 1 |
| 2 | 111726400 | 111729199 | 46.54937 | 74.94913 | 87.9322 | 37.36905 | 0.124584 | 1.837312 | 1 |
| 2 | 111829800 | 111833199 | 45.33973 | 72.72313 | 95.27681 | 40.26259 | -0.04105 | 1.9556 | 1 |
| 2 | 111882800 | 111884799 | 28.20365 | 46.56028 | 62.56751 | 27.01474 | -0.04463 | 1.350408 | 1 |
| 2 | 111917200 | 111919799 | 43.41125 | 70.09883 | 78.7417 | 33.59328 | 0.192114 | 1.74324 | 1 |
| 2 | 111946200 | 111947399 | 22.04731 | 38.07236 | 40.00656 | 17.55083 | 0.31421 | 0.87017 | 1 |
| 2 | 111979400 | 111980999 | 14.67569 | 26.20391 | 31.85396 | 13.99783 | 0.117757 | 0.443277 | 1 |
| 2 | 111989600 | 111992399 | 45.15343 | 72.21331 | 105.2266 | 44.27514 | -0.19904 | 2.050949 | 1 |
| 2 | 112376800 | 112381399 | 63.0116 | 100.2731 | 105.6669 | 44.15171 | 0.259904 | 2.22319 | 1 |
| 2 | 113688000 | 113689599 | 22.65845 | 38.96632 | 41.05773 | 18.00971 | 0.309055 | 0.910247 | 1 |
| 2 | 114310000 | 114312199 | 17.53862 | 30.49618 | 47.91101 | 21.04593 | -0.26178 | 0.790837 | 1 |
| 2 | 114991800 | 114995399 | 74.68024 | 117.4861 | 144.1744 | 59.81299 | 0.026248 | 2.525881 | 1 |
| 2 | 115618600 | 115620999 | 21.01362 | 36.27713 | 41.5689 | 18.23607 | 0.18962 | 0.868886 | 1 |
| 2 | 116307400 | 116309799 | 21.0348 | 36.7025 | 38.14739 | 16.79364 | 0.33199 | 0.74293 | 1 |
| 2 | 116323600 | 116325399 | 45.59446 | 73.05143 | 98.61559 | 41.60398 | -0.08601 | 1.992778 | 1 |
| 2 | 116331000 | 116333999 | 36.12946 | 58.80169 | 73.89541 | 31.72147 | 0.041219 | 1.5624 | 1 |
| 2 | 118387200 | 118389399 | 22.9642 | 39.17448 | 49.55675 | 21.72527 | 0.044468 | 0.994766 | 1 |
| 2 | 119376800 | 119378599 | 13.30787 | 23.37989 | 38.36467 | 16.98202 | -0.32325 | 0.596067 | 1 |
| 2 | 120463800 | 120471399 | 58.25914 | 92.37512 | 128.6742 | 53.55687 | -0.1472 | 2.332587 | 1 |
| 2 | 120969000 | 120971199 | 21.81483 | 36.6316 | 55.37279 | 24.17826 | -0.21299 | 1.136448 | 1 |
| 2 | 122220200 | 122220999 | 9.167979 | 17.56783 | 24.03459 | 9.470386 | 0.070817 | -0.05838 | 1 |
| 2 | 122334400 | 122337599 | 32.84392 | 54.13043 | 66.87435 | 28.85086 | 0.075516 | 1.38644 | 1 |
| 2 | 122484400 | 122485999 | 16.4089 | 28.1696 | 48.47694 | 21.26273 | -0.39619 | 0.909428 | 1 |
| 2 | 123053200 | 123053799 | 10.13491 | 18.78898 | 27.32299 | 11.47482 | -0.09391 | 0.184863 | 1 |
| 2 | 123066800 | 123071399 | 77.39705 | 121.7054 | 130.1395 | 53.98724 | 0.224156 | 2.53358 | 1 |
| 2 | 123364200 | 123364599 | 11.30016 | 20.53228 | 28.783 | 12.41442 | -0.06824 | 0.312144 | 1 |
| 2 | 123410600 | 123412599 | 12.47789 | 22.36966 | 37.2719 | 16.32311 | -0.33096 | 0.416453 | 1 |
| 2 | 123440200 | 123444999 | 70.25023 | 111.2368 | 117.5185 | 48.86462 | 0.24937 | 2.37397 | 1 |
| 2 | 123745400 | 123749399 | 66.98168 | 106.1312 | 124.3714 | 51.73132 | 0.100805 | 2.359288 | 1 |
| 2 | 124096800 | 124098999 | 19.16052 | 32.75275 | 49.53407 | 21.71891 | -0.21157 | 0.962838 | 1 |
| 2 | 124191000 | 124197399 | 65.04345 | 103.1954 | 119.7349 | 49.86671 | 0.116935 | 2.317587 | 1 |
| 2 | 124640200 | 124643399 | 28.0895 | 46.27793 | 67.50478 | 29.1184 | -0.16364 | 1.378597 | 1 |
| 2 | 124671000 | 124673599 | 52.69824 | 84.34346 | 87.84418 | 37.00096 | 0.285738 | 2.029857 | 1 |
| 2 | 125095400 | 125097199 | 24.13059 | 41.60239 | 46.28866 | 20.30578 | 0.231533 | 0.901664 | 1 |
| 2 | 125472200 | 125474399 | 25.53994 | 42.42295 | 68.03669 | 29.50114 | -0.29829 | 1.278857 | 1 |

| | | | | | | | | | |
|---|---|---|---|---|---|---|---|---|---|
| 2 | 128401600 | 128404599 | 22.41713 | 37.62985 | 55.96433 | 24.43117 | -0.18968 | 1.141489 | 1 |
| 2 | 128980800 | 128982799 | 8.385035 | 15.85421 | 28.69541 | 11.65408 | -0.36855 | 0.046083 | 1 |
| 2 | 129191000 | 129193599 | 36.57677 | 59.65004 | 69.53871 | 29.87323 | 0.151329 | 1.531807 | 1 |
| 2 | 129220200 | 129222999 | 23.43465 | 39.59022 | 50.11919 | 21.92722 | 0.04222 | 1.08184 | 1 |
| 2 | 131487200 | 131493399 | 66.38819 | 104.8127 | 137.4184 | 57.05404 | -0.06508 | 2.445842 | 1 |
| 2 | 131620400 | 131623399 | 42.23951 | 68.22701 | 90.91003 | 38.79946 | -0.05327 | 1.736692 | 1 |
| 2 | 131625200 | 131629799 | 44.7246 | 72.14726 | 99.08737 | 42.21868 | -0.09899 | 1.784188 | 1 |
| 2 | 132909000 | 132911999 | 23.41434 | 40.20579 | 44.19508 | 19.385 | 0.247827 | 0.931552 | 1 |
| 2 | 133075400 | 133077599 | 15.84245 | 27.69257 | 40.48391 | 17.8438 | -0.15751 | 0.695627 | 1 |
| 2 | 133147000 | 133148199 | 12.89848 | 23.30615 | 35.24269 | 15.30688 | -0.18394 | 0.367716 | 1 |
| 2 | 134163800 | 134164799 | 14.16037 | 26.02827 | 25.99485 | 11.04476 | 0.432415 | 0.262042 | 1 |
| 2 | 134431200 | 134433599 | 41.16584 | 66.74531 | 71.2307 | 30.50682 | 0.271782 | 1.638751 | 1 |
| 2 | 134508600 | 134509399 | 15.80423 | 27.74485 | 34.76619 | 15.37944 | 0.063847 | 0.616721 | 1 |
| 2 | 134588400 | 134589199 | 12.5307 | 22.6025 | 27.73096 | 12.0704 | 0.114269 | 0.360423 | 1 |
| 2 | 134647800 | 134648799 | 12.99215 | 23.59379 | 28.58135 | 12.33583 | 0.140215 | 0.326361 | 1 |
| 2 | 135354000 | 135354599 | 12.64634 | 22.38161 | 34.7454 | 15.36542 | -0.23967 | 0.5019 | 1 |
| 2 | 135789600 | 135790399 | 12.22807 | 22.85166 | 26.69895 | 11.07196 | 0.233345 | 0.150122 | 1 |
| 2 | 136116000 | 136116999 | 13.92872 | 24.56033 | 38.23988 | 16.92326 | -0.24616 | 0.556652 | 1 |
| 2 | 136391600 | 136394999 | 28.90209 | 47.42439 | 73.78268 | 31.77195 | -0.25877 | 1.433188 | 1 |
| 2 | 136524000 | 136525999 | 22.84243 | 38.69549 | 49.56972 | 21.70483 | 0.025418 | 1.056914 | 1 |
| 2 | 136651400 | 136654199 | 37.3553 | 61.14633 | 62.2519 | 26.77911 | 0.350085 | 1.479322 | 1 |
| 2 | 136704800 | 136705799 | 12.48564 | 23.03265 | 29.67713 | 12.54773 | 0.071685 | 0.232961 | 1 |
| 2 | 136807200 | 136810399 | 32.61959 | 53.21522 | 78.1343 | 33.57011 | -0.18027 | 1.525281 | 1 |
| 2 | 137108000 | 137108599 | 13.2247 | 23.91039 | 28.97569 | 12.57521 | 0.134315 | 0.352437 | 1 |
| 2 | 137193800 | 137194999 | 30.1901 | 49.27315 | 69.05825 | 29.67531 | -0.11276 | 1.509835 | 1 |
| 2 | 137236200 | 137240999 | 71.53496 | 112.2776 | 163.5023 | 67.81104 | -0.22294 | 2.573699 | 1 |
| 2 | 137730400 | 137731199 | 9.828681 | 18.15798 | 30.92404 | 13.04878 | -0.32555 | 0.208919 | 1 |
| 2 | 137876800 | 137879399 | 36.06419 | 58.76728 | 66.54556 | 28.5791 | 0.192391 | 1.540116 | 1 |
| 2 | 138359200 | 138366599 | 156.7277 | 233.8139 | 269.6784 | 111.1516 | 0.05356 | 3.569805 | 1 |
| 2 | 138403600 | 138415799 | 334.9215 | 465.5322 | 622.0958 | 258.1933 | -0.25863 | 4.671015 | 1 |
| 2 | 138416600 | 138418799 | 19.98773 | 34.86178 | 40.80323 | 17.95838 | 0.161271 | 0.745483 | 1 |
| 2 | 138420200 | 138422399 | 15.88002 | 27.64101 | 40.164 | 17.65446 | -0.1492 | 0.757497 | 1 |
| 2 | 138686400 | 138687599 | 14.51444 | 25.63878 | 36.61442 | 16.19813 | -0.12122 | 0.536982 | 1 |
| 2 | 138774800 | 138778799 | 53.45317 | 85.58803 | 90.95644 | 38.33551 | 0.257358 | 2.017623 | 1 |
| 2 | 139000200 | 139001799 | 15.68936 | 27.6143 | 40.22895 | 17.81005 | -0.1519 | 0.590062 | 1 |
| 2 | 139370200 | 139370999 | 14.31636 | 25.50338 | 30.0865 | 13.24689 | 0.159215 | 0.454457 | 1 |
| 2 | 139738400 | 139740999 | 39.44073 | 63.99224 | 72.68991 | 31.14756 | 0.182915 | 1.623787 | 1 |
| 2 | 140303400 | 140305799 | 21.27029 | 35.90881 | 53.26055 | 23.29666 | -0.18546 | 1.084517 | 1 |
| 2 | 142043200 | 142044799 | 11.30423 | 21.10895 | 26.74895 | 11.10715 | 0.115628 | 0.147779 | 1 |
| 2 | 142622400 | 142623599 | 12.07362 | 22.03512 | 34.39977 | 14.75495 | -0.21793 | 0.302474 | 1 |
| 2 | 142884400 | 142885599 | 14.88551 | 26.46197 | 29.4909 | 13.00753 | 0.238969 | 0.46888 | 1 |
| 2 | 143136000 | 143140399 | 56.15459 | 89.28653 | 128.2634 | 53.53496 | -0.18767 | 2.250195 | 1 |
| 2 | 143471600 | 143474199 | 24.48682 | 40.7857 | 58.59872 | 25.45602 | -0.13981 | 1.236364 | 1 |
| 2 | 144878200 | 144880799 | 30.3584 | 50.86005 | 54.57732 | 23.79611 | 0.279674 | 1.189055 | 1 |
| 2 | 144981600 | 144982599 | 15.14766 | 26.76873 | 32.60057 | 14.42119 | 0.107714 | 0.529896 | 1 |
| 2 | 145632400 | 145634199 | 13.27232 | 23.57687 | 36.14462 | 15.95217 | -0.21918 | 0.47687 | 1 |
| 2 | 145644800 | 145649999 | 35.04739 | 57.55708 | 60.69747 | 26.14669 | 0.301233 | 1.43008 | 1 |

| | | | | | | | | | |
|---|---|---|---|---|---|---|---|---|---|
| 2 | 147244200 | 147246199 | 10.75633 | 19.70059 | 31.8177 | 13.58483 | -0.26201 | 0.266174 | 1 |
| 2 | 148454200 | 148458599 | 91.42894 | 140.8774 | 186.7027 | 77.55334 | -0.10778 | 2.911901 | 1 |
| 2 | 148827400 | 148828399 | 15.79498 | 27.75769 | 35.66472 | 15.7909 | 0.027781 | 0.600814 | 1 |
| 2 | 149026400 | 149027599 | 14.96016 | 26.63843 | 32.94423 | 14.51057 | 0.090953 | 0.466338 | 1 |
| 2 | 149105600 | 149108399 | 32.60416 | 53.3206 | 74.88617 | 32.20333 | -0.11382 | 1.488492 | 1 |
| 2 | 149331600 | 149333199 | 30.73068 | 50.87169 | 58.4319 | 25.27703 | 0.181392 | 1.321789 | 1 |
| 2 | 149361200 | 149362199 | 16.98727 | 29.66107 | 36.87632 | 16.24127 | 0.074928 | 0.71044 | 1 |
| 2 | 149916200 | 149916999 | 12.93921 | 23.39457 | 29.18716 | 12.66675 | 0.093106 | 0.354428 | 1 |
| 2 | 150306800 | 150310399 | 29.35433 | 49.00257 | 53.49828 | 23.27314 | 0.254921 | 1.220135 | 1 |
| 2 | 150326000 | 150329399 | 33.16782 | 54.57456 | 64.1506 | 27.65656 | 0.146108 | 1.402608 | 1 |
| 2 | 150449800 | 150450799 | 9.711745 | 18.00214 | 25.26341 | 10.61174 | -0.04345 | 0.173917 | 1 |
| 2 | 150619400 | 150622399 | 25.43245 | 42.21862 | 64.77518 | 28.0751 | -0.2347 | 1.280248 | 1 |
| 2 | 150628200 | 150631399 | 13.33267 | 23.32285 | 40.56006 | 17.88862 | -0.40665 | 0.669907 | 1 |
| 2 | 150895000 | 150897199 | 18.01177 | 31.35881 | 36.87773 | 16.2123 | 0.154298 | 0.755277 | 1 |
| 2 | 150936200 | 150939599 | 26.77069 | 44.29068 | 61.90692 | 26.7696 | -0.10096 | 1.321953 | 1 |
| 2 | 151007800 | 151017199 | 18.59293 | 32.64432 | 36.4172 | 16.11139 | 0.230407 | 0.630944 | 1 |
| 2 | 151040600 | 151043599 | 26.81925 | 45.5522 | 47.95592 | 21.01499 | 0.307635 | 1.04809 | 1 |
| 2 | 151045000 | 151048399 | 50.28182 | 80.46473 | 96.88654 | 40.80162 | 0.076627 | 2.034177 | 1 |
| 2 | 151052000 | 151052999 | 12.54962 | 23.33118 | 27.66852 | 11.57531 | 0.201807 | 0.187588 | 1 |
| 2 | 151183000 | 151187799 | 78.34448 | 123.078 | 138.8909 | 57.61055 | 0.145494 | 2.555426 | 1 |
| 2 | 151189400 | 151195999 | 80.32725 | 124.8978 | 167.2507 | 69.43778 | -0.11265 | 2.742252 | 1 |
| 2 | 151212800 | 151222799 | 13.81456 | 25.06572 | 27.54207 | 11.90006 | 0.279179 | 0.335636 | 1 |
| 2 | 152001200 | 152003199 | 28.27166 | 47.26914 | 49.16235 | 21.40802 | 0.324612 | 1.195297 | 1 |
| 2 | 152051600 | 152056599 | 50.07459 | 80.14937 | 107.1117 | 45.12068 | -0.0732 | 2.032322 | 1 |
| 2 | 152997800 | 152999999 | 16.38842 | 28.6124 | 45.71669 | 20.13346 | -0.28538 | 0.724423 | 1 |
| 2 | 153926800 | 153929999 | 47.67335 | 76.39699 | 96.37091 | 40.66893 | 0.011939 | 1.98792 | 1 |
| 2 | 154588200 | 154591999 | 25.21051 | 41.94911 | 64.33947 | 27.93175 | -0.23397 | 1.250837 | 1 |
| 2 | 154961200 | 154973599 | 148.981 | 224.4344 | 240.182 | 99.21618 | 0.174135 | 3.382669 | 1 |
| 2 | 155956200 | 155961799 | 71.16108 | 111.216 | 166.1289 | 68.92582 | -0.26436 | 2.64937 | 1 |
| 2 | 157427400 | 157428399 | 12.2192 | 22.09188 | 32.14652 | 13.95176 | -0.12799 | 0.35305 | 1 |
| 2 | 157494000 | 157494799 | 14.25558 | 25.05618 | 35.82975 | 15.86599 | -0.12565 | 0.590895 | 1 |
| 2 | 159296200 | 159298599 | 29.93801 | 49.73771 | 56.752 | 24.61124 | 0.191332 | 1.277998 | 1 |
| 2 | 159779200 | 159781199 | 14.09347 | 24.76412 | 38.31347 | 16.9617 | -0.23881 | 0.598091 | 1 |
| 2 | 161208000 | 161213999 | 54.49758 | 86.64459 | 129.0688 | 53.86657 | -0.24007 | 2.250938 | 1 |
| 2 | 161278800 | 161281199 | 25.14851 | 41.7177 | 62.97104 | 27.2798 | -0.21124 | 1.287495 | 1 |
| 2 | 161597000 | 161597799 | 8.646193 | 16.25398 | 26.40495 | 10.84065 | -0.22708 | 0.079484 | 1 |
| 2 | 161832600 | 161833599 | 17.56889 | 30.38417 | 42.21993 | 18.52268 | -0.08671 | 0.839577 | 1 |
| 2 | 162021200 | 162022399 | 13.32336 | 23.42552 | 37.36283 | 16.54461 | -0.2822 | 0.58409 | 1 |
| 2 | 162432200 | 162434399 | 23.33151 | 39.59182 | 50.62983 | 22.18055 | 0.027977 | 1.044988 | 1 |
| 2 | 164012000 | 164014799 | 20.1936 | 34.02209 | 55.88405 | 24.42918 | -0.33227 | 1.101121 | 1 |
| 2 | 164517800 | 164521599 | 26.83592 | 44.12401 | 70.62289 | 30.43755 | -0.29831 | 1.402522 | 1 |
| 2 | 165191600 | 165198199 | 115.1777 | 174.3679 | 250.219 | 103.5187 | -0.24267 | 3.277817 | 1 |
| 2 | 167124600 | 167131799 | 131.0973 | 198.402 | 230.168 | 95.21327 | 0.063477 | 3.286704 | 1 |
| 2 | 168546800 | 168548799 | 17.7814 | 30.33396 | 52.1335 | 22.85494 | -0.39569 | 0.979962 | 1 |
| 2 | 168868200 | 168874799 | 76.56137 | 119.0962 | 177.0778 | 73.51276 | -0.26297 | 2.734316 | 1 |
| 2 | 168978200 | 168979999 | 17.31347 | 30.25459 | 38.32004 | 16.88565 | 0.048275 | 0.702919 | 1 |
| 2 | 170112800 | 170115199 | 11.23934 | 20.50125 | 29.70962 | 12.75198 | -0.11161 | 0.289577 | 1 |

| | | | | | | | | | |
|---|---|---|---|---|---|---|---|---|---|
| 2 | 170436200 | 170438399 | 29.80941 | 49.55556 | 54.61784 | 23.69613 | 0.241198 | 1.266661 | 1 |
| 2 | 170553800 | 170555799 | 17.76951 | 31.08396 | 37.20508 | 16.40883 | 0.129469 | 0.686504 | 1 |
| 2 | 170838200 | 170842199 | 32.04301 | 52.30309 | 70.71435 | 30.38819 | -0.06099 | 1.510306 | 1 |
| 2 | 171761800 | 171766399 | 21.99979 | 38.16115 | 44.11734 | 19.35687 | 0.178198 | 0.833902 | 1 |
| 2 | 171848000 | 171850599 | 50.74702 | 80.79494 | 112.1364 | 46.88619 | -0.13598 | 2.20008 | 1 |
| 2 | 172584600 | 172588599 | 28.48072 | 46.79219 | 66.11356 | 28.48476 | -0.11931 | 1.414685 | 1 |
| 2 | 173239000 | 173245199 | 70.97832 | 111.549 | 153.3546 | 63.61315 | -0.13854 | 2.545682 | 1 |
| 2 | 173346400 | 173355199 | 70.24897 | 110.3776 | 152.325 | 63.18419 | -0.1443 | 2.548329 | 1 |
| 2 | 173488600 | 173490199 | 18.83872 | 32.05702 | 50.58134 | 22.17 | -0.27354 | 1.003282 | 1 |
| 2 | 173492800 | 173495999 | 33.31626 | 54.34516 | 73.23131 | 31.46216 | -0.05693 | 1.523148 | 1 |
| 2 | 174323600 | 174325799 | 18.79607 | 32.63169 | 43.6625 | 19.16088 | -0.03151 | 0.810213 | 1 |
| 2 | 174642600 | 174647999 | 40.28034 | 64.94871 | 88.59091 | 37.72443 | -0.08984 | 1.783232 | 1 |
| 2 | 176481800 | 176482199 | 13.25144 | 23.90641 | 28.68396 | 12.48269 | 0.146523 | 0.368183 | 1 |
| 2 | 176848200 | 176848999 | 13.07456 | 24.08488 | 25.52352 | 10.81239 | 0.350326 | 0.239134 | 1 |
| 2 | 176873200 | 176876399 | 34.62917 | 56.72453 | 64.66597 | 27.82598 | 0.187633 | 1.460783 | 1 |
| 2 | 176942400 | 176947199 | 46.32628 | 74.26265 | 90.85663 | 38.36101 | 0.056555 | 1.973756 | 1 |
| 2 | 176985600 | 176986799 | 14.12352 | 25.46393 | 27.85 | 12.12915 | 0.278085 | 0.37205 | 1 |
| 2 | 177088200 | 177090799 | 26.092 | 43.76123 | 51.23346 | 22.35388 | 0.154372 | 1.158517 | 1 |
| 2 | 177120000 | 177121599 | 19.89466 | 34.59299 | 37.39342 | 16.42589 | 0.275282 | 0.785409 | 1 |
| 2 | 177237200 | 177245399 | 180.4667 | 268.0166 | 274.3036 | 112.9972 | 0.219904 | 3.653158 | 1 |
| 2 | 177291600 | 177292599 | 15.63141 | 27.59391 | 32.42208 | 14.34532 | 0.158283 | 0.541362 | 1 |
| 2 | 177400600 | 177402799 | 15.4746 | 26.99775 | 43.09811 | 18.97175 | -0.28426 | 0.723418 | 1 |
| 2 | 177416600 | 177417799 | 14.08744 | 25.14187 | 34.08759 | 14.98525 | -0.0389 | 0.450118 | 1 |
| 2 | 177466000 | 177472199 | 57.45874 | 91.89675 | 99.26586 | 41.76947 | 0.232071 | 2.05997 | 1 |
| 2 | 177488000 | 177490199 | 21.15995 | 36.91933 | 38.11983 | 16.78107 | 0.34145 | 0.743604 | 1 |
| 2 | 177626400 | 177630799 | 48.32918 | 77.35435 | 109.0731 | 45.94583 | -0.1505 | 2.029374 | 1 |
| 2 | 177676200 | 177690999 | 255.5394 | 371.7567 | 389.1847 | 160.4099 | 0.159103 | 3.994164 | 1 |
| 2 | 178048400 | 178050599 | 30.86442 | 50.11702 | 79.57236 | 34.11665 | -0.2979 | 1.595152 | 1 |
| 2 | 178939000 | 178944799 | 48.75191 | 77.57681 | 126.2862 | 52.76322 | -0.36657 | 2.217055 | 1 |
| 2 | 180282600 | 180287399 | 91.0304 | 141.0312 | 173.2161 | 71.94917 | 0.007889 | 2.811197 | 1 |
| 2 | 181171600 | 181174799 | 38.84561 | 63.02887 | 83.22816 | 35.66412 | -0.03376 | 1.62758 | 1 |
| 2 | 181533800 | 181555399 | 512.9829 | 694.8163 | 776.7727 | 323.7014 | -0.03901 | 5.07581 | 1 |
| 2 | 181582000 | 181583799 | 23.47933 | 40.06209 | 50.09711 | 21.96255 | 0.061133 | 0.995682 | 1 |
| 3 | 3310800 | 3318599 | 42.6891 | 68.94439 | 72.91245 | 31.11315 | 0.279206 | 1.733776 | 1 |
| 3 | 3319600 | 3331799 | 400.4603 | 554.9333 | 551.0255 | 228.8054 | 0.165254 | 4.718491 | 1 |
| 3 | 3443200 | 3443999 | 12.30652 | 22.25403 | 36.31099 | 15.75538 | -0.29243 | 0.358021 | 1 |
| 3 | 3909200 | 3910799 | 11.27699 | 20.89609 | 28.81773 | 12.11058 | -0.01875 | 0.20081 | 1 |
| 3 | 3980400 | 3981599 | 14.82633 | 26.42517 | 31.01168 | 13.64845 | 0.167096 | 0.45682 | 1 |
| 3 | 4145800 | 4147799 | 22.60379 | 38.50534 | 46.90825 | 20.56185 | 0.098376 | 1.003032 | 1 |
| 3 | 4255000 | 4255799 | 16.58421 | 28.88698 | 41.56758 | 18.27947 | -0.13539 | 0.750384 | 1 |
| 3 | 4374200 | 4377799 | 71.26116 | 112.3504 | 127.3329 | 52.84095 | 0.14323 | 2.477328 | 1 |
| 3 | 4455400 | 4459799 | 65.91566 | 104.4992 | 120.3603 | 50.08878 | 0.126453 | 2.341273 | 1 |
| 3 | 4506600 | 4508799 | 16.98968 | 30.13298 | 35.36352 | 15.62813 | 0.162838 | 0.508901 | 1 |
| 3 | 5134800 | 5135599 | 12.42734 | 21.87599 | 38.42666 | 17.01166 | -0.41983 | 0.569271 | 1 |
| 3 | 5584600 | 5586199 | 16.6955 | 29.48081 | 30.95881 | 13.69709 | 0.319858 | 0.544328 | 1 |
| 3 | 5590000 | 5592599 | 30.3862 | 50.3166 | 57.87071 | 25.03948 | 0.179521 | 1.316704 | 1 |
| 3 | 5968000 | 5968799 | 10.43165 | 19.71851 | 22.18987 | 9.017184 | 0.311256 | 0.050046 | 1 |

| | | | | | | | | | |
|---|---|---|---|---|---|---|---|---|---|
| 3 | 6328000 | 6336999 | 176.398 | 264.0234 | 265.6071 | 109.5433 | 0.254974 | 3.512369 | 1 |
| 3 | 6361600 | 6373999 | 176.1379 | 259.219 | 324.4564 | 133.6126 | -0.08256 | 3.807181 | 1 |
| 3 | 6440200 | 6445799 | 76.87932 | 120.6651 | 153.6779 | 63.73641 | -0.02984 | 2.574574 | 1 |
| 3 | 6515200 | 6517399 | 29.44963 | 48.36923 | 75.51314 | 32.53031 | -0.26317 | 1.425715 | 1 |
| 3 | 6803200 | 6805599 | 13.00894 | 23.08444 | 35.21597 | 15.55625 | -0.21278 | 0.481903 | 1 |
| 3 | 6863600 | 6865999 | 52.40885 | 83.74468 | 99.04424 | 41.62305 | 0.100113 | 2.082229 | 1 |
| 3 | 6914400 | 6915799 | 29.27754 | 48.66271 | 56.17207 | 24.36744 | 0.174734 | 1.270377 | 1 |
| 3 | 7010400 | 7013199 | 19.49305 | 33.24117 | 48.49539 | 21.25937 | -0.16034 | 0.985286 | 1 |
| 3 | 7042400 | 7043999 | 16.57252 | 29.10231 | 34.15835 | 15.11437 | 0.157667 | 0.614714 | 1 |
| 3 | 7191800 | 7194599 | 21.19074 | 36.71724 | 46.85274 | 20.55636 | 0.036116 | 0.845493 | 1 |
| 3 | 7199400 | 7200799 | 10.06014 | 18.76516 | 28.91909 | 12.03066 | -0.16701 | 0.148625 | 1 |
| 3 | 7263800 | 7268399 | 43.66322 | 69.99205 | 100.7385 | 42.53179 | -0.17743 | 1.976042 | 1 |
| 3 | 7366400 | 7369799 | 46.88149 | 75.29397 | 86.78698 | 36.74757 | 0.145615 | 1.91627 | 1 |
| 3 | 7548200 | 7550399 | 26.21673 | 44.10991 | 51.77365 | 22.62304 | 0.150703 | 1.128369 | 1 |
| 3 | 7606200 | 7615599 | 135.8985 | 207.6107 | 202.7354 | 84.09047 | 0.322877 | 3.097499 | 1 |
| 3 | 7858600 | 7861599 | 31.28663 | 52.22941 | 53.58087 | 23.30939 | 0.344388 | 1.225292 | 1 |
| 3 | 8224800 | 8225599 | 12.12305 | 22.08458 | 29.5961 | 12.72383 | -0.00118 | 0.302277 | 1 |
| 3 | 8288600 | 8296199 | 201.3839 | 296.8093 | 307.1042 | 126.4689 | 0.193933 | 3.785214 | 1 |
| 3 | 8359000 | 8360999 | 11.16636 | 20.43629 | 31.31717 | 13.38474 | -0.18743 | 0.2753 | 1 |
| 3 | 8391200 | 8394399 | 39.35535 | 63.34203 | 89.85302 | 38.1644 | -0.14941 | 1.839011 | 1 |
| 3 | 8498400 | 8499599 | 19.25977 | 33.5911 | 39.31794 | 17.30434 | 0.161318 | 0.741711 | 1 |
| 3 | 8516200 | 8520399 | 56.23396 | 89.821 | 100.2387 | 42.10122 | 0.183098 | 2.099629 | 1 |
| 3 | 8526600 | 8530799 | 56.4008 | 89.33059 | 136.0735 | 56.58737 | -0.27744 | 2.363395 | 1 |
| 3 | 8636200 | 8643799 | 106.5167 | 164.2123 | 175.0767 | 72.72387 | 0.206366 | 2.906409 | 1 |
| 3 | 8652400 | 8656399 | 53.22695 | 85.55959 | 92.77868 | 39.33463 | 0.235042 | 1.900602 | 1 |
| 3 | 8848400 | 8852599 | 88.91835 | 138.1049 | 157.4766 | 65.39049 | 0.117765 | 2.764354 | 1 |
| 3 | 8988200 | 8989199 | 9.929043 | 18.77559 | 23.2236 | 9.429674 | 0.176643 | 0.044248 | 1 |
| 3 | 9108000 | 9116599 | 36.88368 | 60.44512 | 63.59469 | 27.37029 | 0.303387 | 1.462228 | 1 |
| 3 | 9379600 | 9380799 | 11.77534 | 21.61705 | 28.12092 | 11.96861 | 0.051412 | 0.252666 | 1 |
| 3 | 9481800 | 9484599 | 38.86346 | 63.57516 | 64.84536 | 27.88837 | 0.346705 | 1.490608 | 1 |
| 3 | 9594800 | 9595599 | 11.54689 | 21.37248 | 26.09737 | 10.98394 | 0.153691 | 0.199752 | 1 |
| 3 | 9972600 | 9975399 | 33.23068 | 54.93713 | 61.393 | 26.53361 | 0.220725 | 1.34621 | 1 |
| 3 | 10060400 | 10061399 | 10.63361 | 19.36635 | 30.75779 | 13.22746 | -0.24477 | 0.296009 | 1 |
| 3 | 10593800 | 10598199 | 70.97091 | 111.9576 | 130.6311 | 54.21395 | 0.101983 | 2.466731 | 1 |
| 3 | 10854800 | 10856999 | 31.18605 | 51.12363 | 68.70996 | 29.57571 | -0.04873 | 1.448219 | 1 |
| 3 | 11163400 | 11165199 | 17.1871 | 30.13665 | 38.16346 | 16.86531 | 0.048482 | 0.647308 | 1 |
| 3 | 11310000 | 11311599 | 22.94416 | 38.71578 | 55.442 | 24.24725 | -0.13501 | 1.098054 | 1 |
| 3 | 11344400 | 11346399 | 21.40248 | 36.93736 | 44.71288 | 19.61514 | 0.110678 | 0.877658 | 1 |
| 3 | 11450400 | 11455799 | 54.16885 | 86.31268 | 120.4767 | 50.42985 | -0.14296 | 2.178917 | 1 |
| 3 | 11653000 | 11656999 | 46.33232 | 74.87415 | 77.80812 | 33.22846 | 0.305422 | 1.724586 | 1 |
| 3 | 11802800 | 11805399 | 28.01586 | 46.70656 | 54.47203 | 23.67703 | 0.160087 | 1.233151 | 1 |
| 3 | 11855600 | 11857199 | 23.92646 | 40.80612 | 43.76065 | 19.18393 | 0.281923 | 0.994988 | 1 |
| 3 | 11915800 | 11919799 | 35.76005 | 58.6234 | 71.66095 | 30.84629 | 0.087693 | 1.460151 | 1 |
| 3 | 11945200 | 11945999 | 12.45894 | 23.06737 | 26.61618 | 11.19783 | 0.235501 | 0.208672 | 1 |
| 3 | 12036000 | 12038599 | 30.40862 | 50.46182 | 56.22112 | 24.36177 | 0.225512 | 1.291669 | 1 |
| 3 | 12065200 | 12068199 | 27.31016 | 45.71909 | 53.40152 | 23.27248 | 0.157772 | 1.185249 | 1 |
| 3 | 12268600 | 12270599 | 20.52176 | 35.18948 | 53.35135 | 23.39854 | -0.21469 | 0.941766 | 1 |

| | | | | | | | | | |
|---|---|---|---|---|---|---|---|---|---|
| 3 | 12295400 | 12296399 | 25.40645 | 42.54478 | 54.39575 | 23.71021 | 0.027816 | 1.177895 | 1 |
| 3 | 12545200 | 12546199 | 19.04164 | 33.06353 | 41.09402 | 18.03562 | 0.07465 | 0.807629 | 1 |
| 3 | 12684600 | 12687399 | 39.68733 | 64.25377 | 78.05294 | 33.39253 | 0.083457 | 1.671231 | 1 |
| 3 | 12858200 | 12860199 | 26.70532 | 44.7159 | 52.33968 | 22.81274 | 0.154685 | 1.178139 | 1 |
| 3 | 12957200 | 12959599 | 35.83176 | 58.71755 | 62.41939 | 26.86374 | 0.288451 | 1.458775 | 1 |
| 3 | 12973200 | 12974799 | 20.65773 | 35.32856 | 43.30999 | 18.99059 | 0.09028 | 0.956074 | 1 |
| 3 | 13063600 | 13064599 | 9.597399 | 17.93915 | 25.82408 | 10.70546 | -0.06491 | 0.124899 | 1 |
| 3 | 13080400 | 13081599 | 16.67813 | 29.50919 | 30.37446 | 13.43318 | 0.349645 | 0.521142 | 1 |
| 3 | 13290000 | 13292399 | 41.06659 | 65.78619 | 104.0106 | 43.86916 | -0.31381 | 1.99421 | 1 |
| 3 | 13923200 | 13924199 | 13.90138 | 24.99788 | 29.9348 | 13.07613 | 0.145347 | 0.391159 | 1 |
| 3 | 14036600 | 14037199 | 11.39801 | 20.38756 | 33.79257 | 14.8276 | -0.32495 | 0.418024 | 1 |
| 3 | 14277600 | 14280199 | 18.03171 | 31.12009 | 42.22674 | 18.52449 | -0.05321 | 0.859882 | 1 |
| 3 | 14457400 | 14463999 | 138.0703 | 208.9241 | 232.6416 | 96.23201 | 0.122379 | 3.291112 | 1 |
| 3 | 14576000 | 14577399 | 12.80639 | 22.66961 | 37.20974 | 16.45453 | -0.31969 | 0.504696 | 1 |
| 3 | 14676000 | 14677599 | 12.44199 | 22.21335 | 35.16503 | 15.45415 | -0.261 | 0.437262 | 1 |
| 3 | 14732600 | 14738999 | 91.43446 | 140.866 | 192.4531 | 79.94065 | -0.15178 | 2.915329 | 1 |
| 3 | 14770000 | 14778399 | 245.0439 | 354.7673 | 377.993 | 155.8835 | 0.126922 | 4.064811 | 1 |
| 3 | 14779200 | 14785599 | 55.38254 | 88.53617 | 105.8028 | 44.49164 | 0.085861 | 2.073736 | 1 |
| 3 | 14786600 | 14788399 | 35.36251 | 57.30884 | 82.53635 | 35.33652 | -0.16027 | 1.643193 | 1 |
| 3 | 15093400 | 15093999 | 9.599057 | 17.61801 | 30.89907 | 13.15493 | -0.37653 | 0.246455 | 1 |
| 3 | 15319000 | 15321999 | 24.84665 | 41.13918 | 63.41935 | 27.43821 | -0.24173 | 1.309023 | 1 |
| 3 | 15561400 | 15562599 | 22.55207 | 38.24206 | 52.76207 | 23.1097 | -0.08122 | 1.050683 | 1 |
| 3 | 15812400 | 15813599 | 15.29462 | 27.2781 | 31.92138 | 14.04087 | 0.171651 | 0.456958 | 1 |
| 3 | 15928000 | 15928999 | 22.82776 | 39.27062 | 42.59366 | 18.68363 | 0.26753 | 0.909668 | 1 |
| 3 | 16123400 | 16129799 | 58.60647 | 93.3305 | 108.0151 | 45.17801 | 0.126015 | 2.196647 | 1 |
| 3 | 16131200 | 16135199 | 54.90429 | 87.67416 | 104.187 | 43.7433 | 0.092197 | 2.106894 | 1 |
| 3 | 16281600 | 16296399 | 449.3425 | 612.4729 | 711.6623 | 296.2793 | -0.0855 | 4.975216 | 1 |
| 3 | 16302400 | 16307799 | 28.93002 | 47.81752 | 59.4021 | 25.66955 | 0.068478 | 1.334629 | 1 |
| 3 | 16321400 | 16323199 | 13.00108 | 23.09994 | 36.20179 | 15.97454 | -0.25044 | 0.475243 | 1 |
| 3 | 16337200 | 16339799 | 21.29653 | 36.08373 | 57.06079 | 24.98638 | -0.27739 | 1.057555 | 1 |
| 3 | 16410000 | 16412399 | 33.63947 | 55.3578 | 62.29416 | 26.85769 | 0.209005 | 1.403822 | 1 |
| 3 | 16544200 | 16547599 | 30.84126 | 51.24608 | 56.86052 | 24.66087 | 0.231523 | 1.277563 | 1 |
| 3 | 16562600 | 16563399 | 15.10984 | 26.99005 | 28.99473 | 12.73703 | 0.295769 | 0.440745 | 1 |
| 3 | 16631600 | 16633999 | 33.26139 | 53.60566 | 91.66119 | 39.00783 | -0.41585 | 1.790807 | 1 |
| 3 | 16752200 | 16754999 | 46.10539 | 73.99319 | 108.4788 | 45.88115 | -0.20204 | 1.944884 | 1 |
| 3 | 16846600 | 16849599 | 81.34707 | 127.309 | 132.0372 | 54.77371 | 0.262867 | 2.625154 | 1 |
| 3 | 16850400 | 16856599 | 124.6703 | 189.5719 | 209.2028 | 86.66588 | 0.141193 | 3.187588 | 1 |
| 3 | 17019600 | 17021799 | 39.80847 | 64.65385 | 69.30913 | 29.72092 | 0.267629 | 1.603673 | 1 |
| 3 | 17309000 | 17309999 | 17.35628 | 30.60009 | 31.44112 | 13.9148 | 0.350306 | 0.566243 | 1 |
| 3 | 17389800 | 17390999 | 11.06659 | 20.32672 | 29.24258 | 12.43703 | -0.09194 | 0.247893 | 1 |
| 3 | 17401000 | 17403199 | 23.702 | 40.60643 | 42.64664 | 18.70238 | 0.312845 | 0.951255 | 1 |
| 3 | 17478200 | 17480599 | 19.41002 | 33.47288 | 46.87951 | 20.56514 | -0.09895 | 0.880175 | 1 |
| 3 | 17495200 | 17495999 | 15.41287 | 27.02068 | 36.91112 | 16.30935 | -0.05993 | 0.641995 | 1 |
| 3 | 17703000 | 17703999 | 10.08841 | 18.7313 | 29.38097 | 12.31354 | -0.20062 | 0.178841 | 1 |
| 3 | 17799400 | 17800599 | 14.92606 | 26.69905 | 30.41946 | 13.34716 | 0.212443 | 0.432941 | 1 |
| 3 | 17818600 | 17821599 | 27.56367 | 46.35293 | 52.33711 | 22.86577 | 0.206439 | 1.138293 | 1 |
| 3 | 18034800 | 18040199 | 60.08986 | 95.29903 | 127.9873 | 53.28279 | -0.09429 | 2.328016 | 1 |

| | | | | | | | | | |
|---|---|---|---|---|---|---|---|---|---|
| 3 | 18248800 | 18256399 | 62.22331 | 98.70482 | 153.414 | 63.88268 | -0.30449 | 2.329682 | 1 |
| 3 | 18447800 | 18450399 | 28.97016 | 48.35314 | 61.82916 | 26.89443 | 0.027578 | 1.228348 | 1 |
| 3 | 18661800 | 18667999 | 81.43924 | 127.4284 | 145.3397 | 60.29326 | 0.125666 | 2.629045 | 1 |
| 3 | 18777000 | 18779599 | 29.74529 | 49.60528 | 54.03293 | 23.49081 | 0.258256 | 1.233397 | 1 |
| 3 | 19026000 | 19027199 | 19.29323 | 33.00534 | 45.90258 | 20.12789 | -0.09088 | 0.952053 | 1 |
| 3 | 19031400 | 19037199 | 26.99792 | 44.5294 | 68.49355 | 29.55891 | -0.23928 | 1.363953 | 1 |
| 3 | 19148600 | 19151199 | 32.80706 | 53.4453 | 77.33481 | 33.20946 | -0.16068 | 1.546099 | 1 |
| 3 | 19324200 | 19334199 | 60.38268 | 95.80527 | 133.399 | 55.56008 | -0.14572 | 2.315946 | 1 |
| 3 | 19516200 | 19517999 | 15.55039 | 27.90165 | 29.32114 | 12.83187 | 0.330927 | 0.415788 | 1 |
| 3 | 19867600 | 19871199 | 47.28387 | 75.92415 | 93.88348 | 39.73959 | 0.044037 | 1.923875 | 1 |
| 3 | 19890400 | 19893199 | 17.52263 | 30.54762 | 43.10852 | 18.96727 | -0.10716 | 0.747827 | 1 |
| 3 | 19895000 | 19898999 | 45.74443 | 73.66986 | 76.21039 | 32.39826 | 0.306173 | 1.822663 | 1 |
| 3 | 19971800 | 19979199 | 47.50868 | 76.48858 | 83.2934 | 35.39423 | 0.231771 | 1.83888 | 1 |
| 3 | 20538400 | 20539399 | 12.35786 | 22.59827 | 33.888 | 14.49974 | -0.15803 | 0.291744 | 1 |
| 3 | 20627400 | 20631199 | 48.33888 | 77.53808 | 105.9001 | 44.75575 | -0.10089 | 1.9617 | 1 |
| 3 | 20676600 | 20678599 | 16.1213 | 28.07538 | 40.58417 | 17.84489 | -0.14188 | 0.749804 | 1 |
| 3 | 20757600 | 20759199 | 20.45767 | 35.6933 | 37.11407 | 16.33811 | 0.331323 | 0.738438 | 1 |
| 3 | 20794600 | 20795799 | 16.5168 | 28.65839 | 46.34948 | 20.33642 | -0.30379 | 0.811611 | 1 |
| 3 | 20886800 | 20888599 | 18.07464 | 31.23544 | 43.24169 | 18.97051 | -0.08159 | 0.85044 | 1 |
| 3 | 21254200 | 21255399 | 16.30855 | 28.64934 | 36.1687 | 16.0089 | 0.053051 | 0.611272 | 1 |
| 3 | 21435200 | 21437799 | 23.98108 | 40.90139 | 40.67048 | 17.82933 | 0.390504 | 0.990061 | 1 |
| 3 | 21504200 | 21506799 | 34.66357 | 56.35369 | 77.71587 | 33.34809 | -0.09366 | 1.576658 | 1 |
| 3 | 21744800 | 21747799 | 25.04392 | 42.62876 | 45.30182 | 19.85644 | 0.294513 | 1.017916 | 1 |
| 3 | 21968600 | 21970199 | 16.68531 | 29.68033 | 33.76076 | 14.88287 | 0.209989 | 0.484149 | 1 |
| 3 | 22052000 | 22053799 | 24.74435 | 42.40252 | 45.74813 | 20.06291 | 0.274084 | 0.954867 | 1 |
| 3 | 22288400 | 22296599 | 119.7895 | 183.2884 | 212.7926 | 88.294 | 0.075244 | 3.061172 | 1 |
| 3 | 22428000 | 22433599 | 44.62378 | 71.42284 | 102.7062 | 43.26318 | -0.17873 | 2.024543 | 1 |
| 3 | 22436200 | 22437599 | 10.72832 | 20.24031 | 25.82462 | 10.53702 | 0.125864 | 0.073794 | 1 |
| 3 | 22467200 | 22469199 | 21.99707 | 38.00461 | 42.0377 | 18.44217 | 0.240557 | 0.86696 | 1 |
| 3 | 22514600 | 22519199 | 14.17661 | 25.41176 | 28.43395 | 12.45391 | 0.240028 | 0.409363 | 1 |
| 3 | 22578000 | 22579999 | 19.48707 | 33.94087 | 36.27871 | 15.95295 | 0.291549 | 0.757026 | 1 |
| 3 | 22671200 | 22674399 | 10.91236 | 19.90736 | 30.72902 | 13.18742 | -0.2021 | 0.287706 | 1 |
| 3 | 22675200 | 22677199 | 31.19639 | 51.30495 | 62.57785 | 26.97278 | 0.092754 | 1.405973 | 1 |
| 3 | 22712600 | 22715999 | 50.40147 | 80.79715 | 93.74313 | 39.5811 | 0.132968 | 1.978871 | 1 |
| 3 | 22728000 | 22730399 | 10.69675 | 19.37915 | 32.79168 | 14.19522 | -0.34172 | 0.332408 | 1 |
| 3 | 22752800 | 22765199 | 213.0141 | 311.2012 | 364.4798 | 150.1615 | 0.003175 | 3.92629 | 1 |
| 3 | 22783800 | 22786599 | 67.91113 | 106.1015 | 163.4254 | 67.80647 | -0.3089 | 2.653593 | 1 |
| 3 | 22787400 | 22789599 | 20.25077 | 35.34924 | 38.19856 | 16.82058 | 0.276196 | 0.733664 | 1 |
| 3 | 22954000 | 22966399 | 44.32283 | 71.6483 | 76.4231 | 32.64963 | 0.268373 | 1.711441 | 1 |
| 3 | 22967200 | 22971599 | 78.95624 | 124.1549 | 132.8206 | 55.09936 | 0.22347 | 2.536191 | 1 |
| 3 | 23008800 | 23009799 | 16.54551 | 28.91771 | 35.90992 | 15.82599 | 0.076812 | 0.693361 | 1 |
| 3 | 23324800 | 23327599 | 51.22336 | 82.01632 | 95.12857 | 40.09291 | 0.131368 | 2.017467 | 1 |
| 3 | 23328400 | 23331599 | 86.4546 | 134.2258 | 164.4354 | 68.28406 | 0.013948 | 2.770204 | 1 |
| 3 | 23361400 | 23363199 | 41.35815 | 66.45552 | 94.15007 | 39.89379 | -0.15056 | 1.896412 | 1 |
| 3 | 23562200 | 23564199 | 21.5166 | 37.34189 | 39.91419 | 17.51297 | 0.291085 | 0.822655 | 1 |
| 3 | 23963800 | 23966399 | 36.33215 | 59.15509 | 68.03675 | 29.21073 | 0.16917 | 1.555307 | 1 |
| 3 | 23968600 | 23971599 | 42.11015 | 67.78243 | 96.75237 | 41.10062 | -0.15827 | 1.84203 | 1 |

| | | | | | | | | | |
|---|---|---|---|---|---|---|---|---|---|
| 3 | 24076200 | 24078999 | 29.69357 | 48.46555 | 74.20236 | 31.88647 | -0.23992 | 1.510062 | 1 |
| 3 | 24292600 | 24294199 | 26.95597 | 44.98041 | 53.93397 | 23.4566 | 0.120147 | 1.218279 | 1 |
| 3 | 24392400 | 24393199 | 9.708242 | 18.66455 | 22.44804 | 8.761869 | 0.26844 | -0.0855 | 1 |
| 3 | 24708400 | 24709999 | 23.67724 | 40.16545 | 50.63213 | 22.17937 | 0.048584 | 1.049413 | 1 |
| 3 | 25022400 | 25023599 | 27.03099 | 44.58451 | 67.27588 | 29.03357 | -0.21164 | 1.364239 | 1 |
| 3 | 25087800 | 25088999 | 23.29374 | 38.9025 | 58.08862 | 25.27708 | -0.1953 | 1.201927 | 1 |
| 3 | 25154600 | 25156199 | 14.40735 | 25.65077 | 31.44721 | 13.8527 | 0.103313 | 0.459576 | 1 |
| 3 | 25276400 | 25277399 | 19.20376 | 33.03414 | 44.29165 | 19.42841 | -0.03703 | 0.898075 | 1 |
| 3 | 25303800 | 25305599 | 35.69899 | 57.88456 | 76.09095 | 32.58965 | -0.02811 | 1.631032 | 1 |
| 3 | 25317000 | 25320799 | 73.34076 | 115.3676 | 137.7311 | 57.13916 | 0.065804 | 2.525776 | 1 |
| 3 | 25468000 | 25471799 | 45.88923 | 74.06284 | 80.20757 | 34.19642 | 0.243909 | 1.762552 | 1 |
| 3 | 25532600 | 25533599 | 16.92544 | 29.05411 | 45.91834 | 20.14036 | -0.27368 | 0.911796 | 1 |
| 3 | 25903600 | 25904199 | 9.116894 | 17.24926 | 29.88243 | 12.1194 | -0.30446 | 0.049888 | 1 |
| 3 | 26167000 | 26169199 | 18.64311 | 32.36007 | 47.93669 | 21.03409 | -0.17764 | 0.820198 | 1 |
| 3 | 26629800 | 26633799 | 82.85823 | 129.5138 | 137.6311 | 57.10152 | 0.226433 | 2.64822 | 1 |
| 3 | 26821200 | 26822599 | 20.58782 | 35.40858 | 44.2391 | 19.40525 | 0.064261 | 0.90441 | 1 |
| 3 | 26998000 | 27000199 | 25.76345 | 42.8293 | 63.69261 | 27.63315 | -0.18962 | 1.264535 | 1 |
| 3 | 27071000 | 27082399 | 200.5872 | 291.3497 | 409.311 | 168.7333 | -0.26727 | 4.013466 | 1 |
| 3 | 28251600 | 28255999 | 66.02245 | 104.4434 | 127.2114 | 52.86961 | 0.043337 | 2.396906 | 1 |
| 3 | 28381200 | 28382799 | 16.28963 | 28.5478 | 36.79629 | 16.25383 | 0.023294 | 0.65038 | 1 |
| 3 | 28539400 | 28543399 | 31.78103 | 51.81012 | 82.79297 | 35.56241 | -0.30289 | 1.535028 | 1 |
| 3 | 28834800 | 28836399 | 19.82008 | 33.55893 | 52.17133 | 22.84055 | -0.25282 | 1.052693 | 1 |
| 3 | 28903800 | 28910199 | 89.68473 | 140.2753 | 150.9553 | 62.6254 | 0.209021 | 2.643214 | 1 |
| 3 | 28920200 | 28921399 | 18.27513 | 31.57338 | 43.25496 | 18.97617 | -0.0666 | 0.854739 | 1 |
| 3 | 28984400 | 28987199 | 67.88632 | 107.5927 | 112.3016 | 46.72173 | 0.267922 | 2.349803 | 1 |
| 3 | 29147600 | 29148599 | 12.23361 | 22.32076 | 26.09204 | 11.19553 | 0.196678 | 0.286573 | 1 |
| 3 | 29725600 | 29727599 | 27.18001 | 45.58678 | 52.84465 | 23.05711 | 0.168653 | 1.165112 | 1 |
| 3 | 29835600 | 29838599 | 37.16367 | 60.48734 | 71.57497 | 30.72582 | 0.12788 | 1.563999 | 1 |
| 3 | 29963000 | 29967399 | 50.63117 | 81.57449 | 83.04196 | 35.32424 | 0.330076 | 1.820921 | 1 |
| 3 | 30423600 | 30424599 | 13.50581 | 24.60255 | 25.00936 | 10.75248 | 0.395172 | 0.303531 | 1 |
| 3 | 30632600 | 30634199 | 19.09082 | 33.28486 | 38.31975 | 16.86134 | 0.185078 | 0.744429 | 1 |
| 3 | 31275000 | 31281999 | 49.43594 | 78.78492 | 117.0496 | 49.00708 | -0.23253 | 2.168193 | 1 |
| 3 | 31524000 | 31525199 | 13.03015 | 22.95361 | 37.35974 | 16.53696 | -0.31012 | 0.562219 | 1 |
| 3 | 31593800 | 31599399 | 40.05944 | 64.79026 | 68.86744 | 29.44071 | 0.274276 | 1.686198 | 1 |
| 3 | 31690400 | 31695399 | 11.60377 | 21.63263 | 24.63925 | 10.25971 | 0.26535 | 0.156059 | 1 |
| 3 | 31880000 | 31881799 | 26.73731 | 44.85635 | 50.7082 | 22.12869 | 0.204717 | 1.157123 | 1 |
| 3 | 32056000 | 32061399 | 55.26653 | 88.5236 | 95.98468 | 40.47706 | 0.229042 | 2.010005 | 1 |
| 3 | 32195200 | 32197399 | 10.28655 | 19.1203 | 31.02105 | 12.98137 | -0.24743 | 0.178783 | 1 |
| 3 | 32221800 | 32222999 | 11.59116 | 21.07567 | 30.70804 | 13.2332 | -0.12308 | 0.313377 | 1 |
| 3 | 32693200 | 32695199 | 27.16943 | 45.73606 | 49.62845 | 21.68901 | 0.263623 | 1.125121 | 1 |
| 3 | 32854000 | 32860599 | 110.8649 | 169.7141 | 203.0161 | 84.24675 | 0.032673 | 3.047954 | 1 |
| 3 | 33254600 | 33255599 | 16.90994 | 29.57813 | 36.35899 | 16.03466 | 0.091403 | 0.683902 | 1 |
| 3 | 33303400 | 33305599 | 19.71555 | 34.31275 | 38.94969 | 17.1193 | 0.205249 | 0.775778 | 1 |
| 3 | 33336400 | 33341999 | 93.38394 | 144.3865 | 172.286 | 71.56361 | 0.047307 | 2.847821 | 1 |
| 3 | 33373400 | 33382199 | 115.5085 | 176.6904 | 202.8189 | 84.15037 | 0.091293 | 3.064943 | 1 |
| 3 | 33624000 | 33627999 | 41.37022 | 67.4126 | 72.44503 | 31.11257 | 0.26763 | 1.555205 | 1 |
| 3 | 33672600 | 33673799 | 18.10456 | 31.68912 | 35.65121 | 15.73151 | 0.21845 | 0.676737 | 1 |

| | | | | | | | | | |
|---|---|---|---|---|---|---|---|---|---|
| 3 | 34566600 | 34573599 | 79.88717 | 125.562 | 134.6066 | 55.83698 | 0.219955 | 2.545172 | 1 |
| 3 | 35184400 | 35188599 | 40.3556 | 64.89944 | 96.32045 | 40.86348 | -0.21599 | 1.868481 | 1 |
| 3 | 35242600 | 35244399 | 14.89246 | 26.59073 | 30.89488 | 13.57648 | 0.18278 | 0.444207 | 1 |
| 3 | 35345000 | 35347399 | 14.46781 | 25.69967 | 32.49643 | 14.34247 | 0.057126 | 0.477147 | 1 |
| 3 | 35357600 | 35361799 | 31.81066 | 53.24616 | 56.40914 | 24.58114 | 0.298059 | 1.20127 | 1 |
| 3 | 35633800 | 35638799 | 66.849 | 106.0132 | 127.2615 | 52.97794 | 0.067291 | 2.333132 | 1 |
| 3 | 35666000 | 35667799 | 19.76853 | 34.36369 | 39.44828 | 17.32537 | 0.188839 | 0.791955 | 1 |
| 3 | 35683800 | 35691599 | 27.89525 | 46.83244 | 49.11 | 21.44091 | 0.312746 | 1.152378 | 1 |
| 3 | 35819200 | 35827999 | 73.11799 | 115.7516 | 129.0412 | 53.64967 | 0.171686 | 2.385761 | 1 |
| 3 | 36176400 | 36183199 | 92.39515 | 142.901 | 173.8242 | 72.20241 | 0.019929 | 2.842454 | 1 |
| 3 | 36467800 | 36478799 | 188.1266 | 277.0918 | 343.1477 | 141.311 | -0.06607 | 3.795285 | 1 |
| 3 | 36803600 | 36806599 | 21.04312 | 36.68441 | 39.75145 | 17.49003 | 0.271998 | 0.757503 | 1 |
| 3 | 37441200 | 37445599 | 30.65253 | 50.61528 | 65.08076 | 28.10458 | 0.018822 | 1.355868 | 1 |
| 3 | 37535200 | 37535999 | 15.25538 | 26.60617 | 41.12359 | 18.09868 | -0.23763 | 0.727654 | 1 |
| 3 | 38504000 | 38506599 | 33.58812 | 54.46676 | 79.91463 | 34.2296 | -0.18615 | 1.62659 | 1 |
| 3 | 38816400 | 38817199 | 9.538701 | 18.00437 | 23.79983 | 9.701261 | 0.076867 | 0.05605 | 1 |
| 3 | 38857200 | 38858199 | 14.45861 | 25.51311 | 40.80232 | 18.05467 | -0.28427 | 0.557044 | 1 |
| 3 | 39116600 | 39119199 | 19.71022 | 34.30911 | 38.8495 | 17.07703 | 0.208643 | 0.770666 | 1 |
| 3 | 39153800 | 39154999 | 19.08711 | 32.86495 | 55.00569 | 24.12879 | -0.35565 | 0.900772 | 1 |
| 3 | 39228200 | 39235599 | 172.4642 | 256.5854 | 296.0514 | 121.978 | 0.049399 | 3.62252 | 1 |
| 3 | 39499600 | 39505399 | 122.2078 | 186.1658 | 212.15 | 87.93472 | 0.097035 | 3.150468 | 1 |
| 3 | 39717200 | 39717999 | 13.22312 | 23.65123 | 32.10907 | 14.08935 | -0.03894 | 0.425899 | 1 |
| 3 | 39911600 | 39912599 | 10.0323 | 18.25632 | 30.30266 | 13.04503 | -0.30884 | 0.295313 | 1 |
| 3 | 40205000 | 40205999 | 16.63575 | 29.13176 | 36.83822 | 16.26168 | 0.050787 | 0.665458 | 1 |
| 3 | 40406000 | 40406799 | 9.130077 | 17.03978 | 26.33926 | 10.94711 | -0.16999 | 0.130876 | 1 |
| 3 | 40544800 | 40546799 | 24.08251 | 41.16701 | 44.71874 | 19.60758 | 0.263786 | 0.973127 | 1 |
| 3 | 40662800 | 40664599 | 30.82249 | 50.36902 | 68.15334 | 29.29998 | -0.06086 | 1.492199 | 1 |
| 3 | 40812800 | 40820799 | 177.5108 | 262.9551 | 309.1276 | 127.3097 | 0.016698 | 3.698684 | 1 |
| 3 | 40880000 | 40884799 | 98.78166 | 152.3583 | 180.4478 | 74.95479 | 0.055345 | 2.896634 | 1 |
| 3 | 41509000 | 41512799 | 48.78608 | 77.71169 | 116.4329 | 48.71627 | -0.24541 | 2.182925 | 1 |
| 3 | 41723200 | 41724999 | 34.29453 | 55.55235 | 84.37368 | 36.11081 | -0.23735 | 1.653185 | 1 |
| 3 | 41779800 | 41781199 | 17.53689 | 30.89592 | 33.90795 | 15.00873 | 0.255241 | 0.579337 | 1 |
| 3 | 41792000 | 41798599 | 142.4814 | 215.3817 | 229.3949 | 94.85887 | 0.18529 | 3.313109 | 1 |
| 3 | 41832400 | 41834799 | 29.13491 | 48.33444 | 55.57416 | 24.07672 | 0.180216 | 1.291505 | 1 |
| 3 | 41947600 | 41949799 | 16.35351 | 28.74195 | 33.57461 | 14.86667 | 0.16442 | 0.597856 | 1 |
| 3 | 42069800 | 42071399 | 19.75794 | 34.24678 | 40.04329 | 17.56895 | 0.161845 | 0.82624 | 1 |
| 3 | 42278600 | 42279999 | 25.82967 | 42.78873 | 62.37339 | 26.99523 | -0.16119 | 1.30584 | 1 |
| 3 | 42442400 | 42444199 | 18.17542 | 31.87873 | 36.56446 | 16.16259 | 0.190666 | 0.646277 | 1 |
| 3 | 42661400 | 42664199 | 30.03992 | 49.93818 | 52.95005 | 22.97237 | 0.296912 | 1.267541 | 1 |
| 3 | 42780000 | 42782799 | 28.56103 | 46.63313 | 71.76802 | 30.84427 | -0.24697 | 1.501396 | 1 |
| 3 | 42966400 | 42967199 | 10.83721 | 20.1017 | 28.79828 | 12.08561 | -0.07224 | 0.188833 | 1 |
| 3 | 43036400 | 43039799 | 52.58623 | 84.1621 | 93.76868 | 39.49503 | 0.188611 | 2.032764 | 1 |
| 3 | 43141000 | 43141399 | 12.14037 | 22.58094 | 23.16766 | 9.68463 | 0.410861 | 0.172873 | 1 |
| 3 | 43260200 | 43260999 | 19.13474 | 32.64359 | 48.49343 | 21.25921 | -0.18631 | 0.978994 | 1 |
| 3 | 43511800 | 43513399 | 18.16713 | 31.89563 | 33.99872 | 15.04098 | 0.295903 | 0.626637 | 1 |
| 3 | 43567200 | 43568999 | 15.20516 | 27.17407 | 39.79352 | 17.47333 | -0.14864 | 0.45757 | 1 |
| 3 | 43960800 | 43961199 | 10.4452 | 18.85189 | 32.84444 | 14.28556 | -0.38769 | 0.35475 | 1 |

| | | | | | | | | | |
|---|---|---|---|---|---|---|---|---|---|
| 3 | 44033600 | 44035799 | 24.62724 | 41.68878 | 47.98627 | 21.00688 | 0.179034 | 1.065332 | 1 |
| 3 | 44053200 | 44054399 | 20.51163 | 34.83021 | 49.54346 | 21.70906 | -0.1248 | 1.029915 | 1 |
| 3 | 44096000 | 44097399 | 13.69054 | 24.25277 | 42.39335 | 18.74399 | -0.40965 | 0.512857 | 1 |
| 3 | 44201200 | 44202599 | 22.32162 | 38.36163 | 38.08384 | 16.70489 | 0.394492 | 0.9114 | 1 |
| 3 | 44232600 | 44233199 | 9.358056 | 18.04955 | 19.68767 | 7.613468 | 0.419976 | -0.1298 | 1 |
| 3 | 44268200 | 44270599 | 20.32484 | 35.33813 | 40.3935 | 17.74284 | 0.194894 | 0.791324 | 1 |
| 3 | 44354800 | 44355999 | 22.65074 | 38.54279 | 44.35032 | 19.43888 | 0.180231 | 1.011712 | 1 |
| 3 | 44601000 | 44610599 | 161.3078 | 241.5333 | 254.4806 | 104.9626 | 0.188828 | 3.501527 | 1 |
| 3 | 44672000 | 44673599 | 11.67032 | 21.24992 | 32.20772 | 13.85471 | -0.17833 | 0.30659 | 1 |
| 3 | 44693600 | 44694399 | 10.58128 | 20.0696 | 23.76459 | 9.587773 | 0.247216 | 0.031635 | 1 |
| 3 | 44761600 | 44763799 | 15.15301 | 26.5584 | 40.36588 | 17.83205 | -0.2135 | 0.646781 | 1 |
| 3 | 44887200 | 44887599 | 8.76319 | 16.79881 | 24.05802 | 9.468856 | 0.007218 | -0.06472 | 1 |
| 3 | 45451600 | 45454199 | 39.07844 | 63.56448 | 68.37063 | 29.34364 | 0.264566 | 1.574201 | 1 |
| 3 | 45531600 | 45533999 | 17.50301 | 30.22337 | 44.30874 | 19.43816 | -0.16408 | 0.857878 | 1 |
| 3 | 45828800 | 45833399 | 55.66371 | 88.92576 | 99.8934 | 41.96661 | 0.173905 | 2.09251 | 1 |
| 3 | 45946000 | 45946599 | 11.75951 | 21.52947 | 28.71797 | 12.26796 | 0.012013 | 0.271845 | 1 |
| 3 | 46055600 | 46056399 | 16.02636 | 27.95914 | 40.56422 | 17.85584 | -0.14677 | 0.725945 | 1 |
| 3 | 46069000 | 46070199 | 9.257165 | 17.76575 | 24.09166 | 9.451482 | 0.089782 | -0.0693 | 1 |
| 3 | 46427200 | 46429599 | 20.86974 | 36.20509 | 36.44989 | 15.99277 | 0.377088 | 0.81919 | 1 |
| 3 | 46609000 | 46609999 | 13.06986 | 23.88884 | 28.01311 | 11.99321 | 0.194158 | 0.287519 | 1 |
| 3 | 47031400 | 47033199 | 5.657138 | 11.01934 | 21.96996 | 8.014632 | -0.36213 | -0.36268 | 1 |
| 3 | 47332000 | 47333199 | 16.64056 | 28.91399 | 45.27311 | 19.87825 | -0.25707 | 0.79144 | 1 |
| 3 | 47336200 | 47338999 | 14.69392 | 25.68414 | 41.52289 | 18.30126 | -0.3021 | 0.689974 | 1 |
| 3 | 47340600 | 47342399 | 19.52291 | 33.90792 | 42.55669 | 18.68059 | 0.06044 | 0.805021 | 1 |
| 3 | 47361600 | 47362799 | 11.50519 | 20.89325 | 33.30159 | 14.37329 | -0.25322 | 0.326563 | 1 |
| 3 | 47404600 | 47407199 | 29.77835 | 49.68429 | 52.43869 | 22.80463 | 0.303543 | 1.225143 | 1 |
| 3 | 47433400 | 47441399 | 131.1737 | 199.0162 | 227.3426 | 94.11772 | 0.08878 | 3.236547 | 1 |
| 3 | 47546000 | 47547399 | 27.58043 | 45.64339 | 63.67684 | 27.54037 | -0.09824 | 1.322006 | 1 |
| 3 | 47767200 | 47769599 | 42.91916 | 69.24564 | 83.17834 | 35.44737 | 0.094167 | 1.768516 | 1 |
| 3 | 48098400 | 48100799 | 24.95395 | 41.38451 | 63.009 | 27.29149 | -0.22371 | 1.288688 | 1 |
| 3 | 48126800 | 48127599 | 14.01288 | 24.61089 | 40.9195 | 18.10819 | -0.34209 | 0.611484 | 1 |
| 3 | 48304600 | 48305999 | 14.18199 | 25.05028 | 38.15035 | 16.87779 | -0.21364 | 0.537367 | 1 |
| 3 | 48341600 | 48343599 | 20.88501 | 36.04777 | 39.0665 | 17.13817 | 0.269749 | 0.867931 | 1 |
| 3 | 48387800 | 48390999 | 42.31958 | 68.27579 | 84.68087 | 36.08599 | 0.048043 | 1.77022 | 1 |
| 3 | 48404800 | 48405799 | 12.37068 | 22.07203 | 36.25048 | 15.9397 | -0.31409 | 0.445289 | 1 |
| 3 | 48440200 | 48441799 | 17.93481 | 31.9046 | 33.2043 | 14.63693 | 0.337705 | 0.492207 | 1 |
| 3 | 49095400 | 49097199 | 17.1847 | 30.28802 | 33.40955 | 14.78689 | 0.248183 | 0.571545 | 1 |
| 3 | 49246400 | 49252599 | 67.52605 | 107.0856 | 120.7426 | 50.26357 | 0.157534 | 2.333186 | 1 |
| 3 | 49335400 | 49338599 | 56.95187 | 90.64663 | 110.739 | 46.28276 | 0.047307 | 2.215036 | 1 |
| 3 | 49418600 | 49419199 | 9.739748 | 18.41118 | 24.34244 | 9.891305 | 0.080695 | 0.050118 | 1 |
| 3 | 49429800 | 49431599 | 29.40904 | 48.51099 | 59.19922 | 25.54894 | 0.09381 | 1.362846 | 1 |
| 3 | 49554600 | 49556599 | 20.75368 | 35.3809 | 52.75 | 23.12397 | -0.19179 | 0.996055 | 1 |
| 3 | 49698200 | 49702799 | 21.09848 | 35.76776 | 52.19515 | 22.85956 | -0.16181 | 1.047056 | 1 |
| 3 | 49774800 | 49777799 | 25.58437 | 42.02644 | 68.6115 | 29.55884 | -0.32726 | 1.410849 | 1 |
| 3 | 49952600 | 49954599 | 27.8704 | 47.38081 | 47.93729 | 21.0095 | 0.364859 | 1.042365 | 1 |
| 3 | 50338400 | 50339199 | 9.462731 | 17.61325 | 26.33146 | 10.99376 | -0.12727 | 0.151979 | 1 |
| 3 | 50656400 | 50658999 | 37.71472 | 61.43837 | 69.60064 | 29.88788 | 0.191441 | 1.549674 | 1 |

| | | | | | | | | | |
|---|---|---|---|---|---|---|---|---|---|
| 3 | 50918400 | 50920999 | 24.77124 | 41.17469 | 61.95505 | 26.87699 | -0.2065 | 1.26113 | 1 |
| 3 | 51888200 | 51890799 | 21.80623 | 36.92653 | 53.7971 | 23.55322 | -0.15948 | 1.062112 | 1 |
| 3 | 51967200 | 51968599 | 13.85204 | 24.66744 | 36.59254 | 16.11631 | -0.17008 | 0.464659 | 1 |
| 3 | 52594600 | 52597199 | 10.19483 | 18.98223 | 27.81146 | 11.60515 | -0.0981 | 0.158446 | 1 |
| 3 | 52825200 | 52828999 | 48.73421 | 78.15971 | 103.0596 | 43.54468 | -0.05049 | 1.96752 | 1 |
| 3 | 52866600 | 52868799 | 16.86406 | 29.5209 | 35.29523 | 15.57579 | 0.131198 | 0.668605 | 1 |
| 3 | 53042400 | 53045399 | 30.86631 | 50.23995 | 79.83561 | 34.2725 | -0.29635 | 1.555909 | 1 |
| 3 | 53265000 | 53267399 | 17.49959 | 30.4489 | 47.90285 | 21.05129 | -0.26372 | 0.780155 | 1 |
| 3 | 53291600 | 53295199 | 34.67337 | 57.38922 | 62.17075 | 26.89144 | 0.265532 | 1.336584 | 1 |
| 3 | 53632600 | 53635599 | 25.87207 | 42.9958 | 63.20574 | 27.41574 | -0.17308 | 1.26682 | 1 |
| 3 | 54034600 | 54037199 | 28.3113 | 47.40822 | 49.67922 | 21.65409 | 0.313772 | 1.180252 | 1 |
| 3 | 54425400 | 54427399 | 26.95391 | 46.67976 | 44.01829 | 19.31247 | 0.470565 | 0.871537 | 1 |
| 3 | 54484200 | 54496199 | 101.6787 | 156.006 | 213.6519 | 88.69616 | -0.15979 | 3.000853 | 1 |
| 3 | 54611200 | 54611799 | 9.224532 | 17.50088 | 22.82594 | 9.203915 | 0.10991 | 0.015368 | 1 |
| 3 | 54701000 | 54705599 | 45.12335 | 72.15435 | 111.674 | 46.97792 | -0.28614 | 2.056304 | 1 |
| 3 | 54717800 | 54719799 | 16.53125 | 28.83494 | 47.33047 | 20.83198 | -0.32429 | 0.739799 | 1 |
| 3 | 54929000 | 54930799 | 20.95522 | 36.06993 | 50.10128 | 21.97721 | -0.08781 | 0.905416 | 1 |
| 3 | 55128600 | 55131999 | 34.48545 | 55.78109 | 84.59263 | 36.16603 | -0.23709 | 1.683392 | 1 |
| 3 | 55408800 | 55411399 | 15.37985 | 27.2567 | 31.89327 | 14.10026 | 0.166512 | 0.497333 | 1 |
| 3 | 55487400 | 55494199 | 128.4293 | 193.2319 | 258.7287 | 106.8418 | -0.1505 | 3.403993 | 1 |
| 3 | 55693200 | 55695599 | 18.07642 | 31.59872 | 37.21523 | 16.40361 | 0.152666 | 0.699251 | 1 |
| 3 | 56041800 | 56044399 | 52.52988 | 83.86576 | 109.9166 | 46.13575 | -0.04917 | 2.113143 | 1 |
| 3 | 56048200 | 56049199 | 14.86082 | 26.63625 | 34.43497 | 15.08327 | 0.032578 | 0.425594 | 1 |
| 3 | 56231400 | 56233799 | 33.283 | 54.56522 | 67.26337 | 28.95393 | 0.075654 | 1.451128 | 1 |
| 3 | 56420800 | 56421799 | 15.6132 | 27.23483 | 42.13488 | 18.54571 | -0.23894 | 0.729492 | 1 |
| 3 | 56511000 | 56511599 | 10.04011 | 18.492 | 31.4227 | 13.31443 | -0.3265 | 0.230551 | 1 |
| 3 | 56573800 | 56575399 | 14.42008 | 25.53401 | 34.97368 | 15.46477 | -0.06008 | 0.505135 | 1 |
| 3 | 56810800 | 56811799 | 30.8218 | 50.81227 | 56.04732 | 24.18279 | 0.238841 | 1.370412 | 1 |
| 3 | 57162400 | 57165599 | 42.15144 | 67.68771 | 95.98575 | 40.63381 | -0.15303 | 1.917088 | 1 |
| 3 | 57235000 | 57238799 | 28.26764 | 46.53534 | 65.73537 | 28.34586 | -0.11786 | 1.387955 | 1 |
| 3 | 57606800 | 57608399 | 10.83041 | 19.71058 | 31.28133 | 13.46473 | -0.24443 | 0.304044 | 1 |
| 3 | 57919800 | 57920999 | 14.724 | 25.79013 | 41.97537 | 18.53312 | -0.31172 | 0.658058 | 1 |
| 3 | 57958600 | 57961599 | 31.42056 | 51.7389 | 63.94727 | 27.57776 | 0.074374 | 1.389705 | 1 |
| 3 | 58759400 | 58760599 | 13.96711 | 24.44247 | 39.0381 | 17.2232 | -0.28433 | 0.665522 | 1 |
| 3 | 59279600 | 59281399 | 27.29385 | 45.46214 | 53.3905 | 23.19383 | 0.150091 | 1.23834 | 1 |
| 3 | 59472600 | 59479799 | 140.8664 | 211.302 | 267.0364 | 110.1828 | -0.07107 | 3.465973 | 1 |
| 3 | 59516400 | 59517999 | 22.01091 | 36.92358 | 56.42673 | 24.62277 | -0.22864 | 1.149737 | 1 |
| 3 | 59596200 | 59598599 | 24.63012 | 42.15367 | 44.17622 | 19.37158 | 0.315663 | 0.964862 | 1 |
| 3 | 59699400 | 59706599 | 99.00004 | 152.4877 | 181.8484 | 75.53341 | 0.043922 | 2.921149 | 1 |
| 3 | 59709800 | 59713399 | 37.5245 | 61.18148 | 72.33894 | 31.07357 | 0.130793 | 1.540841 | 1 |
| 3 | 60589000 | 60593399 | 51.37914 | 82.3583 | 92.09633 | 38.88137 | 0.185922 | 1.98188 | 1 |
| 3 | 60989600 | 60991999 | 31.18741 | 51.40606 | 65.56072 | 28.28547 | 0.029843 | 1.379855 | 1 |
| 3 | 61619600 | 61620799 | 29.3445 | 47.88714 | 68.49956 | 29.43391 | -0.14227 | 1.507974 | 1 |
| 3 | 61626000 | 61628599 | 28.08776 | 46.33534 | 71.06051 | 30.66905 | -0.235 | 1.364769 | 1 |
| 3 | 61732800 | 61733599 | 14.98742 | 26.59041 | 33.47655 | 14.79016 | 0.062378 | 0.490466 | 1 |
| 3 | 62058400 | 62060999 | 26.20596 | 44.89906 | 44.13777 | 19.3564 | 0.407866 | 0.962662 | 1 |
| 3 | 62064600 | 62070599 | 153.1649 | 230.1917 | 243.7084 | 100.6091 | 0.186618 | 3.429543 | 1 |

| | | | | | | | | | |
|---|---|---|---|---|---|---|---|---|---|
| 3 | 62110400 | 62112199 | 20.58365 | 35.77553 | 38.30119 | 16.82008 | 0.288902 | 0.794149 | 1 |
| 3 | 62112400 | 62114199 | 14.56113 | 25.5023 | 39.43933 | 17.41193 | -0.23825 | 0.655233 | 1 |
| 3 | 62480600 | 62481799 | 18.40411 | 32.06559 | 40.48712 | 17.80735 | 0.052467 | 0.755498 | 1 |
| 3 | 62649200 | 62650999 | 30.9908 | 50.49357 | 77.15842 | 33.13542 | -0.23899 | 1.539294 | 1 |
| 3 | 62729800 | 62731599 | 17.11242 | 29.64196 | 44.26369 | 19.42026 | -0.18969 | 0.825992 | 1 |
| 3 | 62786200 | 62789199 | 26.99638 | 44.39641 | 68.35073 | 29.46068 | -0.24227 | 1.399359 | 1 |
| 3 | 62838800 | 62841199 | 20.40848 | 35.35889 | 39.51426 | 17.3366 | 0.22674 | 0.832603 | 1 |
| 3 | 62914400 | 62917599 | 49.18805 | 78.82192 | 102.8462 | 43.39952 | -0.03678 | 1.993157 | 1 |
| 3 | 63061200 | 63067799 | 102.0514 | 156.9229 | 197.5943 | 82.05474 | -0.0363 | 2.956918 | 1 |
| 3 | 63075800 | 63083799 | 93.40468 | 144.9177 | 166.4815 | 69.14075 | 0.105815 | 2.785572 | 1 |
| 3 | 63125600 | 63127399 | 13.21687 | 23.45596 | 35.09064 | 15.49947 | -0.18499 | 0.479127 | 1 |
| 3 | 63153800 | 63155199 | 27.57517 | 45.93804 | 58.94392 | 25.60891 | 0.022654 | 1.240427 | 1 |
| 3 | 63168000 | 63168599 | 9.95336 | 18.31457 | 29.80847 | 12.64702 | -0.2664 | 0.228969 | 1 |
| 3 | 63436800 | 63438399 | 16.30055 | 28.43036 | 41.55183 | 18.28769 | -0.15748 | 0.731012 | 1 |
| 3 | 63523600 | 63527999 | 27.77363 | 45.94789 | 68.21372 | 29.49583 | -0.18775 | 1.329843 | 1 |
| 3 | 63945600 | 63959599 | 306.1799 | 434.6549 | 470.0585 | 194.4359 | 0.077191 | 4.356393 | 1 |
| 3 | 64233400 | 64235599 | 23.1516 | 39.43836 | 47.96138 | 21.02348 | 0.100896 | 1.007245 | 1 |
| 3 | 64541800 | 64545599 | 26.10328 | 44.21658 | 52.30314 | 22.90125 | 0.139903 | 1.071495 | 1 |
| 3 | 64564800 | 64567199 | 49.08104 | 78.61279 | 88.08468 | 37.14334 | 0.181714 | 2.002193 | 1 |
| 3 | 64667200 | 64673199 | 10.6791 | 19.62762 | 33.50692 | 14.2382 | -0.33634 | 0.251556 | 1 |
| 3 | 64720000 | 64722199 | 34.84441 | 56.56405 | 70.44375 | 30.19661 | 0.051231 | 1.602749 | 1 |
| 3 | 65390600 | 65393199 | 22.87495 | 38.74181 | 61.08229 | 26.74394 | -0.27329 | 1.069234 | 1 |
| 3 | 66543400 | 66545199 | 19.10548 | 33.41198 | 34.17465 | 15.06887 | 0.35531 | 0.694645 | 1 |
| 3 | 66576000 | 66578799 | 21.86522 | 37.32628 | 44.2859 | 19.41568 | 0.136789 | 0.976701 | 1 |
| 3 | 66741800 | 66743999 | 19.77281 | 33.72645 | 46.94093 | 20.57833 | -0.09266 | 0.981616 | 1 |
| 3 | 66821400 | 66821999 | 7.970853 | 15.47758 | 18.88822 | 7.078224 | 0.303036 | -0.25074 | 1 |
| 3 | 66877200 | 66878599 | 17.00596 | 29.25364 | 48.47239 | 21.26228 | -0.34128 | 0.893204 | 1 |
| 3 | 66880600 | 66881799 | 10.47815 | 19.15283 | 32.33784 | 13.84378 | -0.32831 | 0.277043 | 1 |
| 3 | 67003800 | 67004599 | 21.95079 | 36.52238 | 60.54638 | 26.2837 | -0.34555 | 1.242881 | 1 |
| 3 | 67158000 | 67160199 | 14.94474 | 26.30036 | 40.20988 | 17.80206 | -0.22143 | 0.584585 | 1 |
| 3 | 67631000 | 67633399 | 29.387 | 50.05292 | 47.38375 | 20.77003 | 0.460898 | 1.032759 | 1 |
| 3 | 68021600 | 68023399 | 45.8574 | 73.88108 | 79.0499 | 33.62469 | 0.258356 | 1.813315 | 1 |
| 3 | 68585600 | 68590799 | 47.54942 | 75.96847 | 108.7444 | 45.68992 | -0.17503 | 2.084242 | 1 |
| 3 | 68639000 | 68641799 | 60.73125 | 96.56912 | 105.3405 | 43.96989 | 0.208998 | 2.247975 | 1 |
| 3 | 68644600 | 68645799 | 15.03073 | 26.51419 | 36.63928 | 16.2136 | -0.07471 | 0.556146 | 1 |
| 3 | 68646600 | 68648199 | 21.90605 | 37.71494 | 42.12324 | 18.47772 | 0.225703 | 0.898998 | 1 |
| 3 | 68659200 | 68662199 | 27.07445 | 44.95916 | 62.62746 | 27.15039 | -0.09568 | 1.280526 | 1 |
| 3 | 68752000 | 68755799 | 31.15499 | 51.75962 | 56.88609 | 24.66941 | 0.245194 | 1.279953 | 1 |
| 3 | 68911600 | 68913999 | 21.07276 | 35.24592 | 56.88542 | 24.77687 | -0.30706 | 1.178472 | 1 |
| 3 | 69033200 | 69034999 | 12.65414 | 22.37439 | 34.77906 | 15.38319 | -0.24203 | 0.512577 | 1 |
| 3 | 69469000 | 69475799 | 71.37718 | 112.3885 | 141.5817 | 58.74409 | -0.01054 | 2.506045 | 1 |
| 3 | 69544800 | 69545999 | 18.19656 | 31.32938 | 46.38817 | 20.34859 | -0.17921 | 0.890764 | 1 |
| 3 | 69549000 | 69555399 | 76.0955 | 119.5826 | 136.443 | 56.59751 | 0.129944 | 2.546151 | 1 |
| 3 | 69687600 | 69689199 | 23.43365 | 39.59144 | 53.35041 | 23.3414 | -0.04761 | 1.083854 | 1 |
| 3 | 69959000 | 69961599 | 32.99813 | 54.29634 | 57.88418 | 24.95518 | 0.28668 | 1.398317 | 1 |
| 3 | 70409800 | 70410799 | 13.61647 | 24.45583 | 29.52032 | 12.90828 | 0.133006 | 0.396559 | 1 |
| 3 | 70586000 | 70587999 | 19.74721 | 33.94467 | 48.47478 | 21.26286 | -0.12791 | 0.912451 | 1 |

| | | | | | | | | | |
|---|---|---|---|---|---|---|---|---|---|
| 3 | 70595000 | 70596599 | 17.12541 | 29.33687 | 50.03827 | 21.94377 | -0.38376 | 0.937001 | 1 |
| 3 | 70651600 | 70654799 | 38.92076 | 62.49126 | 95.66199 | 40.48832 | -0.26336 | 1.917165 | 1 |
| 3 | 70971600 | 70972799 | 16.98799 | 29.471 | 41.71869 | 18.30451 | -0.11233 | 0.809917 | 1 |
| 3 | 71302600 | 71303999 | 12.31728 | 21.82141 | 35.70318 | 15.78577 | -0.3146 | 0.489415 | 1 |
| 3 | 71431600 | 71433199 | 32.2364 | 52.54154 | 67.10387 | 28.82122 | 0.019525 | 1.530655 | 1 |
| 3 | 72063800 | 72065799 | 17.70668 | 29.92995 | 54.68612 | 23.92926 | -0.48482 | 1.06601 | 1 |
| 3 | 72193400 | 72194999 | 13.1164 | 23.18613 | 38.22385 | 16.90768 | -0.32681 | 0.523295 | 1 |
| 3 | 72359400 | 72360199 | 9.388486 | 17.81098 | 22.31746 | 8.999905 | 0.167286 | 0.016404 | 1 |
| 3 | 72605400 | 72606599 | 25.50482 | 42.28512 | 61.25932 | 26.52801 | -0.15219 | 1.294852 | 1 |
| 3 | 72800400 | 72801999 | 23.15485 | 39.51443 | 41.1937 | 18.05952 | 0.32266 | 0.981959 | 1 |
| 3 | 72922200 | 72925199 | 40.88772 | 65.77886 | 102.6339 | 43.56465 | -0.2874 | 1.861019 | 1 |
| 3 | 73640200 | 73642199 | 23.3608 | 40.00164 | 39.13854 | 17.16323 | 0.414598 | 0.950656 | 1 |
| 3 | 73763400 | 73764999 | 25.78293 | 42.80101 | 59.16571 | 25.64406 | -0.08461 | 1.278307 | 1 |
| 3 | 73779200 | 73780199 | 14.42967 | 26.09327 | 28.4365 | 12.33868 | 0.286899 | 0.360645 | 1 |
| 3 | 73865400 | 73866799 | 22.50511 | 38.64196 | 44.72262 | 19.6163 | 0.173678 | 0.927484 | 1 |
| 3 | 74124600 | 74125599 | 11.54793 | 21.24213 | 25.10404 | 10.65548 | 0.191867 | 0.232171 | 1 |
| 3 | 74365000 | 74365999 | 19.32619 | 33.49018 | 38.60548 | 16.93798 | 0.182329 | 0.825693 | 1 |
| 3 | 74496600 | 74498799 | 17.75789 | 30.72771 | 50.08916 | 21.97542 | -0.31614 | 0.844854 | 1 |
| 3 | 74499600 | 74500999 | 19.29903 | 33.6801 | 35.29966 | 15.5418 | 0.320177 | 0.728809 | 1 |
| 3 | 74526200 | 74531399 | 14.37083 | 26.21781 | 27.39171 | 11.75455 | 0.357559 | 0.306943 | 1 |
| 3 | 74625000 | 74626799 | 21.64399 | 37.42666 | 42.60693 | 18.69238 | 0.199567 | 0.860394 | 1 |
| 3 | 74741600 | 74743799 | 60.95535 | 96.77847 | 109.2015 | 45.51233 | 0.158126 | 2.291647 | 1 |
| 3 | 74747600 | 74748599 | 11.74117 | 21.4062 | 33.31216 | 14.30696 | -0.21455 | 0.303616 | 1 |
| 3 | 75322200 | 75323999 | 21.00562 | 36.25859 | 41.13002 | 18.04347 | 0.204258 | 0.872054 | 1 |
| 3 | 75325800 | 75326599 | 13.53334 | 24.56578 | 27.94929 | 12.06966 | 0.229453 | 0.330004 | 1 |
| 3 | 75371600 | 75374199 | 18.47464 | 31.81244 | 49.00219 | 21.49534 | -0.23612 | 0.89233 | 1 |
| 3 | 75748200 | 75750199 | 17.16557 | 30.32118 | 31.83761 | 14.08494 | 0.320133 | 0.544628 | 1 |
| 3 | 75786800 | 75787999 | 12.07058 | 21.8894 | 34.80121 | 15.04679 | -0.25158 | 0.339534 | 1 |
| 3 | 75875400 | 75878799 | 40.51396 | 65.67719 | 72.8735 | 31.20649 | 0.215516 | 1.641648 | 1 |
| 3 | 76164600 | 76166199 | 25.34038 | 43.07612 | 44.73768 | 19.60707 | 0.327121 | 1.027631 | 1 |
| 3 | 76411600 | 76413599 | 25.817 | 43.85254 | 49.02987 | 21.48601 | 0.221026 | 1.042301 | 1 |
| 3 | 76462600 | 76465399 | 59.9265 | 94.92007 | 130.783 | 54.38878 | -0.13283 | 2.362752 | 1 |
| 3 | 76511800 | 76512399 | 10.2824 | 18.99596 | 29.33095 | 12.37677 | -0.18505 | 0.209345 | 1 |
| 3 | 78401600 | 78403599 | 30.78387 | 50.32964 | 68.28374 | 29.36109 | -0.06429 | 1.484322 | 1 |
| 3 | 78464000 | 78468199 | 90.63805 | 140.9424 | 147.2143 | 61.11842 | 0.245503 | 2.743128 | 1 |
| 3 | 78647600 | 78650999 | 56.25988 | 89.09123 | 136.9812 | 56.95963 | -0.29109 | 2.368118 | 1 |
| 3 | 79538400 | 79541799 | 40.24206 | 65.39093 | 71.9098 | 30.84756 | 0.231565 | 1.594577 | 1 |
| 3 | 79676600 | 79678399 | 24.30955 | 40.52591 | 57.52602 | 25.00384 | -0.12244 | 1.225318 | 1 |
| 3 | 79872800 | 79873999 | 12.05804 | 21.31686 | 37.30957 | 16.50301 | -0.41239 | 0.515585 | 1 |
| 3 | 79925000 | 79927999 | 17.21605 | 29.99248 | 42.05186 | 18.49434 | -0.0981 | 0.751735 | 1 |
| 3 | 80144600 | 80146799 | 26.83776 | 44.79404 | 62.81006 | 27.32111 | -0.10502 | 1.222443 | 1 |
| 3 | 80262600 | 80263599 | 12.90515 | 23.03508 | 34.12041 | 14.9979 | -0.16581 | 0.43986 | 1 |
| 3 | 80351600 | 80353199 | 15.60408 | 27.48865 | 33.12705 | 14.6633 | 0.121017 | 0.567597 | 1 |
| 3 | 80372600 | 80373399 | 12.60562 | 22.42012 | 31.741 | 13.99414 | -0.10371 | 0.456379 | 1 |
| 3 | 80649400 | 80650799 | 16.54018 | 29.02167 | 32.28224 | 14.27431 | 0.23485 | 0.624463 | 1 |
| 3 | 81166000 | 81166599 | 11.70303 | 21.21775 | 28.19975 | 12.19623 | 0.005405 | 0.324814 | 1 |
| 3 | 81221600 | 81222399 | 13.14297 | 23.32946 | 34.19683 | 15.10214 | -0.15525 | 0.478427 | 1 |

| | | | | | | | | | |
|---|---|---|---|---|---|---|---|---|---|
| 3 | 81618800 | 81619799 | 20.96218 | 35.10046 | 58.97823 | 25.70666 | -0.3649 | 1.170235 | 1 |
| 3 | 81704800 | 81705599 | 10.90793 | 20.14486 | 27.75739 | 11.71809 | -0.02224 | 0.213225 | 1 |
| 3 | 82499600 | 82502999 | 36.01943 | 59.15234 | 64.25606 | 27.67941 | 0.258559 | 1.434344 | 1 |
| 3 | 82518400 | 82519399 | 8.905628 | 16.5253 | 29.95079 | 12.5626 | -0.40826 | 0.174483 | 1 |
| 3 | 82783000 | 82785399 | 36.77615 | 59.69717 | 70.74378 | 30.3242 | 0.122613 | 1.607248 | 1 |
| 3 | 83304000 | 83307199 | 45.99376 | 74.00955 | 87.74243 | 37.25557 | 0.10844 | 1.854328 | 1 |
| 3 | 83959400 | 83967199 | 176.3333 | 260.7991 | 319.2605 | 131.48 | -0.04381 | 3.724901 | 1 |
| 3 | 83989400 | 83990399 | 9.498809 | 18.20645 | 21.83283 | 8.596477 | 0.260774 | -0.06201 | 1 |
| 3 | 84271200 | 84275799 | 54.31801 | 86.43189 | 121.3549 | 50.70127 | -0.15356 | 2.222832 | 1 |
| 3 | 84525600 | 84527599 | 21.05013 | 36.60202 | 37.72613 | 16.57201 | 0.343477 | 0.789386 | 1 |
| 3 | 85843000 | 85844799 | 26.60718 | 45.05014 | 50.14311 | 21.95251 | 0.227299 | 1.072973 | 1 |
| 3 | 85854600 | 85856999 | 28.13425 | 46.88443 | 57.24951 | 24.87778 | 0.09398 | 1.238985 | 1 |
| 3 | 86403800 | 86404599 | 9.889299 | 18.4687 | 26.81803 | 11.13388 | -0.07926 | 0.134717 | 1 |
| 3 | 86808000 | 86810599 | 25.76123 | 42.90293 | 62.04377 | 26.95043 | -0.14928 | 1.243308 | 1 |
| 3 | 86849200 | 86851799 | 47.52559 | 76.45153 | 78.08554 | 33.14085 | 0.322561 | 1.863753 | 1 |
| 3 | 87381600 | 87386799 | 37.97082 | 61.96504 | 65.81636 | 28.28106 | 0.285946 | 1.522095 | 1 |
| 3 | 87970200 | 87973999 | 43.77439 | 70.396 | 93.45125 | 39.64502 | -0.05545 | 1.873019 | 1 |
| 3 | 88190000 | 88193799 | 3.982509 | 7.785355 | 17.74603 | 6.09637 | -0.473 | -0.61484 | 1 |
| 3 | 88571000 | 88576999 | 89.64733 | 138.9837 | 161.4258 | 67.04862 | 0.08917 | 2.79616 | 1 |
| 3 | 88630400 | 88635799 | 51.51819 | 82.16771 | 114.1326 | 47.83762 | -0.13443 | 2.143819 | 1 |
| 3 | 89597400 | 89601999 | 43.1854 | 69.37875 | 100.5443 | 42.59173 | -0.18355 | 1.907335 | 1 |
| 3 | 91001400 | 91002599 | 13.25857 | 23.59757 | 35.63377 | 15.70135 | -0.19589 | 0.463685 | 1 |
| 3 | 91004000 | 91007799 | 58.28589 | 92.52265 | 114.5269 | 47.72274 | 0.024511 | 2.29752 | 1 |
| 3 | 91095600 | 91096999 | 16.80878 | 30.12349 | 31.78823 | 13.92559 | 0.323997 | 0.437309 | 1 |
| 3 | 91100800 | 91106999 | 103.6239 | 160.0598 | 167.1192 | 69.4178 | 0.238613 | 2.869519 | 1 |
| 3 | 91116000 | 91117599 | 21.70899 | 37.71515 | 40.85029 | 17.93465 | 0.272079 | 0.808421 | 1 |
| 3 | 91253400 | 91259199 | 110.4469 | 169.8359 | 184.8699 | 76.77088 | 0.173638 | 2.958096 | 1 |
| 3 | 91508000 | 91516999 | 165.8265 | 249.0787 | 255.0856 | 105.2898 | 0.233799 | 3.443933 | 1 |
| 3 | 91589400 | 91591399 | 32.82916 | 53.25946 | 77.28543 | 33.11419 | -0.16961 | 1.616985 | 1 |
| 3 | 91698600 | 91701199 | 34.06622 | 55.48281 | 76.4633 | 32.83224 | -0.09081 | 1.550625 | 1 |
| 3 | 91902400 | 91903999 | 21.3473 | 37.0293 | 38.90769 | 17.07109 | 0.315573 | 0.82514 | 1 |
| 3 | 92092600 | 92094399 | 52.38505 | 83.34117 | 120.9841 | 50.53382 | -0.20186 | 2.227229 | 1 |
| 3 | 92335600 | 92338399 | 21.48479 | 37.28204 | 43.62306 | 19.14022 | 0.161128 | 0.829614 | 1 |
| 3 | 92460000 | 92461799 | 16.94054 | 29.09102 | 47.44679 | 20.81131 | -0.31884 | 0.910634 | 1 |
| 3 | 92585800 | 92588199 | 16.71727 | 29.20685 | 41.45156 | 18.26309 | -0.1152 | 0.707514 | 1 |
| 3 | 93078800 | 93081399 | 41.16577 | 66.27088 | 89.03744 | 37.83038 | -0.07081 | 1.834909 | 1 |
| 3 | 93113000 | 93114199 | 41.46735 | 66.34299 | 108.3088 | 45.59613 | -0.36208 | 2.037766 | 1 |
| 3 | 93118800 | 93123799 | 56.61334 | 89.94166 | 109.1586 | 45.52328 | 0.054073 | 2.270536 | 1 |
| 3 | 93185400 | 93187599 | 9.758495 | 17.34902 | 37.90223 | 16.71642 | -0.72607 | 0.453476 | 1 |
| 3 | 93188600 | 93189599 | 19.41357 | 32.79415 | 52.6594 | 23.03794 | -0.29936 | 1.07395 | 1 |
| 3 | 93190400 | 93191599 | 36.00258 | 58.46798 | 66.03165 | 28.31382 | 0.192535 | 1.59547 | 1 |
| 3 | 93197800 | 93200599 | 38.76247 | 62.36049 | 91.03135 | 38.64035 | -0.19141 | 1.853856 | 1 |
| 3 | 93527000 | 93527799 | 13.62075 | 24.03896 | 35.26912 | 15.60572 | -0.16018 | 0.540618 | 1 |
| 3 | 93605600 | 93611999 | 82.51956 | 129.1435 | 154.445 | 64.06931 | 0.057586 | 2.628755 | 1 |
| 3 | 93649000 | 93650599 | 9.151481 | 17.58783 | 20.2751 | 7.921902 | 0.327581 | -0.09529 | 1 |
| 3 | 93655200 | 93658399 | 16.69957 | 28.92107 | 45.83858 | 20.1111 | -0.27527 | 0.829362 | 1 |
| 3 | 93660600 | 93661799 | 15.38139 | 26.96096 | 38.37601 | 16.95421 | -0.11911 | 0.645812 | 1 |

| | | | | | | | | | | |
|---|---|---|---|---|---|---|---|---|---|---|
| 3 | 93771000 | 93775199 | 47.6166 | 76.65505 | 90.22455 | 38.33424 | 0.119619 | 1.843321 | | 1 |
| 3 | 93788800 | 93791799 | 25.45418 | 42.95848 | 50.13352 | 21.92675 | 0.159174 | 1.099067 | | 1 |
| 3 | 93875400 | 93876599 | 17.13093 | 29.70191 | 43.7457 | 19.19354 | -0.16955 | 0.817612 | | 1 |
| 3 | 93941400 | 93943999 | 24.34537 | 40.42172 | 59.59946 | 25.83521 | -0.17731 | 1.272808 | | 1 |
| 3 | 94059000 | 94061799 | 40.77103 | 65.87954 | 77.93705 | 33.27868 | 0.118649 | 1.718814 | | 1 |
| 3 | 94108400 | 94111399 | 19.41932 | 33.39151 | 42.71667 | 18.73732 | 0.030241 | 0.898659 | | 1 |
| 3 | 94132000 | 94142799 | 169.5024 | 253.7835 | 260.0098 | 107.2415 | 0.229066 | 3.504818 | | 1 |
| 3 | 94161200 | 94162999 | 18.25277 | 31.89155 | 39.79422 | 17.53358 | 0.069635 | 0.711366 | | 1 |
| 3 | 94196200 | 94197399 | 20.36933 | 35.41807 | 35.86745 | 15.7555 | 0.369002 | 0.785772 | | 1 |
| 3 | 94575400 | 94581199 | 61.97068 | 97.93995 | 137.1495 | 56.97231 | -0.15876 | 2.4185 | | 1 |
| 3 | 94614600 | 94617399 | 32.42548 | 53.34497 | 56.65697 | 24.42434 | 0.292039 | 1.399772 | | 1 |
| 3 | 95302400 | 95304199 | 54.86953 | 87.79422 | 92.39469 | 38.90238 | 0.270312 | 2.042543 | | 1 |
| 3 | 96724400 | 96726199 | 22.24531 | 37.82255 | 52.21687 | 22.88533 | -0.08184 | 1.026219 | | 1 |
| 3 | 97165200 | 97167999 | 41.83478 | 67.59686 | 76.55019 | 32.68554 | 0.181472 | 1.719201 | | 1 |
| 3 | 97420600 | 97434999 | 227.1725 | 328.2868 | 426.054 | 175.7466 | -0.16017 | 4.091481 | | 1 |
| 3 | 97622200 | 97623399 | 15.11987 | 26.68472 | 34.56674 | 15.29494 | 0.018295 | 0.546537 | | 1 |
| 3 | 98549600 | 98550999 | 17.09435 | 29.27239 | 49.00171 | 21.48854 | -0.35697 | 0.937991 | | 1 |
| 3 | 98705200 | 98706199 | 11.70806 | 21.46011 | 31.26984 | 13.33705 | -0.11336 | 0.268701 | | 1 |
| 3 | 98740600 | 98741599 | 11.97716 | 22.24387 | 22.56372 | 9.453146 | 0.424353 | 0.176524 | | 1 |
| 3 | 98910800 | 98911999 | 13.7768 | 24.85947 | 27.37201 | 11.90859 | 0.269389 | 0.363382 | | 1 |
| 3 | 98969400 | 98972199 | 47.4985 | 76.36051 | 82.07543 | 34.80288 | 0.24798 | 1.884009 | | 1 |
| 3 | 98989000 | 98993799 | 52.08781 | 83.60224 | 91.53882 | 38.72203 | 0.218453 | 1.943437 | | 1 |
| 3 | 99319000 | 99322799 | 14.91555 | 26.04748 | 42.09194 | 18.54005 | -0.30143 | 0.707778 | | 1 |
| 3 | 99323600 | 99325999 | 35.35417 | 57.27895 | 77.15146 | 33.02411 | -0.06427 | 1.646439 | | 1 |
| 3 | 99414400 | 99417599 | 42.97188 | 69.38353 | 75.35507 | 32.14497 | 0.240472 | 1.741682 | | 1 |
| 3 | 99630200 | 99630599 | 12.02907 | 22.64951 | 25.25274 | 10.33966 | 0.31544 | 0.099633 | | 1 |
| 3 | 100135400 | 100141799 | 63.41299 | 100.7848 | 117.3639 | 48.96491 | 0.114148 | 2.265158 | | 1 |
| 3 | 101417200 | 101419399 | 20.97429 | 35.58007 | 52.71546 | 23.09193 | -0.18356 | 1.042509 | | 1 |
| 3 | 101754000 | 101756399 | 18.42548 | 31.63593 | 51.68099 | 22.66833 | -0.3216 | 0.919319 | | 1 |
| 3 | 101909400 | 101910199 | 9.576468 | 18.09162 | 26.46271 | 10.76658 | -0.06586 | 0.057229 | | 1 |
| 3 | 102273400 | 102274999 | 20.83641 | 35.81395 | 45.30219 | 19.87119 | 0.046374 | 0.913736 | | 1 |
| 3 | 102343000 | 102345199 | 26.14596 | 43.77439 | 53.91497 | 23.49768 | 0.081497 | 1.180465 | | 1 |
| 3 | 102772800 | 102774199 | 17.56042 | 30.32225 | 41.73536 | 18.30922 | -0.07351 | 0.852655 | | 1 |
| 3 | 104360400 | 104361799 | 15.9211 | 28.2882 | 30.88773 | 13.62962 | 0.267917 | 0.482486 | | 1 |
| 3 | 106236800 | 106244399 | 147.8857 | 222.6887 | 240.2443 | 99.2305 | 0.161955 | 3.390679 | | 1 |
| 3 | 106495000 | 106497999 | 46.41489 | 74.60444 | 91.27586 | 38.69366 | 0.061121 | 1.889433 | | 1 |
| 3 | 106567400 | 106568599 | 14.61267 | 25.85686 | 32.24919 | 14.26205 | 0.074506 | 0.514042 | | 1 |
| 3 | 106628800 | 106630199 | 9.349375 | 17.21751 | 30.35911 | 12.86608 | -0.37999 | 0.22021 | | 1 |
| 3 | 107250000 | 107251399 | 10.6762 | 19.21173 | 31.24348 | 13.63972 | -0.29283 | 0.366512 | | 1 |
| 3 | 107302000 | 107305399 | 51.46845 | 81.97699 | 107.0029 | 44.76508 | -0.04695 | 2.184784 | | 1 |
| 3 | 107550200 | 107552199 | 22.3958 | 38.72262 | 47.39002 | 20.79074 | 0.095537 | 0.871041 | | 1 |
| 3 | 108146600 | 108152599 | 78.50983 | 123.3933 | 141.9813 | 58.89589 | 0.117967 | 2.547111 | | 1 |
| 3 | 108388000 | 108390599 | 23.69444 | 39.41503 | 62.92307 | 27.31138 | -0.29146 | 1.251451 | | 1 |
| 3 | 108847200 | 108849999 | 35.37216 | 57.75321 | 63.81765 | 27.42723 | 0.229563 | 1.508531 | | 1 |
| 3 | 109037400 | 109042799 | 77.1121 | 121.3515 | 136.9215 | 56.80645 | 0.147624 | 2.519641 | | 1 |
| 3 | 109052000 | 109055999 | 56.30498 | 89.68499 | 126.7893 | 53.04651 | -0.16188 | 2.193462 | | 1 |
| 3 | 109963200 | 109965599 | 21.46825 | 36.45556 | 53.26172 | 23.33852 | -0.1634 | 1.036186 | | 1 |

| | | | | | | | | | |
|---|---|---|---|---|---|---|---|---|---|
| 3 | 110049600 | 110052599 | 33.17904 | 53.93131 | 72.85068 | 31.25854 | -0.06399 | 1.572874 | 1 |
| 3 | 110232600 | 110234799 | 19.36355 | 33.05973 | 53.78512 | 23.58054 | -0.31683 | 0.97977 | 1 |
| 3 | 110831200 | 110832799 | 18.51714 | 32.51051 | 36.48411 | 16.14067 | 0.221971 | 0.632278 | 1 |
| 3 | 111042800 | 111043599 | 15.13839 | 27.0354 | 29.81219 | 13.09843 | 0.257826 | 0.441487 | 1 |
| 3 | 111090800 | 111094399 | 35.09006 | 57.14864 | 65.81438 | 28.25945 | 0.167557 | 1.545475 | 1 |
| 3 | 111390600 | 111391199 | 14.46928 | 26.07788 | 27.35601 | 11.9194 | 0.337576 | 0.376142 | 1 |
| 3 | 111510600 | 111512799 | 42.71698 | 68.55184 | 97.82982 | 41.37404 | -0.16324 | 1.940118 | 1 |
| 3 | 111949600 | 111951399 | 16.03401 | 28.05891 | 39.43177 | 17.39655 | -0.10077 | 0.677725 | 1 |
| 3 | 112098800 | 112100999 | 28.09464 | 46.38677 | 64.17048 | 27.70952 | -0.08635 | 1.350877 | 1 |
| 3 | 112228200 | 112228999 | 11.00538 | 20.11397 | 30.29548 | 12.97154 | -0.16437 | 0.279868 | 1 |
| 3 | 112623600 | 112625399 | 15.69095 | 27.61072 | 41.2759 | 18.27439 | -0.18914 | 0.594105 | 1 |
| 3 | 112678400 | 112680199 | 21.00409 | 36.28818 | 38.51079 | 16.89489 | 0.300122 | 0.858569 | 1 |
| 3 | 112698600 | 112699599 | 15.82776 | 28.27364 | 30.76428 | 13.51388 | 0.277206 | 0.446921 | 1 |
| 3 | 112746800 | 112748799 | 21.34682 | 36.28118 | 52.71345 | 23.10227 | -0.15525 | 1.028468 | 1 |
| 3 | 112774200 | 112778599 | 35.18924 | 58.43443 | 58.21709 | 25.23841 | 0.386367 | 1.298679 | 1 |
| 3 | 112820000 | 112826199 | 76.4418 | 120.4699 | 141.1681 | 58.57924 | 0.094698 | 2.491528 | 1 |
| 3 | 112972400 | 112976799 | 16.5136 | 28.75405 | 43.15684 | 18.97423 | -0.19585 | 0.759075 | 1 |
| 3 | 112997000 | 113005599 | 253.8772 | 365.359 | 431.0431 | 177.9036 | -0.02857 | 4.154426 | 1 |
| 3 | 113033200 | 113035399 | 24.19115 | 40.62152 | 57.68915 | 25.18953 | -0.12339 | 1.145454 | 1 |
| 3 | 113079000 | 113080399 | 16.46011 | 28.99478 | 34.06773 | 15.07988 | 0.157368 | 0.575798 | 1 |
| 3 | 113108000 | 113113999 | 80.38398 | 126.4622 | 145.6226 | 60.41401 | 0.117944 | 2.528493 | 1 |
| 3 | 113118800 | 113127199 | 211.1482 | 310.6051 | 311.5977 | 128.3191 | 0.235906 | 3.817382 | 1 |
| 3 | 113813200 | 113815799 | 41.72847 | 67.30316 | 79.97529 | 34.06837 | 0.109217 | 1.7738 | 1 |
| 3 | 114072000 | 114073599 | 40.16165 | 64.37762 | 103.5124 | 43.70149 | -0.33692 | 1.97557 | 1 |
| 3 | 114657200 | 114660799 | 40.92441 | 65.86015 | 101.997 | 43.31554 | -0.27604 | 1.851206 | 1 |
| 3 | 115706000 | 115712199 | 45.47646 | 72.88622 | 104.1658 | 43.96736 | -0.16757 | 1.98489 | 1 |
| 3 | 115880600 | 115882599 | 32.42529 | 53.62714 | 61.3183 | 26.50866 | 0.187811 | 1.339742 | 1 |
| 3 | 116527400 | 116528599 | 22.43733 | 37.78949 | 55.96742 | 24.4631 | -0.18351 | 1.112281 | 1 |
| 3 | 117115400 | 117116999 | 23.21238 | 38.67595 | 57.42219 | 24.9502 | -0.18695 | 1.228612 | 1 |
| 3 | 117174600 | 117175599 | 12.73152 | 22.99201 | 32.63855 | 14.18544 | -0.09379 | 0.364982 | 1 |
| 3 | 118371400 | 118373399 | 21.17015 | 36.09005 | 48.49164 | 21.2572 | -0.04228 | 0.992506 | 1 |
| 3 | 119105800 | 119106599 | 12.52602 | 22.27933 | 35.2371 | 15.53505 | -0.26259 | 0.464057 | 1 |
| 3 | 119898200 | 119899399 | 27.6537 | 45.47934 | 64.98684 | 28.01128 | -0.13493 | 1.399098 | 1 |
| 3 | 120295400 | 120296399 | 25.6011 | 43.49367 | 45.81498 | 20.07826 | 0.306715 | 1.036356 | 1 |
| 3 | 120723600 | 120724199 | 10.88901 | 20.58712 | 24.83082 | 10.08615 | 0.212659 | 0.061355 | 1 |
| 3 | 121115800 | 121118799 | 25.23011 | 42.27417 | 58.78925 | 25.63471 | -0.09314 | 1.174095 | 1 |
| 3 | 121653600 | 121655799 | 6.689622 | 12.94877 | 23.25302 | 8.853895 | -0.27063 | -0.1969 | 1 |
| 3 | 122447800 | 122451799 | 40.79301 | 65.58976 | 99.10071 | 42.03081 | -0.24212 | 1.874904 | 1 |
| 3 | 123095600 | 123097199 | 17.95969 | 31.36152 | 40.36294 | 17.77777 | 0.025137 | 0.719118 | 1 |
| 3 | 123726600 | 123727999 | 20.97355 | 36.32434 | 35.96168 | 15.77776 | 0.400664 | 0.833166 | 1 |
| 3 | 123731400 | 123733799 | 22.45846 | 37.76056 | 60.34761 | 26.36401 | -0.29311 | 1.130904 | 1 |
| 3 | 123824600 | 123826999 | 24.17907 | 40.10933 | 63.45781 | 27.49057 | -0.27886 | 1.284838 | 1 |
| 3 | 123832600 | 123833799 | 17.9625 | 30.42438 | 53.16891 | 23.27976 | -0.42085 | 1.044906 | 1 |
| 3 | 123928000 | 123930599 | 40.66142 | 65.91593 | 69.40114 | 29.71942 | 0.291006 | 1.640275 | 1 |
| 3 | 124679200 | 124685199 | 133.9499 | 202.9695 | 226.9142 | 93.90383 | 0.118329 | 3.262554 | 1 |
| 3 | 125823600 | 125824399 | 11.54075 | 20.62761 | 35.25611 | 15.47942 | -0.3698 | 0.425496 | 1 |
| 3 | 125860200 | 125860599 | 7.992696 | 15.4484 | 20.43554 | 7.834584 | 0.156863 | -0.1683 | 1 |

| | | | | | | | | | |
|---|---|---|---|---|---|---|---|---|---|
| 3 | 126302400 | 126306599 | 88.75744 | 137.8754 | 159.4178 | 66.19512 | 0.097876 | 2.762146 | 1 |
| 3 | 127564600 | 127566799 | 22.99279 | 37.98684 | 66.91545 | 28.90726 | -0.43373 | 1.333733 | 1 |
| 3 | 127638800 | 127640199 | 13.71876 | 24.85913 | 30.97771 | 13.40656 | 0.096813 | 0.347225 | 1 |
| 3 | 127650800 | 127653399 | 21.77502 | 37.77583 | 40.91847 | 17.9534 | 0.271797 | 0.827452 | 1 |
| 3 | 128490600 | 128492199 | 17.57318 | 30.6672 | 37.43942 | 16.48356 | 0.101124 | 0.724192 | 1 |
| 3 | 128600600 | 128602199 | 14.24591 | 26.02386 | 29.48657 | 12.63286 | 0.242722 | 0.303 | 1 |
| 3 | 129213600 | 129215799 | 19.41452 | 33.88585 | 35.11178 | 15.46028 | 0.336357 | 0.724274 | 1 |
| 3 | 129265400 | 129270599 | 53.7322 | 85.82756 | 100.994 | 42.42024 | 0.106833 | 2.095627 | 1 |
| 3 | 129445000 | 129448999 | 48.85902 | 78.61341 | 84.28916 | 35.7849 | 0.252968 | 1.858711 | 1 |
| 3 | 129649600 | 129655399 | 43.46616 | 70.21791 | 74.98351 | 32.00781 | 0.265646 | 1.727503 | 1 |
| 3 | 129750400 | 129752399 | 18.23638 | 31.75817 | 44.14034 | 19.4083 | -0.08563 | 0.765713 | 1 |
| 3 | 130736600 | 130738199 | 10.28563 | 18.89027 | 32.40657 | 13.78468 | -0.34464 | 0.248005 | 1 |
| 3 | 131754800 | 131756999 | 17.40123 | 30.26554 | 47.91082 | 21.04945 | -0.27282 | 0.783197 | 1 |
| 3 | 132118200 | 132120199 | 9.826467 | 18.68697 | 20.28313 | 8.137838 | 0.379423 | -0.00144 | 1 |
| 3 | 132223200 | 132229399 | 62.30907 | 98.39948 | 145.2698 | 60.32217 | -0.23578 | 2.43796 | 1 |
| 3 | 132277600 | 132279599 | 32.67536 | 53.56756 | 72.36748 | 31.15067 | -0.05616 | 1.453643 | 1 |
| 3 | 132326800 | 132328399 | 19.02567 | 32.4255 | 52.69802 | 23.10064 | -0.3157 | 0.992911 | 1 |
| 3 | 132534400 | 132535799 | 10.40088 | 18.92576 | 32.85414 | 14.14463 | -0.37311 | 0.305999 | 1 |
| 3 | 132662400 | 132663799 | 18.49015 | 32.38074 | 38.15472 | 16.84356 | 0.151974 | 0.674813 | 1 |
| 3 | 132898000 | 132898799 | 13.18092 | 24.31502 | 27.00358 | 11.41749 | 0.284913 | 0.234341 | 1 |
| 3 | 133347600 | 133348999 | 21.59942 | 36.40619 | 49.06051 | 21.44911 | -0.04759 | 1.096131 | 1 |
| 3 | 133565200 | 133566399 | 13.17521 | 23.13034 | 39.42356 | 17.44023 | -0.37769 | 0.609266 | 1 |
| 3 | 133680400 | 133686999 | 103.2135 | 159.1833 | 183.2959 | 76.13786 | 0.095853 | 2.898884 | 1 |
| 3 | 134303800 | 134306599 | 24.59005 | 42.07312 | 48.47578 | 21.25648 | 0.179391 | 0.973093 | 1 |
| 3 | 134533600 | 134535999 | 17.97954 | 30.76369 | 50.58949 | 22.18339 | -0.33179 | 0.948103 | 1 |
| 3 | 134798600 | 134799399 | 11.18237 | 20.71157 | 25.71157 | 10.81176 | 0.131359 | 0.194264 | 1 |
| 3 | 134800800 | 134802599 | 23.70186 | 40.77144 | 43.10261 | 18.90684 | 0.304361 | 0.915306 | 1 |
| 3 | 134867600 | 134870199 | 29.98879 | 49.29792 | 58.65579 | 25.2777 | 0.128226 | 1.404265 | 1 |
| 3 | 135479800 | 135481399 | 36.28113 | 59.10243 | 67.59823 | 29.0279 | 0.177692 | 1.546604 | 1 |
| 3 | 135827400 | 135828399 | 11.00983 | 19.98474 | 31.77903 | 13.72381 | -0.25029 | 0.322027 | 1 |
| 3 | 136506600 | 136508999 | 20.89799 | 36.16017 | 43.11736 | 18.91669 | 0.133102 | 0.850406 | 1 |
| 3 | 137113000 | 137116799 | 32.26629 | 53.89372 | 55.79926 | 24.28267 | 0.331193 | 1.226336 | 1 |
| 3 | 137155000 | 137164399 | 109.5268 | 168.2111 | 199.6824 | 82.9079 | 0.047317 | 2.983392 | 1 |
| 3 | 137741800 | 137746399 | 48.5659 | 78.13819 | 94.74371 | 40.22069 | 0.075827 | 1.86601 | 1 |
| 3 | 137801000 | 137804799 | 26.99665 | 44.36099 | 71.77375 | 30.92551 | -0.31423 | 1.410954 | 1 |
| 3 | 137911000 | 137913999 | 69.27044 | 108.975 | 137.2514 | 56.94087 | -0.01119 | 2.521757 | 1 |
| 3 | 137915000 | 137915799 | 9.201815 | 17.52759 | 23.90249 | 9.556997 | 0.056516 | -0.01022 | 1 |
| 3 | 137999000 | 138001999 | 31.36953 | 52.36391 | 61.94498 | 26.94678 | 0.139519 | 1.235733 | 1 |
| 3 | 138101800 | 138103799 | 17.56229 | 30.30446 | 50.05654 | 21.95924 | -0.33577 | 0.872174 | 1 |
| 3 | 138194600 | 138195599 | 10.51644 | 19.04079 | 30.77907 | 13.33421 | -0.27683 | 0.329707 | 1 |
| 3 | 138742800 | 138746799 | 35.10957 | 56.83338 | 84.98327 | 36.35322 | -0.21584 | 1.668794 | 1 |
| 3 | 138759400 | 138761799 | 19.44233 | 33.53407 | 53.36287 | 23.40938 | -0.28254 | 0.886122 | 1 |
| 3 | 139070800 | 139073599 | 41.69891 | 67.46159 | 74.39724 | 31.81154 | 0.221622 | 1.685593 | 1 |
| 3 | 139265200 | 139271799 | 143.7754 | 215.5933 | 268.3835 | 110.7288 | -0.04977 | 3.472822 | 1 |
| 3 | 139466800 | 139467799 | 10.76952 | 19.68641 | 31.27673 | 13.38868 | -0.24104 | 0.276522 | 1 |
| 3 | 139608200 | 139609799 | 23.69387 | 40.62143 | 46.85021 | 20.547 | 0.178352 | 0.94881 | 1 |
| 3 | 139825200 | 139829799 | 18.55164 | 31.68826 | 51.10281 | 22.40539 | -0.304 | 0.969578 | 1 |

| | | | | | | | | | |
|---|---|---|---|---|---|---|---|---|---|
| 3 | 139861600 | 139863799 | 29.38042 | 48.56611 | 61.53109 | 26.59112 | 0.040234 | 1.339001 | 1 |
| 3 | 140075400 | 140075799 | 8.77248 | 16.94264 | 21.4783 | 8.263789 | 0.212714 | -0.14445 | 1 |
| 3 | 140377800 | 140378799 | 14.28569 | 25.55001 | 32.96778 | 14.46698 | 0.033704 | 0.433857 | 1 |
| 3 | 140507000 | 140508399 | 17.6275 | 31.18113 | 32.1997 | 14.24111 | 0.344736 | 0.532299 | 1 |
| 3 | 140852800 | 140857799 | 89.0964 | 137.8344 | 174.0135 | 72.28094 | -0.03338 | 2.83636 | 1 |
| 3 | 140858800 | 140860999 | 35.65426 | 57.94315 | 67.83134 | 29.10064 | 0.141779 | 1.580942 | 1 |
| 3 | 140916400 | 140917999 | 38.62142 | 62.09 | 96.01632 | 40.71492 | -0.27569 | 1.877864 | 1 |
| 3 | 142333600 | 142334799 | 15.88476 | 28.03837 | 31.53979 | 13.95522 | 0.220963 | 0.544412 | 1 |
| 3 | 142658800 | 142659599 | 9.955041 | 18.8258 | 27.59261 | 11.20234 | -0.06594 | 0.057756 | 1 |
| 3 | 143840000 | 143841799 | 28.82432 | 47.66051 | 56.03505 | 24.22075 | 0.147801 | 1.329303 | 1 |
| 3 | 144322800 | 144338799 | 213.3836 | 310.0774 | 422.3325 | 174.093 | -0.22197 | 4.008072 | 1 |
| 3 | 144367200 | 144370999 | 39.1284 | 63.17906 | 84.63893 | 36.10625 | -0.06159 | 1.739403 | 1 |
| 3 | 144443800 | 144446799 | 39.34359 | 63.89075 | 75.63434 | 32.43025 | 0.124663 | 1.608593 | 1 |
| 3 | 144562200 | 144565599 | 37.45671 | 60.89889 | 72.16667 | 30.96771 | 0.124686 | 1.579431 | 1 |
| 3 | 144645200 | 144648399 | 36.56538 | 59.05938 | 89.5422 | 38.21356 | -0.23918 | 1.728589 | 1 |
| 3 | 145662400 | 145664999 | 41.02002 | 66.13261 | 84.71682 | 36.06857 | 0.000308 | 1.78531 | 1 |
| 3 | 146818600 | 146819799 | 13.05846 | 23.34819 | 33.60422 | 14.74995 | -0.12314 | 0.429431 | 1 |
| 3 | 146930200 | 146931599 | 14.49456 | 26.21202 | 28.34417 | 12.29778 | 0.297955 | 0.359203 | 1 |
| 3 | 147400000 | 147401799 | 17.70364 | 31.14597 | 32.31958 | 14.30968 | 0.33476 | 0.590875 | 1 |
| 3 | 147822200 | 147826599 | 89.68629 | 139.4519 | 159.0964 | 66.0522 | 0.118237 | 2.74573 | 1 |
| 3 | 147873600 | 147875399 | 30.42998 | 50.7757 | 53.01028 | 23.05439 | 0.319191 | 1.226635 | 1 |
| 3 | 148454800 | 148457399 | 43.03112 | 69.22431 | 84.04363 | 35.67162 | 0.073772 | 1.856492 | 1 |
| 3 | 149060600 | 149062399 | 22.13056 | 37.6918 | 45.87867 | 20.11023 | 0.09958 | 1.002243 | 1 |
| 3 | 150220800 | 150222399 | 18.9168 | 32.49166 | 47.94842 | 21.03148 | -0.17531 | 0.914796 | 1 |
| 3 | 150813800 | 150815399 | 17.64493 | 31.14931 | 32.78509 | 14.50584 | 0.316348 | 0.555377 | 1 |
| 3 | 150938800 | 150942799 | 72.30753 | 113.4465 | 149.2007 | 61.87775 | -0.07653 | 2.578908 | 1 |
| 3 | 150951800 | 150955199 | 28.89508 | 48.16251 | 57.33389 | 24.91777 | 0.130498 | 1.23805 | 1 |
| 3 | 151328200 | 151330399 | 30.38057 | 49.42295 | 73.86497 | 31.7032 | -0.20856 | 1.559683 | 1 |
| 3 | 151384800 | 151386999 | 18.3385 | 32.21019 | 34.45093 | 15.24652 | 0.290994 | 0.621542 | 1 |
| 3 | 152029200 | 152030399 | 16.31005 | 27.95912 | 46.95025 | 20.59068 | -0.36109 | 0.923219 | 1 |
| 3 | 152036000 | 152036599 | 11.8318 | 21.26098 | 29.83226 | 13.03924 | -0.082 | 0.384151 | 1 |
| 3 | 152512600 | 152517599 | 17.18066 | 30.29529 | 27.73576 | 12.27454 | 0.516151 | 0.557262 | 1 |
| 3 | 152807800 | 152808999 | 18.75041 | 32.81614 | 34.12322 | 15.05593 | 0.331567 | 0.679828 | 1 |
| 3 | 153220200 | 153227199 | 78.40939 | 122.4269 | 166.379 | 69.03641 | -0.12922 | 2.670059 | 1 |
| 3 | 153229600 | 153230999 | 20.86613 | 34.97792 | 59.12403 | 25.78803 | -0.37368 | 1.154198 | 1 |
| 3 | 153329600 | 153331599 | 22.78711 | 38.77618 | 47.43933 | 20.79286 | 0.092046 | 1.014296 | 1 |
| 3 | 153422400 | 153424199 | 18.86072 | 32.14559 | 53.2315 | 23.33457 | -0.34264 | 0.992227 | 1 |
| 3 | 153735000 | 153740999 | 59.28976 | 94.22771 | 122.9266 | 51.28361 | -0.04951 | 2.26572 | 1 |
| 3 | 155966400 | 155968799 | 43.03969 | 69.47068 | 73.91851 | 31.51908 | 0.269485 | 1.751817 | 1 |
| 3 | 156132200 | 156132799 | 10.14343 | 18.77589 | 30.41611 | 12.80057 | -0.25105 | 0.199941 | 1 |
| 3 | 156672200 | 156673599 | 15.83669 | 27.61225 | 45.21088 | 19.89399 | -0.32076 | 0.738051 | 1 |
| 3 | 156711400 | 156712999 | 19.26999 | 33.57548 | 37.84611 | 16.64615 | 0.21538 | 0.75419 | 1 |
| 3 | 157378800 | 157380999 | 29.11449 | 48.36774 | 56.70847 | 24.59172 | 0.152264 | 1.275918 | 1 |
| 3 | 157670400 | 157673599 | 39.51091 | 64.57233 | 71.96883 | 30.94118 | 0.218127 | 1.508877 | 1 |
| 3 | 157806800 | 157808199 | 27.00711 | 44.38478 | 66.58546 | 28.69181 | -0.20535 | 1.406775 | 1 |
| 3 | 158038000 | 158040399 | 15.14572 | 26.51675 | 42.54108 | 18.77568 | -0.29087 | 0.671509 | 1 |
| 3 | 158050200 | 158050999 | 10.18246 | 19.30174 | 26.53671 | 10.71956 | 0.031812 | 0.039358 | 1 |

| | | | | | | | | | |
|---|---|---|---|---|---|---|---|---|---|
| 3 | 158442800 | 158445399 | 45.35794 | 72.37344 | 110.3906 | 46.2966 | -0.26849 | 2.123797 | 1 |
| 3 | 158446200 | 158447199 | 9.204467 | 17.67564 | 22.428 | 8.783057 | 0.187252 | -0.08081 | 1 |
| 3 | 158451800 | 158452799 | 18.03535 | 31.68464 | 33.99022 | 15.0454 | 0.286744 | 0.616004 | 1 |
| 3 | 158666000 | 158667199 | 15.29603 | 26.97275 | 37.60248 | 16.641 | -0.08769 | 0.56179 | 1 |
| 3 | 158876800 | 158878199 | 22.91838 | 38.01747 | 64.42618 | 27.90979 | -0.37764 | 1.282616 | 1 |
| 3 | 158955000 | 158955999 | 11.35644 | 20.86525 | 32.3835 | 13.76736 | -0.20062 | 0.253899 | 1 |
| 3 | 159053800 | 159064199 | 129.0721 | 196.4318 | 212.6295 | 88.10785 | 0.17001 | 3.171473 | 1 |
| 3 | 159151200 | 159152999 | 34.0929 | 55.29449 | 84.7446 | 36.30285 | -0.24871 | 1.626911 | 1 |
| 3 | 159264600 | 159269799 | 43.53145 | 70.68538 | 76.51005 | 32.80364 | 0.253399 | 1.617086 | 1 |
| 3 | 159505800 | 159507399 | 15.59981 | 27.62988 | 33.42509 | 14.77927 | 0.118413 | 0.508862 | 1 |
| 4 | 3050000 | 3057599 | 451.2753 | 612.0198 | 751.8336 | 313.2471 | -0.17309 | 5.05422 | 1 |
| 4 | 3114800 | 3116399 | 17.49519 | 30.7977 | 30.05661 | 13.30614 | 0.42339 | 0.581355 | 1 |
| 4 | 3173600 | 3183999 | 313.0203 | 438.4916 | 540.3479 | 223.9935 | -0.13043 | 4.553212 | 1 |
| 4 | 3284600 | 3287199 | 14.53188 | 25.61622 | 37.67996 | 16.67646 | -0.16482 | 0.56191 | 1 |
| 4 | 3310600 | 3317799 | 133.3429 | 201.7616 | 221.7088 | 91.70836 | 0.14143 | 3.291259 | 1 |
| 4 | 3330400 | 3345399 | 265.9499 | 383.2132 | 401.0956 | 165.5147 | 0.145865 | 4.136507 | 1 |
| 4 | 3398800 | 3400799 | 11.67218 | 21.06452 | 33.24301 | 14.46051 | -0.24628 | 0.36375 | 1 |
| 4 | 3812600 | 3814599 | 22.67047 | 38.9484 | 41.59237 | 18.24368 | 0.289521 | 0.919882 | 1 |
| 4 | 4048200 | 4051399 | 44.43887 | 71.17688 | 106.9964 | 45.11745 | -0.24149 | 2.003221 | 1 |
| 4 | 4119600 | 4123599 | 71.16304 | 112.4991 | 118.0984 | 49.06537 | 0.256678 | 2.413739 | 1 |
| 4 | 4164800 | 4167399 | 31.76117 | 51.95112 | 73.66873 | 31.68182 | -0.12749 | 1.482834 | 1 |
| 4 | 4360200 | 4361999 | 23.56605 | 40.00912 | 48.50054 | 21.25083 | 0.104798 | 1.038195 | 1 |
| 4 | 4744600 | 4746399 | 35.60479 | 58.35907 | 62.48971 | 26.89664 | 0.278123 | 1.45434 | 1 |
| 4 | 5044000 | 5046999 | 42.35042 | 68.136 | 95.59024 | 40.57831 | -0.13422 | 1.8589 | 1 |
| 4 | 5189200 | 5192199 | 59.32665 | 94.11998 | 115.7189 | 48.19137 | 0.033414 | 2.31393 | 1 |
| 4 | 5210800 | 5214599 | 94.87928 | 147.2235 | 154.4057 | 64.12444 | 0.237225 | 2.782954 | 1 |
| 4 | 5248600 | 5249599 | 16.68533 | 29.49743 | 29.94616 | 13.24596 | 0.369043 | 0.529653 | 1 |
| 4 | 5312600 | 5315199 | 23.87112 | 40.2721 | 48.54278 | 21.22867 | 0.112622 | 1.092897 | 1 |
| 4 | 5344200 | 5346999 | 25.1883 | 42.16806 | 57.70096 | 25.1466 | -0.06988 | 1.181265 | 1 |
| 4 | 5460000 | 5466199 | 94.79093 | 145.8369 | 182.6053 | 75.83584 | -0.02759 | 2.942586 | 1 |
| 4 | 5607400 | 5610999 | 55.39537 | 88.4524 | 104.2893 | 43.78205 | 0.103405 | 2.109474 | 1 |
| 4 | 5714600 | 5718399 | 47.2604 | 75.56286 | 101.9705 | 42.88895 | -0.08891 | 2.060417 | 1 |
| 4 | 5761000 | 5767999 | 192.2449 | 283.1891 | 320.5884 | 132.0211 | 0.06352 | 3.793394 | 1 |
| 4 | 5873600 | 5885999 | 144.1206 | 216.7334 | 283.0035 | 116.8505 | -0.11497 | 3.417669 | 1 |
| 4 | 5914800 | 5920199 | 68.08667 | 108.109 | 113.8448 | 47.45108 | 0.257725 | 2.297136 | 1 |
| 4 | 6422200 | 6424199 | 18.77543 | 32.18 | 50.59957 | 22.19084 | -0.26703 | 0.936464 | 1 |
| 4 | 6486800 | 6489799 | 27.4899 | 45.88053 | 60.58625 | 26.35309 | -0.01869 | 1.223361 | 1 |
| 4 | 6632600 | 6634999 | 15.13718 | 27.19129 | 28.2947 | 12.37075 | 0.34589 | 0.402749 | 1 |
| 4 | 6648600 | 6650199 | 20.96424 | 35.44763 | 53.78442 | 23.53698 | -0.218 | 1.07089 | 1 |
| 4 | 7163000 | 7171399 | 120.416 | 183.9021 | 191.2615 | 79.32572 | 0.231707 | 3.097006 | 1 |
| 4 | 7278200 | 7280599 | 33.22159 | 54.74612 | 61.28546 | 26.43833 | 0.217259 | 1.383647 | 1 |
| 4 | 7297600 | 7298599 | 17.65007 | 30.37944 | 46.9022 | 20.57389 | -0.23948 | 0.889696 | 1 |
| 4 | 7322400 | 7329799 | 141.5142 | 214.5967 | 223.0074 | 92.30906 | 0.224598 | 3.248373 | 1 |
| 4 | 7354200 | 7355799 | 23.54435 | 40.28037 | 42.16767 | 18.49026 | 0.317219 | 0.962754 | 1 |
| 4 | 7627000 | 7631999 | 39.57613 | 63.49909 | 99.11966 | 41.90715 | -0.29267 | 1.940896 | 1 |
| 4 | 7702600 | 7703599 | 28.65441 | 47.05245 | 66.07798 | 28.46301 | -0.11085 | 1.421972 | 1 |
| 4 | 7821000 | 7822599 | 12.82316 | 22.59121 | 38.30883 | 16.95673 | -0.36914 | 0.559199 | 1 |

| | | | | | | | | | |
|---|---|---|---|---|---|---|---|---|---|
| 4 | 8039600 | 8043399 | 32.01387 | 52.33602 | 73.25607 | 31.49812 | -0.10938 | 1.488977 | 1 |
| 4 | 8114400 | 8120599 | 81.30701 | 126.5369 | 182.2753 | 75.66524 | -0.21682 | 2.729457 | 1 |
| 4 | 8189200 | 8192199 | 43.67257 | 70.23988 | 84.71716 | 35.94538 | 0.082826 | 1.864735 | 1 |
| 4 | 8209600 | 8211199 | 14.92923 | 26.69336 | 31.41009 | 13.78676 | 0.165767 | 0.437356 | 1 |
| 4 | 8309800 | 8316799 | 85.70192 | 132.8911 | 178.826 | 74.27383 | -0.12277 | 2.792993 | 1 |
| 4 | 8398000 | 8407199 | 113.9601 | 173.527 | 242.1848 | 100.371 | -0.19569 | 3.159181 | 1 |
| 4 | 8816000 | 8819599 | 35.82136 | 57.72673 | 92.44503 | 39.33703 | -0.32171 | 1.796188 | 1 |
| 4 | 8970000 | 8971599 | 17.79153 | 31.32996 | 35.3504 | 15.64866 | 0.215079 | 0.58695 | 1 |
| 4 | 9138000 | 9139999 | 15.64317 | 27.35056 | 39.95152 | 17.61259 | -0.1564 | 0.689054 | 1 |
| 4 | 9493800 | 9494999 | 8.716064 | 16.26832 | 27.40821 | 11.38994 | -0.29301 | 0.1304 | 1 |
| 4 | 9584400 | 9586999 | 65.4275 | 102.7184 | 153.5284 | 63.67662 | -0.26014 | 2.562862 | 1 |
| 4 | 9774600 | 9776199 | 13.05344 | 23.16726 | 35.59639 | 15.7221 | -0.22328 | 0.478261 | 1 |
| 4 | 10110200 | 10111799 | 14.44028 | 25.24486 | 40.55369 | 17.87652 | -0.29273 | 0.686782 | 1 |
| 4 | 10163600 | 10166599 | 29.18093 | 48.71913 | 53.49625 | 23.27407 | 0.246625 | 1.21813 | 1 |
| 4 | 10248600 | 10249399 | 12.34606 | 22.84837 | 23.55311 | 9.91506 | 0.396393 | 0.20212 | 1 |
| 4 | 10477000 | 10478799 | 25.94534 | 43.88686 | 48.52525 | 21.23796 | 0.236913 | 1.079971 | 1 |
| 4 | 10663200 | 10664199 | 16.61817 | 29.1655 | 33.76121 | 14.9313 | 0.177688 | 0.624671 | 1 |
| 4 | 10869000 | 10870399 | 27.58598 | 45.57211 | 59.15106 | 25.55182 | 0.005436 | 1.339562 | 1 |
| 4 | 10885000 | 10890799 | 132.3783 | 200.3474 | 226.3216 | 93.62311 | 0.101881 | 3.286388 | 1 |
| 4 | 11431200 | 11432399 | 18.79273 | 33.10044 | 34.7619 | 15.38746 | 0.318242 | 0.58982 | 1 |
| 4 | 11628400 | 11630599 | 31.4353 | 51.28891 | 72.80641 | 31.28228 | -0.13155 | 1.517262 | 1 |
| 4 | 11840400 | 11847199 | 108.675 | 166.987 | 184.9873 | 76.81181 | 0.147526 | 2.972003 | 1 |
| 4 | 12200400 | 12207799 | 81.63277 | 127.1623 | 189.1155 | 78.49379 | -0.26177 | 2.715049 | 1 |
| 4 | 12325200 | 12326399 | 11.37177 | 20.42591 | 32.27997 | 14.11401 | -0.25303 | 0.388248 | 1 |
| 4 | 12443800 | 12450399 | 97.93586 | 151.063 | 178.5362 | 74.1607 | 0.058452 | 2.89382 | 1 |
| 4 | 12719000 | 12719799 | 15.05038 | 26.62104 | 32.64431 | 14.43791 | 0.098468 | 0.520679 | 1 |
| 4 | 12959400 | 12960199 | 16.36692 | 28.43318 | 43.22826 | 18.97817 | -0.21462 | 0.791676 | 1 |
| 4 | 13002000 | 13004399 | 37.05056 | 60.33892 | 75.05332 | 32.22596 | 0.056651 | 1.557964 | 1 |
| 4 | 13180000 | 13185199 | 85.78537 | 132.8023 | 186.1483 | 77.32116 | -0.18339 | 2.823704 | 1 |
| 4 | 13431400 | 13432799 | 10.22151 | 18.51459 | 30.81457 | 13.34257 | -0.31808 | 0.325569 | 1 |
| 4 | 13664200 | 13664799 | 12.3099 | 22.62979 | 27.23338 | 11.56899 | 0.165594 | 0.251335 | 1 |
| 4 | 13680400 | 13680999 | 8.952145 | 16.84531 | 28.02276 | 11.48615 | -0.25931 | 0.08015 | 1 |
| 4 | 14136200 | 14137599 | 25.73688 | 43.40922 | 45.41579 | 19.85972 | 0.316272 | 1.100562 | 1 |
| 4 | 14423600 | 14424599 | 11.8237 | 22.28457 | 24.11555 | 9.857724 | 0.359627 | 0.086437 | 1 |
| 4 | 14638200 | 14642599 | 18.64982 | 31.99787 | 49.53187 | 21.72441 | -0.24423 | 0.926509 | 1 |
| 4 | 15020000 | 15020599 | 13.28444 | 23.4687 | 36.22815 | 16.02689 | -0.23268 | 0.528443 | 1 |
| 4 | 15054400 | 15055599 | 14.37085 | 25.47632 | 30.48058 | 13.47488 | 0.134732 | 0.483836 | 1 |
| 4 | 15240800 | 15244199 | 35.00212 | 56.80051 | 75.0856 | 32.17823 | -0.0351 | 1.611827 | 1 |
| 4 | 15577200 | 15579799 | 30.34655 | 50.36952 | 52.41931 | 22.7177 | 0.323604 | 1.285459 | 1 |
| 4 | 16308800 | 16312199 | 51.31656 | 82.04284 | 101.9747 | 42.88681 | 0.029392 | 2.066368 | 1 |
| 4 | 16782600 | 16783999 | 11.51192 | 20.85657 | 30.62479 | 13.25672 | -0.13865 | 0.330855 | 1 |
| 4 | 16810200 | 16810999 | 13.83594 | 24.83388 | 30.04905 | 13.14667 | 0.129101 | 0.404226 | 1 |
| 4 | 16844000 | 16846599 | 27.15475 | 45.22001 | 54.9945 | 23.88715 | 0.099845 | 1.241987 | 1 |
| 4 | 16885600 | 16888199 | 22.68163 | 39.34285 | 44.67261 | 19.60049 | 0.204151 | 0.84031 | 1 |
| 4 | 17133800 | 17134799 | 17.99535 | 31.32251 | 37.49612 | 16.48161 | 0.128905 | 0.76222 | 1 |
| 4 | 17217200 | 17218399 | 27.0038 | 44.57693 | 61.74189 | 26.65721 | -0.08803 | 1.350072 | 1 |
| 4 | 17407400 | 17409599 | 41.24108 | 66.63114 | 72.73341 | 31.05198 | 0.234262 | 1.720674 | 1 |

| | | | | | | | | | |
|---|---|---|---|---|---|---|---|---|---|
| 4 | 17570600 | 17571999 | 25.15164 | 41.83106 | 58.59442 | 25.42921 | -0.10338 | 1.25521 | 1 |
| 4 | 18021400 | 18023799 | 25.62489 | 42.57468 | 64.19584 | 27.84053 | -0.20956 | 1.271579 | 1 |
| 4 | 18221600 | 18222399 | 10.28486 | 18.80934 | 27.78399 | 11.88642 | -0.13581 | 0.262415 | 1 |
| 4 | 18283200 | 18284599 | 16.74638 | 29.39783 | 35.6455 | 15.76797 | 0.110959 | 0.623464 | 1 |
| 4 | 18383400 | 18385399 | 20.62261 | 35.73467 | 41.5655 | 18.23664 | 0.169254 | 0.835519 | 1 |
| 4 | 18427600 | 18428599 | 13.77986 | 24.3415 | 34.74547 | 15.37128 | -0.12025 | 0.530983 | 1 |
| 4 | 18435800 | 18438399 | 28.82126 | 48.28595 | 53.43892 | 23.3001 | 0.235296 | 1.182442 | 1 |
| 4 | 18508800 | 18509999 | 11.95593 | 21.51349 | 34.7041 | 15.15114 | -0.28138 | 0.384165 | 1 |
| 4 | 18562400 | 18563399 | 16.37413 | 29.17474 | 30.77638 | 13.54845 | 0.319621 | 0.467302 | 1 |
| 4 | 18606600 | 18607799 | 24.16883 | 40.41911 | 55.37572 | 24.11734 | -0.07146 | 1.190241 | 1 |
| 4 | 18677600 | 18679399 | 24.89314 | 41.90962 | 55.50504 | 24.25819 | -0.02284 | 1.124069 | 1 |
| 4 | 18695400 | 18696999 | 17.58647 | 30.91745 | 31.92408 | 14.13662 | 0.341555 | 0.598188 | 1 |
| 4 | 18807800 | 18808799 | 15.8485 | 27.85453 | 31.68861 | 14.03163 | 0.20244 | 0.59138 | 1 |
| 4 | 19061200 | 19063599 | 27.28905 | 46.43909 | 48.48239 | 21.25012 | 0.319994 | 1.032729 | 1 |
| 4 | 19149200 | 19153199 | 57.84048 | 92.18822 | 100.5548 | 42.10671 | 0.212676 | 2.168161 | 1 |
| 4 | 19165400 | 19167999 | 19.78829 | 33.78396 | 49.53889 | 21.71885 | -0.16748 | 0.976208 | 1 |
| 4 | 19292600 | 19299199 | 93.41172 | 143.8929 | 186.4804 | 77.45831 | -0.07584 | 2.91809 | 1 |
| 4 | 19386200 | 19387399 | 14.95899 | 26.64068 | 28.61012 | 12.59985 | 0.293823 | 0.457909 | 1 |
| 4 | 19720000 | 19721399 | 13.01749 | 23.972 | 26.49069 | 11.22705 | 0.289633 | 0.241602 | 1 |
| 4 | 19765000 | 19767399 | 19.72169 | 33.27324 | 54.2589 | 23.72876 | -0.32169 | 1.086521 | 1 |
| 4 | 19812800 | 19816999 | 54.4486 | 87.34466 | 95.42138 | 40.33159 | 0.220789 | 1.964723 | 1 |
| 4 | 19959200 | 19965599 | 66.1462 | 105.1763 | 118.7169 | 49.55181 | 0.159732 | 2.257713 | 1 |
| 4 | 20360400 | 20360999 | 11.23409 | 20.79285 | 26.7409 | 11.25527 | 0.079801 | 0.201768 | 1 |
| 4 | 20419000 | 20421799 | 28.8586 | 47.21498 | 74.96502 | 32.24217 | -0.28983 | 1.474011 | 1 |
| 4 | 20626600 | 20628799 | 19.56933 | 33.936 | 45.26579 | 19.8606 | -0.02723 | 0.829328 | 1 |
| 4 | 20855800 | 20861399 | 32.27845 | 53.29258 | 60.10029 | 25.95128 | 0.20751 | 1.358583 | 1 |
| 4 | 20896800 | 20898399 | 19.64358 | 34.25625 | 39.24851 | 17.27244 | 0.191896 | 0.743726 | 1 |
| 4 | 20975200 | 20977199 | 22.3212 | 38.1787 | 50.63427 | 22.20239 | -0.02289 | 0.969472 | 1 |
| 4 | 21287000 | 21288399 | 23.13507 | 38.79268 | 54.856 | 23.93086 | -0.11704 | 1.156767 | 1 |
| 4 | 21379200 | 21381599 | 35.7996 | 58.95906 | 60.3894 | 26.04516 | 0.344743 | 1.396146 | 1 |
| 4 | 21382600 | 21383799 | 29.71754 | 49.43746 | 46.08197 | 20.00249 | 0.482368 | 1.253707 | 1 |
| 4 | 21421800 | 21424999 | 34.91049 | 57.21772 | 66.86901 | 28.78085 | 0.152247 | 1.457946 | 1 |
| 4 | 21470000 | 21470599 | 11.2184 | 20.82362 | 25.73486 | 10.78952 | 0.140974 | 0.183181 | 1 |
| 4 | 21504000 | 21506199 | 19.43008 | 33.6416 | 42.11531 | 18.47742 | 0.063383 | 0.836714 | 1 |
| 4 | 21577600 | 21581799 | 16.9974 | 30.00411 | 31.3936 | 13.89037 | 0.32486 | 0.550086 | 1 |
| 4 | 22223800 | 22226599 | 35.70578 | 58.53834 | 61.81146 | 26.60734 | 0.298417 | 1.453141 | 1 |
| 4 | 22348600 | 22355599 | 90.52488 | 140.1771 | 180.8408 | 75.11649 | -0.06352 | 2.819878 | 1 |
| 4 | 22356800 | 22359199 | 9.124558 | 17.13037 | 27.44293 | 11.2923 | -0.20968 | 0.095764 | 1 |
| 4 | 22587600 | 22589199 | 14.6938 | 25.91856 | 38.16869 | 16.89056 | -0.16614 | 0.556523 | 1 |
| 4 | 23099400 | 23101799 | 19.98268 | 34.74707 | 36.46115 | 16.01667 | 0.318057 | 0.785623 | 1 |
| 4 | 23197200 | 23198199 | 16.2226 | 28.64938 | 31.97853 | 14.14789 | 0.232347 | 0.541862 | 1 |
| 4 | 23288400 | 23288999 | 13.02828 | 23.77763 | 26.98324 | 11.57368 | 0.23943 | 0.292961 | 1 |
| 4 | 23411200 | 23412799 | 14.81765 | 26.28603 | 38.59085 | 17.05144 | -0.1585 | 0.497957 | 1 |
| 4 | 23652200 | 23652599 | 8.637249 | 16.57009 | 21.92191 | 8.60914 | 0.124138 | -0.0775 | 1 |
| 4 | 24047800 | 24049199 | 18.45677 | 32.04623 | 40.56504 | 17.80285 | 0.048271 | 0.803105 | 1 |
| 4 | 24106600 | 24107999 | 22.76439 | 38.80946 | 42.78292 | 18.75474 | 0.242239 | 0.992606 | 1 |
| 4 | 24120200 | 24122999 | 48.2894 | 77.07942 | 106.4824 | 44.68097 | -0.12528 | 2.114686 | 1 |

| | | | | | | | | | |
|---|---|---|---|---|---|---|---|---|---|
| 4 | 24138000 | 24139399 | 20.0632 | 34.90049 | 37.86422 | 16.63703 | 0.27006 | 0.781008 | 1 |
| 4 | 24193600 | 24194999 | 43.3441 | 69.47724 | 93.19716 | 39.34503 | -0.07605 | 1.975223 | 1 |
| 4 | 25188600 | 25189599 | 11.60661 | 20.46893 | 37.88046 | 16.76368 | -0.49378 | 0.54041 | 1 |
| 4 | 25432000 | 25433399 | 9.06094 | 16.75004 | 27.8757 | 11.75253 | -0.29124 | 0.192533 | 1 |
| 4 | 25445000 | 25447999 | 33.67937 | 55.49544 | 60.78766 | 26.22257 | 0.248484 | 1.385046 | 1 |
| 4 | 25541200 | 25544199 | 34.32538 | 56.18196 | 72.44484 | 31.16288 | 0.009899 | 1.477657 | 1 |
| 4 | 25580000 | 25580999 | 16.47274 | 28.79825 | 34.8591 | 15.36637 | 0.11342 | 0.68559 | 1 |
| 4 | 25727600 | 25730799 | 42.41191 | 68.4946 | 73.32261 | 31.28693 | 0.261676 | 1.735288 | 1 |
| 4 | 25851400 | 25855199 | 50.10035 | 80.26251 | 89.5043 | 37.7542 | 0.188961 | 1.997843 | 1 |
| 4 | 25896000 | 25898199 | 24.35229 | 41.73788 | 45.77089 | 20.07317 | 0.250765 | 0.953137 | 1 |
| 4 | 26057600 | 26058599 | 23.51082 | 39.219 | 56.40078 | 24.52446 | -0.14113 | 1.215626 | 1 |
| 4 | 26137200 | 26140199 | 47.496 | 76.30046 | 85.36926 | 36.16108 | 0.188883 | 1.907997 | 1 |
| 4 | 26180800 | 26183599 | 50.9531 | 81.66117 | 96.63292 | 40.78584 | 0.104147 | 1.988947 | 1 |
| 4 | 26369000 | 26370399 | 14.34159 | 24.94971 | 44.779 | 19.67634 | -0.45259 | 0.764608 | 1 |
| 4 | 26555200 | 26557799 | 24.79951 | 42.28507 | 48.48449 | 21.25438 | 0.185654 | 1.005685 | 1 |
| 4 | 26568200 | 26570199 | 29.89216 | 49.82378 | 53.495 | 23.24916 | 0.278976 | 1.238799 | 1 |
| 4 | 26663200 | 26665399 | 23.43842 | 40.17447 | 42.09387 | 18.46069 | 0.316258 | 0.943747 | 1 |
| 4 | 26718200 | 26720799 | 25.73629 | 43.21433 | 49.62844 | 21.66932 | 0.182025 | 1.143318 | 1 |
| 4 | 27070800 | 27071399 | 9.202963 | 17.40576 | 25.39435 | 10.30712 | -0.05935 | 0.041677 | 1 |
| 4 | 27145000 | 27148199 | 37.20803 | 60.46639 | 83.83794 | 35.96689 | -0.1019 | 1.593604 | 1 |
| 4 | 27329400 | 27331999 | 31.10038 | 51.38756 | 62.90382 | 27.17645 | 0.089611 | 1.347646 | 1 |
| 4 | 27549400 | 27550999 | 17.52868 | 30.74129 | 35.62407 | 15.74557 | 0.175963 | 0.643055 | 1 |
| 4 | 27886600 | 27888599 | 22.76383 | 38.93107 | 42.72927 | 18.73598 | 0.249198 | 0.963966 | 1 |
| 4 | 28416000 | 28419399 | 48.97241 | 78.77127 | 86.79745 | 36.83313 | 0.213067 | 1.868611 | 1 |
| 4 | 28626600 | 28627399 | 12.00805 | 22.40248 | 26.20126 | 10.89681 | 0.228855 | 0.159495 | 1 |
| 4 | 28745600 | 28747399 | 30.76481 | 50.55924 | 68.67645 | 29.59254 | -0.06253 | 1.418326 | 1 |
| 4 | 28765600 | 28768599 | 35.77811 | 58.34036 | 73.50377 | 31.57517 | 0.039445 | 1.534087 | 1 |
| 4 | 29013000 | 29015799 | 59.19048 | 94.13361 | 105.9479 | 44.23087 | 0.164135 | 2.241271 | 1 |
| 4 | 29027000 | 29030399 | 94.30365 | 146.4305 | 151.5884 | 62.94797 | 0.256738 | 2.769745 | 1 |
| 4 | 29709600 | 29711199 | 38.77962 | 62.89346 | 72.3336 | 30.98539 | 0.164443 | 1.627223 | 1 |
| 4 | 29716200 | 29717999 | 28.05849 | 45.99345 | 70.89032 | 30.51334 | -0.24574 | 1.446285 | 1 |
| 4 | 29751200 | 29751999 | 13.53162 | 23.96445 | 33.13033 | 14.64922 | -0.07307 | 0.496111 | 1 |
| 4 | 30188400 | 30188999 | 8.791642 | 16.64361 | 24.39076 | 9.879921 | -0.06276 | 0.030245 | 1 |
| 4 | 30233800 | 30236599 | 50.31029 | 80.58176 | 89.87041 | 37.8964 | 0.188449 | 2.005531 | 1 |
| 4 | 30325800 | 30327999 | 23.8651 | 40.55925 | 48.49752 | 21.2545 | 0.124731 | 1.032572 | 1 |
| 4 | 30349800 | 30351399 | 23.54406 | 39.80429 | 48.53115 | 21.23718 | 0.096424 | 1.076834 | 1 |
| 4 | 30431400 | 30433799 | 19.19123 | 32.87968 | 53.27031 | 23.36138 | -0.31009 | 0.946854 | 1 |
| 4 | 30642400 | 30643799 | 16.28787 | 28.55048 | 39.29462 | 17.36037 | -0.071 | 0.65046 | 1 |
| 4 | 30650600 | 30651199 | 11.44895 | 20.8971 | 28.10773 | 12.05466 | -0.00421 | 0.281897 | 1 |
| 4 | 30989000 | 30999399 | 228.1299 | 327.7292 | 445.5045 | 183.9228 | -0.23543 | 4.178659 | 1 |
| 4 | 31362800 | 31364799 | 19.21101 | 33.02623 | 46.89072 | 20.56809 | -0.11956 | 0.906463 | 1 |
| 4 | 31631000 | 31632999 | 15.82461 | 27.71694 | 38.36206 | 16.93726 | -0.07895 | 0.659615 | 1 |
| 4 | 31635800 | 31638999 | 30.50572 | 50.76694 | 54.63338 | 23.72024 | 0.275652 | 1.257409 | 1 |
| 4 | 31885400 | 31886399 | 11.86642 | 22.21454 | 24.08231 | 9.957382 | 0.344098 | 0.129908 | 1 |
| 4 | 31949200 | 31951999 | 30.38503 | 50.4046 | 55.6773 | 24.12015 | 0.237815 | 1.294917 | 1 |
| 4 | 32593600 | 32596799 | 37.11314 | 60.6629 | 64.15974 | 27.58539 | 0.293682 | 1.495934 | 1 |
| 4 | 32791200 | 32792599 | 21.40823 | 35.90319 | 55.82623 | 24.35672 | -0.25346 | 1.151413 | 1 |

| | | | | | | | | | |
|---|---|---|---|---|---|---|---|---|---|
| 4 | 32794400 | 32796399 | 41.50745 | 66.7971 | 86.46326 | 36.71768 | -0.0178 | 1.846105 | 1 |
| 4 | 33211800 | 33217799 | 19.53118 | 34.03471 | 39.39748 | 17.32983 | 0.177261 | 0.757225 | 1 |
| 4 | 33702000 | 33705199 | 43.16356 | 69.74917 | 79.15088 | 33.79928 | 0.178617 | 1.720666 | 1 |
| 4 | 33800600 | 33802999 | 28.2472 | 47.27475 | 55.65404 | 24.24992 | 0.146423 | 1.191967 | 1 |
| 4 | 34049400 | 34052599 | 18.34259 | 31.55936 | 49.00063 | 21.49413 | -0.24774 | 0.899791 | 1 |
| 4 | 34191200 | 34197999 | 96.31525 | 150.2241 | 151.5943 | 62.90864 | 0.298538 | 2.695122 | 1 |
| 4 | 34540000 | 34543999 | 56.38177 | 89.79346 | 108.9481 | 45.57234 | 0.058092 | 2.193596 | 1 |
| 4 | 35028400 | 35032199 | 45.45094 | 73.24425 | 79.88798 | 33.9933 | 0.231123 | 1.803988 | 1 |
| 4 | 35292800 | 35293799 | 12.41541 | 23.16068 | 25.09133 | 10.43655 | 0.33829 | 0.159963 | 1 |
| 4 | 35379000 | 35380599 | 42.25105 | 68.14498 | 80.56244 | 34.31781 | 0.116562 | 1.775942 | 1 |
| 4 | 35541400 | 35543799 | 22.42927 | 37.30878 | 60.57796 | 26.29268 | -0.31571 | 1.247267 | 1 |
| 4 | 35844400 | 35846799 | 20.21063 | 35.2472 | 40.29537 | 17.7338 | 0.195105 | 0.74821 | 1 |
| 4 | 36133600 | 36135799 | 13.16036 | 24.14614 | 29.02856 | 12.36432 | 0.163506 | 0.26776 | 1 |
| 4 | 37224000 | 37226799 | 47.13181 | 75.68525 | 88.89065 | 37.63018 | 0.118324 | 1.920278 | 1 |
| 4 | 37342400 | 37343599 | 13.51587 | 24.19127 | 29.63622 | 12.99683 | 0.109054 | 0.418775 | 1 |
| 4 | 37600600 | 37601999 | 20.4484 | 35.60689 | 37.31906 | 16.40802 | 0.319855 | 0.768603 | 1 |
| 4 | 38111400 | 38112599 | 17.86589 | 31.2102 | 37.8445 | 16.67271 | 0.110931 | 0.711022 | 1 |
| 4 | 38213400 | 38215399 | 31.26812 | 51.58694 | 59.46266 | 25.6646 | 0.175878 | 1.363524 | 1 |
| 4 | 38546600 | 38551199 | 83.60643 | 130.4865 | 143.5189 | 59.55353 | 0.17517 | 2.676364 | 1 |
| 4 | 39040400 | 39047999 | 137.9175 | 209.0997 | 214.6552 | 88.8464 | 0.241977 | 3.251037 | 1 |
| 4 | 39113000 | 39113999 | 15.00816 | 26.9219 | 31.45013 | 13.76653 | 0.178987 | 0.419036 | 1 |
| 4 | 39266200 | 39266999 | 13.95344 | 25.47695 | 26.98916 | 11.57008 | 0.338798 | 0.29939 | 1 |
| 4 | 39353400 | 39355199 | 23.3508 | 39.51178 | 47.47017 | 20.77835 | 0.117489 | 1.063893 | 1 |
| 4 | 39603800 | 39605199 | 20.39842 | 35.61809 | 36.52604 | 16.08725 | 0.351272 | 0.725484 | 1 |
| 4 | 39760800 | 39761199 | 9.22025 | 17.83993 | 22.12074 | 8.43877 | 0.255713 | -0.16716 | 1 |
| 4 | 39849600 | 39857199 | 21.72622 | 36.97344 | 53.84673 | 23.60138 | -0.15852 | 1.016819 | 1 |
| 4 | 39914400 | 39916999 | 28.10865 | 47.32095 | 51.7915 | 22.63488 | 0.251308 | 1.131147 | 1 |
| 4 | 40433400 | 40445799 | 214.1643 | 313.0887 | 341.896 | 140.851 | 0.105042 | 3.915566 | 1 |
| 4 | 40595600 | 40597599 | 61.06901 | 95.84619 | 155.3705 | 64.43805 | -0.37741 | 2.567766 | 1 |
| 4 | 40607400 | 40612599 | 52.37156 | 83.84292 | 97.84642 | 41.22955 | 0.122274 | 2.023933 | 1 |
| 4 | 40652800 | 40661999 | 194.2447 | 284.0839 | 356.0576 | 146.6814 | -0.0931 | 3.907228 | 1 |
| 4 | 41380800 | 41382599 | 12.23906 | 22.61866 | 26.58668 | 11.21343 | 0.206219 | 0.216116 | 1 |
| 4 | 41841800 | 41842799 | 15.68441 | 28.3398 | 29.88883 | 12.98167 | 0.333057 | 0.378069 | 1 |
| 4 | 43748200 | 43754799 | 74.94531 | 117.6338 | 145.0862 | 60.17336 | 0.016451 | 2.572234 | 1 |
| 4 | 43872800 | 43876399 | 32.22938 | 52.11774 | 84.64021 | 36.17684 | -0.33603 | 1.686143 | 1 |
| 4 | 46285800 | 46288799 | 41.39847 | 67.06172 | 72.15065 | 30.88019 | 0.258932 | 1.658897 | 1 |
| 4 | 46509400 | 46512399 | 27.75037 | 45.45231 | 71.31835 | 30.68728 | -0.27187 | 1.456109 | 1 |
| 4 | 47061000 | 47074999 | 137.0161 | 207.5412 | 220.9856 | 91.44005 | 0.188163 | 3.269579 | 1 |
| 4 | 47473800 | 47475599 | 14.40896 | 25.20952 | 39.49729 | 17.42218 | -0.2569 | 0.671559 | 1 |
| 4 | 48442200 | 48443199 | 17.99482 | 31.43948 | 36.80969 | 16.21813 | 0.161261 | 0.707945 | 1 |
| 4 | 48472600 | 48473799 | 10.0522 | 18.24188 | 31.80089 | 13.73757 | -0.38249 | 0.314741 | 1 |
| 4 | 48501400 | 48516599 | 26.75065 | 44.83667 | 49.11999 | 21.42293 | 0.249899 | 1.166102 | 1 |
| 4 | 49264200 | 49266599 | 26.05293 | 43.31455 | 62.63609 | 27.17653 | -0.14937 | 1.262746 | 1 |
| 4 | 49348000 | 49349399 | 15.85752 | 28.11425 | 31.98551 | 14.13984 | 0.20728 | 0.499056 | 1 |
| 4 | 49360400 | 49362799 | 40.503 | 65.46554 | 80.79137 | 34.51044 | 0.058289 | 1.712214 | 1 |
| 4 | 50133600 | 50134999 | 11.63841 | 21.41145 | 33.50443 | 14.21844 | -0.21066 | 0.250142 | 1 |
| 4 | 50319200 | 50320399 | 11.9413 | 21.43599 | 32.17548 | 14.07546 | -0.17993 | 0.393978 | 1 |

| | | | | | | | | | |
|---|---|---|---|---|---|---|---|---|---|
| 4 | 50337600 | 50347999 | 71.53983 | 113.0759 | 129.0432 | 53.60794 | 0.136162 | 2.420129 | 1 |
| 4 | 50524400 | 50525399 | 16.77431 | 29.24168 | 43.09725 | 18.96175 | -0.16966 | 0.742481 | 1 |
| 4 | 50526400 | 50529999 | 48.84321 | 78.67633 | 84.96333 | 36.12991 | 0.244673 | 1.826162 | 1 |
| 4 | 50700000 | 50700999 | 18.7193 | 32.4375 | 44.22932 | 19.4054 | -0.05904 | 0.830275 | 1 |
| 4 | 50708400 | 50717799 | 148.4629 | 225.0531 | 222.1774 | 91.95551 | 0.298135 | 3.257458 | 1 |
| 4 | 51037600 | 51042399 | 76.12787 | 119.8054 | 130.5275 | 54.15386 | 0.198087 | 2.51775 | 1 |
| 4 | 51059200 | 51062999 | 33.56027 | 55.54389 | 59.16224 | 25.58918 | 0.289895 | 1.33304 | 1 |
| 4 | 51313600 | 51326799 | 186.8341 | 277.7135 | 295.4898 | 121.7355 | 0.165029 | 3.6402 | 1 |
| 4 | 51570200 | 51570999 | 13.41663 | 24.33952 | 29.49187 | 12.74525 | 0.138465 | 0.336102 | 1 |
| 4 | 51589400 | 51590999 | 12.72681 | 23.65918 | 27.12815 | 11.35035 | 0.250029 | 0.18857 | 1 |
| 4 | 51654200 | 51657999 | 11.42356 | 20.77722 | 30.71224 | 13.22999 | -0.1434 | 0.310095 | 1 |
| 4 | 51830600 | 51832399 | 27.12137 | 44.87721 | 65.85469 | 28.47946 | -0.17098 | 1.324462 | 1 |
| 4 | 51871800 | 51874599 | 56.08298 | 89.34828 | 100.7497 | 42.16302 | 0.164167 | 2.18104 | 1 |
| 4 | 51875800 | 51876199 | 7.693299 | 14.46771 | 28.05081 | 11.50952 | -0.4787 | 0.077347 | 1 |
| 4 | 51877400 | 51877999 | 14.07858 | 25.16047 | 27.63304 | 12.13376 | 0.264475 | 0.426952 | 1 |
| 4 | 51922600 | 51923399 | 15.05011 | 26.77081 | 31.3924 | 13.83915 | 0.166376 | 0.469512 | 1 |
| 4 | 51953800 | 51962399 | 112.1338 | 172.3009 | 184.7361 | 76.70746 | 0.194631 | 2.973109 | 1 |
| 4 | 53057800 | 53060399 | 40.03193 | 64.39205 | 92.02906 | 39.05443 | -0.16128 | 1.860343 | 1 |
| 4 | 53365800 | 53366999 | 11.66574 | 21.37628 | 27.65763 | 11.80109 | 0.056741 | 0.262183 | 1 |
| 4 | 53392600 | 53393199 | 9.234063 | 17.02139 | 28.37482 | 12.00955 | -0.29766 | 0.211817 | 1 |
| 4 | 53403400 | 53404599 | 17.77332 | 31.20335 | 32.03752 | 14.17301 | 0.349682 | 0.620919 | 1 |
| 4 | 53640400 | 53642799 | 40.87159 | 66.1969 | 81.09984 | 34.70603 | 0.07168 | 1.664301 | 1 |
| 4 | 53680200 | 53681599 | 15.10123 | 26.84061 | 32.92023 | 14.52232 | 0.101295 | 0.478195 | 1 |
| 4 | 53943800 | 53944999 | 39.452 | 63.68448 | 76.27017 | 32.52495 | 0.099353 | 1.744618 | 1 |
| 4 | 54008800 | 54016599 | 132.1986 | 200.7253 | 221.5947 | 91.75925 | 0.138924 | 3.21993 | 1 |
| 4 | 54220200 | 54222599 | 25.85079 | 43.39189 | 51.78155 | 22.6046 | 0.1268 | 1.148035 | 1 |
| 4 | 54241200 | 54244199 | 23.06179 | 39.34363 | 45.83862 | 20.09538 | 0.162787 | 0.989962 | 1 |
| 4 | 55612800 | 55614399 | 16.66784 | 29.07096 | 43.62367 | 19.19955 | -0.19537 | 0.736565 | 1 |
| 4 | 57333400 | 57335599 | 22.36473 | 38.30556 | 41.67283 | 18.27494 | 0.26217 | 0.946578 | 1 |
| 4 | 58260400 | 58261799 | 23.84728 | 39.83087 | 60.79907 | 26.45817 | -0.22696 | 1.206718 | 1 |
| 4 | 58560400 | 58561799 | 15.20704 | 26.84881 | 33.00265 | 14.60175 | 0.093688 | 0.538967 | 1 |
| 4 | 58680800 | 58684799 | 39.71973 | 64.25265 | 77.67282 | 33.20949 | 0.089297 | 1.68511 | 1 |
| 4 | 58768200 | 58775199 | 111.9426 | 171.0734 | 222.6713 | 92.36695 | -0.09097 | 3.084027 | 1 |
| 4 | 59043200 | 59043599 | 7.256967 | 14.05919 | 22.75134 | 8.627684 | -0.11666 | -0.20711 | 1 |
| 4 | 59117400 | 59118399 | 12.77889 | 22.69882 | 34.16799 | 15.08065 | -0.19291 | 0.470194 | 1 |
| 4 | 59264600 | 59271199 | 97.5025 | 149.7781 | 202.9446 | 84.26498 | -0.14307 | 2.975214 | 1 |
| 4 | 59307200 | 59308599 | 22.83956 | 39.25186 | 43.13225 | 18.9193 | 0.248503 | 0.919318 | 1 |
| 4 | 59454000 | 59456399 | 12.7767 | 23.55845 | 24.94258 | 10.55271 | 0.352694 | 0.228327 | 1 |
| 4 | 59501800 | 59503599 | 17.59877 | 30.90417 | 34.9569 | 15.46759 | 0.210532 | 0.620137 | 1 |
| 4 | 59969400 | 59974399 | 46.30992 | 73.89084 | 115.3239 | 48.3641 | -0.30164 | 2.125919 | 1 |
| 4 | 60028800 | 60042999 | 394.8636 | 548.2368 | 574.0379 | 238.2889 | 0.091547 | 4.689312 | 1 |
| 4 | 60176600 | 60177599 | 15.49868 | 27.04602 | 37.57498 | 16.54288 | -0.08443 | 0.714023 | 1 |
| 4 | 60179200 | 60184399 | 54.2086 | 86.62433 | 97.0603 | 40.79184 | 0.17801 | 2.082385 | 1 |
| 4 | 60197400 | 60199199 | 21.4969 | 37.21755 | 38.93476 | 17.08197 | 0.321304 | 0.842682 | 1 |
| 4 | 60263000 | 60264999 | 26.39381 | 44.28924 | 47.53604 | 20.74709 | 0.279335 | 1.150979 | 1 |
| 4 | 60279400 | 60280999 | 30.10606 | 49.26452 | 66.63309 | 28.66046 | -0.05896 | 1.468624 | 1 |
| 4 | 60281800 | 60282999 | 26.08079 | 43.54482 | 51.25017 | 22.29739 | 0.146884 | 1.20615 | 1 |

| | | | | | | | | | |
|---|---|---|---|---|---|---|---|---|---|
| 4 | 60300000 | 60302199 | 15.93237 | 28.2661 | 30.43634 | 13.44731 | 0.286731 | 0.491265 | 1 |
| 4 | 60333600 | 60333999 | 8.987195 | 16.63689 | 28.95618 | 12.18494 | -0.35285 | 0.190509 | 1 |
| 4 | 60345600 | 60346999 | 29.17385 | 48.63059 | 52.94894 | 23.01306 | 0.258843 | 1.234274 | 1 |
| 4 | 60362600 | 60365199 | 71.23485 | 111.1088 | 160.7092 | 66.69094 | -0.22018 | 2.684351 | 1 |
| 4 | 60368000 | 60370399 | 35.80496 | 57.77884 | 87.58649 | 37.33531 | -0.2404 | 1.754409 | 1 |
| 4 | 60387400 | 60389399 | 17.06033 | 30.02292 | 29.5269 | 13.07245 | 0.412032 | 0.579574 | 1 |
| 4 | 60421600 | 60422799 | 21.21973 | 35.89661 | 51.67709 | 22.61802 | -0.14246 | 1.064217 | 1 |
| 4 | 60423600 | 60430999 | 111.6321 | 171.5488 | 184.0752 | 76.43418 | 0.193611 | 2.970648 | 1 |
| 4 | 60500200 | 60501999 | 8.693001 | 16.75102 | 19.29138 | 7.480423 | 0.339379 | -0.12746 | 1 |
| 4 | 60565600 | 60568199 | 29.96617 | 49.05158 | 73.49961 | 31.61767 | -0.20604 | 1.467897 | 1 |
| 4 | 60599200 | 60600999 | 19.09415 | 33.15787 | 41.04026 | 18.01311 | 0.080512 | 0.804492 | 1 |
| 4 | 61300400 | 61301799 | 30.50544 | 50.29777 | 59.32067 | 25.59667 | 0.142604 | 1.370042 | 1 |
| 4 | 61308000 | 61313599 | 108.5217 | 166.7177 | 179.0313 | 74.33663 | 0.19215 | 2.97556 | 1 |
| 4 | 61313800 | 61320399 | 72.17752 | 113.1153 | 150.8521 | 62.56987 | -0.09787 | 2.598782 | 1 |
| 4 | 61342600 | 61345399 | 11.71873 | 21.4427 | 30.21116 | 12.91574 | -0.06748 | 0.276659 | 1 |
| 4 | 61393000 | 61393599 | 11.24611 | 20.85733 | 23.19549 | 9.736122 | 0.290789 | 0.180181 | 1 |
| 4 | 61521600 | 61522599 | 15.52668 | 27.0247 | 42.15981 | 18.53118 | -0.2514 | 0.757689 | 1 |
| 4 | 61548800 | 61549599 | 22.5832 | 37.9608 | 51.15546 | 22.34655 | -0.04779 | 1.125287 | 1 |
| 4 | 61603200 | 61607999 | 91.16711 | 140.2537 | 192.994 | 80.14963 | -0.16366 | 2.94398 | 1 |
| 4 | 61648800 | 61651399 | 76.76185 | 120.3287 | 142.255 | 59.00255 | 0.076032 | 2.596134 | 1 |
| 4 | 61656000 | 61656599 | 15.03899 | 26.30213 | 36.06119 | 15.90159 | -0.06515 | 0.676435 | 1 |
| 4 | 61677600 | 61677999 | 9.612902 | 18.42961 | 20.74563 | 8.162706 | 0.352166 | -0.06659 | 1 |
| 4 | 61681400 | 61681999 | 10.73565 | 20.42488 | 26.65894 | 10.68602 | 0.11571 | 0.018419 | 1 |
| 4 | 61690800 | 61693199 | 76.62986 | 120.4808 | 128.0216 | 53.1076 | 0.233068 | 2.536448 | 1 |
| 4 | 61806800 | 61834399 | 450.0165 | 615.8778 | 723.8326 | 301.1561 | -0.09607 | 4.912069 | 1 |
| 4 | 62084000 | 62084599 | 13.27879 | 23.85213 | 30.50541 | 13.33771 | 0.050071 | 0.394764 | 1 |
| 4 | 62085800 | 62086399 | 10.06516 | 18.89717 | 23.79953 | 9.792296 | 0.135475 | 0.096576 | 1 |
| 4 | 62097200 | 62099599 | 21.69065 | 36.92966 | 52.22929 | 22.89329 | -0.11628 | 1.010392 | 1 |
| 4 | 62126200 | 62127199 | 9.62723 | 17.81742 | 28.32211 | 11.92202 | -0.22419 | 0.190488 | 1 |
| 4 | 62155400 | 62161399 | 42.72524 | 68.78785 | 89.72532 | 38.12784 | -0.02809 | 1.832974 | 1 |
| 4 | 62228000 | 62230799 | 15.97937 | 28.30038 | 32.79837 | 14.50234 | 0.179798 | 0.509135 | 1 |
| 4 | 62365600 | 62370599 | 57.78868 | 92.10178 | 97.60083 | 40.86738 | 0.254201 | 2.168119 | 1 |
| 4 | 63657200 | 63659199 | 19.23143 | 33.09062 | 46.36542 | 20.33823 | -0.10027 | 0.898566 | 1 |
| 4 | 64107600 | 64111599 | 46.31798 | 75.01541 | 73.5055 | 31.45528 | 0.393185 | 1.672178 | 1 |
| 4 | 64344600 | 64345999 | 19.88427 | 33.71049 | 55.34961 | 24.24135 | -0.33125 | 1.047 | 1 |
| 4 | 64785400 | 64787399 | 27.00439 | 44.42604 | 64.94393 | 27.99667 | -0.16749 | 1.393289 | 1 |
| 4 | 64949400 | 64951999 | 34.07031 | 55.4187 | 82.89265 | 35.57636 | -0.21001 | 1.571705 | 1 |
| 4 | 65556600 | 65557599 | 14.14021 | 25.35608 | 35.57726 | 15.57768 | -0.08443 | 0.42126 | 1 |
| 4 | 65988400 | 65989799 | 16.80163 | 29.20816 | 45.26129 | 19.87918 | -0.24209 | 0.785276 | 1 |
| 4 | 65996400 | 65996799 | 8.175645 | 15.72558 | 20.92906 | 8.157782 | 0.125942 | -0.1093 | 1 |
| 4 | 66272200 | 66274199 | 20.02466 | 34.23925 | 51.65537 | 22.64952 | -0.20801 | 0.96729 | 1 |
| 4 | 66636400 | 66637199 | 13.77288 | 24.78024 | 29.00026 | 12.6621 | 0.178875 | 0.385772 | 1 |
| 4 | 66820400 | 66825999 | 70.91202 | 111.5463 | 149.2166 | 61.90399 | -0.09815 | 2.526346 | 1 |
| 4 | 67150800 | 67152399 | 29.11211 | 47.6532 | 64.91593 | 27.92511 | -0.06891 | 1.460013 | 1 |
| 4 | 67858600 | 67862199 | 47.09539 | 75.60326 | 93.14891 | 39.4142 | 0.048878 | 1.933301 | 1 |
| 4 | 68030400 | 68035399 | 54.19902 | 86.53476 | 97.17398 | 40.79003 | 0.173488 | 2.108466 | 1 |
| 4 | 68208200 | 68209199 | 10.17256 | 19.14365 | 23.25327 | 9.529607 | 0.191712 | 0.080382 | 1 |

| | | | | | | | | | |
|---|---|---|---|---|---|---|---|---|---|
| 4 | 68618200 | 68620599 | 24.4155 | 41.60899 | 49.02336 | 21.48972 | 0.146451 | 1.008487 | 1 |
| 4 | 69081800 | 69082799 | 16.78734 | 29.60268 | 32.86993 | 14.54659 | 0.238896 | 0.561605 | 1 |
| 4 | 69095000 | 69095799 | 9.96663 | 18.38663 | 28.81557 | 12.18267 | -0.20834 | 0.213285 | 1 |
| 4 | 69134200 | 69136599 | 29.63846 | 49.19299 | 60.61495 | 26.2691 | 0.080659 | 1.291073 | 1 |
| 4 | 69328200 | 69329399 | 27.72625 | 45.38096 | 71.94252 | 30.94683 | -0.28715 | 1.463975 | 1 |
| 4 | 69331800 | 69334199 | 35.8113 | 58.83756 | 61.99623 | 26.71114 | 0.302646 | 1.426307 | 1 |
| 4 | 69486200 | 69488599 | 22.73858 | 39.30596 | 40.47589 | 17.75728 | 0.343565 | 0.865175 | 1 |
| 4 | 69825000 | 69827399 | 13.86925 | 25.4594 | 29.62482 | 12.6098 | 0.211348 | 0.274745 | 1 |
| 4 | 70287400 | 70290199 | 17.0911 | 29.99802 | 43.38957 | 19.19091 | -0.14253 | 0.639028 | 1 |
| 4 | 70506600 | 70513599 | 82.38911 | 128.458 | 177.1995 | 73.53792 | -0.15234 | 2.697857 | 1 |
| 4 | 70730800 | 70738199 | 121.1773 | 185.2245 | 191.3862 | 79.39327 | 0.242074 | 3.080123 | 1 |
| 4 | 70750200 | 70753999 | 44.57552 | 71.40537 | 110.4696 | 46.59151 | -0.28267 | 1.9995 | 1 |
| 4 | 70789000 | 70790399 | 17.72298 | 30.76729 | 43.70244 | 19.17786 | -0.11726 | 0.806797 | 1 |
| 4 | 70792200 | 70795599 | 24.72054 | 43.06659 | 38.51846 | 16.94045 | 0.547595 | 0.78354 | 1 |
| 4 | 70863800 | 70865599 | 18.8177 | 33.25377 | 31.79076 | 14.06301 | 0.45516 | 0.55215 | 1 |
| 4 | 71063000 | 71065599 | 42.48164 | 68.98682 | 73.73665 | 31.61661 | 0.271686 | 1.613815 | 1 |
| 4 | 71287600 | 71289599 | 5.970836 | 11.54666 | 24.87329 | 9.512247 | -0.5365 | -0.18486 | 1 |
| 4 | 71417400 | 71418599 | 11.76882 | 21.77777 | 25.62438 | 10.78856 | 0.206695 | 0.203206 | 1 |
| 4 | 71426600 | 71428799 | 15.85916 | 27.77494 | 42.46983 | 18.74945 | -0.22211 | 0.667536 | 1 |
| 4 | 71553200 | 71556599 | 44.24168 | 71.52573 | 82.29976 | 35.16574 | 0.159471 | 1.711431 | 1 |
| 4 | 71733200 | 71733799 | 7.600688 | 14.53461 | 25.10475 | 9.950704 | -0.27008 | -0.04962 | 1 |
| 4 | 71997000 | 71999799 | 19.59563 | 33.3904 | 49.01743 | 21.48689 | -0.16956 | 0.991513 | 1 |
| 4 | 72176000 | 72178799 | 18.08465 | 31.42686 | 46.84708 | 20.57086 | -0.18653 | 0.798541 | 1 |
| 4 | 72420000 | 72422599 | 26.56392 | 44.40426 | 59.51183 | 25.91123 | -0.04007 | 1.201012 | 1 |
| 4 | 72460800 | 72461999 | 26.06663 | 43.29741 | 54.88404 | 23.79793 | 0.040104 | 1.268486 | 1 |
| 4 | 72497200 | 72498399 | 29.89662 | 49.22544 | 58.07129 | 25.04283 | 0.141488 | 1.385753 | 1 |
| 4 | 72630400 | 72632199 | 22.77705 | 38.27021 | 55.41261 | 24.20307 | -0.15112 | 1.133907 | 1 |
| 4 | 72645600 | 72646599 | 11.20312 | 20.5438 | 32.34761 | 13.78832 | -0.22401 | 0.263435 | 1 |
| 4 | 73102200 | 73112399 | 138.0815 | 208.1905 | 257.0915 | 106.2253 | -0.03119 | 3.364445 | 1 |
| 4 | 73491000 | 73492799 | 13.07098 | 23.59195 | 28.99515 | 12.61069 | 0.11214 | 0.361207 | 1 |
| 4 | 73668600 | 73669799 | 20.05954 | 34.27172 | 45.89226 | 20.12126 | -0.03681 | 0.966072 | 1 |
| 4 | 73915000 | 73918399 | 16.71997 | 29.23686 | 42.00357 | 18.51917 | -0.13249 | 0.698062 | 1 |
| 4 | 74523200 | 74524199 | 15.07059 | 26.44281 | 40.36809 | 17.84948 | -0.21971 | 0.630685 | 1 |
| 4 | 74561200 | 74564799 | 26.24901 | 43.49685 | 63.65166 | 27.55435 | -0.16682 | 1.302358 | 1 |
| 4 | 74675400 | 74678199 | 35.0883 | 57.48303 | 66.25758 | 28.51221 | 0.171872 | 1.463849 | 1 |
| 4 | 74913200 | 74916399 | 13.31139 | 23.62217 | 34.19477 | 15.1046 | -0.13748 | 0.481437 | 1 |
| 4 | 74984600 | 74992799 | 132.6523 | 198.7973 | 279.8956 | 115.4646 | -0.22802 | 3.482202 | 1 |
| 4 | 75065600 | 75067199 | 17.37952 | 30.29015 | 42.55333 | 18.7199 | -0.10091 | 0.747205 | 1 |
| 4 | 75272400 | 75274199 | 33.5138 | 54.12019 | 85.13928 | 36.32557 | -0.29255 | 1.727863 | 1 |
| 4 | 75298800 | 75299999 | 12.50868 | 22.44044 | 35.22542 | 15.41783 | -0.24463 | 0.409795 | 1 |
| 4 | 75319000 | 75322599 | 74.39604 | 117.3036 | 130.4561 | 54.13764 | 0.170574 | 2.478275 | 1 |
| 4 | 75428400 | 75429999 | 34.92041 | 56.48696 | 81.34634 | 34.7792 | -0.16263 | 1.680344 | 1 |
| 4 | 75523000 | 75525799 | 19.73794 | 34.41314 | 39.81803 | 17.52062 | 0.177821 | 0.749473 | 1 |
| 4 | 75628800 | 75630999 | 26.64969 | 44.88506 | 51.21953 | 22.38797 | 0.191343 | 1.12228 | 1 |
| 4 | 75849400 | 75849999 | 6.439516 | 12.33015 | 25.13335 | 9.924922 | -0.50142 | -0.07279 | 1 |
| 4 | 76132000 | 76133599 | 22.95239 | 39.25775 | 39.1405 | 17.16252 | 0.387247 | 0.957945 | 1 |
| 4 | 76664600 | 76666399 | 15.9867 | 28.43713 | 32.26211 | 14.22241 | 0.213716 | 0.476563 | 1 |

| | | | | | | | | | |
|---|---|---|---|---|---|---|---|---|---|
| 4 | 77222800 | 77223399 | 13.43919 | 24.51975 | 26.3854 | 11.32178 | 0.315114 | 0.295754 | 1 |
| 4 | 77332400 | 77333399 | 12.92324 | 23.01305 | 35.64084 | 15.69739 | -0.23175 | 0.457598 | 1 |
| 4 | 77506600 | 77507399 | 18.86315 | 32.21032 | 47.97635 | 21.03411 | -0.18979 | 0.970405 | 1 |
| 4 | 78046400 | 78049799 | 51.63581 | 82.89555 | 92.65747 | 39.20874 | 0.18913 | 1.936775 | 1 |
| 4 | 78575800 | 78580199 | 54.27238 | 86.49021 | 111.2856 | 46.59195 | -0.02546 | 2.171504 | 1 |
| 4 | 78775600 | 78778599 | 16.51099 | 28.59134 | 47.40652 | 20.79895 | -0.34019 | 0.83026 | 1 |
| 4 | 78898600 | 78899999 | 13.75412 | 24.38269 | 35.61316 | 15.74393 | -0.15138 | 0.492462 | 1 |
| 4 | 79060800 | 79061999 | 19.79943 | 33.23187 | 57.38101 | 25.04779 | -0.40411 | 1.138119 | 1 |
| 4 | 79146200 | 79150999 | 107.0603 | 164.943 | 175.2666 | 72.80199 | 0.21049 | 2.919283 | 1 |
| 4 | 79478800 | 79480799 | 50.79672 | 81.36817 | 88.83439 | 37.46465 | 0.219282 | 2.002967 | 1 |
| 4 | 79485400 | 79485999 | 8.071822 | 15.62051 | 19.87297 | 7.5753 | 0.219851 | -0.19079 | 1 |
| 4 | 79642800 | 79644999 | 24.75178 | 41.94469 | 48.51295 | 21.24437 | 0.172276 | 1.059207 | 1 |
| 4 | 79869200 | 79872399 | 35.39592 | 58.21261 | 61.34479 | 26.44152 | 0.303073 | 1.41417 | 1 |
| 4 | 80326200 | 80333599 | 60.82629 | 96.43951 | 119.3899 | 49.68943 | 0.022625 | 2.334231 | 1 |
| 4 | 80348800 | 80349599 | 12.89705 | 23.04485 | 31.6474 | 13.89854 | -0.05624 | 0.42887 | 1 |
| 4 | 80885200 | 80886999 | 36.77681 | 59.37948 | 88.00053 | 37.53819 | -0.20697 | 1.740272 | 1 |
| 4 | 81067000 | 81069999 | 48.83003 | 78.15998 | 98.42606 | 41.46387 | 0.012451 | 2.025967 | 1 |
| 4 | 81716600 | 81719399 | 30.46835 | 50.27942 | 67.38827 | 29.08838 | -0.04098 | 1.363477 | 1 |
| 4 | 81727200 | 81728199 | 15.09406 | 26.25001 | 44.76432 | 19.66577 | -0.37937 | 0.771713 | 1 |
| 4 | 82128600 | 82129599 | 13.23608 | 23.68328 | 31.96128 | 14.02007 | -0.03043 | 0.419978 | 1 |
| 4 | 83457800 | 83460399 | 18.11919 | 31.55867 | 36.9351 | 16.24167 | 0.161388 | 0.752148 | 1 |
| 4 | 83561600 | 83563799 | 23.98268 | 40.02653 | 58.6194 | 25.49738 | -0.16739 | 1.212897 | 1 |
| 4 | 85849200 | 85849999 | 20.98808 | 36.3071 | 36.98372 | 16.22559 | 0.359384 | 0.845053 | 1 |
| 4 | 87421200 | 87421799 | 9.278253 | 17.34064 | 27.37481 | 11.3505 | -0.19752 | 0.125509 | 1 |
| 4 | 87529400 | 87531799 | 13.74558 | 24.29163 | 37.17918 | 16.44649 | -0.22033 | 0.528714 | 1 |
| 4 | 88576000 | 88578599 | 18.76838 | 32.26629 | 51.68721 | 22.67204 | -0.29301 | 0.909964 | 1 |
| 4 | 89103200 | 89106799 | 61.50882 | 97.19476 | 138.5713 | 57.55837 | -0.1848 | 2.422549 | 1 |
| 4 | 89108200 | 89111999 | 24.73954 | 42.09967 | 47.42149 | 20.78508 | 0.210765 | 1.021405 | 1 |
| 4 | 89156600 | 89157799 | 13.50554 | 24.35887 | 28.58927 | 12.44528 | 0.1777 | 0.369942 | 1 |
| 4 | 89212000 | 89214999 | 25.89683 | 42.49708 | 69.79319 | 30.05518 | -0.33644 | 1.42374 | 1 |
| 4 | 89228200 | 89229599 | 10.92445 | 19.47933 | 35.28578 | 15.52377 | -0.45517 | 0.435788 | 1 |
| 4 | 89489800 | 89490599 | 12.11733 | 22.2308 | 28.11071 | 11.97458 | 0.091233 | 0.259429 | 1 |
| 4 | 89596400 | 89597199 | 9.196922 | 17.058 | 29.93211 | 12.56261 | -0.36269 | 0.178989 | 1 |
| 4 | 89680800 | 89681599 | 23.35924 | 39.44448 | 48.01863 | 21.00526 | 0.098566 | 1.085598 | 1 |
| 4 | 89690400 | 89693199 | 46.26217 | 74.20103 | 87.73422 | 37.07394 | 0.106648 | 1.954371 | 1 |
| 4 | 90279600 | 90286399 | 84.90747 | 133.1408 | 143.8094 | 59.64526 | 0.206516 | 2.594369 | 1 |
| 4 | 90291600 | 90293199 | 9.66132 | 18.09825 | 30.12145 | 12.44069 | -0.26862 | 0.122408 | 1 |
| 4 | 90448000 | 90448599 | 14.97408 | 26.27888 | 36.27835 | 16.04392 | -0.0753 | 0.618555 | 1 |
| 4 | 90649600 | 90651199 | 14.97441 | 27.04084 | 29.79507 | 12.95058 | 0.269259 | 0.373398 | 1 |
| 4 | 90761600 | 90773799 | 351.2118 | 487.7226 | 604.859 | 251.0756 | -0.15229 | 4.685411 | 1 |
| 4 | 90866000 | 90867799 | 23.97883 | 40.18402 | 58.12662 | 25.34832 | -0.14968 | 1.169077 | 1 |
| 4 | 90884200 | 90886399 | 23.22932 | 39.10541 | 55.45972 | 24.23781 | -0.12124 | 1.11819 | 1 |
| 4 | 90887400 | 90888199 | 11.06664 | 20.21185 | 27.73559 | 11.88583 | -0.03181 | 0.276708 | 1 |
| 4 | 91528000 | 91529199 | 16.875 | 29.50996 | 38.33488 | 16.90263 | 0.01198 | 0.689523 | 1 |
| 4 | 91542600 | 91544399 | 42.77142 | 68.77614 | 87.75869 | 37.22461 | 0.001368 | 1.87252 | 1 |
| 4 | 91676800 | 91677799 | 16.24383 | 28.39908 | 41.00993 | 18.07828 | -0.13978 | 0.69754 | 1 |
| 4 | 91694200 | 91696599 | 33.51324 | 54.81288 | 65.28971 | 28.0776 | 0.123362 | 1.482608 | 1 |

| | | | | | | | | | |
|---|---|---|---|---|---|---|---|---|---|
| 4 | 91762600 | 91763599 | 13.62009 | 23.95821 | 38.30847 | 16.9618 | -0.28568 | 0.581607 | 1 |
| 4 | 91860800 | 91862799 | 27.96443 | 45.95734 | 70.1549 | 30.22968 | -0.23047 | 1.41105 | 1 |
| 4 | 92123600 | 92125799 | 19.5194 | 33.13822 | 51.10438 | 22.39157 | -0.24102 | 1.028523 | 1 |
| 4 | 92165200 | 92166199 | 15.87109 | 28.20387 | 30.84074 | 13.60704 | 0.26587 | 0.480279 | 1 |
| 4 | 92290400 | 92293999 | 49.89849 | 80.04792 | 94.04786 | 39.75286 | 0.116123 | 1.95864 | 1 |
| 4 | 92310400 | 92312399 | 22.53747 | 38.26737 | 50.61759 | 22.17922 | -0.02052 | 1.037256 | 1 |
| 4 | 92395000 | 92396199 | 14.12733 | 24.80337 | 39.3808 | 17.4222 | -0.27593 | 0.614103 | 1 |
| 4 | 92411000 | 92412799 | 30.98439 | 51.10262 | 58.82469 | 25.38584 | 0.177679 | 1.367143 | 1 |
| 4 | 92598400 | 92599199 | 15.22384 | 27.1204 | 30.56846 | 13.45893 | 0.224747 | 0.462976 | 1 |
| 4 | 92872200 | 92875799 | 59.75921 | 94.5679 | 132.7477 | 55.18014 | -0.16073 | 2.384325 | 1 |
| 4 | 92899800 | 92903599 | 39.56111 | 63.9722 | 88.48322 | 37.81821 | -0.10516 | 1.697724 | 1 |
| 4 | 93063800 | 93064399 | 11.60763 | 20.94786 | 29.73556 | 12.93492 | -0.09431 | 0.355024 | 1 |
| 4 | 93222600 | 93224199 | 15.08392 | 26.20509 | 44.26895 | 19.43543 | -0.36601 | 0.785503 | 1 |
| 4 | 93467400 | 93469599 | 25.56204 | 42.78801 | 51.77765 | 22.56326 | 0.106855 | 1.177071 | 1 |
| 4 | 93591800 | 93592399 | 11.67283 | 21.20139 | 32.27296 | 13.92627 | -0.18694 | 0.325146 | 1 |
| 4 | 93791200 | 93794199 | 35.55218 | 57.55077 | 78.09107 | 33.40533 | -0.07601 | 1.665221 | 1 |
| 4 | 93839200 | 93841999 | 41.73453 | 67.40625 | 78.01904 | 33.29447 | 0.149365 | 1.733073 | 1 |
| 4 | 93924800 | 93927199 | 20.3736 | 35.41168 | 41.9904 | 18.44061 | 0.142157 | 0.798615 | 1 |
| 4 | 93947000 | 93955399 | 130.3952 | 198.4408 | 214.7343 | 88.9794 | 0.170452 | 3.172367 | 1 |
| 4 | 94090600 | 94091599 | 13.78895 | 24.12266 | 38.54741 | 17.00194 | -0.28512 | 0.668368 | 1 |
| 4 | 94183000 | 94186399 | 22.75024 | 39.27593 | 45.75825 | 20.07393 | 0.165769 | 0.882359 | 1 |
| 4 | 94700800 | 94701999 | 13.09762 | 23.20513 | 36.65185 | 16.20504 | -0.26408 | 0.495588 | 1 |
| 4 | 94799400 | 94801799 | 41.5895 | 66.95581 | 88.82146 | 37.7409 | -0.05246 | 1.832693 | 1 |
| 4 | 94972000 | 94973399 | 13.5268 | 24.74758 | 28.57579 | 12.21895 | 0.217718 | 0.288739 | 1 |
| 4 | 95868600 | 95871399 | 55.26314 | 88.37516 | 98.16955 | 41.30152 | 0.191635 | 2.062118 | 1 |
| 4 | 95943000 | 95945199 | 29.59831 | 49.40984 | 53.53988 | 23.29123 | 0.265659 | 1.219549 | 1 |
| 4 | 96275000 | 96282199 | 110.1219 | 169.3547 | 183.4916 | 76.19968 | 0.180458 | 2.956415 | 1 |
| 4 | 96282800 | 96285799 | 33.79286 | 54.69328 | 83.60638 | 35.76004 | -0.24777 | 1.668265 | 1 |
| 4 | 96350600 | 96356199 | 76.34767 | 119.5012 | 167.4163 | 69.44925 | -0.17028 | 2.625551 | 1 |
| 4 | 96511800 | 96512999 | 15.36046 | 27.8436 | 28.8048 | 12.46078 | 0.364216 | 0.353546 | 1 |
| 4 | 96882600 | 96883799 | 12.71606 | 23.27397 | 26.52332 | 11.33491 | 0.236951 | 0.273625 | 1 |
| 4 | 97238200 | 97241199 | 51.18486 | 81.6741 | 110.5804 | 46.37878 | -0.09684 | 2.127872 | 1 |
| 4 | 98747400 | 98749399 | 13.95991 | 24.73582 | 40.77858 | 18.02917 | -0.32554 | 0.508093 | 1 |
| 4 | 98783600 | 98784799 | 15.16829 | 26.87646 | 37.02873 | 16.37134 | -0.06803 | 0.508365 | 1 |
| 4 | 98793400 | 98797799 | 45.30014 | 72.89377 | 81.64825 | 34.66821 | 0.190228 | 1.85054 | 1 |
| 4 | 99189600 | 99193799 | 25.43812 | 43.28186 | 45.31176 | 19.86014 | 0.316007 | 1.024115 | 1 |
| 4 | 99320600 | 99322599 | 27.63708 | 46.107 | 52.32521 | 22.75382 | 0.19931 | 1.222112 | 1 |
| 4 | 100426200 | 100428799 | 24.43966 | 41.5602 | 47.96012 | 21.01998 | 0.175858 | 1.027547 | 1 |
| 4 | 100488400 | 100491199 | 45.88933 | 73.69555 | 99.53967 | 42.14302 | -0.08285 | 1.920552 | 1 |
| 4 | 100558800 | 100560999 | 18.29587 | 31.77523 | 49.55259 | 21.75083 | -0.25146 | 0.811895 | 1 |
| 4 | 100793200 | 100798999 | 75.14033 | 117.2793 | 163.1501 | 67.69917 | -0.16328 | 2.674612 | 1 |
| 4 | 101525000 | 101527399 | 19.94661 | 34.6552 | 42.04105 | 18.45805 | 0.109415 | 0.803455 | 1 |
| 4 | 101724800 | 101726799 | 38.6402 | 61.96381 | 99.55442 | 42.05628 | -0.33511 | 1.957761 | 1 |
| 4 | 102562600 | 102563599 | 14.91655 | 26.71614 | 27.52928 | 12.06603 | 0.35777 | 0.419647 | 1 |
| 4 | 103023600 | 103028399 | 60.89744 | 96.47297 | 129.3527 | 53.79822 | -0.09343 | 2.359501 | 1 |
| 4 | 103128200 | 103130799 | 27.05241 | 44.79051 | 59.78093 | 25.86374 | -0.03448 | 1.311676 | 1 |
| 4 | 105296200 | 105297199 | 13.22141 | 23.19314 | 39.99896 | 17.68298 | -0.39443 | 0.625212 | 1 |

| | | | | | | | | | |
|---|---|---|---|---|---|---|---|---|---|
| 4 | 105433600 | 105436199 | 25.82477 | 42.69163 | 67.40511 | 29.13105 | -0.27637 | 1.332447 | 1 |
| 4 | 105536800 | 105538599 | 22.31232 | 37.59539 | 60.90789 | 26.62609 | -0.31256 | 1.112951 | 1 |
| 4 | 106248600 | 106254799 | 78.6428 | 123.8888 | 133.2796 | 55.30292 | 0.217437 | 2.500374 | 1 |
| 4 | 106665600 | 106671799 | 57.25694 | 91.03978 | 122.3812 | 51.0801 | -0.09202 | 2.24923 | 1 |
| 4 | 106749800 | 106752199 | 28.83857 | 47.64246 | 65.52057 | 28.30373 | -0.07789 | 1.345405 | 1 |
| 4 | 108148200 | 108153599 | 73.02751 | 115.2882 | 125.9191 | 52.27822 | 0.198004 | 2.448987 | 1 |
| 4 | 108160400 | 108166799 | 79.74693 | 125.2595 | 152.1702 | 63.11725 | 0.039027 | 2.560342 | 1 |
| 4 | 108272000 | 108274799 | 20.76828 | 36.3081 | 38.09578 | 16.79323 | 0.318621 | 0.714034 | 1 |
| 4 | 110043000 | 110044999 | 30.22521 | 49.8814 | 59.28184 | 25.59035 | 0.132103 | 1.359132 | 1 |
| 4 | 110178600 | 110179999 | 15.35317 | 27.15408 | 32.1171 | 14.205 | 0.150209 | 0.52232 | 1 |
| 4 | 110332200 | 110333599 | 13.42678 | 24.1772 | 28.52312 | 12.44194 | 0.168205 | 0.377165 | 1 |
| 4 | 110350200 | 110351799 | 22.86069 | 38.66898 | 49.05025 | 21.46771 | 0.039528 | 1.069653 | 1 |
| 4 | 110357000 | 110360399 | 30.09334 | 49.60635 | 75.30472 | 32.49042 | -0.22079 | 1.382772 | 1 |
| 4 | 110479600 | 110482399 | 29.15401 | 48.30326 | 67.9887 | 29.4287 | -0.11098 | 1.314569 | 1 |
| 4 | 110545200 | 110545999 | 13.29994 | 24.30344 | 27.54336 | 11.7957 | 0.242966 | 0.290861 | 1 |
| 4 | 110566600 | 110567999 | 20.76095 | 35.28469 | 52.73027 | 23.1096 | -0.19583 | 1.024023 | 1 |
| 4 | 110720400 | 110725999 | 46.43743 | 74.53974 | 91.15677 | 38.56554 | 0.059407 | 1.931878 | 1 |
| 4 | 111192200 | 111193399 | 14.0047 | 24.76992 | 37.1017 | 16.40952 | -0.18905 | 0.519696 | 1 |
| 4 | 111428000 | 111430599 | 6.546561 | 12.68967 | 22.18073 | 8.390323 | -0.22253 | -0.22265 | 1 |
| 4 | 111712400 | 111713399 | 16.45335 | 29.12203 | 29.83725 | 13.19464 | 0.356416 | 0.51391 | 1 |
| 4 | 111792800 | 111794599 | 26.90714 | 45.14411 | 49.64211 | 21.66432 | 0.244478 | 1.156311 | 1 |
| 4 | 111816400 | 111818399 | 27.40509 | 46.03647 | 54.5278 | 23.81507 | 0.137555 | 1.148791 | 1 |
| 4 | 111896000 | 111902199 | 75.87931 | 119.9371 | 127.6838 | 53.04375 | 0.236316 | 2.423221 | 1 |
| 4 | 112248200 | 112253799 | 93.19858 | 145.6624 | 146.774 | 60.89542 | 0.302972 | 2.657949 | 1 |
| 4 | 112396800 | 112397399 | 9.624692 | 18.41422 | 21.78562 | 8.622782 | 0.272959 | -0.04453 | 1 |
| 4 | 112398200 | 112404399 | 71.71753 | 114.1526 | 113.3371 | 47.37114 | 0.346069 | 2.228559 | 1 |
| 4 | 112537800 | 112539599 | 25.54327 | 42.70382 | 64.44072 | 28.0596 | -0.21064 | 1.202305 | 1 |
| 4 | 112644000 | 112650799 | 91.20708 | 143.0234 | 151.0621 | 62.65312 | 0.238828 | 2.598605 | 1 |
| 4 | 112651600 | 112655599 | 52.24755 | 83.62151 | 94.97969 | 40.00609 | 0.160875 | 2.031984 | 1 |
| 4 | 112849800 | 112859599 | 122.0782 | 188.6582 | 166.6633 | 69.22826 | 0.480042 | 2.869821 | 1 |
| 4 | 112965600 | 112978999 | 172.947 | 259.0855 | 258.7744 | 106.7436 | 0.266475 | 3.495528 | 1 |
| 4 | 113228600 | 113240799 | 220.8714 | 323.0346 | 368.6271 | 151.8593 | 0.042184 | 3.91048 | 1 |
| 4 | 113267400 | 113273799 | 158.8771 | 238.2794 | 239.9786 | 99.01412 | 0.255978 | 3.471352 | 1 |
| 4 | 113789200 | 113792599 | 47.12648 | 76.21468 | 75.81179 | 32.40725 | 0.36961 | 1.703233 | 1 |
| 4 | 113865600 | 113876799 | 183.4786 | 274.7309 | 265.4016 | 109.4671 | 0.313933 | 3.507408 | 1 |
| 4 | 114031600 | 114034599 | 40.20761 | 65.29029 | 72.45948 | 31.0677 | 0.217465 | 1.607741 | 1 |
| 4 | 114074800 | 114075999 | 22.95847 | 38.46523 | 53.78599 | 23.45202 | -0.10088 | 1.16353 | 1 |
| 4 | 114346000 | 114347799 | 22.5381 | 38.87503 | 43.10325 | 18.90868 | 0.236672 | 0.885915 | 1 |
| 4 | 114990200 | 114992599 | 18.47839 | 31.80184 | 45.82977 | 20.1034 | -0.14071 | 0.889919 | 1 |
| 4 | 115057000 | 115058399 | 22.00224 | 37.44556 | 45.89488 | 20.11614 | 0.08953 | 1.009127 | 1 |
| 4 | 115176400 | 115179399 | 30.02903 | 50.4784 | 52.39356 | 22.88763 | 0.327452 | 1.149868 | 1 |
| 4 | 115207600 | 115212799 | 72.76588 | 114.7878 | 127.4126 | 52.87793 | 0.173881 | 2.467611 | 1 |
| 4 | 116275200 | 116278999 | 33.3073 | 54.81422 | 59.05267 | 25.46074 | 0.271662 | 1.39742 | 1 |
| 4 | 117050400 | 117052399 | 25.61643 | 42.87597 | 51.22927 | 22.32328 | 0.125172 | 1.178768 | 1 |
| 4 | 117311400 | 117313199 | 23.39516 | 39.82579 | 47.42811 | 20.78857 | 0.130813 | 1.012494 | 1 |
| 4 | 117654400 | 117657399 | 14.86037 | 26.10991 | 36.26226 | 16.0527 | -0.08386 | 0.60004 | 1 |
| 4 | 118601000 | 118601599 | 10.51454 | 19.31685 | 30.34258 | 12.90111 | -0.21736 | 0.246507 | 1 |

| | | | | | | | | | |
|---|---|---|---|---|---|---|---|---|---|
| 4 | 118602400 | 118606999 | 60.38699 | 96.54833 | 98.07756 | 41.25026 | 0.320009 | 2.072496 | 1 |
| 4 | 119420200 | 119421399 | 16.11101 | 28.14485 | 42.5892 | 18.76315 | -0.20698 | 0.711803 | 1 |
| 4 | 120966200 | 120968399 | 21.12972 | 36.13852 | 48.48264 | 21.25882 | -0.03942 | 0.962126 | 1 |
| 4 | 121027800 | 121036799 | 311.7079 | 436.4281 | 561.5745 | 232.8113 | -0.19355 | 4.560761 | 1 |
| 4 | 121150000 | 121152999 | 54.14381 | 86.89028 | 91.71357 | 38.78788 | 0.271034 | 1.951397 | 1 |
| 4 | 121460400 | 121463799 | 66.22845 | 105.3058 | 110.7388 | 46.22122 | 0.261679 | 2.2539 | 1 |
| 4 | 121750000 | 121750799 | 15.72167 | 27.4307 | 37.14792 | 16.3532 | -0.0476 | 0.718726 | 1 |
| 4 | 121780400 | 121781599 | 10.00736 | 18.91438 | 23.37184 | 9.499738 | 0.17779 | 0.05371 | 1 |
| 4 | 122174200 | 122174599 | 9.571371 | 18.26813 | 22.87149 | 9.106058 | 0.184629 | -0.02124 | 1 |
| 4 | 122401400 | 122402799 | 11.71811 | 21.61632 | 31.93086 | 13.501 | -0.12392 | 0.23455 | 1 |
| 4 | 122579200 | 122589399 | 37.246 | 61.07982 | 70.09266 | 30.17549 | 0.178794 | 1.458784 | 1 |
| 4 | 123195600 | 123198399 | 23.53747 | 40.29237 | 43.18152 | 18.93573 | 0.283601 | 0.958299 | 1 |
| 4 | 123668400 | 123676799 | 119.2581 | 181.7315 | 205.7186 | 85.27724 | 0.10704 | 3.142083 | 1 |
| 4 | 123761200 | 123767599 | 89.28622 | 139.1997 | 155.807 | 64.66085 | 0.148713 | 2.700211 | 1 |
| 4 | 125888000 | 125898799 | 114.1998 | 173.4727 | 235.7845 | 97.6479 | -0.16038 | 3.207394 | 1 |
| 4 | 126956200 | 126958199 | 24.53606 | 41.86102 | 47.40642 | 20.78279 | 0.203568 | 0.996785 | 1 |
| 4 | 129706000 | 129707999 | 18.84817 | 32.97075 | 36.69914 | 16.18565 | 0.233876 | 0.693395 | 1 |
| 4 | 131349200 | 131352599 | 26.02101 | 44.09371 | 47.97508 | 21.00857 | 0.260098 | 1.063084 | 1 |
| 4 | 131812400 | 131814999 | 22.05003 | 36.87298 | 60.72627 | 26.44639 | -0.33619 | 1.187549 | 1 |
| 4 | 133771800 | 133773199 | 12.97684 | 22.87932 | 38.28679 | 16.9444 | -0.34971 | 0.551771 | 1 |
| 4 | 135722600 | 135723799 | 17.31968 | 30.669 | 32.70988 | 14.4637 | 0.298986 | 0.520012 | 1 |
| 4 | 136120800 | 136122999 | 16.04783 | 27.98331 | 41.00753 | 18.04559 | -0.16148 | 0.729749 | 1 |
| 4 | 138770200 | 138772599 | 22.40002 | 37.92725 | 56.56963 | 24.76612 | -0.19341 | 1.066384 | 1 |
| 4 | 138853800 | 138855399 | 19.87872 | 34.0035 | 48.48442 | 21.25983 | -0.12689 | 0.95651 | 1 |
| 4 | 138863000 | 138865999 | 29.77619 | 48.72352 | 75.29408 | 32.38547 | -0.2508 | 1.472387 | 1 |
| 4 | 141248200 | 141249799 | 18.49149 | 32.22955 | 38.40151 | 16.8939 | 0.135771 | 0.74687 | 1 |
| 4 | 141574600 | 141582799 | 96.15109 | 149.3072 | 157.3092 | 65.32306 | 0.231453 | 2.77088 | 1 |
| 4 | 141688200 | 141690599 | 24.19231 | 41.78579 | 42.49959 | 18.64465 | 0.361081 | 0.880349 | 1 |
| 4 | 141887400 | 141891399 | 37.63272 | 60.47601 | 95.70223 | 40.55791 | -0.30962 | 1.88848 | 1 |
| 4 | 143189000 | 143191999 | 37.72024 | 61.42977 | 68.97865 | 29.61769 | 0.203874 | 1.553562 | 1 |
| 4 | 143207800 | 143220999 | 195.6273 | 289.3207 | 298.1687 | 122.7937 | 0.204319 | 3.727407 | 1 |
| 4 | 144262000 | 144263599 | 20.52532 | 35.60216 | 42.03859 | 18.44478 | 0.147848 | 0.825055 | 1 |
| 4 | 144264400 | 144266799 | 37.55588 | 60.96477 | 71.52105 | 30.65808 | 0.137143 | 1.607765 | 1 |
| 4 | 144324200 | 144328399 | 96.04709 | 148.8738 | 159.7117 | 66.33879 | 0.203379 | 2.801933 | 1 |
| 4 | 144378800 | 144393999 | 316.7457 | 449.6724 | 450.4242 | 186.3131 | 0.18777 | 4.356051 | 1 |
| 4 | 144543000 | 144544399 | 18.65767 | 31.79669 | 51.64249 | 22.63784 | -0.31482 | 0.989254 | 1 |
| 4 | 144563400 | 144571799 | 78.75894 | 123.6161 | 141.7358 | 58.78358 | 0.121541 | 2.572883 | 1 |
| 4 | 144573000 | 144574999 | 30.49874 | 50.50902 | 57.36383 | 24.82222 | 0.197653 | 1.314381 | 1 |
| 4 | 144575800 | 144578599 | 16.01137 | 28.4019 | 30.8202 | 13.61867 | 0.275338 | 0.492013 | 1 |
| 4 | 144920600 | 144922199 | 15.86672 | 27.38002 | 46.89359 | 20.57172 | -0.38784 | 0.856688 | 1 |
| 4 | 145201400 | 145214199 | 221.3565 | 323.9043 | 358.6053 | 147.7262 | 0.086486 | 3.901885 | 1 |
| 4 | 145233000 | 145233999 | 13.84071 | 25.47786 | 29.05415 | 12.32012 | 0.243883 | 0.255429 | 1 |
| 4 | 145471400 | 145476799 | 77.96402 | 122.4556 | 132.9968 | 55.16429 | 0.200446 | 2.557759 | 1 |
| 4 | 145500800 | 145503199 | 35.05984 | 56.78046 | 76.34779 | 32.67092 | -0.06226 | 1.654013 | 1 |
| 4 | 145552200 | 145552799 | 9.867952 | 18.62788 | 22.29346 | 9.085118 | 0.219828 | 0.055393 | 1 |
| 4 | 146065000 | 146075599 | 241.3274 | 349.7773 | 345.128 | 142.309 | 0.239248 | 4.047199 | 1 |
| 4 | 146102400 | 146115199 | 631.5803 | 850.5181 | 776.653 | 323.9379 | 0.244465 | 5.169019 | 1 |

| | | | | | | | | | |
|---|---|---|---|---|---|---|---|---|---|
| 4 | 146135600 | 146147599 | 237.5556 | 348.3105 | 334.1309 | 137.6266 | 0.295952 | 3.87221 | 1 |
| 4 | 146172600 | 146174599 | 38.45625 | 62.50748 | 77.60613 | 33.29837 | 0.057323 | 1.588053 | 1 |
| 4 | 146176000 | 146177399 | 62.15946 | 98.09146 | 131.903 | 54.75144 | -0.10204 | 2.453233 | 1 |
| 4 | 146178400 | 146180799 | 70.19546 | 110.4889 | 129.2236 | 53.61411 | 0.096117 | 2.510025 | 1 |
| 4 | 146182200 | 146182999 | 15.38809 | 27.38715 | 32.85969 | 14.47924 | 0.133909 | 0.47102 | 1 |
| 4 | 146249400 | 146251599 | 61.65575 | 97.93257 | 112.4779 | 46.89801 | 0.133242 | 2.283136 | 1 |
| 4 | 146254000 | 146256799 | 53.48484 | 85.71569 | 88.9298 | 37.53258 | 0.293516 | 1.992013 | 1 |
| 4 | 146326200 | 146336999 | 439.6132 | 603.0316 | 630.884 | 262.3885 | 0.075131 | 4.87575 | 1 |
| 4 | 146339200 | 146341799 | 36.51842 | 59.66665 | 69.51539 | 29.88363 | 0.154147 | 1.507076 | 1 |
| 4 | 146370000 | 146371199 | 15.91755 | 28.12972 | 33.95818 | 15.02169 | 0.120189 | 0.535974 | 1 |
| 4 | 146415600 | 146427799 | 1951.003 | 2357.494 | 2514.33 | 1061.954 | -0.1369 | 6.911796 | 1 |
| 4 | 146456800 | 146458999 | 9.313954 | 17.5089 | 28.55263 | 11.72284 | -0.2325 | 0.092759 | 1 |
| 4 | 146557600 | 146628199 | 5606.722 | 6221.216 | 6681.086 | 2860.299 | -0.27229 | 8.25447 | 1 |
| 4 | 146643200 | 146643599 | 8.718758 | 16.86221 | 20.39799 | 7.79754 | 0.288317 | -0.17103 | 1 |
| 4 | 146649200 | 146649799 | 10.3063 | 19.34952 | 23.22561 | 9.556478 | 0.204024 | 0.096062 | 1 |
| 4 | 146659600 | 146661599 | 13.61567 | 25.04475 | 31.25655 | 13.2676 | 0.113566 | 0.263305 | 1 |
| 4 | 146687800 | 146690199 | 45.93608 | 73.81824 | 82.47234 | 34.95028 | 0.191531 | 1.893332 | 1 |
| 4 | 146985800 | 147040399 | 1202.726 | 1523.721 | 1621.13 | 680.8381 | -0.06415 | 6.156194 | 1 |
| 4 | 147260600 | 147261199 | 52.97579 | 84.51283 | 94.66942 | 39.69301 | 0.175801 | 2.13309 | 1 |
| 4 | 147263800 | 147266599 | 57.81419 | 93.21312 | 78.10823 | 33.25718 | 0.611507 | 1.8118 | 1 |
| 4 | 147302600 | 147313599 | 1042.549 | 1318.246 | 1630.009 | 684.7038 | -0.28379 | 6.186497 | 1 |
| 4 | 147314400 | 147314999 | 13.64858 | 24.55405 | 29.53415 | 12.89634 | 0.139524 | 0.387166 | 1 |
| 4 | 147419200 | 147438999 | 3365.827 | 3921.137 | 3461.055 | 1469.804 | 0.082308 | 7.487437 | 1 |
| 4 | 147472000 | 147492599 | 2074.663 | 2507.144 | 2614.55 | 1104.269 | -0.10437 | 6.910762 | 1 |
| 4 | 147501600 | 147506999 | 196.072 | 292.9494 | 262.5393 | 108.2407 | 0.419298 | 3.548403 | 1 |
| 4 | 147742000 | 147743199 | 17.69227 | 31.03514 | 30.61087 | 13.53233 | 0.407298 | 0.630534 | 1 |
| 4 | 147745600 | 147762399 | 627.1901 | 839.0989 | 831.4834 | 347.1436 | 0.1171 | 5.272336 | 1 |
| 4 | 147788400 | 147798999 | 125.8881 | 193.3931 | 190.3492 | 79.03581 | 0.317711 | 2.983826 | 1 |
| 4 | 155768200 | 155769199 | 11.76927 | 21.24588 | 32.68654 | 14.21331 | -0.20962 | 0.360025 | 1 |
| 4 | 156345800 | 156347599 | 26.85206 | 44.07663 | 67.07211 | 28.88672 | -0.22655 | 1.421402 | 1 |
| 4 | 156510200 | 156510799 | 9.513037 | 17.90429 | 24.89087 | 10.199 | -0.00098 | 0.080395 | 1 |
| 4 | 156600000 | 156600599 | 11.32994 | 21.10934 | 22.59823 | 9.419493 | 0.352812 | 0.1511 | 1 |
| 5 | 3235200 | 3239199 | 22.36682 | 37.73452 | 52.76116 | 23.07349 | -0.1007 | 1.093178 | 1 |
| 5 | 3322800 | 3330199 | 87.66929 | 136.4533 | 164.7338 | 68.38238 | 0.037857 | 2.728008 | 1 |
| 5 | 3593600 | 3594599 | 9.917272 | 17.995 | 32.32814 | 13.9673 | -0.42565 | 0.316171 | 1 |
| 5 | 3924800 | 3926199 | 17.12845 | 30.21598 | 33.83263 | 14.97146 | 0.227095 | 0.561165 | 1 |
| 5 | 3943400 | 3950999 | 73.26003 | 116.1717 | 120.7415 | 50.26003 | 0.274799 | 2.340892 | 1 |
| 5 | 4176600 | 4180799 | 16.27811 | 29.44074 | 30.30744 | 13.14773 | 0.368752 | 0.376622 | 1 |
| 5 | 4249800 | 4250999 | 15.72035 | 27.43273 | 43.63666 | 19.21157 | -0.27896 | 0.724674 | 1 |
| 5 | 4423000 | 4427199 | 6.248671 | 12.11407 | 22.74623 | 8.598208 | -0.32418 | -0.22802 | 1 |
| 5 | 4512000 | 4514399 | 27.20783 | 46.46822 | 47.37797 | 20.77208 | 0.354739 | 0.998037 | 1 |
| 5 | 4564600 | 4568599 | 29.68869 | 48.565 | 69.22052 | 29.76968 | -0.13466 | 1.476044 | 1 |
| 5 | 4685000 | 4686199 | 23.74688 | 39.31037 | 64.94041 | 28.09239 | -0.34121 | 1.310017 | 1 |
| 5 | 5168400 | 5169799 | 18.66327 | 32.68818 | 36.61705 | 16.16652 | 0.224657 | 0.671754 | 1 |
| 5 | 5281600 | 5288399 | 144.8407 | 218.2358 | 246.7925 | 101.9517 | 0.09484 | 3.378677 | 1 |
| 5 | 5564200 | 5566399 | 17.27524 | 29.93112 | 46.34599 | 20.33399 | -0.24178 | 0.827033 | 1 |
| 5 | 5647000 | 5647999 | 11.79244 | 21.46078 | 32.26099 | 13.88692 | -0.16681 | 0.313714 | 1 |

| | | | | | | | | | |
|---|---|---|---|---|---|---|---|---|---|
| 5 | 5679600 | 5681199 | 17.38352 | 30.28505 | 37.51394 | 16.49885 | 0.080019 | 0.746783 | 1 |
| 5 | 5826200 | 5830599 | 62.12875 | 98.80327 | 121.891 | 50.88446 | 0.031866 | 2.24775 | 1 |
| 5 | 5963800 | 5965399 | 14.67122 | 25.84277 | 36.66491 | 16.22961 | -0.11339 | 0.568858 | 1 |
| 5 | 6057400 | 6060199 | 36.25658 | 59.21829 | 69.03792 | 29.67462 | 0.152827 | 1.509353 | 1 |
| 5 | 6135400 | 6137199 | 30.86346 | 51.21139 | 55.71395 | 24.14031 | 0.259725 | 1.293154 | 1 |
| 5 | 6192000 | 6193199 | 17.46004 | 30.28646 | 44.25275 | 19.41629 | -0.15799 | 0.815864 | 1 |
| 5 | 6235600 | 6236599 | 16.40984 | 28.92893 | 31.10894 | 13.76803 | 0.284987 | 0.561422 | 1 |
| 5 | 6265800 | 6269799 | 15.29265 | 27.36327 | 28.79767 | 12.63226 | 0.326401 | 0.428836 | 1 |
| 5 | 6277400 | 6287399 | 133.8926 | 202.9287 | 219.119 | 90.68445 | 0.168707 | 3.256483 | 1 |
| 5 | 6305200 | 6312399 | 99.7439 | 154.3228 | 165.5725 | 68.77474 | 0.201237 | 2.837053 | 1 |
| 5 | 6441400 | 6441799 | 6.777408 | 13.20217 | 17.38518 | 6.337738 | 0.230508 | -0.36862 | 1 |
| 5 | 6519800 | 6522199 | 30.62899 | 50.28619 | 73.19387 | 31.52587 | -0.16257 | 1.43322 | 1 |
| 5 | 6532800 | 6540799 | 167.4398 | 247.1714 | 326.5942 | 134.4974 | -0.15653 | 3.755913 | 1 |
| 5 | 6605800 | 6607599 | 15.70654 | 27.45501 | 44.63492 | 19.67342 | -0.31017 | 0.700763 | 1 |
| 5 | 6664800 | 6666599 | 20.46274 | 35.57355 | 39.93842 | 17.5416 | 0.220832 | 0.795244 | 1 |
| 5 | 6730200 | 6732399 | 35.26789 | 57.52292 | 69.96845 | 30.05926 | 0.090344 | 1.527298 | 1 |
| 5 | 6968200 | 6969799 | 14.2549 | 25.45422 | 32.94537 | 14.47666 | 0.0279 | 0.44292 | 1 |
| 5 | 7134400 | 7136199 | 24.47248 | 40.96519 | 55.95241 | 24.38287 | -0.06712 | 1.181125 | 1 |
| 5 | 7394000 | 7395799 | 14.00169 | 24.57245 | 36.30823 | 16.05718 | -0.17293 | 0.61167 | 1 |
| 5 | 7449200 | 7450599 | 21.66959 | 37.52063 | 41.49719 | 18.20626 | 0.241372 | 0.845484 | 1 |
| 5 | 7555200 | 7555999 | 17.66974 | 31.10341 | 33.40546 | 14.78884 | 0.285797 | 0.588242 | 1 |
| 5 | 7798600 | 7807599 | 74.51935 | 117.6327 | 131.6593 | 54.65888 | 0.162673 | 2.453116 | 1 |
| 5 | 7810600 | 7813399 | 41.76242 | 67.64579 | 74.47854 | 31.87461 | 0.225602 | 1.662772 | 1 |
| 5 | 7816000 | 7816999 | 14.76151 | 26.55661 | 29.896 | 13.05419 | 0.234426 | 0.395259 | 1 |
| 5 | 7854800 | 7855799 | 18.56308 | 32.29446 | 35.47437 | 15.58801 | 0.252299 | 0.768355 | 1 |
| 5 | 7906800 | 7907999 | 18.52672 | 32.3116 | 37.8907 | 16.67588 | 0.15879 | 0.737941 | 1 |
| 5 | 7987000 | 7988999 | 31.0295 | 51.32296 | 53.48763 | 23.12469 | 0.321204 | 1.328657 | 1 |
| 5 | 8065200 | 8080999 | 354.265 | 496.4162 | 534.2689 | 221.4633 | 0.065262 | 4.550052 | 1 |
| 5 | 8218400 | 8220999 | 46.38061 | 74.67286 | 83.60384 | 35.52648 | 0.191846 | 1.837484 | 1 |
| 5 | 8346200 | 8348599 | 16.76072 | 29.16622 | 43.14201 | 18.9603 | -0.17513 | 0.767343 | 1 |
| 5 | 8514400 | 8515199 | 14.54234 | 25.60293 | 35.23558 | 15.59849 | -0.06983 | 0.572217 | 1 |
| 5 | 8672200 | 8673599 | 12.86187 | 22.48122 | 44.66999 | 19.6879 | -0.59826 | 0.684465 | 1 |
| 5 | 8760600 | 8762399 | 25.03605 | 40.92545 | 73.03841 | 31.40361 | -0.45877 | 1.478042 | 1 |
| 5 | 8813000 | 8815199 | 45.17623 | 72.67318 | 81.97464 | 34.79208 | 0.179616 | 1.860304 | 1 |
| 5 | 8839000 | 8840999 | 19.59721 | 34.21405 | 35.71968 | 15.73065 | 0.325771 | 0.724855 | 1 |
| 5 | 8971600 | 8976199 | 19.98315 | 34.73555 | 35.87016 | 15.75354 | 0.340848 | 0.787491 | 1 |
| 5 | 9046400 | 9047799 | 9.859628 | 18.63964 | 23.79833 | 9.668185 | 0.130882 | 0.048504 | 1 |
| 5 | 9864200 | 9865799 | 15.56998 | 27.67912 | 30.89331 | 13.62594 | 0.236832 | 0.476442 | 1 |
| 5 | 9897000 | 9908399 | 331.1549 | 460.6504 | 627.2326 | 260.2943 | -0.28458 | 4.659332 | 1 |
| 5 | 10020800 | 10023399 | 32.51181 | 53.54092 | 63.90782 | 27.56186 | 0.124618 | 1.393487 | 1 |
| 5 | 10029000 | 10029599 | 10.49981 | 20.01277 | 22.79819 | 9.103185 | 0.315993 | -0.00345 | 1 |
| 5 | 10313200 | 10315799 | 27.27364 | 46.01122 | 50.16532 | 21.93807 | 0.25684 | 1.106447 | 1 |
| 5 | 10575800 | 10576599 | 15.70881 | 27.88133 | 30.38881 | 13.42149 | 0.26956 | 0.486062 | 1 |
| 5 | 10621200 | 10622599 | 14.61886 | 26.02709 | 30.51556 | 13.44241 | 0.167501 | 0.460502 | 1 |
| 5 | 10670800 | 10680199 | 194.8339 | 284.0344 | 382.355 | 157.5608 | -0.20056 | 3.957053 | 1 |
| 5 | 10824600 | 10827599 | 28.61108 | 47.83989 | 50.77559 | 22.11146 | 0.295432 | 1.197097 | 1 |
| 5 | 11007000 | 11008399 | 14.65264 | 25.86336 | 35.60184 | 15.75255 | -0.06887 | 0.544078 | 1 |

| | | | | | | | | | |
|---|---|---|---|---|---|---|---|---|---|
| 5 | 11113200 | 11115199 | 24.89862 | 41.04922 | 63.19196 | 27.26601 | -0.24058 | 1.360158 | 1 |
| 5 | 11475600 | 11477599 | 12.81168 | 22.74405 | 35.18869 | 15.53865 | -0.23273 | 0.476694 | 1 |
| 5 | 11668800 | 11669599 | 20.64085 | 35.84085 | 33.01661 | 14.49067 | 0.504881 | 0.802376 | 1 |
| 5 | 11977200 | 11979199 | 18.3916 | 32.1055 | 37.26139 | 16.40842 | 0.173616 | 0.721084 | 1 |
| 5 | 12013200 | 12013999 | 13.85421 | 24.22346 | 41.10567 | 18.12185 | -0.37129 | 0.683905 | 1 |
| 5 | 12139200 | 12139799 | 11.29904 | 20.94188 | 28.76422 | 12.08423 | -0.01302 | 0.196819 | 1 |
| 5 | 12144600 | 12146999 | 19.01998 | 33.28457 | 37.16912 | 16.39845 | 0.22918 | 0.6886 | 1 |
| 5 | 12302200 | 12303999 | 18.86057 | 32.94223 | 35.32701 | 15.56195 | 0.287415 | 0.715624 | 1 |
| 5 | 12406200 | 12406999 | 14.52276 | 26.36672 | 28.75382 | 12.41418 | 0.290112 | 0.335838 | 1 |
| 5 | 12531000 | 12531799 | 17.59337 | 30.03859 | 49.52478 | 21.71278 | -0.33576 | 0.96792 | 1 |
| 5 | 12570600 | 12571399 | 13.82395 | 24.70716 | 32.02555 | 14.06168 | 0.026709 | 0.433767 | 1 |
| 5 | 12831600 | 12833199 | 19.8387 | 34.53136 | 38.82504 | 17.06585 | 0.218781 | 0.771323 | 1 |
| 5 | 12844000 | 12847399 | 40.56319 | 65.38829 | 106.1077 | 45.16886 | -0.34036 | 1.799562 | 1 |
| 5 | 13092600 | 13094199 | 19.79625 | 34.31919 | 42.61915 | 18.6992 | 0.075339 | 0.828413 | 1 |
| 5 | 13920600 | 13922199 | 74.14372 | 115.2813 | 173.252 | 71.92954 | -0.27891 | 2.741964 | 1 |
| 5 | 13926400 | 13932999 | 92.55102 | 143.1522 | 178.9606 | 74.33592 | -0.01944 | 2.84145 | 1 |
| 5 | 14229800 | 14232999 | 43.80756 | 70.29286 | 99.57393 | 42.10333 | -0.15283 | 1.944833 | 1 |
| 5 | 14401800 | 14404399 | 22.94173 | 39.67239 | 42.50278 | 18.64672 | 0.286672 | 0.863177 | 1 |
| 5 | 14451400 | 14452399 | 21.28014 | 36.02926 | 50.08214 | 21.92557 | -0.09194 | 1.057482 | 1 |
| 5 | 14653400 | 14655999 | 22.66999 | 38.25688 | 57.70132 | 25.2361 | -0.20971 | 1.095369 | 1 |
| 5 | 15652800 | 15654799 | 31.08119 | 50.77659 | 68.90243 | 29.6188 | -0.06536 | 1.495414 | 1 |
| 5 | 15692800 | 15707199 | 183.1878 | 270.0832 | 343.3253 | 141.3857 | -0.10243 | 3.778305 | 1 |
| 5 | 15707800 | 15710799 | 39.69411 | 64.72371 | 69.34635 | 29.7889 | 0.272624 | 1.540574 | 1 |
| 5 | 16011800 | 16012399 | 8.519269 | 16.32054 | 26.87358 | 10.59691 | -0.19498 | -0.05269 | 1 |
| 5 | 16032400 | 16034599 | 23.30961 | 39.42375 | 55.52785 | 24.30189 | -0.11117 | 1.076915 | 1 |
| 5 | 16226800 | 16228599 | 17.80614 | 31.04969 | 41.98143 | 18.47429 | -0.0459 | 0.740298 | 1 |
| 5 | 16277200 | 16279399 | 28.44996 | 46.78615 | 73.16877 | 31.53716 | -0.26493 | 1.405934 | 1 |
| 5 | 16373200 | 16375599 | 40.74809 | 65.55441 | 91.94169 | 39.02633 | -0.13385 | 1.858015 | 1 |
| 5 | 16922200 | 16928799 | 84.02642 | 131.4427 | 151.8313 | 62.98793 | 0.107112 | 2.637316 | 1 |
| 5 | 17244000 | 17244999 | 12.99641 | 23.31193 | 30.576 | 13.38452 | 0.01281 | 0.402373 | 1 |
| 5 | 17306200 | 17309199 | 24.74101 | 41.32522 | 58.72607 | 25.55668 | -0.12417 | 1.205516 | 1 |
| 5 | 17353200 | 17354199 | 14.06584 | 24.7952 | 35.23506 | 15.59433 | -0.1151 | 0.55544 | 1 |
| 5 | 17474400 | 17475399 | 15.95925 | 27.9496 | 38.84453 | 17.14863 | -0.08496 | 0.662259 | 1 |
| 5 | 17886600 | 17889999 | 31.12565 | 50.65649 | 79.55977 | 34.15271 | -0.27966 | 1.555428 | 1 |
| 5 | 18326000 | 18330399 | 70.46665 | 111.3366 | 118.6389 | 49.27609 | 0.234514 | 2.427763 | 1 |
| 5 | 18333400 | 18335599 | 24.61522 | 42.77149 | 39.74012 | 17.44913 | 0.492608 | 0.821441 | 1 |
| 5 | 18419800 | 18422199 | 19.67053 | 34.11656 | 42.05399 | 18.45149 | 0.086026 | 0.821034 | 1 |
| 5 | 18479800 | 18482799 | 31.83941 | 52.49455 | 61.07995 | 26.35522 | 0.162012 | 1.375654 | 1 |
| 5 | 18502800 | 18503799 | 11.04534 | 19.85036 | 32.28332 | 14.10902 | -0.29375 | 0.381946 | 1 |
| 5 | 18558400 | 18561799 | 29.0882 | 47.71504 | 78.46622 | 33.78434 | -0.33874 | 1.440578 | 1 |
| 5 | 18779200 | 18780799 | 16.92234 | 29.30887 | 42.21446 | 18.5211 | -0.13785 | 0.823089 | 1 |
| 5 | 18857200 | 18865799 | 130.3523 | 197.7714 | 218.5581 | 90.4812 | 0.136551 | 3.235716 | 1 |
| 5 | 19051200 | 19055599 | 66.6599 | 105.8189 | 115.0123 | 47.92655 | 0.211847 | 2.301657 | 1 |
| 5 | 19136000 | 19138399 | 21.91449 | 36.97946 | 56.52998 | 24.72331 | -0.22897 | 1.093501 | 1 |
| 5 | 19191800 | 19192599 | 15.97806 | 28.03425 | 37.75347 | 16.69298 | -0.03969 | 0.631365 | 1 |
| 5 | 19351400 | 19357599 | 89.25724 | 139.0173 | 150.8508 | 62.61323 | 0.192302 | 2.715525 | 1 |
| 5 | 19546400 | 19547399 | 18.89429 | 32.89257 | 37.37989 | 16.43221 | 0.203528 | 0.763076 | 1 |

| | | | | | | | | | |
|---|---|---|---|---|---|---|---|---|---|
| 5 | 19680600 | 19681999 | 17.31049 | 30.36819 | 34.58571 | 15.29069 | 0.200849 | 0.634308 | 1 |
| 5 | 19755400 | 19757799 | 19.18685 | 33.1521 | 47.40233 | 20.79572 | -0.12823 | 0.86117 | 1 |
| 5 | 20167200 | 20168199 | 16.54042 | 28.62779 | 45.32733 | 19.88642 | -0.27395 | 0.833559 | 1 |
| 5 | 20178400 | 20180999 | 21.47875 | 37.30195 | 38.92522 | 17.08569 | 0.325814 | 0.81367 | 1 |
| 5 | 20331800 | 20333199 | 15.62943 | 27.58128 | 33.0865 | 14.64026 | 0.128591 | 0.549064 | 1 |
| 5 | 20390400 | 20396599 | 113.8216 | 174.308 | 172.3185 | 71.51444 | 0.307915 | 3.038149 | 1 |
| 5 | 20892200 | 20892999 | 12.84301 | 23.65225 | 27.0077 | 11.44513 | 0.242713 | 0.240643 | 1 |
| 5 | 20938000 | 20939999 | 12.34961 | 22.31964 | 32.62618 | 14.1663 | -0.13506 | 0.35624 | 1 |
| 5 | 21032400 | 21032799 | 7.423645 | 14.345 | 23.75919 | 9.119284 | -0.16637 | -0.16092 | 1 |
| 5 | 21086400 | 21087999 | 15.04419 | 26.2333 | 43.65515 | 19.21073 | -0.34394 | 0.730389 | 1 |
| 5 | 21130600 | 21131199 | 12.37471 | 22.56614 | 26.23503 | 11.26442 | 0.204295 | 0.293737 | 1 |
| 5 | 21537400 | 21542399 | 62.02425 | 99.06218 | 101.7649 | 42.73763 | 0.302252 | 2.108665 | 1 |
| 5 | 22418600 | 22420999 | 12.75392 | 22.6714 | 35.2913 | 15.56664 | -0.24006 | 0.471512 | 1 |
| 5 | 22482800 | 22483799 | 15.25211 | 27.03196 | 31.93914 | 14.12037 | 0.152663 | 0.496531 | 1 |
| 5 | 22531000 | 22532799 | 20.29203 | 34.78774 | 48.47789 | 21.26078 | -0.09353 | 0.93671 | 1 |
| 5 | 22537400 | 22539199 | 24.97177 | 42.88674 | 43.01223 | 18.86626 | 0.379456 | 0.932134 | 1 |
| 5 | 22568400 | 22570199 | 18.56361 | 32.39684 | 37.38269 | 16.45898 | 0.182095 | 0.728946 | 1 |
| 5 | 22808600 | 22812199 | 45.4768 | 73.06714 | 95.59334 | 40.50025 | -0.03608 | 1.905801 | 1 |
| 5 | 23574400 | 23576399 | 32.11704 | 52.1565 | 78.19399 | 33.52828 | -0.21524 | 1.593584 | 1 |
| 5 | 23584200 | 23587399 | 38.04855 | 61.4493 | 84.86907 | 36.21494 | -0.10505 | 1.732622 | 1 |
| 5 | 23927200 | 23941399 | 176.5933 | 262.7922 | 292.1785 | 120.3853 | 0.10317 | 3.618332 | 1 |
| 5 | 24035400 | 24036799 | 16.19381 | 28.25055 | 40.06078 | 17.63395 | -0.11392 | 0.726721 | 1 |
| 5 | 24135000 | 24138599 | 53.35306 | 85.8953 | 83.09183 | 35.30699 | 0.402205 | 1.849194 | 1 |
| 5 | 24428200 | 24430399 | 16.44558 | 28.74886 | 41.97881 | 18.50379 | -0.15597 | 0.698734 | 1 |
| 5 | 25438000 | 25438999 | 29.03529 | 48.08821 | 52.36556 | 22.6603 | 0.258198 | 1.306306 | 1 |
| 5 | 25439800 | 25443599 | 66.55599 | 104.1761 | 153.5578 | 63.70024 | -0.24346 | 2.619561 | 1 |
| 5 | 25782600 | 25790799 | 122.0022 | 184.802 | 250.3799 | 103.6034 | -0.15914 | 3.268577 | 1 |
| 5 | 25973600 | 25975599 | 32.06086 | 52.28307 | 72.9172 | 31.32409 | -0.10665 | 1.525182 | 1 |
| 5 | 26274800 | 26281599 | 103.7326 | 158.8834 | 219.0208 | 90.89989 | -0.17113 | 3.036372 | 1 |
| 5 | 26452600 | 26454999 | 29.08109 | 48.20205 | 67.27639 | 29.12856 | -0.09875 | 1.311213 | 1 |
| 5 | 26590600 | 26597199 | 76.86687 | 120.7213 | 156.3119 | 64.83412 | -0.05297 | 2.56248 | 1 |
| 5 | 27729800 | 27731599 | 13.40047 | 24.55051 | 26.13307 | 11.15336 | 0.33658 | 0.275755 | 1 |
| 5 | 28014000 | 28015799 | 16.82397 | 29.27308 | 46.83321 | 20.58111 | -0.28784 | 0.775051 | 1 |
| 5 | 28798400 | 28802599 | 53.21898 | 84.56335 | 128.5557 | 53.62169 | -0.27014 | 2.266824 | 1 |
| 5 | 28807400 | 28809999 | 22.57674 | 37.64494 | 61.75326 | 26.84471 | -0.33044 | 1.222083 | 1 |
| 5 | 29308800 | 29310599 | 17.16875 | 29.65596 | 44.30706 | 19.43762 | -0.19116 | 0.852973 | 1 |
| 5 | 29560400 | 29563399 | 86.68174 | 133.4669 | 192.9925 | 80.15839 | -0.23419 | 2.926094 | 1 |
| 5 | 29795800 | 29797599 | 15.13718 | 27.19129 | 28.2947 | 12.37075 | 0.34589 | 0.402749 | 1 |
| 5 | 30188000 | 30188799 | 10.45211 | 19.43609 | 29.40915 | 12.29878 | -0.14659 | 0.174679 | 1 |
| 5 | 30249000 | 30257999 | 161.988 | 241.9007 | 285.9399 | 117.8767 | 0.019457 | 3.553865 | 1 |
| 5 | 30291200 | 30294199 | 32.14355 | 52.94054 | 61.65187 | 26.59022 | 0.160164 | 1.389096 | 1 |
| 5 | 31272600 | 31274799 | 19.72624 | 33.63795 | 49.01689 | 21.48794 | -0.15881 | 0.984556 | 1 |
| 5 | 31393200 | 31394799 | 14.76979 | 26.03271 | 40.79626 | 18.05608 | -0.25579 | 0.571024 | 1 |
| 5 | 31740400 | 31742999 | 27.04287 | 44.80167 | 62.60621 | 27.09732 | -0.10052 | 1.306313 | 1 |
| 5 | 31755800 | 31757799 | 18.2281 | 31.54036 | 44.76965 | 19.64159 | -0.11722 | 0.841412 | 1 |
| 5 | 31911600 | 31914199 | 22.59381 | 38.08555 | 54.89384 | 24.00006 | -0.14438 | 1.104541 | 1 |
| 5 | 32954200 | 32956799 | 28.74751 | 47.75937 | 55.06644 | 23.8801 | 0.176318 | 1.272653 | 1 |

| | | | | | | | | | |
|---|---|---|---|---|---|---|---|---|---|
| 5 | 33356000 | 33356999 | 44.37111 | 71.37817 | 78.21359 | 33.19591 | 0.221374 | 1.858079 | 1 |
| 5 | 33916400 | 33918599 | 26.36863 | 43.39817 | 70.56235 | 30.42406 | -0.32034 | 1.390093 | 1 |
| 5 | 34542200 | 34546599 | 53.09295 | 85.40263 | 88.47292 | 37.54639 | 0.30202 | 1.876114 | 1 |
| 5 | 36069600 | 36071199 | 15.09619 | 26.84847 | 31.43604 | 13.86027 | 0.168489 | 0.471726 | 1 |
| 5 | 38083200 | 38085599 | 17.53139 | 30.90745 | 34.88278 | 15.4381 | 0.215423 | 0.573174 | 1 |
| 5 | 38115800 | 38118199 | 25.09891 | 42.71357 | 46.8705 | 20.54367 | 0.248301 | 1.019777 | 1 |
| 5 | 38559600 | 38566799 | 78.19044 | 121.6519 | 169.1662 | 70.22644 | -0.1663 | 2.731247 | 1 |
| 5 | 38754800 | 38756199 | 13.00342 | 23.30464 | 32.5569 | 14.26189 | -0.0785 | 0.411303 | 1 |
| 5 | 39771600 | 39772799 | 20.13142 | 34.12587 | 52.71095 | 23.08496 | -0.24349 | 1.044247 | 1 |
| 5 | 40231800 | 40235799 | 55.10753 | 88.00535 | 99.68975 | 41.85957 | 0.161312 | 2.104343 | 1 |
| 5 | 40564000 | 40566399 | 23.06151 | 38.82442 | 56.00318 | 24.47561 | -0.14564 | 1.117546 | 1 |
| 5 | 40626800 | 40628999 | 22.04594 | 37.80364 | 44.75496 | 19.62841 | 0.140869 | 0.937907 | 1 |
| 5 | 40635400 | 40636399 | 17.55915 | 30.8488 | 31.64774 | 14.00881 | 0.350994 | 0.609612 | 1 |
| 5 | 40718000 | 40720199 | 28.01683 | 46.51397 | 57.70166 | 25.0112 | 0.071045 | 1.281725 | 1 |
| 5 | 40732400 | 40733599 | 20.76552 | 35.93776 | 42.60654 | 18.69269 | 0.141456 | 0.848047 | 1 |
| 5 | 40978600 | 40982199 | 50.52615 | 80.82651 | 116.7354 | 49.13536 | -0.18593 | 2.051637 | 1 |
| 5 | 41556600 | 41561399 | 39.12438 | 62.75999 | 101.6978 | 42.98228 | -0.34688 | 1.949249 | 1 |
| 5 | 41646200 | 41647799 | 14.06326 | 24.90143 | 35.6101 | 15.74628 | -0.12173 | 0.507037 | 1 |
| 5 | 42169000 | 42171199 | 21.36866 | 36.33729 | 44.90103 | 19.67922 | 0.077692 | 1.013483 | 1 |
| 5 | 42320200 | 42320999 | 12.00243 | 22.21536 | 24.54035 | 10.3288 | 0.297325 | 0.200179 | 1 |
| 5 | 42985200 | 42993799 | 147.2882 | 221.7712 | 231.7164 | 95.70608 | 0.208006 | 3.391479 | 1 |
| 5 | 42994600 | 42996599 | 35.16984 | 57.82865 | 61.38182 | 26.45514 | 0.292541 | 1.415172 | 1 |
| 5 | 43369000 | 43369599 | 10.03686 | 18.94161 | 22.29034 | 9.089051 | 0.243267 | 0.059115 | 1 |
| 5 | 43589200 | 43590399 | 18.09087 | 31.58533 | 37.28929 | 16.42227 | 0.149132 | 0.717753 | 1 |
| 5 | 43987600 | 43989199 | 23.65615 | 40.692 | 43.61689 | 19.13241 | 0.284469 | 0.915185 | 1 |
| 5 | 44262400 | 44265399 | 17.06924 | 30.47401 | 31.34814 | 13.77695 | 0.357602 | 0.463058 | 1 |
| 5 | 44293200 | 44293799 | 9.797257 | 17.84696 | 32.39477 | 13.92774 | -0.43549 | 0.295201 | 1 |
| 5 | 44298800 | 44299199 | 9.495728 | 17.94241 | 25.95127 | 10.55435 | -0.04884 | 0.056057 | 1 |
| 5 | 45815600 | 45821799 | 127.8791 | 194.0023 | 222.643 | 92.16996 | 0.082017 | 3.23711 | 1 |
| 5 | 45969800 | 45984799 | 476.7615 | 650.1766 | 738.6544 | 307.4925 | -0.05229 | 4.968026 | 1 |
| 5 | 46003000 | 46005199 | 10.06077 | 19.3242 | 24.12034 | 9.439942 | 0.211648 | -0.06871 | 1 |
| 5 | 46021800 | 46024199 | 13.71631 | 24.32804 | 34.58192 | 15.28234 | -0.11207 | 0.486002 | 1 |
| 5 | 46615200 | 46617799 | 31.25298 | 50.59584 | 83.51604 | 35.72833 | -0.35786 | 1.660156 | 1 |
| 5 | 47320600 | 47321599 | 11.79325 | 21.59694 | 24.27462 | 10.36609 | 0.257861 | 0.259047 | 1 |
| 5 | 47685800 | 47687599 | 22.22404 | 37.75144 | 53.29816 | 23.35682 | -0.11423 | 1.034969 | 1 |
| 5 | 48127400 | 48131199 | 18.13343 | 31.65478 | 37.82491 | 16.65655 | 0.131824 | 0.721592 | 1 |
| 5 | 49701200 | 49703399 | 24.5863 | 41.44335 | 49.07495 | 21.45585 | 0.13816 | 1.10458 | 1 |
| 5 | 49825200 | 49825599 | 7.778289 | 15.09777 | 17.28748 | 6.497875 | 0.390058 | -0.24867 | 1 |
| 5 | 49857000 | 49859599 | 10.02733 | 18.32532 | 29.81774 | 12.76795 | -0.27521 | 0.270142 | 1 |
| 5 | 49910600 | 49912799 | 21.03223 | 35.42035 | 53.74097 | 23.4894 | -0.21809 | 1.107454 | 1 |
| 5 | 50233000 | 50235799 | 34.89119 | 57.57066 | 62.04249 | 26.77931 | 0.272689 | 1.373293 | 1 |
| 5 | 50271000 | 50271799 | 19.79878 | 34.19307 | 40.13688 | 17.60806 | 0.155237 | 0.859744 | 1 |
| 5 | 50343800 | 50346399 | 33.32189 | 54.79753 | 57.89039 | 24.95178 | 0.299416 | 1.406108 | 1 |
| 5 | 50475400 | 50477199 | 16.88491 | 29.26835 | 44.8016 | 19.65668 | -0.22491 | 0.821741 | 1 |
| 5 | 50851000 | 50852199 | 10.24342 | 19.04791 | 30.47954 | 12.74654 | -0.22702 | 0.173726 | 1 |
| 5 | 51132400 | 51133799 | 21.31974 | 36.58561 | 44.74372 | 19.62465 | 0.094258 | 0.926705 | 1 |
| 5 | 51158000 | 51158799 | 9.768585 | 18.60057 | 22.7774 | 9.113986 | 0.209516 | -0.00583 | 1 |

| | | | | | | | | | |
|---|---|---|---|---|---|---|---|---|---|
| 5 | 51630000 | 51632599 | 33.07557 | 54.34443 | 61.1197 | 26.33383 | 0.208898 | 1.41961 | 1 |
| 5 | 52778800 | 52780599 | 15.5284 | 27.18002 | 39.93478 | 17.62171 | -0.16471 | 0.671501 | 1 |
| 5 | 52988600 | 52991799 | 24.2999 | 40.42768 | 64.16868 | 27.85464 | -0.28323 | 1.251294 | 1 |
| 5 | 54606200 | 54608399 | 22.44919 | 38.03941 | 52.20639 | 22.86117 | -0.07361 | 1.057473 | 1 |
| 5 | 54708600 | 54710599 | 31.57525 | 52.0347 | 57.23738 | 24.69232 | 0.242544 | 1.376448 | 1 |
| 5 | 54724400 | 54725199 | 13.55543 | 23.94462 | 36.16822 | 16.00075 | -0.2017 | 0.530005 | 1 |
| 5 | 55168000 | 55171399 | 28.10619 | 46.25715 | 68.56622 | 29.5632 | -0.18733 | 1.392392 | 1 |
| 5 | 55319800 | 55320599 | 16.44726 | 28.89215 | 33.04817 | 14.62733 | 0.194519 | 0.604515 | 1 |
| 5 | 55803200 | 55806799 | 47.86209 | 77.00724 | 84.8314 | 36.01366 | 0.214002 | 1.860656 | 1 |
| 5 | 55883200 | 55884999 | 21.1001 | 36.4099 | 46.31893 | 20.31961 | 0.039251 | 0.879742 | 1 |
| 5 | 56104400 | 56111999 | 113.6437 | 174.0509 | 193.8362 | 80.4462 | 0.136281 | 3.038214 | 1 |
| 5 | 56286200 | 56288999 | 34.82263 | 56.9757 | 63.42286 | 27.27847 | 0.221087 | 1.476097 | 1 |
| 5 | 56327200 | 56328999 | 17.13505 | 30.20678 | 32.86891 | 14.54701 | 0.267806 | 0.56738 | 1 |
| 5 | 56346200 | 56347399 | 21.58206 | 36.48702 | 49.0487 | 21.46361 | -0.0439 | 1.069726 | 1 |
| 5 | 56683800 | 56685799 | 33.46074 | 54.29934 | 78.09251 | 33.46693 | -0.15638 | 1.612368 | 1 |
| 5 | 57164200 | 57166399 | 29.45122 | 49.17203 | 54.56156 | 23.73804 | 0.23162 | 1.221129 | 1 |
| 5 | 57307800 | 57310199 | 24.1306 | 41.72177 | 42.47017 | 18.63232 | 0.360122 | 0.870263 | 1 |
| 5 | 57820200 | 57822799 | 20.18585 | 34.55416 | 53.28217 | 23.36497 | -0.23922 | 0.959097 | 1 |
| 5 | 57829400 | 57830599 | 26.07886 | 42.81692 | 70.30512 | 30.28196 | -0.33585 | 1.419858 | 1 |
| 5 | 58157800 | 58158399 | 12.63508 | 23.17028 | 27.09118 | 11.54733 | 0.203048 | 0.264703 | 1 |
| 5 | 58402400 | 58403199 | 9.308277 | 17.31502 | 28.37405 | 11.85902 | -0.26024 | 0.158659 | 1 |
| 5 | 58727800 | 58731199 | 53.43155 | 85.37477 | 91.10434 | 38.28373 | 0.248129 | 2.081079 | 1 |
| 5 | 58804400 | 58808199 | 57.71629 | 91.96028 | 107.0535 | 44.80777 | 0.118403 | 2.180389 | 1 |
| 5 | 59036600 | 59038199 | 11.33731 | 20.74263 | 31.31198 | 13.38826 | -0.16628 | 0.27865 | 1 |
| 5 | 59083200 | 59084199 | 20.35763 | 35.29722 | 36.97727 | 16.22389 | 0.319788 | 0.821951 | 1 |
| 5 | 59096200 | 59097999 | 15.61333 | 27.60766 | 32.51456 | 14.38147 | 0.156058 | 0.525988 | 1 |
| 5 | 59268200 | 59269799 | 11.97136 | 22.43689 | 26.1839 | 10.80518 | 0.24046 | 0.127152 | 1 |
| 5 | 59282400 | 59284999 | 26.05856 | 43.30747 | 58.15918 | 25.22747 | -0.04292 | 1.264273 | 1 |
| 5 | 59378200 | 59378999 | 10.46557 | 19.41037 | 29.90713 | 12.55269 | -0.1764 | 0.190878 | 1 |
| 5 | 59467800 | 59468999 | 18.80912 | 32.39393 | 46.89506 | 20.57113 | -0.14682 | 0.890387 | 1 |
| 5 | 59807000 | 59807599 | 9.318296 | 17.5024 | 25.37636 | 10.43354 | -0.06536 | 0.09142 | 1 |
| 5 | 60022000 | 60026199 | 66.22614 | 105.1834 | 117.6302 | 49.04221 | 0.171458 | 2.2878 | 1 |
| 5 | 60110600 | 60114199 | 28.06769 | 45.75283 | 75.14282 | 32.27546 | -0.34225 | 1.523789 | 1 |
| 5 | 60203600 | 60205199 | 16.51734 | 28.52448 | 47.93866 | 21.03067 | -0.36043 | 0.855221 | 1 |
| 5 | 60656400 | 60658199 | 13.60726 | 24.46788 | 30.97565 | 13.53131 | 0.065549 | 0.39165 | 1 |
| 5 | 60702200 | 60704799 | 32.07977 | 52.79882 | 61.69429 | 26.6007 | 0.154892 | 1.39617 | 1 |
| 5 | 61121000 | 61122199 | 13.65844 | 24.77791 | 28.90432 | 12.4915 | 0.192771 | 0.335575 | 1 |
| 5 | 61751600 | 61754599 | 39.7988 | 65.27461 | 64.01265 | 27.5595 | 0.40499 | 1.458644 | 1 |
| 5 | 61758000 | 61758999 | 20.07182 | 33.79327 | 55.30783 | 24.17216 | -0.32703 | 1.107643 | 1 |
| 5 | 61820400 | 61823999 | 62.95021 | 100.0761 | 106.263 | 44.34546 | 0.247505 | 2.255287 | 1 |
| 5 | 61966000 | 61969999 | 60.66956 | 95.94938 | 141.0987 | 58.63317 | -0.22845 | 2.402285 | 1 |
| 5 | 62381400 | 62384199 | 35.41241 | 57.22531 | 87.13582 | 37.21023 | -0.24455 | 1.710867 | 1 |
| 5 | 62385000 | 62385399 | 8.913532 | 17.13668 | 19.83947 | 7.743702 | 0.323411 | -0.10067 | 1 |
| 5 | 62458200 | 62461799 | 19.93517 | 33.94166 | 54.89751 | 24.06287 | -0.30917 | 1.005994 | 1 |
| 5 | 62471000 | 62471999 | 19.85633 | 34.39526 | 35.57863 | 15.60978 | 0.33776 | 0.826155 | 1 |
| 5 | 62930600 | 62932399 | 27.00623 | 45.3315 | 50.71788 | 22.14031 | 0.219537 | 1.151427 | 1 |
| 5 | 63025200 | 63028599 | 43.20694 | 69.33159 | 98.52522 | 41.66215 | -0.15732 | 1.943864 | 1 |

| | | | | | | | | | |
|---|---|---|---|---|---|---|---|---|---|
| 5 | 63306800 | 63309399 | 23.03629 | 39.51091 | 43.68494 | 19.15947 | 0.23919 | 0.939124 | 1 |
| 5 | 63525800 | 63526399 | 10.1454 | 19.18077 | 22.73874 | 9.236832 | 0.237199 | 0.047917 | 1 |
| 5 | 63754200 | 63755799 | 19.50094 | 33.1023 | 52.16386 | 22.85483 | -0.27214 | 1.029946 | 1 |
| 5 | 63969000 | 63970399 | 11.5518 | 20.55284 | 34.27974 | 15.10908 | -0.33852 | 0.452591 | 1 |
| 5 | 64167600 | 64176799 | 107.8617 | 164.7916 | 224.6735 | 93.19156 | -0.15811 | 3.088142 | 1 |
| 5 | 64212600 | 64213999 | 13.77052 | 24.35641 | 37.18168 | 16.44481 | -0.21608 | 0.520282 | 1 |
| 5 | 64582400 | 64587799 | 63.03686 | 99.42893 | 137.8813 | 57.22172 | -0.14694 | 2.466427 | 1 |
| 5 | 65166200 | 65169199 | 29.92658 | 49.01077 | 67.2722 | 28.94479 | -0.07954 | 1.457423 | 1 |
| 5 | 65989600 | 65992399 | 24.44679 | 41.73219 | 45.79201 | 20.07594 | 0.249018 | 0.98854 | 1 |
| 5 | 66116400 | 66118399 | 16.12197 | 28.25029 | 38.30466 | 16.91825 | -0.04949 | 0.653465 | 1 |
| 5 | 67117400 | 67120199 | 22.11517 | 37.10822 | 59.22395 | 25.84767 | -0.29117 | 1.147607 | 1 |
| 5 | 68156000 | 68157799 | 14.0732 | 25.01279 | 36.09694 | 15.92401 | -0.13204 | 0.478787 | 1 |
| 5 | 68469000 | 68470799 | 13.72368 | 24.7015 | 28.46877 | 12.42472 | 0.201168 | 0.381209 | 1 |
| 5 | 68961000 | 68961799 | 14.96994 | 26.62121 | 31.97219 | 14.09787 | 0.131985 | 0.473068 | 1 |
| 5 | 68980800 | 68981599 | 12.75192 | 22.82807 | 31.66133 | 13.88281 | -0.06882 | 0.416633 | 1 |
| 5 | 69280200 | 69282399 | 26.50372 | 44.31178 | 50.17394 | 21.84808 | 0.202562 | 1.194193 | 1 |
| 5 | 69688600 | 69689799 | 17.58858 | 30.78358 | 37.28655 | 16.45203 | 0.112553 | 0.679679 | 1 |
| 5 | 69795600 | 69798599 | 27.24239 | 44.98464 | 60.80024 | 26.25623 | -0.05287 | 1.344094 | 1 |
| 5 | 69831800 | 69833999 | 14.96718 | 26.80451 | 31.42998 | 13.7767 | 0.172221 | 0.428235 | 1 |
| 5 | 70367200 | 70367999 | 15.63644 | 27.44878 | 35.7482 | 15.81312 | 0.008406 | 0.617887 | 1 |
| 5 | 70555800 | 70557399 | 26.41416 | 43.77212 | 59.1912 | 25.6241 | -0.05318 | 1.299707 | 1 |
| 5 | 70587000 | 70587999 | 16.98845 | 29.70519 | 37.9129 | 16.71507 | 0.037531 | 0.693608 | 1 |
| 5 | 70701200 | 70702799 | 24.94015 | 41.99408 | 49.08052 | 21.45121 | 0.15694 | 1.115733 | 1 |
| 5 | 70738600 | 70740399 | 21.5558 | 36.59645 | 54.92041 | 24.06378 | -0.20206 | 1.038016 | 1 |
| 5 | 70803800 | 70808599 | 76.98564 | 120.8391 | 132.7769 | 55.06835 | 0.182955 | 2.570131 | 1 |
| 5 | 71499600 | 71500599 | 14.2951 | 24.94602 | 41.1309 | 18.10857 | -0.33029 | 0.713788 | 1 |
| 5 | 71556600 | 71562799 | 105.7171 | 162.9387 | 173.3014 | 71.98656 | 0.20956 | 2.910978 | 1 |
| 5 | 72648800 | 72649599 | 10.43538 | 19.31734 | 29.34495 | 12.34888 | -0.15887 | 0.199398 | 1 |
| 5 | 72939600 | 72942199 | 19.94174 | 34.66391 | 38.93898 | 17.10142 | 0.220018 | 0.79271 | 1 |
| 5 | 73422200 | 73424399 | 16.54565 | 29.30113 | 30.86231 | 13.64648 | 0.317133 | 0.511772 | 1 |
| 5 | 73992000 | 73994799 | 25.7405 | 43.52229 | 47.99767 | 21.00441 | 0.240599 | 1.081702 | 1 |
| 5 | 74050400 | 74053199 | 26.55951 | 44.40602 | 66.79293 | 29.08517 | -0.20601 | 1.207149 | 1 |
| 5 | 74196400 | 74198999 | 23.75052 | 40.18281 | 52.81586 | 23.11731 | -0.01174 | 1.072555 | 1 |
| 5 | 75429400 | 75431399 | 16.42853 | 28.77042 | 41.38047 | 18.26761 | -0.13433 | 0.668532 | 1 |
| 5 | 75469800 | 75470599 | 15.85545 | 28.07568 | 29.94756 | 13.2423 | 0.298931 | 0.506386 | 1 |
| 5 | 75803800 | 75808599 | 55.83813 | 88.65117 | 122.1808 | 50.92016 | -0.13004 | 2.292288 | 1 |
| 5 | 76425600 | 76426199 | 8.47123 | 16.05278 | 23.37667 | 9.449511 | -0.05105 | 0.017224 | 1 |
| 5 | 76582400 | 76583599 | 14.38355 | 25.81222 | 28.92947 | 12.65873 | 0.238688 | 0.405453 | 1 |
| 5 | 77070800 | 77078199 | 97.18823 | 150.1905 | 172.0921 | 71.48324 | 0.105154 | 2.858834 | 1 |
| 5 | 77087200 | 77089399 | 18.14774 | 31.57646 | 49.5328 | 21.76749 | -0.25963 | 0.785156 | 1 |
| 5 | 77324200 | 77324999 | 11.42703 | 21.31902 | 23.64576 | 9.833601 | 0.304899 | 0.147344 | 1 |
| 5 | 77842800 | 77843799 | 16.22057 | 28.6525 | 31.00971 | 13.71863 | 0.276821 | 0.537709 | 1 |
| 5 | 78276000 | 78276799 | 13.06274 | 23.73458 | 27.51204 | 11.86642 | 0.203892 | 0.320222 | 1 |
| 5 | 78289600 | 78294399 | 50.05511 | 80.4493 | 87.5992 | 37.13089 | 0.228797 | 1.89646 | 1 |
| 5 | 78300400 | 78302799 | 31.80586 | 51.77405 | 76.5914 | 32.88092 | -0.19328 | 1.553891 | 1 |
| 5 | 78346800 | 78348799 | 13.96085 | 24.5173 | 38.3717 | 16.97941 | -0.25528 | 0.607324 | 1 |
| 5 | 78402600 | 78406199 | 45.34867 | 72.83741 | 89.75971 | 38.01043 | 0.049462 | 1.911742 | 1 |

| | | | | | | | | | |
|---|---|---|---|---|---|---|---|---|---|
| 5 | 78517600 | 78518199 | 9.624192 | 18.36196 | 23.41536 | 9.330096 | 0.157476 | -0.0157 | 1 |
| 5 | 78621000 | 78623199 | 23.04193 | 39.16271 | 42.30218 | 18.53943 | 0.270823 | 1.020139 | 1 |
| 5 | 79194400 | 79198799 | 57.99995 | 91.89386 | 135.2621 | 56.25938 | -0.2277 | 2.356129 | 1 |
| 5 | 79397000 | 79397799 | 19.82817 | 34.40636 | 38.55168 | 16.91507 | 0.22349 | 0.816181 | 1 |
| 5 | 79471200 | 79473199 | 10.90243 | 20.76915 | 22.73501 | 9.088126 | 0.371384 | 0.002706 | 1 |
| 5 | 79890400 | 79891199 | 13.423 | 24.32989 | 27.0442 | 11.70018 | 0.261308 | 0.336649 | 1 |
| 5 | 79957800 | 79967199 | 125.6267 | 190.68 | 218.7477 | 90.57088 | 0.083145 | 3.225341 | 1 |
| 5 | 79976600 | 79978999 | 37.98297 | 61.44798 | 85.03127 | 36.35774 | -0.10514 | 1.684492 | 1 |
| 5 | 80140800 | 80141599 | 13.55485 | 24.28162 | 28.05801 | 12.29592 | 0.193402 | 0.408435 | 1 |
| 5 | 80605200 | 80605999 | 9.139152 | 16.78668 | 30.92494 | 13.15231 | -0.4458 | 0.238719 | 1 |
| 5 | 80624800 | 80626199 | 23.21065 | 38.5892 | 61.76554 | 26.79886 | -0.29524 | 1.255272 | 1 |
| 5 | 80922000 | 80924799 | 33.13951 | 54.13473 | 73.37271 | 31.53946 | -0.06383 | 1.502149 | 1 |
| 5 | 81352600 | 81353599 | 25.64785 | 42.04421 | 72.23853 | 31.09437 | -0.40191 | 1.441232 | 1 |
| 5 | 81573600 | 81574199 | 14.61931 | 26.1826 | 28.5339 | 12.50682 | 0.277385 | 0.42013 | 1 |
| 5 | 81780400 | 81782399 | 15.80082 | 27.47106 | 46.85506 | 20.58047 | -0.37963 | 0.784223 | 1 |
| 5 | 81999600 | 82004799 | 37.51518 | 61.2876 | 69.90549 | 30.04997 | 0.184478 | 1.508442 | 1 |
| 5 | 82348200 | 82348599 | 6.984251 | 13.40081 | 22.50282 | 8.833666 | -0.21655 | -0.09619 | 1 |
| 5 | 82454800 | 82456399 | 20.7647 | 35.46262 | 47.96591 | 21.02978 | -0.05132 | 0.975838 | 1 |
| 5 | 82787400 | 82789599 | 11.76804 | 21.15525 | 33.66689 | 14.71012 | -0.26278 | 0.385916 | 1 |
| 5 | 82797600 | 82798199 | 9.277255 | 17.72592 | 21.3636 | 8.486282 | 0.242376 | -0.03479 | 1 |
| 5 | 82872600 | 82875399 | 30.34655 | 50.36952 | 52.41931 | 22.7177 | 0.323604 | 1.285459 | 1 |
| 5 | 83014400 | 83015199 | 10.69683 | 19.89988 | 27.794 | 11.6147 | -0.03112 | 0.169062 | 1 |
| 5 | 83029200 | 83031599 | 29.04817 | 47.80621 | 65.86159 | 28.39675 | -0.08206 | 1.392815 | 1 |
| 5 | 83054200 | 83055399 | 21.92379 | 36.44122 | 62.25527 | 27.007 | -0.389 | 1.253386 | 1 |
| 5 | 83329400 | 83331999 | 36.29397 | 59.31852 | 71.98409 | 30.94843 | 0.095791 | 1.502226 | 1 |
| 5 | 83415200 | 83416599 | 10.57476 | 19.90365 | 21.68215 | 8.8834 | 0.347855 | 0.078213 | 1 |
| 5 | 83704000 | 83709999 | 25.67307 | 43.39318 | 46.94097 | 20.53981 | 0.268284 | 1.08244 | 1 |
| 5 | 83930600 | 83932599 | 32.32078 | 52.91363 | 65.31993 | 28.10073 | 0.072849 | 1.465257 | 1 |
| 5 | 84525400 | 84526999 | 31.16789 | 51.2085 | 60.88735 | 26.23338 | 0.128896 | 1.416152 | 1 |
| 5 | 84808000 | 84808599 | 10.41725 | 19.34431 | 27.80674 | 11.65179 | -0.07519 | 0.178264 | 1 |
| 5 | 84901600 | 84902399 | 18.14083 | 31.44881 | 41.62869 | 18.26463 | -0.01636 | 0.818739 | 1 |
| 5 | 85320200 | 85323199 | 23.63047 | 39.56723 | 61.44699 | 26.78221 | -0.25186 | 1.17856 | 1 |
| 5 | 85628800 | 85634199 | 73.95825 | 116.5862 | 129.088 | 53.5683 | 0.176675 | 2.48299 | 1 |
| 5 | 85661600 | 85665399 | 103.9223 | 159.6335 | 174.4663 | 72.44007 | 0.16667 | 2.975395 | 1 |
| 5 | 85989000 | 85990799 | 27.53571 | 46.16794 | 48.06699 | 20.96811 | 0.323102 | 1.163776 | 1 |
| 5 | 86027000 | 86029599 | 9.478132 | 18.23745 | 20.80378 | 8.10054 | 0.347255 | -0.10116 | 1 |
| 5 | 86108800 | 86111799 | 30.35849 | 50.66474 | 53.03337 | 23.06683 | 0.315377 | 1.22358 | 1 |
| 5 | 86145000 | 86147199 | 27.15458 | 46.03041 | 49.59021 | 21.7181 | 0.274275 | 1.065399 | 1 |
| 5 | 86156800 | 86159599 | 21.42268 | 37.51636 | 36.27784 | 16.01502 | 0.435645 | 0.685648 | 1 |
| 5 | 86516200 | 86518799 | 28.46953 | 47.52327 | 47.59289 | 20.70352 | 0.379065 | 1.212435 | 1 |
| 5 | 86716600 | 86722399 | 129.6689 | 196.6921 | 216.6762 | 89.69632 | 0.140872 | 3.238643 | 1 |
| 5 | 86741800 | 86742799 | 21.87624 | 36.97089 | 51.11952 | 22.36732 | -0.08433 | 1.079114 | 1 |
| 5 | 87179600 | 87181599 | 18.27843 | 32.33904 | 33.12779 | 14.65099 | 0.356359 | 0.535531 | 1 |
| 5 | 87216600 | 87217399 | 10.43567 | 19.71316 | 23.72569 | 9.654685 | 0.213127 | 0.058181 | 1 |
| 5 | 87370200 | 87381399 | 307.0041 | 437.4367 | 437.4822 | 180.8487 | 0.195327 | 4.30236 | 1 |
| 5 | 87998000 | 88000599 | 21.20745 | 35.97169 | 50.08379 | 21.93841 | -0.09427 | 1.039753 | 1 |
| 5 | 88208000 | 88213199 | 30.21255 | 50.32658 | 56.27624 | 24.44833 | 0.220513 | 1.248104 | 1 |

| | | | | | | | | | |
|---|---|---|---|---|---|---|---|---|---|
| 5 | 88259800 | 88262199 | 32.68865 | 53.47683 | 66.37311 | 28.54597 | 0.064408 | 1.477722 | 1 |
| 5 | 88326800 | 88329799 | 31.26633 | 52.54701 | 51.85068 | 22.649 | 0.400205 | 1.156333 | 1 |
| 5 | 88391000 | 88395599 | 41.52191 | 67.49357 | 69.53455 | 29.83586 | 0.32584 | 1.586139 | 1 |
| 5 | 88692800 | 88695599 | 17.50777 | 30.31026 | 44.28314 | 19.42847 | -0.15835 | 0.834692 | 1 |
| 5 | 88961600 | 88964599 | 11.81365 | 21.19112 | 34.20434 | 14.97237 | -0.28477 | 0.402083 | 1 |
| 5 | 89725800 | 89726399 | 8.945899 | 17.25632 | 20.38027 | 7.879032 | 0.307803 | -0.12898 | 1 |
| 5 | 90095200 | 90097999 | 11.6332 | 21.61907 | 22.5697 | 9.447326 | 0.383978 | 0.16949 | 1 |
| 5 | 90099400 | 90103599 | 42.12577 | 68.23616 | 78.65351 | 33.66204 | 0.159633 | 1.66518 | 1 |
| 5 | 90518200 | 90520999 | 34.31199 | 56.05504 | 67.04174 | 28.81902 | 0.116317 | 1.501167 | 1 |
| 5 | 90895400 | 90896399 | 12.34425 | 22.85087 | 23.07144 | 9.70891 | 0.426558 | 0.199615 | 1 |
| 5 | 90995400 | 91000999 | 129.6295 | 195.4856 | 262.4073 | 108.4282 | -0.15122 | 3.359181 | 1 |
| 5 | 91379800 | 91382799 | 46.99065 | 75.60975 | 78.20286 | 33.20257 | 0.304888 | 1.854573 | 1 |
| 5 | 91954400 | 91956399 | 16.96145 | 29.67739 | 34.77407 | 15.3397 | 0.160086 | 0.674863 | 1 |
| 5 | 92145800 | 92151199 | 62.52884 | 99.35351 | 118.9014 | 49.59336 | 0.074441 | 2.273553 | 1 |
| 5 | 92657000 | 92662199 | 66.53229 | 105.035 | 136.2626 | 56.57276 | -0.04991 | 2.446652 | 1 |
| 5 | 92665200 | 92668199 | 25.97908 | 44.0327 | 47.43693 | 20.77434 | 0.274314 | 1.059886 | 1 |
| 5 | 92691800 | 92699399 | 213.3476 | 307.7165 | 430.9686 | 177.8191 | -0.27275 | 4.118933 | 1 |
| 5 | 92762000 | 92764199 | 26.19592 | 43.22534 | 64.43546 | 27.81271 | -0.19385 | 1.354423 | 1 |
| 5 | 93153200 | 93155399 | 16.65389 | 29.26352 | 34.02834 | 15.06481 | 0.170919 | 0.603426 | 1 |
| 5 | 93157800 | 93160399 | 19.34891 | 32.81487 | 53.22087 | 23.31162 | -0.31358 | 1.038091 | 1 |
| 5 | 93207000 | 93208999 | 10.7504 | 19.97186 | 30.45803 | 12.75499 | -0.15951 | 0.18429 | 1 |
| 5 | 93213400 | 93220199 | 101.0953 | 155.3526 | 195.1826 | 81.04608 | -0.0338 | 2.969269 | 1 |
| 5 | 93484200 | 93486799 | 21.63121 | 37.45892 | 44.16383 | 19.37628 | 0.149612 | 0.848902 | 1 |
| 5 | 93593800 | 93596799 | 41.52524 | 66.93583 | 93.86718 | 39.95568 | -0.13024 | 1.797955 | 1 |
| 5 | 93601200 | 93615399 | 22.63799 | 39.33532 | 39.78918 | 17.47052 | 0.370426 | 0.810794 | 1 |
| 5 | 93660400 | 93661199 | 18.34107 | 31.90416 | 40.01462 | 17.58167 | 0.061821 | 0.776446 | 1 |
| 5 | 93822200 | 93825599 | 37.47323 | 61.63372 | 72.65611 | 31.31808 | 0.141775 | 1.425832 | 1 |
| 5 | 93972000 | 93972999 | 12.67732 | 22.4986 | 36.21592 | 15.99641 | -0.28982 | 0.480172 | 1 |
| 5 | 94154800 | 94155599 | 32.48355 | 52.87963 | 69.08498 | 29.65888 | -0.01431 | 1.552051 | 1 |
| 5 | 94193600 | 94194199 | 16.36184 | 28.44061 | 39.70491 | 17.43761 | -0.09191 | 0.779118 | 1 |
| 5 | 94202800 | 94203999 | 27.46642 | 45.6908 | 54.90336 | 23.8309 | 0.117186 | 1.257706 | 1 |
| 5 | 94331000 | 94331999 | 25.06899 | 42.33836 | 45.40831 | 19.86437 | 0.280658 | 1.086611 | 1 |
| 5 | 94333400 | 94334599 | 13.88958 | 25.01303 | 30.9212 | 13.48632 | 0.101094 | 0.383949 | 1 |
| 5 | 94342600 | 94343599 | 41.30816 | 66.48405 | 82.43036 | 35.01091 | 0.044414 | 1.840057 | 1 |
| 5 | 94377000 | 94377799 | 12.97902 | 24.08896 | 24.57463 | 10.30467 | 0.415596 | 0.195564 | 1 |
| 5 | 94402200 | 94403199 | 17.98597 | 31.60233 | 33.44981 | 14.80795 | 0.305965 | 0.611931 | 1 |
| 5 | 94477000 | 94478599 | 45.8691 | 73.62293 | 98.70298 | 41.75417 | -0.07304 | 1.936733 | 1 |
| 5 | 94489600 | 94493399 | 61.15996 | 98.30655 | 83.76721 | 35.51152 | 0.582023 | 1.907979 | 1 |
| 5 | 94839600 | 94839999 | 11.32793 | 21.27222 | 24.10429 | 9.914344 | 0.28655 | 0.104645 | 1 |
| 5 | 94861200 | 94865199 | 122.1969 | 184.9503 | 222.8073 | 92.17123 | 0.009248 | 3.282335 | 1 |
| 5 | 94866800 | 94868599 | 69.27183 | 109.4074 | 120.8178 | 50.17151 | 0.182674 | 2.435522 | 1 |
| 5 | 94871200 | 94875799 | 218.9385 | 315.6748 | 417.7828 | 172.3862 | -0.19159 | 4.124408 | 1 |
| 5 | 95361200 | 95361599 | 9.361205 | 18.01728 | 20.3061 | 7.9007 | 0.36584 | -0.10534 | 1 |
| 5 | 95442800 | 95446999 | 45.19688 | 73.1762 | 78.63568 | 33.64268 | 0.259917 | 1.679105 | 1 |
| 5 | 95518000 | 95528599 | 321.9004 | 452.0088 | 496.1986 | 205.6157 | 0.039753 | 4.517466 | 1 |
| 5 | 95820600 | 95821399 | 32.10956 | 52.22839 | 70.82822 | 30.39827 | -0.06888 | 1.562033 | 1 |
| 5 | 95859600 | 95860599 | 13.33348 | 23.48582 | 36.83775 | 16.3062 | -0.25712 | 0.563814 | 1 |

| | | | | | | | | | |
|---|---|---|---|---|---|---|---|---|---|
| 5 | 95969000 | 95969999 | 23.36368 | 39.28013 | 51.17381 | 22.35584 | 0.000785 | 1.12581 | 1 |
| 5 | 96008800 | 96009999 | 11.72592 | 21.82312 | 27.78178 | 11.60169 | 0.102644 | 0.176486 | 1 |
| 5 | 96288800 | 96289599 | 9.68645 | 18.71115 | 23.20102 | 8.919186 | 0.245095 | -0.13384 | 1 |
| 5 | 96708400 | 96710999 | 24.00088 | 41.3862 | 42.05393 | 18.4482 | 0.362054 | 0.895596 | 1 |
| 5 | 97505400 | 97506599 | 15.13277 | 27.73715 | 28.44286 | 12.13144 | 0.3904 | 0.289598 | 1 |
| 5 | 97539400 | 97541199 | 25.13519 | 41.73512 | 63.57874 | 27.56108 | -0.22437 | 1.27667 | 1 |
| 5 | 98262800 | 98264599 | 8.370952 | 15.87229 | 27.11026 | 10.94481 | -0.27786 | 0.022663 | 1 |
| 5 | 98336000 | 98338399 | 18.14856 | 31.75546 | 34.58159 | 15.25519 | 0.265084 | 0.679051 | 1 |
| 5 | 98835400 | 98837799 | 15.45695 | 27.28388 | 40.70549 | 18.01062 | -0.18461 | 0.556321 | 1 |
| 5 | 99719600 | 99721199 | 27.49304 | 44.76464 | 74.55817 | 32.01093 | -0.36346 | 1.542606 | 1 |
| 5 | 100766800 | 100767999 | 12.80403 | 22.75479 | 36.68572 | 16.18511 | -0.29104 | 0.472166 | 1 |
| 5 | 100892400 | 100894399 | 22.20656 | 37.81004 | 44.31424 | 19.42402 | 0.153832 | 1.002086 | 1 |
| 5 | 101459600 | 101460799 | 12.97302 | 22.95287 | 34.74559 | 15.36643 | -0.20366 | 0.507823 | 1 |
| 5 | 102722800 | 102724799 | 25.17204 | 43.01882 | 48.47558 | 21.2544 | 0.211053 | 0.984739 | 1 |
| 5 | 102815200 | 102816799 | 13.8184 | 24.59491 | 34.01989 | 14.98973 | -0.07008 | 0.462689 | 1 |
| 5 | 103081400 | 103083999 | 17.77867 | 31.4097 | 35.98284 | 15.91823 | 0.195378 | 0.556886 | 1 |
| 5 | 103555200 | 103558399 | 33.204 | 54.21776 | 75.5459 | 32.46874 | -0.10403 | 1.510731 | 1 |
| 5 | 104297200 | 104299999 | 26.57561 | 44.39184 | 56.69925 | 24.67543 | 0.029205 | 1.206895 | 1 |
| 5 | 104431000 | 104433199 | 9.114287 | 16.70582 | 31.39381 | 13.39001 | -0.47765 | 0.249261 | 1 |
| 5 | 104603600 | 104608399 | 55.02661 | 87.47985 | 130.2274 | 54.34648 | -0.23921 | 2.254235 | 1 |
| 5 | 104807200 | 104811599 | 92.32465 | 143.5332 | 151.9836 | 63.10054 | 0.225559 | 2.748756 | 1 |
| 5 | 105040200 | 105045199 | 39.43705 | 63.45051 | 98.45229 | 41.79489 | -0.2792 | 1.855475 | 1 |
| 5 | 105067000 | 105083599 | 447.8197 | 612.7479 | 693.6581 | 288.611 | -0.04229 | 4.915148 | 1 |
| 5 | 105240800 | 105243999 | 42.70581 | 68.81069 | 83.41909 | 35.48852 | 0.07872 | 1.805936 | 1 |
| 5 | 105572000 | 105579199 | 70.13833 | 110.2135 | 155.8233 | 64.63602 | -0.17906 | 2.547872 | 1 |
| 5 | 106735800 | 106738799 | 30.63232 | 50.08546 | 78.16244 | 33.60967 | -0.26572 | 1.4882 | 1 |
| 5 | 107079400 | 107081199 | 18.7189 | 32.71842 | 35.67077 | 15.72147 | 0.263575 | 0.702304 | 1 |
| 5 | 107105800 | 107108599 | 24.1953 | 41.25913 | 47.41086 | 20.7839 | 0.182505 | 0.999738 | 1 |
| 5 | 107345800 | 107346399 | 9.829378 | 18.53333 | 25.32378 | 10.3445 | 0.026526 | 0.068935 | 1 |
| 5 | 107401600 | 107408199 | 110.7121 | 168.5703 | 225.4454 | 93.43171 | -0.13429 | 3.158255 | 1 |
| 5 | 107477600 | 107480399 | 34.24954 | 55.57489 | 77.11428 | 33.04573 | -0.10495 | 1.614414 | 1 |
| 5 | 107682400 | 107683399 | 15.36051 | 26.99687 | 36.22661 | 16.04104 | -0.03448 | 0.598503 | 1 |
| 5 | 107976800 | 107978999 | 39.47718 | 63.74611 | 81.03048 | 34.56963 | 0.014122 | 1.737451 | 1 |
| 5 | 108066400 | 108067599 | 10.18697 | 19.29375 | 23.28354 | 9.422496 | 0.216566 | 0.038147 | 1 |
| 5 | 108267400 | 108269399 | 9.463602 | 17.71343 | 27.93123 | 11.55203 | -0.19283 | 0.120048 | 1 |
| 5 | 108922000 | 108924799 | 18.19038 | 31.34102 | 45.31265 | 19.87723 | -0.14498 | 0.87892 | 1 |
| 5 | 109047000 | 109048399 | 14.36211 | 25.69897 | 31.88397 | 13.98575 | 0.090285 | 0.428024 | 1 |
| 5 | 109387400 | 109389199 | 15.30061 | 27.72675 | 29.79363 | 12.8934 | 0.309302 | 0.35584 | 1 |
| 5 | 109424400 | 109434999 | 126.7003 | 194.642 | 184.2663 | 76.51015 | 0.373808 | 2.983084 | 1 |
| 5 | 109789600 | 109796199 | 66.41517 | 105.6904 | 117.3677 | 49.04138 | 0.184434 | 2.229541 | 1 |
| 5 | 109947000 | 109949199 | 20.62603 | 35.46621 | 44.7709 | 19.63839 | 0.049387 | 0.907883 | 1 |
| 5 | 110014200 | 110016199 | 14.13403 | 24.7211 | 42.06875 | 18.55175 | -0.37554 | 0.679712 | 1 |
| 5 | 110275600 | 110284999 | 179.9711 | 268.2746 | 281.4437 | 115.9828 | 0.189169 | 3.588961 | 1 |
| 5 | 110339800 | 110347799 | 59.93654 | 95.40434 | 114.4221 | 47.82707 | 0.073945 | 2.214279 | 1 |
| 5 | 110789600 | 110790999 | 18.33757 | 32.04077 | 38.8068 | 17.09887 | 0.112554 | 0.711561 | 1 |
| 5 | 112999600 | 113003799 | 28.23587 | 46.02888 | 73.37993 | 31.51873 | -0.29936 | 1.523304 | 1 |
| 5 | 113596200 | 113599799 | 45.66177 | 72.93116 | 111.9661 | 47.02393 | -0.27628 | 2.093372 | 1 |

| | | | | | | | | | |
|---|---|---|---|---|---|---|---|---|---|
| 5 | 115024800 | 115027999 | 32.96711 | 53.86795 | 64.69688 | 27.81293 | 0.110473 | 1.493331 | 1 |
| 5 | 115489200 | 115489999 | 9.470078 | 17.70511 | 30.62379 | 12.69041 | -0.32786 | 0.133079 | 1 |
| 5 | 115945800 | 115947599 | 18.49118 | 32.23008 | 41.98655 | 18.47144 | 0.0075 | 0.750766 | 1 |
| 5 | 117201000 | 117208999 | 77.67028 | 120.8991 | 171.0269 | 70.99392 | -0.19049 | 2.722828 | 1 |
| 5 | 118305600 | 118308399 | 20.9933 | 35.31164 | 53.75972 | 23.48889 | -0.22315 | 1.116349 | 1 |
| 5 | 118474000 | 118477199 | 29.91123 | 49.27274 | 67.63751 | 29.17401 | -0.07633 | 1.38587 | 1 |
| 5 | 119038200 | 119043599 | 74.57018 | 117.0671 | 154.3063 | 63.99893 | -0.07909 | 2.568781 | 1 |
| 5 | 119089600 | 119092399 | 29.11924 | 47.64725 | 71.72927 | 30.85187 | -0.21314 | 1.46898 | 1 |
| 5 | 119624000 | 119626199 | 16.73223 | 29.11139 | 44.70519 | 19.64507 | -0.22908 | 0.770867 | 1 |
| 5 | 119688800 | 119692399 | 46.35021 | 74.17463 | 106.6364 | 44.90888 | -0.17866 | 2.030908 | 1 |
| 5 | 119717400 | 119720199 | 17.24741 | 30.30978 | 31.49161 | 13.94307 | 0.332558 | 0.600551 | 1 |
| 5 | 122365000 | 122366999 | 26.20395 | 44.24064 | 53.42017 | 23.36692 | 0.11009 | 1.099959 | 1 |
| 5 | 123521800 | 123525599 | 37.74567 | 61.6341 | 65.6821 | 28.22946 | 0.281816 | 1.514951 | 1 |
| 5 | 123564400 | 123566599 | 28.07162 | 47.25485 | 49.09209 | 21.45461 | 0.326265 | 1.129003 | 1 |
| 5 | 124407000 | 124414999 | 71.5222 | 112.6859 | 132.2128 | 54.86121 | 0.092601 | 2.492637 | 1 |
| 5 | 125745200 | 125747399 | 30.45698 | 50.2964 | 58.86405 | 25.42135 | 0.154207 | 1.350084 | 1 |
| 5 | 125791000 | 125792999 | 22.40076 | 37.7655 | 55.44857 | 24.24364 | -0.171 | 1.102269 | 1 |
| 5 | 126122600 | 126134399 | 241.9464 | 349.1999 | 398.5607 | 164.4329 | 0.023307 | 4.110391 | 1 |
| 5 | 126443600 | 126444199 | 16.6427 | 29.11843 | 38.88343 | 17.15182 | -0.0275 | 0.682631 | 1 |
| 5 | 126969000 | 126971999 | 34.73177 | 57.04202 | 61.22596 | 26.37499 | 0.275864 | 1.429127 | 1 |
| 5 | 127023600 | 127029199 | 46.45165 | 74.45484 | 118.6377 | 50.08161 | -0.32419 | 1.987341 | 1 |
| 5 | 127816400 | 127822599 | 72.41872 | 113.6667 | 144.6973 | 60.012 | -0.02913 | 2.569911 | 1 |
| 5 | 127905800 | 127913199 | 54.01107 | 85.98316 | 132.0595 | 55.20882 | -0.28213 | 2.210182 | 1 |
| 5 | 129096600 | 129100199 | 29.23758 | 49.12701 | 53.47185 | 23.35564 | 0.25913 | 1.15224 | 1 |
| 5 | 130129400 | 130131999 | 25.20779 | 42.7481 | 45.30746 | 19.84496 | 0.297748 | 1.048785 | 1 |
| 5 | 131343800 | 131345199 | 14.51081 | 25.76659 | 33.02788 | 14.58155 | 0.037335 | 0.481499 | 1 |
| 5 | 131467000 | 131474399 | 90.10864 | 137.9299 | 212.6898 | 88.2596 | -0.33338 | 3.044654 | 1 |
| 5 | 133053400 | 133057599 | 48.91044 | 78.67109 | 94.57261 | 40.13221 | 0.087703 | 1.872636 | 1 |
| 5 | 133172400 | 133173799 | 13.31203 | 25.01924 | 25.08167 | 10.30158 | 0.463705 | 0.11967 | 1 |
| 5 | 133284000 | 133286199 | 26.50966 | 43.74788 | 69.62339 | 30.05374 | -0.288 | 1.356643 | 1 |
| 5 | 133349600 | 133352199 | 55.20406 | 87.53027 | 125.8773 | 52.3923 | -0.19313 | 2.33069 | 1 |
| 5 | 133509800 | 133512999 | 30.99694 | 51.36506 | 57.95934 | 25.09035 | 0.207033 | 1.309455 | 1 |
| 5 | 133745600 | 133746599 | 14.31843 | 25.90132 | 28.34926 | 12.29535 | 0.280921 | 0.355982 | 1 |
| 5 | 134041800 | 134047599 | 51.94719 | 82.7253 | 117.9707 | 49.34177 | -0.17472 | 2.192796 | 1 |
| 5 | 134064000 | 134068799 | 38.40267 | 62.31931 | 72.10651 | 30.90171 | 0.156614 | 1.616779 | 1 |
| 5 | 134502400 | 134507999 | 47.2893 | 75.6953 | 113.3402 | 47.74808 | -0.23687 | 2.029058 | 1 |
| 5 | 134855200 | 134857999 | 14.55195 | 25.44736 | 42.54657 | 18.75953 | -0.35037 | 0.683713 | 1 |
| 5 | 136232000 | 136235799 | 52.38371 | 83.45839 | 117.72 | 49.26788 | -0.15822 | 2.180689 | 1 |
| 5 | 136968000 | 136968799 | 19.44855 | 33.62102 | 41.1251 | 18.04213 | 0.096281 | 0.849983 | 1 |
| 5 | 137808200 | 137815399 | 92.51551 | 142.7175 | 181.8766 | 75.54819 | -0.05005 | 2.89047 | 1 |
| 5 | 138200400 | 138205799 | 48.35753 | 77.62979 | 93.04278 | 39.36959 | 0.088599 | 1.933899 | 1 |
| 5 | 138331800 | 138334399 | 24.80727 | 41.87807 | 45.91281 | 20.08241 | 0.248952 | 1.088055 | 1 |
| 5 | 138372000 | 138373199 | 22.31963 | 37.8984 | 50.07533 | 21.94178 | -0.01885 | 1.037989 | 1 |
| 5 | 138442800 | 138445199 | 26.64098 | 45.27365 | 47.40433 | 20.77528 | 0.315425 | 1.039797 | 1 |
| 5 | 138590200 | 138597399 | 76.32769 | 120.1703 | 170.222 | 70.62633 | -0.17968 | 2.517242 | 1 |
| 5 | 138628200 | 138629399 | 14.66242 | 25.96364 | 33.54313 | 14.83215 | 0.024044 | 0.506115 | 1 |
| 5 | 140456800 | 140459399 | 28.84927 | 47.96334 | 54.01876 | 23.43754 | 0.210121 | 1.262771 | 1 |

| | | | | | | | | | |
|---|---|---|---|---|---|---|---|---|---|
| 5 | 140941800 | 140944199 | 26.2321 | 43.51594 | 59.7791 | 25.89731 | -0.07574 | 1.287065 | 1 |
| 5 | 141072800 | 141074599 | 17.03358 | 29.81697 | 34.80623 | 15.35971 | 0.165589 | 0.669402 | 1 |
| 5 | 141590400 | 141592599 | 25.57966 | 43.88268 | 44.14848 | 19.36311 | 0.374902 | 0.949044 | 1 |
| 5 | 141624000 | 141629199 | 63.58594 | 100.5501 | 127.8129 | 53.10918 | -0.01862 | 2.404574 | 1 |
| 5 | 143780200 | 143781599 | 17.03404 | 29.81684 | 38.77971 | 17.11272 | 0.010215 | 0.675279 | 1 |
| 5 | 144331000 | 144334399 | 31.89745 | 52.15675 | 73.72654 | 31.70285 | -0.12328 | 1.487161 | 1 |
| 5 | 144450400 | 144452599 | 18.54011 | 32.15803 | 44.74453 | 19.63202 | -0.08786 | 0.821038 | 1 |
| 5 | 144506600 | 144539399 | 564.2332 | 752.7265 | 975.7243 | 407.5224 | -0.27443 | 5.316816 | 1 |
| 5 | 144659600 | 144660599 | 11.94399 | 22.08914 | 26.12198 | 11.00672 | 0.198658 | 0.209175 | 1 |
| 5 | 145226600 | 145231599 | 43.28939 | 69.60851 | 91.41164 | 38.77366 | -0.04005 | 1.875212 | 1 |
| 5 | 145417000 | 145419399 | 68.55291 | 108.2006 | 128.4983 | 53.34314 | 0.077148 | 2.452933 | 1 |
| 5 | 145593200 | 145630799 | 1712.511 | 2108.016 | 2158.44 | 909.4242 | -0.05089 | 6.618364 | 1 |
| 5 | 145887400 | 145906199 | 523.8877 | 713.0791 | 681.4633 | 283.7685 | 0.194367 | 4.998811 | 1 |
| 5 | 145944000 | 145962599 | 275.7778 | 397.1575 | 429.2059 | 177.1274 | 0.098949 | 4.145881 | 1 |
| 5 | 146031200 | 146034199 | 38.41921 | 63.04247 | 63.76972 | 27.46054 | 0.360706 | 1.450995 | 1 |
| 5 | 146321800 | 146324799 | 13.03073 | 22.83877 | 39.51105 | 17.45445 | -0.39902 | 0.635894 | 1 |
| 5 | 146655800 | 146662199 | 60.58529 | 95.98902 | 136.5034 | 56.77536 | -0.17808 | 2.358075 | 1 |
| 5 | 146810200 | 146813199 | 23.93994 | 39.9644 | 65.40102 | 28.45128 | -0.32721 | 1.213756 | 1 |
| 5 | 146937600 | 146945999 | 103.0553 | 157.2329 | 218.1028 | 90.43748 | -0.18455 | 3.113691 | 1 |
| 5 | 147604000 | 147605999 | 19.03235 | 32.35577 | 53.75115 | 23.55754 | -0.34781 | 1.016652 | 1 |
| 5 | 147827600 | 147828799 | 13.11331 | 23.55123 | 31.50264 | 13.77553 | -0.01448 | 0.396115 | 1 |
| 5 | 148539000 | 148543999 | 32.03017 | 53.05075 | 56.85465 | 24.60366 | 0.281138 | 1.318209 | 1 |
| 5 | 148591200 | 148593599 | 21.0404 | 35.93735 | 50.61533 | 22.19157 | -0.10957 | 0.977403 | 1 |
| 5 | 148885400 | 148893199 | 61.68676 | 97.23814 | 148.1347 | 61.4719 | -0.28311 | 2.478305 | 1 |
| 5 | 148960400 | 148967199 | 24.0963 | 40.74674 | 50.1199 | 21.93385 | 0.083647 | 1.076405 | 1 |
| 5 | 149198200 | 149199399 | 12.57578 | 22.5445 | 33.1296 | 14.50936 | -0.15074 | 0.408706 | 1 |
| 5 | 149218600 | 149220199 | 26.45198 | 44.65042 | 49.0709 | 21.46316 | 0.245534 | 1.100364 | 1 |
| 5 | 149230600 | 149235399 | 49.49959 | 79.23773 | 106.1581 | 44.72593 | -0.07664 | 2.02666 | 1 |
| 5 | 149396800 | 149399799 | 33.5297 | 54.22813 | 86.08103 | 36.79887 | -0.30299 | 1.685513 | 1 |
| 5 | 149756600 | 149760599 | 52.74605 | 84.11041 | 109.0712 | 45.70438 | -0.03574 | 2.148566 | 1 |
| 5 | 149820400 | 149823199 | 29.33104 | 48.93453 | 54.05658 | 23.50717 | 0.237991 | 1.226364 | 1 |
| 5 | 150011200 | 150011799 | 9.327934 | 17.62631 | 26.48736 | 10.77115 | -0.10353 | 0.055032 | 1 |
| 5 | 150314600 | 150319999 | 70.53095 | 111.0459 | 142.7438 | 59.22555 | -0.03974 | 2.507083 | 1 |
| 5 | 150759000 | 150762399 | 49.16654 | 78.8007 | 105.5899 | 44.56873 | -0.07481 | 1.988272 | 1 |
| 5 | 150847800 | 150851599 | 44.58508 | 71.80558 | 86.93861 | 36.96109 | 0.079697 | 1.822909 | 1 |
| 5 | 150905400 | 150906999 | 23.57761 | 40.5348 | 42.09024 | 18.46245 | 0.330037 | 0.919344 | 1 |
| 5 | 150917600 | 150918799 | 16.95272 | 29.99577 | 32.8807 | 14.5415 | 0.259408 | 0.528298 | 1 |
| 5 | 151390400 | 151395199 | 31.04242 | 51.30638 | 55.10656 | 23.81246 | 0.277766 | 1.338407 | 1 |
| 5 | 151396600 | 151399199 | 26.77772 | 45.65159 | 43.11277 | 18.89934 | 0.464459 | 1.007325 | 1 |
| 5 | 151536400 | 151539199 | 32.11796 | 53.70462 | 60.35473 | 26.28323 | 0.213219 | 1.217807 | 1 |
| 5 | 151620400 | 151622399 | 16.62098 | 28.7898 | 42.73513 | 18.7496 | -0.18108 | 0.82172 | 1 |
| 6 | 3093400 | 3094999 | 25.85099 | 43.52591 | 50.11998 | 21.90524 | 0.178475 | 1.124745 | 1 |
| 6 | 3176600 | 3184799 | 184.2568 | 273.5246 | 289.4446 | 119.2291 | 0.171182 | 3.662658 | 1 |
| 6 | 3284600 | 3295399 | 126.2561 | 192.0538 | 210.7678 | 87.32384 | 0.149633 | 3.181086 | 1 |
| 6 | 3655600 | 3658599 | 50.09519 | 80.26937 | 90.85421 | 38.33459 | 0.167842 | 1.992261 | 1 |
| 6 | 3695800 | 3699399 | 53.63076 | 85.56305 | 106.152 | 44.51125 | 0.028771 | 2.134317 | 1 |
| 6 | 3763200 | 3765599 | 21.34373 | 36.21269 | 52.22036 | 22.87619 | -0.14478 | 1.039224 | 1 |

| | | | | | | | | | | |
|---|---|---|---|---|---|---|---|---|---|---|
| 6 | 3820600 | 3821799 | 12.2802 | 22.74968 | 26.11039 | 10.97634 | 0.243945 | 0.202683 | | 1 |
| 6 | 3852200 | 3855199 | 44.14954 | 70.81592 | 94.4278 | 39.90536 | -0.06633 | 1.95543 | | 1 |
| 6 | 4314200 | 4315199 | 11.70965 | 21.86435 | 25.63462 | 10.64582 | 0.226692 | 0.148251 | | 1 |
| 6 | 4472400 | 4474199 | 16.57405 | 28.7442 | 47.40466 | 20.7991 | -0.3319 | 0.818899 | | 1 |
| 6 | 4653000 | 4654799 | 17.95872 | 31.17666 | 42.17979 | 18.50972 | -0.04718 | 0.807765 | | 1 |
| 6 | 4891800 | 4893999 | 16.12552 | 28.24576 | 40.87374 | 18.04715 | -0.14302 | 0.663158 | | 1 |
| 6 | 5064600 | 5065999 | 14.18305 | 25.28179 | 37.1099 | 16.3306 | -0.15412 | 0.461812 | | 1 |
| 6 | 5330400 | 5336199 | 65.58456 | 104.1069 | 121.435 | 50.60057 | 0.110007 | 2.304123 | | 1 |
| 6 | 5476400 | 5482399 | 59.6046 | 95.20586 | 102.1893 | 42.92112 | 0.239193 | 2.102263 | | 1 |
| 6 | 5525000 | 5526399 | 13.82664 | 24.49477 | 33.73469 | 14.9155 | -0.06712 | 0.499147 | | 1 |
| 6 | 5613400 | 5615199 | 25.0089 | 42.08361 | 47.52117 | 20.76575 | 0.206474 | 1.121382 | | 1 |
| 6 | 5845400 | 5848599 | 21.50129 | 36.77572 | 50.08713 | 21.96245 | -0.06107 | 0.96569 | | 1 |
| 6 | 6489400 | 6490599 | 11.56458 | 20.54089 | 34.30161 | 15.14024 | -0.34165 | 0.463651 | | 1 |
| 6 | 6520000 | 6521599 | 18.83915 | 32.68936 | 38.60043 | 16.93645 | 0.148138 | 0.811526 | | 1 |
| 6 | 6854200 | 6856999 | 24.08945 | 40.10557 | 61.43249 | 26.67938 | -0.23202 | 1.239499 | | 1 |
| 6 | 6971800 | 6983399 | 128.9123 | 197.9231 | 188.8779 | 78.4186 | 0.361632 | 2.996682 | | 1 |
| 6 | 6988800 | 6992999 | 94.65738 | 146.5738 | 169.8977 | 70.57102 | 0.090741 | 2.821533 | | 1 |
| 6 | 7113000 | 7113999 | 18.79372 | 32.75416 | 37.87083 | 16.65975 | 0.178923 | 0.748308 | | 1 |
| 6 | 7332200 | 7333799 | 17.26825 | 30.49335 | 32.42323 | 14.34489 | 0.302061 | 0.550988 | | 1 |
| 6 | 7716200 | 7731599 | 217.2176 | 317.4421 | 323.4625 | 133.2598 | 0.204396 | 3.920566 | | 1 |
| 6 | 7781600 | 7783999 | 17.929 | 31.0107 | 51.17532 | 22.45166 | -0.33394 | 0.8508 | | 1 |
| 6 | 7827000 | 7828799 | 14.5392 | 25.71955 | 38.13314 | 16.86515 | -0.17426 | 0.524281 | | 1 |
| 6 | 7840000 | 7842199 | 24.34744 | 42.0564 | 40.97376 | 17.9753 | 0.423032 | 0.880706 | | 1 |
| 6 | 7923000 | 7925399 | 21.58613 | 37.08413 | 41.66698 | 18.27638 | 0.21652 | 0.916546 | | 1 |
| 6 | 8129600 | 8131199 | 21.87144 | 37.77493 | 40.00628 | 17.5508 | 0.302991 | 0.867723 | | 1 |
| 6 | 8613000 | 8617599 | 95.84302 | 147.2706 | 193.5104 | 80.3509 | -0.09848 | 2.968475 | | 1 |
| 6 | 8947400 | 8952599 | 52.12501 | 82.55658 | 136.3573 | 56.70476 | -0.39406 | 2.359353 | | 1 |
| 6 | 9145200 | 9146199 | 13.81528 | 24.38763 | 37.25648 | 16.48436 | -0.2181 | 0.543501 | | 1 |
| 6 | 9288600 | 9290999 | 18.2724 | 31.99574 | 36.11402 | 15.94114 | 0.21373 | 0.67188 | | 1 |
| 6 | 9294000 | 9306599 | 253.7568 | 361.0385 | 502.1826 | 207.646 | -0.28253 | 4.321789 | | 1 |
| 6 | 9586000 | 9586999 | 13.74176 | 24.29627 | 35.17156 | 15.55695 | -0.14004 | 0.520356 | | 1 |
| 6 | 9716200 | 9716999 | 13.16358 | 23.74249 | 34.12018 | 14.85266 | -0.11307 | 0.376801 | | 1 |
| 6 | 9724800 | 9727799 | 27.71372 | 46.34402 | 49.1471 | 21.40363 | 0.2967 | 1.193375 | | 1 |
| 6 | 9845400 | 9846599 | 18.90221 | 33.0296 | 39.83583 | 17.55303 | 0.11867 | 0.716067 | | 1 |
| 6 | 9858200 | 9870399 | 289.3371 | 410.667 | 488.1512 | 201.9262 | -0.05942 | 4.358656 | | 1 |
| 6 | 10327600 | 10329799 | 20.9213 | 35.89517 | 42.72193 | 18.73761 | 0.133422 | 0.92621 | | 1 |
| 6 | 10486800 | 10491599 | 80.05085 | 125.3407 | 145.5961 | 60.39544 | 0.100091 | 2.618045 | | 1 |
| 6 | 10629000 | 10629999 | 18.38695 | 31.9573 | 41.08236 | 18.04134 | 0.026304 | 0.790712 | | 1 |
| 6 | 10688200 | 10690999 | 45.08996 | 72.93639 | 74.68665 | 31.93163 | 0.328163 | 1.696521 | | 1 |
| 6 | 10705200 | 10706199 | 9.805145 | 17.83262 | 31.39836 | 13.52652 | -0.39361 | 0.302935 | | 1 |
| 6 | 10711000 | 10713199 | 26.49234 | 44.57128 | 48.03137 | 20.98747 | 0.273542 | 1.125909 | | 1 |
| 6 | 10738400 | 10740799 | 36.47298 | 59.40978 | 70.46226 | 30.25681 | 0.125349 | 1.550175 | | 1 |
| 6 | 10876000 | 10876799 | 9.854629 | 18.50601 | 24.27152 | 9.982735 | 0.077522 | 0.092981 | | 1 |
| 6 | 10906600 | 10907799 | 20.12218 | 35.07163 | 37.18747 | 16.35978 | 0.303122 | 0.751262 | | 1 |
| 6 | 11161400 | 11162599 | 20.22649 | 34.95359 | 43.66753 | 19.15732 | 0.065732 | 0.858928 | | 1 |
| 6 | 11204600 | 11205999 | 12.25797 | 21.60541 | 37.41157 | 16.55806 | -0.39852 | 0.552049 | | 1 |
| 6 | 11462000 | 11466199 | 35.95844 | 59.161 | 60.94022 | 26.27151 | 0.336027 | 1.40901 | | 1 |

| | | | | | | | | | |
|---|---|---|---|---|---|---|---|---|---|
| 6 | 11677000 | 11678799 | 12.43098 | 22.90696 | 27.0891 | 11.4704 | 0.193507 | 0.235284 | 1 |
| 6 | 11679000 | 11680599 | 19.44552 | 33.40876 | 47.94832 | 21.03161 | -0.13519 | 0.91577 | 1 |
| 6 | 11702800 | 11703799 | 14.702 | 26.1765 | 30.4192 | 13.39936 | 0.180139 | 0.459104 | 1 |
| 6 | 11822600 | 11823999 | 18.41791 | 32.28806 | 32.39819 | 14.31498 | 0.382497 | 0.646072 | 1 |
| 6 | 11829400 | 11833399 | 37.23678 | 60.51014 | 64.11713 | 27.50557 | 0.285081 | 1.583044 | 1 |
| 6 | 11906800 | 11908599 | 11.22093 | 20.74994 | 24.62222 | 10.37612 | 0.193501 | 0.198753 | 1 |
| 6 | 12671400 | 12671999 | 9.119054 | 17.13596 | 25.35838 | 10.41813 | -0.09402 | 0.085168 | 1 |
| 6 | 12838800 | 12840999 | 57.36526 | 91.3895 | 103.7757 | 43.42828 | 0.154037 | 2.183067 | 1 |
| 6 | 13277800 | 13278399 | 12.44848 | 23.07721 | 24.46929 | 10.27687 | 0.358141 | 0.194637 | 1 |
| 6 | 13395000 | 13396799 | 18.33728 | 32.2131 | 33.45251 | 14.80662 | 0.333365 | 0.617561 | 1 |
| 6 | 13450200 | 13464199 | 315.6863 | 444.859 | 498.685 | 206.5291 | 0.014698 | 4.464702 | 1 |
| 6 | 13569800 | 13574799 | 96.78856 | 149.3402 | 180.6777 | 75.05012 | 0.025089 | 2.888987 | 1 |
| 6 | 13741000 | 13743799 | 35.33331 | 57.38456 | 68.27026 | 29.27503 | 0.117698 | 1.593471 | 1 |
| 6 | 13959800 | 13961199 | 17.13867 | 30.28055 | 34.42228 | 15.22771 | 0.206395 | 0.548053 | 1 |
| 6 | 14162000 | 14164199 | 21.33211 | 37.05485 | 38.84797 | 17.05396 | 0.319017 | 0.807997 | 1 |
| 6 | 14459000 | 14461599 | 33.62812 | 55.48117 | 51.95103 | 22.42262 | 0.474823 | 1.363402 | 1 |
| 6 | 14546400 | 14547199 | 14.55684 | 25.69059 | 32.29414 | 14.28942 | 0.06171 | 0.541402 | 1 |
| 6 | 14670200 | 14682999 | 206.6846 | 304.6829 | 322.8173 | 132.94 | 0.160006 | 3.783107 | 1 |
| 6 | 14793600 | 14795199 | 26.30448 | 43.74976 | 64.24428 | 27.88176 | -0.17136 | 1.262376 | 1 |
| 6 | 14800400 | 14802399 | 30.49225 | 50.59694 | 54.06704 | 23.42719 | 0.285515 | 1.289905 | 1 |
| 6 | 15550800 | 15551999 | 16.44374 | 28.51724 | 42.20974 | 18.51977 | -0.17662 | 0.808947 | 1 |
| 6 | 15996600 | 15998399 | 17.95041 | 31.19344 | 39.62678 | 17.40033 | 0.043482 | 0.790195 | 1 |
| 6 | 16252400 | 16254399 | 17.50544 | 29.97757 | 52.18619 | 22.88509 | -0.41349 | 0.940006 | 1 |
| 6 | 16264000 | 16269199 | 69.02354 | 109.4933 | 120.1401 | 50.02777 | 0.197194 | 2.325765 | 1 |
| 6 | 17003200 | 17005999 | 32.49882 | 53.17374 | 66.41425 | 28.56512 | 0.055465 | 1.474674 | 1 |
| 6 | 17902000 | 17902799 | 11.51546 | 20.75133 | 32.74471 | 14.26983 | -0.24776 | 0.370161 | 1 |
| 6 | 18062800 | 18065199 | 28.49409 | 46.884 | 74.26647 | 32.0176 | -0.28302 | 1.401409 | 1 |
| 6 | 18120200 | 18121399 | 15.51082 | 27.03773 | 44.17111 | 19.4336 | -0.31754 | 0.738154 | 1 |
| 6 | 18899200 | 18900999 | 28.09094 | 45.89331 | 73.40916 | 31.55654 | -0.30189 | 1.49335 | 1 |
| 6 | 19096400 | 19097399 | 16.31953 | 28.66665 | 33.58152 | 14.8629 | 0.160326 | 0.60577 | 1 |
| 6 | 19143000 | 19145799 | 34.76512 | 56.35755 | 81.67804 | 34.97675 | -0.16888 | 1.637253 | 1 |
| 6 | 19221000 | 19224199 | 43.81316 | 70.32957 | 101.6429 | 43.00393 | -0.18102 | 1.933332 | 1 |
| 6 | 19324000 | 19327599 | 69.6761 | 110.115 | 127.2831 | 52.87312 | 0.117561 | 2.421004 | 1 |
| 6 | 19390600 | 19392599 | 13.77493 | 25.43414 | 27.54823 | 11.63323 | 0.322241 | 0.236391 | 1 |
| 6 | 19463800 | 19465799 | 25.42543 | 42.89302 | 45.42249 | 19.86393 | 0.298888 | 1.097902 | 1 |
| 6 | 19582200 | 19591399 | 176.1754 | 262.4303 | 279.2105 | 115.0537 | 0.167959 | 3.600905 | 1 |
| 6 | 19643200 | 19654799 | 205.3973 | 297.3388 | 416.4165 | 171.7315 | -0.26747 | 4.063924 | 1 |
| 6 | 19773000 | 19775999 | 49.32059 | 79.03785 | 95.50594 | 40.3048 | 0.073882 | 1.989474 | 1 |
| 6 | 19958800 | 19961599 | 49.96229 | 80.05762 | 93.17821 | 39.31606 | 0.127694 | 1.992751 | 1 |
| 6 | 20023200 | 20024399 | 14.48007 | 25.6834 | 32.0336 | 14.15585 | 0.075619 | 0.485935 | 1 |
| 6 | 20079600 | 20080999 | 11.38852 | 20.62092 | 32.17252 | 13.93689 | -0.22633 | 0.336918 | 1 |
| 6 | 20148800 | 20157199 | 189.7699 | 280.7462 | 298.1254 | 122.7763 | 0.161573 | 3.721278 | 1 |
| 6 | 20186400 | 20189199 | 48.76713 | 77.83436 | 118.1327 | 49.56264 | -0.26098 | 2.120506 | 1 |
| 6 | 20321200 | 20324599 | 68.28376 | 108.2013 | 127.64 | 53.09266 | 0.091328 | 2.359135 | 1 |
| 6 | 20521000 | 20521399 | 8.418512 | 16.09666 | 22.90673 | 9.082489 | 0.006219 | -0.0485 | 1 |
| 6 | 20528000 | 20528799 | 13.9047 | 25.15124 | 25.59985 | 11.10382 | 0.385337 | 0.349177 | 1 |
| 6 | 20532800 | 20538399 | 50.19766 | 80.69671 | 91.54931 | 38.818 | 0.170102 | 1.890439 | 1 |

| | | | | | | | | | |
|---|---|---|---|---|---|---|---|---|---|
| 6 | 20550000 | 20551799 | 22.30934 | 37.4602 | 57.00811 | 24.89156 | -0.2227 | 1.140345 | 1 |
| 6 | 20710400 | 20711799 | 22.11784 | 37.21456 | 54.86465 | 23.97389 | -0.17696 | 1.118809 | 1 |
| 6 | 20793000 | 20793999 | 15.97497 | 28.18233 | 30.05358 | 13.29906 | 0.297236 | 0.547271 | 1 |
| 6 | 21014200 | 21015199 | 15.95708 | 28.06721 | 34.63668 | 15.33625 | 0.086088 | 0.589574 | 1 |
| 6 | 21026400 | 21028199 | 14.73278 | 25.86374 | 34.43147 | 15.2313 | -0.02288 | 0.611231 | 1 |
| 6 | 21393800 | 21394999 | 12.84709 | 23.05471 | 33.20225 | 14.5285 | -0.12064 | 0.40711 | 1 |
| 6 | 22287200 | 22289999 | 20.24486 | 34.01525 | 57.44619 | 25.09122 | -0.37233 | 1.127701 | 1 |
| 6 | 22611400 | 22612599 | 14.54923 | 25.86181 | 30.93975 | 13.64763 | 0.13706 | 0.469296 | 1 |
| 6 | 22680200 | 22686599 | 95.30055 | 147.5617 | 171.8983 | 71.40202 | 0.083487 | 2.822515 | 1 |
| 6 | 23085000 | 23086399 | 22.99386 | 38.91383 | 50.10448 | 21.93246 | 0.018008 | 1.066114 | 1 |
| 6 | 23223600 | 23225199 | 18.88863 | 32.61899 | 46.87501 | 20.56404 | -0.13556 | 0.864651 | 1 |
| 6 | 23364800 | 23365599 | 18.53986 | 32.45546 | 32.79241 | 14.47169 | 0.373008 | 0.674673 | 1 |
| 6 | 23420600 | 23422199 | 14.71004 | 25.80361 | 39.85987 | 17.62097 | -0.23669 | 0.630895 | 1 |
| 6 | 23480200 | 23483999 | 34.75608 | 56.91336 | 67.94455 | 29.23263 | 0.121101 | 1.469309 | 1 |
| 6 | 23557000 | 23558199 | 16.84932 | 29.23302 | 45.30467 | 19.87795 | -0.2425 | 0.813103 | 1 |
| 6 | 23611000 | 23612799 | 20.27051 | 34.58219 | 51.13271 | 22.4163 | -0.17964 | 0.986156 | 1 |
| 6 | 24068600 | 24069799 | 14.32578 | 25.36923 | 34.53921 | 15.27239 | -0.05122 | 0.5044 | 1 |
| 6 | 24923800 | 24931999 | 146.213 | 220.2043 | 245.97 | 101.5997 | 0.112024 | 3.387588 | 1 |
| 6 | 24947200 | 24947999 | 11.04637 | 20.58004 | 23.62774 | 9.849406 | 0.252518 | 0.151656 | 1 |
| 6 | 25076400 | 25079599 | 31.60338 | 51.76974 | 61.43758 | 26.43715 | 0.129899 | 1.452614 | 1 |
| 6 | 25556800 | 25557999 | 12.89259 | 22.63416 | 39.96679 | 17.68054 | -0.42833 | 0.60978 | 1 |
| 6 | 25988200 | 25989999 | 20.68762 | 35.48576 | 47.41713 | 20.7965 | -0.03297 | 0.933825 | 1 |
| 6 | 26164800 | 26169399 | 95.86859 | 148.7087 | 157.3911 | 65.36755 | 0.223714 | 2.78918 | 1 |
| 6 | 26246000 | 26248799 | 28.72757 | 47.97379 | 56.78474 | 24.70852 | 0.138647 | 1.217142 | 1 |
| 6 | 26273600 | 26274799 | 18.48496 | 31.89878 | 43.28357 | 18.98805 | -0.0531 | 0.867281 | 1 |
| 6 | 26834400 | 26835599 | 12.76358 | 23.58331 | 25.06275 | 10.57354 | 0.350596 | 0.219017 | 1 |
| 6 | 26959200 | 26961199 | 22.41752 | 37.81671 | 52.2111 | 22.83233 | -0.08245 | 1.094941 | 1 |
| 6 | 27126800 | 27132799 | 37.86819 | 61.74104 | 67.37644 | 28.94177 | 0.246118 | 1.536667 | 1 |
| 6 | 27139800 | 27143199 | 61.98534 | 98.66978 | 108.2508 | 45.25026 | 0.202366 | 2.214594 | 1 |
| 6 | 27228200 | 27228799 | 9.055581 | 17.03152 | 28.02952 | 11.49855 | -0.24426 | 0.087956 | 1 |
| 6 | 27330600 | 27331399 | 13.07639 | 23.87853 | 25.56663 | 10.95856 | 0.323671 | 0.288197 | 1 |
| 6 | 27553600 | 27554999 | 14.43846 | 25.56193 | 38.67795 | 17.10335 | -0.2029 | 0.517106 | 1 |
| 6 | 27637600 | 27638799 | 11.91607 | 21.3084 | 33.63688 | 14.76267 | -0.25567 | 0.419149 | 1 |
| 6 | 29628400 | 29629999 | 16.34489 | 28.57644 | 40.97465 | 18.06309 | -0.12971 | 0.696241 | 1 |
| 6 | 30726400 | 30728799 | 14.2189 | 25.10437 | 39.72585 | 17.57635 | -0.26887 | 0.545731 | 1 |
| 6 | 30931000 | 30936599 | 169.2005 | 250.1985 | 325.199 | 133.9253 | -0.13049 | 3.728398 | 1 |
| 6 | 31828600 | 31832999 | 59.91978 | 95.16976 | 118.8353 | 49.54591 | 0.012718 | 2.283223 | 1 |
| 6 | 32663000 | 32673599 | 95.41344 | 148.0065 | 159.9659 | 66.43667 | 0.19354 | 2.78955 | 1 |
| 6 | 33048200 | 33050999 | 35.75226 | 58.64037 | 67.5685 | 29.09094 | 0.173029 | 1.451712 | 1 |
| 6 | 33247600 | 33247999 | 8.622192 | 16.04387 | 29.47916 | 12.31429 | -0.42307 | 0.15616 | 1 |
| 6 | 33724400 | 33727399 | 25.98419 | 43.94603 | 52.29184 | 22.88544 | 0.131342 | 1.085157 | 1 |
| 6 | 33753000 | 33754599 | 20.14375 | 35.08167 | 37.3472 | 16.42194 | 0.297474 | 0.765352 | 1 |
| 6 | 34408400 | 34410399 | 38.64378 | 62.60409 | 77.57203 | 33.20237 | 0.055751 | 1.655113 | 1 |
| 6 | 35228200 | 35230199 | 16.92575 | 29.19804 | 48.47078 | 21.26341 | -0.34325 | 0.865189 | 1 |
| 6 | 35397200 | 35400599 | 32.78119 | 53.67039 | 63.71104 | 27.40923 | 0.129189 | 1.464194 | 1 |
| 6 | 35421000 | 35423999 | 33.28075 | 54.14511 | 76.13424 | 32.6781 | -0.12079 | 1.564945 | 1 |
| 6 | 35566000 | 35566999 | 10.38577 | 18.79326 | 32.29504 | 14.00115 | -0.36521 | 0.33576 | 1 |

| | | | | | | | | | |
|---|---|---|---|---|---|---|---|---|---|
| 6 | 35754200 | 35755399 | 19.98122 | 34.27504 | 43.27529 | 18.97999 | 0.048549 | 0.925127 | 1 |
| 6 | 35957200 | 35958799 | 15.65953 | 28.1394 | 29.7455 | 13.00123 | 0.323604 | 0.407312 | 1 |
| 6 | 36118400 | 36123399 | 38.56213 | 62.35227 | 85.8931 | 36.70773 | -0.09937 | 1.700343 | 1 |
| 6 | 36473800 | 36474599 | 13.27873 | 23.47503 | 33.28907 | 14.72462 | -0.11033 | 0.515916 | 1 |
| 6 | 36500400 | 36501599 | 16.50702 | 28.67076 | 44.2499 | 19.42422 | -0.23633 | 0.795894 | 1 |
| 6 | 36687800 | 36688599 | 15.23004 | 26.56157 | 41.61331 | 18.31402 | -0.25712 | 0.727555 | 1 |
| 6 | 36704400 | 36713399 | 167.36 | 247.9001 | 329.8312 | 135.8354 | -0.16188 | 3.699348 | 1 |
| 6 | 37524200 | 37529999 | 27.24889 | 44.97791 | 66.99638 | 28.92424 | -0.1928 | 1.35239 | 1 |
| 6 | 37662800 | 37665999 | 56.38738 | 89.48692 | 130.051 | 54.17821 | -0.20702 | 2.306954 | 1 |
| 6 | 37682800 | 37684999 | 23.64177 | 40.71877 | 49.57718 | 21.74771 | 0.101624 | 0.909621 | 1 |
| 6 | 37699800 | 37702599 | 24.32241 | 41.63553 | 47.92984 | 21.01791 | 0.18069 | 0.965647 | 1 |
| 6 | 37789400 | 37791399 | 23.59493 | 40.19236 | 46.89886 | 20.55768 | 0.160288 | 1.007451 | 1 |
| 6 | 37847200 | 37861199 | 286.3518 | 411.3739 | 433.7194 | 179.0572 | 0.131051 | 4.182379 | 1 |
| 6 | 37861200 | 37873399 | 170.9257 | 254.8228 | 331.9954 | 136.8184 | -0.12302 | 3.587994 | 1 |
| 6 | 37883200 | 37883999 | 13.26077 | 24.38902 | 28.05724 | 11.91152 | 0.230288 | 0.254819 | 1 |
| 6 | 37953400 | 37954599 | 14.96415 | 26.48917 | 34.00128 | 15.03572 | 0.033083 | 0.511991 | 1 |
| 6 | 38255000 | 38255799 | 15.40609 | 27.35989 | 29.96265 | 13.22681 | 0.263281 | 0.480096 | 1 |
| 6 | 38259600 | 38264199 | 101.6945 | 156.4592 | 183.6266 | 76.26006 | 0.065704 | 2.94522 | 1 |
| 6 | 39094400 | 39097599 | 32.36277 | 52.34957 | 83.73518 | 35.80044 | -0.31374 | 1.677487 | 1 |
| 6 | 39104000 | 39106999 | 33.67398 | 54.80857 | 88.64368 | 38.05336 | -0.32183 | 1.567041 | 1 |
| 6 | 39250200 | 39251799 | 17.87768 | 31.54139 | 34.33976 | 15.19552 | 0.267521 | 0.566598 | 1 |
| 6 | 39349400 | 39352199 | 16.93345 | 29.30208 | 45.82623 | 20.10518 | -0.25649 | 0.83532 | 1 |
| 6 | 40698600 | 40701199 | 35.06718 | 57.00745 | 77.53872 | 33.27156 | -0.07378 | 1.58 | 1 |
| 6 | 41177200 | 41178799 | 29.85207 | 48.61076 | 71.18222 | 30.56266 | -0.17818 | 1.541579 | 1 |
| 6 | 41451800 | 41456399 | 64.25571 | 101.535 | 146.031 | 60.65678 | -0.19743 | 2.425084 | 1 |
| 6 | 41783200 | 41784999 | 27.15425 | 45.14564 | 55.52212 | 24.08995 | 0.083631 | 1.260512 | 1 |
| 6 | 41902000 | 41909599 | 76.68166 | 119.8722 | 155.2273 | 64.40125 | -0.05829 | 2.64601 | 1 |
| 6 | 41966000 | 41967399 | 20.7025 | 35.36293 | 51.69054 | 22.66312 | -0.16296 | 0.976021 | 1 |
| 6 | 41992800 | 41998799 | 120.8646 | 184.6745 | 201.3888 | 83.53516 | 0.163913 | 3.088753 | 1 |
| 6 | 42133200 | 42135599 | 22.51284 | 39.11704 | 44.07298 | 19.35119 | 0.215505 | 0.815795 | 1 |
| 6 | 42475400 | 42476399 | 14.52793 | 25.88674 | 32.52972 | 14.31962 | 0.068761 | 0.459471 | 1 |
| 6 | 42669000 | 42670199 | 11.77011 | 21.2447 | 28.77657 | 12.5151 | -0.02682 | 0.353641 | 1 |
| 6 | 42950400 | 42953599 | 54.25109 | 86.26248 | 124.4518 | 51.94557 | -0.19381 | 2.245078 | 1 |
| 6 | 42959800 | 42967999 | 152.4354 | 229.5782 | 234.8907 | 97.02358 | 0.238583 | 3.387541 | 1 |
| 6 | 43058400 | 43059599 | 11.97088 | 21.76832 | 29.60192 | 12.7549 | -0.02422 | 0.311413 | 1 |
| 6 | 43158800 | 43160199 | 13.87558 | 24.76255 | 31.11016 | 13.67691 | 0.070459 | 0.442906 | 1 |
| 6 | 43600600 | 43603399 | 32.49004 | 53.6105 | 58.52054 | 25.25949 | 0.254237 | 1.362875 | 1 |
| 6 | 43735000 | 43740799 | 78.90871 | 123.1379 | 163.4909 | 67.84191 | -0.09627 | 2.679593 | 1 |
| 6 | 43814800 | 43821999 | 16.00767 | 28.40713 | 29.84955 | 13.18514 | 0.321897 | 0.487737 | 1 |
| 6 | 43833000 | 43833999 | 15.91023 | 28.00656 | 31.68492 | 14.02711 | 0.211315 | 0.575337 | 1 |
| 6 | 43896400 | 43903799 | 125.9798 | 189.9459 | 260.9551 | 107.8219 | -0.18489 | 3.362377 | 1 |
| 6 | 43955200 | 43956999 | 20.13735 | 35.09637 | 40.36462 | 17.75695 | 0.186482 | 0.758484 | 1 |
| 6 | 44246000 | 44248199 | 21.18149 | 36.87812 | 39.78927 | 17.49251 | 0.278023 | 0.775995 | 1 |
| 6 | 44935800 | 44936799 | 13.09722 | 23.40876 | 34.13581 | 14.988 | -0.14218 | 0.433887 | 1 |
| 6 | 45462000 | 45463399 | 12.76914 | 23.36683 | 26.56565 | 11.35584 | 0.240245 | 0.276036 | 1 |
| 6 | 46129800 | 46131199 | 16.58264 | 29.0877 | 33.76782 | 14.92752 | 0.173539 | 0.63242 | 1 |
| 6 | 46263800 | 46264999 | 21.8095 | 37.42143 | 42.213 | 18.51449 | 0.210567 | 0.930263 | 1 |

| | | | | | | | | | |
|---|---|---|---|---|---|---|---|---|---|
| 6 | 46553400 | 46554599 | 12.72274 | 22.76022 | 33.61038 | 14.74594 | -0.15958 | 0.423239 | 1 |
| 6 | 46573400 | 46575999 | 21.02017 | 35.38069 | 53.21592 | 23.25605 | -0.20562 | 1.11124 | 1 |
| 6 | 46660800 | 46662999 | 38.46266 | 62.41959 | 69.65638 | 29.85272 | 0.208782 | 1.61612 | 1 |
| 6 | 47976800 | 47987999 | 150.0331 | 223.1507 | 318.457 | 131.2058 | -0.25746 | 3.624883 | 1 |
| 6 | 48217200 | 48218999 | 14.68819 | 25.92663 | 39.18343 | 17.3372 | -0.20302 | 0.550261 | 1 |
| 6 | 48732800 | 48734799 | 18.70687 | 32.54274 | 52.2835 | 22.97338 | -0.29402 | 0.795034 | 1 |
| 6 | 48751800 | 48759999 | 136.425 | 206.1723 | 237.9255 | 98.37829 | 0.06933 | 3.317211 | 1 |
| 6 | 48915400 | 48921999 | 121.0268 | 184.0261 | 219.4694 | 90.91817 | 0.029295 | 3.188082 | 1 |
| 6 | 49372200 | 49373199 | 20.92818 | 35.99116 | 39.67473 | 17.40307 | 0.244287 | 0.902109 | 1 |
| 6 | 49566200 | 49568599 | 30.76607 | 50.41503 | 65.06591 | 28.00242 | 0.009453 | 1.451928 | 1 |
| 6 | 49947400 | 49949599 | 21.25085 | 36.38691 | 46.88466 | 20.56001 | 0.018834 | 0.950099 | 1 |
| 6 | 50086200 | 50088199 | 16.04039 | 27.6126 | 45.8748 | 20.1233 | -0.34482 | 0.878419 | 1 |
| 6 | 50586600 | 50593199 | 98.51765 | 152.1996 | 172.37 | 71.5988 | 0.12165 | 2.864826 | 1 |
| 6 | 51281400 | 51282999 | 22.22009 | 38.56384 | 44.65848 | 19.59462 | 0.175946 | 0.831546 | 1 |
| 6 | 51394200 | 51396199 | 19.13922 | 32.23202 | 56.85735 | 24.85145 | -0.43469 | 1.104176 | 1 |
| 6 | 52065000 | 52067399 | 17.97088 | 31.15285 | 44.77402 | 19.64462 | -0.13448 | 0.825145 | 1 |
| 6 | 53157800 | 53159799 | 22.0552 | 37.04277 | 58.57385 | 25.57927 | -0.27772 | 1.140154 | 1 |
| 6 | 53701800 | 53703999 | 17.69333 | 30.80056 | 45.20825 | 19.87303 | -0.16409 | 0.768546 | 1 |
| 6 | 54216800 | 54219799 | 25.26371 | 42.26587 | 59.44532 | 25.89581 | -0.10945 | 1.18787 | 1 |
| 6 | 54332000 | 54339199 | 89.98747 | 138.5629 | 193.7776 | 80.48496 | -0.186 | 2.925297 | 1 |
| 6 | 54646600 | 54653999 | 47.03999 | 75.24843 | 103.82 | 43.69805 | -0.12003 | 2.044018 | 1 |
| 6 | 55159400 | 55167999 | 133.5008 | 201.3386 | 265.5788 | 109.7412 | -0.12592 | 3.358988 | 1 |
| 6 | 56891400 | 56891999 | 12.84907 | 23.23755 | 33.65812 | 14.60239 | -0.12117 | 0.358488 | 1 |
| 6 | 57351000 | 57352199 | 20.13517 | 34.25229 | 46.9612 | 20.58275 | -0.07159 | 1.00858 | 1 |
| 6 | 57610600 | 57611599 | 10.61427 | 19.4815 | 32.39283 | 13.79072 | -0.3007 | 0.254777 | 1 |
| 6 | 57994400 | 57994999 | 10.86153 | 19.76844 | 30.80403 | 13.25826 | -0.21793 | 0.304055 | 1 |
| 6 | 58044200 | 58050399 | 92.84431 | 143.7407 | 168.4195 | 69.95708 | 0.07501 | 2.823636 | 1 |
| 6 | 58124200 | 58126599 | 27.43028 | 45.28359 | 64.6956 | 27.9336 | -0.13273 | 1.350544 | 1 |
| 6 | 58150400 | 58154399 | 40.13879 | 65.05233 | 71.928 | 30.79578 | 0.220259 | 1.647872 | 1 |
| 6 | 58170600 | 58179199 | 106.5523 | 164.1953 | 184.3853 | 76.59069 | 0.131069 | 2.915688 | 1 |
| 6 | 58217000 | 58229799 | 146.3426 | 220.1012 | 258.7231 | 106.8287 | 0.036742 | 3.414415 | 1 |
| 6 | 58230600 | 58232799 | 22.86209 | 39.53796 | 42.00449 | 18.42815 | 0.298832 | 0.862677 | 1 |
| 6 | 58258600 | 58262199 | 53.24447 | 84.99942 | 110.6396 | 46.43368 | -0.03943 | 2.114954 | 1 |
| 6 | 58274600 | 58275999 | 15.36314 | 26.80707 | 42.0884 | 18.52892 | -0.26023 | 0.720977 | 1 |
| 6 | 58322400 | 58323799 | 25.46172 | 42.5246 | 55.42609 | 24.11899 | 0.000247 | 1.208255 | 1 |
| 6 | 58352600 | 58358799 | 18.45673 | 32.04678 | 44.18808 | 19.39313 | -0.0747 | 0.806939 | 1 |
| 6 | 58360000 | 58363999 | 28.06961 | 47.11917 | 49.11175 | 21.43996 | 0.32148 | 1.154461 | 1 |
| 6 | 58414800 | 58416999 | 24.4766 | 41.57582 | 50.65477 | 22.19809 | 0.097482 | 1.038131 | 1 |
| 6 | 58440800 | 58443799 | 37.38129 | 60.37572 | 86.64107 | 36.97429 | -0.16007 | 1.729758 | 1 |
| 6 | 58689400 | 58690199 | 10.31906 | 18.94818 | 32.41357 | 13.79111 | -0.34067 | 0.250437 | 1 |
| 6 | 58768600 | 58769399 | 15.12687 | 26.50157 | 35.87475 | 15.84248 | -0.04707 | 0.646545 | 1 |
| 6 | 58798200 | 58799399 | 15.26081 | 26.42475 | 44.82276 | 19.66532 | -0.37273 | 0.822219 | 1 |
| 6 | 58801000 | 58804599 | 27.99083 | 45.96751 | 68.3226 | 29.43115 | -0.19244 | 1.419775 | 1 |
| 6 | 59269400 | 59270399 | 11.53142 | 21.25463 | 25.57461 | 10.82543 | 0.168849 | 0.221474 | 1 |
| 6 | 59443600 | 59445599 | 29.59379 | 49.00259 | 53.95955 | 23.34659 | 0.242166 | 1.312874 | 1 |
| 6 | 59450800 | 59452799 | 20.55296 | 35.70403 | 40.96412 | 17.98376 | 0.189483 | 0.806243 | 1 |
| 6 | 59556400 | 59561999 | 76.0505 | 120.1412 | 126.5763 | 52.56892 | 0.250683 | 2.436168 | 1 |

| | | | | | | | | | |
|---|---|---|---|---|---|---|---|---|---|
| 6 | 59562600 | 59563999 | 30.39389 | 49.81438 | 62.88013 | 27.06385 | 0.041471 | 1.446316 | 1 |
| 6 | 59566600 | 59567399 | 10.27966 | 19.55304 | 23.29133 | 9.340489 | 0.246361 | 0.008474 | 1 |
| 6 | 59620400 | 59622199 | 24.47501 | 40.64271 | 62.43126 | 27.06538 | -0.2363 | 1.272367 | 1 |
| 6 | 59661800 | 59668999 | 121.6065 | 184.7542 | 228.3603 | 94.5779 | -0.02324 | 3.205345 | 1 |
| 6 | 60009800 | 60019599 | 205.1541 | 304.1404 | 293.9029 | 121.0492 | 0.300354 | 3.688413 | 1 |
| 6 | 60165600 | 60168599 | 58.90948 | 93.71464 | 107.7799 | 45.01423 | 0.133479 | 2.232455 | 1 |
| 6 | 60288400 | 60291199 | 26.37519 | 43.38972 | 70.52012 | 30.40015 | -0.32001 | 1.395703 | 1 |
| 6 | 60854800 | 60857799 | 42.45508 | 68.35751 | 87.02889 | 36.98564 | 0.006952 | 1.830756 | 1 |
| 6 | 60977000 | 60979399 | 35.04925 | 57.31412 | 66.76747 | 28.7111 | 0.155102 | 1.487555 | 1 |
| 6 | 61314000 | 61317799 | 91.06867 | 141.3508 | 161.3806 | 67.01826 | 0.115202 | 2.776765 | 1 |
| 6 | 61484800 | 61486799 | 19.09472 | 32.85324 | 46.89332 | 20.56975 | -0.12688 | 0.899651 | 1 |
| 6 | 61488800 | 61490399 | 33.69338 | 54.8377 | 72.94102 | 31.31194 | -0.04047 | 1.556764 | 1 |
| 6 | 61860600 | 61863399 | 28.10071 | 46.98655 | 52.94985 | 23.05831 | 0.209224 | 1.196649 | 1 |
| 6 | 62018600 | 62019399 | 11.76207 | 21.78759 | 23.66506 | 9.94975 | 0.322942 | 0.193175 | 1 |
| 6 | 62380600 | 62384599 | 80.74747 | 126.5675 | 147.5505 | 61.19938 | 0.09612 | 2.600739 | 1 |
| 6 | 62730400 | 62731599 | 22.78167 | 39.20034 | 42.05475 | 18.44736 | 0.283291 | 0.906189 | 1 |
| 6 | 62944000 | 62946199 | 22.52469 | 38.61469 | 44.22092 | 19.39379 | 0.188569 | 0.941541 | 1 |
| 6 | 62987600 | 62988399 | 14.23114 | 25.3597 | 29.98989 | 13.20079 | 0.155806 | 0.450005 | 1 |
| 6 | 63167200 | 63167599 | 10.05551 | 18.65823 | 26.26593 | 11.01766 | -0.04638 | 0.17482 | 1 |
| 6 | 63366600 | 63367599 | 13.46767 | 24.64395 | 28.54488 | 12.20266 | 0.2134 | 0.286329 | 1 |
| 6 | 63562000 | 63564199 | 34.56412 | 56.0933 | 70.8171 | 30.35038 | 0.031249 | 1.609177 | 1 |
| 6 | 63677600 | 63678399 | 12.69077 | 23.26227 | 27.62562 | 11.78211 | 0.180202 | 0.269547 | 1 |
| 6 | 64388800 | 64390599 | 17.18829 | 29.92843 | 52.25991 | 22.97626 | -0.41355 | 0.773784 | 1 |
| 6 | 64488800 | 64489799 | 11.5947 | 21.46798 | 25.13079 | 10.57242 | 0.214856 | 0.197224 | 1 |
| 6 | 64764200 | 64765999 | 28.98669 | 47.81956 | 65.5089 | 28.27335 | -0.0726 | 1.361755 | 1 |
| 6 | 64857600 | 64860399 | 32.75923 | 53.27671 | 73.2068 | 31.41731 | -0.088 | 1.566803 | 1 |
| 6 | 64899600 | 64900999 | 10.77886 | 20.09944 | 31.66561 | 13.18028 | -0.19918 | 0.164332 | 1 |
| 6 | 65082800 | 65083599 | 12.3547 | 22.6046 | 28.13919 | 12.03125 | 0.110033 | 0.278024 | 1 |
| 6 | 65185400 | 65185999 | 15.58913 | 27.02725 | 45.32662 | 19.88715 | -0.35601 | 0.813854 | 1 |
| 6 | 65191400 | 65193799 | 33.84909 | 55.23095 | 66.83638 | 28.71788 | 0.098198 | 1.518081 | 1 |
| 6 | 65366000 | 65369599 | 59.12488 | 94.17011 | 99.88596 | 41.78658 | 0.251855 | 2.191079 | 1 |
| 6 | 65444400 | 65445199 | 6.964186 | 13.45948 | 24.93129 | 9.561477 | -0.32566 | -0.16727 | 1 |
| 6 | 66052400 | 66053399 | 16.84932 | 29.23302 | 45.30467 | 19.87795 | -0.2425 | 0.813103 | 1 |
| 6 | 66553200 | 66555399 | 25.63975 | 42.72443 | 57.71165 | 25.07748 | -0.05123 | 1.231047 | 1 |
| 6 | 66557600 | 66558799 | 16.37413 | 29.17474 | 30.77638 | 13.54845 | 0.319621 | 0.467302 | 1 |
| 6 | 66566400 | 66571999 | 56.42151 | 90.28706 | 95.70178 | 40.30087 | 0.260131 | 2.040724 | 1 |
| 6 | 66804600 | 66806199 | 18.26393 | 31.85435 | 42.0086 | 18.48857 | -0.0099 | 0.742355 | 1 |
| 6 | 67468800 | 67471999 | 44.35126 | 71.35315 | 92.86905 | 39.4216 | -0.0263 | 1.85927 | 1 |
| 6 | 67478400 | 67479399 | 5.64374 | 11.03676 | 16.67052 | 5.675192 | 0.124009 | -0.62489 | 1 |
| 6 | 67484800 | 67485799 | 14.92942 | 26.109 | 40.02745 | 17.64969 | -0.22563 | 0.683576 | 1 |
| 6 | 67559000 | 67565599 | 95.78087 | 148.7707 | 158.503 | 65.81622 | 0.215656 | 2.765917 | 1 |
| 6 | 67567600 | 67573999 | 125.9485 | 191.1024 | 218.4945 | 90.45666 | 0.087599 | 3.232782 | 1 |
| 6 | 67592600 | 67594399 | 25.36342 | 43.12794 | 49.027 | 21.48745 | 0.197381 | 1.031681 | 1 |
| 6 | 67602800 | 67604199 | 23.15079 | 39.63277 | 44.73738 | 19.61809 | 0.208952 | 0.956777 | 1 |
| 6 | 67718200 | 67726599 | 133.612 | 203.1045 | 212.1566 | 87.88129 | 0.220006 | 3.196242 | 1 |
| 6 | 67770800 | 67771599 | 11.53776 | 21.02638 | 30.68651 | 13.18633 | -0.12294 | 0.298566 | 1 |
| 6 | 67859600 | 67866599 | 70.55969 | 110.9805 | 155.3451 | 64.44557 | -0.1636 | 2.528567 | 1 |

| | | | | | | | | | |
|---|---|---|---|---|---|---|---|---|---|
| 6 | 68038200 | 68040199 | 15.2054 | 26.96565 | 33.51031 | 14.8104 | 0.080778 | 0.495661 | 1 |
| 6 | 68253800 | 68260199 | 87.01117 | 135.1942 | 185.5494 | 77.04305 | -0.14893 | 2.761516 | 1 |
| 6 | 68265000 | 68267599 | 44.73203 | 72.09477 | 71.67519 | 30.50383 | 0.364736 | 1.795161 | 1 |
| 6 | 68466200 | 68469199 | 29.3542 | 49.02269 | 56.36583 | 24.52702 | 0.180302 | 1.215047 | 1 |
| 6 | 68615400 | 68620599 | 63.82551 | 101.6505 | 112.4053 | 47.0198 | 0.191764 | 2.201289 | 1 |
| 6 | 68710800 | 68711799 | 11.69081 | 21.89216 | 24.0876 | 9.954662 | 0.323367 | 0.126294 | 1 |
| 6 | 68718600 | 68719799 | 13.00631 | 22.91369 | 37.33307 | 16.52486 | -0.31163 | 0.560105 | 1 |
| 6 | 68861800 | 68864599 | 41.03789 | 66.34938 | 72.40404 | 30.93936 | 0.235836 | 1.699104 | 1 |
| 6 | 68925200 | 68927399 | 32.11689 | 52.52789 | 68.98367 | 29.66593 | -0.01709 | 1.483406 | 1 |
| 6 | 68946800 | 68947999 | 17.10052 | 30.16296 | 33.95227 | 15.02478 | 0.21962 | 0.564655 | 1 |
| 6 | 69012400 | 69014799 | 18.56618 | 32.2382 | 42.65134 | 18.7192 | -0.01512 | 0.807269 | 1 |
| 6 | 69095200 | 69104199 | 198.5953 | 293.2592 | 305.98 | 126.0085 | 0.184452 | 3.753543 | 1 |
| 6 | 69129000 | 69130199 | 18.67535 | 32.66291 | 34.74397 | 15.32115 | 0.299086 | 0.692462 | 1 |
| 6 | 69135400 | 69138199 | 15.40563 | 26.9195 | 44.13161 | 19.44581 | -0.32209 | 0.704801 | 1 |
| 6 | 69221600 | 69224999 | 43.98875 | 70.8309 | 97.67599 | 41.51386 | -0.10798 | 1.835842 | 1 |
| 6 | 69297000 | 69309999 | 104.5823 | 161.7585 | 169.9526 | 70.59422 | 0.23109 | 2.846085 | 1 |
| 6 | 69360600 | 69367599 | 109.7142 | 168.9517 | 175.4589 | 72.87712 | 0.242998 | 2.929201 | 1 |
| 6 | 69435800 | 69437599 | 21.21868 | 35.84185 | 51.67924 | 22.60884 | -0.14481 | 1.076723 | 1 |
| 6 | 69763000 | 69764599 | 15.64676 | 27.44229 | 52.71565 | 23.30095 | -0.54998 | 0.659018 | 1 |
| 6 | 69766000 | 69767199 | 15.49901 | 27.35256 | 37.09255 | 16.4127 | -0.04749 | 0.554403 | 1 |
| 6 | 69924200 | 69925799 | 13.30927 | 23.94129 | 31.0186 | 13.54566 | 0.03273 | 0.387624 | 1 |
| 6 | 69927600 | 69929799 | 17.52599 | 30.11931 | 50.05499 | 21.95584 | -0.34581 | 0.907205 | 1 |
| 6 | 69995400 | 69997999 | 38.0782 | 61.75882 | 71.38722 | 30.58107 | 0.157313 | 1.626843 | 1 |
| 6 | 70186000 | 70192199 | 88.73261 | 137.3844 | 176.807 | 73.44098 | -0.06014 | 2.821431 | 1 |
| 6 | 70249000 | 70250799 | 23.29823 | 39.1581 | 62.04647 | 27.10316 | -0.28081 | 1.139023 | 1 |
| 6 | 70285000 | 70287199 | 23.43919 | 40.15056 | 47.40037 | 20.78686 | 0.144681 | 0.957065 | 1 |
| 6 | 70454000 | 70455399 | 28.10328 | 46.67726 | 52.33105 | 22.6899 | 0.216652 | 1.271924 | 1 |
| 6 | 70466000 | 70468399 | 42.71578 | 68.79883 | 80.22824 | 34.11098 | 0.133917 | 1.817757 | 1 |
| 6 | 70596400 | 70599599 | 23.3264 | 39.92274 | 41.63679 | 18.25804 | 0.322657 | 0.957054 | 1 |
| 6 | 70666600 | 70673399 | 143.9917 | 216.62 | 259.2376 | 107.0484 | 0.011216 | 3.409173 | 1 |
| 6 | 71011400 | 71013999 | 48.91036 | 78.23072 | 106.3801 | 44.76775 | -0.09933 | 2.052022 | 1 |
| 6 | 71609000 | 71611599 | 17.44288 | 30.07221 | 48.47076 | 21.26301 | -0.30098 | 0.874308 | 1 |
| 6 | 72054800 | 72062599 | 82.14758 | 128.6343 | 137.4886 | 57.03183 | 0.220118 | 2.616315 | 1 |
| 6 | 72708800 | 72711599 | 21.16375 | 36.60086 | 41.01323 | 17.99327 | 0.222225 | 0.853053 | 1 |
| 6 | 72910400 | 72915199 | 49.53216 | 79.14468 | 109.239 | 45.90466 | -0.12242 | 2.083838 | 1 |
| 6 | 73321200 | 73322599 | 14.98417 | 26.20355 | 39.56978 | 17.44721 | -0.20377 | 0.685666 | 1 |
| 6 | 73430800 | 73434999 | 20.55493 | 35.70083 | 38.4151 | 16.86274 | 0.281684 | 0.805962 | 1 |
| 6 | 73715200 | 73719799 | 51.73478 | 82.87583 | 97.97992 | 41.32546 | 0.10468 | 2.003909 | 1 |
| 6 | 74046000 | 74047799 | 20.62774 | 35.90321 | 39.36462 | 17.30255 | 0.255021 | 0.778466 | 1 |
| 6 | 74340200 | 74341799 | 40.3566 | 64.86357 | 95.41686 | 40.44627 | -0.20424 | 1.88631 | 1 |
| 6 | 74974400 | 74976199 | 18.44462 | 32.29903 | 35.10857 | 15.49802 | 0.267857 | 0.670752 | 1 |
| 6 | 75030800 | 75034999 | 79.90675 | 125.6197 | 137.3104 | 56.96013 | 0.192199 | 2.541571 | 1 |
| 6 | 75620400 | 75624799 | 49.58192 | 80.17338 | 76.84415 | 32.84205 | 0.422775 | 1.711943 | 1 |
| 6 | 75981400 | 75983799 | 23.03691 | 39.3795 | 44.24911 | 19.40166 | 0.215248 | 0.969959 | 1 |
| 6 | 76005000 | 76006199 | 14.22166 | 25.10058 | 37.17549 | 16.4491 | -0.17391 | 0.545823 | 1 |
| 6 | 78130600 | 78133199 | 26.1246 | 44.16696 | 50.13161 | 21.93753 | 0.1992 | 1.086682 | 1 |
| 6 | 78460200 | 78463199 | 40.57322 | 65.04431 | 104.3392 | 44.05768 | -0.33341 | 1.970393 | 1 |

| | | | | | | | | | |
|---|---|---|---|---|---|---|---|---|---|
| 6 | 78692800 | 78693999 | 10.38691 | 18.94925 | 31.84496 | 13.6659 | -0.32353 | 0.288754 | 1 |
| 6 | 79102400 | 79104999 | 46.30126 | 74.60478 | 82.33573 | 35.0282 | 0.213939 | 1.81071 | 1 |
| 6 | 79259800 | 79264999 | 40.94071 | 66.21593 | 73.48875 | 31.41643 | 0.212126 | 1.690799 | 1 |
| 6 | 79273000 | 79275799 | 10.47295 | 19.8531 | 20.58583 | 8.315059 | 0.435712 | 0.025124 | 1 |
| 6 | 79348400 | 79349999 | 9.378744 | 17.8238 | 23.34496 | 9.37933 | 0.108368 | 0.006466 | 1 |
| 6 | 79401800 | 79402999 | 14.62908 | 25.67067 | 40.91603 | 18.09328 | -0.28153 | 0.628416 | 1 |
| 6 | 79499600 | 79500599 | 17.1964 | 29.69723 | 48.46654 | 21.26225 | -0.31824 | 0.860309 | 1 |
| 6 | 79945200 | 79946199 | 21.66699 | 37.07558 | 47.42179 | 20.79448 | 0.029339 | 0.959893 | 1 |
| 6 | 79986200 | 79987999 | 13.02226 | 23.54045 | 29.59444 | 12.84643 | 0.081706 | 0.355291 | 1 |
| 6 | 80376400 | 80377799 | 8.826303 | 16.70975 | 29.31614 | 11.87425 | -0.32074 | 0.042294 | 1 |
| 6 | 80467200 | 80469999 | 26.16836 | 43.17286 | 66.75182 | 28.81056 | -0.24655 | 1.356687 | 1 |
| 6 | 80528800 | 80530599 | 9.237655 | 17.80439 | 21.96517 | 8.511927 | 0.241808 | -0.11471 | 1 |
| 6 | 80633200 | 80635399 | 54.49432 | 87.12515 | 88.92831 | 37.40109 | 0.313128 | 2.064632 | 1 |
| 6 | 81041400 | 81041999 | 7.695736 | 14.82076 | 24.77802 | 9.623887 | -0.19644 | -0.11773 | 1 |
| 6 | 81209600 | 81217399 | 101.155 | 155.2647 | 205.2353 | 85.20673 | -0.10828 | 2.992637 | 1 |
| 6 | 81257600 | 81261599 | 45.63788 | 73.55508 | 85.38289 | 36.33783 | 0.141664 | 1.802639 | 1 |
| 6 | 81311800 | 81318999 | 80.85293 | 125.6776 | 174.7879 | 72.57025 | -0.16754 | 2.749064 | 1 |
| 6 | 81811800 | 81814799 | 10.97116 | 20.65294 | 20.12576 | 8.243563 | 0.507636 | 0.076044 | 1 |
| 6 | 81838000 | 81847999 | 196.4086 | 289.397 | 329.8189 | 135.8227 | 0.054204 | 3.789542 | 1 |
| 6 | 82037800 | 82042399 | 45.59533 | 72.91194 | 109.4017 | 46.0246 | -0.24143 | 2.054736 | 1 |
| 6 | 82076800 | 82079199 | 31.33037 | 50.91215 | 73.98621 | 31.7393 | -0.16945 | 1.577934 | 1 |
| 6 | 82405000 | 82406999 | 23.06433 | 39.61813 | 41.53506 | 18.21844 | 0.315869 | 0.921649 | 1 |
| 6 | 82528000 | 82530199 | 33.67663 | 55.09868 | 68.20978 | 29.33697 | 0.068205 | 1.479683 | 1 |
| 6 | 82601200 | 82603199 | 24.25726 | 40.42171 | 58.11957 | 25.25511 | -0.14096 | 1.228465 | 1 |
| 6 | 82629000 | 82631199 | 17.29825 | 29.7793 | 47.94501 | 21.03144 | -0.29977 | 0.886453 | 1 |
| 6 | 82809800 | 82811199 | 16.78204 | 29.3401 | 39.91227 | 17.59445 | -0.05416 | 0.69652 | 1 |
| 6 | 82922800 | 82925599 | 29.97357 | 48.91971 | 75.57217 | 32.47442 | -0.25294 | 1.510535 | 1 |
| 6 | 85232600 | 85234599 | 15.72201 | 27.34566 | 41.69063 | 18.31695 | -0.21837 | 0.769218 | 1 |
| 6 | 85562400 | 85565399 | 20.17771 | 34.19306 | 53.22627 | 23.30831 | -0.25462 | 1.049122 | 1 |
| 6 | 85708800 | 85717399 | 148.0689 | 222.7499 | 277.705 | 114.6737 | -0.04779 | 3.412012 | 1 |
| 6 | 85980200 | 85982799 | 19.10693 | 32.60963 | 54.36172 | 23.83263 | -0.35208 | 0.979741 | 1 |
| 6 | 87385800 | 87387599 | 20.2211 | 34.41453 | 49.02085 | 21.48629 | -0.12655 | 1.004608 | 1 |
| 6 | 89510600 | 89511999 | 17.34561 | 30.01159 | 47.40841 | 20.79925 | -0.27082 | 0.842668 | 1 |
| 6 | 89519200 | 89521399 | 13.45617 | 24.13122 | 37.24382 | 16.3064 | -0.2209 | 0.421422 | 1 |
| 6 | 89705600 | 89706999 | 24.61819 | 41.4928 | 49.07438 | 21.45493 | 0.139912 | 1.106202 | 1 |
| 6 | 89915000 | 89925799 | 195.6238 | 285.8941 | 377.7918 | 155.6421 | -0.17038 | 3.919485 | 1 |
| 6 | 89930000 | 89935799 | 29.00429 | 49.14146 | 49.60628 | 21.72187 | 0.367744 | 1.074592 | 1 |
| 6 | 90010000 | 90010999 | 15.13642 | 26.54892 | 41.95327 | 18.54486 | -0.26919 | 0.640685 | 1 |
| 6 | 90094400 | 90095399 | 15.25517 | 27.17633 | 30.91547 | 13.61166 | 0.211254 | 0.461523 | 1 |
| 6 | 90265200 | 90275599 | 161.7056 | 240.8991 | 315.8205 | 130.1409 | -0.13299 | 3.598421 | 1 |
| 6 | 91097600 | 91100199 | 28.51592 | 47.20694 | 65.55271 | 28.35807 | -0.09157 | 1.321049 | 1 |
| 6 | 91105000 | 91106399 | 17.00999 | 29.65369 | 45.20677 | 19.89038 | -0.21817 | 0.747356 | 1 |
| 6 | 92091400 | 92093999 | 17.66589 | 31.87832 | 30.61687 | 13.31968 | 0.465518 | 0.398434 | 1 |
| 6 | 92304800 | 92306199 | 15.25458 | 26.8119 | 33.21458 | 14.70788 | 0.080191 | 0.590249 | 1 |
| 6 | 92344800 | 92347999 | 48.90257 | 78.96379 | 83.92928 | 35.80514 | 0.271638 | 1.75563 | 1 |
| 6 | 93045400 | 93045999 | 7.54727 | 14.45555 | 26.28792 | 10.37196 | -0.33839 | -0.06564 | 1 |
| 6 | 93353000 | 93354599 | 13.83308 | 24.13996 | 42.66052 | 18.7821 | -0.42991 | 0.713418 | 1 |

| | | | | | | | | | |
|---|---|---|---|---|---|---|---|---|---|
| 6 | 93585800 | 93586399 | 10.05734 | 18.76853 | 27.85751 | 11.58004 | -0.11218 | 0.143467 | 1 |
| 6 | 93800800 | 93801399 | 10.92125 | 20.50118 | 25.81612 | 10.62491 | 0.135272 | 0.110913 | 1 |
| 6 | 94004800 | 94005799 | 9.39092 | 17.80775 | 23.33258 | 9.418045 | 0.10203 | 0.021914 | 1 |
| 6 | 95011200 | 95012999 | 17.10236 | 29.65173 | 44.25326 | 19.41623 | -0.18862 | 0.817502 | 1 |
| 6 | 95455800 | 95457799 | 35.51344 | 57.80588 | 67.33869 | 28.90807 | 0.150537 | 1.557248 | 1 |
| 6 | 95631800 | 95635799 | 50.04321 | 79.78063 | 118.5177 | 49.64578 | -0.23187 | 2.156975 | 1 |
| 6 | 95777200 | 95778399 | 8.858896 | 16.57332 | 26.39374 | 10.92534 | -0.20773 | 0.114613 | 1 |
| 6 | 95924800 | 95926999 | 27.61724 | 45.4356 | 66.78479 | 28.7906 | -0.17542 | 1.394463 | 1 |
| 6 | 96127800 | 96130799 | 81.92738 | 128.0227 | 138.5419 | 57.48105 | 0.199933 | 2.653474 | 1 |
| 6 | 96572200 | 96573199 | 12.0857 | 21.75201 | 29.71227 | 12.96837 | -0.04232 | 0.374144 | 1 |
| 6 | 96762800 | 96764599 | 16.52677 | 28.64552 | 46.87866 | 20.56795 | -0.32106 | 0.821904 | 1 |
| 6 | 96853000 | 96855999 | 34.37479 | 55.56397 | 86.48192 | 36.94422 | -0.27556 | 1.704279 | 1 |
| 6 | 97035800 | 97039199 | 10.68951 | 19.31404 | 30.30556 | 13.16317 | -0.23649 | 0.342718 | 1 |
| 6 | 97631000 | 97633599 | 29.61217 | 48.59235 | 68.46781 | 29.484 | -0.11608 | 1.433369 | 1 |
| 6 | 98893400 | 98896199 | 21.66967 | 37.37944 | 42.59314 | 18.68494 | 0.197401 | 0.880227 | 1 |
| 6 | 99004000 | 99005999 | 16.838 | 29.03124 | 46.37753 | 20.3448 | -0.28801 | 0.869595 | 1 |
| 6 | 99503200 | 99505999 | 34.65555 | 56.412 | 71.36986 | 30.63913 | 0.031745 | 1.555677 | 1 |
| 6 | 100681600 | 100682799 | 11.29607 | 20.23093 | 32.19127 | 14.10983 | -0.26568 | 0.402331 | 1 |
| 6 | 101562600 | 101563999 | 21.9303 | 36.96164 | 54.88075 | 23.99326 | -0.18712 | 1.102921 | 1 |
| 6 | 101619600 | 101621599 | 21.80679 | 36.23398 | 63.22919 | 27.42283 | -0.41948 | 1.260105 | 1 |
| 6 | 101738000 | 101739999 | 27.77467 | 45.76191 | 68.02814 | 29.34487 | -0.19084 | 1.377616 | 1 |
| 6 | 102017800 | 102018999 | 36.78164 | 59.26521 | 86.62636 | 36.85295 | -0.19051 | 1.80006 | 1 |
| 6 | 102374400 | 102375399 | 9.675896 | 18.51626 | 20.7771 | 8.218416 | 0.349946 | -0.04751 | 1 |
| 6 | 102540000 | 102541799 | 17.3071 | 30.36666 | 34.42535 | 15.22166 | 0.20721 | 0.628876 | 1 |
| 6 | 102638400 | 102642599 | 9.231121 | 17.81745 | 23.14506 | 8.93073 | 0.173591 | -0.12595 | 1 |
| 6 | 102643800 | 102644599 | 14.93267 | 26.10393 | 39.05606 | 17.21538 | -0.19057 | 0.689147 | 1 |
| 6 | 102799800 | 102804599 | 92.98651 | 143.5257 | 182.8135 | 75.93716 | -0.04868 | 2.880998 | 1 |
| 6 | 103110600 | 103113399 | 26.65226 | 43.8748 | 68.99491 | 29.75106 | -0.27221 | 1.385783 | 1 |
| 6 | 103225400 | 103226599 | 12.32072 | 21.88079 | 36.19595 | 15.97831 | -0.32852 | 0.471945 | 1 |
| 6 | 103823600 | 103825799 | 26.78081 | 44.88965 | 51.77713 | 22.58257 | 0.175829 | 1.168224 | 1 |
| 6 | 104040400 | 104042999 | 17.62211 | 30.58691 | 48.47347 | 21.26921 | -0.27464 | 0.815422 | 1 |
| 6 | 104681000 | 104685399 | 66.67631 | 105.4053 | 141.6041 | 58.8308 | -0.09869 | 2.41557 | 1 |
| 6 | 105176200 | 105179199 | 27.25674 | 45.70612 | 51.25058 | 22.3587 | 0.216425 | 1.164779 | 1 |
| 6 | 105229200 | 105231199 | 10.30483 | 19.35126 | 22.73266 | 9.349968 | 0.235389 | 0.093416 | 1 |
| 6 | 105249200 | 105249999 | 11.2358 | 21.02827 | 22.16164 | 9.167455 | 0.384264 | 0.121831 | 1 |
| 6 | 105439000 | 105439999 | 18.52109 | 32.32213 | 37.85617 | 16.66728 | 0.160635 | 0.728846 | 1 |
| 6 | 105468400 | 105470999 | 31.86175 | 52.44477 | 66.67905 | 28.75108 | 0.033496 | 1.399513 | 1 |
| 6 | 105494200 | 105494999 | 10.17423 | 19.31618 | 22.77698 | 9.17059 | 0.256158 | 0.020073 | 1 |
| 6 | 105674000 | 105677599 | 14.22637 | 24.93907 | 38.39745 | 16.96522 | -0.23188 | 0.636199 | 1 |
| 6 | 105926200 | 105927399 | 18.97619 | 33.15817 | 37.13001 | 16.36054 | 0.225061 | 0.709346 | 1 |
| 6 | 106292200 | 106293399 | 14.75953 | 26.23535 | 30.02117 | 13.24255 | 0.201022 | 0.47022 | 1 |
| 6 | 106340800 | 106344999 | 36.68682 | 60.21538 | 62.54425 | 26.93578 | 0.322736 | 1.441847 | 1 |
| 6 | 106373800 | 106376399 | 36.56264 | 59.19071 | 81.71552 | 34.95763 | -0.10078 | 1.668084 | 1 |
| 6 | 107078200 | 107080799 | 19.14278 | 32.98693 | 50.61291 | 22.2024 | -0.23032 | 0.890865 | 1 |
| 6 | 107116800 | 107117999 | 15.73124 | 27.52242 | 42.0092 | 18.52988 | -0.21946 | 0.683884 | 1 |
| 6 | 107276800 | 107278399 | 18.37839 | 31.41915 | 51.63547 | 22.64049 | -0.331 | 0.961235 | 1 |
| 6 | 107567800 | 107570399 | 24.27105 | 41.40361 | 48.48384 | 21.25491 | 0.155476 | 0.998858 | 1 |

| | | | | | | | | | |
|---|---|---|---|---|---|---|---|---|---|
| 6 | 107653600 | 107655999 | 18.18733 | 31.85479 | 38.17988 | 16.85661 | 0.127536 | 0.671376 | 1 |
| 6 | 108540800 | 108542999 | 21.44898 | 36.17226 | 53.7904 | 23.52088 | -0.18923 | 1.095147 | 1 |
| 6 | 109043600 | 109044999 | 19.421 | 33.76964 | 38.93505 | 17.10317 | 0.182632 | 0.785046 | 1 |
| 6 | 109219400 | 109220999 | 21.91056 | 38.0518 | 41.4557 | 18.19638 | 0.263817 | 0.817767 | 1 |
| 6 | 109684800 | 109685999 | 13.80906 | 25.68137 | 27.11252 | 11.33616 | 0.369334 | 0.196036 | 1 |
| 6 | 109689000 | 109691199 | 37.48427 | 60.73286 | 77.44865 | 33.15246 | 0.014524 | 1.650151 | 1 |
| 6 | 109692600 | 109693599 | 23.49999 | 39.77526 | 47.48082 | 20.78476 | 0.126867 | 1.065049 | 1 |
| 6 | 109986400 | 109987399 | 19.72154 | 34.17416 | 41.56524 | 18.23655 | 0.105047 | 0.829639 | 1 |
| 6 | 110017600 | 110018399 | 12.12019 | 22.40203 | 26.62341 | 11.22684 | 0.190748 | 0.215132 | 1 |
| 6 | 110021600 | 110022199 | 10.20567 | 19.09871 | 25.78869 | 10.66938 | 0.02901 | 0.122528 | 1 |
| 6 | 110195200 | 110198199 | 34.00962 | 55.51669 | 75.66283 | 32.51547 | -0.0725 | 1.516542 | 1 |
| 6 | 111047600 | 111049399 | 18.22536 | 31.54365 | 43.73039 | 19.18579 | -0.08322 | 0.837929 | 1 |
| 6 | 111129400 | 111131599 | 26.00575 | 43.04702 | 63.45928 | 27.4505 | -0.17746 | 1.316312 | 1 |
| 6 | 111522800 | 111524799 | 19.65281 | 34.11198 | 41.57923 | 18.24428 | 0.102517 | 0.815725 | 1 |
| 6 | 111872600 | 111877999 | 70.77651 | 111.375 | 150.5115 | 62.44424 | -0.1124 | 2.519509 | 1 |
| 6 | 112044200 | 112044999 | 12.97661 | 23.33329 | 29.08952 | 12.70765 | 0.08769 | 0.383985 | 1 |
| 6 | 112169400 | 112172399 | 23.09853 | 38.84834 | 56.52154 | 24.69475 | -0.15803 | 1.127462 | 1 |
| 6 | 112388200 | 112389199 | 10.7218 | 19.86385 | 26.75242 | 11.24532 | 0.014954 | 0.191324 | 1 |
| 6 | 112687000 | 112694199 | 97.265 | 149.1574 | 203.4456 | 84.45314 | -0.15449 | 3.008969 | 1 |
| 6 | 113370600 | 113372799 | 11.04823 | 20.23123 | 31.26886 | 13.35409 | -0.19929 | 0.268064 | 1 |
| 6 | 115303800 | 115305799 | 25.83987 | 44.1214 | 46.31183 | 20.30426 | 0.312903 | 0.993347 | 1 |
| 6 | 116555400 | 116555999 | 9.032879 | 17.36448 | 19.80007 | 7.73043 | 0.344545 | -0.09945 | 1 |
| 6 | 116844600 | 116851199 | 51.39918 | 81.83768 | 124.0059 | 51.85291 | -0.26239 | 2.201325 | 1 |
| 6 | 116859600 | 116861799 | 15.72432 | 27.34295 | 43.21595 | 18.98417 | -0.27021 | 0.77464 | 1 |
| 6 | 116901000 | 116902199 | 15.54512 | 27.44054 | 36.49341 | 16.14681 | -0.01955 | 0.548838 | 1 |
| 6 | 117310800 | 117312599 | 17.57202 | 30.81939 | 38.67444 | 17.09479 | 0.061595 | 0.647132 | 1 |
| 6 | 117315200 | 117319799 | 30.58693 | 50.29741 | 61.56715 | 26.53597 | 0.087526 | 1.402345 | 1 |
| 6 | 118192000 | 118193799 | 28.6412 | 46.49594 | 80.1793 | 34.37085 | -0.41701 | 1.596966 | 1 |
| 6 | 118404800 | 118408599 | 49.26932 | 79.18557 | 86.69097 | 36.74539 | 0.220999 | 1.895941 | 1 |
| 6 | 118772000 | 118774799 | 26.61891 | 44.0531 | 66.98274 | 28.97061 | -0.22207 | 1.321421 | 1 |
| 6 | 120831400 | 120833399 | 21.24101 | 36.6225 | 42.12122 | 18.47765 | 0.183946 | 0.883045 | 1 |
| 6 | 120859200 | 120861599 | 36.16865 | 58.80227 | 73.69225 | 31.62188 | 0.044126 | 1.578511 | 1 |
| 6 | 121199400 | 121200399 | 12.10122 | 21.24439 | 39.48517 | 17.46717 | -0.50187 | 0.603702 | 1 |
| 6 | 121544800 | 121558199 | 303.7915 | 432.0026 | 478.0451 | 197.6826 | 0.046549 | 4.331528 | 1 |
| 6 | 121665000 | 121667599 | 19.64383 | 34.52625 | 37.34359 | 16.53613 | 0.27481 | 0.622073 | 1 |
| 6 | 121931800 | 121940799 | 150.1927 | 225.7586 | 250.2948 | 103.3323 | 0.120326 | 3.426281 | 1 |
| 6 | 121996400 | 121997399 | 8.94094 | 16.63507 | 28.4446 | 11.88444 | -0.32011 | 0.158126 | 1 |
| 6 | 122937000 | 122940599 | 42.70873 | 68.80716 | 79.76318 | 33.92732 | 0.142968 | 1.8085 | 1 |
| 6 | 122951400 | 122953799 | 32.12972 | 52.28103 | 76.01858 | 32.63041 | -0.16885 | 1.559757 | 1 |
| 6 | 122988800 | 122995999 | 39.79811 | 64.63088 | 67.87271 | 29.10292 | 0.297071 | 1.601732 | 1 |
| 6 | 123018800 | 123024599 | 81.34014 | 127.9033 | 138.6015 | 57.49747 | 0.204944 | 2.539267 | 1 |
| 6 | 123028600 | 123030399 | 16.45965 | 28.73108 | 43.59306 | 19.19583 | -0.21121 | 0.722854 | 1 |
| 6 | 123220400 | 123221999 | 21.09547 | 36.57335 | 41.53151 | 18.22203 | 0.203885 | 0.832678 | 1 |
| 6 | 123346000 | 123351999 | 74.9997 | 118.7764 | 127.2552 | 52.91746 | 0.229367 | 2.37687 | 1 |
| 6 | 123887800 | 123894599 | 58.8414 | 93.66562 | 138.6834 | 57.97185 | -0.2295 | 2.21937 | 1 |
| 6 | 123917200 | 123919199 | 27.88062 | 46.34983 | 55.5045 | 24.08107 | 0.121984 | 1.265123 | 1 |
| 6 | 123922800 | 123930799 | 165.8856 | 247.8647 | 269.2836 | 111.0227 | 0.141862 | 3.542261 | 1 |

| | | | | | | | | | |
|---|---|---|---|---|---|---|---|---|---|
| 6 | 124134000 | 124135599 | 14.73162 | 25.97686 | 39.22086 | 17.35729 | -0.2021 | 0.562758 | 1 |
| 6 | 124214400 | 124216799 | 32.47594 | 53.42529 | 66.19975 | 28.53754 | 0.070181 | 1.406899 | 1 |
| 6 | 124529200 | 124530799 | 29.86307 | 49.02541 | 71.7869 | 30.91899 | -0.17115 | 1.432868 | 1 |
| 6 | 124581600 | 124589999 | 238.2458 | 345.0817 | 385.419 | 158.9364 | 0.059593 | 4.057893 | 1 |
| 6 | 124926000 | 124934199 | 89.63541 | 138.5384 | 178.3845 | 74.09694 | -0.06283 | 2.854406 | 1 |
| 6 | 125275000 | 125276399 | 18.71375 | 32.58283 | 39.98364 | 17.57836 | 0.093331 | 0.765844 | 1 |
| 6 | 125772000 | 125776199 | 38.86223 | 62.64499 | 92.26056 | 39.27321 | -0.2007 | 1.791269 | 1 |
| 6 | 127859800 | 127860999 | 14.17373 | 25.16495 | 30.64082 | 13.52908 | 0.111088 | 0.475691 | 1 |
| 6 | 128642400 | 128645399 | 9.163941 | 17.30978 | 28.11553 | 11.44254 | -0.21615 | 0.059425 | 1 |
| 6 | 128694200 | 128695799 | 20.63889 | 35.7025 | 44.72203 | 19.62055 | 0.062253 | 0.854373 | 1 |
| 6 | 128781600 | 128788799 | 73.45454 | 115.7026 | 142.4778 | 59.11876 | 0.022647 | 2.499461 | 1 |
| 6 | 128836400 | 128840399 | 46.0957 | 74.25542 | 80.49981 | 34.23503 | 0.239208 | 1.816699 | 1 |
| 6 | 128886000 | 128886599 | 30.35739 | 49.65128 | 70.96583 | 30.51859 | -0.13884 | 1.481177 | 1 |
| 6 | 128917400 | 128919999 | 32.93326 | 53.96372 | 63.21603 | 27.20573 | 0.148718 | 1.454023 | 1 |
| 6 | 129043000 | 129044999 | 14.20655 | 25.12084 | 37.12759 | 16.42177 | -0.1699 | 0.524803 | 1 |
| 6 | 129065400 | 129066399 | 11.75132 | 21.38977 | 32.243 | 13.87608 | -0.17068 | 0.31134 | 1 |
| 6 | 129276800 | 129278999 | 21.6441 | 37.18 | 46.31758 | 20.31621 | 0.067957 | 0.920612 | 1 |
| 6 | 129781800 | 129783799 | 35.09304 | 57.47451 | 65.0763 | 28.00052 | 0.197391 | 1.466802 | 1 |
| 6 | 129996400 | 129999199 | 30.91567 | 51.74758 | 62.62866 | 27.29123 | 0.10658 | 1.204327 | 1 |
| 6 | 130118800 | 130125799 | 77.22622 | 121.9887 | 143.0454 | 59.40635 | 0.096336 | 2.441464 | 1 |
| 6 | 130238400 | 130240999 | 16.34981 | 28.71448 | 45.55445 | 20.15884 | -0.27521 | 0.629327 | 1 |
| 6 | 130294600 | 130300399 | 65.48262 | 104.4768 | 111.1911 | 46.62746 | 0.249654 | 2.143963 | 1 |
| 6 | 130368800 | 130381799 | 227.9225 | 334.6794 | 335.1179 | 138.0204 | 0.236157 | 3.848312 | 1 |
| 6 | 130397400 | 130401799 | 66.55573 | 106.3113 | 102.8788 | 43.21215 | 0.388451 | 2.109303 | 1 |
| 6 | 130500800 | 130513599 | 284.9038 | 408.8782 | 411.9548 | 170.0989 | 0.195059 | 4.196624 | 1 |
| 6 | 130612800 | 130616999 | 35.92405 | 58.21237 | 100.1847 | 42.8888 | -0.41692 | 1.655502 | 1 |
| 6 | 130656000 | 130662999 | 91.41958 | 143.5773 | 146.5836 | 60.79911 | 0.289478 | 2.570091 | 1 |
| 6 | 130881800 | 130884199 | 24.29672 | 40.79479 | 67.2188 | 29.34876 | -0.33718 | 1.15634 | 1 |
| 6 | 131084200 | 131087399 | 29.21097 | 48.89709 | 61.96378 | 27.0025 | 0.040423 | 1.197112 | 1 |
| 6 | 131159400 | 131173799 | 169.2918 | 254.7687 | 272.8101 | 112.6633 | 0.171946 | 3.40858 | 1 |
| 6 | 131183600 | 131195999 | 186.2791 | 277.2439 | 303.069 | 124.8742 | 0.127776 | 3.618192 | 1 |
| 6 | 131559000 | 131560399 | 14.33114 | 25.62892 | 37.13381 | 16.29641 | -0.13279 | 0.442423 | 1 |
| 6 | 131567400 | 131575599 | 100.0494 | 155.0277 | 172.8669 | 71.80407 | 0.147325 | 2.811299 | 1 |
| 6 | 131589800 | 131596999 | 113.8848 | 176.0732 | 178.1133 | 73.98416 | 0.28525 | 2.858704 | 1 |
| 6 | 131717400 | 131720199 | 36.21291 | 60.10309 | 57.68434 | 24.99849 | 0.44005 | 1.307398 | 1 |
| 6 | 131741800 | 131743399 | 31.27533 | 51.50115 | 60.5034 | 26.09286 | 0.147445 | 1.387415 | 1 |
| 6 | 131801200 | 131812799 | 53.40538 | 85.99919 | 86.40086 | 36.7252 | 0.348075 | 1.83974 | 1 |
| 6 | 131907400 | 131912999 | 79.04528 | 124.7599 | 129.8547 | 53.90592 | 0.267172 | 2.461652 | 1 |
| 6 | 132054600 | 132055799 | 19.65281 | 34.11198 | 41.57923 | 18.24428 | 0.102517 | 0.815725 | 1 |
| 6 | 132091400 | 132093199 | 19.50442 | 33.40131 | 50.59262 | 22.18629 | -0.2134 | 0.949354 | 1 |
| 6 | 132184000 | 132185599 | 34.53868 | 55.67335 | 87.34236 | 37.17777 | -0.29162 | 1.785732 | 1 |
| 6 | 132279200 | 132281199 | 34.45132 | 56.55198 | 58.87435 | 25.35638 | 0.319544 | 1.434384 | 1 |
| 6 | 132295200 | 132296599 | 14.52093 | 25.92521 | 44.18001 | 19.41586 | -0.36702 | 0.4687 | 1 |
| 6 | 132374400 | 132389999 | 208.8346 | 308.5774 | 318.0246 | 130.9696 | 0.203068 | 3.745005 | 1 |
| 6 | 132390600 | 132392999 | 29.47724 | 49.24686 | 57.33261 | 24.95368 | 0.16253 | 1.216837 | 1 |
| 6 | 132414800 | 132419199 | 52.25874 | 83.30179 | 125.6834 | 52.63785 | -0.25444 | 2.166568 | 1 |
| 6 | 132428600 | 132432399 | 37.14158 | 61.3495 | 64.11846 | 27.69498 | 0.316936 | 1.367075 | 1 |

| | | | | | | | | | |
|---|---|---|---|---|---|---|---|---|---|
| 6 | 132574000 | 132584799 | 132.5464 | 202.6701 | 193.8762 | 80.43242 | 0.353572 | 3.077771 | 1 |
| 6 | 132618800 | 132620199 | 15.17104 | 26.73284 | 33.21017 | 14.69932 | 0.077653 | 0.563197 | 1 |
| 6 | 132851800 | 132853199 | 20.57064 | 35.54051 | 39.56631 | 17.35796 | 0.23141 | 0.858332 | 1 |
| 6 | 133046800 | 133048399 | 21.12638 | 36.68884 | 40.46703 | 17.76209 | 0.246149 | 0.813948 | 1 |
| 6 | 133061200 | 133062599 | 25.19796 | 42.58051 | 49.59872 | 21.70127 | 0.161896 | 1.085135 | 1 |
| 6 | 133172000 | 133174599 | 72.78455 | 114.9004 | 127.2349 | 52.82351 | 0.178139 | 2.450872 | 1 |
| 6 | 133410000 | 133413799 | 52.23552 | 84.16056 | 80.63626 | 34.30163 | 0.417383 | 1.823597 | 1 |
| 6 | 133414600 | 133416799 | 28.17649 | 48.14294 | 49.57884 | 21.73775 | 0.340425 | 0.999627 | 1 |
| 6 | 133598400 | 133600599 | 10.92435 | 20.52558 | 24.18178 | 9.93701 | 0.232071 | 0.098858 | 1 |
| 6 | 133691400 | 133704999 | 248.8195 | 359.5328 | 413.6974 | 170.6518 | 0.013237 | 4.094359 | 1 |
| 6 | 133790800 | 133793599 | 12.98062 | 22.87573 | 36.3147 | 16.07306 | -0.27417 | 0.553899 | 1 |
| 6 | 133814000 | 133816799 | 38.40526 | 62.31584 | 73.84973 | 31.64591 | 0.121982 | 1.620035 | 1 |
| 6 | 133819800 | 133821199 | 22.66468 | 38.22103 | 58.68861 | 25.66243 | -0.23536 | 1.105983 | 1 |
| 6 | 134330200 | 134331599 | 10.58391 | 20.06567 | 24.84721 | 10.03398 | 0.181874 | 0.037135 | 1 |
| 6 | 134980400 | 134983399 | 26.31072 | 43.47302 | 65.72408 | 28.39466 | -0.21399 | 1.338587 | 1 |
| 6 | 136462600 | 136468199 | 51.07674 | 81.18441 | 122.4122 | 51.07496 | -0.25788 | 2.254312 | 1 |
| 6 | 136657200 | 136661799 | 36.28126 | 58.7572 | 77.98374 | 33.37041 | -0.04362 | 1.655415 | 1 |
| 6 | 137236800 | 137238399 | 8.191706 | 15.84996 | 20.39848 | 7.781471 | 0.202016 | -0.18642 | 1 |
| 6 | 137309000 | 137310799 | 22.96808 | 39.80251 | 40.89713 | 17.94345 | 0.347349 | 0.843079 | 1 |
| 6 | 137444600 | 137446999 | 22.99574 | 39.21188 | 45.30752 | 19.86177 | 0.174568 | 0.992449 | 1 |
| 6 | 137452000 | 137454599 | 29.95491 | 49.4561 | 66.22766 | 28.59697 | -0.03968 | 1.355388 | 1 |
| 6 | 137571800 | 137573799 | 11.58059 | 22.21427 | 22.88222 | 8.989126 | 0.481227 | -0.04581 | 1 |
| 6 | 137816200 | 137821999 | 35.16048 | 57.22502 | 80.89572 | 34.72662 | -0.12818 | 1.563646 | 1 |
| 6 | 138848600 | 138851399 | 15.97695 | 28.03599 | 40.80802 | 18.04559 | -0.15143 | 0.633388 | 1 |
| 6 | 139763400 | 139766199 | 9.201857 | 16.98678 | 30.39677 | 12.83982 | -0.39758 | 0.205972 | 1 |
| 6 | 139980400 | 139982199 | 17.67458 | 31.09458 | 31.97442 | 14.15688 | 0.347997 | 0.5923 | 1 |
| 6 | 140022800 | 140024799 | 34.40295 | 56.2424 | 64.93399 | 27.91998 | 0.167757 | 1.48901 | 1 |
| 6 | 140361600 | 140368399 | 98.51468 | 151.23 | 208.1568 | 86.42109 | -0.16647 | 2.989719 | 1 |
| 6 | 140792400 | 140794399 | 9.804475 | 17.72062 | 32.36176 | 14.05065 | -0.45355 | 0.339613 | 1 |
| 6 | 140934600 | 140941799 | 69.92135 | 109.8506 | 144.0468 | 59.74955 | -0.0708 | 2.549252 | 1 |
| 6 | 141020400 | 141022399 | 16.26932 | 28.70543 | 32.37538 | 14.32611 | 0.21675 | 0.550858 | 1 |
| 6 | 141195000 | 141196599 | 14.92603 | 26.16703 | 36.29251 | 16.03593 | -0.08207 | 0.634027 | 1 |
| 6 | 141699000 | 141700399 | 11.34644 | 20.31036 | 32.7486 | 14.36054 | -0.28492 | 0.409077 | 1 |
| 6 | 142022800 | 142025799 | 21.92244 | 37.68676 | 44.74198 | 19.62556 | 0.137493 | 0.915407 | 1 |
| 6 | 142071200 | 142072799 | 25.45503 | 42.58902 | 57.75409 | 25.16022 | -0.05695 | 1.187214 | 1 |
| 6 | 142515800 | 142517799 | 6.717946 | 12.8815 | 25.16931 | 9.898802 | -0.43572 | -0.0841 | 1 |
| 6 | 142630000 | 142631999 | 23.31903 | 39.93328 | 46.32111 | 20.31309 | 0.169855 | 0.956267 | 1 |
| 6 | 142842000 | 142843399 | 16.31042 | 28.51678 | 37.44346 | 16.50671 | -0.00321 | 0.690311 | 1 |
| 6 | 142923200 | 142924999 | 14.18143 | 24.9034 | 42.91215 | 18.98773 | -0.39387 | 0.612631 | 1 |
| 6 | 143074800 | 143077799 | 41.27729 | 66.69663 | 79.17132 | 33.80508 | 0.113748 | 1.720863 | 1 |
| 6 | 143171600 | 143173399 | 17.01963 | 29.35506 | 47.41814 | 20.80152 | -0.30384 | 0.869072 | 1 |
| 6 | 143360200 | 143362199 | 17.6782 | 30.63932 | 44.22858 | 19.40519 | -0.14097 | 0.8219 | 1 |
| 6 | 143532400 | 143534999 | 33.56932 | 55.65946 | 60.34162 | 26.1319 | 0.264674 | 1.313347 | 1 |
| 6 | 143726400 | 143728199 | 22.87574 | 38.05766 | 62.3363 | 27.05891 | -0.32836 | 1.246507 | 1 |
| 6 | 143827400 | 143830799 | 26.33711 | 43.93941 | 60.5114 | 26.31406 | -0.07906 | 1.223625 | 1 |
| 6 | 144135000 | 144138799 | 46.89919 | 75.10756 | 110.0705 | 46.40433 | -0.20512 | 2.011468 | 1 |
| 6 | 144381200 | 144382399 | 17.00141 | 29.8548 | 33.09803 | 14.64492 | 0.239528 | 0.615016 | 1 |

| | | | | | | | | | |
|---|---|---|---|---|---|---|---|---|---|
| 6 | 145529600 | 145531999 | 19.06479 | 32.80111 | 47.41556 | 20.79882 | -0.14514 | 0.899531 | 1 |
| 6 | 145730800 | 145733399 | 28.85266 | 47.94871 | 54.46657 | 23.62494 | 0.197867 | 1.271117 | 1 |
| 6 | 146152200 | 146153399 | 14.5971 | 25.62402 | 38.85637 | 17.18786 | -0.20997 | 0.618451 | 1 |
| 6 | 147070800 | 147072999 | 40.70522 | 65.49152 | 91.57876 | 38.87749 | -0.12939 | 1.853872 | 1 |
| 6 | 147365600 | 147366399 | 11.92972 | 21.943 | 28.12171 | 11.93739 | 0.075554 | 0.243059 | 1 |
| 6 | 147804400 | 147806999 | 40.21541 | 64.81446 | 89.20344 | 37.96293 | -0.10346 | 1.798866 | 1 |
| 6 | 147807800 | 147812999 | 39.38526 | 63.94338 | 68.21992 | 29.24605 | 0.274012 | 1.607829 | 1 |
| 6 | 148070800 | 148073399 | 25.97486 | 43.82157 | 49.60319 | 21.69245 | 0.203067 | 1.104009 | 1 |
| 6 | 148613200 | 148614999 | 10.07737 | 18.74448 | 30.48715 | 12.74355 | -0.24986 | 0.170198 | 1 |
| 6 | 149176000 | 149178399 | 36.94736 | 60.64244 | 63.61207 | 27.39555 | 0.308572 | 1.444675 | 1 |
| 6 | 149198200 | 149211399 | 184.6054 | 273.0092 | 338.7126 | 139.4907 | -0.06324 | 3.727655 | 1 |
| 6 | 149248800 | 149250799 | 8.292681 | 15.97404 | 27.83219 | 10.80303 | -0.25505 | -0.10656 | 1 |
| 6 | 149367400 | 149368599 | 12.20495 | 22.11114 | 32.15569 | 13.92021 | -0.12481 | 0.34097 | 1 |
| 6 | 149383400 | 149385199 | 13.29654 | 23.54064 | 38.72091 | 17.1223 | -0.32269 | 0.509213 | 1 |
| 7 | 4016200 | 4016999 | 13.31242 | 24.08661 | 31.04531 | 13.46108 | 0.046827 | 0.354289 | 1 |
| 7 | 4020200 | 4021999 | 20.53327 | 35.81372 | 37.77871 | 16.62735 | 0.310844 | 0.747443 | 1 |
| 7 | 4195200 | 4198399 | 13.94056 | 24.46671 | 38.88172 | 17.19621 | -0.27727 | 0.617141 | 1 |
| 7 | 4290600 | 4294399 | 78.57585 | 123.558 | 130.3837 | 54.08847 | 0.243216 | 2.534841 | 1 |
| 7 | 4306800 | 4310399 | 48.93562 | 78.41042 | 93.09517 | 39.27942 | 0.098988 | 1.991542 | 1 |
| 7 | 5019800 | 5022199 | 14.1289 | 24.67001 | 42.11993 | 18.55063 | -0.38038 | 0.705685 | 1 |
| 7 | 5203800 | 5206799 | 57.408 | 91.48505 | 108.2826 | 45.33316 | 0.094768 | 2.17504 | 1 |
| 7 | 5946400 | 5965799 | 279.1124 | 406.6029 | 342.7301 | 141.2429 | 0.473505 | 3.975587 | 1 |
| 7 | 6365200 | 6366599 | 19.82156 | 33.90907 | 45.3517 | 19.88633 | -0.03487 | 0.950968 | 1 |
| 7 | 6710600 | 6715599 | 74.35229 | 115.8976 | 166.5852 | 69.1356 | -0.21186 | 2.696966 | 1 |
| 7 | 6729600 | 6735399 | 153.7224 | 227.4507 | 325.4837 | 134.0434 | -0.26835 | 3.715887 | 1 |
| 7 | 7107200 | 7113599 | 79.59739 | 124.8653 | 138.8996 | 57.60649 | 0.164562 | 2.583119 | 1 |
| 7 | 7148400 | 7149999 | 22.92095 | 39.12332 | 47.41981 | 20.78942 | 0.106073 | 0.986649 | 1 |
| 7 | 7153000 | 7155399 | 18.71333 | 32.7247 | 36.13255 | 15.93157 | 0.245338 | 0.69487 | 1 |
| 7 | 7225000 | 7225999 | 31.84439 | 52.17512 | 63.5882 | 27.36492 | 0.091765 | 1.453789 | 1 |
| 7 | 7279600 | 7281599 | 70.62021 | 110.0598 | 164.6354 | 68.32828 | -0.26955 | 2.699132 | 1 |
| 7 | 7365400 | 7367399 | 33.2607 | 54.67957 | 58.88763 | 25.3782 | 0.271612 | 1.412311 | 1 |
| 7 | 7376400 | 7381999 | 70.51413 | 111.5359 | 120.2085 | 49.95621 | 0.219422 | 2.400555 | 1 |
| 7 | 7798400 | 7800399 | 50.86223 | 81.36914 | 93.366 | 39.30233 | 0.145584 | 2.044587 | 1 |
| 7 | 8597000 | 8599799 | 26.47847 | 44.17865 | 52.83407 | 22.9766 | 0.123945 | 1.217403 | 1 |
| 7 | 8983400 | 8984599 | 9.437629 | 18.0697 | 19.86477 | 7.845822 | 0.381873 | -0.05561 | 1 |
| 7 | 11126600 | 11128799 | 28.7601 | 47.66972 | 57.22758 | 24.77763 | 0.117989 | 1.300774 | 1 |
| 7 | 12201800 | 12204199 | 43.38882 | 70.06822 | 77.62608 | 33.12074 | 0.212155 | 1.740083 | 1 |
| 7 | 12511800 | 12517199 | 48.85966 | 78.34861 | 90.69859 | 38.31277 | 0.136694 | 1.965815 | 1 |
| 7 | 12535400 | 12537999 | 20.15837 | 34.21998 | 52.70206 | 23.09098 | -0.23917 | 1.033101 | 1 |
| 7 | 12965200 | 12968799 | 45.74487 | 73.63645 | 83.33011 | 35.40119 | 0.176145 | 1.842285 | 1 |
| 7 | 13583600 | 13594399 | 144.8283 | 221.3359 | 191.5118 | 79.44208 | 0.497679 | 3.092091 | 1 |
| 7 | 13635400 | 13637399 | 34.93741 | 57.45947 | 62.40876 | 26.90031 | 0.259619 | 1.414033 | 1 |
| 7 | 13932800 | 13934599 | 30.91989 | 52.13658 | 49.64357 | 21.70662 | 0.451554 | 1.123101 | 1 |
| 7 | 15794800 | 15798199 | 53.51614 | 85.30684 | 109.6164 | 45.90934 | -0.02313 | 2.163251 | 1 |
| 7 | 16921800 | 16925199 | 43.73158 | 70.41015 | 80.76213 | 34.3205 | 0.157079 | 1.831172 | 1 |
| 7 | 18478600 | 18480799 | 28.77983 | 48.27025 | 52.8553 | 23.06182 | 0.250657 | 1.172027 | 1 |
| 7 | 20180400 | 20180799 | 8.176417 | 15.7244 | 21.44436 | 8.362706 | 0.090286 | -0.10636 | 1 |

| | | | | | | | | | | |
|---|---|---|---|---|---|---|---|---|---|---|
| 7 | 20182000 | 20182599 | 9.385647 | 17.97521 | 19.33697 | 7.631207 | 0.413802 | -0.06123 | | 1 |
| 7 | 20296600 | 20297199 | 24.43093 | 41.73538 | 38.21679 | 16.75583 | 0.50942 | 0.97668 | | 1 |
| 7 | 20494600 | 20495799 | 13.55957 | 23.94088 | 34.21996 | 15.14024 | -0.12259 | 0.5322 | | 1 |
| 7 | 20595600 | 20597999 | 30.95252 | 51.07311 | 59.43133 | 25.65253 | 0.16232 | 1.361259 | | 1 |
| 7 | 20718400 | 20727399 | 209.5056 | 305.3043 | 346.6866 | 142.8683 | 0.04424 | 3.963669 | | 1 |
| 7 | 20763800 | 20764199 | 12.30129 | 22.22422 | 28.78334 | 12.50343 | 0.038711 | 0.353602 | | 1 |
| 7 | 21960800 | 21961399 | 16.96739 | 29.89563 | 28.19218 | 12.4785 | 0.47315 | 0.56461 | | 1 |
| 7 | 21969600 | 21970399 | 14.0939 | 24.69533 | 37.95722 | 16.76411 | -0.22931 | 0.643215 | | 1 |
| 7 | 21986800 | 21987199 | 21.4369 | 37.02519 | 37.57535 | 16.48436 | 0.364382 | 0.863706 | | 1 |
| 7 | 22094400 | 22095199 | 33.13766 | 54.49131 | 56.63707 | 24.41096 | 0.322896 | 1.408337 | | 1 |
| 7 | 22141400 | 22141999 | 17.90498 | 31.04045 | 41.67595 | 18.28537 | -0.03663 | 0.819346 | | 1 |
| 7 | 22202800 | 22203199 | 10.63768 | 20.18244 | 22.7337 | 9.166233 | 0.319823 | 0.028797 | | 1 |
| 7 | 22251200 | 22251999 | 42.30127 | 68.06524 | 81.27955 | 34.50949 | 0.098049 | 1.849394 | | 1 |
| 7 | 22306200 | 22306999 | 10.72074 | 20.0267 | 27.86279 | 11.56249 | -0.01742 | 0.143723 | | 1 |
| 7 | 22451200 | 22451799 | 17.91818 | 31.43555 | 31.7652 | 14.0442 | 0.372839 | 0.634946 | | 1 |
| 7 | 22861400 | 22866199 | 84.74167 | 131.3796 | 170.1186 | 70.65908 | -0.06749 | 2.795401 | | 1 |
| 7 | 23725600 | 23728199 | 22.89684 | 39.29801 | 41.12143 | 18.03613 | 0.318538 | 0.930282 | | 1 |
| 7 | 23732800 | 23734799 | 26.91964 | 44.65801 | 66.56334 | 28.83703 | -0.19317 | 1.294008 | | 1 |
| 7 | 23735600 | 23740199 | 66.03822 | 105.0696 | 110.7012 | 46.2431 | 0.259879 | 2.23375 | | 1 |
| 7 | 23886200 | 23894799 | 114.2061 | 175.7163 | 183.4876 | 76.20278 | 0.234141 | 2.948105 | | 1 |
| 7 | 24568000 | 24570399 | 31.88412 | 51.76207 | 75.95408 | 32.56033 | -0.18466 | 1.601858 | | 1 |
| 7 | 25564400 | 25575599 | 128.1015 | 194.8487 | 219.1207 | 90.78285 | 0.114357 | 3.182926 | | 1 |
| 7 | 25657200 | 25659399 | 24.60344 | 41.92427 | 46.33455 | 20.31087 | 0.238298 | 1.005709 | | 1 |
| 7 | 25769000 | 25771799 | 56.67474 | 90.62761 | 96.26415 | 40.49676 | 0.256012 | 2.063971 | | 1 |
| 7 | 26187200 | 26197599 | 364.6103 | 509.6573 | 523.0289 | 216.8872 | 0.130324 | 4.586858 | | 1 |
| 7 | 26630000 | 26641399 | 162.7879 | 244.1757 | 269.132 | 111.0464 | 0.126082 | 3.470197 | | 1 |
| 7 | 26670400 | 26677599 | 67.58111 | 107.4916 | 131.134 | 54.75538 | 0.048233 | 2.251077 | | 1 |
| 7 | 26687400 | 26691599 | 61.56927 | 98.01265 | 104.7174 | 43.77642 | 0.240616 | 2.210735 | | 1 |
| 7 | 26786000 | 26788399 | 25.52947 | 43.60519 | 49.56946 | 21.73304 | 0.198323 | 0.992547 | | 1 |
| 7 | 27712400 | 27713799 | 10.91118 | 20.37885 | 23.16498 | 9.615879 | 0.271767 | 0.134027 | | 1 |
| 7 | 27720200 | 27729399 | 236.3781 | 340.3867 | 408.9421 | 168.7668 | -0.05387 | 4.143858 | | 1 |
| 7 | 29317600 | 29318399 | 18.59103 | 32.08328 | 43.25853 | 18.9771 | -0.04405 | 0.866062 | | 1 |
| 7 | 29326600 | 29329599 | 53.35073 | 85.46274 | 92.83174 | 39.15289 | 0.226637 | 2.004799 | | 1 |
| 7 | 29732200 | 29735199 | 15.55877 | 27.13674 | 43.65289 | 19.21247 | -0.29514 | 0.731148 | | 1 |
| 7 | 30630400 | 30637599 | 72.36916 | 112.9236 | 167.6145 | 69.55362 | -0.25702 | 2.678621 | | 1 |
| 7 | 30909800 | 30922999 | 166.0519 | 248.9341 | 258.9601 | 106.835 | 0.208748 | 3.48131 | | 1 |
| 7 | 31457200 | 31458599 | 13.1093 | 24.01546 | 27.54178 | 11.75566 | 0.229254 | 0.274802 | | 1 |
| 7 | 31554800 | 31566599 | 184.3752 | 273.8882 | 287.7441 | 118.5369 | 0.182506 | 3.650925 | | 1 |
| 7 | 31682800 | 31684199 | 18.50047 | 31.77051 | 45.8864 | 20.1269 | -0.14428 | 0.914307 | | 1 |
| 7 | 31885000 | 31885999 | 12.68163 | 22.4917 | 33.77274 | 14.9254 | -0.19041 | 0.480387 | | 1 |
| 7 | 32793000 | 32795999 | 51.172 | 82.04457 | 96.0818 | 40.57772 | 0.119814 | 1.977606 | | 1 |
| 7 | 33691800 | 33700399 | 132.1742 | 200.78 | 217.008 | 89.87227 | 0.170007 | 3.210261 | | 1 |
| 7 | 34488200 | 34490599 | 23.21699 | 38.98063 | 58.75254 | 25.65173 | -0.20895 | 1.144148 | | 1 |
| 7 | 34758600 | 34759999 | 12.54276 | 23.13335 | 25.04026 | 10.59013 | 0.321595 | 0.226379 | | 1 |
| 7 | 34881200 | 34885799 | 22.87117 | 39.5176 | 42.02066 | 18.43474 | 0.297287 | 0.870959 | | 1 |
| 7 | 35268800 | 35272199 | 45.65147 | 72.97549 | 106.891 | 44.9455 | -0.20727 | 2.066493 | | 1 |
| 7 | 35498600 | 35499599 | 13.27567 | 23.98696 | 30.0073 | 13.0335 | 0.088029 | 0.359365 | | 1 |

| | | | | | | | | | |
|---|---|---|---|---|---|---|---|---|---|
| 7 | 37283800 | 37286799 | 24.86135 | 41.75953 | 56.60297 | 24.71997 | -0.05631 | 1.144173 | 1 |
| 7 | 38000000 | 38001399 | 11.56787 | 21.25911 | 30.79518 | 13.08758 | -0.1011 | 0.252408 | 1 |
| 7 | 38098400 | 38105199 | 110.6025 | 170.1238 | 175.3799 | 72.83283 | 0.252352 | 2.952188 | 1 |
| 7 | 38186400 | 38193399 | 152.1031 | 229.1469 | 236.9267 | 97.87251 | 0.223817 | 3.382446 | 1 |
| 7 | 38281200 | 38300999 | 292.1093 | 419.7385 | 413.5621 | 170.7295 | 0.229033 | 4.179773 | 1 |
| 7 | 38479800 | 38483999 | 111.5744 | 170.701 | 178.9059 | 74.23162 | 0.222605 | 3.057809 | 1 |
| 7 | 38699000 | 38699399 | 9.299895 | 17.68125 | 22.37672 | 8.982473 | 0.159187 | 0.002455 | 1 |
| 7 | 39080200 | 39081599 | 22.59766 | 37.57216 | 62.76921 | 27.23539 | -0.35678 | 1.254654 | 1 |
| 7 | 39424600 | 39434399 | 22.62504 | 38.70984 | 41.22851 | 18.07853 | 0.292536 | 0.958525 | 1 |
| 7 | 39756800 | 39758799 | 15.73813 | 27.50752 | 41.03064 | 18.08328 | -0.18631 | 0.698816 | 1 |
| 7 | 41058800 | 41059799 | 13.13547 | 23.96349 | 31.65515 | 13.5845 | 0.021113 | 0.306619 | 1 |
| 7 | 41220800 | 41223799 | 40.8661 | 66.5425 | 68.15281 | 29.26142 | 0.336102 | 1.562394 | 1 |
| 7 | 41286400 | 41293599 | 91.06931 | 143.4851 | 130.1039 | 53.98623 | 0.463928 | 2.505265 | 1 |
| 7 | 41331800 | 41333799 | 19.47514 | 33.26763 | 51.65722 | 22.64858 | -0.24971 | 0.97217 | 1 |
| 7 | 41389000 | 41390399 | 22.36908 | 37.80278 | 51.69081 | 22.61711 | -0.06852 | 1.077785 | 1 |
| 7 | 41447000 | 41448999 | 52.52969 | 84.12833 | 86.33969 | 36.40216 | 0.308004 | 2.009167 | 1 |
| 7 | 41633400 | 41655999 | 121.6659 | 186.5588 | 185.1355 | 76.85138 | 0.30384 | 3.018545 | 1 |
| 7 | 42156800 | 42184599 | 332.5325 | 464.5625 | 529.2306 | 219.4776 | -0.02109 | 4.593826 | 1 |
| 7 | 42522000 | 42556799 | 916.1542 | 1185.473 | 1181.744 | 495.2801 | 0.060449 | 5.811479 | 1 |
| 7 | 43732000 | 43732799 | 16.6174 | 28.65625 | 48.46498 | 21.26064 | -0.36973 | 0.872486 | 1 |
| 7 | 43778000 | 43782799 | 59.3403 | 94.37872 | 119.7003 | 49.97722 | -0.00783 | 2.241492 | 1 |
| 7 | 45579600 | 45582199 | 27.18322 | 45.43375 | 51.28008 | 22.32538 | 0.207047 | 1.197861 | 1 |
| 7 | 46616800 | 46619199 | 14.52127 | 25.4066 | 39.45406 | 17.40346 | -0.24427 | 0.670256 | 1 |
| 7 | 46979200 | 46986999 | 118.9491 | 181.5933 | 202.7804 | 84.09585 | 0.128769 | 3.105126 | 1 |
| 7 | 46995200 | 47002199 | 103.8721 | 160.2019 | 176.718 | 73.40552 | 0.157741 | 2.898322 | 1 |
| 7 | 47226200 | 47227799 | 21.04225 | 36.176 | 43.24466 | 18.96882 | 0.127775 | 0.911226 | 1 |
| 7 | 47436000 | 47452999 | 178.3474 | 265.3598 | 285.1818 | 117.5006 | 0.151924 | 3.621369 | 1 |
| 7 | 47560400 | 47562599 | 32.28124 | 52.57158 | 73.81696 | 31.69504 | -0.11772 | 1.54688 | 1 |
| 7 | 47753000 | 47756999 | 51.36137 | 82.02088 | 111.1406 | 46.66618 | -0.09676 | 2.104122 | 1 |
| 7 | 48015000 | 48018199 | 24.08157 | 40.87298 | 51.18511 | 22.4252 | 0.058021 | 1.045331 | 1 |
| 7 | 48019400 | 48021599 | 30.67939 | 50.49218 | 62.60512 | 26.99301 | 0.069564 | 1.394986 | 1 |
| 7 | 48058400 | 48063999 | 96.31214 | 149.1304 | 175.9718 | 73.09406 | 0.064991 | 2.823605 | 1 |
| 7 | 48215000 | 48216599 | 29.93332 | 49.74656 | 54.55274 | 23.66296 | 0.248486 | 1.27262 | 1 |
| 7 | 48960000 | 48962399 | 20.05967 | 34.13217 | 54.36776 | 23.82946 | -0.28736 | 1.010684 | 1 |
| 7 | 49386400 | 49388199 | 17.93343 | 31.39188 | 38.7794 | 17.11291 | 0.084238 | 0.681395 | 1 |
| 7 | 49851200 | 49852599 | 22.53886 | 37.60497 | 59.07022 | 25.68836 | -0.2681 | 1.212097 | 1 |
| 7 | 49885200 | 49891999 | 110.9896 | 168.6473 | 243.0353 | 100.6598 | -0.24435 | 3.201338 | 1 |
| 7 | 49975800 | 49977799 | 14.12885 | 25.10662 | 37.04983 | 16.34717 | -0.16449 | 0.47949 | 1 |
| 7 | 50179400 | 50180399 | 25.00662 | 41.51835 | 56.90446 | 24.66645 | -0.07222 | 1.273839 | 1 |
| 7 | 50522200 | 50523399 | 12.44995 | 22.70884 | 30.10118 | 12.9205 | 0.015984 | 0.298757 | 1 |
| 7 | 50775800 | 50778799 | 43.88814 | 70.83493 | 78.11363 | 33.30472 | 0.217886 | 1.757392 | 1 |
| 7 | 51140800 | 51141399 | 13.1267 | 24.01766 | 25.92731 | 11.08516 | 0.314023 | 0.27645 | 1 |
| 7 | 51975400 | 51986199 | 188.8385 | 276.9311 | 344.899 | 142.0603 | -0.08021 | 3.864181 | 1 |
| 7 | 52211600 | 52212399 | 13.33329 | 24.28984 | 25.48971 | 10.95812 | 0.34957 | 0.302598 | 1 |
| 7 | 52463600 | 52465799 | 24.88105 | 42.18547 | 49.57862 | 21.71443 | 0.149252 | 1.055626 | 1 |
| 7 | 52786400 | 52787599 | 15.36893 | 27.26995 | 32.35574 | 14.2938 | 0.147394 | 0.488611 | 1 |
| 7 | 53027800 | 53028799 | 16.11047 | 28.26548 | 37.72638 | 16.68046 | -0.02692 | 0.632092 | 1 |

| | | | | | | | | | |
|---|---|---|---|---|---|---|---|---|---|
| 7 | 53207000 | 53208199 | 23.88353 | 39.57425 | 61.68957 | 26.70367 | -0.25759 | 1.296598 | 1 |
| 7 | 53326200 | 53327799 | 29.43286 | 48.71372 | 57.77705 | 24.99037 | 0.135254 | 1.320139 | 1 |
| 7 | 53473400 | 53476199 | 25.48491 | 43.02348 | 46.42829 | 20.30809 | 0.271704 | 1.091175 | 1 |
| 7 | 53504200 | 53505799 | 14.86577 | 26.10372 | 36.30249 | 16.06235 | -0.08576 | 0.61 | 1 |
| 7 | 53530000 | 53531599 | 10.85929 | 20.07404 | 26.22536 | 11.05706 | 0.055392 | 0.20328 | 1 |
| 7 | 53534600 | 53535599 | 15.32633 | 27.05506 | 36.58287 | 16.18631 | -0.04318 | 0.549429 | 1 |
| 7 | 53621000 | 53623199 | 25.27359 | 42.81169 | 51.74293 | 22.65605 | 0.108974 | 1.067815 | 1 |
| 7 | 53676600 | 53677599 | 9.655486 | 17.87552 | 27.33572 | 11.50173 | -0.16799 | 0.187956 | 1 |
| 7 | 53698200 | 53700799 | 27.70702 | 45.91411 | 68.3886 | 29.60548 | -0.19235 | 1.309137 | 1 |
| 7 | 53995400 | 53999799 | 58.01997 | 92.89742 | 94.07615 | 39.64852 | 0.326743 | 2.024439 | 1 |
| 7 | 54117000 | 54120199 | 38.19208 | 61.95224 | 75.84657 | 32.49474 | 0.074815 | 1.628562 | 1 |
| 7 | 54571800 | 54573999 | 23.12253 | 39.49483 | 43.25788 | 18.96584 | 0.251938 | 0.977838 | 1 |
| 7 | 54592000 | 54592399 | 19.14898 | 33.33613 | 35.95694 | 15.80673 | 0.278665 | 0.764049 | 1 |
| 7 | 54595200 | 54596799 | 20.51138 | 35.46536 | 43.16854 | 18.93873 | 0.103412 | 0.856887 | 1 |
| 7 | 54620800 | 54625399 | 54.15061 | 86.24813 | 125.4645 | 52.48779 | -0.20313 | 2.192745 | 1 |
| 7 | 54727000 | 54729799 | 25.80329 | 44.02733 | 46.31288 | 20.30354 | 0.309632 | 0.999157 | 1 |
| 7 | 54735600 | 54736399 | 15.85762 | 27.59022 | 43.18764 | 18.97843 | -0.25633 | 0.766485 | 1 |
| 7 | 55214600 | 55217199 | 21.396 | 36.30258 | 51.13971 | 22.40301 | -0.11108 | 1.039485 | 1 |
| 7 | 55335000 | 55337399 | 19.73248 | 33.86477 | 48.477 | 21.26222 | -0.13181 | 0.926523 | 1 |
| 7 | 55730600 | 55731999 | 24.02663 | 40.50594 | 45.97849 | 20.10297 | 0.199121 | 1.10052 | 1 |
| 7 | 56112200 | 56116999 | 43.89969 | 70.57648 | 99.11171 | 42.02822 | -0.137 | 1.883445 | 1 |
| 7 | 56227200 | 56230799 | 10.38242 | 19.80041 | 21.78322 | 8.687323 | 0.367744 | -0.01022 | 1 |
| 7 | 56485400 | 56486399 | 13.28732 | 23.97105 | 29.03363 | 12.63492 | 0.132683 | 0.36672 | 1 |
| 7 | 56575400 | 56577399 | 47.85233 | 76.6994 | 95.83971 | 40.45727 | 0.025938 | 1.981971 | 1 |
| 7 | 56762400 | 56768399 | 48.31701 | 77.3264 | 99.56119 | 41.93252 | -0.01976 | 2.030109 | 1 |
| 7 | 56786800 | 56790799 | 38.69366 | 62.38036 | 90.45733 | 38.51127 | -0.17818 | 1.786927 | 1 |
| 7 | 56870200 | 56874999 | 61.40184 | 97.98107 | 100.159 | 42.01035 | 0.307974 | 2.135127 | 1 |
| 7 | 57023000 | 57026799 | 37.74462 | 60.95651 | 85.91097 | 36.65803 | -0.13425 | 1.733994 | 1 |
| 7 | 57035200 | 57037999 | 9.06717 | 16.74463 | 29.89778 | 12.6224 | -0.39369 | 0.202503 | 1 |
| 7 | 57510600 | 57512399 | 18.6135 | 32.79351 | 34.25129 | 15.16062 | 0.32625 | 0.584919 | 1 |
| 7 | 57517800 | 57520799 | 13.72846 | 24.22423 | 36.19629 | 16.01661 | -0.18671 | 0.54339 | 1 |
| 7 | 57686000 | 57688999 | 35.27523 | 57.06167 | 78.32103 | 33.48606 | -0.09345 | 1.680539 | 1 |
| 7 | 57749800 | 57751199 | 21.92108 | 37.68913 | 40.63262 | 17.82311 | 0.276146 | 0.910005 | 1 |
| 7 | 57772000 | 57776599 | 92.824 | 143.7331 | 160.1322 | 66.51469 | 0.147839 | 2.819445 | 1 |
| 7 | 57826600 | 57828399 | 22.14778 | 37.23934 | 57.55705 | 25.14513 | -0.24494 | 1.127764 | 1 |
| 7 | 58593800 | 58600399 | 91.63251 | 143.5996 | 147.3459 | 61.11581 | 0.279813 | 2.610216 | 1 |
| 7 | 58734000 | 58743999 | 203.9881 | 300.6689 | 313.1432 | 128.9559 | 0.18458 | 3.784542 | 1 |
| 7 | 58855200 | 58855999 | 11.38969 | 21.19791 | 24.613 | 10.27796 | 0.23436 | 0.163867 | 1 |
| 7 | 59147600 | 59153799 | 167.5884 | 249.471 | 283.5889 | 116.85 | 0.071612 | 3.611699 | 1 |
| 7 | 59335600 | 59336799 | 18.23174 | 31.76774 | 36.99197 | 16.27088 | 0.168858 | 0.749002 | 1 |
| 7 | 59803800 | 59806799 | 30.84706 | 50.77057 | 61.48983 | 26.51268 | 0.103373 | 1.394297 | 1 |
| 7 | 59859000 | 59859799 | 21.18149 | 36.87812 | 39.78927 | 17.49251 | 0.278023 | 0.775995 | 1 |
| 7 | 59887000 | 59888599 | 19.07415 | 33.14709 | 46.84241 | 20.56908 | -0.10983 | 0.803228 | 1 |
| 7 | 60422600 | 60425999 | 43.12989 | 69.57595 | 90.98223 | 38.76614 | -0.02833 | 1.776458 | 1 |
| 7 | 60572600 | 60573599 | 11.01342 | 20.39999 | 26.21298 | 11.02161 | 0.081975 | 0.193292 | 1 |
| 7 | 61940800 | 61944599 | 35.59809 | 58.67676 | 59.71616 | 25.76401 | 0.354495 | 1.386953 | 1 |
| 7 | 61984200 | 61986399 | 30.4866 | 50.32953 | 60.50435 | 26.12408 | 0.115585 | 1.355348 | 1 |

| | | | | | | | | | | |
|---|---|---|---|---|---|---|---|---|---|---|
| 7 | 61997000 | 62001399 | 65.37759 | 103.5184 | 128.9365 | 53.6115 | 0.012173 | 2.375359 | | 1 |
| 7 | 62025600 | 62033599 | 159.0362 | 236.1643 | 305.7231 | 125.9389 | -0.11902 | 3.653604 | | 1 |
| 7 | 62068000 | 62070799 | 72.00634 | 113.4695 | 122.7247 | 50.92534 | 0.210091 | 2.487933 | | 1 |
| 7 | 62331800 | 62337399 | 128.0834 | 194.764 | 203.1898 | 84.1752 | 0.2222 | 3.188531 | | 1 |
| 7 | 62461800 | 62465199 | 42.25748 | 68.51821 | 74.1461 | 31.75644 | 0.251857 | 1.642988 | | 1 |
| 7 | 62616200 | 62618199 | 22.38018 | 38.1821 | 48.48568 | 21.2559 | 0.038963 | 0.988811 | | 1 |
| 7 | 62620000 | 62620599 | 8.493326 | 16.12013 | 24.43368 | 9.843797 | -0.10409 | 0.010003 | | 1 |
| 7 | 63044200 | 63045999 | 31.83989 | 52.40002 | 58.21301 | 25.09887 | 0.22758 | 1.396252 | | 1 |
| 7 | 63457000 | 63459399 | 19.51687 | 34.03499 | 43.49666 | 19.14186 | 0.034923 | 0.748812 | | 1 |
| 7 | 63771200 | 63777199 | 72.57312 | 114.651 | 118.1792 | 49.0822 | 0.282226 | 2.431621 | | 1 |
| 7 | 64084600 | 64086799 | 19.41525 | 33.26721 | 45.87851 | 20.11993 | -0.07848 | 0.937106 | | 1 |
| 7 | 64319800 | 64323599 | 37.28672 | 60.28151 | 85.51746 | 36.54054 | -0.14195 | 1.698613 | | 1 |
| 7 | 64836600 | 64837599 | 19.37234 | 32.89728 | 53.74503 | 23.55047 | -0.32395 | 1.028453 | | 1 |
| 7 | 65863800 | 65865999 | 16.40544 | 28.60927 | 42.55903 | 18.72932 | -0.18302 | 0.73344 | | 1 |
| 7 | 66275600 | 66278599 | 29.83542 | 49.25299 | 66.71953 | 28.807 | -0.05624 | 1.357661 | | 1 |
| 7 | 66809800 | 66810199 | 7.532907 | 14.61712 | 19.4436 | 7.318493 | 0.17409 | -0.23875 | | 1 |
| 7 | 66923600 | 66924399 | 15.00814 | 26.25669 | 37.95458 | 16.74109 | -0.14129 | 0.672637 | | 1 |
| 7 | 66985400 | 66988399 | 21.70667 | 37.44523 | 56.25136 | 24.67663 | -0.19985 | 0.896887 | | 1 |
| 7 | 67319200 | 67321199 | 14.86321 | 26.63269 | 30.8506 | 13.51659 | 0.18998 | 0.421484 | | 1 |
| 7 | 69015200 | 69019599 | 59.66845 | 95.30775 | 100.0392 | 42.01799 | 0.271369 | 2.10265 | | 1 |
| 7 | 69373800 | 69376199 | 29.78027 | 48.86344 | 65.05147 | 28.01168 | -0.03445 | 1.433286 | | 1 |
| 7 | 69654800 | 69656599 | 27.51574 | 46.19299 | 49.63169 | 21.668 | 0.277856 | 1.153925 | | 1 |
| 7 | 69888600 | 69892199 | 32.92019 | 54.12288 | 66.868 | 28.81812 | 0.074017 | 1.414927 | | 1 |
| 7 | 70333600 | 70335199 | 15.74265 | 27.39695 | 45.26342 | 19.89369 | -0.3338 | 0.766166 | | 1 |
| 7 | 70413000 | 70415799 | 28.79001 | 48.21683 | 59.59391 | 25.97806 | 0.076401 | 1.189985 | | 1 |
| 7 | 70574800 | 70578799 | 44.62449 | 72.19079 | 77.94152 | 33.32572 | 0.252062 | 1.692785 | | 1 |
| 7 | 71104000 | 71106599 | 38.63476 | 62.81292 | 70.83631 | 30.3966 | 0.19593 | 1.578987 | | 1 |
| 7 | 72291400 | 72293399 | 17.37559 | 30.05646 | 47.93664 | 21.03084 | -0.28464 | 0.846258 | | 1 |
| 7 | 72302200 | 72304799 | 69.41185 | 108.8652 | 139.8532 | 58.00319 | -0.04307 | 2.583075 | | 1 |
| 7 | 72492800 | 72494999 | 27.58043 | 45.64339 | 63.67684 | 27.54037 | -0.09824 | 1.322006 | | 1 |
| 7 | 72596200 | 72597599 | 22.27318 | 37.44312 | 51.67245 | 22.57302 | -0.08197 | 1.124143 | | 1 |
| 7 | 72731800 | 72733599 | 35.79961 | 58.0948 | 72.31489 | 30.9909 | 0.051443 | 1.611841 | | 1 |
| 7 | 72822400 | 72825399 | 23.32017 | 39.11677 | 58.65868 | 25.59544 | -0.20151 | 1.157182 | | 1 |
| 7 | 72930200 | 72931799 | 18.69006 | 32.48467 | 40.0448 | 17.58589 | 0.086561 | 0.790319 | | 1 |
| 7 | 73334600 | 73336999 | 20.2951 | 34.40482 | 55.40675 | 24.26591 | -0.30355 | 1.046935 | | 1 |
| 7 | 73707600 | 73711999 | 28.92832 | 47.48162 | 70.21953 | 30.24142 | -0.18559 | 1.428604 | | 1 |
| 7 | 73848200 | 73851199 | 21.20071 | 36.44203 | 44.74207 | 19.62554 | 0.089139 | 0.912048 | | 1 |
| 7 | 73892200 | 73892999 | 13.55621 | 25.35146 | 25.49063 | 10.55831 | 0.449578 | 0.152951 | | 1 |
| 7 | 74255400 | 74257799 | 12.58634 | 22.43916 | 34.13837 | 15.02094 | -0.20517 | 0.444959 | | 1 |
| 7 | 74368600 | 74369999 | 15.28635 | 26.79313 | 39.8308 | 17.59624 | -0.18174 | 0.645551 | | 1 |
| 7 | 74449000 | 74449799 | 11.52109 | 20.58149 | 32.77021 | 14.39444 | -0.26848 | 0.423583 | | 1 |
| 7 | 74483600 | 74484599 | 15.22158 | 26.57114 | 43.63211 | 19.21355 | -0.32462 | 0.716738 | | 1 |
| 7 | 74810400 | 74816599 | 89.20369 | 138.6316 | 168.7429 | 70.06241 | 0.024331 | 2.754874 | | 1 |
| 7 | 75196800 | 75198599 | 10.75001 | 19.38485 | 33.81068 | 14.7221 | -0.39051 | 0.363404 | | 1 |
| 7 | 75460600 | 75461799 | 13.15089 | 23.22339 | 34.32895 | 15.18793 | -0.17051 | 0.529864 | | 1 |
| 7 | 75622800 | 75623999 | 17.55154 | 30.70308 | 37.29373 | 16.44811 | 0.108488 | 0.687108 | | 1 |
| 7 | 75645600 | 75646799 | 14.96343 | 26.26633 | 37.38577 | 16.53694 | -0.11889 | 0.620165 | | 1 |

| | | | | | | | | | |
|---|---|---|---|---|---|---|---|---|---|
| 7 | 75715000 | 75716599 | 11.98337 | 21.476 | 35.72586 | 15.6504 | -0.32888 | 0.411906 | 1 |
| 7 | 75717400 | 75718199 | 14.09038 | 25.44337 | 28.46277 | 12.37215 | 0.247732 | 0.366192 | 1 |
| 7 | 75899000 | 75900799 | 18.41444 | 32.21897 | 40.31364 | 17.783 | 0.065813 | 0.693257 | 1 |
| 7 | 76216200 | 76216999 | 11.0064 | 20.58031 | 24.72038 | 10.24159 | 0.195162 | 0.132776 | 1 |
| 7 | 76984200 | 76985399 | 18.80918 | 32.14313 | 49.01058 | 21.48889 | -0.22339 | 0.962519 | 1 |
| 7 | 77175800 | 77177199 | 20.32056 | 34.43737 | 52.70397 | 23.08005 | -0.23022 | 1.04812 | 1 |
| 7 | 77304600 | 77306399 | 20.2805 | 34.71775 | 47.94987 | 21.02676 | -0.081 | 0.948878 | 1 |
| 7 | 77411200 | 77412399 | 21.42332 | 36.90293 | 45.25992 | 19.85397 | 0.09132 | 0.895642 | 1 |
| 7 | 77708400 | 77710599 | 16.16512 | 28.30048 | 44.04463 | 19.4387 | -0.24757 | 0.676749 | 1 |
| 7 | 78874000 | 78875399 | 15.17546 | 26.8752 | 31.89142 | 14.10147 | 0.1459 | 0.502955 | 1 |
| 7 | 79786000 | 79787799 | 35.83311 | 58.71556 | 62.98853 | 27.10787 | 0.2753 | 1.459975 | 1 |
| 7 | 80795800 | 80797999 | 33.33129 | 54.82103 | 60.5171 | 26.08556 | 0.236345 | 1.409048 | 1 |
| 7 | 82604400 | 82606999 | 43.09756 | 69.1009 | 100.5691 | 42.47484 | -0.19309 | 1.966374 | 1 |
| 7 | 82628600 | 82632199 | 38.94924 | 63.21008 | 74.42714 | 31.89765 | 0.131539 | 1.618757 | 1 |
| 7 | 82834800 | 82836399 | 19.23661 | 32.63798 | 54.2611 | 23.77025 | -0.34908 | 1.036745 | 1 |
| 7 | 82921000 | 82923999 | 24.44651 | 41.12641 | 51.73638 | 22.6059 | 0.051068 | 1.123601 | 1 |
| 7 | 82980400 | 82986399 | 115.4392 | 176.8369 | 185.7062 | 77.0757 | 0.221166 | 3.034443 | 1 |
| 7 | 83404000 | 83406999 | 21.05703 | 36.34364 | 42.10364 | 18.47053 | 0.173849 | 0.87243 | 1 |
| 7 | 84122200 | 84124799 | 17.50301 | 30.22337 | 44.30874 | 19.43816 | -0.16408 | 0.857878 | 1 |
| 7 | 84368600 | 84370599 | 22.13855 | 37.25229 | 57.57049 | 25.15684 | -0.24475 | 1.120943 | 1 |
| 7 | 84748400 | 84749199 | 9.439859 | 18.06491 | 21.35661 | 8.44719 | 0.275666 | -0.04701 | 1 |
| 7 | 85304200 | 85306799 | 36.93996 | 60.42378 | 68.6291 | 29.51481 | 0.191874 | 1.490205 | 1 |
| 7 | 85358200 | 85360799 | 21.07185 | 36.11123 | 57.23345 | 25.09992 | -0.27864 | 0.954264 | 1 |
| 7 | 85452400 | 85457799 | 52.36665 | 84.30995 | 85.96999 | 36.53166 | 0.326411 | 1.847796 | 1 |
| 7 | 85805600 | 85807999 | 37.46199 | 61.37629 | 68.44467 | 29.45429 | 0.219432 | 1.468039 | 1 |
| 7 | 86437400 | 86439399 | 16.84768 | 29.49056 | 42.46493 | 18.73889 | -0.13588 | 0.681857 | 1 |
| 7 | 86477400 | 86484199 | 25.7785 | 42.87413 | 53.26481 | 23.11641 | 0.069247 | 1.252672 | 1 |
| 7 | 86486800 | 86489599 | 32.33968 | 53.38638 | 62.49842 | 26.98425 | 0.153674 | 1.359857 | 1 |
| 7 | 86527200 | 86528199 | 16.43635 | 28.88397 | 33.66084 | 14.9033 | 0.167951 | 0.602022 | 1 |
| 7 | 87816200 | 87816999 | 9.728517 | 18.16533 | 27.86926 | 11.57366 | -0.15854 | 0.136313 | 1 |
| 7 | 88054000 | 88056399 | 23.48683 | 40.40215 | 50.66488 | 22.22401 | 0.058947 | 0.9234 | 1 |
| 7 | 88059200 | 88063599 | 49.92582 | 79.95071 | 92.35865 | 38.93435 | 0.137457 | 2.010718 | 1 |
| 7 | 88368000 | 88368999 | 13.87433 | 25.0325 | 26.99619 | 11.74682 | 0.299275 | 0.365726 | 1 |
| 7 | 88945200 | 88945799 | 8.571829 | 16.40274 | 19.85897 | 7.856253 | 0.241103 | -0.06172 | 1 |
| 7 | 89221400 | 89223799 | 11.68333 | 20.47853 | 42.6174 | 18.82774 | -0.6639 | 0.635011 | 1 |
| 7 | 89236000 | 89237999 | 9.770059 | 18.04215 | 27.78445 | 11.73065 | -0.1819 | 0.202409 | 1 |
| 7 | 89302400 | 89304799 | 36.19173 | 58.53394 | 79.85644 | 34.13811 | -0.08502 | 1.684625 | 1 |
| 7 | 89348800 | 89356799 | 99.9226 | 153.2261 | 197.9809 | 82.18417 | -0.07651 | 3.009524 | 1 |
| 7 | 89406800 | 89408799 | 15.90336 | 27.48686 | 46.37355 | 20.34457 | -0.3656 | 0.843894 | 1 |
| 7 | 89466400 | 89468399 | 18.50606 | 32.20811 | 43.61 | 19.16774 | -0.04819 | 0.774073 | 1 |
| 7 | 90175000 | 90177999 | 24.67532 | 41.0256 | 62.41058 | 27.07932 | -0.22216 | 1.260329 | 1 |
| 7 | 90267800 | 90268599 | 13.25871 | 23.49811 | 34.64506 | 15.31678 | -0.16498 | 0.49007 | 1 |
| 7 | 90932000 | 90933799 | 9.286436 | 17.42922 | 27.43739 | 11.29557 | -0.18515 | 0.099412 | 1 |
| 7 | 91395400 | 91397799 | 23.10391 | 39.39744 | 47.42498 | 20.79007 | 0.11576 | 0.996466 | 1 |
| 7 | 91879000 | 91880399 | 19.34335 | 33.35601 | 40.70418 | 17.85611 | 0.099057 | 0.872418 | 1 |
| 7 | 92433800 | 92436999 | 37.42861 | 60.31336 | 91.65024 | 38.99557 | -0.2463 | 1.799766 | 1 |
| 7 | 92727800 | 92731199 | 24.74187 | 40.73094 | 69.33131 | 29.89643 | -0.38611 | 1.380768 | 1 |

| | | | | | | | | | |
|---|---|---|---|---|---|---|---|---|---|
| 7 | 92804600 | 92807599 | 42.07273 | 67.95033 | 80.43557 | 34.32432 | 0.116967 | 1.735656 | 1 |
| 7 | 92816200 | 92817799 | 13.22692 | 23.44668 | 34.11461 | 15.08163 | -0.14593 | 0.485751 | 1 |
| 7 | 92884400 | 92886599 | 30.58091 | 50.48681 | 61.15955 | 26.40755 | 0.104537 | 1.354211 | 1 |
| 7 | 93066800 | 93069399 | 20.97539 | 35.43421 | 54.31381 | 23.76209 | -0.23268 | 1.079463 | 1 |
| 7 | 93213800 | 93214999 | 12.2978 | 21.98749 | 36.2086 | 15.8935 | -0.31642 | 0.429506 | 1 |
| 7 | 93642800 | 93644599 | 27.62672 | 46.49622 | 51.23746 | 22.39079 | 0.241495 | 1.131765 | 1 |
| 7 | 93942000 | 93942999 | 15.36818 | 26.98357 | 38.78187 | 17.15811 | -0.1331 | 0.618433 | 1 |
| 7 | 94357200 | 94359399 | 31.01205 | 51.34254 | 58.35033 | 25.24347 | 0.196725 | 1.323547 | 1 |
| 7 | 94444000 | 94445199 | 13.22718 | 23.78447 | 36.20043 | 15.81366 | -0.19873 | 0.399935 | 1 |
| 7 | 94636200 | 94638199 | 21.8006 | 37.438 | 42.1944 | 18.50751 | 0.212013 | 0.92237 | 1 |
| 7 | 94686800 | 94687799 | 11.88211 | 21.40558 | 31.77974 | 13.85455 | -0.16048 | 0.374279 | 1 |
| 7 | 94725800 | 94728199 | 15.65097 | 27.43142 | 38.3265 | 16.93126 | -0.09252 | 0.647248 | 1 |
| 7 | 94934400 | 94935599 | 24.93925 | 42.34104 | 46.38844 | 20.32548 | 0.250303 | 1.042488 | 1 |
| 7 | 95039000 | 95039799 | 10.00496 | 19.19819 | 21.84004 | 8.571573 | 0.340663 | -0.06657 | 1 |
| 7 | 95182000 | 95182799 | 15.15659 | 26.55531 | 38.86126 | 17.16225 | -0.15905 | 0.650594 | 1 |
| 7 | 95561600 | 95564199 | 27.84964 | 45.63203 | 70.42527 | 30.30781 | -0.24791 | 1.448728 | 1 |
| 7 | 95717400 | 95721199 | 38.05159 | 61.35972 | 83.72187 | 35.65457 | -0.0901 | 1.776946 | 1 |
| 7 | 96476200 | 96477799 | 19.97067 | 34.48463 | 43.70223 | 19.17209 | 0.045029 | 0.865969 | 1 |
| 7 | 96796000 | 96798399 | 26.58892 | 43.70245 | 68.04663 | 29.32283 | -0.25873 | 1.407833 | 1 |
| 7 | 96885200 | 96888999 | 39.93368 | 65.12605 | 76.13124 | 32.7056 | 0.147433 | 1.545601 | 1 |
| 7 | 96926800 | 96930199 | 47.51851 | 76.39164 | 85.03311 | 36.05634 | 0.197568 | 1.885699 | 1 |
| 7 | 97569000 | 97569799 | 10.72926 | 19.72951 | 28.23838 | 11.9914 | -0.08297 | 0.236208 | 1 |
| 7 | 97579000 | 97580999 | 23.61716 | 40.62896 | 42.60957 | 18.69061 | 0.315952 | 0.914835 | 1 |
| 7 | 97659200 | 97665599 | 111.4917 | 170.7416 | 189.0239 | 78.44754 | 0.144765 | 3.039579 | 1 |
| 7 | 97916200 | 97920399 | 69.80282 | 110.4556 | 122.9042 | 51.087 | 0.173925 | 2.392253 | 1 |
| 7 | 98452400 | 98455199 | 15.31526 | 26.85035 | 37.81873 | 16.71081 | -0.10412 | 0.639782 | 1 |
| 7 | 101462000 | 101464399 | 26.22771 | 43.5211 | 64.85721 | 28.10277 | -0.19288 | 1.286922 | 1 |
| 7 | 102652000 | 102654399 | 26.49106 | 44.93015 | 49.58424 | 21.71908 | 0.239693 | 1.058263 | 1 |
| 7 | 102721000 | 102729199 | 174.0765 | 259.772 | 264.2213 | 108.9086 | 0.235208 | 3.567174 | 1 |
| 7 | 102754000 | 102756199 | 37.44823 | 60.28777 | 88.7159 | 37.6952 | -0.20173 | 1.829071 | 1 |
| 7 | 102850200 | 102852999 | 23.74943 | 39.38112 | 62.74335 | 27.17367 | -0.28888 | 1.289686 | 1 |
| 7 | 102897800 | 102900199 | 26.78071 | 44.70725 | 55.57346 | 24.17621 | 0.068284 | 1.214063 | 1 |
| 7 | 102937600 | 102954999 | 410.3106 | 563.4331 | 693.2554 | 288.2809 | -0.15727 | 4.858899 | 1 |
| 7 | 103229200 | 103231799 | 21.45162 | 36.48974 | 52.73717 | 23.11422 | -0.14754 | 1.019967 | 1 |
| 7 | 103613800 | 103616599 | 37.48569 | 61.43221 | 62.949 | 27.09249 | 0.341423 | 1.464916 | 1 |
| 7 | 103681600 | 103684599 | 31.32607 | 51.07061 | 72.55608 | 31.1658 | -0.13348 | 1.529691 | 1 |
| 7 | 103689000 | 103690199 | 15.28586 | 27.29708 | 28.99287 | 12.73911 | 0.311885 | 0.443824 | 1 |
| 7 | 103769000 | 103770199 | 18.51262 | 31.84277 | 49.53032 | 21.72628 | -0.25044 | 0.903868 | 1 |
| 7 | 104064600 | 104070199 | 78.92903 | 123.9502 | 141.4761 | 58.68005 | 0.128689 | 2.564512 | 1 |
| 7 | 104151000 | 104152399 | 21.98544 | 37.82455 | 43.24431 | 18.96908 | 0.192111 | 0.910782 | 1 |
| 7 | 104295800 | 104297399 | 12.59865 | 22.51539 | 32.17653 | 14.12894 | -0.1132 | 0.427146 | 1 |
| 7 | 104308200 | 104310799 | 20.5505 | 36.06865 | 35.35533 | 15.63696 | 0.416285 | 0.64787 | 1 |
| 7 | 104368800 | 104371799 | 49.04315 | 78.3288 | 108.1499 | 45.41862 | -0.1236 | 2.096516 | 1 |
| 7 | 104434000 | 104439199 | 57.69721 | 91.93528 | 105.9124 | 44.33378 | 0.133536 | 2.177989 | 1 |
| 7 | 104911600 | 104912999 | 20.75264 | 35.82184 | 42.63728 | 18.70469 | 0.135015 | 0.871093 | 1 |
| 7 | 104984800 | 104986999 | 16.28381 | 28.69845 | 32.59894 | 14.4283 | 0.206196 | 0.5668 | 1 |
| 7 | 105246600 | 105247399 | 12.10845 | 22.61247 | 23.99651 | 9.96292 | 0.369697 | 0.146137 | 1 |

| | | | | | | | | | | |
|---|---|---|---|---|---|---|---|---|---|---|
| 7 | 105892200 | 105892999 | 11.70965 | 21.86435 | 25.63462 | 10.64582 | 0.226692 | 0.148251 | | 1 |
| 7 | 105955400 | 105959599 | 51.4921 | 82.73541 | 93.00962 | 39.40803 | 0.182357 | 1.910589 | | 1 |
| 7 | 106060600 | 106061999 | 16.99835 | 29.68088 | 43.06503 | 18.96712 | -0.14688 | 0.721734 | | 1 |
| 7 | 106303000 | 106308599 | 51.83274 | 82.77954 | 117.9226 | 49.51885 | -0.16873 | 2.103623 | | 1 |
| 7 | 106445400 | 106446599 | 17.81184 | 31.15612 | 37.23579 | 16.42153 | 0.13167 | 0.688505 | | 1 |
| 7 | 106538000 | 106539999 | 27.73823 | 46.51505 | 52.89015 | 23.07446 | 0.196341 | 1.16708 | | 1 |
| 7 | 106564800 | 106566199 | 16.35154 | 29.4687 | 29.19568 | 12.72181 | 0.419532 | 0.394322 | | 1 |
| 7 | 106583000 | 106586199 | 45.29237 | 73.24092 | 75.52125 | 32.27876 | 0.317673 | 1.702559 | | 1 |
| 7 | 106651000 | 106653199 | 27.88831 | 46.98224 | 53.44942 | 23.36432 | 0.195661 | 1.124997 | | 1 |
| 7 | 106663400 | 106669599 | 79.16702 | 123.6489 | 170.54 | 70.76066 | -0.15017 | 2.664331 | | 1 |
| 7 | 106683400 | 106686999 | 18.16643 | 31.25469 | 51.62992 | 22.64743 | -0.33665 | 0.906179 | | 1 |
| 7 | 107186600 | 107188399 | 16.39106 | 28.81843 | 43.34379 | 19.19315 | -0.19856 | 0.607984 | | 1 |
| 7 | 107453000 | 107454399 | 23.10475 | 39.13547 | 48.00264 | 21.01794 | 0.087963 | 1.059504 | | 1 |
| 7 | 107464400 | 107470199 | 69.90079 | 110.7346 | 129.5408 | 53.87612 | 0.103065 | 2.366471 | | 1 |
| 7 | 107555800 | 107558399 | 40.62046 | 65.95295 | 74.10782 | 31.77169 | 0.199489 | 1.613095 | | 1 |
| 7 | 107611800 | 107614799 | 54.34458 | 86.80773 | 100.0072 | 42.00713 | 0.137387 | 2.097144 | | 1 |
| 7 | 107625200 | 107628799 | 67.87136 | 106.1656 | 163.2278 | 67.71622 | -0.30496 | 2.632475 | | 1 |
| 7 | 107641600 | 107643999 | 20.43979 | 35.57294 | 37.74527 | 16.58985 | 0.301932 | 0.774162 | | 1 |
| 7 | 107801000 | 107802399 | 21.86394 | 37.52477 | 45.27015 | 19.85577 | 0.113943 | 0.929347 | | 1 |
| 7 | 108027600 | 108029799 | 16.91097 | 29.70884 | 31.06715 | 13.75134 | 0.323308 | 0.602903 | | 1 |
| 7 | 108071600 | 108073399 | 16.58879 | 29.40323 | 30.86426 | 13.64496 | 0.322564 | 0.503189 | | 1 |
| 7 | 108286800 | 108288599 | 22.42327 | 38.22993 | 47.42869 | 20.79139 | 0.072355 | 0.994867 | | 1 |
| 7 | 108303600 | 108305799 | 21.62548 | 37.14785 | 41.68561 | 18.2845 | 0.218352 | 0.918333 | | 1 |
| 7 | 109711200 | 109713599 | 14.83945 | 26.25123 | 33.08939 | 14.63439 | 0.058912 | 0.517663 | | 1 |
| 7 | 109766200 | 109775799 | 106.9088 | 162.9078 | 226.7526 | 93.99564 | -0.19095 | 3.140858 | | 1 |
| 7 | 110081000 | 110082599 | 15.93344 | 28.08921 | 39.71686 | 17.57766 | -0.10801 | 0.574718 | | 1 |
| 7 | 112028000 | 112030399 | 22.99091 | 38.62472 | 61.43925 | 26.83437 | -0.28638 | 1.141725 | | 1 |
| 7 | 112250600 | 112252199 | 16.22205 | 28.41551 | 40.83722 | 18.03138 | -0.13323 | 0.661845 | | 1 |
| 7 | 113302800 | 113305399 | 27.25249 | 45.83388 | 52.87252 | 23.10034 | 0.175623 | 1.13715 | | 1 |
| 7 | 113361600 | 113362799 | 18.65245 | 32.41689 | 44.69755 | 19.62825 | -0.07453 | 0.795857 | | 1 |
| 7 | 115204000 | 115206599 | 24.94075 | 41.49673 | 64.79933 | 28.12974 | -0.25978 | 1.252779 | | 1 |
| 7 | 116580400 | 116581599 | 17.98591 | 31.02178 | 47.41551 | 20.80037 | -0.22444 | 0.873653 | | 1 |
| 7 | 117419200 | 117422999 | 36.73781 | 59.50054 | 90.91495 | 38.90577 | -0.24629 | 1.663518 | | 1 |
| 7 | 119102000 | 119104399 | 27.34049 | 45.47554 | 58.26839 | 25.28929 | 0.024695 | 1.258359 | | 1 |
| 7 | 119603600 | 119605999 | 31.34323 | 51.35412 | 69.28952 | 29.8185 | -0.05465 | 1.455204 | | 1 |
| 7 | 120047400 | 120047999 | 9.949005 | 18.90089 | 22.23699 | 8.940838 | 0.260912 | 0.010817 | | 1 |
| 7 | 120103600 | 120106799 | 36.35574 | 58.42089 | 97.66881 | 41.38826 | -0.38875 | 1.889564 | | 1 |
| 7 | 120112200 | 120113199 | 11.8716 | 21.41406 | 30.13967 | 13.11851 | -0.08288 | 0.359078 | | 1 |
| 7 | 120158400 | 120161599 | 5.668924 | 11.02129 | 22.35789 | 8.29573 | -0.40971 | -0.30244 | | 1 |
| 7 | 121119200 | 121125399 | 63.78942 | 101.172 | 130.5468 | 54.35145 | -0.03661 | 2.328802 | | 1 |
| 7 | 122540800 | 122543799 | 10.08841 | 18.7313 | 29.38097 | 12.31354 | -0.20062 | 0.178841 | | 1 |
| 7 | 122617000 | 122624599 | 86.54552 | 133.1738 | 194.7407 | 80.87756 | -0.25105 | 2.938488 | | 1 |
| 7 | 124079600 | 124082399 | 30.89345 | 51.26156 | 55.1439 | 23.89344 | 0.275947 | 1.293296 | | 1 |
| 7 | 124126000 | 124128199 | 23.07189 | 39.32911 | 45.84759 | 20.09801 | 0.161813 | 0.997477 | | 1 |
| 7 | 124138800 | 124140399 | 29.07075 | 47.95907 | 59.74539 | 25.78604 | 0.064201 | 1.36044 | | 1 |
| 7 | 124404000 | 124406799 | 20.4696 | 34.56435 | 55.39172 | 24.23039 | -0.29674 | 1.081383 | | 1 |
| 7 | 124409600 | 124411399 | 18.35836 | 32.16592 | 35.13844 | 15.51852 | 0.260775 | 0.662503 | | 1 |

| | | | | | | | | | |
|---|---|---|---|---|---|---|---|---|---|
| 7 | 124608200 | 124610599 | 24.67856 | 42.88374 | 43.46545 | 19.08547 | 0.367558 | 0.82405 | 1 |
| 7 | 125132800 | 125134799 | 36.04952 | 58.92382 | 66.81651 | 28.72757 | 0.19346 | 1.49647 | 1 |
| 7 | 126440400 | 126441399 | 11.34759 | 20.71663 | 31.30058 | 13.42031 | -0.17017 | 0.291315 | 1 |
| 7 | 127217000 | 127220599 | 17.17414 | 30.15338 | 29.83271 | 13.19836 | 0.403091 | 0.613822 | 1 |
| 7 | 127223000 | 127223599 | 10.01704 | 18.97294 | 23.80742 | 9.633405 | 0.160736 | 0.037114 | 1 |
| 7 | 127329800 | 127330999 | 13.97763 | 25.03521 | 30.52118 | 13.37706 | 0.116492 | 0.418834 | 1 |
| 7 | 128197800 | 128206399 | 104.3638 | 159.976 | 203.121 | 84.31351 | -0.05174 | 3.019304 | 1 |
| 7 | 128210400 | 128213799 | 29.26101 | 47.97685 | 67.3021 | 28.97199 | -0.11019 | 1.441023 | 1 |
| 7 | 128590800 | 128591999 | 14.48868 | 25.67206 | 31.58958 | 13.96658 | 0.094365 | 0.494656 | 1 |
| 7 | 129162800 | 129163199 | 29.72007 | 49.25979 | 53.99829 | 23.37911 | 0.248738 | 1.301364 | 1 |
| 7 | 131625600 | 131627799 | 21.79333 | 37.29883 | 44.23522 | 19.39746 | 0.137857 | 0.951982 | 1 |
| 7 | 131735000 | 131736199 | 11.66324 | 21.08092 | 34.27287 | 14.87974 | -0.28722 | 0.356379 | 1 |
| 7 | 132508000 | 132510599 | 55.50823 | 88.75285 | 94.69358 | 39.83006 | 0.249434 | 2.065299 | 1 |
| 7 | 132746600 | 132747999 | 13.39656 | 23.82568 | 37.1378 | 16.37419 | -0.24217 | 0.470993 | 1 |
| 7 | 133603400 | 133604599 | 15.54628 | 27.07344 | 42.1537 | 18.53479 | -0.24848 | 0.750707 | 1 |
| 7 | 134583400 | 134585799 | 42.10218 | 68.20146 | 75.1181 | 32.15018 | 0.225174 | 1.662272 | 1 |
| 7 | 135846400 | 135848399 | 34.501 | 56.23332 | 68.54686 | 29.44113 | 0.086596 | 1.535814 | 1 |
| 7 | 136090800 | 136093399 | 23.51879 | 39.57451 | 57.13679 | 24.96726 | -0.14702 | 1.123627 | 1 |
| 7 | 137016200 | 137017999 | 13.83254 | 24.69491 | 30.58326 | 13.44106 | 0.091283 | 0.438474 | 1 |
| 7 | 137420400 | 137423399 | 36.46371 | 59.46608 | 62.21749 | 26.72817 | 0.306962 | 1.525719 | 1 |
| 7 | 139001800 | 139011999 | 130.6891 | 197.3869 | 268.198 | 110.8747 | -0.16685 | 3.329327 | 1 |
| 7 | 140108400 | 140109399 | 13.43591 | 24.1664 | 30.9788 | 13.52955 | 0.047825 | 0.388491 | 1 |
| 7 | 140118400 | 140119399 | 9.702165 | 18.31486 | 28.09651 | 11.45008 | -0.13647 | 0.066435 | 1 |
| 7 | 140165600 | 140166799 | 15.74197 | 27.39804 | 44.74208 | 19.66559 | -0.31707 | 0.764348 | 1 |
| 7 | 140757800 | 140758399 | 13.96828 | 24.71348 | 32.11602 | 14.20351 | 0.015119 | 0.507178 | 1 |
| 7 | 142334800 | 142337999 | 46.43314 | 74.90448 | 87.96218 | 37.47922 | 0.126505 | 1.780857 | 1 |
| 7 | 142391400 | 142398799 | 87.82188 | 137.5659 | 152.2978 | 63.17267 | 0.169817 | 2.616312 | 1 |
| 7 | 142773400 | 142783399 | 146.4062 | 222.0929 | 250.974 | 103.8962 | 0.104195 | 3.24645 | 1 |
| 7 | 142803000 | 142809599 | 90.67583 | 140.5289 | 186.8294 | 77.60364 | -0.10593 | 2.805765 | 1 |
| 7 | 143164800 | 143167799 | 17.14263 | 29.688 | 48.46887 | 21.2651 | -0.318 | 0.833411 | 1 |
| 7 | 143229400 | 143230399 | 10.76995 | 20.18693 | 25.21393 | 10.40339 | 0.143603 | 0.114942 | 1 |
| 7 | 143409400 | 143410799 | 28.99119 | 46.97906 | 81.56636 | 34.91885 | -0.42925 | 1.63549 | 1 |
| 7 | 143428600 | 143433399 | 55.07653 | 87.93886 | 103.4805 | 43.43943 | 0.106157 | 2.11063 | 1 |
| 7 | 143943200 | 143944199 | 12.25366 | 22.43355 | 27.57019 | 11.77808 | 0.129078 | 0.270795 | 1 |
| 8 | 3195000 | 3197999 | 24.28284 | 40.56598 | 62.57536 | 27.23439 | -0.24216 | 1.205913 | 1 |
| 8 | 3544600 | 3547799 | 50.96481 | 80.99367 | 121.1195 | 50.52757 | -0.24618 | 2.259756 | 1 |
| 8 | 3963200 | 3968999 | 33.88022 | 55.08335 | 79.2942 | 34.02607 | -0.15534 | 1.575674 | 1 |
| 8 | 4026600 | 4035599 | 115.6917 | 176.5596 | 203.6404 | 84.44581 | 0.081757 | 3.110912 | 1 |
| 8 | 4131400 | 4134399 | 46.28687 | 74.49815 | 84.95748 | 36.08498 | 0.164755 | 1.846185 | 1 |
| 8 | 4447400 | 4455399 | 71.25048 | 112.5696 | 125.0214 | 51.92578 | 0.174822 | 2.429935 | 1 |
| 8 | 4595600 | 4597399 | 17.35515 | 30.54543 | 31.82723 | 14.09053 | 0.329049 | 0.583153 | 1 |
| 8 | 4626200 | 4629399 | 73.87709 | 116.3577 | 121.9853 | 50.61517 | 0.254447 | 2.499913 | 1 |
| 8 | 4666200 | 4668799 | 27.3099 | 46.15871 | 53.97258 | 23.61645 | 0.156427 | 1.09669 | 1 |
| 8 | 4669800 | 4680399 | 163.2021 | 247.7177 | 219.1824 | 90.75131 | 0.457571 | 3.236041 | 1 |
| 8 | 4858200 | 4859199 | 24.56804 | 41.9514 | 43.20075 | 18.94034 | 0.340455 | 0.985718 | 1 |
| 8 | 4903200 | 4904999 | 35.84496 | 58.13441 | 72.68987 | 31.13817 | 0.044221 | 1.624918 | 1 |
| 8 | 5128200 | 5131199 | 34.5273 | 56.0522 | 77.39684 | 33.17843 | -0.0973 | 1.604281 | 1 |

| | | | | | | | | | |
|---|---|---|---|---|---|---|---|---|---|
| 8 | 5290200 | 5292599 | 28.99706 | 47.64065 | 66.67402 | 28.72627 | -0.10559 | 1.417107 | 1 |
| 8 | 5559000 | 5560599 | 28.18256 | 46.71919 | 57.13064 | 24.73813 | 0.09153 | 1.298997 | 1 |
| 8 | 5577600 | 5581999 | 60.88256 | 96.178 | 141.1479 | 58.62021 | -0.22681 | 2.429599 | 1 |
| 8 | 5891000 | 5891799 | 11.88844 | 21.3235 | 30.65757 | 13.42078 | -0.11894 | 0.39431 | 1 |
| 8 | 6149800 | 6152199 | 16.66343 | 29.0791 | 44.13135 | 19.42964 | -0.21154 | 0.729327 | 1 |
| 8 | 6201200 | 6203599 | 26.712 | 44.37052 | 62.65431 | 27.16762 | -0.11524 | 1.275176 | 1 |
| 8 | 6323800 | 6324999 | 13.83074 | 25.08888 | 28.39624 | 12.27291 | 0.236063 | 0.336498 | 1 |
| 8 | 6816600 | 6819399 | 18.13671 | 31.83635 | 38.6354 | 17.08958 | 0.109701 | 0.637995 | 1 |
| 8 | 7071800 | 7073799 | 15.29009 | 27.53709 | 30.25011 | 13.19245 | 0.270469 | 0.391567 | 1 |
| 8 | 7380000 | 7380799 | 14.21539 | 25.37671 | 29.19792 | 12.83294 | 0.197015 | 0.440283 | 1 |
| 8 | 7399600 | 7401599 | 16.64982 | 29.39597 | 31.88393 | 14.1068 | 0.273278 | 0.54633 | 1 |
| 8 | 7591200 | 7592599 | 24.05494 | 40.55703 | 48.54704 | 21.22654 | 0.122761 | 1.10266 | 1 |
| 8 | 7730800 | 7733399 | 31.03587 | 51.39878 | 54.59014 | 23.62207 | 0.294105 | 1.314959 | 1 |
| 8 | 8122200 | 8123599 | 26.77288 | 45.01701 | 48.56459 | 21.21671 | 0.272019 | 1.134296 | 1 |
| 8 | 8185600 | 8188999 | 33.77157 | 54.82554 | 78.26231 | 33.54952 | -0.14511 | 1.604695 | 1 |
| 8 | 9048800 | 9051199 | 23.79764 | 40.56555 | 42.76813 | 18.74805 | 0.306342 | 0.998753 | 1 |
| 8 | 9407600 | 9409999 | 36.39088 | 58.9806 | 71.92482 | 30.79536 | 0.079391 | 1.640413 | 1 |
| 8 | 9600200 | 9602399 | 21.16572 | 36.09528 | 46.9196 | 20.56865 | 0.005407 | 0.987727 | 1 |
| 8 | 10198600 | 10200999 | 24.23148 | 40.72971 | 58.78123 | 25.67742 | -0.14649 | 1.138059 | 1 |
| 8 | 11043200 | 11046599 | 25.51585 | 42.09063 | 63.88359 | 27.57103 | -0.21993 | 1.354466 | 1 |
| 8 | 11516600 | 11519399 | 27.76718 | 45.82625 | 65.26309 | 28.17289 | -0.12819 | 1.355518 | 1 |
| 8 | 14236800 | 14238199 | 17.0861 | 29.86547 | 40.40889 | 17.81023 | -0.04648 | 0.701789 | 1 |
| 8 | 14500400 | 14507799 | 90.68995 | 140.4669 | 180.1842 | 74.84363 | -0.05503 | 2.815807 | 1 |
| 8 | 15220400 | 15221799 | 18.2143 | 31.93075 | 36.50651 | 16.13007 | 0.195159 | 0.652604 | 1 |
| 8 | 15568400 | 15570999 | 280.9365 | 402.6187 | 367.0775 | 151.6022 | 0.337118 | 4.216047 | 1 |
| 8 | 15748200 | 15749999 | 20.42267 | 35.3375 | 41.06897 | 18.01801 | 0.170032 | 0.84625 | 1 |
| 8 | 16186000 | 16189799 | 44.01671 | 71.1933 | 78.88873 | 33.72575 | 0.21439 | 1.696885 | 1 |
| 8 | 17238000 | 17240999 | 41.65246 | 67.05398 | 94.06917 | 39.96793 | -0.13309 | 1.837362 | 1 |
| 8 | 17548200 | 17550399 | 34.50428 | 56.05809 | 78.24336 | 33.5604 | -0.1117 | 1.593065 | 1 |
| 8 | 18481600 | 18483799 | 33.0614 | 54.02933 | 63.98085 | 27.50647 | 0.130989 | 1.493514 | 1 |
| 8 | 18730200 | 18732799 | 26.15745 | 43.75683 | 58.86566 | 25.64172 | -0.04545 | 1.193226 | 1 |
| 8 | 19125400 | 19130799 | 72.42521 | 115.0202 | 121.2702 | 50.55589 | 0.256243 | 2.296873 | 1 |
| 8 | 19381400 | 19383599 | 18.55691 | 32.70445 | 33.19183 | 14.69072 | 0.36758 | 0.578728 | 1 |
| 8 | 19452000 | 19454199 | 39.46181 | 64.2107 | 68.94146 | 29.59243 | 0.267529 | 1.571201 | 1 |
| 8 | 19539800 | 19541199 | 17.09222 | 30.18363 | 29.11507 | 12.88117 | 0.442019 | 0.544014 | 1 |
| 8 | 19809000 | 19816999 | 155.1652 | 232.2295 | 302.1256 | 124.6031 | -0.11584 | 3.511466 | 1 |
| 8 | 20045800 | 20049799 | 60.43652 | 95.58075 | 136.9879 | 56.92489 | -0.1914 | 2.400284 | 1 |
| 8 | 20056200 | 20057399 | 27.11006 | 44.83153 | 63.58264 | 27.48548 | -0.12198 | 1.329606 | 1 |
| 8 | 20516400 | 20518999 | 28.31224 | 47.53507 | 53.4265 | 23.32624 | 0.213024 | 1.157912 | 1 |
| 8 | 20547200 | 20572999 | 201.2003 | 298.7891 | 309.6983 | 127.577 | 0.201549 | 3.659864 | 1 |
| 8 | 20645800 | 20648799 | 21.84451 | 37.35655 | 42.28766 | 18.54225 | 0.20487 | 0.961365 | 1 |
| 8 | 20726800 | 20728599 | 29.36707 | 48.79555 | 55.6692 | 24.14384 | 0.191515 | 1.272651 | 1 |
| 8 | 21077800 | 21079999 | 34.14561 | 55.14785 | 83.59099 | 35.67113 | -0.23885 | 1.724019 | 1 |
| 8 | 21192400 | 21193999 | 39.14503 | 63.46007 | 69.30246 | 29.67797 | 0.238443 | 1.635654 | 1 |
| 8 | 21201000 | 21205399 | 37.08095 | 59.77006 | 83.97435 | 35.74322 | -0.13289 | 1.787432 | 1 |
| 8 | 21236000 | 21237599 | 18.63587 | 32.11283 | 45.33514 | 19.88718 | -0.11054 | 0.882887 | 1 |
| 8 | 21241000 | 21243399 | 24.41149 | 40.27699 | 65.98544 | 28.48057 | -0.32972 | 1.350916 | 1 |

| | | | | | | | | | |
|---|---|---|---|---|---|---|---|---|---|
| 8 | 21312400 | 21314399 | 70.68978 | 111.2236 | 134.4227 | 55.76836 | 0.048396 | 2.518672 | 1 |
| 8 | 21441600 | 21448199 | 80.16702 | 126.4351 | 132.1561 | 54.84493 | 0.260206 | 2.477467 | 1 |
| 8 | 21521800 | 21533599 | 134.9897 | 206.5986 | 200.7667 | 83.30901 | 0.331965 | 3.06099 | 1 |
| 8 | 21629800 | 21636599 | 69.14628 | 110.0958 | 108.5335 | 45.38386 | 0.356821 | 2.214099 | 1 |
| 8 | 21713600 | 21716999 | 22.7078 | 39.02771 | 43.14667 | 18.92566 | 0.239836 | 0.918728 | 1 |
| 8 | 21933200 | 21934799 | 9.152264 | 17.20192 | 29.65277 | 12.17815 | -0.31247 | 0.094403 | 1 |
| 8 | 22072000 | 22072799 | 10.02081 | 19.16777 | 21.18326 | 8.389794 | 0.369351 | -0.04111 | 1 |
| 8 | 22355200 | 22367399 | 315.2252 | 446.8313 | 471.7447 | 195.182 | 0.109716 | 4.37846 | 1 |
| 8 | 22478600 | 22480999 | 24.84417 | 41.78844 | 50.13262 | 21.90405 | 0.119374 | 1.126625 | 1 |
| 8 | 23755400 | 23765799 | 98.33435 | 151.4832 | 190.8174 | 79.2603 | -0.03485 | 2.918763 | 1 |
| 8 | 23849800 | 23852199 | 21.52238 | 36.48167 | 48.51576 | 21.24729 | -0.02832 | 1.045101 | 1 |
| 8 | 23949600 | 23953399 | 19.20183 | 33.387 | 43.1065 | 18.93503 | 0.020057 | 0.791316 | 1 |
| 8 | 24021600 | 24023399 | 15.27684 | 26.99859 | 36.01666 | 15.93214 | -0.02322 | 0.534731 | 1 |
| 8 | 24628000 | 24630199 | 20.29836 | 35.09825 | 47.93658 | 21.03056 | -0.06221 | 0.859832 | 1 |
| 8 | 24639400 | 24640799 | 10.68999 | 19.9098 | 25.23961 | 10.52695 | 0.11018 | 0.156281 | 1 |
| 8 | 25048800 | 25052399 | 34.10995 | 55.35502 | 81.77709 | 35.04697 | -0.19501 | 1.61441 | 1 |
| 8 | 25056000 | 25060399 | 51.29679 | 82.52678 | 87.46476 | 37.12725 | 0.269494 | 1.868514 | 1 |
| 8 | 25258800 | 25260399 | 11.01178 | 20.7773 | 19.67309 | 8.025254 | 0.55381 | 0.061155 | 1 |
| 8 | 25315400 | 25320799 | 45.72688 | 73.41674 | 106.9617 | 45.26853 | -0.19231 | 1.929968 | 1 |
| 8 | 25333000 | 25333799 | 10.31304 | 18.47567 | 33.85874 | 14.83711 | -0.46752 | 0.40078 | 1 |
| 8 | 26696200 | 26699799 | 11.23659 | 21.28354 | 24.31645 | 9.838701 | 0.295291 | 0.051049 | 1 |
| 8 | 26747200 | 26748999 | 22.64327 | 38.98895 | 43.10297 | 18.90756 | 0.240345 | 0.900918 | 1 |
| 8 | 26774000 | 26774999 | 15.10472 | 26.99982 | 31.40675 | 13.78845 | 0.182091 | 0.440445 | 1 |
| 8 | 27056400 | 27062399 | 57.96231 | 92.41522 | 102.9565 | 43.1336 | 0.182756 | 2.156863 | 1 |
| 8 | 27096000 | 27097199 | 9.143143 | 17.52179 | 29.33014 | 11.55407 | -0.21751 | -0.04445 | 1 |
| 8 | 27105400 | 27107599 | 19.34763 | 33.35113 | 45.82201 | 20.10097 | -0.07157 | 0.881286 | 1 |
| 8 | 27509800 | 27511799 | 16.13469 | 28.62438 | 34.26501 | 15.13918 | 0.134416 | 0.496901 | 1 |
| 8 | 27793000 | 27803799 | 156.8721 | 233.5326 | 301.8755 | 124.3855 | -0.1136 | 3.609595 | 1 |
| 8 | 27809200 | 27810799 | 43.70332 | 69.90219 | 103.5312 | 43.56887 | -0.22229 | 2.045739 | 1 |
| 8 | 27937200 | 27938599 | 22.02853 | 37.50748 | 46.928 | 20.56972 | 0.060008 | 1.00656 | 1 |
| 8 | 27952800 | 27958199 | 103.1435 | 158.7881 | 184.0202 | 76.43008 | 0.0846 | 2.93333 | 1 |
| 8 | 28698200 | 28705399 | 109.1125 | 166.4021 | 209.267 | 86.76498 | -0.04373 | 3.12431 | 1 |
| 8 | 28882600 | 28883399 | 19.11695 | 33.24863 | 38.45111 | 16.89304 | 0.178333 | 0.780029 | 1 |
| 8 | 28929800 | 28932399 | 26.52913 | 44.62333 | 48.56254 | 21.21815 | 0.259416 | 1.128983 | 1 |
| 8 | 29109400 | 29110799 | 17.5324 | 30.90596 | 31.44112 | 13.91522 | 0.364517 | 0.569128 | 1 |
| 8 | 29394200 | 29394999 | 13.93588 | 24.55055 | 38.2617 | 16.936 | -0.24804 | 0.566948 | 1 |
| 8 | 29708600 | 29711199 | 18.43625 | 31.2077 | 54.23881 | 23.74381 | -0.41304 | 1.052888 | 1 |
| 8 | 29837800 | 29838999 | 15.62162 | 27.11813 | 46.35657 | 20.34201 | -0.38306 | 0.804021 | 1 |
| 8 | 29850000 | 29850999 | 17.21709 | 30.09667 | 35.87547 | 15.81312 | 0.135595 | 0.694759 | 1 |
| 8 | 29905800 | 29908799 | 75.57982 | 118.5396 | 148.445 | 61.56653 | -0.00633 | 2.58649 | 1 |
| 8 | 29973600 | 29974799 | 11.75041 | 22.00524 | 23.61436 | 9.757975 | 0.359518 | 0.126247 | 1 |
| 8 | 30157600 | 30160399 | 30.99934 | 51.49664 | 53.06243 | 23.00987 | 0.33788 | 1.275375 | 1 |
| 8 | 30474400 | 30475799 | 14.32451 | 25.61232 | 29.87436 | 13.1126 | 0.178226 | 0.428307 | 1 |
| 8 | 30589400 | 30590799 | 24.57493 | 40.54258 | 65.98658 | 28.47986 | -0.32034 | 1.35276 | 1 |
| 8 | 30886600 | 30891399 | 17.20718 | 30.24891 | 33.04144 | 14.63157 | 0.260884 | 0.599969 | 1 |
| 8 | 31570200 | 31572799 | 32.72226 | 53.32202 | 77.62428 | 33.33715 | -0.16915 | 1.539693 | 1 |
| 8 | 32966200 | 32967999 | 15.60799 | 27.40433 | 41.3712 | 18.30353 | -0.20366 | 0.625403 | 1 |

| | | | | | | | | | |
|---|---|---|---|---|---|---|---|---|---|
| 8 | 33071000 | 33071999 | 14.45438 | 26.273 | 26.00376 | 11.21061 | 0.431128 | 0.327057 | 1 |
| 8 | 33376000 | 33376999 | 13.76705 | 25.17417 | 29.01558 | 12.4156 | 0.219417 | 0.29301 | 1 |
| 8 | 34524000 | 34530999 | 63.91626 | 100.9468 | 140.8824 | 58.50301 | -0.15467 | 2.435871 | 1 |
| 8 | 35026200 | 35031999 | 35.55707 | 57.47847 | 89.79469 | 38.36236 | -0.28093 | 1.701861 | 1 |
| 8 | 35096000 | 35097599 | 27.28557 | 46.051 | 50.14744 | 21.93309 | 0.258717 | 1.1055 | 1 |
| 8 | 35120400 | 35130599 | 85.82977 | 134.2954 | 151.4641 | 62.83406 | 0.141805 | 2.634109 | 1 |
| 8 | 35203400 | 35204799 | 20.40752 | 35.70217 | 38.58136 | 17.01726 | 0.276198 | 0.701231 | 1 |
| 8 | 36232000 | 36233799 | 12.67846 | 22.30215 | 37.85685 | 16.76179 | -0.37139 | 0.574731 | 1 |
| 8 | 36299600 | 36301799 | 21.22295 | 36.31738 | 44.83774 | 19.66144 | 0.080293 | 0.956002 | 1 |
| 8 | 36441800 | 36443399 | 19.11465 | 33.58107 | 34.32506 | 15.19373 | 0.356016 | 0.621529 | 1 |
| 8 | 36742000 | 36742799 | 12.07719 | 21.87913 | 28.23728 | 12.22432 | 0.046782 | 0.33365 | 1 |
| 8 | 37555200 | 37558599 | 33.82206 | 55.42138 | 61.50411 | 26.46925 | 0.226625 | 1.456442 | 1 |
| 8 | 38020600 | 38021799 | 21.08063 | 35.35268 | 57.43047 | 25.05615 | -0.31644 | 1.151593 | 1 |
| 8 | 38093400 | 38095199 | 18.31815 | 31.50887 | 46.90465 | 20.57456 | -0.18728 | 0.8993 | 1 |
| 8 | 38312200 | 38313799 | 19.4992 | 33.23212 | 51.11358 | 22.40607 | -0.2365 | 0.993246 | 1 |
| 8 | 38440000 | 38441199 | 13.3738 | 23.67179 | 36.11362 | 15.97012 | -0.2149 | 0.505826 | 1 |
| 8 | 38894800 | 38895999 | 16.83783 | 29.67865 | 34.4131 | 15.23086 | 0.176505 | 0.569872 | 1 |
| 8 | 39218800 | 39222399 | 46.54113 | 74.52115 | 108.1253 | 45.57218 | -0.19103 | 2.015302 | 1 |
| 8 | 39286200 | 39287199 | 15.81497 | 27.73022 | 38.80246 | 17.14746 | -0.09472 | 0.64227 | 1 |
| 8 | 39699600 | 39700399 | 11.46918 | 20.82814 | 29.26569 | 12.63113 | -0.07222 | 0.317761 | 1 |
| 8 | 39786200 | 39790599 | 58.75264 | 93.74449 | 114.1384 | 47.86722 | 0.055956 | 2.138347 | 1 |
| 8 | 39835400 | 39838199 | 45.28233 | 72.49112 | 100.4536 | 42.32609 | -0.1251 | 2.018969 | 1 |
| 8 | 39854600 | 39857199 | 29.32247 | 49.05413 | 58.56123 | 25.51003 | 0.126277 | 1.200563 | 1 |
| 8 | 39975200 | 39977399 | 8.313733 | 15.76168 | 28.23158 | 11.40075 | -0.34686 | 0.022836 | 1 |
| 8 | 39995600 | 39997999 | 16.59797 | 28.45947 | 48.99896 | 21.48959 | -0.39709 | 0.923127 | 1 |
| 8 | 40440600 | 40443599 | 48.6249 | 77.93778 | 94.36828 | 39.83618 | 0.071274 | 1.981725 | 1 |
| 8 | 40509200 | 40509799 | 10.19402 | 19.11447 | 26.32308 | 10.85426 | 0.004557 | 0.110823 | 1 |
| 8 | 40821200 | 40823599 | 29.53465 | 48.27027 | 74.86861 | 32.18837 | -0.25723 | 1.491382 | 1 |
| 8 | 41296000 | 41298399 | 27.21331 | 45.20986 | 70.18196 | 30.43478 | -0.25156 | 1.280078 | 1 |
| 8 | 41335400 | 41337599 | 35.03057 | 57.41269 | 60.6223 | 26.09187 | 0.298248 | 1.454526 | 1 |
| 8 | 41973200 | 41980199 | 115.5474 | 176.2749 | 207.6789 | 86.11319 | 0.050709 | 3.118601 | 1 |
| 8 | 42013800 | 42015799 | 19.29782 | 33.68214 | 38.2812 | 16.85582 | 0.203737 | 0.730863 | 1 |
| 8 | 42060800 | 42063199 | 30.76377 | 50.26345 | 68.85483 | 29.5995 | -0.07886 | 1.493332 | 1 |
| 8 | 42126600 | 42128799 | 14.6991 | 25.97577 | 34.96602 | 15.4676 | -0.03634 | 0.528001 | 1 |
| 8 | 42699400 | 42700999 | 17.62301 | 30.48785 | 47.40662 | 20.79839 | -0.24826 | 0.843912 | 1 |
| 8 | 43071600 | 43072799 | 11.35109 | 20.45644 | 33.24811 | 14.48805 | -0.29015 | 0.369255 | 1 |
| 8 | 43269400 | 43270399 | 15.99297 | 27.83149 | 41.63958 | 18.30053 | -0.19136 | 0.761693 | 1 |
| 8 | 43310600 | 43311599 | 13.6835 | 24.49686 | 32.54451 | 14.26951 | -0.00721 | 0.423757 | 1 |
| 8 | 43405400 | 43411799 | 74.20853 | 116.1243 | 160.9401 | 66.76443 | -0.155 | 2.627753 | 1 |
| 8 | 43600800 | 43604599 | 48.96134 | 78.65378 | 90.02183 | 38.13136 | 0.156177 | 1.91146 | 1 |
| 8 | 43712000 | 43715799 | 31.64599 | 51.78064 | 70.45654 | 30.30422 | -0.06776 | 1.473046 | 1 |
| 8 | 43766800 | 43769199 | 44.18049 | 70.99725 | 78.44218 | 33.24251 | 0.207608 | 1.890194 | 1 |
| 8 | 43771600 | 43775199 | 57.92868 | 92.06292 | 111.7274 | 46.61058 | 0.054615 | 2.26228 | 1 |
| 8 | 43995400 | 43996399 | 18.92384 | 32.97852 | 34.4133 | 15.13805 | 0.32629 | 0.745084 | 1 |
| 8 | 44008800 | 44012599 | 23.49997 | 39.59671 | 54.40972 | 23.78543 | -0.07579 | 1.109371 | 1 |
| 8 | 44047600 | 44049799 | 20.45776 | 35.58557 | 36.39274 | 15.99012 | 0.355015 | 0.782634 | 1 |
| 8 | 44054600 | 44057199 | 23.91623 | 39.99107 | 55.42731 | 24.13767 | -0.08819 | 1.188509 | 1 |

| | | | | | | | | | |
|---|---|---|---|---|---|---|---|---|---|
| 8 | 44136200 | 44138999 | 11.45859 | 20.64166 | 34.21006 | 14.91491 | -0.31889 | 0.372378 | 1 |
| 8 | 44144200 | 44144999 | 12.30251 | 23.26305 | 24.75251 | 10.05113 | 0.392392 | 0.072117 | 1 |
| 8 | 44413000 | 44415199 | 21.40466 | 36.37463 | 58.2314 | 25.5203 | -0.29491 | 1.032212 | 1 |
| 8 | 44430200 | 44431199 | 20.14614 | 34.56476 | 45.32107 | 19.87756 | -0.00574 | 0.926228 | 1 |
| 8 | 44686600 | 44687799 | 22.77778 | 38.26916 | 52.20223 | 22.80043 | -0.06529 | 1.131914 | 1 |
| 8 | 45462800 | 45465199 | 28.28447 | 46.46714 | 71.92734 | 30.98887 | -0.25072 | 1.416932 | 1 |
| 8 | 45499600 | 45510199 | 95.76935 | 146.9419 | 203.8838 | 84.6412 | -0.17858 | 2.997728 | 1 |
| 8 | 45632000 | 45632799 | 14.14019 | 25.09384 | 32.03817 | 14.15156 | 0.042497 | 0.479925 | 1 |
| 8 | 45881200 | 45882999 | 21.54498 | 36.70525 | 49.54906 | 21.7196 | -0.04911 | 1.001937 | 1 |
| 8 | 46143400 | 46144799 | 18.08132 | 31.57673 | 42.48768 | 18.71465 | -0.03863 | 0.723341 | 1 |
| 8 | 46382400 | 46384599 | 38.59131 | 62.69523 | 64.72184 | 27.75975 | 0.322294 | 1.592131 | 1 |
| 8 | 46394600 | 46396799 | 18.20973 | 31.67252 | 44.69442 | 19.63664 | -0.10764 | 0.783273 | 1 |
| 8 | 46553800 | 46556999 | 32.45668 | 53.56003 | 66.40658 | 28.66442 | 0.071072 | 1.368695 | 1 |
| 8 | 46807000 | 46813599 | 120.9608 | 184.0781 | 220.168 | 91.23026 | 0.026097 | 3.171315 | 1 |
| 8 | 47873000 | 47880999 | 70.07456 | 110.8862 | 130.657 | 54.30969 | 0.091395 | 2.392864 | 1 |
| 8 | 47901600 | 47909599 | 134.4591 | 203.5218 | 223.3356 | 92.39148 | 0.143838 | 3.283393 | 1 |
| 8 | 47937200 | 47944799 | 64.69804 | 102.6249 | 116.2464 | 48.40322 | 0.151242 | 2.322841 | 1 |
| 8 | 48090400 | 48102799 | 144.3435 | 216.594 | 279.525 | 115.3467 | -0.10086 | 3.460041 | 1 |
| 8 | 48109600 | 48111799 | 17.47683 | 30.82849 | 32.82668 | 14.52651 | 0.299246 | 0.562846 | 1 |
| 8 | 48292000 | 48293199 | 18.77009 | 32.6605 | 36.94621 | 16.23661 | 0.210125 | 0.769458 | 1 |
| 8 | 48476000 | 48477999 | 16.30644 | 28.52279 | 41.96885 | 18.50881 | -0.16694 | 0.688681 | 1 |
| 8 | 48641200 | 48641999 | 16.20113 | 28.68245 | 29.47069 | 13.03192 | 0.352602 | 0.510214 | 1 |
| 8 | 48800600 | 48808999 | 76.16616 | 119.2346 | 169.0126 | 70.11038 | -0.18702 | 2.622842 | 1 |
| 8 | 48963600 | 48967599 | 36.07991 | 58.3172 | 88.45812 | 37.78597 | -0.23865 | 1.706635 | 1 |
| 8 | 49299200 | 49300799 | 11.90022 | 21.3905 | 33.63806 | 14.6989 | -0.24587 | 0.386782 | 1 |
| 8 | 50287200 | 50293799 | 54.12949 | 86.27882 | 114.496 | 47.9486 | -0.06966 | 2.166316 | 1 |
| 8 | 50434000 | 50436599 | 37.01407 | 59.88224 | 74.70491 | 31.94307 | 0.044186 | 1.678284 | 1 |
| 8 | 51216400 | 51217199 | 15.1427 | 27.18363 | 29.77885 | 13.02673 | 0.271564 | 0.40948 | 1 |
| 8 | 51391200 | 51392599 | 18.38285 | 31.69354 | 50.59667 | 22.19562 | -0.28709 | 0.884212 | 1 |
| 8 | 51439000 | 51445399 | 137.8694 | 208.6517 | 218.4759 | 90.37672 | 0.211253 | 3.287523 | 1 |
| 8 | 51503600 | 51507999 | 34.20534 | 55.94052 | 66.15071 | 28.44706 | 0.133659 | 1.483081 | 1 |
| 8 | 51798200 | 51800799 | 38.50867 | 62.13506 | 85.36217 | 36.38203 | -0.09896 | 1.759569 | 1 |
| 8 | 51847000 | 51851199 | 19.77418 | 34.62072 | 38.67433 | 17.06933 | 0.228714 | 0.688317 | 1 |
| 8 | 51877800 | 51880599 | 27.48275 | 46.28386 | 48.55493 | 21.22268 | 0.312306 | 1.123888 | 1 |
| 8 | 52164000 | 52165999 | 17.70131 | 30.5502 | 48.99698 | 21.49455 | -0.29338 | 0.86936 | 1 |
| 8 | 52184400 | 52186199 | 17.58858 | 30.78358 | 37.28655 | 16.45203 | 0.112553 | 0.679679 | 1 |
| 8 | 52187000 | 52189199 | 41.37926 | 66.83088 | 77.43536 | 33.04439 | 0.14779 | 1.732369 | 1 |
| 8 | 52205200 | 52206399 | 16.38579 | 28.62194 | 36.43908 | 16.05261 | 0.040987 | 0.701867 | 1 |
| 8 | 52381000 | 52385199 | 53.13638 | 84.79207 | 101.7509 | 42.67843 | 0.077061 | 2.126301 | 1 |
| 8 | 52761200 | 52761999 | 14.53218 | 25.73128 | 30.26344 | 13.38211 | 0.159019 | 0.501258 | 1 |
| 8 | 52862200 | 52864199 | 28.7623 | 48.02173 | 53.99943 | 23.49345 | 0.212465 | 1.218103 | 1 |
| 8 | 53356800 | 53357599 | 12.66959 | 22.62062 | 33.7179 | 14.81766 | -0.17432 | 0.438236 | 1 |
| 8 | 53716000 | 53721599 | 7.558627 | 14.6872 | 20.05619 | 7.474381 | 0.149908 | -0.27045 | 1 |
| 8 | 54375400 | 54377599 | 11.61959 | 21.60622 | 25.65252 | 10.72847 | 0.200776 | 0.175018 | 1 |
| 8 | 54769600 | 54770599 | 10.76484 | 20.37661 | 22.68925 | 9.193078 | 0.330082 | 0.045166 | 1 |
| 8 | 54982400 | 54983399 | 12.31743 | 21.88397 | 35.19628 | 15.53161 | -0.28772 | 0.467642 | 1 |
| 8 | 55301600 | 55302999 | 14.1013 | 24.95761 | 37.15737 | 16.43193 | -0.17978 | 0.51554 | 1 |

| | | | | | | | | | |
|---|---|---|---|---|---|---|---|---|---|
| 8 | 55320800 | 55321999 | 20.29767 | 34.0283 | 58.4474 | 25.49446 | -0.39653 | 1.153938 | 1 |
| 8 | 55818000 | 55820999 | 21.97816 | 36.90818 | 54.30355 | 23.71232 | -0.17412 | 1.135628 | 1 |
| 8 | 55947800 | 55951599 | 26.14226 | 43.37834 | 63.5914 | 27.55387 | -0.16918 | 1.28823 | 1 |
| 8 | 56016000 | 56022999 | 132.9501 | 201.9357 | 213.4191 | 88.38295 | 0.202189 | 3.213283 | 1 |
| 8 | 56031600 | 56033599 | 12.72879 | 23.43306 | 23.61461 | 10.01237 | 0.421654 | 0.23558 | 1 |
| 8 | 56196000 | 56197199 | 27.28194 | 45.04027 | 64.65654 | 27.9174 | -0.13959 | 1.350116 | 1 |
| 8 | 56433400 | 56434599 | 14.27584 | 25.56501 | 29.98438 | 13.14365 | 0.17182 | 0.420572 | 1 |
| 8 | 56625200 | 56627999 | 25.69262 | 44.24607 | 42.96688 | 18.84795 | 0.426633 | 0.911616 | 1 |
| 8 | 56632800 | 56633599 | 7.894762 | 15.0696 | 23.47484 | 9.344321 | -0.12767 | -0.037 | 1 |
| 8 | 56803200 | 56804199 | 17.72025 | 30.61867 | 47.41362 | 20.80068 | -0.24254 | 0.857782 | 1 |
| 8 | 56824400 | 56825199 | 13.51505 | 24.20909 | 28.4811 | 12.48186 | 0.167469 | 0.408396 | 1 |
| 8 | 57200800 | 57202599 | 12.70699 | 23.28581 | 28.5392 | 12.17615 | 0.13423 | 0.270491 | 1 |
| 8 | 57471400 | 57472599 | 15.30669 | 27.34592 | 28.35125 | 12.45272 | 0.346672 | 0.437975 | 1 |
| 8 | 57493600 | 57494599 | 14.58636 | 25.92571 | 34.07174 | 15.03018 | 0.002412 | 0.479244 | 1 |
| 8 | 57557200 | 57559999 | 38.38735 | 62.33995 | 65.29061 | 27.99578 | 0.300994 | 1.598858 | 1 |
| 8 | 57959400 | 57961399 | 14.93877 | 26.52484 | 31.95572 | 14.10894 | 0.126185 | 0.481842 | 1 |
| 8 | 58127800 | 58129799 | 18.49837 | 32.3652 | 36.69625 | 16.18706 | 0.207341 | 0.687994 | 1 |
| 8 | 58440600 | 58442199 | 15.46049 | 27.07588 | 40.89465 | 18.05356 | -0.20451 | 0.661792 | 1 |
| 8 | 58568000 | 58569599 | 14.94491 | 26.01373 | 43.21366 | 18.99515 | -0.34149 | 0.756767 | 1 |
| 8 | 58750800 | 58751999 | 15.695 | 27.60308 | 35.69676 | 15.80599 | 0.018883 | 0.592107 | 1 |
| 8 | 58840600 | 58842799 | 29.7577 | 48.90073 | 63.48096 | 27.35301 | 0.002676 | 1.413138 | 1 |
| 8 | 59589400 | 59592199 | 34.91821 | 56.56041 | 76.86164 | 32.89481 | -0.07727 | 1.651281 | 1 |
| 8 | 59723400 | 59725399 | 26.37846 | 44.1443 | 58.28075 | 25.39339 | -0.01838 | 1.190441 | 1 |
| 8 | 60027800 | 60029799 | 18.23647 | 31.37429 | 47.41764 | 20.7997 | -0.20914 | 0.895579 | 1 |
| 8 | 60128200 | 60134799 | 122.5222 | 186.1273 | 219.1272 | 90.75125 | 0.046823 | 3.207879 | 1 |
| 8 | 60286800 | 60287599 | 17.03949 | 29.93788 | 33.09127 | 14.64863 | 0.243841 | 0.607149 | 1 |
| 8 | 60527400 | 60528999 | 16.05823 | 28.67898 | 30.308 | 13.31593 | 0.319004 | 0.450048 | 1 |
| 8 | 60597800 | 60599199 | 20.7369 | 34.87503 | 57.45917 | 25.10411 | -0.33669 | 1.122013 | 1 |
| 8 | 60771200 | 60772799 | 21.97573 | 37.59632 | 43.77948 | 19.19701 | 0.164255 | 0.959113 | 1 |
| 8 | 60850200 | 60851599 | 17.27533 | 29.93061 | 42.72289 | 18.74437 | -0.12476 | 0.823225 | 1 |
| 8 | 61182000 | 61185999 | 23.29596 | 39.96607 | 45.77001 | 20.07435 | 0.18861 | 0.939047 | 1 |
| 8 | 61269400 | 61270999 | 18.80355 | 32.03604 | 52.71597 | 23.10788 | -0.33373 | 0.992987 | 1 |
| 8 | 61272600 | 61272999 | 8.401808 | 16.2224 | 18.81806 | 7.249324 | 0.338053 | -0.15311 | 1 |
| 8 | 61455800 | 61457599 | 35.22788 | 57.12874 | 71.91304 | 30.80347 | 0.034487 | 1.625969 | 1 |
| 8 | 62555400 | 62555799 | 9.814901 | 18.17483 | 30.95737 | 13.02149 | -0.32251 | 0.195492 | 1 |
| 8 | 62615400 | 62616599 | 13.86883 | 24.37737 | 39.8904 | 17.6647 | -0.31912 | 0.596759 | 1 |
| 8 | 62703000 | 62703999 | 21.36044 | 35.86599 | 56.9527 | 24.86699 | -0.28379 | 1.137724 | 1 |
| 8 | 62779200 | 62784799 | 49.31929 | 78.93715 | 108.6522 | 45.76692 | -0.1158 | 2.031369 | 1 |
| 8 | 63258000 | 63260999 | 57.52583 | 91.57184 | 114.0734 | 47.68475 | 0.019377 | 2.210937 | 1 |
| 8 | 63309600 | 63311799 | 20.9303 | 36.25402 | 38.50546 | 16.89387 | 0.299788 | 0.836574 | 1 |
| 8 | 63318600 | 63320999 | 25.40523 | 43.04972 | 46.40008 | 20.31879 | 0.273691 | 1.060199 | 1 |
| 8 | 63337200 | 63339399 | 25.74943 | 43.50592 | 51.21439 | 22.40717 | 0.146729 | 1.090758 | 1 |
| 8 | 63527400 | 63529799 | 11.98531 | 21.47323 | 32.23021 | 14.12253 | -0.18136 | 0.407818 | 1 |
| 8 | 63824400 | 63825599 | 17.32776 | 30.48295 | 33.44686 | 14.80896 | 0.254752 | 0.592662 | 1 |
| 8 | 64190600 | 64192599 | 18.50355 | 32.2118 | 42.03169 | 18.47694 | 0.004998 | 0.768619 | 1 |
| 8 | 64197800 | 64198399 | 9.360499 | 17.56245 | 27.45575 | 11.30943 | -0.17544 | 0.104579 | 1 |
| 8 | 64403400 | 64405599 | 8.899753 | 16.53132 | 27.89785 | 11.68522 | -0.30434 | 0.164348 | 1 |

| | | | | | | | | | |
|---|---|---|---|---|---|---|---|---|---|
| 8 | 65129400 | 65131599 | 18.65658 | 32.53906 | 45.14957 | 19.87091 | -0.08312 | 0.74643 | 1 |
| 8 | 65637200 | 65638399 | 12.66584 | 23.33939 | 26.50169 | 11.22192 | 0.251539 | 0.234759 | 1 |
| 8 | 65749200 | 65751199 | 30.44702 | 50.60439 | 51.85093 | 22.49228 | 0.346173 | 1.271714 | 1 |
| 8 | 65812400 | 65813199 | 25.75257 | 43.28353 | 51.22151 | 22.37466 | 0.138902 | 1.135781 | 1 |
| 8 | 65856000 | 65856799 | 10.02542 | 18.26118 | 28.35308 | 12.19124 | -0.21184 | 0.285846 | 1 |
| 8 | 65995200 | 65996599 | 18.55554 | 32.41576 | 38.83353 | 17.10871 | 0.128312 | 0.716207 | 1 |
| 8 | 66289000 | 66289799 | 12.46347 | 22.18899 | 33.69359 | 14.84276 | -0.20347 | 0.453121 | 1 |
| 8 | 66476600 | 66477799 | 10.55391 | 20.14977 | 23.92647 | 9.519744 | 0.261149 | -0.01091 | 1 |
| 8 | 66678000 | 66680399 | 27.7409 | 46.63529 | 49.63257 | 21.68196 | 0.29148 | 1.139857 | 1 |
| 8 | 66755200 | 66755799 | 9.188913 | 17.15333 | 30.04124 | 12.48112 | -0.34847 | 0.139978 | 1 |
| 8 | 67009200 | 67010999 | 18.99604 | 33.0062 | 41.05123 | 18.02434 | 0.073686 | 0.797095 | 1 |
| 8 | 67109000 | 67111799 | 29.32485 | 49.05113 | 53.99837 | 23.52028 | 0.242973 | 1.201486 | 1 |
| 8 | 67487000 | 67489999 | 49.50887 | 78.93016 | 112.9066 | 47.2965 | -0.17741 | 2.153718 | 1 |
| 8 | 67817600 | 67830799 | 327.3812 | 461.2811 | 495.324 | 205.1414 | 0.076548 | 4.466952 | 1 |
| 8 | 67966600 | 67969399 | 17.30429 | 29.89998 | 46.3609 | 20.33885 | -0.24451 | 0.852192 | 1 |
| 8 | 68054600 | 68059199 | 101.5856 | 156.1291 | 186.5665 | 77.46996 | 0.038622 | 2.965377 | 1 |
| 8 | 68243600 | 68245999 | 23.69714 | 39.92325 | 53.857 | 23.54278 | -0.04927 | 1.111636 | 1 |
| 8 | 68246800 | 68248599 | 45.27896 | 72.49519 | 96.83055 | 40.80744 | -0.07189 | 2.013925 | 1 |
| 8 | 68250400 | 68256199 | 28.01254 | 47.23406 | 49.08369 | 21.46179 | 0.325921 | 1.113646 | 1 |
| 8 | 68612200 | 68613999 | 20.24166 | 35.17633 | 36.80491 | 16.16159 | 0.321881 | 0.792374 | 1 |
| 8 | 68691400 | 68693199 | 16.96617 | 29.75886 | 36.22357 | 16.0124 | 0.105478 | 0.640076 | 1 |
| 8 | 69417600 | 69424799 | 101.9704 | 157.0367 | 176.6724 | 73.3818 | 0.127726 | 2.925295 | 1 |
| 8 | 69699000 | 69705799 | 77.00496 | 121.1645 | 134.2036 | 55.67771 | 0.174114 | 2.520905 | 1 |
| 8 | 69851200 | 69855399 | 62.89128 | 99.48739 | 135.4149 | 56.27822 | -0.11677 | 2.396225 | 1 |
| 8 | 69926800 | 69927999 | 41.69279 | 66.77279 | 100.4883 | 42.36762 | -0.24319 | 2.001055 | 1 |
| 8 | 69988000 | 69988599 | 11.69875 | 21.47537 | 28.18493 | 11.99669 | 0.038894 | 0.254141 | 1 |
| 8 | 70116600 | 70123599 | 93.73873 | 146.0727 | 153.3646 | 63.65167 | 0.24041 | 2.709925 | 1 |
| 8 | 70151600 | 70153399 | 22.84379 | 39.11567 | 44.74728 | 19.62277 | 0.189823 | 0.953873 | 1 |
| 8 | 71628200 | 71630999 | 36.23973 | 59.53584 | 62.10491 | 26.75689 | 0.31702 | 1.427525 | 1 |
| 8 | 72219000 | 72225599 | 103.5692 | 159.7309 | 171.5371 | 71.25346 | 0.196365 | 2.899168 | 1 |
| 8 | 72950200 | 72961399 | 154.0244 | 228.8659 | 322.4122 | 132.8223 | -0.24008 | 3.642336 | 1 |
| 8 | 73345800 | 73373199 | 500.9435 | 678.7126 | 791.3556 | 329.7633 | -0.09923 | 5.070659 | 1 |
| 8 | 73585800 | 73599999 | 326.7566 | 457.3141 | 557.6128 | 231.1834 | -0.11651 | 4.567235 | 1 |
| 8 | 73615200 | 73616999 | 20.18714 | 34.24852 | 54.83418 | 24.02083 | -0.29503 | 1.040974 | 1 |
| 8 | 73692200 | 73695199 | 41.52729 | 66.67967 | 96.25204 | 40.74117 | -0.17875 | 1.919675 | 1 |
| 8 | 73809600 | 73811199 | 16.84044 | 29.83848 | 31.75242 | 14.03862 | 0.302596 | 0.509036 | 1 |
| 8 | 74254600 | 74258799 | 68.16581 | 107.0983 | 146.223 | 60.65265 | -0.12889 | 2.548169 | 1 |
| 8 | 74393600 | 74395599 | 29.84544 | 49.10796 | 68.2694 | 29.43227 | -0.09523 | 1.398428 | 1 |
| 8 | 74903000 | 74905999 | 36.66004 | 60.33115 | 60.23803 | 25.97168 | 0.381106 | 1.410315 | 1 |
| 8 | 74917400 | 74918399 | 11.04602 | 19.9289 | 31.79398 | 13.83505 | -0.26197 | 0.358335 | 1 |
| 8 | 74975600 | 74976999 | 19.52242 | 34.02795 | 38.78433 | 17.0628 | 0.199372 | 0.750226 | 1 |
| 8 | 75075800 | 75076799 | 13.16676 | 23.50191 | 33.55112 | 14.74773 | -0.11301 | 0.439336 | 1 |
| 8 | 75158800 | 75161399 | 25.91709 | 43.29276 | 63.82673 | 27.77766 | -0.17712 | 1.212925 | 1 |
| 8 | 75211200 | 75212999 | 17.73983 | 30.74823 | 46.87011 | 20.56419 | -0.21918 | 0.825937 | 1 |
| 8 | 75273400 | 75274999 | 18.94839 | 32.94812 | 40.4911 | 17.78679 | 0.090946 | 0.784682 | 1 |
| 8 | 75310600 | 75316999 | 14.81879 | 26.53834 | 28.37262 | 12.43673 | 0.30458 | 0.419563 | 1 |
| 8 | 75472000 | 75473999 | 31.17831 | 50.72918 | 72.88687 | 31.2853 | -0.15168 | 1.558092 | 1 |

| | | | | | | | | | |
|---|---|---|---|---|---|---|---|---|---|
| 8 | 75872400 | 75873399 | 16.60378 | 28.68373 | 48.46216 | 21.26055 | -0.36777 | 0.856012 | 1 |
| 8 | 75968800 | 75974999 | 81.6075 | 127.9606 | 139.796 | 57.98094 | 0.190071 | 2.59264 | 1 |
| 8 | 76065400 | 76066799 | 17.01339 | 29.24845 | 47.43026 | 20.80477 | -0.31045 | 0.897875 | 1 |
| 8 | 76109800 | 76113599 | 59.94271 | 94.87216 | 139.3895 | 57.94527 | -0.22628 | 2.382448 | 1 |
| 8 | 76264800 | 76268799 | 62.00686 | 98.3708 | 114.912 | 47.85472 | 0.107114 | 2.31731 | 1 |
| 8 | 76299000 | 76301799 | 31.51938 | 51.89137 | 63.23471 | 27.26818 | 0.094675 | 1.39403 | 1 |
| 8 | 76312200 | 76314999 | 15.42485 | 26.99204 | 41.50627 | 18.31129 | -0.23004 | 0.678562 | 1 |
| 8 | 76415200 | 76417399 | 31.51553 | 51.29758 | 80.54176 | 34.57587 | -0.27904 | 1.557325 | 1 |
| 8 | 76469200 | 76470799 | 18.97805 | 33.28451 | 34.8569 | 15.40702 | 0.321127 | 0.646777 | 1 |
| 8 | 76595400 | 76601599 | 127.2769 | 193.5871 | 201.8011 | 83.6063 | 0.223646 | 3.182571 | 1 |
| 8 | 76823600 | 76825399 | 27.68665 | 45.4345 | 68.19655 | 29.3677 | -0.20696 | 1.428567 | 1 |
| 8 | 77020400 | 77021799 | 12.65994 | 22.26905 | 38.33146 | 16.97201 | -0.39153 | 0.574657 | 1 |
| 8 | 77037600 | 77041399 | 45.22147 | 72.52383 | 94.34213 | 39.85957 | -0.03101 | 1.960207 | 1 |
| 8 | 77138400 | 77139199 | 12.00413 | 21.85104 | 27.17967 | 11.69629 | 0.105105 | 0.301992 | 1 |
| 8 | 77288600 | 77289999 | 24.13875 | 41.35256 | 44.22215 | 19.39323 | 0.286914 | 0.956598 | 1 |
| 8 | 77859600 | 77865199 | 55.19813 | 88.31545 | 94.3077 | 39.70494 | 0.24921 | 2.042739 | 1 |
| 8 | 77867000 | 77875999 | 98.82428 | 150.9305 | 224.4477 | 93.09027 | -0.2837 | 3.093046 | 1 |
| 8 | 78025800 | 78032599 | 139.2161 | 210.6789 | 223.575 | 92.48456 | 0.191867 | 3.28894 | 1 |
| 8 | 78375800 | 78377399 | 16.91119 | 29.71129 | 33.62248 | 14.88329 | 0.210108 | 0.608405 | 1 |
| 8 | 80492000 | 80496399 | 86.32662 | 134.1153 | 179.4729 | 74.52129 | -0.11262 | 2.761894 | 1 |
| 8 | 81072200 | 81073199 | 10.31296 | 18.95601 | 30.33022 | 12.88709 | -0.24327 | 0.240673 | 1 |
| 8 | 81237800 | 81245399 | 93.36922 | 143.948 | 201.1616 | 83.55912 | -0.1836 | 2.904442 | 1 |
| 8 | 81355800 | 81360999 | 51.91426 | 82.74782 | 118.2608 | 49.52514 | -0.17647 | 2.163286 | 1 |
| 8 | 81464800 | 81467599 | 21.58704 | 35.96809 | 61.18587 | 26.58565 | -0.3827 | 1.224727 | 1 |
| 8 | 81680200 | 81683399 | 42.07572 | 67.94523 | 87.07102 | 37.14887 | 0.002482 | 1.742344 | 1 |
| 8 | 81799400 | 81800199 | 8.878413 | 16.48628 | 28.90219 | 12.11129 | -0.35969 | 0.166917 | 1 |
| 8 | 81845400 | 81852799 | 124.5588 | 188.8552 | 227.249 | 94.06034 | 0.013048 | 3.247552 | 1 |
| 8 | 81916400 | 81916999 | 12.76746 | 23.36958 | 26.07662 | 11.14302 | 0.267434 | 0.273628 | 1 |
| 8 | 82380600 | 82383199 | 23.16422 | 39.74236 | 45.77736 | 20.07767 | 0.180348 | 0.938465 | 1 |
| 8 | 82879400 | 82880999 | 31.12807 | 51.42953 | 62.86976 | 27.16036 | 0.09157 | 1.349075 | 1 |
| 8 | 83358600 | 83359799 | 8.984654 | 16.90765 | 27.48814 | 11.26569 | -0.22603 | 0.080086 | 1 |
| 8 | 83445200 | 83447999 | 14.6682 | 25.95527 | 32.60158 | 14.41788 | 0.064111 | 0.512604 | 1 |
| 8 | 83743600 | 83749799 | 57.84577 | 92.63837 | 113.6508 | 47.91391 | 0.050873 | 2.026411 | 1 |
| 8 | 83752800 | 83753599 | 15.68466 | 27.39088 | 39.52477 | 17.40942 | -0.13883 | 0.70774 | 1 |
| 8 | 84490800 | 84496199 | 43.28007 | 69.41344 | 103.7845 | 43.85199 | -0.23143 | 1.959109 | 1 |
| 8 | 85117000 | 85118999 | 19.79942 | 34.09018 | 42.20426 | 18.5133 | 0.077806 | 0.889276 | 1 |
| 8 | 85849200 | 85859799 | 144.8292 | 215.5242 | 311.4056 | 128.3075 | -0.2746 | 3.614748 | 1 |
| 8 | 85900000 | 85907199 | 175.1502 | 259.9268 | 297.375 | 122.492 | 0.058282 | 3.666734 | 1 |
| 8 | 86698800 | 86700999 | 19.40368 | 34.05972 | 38.5293 | 17.04239 | 0.210512 | 0.644318 | 1 |
| 8 | 87010800 | 87012399 | 26.97934 | 45.73518 | 50.13579 | 21.95731 | 0.249287 | 1.064734 | 1 |
| 8 | 87104600 | 87110599 | 53.54562 | 85.06721 | 129.8333 | 54.14495 | -0.27608 | 2.272515 | 1 |
| 8 | 87620200 | 87624199 | 23.05304 | 38.83559 | 56.56069 | 24.72417 | -0.15945 | 1.112152 | 1 |
| 8 | 89192800 | 89194399 | 15.65908 | 27.95447 | 28.82856 | 12.67039 | 0.3537 | 0.446326 | 1 |
| 8 | 89218600 | 89221199 | 29.65612 | 49.15932 | 62.26728 | 26.96135 | 0.040847 | 1.307107 | 1 |
| 8 | 90093400 | 90094799 | 22.08085 | 37.79083 | 47.95238 | 21.02746 | 0.040905 | 0.961963 | 1 |
| 8 | 90649200 | 90652199 | 53.8021 | 85.8267 | 102.2831 | 42.88226 | 0.086513 | 2.137379 | 1 |
| 8 | 90811400 | 90822599 | 225.2922 | 326.8658 | 375.5091 | 154.817 | 0.021322 | 4.031756 | 1 |

| | | | | | | | | | |
|---|---|---|---|---|---|---|---|---|---|
| 8 | 90941400 | 90942599 | 21.51786 | 37.42483 | 36.86077 | 16.19447 | 0.409065 | 0.791057 | 1 |
| 8 | 92300600 | 92303399 | 13.13041 | 24.01334 | 22.63786 | 9.684853 | 0.507881 | 0.272356 | 1 |
| 8 | 93809400 | 93812999 | 62.67133 | 99.01631 | 134.1143 | 55.70268 | -0.11112 | 2.426629 | 1 |
| 8 | 93844200 | 93844599 | 9.220538 | 17.65086 | 21.33211 | 8.429735 | 0.245046 | -0.05368 | 1 |
| 8 | 93952800 | 93954399 | 14.7415 | 26.09908 | 37.61992 | 16.63541 | -0.13324 | 0.516646 | 1 |
| 8 | 94025000 | 94026999 | 19.40103 | 33.80931 | 39.41992 | 17.34015 | 0.166941 | 0.75656 | 1 |
| 8 | 94067400 | 94068599 | 17.23428 | 30.36911 | 33.94922 | 15.02637 | 0.228957 | 0.575704 | 1 |
| 8 | 94211800 | 94213199 | 14.14985 | 25.19769 | 31.53693 | 13.88972 | 0.073971 | 0.457929 | 1 |
| 8 | 94717200 | 94722399 | 23.82883 | 41.35892 | 45.73134 | 20.0657 | 0.242445 | 0.840652 | 1 |
| 8 | 94940200 | 94941799 | 13.14036 | 23.05833 | 39.40861 | 17.4267 | -0.38176 | 0.615023 | 1 |
| 8 | 95933400 | 95934999 | 12.95092 | 23.20065 | 33.15437 | 14.52884 | -0.11112 | 0.417213 | 1 |
| 8 | 96989600 | 96992199 | 36.11739 | 59.05292 | 69.07223 | 29.70073 | 0.149158 | 1.494658 | 1 |
| 8 | 97065600 | 97066799 | 16.41564 | 28.60211 | 44.15162 | 19.41955 | -0.2363 | 0.747995 | 1 |
| 8 | 97259600 | 97260999 | 20.36359 | 35.11644 | 43.72745 | 19.18238 | 0.069964 | 0.880877 | 1 |
| 8 | 97500000 | 97501799 | 28.67053 | 47.8594 | 49.70237 | 21.62134 | 0.326835 | 1.215149 | 1 |
| 8 | 97629800 | 97632799 | 50.03936 | 80.23305 | 84.06126 | 35.50416 | 0.280216 | 1.968891 | 1 |
| 8 | 97665800 | 97667599 | 20.56907 | 35.5428 | 42.62177 | 18.69849 | 0.124591 | 0.860391 | 1 |
| 8 | 97841600 | 97843399 | 23.93045 | 39.5873 | 64.87596 | 28.05133 | -0.32973 | 1.319279 | 1 |
| 8 | 97890400 | 97891799 | 15.68199 | 27.3943 | 41.53254 | 18.29717 | -0.20985 | 0.706019 | 1 |
| 8 | 97909600 | 97913199 | 29.13961 | 48.64796 | 59.56293 | 25.9127 | 0.090043 | 1.224793 | 1 |
| 8 | 97965000 | 97967399 | 16.07545 | 28.09691 | 43.59576 | 19.21369 | -0.24305 | 0.704325 | 1 |
| 8 | 98131800 | 98134599 | 21.8768 | 37.29677 | 52.23438 | 22.89799 | -0.10189 | 0.999452 | 1 |
| 8 | 98153400 | 98155999 | 44.50552 | 71.13811 | 107.824 | 45.33278 | -0.25663 | 2.067611 | 1 |
| 8 | 98174000 | 98175799 | 17.02831 | 29.95542 | 34.48346 | 15.268 | 0.18587 | 0.593102 | 1 |
| 8 | 98278200 | 98279999 | 22.06213 | 38.04461 | 42.08806 | 18.4632 | 0.239859 | 0.885203 | 1 |
| 8 | 98291400 | 98293599 | 29.19036 | 48.17314 | 60.93799 | 26.30511 | 0.042296 | 1.356238 | 1 |
| 8 | 98528400 | 98528999 | 9.917444 | 18.55843 | 26.85316 | 11.11068 | -0.07013 | 0.123093 | 1 |
| 8 | 99018200 | 99019599 | 16.2133 | 28.52767 | 32.59916 | 14.43309 | 0.196627 | 0.582998 | 1 |
| 8 | 99154800 | 99156399 | 21.89441 | 37.48692 | 44.77763 | 19.63595 | 0.12774 | 0.95114 | 1 |
| 8 | 99425200 | 99426599 | 33.63731 | 54.55136 | 76.20276 | 32.64184 | -0.11528 | 1.625003 | 1 |
| 8 | 100192800 | 100194199 | 47.85802 | 76.52208 | 97.49661 | 41.01002 | -0.00606 | 2.057415 | 1 |
| 8 | 100717000 | 100719799 | 28.17127 | 47.31178 | 55.62154 | 24.28895 | 0.148338 | 1.156513 | 1 |
| 8 | 100745600 | 100751599 | 63.74968 | 101.58 | 111.8439 | 46.81705 | 0.198731 | 2.184453 | 1 |
| 8 | 100836400 | 100839199 | 25.3204 | 43.20913 | 47.39502 | 20.77818 | 0.249494 | 0.997225 | 1 |
| 8 | 100842600 | 100844999 | 38.25688 | 61.51632 | 94.46175 | 40.06741 | -0.26521 | 1.869233 | 1 |
| 8 | 100976400 | 100980399 | 57.94818 | 92.43304 | 92.65119 | 38.83967 | 0.335614 | 2.140453 | 1 |
| 8 | 101055400 | 101055999 | 9.865798 | 18.6309 | 26.4422 | 10.76716 | -0.02384 | 0.062056 | 1 |
| 8 | 101086000 | 101087599 | 29.89613 | 49.70931 | 51.85639 | 22.50106 | 0.320366 | 1.264382 | 1 |
| 8 | 101249000 | 101250599 | 18.16722 | 31.28614 | 49.53098 | 21.72737 | -0.27546 | 0.89101 | 1 |
| 8 | 101833000 | 101836199 | 40.53444 | 65.73888 | 75.49861 | 32.34097 | 0.166561 | 1.637528 | 1 |
| 8 | 101934800 | 101936199 | 24.81756 | 41.34525 | 55.89992 | 24.28684 | -0.05237 | 1.233455 | 1 |
| 8 | 102272800 | 102278799 | 91.18651 | 141.1607 | 169.4453 | 70.38319 | 0.040048 | 2.824162 | 1 |
| 8 | 102437200 | 102437999 | 11.78619 | 21.60492 | 26.62901 | 11.35652 | 0.126807 | 0.25829 | 1 |
| 8 | 102706800 | 102709399 | 41.59886 | 67.03398 | 80.72198 | 34.34621 | 0.088589 | 1.800806 | 1 |
| 8 | 102874600 | 102875399 | 12.16387 | 21.85719 | 32.71381 | 14.29901 | -0.17482 | 0.392292 | 1 |
| 8 | 102906200 | 102908399 | 25.22474 | 43.07537 | 45.23224 | 19.8311 | 0.312278 | 0.98737 | 1 |
| 8 | 104358400 | 104362199 | 6.734141 | 12.86528 | 25.62644 | 10.1822 | -0.477 | -0.04861 | 1 |

| | | | | | | | | | |
|---|---|---|---|---|---|---|---|---|---|
| 8 | 104531200 | 104535999 | 38.46733 | 62.10066 | 85.0998 | 36.29456 | -0.09448 | 1.74329 | 1 |
| 8 | 104988600 | 105001999 | 138.8851 | 207.4685 | 307.2416 | 126.6662 | -0.30509 | 3.543862 | 1 |
| 8 | 105304200 | 105305999 | 16.91656 | 29.38401 | 46.85703 | 20.56951 | -0.28348 | 0.798092 | 1 |
| 8 | 105503800 | 105505799 | 29.45492 | 48.21492 | 72.92361 | 31.36995 | -0.21948 | 1.465511 | 1 |
| 8 | 107141800 | 107156799 | 263.0602 | 375.0588 | 510.6604 | 211.0772 | -0.24869 | 4.291968 | 1 |
| 8 | 107214800 | 107215199 | 6.368331 | 12.40048 | 17.96653 | 6.582454 | 0.088267 | -0.35229 | 1 |
| 8 | 108342000 | 108345399 | 54.10633 | 86.14179 | 112.3723 | 46.98366 | -0.04678 | 2.202657 | 1 |
| 8 | 108740800 | 108743199 | 8.207888 | 15.44449 | 25.91394 | 10.62217 | -0.27137 | 0.068396 | 1 |
| 8 | 110017200 | 110020799 | 47.59208 | 76.62566 | 84.17501 | 35.77076 | 0.219234 | 1.836873 | 1 |
| 8 | 111063200 | 111064799 | 26.02101 | 43.96724 | 46.94305 | 20.53874 | 0.287111 | 1.08677 | 1 |
| 8 | 113523400 | 113525399 | 16.02534 | 27.96073 | 43.62879 | 19.20628 | -0.25139 | 0.727991 | 1 |
| 8 | 113761800 | 113762599 | 9.792882 | 18.73527 | 21.25559 | 8.413955 | 0.332916 | -0.04347 | 1 |
| 8 | 114282800 | 114287799 | 61.97 | 98.28931 | 128.0197 | 53.3009 | -0.05004 | 2.325808 | 1 |
| 8 | 114288800 | 114291799 | 21.35757 | 35.98472 | 53.75005 | 23.49663 | -0.19559 | 1.104813 | 1 |
| 8 | 114328200 | 114328999 | 15.06343 | 26.3553 | 40.4876 | 17.85943 | -0.22866 | 0.676762 | 1 |
| 8 | 115047200 | 115053799 | 18.61341 | 32.61032 | 35.06439 | 15.48482 | 0.283419 | 0.663927 | 1 |
| 8 | 115463600 | 115469799 | 79.92271 | 125.7853 | 138.7116 | 57.54977 | 0.180706 | 2.520101 | 1 |
| 8 | 115703400 | 115704599 | 12.7638 | 22.71522 | 33.65596 | 14.83084 | -0.16862 | 0.456988 | 1 |
| 8 | 116009800 | 116011599 | 28.79667 | 47.17598 | 65.4628 | 28.17023 | -0.09498 | 1.449707 | 1 |
| 8 | 116459000 | 116466799 | 69.32732 | 108.8313 | 158.1397 | 65.58841 | -0.21961 | 2.567027 | 1 |
| 8 | 117390600 | 117399599 | 111.1592 | 169.0103 | 244.2586 | 101.1854 | -0.24776 | 3.188295 | 1 |
| 8 | 117438000 | 117448599 | 181.7441 | 271.3369 | 276.4552 | 113.9613 | 0.233342 | 3.561058 | 1 |
| 8 | 117520000 | 117524199 | 48.37291 | 77.38571 | 111.7208 | 47.02787 | -0.18525 | 2.04606 | 1 |
| 8 | 117584200 | 117584999 | 12.94284 | 23.08893 | 35.1551 | 15.46028 | -0.20576 | 0.446476 | 1 |
| 8 | 117608400 | 117616999 | 98.81817 | 151.3163 | 218.2725 | 90.58384 | -0.23683 | 3.039963 | 1 |
| 8 | 117615200 | 117626199 | 148.5765 | 221.6159 | 293.1704 | 120.8302 | -0.14433 | 3.573097 | 1 |
| 8 | 117628000 | 117634399 | 60.35244 | 95.72368 | 130.6149 | 54.38136 | -0.11706 | 2.325199 | 1 |
| 8 | 117733600 | 117736599 | 27.23703 | 44.94309 | 63.02606 | 27.20432 | -0.10607 | 1.353578 | 1 |
| 8 | 118203800 | 118208799 | 52.47687 | 83.58801 | 115.4531 | 48.3041 | -0.12833 | 2.186233 | 1 |
| 8 | 118224800 | 118226399 | 14.40782 | 25.15707 | 41.58248 | 18.31395 | -0.33396 | 0.705179 | 1 |
| 8 | 119778600 | 119780599 | 25.61525 | 43.27001 | 45.40761 | 19.86532 | 0.312016 | 1.087711 | 1 |
| 8 | 122138200 | 122141599 | 34.14541 | 55.6086 | 65.64716 | 28.18736 | 0.131905 | 1.54445 | 1 |
| 8 | 123895800 | 123896799 | 17.69796 | 30.6749 | 42.24236 | 18.53376 | -0.07295 | 0.823584 | 1 |
| 8 | 124839800 | 124847599 | 128.6942 | 193.6515 | 262.7329 | 108.4984 | -0.16938 | 3.401645 | 1 |
| 8 | 125579200 | 125586599 | 131.4753 | 200.718 | 198.7157 | 82.41033 | 0.302305 | 3.108859 | 1 |
| 8 | 126030200 | 126034999 | 42.15184 | 67.62023 | 101.0523 | 42.71316 | -0.23027 | 1.949657 | 1 |
| 8 | 126039600 | 126042999 | 32.45917 | 52.74135 | 81.21907 | 34.83962 | -0.25309 | 1.585319 | 1 |
| 8 | 128624800 | 128627399 | 25.30488 | 42.42857 | 60.41577 | 26.35535 | -0.12713 | 1.169884 | 1 |
| 8 | 128814800 | 128819399 | 62.01457 | 98.27996 | 121.0692 | 50.36602 | 0.029118 | 2.349002 | 1 |
| 8 | 129283800 | 129284799 | 19.17225 | 33.4517 | 36.8253 | 16.21134 | 0.249612 | 0.733716 | 1 |
| 8 | 129870600 | 129875799 | 60.52104 | 95.88941 | 137.0379 | 56.99833 | -0.18516 | 2.359139 | 1 |
| 8 | 129991600 | 129999599 | 160.5564 | 239.4351 | 290.8834 | 119.8827 | -0.02182 | 3.578554 | 1 |
| 8 | 130008600 | 130024199 | 62.72926 | 100.0033 | 108.7525 | 45.55404 | 0.217348 | 2.168704 | 1 |
| 9 | 3198400 | 3201199 | 62.51583 | 99.441 | 97.46729 | 40.70112 | 0.363615 | 2.237012 | 1 |
| 9 | 3387600 | 3391199 | 23.41434 | 40.20579 | 44.19508 | 19.385 | 0.247827 | 0.931552 | 1 |
| 9 | 3464400 | 3467399 | 30.24001 | 49.7667 | 67.03739 | 28.90347 | -0.0497 | 1.398105 | 1 |
| 9 | 3531200 | 3534199 | 26.53631 | 43.89162 | 63.1266 | 27.29216 | -0.14215 | 1.322805 | 1 |

| | | | | | | | | | |
|---|---|---|---|---|---|---|---|---|---|
| 9 | 3554800 | 3555999 | 14.71954 | 26.64158 | 29.88312 | 12.95168 | 0.246462 | 0.359655 | 1 |
| 9 | 3597000 | 3597999 | 20.73492 | 35.99186 | 41.51328 | 18.21586 | 0.181795 | 0.819144 | 1 |
| 9 | 3674600 | 3676599 | 35.11921 | 57.02357 | 78.96084 | 33.85345 | -0.10108 | 1.605288 | 1 |
| 9 | 3697600 | 3701199 | 42.84599 | 69.17287 | 80.58637 | 34.3718 | 0.139295 | 1.747161 | 1 |
| 9 | 4020200 | 4021799 | 17.67818 | 30.81743 | 44.1094 | 19.40711 | -0.12776 | 0.747014 | 1 |
| 9 | 4104200 | 4110799 | 72.18384 | 114.8361 | 114.4716 | 47.81148 | 0.339518 | 2.243545 | 1 |
| 9 | 4154800 | 4156799 | 29.41157 | 49.01285 | 52.92713 | 22.99943 | 0.270746 | 1.23929 | 1 |
| 9 | 4205400 | 4210999 | 32.17599 | 52.47805 | 76.03279 | 32.66414 | -0.16142 | 1.523613 | 1 |
| 9 | 4242600 | 4243799 | 16.78151 | 29.4846 | 33.66403 | 14.90217 | 0.197392 | 0.607665 | 1 |
| 9 | 4258400 | 4261799 | 119.4424 | 181.3424 | 210.2874 | 87.07498 | 0.067952 | 3.218492 | 1 |
| 9 | 4490400 | 4494999 | 80.41597 | 126.0308 | 133.3096 | 55.29345 | 0.236105 | 2.601164 | 1 |
| 9 | 4623000 | 4625199 | 26.80595 | 45.59724 | 48.48561 | 21.25078 | 0.293521 | 1.035047 | 1 |
| 9 | 4771200 | 4772199 | 23.93283 | 39.55138 | 62.07371 | 26.82103 | -0.2676 | 1.329005 | 1 |
| 9 | 4871400 | 4873799 | 31.27471 | 51.86586 | 56.78438 | 24.59497 | 0.250594 | 1.302575 | 1 |
| 9 | 5207800 | 5212999 | 72.37696 | 114.4247 | 128.1156 | 53.22866 | 0.163898 | 2.415529 | 1 |
| 9 | 5231200 | 5232199 | 13.00729 | 23.56694 | 29.08835 | 12.59086 | 0.110697 | 0.340902 | 1 |
| 9 | 5395800 | 5401599 | 46.00702 | 73.75362 | 100.6974 | 42.51833 | -0.1014 | 1.974958 | 1 |
| 9 | 5527000 | 5529599 | 19.73713 | 34.81111 | 33.19461 | 14.6897 | 0.457671 | 0.57975 | 1 |
| 9 | 5641200 | 5644999 | 86.98346 | 136.3255 | 142.8938 | 59.26868 | 0.24921 | 2.605408 | 1 |
| 9 | 5731400 | 5732999 | 14.19404 | 25.26289 | 36.58291 | 16.11937 | -0.1359 | 0.470741 | 1 |
| 9 | 5798000 | 5798999 | 17.59492 | 30.90761 | 29.19036 | 12.91963 | 0.469674 | 0.606184 | 1 |
| 9 | 5934000 | 5936599 | 31.9694 | 51.83117 | 76.72396 | 32.85796 | -0.19914 | 1.625705 | 1 |
| 9 | 6079600 | 6081799 | 19.174 | 33.56334 | 35.50618 | 15.66843 | 0.30672 | 0.680235 | 1 |
| 9 | 6129600 | 6134199 | 87.35185 | 135.9567 | 158.9721 | 65.99085 | 0.083893 | 2.727795 | 1 |
| 9 | 6220400 | 6222799 | 22.43397 | 37.6158 | 55.40761 | 24.17098 | -0.17579 | 1.15343 | 1 |
| 9 | 6441600 | 6442999 | 15.90628 | 27.82214 | 40.39794 | 17.81564 | -0.14794 | 0.682966 | 1 |
| 9 | 6500600 | 6501999 | 17.62364 | 30.7672 | 39.35323 | 17.33039 | 0.034029 | 0.719252 | 1 |
| 9 | 6663200 | 6665399 | 42.59926 | 68.66919 | 84.58131 | 36.00447 | 0.056602 | 1.794111 | 1 |
| 9 | 6692400 | 6704599 | 297.8208 | 420.2326 | 499.3682 | 206.7673 | -0.06751 | 4.444809 | 1 |
| 9 | 6715800 | 6720799 | 43.84076 | 71.48449 | 70.47831 | 30.27506 | 0.392297 | 1.55016 | 1 |
| 9 | 6902200 | 6903199 | 11.62904 | 21.78514 | 23.53758 | 9.720811 | 0.350069 | 0.121106 | 1 |
| 9 | 6951000 | 6952599 | 20.87892 | 36.05587 | 40.55996 | 17.79363 | 0.216274 | 0.86484 | 1 |
| 9 | 6978000 | 6980199 | 22.66677 | 39.09957 | 42.56219 | 18.67136 | 0.263098 | 0.884817 | 1 |
| 9 | 7190800 | 7192599 | 13.52815 | 24.31623 | 36.17441 | 15.80747 | -0.16635 | 0.40395 | 1 |
| 9 | 7403400 | 7418599 | 203.5025 | 295.7464 | 385.2248 | 158.795 | -0.15744 | 4.00497 | 1 |
| 9 | 7611200 | 7612799 | 23.35034 | 39.23289 | 55.38274 | 24.18992 | -0.11455 | 1.138223 | 1 |
| 9 | 7761200 | 7764799 | 32.47819 | 52.60801 | 84.23908 | 36.05213 | -0.31349 | 1.64896 | 1 |
| 9 | 7924200 | 7936399 | 142.8849 | 213.98 | 300.4813 | 123.9396 | -0.22517 | 3.498101 | 1 |
| 9 | 8063800 | 8067199 | 40.74321 | 65.756 | 92.85134 | 39.58596 | -0.13817 | 1.758073 | 1 |
| 9 | 8114600 | 8119999 | 22.50689 | 38.41972 | 42.22365 | 18.51149 | 0.246839 | 0.977728 | 1 |
| 9 | 8216800 | 8220799 | 27.02839 | 44.7102 | 67.15931 | 29.03776 | -0.20468 | 1.325748 | 1 |
| 9 | 8375600 | 8378199 | 21.22403 | 36.5236 | 43.17388 | 18.93833 | 0.144005 | 0.900808 | 1 |
| 9 | 9061400 | 9064999 | 30.05749 | 49.51576 | 67.11017 | 28.94707 | -0.05798 | 1.385032 | 1 |
| 9 | 9278000 | 9278799 | 11.3584 | 21.23545 | 24.09793 | 9.986135 | 0.27595 | 0.133298 | 1 |
| 9 | 9292400 | 9294199 | 31.89715 | 52.53872 | 60.03814 | 25.89506 | 0.187374 | 1.384976 | 1 |
| 9 | 9325000 | 9327399 | 17.91319 | 30.8711 | 48.47197 | 21.26329 | -0.26348 | 0.879763 | 1 |
| 9 | 9510000 | 9510799 | 12.19636 | 22.26814 | 28.64973 | 12.28239 | 0.060169 | 0.289752 | 1 |

| | | | | | | | | | | |
|---|---|---|---|---|---|---|---|---|---|---|
| 9 | 9887400 | 9889199 | 13.37876 | 24.77559 | 23.34466 | 9.818646 | 0.525817 | 0.205544 | | 1 |
| 9 | 9920600 | 9928399 | 197.1613 | 289.6773 | 327.3434 | 134.8136 | 0.06257 | 3.8351 | | 1 |
| 9 | 10158400 | 10161399 | 13.54132 | 24.73212 | 27.96203 | 11.98251 | 0.245617 | 0.295115 | | 1 |
| 9 | 10203600 | 10204399 | 9.87562 | 18.37625 | 25.7638 | 10.76223 | -0.03609 | 0.155001 | | 1 |
| 9 | 10557000 | 10559199 | 14.30371 | 25.32812 | 42.36926 | 18.73494 | -0.34692 | 0.520391 | | 1 |
| 9 | 10705600 | 10707799 | 16.35439 | 28.5574 | 45.17907 | 19.89862 | -0.27116 | 0.720582 | | 1 |
| 9 | 11140000 | 11141199 | 26.04409 | 42.82483 | 69.34543 | 29.8878 | -0.31496 | 1.399415 | | 1 |
| 9 | 11178400 | 11182999 | 50.28602 | 80.68631 | 103.5215 | 43.77102 | -0.01034 | 1.953683 | | 1 |
| 9 | 11454400 | 11456799 | 41.58895 | 67.22369 | 74.03361 | 31.6256 | 0.222249 | 1.709025 | | 1 |
| 9 | 11523800 | 11524599 | 9.936951 | 18.91591 | 22.77158 | 9.117011 | 0.233265 | -0.00199 | | 1 |
| 9 | 11632400 | 11634199 | 30.0528 | 48.83829 | 77.82224 | 33.38602 | -0.30198 | 1.576552 | | 1 |
| 9 | 11651800 | 11652599 | 13.5273 | 23.96838 | 35.07657 | 15.50829 | -0.1546 | 0.493802 | | 1 |
| 9 | 12021000 | 12026599 | 71.22966 | 112.7865 | 117.2249 | 48.74229 | 0.272891 | 2.374866 | | 1 |
| 9 | 12159800 | 12166999 | 83.02631 | 130.0338 | 154.3384 | 64.02013 | 0.069323 | 2.616386 | | 1 |
| 9 | 12220200 | 12227799 | 103.2495 | 159.5929 | 165.9833 | 68.94571 | 0.245 | 2.8562 | | 1 |
| 9 | 12286000 | 12287999 | 19.01721 | 33.29133 | 38.67707 | 17.06876 | 0.172305 | 0.684901 | | 1 |
| 9 | 12372400 | 12375199 | 22.10766 | 37.37577 | 57.07908 | 24.97772 | -0.22742 | 1.07939 | | 1 |
| 9 | 12442200 | 12447999 | 42.73127 | 69.00704 | 73.38022 | 31.30957 | 0.271192 | 1.737081 | | 1 |
| 9 | 12471200 | 12474199 | 29.27933 | 48.1012 | 70.8284 | 30.51531 | -0.17883 | 1.419633 | | 1 |
| 9 | 12476400 | 12477799 | 21.83517 | 37.2561 | 49.0189 | 21.48984 | -0.01193 | 0.988169 | | 1 |
| 9 | 12610000 | 12614599 | 66.52273 | 105.3772 | 126.1298 | 52.45594 | 0.069998 | 2.363553 | | 1 |
| 9 | 12737200 | 12740799 | 14.87467 | 26.09381 | 38.84269 | 17.17209 | -0.18347 | 0.629739 | | 1 |
| 9 | 12753400 | 12755999 | 19.95131 | 34.81563 | 37.23557 | 16.39331 | 0.291041 | 0.735736 | | 1 |
| 9 | 12806000 | 12808999 | 14.74112 | 25.85258 | 40.90734 | 18.08075 | -0.27119 | 0.636869 | | 1 |
| 9 | 13080200 | 13081199 | 11.16875 | 20.58406 | 26.69075 | 11.29945 | 0.061806 | 0.223492 | | 1 |
| 9 | 13082600 | 13086599 | 58.55606 | 92.94112 | 126.6265 | 52.75867 | -0.11386 | 2.302227 | | 1 |
| 9 | 13444600 | 13446799 | 30.53659 | 50.92141 | 73.39652 | 31.90774 | -0.14467 | 1.254217 | | 1 |
| 9 | 13803000 | 13804599 | 23.87264 | 40.63809 | 45.84272 | 20.09364 | 0.208714 | 1.012828 | | 1 |
| 9 | 13839600 | 13841199 | 10.28427 | 19.76323 | 21.84272 | 8.53645 | 0.387481 | -0.07798 | | 1 |
| 9 | 14030800 | 14037399 | 61.38748 | 97.87479 | 103.4178 | 43.32592 | 0.258955 | 2.160597 | | 1 |
| 9 | 15045400 | 15047799 | 62.38865 | 97.97138 | 153.2004 | 63.54261 | -0.32493 | 2.558168 | | 1 |
| 9 | 15133800 | 15146599 | 153.5977 | 229.7059 | 303.6477 | 125.2121 | -0.13985 | 3.525566 | | 1 |
| 9 | 15459000 | 15462599 | 30.32111 | 49.90505 | 67.78169 | 29.22558 | -0.06159 | 1.395596 | | 1 |
| 9 | 15759000 | 15761599 | 17.66969 | 30.51035 | 47.41329 | 20.8001 | -0.2479 | 0.862381 | | 1 |
| 9 | 15767400 | 15768999 | 18.46334 | 32.58328 | 39.00126 | 17.25734 | 0.131559 | 0.575271 | | 1 |
| 9 | 16517400 | 16519399 | 25.30985 | 42.05658 | 60.795 | 26.36795 | -0.14881 | 1.267249 | | 1 |
| 9 | 16527000 | 16527799 | 11.57522 | 21.49366 | 24.12728 | 10.11059 | 0.279044 | 0.178601 | | 1 |
| 9 | 16854800 | 16863199 | 91.96322 | 143.1667 | 165.2215 | 68.58268 | 0.103043 | 2.726889 | | 1 |
| 9 | 16995000 | 17001399 | 74.11462 | 116.7393 | 127.3397 | 52.83729 | 0.197327 | 2.498409 | | 1 |
| 9 | 17107800 | 17110599 | 23.40482 | 39.37615 | 57.64968 | 25.19011 | -0.16707 | 1.126199 | | 1 |
| 9 | 17299800 | 17302799 | 32.04338 | 52.60994 | 65.36719 | 28.15515 | 0.065185 | 1.430697 | | 1 |
| 9 | 17542200 | 17548999 | 110.976 | 170.0731 | 185.6368 | 77.0536 | 0.165971 | 3.023756 | | 1 |
| 9 | 17666000 | 17667799 | 17.85374 | 31.34283 | 32.4287 | 14.34541 | 0.338534 | 0.621569 | | 1 |
| 9 | 17753200 | 17754799 | 13.42313 | 24.53705 | 26.87186 | 11.50314 | 0.292067 | 0.285532 | | 1 |
| 9 | 17767200 | 17767999 | 18.58839 | 31.69664 | 51.0912 | 22.39721 | -0.30361 | 0.986462 | | 1 |
| 9 | 17883400 | 17885199 | 21.16064 | 36.60481 | 39.98189 | 17.54094 | 0.259061 | 0.849585 | | 1 |
| 9 | 17919800 | 17922799 | 48.51444 | 77.28338 | 121.7033 | 50.92542 | -0.31705 | 2.18323 | | 1 |

| | | | | | | | | | |
|---|---|---|---|---|---|---|---|---|---|
| 9 | 17944800 | 17945999 | 11.84765 | 21.66496 | 31.2196 | 13.35825 | -0.10082 | 0.282403 | 1 |
| 9 | 18011000 | 18012799 | 23.102 | 38.92641 | 58.18676 | 25.43663 | -0.19682 | 1.112945 | 1 |
| 9 | 18027600 | 18041999 | 282.8396 | 401.6837 | 471.6507 | 195.0812 | -0.04087 | 4.349442 | 1 |
| 9 | 18133600 | 18136199 | 38.99817 | 63.3496 | 76.26939 | 32.71008 | 0.100796 | 1.601659 | 1 |
| 9 | 18172000 | 18174399 | 27.60714 | 45.90155 | 53.93677 | 23.40313 | 0.149173 | 1.259318 | 1 |
| 9 | 18372400 | 18377399 | 44.71235 | 71.96881 | 94.29624 | 40.05544 | -0.03502 | 1.845824 | 1 |
| 9 | 18384400 | 18386999 | 17.55249 | 30.70138 | 41.90874 | 18.48182 | -0.05928 | 0.69487 | 1 |
| 9 | 18690000 | 18693599 | 47.74205 | 76.59202 | 91.19068 | 38.54784 | 0.097048 | 1.950534 | 1 |
| 9 | 18724600 | 18725799 | 15.68201 | 27.77635 | 33.34445 | 14.74352 | 0.129353 | 0.50748 | 1 |
| 9 | 18804000 | 18804999 | 11.41762 | 20.78503 | 33.29559 | 14.32677 | -0.25782 | 0.309479 | 1 |
| 9 | 18836400 | 18842599 | 89.53217 | 139.9336 | 160.5256 | 66.60076 | 0.116097 | 2.656973 | 1 |
| 9 | 18890200 | 18892999 | 18.44389 | 31.76663 | 44.82093 | 19.66132 | -0.10988 | 0.885082 | 1 |
| 9 | 18926400 | 18934399 | 148.3049 | 223.4359 | 238.5109 | 98.52824 | 0.177887 | 3.380769 | 1 |
| 9 | 18975000 | 18977999 | 34.13429 | 56.55885 | 62.03995 | 26.85515 | 0.247753 | 1.324448 | 1 |
| 9 | 19013800 | 19015999 | 29.92909 | 49.75434 | 59.57274 | 25.84577 | 0.12208 | 1.272456 | 1 |
| 9 | 19414400 | 19416599 | 31.39867 | 52.26187 | 56.31097 | 24.45146 | 0.273929 | 1.260372 | 1 |
| 9 | 19605600 | 19617799 | 60.23513 | 96.24224 | 104.6274 | 43.96409 | 0.221296 | 2.093933 | 1 |
| 9 | 19668400 | 19688599 | 267.9486 | 384.5751 | 410.0675 | 169.3176 | 0.113422 | 4.194099 | 1 |
| 9 | 19884600 | 19891399 | 78.59334 | 124.4822 | 131.4825 | 54.67857 | 0.250027 | 2.378661 | 1 |
| 9 | 19979400 | 19980199 | 19.56845 | 34.10472 | 35.31348 | 15.53484 | 0.337389 | 0.748273 | 1 |
| 9 | 19981000 | 19994199 | 213.0501 | 313.5744 | 311.6391 | 128.3345 | 0.25018 | 3.809269 | 1 |
| 9 | 20001600 | 20006799 | 66.58621 | 105.8161 | 116.1371 | 48.44755 | 0.199305 | 2.271546 | 1 |
| 9 | 20280200 | 20284199 | 42.99596 | 69.473 | 74.9156 | 31.98754 | 0.251954 | 1.72094 | 1 |
| 9 | 21429200 | 21433799 | 49.27668 | 78.54427 | 118.3933 | 49.58112 | -0.25312 | 2.163039 | 1 |
| 9 | 21790400 | 21793799 | 48.98178 | 78.24849 | 97.96295 | 41.15371 | 0.017776 | 2.085972 | 1 |
| 9 | 22093800 | 22101599 | 108.1676 | 166.2379 | 169.8089 | 70.51107 | 0.264644 | 2.966592 | 1 |
| 9 | 22135000 | 22143599 | 100.4681 | 154.2991 | 205.1623 | 85.18316 | -0.11613 | 2.981055 | 1 |
| 9 | 22144800 | 22168399 | 379.4323 | 531.8673 | 522.3386 | 216.5063 | 0.197823 | 4.546025 | 1 |
| 9 | 22169400 | 22171399 | 18.50799 | 32.20522 | 41.01618 | 18.0255 | 0.039814 | 0.773829 | 1 |
| 9 | 22206000 | 22217599 | 126.6048 | 193.0742 | 205.5741 | 85.23336 | 0.196202 | 3.128971 | 1 |
| 9 | 22397000 | 22399399 | 14.77134 | 26.3606 | 32.43117 | 14.25791 | 0.100251 | 0.449947 | 1 |
| 9 | 22404800 | 22407799 | 32.50508 | 53.98638 | 56.93752 | 24.6866 | 0.304461 | 1.287834 | 1 |
| 9 | 22791200 | 22794199 | 24.6332 | 41.09678 | 61.98194 | 26.95357 | -0.20971 | 1.221334 | 1 |
| 9 | 22814400 | 22815599 | 13.17521 | 23.13034 | 39.42356 | 17.44023 | -0.37769 | 0.609266 | 1 |
| 9 | 23202000 | 23204999 | 16.73262 | 29.58535 | 32.87846 | 14.54257 | 0.23928 | 0.533783 | 1 |
| 9 | 23927800 | 23929199 | 18.17636 | 31.84356 | 33.53133 | 14.80739 | 0.313354 | 0.658172 | 1 |
| 9 | 24465400 | 24466399 | 9.811878 | 18.17854 | 29.90154 | 12.56862 | -0.27167 | 0.190468 | 1 |
| 9 | 24482800 | 24485399 | 35.34473 | 57.7225 | 58.80619 | 25.27568 | 0.346734 | 1.501931 | 1 |
| 9 | 24513000 | 24515599 | 25.50365 | 42.51946 | 61.9642 | 26.93406 | -0.16031 | 1.232821 | 1 |
| 9 | 25506800 | 25509199 | 19.06199 | 33.17732 | 44.66982 | 19.63418 | -0.04005 | 0.778529 | 1 |
| 9 | 25562200 | 25564399 | 18.74773 | 32.21034 | 51.1476 | 22.43492 | -0.28063 | 0.91447 | 1 |
| 9 | 25687400 | 25693199 | 64.52423 | 102.3 | 120.0735 | 49.97271 | 0.099355 | 2.339297 | 1 |
| 9 | 25705200 | 25707799 | 30.36724 | 50.4315 | 55.14834 | 23.90947 | 0.252422 | 1.281442 | 1 |
| 9 | 25736000 | 25736999 | 16.60092 | 29.19844 | 32.64564 | 14.45479 | 0.22783 | 0.590788 | 1 |
| 9 | 25943000 | 25944999 | 13.16379 | 23.97681 | 26.46189 | 11.37858 | 0.277014 | 0.304034 | 1 |
| 9 | 26474600 | 26479799 | 31.39191 | 51.34604 | 73.14339 | 31.45565 | -0.13409 | 1.480845 | 1 |
| 9 | 27031800 | 27034599 | 20.53209 | 34.88025 | 54.30011 | 23.7965 | -0.25458 | 1.030992 | 1 |

| | | | | | | | | | |
|---|---|---|---|---|---|---|---|---|---|
| 9 | 29425200 | 29426799 | 17.63849 | 30.86055 | 36.22573 | 15.97943 | 0.157483 | 0.681985 | 1 |
| 9 | 29964600 | 29967599 | 27.7343 | 46.79572 | 50.15653 | 21.93527 | 0.28148 | 1.108675 | 1 |
| 9 | 30241400 | 30244199 | 23.67532 | 40.66877 | 48.471 | 21.26075 | 0.131658 | 0.934975 | 1 |
| 9 | 31104800 | 31106199 | 10.90739 | 20.54979 | 23.15651 | 9.471767 | 0.301171 | 0.078878 | 1 |
| 9 | 31678800 | 31680399 | 17.2104 | 30.34413 | 32.28035 | 14.2861 | 0.300225 | 0.563974 | 1 |
| 9 | 33070800 | 33074199 | 33.27035 | 54.02485 | 80.65081 | 34.57956 | -0.20925 | 1.599467 | 1 |
| 9 | 33818200 | 33819599 | 26.90814 | 44.546 | 60.85496 | 26.3261 | -0.06798 | 1.313856 | 1 |
| 9 | 33875200 | 33876399 | 13.02076 | 23.78698 | 25.03569 | 10.72533 | 0.348637 | 0.283409 | 1 |
| 9 | 34329200 | 34330599 | 12.49134 | 22.23968 | 33.2381 | 14.64151 | -0.1805 | 0.45317 | 1 |
| 9 | 34768800 | 34770399 | 13.72173 | 24.70375 | 27.98706 | 12.21092 | 0.226071 | 0.378942 | 1 |
| 9 | 35647800 | 35649999 | 35.40298 | 57.91837 | 63.91336 | 27.48825 | 0.23353 | 1.481134 | 1 |
| 9 | 35682600 | 35685599 | 13.79836 | 24.22875 | 39.88492 | 17.64704 | -0.32788 | 0.610484 | 1 |
| 9 | 36023600 | 36024399 | 14.98663 | 26.49693 | 31.93379 | 14.12487 | 0.122696 | 0.519189 | 1 |
| 9 | 36234000 | 36235799 | 27.49734 | 46.34552 | 49.62443 | 21.69542 | 0.28283 | 1.115635 | 1 |
| 9 | 36517000 | 36519999 | 37.34051 | 61.36215 | 68.93131 | 29.70161 | 0.21068 | 1.434589 | 1 |
| 9 | 36532400 | 36540999 | 84.70319 | 132.8336 | 144.8469 | 60.07493 | 0.192951 | 2.594096 | 1 |
| 9 | 36569600 | 36573399 | 33.26895 | 54.22503 | 78.14697 | 33.56438 | -0.15439 | 1.539536 | 1 |
| 9 | 37242000 | 37244599 | 32.81633 | 53.15068 | 80.02581 | 34.24657 | -0.22496 | 1.650485 | 1 |
| 9 | 38082400 | 38084599 | 33.02796 | 54.60245 | 59.12916 | 25.55534 | 0.265981 | 1.343588 | 1 |
| 9 | 38188600 | 38190399 | 27.35788 | 46.12396 | 48.54714 | 21.22632 | 0.307535 | 1.112153 | 1 |
| 9 | 38462200 | 38463999 | 36.95127 | 60.01401 | 68.50148 | 29.3748 | 0.177328 | 1.596033 | 1 |
| 9 | 38604400 | 38605999 | 23.32378 | 39.11001 | 57.00929 | 24.8706 | -0.16071 | 1.159519 | 1 |
| 9 | 39096800 | 39101999 | 27.56536 | 46.33405 | 49.10563 | 21.45091 | 0.2974 | 1.137175 | 1 |
| 9 | 39311600 | 39314399 | 30.83432 | 51.36581 | 57.45691 | 24.96233 | 0.220018 | 1.249455 | 1 |
| 9 | 39442200 | 39444199 | 22.07327 | 37.26808 | 53.28116 | 23.30636 | -0.13261 | 1.086652 | 1 |
| 9 | 39975000 | 39975999 | 13.25636 | 23.42122 | 36.1985 | 16.01348 | -0.23445 | 0.526295 | 1 |
| 9 | 40078800 | 40081199 | 27.06106 | 45.4594 | 52.32764 | 22.85428 | 0.178656 | 1.145522 | 1 |
| 9 | 40164400 | 40166199 | 23.14584 | 38.77635 | 56.46234 | 24.61786 | -0.15915 | 1.167705 | 1 |
| 9 | 41201400 | 41202999 | 16.91576 | 29.87642 | 32.91832 | 14.56339 | 0.251028 | 0.547225 | 1 |
| 9 | 41817800 | 41820999 | 43.56275 | 69.66253 | 105.6813 | 44.46028 | -0.25717 | 2.053227 | 1 |
| 9 | 41822600 | 41827599 | 19.81086 | 33.33115 | 57.91696 | 25.30724 | -0.41307 | 1.117233 | 1 |
| 9 | 45210800 | 45213399 | 14.83391 | 26.86449 | 28.32269 | 12.26628 | 0.335996 | 0.353611 | 1 |
| 9 | 45726600 | 45728799 | 16.13836 | 28.00194 | 44.77741 | 19.64666 | -0.28755 | 0.806077 | 1 |
| 9 | 46274000 | 46275599 | 19.09619 | 33.15536 | 38.52079 | 16.90534 | 0.171466 | 0.804214 | 1 |
| 9 | 50332000 | 50335799 | 33.31744 | 53.93634 | 84.70695 | 36.23697 | -0.28634 | 1.663551 | 1 |
| 9 | 50471400 | 50472999 | 22.36615 | 37.44034 | 60.21476 | 26.24075 | -0.30209 | 1.177144 | 1 |
| 9 | 51105400 | 51107599 | 16.59639 | 29.06815 | 39.82111 | 17.58115 | -0.06433 | 0.66547 | 1 |
| 9 | 51354800 | 51357799 | 26.35096 | 44.71707 | 47.42465 | 20.77659 | 0.296938 | 1.051564 | 1 |
| 9 | 52295800 | 52296999 | 12.45707 | 22.01015 | 35.2357 | 15.5874 | -0.2849 | 0.518022 | 1 |
| 9 | 52788400 | 52791199 | 12.79508 | 22.76725 | 33.68945 | 14.84759 | -0.1668 | 0.459205 | 1 |
| 9 | 52918600 | 52925399 | 117.172 | 178.3585 | 221.7419 | 91.88954 | -0.02937 | 3.164912 | 1 |
| 9 | 53240800 | 53244799 | 23.28713 | 39.28822 | 53.33984 | 23.32676 | -0.05844 | 1.095695 | 1 |
| 9 | 53272000 | 53274799 | 30.84555 | 50.64111 | 71.71287 | 30.88782 | -0.12309 | 1.430382 | 1 |
| 9 | 53641000 | 53643399 | 48.17563 | 77.25912 | 89.44899 | 37.79041 | 0.136709 | 1.963878 | 1 |
| 9 | 53908800 | 53913399 | 83.95047 | 130.088 | 180.3408 | 74.9085 | -0.16638 | 2.805291 | 1 |
| 9 | 54264200 | 54264799 | 10.4475 | 19.69746 | 23.20393 | 9.479647 | 0.23903 | 0.070392 | 1 |
| 9 | 55578200 | 55578999 | 10.46485 | 19.8556 | 23.26206 | 9.378194 | 0.263594 | 0.027759 | 1 |

| | | | | | | | | | |
|---|---|---|---|---|---|---|---|---|---|
| 9 | 55686800 | 55690399 | 32.01075 | 51.91934 | 81.27955 | 34.8194 | -0.27915 | 1.62145 | 1 |
| 9 | 55733800 | 55736399 | 39.52664 | 63.82567 | 89.33408 | 38.11192 | -0.12455 | 1.741823 | 1 |
| 9 | 57127400 | 57130799 | 45.76102 | 73.47333 | 87.54853 | 37.05383 | 0.097267 | 1.921311 | 1 |
| 9 | 57240600 | 57249599 | 66.55507 | 105.6236 | 115.737 | 48.21535 | 0.199762 | 2.310326 | 1 |
| 9 | 57258800 | 57264399 | 60.34297 | 96.47336 | 97.55192 | 41.02642 | 0.32655 | 2.073251 | 1 |
| 9 | 57271600 | 57282999 | 168.0361 | 251.2596 | 259.5907 | 107.0415 | 0.215275 | 3.529133 | 1 |
| 9 | 58690600 | 58690999 | 10.01971 | 18.33308 | 31.84985 | 13.6171 | -0.36781 | 0.267012 | 1 |
| 9 | 59276600 | 59278599 | 29.24928 | 48.52911 | 51.8253 | 22.45415 | 0.286465 | 1.287225 | 1 |
| 9 | 62351400 | 62353599 | 16.74931 | 28.3445 | 56.79247 | 24.85968 | -0.61733 | 1.053549 | 1 |
| 9 | 62696600 | 62697799 | 11.68928 | 21.34642 | 30.17821 | 12.93454 | -0.07491 | 0.286937 | 1 |
| 9 | 62923000 | 62928199 | 54.14381 | 87.00755 | 95.60651 | 40.51608 | 0.21543 | 1.9109 | 1 |
| 9 | 62968000 | 62972599 | 37.43757 | 61.01076 | 72.45029 | 31.11602 | 0.124046 | 1.545819 | 1 |
| 9 | 63377800 | 63380399 | 26.41268 | 43.77398 | 61.97572 | 26.83135 | -0.11927 | 1.300305 | 1 |
| 9 | 63750000 | 63751599 | 13.99165 | 24.66395 | 38.79389 | 17.16946 | -0.26081 | 0.562948 | 1 |
| 9 | 65246000 | 65248399 | 29.65693 | 49.53331 | 52.96839 | 23.05014 | 0.284702 | 1.214825 | 1 |
| 9 | 65616800 | 65621399 | 63.99381 | 101.4435 | 134.7718 | 56.08128 | -0.07934 | 2.347454 | 1 |
| 9 | 66941000 | 66943799 | 28.49344 | 46.68512 | 71.66215 | 30.83961 | -0.24018 | 1.451376 | 1 |
| 9 | 67274800 | 67275799 | 16.08207 | 28.6276 | 29.87981 | 13.16413 | 0.334207 | 0.470185 | 1 |
| 9 | 67459000 | 67459999 | 18.23826 | 31.00586 | 53.19853 | 23.31535 | -0.39417 | 1.012942 | 1 |
| 9 | 67502600 | 67506199 | 28.74445 | 48.05593 | 49.68354 | 21.63393 | 0.333273 | 1.20047 | 1 |
| 9 | 67732600 | 67733199 | 10.61222 | 20.02366 | 25.84453 | 10.54248 | 0.109802 | 0.072706 | 1 |
| 9 | 68084000 | 68086399 | 33.37949 | 55.05547 | 65.18187 | 28.12985 | 0.137235 | 1.37816 | 1 |
| 9 | 68090800 | 68093599 | 9.370636 | 17.33401 | 29.90546 | 12.59694 | -0.3421 | 0.196015 | 1 |
| 9 | 68434200 | 68436199 | 11.57455 | 21.09982 | 29.70188 | 12.75839 | -0.07063 | 0.296202 | 1 |
| 9 | 69866600 | 69868999 | 17.96267 | 31.16837 | 48.47405 | 21.26863 | -0.24768 | 0.820453 | 1 |
| 9 | 70585800 | 70587399 | 13.66201 | 23.91816 | 41.03609 | 18.11085 | -0.38719 | 0.66027 | 1 |
| 9 | 70749600 | 70751799 | 16.19009 | 28.44038 | 33.62318 | 14.88183 | 0.147222 | 0.60503 | 1 |
| 9 | 70992000 | 70994199 | 26.90811 | 44.75762 | 58.74339 | 25.49593 | -0.00983 | 1.25834 | 1 |
| 9 | 71320200 | 71322199 | 17.30758 | 30.37176 | 36.5577 | 16.16643 | 0.121324 | 0.632394 | 1 |
| 9 | 71331400 | 71332199 | 10.00993 | 18.54556 | 28.87215 | 12.13585 | -0.19253 | 0.191417 | 1 |
| 9 | 71336800 | 71338999 | 24.63756 | 41.80206 | 50.1057 | 21.94997 | 0.12101 | 1.050558 | 1 |
| 9 | 71581800 | 71586999 | 20.55652 | 35.44318 | 41.11252 | 18.03517 | 0.171749 | 0.878092 | 1 |
| 9 | 71967800 | 71968999 | 17.82392 | 30.72819 | 50.06633 | 21.96293 | -0.31648 | 0.879747 | 1 |
| 9 | 72146000 | 72157799 | 245.3391 | 352.9419 | 453.0772 | 187.0084 | -0.15086 | 4.159847 | 1 |
| 9 | 72178800 | 72183199 | 57.18386 | 91.37614 | 98.40398 | 41.35489 | 0.23507 | 2.087089 | 1 |
| 9 | 72884000 | 72886199 | 21.71865 | 37.24114 | 44.80451 | 19.65008 | 0.117876 | 0.939704 | 1 |
| 9 | 73073000 | 73075799 | 19.87375 | 34.62288 | 36.77859 | 16.17527 | 0.300548 | 0.757056 | 1 |
| 9 | 73076600 | 73081199 | 59.49667 | 94.96171 | 103.9603 | 43.61829 | 0.209569 | 2.126237 | 1 |
| 9 | 73247800 | 73250799 | 22.23045 | 38.77589 | 39.67408 | 17.46211 | 0.354525 | 0.755879 | 1 |
| 9 | 73779000 | 73781799 | 20.72585 | 36.19045 | 42.35805 | 18.65606 | 0.161464 | 0.736297 | 1 |
| 9 | 73807000 | 73808799 | 23.96208 | 40.32217 | 57.08089 | 24.94317 | -0.11861 | 1.127456 | 1 |
| 9 | 74534200 | 74535999 | 19.07568 | 32.78739 | 47.94628 | 21.03098 | -0.16202 | 0.909216 | 1 |
| 9 | 74542400 | 74542999 | 16.20834 | 28.67131 | 29.05571 | 12.85056 | 0.371989 | 0.518785 | 1 |
| 9 | 75006000 | 75007199 | 12.20329 | 22.11338 | 31.6457 | 13.69537 | -0.10143 | 0.338648 | 1 |
| 9 | 75404000 | 75406599 | 9.961826 | 18.3917 | 27.30827 | 11.53351 | -0.12973 | 0.20584 | 1 |
| 9 | 75831200 | 75832599 | 15.56926 | 27.21703 | 40.51993 | 17.86078 | -0.18356 | 0.694241 | 1 |
| 9 | 75885800 | 75886599 | 11.75041 | 22.00524 | 23.61436 | 9.757975 | 0.359518 | 0.126247 | 1 |

| | | | | | | | | | |
|---|---|---|---|---|---|---|---|---|---|
| 9 | 76347400 | 76348199 | 8.129859 | 15.67012 | 20.39339 | 7.901587 | 0.165721 | -0.13213 | 1 |
| 9 | 76351000 | 76355399 | 41.93312 | 67.57543 | 88.02384 | 37.45533 | -0.02446 | 1.802659 | 1 |
| 9 | 76578600 | 76582799 | 72.38596 | 114.3613 | 124.8594 | 51.85796 | 0.199416 | 2.431002 | 1 |
| 9 | 76593800 | 76597799 | 12.42442 | 22.9182 | 25.08772 | 10.60825 | 0.305791 | 0.225404 | 1 |
| 9 | 76645400 | 76646599 | 12.34237 | 21.92932 | 34.73093 | 15.32564 | -0.26549 | 0.467695 | 1 |
| 9 | 76728000 | 76729599 | 19.14019 | 33.52327 | 35.0685 | 15.48275 | 0.322945 | 0.672133 | 1 |
| 9 | 77064400 | 77066199 | 21.01727 | 36.74066 | 38.04022 | 16.76768 | 0.337659 | 0.715412 | 1 |
| 9 | 77108200 | 77108999 | 11.07139 | 20.32022 | 30.78818 | 13.1078 | -0.16741 | 0.2552 | 1 |
| 9 | 77516600 | 77518399 | 18.74905 | 32.20874 | 47.9482 | 21.03148 | -0.18783 | 0.912418 | 1 |
| 9 | 77770800 | 77783599 | 158.2261 | 237.8143 | 265.6249 | 109.6588 | 0.10952 | 3.430281 | 1 |
| 9 | 78087400 | 78092199 | 57.71985 | 91.76405 | 116.0938 | 48.44959 | -0.00481 | 2.251767 | 1 |
| 9 | 78161400 | 78163599 | 18.26744 | 31.58501 | 47.40618 | 20.79784 | -0.19764 | 0.85194 | 1 |
| 9 | 78714200 | 78717199 | 49.81098 | 79.66512 | 105.1194 | 44.23242 | -0.05614 | 2.053628 | 1 |
| 9 | 78971600 | 78976199 | 53.92748 | 86.05926 | 111.0848 | 46.59709 | -0.02793 | 2.127037 | 1 |
| 9 | 79309800 | 79310999 | 20.57685 | 34.64631 | 55.8411 | 24.40581 | -0.30497 | 1.110231 | 1 |
| 9 | 79390400 | 79392799 | 9.974601 | 19.04054 | 21.24723 | 8.456738 | 0.349613 | -0.02352 | 1 |
| 9 | 79968200 | 79975599 | 100.893 | 155.1614 | 188.9579 | 78.46966 | 0.011989 | 2.953342 | 1 |
| 9 | 80469600 | 80471399 | 23.18093 | 39.86014 | 43.11744 | 18.91312 | 0.271258 | 0.916123 | 1 |
| 9 | 80837800 | 80838799 | 8.686624 | 16.29497 | 25.8944 | 10.66976 | -0.19937 | 0.093988 | 1 |
| 9 | 80839600 | 80841199 | 19.76524 | 33.91359 | 45.35363 | 19.89195 | -0.03396 | 0.92625 | 1 |
| 9 | 80945400 | 80948199 | 21.84473 | 36.94319 | 59.2365 | 25.92435 | -0.29745 | 1.07898 | 1 |
| 9 | 81016600 | 81017799 | 14.70833 | 25.7337 | 38.47555 | 16.97172 | -0.18972 | 0.671141 | 1 |
| 9 | 81069600 | 81072199 | 40.94718 | 67.03851 | 64.04029 | 27.55062 | 0.441729 | 1.482807 | 1 |
| 9 | 81141000 | 81142799 | 31.21048 | 51.67463 | 59.04168 | 25.54378 | 0.18899 | 1.321382 | 1 |
| 9 | 81501000 | 81501599 | 10.51608 | 20.25468 | 25.46475 | 9.887264 | 0.211782 | -0.08709 | 1 |
| 9 | 81980800 | 81981799 | 11.73929 | 21.82083 | 26.12848 | 10.93383 | 0.187794 | 0.178571 | 1 |
| 9 | 82015200 | 82016399 | 13.79312 | 24.91125 | 29.00411 | 12.60451 | 0.190647 | 0.364397 | 1 |
| 9 | 82261600 | 82263599 | 20.17394 | 34.35577 | 55.41341 | 24.28943 | -0.30506 | 1.007518 | 1 |
| 9 | 82447000 | 82447999 | 18.03423 | 31.1925 | 40.19222 | 17.63314 | 0.021793 | 0.838194 | 1 |
| 9 | 82646400 | 82654399 | 79.54766 | 123.6477 | 163.1612 | 67.74305 | -0.0918 | 2.747673 | 1 |
| 9 | 82969800 | 82977599 | 65.4625 | 103.1702 | 155.4361 | 64.50019 | -0.26735 | 2.485925 | 1 |
| 9 | 83046000 | 83048599 | 42.06846 | 68.02669 | 80.23483 | 34.29279 | 0.124201 | 1.700162 | 1 |
| 9 | 83050800 | 83057799 | 105.3895 | 162.4492 | 173.0228 | 71.87078 | 0.207647 | 2.909361 | 1 |
| 9 | 83058600 | 83062599 | 45.50537 | 73.43379 | 82.66703 | 35.23911 | 0.187915 | 1.765672 | 1 |
| 9 | 83291200 | 83293399 | 18.10862 | 31.38565 | 45.29016 | 19.87096 | -0.14036 | 0.827533 | 1 |
| 9 | 83341800 | 83343199 | 13.44976 | 24.46379 | 28.02195 | 12.07029 | 0.222287 | 0.319153 | 1 |
| 9 | 83966600 | 83967399 | 12.09763 | 22.44188 | 24.11709 | 10.1197 | 0.340252 | 0.189119 | 1 |
| 9 | 84104400 | 84107199 | 27.15605 | 46.19531 | 47.93685 | 21.01036 | 0.328552 | 1.033422 | 1 |
| 9 | 84300200 | 84300999 | 15.25211 | 27.03196 | 31.93914 | 14.12037 | 0.152663 | 0.496531 | 1 |
| 9 | 85798600 | 85801999 | 52.26012 | 83.45387 | 111.8646 | 46.9684 | -0.08124 | 2.10467 | 1 |
| 9 | 85821000 | 85823999 | 58.83364 | 93.67329 | 108.2391 | 45.25903 | 0.128008 | 2.20426 | 1 |
| 9 | 86230000 | 86231199 | 12.05954 | 22.31121 | 25.079 | 10.56185 | 0.271878 | 0.205165 | 1 |
| 9 | 86829200 | 86831599 | 19.32761 | 33.8952 | 37.93634 | 16.76716 | 0.225649 | 0.654344 | 1 |
| 9 | 86984600 | 86988599 | 89.30477 | 139.3033 | 139.8004 | 58.0138 | 0.306608 | 2.689794 | 1 |
| 9 | 87043200 | 87046599 | 40.74026 | 65.67014 | 78.44806 | 33.39212 | 0.100639 | 1.788743 | 1 |
| 9 | 87696800 | 87697999 | 15.39713 | 27.37345 | 31.38633 | 13.84271 | 0.19819 | 0.475571 | 1 |
| 9 | 87720600 | 87722799 | 38.79872 | 62.33547 | 91.99439 | 38.9736 | -0.2095 | 1.8973 | 1 |

| | | | | | | | | | |
|---|---|---|---|---|---|---|---|---|---|
| 9 | 88507800 | 88509199 | 18.63432 | 32.43993 | 37.93777 | 16.67742 | 0.162413 | 0.763579 | 1 |
| 9 | 88553400 | 88560399 | 137.9602 | 207.828 | 245.5812 | 101.446 | 0.031238 | 3.380907 | 1 |
| 9 | 88631000 | 88633399 | 36.37046 | 58.72473 | 82.9092 | 35.36742 | -0.13711 | 1.734562 | 1 |
| 9 | 88702400 | 88704999 | 28.4856 | 47.67583 | 50.19587 | 21.87226 | 0.307105 | 1.189471 | 1 |
| 9 | 88771800 | 88774199 | 39.60094 | 64.02529 | 77.38942 | 33.06911 | 0.088641 | 1.699543 | 1 |
| 9 | 88889600 | 88892999 | 73.18929 | 114.8354 | 152.3002 | 63.16291 | -0.08857 | 2.579981 | 1 |
| 9 | 89219600 | 89221999 | 28.11771 | 47.29689 | 55.65217 | 24.31601 | 0.147126 | 1.141427 | 1 |
| 9 | 89482800 | 89485399 | 30 | 48.86376 | 75.3184 | 32.34149 | -0.25179 | 1.539825 | 1 |
| 9 | 89995000 | 89997199 | 13.8084 | 24.11365 | 42.6276 | 18.77581 | -0.43035 | 0.702244 | 1 |
| 9 | 90799800 | 90805399 | 100.6139 | 155.1004 | 187.7292 | 77.97955 | 0.023437 | 2.907595 | 1 |
| 9 | 90815800 | 90824399 | 161.5318 | 238.9652 | 320.5308 | 132.0034 | -0.17527 | 3.719222 | 1 |
| 9 | 91052800 | 91055199 | 20.19641 | 34.51065 | 52.2303 | 22.90042 | -0.2129 | 0.970723 | 1 |
| 9 | 91105600 | 91106799 | 12.47733 | 22.10423 | 35.20616 | 15.56612 | -0.27599 | 0.490274 | 1 |
| 9 | 91500800 | 91501799 | 10.54442 | 19.15843 | 31.31292 | 13.5062 | -0.2884 | 0.312238 | 1 |
| 9 | 92290400 | 92291599 | 12.40613 | 22.50577 | 32.6997 | 14.13199 | -0.12186 | 0.33672 | 1 |
| 9 | 92396600 | 92399799 | 57.01857 | 90.73746 | 123.1778 | 51.46944 | -0.10487 | 2.223941 | 1 |
| 9 | 92400600 | 92403199 | 41.01307 | 67.00152 | 66.14666 | 28.4341 | 0.392294 | 1.515015 | 1 |
| 9 | 92510400 | 92515399 | 64.30732 | 102.1124 | 114.525 | 47.73667 | 0.167003 | 2.292932 | 1 |
| 9 | 92534200 | 92538199 | 59.98703 | 95.30124 | 114.4698 | 47.73816 | 0.068976 | 2.273596 | 1 |
| 9 | 92750400 | 92751599 | 17.32247 | 30.33111 | 46.7083 | 20.61685 | -0.23247 | 0.684411 | 1 |
| 9 | 92940800 | 92943599 | 29.47259 | 48.90185 | 60.03227 | 26.01077 | 0.085986 | 1.293877 | 1 |
| 9 | 93023400 | 93025799 | 25.28305 | 43.58087 | 43.04324 | 18.88197 | 0.402729 | 0.904782 | 1 |
| 9 | 93248000 | 93249599 | 17.12509 | 30.22066 | 32.32994 | 14.30549 | 0.292757 | 0.554987 | 1 |
| 9 | 93268200 | 93276599 | 176.0756 | 262.2315 | 290.4718 | 119.6919 | 0.109601 | 3.604464 | 1 |
| 9 | 93316800 | 93318399 | 18.81945 | 32.85734 | 34.35484 | 15.12992 | 0.323725 | 0.719101 | 1 |
| 9 | 93510400 | 93512599 | 41.60368 | 67.34458 | 71.38103 | 30.53423 | 0.279463 | 1.67444 | 1 |
| 9 | 93550200 | 93551399 | 10.5283 | 19.09772 | 30.79749 | 13.31121 | -0.27114 | 0.319844 | 1 |
| 9 | 93570800 | 93571399 | 11.21258 | 20.89325 | 22.65241 | 9.44012 | 0.33502 | 0.150076 | 1 |
| 9 | 94009800 | 94010799 | 15.42405 | 27.49291 | 28.97925 | 12.75302 | 0.321243 | 0.455871 | 1 |
| 9 | 94536800 | 94539599 | 25.65864 | 42.4216 | 67.4029 | 29.13228 | -0.28544 | 1.330577 | 1 |
| 9 | 95551600 | 95555399 | 17.77996 | 30.70214 | 47.41019 | 20.79877 | -0.23888 | 0.861159 | 1 |
| 9 | 96628800 | 96632999 | 39.14436 | 63.46079 | 72.23692 | 30.93527 | 0.178798 | 1.636301 | 1 |
| 9 | 96695600 | 96697399 | 24.02784 | 40.16269 | 58.59864 | 25.51272 | -0.16198 | 1.198218 | 1 |
| 9 | 96929400 | 96931199 | 25.05734 | 41.83758 | 56.54671 | 24.60232 | -0.05203 | 1.208836 | 1 |
| 9 | 97212400 | 97213799 | 17.63122 | 29.99368 | 51.59612 | 22.61426 | -0.39761 | 1.004339 | 1 |
| 9 | 97246400 | 97249399 | 25.89089 | 42.92734 | 65.40254 | 28.32319 | -0.22479 | 1.295036 | 1 |
| 9 | 98094400 | 98095999 | 16.48471 | 29.41374 | 31.20712 | 13.72154 | 0.312467 | 0.45916 | 1 |
| 9 | 98339200 | 98343399 | 41.55345 | 66.58373 | 97.46361 | 41.12436 | -0.20236 | 1.98492 | 1 |
| 9 | 98737200 | 98739599 | 25.87954 | 42.70617 | 68.35415 | 29.50501 | -0.29616 | 1.355316 | 1 |
| 9 | 99477400 | 99486399 | 121.175 | 184.2054 | 227.6736 | 94.30983 | -0.0225 | 3.194026 | 1 |
| 9 | 99549400 | 99553999 | 45.15722 | 72.69926 | 85.39273 | 36.28379 | 0.122684 | 1.836647 | 1 |
| 9 | 99785000 | 99791599 | 133.0433 | 201.3609 | 235.2906 | 97.33441 | 0.053168 | 3.286217 | 1 |
| 9 | 99955600 | 99956799 | 20.32099 | 34.07134 | 56.8037 | 24.77929 | -0.3536 | 1.152771 | 1 |
| 9 | 100430400 | 100435599 | 71.76009 | 113.3657 | 131.0179 | 54.41416 | 0.117382 | 2.433233 | 1 |
| 9 | 100855200 | 100862599 | 85.38228 | 132.414 | 172.9004 | 71.81119 | -0.07922 | 2.790454 | 1 |
| 9 | 100902400 | 100909999 | 65.25407 | 103.1765 | 148.8757 | 61.8578 | -0.20142 | 2.411969 | 1 |
| 9 | 100967600 | 100970199 | 38.90567 | 62.62435 | 93.86293 | 39.87318 | -0.22851 | 1.837649 | 1 |

| | | | | | | | | | |
|---|---|---|---|---|---|---|---|---|---|
| 9 | 101184600 | 101186799 | 22.12784 | 37.26731 | 57.03741 | 24.9306 | -0.23074 | 1.112606 | 1 |
| 9 | 101303600 | 101305799 | 21.61752 | 36.44646 | 54.32751 | 23.75376 | -0.19268 | 1.09878 | 1 |
| 9 | 102818200 | 102822199 | 30.65106 | 50.00935 | 74.94413 | 32.20077 | -0.20953 | 1.517393 | 1 |
| 9 | 103241600 | 103243399 | 13.51379 | 23.52925 | 43.72752 | 19.2244 | -0.50265 | 0.744194 | 1 |
| 9 | 103673200 | 103674999 | 10.76357 | 20.3787 | 22.17694 | 8.981313 | 0.36357 | 0.042435 | 1 |
| 9 | 103850200 | 103852799 | 14.01829 | 25.38122 | 27.41803 | 11.87808 | 0.300874 | 0.347241 | 1 |
| 9 | 105344600 | 105346199 | 16.79396 | 29.45675 | 40.79608 | 18.03362 | -0.08003 | 0.645219 | 1 |
| 9 | 106467200 | 106468399 | 16.57323 | 29.10131 | 34.65055 | 15.33115 | 0.137063 | 0.616701 | 1 |
| 9 | 107011200 | 107011799 | 21.66411 | 36.44986 | 52.7019 | 23.0286 | -0.1489 | 1.116628 | 1 |
| 9 | 107089400 | 107092199 | 18.40012 | 31.73451 | 46.88228 | 20.56642 | -0.17566 | 0.87377 | 1 |
| 9 | 109080200 | 109081199 | 19.52606 | 33.02998 | 53.74737 | 23.52382 | -0.31849 | 1.061693 | 1 |
| 9 | 109138400 | 109139999 | 22.73221 | 37.8843 | 61.21245 | 26.60081 | -0.30867 | 1.227504 | 1 |
| 9 | 109236000 | 109251999 | 172.951 | 257.1984 | 326.9294 | 134.6944 | -0.09079 | 3.63118 | 1 |
| 9 | 109297000 | 109305599 | 155.744 | 233.7049 | 258.1138 | 106.5118 | 0.123677 | 3.460267 | 1 |
| 9 | 109403000 | 109411999 | 130.0436 | 197.8363 | 215.7249 | 89.38052 | 0.159009 | 3.180185 | 1 |
| 9 | 109459000 | 109469199 | 112.607 | 173.944 | 172.0067 | 71.44795 | 0.317023 | 2.874018 | 1 |
| 9 | 109535000 | 109537799 | 18.28319 | 32.13463 | 34.36087 | 15.2153 | 0.291283 | 0.609514 | 1 |
| 9 | 110110200 | 110111999 | 53.34949 | 85.43368 | 87.14457 | 36.73654 | 0.316694 | 2.015448 | 1 |
| 9 | 111349400 | 111356799 | 69.14349 | 108.8161 | 157.2736 | 65.25057 | -0.20911 | 2.517562 | 1 |
| 9 | 111393200 | 111395799 | 30.7481 | 50.846 | 62.35429 | 26.95387 | 0.087071 | 1.336785 | 1 |
| 9 | 111675000 | 111676799 | 17.94482 | 31.2039 | 46.838 | 20.57518 | -0.19636 | 0.789184 | 1 |
| 9 | 112787200 | 112788599 | 31.46864 | 51.38626 | 69.89829 | 30.04192 | -0.06926 | 1.505063 | 1 |
| 9 | 113340400 | 113343399 | 15.77257 | 27.58928 | 40.41645 | 17.82436 | -0.16062 | 0.682267 | 1 |
| 9 | 113373600 | 113375999 | 21.04704 | 36.373 | 41.49734 | 18.20526 | 0.19607 | 0.857563 | 1 |
| 9 | 114110600 | 114111199 | 8.727685 | 16.32987 | 26.41691 | 10.93264 | -0.22975 | 0.113565 | 1 |
| 9 | 114112400 | 114113199 | 29.52397 | 48.29072 | 70.91574 | 30.49763 | -0.17774 | 1.477879 | 1 |
| 9 | 114395200 | 114398199 | 25.61456 | 42.30918 | 69.21582 | 29.89656 | -0.32762 | 1.340409 | 1 |
| 9 | 114699200 | 114706999 | 97.09666 | 150.3304 | 163.4504 | 67.89299 | 0.182816 | 2.823577 | 1 |
| 9 | 114707600 | 114709599 | 17.91958 | 30.80491 | 48.47441 | 21.26274 | -0.26739 | 0.902172 | 1 |
| 9 | 114843400 | 114849399 | 60.86414 | 96.43868 | 132.3837 | 55.06413 | -0.1271 | 2.354988 | 1 |
| 9 | 115179600 | 115182399 | 30.65624 | 52.53285 | 40.03734 | 17.55842 | 0.773459 | 0.971074 | 1 |
| 9 | 115190200 | 115195399 | 53.28249 | 84.80579 | 119.6724 | 50.01598 | -0.16041 | 2.212362 | 1 |
| 9 | 115405800 | 115409399 | 25.81692 | 43.28313 | 53.90789 | 23.51504 | 0.065422 | 1.164727 | 1 |
| 9 | 115625400 | 115626999 | 21.5135 | 36.68973 | 46.89647 | 20.55857 | 0.029456 | 0.985961 | 1 |
| 9 | 116270200 | 116272199 | 19.0909 | 32.69033 | 50.06884 | 21.95645 | -0.22948 | 0.945534 | 1 |
| 9 | 116680200 | 116681199 | 14.3358 | 25.47409 | 36.03776 | 15.90076 | -0.10377 | 0.480415 | 1 |
| 9 | 117503400 | 117505799 | 24.43618 | 41.72195 | 51.2013 | 22.4478 | 0.088244 | 0.994063 | 1 |
| 9 | 117807400 | 117814199 | 131.542 | 198.3479 | 264.1171 | 109.1309 | -0.13976 | 3.363049 | 1 |
| 9 | 117972200 | 117974799 | 35.09645 | 56.74158 | 80.58975 | 34.43583 | -0.14351 | 1.694427 | 1 |
| 9 | 117978800 | 117981999 | 24.83329 | 41.11342 | 64.56861 | 27.93372 | -0.26851 | 1.310192 | 1 |
| 9 | 118032800 | 118041599 | 109.0239 | 166.9936 | 197.7161 | 82.0585 | 0.048175 | 3.035042 | 1 |
| 9 | 119037800 | 119044599 | 84.32546 | 130.9747 | 177.7299 | 73.80022 | -0.13297 | 2.763975 | 1 |
| 9 | 119624600 | 119627599 | 25.94295 | 43.53979 | 51.22465 | 22.35929 | 0.147318 | 1.151343 | 1 |
| 9 | 119685400 | 119692399 | 118.2075 | 180.2294 | 209.8307 | 86.9939 | 0.067178 | 3.131311 | 1 |
| 9 | 120130200 | 120134199 | 51.56769 | 82.17772 | 119.5818 | 50.06345 | -0.20273 | 2.172747 | 1 |
| 9 | 120225800 | 120230599 | 42.58134 | 68.53661 | 93.80455 | 39.84478 | -0.09789 | 1.843363 | 1 |
| 9 | 121004400 | 121007399 | 32.92256 | 53.97687 | 65.54947 | 28.21674 | 0.097217 | 1.448062 | 1 |

| | | | | | | | | | |
|---|---|---|---|---|---|---|---|---|---|
| 9 | 121008800 | 121009399 | 10.02065 | 18.96723 | 25.39855 | 10.29163 | 0.065738 | 0.045316 | 1 |
| 9 | 121010200 | 121010399 | 7.730487 | 14.97862 | 19.97471 | 7.569883 | 0.161618 | -0.20996 | 1 |
| 9 | 121374000 | 121375599 | 21.42233 | 37.05353 | 42.59243 | 18.68618 | 0.185693 | 0.856082 | 1 |
| 9 | 121830200 | 121831799 | 28.28334 | 47.20793 | 53.97457 | 23.4782 | 0.188568 | 1.220008 | 1 |
| 9 | 122694000 | 122702199 | 123.477 | 187.263 | 235.4476 | 97.46134 | -0.04995 | 3.243068 | 1 |
| 9 | 122732000 | 122744199 | 176.5215 | 260.8672 | 349.7524 | 144.0361 | -0.17605 | 3.739357 | 1 |
| 9 | 123021600 | 123022399 | 14.50983 | 26.38608 | 29.38886 | 12.6626 | 0.261956 | 0.329727 | 1 |
| 9 | 123088400 | 123090199 | 15.73973 | 27.13191 | 47.94246 | 21.03116 | -0.43299 | 0.867713 | 1 |
| 9 | 123288200 | 123291199 | 773.9717 | 1012.644 | 985.7308 | 412.6686 | 0.110716 | 5.630665 | 1 |
| 9 | 123447600 | 123449599 | 20.64071 | 35.94307 | 43.05004 | 18.92818 | 0.12818 | 0.778631 | 1 |
| 9 | 123576400 | 123581599 | 59.31958 | 94.51151 | 106.4225 | 44.54151 | 0.16627 | 2.182414 | 1 |
| 9 | 123616600 | 123618599 | 29.70944 | 49.05835 | 60.96071 | 26.3255 | 0.06803 | 1.351645 | 1 |
| 9 | 123701800 | 123702799 | 224.8641 | 323.7686 | 429.2437 | 177.1478 | -0.19609 | 4.145575 | 1 |
| 9 | 124017800 | 124020799 | 1065.016 | 1356.444 | 1423.655 | 597.5862 | -0.0368 | 6.069374 | 1 |
| 9 | 124110600 | 124112999 | 29.74578 | 49.88484 | 51.83704 | 22.61595 | 0.325848 | 1.172591 | 1 |
| 10 | 3182400 | 3189999 | 93.09722 | 144.7298 | 151.6109 | 62.94611 | 0.241021 | 2.748748 | 1 |
| 10 | 3261200 | 3268199 | 54.0744 | 86.61327 | 90.85125 | 38.31173 | 0.276793 | 2.009194 | 1 |
| 10 | 3569400 | 3571999 | 45.03828 | 72.36289 | 87.95294 | 37.26345 | 0.069926 | 1.900701 | 1 |
| 10 | 3922400 | 3924799 | 20.68825 | 35.29396 | 46.92316 | 20.57082 | -0.02687 | 0.982601 | 1 |
| 10 | 4310200 | 4321199 | 124.7823 | 188.7343 | 254.1634 | 105.12 | -0.15222 | 3.298856 | 1 |
| 10 | 4742400 | 4744999 | 20.76373 | 35.72815 | 55.03188 | 24.1398 | -0.23648 | 0.914934 | 1 |
| 10 | 5012600 | 5014199 | 15.66023 | 27.33516 | 39.0297 | 17.1862 | -0.12374 | 0.713101 | 1 |
| 10 | 5039600 | 5040199 | 10.74877 | 19.48031 | 31.34287 | 13.56201 | -0.26865 | 0.330121 | 1 |
| 10 | 5168000 | 5172399 | 25.5029 | 43.34373 | 47.41605 | 20.78058 | 0.252481 | 1.034867 | 1 |
| 10 | 5665600 | 5677799 | 212.7528 | 308.6195 | 389.8588 | 160.736 | -0.11583 | 4.033694 | 1 |
| 10 | 6026800 | 6045799 | 363.0672 | 504.0477 | 625.6418 | 259.7136 | -0.15393 | 4.690188 | 1 |
| 10 | 6185800 | 6187999 | 13.2567 | 24.01287 | 29.00327 | 12.55793 | 0.141318 | 0.342696 | 1 |
| 10 | 6298600 | 6302399 | 40.11294 | 65.31175 | 73.84974 | 31.70662 | 0.193638 | 1.56557 | 1 |
| 10 | 6348200 | 6350399 | 40.81518 | 65.63545 | 90.41412 | 38.35513 | -0.10871 | 1.868082 | 1 |
| 10 | 6351200 | 6356399 | 87.77322 | 135.3634 | 185.8465 | 77.19733 | -0.15772 | 2.894493 | 1 |
| 10 | 6363200 | 6365199 | 25.45941 | 43.07779 | 45.3905 | 19.86778 | 0.30624 | 1.073071 | 1 |
| 10 | 6816000 | 6822799 | 88.22026 | 137.9234 | 151.1894 | 62.72535 | 0.181927 | 2.65035 | 1 |
| 10 | 7114600 | 7117199 | 20.17001 | 34.73514 | 46.87358 | 20.56166 | -0.04606 | 0.894945 | 1 |
| 10 | 7298400 | 7303199 | 62.50729 | 99.17787 | 121.0067 | 50.39956 | 0.044612 | 2.315022 | 1 |
| 10 | 7314800 | 7318199 | 44.60857 | 71.64658 | 97.26408 | 41.18646 | -0.08994 | 1.91533 | 1 |
| 10 | 7345600 | 7347599 | 21.07518 | 36.10447 | 51.15765 | 22.43474 | -0.11756 | 0.951788 | 1 |
| 10 | 7363800 | 7368599 | 76.95067 | 121.1197 | 138.9843 | 57.66365 | 0.123526 | 2.516255 | 1 |
| 10 | 7489400 | 7491399 | 18.78768 | 32.7531 | 38.95454 | 17.1396 | 0.138536 | 0.748161 | 1 |
| 10 | 7851600 | 7852599 | 15.19333 | 26.4663 | 42.13591 | 18.53027 | -0.28048 | 0.743572 | 1 |
| 10 | 8137600 | 8140799 | 23.07757 | 39.58246 | 43.17808 | 18.93721 | 0.258607 | 0.939248 | 1 |
| 10 | 8194800 | 8196799 | 18.60035 | 31.51369 | 52.6524 | 23.05559 | -0.35623 | 1.044973 | 1 |
| 10 | 8353400 | 8354599 | 9.815443 | 18.0125 | 31.34548 | 13.34768 | -0.36621 | 0.245945 | 1 |
| 10 | 8413000 | 8415599 | 21.05446 | 35.82509 | 50.07634 | 21.94854 | -0.09948 | 1.011608 | 1 |
| 10 | 8476000 | 8476399 | 12.33949 | 22.14105 | 31.26473 | 13.68212 | -0.09237 | 0.400326 | 1 |
| 10 | 8984600 | 8986599 | 20.04311 | 34.382 | 49.54308 | 21.729 | -0.14143 | 0.932775 | 1 |
| 10 | 9573400 | 9575399 | 20.49886 | 35.22353 | 44.27319 | 19.41967 | 0.055438 | 0.913596 | 1 |
| 10 | 9775600 | 9777999 | 16.47287 | 28.45007 | 48.46824 | 21.26305 | -0.38005 | 0.85327 | 1 |

| | | | | | | | | | |
|---|---|---|---|---|---|---|---|---|---|
| 10 | 10038000 | 10039399 | 18.16449 | 31.74172 | 38.80546 | 17.09955 | 0.099135 | 0.708892 | 1 |
| 10 | 10290800 | 10292999 | 19.77853 | 34.71169 | 34.84645 | 15.41113 | 0.381956 | 0.642485 | 1 |
| 10 | 10468000 | 10472399 | 44.90981 | 72.25157 | 88.41727 | 37.53175 | 0.062477 | 1.860045 | 1 |
| 10 | 10542400 | 10543799 | 16.1673 | 28.108 | 43.19694 | 18.97359 | -0.23006 | 0.778504 | 1 |
| 10 | 10594800 | 10595999 | 13.43733 | 24.04828 | 30.51361 | 13.38274 | 0.058361 | 0.418692 | 1 |
| 10 | 11251800 | 11253999 | 33.45479 | 54.64785 | 65.86678 | 28.31265 | 0.105042 | 1.499448 | 1 |
| 10 | 11332600 | 11334399 | 12.00638 | 21.8485 | 32.19496 | 13.85978 | -0.13859 | 0.313173 | 1 |
| 10 | 11336000 | 11348399 | 194.991 | 288.2002 | 315.4552 | 129.9119 | 0.116621 | 3.737827 | 1 |
| 10 | 11384600 | 11386199 | 17.64967 | 30.31844 | 48.47589 | 21.26291 | -0.29044 | 0.908967 | 1 |
| 10 | 11510400 | 11512999 | 52.26544 | 83.78465 | 84.73879 | 35.77883 | 0.330683 | 1.979847 | 1 |
| 10 | 11552200 | 11566399 | 222.4113 | 325.1109 | 362.5569 | 149.3637 | 0.074622 | 3.918798 | 1 |
| 10 | 11568000 | 11572199 | 132.1589 | 199.0724 | 251.7209 | 103.9799 | -0.06652 | 3.381808 | 1 |
| 10 | 11647400 | 11653199 | 87.77441 | 137.0177 | 154.9194 | 64.28292 | 0.13561 | 2.677271 | 1 |
| 10 | 11667600 | 11673999 | 43.89784 | 70.68459 | 87.47438 | 37.17806 | 0.04788 | 1.830688 | 1 |
| 10 | 11676600 | 11677799 | 17.03533 | 29.94439 | 34.02441 | 15.0671 | 0.204186 | 0.601151 | 1 |
| 10 | 11832000 | 11834399 | 18.31006 | 32.08817 | 34.10548 | 15.06505 | 0.300105 | 0.656589 | 1 |
| 10 | 12879000 | 12880799 | 22.1392 | 37.40891 | 55.41677 | 24.24633 | -0.18362 | 1.08355 | 1 |
| 10 | 12963000 | 12972799 | 123.8862 | 187.7732 | 239.8348 | 99.25959 | -0.07334 | 3.25504 | 1 |
| 10 | 14458800 | 14460999 | 29.38522 | 49.06084 | 55.69845 | 24.23236 | 0.198663 | 1.220849 | 1 |
| 10 | 14621000 | 14623199 | 16.47415 | 29.09699 | 30.90792 | 13.67392 | 0.30345 | 0.539265 | 1 |
| 10 | 14797800 | 14801599 | 62.11615 | 98.6577 | 120.5471 | 50.25935 | 0.043957 | 2.286347 | 1 |
| 10 | 14879000 | 14880999 | 20.31778 | 34.94824 | 44.26068 | 19.41479 | 0.044851 | 0.903219 | 1 |
| 10 | 15035600 | 15038399 | 30.23232 | 50.29021 | 57.93032 | 25.14417 | 0.177765 | 1.264638 | 1 |
| 10 | 15057400 | 15061799 | 42.65129 | 69.15098 | 72.61006 | 31.09665 | 0.295081 | 1.643091 | 1 |
| 10 | 15217800 | 15219199 | 9.028024 | 16.95637 | 26.42506 | 10.86561 | -0.16884 | 0.091993 | 1 |
| 10 | 15262400 | 15263399 | 17.85366 | 31.22974 | 36.76236 | 16.21294 | 0.153503 | 0.688599 | 1 |
| 10 | 15409800 | 15410399 | 9.236329 | 17.62659 | 24.49013 | 9.752924 | 0.034968 | -0.02052 | 1 |
| 10 | 15501000 | 15502999 | 29.07766 | 47.72525 | 63.41837 | 27.31201 | -0.03162 | 1.425314 | 1 |
| 10 | 15669800 | 15670399 | 10.99149 | 19.9141 | 31.75446 | 13.74537 | -0.25654 | 0.331903 | 1 |
| 10 | 15891000 | 15891799 | 12.66333 | 22.52116 | 35.20665 | 15.52302 | -0.24603 | 0.464876 | 1 |
| 10 | 16179400 | 16180599 | 17.63566 | 30.86391 | 38.21896 | 16.86249 | 0.080648 | 0.680196 | 1 |
| 10 | 16192000 | 16193199 | 16.69891 | 29.13382 | 38.94351 | 17.14293 | -0.02928 | 0.723751 | 1 |
| 10 | 16378000 | 16380599 | 17.94947 | 31.50213 | 33.50825 | 14.81948 | 0.29891 | 0.629645 | 1 |
| 10 | 16567600 | 16568599 | 16.06079 | 28.21215 | 34.15366 | 15.11612 | 0.11322 | 0.606286 | 1 |
| 10 | 16956800 | 16958199 | 13.30514 | 23.94696 | 34.06868 | 14.86963 | -0.10112 | 0.389468 | 1 |
| 10 | 17012000 | 17013399 | 20.29904 | 35.44014 | 36.69168 | 16.15894 | 0.33779 | 0.730032 | 1 |
| 10 | 17017600 | 17019799 | 45.24964 | 73.03538 | 74.09183 | 31.59166 | 0.338105 | 1.75772 | 1 |
| 10 | 17083400 | 17084799 | 15.26451 | 26.81499 | 39.30663 | 17.39765 | -0.16132 | 0.611839 | 1 |
| 10 | 17157800 | 17158999 | 11.00651 | 19.89626 | 32.25565 | 14.00186 | -0.28292 | 0.346324 | 1 |
| 10 | 17196200 | 17196599 | 10.71329 | 20.04053 | 24.75968 | 10.25069 | 0.155723 | 0.128132 | 1 |
| 10 | 17200000 | 17200999 | 10.79645 | 20.1532 | 24.69569 | 10.26108 | 0.163042 | 0.140756 | 1 |
| 10 | 17372600 | 17384799 | 195.9606 | 287.6575 | 360.1832 | 148.3466 | -0.0866 | 3.84997 | 1 |
| 10 | 17573000 | 17573999 | 13.38081 | 24.85348 | 26.54663 | 11.12011 | 0.350473 | 0.197589 | 1 |
| 10 | 17674800 | 17675599 | 11.04302 | 20.13831 | 31.21413 | 13.40177 | -0.20859 | 0.291014 | 1 |
| 10 | 17692800 | 17697999 | 27.91237 | 46.93012 | 51.7941 | 22.62719 | 0.239234 | 1.141035 | 1 |
| 10 | 18013600 | 18016399 | 22.90589 | 38.35802 | 58.15923 | 25.35089 | -0.21743 | 1.170267 | 1 |
| 10 | 18184600 | 18191399 | 110.6048 | 169.5358 | 202.7736 | 84.16885 | 0.034348 | 3.023265 | 1 |

| | | | | | | | | | |
|---|---|---|---|---|---|---|---|---|---|
| 10 | 18535400 | 18537399 | 19.93809 | 34.53107 | 42.61993 | 18.69898 | 0.083848 | 0.837496 | 1 |
| 10 | 19412400 | 19415199 | 30.04344 | 50.15178 | 61.40379 | 26.71185 | 0.089997 | 1.228481 | 1 |
| 10 | 19823800 | 19827599 | 10.08841 | 18.7313 | 29.38097 | 12.31354 | -0.20062 | 0.178841 | 1 |
| 10 | 19860800 | 19865599 | 57.61742 | 91.42288 | 125.7045 | 52.35823 | -0.12748 | 2.310992 | 1 |
| 10 | 21574800 | 21575799 | 14.70151 | 26.47446 | 32.39683 | 14.13041 | 0.115579 | 0.392554 | 1 |
| 10 | 21610800 | 21615399 | 20.20808 | 35.10018 | 39.43448 | 17.31076 | 0.219653 | 0.80446 | 1 |
| 10 | 21907200 | 21909199 | 21.73009 | 37.91598 | 38.71962 | 17.04565 | 0.357474 | 0.749491 | 1 |
| 10 | 21942600 | 21945199 | 37.79565 | 61.55832 | 66.24105 | 28.4432 | 0.265229 | 1.549459 | 1 |
| 10 | 21964800 | 21969399 | 72.34997 | 114.1756 | 136.3687 | 56.60582 | 0.068741 | 2.460552 | 1 |
| 10 | 22338200 | 22340799 | 27.67313 | 46.40046 | 54.56263 | 23.80236 | 0.147977 | 1.168241 | 1 |
| 10 | 22399000 | 22401399 | 52.80033 | 84.58647 | 91.69618 | 38.67771 | 0.22962 | 2.00102 | 1 |
| 10 | 22402600 | 22407999 | 120.4155 | 182.8947 | 215.9136 | 89.41563 | 0.042658 | 3.21054 | 1 |
| 10 | 22432800 | 22439799 | 81.50052 | 127.7977 | 161.7462 | 67.08464 | -0.02189 | 2.59524 | 1 |
| 10 | 22497200 | 22497999 | 15.1053 | 26.38083 | 40.58667 | 17.87788 | -0.23073 | 0.706779 | 1 |
| 10 | 22566000 | 22569199 | 43.33333 | 69.89732 | 72.68085 | 30.9643 | 0.301484 | 1.770884 | 1 |
| 10 | 22677800 | 22679799 | 11.19482 | 20.08084 | 32.27124 | 14.12633 | -0.27831 | 0.394461 | 1 |
| 10 | 22732600 | 22735599 | 23.30176 | 38.89453 | 61.38933 | 26.70412 | -0.27511 | 1.210307 | 1 |
| 10 | 22817200 | 22822199 | 51.28723 | 81.93909 | 101.0292 | 42.44744 | 0.039892 | 2.086806 | 1 |
| 10 | 23338200 | 23339399 | 19.1938 | 33.1423 | 42.70719 | 18.73555 | 0.021149 | 0.862217 | 1 |
| 10 | 23533200 | 23539399 | 99.1135 | 152.8044 | 178.3951 | 74.10231 | 0.075569 | 2.903363 | 1 |
| 10 | 23602200 | 23604599 | 29.95069 | 49.02604 | 74.13052 | 31.88903 | -0.21911 | 1.467754 | 1 |
| 10 | 23735400 | 23750399 | 212.9691 | 312.7845 | 336.5776 | 138.6199 | 0.1326 | 3.843064 | 1 |
| 10 | 23753000 | 23760599 | 74.35006 | 115.9119 | 165.2899 | 68.59648 | -0.20026 | 2.694315 | 1 |
| 10 | 23816600 | 23817399 | 11.98966 | 21.46738 | 33.71114 | 14.77934 | -0.24682 | 0.41449 | 1 |
| 10 | 24034000 | 24036599 | 27.15142 | 45.08232 | 57.69949 | 25.01222 | 0.026119 | 1.276691 | 1 |
| 10 | 24116200 | 24117599 | 17.52024 | 30.93103 | 32.82964 | 14.52538 | 0.304388 | 0.554604 | 1 |
| 10 | 24386200 | 24388799 | 47.77194 | 76.81762 | 89.75103 | 38.07019 | 0.128222 | 1.878728 | 1 |
| 10 | 24560800 | 24565599 | 44.533 | 71.60562 | 86.43354 | 36.66019 | 0.081227 | 1.875329 | 1 |
| 10 | 24670800 | 24672199 | 10.47876 | 19.07186 | 32.8022 | 14.11821 | -0.35994 | 0.302173 | 1 |
| 10 | 24730000 | 24731399 | 18.68075 | 32.65144 | 35.28209 | 15.55004 | 0.276532 | 0.703697 | 1 |
| 10 | 24868400 | 24870599 | 16.98237 | 29.39602 | 47.40661 | 20.79874 | -0.30047 | 0.835856 | 1 |
| 10 | 24925200 | 24926599 | 13.7095 | 24.33681 | 36.59137 | 16.15866 | -0.192 | 0.483654 | 1 |
| 10 | 24976200 | 24978799 | 18.50271 | 32.21302 | 41.51023 | 18.24867 | 0.023009 | 0.766798 | 1 |
| 10 | 25046600 | 25049199 | 25.16687 | 41.74507 | 62.93878 | 27.26443 | -0.20955 | 1.288966 | 1 |
| 10 | 25769200 | 25778199 | 98.57171 | 152.1251 | 170.3335 | 70.75325 | 0.136931 | 2.883872 | 1 |
| 10 | 25818200 | 25820799 | 15.03178 | 26.24127 | 40.5546 | 17.85898 | -0.23744 | 0.710079 | 1 |
| 10 | 25843800 | 25847199 | 38.7711 | 62.82011 | 72.40319 | 30.99355 | 0.160178 | 1.649353 | 1 |
| 10 | 26470200 | 26471399 | 20.69357 | 36.06652 | 38.73367 | 17.03969 | 0.284829 | 0.757576 | 1 |
| 10 | 26516800 | 26521199 | 64.17212 | 101.7376 | 120.7065 | 50.23408 | 0.083778 | 2.340015 | 1 |
| 10 | 26532400 | 26534399 | 25.02725 | 43.01988 | 45.19787 | 19.82547 | 0.313052 | 0.929519 | 1 |
| 10 | 26603000 | 26611599 | 141.4004 | 213.9464 | 218.3513 | 90.31865 | 0.247916 | 3.292561 | 1 |
| 10 | 26805200 | 26812399 | 124.457 | 188.6193 | 214.0741 | 88.59526 | 0.096822 | 3.255387 | 1 |
| 10 | 27538000 | 27539199 | 8.543279 | 16.15609 | 27.59519 | 11.20333 | -0.28495 | 0.04439 | 1 |
| 10 | 27600000 | 27601799 | 19.48678 | 33.11686 | 50.58478 | 22.16919 | -0.22718 | 1.017955 | 1 |
| 10 | 27668000 | 27672199 | 28.64873 | 47.99136 | 58.43949 | 25.47856 | 0.097809 | 1.186653 | 1 |
| 10 | 27694200 | 27696399 | 25.67157 | 42.88452 | 51.74628 | 22.52083 | 0.111108 | 1.201226 | 1 |
| 10 | 27700600 | 27702399 | 26.41465 | 44.26192 | 48.596 | 21.19091 | 0.246781 | 1.167165 | 1 |

| | | | | | | | | | |
|---|---|---|---|---|---|---|---|---|---|
| 10 | 27811800 | 27813599 | 14.22838 | 25.63792 | 28.43369 | 12.39246 | 0.257608 | 0.378944 | 1 |
| 10 | 28402600 | 28404599 | 22.832 | 38.71063 | 45.92388 | 20.11362 | 0.135906 | 1.046163 | 1 |
| 10 | 28479600 | 28481999 | 31.46468 | 52.38773 | 55.24999 | 23.99538 | 0.304698 | 1.2551 | 1 |
| 10 | 28808000 | 28810799 | 21.27513 | 36.62149 | 39.04714 | 17.12826 | 0.292267 | 0.892373 | 1 |
| 10 | 28971800 | 28974599 | 22.10812 | 37.58026 | 61.04713 | 26.75489 | -0.31588 | 1.035993 | 1 |
| 10 | 29019400 | 29020999 | 9.540682 | 17.42978 | 32.86812 | 14.08036 | -0.48757 | 0.272116 | 1 |
| 10 | 29072400 | 29074599 | 44.07457 | 70.7483 | 87.23505 | 36.90745 | 0.047644 | 1.928555 | 1 |
| 10 | 29162600 | 29166599 | 46.89648 | 75.20905 | 99.53798 | 42.0539 | -0.0559 | 1.965033 | 1 |
| 10 | 29829400 | 29830399 | 15.79195 | 27.65722 | 39.35286 | 17.37337 | -0.11875 | 0.661459 | 1 |
| 10 | 30357000 | 30361799 | 92.66417 | 143.4688 | 167.3934 | 69.53084 | 0.081142 | 2.82215 | 1 |
| 10 | 30669800 | 30675399 | 56.78721 | 90.61046 | 127.9368 | 53.65563 | -0.1574 | 2.138775 | 1 |
| 10 | 30689400 | 30691999 | 47.76828 | 76.31253 | 107.1229 | 45.00394 | -0.14698 | 2.087549 | 1 |
| 10 | 30694800 | 30699199 | 42.98786 | 69.3123 | 83.87648 | 35.71554 | 0.082436 | 1.785528 | 1 |
| 10 | 30930600 | 30931599 | 11.37002 | 20.52785 | 33.2741 | 14.46643 | -0.28406 | 0.359666 | 1 |
| 10 | 31355800 | 31361999 | 82.3641 | 129.0802 | 147.0405 | 60.98889 | 0.129248 | 2.60402 | 1 |
| 10 | 31483000 | 31485999 | 21.17626 | 35.66751 | 56.44153 | 24.67074 | -0.27866 | 1.108483 | 1 |
| 10 | 31575000 | 31582999 | 117.1292 | 179.3249 | 185.6361 | 77.03658 | 0.241207 | 3.04568 | 1 |
| 10 | 31760800 | 31762199 | 12.33396 | 21.8097 | 35.76198 | 15.81774 | -0.31875 | 0.510875 | 1 |
| 10 | 31921200 | 31921999 | 9.660435 | 18.42088 | 20.12267 | 8.027501 | 0.376575 | -0.02527 | 1 |
| 10 | 32374800 | 32375999 | 18.12534 | 31.35716 | 45.3054 | 19.87654 | -0.14266 | 0.844381 | 1 |
| 10 | 32508200 | 32509799 | 15.49976 | 27.48778 | 33.31239 | 14.72288 | 0.116475 | 0.49369 | 1 |
| 10 | 32565600 | 32567799 | 26.26018 | 44.74184 | 46.8683 | 20.54464 | 0.315399 | 1.016196 | 1 |
| 10 | 32800400 | 32800999 | 9.018621 | 16.7868 | 27.88365 | 11.64161 | -0.27824 | 0.151549 | 1 |
| 10 | 32809200 | 32816199 | 103.4695 | 159.3371 | 192.0637 | 79.77408 | 0.028237 | 2.929119 | 1 |
| 10 | 33144600 | 33146399 | 18.37953 | 32.24123 | 35.44024 | 15.6675 | 0.251616 | 0.641752 | 1 |
| 10 | 33215800 | 33216599 | 10.963 | 19.86129 | 32.23369 | 13.95385 | -0.28206 | 0.331884 | 1 |
| 10 | 33367600 | 33368599 | 13.17685 | 24.12302 | 24.52027 | 10.47554 | 0.401303 | 0.271637 | 1 |
| 10 | 33384600 | 33385599 | 16.6827 | 29.35333 | 31.1115 | 13.77447 | 0.304652 | 0.583385 | 1 |
| 10 | 33517400 | 33519999 | 26.12921 | 44.01885 | 52.29714 | 22.86063 | 0.133373 | 1.118865 | 1 |
| 10 | 33551200 | 33558399 | 79.47082 | 123.6688 | 169.1118 | 70.20175 | -0.14192 | 2.727739 | 1 |
| 10 | 33580000 | 33581999 | 27.17105 | 45.44628 | 53.99941 | 23.52 | 0.133083 | 1.191937 | 1 |
| 10 | 33684200 | 33688399 | 55.15969 | 87.60679 | 123.7624 | 51.59862 | -0.16512 | 2.280663 | 1 |
| 10 | 33696800 | 33697999 | 21.0184 | 36.27127 | 39.54542 | 17.34817 | 0.261026 | 0.870255 | 1 |
| 10 | 33711800 | 33712799 | 19.41781 | 33.65477 | 38.05613 | 16.69704 | 0.20994 | 0.822698 | 1 |
| 10 | 33766600 | 33768599 | 24.30625 | 40.67844 | 61.45618 | 26.77777 | -0.21223 | 1.186739 | 1 |
| 10 | 34767400 | 34769399 | 24.40129 | 40.84473 | 57.64017 | 25.11786 | -0.11418 | 1.180711 | 1 |
| 10 | 34913400 | 34914599 | 10.7518 | 19.96928 | 30.99868 | 12.98599 | -0.18532 | 0.186825 | 1 |
| 10 | 34955400 | 34956999 | 14.39571 | 25.50683 | 35.58056 | 15.73115 | -0.0857 | 0.502231 | 1 |
| 10 | 35026600 | 35029399 | 34.16058 | 56.86761 | 58.10474 | 25.23278 | 0.350328 | 1.269495 | 1 |
| 10 | 35186000 | 35187199 | 27.32066 | 44.49993 | 75.79439 | 32.54594 | -0.39533 | 1.536914 | 1 |
| 10 | 35258800 | 35261599 | 28.91597 | 48.23026 | 56.21252 | 24.44094 | 0.160931 | 1.230492 | 1 |
| 10 | 35509400 | 35511199 | 9.332084 | 17.37733 | 30.52585 | 12.73596 | -0.35785 | 0.15755 | 1 |
| 10 | 35644400 | 35647799 | 46.96786 | 75.72639 | 91.35467 | 38.89775 | 0.086809 | 1.796368 | 1 |
| 10 | 35816200 | 35817599 | 16.37344 | 28.99022 | 32.34068 | 14.30076 | 0.2345 | 0.51514 | 1 |
| 10 | 36036600 | 36037199 | 11.44148 | 21.13637 | 25.64364 | 10.82147 | 0.16038 | 0.209675 | 1 |
| 10 | 36101000 | 36102199 | 9.727052 | 18.49948 | 20.68311 | 8.296828 | 0.336681 | -0.00403 | 1 |
| 10 | 36151600 | 36153199 | 23.10432 | 39.0199 | 54.36673 | 23.78325 | -0.09554 | 1.090221 | 1 |

| | | | | | | | | | |
|---|---|---|---|---|---|---|---|---|---|
| 10 | 36368400 | 36370199 | 21.66633 | 37.36912 | 45.79039 | 20.08745 | 0.093035 | 0.887153 | 1 |
| 10 | 36724200 | 36725599 | 7.696409 | 14.91514 | 19.42791 | 7.35676 | 0.196059 | -0.21599 | 1 |
| 10 | 36849400 | 36852799 | 18.48327 | 31.93584 | 43.20575 | 18.95464 | -0.0487 | 0.852291 | 1 |
| 10 | 37846000 | 37847199 | 15.74917 | 27.38656 | 42.71819 | 18.76568 | -0.25121 | 0.774746 | 1 |
| 10 | 37921200 | 37923999 | 37.68857 | 60.86227 | 83.10652 | 35.45858 | -0.08881 | 1.733148 | 1 |
| 10 | 38140000 | 38140799 | 16.10769 | 28.16675 | 37.41246 | 16.49523 | -0.01982 | 0.685626 | 1 |
| 10 | 38466000 | 38471599 | 98.79791 | 152.8741 | 171.8003 | 71.36157 | 0.134541 | 2.836525 | 1 |
| 10 | 38475600 | 38476399 | 14.91972 | 26.71044 | 28.47822 | 12.48643 | 0.308449 | 0.424087 | 1 |
| 10 | 38648200 | 38649399 | 28.9853 | 46.97506 | 80.47829 | 34.45604 | -0.40989 | 1.632139 | 1 |
| 10 | 38771600 | 38773999 | 25.42297 | 42.89699 | 48.01542 | 20.99906 | 0.219176 | 1.098417 | 1 |
| 10 | 38808400 | 38809599 | 17.17858 | 29.8239 | 40.712 | 17.86621 | -0.05989 | 0.802186 | 1 |
| 10 | 39627800 | 39630199 | 41.45142 | 66.69745 | 92.73273 | 39.37196 | -0.12104 | 1.851176 | 1 |
| 10 | 39635800 | 39640999 | 57.72942 | 91.99525 | 98.26653 | 41.13865 | 0.242561 | 2.173847 | 1 |
| 10 | 39752800 | 39760199 | 79.10287 | 124.5614 | 140.784 | 58.41364 | 0.145838 | 2.509492 | 1 |
| 10 | 40139800 | 40141799 | 22.32094 | 37.78098 | 52.23733 | 22.86717 | -0.08444 | 1.063471 | 1 |
| 10 | 40437000 | 40443399 | 153.8275 | 231.2967 | 235.5845 | 97.26765 | 0.243001 | 3.419965 | 1 |
| 10 | 40520600 | 40524999 | 83.46942 | 128.8002 | 182.283 | 75.71692 | -0.2008 | 2.882908 | 1 |
| 10 | 41433800 | 41440399 | 45.69898 | 73.31231 | 94.05967 | 39.75893 | -0.0106 | 1.950932 | 1 |
| 10 | 41602800 | 41604999 | 14.75137 | 27.11733 | 28.48102 | 12.09944 | 0.360107 | 0.269999 | 1 |
| 10 | 42013200 | 42015799 | 23.29208 | 39.38276 | 58.77056 | 25.71882 | -0.19422 | 1.085002 | 1 |
| 10 | 42140400 | 42143399 | 22.46571 | 37.75056 | 55.93543 | 24.43211 | -0.18427 | 1.132767 | 1 |
| 10 | 42536800 | 42546399 | 82.05777 | 128.6086 | 139.4029 | 57.82062 | 0.200897 | 2.600033 | 1 |
| 10 | 42544400 | 42551999 | 49.51985 | 79.43759 | 93.51232 | 39.52429 | 0.113219 | 1.955746 | 1 |
| 10 | 42724200 | 42733799 | 97.07869 | 148.5068 | 215.2029 | 89.28967 | -0.24454 | 3.058921 | 1 |
| 10 | 43647600 | 43648599 | 12.1905 | 21.93661 | 33.2273 | 14.50583 | -0.19071 | 0.385134 | 1 |
| 10 | 43649200 | 43650799 | 11.7503 | 21.26953 | 31.1802 | 13.51216 | -0.13712 | 0.34109 | 1 |
| 10 | 43749200 | 43758399 | 30.11222 | 49.30331 | 67.79423 | 29.16653 | -0.08222 | 1.461692 | 1 |
| 10 | 44516800 | 44517799 | 10.39855 | 19.13832 | 32.43613 | 13.75738 | -0.32438 | 0.235924 | 1 |
| 10 | 44667400 | 44669799 | 11.74421 | 22.01676 | 26.23478 | 10.82152 | 0.211329 | 0.12505 | 1 |
| 10 | 44779200 | 44783999 | 51.19726 | 82.14856 | 97.07323 | 41.04467 | 0.108169 | 1.954376 | 1 |
| 10 | 45713200 | 45715799 | 34.24379 | 56.24944 | 65.19988 | 28.08875 | 0.165324 | 1.428254 | 1 |
| 10 | 46256200 | 46256999 | 17.97842 | 30.81701 | 49.53145 | 21.72258 | -0.29849 | 0.931023 | 1 |
| 10 | 46462200 | 46462799 | 10.25014 | 19.1832 | 25.30394 | 10.46765 | 0.062883 | 0.12248 | 1 |
| 10 | 46848400 | 46851999 | 42.67512 | 68.45482 | 99.06076 | 41.86686 | -0.18405 | 1.952904 | 1 |
| 10 | 47015400 | 47016599 | 16.73518 | 29.41608 | 35.05702 | 15.52445 | 0.135703 | 0.601589 | 1 |
| 10 | 47203200 | 47204999 | 34.25886 | 56.13999 | 60.49247 | 26.03444 | 0.269028 | 1.455228 | 1 |
| 10 | 47301800 | 47303199 | 14.866 | 26.36508 | 31.40614 | 13.87984 | 0.141327 | 0.48622 | 1 |
| 10 | 47313800 | 47315399 | 38.25629 | 62.018 | 70.72277 | 30.28574 | 0.17617 | 1.636792 | 1 |
| 10 | 47545400 | 47547599 | 42.39564 | 68.60021 | 71.49438 | 30.57407 | 0.303244 | 1.68308 | 1 |
| 10 | 47621800 | 47635599 | 305.5524 | 430.3115 | 507.0975 | 210.0346 | -0.05828 | 4.473355 | 1 |
| 10 | 47741800 | 47742599 | 11.73942 | 21.28322 | 32.1946 | 13.92439 | -0.18037 | 0.333583 | 1 |
| 10 | 48104400 | 48108199 | 18.25813 | 31.71639 | 44.73285 | 19.63784 | -0.10699 | 0.802594 | 1 |
| 10 | 48391000 | 48391599 | 11.06753 | 20.89172 | 23.15453 | 9.436605 | 0.329299 | 0.067753 | 1 |
| 10 | 48557800 | 48559999 | 28.53923 | 47.53993 | 57.21247 | 24.85528 | 0.114932 | 1.245966 | 1 |
| 10 | 48791600 | 48792399 | 11.44148 | 21.13637 | 25.64364 | 10.82147 | 0.16038 | 0.209675 | 1 |
| 10 | 48865400 | 48873399 | 105.3906 | 162.4381 | 173.0227 | 71.87079 | 0.207462 | 2.910705 | 1 |
| 10 | 49711400 | 49712799 | 18.10081 | 31.71618 | 39.2304 | 17.32639 | 0.08234 | 0.666996 | 1 |

| | | | | | | | | | |
|---|---|---|---|---|---|---|---|---|---|
| 10 | 49849000 | 49849799 | 16.56632 | 29.11125 | 33.6194 | 14.88446 | 0.180919 | 0.602764 | 1 |
| 10 | 50060400 | 50060999 | 13.42433 | 23.70349 | 34.25661 | 15.15628 | -0.13833 | 0.531421 | 1 |
| 10 | 50128800 | 50132399 | 77.90428 | 122.4958 | 128.8648 | 53.45804 | 0.247603 | 2.535047 | 1 |
| 10 | 50296200 | 50299199 | 49.77274 | 79.42795 | 107.3041 | 45.01036 | -0.09351 | 2.124059 | 1 |
| 10 | 50301600 | 50302199 | 12.9367 | 23.40108 | 28.22643 | 12.24257 | 0.142125 | 0.349815 | 1 |
| 10 | 50430600 | 50431799 | 18.16693 | 31.73798 | 40.35737 | 17.78045 | 0.042582 | 0.714541 | 1 |
| 10 | 51008000 | 51010999 | 42.6909 | 68.94173 | 74.64948 | 31.85106 | 0.245122 | 1.736799 | 1 |
| 10 | 51243200 | 51243999 | 12.968 | 23.04963 | 35.67358 | 15.73658 | -0.23223 | 0.470881 | 1 |
| 10 | 51254200 | 51255599 | 11.62899 | 21.1443 | 34.87759 | 15.03014 | -0.30118 | 0.322203 | 1 |
| 10 | 51696400 | 51697799 | 10.93677 | 20.35054 | 26.70029 | 11.15367 | 0.058737 | 0.16599 | 1 |
| 10 | 51867000 | 51867799 | 10.9491 | 19.87364 | 28.29268 | 12.21889 | -0.09167 | 0.313231 | 1 |
| 10 | 51874600 | 51877999 | 30.49448 | 50.51353 | 52.42014 | 22.68659 | 0.327473 | 1.308529 | 1 |
| 10 | 52428800 | 52431399 | 22.34635 | 38.26867 | 44.79735 | 19.64492 | 0.156916 | 0.95333 | 1 |
| 10 | 52447000 | 52448799 | 23.46666 | 39.22208 | 59.19231 | 25.77012 | -0.21063 | 1.195386 | 1 |
| 10 | 52524600 | 52526199 | 20.59167 | 34.53442 | 55.80876 | 24.34939 | -0.30889 | 1.147717 | 1 |
| 10 | 52566200 | 52567999 | 18.43472 | 31.33605 | 52.12808 | 22.84598 | -0.34965 | 1.01544 | 1 |
| 10 | 53175200 | 53176999 | 32.51196 | 52.7407 | 76.02577 | 32.57257 | -0.16012 | 1.617662 | 1 |
| 10 | 53345800 | 53354199 | 117.6488 | 179.5871 | 208.4513 | 86.44534 | 0.072861 | 3.107242 | 1 |
| 10 | 53776000 | 53777199 | 29.59465 | 48.93538 | 57.6143 | 24.90377 | 0.145846 | 1.334688 | 1 |
| 10 | 54024600 | 54025799 | 10.48293 | 19.54959 | 27.35666 | 11.38481 | -0.02864 | 0.153106 | 1 |
| 10 | 54467600 | 54467999 | 8.444533 | 16.30049 | 19.87168 | 7.664872 | 0.265423 | -0.14426 | 1 |
| 10 | 54494400 | 54496799 | 16.4989 | 28.77846 | 40.04292 | 17.62442 | -0.08675 | 0.730031 | 1 |
| 10 | 54783600 | 54786199 | 16.55442 | 28.78586 | 43.72025 | 19.20547 | -0.21302 | 0.780481 | 1 |
| 10 | 54955400 | 54959199 | 56.61204 | 89.98187 | 116.5872 | 48.64419 | -0.03948 | 2.258222 | 1 |
| 10 | 55346600 | 55349199 | 24.54239 | 41.16382 | 59.34755 | 25.89481 | -0.14506 | 1.160647 | 1 |
| 10 | 55363800 | 55366199 | 19.05494 | 33.33453 | 36.09311 | 15.91921 | 0.27342 | 0.690883 | 1 |
| 10 | 55380400 | 55382799 | 17.69675 | 30.70442 | 43.18627 | 18.9485 | -0.10328 | 0.811721 | 1 |
| 10 | 55450600 | 55451399 | 13.34049 | 23.56617 | 35.79324 | 15.8347 | -0.20924 | 0.530657 | 1 |
| 10 | 55456800 | 55457199 | 14.19753 | 25.6858 | 28.39768 | 12.31445 | 0.266713 | 0.355092 | 1 |
| 10 | 55544800 | 55546399 | 8.974823 | 16.45642 | 30.90599 | 13.17515 | -0.47601 | 0.245883 | 1 |
| 10 | 55620200 | 55621799 | 25.52042 | 42.52128 | 61.56777 | 26.75136 | -0.15112 | 1.235423 | 1 |
| 10 | 55717600 | 55718399 | 10.85899 | 19.59158 | 32.76003 | 14.25533 | -0.32959 | 0.3574 | 1 |
| 10 | 55861800 | 55862599 | 11.85577 | 22.28947 | 24.19229 | 9.930867 | 0.351082 | 0.10523 | 1 |
| 10 | 56103800 | 56105799 | 10.42893 | 19.11298 | 31.83048 | 13.57866 | -0.30516 | 0.259447 | 1 |
| 10 | 56189800 | 56190999 | 12.62896 | 22.57693 | 35.18157 | 15.44422 | -0.2372 | 0.431277 | 1 |
| 10 | 56379000 | 56380999 | 21.46505 | 36.89671 | 40.58926 | 17.80396 | 0.246961 | 0.907836 | 1 |
| 10 | 56785800 | 56786999 | 13.28961 | 24.32584 | 28.55077 | 12.19992 | 0.194897 | 0.28298 | 1 |
| 10 | 56905400 | 56906599 | 17.14548 | 29.77445 | 46.86332 | 20.56915 | -0.26464 | 0.804409 | 1 |
| 10 | 56974400 | 56976199 | 24.11762 | 40.63197 | 49.60162 | 21.68266 | 0.094372 | 1.108199 | 1 |
| 10 | 57131000 | 57132199 | 11.25544 | 20.59974 | 36.12724 | 15.4401 | -0.38141 | 0.282753 | 1 |
| 10 | 57208800 | 57210199 | 28.38669 | 46.41308 | 69.03938 | 29.68663 | -0.19656 | 1.479482 | 1 |
| 10 | 57657800 | 57658799 | 11.63947 | 21.2799 | 28.12875 | 12.03884 | 0.022782 | 0.27496 | 1 |
| 10 | 57748600 | 57750599 | 422.1037 | 573.6376 | 756.1044 | 314.9263 | -0.27168 | 5.021532 | 1 |
| 10 | 57976200 | 57977999 | 10.57796 | 19.41869 | 27.74892 | 11.81047 | -0.08298 | 0.243397 | 1 |
| 10 | 58868800 | 58874599 | 45.95682 | 73.41211 | 108.5711 | 45.60663 | -0.22226 | 2.08813 | 1 |
| 10 | 59323600 | 59328999 | 60.79068 | 96.63011 | 133.4465 | 55.68223 | -0.1313 | 2.264095 | 1 |
| 10 | 59588200 | 59589599 | 14.2896 | 25.09884 | 39.86041 | 17.64066 | -0.27634 | 0.608977 | 1 |

| | | | | | | | | | |
|---|---|---|---|---|---|---|---|---|---|
| 10 | 59997000 | 59998999 | 19.3958 | 33.08463 | 53.76529 | 23.57011 | -0.31538 | 0.989058 | 1 |
| 10 | 60075600 | 60076999 | 14.07075 | 24.78958 | 37.73027 | 16.70058 | -0.21407 | 0.565668 | 1 |
| 10 | 61050800 | 61055799 | 47.48078 | 75.75357 | 117.2989 | 49.18739 | -0.29036 | 2.130609 | 1 |
| 10 | 61750200 | 61758599 | 92.0957 | 141.4437 | 206.0615 | 85.55675 | -0.24778 | 2.977671 | 1 |
| 10 | 62443000 | 62445399 | 20.30406 | 34.45819 | 50.06878 | 21.93548 | -0.1555 | 1.033167 | 1 |
| 10 | 62510400 | 62515999 | 30.49874 | 49.59893 | 79.32584 | 34.04297 | -0.30652 | 1.56508 | 1 |
| 10 | 63189000 | 63196199 | 92.68965 | 144.5031 | 153.1193 | 63.54559 | 0.227649 | 2.701121 | 1 |
| 10 | 63279000 | 63281199 | 33.18773 | 54.41976 | 61.10884 | 26.30688 | 0.209985 | 1.443253 | 1 |
| 10 | 63347400 | 63348999 | 20.4721 | 35.53779 | 38.90514 | 17.07206 | 0.256744 | 0.812568 | 1 |
| 10 | 63484200 | 63485999 | 13.37325 | 23.98318 | 33.57979 | 14.70184 | -0.08111 | 0.412766 | 1 |
| 10 | 63722200 | 63722799 | 8.859214 | 16.9304 | 22.87424 | 9.08269 | 0.078523 | -0.03952 | 1 |
| 10 | 64457400 | 64460199 | 45.39291 | 72.74722 | 97.0517 | 40.96021 | -0.06857 | 1.983314 | 1 |
| 10 | 64620000 | 64622599 | 51.24941 | 82.08942 | 89.67528 | 37.81655 | 0.21833 | 2.004609 | 1 |
| 10 | 64935600 | 64936399 | 8.979457 | 17.0322 | 29.39536 | 11.85834 | -0.29212 | 0.03082 | 1 |
| 10 | 65096400 | 65099999 | 61.96004 | 98.58984 | 113.1904 | 47.28873 | 0.136298 | 2.228093 | 1 |
| 10 | 65476600 | 65482999 | 59.78885 | 94.76078 | 120.5517 | 50.15504 | -0.0171 | 2.345659 | 1 |
| 10 | 65497400 | 65500599 | 27.14393 | 45.42341 | 52.85518 | 23.02882 | 0.163247 | 1.188324 | 1 |
| 10 | 65664000 | 65665799 | 24.90092 | 42.17448 | 49.04626 | 21.47437 | 0.164482 | 1.067388 | 1 |
| 10 | 65736400 | 65738599 | 23.96115 | 40.05961 | 55.39572 | 24.12142 | -0.08486 | 1.191608 | 1 |
| 10 | 65772000 | 65774999 | 29.63286 | 48.51189 | 74.07483 | 31.86676 | -0.23311 | 1.465633 | 1 |
| 10 | 66143000 | 66146199 | 25.94765 | 42.91999 | 60.196 | 26.02599 | -0.10586 | 1.320457 | 1 |
| 10 | 66334400 | 66335999 | 20.84701 | 36.09822 | 37.86296 | 16.61182 | 0.317449 | 0.834575 | 1 |
| 10 | 66544600 | 66558399 | 225.9707 | 327.8651 | 396.2909 | 163.3842 | -0.05189 | 4.031116 | 1 |
| 10 | 66562800 | 66566999 | 62.91133 | 99.97125 | 105.4592 | 43.99106 | 0.256328 | 2.267122 | 1 |
| 10 | 67183200 | 67185399 | 24.65968 | 41.90036 | 46.91367 | 20.56 | 0.21917 | 1.035677 | 1 |
| 10 | 67333600 | 67335999 | 30.87349 | 51.10627 | 58.45946 | 25.28833 | 0.187316 | 1.322227 | 1 |
| 10 | 67589800 | 67591999 | 21.83471 | 37.15647 | 50.61836 | 22.18629 | -0.06254 | 1.01363 | 1 |
| 10 | 67826200 | 67828799 | 38.9661 | 63.17825 | 74.22883 | 31.7907 | 0.133412 | 1.637759 | 1 |
| 10 | 67844600 | 67853599 | 84.52844 | 131.1126 | 178.3162 | 74.0586 | -0.13772 | 2.786605 | 1 |
| 10 | 67854400 | 67855999 | 27.64711 | 46.09 | 56.67142 | 24.63242 | 0.084002 | 1.233174 | 1 |
| 10 | 67886600 | 67889599 | 31.84679 | 51.71495 | 80.85691 | 34.66877 | -0.27572 | 1.59724 | 1 |
| 10 | 67913400 | 67916399 | 17.25698 | 30.04416 | 45.75035 | 20.11268 | -0.21686 | 0.766465 | 1 |
| 10 | 68078800 | 68082399 | 34.7091 | 55.92795 | 88.37769 | 37.60011 | -0.30264 | 1.797497 | 1 |
| 10 | 68177800 | 68182399 | 47.3854 | 76.11459 | 103.7625 | 43.94532 | -0.09582 | 1.917464 | 1 |
| 10 | 68209800 | 68212199 | 22.1119 | 36.79339 | 62.75746 | 27.24505 | -0.38653 | 1.244061 | 1 |
| 10 | 68219000 | 68221599 | 29.9893 | 50.25963 | 55.76385 | 24.31892 | 0.231557 | 1.182026 | 1 |
| 10 | 68449600 | 68450799 | 15.44609 | 27.29999 | 35.00127 | 15.48272 | 0.033854 | 0.533495 | 1 |
| 10 | 68929000 | 68934199 | 70.12994 | 110.196 | 151.143 | 62.69431 | -0.13539 | 2.547916 | 1 |
| 10 | 68935200 | 68938199 | 24.64434 | 41.23793 | 56.61098 | 24.66423 | -0.07455 | 1.182269 | 1 |
| 10 | 68945800 | 68948799 | 35.79426 | 58.25307 | 69.18044 | 29.69688 | 0.122587 | 1.560091 | 1 |
| 10 | 69443400 | 69445199 | 16.65872 | 29.08817 | 40.49861 | 17.83692 | -0.08752 | 0.71625 | 1 |
| 10 | 69837800 | 69843199 | 36.76663 | 59.39708 | 88.86 | 37.93284 | -0.21959 | 1.723678 | 1 |
| 10 | 70306800 | 70315199 | 47.42405 | 76.49589 | 79.97447 | 34.07171 | 0.293653 | 1.779843 | 1 |
| 10 | 70492200 | 70499999 | 77.83483 | 121.6612 | 156.2767 | 64.83739 | -0.04678 | 2.64874 | 1 |
| 10 | 70726000 | 70726999 | 21.90913 | 36.47492 | 61.74469 | 26.81462 | -0.37566 | 1.234841 | 1 |
| 10 | 71283400 | 71288399 | 58.66584 | 93.26755 | 120.541 | 50.30533 | -0.03557 | 2.253046 | 1 |
| 10 | 71461200 | 71462999 | 18.49585 | 32.56153 | 37.91668 | 16.78558 | 0.169668 | 0.598371 | 1 |

| | | | | | | | | | |
|---|---|---|---|---|---|---|---|---|---|
| 10 | 71471600 | 71474199 | 29.01632 | 48.89656 | 53.99869 | 23.60647 | 0.238455 | 1.12873 | 1 |
| 10 | 71529200 | 71529999 | 8.826728 | 16.84242 | 26.16054 | 10.42139 | -0.12489 | -0.01825 | 1 |
| 10 | 71534200 | 71536199 | 19.93936 | 34.31634 | 43.73029 | 19.18246 | 0.036075 | 0.894938 | 1 |
| 10 | 71539000 | 71545599 | 61.00448 | 96.3812 | 140.3703 | 58.30055 | -0.21569 | 2.426229 | 1 |
| 10 | 71843400 | 71845399 | 37.40208 | 60.43319 | 80.09691 | 34.19899 | -0.04494 | 1.71648 | 1 |
| 10 | 71883800 | 71885599 | 16.23013 | 28.29292 | 44.17949 | 19.43786 | -0.25251 | 0.743492 | 1 |
| 10 | 71955400 | 71960199 | 59.57986 | 95.39267 | 109.1127 | 45.984 | 0.151349 | 2.033598 | 1 |
| 10 | 72291600 | 72293399 | 16.85919 | 29.45797 | 43.05644 | 18.97147 | -0.15729 | 0.711834 | 1 |
| 10 | 72433200 | 72434599 | 11.44852 | 20.74494 | 34.28584 | 14.83861 | -0.30803 | 0.338738 | 1 |
| 10 | 72772400 | 72777999 | 40.26895 | 64.88009 | 84.69905 | 36.02969 | -0.02798 | 1.804641 | 1 |
| 10 | 73272400 | 73274199 | 16.29168 | 28.68293 | 33.12848 | 14.66567 | 0.18173 | 0.579068 | 1 |
| 10 | 73437200 | 73438199 | 19.91531 | 34.56357 | 37.99894 | 16.67264 | 0.250764 | 0.813167 | 1 |
| 10 | 73955000 | 73957799 | 34.72295 | 56.88743 | 61.11915 | 26.30159 | 0.273078 | 1.458961 | 1 |
| 10 | 74549400 | 74556599 | 68.43271 | 107.825 | 144.8016 | 60.08538 | -0.10191 | 2.491525 | 1 |
| 10 | 74603200 | 74604599 | 17.64462 | 31.04159 | 31.64085 | 14.00922 | 0.360803 | 0.593725 | 1 |
| 10 | 74676000 | 74677199 | 8.164928 | 15.89328 | 20.80613 | 7.64434 | 0.228643 | -0.31121 | 1 |
| 10 | 74713000 | 74715599 | 27.01131 | 45.32502 | 49.11479 | 21.43602 | 0.265545 | 1.153779 | 1 |
| 10 | 75352400 | 75355399 | 18.95745 | 31.89937 | 55.81198 | 24.38839 | -0.42303 | 1.108016 | 1 |
| 10 | 75667800 | 75671399 | 22.27422 | 37.00078 | 63.85089 | 27.68737 | -0.40353 | 1.264306 | 1 |
| 10 | 75886800 | 75891399 | 63.80021 | 101.253 | 121.7976 | 50.74087 | 0.065379 | 2.309149 | 1 |
| 10 | 76046400 | 76054599 | 62.24207 | 98.51185 | 135.5223 | 56.33776 | -0.13151 | 2.383653 | 1 |
| 10 | 76752400 | 76759799 | 100.6188 | 154.5793 | 205.2418 | 85.21987 | -0.11373 | 2.974754 | 1 |
| 10 | 77555000 | 77558399 | 45.66844 | 73.50746 | 78.12285 | 33.18508 | 0.266373 | 1.841696 | 1 |
| 10 | 77930600 | 77937799 | 73.5332 | 115.0977 | 161.7301 | 67.09031 | -0.17456 | 2.622637 | 1 |
| 10 | 78645600 | 78647399 | 18.01975 | 31.54132 | 39.24685 | 17.31843 | 0.073746 | 0.682005 | 1 |
| 10 | 78874400 | 78895799 | 55.49685 | 89.19861 | 94.69341 | 40.14006 | 0.265401 | 1.906974 | 1 |
| 10 | 78897600 | 78908999 | 248.4351 | 365.6491 | 309.568 | 127.4816 | 0.481318 | 3.814281 | 1 |
| 10 | 79081000 | 79082599 | 26.43675 | 44.68504 | 50.13653 | 21.93825 | 0.215801 | 1.089369 | 1 |
| 10 | 79172400 | 79175399 | 44.44939 | 71.36297 | 95.13325 | 40.26017 | -0.06441 | 1.926708 | 1 |
| 10 | 81563600 | 81565999 | 67.52575 | 106.9881 | 108.878 | 45.28586 | 0.303999 | 2.356949 | 1 |
| 10 | 81672000 | 81672599 | 10.86608 | 20.42904 | 23.12419 | 9.49223 | 0.290505 | 0.090955 | 1 |
| 10 | 81796800 | 81797799 | 14.65681 | 25.70024 | 39.33385 | 17.38272 | -0.22348 | 0.634595 | 1 |
| 10 | 81813400 | 81814999 | 35.2218 | 57.58533 | 57.70202 | 24.81109 | 0.371725 | 1.486175 | 1 |
| 10 | 81875400 | 81877599 | 18.11065 | 31.68238 | 50.58747 | 22.31195 | -0.2846 | 0.707314 | 1 |
| 10 | 83089400 | 83092599 | 18.32 | 32.0523 | 34.58295 | 15.25453 | 0.278405 | 0.681757 | 1 |
| 10 | 83305400 | 83313999 | 231.9042 | 334.151 | 432.646 | 178.5345 | -0.16091 | 4.135068 | 1 |
| 10 | 83318000 | 83322599 | 51.64589 | 82.94024 | 90.85129 | 38.46455 | 0.218809 | 1.923495 | 1 |
| 10 | 83538600 | 83541399 | 14.53783 | 25.52259 | 42.92831 | 18.99069 | -0.35906 | 0.6203 | 1 |
| 10 | 83551600 | 83555399 | 17.05572 | 29.79155 | 41.45419 | 18.2617 | -0.08683 | 0.712844 | 1 |
| 10 | 83912600 | 83914399 | 17.57481 | 31.01356 | 34.37951 | 15.21247 | 0.241784 | 0.562931 | 1 |
| 10 | 84073000 | 84081199 | 95.18769 | 147.7838 | 167.355 | 69.49652 | 0.12725 | 2.774877 | 1 |
| 10 | 84087200 | 84088399 | 19.12149 | 33.38377 | 37.26034 | 16.40922 | 0.229827 | 0.724484 | 1 |
| 10 | 84108400 | 84110599 | 19.52948 | 33.4566 | 52.19368 | 22.88908 | -0.25573 | 0.948257 | 1 |
| 10 | 84893600 | 84898999 | 64.14598 | 101.8305 | 123.4004 | 51.42311 | 0.055132 | 2.301732 | 1 |
| 10 | 85045800 | 85049399 | 49.1054 | 78.78018 | 90.06776 | 38.07387 | 0.155483 | 1.952562 | 1 |
| 10 | 85552400 | 85553199 | 17.66416 | 30.9556 | 32.73139 | 14.45784 | 0.307631 | 0.65143 | 1 |
| 10 | 85603800 | 85606799 | 25.51042 | 43.33065 | 50.11392 | 21.96089 | 0.172379 | 1.042642 | 1 |

| | | | | | | | | | |
|---|---|---|---|---|---|---|---|---|---|
| 10 | 86068800 | 86071399 | 43.73157 | 70.30809 | 89.18707 | 37.82117 | 0.00952 | 1.879274 | 1 |
| 10 | 86511800 | 86519399 | 101.175 | 154.348 | 219.9349 | 91.195 | -0.22341 | 3.116325 | 1 |
| 10 | 86520200 | 86521999 | 14.37334 | 25.23489 | 39.3147 | 17.39273 | -0.24895 | 0.613507 | 1 |
| 10 | 86637200 | 86639799 | 23.65332 | 40.69733 | 42.56128 | 18.66951 | 0.319947 | 0.911831 | 1 |
| 10 | 86978000 | 86983599 | 109.9001 | 168.6543 | 177.8259 | 73.82528 | 0.217357 | 2.996533 | 1 |
| 10 | 87015800 | 87017599 | 21.8271 | 36.80197 | 55.92726 | 24.45366 | -0.22043 | 1.101703 | 1 |
| 10 | 87240200 | 87243799 | 42.87669 | 69.07713 | 79.18059 | 33.67907 | 0.159146 | 1.808944 | 1 |
| 10 | 88192400 | 88195999 | 38.00189 | 61.38039 | 84.48248 | 36.05486 | -0.09994 | 1.728159 | 1 |
| 10 | 88339200 | 88340399 | 12.33925 | 21.80634 | 38.26878 | 16.92848 | -0.41665 | 0.521337 | 1 |
| 10 | 88803600 | 88805399 | 32.27352 | 53.04193 | 65.03029 | 28.02214 | 0.084915 | 1.416538 | 1 |
| 10 | 89678200 | 89683199 | 71.51338 | 112.7402 | 142.7299 | 59.22989 | -0.01633 | 2.482127 | 1 |
| 10 | 90015600 | 90016799 | 12.77732 | 23.55434 | 29.6998 | 12.5692 | 0.101991 | 0.241314 | 1 |
| 10 | 90307000 | 90310399 | 7.505707 | 14.29053 | 25.05238 | 10.02613 | -0.30297 | -0.01616 | 1 |
| 10 | 90585000 | 90586999 | 19.59737 | 33.93857 | 40.57441 | 17.8015 | 0.129665 | 0.833958 | 1 |
| 10 | 90917600 | 90919399 | 15.53109 | 26.96551 | 43.24406 | 18.9782 | -0.29141 | 0.794424 | 1 |
| 10 | 93209800 | 93217199 | 90.31528 | 140.3918 | 158.3322 | 65.73759 | 0.134549 | 2.749839 | 1 |
| 10 | 94196800 | 94198599 | 19.72863 | 33.87039 | 50.61469 | 22.20047 | -0.19356 | 0.925262 | 1 |
| 10 | 94527000 | 94533399 | 84.58103 | 131.6972 | 171.8646 | 71.33835 | -0.07391 | 2.721643 | 1 |
| 10 | 94816000 | 94822199 | 59.61634 | 94.50259 | 125.9184 | 52.39626 | -0.08357 | 2.341577 | 1 |
| 10 | 95204400 | 95208199 | 41.15258 | 66.20165 | 95.51358 | 40.53939 | -0.17473 | 1.859279 | 1 |
| 10 | 96722800 | 96725999 | 53.89691 | 86.33875 | 100.3484 | 42.32395 | 0.129242 | 2.010031 | 1 |
| 10 | 96748000 | 96756199 | 188.2321 | 277.4633 | 340.1897 | 140.0941 | -0.0506 | 3.781888 | 1 |
| 10 | 97155400 | 97156799 | 25.75821 | 43.37546 | 49.08294 | 21.45286 | 0.203399 | 1.117866 | 1 |
| 10 | 97396800 | 97399399 | 39.11676 | 63.4163 | 75.45126 | 32.31191 | 0.115172 | 1.640249 | 1 |
| 10 | 97528600 | 97530399 | 40.78121 | 65.30519 | 106.5016 | 44.89491 | -0.35907 | 2.007131 | 1 |
| 10 | 97720200 | 97721599 | 12.839 | 23.95726 | 27.76948 | 11.5453 | 0.241964 | 0.168576 | 1 |
| 10 | 97749400 | 97750599 | 14.13403 | 24.7211 | 42.06875 | 18.55175 | -0.37554 | 0.679712 | 1 |
| 10 | 97752600 | 97752999 | 11.53325 | 21.75159 | 23.62625 | 9.646868 | 0.355731 | 0.078966 | 1 |
| 10 | 97833800 | 97834399 | 7.085604 | 13.78119 | 19.54594 | 7.228388 | 0.10565 | -0.30626 | 1 |
| 10 | 97881200 | 97882199 | 12.80158 | 22.67446 | 34.71219 | 15.34828 | -0.21921 | 0.494115 | 1 |
| 10 | 97925600 | 97927999 | 19.83259 | 33.65443 | 52.16598 | 22.85352 | -0.24843 | 1.034387 | 1 |
| 10 | 98057800 | 98060799 | 34.83538 | 57.52498 | 65.43777 | 28.25967 | 0.195055 | 1.367522 | 1 |
| 10 | 98173000 | 98173799 | 11.90227 | 21.08499 | 36.78621 | 16.2648 | -0.40657 | 0.491257 | 1 |
| 10 | 98263800 | 98266199 | 24.35777 | 41.90808 | 43.60334 | 19.12659 | 0.327319 | 0.916759 | 1 |
| 10 | 98534600 | 98536599 | 15.83511 | 27.70271 | 42.50141 | 18.74617 | -0.22694 | 0.68449 | 1 |
| 10 | 99166400 | 99171799 | 47.67834 | 76.81791 | 84.70985 | 36.03432 | 0.214936 | 1.814374 | 1 |
| 10 | 99173200 | 99177399 | 61.11855 | 97.23567 | 106.4272 | 44.45357 | 0.204888 | 2.230773 | 1 |
| 10 | 99279600 | 99282199 | 45.48681 | 72.62379 | 112.401 | 47.18091 | -0.28856 | 2.10466 | 1 |
| 10 | 99322000 | 99323199 | 17.93204 | 31.10564 | 46.33877 | 20.33155 | -0.18592 | 0.820046 | 1 |
| 10 | 99381200 | 99382799 | 36.83595 | 59.51389 | 77.33015 | 33.0144 | -0.01655 | 1.714475 | 1 |
| 10 | 99383600 | 99388599 | 109.1933 | 167.1423 | 197.4989 | 81.95579 | 0.050299 | 3.049578 | 1 |
| 10 | 99447800 | 99449599 | 26.95189 | 45.40082 | 51.25627 | 22.40505 | 0.206628 | 1.118962 | 1 |
| 10 | 99560600 | 99561999 | 16.16375 | 28.11175 | 44.73769 | 19.65471 | -0.28023 | 0.775142 | 1 |
| 10 | 100012200 | 100015199 | 37.8096 | 61.2359 | 82.22806 | 35.18812 | -0.06035 | 1.66052 | 1 |
| 10 | 100221400 | 100223999 | 25.8533 | 44.39731 | 43.53406 | 19.09511 | 0.411885 | 0.937742 | 1 |
| 10 | 100414000 | 100416999 | 34.82249 | 57.59921 | 59.29072 | 25.63439 | 0.338958 | 1.34228 | 1 |
| 10 | 100448400 | 100449599 | 10.42305 | 19.21406 | 33.57142 | 14.20801 | -0.36574 | 0.230154 | 1 |

| | | | | | | | | | |
|---|---|---|---|---|---|---|---|---|---|
| 10 | 101402400 | 101403599 | 22.91668 | 38.51623 | 53.81351 | 23.50673 | -0.09979 | 1.130023 | 1 |
| 10 | 102379000 | 102381799 | 9.790517 | 18.20481 | 27.33282 | 11.43156 | -0.13545 | 0.164153 | 1 |
| 10 | 103411000 | 103411999 | 12.85137 | 23.47334 | 26.0042 | 11.14272 | 0.274895 | 0.284862 | 1 |
| 10 | 103496800 | 103499399 | 40.34041 | 64.88165 | 97.69272 | 41.45181 | -0.23658 | 1.865278 | 1 |
| 10 | 103589600 | 103590399 | 11.84645 | 21.35006 | 31.25938 | 13.62072 | -0.14013 | 0.369695 | 1 |
| 10 | 104376400 | 104377399 | 16.26537 | 29.48604 | 29.80195 | 12.89098 | 0.397756 | 0.362296 | 1 |
| 10 | 104381200 | 104381999 | 15.74623 | 27.63217 | 36.83829 | 16.29066 | -0.025 | 0.628022 | 1 |
| 10 | 104409800 | 104415999 | 62.9341 | 100.0447 | 121.9173 | 50.87534 | 0.04899 | 2.259647 | 1 |
| 10 | 104578400 | 104580999 | 18.92624 | 33.52439 | 33.64924 | 14.87813 | 0.386385 | 0.529586 | 1 |
| 10 | 104589800 | 104591399 | 22.28019 | 37.056 | 61.63882 | 26.75073 | -0.35046 | 1.249526 | 1 |
| 10 | 104600800 | 104601599 | 14.92045 | 26.04181 | 41.60357 | 18.31866 | -0.28502 | 0.715106 | 1 |
| 10 | 104705600 | 104712199 | 85.69075 | 133.4013 | 180.5352 | 74.93937 | -0.12652 | 2.72514 | 1 |
| 10 | 104725800 | 104726599 | 12.24603 | 22.60787 | 24.11788 | 10.18666 | 0.343967 | 0.217004 | 1 |
| 10 | 104811000 | 104812799 | 21.4977 | 37.19925 | 38.34501 | 16.82296 | 0.342233 | 0.844384 | 1 |
| 10 | 105016000 | 105017799 | 21.49695 | 36.50676 | 47.46175 | 20.79748 | 0.004333 | 1.027489 | 1 |
| 10 | 105073200 | 105074599 | 174.5609 | 257.1116 | 331.823 | 136.6474 | -0.12558 | 3.79366 | 1 |
| 10 | 105229000 | 105231399 | 18.39902 | 32.36748 | 37.88205 | 16.77269 | 0.161904 | 0.604261 | 1 |
| 10 | 105496200 | 105496999 | 10.30473 | 19.51444 | 23.25185 | 9.412077 | 0.234314 | 0.03926 | 1 |
| 10 | 105512200 | 105515799 | 38.77881 | 62.61722 | 79.27767 | 33.82101 | 0.019844 | 1.735037 | 1 |
| 10 | 105643400 | 105647199 | 41.64212 | 67.25852 | 89.31964 | 38.1181 | -0.04851 | 1.737781 | 1 |
| 10 | 105716600 | 105722199 | 114.5378 | 175.6931 | 187.0474 | 77.64889 | 0.202928 | 3.00734 | 1 |
| 10 | 105928400 | 105930199 | 24.99776 | 41.91686 | 56.06646 | 24.45923 | -0.03718 | 1.161716 | 1 |
| 10 | 106017400 | 106024799 | 97.03449 | 149.9101 | 172.4669 | 71.63905 | 0.099008 | 2.863932 | 1 |
| 10 | 106232200 | 106233199 | 27.38698 | 45.67854 | 49.16052 | 21.37434 | 0.275588 | 1.21994 | 1 |
| 10 | 106277600 | 106279199 | 24.6574 | 40.89567 | 62.94025 | 27.26307 | -0.23918 | 1.287793 | 1 |
| 10 | 106371000 | 106372399 | 20.98346 | 35.48917 | 52.17851 | 22.83597 | -0.17267 | 1.067987 | 1 |
| 10 | 106381000 | 106382599 | 21.94916 | 37.9554 | 41.55005 | 18.22873 | 0.255818 | 0.860016 | 1 |
| 10 | 106935800 | 106941799 | 104.3712 | 159.481 | 213.4349 | 88.53279 | -0.13118 | 3.084102 | 1 |
| 10 | 107201000 | 107203199 | 33.62099 | 54.62333 | 77.14232 | 33.08828 | -0.12862 | 1.588835 | 1 |
| 10 | 107396200 | 107396799 | 10.78169 | 19.91442 | 27.78684 | 11.72836 | -0.03994 | 0.212243 | 1 |
| 10 | 107930200 | 107934399 | 44.24997 | 70.96793 | 97.18197 | 41.06129 | -0.1049 | 1.958909 | 1 |
| 10 | 108324800 | 108329599 | 36.57439 | 59.73091 | 69.08677 | 29.69445 | 0.164097 | 1.511423 | 1 |
| 10 | 108832000 | 108833799 | 20.1105 | 34.97458 | 38.90906 | 17.0936 | 0.234 | 0.786532 | 1 |
| 10 | 108919200 | 108919999 | 12.62597 | 22.88457 | 29.21099 | 12.63806 | 0.063236 | 0.338206 | 1 |
| 10 | 109034800 | 109036799 | 28.65255 | 47.98509 | 54.59984 | 23.80058 | 0.195314 | 1.18404 | 1 |
| 10 | 109147600 | 109151799 | 44.13859 | 71.25404 | 84.51595 | 36.04415 | 0.113276 | 1.75271 | 1 |
| 10 | 109156000 | 109158799 | 37.21433 | 60.60089 | 62.05277 | 26.6431 | 0.336539 | 1.550092 | 1 |
| 10 | 109242600 | 109244799 | 14.77625 | 26.35265 | 33.94761 | 14.93257 | 0.033664 | 0.456503 | 1 |
| 10 | 109419000 | 109419999 | 14.09971 | 25.42707 | 27.04661 | 11.77575 | 0.318612 | 0.371232 | 1 |
| 10 | 109850000 | 109852399 | 18.46063 | 31.9475 | 46.34792 | 20.33412 | -0.14848 | 0.844435 | 1 |
| 10 | 110580600 | 110581799 | 11.55505 | 20.50133 | 33.32707 | 14.72377 | -0.30401 | 0.46814 | 1 |
| 10 | 110682200 | 110684199 | 35.00262 | 56.93914 | 69.56454 | 29.85687 | 0.081349 | 1.565527 | 1 |
| 10 | 111229400 | 111232399 | 32.19905 | 52.69531 | 71.55762 | 30.77987 | -0.06466 | 1.473977 | 1 |
| 10 | 111298400 | 111300799 | 20.33972 | 35.33183 | 45.22333 | 19.85298 | 0.032327 | 0.812486 | 1 |
| 10 | 111935000 | 111941999 | 115.8671 | 177.702 | 207.2637 | 86.03927 | 0.071198 | 3.013445 | 1 |
| 10 | 112141600 | 112143799 | 22.46784 | 38.89908 | 42.03753 | 18.44327 | 0.274688 | 0.852477 | 1 |
| 10 | 112159000 | 112162399 | 18.49348 | 31.7818 | 46.3937 | 20.34987 | -0.15938 | 0.908017 | 1 |

| | | | | | | | | | |
|---|---|---|---|---|---|---|---|---|---|
| 10 | 112260200 | 112261999 | 26.08468 | 43.22151 | 61.39274 | 26.57519 | -0.12396 | 1.301175 | 1 |
| 10 | 112422800 | 112424199 | 23.88207 | 39.70551 | 61.76603 | 26.79909 | -0.25422 | 1.258389 | 1 |
| 10 | 112534000 | 112535999 | 32.77388 | 53.88769 | 59.40959 | 25.60484 | 0.23802 | 1.409559 | 1 |
| 10 | 112652000 | 112654799 | 31.25247 | 51.79948 | 64.76183 | 28.03922 | 0.059531 | 1.313595 | 1 |
| 10 | 112707800 | 112708799 | 10.86019 | 19.66934 | 32.78746 | 14.19872 | -0.32055 | 0.335648 | 1 |
| 10 | 113075000 | 113078599 | 26.38658 | 44.39917 | 50.69449 | 22.15201 | 0.190407 | 1.12754 | 1 |
| 10 | 113224800 | 113225799 | 11.77913 | 21.7628 | 24.17477 | 10.19991 | 0.287141 | 0.209118 | 1 |
| 10 | 113334200 | 113340399 | 112.7249 | 172.9079 | 194.9847 | 80.94362 | 0.119968 | 3.008541 | 1 |
| 10 | 113827000 | 113828599 | 12.73721 | 22.84907 | 35.15453 | 15.38757 | -0.21616 | 0.40923 | 1 |
| 10 | 113891400 | 113892399 | 15.53095 | 27.09008 | 43.64212 | 19.20856 | -0.29731 | 0.729193 | 1 |
| 10 | 114001000 | 114003399 | 24.63121 | 41.8155 | 51.18316 | 22.42591 | 0.090895 | 1.046409 | 1 |
| 10 | 114366600 | 114368999 | 28.07162 | 47.25485 | 49.09209 | 21.45461 | 0.326265 | 1.129003 | 1 |
| 10 | 114694400 | 114697199 | 26.94491 | 44.99549 | 60.04144 | 26.1251 | -0.03374 | 1.214421 | 1 |
| 10 | 114869800 | 114872799 | 36.06813 | 58.33205 | 82.65334 | 35.33286 | -0.13958 | 1.687297 | 1 |
| 10 | 115139000 | 115141599 | 30.12594 | 49.66257 | 59.90483 | 25.84664 | 0.11008 | 1.370541 | 1 |
| 10 | 115408400 | 115410999 | 25.38909 | 42.2112 | 62.04495 | 26.92018 | -0.17284 | 1.259769 | 1 |
| 10 | 115552200 | 115554399 | 22.75166 | 38.47467 | 57.11266 | 24.99441 | -0.18661 | 1.079363 | 1 |
| 10 | 115638200 | 115639799 | 15.04425 | 26.77834 | 33.88177 | 14.92813 | 0.057725 | 0.469273 | 1 |
| 10 | 115872400 | 115874399 | 10.67247 | 19.79234 | 29.86763 | 12.53765 | -0.1471 | 0.190379 | 1 |
| 10 | 116249000 | 116261199 | 37.39242 | 60.36093 | 85.19985 | 36.3338 | -0.13716 | 1.745271 | 1 |
| 10 | 116456400 | 116458799 | 36.47669 | 59.48113 | 72.91972 | 31.3246 | 0.078839 | 1.535801 | 1 |
| 10 | 116469400 | 116471599 | 23.20884 | 39.13425 | 58.79611 | 25.70855 | -0.20416 | 1.106413 | 1 |
| 10 | 117075400 | 117077399 | 13.53729 | 23.8871 | 37.17466 | 16.4495 | -0.24524 | 0.54248 | 1 |
| 10 | 117134000 | 117136599 | 18.5191 | 32.74111 | 35.2348 | 15.58509 | 0.285243 | 0.548994 | 1 |
| 10 | 117275800 | 117276599 | 8.074494 | 15.57107 | 24.23811 | 9.377984 | -0.08833 | -0.12407 | 1 |
| 10 | 117325600 | 117328999 | 24.73166 | 40.93096 | 65.53876 | 28.34664 | -0.29633 | 1.316718 | 1 |
| 10 | 117371600 | 117373399 | 16.26375 | 28.83752 | 31.20954 | 13.79557 | 0.278685 | 0.495757 | 1 |
| 10 | 117864400 | 117871999 | 89.8061 | 138.3351 | 190.2785 | 79.03564 | -0.1617 | 2.917486 | 1 |
| 10 | 117960600 | 117963399 | 31.42434 | 51.22016 | 73.33919 | 31.49979 | -0.14499 | 1.53114 | 1 |
| 10 | 118570800 | 118572599 | 18.3839 | 31.84729 | 50.652 | 22.22313 | -0.28049 | 0.837863 | 1 |
| 10 | 118745200 | 118747199 | 19.95517 | 34.64024 | 40.01853 | 17.56056 | 0.179577 | 0.813565 | 1 |
| 10 | 118995400 | 118997399 | 17.13374 | 29.62186 | 48.46871 | 21.26392 | -0.32173 | 0.848327 | 1 |
| 10 | 119178600 | 119185799 | 90.12821 | 138.8179 | 201.9549 | 83.88455 | -0.24263 | 2.921146 | 1 |
| 10 | 119488800 | 119496199 | 78.33978 | 122.3885 | 160.8231 | 66.72712 | -0.08009 | 2.658344 | 1 |
| 10 | 119497400 | 119499999 | 27.90245 | 46.03017 | 68.76056 | 29.67656 | -0.19715 | 1.362172 | 1 |
| 10 | 119531400 | 119538599 | 83.0062 | 128.4841 | 189.1594 | 78.57198 | -0.25429 | 2.82545 | 1 |
| 10 | 119605400 | 119607799 | 33.02506 | 53.90416 | 77.30893 | 33.22166 | -0.146 | 1.51638 | 1 |
| 10 | 119685000 | 119685999 | 11.58962 | 21.07791 | 30.20629 | 13.01308 | -0.09899 | 0.311023 | 1 |
| 10 | 119926800 | 119933799 | 95.42992 | 146.8737 | 191.8764 | 79.6901 | -0.08837 | 2.936138 | 1 |
| 10 | 120096200 | 120097199 | 10.46372 | 19.2755 | 29.82603 | 12.63495 | -0.19214 | 0.228524 | 1 |
| 10 | 120155200 | 120157599 | 28.17216 | 46.37386 | 68.42932 | 29.50637 | -0.18067 | 1.392338 | 1 |
| 10 | 120168800 | 120177399 | 62.3865 | 98.66668 | 141.8568 | 58.94877 | -0.19594 | 2.401842 | 1 |
| 10 | 120354800 | 120357199 | 29.01299 | 47.91002 | 67.81571 | 29.28651 | -0.11943 | 1.352394 | 1 |
| 10 | 120720000 | 120727599 | 110.358 | 167.3119 | 246.1411 | 101.8778 | -0.27676 | 3.247049 | 1 |
| 10 | 120802200 | 120804399 | 18.13398 | 31.24775 | 47.41727 | 20.80054 | -0.21437 | 0.882419 | 1 |
| 10 | 121106800 | 121109199 | 26.91561 | 44.53671 | 64.2041 | 27.76563 | -0.14543 | 1.321691 | 1 |
| 10 | 121532600 | 121534599 | 22.06004 | 37.28403 | 52.21511 | 22.84717 | -0.10288 | 1.076605 | 1 |

| | | | | | | | | | |
|---|---|---|---|---|---|---|---|---|---|
| 10 | 121553800 | 121560799 | 97.16295 | 150.2249 | 158.4649 | 65.82265 | 0.224923 | 2.848612 | 1 |
| 10 | 121616400 | 121622999 | 74.17581 | 116.1203 | 159.536 | 66.17916 | -0.14197 | 2.620051 | 1 |
| 10 | 121910800 | 121924399 | 14.66913 | 25.95412 | 41.2567 | 18.24595 | -0.27432 | 0.527039 | 1 |
| 10 | 121949200 | 121950999 | 15.62067 | 26.94631 | 47.94127 | 21.03111 | -0.44245 | 0.862549 | 1 |
| 10 | 121994600 | 121995799 | 15.04975 | 26.29296 | 43.09186 | 18.98554 | -0.32172 | 0.702979 | 1 |
| 10 | 122226000 | 122228199 | 34.02157 | 54.9991 | 83.6229 | 35.72797 | -0.24169 | 1.698296 | 1 |
| 10 | 122260800 | 122261599 | 11.70561 | 21.32576 | 30.68159 | 13.18925 | -0.10277 | 0.301869 | 1 |
| 10 | 122467200 | 122468399 | 16.57029 | 28.66143 | 43.81201 | 19.22124 | -0.22349 | 0.838585 | 1 |
| 10 | 122651000 | 122652999 | 14.89925 | 26.15634 | 41.3736 | 18.30246 | -0.27074 | 0.622014 | 1 |
| 10 | 122674200 | 122679799 | 146.2004 | 219.5157 | 249.9322 | 103.1522 | 0.080583 | 3.446857 | 1 |
| 10 | 123192000 | 123194599 | 19.18042 | 32.59541 | 51.63034 | 22.62738 | -0.2794 | 1.018298 | 1 |
| 10 | 123465200 | 123466799 | 15.85408 | 27.78434 | 41.92241 | 18.51807 | -0.203 | 0.656009 | 1 |
| 10 | 123650000 | 123651999 | 16.35934 | 29.20564 | 30.25483 | 13.29684 | 0.347274 | 0.453955 | 1 |
| 10 | 123692000 | 123702799 | 213.8292 | 312.2537 | 374.6564 | 154.3612 | -0.03229 | 3.933001 | 1 |
| 10 | 123923000 | 123923799 | 14.80496 | 26.30643 | 30.54879 | 13.47947 | 0.179665 | 0.474582 | 1 |
| 10 | 124302600 | 124306399 | 48.75095 | 78.31869 | 84.62298 | 35.84644 | 0.239149 | 1.907696 | 1 |
| 10 | 125081800 | 125086399 | 40.86046 | 65.83458 | 97.8989 | 41.64606 | -0.21546 | 1.811622 | 1 |
| 10 | 125291400 | 125291999 | 9.042914 | 17.21429 | 26.09873 | 10.44876 | -0.09708 | -0.00287 | 1 |
| 10 | 125321000 | 125321999 | 11.79906 | 21.44698 | 34.87039 | 15.03277 | -0.28084 | 0.325479 | 1 |
| 10 | 125337400 | 125344399 | 76.69871 | 119.9185 | 171.4431 | 71.12774 | -0.20075 | 2.645342 | 1 |
| 10 | 125543200 | 125544999 | 23.30594 | 39.83014 | 44.75046 | 19.62112 | 0.215303 | 0.972708 | 1 |
| 10 | 125876000 | 125883599 | 111.3732 | 170.0143 | 229.3277 | 95.10371 | -0.14364 | 3.106267 | 1 |
| 10 | 126017000 | 126026599 | 128.015 | 194.9182 | 213.6399 | 88.53949 | 0.152603 | 3.161883 | 1 |
| 10 | 126039600 | 126041799 | 33.66185 | 55.28053 | 57.29866 | 24.68237 | 0.326055 | 1.42366 | 1 |
| 10 | 126120000 | 126122799 | 21.24344 | 35.93574 | 54.89516 | 24.02631 | -0.22778 | 1.067885 | 1 |
| 10 | 126419600 | 126425599 | 87.3712 | 134.9423 | 193.9799 | 80.5752 | -0.22232 | 2.868271 | 1 |
| 10 | 126426400 | 126430399 | 47.40433 | 76.14748 | 97.42343 | 41.26256 | -0.00438 | 1.912194 | 1 |
| 10 | 126444800 | 126449799 | 63.32915 | 100.8991 | 106.8034 | 44.7005 | 0.255294 | 2.185401 | 1 |
| 10 | 126659400 | 126663999 | 52.69509 | 84.24552 | 97.12224 | 40.84517 | 0.137726 | 2.065709 | 1 |
| 10 | 127054800 | 127057799 | 37.90879 | 61.67501 | 71.20535 | 30.56262 | 0.162872 | 1.569671 | 1 |
| 10 | 127662800 | 127665999 | 46.70641 | 74.8299 | 106.6119 | 44.97463 | -0.16377 | 1.996904 | 1 |
| 10 | 127764400 | 127768599 | 32.07052 | 52.31645 | 79.42957 | 34.1259 | -0.22854 | 1.523798 | 1 |
| 10 | 127774800 | 127776999 | 22.66041 | 38.34167 | 56.05873 | 24.53732 | -0.16484 | 1.071377 | 1 |
| 10 | 128618000 | 128622999 | 77.26767 | 121.5925 | 134.3465 | 55.73789 | 0.177805 | 2.519903 | 1 |
| 10 | 128894600 | 128896199 | 15.93676 | 27.88003 | 38.88434 | 17.15052 | -0.09002 | 0.679241 | 1 |
| 10 | 129025200 | 129025999 | 10.24704 | 18.54885 | 31.82655 | 13.79187 | -0.36245 | 0.332548 | 1 |
| 10 | 129055200 | 129055999 | 17.80068 | 31.16995 | 37.67081 | 16.62841 | 0.115559 | 0.671503 | 1 |
| 10 | 129170400 | 129172599 | 32.63407 | 54.23625 | 55.84604 | 24.22435 | 0.338999 | 1.279495 | 1 |
| 10 | 129230400 | 129233399 | 25.09477 | 41.85446 | 62.60613 | 27.21993 | -0.19788 | 1.225656 | 1 |
| 10 | 129248200 | 129249599 | 20.4574 | 35.1733 | 42.20855 | 18.51438 | 0.122193 | 0.905115 | 1 |
| 10 | 129429200 | 129430199 | 24.72772 | 41.41946 | 52.24516 | 22.7767 | 0.047293 | 1.17056 | 1 |
| 10 | 129531800 | 129534399 | 33.81923 | 55.21433 | 68.66709 | 29.51075 | 0.05944 | 1.509956 | 1 |
| 10 | 129760800 | 129763599 | 25.20913 | 42.56564 | 49.60198 | 21.69748 | 0.161267 | 1.09215 | 1 |
| 10 | 129861600 | 129863399 | 11.69557 | 21.90416 | 28.4353 | 11.74862 | 0.086806 | 0.137055 | 1 |
| 10 | 129952000 | 129953599 | 20.04419 | 34.21241 | 50.60806 | 22.18715 | -0.18023 | 0.978812 | 1 |
| 10 | 130036600 | 130038399 | 28.41935 | 47.35138 | 57.85165 | 25.1367 | 0.093156 | 1.241289 | 1 |
| 10 | 130277200 | 130282999 | 59.17683 | 94.25773 | 122.3111 | 51.17164 | -0.03849 | 2.194684 | 1 |

| | | | | | | | | | |
|---|---|---|---|---|---|---|---|---|---|
| 10 | 130318400 | 130324799 | 97.37329 | 149.9397 | 193.1774 | 80.23595 | -0.06778 | 2.927514 | 1 |
| 10 | 130358600 | 130360599 | 24.53566 | 41.72539 | 47.95532 | 21.01796 | 0.181666 | 1.026397 | 1 |
| 10 | 130400400 | 130409399 | 215.0188 | 315.0112 | 348.1874 | 143.4237 | 0.090516 | 3.882565 | 1 |
| 11 | 3078800 | 3093799 | 1238.33 | 1565.407 | 1345.812 | 565.3384 | 0.240104 | 6.191135 | 1 |
| 11 | 3311600 | 3314599 | 24.8455 | 41.69471 | 58.82792 | 25.67682 | -0.11401 | 1.155102 | 1 |
| 11 | 3574800 | 3580399 | 61.4549 | 97.30854 | 144.0112 | 59.87994 | -0.23625 | 2.374668 | 1 |
| 11 | 3899400 | 3909199 | 136.2202 | 206.4771 | 220.8064 | 91.38566 | 0.182764 | 3.256394 | 1 |
| 11 | 5311000 | 5315399 | 96.75785 | 148.3974 | 199.7174 | 82.90687 | -0.13506 | 3.006393 | 1 |
| 11 | 6184200 | 6190999 | 141.5917 | 213.9659 | 238.9655 | 98.80617 | 0.116446 | 3.319476 | 1 |
| 11 | 6192200 | 6199199 | 27.6253 | 46.12697 | 50.72847 | 22.07144 | 0.244487 | 1.211552 | 1 |
| 11 | 6397200 | 6401999 | 57.74672 | 92.14958 | 101.2031 | 42.44901 | 0.204653 | 2.128743 | 1 |
| 11 | 6483600 | 6483999 | 8.607525 | 16.61935 | 23.7438 | 9.147532 | 0.039545 | -0.13651 | 1 |
| 11 | 6525200 | 6527999 | 22.56474 | 37.92044 | 52.74801 | 23.04051 | -0.09346 | 1.128081 | 1 |
| 11 | 6703600 | 6705799 | 22.67399 | 37.91298 | 55.87656 | 24.33295 | -0.17646 | 1.186371 | 1 |
| 11 | 6735200 | 6743399 | 118.5043 | 180.8073 | 210.1534 | 87.14163 | 0.070363 | 3.117144 | 1 |
| 11 | 6747400 | 6750999 | 60.25026 | 95.46021 | 134.4544 | 55.92374 | -0.16422 | 2.356873 | 1 |
| 11 | 6794600 | 6804199 | 108.3266 | 167.1693 | 189.2113 | 78.59477 | 0.121288 | 2.891146 | 1 |
| 11 | 6983200 | 6985799 | 22.21028 | 37.67467 | 52.75968 | 23.11092 | -0.10263 | 1.046266 | 1 |
| 11 | 7530400 | 7535999 | 125.6622 | 190.6587 | 216.6586 | 89.69529 | 0.096353 | 3.233247 | 1 |
| 11 | 7670400 | 7671799 | 12.14816 | 21.77006 | 35.21174 | 15.42594 | -0.28859 | 0.412798 | 1 |
| 11 | 7789800 | 7802999 | 212.1831 | 309.1266 | 395.4434 | 162.9652 | -0.12793 | 3.968589 | 1 |
| 11 | 8785800 | 8788999 | 47.21104 | 75.54683 | 112.9468 | 47.56165 | -0.23519 | 2.036633 | 1 |
| 11 | 8872800 | 8873799 | 20.92661 | 35.57012 | 47.98471 | 21.02999 | -0.04856 | 1.019543 | 1 |
| 11 | 8876200 | 8876799 | 9.836301 | 18.06548 | 29.31728 | 12.47019 | -0.26463 | 0.238628 | 1 |
| 11 | 9664600 | 9665999 | 10.46814 | 19.26904 | 31.38729 | 13.31005 | -0.26695 | 0.235906 | 1 |
| 11 | 9699600 | 9700599 | 17.44709 | 30.2116 | 44.26484 | 19.42056 | -0.16247 | 0.830972 | 1 |
| 11 | 9832200 | 9840799 | 182.8077 | 268.0936 | 366.3639 | 150.9012 | -0.21401 | 3.863722 | 1 |
| 11 | 10056600 | 10059199 | 25.3039 | 43.05463 | 49.02548 | 21.4879 | 0.195182 | 1.027178 | 1 |
| 11 | 10147400 | 10151399 | 47.61525 | 76.52769 | 80.3267 | 34.04813 | 0.281654 | 1.891264 | 1 |
| 11 | 10151000 | 10156199 | 50.20638 | 80.51383 | 96.50171 | 40.7685 | 0.086787 | 1.97014 | 1 |
| 11 | 10754200 | 10757999 | 45.35816 | 73.11444 | 85.22939 | 36.27903 | 0.135848 | 1.798934 | 1 |
| 11 | 10998600 | 11002599 | 86.43272 | 134.9884 | 142.7486 | 59.22979 | 0.232557 | 2.666676 | 1 |
| 11 | 11003600 | 11009799 | 94.22254 | 146.9003 | 154.6961 | 64.19948 | 0.236654 | 2.701842 | 1 |
| 11 | 11327800 | 11328399 | 11.62098 | 21.6039 | 26.16264 | 10.94603 | 0.171922 | 0.177562 | 1 |
| 11 | 12706000 | 12706999 | 15.1792 | 26.61688 | 36.8474 | 16.28423 | -0.07921 | 0.635146 | 1 |
| 11 | 12721600 | 12724599 | 29.73342 | 49.91638 | 60.27678 | 26.31552 | 0.109741 | 1.167605 | 1 |
| 11 | 12855400 | 12856599 | 12.74326 | 22.73788 | 35.64156 | 15.67131 | -0.24741 | 0.443361 | 1 |
| 11 | 12886200 | 12887999 | 22.20827 | 37.44818 | 52.22189 | 22.83418 | -0.09689 | 1.096499 | 1 |
| 11 | 12964400 | 12965399 | 19.53116 | 33.86382 | 42.64567 | 18.71091 | 0.055358 | 0.827151 | 1 |
| 11 | 13007000 | 13009399 | 14.22536 | 25.20915 | 39.69023 | 17.54757 | -0.25953 | 0.505446 | 1 |
| 11 | 13090800 | 13091599 | 11.12651 | 20.24564 | 30.74617 | 13.23751 | -0.18156 | 0.305892 | 1 |
| 11 | 13265000 | 13267199 | 16.05539 | 28.21753 | 34.59946 | 15.32025 | 0.094827 | 0.598293 | 1 |
| 11 | 13329000 | 13329799 | 20.22689 | 34.55633 | 45.37992 | 19.89657 | -0.00878 | 0.966764 | 1 |
| 11 | 13495600 | 13497599 | 25.28269 | 41.88451 | 62.43855 | 27.02428 | -0.19347 | 1.301749 | 1 |
| 11 | 13760000 | 13763199 | 22.7447 | 38.39387 | 54.91125 | 24.01791 | -0.13318 | 1.092775 | 1 |
| 11 | 13862400 | 13865399 | 35.25913 | 57.88334 | 61.4028 | 26.44638 | 0.292478 | 1.433489 | 1 |
| 11 | 14045600 | 14046799 | 15.97584 | 28.30424 | 31.30423 | 13.8407 | 0.247145 | 0.50278 | 1 |

| | | | | | | | | | |
|---|---|---|---|---|---|---|---|---|---|
| 11 | 14077200 | 14078799 | 25.9925 | 43.3896 | 54.97817 | 23.91661 | 0.040786 | 1.2121 | 1 |
| 11 | 14082200 | 14084599 | 23.76896 | 40.48248 | 46.89916 | 20.55756 | 0.170589 | 1.009713 | 1 |
| 11 | 14092400 | 14098199 | 29.32545 | 48.58032 | 61.09208 | 26.4409 | 0.051157 | 1.312631 | 1 |
| 11 | 14220800 | 14221999 | 12.61521 | 23.4412 | 23.5949 | 9.878077 | 0.436491 | 0.183476 | 1 |
| 11 | 14518800 | 14524199 | 65.04041 | 103.1988 | 113.9357 | 47.45525 | 0.188617 | 2.314244 | 1 |
| 11 | 14548000 | 14550199 | 20.81797 | 36.13639 | 37.88764 | 16.62505 | 0.318842 | 0.814063 | 1 |
| 11 | 14588600 | 14589199 | 10.74143 | 20.18843 | 23.77643 | 9.765093 | 0.233723 | 0.096666 | 1 |
| 11 | 14711200 | 14714199 | 31.53977 | 52.08175 | 59.56734 | 25.72413 | 0.187246 | 1.352841 | 1 |
| 11 | 14846400 | 14850399 | 71.09312 | 111.3263 | 161.2893 | 66.90452 | -0.21814 | 2.614299 | 1 |
| 11 | 14999000 | 15001399 | 30.34218 | 49.40997 | 78.78762 | 33.82838 | -0.30084 | 1.546176 | 1 |
| 11 | 15142600 | 15143399 | 11.15807 | 20.11808 | 33.22863 | 14.47102 | -0.31281 | 0.36376 | 1 |
| 11 | 15728000 | 15728999 | 12.15166 | 21.67463 | 34.20577 | 15.04449 | -0.25718 | 0.440233 | 1 |
| 11 | 15975200 | 15976799 | 27.85382 | 46.51554 | 55.02285 | 23.94215 | 0.139672 | 1.21498 | 1 |
| 11 | 16054600 | 16056599 | 15.79729 | 27.75477 | 41.27578 | 18.27136 | -0.18234 | 0.612821 | 1 |
| 11 | 16154200 | 16157799 | 86.05311 | 134.4694 | 142.749 | 59.22659 | 0.227614 | 2.656357 | 1 |
| 11 | 16207000 | 16208999 | 10.36252 | 19.16927 | 26.73453 | 11.26091 | -0.03741 | 0.195379 | 1 |
| 11 | 16521200 | 16521399 | 9.964577 | 19.05662 | 21.77488 | 8.628772 | 0.321264 | -0.0367 | 1 |
| 11 | 17048800 | 17049599 | 13.59295 | 24.8462 | 26.01287 | 11.13615 | 0.356961 | 0.287947 | 1 |
| 11 | 17080800 | 17083199 | 7.731366 | 14.71781 | 25.55653 | 10.23137 | -0.29092 | -0.01645 | 1 |
| 11 | 17357000 | 17357799 | 12.85447 | 23.39556 | 28.60955 | 12.31381 | 0.129333 | 0.313107 | 1 |
| 11 | 17468000 | 17469799 | 9.164219 | 17.18602 | 29.59692 | 12.20121 | -0.31542 | 0.108896 | 1 |
| 11 | 17639600 | 17641199 | 17.67038 | 30.72714 | 41.05852 | 18.03651 | -0.02926 | 0.778583 | 1 |
| 11 | 17922800 | 17924999 | 27.95299 | 46.07186 | 64.04293 | 27.6299 | -0.09402 | 1.372017 | 1 |
| 11 | 18001600 | 18003999 | 21.24391 | 36.61804 | 39.59634 | 17.3699 | 0.272578 | 0.882687 | 1 |
| 11 | 18056400 | 18061399 | 49.90754 | 79.48462 | 115.5246 | 48.32491 | -0.20191 | 2.188843 | 1 |
| 11 | 18631200 | 18633799 | 20.26991 | 35.30804 | 43.53274 | 19.14405 | 0.086419 | 0.770142 | 1 |
| 11 | 18729200 | 18731999 | 42.28143 | 68.00061 | 98.21202 | 41.66958 | -0.17671 | 1.869301 | 1 |
| 11 | 18754800 | 18758799 | 73.82681 | 116.2634 | 131.4314 | 54.53369 | 0.145662 | 2.50307 | 1 |
| 11 | 18762000 | 18767599 | 69.60729 | 109.5279 | 148.8176 | 61.74098 | -0.12026 | 2.519537 | 1 |
| 11 | 19031000 | 19033199 | 18.04307 | 30.86667 | 50.05143 | 21.94713 | -0.31153 | 0.951729 | 1 |
| 11 | 19539000 | 19540599 | 12.98398 | 23.0248 | 37.19155 | 16.43818 | -0.29556 | 0.488262 | 1 |
| 11 | 19732600 | 19741199 | 69.09535 | 108.7533 | 151.0261 | 62.65954 | -0.1514 | 2.513633 | 1 |
| 11 | 19853000 | 19860399 | 81.46477 | 127.3875 | 177.1735 | 73.50412 | -0.16092 | 2.64619 | 1 |
| 11 | 20132800 | 20135999 | 35.50576 | 57.64835 | 78.15646 | 33.50736 | -0.07078 | 1.60404 | 1 |
| 11 | 20281800 | 20283399 | 14.55312 | 25.50878 | 37.90087 | 16.74411 | -0.18072 | 0.639764 | 1 |
| 11 | 20356800 | 20375599 | 321.4209 | 452.6289 | 543.1287 | 224.9608 | -0.08445 | 4.475558 | 1 |
| 11 | 20467200 | 20473399 | 31.35721 | 52.2155 | 52.39756 | 22.7584 | 0.376324 | 1.254884 | 1 |
| 11 | 20523800 | 20526399 | 25.68295 | 43.27811 | 46.97193 | 20.53442 | 0.26338 | 1.108707 | 1 |
| 11 | 20897000 | 20899199 | 21.95077 | 37.77862 | 46.85708 | 20.55404 | 0.074927 | 0.907916 | 1 |
| 11 | 21135200 | 21146999 | 146.0591 | 218.7188 | 295.22 | 121.7679 | -0.16821 | 3.499592 | 1 |
| 11 | 21159000 | 21162799 | 25.15322 | 42.5349 | 46.93788 | 20.54144 | 0.239664 | 1.075931 | 1 |
| 11 | 21429800 | 21432399 | 22.26417 | 38.14964 | 43.22272 | 18.95527 | 0.203946 | 0.944899 | 1 |
| 11 | 21786800 | 21789799 | 38.87862 | 63.02864 | 81.14408 | 34.7492 | 0.001621 | 1.644496 | 1 |
| 11 | 21855400 | 21857599 | 18.40905 | 31.92032 | 46.3359 | 20.33004 | -0.14882 | 0.825692 | 1 |
| 11 | 22526200 | 22529199 | 20.79034 | 35.4147 | 56.00279 | 24.54829 | -0.27664 | 1.007283 | 1 |
| 11 | 22561600 | 22564599 | 26.56881 | 44.32814 | 57.25094 | 24.89698 | 0.01323 | 1.219554 | 1 |
| 11 | 22591400 | 22592799 | 10.68984 | 19.77284 | 30.36484 | 12.79174 | -0.17586 | 0.206382 | 1 |

| | | | | | | | | | |
|---|---|---|---|---|---|---|---|---|---|
| 11 | 22699000 | 22701199 | 10.39336 | 19.38002 | 28.37365 | 11.81065 | -0.0941 | 0.153179 | 1 |
| 11 | 22966200 | 22969199 | 30.60925 | 50.60242 | 68.09297 | 29.42891 | -0.04655 | 1.343427 | 1 |
| 11 | 23134800 | 23136599 | 34.73721 | 56.86637 | 63.90609 | 27.49182 | 0.207937 | 1.470278 | 1 |
| 11 | 23166800 | 23171599 | 24.73549 | 41.33271 | 59.84679 | 26.05104 | -0.15112 | 1.201876 | 1 |
| 11 | 23194400 | 23195599 | 20.05236 | 34.91747 | 35.32285 | 15.53041 | 0.370722 | 0.763084 | 1 |
| 11 | 23474200 | 23487399 | 60.77106 | 96.3931 | 128.8537 | 53.65116 | -0.08736 | 2.324943 | 1 |
| 11 | 23801000 | 23801799 | 8.741998 | 16.82912 | 21.94678 | 8.533781 | 0.15771 | -0.10983 | 1 |
| 11 | 26458400 | 26459999 | 19.38549 | 33.30444 | 45.33482 | 19.88522 | -0.05915 | 0.911808 | 1 |
| 11 | 26502000 | 26505999 | 56.68727 | 90.47816 | 105.7563 | 44.37215 | 0.115248 | 2.123366 | 1 |
| 11 | 26682600 | 26683399 | 13.49455 | 23.64262 | 40.50146 | 17.88617 | -0.385 | 0.646008 | 1 |
| 11 | 27395400 | 27396799 | 19.15926 | 33.19023 | 44.21173 | 19.39751 | -0.02566 | 0.834024 | 1 |
| 11 | 27483600 | 27485799 | 34.71248 | 57.07756 | 62.44762 | 26.91467 | 0.248985 | 1.415035 | 1 |
| 11 | 27528400 | 27529999 | 24.32263 | 40.6764 | 54.41269 | 23.69795 | -0.03726 | 1.185579 | 1 |
| 11 | 27723200 | 27724599 | 11.35644 | 20.86525 | 32.3835 | 13.76736 | -0.20062 | 0.253899 | 1 |
| 11 | 27988000 | 27989799 | 23.13306 | 39.8016 | 40.0823 | 17.5821 | 0.374353 | 0.909068 | 1 |
| 11 | 28047200 | 28049399 | 33.84885 | 55.29612 | 65.22036 | 28.03564 | 0.136325 | 1.500403 | 1 |
| 11 | 28179800 | 28182599 | 15.52296 | 27.02792 | 43.68359 | 19.20529 | -0.30224 | 0.754261 | 1 |
| 11 | 28189000 | 28190399 | 12.3933 | 22.38407 | 33.67477 | 14.63347 | -0.177 | 0.363159 | 1 |
| 11 | 28303600 | 28304199 | 12.10722 | 22.42493 | 27.1424 | 11.41314 | 0.167414 | 0.20422 | 1 |
| 11 | 28728000 | 28730999 | 17.51516 | 30.62625 | 36.79835 | 16.22372 | 0.124079 | 0.692589 | 1 |
| 11 | 29018400 | 29020799 | 13.06088 | 23.73688 | 27.02397 | 11.65242 | 0.230004 | 0.317875 | 1 |
| 11 | 29209200 | 29210399 | 11.82163 | 22.08087 | 24.55913 | 10.19335 | 0.30282 | 0.144109 | 1 |
| 11 | 29281400 | 29284199 | 14.31954 | 25.89969 | 24.54345 | 10.6466 | 0.487631 | 0.349566 | 1 |
| 11 | 30021000 | 30022399 | 22.04889 | 37.29767 | 55.4671 | 24.27652 | -0.18923 | 1.071545 | 1 |
| 11 | 30203400 | 30205799 | 21.09545 | 37.01384 | 35.35883 | 15.63449 | 0.453335 | 0.656205 | 1 |
| 11 | 30247200 | 30248199 | 13.03218 | 23.38428 | 33.58706 | 14.69825 | -0.11728 | 0.406506 | 1 |
| 11 | 30260800 | 30273399 | 215.0525 | 314.863 | 358.691 | 147.7561 | 0.046118 | 3.892101 | 1 |
| 11 | 30983800 | 30986599 | 81.52034 | 127.1891 | 147.2278 | 61.09463 | 0.101159 | 2.682117 | 1 |
| 11 | 31286200 | 31287799 | 16.60777 | 29.35941 | 33.92035 | 15.00394 | 0.183426 | 0.537177 | 1 |
| 11 | 31401000 | 31407399 | 125.8716 | 190.2636 | 246.1508 | 101.7861 | -0.09517 | 3.310931 | 1 |
| 11 | 31532600 | 31534599 | 19.39907 | 33.49267 | 49.55093 | 21.73781 | -0.17748 | 0.872332 | 1 |
| 11 | 31722000 | 31729399 | 82.69361 | 128.9513 | 164.7223 | 68.3584 | -0.04142 | 2.693907 | 1 |
| 11 | 32934400 | 32935999 | 17.35678 | 30.09514 | 49.53951 | 21.73563 | -0.32942 | 0.826217 | 1 |
| 11 | 34637600 | 34638999 | 20.39457 | 35.1975 | 43.72623 | 19.1823 | 0.073573 | 0.874517 | 1 |
| 11 | 35601200 | 35603799 | 27.38389 | 45.48615 | 63.24968 | 27.4256 | -0.09313 | 1.278144 | 1 |
| 11 | 38260800 | 38263199 | 25.5491 | 42.69457 | 59.89727 | 26.07392 | -0.10573 | 1.203983 | 1 |
| 11 | 38870600 | 38871399 | 14.93443 | 26.87804 | 28.41755 | 12.40393 | 0.324882 | 0.391638 | 1 |
| 11 | 39882200 | 39882999 | 11.04937 | 20.70263 | 24.74971 | 10.21857 | 0.20604 | 0.121093 | 1 |
| 11 | 40309600 | 40311399 | 26.66501 | 44.09965 | 60.13945 | 25.99872 | -0.06547 | 1.326398 | 1 |
| 11 | 40551400 | 40552599 | 13.98099 | 24.89844 | 32.02879 | 14.10566 | 0.034632 | 0.457054 | 1 |
| 11 | 40736200 | 40739199 | 22.35826 | 37.89898 | 59.38929 | 26.00941 | -0.26444 | 1.05806 | 1 |
| 11 | 40809200 | 40810999 | 31.6016 | 52.25001 | 59.54267 | 25.73598 | 0.192646 | 1.339664 | 1 |
| 11 | 40862000 | 40862999 | 18.23933 | 31.90429 | 32.2769 | 14.23474 | 0.371049 | 0.682934 | 1 |
| 11 | 40994800 | 40996399 | 13.5189 | 24.52217 | 29.06388 | 12.56213 | 0.170256 | 0.338474 | 1 |
| 11 | 41022600 | 41025999 | 59.04115 | 93.35992 | 132.3344 | 54.98698 | -0.17562 | 2.40341 | 1 |
| 11 | 41047800 | 41048799 | 12.93615 | 23.51876 | 26.59149 | 11.46071 | 0.240449 | 0.314613 | 1 |
| 11 | 41109800 | 41115799 | 10.03185 | 18.4017 | 30.85467 | 13.14638 | -0.31317 | 0.251745 | 1 |

| | | | | | | | | | |
|---|---|---|---|---|---|---|---|---|---|
| 11 | 41780200 | 41780999 | 17.10836 | 29.72901 | 43.18199 | 18.96098 | -0.14916 | 0.79001 | 1 |
| 11 | 42228000 | 42229999 | 13.33651 | 23.86102 | 38.8195 | 17.02967 | -0.29844 | 0.438658 | 1 |
| 11 | 42669000 | 42670399 | 11.23881 | 20.40376 | 31.19345 | 13.46813 | -0.19398 | 0.319147 | 1 |
| 11 | 42705600 | 42706599 | 12.55105 | 22.4734 | 32.11842 | 14.08063 | -0.11181 | 0.413483 | 1 |
| 11 | 43193000 | 43195999 | 43.41912 | 69.887 | 82.32626 | 34.97063 | 0.118218 | 1.840494 | 1 |
| 11 | 43294800 | 43295999 | 21.39648 | 36.30206 | 54.91252 | 24.05552 | -0.2135 | 1.043057 | 1 |
| 11 | 43345400 | 43348199 | 36.67936 | 59.81233 | 67.9045 | 29.1703 | 0.189359 | 1.531252 | 1 |
| 11 | 45629000 | 45631199 | 14.09386 | 25.27497 | 31.44683 | 13.76823 | 0.088195 | 0.412647 | 1 |
| 11 | 45652000 | 45656799 | 39.67268 | 64.21111 | 86.49292 | 36.99527 | -0.06574 | 1.680128 | 1 |
| 11 | 45678800 | 45679399 | 13.71673 | 25.05087 | 27.45205 | 11.76492 | 0.290347 | 0.296054 | 1 |
| 11 | 46303400 | 46306999 | 58.10621 | 92.27331 | 122.684 | 51.14167 | -0.07796 | 2.287248 | 1 |
| 11 | 46394000 | 46396399 | 48.78742 | 77.97821 | 103.1107 | 43.34792 | -0.06013 | 2.073755 | 1 |
| 11 | 46463200 | 46465599 | 20.51014 | 35.62253 | 38.89927 | 17.07512 | 0.260481 | 0.805852 | 1 |
| 11 | 46657000 | 46657799 | 13.7775 | 24.93959 | 28.49637 | 12.34875 | 0.220233 | 0.350082 | 1 |
| 11 | 47309200 | 47319799 | 374.6067 | 515.9491 | 661.8063 | 275.0728 | -0.21291 | 4.811622 | 1 |
| 11 | 47358400 | 47359999 | 22.16051 | 37.11375 | 59.1614 | 25.78945 | -0.28935 | 1.169116 | 1 |
| 11 | 47559800 | 47561199 | 15.95353 | 28.22407 | 32.96525 | 14.57928 | 0.168272 | 0.523369 | 1 |
| 11 | 47685000 | 47685999 | 15.17529 | 27.03827 | 30.52226 | 13.43676 | 0.222615 | 0.46075 | 1 |
| 11 | 47740600 | 47742399 | 39.68505 | 64.00413 | 79.90948 | 34.03812 | 0.038013 | 1.773643 | 1 |
| 11 | 48235800 | 48238799 | 16.53855 | 28.96796 | 42.97917 | 18.97606 | -0.17884 | 0.670845 | 1 |
| 11 | 48342200 | 48343599 | 17.51662 | 30.42737 | 44.72617 | 19.63453 | -0.16649 | 0.797311 | 1 |
| 11 | 48414400 | 48416599 | 16.82303 | 29.10602 | 46.358 | 20.33837 | -0.28269 | 0.838291 | 1 |
| 11 | 48517400 | 48519199 | 18.57349 | 32.51854 | 36.08182 | 15.92509 | 0.238257 | 0.67531 | 1 |
| 11 | 48776400 | 48777999 | 17.72484 | 30.76497 | 41.14834 | 18.055 | -0.0308 | 0.806517 | 1 |
| 11 | 49334200 | 49340999 | 106.4746 | 163.8132 | 184.314 | 76.54553 | 0.126516 | 2.947221 | 1 |
| 11 | 49493400 | 49494399 | 14.11848 | 25.58782 | 28.97036 | 12.53514 | 0.234769 | 0.346768 | 1 |
| 11 | 50335200 | 50339999 | 53.16421 | 84.62775 | 116.8277 | 48.83523 | -0.12858 | 2.208643 | 1 |
| 11 | 51033400 | 51042999 | 274.5208 | 392.4265 | 459.66 | 189.9141 | -0.02787 | 4.253895 | 1 |
| 11 | 52758600 | 52760799 | 29.30061 | 48.62289 | 63.44314 | 27.49113 | -0.00175 | 1.293886 | 1 |
| 11 | 53086000 | 53088999 | 35.73644 | 57.87843 | 88.96314 | 38.07038 | -0.25483 | 1.660381 | 1 |
| 11 | 53126800 | 53130999 | 28.55581 | 46.67421 | 74.75841 | 32.14192 | -0.3034 | 1.487712 | 1 |
| 11 | 53816000 | 53818599 | 23.05647 | 39.62386 | 42.10796 | 18.47001 | 0.296647 | 0.920152 | 1 |
| 11 | 54278200 | 54279599 | 14.44946 | 25.40365 | 41.35425 | 18.31231 | -0.31181 | 0.597809 | 1 |
| 11 | 54469000 | 54475399 | 136.0751 | 204.2038 | 278.005 | 114.722 | -0.17781 | 3.457503 | 1 |
| 11 | 54745600 | 54748399 | 30.22535 | 49.09255 | 77.71669 | 33.32766 | -0.29326 | 1.587937 | 1 |
| 11 | 54889600 | 54892599 | 19.17902 | 33.43686 | 37.86757 | 16.66182 | 0.208774 | 0.746479 | 1 |
| 11 | 55704600 | 55708999 | 28.18234 | 47.39572 | 50.17762 | 21.92264 | 0.299011 | 1.137649 | 1 |
| 11 | 55874400 | 55875399 | 11.90885 | 22.369 | 26.26312 | 10.79749 | 0.236324 | 0.114297 | 1 |
| 11 | 56184200 | 56191199 | 99.05111 | 151.952 | 211.839 | 87.94178 | -0.18562 | 3.002975 | 1 |
| 11 | 56579600 | 56581799 | 22.21713 | 38.22772 | 47.38773 | 20.78666 | 0.075598 | 0.910787 | 1 |
| 11 | 56697200 | 56698799 | 13.10493 | 23.12765 | 37.73416 | 16.69658 | -0.31287 | 0.540091 | 1 |
| 11 | 57427600 | 57431199 | 41.37383 | 66.83792 | 82.33342 | 35.14529 | 0.06002 | 1.727228 | 1 |
| 11 | 58280400 | 58283799 | 41.86072 | 67.68107 | 71.935 | 30.73971 | 0.273827 | 1.699909 | 1 |
| 11 | 58365600 | 58367199 | 24.12015 | 40.92706 | 44.86098 | 19.65282 | 0.249621 | 1.04295 | 1 |
| 11 | 58404600 | 58413199 | 134.9829 | 202.6046 | 281.2119 | 116.0515 | -0.20543 | 3.453316 | 1 |
| 11 | 58579600 | 58589399 | 155.3575 | 231.6693 | 310.97 | 128.1597 | -0.16571 | 3.579643 | 1 |
| 11 | 60855200 | 60856199 | 10.95673 | 20.06764 | 30.79128 | 13.14659 | -0.1883 | 0.26712 | 1 |

| | | | | | | | | | |
|---|---|---|---|---|---|---|---|---|---|
| 11 | 62398000 | 62399999 | 10.54315 | 19.91883 | 22.76403 | 9.260776 | 0.288113 | 0.058035 | 1 |
| 11 | 62664000 | 62676799 | 217.7546 | 320.576 | 317.9215 | 130.922 | 0.253569 | 3.806074 | 1 |
| 11 | 62683600 | 62690399 | 139.4917 | 209.7508 | 271.3093 | 112.019 | -0.10147 | 3.418394 | 1 |
| 11 | 62879600 | 62881399 | 16.25611 | 28.47994 | 39.30263 | 17.35625 | -0.07488 | 0.658061 | 1 |
| 11 | 63094600 | 63097599 | 20.97511 | 36.16865 | 45.79704 | 20.09023 | 0.045821 | 0.885598 | 1 |
| 11 | 63099000 | 63100599 | 20.11018 | 34.70591 | 45.79718 | 20.09079 | -0.0133 | 0.873438 | 1 |
| 11 | 63696000 | 63696599 | 9.790517 | 18.20481 | 27.33282 | 11.43156 | -0.13545 | 0.164153 | 1 |
| 11 | 64182800 | 64184399 | 15.27161 | 26.89193 | 40.19438 | 17.79303 | -0.1886 | 0.580137 | 1 |
| 11 | 64312800 | 64314399 | 21.66042 | 37.3918 | 39.49708 | 17.32715 | 0.306577 | 0.869966 | 1 |
| 11 | 65347200 | 65347999 | 18.0561 | 31.63671 | 32.30367 | 14.26671 | 0.357876 | 0.657012 | 1 |
| 11 | 66351600 | 66352199 | 10.06961 | 18.64538 | 25.78291 | 10.84584 | -0.02362 | 0.185897 | 1 |
| 11 | 67585800 | 67586799 | 13.03873 | 23.0245 | 38.31332 | 16.95083 | -0.34054 | 0.537343 | 1 |
| 11 | 69604400 | 69606599 | 14.70812 | 26.16989 | 28.07934 | 12.37566 | 0.294201 | 0.459363 | 1 |
| 11 | 70251400 | 70255599 | 35.45884 | 57.90182 | 67.13527 | 28.85435 | 0.160479 | 1.510385 | 1 |
| 11 | 71098800 | 71099999 | 13.34183 | 23.68293 | 37.20491 | 16.4303 | -0.25472 | 0.485568 | 1 |
| 11 | 71195000 | 71199399 | 82.42465 | 128.9734 | 154.288 | 64.00526 | 0.056973 | 2.632502 | 1 |
| 11 | 71273600 | 71275799 | 28.31669 | 46.5171 | 64.40433 | 27.74693 | -0.09015 | 1.414739 | 1 |
| 11 | 72167800 | 72170799 | 39.7951 | 64.12923 | 81.03648 | 34.48152 | 0.019276 | 1.796829 | 1 |
| 11 | 73218200 | 73221199 | 29.04793 | 47.71063 | 65.19988 | 28.08775 | -0.07155 | 1.416631 | 1 |
| 11 | 73260000 | 73260599 | 9.183932 | 17.27476 | 27.48838 | 11.27452 | -0.19593 | 0.086367 | 1 |
| 11 | 73305800 | 73306799 | 11.89839 | 22.1626 | 25.61635 | 10.68229 | 0.242447 | 0.165767 | 1 |
| 11 | 73319800 | 73321199 | 17.33887 | 30.84062 | 31.66416 | 13.95948 | 0.357009 | 0.485423 | 1 |
| 11 | 73357800 | 73361399 | 51.61205 | 82.71621 | 94.2831 | 39.79329 | 0.158066 | 1.989558 | 1 |
| 11 | 73452200 | 73454999 | 59.61035 | 94.9024 | 99.71844 | 41.69162 | 0.264766 | 2.205247 | 1 |
| 11 | 73511800 | 73522199 | 150.3727 | 227.449 | 242.0426 | 100.1082 | 0.187256 | 3.303272 | 1 |
| 11 | 73584400 | 73588799 | 55.0693 | 88.15342 | 100.7769 | 42.45864 | 0.151826 | 2.031708 | 1 |
| 11 | 73590400 | 73595999 | 46.91142 | 75.62838 | 84.16707 | 35.83306 | 0.202813 | 1.798242 | 1 |
| 11 | 73644600 | 73645599 | 22.28985 | 38.10722 | 45.84196 | 20.10027 | 0.117286 | 0.968484 | 1 |
| 11 | 73720600 | 73733599 | 181.2325 | 267.4863 | 338.0103 | 139.1985 | -0.09241 | 3.760816 | 1 |
| 11 | 73916200 | 73917799 | 24.07945 | 41.3152 | 45.24398 | 19.84383 | 0.253052 | 0.942528 | 1 |
| 11 | 73981000 | 73983599 | 30.69752 | 50.68477 | 57.29142 | 24.7392 | 0.204168 | 1.350043 | 1 |
| 11 | 74010800 | 74021199 | 150.8354 | 227.0475 | 249.1556 | 102.9011 | 0.136928 | 3.398982 | 1 |
| 11 | 74096800 | 74098199 | 19.26707 | 33.57996 | 39.86602 | 17.53773 | 0.140866 | 0.752572 | 1 |
| 11 | 75245200 | 75257399 | 373.601 | 514.5075 | 688.6181 | 286.2218 | -0.27439 | 4.813529 | 1 |
| 11 | 76599800 | 76602799 | 32.13901 | 52.22983 | 76.44237 | 32.79464 | -0.17961 | 1.579378 | 1 |
| 11 | 78879600 | 78889799 | 117.6393 | 180.5948 | 187.0184 | 77.64535 | 0.243695 | 2.994054 | 1 |
| 11 | 79638600 | 79641399 | 32.21665 | 52.48112 | 73.45257 | 31.54181 | -0.11283 | 1.540631 | 1 |
| 11 | 79872400 | 79874399 | 17.80863 | 31.1599 | 35.23025 | 15.54101 | 0.211396 | 0.680867 | 1 |
| 11 | 80695400 | 80696399 | 10.52578 | 19.80545 | 22.62735 | 9.275419 | 0.278756 | 0.080946 | 1 |
| 11 | 83145400 | 83146599 | 11.49566 | 21.63029 | 23.68772 | 9.710158 | 0.339849 | 0.095222 | 1 |
| 11 | 86713600 | 86716399 | 27.38155 | 45.96125 | 52.34043 | 22.84854 | 0.194106 | 1.154823 | 1 |
| 11 | 86734400 | 86741799 | 83.04785 | 128.8964 | 172.1183 | 71.47783 | -0.11059 | 2.775531 | 1 |
| 11 | 87372600 | 87374199 | 17.70486 | 30.6561 | 50.07289 | 21.96872 | -0.31868 | 0.841408 | 1 |
| 11 | 88686200 | 88688799 | 28.41842 | 46.68542 | 69.11355 | 29.7761 | -0.18658 | 1.415342 | 1 |
| 11 | 88799000 | 88801999 | 34.93829 | 56.4692 | 81.05316 | 34.62067 | -0.15918 | 1.702953 | 1 |
| 11 | 88934800 | 88938999 | 26.46047 | 44.12679 | 63.39601 | 27.5612 | -0.13997 | 1.229492 | 1 |
| 11 | 89123200 | 89125999 | 18.33216 | 31.91599 | 38.46494 | 16.90987 | 0.119229 | 0.762155 | 1 |

| | | | | | | | | | |
|---|---|---|---|---|---|---|---|---|---|
| 11 | 90659400 | 90660399 | 8.366058 | 15.98941 | 29.69467 | 11.78861 | -0.37665 | -0.02615 | 1 |
| 11 | 90682200 | 90684199 | 22.1285 | 38.24932 | 42.59244 | 18.6858 | 0.231168 | 0.865931 | 1 |
| 11 | 90855200 | 90855599 | 10.47603 | 19.8502 | 21.59507 | 8.73085 | 0.365705 | 0.030614 | 1 |
| 11 | 90856200 | 90865199 | 191.5351 | 280.0278 | 358.839 | 147.8303 | -0.12552 | 3.91234 | 1 |
| 11 | 90892600 | 90894399 | 9.282617 | 17.33688 | 28.95827 | 12.02104 | -0.27989 | 0.133292 | 1 |
| 11 | 90911000 | 90911999 | 12.55568 | 22.22381 | 37.78935 | 16.71114 | -0.3702 | 0.507389 | 1 |
| 11 | 91533600 | 91536599 | 35.00769 | 57.20036 | 65.34158 | 28.08974 | 0.182564 | 1.500553 | 1 |
| 11 | 91811200 | 91818599 | 137.1038 | 207.0537 | 238.781 | 98.71026 | 0.069446 | 3.330645 | 1 |
| 11 | 92063600 | 92073599 | 144.454 | 218.0237 | 251.6966 | 104.0299 | 0.067195 | 3.345564 | 1 |
| 11 | 92585000 | 92585799 | 14.27505 | 25.03215 | 39.42085 | 17.42194 | -0.26422 | 0.63447 | 1 |
| 11 | 92752400 | 92753999 | 22.43509 | 38.89362 | 41.42725 | 18.17625 | 0.295732 | 0.837349 | 1 |
| 11 | 92979000 | 92980599 | 19.87362 | 34.66854 | 42.99026 | 18.92292 | 0.078457 | 0.748363 | 1 |
| 11 | 96500400 | 96501399 | 11.30656 | 20.65042 | 30.77098 | 13.18629 | -0.14989 | 0.28653 | 1 |
| 11 | 96591800 | 96593799 | 9.725331 | 18.62834 | 22.9188 | 9.041173 | 0.221719 | -0.04954 | 1 |
| 11 | 98949600 | 98951599 | 18.46581 | 31.73755 | 46.38799 | 20.34743 | -0.16122 | 0.906225 | 1 |
| 11 | 99145800 | 99148399 | 18.44466 | 31.97946 | 50.10052 | 21.98173 | -0.25848 | 0.831386 | 1 |
| 11 | 99629600 | 99631199 | 13.1056 | 23.39405 | 37.19937 | 16.35059 | -0.26761 | 0.44707 | 1 |
| 11 | 99741800 | 99742399 | 10.23124 | 19.06427 | 25.78219 | 10.74477 | 0.01846 | 0.150719 | 1 |
| 11 | 99779200 | 99784799 | 62.71673 | 98.97657 | 145.416 | 60.36228 | -0.22957 | 2.455428 | 1 |
| 11 | 100694800 | 100696799 | 18.82673 | 32.12422 | 52.17863 | 22.87513 | -0.31466 | 0.979781 | 1 |
| 11 | 101538800 | 101540799 | 18.71245 | 32.72562 | 35.6274 | 15.70984 | 0.265605 | 0.692969 | 1 |
| 11 | 102230200 | 102230999 | 6.535189 | 12.66587 | 23.32069 | 8.82728 | -0.2984 | -0.21933 | 1 |
| 11 | 102976200 | 102976999 | 16.97444 | 29.4896 | 41.1705 | 18.07516 | -0.09216 | 0.790893 | 1 |
| 11 | 102977800 | 102978199 | 9.674162 | 18.51913 | 19.77393 | 7.814111 | 0.422432 | -0.05324 | 1 |
| 11 | 104381400 | 104385999 | 49.38984 | 78.92903 | 114.3054 | 48.04379 | -0.19142 | 2.080706 | 1 |
| 11 | 105740000 | 105745999 | 59.96178 | 95.79598 | 100.402 | 42.18265 | 0.27383 | 2.09636 | 1 |
| 11 | 106903600 | 106906599 | 17.3721 | 30.41644 | 35.75144 | 15.77984 | 0.155665 | 0.66918 | 1 |
| 11 | 107044200 | 107047199 | 25.87587 | 42.89355 | 60.34153 | 26.12774 | -0.11005 | 1.293703 | 1 |
| 11 | 107049400 | 107050599 | 21.29319 | 36.43422 | 45.8473 | 20.10399 | 0.052771 | 0.956325 | 1 |
| 11 | 107266000 | 107267799 | 14.09468 | 24.69485 | 38.45435 | 16.98258 | -0.24805 | 0.645159 | 1 |
| 11 | 107612400 | 107614199 | 15.73873 | 27.50551 | 45.70279 | 20.14052 | -0.34146 | 0.70653 | 1 |
| 11 | 108365400 | 108366799 | 16.4715 | 28.63099 | 47.38161 | 20.80895 | -0.33661 | 0.78736 | 1 |
| 11 | 109501000 | 109502999 | 14.8468 | 25.97113 | 39.45748 | 17.40212 | -0.21282 | 0.675668 | 1 |
| 11 | 109515000 | 109516199 | 10.30624 | 19.21946 | 28.89004 | 12.0236 | -0.13204 | 0.150681 | 1 |
| 11 | 109801200 | 109804599 | 39.61912 | 64.11231 | 77.54175 | 33.162 | 0.089073 | 1.679198 | 1 |
| 11 | 111072400 | 111074999 | 29.38223 | 48.95756 | 50.77771 | 22.06358 | 0.328754 | 1.23857 | 1 |
| 11 | 111078200 | 111083999 | 60.0492 | 95.8594 | 106.6215 | 44.7453 | 0.186956 | 2.121074 | 1 |
| 11 | 111110200 | 111115799 | 69.40859 | 109.5986 | 132.9119 | 55.18745 | 0.047479 | 2.444473 | 1 |
| 11 | 111184600 | 111187399 | 33.35491 | 55.09476 | 64.31563 | 27.78013 | 0.157819 | 1.357453 | 1 |
| 11 | 112321000 | 112323599 | 31.27542 | 51.01172 | 69.86254 | 30.0139 | -0.08025 | 1.520073 | 1 |
| 11 | 112746800 | 112753399 | 60.09168 | 95.25746 | 134.155 | 55.82443 | -0.16339 | 2.342637 | 1 |
| 11 | 113343600 | 113345599 | 15.48132 | 27.14225 | 40.85904 | 18.05465 | -0.19973 | 0.644565 | 1 |
| 11 | 114553000 | 114555199 | 13.84104 | 24.68279 | 29.19273 | 12.84189 | 0.156533 | 0.443152 | 1 |
| 11 | 114858800 | 114876599 | 333.3684 | 466.6941 | 584.2118 | 242.201 | -0.15404 | 4.563958 | 1 |
| 11 | 115650400 | 115655199 | 42.73653 | 68.62263 | 95.20853 | 40.30064 | -0.12171 | 1.918206 | 1 |
| 11 | 116648000 | 116652399 | 48.56514 | 77.90846 | 95.6922 | 40.44753 | 0.052102 | 1.954576 | 1 |
| 11 | 120094600 | 120096799 | 22.00151 | 37.07018 | 54.84147 | 23.97384 | -0.18181 | 1.107783 | 1 |

| | | | | | | | | | |
|---|---|---|---|---|---|---|---|---|---|
| 11 | 120216400 | 120233799 | 255.6304 | 370.0446 | 396.4335 | 163.4912 | 0.118842 | 4.068063 | 1 |
| 11 | 120644600 | 120653399 | 164.9889 | 244.5558 | 313.0225 | 128.9218 | -0.10513 | 3.687088 | 1 |
| 11 | 121198400 | 121214999 | 526.3417 | 705.1336 | 871.6154 | 363.8255 | -0.19982 | 5.249367 | 1 |
| 11 | 121719200 | 121724599 | 44.69191 | 71.46738 | 105.458 | 44.36563 | -0.21732 | 2.0545 | 1 |
| 12 | 3221200 | 3223599 | 19.62074 | 34.17055 | 39.38117 | 17.31621 | 0.18343 | 0.764881 | 1 |
| 12 | 3283200 | 3290999 | 110.4727 | 168.1343 | 209.3265 | 86.7405 | -0.03158 | 3.165336 | 1 |
| 12 | 3411800 | 3418999 | 76.61078 | 119.0815 | 174.8739 | 72.60614 | -0.24595 | 2.748249 | 1 |
| 12 | 3575200 | 3588399 | 34.07982 | 55.76602 | 77.56504 | 33.36233 | -0.09932 | 1.484919 | 1 |
| 12 | 4928200 | 4929999 | 16.33112 | 28.76589 | 34.60075 | 15.316 | 0.123709 | 0.577076 | 1 |
| 12 | 4964600 | 4965599 | 12.88522 | 23.78654 | 26.5326 | 11.20744 | 0.279736 | 0.227449 | 1 |
| 12 | 6284600 | 6287399 | 20.85306 | 35.29928 | 51.63977 | 22.60615 | -0.16535 | 1.060473 | 1 |
| 12 | 6337000 | 6337799 | 10.88801 | 20.58977 | 24.30285 | 9.867083 | 0.244262 | 0.058638 | 1 |
| 12 | 6371800 | 6372999 | 12.90934 | 23.75144 | 24.41352 | 10.3594 | 0.392037 | 0.239976 | 1 |
| 12 | 6486000 | 6488599 | 27.97772 | 46.31646 | 68.76963 | 29.74992 | -0.18784 | 1.324106 | 1 |
| 12 | 6596800 | 6598999 | 27.62941 | 47.10781 | 45.77691 | 20.06736 | 0.423507 | 1.014548 | 1 |
| 12 | 6624200 | 6625799 | 17.46585 | 31.34904 | 30.18926 | 13.20805 | 0.456535 | 0.429383 | 1 |
| 12 | 6722400 | 6723599 | 11.46743 | 21.25805 | 27.70827 | 11.63758 | 0.062434 | 0.195277 | 1 |
| 12 | 6821000 | 6821799 | 18.00542 | 30.8493 | 47.44471 | 20.80666 | -0.23513 | 0.934035 | 1 |
| 12 | 6851800 | 6855599 | 12.97354 | 23.30618 | 34.68213 | 15.16259 | -0.16695 | 0.403102 | 1 |
| 12 | 7676800 | 7682399 | 102.1165 | 157.3127 | 175.9232 | 73.07384 | 0.136739 | 2.918836 | 1 |
| 12 | 7686400 | 7689799 | 33.67305 | 55.41793 | 60.17409 | 25.9446 | 0.260385 | 1.399302 | 1 |
| 12 | 7851800 | 7852399 | 9.046227 | 16.83545 | 27.90051 | 11.65185 | -0.27508 | 0.15407 | 1 |
| 12 | 9461400 | 9463999 | 13.8333 | 25.26668 | 26.90416 | 11.52845 | 0.331618 | 0.294597 | 1 |
| 12 | 9493400 | 9495999 | 28.10814 | 46.20887 | 67.35769 | 29.02845 | -0.16387 | 1.405286 | 1 |
| 12 | 9534200 | 9537199 | 27.01995 | 45.52762 | 49.08548 | 21.45789 | 0.272934 | 1.116272 | 1 |
| 12 | 9675000 | 9677199 | 12.99551 | 23.00694 | 35.73328 | 15.80128 | -0.24015 | 0.503501 | 1 |
| 12 | 10747800 | 10750999 | 33.62607 | 55.64906 | 58.54524 | 25.32114 | 0.307721 | 1.334627 | 1 |
| 12 | 10879200 | 10879999 | 16.41783 | 28.68199 | 38.92808 | 17.15063 | -0.05091 | 0.704001 | 1 |
| 12 | 11010600 | 11012999 | 42.81265 | 68.68511 | 92.5453 | 39.12156 | -0.08098 | 1.946273 | 1 |
| 12 | 11045600 | 11051199 | 62.22301 | 98.66035 | 133.1983 | 55.44001 | -0.10217 | 2.337029 | 1 |
| 12 | 11080600 | 11082399 | 15.37683 | 26.96876 | 38.34218 | 16.94766 | -0.11745 | 0.636099 | 1 |
| 12 | 11158800 | 11159399 | 7.556566 | 14.51141 | 23.02284 | 9.015479 | -0.13221 | -0.0968 | 1 |
| 12 | 11557800 | 11560799 | 27.08576 | 45.41856 | 55.61863 | 24.26403 | 0.089685 | 1.167574 | 1 |
| 12 | 11597600 | 11598999 | 21.94524 | 37.42426 | 47.95914 | 21.02435 | 0.025812 | 0.991347 | 1 |
| 12 | 12706800 | 12714399 | 138.6366 | 208.4907 | 268.097 | 110.6964 | -0.09282 | 3.415777 | 1 |
| 12 | 13475400 | 13476599 | 26.29196 | 43.44816 | 61.82562 | 26.71302 | -0.12681 | 1.333967 | 1 |
| 12 | 13946400 | 13947599 | 17.75626 | 31.22495 | 33.85899 | 14.99257 | 0.271394 | 0.599013 | 1 |
| 12 | 14093800 | 14094799 | 14.88142 | 26.99511 | 28.86172 | 12.47391 | 0.31788 | 0.346155 | 1 |
| 12 | 14444200 | 14444999 | 15.21016 | 26.67363 | 35.89545 | 15.86475 | -0.03846 | 0.633271 | 1 |
| 12 | 14873000 | 14873799 | 9.996343 | 19.21583 | 21.87168 | 8.540236 | 0.346709 | -0.0832 | 1 |
| 12 | 14986800 | 14988999 | 22.37284 | 38.07982 | 54.41534 | 23.85091 | -0.13122 | 1.018297 | 1 |
| 12 | 15075800 | 15076799 | 13.61605 | 23.96202 | 36.31638 | 16.0783 | -0.20844 | 0.573514 | 1 |
| 12 | 15110600 | 15111799 | 11.15511 | 20.31718 | 32.81467 | 14.11112 | -0.26879 | 0.305277 | 1 |
| 12 | 15206000 | 15208199 | 12.89503 | 23.04758 | 35.12616 | 15.42272 | -0.20568 | 0.432957 | 1 |
| 12 | 15350400 | 15352199 | 22.36732 | 38.56736 | 45.22759 | 19.84039 | 0.155989 | 0.890296 | 1 |
| 12 | 15426800 | 15429599 | 38.27426 | 61.61971 | 93.69041 | 39.81086 | -0.24878 | 1.831284 | 1 |
| 12 | 15472800 | 15473999 | 9.948445 | 18.51537 | 29.42813 | 12.28887 | -0.21547 | 0.16408 | 1 |

| | | | | | | | | | |
|---|---|---|---|---|---|---|---|---|---|
| 12 | 16465000 | 16465999 | 12.383 | 22.04562 | 33.30895 | 14.67338 | -0.19599 | 0.454567 | 1 |
| 12 | 17168000 | 17175999 | 84.19287 | 131.1743 | 165.365 | 68.6341 | -0.02339 | 2.709464 | 1 |
| 12 | 17512200 | 17517599 | 28.20831 | 47.33457 | 49.12671 | 21.44151 | 0.327495 | 1.155805 | 1 |
| 12 | 17696200 | 17709799 | 53.20207 | 84.58329 | 127.8457 | 53.3551 | -0.26105 | 2.250297 | 1 |
| 12 | 18927000 | 18965399 | 1075.968 | 1379.243 | 1205.452 | 505.6761 | 0.236584 | 5.964933 | 1 |
| 12 | 19410400 | 19411799 | 59.09421 | 93.90406 | 106.8061 | 44.55219 | 0.147869 | 2.266109 | 1 |
| 12 | 19428200 | 19429799 | 14.59829 | 26.05728 | 28.56669 | 12.55671 | 0.266042 | 0.440593 | 1 |
| 12 | 19437200 | 19441799 | 36.95756 | 59.46483 | 92.64375 | 39.33249 | -0.2849 | 1.846729 | 1 |
| 12 | 20169400 | 20171799 | 28.77768 | 47.88984 | 53.43176 | 23.19808 | 0.223792 | 1.253053 | 1 |
| 12 | 20596800 | 20598999 | 39.08677 | 63.22696 | 73.37483 | 31.37121 | 0.148064 | 1.683235 | 1 |
| 12 | 20664400 | 20666199 | 28.2978 | 45.99284 | 78.67266 | 33.75441 | -0.40377 | 1.570931 | 1 |
| 12 | 21466400 | 21468399 | 22.64476 | 38.74396 | 42.16722 | 18.49016 | 0.261298 | 0.957097 | 1 |
| 12 | 21628600 | 21632999 | 42.6244 | 69.04055 | 74.28942 | 31.79318 | 0.258569 | 1.662559 | 1 |
| 12 | 21644400 | 21662799 | 336.8934 | 472.4515 | 548.0703 | 227.156 | -0.04172 | 4.53735 | 1 |
| 12 | 22280800 | 22282599 | 36.32045 | 59.2608 | 60.85824 | 26.14881 | 0.334351 | 1.521497 | 1 |
| 12 | 22518800 | 22525799 | 234.9635 | 340.2465 | 352.6986 | 145.448 | 0.166841 | 4.060371 | 1 |
| 12 | 22672800 | 22673799 | 23.39209 | 38.77267 | 61.68477 | 26.70764 | -0.28688 | 1.290876 | 1 |
| 12 | 23472400 | 23473599 | 30.37818 | 50.32718 | 51.32409 | 22.21416 | 0.352604 | 1.306915 | 1 |
| 12 | 23475800 | 23477999 | 58.57958 | 93.40588 | 90.78532 | 38.03814 | 0.379458 | 2.154263 | 1 |
| 12 | 23689600 | 23693599 | 159.017 | 237.2578 | 259.5791 | 106.9916 | 0.129833 | 3.568441 | 1 |
| 12 | 24003400 | 24006599 | 38.92309 | 63.49477 | 66.52779 | 28.58277 | 0.305167 | 1.529397 | 1 |
| 12 | 24012200 | 24013999 | 34.16329 | 55.92276 | 61.68909 | 26.53727 | 0.234596 | 1.466969 | 1 |
| 12 | 24018600 | 24019599 | 20.61616 | 35.28633 | 40.2963 | 17.67031 | 0.192417 | 0.944557 | 1 |
| 12 | 25135000 | 25136599 | 15.02936 | 26.79683 | 29.89519 | 13.15301 | 0.239865 | 0.451885 | 1 |
| 12 | 27003400 | 27006399 | 36.41089 | 58.94522 | 85.25676 | 36.47264 | -0.16784 | 1.669445 | 1 |
| 12 | 27282000 | 27286399 | 51.0261 | 81.89498 | 89.03439 | 37.6604 | 0.228614 | 1.941459 | 1 |
| 12 | 27473000 | 27474799 | 24.44038 | 41.74122 | 43.67852 | 19.15009 | 0.317368 | 0.982126 | 1 |
| 12 | 28386400 | 28389999 | 55.8007 | 88.73819 | 125.8279 | 52.52691 | -0.1687 | 2.244309 | 1 |
| 12 | 28541200 | 28543399 | 23.02783 | 39.52517 | 44.19578 | 19.38466 | 0.223138 | 0.932923 | 1 |
| 12 | 28783200 | 28784799 | 14.69569 | 25.91586 | 39.20011 | 17.3478 | -0.20473 | 0.560633 | 1 |
| 12 | 30017800 | 30020399 | 23.70899 | 40.59324 | 49.01852 | 21.49565 | 0.112027 | 0.963106 | 1 |
| 12 | 30140800 | 30143599 | 28.4328 | 46.67185 | 65.04613 | 28.0143 | -0.10014 | 1.423267 | 1 |
| 12 | 31017800 | 31019199 | 14.87263 | 26.61911 | 29.39361 | 12.89065 | 0.257793 | 0.42626 | 1 |
| 12 | 31078400 | 31079799 | 9.845083 | 18.24135 | 29.39225 | 12.35335 | -0.24179 | 0.190444 | 1 |
| 12 | 32023000 | 32024999 | 17.46499 | 30.46554 | 37.38605 | 16.4557 | 0.093494 | 0.727432 | 1 |
| 12 | 32139400 | 32142799 | 76.12425 | 118.794 | 167.5994 | 69.54671 | -0.18377 | 2.678568 | 1 |
| 12 | 32308000 | 32310199 | 40.93559 | 66.29994 | 72.57328 | 31.05695 | 0.233853 | 1.662374 | 1 |
| 12 | 32467800 | 32468999 | 11.65606 | 20.89462 | 32.23344 | 14.11751 | -0.22026 | 0.401575 | 1 |
| 12 | 32645400 | 32647199 | 17.41479 | 30.24964 | 44.74048 | 19.64046 | -0.17529 | 0.798562 | 1 |
| 12 | 33285200 | 33288399 | 12.80105 | 22.75894 | 35.67661 | 15.73439 | -0.25036 | 0.467854 | 1 |
| 12 | 33794600 | 33800599 | 102.2086 | 157.1871 | 190.1141 | 78.94999 | 0.021905 | 2.953097 | 1 |
| 12 | 33816400 | 33817399 | 11.70593 | 20.78549 | 34.75715 | 15.34532 | -0.34398 | 0.466618 | 1 |
| 12 | 33831400 | 33833199 | 11.54741 | 20.69879 | 32.30196 | 14.14814 | -0.23664 | 0.40299 | 1 |
| 12 | 33930000 | 33932599 | 20.89807 | 36.14294 | 51.76926 | 22.71217 | -0.13053 | 0.866404 | 1 |
| 12 | 34049600 | 34051199 | 11.62935 | 21.02052 | 30.23742 | 13.12705 | -0.11143 | 0.34765 | 1 |
| 12 | 34059000 | 34060199 | 13.17711 | 24.12277 | 29.00789 | 12.39325 | 0.160015 | 0.280563 | 1 |
| 12 | 34783400 | 34784399 | 18.59462 | 32.37032 | 38.41467 | 16.88691 | 0.141319 | 0.763475 | 1 |

| | | | | | | | | | |
|---|---|---|---|---|---|---|---|---|---|
| 12 | 35302000 | 35304199 | 42.5764 | 68.49769 | 89.30382 | 37.90819 | -0.02871 | 1.855953 | 1 |
| 12 | 35402800 | 35403799 | 11.26027 | 20.99469 | 25.66763 | 10.68582 | 0.163946 | 0.15408 | 1 |
| 12 | 35802800 | 35804799 | 20.80947 | 36.49292 | 36.78495 | 16.25787 | 0.375965 | 0.661531 | 1 |
| 12 | 36148200 | 36154999 | 102.3129 | 156.6015 | 203.837 | 84.5848 | -0.08919 | 3.048233 | 1 |
| 12 | 36178800 | 36179799 | 11.25982 | 21.1863 | 24.70336 | 10.12668 | 0.249678 | 0.094266 | 1 |
| 12 | 36243200 | 36249599 | 90.03244 | 139.951 | 170.4763 | 70.77978 | 0.023504 | 2.751886 | 1 |
| 12 | 36335000 | 36335599 | 10.22271 | 18.57626 | 32.32408 | 13.94013 | -0.37819 | 0.310386 | 1 |
| 12 | 36468800 | 36471599 | 25.59551 | 42.78726 | 71.33826 | 31.06123 | -0.35414 | 1.20934 | 1 |
| 12 | 37208800 | 37211599 | 20.65863 | 35.06503 | 51.11643 | 22.39544 | -0.16015 | 1.035112 | 1 |
| 12 | 37893400 | 37894799 | 12.37956 | 22.28446 | 32.60983 | 14.23021 | -0.1412 | 0.380011 | 1 |
| 12 | 38124600 | 38125199 | 11.57365 | 20.65581 | 34.30118 | 15.07896 | -0.32948 | 0.434661 | 1 |
| 12 | 38522000 | 38523999 | 26.019 | 43.85949 | 47.49143 | 20.76369 | 0.266765 | 1.107549 | 1 |
| 12 | 39029400 | 39034399 | 47.43144 | 75.75078 | 108.7842 | 45.68171 | -0.18028 | 2.096857 | 1 |
| 12 | 39480800 | 39486999 | 55.17689 | 87.62 | 131.3695 | 54.76133 | -0.25107 | 2.287017 | 1 |
| 12 | 39949000 | 39950799 | 20.45362 | 35.29103 | 41.63997 | 18.26694 | 0.147509 | 0.872832 | 1 |
| 12 | 40919400 | 40929799 | 122.845 | 186.3156 | 263.9925 | 109.2791 | -0.22217 | 3.242891 | 1 |
| 12 | 41049600 | 41051199 | 15.01403 | 26.81621 | 29.86306 | 13.11958 | 0.243859 | 0.440651 | 1 |
| 12 | 41061800 | 41063799 | 26.03966 | 44.13977 | 48.51115 | 21.24542 | 0.245747 | 1.063581 | 1 |
| 12 | 41412800 | 41416799 | 64.5439 | 102.5281 | 104.4424 | 43.55102 | 0.30615 | 2.278849 | 1 |
| 12 | 42287400 | 42288999 | 22.03655 | 37.39651 | 47.98806 | 21.02357 | 0.023093 | 1.037389 | 1 |
| 12 | 42675600 | 42676999 | 27.67132 | 45.37961 | 71.07994 | 30.6009 | -0.26871 | 1.437373 | 1 |
| 12 | 42791800 | 42792999 | 22.666 | 37.98001 | 56.98056 | 24.84691 | -0.20212 | 1.164529 | 1 |
| 12 | 42814000 | 42814999 | 12.1282 | 22.60814 | 27.76087 | 11.56142 | 0.157284 | 0.168234 | 1 |
| 12 | 42916800 | 42922399 | 119.8317 | 182.8879 | 187.9189 | 77.92741 | 0.248393 | 3.109369 | 1 |
| 12 | 43395800 | 43396999 | 23.37097 | 38.79053 | 62.86107 | 27.24258 | -0.31327 | 1.275358 | 1 |
| 12 | 43704200 | 43707399 | 19.91315 | 34.78561 | 41.89554 | 18.45711 | 0.120577 | 0.728369 | 1 |
| 12 | 43715400 | 43725799 | 293.2895 | 411.4671 | 570.2331 | 236.327 | -0.29764 | 4.530196 | 1 |
| 12 | 43794400 | 43796199 | 39.50626 | 63.66274 | 91.07943 | 38.75393 | -0.1595 | 1.80118 | 1 |
| 12 | 44783000 | 44785199 | 39.06376 | 63.43656 | 65.89718 | 28.25526 | 0.312746 | 1.6015 | 1 |
| 12 | 44815800 | 44819799 | 67.32874 | 105.8487 | 146.4779 | 60.76327 | -0.14762 | 2.53529 | 1 |
| 12 | 45342200 | 45344799 | 28.6637 | 47.31563 | 65.49814 | 28.27839 | -0.0874 | 1.353724 | 1 |
| 12 | 45431800 | 45434199 | 35.44811 | 58.46761 | 62.01254 | 26.76213 | 0.295435 | 1.380205 | 1 |
| 12 | 46090800 | 46092799 | 37.00005 | 60.12488 | 66.191 | 28.39504 | 0.2301 | 1.585614 | 1 |
| 12 | 46298200 | 46302599 | 49.03563 | 78.26637 | 115.4321 | 48.43267 | -0.21964 | 2.117641 | 1 |
| 12 | 46317200 | 46322599 | 53.58039 | 85.58678 | 110.4812 | 46.40642 | -0.02651 | 2.097205 | 1 |
| 12 | 46864600 | 46866799 | 16.62253 | 28.78882 | 43.24529 | 18.97335 | -0.19823 | 0.823473 | 1 |
| 12 | 47406800 | 47408799 | 26.22595 | 44.09673 | 51.23171 | 22.38182 | 0.165383 | 1.133706 | 1 |
| 12 | 47858600 | 47860199 | 19.57033 | 33.66698 | 49.54241 | 21.73168 | -0.17082 | 0.905416 | 1 |
| 12 | 47912000 | 47913199 | 19.48723 | 33.17785 | 49.01799 | 21.48568 | -0.17886 | 1.000142 | 1 |
| 12 | 48040600 | 48044199 | 34.85641 | 56.88549 | 77.48694 | 33.29649 | -0.07198 | 1.523265 | 1 |
| 12 | 48301400 | 48302399 | 15.34965 | 27.29342 | 31.3421 | 13.82133 | 0.196064 | 0.47336 | 1 |
| 12 | 48362000 | 48362999 | 10.45262 | 19.8765 | 23.28502 | 9.343452 | 0.269551 | 0.012302 | 1 |
| 12 | 48606200 | 48608399 | 19.66763 | 34.23065 | 36.31967 | 15.96373 | 0.302075 | 0.768504 | 1 |
| 12 | 48717000 | 48718599 | 22.81001 | 39.13283 | 42.69177 | 18.72427 | 0.258579 | 0.936434 | 1 |
| 12 | 48896000 | 48897999 | 18.14024 | 31.77911 | 37.12325 | 16.39307 | 0.164388 | 0.665485 | 1 |
| 12 | 49042600 | 49043799 | 19.45093 | 32.79116 | 54.21369 | 23.70345 | -0.34141 | 1.094084 | 1 |
| 12 | 49056000 | 49058199 | 19.92534 | 34.02753 | 51.67103 | 22.65414 | -0.21774 | 0.9756 | 1 |

| | | | | | | | | | |
|---|---|---|---|---|---|---|---|---|---|
| 12 | 49103600 | 49104999 | 19.3688 | 33.42598 | 40.70175 | 17.85547 | 0.102411 | 0.866038 | 1 |
| 12 | 49170800 | 49173799 | 30.28499 | 49.83069 | 74.5341 | 32.13296 | -0.20053 | 1.405861 | 1 |
| 12 | 49266200 | 49270199 | 15.00906 | 26.19677 | 42.0704 | 18.52435 | -0.29272 | 0.713724 | 1 |
| 12 | 49297400 | 49299399 | 33.05654 | 53.97221 | 73.11317 | 31.42217 | -0.06352 | 1.510871 | 1 |
| 12 | 49651000 | 49653799 | 40.48287 | 65.68589 | 76.89127 | 32.94887 | 0.139692 | 1.627753 | 1 |
| 12 | 49747000 | 49749599 | 36.06726 | 58.97437 | 66.23862 | 28.4829 | 0.207572 | 1.491379 | 1 |
| 12 | 50448800 | 50452599 | 45.36458 | 72.95102 | 100.8402 | 42.77772 | -0.11375 | 1.880605 | 1 |
| 12 | 50507600 | 50513799 | 129.0485 | 195.9502 | 215.5762 | 89.26815 | 0.143959 | 3.218168 | 1 |
| 12 | 50826800 | 50829199 | 32.97068 | 54.53825 | 57.3609 | 24.80069 | 0.30808 | 1.337728 | 1 |
| 12 | 51347600 | 51355599 | 98.24359 | 151.612 | 179.7567 | 74.66747 | 0.054404 | 2.885255 | 1 |
| 12 | 51518000 | 51520599 | 30.62358 | 50.76049 | 54.63219 | 23.65459 | 0.275067 | 1.302269 | 1 |
| 12 | 51524200 | 51544799 | 355.4718 | 493.7488 | 658.2039 | 273.2095 | -0.25616 | 4.682262 | 1 |
| 12 | 51548200 | 51551799 | 14.14708 | 25.58692 | 29.35726 | 12.73529 | 0.213031 | 0.357373 | 1 |
| 12 | 52019200 | 52022599 | 24.96367 | 42.37277 | 48.4972 | 21.24797 | 0.187336 | 1.044181 | 1 |
| 12 | 52269200 | 52271599 | 22.29364 | 37.42105 | 55.93161 | 24.41627 | -0.19679 | 1.142579 | 1 |
| 12 | 54232800 | 54235399 | 23.08216 | 38.67367 | 60.93374 | 26.56914 | -0.2727 | 1.167841 | 1 |
| 12 | 54413400 | 54414599 | 12.73263 | 23.1119 | 26.57631 | 11.47702 | 0.214289 | 0.321301 | 1 |
| 12 | 54435800 | 54443599 | 107.6171 | 165.0674 | 185.5044 | 77.00773 | 0.124889 | 3.006917 | 1 |
| 12 | 54624600 | 54627199 | 37.19489 | 60.13335 | 86.42478 | 36.92847 | -0.16065 | 1.700524 | 1 |
| 12 | 54663400 | 54677799 | 193.1125 | 283.7548 | 343.731 | 141.5618 | -0.03753 | 3.833676 | 1 |
| 12 | 54819400 | 54822199 | 18.23437 | 31.05571 | 51.61842 | 22.62615 | -0.34806 | 0.995075 | 1 |
| 12 | 55316200 | 55319799 | 14.54649 | 26.31748 | 31.43178 | 13.62993 | 0.15588 | 0.363354 | 1 |
| 12 | 55597000 | 55602199 | 59.14815 | 93.66688 | 137.9615 | 57.36783 | -0.22921 | 2.369002 | 1 |
| 12 | 55678800 | 55680599 | 24.97062 | 41.41173 | 59.09856 | 25.59755 | -0.13053 | 1.286907 | 1 |
| 12 | 56630600 | 56632399 | 15.23204 | 26.49543 | 43.21125 | 18.98594 | -0.31525 | 0.766939 | 1 |
| 12 | 56797200 | 56801599 | 45.03663 | 72.53608 | 76.99718 | 32.73673 | 0.269242 | 1.821117 | 1 |
| 12 | 57014000 | 57015199 | 12.36428 | 22.82355 | 28.63499 | 12.09643 | 0.11091 | 0.227034 | 1 |
| 12 | 58164600 | 58168599 | 66.41667 | 104.7428 | 132.3761 | 54.93584 | -0.01342 | 2.470126 | 1 |
| 12 | 58203000 | 58205199 | 9.628512 | 17.57011 | 29.43001 | 12.62557 | -0.318 | 0.277618 | 1 |
| 12 | 58705800 | 58707999 | 25.0668 | 41.98037 | 58.77498 | 25.62072 | -0.10281 | 1.178625 | 1 |
| 12 | 58802800 | 58804399 | 17.10236 | 29.65173 | 44.25326 | 19.41623 | -0.18862 | 0.817502 | 1 |
| 12 | 58816000 | 58817599 | 24.17302 | 40.50253 | 57.13737 | 24.91424 | -0.11375 | 1.167524 | 1 |
| 12 | 58903800 | 58905799 | 22.24644 | 38.17902 | 43.70975 | 19.17128 | 0.189252 | 0.930885 | 1 |
| 12 | 59286400 | 59288999 | 25.33846 | 42.39557 | 64.48856 | 28.09471 | -0.22217 | 1.192236 | 1 |
| 12 | 59432600 | 59435399 | 24.66898 | 41.27736 | 59.92442 | 26.10711 | -0.15499 | 1.185799 | 1 |
| 12 | 59699000 | 59701199 | 22.50503 | 38.93928 | 45.20154 | 19.83105 | 0.171584 | 0.859815 | 1 |
| 12 | 60253600 | 60255199 | 20.9522 | 36.60115 | 38.65508 | 17.02965 | 0.309227 | 0.727821 | 1 |
| 12 | 60318400 | 60320399 | 29.63022 | 48.68259 | 71.40113 | 30.76338 | -0.17307 | 1.420276 | 1 |
| 12 | 60358200 | 60360199 | 23.03096 | 40.01368 | 41.96174 | 18.42315 | 0.318731 | 0.819417 | 1 |
| 12 | 60515400 | 60518599 | 73.38007 | 114.3508 | 161.8871 | 67.18855 | -0.19029 | 2.700617 | 1 |
| 12 | 60804000 | 60805399 | 15.18974 | 26.60574 | 40.41422 | 17.84374 | -0.21258 | 0.658436 | 1 |
| 12 | 61115600 | 61118799 | 44.87152 | 72.39921 | 78.19144 | 33.32431 | 0.247372 | 1.76813 | 1 |
| 12 | 61200800 | 61201999 | 19.88501 | 34.44275 | 41.63219 | 18.26568 | 0.114027 | 0.837097 | 1 |
| 12 | 61337200 | 61340999 | 48.33069 | 77.24112 | 110.6663 | 46.51823 | -0.17586 | 2.07527 | 1 |
| 12 | 61416200 | 61417199 | 11.95659 | 21.51251 | 30.71841 | 13.41221 | -0.10626 | 0.377806 | 1 |
| 12 | 61418600 | 61421199 | 29.71242 | 49.05249 | 69.54936 | 30.0299 | -0.12187 | 1.359271 | 1 |
| 12 | 61517600 | 61519799 | 24.01319 | 40.76561 | 49.56309 | 21.71597 | 0.100637 | 1.043508 | 1 |

| | | | | | | | | | |
|---|---|---|---|---|---|---|---|---|---|
| 12 | 61751800 | 61752599 | 11.22375 | 20.1167 | 33.77767 | 14.79568 | -0.34185 | 0.403438 | 1 |
| 12 | 61939200 | 61939999 | 8.875079 | 16.55662 | 27.91741 | 11.61162 | -0.29525 | 0.136557 | 1 |
| 12 | 62016200 | 62019399 | 52.75258 | 84.22598 | 98.53822 | 41.36313 | 0.114482 | 2.107398 | 1 |
| 12 | 62354600 | 62358399 | 37.4345 | 61.78135 | 60.48691 | 26.11121 | 0.41087 | 1.377027 | 1 |
| 12 | 63067000 | 63068599 | 16.63309 | 29.01585 | 46.78929 | 20.59524 | -0.29885 | 0.736663 | 1 |
| 12 | 63343200 | 63348799 | 109.5404 | 168.1894 | 182.682 | 75.84671 | 0.175115 | 2.987134 | 1 |
| 12 | 63396800 | 63399599 | 38.34507 | 61.9495 | 80.35831 | 34.30535 | -0.01418 | 1.720752 | 1 |
| 12 | 63439800 | 63441199 | 19.37885 | 33.82606 | 34.23717 | 15.07588 | 0.37023 | 0.723793 | 1 |
| 12 | 63737400 | 63739199 | 27.74204 | 45.86284 | 60.29512 | 26.05894 | -0.01292 | 1.332315 | 1 |
| 12 | 63744800 | 63746799 | 30.14631 | 51.14826 | 49.03869 | 21.48249 | 0.441965 | 1.068104 | 1 |
| 12 | 64440600 | 64444199 | 48.75633 | 78.10911 | 93.02137 | 39.23757 | 0.094278 | 1.996594 | 1 |
| 12 | 64781600 | 64789599 | 106.2735 | 163.0885 | 190.5198 | 79.09519 | 0.069604 | 2.996888 | 1 |
| 12 | 64888800 | 64891399 | 35.80017 | 58.53854 | 62.33528 | 26.80459 | 0.284282 | 1.486641 | 1 |
| 12 | 65072800 | 65073999 | 14.37225 | 25.82755 | 29.88326 | 13.06088 | 0.194077 | 0.398331 | 1 |
| 12 | 65271600 | 65274199 | 50.29078 | 80.36659 | 98.4905 | 41.395 | 0.049026 | 2.077407 | 1 |
| 12 | 65678600 | 65681999 | 36.05423 | 58.44238 | 78.36139 | 33.55438 | -0.05704 | 1.639662 | 1 |
| 12 | 65684800 | 65687399 | 17.29962 | 30.38073 | 35.49812 | 15.70788 | 0.163967 | 0.618585 | 1 |
| 12 | 65852800 | 65855199 | 38.63024 | 62.62764 | 67.2255 | 28.78926 | 0.26335 | 1.63591 | 1 |
| 12 | 66150000 | 66150799 | 12.58351 | 22.65145 | 31.63612 | 13.80541 | -0.07411 | 0.380947 | 1 |
| 12 | 66330000 | 66331599 | 44.69324 | 71.79096 | 83.61165 | 35.41161 | 0.130963 | 1.906235 | 1 |
| 12 | 66459400 | 66461199 | 23.82538 | 40.27065 | 49.05675 | 21.46568 | 0.097618 | 1.079394 | 1 |
| 12 | 67795800 | 67797999 | 24.37248 | 40.34798 | 66.17037 | 28.62543 | -0.33078 | 1.31135 | 1 |
| 12 | 67867200 | 67870199 | 39.91668 | 64.36178 | 85.01819 | 36.20266 | -0.04365 | 1.780766 | 1 |
| 12 | 67881400 | 67884199 | 21.81216 | 36.20529 | 63.75906 | 27.63283 | -0.43276 | 1.272229 | 1 |
| 12 | 68670000 | 68670999 | 15.26612 | 27.39624 | 30.23389 | 13.22942 | 0.260633 | 0.412575 | 1 |
| 12 | 68673800 | 68676399 | 23.94827 | 40.52792 | 46.95486 | 20.55359 | 0.169788 | 1.066766 | 1 |
| 12 | 68960600 | 68972599 | 186.6272 | 272.7895 | 370.4451 | 152.6139 | -0.2095 | 3.916169 | 1 |
| 12 | 69056000 | 69061599 | 62.21563 | 99.18535 | 117.8272 | 49.35587 | 0.090078 | 2.169581 | 1 |
| 12 | 69478800 | 69481199 | 14.61475 | 26.39141 | 26.52846 | 11.5307 | 0.401362 | 0.366633 | 1 |
| 12 | 69594000 | 69594599 | 9.539743 | 17.71546 | 25.90772 | 10.85551 | -0.0992 | 0.168259 | 1 |
| 12 | 69606400 | 69610999 | 51.1853 | 81.76596 | 115.9065 | 48.68945 | -0.16123 | 2.094998 | 1 |
| 12 | 70055800 | 70058799 | 32.94778 | 53.65453 | 77.98082 | 33.48232 | -0.16742 | 1.551351 | 1 |
| 12 | 70235400 | 70237399 | 15.22019 | 26.65869 | 44.05507 | 19.45102 | -0.33367 | 0.664425 | 1 |
| 12 | 70969400 | 70974199 | 72.86635 | 113.6558 | 157.5249 | 65.36939 | -0.15866 | 2.684338 | 1 |
| 12 | 71642600 | 71643399 | 11.33379 | 20.75172 | 29.78625 | 12.72293 | -0.09286 | 0.271425 | 1 |
| 12 | 72335400 | 72342199 | 65.08656 | 103.9286 | 105.3816 | 44.24167 | 0.320643 | 2.118989 | 1 |
| 12 | 72645000 | 72649999 | 59.85754 | 95.44395 | 105.1833 | 44.07098 | 0.198478 | 2.155967 | 1 |
| 12 | 72807200 | 72808999 | 13.96512 | 24.35183 | 43.68231 | 19.22276 | -0.45162 | 0.724986 | 1 |
| 12 | 73149800 | 73151599 | 24.68489 | 41.26106 | 55.48376 | 24.1566 | -0.04474 | 1.194009 | 1 |
| 12 | 73303000 | 73311399 | 80.09149 | 126.508 | 140.4168 | 58.31312 | 0.175417 | 2.444868 | 1 |
| 12 | 74611000 | 74612199 | 10.26779 | 19.15263 | 27.86542 | 11.59218 | -0.08424 | 0.149577 | 1 |
| 12 | 74613400 | 74617199 | 57.14135 | 91.25322 | 95.02084 | 39.89905 | 0.282572 | 2.104068 | 1 |
| 12 | 74688400 | 74688999 | 7.508191 | 14.48156 | 20.93016 | 8.093987 | 0.017916 | -0.14544 | 1 |
| 12 | 74745000 | 74749799 | 51.51918 | 82.55431 | 88.93715 | 37.52793 | 0.239025 | 1.991538 | 1 |
| 12 | 74853800 | 74862199 | 111.5004 | 170.206 | 238.3995 | 98.8655 | -0.19796 | 3.107812 | 1 |
| 12 | 74906800 | 74909599 | 9.255586 | 17.26421 | 31.11232 | 12.94333 | -0.3907 | 0.150223 | 1 |
| 12 | 74972400 | 74973799 | 13.59697 | 24.07313 | 37.66966 | 16.65735 | -0.25108 | 0.508753 | 1 |

| | | | | | | | | | |
|---|---|---|---|---|---|---|---|---|---|
| 12 | 75948000 | 75949199 | 18.9185 | 32.48733 | 54.34939 | 23.83896 | -0.3556 | 0.927239 | 1 |
| 12 | 77435600 | 77436599 | 15.51616 | 27.32028 | 35.16729 | 15.56793 | 0.026184 | 0.577091 | 1 |
| 12 | 77848200 | 77850999 | 36.27684 | 58.45622 | 87.86968 | 37.38594 | -0.23069 | 1.796654 | 1 |
| 12 | 78349600 | 78352799 | 48.52248 | 77.78327 | 102.2955 | 43.1907 | -0.04754 | 1.98091 | 1 |
| 12 | 78397600 | 78399399 | 702.4084 | 920.2122 | 1070.87 | 448.2509 | -0.14497 | 5.609271 | 1 |
| 12 | 78514400 | 78515599 | 14.31456 | 25.90663 | 27.36129 | 11.85981 | 0.332677 | 0.351365 | 1 |
| 12 | 79153200 | 79154799 | 12.15166 | 21.67463 | 34.20577 | 15.04449 | -0.25718 | 0.440233 | 1 |
| 12 | 79534200 | 79535999 | 14.54009 | 25.71808 | 38.65177 | 17.09493 | -0.19382 | 0.526378 | 1 |
| 12 | 79613400 | 79616199 | 25.08662 | 41.31851 | 65.88246 | 28.41507 | -0.29173 | 1.375403 | 1 |
| 12 | 80198600 | 80200199 | 17.47692 | 30.55557 | 41.90145 | 18.47164 | -0.06594 | 0.700229 | 1 |
| 12 | 81440200 | 81442799 | 24.25054 | 41.04019 | 54.95982 | 24.05774 | -0.03849 | 1.077065 | 1 |
| 12 | 81500400 | 81502799 | 28.74116 | 47.33583 | 67.55268 | 29.13496 | -0.13234 | 1.384468 | 1 |
| 12 | 82596600 | 82599599 | 44.55074 | 71.58311 | 88.46804 | 37.48438 | 0.046008 | 1.898667 | 1 |
| 12 | 82882400 | 82885399 | 30.75933 | 50.75284 | 71.59902 | 30.90413 | -0.11482 | 1.367488 | 1 |
| 12 | 82945400 | 82950799 | 34.39937 | 56.77764 | 61.43056 | 26.52033 | 0.267087 | 1.367918 | 1 |
| 12 | 84109000 | 84110399 | 15.7853 | 27.91081 | 32.02646 | 14.16568 | 0.193455 | 0.526839 | 1 |
| 12 | 84464000 | 84465799 | 14.19021 | 24.8718 | 42.02199 | 18.56421 | -0.36509 | 0.647602 | 1 |
| 12 | 84628800 | 84631399 | 35.07583 | 57.05497 | 71.12692 | 30.52681 | 0.052279 | 1.568984 | 1 |
| 12 | 85049000 | 85050799 | 26.75571 | 44.37467 | 61.99532 | 26.85328 | -0.10003 | 1.293552 | 1 |
| 12 | 85409400 | 85411999 | 19.31956 | 33.14567 | 50.59767 | 22.1918 | -0.22421 | 0.931387 | 1 |
| 12 | 85551400 | 85553599 | 32.58142 | 53.11295 | 73.44415 | 31.54634 | -0.0947 | 1.532612 | 1 |
| 12 | 85715800 | 85718599 | 26.6227 | 43.99331 | 67.49803 | 29.16325 | -0.23522 | 1.338911 | 1 |
| 12 | 85819600 | 85822599 | 25.7037 | 43.63939 | 45.28151 | 19.84054 | 0.328292 | 1.040943 | 1 |
| 12 | 86687600 | 86689399 | 14.32722 | 25.88913 | 26.42106 | 11.47463 | 0.380134 | 0.358779 | 1 |
| 12 | 86751400 | 86752999 | 12.04138 | 21.80583 | 33.70961 | 14.60042 | -0.21302 | 0.344689 | 1 |
| 12 | 86803200 | 86811399 | 79.83933 | 123.3165 | 192.2465 | 79.85518 | -0.33915 | 2.863608 | 1 |
| 12 | 87193600 | 87207399 | 200.3081 | 292.5756 | 384.9276 | 158.5893 | -0.16483 | 3.928719 | 1 |
| 12 | 87630400 | 87630999 | 10.85007 | 19.78918 | 30.80201 | 13.22254 | -0.21392 | 0.291554 | 1 |
| 12 | 87655800 | 87656999 | 19.59514 | 34.05193 | 41.06772 | 18.03317 | 0.118035 | 0.799845 | 1 |
| 12 | 87875600 | 87876999 | 20.39601 | 35.24962 | 42.62162 | 18.69852 | 0.112735 | 0.857934 | 1 |
| 12 | 87878000 | 87879799 | 10.50074 | 19.79457 | 23.23754 | 9.496042 | 0.24388 | 0.073011 | 1 |
| 12 | 87895600 | 87914599 | 306.0138 | 430.029 | 570.4626 | 236.3613 | -0.23218 | 4.506285 | 1 |
| 12 | 87926000 | 87930799 | 43.33449 | 70.02938 | 82.97689 | 35.43569 | 0.116563 | 1.723742 | 1 |
| 12 | 87995600 | 87998799 | 63.4109 | 100.9014 | 110.5771 | 46.19881 | 0.203369 | 2.227638 | 1 |
| 12 | 88121000 | 88122799 | 13.21226 | 24.49592 | 26.01662 | 10.92504 | 0.35609 | 0.205124 | 1 |
| 12 | 88128200 | 88128399 | 6.521351 | 12.73163 | 15.87353 | 5.605134 | 0.350806 | -0.48653 | 1 |
| 12 | 88132200 | 88132999 | 11.1444 | 20.22168 | 32.25775 | 13.93686 | -0.2554 | 0.325271 | 1 |
| 12 | 88152800 | 88166799 | 164.6215 | 245.4027 | 279.6844 | 115.2586 | 0.069778 | 3.585841 | 1 |
| 12 | 88171000 | 88190399 | 247.1079 | 358.9945 | 363.1557 | 149.7031 | 0.206619 | 4.01282 | 1 |
| 12 | 88305600 | 88321399 | 73.05304 | 115.6172 | 123.15 | 51.19328 | 0.237019 | 2.390478 | 1 |
| 12 | 90331200 | 90332999 | 15.71954 | 27.34881 | 43.71255 | 19.20858 | -0.28628 | 0.767642 | 1 |
| 12 | 90442600 | 90445399 | 29.30889 | 49.20105 | 55.74437 | 24.33594 | 0.201392 | 1.162067 | 1 |
| 12 | 90807400 | 90809199 | 25.49408 | 42.37082 | 65.27694 | 28.3156 | -0.24044 | 1.269163 | 1 |
| 12 | 91125800 | 91127199 | 19.6177 | 33.43248 | 47.97773 | 21.03136 | -0.13685 | 0.990051 | 1 |
| 12 | 91130200 | 91131599 | 14.22391 | 25.00785 | 34.81983 | 15.41868 | -0.08727 | 0.584785 | 1 |
| 12 | 91191600 | 91198199 | 68.03834 | 107.7836 | 123.1714 | 51.22696 | 0.136786 | 2.363517 | 1 |
| 12 | 91572200 | 91574799 | 23.26692 | 39.70608 | 46.89071 | 20.55713 | 0.143431 | 0.989626 | 1 |

| | | | | | | | | | |
|---|---|---|---|---|---|---|---|---|---|
| 12 | 91908800 | 91909399 | 8.692577 | 16.37254 | 26.44988 | 10.8239 | -0.21496 | 0.069973 | 1 |
| 12 | 92138400 | 92139999 | 24.54954 | 40.6022 | 61.65028 | 26.65241 | -0.22 | 1.320564 | 1 |
| 12 | 92714200 | 92714799 | 7.896931 | 15.06701 | 24.52197 | 9.770374 | -0.19173 | -0.03135 | 1 |
| 12 | 93068600 | 93072599 | 15.36955 | 26.87923 | 42.53297 | 18.75478 | -0.27128 | 0.687796 | 1 |
| 12 | 93072800 | 93076799 | 64.35115 | 101.8974 | 120.98 | 50.30426 | 0.081255 | 2.371775 | 1 |
| 12 | 93224600 | 93226199 | 20.29882 | 34.46487 | 51.6476 | 22.63026 | -0.19986 | 1.030538 | 1 |
| 12 | 93505400 | 93510399 | 83.29337 | 129.4674 | 172.2665 | 71.5237 | -0.10383 | 2.749824 | 1 |
| 12 | 93916000 | 93917399 | 9.194498 | 17.69107 | 21.91687 | 8.534817 | 0.229042 | -0.10052 | 1 |
| 12 | 94239800 | 94243799 | 41.0585 | 66.14119 | 80.08754 | 34.05983 | 0.080041 | 1.809005 | 1 |
| 12 | 94277800 | 94286599 | 166.9735 | 250.1798 | 266.452 | 109.9142 | 0.174114 | 3.491619 | 1 |
| 12 | 94516600 | 94517199 | 7.89014 | 15.17074 | 23.05046 | 8.994317 | -0.0652 | -0.10279 | 1 |
| 12 | 94821400 | 94822199 | 10.75739 | 20.0544 | 23.19158 | 9.657196 | 0.24353 | 0.144504 | 1 |
| 12 | 95307200 | 95307799 | 9.543832 | 17.53616 | 29.34681 | 12.47517 | -0.30795 | 0.234272 | 1 |
| 12 | 95381800 | 95382399 | 12.51482 | 22.94702 | 27.62995 | 11.77887 | 0.160838 | 0.266177 | 1 |
| 12 | 95551600 | 95552799 | 22.20342 | 37.51982 | 52.75939 | 23.0841 | -0.10876 | 1.078639 | 1 |
| 12 | 95584200 | 95584999 | 16.82663 | 29.26891 | 41.13965 | 18.07564 | -0.10169 | 0.772753 | 1 |
| 12 | 95745400 | 95747799 | 36.5281 | 58.93919 | 85.14892 | 36.29102 | -0.17138 | 1.755138 | 1 |
| 12 | 95788800 | 95789599 | 12.54931 | 22.47505 | 31.62612 | 13.86233 | -0.08935 | 0.411261 | 1 |
| 12 | 96105200 | 96108199 | 43.30698 | 69.49094 | 98.27238 | 41.55418 | -0.15037 | 1.942973 | 1 |
| 12 | 96248600 | 96250399 | 19.85636 | 34.72412 | 42.90225 | 18.91673 | 0.083872 | 0.711658 | 1 |
| 12 | 96477400 | 96478599 | 14.50921 | 25.1732 | 45.85073 | 20.11754 | -0.47425 | 0.803668 | 1 |
| 12 | 96813800 | 96817599 | 66.5793 | 104.5431 | 156.0003 | 64.70316 | -0.25761 | 2.561013 | 1 |
| 12 | 96829600 | 96832599 | 43.83182 | 70.12343 | 108.3612 | 45.61593 | -0.28308 | 2.039315 | 1 |
| 12 | 97003800 | 97006399 | 33.48365 | 54.26549 | 76.617 | 32.80294 | -0.13155 | 1.638681 | 1 |
| 12 | 98087200 | 98089399 | 24.89501 | 42.03508 | 56.07619 | 24.52947 | -0.03312 | 1.09953 | 1 |
| 12 | 98319200 | 98324799 | 58.13884 | 92.31019 | 132.8632 | 55.37598 | -0.19246 | 2.294274 | 1 |
| 12 | 98895600 | 98896599 | 13.95813 | 25.24093 | 29.98231 | 13.00917 | 0.163328 | 0.360231 | 1 |
| 12 | 99471000 | 99477399 | 86.22981 | 133.1645 | 180.4644 | 74.96117 | -0.13746 | 2.867805 | 1 |
| 12 | 99608400 | 99616399 | 163.7233 | 242.361 | 330.0052 | 135.9062 | -0.19607 | 3.709625 | 1 |
| 12 | 99660400 | 99667199 | 51.29765 | 81.88533 | 113.7698 | 47.74224 | -0.13349 | 2.116119 | 1 |
| 12 | 101175000 | 101176599 | 19.39908 | 33.79903 | 37.81854 | 16.6336 | 0.225914 | 0.754855 | 1 |
| 12 | 101621600 | 101622999 | 17.82641 | 31.27357 | 32.62602 | 14.4244 | 0.326872 | 0.634775 | 1 |
| 12 | 101701600 | 101703199 | 27.28364 | 44.52687 | 74.39694 | 31.96873 | -0.36571 | 1.506668 | 1 |
| 12 | 102519600 | 102522599 | 26.04499 | 42.74809 | 71.58903 | 30.83093 | -0.36445 | 1.422329 | 1 |
| 12 | 102833000 | 102834999 | 22.58122 | 37.49487 | 62.78867 | 27.2188 | -0.36039 | 1.268658 | 1 |
| 12 | 103078600 | 103079599 | 11.05545 | 20.34197 | 30.29077 | 12.85206 | -0.13897 | 0.239751 | 1 |
| 12 | 103582200 | 103584399 | 25.74946 | 42.78589 | 60.33114 | 26.16627 | -0.11311 | 1.267266 | 1 |
| 12 | 103717200 | 103720599 | 60.56842 | 96.3829 | 106.0978 | 44.33027 | 0.197016 | 2.221981 | 1 |
| 12 | 103747200 | 103749599 | 39.43681 | 63.8031 | 72.22399 | 30.88271 | 0.184096 | 1.680811 | 1 |
| 12 | 103750600 | 103755799 | 64.62864 | 102.5151 | 122.232 | 50.89564 | 0.077376 | 2.324928 | 1 |
| 12 | 104272200 | 104281599 | 126.0439 | 191.1491 | 243.6197 | 100.843 | -0.06957 | 3.245468 | 1 |
| 12 | 104346000 | 104349799 | 44.85801 | 72.3495 | 84.14309 | 35.843 | 0.140118 | 1.78113 | 1 |
| 12 | 105963400 | 105965399 | 14.35504 | 25.24791 | 36.7223 | 16.2599 | -0.14988 | 0.583495 | 1 |
| 12 | 107022800 | 107024999 | 21.43251 | 36.74088 | 45.79376 | 20.08352 | 0.066719 | 0.936666 | 1 |
| 12 | 109202200 | 109212599 | 45.25927 | 73.00812 | 87.13649 | 37.12563 | 0.103105 | 1.779483 | 1 |
| 12 | 109784800 | 109792399 | 77.84803 | 121.7169 | 163.0606 | 67.64995 | -0.10704 | 2.644845 | 1 |
| 12 | 109958400 | 109964999 | 59.59945 | 95.08217 | 103.2746 | 43.30257 | 0.220202 | 2.139511 | 1 |

| | | | | | | | | | |
|---|---|---|---|---|---|---|---|---|---|
| 12 | 112743400 | 112743799 | 16.34192 | 28.47006 | 40.59905 | 17.85556 | -0.1222 | 0.747321 | 1 |
| 12 | 112744600 | 112746999 | 52.53285 | 83.40508 | 126.3757 | 52.66942 | -0.26647 | 2.292369 | 1 |
| 12 | 113344800 | 113345599 | 8.840326 | 16.69028 | 25.42477 | 10.35823 | -0.12533 | 0.053421 | 1 |
| 12 | 113359400 | 113363799 | 63.40801 | 100.6908 | 112.7485 | 46.99915 | 0.169409 | 2.289045 | 1 |
| 12 | 113544200 | 113546799 | 33.81059 | 55.36237 | 74.83613 | 32.197 | -0.05772 | 1.471927 | 1 |
| 12 | 113614400 | 113620599 | 85.92152 | 134.2154 | 157.0183 | 65.14936 | 0.087187 | 2.664486 | 1 |
| 12 | 113643600 | 113649999 | 84.96333 | 133.3945 | 138.7262 | 57.53765 | 0.262494 | 2.569812 | 1 |
| 12 | 113669200 | 113675399 | 65.54174 | 104.2324 | 112.0975 | 46.79702 | 0.2296 | 2.248036 | 1 |
| 12 | 113688200 | 113695399 | 80.53583 | 127.1791 | 131.1611 | 54.4627 | 0.281008 | 2.449001 | 1 |
| 12 | 113757800 | 113760599 | 18.45415 | 32.28491 | 37.68405 | 16.62152 | 0.165555 | 0.689809 | 1 |
| 12 | 113881600 | 113892799 | 185.2404 | 274.6702 | 319.0656 | 131.4154 | 0.035201 | 3.683557 | 1 |
| 12 | 113895400 | 113896199 | 7.705471 | 14.81042 | 25.27579 | 9.872689 | -0.23351 | -0.09785 | 1 |
| 12 | 114022600 | 114023999 | 16.8753 | 29.59964 | 43.98018 | 19.44125 | -0.181 | 0.651351 | 1 |
| 12 | 114059200 | 114061399 | 23.31972 | 40.63472 | 41.82477 | 18.397 | 0.345871 | 0.778891 | 1 |
| 12 | 114305600 | 114308399 | 34.74395 | 57.09107 | 64.4645 | 27.77607 | 0.203284 | 1.427801 | 1 |
| 12 | 114313000 | 114321999 | 29.87449 | 49.85702 | 52.95631 | 23.03332 | 0.294449 | 1.224751 | 1 |
| 12 | 114418400 | 114433199 | 205.3467 | 304.5779 | 298.5438 | 122.9669 | 0.2805 | 3.680361 | 1 |
| 12 | 115008200 | 115015199 | 120.4197 | 184.9745 | 181.7567 | 75.46708 | 0.320028 | 2.982784 | 1 |
| 12 | 115421800 | 115430599 | 101.0301 | 157.2084 | 174.3579 | 72.3802 | 0.159796 | 2.741724 | 1 |
| 12 | 115682800 | 115690199 | 164.0849 | 245.8748 | 249.1856 | 102.7935 | 0.245803 | 3.48841 | 1 |
| 12 | 115996200 | 115997799 | 22.4979 | 38.78989 | 41.53348 | 18.21982 | 0.286708 | 0.887136 | 1 |
| 12 | 116361400 | 116369599 | 159.3062 | 236.7708 | 320.6223 | 132.0881 | -0.18278 | 3.639095 | 1 |
| 12 | 118316800 | 118322199 | 94.81617 | 146.7323 | 160.5341 | 66.68183 | 0.173344 | 2.830798 | 1 |
| 12 | 118974000 | 118976399 | 9.837083 | 17.99893 | 32.88341 | 14.05899 | -0.44022 | 0.266069 | 1 |
| 12 | 119098400 | 119099999 | 11.97117 | 21.49262 | 35.72926 | 15.62825 | -0.32644 | 0.40035 | 1 |
| 12 | 119260800 | 119264399 | 39.78652 | 64.13756 | 82.97828 | 35.32383 | -0.01406 | 1.787113 | 1 |
| 12 | 119314600 | 119317399 | 13.02159 | 23.18186 | 35.21715 | 15.51434 | -0.20403 | 0.462093 | 1 |
| 12 | 119512200 | 119514799 | 18.84244 | 32.10714 | 51.10841 | 22.40348 | -0.28589 | 0.991654 | 1 |
| 12 | 119622200 | 119622799 | 10.03217 | 19.15532 | 20.6567 | 8.217264 | 0.398851 | -0.0279 | 1 |
| 12 | 119653600 | 119655399 | 23.86737 | 40.06196 | 58.64799 | 25.6018 | -0.1669 | 1.153215 | 1 |
| 13 | 3134800 | 3142999 | 114.2244 | 174.3467 | 208.572 | 86.49386 | 0.029252 | 3.107295 | 1 |
| 13 | 3147000 | 3160799 | 221.2709 | 322.9459 | 354.9392 | 146.2458 | 0.093468 | 3.941536 | 1 |
| 13 | 3184800 | 3191199 | 104.5531 | 160.6695 | 173.1657 | 71.90413 | 0.187239 | 2.968191 | 1 |
| 13 | 3247000 | 3250799 | 29.77157 | 50.20625 | 49.63621 | 21.70419 | 0.397533 | 1.118311 | 1 |
| 13 | 3422400 | 3448199 | 613.5933 | 823.1711 | 799.3902 | 333.6087 | 0.150218 | 5.228311 | 1 |
| 13 | 3660000 | 3674399 | 287.9031 | 410.5945 | 450.8347 | 186.3399 | 0.061966 | 4.288251 | 1 |
| 13 | 3725600 | 3728599 | 18.38545 | 31.95954 | 51.7641 | 22.73441 | -0.30589 | 0.800488 | 1 |
| 13 | 3770000 | 3772599 | 23.10208 | 39.69304 | 46.31695 | 20.31607 | 0.16214 | 0.927333 | 1 |
| 13 | 4125200 | 4126999 | 35.70371 | 57.97984 | 67.06157 | 28.75443 | 0.158143 | 1.596954 | 1 |
| 13 | 4174000 | 4178199 | 13.26703 | 23.86775 | 34.65488 | 15.1323 | -0.13055 | 0.395457 | 1 |
| 13 | 4233000 | 4233799 | 14.43796 | 25.56386 | 34.15014 | 15.10083 | -0.02383 | 0.508813 | 1 |
| 13 | 4307200 | 4309799 | 25.61059 | 43.27895 | 50.66835 | 22.16945 | 0.154652 | 1.088731 | 1 |
| 13 | 4404400 | 4408399 | 59.59936 | 94.86716 | 113.0551 | 47.25535 | 0.083118 | 2.213732 | 1 |
| 13 | 4847400 | 4851199 | 28.15718 | 47.31368 | 54.58021 | 23.84007 | 0.175498 | 1.147737 | 1 |
| 13 | 4877400 | 4880599 | 33.35845 | 54.51402 | 77.46748 | 33.30443 | -0.13153 | 1.500352 | 1 |
| 13 | 4951600 | 4956799 | 106.8405 | 163.2537 | 209.8657 | 87.05225 | -0.07318 | 3.084631 | 1 |
| 13 | 4973800 | 4974599 | 11.30812 | 20.68182 | 28.63805 | 12.25092 | -0.04344 | 0.270675 | 1 |

| | | | | | | | | | |
|---|---|---|---|---|---|---|---|---|---|
| 13 | 5113400 | 5114799 | 20.12569 | 34.50034 | 42.26407 | 18.53551 | 0.091805 | 0.930265 | 1 |
| 13 | 5148000 | 5149399 | 16.62919 | 28.92652 | 45.77523 | 20.11284 | -0.27217 | 0.775751 | 1 |
| 13 | 5180600 | 5181999 | 15.87116 | 27.57625 | 44.25279 | 19.42988 | -0.29225 | 0.787627 | 1 |
| 13 | 5293600 | 5300399 | 37.05175 | 59.87674 | 85.57001 | 36.5434 | -0.15318 | 1.712616 | 1 |
| 13 | 5328000 | 5330199 | 14.02111 | 24.50108 | 40.63143 | 17.90401 | -0.33821 | 0.694037 | 1 |
| 13 | 5744400 | 5747799 | 35.37028 | 57.97041 | 65.80506 | 28.32273 | 0.194112 | 1.458077 | 1 |
| 13 | 6011600 | 6012399 | 13.12081 | 23.54453 | 33.52848 | 14.67198 | -0.1051 | 0.405086 | 1 |
| 13 | 6233400 | 6235199 | 13.31983 | 23.73607 | 40.34664 | 17.75937 | -0.36479 | 0.465461 | 1 |
| 13 | 6723600 | 6726199 | 35.32793 | 56.89376 | 92.75737 | 39.43197 | -0.34868 | 1.81572 | 1 |
| 13 | 6778600 | 6781199 | 15.66847 | 27.51714 | 40.27498 | 17.82216 | -0.15935 | 0.618031 | 1 |
| 13 | 6841600 | 6842799 | 18.81392 | 32.61052 | 39.60167 | 17.37519 | 0.107509 | 0.82171 | 1 |
| 13 | 6933600 | 6935599 | 15.092 | 26.45137 | 37.29289 | 16.47447 | -0.10561 | 0.640691 | 1 |
| 13 | 6952600 | 6954599 | 20.78893 | 36.05118 | 45.23399 | 19.84665 | 0.060553 | 0.833373 | 1 |
| 13 | 7103200 | 7105999 | 29.34506 | 48.4249 | 71.58373 | 30.89904 | -0.18186 | 1.36735 | 1 |
| 13 | 7180200 | 7182599 | 50.09591 | 79.93764 | 111.8197 | 46.89951 | -0.1438 | 2.127624 | 1 |
| 13 | 7219400 | 7224199 | 71.39698 | 113.1611 | 127.5829 | 53.08144 | 0.156882 | 2.356244 | 1 |
| 13 | 7327400 | 7330199 | 23.32651 | 38.9701 | 58.60538 | 25.50727 | -0.20552 | 1.199965 | 1 |
| 13 | 7338200 | 7340599 | 6.090579 | 11.84703 | 22.44441 | 8.293956 | -0.3069 | -0.31184 | 1 |
| 13 | 7388200 | 7390199 | 25.39422 | 42.73729 | 50.14874 | 21.91477 | 0.151134 | 1.121821 | 1 |
| 13 | 7536600 | 7543199 | 93.14096 | 143.7841 | 186.8041 | 77.5947 | -0.07707 | 2.878496 | 1 |
| 13 | 7763800 | 7766199 | 19.95333 | 34.7665 | 39.78151 | 17.49754 | 0.193627 | 0.757842 | 1 |
| 13 | 7876400 | 7881399 | 8.525899 | 16.08439 | 28.63289 | 11.68405 | -0.35094 | 0.062172 | 1 |
| 13 | 7958600 | 7960399 | 20.23805 | 35.37679 | 37.0924 | 16.34923 | 0.319632 | 0.711502 | 1 |
| 13 | 8078000 | 8082199 | 41.09851 | 66.2247 | 97.7912 | 41.60623 | -0.20522 | 1.806865 | 1 |
| 13 | 8087200 | 8088199 | 12.71139 | 23.46166 | 27.14797 | 11.47002 | 0.227227 | 0.230443 | 1 |
| 13 | 8349800 | 8351599 | 20.6653 | 35.81027 | 44.71401 | 19.61806 | 0.067365 | 0.839451 | 1 |
| 13 | 8859800 | 8860399 | 10.2748 | 19.00607 | 26.79781 | 11.28831 | -0.05282 | 0.196868 | 1 |
| 13 | 8950000 | 8959799 | 182.6117 | 270.9603 | 295.0603 | 121.5368 | 0.129296 | 3.670217 | 1 |
| 13 | 9061200 | 9062399 | 30.10314 | 49.68964 | 57.19406 | 24.69244 | 0.178103 | 1.352697 | 1 |
| 13 | 9436000 | 9439999 | 51.3445 | 81.73812 | 126.2582 | 52.78352 | -0.29033 | 2.20549 | 1 |
| 13 | 9452000 | 9455599 | 54.96248 | 87.63314 | 113.648 | 47.61344 | -0.03605 | 2.157475 | 1 |
| 13 | 9899000 | 9912599 | 201.8122 | 297.5926 | 308.5441 | 127.0626 | 0.191678 | 3.776418 | 1 |
| 13 | 10351800 | 10352799 | 15.04019 | 26.38561 | 37.83601 | 16.72781 | -0.12976 | 0.628583 | 1 |
| 13 | 10652600 | 10653599 | 21.32537 | 36.67686 | 38.60452 | 16.93367 | 0.31064 | 0.900538 | 1 |
| 13 | 11149200 | 11152399 | 7.26213 | 14.1442 | 17.39953 | 6.355862 | 0.32598 | -0.35199 | 1 |
| 13 | 11184200 | 11186599 | 32.37548 | 53.61609 | 59.68458 | 25.82611 | 0.226488 | 1.322597 | 1 |
| 13 | 11210200 | 11211399 | 23.09975 | 39.40439 | 45.84629 | 20.09858 | 0.164774 | 0.991686 | 1 |
| 13 | 11266600 | 11271599 | 75.53773 | 117.6965 | 167.4665 | 69.50569 | -0.19773 | 2.705546 | 1 |
| 13 | 11514800 | 11517999 | 18.39158 | 32.54114 | 36.29091 | 16.04972 | 0.234497 | 0.54185 | 1 |
| 13 | 11522800 | 11524599 | 11.15511 | 20.31718 | 32.81467 | 14.11112 | -0.26879 | 0.305277 | 1 |
| 13 | 12509400 | 12512199 | 21.55869 | 36.35208 | 59.15229 | 25.86435 | -0.3187 | 1.104307 | 1 |
| 13 | 12698400 | 12702199 | 65.82327 | 103.6608 | 142.616 | 59.17452 | -0.1373 | 2.499642 | 1 |
| 13 | 12704600 | 12706999 | 14.56578 | 25.4989 | 42.51625 | 18.76318 | -0.34642 | 0.666811 | 1 |
| 13 | 13100600 | 13100999 | 13.85207 | 25.23616 | 23.60205 | 10.14599 | 0.514949 | 0.303028 | 1 |
| 13 | 13593400 | 13597199 | 83.3151 | 129.8924 | 146.4589 | 60.7812 | 0.138171 | 2.695205 | 1 |
| 13 | 14068600 | 14073799 | 87.72706 | 137.2397 | 149.3726 | 61.96755 | 0.192901 | 2.638972 | 1 |
| 13 | 14124000 | 14127399 | 14.66062 | 25.76669 | 39.2308 | 17.37123 | -0.21603 | 0.597136 | 1 |

| | | | | | | | | | |
|---|---|---|---|---|---|---|---|---|---|
| 13 | 14139000 | 14140599 | 16.62937 | 28.43274 | 50.56021 | 22.16925 | -0.44415 | 0.951382 | 1 |
| 13 | 14870800 | 14871599 | 16.00243 | 27.89957 | 41.07825 | 18.07485 | -0.16806 | 0.734928 | 1 |
| 13 | 14915800 | 14919599 | 16.61867 | 29.03628 | 40.96699 | 18.05026 | -0.1066 | 0.706923 | 1 |
| 13 | 14946000 | 14948199 | 12.9344 | 23.52123 | 26.11383 | 11.25148 | 0.2669 | 0.31226 | 1 |
| 13 | 15025400 | 15035999 | 131.6381 | 199.086 | 255.8841 | 105.8287 | -0.08509 | 3.301674 | 1 |
| 13 | 15201200 | 15202599 | 19.28699 | 33.01395 | 46.92386 | 20.57668 | -0.12204 | 0.947579 | 1 |
| 13 | 15421200 | 15422999 | 21.26046 | 36.24416 | 54.38814 | 23.84204 | -0.20113 | 1.000149 | 1 |
| 13 | 15425000 | 15428799 | 21.75079 | 36.49623 | 57.9828 | 25.30573 | -0.28444 | 1.149559 | 1 |
| 13 | 15607000 | 15607399 | 11.17905 | 20.57181 | 25.24735 | 10.71048 | 0.138676 | 0.229272 | 1 |
| 13 | 16142400 | 16145199 | 29.1657 | 48.37085 | 60.54117 | 26.22311 | 0.058102 | 1.29888 | 1 |
| 13 | 16272000 | 16278399 | 112.8107 | 171.7928 | 219.4194 | 90.93868 | -0.06775 | 3.155585 | 1 |
| 13 | 16565600 | 16566799 | 14.63535 | 25.49208 | 43.71607 | 19.22413 | -0.38707 | 0.746412 | 1 |
| 13 | 16721800 | 16722599 | 16.41428 | 29.09141 | 31.84728 | 14.07979 | 0.262174 | 0.504453 | 1 |
| 13 | 17377600 | 17378999 | 25.89157 | 43.0948 | 55.44954 | 24.07715 | 0.018818 | 1.244625 | 1 |
| 13 | 18225000 | 18230199 | 89.36515 | 137.9509 | 198.5194 | 82.46104 | -0.22446 | 2.878801 | 1 |
| 13 | 18783400 | 18785399 | 12.84429 | 23.11016 | 32.0636 | 13.9993 | -0.0653 | 0.384952 | 1 |
| 13 | 19175400 | 19180399 | 23.2741 | 39.57896 | 54.4046 | 23.84451 | -0.07558 | 1.031219 | 1 |
| 13 | 19622000 | 19623999 | 18.14289 | 30.99084 | 50.58883 | 22.18009 | -0.32159 | 0.964043 | 1 |
| 13 | 20098600 | 20105999 | 120.9189 | 182.9309 | 257.0296 | 106.3127 | -0.21314 | 3.29306 | 1 |
| 13 | 20178200 | 20182399 | 24.83371 | 41.85525 | 49.61203 | 21.68986 | 0.136652 | 1.105472 | 1 |
| 13 | 20888400 | 20889599 | 13.32513 | 24.03059 | 26.0439 | 11.33896 | 0.291569 | 0.361165 | 1 |
| 13 | 21818200 | 21820199 | 11.21897 | 20.31777 | 31.3073 | 13.55888 | -0.2076 | 0.337505 | 1 |
| 13 | 21834200 | 21837799 | 72.806 | 114.6907 | 129.5188 | 53.74222 | 0.147495 | 2.495765 | 1 |
| 13 | 21997400 | 21998599 | 15.39635 | 26.77552 | 40.21073 | 17.66521 | -0.19667 | 0.767464 | 1 |
| 13 | 22304800 | 22310599 | 22.88287 | 38.63699 | 57.12594 | 24.98855 | -0.181 | 1.092304 | 1 |
| 13 | 22349400 | 22351399 | 20.23871 | 35.37527 | 37.60921 | 16.57596 | 0.299692 | 0.713391 | 1 |
| 13 | 22394800 | 22397199 | 24.76298 | 41.2357 | 63.6923 | 27.66451 | -0.24403 | 1.241842 | 1 |
| 13 | 22465600 | 22467999 | 27.65043 | 46.24325 | 52.92245 | 23.0494 | 0.187058 | 1.193658 | 1 |
| 13 | 22490400 | 22500399 | 138.5308 | 209.6973 | 233.5837 | 96.63284 | 0.122325 | 3.283983 | 1 |
| 13 | 22552600 | 22554399 | 25.90488 | 43.70432 | 46.96497 | 20.53882 | 0.27778 | 1.100234 | 1 |
| 13 | 22732600 | 22734599 | 35.85063 | 57.76982 | 92.21357 | 39.23453 | -0.31713 | 1.799504 | 1 |
| 13 | 23360400 | 23362999 | 31.78933 | 52.72888 | 56.8039 | 24.60651 | 0.273862 | 1.30229 | 1 |
| 13 | 23822800 | 23834199 | 153.9752 | 230.1493 | 284.7213 | 117.3956 | -0.04497 | 3.534297 | 1 |
| 13 | 24659800 | 24660799 | 15.43414 | 27.31785 | 36.99565 | 16.36026 | -0.04406 | 0.520671 | 1 |
| 13 | 25084400 | 25085999 | 10.07104 | 19.06388 | 24.87708 | 10.0789 | 0.103157 | 0.045238 | 1 |
| 13 | 25190000 | 25192599 | 35.5761 | 57.98566 | 69.40126 | 29.8081 | 0.112883 | 1.537072 | 1 |
| 13 | 25409600 | 25410399 | 10.08067 | 19.1025 | 22.71044 | 9.180328 | 0.239013 | 0.03009 | 1 |
| 13 | 25729200 | 25730199 | 17.13279 | 30.20982 | 31.88077 | 14.10902 | 0.311918 | 0.563281 | 1 |
| 13 | 25864200 | 25865999 | 21.96695 | 38.11824 | 42.0225 | 18.43795 | 0.246712 | 0.829973 | 1 |
| 13 | 26360000 | 26360599 | 11.20347 | 20.85302 | 24.73284 | 10.32659 | 0.204483 | 0.164355 | 1 |
| 13 | 26625600 | 26626999 | 22.44594 | 38.3055 | 42.78053 | 18.75523 | 0.22376 | 0.982227 | 1 |
| 13 | 26863200 | 26863999 | 13.19151 | 23.81777 | 27.97098 | 12.15997 | 0.177747 | 0.357507 | 1 |
| 13 | 26874000 | 26876799 | 38.75191 | 63.26051 | 67.49657 | 29.00652 | 0.279814 | 1.522067 | 1 |
| 13 | 26952600 | 26954399 | 24.41005 | 40.25282 | 63.73864 | 27.50451 | -0.28103 | 1.355921 | 1 |
| 13 | 27007400 | 27009599 | 13.23754 | 23.43752 | 35.1407 | 15.53896 | -0.1897 | 0.500806 | 1 |
| 13 | 27021000 | 27021799 | 11.66518 | 20.96927 | 33.72063 | 14.73424 | -0.27757 | 0.38734 | 1 |
| 13 | 27444600 | 27446199 | 11.62853 | 20.9247 | 31.70009 | 13.83946 | -0.19112 | 0.376002 | 1 |

| | | | | | | | | | | |
|---|---|---|---|---|---|---|---|---|---|---|
| 13 | 27548800 | 27552199 | 47.03956 | 75.8261 | 80.59836 | 34.30856 | 0.268754 | 1.800033 | | 1 |
| 13 | 27629800 | 27636799 | 168.484 | 251.9718 | 254.6637 | 105.0133 | 0.247202 | 3.525492 | | 1 |
| 13 | 27725800 | 27726599 | 13.32502 | 23.80452 | 35.06485 | 15.40213 | -0.15734 | 0.437718 | | 1 |
| 13 | 28082600 | 28083999 | 18.03808 | 31.37998 | 44.16303 | 19.40551 | -0.10375 | 0.778899 | | 1 |
| 13 | 28152400 | 28152999 | 12.91147 | 23.74935 | 24.90738 | 10.57259 | 0.362824 | 0.242427 | | 1 |
| 13 | 28167000 | 28167999 | 20.93186 | 36.10666 | 41.11208 | 18.03522 | 0.198588 | 0.876601 | | 1 |
| 13 | 28313000 | 28313399 | 7.908562 | 15.25957 | 23.1295 | 8.934757 | -0.04799 | -0.13413 | | 1 |
| 13 | 28519200 | 28527599 | 117.3054 | 179.4627 | 217.341 | 90.17841 | 0.014197 | 3.064274 | | 1 |
| 13 | 28568000 | 28570199 | 42.56615 | 68.60679 | 82.22275 | 34.99453 | 0.095789 | 1.796426 | | 1 |
| 13 | 28659200 | 28662599 | 43.17705 | 69.25087 | 103.4106 | 43.69654 | -0.22952 | 1.959126 | | 1 |
| 13 | 29035600 | 29038999 | 41.38181 | 66.60539 | 97.98553 | 41.62024 | -0.20182 | 1.845524 | | 1 |
| 13 | 29076600 | 29077999 | 10.48172 | 19.37965 | 31.46434 | 13.26258 | -0.25592 | 0.211881 | | 1 |
| 13 | 30306600 | 30308399 | 23.12651 | 38.87887 | 50.13066 | 21.8997 | 0.015571 | 1.12277 | | 1 |
| 13 | 30560800 | 30566399 | 53.90271 | 85.91315 | 119.968 | 50.23675 | -0.14313 | 2.167744 | | 1 |
| 13 | 31422800 | 31424599 | 43.2409 | 69.82606 | 72.2337 | 30.8181 | 0.310758 | 1.739474 | | 1 |
| 13 | 31775800 | 31784799 | 82.12834 | 127.7291 | 164.6914 | 68.37278 | -0.05785 | 2.738988 | | 1 |
| 13 | 32068200 | 32069999 | 14.10052 | 24.95919 | 36.64933 | 16.20688 | -0.15984 | 0.513432 | | 1 |
| 13 | 32378200 | 32379199 | 17.21471 | 30.23562 | 34.57008 | 15.29894 | 0.19526 | 0.615943 | | 1 |
| 13 | 32530800 | 32537799 | 109.1476 | 167.3024 | 179.7782 | 74.62377 | 0.188728 | 3.019521 | | 1 |
| 13 | 32820600 | 32826799 | 51.96304 | 82.77329 | 120.6853 | 50.49635 | -0.20623 | 2.184878 | | 1 |
| 13 | 33304400 | 33305399 | 13.42379 | 23.78987 | 34.15055 | 15.09825 | -0.12662 | 0.490931 | | 1 |
| 13 | 33557000 | 33558399 | 31.88013 | 51.73121 | 76.47669 | 32.77298 | -0.19603 | 1.611333 | | 1 |
| 13 | 33608200 | 33608599 | 8.873616 | 16.19199 | 31.94422 | 13.7048 | -0.55259 | 0.277718 | | 1 |
| 13 | 33626800 | 33633399 | 117.0117 | 178.49 | 210.4833 | 87.2739 | 0.049256 | 3.121752 | | 1 |
| 13 | 33651800 | 33654399 | 27.88776 | 46.87115 | 52.30603 | 22.84824 | 0.223339 | 1.146787 | | 1 |
| 13 | 33723400 | 33725799 | 22.54716 | 38.15439 | 51.68914 | 22.6255 | -0.05508 | 1.067443 | | 1 |
| 13 | 33770800 | 33783199 | 201.2708 | 297.3621 | 323.9531 | 133.4109 | 0.122874 | 3.745783 | | 1 |
| 13 | 33829400 | 33835199 | 83.20959 | 131.1209 | 132.2888 | 54.89389 | 0.310247 | 2.496958 | | 1 |
| 13 | 33865000 | 33872799 | 110.6865 | 170.5616 | 178.4137 | 74.1102 | 0.233322 | 2.917536 | | 1 |
| 13 | 33884000 | 33891599 | 129.0128 | 196.4452 | 213.0705 | 88.30455 | 0.167745 | 3.161131 | | 1 |
| 13 | 33916600 | 33922599 | 41.00888 | 66.08665 | 89.43726 | 38.05729 | -0.07943 | 1.802613 | | 1 |
| 13 | 33957000 | 33963599 | 100.2243 | 154.1604 | 200.8198 | 83.39767 | -0.0849 | 2.951023 | | 1 |
| 13 | 34348400 | 34351399 | 37.95308 | 61.54204 | 74.17069 | 31.76819 | 0.096903 | 1.633099 | | 1 |
| 13 | 35180800 | 35184799 | 45.15657 | 73.17112 | 74.02598 | 31.68973 | 0.347865 | 1.660211 | | 1 |
| 13 | 35239600 | 35240999 | 15.67731 | 27.64073 | 36.6082 | 16.20149 | -0.01407 | 0.56559 | | 1 |
| 13 | 35750000 | 35752199 | 25.33205 | 42.3308 | 58.2615 | 25.36144 | -0.07823 | 1.20112 | | 1 |
| 13 | 35765200 | 35766999 | 13.95409 | 24.81822 | 35.50879 | 15.65544 | -0.11932 | 0.469998 | | 1 |
| 13 | 35928000 | 35935199 | 103.1205 | 158.8643 | 193.9645 | 80.56806 | 0.010203 | 2.921123 | | 1 |
| 13 | 36108200 | 36116999 | 123.6332 | 189.0115 | 202.1398 | 83.85764 | 0.192648 | 3.078446 | | 1 |
| 13 | 36365400 | 36373999 | 82.48996 | 130.2961 | 137.525 | 57.11208 | 0.247912 | 2.446941 | | 1 |
| 13 | 36796600 | 36797999 | 15.21101 | 26.67241 | 40.37795 | 17.84436 | -0.20768 | 0.641168 | | 1 |
| 13 | 37037000 | 37037999 | 12.00241 | 21.85313 | 26.70075 | 11.48676 | 0.131047 | 0.299628 | | 1 |
| 13 | 37162200 | 37163599 | 19.00996 | 32.98444 | 38.10593 | 16.71946 | 0.179581 | 0.812272 | | 1 |
| 13 | 37351000 | 37352199 | 16.48325 | 28.61863 | 39.71724 | 17.43039 | -0.08363 | 0.795251 | | 1 |
| 13 | 39150800 | 39152999 | 27.24717 | 45.61693 | 53.96987 | 23.52124 | 0.139283 | 1.183551 | | 1 |
| 13 | 39334400 | 39336599 | 26.89119 | 44.63171 | 60.37982 | 26.16662 | -0.05366 | 1.283952 | | 1 |
| 13 | 39688400 | 39693599 | 60.64669 | 96.01762 | 143.5004 | 59.66401 | -0.2505 | 2.37687 | | 1 |

| | | | | | | | | | |
|---|---|---|---|---|---|---|---|---|---|
| 13 | 39728400 | 39729399 | 18.76965 | 32.26453 | 48.47499 | 21.26296 | -0.20084 | 0.90791 | 1 |
| 13 | 39862000 | 39866199 | 47.17579 | 75.85371 | 95.59529 | 40.54508 | 0.019082 | 1.882806 | 1 |
| 13 | 40428800 | 40431599 | 20.77183 | 35.19972 | 55.424 | 24.27069 | -0.2712 | 1.051873 | 1 |
| 13 | 40524400 | 40526199 | 21.30317 | 37.24782 | 39.11505 | 17.24434 | 0.317405 | 0.716571 | 1 |
| 13 | 41946000 | 41947799 | 16.97783 | 29.76381 | 36.63594 | 16.18757 | 0.089253 | 0.646338 | 1 |
| 13 | 41970600 | 41973599 | 23.9677 | 39.93276 | 63.05725 | 27.39793 | -0.27577 | 1.23361 | 1 |
| 13 | 42053600 | 42055399 | 16.95163 | 29.63711 | 43.01095 | 18.9607 | -0.14717 | 0.70125 | 1 |
| 13 | 42133200 | 42137999 | 91.16197 | 142.3437 | 151.8449 | 63.00548 | 0.219742 | 2.673141 | 1 |
| 13 | 42347400 | 42348599 | 13.12707 | 23.25517 | 36.67906 | 16.21736 | -0.26202 | 0.49777 | 1 |
| 13 | 42352600 | 42354799 | 35.23819 | 57.41922 | 73.0854 | 31.38733 | 0.024096 | 1.545295 | 1 |
| 13 | 42481400 | 42482999 | 19.57898 | 34.24779 | 38.63738 | 17.03991 | 0.21439 | 0.699119 | 1 |
| 13 | 42619000 | 42619599 | 11.73064 | 21.42331 | 29.69601 | 12.72686 | -0.04643 | 0.286944 | 1 |
| 13 | 43047000 | 43050199 | 18.72272 | 32.09595 | 54.41594 | 23.86492 | -0.37542 | 0.936623 | 1 |
| 13 | 43433000 | 43433599 | 13.89151 | 25.37463 | 26.45031 | 11.33307 | 0.362342 | 0.294605 | 1 |
| 13 | 43537600 | 43540399 | 31.87738 | 52.13008 | 73.19872 | 31.47724 | -0.11357 | 1.484659 | 1 |
| 13 | 43541200 | 43541799 | 10.17376 | 18.85752 | 27.3475 | 11.488 | -0.09007 | 0.187358 | 1 |
| 13 | 43723400 | 43726799 | 38.75001 | 62.79186 | 78.39549 | 33.56099 | 0.045169 | 1.649442 | 1 |
| 13 | 44411800 | 44420599 | 114.3403 | 173.9308 | 225.6947 | 93.5087 | -0.09187 | 3.177291 | 1 |
| 13 | 44443000 | 44445199 | 22.03142 | 37.50259 | 51.69056 | 22.65679 | -0.07927 | 1.013327 | 1 |
| 13 | 45974000 | 45975599 | 13.84356 | 24.56709 | 37.07589 | 16.37695 | -0.19771 | 0.486605 | 1 |
| 13 | 46231800 | 46233199 | 12.93796 | 22.99513 | 36.15165 | 15.94812 | -0.25489 | 0.470827 | 1 |
| 13 | 46528200 | 46531999 | 7.759568 | 14.82722 | 26.74499 | 10.62224 | -0.335 | -0.03718 | 1 |
| 13 | 46533400 | 46538399 | 50.86114 | 81.21149 | 110.9448 | 46.57541 | -0.10871 | 2.107353 | 1 |
| 13 | 46728400 | 46730199 | 16.81507 | 29.19148 | 42.71464 | 18.74452 | -0.15979 | 0.802357 | 1 |
| 13 | 46983000 | 46985599 | 38.41312 | 62.24934 | 72.4528 | 31.01819 | 0.146242 | 1.645043 | 1 |
| 13 | 47117600 | 47120199 | 27.3692 | 45.8749 | 53.99091 | 23.54779 | 0.146773 | 1.170874 | 1 |
| 13 | 47157200 | 47169399 | 207.9695 | 300.5954 | 427.3873 | 176.2921 | -0.29147 | 4.087329 | 1 |
| 13 | 47359600 | 47360999 | 19.44227 | 33.87766 | 36.83949 | 16.204 | 0.26701 | 0.753123 | 1 |
| 13 | 47680800 | 47681999 | 14.88526 | 26.1785 | 34.78434 | 15.40204 | -0.01995 | 0.586052 | 1 |
| 13 | 47892600 | 47893799 | 16.22599 | 28.51275 | 31.2695 | 13.84389 | 0.255206 | 0.597114 | 1 |
| 13 | 48407200 | 48408999 | 24.11207 | 40.44725 | 59.27522 | 25.86534 | -0.1685 | 1.159167 | 1 |
| 13 | 50056800 | 50062799 | 47.29267 | 76.08358 | 81.8672 | 34.7503 | 0.247653 | 1.860758 | 1 |
| 13 | 50174800 | 50175799 | 16.51143 | 28.87995 | 37.43838 | 16.51014 | 0.015204 | 0.685621 | 1 |
| 13 | 50237200 | 50239199 | 22.31763 | 38.06156 | 45.86868 | 20.10803 | 0.114231 | 0.991252 | 1 |
| 13 | 50287800 | 50291799 | 36.23441 | 58.59979 | 85.67197 | 36.62232 | -0.18468 | 1.689456 | 1 |
| 13 | 50363200 | 50364799 | 16.8192 | 29.87639 | 32.70549 | 14.43479 | 0.263834 | 0.491617 | 1 |
| 13 | 50692000 | 50693399 | 10.4944 | 19.23072 | 31.33797 | 13.37068 | -0.27384 | 0.261864 | 1 |
| 13 | 50842200 | 50844799 | 21.8265 | 36.67194 | 55.36679 | 24.18283 | -0.21123 | 1.131281 | 1 |
| 13 | 50875400 | 50877399 | 19.60397 | 34.0525 | 41.01945 | 18.00706 | 0.119537 | 0.803458 | 1 |
| 13 | 50883600 | 50886199 | 34.81893 | 56.54751 | 68.27044 | 29.27447 | 0.096497 | 1.592284 | 1 |
| 13 | 51032000 | 51035599 | 32.99647 | 54.07537 | 74.17914 | 31.92585 | -0.07838 | 1.459392 | 1 |
| 13 | 51239000 | 51241199 | 26.21939 | 44.01035 | 53.40789 | 23.31448 | 0.102691 | 1.15143 | 1 |
| 13 | 51262000 | 51272399 | 126.3844 | 191.7896 | 255.7819 | 105.8979 | -0.13422 | 3.232517 | 1 |
| 13 | 51536200 | 51537399 | 20.20429 | 34.68269 | 42.27773 | 18.54355 | 0.099303 | 0.919843 | 1 |
| 13 | 51791200 | 51793999 | 22.82103 | 38.10997 | 60.25148 | 26.21692 | -0.27749 | 1.203019 | 1 |
| 13 | 52246200 | 52248199 | 16.98512 | 29.47522 | 47.93531 | 21.03231 | -0.31179 | 0.814027 | 1 |
| 13 | 52284600 | 52289999 | 47.10861 | 76.25694 | 75.42121 | 32.26292 | 0.37909 | 1.684002 | 1 |

| | | | | | | | | | |
|---|---|---|---|---|---|---|---|---|---|
| 13 | 52295600 | 52300999 | 77.83103 | 121.7156 | 165.4402 | 68.63572 | -0.1277 | 2.64126 | 1 |
| 13 | 52416800 | 52418599 | 11.47926 | 20.94927 | 31.78946 | 13.63624 | -0.17695 | 0.294629 | 1 |
| 13 | 52532400 | 52537799 | 91.4929 | 142.2096 | 165.9217 | 68.88958 | 0.085512 | 2.753084 | 1 |
| 13 | 53080600 | 53081799 | 11.6157 | 21.44042 | 26.63865 | 11.25404 | 0.12508 | 0.218095 | 1 |
| 13 | 53237400 | 53240799 | 16.27008 | 28.52014 | 32.72493 | 14.45824 | 0.190102 | 0.637035 | 1 |
| 13 | 53418600 | 53421599 | 22.6354 | 37.85751 | 62.52353 | 27.23166 | -0.34036 | 1.187208 | 1 |
| 13 | 53559400 | 53560599 | 10.72152 | 19.86423 | 31.49021 | 13.23603 | -0.21904 | 0.200423 | 1 |
| 13 | 54051400 | 54053199 | 22.9349 | 39.10146 | 49.02177 | 21.48972 | 0.057249 | 0.998962 | 1 |
| 13 | 54270000 | 54272399 | 20.15103 | 35.064 | 38.90306 | 17.09677 | 0.237991 | 0.779664 | 1 |
| 13 | 55173600 | 55175799 | 37.09584 | 60.46119 | 66.01836 | 28.35488 | 0.244963 | 1.540081 | 1 |
| 13 | 56393800 | 56396399 | 28.80263 | 48.01283 | 51.32544 | 22.30774 | 0.285232 | 1.230964 | 1 |
| 13 | 56680800 | 56683799 | 46.94836 | 75.1216 | 102.9106 | 43.33144 | -0.10935 | 2.037153 | 1 |
| 13 | 56864600 | 56866199 | 14.1744 | 25.29695 | 30.09868 | 13.23174 | 0.148544 | 0.442429 | 1 |
| 13 | 57123200 | 57127599 | 44.12875 | 71.41487 | 74.31471 | 31.78354 | 0.30559 | 1.683165 | 1 |
| 13 | 57344200 | 57345599 | 11.6659 | 21.98396 | 24.73361 | 10.11286 | 0.304053 | 0.089574 | 1 |
| 13 | 58515400 | 58533599 | 322.2267 | 450.468 | 601.1092 | 249.2604 | -0.24822 | 4.584167 | 1 |
| 13 | 58610400 | 58612599 | 21.28797 | 37.03563 | 39.86452 | 17.5173 | 0.28143 | 0.788648 | 1 |
| 13 | 59259600 | 59266799 | 73.28793 | 114.447 | 159.4049 | 66.14161 | -0.16447 | 2.663301 | 1 |
| 13 | 59431600 | 59434599 | 26.44861 | 44.52609 | 49.09163 | 21.45491 | 0.240681 | 1.120409 | 1 |
| 13 | 60033200 | 60036399 | 29.37235 | 48.42888 | 70.63589 | 30.47898 | -0.16312 | 1.373937 | 1 |
| 13 | 60348400 | 60356999 | 115.7214 | 176.043 | 218.9367 | 90.71047 | -0.03057 | 3.176968 | 1 |
| 13 | 60748800 | 60750999 | 14.3419 | 25.18706 | 37.76585 | 16.71165 | -0.19398 | 0.605476 | 1 |
| 13 | 61127200 | 61129599 | 20.07693 | 34.10387 | 55.88898 | 24.49213 | -0.32838 | 1.032347 | 1 |
| 13 | 61142200 | 61145799 | 45.77565 | 73.82127 | 90.78576 | 38.66705 | 0.059533 | 1.789659 | 1 |
| 13 | 61276200 | 61293999 | 313.2732 | 446.6608 | 444.8881 | 183.89 | 0.202161 | 4.293702 | 1 |
| 13 | 62074200 | 62075999 | 27.27371 | 46.44673 | 46.30159 | 20.29544 | 0.386462 | 1.021887 | 1 |
| 13 | 62462400 | 62482399 | 185.1588 | 276.5514 | 278.6725 | 114.8847 | 0.249823 | 3.553953 | 1 |
| 13 | 62754200 | 62768399 | 226.1161 | 331.3234 | 350.9032 | 144.5409 | 0.152339 | 3.881358 | 1 |
| 13 | 63150800 | 63152799 | 27.03771 | 45.51407 | 51.75484 | 22.61826 | 0.196402 | 1.129605 | 1 |
| 13 | 63302400 | 63305199 | 31.90673 | 52.60245 | 59.50084 | 25.67321 | 0.202575 | 1.373762 | 1 |
| 13 | 63369600 | 63373399 | 26.56357 | 44.48459 | 51.78978 | 22.57499 | 0.162433 | 1.176227 | 1 |
| 13 | 63570800 | 63573199 | 30.40746 | 50.69036 | 57.39746 | 24.94757 | 0.202482 | 1.241782 | 1 |
| 13 | 63621600 | 63623599 | 18.73039 | 32.70619 | 41.88729 | 18.44876 | 0.0321 | 0.724877 | 1 |
| 13 | 64459800 | 64461399 | 10.71885 | 19.23448 | 33.25986 | 14.55435 | -0.38316 | 0.389537 | 1 |
| 13 | 64735800 | 64738199 | 31.38076 | 51.7311 | 59.87486 | 25.8337 | 0.169557 | 1.376156 | 1 |
| 13 | 64768800 | 64770799 | 22.3177 | 38.25131 | 46.3261 | 20.31668 | 0.108025 | 0.944 | 1 |
| 13 | 64771400 | 64773799 | 26.17027 | 44.20748 | 53.38315 | 23.35452 | 0.110112 | 1.097362 | 1 |
| 13 | 64796400 | 64797399 | 19.39444 | 33.80514 | 35.31216 | 15.53533 | 0.324773 | 0.745661 | 1 |
| 13 | 64821400 | 64825599 | 31.19978 | 51.45353 | 60.64694 | 26.1713 | 0.143509 | 1.367559 | 1 |
| 13 | 64851000 | 64852199 | 10.85742 | 20.07635 | 30.35799 | 12.79446 | -0.15413 | 0.209848 | 1 |
| 13 | 65120200 | 65122199 | 29.75478 | 49.58625 | 51.31069 | 22.29742 | 0.332101 | 1.239128 | 1 |
| 13 | 65143600 | 65148599 | 25.21572 | 42.12928 | 55.47291 | 24.14499 | -0.01447 | 1.201621 | 1 |
| 13 | 65203200 | 65212799 | 120.913 | 184.6446 | 194.7053 | 80.7524 | 0.211691 | 3.099256 | 1 |
| 13 | 65243800 | 65256199 | 206.7831 | 305.5497 | 335.58 | 138.1994 | 0.111388 | 3.745382 | 1 |
| 13 | 65259800 | 65261399 | 19.06693 | 32.71586 | 48.47759 | 21.26213 | -0.18148 | 0.924955 | 1 |
| 13 | 65564600 | 65569599 | 55.91264 | 88.61636 | 130.4633 | 54.27813 | -0.22751 | 2.345432 | 1 |
| 13 | 65598200 | 65602999 | 103.2983 | 158.5982 | 173.7775 | 72.14925 | 0.162462 | 2.984339 | 1 |

| | | | | | | | | | |
|---|---|---|---|---|---|---|---|---|---|
| 13 | 65709200 | 65712799 | 87.55779 | 136.1785 | 149.3842 | 62.01751 | 0.175086 | 2.739604 | 1 |
| 13 | 65807400 | 65813399 | 173.3287 | 258.0498 | 257.2123 | 105.9832 | 0.261288 | 3.609113 | 1 |
| 13 | 66069000 | 66073799 | 34.29034 | 56.5904 | 58.51791 | 25.26065 | 0.332239 | 1.368291 | 1 |
| 13 | 66159800 | 66162599 | 43.96033 | 70.87497 | 72.12568 | 30.70894 | 0.331725 | 1.78504 | 1 |
| 13 | 66347000 | 66349199 | 38.20827 | 61.91499 | 70.38098 | 30.13035 | 0.180264 | 1.647199 | 1 |
| 13 | 67208400 | 67211399 | 37.71832 | 60.76441 | 92.4195 | 39.30807 | -0.2481 | 1.807487 | 1 |
| 13 | 67755800 | 67758199 | 10.04789 | 17.85075 | 39.86191 | 17.59133 | -0.75876 | 0.45933 | 1 |
| 13 | 67789400 | 67797199 | 194.0606 | 284.5042 | 330.1102 | 135.9715 | 0.021472 | 3.869019 | 1 |
| 13 | 67936400 | 67941799 | 71.03925 | 112.6511 | 122.3149 | 50.91544 | 0.211817 | 2.338388 | 1 |
| 13 | 67961800 | 67975599 | 234.978 | 344.3473 | 345.4495 | 142.2934 | 0.230679 | 3.88029 | 1 |
| 13 | 68006000 | 68012599 | 65.73509 | 103.8058 | 142.6885 | 59.24912 | -0.13294 | 2.440733 | 1 |
| 13 | 68638600 | 68643599 | 63.27452 | 100.7008 | 117.7417 | 49.20329 | 0.110319 | 2.223989 | 1 |
| 13 | 68662400 | 68665199 | 40.3649 | 65.57478 | 72.23618 | 30.98208 | 0.228808 | 1.60191 | 1 |
| 13 | 68744800 | 68747599 | 20.56355 | 35.12753 | 51.68376 | 22.66025 | -0.17233 | 0.975456 | 1 |
| 13 | 69603000 | 69605199 | 21.35012 | 36.27881 | 52.21801 | 22.88473 | -0.14197 | 1.026329 | 1 |
| 13 | 70412000 | 70412799 | 11.05517 | 20.68942 | 27.35054 | 11.31465 | 0.059471 | 0.13417 | 1 |
| 13 | 70961600 | 70962799 | 17.28453 | 30.44779 | 34.98957 | 15.48785 | 0.189198 | 0.581894 | 1 |
| 13 | 71076200 | 71077799 | 18.15582 | 31.86062 | 37.95761 | 16.78564 | 0.135896 | 0.636985 | 1 |
| 13 | 71167800 | 71169999 | 24.33605 | 41.41009 | 48.49208 | 21.25421 | 0.154969 | 1.022902 | 1 |
| 13 | 71460800 | 71464599 | 40.15662 | 65.57438 | 67.22883 | 28.89479 | 0.337519 | 1.519236 | 1 |
| 13 | 71529400 | 71533399 | 13.36686 | 24.37499 | 27.10771 | 11.63936 | 0.267818 | 0.303446 | 1 |
| 13 | 72027000 | 72028399 | 19.00495 | 32.45451 | 47.4593 | 20.80752 | -0.16337 | 0.969361 | 1 |
| 13 | 72153400 | 72157799 | 62.12521 | 98.80866 | 114.79 | 47.9275 | 0.118589 | 2.242806 | 1 |
| 13 | 72183600 | 72185199 | 29.4699 | 48.81706 | 57.76292 | 24.99912 | 0.138767 | 1.311156 | 1 |
| 13 | 73138800 | 73139599 | 10.71453 | 20.03822 | 25.26469 | 10.46386 | 0.126129 | 0.130737 | 1 |
| 13 | 73277400 | 73278999 | 13.25741 | 23.42035 | 36.70009 | 16.23575 | -0.25434 | 0.528377 | 1 |
| 13 | 73290000 | 73296999 | 136.242 | 205.8117 | 238.0999 | 98.43772 | 0.065246 | 3.324673 | 1 |
| 13 | 74564000 | 74611399 | 457.3554 | 631.2638 | 638.276 | 265.2026 | 0.133344 | 4.781851 | 1 |
| 13 | 74743000 | 74766199 | 134.1868 | 205.2378 | 204.2099 | 84.7252 | 0.297196 | 3.073216 | 1 |
| 13 | 74837600 | 74839399 | 17.46 | 30.57106 | 35.6815 | 15.74929 | 0.165651 | 0.66787 | 1 |
| 13 | 75585600 | 75591999 | 53.16659 | 84.70271 | 120.7854 | 50.54835 | -0.17401 | 2.182566 | 1 |
| 13 | 75596000 | 75602599 | 149.2727 | 224.5126 | 252.2826 | 104.1699 | 0.101721 | 3.41451 | 1 |
| 13 | 75779400 | 75780199 | 10.27054 | 18.81971 | 28.25969 | 12.05799 | -0.15669 | 0.252004 | 1 |
| 13 | 75844200 | 75848799 | 50.0378 | 80.05797 | 97.07655 | 40.86988 | 0.066169 | 2.038758 | 1 |
| 13 | 76003400 | 76015999 | 144.8897 | 216.6978 | 301.6877 | 124.4055 | -0.21447 | 3.522676 | 1 |
| 13 | 76178600 | 76185799 | 118.3903 | 180.4007 | 225.2124 | 93.35677 | -0.03412 | 3.144656 | 1 |
| 13 | 76233000 | 76233999 | 10.59863 | 19.87621 | 24.70136 | 10.18288 | 0.151825 | 0.10868 | 1 |
| 13 | 76350600 | 76352599 | 26.84784 | 44.14709 | 69.438 | 29.92787 | -0.27314 | 1.401427 | 1 |
| 13 | 77189800 | 77190999 | 18.54343 | 32.04969 | 44.77155 | 19.64175 | -0.09461 | 0.852848 | 1 |
| 13 | 77375200 | 77385999 | 258.8437 | 372.1951 | 391.3975 | 161.5619 | 0.136099 | 4.166092 | 1 |
| 13 | 77503600 | 77504599 | 10.74655 | 19.94095 | 28.76893 | 12.06719 | -0.08206 | 0.183834 | 1 |
| 13 | 77520000 | 77521599 | 31.67002 | 52.1882 | 55.10056 | 23.76998 | 0.301461 | 1.374703 | 1 |
| 13 | 77564600 | 77565399 | 12.23079 | 22.07633 | 35.76378 | 15.55378 | -0.28406 | 0.369122 | 1 |
| 13 | 77574600 | 77579199 | 56.87827 | 90.83747 | 102.2867 | 42.9529 | 0.16995 | 2.104859 | 1 |
| 13 | 77581800 | 77585999 | 16.56646 | 29.69588 | 32.27177 | 14.13474 | 0.2819 | 0.434148 | 1 |
| 13 | 77701000 | 77703199 | 20.41573 | 35.22527 | 45.79108 | 20.08798 | 0.008149 | 0.876484 | 1 |
| 13 | 77717200 | 77717999 | 19.19657 | 33.13837 | 40.20158 | 17.6362 | 0.107851 | 0.86188 | 1 |

| | | | | | | | | | |
|---|---|---|---|---|---|---|---|---|---|
| 13 | 77751200 | 77755399 | 39.75797 | 64.31145 | 70.51611 | 30.14854 | 0.229734 | 1.684099 | 1 |
| 13 | 77762600 | 77767599 | 55.64395 | 88.89683 | 106.3798 | 44.69352 | 0.08286 | 2.092796 | 1 |
| 13 | 78197400 | 78199599 | 21.0197 | 35.50429 | 57.54089 | 25.1729 | -0.31284 | 1.084755 | 1 |
| 13 | 78222400 | 78223199 | 13.40825 | 23.8101 | 32.65658 | 14.41695 | -0.0596 | 0.473612 | 1 |
| 13 | 78379600 | 78381999 | 49.27443 | 79.11076 | 91.71397 | 38.81408 | 0.136702 | 1.930109 | 1 |
| 13 | 78985600 | 78986199 | 10.49981 | 20.01277 | 22.79819 | 9.103185 | 0.315993 | -0.00345 | 1 |
| 13 | 79411600 | 79414199 | 44.12104 | 71.15575 | 71.08065 | 30.2759 | 0.358929 | 1.776016 | 1 |
| 13 | 79478600 | 79479999 | 22.0429 | 36.93444 | 59.10791 | 25.77392 | -0.29496 | 1.165159 | 1 |
| 13 | 80171800 | 80172599 | 11.36743 | 20.53283 | 32.27394 | 14.02335 | -0.23932 | 0.355076 | 1 |
| 13 | 80425200 | 80426599 | 17.76318 | 30.98625 | 36.35978 | 16.00416 | 0.157806 | 0.727026 | 1 |
| 13 | 80577000 | 80577999 | 11.92118 | 21.70255 | 27.57994 | 11.86682 | 0.074206 | 0.299607 | 1 |
| 13 | 80712000 | 80712799 | 10.98902 | 19.83765 | 31.24232 | 13.58508 | -0.24299 | 0.35141 | 1 |
| 13 | 80743600 | 80745799 | 16.28294 | 28.70023 | 32.11552 | 14.21401 | 0.227813 | 0.564754 | 1 |
| 13 | 81218800 | 81220599 | 23.23612 | 40.20915 | 42.47325 | 18.63415 | 0.307151 | 0.85794 | 1 |
| 13 | 81230600 | 81238999 | 118.6947 | 181.4715 | 194.9022 | 80.85614 | 0.186588 | 3.074779 | 1 |
| 13 | 82337800 | 82340199 | 62.78168 | 99.86269 | 102.2172 | 42.68392 | 0.301139 | 2.237794 | 1 |
| 13 | 82482400 | 82485399 | 44.49853 | 71.42941 | 101.033 | 42.74561 | -0.15002 | 1.934863 | 1 |
| 13 | 82968400 | 82969199 | 15.57797 | 27.40781 | 35.09494 | 15.53831 | 0.033129 | 0.583804 | 1 |
| 13 | 83334400 | 83335999 | 19.02509 | 32.49982 | 52.71036 | 23.11037 | -0.31223 | 0.971958 | 1 |
| 13 | 83571800 | 83574199 | 9.958279 | 18.50178 | 28.35459 | 11.87282 | -0.16618 | 0.172762 | 1 |
| 13 | 83833400 | 83837199 | 36.56239 | 59.67266 | 65.56157 | 28.17262 | 0.237418 | 1.51863 | 1 |
| 13 | 83872400 | 83875999 | 25.85701 | 43.14147 | 58.24118 | 25.32795 | -0.05039 | 1.219973 | 1 |
| 13 | 84429200 | 84430199 | 12.3007 | 22.94954 | 25.12847 | 10.4498 | 0.323411 | 0.158936 | 1 |
| 13 | 84560000 | 84562799 | 30.28041 | 49.62657 | 72.48645 | 31.19796 | -0.1686 | 1.452917 | 1 |
| 13 | 84637400 | 84639399 | 13.82484 | 24.36789 | 39.81508 | 17.62172 | -0.31554 | 0.564159 | 1 |
| 13 | 84683800 | 84689399 | 82.96719 | 129.4126 | 158.2352 | 65.66408 | 0.021925 | 2.687906 | 1 |
| 13 | 84957200 | 84959999 | 26.29284 | 44.20281 | 54.48057 | 23.80012 | 0.080425 | 1.138905 | 1 |
| 13 | 84982000 | 84983599 | 21.42421 | 35.9965 | 58.52743 | 25.56342 | -0.31772 | 1.134962 | 1 |
| 13 | 84996000 | 84996999 | 10.71565 | 19.74426 | 28.75018 | 12.17528 | -0.10487 | 0.225535 | 1 |
| 13 | 85060800 | 85061799 | 14.57153 | 25.66634 | 36.72402 | 16.25588 | -0.12542 | 0.57021 | 1 |
| 13 | 85084200 | 85085999 | 21.99895 | 38.02284 | 43.1302 | 18.92168 | 0.204608 | 0.867041 | 1 |
| 13 | 85184000 | 85188399 | 31.23701 | 51.82969 | 57.37615 | 24.86001 | 0.234707 | 1.296402 | 1 |
| 13 | 85260000 | 85261799 | 28.14591 | 46.86409 | 58.90281 | 25.58252 | 0.05242 | 1.249366 | 1 |
| 13 | 85332200 | 85333599 | 21.61586 | 37.31437 | 42.62269 | 18.69832 | 0.194209 | 0.87508 | 1 |
| 13 | 85436400 | 85437999 | 19.15346 | 32.43999 | 50.5936 | 22.15184 | -0.25741 | 1.044832 | 1 |
| 13 | 85550000 | 85556199 | 91.07539 | 140.8704 | 187.073 | 77.70564 | -0.10649 | 2.841614 | 1 |
| 13 | 85666000 | 85667399 | 17.70486 | 30.6561 | 50.07289 | 21.96872 | -0.31868 | 0.841408 | 1 |
| 13 | 85819600 | 85821799 | 28.32504 | 46.46016 | 73.47043 | 31.63248 | -0.28237 | 1.438383 | 1 |
| 13 | 85880200 | 85881199 | 15.90602 | 27.90547 | 35.22823 | 15.57531 | 0.053074 | 0.627209 | 1 |
| 13 | 86029800 | 86030399 | 11.64197 | 21.56903 | 28.22501 | 11.864 | 0.055943 | 0.20128 | 1 |
| 13 | 86313000 | 86315599 | 27.70215 | 47.39228 | 46.24922 | 20.27995 | 0.41797 | 0.981882 | 1 |
| 13 | 86436000 | 86438199 | 30.52776 | 50.53305 | 61.10729 | 26.4334 | 0.107425 | 1.325865 | 1 |
| 13 | 86779400 | 86780399 | 19.23668 | 32.53628 | 54.78828 | 23.97846 | -0.36768 | 1.06368 | 1 |
| 13 | 86855600 | 86857399 | 17.74273 | 31.00146 | 37.75885 | 16.63715 | 0.104415 | 0.705082 | 1 |
| 13 | 86906200 | 86908999 | 33.684 | 55.58388 | 59.70232 | 25.77023 | 0.277522 | 1.366811 | 1 |
| 13 | 87068600 | 87076599 | 78.97104 | 124.3292 | 133.388 | 55.34348 | 0.220681 | 2.510829 | 1 |
| 13 | 87217800 | 87220199 | 23.9755 | 40.39946 | 49.073 | 21.4527 | 0.101563 | 1.106202 | 1 |

| | | | | | | | | | |
|---|---|---|---|---|---|---|---|---|---|
| 13 | 87263600 | 87265999 | 23.42818 | 40.59407 | 43.03843 | 18.88286 | 0.302338 | 0.850362 | 1 |
| 13 | 87294200 | 87294999 | 10.51454 | 19.31685 | 30.34258 | 12.90111 | -0.21736 | 0.246507 | 1 |
| 13 | 87316600 | 87318799 | 13.75231 | 25.02115 | 26.84325 | 11.5585 | 0.316202 | 0.314815 | 1 |
| 13 | 87320600 | 87322799 | 9.733287 | 18.61152 | 22.88875 | 9.074043 | 0.215708 | -0.03335 | 1 |
| 13 | 87333000 | 87339399 | 80.45841 | 126.2633 | 152.027 | 63.05048 | 0.05089 | 2.578959 | 1 |
| 13 | 87508600 | 87510599 | 22.99334 | 38.81941 | 59.2734 | 25.92708 | -0.22724 | 1.097857 | 1 |
| 13 | 87542800 | 87545599 | 54.36904 | 86.80152 | 94.66755 | 39.73488 | 0.215477 | 2.109819 | 1 |
| 13 | 87703600 | 87707599 | 69.36538 | 109.3295 | 129.1837 | 53.60694 | 0.082856 | 2.483006 | 1 |
| 13 | 87774600 | 87782199 | 140.9036 | 213.0228 | 226.2028 | 93.54254 | 0.189765 | 3.309538 | 1 |
| 13 | 87814000 | 87821199 | 102.4017 | 157.8705 | 188.2893 | 78.21219 | 0.044749 | 2.905685 | 1 |
| 13 | 89005200 | 89006399 | 17.87545 | 31.19494 | 44.08558 | 19.40978 | -0.10931 | 0.733477 | 1 |
| 13 | 89304800 | 89306399 | 15.70102 | 27.3713 | 42.62676 | 18.75502 | -0.24869 | 0.73718 | 1 |
| 13 | 90008000 | 90010599 | 18.4413 | 31.77021 | 47.41572 | 20.79972 | -0.19059 | 0.885441 | 1 |
| 13 | 90364200 | 90371599 | 123.8433 | 188.6315 | 219.2754 | 90.88443 | 0.068219 | 3.154464 | 1 |
| 13 | 90811200 | 90812399 | 20.26051 | 34.69633 | 44.85476 | 19.67009 | 0.014326 | 0.94553 | 1 |
| 13 | 91407800 | 91408599 | 15.11744 | 26.97508 | 31.42836 | 13.8184 | 0.178357 | 0.451708 | 1 |
| 13 | 91771600 | 91774199 | 29.32909 | 49.67506 | 55.15762 | 24.15025 | 0.230855 | 1.08722 | 1 |
| 13 | 92204800 | 92206199 | 13.39872 | 23.94009 | 38.23252 | 16.7912 | -0.27314 | 0.444108 | 1 |
| 13 | 93334200 | 93340199 | 70.11232 | 110.6549 | 131.623 | 54.63854 | 0.074756 | 2.45485 | 1 |
| 13 | 93355600 | 93357999 | 32.94412 | 54.35878 | 58.43634 | 25.22295 | 0.276299 | 1.366473 | 1 |
| 13 | 93898000 | 93899799 | 18.25466 | 31.60436 | 49.55245 | 21.74031 | -0.26007 | 0.842161 | 1 |
| 13 | 94120600 | 94123399 | 25.22329 | 41.93773 | 66.02031 | 28.64605 | -0.27156 | 1.261109 | 1 |
| 13 | 94423800 | 94425999 | 16.31076 | 28.51635 | 41.47328 | 18.28268 | -0.15017 | 0.696146 | 1 |
| 13 | 94454800 | 94456599 | 12.01082 | 21.73062 | 31.6186 | 13.71041 | -0.12716 | 0.34519 | 1 |
| 13 | 94618600 | 94622199 | 48.22813 | 77.67916 | 88.36646 | 37.57257 | 0.169622 | 1.829583 | 1 |
| 13 | 94867000 | 94875799 | 144.1553 | 216.9388 | 248.9298 | 102.8011 | 0.072261 | 3.402134 | 1 |
| 13 | 94877600 | 94879199 | 12.05833 | 22.1542 | 30.17815 | 12.83029 | -0.01372 | 0.253797 | 1 |
| 13 | 95767600 | 95769199 | 16.3884 | 28.96496 | 31.93348 | 14.12574 | 0.250518 | 0.534284 | 1 |
| 13 | 95961200 | 95963399 | 12.97195 | 23.15797 | 37.22136 | 16.35876 | -0.28284 | 0.446237 | 1 |
| 13 | 96954400 | 96956799 | 15.58705 | 27.30251 | 41.97532 | 18.5335 | -0.22981 | 0.664184 | 1 |
| 13 | 98502000 | 98503799 | 16.34865 | 28.30716 | 44.80357 | 19.65688 | -0.27336 | 0.827468 | 1 |
| 13 | 99123600 | 99124199 | 9.063926 | 17.08941 | 23.31008 | 9.522539 | 0.029211 | 0.057133 | 1 |
| 13 | 99125200 | 99129799 | 48.84817 | 77.64277 | 124.4001 | 51.90476 | -0.3453 | 2.253656 | 1 |
| 13 | 99130600 | 99133799 | 34.12894 | 55.18782 | 83.43072 | 35.65849 | -0.23304 | 1.688382 | 1 |
| 13 | 99135000 | 99145799 | 128.2009 | 195.1457 | 211.8572 | 87.79319 | 0.166029 | 3.167573 | 1 |
| 13 | 99263200 | 99271599 | 95.25627 | 147.5807 | 170.9088 | 70.99077 | 0.092645 | 2.811094 | 1 |
| 13 | 99411400 | 99414599 | 26.25592 | 43.60744 | 68.2884 | 29.60805 | -0.2641 | 1.279635 | 1 |
| 13 | 100220800 | 100222599 | 15.7521 | 27.81863 | 37.53016 | 16.60402 | -0.03983 | 0.547596 | 1 |
| 13 | 100710000 | 100712399 | 25.13033 | 42.25042 | 51.74083 | 22.60357 | 0.089699 | 1.132034 | 1 |
| 13 | 100746600 | 100749999 | 45.55153 | 73.06386 | 92.01971 | 38.887 | 0.01582 | 1.955535 | 1 |
| 13 | 101082800 | 101085199 | 23.58778 | 39.72296 | 55.4571 | 24.23927 | -0.0986 | 1.117184 | 1 |
| 13 | 101415400 | 101416199 | 14.24647 | 26.32555 | 29.1329 | 12.28921 | 0.292436 | 0.23847 | 1 |
| 13 | 101516800 | 101518599 | 23.88157 | 39.91387 | 59.29074 | 25.81228 | -0.18795 | 1.195279 | 1 |
| 13 | 102095800 | 102100199 | 58.10265 | 92.23814 | 127.0153 | 52.93026 | -0.12895 | 2.297247 | 1 |
| 13 | 102530000 | 102531999 | 13.42375 | 23.79011 | 38.16488 | 16.87298 | -0.2863 | 0.497125 | 1 |
| 13 | 102627200 | 102629399 | 23.45329 | 39.37932 | 59.81071 | 26.11455 | -0.21993 | 1.147184 | 1 |
| 13 | 104124000 | 104128599 | 54.68624 | 86.99837 | 125.5654 | 52.44416 | -0.19363 | 2.231853 | 1 |

| | | | | | | | | | | |
|---|---|---|---|---|---|---|---|---|---|---|
| 13 | 104131000 | 104132799 | 12.43056 | 23.09903 | 24.46023 | 10.23977 | 0.363322 | 0.181018 | | 1 |
| 13 | 104904000 | 104904799 | 14.10978 | 24.74667 | 38.91534 | 17.20108 | -0.26214 | 0.629738 | | 1 |
| 13 | 104963600 | 104967599 | 44.42091 | 71.68772 | 81.53697 | 34.75987 | 0.173157 | 1.760228 | | 1 |
| 13 | 105127400 | 105128199 | 11.03235 | 20.24752 | 26.723 | 11.37789 | 0.030585 | 0.246254 | | 1 |
| 13 | 105130000 | 105135399 | 81.97329 | 128.2886 | 141.2113 | 58.57946 | 0.177106 | 2.626199 | | 1 |
| 13 | 105874800 | 105880399 | 96.65872 | 149.4555 | 172.7882 | 71.77218 | 0.092878 | 2.848155 | | 1 |
| 13 | 106431600 | 106433799 | 28.9428 | 47.93118 | 58.87116 | 25.47408 | 0.085031 | 1.3126 | | 1 |
| 13 | 106634200 | 106637199 | 33.21557 | 54.85646 | 58.53189 | 25.27935 | 0.28717 | 1.355793 | | 1 |
| 13 | 106768600 | 106770399 | 27.83357 | 46.25642 | 62.21197 | 26.98504 | -0.04515 | 1.272173 | | 1 |
| 13 | 106786600 | 106787599 | 17.73346 | 30.48711 | 46.92086 | 20.58134 | -0.23517 | 0.903137 | | 1 |
| 13 | 106815800 | 106817199 | 22.42165 | 38.62241 | 41.58975 | 18.24399 | 0.278353 | 0.896461 | | 1 |
| 13 | 106920200 | 106928199 | 164.2003 | 244.8029 | 289.554 | 119.3285 | 0.016397 | 3.58374 | | 1 |
| 13 | 107182400 | 107185199 | 24.08672 | 41.29756 | 44.72456 | 19.61486 | 0.268893 | 0.948699 | | 1 |
| 13 | 108146400 | 108149799 | 34.652 | 56.53316 | 75.72746 | 32.5365 | -0.04821 | 1.525783 | | 1 |
| 13 | 108748400 | 108748999 | 9.355829 | 17.44648 | 26.88508 | 11.18984 | -0.16684 | 0.142103 | | 1 |
| 13 | 109053600 | 109055399 | 14.86052 | 26.21767 | 40.7523 | 18.03318 | -0.24368 | 0.559351 | | 1 |
| 13 | 109508800 | 109509799 | 16.12609 | 28.37005 | 33.17172 | 14.68703 | 0.163707 | 0.586365 | | 1 |
| 13 | 109904400 | 109905199 | 16.54906 | 28.69581 | 44.80013 | 19.65623 | -0.25315 | 0.816732 | | 1 |
| 13 | 110445400 | 110447199 | 30.81168 | 50.24401 | 73.17568 | 31.43468 | -0.16898 | 1.52455 | | 1 |
| 13 | 111411200 | 111412799 | 11.39504 | 20.2275 | 36.74738 | 16.22805 | -0.46331 | 0.469181 | | 1 |
| 13 | 112149600 | 112151999 | 29.83602 | 48.72863 | 71.4495 | 30.71053 | -0.17708 | 1.499455 | | 1 |
| 13 | 112484600 | 112487599 | 34.78658 | 56.22007 | 82.73603 | 35.33629 | -0.19524 | 1.705699 | | 1 |
| 13 | 112662000 | 112662799 | 10.2559 | 19.03122 | 29.9044 | 12.54284 | -0.204 | 0.184877 | | 1 |
| 13 | 112788000 | 112790399 | 56.61243 | 90.16685 | 103.0069 | 43.09114 | 0.144966 | 2.191445 | | 1 |
| 13 | 112963400 | 112964399 | 16.00942 | 28.11082 | 36.24046 | 16.03438 | 0.023031 | 0.61765 | | 1 |
| 13 | 113313200 | 113314399 | 13.56799 | 23.93375 | 34.25645 | 15.15917 | -0.12504 | 0.542645 | | 1 |
| 13 | 113465800 | 113467999 | 28.37684 | 47.78747 | 53.4637 | 23.36793 | 0.219689 | 1.129747 | | 1 |
| 13 | 113693200 | 113700599 | 75.25165 | 117.4726 | 164.8969 | 68.42287 | -0.17608 | 2.670763 | | 1 |
| 13 | 113702800 | 113705199 | 24.46615 | 41.39451 | 47.462 | 20.77408 | 0.184626 | 1.068743 | | 1 |
| 13 | 113784000 | 113795199 | 73.47379 | 115.1215 | 149.4439 | 61.98717 | -0.05927 | 2.602831 | | 1 |
| 13 | 113930000 | 113932199 | 16.02735 | 27.87036 | 42.18512 | 18.53154 | -0.2081 | 0.774245 | | 1 |
| 13 | 113942200 | 113944599 | 19.23676 | 33.08339 | 48.47428 | 21.26296 | -0.1647 | 0.905286 | | 1 |
| 13 | 114618800 | 114620199 | 30.88477 | 50.17465 | 74.29043 | 31.86353 | -0.19666 | 1.584106 | | 1 |
| 13 | 114678000 | 114678599 | 9.595356 | 18.06413 | 24.33318 | 9.965909 | 0.044418 | 0.076196 | | 1 |
| 13 | 114679200 | 114680599 | 18.47406 | 32.53809 | 35.42229 | 15.67988 | 0.266788 | 0.591696 | | 1 |
| 13 | 114949400 | 114950799 | 20.19857 | 34.59277 | 49.01158 | 21.4931 | -0.1175 | 0.94733 | | 1 |
| 13 | 115300200 | 115301799 | 16.28922 | 28.54848 | 40.3276 | 17.81443 | -0.10839 | 0.654355 | | 1 |
| 13 | 115727400 | 115728199 | 10.81307 | 19.73471 | 29.77587 | 12.7715 | -0.16858 | 0.284399 | | 1 |
| 13 | 116233000 | 116234399 | 10.74593 | 19.26211 | 31.37088 | 13.74033 | -0.29783 | 0.394455 | | 1 |
| 13 | 116450200 | 116452199 | 24.07632 | 40.77345 | 50.65276 | 22.17703 | 0.069438 | 1.063711 | | 1 |
| 13 | 116713400 | 116715799 | 28.75189 | 48.04161 | 52.91266 | 23.0319 | 0.24228 | 1.208894 | | 1 |
| 13 | 116718400 | 116720799 | 18.96078 | 32.9323 | 36.56701 | 16.05158 | 0.236683 | 0.796447 | | 1 |
| 13 | 116930000 | 116932199 | 23.07393 | 39.44757 | 46.87814 | 20.5546 | 0.134788 | 0.972134 | | 1 |
| 13 | 117301800 | 117312199 | 269.423 | 382.4637 | 504.8837 | 208.8421 | -0.2104 | 4.355265 | | 1 |
| 13 | 117500600 | 117502399 | 43.76879 | 70.46496 | 87.85547 | 37.33096 | 0.036852 | 1.836516 | | 1 |
| 13 | 117576600 | 117583599 | 103.1163 | 158.3237 | 216.6469 | 89.94825 | -0.15783 | 2.987939 | | 1 |
| 13 | 117683600 | 117684399 | 18.01556 | 31.41084 | 37.92069 | 16.68598 | 0.116993 | 0.739121 | | 1 |

| | | | | | | | | | |
|---|---|---|---|---|---|---|---|---|---|
| 13 | 118207000 | 118210999 | 91.35068 | 141.3899 | 169.5831 | 70.44049 | 0.04102 | 2.827164 | 1 |
| 13 | 118404800 | 118406399 | 20.17902 | 35.00786 | 38.9565 | 17.09258 | 0.233156 | 0.817974 | 1 |
| 13 | 118459400 | 118460599 | 13.97998 | 24.57623 | 39.3416 | 17.42137 | -0.28761 | 0.593409 | 1 |
| 13 | 118658800 | 118659599 | 14.36992 | 25.31759 | 32.81386 | 14.52432 | 0.016794 | 0.559158 | 1 |
| 13 | 119271600 | 119274199 | 33.01477 | 54.62661 | 60.30063 | 26.0768 | 0.238471 | 1.334451 | 1 |
| 13 | 119353600 | 119355199 | 20.77091 | 36.09119 | 38.41062 | 16.86525 | 0.297569 | 0.801748 | 1 |
| 13 | 119631000 | 119631999 | 42.19991 | 67.66113 | 103.5791 | 43.74561 | -0.26587 | 1.968244 | 1 |
| 13 | 120674400 | 120675999 | 16.89703 | 29.83937 | 30.31674 | 13.41278 | 0.367158 | 0.541665 | 1 |
| 13 | 120680600 | 120686399 | 105.9307 | 161.7418 | 208.6782 | 86.54464 | -0.07926 | 3.099012 | 1 |
| 14 | 3164800 | 3167999 | 25.01157 | 42.42151 | 44.30548 | 19.40695 | 0.318991 | 1.047976 | 1 |
| 14 | 3377400 | 3379199 | 21.80623 | 38.01597 | 40.27673 | 17.72151 | 0.304428 | 0.762763 | 1 |
| 14 | 3591600 | 3603599 | 37.22912 | 61.26004 | 63.84495 | 27.52536 | 0.319304 | 1.413832 | 1 |
| 14 | 4031200 | 4033599 | 29.83846 | 48.87211 | 71.09668 | 30.59182 | -0.16308 | 1.460065 | 1 |
| 14 | 4237000 | 4245599 | 181.8164 | 268.6869 | 307.1503 | 126.4915 | 0.053797 | 3.738965 | 1 |
| 14 | 4701200 | 4702399 | 17.03754 | 29.94095 | 35.5246 | 15.72869 | 0.142022 | 0.607162 | 1 |
| 14 | 4870800 | 4875399 | 86.93213 | 135.6175 | 151.6467 | 62.92903 | 0.15087 | 2.687002 | 1 |
| 14 | 4919600 | 4922199 | 22.54612 | 38.87394 | 43.68041 | 19.16165 | 0.217531 | 0.892474 | 1 |
| 14 | 4971200 | 4973399 | 16.02334 | 29.01495 | 28.73916 | 12.44859 | 0.425386 | 0.364147 | 1 |
| 14 | 5142600 | 5143799 | 11.80557 | 21.30265 | 33.21564 | 14.45081 | -0.22922 | 0.364632 | 1 |
| 14 | 5231200 | 5231999 | 9.030678 | 16.85359 | 26.38867 | 10.96839 | -0.18811 | 0.132382 | 1 |
| 14 | 5257000 | 5264999 | 78.65276 | 122.8876 | 161.8861 | 67.1676 | -0.08363 | 2.656951 | 1 |
| 14 | 5443000 | 5445199 | 28.37693 | 46.42753 | 73.13713 | 31.45615 | -0.27847 | 1.471918 | 1 |
| 14 | 5494800 | 5496399 | 17.95351 | 31.18669 | 45.79395 | 20.10345 | -0.16487 | 0.802984 | 1 |
| 14 | 5560600 | 5562999 | 13.25407 | 23.60435 | 34.12718 | 15.02955 | -0.1329 | 0.457176 | 1 |
| 14 | 5843800 | 5853199 | 126.6051 | 192.9565 | 207.6148 | 86.06719 | 0.180391 | 3.142267 | 1 |
| 14 | 6126600 | 6127599 | 15.23604 | 26.74223 | 41.37682 | 18.30079 | -0.23903 | 0.627601 | 1 |
| 14 | 6650600 | 6652999 | 48.32596 | 77.31711 | 94.98852 | 39.98933 | 0.04731 | 2.03845 | 1 |
| 14 | 6909400 | 6910199 | 12.16895 | 22.51763 | 28.73209 | 12.0974 | 0.090432 | 0.21422 | 1 |
| 14 | 6913000 | 6925199 | 170.7732 | 254.5989 | 263.2456 | 108.4863 | 0.210289 | 3.583799 | 1 |
| 14 | 6993800 | 6998399 | 31.74909 | 52.57954 | 54.59049 | 23.62209 | 0.32679 | 1.318034 | 1 |
| 14 | 7005400 | 7010599 | 80.63805 | 125.0582 | 173.3735 | 72.00752 | -0.16549 | 2.789103 | 1 |
| 14 | 7167200 | 7173799 | 70.42907 | 110.9861 | 147.5267 | 61.21725 | -0.08699 | 2.490824 | 1 |
| 14 | 7433800 | 7436399 | 21.11125 | 36.66857 | 40.38489 | 17.72786 | 0.24814 | 0.808917 | 1 |
| 14 | 7437200 | 7444999 | 131.1919 | 199.1592 | 235.7322 | 97.60786 | 0.038252 | 3.224598 | 1 |
| 14 | 7740800 | 7741599 | 11.09518 | 20.18255 | 31.73618 | 13.66889 | -0.23203 | 0.308238 | 1 |
| 14 | 7765800 | 7767999 | 19.65389 | 33.69954 | 50.08294 | 21.965 | -0.186 | 0.934016 | 1 |
| 14 | 7847800 | 7850399 | 28.98597 | 47.38473 | 73.87809 | 31.76525 | -0.26451 | 1.483881 | 1 |
| 14 | 7886800 | 7892199 | 71.29616 | 112.4621 | 134.7198 | 55.91001 | 0.063968 | 2.469012 | 1 |
| 14 | 8172000 | 8179199 | 85.26874 | 132.06 | 187.3809 | 77.83299 | -0.2005 | 2.815304 | 1 |
| 14 | 8220600 | 8221999 | 25.25321 | 42.41978 | 49.09382 | 21.44118 | 0.170922 | 1.137179 | 1 |
| 14 | 8225200 | 8230399 | 79.94723 | 125.3747 | 147.441 | 61.15089 | 0.084082 | 2.589998 | 1 |
| 14 | 8338400 | 8339199 | 12.38062 | 22.99319 | 25.56205 | 10.70929 | 0.293016 | 0.187592 | 1 |
| 14 | 8530000 | 8538799 | 63.08588 | 100.4619 | 115.0582 | 48.12246 | 0.141046 | 2.203557 | 1 |
| 14 | 8614400 | 8615999 | 17.94518 | 30.91129 | 46.37207 | 20.34182 | -0.19801 | 0.88292 | 1 |
| 14 | 8715000 | 8721399 | 80.16336 | 124.6856 | 169.2044 | 70.24531 | -0.13145 | 2.736963 | 1 |
| 14 | 8775000 | 8776999 | 36.84419 | 59.36972 | 91.10844 | 38.76326 | -0.26047 | 1.801853 | 1 |
| 14 | 8829000 | 8830999 | 44.11513 | 71.28811 | 75.79147 | 32.36547 | 0.27249 | 1.71999 | 1 |

| | | | | | | | | | |
|---|---|---|---|---|---|---|---|---|---|
| 14 | 9721600 | 9723799 | 27.17392 | 44.61551 | 67.11275 | 28.90712 | -0.20981 | 1.419454 | 1 |
| 14 | 10205200 | 10207999 | 14.94936 | 26.30651 | 39.27801 | 17.38975 | -0.18713 | 0.588187 | 1 |
| 14 | 10214800 | 10216999 | 31.90292 | 51.89948 | 68.66874 | 29.47295 | -0.03329 | 1.557087 | 1 |
| 14 | 10490600 | 10491999 | 21.44525 | 36.86693 | 42.68835 | 18.72477 | 0.173473 | 0.911331 | 1 |
| 14 | 10997600 | 11004599 | 150.5141 | 225.4047 | 277.2075 | 114.3398 | -0.03395 | 3.498475 | 1 |
| 14 | 11141400 | 11147399 | 116.4981 | 177.3988 | 214.5754 | 88.92967 | 0.010655 | 3.156591 | 1 |
| 14 | 11171400 | 11172399 | 16.25243 | 28.60376 | 31.21212 | 13.81829 | 0.263132 | 0.579025 | 1 |
| 14 | 11183200 | 11183999 | 21.91245 | 37.1352 | 50.61319 | 22.16446 | -0.06363 | 1.05179 | 1 |
| 14 | 11299600 | 11301799 | 41.54875 | 67.37652 | 72.97303 | 31.25658 | 0.250764 | 1.638092 | 1 |
| 14 | 11860600 | 11861399 | 20.55602 | 35.69908 | 35.45981 | 15.56449 | 0.396674 | 0.803902 | 1 |
| 14 | 12356400 | 12359399 | 17.42431 | 30.6402 | 34.52171 | 15.28612 | 0.216604 | 0.600876 | 1 |
| 14 | 12382600 | 12383599 | 16.68383 | 29.35196 | 31.58908 | 13.98623 | 0.282638 | 0.585406 | 1 |
| 14 | 12553000 | 12554999 | 12.73677 | 23.38242 | 38.53855 | 16.40213 | -0.28814 | 0.281942 | 1 |
| 14 | 12609200 | 12609999 | 17.85814 | 31.09872 | 38.48696 | 16.92228 | 0.081205 | 0.756311 | 1 |
| 14 | 13044600 | 13055399 | 316.4374 | 443.1827 | 550.1464 | 228.063 | -0.14131 | 4.556376 | 1 |
| 14 | 13121800 | 13122599 | 11.87678 | 21.77338 | 30.22167 | 12.88675 | -0.04335 | 0.267228 | 1 |
| 14 | 13577200 | 13584399 | 109.0376 | 166.9681 | 209.641 | 87.00211 | -0.03678 | 3.041569 | 1 |
| 14 | 13929000 | 13931599 | 28.92217 | 47.79906 | 71.2092 | 30.76934 | -0.19291 | 1.346526 | 1 |
| 14 | 14180800 | 14185799 | 60.2537 | 96.14335 | 104.9031 | 43.99658 | 0.213931 | 2.134857 | 1 |
| 14 | 14284000 | 14284999 | 11.20828 | 20.53781 | 33.95514 | 14.48836 | -0.29513 | 0.270695 | 1 |
| 14 | 14392800 | 14395599 | 31.34234 | 51.41166 | 66.46878 | 28.6185 | 0.007438 | 1.439719 | 1 |
| 14 | 14772200 | 14775199 | 38.02249 | 61.82769 | 67.18309 | 28.8307 | 0.24958 | 1.57452 | 1 |
| 14 | 15068600 | 15075799 | 45.70522 | 73.22783 | 97.67237 | 41.20513 | -0.06874 | 1.992023 | 1 |
| 14 | 15085000 | 15086199 | 20.34084 | 35.16236 | 41.12625 | 18.04262 | 0.160701 | 0.855821 | 1 |
| 14 | 15569000 | 15571399 | 29.9017 | 48.95239 | 67.88487 | 29.20395 | -0.09453 | 1.461287 | 1 |
| 14 | 15792800 | 15795199 | 22.40139 | 38.02583 | 48.5077 | 21.25295 | 0.031633 | 1.037789 | 1 |
| 14 | 15800200 | 15802199 | 13.4915 | 23.99983 | 32.55147 | 14.34939 | -0.04232 | 0.461197 | 1 |
| 14 | 15805200 | 15808399 | 17.77447 | 30.70857 | 49.0049 | 21.4987 | -0.28601 | 0.858051 | 1 |
| 14 | 15810000 | 15811999 | 23.92725 | 39.97854 | 56.49779 | 24.59184 | -0.11614 | 1.197805 | 1 |
| 14 | 15911200 | 15914199 | 49.52732 | 78.77358 | 120.3518 | 50.25102 | -0.27587 | 2.234926 | 1 |
| 14 | 15950800 | 15952599 | 17.45538 | 30.77243 | 34.95594 | 15.47058 | 0.206222 | 0.574533 | 1 |
| 14 | 16145400 | 16147599 | 42.18198 | 68.0731 | 76.18225 | 32.47627 | 0.196471 | 1.756062 | 1 |
| 14 | 16186200 | 16190199 | 54.65734 | 86.6044 | 130.5385 | 54.29674 | -0.26175 | 2.35284 | 1 |
| 14 | 16301200 | 16302399 | 12.4026 | 22.9555 | 27.62654 | 11.62811 | 0.174472 | 0.211193 | 1 |
| 14 | 16474600 | 16476399 | 18.56248 | 32.39886 | 40.40782 | 17.79227 | 0.070343 | 0.731001 | 1 |
| 14 | 17521600 | 17523199 | 13.87171 | 24.53608 | 33.16987 | 14.67037 | -0.04159 | 0.512756 | 1 |
| 14 | 17529200 | 17532999 | 44.62095 | 71.7739 | 75.52733 | 32.05179 | 0.279479 | 1.85884 | 1 |
| 14 | 17613000 | 17615599 | 20.28989 | 33.9456 | 57.91198 | 25.22826 | -0.3868 | 1.17422 | 1 |
| 14 | 17872600 | 17879199 | 29.63384 | 48.88458 | 56.58225 | 24.42011 | 0.169873 | 1.358837 | 1 |
| 14 | 17969400 | 17971399 | 18.61305 | 31.40594 | 54.76189 | 23.94945 | -0.41796 | 1.08159 | 1 |
| 14 | 18002400 | 18002999 | 11.293 | 20.08096 | 35.8429 | 15.80562 | -0.43607 | 0.459656 | 1 |
| 14 | 18029400 | 18031999 | 39.411 | 63.73101 | 74.53705 | 31.85854 | 0.1364 | 1.691699 | 1 |
| 14 | 18225600 | 18226999 | 10.40966 | 19.73269 | 25.43565 | 10.27449 | 0.124487 | 0.040313 | 1 |
| 14 | 18229000 | 18229999 | 15.77006 | 27.18867 | 46.39296 | 20.35154 | -0.38258 | 0.866239 | 1 |
| 14 | 18235000 | 18235599 | 8.026866 | 15.40238 | 23.54552 | 9.241118 | -0.08203 | -0.08193 | 1 |
| 14 | 18259000 | 18261999 | 14.4075 | 25.27832 | 39.36898 | 17.40711 | -0.24836 | 0.62542 | 1 |
| 14 | 18311800 | 18313999 | 19.26808 | 32.89185 | 49.01382 | 21.48844 | -0.19068 | 0.97358 | 1 |

| | | | | | | | | | |
|---|---|---|---|---|---|---|---|---|---|
| 14 | 18358000 | 18361599 | 38.70475 | 62.33164 | 86.94303 | 36.95997 | -0.12409 | 1.818606 | 1 |
| 14 | 18397000 | 18400999 | 15.22317 | 26.50253 | 41.66401 | 18.31591 | -0.26233 | 0.752616 | 1 |
| 14 | 18404400 | 18405199 | 9.675801 | 17.45135 | 32.89047 | 14.31778 | -0.50107 | 0.353021 | 1 |
| 14 | 18575600 | 18579399 | 40.0706 | 64.72684 | 76.08426 | 32.47402 | 0.127335 | 1.725256 | 1 |
| 14 | 19181400 | 19182199 | 24.93198 | 42.00505 | 46.47131 | 20.31445 | 0.235919 | 1.108328 | 1 |
| 14 | 19352000 | 19354199 | 26.43746 | 44.23217 | 53.93873 | 23.4979 | 0.095813 | 1.187973 | 1 |
| 14 | 19520000 | 19520799 | 19.1863 | 33.27197 | 39.60106 | 17.37517 | 0.136567 | 0.820082 | 1 |
| 14 | 19675600 | 19687799 | 349.5392 | 489.8614 | 515.8733 | 213.8335 | 0.096823 | 4.547443 | 1 |
| 14 | 19792600 | 19794599 | 11.14661 | 20.96608 | 25.79086 | 10.57864 | 0.172384 | 0.098519 | 1 |
| 14 | 20324200 | 20331399 | 50.16873 | 80.12986 | 116.3344 | 48.8587 | -0.19589 | 2.09799 | 1 |
| 14 | 21415400 | 21417399 | 16.19369 | 28.55111 | 35.4643 | 15.69525 | 0.077844 | 0.56455 | 1 |
| 14 | 21565600 | 21567799 | 24.46671 | 40.89117 | 58.66991 | 25.54174 | -0.13793 | 1.199655 | 1 |
| 14 | 21591400 | 21593199 | 13.89893 | 24.87354 | 36.65897 | 16.07844 | -0.15653 | 0.435439 | 1 |
| 14 | 21644400 | 21649399 | 88.0124 | 135.5365 | 194.2165 | 80.66847 | -0.22097 | 2.922604 | 1 |
| 14 | 21932400 | 21934799 | 20.84557 | 36.24374 | 38.32398 | 16.83388 | 0.30686 | 0.791796 | 1 |
| 14 | 22268800 | 22269999 | 19.99268 | 34.7243 | 43.63596 | 19.1545 | 0.05875 | 0.810697 | 1 |
| 14 | 22885200 | 22887599 | 37.25125 | 60.1843 | 78.81644 | 33.64853 | -0.02781 | 1.718562 | 1 |
| 14 | 23025800 | 23029399 | 34.08388 | 56.04217 | 64.65383 | 27.86517 | 0.17268 | 1.415579 | 1 |
| 14 | 23587600 | 23592799 | 78.05417 | 123.2505 | 127.4752 | 52.93007 | 0.276781 | 2.448548 | 1 |
| 14 | 24123000 | 24126599 | 16.7302 | 28.87765 | 48.4651 | 21.2613 | -0.35851 | 0.863056 | 1 |
| 14 | 24373000 | 24378399 | 70.12969 | 110.8508 | 127.4362 | 52.9413 | 0.125633 | 2.418896 | 1 |
| 14 | 24431200 | 24432799 | 14.61005 | 25.52217 | 41.08641 | 18.1008 | -0.29582 | 0.700666 | 1 |
| 14 | 24462400 | 24467999 | 36.0101 | 59.07216 | 63.04856 | 27.14665 | 0.283284 | 1.446894 | 1 |
| 14 | 27250200 | 27251199 | 23.32165 | 39.81092 | 42.70121 | 18.72087 | 0.281783 | 0.981485 | 1 |
| 14 | 27296200 | 27302799 | 93.71559 | 144.9432 | 189.4056 | 78.67456 | -0.08336 | 2.844509 | 1 |
| 14 | 27675800 | 27676999 | 18.38174 | 32.11436 | 35.35963 | 15.5789 | 0.249595 | 0.70961 | 1 |
| 14 | 27679800 | 27681999 | 20.09516 | 35.07867 | 37.79809 | 16.64456 | 0.280479 | 0.734374 | 1 |
| 14 | 28332200 | 28336199 | 40.33495 | 64.88776 | 98.45583 | 41.78951 | -0.24725 | 1.858192 | 1 |
| 14 | 28831800 | 28834799 | 35.86862 | 58.25647 | 77.56232 | 33.26062 | -0.04423 | 1.598939 | 1 |
| 14 | 28968800 | 28976599 | 244.0619 | 348.7669 | 453.1742 | 187.2444 | -0.17795 | 4.257047 | 1 |
| 14 | 29275400 | 29277599 | 16.23665 | 28.50115 | 34.20297 | 15.12902 | 0.12583 | 0.619003 | 1 |
| 14 | 31837000 | 31839599 | 14.61931 | 26.1826 | 28.5339 | 12.50682 | 0.277385 | 0.42013 | 1 |
| 14 | 33151800 | 33159199 | 76.11615 | 119.7167 | 144.304 | 59.86495 | 0.0518 | 2.532924 | 1 |
| 14 | 33369200 | 33372999 | 67.07087 | 105.8534 | 137.1785 | 56.94436 | -0.04873 | 2.454867 | 1 |
| 14 | 33388400 | 33394199 | 49.37593 | 79.24268 | 87.44053 | 36.98349 | 0.20714 | 1.940143 | 1 |
| 14 | 33585000 | 33586199 | 17.56991 | 30.1261 | 48.4811 | 21.26247 | -0.30002 | 0.927751 | 1 |
| 14 | 33645000 | 33649599 | 36.27773 | 58.48136 | 91.03512 | 38.75373 | -0.28035 | 1.78578 | 1 |
| 14 | 34511000 | 34512999 | 25.98716 | 43.56687 | 60.57252 | 26.42156 | -0.09289 | 1.170764 | 1 |
| 14 | 34515400 | 34522199 | 72.35127 | 114.3609 | 128.7647 | 53.49283 | 0.155578 | 2.420049 | 1 |
| 14 | 34524200 | 34530399 | 71.02425 | 112.9095 | 113.429 | 47.33492 | 0.327262 | 2.264294 | 1 |
| 14 | 34916800 | 34919599 | 18.00113 | 30.7975 | 50.58535 | 22.1814 | -0.33008 | 0.949854 | 1 |
| 14 | 35234600 | 35236999 | 29.84922 | 49.59201 | 54.59282 | 23.67547 | 0.242876 | 1.274295 | 1 |
| 14 | 35436800 | 35438199 | 16.86951 | 29.12368 | 47.41589 | 20.80114 | -0.31483 | 0.860234 | 1 |
| 14 | 35459200 | 35460999 | 15.50586 | 27.11661 | 42.52066 | 18.74877 | -0.25829 | 0.688493 | 1 |
| 14 | 35674000 | 35675599 | 17.23143 | 30.20634 | 35.70028 | 15.77324 | 0.147752 | 0.649271 | 1 |
| 14 | 35686800 | 35688399 | 17.9718 | 31.47128 | 36.65189 | 16.17938 | 0.168785 | 0.670374 | 1 |
| 14 | 35739200 | 35741599 | 36.1672 | 58.64158 | 72.12425 | 30.88988 | 0.067613 | 1.630496 | 1 |

| | | | | | | | | | |
|---|---|---|---|---|---|---|---|---|---|
| 14 | 35917400 | 35924199 | 63.8319 | 101.078 | 130.2112 | 54.14341 | -0.0363 | 2.371816 | 1 |
| 14 | 36028800 | 36029799 | 14.19102 | 25.26812 | 35.55573 | 15.66122 | -0.09434 | 0.466408 | 1 |
| 14 | 36098200 | 36099399 | 15.67526 | 28.11046 | 30.2686 | 13.25257 | 0.295452 | 0.421106 | 1 |
| 14 | 36154800 | 36157199 | 23.56375 | 40.24649 | 45.82164 | 20.08981 | 0.196178 | 0.981704 | 1 |
| 14 | 36267000 | 36267799 | 12.48571 | 22.81419 | 26.15395 | 11.20152 | 0.226579 | 0.282059 | 1 |
| 14 | 36317600 | 36318999 | 11.91049 | 21.5766 | 33.21531 | 14.37982 | -0.20645 | 0.341474 | 1 |
| 14 | 36919200 | 36922399 | 29.43457 | 47.83143 | 75.82224 | 32.52692 | -0.29448 | 1.576685 | 1 |
| 14 | 36956600 | 36958599 | 16.73655 | 29.41382 | 36.06256 | 15.96771 | 0.094963 | 0.605597 | 1 |
| 14 | 37312400 | 37313999 | 22.06967 | 37.89102 | 42.6744 | 18.71778 | 0.212917 | 0.925206 | 1 |
| 14 | 37664200 | 37665399 | 15.36997 | 26.98044 | 40.32199 | 17.836 | -0.18924 | 0.624377 | 1 |
| 14 | 38051200 | 38052799 | 18.27041 | 31.99807 | 31.26373 | 13.80228 | 0.421067 | 0.662127 | 1 |
| 14 | 38183600 | 38185199 | 17.31491 | 30.59449 | 37.47852 | 16.57945 | 0.099057 | 0.552972 | 1 |
| 14 | 38190400 | 38198399 | 159.5379 | 239.8414 | 250.5001 | 103.4212 | 0.206575 | 3.42541 | 1 |
| 14 | 39005000 | 39005599 | 9.644067 | 17.54496 | 31.4006 | 13.52239 | -0.41668 | 0.299648 | 1 |
| 14 | 39234600 | 39236399 | 32.67674 | 53.7344 | 61.01048 | 26.29614 | 0.195758 | 1.410062 | 1 |
| 14 | 39274400 | 39277199 | 32.96141 | 53.53776 | 80.58058 | 34.55638 | -0.22065 | 1.593904 | 1 |
| 14 | 39452800 | 39456599 | 46.68764 | 75.068 | 84.35842 | 35.77859 | 0.184159 | 1.879115 | 1 |
| 14 | 39626600 | 39628799 | 7.875864 | 15.09208 | 22.9886 | 9.057846 | -0.08208 | -0.07228 | 1 |
| 14 | 39818600 | 39820399 | 33.95142 | 55.03353 | 76.2132 | 32.63441 | -0.10349 | 1.635156 | 1 |
| 14 | 39848200 | 39859599 | 252.7792 | 361.1779 | 479.6576 | 198.1909 | -0.20962 | 4.259416 | 1 |
| 14 | 40007000 | 40007799 | 12.98174 | 23.81436 | 29.63631 | 12.62611 | 0.113989 | 0.270714 | 1 |
| 14 | 40058400 | 40060799 | 26.41174 | 44.59573 | 51.76846 | 22.64509 | 0.166789 | 1.099248 | 1 |
| 14 | 40308800 | 40311599 | 23.78123 | 40.53163 | 48.48374 | 21.25333 | 0.124583 | 1.002423 | 1 |
| 14 | 40359200 | 40367199 | 132.7618 | 201.1366 | 233.8471 | 96.76791 | 0.061667 | 3.266258 | 1 |
| 14 | 40385000 | 40386399 | 19.06793 | 33.36898 | 36.69335 | 16.18875 | 0.251381 | 0.688694 | 1 |
| 14 | 40447800 | 40449399 | 23.00991 | 39.09165 | 50.08814 | 21.95097 | 0.025242 | 1.033403 | 1 |
| 14 | 40487800 | 40490199 | 19.83719 | 34.14327 | 46.87775 | 20.56315 | -0.07105 | 0.898199 | 1 |
| 14 | 40503000 | 40504799 | 14.57928 | 25.65341 | 41.32574 | 18.29662 | -0.29638 | 0.588473 | 1 |
| 14 | 40910400 | 40916599 | 113.334 | 173.5124 | 194.2847 | 80.62543 | 0.128065 | 3.045766 | 1 |
| 14 | 41064400 | 41066399 | 28.48673 | 47.79952 | 50.74059 | 22.14523 | 0.295128 | 1.160285 | 1 |
| 14 | 41152200 | 41154199 | 21.37918 | 36.84002 | 44.19418 | 19.38667 | 0.123196 | 0.890443 | 1 |
| 14 | 41155200 | 41157199 | 15.34305 | 27.1743 | 30.9126 | 13.66843 | 0.206618 | 0.501696 | 1 |
| 14 | 41188800 | 41193199 | 85.00598 | 134.0538 | 132.3572 | 54.9277 | 0.342218 | 2.483057 | 1 |
| 14 | 41200200 | 41217399 | 299.4681 | 432.2582 | 376.4429 | 155.306 | 0.413477 | 4.11337 | 1 |
| 14 | 41436200 | 41437199 | 26.90605 | 44.49418 | 61.94742 | 26.77861 | -0.09538 | 1.326842 | 1 |
| 14 | 41444600 | 41448399 | 94.96169 | 145.579 | 200.5819 | 83.26074 | -0.1694 | 3.014758 | 1 |
| 14 | 41876400 | 41878199 | 31.90948 | 52.59839 | 53.97714 | 23.28826 | 0.342576 | 1.372526 | 1 |
| 14 | 42048400 | 42049199 | 16.41978 | 29.07997 | 30.91298 | 13.66872 | 0.303999 | 0.510977 | 1 |
| 14 | 42050000 | 42056399 | 114.1503 | 174.7957 | 167.8894 | 69.67482 | 0.349373 | 3.040135 | 1 |
| 14 | 42061000 | 42063199 | 31.63299 | 52.19087 | 60.57757 | 26.14622 | 0.165928 | 1.365419 | 1 |
| 14 | 42130600 | 42131599 | 20.0231 | 33.96455 | 52.68962 | 23.08021 | -0.2496 | 1.039826 | 1 |
| 14 | 42140000 | 42143999 | 82.96955 | 128.5484 | 170.2733 | 70.72682 | -0.10091 | 2.806672 | 1 |
| 14 | 42156000 | 42158799 | 39.72007 | 64.36456 | 67.98147 | 29.10303 | 0.285975 | 1.646518 | 1 |
| 14 | 42177000 | 42178599 | 7.911136 | 15.05024 | 26.08989 | 10.45928 | -0.29 | -0.00713 | 1 |
| 14 | 42181000 | 42182799 | 15.83089 | 28.10761 | 33.29074 | 14.69892 | 0.150385 | 0.488811 | 1 |
| 14 | 42241200 | 42241799 | 10.09315 | 19.08906 | 22.19862 | 9.010346 | 0.265589 | 0.042555 | 1 |
| 14 | 42388200 | 42396999 | 147.3435 | 222.0466 | 222.7898 | 92.04042 | 0.267535 | 3.374458 | 1 |

| | | | | | | | | | |
|---|---|---|---|---|---|---|---|---|---|
| 14 | 42408000 | 42410599 | 34.65765 | 56.75486 | 59.85066 | 25.7508 | 0.299701 | 1.465545 | 1 |
| 14 | 42476400 | 42477199 | 17.17826 | 29.57872 | 46.90411 | 20.57497 | -0.27777 | 0.884279 | 1 |
| 14 | 42536600 | 42537799 | 25.12864 | 41.80293 | 57.53066 | 24.97173 | -0.07797 | 1.250995 | 1 |
| 14 | 42558000 | 42560599 | 43.79395 | 70.4784 | 80.38188 | 34.13632 | 0.164483 | 1.846641 | 1 |
| 14 | 42633800 | 42636399 | 51.74955 | 82.31062 | 117.8374 | 49.20331 | -0.18217 | 2.234006 | 1 |
| 14 | 42664200 | 42664799 | 11.61273 | 22.12783 | 24.96825 | 9.975551 | 0.328632 | 0.01365 | 1 |
| 14 | 42841200 | 42841999 | 21.10929 | 36.73284 | 43.00934 | 18.90182 | 0.160518 | 0.788149 | 1 |
| 14 | 42856600 | 42857799 | 41.27394 | 66.854 | 69.69833 | 29.82864 | 0.304143 | 1.660167 | 1 |
| 14 | 42887000 | 42888399 | 60.73856 | 96.69932 | 102.04 | 42.66273 | 0.258638 | 2.206281 | 1 |
| 14 | 43068800 | 43071599 | 24.16662 | 41.85692 | 42.45806 | 18.628 | 0.365695 | 0.855456 | 1 |
| 14 | 43075600 | 43076199 | 11.39628 | 21.13058 | 27.2441 | 11.43936 | 0.078624 | 0.194257 | 1 |
| 14 | 43905600 | 43906999 | 29.17464 | 47.77996 | 64.79278 | 27.87814 | -0.062 | 1.456014 | 1 |
| 14 | 43929600 | 43932199 | 20.10516 | 34.62271 | 50.08279 | 21.96941 | -0.14586 | 0.900023 | 1 |
| 14 | 43972600 | 43974999 | 29.16766 | 47.60471 | 71.97389 | 30.92827 | -0.22189 | 1.506786 | 1 |
| 14 | 44083200 | 44087999 | 50.07731 | 80.48275 | 94.38414 | 40.00523 | 0.121919 | 1.899689 | 1 |
| 14 | 44164200 | 44166399 | 24.46432 | 41.83053 | 47.93968 | 21.02029 | 0.186959 | 0.978559 | 1 |
| 14 | 44587400 | 44588199 | 12.63451 | 22.39278 | 36.68007 | 16.21633 | -0.31607 | 0.488811 | 1 |
| 14 | 44621200 | 44622999 | 15.88162 | 28.0425 | 33.95554 | 15.02306 | 0.115328 | 0.544318 | 1 |
| 14 | 44906000 | 44915399 | 154.592 | 231.7512 | 260.9007 | 107.6354 | 0.094847 | 3.479225 | 1 |
| 14 | 45257000 | 45259399 | 24.85225 | 41.91978 | 49.07438 | 21.45997 | 0.154595 | 1.098672 | 1 |
| 14 | 45793600 | 45796999 | 43.86237 | 70.68072 | 79.43049 | 33.79617 | 0.188034 | 1.80377 | 1 |
| 14 | 46000000 | 46001599 | 15.33279 | 26.92517 | 38.24673 | 16.92334 | -0.11631 | 0.61438 | 1 |
| 14 | 46132400 | 46134199 | 21.37434 | 36.108 | 58.63228 | 25.65137 | -0.3156 | 1.086705 | 1 |
| 14 | 46284200 | 46288199 | 64.14403 | 101.5581 | 140.3708 | 58.3638 | -0.13789 | 2.376669 | 1 |
| 14 | 46448200 | 46449999 | 14.70301 | 25.70607 | 41.61233 | 18.34433 | -0.30353 | 0.691879 | 1 |
| 14 | 46991600 | 46998399 | 88.53924 | 136.6886 | 190.472 | 79.11828 | -0.17796 | 2.875566 | 1 |
| 14 | 47015200 | 47018399 | 35.24373 | 57.70306 | 67.41564 | 29.00319 | 0.152086 | 1.471967 | 1 |
| 14 | 47034200 | 47039599 | 53.6217 | 85.4823 | 125.6633 | 52.63621 | -0.21687 | 2.163026 | 1 |
| 14 | 47462400 | 47465199 | 20.3513 | 35.08562 | 42.63839 | 18.70447 | 0.104709 | 0.878954 | 1 |
| 14 | 47893000 | 47894199 | 9.025259 | 17.37504 | 20.84313 | 8.103818 | 0.277326 | -0.11046 | 1 |
| 14 | 48023000 | 48025599 | 19.8889 | 33.99244 | 52.22201 | 22.89718 | -0.23435 | 0.967849 | 1 |
| 14 | 48261000 | 48265199 | 43.96941 | 70.98056 | 82.04882 | 34.99174 | 0.150387 | 1.751604 | 1 |
| 14 | 48348400 | 48349799 | 21.42612 | 36.89836 | 42.65514 | 18.7112 | 0.176299 | 0.895273 | 1 |
| 14 | 48467200 | 48468799 | 18.49077 | 31.63491 | 52.18838 | 22.8843 | -0.3364 | 0.953988 | 1 |
| 14 | 48712800 | 48713999 | 12.1126 | 21.82445 | 31.18694 | 13.59328 | -0.10576 | 0.371448 | 1 |
| 14 | 49094400 | 49095999 | 11.28857 | 20.95608 | 30.41554 | 12.75009 | -0.08992 | 0.190813 | 1 |
| 14 | 49367600 | 49368799 | 19.50757 | 33.76927 | 45.79675 | 20.09248 | -0.05164 | 0.844234 | 1 |
| 14 | 50090800 | 50093599 | 27.10555 | 45.77099 | 48.53647 | 21.23184 | 0.296928 | 1.098859 | 1 |
| 14 | 50121600 | 50123799 | 24.04641 | 39.88375 | 60.61583 | 26.25688 | -0.221 | 1.285269 | 1 |
| 14 | 50224200 | 50234999 | 172.4202 | 257.748 | 259.7413 | 107.0982 | 0.250857 | 3.534493 | 1 |
| 14 | 50245200 | 50247199 | 33.46313 | 54.69413 | 66.46871 | 28.57762 | 0.093775 | 1.491501 | 1 |
| 14 | 50826000 | 50828799 | 56.0312 | 89.27547 | 115.7989 | 48.46827 | -0.0374 | 2.179844 | 1 |
| 14 | 50861600 | 50863599 | 21.72058 | 36.37478 | 60.18299 | 26.23378 | -0.34284 | 1.170507 | 1 |
| 14 | 50895200 | 50901599 | 108.1361 | 165.7687 | 193.4585 | 80.30342 | 0.069857 | 3.019157 | 1 |
| 14 | 50949600 | 50966399 | 78.7019 | 123.2882 | 140.966 | 58.4719 | 0.123412 | 2.607962 | 1 |
| 14 | 51625200 | 51626199 | 20.04957 | 34.66682 | 42.10164 | 18.47066 | 0.106545 | 0.851282 | 1 |
| 14 | 51630000 | 51630999 | 13.1162 | 23.81608 | 28.5387 | 12.31935 | 0.155569 | 0.327268 | 1 |

| | | | | | | | | | |
|---|---|---|---|---|---|---|---|---|---|
| 14 | 51725200 | 51727199 | 26.56373 | 44.57499 | 50.17157 | 21.89757 | 0.210962 | 1.153601 | 1 |
| 14 | 52103600 | 52105399 | 13.5391 | 24.41358 | 29.43307 | 12.81641 | 0.138577 | 0.370761 | 1 |
| 14 | 52740800 | 52741999 | 17.12148 | 29.81108 | 41.58851 | 18.28415 | -0.09086 | 0.760103 | 1 |
| 14 | 52837800 | 52838999 | 22.19144 | 37.77709 | 52.7357 | 23.115 | -0.09744 | 1.017588 | 1 |
| 14 | 52907200 | 52909399 | 29.16454 | 47.74949 | 71.052 | 30.5675 | -0.19585 | 1.462483 | 1 |
| 14 | 52914800 | 52916999 | 28.13596 | 47.24966 | 52.87679 | 23.0895 | 0.21923 | 1.154377 | 1 |
| 14 | 52930800 | 52935999 | 22.17789 | 37.14325 | 58.5929 | 25.5418 | -0.27429 | 1.169254 | 1 |
| 14 | 53106200 | 53108599 | 10.72926 | 19.72951 | 28.23838 | 11.9914 | -0.08297 | 0.236208 | 1 |
| 14 | 53111600 | 53112399 | 13.04279 | 23.75943 | 26.51292 | 11.39872 | 0.261436 | 0.303113 | 1 |
| 14 | 53249400 | 53251199 | 22.71153 | 38.03423 | 55.8597 | 24.34923 | -0.17141 | 1.171406 | 1 |
| 14 | 53422600 | 53423199 | 9.042137 | 17.35166 | 19.81123 | 7.774047 | 0.335961 | -0.08273 | 1 |
| 14 | 53637400 | 53642599 | 55.32502 | 88.71584 | 95.80436 | 40.46936 | 0.236769 | 1.976516 | 1 |
| 14 | 53715800 | 53717199 | 22.43058 | 38.32863 | 40.75255 | 17.868 | 0.294726 | 0.968193 | 1 |
| 14 | 54172600 | 54176199 | 56.06922 | 88.83247 | 128.9769 | 53.64443 | -0.20796 | 2.354791 | 1 |
| 14 | 54528600 | 54535599 | 108.8997 | 165.3742 | 232.3687 | 96.22496 | -0.20859 | 3.212275 | 1 |
| 14 | 54551400 | 54560999 | 108.9978 | 166.5535 | 210.0491 | 87.12874 | -0.04557 | 3.084498 | 1 |
| 14 | 54605800 | 54612199 | 61.47592 | 97.44763 | 121.4023 | 50.51565 | 0.013206 | 2.34156 | 1 |
| 14 | 54771800 | 54773599 | 11.35761 | 20.44679 | 31.2837 | 13.64989 | -0.20452 | 0.371989 | 1 |
| 14 | 54775000 | 54776599 | 17.58684 | 30.53464 | 43.19267 | 18.95559 | -0.11124 | 0.804381 | 1 |
| 14 | 55022800 | 55026999 | 41.94953 | 67.50484 | 94.70014 | 40.21238 | -0.1338 | 1.850009 | 1 |
| 14 | 55527200 | 55528999 | 22.54237 | 38.16153 | 57.13915 | 25.01412 | -0.199 | 1.068438 | 1 |
| 14 | 56211800 | 56225799 | 274.5766 | 389.8798 | 498.0168 | 205.9927 | -0.16258 | 4.35198 | 1 |
| 14 | 57047800 | 57050599 | 18.12417 | 31.05387 | 51.12894 | 22.42243 | -0.33327 | 0.936536 | 1 |
| 14 | 57070400 | 57073199 | 31.28212 | 51.00358 | 80.36705 | 34.52207 | -0.28252 | 1.530543 | 1 |
| 14 | 57235200 | 57237799 | 41.17207 | 66.26083 | 90.09008 | 38.26074 | -0.0885 | 1.844937 | 1 |
| 14 | 57465200 | 57467999 | 6.803432 | 13.18283 | 23.93554 | 9.069396 | -0.28056 | -0.21358 | 1 |
| 14 | 58644600 | 58646399 | 21.55995 | 37.26648 | 51.7754 | 22.71447 | -0.08688 | 0.874393 | 1 |
| 14 | 59401600 | 59403399 | 18.95356 | 32.73663 | 45.30904 | 19.87715 | -0.0814 | 0.863164 | 1 |
| 14 | 60011600 | 60014199 | 25.58568 | 42.91014 | 50.16444 | 21.88685 | 0.156459 | 1.15608 | 1 |
| 14 | 60582200 | 60583999 | 13.79708 | 24.90395 | 34.62586 | 15.05851 | -0.06485 | 0.377677 | 1 |
| 14 | 60678800 | 60679799 | 12.00951 | 22.38222 | 24.04501 | 10.01745 | 0.348363 | 0.159095 | 1 |
| 14 | 60801200 | 60803399 | 30.7405 | 51.09879 | 55.80211 | 24.20842 | 0.254405 | 1.270408 | 1 |
| 14 | 61716000 | 61717799 | 20.78674 | 35.77224 | 44.23847 | 19.40533 | 0.079174 | 0.90058 | 1 |
| 14 | 61971600 | 61974199 | 32.85899 | 53.98543 | 63.44709 | 27.33592 | 0.14552 | 1.420001 | 1 |
| 14 | 62292000 | 62293399 | 17.45753 | 31.38506 | 30.68604 | 13.40702 | 0.436073 | 0.420081 | 1 |
| 14 | 62882600 | 62887199 | 38.66724 | 62.41289 | 86.74601 | 36.98863 | -0.11512 | 1.750028 | 1 |
| 14 | 63107200 | 63110599 | 34.70363 | 56.51178 | 79.57808 | 34.16766 | -0.12203 | 1.555297 | 1 |
| 14 | 65172200 | 65178399 | 71.3508 | 112.8846 | 125.8872 | 52.32211 | 0.170491 | 2.396991 | 1 |
| 14 | 65861600 | 65870799 | 86.67122 | 134.423 | 173.5029 | 72.06061 | -0.06246 | 2.789063 | 1 |
| 14 | 65898600 | 65902799 | 41.82411 | 67.50505 | 78.63603 | 33.52837 | 0.138985 | 1.751671 | 1 |
| 14 | 66037600 | 66039599 | 22.5268 | 38.18463 | 54.94665 | 24.06375 | -0.14179 | 1.055163 | 1 |
| 14 | 66847600 | 66855599 | 125.0866 | 188.7307 | 261.2476 | 107.9663 | -0.1949 | 3.348179 | 1 |
| 14 | 66863800 | 66865599 | 22.58492 | 38.78526 | 43.17599 | 18.93806 | 0.229709 | 0.926069 | 1 |
| 14 | 67400400 | 67402399 | 19.05211 | 33.33857 | 38.1117 | 16.81335 | 0.195384 | 0.689111 | 1 |
| 14 | 67431000 | 67432999 | 24.15976 | 40.66264 | 50.12844 | 21.90637 | 0.080243 | 1.118168 | 1 |
| 14 | 67737200 | 67740199 | 32.93606 | 53.84851 | 73.52227 | 31.61362 | -0.07354 | 1.488692 | 1 |
| 14 | 68238600 | 68241599 | 43.42745 | 69.99342 | 84.66043 | 36.03097 | 0.082497 | 1.800086 | 1 |

| | | | | | | | | | |
|---|---|---|---|---|---|---|---|---|---|
| 14 | 68713000 | 68714399 | 23.49014 | 39.24946 | 56.97811 | 24.80141 | -0.15473 | 1.197775 | 1 |
| 14 | 68918200 | 68921199 | 38.93583 | 62.69581 | 92.64735 | 39.37766 | -0.20742 | 1.82673 | 1 |
| 14 | 71455200 | 71458799 | 35.30307 | 57.14445 | 80.10315 | 34.2647 | -0.12286 | 1.668911 | 1 |
| 14 | 71669600 | 71671999 | 8.782326 | 16.77163 | 24.514 | 9.748323 | -0.03569 | -0.02945 | 1 |
| 14 | 73424000 | 73426999 | 20.52481 | 35.08894 | 56.08988 | 24.59362 | -0.29146 | 0.971359 | 1 |
| 14 | 73431800 | 73432799 | 12.17331 | 21.58402 | 36.70863 | 16.22676 | -0.36966 | 0.482015 | 1 |
| 14 | 73502200 | 73503999 | 31.42124 | 52.08971 | 59.04325 | 25.56687 | 0.200614 | 1.308305 | 1 |
| 14 | 73837200 | 73845399 | 104.2035 | 160.6937 | 176.8588 | 73.46405 | 0.160878 | 2.900516 | 1 |
| 14 | 74026200 | 74027599 | 13.69426 | 24.3562 | 36.08037 | 15.90738 | -0.16908 | 0.470522 | 1 |
| 14 | 74374600 | 74376199 | 15.85483 | 27.67563 | 41.55731 | 18.29953 | -0.19612 | 0.717888 | 1 |
| 14 | 74545600 | 74547799 | 26.17628 | 43.20702 | 69.61877 | 30.0562 | -0.30578 | 1.35295 | 1 |
| 14 | 74677600 | 74678999 | 23.4517 | 38.79886 | 65.9448 | 28.51528 | -0.38216 | 1.317684 | 1 |
| 14 | 75649600 | 75651799 | 33.81937 | 55.21418 | 72.16613 | 31.01444 | -0.01212 | 1.511756 | 1 |
| 14 | 75664400 | 75664799 | 11.32266 | 21.07709 | 24.1993 | 10.10202 | 0.251165 | 0.162816 | 1 |
| 14 | 76476000 | 76483999 | 73.98524 | 115.9131 | 157.4821 | 65.3219 | -0.12499 | 2.60586 | 1 |
| 14 | 77079000 | 77080399 | 21.9096 | 37.37169 | 47.42774 | 20.79176 | 0.039831 | 0.98803 | 1 |
| 14 | 77314000 | 77315199 | 13.62471 | 24.13719 | 34.67905 | 15.33302 | -0.12799 | 0.498227 | 1 |
| 14 | 77470600 | 77475199 | 71.21136 | 111.9595 | 135.2063 | 56.08792 | 0.048685 | 2.53432 | 1 |
| 14 | 77536600 | 77537599 | 19.52189 | 33.45953 | 44.83318 | 19.66196 | -0.03706 | 0.932633 | 1 |
| 14 | 78137000 | 78138399 | 15.75329 | 27.60917 | 41.43379 | 18.30306 | -0.19513 | 0.655382 | 1 |
| 14 | 79023800 | 79025999 | 5.936851 | 11.60281 | 17.14526 | 5.930226 | 0.135125 | -0.56201 | 1 |
| 14 | 79331000 | 79332799 | 12.18462 | 22.48236 | 29.75984 | 12.57889 | 0.033751 | 0.232503 | 1 |
| 14 | 79663800 | 79666599 | 48.92868 | 78.10842 | 117.7062 | 49.39827 | -0.25038 | 2.114198 | 1 |
| 14 | 79689000 | 79691199 | 29.02224 | 47.89749 | 64.89585 | 28.01432 | -0.05652 | 1.357559 | 1 |
| 14 | 80592400 | 80594399 | 25.24656 | 42.24344 | 52.777 | 22.99309 | 0.06109 | 1.184688 | 1 |
| 14 | 80863600 | 80866399 | 15.85655 | 28.0743 | 30.42802 | 13.45507 | 0.275937 | 0.5085 | 1 |
| 14 | 80972400 | 80980399 | 93.26951 | 144.1271 | 191.1074 | 79.38181 | -0.10535 | 2.860987 | 1 |
| 14 | 81282400 | 81284599 | 31.77069 | 51.70286 | 70.87209 | 30.42261 | -0.08383 | 1.55379 | 1 |
| 14 | 81633600 | 81634399 | 7.886807 | 15.17601 | 26.48373 | 10.31117 | -0.26091 | -0.10201 | 1 |
| 14 | 81796600 | 81799999 | 34.45725 | 56.04296 | 80.33978 | 34.47955 | -0.14891 | 1.574267 | 1 |
| 14 | 81917000 | 81917799 | 15.55613 | 27.70205 | 30.86458 | 13.59356 | 0.240752 | 0.465393 | 1 |
| 14 | 82034200 | 82035799 | 30.95997 | 51.06161 | 56.08974 | 24.20542 | 0.245014 | 1.365114 | 1 |
| 14 | 82284400 | 82285399 | 11.97008 | 21.60595 | 36.32457 | 15.79748 | -0.33712 | 0.369649 | 1 |
| 14 | 82320800 | 82323999 | 48.36527 | 77.35305 | 103.3155 | 43.47346 | -0.0736 | 2.051921 | 1 |
| 14 | 82418400 | 82421199 | 25.9536 | 44.2148 | 45.78675 | 20.07037 | 0.331832 | 1.013079 | 1 |
| 14 | 82601400 | 82602999 | 32.3186 | 52.86111 | 60.76712 | 26.13306 | 0.174539 | 1.475648 | 1 |
| 14 | 82646400 | 82648399 | 18.84933 | 32.96781 | 37.71629 | 16.63212 | 0.194467 | 0.697203 | 1 |
| 14 | 82710800 | 82712799 | 27.37093 | 45.97849 | 48.04254 | 20.98245 | 0.317862 | 1.143206 | 1 |
| 14 | 82778400 | 82780799 | 28.81316 | 47.92507 | 56.13937 | 24.36527 | 0.153697 | 1.261345 | 1 |
| 14 | 82936000 | 82938799 | 26.04817 | 43.8201 | 49.62218 | 21.682 | 0.202268 | 1.127196 | 1 |
| 14 | 83129600 | 83131999 | 24.06865 | 40.4362 | 58.21183 | 25.42492 | -0.14291 | 1.141263 | 1 |
| 14 | 83536000 | 83539399 | 37.06363 | 59.86071 | 88.3528 | 37.70319 | -0.20062 | 1.730991 | 1 |
| 14 | 83768000 | 83769199 | 12.95089 | 23.88496 | 28.11185 | 11.88995 | 0.201481 | 0.23734 | 1 |
| 14 | 84286400 | 84287599 | 10.74303 | 20.18474 | 29.65095 | 12.18462 | -0.08404 | 0.111295 | 1 |
| 14 | 84473400 | 84473999 | 12.94271 | 23.71801 | 23.51692 | 10.03327 | 0.438213 | 0.261031 | 1 |
| 14 | 84646600 | 84647799 | 10.77663 | 20.17843 | 22.70245 | 9.383674 | 0.291565 | 0.116169 | 1 |
| 14 | 84770400 | 84770999 | 10.65054 | 20.15823 | 23.24114 | 9.418676 | 0.280081 | 0.046788 | 1 |

| | | | | | | | | | |
|---|---|---|---|---|---|---|---|---|---|
| 14 | 84987600 | 84989199 | 10.99649 | 20.17499 | 32.34738 | 13.77883 | -0.24942 | 0.257636 | 1 |
| 14 | 85434400 | 85435399 | 13.98337 | 24.48815 | 41.524 | 18.33086 | -0.37037 | 0.6581 | 1 |
| 14 | 85677200 | 85678999 | 15.86728 | 28.20997 | 33.82522 | 14.91806 | 0.133894 | 0.482217 | 1 |
| 14 | 85759800 | 85763399 | 44.03672 | 70.88756 | 82.01108 | 34.84136 | 0.144354 | 1.83775 | 1 |
| 14 | 86118600 | 86120999 | 17.59921 | 30.51892 | 49.01291 | 21.50466 | -0.29405 | 0.823846 | 1 |
| 14 | 86367600 | 86373999 | 57.36448 | 91.50003 | 102.8326 | 43.10649 | 0.170805 | 2.143757 | 1 |
| 14 | 86511600 | 86512399 | 12.2943 | 22.72233 | 26.11306 | 11.01262 | 0.23881 | 0.2161 | 1 |
| 14 | 86829600 | 86830399 | 24.14901 | 40.59618 | 49.08375 | 21.44209 | 0.108118 | 1.127271 | 1 |
| 14 | 87151200 | 87154199 | 37.24915 | 60.06491 | 90.13875 | 38.38664 | -0.22716 | 1.778551 | 1 |
| 14 | 87437800 | 87443399 | 64.78508 | 102.9756 | 113.0456 | 47.16791 | 0.199273 | 2.263415 | 1 |
| 14 | 88033400 | 88034399 | 13.89585 | 24.60582 | 36.11492 | 15.96939 | -0.15909 | 0.506179 | 1 |
| 14 | 88150800 | 88153399 | 20.14683 | 34.40911 | 53.27667 | 23.35828 | -0.24565 | 0.978588 | 1 |
| 14 | 88178000 | 88178799 | 13.04913 | 23.92163 | 26.50172 | 11.30118 | 0.279728 | 0.267548 | 1 |
| 14 | 88412200 | 88412999 | 7.723498 | 14.8776 | 23.64294 | 9.177035 | -0.12253 | -0.12085 | 1 |
| 14 | 89091800 | 89095199 | 39.62098 | 63.90821 | 83.15888 | 35.42753 | -0.02138 | 1.769794 | 1 |
| 14 | 89500200 | 89501199 | 10.00873 | 18.98612 | 25.44262 | 10.26195 | 0.070468 | 0.030072 | 1 |
| 14 | 89731200 | 89733199 | 13.25871 | 23.4984 | 38.6834 | 17.10211 | -0.32339 | 0.496264 | 1 |
| 14 | 89818600 | 89822199 | 88.98905 | 137.8996 | 159.2198 | 66.13542 | 0.0972 | 2.803045 | 1 |
| 14 | 90134000 | 90142999 | 168.6492 | 250.8062 | 299.7804 | 123.5093 | -0.00204 | 3.628693 | 1 |
| 14 | 90320600 | 90322799 | 36.04666 | 59.00956 | 65.77928 | 28.29756 | 0.219579 | 1.474359 | 1 |
| 14 | 90511400 | 90517999 | 115.989 | 177.5303 | 185.0041 | 76.76943 | 0.231314 | 3.051127 | 1 |
| 14 | 90531600 | 90533799 | 38.09866 | 61.36881 | 91.51573 | 38.91602 | -0.21991 | 1.813722 | 1 |
| 14 | 90753600 | 90754599 | 21.24565 | 36.75813 | 38.50817 | 16.89447 | 0.319185 | 0.848078 | 1 |
| 14 | 90919800 | 90922599 | 40.80577 | 66.04055 | 70.96524 | 30.35215 | 0.259429 | 1.67392 | 1 |
| 14 | 91160400 | 91162599 | 28.89613 | 48.26548 | 51.29782 | 22.32399 | 0.293552 | 1.2116 | 1 |
| 14 | 91210200 | 91213199 | 35.65886 | 58.28674 | 66.02894 | 28.38921 | 0.194954 | 1.496775 | 1 |
| 14 | 91362800 | 91364199 | 16.87979 | 29.41307 | 42.04911 | 18.49566 | -0.12602 | 0.746524 | 1 |
| 14 | 91496400 | 91498799 | 36.8742 | 59.99499 | 62.54497 | 26.84595 | 0.309829 | 1.561654 | 1 |
| 14 | 91516800 | 91517999 | 20.79673 | 35.17151 | 50.08121 | 21.91774 | -0.1266 | 1.065182 | 1 |
| 14 | 91651400 | 91653199 | 25.14538 | 42.22891 | 50.14376 | 21.89654 | 0.134049 | 1.141185 | 1 |
| 14 | 91884000 | 91886399 | 32.88278 | 54.62668 | 58.75314 | 25.47798 | 0.276237 | 1.285626 | 1 |
| 14 | 91914000 | 91915199 | 22.20555 | 37.99876 | 39.21797 | 17.19716 | 0.337742 | 0.951581 | 1 |
| 14 | 92170200 | 92171999 | 18.43201 | 32.16822 | 45.15325 | 19.88063 | -0.09951 | 0.73797 | 1 |
| 14 | 92212200 | 92214599 | 26.97399 | 44.56856 | 69.78058 | 30.14701 | -0.2644 | 1.343388 | 1 |
| 14 | 92228800 | 92233799 | 95.18506 | 146.7154 | 186.23 | 77.35687 | -0.04523 | 2.906903 | 1 |
| 14 | 92464600 | 92467599 | 25.73493 | 44.02912 | 44.70642 | 19.60352 | 0.361067 | 0.97447 | 1 |
| 14 | 92620600 | 92622799 | 16.28603 | 28.36237 | 43.6524 | 19.19405 | -0.23205 | 0.754447 | 1 |
| 14 | 92669600 | 92670599 | 13.94146 | 24.40741 | 40.49495 | 17.87204 | -0.33916 | 0.65976 | 1 |
| 14 | 92839200 | 92841999 | 36.10568 | 58.61656 | 75.41406 | 32.32926 | 0.004459 | 1.604813 | 1 |
| 14 | 93166200 | 93168199 | 30.04233 | 48.93578 | 75.72479 | 32.51673 | -0.25731 | 1.541039 | 1 |
| 14 | 93181000 | 93181999 | 15.08662 | 26.25718 | 43.71675 | 19.21481 | -0.34483 | 0.759248 | 1 |
| 14 | 93286800 | 93287799 | 14.56325 | 25.39061 | 39.13057 | 17.21794 | -0.23353 | 0.723994 | 1 |
| 14 | 93431600 | 93433399 | 25.51828 | 42.08839 | 64.9952 | 28.04902 | -0.24492 | 1.357012 | 1 |
| 14 | 93451800 | 93454199 | 36.19612 | 58.81779 | 72.38876 | 31.05352 | 0.069614 | 1.587353 | 1 |
| 14 | 94123000 | 94124399 | 9.667416 | 17.96042 | 29.41811 | 12.31904 | -0.26156 | 0.173312 | 1 |
| 14 | 95049200 | 95051399 | 40.0641 | 64.84037 | 71.8547 | 30.73264 | 0.215135 | 1.674329 | 1 |
| 14 | 95078400 | 95081799 | 25.70126 | 43.46575 | 47.46487 | 20.77267 | 0.254769 | 1.078541 | 1 |

| | | | | | | | | | |
|---|---|---|---|---|---|---|---|---|---|
| 14 | 95218800 | 95219799 | 14.46083 | 25.42668 | 34.22113 | 15.15329 | -0.03866 | 0.58218 | 1 |
| 14 | 95276000 | 95276599 | 8.718239 | 16.86294 | 19.854 | 7.585941 | 0.32777 | -0.17408 | 1 |
| 14 | 95328000 | 95331599 | 61.56798 | 98.13025 | 103.1454 | 43.19211 | 0.266005 | 2.171416 | 1 |
| 14 | 95440400 | 95441399 | 12.87594 | 23.4457 | 27.99529 | 12.04232 | 0.163773 | 0.306913 | 1 |
| 14 | 95450800 | 95452999 | 23.63085 | 40.25796 | 49.55901 | 21.72392 | 0.083335 | 1.009621 | 1 |
| 14 | 95600000 | 95602399 | 22.53224 | 39.23626 | 40.83865 | 17.95569 | 0.329806 | 0.783885 | 1 |
| 14 | 95775000 | 95775799 | 13.6853 | 24.8622 | 29.55221 | 12.74865 | 0.167857 | 0.331839 | 1 |
| 14 | 95798800 | 95801399 | 42.81848 | 69.15019 | 84.4209 | 36.02191 | 0.072402 | 1.738921 | 1 |
| 14 | 96346200 | 96348599 | 25.13735 | 41.8559 | 68.79644 | 29.88003 | -0.33347 | 1.248595 | 1 |
| 14 | 96521200 | 96522999 | 34.95175 | 57.14674 | 60.38518 | 25.96529 | 0.295393 | 1.487359 | 1 |
| 14 | 96527000 | 96528599 | 24.3308 | 41.62076 | 43.67598 | 19.15145 | 0.313672 | 0.968035 | 1 |
| 14 | 96642200 | 96643199 | 14.55671 | 25.58548 | 38.7658 | 17.16613 | -0.2089 | 0.59647 | 1 |
| 14 | 97031200 | 97032799 | 16.57506 | 28.98499 | 39.37612 | 17.3615 | -0.05228 | 0.689939 | 1 |
| 14 | 97382200 | 97383999 | 18.94088 | 32.84622 | 41.08282 | 18.0253 | 0.065154 | 0.819115 | 1 |
| 14 | 97560800 | 97563599 | 50.30946 | 80.52753 | 95.6616 | 40.29892 | 0.096444 | 2.027335 | 1 |
| 14 | 97951600 | 97952999 | 12.14816 | 21.77006 | 35.21174 | 15.42594 | -0.28859 | 0.412798 | 1 |
| 14 | 98474600 | 98475999 | 25.25492 | 42.01212 | 61.44701 | 26.67127 | -0.16561 | 1.253256 | 1 |
| 14 | 100622000 | 100623399 | 20.29802 | 34.62897 | 54.3624 | 23.83218 | -0.26578 | 0.989922 | 1 |
| 14 | 100761400 | 100762599 | 14.93557 | 26.68199 | 33.42737 | 14.68273 | 0.075066 | 0.446159 | 1 |
| 14 | 100930400 | 100933999 | 61.47395 | 97.56252 | 115.9594 | 48.30934 | 0.082683 | 2.305391 | 1 |
| 14 | 101093000 | 101096599 | 43.76227 | 70.22969 | 98.00989 | 41.45053 | -0.1311 | 1.939134 | 1 |
| 14 | 101245400 | 101251199 | 107.8589 | 164.7188 | 222.6281 | 92.33428 | -0.14603 | 3.096918 | 1 |
| 14 | 101815800 | 101821999 | 101.8796 | 156.1985 | 198.4936 | 82.39494 | -0.05274 | 3.01457 | 1 |
| 14 | 102056800 | 102057999 | 10.53252 | 19.46266 | 28.25241 | 11.91781 | -0.0964 | 0.206333 | 1 |
| 14 | 102135800 | 102138199 | 18.22408 | 31.92036 | 34.56792 | 15.2624 | 0.273141 | 0.664069 | 1 |
| 14 | 102368200 | 102369199 | 11.2177 | 20.42238 | 29.68476 | 12.77205 | -0.11827 | 0.299714 | 1 |
| 14 | 102380600 | 102381599 | 12.49743 | 22.15088 | 32.80707 | 14.50393 | -0.17117 | 0.481818 | 1 |
| 14 | 103050200 | 103055399 | 64.69281 | 102.2313 | 132.6423 | 55.09707 | -0.04896 | 2.421314 | 1 |
| 14 | 103872200 | 103877799 | 96.62111 | 149.6367 | 158.8845 | 65.99637 | 0.2173 | 2.818474 | 1 |
| 14 | 104247000 | 104248599 | 14.91078 | 26.13821 | 39.40483 | 17.40995 | -0.20159 | 0.643481 | 1 |
| 14 | 104262400 | 104263199 | 12.28931 | 21.84201 | 34.66983 | 15.29458 | -0.26862 | 0.46328 | 1 |
| 14 | 104309200 | 104309799 | 14.52829 | 25.52956 | 36.75642 | 16.27314 | -0.1356 | 0.596396 | 1 |
| 14 | 104346800 | 104348399 | 17.7193 | 31.12971 | 28.20779 | 12.48597 | 0.529073 | 0.602932 | 1 |
| 14 | 104819400 | 104822399 | 29.68862 | 49.69517 | 51.34796 | 22.37594 | 0.333948 | 1.188573 | 1 |
| 14 | 104948600 | 104953999 | 168.7522 | 250.4242 | 287.1155 | 118.2657 | 0.055165 | 3.66604 | 1 |
| 14 | 105222600 | 105225399 | 28.03516 | 46.94539 | 50.20501 | 21.88314 | 0.284555 | 1.180704 | 1 |
| 14 | 105405800 | 105407399 | 18.96028 | 33.22578 | 36.0344 | 15.91633 | 0.271065 | 0.663695 | 1 |
| 14 | 105449000 | 105452399 | 71.08508 | 112.3456 | 116.0609 | 48.21224 | 0.279485 | 2.420699 | 1 |
| 14 | 105688600 | 105691199 | 21.42924 | 37.16829 | 39.37365 | 17.27549 | 0.303745 | 0.82566 | 1 |
| 14 | 106098200 | 106101399 | 31.8961 | 52.10454 | 73.05385 | 31.40261 | -0.11251 | 1.50035 | 1 |
| 14 | 106207600 | 106214399 | 103.2071 | 158.243 | 205.3993 | 85.26217 | -0.08324 | 3.013774 | 1 |
| 14 | 106215200 | 106217399 | 28.04756 | 46.66642 | 55.07898 | 23.91028 | 0.142825 | 1.252505 | 1 |
| 14 | 106240200 | 106243599 | 44.77649 | 71.77544 | 98.07088 | 41.40449 | -0.10258 | 1.976601 | 1 |
| 14 | 106343000 | 106345599 | 57.63228 | 91.01972 | 141.4251 | 58.72763 | -0.30934 | 2.433844 | 1 |
| 14 | 106544200 | 106544999 | 19.14125 | 32.70365 | 52.19133 | 22.88311 | -0.28904 | 0.969175 | 1 |
| 14 | 106596800 | 106598399 | 21.30493 | 37.10088 | 38.19725 | 16.79484 | 0.345342 | 0.771188 | 1 |
| 14 | 106684400 | 106686999 | 27.76512 | 46.46663 | 53.97593 | 23.5183 | 0.165684 | 1.189678 | 1 |

| | | | | | | | | | |
|---|---|---|---|---|---|---|---|---|---|
| 14 | 107201200 | 107202399 | 17.79422 | 30.26633 | 51.59668 | 22.61422 | -0.38465 | 1.006589 | 1 |
| 14 | 107402800 | 107405999 | 23.46855 | 39.1189 | 62.49235 | 27.16028 | -0.29245 | 1.226279 | 1 |
| 14 | 107484800 | 107486599 | 17.81904 | 31.14764 | 41.90435 | 18.46987 | -0.03846 | 0.705591 | 1 |
| 14 | 107994000 | 107994999 | 24.45863 | 41.39668 | 44.38396 | 19.42895 | 0.281287 | 1.065701 | 1 |
| 14 | 109193400 | 109194199 | 11.61099 | 20.94416 | 30.70674 | 13.36518 | -0.14126 | 0.359601 | 1 |
| 14 | 109248400 | 109252799 | 51.02031 | 81.40348 | 111.1039 | 46.59183 | -0.10858 | 2.130814 | 1 |
| 14 | 109292000 | 109292999 | 10.70844 | 20.10295 | 22.14479 | 9.112962 | 0.326678 | 0.096554 | 1 |
| 14 | 109451800 | 109453999 | 26.39175 | 44.29201 | 50.16984 | 21.89856 | 0.201846 | 1.151521 | 1 |
| 14 | 109579000 | 109586399 | 69.98419 | 110.2079 | 155.1972 | 64.3943 | -0.17095 | 2.505929 | 1 |
| 14 | 109674600 | 109676199 | 15.09189 | 27.02487 | 31.38794 | 13.75955 | 0.18573 | 0.429081 | 1 |
| 14 | 110092200 | 110094599 | 32.02648 | 52.87162 | 56.24173 | 24.28358 | 0.291531 | 1.355075 | 1 |
| 14 | 110290000 | 110295999 | 116.9739 | 178.1551 | 208.103 | 86.25183 | 0.061149 | 3.152788 | 1 |
| 14 | 110315800 | 110318599 | 28.51709 | 47.13651 | 59.2861 | 25.61999 | 0.050718 | 1.334765 | 1 |
| 14 | 110595400 | 110596399 | 13.67614 | 24.92182 | 27.92374 | 12.00005 | 0.255863 | 0.308574 | 1 |
| 14 | 110598200 | 110600599 | 54.42123 | 86.62 | 107.8549 | 45.07825 | 0.019894 | 2.210428 | 1 |
| 14 | 110602200 | 110602999 | 19.38464 | 32.88054 | 51.62136 | 22.61206 | -0.26673 | 1.036806 | 1 |
| 14 | 110790200 | 110790999 | 8.933082 | 16.8623 | 25.91646 | 10.56265 | -0.13903 | 0.054591 | 1 |
| 14 | 111141600 | 111143399 | 11.87814 | 21.11353 | 35.26029 | 15.55362 | -0.34119 | 0.462983 | 1 |
| 14 | 111207800 | 111209799 | 36.58801 | 59.0331 | 89.66684 | 38.21429 | -0.24358 | 1.760099 | 1 |
| 14 | 111243000 | 111245999 | 31.46985 | 52.22177 | 61.87259 | 26.81028 | 0.13703 | 1.299806 | 1 |
| 14 | 111610800 | 111613399 | 26.97623 | 45.74115 | 49.04392 | 21.48072 | 0.281149 | 1.061665 | 1 |
| 14 | 111616800 | 111619599 | 41.81057 | 67.13078 | 102.1456 | 43.23211 | -0.25476 | 1.922385 | 1 |
| 14 | 112017800 | 112019799 | 10.90885 | 20.5477 | 23.67036 | 9.685816 | 0.269086 | 0.081555 | 1 |
| 14 | 112109400 | 112114999 | 77.88729 | 121.984 | 140.4565 | 58.26194 | 0.113049 | 2.611463 | 1 |
| 14 | 112185800 | 112187999 | 24.91635 | 41.41877 | 64.81881 | 28.12068 | -0.26304 | 1.262173 | 1 |
| 14 | 112296200 | 112298999 | 22.34973 | 38.32524 | 47.39591 | 20.78669 | 0.078103 | 0.941246 | 1 |
| 14 | 112500000 | 112502399 | 50.44606 | 80.90891 | 91.37128 | 38.60885 | 0.172721 | 1.963561 | 1 |
| 14 | 112537000 | 112538799 | 15.06591 | 26.58603 | 38.68537 | 17.11778 | -0.14865 | 0.556144 | 1 |
| 14 | 112632800 | 112634199 | 20.76446 | 35.93982 | 38.53329 | 16.9057 | 0.286034 | 0.842503 | 1 |
| 14 | 112870400 | 112872999 | 28.76354 | 48.14147 | 49.67382 | 21.64491 | 0.336087 | 1.18902 | 1 |
| 14 | 112885400 | 112885999 | 10.9873 | 19.83917 | 30.75245 | 13.36816 | -0.21991 | 0.349115 | 1 |
| 14 | 113529800 | 113531599 | 19.72803 | 34.42926 | 37.715 | 16.60538 | 0.256581 | 0.732997 | 1 |
| 14 | 113542600 | 113544199 | 22.27645 | 37.57447 | 56.51625 | 24.71415 | -0.20566 | 1.099422 | 1 |
| 14 | 113550200 | 113554199 | 51.77294 | 82.56088 | 111.9676 | 46.91981 | -0.10023 | 2.147408 | 1 |
| 14 | 113610400 | 113612599 | 18.22937 | 30.97331 | 52.67064 | 23.08294 | -0.38148 | 1.017044 | 1 |
| 14 | 113641200 | 113642799 | 22.56108 | 38.6869 | 46.33117 | 20.31969 | 0.124304 | 0.942075 | 1 |
| 14 | 113773000 | 113780599 | 91.35869 | 142.0151 | 173.0119 | 71.83231 | 0.023346 | 2.752626 | 1 |
| 14 | 113821600 | 113824599 | 28.77254 | 47.80745 | 60.62177 | 26.29169 | 0.039498 | 1.275321 | 1 |
| 14 | 113866800 | 113869999 | 17.21915 | 30.47818 | 31.15104 | 13.77556 | 0.359786 | 0.520023 | 1 |
| 14 | 114211800 | 114212999 | 13.51472 | 24.53218 | 27.59327 | 11.9158 | 0.246225 | 0.331474 | 1 |
| 14 | 114335800 | 114340399 | 133.7077 | 200.7461 | 247.098 | 101.9808 | -0.03191 | 3.445566 | 1 |
| 14 | 114371400 | 114372599 | 15.50081 | 27.2196 | 37.74104 | 16.69926 | -0.08156 | 0.615198 | 1 |
| 14 | 114387400 | 114390799 | 54.05562 | 86.65154 | 98.91973 | 41.76425 | 0.156222 | 1.986629 | 1 |
| 14 | 114445000 | 114449199 | 49.73038 | 79.81983 | 85.64167 | 36.22838 | 0.247706 | 1.937306 | 1 |
| 14 | 114661200 | 114662799 | 15.58352 | 27.51548 | 34.99786 | 15.4843 | 0.044692 | 0.54477 | 1 |
| 14 | 114748400 | 114751399 | 37.75022 | 61.25998 | 71.42603 | 30.60988 | 0.145581 | 1.615382 | 1 |
| 14 | 114940800 | 114942799 | 20.63156 | 35.89388 | 38.37042 | 16.86084 | 0.291368 | 0.783649 | 1 |

| | | | | | | | | | |
|---|---|---|---|---|---|---|---|---|---|
| 14 | 115140000 | 115142599 | 30.81139 | 50.68515 | 66.64482 | 28.72896 | -0.01524 | 1.40272 | 1 |
| 14 | 115540800 | 115542799 | 19.68407 | 33.83482 | 49.54518 | 21.73235 | -0.16406 | 0.912209 | 1 |
| 14 | 116202200 | 116202799 | 10.02578 | 18.51748 | 30.40493 | 12.83411 | -0.27331 | 0.212356 | 1 |
| 14 | 116203600 | 116205399 | 22.0092 | 37.80921 | 46.87061 | 20.55882 | 0.075186 | 0.925655 | 1 |
| 14 | 116310400 | 116315399 | 77.92338 | 121.0899 | 178.9741 | 74.31145 | -0.25555 | 2.753316 | 1 |
| 14 | 116702200 | 116703599 | 17.72208 | 30.76824 | 43.18207 | 18.95048 | -0.09998 | 0.805021 | 1 |
| 14 | 116750400 | 116752999 | 33.94731 | 55.00245 | 83.25743 | 35.63907 | -0.2322 | 1.646761 | 1 |
| 14 | 116878400 | 116879799 | 7.586878 | 14.48346 | 24.53144 | 9.7627 | -0.24745 | -0.03911 | 1 |
| 14 | 116964800 | 116969399 | 76.29263 | 119.9926 | 142.8814 | 59.27464 | 0.069363 | 2.531218 | 1 |
| 14 | 117025800 | 117026599 | 20.46056 | 35.54973 | 33.44176 | 14.68296 | 0.47482 | 0.792971 | 1 |
| 14 | 117143200 | 117145599 | 16.21779 | 28.48204 | 41.84029 | 18.51493 | -0.16459 | 0.624578 | 1 |
| 14 | 117315000 | 117323799 | 167.8558 | 248.2603 | 337.0568 | 138.8089 | -0.19307 | 3.724695 | 1 |
| 14 | 117415400 | 117417799 | 21.95143 | 37.51867 | 44.81095 | 19.64779 | 0.127567 | 0.968283 | 1 |
| 14 | 117459000 | 117460599 | 10.90029 | 20.15703 | 29.83883 | 12.57445 | -0.12337 | 0.20983 | 1 |
| 14 | 117493800 | 117496399 | 48.08029 | 76.92351 | 105.7472 | 44.51818 | -0.11458 | 2.042581 | 1 |
| 14 | 117520000 | 117523599 | 8.091555 | 15.40754 | 28.33923 | 11.33909 | -0.37237 | -0.0051 | 1 |
| 14 | 117524400 | 117526599 | 31.22088 | 52.35015 | 52.46536 | 22.88774 | 0.377804 | 1.177031 | 1 |
| 14 | 117700800 | 117702999 | 22.04333 | 37.72899 | 59.52825 | 26.10369 | -0.2725 | 0.971254 | 1 |
| 14 | 118098800 | 118104399 | 82.92455 | 128.6014 | 175.6923 | 72.97083 | -0.14442 | 2.789968 | 1 |
| 14 | 118113600 | 118117399 | 28.10953 | 46.67023 | 61.70175 | 26.74601 | -0.02062 | 1.282486 | 1 |
| 14 | 118265000 | 118272999 | 120.9818 | 183.8884 | 241.5842 | 100.0683 | -0.11063 | 3.197693 | 1 |
| 14 | 118515200 | 118520599 | 39.3675 | 63.90191 | 68.22435 | 29.24368 | 0.272715 | 1.611405 | 1 |
| 14 | 118708800 | 118713199 | 38.38084 | 61.79354 | 95.2955 | 40.49501 | -0.26916 | 1.829138 | 1 |
| 14 | 119612400 | 119616199 | 54.83995 | 87.71425 | 104.2829 | 43.88471 | 0.094079 | 2.056313 | 1 |
| 14 | 119678600 | 119685199 | 76.11909 | 120.0041 | 130.9204 | 54.32935 | 0.198088 | 2.481172 | 1 |
| 14 | 119934400 | 119935399 | 15.21937 | 27.0771 | 34.38787 | 15.15723 | 0.051974 | 0.474472 | 1 |
| 14 | 120170800 | 120173199 | 38.85069 | 63.10649 | 73.82128 | 31.65883 | 0.142161 | 1.601754 | 1 |
| 14 | 120236200 | 120237999 | 19.60159 | 34.20548 | 42.42649 | 18.67871 | 0.078065 | 0.739961 | 1 |
| 14 | 120361800 | 120364399 | 24.58207 | 41.03697 | 59.26041 | 25.78019 | -0.14725 | 1.21132 | 1 |
| 14 | 120416000 | 120417399 | 32.2326 | 52.136 | 82.88952 | 35.43738 | -0.30505 | 1.679659 | 1 |
| 14 | 120626200 | 120628599 | 23.39968 | 38.86958 | 62.81592 | 27.2382 | -0.30917 | 1.26757 | 1 |
| 14 | 120673400 | 120681399 | 99.51406 | 152.3261 | 219.0968 | 90.9182 | -0.2331 | 3.047299 | 1 |
| 14 | 120689800 | 120698599 | 81.62941 | 128.0839 | 140.0565 | 58.08491 | 0.189556 | 2.581029 | 1 |
| 14 | 121241200 | 121249599 | 54.94125 | 87.70586 | 108.4391 | 45.50848 | 0.034586 | 2.115517 | 1 |
| 14 | 121746400 | 121751799 | 73.96821 | 117.0486 | 123.1039 | 51.17038 | 0.255112 | 2.393762 | 1 |
| 14 | 122332600 | 122334199 | 28.30051 | 46.9212 | 59.96032 | 25.96588 | 0.028152 | 1.298576 | 1 |
| 14 | 123011600 | 123014999 | 56.63922 | 90.22977 | 117.155 | 49.02501 | -0.03909 | 2.187554 | 1 |
| 14 | 123320800 | 123322599 | 24.72437 | 41.34782 | 62.11446 | 27.05206 | -0.20414 | 1.19456 | 1 |
| 14 | 123583600 | 123585799 | 19.92263 | 34.35336 | 52.82231 | 23.17206 | -0.23331 | 0.889876 | 1 |
| 14 | 123717600 | 123720399 | 39.44877 | 63.73475 | 76.53651 | 32.67522 | 0.096905 | 1.719387 | 1 |
| 14 | 123781600 | 123783999 | 35.16118 | 57.28439 | 66.85282 | 28.70898 | 0.148825 | 1.54279 | 1 |
| 14 | 123897200 | 123898999 | 7.08452 | 13.78233 | 17.37502 | 6.413322 | 0.276992 | -0.31933 | 1 |
| 14 | 123932600 | 123934599 | 25.05134 | 41.36126 | 64.94311 | 28.04279 | -0.26844 | 1.34418 | 1 |
| 14 | 124047400 | 124049199 | 11.53095 | 20.72583 | 34.25127 | 14.9674 | -0.31673 | 0.38873 | 1 |
| 14 | 124057800 | 124059199 | 16.93642 | 29.54196 | 42.55631 | 18.7308 | -0.13675 | 0.734204 | 1 |
| 14 | 124067800 | 124069999 | 23.48654 | 40.38164 | 46.29336 | 20.30615 | 0.187712 | 0.921162 | 1 |
| 14 | 124234600 | 124237599 | 16.76225 | 29.50388 | 33.4053 | 14.78897 | 0.209855 | 0.582011 | 1 |

| | | | | | | | | | |
|---|---|---|---|---|---|---|---|---|---|
| 14 | 124267000 | 124269799 | 39.58558 | 64.31748 | 75.96406 | 32.58424 | 0.128652 | 1.597129 | 1 |
| 14 | 124571200 | 124571799 | 14.5784 | 25.55931 | 35.96272 | 15.89112 | -0.10236 | 0.633987 | 1 |
| 14 | 124628800 | 124633599 | 74.86589 | 118.3727 | 120.3936 | 50.02314 | 0.302465 | 2.41378 | 1 |
| 14 | 124661600 | 124663199 | 32.33872 | 53.08762 | 59.24539 | 25.51689 | 0.219619 | 1.429026 | 1 |
| 14 | 124727800 | 124729199 | 16.76325 | 29.5775 | 33.12913 | 14.65958 | 0.227149 | 0.560368 | 1 |
| 14 | 124972000 | 124975399 | 43.97431 | 70.96136 | 93.09905 | 39.68483 | -0.03253 | 1.769473 | 1 |
| 14 | 125015200 | 125018199 | 36.64063 | 60.09438 | 63.50731 | 27.34184 | 0.297473 | 1.453363 | 1 |
| 15 | 3114600 | 3116999 | 25.11316 | 42.82247 | 44.73289 | 19.60987 | 0.319507 | 1.000282 | 1 |
| 15 | 3399400 | 3403399 | 77.78201 | 122.9201 | 128.6648 | 53.44589 | 0.260447 | 2.428598 | 1 |
| 15 | 3576000 | 3580999 | 77.25947 | 120.6294 | 165.1737 | 68.53641 | -0.14008 | 2.669622 | 1 |
| 15 | 3834800 | 3837799 | 37.41127 | 60.33594 | 82.41359 | 35.104 | -0.09138 | 1.771493 | 1 |
| 15 | 3944200 | 3944999 | 14.37209 | 25.99994 | 27.89957 | 12.09941 | 0.309498 | 0.356008 | 1 |
| 15 | 3977400 | 3983799 | 63.39127 | 100.8863 | 113.2831 | 47.33987 | 0.16859 | 2.224732 | 1 |
| 15 | 4315400 | 4316599 | 17.27949 | 30.26646 | 36.77405 | 16.23672 | 0.108118 | 0.664806 | 1 |
| 15 | 4636800 | 4637999 | 17.95479 | 31.17445 | 42.62616 | 18.70734 | -0.06263 | 0.802635 | 1 |
| 15 | 4700600 | 4702999 | 25.12468 | 41.80895 | 62.52204 | 27.14393 | -0.19747 | 1.250798 | 1 |
| 15 | 4710200 | 4716399 | 115.0391 | 176.4413 | 183.6952 | 76.25607 | 0.234994 | 3.010179 | 1 |
| 15 | 4804600 | 4806199 | 18.37748 | 31.49658 | 47.44725 | 20.8083 | -0.20522 | 0.934443 | 1 |
| 15 | 4863000 | 4867399 | 42.75882 | 68.9573 | 77.00455 | 32.79816 | 0.198503 | 1.778615 | 1 |
| 15 | 4891400 | 4891999 | 8.3959 | 16.22878 | 18.80591 | 7.207615 | 0.34643 | -0.17071 | 1 |
| 15 | 4894400 | 4895599 | 10.97922 | 19.50843 | 34.88583 | 15.39327 | -0.43928 | 0.460332 | 1 |
| 15 | 5053000 | 5054799 | 13.17415 | 23.49535 | 35.58158 | 15.65133 | -0.19851 | 0.448089 | 1 |
| 15 | 5094200 | 5096799 | 26.03592 | 44.06708 | 49.59329 | 21.70944 | 0.211473 | 1.074794 | 1 |
| 15 | 5121000 | 5125799 | 8.393152 | 15.9434 | 27.71022 | 11.144 | -0.29779 | 0.012678 | 1 |
| 15 | 5295800 | 5296999 | 14.94356 | 26.29923 | 32.25295 | 14.27919 | 0.095399 | 0.572501 | 1 |
| 15 | 5404200 | 5415799 | 158.6412 | 238.1421 | 248.2618 | 102.4566 | 0.207382 | 3.453231 | 1 |
| 15 | 5483800 | 5484799 | 11.58541 | 21.08406 | 33.28917 | 14.32947 | -0.2374 | 0.312769 | 1 |
| 15 | 5714200 | 5716399 | 36.44432 | 59.31903 | 68.70221 | 29.49305 | 0.158799 | 1.559256 | 1 |
| 15 | 5858800 | 5859799 | 9.205112 | 17.23788 | 27.43902 | 11.33922 | -0.20502 | 0.115334 | 1 |
| 15 | 6050800 | 6052599 | 20.04285 | 34.93239 | 37.24882 | 16.38648 | 0.295124 | 0.752538 | 1 |
| 15 | 6164400 | 6166399 | 31.59671 | 53.53846 | 49.0496 | 21.47868 | 0.5073 | 1.085663 | 1 |
| 15 | 6221800 | 6223599 | 28.37748 | 47.32002 | 53.45989 | 23.24037 | 0.205746 | 1.228754 | 1 |
| 15 | 6228800 | 6230799 | 24.38835 | 41.27787 | 50.63926 | 22.16721 | 0.087574 | 1.072527 | 1 |
| 15 | 6255200 | 6256399 | 17.63597 | 30.86312 | 34.73473 | 15.32467 | 0.217992 | 0.676242 | 1 |
| 15 | 6464600 | 6465599 | 11.41851 | 20.68134 | 31.18853 | 13.50564 | -0.17706 | 0.334619 | 1 |
| 15 | 6558800 | 6560399 | 14.26753 | 25.04137 | 37.38584 | 16.53508 | -0.18754 | 0.616792 | 1 |
| 15 | 6561200 | 6563599 | 22.51912 | 37.67788 | 55.86946 | 24.33958 | -0.18522 | 1.179364 | 1 |
| 15 | 6608600 | 6610799 | 24.9859 | 41.93261 | 51.20416 | 22.35002 | 0.093842 | 1.148724 | 1 |
| 15 | 6637400 | 6638199 | 13.22141 | 23.19314 | 39.99896 | 17.68298 | -0.39443 | 0.625212 | 1 |
| 15 | 6722000 | 6724999 | 21.55139 | 37.27926 | 42.05397 | 18.44998 | 0.212698 | 0.854968 | 1 |
| 15 | 7158400 | 7161199 | 9.59197 | 17.49287 | 31.35323 | 13.46079 | -0.4163 | 0.282377 | 1 |
| 15 | 7484600 | 7485599 | 9.890853 | 18.97393 | 23.48444 | 9.224345 | 0.21824 | -0.06342 | 1 |
| 15 | 7939000 | 7941999 | 43.84899 | 70.76388 | 75.19939 | 32.05761 | 0.270989 | 1.758802 | 1 |
| 15 | 7998800 | 7999999 | 4.773504 | 9.329711 | 20.20934 | 6.980103 | -0.40821 | -0.57327 | 1 |
| 15 | 8004000 | 8005599 | 19.02388 | 32.61363 | 45.86344 | 20.11416 | -0.1065 | 0.928875 | 1 |
| 15 | 8501600 | 8503199 | 23.03487 | 39.5133 | 43.16341 | 18.93093 | 0.25657 | 0.937476 | 1 |
| 15 | 8566000 | 8566799 | 9.47829 | 17.83586 | 26.35861 | 10.80353 | -0.09 | 0.07906 | 1 |

| | | | | | | | | | |
|---|---|---|---|---|---|---|---|---|---|
| 15 | 8942400 | 8944999 | 17.39946 | 30.26748 | 46.83111 | 20.57723 | -0.23993 | 0.779584 | 1 |
| 15 | 9223200 | 9227799 | 30.94318 | 51.43464 | 54.67312 | 23.7183 | 0.293249 | 1.271146 | 1 |
| 15 | 9328800 | 9333599 | 76.77808 | 120.8159 | 140.8007 | 58.41522 | 0.100902 | 2.521487 | 1 |
| 15 | 9442600 | 9443799 | 16.78744 | 29.22525 | 40.08494 | 17.62151 | -0.06645 | 0.758347 | 1 |
| 15 | 9874600 | 9875399 | 13.16097 | 24.12988 | 25.60661 | 10.91749 | 0.341922 | 0.267549 | 1 |
| 15 | 9902600 | 9903399 | 11.26752 | 20.72699 | 29.78324 | 12.64102 | -0.08803 | 0.243153 | 1 |
| 15 | 10063000 | 10063799 | 8.870117 | 16.64915 | 28.52975 | 11.74373 | -0.30631 | 0.098921 | 1 |
| 15 | 10349800 | 10351799 | 20.48438 | 34.79375 | 55.94919 | 24.51803 | -0.30126 | 1.032287 | 1 |
| 15 | 10363400 | 10364399 | 16.46843 | 28.97647 | 35.09578 | 15.53847 | 0.113141 | 0.590024 | 1 |
| 15 | 10368200 | 10368999 | 10.66084 | 19.68709 | 29.2648 | 12.35569 | -0.13154 | 0.212293 | 1 |
| 15 | 10425400 | 10427799 | 15.84399 | 27.53538 | 43.21186 | 18.97559 | -0.26018 | 0.782139 | 1 |
| 15 | 10603800 | 10607799 | 49.06294 | 78.34219 | 114.2597 | 47.9687 | -0.20288 | 2.105409 | 1 |
| 15 | 10785600 | 10788999 | 44.10701 | 71.37288 | 79.78992 | 34.12283 | 0.202272 | 1.686803 | 1 |
| 15 | 12068200 | 12075199 | 62.01768 | 98.44829 | 136.5326 | 56.89346 | -0.13928 | 2.303364 | 1 |
| 15 | 12092800 | 12100399 | 78.4052 | 122.3663 | 163.4917 | 67.84144 | -0.1052 | 2.677161 | 1 |
| 15 | 12993200 | 12994799 | 28.81629 | 47.39363 | 69.62208 | 30.00982 | -0.17476 | 1.404852 | 1 |
| 15 | 13004600 | 13007599 | 33.82507 | 55.10078 | 74.30145 | 31.90624 | -0.0593 | 1.544357 | 1 |
| 15 | 13931400 | 13934199 | 22.9726 | 39.6646 | 47.92875 | 21.0262 | 0.113362 | 0.885277 | 1 |
| 15 | 14479800 | 14489599 | 162.4583 | 242.74 | 283.7307 | 116.9788 | 0.03637 | 3.542851 | 1 |
| 15 | 14693000 | 14698599 | 93.70087 | 145.3402 | 165.0679 | 68.5563 | 0.12201 | 2.789012 | 1 |
| 15 | 14726200 | 14728799 | 43.63312 | 70.43215 | 75.36687 | 32.13881 | 0.261395 | 1.750744 | 1 |
| 15 | 14840800 | 14854999 | 344.8624 | 486.3716 | 485.5462 | 201.0605 | 0.183061 | 4.453783 | 1 |
| 15 | 14871800 | 14879399 | 162.0575 | 240.816 | 299.8274 | 123.5189 | -0.06191 | 3.641897 | 1 |
| 15 | 14980200 | 14982399 | 20.12893 | 35.35688 | 35.78555 | 15.83789 | 0.370173 | 0.632374 | 1 |
| 15 | 15047800 | 15061799 | 408.9829 | 561.7913 | 661.0269 | 274.8651 | -0.09234 | 4.853729 | 1 |
| 15 | 15077000 | 15085199 | 22.72855 | 38.99188 | 40.13301 | 17.60198 | 0.342192 | 0.932482 | 1 |
| 15 | 15265400 | 15266199 | 10.66557 | 19.94533 | 25.75042 | 10.66623 | 0.091838 | 0.130782 | 1 |
| 15 | 15441600 | 15443799 | 33.48612 | 54.78817 | 65.93352 | 28.35817 | 0.108937 | 1.477184 | 1 |
| 15 | 15529600 | 15531999 | 32.28574 | 53.27793 | 56.72577 | 24.48569 | 0.29022 | 1.362455 | 1 |
| 15 | 15784000 | 15790199 | 100.6257 | 155.5112 | 162.5288 | 67.51083 | 0.237789 | 2.857956 | 1 |
| 15 | 15938000 | 15938799 | 11.21385 | 20.8326 | 28.82737 | 12.07232 | -0.02014 | 0.184721 | 1 |
| 15 | 16180200 | 16182999 | 35.66373 | 57.98508 | 75.43443 | 32.37021 | -0.00959 | 1.576995 | 1 |
| 15 | 16350400 | 16353399 | 18.40942 | 32.23369 | 39.24153 | 17.32075 | 0.105175 | 0.679976 | 1 |
| 15 | 16361800 | 16363399 | 8.563612 | 16.044 | 29.03713 | 11.99025 | -0.38832 | 0.10871 | 1 |
| 15 | 16391800 | 16396599 | 40.64304 | 66.37575 | 67.90316 | 29.18575 | 0.34074 | 1.518988 | 1 |
| 15 | 16906800 | 16907999 | 15.66499 | 27.67316 | 32.91881 | 14.56299 | 0.141009 | 0.535102 | 1 |
| 15 | 16908800 | 16912999 | 45.58322 | 73.51152 | 85.27503 | 36.32032 | 0.143659 | 1.786799 | 1 |
| 15 | 16914200 | 16915599 | 15.78445 | 28.02294 | 29.79956 | 13.15834 | 0.305012 | 0.482529 | 1 |
| 15 | 17010400 | 17013199 | 34.08877 | 55.9432 | 66.276 | 28.54122 | 0.133392 | 1.439864 | 1 |
| 15 | 17038800 | 17039799 | 12.64507 | 22.55077 | 32.72495 | 14.39531 | -0.13685 | 0.44287 | 1 |
| 15 | 17093200 | 17096599 | 43.66178 | 70.66817 | 73.61062 | 31.48538 | 0.304343 | 1.679924 | 1 |
| 15 | 17162000 | 17162599 | 8.462975 | 16.2733 | 20.89778 | 8.152703 | 0.175703 | -0.10405 | 1 |
| 15 | 17168600 | 17169799 | 25.03538 | 41.31111 | 66.66597 | 28.7757 | -0.30799 | 1.350964 | 1 |
| 15 | 17266200 | 17268199 | 14.53893 | 25.52111 | 39.82838 | 17.61735 | -0.2514 | 0.618286 | 1 |
| 15 | 17918800 | 17920999 | 33.87674 | 55.57949 | 63.76376 | 27.45605 | 0.179517 | 1.443717 | 1 |
| 15 | 18125400 | 18127999 | 42.95645 | 68.97892 | 93.75265 | 39.68715 | -0.09196 | 1.918003 | 1 |
| 15 | 18163000 | 18165399 | 28.30637 | 46.98734 | 58.30366 | 25.26942 | 0.070629 | 1.283961 | 1 |

| | | | | | | | | | |
|---|---|---|---|---|---|---|---|---|---|
| 15 | 18313800 | 18315199 | 18.19327 | 31.69757 | 37.95592 | 16.69383 | 0.128639 | 0.750736 | 1 |
| 15 | 18524200 | 18525799 | 38.10756 | 61.77616 | 67.78878 | 29.02906 | 0.231537 | 1.635799 | 1 |
| 15 | 18526600 | 18527799 | 14.8261 | 26.68625 | 28.81272 | 12.57454 | 0.294724 | 0.389305 | 1 |
| 15 | 18682000 | 18684399 | 15.42998 | 27.19788 | 39.13215 | 17.31936 | -0.13333 | 0.568701 | 1 |
| 15 | 18731000 | 18731599 | 12.68369 | 22.92592 | 31.09411 | 13.49924 | -0.02738 | 0.355766 | 1 |
| 15 | 19049200 | 19051399 | 31.6816 | 51.89509 | 62.99649 | 27.10734 | 0.097334 | 1.456218 | 1 |
| 15 | 19402600 | 19405799 | 44.89123 | 72.11454 | 85.9583 | 36.40952 | 0.097721 | 1.905483 | 1 |
| 15 | 19519600 | 19521399 | 21.74084 | 37.55164 | 40.02867 | 17.56065 | 0.293718 | 0.86711 | 1 |
| 15 | 19580000 | 19580599 | 10.38923 | 19.60813 | 21.17254 | 8.631823 | 0.366281 | 0.056999 | 1 |
| 15 | 19639600 | 19640999 | 14.10686 | 24.9467 | 36.67476 | 16.2212 | -0.16207 | 0.524082 | 1 |
| 15 | 19646600 | 19651199 | 66.0681 | 104.7401 | 123.7717 | 51.50807 | 0.089489 | 2.342042 | 1 |
| 15 | 19678000 | 19680599 | 48.66532 | 77.44943 | 115.9181 | 48.44038 | -0.24528 | 2.212645 | 1 |
| 15 | 19892000 | 19894799 | 33.73301 | 55.977 | 57.5181 | 24.92332 | 0.341887 | 1.30263 | 1 |
| 15 | 19967800 | 19968999 | 13.48303 | 24.47298 | 28.4065 | 12.26804 | 0.20055 | 0.329985 | 1 |
| 15 | 20159000 | 20161599 | 39.89909 | 64.51007 | 72.81519 | 31.11622 | 0.187274 | 1.696464 | 1 |
| 15 | 20329800 | 20330799 | 8.876378 | 16.55528 | 28.43493 | 11.83143 | -0.32218 | 0.139136 | 1 |
| 15 | 20480600 | 20483399 | 41.75141 | 67.51009 | 72.40022 | 30.94192 | 0.261066 | 1.698446 | 1 |
| 15 | 20498200 | 20499799 | 20.6677 | 35.69177 | 41.56852 | 18.23609 | 0.166339 | 0.863995 | 1 |
| 15 | 20827400 | 20829599 | 16.38525 | 28.49782 | 46.84394 | 20.58043 | -0.32669 | 0.779973 | 1 |
| 15 | 21443600 | 21444799 | 22.60396 | 38.36411 | 47.98994 | 21.02517 | 0.059827 | 1.040138 | 1 |
| 15 | 21750400 | 21752599 | 19.96935 | 34.7759 | 38.35937 | 16.86638 | 0.24645 | 0.766412 | 1 |
| 15 | 21796000 | 21797799 | 24.92084 | 42.42714 | 45.80383 | 20.07678 | 0.271805 | 1.014379 | 1 |
| 15 | 21977000 | 21978799 | 17.0016 | 29.99748 | 33.37115 | 14.76673 | 0.23655 | 0.558339 | 1 |
| 15 | 21992600 | 21994199 | 25.67301 | 43.05182 | 52.81501 | 23.04168 | 0.087194 | 1.161359 | 1 |
| 15 | 22113800 | 22114799 | 11.12819 | 21.00302 | 23.18666 | 9.452765 | 0.334709 | 0.070383 | 1 |
| 15 | 22272600 | 22275199 | 25.82009 | 43.37796 | 52.28242 | 22.83504 | 0.112569 | 1.141893 | 1 |
| 15 | 22362600 | 22371999 | 177.8203 | 264.1228 | 301.8771 | 124.3577 | 0.06088 | 3.651837 | 1 |
| 15 | 22487800 | 22489799 | 23.57705 | 39.57376 | 54.90166 | 23.96605 | -0.08966 | 1.146466 | 1 |
| 15 | 22621600 | 22623999 | 29.55948 | 49.62126 | 54.04094 | 23.59215 | 0.2583 | 1.162977 | 1 |
| 15 | 22699000 | 22703399 | 75.03962 | 118.1957 | 126.7875 | 52.60812 | 0.221441 | 2.49828 | 1 |
| 15 | 22704200 | 22705199 | 17.41371 | 30.32907 | 36.43998 | 16.02368 | 0.123622 | 0.746932 | 1 |
| 15 | 22713800 | 22716399 | 26.54925 | 45.1375 | 46.31767 | 20.29976 | 0.344478 | 1.033302 | 1 |
| 15 | 22751200 | 22753399 | 19.96224 | 34.1518 | 50.06772 | 21.95437 | -0.16673 | 0.95893 | 1 |
| 15 | 23467400 | 23468999 | 16.89908 | 29.73321 | 34.02615 | 15.065 | 0.194507 | 0.590334 | 1 |
| 15 | 23738000 | 23744799 | 127.9087 | 194.6445 | 215.0474 | 89.10747 | 0.140434 | 3.173287 | 1 |
| 15 | 23800600 | 23803399 | 26.90653 | 45.53517 | 50.13704 | 21.94667 | 0.242922 | 1.080118 | 1 |
| 15 | 23821600 | 23829399 | 98.56186 | 152.1634 | 176.6225 | 73.36545 | 0.085427 | 2.878228 | 1 |
| 15 | 23952600 | 23953799 | 17.55268 | 31.33005 | 30.65799 | 13.4762 | 0.42889 | 0.464187 | 1 |
| 15 | 23956200 | 23965799 | 145.7072 | 219.4725 | 260.9797 | 107.8035 | 0.022 | 3.385463 | 1 |
| 15 | 24064200 | 24066799 | 21.26347 | 36.13777 | 54.35808 | 23.82295 | -0.20523 | 1.028077 | 1 |
| 15 | 24214800 | 24215599 | 11.13768 | 20.98392 | 22.18583 | 9.074547 | 0.392808 | 0.079831 | 1 |
| 15 | 24411400 | 24412599 | 16.56245 | 28.90422 | 37.46547 | 16.49534 | 0.01506 | 0.717481 | 1 |
| 15 | 24668600 | 24669399 | 12.09755 | 22.11173 | 27.08928 | 11.59766 | 0.131464 | 0.277778 | 1 |
| 15 | 25053400 | 25053999 | 11.62484 | 21.54335 | 27.26456 | 11.45591 | 0.104727 | 0.200245 | 1 |
| 15 | 25055000 | 25056999 | 18.73442 | 32.93973 | 38.48467 | 17.04134 | 0.164271 | 0.615378 | 1 |
| 15 | 25116400 | 25117599 | 15.5092 | 27.20759 | 40.33204 | 17.83089 | -0.17758 | 0.634917 | 1 |
| 15 | 25364200 | 25365799 | 21.78691 | 36.66058 | 55.94156 | 24.44503 | -0.22649 | 1.117524 | 1 |

| | | | | | | | | | |
|---|---|---|---|---|---|---|---|---|---|
| 15 | 25699800 | 25703599 | 35.72365 | 57.81215 | 88.24563 | 37.74141 | -0.24589 | 1.675386 | 1 |
| 15 | 26119200 | 26121999 | 35.88674 | 58.1727 | 85.56192 | 36.6385 | -0.19023 | 1.641539 | 1 |
| 15 | 26177400 | 26180999 | 46.38973 | 74.45778 | 102.4726 | 43.34814 | -0.11081 | 1.939366 | 1 |
| 15 | 26248200 | 26249799 | 19.47082 | 33.989 | 36.75722 | 16.18626 | 0.275172 | 0.727911 | 1 |
| 15 | 27480200 | 27482799 | 19.14582 | 32.85989 | 51.71005 | 22.68036 | -0.26806 | 0.924034 | 1 |
| 15 | 29394800 | 29395799 | 11.69062 | 20.79762 | 34.24673 | 15.09588 | -0.32034 | 0.453414 | 1 |
| 15 | 29469800 | 29471599 | 27.35676 | 45.45108 | 58.78851 | 25.49507 | 0.011024 | 1.272039 | 1 |
| 15 | 29798600 | 29805799 | 123.2372 | 187.663 | 209.5049 | 86.8285 | 0.12622 | 3.158055 | 1 |
| 15 | 29983600 | 29984999 | 16.49484 | 28.60713 | 41.23259 | 18.09106 | -0.1383 | 0.809166 | 1 |
| 15 | 30042600 | 30043999 | 15.78538 | 27.5792 | 36.98732 | 16.29637 | -0.03366 | 0.697037 | 1 |
| 15 | 30447400 | 30449999 | 40.03556 | 64.63039 | 82.60262 | 35.22814 | 0.005863 | 1.748281 | 1 |
| 15 | 30753800 | 30755599 | 51.05638 | 81.64801 | 96.12801 | 40.44272 | 0.107916 | 2.057871 | 1 |
| 15 | 30932200 | 30936799 | 49.1407 | 78.68227 | 105.0951 | 44.29391 | -0.07188 | 2.017971 | 1 |
| 15 | 31893600 | 31894399 | 11.27159 | 20.57528 | 32.29622 | 13.84939 | -0.22535 | 0.291299 | 1 |
| 15 | 31931400 | 31935599 | 76.22023 | 120.1404 | 127.5031 | 52.90992 | 0.23763 | 2.485664 | 1 |
| 15 | 31946200 | 31947399 | 16.3031 | 28.42763 | 43.10535 | 18.96841 | -0.21042 | 0.73657 | 1 |
| 15 | 32741800 | 32743199 | 14.25317 | 26.00649 | 27.43745 | 11.77229 | 0.343798 | 0.306018 | 1 |
| 15 | 32778200 | 32781199 | 20.15532 | 34.05302 | 53.76638 | 23.52347 | -0.27526 | 1.074176 | 1 |
| 15 | 32782800 | 32787199 | 74.8654 | 117.8079 | 144.9506 | 60.13704 | 0.02274 | 2.521303 | 1 |
| 15 | 32855800 | 32857599 | 19.25387 | 33.05978 | 51.68784 | 22.67142 | -0.2585 | 0.923257 | 1 |
| 15 | 32879600 | 32881399 | 36.09878 | 59.00154 | 62.71976 | 26.96571 | 0.286398 | 1.496118 | 1 |
| 15 | 32912000 | 32912799 | 10.98587 | 20.43856 | 26.72438 | 11.16683 | 0.063469 | 0.168509 | 1 |
| 15 | 33117000 | 33121999 | 139.8583 | 211.3379 | 223.9371 | 92.59112 | 0.192221 | 3.318679 | 1 |
| 15 | 33525000 | 33527599 | 19.84325 | 34.77085 | 37.02531 | 16.35312 | 0.29741 | 0.671696 | 1 |
| 15 | 33565800 | 33570799 | 56.06039 | 89.87911 | 92.48148 | 39.0552 | 0.306076 | 1.98296 | 1 |
| 15 | 33612400 | 33613399 | 13.32447 | 23.49916 | 39.30452 | 17.39379 | -0.3488 | 0.553439 | 1 |
| 15 | 33659000 | 33663199 | 54.77559 | 87.80967 | 97.59908 | 41.20954 | 0.19476 | 1.984393 | 1 |
| 15 | 33744800 | 33746199 | 22.41127 | 37.37359 | 61.75145 | 26.84598 | -0.3408 | 1.22009 | 1 |
| 15 | 34051200 | 34053999 | 24.72463 | 41.22124 | 60.38607 | 26.24831 | -0.16788 | 1.225777 | 1 |
| 15 | 34221800 | 34233999 | 261.1934 | 375.1668 | 419.175 | 173.0574 | 0.047128 | 4.182449 | 1 |
| 15 | 34265800 | 34267399 | 23.67362 | 40.47313 | 51.18845 | 22.44455 | 0.045248 | 0.980188 | 1 |
| 15 | 34313200 | 34314999 | 11.96066 | 22.52447 | 25.23338 | 10.32881 | 0.308768 | 0.09702 | 1 |
| 15 | 34762400 | 34764199 | 17.19324 | 30.12825 | 34.7167 | 15.33387 | 0.184147 | 0.652995 | 1 |
| 15 | 34872000 | 34873599 | 36.57399 | 59.91511 | 67.58697 | 29.0839 | 0.202887 | 1.468482 | 1 |
| 15 | 34922800 | 34923599 | 11.63163 | 21.79826 | 27.33661 | 11.28254 | 0.13748 | 0.129246 | 1 |
| 15 | 34992000 | 34994199 | 24.99425 | 42.22702 | 50.12277 | 21.9291 | 0.134872 | 1.088882 | 1 |
| 15 | 35116600 | 35120199 | 40.97464 | 65.5075 | 109.6802 | 46.12636 | -0.39959 | 2.059207 | 1 |
| 15 | 35589600 | 35590399 | 17.43224 | 30.22902 | 42.19784 | 18.51451 | -0.09247 | 0.815464 | 1 |
| 15 | 36222600 | 36223999 | 16.3277 | 28.32635 | 44.26755 | 19.42286 | -0.25452 | 0.808703 | 1 |
| 15 | 37077000 | 37092799 | 221.3288 | 321.4128 | 401.0499 | 165.3311 | -0.09654 | 4.018045 | 1 |
| 15 | 37402600 | 37405199 | 29.37131 | 48.43043 | 66.5706 | 28.72564 | -0.07772 | 1.370788 | 1 |
| 15 | 37845600 | 37850399 | 101.4118 | 155.8911 | 185.5411 | 77.04613 | 0.044572 | 2.961424 | 1 |
| 15 | 37960600 | 37961799 | 12.17793 | 21.48668 | 38.28795 | 16.94241 | -0.43953 | 0.537278 | 1 |
| 15 | 38299200 | 38303399 | 28.92683 | 47.02506 | 77.6985 | 33.34005 | -0.35368 | 1.567738 | 1 |
| 15 | 38442000 | 38443999 | 21.36655 | 36.66681 | 46.3588 | 20.33307 | 0.046651 | 0.932096 | 1 |
| 15 | 38461600 | 38467199 | 79.76149 | 125.4079 | 136.537 | 56.64027 | 0.198055 | 2.539026 | 1 |
| 15 | 38531400 | 38533999 | 30.886 | 50.45538 | 71.27693 | 30.63899 | -0.12333 | 1.495769 | 1 |

| | | | | | | | | | |
|---|---|---|---|---|---|---|---|---|---|
| 15 | 39039600 | 39045999 | 78.17662 | 122.6941 | 138.1385 | 57.29118 | 0.147736 | 2.57434 | 1 |
| 15 | 39047600 | 39049199 | 16.13125 | 28.52069 | 39.08655 | 17.28866 | -0.06174 | 0.540852 | 1 |
| 15 | 39852200 | 39857999 | 19.63681 | 33.41122 | 51.64651 | 22.63666 | -0.2441 | 1.008577 | 1 |
| 15 | 39912600 | 39913399 | 10.49944 | 19.79678 | 22.73592 | 9.287345 | 0.27581 | 0.070325 | 1 |
| 15 | 39955800 | 39957199 | 18.60781 | 32.35675 | 37.97711 | 16.68272 | 0.157066 | 0.779309 | 1 |
| 15 | 39958200 | 39960999 | 19.40589 | 32.85177 | 53.19249 | 23.28635 | -0.31121 | 1.055818 | 1 |
| 15 | 40086800 | 40088999 | 18.58799 | 31.99713 | 50.61345 | 22.20192 | -0.2745 | 0.896186 | 1 |
| 15 | 41795400 | 41802599 | 124.3877 | 187.6419 | 262.1195 | 108.3204 | -0.20827 | 3.352044 | 1 |
| 15 | 41999800 | 42002399 | 12.88215 | 23.18476 | 31.09993 | 13.57456 | -0.01647 | 0.38272 | 1 |
| 15 | 42021400 | 42022599 | 21.94944 | 37.41902 | 45.86929 | 20.10766 | 0.089623 | 0.992492 | 1 |
| 15 | 42063800 | 42067399 | 42.36666 | 68.32439 | 87.38257 | 37.21808 | 0.003144 | 1.780543 | 1 |
| 15 | 42459800 | 42462399 | 33.15328 | 54.66661 | 69.31879 | 29.91153 | 0.038291 | 1.382365 | 1 |
| 15 | 42801200 | 42802399 | 17.48308 | 30.1694 | 47.4142 | 20.80011 | -0.2643 | 0.866345 | 1 |
| 15 | 43195200 | 43195799 | 12.47333 | 22.37874 | 31.22612 | 13.66649 | -0.07547 | 0.401184 | 1 |
| 15 | 43759800 | 43761199 | 22.94444 | 38.42497 | 60.33552 | 26.30054 | -0.26775 | 1.172519 | 1 |
| 15 | 43818600 | 43819799 | 13.08551 | 23.42921 | 34.12757 | 14.96206 | -0.13914 | 0.422504 | 1 |
| 15 | 43880200 | 43880799 | 10.32045 | 18.94624 | 28.31012 | 12.04909 | -0.14675 | 0.243919 | 1 |
| 15 | 43987000 | 43989199 | 13.70981 | 24.45406 | 37.14415 | 16.33674 | -0.20263 | 0.454867 | 1 |
| 15 | 44338400 | 44340799 | 32.70541 | 52.86007 | 85.24429 | 36.41065 | -0.3268 | 1.701702 | 1 |
| 15 | 44522200 | 44528399 | 92.67102 | 144.0327 | 159.9025 | 66.39099 | 0.157064 | 2.754427 | 1 |
| 15 | 44585400 | 44587799 | 19.74969 | 34.42792 | 39.90673 | 17.55782 | 0.17539 | 0.754645 | 1 |
| 15 | 44611000 | 44613999 | 16.08038 | 28.31765 | 30.05322 | 13.30366 | 0.303067 | 0.566542 | 1 |
| 15 | 44658000 | 44659599 | 13.12417 | 23.17823 | 38.2442 | 16.91984 | -0.32857 | 0.533835 | 1 |
| 15 | 44853000 | 44854399 | 22.52871 | 38.09504 | 55.48493 | 24.28157 | -0.15932 | 1.07642 | 1 |
| 15 | 44897800 | 44899999 | 23.14255 | 39.93027 | 43.59357 | 19.12396 | 0.258944 | 0.885529 | 1 |
| 15 | 45058200 | 45060199 | 25.17203 | 42.61516 | 48.51833 | 21.24042 | 0.194813 | 1.067945 | 1 |
| 15 | 45439000 | 45442199 | 34.31949 | 56.11381 | 75.92345 | 32.64684 | -0.06042 | 1.493961 | 1 |
| 15 | 45552600 | 45554199 | 26.29216 | 45.11966 | 47.36919 | 20.77623 | 0.313714 | 0.952706 | 1 |
| 15 | 45555000 | 45560199 | 71.67301 | 112.8603 | 147.411 | 61.16316 | -0.06259 | 2.505519 | 1 |
| 15 | 45860400 | 45861599 | 11.07619 | 20.31373 | 32.36358 | 13.79263 | -0.24058 | 0.262486 | 1 |
| 15 | 46041400 | 46043999 | 19.14365 | 33.34719 | 36.7794 | 16.17449 | 0.2465 | 0.75366 | 1 |
| 15 | 46071800 | 46073599 | 9.877945 | 18.30823 | 28.38325 | 11.92397 | -0.18576 | 0.187903 | 1 |
| 15 | 46751600 | 46752399 | 14.5259 | 25.5313 | 35.27259 | 15.61928 | -0.07629 | 0.590379 | 1 |
| 15 | 46847600 | 46849399 | 18.8989 | 32.85425 | 45.23207 | 19.86642 | -0.07224 | 0.795305 | 1 |
| 15 | 47043400 | 47044599 | 13.68524 | 24.12882 | 38.81639 | 17.17871 | -0.29295 | 0.559295 | 1 |
| 15 | 47113000 | 47113799 | 15.88892 | 27.91953 | 32.69414 | 14.47422 | 0.160983 | 0.597472 | 1 |
| 15 | 47425400 | 47427199 | 20.25647 | 35.03283 | 40.06928 | 17.57916 | 0.192881 | 0.848664 | 1 |
| 15 | 48336200 | 48338599 | 36.59195 | 59.0624 | 90.39432 | 38.54364 | -0.25386 | 1.74925 | 1 |
| 15 | 48715800 | 48716799 | 13.23934 | 24.03667 | 28.49535 | 12.30256 | 0.170736 | 0.328174 | 1 |
| 15 | 48903200 | 48903999 | 14.29285 | 25.70268 | 32.46693 | 14.18182 | 0.068777 | 0.399364 | 1 |
| 15 | 49030400 | 49032599 | 16.11753 | 28.81284 | 32.22286 | 14.14566 | 0.238365 | 0.446155 | 1 |
| 15 | 49033600 | 49043199 | 184.3958 | 269.7689 | 379.8088 | 156.4627 | -0.26032 | 3.902104 | 1 |
| 15 | 49050200 | 49052799 | 27.44954 | 45.99161 | 53.47509 | 23.31708 | 0.164196 | 1.173398 | 1 |
| 15 | 49105200 | 49108199 | 40.94516 | 66.20746 | 78.15166 | 33.40061 | 0.123 | 1.699077 | 1 |
| 15 | 49112000 | 49113199 | 29.4327 | 48.16116 | 63.81319 | 27.44725 | -0.02935 | 1.465076 | 1 |
| 15 | 49144800 | 49146199 | 12.85447 | 23.39556 | 28.60955 | 12.31381 | 0.129333 | 0.313107 | 1 |
| 15 | 49185400 | 49188799 | 42.15254 | 67.65134 | 95.07907 | 40.21549 | -0.14109 | 1.934296 | 1 |

| | | | | | | | | | |
|---|---|---|---|---|---|---|---|---|---|
| 15 | 49277800 | 49278399 | 11.59327 | 21.6403 | 26.26467 | 10.91323 | 0.177055 | 0.153866 | 1 |
| 15 | 49503600 | 49508799 | 25.48284 | 42.70963 | 57.19159 | 24.94248 | -0.03882 | 1.169209 | 1 |
| 15 | 49568000 | 49569599 | 21.86064 | 37.61699 | 43.71949 | 19.17763 | 0.168439 | 0.90706 | 1 |
| 15 | 49725400 | 49728199 | 30.41078 | 50.56306 | 56.80332 | 24.64693 | 0.213771 | 1.270057 | 1 |
| 15 | 49782200 | 49784399 | 7.892358 | 15.07228 | 27.88159 | 11.08715 | -0.37302 | -0.03034 | 1 |
| 15 | 49920400 | 49921599 | 16.49112 | 29.07652 | 30.52896 | 13.51094 | 0.319268 | 0.558014 | 1 |
| 15 | 49950400 | 49952999 | 27.09984 | 45.39537 | 54.50687 | 23.76356 | 0.118034 | 1.177924 | 1 |
| 15 | 50227200 | 50228199 | 18.90011 | 32.51096 | 45.3319 | 19.88469 | -0.09333 | 0.898444 | 1 |
| 15 | 50306600 | 50308799 | 8.732517 | 16.25349 | 28.93739 | 12.08252 | -0.37771 | 0.152103 | 1 |
| 15 | 50911600 | 50912799 | 66.10848 | 103.3148 | 159.4005 | 66.13459 | -0.31098 | 2.647332 | 1 |
| 15 | 50974600 | 50975599 | 11.17086 | 19.7406 | 37.43464 | 16.55965 | -0.52724 | 0.51716 | 1 |
| 15 | 51891000 | 51891799 | 16.42447 | 28.77507 | 34.83149 | 15.38184 | 0.113516 | 0.652956 | 1 |
| 15 | 52094800 | 52097199 | 21.22879 | 36.02029 | 51.67107 | 22.63606 | -0.13717 | 1.038817 | 1 |
| 15 | 53437400 | 53438399 | 17.67561 | 30.92147 | 35.9005 | 15.83423 | 0.173557 | 0.68739 | 1 |
| 15 | 53846200 | 53847799 | 14.49308 | 25.42186 | 39.94258 | 17.65685 | -0.26081 | 0.633544 | 1 |
| 15 | 54049200 | 54051399 | 26.04491 | 43.73137 | 45.46432 | 19.8508 | 0.325174 | 1.141008 | 1 |
| 15 | 54074000 | 54075999 | 45.54977 | 72.51382 | 117.4287 | 49.09289 | -0.35827 | 2.198949 | 1 |
| 15 | 55020800 | 55022599 | 22.45588 | 37.83719 | 57.05208 | 24.94049 | -0.20927 | 1.109973 | 1 |
| 15 | 55028000 | 55032199 | 55.17963 | 88.08708 | 101.1262 | 42.43994 | 0.14143 | 2.115006 | 1 |
| 15 | 55473000 | 55475399 | 7.918353 | 15.1264 | 26.73825 | 10.62584 | -0.30675 | -0.03329 | 1 |
| 15 | 55546800 | 55548999 | 37.13621 | 59.79353 | 92.2543 | 39.20635 | -0.26965 | 1.825956 | 1 |
| 15 | 55896200 | 55902399 | 172.3628 | 255.6092 | 294.9557 | 121.4867 | 0.044968 | 3.678192 | 1 |
| 15 | 56075800 | 56077599 | 16.39678 | 28.70219 | 40.86625 | 18.03391 | -0.1198 | 0.674106 | 1 |
| 15 | 56224200 | 56225999 | 17.11691 | 29.51253 | 49.53534 | 21.73007 | -0.35921 | 0.872861 | 1 |
| 15 | 56383200 | 56385399 | 24.24632 | 40.54951 | 56.5236 | 24.61758 | -0.09644 | 1.188895 | 1 |
| 15 | 56770800 | 56772399 | 32.07767 | 52.47019 | 62.27726 | 26.78308 | 0.128646 | 1.477117 | 1 |
| 15 | 56850800 | 56851799 | 16.90861 | 28.97544 | 50.57241 | 22.17863 | -0.4169 | 0.931004 | 1 |
| 15 | 56852600 | 56853399 | 11.58917 | 21.65708 | 24.12986 | 10.00704 | 0.30155 | 0.139446 | 1 |
| 15 | 57142000 | 57143399 | 14.84827 | 26.03777 | 37.84171 | 16.72428 | -0.14908 | 0.63353 | 1 |
| 15 | 57154400 | 57157599 | 40.66899 | 65.82737 | 74.62628 | 31.92092 | 0.182594 | 1.676325 | 1 |
| 15 | 57286800 | 57287999 | 15.28522 | 26.75676 | 39.4918 | 17.42755 | -0.17104 | 0.670181 | 1 |
| 15 | 57292000 | 57292999 | 12.02302 | 21.71903 | 30.61692 | 13.3018 | -0.08347 | 0.352616 | 1 |
| 15 | 58565200 | 58570999 | 80.37169 | 125.4623 | 162.4984 | 67.42778 | -0.06018 | 2.674324 | 1 |
| 15 | 58694400 | 58695999 | 14.29708 | 24.89682 | 43.20725 | 18.99747 | -0.40426 | 0.7464 | 1 |
| 15 | 58878200 | 58880999 | 31.53565 | 51.22151 | 81.37067 | 34.89499 | -0.29831 | 1.592773 | 1 |
| 15 | 59035800 | 59041999 | 64.82645 | 102.2633 | 138.1198 | 57.32151 | -0.10882 | 2.465239 | 1 |
| 15 | 59848600 | 59849999 | 15.7085 | 27.29207 | 45.28971 | 19.88445 | -0.34045 | 0.788665 | 1 |
| 15 | 59886800 | 59888599 | 15.53032 | 27.02211 | 44.21887 | 19.4322 | -0.32015 | 0.764962 | 1 |
| 15 | 60351200 | 60352799 | 24.58023 | 41.45067 | 50.66879 | 22.15554 | 0.092455 | 1.102119 | 1 |
| 15 | 60463600 | 60467799 | 72.24129 | 114.016 | 132.4843 | 54.99635 | 0.108477 | 2.456153 | 1 |
| 15 | 60672600 | 60675199 | 31.17908 | 50.86489 | 72.54874 | 31.17023 | -0.13845 | 1.520397 | 1 |
| 15 | 60819200 | 60819999 | 15.33349 | 26.92432 | 38.75661 | 17.14783 | -0.13539 | 0.616366 | 1 |
| 15 | 62393400 | 62395399 | 18.03746 | 31.38067 | 40.03324 | 17.59148 | 0.037439 | 0.773222 | 1 |
| 15 | 62504400 | 62509199 | 48.21261 | 77.25874 | 89.5698 | 37.79703 | 0.133414 | 1.985729 | 1 |
| 15 | 63068000 | 63068599 | 11.55495 | 21.52048 | 27.19934 | 11.34458 | 0.114001 | 0.166354 | 1 |
| 15 | 63205400 | 63207399 | 20.45912 | 34.83445 | 52.19128 | 22.8767 | -0.19908 | 1.006585 | 1 |
| 15 | 63296400 | 63297799 | 19.51249 | 33.64662 | 43.73848 | 19.18718 | 0.008125 | 0.876902 | 1 |

| | | | | | | | | | |
|---|---|---|---|---|---|---|---|---|---|
| 15 | 63432600 | 63433199 | 8.107076 | 14.82277 | 30.96066 | 13.24976 | -0.63203 | 0.256259 | 1 |
| 15 | 63657200 | 63663399 | 100.8742 | 156.4825 | 173.209 | 71.93637 | 0.159202 | 2.79285 | 1 |
| 15 | 65139400 | 65141799 | 25.58612 | 43.09018 | 52.84205 | 23.0966 | 0.087759 | 1.119226 | 1 |
| 15 | 65142600 | 65145999 | 35.63414 | 57.6391 | 82.96953 | 35.4721 | -0.16204 | 1.685148 | 1 |
| 15 | 65245600 | 65248399 | 32.85849 | 54.85184 | 54.16632 | 23.56365 | 0.399248 | 1.231956 | 1 |
| 15 | 65277200 | 65278399 | 14.32052 | 25.04521 | 41.57584 | 18.33311 | -0.33981 | 0.682503 | 1 |
| 15 | 65334200 | 65336399 | 32.27844 | 52.57578 | 75.59759 | 32.46157 | -0.15175 | 1.545254 | 1 |
| 15 | 65442000 | 65444999 | 17.15449 | 29.67338 | 45.83693 | 20.10964 | -0.23874 | 0.841502 | 1 |
| 15 | 65928800 | 65930399 | 26.41714 | 43.33589 | 71.96522 | 30.98598 | -0.35259 | 1.43252 | 1 |
| 15 | 67568200 | 67569199 | 13.57012 | 24.11194 | 34.60268 | 15.26841 | -0.12419 | 0.474251 | 1 |
| 15 | 67706600 | 67708199 | 15.23403 | 26.55741 | 44.18667 | 19.44144 | -0.34368 | 0.736812 | 1 |
| 15 | 67781200 | 67783999 | 27.70549 | 45.37481 | 70.90032 | 30.50627 | -0.26596 | 1.454904 | 1 |
| 15 | 68756600 | 68757999 | 13.00799 | 22.85444 | 38.88784 | 17.21449 | -0.37528 | 0.594503 | 1 |
| 15 | 69015200 | 69017599 | 15.66534 | 27.52234 | 37.70615 | 16.69091 | -0.06437 | 0.608071 | 1 |
| 15 | 69065800 | 69066399 | 9.494329 | 18.21319 | 19.26325 | 7.566426 | 0.444097 | -0.07648 | 1 |
| 15 | 69624200 | 69625199 | 11.10845 | 20.26994 | 29.24361 | 12.54619 | -0.10463 | 0.286266 | 1 |
| 15 | 69862000 | 69863399 | 15.48404 | 27.38068 | 32.55823 | 14.40062 | 0.142376 | 0.525202 | 1 |
| 15 | 69954200 | 69955599 | 21.06153 | 35.7392 | 48.51042 | 21.25193 | -0.05763 | 1.034576 | 1 |
| 15 | 70483000 | 70484999 | 24.55842 | 41.22433 | 57.67506 | 25.17801 | -0.10182 | 1.151214 | 1 |
| 15 | 70782600 | 70786599 | 42.48564 | 68.65798 | 74.32661 | 31.74176 | 0.246556 | 1.71546 | 1 |
| 15 | 70976000 | 70976799 | 14.33662 | 25.61894 | 30.54584 | 13.41396 | 0.146548 | 0.43642 | 1 |
| 15 | 71134400 | 71135599 | 8.930003 | 16.56747 | 29.95968 | 12.56949 | -0.4051 | 0.176983 | 1 |
| 15 | 71262400 | 71263599 | 14.10365 | 25.62029 | 28.46221 | 12.2796 | 0.264645 | 0.332237 | 1 |
| 15 | 71467800 | 71473799 | 92.34818 | 143.7168 | 152.2571 | 63.20446 | 0.225958 | 2.732279 | 1 |
| 15 | 71965800 | 71968999 | 30.1486 | 49.17575 | 73.74452 | 31.6821 | -0.21078 | 1.51902 | 1 |
| 15 | 72300800 | 72301599 | 12.75882 | 22.72052 | 32.18987 | 14.17733 | -0.10378 | 0.450475 | 1 |
| 15 | 73063600 | 73069599 | 82.77483 | 129.7447 | 139.9213 | 58.03504 | 0.208333 | 2.599637 | 1 |
| 15 | 73750000 | 73750799 | 17.70713 | 31.03375 | 31.82615 | 14.05907 | 0.351615 | 0.649572 | 1 |
| 15 | 74115000 | 74121399 | 49.864 | 80.07969 | 90.36345 | 38.25876 | 0.176053 | 1.920877 | 1 |
| 15 | 74946400 | 74950599 | 44.81095 | 72.36014 | 81.63775 | 34.82951 | 0.185814 | 1.744831 | 1 |
| 15 | 74975400 | 74975999 | 7.479527 | 14.20729 | 26.06723 | 10.48421 | -0.37518 | -0.00031 | 1 |
| 15 | 75082000 | 75086999 | 72.67809 | 113.8711 | 151.4957 | 62.83846 | -0.09469 | 2.603596 | 1 |
| 15 | 75476400 | 75478799 | 11.59116 | 21.07567 | 30.70804 | 13.2332 | -0.12308 | 0.313377 | 1 |
| 15 | 76444800 | 76447199 | 16.27224 | 28.05194 | 47.94438 | 21.03205 | -0.38496 | 0.87228 | 1 |
| 15 | 77017800 | 77021399 | 43.14978 | 69.53999 | 81.32085 | 34.60588 | 0.130919 | 1.798811 | 1 |
| 15 | 79861600 | 79862999 | 25.64249 | 42.92175 | 53.90454 | 23.48978 | 0.05349 | 1.180882 | 1 |
| 15 | 80025200 | 80026999 | 14.75843 | 26.56298 | 28.90906 | 12.61746 | 0.283402 | 0.390735 | 1 |
| 15 | 81084200 | 81092199 | 65.0295 | 103.1652 | 116.4621 | 48.4999 | 0.156331 | 2.319103 | 1 |
| 15 | 82437000 | 82446199 | 139.5367 | 209.0229 | 285.3244 | 117.6941 | -0.18398 | 3.491993 | 1 |
| 15 | 82749800 | 82752399 | 23.96324 | 40.21642 | 60.37318 | 26.35223 | -0.20314 | 1.154884 | 1 |
| 15 | 83080800 | 83081599 | 17.93722 | 30.8988 | 47.9485 | 21.03338 | -0.24642 | 0.888352 | 1 |
| 15 | 83167400 | 83172199 | 52.62256 | 83.71649 | 123.0352 | 51.38835 | -0.21968 | 2.227087 | 1 |
| 15 | 84037000 | 84037799 | 11.54858 | 21.72556 | 24.13944 | 9.898652 | 0.318338 | 0.096353 | 1 |
| 15 | 85015200 | 85017999 | 25.57827 | 42.12191 | 64.7784 | 27.93711 | -0.23979 | 1.378343 | 1 |
| 15 | 85698800 | 85706199 | 114.1912 | 172.7883 | 250.4182 | 103.5848 | -0.25755 | 3.286935 | 1 |
| 15 | 86646200 | 86648599 | 33.56494 | 54.4889 | 77.80109 | 33.35127 | -0.14552 | 1.603124 | 1 |
| 15 | 86769200 | 86770199 | 11.46743 | 21.25805 | 27.70827 | 11.63758 | 0.062434 | 0.195277 | 1 |

| | | | | | | | | | |
|---|---|---|---|---|---|---|---|---|---|
| 15 | 87173600 | 87179999 | 64.82756 | 102.5301 | 124.9612 | 51.92851 | 0.042155 | 2.400265 | 1 |
| 15 | 89491800 | 89501199 | 31.73013 | 52.41664 | 59.00085 | 25.48639 | 0.21027 | 1.349038 | 1 |
| 15 | 89539800 | 89541599 | 27.92033 | 48.07429 | 42.90168 | 18.81925 | 0.548054 | 0.923169 | 1 |
| 15 | 89592600 | 89600999 | 119.6984 | 182.3459 | 225.5788 | 93.50129 | -0.02131 | 3.150793 | 1 |
| 15 | 89741800 | 89744599 | 16.33787 | 28.59017 | 42.48339 | 18.74261 | -0.18104 | 0.683176 | 1 |
| 15 | 89825000 | 89827999 | 31.42779 | 51.80177 | 62.83879 | 27.11101 | 0.10185 | 1.377265 | 1 |
| 15 | 89835400 | 89836999 | 12.38462 | 22.27859 | 34.13579 | 14.90657 | -0.20793 | 0.386804 | 1 |
| 15 | 90986200 | 90988199 | 22.53966 | 38.59392 | 45.81001 | 20.08876 | 0.136787 | 0.954231 | 1 |
| 15 | 91269600 | 91271999 | 17.78511 | 30.88987 | 47.39214 | 20.80324 | -0.22795 | 0.805799 | 1 |
| 15 | 91303000 | 91308399 | 51.57 | 82.54407 | 99.33548 | 41.84579 | 0.077745 | 2.02937 | 1 |
| 15 | 91460200 | 91461399 | 13.67151 | 24.68299 | 30.03451 | 13.058 | 0.126997 | 0.368099 | 1 |
| 15 | 91765400 | 91770199 | 56.05339 | 89.80252 | 106.7515 | 45.03162 | 0.096945 | 2.008979 | 1 |
| 15 | 91876600 | 91884599 | 93.83976 | 145.3771 | 187.4081 | 77.84418 | -0.06193 | 2.814631 | 1 |
| 15 | 91888200 | 91890999 | 28.04058 | 46.23176 | 62.23695 | 26.85456 | -0.0476 | 1.369554 | 1 |
| 15 | 92430600 | 92432199 | 14.97982 | 26.20983 | 41.46031 | 18.28898 | -0.27093 | 0.674573 | 1 |
| 15 | 92514000 | 92517399 | 31.12659 | 50.62654 | 79.50703 | 34.12234 | -0.28019 | 1.564109 | 1 |
| 15 | 93062000 | 93068199 | 67.91494 | 106.5092 | 160.2723 | 66.47239 | -0.27108 | 2.585472 | 1 |
| 15 | 93512200 | 93514399 | 35.93929 | 58.34612 | 69.55084 | 29.81563 | 0.114313 | 1.60193 | 1 |
| 15 | 93933000 | 93936399 | 41.02292 | 66.67121 | 74.19014 | 31.82979 | 0.214796 | 1.595791 | 1 |
| 15 | 93993800 | 94000599 | 88.38943 | 136.8995 | 190.8078 | 79.2564 | -0.17471 | 2.816296 | 1 |
| 15 | 94241600 | 94245799 | 41.6712 | 67.02322 | 97.25665 | 41.26785 | -0.18335 | 1.867207 | 1 |
| 15 | 94513000 | 94514799 | 18.65191 | 31.75373 | 53.24363 | 23.33776 | -0.36094 | 0.999772 | 1 |
| 15 | 94630800 | 94631799 | 14.29284 | 25.77361 | 26.87814 | 11.70338 | 0.346632 | 0.370677 | 1 |
| 15 | 94712200 | 94715799 | 40.00277 | 64.99574 | 69.06007 | 29.62296 | 0.28089 | 1.593454 | 1 |
| 15 | 95205400 | 95206999 | 13.03042 | 23.27574 | 32.09684 | 14.0997 | -0.06254 | 0.431911 | 1 |
| 15 | 95490800 | 95492799 | 18.48932 | 32.38273 | 33.66457 | 14.86269 | 0.331972 | 0.666983 | 1 |
| 15 | 96248800 | 96251199 | 30.33784 | 50.38119 | 52.42495 | 22.72828 | 0.323829 | 1.279368 | 1 |
| 15 | 96395600 | 96398199 | 42.44113 | 68.38065 | 90.43533 | 38.47049 | -0.04667 | 1.810738 | 1 |
| 15 | 96399800 | 96401999 | 39.34009 | 63.94119 | 70.20934 | 30.12363 | 0.234052 | 1.585594 | 1 |
| 15 | 96644800 | 96647599 | 24.75389 | 41.37839 | 64.344 | 28.01504 | -0.25379 | 1.20174 | 1 |
| 15 | 96684000 | 96686399 | 16.84679 | 29.23698 | 47.93414 | 21.03184 | -0.32336 | 0.813391 | 1 |
| 15 | 96802200 | 96806199 | 31.51741 | 51.25191 | 76.89394 | 32.99845 | -0.21455 | 1.569195 | 1 |
| 15 | 97200400 | 97203599 | 29.29486 | 47.9046 | 73.40585 | 31.56578 | -0.23928 | 1.47954 | 1 |
| 15 | 97376000 | 97380399 | 75.74237 | 119.3018 | 131.3779 | 54.51281 | 0.183611 | 2.500042 | 1 |
| 15 | 97392200 | 97394399 | 15.33169 | 27.18497 | 30.37893 | 13.42837 | 0.232817 | 0.488753 | 1 |
| 15 | 97407400 | 97411999 | 63.70719 | 101.1766 | 115.6968 | 48.23325 | 0.1393 | 2.286382 | 1 |
| 15 | 97451200 | 97453599 | 33.04967 | 54.382 | 60.08124 | 25.90255 | 0.235374 | 1.400471 | 1 |
| 15 | 98216400 | 98224199 | 72.30029 | 114.0655 | 134.4472 | 55.80039 | 0.087475 | 2.466143 | 1 |
| 15 | 100725400 | 100730599 | 43.69982 | 69.87815 | 108.4128 | 45.60614 | -0.28957 | 2.054861 | 1 |
| 15 | 101054200 | 101055599 | 25.60538 | 43.1671 | 51.21265 | 22.39141 | 0.135442 | 1.109507 | 1 |
| 15 | 101927800 | 101929199 | 13.51325 | 24.07706 | 36.60217 | 16.11326 | -0.2048 | 0.458563 | 1 |
| 15 | 103318800 | 103321199 | 31.28135 | 50.54443 | 85.28421 | 36.41468 | -0.39235 | 1.708533 | 1 |
| 15 | 103338600 | 103339199 | 8.127219 | 15.67305 | 18.88195 | 7.305221 | 0.278416 | -0.14108 | 1 |
| 15 | 103503400 | 103506399 | 28.4231 | 47.03017 | 75.36131 | 32.59021 | -0.29768 | 1.333839 | 1 |
| 15 | 103514000 | 103516399 | 33.41323 | 54.17856 | 75.80342 | 32.46663 | -0.11786 | 1.625743 | 1 |
| 15 | 103517200 | 103521199 | 41.42368 | 67.01312 | 85.06192 | 36.37038 | 0.019105 | 1.691173 | 1 |
| 15 | 103558600 | 103559599 | 12.90366 | 23.29588 | 32.63299 | 14.18835 | -0.07505 | 0.368185 | 1 |

| | | | | | | | | | |
|---|---|---|---|---|---|---|---|---|---|
| 15 | 103590800 | 103591599 | 9.471933 | 17.70276 | 26.35828 | 10.92887 | -0.11335 | 0.126457 | 1 |
| 15 | 103646600 | 103649399 | 31.91319 | 52.51783 | 56.65393 | 24.4261 | 0.269794 | 1.394368 | 1 |
| 15 | 103706600 | 103715199 | 168.9062 | 251.2194 | 271.908 | 112.0273 | 0.1412 | 3.625862 | 1 |
| 15 | 103857400 | 103861999 | 21.07298 | 36.20677 | 43.20357 | 18.95044 | 0.130056 | 0.914411 | 1 |
| 15 | 103946800 | 103951599 | 98.43118 | 152.5 | 163.995 | 68.11899 | 0.199425 | 2.812744 | 1 |
| 15 | 103953800 | 103961599 | 122.4249 | 186.8491 | 203.9855 | 84.59038 | 0.16105 | 3.111707 | 1 |
| 16 | 3203400 | 3203799 | 14.33371 | 25.62457 | 29.58144 | 12.98582 | 0.193291 | 0.432012 | 1 |
| 16 | 3560600 | 3563999 | 25.17398 | 41.60223 | 62.27484 | 26.90781 | -0.19964 | 1.330914 | 1 |
| 16 | 3585000 | 3587199 | 16.71632 | 29.31208 | 35.69568 | 15.77517 | 0.104745 | 0.640988 | 1 |
| 16 | 3610400 | 3634799 | 360.0765 | 505.4246 | 517.207 | 214.3337 | 0.140493 | 4.524794 | 1 |
| 16 | 3638200 | 3641799 | 46.66972 | 74.87408 | 88.8319 | 37.55257 | 0.102205 | 1.948727 | 1 |
| 16 | 3845000 | 3851199 | 55.68361 | 88.3401 | 127.9322 | 53.27744 | -0.20242 | 2.314952 | 1 |
| 16 | 3983400 | 3985799 | 32.28503 | 52.66898 | 76.32682 | 32.79343 | -0.16143 | 1.523598 | 1 |
| 16 | 4562400 | 4563599 | 16.03479 | 28.75181 | 29.6181 | 12.96905 | 0.358699 | 0.421429 | 1 |
| 16 | 4566200 | 4570799 | 56.7665 | 90.52158 | 105.3459 | 44.14338 | 0.120131 | 2.153213 | 1 |
| 16 | 4633800 | 4647599 | 192.3475 | 282.7846 | 349.4094 | 143.8955 | -0.06535 | 3.82567 | 1 |
| 16 | 5748400 | 5749799 | 14.11254 | 24.74396 | 40.94573 | 18.09387 | -0.33541 | 0.637579 | 1 |
| 16 | 5888400 | 5890799 | 19.7142 | 33.89144 | 52.25385 | 22.92058 | -0.23814 | 0.914386 | 1 |
| 16 | 7278400 | 7282599 | 54.32775 | 86.41945 | 122.2996 | 51.07312 | -0.16543 | 2.233554 | 1 |
| 16 | 7726200 | 7728799 | 27.92657 | 47.22063 | 50.14674 | 21.94516 | 0.294868 | 1.091503 | 1 |
| 16 | 8071800 | 8073799 | 15.06429 | 27.07907 | 30.85192 | 13.48094 | 0.216456 | 0.403791 | 1 |
| 16 | 8885400 | 8895799 | 102.9547 | 158.5522 | 180.7903 | 75.09215 | 0.108357 | 2.925769 | 1 |
| 16 | 9960600 | 9971399 | 103.6157 | 160.0449 | 175.5474 | 72.91868 | 0.167549 | 2.87023 | 1 |
| 16 | 10923800 | 10924799 | 22.82883 | 39.02058 | 42.18655 | 18.49718 | 0.270675 | 0.967094 | 1 |
| 16 | 10996600 | 10998999 | 28.5207 | 46.96913 | 63.64319 | 27.44793 | -0.05781 | 1.380986 | 1 |
| 16 | 11104400 | 11115599 | 66.53525 | 105.3251 | 128.8728 | 53.57824 | 0.037504 | 2.380045 | 1 |
| 16 | 11676400 | 11678599 | 19.96311 | 34.0569 | 52.1946 | 22.88179 | -0.23119 | 0.986506 | 1 |
| 16 | 12139600 | 12140599 | 11.9422 | 21.92744 | 27.1084 | 11.53476 | 0.124775 | 0.251182 | 1 |
| 16 | 12280000 | 12281199 | 14.41529 | 25.27007 | 37.4308 | 16.53802 | -0.17622 | 0.637091 | 1 |
| 16 | 12657200 | 12658199 | 12.99372 | 23.591 | 29.07633 | 12.55321 | 0.115115 | 0.328701 | 1 |
| 16 | 12947800 | 12948599 | 10.22271 | 18.57626 | 32.32408 | 13.94013 | -0.37819 | 0.310386 | 1 |
| 16 | 13358600 | 13367399 | 146.1576 | 220.2094 | 245.9358 | 101.5965 | 0.11277 | 3.379887 | 1 |
| 16 | 14491600 | 14493999 | 25.23361 | 42.03794 | 59.26778 | 25.7499 | -0.11274 | 1.235289 | 1 |
| 16 | 14547400 | 14549199 | 9.205112 | 17.23788 | 27.43902 | 11.33922 | -0.20502 | 0.115334 | 1 |
| 16 | 14633000 | 14634399 | 23.68686 | 39.30261 | 64.0089 | 27.73408 | -0.3205 | 1.280404 | 1 |
| 16 | 14910200 | 14912999 | 16.66205 | 28.97453 | 47.37556 | 20.81186 | -0.31919 | 0.783081 | 1 |
| 16 | 14957800 | 14959999 | 24.95187 | 42.38775 | 48.48994 | 21.25012 | 0.188093 | 1.036908 | 1 |
| 16 | 15044600 | 15045399 | 42.66271 | 68.51319 | 91.11827 | 38.57707 | -0.0605 | 1.914886 | 1 |
| 16 | 15251800 | 15252799 | 13.31797 | 23.74154 | 35.17481 | 15.47806 | -0.16704 | 0.454834 | 1 |
| 16 | 15256200 | 15260799 | 57.08356 | 91.17913 | 105.4738 | 44.30075 | 0.131403 | 2.101732 | 1 |
| 16 | 15273400 | 15281799 | 163.7618 | 244.984 | 272.304 | 112.2941 | 0.110488 | 3.519899 | 1 |
| 16 | 15641600 | 15644199 | 37.38277 | 60.87685 | 64.33039 | 27.6213 | 0.291222 | 1.551248 | 1 |
| 16 | 15844400 | 15847799 | 52.81323 | 84.88062 | 94.10507 | 39.88748 | 0.202823 | 1.904444 | 1 |
| 16 | 16030400 | 16032199 | 10.83386 | 19.49015 | 31.24182 | 13.64211 | -0.27233 | 0.36968 | 1 |
| 16 | 16191400 | 16194599 | 79.08112 | 123.434 | 158.8104 | 65.89823 | -0.0507 | 2.675631 | 1 |
| 16 | 16425000 | 16442999 | 58.85771 | 93.93509 | 108.1379 | 45.36645 | 0.137052 | 2.128942 | 1 |
| 16 | 16636000 | 16638599 | 12.20933 | 22.48687 | 24.58064 | 10.41557 | 0.305572 | 0.230245 | 1 |

| | | | | | | | | | |
|---|---|---|---|---|---|---|---|---|---|
| 16 | 16755000 | 16761599 | 82.66403 | 127.9753 | 189.3689 | 78.65893 | -0.26143 | 2.821356 | 1 |
| 16 | 17087600 | 17090199 | 28.72414 | 46.84333 | 73.42862 | 31.54418 | -0.27467 | 1.51939 | 1 |
| 16 | 17431600 | 17433199 | 16.51781 | 28.74703 | 46.30131 | 20.35033 | -0.29739 | 0.770005 | 1 |
| 16 | 17498400 | 17500199 | 27.22882 | 44.99921 | 61.95808 | 26.77142 | -0.07947 | 1.335012 | 1 |
| 16 | 17828800 | 17830399 | 11.08396 | 19.806 | 33.32638 | 14.63537 | -0.34733 | 0.421017 | 1 |
| 16 | 17923800 | 17931799 | 78.42711 | 121.8532 | 176.081 | 73.11193 | -0.22318 | 2.756292 | 1 |
| 16 | 18662200 | 18665999 | 53.94437 | 86.26088 | 105.3885 | 44.33477 | 0.05446 | 2.06422 | 1 |
| 16 | 18797000 | 18798999 | 31.33349 | 52.99909 | 51.29605 | 22.45041 | 0.428237 | 1.100378 | 1 |
| 16 | 18854400 | 18855399 | 14.5896 | 26.23751 | 31.40697 | 13.71829 | 0.145966 | 0.398873 | 1 |
| 16 | 18899400 | 18904199 | 17.17139 | 30.14827 | 30.54445 | 13.51314 | 0.368691 | 0.612328 | 1 |
| 16 | 18989600 | 18991399 | 24.15636 | 40.86256 | 51.18243 | 22.40122 | 0.057615 | 1.075658 | 1 |
| 16 | 19069600 | 19071599 | 21.35099 | 37.19849 | 38.71156 | 17.02599 | 0.32999 | 0.766051 | 1 |
| 16 | 19115200 | 19118799 | 47.20435 | 76.22385 | 80.64265 | 34.40431 | 0.278351 | 1.75022 | 1 |
| 16 | 19220800 | 19222399 | 20.83511 | 35.81606 | 52.27866 | 22.93141 | -0.1595 | 0.919523 | 1 |
| 16 | 19367200 | 19374199 | 87.59914 | 136.0304 | 185.6659 | 77.09797 | -0.1416 | 2.769357 | 1 |
| 16 | 19713200 | 19714999 | 14.87305 | 26.0185 | 42.52795 | 18.75719 | -0.31788 | 0.6797 | 1 |
| 16 | 19925600 | 19929199 | 29.91227 | 48.98331 | 75.45011 | 32.46232 | -0.24555 | 1.462134 | 1 |
| 16 | 20397000 | 20399799 | 45.1636 | 72.63688 | 84.93665 | 36.03774 | 0.127387 | 1.867882 | 1 |
| 16 | 20801000 | 20804199 | 38.55345 | 62.73295 | 77.52975 | 33.27893 | 0.065051 | 1.574645 | 1 |
| 16 | 20821800 | 20823799 | 24.23756 | 41.32206 | 47.94883 | 21.01936 | 0.168453 | 1.003046 | 1 |
| 16 | 20913800 | 20916399 | 18.41928 | 32.44792 | 39.00643 | 17.26575 | 0.124386 | 0.593638 | 1 |
| 16 | 21567600 | 21570599 | 40.59958 | 65.82137 | 82.92623 | 35.51349 | 0.032767 | 1.645885 | 1 |
| 16 | 21839200 | 21841199 | 23.89811 | 40.86147 | 50.65621 | 22.21138 | 0.07396 | 0.977965 | 1 |
| 16 | 22090000 | 22104199 | 260.9929 | 374.5198 | 433.073 | 178.822 | -0.00378 | 4.196852 | 1 |
| 16 | 22734000 | 22736999 | 15.65953 | 28.1394 | 29.7455 | 13.00123 | 0.323604 | 0.407312 | 1 |
| 16 | 22849600 | 22851599 | 29.74549 | 49.49223 | 54.56801 | 23.6887 | 0.240818 | 1.258321 | 1 |
| 16 | 24005000 | 24007199 | 17.14161 | 30.19771 | 31.92717 | 14.13215 | 0.308777 | 0.573556 | 1 |
| 16 | 25735200 | 25737599 | 16.50223 | 29.38118 | 32.23401 | 14.19901 | 0.262626 | 0.474631 | 1 |
| 16 | 26046400 | 26046999 | 8.972303 | 17.28983 | 18.23907 | 7.071666 | 0.465035 | -0.1281 | 1 |
| 16 | 26684600 | 26686399 | 24.19367 | 40.61808 | 51.71492 | 22.57823 | 0.033785 | 1.142925 | 1 |
| 16 | 26691800 | 26692199 | 12.92322 | 23.7174 | 29.09226 | 12.38695 | 0.135232 | 0.26586 | 1 |
| 16 | 26896400 | 26899199 | 51.17774 | 81.83256 | 100.6496 | 42.33804 | 0.044767 | 2.061958 | 1 |
| 16 | 27282600 | 27286399 | 63.73038 | 100.8471 | 131.1761 | 54.52544 | -0.05102 | 2.389019 | 1 |
| 16 | 27815600 | 27816399 | 13.74435 | 24.82171 | 29.45746 | 12.80247 | 0.163041 | 0.364369 | 1 |
| 16 | 27965200 | 27966399 | 12.04481 | 21.80169 | 34.75522 | 15.06224 | -0.25771 | 0.349318 | 1 |
| 16 | 27994800 | 27995799 | 16.56968 | 29.10635 | 32.22135 | 14.26102 | 0.241737 | 0.606751 | 1 |
| 16 | 28111600 | 28114399 | 23.87354 | 39.55033 | 60.05766 | 25.99392 | -0.21991 | 1.29718 | 1 |
| 16 | 28938000 | 28940599 | 12.6239 | 22.48371 | 36.17934 | 15.93235 | -0.28648 | 0.455899 | 1 |
| 16 | 29092000 | 29092799 | 11.76271 | 21.25399 | 30.69064 | 13.32976 | -0.11748 | 0.350841 | 1 |
| 16 | 29514800 | 29523199 | 97.0124 | 150.0181 | 187.4056 | 77.84387 | -0.01864 | 2.848529 | 1 |
| 16 | 29603200 | 29605599 | 5.819114 | 11.27 | 23.23482 | 8.824143 | -0.4635 | -0.21343 | 1 |
| 16 | 29729800 | 29731399 | 20.19492 | 34.88016 | 42.62208 | 18.69836 | 0.097362 | 0.861922 | 1 |
| 16 | 29782800 | 29785799 | 18.53837 | 31.89455 | 50.07977 | 21.96746 | -0.26409 | 0.898945 | 1 |
| 16 | 29895400 | 29898999 | 19.48586 | 33.68248 | 44.74733 | 19.6312 | -0.02246 | 0.855615 | 1 |
| 16 | 30707600 | 30715799 | 72.99121 | 113.9232 | 165.9833 | 68.87492 | -0.22995 | 2.673535 | 1 |
| 16 | 31959400 | 31966999 | 105.5595 | 160.0956 | 246.9366 | 102.2182 | -0.34452 | 3.239529 | 1 |
| 16 | 32403200 | 32404199 | 11.17206 | 20.91527 | 26.27084 | 10.86174 | 0.133352 | 0.129994 | 1 |

| | | | | | | | | | |
|---|---|---|---|---|---|---|---|---|---|
| 16 | 33110400 | 33114399 | 42.27843 | 68.05035 | 93.63965 | 39.77583 | -0.10557 | 1.842768 | 1 |
| 16 | 33543400 | 33544799 | 14.91115 | 26.72568 | 29.90008 | 13.09721 | 0.240225 | 0.4193 | 1 |
| 16 | 33669200 | 33669999 | 77.83957 | 122.3261 | 132.3547 | 54.9018 | 0.206498 | 2.547541 | 1 |
| 16 | 33671400 | 33673999 | 36.72765 | 59.37039 | 80.20612 | 34.26566 | -0.0717 | 1.701568 | 1 |
| 16 | 35515400 | 35517999 | 29.03695 | 47.87767 | 64.82979 | 27.97412 | -0.05612 | 1.369041 | 1 |
| 16 | 35769200 | 35773199 | 35.27711 | 57.79209 | 61.29095 | 26.37489 | 0.291658 | 1.4592 | 1 |
| 16 | 36047200 | 36051799 | 57.39448 | 91.57101 | 109.951 | 46.10653 | 0.075899 | 2.137681 | 1 |
| 16 | 37518600 | 37521399 | 25.76906 | 43.47023 | 51.21422 | 22.39634 | 0.145489 | 1.104717 | 1 |
| 16 | 37955000 | 37957999 | 32.01123 | 52.24023 | 79.11993 | 33.99773 | -0.22455 | 1.520954 | 1 |
| 16 | 39638800 | 39639799 | 15.199 | 26.5955 | 39.45359 | 17.40498 | -0.1785 | 0.673635 | 1 |
| 16 | 40537800 | 40539399 | 33.6363 | 55.17361 | 64.40816 | 27.7311 | 0.154348 | 1.444532 | 1 |
| 16 | 40632400 | 40634999 | 26.21709 | 44.34504 | 51.76861 | 22.65724 | 0.158749 | 1.083042 | 1 |
| 16 | 40880600 | 40881399 | 13.67332 | 23.96588 | 40.49572 | 17.88998 | -0.36524 | 0.641128 | 1 |
| 16 | 41313200 | 41313999 | 8.644127 | 16.33385 | 24.38343 | 9.915992 | -0.09421 | 0.041642 | 1 |
| 16 | 41442400 | 41443399 | 12.05344 | 21.32116 | 34.8458 | 15.41142 | -0.31359 | 0.505094 | 1 |
| 16 | 41681000 | 41681599 | 11.89715 | 21.73655 | 28.19427 | 12.0769 | 0.049404 | 0.283099 | 1 |
| 16 | 41766600 | 41768599 | 19.44887 | 33.51971 | 52.80197 | 23.16279 | -0.26818 | 0.892556 | 1 |
| 16 | 41823200 | 41824199 | 15.61844 | 27.9688 | 29.39906 | 12.88728 | 0.329064 | 0.429532 | 1 |
| 16 | 42292800 | 42296799 | 61.11353 | 96.97688 | 120.4831 | 50.18685 | 0.018673 | 2.309495 | 1 |
| 16 | 42476600 | 42478199 | 20.23741 | 35.0594 | 40.53485 | 17.78433 | 0.177828 | 0.833614 | 1 |
| 16 | 42599200 | 42600599 | 16.66099 | 28.89118 | 44.7862 | 19.65015 | -0.24304 | 0.815492 | 1 |
| 16 | 42655400 | 42656399 | 12.05114 | 22.03083 | 27.04872 | 11.57771 | 0.128432 | 0.275377 | 1 |
| 16 | 42671600 | 42676199 | 9.098892 | 17.06308 | 25.33101 | 10.44305 | -0.10274 | 0.097115 | 1 |
| 16 | 42761800 | 42763199 | 24.88932 | 42.06293 | 50.13092 | 21.93475 | 0.128982 | 1.083958 | 1 |
| 16 | 42805000 | 42806199 | 20.77909 | 34.85976 | 55.78959 | 24.34588 | -0.29487 | 1.146268 | 1 |
| 16 | 42962800 | 42963799 | 34.3535 | 55.46462 | 84.42653 | 36.0118 | -0.24546 | 1.736495 | 1 |
| 16 | 43588400 | 43590399 | 48.54983 | 77.92388 | 88.55652 | 37.46062 | 0.164864 | 1.936447 | 1 |
| 16 | 44223400 | 44226199 | 30.27639 | 49.52944 | 78.86103 | 33.91654 | -0.29408 | 1.479366 | 1 |
| 16 | 44414800 | 44418599 | 46.06979 | 74.21989 | 88.04619 | 37.4489 | 0.109674 | 1.818842 | 1 |
| 16 | 44739600 | 44741999 | 26.44582 | 44.82439 | 51.75786 | 22.66673 | 0.174639 | 1.067206 | 1 |
| 16 | 45287400 | 45290799 | 17.29404 | 30.09825 | 43.66102 | 19.18999 | -0.14706 | 0.769972 | 1 |
| 16 | 46566000 | 46566599 | 8.46091 | 16.2763 | 25.33112 | 9.870102 | -0.09832 | -0.09739 | 1 |
| 16 | 46939600 | 46942799 | 34.43389 | 56.26615 | 65.42325 | 28.12546 | 0.157094 | 1.49673 | 1 |
| 16 | 47032600 | 47034399 | 19.27062 | 33.03661 | 51.13937 | 22.42801 | -0.24449 | 0.937335 | 1 |
| 16 | 47121200 | 47122999 | 23.84995 | 39.90448 | 64.1932 | 27.96579 | -0.30251 | 1.190817 | 1 |
| 16 | 47124000 | 47126199 | 34.09696 | 55.48579 | 73.93406 | 31.73641 | -0.04315 | 1.562377 | 1 |
| 16 | 47195600 | 47196799 | 12.91841 | 23.01886 | 34.13278 | 15.02525 | -0.16875 | 0.451074 | 1 |
| 16 | 47230000 | 47231799 | 17.52612 | 30.74554 | 37.61494 | 16.62951 | 0.098003 | 0.641161 | 1 |
| 16 | 47516800 | 47523599 | 131.6613 | 199.1477 | 243.7477 | 100.8135 | -0.01442 | 3.298201 | 1 |
| 16 | 47553400 | 47554199 | 11.94538 | 22.29437 | 25.65579 | 10.6628 | 0.252725 | 0.154373 | 1 |
| 16 | 47621600 | 47622399 | 9.370042 | 18.00204 | 20.30293 | 7.940152 | 0.35803 | -0.0885 | 1 |
| 16 | 47841800 | 47844599 | 28.67012 | 46.87086 | 70.92044 | 30.49414 | -0.22131 | 1.48115 | 1 |
| 16 | 47975200 | 47976199 | 8.645086 | 16.55723 | 27.46604 | 10.83868 | -0.20686 | -0.04863 | 1 |
| 16 | 47989000 | 47990999 | 13.28629 | 24.13132 | 36.42395 | 15.71797 | -0.17555 | 0.341116 | 1 |
| 16 | 48004000 | 48013599 | 150.0542 | 226.202 | 237.1748 | 97.99144 | 0.204458 | 3.369022 | 1 |
| 16 | 48045600 | 48046599 | 11.51826 | 20.7467 | 33.74601 | 14.71477 | -0.29189 | 0.374712 | 1 |
| 16 | 48308400 | 48310799 | 28.04141 | 46.93572 | 56.22397 | 24.50003 | 0.121391 | 1.189788 | 1 |

| | | | | | | | | | |
|---|---|---|---|---|---|---|---|---|---|
| 16 | 48475400 | 48477199 | 11.79433 | 21.21481 | 36.25994 | 15.83536 | -0.36476 | 0.387788 | 1 |
| 16 | 48675600 | 48676999 | 13.7997 | 24.56379 | 37.17471 | 16.37858 | -0.19893 | 0.470447 | 1 |
| 16 | 48783000 | 48784199 | 10.08895 | 19.09366 | 20.71769 | 8.397576 | 0.366723 | 0.034376 | 1 |
| 16 | 49048800 | 49050199 | 22.15304 | 38.18841 | 40.0087 | 17.55084 | 0.317961 | 0.885328 | 1 |
| 16 | 49306600 | 49311999 | 55.29107 | 88.4893 | 96.04049 | 40.45076 | 0.226281 | 2.035899 | 1 |
| 16 | 49906600 | 49908999 | 31.3025 | 51.70317 | 60.58958 | 26.17021 | 0.152341 | 1.351602 | 1 |
| 16 | 50586400 | 50587799 | 13.78373 | 24.24525 | 37.80682 | 16.7398 | -0.24965 | 0.582434 | 1 |
| 16 | 50593600 | 50596799 | 53.63991 | 85.3455 | 115.3932 | 48.20475 | -0.09933 | 2.223391 | 1 |
| 16 | 50814600 | 50816799 | 62.30278 | 98.43695 | 127.7894 | 53.07648 | -0.04993 | 2.424035 | 1 |
| 16 | 50949400 | 50950399 | 9.913979 | 18.94482 | 19.20619 | 7.627317 | 0.489772 | -0.03758 | 1 |
| 16 | 51486800 | 51489199 | 22.39552 | 38.87517 | 46.83603 | 20.55173 | 0.1191 | 0.832603 | 1 |
| 16 | 51622600 | 51625199 | 25.47755 | 42.26972 | 67.53646 | 29.26054 | -0.29306 | 1.289863 | 1 |
| 16 | 51765600 | 51768399 | 24.80967 | 42.17195 | 49.56882 | 21.72432 | 0.149242 | 1.032172 | 1 |
| 16 | 51770400 | 51777399 | 87.12921 | 134.7815 | 184.6929 | 76.71692 | -0.15159 | 2.837715 | 1 |
| 16 | 52074400 | 52076599 | 32.49859 | 53.47097 | 67.91992 | 29.28102 | 0.03459 | 1.406428 | 1 |
| 16 | 52100400 | 52101799 | 15.37893 | 27.57779 | 31.32628 | 13.71618 | 0.218707 | 0.421591 | 1 |
| 16 | 52483200 | 52489799 | 120.0775 | 183.88 | 189.7179 | 78.73459 | 0.246269 | 3.042782 | 1 |
| 16 | 52526600 | 52528599 | 25.59143 | 42.98726 | 51.21943 | 22.3693 | 0.129062 | 1.140445 | 1 |
| 16 | 52832400 | 52839599 | 57.18344 | 91.04369 | 106.4917 | 44.52688 | 0.110408 | 2.202947 | 1 |
| 16 | 53168000 | 53168999 | 22.04133 | 37.7011 | 41.71266 | 18.29044 | 0.237664 | 0.957389 | 1 |
| 16 | 53182000 | 53183399 | 14.70401 | 25.73687 | 39.44845 | 17.407 | -0.22547 | 0.665491 | 1 |
| 16 | 53304200 | 53305999 | 17.67942 | 30.93393 | 39.27625 | 17.32587 | 0.044747 | 0.686031 | 1 |
| 16 | 53350400 | 53351399 | 20.2603 | 34.51372 | 47.4596 | 20.80356 | -0.07559 | 0.995831 | 1 |
| 16 | 53411000 | 53411599 | 8.045417 | 15.37733 | 25.08972 | 9.959097 | -0.19064 | -0.04064 | 1 |
| 16 | 54284800 | 54286199 | 29.64542 | 48.41801 | 73.26234 | 31.48989 | -0.22236 | 1.498642 | 1 |
| 16 | 54327400 | 54328999 | 8.008238 | 15.32897 | 26.26494 | 10.38133 | -0.25539 | -0.05384 | 1 |
| 16 | 54710600 | 54713399 | 21.91199 | 37.48403 | 58.3576 | 25.58924 | -0.25338 | 0.975113 | 1 |
| 16 | 54778600 | 54781199 | 39.40249 | 64.15473 | 67.34752 | 28.91491 | 0.300559 | 1.560314 | 1 |
| 16 | 54882000 | 54891799 | 152.9047 | 230.0199 | 257.7931 | 106.4508 | 0.105754 | 3.411583 | 1 |
| 16 | 54939400 | 54941999 | 31.14659 | 51.59894 | 60.65427 | 26.25202 | 0.148218 | 1.317343 | 1 |
| 16 | 54974800 | 54975999 | 20.63088 | 35.8957 | 37.86223 | 16.6384 | 0.310613 | 0.781839 | 1 |
| 16 | 55142200 | 55146999 | 82.52747 | 128.6054 | 156.2331 | 64.84198 | 0.030208 | 2.70396 | 1 |
| 16 | 55474200 | 55475999 | 23.88304 | 40.18049 | 63.30408 | 27.66076 | -0.27263 | 1.13199 | 1 |
| 16 | 55500200 | 55502799 | 40.27641 | 64.99912 | 81.64322 | 34.8051 | 0.03033 | 1.755393 | 1 |
| 16 | 55664200 | 55665199 | 17.64496 | 30.98338 | 34.99578 | 15.48399 | 0.212673 | 0.622205 | 1 |
| 16 | 55714600 | 55716399 | 16.09852 | 28.02366 | 43.65773 | 19.19116 | -0.24959 | 0.758834 | 1 |
| 16 | 55933200 | 55934599 | 18.86461 | 32.9324 | 38.33197 | 16.88024 | 0.169596 | 0.726861 | 1 |
| 16 | 56149200 | 56160599 | 134.2628 | 203.2931 | 225.2878 | 93.20901 | 0.130083 | 3.276686 | 1 |
| 16 | 56758800 | 56763599 | 13.99763 | 24.46969 | 42.10433 | 18.55922 | -0.39134 | 0.6884 | 1 |
| 16 | 57084400 | 57086399 | 33.61107 | 54.69559 | 75.60939 | 32.45565 | -0.09603 | 1.562116 | 1 |
| 16 | 57104200 | 57105999 | 24.15731 | 41.1269 | 46.90489 | 20.55938 | 0.193067 | 1.015934 | 1 |
| 16 | 57516400 | 57517599 | 14.81427 | 26.166 | 36.07667 | 15.96053 | -0.07084 | 0.538607 | 1 |
| 16 | 57788000 | 57788599 | 11.31557 | 20.63293 | 29.76625 | 12.78199 | -0.10531 | 0.294351 | 1 |
| 16 | 58026400 | 58028599 | 24.17227 | 40.15953 | 59.61805 | 25.85448 | -0.18706 | 1.26474 | 1 |
| 16 | 58160800 | 58164399 | 96.85629 | 149.2112 | 167.1937 | 69.44798 | 0.13391 | 2.916182 | 1 |
| 16 | 58628200 | 58631199 | 34.88027 | 57.18443 | 61.74123 | 26.57701 | 0.266381 | 1.451148 | 1 |
| 16 | 58702000 | 58704399 | 18.23531 | 31.92809 | 36.21614 | 15.98507 | 0.206792 | 0.675209 | 1 |

| | | | | | | | | | |
|---|---|---|---|---|---|---|---|---|---|
| 16 | 58997800 | 59000199 | 65.5432 | 103.7416 | 113.0601 | 47.00119 | 0.204168 | 2.381769 | 1 |
| 16 | 59058800 | 59060399 | 29.65649 | 48.47952 | 73.87864 | 31.76516 | -0.23167 | 1.486781 | 1 |
| 16 | 59062600 | 59076399 | 55.08813 | 87.81543 | 118.2804 | 49.54015 | -0.09094 | 2.164535 | 1 |
| 16 | 59172800 | 59178799 | 37.73538 | 60.72408 | 91.25275 | 38.74909 | -0.23274 | 1.842424 | 1 |
| 16 | 59403600 | 59405199 | 9.609311 | 18.04387 | 30.19101 | 12.41908 | -0.27177 | 0.105434 | 1 |
| 16 | 59420800 | 59423399 | 27.86858 | 46.13196 | 61.50556 | 26.60595 | -0.03299 | 1.31766 | 1 |
| 16 | 59604000 | 59605199 | 10.70599 | 19.45536 | 30.32027 | 13.0752 | -0.21986 | 0.310815 | 1 |
| 16 | 59718800 | 59721799 | 28.83072 | 47.14096 | 68.66204 | 29.5248 | -0.1663 | 1.478143 | 1 |
| 16 | 59807000 | 59808599 | 12.12593 | 21.804 | 35.7317 | 15.60905 | -0.30463 | 0.391848 | 1 |
| 16 | 60377400 | 60379399 | 22.30037 | 38.37248 | 46.32255 | 20.31944 | 0.113792 | 0.911029 | 1 |
| 16 | 60549800 | 60551999 | 62.34245 | 98.44782 | 137.5767 | 57.12639 | -0.15671 | 2.438497 | 1 |
| 16 | 60567200 | 60568799 | 25.02328 | 41.96407 | 54.95926 | 23.97781 | -0.00685 | 1.160414 | 1 |
| 16 | 60593000 | 60594599 | 32.58907 | 53.05522 | 73.30056 | 31.4695 | -0.09479 | 1.551503 | 1 |
| 16 | 60751000 | 60753199 | 15.87366 | 27.45913 | 43.29307 | 18.99363 | -0.268 | 0.831629 | 1 |
| 16 | 60752800 | 60756999 | 43.28789 | 69.6666 | 85.13043 | 36.15489 | 0.065214 | 1.845978 | 1 |
| 16 | 60880400 | 60884599 | 58.97724 | 93.9542 | 123.0018 | 51.47151 | -0.051 | 2.191482 | 1 |
| 16 | 60928800 | 60930799 | 21.33022 | 37.14926 | 40.35178 | 17.74347 | 0.26853 | 0.77474 | 1 |
| 16 | 61071000 | 61080999 | 304.6967 | 427.2458 | 550.0803 | 227.9935 | -0.19224 | 4.538085 | 1 |
| 16 | 61143600 | 61156999 | 251.9223 | 364.3219 | 392.2084 | 161.7696 | 0.110435 | 4.081625 | 1 |
| 16 | 61408200 | 61409399 | 12.22516 | 22.64027 | 27.11024 | 11.40176 | 0.182509 | 0.205212 | 1 |
| 16 | 61421000 | 61423199 | 25.38841 | 43.08112 | 47.43608 | 20.78122 | 0.243034 | 1.048873 | 1 |
| 16 | 61535800 | 61537799 | 36.36035 | 58.97665 | 71.54688 | 30.65099 | 0.087895 | 1.623943 | 1 |
| 16 | 62165400 | 62167599 | 21.73672 | 37.55937 | 42.0704 | 18.45659 | 0.222601 | 0.865732 | 1 |
| 16 | 62208800 | 62213599 | 69.32563 | 110.1437 | 122.3558 | 51.0298 | 0.181581 | 2.282831 | 1 |
| 16 | 62215000 | 62216199 | 25.8028 | 43.52993 | 46.9734 | 20.5422 | 0.271818 | 1.101347 | 1 |
| 16 | 62530400 | 62532599 | 35.21865 | 57.1925 | 68.69613 | 29.45542 | 0.103814 | 1.596697 | 1 |
| 16 | 62577800 | 62579599 | 22.81158 | 38.84935 | 46.9136 | 20.56378 | 0.110977 | 1.007005 | 1 |
| 16 | 62666800 | 62667599 | 9.326315 | 17.23162 | 28.32443 | 11.95021 | -0.2746 | 0.197004 | 1 |
| 16 | 62694600 | 62699199 | 60.70711 | 96.35287 | 120.6467 | 50.26596 | 0.007735 | 2.304076 | 1 |
| 16 | 62700200 | 62701599 | 23.5365 | 40.16167 | 46.88132 | 20.55283 | 0.160046 | 0.990745 | 1 |
| 16 | 63259400 | 63260799 | 19.99202 | 34.19926 | 50.06486 | 21.95294 | -0.16464 | 0.960679 | 1 |
| 16 | 63280600 | 63283399 | 32.67303 | 52.97872 | 75.40782 | 32.29735 | -0.14249 | 1.626439 | 1 |
| 16 | 63653200 | 63654999 | 55.63742 | 88.42793 | 115.5294 | 48.20021 | -0.05151 | 2.258861 | 1 |
| 16 | 64122600 | 64123799 | 14.79796 | 26.18656 | 36.02213 | 15.93039 | -0.06654 | 0.517456 | 1 |
| 16 | 64228800 | 64231399 | 18.38521 | 32.30484 | 35.45353 | 15.6954 | 0.254028 | 0.617675 | 1 |
| 16 | 64270400 | 64271199 | 12.34527 | 22.32452 | 27.22582 | 11.81303 | 0.125806 | 0.342927 | 1 |
| 16 | 64392400 | 64393399 | 18.73146 | 32.5901 | 37.91662 | 16.66205 | 0.169696 | 0.771166 | 1 |
| 16 | 64597600 | 64600599 | 31.66217 | 52.59931 | 60.17838 | 26.09616 | 0.187466 | 1.287066 | 1 |
| 16 | 64784800 | 64787199 | 26.97455 | 44.46314 | 67.36878 | 29.06585 | -0.21797 | 1.370039 | 1 |
| 16 | 64788000 | 64793599 | 56.65972 | 90.46148 | 104.1664 | 43.7238 | 0.13727 | 2.113399 | 1 |
| 16 | 64849200 | 64855999 | 88.14298 | 137.1573 | 152.8892 | 63.46791 | 0.152534 | 2.73068 | 1 |
| 16 | 64881800 | 64883999 | 29.93044 | 49.48728 | 56.12549 | 24.25911 | 0.199543 | 1.332221 | 1 |
| 16 | 64996400 | 64997799 | 20.83778 | 36.3455 | 36.73437 | 16.16789 | 0.372274 | 0.747038 | 1 |
| 16 | 64999800 | 65000399 | 6.984266 | 13.49215 | 21.01961 | 8.078444 | -0.07988 | -0.16797 | 1 |
| 16 | 65358600 | 65360199 | 12.04346 | 21.28063 | 35.82251 | 15.8468 | -0.35672 | 0.517756 | 1 |
| 16 | 65706800 | 65707799 | 13.85221 | 24.557 | 31.63527 | 13.98533 | 0.028878 | 0.484879 | 1 |
| 16 | 65846200 | 65848599 | 24.99883 | 42.71611 | 43.62332 | 19.12666 | 0.352399 | 0.978516 | 1 |

| | | | | | | | | | |
|---|---|---|---|---|---|---|---|---|---|
| 16 | 65936200 | 65938199 | 32.34886 | 53.22579 | 57.27908 | 24.69387 | 0.273136 | 1.396855 | 1 |
| 16 | 65958600 | 65961799 | 53.0116 | 84.80776 | 96.39977 | 40.57943 | 0.159096 | 2.045535 | 1 |
| 16 | 66025400 | 66026599 | 8.527161 | 15.6197 | 30.4125 | 12.98272 | -0.52915 | 0.246259 | 1 |
| 16 | 66108600 | 66109199 | 15.172 | 26.86965 | 31.10982 | 13.75581 | 0.18157 | 0.504457 | 1 |
| 16 | 66286400 | 66287399 | 13.01785 | 23.2892 | 32.57216 | 14.28991 | -0.08157 | 0.422758 | 1 |
| 16 | 66320600 | 66322199 | 12.37309 | 22.41431 | 31.64048 | 13.69828 | -0.08214 | 0.341889 | 1 |
| 16 | 66349800 | 66350599 | 11.43977 | 21.50332 | 26.29964 | 10.79937 | 0.179233 | 0.105913 | 1 |
| 16 | 66551000 | 66552399 | 18.96235 | 32.85957 | 46.33958 | 20.33128 | -0.1074 | 0.835058 | 1 |
| 16 | 67130800 | 67131599 | 14.55124 | 25.86001 | 31.42794 | 13.8654 | 0.114304 | 0.471452 | 1 |
| 16 | 67187400 | 67188999 | 20.34976 | 35.52622 | 36.72472 | 16.17278 | 0.340021 | 0.732 | 1 |
| 16 | 67359000 | 67360199 | 15.69669 | 27.75922 | 31.93456 | 14.12451 | 0.189712 | 0.522521 | 1 |
| 16 | 67896400 | 67899599 | 36.21857 | 59.20244 | 65.67019 | 28.23511 | 0.225196 | 1.495826 | 1 |
| 16 | 68305400 | 68308599 | 39.97226 | 64.69562 | 79.35869 | 33.94363 | 0.069015 | 1.67741 | 1 |
| 16 | 68344000 | 68346199 | 24.4093 | 40.90964 | 60.42405 | 26.35193 | -0.17979 | 1.169665 | 1 |
| 16 | 68584000 | 68584999 | 18.38319 | 32.31146 | 33.44543 | 14.81034 | 0.338072 | 0.609677 | 1 |
| 16 | 68959200 | 68963799 | 40.63953 | 65.6709 | 77.71919 | 33.18816 | 0.118162 | 1.714121 | 1 |
| 16 | 69211000 | 69211799 | 15.31903 | 27.06751 | 32.60055 | 14.42161 | 0.123576 | 0.532828 | 1 |
| 16 | 69502400 | 69504199 | 27.13944 | 45.33034 | 53.93513 | 23.4714 | 0.131247 | 1.209019 | 1 |
| 16 | 69585000 | 69585599 | 10.51849 | 19.76457 | 25.28479 | 10.38889 | 0.114368 | 0.098365 | 1 |
| 16 | 69660000 | 69666399 | 121.4618 | 183.9938 | 248.6615 | 102.8942 | -0.15546 | 3.26636 | 1 |
| 16 | 70339000 | 70341799 | 39.73832 | 64.33901 | 73.62909 | 31.5009 | 0.169381 | 1.666407 | 1 |
| 16 | 70614400 | 70615399 | 17.31964 | 29.51331 | 51.59164 | 22.61536 | -0.42026 | 0.988313 | 1 |
| 16 | 70743800 | 70745799 | 21.00706 | 35.39639 | 55.36236 | 24.20191 | -0.26184 | 1.103385 | 1 |
| 16 | 70779600 | 70782999 | 48.81281 | 78.41163 | 88.06158 | 37.2986 | 0.183367 | 1.912346 | 1 |
| 16 | 71266800 | 71268399 | 19.27913 | 33.11601 | 49.53654 | 21.72811 | -0.19477 | 0.91839 | 1 |
| 16 | 71278200 | 71279399 | 23.51354 | 39.3468 | 54.28914 | 23.65305 | -0.08161 | 1.180815 | 1 |
| 16 | 71589400 | 71590799 | 20.42267 | 35.3375 | 41.06897 | 18.01801 | 0.170032 | 0.84625 | 1 |
| 16 | 71772800 | 71773799 | 16.85156 | 29.65911 | 32.99793 | 14.60882 | 0.235113 | 0.58419 | 1 |
| 16 | 71818400 | 71820599 | 33.94303 | 55.55207 | 62.07378 | 26.70095 | 0.21596 | 1.471377 | 1 |
| 16 | 72419800 | 72421599 | 14.69273 | 25.74472 | 39.89727 | 17.62101 | -0.24134 | 0.648272 | 1 |
| 16 | 72713600 | 72715599 | 19.23823 | 33.7002 | 37.12382 | 16.39231 | 0.24876 | 0.674363 | 1 |
| 16 | 73151000 | 73152799 | 20.57931 | 35.31969 | 48.47665 | 21.26215 | -0.07132 | 0.930459 | 1 |
| 16 | 73349800 | 73351199 | 13.4401 | 23.68797 | 34.33002 | 15.19432 | -0.14331 | 0.55224 | 1 |
| 16 | 73477000 | 73478799 | 17.79681 | 30.869 | 46.8714 | 20.56521 | -0.2133 | 0.821177 | 1 |
| 16 | 73794600 | 73795599 | 12.78726 | 22.68958 | 32.73116 | 14.45941 | -0.13262 | 0.47472 | 1 |
| 16 | 73901200 | 73901999 | 10.23665 | 19.05699 | 27.82793 | 11.61523 | -0.09347 | 0.16097 | 1 |
| 16 | 74002600 | 74003399 | 20.0363 | 33.94435 | 51.10553 | 22.37781 | -0.20659 | 1.049639 | 1 |
| 16 | 74375000 | 74375599 | 11.67402 | 21.70884 | 25.18203 | 10.53055 | 0.234414 | 0.174989 | 1 |
| 16 | 74573600 | 74575999 | 30.62622 | 50.31447 | 61.47522 | 26.48576 | 0.089652 | 1.414817 | 1 |
| 16 | 75436800 | 75439799 | 97.70667 | 150.9089 | 158.1808 | 65.70498 | 0.232914 | 2.867869 | 1 |
| 16 | 75533000 | 75534599 | 16.69227 | 28.74169 | 48.99637 | 21.49276 | -0.3818 | 0.883812 | 1 |
| 16 | 75595600 | 75597399 | 9.839313 | 18.14502 | 29.86085 | 12.63096 | -0.27884 | 0.21726 | 1 |
| 16 | 75939200 | 75940399 | 9.889862 | 18.0851 | 28.86277 | 12.34963 | -0.24653 | 0.264399 | 1 |
| 16 | 75971000 | 75974199 | 43.57152 | 70.07644 | 83.25517 | 35.3244 | 0.104539 | 1.86578 | 1 |
| 16 | 75985400 | 75990399 | 12.2993 | 22.91289 | 23.99397 | 10.00314 | 0.384096 | 0.16367 | 1 |
| 16 | 76689800 | 76690799 | 11.07368 | 20.46058 | 30.89273 | 13.03337 | -0.15293 | 0.218275 | 1 |
| 16 | 76911800 | 76912799 | 15.21438 | 26.95122 | 30.48033 | 13.47681 | 0.215179 | 0.498778 | 1 |

| | | | | | | | | | |
|---|---|---|---|---|---|---|---|---|---|
| 16 | 76934400 | 76935599 | 14.45326 | 26.27545 | 29.97328 | 12.91917 | 0.227508 | 0.333532 | 1 |
| 16 | 76989400 | 76991599 | 30.23938 | 49.48439 | 63.33678 | 27.24298 | 0.020485 | 1.465861 | 1 |
| 16 | 77055400 | 77056999 | 27.52519 | 45.35858 | 72.45864 | 31.25828 | -0.29423 | 1.379703 | 1 |
| 16 | 77516200 | 77519799 | 45.41507 | 73.00416 | 97.98429 | 41.54307 | -0.07203 | 1.889737 | 1 |
| 16 | 77734600 | 77736599 | 18.79726 | 32.15623 | 51.6607 | 22.65274 | -0.29834 | 0.955258 | 1 |
| 16 | 77925200 | 77927799 | 21.61483 | 36.68441 | 55.44056 | 24.28923 | -0.21214 | 1.042924 | 1 |
| 16 | 78148000 | 78148399 | 9.374263 | 17.5356 | 24.43315 | 10.11562 | -0.01596 | 0.117554 | 1 |
| 16 | 78242600 | 78248999 | 114.3226 | 172.9931 | 252.4256 | 104.4163 | -0.26731 | 3.286533 | 1 |
| 16 | 78287200 | 78291199 | 44.2585 | 70.99639 | 104.1434 | 44.01656 | -0.20361 | 1.954559 | 1 |
| 16 | 78398400 | 78400999 | 48.15061 | 77.24204 | 95.26242 | 40.26474 | 0.046203 | 1.955775 | 1 |
| 16 | 78681600 | 78682199 | 11.86809 | 21.63602 | 28.66676 | 12.31298 | 0.016257 | 0.295686 | 1 |
| 16 | 78684200 | 78686799 | 52.9251 | 84.53584 | 98.18372 | 41.23809 | 0.125601 | 2.095191 | 1 |
| 16 | 78803400 | 78805599 | 19.1022 | 33.26902 | 37.3755 | 16.43478 | 0.220143 | 0.758697 | 1 |
| 16 | 78991600 | 78994999 | 42.85482 | 68.88944 | 85.55743 | 36.27503 | 0.039756 | 1.879702 | 1 |
| 16 | 79339400 | 79344799 | 73.56232 | 115.5934 | 139.9342 | 58.04516 | 0.044584 | 2.547723 | 1 |
| 16 | 79457600 | 79459199 | 24.53902 | 40.57852 | 66.71109 | 28.8372 | -0.33443 | 1.324923 | 1 |
| 16 | 79577400 | 79579199 | 10.83303 | 19.61898 | 33.27523 | 14.41103 | -0.34564 | 0.335628 | 1 |
| 16 | 79649000 | 79655999 | 86.17902 | 134.1314 | 156.7567 | 65.0712 | 0.084638 | 2.726578 | 1 |
| 16 | 79687000 | 79688999 | 14.27209 | 25.25306 | 37.66743 | 16.65839 | -0.18262 | 0.520603 | 1 |
| 16 | 79711200 | 79713199 | 20.37121 | 35.25346 | 47.39771 | 20.79468 | -0.03931 | 0.853464 | 1 |
| 16 | 79714400 | 79715399 | 15.35488 | 27.62342 | 32.33917 | 14.12131 | 0.178012 | 0.402916 | 1 |
| 16 | 79739200 | 79741599 | 18.48032 | 32.39987 | 34.55539 | 15.26908 | 0.295138 | 0.651637 | 1 |
| 16 | 79903800 | 79904999 | 19.29722 | 33.41292 | 38.60788 | 16.93862 | 0.178656 | 0.832284 | 1 |
| 16 | 79906400 | 79909199 | 27.24241 | 45.1628 | 63.15299 | 27.34676 | -0.10139 | 1.300469 | 1 |
| 16 | 80554200 | 80555399 | 21.92994 | 37.10803 | 53.81356 | 23.55504 | -0.15296 | 1.068111 | 1 |
| 16 | 80609000 | 80612399 | 21.60671 | 37.46493 | 39.89231 | 17.5029 | 0.296282 | 0.829925 | 1 |
| 16 | 80819200 | 80826599 | 139.2085 | 210.5063 | 230.8124 | 95.45536 | 0.143817 | 3.303965 | 1 |
| 16 | 80868000 | 80870599 | 24.57205 | 41.79408 | 43.26695 | 18.96338 | 0.332046 | 1.021359 | 1 |
| 16 | 81475200 | 81476999 | 16.16152 | 28.9322 | 32.76933 | 14.36848 | 0.221378 | 0.439285 | 1 |
| 16 | 81537000 | 81539399 | 24.53964 | 42.00748 | 44.1519 | 19.36124 | 0.311443 | 0.961357 | 1 |
| 16 | 81540400 | 81542999 | 13.53169 | 24.30978 | 33.06163 | 14.45367 | -0.03792 | 0.402086 | 1 |
| 16 | 81544200 | 81545399 | 11.35739 | 21.08362 | 25.59221 | 10.72832 | 0.166289 | 0.180183 | 1 |
| 16 | 81656000 | 81656599 | 12.52475 | 23.16022 | 24.18194 | 10.19104 | 0.377341 | 0.212065 | 1 |
| 16 | 81843000 | 81845199 | 9.209486 | 17.51863 | 21.81384 | 8.746734 | 0.18342 | -0.00571 | 1 |
| 16 | 82224400 | 82225399 | 8.799329 | 17.09053 | 19.43949 | 7.270232 | 0.406488 | -0.24485 | 1 |
| 16 | 82379800 | 82383399 | 51.25529 | 81.97769 | 99.35546 | 41.80894 | 0.066344 | 2.052462 | 1 |
| 16 | 82384600 | 82386399 | 26.26244 | 44.25966 | 53.95052 | 23.58602 | 0.096442 | 1.116939 | 1 |
| 16 | 82429400 | 82433999 | 106.6659 | 163.5192 | 192.1441 | 79.75806 | 0.060017 | 3.017402 | 1 |
| 16 | 82464000 | 82465599 | 14.80341 | 26.47129 | 31.45287 | 13.80426 | 0.151987 | 0.436516 | 1 |
| 16 | 82692800 | 82694999 | 18.28534 | 32.13083 | 35.89105 | 15.88971 | 0.228538 | 0.615513 | 1 |
| 16 | 82714400 | 82716199 | 24.75419 | 40.93822 | 66.58805 | 28.7859 | -0.319 | 1.326673 | 1 |
| 16 | 82751800 | 82752799 | 11.69522 | 21.70789 | 24.57565 | 10.30403 | 0.266148 | 0.182138 | 1 |
| 16 | 82753800 | 82759799 | 96.78553 | 149.2851 | 184.9534 | 76.82629 | -0.00953 | 2.894194 | 1 |
| 16 | 83057000 | 83059599 | 22.32055 | 37.77014 | 54.87944 | 24.02181 | -0.15569 | 1.071717 | 1 |
| 16 | 83128600 | 83130999 | 12.10302 | 21.73161 | 35.19133 | 15.39161 | -0.28879 | 0.398966 | 1 |
| 16 | 83246200 | 83247599 | 33.95799 | 55.35901 | 63.4872 | 27.26992 | 0.174594 | 1.528118 | 1 |
| 16 | 83280400 | 83281599 | 13.98091 | 24.49123 | 39.49989 | 17.44176 | -0.29835 | 0.650356 | 1 |

| | | | | | | | | | |
|---|---|---|---|---|---|---|---|---|---|
| 16 | 83600400 | 83602199 | 19.35525 | 33.73469 | 38.8844 | 17.10623 | 0.1834 | 0.752779 | 1 |
| 16 | 83741400 | 83743599 | 24.6457 | 41.65771 | 56.66984 | 24.79703 | -0.06123 | 1.088593 | 1 |
| 16 | 84123600 | 84125199 | 14.76299 | 26.55352 | 30.39512 | 13.27456 | 0.210308 | 0.397518 | 1 |
| 16 | 84183400 | 84184199 | 11.54229 | 20.5614 | 31.39614 | 13.82364 | -0.21062 | 0.439553 | 1 |
| 16 | 84370800 | 84376399 | 59.09144 | 94.16028 | 111.9354 | 46.85727 | 0.088345 | 2.17832 | 1 |
| 16 | 84468200 | 84470799 | 28.62102 | 47.24658 | 76.98308 | 33.23762 | -0.32197 | 1.362859 | 1 |
| 16 | 84575400 | 84577399 | 22.27537 | 38.42147 | 40.99476 | 17.98372 | 0.291829 | 0.881185 | 1 |
| 16 | 84603800 | 84605199 | 15.85213 | 28.08007 | 32.34222 | 14.30013 | 0.188853 | 0.506221 | 1 |
| 16 | 84857200 | 84859399 | 16.9213 | 30.12539 | 31.12482 | 13.71115 | 0.34864 | 0.474979 | 1 |
| 16 | 85037400 | 85038199 | 10.1133 | 18.94078 | 24.71142 | 10.21004 | 0.079507 | 0.112168 | 1 |
| 16 | 85446800 | 85449799 | 33.23788 | 54.55777 | 61.65705 | 26.55436 | 0.201382 | 1.43222 | 1 |
| 16 | 85768200 | 85769999 | 9.73102 | 18.27865 | 29.08146 | 11.95576 | -0.199 | 0.101207 | 1 |
| 16 | 85871200 | 85873599 | 70.30419 | 110.507 | 127.4447 | 52.86655 | 0.114811 | 2.538577 | 1 |
| 16 | 85878000 | 85878799 | 22.18999 | 38.09044 | 42.2124 | 18.51486 | 0.236179 | 0.928889 | 1 |
| 16 | 85909400 | 85911599 | 18.36297 | 31.87528 | 45.79489 | 20.09487 | -0.13372 | 0.813511 | 1 |
| 16 | 86552600 | 86553999 | 9.259125 | 17.46052 | 26.97081 | 11.01472 | -0.14878 | 0.06741 | 1 |
| 16 | 86746600 | 86748599 | 13.06469 | 23.897 | 26.51483 | 11.341 | 0.274454 | 0.280323 | 1 |
| 16 | 86945400 | 86947199 | 29.06222 | 47.84369 | 63.01568 | 27.17316 | -0.01717 | 1.387962 | 1 |
| 16 | 87173000 | 87174399 | 20.67452 | 35.99121 | 37.85638 | 16.64152 | 0.314754 | 0.774938 | 1 |
| 16 | 88751000 | 88751799 | 11.57871 | 21.48905 | 25.11865 | 10.53342 | 0.220228 | 0.183663 | 1 |
| 16 | 88943600 | 88948199 | 18.03696 | 30.81631 | 49.00863 | 21.48763 | -0.28389 | 0.962111 | 1 |
| 16 | 88980000 | 88986799 | 93.74124 | 145.7063 | 155.3871 | 64.51554 | 0.215064 | 2.751967 | 1 |
| 16 | 88992200 | 88994399 | 35.36362 | 57.50135 | 68.34268 | 29.3278 | 0.120574 | 1.57275 | 1 |
| 16 | 89209600 | 89211799 | 21.08246 | 36.44004 | 43.13038 | 18.92178 | 0.143452 | 0.861198 | 1 |
| 16 | 89735200 | 89743399 | 91.92999 | 141.2067 | 207.0169 | 85.95489 | -0.25673 | 2.9751 | 1 |
| 16 | 89857400 | 89864399 | 111.6173 | 169.3496 | 231.5678 | 95.86763 | -0.17042 | 3.231045 | 1 |
| 16 | 90369600 | 90372399 | 52.76978 | 84.16377 | 111.6469 | 46.79563 | -0.06816 | 2.143652 | 1 |
| 16 | 90373800 | 90383799 | 128.6841 | 195.0652 | 230.9906 | 95.60234 | 0.035869 | 3.253454 | 1 |
| 16 | 90449000 | 90457399 | 30.10158 | 49.56975 | 62.14472 | 26.80103 | 0.053939 | 1.384128 | 1 |
| 16 | 90904000 | 90906999 | 33.34115 | 54.36648 | 76.0927 | 32.68727 | -0.11189 | 1.530185 | 1 |
| 16 | 91184800 | 91187399 | 19.13679 | 33.35708 | 39.37108 | 17.32136 | 0.14931 | 0.750064 | 1 |
| 16 | 91750400 | 91753199 | 30.03358 | 49.77299 | 57.32663 | 24.81794 | 0.177561 | 1.305631 | 1 |
| 16 | 93080800 | 93082999 | 23.68176 | 40.49854 | 43.70302 | 19.16286 | 0.273449 | 0.968189 | 1 |
| 16 | 93500400 | 93500799 | 7.735309 | 14.96936 | 20.48988 | 7.810296 | 0.116372 | -0.18873 | 1 |
| 16 | 93615800 | 93623199 | 102.3142 | 156.5871 | 222.3202 | 92.25242 | -0.21457 | 3.051653 | 1 |
| 16 | 93628800 | 93630799 | 9.278181 | 16.93492 | 30.85816 | 13.23425 | -0.43896 | 0.273352 | 1 |
| 16 | 94746600 | 94748599 | 24.58797 | 41.53457 | 49.59442 | 21.69698 | 0.126283 | 1.086796 | 1 |
| 16 | 95695800 | 95698799 | 20.162 | 34.74741 | 43.72827 | 19.18238 | 0.054544 | 0.884781 | 1 |
| 16 | 95995400 | 95996999 | 20.4539 | 35.08091 | 45.8355 | 20.10264 | -0.00077 | 0.930823 | 1 |
| 16 | 96781000 | 96781999 | 11.17892 | 20.27589 | 32.78619 | 14.17252 | -0.27526 | 0.329956 | 1 |
| 16 | 97274600 | 97278599 | 24.22255 | 40.45516 | 62.08148 | 27.01518 | -0.23474 | 1.20633 | 1 |
| 16 | 97521400 | 97526199 | 81.35692 | 126.9982 | 149.4147 | 61.99887 | 0.078314 | 2.673682 | 1 |
| 16 | 97576200 | 97580199 | 33.84491 | 54.71941 | 86.09559 | 36.78955 | -0.29082 | 1.695236 | 1 |
| 16 | 97836000 | 97878399 | 213.2745 | 316.9077 | 291.7395 | 120.1863 | 0.37336 | 3.650669 | 1 |
| 16 | 97890200 | 97898399 | 129.4596 | 196.8596 | 205.6589 | 85.19846 | 0.22023 | 3.187935 | 1 |
| 17 | 3957000 | 3957999 | 17.15232 | 30.23742 | 36.00542 | 15.93513 | 0.138572 | 0.574979 | 1 |
| 17 | 5417800 | 5420799 | 36.68468 | 59.48149 | 75.62783 | 32.39077 | 0.019818 | 1.633401 | 1 |

| | | | | | | | | | |
|---|---|---|---|---|---|---|---|---|---|
| 17 | 5819200 | 5832599 | 162.2124 | 239.8423 | 344.0895 | 141.7047 | -0.27299 | 3.729175 | 1 |
| 17 | 7759800 | 7761599 | 24.90523 | 42.04137 | 51.74896 | 22.63482 | 0.082505 | 1.09548 | 1 |
| 17 | 8140400 | 8141799 | 13.12781 | 23.97874 | 29.06167 | 12.45253 | 0.146143 | 0.294703 | 1 |
| 17 | 8209000 | 8211199 | 19.54355 | 34.14163 | 36.14598 | 15.92462 | 0.305689 | 0.715561 | 1 |
| 17 | 10004000 | 10011399 | 92.74171 | 144.3098 | 150.8517 | 62.6223 | 0.245097 | 2.73299 | 1 |
| 17 | 10169400 | 10170599 | 14.7006 | 26.47599 | 27.88741 | 12.16173 | 0.330998 | 0.3839 | 1 |
| 17 | 10536800 | 10539399 | 20.7306 | 34.75581 | 57.99257 | 25.29686 | -0.35498 | 1.152402 | 1 |
| 17 | 10663200 | 10667199 | 19.33223 | 33.79046 | 36.71064 | 16.17978 | 0.268711 | 0.708797 | 1 |
| 17 | 11321000 | 11324799 | 57.6852 | 92.01018 | 102.3141 | 42.8882 | 0.186069 | 2.14436 | 1 |
| 17 | 11506400 | 11507799 | 9.057718 | 16.91051 | 30.04775 | 12.48122 | -0.3688 | 0.138944 | 1 |
| 17 | 11718800 | 11719999 | 14.88873 | 26.46431 | 33.03886 | 14.574 | 0.076122 | 0.478775 | 1 |
| 17 | 12034200 | 12035999 | 17.7434 | 31.24131 | 32.32231 | 14.30856 | 0.339515 | 0.582837 | 1 |
| 17 | 12772800 | 12779599 | 100.9494 | 155.0578 | 211.4299 | 87.78696 | -0.1523 | 2.97963 | 1 |
| 17 | 13031400 | 13043399 | 206.882 | 302.2844 | 344.4259 | 141.8979 | 0.043053 | 3.922849 | 1 |
| 17 | 13205400 | 13207399 | 16.07566 | 28.31219 | 36.5432 | 16.17624 | 0.022198 | 0.578126 | 1 |
| 17 | 13285000 | 13286999 | 12.55329 | 22.3885 | 33.60862 | 14.78325 | -0.18574 | 0.440543 | 1 |
| 17 | 13346200 | 13348999 | 13.80426 | 25.12326 | 25.90976 | 11.15232 | 0.373309 | 0.312472 | 1 |
| 17 | 13376800 | 13383199 | 96.87152 | 148.2629 | 208.7553 | 86.62436 | -0.2025 | 3.04869 | 1 |
| 17 | 13385800 | 13394199 | 96.88834 | 148.7101 | 210.2061 | 87.27 | -0.20498 | 2.992078 | 1 |
| 17 | 13468200 | 13471399 | 122.2125 | 184.6458 | 263.7143 | 109.0326 | -0.23832 | 3.320819 | 1 |
| 17 | 14239400 | 14242199 | 21.63445 | 36.91 | 50.62596 | 22.19419 | -0.0718 | 0.988952 | 1 |
| 17 | 14409400 | 14410799 | 15.28155 | 27.13119 | 34.93345 | 15.42475 | 0.030597 | 0.489803 | 1 |
| 17 | 15086400 | 15088799 | 38.82979 | 62.91125 | 68.2241 | 29.20326 | 0.247797 | 1.650196 | 1 |
| 17 | 15090400 | 15092599 | 29.93113 | 49.56397 | 58.92765 | 25.49712 | 0.131824 | 1.316553 | 1 |
| 17 | 15760400 | 15767199 | 55.35681 | 88.55221 | 102.4559 | 43.12364 | 0.133459 | 2.055171 | 1 |
| 17 | 15826600 | 15831199 | 79.76647 | 124.9032 | 145.0775 | 60.17994 | 0.100271 | 2.617391 | 1 |
| 17 | 16349200 | 16351999 | 51.65498 | 82.52617 | 107.8916 | 45.3303 | -0.04449 | 2.089646 | 1 |
| 17 | 16380800 | 16382199 | 12.13663 | 21.91035 | 35.27304 | 15.33688 | -0.27464 | 0.36827 | 1 |
| 17 | 16844600 | 16847199 | 27.28696 | 44.98247 | 69.15979 | 29.83979 | -0.23899 | 1.370377 | 1 |
| 17 | 17091600 | 17097999 | 140.5385 | 212.57 | 229.8147 | 95.05014 | 0.164432 | 3.300242 | 1 |
| 17 | 17170200 | 17172199 | 23.15056 | 39.51958 | 46.87589 | 20.55114 | 0.137093 | 0.984442 | 1 |
| 17 | 17311800 | 17312999 | 8.271073 | 15.50949 | 27.99543 | 11.54299 | -0.38253 | 0.098767 | 1 |
| 17 | 17391400 | 17391999 | 11.10999 | 20.26787 | 29.74142 | 12.76361 | -0.1293 | 0.288649 | 1 |
| 17 | 17450000 | 17457199 | 77.88445 | 122.9806 | 132.1173 | 54.85712 | 0.222067 | 2.449423 | 1 |
| 17 | 17692000 | 17694199 | 21.98773 | 37.37224 | 52.23254 | 22.89159 | -0.09964 | 1.026245 | 1 |
| 17 | 18148200 | 18149399 | 16.72896 | 29.59289 | 30.91093 | 13.67101 | 0.32853 | 0.525407 | 1 |
| 17 | 18301600 | 18303599 | 34.73358 | 57.25032 | 60.2503 | 25.99418 | 0.306249 | 1.384775 | 1 |
| 17 | 18827600 | 18831799 | 28.10679 | 45.83844 | 73.78457 | 31.69757 | -0.31274 | 1.519033 | 1 |
| 17 | 18957000 | 18958799 | 29.38399 | 48.88059 | 55.5719 | 24.12104 | 0.196761 | 1.262657 | 1 |
| 17 | 19215400 | 19217399 | 72.40631 | 113.9015 | 134.3006 | 55.71626 | 0.083768 | 2.524775 | 1 |
| 17 | 19322800 | 19324799 | 34.24712 | 55.37731 | 90.59568 | 38.71935 | -0.34671 | 1.696662 | 1 |
| 17 | 19538000 | 19551199 | 338.6737 | 478.2635 | 458.1608 | 189.6797 | 0.244425 | 4.434471 | 1 |
| 17 | 19705200 | 19705599 | 9.087973 | 17.51105 | 18.18548 | 7.052827 | 0.4868 | -0.12686 | 1 |
| 17 | 19711800 | 19721399 | 53.0168 | 85.61761 | 79.00492 | 33.70969 | 0.475323 | 1.754559 | 1 |
| 17 | 19872400 | 19874199 | 26.2408 | 44.79791 | 45.25078 | 19.83878 | 0.368102 | 0.996453 | 1 |
| 17 | 20335200 | 20336799 | 16.05604 | 28.92785 | 30.77452 | 13.41459 | 0.317085 | 0.397086 | 1 |
| 17 | 20518600 | 20522199 | 52.79141 | 84.73764 | 92.2254 | 39.01674 | 0.227246 | 1.942943 | 1 |

| | | | | | | | | | |
|---|---|---|---|---|---|---|---|---|---|
| 17 | 20790400 | 20791799 | 39.68585 | 64.00285 | 77.54095 | 33.02752 | 0.081246 | 1.773981 | 1 |
| 17 | 20859000 | 20860399 | 20.99576 | 35.47179 | 57.01307 | 24.94368 | -0.30091 | 1.081608 | 1 |
| 17 | 21083800 | 21089999 | 88.9478 | 138.7492 | 145.9587 | 60.56925 | 0.238762 | 2.689554 | 1 |
| 17 | 21164800 | 21166799 | 22.29834 | 38.16794 | 41.65515 | 18.26627 | 0.257415 | 0.950817 | 1 |
| 17 | 21202200 | 21211399 | 181.131 | 268.7098 | 289.2509 | 119.1414 | 0.145678 | 3.673044 | 1 |
| 17 | 21259400 | 21265999 | 111.4897 | 170.7698 | 198.7494 | 82.48725 | 0.072888 | 3.036149 | 1 |
| 17 | 21308200 | 21315199 | 70.58047 | 111.3762 | 132.5143 | 55.00415 | 0.074208 | 2.458674 | 1 |
| 17 | 21316000 | 21329799 | 194.1078 | 285.0365 | 348.1397 | 143.3832 | -0.05028 | 3.843888 | 1 |
| 17 | 21433000 | 21442799 | 149.9242 | 224.6386 | 278.892 | 115.0463 | -0.04691 | 3.488342 | 1 |
| 17 | 21519000 | 21521199 | 36.13843 | 58.34563 | 87.78364 | 37.44308 | -0.22877 | 1.741073 | 1 |
| 17 | 21818200 | 21818999 | 9.904957 | 18.57587 | 26.87539 | 11.07855 | -0.06572 | 0.108762 | 1 |
| 17 | 22034200 | 22036199 | 52.63113 | 83.75262 | 120.0091 | 50.14307 | -0.18272 | 2.221218 | 1 |
| 17 | 22077000 | 22079199 | 28.91901 | 48.22494 | 54.00336 | 23.47722 | 0.218462 | 1.231297 | 1 |
| 17 | 22384400 | 22385799 | 16.18279 | 28.55152 | 29.40963 | 13.01387 | 0.346861 | 0.544621 | 1 |
| 17 | 22402800 | 22407999 | 73.79909 | 116.7592 | 136.063 | 56.5517 | 0.107134 | 2.401659 | 1 |
| 17 | 22642800 | 22644799 | 15.59199 | 27.64278 | 33.38828 | 14.76014 | 0.121175 | 0.498073 | 1 |
| 17 | 22893000 | 22893799 | 11.20986 | 20.33809 | 31.7988 | 13.74057 | -0.22647 | 0.327641 | 1 |
| 17 | 23002800 | 23016799 | 96.79891 | 151.1637 | 162.4219 | 67.39339 | 0.209484 | 2.677169 | 1 |
| 17 | 23040800 | 23045799 | 50.12191 | 80.0355 | 121.8882 | 51.17349 | -0.26514 | 2.108811 | 1 |
| 17 | 23085600 | 23087199 | 10.34137 | 19.30834 | 25.7521 | 10.69568 | 0.042109 | 0.137738 | 1 |
| 17 | 23094400 | 23097199 | 60.24524 | 95.58224 | 121.2427 | 50.4944 | -0.01148 | 2.315665 | 1 |
| 17 | 23229400 | 23230999 | 13.76489 | 24.49035 | 38.71102 | 17.0622 | -0.26187 | 0.474719 | 1 |
| 17 | 23419400 | 23432799 | 145.0472 | 219.1982 | 231.4064 | 95.68212 | 0.197671 | 3.317789 | 1 |
| 17 | 23720800 | 23721999 | 26.83499 | 44.89716 | 55.01075 | 23.96487 | 0.089002 | 1.191468 | 1 |
| 17 | 24226400 | 24228799 | 21.86721 | 36.57817 | 60.20004 | 26.22214 | -0.33524 | 1.182547 | 1 |
| 17 | 28882000 | 28883799 | 19.54264 | 34.14297 | 35.64092 | 15.70296 | 0.325975 | 0.71368 | 1 |
| 17 | 29126600 | 29129599 | 20.72303 | 34.6733 | 60.74381 | 26.46336 | -0.42512 | 1.174668 | 1 |
| 17 | 31461800 | 31464399 | 19.35748 | 33.25148 | 50.0802 | 21.96659 | -0.20469 | 0.917072 | 1 |
| 17 | 31767800 | 31774399 | 75.35126 | 117.4054 | 172.7143 | 71.68378 | -0.24578 | 2.705317 | 1 |
| 17 | 32778600 | 32786399 | 95.77518 | 148.3553 | 187.826 | 78.01782 | -0.03607 | 2.816825 | 1 |
| 17 | 32794600 | 32806999 | 137.0255 | 206.7754 | 255.6709 | 105.6667 | -0.03198 | 3.346866 | 1 |
| 17 | 32811200 | 32819599 | 125.8897 | 191.6564 | 207.6149 | 86.03905 | 0.169356 | 3.163542 | 1 |
| 17 | 32933000 | 32940199 | 162.0002 | 242.6258 | 259.7103 | 107.1244 | 0.166306 | 3.497955 | 1 |
| 17 | 33231400 | 33250599 | 434.126 | 599.6818 | 614.9866 | 255.4933 | 0.114096 | 4.768582 | 1 |
| 17 | 33261800 | 33273399 | 192.9901 | 282.0328 | 361.6849 | 149.0068 | -0.12722 | 3.919731 | 1 |
| 17 | 33351000 | 33353999 | 28.01509 | 47.09407 | 52.90676 | 23.11197 | 0.213632 | 1.142861 | 1 |
| 17 | 33497200 | 33498399 | 16.16714 | 28.29723 | 37.89746 | 16.72231 | -0.03163 | 0.672674 | 1 |
| 17 | 33585800 | 33593399 | 106.0717 | 164.2245 | 165.6699 | 68.81496 | 0.290772 | 2.829003 | 1 |
| 17 | 33632200 | 33633999 | 30.92907 | 51.19906 | 57.29805 | 24.78619 | 0.218831 | 1.322512 | 1 |
| 17 | 33642400 | 33643399 | 19.98479 | 34.94193 | 36.13502 | 15.93027 | 0.339512 | 0.706337 | 1 |
| 17 | 34322600 | 34327199 | 38.17222 | 61.68827 | 88.31172 | 37.71491 | -0.15569 | 1.714722 | 1 |
| 17 | 34460000 | 34461799 | 26.06094 | 43.44504 | 62.11716 | 26.99916 | -0.13297 | 1.233172 | 1 |
| 17 | 34777800 | 34780799 | 19.83044 | 33.59988 | 52.16732 | 22.84362 | -0.2509 | 1.047178 | 1 |
| 17 | 34985400 | 34987399 | 10.54563 | 19.71895 | 26.29489 | 10.89372 | 0.045391 | 0.132342 | 1 |
| 17 | 35591000 | 35602999 | 85.17448 | 133.09 | 142.2332 | 59.0129 | 0.217913 | 2.65589 | 1 |
| 17 | 36248000 | 36256199 | 126.9517 | 190.9481 | 271.1906 | 111.9784 | -0.23587 | 3.410503 | 1 |
| 17 | 36310400 | 36318599 | 101.6653 | 156.9355 | 162.5021 | 67.50016 | 0.249884 | 2.87946 | 1 |

| | | | | | | | | | |
|---|---|---|---|---|---|---|---|---|---|
| 17 | 36319800 | 36320599 | 13.19328 | 23.5752 | 30.6339 | 13.45294 | 0.023123 | 0.428391 | 1 |
| 17 | 36365200 | 36369599 | 47.32735 | 75.53828 | 114.8946 | 48.20582 | -0.26401 | 2.11818 | 1 |
| 17 | 36797200 | 36799599 | 25.38221 | 42.41156 | 57.66828 | 25.10207 | -0.06072 | 1.202801 | 1 |
| 17 | 36882000 | 36884399 | 20.30807 | 35.08221 | 40.62014 | 17.82021 | 0.175003 | 0.860548 | 1 |
| 17 | 37419600 | 37420799 | 20.49434 | 35.64374 | 37.80991 | 16.6112 | 0.302305 | 0.784877 | 1 |
| 17 | 37607800 | 37608399 | 12.45413 | 22.29399 | 32.68229 | 14.33101 | -0.14834 | 0.417116 | 1 |
| 17 | 37665000 | 37666599 | 19.92613 | 34.69139 | 40.39764 | 17.75962 | 0.168466 | 0.771776 | 1 |
| 17 | 37970400 | 37971199 | 15.90028 | 28.32945 | 28.44346 | 12.52182 | 0.390652 | 0.460628 | 1 |
| 17 | 38054600 | 38055599 | 16.03503 | 28.41764 | 32.96956 | 14.57612 | 0.178952 | 0.506302 | 1 |
| 17 | 38164200 | 38164799 | 8.385459 | 16.24011 | 20.44119 | 7.763578 | 0.240131 | -0.19738 | 1 |
| 17 | 38355400 | 38362599 | 121.6141 | 186.0426 | 204.7861 | 84.96791 | 0.151798 | 3.065433 | 1 |
| 17 | 38501200 | 38503599 | 29.92038 | 51.18769 | 44.59324 | 19.55383 | 0.58122 | 0.991245 | 1 |
| 17 | 39200200 | 39202399 | 17.75785 | 30.44039 | 48.48231 | 21.26256 | -0.28516 | 0.931873 | 1 |
| 17 | 39368000 | 39383399 | 224.2694 | 332.1399 | 288.6927 | 118.893 | 0.451775 | 3.709067 | 1 |
| 17 | 39871600 | 39872999 | 18.2435 | 32.13276 | 32.41768 | 14.34981 | 0.376067 | 0.584707 | 1 |
| 17 | 40302600 | 40303999 | 39.12634 | 63.47001 | 69.66561 | 29.84744 | 0.232049 | 1.625976 | 1 |
| 17 | 40394000 | 40394599 | 10.55374 | 19.89587 | 22.77302 | 9.305109 | 0.280506 | 0.072944 | 1 |
| 17 | 40489600 | 40494999 | 47.35423 | 76.38203 | 78.91374 | 33.6191 | 0.310688 | 1.778778 | 1 |
| 17 | 40680800 | 40683399 | 30.62834 | 50.65837 | 60.59459 | 26.19691 | 0.122993 | 1.333903 | 1 |
| 17 | 40794200 | 40794799 | 10.48043 | 20.055 | 23.41181 | 9.261721 | 0.292836 | -0.03251 | 1 |
| 17 | 40929600 | 40942999 | 80.79044 | 126.8827 | 135.1484 | 56.05604 | 0.228388 | 2.561967 | 1 |
| 17 | 40956000 | 40957999 | 27.05984 | 45.88443 | 47.96572 | 21.00869 | 0.317661 | 1.061931 | 1 |
| 17 | 41007400 | 41008799 | 16.93861 | 29.3645 | 41.70541 | 18.29829 | -0.11737 | 0.814707 | 1 |
| 17 | 41015400 | 41017199 | 17.64289 | 31.15291 | 35.81775 | 15.84691 | 0.189569 | 0.557371 | 1 |
| 17 | 41363200 | 41365199 | 30.32172 | 49.43356 | 77.78882 | 33.41356 | -0.28063 | 1.52873 | 1 |
| 17 | 41501400 | 41504999 | 51.34975 | 82.09561 | 113.9583 | 47.9264 | -0.12973 | 2.069894 | 1 |
| 17 | 42027400 | 42030199 | 44.00854 | 70.40365 | 104.5057 | 43.99067 | -0.2252 | 2.040046 | 1 |
| 17 | 42317600 | 42318799 | 26.05198 | 42.8474 | 67.05155 | 28.90188 | -0.26563 | 1.395835 | 1 |
| 17 | 42382800 | 42383999 | 16.67167 | 28.96074 | 42.70715 | 18.74864 | -0.17082 | 0.793073 | 1 |
| 17 | 43208600 | 43211199 | 48.55499 | 78.08334 | 82.09818 | 34.82829 | 0.28023 | 1.874284 | 1 |
| 17 | 43255000 | 43259999 | 63.71759 | 101.3155 | 115.3573 | 48.14988 | 0.147196 | 2.25146 | 1 |
| 17 | 43572600 | 43579199 | 70.45664 | 110.6216 | 166.4425 | 69.03341 | -0.26968 | 2.565375 | 1 |
| 17 | 43632200 | 43636199 | 37.64491 | 61.21226 | 79.91015 | 34.29205 | -0.01453 | 1.58202 | 1 |
| 17 | 43772800 | 43774199 | 14.9009 | 26.74399 | 30.87207 | 13.50733 | 0.196384 | 0.412247 | 1 |
| 17 | 44034800 | 44036399 | 19.50795 | 34.09557 | 40.28954 | 17.75624 | 0.147943 | 0.715415 | 1 |
| 17 | 44725600 | 44728599 | 31.38881 | 52.99815 | 49.08247 | 21.47 | 0.49151 | 1.112567 | 1 |
| 17 | 45419800 | 45422999 | 28.8421 | 47.16214 | 74.39603 | 31.99106 | -0.28105 | 1.480736 | 1 |
| 17 | 45495200 | 45496199 | 12.02014 | 22.58046 | 25.6959 | 10.56236 | 0.281202 | 0.112714 | 1 |
| 17 | 45773400 | 45774199 | 10.95488 | 20.21754 | 28.79782 | 12.1688 | -0.07074 | 0.220658 | 1 |
| 17 | 45954800 | 45955999 | 13.2322 | 23.44189 | 32.68529 | 14.45149 | -0.08504 | 0.490208 | 1 |
| 17 | 47339800 | 47341399 | 17.96402 | 31.51817 | 34.66287 | 15.32623 | 0.251212 | 0.639027 | 1 |
| 17 | 47394200 | 47411599 | 185.5358 | 273.5697 | 352.5444 | 145.1823 | -0.12202 | 3.777254 | 1 |
| 17 | 48518600 | 48521399 | 35.29418 | 57.03504 | 85.08163 | 36.33367 | -0.21493 | 1.710962 | 1 |
| 17 | 48939200 | 48940199 | 15.29873 | 27.1021 | 29.63678 | 13.1037 | 0.263704 | 0.498897 | 1 |
| 17 | 48952800 | 48955799 | 19.59153 | 33.39436 | 51.12902 | 22.4131 | -0.23 | 0.990369 | 1 |
| 17 | 50382400 | 50382999 | 13.67154 | 24.76246 | 27.91719 | 12.0888 | 0.239953 | 0.343089 | 1 |
| 17 | 50384000 | 50388799 | 88.30235 | 137.7562 | 141.5928 | 58.75688 | 0.27229 | 2.687235 | 1 |

| | | | | | | | | | |
|---|---|---|---|---|---|---|---|---|---|
| 17 | 50724200 | 50732399 | 120.4401 | 182.565 | 252.4594 | 104.4872 | -0.18775 | 3.253068 | 1 |
| 17 | 50965400 | 50974599 | 167.6729 | 250.7979 | 268.0541 | 110.5383 | 0.166791 | 3.523353 | 1 |
| 17 | 51266600 | 51267999 | 40.10372 | 64.35843 | 98.42184 | 41.62474 | -0.26278 | 1.936605 | 1 |
| 17 | 51521400 | 51523799 | 30.62779 | 51.13649 | 55.23138 | 24.02947 | 0.27033 | 1.222982 | 1 |
| 17 | 52009600 | 52012599 | 27.96703 | 47.05143 | 49.63796 | 21.68935 | 0.304071 | 1.132472 | 1 |
| 17 | 52138200 | 52142599 | 36.90346 | 59.46223 | 94.80811 | 40.33438 | -0.31571 | 1.801233 | 1 |
| 17 | 52169800 | 52174199 | 39.4004 | 63.91941 | 72.16659 | 30.92068 | 0.191508 | 1.625018 | 1 |
| 17 | 52687400 | 52689999 | 29.99979 | 49.90154 | 52.96829 | 22.98985 | 0.295399 | 1.259869 | 1 |
| 17 | 52792000 | 52798799 | 104.3684 | 160.4469 | 185.5179 | 77.03722 | 0.086343 | 2.961299 | 1 |
| 17 | 53268400 | 53269799 | 13.37009 | 24.11037 | 29.93484 | 13.03387 | 0.096432 | 0.369854 | 1 |
| 17 | 53666400 | 53670399 | 26.56147 | 43.83619 | 65.30245 | 28.1898 | -0.19297 | 1.349882 | 1 |
| 17 | 53671200 | 53675999 | 10.49243 | 19.80868 | 24.79738 | 10.10341 | 0.155302 | 0.066221 | 1 |
| 17 | 53785600 | 53787199 | 15.65005 | 27.27387 | 41.59175 | 18.29602 | -0.21862 | 0.738624 | 1 |
| 17 | 54127200 | 54132999 | 147.5558 | 221.7223 | 238.4313 | 98.4259 | 0.163906 | 3.431114 | 1 |
| 17 | 54288600 | 54292999 | 59.32971 | 94.55833 | 97.82583 | 40.96189 | 0.288785 | 2.168254 | 1 |
| 17 | 54338000 | 54339399 | 9.081418 | 17.17417 | 25.94538 | 10.53298 | -0.10942 | 0.043292 | 1 |
| 17 | 54346800 | 54348199 | 23.34671 | 39.59302 | 44.33995 | 19.42134 | 0.218665 | 1.042011 | 1 |
| 17 | 54356400 | 54358199 | 18.39635 | 31.83023 | 44.24934 | 19.4133 | -0.08727 | 0.842148 | 1 |
| 17 | 54413600 | 54416999 | 40.81848 | 65.75268 | 91.78104 | 39.03127 | -0.12472 | 1.815351 | 1 |
| 17 | 54419200 | 54420599 | 22.10905 | 36.79694 | 57.85497 | 25.12063 | -0.2693 | 1.238024 | 1 |
| 17 | 54439400 | 54444199 | 69.6729 | 110.3887 | 119.2915 | 49.61685 | 0.217465 | 2.360755 | 1 |
| 17 | 54726400 | 54731199 | 69.2208 | 108.6401 | 156.5433 | 64.92447 | -0.20776 | 2.57216 | 1 |
| 17 | 54738600 | 54742399 | 64.79441 | 102.9586 | 112.7373 | 47.02474 | 0.202541 | 2.272428 | 1 |
| 17 | 54810400 | 54818199 | 123.7797 | 188.3091 | 213.986 | 88.66033 | 0.099543 | 3.178138 | 1 |
| 17 | 55076600 | 55078399 | 12.55809 | 23.31518 | 25.55687 | 10.71187 | 0.312812 | 0.191102 | 1 |
| 17 | 55115800 | 55116599 | 10.80092 | 19.87897 | 30.3325 | 12.86507 | -0.17334 | 0.237821 | 1 |
| 17 | 55311000 | 55312199 | 15.31185 | 26.82171 | 46.18696 | 20.39388 | -0.39269 | 0.668669 | 1 |
| 17 | 55479400 | 55481599 | 19.98621 | 33.88571 | 52.17121 | 22.8498 | -0.2388 | 1.042171 | 1 |
| 17 | 55683400 | 55685399 | 33.2869 | 54.81702 | 58.3737 | 25.17507 | 0.288822 | 1.391476 | 1 |
| 17 | 55881000 | 55882999 | 15.05707 | 27.36291 | 28.46842 | 12.27629 | 0.359339 | 0.338681 | 1 |
| 17 | 55895400 | 55896599 | 13.6536 | 24.29688 | 31.08845 | 13.70033 | 0.041703 | 0.457739 | 1 |
| 17 | 55982800 | 55990199 | 132.6875 | 200.6279 | 227.404 | 94.04318 | 0.095872 | 3.304829 | 1 |
| 17 | 56179600 | 56180999 | 25.72198 | 43.44827 | 51.19277 | 22.39595 | 0.145505 | 1.095155 | 1 |
| 17 | 57741200 | 57742599 | 22.62692 | 38.58088 | 45.86185 | 20.10469 | 0.133858 | 0.994088 | 1 |
| 17 | 58038600 | 58041399 | 38.92806 | 63.11016 | 70.20037 | 30.06314 | 0.212039 | 1.637251 | 1 |
| 17 | 58044000 | 58045199 | 15.13618 | 26.78644 | 34.50163 | 15.25781 | 0.02798 | 0.517061 | 1 |
| 17 | 58093200 | 58094599 | 22.84053 | 39.11999 | 40.13626 | 17.60109 | 0.346404 | 0.94692 | 1 |
| 17 | 58166200 | 58168399 | 28.63401 | 47.41126 | 61.07034 | 26.42239 | 0.01656 | 1.315509 | 1 |
| 17 | 58262000 | 58264199 | 33.64789 | 55.45163 | 64.66114 | 27.89522 | 0.158592 | 1.387705 | 1 |
| 17 | 58631600 | 58633199 | 20.39734 | 34.6849 | 52.20142 | 22.87879 | -0.2059 | 1.016287 | 1 |
| 17 | 58856200 | 58858999 | 31.72936 | 51.83328 | 73.63138 | 31.65112 | -0.13133 | 1.499823 | 1 |
| 17 | 59037200 | 59040399 | 23.03583 | 38.61395 | 63.03858 | 27.49497 | -0.32368 | 1.167706 | 1 |
| 17 | 59145400 | 59158199 | 264.5254 | 375.0561 | 502.7418 | 207.9985 | -0.23424 | 4.372814 | 1 |
| 17 | 59351800 | 59353599 | 21.66693 | 36.51116 | 52.17601 | 22.80953 | -0.13196 | 1.103 | 1 |
| 17 | 59809800 | 59811199 | 16.95869 | 29.77667 | 37.15093 | 16.43676 | 0.069954 | 0.624405 | 1 |
| 17 | 60581200 | 60583999 | 30.64406 | 51.28144 | 53.49574 | 23.30814 | 0.320346 | 1.200245 | 1 |
| 17 | 60615400 | 60621399 | 109.5009 | 167.3223 | 203.5923 | 84.45046 | 0.006072 | 3.084051 | 1 |

| | | | | | | | | | |
|---|---|---|---|---|---|---|---|---|---|
| 17 | 61051800 | 61053199 | 18.84714 | 32.678 | 41.6292 | 18.26493 | 0.038815 | 0.822118 | 1 |
| 17 | 61096200 | 61105399 | 102.7029 | 159.358 | 176.8562 | 73.44846 | 0.155685 | 2.790132 | 1 |
| 17 | 61681000 | 61682199 | 16.22734 | 29.01208 | 31.28149 | 13.73123 | 0.291063 | 0.446251 | 1 |
| 17 | 61757000 | 61757799 | 8.419066 | 16.2032 | 24.19234 | 9.414031 | -0.03732 | -0.10609 | 1 |
| 17 | 61833800 | 61837599 | 41.24749 | 66.39067 | 96.75608 | 41.09935 | -0.18829 | 1.841122 | 1 |
| 17 | 62049200 | 62049999 | 12.27434 | 22.60773 | 24.47969 | 10.3721 | 0.318956 | 0.228763 | 1 |
| 17 | 62066200 | 62068399 | 17.62433 | 30.70667 | 43.10426 | 18.95832 | -0.09972 | 0.755532 | 1 |
| 17 | 62699000 | 62700999 | 43.23134 | 69.46048 | 88.83199 | 37.63652 | -0.00331 | 1.897795 | 1 |
| 17 | 62709000 | 62709799 | 12.12119 | 22.28779 | 24.02636 | 10.20357 | 0.322982 | 0.23601 | 1 |
| 17 | 63065400 | 63067799 | 23.98009 | 39.79686 | 65.7943 | 28.51147 | -0.34214 | 1.279226 | 1 |
| 17 | 63396000 | 63398199 | 26.6541 | 44.63108 | 54.46931 | 23.74127 | 0.094653 | 1.18147 | 1 |
| 17 | 63427800 | 63436599 | 182.4965 | 268.5888 | 344.8128 | 141.9955 | -0.11905 | 3.807534 | 1 |
| 17 | 63438000 | 63441999 | 74.33099 | 117.3446 | 131.1355 | 54.44361 | 0.164973 | 2.450793 | 1 |
| 17 | 63514400 | 63516799 | 26.47184 | 43.62118 | 69.07351 | 29.79749 | -0.28163 | 1.37264 | 1 |
| 17 | 63978000 | 63979799 | 18.60499 | 32.06297 | 45.32743 | 19.88386 | -0.11253 | 0.881071 | 1 |
| 17 | 64008600 | 64009599 | 13.55454 | 23.94579 | 39.72912 | 17.57568 | -0.33651 | 0.534049 | 1 |
| 17 | 64469200 | 64472799 | 51.52252 | 82.02071 | 121.6129 | 50.84086 | -0.23136 | 2.205378 | 1 |
| 17 | 64696200 | 64698999 | 24.15413 | 41.19405 | 43.72103 | 19.16655 | 0.296742 | 0.994383 | 1 |
| 17 | 64729200 | 64731399 | 23.13281 | 40.27958 | 38.76858 | 17.04487 | 0.44232 | 0.786286 | 1 |
| 17 | 65052000 | 65052799 | 11.45786 | 21.46572 | 24.05842 | 9.934932 | 0.297591 | 0.120104 | 1 |
| 17 | 65390400 | 65393199 | 38.66947 | 62.72123 | 69.86437 | 29.92991 | 0.210654 | 1.625001 | 1 |
| 17 | 65597800 | 65598599 | 10.24811 | 19.41876 | 22.19145 | 8.971557 | 0.295587 | 0.031123 | 1 |
| 17 | 65637800 | 65638799 | 15.49197 | 27.05126 | 37.54056 | 16.53424 | -0.08277 | 0.704825 | 1 |
| 17 | 65762600 | 65765999 | 30.37018 | 49.90131 | 66.44747 | 28.6298 | -0.03402 | 1.416769 | 1 |
| 17 | 65775200 | 65778199 | 21.23272 | 35.51698 | 59.06047 | 25.72584 | -0.35005 | 1.179646 | 1 |
| 17 | 65930400 | 65932399 | 19.6066 | 34.04868 | 42.58492 | 18.69139 | 0.065504 | 0.808802 | 1 |
| 17 | 66472400 | 66491999 | 302.8923 | 431.0759 | 465.7857 | 192.5867 | 0.082104 | 4.319314 | 1 |
| 17 | 66918600 | 66920799 | 26.03606 | 42.86562 | 69.4227 | 29.93706 | -0.31447 | 1.383975 | 1 |
| 17 | 67312000 | 67315399 | 28.0593 | 46.44431 | 67.7582 | 29.30924 | -0.16262 | 1.323509 | 1 |
| 17 | 67572600 | 67575199 | 22.44998 | 37.65178 | 57.02521 | 24.88045 | -0.21581 | 1.152666 | 1 |
| 17 | 67613600 | 67614999 | 12.08565 | 22.46317 | 29.87327 | 12.50201 | 0.037143 | 0.189695 | 1 |
| 17 | 67731800 | 67738399 | 84.44523 | 131.3109 | 170.0751 | 70.60959 | -0.06454 | 2.743724 | 1 |
| 17 | 67744400 | 67751799 | 41.89988 | 67.56924 | 85.96619 | 36.61315 | 0.010628 | 1.780783 | 1 |
| 17 | 67882800 | 67888999 | 73.04804 | 114.3671 | 156.1286 | 64.76499 | -0.13265 | 2.61759 | 1 |
| 17 | 68601200 | 68610199 | 178.8221 | 264.0622 | 350.7701 | 144.454 | -0.16374 | 3.752339 | 1 |
| 17 | 68882000 | 68883599 | 17.83265 | 31.30034 | 36.14812 | 15.9884 | 0.180988 | 0.634418 | 1 |
| 17 | 68896200 | 68898399 | 22.0996 | 37.38913 | 53.81473 | 23.5544 | -0.14214 | 1.070292 | 1 |
| 17 | 69143200 | 69144999 | 17.0108 | 29.30238 | 45.86888 | 20.1211 | -0.25904 | 0.878476 | 1 |
| 17 | 70015200 | 70016999 | 32.63881 | 53.02865 | 77.60346 | 33.28656 | -0.17981 | 1.586425 | 1 |
| 17 | 70209200 | 70210199 | 11.86686 | 21.99149 | 26.69413 | 11.21651 | 0.164318 | 0.19972 | 1 |
| 17 | 70456000 | 70457799 | 42.73092 | 68.781 | 82.32468 | 34.97123 | 0.095316 | 1.837973 | 1 |
| 17 | 70535200 | 70537799 | 21.62941 | 37.09566 | 42.66038 | 18.71003 | 0.182576 | 0.930903 | 1 |
| 17 | 71093600 | 71095999 | 19.9929 | 34.45322 | 48.474 | 21.26414 | -0.10587 | 0.889512 | 1 |
| 17 | 71100000 | 71101999 | 14.53673 | 25.4024 | 44.12216 | 19.44238 | -0.40523 | 0.696756 | 1 |
| 17 | 71297000 | 71299999 | 21.11451 | 35.62359 | 53.25405 | 23.28941 | -0.19677 | 1.089359 | 1 |
| 17 | 71727800 | 71729999 | 60.35881 | 95.49069 | 131.1496 | 54.5083 | -0.12959 | 2.392805 | 1 |
| 17 | 72442200 | 72445399 | 48.15873 | 77.28597 | 90.48834 | 38.27013 | 0.121751 | 1.942214 | 1 |

| | | | | | | | | | |
|---|---|---|---|---|---|---|---|---|---|
| 17 | 72752400 | 72758999 | 61.53918 | 97.45624 | 127.4573 | 53.00073 | -0.05795 | 2.367489 | 1 |
| 17 | 72941800 | 72944399 | 29.35448 | 47.94228 | 76.6067 | 32.92723 | -0.30081 | 1.50099 | 1 |
| 17 | 73076600 | 73078199 | 11.39525 | 20.61561 | 29.68129 | 12.87136 | -0.11191 | 0.33746 | 1 |
| 17 | 74168200 | 74170199 | 24.53517 | 40.76508 | 60.13464 | 26.07966 | -0.1779 | 1.266799 | 1 |
| 17 | 74391600 | 74395599 | 49.84716 | 79.45632 | 113.5158 | 47.54077 | -0.17581 | 2.163178 | 1 |
| 17 | 74416800 | 74418599 | 10.51583 | 19.31437 | 30.85565 | 13.12375 | -0.24198 | 0.248947 | 1 |
| 17 | 74578800 | 74580199 | 19.3924 | 33.29587 | 47.94747 | 21.03072 | -0.14027 | 0.920131 | 1 |
| 17 | 75071000 | 75074799 | 45.04032 | 72.59883 | 83.89298 | 35.70809 | 0.148346 | 1.799473 | 1 |
| 17 | 75121600 | 75124599 | 19.51804 | 34.31805 | 35.81538 | 15.85931 | 0.326347 | 0.615328 | 1 |
| 17 | 75653800 | 75654999 | 14.48403 | 26.22796 | 29.85467 | 12.93155 | 0.225823 | 0.354088 | 1 |
| 17 | 75658800 | 75660399 | 20.0354 | 34.49185 | 46.32908 | 20.32261 | -0.03959 | 0.894099 | 1 |
| 17 | 75705400 | 75706799 | 20.31063 | 34.38817 | 53.77546 | 23.54256 | -0.2613 | 1.056697 | 1 |
| 17 | 75864600 | 75870599 | 74.24423 | 117.0587 | 134.9552 | 56.00438 | 0.118655 | 2.47987 | 1 |
| 17 | 75975400 | 75976799 | 19.41135 | 33.66253 | 43.12958 | 18.92333 | 0.030502 | 0.823294 | 1 |
| 17 | 76523800 | 76525999 | 33.90133 | 55.4177 | 78.05245 | 33.55956 | -0.11838 | 1.49882 | 1 |
| 17 | 76698200 | 76699199 | 17.41664 | 30.24729 | 42.16045 | 18.50573 | -0.09001 | 0.798291 | 1 |
| 17 | 76873000 | 76878199 | 81.29953 | 127.5447 | 142.5135 | 59.10484 | 0.158229 | 2.584036 | 1 |
| 17 | 77206400 | 77207399 | 15.81091 | 27.56041 | 39.56737 | 17.40796 | -0.13183 | 0.73386 | 1 |
| 17 | 77495400 | 77496199 | 14.74233 | 26.40769 | 31.89885 | 13.9796 | 0.129638 | 0.425081 | 1 |
| 17 | 77539400 | 77540199 | 19.55405 | 33.69347 | 42.74144 | 18.74938 | 0.04312 | 0.883469 | 1 |
| 17 | 77669200 | 77670999 | 12.93566 | 22.91624 | 37.67189 | 16.65631 | -0.32156 | 0.496851 | 1 |
| 17 | 78083200 | 78089399 | 136.5635 | 207.6201 | 211.6834 | 87.68895 | 0.255088 | 3.194268 | 1 |
| 17 | 78100400 | 78108399 | 126.5541 | 191.6305 | 240.2492 | 99.40059 | -0.04771 | 3.275572 | 1 |
| 17 | 78132200 | 78132999 | 14.26478 | 25.75665 | 27.97181 | 12.15971 | 0.290213 | 0.367103 | 1 |
| 17 | 78227600 | 78230399 | 32.6771 | 53.83641 | 66.83677 | 28.83032 | 0.068288 | 1.390282 | 1 |
| 17 | 79355600 | 79358199 | 12.66292 | 23.3285 | 26.61253 | 11.27246 | 0.244929 | 0.2387 | 1 |
| 17 | 79441200 | 79442599 | 14.11002 | 25.8621 | 27.47355 | 11.71827 | 0.339758 | 0.279773 | 1 |
| 17 | 79862600 | 79869799 | 113.9068 | 174.1075 | 219.7386 | 91.15434 | -0.0463 | 3.080737 | 1 |
| 17 | 80870200 | 80879399 | 126.9666 | 191.5284 | 263.7838 | 109.0075 | -0.18787 | 3.35333 | 1 |
| 17 | 81161200 | 81162799 | 23.76704 | 39.35387 | 62.11665 | 26.87565 | -0.27556 | 1.306368 | 1 |
| 17 | 81537800 | 81540799 | 51.6657 | 82.85304 | 90.05751 | 38.04534 | 0.227502 | 1.967855 | 1 |
| 17 | 81541800 | 81545399 | 78.89556 | 124.1137 | 130.8701 | 54.29328 | 0.244786 | 2.526662 | 1 |
| 17 | 81990600 | 81992399 | 17.61127 | 30.41971 | 46.88274 | 20.56756 | -0.23598 | 0.856988 | 1 |
| 17 | 82023400 | 82024199 | 11.9967 | 22.22182 | 23.07744 | 9.702926 | 0.387013 | 0.192654 | 1 |
| 17 | 82279200 | 82282599 | 87.88826 | 135.6712 | 177.0768 | 73.55421 | -0.08372 | 2.875876 | 1 |
| 17 | 82722200 | 82723799 | 11.13357 | 20.35475 | 33.9209 | 14.51802 | -0.30941 | 0.284907 | 1 |
| 17 | 82768400 | 82769999 | 14.31853 | 25.37777 | 35.01088 | 15.47834 | -0.06978 | 0.495735 | 1 |
| 17 | 83057200 | 83060399 | 64.37529 | 102.4255 | 110.9043 | 46.32901 | 0.220549 | 2.231557 | 1 |
| 17 | 83075000 | 83076599 | 18.64683 | 32.55676 | 41.45911 | 18.25893 | 0.040534 | 0.729526 | 1 |
| 17 | 83156400 | 83163999 | 95.20161 | 147.1208 | 177.8413 | 73.87132 | 0.028029 | 2.857918 | 1 |
| 17 | 83288000 | 83290599 | 32.83926 | 53.28187 | 78.43506 | 33.60957 | -0.1901 | 1.615604 | 1 |
| 17 | 83454200 | 83458999 | 9.024158 | 16.78193 | 29.96574 | 12.53006 | -0.38382 | 0.161761 | 1 |
| 17 | 83709800 | 83711599 | 22.50322 | 38.45344 | 43.28516 | 18.97846 | 0.21276 | 0.972102 | 1 |
| 17 | 83790800 | 83810999 | 265.7743 | 378.4907 | 513.2168 | 212.1751 | -0.24444 | 4.308952 | 1 |
| 17 | 84576000 | 84578999 | 31.08993 | 50.33261 | 84.09693 | 35.97723 | -0.37533 | 1.659675 | 1 |
| 17 | 85282600 | 85285599 | 33.22575 | 54.58541 | 72.48762 | 31.23037 | -0.03024 | 1.430352 | 1 |
| 17 | 85493000 | 85494399 | 16.26092 | 28.38337 | 40.99585 | 18.05184 | -0.14045 | 0.718063 | 1 |

| | | | | | | | | | |
|---|---|---|---|---|---|---|---|---|---|
| 17 | 85727000 | 85728799 | 23.17009 | 39.73327 | 44.19719 | 19.38386 | 0.230431 | 0.941353 | 1 |
| 17 | 86896200 | 86897599 | 13.20228 | 23.68303 | 33.58342 | 14.70005 | -0.09912 | 0.409638 | 1 |
| 17 | 86996600 | 86999199 | 37.53271 | 60.62193 | 82.51143 | 35.21319 | -0.08385 | 1.727413 | 1 |
| 17 | 88123800 | 88127999 | 48.24922 | 77.14263 | 107.8618 | 45.36582 | -0.14005 | 2.063606 | 1 |
| 17 | 88212400 | 88219599 | 14.32229 | 25.89601 | 29.35145 | 12.73757 | 0.230212 | 0.360591 | 1 |
| 17 | 88268200 | 88273399 | 58.11439 | 92.60445 | 122.0855 | 51.10702 | -0.06068 | 2.17913 | 1 |
| 17 | 88542400 | 88544999 | 55.1975 | 87.97447 | 109.4131 | 45.81518 | 0.02365 | 2.167528 | 1 |
| 17 | 88619400 | 88621599 | 21.01118 | 36.5631 | 39.81774 | 17.49946 | 0.264607 | 0.782244 | 1 |
| 17 | 88783800 | 88785199 | 15.25057 | 26.7123 | 43.53851 | 19.2232 | -0.31377 | 0.664516 | 1 |
| 17 | 88837200 | 88839199 | 15.45522 | 27.16916 | 35.16622 | 15.57247 | 0.017276 | 0.593148 | 1 |
| 17 | 89230600 | 89231599 | 11.93511 | 22.59087 | 23.68678 | 9.598075 | 0.416072 | 0.059171 | 1 |
| 17 | 89340200 | 89341999 | 22.94577 | 38.42361 | 60.89067 | 26.54097 | -0.28097 | 1.173945 | 1 |
| 17 | 89460600 | 89464799 | 58.2329 | 92.64484 | 115.4533 | 48.22376 | 0.017967 | 2.230771 | 1 |
| 17 | 89980200 | 89981599 | 12.29536 | 22.09784 | 36.24669 | 15.84056 | -0.30638 | 0.397257 | 1 |
| 17 | 90409400 | 90412399 | 73.86077 | 116.0675 | 133.036 | 55.18407 | 0.123372 | 2.545738 | 1 |
| 17 | 91579000 | 91582199 | 37.15602 | 60.23405 | 84.11537 | 36.02076 | -0.1155 | 1.641241 | 1 |
| 17 | 91782200 | 91785599 | 64.44578 | 102.375 | 111.1754 | 46.35983 | 0.214045 | 2.280012 | 1 |
| 17 | 91954400 | 91955799 | 23.81948 | 40.51897 | 44.84164 | 19.65374 | 0.236189 | 1.019932 | 1 |
| 17 | 92037800 | 92039599 | 8.878764 | 17.03596 | 20.82873 | 8.176338 | 0.237297 | -0.0809 | 1 |
| 17 | 92136000 | 92139799 | 43.68814 | 70.59887 | 72.97393 | 31.16273 | 0.313031 | 1.716175 | 1 |
| 17 | 92141800 | 92145399 | 65.80819 | 104.4008 | 111.5237 | 46.44341 | 0.235921 | 2.320318 | 1 |
| 17 | 92395200 | 92397599 | 20.38945 | 34.94756 | 55.54636 | 24.36037 | -0.28266 | 0.947759 | 1 |
| 17 | 92410800 | 92412199 | 17.15738 | 30.33533 | 31.79266 | 14.06241 | 0.323323 | 0.534111 | 1 |
| 17 | 92515200 | 92515799 | 11.83961 | 22.2584 | 24.62813 | 10.11031 | 0.322931 | 0.103752 | 1 |
| 17 | 92620400 | 92622999 | 61.51047 | 97.99263 | 105.4891 | 44.14489 | 0.230914 | 2.188114 | 1 |
| 17 | 92763800 | 92769399 | 93.76215 | 145.9316 | 153.0671 | 63.53975 | 0.240454 | 2.730357 | 1 |
| 17 | 92829800 | 92830999 | 16.38544 | 28.40594 | 46.35683 | 20.33911 | -0.31692 | 0.819513 | 1 |
| 17 | 93156000 | 93158799 | 22.08855 | 38.35676 | 41.45689 | 18.19566 | 0.275239 | 0.820292 | 1 |
| 17 | 94096200 | 94101399 | 53.72883 | 86.1543 | 91.44113 | 38.62491 | 0.261679 | 1.974728 | 1 |
| 17 | 94240000 | 94244199 | 51.93969 | 83.03885 | 107.396 | 45.16664 | -0.02786 | 2.067917 | 1 |
| 17 | 94441400 | 94442599 | 14.04376 | 25.25245 | 29.09031 | 12.70785 | 0.201228 | 0.393522 | 1 |
| 17 | 94655400 | 94656599 | 16.1271 | 28.36801 | 37.618 | 16.6561 | -0.01722 | 0.594422 | 1 |
| 17 | 94764000 | 94766199 | 23.12305 | 38.98093 | 55.43416 | 24.23691 | -0.12503 | 1.107138 | 1 |
| 17 | 94842200 | 94844399 | 36.90213 | 59.91281 | 68.5555 | 29.39023 | 0.173265 | 1.601993 | 1 |
| 17 | 94881600 | 94883199 | 30.64087 | 50.63618 | 56.11062 | 24.24417 | 0.232914 | 1.341125 | 1 |
| 17 | 94989800 | 94993599 | 54.59433 | 87.26746 | 96.54197 | 40.59162 | 0.196835 | 2.071822 | 1 |
| 18 | 3382600 | 3383799 | 13.09465 | 23.69495 | 27.55984 | 11.94831 | 0.194315 | 0.344622 | 1 |
| 18 | 3585400 | 3588999 | 50.05526 | 80.35986 | 96.41934 | 40.80192 | 0.087131 | 1.936117 | 1 |
| 18 | 4177200 | 4178199 | 16.53558 | 28.97826 | 39.34364 | 17.37877 | -0.05131 | 0.657282 | 1 |
| 18 | 4222400 | 4223799 | 22.61278 | 38.2315 | 51.14562 | 22.38156 | -0.03695 | 1.076318 | 1 |
| 18 | 4274600 | 4276799 | 14.24206 | 24.99243 | 38.86066 | 17.18491 | -0.24603 | 0.620662 | 1 |
| 18 | 4426800 | 4427799 | 14.43232 | 25.89778 | 32.94946 | 14.41869 | 0.056614 | 0.414075 | 1 |
| 18 | 4472600 | 4474599 | 12.9052 | 23.75024 | 23.61222 | 10.0157 | 0.440654 | 0.238995 | 1 |
| 18 | 4483200 | 4485599 | 21.4095 | 36.92578 | 43.66815 | 19.15645 | 0.144041 | 0.882424 | 1 |
| 18 | 4560600 | 4563599 | 28.89281 | 48.16605 | 52.92174 | 23.00242 | 0.245806 | 1.233339 | 1 |
| 18 | 4931800 | 4932999 | 13.27475 | 24.15125 | 32.6566 | 14.06242 | -0.01565 | 0.324698 | 1 |
| 18 | 5434400 | 5441799 | 68.87731 | 107.9852 | 163.5612 | 67.83862 | -0.28088 | 2.59121 | 1 |

| | | | | | | | | | |
|---|---|---|---|---|---|---|---|---|---|
| 18 | 5499400 | 5503199 | 44.51387 | 71.5805 | 88.36778 | 37.48487 | 0.048984 | 1.873063 | 1 |
| 18 | 5573400 | 5577599 | 57.94762 | 92.59686 | 105.2135 | 44.21682 | 0.157826 | 2.090756 | 1 |
| 18 | 5658200 | 5661399 | 45.875 | 74.32298 | 79.46518 | 34.01405 | 0.268075 | 1.667832 | 1 |
| 18 | 5672000 | 5679999 | 36.1023 | 58.46163 | 91.89991 | 39.32249 | -0.28737 | 1.666501 | 1 |
| 18 | 5868200 | 5872999 | 62.54654 | 99.41775 | 106.3728 | 44.38384 | 0.236267 | 2.258863 | 1 |
| 18 | 6042200 | 6047399 | 87.04519 | 134.432 | 181.4841 | 75.38471 | -0.13186 | 2.867964 | 1 |
| 18 | 6081000 | 6082799 | 28.05273 | 45.85497 | 73.61331 | 31.65036 | -0.3066 | 1.484168 | 1 |
| 18 | 6294200 | 6295599 | 14.33637 | 25.72622 | 29.37089 | 12.85245 | 0.212016 | 0.405415 | 1 |
| 18 | 6542200 | 6551399 | 143.7517 | 217.2157 | 242.9565 | 100.4542 | 0.114219 | 3.321387 | 1 |
| 18 | 6553000 | 6557999 | 56.08844 | 89.7194 | 93.53869 | 39.36842 | 0.283398 | 2.053645 | 1 |
| 18 | 6921400 | 6922199 | 12.4946 | 22.99066 | 25.15505 | 10.67261 | 0.30318 | 0.241017 | 1 |
| 18 | 7769800 | 7774999 | 66.24639 | 105.0444 | 115.4128 | 48.03938 | 0.1947 | 2.334965 | 1 |
| 18 | 8243400 | 8252399 | 54.43171 | 86.34555 | 129.5151 | 53.93138 | -0.25315 | 2.318128 | 1 |
| 18 | 8267800 | 8273999 | 62.07435 | 98.7709 | 108.2798 | 45.23665 | 0.202814 | 2.226899 | 1 |
| 18 | 8415000 | 8420799 | 88.91728 | 138.1249 | 162.2817 | 67.38421 | 0.074829 | 2.762253 | 1 |
| 18 | 9461800 | 9463199 | 17.23038 | 30.59182 | 32.23208 | 14.23171 | 0.318432 | 0.497697 | 1 |
| 18 | 9478000 | 9479599 | 17.29544 | 30.38521 | 33.0028 | 14.60851 | 0.268875 | 0.608629 | 1 |
| 18 | 9591000 | 9597199 | 20.31069 | 34.04732 | 58.42696 | 25.48422 | -0.39519 | 1.155514 | 1 |
| 18 | 9678600 | 9680999 | 36.10726 | 58.90626 | 67.14578 | 28.84917 | 0.184064 | 1.524788 | 1 |
| 18 | 9904600 | 9912599 | 51.08888 | 81.95226 | 86.65704 | 36.62526 | 0.267675 | 1.955765 | 1 |
| 18 | 10181200 | 10182799 | 18.70488 | 32.10837 | 44.82933 | 19.66269 | -0.09584 | 0.914738 | 1 |
| 18 | 10274200 | 10277399 | 36.45324 | 59.30569 | 73.3187 | 31.46947 | 0.064386 | 1.568243 | 1 |
| 18 | 10600800 | 10603999 | 30.40807 | 49.84818 | 74.37851 | 32.01567 | -0.19916 | 1.45019 | 1 |
| 18 | 10669200 | 10675999 | 30.22195 | 49.82972 | 59.43935 | 25.64774 | 0.126197 | 1.36688 | 1 |
| 18 | 10828600 | 10834999 | 54.60233 | 87.30862 | 103.0618 | 43.35289 | 0.103896 | 2.063606 | 1 |
| 18 | 10896400 | 10898999 | 23.86567 | 40.7764 | 50.65735 | 22.21058 | 0.070757 | 0.983843 | 1 |
| 18 | 11341200 | 11345599 | 56.71143 | 90.78529 | 99.99976 | 42.13332 | 0.205395 | 2.032284 | 1 |
| 18 | 11386600 | 11389599 | 33.26482 | 54.45739 | 69.53928 | 29.91549 | 0.024087 | 1.469314 | 1 |
| 18 | 11393200 | 11399399 | 88.90541 | 138.4562 | 167.2222 | 69.4094 | 0.037895 | 2.720865 | 1 |
| 18 | 11879000 | 11883199 | 57.50741 | 91.69651 | 114.2941 | 47.88826 | 0.021138 | 2.157008 | 1 |
| 18 | 12130200 | 12131799 | 18.5303 | 32.47712 | 40.20326 | 17.75993 | 0.0811 | 0.665449 | 1 |
| 18 | 12152800 | 12156999 | 27.05294 | 44.73668 | 68.84072 | 29.75826 | -0.23943 | 1.331083 | 1 |
| 18 | 12736800 | 12739199 | 21.98594 | 37.02159 | 58.111 | 25.39846 | -0.26707 | 1.115056 | 1 |
| 18 | 13220600 | 13222799 | 28.89959 | 48.37814 | 51.83152 | 22.58775 | 0.281949 | 1.188646 | 1 |
| 18 | 13299200 | 13299999 | 9.527536 | 17.63206 | 28.36651 | 11.9415 | -0.24116 | 0.191977 | 1 |
| 18 | 13410800 | 13412399 | 23.49755 | 39.52644 | 55.49641 | 24.24817 | -0.10689 | 1.124434 | 1 |
| 18 | 13714600 | 13717599 | 29.41743 | 49.2516 | 52.38226 | 22.82968 | 0.292483 | 1.189657 | 1 |
| 18 | 14141800 | 14143399 | 27.45545 | 45.75337 | 56.67742 | 24.62922 | 0.073299 | 1.235981 | 1 |
| 18 | 14803600 | 14804599 | 13.80814 | 25.11823 | 26.8899 | 11.58117 | 0.319177 | 0.317186 | 1 |
| 18 | 15018000 | 15018799 | 11.61093 | 21.10153 | 28.09558 | 12.11463 | 0.005337 | 0.307809 | 1 |
| 18 | 15621600 | 15624599 | 30.07502 | 49.9802 | 54.57085 | 23.67021 | 0.25473 | 1.273074 | 1 |
| 18 | 15646400 | 15648999 | 18.29665 | 31.64423 | 45.82818 | 20.10571 | -0.14615 | 0.848589 | 1 |
| 18 | 15828200 | 15828999 | 32.56313 | 52.75694 | 75.59782 | 32.35903 | -0.1532 | 1.644923 | 1 |
| 18 | 16120200 | 16122799 | 24.75622 | 42.12486 | 45.284 | 19.84809 | 0.27783 | 1.017955 | 1 |
| 18 | 16125000 | 16127199 | 22.58324 | 37.90392 | 55.9662 | 24.43005 | -0.17928 | 1.143574 | 1 |
| 18 | 16654400 | 16656399 | 24.49951 | 41.54784 | 46.91523 | 20.54931 | 0.206747 | 1.049057 | 1 |
| 18 | 16982200 | 16988199 | 40.41842 | 65.2521 | 80.0636 | 34.14777 | 0.064648 | 1.744029 | 1 |

| | | | | | | | | | |
|---|---|---|---|---|---|---|---|---|---|
| 18 | 17127400 | 17128399 | 10.12926 | 18.94621 | 26.86489 | 11.12407 | -0.0418 | 0.12926 | 1 |
| 18 | 17445600 | 17455599 | 103.1971 | 159.2512 | 178.8302 | 74.28267 | 0.132665 | 2.888192 | 1 |
| 18 | 17572800 | 17574199 | 20.4806 | 34.79984 | 54.32287 | 23.80711 | -0.25853 | 1.027616 | 1 |
| 18 | 17587000 | 17587799 | 11.72769 | 21.10912 | 31.13785 | 13.5894 | -0.15273 | 0.372309 | 1 |
| 18 | 18071200 | 18075799 | 79.38663 | 124.6499 | 129.7853 | 53.82996 | 0.260895 | 2.563805 | 1 |
| 18 | 18455400 | 18456199 | 15.11673 | 27.18067 | 29.88969 | 13.05751 | 0.267633 | 0.401576 | 1 |
| 18 | 18848200 | 18848799 | 11.35203 | 21.24314 | 22.10824 | 9.1473 | 0.40189 | 0.122878 | 1 |
| 18 | 18863000 | 18863599 | 8.498358 | 16.22127 | 26.73411 | 10.64237 | -0.20899 | -0.02318 | 1 |
| 18 | 18923600 | 18924399 | 11.12403 | 20.80247 | 24.17893 | 10.01547 | 0.242388 | 0.131215 | 1 |
| 18 | 18979200 | 18992799 | 239.7961 | 348.5124 | 356.0912 | 146.7848 | 0.192777 | 4.006221 | 1 |
| 18 | 19130600 | 19132199 | 28.48401 | 48.01924 | 50.70026 | 22.16726 | 0.303084 | 1.120333 | 1 |
| 18 | 19393000 | 19394799 | 22.18624 | 37.94656 | 43.79286 | 19.20247 | 0.177127 | 0.963182 | 1 |
| 18 | 19566600 | 19571799 | 19.80048 | 34.31245 | 40.57348 | 17.80144 | 0.145664 | 0.829864 | 1 |
| 18 | 19682800 | 19683399 | 10.41713 | 19.73797 | 21.67984 | 8.765719 | 0.352292 | 0.032157 | 1 |
| 18 | 19745000 | 19745999 | 11.81715 | 21.18665 | 31.25962 | 13.68923 | -0.15614 | 0.400234 | 1 |
| 18 | 20747800 | 20755999 | 86.41862 | 132.9063 | 193.5166 | 80.36318 | -0.24545 | 2.948704 | 1 |
| 18 | 20758400 | 20762999 | 48.936 | 78.05968 | 105.6117 | 44.27426 | -0.09629 | 2.135679 | 1 |
| 18 | 21161600 | 21164399 | 29.42407 | 48.89026 | 65.27022 | 28.30782 | -0.03461 | 1.280088 | 1 |
| 18 | 21392200 | 21396399 | 51.5011 | 82.39909 | 101.7834 | 42.85142 | 0.039507 | 2.044203 | 1 |
| 18 | 21432000 | 21434199 | 16.29306 | 27.84941 | 50.5648 | 22.17073 | -0.47422 | 0.950953 | 1 |
| 18 | 21784000 | 21787199 | 27.5576 | 45.01915 | 72.42893 | 31.13445 | -0.31033 | 1.489779 | 1 |
| 18 | 22333600 | 22334599 | 11.70394 | 21.32785 | 30.1735 | 12.96693 | -0.07837 | 0.299499 | 1 |
| 18 | 22570800 | 22573399 | 29.30916 | 48.70454 | 60.61922 | 26.29262 | 0.066332 | 1.276496 | 1 |
| 18 | 23010600 | 23011599 | 11.99179 | 21.56773 | 31.24093 | 13.64468 | -0.12701 | 0.382365 | 1 |
| 18 | 23366200 | 23374599 | 29.04983 | 48.47382 | 53.47749 | 23.2579 | 0.239905 | 1.224036 | 1 |
| 18 | 23377000 | 23377999 | 16.8621 | 29.64426 | 34.02458 | 15.06673 | 0.189784 | 0.598319 | 1 |
| 18 | 23669800 | 23672199 | 32.37425 | 53.61846 | 59.11894 | 25.5828 | 0.240261 | 1.321289 | 1 |
| 18 | 24076800 | 24082799 | 62.13656 | 99.04656 | 108.2377 | 45.3309 | 0.210142 | 2.170679 | 1 |
| 18 | 24614400 | 24618799 | 75.39935 | 118.8223 | 129.6979 | 53.81929 | 0.196918 | 2.489427 | 1 |
| 18 | 24670600 | 24674399 | 47.93076 | 76.54687 | 113.3131 | 47.58234 | -0.22402 | 2.099196 | 1 |
| 18 | 24708800 | 24709999 | 15.58976 | 27.29564 | 40.96909 | 18.08271 | -0.19527 | 0.66987 | 1 |
| 18 | 24838000 | 24840599 | 28.42836 | 47.33781 | 54.52172 | 23.68082 | 0.17806 | 1.245608 | 1 |
| 18 | 25301000 | 25303199 | 17.85957 | 31.06474 | 35.96287 | 15.80081 | 0.176774 | 0.766787 | 1 |
| 18 | 25345200 | 25347199 | 35.84891 | 58.17752 | 69.88369 | 29.95003 | 0.102814 | 1.611181 | 1 |
| 18 | 25572800 | 25576399 | 50.0834 | 80.28509 | 95.48448 | 40.31469 | 0.097262 | 1.979925 | 1 |
| 18 | 26290000 | 26290999 | 11.96386 | 21.50252 | 33.18644 | 14.5029 | -0.21883 | 0.389105 | 1 |
| 18 | 26967400 | 26972399 | 73.16878 | 115.2644 | 132.0669 | 54.79969 | 0.126635 | 2.496407 | 1 |
| 18 | 27137200 | 27138599 | 15.14101 | 26.92947 | 30.99287 | 13.66431 | 0.193328 | 0.47178 | 1 |
| 18 | 28147400 | 28149799 | 12.01932 | 21.64003 | 32.73322 | 14.28071 | -0.18811 | 0.379697 | 1 |
| 18 | 28184600 | 28185199 | 10.05931 | 19.0876 | 24.37581 | 9.825626 | 0.140521 | 0.027231 | 1 |
| 18 | 28220200 | 28221199 | 14.55161 | 25.69917 | 33.21035 | 14.69278 | 0.022348 | 0.535081 | 1 |
| 18 | 28487000 | 28488199 | 21.45186 | 36.23167 | 50.08837 | 21.91211 | -0.08419 | 1.079284 | 1 |
| 18 | 28576400 | 28579399 | 10.70595 | 20.6135 | 20.08214 | 7.804828 | 0.574665 | -0.09873 | 1 |
| 18 | 28593200 | 28594799 | 18.65771 | 32.70006 | 36.06704 | 15.93259 | 0.246889 | 0.660234 | 1 |
| 18 | 29152800 | 29153999 | 21.47867 | 36.81248 | 41.24368 | 18.08772 | 0.220166 | 0.937941 | 1 |
| 18 | 30926200 | 30926999 | 10.25283 | 19.03533 | 28.84989 | 12.09205 | -0.15143 | 0.179818 | 1 |
| 18 | 31119800 | 31122399 | 16.15682 | 28.59647 | 30.86095 | 13.64741 | 0.281868 | 0.51437 | 1 |

| | | | | | | | | | |
|---|---|---|---|---|---|---|---|---|---|
| 18 | 31162600 | 31163799 | 18.99686 | 33.00491 | 41.56759 | 18.25011 | 0.05565 | 0.798883 | 1 |
| 18 | 31765600 | 31768599 | 17.43499 | 29.75878 | 51.61272 | 22.6276 | -0.40869 | 0.972051 | 1 |
| 18 | 32254400 | 32260999 | 67.85869 | 106.6994 | 155.7926 | 64.62865 | -0.22475 | 2.534319 | 1 |
| 18 | 32300600 | 32303399 | 28.26999 | 47.23256 | 55.08218 | 23.97503 | 0.160057 | 1.209987 | 1 |
| 18 | 33018400 | 33019199 | 13.14574 | 24.37717 | 25.98406 | 10.90858 | 0.351044 | 0.202626 | 1 |
| 18 | 33373400 | 33376399 | 42.19414 | 68.48457 | 71.15208 | 30.49679 | 0.311858 | 1.623149 | 1 |
| 18 | 33711000 | 33713399 | 28.18818 | 47.43932 | 52.32201 | 22.86447 | 0.240267 | 1.135833 | 1 |
| 18 | 34059600 | 34061999 | 24.85356 | 42.22879 | 47.42413 | 20.78367 | 0.214728 | 1.034942 | 1 |
| 18 | 34164800 | 34169199 | 55.58437 | 88.45203 | 122.7757 | 51.29893 | -0.13697 | 2.221752 | 1 |
| 18 | 34689400 | 34691799 | 20.15554 | 34.30062 | 55.41488 | 24.28872 | -0.30758 | 1.013234 | 1 |
| 18 | 34765000 | 34767599 | 16.61102 | 29.04728 | 43.00781 | 18.96244 | -0.17596 | 0.695877 | 1 |
| 18 | 34908800 | 34911199 | 20.39201 | 34.4629 | 56.95758 | 24.92185 | -0.34089 | 1.077875 | 1 |
| 18 | 35027000 | 35029199 | 19.13614 | 32.64213 | 49.01428 | 21.48735 | -0.20177 | 0.980593 | 1 |
| 18 | 36611000 | 36622199 | 109.6641 | 168.4867 | 173.8996 | 72.20671 | 0.249366 | 2.972924 | 1 |
| 18 | 37100600 | 37103199 | 91.72591 | 142.3759 | 148.2204 | 61.55276 | 0.24825 | 2.775226 | 1 |
| 18 | 37106800 | 37110599 | 100.4975 | 154.2486 | 193.0125 | 80.13178 | -0.02928 | 2.992495 | 1 |
| 18 | 37128800 | 37135199 | 90.06601 | 140.0606 | 154.5421 | 64.16011 | 0.166524 | 2.744224 | 1 |
| 18 | 37138200 | 37141799 | 77.3769 | 121.4278 | 136.2309 | 56.49932 | 0.152727 | 2.57574 | 1 |
| 18 | 37359800 | 37361999 | 38.36877 | 62.20891 | 75.16411 | 32.19094 | 0.09314 | 1.638518 | 1 |
| 18 | 37395000 | 37400999 | 15.31469 | 27.49917 | 32.82838 | 14.35775 | 0.148367 | 0.414493 | 1 |
| 18 | 37450600 | 37458599 | 136.4239 | 206.5531 | 228.0024 | 94.33036 | 0.135682 | 3.279738 | 1 |
| 18 | 37462200 | 37470999 | 135.7555 | 205.5765 | 236.5149 | 97.85752 | 0.076217 | 3.276513 | 1 |
| 18 | 37473800 | 37479999 | 75.51642 | 117.7143 | 170.7626 | 70.86911 | -0.22512 | 2.698243 | 1 |
| 18 | 39905600 | 39908599 | 38.46185 | 61.71599 | 96.88686 | 40.96791 | -0.30071 | 1.937049 | 1 |
| 18 | 39953600 | 39954599 | 14.80194 | 26.4743 | 30.9578 | 13.58455 | 0.175109 | 0.434309 | 1 |
| 18 | 41513200 | 41515199 | 15.55128 | 27.2519 | 39.38847 | 17.39781 | -0.14111 | 0.652459 | 1 |
| 18 | 41596200 | 41599199 | 38.08893 | 61.34654 | 91.52144 | 38.91239 | -0.22072 | 1.816768 | 1 |
| 18 | 41642800 | 41644199 | 24.59622 | 40.9618 | 60.32148 | 26.20194 | -0.17537 | 1.23792 | 1 |
| 18 | 42007200 | 42009999 | 42.02211 | 67.53142 | 90.64932 | 38.41787 | -0.07262 | 1.892367 | 1 |
| 18 | 42314200 | 42321599 | 81.60547 | 127.436 | 148.017 | 61.41644 | 0.097245 | 2.665634 | 1 |
| 18 | 42605000 | 42622999 | 173.9999 | 255.6331 | 365.1508 | 150.3847 | -0.27539 | 3.83478 | 1 |
| 18 | 42718600 | 42720599 | 16.44203 | 28.87428 | 34.19774 | 15.13229 | 0.144771 | 0.613978 | 1 |
| 18 | 43135600 | 43138199 | 29.44429 | 48.32157 | 65.04917 | 28.013 | -0.05033 | 1.42976 | 1 |
| 18 | 43968400 | 43969799 | 18.59374 | 32.51205 | 36.30128 | 16.00451 | 0.229535 | 0.699465 | 1 |
| 18 | 44032600 | 44034599 | 35.47795 | 57.86816 | 67.58382 | 29.03528 | 0.148935 | 1.526956 | 1 |
| 18 | 44840200 | 44842999 | 10.02969 | 18.51056 | 31.98045 | 13.51317 | -0.34747 | 0.219806 | 1 |
| 18 | 44855000 | 44855999 | 12.46639 | 23.05142 | 29.23193 | 12.31991 | 0.098177 | 0.221168 | 1 |
| 18 | 44931600 | 44933599 | 31.01977 | 51.62933 | 58.57761 | 25.43506 | 0.199616 | 1.260388 | 1 |
| 18 | 45165400 | 45165999 | 10.35297 | 18.90435 | 30.31057 | 12.99279 | -0.25511 | 0.279203 | 1 |
| 18 | 45291600 | 45293399 | 18.93436 | 33.10349 | 36.67492 | 16.16753 | 0.24053 | 0.701448 | 1 |
| 18 | 45345200 | 45354399 | 181.8645 | 270.3289 | 284.5069 | 117.2109 | 0.180795 | 3.63903 | 1 |
| 18 | 45365600 | 45366799 | 9.738862 | 18.63404 | 21.22525 | 8.399197 | 0.327388 | -0.04621 | 1 |
| 18 | 45428800 | 45429399 | 10.06688 | 18.44464 | 28.38798 | 12.11441 | -0.19159 | 0.254939 | 1 |
| 18 | 45496000 | 45498399 | 56.83411 | 90.5683 | 98.35895 | 41.17719 | 0.218676 | 2.172008 | 1 |
| 18 | 46043400 | 46044599 | 15.14536 | 27.1104 | 31.92766 | 14.0006 | 0.165555 | 0.433558 | 1 |
| 18 | 46449600 | 46453399 | 34.74152 | 56.30612 | 82.92597 | 35.50503 | -0.19235 | 1.641672 | 1 |
| 18 | 46701600 | 46704199 | 27.90565 | 46.5097 | 61.80234 | 26.85847 | -0.02771 | 1.238677 | 1 |

| | | | | | | | | | |
|---|---|---|---|---|---|---|---|---|---|
| 18 | 47071800 | 47079799 | 84.84141 | 131.2224 | 192.3671 | 79.90466 | -0.249 | 2.840231 | 1 |
| 18 | 47517600 | 47520599 | 36.93709 | 60.03357 | 72.12016 | 30.94275 | 0.104632 | 1.58315 | 1 |
| 18 | 47649600 | 47652399 | 20.96346 | 36.05703 | 49.55406 | 21.73667 | -0.07275 | 0.911752 | 1 |
| 18 | 47876200 | 47879399 | 47.91084 | 77.01164 | 83.54952 | 35.4201 | 0.234279 | 1.888543 | 1 |
| 18 | 47969400 | 47969999 | 8.732871 | 16.8422 | 21.98325 | 8.503375 | 0.163411 | -0.12693 | 1 |
| 18 | 48176800 | 48180999 | 40.34847 | 64.78388 | 98.05922 | 41.5032 | -0.24716 | 1.91881 | 1 |
| 18 | 48674400 | 48675999 | 15.12441 | 26.34711 | 44.73704 | 19.6746 | -0.37287 | 0.748927 | 1 |
| 18 | 49523600 | 49525199 | 23.59411 | 39.48606 | 54.87974 | 23.91206 | -0.09219 | 1.178046 | 1 |
| 18 | 49752200 | 49754599 | 35.20116 | 57.61358 | 68.04322 | 29.26914 | 0.136249 | 1.475801 | 1 |
| 18 | 49825200 | 49831599 | 125.607 | 190.0995 | 247.1899 | 102.2558 | -0.10096 | 3.285927 | 1 |
| 18 | 50400800 | 50401999 | 24.69554 | 41.19351 | 54.38414 | 23.64695 | -0.01824 | 1.215375 | 1 |
| 18 | 50671200 | 50672599 | 13.34733 | 24.02722 | 29.09908 | 12.69775 | 0.130432 | 0.380864 | 1 |
| 18 | 51750600 | 51752799 | 24.80599 | 42.82614 | 42.46644 | 18.62984 | 0.397376 | 0.886449 | 1 |
| 18 | 51831200 | 51832399 | 15.88259 | 28.04107 | 30.58186 | 13.53074 | 0.265534 | 0.540272 | 1 |
| 18 | 52519400 | 52523199 | 42.72662 | 68.88903 | 83.59641 | 35.59491 | 0.078406 | 1.787138 | 1 |
| 18 | 52527000 | 52528199 | 16.29403 | 28.85558 | 31.44866 | 13.90575 | 0.268237 | 0.512296 | 1 |
| 18 | 52574200 | 52575599 | 14.41598 | 26.12206 | 28.42252 | 12.301 | 0.29153 | 0.348584 | 1 |
| 18 | 52672800 | 52675199 | 24.13447 | 40.90144 | 50.10797 | 21.94331 | 0.089576 | 1.058347 | 1 |
| 18 | 53468600 | 53470599 | 28.63569 | 47.0836 | 69.62562 | 30.00784 | -0.18444 | 1.407245 | 1 |
| 18 | 53604600 | 53608599 | 38.86293 | 62.48852 | 95.4764 | 40.49561 | -0.25812 | 1.87026 | 1 |
| 18 | 53968600 | 53968999 | 11.46267 | 21.71131 | 23.73083 | 9.602192 | 0.358394 | 0.050534 | 1 |
| 18 | 54008400 | 54009599 | 14.24538 | 24.98927 | 37.39299 | 16.53085 | -0.19085 | 0.624567 | 1 |
| 18 | 54101600 | 54104399 | 35.69238 | 58.39544 | 69.83754 | 30.03684 | 0.117798 | 1.483188 | 1 |
| 18 | 54156000 | 54157599 | 23.54981 | 39.16186 | 61.76242 | 26.80167 | -0.27394 | 1.254482 | 1 |
| 18 | 54376200 | 54379999 | 35.14048 | 57.03643 | 87.23738 | 37.39144 | -0.24478 | 1.616994 | 1 |
| 18 | 54459400 | 54468399 | 81.21914 | 126.1246 | 169.8965 | 70.54962 | -0.1226 | 2.765332 | 1 |
| 18 | 54608800 | 54610199 | 11.97819 | 21.27392 | 37.70135 | 16.64189 | -0.42749 | 0.470157 | 1 |
| 18 | 55448000 | 55451599 | 50.42484 | 80.82097 | 91.08367 | 38.44832 | 0.174561 | 1.983785 | 1 |
| 18 | 55453600 | 55455999 | 31.19031 | 51.24199 | 64.23246 | 27.67386 | 0.052836 | 1.41938 | 1 |
| 18 | 56472800 | 56475999 | 38.49167 | 62.51271 | 74.99074 | 32.15634 | 0.105692 | 1.604356 | 1 |
| 18 | 56535600 | 56536399 | 21.90565 | 37.36072 | 46.40624 | 20.34376 | 0.070852 | 0.991171 | 1 |
| 18 | 56630200 | 56633799 | 37.31659 | 60.10405 | 91.72298 | 38.99944 | -0.25327 | 1.816209 | 1 |
| 18 | 56644200 | 56646199 | 24.26394 | 41.73974 | 45.18981 | 19.8224 | 0.270052 | 0.91819 | 1 |
| 18 | 58419600 | 58421799 | 19.44317 | 33.41297 | 46.89944 | 20.57169 | -0.10312 | 0.912433 | 1 |
| 18 | 58454400 | 58455999 | 22.66795 | 38.62958 | 44.3357 | 19.4348 | 0.184267 | 0.998534 | 1 |
| 18 | 58985400 | 58987799 | 26.1323 | 43.39686 | 68.77944 | 29.81845 | -0.28136 | 1.282004 | 1 |
| 18 | 59235000 | 59235799 | 11.56228 | 21.70237 | 24.13031 | 9.932749 | 0.313011 | 0.110883 | 1 |
| 18 | 59246400 | 59248799 | 26.70784 | 45.04411 | 51.20974 | 22.39291 | 0.196802 | 1.111737 | 1 |
| 18 | 59533400 | 59534599 | 15.68081 | 28.03989 | 32.42005 | 14.2289 | 0.19114 | 0.445056 | 1 |
| 18 | 59542000 | 59543399 | 15.35479 | 27.15054 | 33.08912 | 14.6356 | 0.107057 | 0.526509 | 1 |
| 18 | 59637600 | 59638199 | 10.2411 | 19.05104 | 24.80403 | 10.36618 | 0.069796 | 0.159484 | 1 |
| 18 | 59662200 | 59663599 | 13.81399 | 24.52657 | 39.23237 | 17.32237 | -0.28062 | 0.490052 | 1 |
| 18 | 60062000 | 60065199 | 30.08047 | 49.97054 | 64.16345 | 27.82383 | 0.021417 | 1.283664 | 1 |
| 18 | 60086600 | 60088399 | 20.08523 | 34.53157 | 46.36724 | 20.33852 | -0.03929 | 0.910532 | 1 |
| 18 | 60434600 | 60435999 | 22.66449 | 38.95734 | 43.13218 | 18.91937 | 0.237721 | 0.916937 | 1 |
| 18 | 60545000 | 60546599 | 11.63353 | 21.79418 | 23.15736 | 9.563324 | 0.374588 | 0.12258 | 1 |
| 18 | 60589400 | 60590199 | 9.550847 | 18.30718 | 21.3295 | 8.396993 | 0.302586 | -0.06213 | 1 |

| | | | | | | | | | |
|---|---|---|---|---|---|---|---|---|---|
| 18 | 60665800 | 60668199 | 29.61158 | 48.72017 | 70.95354 | 30.58866 | -0.16212 | 1.402386 | 1 |
| 18 | 61348800 | 61350799 | 11.60911 | 21.84611 | 23.13834 | 9.483228 | 0.387792 | 0.093631 | 1 |
| 18 | 61731600 | 61740199 | 146.6526 | 221.4847 | 236.2083 | 97.64821 | 0.182252 | 3.331718 | 1 |
| 18 | 62251000 | 62253399 | 17.10594 | 30.14784 | 37.51624 | 16.60464 | 0.07506 | 0.57899 | 1 |
| 18 | 63007600 | 63010199 | 23.08086 | 39.72621 | 43.60415 | 19.12717 | 0.250484 | 0.9062 | 1 |
| 18 | 64572800 | 64574999 | 20.73515 | 35.99161 | 37.94376 | 16.64938 | 0.311026 | 0.815309 | 1 |
| 18 | 65489400 | 65493799 | 43.58922 | 70.05308 | 102.5518 | 43.46515 | -0.19748 | 1.895172 | 1 |
| 18 | 66881800 | 66884599 | 8.891396 | 16.61998 | 27.94737 | 11.58512 | -0.28726 | 0.124983 | 1 |
| 18 | 67012600 | 67014799 | 28.82257 | 47.73416 | 58.26525 | 25.21265 | 0.094027 | 1.312302 | 1 |
| 18 | 67436800 | 67439799 | 30.92347 | 50.70767 | 69.1379 | 29.76351 | -0.06921 | 1.445926 | 1 |
| 18 | 68054800 | 68056999 | 17.98484 | 31.4533 | 37.74573 | 16.64399 | 0.125772 | 0.693104 | 1 |
| 18 | 68825000 | 68826599 | 136.8044 | 206.25 | 231.0443 | 95.46124 | 0.109231 | 3.365148 | 1 |
| 18 | 69200600 | 69202399 | 15.88827 | 27.84604 | 45.08919 | 19.918 | -0.30445 | 0.660078 | 1 |
| 18 | 69284200 | 69286399 | 18.86901 | 32.54688 | 44.78335 | 19.64572 | -0.07347 | 0.872414 | 1 |
| 18 | 69376400 | 69378599 | 16.83075 | 29.26232 | 47.91278 | 21.04534 | -0.32123 | 0.78744 | 1 |
| 18 | 69859000 | 69863199 | 29.51003 | 48.56218 | 68.84116 | 29.68034 | -0.12325 | 1.396069 | 1 |
| 18 | 70327800 | 70328799 | 12.56215 | 23.10443 | 26.04657 | 11.05771 | 0.259373 | 0.244439 | 1 |
| 18 | 70490000 | 70491799 | 27.87304 | 46.75241 | 51.26805 | 22.3702 | 0.248443 | 1.163255 | 1 |
| 18 | 70492600 | 70493999 | 19.7428 | 34.57881 | 37.00092 | 16.33595 | 0.290329 | 0.677195 | 1 |
| 18 | 71213600 | 71216599 | 39.78807 | 64.22058 | 76.8501 | 32.76805 | 0.100351 | 1.748534 | 1 |
| 18 | 71259000 | 71260999 | 25.32151 | 43.06232 | 45.29416 | 19.85164 | 0.309075 | 1.026386 | 1 |
| 18 | 71272000 | 71272599 | 7.692332 | 14.46925 | 27.53207 | 11.29223 | -0.45128 | 0.074683 | 1 |
| 18 | 71341600 | 71341999 | 9.621838 | 18.41799 | 20.23253 | 7.996558 | 0.381228 | -0.05312 | 1 |
| 18 | 71763400 | 71764999 | 19.73048 | 33.35561 | 54.79149 | 23.97641 | -0.33204 | 1.070208 | 1 |
| 18 | 72623400 | 72623999 | 10.32923 | 19.47698 | 23.24137 | 9.492875 | 0.22101 | 0.069305 | 1 |
| 18 | 72638600 | 72640399 | 19.38363 | 33.8633 | 35.78324 | 15.76512 | 0.308621 | 0.716267 | 1 |
| 18 | 72840600 | 72841199 | 7.34605 | 14.26509 | 21.15873 | 7.925789 | 0.023982 | -0.25528 | 1 |
| 18 | 72890000 | 72893599 | 29.98118 | 49.47005 | 81.26487 | 35.07653 | -0.33382 | 1.375005 | 1 |
| 18 | 73065000 | 73067199 | 21.93027 | 38.00238 | 40.47467 | 17.75808 | 0.295869 | 0.839782 | 1 |
| 18 | 73741800 | 73744399 | 30.07296 | 49.21595 | 66.71747 | 28.69795 | -0.06212 | 1.465958 | 1 |
| 18 | 73820200 | 73822599 | 34.83522 | 56.85198 | 67.53266 | 29.01928 | 0.125114 | 1.517729 | 1 |
| 18 | 74014000 | 74021399 | 82.24262 | 129.4032 | 138.7047 | 57.54498 | 0.221352 | 2.526686 | 1 |
| 18 | 74487800 | 74511999 | 584.2685 | 782.5477 | 872.4889 | 364.2032 | -0.05136 | 5.253928 | 1 |
| 18 | 75065400 | 75066799 | 27.80723 | 46.29753 | 57.22043 | 24.85093 | 0.07662 | 1.247577 | 1 |
| 18 | 75067800 | 75069999 | 15.39286 | 27.76815 | 30.32762 | 13.19887 | 0.28086 | 0.384392 | 1 |
| 18 | 76324200 | 76326999 | 20.32358 | 34.37441 | 54.84205 | 24.00195 | -0.29018 | 1.066871 | 1 |
| 18 | 77147800 | 77152799 | 48.24531 | 77.06618 | 109.1726 | 45.85793 | -0.16031 | 2.09163 | 1 |
| 18 | 78739000 | 78740799 | 13.45462 | 24.67575 | 24.11635 | 10.27787 | 0.460403 | 0.264599 | 1 |
| 18 | 79548600 | 79551399 | 27.60166 | 45.49828 | 65.19865 | 28.12994 | -0.13774 | 1.369482 | 1 |
| 18 | 79751200 | 79755599 | 98.55289 | 152.1966 | 169.3082 | 70.32711 | 0.147042 | 2.872143 | 1 |
| 18 | 79766800 | 79768599 | 13.79961 | 24.16542 | 39.00557 | 17.21853 | -0.29954 | 0.653258 | 1 |
| 18 | 79870800 | 79875399 | 65.59414 | 103.6168 | 135.3543 | 56.21293 | -0.05916 | 2.432339 | 1 |
| 18 | 79949600 | 79952199 | 24.06104 | 41.36017 | 44.69127 | 19.60327 | 0.272744 | 0.925092 | 1 |
| 18 | 80929000 | 80930999 | 45.21255 | 72.57658 | 88.72471 | 37.53895 | 0.060022 | 1.929773 | 1 |
| 18 | 81425000 | 81426799 | 18.38277 | 32.00984 | 37.87991 | 16.65455 | 0.145355 | 0.757133 | 1 |
| 18 | 82712600 | 82713799 | 22.52474 | 37.20788 | 65.69168 | 28.37561 | -0.43693 | 1.334268 | 1 |
| 18 | 82718800 | 82719599 | 21.92167 | 37.12088 | 50.61374 | 22.15929 | -0.06424 | 1.058961 | 1 |

| | | | | | | | | | |
|---|---|---|---|---|---|---|---|---|---|
| 18 | 83061600 | 83064799 | 40.77965 | 65.75872 | 84.42225 | 35.95277 | -0.00246 | 1.780672 | 1 |
| 18 | 83117600 | 83120599 | 28.25489 | 46.97779 | 55.05402 | 23.88765 | 0.152995 | 1.262155 | 1 |
| 18 | 83224800 | 83227599 | 21.13164 | 35.88243 | 54.87981 | 24.04723 | -0.22924 | 1.036396 | 1 |
| 18 | 85050800 | 85053799 | 15.44899 | 26.77909 | 46.36474 | 20.3425 | -0.40183 | 0.816745 | 1 |
| 18 | 85132600 | 85133399 | 13.29136 | 23.80773 | 32.15862 | 14.09388 | -0.03033 | 0.418994 | 1 |
| 18 | 85343000 | 85347199 | 16.94219 | 29.81666 | 32.60038 | 14.43301 | 0.260233 | 0.586368 | 1 |
| 18 | 85801400 | 85802599 | 16.33533 | 29.23315 | 28.42191 | 12.46592 | 0.440558 | 0.437519 | 1 |
| 18 | 85914600 | 85916799 | 28.18323 | 47.16296 | 50.21382 | 21.87809 | 0.29096 | 1.187734 | 1 |
| 18 | 85935200 | 85936999 | 35.98282 | 58.49657 | 65.62423 | 28.1592 | 0.203475 | 1.574814 | 1 |
| 18 | 85982200 | 85983999 | 16.60349 | 28.95436 | 41.53805 | 18.2799 | -0.13083 | 0.734314 | 1 |
| 18 | 86225400 | 86226599 | 23.12024 | 38.74754 | 54.86233 | 23.92573 | -0.11889 | 1.161836 | 1 |
| 18 | 87168000 | 87168799 | 13.76626 | 24.36324 | 34.67782 | 15.33559 | -0.11517 | 0.509768 | 1 |
| 18 | 87178600 | 87181599 | 37.68382 | 61.17378 | 71.62392 | 30.70267 | 0.140014 | 1.606885 | 1 |
| 18 | 87250000 | 87252399 | 26.27851 | 43.93736 | 55.02079 | 23.95935 | 0.057641 | 1.195267 | 1 |
| 18 | 87591800 | 87592599 | 14.53201 | 26.15563 | 29.8586 | 13.03179 | 0.214726 | 0.389797 | 1 |
| 18 | 87901200 | 87903799 | 30.06934 | 50.04836 | 59.4927 | 25.83273 | 0.132683 | 1.261393 | 1 |
| 18 | 87971400 | 87972199 | 12.66083 | 22.82912 | 28.1532 | 12.25963 | 0.106364 | 0.363589 | 1 |
| 18 | 88430200 | 88433199 | 3.986897 | 7.78276 | 21.02163 | 7.437037 | -0.75409 | -0.49874 | 1 |
| 18 | 88637400 | 88638199 | 12.2112 | 22.48464 | 25.06747 | 10.62551 | 0.276927 | 0.232706 | 1 |
| 18 | 89388000 | 89388999 | 14.91356 | 26.13385 | 37.93186 | 16.75416 | -0.14705 | 0.647306 | 1 |
| 18 | 89425600 | 89426999 | 14.32229 | 25.89601 | 29.35145 | 12.73757 | 0.230212 | 0.360591 | 1 |
| 18 | 89686800 | 89689799 | 25.69745 | 43.72039 | 51.19014 | 22.43651 | 0.155115 | 1.032272 | 1 |
| 18 | 89706600 | 89707199 | 9.622154 | 17.82162 | 26.81049 | 11.27401 | -0.14402 | 0.182958 | 1 |
| 18 | 89934800 | 89936399 | 16.06566 | 28.11502 | 36.35074 | 16.03762 | 0.018817 | 0.67037 | 1 |
| 18 | 90002600 | 90004199 | 15.9365 | 28.21673 | 30.32243 | 13.40829 | 0.288062 | 0.507121 | 1 |
| 18 | 90022000 | 90027399 | 79.9871 | 126.6109 | 123.1762 | 51.20819 | 0.367754 | 2.391993 | 1 |
| 18 | 90287200 | 90289399 | 43.80065 | 70.26068 | 95.10741 | 40.19638 | -0.0879 | 1.952729 | 1 |
| 18 | 90315800 | 90317199 | 10.32792 | 19.67732 | 24.42797 | 9.762143 | 0.191682 | 0.001114 | 1 |
| 18 | 90488400 | 90494399 | 69.25122 | 109.7867 | 117.9908 | 49.10245 | 0.226202 | 2.344203 | 1 |
| 18 | 90575800 | 90582599 | 112.7459 | 172.8757 | 180.8191 | 75.05927 | 0.228007 | 3.01468 | 1 |
| 18 | 90618000 | 90620399 | 33.16815 | 54.52206 | 65.59739 | 28.26866 | 0.112149 | 1.418377 | 1 |
| 19 | 3217800 | 3223199 | 76.13232 | 120.1206 | 128.0601 | 53.15161 | 0.232204 | 2.464685 | 1 |
| 19 | 4453000 | 4453799 | 12.22003 | 22.64828 | 25.56822 | 10.74192 | 0.268127 | 0.197677 | 1 |
| 19 | 6670600 | 6675799 | 54.046 | 85.85829 | 128.5681 | 53.61492 | -0.2487 | 2.274241 | 1 |
| 19 | 7361600 | 7364399 | 28.33776 | 47.47485 | 53.47666 | 23.31622 | 0.209839 | 1.178651 | 1 |
| 19 | 7517600 | 7521999 | 43.05495 | 69.36283 | 99.99769 | 42.5322 | -0.17104 | 1.82027 | 1 |
| 19 | 7751200 | 7752399 | 26.41853 | 43.28252 | 69.62412 | 29.96168 | -0.3075 | 1.446644 | 1 |
| 19 | 7827000 | 7829199 | 26.49042 | 44.56288 | 53.92879 | 23.56355 | 0.106785 | 1.134359 | 1 |
| 19 | 7848400 | 7855999 | 90.44205 | 140.7668 | 165.598 | 68.74126 | 0.075145 | 2.729293 | 1 |
| 19 | 7860200 | 7863799 | 42.42992 | 68.34367 | 94.68773 | 40.26327 | -0.11412 | 1.820499 | 1 |
| 19 | 7916200 | 7922799 | 40.29424 | 65.45682 | 74.73515 | 32.05275 | 0.177171 | 1.602579 | 1 |
| 19 | 8005400 | 8018599 | 233.9472 | 341.3002 | 344.0336 | 141.7587 | 0.217578 | 3.9488 | 1 |
| 19 | 8095800 | 8103999 | 96.197 | 150.7745 | 143.286 | 59.43099 | 0.390501 | 2.610688 | 1 |
| 19 | 8125400 | 8133999 | 158.1668 | 236.8509 | 260.7506 | 107.5506 | 0.125626 | 3.499587 | 1 |
| 19 | 8332600 | 8334399 | 22.90015 | 39.65131 | 40.43328 | 17.73948 | 0.358112 | 0.850977 | 1 |
| 19 | 8394400 | 8395999 | 24.36558 | 41.42924 | 45.84489 | 20.09272 | 0.236113 | 1.02525 | 1 |
| 19 | 8399000 | 8403999 | 31.78384 | 52.74272 | 57.37395 | 24.86092 | 0.259914 | 1.297584 | 1 |

| | | | | | | | | | |
|---|---|---|---|---|---|---|---|---|---|
| 19 | 8504600 | 8507399 | 20.07948 | 34.54312 | 47.41347 | 20.79783 | -0.07072 | 0.905884 | 1 |
| 19 | 8521400 | 8530999 | 133.4464 | 201.585 | 257.1417 | 106.3096 | -0.07561 | 3.325264 | 1 |
| 19 | 8795800 | 8798599 | 18.12544 | 31.05264 | 47.95944 | 21.03227 | -0.2413 | 0.934492 | 1 |
| 19 | 9283400 | 9285399 | 16.40893 | 28.93036 | 34.49015 | 15.26414 | 0.136873 | 0.565461 | 1 |
| 19 | 9286400 | 9290999 | 65.25679 | 102.8947 | 139.9071 | 58.05978 | -0.119 | 2.473751 | 1 |
| 19 | 9398200 | 9402199 | 51.40062 | 82.32879 | 100.3596 | 42.31999 | 0.060336 | 2.008433 | 1 |
| 19 | 9411000 | 9412599 | 21.40292 | 36.80712 | 41.64704 | 18.26821 | 0.206711 | 0.906187 | 1 |
| 19 | 9454800 | 9472799 | 326.4334 | 460.9769 | 485.8494 | 201.143 | 0.106702 | 4.434236 | 1 |
| 19 | 9507200 | 9509399 | 20.80637 | 36.20314 | 36.85558 | 16.19662 | 0.36161 | 0.78079 | 1 |
| 19 | 11101200 | 11103399 | 32.74892 | 54.64006 | 55.33396 | 24.06357 | 0.362978 | 1.236493 | 1 |
| 19 | 11845200 | 11846799 | 15.84446 | 27.68989 | 42.02702 | 18.52045 | -0.21142 | 0.701302 | 1 |
| 19 | 12359800 | 12360999 | 13.21041 | 23.81711 | 34.65829 | 15.0947 | -0.13143 | 0.381392 | 1 |
| 19 | 12429400 | 12431999 | 45.50303 | 73.17395 | 83.09774 | 35.25135 | 0.169301 | 1.8718 | 1 |
| 19 | 12636000 | 12643799 | 105.9812 | 161.9647 | 237.7715 | 98.63095 | -0.26441 | 3.083628 | 1 |
| 19 | 12881400 | 12884199 | 33.02773 | 53.84925 | 71.30012 | 30.62723 | -0.0321 | 1.529155 | 1 |
| 19 | 13059800 | 13063399 | 28.20577 | 47.23401 | 49.13882 | 21.41925 | 0.324247 | 1.179904 | 1 |
| 19 | 13095200 | 13100999 | 89.32538 | 139.4186 | 150.7503 | 62.55393 | 0.199798 | 2.680995 | 1 |
| 19 | 13532200 | 13532599 | 10.54844 | 19.78533 | 24.67292 | 10.16832 | 0.147047 | 0.106101 | 1 |
| 19 | 13554600 | 13555999 | 23.02641 | 39.39463 | 40.66382 | 17.83084 | 0.33747 | 0.958616 | 1 |
| 19 | 13634400 | 13635399 | 22.97507 | 39.34345 | 40.61804 | 17.81211 | 0.337368 | 0.949132 | 1 |
| 19 | 13660600 | 13664399 | 40.15857 | 64.93374 | 80.90258 | 34.57505 | 0.045197 | 1.700234 | 1 |
| 19 | 13876600 | 13877599 | 11.82025 | 22.09929 | 22.20332 | 9.200755 | 0.451032 | 0.135204 | 1 |
| 19 | 14244800 | 14245799 | 13.99245 | 24.4195 | 41.64811 | 18.33704 | -0.37882 | 0.712329 | 1 |
| 19 | 14665000 | 14668399 | 13.13 | 23.35172 | 37.70059 | 16.62226 | -0.29246 | 0.471517 | 1 |
| 19 | 15500400 | 15501999 | 21.51029 | 37.54616 | 38.08637 | 16.77071 | 0.366955 | 0.739788 | 1 |
| 19 | 15930400 | 15934999 | 10.31218 | 19.34258 | 25.25427 | 10.40757 | 0.081543 | 0.106617 | 1 |
| 19 | 16056400 | 16063599 | 153.5745 | 230.6053 | 242.4086 | 100.0491 | 0.195796 | 3.445931 | 1 |
| 19 | 16412600 | 16415199 | 20.54356 | 34.47195 | 54.25814 | 23.68075 | -0.27105 | 1.138626 | 1 |
| 19 | 18225800 | 18227999 | 18.8228 | 32.48433 | 52.79308 | 23.15983 | -0.31258 | 0.878336 | 1 |
| 19 | 18334800 | 18336999 | 12.55213 | 22.23063 | 35.30187 | 15.60929 | -0.27157 | 0.496815 | 1 |
| 19 | 18480400 | 18483599 | 35.81855 | 58.11963 | 79.93692 | 34.25479 | -0.09234 | 1.617622 | 1 |
| 19 | 19216400 | 19218999 | 14.30205 | 25.53158 | 36.65781 | 16.11158 | -0.12096 | 0.455153 | 1 |
| 19 | 19575600 | 19577799 | 24.09487 | 40.14626 | 63.12108 | 27.42636 | -0.26959 | 1.232561 | 1 |
| 19 | 20315000 | 20317999 | 21.4119 | 36.36384 | 65.02161 | 28.49247 | -0.4541 | 1.043572 | 1 |
| 19 | 20432600 | 20433999 | 18.16938 | 31.61153 | 39.50465 | 17.35967 | 0.06704 | 0.772065 | 1 |
| 19 | 20550600 | 20551399 | 16.57725 | 29.09547 | 33.71873 | 14.91329 | 0.175976 | 0.622611 | 1 |
| 19 | 21194600 | 21196599 | 25.10186 | 42.19937 | 49.08872 | 21.44448 | 0.163629 | 1.129901 | 1 |
| 19 | 21249000 | 21250599 | 17.80034 | 30.60982 | 46.90456 | 20.57435 | -0.229 | 0.898343 | 1 |
| 19 | 21909800 | 21912399 | 36.60469 | 59.16632 | 87.7893 | 37.50102 | -0.20687 | 1.707949 | 1 |
| 19 | 21914600 | 21915999 | 22.38085 | 38.01016 | 44.85533 | 19.65573 | 0.143485 | 1.026334 | 1 |
| 19 | 22627600 | 22629599 | 22.64158 | 37.57807 | 63.17915 | 27.37951 | -0.36606 | 1.277395 | 1 |
| 19 | 22991800 | 22995999 | 39.83202 | 64.72138 | 72.60156 | 31.14316 | 0.202878 | 1.5936 | 1 |
| 19 | 23109400 | 23114199 | 41.65815 | 66.88404 | 96.42718 | 40.80997 | -0.17712 | 1.921736 | 1 |
| 19 | 23179200 | 23181799 | 35.56189 | 57.72081 | 80.91621 | 34.68225 | -0.11934 | 1.613204 | 1 |
| 19 | 23834200 | 23836999 | 26.54814 | 45.28717 | 45.2142 | 19.82149 | 0.384602 | 1.002202 | 1 |
| 19 | 23906800 | 23912999 | 68.19454 | 107.0889 | 158.3793 | 65.69066 | -0.24467 | 2.560134 | 1 |
| 19 | 24063800 | 24064999 | 11.72182 | 21.43766 | 31.24341 | 13.36611 | -0.11676 | 0.281466 | 1 |

| | | | | | | | | | |
|---|---|---|---|---|---|---|---|---|---|
| 19 | 24663400 | 24664399 | 11.03771 | 20.36623 | 29.27042 | 12.37195 | -0.08416 | 0.22165 | 1 |
| 19 | 25308000 | 25309199 | 13.84967 | 24.47067 | 34.14683 | 15.10572 | -0.08779 | 0.525386 | 1 |
| 19 | 25452800 | 25456599 | 13.88372 | 24.44381 | 37.7288 | 16.70216 | -0.23461 | 0.570923 | 1 |
| 19 | 25795600 | 25797599 | 16.16166 | 28.19846 | 47.32608 | 20.83384 | -0.35624 | 0.732578 | 1 |
| 19 | 26054200 | 26058599 | 29.3292 | 49.04324 | 54.06077 | 23.54286 | 0.241006 | 1.20225 | 1 |
| 19 | 27728600 | 27729399 | 11.23524 | 20.10181 | 33.79418 | 14.82494 | -0.34505 | 0.414923 | 1 |
| 19 | 28001800 | 28005199 | 38.96075 | 63.12069 | 75.87877 | 32.47879 | 0.099414 | 1.653817 | 1 |
| 19 | 28169400 | 28172399 | 26.14858 | 44.11072 | 52.83863 | 23.10675 | 0.121631 | 1.108164 | 1 |
| 19 | 28292600 | 28297199 | 30.96881 | 51.12967 | 61.68701 | 26.63618 | 0.110417 | 1.356251 | 1 |
| 19 | 29176800 | 29178799 | 13.99463 | 25.12131 | 29.87068 | 13.06749 | 0.153887 | 0.401384 | 1 |
| 19 | 29973200 | 29975599 | 31.07702 | 51.33915 | 57.83448 | 24.98303 | 0.209131 | 1.346279 | 1 |
| 19 | 30165000 | 30167999 | 42.4915 | 68.9387 | 73.28668 | 31.40214 | 0.278211 | 1.629628 | 1 |
| 19 | 30198000 | 30201799 | 43.58817 | 69.97115 | 107.6432 | 45.54531 | -0.27095 | 1.932121 | 1 |
| 19 | 30582200 | 30583199 | 16.09435 | 28.17961 | 36.34393 | 16.04152 | 0.022415 | 0.662871 | 1 |
| 19 | 30591200 | 30593599 | 27.76365 | 45.83138 | 63.5773 | 27.44969 | -0.09034 | 1.351684 | 1 |
| 19 | 30638000 | 30638599 | 17.78392 | 30.89224 | 42.67139 | 18.7327 | -0.07694 | 0.798388 | 1 |
| 19 | 30740400 | 30741199 | 14.56895 | 25.67066 | 35.2352 | 15.59578 | -0.06552 | 0.564089 | 1 |
| 19 | 31007200 | 31008999 | 16.14564 | 28.21989 | 52.24627 | 23.02639 | -0.49718 | 0.7127 | 1 |
| 19 | 31837600 | 31840399 | 17.11428 | 29.72226 | 46.86088 | 20.56895 | -0.26712 | 0.802518 | 1 |
| 19 | 32413000 | 32420799 | 83.46981 | 129.7249 | 167.2838 | 69.45623 | -0.0588 | 2.75183 | 1 |
| 19 | 32722400 | 32723399 | 11.64038 | 21.57166 | 27.70263 | 11.64035 | 0.08329 | 0.198767 | 1 |
| 19 | 32853200 | 32860399 | 91.5336 | 141.643 | 175.9689 | 73.09309 | -0.00991 | 2.832009 | 1 |
| 19 | 32996600 | 32999399 | 13.69391 | 24.8956 | 28.43358 | 12.25515 | 0.225696 | 0.323329 | 1 |
| 19 | 33383600 | 33386599 | 60.52572 | 96.36758 | 105.9617 | 44.3069 | 0.199494 | 2.205979 | 1 |
| 19 | 33421200 | 33421999 | 13.36868 | 24.41243 | 28.98943 | 12.42556 | 0.174942 | 0.296564 | 1 |
| 19 | 33459000 | 33461199 | 18.06036 | 31.46927 | 45.18874 | 19.87628 | -0.13244 | 0.757854 | 1 |
| 19 | 33569200 | 33573199 | 91.82675 | 141.1911 | 200.4931 | 83.25775 | -0.20961 | 2.956452 | 1 |
| 19 | 33706800 | 33708199 | 29.73194 | 49.41477 | 48.67146 | 21.11273 | 0.402971 | 1.26666 | 1 |
| 19 | 33738600 | 33740399 | 22.17317 | 37.10098 | 57.49934 | 25.05046 | -0.24881 | 1.177982 | 1 |
| 19 | 33810800 | 33812799 | 18.31707 | 32.07591 | 39.1924 | 17.30086 | 0.099824 | 0.675941 | 1 |
| 19 | 35133800 | 35135799 | 17.35353 | 30.44417 | 34.13157 | 15.09011 | 0.22349 | 0.634393 | 1 |
| 19 | 35210000 | 35212599 | 27.91135 | 46.58949 | 55.6296 | 24.19892 | 0.126125 | 1.218644 | 1 |
| 19 | 35433400 | 35434999 | 14.93592 | 26.87463 | 28.90531 | 12.61993 | 0.300035 | 0.393901 | 1 |
| 19 | 35643400 | 35644999 | 26.00072 | 42.9989 | 63.43871 | 27.42409 | -0.17871 | 1.326686 | 1 |
| 19 | 35810400 | 35811999 | 18.97228 | 33.20169 | 33.67558 | 14.85738 | 0.367387 | 0.682632 | 1 |
| 19 | 35819200 | 35823599 | 7.904369 | 15.14834 | 28.03996 | 11.04473 | -0.36105 | -0.05997 | 1 |
| 19 | 35879600 | 35882399 | 35.32342 | 57.72051 | 63.69362 | 27.38204 | 0.232423 | 1.499126 | 1 |
| 19 | 36046800 | 36050399 | 42.24638 | 67.99902 | 81.60433 | 34.66371 | 0.091461 | 1.838049 | 1 |
| 19 | 36990400 | 36996999 | 70.27435 | 110.5837 | 147.0069 | 60.99017 | -0.08873 | 2.518798 | 1 |
| 19 | 38327600 | 38336999 | 27.80722 | 45.76823 | 71.95089 | 31.02344 | -0.27191 | 1.393432 | 1 |
| 19 | 38364600 | 38369799 | 74.67567 | 115.9694 | 174.1233 | 72.30421 | -0.27891 | 2.762241 | 1 |
| 19 | 38481600 | 38485399 | 22.53028 | 37.5732 | 62.39165 | 27.12483 | -0.34806 | 1.218897 | 1 |
| 19 | 39083800 | 39090199 | 111.6241 | 171.5647 | 184.2167 | 76.49465 | 0.192818 | 2.967534 | 1 |
| 19 | 39134000 | 39136599 | 55.1657 | 87.95054 | 103.8583 | 43.50748 | 0.09881 | 2.158604 | 1 |
| 19 | 39144400 | 39148399 | 58.49111 | 93.24919 | 109.2813 | 45.77716 | 0.109669 | 2.162344 | 1 |
| 19 | 39224400 | 39225399 | 16.70242 | 29.64633 | 30.77725 | 13.59249 | 0.339442 | 0.493144 | 1 |
| 19 | 39255200 | 39255999 | 24.0647 | 40.69265 | 50.65909 | 22.16971 | 0.06632 | 1.076272 | 1 |

| | | | | | | | | | |
|---|---|---|---|---|---|---|---|---|---|
| 19 | 39289000 | 39296999 | 42.65458 | 69.14503 | 74.96896 | 32.10303 | 0.248719 | 1.647371 | 1 |
| 19 | 39329400 | 39339399 | 154.1182 | 233.4112 | 222.8511 | 92.20763 | 0.345214 | 3.277545 | 1 |
| 19 | 39441800 | 39444399 | 32.53167 | 53.93287 | 58.58883 | 25.37084 | 0.261729 | 1.309942 | 1 |
| 19 | 39644400 | 39647199 | 42.93908 | 69.35201 | 81.68327 | 34.85836 | 0.124266 | 1.737391 | 1 |
| 19 | 39701800 | 39703999 | 22.49512 | 38.46944 | 54.48175 | 23.88911 | -0.11736 | 0.975457 | 1 |
| 19 | 39846200 | 39847799 | 15.05257 | 26.61555 | 38.13079 | 16.86565 | -0.12524 | 0.53312 | 1 |
| 19 | 39889200 | 39894999 | 46.78296 | 75.33269 | 87.09966 | 37.02055 | 0.145785 | 1.832759 | 1 |
| 19 | 39903000 | 39905599 | 40.91658 | 66.26106 | 74.9545 | 32.07307 | 0.186361 | 1.665399 | 1 |
| 19 | 40052800 | 40057799 | 30.18025 | 50.2723 | 56.85431 | 24.69931 | 0.204242 | 1.247963 | 1 |
| 19 | 40268200 | 40272399 | 51.1923 | 82.05404 | 95.2292 | 40.20046 | 0.132312 | 1.983928 | 1 |
| 19 | 40861400 | 40863799 | 31.69701 | 52.07064 | 72.96911 | 31.43698 | -0.10768 | 1.426596 | 1 |
| 19 | 41130800 | 41132599 | 22.05818 | 37.15658 | 50.63011 | 22.13121 | -0.06364 | 1.103753 | 1 |
| 19 | 42331000 | 42337399 | 91.62451 | 141.0648 | 198.5207 | 82.45345 | -0.19522 | 2.929573 | 1 |
| 19 | 42925000 | 42933399 | 88.16496 | 138.2503 | 142.8984 | 59.26734 | 0.269961 | 2.597254 | 1 |
| 19 | 43034400 | 43035999 | 27.73128 | 45.94857 | 58.15457 | 25.17299 | 0.042025 | 1.304843 | 1 |
| 19 | 43063000 | 43064199 | 13.39785 | 24.08396 | 28.50736 | 12.45627 | 0.161753 | 0.386619 | 1 |
| 19 | 45118000 | 45119399 | 14.57867 | 26.08628 | 30.91939 | 13.56273 | 0.15596 | 0.428956 | 1 |
| 19 | 45692600 | 45702999 | 84.11633 | 129.8602 | 192.3831 | 79.91208 | -0.2662 | 2.874108 | 1 |
| 19 | 47169800 | 47170999 | 18.12044 | 31.48951 | 46.3153 | 20.33754 | -0.16721 | 0.79865 | 1 |
| 19 | 47579400 | 47583599 | 52.55486 | 83.63773 | 118.6835 | 49.59455 | -0.16859 | 2.216937 | 1 |
| 19 | 47622400 | 47623199 | 17.33545 | 30.52455 | 32.60095 | 14.43177 | 0.294246 | 0.58403 | 1 |
| 19 | 48220000 | 48221599 | 27.07961 | 44.87789 | 59.218 | 25.63689 | -0.01793 | 1.301327 | 1 |
| 19 | 48890800 | 48892399 | 16.79207 | 29.64577 | 34.96191 | 15.46876 | 0.153218 | 0.554733 | 1 |
| 19 | 48963800 | 48966399 | 18.06247 | 30.84217 | 51.09797 | 22.40266 | -0.34283 | 0.970294 | 1 |
| 19 | 49010600 | 49011599 | 25.35873 | 42.36887 | 51.7349 | 22.51821 | 0.09404 | 1.19873 | 1 |
| 19 | 49058400 | 49059199 | 13.82201 | 24.2744 | 38.39588 | 16.99066 | -0.27041 | 0.606586 | 1 |
| 19 | 49164800 | 49166199 | 22.68832 | 38.87444 | 44.7865 | 19.64099 | 0.179949 | 0.950243 | 1 |
| 19 | 49448600 | 49449799 | 18.75844 | 32.37499 | 42.24349 | 18.53186 | 0.003131 | 0.865075 | 1 |
| 19 | 49511600 | 49512199 | 11.36249 | 20.64044 | 29.65644 | 12.79508 | -0.1043 | 0.313028 | 1 |
| 19 | 49696400 | 49698999 | 26.71161 | 45.1358 | 47.46751 | 20.76843 | 0.30886 | 1.0899 | 1 |
| 19 | 50094800 | 50097199 | 29.28746 | 47.98963 | 71.8123 | 30.9052 | -0.20352 | 1.452018 | 1 |
| 19 | 50528400 | 50530199 | 20.95461 | 36.20742 | 44.72198 | 19.61984 | 0.08209 | 0.865756 | 1 |
| 19 | 51127800 | 51129599 | 29.20369 | 48.3007 | 69.01515 | 29.83681 | -0.13279 | 1.336374 | 1 |
| 19 | 51130400 | 51131399 | 12.73153 | 22.75041 | 32.1381 | 14.11335 | -0.09676 | 0.427987 | 1 |
| 19 | 51236600 | 51240799 | 45.19678 | 72.69922 | 90.38486 | 38.35623 | 0.039318 | 1.866363 | 1 |
| 19 | 51418800 | 51419799 | 20.97386 | 36.17105 | 45.26758 | 19.85804 | 0.062672 | 0.883895 | 1 |
| 19 | 51447400 | 51449199 | 16.90637 | 29.58328 | 41.39366 | 18.26094 | -0.09478 | 0.68416 | 1 |
| 19 | 51537800 | 51538599 | 10.88584 | 20.04133 | 29.77606 | 12.62386 | -0.13493 | 0.233882 | 1 |
| 19 | 51638000 | 51638799 | 10.71124 | 20.26107 | 24.30872 | 9.863729 | 0.221562 | 0.054889 | 1 |
| 19 | 51668600 | 51671599 | 35.54854 | 59.45631 | 52.51471 | 22.87281 | 0.559716 | 1.221071 | 1 |
| 19 | 51886200 | 51888799 | 17.36538 | 30.07772 | 46.35651 | 20.33841 | -0.23505 | 0.832627 | 1 |
| 19 | 52251800 | 52254199 | 17.84651 | 31.24117 | 39.24543 | 17.31924 | 0.060054 | 0.67929 | 1 |
| 19 | 52762400 | 52764399 | 36.14466 | 58.72082 | 67.27458 | 28.85479 | 0.172388 | 1.588304 | 1 |
| 19 | 53997200 | 53998199 | 15.86785 | 27.75922 | 38.45504 | 16.96104 | -0.08012 | 0.680636 | 1 |
| 19 | 54047800 | 54050199 | 27.35796 | 45.44928 | 59.34174 | 25.73351 | -0.00252 | 1.273383 | 1 |
| 19 | 54260400 | 54261399 | 12.04926 | 21.92261 | 32.2132 | 13.87073 | -0.13464 | 0.315545 | 1 |
| 19 | 55900800 | 55903599 | 17.50234 | 29.97946 | 51.11805 | 22.41711 | -0.3836 | 0.93673 | 1 |

| | | | | | | | | | |
|---|---|---|---|---|---|---|---|---|---|
| 19 | 55970200 | 55973199 | 30.88816 | 51.27084 | 56.25951 | 24.38276 | 0.247423 | 1.289899 | 1 |
| 19 | 56166800 | 56174399 | 82.86441 | 128.7823 | 193.2797 | 80.24993 | -0.27751 | 2.75572 | 1 |
| 19 | 56598600 | 56600399 | 19.24715 | 33.2921 | 45.81279 | 20.09904 | -0.07275 | 0.851359 | 1 |
| 19 | 56841600 | 56843199 | 12.63862 | 22.32814 | 38.75596 | 17.14511 | -0.40114 | 0.527138 | 1 |
| 19 | 57554200 | 57556199 | 21.05839 | 36.65193 | 38.2854 | 16.82794 | 0.324556 | 0.778724 | 1 |
| 19 | 57623800 | 57625199 | 23.87671 | 40.10798 | 56.56651 | 24.70495 | -0.1134 | 1.139953 | 1 |
| 19 | 57820800 | 57821799 | 11.97756 | 22.22921 | 28.2362 | 11.84081 | 0.101046 | 0.194726 | 1 |
| 19 | 57850000 | 57852199 | 27.87151 | 47.019 | 50.70492 | 22.17392 | 0.27266 | 1.11068 | 1 |
| 19 | 57941400 | 57942999 | 12.16226 | 22.04294 | 31.6205 | 13.68143 | -0.10477 | 0.336304 | 1 |
| 19 | 58069600 | 58076599 | 78.72807 | 123.346 | 148.1855 | 61.46562 | 0.052268 | 2.606869 | 1 |
| 19 | 58606200 | 58608799 | 29.26534 | 48.79049 | 54.00716 | 23.47523 | 0.235143 | 1.235273 | 1 |
| 19 | 58880400 | 58889999 | 64.22889 | 101.1133 | 149.2925 | 61.94184 | -0.23947 | 2.507345 | 1 |
| 19 | 59142200 | 59143799 | 15.68519 | 27.91593 | 27.96221 | 12.32122 | 0.393084 | 0.464295 | 1 |
| 19 | 59182000 | 59183199 | 12.50944 | 22.13804 | 35.26954 | 15.59739 | -0.27684 | 0.503215 | 1 |
| 19 | 59184200 | 59189999 | 25.87097 | 42.8975 | 65.43911 | 28.34048 | -0.22661 | 1.29357 | 1 |
| 19 | 59267200 | 59270599 | 37.51948 | 61.28049 | 65.19088 | 28.02096 | 0.284576 | 1.509489 | 1 |
| 19 | 60913000 | 60913799 | 21.84751 | 38.17174 | 38.10299 | 16.78825 | 0.390307 | 0.730052 | 1 |
| 19 | 60914800 | 60916199 | 35.43405 | 57.26564 | 80.47653 | 34.37062 | -0.12888 | 1.706411 | 1 |
| 19 | 61092600 | 61097399 | 62.50246 | 100.0108 | 102.0372 | 42.96444 | 0.315045 | 2.054865 | 1 |
| 19 | 61114600 | 61121399 | 70.32402 | 111.9339 | 109.1972 | 45.64045 | 0.371366 | 2.224398 | 1 |
| 19 | 61142400 | 61144999 | 36.00718 | 59.28647 | 60.92959 | 26.27542 | 0.339775 | 1.400597 | 1 |
| 19 | 61154400 | 61156599 | 17.66974 | 31.10341 | 33.40546 | 14.78884 | 0.285797 | 0.588242 | 1 |
| X | 3953800 | 3956999 | 50.89732 | 81.66688 | 87.58615 | 37.03305 | 0.247814 | 1.951937 | 1 |
| X | 4913600 | 4927199 | 434.0493 | 604.8874 | 527.5898 | 218.8814 | 0.360475 | 4.635497 | 1 |
| X | 5174200 | 5176999 | 26.64763 | 45.46785 | 45.79207 | 20.07522 | 0.372269 | 1.004341 | 1 |
| X | 6936800 | 6938999 | 45.59897 | 73.67261 | 76.79651 | 32.78701 | 0.300632 | 1.730809 | 1 |
| X | 8432600 | 8445199 | 20.53645 | 35.80281 | 36.80338 | 16.1934 | 0.347952 | 0.752768 | 1 |
| X | 8868200 | 8873599 | 26.20172 | 44.486 | 47.96611 | 21.01698 | 0.273297 | 1.04983 | 1 |
| X | 8926600 | 8930999 | 66.4797 | 105.7648 | 112.1586 | 46.84743 | 0.250461 | 2.239651 | 1 |
| X | 9340000 | 9344199 | 39.07318 | 64.35291 | 68.62887 | 29.59923 | 0.286761 | 1.412432 | 1 |
| X | 9538200 | 9551999 | 171.6857 | 256.9711 | 278.7602 | 114.9681 | 0.146258 | 3.513399 | 1 |
| X | 9761800 | 9762999 | 15.86233 | 28.3446 | 33.43071 | 14.68115 | 0.162002 | 0.452542 | 1 |
| X | 9790800 | 9792399 | 29.71855 | 49.22869 | 55.5146 | 24.02598 | 0.208024 | 1.312166 | 1 |
| X | 10162000 | 10163999 | 28.54469 | 47.81512 | 58.39557 | 25.45875 | 0.093631 | 1.187731 | 1 |
| X | 10389000 | 10391799 | 9.505193 | 17.46767 | 32.48808 | 13.8078 | -0.45858 | 0.244688 | 1 |
| X | 11148200 | 11150999 | 31.36968 | 53.30724 | 47.37273 | 20.76136 | 0.551296 | 1.060134 | 1 |
| X | 11474400 | 11477799 | 45.66563 | 73.80766 | 80.39099 | 34.33803 | 0.238037 | 1.722638 | 1 |
| X | 14015800 | 14018199 | 16.43911 | 29.51832 | 32.40901 | 14.17428 | 0.268878 | 0.425518 | 1 |
| X | 14044800 | 14046199 | 17.07528 | 30.4392 | 34.90889 | 15.36121 | 0.200231 | 0.478322 | 1 |
| X | 14683200 | 14691399 | 91.56191 | 143.5244 | 140.0304 | 58.08005 | 0.352721 | 2.606022 | 1 |
| X | 15666400 | 15669799 | 24.61384 | 42.70037 | 48.49091 | 21.27589 | 0.204008 | 0.852868 | 1 |
| X | 17291200 | 17293199 | 20.43752 | 35.78655 | 35.57436 | 15.70277 | 0.396314 | 0.685651 | 1 |
| X | 17359400 | 17362199 | 19.76485 | 34.67119 | 37.52538 | 16.58924 | 0.274046 | 0.655689 | 1 |
| X | 17977800 | 17980599 | 26.11552 | 44.80191 | 45.74572 | 20.06365 | 0.353621 | 0.953225 | 1 |
| X | 18491200 | 18492399 | 16.55747 | 29.74695 | 30.84363 | 13.4835 | 0.351412 | 0.419651 | 1 |
| X | 18661600 | 18663599 | 16.86705 | 29.53905 | 45.65303 | 20.15415 | -0.23737 | 0.681779 | 1 |
| X | 19494000 | 19495799 | 10.51323 | 19.47504 | 32.0461 | 13.47277 | -0.27237 | 0.203438 | 1 |

| | | | | | | | | | |
|---|---|---|---|---|---|---|---|---|---|
| X | 19527600 | 19529999 | 31.02575 | 51.4316 | 59.49204 | 25.76035 | 0.171474 | 1.309756 | 1 |
| X | 21017000 | 21021399 | 53.50823 | 86.43528 | 78.24529 | 33.39756 | 0.503401 | 1.749098 | 1 |
| X | 21912000 | 21917799 | 72.12204 | 114.912 | 118.8055 | 49.7259 | 0.289305 | 2.200413 | 1 |
| X | 23456000 | 23464599 | 131.8746 | 199.6143 | 237.9021 | 98.41818 | 0.024838 | 3.285105 | 1 |
| X | 23776600 | 23786199 | 49.49873 | 79.94411 | 81.43065 | 34.74911 | 0.333297 | 1.750715 | 1 |
| X | 25398800 | 25416999 | 220.5828 | 328.2047 | 272.2882 | 112.1883 | 0.525124 | 3.628131 | 1 |
| X | 26065800 | 26075799 | 136.304 | 208.7419 | 191.0089 | 79.27151 | 0.419378 | 3.048953 | 1 |
| X | 26159600 | 26160199 | 14.99005 | 26.97345 | 28.46597 | 12.42755 | 0.327432 | 0.393937 | 1 |
| X | 26368200 | 26374799 | 49.55822 | 79.97798 | 87.17008 | 37.16103 | 0.234573 | 1.77596 | 1 |
| X | 27125400 | 27125799 | 11.91058 | 22.456 | 22.69548 | 9.27187 | 0.459008 | 0.085169 | 1 |
| X | 28647600 | 28647999 | 13.8608 | 25.18547 | 28.54836 | 12.31289 | 0.236224 | 0.330422 | 1 |
| X | 29766200 | 29767199 | 16.33576 | 28.96174 | 30.48212 | 13.47536 | 0.319005 | 0.499455 | 1 |
| X | 29996800 | 29997999 | 29.24064 | 48.9958 | 50.1867 | 21.88412 | 0.346646 | 1.181118 | 1 |
| X | 30621200 | 30621599 | 8.339953 | 16.19927 | 20.59683 | 7.699361 | 0.247846 | -0.2485 | 1 |
| X | 31947800 | 31948199 | 15.71786 | 28.20653 | 28.01702 | 12.25954 | 0.412251 | 0.415911 | 1 |
| X | 32135000 | 32135799 | 20.20815 | 35.11187 | 36.53418 | 16.04092 | 0.330158 | 0.798789 | 1 |
| X | 32489200 | 32522199 | 921.6011 | 1204.138 | 1120.947 | 469.3427 | 0.173148 | 5.65545 | 1 |
| X | 32894000 | 32896799 | 24.55121 | 42.39687 | 41.96777 | 18.41122 | 0.399999 | 0.883521 | 1 |
| X | 33743600 | 33753199 | 178.96 | 270.5059 | 222.8609 | 92.15494 | 0.555407 | 3.323462 | 1 |
| X | 34371800 | 34372199 | 10.41942 | 19.49888 | 24.7975 | 10.25915 | 0.115204 | 0.123479 | 1 |
| X | 34562000 | 34562799 | 17.10462 | 30.1529 | 32.51031 | 14.38832 | 0.281189 | 0.568811 | 1 |
| X | 34563800 | 34564799 | 12.44217 | 23.10319 | 25.59591 | 10.72621 | 0.297843 | 0.190099 | 1 |
| X | 34861000 | 34865799 | 28.52244 | 48.807 | 45.16102 | 19.80315 | 0.4946 | 0.983747 | 1 |
| X | 35179800 | 35184799 | 55.48908 | 89.07513 | 87.29031 | 36.93354 | 0.378501 | 1.945245 | 1 |
| X | 35288400 | 35289999 | 23.86237 | 40.93846 | 42.16272 | 18.49224 | 0.341365 | 0.941078 | 1 |
| X | 35470200 | 35471999 | 20.64688 | 35.85568 | 38.46893 | 16.88494 | 0.285961 | 0.809744 | 1 |
| X | 35861400 | 35863399 | 21.27474 | 37.23192 | 39.72037 | 17.52465 | 0.294921 | 0.706415 | 1 |
| X | 36394000 | 36394399 | 15.21866 | 27.13542 | 32.49826 | 14.29781 | 0.138331 | 0.460544 | 1 |
| X | 37207600 | 37212599 | 79.905 | 125.8423 | 136.9013 | 56.80369 | 0.201046 | 2.50824 | 1 |
| X | 37592400 | 37594799 | 21.99912 | 38.27315 | 40.88896 | 17.96756 | 0.292209 | 0.793758 | 1 |
| X | 38294200 | 38295199 | 23.66937 | 40.00244 | 51.67741 | 22.61163 | 0.013283 | 1.085431 | 1 |
| X | 38420400 | 38421599 | 19.83943 | 34.12634 | 47.94721 | 21.03189 | -0.1042 | 0.908172 | 1 |
| X | 39336400 | 39337399 | 15.86275 | 28.0995 | 31.54382 | 13.94706 | 0.226054 | 0.507727 | 1 |
| X | 40394800 | 40395199 | 9.704714 | 18.67226 | 25.98509 | 10.11986 | 0.062275 | -0.08466 | 1 |
| X | 40571200 | 40572599 | 25.06788 | 42.0756 | 49.09172 | 21.4297 | 0.159414 | 1.147685 | 1 |
| X | 41132000 | 41134999 | 34.88135 | 57.09909 | 74.78506 | 32.17108 | -0.01243 | 1.479639 | 1 |
| X | 42748800 | 42751399 | 26.61277 | 45.14439 | 47.43457 | 20.77856 | 0.310349 | 1.057122 | 1 |
| X | 43286600 | 43287199 | 10.48126 | 20.05322 | 23.96985 | 9.48707 | 0.258293 | -0.02965 | 1 |
| X | 43504800 | 43506799 | 31.02628 | 52.74198 | 47.91303 | 20.99873 | 0.519664 | 1.054407 | 1 |
| X | 43814600 | 43822399 | 102.1142 | 159.2429 | 156.6197 | 64.99542 | 0.335391 | 2.70319 | 1 |
| X | 44883600 | 44886799 | 21.39209 | 37.19209 | 39.42403 | 17.31652 | 0.303467 | 0.799405 | 1 |
| X | 45610600 | 45612999 | 28.08401 | 48.38895 | 41.90729 | 18.38343 | 0.591495 | 0.920223 | 1 |
| X | 46954800 | 46955999 | 19.71496 | 34.31459 | 38.44361 | 16.89775 | 0.224135 | 0.773962 | 1 |
| X | 48513800 | 48515799 | 25.36402 | 42.89354 | 47.48036 | 20.77917 | 0.23536 | 1.080265 | 1 |
| X | 48781200 | 48784199 | 31.65044 | 52.36808 | 65.03403 | 28.12333 | 0.069116 | 1.338794 | 1 |
| X | 49252600 | 49256399 | 47.81347 | 77.43821 | 74.63286 | 31.93794 | 0.417082 | 1.677503 | 1 |
| X | 49300400 | 49302399 | 15.80612 | 28.54251 | 30.86589 | 13.41615 | 0.296341 | 0.38353 | 1 |

| | | | | | | | | | |
|---|---|---|---|---|---|---|---|---|---|
| X | 49486400 | 49487199 | 15.66115 | 27.66414 | 35.66877 | 15.77981 | 0.025764 | 0.546323 | 1 |
| X | 49835600 | 49837599 | 24.25864 | 41.43117 | 46.87489 | 20.5515 | 0.205204 | 0.986514 | 1 |
| X | 50768800 | 50770799 | 29.33037 | 49.73793 | 48.50204 | 21.24404 | 0.41773 | 1.073355 | 1 |
| X | 50888800 | 50891999 | 33.33565 | 55.0531 | 63.55678 | 27.44897 | 0.17385 | 1.361441 | 1 |
| X | 52755000 | 52762799 | 91.71978 | 142.8597 | 148.1297 | 61.48332 | 0.257896 | 2.717963 | 1 |
| X | 53104000 | 53110399 | 80.03796 | 126.7479 | 121.9854 | 50.72458 | 0.383823 | 2.381939 | 1 |
| X | 53767000 | 53769399 | 33.39754 | 56.84999 | 43.99497 | 19.28368 | 0.750395 | 1.047793 | 1 |
| X | 57839200 | 57841799 | 27.97361 | 47.27436 | 48.53001 | 21.23414 | 0.343634 | 1.095647 | 1 |
| X | 57906600 | 57908199 | 18.45845 | 32.09727 | 44.71221 | 19.64144 | -0.08901 | 0.789655 | 1 |
| X | 57974200 | 57975799 | 22.31724 | 39.13226 | 35.32777 | 15.61222 | 0.534575 | 0.673507 | 1 |
| X | 58328800 | 58330599 | 16.44382 | 29.02963 | 32.51212 | 14.385 | 0.227394 | 0.54899 | 1 |
| X | 59140600 | 59141999 | 8.445632 | 16.29888 | 20.92106 | 8.077339 | 0.190193 | -0.13826 | 1 |
| X | 60751200 | 60756599 | 59.11919 | 95.03287 | 87.68722 | 37.17708 | 0.467569 | 1.907347 | 1 |
| X | 60796600 | 60800199 | 22.31374 | 38.28135 | 46.89013 | 20.56564 | 0.092213 | 0.941237 | 1 |
| X | 61115000 | 61116999 | 34.43261 | 56.28643 | 71.52112 | 30.75139 | 0.029817 | 1.497565 | 1 |
| X | 61934200 | 61935399 | 17.04251 | 29.72088 | 42.59809 | 18.74676 | -0.12937 | 0.740067 | 1 |
| X | 61945400 | 61949999 | 60.27582 | 96.68367 | 93.73738 | 39.61432 | 0.392587 | 1.976578 | 1 |
| X | 62490600 | 62490999 | 9.565156 | 18.50833 | 20.40062 | 7.780501 | 0.424786 | -0.168 | 1 |
| X | 63079600 | 63080999 | 19.16065 | 33.56607 | 36.67867 | 16.19648 | 0.260459 | 0.673746 | 1 |
| X | 63751000 | 63754599 | 35.98219 | 59.68491 | 61.63154 | 26.69819 | 0.334761 | 1.317242 | 1 |
| X | 64003400 | 64004799 | 15.16621 | 28.0531 | 28.58768 | 12.04285 | 0.412174 | 0.240002 | 1 |
| X | 64136800 | 64140399 | 41.47778 | 67.23085 | 74.46006 | 31.88283 | 0.218032 | 1.649304 | 1 |
| X | 64860800 | 64863399 | 31.6318 | 52.53845 | 58.49637 | 25.36328 | 0.226556 | 1.287664 | 1 |
| X | 64868800 | 64872399 | 22.31094 | 38.86499 | 40.22235 | 17.68875 | 0.337884 | 0.774341 | 1 |
| X | 65186600 | 65191399 | 71.61178 | 113.8505 | 114.5644 | 47.81171 | 0.324979 | 2.265763 | 1 |
| X | 66268600 | 66271399 | 28.05503 | 47.42064 | 53.97155 | 23.61638 | 0.195273 | 1.099964 | 1 |
| X | 67226400 | 67228599 | 17.18118 | 30.39994 | 35.92691 | 15.8887 | 0.151422 | 0.535398 | 1 |
| X | 67469000 | 67472199 | 51.43473 | 82.78291 | 82.99911 | 35.25277 | 0.350176 | 1.856331 | 1 |
| X | 68403000 | 68405199 | 26.10361 | 44.3486 | 48.49497 | 21.25281 | 0.253155 | 1.045261 | 1 |
| X | 68679200 | 68679999 | 8.704523 | 16.46013 | 27.0497 | 10.98311 | -0.22971 | 0.045418 | 1 |
| X | 69098600 | 69099799 | 13.89925 | 25.07626 | 30.56216 | 13.29927 | 0.123952 | 0.375902 | 1 |
| X | 69295400 | 69299399 | 49.07588 | 78.97628 | 83.17725 | 35.32188 | 0.279082 | 1.856401 | 1 |
| X | 69654800 | 69658199 | 35.85622 | 58.99019 | 61.83757 | 26.65786 | 0.310896 | 1.413399 | 1 |
| X | 70598800 | 70602199 | 41.79485 | 68.45822 | 66.10979 | 28.44654 | 0.426479 | 1.484862 | 1 |
| X | 71313200 | 71314599 | 19.27208 | 33.87475 | 33.89903 | 15.01155 | 0.386601 | 0.615821 | 1 |
| X | 71621400 | 71622799 | 32.79691 | 53.30904 | 70.73784 | 30.35155 | -0.03827 | 1.574728 | 1 |
| X | 72218000 | 72220399 | 10.05416 | 18.9144 | 24.29493 | 9.962245 | 0.111074 | 0.084639 | 1 |
| X | 72396400 | 72398799 | 36.11701 | 59.0659 | 67.07431 | 28.84406 | 0.19202 | 1.493424 | 1 |
| X | 72505600 | 72509999 | 22.15634 | 38.38848 | 39.98069 | 17.54126 | 0.32805 | 0.842388 | 1 |
| X | 73127600 | 73130199 | 28.60691 | 48.99354 | 43.59088 | 19.11596 | 0.551251 | 0.977645 | 1 |
| X | 73228000 | 73234199 | 22.4417 | 39.29947 | 36.3105 | 16.02893 | 0.501189 | 0.693815 | 1 |
| X | 73683200 | 73686599 | 28.97504 | 48.22667 | 54.51158 | 23.66974 | 0.205122 | 1.253017 | 1 |
| X | 73819600 | 73821999 | 19.51194 | 34.24357 | 35.04108 | 15.4971 | 0.354702 | 0.644422 | 1 |
| X | 73833000 | 73840599 | 71.41114 | 113.2102 | 112.6154 | 46.86517 | 0.337623 | 2.347086 | 1 |
| X | 74326200 | 74330399 | 62.71068 | 100.7567 | 84.65431 | 35.86568 | 0.601591 | 1.925594 | 1 |
| X | 74999600 | 75005799 | 82.08116 | 128.9333 | 140.1749 | 58.14226 | 0.199081 | 2.561349 | 1 |
| X | 75008600 | 75010799 | 33.11949 | 54.0467 | 72.9092 | 31.32854 | -0.05801 | 1.519926 | 1 |

| | | | | | | | | | |
|---|---|---|---|---|---|---|---|---|---|
| X | 75895200 | 75899399 | 43.52829 | 70.46934 | 76.19066 | 32.59481 | 0.251076 | 1.679053 | 1 |
| X | 76357800 | 76359799 | 13.39059 | 23.68927 | 36.78597 | 16.26912 | -0.24014 | 0.517979 | 1 |
| X | 77325800 | 77333399 | 93.53313 | 147.4215 | 132.0353 | 54.79026 | 0.482112 | 2.501042 | 1 |
| X | 77397600 | 77403399 | 85.99746 | 135.8432 | 130.6858 | 54.27326 | 0.38154 | 2.447387 | 1 |
| X | 77983000 | 77984999 | 27.58177 | 46.45294 | 48.55448 | 21.22281 | 0.317523 | 1.122783 | 1 |
| X | 77997400 | 78001399 | 55.65824 | 89.699 | 89.33207 | 38.01527 | 0.362177 | 1.824566 | 1 |
| X | 78112200 | 78121799 | 161.8396 | 242.9201 | 248.4463 | 102.5302 | 0.234833 | 3.45683 | 1 |
| X | 78267600 | 78288799 | 289.7775 | 417.1029 | 406.0054 | 167.5648 | 0.249014 | 4.154862 | 1 |
| X | 78327800 | 78336999 | 101.4001 | 158.9381 | 137.6299 | 57.08469 | 0.524584 | 2.612481 | 1 |
| X | 78585200 | 78591599 | 58.38637 | 93.83285 | 94.69996 | 40.13664 | 0.338079 | 1.916813 | 1 |
| X | 78761200 | 78763399 | 36.13905 | 59.00743 | 64.86649 | 27.87836 | 0.237098 | 1.512484 | 1 |
| X | 78904600 | 78906999 | 35.93968 | 58.45614 | 74.04841 | 31.77988 | 0.029188 | 1.574074 | 1 |
| X | 79301400 | 79302199 | 12.0812 | 22.85013 | 24.8005 | 10.06543 | 0.364879 | 0.06992 | 1 |
| X | 79415000 | 79416999 | 17.74755 | 30.84123 | 50.6129 | 22.22454 | -0.32454 | 0.804261 | 1 |
| X | 79639200 | 79641199 | 31.24997 | 51.69278 | 58.83648 | 25.43896 | 0.194514 | 1.335518 | 1 |
| X | 79798000 | 79800199 | 22.71604 | 38.97911 | 45.29625 | 19.86688 | 0.167865 | 0.938032 | 1 |
| X | 79931600 | 79933999 | 27.93828 | 47.18757 | 47.99976 | 20.99848 | 0.356748 | 1.099505 | 1 |
| X | 80807800 | 80809399 | 17.38281 | 30.4005 | 35.83557 | 15.80174 | 0.151599 | 0.688089 | 1 |
| X | 81024600 | 81026999 | 51.6156 | 82.22301 | 116.3477 | 48.68404 | -0.16301 | 2.185149 | 1 |
| X | 81438400 | 81439199 | 17.15434 | 30.85886 | 30.22441 | 13.19851 | 0.434119 | 0.413818 | 1 |
| X | 81698000 | 81698999 | 18.33548 | 32.04356 | 40.41197 | 17.80889 | 0.05461 | 0.71322 | 1 |
| X | 81785800 | 81787799 | 21.25329 | 37.01852 | 43.5429 | 19.14772 | 0.154149 | 0.778084 | 1 |
| X | 82457600 | 82458599 | 12.31456 | 22.91557 | 26.15012 | 10.92279 | 0.259064 | 0.177932 | 1 |
| X | 83179200 | 83180199 | 13.65968 | 24.34784 | 34.16889 | 15.0369 | -0.08925 | 0.456589 | 1 |
| X | 83588800 | 83589799 | 13.74369 | 24.94774 | 26.6792 | 11.52111 | 0.318819 | 0.332417 | 1 |
| X | 83651000 | 83652199 | 13.8121 | 24.72033 | 32.15228 | 14.10079 | 0.023283 | 0.42832 | 1 |
| X | 85008600 | 85013999 | 33.95567 | 55.8749 | 61.76577 | 26.62922 | 0.2346 | 1.404791 | 1 |
| X | 85072000 | 85076399 | 35.19435 | 58.51055 | 59.27495 | 25.71754 | 0.362508 | 1.288559 | 1 |
| X | 85572000 | 85581199 | 143.1375 | 218.0558 | 208.3761 | 86.37124 | 0.350875 | 3.156875 | 1 |
| X | 85865200 | 85866799 | 17.10693 | 30.14646 | 33.98948 | 15.04402 | 0.216775 | 0.574944 | 1 |
| X | 86127600 | 86128399 | 13.16506 | 23.40452 | 34.20978 | 15.08835 | -0.14984 | 0.469648 | 1 |
| X | 86406600 | 86408399 | 11.07182 | 20.65149 | 30.00248 | 12.48213 | -0.08246 | 0.161203 | 1 |
| X | 86881600 | 86882999 | 16.23774 | 28.27601 | 43.17803 | 18.98451 | -0.22053 | 0.757752 | 1 |
| X | 87198000 | 87200399 | 32.46618 | 54.57382 | 53.54647 | 23.39108 | 0.408365 | 1.160298 | 1 |
| X | 87343600 | 87350799 | 74.36261 | 118.761 | 96.00298 | 40.31429 | 0.647331 | 2.122697 | 1 |
| X | 87454600 | 87457399 | 46.2908 | 74.64276 | 80.97359 | 34.48236 | 0.239964 | 1.791025 | 1 |
| X | 87923400 | 87929799 | 95.40388 | 148.2565 | 169.3389 | 70.31066 | 0.11591 | 2.760764 | 1 |
| X | 87996000 | 88003399 | 68.35939 | 109.0685 | 111.9323 | 46.9394 | 0.302101 | 2.150539 | 1 |
| X | 88054400 | 88055399 | 14.97029 | 27.0271 | 29.91395 | 13.00604 | 0.262808 | 0.377158 | 1 |
| X | 88056200 | 88068199 | 163.3692 | 245.1022 | 255.5792 | 105.4607 | 0.206326 | 3.467027 | 1 |
| X | 88465800 | 88466599 | 21.79281 | 37.59913 | 43.19814 | 18.95046 | 0.185839 | 0.884101 | 1 |
| X | 88529400 | 88532399 | 54.38751 | 87.26732 | 86.48656 | 36.56829 | 0.361516 | 1.956228 | 1 |
| X | 88567000 | 88568599 | 21.12049 | 36.83569 | 35.80254 | 15.75698 | 0.428538 | 0.749869 | 1 |
| X | 88814400 | 88822199 | 113.0937 | 174.9055 | 167.0504 | 69.38874 | 0.368433 | 2.852425 | 1 |
| X | 89292200 | 89294599 | 19.19682 | 33.58819 | 37.682 | 16.62228 | 0.222544 | 0.693216 | 1 |
| X | 89427800 | 89428999 | 16.40512 | 28.73179 | 39.90605 | 17.61795 | -0.0839 | 0.668096 | 1 |
| X | 89714600 | 89716599 | 21.12745 | 36.11756 | 49.01339 | 21.49073 | -0.05585 | 0.970155 | 1 |

| | | | | | | | | | |
|---|---|---|---|---|---|---|---|---|---|
| X | 89818000 | 89819599 | 20.86386 | 36.25674 | 36.96288 | 16.23072 | 0.359472 | 0.800177 | 1 |
| X | 89961000 | 89962199 | 12.39558 | 23.32576 | 25.24011 | 10.34489 | 0.357217 | 0.106999 | 1 |
| X | 89993600 | 90002999 | 64.47655 | 103.4315 | 90.74189 | 38.35391 | 0.536657 | 1.976726 | 1 |
| X | 90121800 | 90125599 | 50.5973 | 81.59357 | 82.45314 | 35.11614 | 0.342204 | 1.795862 | 1 |
| X | 90169000 | 90172599 | 57.6088 | 92.04362 | 97.43268 | 40.93942 | 0.259658 | 2.092385 | 1 |
| X | 91755000 | 91757599 | 17.81196 | 31.38428 | 35.96952 | 15.92087 | 0.193157 | 0.58437 | 1 |
| X | 91843600 | 91844999 | 11.86012 | 22.2766 | 26.30014 | 10.81341 | 0.228579 | 0.115829 | 1 |
| X | 92962800 | 92967399 | 71.0483 | 111.7598 | 149.324 | 61.94849 | -0.09642 | 2.528633 | 1 |
| X | 93023000 | 93025399 | 28.59443 | 47.51371 | 62.15115 | 26.95584 | -0.00513 | 1.278262 | 1 |
| X | 93874200 | 93874999 | 12.47431 | 23.29575 | 27.24579 | 11.31194 | 0.230677 | 0.158847 | 1 |
| X | 94377000 | 94378799 | 8.877237 | 16.75014 | 26.47814 | 10.80077 | -0.17981 | 0.061447 | 1 |
| X | 94461200 | 94476799 | 186.4423 | 277.64 | 303.7452 | 125.1599 | 0.127356 | 3.609319 | 1 |
| X | 94603200 | 94609999 | 57.79899 | 92.45348 | 98.79318 | 41.57858 | 0.247883 | 2.058318 | 1 |
| X | 94802200 | 94803599 | 25.47796 | 43.1832 | 47.98192 | 21.013 | 0.229987 | 1.060958 | 1 |
| X | 94823800 | 94824799 | 15.50761 | 27.99841 | 27.92109 | 12.13891 | 0.412538 | 0.377603 | 1 |
| X | 95315000 | 95317999 | 30.50403 | 50.384 | 63.33368 | 27.35534 | 0.051434 | 1.351596 | 1 |
| X | 95582400 | 95584999 | 37.81682 | 61.77648 | 66.7792 | 28.70542 | 0.261748 | 1.510854 | 1 |
| X | 95702800 | 95703599 | 13.89437 | 25.36778 | 27.07716 | 11.60856 | 0.32812 | 0.300857 | 1 |
| X | 96876000 | 96877999 | 11.8792 | 21.41287 | 34.77988 | 15.15136 | -0.28893 | 0.376106 | 1 |
| X | 96975200 | 96979999 | 37.61182 | 60.84187 | 101.1575 | 43.25006 | -0.36956 | 1.694719 | 1 |
| X | 96983000 | 96985199 | 25.32317 | 43.73566 | 41.96694 | 18.41093 | 0.444798 | 0.886916 | 1 |
| X | 97075400 | 97077799 | 16.82056 | 29.79242 | 32.88397 | 14.53997 | 0.250096 | 0.516836 | 1 |
| X | 97861600 | 97863799 | 17.10604 | 29.87657 | 52.22643 | 23.00578 | -0.4146 | 0.731955 | 1 |
| X | 98929800 | 98932199 | 23.01302 | 39.24751 | 56.06147 | 24.57637 | -0.13016 | 1.006907 | 1 |
| X | 99050600 | 99051399 | 14.83137 | 26.05093 | 34.79425 | 15.39883 | -0.02776 | 0.601955 | 1 |
| X | 99471800 | 99473999 | 26.73101 | 44.086 | 68.31727 | 29.48194 | -0.24994 | 1.365932 | 1 |
| X | 100737400 | 100742399 | 124.72 | 188.5803 | 244.7436 | 101.2139 | -0.09933 | 3.306306 | 1 |
| X | 101374800 | 101376399 | 17.60024 | 31.18547 | 32.75626 | 14.48238 | 0.321399 | 0.516627 | 1 |
| X | 101610200 | 101612399 | 15.15543 | 27.08606 | 31.44668 | 13.80787 | 0.184793 | 0.442694 | 1 |
| X | 102079800 | 102082599 | 25.16697 | 42.83412 | 47.95619 | 21.01973 | 0.219596 | 1.023492 | 1 |
| X | 102137000 | 102140999 | 21.29239 | 36.42718 | 56.07216 | 24.58727 | -0.23696 | 0.970528 | 1 |
| X | 102793200 | 102793999 | 18.8845 | 33.08554 | 33.66101 | 14.86518 | 0.363008 | 0.664911 | 1 |
| X | 103092800 | 103095199 | 39.20402 | 63.99971 | 70.30007 | 30.21152 | 0.238014 | 1.526134 | 1 |
| X | 104612800 | 104615199 | 28.6175 | 48.72691 | 48.476 | 21.24827 | 0.389425 | 1.032573 | 1 |
| X | 104625400 | 104627399 | 26.48159 | 44.7312 | 49.06626 | 21.46561 | 0.248319 | 1.095014 | 1 |
| X | 106100600 | 106101399 | 11.41149 | 21.64253 | 20.74095 | 8.369319 | 0.55083 | 0.035619 | 1 |
| X | 106979000 | 106985999 | 101.3128 | 157.65 | 157.0382 | 65.19026 | 0.314627 | 2.739385 | 1 |
| X | 107165000 | 107166399 | 15.83197 | 28.18381 | 34.43094 | 15.16857 | 0.108336 | 0.476975 | 1 |
| X | 107414000 | 107416599 | 28.25065 | 48.31052 | 47.92159 | 21.01256 | 0.394584 | 0.993609 | 1 |
| X | 108874000 | 108877599 | 36.88106 | 60.28555 | 70.1363 | 30.15537 | 0.156649 | 1.502543 | 1 |
| X | 108914400 | 108915199 | 10.59086 | 20.27125 | 22.28904 | 8.81139 | 0.379392 | -0.03705 | 1 |
| X | 109027000 | 109027199 | 7.751107 | 15.08289 | 18.45787 | 6.799302 | 0.322556 | -0.31181 | 1 |
| X | 109460200 | 109462799 | 52.89039 | 84.55465 | 96.92518 | 40.76017 | 0.145908 | 2.069994 | 1 |
| X | 109481200 | 109483199 | 22.40879 | 38.84209 | 41.47954 | 18.19911 | 0.292076 | 0.840457 | 1 |
| X | 109923600 | 109926399 | 37.53339 | 61.05574 | 68.00145 | 29.18597 | 0.214531 | 1.571722 | 1 |
| X | 110032200 | 110034199 | 26.74241 | 45.57806 | 47.39612 | 20.77654 | 0.325969 | 1.015417 | 1 |
| X | 110048000 | 110054599 | 85.25802 | 133.8812 | 143.6769 | 59.59237 | 0.217436 | 2.570216 | 1 |

| | | | | | | | | | |
|---|---|---|---|---|---|---|---|---|---|
| X | 110697200 | 110698599 | 23.59729 | 39.80625 | 54.37533 | 23.77865 | -0.06715 | 1.102016 | 1 |
| X | 111060400 | 111061599 | 11.96976 | 22.50089 | 24.15763 | 9.918796 | 0.366274 | 0.106297 | 1 |
| X | 111344400 | 111345999 | 11.82848 | 21.8617 | 27.68447 | 11.67512 | 0.099673 | 0.215673 | 1 |
| X | 111510000 | 111511799 | 32.09498 | 54.08325 | 49.63609 | 21.69913 | 0.504597 | 1.136572 | 1 |
| X | 111748200 | 111750199 | 20.02617 | 35.06464 | 37.58301 | 16.58875 | 0.288061 | 0.685882 | 1 |
| X | 112518400 | 112520199 | 23.66162 | 40.50659 | 45.2918 | 19.86119 | 0.222713 | 0.962358 | 1 |
| X | 112820200 | 112821799 | 14.42541 | 26.41524 | 28.04431 | 11.97708 | 0.339517 | 0.290382 | 1 |
| X | 113035800 | 113046199 | 218.1094 | 319.5312 | 347.1398 | 142.9924 | 0.115372 | 3.884016 | 1 |
| X | 113323800 | 113328999 | 63.32385 | 100.9787 | 121.2182 | 50.79479 | 0.075431 | 2.16434 | 1 |
| X | 114030000 | 114030999 | 16.46846 | 29.41146 | 32.83648 | 14.4269 | 0.24023 | 0.458169 | 1 |
| X | 115110200 | 115112399 | 20.53438 | 36.14988 | 34.28386 | 15.17738 | 0.4646 | 0.607597 | 1 |
| X | 115114000 | 115116599 | 47.10292 | 75.94943 | 82.36655 | 35.07385 | 0.240321 | 1.793638 | 1 |
| X | 115404200 | 115404999 | 13.78985 | 25.50918 | 28.1715 | 11.86631 | 0.297288 | 0.229573 | 1 |
| X | 115829800 | 115831599 | 22.37856 | 39.29117 | 34.24468 | 15.15033 | 0.585091 | 0.652238 | 1 |
| X | 116763600 | 116765399 | 29.70743 | 49.5229 | 55.06726 | 23.93461 | 0.228665 | 1.240245 | 1 |
| X | 117262800 | 117265399 | 28.58358 | 49.17976 | 42.44156 | 18.61699 | 0.596183 | 0.935272 | 1 |
| X | 117607400 | 117613199 | 25.47444 | 44.21233 | 40.28353 | 17.67505 | 0.520563 | 0.842478 | 1 |
| X | 118418200 | 118419799 | 22.01169 | 37.35622 | 53.25632 | 23.33195 | -0.12817 | 1.044525 | 1 |
| X | 118857800 | 118859799 | 17.62628 | 31.39433 | 31.22508 | 13.7502 | 0.403857 | 0.479712 | 1 |
| X | 118988800 | 119001599 | 146.2645 | 220.3647 | 251.042 | 103.7051 | 0.08413 | 3.381803 | 1 |
| X | 120060200 | 120062599 | 27.84761 | 47.24796 | 56.23652 | 24.63563 | 0.131061 | 1.068397 | 1 |
| X | 120235000 | 120241199 | 69.58206 | 110.5408 | 117.9522 | 49.18864 | 0.239449 | 2.285918 | 1 |
| X | 120244000 | 120246199 | 36.2548 | 58.76374 | 80.63781 | 34.52752 | -0.09045 | 1.643221 | 1 |
| X | 120865600 | 120867999 | 19.24978 | 33.97171 | 36.2999 | 16.06199 | 0.294531 | 0.575216 | 1 |
| X | 121088800 | 121089999 | 22.42951 | 38.88304 | 40.00211 | 17.55095 | 0.345904 | 0.840173 | 1 |
| X | 121196200 | 121199199 | 31.17916 | 53.05365 | 45.7274 | 20.04242 | 0.595521 | 1.045954 | 1 |
| X | 121204600 | 121209399 | 43.94626 | 71.78237 | 65.15222 | 28.00506 | 0.513154 | 1.52457 | 1 |
| X | 121633400 | 121635599 | 37.02547 | 60.5318 | 61.56322 | 26.47096 | 0.350327 | 1.495781 | 1 |
| X | 121851200 | 121855799 | 53.95971 | 86.80476 | 88.15322 | 37.41565 | 0.330876 | 1.87701 | 1 |
| X | 121938600 | 121941599 | 43.33189 | 70.29134 | 75.42229 | 32.31355 | 0.264673 | 1.63807 | 1 |
| X | 122213800 | 122215599 | 18.79108 | 33.01703 | 33.98859 | 15.04649 | 0.346041 | 0.619401 | 1 |
| X | 122250600 | 122253199 | 43.85727 | 70.70999 | 82.14069 | 34.97378 | 0.141253 | 1.791959 | 1 |
| X | 122733000 | 122735399 | 25.34102 | 43.36261 | 44.73787 | 19.61759 | 0.338235 | 0.972762 | 1 |
| X | 122834400 | 122835399 | 18.93777 | 32.57345 | 50.58504 | 22.18892 | -0.24804 | 0.909396 | 1 |
| X | 122886000 | 122888599 | 23.97611 | 41.18699 | 44.68108 | 19.59841 | 0.266797 | 0.929442 | 1 |
| X | 123062200 | 123068599 | 78.05086 | 123.9334 | 125.8207 | 52.44405 | 0.310475 | 2.308597 | 1 |
| X | 124923600 | 124924999 | 19.87205 | 34.32146 | 42.69341 | 18.72964 | 0.071927 | 0.863754 | 1 |
| X | 125045600 | 125052999 | 81.99497 | 129.412 | 139.6843 | 57.98562 | 0.214758 | 2.466909 | 1 |
| X | 125127400 | 125138799 | 135.1565 | 206.8854 | 201.8015 | 83.74136 | 0.326728 | 3.058669 | 1 |
| X | 125454200 | 125456799 | 41.4465 | 67.44041 | 68.02206 | 29.19784 | 0.357478 | 1.574804 | 1 |
| X | 125724600 | 125729799 | 19.72286 | 35.64293 | 28.97855 | 12.5836 | 0.706454 | 0.403109 | 1 |
| X | 125913200 | 125914799 | 22.75964 | 39.46861 | 41.99024 | 18.42344 | 0.297619 | 0.83875 | 1 |
| X | 126284000 | 126285999 | 15.24342 | 27.04374 | 29.05914 | 12.83887 | 0.289757 | 0.483729 | 1 |
| X | 126638800 | 126640599 | 24.88466 | 42.6297 | 45.77133 | 20.07258 | 0.281058 | 0.960149 | 1 |
| X | 126705800 | 126719199 | 190.7546 | 285.631 | 258.011 | 106.4192 | 0.410776 | 3.50809 | 1 |
| X | 126783000 | 126785799 | 25.92323 | 43.6525 | 61.11659 | 26.71229 | -0.10279 | 1.131002 | 1 |
| X | 127629800 | 127631399 | 12.49749 | 22.98435 | 26.13061 | 11.09415 | 0.247458 | 0.24592 | 1 |

| | | | | | | | | | |
|---|---|---|---|---|---|---|---|---|---|
| X | 130244400 | 130248599 | 33.48403 | 55.27271 | 59.57421 | 25.7208 | 0.272641 | 1.364462 | 1 |
| X | 130878400 | 130879199 | 11.01693 | 20.77724 | 25.30771 | 10.33391 | 0.191822 | 0.080064 | 1 |
| X | 131105600 | 131111199 | 69.82764 | 111.2213 | 109.661 | 45.8785 | 0.357149 | 2.203529 | 1 |
| X | 132033000 | 132034399 | 20.77011 | 35.7123 | 46.88695 | 20.56654 | -0.00696 | 0.913772 | 1 |
| X | 132160800 | 132176399 | 228.233 | 332.9675 | 349.6489 | 144.0723 | 0.158605 | 3.949039 | 1 |
| X | 132335000 | 132335999 | 13.24537 | 24.20217 | 28.00822 | 11.99574 | 0.212762 | 0.290854 | 1 |
| X | 132585400 | 132592399 | 86.77075 | 136.5607 | 133.2443 | 55.28134 | 0.357076 | 2.526305 | 1 |
| X | 133864000 | 133867199 | 37.50443 | 60.68479 | 91.62354 | 39.18169 | -0.23033 | 1.686883 | 1 |
| X | 134537200 | 134538999 | 16.75428 | 29.64136 | 30.75685 | 13.60253 | 0.337906 | 0.521867 | 1 |
| X | 136399600 | 136402399 | 25.66731 | 44.46798 | 42.98024 | 18.85725 | 0.435244 | 0.86216 | 1 |
| X | 140681000 | 140687399 | 65.9487 | 104.2969 | 144.315 | 59.96927 | -0.14078 | 2.407362 | 1 |
| X | 141305400 | 141307799 | 26.77347 | 45.14783 | 49.0777 | 21.45961 | 0.261208 | 1.109967 | 1 |
| X | 141952600 | 141955599 | 8.564807 | 16.59513 | 22.76843 | 8.628791 | 0.120722 | -0.19152 | 1 |
| X | 143100800 | 143102999 | 22.84425 | 39.55348 | 43.1294 | 18.92235 | 0.261959 | 0.857493 | 1 |
| X | 143485400 | 143486399 | 14.40565 | 25.48785 | 37.64228 | 16.64747 | -0.16847 | 0.521393 | 1 |
| X | 143897200 | 143898799 | 16.78261 | 29.94195 | 34.7731 | 15.29091 | 0.18253 | 0.467163 | 1 |
| X | 144966000 | 144966599 | 10.55292 | 20.15246 | 23.38165 | 9.298498 | 0.294973 | -0.01374 | 1 |
| X | 145098000 | 145099599 | 17.19567 | 30.34634 | 34.99598 | 15.48506 | 0.185219 | 0.562684 | 1 |
| X | 145483400 | 145483999 | 11.74932 | 22.4535 | 22.81199 | 9.058012 | 0.487118 | -0.00798 | 1 |
| X | 146636800 | 146637799 | 17.50788 | 30.63548 | 35.75984 | 15.77542 | 0.165617 | 0.679371 | 1 |
| X | 147028200 | 147028999 | 17.11806 | 30.11823 | 33.08627 | 14.64889 | 0.253294 | 0.59124 | 1 |
| X | 147396600 | 147398999 | 41.20714 | 66.66558 | 72.1724 | 30.86008 | 0.2483 | 1.687046 | 1 |
| X | 147451200 | 147456199 | 15.41885 | 27.5025 | 31.3805 | 13.80176 | 0.207967 | 0.455603 | 1 |
| X | 147583600 | 147586199 | 44.00026 | 70.88267 | 81.9066 | 34.83529 | 0.147476 | 1.81628 | 1 |
| X | 147662800 | 147668199 | 62.3849 | 100.0014 | 97.88109 | 41.32448 | 0.377771 | 1.99962 | 1 |
| X | 147721400 | 147724199 | 27.809 | 46.49531 | 55.60185 | 24.21205 | 0.123926 | 1.201964 | 1 |
| X | 148438000 | 148439999 | 25.85692 | 44.27248 | 44.68396 | 19.59491 | 0.369845 | 0.967355 | 1 |
| X | 148716800 | 148721199 | 28.40122 | 46.63031 | 72.72776 | 31.32558 | -0.26186 | 1.428962 | 1 |
| X | 148795800 | 148796199 | 9.316383 | 17.90707 | 19.24876 | 7.518086 | 0.428363 | -0.09714 | 1 |
| X | 148892000 | 148894399 | 34.31445 | 57.57005 | 53.60435 | 23.39301 | 0.483689 | 1.185072 | 1 |
| X | 149521800 | 149525199 | 16.41693 | 28.68925 | 39.5185 | 17.41443 | -0.0719 | 0.703852 | 1 |
| X | 149525200 | 149527399 | 15.25517 | 27.0251 | 33.09168 | 14.63124 | 0.101429 | 0.506531 | 1 |
| X | 150302600 | 150305599 | 26.4696 | 45.23454 | 45.80527 | 20.08352 | 0.36487 | 0.991587 | 1 |
| X | 150366400 | 150367399 | 14.80895 | 25.93222 | 42.53758 | 18.77726 | -0.32277 | 0.666057 | 1 |
| X | 151705000 | 151707599 | 9.050358 | 17.20719 | 28.30908 | 11.36333 | -0.21676 | 0.015215 | 1 |
| X | 152280600 | 152283399 | 47.47964 | 77.02057 | 66.88569 | 28.65642 | 0.569212 | 1.64413 | 1 |
| X | 153195200 | 153209399 | 225.48 | 332.1721 | 315.6313 | 129.9798 | 0.316142 | 3.796812 | 1 |
| X | 153439600 | 153443199 | 40.12103 | 65.84996 | 66.66001 | 28.70792 | 0.359797 | 1.4566 | 1 |
| X | 153566800 | 153568199 | 17.99428 | 32.01214 | 33.126 | 14.60174 | 0.345848 | 0.490807 | 1 |
| X | 154010600 | 154013399 | 29.71581 | 49.23468 | 64.472 | 27.90675 | -0.00704 | 1.315291 | 1 |
| X | 154470200 | 154471399 | 20.79158 | 35.8261 | 44.78678 | 19.64661 | 0.06415 | 0.892755 | 1 |
| X | 154512800 | 154515399 | 28.03967 | 47.48929 | 52.31965 | 22.90741 | 0.242148 | 1.081312 | 1 |
| X | 154561600 | 154564799 | 39.06794 | 63.57192 | 64.70148 | 27.7728 | 0.344579 | 1.569746 | 1 |
| X | 154620200 | 154622799 | 33.46223 | 55.76614 | 59.70465 | 25.9455 | 0.283134 | 1.258832 | 1 |
| X | 155100400 | 155107199 | 88.88556 | 139.2662 | 144.1916 | 59.80876 | 0.266615 | 2.61254 | 1 |
| X | 155391200 | 155394199 | 14.66275 | 27.87218 | 25.42265 | 10.20913 | 0.626969 | 0.059174 | 1 |
| X | 156893600 | 156896999 | 32.83271 | 54.37 | 68.66661 | 29.71148 | 0.044949 | 1.334747 | 1 |

| | | | | | | | | | |
|---|---|---|---|---|---|---|---|---|---|
| X | 157382400 | 157384399 | 20.75679 | 36.23695 | 38.76565 | 17.07161 | 0.290806 | 0.737589 | 1 |
| X | 157652200 | 157653399 | 16.84169 | 29.50841 | 39.83676 | 17.59342 | -0.04321 | 0.662682 | 1 |
| X | 158409600 | 158411399 | 18.81817 | 33.11844 | 37.04887 | 16.40245 | 0.227424 | 0.606937 | 1 |
| X | 159857000 | 159864599 | 92.84748 | 145.3966 | 144.8438 | 60.08253 | 0.321596 | 2.623819 | 1 |
| X | 159883400 | 159892199 | 105.5168 | 162.2443 | 195.3924 | 81.13984 | 0.02787 | 2.959879 | 1 |
| X | 160495200 | 160495799 | 10.55001 | 19.38333 | 29.85285 | 12.69173 | -0.18882 | 0.246501 | 1 |
| X | 160499600 | 160501599 | 15.17812 | 27.26041 | 28.02195 | 12.25318 | 0.36368 | 0.406497 | 1 |
| X | 160959000 | 160963999 | 15.91284 | 27.94656 | 38.77866 | 17.16015 | -0.08257 | 0.619015 | 1 |
| X | 161156800 | 161157999 | 14.9567 | 27.06424 | 29.40754 | 12.74943 | 0.291904 | 0.362908 | 1 |
| X | 161542200 | 161544599 | 29.89036 | 50.45771 | 49.61633 | 21.70217 | 0.405473 | 1.113712 | 1 |
| X | 162137600 | 162140999 | 20.56686 | 35.98254 | 37.11051 | 16.36912 | 0.343419 | 0.70152 | 1 |
| X | 163397000 | 163399399 | 19.21615 | 33.35691 | 38.59892 | 16.93748 | 0.177377 | 0.809871 | 1 |
| X | 164589200 | 164599799 | 270.0944 | 387.7218 | 412.0954 | 170.1503 | 0.11831 | 4.192564 | 1 |
| X | 164637200 | 164639399 | 30.63261 | 50.74761 | 62.91443 | 27.23031 | 0.071634 | 1.316907 | 1 |
| X | 164707400 | 164711999 | 48.69542 | 78.77568 | 84.22673 | 36.01284 | 0.266113 | 1.706375 | 1 |
| X | 166958200 | 166960999 | 25.65661 | 43.86642 | 47.39559 | 20.78161 | 0.271675 | 0.98373 | 1 |
| X | 167276400 | 167281199 | 61.47257 | 97.8587 | 119.7759 | 50.07153 | 0.044828 | 2.216509 | 1 |
| X | 168773800 | 168779999 | 1858.97 | 2219.097 | 3106.204 | 1314.21 | -0.54704 | 7.103603 | 1 |
| X | 168785200 | 168791999 | 1345.244 | 1659.793 | 2080.699 | 876.4176 | -0.33942 | 6.580362 | 1 |
| X | 168794400 | 168797599 | 330.9816 | 453.4239 | 744.0064 | 309.5007 | -0.57591 | 4.894259 | 1 |
| X | 168855600 | 168858999 | 185.8587 | 271.9774 | 349.5065 | 143.9774 | -0.12832 | 3.897673 | 1 |
| X | 168909000 | 168910799 | 100.8406 | 153.3048 | 228.5729 | 94.68438 | -0.29277 | 3.188148 | 1 |
| X | 168960800 | 168962199 | 76.37093 | 119.3476 | 147.9967 | 61.40345 | 0.00377 | 2.650892 | 1 |
| X | 169012200 | 169016199 | 314.9683 | 439.1196 | 573.9838 | 238.1134 | -0.22239 | 4.624512 | 1 |
| X | 169121000 | 169122399 | 12.38559 | 21.87413 | 36.34437 | 16.07928 | -0.3382 | 0.527846 | 1 |
| X | 169179200 | 169182799 | 192.3505 | 276.3499 | 457.0269 | 188.6775 | -0.51803 | 4.17513 | 1 |
| X | 169264800 | 169265599 | 20.67792 | 34.38783 | 62.66892 | 27.19599 | -0.48178 | 1.245088 | 1 |
| X | 169271200 | 169271599 | 10.30231 | 18.96973 | 26.82471 | 11.3704 | -0.06339 | 0.223493 | 1 |
| Y | 136400 | 136999 | 7.469656 | 14.4377 | 17.35992 | 6.65493 | 0.293219 | -0.18693 | 1 |
| Y | 171200 | 171799 | 24.93932 | 41.13299 | 54.77632 | 23.63889 | -0.0319 | 1.344762 | 1 |
| Y | 179200 | 180599 | 13.29348 | 24.35751 | 19.04275 | 8.125909 | 0.778129 | 0.251075 | 1 |
| Y | 689600 | 696399 | 60.05316 | 95.02969 | 106.6137 | 44.31604 | 0.162089 | 2.377867 | 1 |
| Y | 700800 | 706399 | 53.7935 | 85.93629 | 74.12791 | 31.14128 | 0.554173 | 2.079016 | 1 |
| Y | 707200 | 712199 | 49.44969 | 79.39693 | 61.71997 | 26.12277 | 0.71216 | 1.912149 | 1 |
| Y | 884800 | 885399 | 11.95728 | 21.77973 | 22.75844 | 9.785824 | 0.355411 | 0.284602 | 1 |
| Y | 953000 | 959199 | 67.75593 | 107.1036 | 95.13175 | 39.52185 | 0.497284 | 2.405911 | 1 |
| Y | 959000 | 965599 | 39.28998 | 63.60242 | 56.54395 | 24.18832 | 0.53251 | 1.655879 | 1 |
| Y | 1276200 | 1279599 | 29.00438 | 47.954 | 45.11855 | 19.50098 | 0.468157 | 1.316952 | 1 |
| Y | 1401200 | 1402199 | 10.20434 | 18.98289 | 21.28726 | 8.896254 | 0.283278 | 0.14642 | 1 |
| Y | 1771400 | 1772199 | 12.858 | 23.30028 | 22.2231 | 9.617266 | 0.480266 | 0.319567 | 1 |
| Y | 1800600 | 1801799 | 12.27658 | 21.74007 | 30.41085 | 13.44697 | -0.08981 | 0.483338 | 1 |
| Y | 1802800 | 1803599 | 11.4026 | 20.42552 | 27.85841 | 12.20877 | -0.04432 | 0.392405 | 1 |
| Y | 1822400 | 1827399 | 33.97171 | 55.75887 | 43.64247 | 18.79538 | 0.729637 | 1.416886 | 1 |
| Y | 1852600 | 1854199 | 11.79749 | 20.68338 | 38.55843 | 17.03785 | -0.50618 | 0.621658 | 1 |
| Y | 2026600 | 2027599 | 13.71301 | 24.93924 | 20.15711 | 8.684025 | 0.72165 | 0.30044 | 1 |
| Y | 2164000 | 2165799 | 16.09957 | 28.64216 | 21.14713 | 9.321325 | 0.829843 | 0.451396 | 1 |
| Y | 2308800 | 2309799 | 11.34488 | 21.13033 | 19.20531 | 8.009635 | 0.58653 | 0.14305 | 1 |

| | | | | | | | | | |
|---|---|---|---|---|---|---|---|---|---|
| Y | 2361000 | 2364399 | 37.16036 | 60.57708 | 46.74769 | 20.07943 | 0.744398 | 1.520059 | 1 |
| Y | 2407800 | 2408999 | 14.79982 | 25.83161 | 36.08351 | 15.88823 | -0.0923 | 0.703078 | 1 |
| Y | 2516000 | 2517999 | 16.92412 | 29.70372 | 26.86453 | 11.88159 | 0.531675 | 0.60933 | 1 |
| Y | 2580800 | 2587599 | 64.68067 | 101.76 | 116.7479 | 48.43499 | 0.123299 | 2.511434 | 1 |
| Y | 2756200 | 2762999 | 74.5155 | 116.5103 | 120.2887 | 49.90472 | 0.268386 | 2.633139 | 1 |
| Y | 3335200 | 3337999 | 19.62758 | 32.8608 | 54.20991 | 23.62608 | -0.33851 | 1.159512 | 1 |
| Y | 3913400 | 3915199 | 13.11908 | 23.81746 | 21.62953 | 9.338744 | 0.552578 | 0.309122 | 1 |
| Y | 3948800 | 3950799 | 17.02086 | 29.63881 | 32.59998 | 14.33328 | 0.250483 | 0.74112 | 1 |
| Y | 3959600 | 3963799 | 39.17103 | 63.33058 | 58.00407 | 24.78578 | 0.487916 | 1.684949 | 1 |
| Y | 3994600 | 4005199 | 85.36153 | 133.3256 | 104.5883 | 43.39585 | 0.662925 | 2.654623 | 1 |
| Y | 4065600 | 4066599 | 11.84184 | 20.91389 | 31.9499 | 14.13509 | -0.2178 | 0.512845 | 1 |
| Y | 4191600 | 4194199 | 23.27763 | 40.06436 | 28.05789 | 12.30772 | 0.895858 | 0.884008 | 1 |
| Y | 4247800 | 4249799 | 13.77204 | 24.08638 | 37.08356 | 16.35245 | -0.23167 | 0.670082 | 1 |
| Y | 4272400 | 4273799 | 12.03906 | 22.17474 | 25.07098 | 10.6224 | 0.257249 | 0.229286 | 1 |
| Y | 4291400 | 4292399 | 8.339613 | 15.97316 | 18.33197 | 7.232912 | 0.320781 | -0.07975 | 1 |
| Y | 4420400 | 4421999 | 16.59164 | 29.3469 | 22.18444 | 9.811745 | 0.791986 | 0.503829 | 1 |
| Y | 4433200 | 4438999 | 12.38377 | 22.50296 | 22.23223 | 9.588054 | 0.432929 | 0.299832 | 1 |
| Y | 4447600 | 4448599 | 15.15249 | 26.52438 | 31.5112 | 13.90574 | 0.140291 | 0.650548 | 1 |
| Y | 4449400 | 4450999 | 17.17378 | 29.86562 | 33.12518 | 14.55211 | 0.238286 | 0.761376 | 1 |
| Y | 4453600 | 4461199 | 77.9189 | 121.7944 | 115.7621 | 48.02827 | 0.387213 | 2.639549 | 1 |
| Y | 4596400 | 4597199 | 9.416518 | 17.77515 | 19.80244 | 8.070498 | 0.321921 | 0.041545 | 1 |
| Y | 4654200 | 4655599 | 10.98213 | 19.84374 | 29.30326 | 12.72705 | -0.14942 | 0.342218 | 1 |
| Y | 4674400 | 4677599 | 21.99986 | 37.49124 | 36.2939 | 15.90964 | 0.428748 | 0.981407 | 1 |
| Y | 4796000 | 4804599 | 108.036 | 162.7623 | 234.793 | 96.99162 | -0.25623 | 3.371103 | 1 |
| Y | 4833000 | 4837799 | 36.19574 | 58.78284 | 53.88998 | 23.10792 | 0.492582 | 1.582982 | 1 |
| Y | 4843600 | 4844799 | 14.7035 | 26.08129 | 24.76315 | 10.94243 | 0.466906 | 0.469329 | 1 |
| Y | 4845600 | 4846399 | 11.09052 | 20.86152 | 17.15282 | 7.035037 | 0.748976 | 0.071707 | 1 |
| Y | 4851000 | 4855999 | 43.95965 | 70.8459 | 58.57722 | 24.92772 | 0.629893 | 1.785774 | 1 |
| Y | 4865400 | 4866599 | 13.6645 | 23.77383 | 40.73159 | 17.89883 | -0.38587 | 0.751729 | 1 |
| Y | 4903200 | 4905399 | 16.19192 | 27.85128 | 41.86355 | 18.36329 | -0.20111 | 0.881693 | 1 |
| Y | 5108000 | 5109399 | 11.46323 | 20.98904 | 22.25803 | 9.507114 | 0.340584 | 0.249221 | 1 |
| Y | 5227600 | 5229199 | 13.77167 | 25.16618 | 18.60571 | 7.967782 | 0.854708 | 0.269107 | 1 |
| Y | 5364000 | 5364599 | 9.911027 | 18.83521 | 17.21607 | 6.917049 | 0.622938 | -0.00829 | 1 |
| Y | 5487400 | 5493999 | 52.10415 | 83.1644 | 79.58221 | 33.39107 | 0.403372 | 2.106591 | 1 |
| Y | 5497800 | 5498999 | 14.89576 | 26.52751 | 22.70221 | 9.998195 | 0.618981 | 0.440212 | 1 |
| Y | 5501800 | 5505199 | 31.25352 | 50.972 | 57.44913 | 24.68022 | 0.200005 | 1.509148 | 1 |
| Y | 5517000 | 5521799 | 34.24677 | 56.05878 | 46.71499 | 20.09565 | 0.638038 | 1.454325 | 1 |
| Y | 5540200 | 5541799 | 14.07578 | 25.31988 | 21.68049 | 9.46783 | 0.626431 | 0.369636 | 1 |
| Y | 5542600 | 5544599 | 20.15655 | 34.8474 | 33.0797 | 14.51253 | 0.460424 | 0.838976 | 1 |
| Y | 5637000 | 5638999 | 16.45339 | 28.90147 | 26.85354 | 11.88504 | 0.492924 | 0.593042 | 1 |
| Y | 5869200 | 5875199 | 49.65206 | 79.43035 | 76.04577 | 32.00981 | 0.406221 | 2.033086 | 1 |
| Y | 5919200 | 5921999 | 17.88892 | 31.16578 | 30.54681 | 13.43473 | 0.416042 | 0.734608 | 1 |
| Y | 6113200 | 6115399 | 18.60015 | 32.40203 | 29.00828 | 12.75737 | 0.546022 | 0.736311 | 1 |
| Y | 6162000 | 6166599 | 43.39567 | 69.60966 | 76.40324 | 32.29023 | 0.213956 | 1.941886 | 1 |
| Y | 6167800 | 6168999 | 15.26897 | 26.57814 | 36.65838 | 16.11381 | -0.0745 | 0.745144 | 1 |
| Y | 6528200 | 6536399 | 72.66123 | 115.0654 | 77.83675 | 32.36635 | 0.89158 | 2.361358 | 1 |
| Y | 6540000 | 6545399 | 41.11701 | 66.31962 | 61.63244 | 26.25594 | 0.46307 | 1.758965 | 1 |

| | | | | | | | | | |
|---|---|---|---|---|---|---|---|---|---|
| Y | 6546200 | 6555199 | 78.38568 | 123.0167 | 100.1431 | 41.53283 | 0.614864 | 2.564687 | 1 |
| Y | 6760400 | 6763799 | 46.30017 | 74.01363 | 90.18529 | 37.92193 | 0.057833 | 2.058916 | 1 |
| Y | 6767600 | 6769199 | 13.68722 | 24.32858 | 26.75903 | 11.8039 | 0.257702 | 0.454613 | 1 |
| Y | 6770600 | 6775999 | 36.55358 | 59.60718 | 48.27688 | 20.7387 | 0.675215 | 1.515601 | 1 |
| Y | 6786000 | 6787199 | 11.92091 | 22.03697 | 19.17142 | 8.082771 | 0.637531 | 0.190653 | 1 |
| Y | 6951200 | 6954199 | 32.44349 | 52.737 | 59.47938 | 25.52145 | 0.195658 | 1.561686 | 1 |
| Y | 7043000 | 7043599 | 10.01623 | 18.89381 | 19.27313 | 7.866544 | 0.446273 | 0.051096 | 1 |
| Y | 7050000 | 7050999 | 11.35175 | 20.20776 | 30.41284 | 13.39864 | -0.1906 | 0.438886 | 1 |
| Y | 8150400 | 8152799 | 17.3243 | 30.35968 | 31.14996 | 13.75917 | 0.350609 | 0.644362 | 1 |
| Y | 8393600 | 8395599 | 15.37136 | 27.08472 | 26.83446 | 11.87743 | 0.402086 | 0.548917 | 1 |
| Y | 8444200 | 8445999 | 15.00117 | 26.29237 | 30.98305 | 13.68713 | 0.151945 | 0.628616 | 1 |
| Y | 9003000 | 9005399 | 17.77799 | 31.15987 | 25.85148 | 11.42038 | 0.655483 | 0.633445 | 1 |
| Y | 9032000 | 9032599 | 7.867855 | 14.90327 | 21.91831 | 8.867712 | -0.06623 | 0.009231 | 1 |
| Y | 9038000 | 9041399 | 27.40351 | 45.73133 | 39.96017 | 17.38019 | 0.575039 | 1.201258 | 1 |
| Y | 9045400 | 9046399 | 11.84878 | 21.21558 | 27.3451 | 11.98819 | 0.036414 | 0.397264 | 1 |
| Y | 9051400 | 9056599 | 50.12806 | 80.17175 | 75.48418 | 31.76215 | 0.429878 | 2.041027 | 1 |
| Y | 9164800 | 9166399 | 11.60219 | 21.28884 | 21.74712 | 9.261963 | 0.397213 | 0.237246 | 1 |
| Y | 9472800 | 9474599 | 17.44202 | 30.57698 | 25.8481 | 11.42077 | 0.628517 | 0.627925 | 1 |
| Y | 9580000 | 9582199 | 18.44888 | 32.18198 | 28.48032 | 12.53583 | 0.562783 | 0.715661 | 1 |
| Y | 9648200 | 9653999 | 55.69683 | 88.94875 | 70.9812 | 29.80599 | 0.665743 | 2.089572 | 1 |
| Y | 9820400 | 9820999 | 7.508451 | 13.77615 | 30.48528 | 12.9849 | -0.70972 | 0.2299 | 1 |
| Y | 9915000 | 9915599 | 8.158559 | 15.41521 | 21.89948 | 8.907213 | -0.02343 | 0.029325 | 1 |
| Y | 9948400 | 9949599 | 9.227933 | 16.88158 | 26.901 | 11.50385 | -0.24398 | 0.249362 | 1 |
| Y | 9956800 | 9959399 | 23.09046 | 39.85889 | 27.01356 | 11.85068 | 0.943726 | 0.855474 | 1 |
| Y | 9989800 | 9990599 | 11.90889 | 21.79402 | 21.73747 | 9.291152 | 0.427733 | 0.254467 | 1 |
| Y | 10173200 | 10180599 | 44.26258 | 70.3601 | 112.0289 | 46.74698 | -0.33612 | 2.241458 | 1 |
| Y | 10359400 | 10360599 | 13.47871 | 24.30725 | 22.2068 | 9.668524 | 0.536364 | 0.351941 | 1 |
| Y | 10374600 | 10377199 | 14.5981 | 25.78344 | 26.82888 | 11.86932 | 0.33303 | 0.517789 | 1 |
| Y | 10379600 | 10387399 | 61.41684 | 97.80906 | 73.58707 | 30.77961 | 0.746268 | 2.185401 | 1 |
| Y | 10454400 | 10456199 | 15.23203 | 26.86504 | 26.83256 | 11.87431 | 0.391059 | 0.537785 | 1 |
| Y | 10545200 | 10547199 | 15.78629 | 27.81492 | 25.80582 | 11.42221 | 0.49626 | 0.549236 | 1 |
| Y | 10635200 | 10636399 | 11.64248 | 21.52657 | 19.69511 | 8.301301 | 0.565685 | 0.188697 | 1 |
| Y | 10643200 | 10644199 | 15.21474 | 26.62619 | 32.53935 | 14.35619 | 0.099805 | 0.658423 | 1 |
| Y | 10705000 | 10709999 | 261.6173 | 376.1211 | 346.6662 | 143.1013 | 0.326002 | 4.167903 | 1 |
| Y | 13416200 | 13417799 | 12.16543 | 22.31359 | 20.70595 | 8.823241 | 0.534258 | 0.241055 | 1 |
| Y | 14089000 | 14089999 | 22.37232 | 37.12232 | 54.22076 | 23.4934 | -0.1638 | 1.26698 | 1 |
| Y | 14109800 | 14110399 | 10.37105 | 18.99596 | 24.84579 | 10.60734 | 0.040711 | 0.246479 | 1 |
| Y | 20224800 | 20231999 | 49.53517 | 78.15088 | 132.6968 | 55.07427 | -0.4383 | 2.449035 | 1 |
| Y | 20304600 | 20306999 | 10.3545 | 19.53764 | 17.69703 | 7.218758 | 0.616791 | 0.046291 | 1 |
| Y | 20353400 | 20354199 | 13.8143 | 24.80392 | 22.71275 | 9.933988 | 0.528773 | 0.381853 | 1 |
| Y | 20535800 | 20538799 | 16.60093 | 29.12289 | 27.38335 | 12.10628 | 0.475868 | 0.615584 | 1 |
| Y | 20569000 | 20569799 | 9.276987 | 17.47455 | 20.3129 | 8.311475 | 0.256236 | 0.055456 | 1 |
| Y | 20663800 | 20673599 | 95.76116 | 144.9682 | 227.7642 | 94.22784 | -0.37331 | 3.270941 | 1 |
| Y | 20907600 | 20908599 | 15.51865 | 26.53157 | 45.95262 | 20.14882 | -0.40624 | 0.94017 | 1 |
| Y | 22711800 | 22713399 | 10.87277 | 20.44286 | 18.65424 | 7.656581 | 0.599154 | 0.077542 | 1 |
| Y | 23461400 | 23464399 | 15.34299 | 27.09013 | 25.80152 | 11.41556 | 0.459783 | 0.523964 | 1 |
| Y | 23480800 | 23481599 | 9.620575 | 17.74902 | 25.83063 | 10.92002 | -0.10203 | 0.20237 | 1 |

| | | | | | | | | | |
|---|---|---|---|---|---|---|---|---|---|
| Y | 29025400 | 29026199 | 10.51279 | 19.29199 | 24.80336 | 10.56206 | 0.067973 | 0.236915 | 1 |
| Y | 29149800 | 29152399 | 24.21456 | 40.75453 | 39.90958 | 17.44063 | 0.411152 | 1.104215 | 1 |
| Y | 29184200 | 29184799 | 8.401554 | 15.94646 | 20.36575 | 8.20435 | 0.141076 | -0.00463 | 1 |
| Y | 29397800 | 29398799 | 10.49024 | 19.30314 | 23.83055 | 10.11024 | 0.130037 | 0.218946 | 1 |
| Y | 32078000 | 32078999 | 13.08658 | 23.30792 | 27.73633 | 12.21421 | 0.146449 | 0.440096 | 1 |
| Y | 33872600 | 33873399 | 9.130673 | 16.91272 | 24.37904 | 10.25041 | -0.08267 | 0.171021 | 1 |
| Y | 37995000 | 37996599 | 10.72787 | 19.4274 | 26.35011 | 11.41269 | -0.02512 | 0.318121 | 1 |
| Y | 39185200 | 39186399 | 8.658394 | 16.47066 | 19.33246 | 7.751433 | 0.267784 | -0.02084 | 1 |
| Y | 39222800 | 39223999 | 11.80501 | 20.68313 | 39.08118 | 17.25937 | -0.52554 | 0.633332 | 1 |
| Y | 39959200 | 39961599 | 29.64428 | 49.06089 | 49.21888 | 21.28787 | 0.376058 | 1.313559 | 1 |
| Y | 39963600 | 39964599 | 12.00698 | 21.76514 | 24.22818 | 10.48076 | 0.259146 | 0.316427 | 1 |
| Y | 40021800 | 40023199 | 11.71896 | 21.60366 | 20.72536 | 8.770946 | 0.493759 | 0.209498 | 1 |
| Y | 40108800 | 40111799 | 5.539743 | 10.72066 | 22.6596 | 8.636774 | -0.50397 | -0.20676 | 1 |
| Y | 40165200 | 40170799 | 46.87806 | 75.40522 | 60.69298 | 25.75683 | 0.665239 | 1.851161 | 1 |
| Y | 40922000 | 40924599 | 17.91347 | 31.16576 | 31.58588 | 13.87975 | 0.36785 | 0.756128 | 1 |
| Y | 41158000 | 41158799 | 10.43043 | 19.24965 | 22.80872 | 9.638049 | 0.192879 | 0.198295 | 1 |
| Y | 41633000 | 41636399 | 21.48026 | 36.65611 | 35.77874 | 15.68555 | 0.41717 | 0.965392 | 1 |
| Y | 41666200 | 41666799 | 9.123471 | 17.26511 | 19.82023 | 8.035126 | 0.285672 | 0.021576 | 1 |
| Y | 41684800 | 41686399 | 11.92937 | 21.19683 | 29.38672 | 12.96675 | -0.07451 | 0.451178 | 1 |
| Y | 41714800 | 41715999 | 8.52669 | 15.94734 | 23.91462 | 9.903863 | -0.12207 | 0.106084 | 1 |
| Y | 41867200 | 41868599 | 10.48602 | 18.90443 | 28.34153 | 12.34427 | -0.17394 | 0.347367 | 1 |
| Y | 41967600 | 41968799 | 15.5395 | 27.13762 | 34.0612 | 15.00297 | 0.061405 | 0.695681 | 1 |
| Y | 42315200 | 42317999 | 16.18937 | 28.3317 | 29.97987 | 13.22692 | 0.306409 | 0.657782 | 1 |
| Y | 42329800 | 42333199 | 30.56415 | 50.55939 | 42.57083 | 18.40649 | 0.627873 | 1.313096 | 1 |
| Y | 42566200 | 42568999 | 16.86984 | 29.41288 | 32.54123 | 14.31879 | 0.242251 | 0.72241 | 1 |
| Y | 43171600 | 43174999 | 37.52635 | 60.58814 | 65.62018 | 27.99054 | 0.244421 | 1.727681 | 1 |
| Y | 43175800 | 43178799 | 19.89114 | 34.66579 | 28.82032 | 12.67812 | 0.652375 | 0.737237 | 1 |
| Y | 43186400 | 43188799 | 15.66214 | 27.5946 | 26.32061 | 11.6502 | 0.456522 | 0.550517 | 1 |
| Y | 43210000 | 43210999 | 10.08574 | 18.7603 | 21.80698 | 9.115058 | 0.231718 | 0.147954 | 1 |
| Y | 43219400 | 43221199 | 12.40329 | 22.24481 | 25.80585 | 11.29779 | 0.188739 | 0.38696 | 1 |
| Y | 43222000 | 43223799 | 15.81884 | 27.81326 | 27.28908 | 12.08385 | 0.414831 | 0.575759 | 1 |
| Y | 43256600 | 43257399 | 14.90097 | 25.99458 | 37.65818 | 16.57615 | -0.14448 | 0.716532 | 1 |
| Y | 43570000 | 43574399 | 13.16441 | 23.54937 | 25.28804 | 11.09505 | 0.297404 | 0.407321 | 1 |
| Y | 43665400 | 43666199 | 10.53098 | 19.2829 | 24.84474 | 10.61139 | 0.061857 | 0.249841 | 1 |
| Y | 43842400 | 43844399 | 12.7781 | 22.73666 | 27.33793 | 12.04852 | 0.130831 | 0.442991 | 1 |
| Y | 43863400 | 43865999 | 19.1127 | 33.19885 | 30.57783 | 13.42324 | 0.504935 | 0.78292 | 1 |
| Y | 43884800 | 43886799 | 17.72042 | 30.65049 | 35.71998 | 15.6711 | 0.165991 | 0.827588 | 1 |
| Y | 44285400 | 44285999 | 9.166987 | 17.51794 | 17.25296 | 6.851171 | 0.530834 | -0.05389 | 1 |
| Y | 44428400 | 44430199 | 14.7336 | 25.58563 | 41.73522 | 18.31816 | -0.31578 | 0.785065 | 1 |
| Y | 44608200 | 44615599 | 72.38396 | 112.698 | 137.7666 | 57.18535 | 0.020228 | 2.709759 | 1 |
| Y | 53870000 | 53870599 | 12.65469 | 22.20484 | 34.54094 | 15.27346 | -0.24647 | 0.604563 | 1 |
| Y | 53874200 | 53874799 | 12.94721 | 22.71676 | 34.03205 | 15.04768 | -0.19247 | 0.606109 | 1 |
| Y | 58856800 | 58858399 | 9.552538 | 17.42383 | 27.40453 | 11.76308 | -0.22872 | 0.271159 | 1 |
| Y | 58860400 | 58863599 | 30.34131 | 48.95327 | 84.06435 | 35.82562 | -0.42008 | 1.74911 | 1 |
| Y | 69822800 | 69823199 | 7.867855 | 14.90327 | 21.91831 | 8.867712 | -0.06623 | 0.009231 | 1 |
| Y | 74585200 | 74586199 | 10.82247 | 19.14917 | 34.42871 | 15.22609 | -0.45023 | 0.492187 | 1 |
| Y | 78024400 | 78025199 | 10.10859 | 18.8392 | 21.81786 | 9.089791 | 0.240927 | 0.136544 | 1 |

| | | | | | | | | | | |
|---|---|---|---|---|---|---|---|---|---|---|
| Y | 84721600 | 84722399 | 8.325622 | 15.40957 | 27.9354 | 11.75812 | -0.41182 | 0.180134 | | 1 |
| Y | 85105200 | 85107199 | 17.39713 | 29.98407 | 39.7975 | 17.45809 | -0.02182 | 0.866229 | | 1 |
| Y | 86580200 | 86581599 | 10.26972 | 18.4016 | 30.42023 | 13.32817 | -0.31964 | 0.389919 | | 1 |
| Y | 86808400 | 86811199 | 24.13421 | 40.96235 | 34.26063 | 15.01026 | 0.638069 | 1.022652 | | 1 |
| Y | 87095200 | 87096599 | 12.94934 | 23.62126 | 20.68728 | 8.878998 | 0.610101 | 0.278545 | | 1 |
| Y | 87317200 | 87322599 | 15.18313 | 27.03486 | 22.18447 | 9.771522 | 0.679066 | 0.441914 | | 1 |
| Y | 88328800 | 88329799 | 9.230787 | 17.58965 | 18.29105 | 7.307315 | 0.445741 | -0.02954 | | 1 |
| Y | 88737600 | 88741199 | 15.58993 | 27.52929 | 24.77079 | 10.95928 | 0.541309 | 0.52137 | | 1 |
| Y | 88863800 | 88864799 | 14.98014 | 26.10694 | 37.12829 | 16.33278 | -0.11827 | 0.727679 | | 1 |
| Y | 89033000 | 89035999 | 18.93224 | 32.91096 | 30.05645 | 13.2008 | 0.517193 | 0.769841 | | 1 |
| Y | 89136200 | 89138799 | 24.76375 | 40.68306 | 65.82477 | 28.35963 | -0.31428 | 1.404102 | | 1 |
| Y | 89785200 | 89787799 | 20.80707 | 36.02157 | 29.57271 | 12.97451 | 0.66928 | 0.823357 | | 1 |
| Y | 90728800 | 90731999 | 21.12476 | 35.96473 | 39.79504 | 17.44301 | 0.236465 | 0.991532 | | 1 |
| Y | 90734600 | 90737599 | 20.96608 | 35.51402 | 47.99732 | 21.01581 | -0.05164 | 1.04636 | | 1 |
| Y | 90738400 | 90740799 | 14.45813 | 25.56134 | 26.82689 | 11.86611 | 0.321297 | 0.506406 | | 1 |
| Y | 90742200 | 90744399 | 18.47279 | 31.86805 | 39.21255 | 17.20198 | 0.087225 | 0.861497 | | 1 |
| Y | 90771200 | 90774199 | 768.5171 | 982.0316 | 1564.78 | 656.612 | -0.6343 | 6.014898 | | 1 |
| Y | 90777600 | 90780599 | 23.03999 | 38.6051 | 54.80542 | 23.8978 | -0.12259 | 1.166058 | | 1 |
| Y | 90797600 | 90800399 | 255.1072 | 358.3598 | 542.4538 | 224.7692 | -0.42303 | 4.509218 | | 1 |
| Y | 90807200 | 90825799 | 8078.719 | 8698.504 | 7557.302 | 3251.872 | -0.01066 | 8.731732 | | 1 |
| Y | 90836400 | 90840599 | 19.20102 | 31.78161 | 62.83667 | 27.18023 | -0.59937 | 1.297116 | | 1 |
